# Analysis of virus-specific B cell epitopes reveals extensive antigen processing prior to recognition

**DOI:** 10.1101/2023.12.15.571861

**Authors:** Alvaro Ras-Carmona, Pedro A. Reche

## Abstract

B cell epitopes must be solvent accessible for recognition by cognate B cells and antibodies. Here, we sought to study such premise for B cell epitopes targeted during infection in humans, available at the Immune Epitope Database. Most of these B cell epitopes were virus-specific linear B cell epitopes and so we focused on them, analyzing first the localization of the relevant antigens. Antigen localization could be unequivocally assigned to 26498 linear B cell epitopes. Of those, 18832 B cell epitopes belonged to antigens that remain enclosed in host cells and/or virus particles, hidden to antibody recognition, while just 7666 lie in ectodomains of viral envelope antigens and/or mature secreted antigens, visible to antibody recognition. Next, we selected B cell epitopes that mapped in antigens with known tertiary (3D-)structures and determined residue relative solvent accessibility (rRSA), comparing them with those of conformational B cell epitopes obtained from available 3D-structures of antigen-antibody complexes. rRSA values computed form linear B cell epitopes had a median value of 23.00%, while that of conformational B cell epitopes was 48.50%. Moreover, considering average rRSA values per entire epitopes (eRSA), only 32.72% of the linear B cell epitopes had eRSA values minimally comparable to those of conformational B cell epitopes. In sum, our results point that most virus-specific B cell epitopes targeted during infection are unreachable to antibody recognition on intact viral particles and/or host cells. Hence, we must conclude that antigen recognition by antibodies must be preceded by degradation/processing of viral particles and infected cells.

## 1. INTRODUCTION

B cells recognize freely accessible antigens through their surface receptor (BCR), encompassing a membrane bound antibody. B cells become activated upon antigen recognition and with the assistance of T helper cells secrete antibodies that mediate humoral immunity (1). B cells and antibodies do not recognize the antigen as a whole, but solvent accessible portions known as B cell epitopes (2). These epitopes can be classified as conformational or linear. Conformational B cell epitopes, also known as discontinuous B cell epitopes, encompass residues that are not sequential but are close in space in the antigen tertiary (3D-) structure (2, 3). Linear B cell epitopes, also known as continuous B cell epitopes, consist of sequential residues and, unlike conformational B cell epitopes, can be recognized by antibodies out of the remaining protein context (2, 3). Given the protein fold, it is often considered that most B cell epitopes are conformational (4).

B cell epitopes in protein antigens can be identified through different experimental methodologies. Arguably, the most accurate approach to identify B cell epitopes is to solve the 3D-structure of the relevant antigen–antibody complex by techniques like X-ray crystallography (5). This approach serves to define *bona-fide* B cell epitopes in native antigens but its reach is limited by the need to have crystals of highly purified protein-antibody complexes. Thereby, researchers often turn to alternative approaches for B cell epitope mapping, like determining antibody binding to overlapping peptides spanning the protein antigen (5–7). Unfortunately, these approaches are poorly suited to identify conformational B cell epitopes, recognizing mostly linear B cell epitopes which may or may not be solvent accessible on intact native antigen. B cell epitopes with reported antibody binding assays are collected in dedicated databases (8). Currently, the mayor B cell epitope repository is the immune epitope database (IEDB) (9, 10), which includes over 220000 B cell epitopes (release 17/07/2023). Unlike other databases, IEDB allows to identify the context, including immunogens, relating antibodies and cognate B cell epitopes. In this work, we examined B cell epitopes targeted during the course of a natural infection in humans.

We found that the vast majority of B cell epitopes recorded at IEDB as recognized during the course of infection in humans are not conformational but linear and derived from viruses. Focusing on these virus-specific linear B cell epitopes, we found that a large portion (64.44 % of them) resides in viral antigens that remain enclosed in host cells and/or viral particles, hidden to antibody recognition. Moreover, we also found that those mapping in visible antigens (e.g. ectodomain of envelop proteins) have solvent accessibilities incompatible with antigen recognition by antibodies occurring on intact cells or viral particles. Collectively, these results support that extensive antigen degradation/processing must occur prior to recognition by antibodies, which shapes B cell epitope and antibodies repertoire. We argue that innate immune processes, such as phagocytosis, are likely involved in exposing hidden antigens and antigen fragments to antibody recognition.

## 2. RESULTS

### 2.1 Characterization of B cell epitopes targeted during the course of a natural infection

B cell epitope identification is of great practical relevance and a matter of intense activity. As a result, thousands of B cell epitopes have already been reported in the literature and are recorded in databases, such as the IEDB. These B cell epitopes have been identified under different experimental contexts and here we sought to characterize those targeted by humans during infection. We identified and selected 55642 of such B cell epitopes from IEDB, all including between 8 and 25 residues. This size range was chosen to limit the chance of considering B cell epitopes with excess residues that are not part of the B cell epitope. Interestingly, the vast majority of these epitopes (54996, 98.84%) were reported as linear B cell epitopes (Figure 1). These linear B cell epitopes covered 4099 distinct antigen proteins and we subsequently classified them by the taxa of the source organisms (Supplementary Dataset 1). As shown in Figure 1, 87.96 % of the linear B cell epitopes (48377 out of 54996) were from viruses and a mere 1.78 % (979 B cell epitopes) and 10.26 % (5640 B cell epitopes) were from bacteria and eukaryotic organisms, respectively. Given the abundance of virus-specific B cell epitopes, we focused on them for further structural and location analyses.

**Figure 1.**
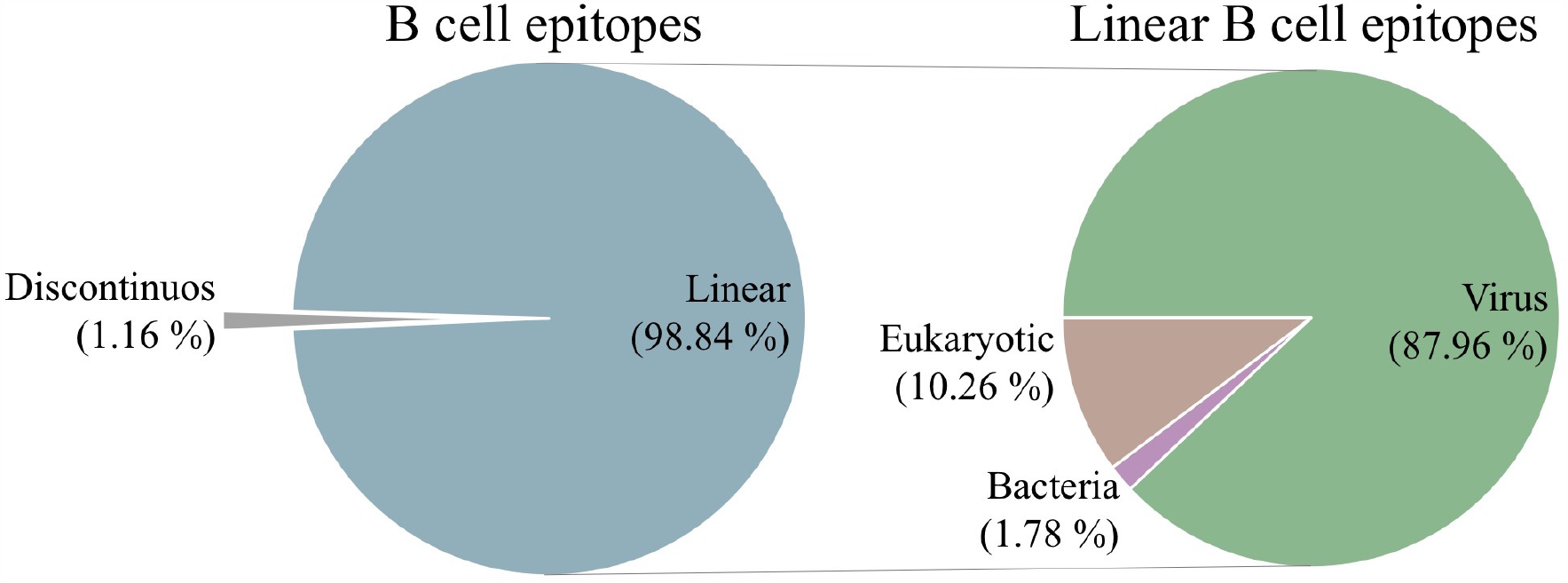
B cell epitopes targeted by humans during infection. A total of 55642 unique B cell epitopes with 8 to 25 residues were identified in IEDB to be targeted during infection in humans. Of those, 98.84 % were linear B cell epitopes (left pie chart) with 87.96 % mapping in virus antigens (right pie chart).

### 2.2 Analysis of the visibility of viral antigens encompassing linear B cell epitopes

Antibodies can only target antigens that are visible and solvent accessible to scrutiny. Thereby, we analyzed the location in host cells and viral particles of the 3010 antigens that encompassed the selected 48377 virus-specific linear B cell epitopes, using the relevant UNIPROT annotations (details in Methods). It is important to remind that all viral proteins are produced by the host cells and hence have sub-cellular location annotations in UNIPROT. Taking in consideration these annotations, we classified viral proteins into four groups attending to their visibility to antibody recognition. Non-structural viral proteins remaining within cells (cytoplasm, nucleus, endoplasmic reticulum, etc) and structural proteins enclosed in viral particles were considered as hidden to antibody recognition, and included into a group that we termed as Cell/Virus Enclosed. In addition, we considered three other major groups of viral antigens: secreted proteins, consisting of non-structural viral proteins that are secreted by cells (e.g. proteins interfering with immune response); envelope proteins, consisting of proteins that are located in the cell plasma membrane and subsequently into the viral envelope; and capsid proteins, which form part of viral capsids. Envelope and secreted proteins can be considered readily visible to antibody recognition, while capsid proteins may or may not be visible for recognition depending on the type of virus. We were able to classify 2187 out of 3010 viral antigens into one of these groups, collectively encompassing 29225 of the 48377 selected linear B cell epitopes (Figure 2A). The B cell epitopes that were not classified into the mentioned antigens groups corresponded to viral polyproteins and/or protein antigens without sub-cellular location annotations in UNIPROT records. There are some overlaps in the classification of viral antigens but most of them are classified as Cell/Virus Enclosed (1195) and envelope (875) antigens (Figure 2A). These two groups of viral antigens encompass 27924 of the 29225 B cell epitopes distributed as follows: 18832 B cell epitopes (64.44 %) in Cell/Virus Enclosed antigens and 9092 epitopes (31.11 %) in envelope antigens (Figure 2B).

**Figure 2.**
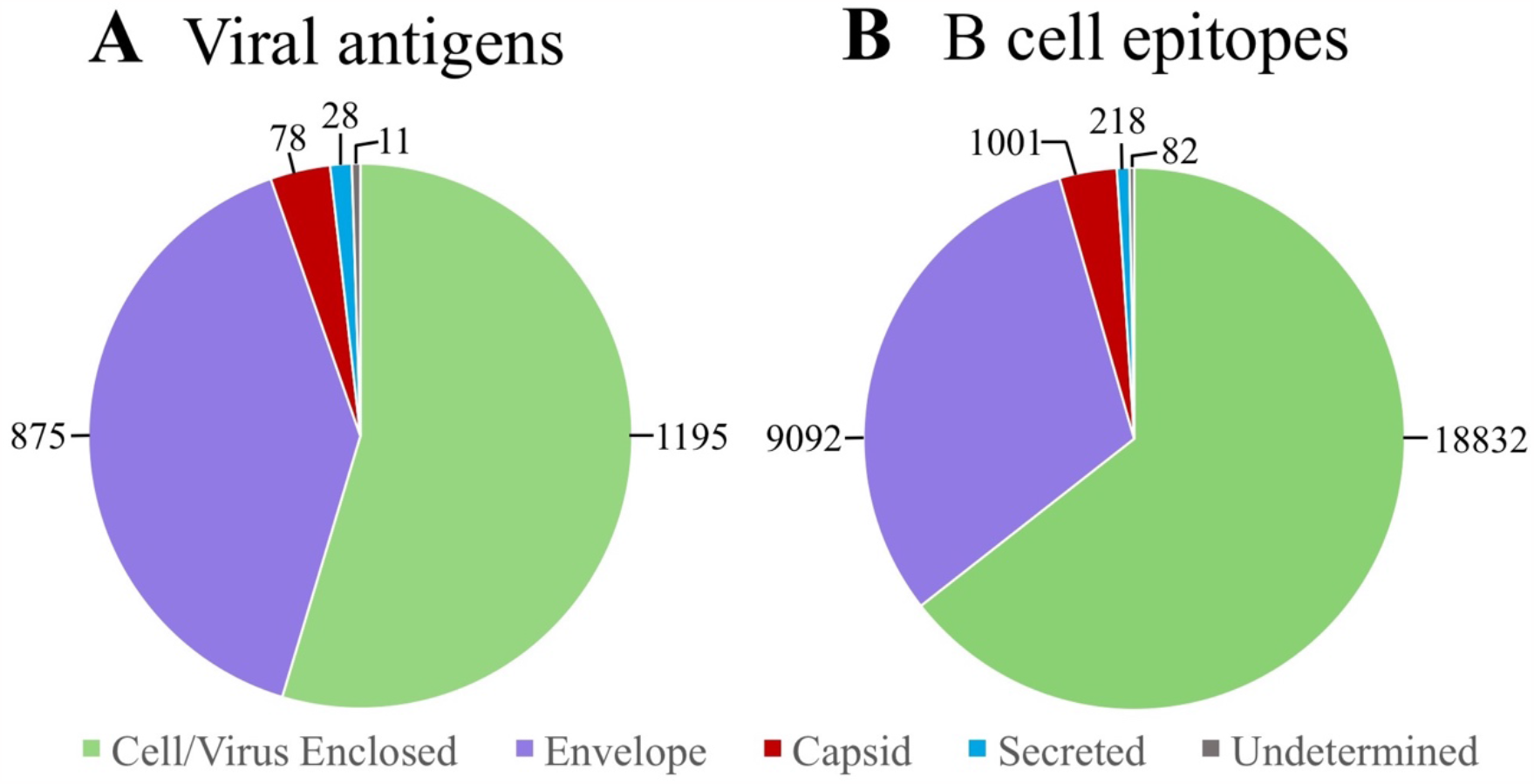
Classification of antigens and linear B cell epitopes from viruses. Pie chart depicting the number of viral antigens (**A**) and B cell epitopes (**B**) belonging to different antigen groups related to their visibility to antibody recognition: Cell/Virus Enclosed (in green, hidden); Envelope (in violet, visible); Capsid (in red, undetermined visibility); and Secreted (in blue, visible). Viral antigens and respective B cell epitopes that could not be assigned to a single group are noted as undetermined (in grey).

As noted, envelope antigens can be visible to antibody recognition, but only the region surfacing the cell and/or the viral particle (ectodomain). Thereby, we determined the topology of viral envelope antigens and accordingly B cell epitopes lying in ectodomains, transmembranes and regions underneath the cell membrane and/or the viral envelope membrane (underneath membrane). Of the 9092 linear B cell epitopes in viral envelope proteins, 7448 of them reside in ectodomain regions, readily visible to antibody recognition (Figure 3A). In this context, the number of B cell epitopes in antigen regions hidden to antibody recognition rise to 19439, corresponding to 68.33 % of all virus-specific linear B cell epitopes that could be classified according to the visibility of their antigens (Figure 3B and Supplementary Dataset 2).

**Figure 3.**
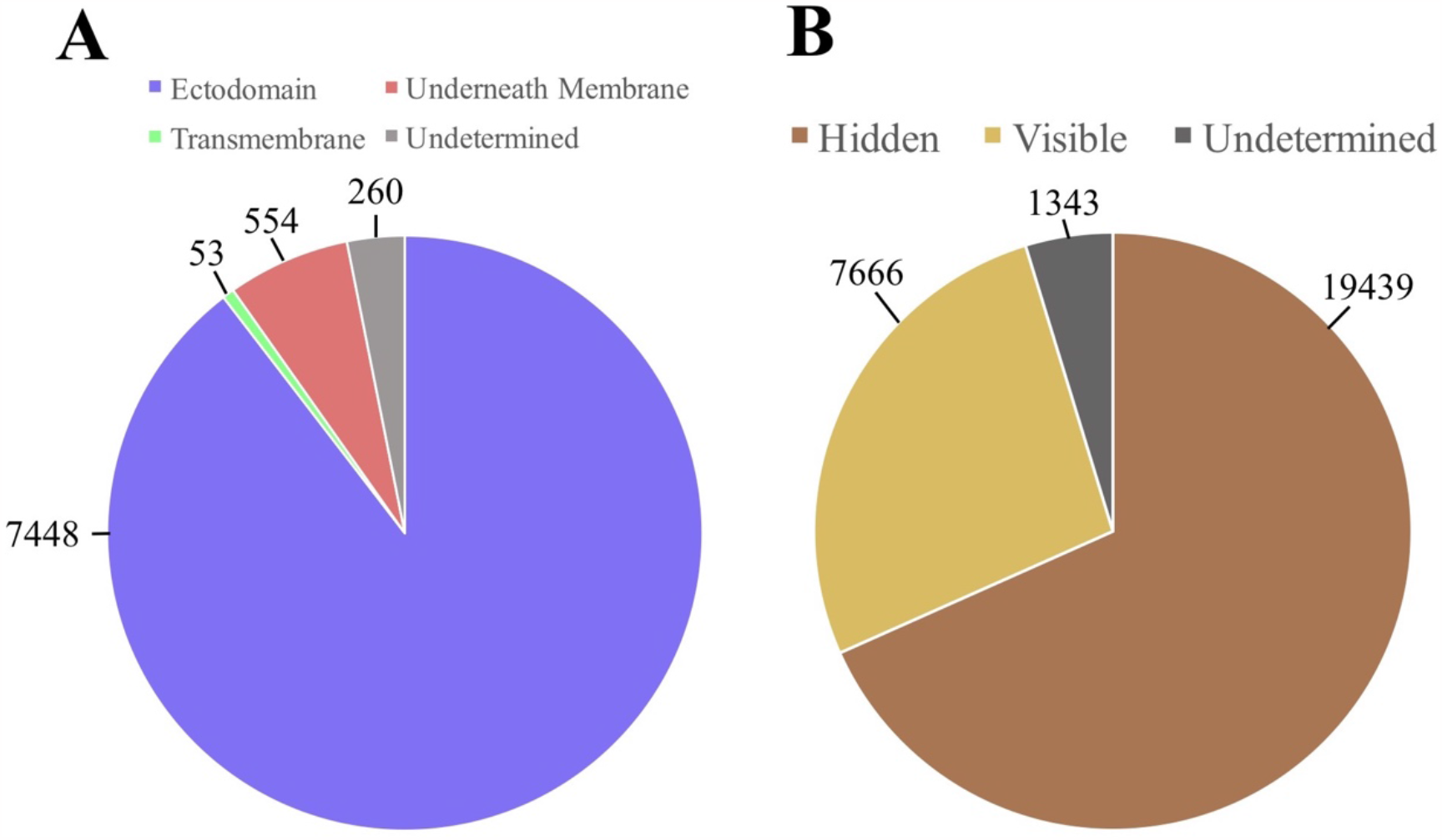
Analysis of B cell epitopes in viral envelope antigens. (**A**) Pie chart depicting the number of viral envelope B cell epitopes mapping in distinct regions: ectodomain (purple), transmembrane (green) or underneath plasmatic and/or envelope membrane (red). B cell epitopes that included segments mapping in more than one region are noted as undetermined (grey). (**B**) Pie chart depicting the number of virus-specific linear B cell epitopes classified according to their visibility. Hidden (brown): B cell epitopes in antigens enclosed in cells/viruses plus B cell epitopes in hidden regions of envelope proteins; Visible (yellow): B cell epitopes in ectodomains of envelope proteins and mature secreted proteins; Undetermined (grey): B cell epitopes that could not be unequivocally classified as visible or hidden.

### 2.3 Most linear B cell epitopes are not solvent accessible in the native antigens

B cell epitopes must also be solvent accessible to be recognized by antibodies. Thereby, we investigated if potentially recognizable linear B cell epitopes (located in ectodomains of viral envelope antigens and secreted antigens) were actually solvent accessible in the native 3D-structure of the relevant antigens. To that end, we selected 220 linear B cell epitopes that mapped entirely in envelope antigens with available 3D-structures. Next, we computed the relative solvent accessibility of residues (rRSA), distinguishing between those in linear B cell epitopes (1429 residues) and the remaining antigen sequence (non-B cell epitopes) (1487 residues). Subsequently, we compared rRSA from linear B cell epitopes and non-B cell epitopes with those from 483 conformational B cell epitopes extracted from the 3D-structure of antigen-antibody complexes (details in Methods) (Figure 4). The rRSAs from linear B cell epitopes and non-B cell epitopes are very similar with no statistical difference between them, as judge by one-sided Mann-Whitney U Test (*p*-value = 0.4177). The median of rRSA values from linear B cell epitopes and non-B cell epitopes is 23.00 % and 22.40 %, respectively. In fact, it is worth noting about 50% of the antigen residues mapped within B cell epitopes. In contrast, rRSA values in conformational B cell epitopes are significantly higher than those of linear B cell epitopes (*p*-value < 2.2 · 10^-16^) (median rRSA = 48.5 %).

**Figure 4.**
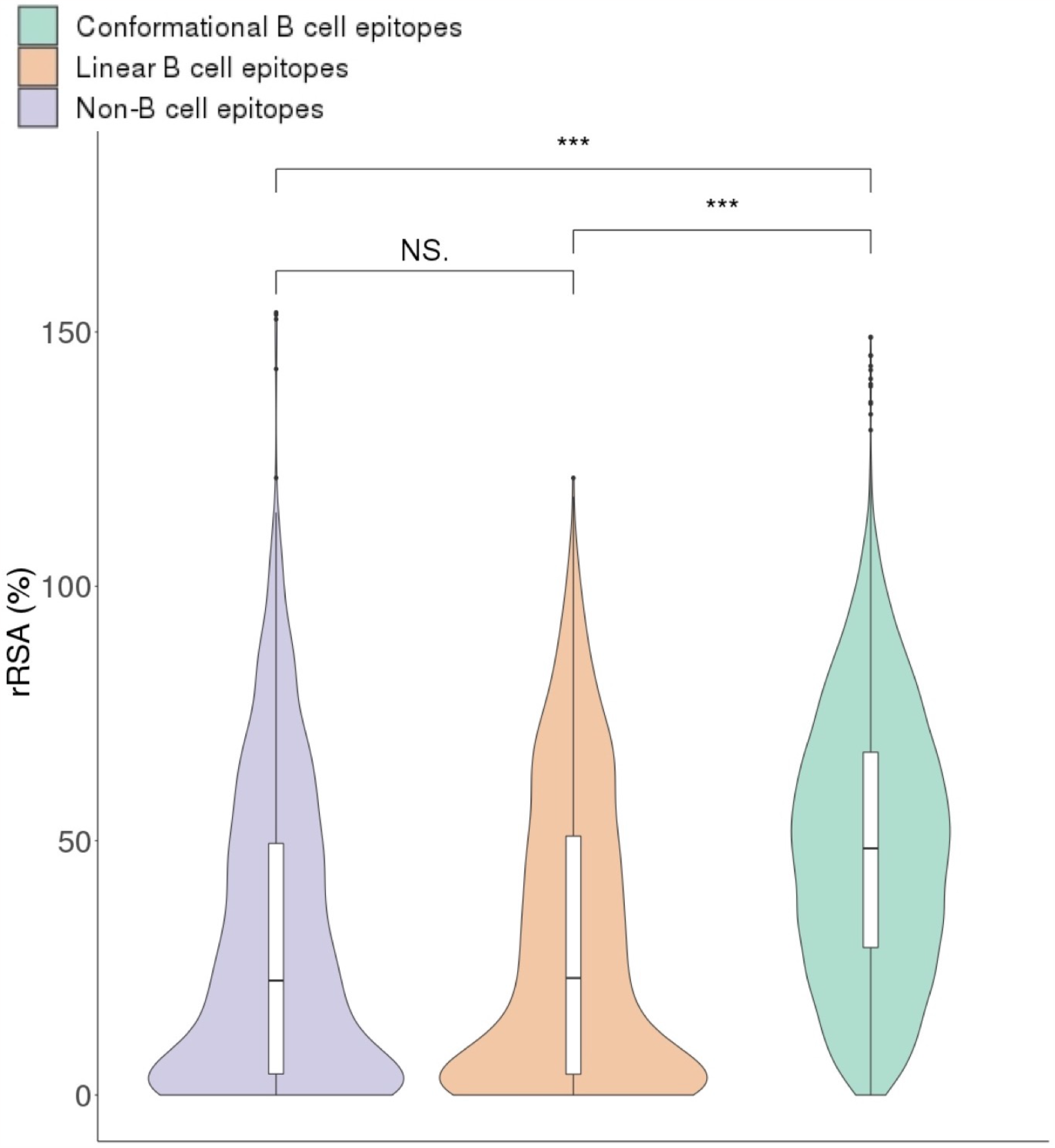
Residue accessibility of B cell and non-B cell epitopes. Violin plots depicting rRSA from conformational B cell epitopes (green), linear B cell epitopes (orange) and non-B cell epitopes (purple). Statistical differences were identified using Mann-Whitney U Tests (*** *p*-value ≤ 0.001, NS *p*-value ≥ 0.05).

To complete the analysis, we also assigned to each B cell epitope an entire RSA (eRSA) that was computed as an average value of the relevant rRSA values of the residues comprising the B cell epitope (Supplementary Dataset 3). To establish a threshold of solvent accessibility, we carried out the same calculations for 483 conformational B cell epitopes (Supplementary Dataset 4). As we show in Figure 5, eRSA values of conformational B cell epitopes were significantly higher than those of linear B cell epitopes (*p*-value < 2.2 · 10^-16^). The median eRSA value of linear B cell epitopes is 29.34 %, while the median eRSA value of conformational B cell epitopes is 49.29% (Figure 5). Using the minimum eRSA value (without outliers) of conformational B cell epitopes as a threshold (33.53 %), we determined that 67.28 % of linear B cell epitopes are not accessible in the antigen 3D-structure.

**Figure 5.**
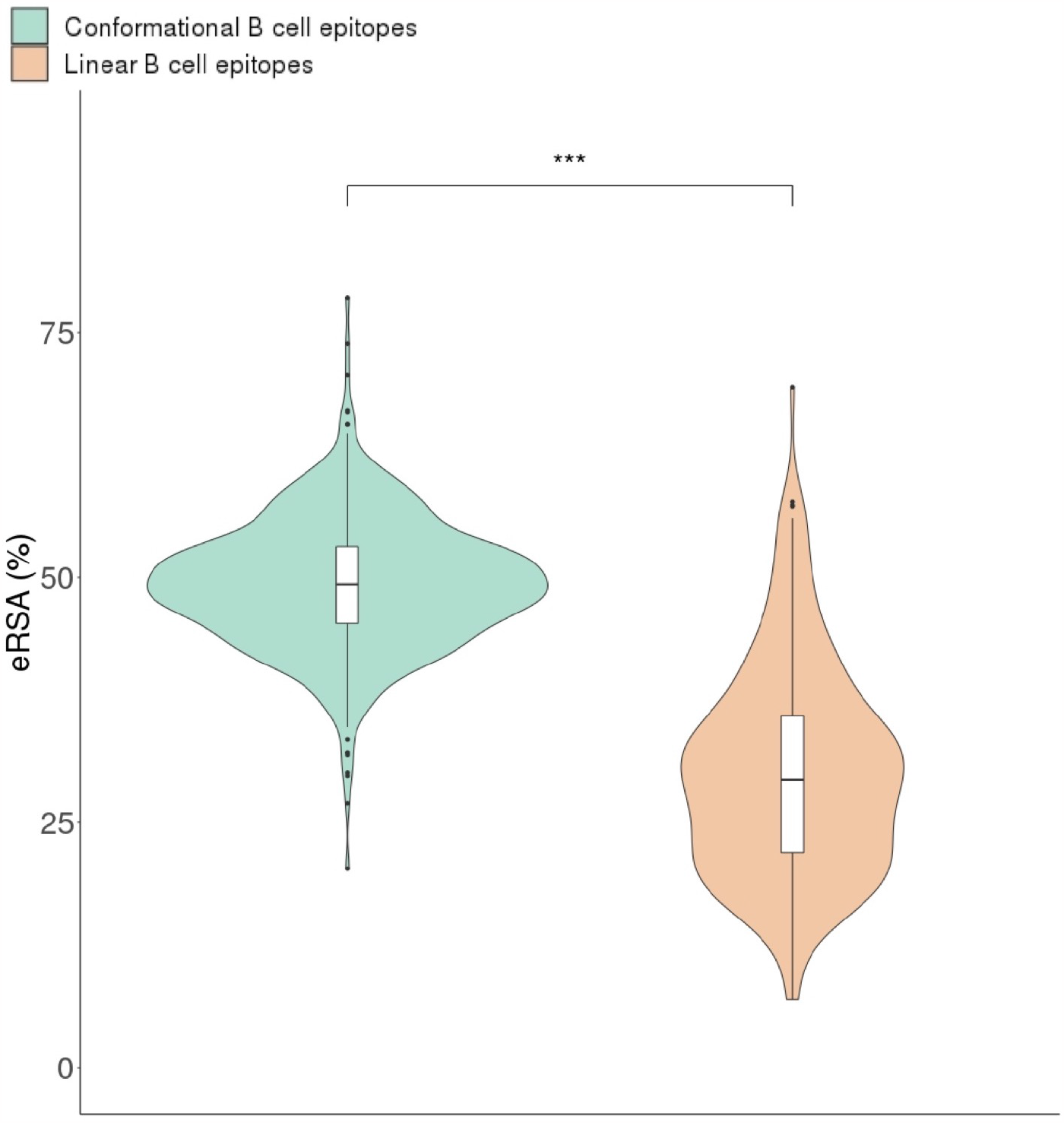
Solvent accessibility of B cell epitopes. Relative solvent accessibility of entire B cell epitopes (eRSA) was computed for linear (orange) and conformational (green) B cell epitopes and plotted using violin representations. Mann-Whitney U Tests were used to identify statistical differences between groups (*** p-value ≤ 0.001).

## 3. DISCUSSION

B cell receptors and antibodies can only recognize visible and solvent accessible antigens. Thereby, given the 3D-structure of proteins, it is often assumed that most B cell epitopes in protein antigens are conformational, encompassing non-sequential residues that are in close proximity in the molecular surface of the antigen (4). Currently, there are numerous known B cell epitopes that have been identified under different experimental settings and are collected in dedicated databases such as the IEDB, the largest repository. In this work, we sought to characterize protein B cell epitopes available at IEDB that are targeted by humans during infection. It is worth noting that the these B cell epitopes reflect what the immune system sees from a pathogen during infection but also the interest of researches for particular antigens. The overwhelming majority (86.94 %) of the selected B cell epitopes are linear and from viruses (Figure 1). This result denotes, on the one hand, that researches have had more success in determining the targets antibody responses in viral infections than in infections caused by other pathogens; virus are very simple organisms and B cell epitope mapping studies are easier to tackle. On the other hand, the result appears to contradict the assumption that most B cell epitopes are conformational. This apparent contradiction could be the result of technological biases. The most common approaches to B cell epitope mapping rely on detecting antibody binding to antigen fragments (peptides) and hence yield only linear B cell epitopes. Nonetheless, the identified linear B cell epitope peptides must have been visible and accessible to antibody recognition during the course of infection; but not necessarily on intact antigens. Thereby, we investigated to what extent the reported virus-specific linear B cell epitopes are visible and accessible to antibody recognition on host cells and/or viral particles. To that end, we focused on a subset of linear B cell epitopes from viral antigens with UNIPROT annotations enabling such investigation.

We were surprised to find that most virus-specific B cell epitopes (64.44 %) lied on viral antigens that remain enclosed in host cells or viral particles, precluding their recognition by antibodies on those contexts (Figure 2B). In comparison, linear B cell epitopes on envelope viral proteins represented only 31.11%, in spite of being visible to antibody recognition on viral particles and of particular interest. Envelope proteins play key roles in viral entry into host cells and spreading infection, and are main targets of neutralizing and anti-viral antibodies. Neutralizing antibodies, such those against SARS-CoV2 Spike protein, preclude binding to cognate host receptors required for viral entry (11, 12). Non-neutralizing anti-viral antibodies prevent the function of envelope proteins required for spreading infection. For example, antibodies targeting influenza A virus envelope protein neuraminidase limit infection because they can inhibit the enzymatic activity of neuraminidase, which is required by the virus to leave infected cells (13, 14). Interestingly, a substantial number of B cell epitopes in envelope proteins map outside of the ectodomain (6.68 %), the protein region visible to antibody recognition (Figure 3A). Overall, we estimated that only 23.43 % of virus-specific linear B cell epitopes targeted during the course of infection are in antigen regions visible to antibody recognition on viral particles and/or host cells (Figure 3B).

We also found evidence indicating that virus-specific linear B cell epitopes in ectodomains may not be solvent accessible (Figure 4 and Figure 5). We came to this evidence after evaluating the solvent accessibility of linear B cell epitopes located in ectodomains of envelope proteins with known 3D-structures. We found out that antigen residues in and out of linear B cell epitopes have very similar RSA values that are nonetheless much lower than those of conformational B cell epitopes defined by antigen-antibody 3D-structures (Figure 4). Moreover, we estimated that 67.28 % of the selected linear B cell epitopes have entire relative solvent accessibility (eRSA) that are lower than the minimum eRSA determined for conformational B cell epitopes. Therefore, the majority of the examined B cell epitopes have solvent accessibilities that are incompatible with their recognition on native antigens, which likely holds true for all remaining linear B cell epitopes on envelope proteins. If this is the case, a mere 8.82 % of all virus-specific B cell epitopes that are known to be targeted during infection could have been recognized on native antigens visible on viral particles and/or host cells.

That the vast majority of linear B cell epitopes have locations and solvent accessibilities unsuitable for antibody recognition on native antigens visible on viral particles and/or host cells is highly revealing. It implies that most B cell epitopes are recognized after the participation of processes that makes them visible and accessible to antibodies/B cells. One of such processes may be phagocytosis. Phagocyte cells such neutrophils are recruited early on to the sites of infection, where they engulf viral particles and dead infected cells. Subsequently, these cells particulate and degrade the engulfed material and antigens in phagolysosomes, releasing their content by exocytosis to keep phagocyting (15–18). Sentinel cells like dendritic cells and tissue-resident macrophages can also contribute to this process. Antigen degradation may further proceed in the extracellular space with the participation of different proteases such as matrix metalloproteases (19, 20) and proteases released by phagocytes (15–18). As a result of all these processes, new antigens and antigen fragments are available for B cell recognition and generation of specific antibodies that would be otherwise hidden.

## 4. CONCLUSIONS AND LIMITATIONS

In sum, our analysis of available B cell epitope data points that antibodies elicited during infections target mostly linear B cell epitopes requiring prior degradation of pathogens and/or infected cells. Likewise, it unravels that B cell epitope and antibody repertoires are largely shaped by antigen degradation processes, unconnected to presentation by major histocompatibility complex (MHC) molecules. We are aware that the extent of this finding could be limited by the quality of the available B cell epitope data. For example, we cannot discard that discriminating B cell epitopes by binding affinity to antibodies will somewhat alter our results. Unfortunately, binding affinity of B cell epitopes to antibodies is seldom reported and so we used B cell epitopes that were recognized by humans in the context of infection. Likewise, we do not know, nor that information is available, whether B cell epitopes used in this study resulted from the engagement of naive B cells or pre-existing cross-reactive memory B cells. In fact, we actually speculate that the recognition of so many, unlikely protective, hidden linear B cell epitopes from pathogens could be an strategy for a prompt elicitation of a network of cross-reactive antibodies and memory B cells that could be actually protective against unrelated infections. Indeed, we have previously identified potentially cross-reactive linear B cell epitopes between bacteria and viruses, including SARS-CoV-2, anticipating this possibility (21, 22).

## 5. MATERIALS AND METHODS

### 5.1 B cell epitope data collection and processing

B cell epitopes were obtained from the Immune Epitope Database (IEDB) (9, 10), after a search for B cell epitopes with positive assays linked to infectious diseases in humans. B cell epitope assay data were downloaded and parsed, selecting the epitope amino acid sequence (only unique sequences were considered), type of epitope (linear or conformational) and NCBI and UNIPROT accessions of source antigens. B cell epitopes that did not include NCBI or UNIPROT accessions, or did not have between 8 and 25 residues were discarded. Amino acid sequences of antigens were obtained from UNIPROT using the relevant accessions and matching of linear B cell epitopes was verified through BLAST searches as described elsewhere in Ras-Carmona et al (23). Linear B cell epitopes that did not match the sequence of source antigens (identity ≥ 90 % over ≥ 90 % of the epitope length) were discarded. CD-HIT (24) was used to cluster overlapping linear B cell epitopes (100 % identity threshold). B cell epitope clusters were processed selecting overlapping regions including 8-25 residues as representative linear B cell epitopes. An additional dataset consisting of conformational B cell epitopes was obtained as described elsewhere (25) from known 3D-structures of antigen-antibody complexes available at the abYbank/AbDb database (26). Conformational B cell epitopes consisted of antigen residues with atoms contacting the antibody (≤ 4 Å distance).

### 5.2 Antigen annotations

Taxonomy, sub-cellular location and transmembrane topology of antigens were collected after protein records in NCBI and UNIPROT and recorded for every antigen. Taxonomy information was obtained from the NCBI taxonomy database (27), using the corresponding NCBI tax identifiers. Sub-cellular location of proteins and transmembrane topology of plasma membrane-proteins were obtained from UNIPROT. Transmembrane topology was predicted using TMHMM (28) for protein antigens with plasma membrane sub-cellular locations but without relevant information on the subject.

### 5.3 Computation of relative solvent accessibility

Solvent accessibility was computed for linear B cell epitopes in antigens with known 3D-structure and conformational B cell epitopes defined by the 3D-structure of antigen-antibody complexes available in abYbank/AbDb database (26). For linear B cell epitopes, 3D-structure coordinates of protein antigens were obtained from the Protein Database Bank (PDB) at Brookhaven (29) after the relevant PDB codes in NCBI protein records. PyMOL Molecular Graphics System (Version 1.8 Schrödinger) was used to remove redundant molecules/chains concurring in crystallization units and to discard linear B cell epitopes that did not map in the relevant 3D-structures. For conformational B cell epitopes, antibody chains were removed from antigen-antibody 3D-structures prior to further analysis using also PyMOL. Relative solvent accessibility (RSA) of protein antigen residues (rRSA) in % values were obtained upon the relevant 3D-structure coordinates using NACCESS (30) and entire B cell epitope RSA (eRSA) values (%) were subsequently computed using the following equation:

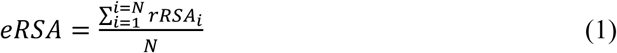

where N is the number of residues of the B cell epitope and rRSA_i_ is the rRSA of residues (i = 1, i = 2, …, i = N) included in the B cell epitope.

### 5.4 Graphics and statistics

All plots were generated in RStudio with the help of specific R packages. Statistical differences between groups of B cell epitopes with regard to rRSA and eRSA values were analyzed in RStudio, using non-parametric one-sided Mann-Whitney U Tests. *p* < 0.05 was considered significant.

## ACKNOWLEDGMENTS

We wish to thank to PAR lab members for comments and critical reading.

## SUPPLEMENTARY MATERIAL

**Supplementary Dataset 1**. Linear B cell epitopes targeted during infection in humans

**Supplementary Dataset 2**. Virus-specific linear B cell epitopes with location

**Supplementary Dataset 3**. Virus-specific linear B cell epitopes mapping in ectodomains of viral envelope proteins with available tertiary structures

**Supplementary Dataset 4**. Discontinuous B cell epitopes extracted from antibody-antigen tertiary structures

## REFERENCES

1. A. K. Abbas, A. H. Lichtman, S. Pillai, D. L. Baker, A. Baker, Cellular and molecular immunology (2018).

2. J. L. Sanchez-Trincado, M. Gomez-Perosanz, P. A. Reche, Fundamentals and Methods for T- and B-Cell Epitope Prediction. J. Immunol. Res. 2017, 2680160 (2017).

3. M. H. V. Van Regenmortel, “What is a B-cell epitope?” in Methods in Molecular Biology, Second, M. Schutkowski, U. Reineke, Eds. (Humana Press, 2009), pp. 3–20.

4. S. Ferdous, S. Kelm, T. S. Baker, J. Shi, A. C. R. Martin, B-cell epitopes: Discontinuity and conformational analysis. Mol. Immunol. 114, 643–650 (2019).

5. J. Nilvebrant, J. Rockberg, An Introduction to Epitope Mapping. Methods Mol. Biol. 1785, 1–10 (2018).

6. L. B. Hansen, S. Buus, C. Schafer-Nielsen, Identification and mapping of linear antibody epitopes in human serum albumin using high-density Peptide arrays. PLoS One 8, e68902 (2013).

7. L. C. Szymczak, H.-Y. Kuo, M. Mrksich, Peptide Arrays: Development and Application. Anal. Chem. 90, 266–282 (2018).

8. N. Salimi, W. Fleri, B. Peters, A. Sette, Design and utilization of epitope-based databases and predictive tools. Immunogenetics 62, 185–196 (2010).

9. R. Vita, et al., The immune epitope database (IEDB) 3.0. Nucleic Acids Res. 43, D405–D412 (2015).

10. R. Vita, et al., The Immune Epitope Database (IEDB): 2018 update. Nucleic Acids Res. 47, D339–D343 (2019).

11. B. Ju, et al., Human neutralizing antibodies elicited by SARS-CoV-2 infection. Nature 584, 115–119 (2020).

12. Y. Huang, C. Yang, X. Xu, W. Xu, S. Liu, Structural and functional properties of SARS-CoV-2 spike protein: potential antivirus drug development for COVID-19. Acta Pharmacol. Sin. 41, 1141–1149 (2020).

13. J. L. McAuley, B. P. Gilbertson, S. Trifkovic, L. E. Brown, J. L. McKimm-Breschkin, Influenza Virus Neuraminidase Structure and Functions . Front. Microbiol. 10 (2019).

14. N. Abbadi, J. J. Mousa, Broadly Protective Neuraminidase-Based Influenza Vaccines and Monoclonal Antibodies: Target Epitopes and Mechanisms of Action. Viruses 15 (2023).

15. S. Gordon, Phagocytosis: An Immunobiologic Process. Immunity 44, 463–475 (2016).

16. D. M. Underhill, A. Ozinsky, Phagocytosis of Microbes: Complexity in Action. Annu. Rev. Immunol. 20, 825–852 (2002).

17. E. Uribe-Querol, C. Rosales, Phagocytosis: Our Current Understanding of a Universal Biological Process. Front. Immunol. 11, 1066 (2020).

18. S. D. Catz, K. R. McLeish, Therapeutic targeting of neutrophil exocytosis. J. Leukoc. Biol. 107, 393–408 (2020).

19. D. Young, N. Das, A. Anowai, A. Dufour, Matrix Metalloproteases as Influencers of the Cells’ Social Media. Int. J. Mol. Sci. 20 (2019).

20. L. Nissinen, V.-M. Kähäri, Matrix metalloproteinases in inflammation. Biochim. Biophys. Acta - Gen. Subj. 1840, 2571–2580 (2014).

21. P. A. Reche, Potential Cross-Reactive Immunity to SARS-CoV-2 From Common Human Pathogens and Vaccines. Front. Immunol. 11, 586984 (2020).

22. A. Bodas-Pinedo, et al., Combining different bacteria in vaccine formulations enhances the chance for antiviral cross-reactive immunity: a detailed in silico analysis for influenza A virus . Front. Immunol. 14 (2023).

23. A. Ras-Carmona, M. Gomez-Perosanz, P. A. Reche, Prediction of unconventional protein secretion by exosomes. BMC Bioinformatics 22, 333 (2021).

24. W. Li, A. Godzik, Cd-hit: a fast program for clustering and comparing large sets of protein or nucleotide sequences. Bioinformatics 22, 1658–1659 (2006).

25. A. Ras-Carmona, H. F. Pelaez-Prestel, E. M. Lafuente, P. A. Reche, BCEPS: A Web Server to Predict Linear B Cell Epitopes with Enhanced Immunogenicity and Cross-Reactivity. Cells 10 (2021).

26. S. Ferdous, A. C. R. Martin, AbDb: antibody structure database-a database of PDB-derived antibody structures. Database (Oxford). 2018, bay040 (2018).

27. C. L. Schoch, et al., NCBI Taxonomy: a comprehensive update on curation, resources and tools. Database (Oxford). 2020 (2020).

28. A. Krogh, B. Larsson, G. von Heijne, E. L. . Sonnhammer, Predicting transmembrane protein topology with a hidden markov model: application to complete genomes. J. Mol. Biol. 305, 567–580 (2001).

29. H. M. Berman, et al., The Protein Data Bank. Nucleic Acids Res. 28, 235–242 (2000).

30. S. Hubbard, M. S. Building, NACCESS, Computer Program (1993).

