## Supplementary Dataset 1 for "Analysis of virus-specific B cell epitopes reveals extensive antigen processing prior to recognition"

**Supplementary Dataset 1.** Linear B cell epitopes targeted during infection in humans

| Sequence | Virus | Bacteria | Eukaryotic | IEDB | NCBI | UNIPROT |
|---|---|---|---|---|---|---|
| AADMIMHTPGCVPC | 1 | 0 | 0 | 92 | AAA45721.1 | P27958 |
| AAECPFLPKPKVA | 1 | 0 | 0 | 101 | AAO86636.1 | O36307 |
| AAEESKLPINPLSNSLLRH | 1 | 0 | 0 | 103 | BAA03375.1 | P27958 |
| AAHIPYLEQGMHLAEQFKQK | 1 | 0 | 0 | 200 | Q81754.3 | P27958 |
| AALVSGTATAGWTFGAG | 1 | 0 | 0 | 307 | NP_828851.1 | P59594 |
| AAQAALGLLQTASR | 1 | 0 | 0 | 384 | SRC279960 | P27958 |
| AAQGESSTQKVRPSHTNNPK | 1 | 0 | 0 | 387 | AAA45411.1 | P03410 |
| AARALAHGVRVLEDGVNYATGNLPA | 1 | 0 | 0 | 406 | SRC279960 | P27958 |
| AAYFVGYLKPTTFMLKY | 1 | 0 | 0 | 525 | P59594.1 | P59594 |
| ACKPLLREDVTFQV | 1 | 0 | 0 | 568 | BAA01583.1 | P27958 |
| ACSLPEEAHTAIHS | 1 | 0 | 0 | 599 | P26660.3 | P27958 |
| ACSLPQEARTAIHS | 1 | 0 | 0 | 600 | AAA45665.1 | P27958 |
| ADDVILHTPGCVPC | 1 | 0 | 0 | 652 | Q81258.3 | P27958 |
| ADIDKLIDYAASGD | 1 | 0 | 0 | 677 | P27313.1 | P27313 |
| ADLVAAQKLATKP | 1 | 0 | 0 | 723 | AAO86636.1 | O36307 |
| AEDAAEISVPAEIL | 1 | 0 | 0 | 848 | SRC279960 | P27958 |
| AEEKLKKKSSFYQS | 1 | 0 | 0 | 881 | P27313.1 | P27313 |
| AEHVDTSYECDIPIGA | 1 | 0 | 0 | 971 | NP_828851.1 | P59594 |
| AERAVEVDPDDVNK | 1 | 0 | 0 | 1146 | P27313.1 | P27313 |
| AETHVTGGNAGRTTAGLVGLLTPG | 1 | 0 | 0 | 1189 | P27958.3 | P27958 |
| AFAFAGNHVSPTHYVPESDA | 1 | 0 | 0 | 1248 | AAA86907.1 | P27958 |
| AFASRGNHDSPTHYVPESD | 1 | 0 | 0 | 1258 | AAA75355.1 | P27958 |
| AFASRGNHVAPTHYVTESDA | 1 | 0 | 0 | 1260 | P26660.3 | P27958 |
| AFASRGNHVAPTHYVVESDA | 1 | 0 | 0 | 1261 | P26661.3 | P27958 |
| AFASRGNHVSPRHYVPESE | 1 | 0 | 0 | 1263 | AAA52748.1 | P27958 |
| AFASRGNHVSPTHYVPESDA | 1 | 0 | 0 | 1265 | BAB18806.1 | P27958 |
| AFDEMEECASHLPYIEQGMQ | 1 | 0 | 0 | 1276 | AAC53957.1 | P27958 |
| AFFSILQDMRNTIM | 1 | 0 | 0 | 1299 | P27313.1 | P27313 |
| AGCLIGAEHVDTSYECD | 1 | 0 | 0 | 1460 | NP_828851.1 | P59594 |
| AGDLESSVDDPRSEEDKRFE | 1 | 0 | 0 | 1470 | P08617.1 | P08617 |
| AGGQLFYSRPVVSANGEPTVKLYTS | 1 | 0 | 0 | 1533 | P29326.1 | Q81871 |
| AGHATLREHLRDIKAENTDA | 1 | 0 | 0 | 1547 | AAA19496.1 | P10211 |
| AGPRVRQPARPLGSAWRDQAQRPAV | 1 | 0 | 0 | 1646 | P29326.1 | Q81871 |
| AHLSLRGLPVCAFSS | 1 | 0 | 0 | 1799 | ABK27207.1 | Q69027 |
| AHWGVLAGIAYFSMVGNWAK | 1 | 0 | 0 | 1827 | AAA45676.1 | P27958 |
| AIAYYRGKDSSIIKDGDLVVC | 1 | 0 | 0 | 1859 | CAA75607.1 | A0A159D718 |
| AICKRIPDKKPGKKT | 1 | 0 | 0 | 1861 | SRC279969 | O36633 |
| AIGIITLYLGAVVQA | 1 | 0 | 0 | 1923 | QCZ25049.1 | P17763 |
| AIILLTCAVHPELI | 1 | 0 | 0 | 1948 | P26662.3 | P27958 |
| AILSSLTVTQLLRR | 1 | 0 | 0 | 2005 | P26664.3 | P27958 |
| AIPTNFSISITTEVMPV | 1 | 0 | 0 | 2053 | NP_828851.1 | P59594 |
| AKKVPETFPELKPGESRHTS | 1 | 0 | 0 | 2226 | P08617.1 | P08617 |
| AKWAVPTTRTDDKLR | 1 | 0 | 0 | 2322 | P08362.1 | Q786F2 |
| ALAALAEKSFPSSS | 1 | 0 | 0 | 2365 | SRC279960 | P27958 |
| ALAELATRSFGSSS | 1 | 0 | 0 | 2380 | P26664.3 | P27958 |
| ALGGWGLHQPPPQIFLKILP | 1 | 0 | 0 | 2558 | P07299.1 | Q9PZT0 |
| ALGLLQQASKQAEAAAPVV | 1 | 0 | 0 | 2566 | AAC53936.1 | P27958 |
| ALGVINTLEWIPRFK | 1 | 0 | 0 | 2598 | P08362.1 | Q786F2 |
| ALHQVFGAIYGAAFSGVSWT | 1 | 0 | 0 | 2612 | ABX25775.1 | P17763 |
| ALQEQCRFPNITNSHVPILQERP | 1 | 0 | 0 | 2838 | AAL05538.1 | P03381 |
| AMDAWNQPSGISSPPSPSLN | 1 | 0 | 0 | 3069 | AAA45412.1 | Q85601 |
| AMTPTVATRDGKLPATQLRR | 1 | 0 | 0 | 3200 | AAA45676.1 | P27958 |
| ANCASILCKCYTTGT | 1 | 0 | 0 | 3236 | P69353.1 | Q786F3 |
| ANKEGIVWVATEGALN | 1 | 0 | 0 | 3325 | AAS48456.1 | P59595 |
| APAHDTPPVPDVDSRGAILRRQYNL | 1 | 0 | 0 | 3476 | P29326.1 | Q81871 |
| APDYVPPTVHGCALPPR | 1 | 0 | 0 | 3518 | Q81258.3 | P27958 |
| APEARQAIKSLT | 1 | 0 | 0 | 3525 | P26663.3 | P27958 |
| APEARQAIRSLT | 1 | 0 | 0 | 3526 | P29846.3 | P27958 |
| APHLPYIEQGMQLAEQFKQK | 1 | 0 | 0 | 3592 | BAA20975.1 | P27958 |
| APLSPLLPLQDGTNTHIMATEASNY | 1 | 0 | 0 | 3648 | P29326.1 | Q81871 |
| APLTAVAPAHDTPPVPDVDSRGAIL | 1 | 0 | 0 | 3651 | P29326.1 | Q81871 |
| APLTRGSCRKRNRSPER | 1 | 0 | 0 | 3652 | YP_001129431.1 | Q9QR71 |
| APPLRAWKSRARAV | 1 | 0 | 0 | 3686 | P26661.3 | P27958 |
| APPLRTWKSRARAV | 1 | 0 | 0 | 3687 | AAF59940.1 | P27958 |
| APPSQPFPWTHCYQPRL | 1 | 0 | 0 | 3702 | P03383.1 | P03383 |
| APRITFGGPTDSTDNNQN | 1 | 0 | 0 | 3779 | P59595.1 | P59595 |
| AQAHTTRGLTNLFSSG | 1 | 0 | 0 | 3938 | SRC279960 | P27958 |
| AQGRNVRTFTSFFSPG | 1 | 0 | 0 | 3973 | SRC279960 | P27958 |
| AQKFNGLTVLPPLLTDD | 1 | 0 | 0 | 3983 | NP_828851.1 | P59594 |
| AQRVEFLLKAWAEK | 1 | 0 | 0 | 4059 | P26660.3 | P27958 |
| AQSRNVRTFTGLFTSG | 1 | 0 | 0 | 4071 | SRC279960 | P27958 |
| AQTFLATCINGVCW | 1 | 0 | 0 | 4080 | P26664.3 | P27958 |
| AQVLSEMVMCGGSLY | 1 | 0 | 0 | 4092 | P0DTD1.1 | P0DTD1 |
| ARATIRYRPLVPNAVGGYAISISFW | 1 | 0 | 0 | 4121 | P29326.1 | Q81871 |
| ARGYISTRVEMGEAA | 1 | 0 | 0 | 4152 | P14340.2 | P17763 |
| ARSLDWTKVTLDGRPLSTIQQYSKT | 1 | 0 | 0 | 4260 | P29326.1 | Q81871 |
| ASHLPYFEQGMQLAEQFKQK | 1 | 0 | 0 | 4450 | AAA45721.1 | P27958 |
| ASHLPYIEQGMHLAEQFKQK | 1 | 0 | 0 | 4451 | AAC53941.1 | P27958 |
| ASHLPYIEQGMQLAAEFKQK | 1 | 0 | 0 | 4452 | AAA79971.1 | P27958 |
| ASHLPYIEQGMQLAEQFKQK | 1 | 0 | 0 | 4453 | AAC53954.1 | P27958 |
| ASHLPYIEQGMQLAEQLKQK | 1 | 0 | 0 | 4454 | AAC53956.1 | P27958 |
| ASHLPYIEQGVQLAEQFKQK | 1 | 0 | 0 | 4455 | AAA75355.1 | P27958 |
| ASICQMFCFWRGDLVFDFQV | 1 | 0 | 0 | 4467 | AAR01187.1 | P08617 |
| ASKAALIEEGQRMAEMLKSK | 1 | 0 | 0 | 4498 | AAF59945.1 | P27958 |
| ASKATLIEEGQRIAEMLKSK | 1 | 0 | 0 | 4499 | BAA01956.1 | P27958 |
| ASQLPYIEQGMQLAEQFKQK | 1 | 0 | 0 | 4718 | BAA01583.1 | P27958 |
| ASRAALIEEGQRIAEMLKSK | 1 | 0 | 0 | 4733 | AAU89634.1 | P27958 |
| ASRAALLEEGQRIAEMLKSK | 1 | 0 | 0 | 4734 | AAC53939.1 | P27958 |
| ASSRSSSRSRGNSRNST | 1 | 0 | 0 | 4782 | AAS48456.1 | P59595 |

| | | | | | | |
|---|---|---|---|---|---|---|
| ASTNRQSGRQPTPISPPLRDSHPQ | 1 | 0 | 0 | 4802 | P03140.3 | Q76R62 |
| ASWAIKWEYVVLLFLLLADA | 1 | 0 | 0 | 4833 | AAA45676.1 | P27958 |
| ASWFTALTQHGKEELRFPRGQ | 1 | 0 | 0 | 4838 | ABI96968.1 | P59595 |
| ATCMQADLEVMTST | 1 | 0 | 0 | 4911 | P26660.3 | P27958 |
| ATCMQVNLEIMTST | 1 | 0 | 0 | 4912 | BAA03177.1 | P27958 |
| ATSRSTMQLASFFTPG | 1 | 0 | 0 | 5147 | SRC279960 | P27958 |
| ATSRSTSQLTSLFSFG | 1 | 0 | 0 | 5148 | SRC279960 | P27958 |
| AVARSAQSLTSLFVSG | 1 | 0 | 0 | 5270 | SRC279960 | P27958 |
| AVASRRRPTTAGAAPLTAVAPAHDT | 1 | 0 | 0 | 5275 | P29326.1 | Q81871 |
| AVDSAARIHDFRYSQLAKLG | 1 | 0 | 0 | 5298 | P07299.1 | Q9PZT0 |
| AVDTGSGGGGQPHDTAPRGARKKQ | 1 | 0 | 0 | 5303 | P14348.1 | P14348 |
| AVEVDPDDVNKNTL | 1 | 0 | 0 | 5312 | P27313.1 | P27313 |
| AVGIMTVTMTFKLGPRKATG | 1 | 0 | 0 | 5346 | P07299.1 | Q9PZT0 |
| AVGYRASRTVSLFSPG | 1 | 0 | 0 | 5356 | SRC279960 | P27958 |
| AVIEPIVATNWQKL | 1 | 0 | 0 | 5368 | CAJ41284.1 | P27958 |
| AVILLMCAVHPTLV | 1 | 0 | 0 | 5380 | P26664.3 | P27958 |
| AVIRRANNVLKNEMKRYKGL | 1 | 0 | 0 | 5387 | P03418.1 | O42053 |
| AVLTSMLTDPSHITAETAKR | 1 | 0 | 0 | 5467 | P26662.3 | P27958 |
| AVSLDFSPGTSGSPI | 1 | 0 | 0 | 5542 | P14340.2 | P17763 |
| AVVAPDKEVLYEAFDEMEEC | 1 | 0 | 0 | 5591 | AAU89634.1 | P27958 |
| AYFPREGVFVFNGTSWF | 1 | 0 | 0 | 5773 | NP_828851.1 | P59594 |
| AYQVRNSTGLYHVTNDCPNS | 1 | 0 | 0 | 5907 | AAA45676.1 | P27958 |
| AYSNNTIAIPTNFSISI | 1 | 0 | 0 | 5927 | NP_828851.1 | P59594 |
| CANLLLQYGSFCTQLNRALSGIA | 1 | 0 | 0 | 6011 | P59594.1 | P59594 |
| CASKAALIEEGQRMAEMLKS | 1 | 0 | 0 | 6034 | AAF59945.1 | P27958 |
| CASYHTVSLLRSTSQKS | 1 | 0 | 0 | 6040 | NP_828851.1 | P59594 |
| CCDLDPQARVAIKS | 1 | 0 | 0 | 6061 | P26664.3 | P27958 |
| CCNLEPEARKVISS | 1 | 0 | 0 | 6067 | Q81258.3 | P27958 |
| CDIPIGAGICASYHTVS | 1 | 0 | 0 | 6110 | NP_828851.1 | P59594 |
| CFKKPPKYPPALPI | 1 | 0 | 0 | 6233 | Q81258.3 | P27958 |
| CFLNISNTHVSV | 1 | 0 | 0 | 6241 | P03383.1 | P03383 |
| CFNPAHTLTLSQP | 1 | 0 | 0 | 6246 | AAO86638.1 | Q9E006 |
| CGPKLSTDLIKNQCVNF | 1 | 0 | 0 | 6332 | P59594.1 | P59594 |
| CGSVFLVGQLFTFSPRRHWT | 1 | 0 | 0 | 6355 | AAA45676.1 | P27958 |
| CHARKAVTHINS | 1 | 0 | 0 | 6375 | P26664.3 | P27958 |
| CHILGPDCCIEPHDW | 1 | 0 | 0 | 6391 | ALG01342.1 | Q05320 |
| CILRQLRHLPSRVRYPHYS | 1 | 0 | 0 | 6429 | P03381.1 | P03381 |
| CNYSKFWYLEHAK | 1 | 0 | 0 | 6742 | AAL01687.1 | P09991 |
| CPECRPLGLQGCAFQSTVAELQRLK | 1 | 0 | 0 | 6757 | P29326.1 | Q81871 |
| CPKDELRPLEKVLESKTRAI | 1 | 0 | 0 | 6777 | P08617.1 | P08617 |
| CPPLRAWRHRARAV | 1 | 0 | 0 | 6824 | Q81258.3 | P27958 |
| CRFPNITNSHVPILQERPPLENR | 1 | 0 | 0 | 6919 | P03381.1 | P03381 |
| CSQHLPYIEQGMQLA | 1 | 0 | 0 | 7063 | SRC279960 | P27958 |
| CSQNPLAELKCSVKSFE | 1 | 0 | 0 | 7064 | P59594.1 | P59594 |
| CSRIIPARARVECFD | 1 | 0 | 0 | 7072 | P0DTD1.1 | P0DTD1 |
| CTIAALGSSDRDTVV | 1 | 0 | 0 | 7155 | AAA96965.1 | A0A159D718 |
| CTINYTIFKIRMYVGGVEHR | 1 | 0 | 0 | 7160 | AAA45676.1 | P27958 |
| CTIPAQGTSMFPSCCCTKPSDGNC | 1 | 0 | 0 | 7161 | BAF42669.1 | Q76R62 |
| CTIPAQGTSVFPSCCCTKPSDGNC | 1 | 0 | 0 | 7162 | AAC06360.1 | Q76R62 |
| CTPPALNCYWPLNDYGF | 1 | 0 | 0 | 7193 | P59594.1 | P59594 |
| CVLADSESGGHITHS | 1 | 0 | 0 | 7279 | P08362.1 | Q786F2 |
| CVLAWNTRNIDATSTGN | 1 | 0 | 0 | 7280 | P59594.1 | P59594 |
| CWGELMTLATWVGVNLEDPASRDLV | 1 | 0 | 0 | 7332 | P03146.1 | Q76R61 |
| CYGVSATKLNDLCFSNV | 1 | 0 | 0 | 7383 | P59594.1 | P59594 |
| CYWPLNDYGFYTTTGIG | 1 | 0 | 0 | 7452 | P59594.1 | P59594 |
| DAEKAVEVDPDDV | 1 | 0 | 0 | 7496 | AAO86636.1 | O36307 |
| DAGESSGTGAEVVPFN | 1 | 0 | 0 | 7517 | P07298.1 | Q9PZT1 |
| DAHFLSQTKQSGENLPYLVA | 1 | 0 | 0 | 7528 | AAY82027.1 | P27958 |
| DAIMTRCLAVHECFV | 1 | 0 | 0 | 7545 | P0DTD1.1 | P0DTD1 |
| DAVSRKKMDTKPTD | 1 | 0 | 0 | 7671 | P27313.1 | P27313 |
| DCHAPTYLPAEVDGD | 1 | 0 | 0 | 7704 | P08362.1 | Q786F2 |
| DDFHTDEDKLDTNSVYEPYY | 1 | 0 | 0 | 7771 | AAK01056.1 | P09259 |
| DDGDKGSDVESYSSMPPLE | 1 | 0 | 0 | 7778 | P26662.3 | P27958 |
| DDKDPQFKDNVILLNKHIDA | 1 | 0 | 0 | 7808 | P59595.1 | P59595 |
| DDKLRMETCFQQACK | 1 | 0 | 0 | 7812 | P08362.1 | Q786F2 |
| DDLPTTMFQPVRAPCIQTAW | 1 | 0 | 0 | 7822 | AAG48731.1 | P03410 |
| DDNGDRPPEPAAQGESSTQK | 1 | 0 | 0 | 7833 | AAA45411.1 | P03410 |
| DECHAQDATSILGI | 1 | 0 | 0 | 7920 | Q81258.3 | P27958 |
| DECHAVDSTTILGI | 1 | 0 | 0 | 7921 | P26660.3 | P27958 |
| DEFDPLVAEEDEREVSVPAEILR | 1 | 0 | 0 | 7979 | SRC279960 | P27958 |
| DEMEECSQHLPYIEQGMMLA | 1 | 0 | 0 | 8095 | P26664.3 | P27958 |
| DEQEQQEEQEQQEEQEQ | 1 | 0 | 0 | 8131 | YP_001129431.1 | Q9QR71 |
| DEVGLRTPQRFTDLV | 1 | 0 | 0 | 8195 | P06830.1 | Q786F2 |
| DFALELEFRNLTPGNTNTRVSRYSS | 1 | 0 | 0 | 8233 | P29326.1 | Q81871 |
| DFCGKGYHLMSFPQAAP | 1 | 0 | 0 | 8239 | NP_828851.1 | P59594 |
| DFDEMEECASHLPYIEQGMQ | 1 | 0 | 0 | 8241 | ABU97067.1 | P27958 |
| DGIMHTRCHCGAEI | 1 | 0 | 0 | 8408 | P26664.3 | P27958 |
| DGIMQTTCPCGAQI | 1 | 0 | 0 | 8409 | P26662.3 | P27958 |
| DGIYFAATEKSNVVRGW | 1 | 0 | 0 | 8413 | P59594.1 | P59594 |
| DGSNVLAALAALAE | 1 | 0 | 0 | 8494 | SRC279960 | P27958 |
| DGTRDTLPQSPGP | 1 | 0 | 0 | 8503 | NP_044666.1 | P06484 |
| DGVMSTRCPCGAAI | 1 | 0 | 0 | 8514 | Q81258.3 | P27958 |
| DGVRLHRYAPVCKPLLRE | 1 | 0 | 0 | 8517 | P26662.3 | P27958 |
| DHCPVVEVNGVTIQV | 1 | 0 | 0 | 8530 | P69353.1 | Q786F3 |
| DHILEPSIPWKSKLLTLVQL | 1 | 0 | 0 | 8544 | AAF76481.1 | P03381 |
| DHMSIYKFMGRSHFLCTFTF | 1 | 0 | 0 | 8554 | P08617.1 | P08617 |
| DKGIAIPHDIDLGESRVVIQDYDNQ | 1 | 0 | 0 | 8875 | P29326.1 | Q81871 |
| DKGSDVESYSSMPPLEGEP | 1 | 0 | 0 | 8877 | P26662.3 | P27958 |
| DKKGKVVGLYGNGVV | 1 | 0 | 0 | 8911 | P14340.2 | P17763 |
| DKLTTREIEQVELLKRIYDKL | 1 | 0 | 0 | 8937 | AAR18269.1 | A0A096XVH2 |
| DKPGSKKTQGEKFFLIHSAD | 1 | 0 | 0 | 8960 | P08617.1 | P08617 |

| | | | | | | |
|---|---|---|---|---|---|---|
| DLDFSAFDASLSPFMIREAG | 1 | 0 | 0 | 9043 | P08617.1 | P08617 |
| DLHPGSGKTRKYLPA | 1 | 0 | 0 | 9110 | AEX65549.1 | P17763 |
| DLLALSADQALQPPCPNLVSYSS | 1 | 0 | 0 | 9157 | AAA96674.1 | P03381 |
| DLPSGFNTLKPIFKLPL | 1 | 0 | 0 | 9242 | P59594.1 | P59594 |
| DLSNCMVALGELKLA | 1 | 0 | 0 | 9281 | P08362.1 | Q786F2 |
| DLVKFISDKIKFLNP | 1 | 0 | 0 | 9319 | P08362.1 | Q786F2 |
| DMIAAYTAALVSGTATA | 1 | 0 | 0 | 9365 | NP_828851.1 | P59594 |
| DNNDSGHSTGADTG | 1 | 0 | 0 | 9511 | P26661.3 | P27958 |
| DPDYVPPVVHGCPLPPT | 1 | 0 | 0 | 9619 | BAA01583.1 | P27958 |
| DPQARVAIKSLT | 1 | 0 | 0 | 9733 | P26664.3 | P27958 |
| DPRRRSRNLGKVIDTLTCGL | 1 | 0 | 0 | 9751 | BAA02670.1 | P27958 |
| DPVIDRLYLSSHRGV | 1 | 0 | 0 | 9804 | P08362.1 | Q786F2 |
| DQRPYCWHYPPKPCGIVPA | 1 | 0 | 0 | 9899 | AAA45676.1 | P27958 |
| DQYCADVAAEELMNA | 1 | 0 | 0 | 9929 | P06830.1 | Q786F2 |
| DRCTTFDDVQAPNYTQH | 1 | 0 | 0 | 9956 | P59594.1 | P59594 |
| DRLNQLESKVSGKGQQQQ | 1 | 0 | 0 | 10010 | AAS48456.1 | P59595 |
| DRSGAPTYSWGENDTDVFVL | 1 | 0 | 0 | 10050 | AAA45676.1 | P27958 |
| DRTAVTGASYFTSVDQSSVH | 1 | 0 | 0 | 10057 | P08617.1 | P08617 |
| DSAVAEFFQSFPSGEPSNSK | 1 | 0 | 0 | 10080 | P08617.1 | P08617 |
| DSGCVVSWKNKELKC | 1 | 0 | 0 | 10125 | AAB33855.1 | P17763 |
| DSGSRTPPDELALS | 1 | 0 | 0 | 10131 | P26660.3 | P27958 |
| DSITWQLQAAVLHV | 1 | 0 | 0 | 10149 | P26660.3 | P27958 |
| DSLNTGWLAGLFYHHKFNSS | 1 | 0 | 0 | 10167 | AAA45676.1 | P27958 |
| DSPDADLIEANLLWRQEMG | 1 | 0 | 0 | 10200 | P26662.3 | P27958 |
| DSRGAILRRQYNLSTSPLTSSVATG | 1 | 0 | 0 | 10221 | P29326.1 | Q81871 |
| DTGCAIDITRKEMRC | 1 | 0 | 0 | 10340 | NP_775667.1 | P0DOH9 |
| DVESYSSMPPLEGEPGDPDL | 1 | 0 | 0 | 10584 | BAB18806.1 | P27958 |
| DVKFPGGGQIVGGVYLLPRRGPRLG | 1 | 0 | 0 | 10640 | P26664.3 | P27958 |
| DVNAIDIEEPSGQT | 1 | 0 | 0 | 10677 | P27313.1 | P27313 |
| DVTRIESDSKVIVL | 1 | 0 | 0 | 10753 | P26661.3 | P27958 |
| DVTRIESESKVVVL | 1 | 0 | 0 | 10754 | P26660.3 | P27958 |
| DVVLMKVPTIPKFRDITQHF | 1 | 0 | 0 | 10774 | P08617.1 | P08617 |
| DVVRDLPSGFNTLKPI | 1 | 0 | 0 | 10779 | NP_828851.1 | P59594 |
| DVVSKLPLAVMG | 1 | 0 | 0 | 10781 | P27958.3 | P27958 |
| DVVSTLPQAVMG | 1 | 0 | 0 | 10783 | P29846.3 | P27958 |
| DVVSTLPQVVMG | 1 | 0 | 0 | 10784 | P26663.3 | P27958 |
| DVVTKLPLAVMG | 1 | 0 | 0 | 10787 | P26664.3 | P27958 |
| DWICEVLSDFKTWL | 1 | 0 | 0 | 10812 | P26664.3 | P27958 |
| DWICTVLTDFKTWL | 1 | 0 | 0 | 10813 | AAA45721.1 | P27958 |
| DWVCSILTDFKNWL | 1 | 0 | 0 | 10832 | P26661.3 | P27958 |
| DWVCSVLADFKAWL | 1 | 0 | 0 | 10833 | Q81258.3 | P27958 |
| DWVCTILTDFKNWL | 1 | 0 | 0 | 10834 | P26660.3 | P27958 |
| EAAAPVVESKWRAL | 1 | 0 | 0 | 10983 | P26662.3 | P27958 |
| EAHFTDPASIAARGY | 1 | 0 | 0 | 11063 | P17763.2 | P17763 |
| EAMTRYSAPPGDPPQPEYD | 1 | 0 | 0 | 11139 | P26662.3 | P27958 |
| EAMYTPHTVLQAVGA | 1 | 0 | 0 | 11142 | P0DTD1.1 | P0DTD1 |
| EASLPYGANKEGIVWVA | 1 | 0 | 0 | 11189 | AAS48456.1 | P59595 |
| EATSGLVMLCIHGSPVNSYTNTPYT | 1 | 0 | 0 | 11204 | BAA01867.1 | Q81871 |
| EDGVNYATGNLPGCSFSIFLLALLS | 1 | 0 | 0 | 11341 | P26664.3 | P27958 |
| EDINGIRRPKHLYV | 1 | 0 | 0 | 11354 | P27313.1 | P27313 |
| EDPVLAKKVPETFPELKPGE | 1 | 0 | 0 | 11420 | P08617.1 | P08617 |
| EDSITIPYQGSGKGV | 1 | 0 | 0 | 11441 | P08362.1 | Q786F2 |
| EEACKLTPPHSAKSKFGYG | 1 | 0 | 0 | 11488 | P26662.3 | P27958 |
| EEDEREISVPAEIL | 1 | 0 | 0 | 11543 | P26664.3 | P27958 |
| EEEEEEEGAGDGE | 1 | 0 | 0 | 11576 | NP_044666.1 | P06484 |
| EEEEGAGDGEHLE | 1 | 0 | 0 | 11579 | NP_044666.1 | P06484 |
| EELGEEIRLKVFVLG | 1 | 0 | 0 | 11748 | AAP79798.1 | Q69027 |
| EEMLDNRATLQAIAS | 1 | 0 | 0 | 11787 | P0DTC1.1 | P0DTD1 |
| EFDEMEACAPHLPYIEQGMQ | 1 | 0 | 0 | 11963 | BAA20975.1 | P27958 |
| EFDEMEECASHLPYIEQGMH | 1 | 0 | 0 | 11966 | AAC53941.1 | P27958 |
| EFDEMEECASHLPYIEQGMQ | 1 | 0 | 0 | 11967 | AAC53954.1 | P27958 |
| EFDEMEECASHLPYIEQGVQ | 1 | 0 | 0 | 11968 | AAA75355.1 | P27958 |
| EFDEMEECASQLPYIEQGMQ | 1 | 0 | 0 | 11969 | BAA01583.1 | P27958 |
| EFDEMEECGSHLPYIEQGMQ | 1 | 0 | 0 | 11970 | AAC53960.1 | P27958 |
| EFDEMEECPSHLPYIEQGMQ | 1 | 0 | 0 | 11971 | AAC53929.1 | P27958 |
| EFDEMEECSQHLPYIEQGMM | 1 | 0 | 0 | 11974 | AAA45677.1 | P27958 |
| EFEPVGSAPPSEGE | 1 | 0 | 0 | 11987 | P26661.3 | P27958 |
| EFYEKVTGTDLELIQILKDH | 1 | 0 | 0 | 12111 | P07299.1 | Q9PZT0 |
| EGAGDGEHLEGGD | 1 | 0 | 0 | 12118 | NP_044666.1 | P06484 |
| EGGDGTRDTLPQS | 1 | 0 | 0 | 12169 | NP_044666.1 | P06484 |
| EGHRMAWDMMMNWS | 1 | 0 | 0 | 12184 | SRC279960 | P27958 |
| EHLEGGDGTRDTL | 1 | 0 | 0 | 12346 | NP_044666.1 | P06484 |
| EIDKGIYQTSNFRVVPS | 1 | 0 | 0 | 12417 | P59594.1 | P59594 |
| EIRQVGDFHYVTGM | 1 | 0 | 0 | 12571 | P26664.3 | P27958 |
| EKLNNLLEGNDSDNDLSLEDF | 1 | 0 | 0 | 12799 | P03421.1 | O42062 |
| EKVTGTDLELIQILKDHYNI | 1 | 0 | 0 | 12901 | P07299.1 | Q9PZT0 |
| ELAAALRGMGINAV | 1 | 0 | 0 | 12921 | BAA03177.1 | P27958 |
| ELAAKLSGLGLNAV | 1 | 0 | 0 | 12925 | Q9WMX2.3 | P27958 |
| ELAAKLVALGINAV | 1 | 0 | 0 | 12927 | P26664.3 | P27958 |
| ELATKTFGSSGSSAVDSGTA | 1 | 0 | 0 | 12955 | P26662.3 | P27958 |
| ELGAFFSILQDMR | 1 | 0 | 0 | 13064 | AAO86636.1 | O36307 |
| ELGLISIEAASDI | 1 | 0 | 0 | 13080 | AAO86638.1 | Q9E006 |
| ELKLAALCHGEDSIT | 1 | 0 | 0 | 13126 | P08362.1 | Q786F2 |
| ELRQLAQSLIDTKVKEISNQEPLKL | 1 | 0 | 0 | 13253 | AAO86636.1 | O36307 |
| ELRYSWKTWGKAKMLSTELH | 1 | 0 | 0 | 13267 | CAA78918.1 | P17763 |
| ELYDTLKKVKKTV | 1 | 0 | 0 | 13326 | AAO86638.1 | Q9E006 |
| EMGEAAGIFMTATPP | 1 | 0 | 0 | 13373 | P14340.2 | P17763 |
| EPEQVELQPPPQGG | 1 | 0 | 0 | 13641 | P26660.3 | P27958 |
| EPEYFNSVCRLMKTI | 1 | 0 | 0 | 13646 | P0DTD1.1 | P0DTD1 |
| EPHDWTKNITDKIDQ | 1 | 0 | 0 | 13658 | ALG01342.1 | Q05320 |
| EPKKDKKKKTDEAQPL | 1 | 0 | 0 | 13680 | NP_828858.1 | P59595 |

| | | | | | | |
|---|---|---|---|---|---|---|
| EPLRTSVDKPGSKKTQGEKF | 1 | 0 | 0 | 13710 | P08617.1 | P08617 |
| ERSDLEPSIPSEYM | 1 | 0 | 0 | 14074 | P26660.3 | P27958 |
| ESAGTQEDAAALRAFTEAMT | 1 | 0 | 0 | 14110 | P26662.3 | P27958 |
| ESLTTTSTALGKLQDVV | 1 | 0 | 0 | 14208 | NP_828851.1 | P59594 |
| ETRTTNQFLAVSKGN | 1 | 0 | 0 | 14536 | P08362.1 | Q786F2 |
| EVDGDVKLSSNLVIL | 1 | 0 | 0 | 14641 | P08362.1 | Q786F2 |
| EVGSHQVEPLRTSVDKPGSK | 1 | 0 | 0 | 14704 | P08617.1 | P08617 |
| EVIAPAVQTNWQKL | 1 | 0 | 0 | 14721 | P26664.3 | P27958 |
| EVKPGTPAQEIEML | 1 | 0 | 0 | 14751 | P27313.1 | P27313 |
| EVLYEAFDEMEECASKAALI | 1 | 0 | 0 | 14774 | Q9IZA3.1 | P27958 |
| EVRRVGDFHYITGA | 1 | 0 | 0 | 14865 | Q81258.3 | P27958 |
| EVTQHGSFSYVTGL | 1 | 0 | 0 | 14887 | P26661.3 | P27958 |
| EVTQHGSYHYITGL | 1 | 0 | 0 | 14888 | P26660.3 | P27958 |
| EVTRVGDFHYVTGM | 1 | 0 | 0 | 14889 | P26662.3 | P27958 |
| FAGNHVSPTHYVPESDAAAR | 1 | 0 | 0 | 15184 | AAA86907.1 | P27958 |
| FCIDPTKGGKKPAR | 1 | 0 | 0 | 15328 | P26661.3 | P27958 |
| FCVDPTKGGKKAAR | 1 | 0 | 0 | 15352 | P26660.3 | P27958 |
| FCVQPEKGGRKPAR | 1 | 0 | 0 | 15356 | P26663.3 | P27958 |
| FDFQVFPTKYHSGRLLFCFV | 1 | 0 | 0 | 15384 | P08617.1 | P08617 |
| FDIIAGAHWGILAGLA | 1 | 0 | 0 | 15407 | AAA45546.1 | P27958 |
| FDPLVAEEDEREIS | 1 | 0 | 0 | 15439 | P26664.3 | P27958 |
| FEKMVSLLSVLLSMQ | 1 | 0 | 0 | 15579 | P0DTC1.1 | P0DTD1 |
| FEPLRAETDDVEPS | 1 | 0 | 0 | 15615 | Q81258.3 | P27958 |
| FFAVSKPMGTQTHTMIF | 1 | 0 | 0 | 15783 | P59594.1 | P59594 |
| FFSTFKCYGVSATKLND | 1 | 0 | 0 | 15903 | P59594.1 | P59594 |
| FFTELDGVRLHRFA | 1 | 0 | 0 | 15913 | P26664.3 | P27958 |
| FFTEVDGVRLHRYA | 1 | 0 | 0 | 15918 | P26662.3 | P27958 |
| FFTPKESEQLKVIKTFEGIL | 1 | 0 | 0 | 15925 | AAO86638.1 | Q9E006 |
| FGCTWMNSTGFTKVCGAPPC | 1 | 0 | 0 | 15959 | AAA45676.1 | P27958 |
| FHPPSSNQCESTPPIAMDAW | 1 | 0 | 0 | 16132 | AAA45412.1 | Q85601 |
| FKDKVLGLLQRASQQEAEIR | 1 | 0 | 0 | 16354 | Q81487.3 | P27958 |
| FKEKILGLLQRATQQQAVIE | 1 | 0 | 0 | 16359 | AAC53966.1 | P27958 |
| FKEKVLGFISTTGQKAETLK | 1 | 0 | 0 | 16360 | AAA65789.1 | P27958 |
| FKEKVLGLLQRATQQQAVIE | 1 | 0 | 0 | 16363 | AAC03058.1 | P27958 |
| FKHLREFVFKNKDGFLY | 1 | 0 | 0 | 16376 | P59594.1 | P59594 |
| FKNKDGFLYVYKGYQPI | 1 | 0 | 0 | 16417 | P59594.1 | P59594 |
| FKQKALGLLQTASR | 1 | 0 | 0 | 16437 | P26664.3 | P27958 |
| FKQKALGLLQTSTKQAEAAA | 1 | 0 | 0 | 16448 | AAC53938.1 | P27958 |
| FKQKARGLLQTATKQAEAAA | 1 | 0 | 0 | 16449 | SRC279960 | P27958 |
| FLIDGPDTSECPNERRA | 1 | 0 | 0 | 16649 | Q6TFL5 | P17763 |
| FLNTEPSQLPPTAPPLLPHSNLDHI | 1 | 0 | 0 | 16790 | AAL05536.1 | P03381 |
| FLVNAWKSKKCP | 1 | 0 | 0 | 16957 | P29846.3 | P27958 |
| FLVNAWKSKKNP | 1 | 0 | 0 | 16958 | AAL91977.1 | P27958 |
| FLVNTWKSKKCP | 1 | 0 | 0 | 16961 | P26662.3 | P27958 |
| FLVNTWKSKKNP | 1 | 0 | 0 | 16962 | P26663.3 | P27958 |
| FLVQAWKSKKTP | 1 | 0 | 0 | 16963 | P26664.3 | P27958 |
| FNASDSVGQQIKVIPVDPYF | 1 | 0 | 0 | 17129 | P08617.1 | P08617 |
| FNMVYMPASWVMRIM | 1 | 0 | 0 | 17203 | P0DTC1.1 | P0DTD1 |
| FNVPIKEAGEDCHAP | 1 | 0 | 0 | 17243 | P08362.1 | Q786F2 |
| FPELKPGESRHTSDHMSIYK | 1 | 0 | 0 | 17278 | P08617.1 | P08617 |
| FPTKYHSGRLLFCFVPGNEL | 1 | 0 | 0 | 17426 | P08617.1 | P08617 |
| FQDLSDNPCKVDL | 1 | 0 | 0 | 17466 | AAO86638.1 | Q9E006 |
| FQQFGRDVSDFTDSVRDPKT | 1 | 0 | 0 | 17524 | P59594.1 | P59594 |
| FQSFPSGEPSNSKLSGFFQS | 1 | 0 | 0 | 17531 | P08617.1 | P08617 |
| FQSMRKHTPYRNGCLEPTLG | 1 | 0 | 0 | 17535 | AAG48731.1 | P03410 |
| FRDLTWCINPPERIK | 1 | 0 | 0 | 17572 | P08362.1 | Q786F2 |
| FRSDTLYLTQDLFLPFY | 1 | 0 | 0 | 17644 | P59594.1 | P59594 |
| FSILMAQDTLDQNL | 1 | 0 | 0 | 17743 | P26660.3 | P27958 |
| FSPGPSQKIQLVNT | 1 | 0 | 0 | 17801 | BAD73977.1 | P27958 |
| FTDSVRDPKTSEILDIS | 1 | 0 | 0 | 17929 | P59594.1 | P59594 |
| FTFSPRRHWTTQGCNCSIYP | 1 | 0 | 0 | 17960 | AAA45676.1 | P27958 |
| FTKVCGAPPCVIGGAGNNTL | 1 | 0 | 0 | 17997 | AAA45676.1 | P27958 |
| FTKWTWETKGDL | 1 | 0 | 0 | 17998 | P27312.1 | P27312 |
| FTPGAKQNIQLINT | 1 | 0 | 0 | 18037 | Q03463.1 | P27958 |
| FTQCSMNVVSKTLFRKSPIY | 1 | 0 | 0 | 18053 | P08617.1 | P08617 |
| FVFNGTSWFITQRNFFS | 1 | 0 | 0 | 18161 | NP_828851.1 | P59594 |
| FVGAGLAGAAIGSV | 1 | 0 | 0 | 18173 | P26664.3 | P27958 |
| FVVSGLAGAAIGGI | 1 | 0 | 0 | 18320 | Q81258.3 | P27958 |
| FWRGDLVFDFQVFPTKYHSG | 1 | 0 | 0 | 18364 | P08617.1 | P08617 |
| GAGGGAGGAGAGGGAGGAGA | 1 | 0 | 0 | 18588 | P03211.1 | P03211 |
| GAGKALTGLSTGTSQNTRIS | 1 | 0 | 0 | 18595 | P07299.1 | Q9PZT0 |
| GALNTPKDHIGTRNPNNNA | 1 | 0 | 0 | 18671 | AAS48456.1 | P59595 |
| GAQAVARSLDWTKVTLDGR | 1 | 0 | 0 | 18736 | AAG16766.1 | Q81871 |
| GAQITGHVKNGSMRIVGPKT | 1 | 0 | 0 | 18742 | P26662.3 | P27958 |
| GARSLTPCTCGSSD | 1 | 0 | 0 | 18753 | AAA45721.1 | P27958 |
| GCPLPSTKAPPIPPPRRK | 1 | 0 | 0 | 18893 | P26662.3 | P27958 |
| GCQSWTCPYTGAVSSPYWKFQHDVN | 1 | 0 | 0 | 18895 | AAA96674.1 | P03381 |
| GDGEHLEGGDGTR | 1 | 0 | 0 | 18985 | NP_044666.1 | P06484 |
| GDIVQQPSDETAAS | 1 | 0 | 0 | 19007 | P26661.3 | P27958 |
| GDSSNTGAGKALTGLSTGTS | 1 | 0 | 0 | 19119 | P07299.1 | Q9PZT0 |
| GDYSPSCCTLTIGVSSYHSKP | 1 | 0 | 0 | 19192 | AAA96674.1 | P03381 |
| GEGAPPAKRARTDQME | 1 | 0 | 0 | 19269 | AAA45723.1 | P29996 |
| GEGAVQWMNRLIAFASRGNH | 1 | 0 | 0 | 19276 | BAB18806.1 | P27958 |
| GEGPVQWMNRLIAFASRGNH | 1 | 0 | 0 | 19289 | Q81495.3 | P27958 |
| GEIRPSAPPLPPVA | 1 | 0 | 0 | 19311 | Q03499.1 | Q81870 |
| GFDEMEECASHLPYIEQGMQ | 1 | 0 | 0 | 19511 | AAC53946.1 | P27958 |
| GFFGAIAGFIE | 1 | 0 | 0 | 19520 | SRC279975 | P03460 |
| GFMKQYGECLGDINARD | 1 | 0 | 0 | 19585 | NP_828851.1 | P59594 |
| GFYTTTGIGYQPYRVVV | 1 | 0 | 0 | 19657 | P59594.1 | P59594 |
| GGAKTPSDAVQNILQKIEKIKNTEE | 1 | 0 | 0 | 19678 | AAA45992.1 | Q6SW99 |
| GGAKTPSDAVQNISQKIEKIKNTEE | 1 | 0 | 0 | 19679 | SRC7948 | Q6SW99 |

| | | | | | | |
|---|---|---|---|---|---|---|
| GGESSEELSESSFFNLIT | 1 | 0 | 0 | 19723 | P07298.1 | Q9PZT1 |
| GGFWGDRVDSQPFAIPYIHPTNPFA | 1 | 0 | 0 | 19736 | P29326.1 | Q81871 |
| GGIKITHFTTWTSIPTLAAQ | 1 | 0 | 0 | 19887 | P08617.1 | P08617 |
| GGITGSMSLASGKSLLHEVDKD | 1 | 0 | 0 | 19891 | AAL05536.1 | P03381 |
| GGTYVTGGTMAKNTLGITSLFSPG | 1 | 0 | 0 | 20062 | Q9WMX2.3 | P27958 |
| GGYYSASYSDPCSLK | 1 | 0 | 0 | 20108 | AAL05536.1 | P03381 |
| GHEWVTDFKGKTVWF | 1 | 0 | 0 | 20130 | P14340.2 | P17763 |
| GHIELEGKPALVPDKEVLYQ | 1 | 0 | 0 | 20141 | Q81258.3 | P27958 |
| GHIELGGKPAIVPDKEVLYQ | 1 | 0 | 0 | 20143 | Q81594 | P27958 |
| GHIELGGKPALVPDKEVLYQ | 1 | 0 | 0 | 20144 | AAC03058.1 | P27958 |
| GHIELGGKPALVPDRQVLYQ | 1 | 0 | 0 | 20145 | Q81487.3 | P27958 |
| GHWMDATFNTKTA | 1 | 0 | 0 | 20197 | AAO86638.1 | Q9E006 |
| GHWSRDCTQPRPPPGPCPLCQDP | 1 | 0 | 0 | 20199 | AAD34841.1 | P03346 |
| GIGSAVVSFAIKWE | 1 | 0 | 0 | 20285 | P26662.3 | P27958 |
| GIGVTQNVLYENQKQIA | 1 | 0 | 0 | 20288 | NP_828851.1 | P59594 |
| GIIWVAVILHPRLV | 1 | 0 | 0 | 20310 | P26661.3 | P27958 |
| GITLKQATTAPCAVMDITGV | 1 | 0 | 0 | 20462 | P08617.1 | P08617 |
| GKNRAINMLKRGLPR | 1 | 0 | 0 | 20650 | NP_775663.1 | P19110 |
| GLDHILSLADIEEEQMIQSV | 1 | 0 | 0 | 20797 | P08617.1 | P08617 |
| GLDLLPWEQGGLCKAIQEQCCFLN | 1 | 0 | 0 | 20803 | SRC276213 | P03383 |
| GLLRYHTYARFG | 1 | 0 | 0 | 21026 | NP_740550.1 | P08617 |
| GLVRKNLVQFGVGEKNGCVR | 1 | 0 | 0 | 21210 | P08617.1 | P08617 |
| GMGVSCTVTREDGTNRR | 1 | 0 | 0 | 21277 | P06830.1 | Q786F2 |
| GMMLAEQFKQKALGLLQTAS | 1 | 0 | 0 | 21313 | P26664.3 | P27958 |
| GNFGDQDLIRQGTDYKHW | 1 | 0 | 0 | 21424 | AAS48456.1 | P59595 |
| GNKTLAGPKGPVTQ | 1 | 0 | 0 | 21459 | P26661.3 | P27958 |
| GNKTLAGSRGPVTQ | 1 | 0 | 0 | 21460 | P26660.3 | P27958 |
| GNQHVDDRPSPAPAPKRALGTL | 1 | 0 | 0 | 21502 | ADE95846.1 | Q913Y7 |
| GPRLGVRAIRKTSERSQPRG | 1 | 0 | 0 | 21756 | BAA02670.1 | P27958 |
| GPSYGFQYSPGQRVEFLVNT | 1 | 0 | 0 | 21804 | P26662.3 | P27958 |
| GPVYCFTPSPVVVGTTDR | 1 | 0 | 0 | 21831 | P27958.3 | P27958 |
| GPVYVSDSVTLVNVATGAQAVARSL | 1 | 0 | 0 | 21834 | P29326.1 | Q81871 |
| GQIFSRSASPIPR | 1 | 0 | 0 | 21888 | AAD50663.1 | P03362 |
| GQLKLNWFKKGSSIG | 1 | 0 | 0 | 21908 | P14340.2 | P17763 |
| GQLSDHPHALSSSSSHAEPR | 1 | 0 | 0 | 21912 | P07299.1 | Q9PZT0 |
| GQRVEFLVQAWKSK | 1 | 0 | 0 | 21953 | P26664.3 | P27958 |
| GRGYNVSSIVTMTSQ | 1 | 0 | 0 | 22063 | P06830.1 | Q786F2 |
| GRIILSGKPAIIPDREALYQ | 1 | 0 | 0 | 22074 | AAC53962.1 | P27958 |
| GRIILSGKPAIIPDREVLYQ | 1 | 0 | 0 | 22075 | AAC53965.1 | P27958 |
| GRIILSGKPATIPDREALYR | 1 | 0 | 0 | 22076 | AAC53955.1 | P27958 |
| GRIILSGKPATIPDREAVLY | 1 | 0 | 0 | 22077 | SRC279960 | P27958 |
| GRIILSGKPATIPDREVLYR | 1 | 0 | 0 | 22078 | AAC53938.1 | P27958 |
| GRIILSGKPATVPDREVLYQ | 1 | 0 | 0 | 22079 | AAC53954.1 | P27958 |
| GRIILSGKPAVIPDRELLYR | 1 | 0 | 0 | 22080 | AAC53929.1 | P27958 |
| GRIILSGKPAVIPDREVLYQ | 1 | 0 | 0 | 22081 | AAC53946.1 | P27958 |
| GRIILSGKPAVIPDREVLYR | 1 | 0 | 0 | 22082 | AAC53960.1 | P27958 |
| GRIILSGKPAVVPDREVLYQ | 1 | 0 | 0 | 22083 | ABU97068.1 | P27958 |
| GRIILSGRPAIIPDREALYQ | 1 | 0 | 0 | 22084 | AAC53969.1 | P27958 |
| GRIILSGRPAIIPDREVLYQ | 1 | 0 | 0 | 22086 | AAC53943.1 | P27958 |
| GRIILSGRPAIIPDREVLYR | 1 | 0 | 0 | 22087 | AAC53967.1 | P27958 |
| GRIILSGRPAIVPDRELLYQ | 1 | 0 | 0 | 22088 | AAA72945.1 | P27958 |
| GRIILSGRPAVIPDREVLYQ | 1 | 0 | 0 | 22089 | BAA06303.1 | P27958 |
| GRIILSGRPAVIPDREVLYR | 1 | 0 | 0 | 22090 | AAA75355.1 | P27958 |
| GRIILSGRPAVVPDREVLYQ | 1 | 0 | 0 | 22091 | AAA79971.1 | P27958 |
| GRIILSGRPAVVPDREVLYR | 1 | 0 | 0 | 22092 | BAA20975.1 | P27958 |
| GRIILSGRPVIVPDREVLYR | 1 | 0 | 0 | 22093 | AAC53963.1 | P27958 |
| GRIILTGKPAVIPDREVLYQ | 1 | 0 | 0 | 22094 | AAC53934.1 | P27958 |
| GRIVLSGKPAIIPDREVLYR | 1 | 0 | 0 | 22104 | CAB53095.1 | P27958 |
| GRIVLSGRPAIIPDREVLYR | 1 | 0 | 0 | 22105 | BAA01582.1 | P27958 |
| GRIVLSGRPAVIPDREVLYQ | 1 | 0 | 0 | 22106 | BAA01728.1 | P27958 |
| GRIVLSGSPAIVPDREVLYQ | 1 | 0 | 0 | 22107 | ABU97067.1 | P27958 |
| GRLGIYRYVSTGER | 1 | 0 | 0 | 22121 | P26660.3 | P27958 |
| GRLGTYRYVASGER | 1 | 0 | 0 | 22122 | Q81258.3 | P27958 |
| GRLGTYRYVSSGER | 1 | 0 | 0 | 22123 | BAA03177.1 | P27958 |
| GRLHINQRTVVAPDKEVLYE | 1 | 0 | 0 | 22124 | AAC53939.1 | P27958 |
| GRLHLINQRAVVAPDKEVLY | 1 | 0 | 0 | 22125 | SRC279960 | P27958 |
| GRLHLNDQVVVTPDKEILYE | 1 | 0 | 0 | 22126 | AAP55688.1 | P27958 |
| GRLHLNDRVVVAPDKEILYE | 1 | 0 | 0 | 22128 | BAA01761.1 | P27958 |
| GRLHVNERAVVAPDKEVLYE | 1 | 0 | 0 | 22129 | AAC53944.1 | P27958 |
| GRLHVNQRAVVAPDKEVLYE | 1 | 0 | 0 | 22130 | BAA00792.1 | P27958 |
| GRLVLSGKPAIIPDREVLYR | 1 | 0 | 0 | 22145 | AAC53959.1 | P27958 |
| GRRQPIPKARRPEGRTWAQPGYPWP | 1 | 0 | 0 | 22194 | P26664.3 | P27958 |
| GRVASSTQSLVSWLSQGPSQK | 1 | 0 | 0 | 22229 | BAA14233.1 | P27958 |
| GRVILSGKPAIIPDREVLYR | 1 | 0 | 0 | 22236 | BAA01515.1 | P27958 |
| GRVILSGKPATIPDREVLYQ | 1 | 0 | 0 | 22237 | AAC53937.1 | P27958 |
| GRVILSGRPAIVPDREVLYR | 1 | 0 | 0 | 22238 | AAC53968.1 | P27958 |
| GRVVLSGQPAVIPDREVLYQ | 1 | 0 | 0 | 22247 | ABL63009.1 | P27958 |
| GRWNPQPGVYPPHAA | 1 | 0 | 0 | 22253 | NP_050020.1 | Q9PZT0 |
| GSAGHTTAGLVGLLTPGAKQN | 1 | 0 | 0 | 22269 | AAB67037.1 | P27958 |
| GSAGHTVSGFVSLLAPGAKQN | 1 | 0 | 0 | 22271 | AAA45676.1 | P27958 |
| GSGKTRKYLPAIVRE | 1 | 0 | 0 | 22346 | AEX65549.1 | P17763 |
| GSHLPYIEQGMQLAEQFKQK | 1 | 0 | 0 | 22367 | AAC53960.1 | P27958 |
| GSKTLAGPKGPITQ | 1 | 0 | 0 | 22396 | P26662.3 | P27958 |
| GSQLPCEPEPDVAVLTSML | 1 | 0 | 0 | 22469 | P26662.3 | P27958 |
| GSRTLAGAKHPALQ | 1 | 0 | 0 | 22491 | Q81258.3 | P27958 |
| GSSDLYLVTRHADVIPVRRR | 1 | 0 | 0 | 22497 | P26664.3 | P27958 |
| GSSGSSAVDSGTATGPPDQA | 1 | 0 | 0 | 22505 | P26662.3 | P27958 |
| GSTAHNARTLTGMFSLGARQK | 1 | 0 | 0 | 22536 | P26660.3 | P27958 |
| GTKSLEPCTCGAVD | 1 | 0 | 0 | 22732 | P26660.3 | P27958 |
| GTPSMDALSALLSNTLSLAS | 1 | 0 | 0 | 22786 | AAA45412.1 | Q85601 |
| GTRIRFKDDTSFED | 1 | 0 | 0 | 22799 | P27313.1 | P27313 |

| | | | | | | |
|---|---|---|---|---|---|---|
| GTRTIASPKGPVIQ | 1 | 0 | 0 | 22806 | P26664.3 | P27958 |
| GVLSVDLSLTVELKI | 1 | 0 | 0 | 23048 | P06830.1 | Q786F2 |
| GVLTPSSKRFQPFQQFG | 1 | 0 | 0 | 23051 | P59594.1 | P59594 |
| GVMLAAQMFIVSPQ | 1 | 0 | 0 | 23058 | P26660.3 | P27958 |
| GVPIELQVECFTWDQ | 1 | 0 | 0 | 23096 | P06830.1 | Q786F2 |
| GVSGSSITTLSSGPHSLSPA | 1 | 0 | 0 | 23142 | AAC56957.1 | Q2HR63 |
| GVTRPSAPPLPHVV | 1 | 0 | 0 | 23176 | P69616.1 | Q81870 |
| GWGLHQPPPQIFLKILPQSG | 1 | 0 | 0 | 23246 | P07299.1 | Q9PZT0 |
| GWGLNWDLGLSQWAREALQTGIT | 1 | 0 | 0 | 23247 | AAL05536.1 | P03381 |
| GWVFGSTMNNKSQSVII | 1 | 0 | 0 | 23296 | P59594.1 | P59594 |
| GYKVLVLNPSVAATLGFG | 1 | 0 | 0 | 23392 | P29846.3 | P27958 |
| GYLPISGEMEKVT | 1 | 0 | 0 | 23404 | AAO86638.1 | Q9E006 |
| GYNVSSIVTMTSQGM | 1 | 0 | 0 | 23422 | P08362.1 | Q786F2 |
| GYPWPLYGNEGCGWAGWLLSPRGSR | 1 | 0 | 0 | 23432 | P26664.3 | P27958 |
| GYQPYRVVVLSFELLNA | 1 | 0 | 0 | 23437 | P59594.1 | P59594 |
| GYSDRVRSITLGNS | 1 | 0 | 0 | 23455 | AAB59927.1 | Q86887 |
| HASGHRMAWDMMMNWSPTTA | 1 | 0 | 0 | 23564 | BAA02670.1 | P27958 |
| HDSPTHYVPESDAAARVTQI | 1 | 0 | 0 | 23645 | AAA75355.1 | P27958 |
| HEQQLVTARQKLK | 1 | 0 | 0 | 23727 | AAO86636.1 | O36307 |
| HFKDPQYPVWELTIRVWSEL | 1 | 0 | 0 | 23791 | P08617.1 | P08617 |
| HFLCTFTFNSNNKEYTFPIT | 1 | 0 | 0 | 23797 | P08617.1 | P08617 |
| HGKEELRFPRGQGVPI | 1 | 0 | 0 | 23917 | AAS48456.1 | P59595 |
| HGVRGLTNLFS | 1 | 0 | 0 | 23965 | BAA02347.1 | P27958 |
| HGVRGLTSLFS | 1 | 0 | 0 | 23966 | BAA02343.1 | P27958 |
| HGVTKPKQVIKLDADPVESQ | 1 | 0 | 0 | 23972 | P08617.1 | P08617 |
| HHFQSHGQLSDHPHALSSSS | 1 | 0 | 0 | 23985 | P07299.1 | Q9PZT0 |
| HHPSTNQEQTSLYVQAS | 1 | 0 | 0 | 24012 | P04664.1 | P03452 |
| HIDLLVGSATLCSALYVGDL | 1 | 0 | 0 | 24034 | AAA45676.1 | P27958 |
| HIFYRRFLGKDGRRDPK | 1 | 0 | 0 | 24046 | YP_001129431.1 | Q9QR71 |
| HITHSGMEGMGVSCT | 1 | 0 | 0 | 24104 | P08362.1 | Q786F2 |
| HMTNYLEQPVSNDLS | 1 | 0 | 0 | 24352 | P06830.1 | Q786F2 |
| HNASGKEAKVCTISPIMGYS | 1 | 0 | 0 | 24362 | P07299.1 | Q9PZT0 |
| HPTNPFAPDVTAAAGAGPRVRQPAR | 1 | 0 | 0 | 24518 | P29326.1 | Q81871 |
| HRMAWNMMMNWSPT | 1 | 0 | 0 | 24696 | P27958.3 | P27958 |
| HSDHAESSWVNRGESSRKAY | 1 | 0 | 0 | 24730 | AAK01056.1 | P09259 |
| HSVRGFTSLFSA | 1 | 0 | 0 | 24871 | BAA02337.1 | P27958 |
| HSVRGLTSLFS | 1 | 0 | 0 | 24872 | BAA02342.1 | P27958 |
| HTFDDFCPECRPLGLQGCAFQSTVA | 1 | 0 | 0 | 24910 | P29326.1 | Q81871 |
| HTINHTFGNPVIPFKDG | 1 | 0 | 0 | 24923 | P59594.1 | P59594 |
| HTPGCVPCVREGNSSRCWVA | 1 | 0 | 0 | 24954 | BAA02670.1 | P27958 |
| HTPLFSFLTASPA | 1 | 0 | 0 | 24960 | NP_044666.1 | P06484 |
| HTSSMRGVYYPDEIFRS | 1 | 0 | 0 | 24978 | P59594.1 | P59594 |
| HTVMPLSAPTLVPQE | 1 | 0 | 0 | 24998 | P0DTD1.1 | P0DTD1 |
| HTYARFGIEIQVQINPTPFQ | 1 | 0 | 0 | 25004 | P08617.1 | P08617 |
| HTYSPHELSRVAAT | 1 | 0 | 0 | 25009 | P26661.3 | P27958 |
| HTYTPHELTRVASA | 1 | 0 | 0 | 25010 | P26660.3 | P27958 |
| HVAPTHYVTESDASQRVTQL | 1 | 0 | 0 | 25021 | AAU89634.1 | P27958 |
| HVAPTHYVVESDASQRVTQV | 1 | 0 | 0 | 25022 | BAA01761.1 | P27958 |
| HVDTSYECDIPIGAGIC | 1 | 0 | 0 | 25031 | NP_828851.1 | P59594 |
| HVIWDVGLQSS | 1 | 0 | 0 | 25054 | CAA29172.1 | Q86887 |
| HVSPAHYVPESDAAARVTAL | 1 | 0 | 0 | 25092 | Q81495.3 | P27958 |
| HVSPRHYVPESEPAARVTQI | 1 | 0 | 0 | 25093 | AAA45721.1 | P27958 |
| HVSPTHYVPESDAAAKVTAL | 1 | 0 | 0 | 25095 | Q68870.1 | P27958 |
| HVSPTHYVPESDAAARVTQI | 1 | 0 | 0 | 25096 | AAA86907.1 | P27958 |
| HVWIPPLNARGGRDAIILLMCAV | 1 | 0 | 0 | 25119 | AAA72945.1 | P27958 |
| HYTNESKFKFILE | 1 | 0 | 0 | 25200 | AAO86638.1 | Q9E006 |
| IAAEQDRNTREVFAQVK | 1 | 0 | 0 | 25219 | NP_828851.1 | P59594 |
| IADYNYKLPDDFMGCVL | 1 | 0 | 0 | 25250 | P59594.1 | P59594 |
| IAGLIAIVMVTILLCCM | 1 | 0 | 0 | 25293 | NP_828851.1 | P59594 |
| IAHQFKEKILGLLQRATQQQ | 1 | 0 | 0 | 25308 | AAC53966.1 | P27958 |
| IAHQFKEKVLGLLQRATQQQ | 1 | 0 | 0 | 25309 | AAC03058.1 | P27958 |
| IAPGQTGVIADYNYKLP | 1 | 0 | 0 | 25378 | P59594.1 | P59594 |
| IAQQFKDKVLGLLQRASQQE | 1 | 0 | 0 | 25397 | Q81487.3 | P27958 |
| IAYFSMVGNWAKVLVVLLLFAGVDA | 1 | 0 | 0 | 25480 | P26664.3 | P27958 |
| ICESAGVQEDAASL | 1 | 0 | 0 | 25497 | P26664.3 | P27958 |
| IDANDCLVTPSVK | 1 | 0 | 0 | 25538 | AAO86638.1 | Q9E006 |
| IDAVNKRGRKQNKRG | 1 | 0 | 0 | 25541 | NP_775663.1 | P19110 |
| IDMPSWPPVQSTSSPGTPSM | 1 | 0 | 0 | 25631 | AAA45412.1 | Q85601 |
| IDQKVKEISNQEPL | 1 | 0 | 0 | 25653 | P27313.1 | P27313 |
| IDRLITGRLQSLQTY | 1 | 0 | 0 | 25662 | P0DTC2.1 | P0DTC2 |
| IEEVALGHEGEIPF | 1 | 0 | 0 | 25791 | P26661.3 | P27958 |
| IEEVALGQEGEIPF | 1 | 0 | 0 | 25792 | P26660.3 | P27958 |
| IEEVALSNTGEIPF | 1 | 0 | 0 | 25794 | P26662.3 | P27958 |
| IEEVALSTTGEIPF | 1 | 0 | 0 | 25795 | P26664.3 | P27958 |
| IEKTNEKFHQIEK | 1 | 0 | 0 | 25863 | 0704243B | P03452 |
| IEPSWADVKKDLISY | 1 | 0 | 0 | 25929 | P14340.2 | P17763 |
| IGTRNPNNNAATVLQL | 1 | 0 | 0 | 26329 | NP_828858.1 | P59595 |
| IGVTGVIIAVIALFC | 1 | 0 | 0 | 26343 | ALG01342.1 | Q05320 |
| IIHDFVDKTLPDQGD | 1 | 0 | 0 | 26510 | ALG01342.1 | Q05320 |
| IIMDEAHFTDPASIA | 1 | 0 | 0 | 26592 | P17763.2 | P17763 |
| IIMYAPTLWARMIL | 1 | 0 | 0 | 26600 | P26662.3 | P27958 |
| IINNSTNVVIRACNFEL | 1 | 0 | 0 | 26610 | P59594.1 | P59594 |
| IIPDREVLYREFDEMEECSQ | 1 | 0 | 0 | 26623 | P26664.3 | P27958 |
| IKRRLRTLILAPTRV | 1 | 0 | 0 | 26895 | ACA48751.1 | P17763 |
| IKVIPVDPYFFQMTNTNPDQ | 1 | 0 | 0 | 26927 | P08617.1 | P08617 |
| ILAPTRVVAAEMEEA | 1 | 0 | 0 | 26965 | P14340.2 | P17763 |
| ILEENMEVEIWTREGEKKKL | 1 | 0 | 0 | 27016 | NP_740321.1 | P17763 |
| ILRKSRRFAQALPV | 1 | 0 | 0 | 27291 | P26664.3 | P27958 |
| ILRRHVGQGEGAVQWMNRLI | 1 | 0 | 0 | 27307 | AAF59945.1 | P27958 |
| ILYEAFDEMEECASKAALIE | 1 | 0 | 0 | 27422 | AAF59945.1 | P27958 |
| IMASENSSAMVQVHS | 1 | 0 | 0 | 27448 | AGB56758.1 | Q05320 |

| | | | | | | |
|---|---|---|---|---|---|---|
| IMASKTVGTAEEKL | 1 | 0 | 0 | 27451 | P27313.1 | P27313 |
| IMGGECPTAEDMVN | 1 | 0 | 0 | 27494 | Q81258.3 | P27958 |
| IMGYSTPWRYLDFNALNLFF | 1 | 0 | 0 | 27498 | P07299.1 | Q9PZT0 |
| IMSGEKPSMEDVVN | 1 | 0 | 0 | 27554 | P26660.3 | P27958 |
| IMSGEKPTVEDVVN | 1 | 0 | 0 | 27555 | P26661.3 | P27958 |
| IMTVTMTFKLGPRKATGRWN | 1 | 0 | 0 | 27573 | P07299.1 | Q9PZT0 |
| INAYTTGPCTPSPA | 1 | 0 | 0 | 27595 | P26662.3 | P27958 |
| INCYTEGPCVPKPP | 1 | 0 | 0 | 27598 | P26661.3 | P27958 |
| INCYTEGQCVPKPA | 1 | 0 | 0 | 27599 | P26660.3 | P27958 |
| INEYTTGPSTPCPS | 1 | 0 | 0 | 27627 | Q81258.3 | P27958 |
| INLRTNNSATIV | 1 | 0 | 0 | 27685 | AAA74400.1 | Q86887 |
| INNVNLYYVFSKIFGKSPVF | 1 | 0 | 0 | 27703 | P08617.1 | P08617 |
| INPTPFQQGGLICAMVPGDQ | 1 | 0 | 0 | 27722 | P08617.1 | P08617 |
| INQRTVVAPDKEVLYEAFDE | 1 | 0 | 0 | 27730 | AAC53939.1 | P27958 |
| IPHDIDLGESRVVIQDYDNQHEQDR | 1 | 0 | 0 | 27861 | P29326.1 | Q81871 |
| IPKPFFRDEVSFCV | 1 | 0 | 0 | 27903 | P26660.3 | P27958 |
| IPPPRRKRTVVLTE | 1 | 0 | 0 | 27980 | P26662.3 | P27958 |
| IPRFKVSPYLFNVPI | 1 | 0 | 0 | 28041 | P08362.1 | Q786F2 |
| IQQVVDADSKIVQLS | 1 | 0 | 0 | 28211 | P0DTC1.1 | P0DTD1 |
| IRFKDDTSFEDING | 1 | 0 | 0 | 28291 | P27313.1 | P27313 |
| IRSAEVVSRMLVH | 1 | 0 | 0 | 28396 | AAO86638.1 | Q9E006 |
| IRSEELSFTVVSNGA | 1 | 0 | 0 | 28398 | NP_066246.1 | Q05320 |
| IRYQTTATKSEHTGR | 1 | 0 | 0 | 28423 | QTX93491.1 | P17763 |
| ISESQGNEEDERNL | 1 | 0 | 0 | 28482 | P26661.3 | P27958 |
| ISESQGTEEDERNL | 1 | 0 | 0 | 28483 | P26660.3 | P27958 |
| ISFHFQMEKQVIH | 1 | 0 | 0 | 28492 | AAO86638.1 | Q9E006 |
| ISGHRMAWDMMMNWS | 1 | 0 | 0 | 28509 | ABR25251.1 | P27958 |
| ISPCAFGGVSVITPGTN | 1 | 0 | 0 | 28643 | NP_828851.1 | P59594 |
| ISYANGSGLDERPY | 1 | 0 | 0 | 28786 | P27958.3 | P27958 |
| ITGHRMAWDMMLNWS | 1 | 0 | 0 | 28910 | AAP55688.1 | P27958 |
| ITGHRMAWDMMVNWS | 1 | 0 | 0 | 28914 | SRC279960 | P27958 |
| ITGQRMAWDMMMNWS | 1 | 0 | 0 | 28922 | AEI00313.1 | P27958 |
| ITLPQDAVSRTQRRGRTGRG | 1 | 0 | 0 | 28982 | AAA45676.1 | P27958 |
| ITRVESENKVVILDSFDPLV | 1 | 0 | 0 | 29077 | AGN33396.1 | P27958 |
| ITTIEDPVLAKKVPETFPEL | 1 | 0 | 0 | 29116 | P08617.1 | P08617 |
| ITVHTGDQHQVGNET | 1 | 0 | 0 | 29132 | QHR82515.1 | P17763 |
| ITYNESHSSDQRPY | 1 | 0 | 0 | 29171 | Q9WMX2.3 | P27958 |
| IVDLMCHATFTMRLL | 1 | 0 | 0 | 29233 | P17763.2 | P17763 |
| IVWVATEGALNTPKDHI | 1 | 0 | 0 | 29504 | AAS48456.1 | P59595 |
| KAAASKVKANLLSV | 1 | 0 | 0 | 29744 | P26664.3 | P27958 |
| KAGSTVYAVTAVTSIFSVG | 1 | 0 | 0 | 29812 | SRC279960 | P27958 |
| KAISQIQESLTTTSTAL | 1 | 0 | 0 | 29832 | NP_828851.1 | P59594 |
| KAKLMPQLPGIPFV | 1 | 0 | 0 | 29839 | P26664.3 | P27958 |
| KALENKKKQLSAGGKNLS | 1 | 0 | 0 | 29852 | CAA44973.1 | P29996 |
| KALTGLSTGTSQNTRISLRP | 1 | 0 | 0 | 29879 | P07299.1 | Q9PZT0 |
| KANGNGSGDNGDTTW | 1 | 0 | 0 | 29899 | P03277.3 | P03277 |
| KARRSEGRSWAQ | 1 | 0 | 0 | 29938 | Q81258.3 | P27958 |
| KATAACRAAKLQDCTMLVN | 1 | 0 | 0 | 29957 | P26662.3 | P27958 |
| KDISQLTQAIVKNHKNLLKIAQY | 1 | 0 | 0 | 30178 | AAA96674.1 | P03381 |
| KDNRIPSYGVLSVDL | 1 | 0 | 0 | 30220 | P08362.1 | Q786F2 |
| KEFSEVEGRIQDLEKYV | 1 | 0 | 0 | 30386 | 0704243B | P03452 |
| KEGEKKKLRPRWLDA | 1 | 0 | 0 | 30395 | QHR82521.1 | P17763 |
| KEGERKKLRPRWLDA | 1 | 0 | 0 | 30397 | P17763.2 | P17763 |
| KERASRVKARMLTI | 1 | 0 | 0 | 30585 | Q81258.3 | P27958 |
| KERSSLRYGNVLDV | 1 | 0 | 0 | 30597 | P27313.1 | P27313 |
| KEVPALTAVETGAT | 1 | 0 | 0 | 30661 | ACK37265.1 | P03300 |
| KFLNPDREYDFRDLT | 1 | 0 | 0 | 30774 | P08362.1 | Q786F2 |
| KFQPESPARLASAILNA | 1 | 0 | 0 | 30831 | Q6TFL5 | P17763 |
| KGIYQTSNFRVVPSGDVVRF | 1 | 0 | 0 | 30988 | P59594.1 | P59594 |
| KGLPIRYQTTATKSE | 1 | 0 | 0 | 31003 | QTX93489.1 | P17763 |
| KHKKVSTNLCTHSFRKKQVR | 1 | 0 | 0 | 31124 | P59633.1 | P59633 |
| KHLYVSMPTAQSTM | 1 | 0 | 0 | 31133 | P27313.1 | P27313 |
| KHWIGRLMWWNQYT | 1 | 0 | 0 | 31165 | Q81258.3 | P27958 |
| KINLADRMLGLSGVQEIKEQ | 1 | 0 | 0 | 31367 | P08617.1 | P08617 |
| KKISNCVADYSVLYNST | 1 | 0 | 0 | 31581 | P59594.1 | P59594 |
| KKPNRQGLGYYSPSYNDP | 1 | 0 | 0 | 31664 | NP_041006.1 | P03383 |
| KKSAAEASKKPRQKRTATKQYNVTQ | 1 | 0 | 0 | 31693 | ABI96968.1 | P59595 |
| KLDDKDPQFKDNVILL | 1 | 0 | 0 | 31780 | AAS48456.1 | P59595 |
| KLDVVKLLYNEQFAVQGLLR | 1 | 0 | 0 | 31794 | P08617.1 | P08617 |
| KLFIRQEEVQQELYSP | 1 | 0 | 0 | 31847 | P59635.1 | P59635 |
| KLGVWKSPTDMQSWV | 1 | 0 | 0 | 31894 | P08362.1 | Q786F2 |
| KLIAQGGKAKICGL | 1 | 0 | 0 | 31903 | Q81258.3 | P27958 |
| KLLSQGGRAATCGK | 1 | 0 | 0 | 32006 | P26662.3 | P27958 |
| KLNDLCFSNVYADSFVV | 1 | 0 | 0 | 32037 | P59594.1 | P59594 |
| KLNTDHAGSNDNIALLVQ | 1 | 0 | 0 | 32059 | YP_009825055.1 | P59596 |
| KLPDDFMGCVLAWNTRN | 1 | 0 | 0 | 32070 | P59594.1 | P59594 |
| KLSYGIATVREVLSD | 1 | 0 | 0 | 32185 | P0DTD1.1 | P0DTD1 |
| KMDTKPTDPTGIEP | 1 | 0 | 0 | 32296 | P27313.1 | P27313 |
| KNDGTTVDLTVDQAWRGKGE | 1 | 0 | 0 | 32462 | P08617.1 | P08617 |
| KNEQAGQVQVLSSV | 1 | 0 | 0 | 32475 | P26661.3 | P27958 |
| KNGKKVIQLSRKTFD | 1 | 0 | 0 | 32500 | P14340.2 | P17763 |
| KNHKNLLKIAQYAAQNRRGLDLL | 1 | 0 | 0 | 32506 | AAU04921.1 | P03381 |
| KNPHNDMLNVNLAKTNDAIK | 1 | 0 | 0 | 32589 | P08617.1 | P08617 |
| KNQVEGEVQIVSTATQTFLA | 1 | 0 | 0 | 32599 | AAB67037.1 | P27958 |
| KNVVTGEVQVLYTA | 1 | 0 | 0 | 32635 | SRC279960 | P27958 |
| KPAIIPDREVLYREFDEMEE | 1 | 0 | 0 | 32650 | AAA45676.1 | P27958 |
| KPDYEPPVVHGCPLPPP | 1 | 0 | 0 | 32684 | P26664.3 | P27958 |
| KPNRPVVPSPDPN | 1 | 0 | 0 | 32812 | NP_044666.1 | P06484 |
| KPTTFMLKYDENGTITD | 1 | 0 | 0 | 32912 | P59594.1 | P59594 |
| KQGGENFAYLTAYQA | 1 | 0 | 0 | 32984 | P26661.3 | P27958 |
| KQNIQLINTNGSWH | 1 | 0 | 0 | 33043 | P27958.3 | P27958 |

| | | | | | | |
|---|---|---|---|---|---|---|
| KQQGLNFSYLAAYQA | 1 | 0 | 0 | 33055 | ADE10208.1 | P27958 |
| KQSGENFAYLTAYQA | 1 | 0 | 0 | 33065 | P26660.3 | P27958 |
| KQSGENLPYLVAYQA | 1 | 0 | 0 | 33066 | P26664.3 | P27958 |
| KRAASKVSARLLTV | 1 | 0 | 0 | 33107 | P26661.3 | P27958 |
| KRLHQWINEDCSTP | 1 | 0 | 0 | 33183 | P26662.3 | P27958 |
| KRTATKQYNVTQAFGRR | 1 | 0 | 0 | 33220 | AAS48456.1 | P59595 |
| KRYISWCLWWLQYF | 1 | 0 | 0 | 33259 | P26664.3 | P27958 |
| KSEFDRDAAMQRKLE | 1 | 0 | 0 | 33297 | P0DTC1.1 | P0DTD1 |
| KSEHTGREIVDLMCH | 1 | 0 | 0 | 33299 | P27909.2 | P17763 |
| KSKFGYSAKDVRSL | 1 | 0 | 0 | 33366 | SRC279960 | P27958 |
| KSRYGFGAKEVRSL | 1 | 0 | 0 | 33504 | P26661.3 | P27958 |
| KSVCGPVYCFTPSPVVVGTT | 1 | 0 | 0 | 33541 | AAA45676.1 | P27958 |
| KTLLSRFLWWLCYL | 1 | 0 | 0 | 33761 | P26660.3 | P27958 |
| KTMIRFGDVGLDLDFSAFDA | 1 | 0 | 0 | 33776 | P08617.1 | P08617 |
| KTSVDCNMYICGDSTEC | 1 | 0 | 0 | 33874 | NP_828851.1 | P59594 |
| KVKVEYQKTMGSM | 1 | 0 | 0 | 34082 | AAO86638.1 | Q9E006 |
| KVLHKRTLGLSAMSTT | 1 | 0 | 0 | 34103 | ABK27207.1 | Q69027 |
| KVLSPLQDNNDSGH | 1 | 0 | 0 | 34115 | P26661.3 | P27958 |
| KVNFPHGMLDLEEIAANSKD | 1 | 0 | 0 | 34135 | P08617.1 | P08617 |
| KVTAFQHQNSKKTTK | 1 | 0 | 0 | 34210 | P59633.1 | P59633 |
| KVVGLYGNGVVTRSG | 1 | 0 | 0 | 34231 | P14340.2 | P17763 |
| LAAEFKQKALGLLQTATKQA | 1 | 0 | 0 | 34661 | AAA79971.1 | P27958 |
| LAEQFKQKALGLLQQASKQA | 1 | 0 | 0 | 34744 | AAC53936.1 | P27958 |
| LAEQFKQKALGLLQTATRQA | 1 | 0 | 0 | 34752 | AAC53943.1 | P27958 |
| LAEQFKQKALGLLQTSTKQA | 1 | 0 | 0 | 34754 | AAC53938.1 | P27958 |
| LAEQFKQKARGLLQTATKQA | 1 | 0 | 0 | 34755 | SRC279960 | P27958 |
| LAEQLKQKALGLLQTATKQA | 1 | 0 | 0 | 34757 | AAC53956.1 | P27958 |
| LAGHGAFQITSLFTRG | 1 | 0 | 0 | 34779 | SRC279960 | P27958 |
| LALLSCLTVPASAYQVRNSTGLYHV | 1 | 0 | 0 | 34854 | P26664.3 | P27958 |
| LALPAPHLTLPFNWTHCFDPQIQ | 1 | 0 | 0 | 34858 | A45714 | P03381 |
| LALSADQALQPPCPNLVSYSSYH | 1 | 0 | 0 | 34862 | AAA96674.1 | P03381 |
| LAPHSALALLEDTMDYPARAHTFDD | 1 | 0 | 0 | 34904 | BAA01867.1 | Q81871 |
| LAPSTLRSLRKRRLSSP | 1 | 0 | 0 | 34914 | YP_001129431.1 | Q9QR71 |
| LAPVPPPATRRRRAVPI | 1 | 0 | 0 | 34925 | P03383.1 | P03383 |
| LARGSPPSLASSSASQLSA | 1 | 0 | 0 | 34942 | P26662.3 | P27958 |
| LASPGESQDTPGPC | 1 | 0 | 0 | 34968 | SRC279960 | P27958 |
| LATCPEHQLTWDPIDGRVVS | 1 | 0 | 0 | 34981 | AAG48731.1 | P03410 |
| LATVAYFNMVYMPAS | 1 | 0 | 0 | 34992 | P0DTC1.1 | P0DTD1 |
| LAWLYAAVINGDRWF | 1 | 0 | 0 | 35022 | P0DTC1.1 | P0DTD1 |
| LCENPEWAPLKDNRI | 1 | 0 | 0 | 35050 | P08362.1 | Q786F2 |
| LCLFLLPSLATVAYF | 1 | 0 | 0 | 35072 | P0DTC1.1 | P0DTD1 |
| LCPFGEVFNATKFPSVY | 1 | 0 | 0 | 35083 | P59594.1 | P59594 |
| LCQYLNTLTLAVPYN | 1 | 0 | 0 | 35092 | P0DTD1.1 | P0DTD1 |
| LCSALYVGDLCGSVFLVGQL | 1 | 0 | 0 | 35094 | AAA45676.1 | P27958 |
| LDAYNMMISAGFSLW | 1 | 0 | 0 | 35122 | P0DTD1.1 | P0DTD1 |
| LDENGLLLGVHPRLAQRILF | 1 | 0 | 0 | 35143 | P08617.1 | P08617 |
| LDFSLPSSSSYSY | 1 | 0 | 0 | 35161 | AAO86638.1 | Q9E006 |
| LDKNHVADIDKLID | 1 | 0 | 0 | 35199 | P27313.1 | P27313 |
| LDMYSVMLTNDNTSR | 1 | 0 | 0 | 35230 | P0DTD1.1 | P0DTD1 |
| LDNPLENPSSLFDLVARIKN | 1 | 0 | 0 | 35244 | P07299.1 | Q9PZT0 |
| LDPMVEERSDLEPS | 1 | 0 | 0 | 35253 | P26660.3 | P27958 |
| LDSMTEVEDDREPS | 1 | 0 | 0 | 35292 | P26661.3 | P27958 |
| LDTEVAASCGGVVL | 1 | 0 | 0 | 35304 | P26664.3 | P27958 |
| LEGKPALVPDKEVLYQQYDE | 1 | 0 | 0 | 35488 | Q81258.3 | P27958 |
| LEKMADQAMTQMYKQ | 1 | 0 | 0 | 35555 | P0DTD1.1 | P0DTD1 |
| LEWRNTSGLYVLT | 1 | 0 | 0 | 35756 | Q81258.3 | P27958 |
| LFDMSKFPLKLRGTA | 1 | 0 | 0 | 35815 | P0DTD1.1 | P0DTD1 |
| LFPTQIQVRNIMSP | 1 | 0 | 0 | 35947 | P27313.1 | P27313 |
| LFSLMPDVAHSLA | 1 | 0 | 0 | 35982 | AAO86638.1 | Q9E006 |
| LFTYHKFNSSGCPERLASCRSLDD | 1 | 0 | 0 | 36007 | SRC279960 | P27958 |
| LGGKPAIVPDKEVLYQQYDE | 1 | 0 | 0 | 36124 | Q81594 | P27958 |
| LGGKPALVPDKEVLYQQYDE | 1 | 0 | 0 | 36126 | AAC03058.1 | P27958 |
| LGGKPALVPDRQVLYQQYDE | 1 | 0 | 0 | 36127 | Q81487.3 | P27958 |
| LGINITNFRAILTAFSP | 1 | 0 | 0 | 36167 | P59594.1 | P59594 |
| LGMTATSADKNVPQLKPVSE | 1 | 0 | 0 | 36223 | P08617.1 | P08617 |
| LGPADGMVSKGWRL | 1 | 0 | 0 | 36241 | P26664.3 | P27958 |
| LGPADSLEGQGWRL | 1 | 0 | 0 | 36244 | AAA45721.1 | P27958 |
| LGPLYLIQATITTT | 1 | 0 | 0 | 36264 | Q81258.3 | P27958 |
| LGSINQAMVTRCEPVVCYLY | 1 | 0 | 0 | 36319 | P08617.1 | P08617 |
| LGVYDYLVSTQEFRY | 1 | 0 | 0 | 36351 | P0DTC1.1 | P0DTD1 |
| LHDIGNPKAIKCVPQ | 1 | 0 | 0 | 36365 | P0DTD1.1 | P0DTD1 |
| LHQPPPQIFLKILPQSGPIG | 1 | 0 | 0 | 36413 | P07299.1 | Q9PZT0 |
| LHTPGCVPCVREGNASRCWV | 1 | 0 | 0 | 36438 | AAA45676.1 | P27958 |
| LHYRNQGWRSVETSGVAEEEATSGL | 1 | 0 | 0 | 36443 | P29326.1 | Q81871 |
| LIGLLAIAGIRLHRAAIYTAEIHK | 1 | 0 | 0 | 36538 | P08362.1 | Q786F2 |
| LIIMDEAHFTDPASI | 1 | 0 | 0 | 36560 | P14340.2 | P17763 |
| LIKNQCVNFNFNGLTGT | 1 | 0 | 0 | 36579 | P59594.1 | P59594 |
| LINQRAVVAPDKEVLYEAFD | 1 | 0 | 0 | 36645 | SRC279960 | P27958 |
| LINTNGSWHINRTALNCNDSLNTGA | 1 | 0 | 0 | 36649 | SRC279960 | P27958 |
| LKCSVKSFEIDKGIYQT | 1 | 0 | 0 | 36815 | P59594.1 | P59594 |
| LKDAERAVEVDPDD | 1 | 0 | 0 | 36816 | P27313.1 | P27313 |
| LKGACSCGSCCKFDEDD | 1 | 0 | 0 | 36856 | NP_828851.1 | P59594 |
| LKIKIASGFGPLITH | 1 | 0 | 0 | 36890 | P08362.1 | Q786F2 |
| LKILPQSGPIGGIKSMGITT | 1 | 0 | 0 | 36893 | P07299.1 | Q9PZT0 |
| LKLFAAETLKATEET | 1 | 0 | 0 | 36937 | P0DTD1.1 | P0DTD1 |
| LKPIFKLPLGINITNFR | 1 | 0 | 0 | 36991 | P59594.1 | P59594 |
| LKQKALGLLQTATKQAEAAA | 1 | 0 | 0 | 36999 | AAC53956.1 | P27958 |
| LKSKIQGLLQQASK | 1 | 0 | 0 | 37021 | P26660.3 | P27958 |
| LKSKIQGLLQQATR | 1 | 0 | 0 | 37023 | P26661.3 | P27958 |
| LLFVGVGLFLLPAR | 1 | 0 | 0 | 37304 | P26660.3 | P27958 |
| LLISQAEAALENLVILNAAS | 1 | 0 | 0 | 37425 | BAA01582.1 | P27958 |

| | | | | | | |
|---|---|---|---|---|---|---|
| LLKIAQYAAQNRRGLDLLFWEQG | 1 | 0 | 0 | 37441 | AAU04921.1 | P03381 |
| LLLTVGGIFLLPAR | 1 | 0 | 0 | 37560 | SRC279960 | P27958 |
| LLPHSNLDHILEPSIPWKSKLLT | 1 | 0 | 0 | 37661 | AAA96674.1 | P03381 |
| LLRSTSQKSIVAYTMSL | 1 | 0 | 0 | 37758 | NP_828851.1 | P59594 |
| LLSPRGSRPSWG | 1 | 0 | 0 | 37814 | P27958.3 | P27958 |
| LLSVGVGIFLLPAR | 1 | 0 | 0 | 37833 | P26661.3 | P27958 |
| LLVILAGPCILRQLRHLPSRVRY | 1 | 0 | 0 | 37899 | AAU04921.1 | P03381 |
| LLYRLGAVQNEITLTHPVTKYIMTC | 1 | 0 | 0 | 37987 | P26664.3 | P27958 |
| LMESELVIGAVIIRGHLR | 1 | 0 | 0 | 38014 | P59596.1 | P59596 |
| LMSFPQAAPHGVVFLHV | 1 | 0 | 0 | 38118 | NP_828851.1 | P59594 |
| LNDQVVVTPDKEILYEAFDE | 1 | 0 | 0 | 38192 | AAP55688.1 | P27958 |
| LNDRVVVAPDKEILYEAFDE | 1 | 0 | 0 | 38193 | BAA01761.1 | P27958 |
| LNPSVAATLGFGAYMSKAHG | 1 | 0 | 0 | 38311 | P26662.3 | P27958 |
| LNTLTLAVPYNMRVI | 1 | 0 | 0 | 38352 | P0DTD1.1 | P0DTD1 |
| LPAIVREAIKRRLRT | 1 | 0 | 0 | 38401 | ACA48751.1 | P17763 |
| LPGCSFSIFLLALLSCLTVPASAYQ | 1 | 0 | 0 | 38500 | P26664.3 | P27958 |
| LPHSNLDHILEPSIPWKSKLLTLV | 1 | 0 | 0 | 38524 | AAL05539.1 | P03381 |
| LPHVVDLPQLGPRR | 1 | 0 | 0 | 38528 | P69616.1 | Q81870 |
| LPLRGKLSFWEAGTTKAGYPYNYNT | 1 | 0 | 0 | 38586 | P29326.1 | Q81871 |
| LPPVADLPQPGLRR | 1 | 0 | 0 | 38632 | Q03499.1 | Q81870 |
| LPQSGPIGGIKSMGITTLVQ | 1 | 0 | 0 | 38659 | P07299.1 | Q9PZT0 |
| LQAAVLHVPGCVPC | 1 | 0 | 0 | 38799 | P26660.3 | P27958 |
| LQAMRKYSPFRNGYMEPTLG | 1 | 0 | 0 | 38810 | AAG31572.1 | P03409 |
| LQNSMSGASADSTQA | 1 | 0 | 0 | 38925 | NP_828858.1 | P59595 |
| LQTTGRPSHEAPN | 1 | 0 | 0 | 39001 | NP_044666.1 | P06484 |
| LRFTSARRMETTVN | 1 | 0 | 0 | 39089 | ABK27207.1 | Q69027 |
| LRMAGHSLGRCDIKDLPK | 1 | 0 | 0 | 39141 | P59596.1 | P59596 |
| LRPGPVSQPYHHWDTDK | 1 | 0 | 0 | 39164 | P07299.1 | Q9PZT0 |
| LRRTQSMGIQLDQ | 1 | 0 | 0 | 39198 | AAO86636.1 | O36307 |
| LRTLILAPTRVVAAE | 1 | 0 | 0 | 39225 | P14340.2 | P17763 |
| LRYHNKVYCTTSKSA | 1 | 0 | 0 | 39253 | P26660.3 | P27958 |
| LSFSCYLSVTEQSEFYFPRA | 1 | 0 | 0 | 39352 | P08617.1 | P08617 |
| LSGHRMAWDMMMNWS | 1 | 0 | 0 | 39367 | BAA03375.1 | P27958 |
| LSGKPAIIPDREALYQEFDE | 1 | 0 | 0 | 39370 | AAC53962.1 | P27958 |
| LSGKPAIIPDREALYQQFDE | 1 | 0 | 0 | 39371 | AAA65789.1 | P27958 |
| LSGKPAIIPDREVLYQAFDE | 1 | 0 | 0 | 39372 | AAC53957.1 | P27958 |
| LSGKPAIIPDREVLYQEFDE | 1 | 0 | 0 | 39373 | AAC53965.1 | P27958 |
| LSGKPAIIPDREVLYREFDE | 1 | 0 | 0 | 39375 | P26664.3 | P27958 |
| LSGKPATIPDREALYREFDE | 1 | 0 | 0 | 39376 | AAC53955.1 | P27958 |
| LSGKPATIPDREAVLYQEFD | 1 | 0 | 0 | 39377 | SRC279960 | P27958 |
| LSGKPATIPDREVLYQEFDE | 1 | 0 | 0 | 39378 | AAC53937.1 | P27958 |
| LSGKPATIPDREVLYREFDE | 1 | 0 | 0 | 39379 | AAC53938.1 | P27958 |
| LSGKPATVPDREVLYQEFDE | 1 | 0 | 0 | 39380 | AAC53954.1 | P27958 |
| LSGKPAVIPDRELLYREFDE | 1 | 0 | 0 | 39381 | AAC53929.1 | P27958 |
| LSGKPAVIPDREVLYQAFDE | 1 | 0 | 0 | 39382 | AAC53931.1 | P27958 |
| LSGKPAVIPDREVLYQGFDE | 1 | 0 | 0 | 39383 | AAC53946.1 | P27958 |
| LSGKPAVIPDREVLYREFDE | 1 | 0 | 0 | 39384 | AAC53960.1 | P27958 |
| LSGKPAVVPDREVLYQEFDE | 1 | 0 | 0 | 39385 | ABU97068.1 | P27958 |
| LSGQPAVIPDREVLYQQFDE | 1 | 0 | 0 | 39391 | ABL63009.1 | P27958 |
| LSGRPAIIPDREALYQEFDE | 1 | 0 | 0 | 39392 | AAC53969.1 | P27958 |
| LSGRPAIIPDREVLHREFDE | 1 | 0 | 0 | 39393 | AAA45616.1 | P27958 |
| LSGRPAIIPDREVLYQEFDE | 1 | 0 | 0 | 39394 | AAC53943.1 | P27958 |
| LSGRPAIIPDREVLYREFDE | 1 | 0 | 0 | 39395 | AAC53967.1 | P27958 |
| LSGRPAIVPDRELLYQEFDE | 1 | 0 | 0 | 39396 | AAA72945.1 | P27958 |
| LSGRPAIVPDREVLYQEFDE | 1 | 0 | 0 | 39398 | AAA86907.1 | P27958 |
| LSGRPAIVPDREVLYREFDE | 1 | 0 | 0 | 39399 | AAC53968.1 | P27958 |
| LSGRPAVIPDREVLYQEFDE | 1 | 0 | 0 | 39400 | BAA06303.1 | P27958 |
| LSGRPAVIPDREVLYREFDE | 1 | 0 | 0 | 39401 | AAA75355.1 | P27958 |
| LSGRPAVVPDREVLYQEFDE | 1 | 0 | 0 | 39402 | AAA79971.1 | P27958 |
| LSGRPAVVPDREVLYREFDE | 1 | 0 | 0 | 39403 | BAA20975.1 | P27958 |
| LSGRPVIVPDREVLYREFDE | 1 | 0 | 0 | 39404 | AAC53963.1 | P27958 |
| LSGSPAIVPDREVLYQDFDE | 1 | 0 | 0 | 39405 | ABU97067.1 | P27958 |
| LSLASPPSPPREPPRPSRSL | 1 | 0 | 0 | 39473 | AAA45412.1 | Q85601 |
| LSLLDLYLGRGYNVS | 1 | 0 | 0 | 39483 | P08362.1 | Q786F2 |
| LSPVALRQMSCAAGT | 1 | 0 | 0 | 39582 | P0DTC1.1 | P0DTD1 |
| LSTTEWQILPCSFTT | 1 | 0 | 0 | 39719 | P26662.3 | P27958 |
| LSTWHVLYSPNVSVPSSSSTPLL | 1 | 0 | 0 | 39726 | P03381.1 | P03381 |
| LSVDLSLTVELKIKI | 1 | 0 | 0 | 39732 | P08362.1 | Q786F2 |
| LTCGFADLMGYIP | 1 | 0 | 0 | 39825 | P27958.3 | P27958 |
| LTDIQEDITRHEQQ | 1 | 0 | 0 | 39849 | P27313.1 | P27313 |
| LTGHRMAWDMMMNWS | 1 | 0 | 0 | 39914 | BAB18806.1 | P27958 |
| LTGHRMAWDMMQNWS | 1 | 0 | 0 | 39915 | SRC279960 | P27958 |
| LTGKPAVIPDREILYQQFDE | 1 | 0 | 0 | 39917 | SRC279960 | P27958 |
| LTGKPAVIPDREVLYQEFDE | 1 | 0 | 0 | 39918 | AAC53934.1 | P27958 |
| LTGKPAVVPDREILYQQFDE | 1 | 0 | 0 | 39919 | Q5I2N3.3 | P27958 |
| LTIMDLHPGSGKTRK | 1 | 0 | 0 | 39967 | AEX65549.1 | P17763 |
| LTPAWRIYSTGNNVFQ | 1 | 0 | 0 | 40075 | P59594.1 | P59594 |
| LTPTLAARNATIPTTTIRHH | 1 | 0 | 0 | 40129 | BAA02670.1 | P27958 |
| LVCAPAPCNFFTSA | 1 | 0 | 0 | 40283 | AAP79798.1 | Q69027 |
| LVEKPNLSSKRSELS | 1 | 0 | 0 | 40319 | P08362.1 | Q786F2 |
| LVIGAVIIRGHLRMAGHSLGR | 1 | 0 | 0 | 40389 | P59596.1 | P59596 |
| LVNSVSTKEGDSSNTGAGKA | 1 | 0 | 0 | 40495 | P07299.1 | Q9PZT0 |
| LVQLTLQSTNYTCIVCIDRASLST | 1 | 0 | 0 | 40541 | AAL05538.1 | P03381 |
| LYCGGPMFNSKGAQ | 1 | 0 | 0 | 40724 | Q81258.3 | P27958 |
| LYKSNHNNVYWLTIP | 1 | 0 | 0 | 40826 | P08362.1 | Q786F2 |
| LYNEQFAVQGLLRYHTYARF | 1 | 0 | 0 | 40866 | P08617.1 | P08617 |
| LYQDVNCTDVSTAIHAD | 1 | 0 | 0 | 40905 | NP_828851.1 | P59594 |
| LYREFDEMEECSQHLPYIEQ | 1 | 0 | 0 | 40925 | P26664.3 | P27958 |
| LYSHHFQSHGQLSDHPHALS | 1 | 0 | 0 | 40948 | P07299.1 | Q9PZT0 |
| LYVGGPLTNSRGEN | 1 | 0 | 0 | 40980 | P26664.3 | P27958 |
| LYVGGPMTNSKGQS | 1 | 0 | 0 | 40982 | P26661.3 | P27958 |

| | | | | | | |
|---|---|---|---|---|---|---|
| MAARLCCQLDPARDV | 1 | 0 | 0 | 41004 | CAB41697.1 | Q69027 |
| MACMSADLEVTTST | 1 | 0 | 0 | 41011 | Q81258.3 | P27958 |
| MACMSADLEVVTST | 1 | 0 | 0 | 41012 | P26663.3 | P27958 |
| MAHFPGFGQSLLYGYPVYVF | 1 | 0 | 0 | 41048 | AAG48731.1 | P03410 |
| MASGGGETALALLLLDRL | 1 | 0 | 0 | 41120 | AAS48456.1 | P59595 |
| MDIDPYKEFGATVELLSFLP | 1 | 0 | 0 | 41238 | P03146.1 | Q76R61 |
| MDYPARAHTFDDFCPECRPLGLQGC | 1 | 0 | 0 | 41293 | BAA01867.1 | Q81871 |
| MEACAPHLPYIEQGMQLAEQ | 1 | 0 | 0 | 41296 | BAA20975.1 | P27958 |
| MEECASHLPYFEQGMQLAEQ | 1 | 0 | 0 | 41320 | AAA45721.1 | P27958 |
| MEECASHLPYIEQGMHLAEQ | 1 | 0 | 0 | 41322 | AAC53941.1 | P27958 |
| MEECASHLPYIEQGMQLAAE | 1 | 0 | 0 | 41323 | AAA79971.1 | P27958 |
| MEECASHLPYIEQGMQLAEQ | 1 | 0 | 0 | 41324 | AAC53954.1 | P27958 |
| MEECASHLPYIEQGVQLAEQ | 1 | 0 | 0 | 41325 | AAA75355.1 | P27958 |
| MEECASQLPYIEQGMQLAEQ | 1 | 0 | 0 | 41329 | BAA01583.1 | P27958 |
| MEECASRAALLEEGQRIAEM | 1 | 0 | 0 | 41331 | AAC53939.1 | P27958 |
| MEECGSHLPYIEQGMQLAEQ | 1 | 0 | 0 | 41332 | AAC53960.1 | P27958 |
| MEECPSHLPYIEQGMQLAEQ | 1 | 0 | 0 | 41333 | AAC53929.1 | P27958 |
| MEECSASLPYMDETRAIAGQ | 1 | 0 | 0 | 41334 | AAA65789.1 | P27958 |
| MEECSKARPYIEQAQVIAHQ | 1 | 0 | 0 | 41335 | AAC53958.1 | P27958 |
| MEECSKHLPLVEHGLQLAEQ | 1 | 0 | 0 | 41336 | O39929.3 | P27958 |
| MEECSQAAPYIEQAQAIAHQ | 1 | 0 | 0 | 41338 | AAC03058.1 | P27958 |
| MEECSQAAPYIEQAQVIAHQ | 1 | 0 | 0 | 41339 | Q81594 | P27958 |
| MEECSQAGPYIEQAQVIAHQ | 1 | 0 | 0 | 41340 | AAC53966.1 | P27958 |
| MEECSQARPYIEQAQVIAHQ | 1 | 0 | 0 | 41341 | Q81495.3 | P27958 |
| MEECSQHLPYIEQGMMLAEQ | 1 | 0 | 0 | 41343 | AAA45677.1 | P27958 |
| MEECSQSAPYIEQAQAIAQQ | 1 | 0 | 0 | 41344 | Q81487.3 | P27958 |
| MEEKATYVHKKNDGTTVDLT | 1 | 0 | 0 | 41351 | P08617.1 | P08617 |
| MENKAWLVHRQWFLD | 1 | 0 | 0 | 41407 | P14340.2 | P17763 |
| MEVTPSGTWLTYHGAIK | 1 | 0 | 0 | 41461 | AAS48456.1 | P59595 |
| MFIFLLFLTLTSGSDLD | 1 | 0 | 0 | 41504 | P59594.1 | P59594 |
| MFLPMLPAPPPGQPSGRRRGRRSGG | 1 | 0 | 0 | 41513 | P29326.1 | Q81871 |
| MGGWSSKPRKGMGTNLSVPNP | 1 | 0 | 0 | 41584 | P03141.3 | Q76R62 |
| MGTNLSVPNPLGFFPDHQLDP | 1 | 0 | 0 | 41631 | P03141.3 | Q76R62 |
| MGTVNKPVVGVLMGFGIITG | 1 | 0 | 0 | 41636 | AAK01056.1 | P09259 |
| MHGDTPTLHEYMLDLQPETT | 1 | 0 | 0 | 41660 | P03129.1 | P03129 |
| MIDRPYVLLAVLFVM | 1 | 0 | 0 | 41693 | P06830.1 | Q786F2 |
| MISGPCPKAGQPSLVVQSSL | 1 | 0 | 0 | 41775 | AAG48731.1 | P03410 |
| MKTVAFDLSSPQKSGTGPQPGSAGM | 1 | 0 | 0 | 41884 | AAA45992.1 | Q6SW99 |
| MLKYDENGTITDAVDCS | 1 | 0 | 0 | 41984 | P59594.1 | P59594 |
| MLPAPPPGQPSGRRRGRRSGGSGGG | 1 | 0 | 0 | 42026 | P29326.1 | Q81871 |
| MLSKYSLPIVEEPEDYKEAS | 1 | 0 | 0 | 42060 | P08617.1 | P08617 |
| MNALVNSTLLETRTT | 1 | 0 | 0 | 42187 | P08362.1 | Q786F2 |
| MPKTRRQRTRRARRNRPPTP | 1 | 0 | 0 | 42289 | CAA61542.1 | P03410 |
| MPNMLRIMASLVLAR | 1 | 0 | 0 | 42301 | P0DTD1.1 | P0DTD1 |
| MQSWVPLSTDDPVID | 1 | 0 | 0 | 42413 | P08362.1 | Q786F2 |
| MSDLTDIQEDITRH | 1 | 0 | 0 | 42532 | P27313.1 | P27313 |
| MSDNGPQSNQRSAPRIT | 1 | 0 | 0 | 42541 | P59595.1 | P59595 |
| MSLSLLDLYLGRGYN | 1 | 0 | 0 | 42603 | P06830.1 | Q786F2 |
| MSNFKVRDPVIQERLDHDYA | 1 | 0 | 0 | 42614 | AAC56957.1 | Q2HR63 |
| MSRIAAGDLESSVDDPRSEE | 1 | 0 | 0 | 42646 | P08617.1 | P08617 |
| MSTLQELQENITA | 1 | 0 | 0 | 42678 | AAO86636.1 | O36307 |
| MSTNPKPQKKNKRNTNRRPQDV | 1 | 0 | 0 | 42686 | AAA45676.1 | P27958 |
| MSTNPKPQRKTKRNTNRRPQDV | 1 | 0 | 0 | 42700 | AAS15483.1 | P27958 |
| MVTILLCCMTSCCSCLK | 1 | 0 | 0 | 42999 | NP_828851.1 | P59594 |
| MVYMPASWVMRIMTW | 1 | 0 | 0 | 43010 | P0DTC1.1 | P0DTD1 |
| MYRVFEVSVIRNPGL | 1 | 0 | 0 | 43084 | P06830.1 | Q786F2 |
| NAFNCTFEYISDAFSLD | 1 | 0 | 0 | 43145 | P59594.1 | P59594 |
| NAPATVCGPKLSTDLIK | 1 | 0 | 0 | 43264 | P59594.1 | P59594 |
| NASSEVAVLYQDVNCTD | 1 | 0 | 0 | 43313 | NP_828851.1 | P59594 |
| NDHEPQISPGGLEPPSEKHFRETEV | 1 | 0 | 0 | 43449 | P03409.2 | P03409 |
| NDWDFVVTTDISEMG | 1 | 0 | 0 | 43542 | P14340.2 | P17763 |
| NFELCDNPFFAVSKPMG | 1 | 0 | 0 | 43825 | P59594.1 | P59594 |
| NFISGIQYLAGLSTLPGNPA | 1 | 0 | 0 | 43854 | P29846.3 | P27958 |
| NFNGLTGTGVLTPSSKR | 1 | 0 | 0 | 43893 | P59594.1 | P59594 |
| NGIRRPKHLYVSMP | 1 | 0 | 0 | 44002 | P27313.1 | P27313 |
| NHVADIDKLIDYAA | 1 | 0 | 0 | 44172 | P27313.1 | P27313 |
| NIDATSTGNYNYKYRYL | 1 | 0 | 0 | 44198 | P59594.1 | P59594 |
| NIGGEGNHNESPL | 1 | 0 | 0 | 44232 | SRC279960 | P27958 |
| NIGGVGNNTLTCPT | 1 | 0 | 0 | 44233 | ABU97067.1 | P27958 |
| NIGTGTSAYTSLNVLARFTD | 1 | 0 | 0 | 44235 | P08617.1 | P08617 |
| NILKDNQQKIEKAIEEADEF | 1 | 0 | 0 | 44278 | P08617.1 | P08617 |
| NISGHVRMGTMKIT | 1 | 0 | 0 | 44340 | P26661.3 | P27958 |
| NISLDNPLENPSSLFDLVAR | 1 | 0 | 0 | 44350 | P07299.1 | Q9PZT0 |
| NITRVESETKVVVL | 1 | 0 | 0 | 44380 | Q81258.3 | P27958 |
| NKRNTNRRPQDVKFPGGGQIVGGVY | 1 | 0 | 0 | 44590 | P26664.3 | P27958 |
| NKSTLQNRRAAVS | 1 | 0 | 0 | 44602 | AAO86636.1 | O36307 |
| NLDDRDRSELSPLLLSTTEW | 1 | 0 | 0 | 44662 | BAA02670.1 | P27958 |
| NLINTNGSWHIN | 1 | 0 | 0 | 44747 | AAP55702.1 | P27958 |
| NLNFEMYGAVYSVN | 1 | 0 | 0 | 44816 | P26661.3 | P27958 |
| NLNFEMYGAVYSVS | 1 | 0 | 0 | 44817 | P26660.3 | P27958 |
| NLSGKAINHIRS | 1 | 0 | 0 | 44869 | P29846.3 | P27958 |
| NLSGKAVNHIRS | 1 | 0 | 0 | 44870 | BAK78910.1 | P27958 |
| NLSSKAVNHIHS | 1 | 0 | 0 | 44882 | P26663.3 | P27958 |
| NLVILPGQDLQYVLA | 1 | 0 | 0 | 44915 | P08362.1 | Q786F2 |
| NLVLNRDVSFQDL | 1 | 0 | 0 | 44917 | AAO86638.1 | Q9E006 |
| NNKSQSVIIINNSTNVV | 1 | 0 | 0 | 45151 | P59594.1 | P59594 |
| NNSPARPETSRPK | 1 | 0 | 0 | 45247 | NP_044666.1 | P06484 |
| NPLYGSRLGVPDTLGGDPKF | 1 | 0 | 0 | 45398 | P07299.1 | Q9PZT0 |
| NPSPKTVYVKEAIDRRLHFK | 1 | 0 | 0 | 45478 | P08617.1 | P08617 |
| NPSSLFDLVARIKNNLKNSP | 1 | 0 | 0 | 45480 | P07299.1 | Q9PZT0 |
| NPVIPFKDGIYFAATEK | 1 | 0 | 0 | 45506 | P59594.1 | P59594 |

| | | | | | | |
|---|---|---|---|---|---|---|
| NPVRASVLRYDDFHTDEDKL | 1 | 0 | 0 | 45512 | AAK01056.1 | P09259 |
| NREGKIVGLYGNGVV | 1 | 0 | 0 | 45669 | P17763.2 | P17763 |
| NREGKVVGLYGNGVV | 1 | 0 | 0 | 45670 | ACY70710.1 | P17763 |
| NRGKMDVSGVQAPVGAITTI | 1 | 0 | 0 | 45681 | P08617.1 | P08617 |
| NRLIAFAFAGNHVSPTHYVP | 1 | 0 | 0 | 45696 | AAA86907.1 | P27958 |
| NRLIAFASRGNHDSPTHYV | 1 | 0 | 0 | 45697 | AAA75355.1 | P27958 |
| NRLIAFASRGNHVAPTHYVT | 1 | 0 | 0 | 45698 | BAA00792.1 | P27958 |
| NRLIAFASRGNHVAPTHYVV | 1 | 0 | 0 | 45699 | BAA01761.1 | P27958 |
| NRLIAFASRGNHVSPRHYV | 1 | 0 | 0 | 45702 | AAA52748.1 | P27958 |
| NRLIAFASRGNHVSPTHYVP | 1 | 0 | 0 | 45704 | BAB18806.1 | P27958 |
| NRRGLDLLFWEQGGLCKALQEQCRF | 1 | 0 | 0 | 45728 | AAL05538.1 | P03381 |
| NSLILPPFSLSPVPTLGSRSRRA | 1 | 0 | 0 | 45894 | P03381.1 | P03381 |
| NTIMASKSVGTAE | 1 | 0 | 0 | 46127 | AAO86636.1 | O36307 |
| NTNSGPDDQIGYYRRATR | 1 | 0 | 0 | 46198 | AAS48456.1 | P59595 |
| NTTASDQLLVENAAGHRVA | 1 | 0 | 0 | 46276 | AAG16766.1 | Q81871 |
| NVFFLLLFSLTHFPLAQ | 1 | 0 | 0 | 46375 | P03383.1 | P03383 |
| NVFQTQAGCLIGAEHVD | 1 | 0 | 0 | 46379 | NP_828851.1 | P59594 |
| NVLDVNAIDIEEPS | 1 | 0 | 0 | 46417 | P27313.1 | P27313 |
| NVYADSFVVKGDDVRQI | 1 | 0 | 0 | 46522 | P59594.1 | P59594 |
| NYNLIIMDEAHFTDP | 1 | 0 | 0 | 46669 | P14340.2 | P17763 |
| NYNYKYRYLRHGKLRPF | 1 | 0 | 0 | 46681 | P59594.1 | P59594 |
| PAADRAAAPDAVAA | 1 | 0 | 0 | 46738 | SRC279960 | P27958 |
| PAANQVGVGAFGPRLTPPHGGI | 1 | 0 | 0 | 46745 | P03141.3 | Q76R62 |
| PAASSCHNASGKEAKVCTIS | 1 | 0 | 0 | 46753 | P07299.1 | Q9PZT0 |
| PAEILRKPRKFPPALPIW | 1 | 0 | 0 | 46778 | P26662.3 | P27958 |
| PAFGANSNNPDWDFNPVKDDWP | 1 | 0 | 0 | 46793 | P03141.3 | Q76R62 |
| PAFPLAEDVEKDK | 1 | 0 | 0 | 46796 | NP_044666.1 | P06484 |
| PAIIPDREALYQEFDEMEEC | 1 | 0 | 0 | 46823 | AAC53969.1 | P27958 |
| PAIIPDREALYQQFDEMEEC | 1 | 0 | 0 | 46824 | AAA65789.1 | P27958 |
| PAIIPDREVLHREFDEMEEC | 1 | 0 | 0 | 46826 | AAA45616.1 | P27958 |
| PAIIPDREVLYQAFDEMEEC | 1 | 0 | 0 | 46827 | AAC53957.1 | P27958 |
| PAIIPDREVLYQEFDEMEEC | 1 | 0 | 0 | 46828 | AAC53943.1 | P27958 |
| PAIIPDREVLYREFDEMEEC | 1 | 0 | 0 | 46829 | CAB53095.1 | P27958 |
| PAIVPDKEVLYQQYDEMEEC | 1 | 0 | 0 | 46833 | Q81594 | P27958 |
| PAIVPDRELLYQEFDEMEEC | 1 | 0 | 0 | 46834 | AAA72945.1 | P27958 |
| PAIVPDREVLYQDFDEMEEC | 1 | 0 | 0 | 46835 | ABU97067.1 | P27958 |
| PAIVPDREVLYREFDEMEEC | 1 | 0 | 0 | 46836 | AAC53968.1 | P27958 |
| PALTAVETGATNPL | 1 | 0 | 0 | 46859 | NP_740471.1 | P03300 |
| PALVPDKEVLYQQYDEMEEC | 1 | 0 | 0 | 46861 | AAC03058.1 | P27958 |
| PALVPDRQVLYQQYDEMEEC | 1 | 0 | 0 | 46862 | Q81487.3 | P27958 |
| PANKEESISFHFQ | 1 | 0 | 0 | 46871 | AAO86638.1 | Q9E006 |
| PAPQAPYQGYQEPPAPQAPY | 1 | 0 | 0 | 46895 | P03204.1 | P03204 |
| PAPSGCPPDSDAES | 1 | 0 | 0 | 46900 | P26664.3 | P27958 |
| PARDVLCLRPVGAES | 1 | 0 | 0 | 46927 | CAB41697.1 | Q69027 |
| PARPETSRPKTPP | 1 | 0 | 0 | 46933 | NP_044666.1 | P06484 |
| PASTNRQSGRQPTPISPPLRDSHP | 1 | 0 | 0 | 46957 | CAK55122.1 | Q76R62 |
| PATIPDREALYREFDEMEEC | 1 | 0 | 0 | 46970 | AAC53955.1 | P27958 |
| PATIPDREAVLYQEFDEMEE | 1 | 0 | 0 | 46971 | SRC279960 | P27958 |
| PATIPDREVLYQEFDEMEEC | 1 | 0 | 0 | 46972 | AAC53937.1 | P27958 |
| PATIPDREVLYREFDEMEEC | 1 | 0 | 0 | 46973 | AAC53938.1 | P27958 |
| PATVPDREVLYQEFDEMEEC | 1 | 0 | 0 | 46982 | AAC53954.1 | P27958 |
| PAVIPDRELLYREFDEMEEC | 1 | 0 | 0 | 46989 | AAC53929.1 | P27958 |
| PAVIPDREVLYQAFDEMEEC | 1 | 0 | 0 | 46990 | AAC53931.1 | P27958 |
| PAVIPDREVLYQEFDEMEEC | 1 | 0 | 0 | 46991 | BAA06303.1 | P27958 |
| PAVIPDREVLYQGFDEMEEC | 1 | 0 | 0 | 46992 | AAC53946.1 | P27958 |
| PAVIPDREVLYQQFDEMEEC | 1 | 0 | 0 | 46993 | ABL63009.1 | P27958 |
| PAVIPDREVLYREFDEMEEC | 1 | 0 | 0 | 46994 | AAC53960.1 | P27958 |
| PAVVPDREILYQQFDEMEEC | 1 | 0 | 0 | 46998 | Q5I2N3.3 | P27958 |
| PAVVPDREVLYQEFDEMEEC | 1 | 0 | 0 | 46999 | ABU97068.1 | P27958 |
| PAVVPDREVLYREFDEMEAC | 1 | 0 | 0 | 47000 | BAA20975.1 | P27958 |
| PAVVPDREVLYREFDEMEEC | 1 | 0 | 0 | 47001 | AAF65962.1 | P27958 |
| PCKPLLRDDITFMV | 1 | 0 | 0 | 47026 | Q81258.3 | P27958 |
| PCKPLLREEVSFRV | 1 | 0 | 0 | 47027 | P26664.3 | P27958 |
| PCRFVTDTPKGPKVK | 1 | 0 | 0 | 47038 | P0DTC1.1 | P0DTD1 |
| PCSFGGVSVITPGTNA | 1 | 0 | 0 | 47042 | QHR63300.2 | Q3LZX1 |
| PCSLKCPYLGCQSWTCPYTGAVS | 1 | 0 | 0 | 47047 | AAA96674.1 | P03381 |
| PDCCIEPHDWTKNIT | 1 | 0 | 0 | 47064 | ALG01342.1 | Q05320 |
| PDDDPQPGPSREYRYVL | 1 | 0 | 0 | 47068 | YP_001129431.1 | Q9QR71 |
| PDELALSETGMVPL | 1 | 0 | 0 | 47084 | SRC279960 | P27958 |
| PDKEILYEAFDEMEECASKA | 1 | 0 | 0 | 47120 | AAF59945.1 | P27958 |
| PDKEVLYEAFDEMEECASKA | 1 | 0 | 0 | 47122 | Q9IZA3.1 | P27958 |
| PDKEVLYEAFDEMEECASRA | 1 | 0 | 0 | 47123 | AAU89634.1 | P27958 |
| PDKEVLYQQYDEMEECSKAR | 1 | 0 | 0 | 47125 | AAC53958.1 | P27958 |
| PDKEVLYQQYDEMEECSQAA | 1 | 0 | 0 | 47126 | AAC03058.1 | P27958 |
| PDKEVLYQQYDEMEECSQAG | 1 | 0 | 0 | 47127 | AAC53966.1 | P27958 |
| PDKEVLYQQYDEMEECSQAR | 1 | 0 | 0 | 47128 | Q81495.3 | P27958 |
| PDPNNSPARPETS | 1 | 0 | 0 | 47159 | NP_044666.1 | P06484 |
| PDREALYQEFDEMEECASHL | 1 | 0 | 0 | 47174 | AAC53969.1 | P27958 |
| PDREALYQQFDEMEECSASL | 1 | 0 | 0 | 47175 | AAA65789.1 | P27958 |
| PDREALYREFDEMEECASHL | 1 | 0 | 0 | 47176 | AAC53955.1 | P27958 |
| PDRELLYREFDEMEECPSHL | 1 | 0 | 0 | 47179 | AAC53929.1 | P27958 |
| PDREVLHREFDEMEECASHL | 1 | 0 | 0 | 47180 | AAA45616.1 | P27958 |
| PDREVLYQAFDEMEECASHL | 1 | 0 | 0 | 47181 | AAC53957.1 | P27958 |
| PDREVLYQDFDEMEECASHL | 1 | 0 | 0 | 47182 | ABU97067.1 | P27958 |
| PDREVLYQEFDEMEECASHL | 1 | 0 | 0 | 47183 | ABU97068.1 | P27958 |
| PDREVLYQEFDEMEECASQL | 1 | 0 | 0 | 47184 | BAA01583.1 | P27958 |
| PDREVLYQEFDEMEECGSHL | 1 | 0 | 0 | 47185 | BAA01728.1 | P27958 |
| PDREVLYQEFDEMEECSQHL | 1 | 0 | 0 | 47186 | AAA45534.1 | P27958 |
| PDREVLYQGFDEMEECASHL | 1 | 0 | 0 | 47187 | AAC53946.1 | P27958 |
| PDREVLYQQFDEMEECSKHL | 1 | 0 | 0 | 47188 | O39929.3 | P27958 |
| PDREVLYREFDEMEACAPHL | 1 | 0 | 0 | 47190 | BAA20975.1 | P27958 |

| | | | | | | |
|---|---|---|---|---|---|---|
| PDREVLYREFDEMEECAAHI | 1 | 0 | 0 | 47191 | Q81754.3 | P27958 |
| PDREVLYREFDEMEECASHL | 1 | 0 | 0 | 47192 | ABV46292.1 | P27958 |
| PDREVLYREFDEMEECGSHL | 1 | 0 | 0 | 47193 | AAC53960.1 | P27958 |
| PDREVLYREFDEMEECSQHL | 1 | 0 | 0 | 47194 | AAA45677.1 | P27958 |
| PDRQVLYQQYDEMEECSQSA | 1 | 0 | 0 | 47198 | Q81487.3 | P27958 |
| PDSDAESYSSMPPL | 1 | 0 | 0 | 47200 | P26664.3 | P27958 |
| PDVTAAAGAGPRVRQPARPLGSAWR | 1 | 0 | 0 | 47224 | P29326.1 | Q81871 |
| PEEFEGAGDGEPPEDDDS | 1 | 0 | 0 | 47266 | NP_044534.1 | P13290 |
| PERIKLDYDQYCADV | 1 | 0 | 0 | 47398 | P08362.1 | Q786F2 |
| PGLGAPVFHMTNYLE | 1 | 0 | 0 | 47694 | P08362.1 | Q786F2 |
| PGTPAQEIEMLKRN | 1 | 0 | 0 | 47765 | P27313.1 | P27313 |
| PHGVVFLHVTYVPSQER | 1 | 0 | 0 | 47823 | NP_828851.1 | P59594 |
| PHWTKKPNRNGGGYYSASYSDP | 1 | 0 | 0 | 47861 | P03381.1 | P03381 |
| PIAAAGQLDLSGWF | 1 | 0 | 0 | 47865 | P26664.3 | P27958 |
| PIELQVECFTWDQKL | 1 | 0 | 0 | 47890 | P08362.1 | Q786F2 |
| PIGVPSQVVYVPPAYIDMPS | 1 | 0 | 0 | 47910 | AAA45412.1 | Q85601 |
| PIPAASQLDLSGWF | 1 | 0 | 0 | 47941 | P26662.3 | P27958 |
| PISLLFNEKEADDNDHEPQI | 1 | 0 | 0 | 47984 | AAG31572.1 | P03409 |
| PISYANGSGPDQRPYCWHY | 1 | 0 | 0 | 47989 | AAA45676.1 | P27958 |
| PIWFLNTEPSQLPPTAPPLLPHS | 1 | 0 | 0 | 48016 | AAA96674.1 | P03381 |
| PKEITVATSRTLSYYKL | 1 | 0 | 0 | 48052 | P59596.1 | P59596 |
| PKPKVASEAFMST | 1 | 0 | 0 | 48115 | AAO86636.1 | O36307 |
| PKPQRKTKRNTNRRPQDVKFPGG | 1 | 0 | 0 | 48126 | BAB18806.1 | P27958 |
| PKQRRPQGLPNNTASWFT | 1 | 0 | 0 | 48131 | P59595.1 | P59595 |
| PKTSEILDISPCAFGGV | 1 | 0 | 0 | 48155 | NP_828851.1 | P59594 |
| PLDFEMYGATYSVT | 1 | 0 | 0 | 48285 | Q81258.3 | P27958 |
| PLEGEPGDPDLSDGSWST | 1 | 0 | 0 | 48301 | P26662.3 | P27958 |
| PLENPSSLFDLVARIKNNLK | 1 | 0 | 0 | 48307 | P07299.1 | Q9PZT0 |
| PLENRVLTGWGLNWDLGLSQWAR | 1 | 0 | 0 | 48309 | A45714 | P03381 |
| PLGFLPDHQLDPAFGANSTNP | 1 | 0 | 0 | 48339 | P03142.4 | Q76R62 |
| PLGSAWRDQAQRPAVASRRRPTTAG | 1 | 0 | 0 | 48350 | P29326.1 | Q81871 |
| PLITHGSGMDLYKSN | 1 | 0 | 0 | 48370 | P08362.1 | Q786F2 |
| PLLDRWKAPDYVPP | 1 | 0 | 0 | 48390 | Q81258.3 | P27958 |
| PLNSNAMLSTESMMSRIAAG | 1 | 0 | 0 | 48431 | P08617.1 | P08617 |
| PLTDFDQGWGPISYANGSGP | 1 | 0 | 0 | 48498 | AAA45676.1 | P27958 |
| PLVEHGLQLAEQFKQKALGL | 1 | 0 | 0 | 48519 | O39929.3 | P27958 |
| PLVESWKRPDYQPA | 1 | 0 | 0 | 48520 | P26660.3 | P27958 |
| PMGTQTHTMIFDNAFNC | 1 | 0 | 0 | 48554 | P59594.1 | P59594 |
| PMKTDLELDFSLP | 1 | 0 | 0 | 48566 | AAO86638.1 | Q9E006 |
| PNYSRALWRVAAEE | 1 | 0 | 0 | 48728 | P26662.3 | P27958 |
| PPHRPGVRMRRVPVTHP | 1 | 0 | 0 | 48799 | YP_001129431.1 | Q9QR71 |
| PPKPCGIVPAKSVCGPVYCF | 1 | 0 | 0 | 48809 | AAA45676.1 | P27958 |
| PPLLESWKDPDYVPPVVHG | 1 | 0 | 0 | 48823 | P26662.3 | P27958 |
| PPPGRRPFFHPVGE | 1 | 0 | 0 | 48852 | P03211.1 | P03211 |
| PPPQIFLKILPQSGPIGGIK | 1 | 0 | 0 | 48879 | P07299.1 | Q9PZT0 |
| PPQGGVVTPGSGSG | 1 | 0 | 0 | 48894 | P26660.3 | P27958 |
| PPSPGGESDSESCS | 1 | 0 | 0 | 48921 | Q81258.3 | P27958 |
| PPYVEPTAPQVL | 1 | 0 | 0 | 48955 | AAD50663.1 | P03362 |
| PQSAVDSAARIHDFRYSQLA | 1 | 0 | 0 | 49120 | P07299.1 | Q9PZT0 |
| PRGSRPSWGPTDPRRRSRNLGKVID | 1 | 0 | 0 | 49196 | P26664.3 | P27958 |
| PSDTMQTRHVKNYHSRSES | 1 | 0 | 0 | 49349 | BAF95214.1 | Q86887 |
| PSEGECEVIDSDSK | 1 | 0 | 0 | 49352 | P26661.3 | P27958 |
| PSHEAPNMTQTGT | 1 | 0 | 0 | 49395 | NP_044666.1 | P06484 |
| PSHLPYIEQGMQLAEQFKQK | 1 | 0 | 0 | 49398 | AAC53929.1 | P27958 |
| PSRSLPLPPLLSPPRFHPPS | 1 | 0 | 0 | 49498 | AAA45412.1 | Q85601 |
| PSVYAWERKKISNCVAD | 1 | 0 | 0 | 49557 | P59594.1 | P59594 |
| PTDPRRRSRNLGKVIDTLTCGFADL | 1 | 0 | 0 | 49605 | P26664.3 | P27958 |
| PTDSTDNNQNGGRNGARPKQRRPQ | 1 | 0 | 0 | 49608 | P59595.1 | P59595 |
| PTPSDNPFSKLYKETIETFD | 1 | 0 | 0 | 49749 | P03421.1 | O42062 |
| PTPSPAPSRPFSVLRANDVL | 1 | 0 | 0 | 49750 | AAG16766.1 | Q81871 |
| PTSTFLLHTDFEQVMC | 1 | 0 | 0 | 49778 | P07298.1 | Q9PZT1 |
| PTVKLYTSVENAQQDKGIAIPHDID | 1 | 0 | 0 | 49810 | P29326.1 | Q81871 |
| PVIVPDREVLYREFDEMEEC | 1 | 0 | 0 | 49890 | AAC53963.1 | P27958 |
| PVSMAKTSVDCNMYICGDS | 1 | 0 | 0 | 49968 | NP_828851.1 | P59594 |
| PVYVFGDCVQADWCPVSGGL | 1 | 0 | 0 | 50022 | AAG48731.1 | P03410 |
| PWRYLDFNALNLFFSPLEFQ | 1 | 0 | 0 | 50051 | P07299.1 | Q9PZT0 |
| PWYVWLGFIAGLIAIVM | 1 | 0 | 0 | 50058 | NP_828851.1 | P59594 |
| PYFEQGMQLAEQFKQKALGL | 1 | 0 | 0 | 50084 | AAA45721.1 | P27958 |
| PYIEQAQVIAHQFKEKILGL | 1 | 0 | 0 | 50099 | AAC53966.1 | P27958 |
| PYIEQAQVIAHQFKEKVLGL | 1 | 0 | 0 | 50100 | Q81594 | P27958 |
| PYIEQGMHLAEQFKQKALGL | 1 | 0 | 0 | 50101 | AAC53941.1 | P27958 |
| PYIEQGMMLAEQFKQKALGL | 1 | 0 | 0 | 50103 | AAA45677.1 | P27958 |
| PYIEQGMQLAAEFKQKALGL | 1 | 0 | 0 | 50105 | AAA79971.1 | P27958 |
| PYIEQGMQLAEQFKQKALGL | 1 | 0 | 0 | 50106 | ABU97067.1 | P27958 |
| PYIEQGMQLAEQFKQKARGL | 1 | 0 | 0 | 50107 | SRC279960 | P27958 |
| PYIEQGMQLAEQLKQKALGL | 1 | 0 | 0 | 50108 | AAC53956.1 | P27958 |
| PYIEQGVQLAEQFKQKALGL | 1 | 0 | 0 | 50109 | AAA75355.1 | P27958 |
| PYKELRLEVGKQRLKYAQEE | 1 | 0 | 0 | 50119 | P08617.1 | P08617 |
| QAGRTAHAVTNIFSVG | 1 | 0 | 0 | 50273 | SRC279960 | P27958 |
| QALDAHYDSVLKDI | 1 | 0 | 0 | 50300 | P26660.3 | P27958 |
| QALIDQKVKEISNQ | 1 | 0 | 0 | 50307 | P27313.1 | P27313 |
| QAQRPAVASRRRPTTAGAAPLTAVA | 1 | 0 | 0 | 50343 | P29326.1 | Q81871 |
| QASDDGDKGSDVES | 1 | 0 | 0 | 50354 | P26662.3 | P27958 |
| QDCNCSIYPGHASGHRMAWD | 1 | 0 | 0 | 50445 | BAA02670.1 | P27958 |
| QDGTNTHIMATEASNYAQYRVARAT | 1 | 0 | 0 | 50464 | P29326.1 | Q81871 |
| QDIQPAIQSSWPKL | 1 | 0 | 0 | 50476 | P26661.3 | P27958 |
| QDIQPAVQASWPKV | 1 | 0 | 0 | 50477 | P26660.3 | P27958 |
| QDKGIAIPHDIDLGESRVVI | 1 | 0 | 0 | 50481 | AAG16766.1 | Q81871 |
| QDMRNTIMASKTVG | 1 | 0 | 0 | 50494 | P27313.1 | P27313 |
| QDPSKFTEPVKD | 1 | 0 | 0 | 50510 | CAA29172.1 | Q86887 |
| QDRPTPSPAPSRPFSVLRA | 1 | 0 | 0 | 50514 | AAG16766.1 | Q81871 |

| | | | | | | |
|---|---|---|---|---|---|---|
| QDVKFPGGGQIVGGVYLLPRRG | 1 | 0 | 0 | 50526 | AAA45676.1 | P27958 |
| QDVLDKNHVADIDK | 1 | 0 | 0 | 50528 | P27313.1 | P27313 |
| QDYDNQHEQDRPTPSPAPSRPFSVL | 1 | 0 | 0 | 50539 | P29326.1 | Q81871 |
| QEEQEQQEEQEQELEEQ | 1 | 0 | 0 | 50578 | YP_001129431.1 | Q9QR71 |
| QEQEEQELEEVEEQEQE | 1 | 0 | 0 | 50674 | YP_001129431.1 | Q9QR71 |
| QFAALGGWGLHQPPPQIFLK | 1 | 0 | 0 | 50735 | P07299.1 | Q9PZT0 |
| QFAPSASAFFGMSRIGM | 1 | 0 | 0 | 50741 | AAS48456.1 | P59595 |
| QFDEMEECSASLPYMDETRA | 1 | 0 | 0 | 50744 | AAA65789.1 | P27958 |
| QFDEMEECSKHLPLVEHGLQ | 1 | 0 | 0 | 50745 | O39929.3 | P27958 |
| QFKDNVILLNKHIDAYK | 1 | 0 | 0 | 50781 | AAS48456.1 | P59595 |
| QFPFNASDSVGQQIKVIPVD | 1 | 0 | 0 | 50809 | P08617.1 | P08617 |
| QFVEFYEKVTGTDLELIQIL | 1 | 0 | 0 | 50820 | P07299.1 | Q9PZT0 |
| QGCAFQSTVAELQRLKMKVGKTREL | 1 | 0 | 0 | 50837 | P29326.1 | Q81871 |
| QGMHLAEQFKQKALGLLQTA | 1 | 0 | 0 | 50903 | AAC53941.1 | P27958 |
| QGMQLAAEFKQKALGLLQTA | 1 | 0 | 0 | 50911 | AAA79971.1 | P27958 |
| QGMQLAEQFKQKALGLLQQA | 1 | 0 | 0 | 50913 | AAC53936.1 | P27958 |
| QGMQLAEQFKQKALGLLQTA | 1 | 0 | 0 | 50914 | ABU97067.1 | P27958 |
| QGMQLAEQFKQKARGLLQTA | 1 | 0 | 0 | 50916 | SRC279960 | P27958 |
| QGMQLAEQLKQKALGLLQTA | 1 | 0 | 0 | 50917 | AAC53956.1 | P27958 |
| QGSGKGVSFQLVKLG | 1 | 0 | 0 | 50957 | P06830.1 | Q786F2 |
| QGVPESPFSRTGEGLD | 1 | 0 | 0 | 50977 | CAA44973.1 | P29996 |
| QGVQLAEQFKQKALGLLQTA | 1 | 0 | 0 | 50980 | AAA75355.1 | P27958 |
| QHDVNFTQEVSRLNINLHFSKCG | 1 | 0 | 0 | 50995 | AAA96674.1 | P03381 |
| QIANQFNKAISQIQESL | 1 | 0 | 0 | 51043 | NP_828851.1 | P59594 |
| QIFLKILPQSGPIGGIKSMG | 1 | 0 | 0 | 51063 | P07299.1 | Q9PZT0 |
| QIGYYRRATRRVRGGDGK | 1 | 0 | 0 | 51074 | AAS48456.1 | P59595 |
| QLINSNGSWHIN | 1 | 0 | 0 | 51370 | ABL63009.1 | P27958 |
| QLINTNGSWHINSTA | 1 | 0 | 0 | 51372 | AAB02127.1 | P27958 |
| QLINTNGSWHVN | 1 | 0 | 0 | 51374 | AAA16300.1 | P27958 |
| QLLRIPQAIMDM | 1 | 0 | 0 | 51428 | P27958.3 | P27958 |
| QLNRALSGIAAEQDRNT | 1 | 0 | 0 | 51456 | NP_828851.1 | P59594 |
| QLPQGTTLPKGFYAEGSRGGSQ | 1 | 0 | 0 | 51485 | P59595.1 | P59595 |
| QLVNSNGSWHIN | 1 | 0 | 0 | 51574 | AAD56189.1 | P27958 |
| QNVPDPQVGITTMKDLKGKA | 1 | 0 | 0 | 51782 | P08617.1 | P08617 |
| QPELAPEDPED | 1 | 0 | 0 | 51806 | ABM66847.1 | Q69091 |
| QPFAIPYIHPTNPFAPDVTAAAGAG | 1 | 0 | 0 | 51811 | P29326.1 | Q81871 |
| QPGPSREYRYVLRTSPP | 1 | 0 | 0 | 51844 | YP_001129431.1 | Q9QR71 |
| QPSGRRRGRRSGGSGGGFWGDRVDS | 1 | 0 | 0 | 51965 | P29326.1 | Q81871 |
| QQACKGKIQALCENP | 1 | 0 | 0 | 52000 | P08362.1 | Q786F2 |
| QRLKYAQEELSNEVLPPPRK | 1 | 0 | 0 | 52228 | P08617.1 | P08617 |
| QRQKKQPTVTLLPAADMDDFSRQ | 1 | 0 | 0 | 52244 | P59595.1 | P59595 |
| QSHGQLSDHPHALSSSSSHA | 1 | 0 | 0 | 52333 | P07299.1 | Q9PZT0 |
| QSNQRSAPRITFGGPTDS | 1 | 0 | 0 | 52389 | AAS48456.1 | P59595 |
| QSPARTSSDPGTNTT | 1 | 0 | 0 | 52394 | AGB56758.1 | Q05320 |
| QTHRPHPDAELVDA | 1 | 0 | 0 | 52529 | Q81258.3 | P27958 |
| QVKARNIISPVMG | 1 | 0 | 0 | 52720 | AAO86636.1 | O36307 |
| QVLDDHYKTALKEV | 1 | 0 | 0 | 52732 | Q81258.3 | P27958 |
| QVLDSHYQDVLKEV | 1 | 0 | 0 | 52735 | P26664.3 | P27958 |
| QVRNIMSPVMGVIG | 1 | 0 | 0 | 52778 | P27313.1 | P27313 |
| QYDEMEECSQAAPYIEQAQA | 1 | 0 | 0 | 52878 | AAC03058.1 | P27958 |
| QYDEMEECSQAAPYIEQAQV | 1 | 0 | 0 | 52879 | Q81594 | P27958 |
| QYDEMEECSQSAPYIEQAQA | 1 | 0 | 0 | 52882 | Q81487.3 | P27958 |
| QYLYGVGSSIASWAIKWEYV | 1 | 0 | 0 | 53004 | AAA45676.1 | P27958 |
| QYVLATYDTSRVEHA | 1 | 0 | 0 | 53046 | P08362.1 | Q786F2 |
| RAAVCTRGVAKAVDFIPVEN | 1 | 0 | 0 | 53073 | AAY82033.1 | P27958 |
| RAETDDVEPSVAAEC | 1 | 0 | 0 | 53092 | Q81258.3 | P27958 |
| RAILTAFSPAQDIWGTS | 1 | 0 | 0 | 53116 | P59594.1 | P59594 |
| RCEPVVCYLYGKRGGGKSLT | 1 | 0 | 0 | 53267 | P08617.1 | P08617 |
| RCESALHVWVPPLLA | 1 | 0 | 0 | 53269 | Q81258.3 | P27958 |
| RCHARKAVAHINSV | 1 | 0 | 0 | 53276 | P27958.3 | P27958 |
| RDDITFMVGLHSYT | 1 | 0 | 0 | 53331 | Q81258.3 | P27958 |
| RDEVDFCVGLNSFV | 1 | 0 | 0 | 53336 | SRC279960 | P27958 |
| RDEVTFTVGLNSFV | 1 | 0 | 0 | 53337 | P26661.3 | P27958 |
| RDGKLPATQLRRHIDLLVGSATLCS | 1 | 0 | 0 | 53351 | P26664.3 | P27958 |
| RDHVNLVLNRDVS | 1 | 0 | 0 | 53361 | AAO86638.1 | Q9E006 |
| RDTLPQSPGPAFP | 1 | 0 | 0 | 53429 | NP_044666.1 | P06484 |
| RDVSDFTDSVRDPKTSEI | 1 | 0 | 0 | 53435 | P59594.1 | P59594 |
| REEVSFRVGLHEYP | 1 | 0 | 0 | 53494 | P26664.3 | P27958 |
| REVLYQEFDEMEECSQHLPYIE | 1 | 0 | 0 | 53665 | AAB66324.1 | P27958 |
| RFDLELTFVIT | 1 | 0 | 0 | 53691 | BAF95214.1 | Q86887 |
| RFPRGQGVPINTNSGPDDQI | 1 | 0 | 0 | 53758 | AAS48456.1 | P59595 |
| RFQEALKKSLKKP | 1 | 0 | 0 | 53763 | AAO86638.1 | Q9E006 |
| RGADGTAELTTTAATRFMKDLYFTS | 1 | 0 | 0 | 53816 | P29326.1 | Q81871 |
| RGERCDLEDRDR | 1 | 0 | 0 | 53858 | AAB02127.1 | P27958 |
| RGFTSLFSAFSAQ | 1 | 0 | 0 | 53868 | SRC279960 | P27958 |
| RGFTSLFSAGSAQ | 1 | 0 | 0 | 53870 | BAA02337.1 | P27958 |
| RGKGEGLPGMCGGALVSSNQ | 1 | 0 | 0 | 53895 | P08617.1 | P08617 |
| RGLTSLFSAFSAQ | 1 | 0 | 0 | 53923 | SRC279960 | P27958 |
| RGNQGFPWDILFPADPPFSPQSC | 1 | 0 | 0 | 53929 | AAG26087.1 | P29996 |
| RGPRLGVRATRKTSERSQPRGRRQP | 1 | 0 | 0 | 53945 | P26664.3 | P27958 |
| RHLIFCHSKKKCDELAAKLSAL | 1 | 0 | 0 | 54064 | BAA03375.1 | P27958 |
| RHLLLCLLLLTVGG | 1 | 0 | 0 | 54065 | SRC279960 | P27958 |
| RHRLRRGADGTAELTTTAATRFMKD | 1 | 0 | 0 | 54090 | P29326.1 | Q81871 |
| RIHDFRYSQLAKLGINPYTH | 1 | 0 | 0 | 54158 | P07299.1 | Q9PZT0 |
| RIMKVEFTQCSMNVVSKTLF | 1 | 0 | 0 | 54197 | P08617.1 | P08617 |
| RIRADFNASMDLL | 1 | 0 | 0 | 54232 | P26660.3 | P27958 |
| RITFGGPTDSTDNNQNGGR | 1 | 0 | 0 | 54296 | AAS48456.1 | P59595 |
| RKFPPALPIWARPDYNPPLL | 1 | 0 | 0 | 54346 | P26662.3 | P27958 |
| RKPARLIVFPDL | 1 | 0 | 0 | 54402 | P26663.3 | P27958 |
| RKPLVSFKEDPTPSDNPFSK | 1 | 0 | 0 | 54408 | P03421.1 | O42062 |
| RKTSERSQPRGRRQPIPKARRPEGR | 1 | 0 | 0 | 54442 | P26662.3 | P27958 |

| | | | | | | |
|---|---|---|---|---|---|---|
| RLGVPDTLGGDPKFRSLTHE | 1 | 0 | 0 | 54566 | P07299.1 | Q9PZT0 |
| RLLARGGRAAICGK | 1 | 0 | 0 | 54633 | P26664.3 | P27958 |
| RMADAVSRKKMDTK | 1 | 0 | 0 | 54848 | P27313.1 | P27313 |
| RMMATKDSFQSFN | 1 | 0 | 0 | 54923 | AAO86638.1 | Q9E006 |
| RMYVGGVEHRLEAACNWTRG | 1 | 0 | 0 | 54967 | AAA45676.1 | P27958 |
| RNEKWREQLGQKFEE | 1 | 0 | 0 | 54980 | ACP44180.1 | P03508 |
| RNFTTAPAICHEGKAYF | 1 | 0 | 0 | 54989 | NP_828851.1 | P59594 |
| RNLTPGNTNTRVSRYSSTARHRLRR | 1 | 0 | 0 | 55026 | P29326.1 | Q81871 |
| RNRSPERCDLGDDLHLQ | 1 | 0 | 0 | 55055 | YP_001129431.1 | Q9QR71 |
| RNTIMASKTVGTAE | 1 | 0 | 0 | 55066 | P27313.1 | P27313 |
| RPDYQPATVAGCALPPP | 1 | 0 | 0 | 55137 | P26660.3 | P27958 |
| RPEGRTWAQPGYPWPLYGNEGCGWA | 1 | 0 | 0 | 55143 | P26664.3 | P27958 |
| RPFERDISNVPFSPDGK | 1 | 0 | 0 | 55149 | P59594.1 | P59594 |
| RPGYEPPTVLGCALPPT | 1 | 0 | 0 | 55168 | P26661.3 | P27958 |
| RPIDDFDQGWGPITY | 1 | 0 | 0 | 55174 | AAN40662.1 | P27958 |
| RPKQRRPQGLPNNTASWF | 1 | 0 | 0 | 55196 | AAS48456.1 | P59595 |
| RPLGLQGCAFQSTVAELQRLKMKVG | 1 | 0 | 0 | 55207 | P29326.1 | Q81871 |
| RPLIDFDQGWGPISY | 1 | 0 | 0 | 55211 | SRC279960 | P27958 |
| RPLTETRGDLFSGDEDSDSSD | 1 | 0 | 0 | 55227 | AAA45992.1 | Q6SW99 |
| RPPTPWPISQDSDRASYMDT | 1 | 0 | 0 | 55275 | AAA45412.1 | Q85601 |
| RPSWGPTDPRRRSRNLGKVLDTLT | 1 | 0 | 0 | 55351 | SRC279960 | P27958 |
| RPTTAGAAPLTAVAPAHDTPPVPDV | 1 | 0 | 0 | 55362 | P29326.1 | Q81871 |
| RPYCWHYPPQQCGIVPASQV | 1 | 0 | 0 | 55383 | BAA02670.1 | P27958 |
| RQKLKDAERAVEVD | 1 | 0 | 0 | 55436 | P27313.1 | P27313 |
| RQYNLSTSPLTSSVATGTNLVLYAA | 1 | 0 | 0 | 55519 | P29326.1 | Q81871 |
| RRGRRSGGSGGGFWGDRVDSQPFAI | 1 | 0 | 0 | 55587 | P29326.1 | Q81871 |
| RRKLTNPANKEES | 1 | 0 | 0 | 55627 | AAO86638.1 | Q9E006 |
| RRLHAWITEDCPVP | 1 | 0 | 0 | 55643 | P26661.3 | P27958 |
| RRLHNWITEDCPIP | 1 | 0 | 0 | 55644 | P26660.3 | P27958 |
| RRLHQWINEDYPSP | 1 | 0 | 0 | 55645 | Q81258.3 | P27958 |
| RRLHQWISSECTTP | 1 | 0 | 0 | 55646 | P26664.3 | P27958 |
| RRPKHLYVSMPTAQ | 1 | 0 | 0 | 55675 | P27313.1 | P27313 |
| RRQPIPKARRPEGRAWAQPG | 1 | 0 | 0 | 55699 | BAA02670.1 | P27958 |
| RSELSQLSMYRVFEV | 1 | 0 | 0 | 55826 | P08362.1 | Q786F2 |
| RSFGSSSTSGITGD | 1 | 0 | 0 | 55834 | P26664.3 | P27958 |
| RSKFGYSAKDVRSL | 1 | 0 | 0 | 55870 | Q81258.3 | P27958 |
| RSKYGFGAKEVRSL | 1 | 0 | 0 | 55877 | P26660.3 | P27958 |
| RSRGNSRNSTPGSSR | 1 | 0 | 0 | 55940 | AAS48456.1 | P59595 |
| RSRRAVPVAVWLVSALAMGAGVA | 1 | 0 | 0 | 55950 | AAU04921.1 | P03381 |
| RTALNCNDSLQTGFLAALFY | 1 | 0 | 0 | 56017 | BAA02670.1 | P27958 |
| RVCSCLWMMLLISQAEA | 1 | 0 | 0 | 56248 | AAA45676.1 | P27958 |
| RVEAQLHVWIPPLN | 1 | 0 | 0 | 56259 | P26664.3 | P27958 |
| RVEHAVVYYVYSPSR | 1 | 0 | 0 | 56275 | P08362.1 | Q786F2 |
| RVFEVGVIRNPGLGA | 1 | 0 | 0 | 56283 | P08362.1 | Q786F2 |
| RVGTDSGFAAYNRYRI | 1 | 0 | 0 | 56302 | NP_828855.1 | P59596 |
| RVTVSTRRSQQTIIPNIG | 1 | 0 | 0 | 56434 | P04664.1 | P03452 |
| RVVVTPDKEILYEA | 1 | 0 | 0 | 56448 | AAP55696.1 | P27958 |
| RYGNVLDVNAIDIE | 1 | 0 | 0 | 56537 | P27313.1 | P27313 |
| RYSSTARHRLRRGADGTAELTTTAA | 1 | 0 | 0 | 56660 | P29326.1 | Q81871 |
| SAAHTAFGIANLFRMG | 1 | 0 | 0 | 56701 | SRC279960 | P27958 |
| SAARIHDFRYSQLAKLGINP | 1 | 0 | 0 | 56712 | P07299.1 | Q9PZT0 |
| SAARSTGSFAKLFMPG | 1 | 0 | 0 | 56713 | SRC279960 | P27958 |
| SAASCNGFLYFVIF | 1 | 0 | 0 | 56715 | P26660.3 | P27958 |
| SAAYATSSFSSLFTRG | 1 | 0 | 0 | 56718 | SRC279960 | P27958 |
| SAAYTAFGIANLFRVG | 1 | 0 | 0 | 56719 | SRC279960 | P27958 |
| SAGGQLFYSRPVVSANGE | 1 | 0 | 0 | 56796 | P29326.1 | Q81871 |
| SAGRSTGSLVRLFAPG | 1 | 0 | 0 | 56804 | SRC279960 | P27958 |
| SALEWIDLDSSL | 1 | 0 | 0 | 56856 | P27312.1 | P27312 |
| SAMSTTDLEAYFKDC | 1 | 0 | 0 | 56900 | ABK27207.1 | Q69027 |
| SANGEPTVKLYTSVENAQQDKGIAI | 1 | 0 | 0 | 56908 | P29326.1 | Q81871 |
| SARDGIRDAMIGLREEMIEK | 1 | 0 | 0 | 56962 | P03421.1 | O42062 |
| SASLPYMDETRAIAGQFKEK | 1 | 0 | 0 | 56993 | AAA65789.1 | P27958 |
| SASQLSAPSLKATCTTHHDS | 1 | 0 | 0 | 56996 | P26662.3 | P27958 |
| SDPSQGGGIKITHFTTWTSI | 1 | 0 | 0 | 57261 | P08617.1 | P08617 |
| SDSESCSSMPPLEG | 1 | 0 | 0 | 57279 | Q81258.3 | P27958 |
| SDSVGQQIKVIPVDPYFFQM | 1 | 0 | 0 | 57296 | P08617.1 | P08617 |
| SEAFMSTNKMYFL | 1 | 0 | 0 | 57351 | AAO86636.1 | O36307 |
| SEKTIWSLIAWQRSNAEFEQ | 1 | 0 | 0 | 57508 | P08617.1 | P08617 |
| SELLSLINDMPITNDQKKLMS | 1 | 0 | 0 | 57524 | P12568.1 | O36634 |
| SELSPLLLTTTQWQVLPCSF | 1 | 0 | 0 | 57540 | AAA45676.1 | P27958 |
| SERSQPRGRRQPIPKARRPEGRT | 1 | 0 | 0 | 57617 | P26663.3 | P27958 |
| SESTIGDVLQQMAI | 1 | 0 | 0 | 57635 | AAA03612.1 | P27958 |
| SFGVWIRTPPAYRPPNAPIL | 1 | 0 | 0 | 57784 | CAL29866.1 | P0C767 |
| SFHNLHLLFDEYTNIPVSIL | 1 | 0 | 0 | 57789 | AAG48731.1 | P03410 |
| SFQSFNLTEPHIT | 1 | 0 | 0 | 57875 | AAO86638.1 | Q9E006 |
| SFWEAGTTKAGYPYNYNTTA | 1 | 0 | 0 | 57943 | AAG16766.1 | Q81871 |
| SGDVVRFPNITNLCPFG | 1 | 0 | 0 | 58001 | P59594.1 | P59594 |
| SGHMAWDMMNWSPT | 1 | 0 | 0 | 58073 | SRC279960 | P27958 |
| SGKGVSFQLVKLGVW | 1 | 0 | 0 | 58103 | P08362.1 | Q786F2 |
| SGKPAIIPDREVLYREFDEM | 1 | 0 | 0 | 58112 | AAA45676.1 | P27958 |
| SGKSLLHEVDKDISQLTQAIVK | 1 | 0 | 0 | 58115 | AAL05538.1 | P03381 |
| SGNCDVVIGIINNTVYD | 1 | 0 | 0 | 58143 | NP_828851.1 | P59594 |
| SGSLGTLSSPSPSAV | 1 | 0 | 0 | 58234 | CAB41697.1 | Q69027 |
| SGVFTTSMGNTMTC | 1 | 0 | 0 | 58285 | P26661.3 | P27958 |
| SGVLPTSFGNTITC | 1 | 0 | 0 | 58291 | Q81258.3 | P27958 |
| SGVLTTSCGNTLTC | 1 | 0 | 0 | 58293 | P26664.3 | P27958 |
| SGVLTTSMGNTITC | 1 | 0 | 0 | 58294 | P26660.3 | P27958 |
| SGYKRMMATKDSF | 1 | 0 | 0 | 58324 | AAO86638.1 | Q9E006 |
| SHIECRKPYKELRLEVGKQR | 1 | 0 | 0 | 58366 | P08617.1 | P08617 |
| SHRGVIADNQAKWAV | 1 | 0 | 0 | 58403 | P08362.1 | Q786F2 |
| SIEHQVKDVLTPLFK | 1 | 0 | 0 | 58485 | P08362.1 | Q786F2 |

| | | | | | | |
|---|---|---|---|---|---|---|
| SILQDMRNTIMASK | 1 | 0 | 0 | 58614 | P27313.1 | P27313 |
| SIRNGTYDHDVYRDE | 1 | 0 | 0 | 58691 | AFM72565.1 | P03452 |
| SISFWPQTTTTPTSVDMNSITSTDV | 1 | 0 | 0 | 58700 | P29326.1 | Q81871 |
| SIVYEAADAILHTPGCVPCV | 1 | 0 | 0 | 58745 | AAA45676.1 | P27958 |
| SIYPGTITGHRMAW | 1 | 0 | 0 | 58768 | P26660.3 | P27958 |
| SKARPYIEQAQVIAHQFKEK | 1 | 0 | 0 | 58788 | AAC53958.1 | P27958 |
| SKHLPLVEHGLQLAEQFKQK | 1 | 0 | 0 | 58845 | O39929.3 | P27958 |
| SKLLTLVQLTLQSTNYTCIVCID | 1 | 0 | 0 | 58884 | AAA96674.1 | P03381 |
| SKTRAIDACPLDYSILCRMY | 1 | 0 | 0 | 58951 | P08617.1 | P08617 |
| SKTVGTAEEKLKKK | 1 | 0 | 0 | 58954 | P27313.1 | P27313 |
| SLGADSSIAYSNNTIAI | 1 | 0 | 0 | 59120 | NP_828851.1 | P59594 |
| SLGKMVHQIFGSAYT | 1 | 0 | 0 | 59137 | QTX93593.1 | P17763 |
| SLIAWQRSNAEFEQNLENAQ | 1 | 0 | 0 | 59160 | P08617.1 | P08617 |
| SLINTNGSWHIN | 1 | 0 | 0 | 59192 | AAP55688.1 | P27958 |
| SLKLESSCNFDL | 1 | 0 | 0 | 59222 | P27312.1 | P27312 |
| SLLLEVDKDISHLTQAIVKNHQN | 1 | 0 | 0 | 59270 | P03383.1 | P03383 |
| SLNTGWLAGLFYQHKFNS | 1 | 0 | 0 | 59375 | P27958.3 | P27958 |
| SLSPVPTLGSRSRRAVPVAV | 1 | 0 | 0 | 59467 | P03381.1 | P03381 |
| SLSTNLDVTNSIEHQ | 1 | 0 | 0 | 59483 | P08362.1 | Q786F2 |
| SLYPGHLSGHRMAW | 1 | 0 | 0 | 59619 | Q81258.3 | P27958 |
| SNTVKVVGKGGHS | 1 | 0 | 0 | 59939 | AAO86638.1 | Q9E006 |
| SNVPFSPDGKPCTPPAL | 1 | 0 | 0 | 59944 | P59594.1 | P59594 |
| SPAPSRPFSVLRANDVLWLS | 1 | 0 | 0 | 59984 | AAG16766.1 | Q81871 |
| SPAQDIWGTSAAAYFVG | 1 | 0 | 0 | 59985 | P59594.1 | P59594 |
| SPDGKPCTPPALNCYWP | 1 | 0 | 0 | 60014 | P59594.1 | P59594 |
| SPPLVHDSDLEHVLTPS | 1 | 0 | 0 | 60186 | NP_041006.1 | P03383 |
| SPSAVPTDHGAHLSL | 1 | 0 | 0 | 60249 | P03165.2 | Q69027 |
| SPSLNLASVPKTSTPPGEKP | 1 | 0 | 0 | 60259 | AAA45412.1 | Q85601 |
| SQAAPYIEQAQAIAHQFKEK | 1 | 0 | 0 | 60359 | AAC03058.1 | P27958 |
| SQAAPYIEQAQVIAHQFKEK | 1 | 0 | 0 | 60360 | Q81594 | P27958 |
| SQAGPYIEQAQVIAHQFKEK | 1 | 0 | 0 | 60367 | AAC53966.1 | P27958 |
| SQARPYIEQAQVIAHQFKEK | 1 | 0 | 0 | 60377 | Q81495.3 | P27958 |
| SQAWQPGVAMPNLYK | 1 | 0 | 0 | 60394 | P0DTD1.1 | P0DTD1 |
| SQGISDDDNDSAVAEFFQSF | 1 | 0 | 0 | 60441 | P08617.1 | P08617 |
| SQHLPYIEQGMMLAEQFKQK | 1 | 0 | 0 | 60450 | P26664.3 | P27958 |
| SQSAPYIEQAQAIAQQFKDK | 1 | 0 | 0 | 60545 | Q81487.3 | P27958 |
| SQWAREALQTGITLVALLLLVIL | 1 | 0 | 0 | 60591 | P03381.1 | P03381 |
| SRGAPGGGFVPSLQGVP | 1 | 0 | 0 | 60663 | CAA44973.1 | P29996 |
| SRGNHDSPTHYVPESDAAAR | 1 | 0 | 0 | 60674 | AAA75355.1 | P27958 |
| SRGNHVAPTHYVTESDASQR | 1 | 0 | 0 | 60675 | AAU89634.1 | P27958 |
| SRGNHVAPTHYVVESDASQR | 1 | 0 | 0 | 60676 | BAA01761.1 | P27958 |
| SRGNHVSPAHYVPESDAAAR | 1 | 0 | 0 | 60677 | Q81495.3 | P27958 |
| SRGNHVSPRHYVPESEPAAR | 1 | 0 | 0 | 60678 | AAA45721.1 | P27958 |
| SRGNHVSPTHYVPESDAAAK | 1 | 0 | 0 | 60679 | Q68870.1 | P27958 |
| SRGNHVSPTHYVPESDAAAR | 1 | 0 | 0 | 60680 | P26662.3 | P27958 |
| SRGNSPARMASGGGETAL | 1 | 0 | 0 | 60681 | AAS48456.1 | P59595 |
| SRKAYDHNSPYIWPRNDYDG | 1 | 0 | 0 | 60710 | AAK01056.1 | P09259 |
| SRKKMDTKPTDPTG | 1 | 0 | 0 | 60714 | P27313.1 | P27313 |
| SRKSYKKMLKEMGEVAPEYR | 1 | 0 | 0 | 60721 | P03418.1 | O42053 |
| SRNSTPGSSRGNSPARMA | 1 | 0 | 0 | 60749 | AAS48456.1 | P59595 |
| SRSTSLSVSLVLVGVVTLYLGAMV | 1 | 0 | 0 | 60811 | AAS49691.2 | P17763 |
| SRVKNLNSSEGVPDLLV | 1 | 0 | 0 | 60827 | P59637.1 | P59637 |
| SRVVIQDYDNQHEQDRPTPSPAPSR | 1 | 0 | 0 | 60833 | P29326.1 | Q81871 |
| SSAEGDLVGWPSPP | 1 | 0 | 0 | 60880 | P26660.3 | P27958 |
| SSCHNASGKEAKVCTISPIM | 1 | 0 | 0 | 60923 | P07299.1 | Q9PZT0 |
| SSEVAVLYQDVNCTDV | 1 | 0 | 0 | 60978 | P59594.1 | P59594 |
| SSGLYHVTNDC | 1 | 0 | 0 | 61037 | NP_671491.1 | P27958 |
| SSIVYEAADAILHT | 1 | 0 | 0 | 61123 | P26664.3 | P27958 |
| SSIVYEAADMIMHT | 1 | 0 | 0 | 61124 | P26662.3 | P27958 |
| SSIVYEADDVILHT | 1 | 0 | 0 | 61125 | Q81258.3 | P27958 |
| SSKLLPKMPGIPFI | 1 | 0 | 0 | 61139 | P26661.3 | P27958 |
| SSLRDHINVIVS | 1 | 0 | 0 | 61186 | P27312.1 | P27312 |
| SSLRYGNVLDVNAI | 1 | 0 | 0 | 61192 | P27313.1 | P27313 |
| SSPCHNSLILPPFSLSPVPTLGSRS | 1 | 0 | 0 | 61260 | AAU04921.1 | P03381 |
| SSTRGGSRVEGIFAGLFMNA | 1 | 0 | 0 | 61410 | P03418.1 | O42053 |
| SSTTSKVPPSPGGE | 1 | 0 | 0 | 61420 | Q81258.3 | P27958 |
| STCSEEDDSVVCCS | 1 | 0 | 0 | 61541 | P26660.3 | P27958 |
| STGADTGGDIVQQP | 1 | 0 | 0 | 61613 | P26661.3 | P27958 |
| STGTSQNTRISLRPGPVSQP | 1 | 0 | 0 | 61638 | P07299.1 | Q9PZT0 |
| STLQVSDVDKLVCRD | 1 | 0 | 0 | 61752 | AGB56832.1 | P60170 |
| STNPKPQKKNKRNTNRRPQDVKFP | 1 | 0 | 0 | 61789 | P26664.3 | P27958 |
| STVSDQEDSEICCS | 1 | 0 | 0 | 61992 | SRC279960 | P27958 |
| STVSEEASAEDVVC | 1 | 0 | 0 | 61994 | SRC279960 | P27958 |
| STVSSEANAEDVVC | 1 | 0 | 0 | 61996 | P26664.3 | P27958 |
| SVGHMTGRFTSLFSVG | 1 | 0 | 0 | 62141 | SRC279960 | P27958 |
| SVLCPRRGHKKTV | 1 | 0 | 0 | 62199 | AAO86638.1 | Q9E006 |
| SVSHARPRLLLLCL | 1 | 0 | 0 | 62325 | P26661.3 | P27958 |
| SVSRARTRHLLLCL | 1 | 0 | 0 | 62334 | Q81258.3 | P27958 |
| SVSTKEGDSSNTGAGKALTG | 1 | 0 | 0 | 62336 | P07299.1 | Q9PZT0 |
| SWEVGKPRPPLNRNY | 1 | 0 | 0 | 62435 | P0DTD1.1 | P0DTD1 |
| SWGANDTDVFVL | 1 | 0 | 0 | 62447 | P27958.3 | P27958 |
| SYHSKPCNPAQPVCSWTLDLLAL | 1 | 0 | 0 | 62567 | AAA96674.1 | P03381 |
| SYMDTPSTCLAIVYRPIGVP | 1 | 0 | 0 | 62650 | AAA45412.1 | Q85601 |
| SYSDPCSLKCPYLGCQSWTCPYT | 1 | 0 | 0 | 62700 | AAA96674.1 | P03381 |
| TAAAKAAGLRNPDF | 1 | 0 | 0 | 62771 | BAA00972.1 | P27958 |
| TAAVTSPLTTSQTL | 1 | 0 | 0 | 62807 | P26664.3 | P27958 |
| TAEELEAIKHQLNPKDNDVEL | 1 | 0 | 0 | 62828 | P03418.1 | O42053 |
| TAETAKRRLARGSPPSLA | 1 | 0 | 0 | 62837 | P26662.3 | P27958 |
| TAGHGAYGITSLFTRG | 1 | 0 | 0 | 62852 | SRC279960 | P27958 |
| TAGRTMSGFTSFLSPG | 1 | 0 | 0 | 62867 | SRC279960 | P27958 |
| TAGWTFGAGAALQIPFA | 1 | 0 | 0 | 62872 | NP_828851.1 | P59594 |

| | | | | | | |
|---|---|---|---|---|---|---|
| TALGKLQDVVNQNAQAL | 1 | 0 | 0 | 62908 | NP_828851.1 | P59594 |
| TALVVAQLLRIPQAI | 1 | 0 | 0 | 62924 | Q5EG65.3 | P27958 |
| TALVVSQLLRIPQAV | 1 | 0 | 0 | 62925 | P26662.3 | P27958 |
| TANHDSPDAELIEA | 1 | 0 | 0 | 62933 | P26664.3 | P27958 |
| TAQSTMKAEEITP | 1 | 0 | 0 | 62975 | AAO86636.1 | O36307 |
| TASITSPLTTQNTL | 1 | 0 | 0 | 62993 | P26662.3 | P27958 |
| TATALPDQASDDGD | 1 | 0 | 0 | 63008 | P26663.3 | P27958 |
| TAWDFGSLGGVFTSIG | 1 | 0 | 0 | 63044 | ABX25775.1 | P17763 |
| TDPASIAARGYISTR | 1 | 0 | 0 | 63206 | P17763.2 | P17763 |
| TDQMEVDSGPRKRPLRGG | 1 | 0 | 0 | 63217 | CAA44973.1 | P29996 |
| TDWDVKGGGSPLYRHGD | 1 | 0 | 0 | 63273 | AAA96965.1 | A0A159D718 |
| TEASNYAQYRVARATIRYRPLVPNA | 1 | 0 | 0 | 63307 | P29326.1 | Q81871 |
| TECANLLLQYGSFCTQL | 1 | 0 | 0 | 63309 | NP_828851.1 | P59594 |
| TEGRTANAFASLLSPG | 1 | 0 | 0 | 63384 | SRC279960 | P27958 |
| TEGRTANTFASLLSPG | 1 | 0 | 0 | 63385 | SRC279960 | P27958 |
| TESTLSTALAELAT | 1 | 0 | 0 | 63540 | P26664.3 | P27958 |
| TESTVSSALAELATKTFGS | 1 | 0 | 0 | 63543 | P26662.3 | P27958 |
| TETAAPAVADARKPPSGKKK | 1 | 0 | 0 | 63545 | AAC56957.1 | Q2HR63 |
| TEVEDDREPSVPSEY | 1 | 0 | 0 | 63571 | P26661.3 | P27958 |
| TFHTMWHVTRGAVLM | 1 | 0 | 0 | 63672 | QTX93024.1 | P17763 |
| TFKLSYGIATVREVL | 1 | 0 | 0 | 63688 | P0DTD1.1 | P0DTD1 |
| TGHMLDMYSVMLTND | 1 | 0 | 0 | 63847 | P0DTD1.1 | P0DTD1 |
| TGLSTGTSQNTRISLRPGPV | 1 | 0 | 0 | 63887 | P07299.1 | Q9PZT0 |
| TGLTHIDAHFLSQTKQSGEN | 1 | 0 | 0 | 63890 | AAY82033.1 | P27958 |
| TGMQSWVPLSTDDPV | 1 | 0 | 0 | 63895 | P06830.1 | Q786F2 |
| TGREIVDLMCHATFT | 1 | 0 | 0 | 63946 | QUD37222.1 | P17763 |
| TGRPSHEAPNMTQ | 1 | 0 | 0 | 63953 | NP_044666.1 | P06484 |
| TGTDLELIQILKDHYNISLD | 1 | 0 | 0 | 63973 | P07299.1 | Q9PZT0 |
| THYVPESDAAARVTQILSSL | 1 | 0 | 0 | 64105 | AAA86907.1 | P27958 |
| TIRGQFSNMSLSLLD | 1 | 0 | 0 | 64444 | P08362.1 | Q786F2 |
| TITDAVDCSQNPLAELK | 1 | 0 | 0 | 64477 | P59594.1 | P59594 |
| TITTGSPITYSTYGK | 1 | 0 | 0 | 64490 | P26664.3 | P27958 |
| TKEGDSSNTGAGKALTGLST | 1 | 0 | 0 | 64547 | P07299.1 | Q9PZT0 |
| TKKNYSELDDEEPMELDYP | 1 | 0 | 0 | 64600 | P59634.1 | P59634 |
| TKKPNRNGGGYYSASYSDPCSL | 1 | 0 | 0 | 64602 | AAA96674.1 | P03381 |
| TKPTDPTGIEPDDH | 1 | 0 | 0 | 64645 | P27313.1 | P27313 |
| TKPVASDYWDGYSGQLVCII | 1 | 0 | 0 | 64651 | P08617.1 | P08617 |
| TKRNTNRRPQDVKFPGGGQI | 1 | 0 | 0 | 64662 | BAA02670.1 | P27958 |
| TKSGEWLLGILNG | 1 | 0 | 0 | 64671 | AAO86638.1 | Q9E006 |
| TLETKLGELKRQL | 1 | 0 | 0 | 64782 | AAO86636.1 | O36307 |
| TLGGDPKFRSLTHEDHAIQP | 1 | 0 | 0 | 64815 | P07299.1 | Q9PZT0 |
| TLKYFGGFNFSQILPDP | 1 | 0 | 0 | 64888 | NP_828851.1 | P59594 |
| TLNKDQLLSSSKYTIQRSTG | 1 | 0 | 0 | 64978 | P03418.1 | O42053 |
| TLPFNWTHCFDPQIQAIVS | 1 | 0 | 0 | 65003 | AAL05539.1 | P03381 |
| TLTLAMPKTTYELKMECPH | 1 | 0 | 0 | 65099 | AAO86638.1 | Q9E006 |
| TLTSGSDLDRCTTFDDV | 1 | 0 | 0 | 65110 | P59594.1 | P59594 |
| TMTSQGMYGGTYPVE | 1 | 0 | 0 | 65269 | P06830.1 | Q786F2 |
| TNLEDPASRDLVVSY | 1 | 0 | 0 | 65360 | P03147.1 | Q76R61 |
| TNTRVSRYSSTARHRLRRGADGTAE | 1 | 0 | 0 | 65464 | P29326.1 | Q81871 |
| TNVDQDLVGWPAPP | 1 | 0 | 0 | 65473 | Q81258.3 | P27958 |
| TNVDQDLVGWPAPQ | 1 | 0 | 0 | 65474 | P26664.3 | P27958 |
| TNVVIRACNFELCDNPF | 1 | 0 | 0 | 65479 | P59594.1 | P59594 |
| TNYLEQPVSNDLSNC | 1 | 0 | 0 | 65486 | P08362.1 | Q786F2 |
| TPCAAEEQKLPINA | 1 | 0 | 0 | 65515 | P26664.3 | P27958 |
| TPCSAEEEKLPISP | 1 | 0 | 0 | 65520 | Q81258.3 | P27958 |
| TPLFKIIGDEVGLRT | 1 | 0 | 0 | 65635 | P08362.1 | Q786F2 |
| TPLLYPSLALPAPHLTLPFNWTH | 1 | 0 | 0 | 65644 | P03381.1 | P03381 |
| TPPMPSIGLEEEE | 1 | 0 | 0 | 65691 | NP_044666.1 | P06484 |
| TPPPRRRRTVGLSE | 1 | 0 | 0 | 65694 | P26660.3 | P27958 |
| TPPTTPVSPKVPPAFFQSMR | 1 | 0 | 0 | 65699 | AAG48731.1 | P03410 |
| TPSPVVVGTTDRSGAPTYSW | 1 | 0 | 0 | 65774 | AAA45676.1 | P27958 |
| TPVGLAVDTPWVEKESALSI | 1 | 0 | 0 | 65797 | P08617.1 | P08617 |
| TQAHTTMRLANLFTSG | 1 | 0 | 0 | 65820 | SRC279960 | P27958 |
| TQDLFLPFYSNVTGFHT | 1 | 0 | 0 | 65832 | P59594.1 | P59594 |
| TQGCNCSIYPGHITGHRMAW | 1 | 0 | 0 | 65848 | AAA45676.1 | P27958 |
| TQIQVRNIMSPVMG | 1 | 0 | 0 | 65881 | P27313.1 | P27313 |
| TQSFLATCVNGVCW | 1 | 0 | 0 | 65931 | P26662.3 | P27958 |
| TQSFLGTTISGVLW | 1 | 0 | 0 | 65932 | P26660.3 | P27958 |
| TQTFLGTSISGVLW | 1 | 0 | 0 | 65947 | P26661.3 | P27958 |
| TQWQVLPCSFTTLPALSTGL | 1 | 0 | 0 | 65971 | AAA45676.1 | P27958 |
| TRKYLPAIVREAIKR | 1 | 0 | 0 | 66049 | AEX65549.1 | P17763 |
| TRLTSKGRPLVPT | 1 | 0 | 0 | 66066 | NP_044666.1 | P06484 |
| TRRVRGGDGKMKELSPRW | 1 | 0 | 0 | 66094 | AAS48456.1 | P59595 |
| TSAEGDLVGWPSPP | 1 | 0 | 0 | 66144 | P26661.3 | P27958 |
| TSGITGDNTTTSSE | 1 | 0 | 0 | 66219 | P26664.3 | P27958 |
| TSKLFPKMPGLPFI | 1 | 0 | 0 | 66284 | P26660.3 | P27958 |
| TSLGAGPVSISAVAVLAPHSALALL | 1 | 0 | 0 | 66296 | P29326.1 | Q81871 |
| TSLNDRQGNATK | 1 | 0 | 0 | 66304 | P03277.3 | P03277 |
| TSNFRVVPSGDVVRFPN | 1 | 0 | 0 | 66337 | P59594.1 | P59594 |
| TSQGMYGGTYLVEKP | 1 | 0 | 0 | 66385 | P08362.1 | Q786F2 |
| TSQNTRISLRPGPVSQPYHH | 1 | 0 | 0 | 66389 | P07299.1 | Q9PZT0 |
| TSRSASLRQKKVTFDRLQVL | 1 | 0 | 0 | 66406 | P26662.3 | P27958 |
| TSRTLSYYKLGASQRV | 1 | 0 | 0 | 66409 | P59596.1 | P59596 |
| TSRVEHAVVYYVYSP | 1 | 0 | 0 | 66410 | P06830.1 | Q786F2 |
| TSVENAQQDKGIAIPHDIDLGESRV | 1 | 0 | 0 | 66481 | P29326.1 | Q81871 |
| TTAATRFMKDLYFTSTNGVGEIGRG | 1 | 0 | 0 | 66511 | P29326.1 | Q81871 |
| TTATKSEHTGREIVD | 1 | 0 | 0 | 66547 | QTX93593.1 | P17763 |
| TTFHQTLQDPRVRGL | 1 | 0 | 0 | 66607 | P03138.3 | Q76R62 |
| TTHGKAYDVDMVDA | 1 | 0 | 0 | 66633 | P26660.3 | P27958 |
| TTHKTAYDCDMVDA | 1 | 0 | 0 | 66635 | P26661.3 | P27958 |
| TTKAGYPYNYNTTASDQLLVENAAG | 1 | 0 | 0 | 66667 | P29326.1 | Q81871 |

| | | | | | | |
|---|---|---|---|---|---|---|
| TTLPALSTGLIHLHQNIVDV | 1 | 0 | 0 | 66705 | AAA45676.1 | P27958 |
| TTLVQYAVGIMTVTMTFKLG | 1 | 0 | 0 | 66719 | P07299.1 | Q9PZT0 |
| TTVNAHQILPKVLHK | 1 | 0 | 0 | 66895 | AAP79798.1 | Q69027 |
| TTVSTEQNVPDPQVGITTMK | 1 | 0 | 0 | 66906 | P08617.1 | P08617 |
| TVAGCALPPPKKTP | 1 | 0 | 0 | 66938 | P26660.3 | P27958 |
| TVAHATRRFTNLLTRG | 1 | 0 | 0 | 66939 | SRC279960 | P27958 |
| TVARATMQFTGLFTRG | 1 | 0 | 0 | 66947 | SRC279960 | P27958 |
| TVLGCALPPTPQTP | 1 | 0 | 0 | 67069 | P26661.3 | P27958 |
| TVVAPDKEVLYEAFDEMEEC | 1 | 0 | 0 | 67191 | AAC53939.1 | P27958 |
| TVYCNGVKKVIL | 1 | 0 | 0 | 67217 | P27312.1 | P27312 |
| TWGENETDVLLLNNTRPPQ | 1 | 0 | 0 | 67257 | ACE82411.1 | P27958 |
| TWNLDLNSLTTDQRLH | 1 | 0 | 0 | 67283 | P03383.1 | P03383 |
| TYNWGANDTDVFVL | 1 | 0 | 0 | 67404 | Q03463.1 | P27958 |
| TYNWGENETDVFLL | 1 | 0 | 0 | 67405 | AAP55692.1 | P27958 |
| VAAEEYVEVTRVGDFHYVT | 1 | 0 | 0 | 67514 | P26662.3 | P27958 |
| VAAQLAPPSAASAF | 1 | 0 | 0 | 67531 | P26662.3 | P27958 |
| VAESDGVDEDRAAL | 1 | 0 | 0 | 67572 | Q81258.3 | P27958 |
| VAFKIMSGEVPSTEDLVNLL | 1 | 0 | 0 | 67584 | P26664.3 | P27958 |
| VAGRKKLEELERDLRKTK | 1 | 0 | 0 | 67615 | CAA44973.1 | P29996 |
| VAHDASGKRVYYLT | 1 | 0 | 0 | 67624 | P26662.3 | P27958 |
| VAHDGAGKRVYYLT | 1 | 0 | 0 | 67625 | P26664.3 | P27958 |
| VAKLVTQEMFQNIDKKIESQ | 1 | 0 | 0 | 67647 | P08617.1 | P08617 |
| VALDSRGRRRYFLT | 1 | 0 | 0 | 67662 | P26661.3 | P27958 |
| VALGPQGRRRYYLT | 1 | 0 | 0 | 67667 | P26660.3 | P27958 |
| VARALAHGVRVLEDGVNFAT | 1 | 0 | 0 | 67730 | P26660.3 | P27958 |
| VARDDKGRRYYYLT | 1 | 0 | 0 | 67732 | Q81258.3 | P27958 |
| VASHVRVNVYLSAINLECFA | 1 | 0 | 0 | 67746 | P08617.1 | P08617 |
| VATHLAGPQSSSAF | 1 | 0 | 0 | 67768 | Q81258.3 | P27958 |
| VAVRYVGATTASIRS | 1 | 0 | 0 | 67802 | Q81258.3 | P27958 |
| VCEYQGNTISGYK | 1 | 0 | 0 | 67837 | AAO86638.1 | Q9E006 |
| VCGPKLSTDLIKNQCV | 1 | 0 | 0 | 67844 | NP_828851.1 | P59594 |
| VDDPRSEEDKRFESHIECRK | 1 | 0 | 0 | 67922 | P08617.1 | P08617 |
| VDMNSITSTDVRILVQPGIASELVI | 1 | 0 | 0 | 68025 | P29326.1 | Q81871 |
| VDMVAGAHWGVLAGLA | 1 | 0 | 0 | 68027 | P26662.3 | P27958 |
| VDPDDVNKNTLQAR | 1 | 0 | 0 | 68034 | P27313.1 | P27313 |
| VEEEIYQCCNLEPE | 1 | 0 | 0 | 68157 | Q81258.3 | P27958 |
| VEEQEQEQEEQEEQELE | 1 | 0 | 0 | 68167 | YP_001129431.1 | Q9QR71 |
| VEEQEQEQEEQELEEVE | 1 | 0 | 0 | 68168 | YP_001129431.1 | Q9QR71 |
| VELKIKIASGFGPLI | 1 | 0 | 0 | 68272 | P06830.1 | Q786F2 |
| VENLETTMRSPVFT | 1 | 0 | 0 | 68302 | P26664.3 | P27958 |
| VESLDVATRTPSFS | 1 | 0 | 0 | 68346 | P26661.3 | P27958 |
| VESMETTMRSPVFT | 1 | 0 | 0 | 68347 | P26662.3 | P27958 |
| VETLDIVTRSPTFS | 1 | 0 | 0 | 68379 | P26660.3 | P27958 |
| VFHLYLQYIRKLHDE | 1 | 0 | 0 | 68484 | P0DTD1.1 | P0DTD1 |
| VFSPAASSCHNASGKEAKVC | 1 | 0 | 0 | 68567 | P07299.1 | Q9PZT0 |
| VFVLGGCRHKLVCAPAPCNF | 1 | 0 | 0 | 68600 | AAP79798.1 | Q69027 |
| VGAESCGRPFSGSLG | 1 | 0 | 0 | 68618 | ABG23418.1 | Q69027 |
| VGGPLTNSKGQNCGYRRCR | 1 | 0 | 0 | 68664 | P26662.3 | P27958 |
| VGGVYLLPRRGPRLGVRAIR | 1 | 0 | 0 | 68674 | BAA02670.1 | P27958 |
| VGGVYLLPRRGPRLGVRATRKTSER | 1 | 0 | 0 | 68676 | P26664.3 | P27958 |
| VGLRTPQRFTDLVKF | 1 | 0 | 0 | 68723 | P08362.1 | Q786F2 |
| VGMVVAHVLRLPQTL | 1 | 0 | 0 | 68730 | Q81258.3 | P27958 |
| VGTAEEKLKKKSSF | 1 | 0 | 0 | 68800 | P27313.1 | P27313 |
| VHDSDLEHVLTPSTSWTTKILKFI | 1 | 0 | 0 | 68842 | NP_041006.1 | P03383 |
| VIGGAGNNTLHCPTDCFRKH | 1 | 0 | 0 | 68962 | AAA45676.1 | P27958 |
| VIGIINNTVYDPLQPEL | 1 | 0 | 0 | 68971 | NP_828851.1 | P59594 |
| VIPSERLHYRNQGWRSVETSGVAEE | 1 | 0 | 0 | 69074 | P29326.1 | Q81871 |
| VITPGTNASSEVAVLY | 1 | 0 | 0 | 69129 | P59594.1 | P59594 |
| VKEAIDRRLHFKVEVKPASF | 1 | 0 | 0 | 69203 | P08617.1 | P08617 |
| VKNHKNLLKIAQYAAQNRRGLDL | 1 | 0 | 0 | 69279 | AAU04921.1 | P03381 |
| VKPASFFKNPHNDMLNVNLA | 1 | 0 | 0 | 69287 | P08617.1 | P08617 |
| VKQMYKTPTLKYFGGFN | 1 | 0 | 0 | 69299 | NP_828851.1 | P59594 |
| VLFVMFLSLIGLLAI | 1 | 0 | 0 | 69496 | P06830.1 | Q786F2 |
| VLLFLLLADARVCSCLWMML | 1 | 0 | 0 | 69593 | AAA45676.1 | P27958 |
| VLNRDVSFQDLSD | 1 | 0 | 0 | 69673 | AAO86638.1 | Q9E006 |
| VLPPLLTDDMIAAYTAA | 1 | 0 | 0 | 69699 | NP_828851.1 | P59594 |
| VLPPPRKMKGLFSQAKISLF | 1 | 0 | 0 | 69701 | P08617.1 | P08617 |
| VLQQLAVKTFGQPP | 1 | 0 | 0 | 69723 | AAA17048.1 | P27958 |
| VLRTSPPHRPGVRMRRV | 1 | 0 | 0 | 69746 | YP_001129431.1 | Q9QR71 |
| VLSFELLNAPATVCGPK | 1 | 0 | 0 | 69760 | P59594.1 | P59594 |
| VLTLDNQDLNGNWYD | 1 | 0 | 0 | 69808 | P0DTD1.1 | P0DTD1 |
| VLVLFGFFTLSPWY | 1 | 0 | 0 | 69828 | Q81258.3 | P27958 |
| VLVLNPSVAATLGFGAYMSK | 1 | 0 | 0 | 69833 | P26664.3 | P27958 |
| VLVVLLLFAGVDAETH | 1 | 0 | 0 | 69847 | AAA45676.1 | P27958 |
| VLYEAFDEMEECASKATLIE | 1 | 0 | 0 | 69861 | BAA01956.1 | P27958 |
| VLYEAFDEMEECASRAALIE | 1 | 0 | 0 | 69862 | AAU89634.1 | P27958 |
| VLYEAFDEMEECASRAALLE | 1 | 0 | 0 | 69863 | AAC53939.1 | P27958 |
| VLYENQKQIANQFNKAI | 1 | 0 | 0 | 69865 | NP_828851.1 | P59594 |
| VLYQAFDEMEECASHLPYIE | 1 | 0 | 0 | 69880 | AAC53957.1 | P27958 |
| VLYQEFDEMEECASHLPYFE | 1 | 0 | 0 | 69884 | AAA45721.1 | P27958 |
| VLYQEFDEMEECASHLPYIE | 1 | 0 | 0 | 69885 | ABU97068.1 | P27958 |
| VLYQGFDEMEECASHLPYIE | 1 | 0 | 0 | 69889 | AAC53946.1 | P27958 |
| VLYREFDEMEECSQHLPYIE | 1 | 0 | 0 | 69905 | P26664.3 | P27958 |
| VLYSPNVSVPSSSSTPLLYPSLA | 1 | 0 | 0 | 69909 | P03381.1 | P03381 |
| VMSGEMPSTEDLVN | 1 | 0 | 0 | 70016 | P26662.3 | P27958 |
| VMTVEIRKQNMTEFMELWSQ | 1 | 0 | 0 | 70026 | P08617.1 | P08617 |
| VNERAVVAPDKEVLYEAFDE | 1 | 0 | 0 | 70061 | AAC53944.1 | P27958 |
| VNQRAVVAPDKEVLYEAFDE | 1 | 0 | 0 | 70170 | BAA00792.1 | P27958 |
| VPAAYAAQGYK | 1 | 0 | 0 | 70240 | P26664.3 | P27958 |
| VPDTLGGDPKFRSLTHEDHA | 1 | 0 | 0 | 70275 | P07299.1 | Q9PZT0 |
| VPGPFFRDEVTFTV | 1 | 0 | 0 | 70311 | P26661.3 | P27958 |

| | | | | | | |
|---|---|---|---|---|---|---|
| VPNAVGGYAISISFWPQTTTTPTSV | 1 | 0 | 0 | 70378 | P29326.1 | Q81871 |
| VPPLRVWRHRARSV | 1 | 0 | 0 | 70421 | P26662.3 | P27958 |
| VPPPRKKRTVVLTE | 1 | 0 | 0 | 70426 | P26664.3 | P27958 |
| VPPPRRKRTIQLDG | 1 | 0 | 0 | 70427 | Q81258.3 | P27958 |
| VPPPRRRRAKVLTQ | 1 | 0 | 0 | 70428 | P26661.3 | P27958 |
| VPSLEDIRSAEVV | 1 | 0 | 0 | 70478 | AAO86638.1 | Q9E006 |
| VPSPDPNNSPARP | 1 | 0 | 0 | 70481 | NP_044666.1 | P06484 |
| VPTNVSSTTQPQL | 1 | 0 | 0 | 70496 | NP_044666.1 | P06484 |
| VQAPNYTQHTSSMRGVY | 1 | 0 | 0 | 70527 | P59594.1 | P59594 |
| VQMAFMKLAALTGT | 1 | 0 | 0 | 70609 | P26662.3 | P27958 |
| VQMVIIKLGALTGT | 1 | 0 | 0 | 70612 | P26664.3 | P27958 |
| VQWMNRLIAFAFAGNHVSP | 1 | 0 | 0 | 70667 | AAA86907.1 | P27958 |
| VQWMNRLIAFASRGNHDSP | 1 | 0 | 0 | 70669 | AAA75355.1 | P27958 |
| VQWMNRLIAFASRGNHVAPT | 1 | 0 | 0 | 70670 | BAA00792.1 | P27958 |
| VQWMNRLIAFASRGNHVSPA | 1 | 0 | 0 | 70672 | Q81495.3 | P27958 |
| VQWMNRLIAFASRGNHVSPT | 1 | 0 | 0 | 70673 | BAB18806.1 | P27958 |
| VQYAVGIMTVTMTFKLGPRK | 1 | 0 | 0 | 70677 | P07299.1 | Q9PZT0 |
| VREAIKRRLRTLILA | 1 | 0 | 0 | 70708 | ACA48803.1 | P17763 |
| VRQPARPLGSAWRDQAQRPAVASRR | 1 | 0 | 0 | 70808 | P29326.1 | Q81871 |
| VRRRGDSRGSLLSP | 1 | 0 | 0 | 70816 | P26664.3 | P27958 |
| VRVPNYNLIIMDEAH | 1 | 0 | 0 | 70848 | QTX93024.1 | P17763 |
| VSGCPMRLNMASLEEKGRHF | 1 | 0 | 0 | 70945 | P08617.1 | P08617 |
| VSGGLCSTRLHRHALLATCP | 1 | 0 | 0 | 70950 | AAG48731.1 | P03410 |
| VSGHRMAWDMMMNWS | 1 | 0 | 0 | 70954 | AAA86907.1 | P27958 |
| VSKGNCSGPTTIRGQ | 1 | 0 | 0 | 71009 | P08362.1 | Q786F2 |
| VSLLSVLLSMQGAVD | 1 | 0 | 0 | 71036 | P0DTC1.1 | P0DTD1 |
| VSSHRLDTGEVPALQ | 1 | 0 | 0 | 71118 | ABI20542.1 | A0A0M3PZA5 |
| VSVITPGTNASSEVAVL | 1 | 0 | 0 | 71189 | NP_828851.1 | P59594 |
| VSWFASTGGRDSKIDVWSLV | 1 | 0 | 0 | 71216 | AAX68554.1 | A0A159D718 |
| VSWVMKIGIGVLLTW | 1 | 0 | 0 | 71219 | AHG23230.1 | P17763 |
| VSYSSYHATYSLYLFPHWTKKPNR | 1 | 0 | 0 | 71235 | AAA96674.1 | P03381 |
| VTGHRMAWDMMMNWS | 1 | 0 | 0 | 71323 | BAA09919.1 | P27958 |
| VTGHRMAWDMMQNWS | 1 | 0 | 0 | 71324 | BAA13621.1 | P27958 |
| VTMECSPRTGLDFNE | 1 | 0 | 0 | 71418 | QTX93024.1 | P17763 |
| VTYVPSQERNFTTAPAI | 1 | 0 | 0 | 71589 | NP_828851.1 | P59594 |
| VVHGCPLPPPKSPP | 1 | 0 | 0 | 71686 | P26664.3 | P27958 |
| VVHGCPLPPTKAPP | 1 | 0 | 0 | 71687 | BAA01583.1 | P27958 |
| VVPLATYSLTGLWS | 1 | 0 | 0 | 71786 | P26660.3 | P27958 |
| VVVAPDKEILYEAFDEMEEC | 1 | 0 | 0 | 71866 | BAA01761.1 | P27958 |
| VVVTPDKEILYEAFDEMEEC | 1 | 0 | 0 | 71888 | AAP55688.1 | P27958 |
| VYYPDEIFRSDTLYLTQD | 1 | 0 | 0 | 72205 | P59594.1 | P59594 |
| WDEGIIKSTIVEAAK | 1 | 0 | 0 | 72295 | P07298.1 | Q9PZT1 |
| WDQKLWCRHFCVLAD | 1 | 0 | 0 | 72324 | P08362.1 | Q786F2 |
| WGLNWDLGLSQWAREALQTGITL | 1 | 0 | 0 | 72481 | P03381.1 | P03381 |
| WGTSAAAYFVGYLKPTT | 1 | 0 | 0 | 72519 | P59594.1 | P59594 |
| WHVLYTPNISIPQQTSSRTILFP | 1 | 0 | 0 | 72558 | P03383.1 | P03383 |
| WIPAGIGVTGVIIAV | 1 | 0 | 0 | 72638 | ALG01342.1 | Q05320 |
| WLGFIAGLIAIVMVT | 1 | 0 | 0 | 72717 | P0DTC2.1 | P0DTC2 |
| WLTIPPMKNLALGVI | 1 | 0 | 0 | 72808 | P08362.1 | Q786F2 |
| WLTYHGAIKLDDKDPQF | 1 | 0 | 0 | 72816 | AAS48456.1 | P59595 |
| WNRKELLVTFKNAHA | 1 | 0 | 0 | 72905 | QTX93593.1 | P17763 |
| WQRSNAEFEQNLENAQWFAF | 1 | 0 | 0 | 73004 | P08617.1 | P08617 |
| WRIYSTGNNVFQTQAGC | 1 | 0 | 0 | 73036 | NP_828851.1 | P59594 |
| WSGEDSATLGAGGVL | 1 | 0 | 0 | 73072 | SRC279960 | P27958 |
| WTHCFDPQIQAIVSSPCHNSLIL | 1 | 0 | 0 | 73133 | A45714 | P03381 |
| WTHCYQPRLQAITTDNCNNSIIL | 1 | 0 | 0 | 73134 | P03383.1 | P03383 |
| WTWETKGDLAEN | 1 | 0 | 0 | 73171 | P27312.1 | P27312 |
| WYGMEIRPLSEKEENMV | 1 | 0 | 0 | 73272 | Q6TFL5 | P17763 |
| YAAQNRRGLDLLFWEQGGL | 1 | 0 | 0 | 73327 | P03381.1 | P03381 |
| YALADASLKMADPNRFRGKD | 1 | 0 | 0 | 73385 | ABM66847.1 | Q69091 |
| YCADVAAEELMNALV | 1 | 0 | 0 | 73483 | P08362.1 | Q786F2 |
| YDASVHGQIGAALL | 1 | 0 | 0 | 73521 | P26660.3 | P27958 |
| YEAADMIMHTP | 1 | 0 | 0 | 73615 | P26662.3 | P27958 |
| YEPYYHSDHAESSWVNRGES | 1 | 0 | 0 | 73744 | AAK01056.1 | P09259 |
| YEVRNVSGIYHVTNDCSNS | 1 | 0 | 0 | 73787 | BAA09919.1 | P27958 |
| YFKDCLFKDWEELGEE | 1 | 0 | 0 | 73843 | ABK27207.1 | Q69027 |
| YFVLTSHTVMPLSAP | 1 | 0 | 0 | 73959 | P0DTD1.1 | P0DTD1 |
| YGAGVAGALVAFKIMSGEVP | 1 | 0 | 0 | 73975 | AAB67037.1 | P27958 |
| YGSRLGVPDTLGGDPKFRSL | 1 | 0 | 0 | 74084 | P07299.1 | Q9PZT0 |
| YHVTNDCPNSSIVYEAADAI | 1 | 0 | 0 | 74152 | AAA45676.1 | P27958 |
| YICGDSTECANLLLQYG | 1 | 0 | 0 | 74173 | NP_828851.1 | P59594 |
| YIKARAACRAAG | 1 | 0 | 0 | 74272 | P26664.3 | P27958 |
| YISDAFSLDVSEKSGNF | 1 | 0 | 0 | 74367 | P59594.1 | P59594 |
| YKMFLHTGAVIVLPEDDLPT | 1 | 0 | 0 | 74479 | AAG48731.1 | P03410 |
| YKRRMADAVSRKKM | 1 | 0 | 0 | 74503 | P27313.1 | P27313 |
| YLDFNALNLFFSPLEFQHLI | 1 | 0 | 0 | 74601 | P07299.1 | Q9PZT0 |
| YLGTGPEASLPYGANK | 1 | 0 | 0 | 74684 | NP_828858.1 | P59595 |
| YLIKRRKFPPALPP | 1 | 0 | 0 | 74705 | P26661.3 | P27958 |
| YLKASAACRAAK | 1 | 0 | 0 | 74728 | P26663.3 | P27958 |
| YLRRMGGIQRRDHLQTLRDQ | 1 | 0 | 0 | 74902 | AAC56957.1 | Q2HR63 |
| YLVQKRHYLVFLIAHHYYEA | 1 | 0 | 0 | 74975 | AAC56957.1 | Q2HR63 |
| YPFRLPIKGVPIELQ | 1 | 0 | 0 | 75287 | P08362.1 | Q786F2 |
| YPGHITGHRMAWDMMMNWSPTTALV | 1 | 0 | 0 | 75294 | P26664.3 | P27958 |
| YPTFGEHKQEKDLEY | 1 | 0 | 0 | 75418 | ACS12928.1 | A0A0M3PZA5 |
| YQPIDVVRDLPSGFNTL | 1 | 0 | 0 | 75528 | P59594.1 | P59594 |
| YQVRNSSGLYHV | 1 | 0 | 0 | 75567 | P27958.3 | P27958 |
| YSCMVCVDRSSLSSWHVLYTP | 1 | 0 | 0 | 75707 | P03383.1 | P03383 |
| YSPSRSFSYFYPFRL | 1 | 0 | 0 | 75863 | P08362.1 | Q786F2 |
| YSRPVVSANGEPTVKLYTSVENAQQ | 1 | 0 | 0 | 75883 | P29326.1 | Q81871 |
| YSSMPPLEGEPGDPDLSDGS | 1 | 0 | 0 | 75890 | P26662.3 | P27958 |
| YSTPWRYLDFNALNLFFSPL | 1 | 0 | 0 | 75905 | P07299.1 | Q9PZT0 |

| | | | | | | |
|---|---|---|---|---|---|---|
| YSVLYNSTFFSTFKCYG | 1 | 0 | 0 | 75920 | P59594.1 | P59594 |
| YTCIVCIDRASLSTWHVLYSP | 1 | 0 | 0 | 75984 | P03381.1 | P03381 |
| YTGPVSSPSWKFHSDV | 1 | 0 | 0 | 76065 | P03383.1 | P03383 |
| YTNIPVSILFNKEEADDNGD | 1 | 0 | 0 | 76132 | AAG48731.1 | P03410 |
| YVYKGYQPIDVVRDLPS | 1 | 0 | 0 | 76446 | P59594.1 | P59594 |
| AVYVKSKVGNYR | 1 | 0 | 0 | 77840 | BAM16435.1 | Q01455 |
| IQAIVSSPCHNSLILPPFSLSPV | 1 | 0 | 0 | 78425 | P03381.1 | P03381 |
| LPPTAPPLLPHSNLDHILEPSIPW | 1 | 0 | 0 | 78454 | P03381.1 | P03381 |
| QGGLCKALQEQCRFPNITNSHVP | 1 | 0 | 0 | 78492 | P03381.1 | P03381 |
| RASLSTWHVLYSPNVSVPSS | 1 | 0 | 0 | 78500 | P03381.1 | P03381 |
| MPKTRRRPRRSQRKRPPTPW | 1 | 0 | 0 | 78749 | AAF71372.1 | P0C205 |
| PPALPIWARPDYNPPLLESWKDPD | 1 | 0 | 0 | 78764 | P26662.3 | P27958 |
| TGAPSLGDYVRPAYIVTPYW | 1 | 0 | 0 | 78850 | AAF71372.1 | P0C205 |
| TPSMDALSAQLYSSLSLDSP | 1 | 0 | 0 | 78863 | AAF71372.1 | P0C205 |
| TWAQPGYPWPLYGNEGMGWAGWLLS | 1 | 0 | 0 | 78868 | P26662.3 | P27958 |
| VIPDREVLYQEFDEMEECAS | 1 | 0 | 0 | 78872 | P26662.3 | P27958 |
| ATRKTSERSQPRGRRQPI | 1 | 0 | 0 | 78932 | Q68948 | P27958 |
| LGIASFLTRGPKQNI | 1 | 0 | 0 | 79013 | BAC41283.1 | P27958 |
| PKARQPEGRAWAQPG | 1 | 0 | 0 | 79041 | Q68948 | P27958 |
| QLVNTNGSWHINRTALNC | 1 | 0 | 0 | 79053 | Q68948 | P27958 |
| RRVASFFSPGSAQKI | 1 | 0 | 0 | 79075 | Q68948 | P27958 |
| SLYPGHVSGHRMAWDMMM | 1 | 0 | 0 | 79086 | Q68948 | P27958 |
| YEVRNVSGVYHVTNDCS | 1 | 0 | 0 | 79123 | Q68948 | P27958 |
| FHISCLTFGRETVIEYLVSFG | 1 | 0 | 0 | 79546 | CAM58990.1 | P0C767 |
| IPKDRRSTGKSWGKPGY | 1 | 0 | 0 | 79576 | P26660.3 | P27958 |
| IRTPPAYRPPNAPILSTLPETTV | 1 | 0 | 0 | 79577 | P03146.1 | Q76R61 |
| KFRQLLWFHISCLTFGRETVIEY | 1 | 0 | 0 | 79583 | P03146.1 | Q76R61 |
| LCWGELMTLATWVGVNLEDPASRDL | 1 | 0 | 0 | 79594 | P03146.1 | Q76R61 |
| LSTLPETTVVRRRGRSPRRRTPSPR | 1 | 0 | 0 | 79609 | P03146.1 | Q76R61 |
| LYREALESPEHCSPHHTALRQA | 1 | 0 | 0 | 79613 | P03146.1 | Q76R61 |
| VSYVNTNMGLKFRQL | 1 | 0 | 0 | 79677 | P03146.1 | Q76R61 |
| AQQPSTATSVDAP | 1 | 0 | 0 | 80440 | AAO84298.1 | Q86887 |
| ARGACVTIMTVDNPA | 1 | 0 | 0 | 80446 | NP_740471.1 | P03300 |
| DDIEQQADNMITEML | 1 | 0 | 0 | 80922 | P08669.1 | P08669 |
| EASGPTHSKEIPALT | 1 | 0 | 0 | 81394 | NP_740471.1 | P03300 |
| EIPALTAVETGATNP | 1 | 0 | 0 | 81565 | NP_740471.1 | P03300 |
| GPTHSKEIPALTAVE | 1 | 0 | 0 | 82831 | NP_740471.1 | P03300 |
| HSKEIPALTAVETGA | 1 | 0 | 0 | 83234 | NP_740471.1 | P03300 |
| HTSQVVPSDTMQTRH | 1 | 0 | 0 | 83266 | AAO84298.1 | Q86887 |
| IPALTAAETGHTSQV | 1 | 0 | 0 | 83789 | AAO84298.1 | Q86887 |
| IPYCNYSKYWYLNHT | 1 | 0 | 0 | 83829 | P08669.1 | P08669 |
| KLEFFTYLRFDLELT | 1 | 0 | 0 | 84461 | AAO84298.1 | Q86887 |
| LPHSLNDHILEPSIPWKSK | 1 | 0 | 0 | 85672 | SRC276211 | P03381 |
| SDTVQTRHVVQHRSR | 1 | 0 | 0 | 88446 | NP_740471.1 | P03300 |
| SESSIESFFARGACV | 1 | 0 | 0 | 88495 | NP_740471.1 | P03300 |
| SRSESSVENFLCRSA | 1 | 0 | 0 | 89062 | AAO84298.1 | Q86887 |
| SYTNNSKKGYAEWVINTRQV | 1 | 0 | 0 | 89316 | AAO84298.1 | Q86887 |
| VSSRPTNSESIPALT | 1 | 0 | 0 | 90405 | AAO84298.1 | Q86887 |
| GLDLLFWEQGGLCKALQEQCRFPN | 1 | 0 | 0 | 94530 | AAL05538.1 | P03381 |
| LLTLVQLTLQSTNYTCIVCIDRA | 1 | 0 | 0 | 94590 | AAA96674.1 | P03381 |
| QLRHLPSRVRYPHYSLIKPESSL | 1 | 0 | 0 | 94658 | CAA63041.1 | P03381 |
| SLLHEVDKDISQLTQAIVKNHKN | 1 | 0 | 0 | 94690 | A45714 | P03381 |
| VPTIAHGDGRRPSKQRTFI | 1 | 0 | 0 | 94962 | P03226.1 | P03226 |
| YLKDQQLLGIWGCSGKL | 1 | 0 | 0 | 94978 | AAC28452.1 | P03378 |
| ALESPEHCSPHHTALRQAIL | 1 | 0 | 0 | 96088 | P03146.1 | Q76R61 |
| LWFHISCLTFGRETVIEYLV | 1 | 0 | 0 | 96630 | P03146.1 | Q76R61 |
| SDFFPSVRDLLDTASALYRE | 1 | 0 | 0 | 96862 | P03146.1 | Q76R61 |
| WVGVNLEDPASRDLVVSYVN | 1 | 0 | 0 | 97104 | P03146.1 | Q76R61 |
| ADQELGDAPFLDRLRRDQKS | 1 | 0 | 0 | 97170 | ABP35610.1 | P03496 |
| DAVATTHSWIPKRNRSIL | 1 | 0 | 0 | 97236 | ABO21688.1 | P03431 |
| EKANPVNDLCYPGDFNDYEELKH | 1 | 0 | 0 | 97267 | ABW90125.1 | P03452 |
| HKIFKMEKGKVVKSVELDAPNYHY | 1 | 0 | 0 | 97383 | AAT73329.1 | P03468 |
| KAGLLVSDGGPNLY | 1 | 0 | 0 | 97407 | ABO21688.1 | P03431 |
| LFGAIAGFIEGGWQGMVDGWYG | 1 | 0 | 0 | 97441 | ABW90125.1 | P03452 |
| MASQGTKRSYEQMETGGERQNATE | 1 | 0 | 0 | 97491 | AAT70630.1 | P03466 |
| SLLTEVETPTRNEWECRCSDSSDP | 1 | 0 | 0 | 97651 | ABP35634.1 | P06821 |
| TNQEDLLVLWGIHHPNDAAEQT | 1 | 0 | 0 | 97700 | ABW90125.1 | P03452 |
| VNSDTVGWSWPDGAELPFTID | 1 | 0 | 0 | 97748 | AAT73329.1 | P03468 |
| WCCRSCTLPPLRYRGEDGCW | 1 | 0 | 0 | 101410 | AAL58460.1 | P17763 |
| DPSLKMADPNRFRGKNLPVL | 1 | 0 | 0 | 101555 | P03172.2 | Q69467 |
| PELVPEDPEDSALLEDPAGT | 1 | 0 | 0 | 101960 | P03172.2 | Q69467 |
| VSSQIPPNWHIPSIQDVAPH | 1 | 0 | 0 | 102215 | P03172.2 | Q69467 |
| CASHLPYIEQGMQLAEQFKQ | 1 | 0 | 0 | 103822 | BAB18806.1 | P27958 |
| GMQLAEQFKQKALGLLQTAT | 1 | 0 | 0 | 103941 | BAB18806.1 | P27958 |
| HSYSPGEINRVASCLRKLGV | 1 | 0 | 0 | 103977 | BAB18806.1 | P27958 |
| KTKRNTNRRPQDVKFPGGGQ | 1 | 0 | 0 | 104017 | BAB18806.1 | P27958 |
| LPRRGPRLGVRATRKTSERS | 1 | 0 | 0 | 104050 | BAB18806.1 | P27958 |
| NHVSPTHYVPESDAAARVTQ | 1 | 0 | 0 | 104148 | BAB18806.1 | P27958 |
| PPSLASSSASQLSAPSLKA | 1 | 0 | 0 | 104181 | BAB18806.1 | P27958 |
| PRGRRQPIPKARRPEGRTWA | 1 | 0 | 0 | 104184 | BAB18810.1 | P27958 |
| PRRGPRLGVRATRKTSERSQ | 1 | 0 | 0 | 104187 | BAB18806.1 | P27958 |
| RPAVIPDREVLYQEFDEMEE | 1 | 0 | 0 | 104225 | BAA25076.1 | P27958 |
| RPQDVKFPGGGQIVGGVYLL | 1 | 0 | 0 | 104226 | BAB18806.1 | P27958 |
| SHITAETAKRRLARGSPPSL | 1 | 0 | 0 | 104246 | BAB18806.1 | P27958 |
| TRKTSERSQPRGRRQPIPKA | 1 | 0 | 0 | 104315 | BAB18806.1 | P27958 |
| GTTDHVGVPTYDWGK | 1 | 0 | 0 | 108412 | SRC280504 | P27958 |
| NDSLNTGFLAFLFYTKK | 1 | 0 | 0 | 108551 | SRC279960 | P27958 |
| HKSAIVTLTYDSEWQRDQ | 1 | 0 | 0 | 110284 | ACS92695.1 | P03120 |
| IPKPSPWAPKKHRRLSSDQD | 1 | 0 | 0 | 110293 | P06922.1 | P06922 |
| KHRRLSSDQDQSQTPETPAT | 1 | 0 | 0 | 110303 | P06922.1 | P06922 |
| PKRLRLPHIRDDDAPPSHQP | 1 | 0 | 0 | 110364 | P03172.2 | Q69467 |

| | | | | | | |
|---|---|---|---|---|---|---|
| QSQTPETPATPLSCCTETQW | 1 | 0 | 0 | 110388 | P06922.1 | P06922 |
| STWPTTPPRPIPKPSPWAPK | 1 | 0 | 0 | 110427 | P06922.1 | P06922 |
| TKYPLLKLLGSTWPTTPPRP | 1 | 0 | 0 | 110434 | P06922.1 | P06922 |
| VTFCCKCDSTLRLCVQ | 1 | 0 | 0 | 110456 | ABL96585.1 | P03129 |
| YEQLNDSSEEEDEIDG | 1 | 0 | 0 | 110460 | P03129.1 | P03129 |
| YYVLHLCLAATKYPLLKLLG | 1 | 0 | 0 | 110468 | P06922.1 | P06922 |
| EKTGILTVTYHSETQRTK | 1 | 0 | 0 | 110571 | ABP99755.1 | T2A704 |
| GDICNTMHYTNWTHIYICEE | 1 | 0 | 0 | 110589 | ACS92695.1 | P03120 |
| HKHAIVTVTYDSEEQRQQ | 1 | 0 | 0 | 110625 | CAS89355.1 | P04015 |
| KCDSTLRLCVQSTHVDIRTLE | 1 | 0 | 0 | 110640 | ABL96585.1 | P03129 |
| LCLIGCKPPIGEHWGKGSPC | 1 | 0 | 0 | 110652 | ACG75893.1 | P03101 |
| MHQKRTAMFQDPQERPRKLPQLC | 1 | 0 | 0 | 110669 | P03126.1 | P03126 |
| PMDTFIVSTNPNTVTSSTPI | 1 | 0 | 0 | 110685 | ACL12316.1 | P03107 |
| QSQAITCQKPTPEKEKPDPYK | 1 | 0 | 0 | 110696 | CAS89491.1 | P04012 |
| VGENVPDDLYIKGSG | 1 | 0 | 0 | 110733 | ACG75893.1 | P03101 |
| EDTYRFVTSQAIACQKHTPPA | 1 | 0 | 0 | 110825 | ACG75893.1 | P03101 |
| GLKAKPKFTLGKRKATPTTS | 1 | 0 | 0 | 110863 | ACG75893.1 | P03101 |
| GLKAKPKLKRAAPTSTRTSS | 1 | 0 | 0 | 110864 | ACL12333.1 | P03101 |
| GQAEPDRAHYNIVTFCCKCD | 1 | 0 | 0 | 110869 | P03129.1 | P03129 |
| GYRGRSSIRTGVKRPAVSKA | 1 | 0 | 0 | 110880 | NP_040304.1 | P04012 |
| HEYMLDLQPETTDLYCYE | 1 | 0 | 0 | 110884 | P03129.1 | P03129 |
| LYIKGSGSTANLASSNYFPT | 1 | 0 | 0 | 110938 | ACG75893.1 | P03101 |
| PETTDLYCYEQLNDSSEEEDE | 1 | 0 | 0 | 110960 | P03129.1 | P03129 |
| PNNNKILVPKVSGLQYRVFR | 1 | 0 | 0 | 110965 | ACG75893.1 | P03101 |
| RTGYIPLGTRPPT | 1 | 0 | 0 | 111013 | ACL12316.1 | P03107 |
| YMLRKRRKRLPYFF | 1 | 0 | 0 | 111065 | ACL12316.1 | P03107 |
| ACQKHTPPAPKEDDPLKKYT | 1 | 0 | 0 | 111088 | AAM29174.1 | P03101 |
| CQDKILTHYENDSTD | 1 | 0 | 0 | 111167 | ACS92695.1 | P03120 |
| EEASVTVVEGQVDYY | 1 | 0 | 0 | 111233 | ACS92695.1 | P03120 |
| KGSPCTNVAVNPGDCPPLEL | 1 | 0 | 0 | 111450 | ACA14209.1 | P03101 |
| LSKLLCVSPMCMMIEPPKLR | 1 | 0 | 0 | 111521 | P03114.2 | P03114 |
| NGICWGNQLFVTVVDTTRST | 1 | 0 | 0 | 111581 | ACA14209.1 | P03101 |
| NKFGFPDTSFYNPDTQRLVW | 1 | 0 | 0 | 111583 | ACA14209.1 | P03101 |
| NKSEVPLDICTSICKYPDYI | 1 | 0 | 0 | 111585 | ACA14209.1 | P03101 |
| QVILCPTSVFSSNEV | 1 | 0 | 0 | 111693 | ACS92695.1 | P03120 |
| RWTGRCMSCCRSSRTRRETQL | 1 | 0 | 0 | 111750 | ACS92692.1 | P03126 |
| STHNYEEIPMDTFIVST | 1 | 0 | 0 | 111808 | ACL12316.1 | P03107 |
| STHNYEEIRMDTFIVST | 1 | 0 | 0 | 111809 | SRC276209 | P03107 |
| TEETQTTIQRPRSEP | 1 | 0 | 0 | 111821 | ACS92695.1 | P03120 |
| TSHPIEEIPLDTFVI | 1 | 0 | 0 | 111851 | AAF00067.1 | P04013 |
| TSHPIEEIPLDTFVV | 1 | 0 | 0 | 111852 | AAK01849.1 | P04013 |
| TSIFRNPVFTEPSVTQPQPPVE | 1 | 0 | 0 | 111853 | AAF00067.1 | P04013 |
| VDNRECISMDYKQTQLCLIG | 1 | 0 | 0 | 111890 | ACA14209.1 | P03101 |
| VHTGFGAMDFTTLQANKSEV | 1 | 0 | 0 | 111915 | ACG75893.1 | P03101 |
| VRYKCGKNRETIEKLLSKLL | 1 | 0 | 0 | 111921 | P03114.2 | P03114 |
| SGYIPANTTIPF | 1 | 0 | 0 | 112837 | ACL12316.1 | P03107 |
| EFGKAKGSRAIWYMW | 1 | 0 | 0 | 113359 | P17763.2 | P17763 |
| WKSKLLTLVQLT | 1 | 0 | 0 | 118221 | AAT38829.2 | P03381 |
| ALTGATEIQNSGGTS | 1 | 0 | 0 | 119103 | QTX93489.1 | P17763 |
| AYTALFSGVSWVMKI | 1 | 0 | 0 | 119110 | QHJ86727.1 | P17763 |
| CLEPIEGKVVQYENL | 1 | 0 | 0 | 119113 | QTX93500.1 | P17763 |
| EPVNIEAEPPFGESN | 1 | 0 | 0 | 119144 | QTX93489.1 | P17763 |
| GATTETPTWNRKELL | 1 | 0 | 0 | 119169 | QTX93500.1 | P17763 |
| GDNALKINWYKKGSS | 1 | 0 | 0 | 119171 | QHJ86727.1 | P17763 |
| IEGKITNITTDSRCP | 1 | 0 | 0 | 119209 | QTX93491.1 | P17763 |
| KVEYKGEDAPCKIPF | 1 | 0 | 0 | 119238 | QTX93489.1 | P17763 |
| LDIELQKTEATQLAT | 1 | 0 | 0 | 119241 | QTX93474.1 | P17763 |
| LECSPRTGLDFNEMI | 1 | 0 | 0 | 119243 | QTX93489.1 | P17763 |
| LKCRLKMDKLELKGM | 1 | 0 | 0 | 119252 | AAC63321.1 | P17763 |
| LTWIGLNSKNTSMSF | 1 | 0 | 0 | 119265 | QTX93489.1 | P17763 |
| MRCVGVGNRDFVEGL | 1 | 0 | 0 | 119276 | AAW28141.1 | P17763 |
| NKAWMVHRQWFFDLP | 1 | 0 | 0 | 119290 | QTX93489.1 | P17763 |
| QGKAHNGRLITANPV | 1 | 0 | 0 | 119312 | QTX93500.1 | P17763 |
| SYAMCTNTFVLKKEV | 1 | 0 | 0 | 119351 | AHG23230.1 | P17763 |
| TNTFVLKKEVSETQH | 1 | 0 | 0 | 119364 | AHG23230.1 | P17763 |
| VDVVLEHGGCVTTMA | 1 | 0 | 0 | 119377 | QTX93593.1 | P17763 |
| WGNGCGLFGKGSLVT | 1 | 0 | 0 | 119397 | AQV12170.1 | P17763 |
| AGAGGGAGGAGAGGGAGGAGC | 1 | 0 | 0 | 119738 | SRC279943 | P03211 |
| KILPEVKEKHEFLSRL | 1 | 0 | 0 | 120682 | CAA29172.1 | Q86887 |
| GSDDYVWLSRLLDLAPNYRQ | 1 | 0 | 0 | 125428 | AAK06654.1 | F5HF49 |
| KLTPLPEARLLDLSSWFTVG | 1 | 0 | 0 | 125463 | BAA00792.1 | P27958 |
| CPLPPPRSPPVPPPRK | 1 | 0 | 0 | 127939 | AAB66324.1 | P27958 |
| NPPLVETWKKPDYEPP | 1 | 0 | 0 | 128065 | AAB66324.1 | P27958 |
| VTNDCSNSSIVYEAADVI | 1 | 0 | 0 | 128141 | ABF71404.1 | P27958 |
| PDQRPYCWHYPPKPC | 1 | 0 | 0 | 130994 | ABR25251.1 | P27958 |
| PPPPSSPTHDPPDSDPQIPPP | 1 | 0 | 0 | 132011 | AAD50663.1 | P03362 |
| AKIIGADVQNTTFII | 1 | 0 | 0 | 132152 | P17763.2 | P17763 |
| DSGCVINWKGRELKC | 1 | 0 | 0 | 132233 | P17763.2 | P17763 |
| LRFKGEDGCWYGMEI | 1 | 0 | 0 | 132452 | P17763.2 | P17763 |
| MIIPKIYGGPISQHN | 1 | 0 | 0 | 132477 | P17763.2 | P17763 |
| PECPDDQRAWNIWEV | 1 | 0 | 0 | 132520 | QFS19562.1 | P17763 |
| TRLENIMWKQISNEL | 1 | 0 | 0 | 132701 | P17763.2 | P17763 |
| WIESEKNETWKLARA | 1 | 0 | 0 | 132736 | P17763.2 | P17763 |
| EEAIERAVVHVA | 1 | 0 | 0 | 135913 | ACL31285.1 | Q86887 |
| GDVEEAIERAVV | 1 | 0 | 0 | 135931 | ACL31285.1 | Q86887 |
| IERAVVHVADTM | 1 | 0 | 0 | 135936 | ACL31285.1 | Q86887 |
| LTAVETGHTSQV | 1 | 0 | 0 | 135958 | 1D4M_1 | Q86887 |
| PSNSASVPALTA | 1 | 0 | 0 | 135974 | 1D4M_1 | Q86887 |
| QSRRRGDMSTLN | 1 | 0 | 0 | 135980 | BAA00518.1 | Q86887 |
| RRGDMSTLNTHG | 1 | 0 | 0 | 135985 | BAA00518.1 | Q86887 |
| SASVPALTAVET | 1 | 0 | 0 | 135990 | 1D4M_1 | Q86887 |

| | | | | | | |
|---|---|---|---|---|---|---|
| TVAQSRRRGDMS | 1 | 0 | 0 | 136005 | BAA00518.1 | Q86887 |
| VETGHTSQVTPS | 1 | 0 | 0 | 136006 | 1D4M_1 | Q86887 |
| VMIKSLPALNSPTVEEC | 1 | 0 | 0 | 136010 | AAR16005.1 | Q86887 |
| VPALTAVETGHT | 1 | 0 | 0 | 136011 | 1D4M_1 | Q86887 |
| QLEKALDCQIYG | 1 | 0 | 0 | 136573 | BAH79736.1 | P27958 |
| VIPMVTQIAMTDTTP | 1 | 0 | 0 | 136894 | P17763.2 | P17763 |
| GLPIRYQTPAIRAEH | 1 | 0 | 0 | 139762 | P14340.2 | P17763 |
| LVDYPYRLWHYPCTI | 1 | 0 | 0 | 140577 | ABR25251.1 | P27958 |
| KSLSSTEFIPN | 1 | 0 | 0 | 144712 | P13285.1 | P13285 |
| RIEDPPFNSLL | 1 | 0 | 0 | 144763 | P13285.1 | P13285 |
| EPGQLKLNWFKKGSS | 1 | 0 | 0 | 144885 | P14340.2 | P17763 |
| CNTKCQTPMGAINSS | 1 | 0 | 0 | 148521 | ABW06108.1 | P03452 |
| DSLSPRRAGAKAGPGLSPGT | 1 | 0 | 0 | 150173 | SRC280102 | P0C045 |
| EKNEQELLELDKWASLW | 1 | 0 | 0 | 150180 | P03375.1 | P03378 |
| LPQGASQEPSRMAWGPLKTG | 1 | 0 | 0 | 150397 | SRC280102 | P0C045 |
| LTIGNSTITTQEAAN | 1 | 0 | 0 | 150419 | ACY00666.1 | A0A0M3PZA5 |
| MAVGMVSILASSLLK | 1 | 0 | 0 | 150433 | P14340.2 | P17763 |
| NVKFPGGGQIVGGVYVLPRR | 1 | 0 | 0 | 150483 | AAB40040.1 | P27958 |
| PLEGTTNPNGYANWD | 1 | 0 | 0 | 150496 | ABI20542.1 | A0A0M3PZA5 |
| RNLLELCQVETILEV | 1 | 0 | 0 | 150533 | ACY00666.1 | A0A0M3PZA5 |
| TEDSHPPYKQTQPGA | 1 | 0 | 0 | 150616 | ACM47545.1 | A0A0M3PZA5 |
| TFTMRLLSPVRVPNY | 1 | 0 | 0 | 150617 | P17763.2 | P17763 |
| TKRNTIRRPQNVKFPGGGQI | 1 | 0 | 0 | 150626 | AAB40040.1 | P27958 |
| TSWGTSRSSVLPQGASQEPS | 1 | 0 | 0 | 150661 | SRC280102 | P0C045 |
| VGGVYVLPRRGPRLGVRATR | 1 | 0 | 0 | 150678 | ADP37774.1 | P27958 |
| WISNTHYRAHARDGV | 1 | 0 | 0 | 150723 | ACY00666.1 | A0A0M3PZA5 |
| NWFDISKWLWYI | 1 | 0 | 0 | 150874 | P19550.1 | P03378 |
| EKNVTVTHSVNLLED | 1 | 0 | 0 | 150978 | ACP41105.1 | P03452 |
| ILGNPECESLSTASS | 1 | 0 | 0 | 150998 | ACP41105.1 | P03452 |
| IPSIQSRGLFGAIAG | 1 | 0 | 0 | 150999 | ACP41105.1 | P03452 |
| LVKKGNSYPKLSKSY | 1 | 0 | 0 | 151031 | ACP41105.1 | P03452 |
| LYQNADTYVFVGSSR | 1 | 0 | 0 | 151032 | ACP41105.1 | P03452 |
| SQLKNNAKEIGNGCF | 1 | 0 | 0 | 151056 | ACP41105.1 | P03452 |
| TPKGAINTSLPFQNI | 1 | 0 | 0 | 151064 | ACP41105.1 | P03452 |
| VDDGFLDIWTYNAEL | 1 | 0 | 0 | 151067 | ACP41105.1 | P03452 |
| FNPNKDTWPDANKVG | 1 | 0 | 0 | 153257 | P03138.3 | Q76R62 |
| LAPTRVVAAEMEEAL | 1 | 0 | 0 | 156711 | P14340.2 | P17763 |
| IVPAKSVCGPVYCFTPSPVV | 1 | 0 | 0 | 161246 | AAB67036.1 | P27958 |
| SKAFKFARTPADTG | 1 | 0 | 0 | 161696 | P06935.2 | Q9Q6P4 |
| SLTVQTHGESTLA | 1 | 0 | 0 | 161698 | YP_001527879.1 | Q9Q6P4 |
| TLDVRMINIEASQLA | 1 | 0 | 0 | 161711 | AAK31640.1 | P0DOH9 |
| TTVESHGKIGATQAGRF | 1 | 0 | 0 | 161715 | P06935.2 | Q9Q6P4 |
| NNTRKSIHLGPGRAFYATGDIIG | 1 | 0 | 0 | 163040 | ADZ13293.1 | P03378 |
| RKRIRVGPGQTVY | 1 | 0 | 0 | 163244 | AAB01026.1 | P03378 |
| TKKGIAIGPGRTLY | 1 | 0 | 0 | 163621 | P12490.1 | P03378 |
| TRKSIRIGPGQAFY | 1 | 0 | 0 | 163689 | AAL59843.1 | P03378 |
| NPVENYIDSVLNEVLVVPNIQ | 1 | 0 | 0 | 164167 | AAQ19944.1 | P07210 |
| LVCASRELERFAVNPGLLE | 1 | 0 | 0 | 165638 | AAN08336.1 | P03349 |
| PPLLDRWKTPDYVPPTVHG | 1 | 0 | 0 | 176375 | ACZ60117.1 | P27958 |
| PPVIETWKRPDYEPPTVLG | 1 | 0 | 0 | 176376 | AAP55699.1 | P27958 |
| QRIAEMLKSKIQGLL | 1 | 0 | 0 | 176385 | AAF59940.1 | P27958 |
| QVIAHQFKEKVLGLL | 1 | 0 | 0 | 176401 | ACZ60112.1 | P27958 |
| STKDNFNVYKA | 1 | 0 | 0 | 177299 | AEJ18140.1 | P0DOK1 |
| TKDNFNVYKATRPYLAHC | 1 | 0 | 0 | 177328 | AEJ18140.1 | P0DOK1 |
| VPRNAELGDRKGKIHIPF | 1 | 0 | 0 | 177356 | ABD95938.1 | Q8JUX5 |
| VPRNAELGDRQGKIHIPF | 1 | 0 | 0 | 177357 | ADC53733.1 | Q8JUX5 |
| NIQLINTNGSWHINST | 1 | 0 | 0 | 178301 | AAB67036.1 | P27958 |
| NEQELLELDKWASLWN | 1 | 0 | 0 | 179863 | ADZ13338.1 | P03378 |
| GVFTTNIWLKLKEKQ | 1 | 0 | 0 | 180496 | QTX93151.1 | P17763 |
| HKKLAEAIFKLTYQN | 1 | 0 | 0 | 180500 | P14340.2 | P17763 |
| KNPHAKKQDVVVLGS | 1 | 0 | 0 | 180568 | P14340.2 | P17763 |
| KRYLPAIVREAIKRG | 1 | 0 | 0 | 180575 | P14340.2 | P17763 |
| LDFELIKTEAKQPAT | 1 | 0 | 0 | 180590 | QTX92984.1 | P17763 |
| MAMTDTTPFGQQRVF | 1 | 0 | 0 | 180643 | P14340.2 | P17763 |
| PNYNLIIMDEAHFTD | 1 | 0 | 0 | 180698 | P14340.2 | P17763 |
| STRVEMGEAAGIFMT | 1 | 0 | 0 | 180756 | QTX93024.1 | P17763 |
| TLRVLNLVENWLNNN | 1 | 0 | 0 | 180774 | QTX93024.1 | P17763 |
| VTDFKGKTVWFVPSI | 1 | 0 | 0 | 180807 | QTX93024.1 | P17763 |
| WGNGCGLFGKGGIVT | 1 | 0 | 0 | 180816 | BAG84557.1 | P17763 |
| QAQGLLTTVPAAPP | 1 | 0 | 0 | 180917 | AAR99337.1 | Q76R62 |
| SQISSHSPTCCPPICPGYRW | 1 | 0 | 0 | 180928 | AAR99337.1 | Q76R62 |
| SSGTVSPAQNTVSAISSI | 1 | 0 | 0 | 180929 | ADB55510.1 | Q76R62 |
| AKGSRAIWYMWLGAR | 1 | 0 | 0 | 181081 | P17763.2 | P17763 |
| GARYLEFEALGFLNE | 1 | 0 | 0 | 181095 | ACJ09459.1 | P17763 |
| TDTTPFGQQRVFKEK | 1 | 0 | 0 | 181156 | P17763.2 | P17763 |
| GDPIADMIDQTVNNQ | 1 | 0 | 0 | 181321 | ABX55895.1 | A0A0M3PZA5 |
| NNNTRKSINIGPGRALYTT | 1 | 0 | 0 | 187169 | P35961.1 | P03378 |
| SLWNWFDITNWLWYIK | 1 | 0 | 0 | 187194 | Q0ED31 | P03378 |
| DISEATQVEQHHRRT | 1 | 0 | 0 | 187296 | AGB56830.1 | Q05320 |
| EDHKIMASENSSAMV | 1 | 0 | 0 | 187314 | AGB56830.1 | Q05320 |
| RTSSDPGTNTTTEDH | 1 | 0 | 0 | 187509 | AGB56758.1 | Q05320 |
| TGEESASSGKLGLIT | 1 | 0 | 0 | 187541 | AGB56758.1 | Q05320 |
| TIRYQATGFGTNETE | 1 | 0 | 0 | 187549 | AGB56832.1 | P60170 |
| EIPFYGKAIPIETIKG | 1 | 0 | 0 | 187636 | AFC36922.1 | P27958 |
| LHAPTGSGKSTKVPAA | 1 | 0 | 0 | 187678 | AFC36922.1 | P27958 |
| IMKNHLRDIMG | 1 | 0 | 0 | 189683 | ADY11070.1 | P08669 |
| KCNEKHDEEFCDMLRL | 1 | 0 | 0 | 189691 | ADY11070.1 | P08669 |
| NRGGDENPRGPVSR | 1 | 0 | 0 | 189777 | AAB48506.1 | P89522 |
| SDAHKKNLYDHAL | 1 | 0 | 0 | 189839 | ADY11070.1 | P08669 |
| AIALDFKPGTSGSPI | 1 | 0 | 0 | 190703 | P17763.2 | P17763 |
| ANCLRKNGKKVIQLS | 1 | 0 | 0 | 190710 | QTX93489.1 | P17763 |

| | | | | | | |
|---|---|---|---|---|---|---|
| ARGYISTRVGMGEAA | 1 | 0 | 0 | 190711 | P17763.2 | P17763 |
| ASIAARGYISTRVGM | 1 | 0 | 0 | 190712 | P17763.2 | P17763 |
| AVHADMGYWIESQKN | 1 | 0 | 0 | 190715 | QTX93489.1 | P17763 |
| CLKPVILTDGPERVI | 1 | 0 | 0 | 190718 | QTX93489.1 | P17763 |
| CRSCTLPPLRYMGED | 1 | 0 | 0 | 190721 | QTX93489.1 | P17763 |
| EAAAIFMTATPPGTA | 1 | 0 | 0 | 190732 | QTX93489.1 | P17763 |
| EEMFKKRNLTIMDLH | 1 | 0 | 0 | 190735 | QTX93479.1 | P17763 |
| EMGANFKADRVIDPR | 1 | 0 | 0 | 190737 | QTX93489.1 | P17763 |
| EPGKNPKNFQTMPGT | 1 | 0 | 0 | 190739 | QPI11927.1 | P17763 |
| EVYTQLCDHRLMSAA | 1 | 0 | 0 | 190747 | QBA29683.1 | P17763 |
| FIIDGPNTPECPSAS | 1 | 0 | 0 | 190751 | Q6YMS3.1 | P17763 |
| FQTMPGTFQTTTGEI | 1 | 0 | 0 | 190759 | QIS48873.1 | P17763 |
| FVVTTDISEMGANFK | 1 | 0 | 0 | 190761 | P14340.2 | P17763 |
| GEIGAIALDFKPGTS | 1 | 0 | 0 | 190765 | QFS19205.1 | P17763 |
| GKTVWFVPSIKAGND | 1 | 0 | 0 | 190774 | QTX92786.1 | P17763 |
| HLGKLELDFNYCEGT | 1 | 0 | 0 | 190783 | QTX93479.1 | P17763 |
| HRLMSAAVKDERAVH | 1 | 0 | 0 | 190786 | Q6YMS3.1 | P17763 |
| HTLWSNGVLESDMII | 1 | 0 | 0 | 190787 | QTX93489.1 | P17763 |
| HTQTAGPWHLGKLEL | 1 | 0 | 0 | 190788 | QTX93479.1 | P17763 |
| IFMTATPPGTADAFP | 1 | 0 | 0 | 190793 | QHR82515.1 | P17763 |
| ISYGGGWRLSAQWQK | 1 | 0 | 0 | 190798 | QTX93491.1 | P17763 |
| KFQADSPKRLATAIA | 1 | 0 | 0 | 190810 | QTX93479.1 | P17763 |
| KKDLISYGGGWRLSA | 1 | 0 | 0 | 190815 | QTX93491.1 | P17763 |
| KKKLRPRWLDARTYS | 1 | 0 | 0 | 190816 | QHR82521.1 | P17763 |
| KKRNLTIMDLHPGSG | 1 | 0 | 0 | 190817 | P27909.2 | P17763 |
| KKVIQLSRKTFDTEY | 1 | 0 | 0 | 190819 | QTX93491.1 | P17763 |
| KLTVVVGDIIGVLEQ | 1 | 0 | 0 | 190823 | QTX93489.1 | P17763 |
| KQIANELNYILWENN | 1 | 0 | 0 | 190825 | QTX93489.1 | P17763 |
| KTFDTEYQKTKLNDW | 1 | 0 | 0 | 190831 | QIS48873.1 | P17763 |
| LKLREVYTQLCDHRL | 1 | 0 | 0 | 190842 | QBA29683.1 | P17763 |
| MYFHRRDLRLASNAI | 1 | 0 | 0 | 190863 | QTX93491.1 | P17763 |
| NEWITDFAGKTVWFV | 1 | 0 | 0 | 190864 | QFN25407.1 | P17763 |
| NFKADRVIDPRRCLK | 1 | 0 | 0 | 190865 | QTX93489.1 | P17763 |
| NGKRLEPNWASVKKD | 1 | 0 | 0 | 190868 | QTX93474.1 | P17763 |
| NPKNFQTMPGTFQTT | 1 | 0 | 0 | 190879 | QPI11927.1 | P17763 |
| NPQKENDQYIFTGQP | 1 | 0 | 0 | 190880 | QTX93489.1 | P17763 |
| PELEEEMFKKRNLTI | 1 | 0 | 0 | 190887 | QTX93479.1 | P17763 |
| PGTFQTTTGEIGAIA | 1 | 0 | 0 | 190891 | QIS48873.1 | P17763 |
| PLALKEFKDFAAGRK | 1 | 0 | 0 | 190894 | QTX93479.1 | P17763 |
| QNSSFIIDGPNTPEC | 1 | 0 | 0 | 190923 | QTX93489.1 | P17763 |
| RPRWLDARTYSDPLA | 1 | 0 | 0 | 190956 | P17763.2 | P17763 |
| SLRTTTVSGKLIHEW | 1 | 0 | 0 | 190977 | QTX93593.1 | P17763 |
| SNGVLESDMIIPKSL | 1 | 0 | 0 | 190985 | QTX93593.1 | P17763 |
| SPIINREGKVVGLYG | 1 | 0 | 0 | 190986 | QTX93593.1 | P17763 |
| SWKLEKASLIEVKTC | 1 | 0 | 0 | 190991 | QHR82521.1 | P17763 |
| TDFAGKTVWFVPSIK | 1 | 0 | 0 | 190992 | AHG23230.1 | P17763 |
| TDISEMGANFKADRV | 1 | 0 | 0 | 190993 | QTX93593.1 | P17763 |
| TEQYKFQADSPKRLA | 1 | 0 | 0 | 190996 | QTX93474.1 | P17763 |
| TEYQKTKLNDWDFVV | 1 | 0 | 0 | 190997 | QTX93374.1 | P17763 |
| TWGKAKIVTAETQNS | 1 | 0 | 0 | 191008 | QTX93593.1 | P17763 |
| VILTDGPERVILAGP | 1 | 0 | 0 | 191016 | QTX93593.1 | P17763 |
| VINWKGKELKCGSGI | 1 | 0 | 0 | 191017 | QTX93593.1 | P17763 |
| VLTYNGKRLEPNWAS | 1 | 0 | 0 | 191021 | QHD57352.1 | P17763 |
| WASVKKDLISYGGGW | 1 | 0 | 0 | 191025 | P17763.2 | P17763 |
| WNSGNEWITDFAGKT | 1 | 0 | 0 | 191027 | AHG23230.1 | P17763 |
| DRSGAPTYSWGAND | 1 | 0 | 0 | 195013 | AAB02127.1 | P27958 |
| EHAYPNASHPWDEDVMPDLPY | 1 | 0 | 0 | 195018 | AFE88887.1 | Q3YPH4 |
| EHAYPNATHPWDEDVMPELPYE | 1 | 0 | 0 | 195020 | AFE88888.1 | B9UYK7 |
| GEHAYPNATHPWDEDVMPELPYQT | 1 | 0 | 0 | 195023 | ADJ37024.1 | Q3YPH4 |
| HAYPNASHPWDEDVMPDLPYKT | 1 | 0 | 0 | 195026 | AFE88887.1 | Q3YPH4 |
| MSDTDIQDQQPDTVDAPQNT | 1 | 0 | 0 | 195042 | AFE88887.1 | Q3YPH4 |
| MSENEIQDQQPSDSMEERGG | 1 | 0 | 0 | 195043 | AFE88888.1 | B9UYK7 |
| VGPAASDTAAWMVGVDPEGANINS | 1 | 0 | 0 | 195059 | ADJ37024.1 | Q3YPH4 |
| AKFKCVTKLEGKIVQ | 1 | 0 | 0 | 195145 | P17763.2 | P17763 |
| AKLALDNIVMLHTTE | 1 | 0 | 0 | 195146 | Q2YHF0.1 | P17763 |
| AKWLWGFLSRNKKPR | 1 | 0 | 0 | 195150 | QPD98749.1 | P17763 |
| AMVLSIVSLFPLCLS | 1 | 0 | 0 | 195162 | QTX92488.1 | P17763 |
| ANCLRKNGKRVIQLS | 1 | 0 | 0 | 195163 | QFS19562.1 | P17763 |
| ATWAKNIQTAINQVR | 1 | 0 | 0 | 195188 | AAG15002.1 | P17763 |
| CHATFTMRLLSPVRV | 1 | 0 | 0 | 195201 | P17763.2 | P17763 |
| CLGKAYAQMWSLMYF | 1 | 0 | 0 | 195204 | QTX93491.1 | P17763 |
| CTILAVVSVSPLLLT | 1 | 0 | 0 | 195207 | QTX93067.1 | P17763 |
| CVYNMMGKREKKLGE | 1 | 0 | 0 | 195208 | P17763.2 | P17763 |
| DELIGRARISQGAGW | 1 | 0 | 0 | 195213 | P14340.2 | P17763 |
| DTLTILLKATLLAVS | 1 | 0 | 0 | 195228 | QFS19562.1 | P17763 |
| EDHWFSRENSLSGVE | 1 | 0 | 0 | 195233 | P17763.2 | P17763 |
| EEFISKVRSNAAIGA | 1 | 0 | 0 | 195235 | Q2YHF0.1 | P17763 |
| ELIGRARISQGAGWS | 1 | 0 | 0 | 195249 | P14340.2 | P17763 |
| ELLVTFKNAHAKKQE | 1 | 0 | 0 | 195252 | QTX93489.1 | P17763 |
| ENLKYSVIVTVHTGD | 1 | 0 | 0 | 195260 | P17763.2 | P17763 |
| EPDYEVDEDIFRKKR | 1 | 0 | 0 | 195263 | Q2YHF0.1 | P17763 |
| ETLMLVALLGAMTAG | 1 | 0 | 0 | 195274 | Q2YHF0.1 | P17763 |
| ETTILDVDLRPASAW | 1 | 0 | 0 | 195276 | Q2YHF0.1 | P17763 |
| FALLAGFMAYMIGQT | 1 | 0 | 0 | 195283 | Q2YHF0.1 | P17763 |
| FHTMWHVTRGAVLMY | 1 | 0 | 0 | 195296 | P17763.2 | P17763 |
| FLRFLAIPPTAGVLA | 1 | 0 | 0 | 195306 | AYA22932.1 | P17763 |
| FNEMILMKMKKKTWL | 1 | 0 | 0 | 195310 | Q58HT7.1 | P17763 |
| FNMLKRARNRVSTGS | 1 | 0 | 0 | 195314 | AAT76483.1 | P17763 |
| FNMLKRARNRVSTVS | 1 | 0 | 0 | 195315 | P17763.2 | P17763 |
| FQRVLIFILLTAVAP | 1 | 0 | 0 | 195322 | QTX92964.1 | P17763 |
| FTMRLLSPVRVPNYN | 1 | 0 | 0 | 195338 | P17763.2 | P17763 |

| | | | | | | |
|---|---|---|---|---|---|---|
| GGMLVRNPLSRNSTH | 1 | 0 | 0 | 195360 | P17763.2 | P17763 |
| GHLKCRLRMDKLQLK | 1 | 0 | 0 | 195366 | P14340.2 | P17763 |
| GIIFILLMLVTPSMA | 1 | 0 | 0 | 195368 | P17763.2 | P17763 |
| GIYRILQRGLLGRSQ | 1 | 0 | 0 | 195377 | P17763.2 | P17763 |
| GKVRKDIPQWQPSKG | 1 | 0 | 0 | 195382 | QTX93489.1 | P17763 |
| GLAFSLIKNAQTPRR | 1 | 0 | 0 | 195383 | Q2YHF0.1 | P17763 |
| GLLFSIMKNTANTRR | 1 | 0 | 0 | 195393 | QQW40796.1 | P17763 |
| GNIVSSVNMVSRLLL | 1 | 0 | 0 | 195410 | QTX93504.1 | P17763 |
| GTSGSPIINRKGKVI | 1 | 0 | 0 | 195425 | Q2YHF0.1 | P17763 |
| GTSGSPIVNREGKIV | 1 | 0 | 0 | 195426 | P17763.2 | P17763 |
| HATMLDVDLRPASAW | 1 | 0 | 0 | 195441 | QOW96372.1 | P17763 |
| HNQTFLIDGPETAEC | 1 | 0 | 0 | 195458 | P14340.2 | P17763 |
| IGETLGEKWKSRLNA | 1 | 0 | 0 | 195482 | P14340.2 | P17763 |
| IGFRKEIGRMLNILN | 1 | 0 | 0 | 195483 | Q5UCB8.1 | P17763 |
| ILIRTGLLVISGVFP | 1 | 0 | 0 | 195501 | QRG47112.1 | P17763 |
| ILSITISEDGSMSIK | 1 | 0 | 0 | 195506 | P14340.2 | P17763 |
| IMKDGRKLVVPCRPQ | 1 | 0 | 0 | 195514 | QTX93489.1 | P17763 |
| INMLKRVRNRVSTGS | 1 | 0 | 0 | 195518 | QTX93489.1 | P17763 |
| IQRTVFFILMMLVAP | 1 | 0 | 0 | 195526 | Q2YHF0.1 | P17763 |
| IQRTVFFVLMMLVAP | 1 | 0 | 0 | 195527 | Q58HT7.1 | P17763 |
| ITQKGIIFILLMLVT | 1 | 0 | 0 | 195544 | P17763.2 | P17763 |
| KEISNMLNIMNRRKR | 1 | 0 | 0 | 195557 | P17763.2 | P17763 |
| KEISNMLSIINKRKK | 1 | 0 | 0 | 195559 | ADI44327.1 | P17763 |
| KLFMALVAFLRFLTI | 1 | 0 | 0 | 195573 | ABK78722.1 | P17763 |
| KLGYILRDISKIPGG | 1 | 0 | 0 | 195574 | P17763.2 | P17763 |
| KLTIMTGDIKGIMQA | 1 | 0 | 0 | 195576 | P14340.2 | P17763 |
| KNKAIKILIGFRKEI | 1 | 0 | 0 | 195580 | ARJ31953.1 | P17763 |
| LAVFLLLIMGQLTWN | 1 | 0 | 0 | 195613 | P27909.2 | P17763 |
| LCAGQLLLMRTTWAF | 1 | 0 | 0 | 195616 | Q2YHF0.1 | P17763 |
| LCTSQILLMRTTWAL | 1 | 0 | 0 | 195620 | P17763.2 | P17763 |
| LDIELLKTEVTNPAV | 1 | 0 | 0 | 195623 | P17763.2 | P17763 |
| LFTFVLLLSGQITWR | 1 | 0 | 0 | 195630 | QTX93504.1 | P17763 |
| LGFFLRKLTSRENLL | 1 | 0 | 0 | 195632 | QTX93489.1 | P17763 |
| LGQIMLLILCTSQIL | 1 | 0 | 0 | 195637 | P17763.2 | P17763 |
| LIMIGSNASDRMGMG | 1 | 0 | 0 | 195644 | QTX93489.1 | P17763 |
| LISLTCSNTIFTLTV | 1 | 0 | 0 | 195645 | QPI11927.1 | P17763 |
| LKGFKKEISNMLSII | 1 | 0 | 0 | 195649 | ADI44327.1 | P17763 |
| LLLMRTSWALCEALT | 1 | 0 | 0 | 195664 | QTX93491.1 | P17763 |
| LLSLTFIKTTFSLHY | 1 | 0 | 0 | 195670 | QFS19562.1 | P17763 |
| LLSPIRVPNYNLIIM | 1 | 0 | 0 | 195671 | Q9WDA6.1 | P17763 |
| LLSPVRVPNYNLIIM | 1 | 0 | 0 | 195672 | P14340.2 | P17763 |
| LMLVAHYAIIGPGLQ | 1 | 0 | 0 | 195682 | QFS19562.1 | P17763 |
| LQKQSHWVEITALIL | 1 | 0 | 0 | 195699 | Q2YHF0.1 | P17763 |
| LRHPGFTIMAAILAY | 1 | 0 | 0 | 195702 | P14340.2 | P17763 |
| LRTLILAPTRVVASE | 1 | 0 | 0 | 195705 | QFS19562.1 | P17763 |
| LSLTFIRSTMSLVMA | 1 | 0 | 0 | 195711 | AKQ00034.1 | P17763 |
| LTQKVVIFILLMLVT | 1 | 0 | 0 | 195726 | QTX93461.1 | P17763 |
| LTVAWRTATLILAGV | 1 | 0 | 0 | 195729 | QTX93491.1 | P17763 |
| LWTALISLTCSNTIF | 1 | 0 | 0 | 195739 | QPI11927.1 | P17763 |
| LWTALVSLTCSNTIF | 1 | 0 | 0 | 195740 | QTX93504.1 | P17763 |
| MAVFKMSPGYVLGVF | 1 | 0 | 0 | 195746 | Q5UCB8.1 | P17763 |
| MAVGIVSILLSSLLK | 1 | 0 | 0 | 195747 | P17763.2 | P17763 |
| MGKVRKDIPQWEPSK | 1 | 0 | 0 | 195755 | P17763.2 | P17763 |
| MLLILCTSQILLMRT | 1 | 0 | 0 | 195760 | P17763.2 | P17763 |
| MTLGMCCIITASILL | 1 | 0 | 0 | 195767 | P14340.2 | P17763 |
| MYWISNGTGNIVSSV | 1 | 0 | 0 | 195775 | QTX93500.1 | P17763 |
| NEDHWFSRGNSLSGV | 1 | 0 | 0 | 195781 | QRG47115.1 | P17763 |
| NELNYILWENNIKLT | 1 | 0 | 0 | 195782 | QTX93489.1 | P17763 |
| NGTVMDVISRRDQRG | 1 | 0 | 0 | 195786 | P17763.2 | P17763 |
| NILTVLLKTALLIVS | 1 | 0 | 0 | 195789 | QTX93479.1 | P17763 |
| NLLWKQIANELNYIL | 1 | 0 | 0 | 195795 | QTX93479.1 | P17763 |
| NMISRMLINRFTMKH | 1 | 0 | 0 | 195800 | QRG47107.1 | P17763 |
| NMISRMLINRFTMRH | 1 | 0 | 0 | 195801 | P14340.2 | P17763 |
| NVMWKQITNELNYVL | 1 | 0 | 0 | 195822 | Q2YHF0.1 | P17763 |
| NYLDYMTSMKRFKNE | 1 | 0 | 0 | 195824 | ABZ81543.1 | P17763 |
| PETMETLLLLGLMIL | 1 | 0 | 0 | 195828 | QTX93489.1 | P17763 |
| PVRVPNYNMIIMDEA | 1 | 0 | 0 | 195841 | P17763.2 | P17763 |
| QMWSLMYFHRRDLRL | 1 | 0 | 0 | 195858 | P14340.2 | P17763 |
| QTLTILIRTGLLVIS | 1 | 0 | 0 | 195869 | P14340.2 | P17763 |
| RAIWYMWLGARYLEF | 1 | 0 | 0 | 195875 | ACJ09459.1 | P17763 |
| RHTIENSTANVSLAA | 1 | 0 | 0 | 195891 | QTX93489.1 | P17763 |
| RLLSPVRVPNYNMII | 1 | 0 | 0 | 195897 | P17763.2 | P17763 |
| RNPGFALLAGFMAYM | 1 | 0 | 0 | 195901 | Q2YHF0.1 | P17763 |
| RNSTFLIDGPDTSEC | 1 | 0 | 0 | 195902 | Q5UCB8.1 | P17763 |
| RPAKNGTVMDVISRR | 1 | 0 | 0 | 195903 | P17763.2 | P17763 |
| SCVYNMMGKREKKLG | 1 | 0 | 0 | 195939 | QTX93489.1 | P17763 |
| SGNIVSSVNMISRML | 1 | 0 | 0 | 195948 | P14340.2 | P17763 |
| SGVSWTMKILIGVVI | 1 | 0 | 0 | 195953 | QQW40772.1 | P17763 |
| SNILDIDLRPASAWT | 1 | 0 | 0 | 195974 | QTX93024.1 | P17763 |
| SSTPLAENTNSVTNI | 1 | 0 | 0 | 195999 | QTX92319.1 | P17763 |
| TCVYNMMGKREKKLG | 1 | 0 | 0 | 196031 | P17763.2 | P17763 |
| TGNIVSSVNMISRML | 1 | 0 | 0 | 196040 | QTX93073.1 | P17763 |
| TKTDFGFYQVKTETT | 1 | 0 | 0 | 196052 | QTX92372.1 | P17763 |
| TMPLVMAWRTIMAVL | 1 | 0 | 0 | 196060 | QTX92385.1 | P17763 |
| TMSLVMAWRTIMAVL | 1 | 0 | 0 | 196061 | AKQ00032.1 | P17763 |
| TRMVMTTTANWLWAL | 1 | 0 | 0 | 196071 | QDZ58857.1 | P17763 |
| TSREVLLLTIGLSLV | 1 | 0 | 0 | 196079 | QFS19562.1 | P17763 |
| VDYMPVMKRYSAPFE | 1 | 0 | 0 | 196105 | QOV09272.1 | P17763 |
| VIEHLERLQRKHGGM | 1 | 0 | 0 | 196126 | QTX93593.1 | P17763 |
| VILKDGPERVILAGP | 1 | 0 | 0 | 196130 | P17763.2 | P17763 |
| VLDDGIYRILQRGLL | 1 | 0 | 0 | 196138 | P17763.2 | P17763 |

| | | | | | | |
|---|---|---|---|---|---|---|
| VMAWRTIMAVLFVVT | 1 | 0 | 0 | 196153 | QTX92385.1 | P17763 |
| VRNMEKYQLAVTIMA | 1 | 0 | 0 | 196174 | QTX93213.1 | P17763 |
| VSSVNMISRMLINRF | 1 | 0 | 0 | 196187 | QTX93024.1 | P17763 |
| VVIFILLMLVTPSMT | 1 | 0 | 0 | 196203 | QTX93593.1 | P17763 |
| WAFCEVLTLATGPIL | 1 | 0 | 0 | 196208 | QTX92385.1 | P17763 |
| WGTIKKSKAINVLRG | 1 | 0 | 0 | 196213 | QDR24384.1 | P17763 |
| WKTMAMVLSIVSLFP | 1 | 0 | 0 | 196216 | QTX92650.1 | P17763 |
| WKVSCTILAAVSVSP | 1 | 0 | 0 | 196217 | QTX93073.1 | P17763 |
| WNIVKLMSGKDVFYL | 1 | 0 | 0 | 196222 | AAC63341.1 | P17763 |
| YAQMWSLMYFHRRDL | 1 | 0 | 0 | 196230 | QTX93201.1 | P17763 |
| YQNKVVKVLRPTPKG | 1 | 0 | 0 | 196238 | QTX92385.1 | P17763 |
| YQNKVVRVQRPTPRG | 1 | 0 | 0 | 196243 | QTX93024.1 | P17763 |
| NDKHSNGTIKDRSPY | 1 | 0 | 0 | 215815 | ACQ63272.1 | P03468 |
| LKYNPSRVEAFHRYGTTVNC | 1 | 0 | 0 | 226063 | AAA19496.1 | P10211 |
| ATTQNDRMPGPELSG | 1 | 0 | 0 | 227054 | AGB56828.1 | Q05127 |
| CDIENNPGLCYASQM | 1 | 0 | 0 | 227058 | AGB56828.1 | Q05127 |
| DDIPFPGPINDDDNP | 1 | 0 | 0 | 227076 | AGB56827.1 | P18272 |
| DETVPQSVREAFNNL | 1 | 0 | 0 | 227084 | AGB56828.1 | Q05127 |
| DPGTNTTTEDHKIMA | 1 | 0 | 0 | 227099 | AGB56758.1 | Q05320 |
| EHGQPPPGPSLYEES | 1 | 0 | 0 | 227128 | AGB56828.1 | Q05127 |
| ENSSAMVQVHSQGRE | 1 | 0 | 0 | 227141 | AGB56758.1 | Q05320 |
| FGTNETEYLFEVDNL | 1 | 0 | 0 | 227170 | AGB56832.1 | P60170 |
| GDLTMVITQDCDTCH | 1 | 0 | 0 | 227198 | 4LDI_A | Q05128 |
| HLGLDDQEKKILMNF | 1 | 0 | 0 | 227244 | AGB56827.1 | P18272 |
| HNTPVYKLDISEATQ | 1 | 0 | 0 | 227248 | AGB56830.1 | Q05320 |
| IESRDETVPQSVREA | 1 | 0 | 0 | 227263 | AGB56828.1 | Q05127 |
| IEVPETLVHKLTGKK | 1 | 0 | 0 | 227264 | 4LDI_A | Q05128 |
| IGLDPVAPGDLTMVI | 1 | 0 | 0 | 227266 | 4LDI_A | Q05128 |
| KPGPDNSTHNTPVYK | 1 | 0 | 0 | 227307 | AGB56758.1 | Q05320 |
| KSGKKGNSADLTSPE | 1 | 0 | 0 | 227312 | 4LDI_A | Q05128 |
| LFDLDEDDEDTKPVP | 1 | 0 | 0 | 227330 | AGB56827.1 | P18272 |
| LLPNKSGKKGNSADL | 1 | 0 | 0 | 227347 | 4LDI_A | Q05128 |
| LMTGRIPVSDIFCDI | 1 | 0 | 0 | 227350 | AGB56828.1 | Q05127 |
| LPKYIGLDPVAPGDL | 1 | 0 | 0 | 227357 | 4LDI_A | Q05128 |
| LRPILLPNKSGKKGN | 1 | 0 | 0 | 227365 | 4LDI_A | Q05128 |
| LTEENFGKPDISAKD | 1 | 0 | 0 | 227373 | AGL73451.1 | B8XCM8 |
| MVITQDCDTCHSPAS | 1 | 0 | 0 | 227398 | 4LDI_A | Q05128 |
| NISGQSPARTSSDPG | 1 | 0 | 0 | 227409 | AGB56758.1 | Q05320 |
| NNPGLCYASQMQQTK | 1 | 0 | 0 | 227416 | AGB56828.1 | Q05127 |
| NTTTEDHKIMASENS | 1 | 0 | 0 | 227422 | AGB56758.1 | Q05320 |
| PPKVVNYEAGEWAEN | 1 | 0 | 0 | 227450 | AGL73460.1 | B8XCN0 |
| PQSVREAFNNLNSTT | 1 | 0 | 0 | 227458 | AAM76032.1 | Q05127 |
| QKKNEISFQQTNAMV | 1 | 0 | 0 | 227489 | AGL73422.1 | B0LPL4 |
| QRQIQVHAEQGLIQY | 1 | 0 | 0 | 227498 | AGL73450.1 | B8XCM7 |
| RELDHLGLDDQEKKI | 1 | 0 | 0 | 227509 | AGL73450.1 | B8XCM7 |
| RIPVSDIFCDIENNP | 1 | 0 | 0 | 227512 | AGB56828.1 | Q05127 |
| RSTKGGQQKNSQKGQ | 1 | 0 | 0 | 227532 | AGB56773.1 | P18272 |
| SDIFCDIENNPGLCY | 1 | 0 | 0 | 227547 | AGB56828.1 | Q05127 |
| STTSLTEENFGKPDI | 1 | 0 | 0 | 227589 | AGB56828.1 | Q05127 |
| TATAAATEAYWAEHG | 1 | 0 | 0 | 227601 | AGL73451.1 | B8XCM8 |
| TSGHYDDDDDIPFPG | 1 | 0 | 0 | 227631 | AGB56773.1 | P18272 |
| TVVSNGAKNISGQSP | 1 | 0 | 0 | 227646 | NP_066246.1 | Q05320 |
| VCVFQLQDGKTLGLK | 1 | 0 | 0 | 227651 | AGB56828.1 | Q05127 |
| VYKLDISEATQVEQH | 1 | 0 | 0 | 227687 | AGB56830.1 | Q05320 |
| GDVPEAVENAVARVA | 1 | 0 | 0 | 229649 | AAK13386.1 | Q86887 |
| GPQGEAVERAIARVA | 1 | 0 | 0 | 229665 | SRC292625 | A0A0M3PZA5 |
| NDVRQAVEGAIGRVA | 1 | 0 | 0 | 229747 | AAD09601.1 | Q86887 |
| NSWGSQMDLTDPLCI | 1 | 0 | 0 | 229755 | AAB23363.1 | O73556 |
| SPVEESIERSIGRVA | 1 | 0 | 0 | 229799 | ABX60142.1 | Q86887 |
| AMTDTTPFGQQRVFK | 1 | 0 | 0 | 230626 | P17763.2 | P17763 |
| GKAKGSRAIWYMWLG | 1 | 0 | 0 | 230858 | P17763.2 | P17763 |
| GSRAIWYMWLGARYL | 1 | 0 | 0 | 230934 | ACJ09459.1 | P17763 |
| ISRRDQRGSGQVVTY | 1 | 0 | 0 | 231017 | Q9WDA6.1 | P17763 |
| RYLEFEALGFLNEDH | 1 | 0 | 0 | 231420 | ABQ18244.1 | P17763 |
| TTPFGQQRVFKEKVD | 1 | 0 | 0 | 231543 | P17763.2 | P17763 |
| ELDKWASLWNWFNITNWLWYIK | 1 | 0 | 0 | 236082 | NP_057856.1 | P03378 |
| ASFTVSSEKTILTMGEYG | 1 | 0 | 0 | 237645 | ADQ00969.1 | P14336 |
| ECPLERRKTGVFTVAEFG | 1 | 0 | 0 | 237691 | ADQ00969.1 | P14336 |
| ENPAKTREYCLHAKLSDT | 1 | 0 | 0 | 237702 | ADQ00969.1 | P14336 |
| GKALYFLNDMAKTRKDIG | 1 | 0 | 0 | 237732 | ADQ00969.1 | P14336 |
| GLLVISGLFPVSIPITAA | 1 | 0 | 0 | 237736 | AAC59275.1 | P17763 |
| GLNTFTNMEAQLIRQMEG | 1 | 0 | 0 | 237737 | AAA99437.1 | P17763 |
| LALPWKHEGAQNWNNAER | 1 | 0 | 0 | 237808 | ADQ00969.1 | P14336 |
| LNTLTNIKVQLIRMMEGE | 1 | 0 | 0 | 237829 | ADQ00969.1 | P14336 |
| NTLTNIKVQLIRMMEGEG | 1 | 0 | 0 | 237863 | ADQ00969.1 | P14336 |
| PLDDRFGKALYFLNDMAK | 1 | 0 | 0 | 237877 | ADQ00969.1 | P14336 |
| PWLAWHVAANVSSVTDRS | 1 | 0 | 0 | 237885 | ADQ00969.1 | P14336 |
| SEWTNVDIQPARSWGTYV | 1 | 0 | 0 | 237924 | ADQ00969.1 | P14336 |
| WSIHASGAWMTTEDMLDV | 1 | 0 | 0 | 238032 | ADQ00969.1 | P14336 |
| CFKCCRRTRGLFKYRHLKD | 1 | 0 | 0 | 244173 | ABB30034.1 | Q8JSZ3 |
| FMFGWRILFCFKCCRRTRG | 1 | 0 | 0 | 244201 | ABB30034.1 | Q8JSZ3 |
| KIPLLGKMAIYICRMSNHP | 1 | 0 | 0 | 244229 | ABB30034.1 | Q8JSZ3 |
| LHKEWPHSRNWRCNPTWCW | 1 | 0 | 0 | 244240 | ABB30034.1 | Q8JSZ3 |
| MSRSESRKNRGGREEILE | 1 | 0 | 0 | 244254 | AAA45723.1 | P29996 |
| PGGDPRGAPGGGFVPTML | 1 | 0 | 0 | 244264 | CAE51162.1 | P0C6L3 |
| PGGSPRGAPGGGFVPRML | 1 | 0 | 0 | 244265 | CAJ66091.1 | P0C6L3 |
| RCGCTSSTCLHKEWPHSRN | 1 | 0 | 0 | 244270 | AAT81454.1 | Q8JSZ3 |
| RGDQQFPWGQQPPPPPRLPLLEC | 1 | 0 | 0 | 244271 | CAJ66091.1 | P0C6L3 |
| RGTQQFPWGNTPPRPPRLPLLEC | 1 | 0 | 0 | 244272 | AGA20452.1 | P0C6L3 |
| TCTGCYACSSGISCKVRIH | 1 | 0 | 0 | 244287 | ABB30034.1 | Q8JSZ3 |
| VPESPFSRTGEGLDIRGNQGFP | 1 | 0 | 0 | 244302 | AAA45723.1 | P29996 |

| | | | | | | |
|---|---|---|---|---|---|---|
| VPESPFSRTGEGLDVRGTQQFP | 1 | 0 | 0 | 244303 | CAE51162.1 | P0C6L3 |
| VPESPFTRRGDGLDIRGDQQFP | 1 | 0 | 0 | 244304 | CAJ66091.1 | P0C6L3 |
| NWFDITNWLWYIKLFIMIVGGLV | 1 | 0 | 0 | 347770 | ABC47957.1 | P03378 |
| KGKSLLFKTENGVNMC | 1 | 0 | 0 | 423017 | AAK67712.1 | P17763 |
| AAVPVLAFDAARLRLLE | 1 | 0 | 0 | 429189 | AFJ06845.2 | P03189 |
| PGRRPFFHPVGEAD | 1 | 0 | 0 | 431702 | AFY97913.1 | P03211 |
| AETQHGTTVVKVKYE | 1 | 0 | 0 | 489122 | Q2YHF0.1 | P17763 |
| AETQHGTVLVQVKYE | 1 | 0 | 0 | 489123 | P17763.2 | P17763 |
| APCKIPFSTEDGQGK | 1 | 0 | 0 | 489134 | QTX93504.1 | P17763 |
| FGDSYIVIGVGDSAL | 1 | 0 | 0 | 489159 | Q2YHF0.1 | P17763 |
| GRIISSTPFAENTNS | 1 | 0 | 0 | 489214 | Q2YHF0.1 | P17763 |
| GRLITANPIVTDKEK | 1 | 0 | 0 | 489215 | P17763.2 | P17763 |
| GRLITVNPIVTEKDS | 1 | 0 | 0 | 489216 | P14340.2 | P17763 |
| IEAEPPFGESNIVIG | 1 | 0 | 0 | 489229 | QTX93491.1 | P17763 |
| IEIRDVNKEKVVGRI | 1 | 0 | 0 | 489231 | Q2YHF0.1 | P17763 |
| KEKPVNIEAEPPFGE | 1 | 0 | 0 | 489248 | P17763.2 | P17763 |
| KGMSYAMCLNTFVLK | 1 | 0 | 0 | 489249 | AAC63321.1 | P17763 |
| KYEGAGAPCKVPIEI | 1 | 0 | 0 | 489272 | Q58HT7.1 | P17763 |
| MCSGKFSIDKEMAET | 1 | 0 | 0 | 489297 | Q2YHF0.1 | P17763 |
| MCTGKFKVVKEIAET | 1 | 0 | 0 | 489298 | P14340.2 | P17763 |
| MCTGSFKLEKEVAET | 1 | 0 | 0 | 489299 | P17763.2 | P17763 |
| TNSVTNIELEPPFGD | 1 | 0 | 0 | 489394 | QTX92385.1 | P17763 |
| VIGIGDKALKINWYK | 1 | 0 | 0 | 489400 | QTX93593.1 | P17763 |
| VLKKEVSETQHGTIL | 1 | 0 | 0 | 489408 | QTX93593.1 | P17763 |
| EVLPPPRKKKGLFSQ | 1 | 0 | 0 | 489533 | AAA45474.1 | P08617 |
| LEVGKQRLKYAQEEL | 1 | 0 | 0 | 489637 | AAA45474.1 | P08617 |
| RSNTTGTLIWKV | 1 | 0 | 0 | 502007 | ALT19772.1 | B8XCN0 |
| ASPQLEGFSAESRKL | 1 | 0 | 0 | 504100 | ACP41104.1 | P03433 |
| GRDRIMAWTVVNSIC | 1 | 0 | 0 | 504102 | ACP41104.1 | P03433 |
| LAKGEKANVLIGQGD | 1 | 0 | 0 | 504103 | P21428.1 | P03428 |
| NPECELLISRESWSY | 1 | 0 | 0 | 504111 | ABU99069.1 | P03452 |
| PFQNIHPVTIGECPK | 1 | 0 | 0 | 504113 | AAB52905.1 | P03452 |
| QGALLNDKHSNGTIK | 1 | 0 | 0 | 504114 | ACP41107.1 | P03468 |
| PTADSPLTASPPATAPGPSA | 1 | 0 | 0 | 523862 | AIL27778.1 | P13290 |
| ELYHYQECVRGTTVL | 1 | 0 | 0 | 531543 | P0DTC7.1 | P0DTC7 |
| QLRARSVSPKLFIRQ | 1 | 0 | 0 | 533379 | P0DTC7.1 | P0DTC7 |
| AIYGAAFSGVSWTMKILI | 1 | 0 | 0 | 538806 | ABW06583.1 | P17763 |
| ATEIQMSSGNLLFTGHLK | 1 | 0 | 0 | 538813 | ABW06583.1 | P17763 |
| DTQGSNWIQKETLVNFKN | 1 | 0 | 0 | 538824 | ABW06583.1 | P17763 |
| GKVVLPENLEYTIVITPH | 1 | 0 | 0 | 538853 | ABW06583.1 | P17763 |
| GLFGKGGIVTCAMFTCKK | 1 | 0 | 0 | 538855 | ABW06583.1 | P17763 |
| GSPCKIPFEIMDLEKRHV | 1 | 0 | 0 | 538856 | ABW06583.1 | P17763 |
| KAWLVHRQWFLDLPLPWL | 1 | 0 | 0 | 538878 | ABW06583.1 | P17763 |
| KGMSYSMCTGKFKIVKEI | 1 | 0 | 0 | 538883 | ABW06583.1 | P17763 |
| KILIGVIITWIGMNSRST | 1 | 0 | 0 | 538884 | ABW06583.1 | P17763 |
| LGSQEGAMHTALTGATEI | 1 | 0 | 0 | 538914 | ABW06583.1 | P17763 |
| LPWLPGADTQGSNWIQKE | 1 | 0 | 0 | 538916 | ABW06583.1 | P17763 |
| MHTALTGATEIQMSSGNL | 1 | 0 | 0 | 538919 | ABW06583.1 | P17763 |
| MRCIGISNRDFVEGVSGG | 1 | 0 | 0 | 538922 | ABW06583.1 | P17763 |
| NLEYTIVITPHSGEEHAV | 1 | 0 | 0 | 538925 | ABW06583.1 | P17763 |
| NRDFVEGVSGGSWVDIVL | 1 | 0 | 0 | 538926 | ABW06583.1 | P17763 |
| PFGDSYIIIGVEPGQLKL | 1 | 0 | 0 | 538928 | ABW06583.1 | P17763 |
| RVQYEGDGSPCKIPFEIM | 1 | 0 | 0 | 538961 | ABW06583.1 | P17763 |
| SCVTTMAKNKPTLDFELI | 1 | 0 | 0 | 538964 | AAB33854.1 | P17763 |
| SGNLLFTGHLKCRLRMDK | 1 | 0 | 0 | 538965 | ABW06583.1 | P17763 |
| SGVSWTMKILIGVIITWI | 1 | 0 | 0 | 538966 | ABW06583.1 | P17763 |
| SLVLVGVVTLYLGAMVQA | 1 | 0 | 0 | 538968 | ABW06583.1 | P17763 |
| VFTSIGKALHQVFGAIYG | 1 | 0 | 0 | 538990 | ABW06583.1 | P17763 |
| VKEIAETQHGTIVIRVQY | 1 | 0 | 0 | 538991 | ABW06583.1 | P17763 |
| VLLQMEEKAWLVHRQWFL | 1 | 0 | 0 | 538992 | ABW06583.1 | P17763 |
| VNIEAEPPFGDSYIIIGV | 1 | 0 | 0 | 538994 | ABW06583.1 | P17763 |
| VSGGSWVDIVLEHGSCVT | 1 | 0 | 0 | 538995 | ABW06583.1 | P17763 |
| VSLVLVGVVTLYLGAMVQ | 1 | 0 | 0 | 538996 | ABW06583.1 | P17763 |
| VTEKDSPVNIEAEPPFGD | 1 | 0 | 0 | 538997 | ABW06583.1 | P17763 |
| AAAIFMTATPPGATD | 1 | 0 | 0 | 539020 | Q58HT7.1 | P17763 |
| AAAIFMTATPPGSID | 1 | 0 | 0 | 539021 | AHG06382.1 | P17763 |
| AAAIFMTATPPGTTD | 1 | 0 | 0 | 539024 | Q2YHF0.1 | P17763 |
| AAGIFMTATPPGSRD | 1 | 0 | 0 | 539025 | P14340.2 | P17763 |
| AEMEEALRGLPIRYQ | 1 | 0 | 0 | 539041 | P14340.2 | P17763 |
| AFIAFLRFLAIPPTA | 1 | 0 | 0 | 539043 | P17763.2 | P17763 |
| AGLLLRKLTSKELMM | 1 | 0 | 0 | 539054 | P14340.2 | P17763 |
| ALFLAHAIGTSITQK | 1 | 0 | 0 | 539074 | P17763.2 | P17763 |
| ALNDTWKMEKASFIE | 1 | 0 | 0 | 539079 | QRG47112.1 | P17763 |
| AVSFVTLITGNMSFR | 1 | 0 | 0 | 539108 | P14340.2 | P17763 |
| AVSGVYPMSIPATLF | 1 | 0 | 0 | 539109 | P27909.2 | P17763 |
| DIDLRPASAWTLYAV | 1 | 0 | 0 | 539163 | P14340.2 | P17763 |
| DILTEIASLPTYLSS | 1 | 0 | 0 | 539165 | Q2YHF0.1 | P17763 |
| DKRSVALAPHVGMGL | 1 | 0 | 0 | 539168 | QTX93489.1 | P17763 |
| ECLRRRVTRKHMILA | 1 | 0 | 0 | 539198 | AMP43484.1 | P17763 |
| EFTKKVRTNAAMGAV | 1 | 0 | 0 | 539211 | QTX93489.1 | P17763 |
| EMLRTRVGTKHAILL | 1 | 0 | 0 | 539231 | P14340.2 | P17763 |
| ENLEYTIVITPHSGE | 1 | 0 | 0 | 539236 | BAG84557.1 | P17763 |
| ETYQLWTALISLTCS | 1 | 0 | 0 | 539243 | QIS48873.1 | P17763 |
| FEEYKRSGILEVDRT | 1 | 0 | 0 | 539251 | Q2YHF0.1 | P17763 |
| FITFLRVLSIPPTAG | 1 | 0 | 0 | 539255 | Q2YHF0.1 | P17763 |
| FKKEISNMLNIMNRR | 1 | 0 | 0 | 539257 | P17763.2 | P17763 |
| FKKNGAIKVLRGFKK | 1 | 0 | 0 | 539259 | P17763.2 | P17763 |
| FLRFLTIPPTAGILK | 1 | 0 | 0 | 539264 | P14340.2 | P17763 |
| FNTYKRSGIMEVDRS | 1 | 0 | 0 | 539267 | QFS19562.1 | P17763 |
| GAYRIKQRGILGYSQ | 1 | 0 | 0 | 539283 | QRG47115.1 | P17763 |
| GGVSWMVRILIGLLV | 1 | 0 | 0 | 539298 | Q2YHF0.1 | P17763 |

| | | | | | | |
|---|---|---|---|---|---|---|
| GIFLRKLTSRETALM | 1 | 0 | 0 | 539301 | Q2YHF0.1 | P17763 |
| GIYRIMQRGLLGRSQ | 1 | 0 | 0 | 539309 | QFS19562.1 | P17763 |
| GLAFSIMKSVGTGKR | 1 | 0 | 0 | 539321 | QTX93374.1 | P17763 |
| GLAFSLMKSLGGGRR | 1 | 0 | 0 | 539322 | P17763.2 | P17763 |
| GLLFSIMKNTTNTRR | 1 | 0 | 0 | 539327 | P14340.2 | P17763 |
| GTRMVMTTTANWLWA | 1 | 0 | 0 | 539354 | AUN35144.1 | P17763 |
| GTYGLNTFTNMEAQL | 1 | 0 | 0 | 539355 | P17763.2 | P17763 |
| GVFLRKLTSRETALM | 1 | 0 | 0 | 539360 | Q5UCB8.1 | P17763 |
| GVGLAMATTLQLPED | 1 | 0 | 0 | 539363 | QTX93489.1 | P17763 |
| IDDRFATALTALNDM | 1 | 0 | 0 | 539396 | P17763.2 | P17763 |
| IDNFSLGVLGMALFL | 1 | 0 | 0 | 539397 | P14340.2 | P17763 |
| IEGKVVQHENLKYTV | 1 | 0 | 0 | 539399 | QTX93457.1 | P17763 |
| KCGSCVYNMMGKREK | 1 | 0 | 0 | 539458 | QTX93489.1 | P17763 |
| KEISSMLNIMNRRKR | 1 | 0 | 0 | 539459 | P27909.2 | P17763 |
| KKEISSMLNIMNRRK | 1 | 0 | 0 | 539475 | P27909.2 | P17763 |
| KMLLNRFTTRHRKPT | 1 | 0 | 0 | 539493 | CCM44290.1 | P17763 |
| KNKAIKILTGFRKEI | 1 | 0 | 0 | 539496 | Q2YHF0.1 | P17763 |
| KSLLFKTASGINMCT | 1 | 0 | 0 | 539515 | QTX93489.1 | P17763 |
| KSLLFKTSAGVNMCT | 1 | 0 | 0 | 539516 | P17763.2 | P17763 |
| KVIGLYGNGVVTKSG | 1 | 0 | 0 | 539520 | Q2YHF0.1 | P17763 |
| LDNLAVLHTAEAGGR | 1 | 0 | 0 | 539545 | P14340.2 | P17763 |
| LGCYSQVNPLTLTAA | 1 | 0 | 0 | 539551 | P17763.2 | P17763 |
| LGEFGKAKGSRAIWY | 1 | 0 | 0 | 539552 | P17763.2 | P17763 |
| LGMMILKIVRNMEKY | 1 | 0 | 0 | 539561 | QQW40674.1 | P17763 |
| LGMMVLKIVRNMEKY | 1 | 0 | 0 | 539562 | QTX93014.1 | P17763 |
| LIATFKIQPFLALGF | 1 | 0 | 0 | 539568 | QTX93489.1 | P17763 |
| LLAAFKVRPTFAAGL | 1 | 0 | 0 | 539583 | P14340.2 | P17763 |
| LLLNRFTMTHRRPTI | 1 | 0 | 0 | 539589 | QTX93489.1 | P17763 |
| LNTFTNMEVQLIRQM | 1 | 0 | 0 | 539605 | P27909.2 | P17763 |
| LPTFMTQKARDALDN | 1 | 0 | 0 | 539608 | AHG25307.1 | P17763 |
| LPTFMTQKARNALDN | 1 | 0 | 0 | 539609 | QQW40770.1 | P17763 |
| LRGFKKEISNMLNIM | 1 | 0 | 0 | 539616 | P17763.2 | P17763 |
| LRGFKKEISSMLNIM | 1 | 0 | 0 | 539617 | P27909.2 | P17763 |
| LSLTFIRSTTPLVMA | 1 | 0 | 0 | 539628 | AHG06383.1 | P17763 |
| LTMKNKAWMVHRQWF | 1 | 0 | 0 | 539634 | QTX93489.1 | P17763 |
| LVALLGAMTAGIFLF | 1 | 0 | 0 | 539638 | Q2YHF0.1 | P17763 |
| LVTFKNPHAKKQDVV | 1 | 0 | 0 | 539648 | P14340.2 | P17763 |
| MGCYSQVNPTTLTAS | 1 | 0 | 0 | 539662 | Q2YHF0.1 | P17763 |
| MLLNRFTMAHRKPTY | 1 | 0 | 0 | 539678 | P17763.2 | P17763 |
| MTSRMLLNRFTMAHR | 1 | 0 | 0 | 539691 | P17763.2 | P17763 |
| MVSRLLLNRFTMTHR | 1 | 0 | 0 | 539701 | QTX93489.1 | P17763 |
| NAIFKLTYQNKVVKV | 1 | 0 | 0 | 539704 | QTX93479.1 | P17763 |
| NPKNFQTMPGIFQTT | 1 | 0 | 0 | 539742 | QTX93504.1 | P17763 |
| PGLFKTLTGEIGAVT | 1 | 0 | 0 | 539767 | Q2YHF0.1 | P17763 |
| QKETLVTFKNPHAKK | 1 | 0 | 0 | 539806 | P14340.2 | P17763 |
| RGTAKLRWFVERNLV | 1 | 0 | 0 | 539851 | P17763.2 | P17763 |
| RKNGKKVIQLSRKTF | 1 | 0 | 0 | 539857 | P14340.2 | P17763 |
| RKNGKRVIQLSRKTF | 1 | 0 | 0 | 539858 | QFS19562.1 | P17763 |
| RQDLLVTFKTAHAKK | 1 | 0 | 0 | 539869 | P17763.2 | P17763 |
| RRDLRLAANAICSAV | 1 | 0 | 0 | 539871 | P17763.2 | P17763 |
| RRDLRLASNAICSAV | 1 | 0 | 0 | 539872 | QTX93489.1 | P17763 |
| SGNIVSSVNTTSKML | 1 | 0 | 0 | 539901 | Q2YHF0.1 | P17763 |
| SHHFHQLIMKDGREI | 1 | 0 | 0 | 539905 | P17763.2 | P17763 |
| SRETALMVIGMAMTT | 1 | 0 | 0 | 539930 | QTX92385.1 | P17763 |
| STRVEMGEAAAIFMT | 1 | 0 | 0 | 539938 | QTX92385.1 | P17763 |
| SVALVPHVGMGLETR | 1 | 0 | 0 | 539943 | QTX93024.1 | P17763 |
| TSIFKLTYQNKVVRV | 1 | 0 | 0 | 540001 | P17763.2 | P17763 |
| TTFSLHYAWKTMAMV | 1 | 0 | 0 | 540005 | QTX92825.1 | P17763 |
| TYGLNTFTNMEAQLI | 1 | 0 | 0 | 540018 | P17763.2 | P17763 |
| TYGLNTFTNMEVQLI | 1 | 0 | 0 | 540020 | QFS19562.1 | P17763 |
| VGGLFTSLGKAVHQV | 1 | 0 | 0 | 540036 | QTX92219.1 | P17763 |
| VGLLFRRLTSREVLL | 1 | 0 | 0 | 540038 | P17763.2 | P17763 |
| VLMHKGKRIEPSWAD | 1 | 0 | 0 | 540056 | QTX93201.1 | P17763 |
| VNGVVRLLTKPWDVI | 1 | 0 | 0 | 540064 | P17763.2 | P17763 |
| VNGVVRLLTKPWDVV | 1 | 0 | 0 | 540065 | QTX93073.1 | P17763 |
| VREAIKRKLRTLILA | 1 | 0 | 0 | 540074 | QFS19562.1 | P17763 |
| VREAIKRKLRTLVLA | 1 | 0 | 0 | 540075 | QMT58616.1 | P17763 |
| VVQYENLKYTVIITV | 1 | 0 | 0 | 540102 | QTX93593.1 | P17763 |
| WCGSLIGLTARATWA | 1 | 0 | 0 | 540112 | P17763.2 | P17763 |
| WGSFKKNGAIKVLRG | 1 | 0 | 0 | 540117 | P17763.2 | P17763 |
| WGTFKKSGAIKVLKG | 1 | 0 | 0 | 540121 | QBM00871.1 | P17763 |
| WGTFKKSGAIKVLRG | 1 | 0 | 0 | 540123 | QHR82521.1 | P17763 |
| WQLMYFHRRDLRLAA | 1 | 0 | 0 | 540138 | P17763.2 | P17763 |
| WSSFKKNGAIKVLRG | 1 | 0 | 0 | 540141 | QUD37222.1 | P17763 |
| YAQIQPHWIAASIIL | 1 | 0 | 0 | 540148 | QTX93024.1 | P17763 |
| AVAKCNEKHDEEFCD | 1 | 0 | 0 | 540184 | P08669.1 | P08669 |
| DQIEQEADNMITEML | 1 | 0 | 0 | 540192 | AAL01685.1 | P09991 |
| HDEEFCDMLRLFDFN | 1 | 0 | 0 | 540214 | P08669.1 | P08669 |
| SDIEQEANNMITEML | 1 | 0 | 0 | 540266 | Q2A069.1 | Q2A069 |
| ATDHAQAIKVHTAAMKVG | 1 | 0 | 0 | 558229 | AFL65790.1 | P03316 |
| EITVMSSEVLPSTNQEYI | 1 | 0 | 0 | 558247 | P03316.1 | P03316 |
| HVKGTIDHPVLSKLKFTK | 1 | 0 | 0 | 558264 | AFL65790.1 | P03316 |
| KPGKRQRMALKLEADRLF | 1 | 0 | 0 | 558280 | AFL65790.1 | P03316 |
| MNRGFFNMLGRRPFPAAT | 1 | 0 | 0 | 558294 | AFL65790.1 | P03316 |
| NIPISIDIPNAAFIRTSD | 1 | 0 | 0 | 558299 | NP_740677.1 | P03316 |
| PISASFTPFDHKVVIHRG | 1 | 0 | 0 | 558303 | NP_740677.1 | P03316 |
| QTKWVFNSPDLIRHADHT | 1 | 0 | 0 | 558309 | AFL65790.1 | P03316 |
| SVTDDFTLTSPYLGTCSY | 1 | 0 | 0 | 558330 | AFL65790.1 | P03316 |
| TTAVSALVIGQATRPQNP | 1 | 0 | 0 | 558338 | AFL65790.1 | P03316 |
| TWNSKGKTIKTTPEGTEE | 1 | 0 | 0 | 558342 | AFL65790.1 | P03316 |
| YNYDFPEYGAMKPGAFGD | 1 | 0 | 0 | 558349 | NP_740677.1 | P03316 |

| | | | | | | |
|---|---|---|---|---|---|---|
| TDVSTAIHADQLTPAWRIYSTG | 1 | 0 | 0 | 558510 | P59594.1 | P59594 |
| AIPNFENLRRNWLEL | 1 | 0 | 0 | 561863 | BAJ49820.1 | A0A0M3PZA5 |
| AVLTSKDPRLEVDFE | 1 | 0 | 0 | 561868 | AAD22046.3 | A0A0M3PZA5 |
| DAIRRRFYMDCDIEV | 1 | 0 | 0 | 561869 | ACY00666.1 | A0A0M3PZA5 |
| DELRSLDKIKKGKSR | 1 | 0 | 0 | 561870 | ADO85560.1 | A0A0M3PZA5 |
| DVSVNRFYTLDTKLW | 1 | 0 | 0 | 561871 | ACY00666.1 | A0A0M3PZA5 |
| EPDEYVTQAALHYAN | 1 | 0 | 0 | 561872 | AAB39968.1 | A0A0M3PZA5 |
| ESIRWTKDARNTQDH | 1 | 0 | 0 | 561873 | ACY00666.1 | A0A0M3PZA5 |
| GPPKFRPIRISLEEK | 1 | 0 | 0 | 561878 | ACY00666.1 | A0A0M3PZA5 |
| GRLNINGPTRTKLEP | 1 | 0 | 0 | 561879 | ACY00666.1 | A0A0M3PZA5 |
| GRQGFCAGLKRSYFA | 1 | 0 | 0 | 561880 | ACY00666.1 | A0A0M3PZA5 |
| IHIPGEVRNLLELCQ | 1 | 0 | 0 | 561882 | ACY00666.1 | A0A0M3PZA5 |
| IKKRDILDPTTRDVS | 1 | 0 | 0 | 561883 | ADG57604.1 | A0A0M3PZA5 |
| INHTHHVYRNKTYCV | 1 | 0 | 0 | 561884 | AEK98534.1 | A0A0M3PZA5 |
| IPETPTNVERHLNRA | 1 | 0 | 0 | 561885 | ACY00666.1 | A0A0M3PZA5 |
| ITGYAQMRRKVELFT | 1 | 0 | 0 | 561886 | AEI71367.1 | A0A0M3PZA5 |
| KLCSENNTANFKRCS | 1 | 0 | 0 | 561887 | ACY00666.1 | A0A0M3PZA5 |
| KRMNNYMQFKSKHRI | 1 | 0 | 0 | 561888 | ACY00666.1 | A0A0M3PZA5 |
| LAWHNGKDEYEKFVS | 1 | 0 | 0 | 561889 | ACK37353.1 | A0A0M3PZA5 |
| LELAKTGKEYGLTMT | 1 | 0 | 0 | 561890 | ACK37353.1 | A0A0M3PZA5 |
| LPPDPDHFDGYKQQV | 1 | 0 | 0 | 561892 | AEK98149.1 | A0A0M3PZA5 |
| LRDRKSKVRYSVDTV | 1 | 0 | 0 | 561893 | ACY00666.1 | A0A0M3PZA5 |
| LRRNIRQVQTDQGHF | 1 | 0 | 0 | 561894 | AAD22046.3 | A0A0M3PZA5 |
| NLEQSAASQEDLEAM | 1 | 0 | 0 | 561896 | AAD22046.3 | A0A0M3PZA5 |
| NLSGKPTHRTMMYNF | 1 | 0 | 0 | 561897 | ACY00666.1 | A0A0M3PZA5 |
| NQLKQLDINTSKMSM | 1 | 0 | 0 | 561899 | ACK37353.1 | A0A0M3PZA5 |
| NYLFKANPNYAGNSI | 1 | 0 | 0 | 561900 | AEI71367.1 | A0A0M3PZA5 |
| PGGPLPKDRATAMLG | 1 | 0 | 0 | 561902 | ACY00666.1 | A0A0M3PZA5 |
| PRPMRNQNYLFKANP | 1 | 0 | 0 | 561903 | AEI71367.1 | A0A0M3PZA5 |
| PTGEVVPQLLQYMFV | 1 | 0 | 0 | 561904 | ADB77848.1 | A0A0M3PZA5 |
| PYSTYVKDELRSLDK | 1 | 0 | 0 | 561907 | ADO85560.1 | A0A0M3PZA5 |
| QGALLVAILPEYVIG | 1 | 0 | 0 | 561909 | ACK37353.1 | A0A0M3PZA5 |
| QTASIQGDRVADVIE | 1 | 0 | 0 | 561911 | ACK37353.1 | A0A0M3PZA5 |
| RFPVSAQAGKGELCA | 1 | 0 | 0 | 561912 | ACY00666.1 | A0A0M3PZA5 |
| RHSQPGKTIWVEHKL | 1 | 0 | 0 | 561913 | AAD22046.3 | A0A0M3PZA5 |
| SKMKFYMDKYGLDLP | 1 | 0 | 0 | 561914 | AEK98145.1 | A0A0M3PZA5 |
| STQRSGSHENSNSAT | 1 | 0 | 0 | 561915 | ACY00666.1 | A0A0M3PZA5 |
| TEGSTINYTTINYYK | 1 | 0 | 0 | 561916 | ACY00666.1 | A0A0M3PZA5 |
| TGEVPALQAAEIGAS | 1 | 0 | 0 | 561917 | ABI20542.1 | A0A0M3PZA5 |
| TVAGGTGTEDSHPPY | 1 | 0 | 0 | 561918 | ACY00666.1 | A0A0M3PZA5 |
| VDKPTRPDVSVNRFY | 1 | 0 | 0 | 561919 | ACY00666.1 | A0A0M3PZA5 |
| VGSSKSKYPLVVRIY | 1 | 0 | 0 | 561920 | ABI20542.1 | A0A0M3PZA5 |
| VKPNKETGRLNINGP | 1 | 0 | 0 | 561921 | ACY00666.1 | A0A0M3PZA5 |
| VTLDTNEKFRDITKF | 1 | 0 | 0 | 561924 | ACK37353.1 | A0A0M3PZA5 |
| VTQGFPTEPKPGTNQ | 1 | 0 | 0 | 561925 | ACK37353.1 | A0A0M3PZA5 |
| VVSELIREYNSRSAI | 1 | 0 | 0 | 561926 | ACK37353.1 | A0A0M3PZA5 |
| VYYCNSKRKHYPVSF | 1 | 0 | 0 | 561927 | ACK37353.1 | A0A0M3PZA5 |
| YMDCDIEVTDSYKTD | 1 | 0 | 0 | 561928 | ACY00666.1 | A0A0M3PZA5 |
| YTLDTKLWEKSSKGW | 1 | 0 | 0 | 561929 | ACY00666.1 | A0A0M3PZA5 |
| YTTINYYKDSYAATA | 1 | 0 | 0 | 561930 | ACY00666.1 | A0A0M3PZA5 |
| YVEASEYYPARYQSH | 1 | 0 | 0 | 561931 | ACY00666.1 | A0A0M3PZA5 |
| PALQAAETGASSNA | 1 | 0 | 0 | 566609 | ACF21979.1 | A0A0M3PZA5 |
| IWGCSGKLICTTAVP | 1 | 0 | 0 | 566698 | P03378.1 | P03378 |
| LRGFKKEISNMLSII | 1 | 0 | 0 | 591709 | QDG00742.1 | P17763 |
| YGTCHHKKGEARRSR | 1 | 0 | 0 | 591797 | AHZ13508.1 | Q32ZE1 |
| NNTRKSIRIGPGQTFYATGDIIG | 1 | 0 | 0 | 604576 | AGW27143.1 | P03378 |
| YNKRKRIHIGPGRAFYTTKNIIG | 1 | 0 | 0 | 604752 | P05877.1 | P03378 |
| LDAACNWTRGERCD | 1 | 0 | 0 | 607082 | BAA01582.1 | P27958 |
| SQKIQLVNTNGSWHIN | 1 | 0 | 0 | 607134 | BAA01728.1 | P27958 |
| GILKRWGTIKKSKAINVL | 1 | 0 | 0 | 608474 | ACO06168.1 | P17763 |
| GRGPLKLFMALVAFLRFL | 1 | 0 | 0 | 608515 | ACJ04259.1 | P17763 |
| IKKSKAINVLRGFRKEI | 1 | 0 | 0 | 609022 | ACO06168.1 | P17763 |
| KEIGRMLNILNRRRRTA | 1 | 0 | 0 | 609231 | AHG06353.1 | P17763 |
| KRERNRVSTVQQLTKRF | 1 | 0 | 0 | 609471 | ACJ04259.1 | P17763 |
| MALVAFLRFLTIPPTAGI | 1 | 0 | 0 | 609994 | ACO06168.1 | P17763 |
| NILNRRRRTAGMIIMLI | 1 | 0 | 0 | 610172 | AHG25307.1 | P17763 |
| NNQRKKARNTPFNMLKR | 1 | 0 | 0 | 610175 | AHG06351.1 | P17763 |
| NVLRGFRKEIGRMLNIL | 1 | 0 | 0 | 610208 | ACJ04265.1 | P17763 |
| RNTPFNMLKRERNRVSTV | 1 | 0 | 0 | 611061 | AHG06353.1 | P17763 |
| RTAGMIIMLIPTVMA | 1 | 0 | 0 | 612083 | AHG25306.1 | P17763 |
| ELDKWASLWNWFDITNWLWYIR | 1 | 0 | 0 | 688897 | Q73372.2 | P03378 |
| LLNNTRPPRGNWF | 1 | 0 | 0 | 697275 | ABR27415.1 | P27958 |
| NCNDSLNTGFLAALFYTHRF | 1 | 0 | 0 | 697287 | BAA09072.1 | P27958 |
| SRAAHRVTTFITR | 1 | 0 | 0 | 697336 | AMQ48485.1 | P27958 |
| AAILAYTIGTTHFQR | 1 | 0 | 0 | 738424 | P14340.2 | P17763 |
| AGLAFSIMKSVGTGK | 1 | 0 | 0 | 738453 | QTX93374.1 | P17763 |
| AGLLFSIMKNTTNTR | 1 | 0 | 0 | 738457 | P14340.2 | P17763 |
| AINVLRGFRKEIGRM | 1 | 0 | 0 | 738469 | P14340.2 | P17763 |
| ANGILAQGKKMIRPQ | 1 | 0 | 0 | 738503 | P27909.2 | P17763 |
| DLILEIGKLPQHLTL | 1 | 0 | 0 | 738677 | P27909.2 | P17763 |
| EATLGGFGSLGLDC | 1 | 0 | 0 | 738737 | ALL27019.1 | Q32ZE1 |
| ECLRRRVTRKHMILV | 1 | 0 | 0 | 738742 | Q2YHF0.1 | P17763 |
| EEFLDYMPSMKRFRK | 1 | 0 | 0 | 738753 | ABZ90961.1 | P17763 |
| ENLEYTVVVTVHNGD | 1 | 0 | 0 | 738809 | Q2YHF0.1 | P17763 |
| ERELHKLGKCGSCVY | 1 | 0 | 0 | 738829 | QTX93489.1 | P17763 |
| FGFYQVKTETTILDV | 1 | 0 | 0 | 738897 | Q58HT7.1 | P17763 |
| FLRFLAIPPTAGILA | 1 | 0 | 0 | 738927 | P17763.2 | P17763 |
| GLGKTSIGLLCVMAS | 1 | 0 | 0 | 739034 | QFS19562.1 | P17763 |
| GLPIRYQTPAVKSEH | 1 | 0 | 0 | 739036 | Q2YHF0.1 | P17763 |
| GPERVILAGPIPVTP | 1 | 0 | 0 | 739048 | Q2YHF0.1 | P17763 |
| HQVFGSVYTTMFGGV | 1 | 0 | 0 | 739120 | Q2YHF0.1 | P17763 |

| | | | | | | |
|---|---|---|---|---|---|---|
| IKGIMQAGKRSLRPQ | 1 | 0 | 0 | 739192 | QTX92981.1 | P17763 |
| IMIEEVMRSRWSRKM | 1 | 0 | 0 | 739215 | P17763.2 | P17763 |
| KKNGAIKVLRGFKKE | 1 | 0 | 0 | 739322 | P17763.2 | P17763 |
| KVRSNAAIGAVFVDE | 1 | 0 | 0 | 739406 | P17763.2 | P17763 |
| LLVLCAVQLLLMRTS | 1 | 0 | 0 | 739526 | QTX93489.1 | P17763 |
| LPAIVREAIKRKLRT | 1 | 0 | 0 | 739540 | QFS19562.1 | P17763 |
| LPTYLSSRAKLALDN | 1 | 0 | 0 | 739549 | Q2YHF0.1 | P17763 |
| MKIGIGVLLTWIGLN | 1 | 0 | 0 | 739669 | QTX93491.1 | P17763 |
| PGIFQTTTGEIGAIA | 1 | 0 | 0 | 739889 | QTX93504.1 | P17763 |
| QNKVVRVQRPAKNGT | 1 | 0 | 0 | 740024 | P17763.2 | P17763 |
| REAIKRGLRTLILAP | 1 | 0 | 0 | 740081 | P14340.2 | P17763 |
| RGNQFCIKILNPYMP | 1 | 0 | 0 | 740100 | P17763.2 | P17763 |
| RMLINRFTMRHKKAT | 1 | 0 | 0 | 740139 | P14340.2 | P17763 |
| RPPFNMLKRERNRVS | 1 | 0 | 0 | 740156 | Q2YHF0.1 | P17763 |
| RTLILAPTRVVAAEM | 1 | 0 | 0 | 740193 | P14340.2 | P17763 |
| RWGTFKKSGAIKVLK | 1 | 0 | 0 | 740207 | ADI44327.1 | P17763 |
| SGVSWTMKIGIGVLL | 1 | 0 | 0 | 740267 | QFS19562.1 | P17763 |
| SSVNMVSRLLLNRFT | 1 | 0 | 0 | 740392 | QTX93593.1 | P17763 |
| TKGVERLKRMAISGD | 1 | 0 | 0 | 740478 | QTX93593.1 | P17763 |
| TWNDLIRLCIMVGAN | 1 | 0 | 0 | 740566 | P17763.2 | P17763 |
| TYLALLAAFKVRPTF | 1 | 0 | 0 | 740571 | QTX93024.1 | P17763 |
| VVTLIPLCRTSCLQK | 1 | 0 | 0 | 740701 | QTX92385.1 | P17763 |
| WNLVRLQSGVDVFFI | 1 | 0 | 0 | 740726 | QTX93201.1 | P17763 |
| YFHRRDLRLAANAIC | 1 | 0 | 0 | 740761 | P17763.2 | P17763 |
| AANGGPASRPPPVPSPATTK | 1 | 0 | 0 | 741180 | P08666.2 | P08666 |
| AATQAANGGPATPAPPAPGP | 1 | 0 | 0 | 741184 | AMB65625.1 | P10211 |
| ALVSAMERTEHKARKKGTSA | 1 | 0 | 0 | 741209 | P08666.2 | P08666 |
| APKRIRLPHIREDDQPSSHQ | 1 | 0 | 0 | 741216 | GI:1089747699 | Q69091 |
| APRLGPSSVYTPGASRPTPP | 1 | 0 | 0 | 741218 | P13291.1 | P13291 |
| APTGDTKPKKNKKPKNPPPP | 1 | 0 | 0 | 741221 | AMB65625.1 | P10211 |
| ARKRKTKKPPKRPEATPPPD | 1 | 0 | 0 | 741230 | P08666.2 | P08666 |
| ASATGPTYIRVADSELYADW | 1 | 0 | 0 | 741238 | P89475.1 | P89475 |
| ASNAAPSASPRNASAPRTTP | 1 | 0 | 0 | 741243 | Q89730.1 | Q89730 |
| ATPAPPAPGPAPTGDTKPKK | 1 | 0 | 0 | 741256 | AMB65625.1 | P10211 |
| DDHETDMELKPAKVATRTSR | 1 | 0 | 0 | 741286 | P08666.2 | P08666 |
| DGCGPLHPSWVSLMPPKQVP | 1 | 0 | 0 | 741291 | P04488.1 | P04488 |
| DGIIELFHYPLGNHCPRVVH | 1 | 0 | 0 | 741294 | P13291.1 | P13291 |
| DHTPPMPSIGLE | 1 | 0 | 0 | 741300 | P06484.1 | P06484 |
| DPTPAPGDTGTPAPASGERA | 1 | 0 | 0 | 741315 | ATB17382.1 | P13291 |
| DRIFARRYNATHIKVGQPQY | 1 | 0 | 0 | 741318 | AAA19496.1 | P10211 |
| DSPSTNGSGFEILSPTAPSV | 1 | 0 | 0 | 741322 | P89475.1 | P89475 |
| DTLPQSPGPAFPLAEDVEKD | 1 | 0 | 0 | 741329 | P06484.1 | P06484 |
| EEEEEEGAGDGEHL | 1 | 0 | 0 | 741352 | P06484.1 | P06484 |
| EGQLGENNELRLTRDALEPC | 1 | 0 | 0 | 741366 | P08666.2 | P08666 |
| EILSPTAPSVYPHSEGRKSR | 1 | 0 | 0 | 741370 | P89475.1 | P89475 |
| ENIAPYKFKATMYYKDVTVS | 1 | 0 | 0 | 741381 | AAA19496.1 | P10211 |
| FGSGSPGRRHSQASYPSVLW | 1 | 0 | 0 | 741410 | P89475.1 | P89475 |
| FHRYGTTVNCIVEEVDARSV | 1 | 0 | 0 | 741412 | AAA19496.1 | P10211 |
| FIHRCQRRYRRPRGQIYNPG | 1 | 0 | 0 | 741413 | P13291.1 | P13291 |
| FPLAEDVEKDKPNRPVVPSP | 1 | 0 | 0 | 741427 | P06484.1 | P06484 |
| FWVRRRAQMAPKRLRLPHIR | 1 | 0 | 0 | 741438 | P03172.2 | Q69467 |
| GPEEFEGAGDGEPP | 1 | 0 | 0 | 741481 | AIL27778.1 | P13290 |
| GPVVLLSVSPRPRSGPTAPQ | 1 | 0 | 0 | 741485 | P13291.1 | P13291 |
| HGVRSKYALVDASLKMADPN | 1 | 0 | 0 | 741519 | Q69091.1 | Q69091 |
| HPNTPPKPRRRSSSSTTMPS | 1 | 0 | 0 | 741530 | P13291.1 | P13291 |
| IPSIQDAATPYHPPATPNNM | 1 | 0 | 0 | 741563 | P36318.1 | Q69091 |
| IYEACLYHPQLPECLSPADA | 1 | 0 | 0 | 741576 | P89475.1 | P89475 |
| LIIGALAGSTLAVLVIGGIA | 1 | 0 | 0 | 741658 | P03172.2 | Q69467 |
| LLRVRTATRDYAGLYVLRVW | 1 | 0 | 0 | 741670 | P13291.1 | P13291 |
| LRLPHIRDDDAPPSHQPLFY | 1 | 0 | 0 | 741691 | P03172.2 | Q69467 |
| LTSIAEESEPGPVVLLSVSP | 1 | 0 | 0 | 741706 | P13291.1 | P13291 |
| MARLGAELRSHPNTPPKPRR | 1 | 0 | 0 | 741723 | P13291.1 | P13291 |
| NKKPKNPPPPRPAGDNATVA | 1 | 0 | 0 | 741770 | AMB65625.1 | P10211 |
| PAPVPTPTPDDYDEEDDAGV | 1 | 0 | 0 | 741786 | P89475.1 | P89475 |
| PAVAFTLCRSTHHAHSPAYP | 1 | 0 | 0 | 741790 | P13291.1 | P13291 |
| PLARYGSRVQIRCRFPNSTR | 1 | 0 | 0 | 741815 | Q89730.1 | Q89730 |
| PPKTGPPKTSSEPVRCNRHD | 1 | 0 | 0 | 741823 | Q89730.1 | Q89730 |
| PPMPSIGLEEEEEEGAGDG | 1 | 0 | 0 | 741824 | P06484.1 | P06484 |
| PPNSTRSASESRHRLTVAQV | 1 | 0 | 0 | 741825 | P13291.1 | P13291 |
| PPVPSPATTKARKRKTKKPP | 1 | 0 | 0 | 741832 | P08666.2 | P08666 |
| PQASTTPFPTGDPKPQPPGV | 1 | 0 | 0 | 741833 | P06487.1 | P06487 |
| QPPPVPSPATTKARKRKTKK | 1 | 0 | 0 | 741875 | ADG45180.1 | P08666 |
| QSRKPPNPTPPPPGASANAS | 1 | 0 | 0 | 741879 | P06437.2 | P10211 |
| QVASVVLVVEPAPVPTPTPD | 1 | 0 | 0 | 741882 | P89475.1 | P89475 |
| RAPHPAPSARGPLRLGAVLG | 1 | 0 | 0 | 741889 | P89475.1 | P89475 |
| RASGGVAATVAANGGPASRP | 1 | 0 | 0 | 741891 | P08666.2 | P08666 |
| RLTSKGRPLVPTPQHTPLFS | 1 | 0 | 0 | 741919 | P06484.1 | P06484 |
| RNASAPRTTPTPPQPRKATK | 1 | 0 | 0 | 741924 | Q89730.1 | Q89730 |
| RPLTTFGSGSPGRRHSQASY | 1 | 0 | 0 | 741930 | P89475.1 | P89475 |
| RPRGQIYNPGGVSCAVNEAA | 1 | 0 | 0 | 741931 | P13291.1 | P13291 |
| RTITVGPRGNASNAAPSASP | 1 | 0 | 0 | 741947 | Q89730.1 | Q89730 |
| RTTTSPSSPRDPTPAPGDTG | 1 | 0 | 0 | 741952 | ATB17382.1 | P13291 |
| RVYHIQPSLEDPFQPPSIPI | 1 | 0 | 0 | 741959 | P03172.2 | Q69467 |
| SAAPAAPAAPRASGGVAATV | 1 | 0 | 0 | 741963 | P08666.2 | P08666 |
| SCDPAQLPFSAPRLGPSSVY | 1 | 0 | 0 | 741972 | P13291.1 | P13291 |
| SEPVRCNRHDPLARYGSRVQ | 1 | 0 | 0 | 741979 | Q89730.1 | Q89730 |
| SKASTAKPAPPPKTGPPKTS | 1 | 0 | 0 | 741984 | Q89730.1 | Q89730 |
| SLWQDPPERPDSPSTNGSGF | 1 | 0 | 0 | 741988 | P89475.1 | P89475 |
| SVCTMTKWQEVDEMLRSEYG | 1 | 0 | 0 | 742012 | AAA19496.1 | P10211 |
| THHAHSPAYPTLELGLARQP | 1 | 0 | 0 | 742040 | P13291.1 | P13291 |
| TKARATAPTTRNLLTTPKFT | 1 | 0 | 0 | 742045 | AAA19496.1 | P10211 |

| | | | | | | |
|---|---|---|---|---|---|---|
| TLELGLARQPLLRVRTATRD | 1 | 0 | 0 | 742048 | P13291.1 | P13291 |
| TLGVLVASAAPSSPGTPGVA | 1 | 0 | 0 | 742050 | AAA19496.1 | P10211 |
| TPAPASGERAPPNSTRSASE | 1 | 0 | 0 | 742067 | ATB17382.1 | P13291 |
| TPGASRPTPPRTTTSPSSPR | 1 | 0 | 0 | 742069 | ATB17382.1 | P13291 |
| TPPQPRKATKSKASTAKPAP | 1 | 0 | 0 | 742072 | Q89730.1 | Q89730 |
| TVGRRVSARMLGDVMAVSTC | 1 | 0 | 0 | 742098 | P08666.2 | P08666 |
| VADSELYADWSSDSEGERDG | 1 | 0 | 0 | 742103 | P89475.1 | P89475 |
| VAWDWVPKRPSVCTMTKWQE | 1 | 0 | 0 | 742106 | AAA19496.1 | P10211 |
| VCRSTAKYVRNNMETTAFHR | 1 | 0 | 0 | 742108 | P08666.2 | P08666 |
| VGSATNASLFVLGVALSANG | 1 | 0 | 0 | 742127 | ATB17382.1 | P13291 |
| VSDSLVDAGAVGPQGFVEED | 1 | 0 | 0 | 742160 | P13291.1 | P13291 |
| VYWMRRHTQKAPKRIRLPHI | 1 | 0 | 0 | 742182 | Q69091.1 | Q69091 |
| YPHSEGRKSRRPLTTFGSGS | 1 | 0 | 0 | 742228 | P89475.1 | P89475 |
| DTGWLAGLFYYHK | 1 | 0 | 0 | 742498 | ACA50657.1 | P27958 |
| NTGFLASLFYTHK | 1 | 0 | 0 | 742508 | YP_001469632.1 | P27958 |
| EDLALPWKHKDNQDWN | 1 | 0 | 0 | 745102 | ADK37753.1 | Q04538 |
| SLWNWFDITKWLWYIK | 1 | 0 | 0 | 745647 | Q75760 | P03378 |
| FGKAKGSRAIWYMWL | 1 | 0 | 0 | 753348 | P17763.2 | P17763 |
| GARFLEFEALGFLNE | 1 | 0 | 0 | 755186 | P14340.2 | P17763 |
| GSRAIWYMWLGARFL | 1 | 0 | 0 | 755407 | P17763.2 | P17763 |
| TDKIDQIIHDFIDKPL | 1 | 0 | 0 | 769761 | YP_003815435.1 | B8XCN0 |
| TDKINQIIHDFIDNPL | 1 | 0 | 0 | 769763 | ALT19781.1 | B0LPL7 |
| AAAQKNFTMKLCKDASDILQ | 1 | 0 | 0 | 779897 | APU51914.1 | A0A0M3PZA5 |
| APTGQNTQVSSHRLDTGKVP | 1 | 0 | 0 | 779917 | APU51914.1 | A0A0M3PZA5 |
| AREKVEFLNNLKQLPLLENQ | 1 | 0 | 0 | 779918 | APU51914.1 | A0A0M3PZA5 |
| EDTHPPYKQTQPGADGFELQ | 1 | 0 | 0 | 779950 | APU51914.1 | A0A0M3PZA5 |
| EQGVSDYIKGLGDAFGTGFT | 1 | 0 | 0 | 779956 | APU51914.1 | A0A0M3PZA5 |
| GHFTMLGVRDRLAVLPRHSQ | 1 | 0 | 0 | 779982 | APU51914.1 | A0A0M3PZA5 |
| GKGELCAVFRADPGRNGPWQ | 1 | 0 | 0 | 779984 | APU51914.1 | A0A0M3PZA5 |
| HPYVLDAGIPISQLTVCPHQ | 1 | 0 | 0 | 779999 | APU51914.1 | A0A0M3PZA5 |
| ISDLLASVDSEEVRQYCRDQ | 1 | 0 | 0 | 780016 | APU51914.1 | A0A0M3PZA5 |
| LEVDFEQALFSKYVGNTLHE | 1 | 0 | 0 | 780039 | APU51914.1 | A0A0M3PZA5 |
| LGDAFGTGFTDAVSREVEAL | 1 | 0 | 0 | 780041 | APU51914.1 | A0A0M3PZA5 |
| LIREYSNRSAIGNTIEALFQ | 1 | 0 | 0 | 780045 | APU51914.1 | A0A0M3PZA5 |
| PTRTKLEPSVFHDIFEGNKE | 1 | 0 | 0 | 780100 | APT69924.1 | A0A0M3PZA5 |
| SHRLDTGKVPALQAAEIGAS | 1 | 0 | 0 | 780145 | APU51914.1 | A0A0M3PZA5 |
| SLDFALSLLRRNIRQVQTDQ | 1 | 0 | 0 | 780148 | APU51914.1 | A0A0M3PZA5 |
| SLEEKPAPDAISDLLASVDS | 1 | 0 | 0 | 780149 | APU51914.1 | A0A0M3PZA5 |
| YHSSVYSLPPDPDHFDGYKQ | 1 | 0 | 0 | 780225 | APU51914.1 | A0A0M3PZA5 |
| ADREIHQLHLSICKKRKTGS | 1 | 0 | 0 | 839081 | APG38070.1 | Q8JSZ3 |
| AVCKRMCFRATIEASRRALL | 1 | 0 | 0 | 839163 | ABB30036.1 | Q8JSZ3 |
| DYYCNMGDWPSCTYTGVTQH | 1 | 0 | 0 | 839257 | APG38070.1 | Q8JSZ3 |
| EGFFDLMHVQKVLSASTVCK | 1 | 0 | 0 | 839267 | APG38070.1 | Q8JSZ3 |
| EIITLHPKIEEGFFDLMHVQ | 1 | 0 | 0 | 839273 | APG38058.1 | Q8JSZ3 |
| ETAEIHDDNYGGPGDKITIC | 1 | 0 | 0 | 839290 | APG38070.1 | Q8JSZ3 |
| FASLACTGCYACSSGISCKV | 1 | 0 | 0 | 839311 | APG38064.1 | Q8JSZ3 |
| GGPGDKITICNGSTIVDQRL | 1 | 0 | 0 | 839541 | APG38070.1 | Q8JSZ3 |
| HSAMSRIPTPHTATRVSTEN | 1 | 0 | 0 | 839646 | ABB30036.1 | Q8JSZ3 |
| LAVCKRMCFRATIEASRRAL | 1 | 0 | 0 | 839843 | ABB30036.1 | Q8JSZ3 |
| LRTETAEIHDDNYGGPGDKI | 1 | 0 | 0 | 839968 | ABB30036.1 | Q8JSZ3 |
| MSAAPTAVQDIHPSPTNRSK | 1 | 0 | 0 | 840061 | ACT88367.1 | Q8JSZ3 |
| MTSPAQSILLMSAAPTAVQD | 1 | 0 | 0 | 840063 | ACT88367.1 | Q8JSZ3 |
| NSTTFHQALLDPRVRGLYFPAGG | 1 | 0 | 0 | 840115 | AAN03492.1 | Q76R62 |
| NVMLAVCKRMCFRATIEASR | 1 | 0 | 0 | 840124 | APG38070.1 | Q8JSZ3 |
| PQSILIEHKGTIIGKQNDTC | 1 | 0 | 0 | 840154 | APG38070.1 | Q8JSZ3 |
| RKPLFLDSIVKGMKNLLNST | 1 | 0 | 0 | 840222 | APG38061.1 | Q8JSZ3 |
| RLTSDGLARHVTQCPKRKEK | 1 | 0 | 0 | 840226 | APG38058.1 | Q8JSZ3 |
| SICKKRKTGSNVMLAVCKRM | 1 | 0 | 0 | 840289 | APG38070.1 | Q8JSZ3 |
| SLETSLSIEAPWGAINVQST | 1 | 0 | 0 | 840307 | APG38070.1 | Q8JSZ3 |
| SSAQQTTPSPMTSPAQSILL | 1 | 0 | 0 | 840339 | ACT88367.1 | Q8JSZ3 |
| STPQGIYHPARSLLSVSSPK | 1 | 0 | 0 | 840355 | ABB30036.1 | Q8JSZ3 |
| SVLRQYKTEIKIGKASTGFR | 1 | 0 | 0 | 840374 | ACT88367.1 | Q8JSZ3 |
| TSPSSSPSTPSTPQGIYHPA | 1 | 0 | 0 | 840462 | ABB30036.1 | Q8JSZ3 |
| TTFVICILTLTICVVSTSAV | 1 | 0 | 0 | 840469 | APG38070.1 | Q8JSZ3 |
| VTQCPKRKEKVEETELYLNL | 1 | 0 | 0 | 840603 | APG38067.1 | Q8JSZ3 |
| GAFFLYDRLAST | 1 | 0 | 0 | 852598 | ALT19781.1 | B0LPL7 |
| IWYMWLGARYLEFEA | 1 | 0 | 0 | 857703 | ACJ09459.1 | P17763 |
| DLPVWLAYRVAAEGI | 1 | 0 | 0 | 857764 | P14340.2 | P17763 |
| GTIGAVSLDFSPGTS | 1 | 0 | 0 | 857771 | P14340.2 | P17763 |
| KVIQLSRKTFDSEYV | 1 | 0 | 0 | 857782 | P14340.2 | P17763 |
| AWEDGICGISSVSRMEN | 1 | 0 | 0 | 857846 | APO08504.1 | Q32ZE1 |
| CGTGVFVYNDVEAWR | 1 | 0 | 0 | 857851 | ANW07474.1 | Q32ZE1 |
| DVGCSVDFSKKETRCGTG | 1 | 0 | 0 | 857862 | ANW07474.1 | Q32ZE1 |
| EELEIRFEECPGTKV | 1 | 0 | 0 | 857867 | APO08504.1 | Q32ZE1 |
| EENGVQLTVVVGSVK | 1 | 0 | 0 | 857868 | APO08504.1 | Q32ZE1 |
| EIRPRKEPESNLVRSMVT | 1 | 0 | 0 | 857871 | APO08504.1 | Q32ZE1 |
| GPWHSEELEIRFEEC | 1 | 0 | 0 | 857901 | APO08504.1 | Q32ZE1 |
| IEESDLIIPKSLAGP | 1 | 0 | 0 | 857913 | APO08504.1 | Q32ZE1 |
| KDGCWYGMEIRPRKE | 1 | 0 | 0 | 857921 | APO08504.1 | Q32ZE1 |
| KNDTWRLKRAHLIEM | 1 | 0 | 0 | 857933 | APO08504.1 | Q32ZE1 |
| LPHGWKAWGKSYFVR | 1 | 0 | 0 | 857961 | APO08504.1 | Q32ZE1 |
| PAVIGTAVKGKEAVH | 1 | 0 | 0 | 857988 | APO08504.1 | Q32ZE1 |
| PESNLVRSMVTAGST | 1 | 0 | 0 | 857989 | APO08504.1 | Q32ZE1 |
| PLKHRAWNSFLVEDH | 1 | 0 | 0 | 857991 | APO08504.1 | Q32ZE1 |
| PVPVNELPHGWKAWG | 1 | 0 | 0 | 857994 | APO08504.1 | Q32ZE1 |
| QLTVVVGSVKNPMWR | 1 | 0 | 0 | 858011 | APO08504.1 | Q32ZE1 |
| RRLAAVKQAWEDGI | 1 | 0 | 0 | 858032 | APO08504.1 | Q32ZE1 |
| TTASGRVIEEWCCR | 1 | 0 | 0 | 858086 | APO08504.1 | Q32ZE1 |
| VEAWRDRYKYHPDSPRRLAA | 1 | 0 | 0 | 858092 | ANW07474.1 | Q32ZE1 |
| VKNPMWRGPQRLPVPVNELPHG | 1 | 0 | 0 | 858099 | APO08504.1 | Q32ZE1 |
| VSRMENIMWRSVEGE | 1 | 0 | 0 | 858103 | APO08504.1 | Q32ZE1 |

| | | | | | | |
|---|---|---|---|---|---|---|
| YGMEIRPRKEPESNL | 1 | 0 | 0 | 858112 | APO08504.1 | Q32ZE1 |
| DIPYLGKREDLWCGS | 1 | 0 | 0 | 862475 | Q2YHF0.1 | P17763 |
| RAIWYMWLGARFLEF | 1 | 0 | 0 | 862966 | P17763.2 | P17763 |
| YMWLGARFLEFEALG | 1 | 0 | 0 | 863265 | P17763.2 | P17763 |
| ELRDKKQKAYALFYRPDVV | 1 | 0 | 0 | 863965 | Q1PHM9 | P03378 |
| AEALKGMPIRYQTTA | 1 | 0 | 0 | 865944 | P17763.2 | P17763 |
| AGFMAYMIGQTGIQR | 1 | 0 | 0 | 865962 | Q2YHF0.1 | P17763 |
| AGVNMCTLIAMDLGE | 1 | 0 | 0 | 865971 | P17763.2 | P17763 |
| AIWYMWLGARFLEFE | 1 | 0 | 0 | 866015 | P17763.2 | P17763 |
| AKEAVEDERFWDLVH | 1 | 0 | 0 | 866017 | P17763.2 | P17763 |
| ARFLEFEALGFLNED | 1 | 0 | 0 | 866164 | ASK38693.1 | P17763 |
| ATLFVWYFWQKKKQR | 1 | 0 | 0 | 866201 | P17763.2 | P17763 |
| AVLFVVTLIPLCRTS | 1 | 0 | 0 | 866227 | Q2YHF0.1 | P17763 |
| AYAQMWSLMYFHRRD | 1 | 0 | 0 | 866244 | QTX93491.1 | P17763 |
| CTILAAVSVSPLLLT | 1 | 0 | 0 | 866267 | QRG47124.1 | P17763 |
| DGIVAIDLDPVVYDA | 1 | 0 | 0 | 866400 | P17763.2 | P17763 |
| DTAWDFGSIGGVFTS | 1 | 0 | 0 | 866451 | P17763.2 | P17763 |
| DTAWDFGSLGGVFTS | 1 | 0 | 0 | 866452 | P14340.2 | P17763 |
| DVPMAGPLVAGGLLI | 1 | 0 | 0 | 866459 | QTX93489.1 | P17763 |
| EALRGLPIRYQTPAI | 1 | 0 | 0 | 866477 | P14340.2 | P17763 |
| EDKTPVTTWEDVPYL | 1 | 0 | 0 | 866489 | QTX93491.1 | P17763 |
| EEALKGLPIRYQTTA | 1 | 0 | 0 | 866506 | QTX93489.1 | P17763 |
| EGAMHSALAGATEVD | 1 | 0 | 0 | 866549 | Q2YHF0.1 | P17763 |
| EGAMHTALTGATEIQ | 1 | 0 | 0 | 866551 | P17763.2 | P17763 |
| EITALILGAQALPVY | 1 | 0 | 0 | 866577 | Q2YHF0.1 | P17763 |
| EKYQLAVTIMAILCV | 1 | 0 | 0 | 866585 | P14340.2 | P17763 |
| EKYQLAVTIMAISCV | 1 | 0 | 0 | 866586 | QRG47112.1 | P17763 |
| ELIDGISLGLILLKI | 1 | 0 | 0 | 866594 | Q5UCB8.1 | P17763 |
| EPHWIAASIILEFFL | 1 | 0 | 0 | 866618 | P17763.2 | P17763 |
| ESWILRNPGFALLAG | 1 | 0 | 0 | 866646 | Q2YHF0.1 | P17763 |
| ETAWDFGSVGGLLTS | 1 | 0 | 0 | 866647 | Q2YHF0.1 | P17763 |
| FDFCEGTTVVVTEDC | 1 | 0 | 0 | 866681 | P14340.2 | P17763 |
| FDLCEGTTVVVDEHC | 1 | 0 | 0 | 866682 | P17763.2 | P17763 |
| FFLMVLLIPEPDRQR | 1 | 0 | 0 | 866690 | P17763.2 | P17763 |
| FGESYIVIGAGEKAL | 1 | 0 | 0 | 866692 | QFS19562.1 | P17763 |
| FKSIQHLTVTEEIAV | 1 | 0 | 0 | 866719 | P14340.2 | P17763 |
| FLFFMQGKGIGKLSM | 1 | 0 | 0 | 866731 | Q2YHF0.1 | P17763 |
| FTILALFLAHYIGTS | 1 | 0 | 0 | 866788 | QTX93489.1 | P17763 |
| FWNTTIAVSMANIFR | 1 | 0 | 0 | 866802 | P17763.2 | P17763 |
| FWNTTIAVSTANIFR | 1 | 0 | 0 | 866803 | Q2YHF0.1 | P17763 |
| GEGLHRLGYILEDID | 1 | 0 | 0 | 866823 | Q2YHF0.1 | P17763 |
| GFALLAGFMAYMIGQ | 1 | 0 | 0 | 866827 | Q2YHF0.1 | P17763 |
| GGVSWMIRILIGFLV | 1 | 0 | 0 | 866846 | ARM59249.1 | P17763 |
| GGVSWMVRILIGFLV | 1 | 0 | 0 | 866847 | Q58HT7.1 | P17763 |
| GISLGLILLKIVTQF | 1 | 0 | 0 | 866892 | ANK35835.1 | P17763 |
| GLPIRYQTPAIKTEH | 1 | 0 | 0 | 866941 | QNG98968.1 | P17763 |
| GNIVSAVNMTSRMLL | 1 | 0 | 0 | 866989 | P17763.2 | P17763 |
| HDLMEFIDGISLGLI | 1 | 0 | 0 | 867048 | Q2YHF0.1 | P17763 |
| HQIFGTAYGVLFSGV | 1 | 0 | 0 | 867087 | P17763.2 | P17763 |
| HQVFGTAYGVLFSGV | 1 | 0 | 0 | 867089 | P27909.2 | P17763 |
| HTALTGATEIQTSGT | 1 | 0 | 0 | 867101 | P17763.2 | P17763 |
| HTLWSNGVLESEMII | 1 | 0 | 0 | 867105 | P17763.2 | P17763 |
| IAASIILEFFLMVLL | 1 | 0 | 0 | 867112 | P17763.2 | P17763 |
| IAGVFFTFVLLLSGQ | 1 | 0 | 0 | 867116 | QDG00730.1 | P17763 |
| IASAIVLEFFMMVLL | 1 | 0 | 0 | 867122 | QTX93489.1 | P17763 |
| IENTSANLSLAAIAN | 1 | 0 | 0 | 867143 | Q2YHF0.1 | P17763 |
| IFRGSYLAGAGLAFS | 1 | 0 | 0 | 867156 | P17763.2 | P17763 |
| IKRKLRTLILAPTRV | 1 | 0 | 0 | 867205 | QFS19562.1 | P17763 |
| ILKLLTDFQSHQLWA | 1 | 0 | 0 | 867232 | QFS19562.1 | P17763 |
| ILLVAVSFVTLITGN | 1 | 0 | 0 | 867241 | P14340.2 | P17763 |
| ITAAAWYLWEVKKQR | 1 | 0 | 0 | 867327 | P14340.2 | P17763 |
| ITQFETYQLWTALIS | 1 | 0 | 0 | 867340 | Q6YMS3.1 | P17763 |
| IVSILLSSLLKNDVP | 1 | 0 | 0 | 867359 | P17763.2 | P17763 |
| KAYAQMWSLMYFHRR | 1 | 0 | 0 | 867376 | QTX93491.1 | P17763 |
| KGMPIRYQTTAVKSE | 1 | 0 | 0 | 867396 | P17763.2 | P17763 |
| KKHMIAGVFFTFVLL | 1 | 0 | 0 | 867416 | QCZ25049.1 | P17763 |
| KPLDERFGTSLLFLN | 1 | 0 | 0 | 867473 | ARJ31952.1 | P17763 |
| KRSLRPQPTELKYSW | 1 | 0 | 0 | 867488 | QTX92981.1 | P17763 |
| KSGDYVSAITQAERI | 1 | 0 | 0 | 867494 | Q2YHF0.1 | P17763 |
| KTPEGEVGAIALDFK | 1 | 0 | 0 | 867513 | P17763.2 | P17763 |
| KVLNPYMPTVIEELE | 1 | 0 | 0 | 867522 | Q2YHF0.1 | P17763 |
| LCLTLFVEECLRRRV | 1 | 0 | 0 | 867566 | Q2YHF0.1 | P17763 |
| LCVSIMIEEVMRSRW | 1 | 0 | 0 | 867572 | P27909.2 | P17763 |
| LFLEEMLRTRVGTKH | 1 | 0 | 0 | 867630 | P14340.2 | P17763 |
| LGGLTWMDLLRALIM | 1 | 0 | 0 | 867643 | Q2YHF0.1 | P17763 |
| LGLMALKLITQFETY | 1 | 0 | 0 | 867654 | Q6YMS3.1 | P17763 |
| LGMALFLEEMLRTRV | 1 | 0 | 0 | 867657 | P14340.2 | P17763 |
| LHWFRKGSSIGKMFE | 1 | 0 | 0 | 867697 | Q2YHF0.1 | P17763 |
| LIGNEEFLDYMPSMK | 1 | 0 | 0 | 867706 | ABZ90961.1 | P17763 |
| LLCVMASSVLLWMAS | 1 | 0 | 0 | 867760 | QFS19562.1 | P17763 |
| LLLIMGQLTWNDLIR | 1 | 0 | 0 | 867803 | P27909.2 | P17763 |
| LLRNDVPMAGPLVAG | 1 | 0 | 0 | 867834 | QTX93489.1 | P17763 |
| LPVYLMTLMKGASRR | 1 | 0 | 0 | 867894 | Q5UCB8.1 | P17763 |
| LQRKYGGALVRNPLS | 1 | 0 | 0 | 867902 | P14340.2 | P17763 |
| LRHPGFTVIALFLAH | 1 | 0 | 0 | 867912 | P17763.2 | P17763 |
| LRKLTSKELMMATIG | 1 | 0 | 0 | 867914 | QRG47115.1 | P17763 |
| LVSILASSLLRNDVP | 1 | 0 | 0 | 868028 | QTX93489.1 | P17763 |
| LVTFKTAHAKKQEVV | 1 | 0 | 0 | 868031 | P17763.2 | P17763 |
| LYAVATTILTPMLRH | 1 | 0 | 0 | 868036 | Q2YHF0.1 | P17763 |
| MAEIPLQWIASAIVL | 1 | 0 | 0 | 868043 | QPI11927.1 | P17763 |
| MASSVLLWMASVEPH | 1 | 0 | 0 | 868055 | QFS19562.1 | P17763 |

| | | | | | | |
|---|---|---|---|---|---|---|
| MAVFKMSPGYVLGIF | 1 | 0 | 0 | 868056 | Q2YHF0.1 | P17763 |
| MAYMIGQTGIQRTVF | 1 | 0 | 0 | 868057 | Q2YHF0.1 | P17763 |
| MEVTARWLWGFLSRN | 1 | 0 | 0 | 868089 | P17763.2 | P17763 |
| MGQLTWNDLIRLCIM | 1 | 0 | 0 | 868100 | P17763.2 | P17763 |
| MIIMDEAHFTDPASI | 1 | 0 | 0 | 868113 | P17763.2 | P17763 |
| MLIACYVISGSSADL | 1 | 0 | 0 | 868126 | P17763.2 | P17763 |
| MVHRQWFFDLPLPWT | 1 | 0 | 0 | 868175 | QTX93489.1 | P17763 |
| MVSILASSLLKNDIP | 1 | 0 | 0 | 868180 | P14340.2 | P17763 |
| MWHVTRGAVLMHRGK | 1 | 0 | 0 | 868183 | QRG47112.1 | P17763 |
| NEGIMAIGIVSILLS | 1 | 0 | 0 | 868195 | P27909.2 | P17763 |
| NEGIMAVGIVSILLS | 1 | 0 | 0 | 868196 | P17763.2 | P17763 |
| NIQVAINQVRRLIGN | 1 | 0 | 0 | 868238 | P17763.2 | P17763 |
| NMEAQLIRQMESEGI | 1 | 0 | 0 | 868259 | P17763.2 | P17763 |
| NPTVDGITVIDLEPI | 1 | 0 | 0 | 868268 | QRG47106.1 | P17763 |
| NPTVDGIVAIDLDPV | 1 | 0 | 0 | 868269 | P17763.2 | P17763 |
| NSRNTSMAMTCIAVG | 1 | 0 | 0 | 868280 | Q58HT7.1 | P17763 |
| PFLALGFFLRKLTSR | 1 | 0 | 0 | 868303 | QTX93489.1 | P17763 |
| PKWLDARVYADPMAL | 1 | 0 | 0 | 868325 | Q2YHF0.1 | P17763 |
| PLQWIASAIVLEFFM | 1 | 0 | 0 | 868330 | QTX93489.1 | P17763 |
| PMTVAAMGVPPLPLF | 1 | 0 | 0 | 868340 | QTX93489.1 | P17763 |
| PNYNMIIMDEAHFTD | 1 | 0 | 0 | 868343 | P17763.2 | P17763 |
| PRWLDARVYADPMAL | 1 | 0 | 0 | 868371 | AQV12649.1 | P17763 |
| PTTLTASLVMLLVHY | 1 | 0 | 0 | 868413 | Q2YHF0.1 | P17763 |
| PTVDGITVIDLDPIP | 1 | 0 | 0 | 868414 | P14340.2 | P17763 |
| PTVDGITVIDLEPIP | 1 | 0 | 0 | 868415 | QRG47106.1 | P17763 |
| PYMPSVIEKMEALQR | 1 | 0 | 0 | 868423 | P14340.2 | P17763 |
| QNILTAIQQVRSLIG | 1 | 0 | 0 | 868544 | QTX93504.1 | P17763 |
| QPHWIAASIILEFFL | 1 | 0 | 0 | 868554 | P14340.2 | P17763 |
| QPQWIAASIILEFFL | 1 | 0 | 0 | 868560 | Q2YHF0.1 | P17763 |
| QSHQLWATLLSLTFI | 1 | 0 | 0 | 868583 | QFS19562.1 | P17763 |
| QSHQLWATLLSLTFV | 1 | 0 | 0 | 868584 | P17763.2 | P17763 |
| QTTTGEIGAIALDFK | 1 | 0 | 0 | 868601 | QTX93491.1 | P17763 |
| QVPFCSHHFHELIMK | 1 | 0 | 0 | 868619 | P14340.2 | P17763 |
| QVPFCSHHFHELVMK | 1 | 0 | 0 | 868620 | QRG47115.1 | P17763 |
| QVRSLIGNEEFLDYM | 1 | 0 | 0 | 868623 | ABZ90961.1 | P17763 |
| RDLRLAANAICSAVP | 1 | 0 | 0 | 868637 | P17763.2 | P17763 |
| RFLEFEALGFMNEDH | 1 | 0 | 0 | 868641 | P17763.2 | P17763 |
| RHPGFTILALFLAHY | 1 | 0 | 0 | 868649 | QTX93489.1 | P17763 |
| RKMLMTGTLAVFLLL | 1 | 0 | 0 | 868665 | P17763.2 | P17763 |
| RLASNAICSAVPVHW | 1 | 0 | 0 | 868670 | QTX93489.1 | P17763 |
| RLATAIAGAWENGVC | 1 | 0 | 0 | 868671 | QTX93489.1 | P17763 |
| RSQVGVGVFQENVFH | 1 | 0 | 0 | 868711 | QTX92726.1 | P17763 |
| RTIMAVFFVVTLIPL | 1 | 0 | 0 | 868715 | ANW72070.1 | P17763 |
| RTIMAVLFVVTLIPL | 1 | 0 | 0 | 868716 | Q2YHF0.1 | P17763 |
| SANLSLAAIANQAAV | 1 | 0 | 0 | 868753 | Q2YHF0.1 | P17763 |
| SCVPNAVILQNAWKV | 1 | 0 | 0 | 868762 | QRG47120.1 | P17763 |
| SLGILCVSIMIEEVM | 1 | 0 | 0 | 868915 | QPZ88407.1 | P17763 |
| SLGLLCISIMIEEVM | 1 | 0 | 0 | 868916 | P17763.2 | P17763 |
| SLGVLGMALFLEEML | 1 | 0 | 0 | 868918 | QTX93024.1 | P17763 |
| STESHNQTFLIDGPE | 1 | 0 | 0 | 869038 | QTX93189.1 | P17763 |
| SWPLNEAIMAVGMVS | 1 | 0 | 0 | 869079 | QTX93024.1 | P17763 |
| SYAQMWSLMYFHRRD | 1 | 0 | 0 | 869082 | QTX93208.1 | P17763 |
| SYAQMWTLMYFHRRD | 1 | 0 | 0 | 869083 | QTX93073.1 | P17763 |
| TEQYKFQADSPKRLS | 1 | 0 | 0 | 869135 | P17763.2 | P17763 |
| TGLDFNEMVLLQMED | 1 | 0 | 0 | 869152 | QRG47133.1 | P17763 |
| TGLDFNEMVLLQMEN | 1 | 0 | 0 | 869153 | QTX93201.1 | P17763 |
| TIGIVLLSQSTIPET | 1 | 0 | 0 | 869180 | QTX93201.1 | P17763 |
| TIRVLKMVEPWLKNN | 1 | 0 | 0 | 869191 | QTX93593.1 | P17763 |
| TKHAILLVAVSFVTL | 1 | 0 | 0 | 869200 | QTX93201.1 | P17763 |
| TKHAVLLVAVSFMTL | 1 | 0 | 0 | 869201 | QGW51895.1 | P17763 |
| TKKEEPVNIEAEPPF | 1 | 0 | 0 | 869202 | QTX93593.1 | P17763 |
| TLFFLSGKGLGKTSI | 1 | 0 | 0 | 869217 | QFS19562.1 | P17763 |
| TMGVLCLAILFEEVM | 1 | 0 | 0 | 869269 | QTX93474.1 | P17763 |
| TQKGIIFILLMLVTP | 1 | 0 | 0 | 869300 | P17763.2 | P17763 |
| TQKVVIFILLMLVTP | 1 | 0 | 0 | 869302 | QTX93593.1 | P17763 |
| TTMFGGVSWMVRILI | 1 | 0 | 0 | 869337 | QTX92385.1 | P17763 |
| TYLALIATFKIQPFL | 1 | 0 | 0 | 869372 | QTX93593.1 | P17763 |
| VEMGEAAAIFMTATP | 1 | 0 | 0 | 869421 | QTX92385.1 | P17763 |
| VGMGEAAAIFMTATP | 1 | 0 | 0 | 869451 | P17763.2 | P17763 |
| WEPSKGWNDWQQVPF | 1 | 0 | 0 | 869682 | P17763.2 | P17763 |
| WNLVRLQSGVDVFFT | 1 | 0 | 0 | 869694 | QTX93213.1 | P17763 |
| WNTGEEVQVIAVEPG | 1 | 0 | 0 | 869696 | QFS19562.1 | P17763 |
| WQKGEEVQVIAVEPG | 1 | 0 | 0 | 869697 | QTX93593.1 | P17763 |
| WRTATLILAGVSLLP | 1 | 0 | 0 | 869700 | QTX93593.1 | P17763 |
| YGWNIVKLMSGKDVF | 1 | 0 | 0 | 869726 | AAC63341.1 | P17763 |
| YHGSYEAPSTGSASS | 1 | 0 | 0 | 869730 | QTX92385.1 | P17763 |
| FTDPSSIAARGYIST | 1 | 0 | 0 | 870244 | QFS19562.1 | P17763 |
| SIAARGYISTRVEMG | 1 | 0 | 0 | 871207 | QTX93024.1 | P17763 |
| TLVDRGWGNGCGLFGKGS | 1 | 0 | 0 | 886475 | AHZ13508.1 | Q32ZE1 |
| VDRGWGNGCGLFGKGSLV | 1 | 0 | 0 | 886505 | AGR44695.1 | P17763 |
| AGPISQHNHRPGYHTQTA | 1 | 0 | 0 | 919518 | AGH08163.1 | P17763 |
| AGPLSHHNTREGYRTQMK | 1 | 0 | 0 | 919519 | AHZ13508.1 | Q32ZE1 |
| ANPVITESTENSKMMLEL | 1 | 0 | 0 | 919521 | AHZ13508.1 | Q32ZE1 |
| ANPVVTKKEEPVNIEA | 1 | 0 | 0 | 919522 | AHG23217.1 | P17763 |
| AQMAVDMQTLTPV | 1 | 0 | 0 | 919524 | AHZ13508.1 | Q32ZE1 |
| CGIRSTTRMENLLWKQIA | 1 | 0 | 0 | 919526 | AHG23217.1 | P17763 |
| CGISSVSRMENIMWRSVE | 1 | 0 | 0 | 919527 | AHZ13508.1 | Q32ZE1 |
| CNLTSTWVTYGTCNQAG | 1 | 0 | 0 | 919528 | AHG23217.1 | P17763 |
| CNTTSTWVVYGTCHHKKG | 1 | 0 | 0 | 919529 | AHZ13508.1 | Q32ZE1 |
| DKPTVDIELVTTTVSNMA | 1 | 0 | 0 | 919530 | AHZ13508.1 | Q32ZE1 |
| DMGCVINWKGKELKCGSG | 1 | 0 | 0 | 919531 | AHG23217.1 | P17763 |

| | | | | | | |
|---|---|---|---|---|---|---|
| DRNDAGEAISFPTTLGMN | 1 | 0 | 0 | 919534 | AHZ13508.1 | Q32ZE1 |
| ECTMPPLSFRAK | 1 | 0 | 0 | 919535 | AHZ13508.1 | Q32ZE1 |
| EFKDAHAKRQTVVVLGSQ | 1 | 0 | 0 | 919537 | AHZ13508.1 | Q32ZE1 |
| EPIEGKVVQYENLKYTVI | 1 | 0 | 0 | 919539 | AHG23217.1 | P17763 |
| EPRTGLDFSDLYY | 1 | 0 | 0 | 919540 | AHZ13508.1 | Q32ZE1 |
| FKSLFGGMSWFSQILIGT | 1 | 0 | 0 | 919545 | AHZ13508.1 | Q32ZE1 |
| GALEAEMDGAKGRLSSGH | 1 | 0 | 0 | 919548 | AHZ13508.1 | Q32ZE1 |
| GATEIQNSGGTSIFAGH | 1 | 0 | 0 | 919549 | AHG23217.1 | P17763 |
| GDIIGVLEQGKRTLTPQPMELK | 1 | 0 | 0 | 919550 | AHG23217.1 | P17763 |
| GDQHQVGNETQGVTAEIT | 1 | 0 | 0 | 919551 | AHG23217.1 | P17763 |
| GDTLKECPLKHRAWNSFL | 1 | 0 | 0 | 919552 | AHZ13508.1 | Q32ZE1 |
| GEMCDDTVTYKCPHITE | 1 | 0 | 0 | 919553 | AHG23217.1 | P17763 |
| GHETDENRAKVEITPNSP | 1 | 0 | 0 | 919554 | AHZ13508.1 | Q32ZE1 |
| GHMCDATMSYECPMLDEG | 1 | 0 | 0 | 919555 | AHZ13508.1 | Q32ZE1 |
| GKAKIVTAETQNSSFIID | 1 | 0 | 0 | 919556 | AHG23217.1 | P17763 |
| GKNERGKSLLFKTASGIN | 1 | 0 | 0 | 919557 | AHG23217.1 | P17763 |
| GKSYFVRAAKTNNSFVVD | 1 | 0 | 0 | 919558 | AHZ13508.1 | Q32ZE1 |
| GNETQGVTAEITPQASTT | 1 | 0 | 0 | 919560 | AHG23217.1 | P17763 |
| GPNTPECPSASRAWNVWE | 1 | 0 | 0 | 919561 | AHG23217.1 | P17763 |
| GSVKNPMWRGPQRLPVPVNELP | 1 | 0 | 0 | 919566 | AHZ13508.1 | Q32ZE1 |
| HITEVEPEDIDCWCNLT | 1 | 0 | 0 | 919567 | AHG23217.1 | P17763 |
| HSTRKLQTRSQTWLESREY | 1 | 0 | 0 | 919569 | AHZ13508.1 | Q32ZE1 |
| HWNNKEALVEF | 1 | 0 | 0 | 919571 | AHZ13508.1 | Q32ZE1 |
| IFVTNEVHTWTEQYKFQA | 1 | 0 | 0 | 919574 | AHG23217.1 | P17763 |
| IRFEECPGTKVHVEETCG | 1 | 0 | 0 | 919577 | AHZ13508.1 | Q32ZE1 |
| ISEKEENMVKSLVSAG | 1 | 0 | 0 | 919580 | AHG23217.1 | P17763 |
| KCYIQIMDLGHMCDATMS | 1 | 0 | 0 | 919581 | AHZ13508.1 | Q32ZE1 |
| KKGSSIGKMFEATARGAR | 1 | 0 | 0 | 919586 | AHG23217.1 | P17763 |
| KKMTGKSIQPENLEYRIM | 1 | 0 | 0 | 919587 | AHZ13508.1 | Q32ZE1 |
| KVEITPNSPRAEATLGGF | 1 | 0 | 0 | 919592 | AHZ13508.1 | Q32ZE1 |
| KVHVEETCGTRGPSLRST | 1 | 0 | 0 | 919594 | AHZ13508.1 | Q32ZE1 |
| LDEGVEPDDVDCWCNTT | 1 | 0 | 0 | 919600 | AHZ13508.1 | Q32ZE1 |
| LDFNYCEGTTVVITENCG | 1 | 0 | 0 | 919601 | AHG23217.1 | P17763 |
| LFKTASGINMCTLIAMDL | 1 | 0 | 0 | 919603 | AHG23217.1 | P17763 |
| LVTCAKFACSKKMTGKSI | 1 | 0 | 0 | 919606 | AHZ13508.1 | Q32ZE1 |
| LVTCAKFQCLEPIEGKVV | 1 | 0 | 0 | 919607 | AHG23217.1 | P17763 |
| MCTLIAMDLGEMCDDTVT | 1 | 0 | 0 | 919608 | AHG23217.1 | P17763 |
| NEMILLTMKNKAWMVHRQ | 1 | 0 | 0 | 919612 | AHG23217.1 | P17763 |
| NKPTLDIELQKTEATQLA | 1 | 0 | 0 | 919613 | AHG23217.1 | P17763 |
| PENLEYRIMLSVHGSQHS | 1 | 0 | 0 | 919616 | AHZ13508.1 | Q32ZE1 |
| PLRYMGEDGCWYGMEIRP | 1 | 0 | 0 | 919617 | AHG23217.1 | P17763 |
| PLSFRAKDGCWYGMEIRP | 1 | 0 | 0 | 919618 | AHZ13508.1 | Q32ZE1 |
| QIANELNYILWENNIK | 1 | 0 | 0 | 919622 | AHG23217.1 | P17763 |
| RCPTQGEAVLPEEQDQNY | 1 | 0 | 0 | 919626 | AHG23217.1 | P17763 |
| RCPTQGEAYLDKQSDTQY | 1 | 0 | 0 | 919627 | AHZ13508.1 | Q32ZE1 |
| RDGEPRMIVGKNERGKSL | 1 | 0 | 0 | 919628 | AHG23217.1 | P17763 |
| REGYRTQMKGPWHSEELE | 1 | 0 | 0 | 919629 | AHZ13508.1 | Q32ZE1 |
| RGSAYYMYLDRNDAGEAI | 1 | 0 | 0 | 919630 | AHZ13508.1 | Q32ZE1 |
| RKEPESNLVRSMVTAG | 1 | 0 | 0 | 919631 | AHZ13508.1 | Q32ZE1 |
| RPGYHTQTAGPWHLGKLE | 1 | 0 | 0 | 919632 | AHG23217.1 | P17763 |
| RSGSTIGKAFEATVRGAK | 1 | 0 | 0 | 919640 | AHZ13508.1 | Q32ZE1 |
| SDLIIPKSLAGPLSHHNT | 1 | 0 | 0 | 919645 | AHZ13508.1 | Q32ZE1 |
| SDLYYLTMNNKHWLVHKE | 1 | 0 | 0 | 919646 | AHZ13508.1 | Q32ZE1 |
| SDMIIPKSLAGPISQHNH | 1 | 0 | 0 | 919647 | AGH08163.1 | P17763 |
| SEELEIRFEECP | 1 | 0 | 0 | 919648 | AHZ13508.1 | Q32ZE1 |
| SFPTTLGMNKCYIQIMDL | 1 | 0 | 0 | 919649 | AHZ13508.1 | Q32ZE1 |
| SGMIVNDTGHETDENRAK | 1 | 0 | 0 | 919650 | AHZ13508.1 | Q32ZE1 |
| SLCTAAFTFTKIPA | 1 | 0 | 0 | 919651 | AHZ13508.1 | Q32ZE1 |
| SPRTGLDFNEMIL | 1 | 0 | 0 | 919655 | AHG23217.1 | P17763 |
| SVEGELNAILEENGVQ | 1 | 0 | 0 | 919659 | AHZ13508.1 | Q32ZE1 |
| TAEITPQASTTEAILPEY | 1 | 0 | 0 | 919660 | AHG23217.1 | P17763 |
| TASGRVIEEWCCRECTMP | 1 | 0 | 0 | 919661 | AHZ13508.1 | Q32ZE1 |
| TFKNAHAKKQEVVVLGSQ | 1 | 0 | 0 | 919663 | AHG23217.1 | P17763 |
| TILIKVEYKGEDAPCKIP | 1 | 0 | 0 | 919664 | Q98UK6 | P17763 |
| TRGPSLRSTTASGRVIEE | 1 | 0 | 0 | 919668 | AHZ13508.1 | Q32ZE1 |
| TRGPSLRTTTVSGKLIHE | 1 | 0 | 0 | 919669 | AHG23217.1 | P17763 |
| TVSGKLIHEWCCRSCTLP | 1 | 0 | 0 | 919672 | AHG23217.1 | P17763 |
| TVTVEVQYAGTDGPCKVP | 1 | 0 | 0 | 919673 | AHZ13508.1 | Q32ZE1 |
| TVVITENCGTRGPSLRTT | 1 | 0 | 0 | 919674 | AHG23217.1 | P17763 |
| VCKHTYVDRGWGNGCGLF | 1 | 0 | 0 | 919676 | AHG23217.1 | P17763 |
| VCKRTLVDRGWGNGCGLF | 1 | 0 | 0 | 919677 | AHZ13508.1 | Q32ZE1 |
| VEDHGFGVFHTSVWLKVREDYS | 1 | 0 | 0 | 919678 | AHZ13508.1 | Q32ZE1 |
| VEDYGFGVFTTNIWLKLREVYT | 1 | 0 | 0 | 919679 | AHG23217.1 | P17763 |
| VFVYNDVEAWRDRYKYHP | 1 | 0 | 0 | 919680 | AHZ13508.1 | Q32ZE1 |
| VGMGLDTRTQTWMSAEGAW | 1 | 0 | 0 | 919681 | AHG23217.1 | P17763 |
| WFFDLPLPWTSGATTETP | 1 | 0 | 0 | 919685 | AHG23217.1 | P17763 |
| WFHDIPLPWHAGADTGTP | 1 | 0 | 0 | 919686 | AHZ13508.1 | Q32ZE1 |
| YEASISDMASDS | 1 | 0 | 0 | 919689 | AHZ13508.1 | Q32ZE1 |
| YECPMLDEGVEPDDVDCW | 1 | 0 | 0 | 919690 | AHZ13508.1 | Q32ZE1 |
| YENLKYTVIITVHTGDQH | 1 | 0 | 0 | 919692 | AHG23217.1 | P17763 |
| YKCPHITEVEPEDIDCW | 1 | 0 | 0 | 919693 | AHG23217.1 | P17763 |
| YTALFSGVSWVMKIGIGV | 1 | 0 | 0 | 919697 | AHG23217.1 | P17763 |
| AECPNTNRAWNSLEV | 1 | 0 | 0 | 922138 | P14340.2 | P17763 |
| AELTGYGTVTMECSP | 1 | 0 | 0 | 922141 | P14340.2 | P17763 |
| AEPPFGDSYIIIGVE | 1 | 0 | 0 | 922144 | P14340.2 | P17763 |
| AGPWHLGKLEMDFDF | 1 | 0 | 0 | 922149 | P14340.2 | P17763 |
| AHEEGICGIRSVTRL | 1 | 0 | 0 | 922151 | P14340.2 | P17763 |
| AIQKAHEEGICGIRS | 1 | 0 | 0 | 922154 | P14340.2 | P17763 |
| AKKQDVVVLGSQEGA | 1 | 0 | 0 | 922156 | P14340.2 | P17763 |
| AKMLSTESHNQTFLI | 1 | 0 | 0 | 922157 | P14340.2 | P17763 |
| ALNDTWKIEKASFIE | 1 | 0 | 0 | 922158 | P14340.2 | P17763 |

| | | | | | | |
|---|---|---|---|---|---|---|
| AVHADMGYWIESALN | 1 | 0 | 0 | 922170 | P14340.2 | P17763 |
| AWNSLEVEDYGFGVF | 1 | 0 | 0 | 922175 | P14340.2 | P17763 |
| CGLFGKGGIVTCAMF | 1 | 0 | 0 | 922179 | BAG84557.1 | P17763 |
| CRLRMDKLQLKGMSY | 1 | 0 | 0 | 922183 | P14340.2 | P17763 |
| CSPRTGLDFNEMVLL | 1 | 0 | 0 | 922185 | P17763.2 | P17763 |
| CWYGMEIRPLKEKEE | 1 | 0 | 0 | 922186 | P14340.2 | P17763 |
| DGSPCKIPFEIMDLE | 1 | 0 | 0 | 922190 | P14340.2 | P17763 |
| DMGYWIESALNDTWK | 1 | 0 | 0 | 922196 | P14340.2 | P17763 |
| DVVVLGSQEGAMHTA | 1 | 0 | 0 | 922206 | P14340.2 | P17763 |
| EAKQPATLRKYCIEA | 1 | 0 | 0 | 922208 | P14340.2 | P17763 |
| EDCGNRGPSLRTTTA | 1 | 0 | 0 | 922210 | P14340.2 | P17763 |
| EGTTVVVTEDCGNRG | 1 | 0 | 0 | 922218 | P14340.2 | P17763 |
| EGVSGGSWVDIVLEH | 1 | 0 | 0 | 922219 | P14340.2 | P17763 |
| EIKITPQSSITEAEL | 1 | 0 | 0 | 922221 | P14340.2 | P17763 |
| EKASFIEVKSCHWPK | 1 | 0 | 0 | 922223 | P14340.2 | P17763 |
| EKQDVFCDSKLMSAA | 1 | 0 | 0 | 922224 | P14340.2 | P17763 |
| ENEVKLTIMTGDIKG | 1 | 0 | 0 | 922229 | P14340.2 | P17763 |
| EQDKRFVCKHSMVDR | 1 | 0 | 0 | 922230 | P14340.2 | P17763 |
| ESPSKLASAIQKAHE | 1 | 0 | 0 | 922235 | P14340.2 | P17763 |
| FIEVKSCHWPKSHTL | 1 | 0 | 0 | 922242 | P14340.2 | P17763 |
| FLIDGPETAECPNTN | 1 | 0 | 0 | 922245 | P14340.2 | P17763 |
| FNEMVLLQMENKAWL | 1 | 0 | 0 | 922247 | P14340.2 | P17763 |
| GEDGCWYGMEIRPLK | 1 | 0 | 0 | 922253 | P14340.2 | P17763 |
| GICGIRSVTRLENLM | 1 | 0 | 0 | 922264 | P14340.2 | P17763 |
| GISNRDFVEGVSGGS | 1 | 0 | 0 | 922266 | P14340.2 | P17763 |
| GKLITEWCCRSCTLP | 1 | 0 | 0 | 922268 | P14340.2 | P17763 |
| GPETAECPNTNRAWN | 1 | 0 | 0 | 922272 | P14340.2 | P17763 |
| GPVSQHNYRPGYHTQ | 1 | 0 | 0 | 922278 | P14340.2 | P17763 |
| GVEPGQLKLNWFKKG | 1 | 0 | 0 | 922291 | P14340.2 | P17763 |
| GVFTTNIWLKLREKQ | 1 | 0 | 0 | 922292 | P14340.2 | P17763 |
| GYGTVTMECSPRTGL | 1 | 0 | 0 | 922301 | P14340.2 | P17763 |
| HAVGNDTGKHGKEIK | 1 | 0 | 0 | 922302 | P14340.2 | P17763 |
| HLGKLEMDFDFCEGT | 1 | 0 | 0 | 922309 | P14340.2 | P17763 |
| HRQWFLDLPLPWLPG | 1 | 0 | 0 | 922311 | P14340.2 | P17763 |
| HTQTAGPWHLGKLEM | 1 | 0 | 0 | 922312 | P14340.2 | P17763 |
| IRSVTRLENLMWKQI | 1 | 0 | 0 | 922324 | P14340.2 | P17763 |
| ITDNVHTWTEQYKFQ | 1 | 0 | 0 | 922326 | P14340.2 | P17763 |
| ITPHSGEEHAVGNDT | 1 | 0 | 0 | 922328 | BAG84557.1 | P17763 |
| IVTEKDSPVNIEAEP | 1 | 0 | 0 | 922331 | P14340.2 | P17763 |
| KFKVVKEIAETQHGT | 1 | 0 | 0 | 922338 | P14340.2 | P17763 |
| KHGKEIKITPQSSIT | 1 | 0 | 0 | 922342 | P14340.2 | P17763 |
| KLASAIQKAHEEGIC | 1 | 0 | 0 | 922346 | P14340.2 | P17763 |
| KNFAGPVSQHNYRPG | 1 | 0 | 0 | 922347 | P14340.2 | P17763 |
| KNKELKCGSGIFITD | 1 | 0 | 0 | 922348 | AAB33855.1 | P17763 |
| KQITPELNHILSENE | 1 | 0 | 0 | 922350 | P14340.2 | P17763 |
| KSCHWPKSHTLWSNG | 1 | 0 | 0 | 922353 | P14340.2 | P17763 |
| LEHGSCVTTMAKNKP | 1 | 0 | 0 | 922370 | P27909.2 | P17763 |
| LEMDFDFCEGTTVVV | 1 | 0 | 0 | 922371 | P14340.2 | P17763 |
| LESEMIIPKNFAGPV | 1 | 0 | 0 | 922372 | P14340.2 | P17763 |
| LEVEDYGFGVFTTNI | 1 | 0 | 0 | 922373 | P14340.2 | P17763 |
| LKCGSGIFITDNVHT | 1 | 0 | 0 | 922379 | AAB33855.1 | P17763 |
| LKEKEENLVNSLVTA | 1 | 0 | 0 | 922380 | P14340.2 | P17763 |
| LKLREKQDVFCDSKL | 1 | 0 | 0 | 922381 | P14340.2 | P17763 |
| LRYRGEDGCWYGMEI | 1 | 0 | 0 | 922403 | P14340.2 | P17763 |
| MDKLQLKGMSYSMCT | 1 | 0 | 0 | 922414 | P14340.2 | P17763 |
| MEIRPLKEKEENLVN | 1 | 0 | 0 | 922415 | P14340.2 | P17763 |
| MIIPKNFAGPVSQHN | 1 | 0 | 0 | 922416 | P14340.2 | P17763 |
| MQAGKRSLRPQPTEL | 1 | 0 | 0 | 922421 | QTX92981.1 | P17763 |
| MSYSMCTGKFKVVKE | 1 | 0 | 0 | 922423 | P14340.2 | P17763 |
| MTGDIKGIMQAGKRS | 1 | 0 | 0 | 922424 | P14340.2 | P17763 |
| NDTGKHGKEIKITPQ | 1 | 0 | 0 | 922427 | P14340.2 | P17763 |
| NLMWKQITPELNHIL | 1 | 0 | 0 | 922440 | P14340.2 | P17763 |
| NRGPSLRTTTASGKL | 1 | 0 | 0 | 922445 | P14340.2 | P17763 |
| NTNRAWNSLEVEDYG | 1 | 0 | 0 | 922451 | P14340.2 | P17763 |
| PATLRKYCIEAKLTN | 1 | 0 | 0 | 922462 | P14340.2 | P17763 |
| PELNHILSENEVKLT | 1 | 0 | 0 | 922473 | P14340.2 | P17763 |
| PLKEKEENLVNSLVT | 1 | 0 | 0 | 922492 | P14340.2 | P17763 |
| QHNYRPGYHTQTAGP | 1 | 0 | 0 | 922530 | P14340.2 | P17763 |
| QLKGMSYSMCTGKFK | 1 | 0 | 0 | 922532 | P14340.2 | P17763 |
| RPQPTELKYSWKTWG | 1 | 0 | 0 | 922551 | QTX92984.1 | P17763 |
| SGEEHAVGNDTGKHG | 1 | 0 | 0 | 922571 | P14340.2 | P17763 |
| SGIFITDNVHTWTEQ | 1 | 0 | 0 | 922572 | P14340.2 | P17763 |
| SITEAELTGYGTVTM | 1 | 0 | 0 | 922579 | QTX93024.1 | P17763 |
| SLRTTTASGKLITEW | 1 | 0 | 0 | 922590 | QTX93024.1 | P17763 |
| SNWIQKETLVTFKNP | 1 | 0 | 0 | 922592 | QTX93024.1 | P17763 |
| TELKYSWKTWGKAKM | 1 | 0 | 0 | 922614 | QTX93024.1 | P17763 |
| TEQYKFQPESPSKLA | 1 | 0 | 0 | 922615 | QTX93024.1 | P17763 |
| TEWCCRSCTLPPLRY | 1 | 0 | 0 | 922616 | QTX93024.1 | P17763 |
| TGATEIQMSSGNLLF | 1 | 0 | 0 | 922618 | QTX93024.1 | P17763 |
| TLPPLRYRGEDGCWY | 1 | 0 | 0 | 922626 | QFS19562.1 | P17763 |
| TNIWLKLREKQDVFC | 1 | 0 | 0 | 922629 | QRG47112.1 | P17763 |
| TPQSSITEAELTGYG | 1 | 0 | 0 | 922631 | QTX93024.1 | P17763 |
| TRLENLMWKQITPEL | 1 | 0 | 0 | 922634 | QTX93024.1 | P17763 |
| TTASGKLITEWCCRS | 1 | 0 | 0 | 922640 | QTX93024.1 | P17763 |
| TVNPIVTEKDSPVNI | 1 | 0 | 0 | 922642 | QTX93024.1 | P17763 |
| TWGKAKMLSTESHNQ | 1 | 0 | 0 | 922643 | QTX93189.1 | P17763 |
| TWKIEKASFIEVKSC | 1 | 0 | 0 | 922644 | QSV27725.1 | P17763 |
| VDIVLEHGSCVTTMA | 1 | 0 | 0 | 922649 | QTX93024.1 | P17763 |
| VFCDSKLMSAAIKDN | 1 | 0 | 0 | 922650 | QTX93213.1 | P17763 |
| VHTWTEQYKFQPESP | 1 | 0 | 0 | 922651 | QTX93024.1 | P17763 |
| VLLQMENKAWLVHRQ | 1 | 0 | 0 | 922657 | QTX93201.1 | P17763 |

| | | | | | | |
|---|---|---|---|---|---|---|
| VVQPENLEYTIVITP | 1 | 0 | 0 | 922686 | QTX93213.1 | P17763 |
| VVSWKNKELKCGSGI | 1 | 0 | 0 | 922687 | AAB33855.1 | P17763 |
| VVVTEDCGNRGPSLR | 1 | 0 | 0 | 922688 | QTX93189.1 | P17763 |
| WIESALNDTWKIEKA | 1 | 0 | 0 | 922691 | QTX93201.1 | P17763 |
| YIIIGVEPGQLKLNW | 1 | 0 | 0 | 922702 | QTX93189.1 | P17763 |
| KQAIITHGAVIDWNQ | 1 | 0 | 0 | 922866 | YP_009508562.1 | A8HC79 |
| NKDIAVLGPVIDWNV | 1 | 0 | 0 | 922916 | AGM61339.1 | P23073 |
| NKDIQVLGPVIDWNV | 1 | 0 | 0 | 922921 | AAB48113.1 | Q87041 |
| NKDIVTLGPVIDWNV | 1 | 0 | 0 | 922924 | AFX98100.1 | K7YEX3 |
| NQDIQVYGPVIDWNI | 1 | 0 | 0 | 922927 | YP_009508889.1 | Q7SIS5 |
| RHAIITPGPVIDWNS | 1 | 0 | 0 | 922960 | YP_009508583.1 | A0A0G3Y5C1 |
| RNAITVPGIILDWNS | 1 | 0 | 0 | 922968 | YP_009508567.1 | D5JWU7 |
| NSTHNTPVYKLDISE | 1 | 0 | 0 | 929450 | ALG01342.1 | Q05320 |
| TKNITDKIDQIIHDFVDKTL | 1 | 0 | 0 | 931784 | ALG01342.1 | Q05320 |
| VDKTLPDQGDNDNWWTGWRQWIPAG | 1 | 0 | 0 | 932264 | ALG01342.1 | Q05320 |
| IITWIGMNSRSTSLSVSL | 1 | 0 | 0 | 947360 | Q9Q4T1 | P17763 |
| RPYCWHYAPQPCGIVPA | 1 | 0 | 0 | 950827 | CAA03854.1 | P27958 |
| AEPDDVDCWCNATDTWV | 1 | 0 | 0 | 952005 | BAA00395.1 | P17763 |
| AWEEGVCGIRSATRLENI | 1 | 0 | 0 | 952040 | CAA49193.1 | P17763 |
| DDQRAWNIWEVEDYGFGI | 1 | 0 | 0 | 952057 | CAA49193.1 | P17763 |
| EGTTVVVDEHCGNRGPSL | 1 | 0 | 0 | 952073 | CAA49193.1 | P17763 |
| EIQLTDYGALTLDCSPRT | 1 | 0 | 0 | 952075 | BAA00395.1 | P17763 |
| ELCEDTMTYKCPRITEA | 1 | 0 | 0 | 952078 | AXP83898.1 | P17763 |
| EVTNPAVLRKLCIEAKIS | 1 | 0 | 0 | 952088 | BAA00395.1 | P17763 |
| FGTAYGVLFSGVSWTMKI | 1 | 0 | 0 | 952095 | BAA00395.1 | P17763 |
| FSGVSWTMKIGIGILLTW | 1 | 0 | 0 | 952111 | BAA00395.1 | P17763 |
| GIFTTNIWLKLRDSYTQV | 1 | 0 | 0 | 952132 | CAA49193.1 | P17763 |
| GPNTPECPDDQRAWNIWE | 1 | 0 | 0 | 952144 | CAA49193.1 | P17763 |
| HKKGEARRSRRAVTLPSH | 1 | 0 | 0 | 952157 | AHZ13508.1 | Q32ZE1 |
| HMCDATMSYECPMLDEGV | 1 | 0 | 0 | 952158 | AHZ13508.1 | Q32ZE1 |
| IRSATRLENIMWKQISNE | 1 | 0 | 0 | 952178 | CAA49193.1 | P17763 |
| ISDMASDSRCPTQGEAYL | 1 | 0 | 0 | 952180 | AHZ13508.1 | Q32ZE1 |
| ISNTTTDSRCPTQGEATL | 1 | 0 | 0 | 952181 | BAA00395.1 | P17763 |
| IVQYENLKYSVIVTVHTG | 1 | 0 | 0 | 952185 | BAA00395.1 | P17763 |
| KFQADSPKRLSAAIGKAW | 1 | 0 | 0 | 952193 | CAA49193.1 | P17763 |
| KGRELKCGSGIFVTNEVH | 1 | 0 | 0 | 952195 | CAA49193.1 | P17763 |
| KMFEATARGARRMAILGD | 1 | 0 | 0 | 952215 | BAA00395.1 | P17763 |
| KSWLVHKQWFLDLPLPWT | 1 | 0 | 0 | 952223 | BAA00395.1 | P17763 |
| LAQGKKMIRPQPMEHKYS | 1 | 0 | 0 | 952226 | AWV55242.1 | P17763 |
| LELDFDLCEGTTVVVDEH | 1 | 0 | 0 | 952238 | CAA49193.1 | P17763 |
| MVLLTMKEKSWLVHKQWF | 1 | 0 | 0 | 952280 | BAA01388.1 | P17763 |
| NELNHILLENDMKFTVVV | 1 | 0 | 0 | 952283 | CAA49193.1 | P17763 |
| NIMWKQISNELNHILLEN | 1 | 0 | 0 | 952286 | CAA49193.1 | P17763 |
| QETWNRQDLLVTFKTAHA | 1 | 0 | 0 | 952302 | BAA00395.1 | P17763 |
| QNTTFIIDGPNTPECPDD | 1 | 0 | 0 | 952311 | CAA49193.1 | P17763 |
| QRVETWALRHPGFTVIAL | 1 | 0 | 0 | 952315 | AWV55242.1 | P17763 |
| RKLCIEAKISNTTTDSRC | 1 | 0 | 0 | 952341 | BAA00395.1 | P17763 |
| RLSAAIGKAWEEGVCGIR | 1 | 0 | 0 | 952348 | CAA49193.1 | P17763 |
| RPQPMEHKYSWKSWGKAK | 1 | 0 | 0 | 952354 | AWV55228.1 | P17763 |
| RTGLDFNEMVLLTMKEKS | 1 | 0 | 0 | 952360 | BAA01388.1 | P17763 |
| SAAIKDSKAVHADMGYWI | 1 | 0 | 0 | 952367 | CAA49193.1 | P17763 |
| TEHGTTATITPQAPTSEI | 1 | 0 | 0 | 952417 | C32401 | P17763 |
| TGVFVYNDVEAWRDRYKY | 1 | 0 | 0 | 952422 | AHZ13508.1 | Q32ZE1 |
| TVSNMAEVRSYCYEASIS | 1 | 0 | 0 | 952447 | AHZ13508.1 | Q32ZE1 |
| VGKLVHQIFGTAYGVLFS | 1 | 0 | 0 | 952576 | BAJ72472.1 | P17763 |
| WEVEDYGFGIFTTNIWLK | 1 | 0 | 0 | 952595 | CAA49193.1 | P17763 |
| WFLDLPLPWTSGASTSQE | 1 | 0 | 0 | 952596 | BAA00395.1 | P17763 |
| WPKSHTLWSNGVLESEMI | 1 | 0 | 0 | 952603 | CAA49193.1 | P17763 |
| GWVAGLFYYHKFN | 1 | 0 | 0 | 969051 | BAA03732.1 | P27958 |
| AFWETKKNLTRKIRSEEL | 1 | 0 | 0 | 999356 | ALG01342.1 | Q05320 |
| IPLGVIHNSTLQVSDVD | 1 | 0 | 0 | 1021615 | ALG01342.1 | Q05320 |
| LITNTIAGVAGLITGGRR | 1 | 0 | 0 | 1029633 | ALG01342.1 | Q05320 |
| LQLFLRATTELRTFSILN | 1 | 0 | 0 | 1031292 | ALG01342.1 | Q05320 |
| RRTRREVIVNAQPKCNPN | 1 | 0 | 0 | 1045294 | ALG01342.1 | Q05320 |
| VNAQPKCNPNLHYWTTQD | 1 | 0 | 0 | 1058471 | ALG01342.1 | Q05320 |
| PNVDLINYLPK | 1 | 0 | 0 | 1065962 | O80066 | O80066 |
| DITWEKDAEVTGNSPRLDVALD | 1 | 0 | 0 | 1066068 | AMO03410.2 | Q32ZE1 |
| ACLGKAYAQMWSLMY | 1 | 0 | 0 | 1066543 | QTX93491.1 | P17763 |
| LKRWGQLKKNKAIKI | 1 | 0 | 0 | 1067635 | Q2YHF0.1 | P17763 |
| GSQHSGMIVNDTGHETDENRA | 1 | 0 | 0 | 1068729 | AMK49164.2 | Q32ZE1 |
| CLKGCCSCGSCCKFD | 1 | 0 | 0 | 1069288 | YP_009724390.1 | P0DTC2 |
| IAIVMVTIMLCCMTS | 1 | 0 | 0 | 1070307 | P0DTC2.1 | P0DTC2 |
| KPTKRSFIEDLLFNKVTL | 1 | 0 | 0 | 1070805 | NP_828851.1 | P59594 |
| LTDEMIAQYTSALLA | 1 | 0 | 0 | 1071338 | YP_009724390.1 | P0DTC2 |
| PWYIWLGFIAGLIAI | 1 | 0 | 0 | 1071825 | P0DTC2.1 | P0DTC2 |
| QPRTFLLKYNENGTI | 1 | 0 | 0 | 1072010 | YP_009724390.1 | P0DTC2 |
| SFTRGVYYPDKVFRS | 1 | 0 | 0 | 1072624 | YP_009724390.1 | P0DTC2 |
| SLSSTASALGKLQDV | 1 | 0 | 0 | 1072822 | P0DTC2.1 | P0DTC2 |
| VLTPSSKRFQPFQQFGRD | 1 | 0 | 0 | 1073900 | NP_828851.1 | P59594 |
| GDAALALLLLDRLNQL | 1 | 0 | 0 | 1074903 | YP_009724397.2 | P0DTC9 |
| SLLLCRMNSRNYIA | 1 | 0 | 0 | 1075055 | QRA56788.1 | A0A663DJA2 |
| AAEQDRNTREVFAQVKQM | 1 | 0 | 0 | 1087450 | YP_009825051.1 | P59594 |
| GIAVEQDKNTQEVFAQVK | 1 | 0 | 0 | 1087531 | QHD43416.1 | P0DTC2 |
| KKDKKKKADETQALPQRQ | 1 | 0 | 0 | 1087574 | QHD43423.2 | P0DTC9 |
| KTFPPTEPKKDKKKKTDE | 1 | 0 | 0 | 1087598 | YP_009825061.1 | P59595 |
| PINLVRDLPQGFSALEPL | 1 | 0 | 0 | 1087680 | QHD43416.1 | P0DTC2 |
| QMYKTPTLKYFGGFNFSQ | 1 | 0 | 0 | 1087693 | YP_009825051.1 | P59594 |
| ADLIIERREGSDVCYPGKFVNEEAL | 1 | 0 | 0 | 1126060 | AGL44438.1 | P03452 |
| AEEDGTGCFEIFHKCDDDCMASIRN | 1 | 0 | 0 | 1126261 | AGL44438.1 | P03452 |
| ATGMKNVPEIPKGRGLFGAIAGFIE | 1 | 0 | 0 | 1129863 | AGL44438.1 | P03452 |
| ATGNLVVPRYAFAMERNAGSGIIIS | 1 | 0 | 0 | 1129864 | ACP41105.1 | P03452 |

| | | | | | | |
|---|---|---|---|---|---|---|
| AVSNGTKVNTLTERGVEVVNATETV | 1 | 0 | 0 | 1130264 | AGL44438.1 | P03452 |
| CFILLAIVMGLVFICVKNGNMRCTI | 1 | 0 | 0 | 1130442 | AGL44438.1 | P03452 |
| CYPGKFVNEEALRQILRESGGIDKE | 1 | 0 | 0 | 1130598 | AGL44438.1 | P03452 |
| DGFLDIWTYNAELLVLLENERTLDY | 1 | 0 | 0 | 1131855 | ACP41105.1 | P03452 |
| DSNKGVTAACPHAGAKSFYKNLIWL | 1 | 0 | 0 | 1134662 | ACP41105.1 | P03452 |
| DSNVKNLYEKVRSQLKNNAKEIGNG | 1 | 0 | 0 | 1134664 | ACP41105.1 | P03452 |
| DVTDVTQLYLGGMSY | 1 | 0 | 0 | 1135253 | P0DTD1.1 | P0DTD1 |
| EEAKLNREEIDGVKLESTRIYQILA | 1 | 0 | 0 | 1136292 | ACP41105.1 | P03452 |
| EGDCYHSGGTIISNLPFQNIDSRAV | 1 | 0 | 0 | 1136912 | AGL44438.1 | P03452 |
| EITNKVNSVIEKMNTQFTAVGKEFN | 1 | 0 | 0 | 1137428 | ACP41105.1 | P03452 |
| EMKWLLSNTDNAAFPQMTKSYKNTR | 1 | 0 | 0 | 1138304 | AGL44438.1 | P03452 |
| FEFYHKCDNTCMESVKNGTYDYPKY | 1 | 0 | 0 | 1140522 | ACP41105.1 | P03452 |
| FEIFPKTSSWPNHDSNKGVTAACPH | 1 | 0 | 0 | 1140533 | ACP41105.1 | P03452 |
| FPQMTKSYKNTRKSPALIVWGIHHS | 1 | 0 | 0 | 1141159 | AGL44438.1 | P03452 |
| FVPSPGARPQVNGLSGRIDFHWLML | 1 | 0 | 0 | 1141600 | AGL44438.1 | P03452 |
| GAKSFYKNLIWLVKKGNSYPKLSKS | 1 | 0 | 0 | 1141843 | ACP41105.1 | P03452 |
| GFIEGGWTGMVDGWYGYHHQNEQGS | 1 | 0 | 0 | 1142694 | ACP41105.1 | P03452 |
| GNVINWTRDSITEVWSYNAELLVAM | 1 | 0 | 0 | 1144095 | AGL44438.1 | P03452 |
| GTCYPGDFIDYEELREQLSSVSSFE | 1 | 0 | 0 | 1145632 | ACP41105.1 | P03452 |
| GWEGLIDGWYGFRHQNAQGEGTAAD | 1 | 0 | 0 | 1146121 | AGL44438.1 | P03452 |
| HPITIGKCPKYVKSTKLRLATGLRN | 1 | 0 | 0 | 1146809 | ACP41105.1 | P03452 |
| HQNAQGEGTAADYKSTQSAIDQITG | 1 | 0 | 0 | 1146898 | AGL44438.1 | P03452 |
| IAPDRASFLRGKSMGIQSGVQVDAN | 1 | 0 | 0 | 1147379 | AGL44438.1 | P03452 |
| ICVKNGNMRCTICI | 1 | 0 | 0 | 1147484 | AGL44438.1 | P03452 |
| IHHPSTSADQQSLYQNADTYVFVGS | 1 | 0 | 0 | 1148302 | ACP41105.1 | P03452 |
| INDKGKEVLVLWGIHHPSTSADQQS | 1 | 0 | 0 | 1148935 | ACP41105.1 | P03452 |
| IPTNADKICLGHHAVSNGTKVNTLT | 1 | 0 | 0 | 1149354 | AGL44438.1 | P03452 |
| KCDDDCMASIRNNTYDHSKYREEAM | 1 | 0 | 0 | 1150868 | AGL44438.1 | P03452 |
| KCPRYVKQRSLLLATGMKNVPEIPK | 1 | 0 | 0 | 1150881 | AGL44438.1 | P03452 |
| KGAINTSLPFQNIHPITIGKCPKYV | 1 | 0 | 0 | 1151775 | ACP41105.1 | P03452 |
| KHNGKLCKLRGVAPLHLGKCNIAGW | 1 | 0 | 0 | 1152132 | ACP41105.1 | P03452 |
| KKGNSYPKLSKSYINDKGKEVLVLW | 1 | 0 | 0 | 1152423 | ACP41105.1 | P03452 |
| KLESTRIYQILAIYSTVASSLVLVV | 1 | 0 | 0 | 1152749 | ACP41105.1 | P03452 |
| KLVTVGSSNYQQSFVPSPGARPQVN | 1 | 0 | 0 | 1152983 | AGL44438.1 | P03452 |
| KLYERVKRQLRENAEEDGTGCFEIF | 1 | 0 | 0 | 1152987 | AGL44438.1 | P03452 |
| KMVSLLSVLLSMQGA | 1 | 0 | 0 | 1153055 | P0DTC1.1 | P0DTD1 |
| KSTQSAIDQITGKLNRLIEKTNQQF | 1 | 0 | 0 | 1154324 | AGL44438.1 | P03452 |
| LAKDTTEAFEKMVSL | 1 | 0 | 0 | 1155225 | P0DTC1.1 | P0DTD1 |
| LEKRIENLNKKVDDGFLDIWTYNAE | 1 | 0 | 0 | 1156056 | ACP41105.1 | P03452 |
| LGAISFWMCSNGSLQCRICI | 1 | 0 | 0 | 1156351 | ACP41105.1 | P03452 |
| LGNPECESLSTASSWSYIVETPSSD | 1 | 0 | 0 | 1156513 | ACP41105.1 | P03452 |
| LIDNEFNEVEKQIGNVINWTRDSIT | 1 | 0 | 0 | 1156755 | AGL44438.1 | P03452 |
| LNRLIEKTNQQFELIDNEFNEVEKQ | 1 | 0 | 0 | 1157672 | AGL44438.1 | P03452 |
| LREQLSSVSSFERFEIFPKTSSWPN | 1 | 0 | 0 | 1158864 | ACP41105.1 | P03452 |
| LSGRIDFHWLMLNPNDTVTFSFNGA | 1 | 0 | 0 | 1159143 | AGL44438.1 | P03452 |
| LVLLENERTLDYHDSNVKNLYEKVR | 1 | 0 | 0 | 1159730 | ACP41105.1 | P03452 |
| MERNAGSGIIISDTPVHDCNTTCQT | 1 | 0 | 0 | 1160155 | ACP41105.1 | P03452 |
| MGFTYSGIRTNGATSACRRSGSSFY | 1 | 0 | 0 | 1160199 | AGL44438.1 | P03452 |
| MGIQSGVQVDANCEGDCYHSGGTII | 1 | 0 | 0 | 1160208 | AGL44438.1 | P03452 |
| MKAILVVLLYTFATANADTLCIGYH | 1 | 0 | 0 | 1160252 | ACP41105.1 | P03452 |
| MNTQILVFALIAIIPTNADKICLGH | 1 | 0 | 0 | 1160366 | AGL44438.1 | P03452 |
| NLPFQNIDSRAVGKCPRYVKQRSLL | 1 | 0 | 0 | 1161651 | AGL44438.1 | P03452 |
| NNSTDTVDTVLEKNVTVTHSVNLLE | 1 | 0 | 0 | 1161800 | ACP41105.1 | P03452 |
| NPYVCNAPGCDVTDV | 1 | 0 | 0 | 1162137 | P0DTD1.1 | P0DTD1 |
| NQHTIDLADSEMDKLYERVKRQLRE | 1 | 0 | 0 | 1162178 | AGL44438.1 | P03452 |
| NRIQIDPVKLSSGYKDVILWFSFGA | 1 | 0 | 0 | 1162286 | AGL44438.1 | P03452 |
| NTQFTAVGKEFNHLEKRIENLNKKV | 1 | 0 | 0 | 1162644 | ACP41105.1 | P03452 |
| NVTVTHSVNLLEDKHNGKLCKLRGV | 1 | 0 | 0 | 1162859 | ACP41105.1 | P03452 |
| PIDKCSRIIPARARV | 1 | 0 | 0 | 1163184 | P0DTD1.1 | P0DTD1 |
| PKVRDQEGRMNYYWTLVEPGDKITF | 1 | 0 | 0 | 1163235 | ACP41105.1 | P03452 |
| PLHLGKCNIAGWILGNPECESLSTA | 1 | 0 | 0 | 1163247 | ACP41105.1 | P03452 |
| PNDTVTFSFNGAFIAPDRASFLRGK | 1 | 0 | 0 | 1163297 | AGL44438.1 | P03452 |
| PSIQSRGLFGAIAGFIEGGWTGMVD | 1 | 0 | 0 | 1163468 | ACP41105.1 | P03452 |
| QILRESGGIDKEAMGFTYSGIRTNG | 1 | 0 | 0 | 1164116 | AGL44438.1 | P03452 |
| QLKNNAKEIGNGCFEFYHKCDNTCM | 1 | 0 | 0 | 1164264 | ACP41105.1 | P03452 |
| RGLFGAIAGFIENGWEGLIDGWYGF | 1 | 0 | 0 | 1165813 | AGL44438.1 | P03452 |
| RGVEVVNATETVERTNIPRICSKGK | 1 | 0 | 0 | 1165884 | AGL44438.1 | P03452 |
| RTNIPRICSKGKRTVDLGQCGLLGT | 1 | 0 | 0 | 1167503 | AGL44438.1 | P03452 |
| RYSKKFKPEIAIRPKVRDQEGRMNY | 1 | 0 | 0 | 1167778 | ACP41105.1 | P03452 |
| SPALIVWGIHHSVSTAEQTKLYGSG | 1 | 0 | 0 | 1170135 | AGL44438.1 | P03452 |
| STAEQTKLYGSGNKLVTVGSSNYQQ | 1 | 0 | 0 | 1171846 | AGL44438.1 | P03452 |
| STKLRLATGLRNIPSIQSRGLFGAI | 1 | 0 | 0 | 1171943 | ACP41105.1 | P03452 |
| SVKNGTYDYPKYSEEAKLNREEIDG | 1 | 0 | 0 | 1172218 | ACP41105.1 | P03452 |
| SWSYIVETPSSDNGTCYPGDFIDYE | 1 | 0 | 0 | 1172375 | ACP41105.1 | P03452 |
| TANADTLCIGYHANNSTDTVDTVLE | 1 | 0 | 0 | 1172621 | ACP41105.1 | P03452 |
| TGPPQCDQFLEFSADLIIERREGSD | 1 | 0 | 0 | 1173378 | AGL44438.1 | P03452 |
| TPVHDCNTTCQTPKGAINTSLPFQN | 1 | 0 | 0 | 1174631 | ACP41105.1 | P03452 |
| TSACRRSGSSFYAEMKWLLSNTDNA | 1 | 0 | 0 | 1174935 | AGL44438.1 | P03452 |
| TVDLGQCGLLGTITGPPQCDQFLEF | 1 | 0 | 0 | 1175353 | AGL44438.1 | P03452 |
| TYDHSKYREEAMQNRIQIDPVKLSS | 1 | 0 | 0 | 1175592 | AGL44438.1 | P03452 |
| VFCTVNALPETTADI | 1 | 0 | 0 | 1176720 | P0DTD1.1 | P0DTD1 |
| VLTSHTVMPLSAPTL | 1 | 0 | 0 | 1177791 | P0DTD1.1 | P0DTD1 |
| VVLLSVLQQLRVESS | 1 | 0 | 0 | 1179524 | P0DTC1.1 | P0DTD1 |
| VWSYNAELLVAMENQHTIDLADSEM | 1 | 0 | 0 | 1179659 | AGL44438.1 | P03452 |
| WTLVEPGDKITFEATGNLVVPRYAF | 1 | 0 | 0 | 1179982 | ACP41105.1 | P03452 |
| WYGYHHQNEQGSGYAADLKSTQNAI | 1 | 0 | 0 | 1180001 | ACP41105.1 | P03452 |
| YAADLKSTQNAIDEITNKVNSVIEK | 1 | 0 | 0 | 1180011 | ACP41105.1 | P03452 |
| YKDVILWFSFGASCFILLAIVMGLV | 1 | 0 | 0 | 1180540 | AGL44438.1 | P03452 |
| YQNADTYVFVGSSRYSKKFKPEIAI | 1 | 0 | 0 | 1180945 | ACP41105.1 | P03452 |
| YSTVASSLVLVVSLGAISFWMCSNG | 1 | 0 | 0 | 1181119 | ACP41105.1 | P03452 |
| EGDMFPFLDGLINAPIDFDPVPN | 1 | 0 | 0 | 1204102 | AFV31349.1 | P35253 |

| | | | | | | |
|---|---|---|---|---|---|---|
| ILWREGDMFPFLDGLINAP | 1 | 0 | 0 | 1233642 | AFV31349.1 | P35253 |
| EFVFKNIDGYFKIYS | 1 | 0 | 0 | 1309113 | P0DTC2.1 | P0DTC2 |
| AEIRASANLAATKMSECVLG | 1 | 0 | 0 | 1309418 | QII57161.1 | P0DTC2 |
| APHGVVFLHVTYVPAQEKNF | 1 | 0 | 0 | 1309428 | QII57161.1 | P0DTC2 |
| AYSNNSIAIPTNFTISVTTE | 1 | 0 | 0 | 1309441 | QII57161.1 | P0DTC2 |
| AYYVGYLQPRTFLLKYNENG | 1 | 0 | 0 | 1309442 | QII57161.1 | P0DTC2 |
| CMTSCCSCLKGCCSCGSCCK | 1 | 0 | 0 | 1309443 | QII57161.1 | P0DTC2 |
| DAVRDPQTLEILDITPCSFG | 1 | 0 | 0 | 1309444 | QII57161.1 | P0DTC2 |
| DFGGFNFSQILPDPSKPSKR | 1 | 0 | 0 | 1309447 | QII57161.1 | P0DTC2 |
| DPLSETKCTLKSFTVEKGIY | 1 | 0 | 0 | 1309450 | QII57161.1 | P0DTC2 |
| EHVNNSYECDIPIGAGICAS | 1 | 0 | 0 | 1309461 | QII57161.1 | P0DTC2 |
| EPLVDLPIGINITRFQTLLA | 1 | 0 | 0 | 1309462 | QII57161.1 | P0DTC2 |
| ESNKKFLPFQQFGRDIADTT | 1 | 0 | 0 | 1309464 | QII57161.1 | P0DTC2 |
| FKNHTSPDVDLGDISGINAS | 1 | 0 | 0 | 1309469 | QII57161.1 | P0DTC2 |
| FKNIDGYFKIYSKHTPINLV | 1 | 0 | 0 | 1309470 | QII57161.1 | P0DTC2 |
| GCCSCGSCCKFDEDDSEPVL | 1 | 0 | 0 | 1309475 | QII57161.1 | P0DTC2 |
| GVSVITPGTNTSNQVAVLYQ | 1 | 0 | 0 | 1309480 | QII57161.1 | P0DTC2 |
| GYQPYRVVVLSFELLHAPAT | 1 | 0 | 0 | 1309482 | QII57161.1 | P0DTC2 |
| ILDITPCSFGGVSVITPGTN | 1 | 0 | 0 | 1309492 | QII57161.1 | P0DTC2 |
| ILPVSMTKTSVDCTMYICGD | 1 | 0 | 0 | 1309493 | QII57161.1 | P0DTC2 |
| KLPDDFTGCVIAWNSNNLDS | 1 | 0 | 0 | 1309501 | QII57161.1 | P0DTC2 |
| KNTQEVFAQVKQIYKTPPIK | 1 | 0 | 0 | 1309503 | QII57161.1 | P0DTC2 |
| KSFTVEKGIYQTSNFRVQPT | 1 | 0 | 0 | 1309505 | QII57161.1 | P0DTC2 |
| LGDISGINASVVNIQKEIDR | 1 | 0 | 0 | 1309512 | QII57161.1 | P0DTC2 |
| LNEVAKNLNESLIDLQELGK | 1 | 0 | 0 | 1309518 | QII57161.1 | P0DTC2 |
| LPDPSKPSKRSFIEDLLFNK | 1 | 0 | 0 | 1309519 | QII57161.1 | P0DTC2 |
| LPPAYTNSFTRGVYYPDKVF | 1 | 0 | 0 | 1309522 | QII57161.1 | P0DTC2 |
| LSSNFGAISSVLNDILSRLD | 1 | 0 | 0 | 1309523 | QII57161.1 | P0DTC2 |
| LYENQKLIANQFNSAIGKIQ | 1 | 0 | 0 | 1309524 | QII57161.1 | P0DTC2 |
| NGLTGTGVLTESNKKFLPFQ | 1 | 0 | 0 | 1309531 | QII57161.1 | P0DTC2 |
| NGLTVLPPLLTDEMIAQYTS | 1 | 0 | 0 | 1309532 | QII57161.1 | P0DTC2 |
| NGTKRFDNPVLPFNDGVYFA | 1 | 0 | 0 | 1309533 | QII57161.1 | P0DTC2 |
| NITRFQTLLALHRSYLTPGD | 1 | 0 | 0 | 1309534 | QII57161.1 | P0DTC2 |
| NLCPFGEVFNATRFASVYAW | 1 | 0 | 0 | 1309535 | QII57161.1 | P0DTC2 |
| NNCTFEYVSQPFLMDLEGKQ | 1 | 0 | 0 | 1309536 | QII57161.1 | P0DTC2 |
| PFLMDLEGKQGNFKNLREFV | 1 | 0 | 0 | 1309546 | QII57161.1 | P0DTC2 |
| PTKLNDLCFTNVYADSFVIR | 1 | 0 | 0 | 1309553 | QII57161.1 | P0DTC2 |
| QFCNDPFLGVYYHKNNKSWM | 1 | 0 | 0 | 1309556 | QII57161.1 | P0DTC2 |
| QFGRDIADTTDAVRDPQTLE | 1 | 0 | 0 | 1309557 | QII57161.1 | P0DTC2 |
| QRNFYEPQIITTDNTFVSGN | 1 | 0 | 0 | 1309561 | QII57161.1 | P0DTC2 |
| QTGKIADYNYKLPDDFTGCV | 1 | 0 | 0 | 1309563 | QII57161.1 | P0DTC2 |
| QTRAGCLIGAEHVNNSYECD | 1 | 0 | 0 | 1309564 | QII57161.1 | P0DTC2 |
| QYGDCLGDIAARDLICAQKF | 1 | 0 | 0 | 1309566 | QII57161.1 | P0DTC2 |
| RDLPQGFSALEPLVDLPIGI | 1 | 0 | 0 | 1309567 | QII57161.1 | P0DTC2 |
| RGVYYPDKVFRSSVLHSTQD | 1 | 0 | 0 | 1309570 | QII57161.1 | P0DTC2 |
| RLFRKSNLKPFERDISTEIY | 1 | 0 | 0 | 1309573 | QII57161.1 | P0DTC2 |
| RVYSTGSNVFQTRAGCLIGA | 1 | 0 | 0 | 1309576 | QII57161.1 | P0DTC2 |
| SFELLHAPATVCGPKKSTNL | 1 | 0 | 0 | 1309577 | QII57161.1 | P0DTC2 |
| SFIEDLLFNKVTLADAGFIK | 1 | 0 | 0 | 1309578 | QII57161.1 | P0DTC2 |
| SLIDLQELGKYEQYIKWPWY | 1 | 0 | 0 | 1309581 | QII57161.1 | P0DTC2 |
| STECSNLLLQYGSFCTQLNR | 1 | 0 | 0 | 1309586 | QII57161.1 | P0DTC2 |
| STEKSNIIRGWIFGTTLDSK | 1 | 0 | 0 | 1309587 | QII57161.1 | P0DTC2 |
| TFLLKYNENGTITDAVDCAL | 1 | 0 | 0 | 1309592 | QII57161.1 | P0DTC2 |
| TITDAVDCALDPLSETKCTL | 1 | 0 | 0 | 1309593 | QII57161.1 | P0DTC2 |
| TNFTISVTTEILPVSMTKTS | 1 | 0 | 0 | 1309595 | QII57161.1 | P0DTC2 |
| TTAPAICHDGKAHFPREGVF | 1 | 0 | 0 | 1309597 | QII57161.1 | P0DTC2 |
| TVYDPLQPELDSFKEELDKY | 1 | 0 | 0 | 1309598 | QII57161.1 | P0DTC2 |
| TYVPAQEKNFTTAPAICHDG | 1 | 0 | 0 | 1309599 | QII57161.1 | P0DTC2 |
| TYVTQQLIRAAEIRASANLA | 1 | 0 | 0 | 1309600 | QII57161.1 | P0DTC2 |
| VLNDILSRLDKVEAEVQIDR | 1 | 0 | 0 | 1309604 | QII57161.1 | P0DTC2 |
| VSNGTHWFVTQRNFYEPQII | 1 | 0 | 0 | 1309605 | QII57161.1 | P0DTC2 |
| VTLADAGFIKQYGDCLGDIA | 1 | 0 | 0 | 1309606 | QII57161.1 | P0DTC2 |
| VVNIQKEIDRLNEVAKNLNE | 1 | 0 | 0 | 1309608 | QII57161.1 | P0DTC2 |
| YGSFCTQLNRALTGIAVEQD | 1 | 0 | 0 | 1309617 | QII57161.1 | P0DTC2 |
| YSKHTPINLVRDLPQGFSAL | 1 | 0 | 0 | 1309621 | QII57161.1 | P0DTC2 |
| LYVYKGYQPIDVVRDL | 1 | 0 | 0 | 1309876 | NP_828851.1 | P59594 |
| MYSFVSEETGTLIVNSV | 1 | 0 | 0 | 1309883 | QHO62113.1 | P0DTC4 |
| AIGLALYYPSARIVY | 1 | 0 | 0 | 1310263 | P0DTD1.1 | P0DTD1 |
| ASWFTALTQHGKEDL | 1 | 0 | 0 | 1310292 | P0DTC9.1 | P0DTC9 |
| CQPILLLDQALVSDV | 1 | 0 | 0 | 1310314 | P0DTC1.1 | P0DTD1 |
| DNFKFVCDNIKFADD | 1 | 0 | 0 | 1310333 | P0DTC1.1 | P0DTD1 |
| EAFEKMVSLLSVLLS | 1 | 0 | 0 | 1310344 | P0DTC1.1 | P0DTD1 |
| ECVLGQSKRVDFCGK | 1 | 0 | 0 | 1310345 | YP_009724390.1 | P0DTC2 |
| EETFKLSYGIATVRE | 1 | 0 | 0 | 1310350 | P0DTD1.1 | P0DTD1 |
| EFYAYLRKHFSMMIL | 1 | 0 | 0 | 1310354 | P0DTD1.1 | P0DTD1 |
| EIIKSQDLSVVSKVV | 1 | 0 | 0 | 1310358 | P0DTD1.1 | P0DTD1 |
| ELEDFIPMDSTVKNY | 1 | 0 | 0 | 1310364 | P0DTD1.1 | P0DTD1 |
| EWFLAYILFTRFFYV | 1 | 0 | 0 | 1310378 | P0DTC1.1 | P0DTD1 |
| FKIYSKHTPINLVRD | 1 | 0 | 0 | 1310401 | P0DTC2.1 | P0DTC2 |
| FLGIITTVAAFHQEC | 1 | 0 | 0 | 1310404 | YP_009724396.1 | P0DTC8 |
| FMGRIRSVYPVASPN | 1 | 0 | 0 | 1310410 | P0DTC1.1 | P0DTD1 |
| FNGLTVLPPLLTDEM | 1 | 0 | 0 | 1310415 | P0DTC2.1 | P0DTC2 |
| FVDGVPFVVSTGYHF | 1 | 0 | 0 | 1310424 | P0DTD1.1 | P0DTD1 |
| GAAAYYVGYLQPRTF | 1 | 0 | 0 | 1310431 | P0DTC2.1 | P0DTC2 |
| GDYFVLTSHTVMPLS | 1 | 0 | 0 | 1310439 | P0DTD1.1 | P0DTD1 |
| HSLSHFVNLDNLRAN | 1 | 0 | 0 | 1310475 | P0DTC1.1 | P0DTD1 |
| IAGLIAIVMVTIMLC | 1 | 0 | 0 | 1310479 | P0DTC2.1 | P0DTC2 |
| IGFDYVYNPFMIDVQ | 1 | 0 | 0 | 1310486 | P0DTD1.1 | P0DTD1 |
| IGINITRFQTLLALH | 1 | 0 | 0 | 1310487 | P0DTC2.1 | P0DTC2 |
| ILASFSASTSAFVET | 1 | 0 | 0 | 1310495 | P0DTC1.1 | P0DTD1 |
| IMQLFFSYFAVHFIS | 1 | 0 | 0 | 1310500 | P0DTC1.1 | P0DTD1 |

| | | | | | | |
|---|---|---|---|---|---|---|
| INVFAFPFTIYSLLL | 1 | 0 | 0 | 1310502 | QNU10373.1 | A0A663DJA2 |
| IQWMVMFTPLVPFWI | 1 | 0 | 0 | 1310505 | P0DTC1.1 | P0DTD1 |
| IWLGFIAGLIAIVMV | 1 | 0 | 0 | 1310516 | YP_009724390.1 | P0DTC2 |
| IWNLDYIINLIIKNL | 1 | 0 | 0 | 1310517 | P0DTC6.1 | P0DTC6 |
| KIADKYVRNLQHRLY | 1 | 0 | 0 | 1310537 | P0DTD1.1 | P0DTD1 |
| LIIMRTFKVSIWNLD | 1 | 0 | 0 | 1310581 | P0DTC6.1 | P0DTC6 |
| LIISVTSNYSGVVTT | 1 | 0 | 0 | 1310582 | P0DTC1.1 | P0DTD1 |
| LMIERFVSLAIDAYP | 1 | 0 | 0 | 1310604 | P0DTD1.1 | P0DTD1 |
| LMSFTVLCLTPVYSF | 1 | 0 | 0 | 1310605 | P0DTC1.1 | P0DTD1 |
| LNNDYYRSLPGVFCG | 1 | 0 | 0 | 1310607 | P0DTC1.1 | P0DTD1 |
| LQQLRVESSSKLWAQ | 1 | 0 | 0 | 1310615 | P0DTC1.1 | P0DTD1 |
| LYQDVNCTEVPVAIH | 1 | 0 | 0 | 1310635 | P0DTC2.1 | P0DTC2 |
| MGIIAMSAFAMMFVK | 1 | 0 | 0 | 1310640 | P0DTC1.1 | P0DTD1 |
| NMFITREEAIRHVRA | 1 | 0 | 0 | 1310674 | P0DTD1.1 | P0DTD1 |
| NRNRFLYIIKLIFLW | 1 | 0 | 0 | 1310685 | P0DTC5.1 | P0DTC5 |
| NSFSGYLKLTDNVYI | 1 | 0 | 0 | 1310687 | P0DTC1.1 | P0DTD1 |
| NVFQTRAGCLIGAEH | 1 | 0 | 0 | 1310695 | P0DTC2.1 | P0DTC2 |
| NYEDLLIRKSNHNFL | 1 | 0 | 0 | 1310702 | P0DTC1.1 | P0DTD1 |
| PRRARSVASQSIIAY | 1 | 0 | 0 | 1310725 | P0DTC2.1 | P0DTC2 |
| PSGTWLTYTGAIKLD | 1 | 0 | 0 | 1310726 | P0DTC9.1 | P0DTC9 |
| QEEVQELYSPIFLIV | 1 | 0 | 0 | 1310732 | P0DTC7.1 | P0DTC7 |
| QLGGLHLLIGLAKRF | 1 | 0 | 0 | 1310738 | P0DTD1.1 | P0DTD1 |
| QPTESIVRFPNITNL | 1 | 0 | 0 | 1310745 | P0DTC2.1 | P0DTC2 |
| QPYVFIKRSDARTAP | 1 | 0 | 0 | 1310748 | P0DTC1.1 | P0DTD1 |
| RAMPNMLRIMASLVL | 1 | 0 | 0 | 1310760 | P0DTD1.1 | P0DTD1 |
| RFFYVLGLAAIMQLF | 1 | 0 | 0 | 1310766 | P0DTC1.1 | P0DTD1 |
| SHNIALIWNVKDFMS | 1 | 0 | 0 | 1310801 | P0DTC1.1 | P0DTD1 |
| TITSGWTFGAGAALQ | 1 | 0 | 0 | 1310847 | P0DTC2.1 | P0DTC2 |
| TLACFVLAAVYRINW | 1 | 0 | 0 | 1310849 | P0DTC5.1 | P0DTC5 |
| VDDIVKTDGTLMIER | 1 | 0 | 0 | 1310878 | P0DTD1.1 | P0DTD1 |
| VLKKLKKSLNVAKSE | 1 | 0 | 0 | 1310897 | P0DTC1.1 | P0DTD1 |
| VPFWITIAYIICIST | 1 | 0 | 0 | 1310912 | P0DTC1.1 | P0DTD1 |
| WEIVKFISTCACEIV | 1 | 0 | 0 | 1310937 | P0DTC1.1 | P0DTD1 |
| WICLLQFAYANRNRF | 1 | 0 | 0 | 1310941 | YP_009724393.1 | P0DTC5 |
| WLIINLVQMAPISAM | 1 | 0 | 0 | 1310942 | P0DTC1.1 | P0DTD1 |
| YIRVGARKSAPLIEL | 1 | 0 | 0 | 1310959 | P0DTC8.1 | P0DTC8 |
| YRRLISMMGFKMNYQ | 1 | 0 | 0 | 1310976 | P0DTD1.1 | P0DTD1 |
| YSLRLIDAMMFTSDL | 1 | 0 | 0 | 1310977 | P0DTC1.1 | P0DTD1 |
| CCSCGSCCKFDEDDSEPVLKGVKL | 1 | 0 | 0 | 1311657 | QKE11719.1 | P0DTC2 |
| DAYKTFPPTEPKKDKKKKADETQA | 1 | 0 | 0 | 1311663 | QHD43423.2 | P0DTC9 |
| FAQVKQIYKTPPIKDFGGFNFSQI | 1 | 0 | 0 | 1311674 | QKE11719.1 | P0DTC2 |
| KGQQQQGQTVTKKSAAEASKKPRQ | 1 | 0 | 0 | 1311721 | QHD43423.2 | P0DTC9 |
| KSFDLGDELGTDPYEDFQENWNTKH | 1 | 0 | 0 | 1311732 | QHD43415.1 | P0DTD1 |
| LLLLDRLNQLESKMSGKGQQQQGQ | 1 | 0 | 0 | 1311743 | QHD43423.2 | P0DTC9 |
| LPQRQKKQQTVTLLPAADLDDFSK | 1 | 0 | 0 | 1311750 | QHD43423.2 | P0DTC9 |
| NNAAIVLQLPQGTTLPKGFYAEGS | 1 | 0 | 0 | 1311774 | QHD43423.2 | P0DTC9 |
| PDPSKPSKRSFIEDLLFNKVTLAD | 1 | 0 | 0 | 1311782 | QKE11719.1 | P0DTC2 |
| TGTGVLTESNKKFLPFQQFGRDIA | 1 | 0 | 0 | 1311868 | QKE11719.1 | P0DTC2 |
| YDPLQPELDSFKEELDKYFKNHTS | 1 | 0 | 0 | 1311941 | QKE11719.1 | P0DTC2 |
| AIAMACLVGLMWLSY | 1 | 0 | 0 | 1312122 | P0DTC5.1 | P0DTC5 |
| AVYRINWITGGIAIA | 1 | 0 | 0 | 1312246 | P0DTC5.1 | P0DTC5 |
| CLLQFAYANRNRFLY | 1 | 0 | 0 | 1312263 | P0DTC5.1 | P0DTC5 |
| EILPVSMTKTSVDCT | 1 | 0 | 0 | 1312357 | P0DTC2.1 | P0DTC2 |
| ELVIGAVILRGHLRI | 1 | 0 | 0 | 1312367 | P0DTC5.1 | P0DTC5 |
| FLYIIKLIFLWLLWP | 1 | 0 | 0 | 1312469 | P0DTC5.1 | P0DTC5 |
| FPREGVFVSNGTHWF | 1 | 0 | 0 | 1312481 | P0DTC2.1 | P0DTC2 |
| GAVILRGHLRIAGHH | 1 | 0 | 0 | 1312531 | P0DTC5.1 | P0DTC5 |
| GFLFLTWICLLQFAY | 1 | 0 | 0 | 1312541 | P0DTC5.1 | P0DTC5 |
| HGTILTRPLLESELV | 1 | 0 | 0 | 1312642 | P0DTC5.1 | P0DTC5 |
| IKLIFLWLLWPVTLA | 1 | 0 | 0 | 1312710 | YP_009724393.1 | P0DTC5 |
| ILTALRLCAYCCNIV | 1 | 0 | 0 | 1312738 | P0DTC4.1 | P0DTC4 |
| INITRFQTLLALHRS | 1 | 0 | 0 | 1312746 | P0DTC2.1 | P0DTC2 |
| INWITGGIAIAMACL | 1 | 0 | 0 | 1312747 | P0DTC5.1 | P0DTC5 |
| LDPLSETKCTLKSFT | 1 | 0 | 0 | 1312959 | P0DTC2.1 | P0DTC2 |
| LEQWNLVIGFLFLTW | 1 | 0 | 0 | 1312964 | P0DTC5.1 | P0DTC5 |
| LLESELVIGAVILRG | 1 | 0 | 0 | 1312993 | YP_009724393.1 | P0DTC5 |
| LRGHLRIAGHHLGRC | 1 | 0 | 0 | 1313076 | YP_009724393.1 | P0DTC5 |
| LRLCAYCCNIVNVSL | 1 | 0 | 0 | 1313078 | YP_009724392.1 | P0DTC4 |
| LSFELLHAPATVCGP | 1 | 0 | 0 | 1313083 | P0DTC2.1 | P0DTC2 |
| LTRPLLESELVIGAV | 1 | 0 | 0 | 1313109 | P0DTC5.1 | P0DTC5 |
| NSSPDDQIGYYRRAT | 1 | 0 | 0 | 1313250 | QHO62115.1 | P0DTC9 |
| NVPLHGTILTRPLLE | 1 | 0 | 0 | 1313260 | YP_009724393.1 | P0DTC5 |
| NVVIKVCEFQFCNDP | 1 | 0 | 0 | 1313262 | P0DTC2.1 | P0DTC2 |
| RTFLLKYNENGTITD | 1 | 0 | 0 | 1313495 | P0DTC2.1 | P0DTC2 |
| RTRSMWSFNPETNIL | 1 | 0 | 0 | 1313501 | P0DTC5.1 | P0DTC5 |
| TNLCPFGEVFNATRF | 1 | 0 | 0 | 1313768 | P0DTC2.1 | P0DTC2 |
| VSEETGTLIVNSVLL | 1 | 0 | 0 | 1313919 | P0DTC4.1 | P0DTC4 |
| VSLVKPSFYVYSRVK | 1 | 0 | 0 | 1313920 | YP_009724392.1 | P0DTC4 |
| VTYVPAQEKNFTTAP | 1 | 0 | 0 | 1313930 | QHO62112.1 | P0DTC2 |
| AVMSLKEGQINDMIL | 1 | 0 | 0 | 1315054 | P0DTD1.1 | P0DTD1 |
| EDFLEYHDVRVVLDF | 1 | 0 | 0 | 1315736 | P0DTC8.1 | P0DTC8 |
| ELHLSWEVGKPRPPL | 1 | 0 | 0 | 1315881 | P0DTD1.1 | P0DTD1 |
| FSQILPDPSKPSKRSFIE | 1 | 0 | 0 | 1316945 | P0DTC2.1 | P0DTC2 |
| GSLVVRCSFYEDFLE | 1 | 0 | 0 | 1317791 | YP_009724396.1 | P0DTC8 |
| GYKKPASRELKVTFF | 1 | 0 | 0 | 1317914 | P0DTC1.1 | P0DTD1 |
| HTTDPSFLGRYMSAL | 1 | 0 | 0 | 1318193 | P0DTC1.1 | P0DTD1 |
| IDAYPLTKHPNQEYA | 1 | 0 | 0 | 1318319 | P0DTD1.1 | P0DTD1 |
| IGNYTVSCLPFTINC | 1 | 0 | 0 | 1318448 | YP_009724396.1 | P0DTC8 |
| LTQHGKEDLKFPRGQ | 1 | 0 | 0 | 1321406 | P0DTC9.1 | P0DTC9 |
| PCPIHFYSKWYIRVG | 1 | 0 | 0 | 1322874 | YP_009724396.1 | P0DTC8 |
| PRPPLNRNYVFTGYR | 1 | 0 | 0 | 1323024 | P0DTD1.1 | P0DTD1 |

| | | | | | | |
|---|---|---|---|---|---|---|
| QEPKLGSLVVRCSFY | 1 | 0 | 0 | 1323204 | YP_009724396.1 | P0DTC8 |
| RCSFYEDFLEYHDVR | 1 | 0 | 0 | 1323753 | YP_009724396.1 | P0DTC8 |
| RPLLESELVIGAVIL | 1 | 0 | 0 | 1324008 | P0DTC5.1 | P0DTC5 |
| SLQSCTQHQPYVVDD | 1 | 0 | 0 | 1324722 | YP_009724396.1 | P0DTC8 |
| TDTGVEHVTFFIYNK | 1 | 0 | 0 | 1325392 | P0DTC3.1 | P0DTC3 |
| TQHQPYVVDDPCPIH | 1 | 0 | 0 | 1325852 | YP_009724396.1 | P0DTC8 |
| TTVAAFHQECSLQSC | 1 | 0 | 0 | 1325992 | YP_009724396.1 | P0DTC8 |
| WDLTAFGLVAEWFLA | 1 | 0 | 0 | 1327752 | P0DTC1.1 | P0DTD1 |
| WHYTSYTIVGALYVT | 1 | 0 | 0 | 1327764 | APF29071.1 | A0A0P0G321 |
| YDANYFLCWHTNCYD | 1 | 0 | 0 | 1327904 | P0DTC3.1 | P0DTC3 |
| YRINWITGGIAIAMA | 1 | 0 | 0 | 1328509 | P0DTC5.1 | P0DTC5 |
| YVVDDPCPIHFYSKW | 1 | 0 | 0 | 1328882 | YP_009724396.1 | P0DTC8 |
| ASVVNIQKEIDRLNEV | 1 | 0 | 0 | 1329153 | YP_009724390.1 | P0DTC2 |
| LKSFTVEKGIYQTSNF | 1 | 0 | 0 | 1329686 | YP_009724390.1 | P0DTC2 |
| NASVVNIQKEIDRLNE | 1 | 0 | 0 | 1329789 | YP_009724390.1 | P0DTC2 |
| TLKSFTVEKGIYQTSN | 1 | 0 | 0 | 1330210 | YP_009724390.1 | P0DTC2 |
| AGCLIGAEHVNNSYECDIPIGA | 1 | 0 | 0 | 1330816 | YP_009724390.1 | P0DTC2 |
| DNQDLNGNWYDFGDF | 1 | 0 | 0 | 1331320 | P0DTD1.1 | P0DTD1 |
| FKDQVILLNKHIDAYK | 1 | 0 | 0 | 1331955 | YP_009724397.2 | P0DTC9 |
| GLPGTILRTTNGDFL | 1 | 0 | 0 | 1332093 | YP_009724389.1 | P0DTD1 |
| LDSKVGGNYNYLYRLFRKSNLKPF | 1 | 0 | 0 | 1332591 | YP_009724390.1 | P0DTC2 |
| MKFLVFLGIITTVAA | 1 | 0 | 0 | 1332793 | YP_009724396.1 | P0DTC8 |
| QTQGNFGDQELIRQGTDYKHW | 1 | 0 | 0 | 1333233 | YP_009724397.2 | P0DTC9 |
| VATEGALNTPKDHIGTRNPA | 1 | 0 | 0 | 1334138 | YP_009724397.2 | P0DTC9 |
| ANNAAIVLQLPQGTTLPKGFY | 1 | 0 | 0 | 1334455 | QJF11943.1 | P0DTC9 |
| NNNAATVLQLPQGTTLPKGFY | 1 | 0 | 0 | 1334471 | AAP49024.1 | P59595 |
| PIKDFGGFNFSQILPDPSKP | 1 | 0 | 0 | 1334476 | QHO62112.1 | P0DTC2 |
| TWFHAIHVSGTNGTKRFDNPVLP | 1 | 0 | 0 | 1334486 | QHO62112.1 | P0DTC2 |
| YIIKLIFLWLLWPVTL | 1 | 0 | 0 | 1334490 | QHO62114.1 | P0DTC5 |
| IYQAGSTPCNGVEGFNCYFPLQSY | 1 | 0 | 0 | 1337002 | YP_009724390.1 | P0DTC2 |
| DALFAYTKRNVIPTI | 1 | 0 | 0 | 1339900 | QSE17078.1 | P0DTD1 |
| EYYHTTDPSFLGRYM | 1 | 0 | 0 | 1340790 | QSE17078.1 | P0DTD1 |
| FSAVGNICYTPSKLI | 1 | 0 | 0 | 1341076 | QSE17078.1 | P0DTD1 |
| GNCDTLKEILVTYNC | 1 | 0 | 0 | 1341474 | QSE17078.1 | P0DTD1 |
| GVLVPHVGEIPVAYR | 1 | 0 | 0 | 1341587 | QSE17078.1 | P0DTD1 |
| HIQWMVMFTPLVPFW | 1 | 0 | 0 | 1341640 | QSE17078.1 | P0DTD1 |
| HRLYECLYRNRDVDT | 1 | 0 | 0 | 1341683 | QSE17078.1 | P0DTD1 |
| IFTQTTETAHSCNVN | 1 | 0 | 0 | 1341806 | QSE17078.1 | P0DTD1 |
| LQLCTFTRSTNSRIK | 1 | 0 | 0 | 1343153 | QSE17078.1 | P0DTD1 |
| LYQPPQTSITSAVLQ | 1 | 0 | 0 | 1343343 | QSU72953.1 | P0DTD1 |
| PQTSITSAVLQSGFR | 1 | 0 | 0 | 1343953 | QSU72953.1 | P0DTD1 |
| QIELKFNPPALQDAY | 1 | 0 | 0 | 1344287 | QSE17078.1 | P0DTD1 |
| QLREPMLQSADAQSF | 1 | 0 | 0 | 1344330 | QSE17078.1 | P0DTD1 |
| QQLRVESSSKLWAQC | 1 | 0 | 0 | 1344382 | QSE17078.1 | P0DTD1 |
| TTNIVTRCLNRVCTN | 1 | 0 | 0 | 1345919 | QSE17078.1 | P0DTD1 |
| VESSSKLWAQCVQLH | 1 | 0 | 0 | 1346061 | QSE17078.1 | P0DTD1 |
| YVQIPTTCANDPVGF | 1 | 0 | 0 | 1346663 | QSE17078.1 | P0DTD1 |
| AAYSRYRIGNYKLNTD | 1 | 0 | 0 | 1377315 | YP_009724393.1 | P0DTC5 |
| ADSCLQGDKCNIFANF | 1 | 0 | 0 | 1377371 | YP_009555241.1 | P36334 |
| ADSVVEVVTTSLTPCG | 1 | 0 | 0 | 1377374 | YP_009555238.1 | P0C6X6 |
| ADSYYSYIMPMLTMCH | 1 | 0 | 0 | 1377375 | YP_009555238.1 | P0C6X6 |
| AEVAVKMFDAYVNTFS | 1 | 0 | 0 | 1377423 | YP_009724389.1 | P0DTD1 |
| AGDSGFAAYSRYRIGN | 1 | 0 | 0 | 1377468 | YP_009724393.1 | P0DTC5 |
| AGHHLGRCDIKDLPKE | 1 | 0 | 0 | 1377483 | YP_009724393.1 | P0DTC5 |
| AIKLDDKDPNFKDQVI | 1 | 0 | 0 | 1377559 | YP_009724397.2 | P0DTC9 |
| AISASIGDIIQRLDVLEQD | 1 | 0 | 0 | 1377571 | YP_009047204.1 | K9N5Q8 |
| APHGARSLTPCTCGSS | 1 | 0 | 0 | 1377751 | AGN33396.1 | P27958 |
| ARMAGNGGDAALALLL | 1 | 0 | 0 | 1377826 | YP_009724397.2 | P0DTC9 |
| ASQRVAGDSGFAAYSR | 1 | 0 | 0 | 1377882 | YP_009724393.1 | P0DTC5 |
| AYSRYRIGNYKLNTDH | 1 | 0 | 0 | 1378028 | YP_009724393.1 | P0DTC5 |
| CSVEDFNVWALSNGFS | 1 | 0 | 0 | 1378250 | YP_009555238.1 | P0C6X6 |
| CTIAQLTDAALSIKEN | 1 | 0 | 0 | 1378263 | YP_009555239.1 | Q80872 |
| DAALALLLLDRLNQLE | 1 | 0 | 0 | 1378299 | YP_009724397.2 | P0DTC9 |
| DDKDPSFKDNVILLNKH | 1 | 0 | 0 | 1378399 | QIQ54056.1 | P59595 |
| DESQPLPQRQKKQQTVT | 1 | 0 | 0 | 1378496 | QIQ54056.1 | P59595 |
| DFSRVSAKPPPGDQFKH | 1 | 0 | 0 | 1378563 | YP_009724389.1 | P0DTD1 |
| DGVVCTEIDPKLDGYYK | 1 | 0 | 0 | 1378647 | QIQ54047.1 | P0C6X7 |
| DKCVNFNFNGLTGTGVL | 1 | 0 | 0 | 1378736 | QIQ54048.1 | P59594 |
| DMDDFSRQLQNSMSGAS | 1 | 0 | 0 | 1378859 | YP_009825061.1 | P59595 |
| DSGFAAYSRYRIGNYK | 1 | 0 | 0 | 1379081 | YP_009724393.1 | P0DTC5 |
| DTSYECDIPIGAGICA | 1 | 0 | 0 | 1379190 | YP_009825051.1 | P59594 |
| ECSKASSRSAIEDLLFDKVKLSDV | 1 | 0 | 0 | 1379358 | YP_009555241.1 | P36334 |
| EGIIWVATEGALNTPK | 1 | 0 | 0 | 1379562 | QIQ54056.1 | P59595 |
| EGIVWVATEGALNTPKD | 1 | 0 | 0 | 1379563 | YP_009825061.1 | P59595 |
| EVGSKSPIQYIDIGNYTVSC | 1 | 0 | 0 | 1380126 | QHR63307.1 | Q3LZX5 |
| EVPMAIHAEQLTPAWRVYS | 1 | 0 | 0 | 1380140 | QIQ54048.1 | P59594 |
| FAAYSRYRIGNYKLNT | 1 | 0 | 0 | 1380209 | YP_009724393.1 | P0DTC5 |
| FAVRKEGQDVIFSQFD | 1 | 0 | 0 | 1380243 | YP_009555238.1 | P0C6X6 |
| FELFAKRKVGLTPPLS | 1 | 0 | 0 | 1380313 | NP_073549.1 | A0A0P0LKV0 |
| FGAALAMEKVNECVKS | 1 | 0 | 0 | 1380354 | YP_173238.1 | A0A140H1H1 |
| FKKNAWNIGEPKSILS | 1 | 0 | 0 | 1380467 | QIQ54047.1 | P0C6X7 |
| FLQILPDPSKPSKRSFIED | 1 | 0 | 0 | 1380525 | QIQ54048.1 | P59594 |
| FNFLQILPDPSKPSKR | 1 | 0 | 0 | 1380541 | QIQ54048.1 | P59594 |
| FNFNGLTGTGVLTESN | 1 | 0 | 0 | 1380542 | YP_009724390.1 | P0DTC2 |
| GASQRVAGDSGFAAYS | 1 | 0 | 0 | 1380881 | YP_009724393.1 | P0DTC5 |
| GDIILKPANDGLKITE | 1 | 0 | 0 | 1380937 | QHR63299.2 | P0C6W2 |
| GDSGFAAYSRYRIGNY | 1 | 0 | 0 | 1380962 | YP_009724393.1 | P0DTC5 |
| GEIKDATPSDFVRATA | 1 | 0 | 0 | 1381002 | YP_009724391.1 | P0DTC3 |
| GERSGARSKQRRPQGL | 1 | 0 | 0 | 1381027 | YP_009724397.2 | P0DTC9 |
| GETALALLLLDRLNQLESKVSGKG | 1 | 0 | 0 | 1381034 | YP_009825061.1 | P59595 |
| GFAAYSRYRIGNYKLN | 1 | 0 | 0 | 1381044 | YP_009724393.1 | P0DTC5 |

| | | | | | | |
|---|---|---|---|---|---|---|
| GGDAALALLLLDRLNQ | 1 | 0 | 0 | 1381105 | YP_009724397.2 | P0DTC9 |
| GGDGKMKDLSPRWYFYY | 1 | 0 | 0 | 1381108 | YP_009724397.2 | P0DTC9 |
| GGDGKMKELSPRWYFY | 1 | 0 | 0 | 1381109 | YP_009825061.1 | P59595 |
| GGFNFLQILPDPSKPS | 1 | 0 | 0 | 1381121 | QIQ54048.1 | P59594 |
| GHHLGRCDIKDLPKEI | 1 | 0 | 0 | 1381236 | YP_009724393.1 | P0DTC5 |
| GHSLGRCDIKDLPKEITVATSRTLS | 1 | 0 | 0 | 1381252 | YP_009825055.1 | P59596 |
| GNGGDAALALLLLDRL | 1 | 0 | 0 | 1381497 | YP_009724397.2 | P0DTC9 |
| GQGLNGYTVEEARTVL | 1 | 0 | 0 | 1381626 | QHR63299.2 | P0C6W2 |
| GSRSARSAIEDLLFDKVTIAD | 1 | 0 | 0 | 1381810 | YP_009047204.1 | K9N5Q8 |
| GVVNPAMDPIYDEPTTT | 1 | 0 | 0 | 1381942 | QIQ54049.1 | P59632 |
| HHLGRCDIKDLPKEIT | 1 | 0 | 0 | 1382079 | YP_009724393.1 | P0DTC5 |
| HLGRCDIKDLPKEITV | 1 | 0 | 0 | 1382130 | YP_009724393.1 | P0DTC5 |
| IHADQLTPAWRIYSTGNNV | 1 | 0 | 0 | 1382573 | YP_009825051.1 | P59594 |
| IHADQLTPTWRVYSTGSNVFQTR | 1 | 0 | 0 | 1382575 | YP_009724390.1 | P0DTC2 |
| IHAEQLTPAWRVYSAGANVFQT | 1 | 0 | 0 | 1382576 | QIQ54048.1 | P59594 |
| ILLDDFVALVKSLNLNC | 1 | 0 | 0 | 1382727 | YP_009555238.1 | P0C6X6 |
| ISVQELRYMNANGLRP | 1 | 0 | 0 | 1383031 | YP_009555238.1 | P0C6X6 |
| ISYEEQDALFALTKRN | 1 | 0 | 0 | 1383032 | YP_003766.2 | H9EJ42 |
| ITADIELALLRA | 1 | 0 | 0 | 1383036 | ASY99815.1 | K9N7A1 |
| ITFDNLKTLLSLREVR | 1 | 0 | 0 | 1383044 | YP_009724389.1 | P0DTD1 |
| ITIDKFAIPNGRKVDL | 1 | 0 | 0 | 1383054 | YP_009555241.1 | P36334 |
| ITISSFKWDLTAFGLV | 1 | 0 | 0 | 1383059 | YP_009724389.1 | P0DTD1 |
| ITKDEAIKRVRGWVGF | 1 | 0 | 0 | 1383061 | YP_173236.1 | A0A140H1G9 |
| ITREEAIKHVRAWVGF | 1 | 0 | 0 | 1383083 | QIQ54047.1 | P0C6X7 |
| ITREEAIRHVRAWIGF | 1 | 0 | 0 | 1383084 | YP_009724389.1 | P0DTD1 |
| ITREEAVRHVRAWIGF | 1 | 0 | 0 | 1383085 | QHR63299.2 | P0C6W2 |
| ITRVESENKVVVLDSFDPLV | 1 | 0 | 0 | 1383089 | ANN11859.1 | P27958 |
| ITVEAAIELSKELKRP | 1 | 0 | 0 | 1383099 | YP_173236.1 | A0A140H1G9 |
| ITVEAALDLSKELKRP | 1 | 0 | 0 | 1383100 | YP_009555238.1 | P0C6X6 |
| ITVPEEFASHVANFQK | 1 | 0 | 0 | 1383102 | YP_009047202.1 | K9N7C7 |
| ITVSQIQLSLLKVTAF | 1 | 0 | 0 | 1383103 | YP_009825053.1 | P59633 |
| KDATPSDFVRATATIP | 1 | 0 | 0 | 1383237 | YP_009724391.1 | P0DTC3 |
| KDEDDNLIDSYFVVKR | 1 | 0 | 0 | 1383241 | YP_009724389.1 | P0DTD1 |
| KGIYQTSNFRVQPTIS | 1 | 0 | 0 | 1383389 | QIQ54048.1 | P59594 |
| KIVDEPEEHVQIHTIDG | 1 | 0 | 0 | 1383492 | YP_009724391.1 | P0DTC3 |
| KKQPTVTLLPAADMDDFSRQLQNSM | 1 | 0 | 0 | 1383528 | YP_009825061.1 | P59595 |
| KLDDKDPNFKDQVILLNKH | 1 | 0 | 0 | 1383548 | YP_009724397.2 | P0DTC9 |
| KLDDKDPSFKDNVILL | 1 | 0 | 0 | 1383549 | QIQ54056.1 | P59595 |
| KLNTDHSSSSDNIALLVQ | 1 | 0 | 0 | 1383583 | YP_009724393.1 | P0DTC5 |
| KNAWNIGEPKSILSPLY | 1 | 0 | 0 | 1383655 | QIQ54047.1 | P0C6X7 |
| KQGEIKDATPSDFVRA | 1 | 0 | 0 | 1383768 | YP_009724391.1 | P0DTC3 |
| KSAPLIELCVDEAGSK | 1 | 0 | 0 | 1383843 | YP_009724396.1 | P0DTC8 |
| KTIEDALRNSARLESA | 1 | 0 | 0 | 1383944 | NP_073551.1 | A0A0P0K6L9 |
| KTVGELGDVRETMSHL | 1 | 0 | 0 | 1383976 | QIQ54047.1 | P0C6X7 |
| KTVGELGDVRETMSYL | 1 | 0 | 0 | 1383977 | YP_009724389.1 | P0DTD1 |
| KTVGELGDVRETMTHL | 1 | 0 | 0 | 1383978 | NP_828849.7 | P0C6X7 |
| KTVLGEYVFDKSELTN | 1 | 0 | 0 | 1383981 | YP_009555238.1 | P0C6X6 |
| LALLLLDRLNQLESKM | 1 | 0 | 0 | 1384139 | YP_009724397.2 | P0DTC9 |
| LAVIGDFNCTNFAIND | 1 | 0 | 0 | 1384168 | YP_173238.1 | A0A140H1H1 |
| LCVDEAGSKSPIQYIDIGNY | 1 | 0 | 0 | 1384204 | YP_009724396.1 | P0DTC8 |
| LCVDEVGSKSPIQYIDI | 1 | 0 | 0 | 1384205 | QHR63307.1 | Q3LZX5 |
| LDGEVITFDNLKTLLS | 1 | 0 | 0 | 1384227 | YP_009724389.1 | P0DTD1 |
| LEDLLFSKVVTSGLGT | 1 | 0 | 0 | 1384293 | YP_003767.1 | A0A0P0G321 |
| LESKMSGKGQQQQSQTVTKK | 1 | 0 | 0 | 1384356 | QHR63308.1 | Q3LZX4 |
| LFQTTVELQGLIDQIN | 1 | 0 | 0 | 1384395 | YP_003767.1 | A0A0P0G321 |
| LGASQRVAGDSGFAAY | 1 | 0 | 0 | 1384412 | YP_009724393.1 | P0DTC5 |
| LGNSTGIDFQDELDEFFKNVSTSIP | 1 | 0 | 0 | 1384455 | YP_009047204.1 | K9N5Q8 |
| LKSLTVEKGIYQTSNF | 1 | 0 | 0 | 1384613 | QIQ54048.1 | P59594 |
| LKYFGGFNFSQILPDPL | 1 | 0 | 0 | 1384627 | YP_009825051.1 | P59594 |
| LLALHRSYLTPGDSSS | 1 | 0 | 0 | 1384629 | YP_009724390.1 | P0DTC2 |
| LNALESKMSGKGSQQQSQTVTK | 1 | 0 | 0 | 1384735 | QIQ54056.1 | P59595 |
| LNQLESKMSGKGQQQQS | 1 | 0 | 0 | 1384789 | QHR63308.1 | Q3LZX4 |
| LSIADLACAQYYNGIM | 1 | 0 | 0 | 1385058 | NP_073551.1 | A0A0P0K6L9 |
| LSIEEFADVVKEQVSD | 1 | 0 | 0 | 1385059 | YP_009047202.1 | K9N7C7 |
| LTITELLQFVTDPSLI | 1 | 0 | 0 | 1385165 | YP_003766.2 | H9EJ42 |
| LTITELLQFVTDPTLI | 1 | 0 | 0 | 1385166 | NP_073549.1 | A0A0P0LKV0 |
| LTIVDIPQSLTFSYDG | 1 | 0 | 0 | 1385167 | YP_009047202.1 | K9N7C7 |
| LTQTFTSMLTNYFLDDVEM | 1 | 0 | 0 | 1385188 | YP_173236.1 | A0A140H1G9 |
| LTVGDIFVLTSHSVAT | 1 | 0 | 0 | 1385202 | YP_009047202.1 | K9N7C7 |
| LTVSDDDFVSAVANAH | 1 | 0 | 0 | 1385206 | YP_003766.2 | H9EJ42 |
| MADNGTITVEELKKLLEQWNLVI | 1 | 0 | 0 | 1385326 | QHR63303.1 | Q3LZX9 |
| MADSNGTITVEELKKLLEQWNLVI | 1 | 0 | 0 | 1385329 | YP_009724393.1 | P0DTC5 |
| MSANNGTITVEELKKLLEQWNLVI | 1 | 0 | 0 | 1385614 | QIQ54051.1 | P59596 |
| MTLDLLKDIKFVIPAL | 1 | 0 | 0 | 1385637 | YP_009047202.1 | K9N7C7 |
| NAAIVLQLPQGTALPKGFYAEGS | 1 | 0 | 0 | 1385685 | QIQ54056.1 | P59595 |
| NGGDAALALLLLDRLN | 1 | 0 | 0 | 1385897 | YP_009724397.2 | P0DTC9 |
| NGKTVLGEYVFDKSEL | 1 | 0 | 0 | 1385911 | YP_009555238.1 | P0C6X6 |
| NKDGIIWVATEGALNTP | 1 | 0 | 0 | 1386017 | YP_009724397.2 | P0DTC9 |
| NQRSAPRITFGGPTDSTDNN | 1 | 0 | 0 | 1386304 | YP_009825061.1 | P59595 |
| NTVSELVYENKFVPVKE | 1 | 0 | 0 | 1386438 | NP_073549.1 | A0A0P0LKV0 |
| PEAGLPYGANKDGIIWVAT | 1 | 0 | 0 | 1386574 | YP_009724397.2 | P0DTC9 |
| PLQPELDSFKEELDKYFKNHTSPDV | 1 | 0 | 0 | 1386752 | P0DTC2.1 | P0DTC2 |
| PPIKDFGGFNFSQILP | 1 | 0 | 0 | 1386807 | YP_009724390.1 | P0DTC2 |
| QDVNCTDVSTAIHADQLTPAWRIYS | 1 | 0 | 0 | 1387042 | YP_009825051.1 | P59594 |
| QDVNCTEVPVAIHADQLTPTWRVYS | 1 | 0 | 0 | 1387044 | YP_009724390.1 | P0DTC2 |
| QGTALPKGFYAEGSRG | 1 | 0 | 0 | 1387119 | QIQ54056.1 | P59595 |
| QNGERSGARPKQRRPQ | 1 | 0 | 0 | 1387258 | QHR63308.1 | Q3LZX4 |
| QNQRNAPRITFGGPSDSTGSN | 1 | 0 | 0 | 1387279 | YP_009724397.2 | P0DTC9 |
| QPLPQRQKKQQTVTLLPA | 1 | 0 | 0 | 1387305 | QIQ54056.1 | P59595 |
| QRVAGDSGFAAYSRYR | 1 | 0 | 0 | 1387385 | YP_009724393.1 | P0DTC5 |
| QSQTVTKKSAAEASKKPRQ | 1 | 0 | 0 | 1387411 | QIQ54056.1 | P59595 |

| | | | | | | |
|---|---|---|---|---|---|---|
| QTTVELQGLIDQINSTY | 1 | 0 | 0 | 1387449 | YP_003767.1 | A0A0P0G321 |
| RAVKDGIVWVHEDGATD | 1 | 0 | 0 | 1387520 | YP_009047211.1 | K9N4V7 |
| RLGVRTTRKTSERSQPRGRR | 1 | 0 | 0 | 1387738 | AGN33396.1 | P27958 |
| RRPQGLPNNTASWFTA | 1 | 0 | 0 | 1387894 | YP_009724397.2 | P0DTC9 |
| RSKQRRPQGLPNNTAS | 1 | 0 | 0 | 1387924 | YP_009724397.2 | P0DTC9 |
| RSSRIAGRSALEDLLFSKVVTSGL | 1 | 0 | 0 | 1387943 | YP_003767.1 | A0A0P0G321 |
| RSVASQSIIAYTMSLGAENSVAYSN | 1 | 0 | 0 | 1387949 | YP_009724390.1 | P0DTC2 |
| RVAGDSGFAAYSRYRI | 1 | 0 | 0 | 1387989 | YP_009724393.1 | P0DTC5 |
| SACKTIEDALRLSAHLETND | 1 | 0 | 0 | 1388052 | YP_003767.1 | A0A0P0G321 |
| SGFAAYSRYRIGNYKL | 1 | 0 | 0 | 1388327 | YP_009724393.1 | P0DTC5 |
| SGSRVAGRSAIEDILFSKLVTSGL | 1 | 0 | 0 | 1388480 | NP_073551.1 | A0A0P0K6L9 |
| SGVVNPAMEPIYDEPTTTTSV | 1 | 0 | 0 | 1388497 | QHR63301.1 | Q3LZX0 |
| SHITAEAAGRRLARGSPPSL | 1 | 0 | 0 | 1388533 | ANN11856.1 | P27958 |
| SHITAEAAGRRLARGSPPSV | 1 | 0 | 0 | 1388534 | AGN33396.1 | P27958 |
| SKQRRPQGLPNNTASW | 1 | 0 | 0 | 1388658 | YP_009724397.2 | P0DTC9 |
| SKVSGKGQQQQGQTVTKKS | 1 | 0 | 0 | 1388672 | YP_009825061.1 | P59595 |
| SNASSSFDCIVNLLFT | 1 | 0 | 0 | 1388793 | YP_003767.1 | A0A0P0G321 |
| SPASSQTSAKSLARSQ | 1 | 0 | 0 | 1388859 | NP_073556.1 | A0A0N7J131 |
| SPASTVHATATIPLQA | 1 | 0 | 0 | 1388860 | YP_009825052.1 | P59632 |
| SPDVDLGDISGINASVV | 1 | 0 | 0 | 1388866 | P0DTC2.1 | P0DTC2 |
| SPVFTDNSSPPAVPQSF | 1 | 0 | 0 | 1388933 | AGN33396.1 | P27958 |
| SQASSRSSSRSRGNSR | 1 | 0 | 0 | 1388946 | YP_009825061.1 | P59595 |
| SQRVAGDSGFAAYSRY | 1 | 0 | 0 | 1388977 | YP_009724393.1 | P0DTC5 |
| SRASSGNRSGNGILKW | 1 | 0 | 0 | 1388989 | YP_009555245.1 | P33469 |
| SRASSLSRNSSRSSSQ | 1 | 0 | 0 | 1388990 | YP_009047211.1 | K9N4V7 |
| SRIVDEPEDHVQIHTI | 1 | 0 | 0 | 1388999 | QIQ54049.1 | P59632 |
| SRYRIGNYKLNTDHSS | 1 | 0 | 0 | 1389032 | YP_009724393.1 | P0DTC5 |
| SSGVVNPVMEPIYDEPTTTTSVPL | 1 | 0 | 0 | 1389065 | YP_009724391.1 | P0DTC3 |
| STGSRSARSAIEDLLF | 1 | 0 | 0 | 1389156 | YP_009047204.1 | K9N5Q8 |
| STSQKSIVAYTMSLGADSSIAY | 1 | 0 | 0 | 1389187 | YP_009825051.1 | P59594 |
| SVASLSAPTLVPQENY | 1 | 0 | 0 | 1389203 | YP_173236.1 | A0A140H1G9 |
| SVASSFVGMPSFVAYE | 1 | 0 | 0 | 1389205 | NP_073549.1 | A0A0P0LKV0 |
| SVASSFVSMPSYIAYE | 1 | 0 | 0 | 1389206 | YP_003766.2 | H9EJ42 |
| SVNQRSIIAYTMSLGAENSVAYSN | 1 | 0 | 0 | 1389235 | QIQ54048.1 | P59594 |
| TATGDVVLATDDLYVK | 1 | 0 | 0 | 1389324 | YP_009555238.1 | P0C6X6 |
| TDEAQPLPQRQKKQPT | 1 | 0 | 0 | 1389360 | YP_009825061.1 | P59595 |
| TGDEVRQIAPGQTGKIADY | 1 | 0 | 0 | 1389530 | QHR63300.2 | Q3LZX1 |
| TLSYYKLGASQRVAGDSGFA | 1 | 0 | 0 | 1389743 | YP_009724393.1 | P0DTC5 |
| TLSYYKLGASQRVGTDSGFA | 1 | 0 | 0 | 1389744 | YP_009825055.1 | P59596 |
| TPSSKRFQPFQQFGRDVSDFTDSVR | 1 | 0 | 0 | 1389873 | YP_009825051.1 | P59594 |
| TTLEETKFLTNKLLLF | 1 | 0 | 0 | 1390041 | NP_828849.7 | P0C6X7 |
| TTVDNINLHTQVVDMS | 1 | 0 | 0 | 1390061 | YP_009724389.1 | P0DTD1 |
| TVGELGDVRETMNYLF | 1 | 0 | 0 | 1390082 | QHR63299.2 | P0C6W2 |
| TVVGQLEQTNMHSPDVI | 1 | 0 | 0 | 1390125 | YP_009047202.1 | K9N7C7 |
| VAFELFAKRKMGLTPP | 1 | 0 | 0 | 1390173 | YP_003766.2 | H9EJ42 |
| VAGDSGFAAYSRYRIG | 1 | 0 | 0 | 1390174 | YP_009724393.1 | P0DTC5 |
| VAIEQLLYAIQQLYTG | 1 | 0 | 0 | 1390179 | YP_009047202.1 | K9N7C7 |
| VATGDVVLASDDLYVK | 1 | 0 | 0 | 1390205 | YP_173236.1 | A0A140H1G9 |
| VDSSQGSEYQYVIFCQ | 1 | 0 | 0 | 1390290 | YP_009047202.1 | K9N7C7 |
| VQTVEDLRARMTYVCQC | 1 | 0 | 0 | 1390824 | YP_009047202.1 | K9N7C7 |
| VRGGDGKMKDLSPRWY | 1 | 0 | 0 | 1390837 | QIQ54056.1 | P59595 |
| VRGGDGKMKELSPRWY | 1 | 0 | 0 | 1390838 | YP_009825061.1 | P59595 |
| VSVEQLEPCAQSRLLS | 1 | 0 | 0 | 1390917 | NP_073549.1 | A0A0P0LKV0 |
| VSVEQLLASIQHLHEG | 1 | 0 | 0 | 1390918 | YP_003766.2 | H9EJ42 |
| VTIAEILLIIMRTFKV | 1 | 0 | 0 | 1390941 | YP_009724394.1 | P0DTC6 |
| VTLDKLRDYLADYDVA | 1 | 0 | 0 | 1390947 | YP_009047202.1 | K9N7C7 |
| VTLEQYSTCDICKSTV | 1 | 0 | 0 | 1390951 | YP_003766.2 | H9EJ42 |
| VTLGDFTIMSGRMSLT | 1 | 0 | 0 | 1390952 | YP_173236.1 | A0A140H1G9 |
| VTTTLEETKFLTENLLLYA | 1 | 0 | 0 | 1390975 | QIQ54047.1 | P0C6X7 |
| VTVDVLKDMPVLKTIN | 1 | 0 | 0 | 1390978 | YP_173236.1 | A0A140H1G9 |
| VTVELEPPCRFVIDTP | 1 | 0 | 0 | 1390979 | NP_073549.1 | A0A0P0LKV0 |
| VYSTGSNVFQTRAGCLIGAE | 1 | 0 | 0 | 1391065 | YP_009724390.1 | P0DTC2 |
| WTADEAIKFLKEWNFS | 1 | 0 | 0 | 1391194 | YP_009555244.1 | Q01455 |
| YFKNHTSPDVDLGDISGINA | 1 | 0 | 0 | 1391352 | YP_009724390.1 | P0DTC2 |
| YSRYRIGNYKLNTDHS | 1 | 0 | 0 | 1391666 | YP_009724393.1 | P0DTC5 |
| YTLSDLPVYVTVAKVQ | 1 | 0 | 0 | 1391685 | YP_173241.1 | Q0ZJ82 |
| YTMADLVYALRHFDEG | 1 | 0 | 0 | 1391686 | YP_009724389.1 | P0DTD1 |
| YTMMDLCFALRNFDEK | 1 | 0 | 0 | 1391688 | NP_073549.1 | A0A0P0LKV0 |
| YTMMDLVYALRHFDQN | 1 | 0 | 0 | 1391689 | YP_009047202.1 | K9N7C7 |
| YTMMDLVYAMRNFDEQ | 1 | 0 | 0 | 1391690 | YP_003766.2 | H9EJ42 |
| YVVDDIYYPASCNGVL | 1 | 0 | 0 | 1391749 | YP_003766.2 | H9EJ42 |
| GREIVDLMCHATFTM | 1 | 0 | 0 | 1392191 | P27909.2 | P17763 |
| VSCLPFTINCQEPKL | 1 | 0 | 0 | 1392500 | YP_009724396.1 | P0DTC8 |
| YHDVRVVLDFI | 1 | 0 | 0 | 1392528 | YP_009724396.1 | P0DTC8 |
| NLVRDLPQGFSALEPLV | 1 | 0 | 0 | 1393954 | P0DTC2.1 | P0DTC2 |
| CASYQTQTNSHRRAR | 1 | 0 | 0 | 1397261 | QSJ05725.1 | P0DTC2 |
| CVIAWNSKNLDSKVG | 1 | 0 | 0 | 1397263 | QRO03339.1 | P0DTC2 |
| EPQIITTHNTFVSGN | 1 | 0 | 0 | 1397275 | QSJ05725.1 | P0DTC2 |
| FELLHAPVTVCGPKK | 1 | 0 | 0 | 1397279 | QSW39933.1 | P0DTC2 |
| GLTGTGVLTESNKKFLPFQQF | 1 | 0 | 0 | 1397311 | P0DTC2.1 | P0DTC2 |
| GRDIADTTDAVRDPQTLEILDI | 1 | 0 | 0 | 1397316 | P0DTC2.1 | P0DTC2 |
| IAPGQTGNIADYNYK | 1 | 0 | 0 | 1397326 | QTA94406.1 | P0DTC2 |
| KHIDAYKTFPPTEPKKDKKKK | 1 | 0 | 0 | 1397350 | P0DTC9.1 | P0DTC9 |
| MSLNGPQNQRNAPRI | 1 | 0 | 0 | 1397370 | QUF24372.1 | P0DTC9 |
| NCVADYSFLYNSASF | 1 | 0 | 0 | 1397372 | QWO74701.1 | P0DTC2 |
| NDPFLGVYHKNNKS | 1 | 0 | 0 | 1397373 | QSJ05725.1 | P0DTC2 |
| NGVGYQPHRVVVLSF | 1 | 0 | 0 | 1397378 | QQW60654.1 | P0DTC2 |
| NKVTLADVGFIKQYG | 1 | 0 | 0 | 1397381 | QQP64673.1 | P0DTC2 |
| NNSIAIPINFTISVT | 1 | 0 | 0 | 1397389 | QSJ05725.1 | P0DTC2 |
| NQLESKMFGKGQQQQ | 1 | 0 | 0 | 1397390 | QTY61797.1 | P0DTC9 |
| PKKDKKKKTDEAQPLPQRQKKQP | 1 | 0 | 0 | 1397392 | SRC558394 | P0DTC9 |

| | | | | | | |
|---|---|---|---|---|---|---|
| QPLPQRQKKQPTVTLLPAADM | 1 | 0 | 0 | 1397401 | SRC558394 | P0DTC9 |
| QVAVLYQGVNCTEVP | 1 | 0 | 0 | 1397404 | CAD0240757.1 | P0DTC2 |
| RPQGLPNNTASWFTALTQHGK | 1 | 0 | 0 | 1397407 | P0DTC9.1 | P0DTC9 |
| RVYSSANQCTFEYVS | 1 | 0 | 0 | 1397414 | SRC558393 | P0DTC2 |
| SVLNDILARLDKVEA | 1 | 0 | 0 | 1397436 | QSJ05725.1 | P0DTC2 |
| SYGFQPTYGVGYQPY | 1 | 0 | 0 | 1397439 | QSJ05725.1 | P0DTC2 |
| TWFHAISGTNGTK | 1 | 0 | 0 | 1397450 | QSJ05725.1 | P0DTC2 |
| VDLPIGIQITRFQTL | 1 | 0 | 0 | 1397452 | SRC558393 | P0DTC2 |
| VEGFNCYLPLQSYGF | 1 | 0 | 0 | 1397453 | QUL62581.1 | P0DTC2 |
| VRQIAPGETGKIADY | 1 | 0 | 0 | 1397460 | SRC558393 | P0DTC2 |
| YQTQTNSHRRARSVA | 1 | 0 | 0 | 1397473 | QSJ05725.1 | P0DTC2 |
| AAADGIIYPTGK | 1 | 0 | 0 | 1406431 | AVV62526.1 | K9N5Q8 |
| AAAGAPFYLNVQ | 1 | 0 | 0 | 1406433 | ARB66070.1 | P36334 |
| AAAGLEAPFLYL | 1 | 0 | 0 | 1406434 | APO40580.1 | P59632 |
| AAAGVPLYINVQ | 1 | 0 | 0 | 1406437 | AIL49503.1 | P36334 |
| AAAGVPLYLNVQ | 1 | 0 | 0 | 1406438 | ARB66061.1 | P36334 |
| AAALEDEFTPFD | 1 | 0 | 0 | 1406448 | 3M3V_A | P0C6X7 |
| AAALKSLGFEKP | 1 | 0 | 0 | 1406450 | ARB07429.1 | A0A0N7J131 |
| AAANATGSVIIS | 1 | 0 | 0 | 1406454 | AVV62537.1 | K9N5Q8 |
| AAANATGTVIIS | 1 | 0 | 0 | 1406455 | AVK87439.1 | K9N5Q8 |
| AAANIFVVNKDT | 1 | 0 | 0 | 1406456 | AVV62535.1 | K9N7C7 |
| AAANIGSNHTGV | 1 | 0 | 0 | 1406457 | APO40578.1 | P0C6X7 |
| AAANSTGNLIIS | 1 | 0 | 0 | 1406458 | AVV62526.1 | K9N5Q8 |
| AAANSTGTVIIR | 1 | 0 | 0 | 1406459 | QBF80608.1 | K9N5Q8 |
| AAAPFTKPFAVY | 1 | 0 | 0 | 1406460 | 2VRI_A | H9EJ42 |
| AAAQGFQACRFI | 1 | 0 | 0 | 1406461 | AVA26872.1 | H9EJ42 |
| AAASAVKLQNNE | 1 | 0 | 0 | 1406463 | AVV62524.1 | K9N7C7 |
| AAAVDEAFPLNE | 1 | 0 | 0 | 1406465 | AHI48527.1 | K9N638 |
| AAAVDEVFPLDE | 1 | 0 | 0 | 1406466 | ASU90174.1 | K9N638 |
| AAAYMKLSDALK | 1 | 0 | 0 | 1406471 | AXN92247.1 | K9N7C7 |
| AACAVDVNVCHS | 1 | 0 | 0 | 1406477 | ARU07575.1 | A0A140H1G9 |
| AACCHLAKAIND | 1 | 0 | 0 | 1406478 | AVP78030.1 | P0C6W2 |
| AACHDLLSEYGT | 1 | 0 | 0 | 1406480 | BBA20983.1 | A0A140H1H1 |
| AADAERMAMYTG | 1 | 0 | 0 | 1406490 | APT69876.1 | A0A0P0K6L9 |
| AADGIIYPTGKS | 1 | 0 | 0 | 1406492 | AVV62526.1 | K9N5Q8 |
| AADGLKITEELQ | 1 | 0 | 0 | 1406493 | APO40578.1 | P0C6X7 |
| AADINIHDLPIC | 1 | 0 | 0 | 1406494 | AGT51530.1 | P30215 |
| AADINIPDLLIC | 1 | 0 | 0 | 1406495 | AGT51729.1 | P30215 |
| AADINIPHLPIC | 1 | 0 | 0 | 1406497 | AXX83356.1 | P30215 |
| AADINNPDLPIC | 1 | 0 | 0 | 1406498 | P30215.1 | P30215 |
| AADINTPDVPIC | 1 | 0 | 0 | 1406499 | AAA73152.1 | P30215 |
| AADLDEFSKQLQ | 1 | 0 | 0 | 1406502 | AVP78038.1 | Q3LZX4 |
| AADLPCTLHFSL | 1 | 0 | 0 | 1406505 | ARU07575.1 | A0A140H1G9 |
| AAECTIFKDSSG | 1 | 0 | 0 | 1406513 | AVP78041.1 | P0C6W2 |
| AAECTIFKNAQG | 1 | 0 | 0 | 1406514 | APO40578.1 | P0C6X7 |
| AAEELASPMKSD | 1 | 0 | 0 | 1406515 | ARU07575.1 | A0A140H1G9 |
| AAEGVPFYLNVQ | 1 | 0 | 0 | 1406518 | AIL49550.1 | P36334 |
| AAEIHRDCDFNK | 1 | 0 | 0 | 1406519 | AVV62535.1 | K9N7C7 |
| AAEKEVLSELYG | 1 | 0 | 0 | 1406521 | AVV62535.1 | K9N7C7 |
| AAEKLASPMKSD | 1 | 0 | 0 | 1406522 | AZS52616.1 | A0A140H1G9 |
| AAELDDFSKQLQ | 1 | 0 | 0 | 1406523 | APO40586.1 | P59595 |
| AAEQVVAKPKTR | 1 | 0 | 0 | 1406526 | AVV62524.1 | K9N7C7 |
| AAESHMDADFAK | 1 | 0 | 0 | 1406527 | SRC265912 | P0C6X7 |
| AAETLKANEETF | 1 | 0 | 0 | 1406531 | APO40578.1 | P0C6X7 |
| AAEYREYMEAYN | 1 | 0 | 0 | 1406536 | AVV62535.1 | K9N7C7 |
| AAEYREYVEAYN | 1 | 0 | 0 | 1406537 | AVV62524.1 | K9N7C7 |
| AAFATHVCFKDC | 1 | 0 | 0 | 1406539 | ABE97136.1 | H9EJ42 |
| AAFDCGDYAACK | 1 | 0 | 0 | 1406541 | ARB66062.1 | P36334 |
| AAFDFSSDVFKI | 1 | 0 | 0 | 1406543 | QDH43737.1 | H9EJ42 |
| AAFDLSKELKRP | 1 | 0 | 0 | 1406544 | QBQ01836.1 | P0C6X6 |
| AAFHVNSSEPAL | 1 | 0 | 0 | 1406552 | AIX10749.1 | P36334 |
| AAFISPYNSQNF | 1 | 0 | 0 | 1406553 | AXX83330.1 | P0C6X6 |
| AAFLELLPNVFK | 1 | 0 | 0 | 1406554 | AWK59942.1 | H9EJ42 |
| AAFNAVCSVIDL | 1 | 0 | 0 | 1406555 | ANI69909.1 | K9N7C7 |
| AAFNNLLGYDFN | 1 | 0 | 0 | 1406556 | AUM60013.1 | K9N7C7 |
| AAFNWVLLACTL | 1 | 0 | 0 | 1406557 | AVV62524.1 | K9N7C7 |
| AAFYVYKLQPLT | 1 | 0 | 0 | 1406572 | AXN92228.1 | K9N5Q8 |
| AAGNIFVVNKDT | 1 | 0 | 0 | 1406586 | AVV62524.1 | K9N7C7 |
| AAGQQFVKETTD | 1 | 0 | 0 | 1406587 | AVQ05261.1 | P0C6X6 |
| AAGVGNTFICEE | 1 | 0 | 0 | 1406592 | AVN89451.1 | K9N7C7 |
| AAGYGGFGCKDS | 1 | 0 | 0 | 1406597 | AEN19363.1 | P0C6X6 |
| AAHADPNHFTLG | 1 | 0 | 0 | 1406598 | AUM60024.1 | K9N5Q8 |
| AAHADPNRFILG | 1 | 0 | 0 | 1406599 | AUM60014.1 | K9N5Q8 |
| AAHPPPGEQFKH | 1 | 0 | 0 | 1406600 | AVN89289.1 | K9N7C7 |
| AAIASNCYSSLI | 1 | 0 | 0 | 1406602 | AXN92228.1 | K9N5Q8 |
| AAIDLAHTKLQS | 1 | 0 | 0 | 1406604 | AVV62535.1 | K9N7C7 |
| AAIDLAYNKLQS | 1 | 0 | 0 | 1406605 | AVV62524.1 | K9N7C7 |
| AAIGDLKCPLDS | 1 | 0 | 0 | 1406607 | AIL49486.1 | P36334 |
| AAIGDLNCTLDP | 1 | 0 | 0 | 1406608 | BBO58773.1 | P36334 |
| AAIHADQLTPAW | 1 | 0 | 0 | 1406609 | Q3I5J5.1 | Q3LZX1 |
| AAIKLVPTTTGG | 1 | 0 | 0 | 1406614 | AGT51379.1 | H9EJ42 |
| AAIKLVPTTTGI | 1 | 0 | 0 | 1406615 | AWK59942.1 | H9EJ42 |
| AAIKRFKNGFQG | 1 | 0 | 0 | 1406617 | AGT51657.1 | P0C6X6 |
| AAIKRLKNGFQG | 1 | 0 | 0 | 1406618 | AEN19363.1 | P0C6X6 |
| AAIKRLYMGFQG | 1 | 0 | 0 | 1406619 | ARU07575.1 | A0A140H1G9 |
| AAILNGCNWFVK | 1 | 0 | 0 | 1406621 | AUM60013.1 | K9N7C7 |
| AAINFNQPLVVD | 1 | 0 | 0 | 1406629 | AUM60024.1 | K9N5Q8 |
| AAINNASNGAVQ | 1 | 0 | 0 | 1406630 | AUM60013.1 | K9N7C7 |
| AAINSVPWDTIA | 1 | 0 | 0 | 1406631 | AVP78041.1 | P0C6W2 |
| AAINSVPWSKIF | 1 | 0 | 0 | 1406632 | ATO98143.1 | P0C6W2 |
| AAIPDYNYYKYN | 1 | 0 | 0 | 1406634 | ACV87150.1 | A0A140H1G9 |

| | | | | | | |
|---|---|---|---|---|---|---|
| AAIQNCAWDYHI | 1 | 0 | 0 | 1406639 | AST12962.1 | A0A0P0LKV0 |
| AAIQNCAWDYHV | 1 | 0 | 0 | 1406640 | NP_073549.1 | A0A0P0LKV0 |
| AAISSNCYSSLI | 1 | 0 | 0 | 1406647 | AHL18090.1 | K9N5Q8 |
| AAIVLQLPQGTS | 1 | 0 | 0 | 1406655 | APO40586.1 | P59595 |
| AAIVSNCYSSLI | 1 | 0 | 0 | 1406657 | AHI48733.1 | K9N5Q8 |
| AAIVTQFAPGTQ | 1 | 0 | 0 | 1406658 | AVV62533.1 | K9N4V7 |
| AAIYYLNSKIVD | 1 | 0 | 0 | 1406660 | AGT51518.1 | P0C6X6 |
| AAKEGINSLADV | 1 | 0 | 0 | 1406663 | AVV62535.1 | K9N7C7 |
| AAKEGLNSLADV | 1 | 0 | 0 | 1406664 | AVV62524.1 | K9N7C7 |
| AAKICIVDNVYM | 1 | 0 | 0 | 1406667 | YP_009555247.1 | P0C6X6 |
| AAKLQNNEIKPS | 1 | 0 | 0 | 1406679 | ALD51903.1 | K9N7C7 |
| AAKLSPWFMDGQ | 1 | 0 | 0 | 1406681 | AVV62524.1 | K9N7C7 |
| AAKNKMRHKRIS | 1 | 0 | 0 | 1406682 | AID66496.1 | K9N4V7 |
| AAKNKMRHKRVA | 1 | 0 | 0 | 1406684 | AVV62533.1 | K9N4V7 |
| AALAAKEGLNSL | 1 | 0 | 0 | 1406689 | AVV62524.1 | K9N7C7 |
| AALAVNACLIPF | 1 | 0 | 0 | 1406692 | AUM60018.1 | K9N7D2 |
| AALAVNACLIPL | 1 | 0 | 0 | 1406693 | ALB08293.1 | K9N7D2 |
| AALFASLDVKPY | 1 | 0 | 0 | 1406700 | AWK59942.1 | H9EJ42 |
| AALGLTVSDDDF | 1 | 0 | 0 | 1406707 | AGT51337.1 | H9EJ42 |
| AALISGTATAGW | 1 | 0 | 0 | 1406711 | AVP78031.1 | Q3LZX1 |
| AALKSEKHQDHV | 1 | 0 | 0 | 1406719 | AGV08406.1 | K9N7C7 |
| AALNNEKRPDHV | 1 | 0 | 0 | 1406729 | AVV62535.1 | K9N7C7 |
| AALNSEKHQDHV | 1 | 0 | 0 | 1406730 | AXN92247.1 | K9N7C7 |
| AALNSEKRPDHV | 1 | 0 | 0 | 1406731 | AUM60013.1 | K9N7C7 |
| AALNVFVSHTLT | 1 | 0 | 0 | 1406732 | AIW52757.1 | A0A0P0K6L9 |
| AALPQSKESNFL | 1 | 0 | 0 | 1406738 | AQZ41321.1 | K9N7C7 |
| AALPYGANKEGI | 1 | 0 | 0 | 1406740 | APO40586.1 | P59595 |
| AALRAGSFDKVY | 1 | 0 | 0 | 1406743 | AIY60536.1 | K9N7C7 |
| AALRAVSFADNN | 1 | 0 | 0 | 1406744 | QDP16202.1 | K9N4V7 |
| AALSFKENVDFI | 1 | 0 | 0 | 1406747 | AXX83325.1 | Q80872 |
| AAMASQKPQRAP | 1 | 0 | 0 | 1406760 | AVV62533.1 | K9N4V7 |
| AAMLSDSHIVSK | 1 | 0 | 0 | 1406765 | ABB90484.1 | A0A0N7J131 |
| AAMSGQILLNCA | 1 | 0 | 0 | 1406767 | NP_073549.1 | A0A0P0LKV0 |
| AAMTEQLQMGFG | 1 | 0 | 0 | 1406768 | AWF93656.1 | K9N5Q8 |
| AAMYKEARAVNR | 1 | 0 | 0 | 1406771 | NP_073549.1 | A0A0P0LKV0 |
| AANATGSVIIST | 1 | 0 | 0 | 1406774 | AVV62537.1 | K9N5Q8 |
| AANATGTVIISP | 1 | 0 | 0 | 1406775 | AVK87439.1 | K9N5Q8 |
| AANDWKPGLAMP | 1 | 0 | 0 | 1406776 | AVV62524.1 | K9N7C7 |
| AANGGTSFCCRH | 1 | 0 | 0 | 1406781 | AVV62535.1 | K9N7C7 |
| AANHSIPDLQKA | 1 | 0 | 0 | 1406782 | ARO76381.1 | P0C6U8 |
| AANIGSNHTGVV | 1 | 0 | 0 | 1406784 | APO40578.1 | P0C6X7 |
| AANIVNVSLVKP | 1 | 0 | 0 | 1406786 | 5X29_A | P59637 |
| AANNGHKQIVKS | 1 | 0 | 0 | 1406792 | AUM60017.1 | K9N643 |
| AANNTIGIVSQP | 1 | 0 | 0 | 1406793 | AUM60013.1 | K9N7C7 |
| AANPSTPPYTFT | 1 | 0 | 0 | 1406795 | AUM60013.1 | K9N7C7 |
| AANPTNCITPVN | 1 | 0 | 0 | 1406797 | ALJ76278.1 | K9N5Q8 |
| AANSTGNLIISS | 1 | 0 | 0 | 1406800 | AVV62526.1 | K9N5Q8 |
| AANSTGTVIINP | 1 | 0 | 0 | 1406801 | AJD81440.1 | K9N5Q8 |
| AANSTGTVIIRP | 1 | 0 | 0 | 1406802 | QBF80608.1 | K9N5Q8 |
| AANSTGTVIISP | 1 | 0 | 0 | 1406803 | AXN92228.1 | K9N5Q8 |
| AANTVIWDFNRE | 1 | 0 | 0 | 1406807 | APO40578.1 | P0C6X7 |
| AANTVIWDHKRE | 1 | 0 | 0 | 1406808 | AVP78041.1 | P0C6W2 |
| AANVGRWSASIN | 1 | 0 | 0 | 1406812 | ABB90509.1 | A0A0P0K6L9 |
| AANVSVSRFDPS | 1 | 0 | 0 | 1406814 | AIV41831.1 | P36334 |
| AANVSVSRFNPS | 1 | 0 | 0 | 1406815 | CAA79896.1 | P36334 |
| AAPAAPRAVSFA | 1 | 0 | 0 | 1406817 | AQZ41292.1 | K9N4V7 |
| AAPCAVSFADNN | 1 | 0 | 0 | 1406818 | AKK52599.1 | K9N4V7 |
| AAPKSRVQQNKS | 1 | 0 | 0 | 1406823 | AIX10718.1 | P33469 |
| AAPKVITKKDAA | 1 | 0 | 0 | 1406824 | AVV62544.1 | K9N4V7 |
| AAPNNTVSWYTG | 1 | 0 | 0 | 1406826 | AXN92267.1 | K9N4V7 |
| AAPRAISFADNN | 1 | 0 | 0 | 1406829 | ANI69896.1 | K9N4V7 |
| AAPRAVSFAYNN | 1 | 0 | 0 | 1406832 | AVN89381.1 | K9N4V7 |
| AAPRAVTFGDNN | 1 | 0 | 0 | 1406833 | AVV62533.1 | K9N4V7 |
| AAQADLQHEDSD | 1 | 0 | 0 | 1406835 | ATQ39389.1 | K9N7C7 |
| AAQAMEKVNERV | 1 | 0 | 0 | 1406837 | ARB66064.1 | P36334 |
| AAQCDCLCTPDP | 1 | 0 | 0 | 1406838 | CAA83660.1 | P36334 |
| AARGPAGVEFDV | 1 | 0 | 0 | 1406858 | AIL23988.1 | K9N7C7 |
| AARGPAGVESDI | 1 | 0 | 0 | 1406859 | AVV62524.1 | K9N7C7 |
| AARLTPCGSGIS | 1 | 0 | 0 | 1406867 | AVP78041.1 | P0C6W2 |
| AARNGVVDSDVD | 1 | 0 | 0 | 1406870 | APO40578.1 | P0C6X7 |
| AARPPPGEHFKH | 1 | 0 | 0 | 1406871 | ALW82707.1 | K9N7C7 |
| AARSSQLASDKV | 1 | 0 | 0 | 1406880 | AUM60014.1 | K9N5Q8 |
| AASAVNVCLIPL | 1 | 0 | 0 | 1406885 | AHI48543.1 | K9N7D2 |
| AASDITLKCCIV | 1 | 0 | 0 | 1406887 | AGW80946.1 | A0A0P0LKV0 |
| AASDITLKCCTV | 1 | 0 | 0 | 1406888 | NP_073549.1 | A0A0P0LKV0 |
| AASGNLLLDKRTTC | 1 | 0 | 0 | 1406890 | P0DTD1.1 | P0DTD1 |
| AASNGFRACRFV | 1 | 0 | 0 | 1406898 | NP_073549.1 | A0A0P0LKV0 |
| AASNIFLITKDT | 1 | 0 | 0 | 1406899 | AUM60013.1 | K9N7C7 |
| AASNIFVINRDT | 1 | 0 | 0 | 1406901 | ALA49659.1 | K9N7C7 |
| AASQAWQPGVAM | 1 | 0 | 0 | 1406903 | 3R24_A | P0C6X7 |
| AASSAVKLQNNE | 1 | 0 | 0 | 1406904 | AUM60013.1 | K9N7C7 |
| AASVNPANMRNA | 1 | 0 | 0 | 1406906 | AVV62524.1 | K9N7C7 |
| AASVNPASMRNA | 1 | 0 | 0 | 1406907 | AUM60013.1 | K9N7C7 |
| AASVNQTVLRNS | 1 | 0 | 0 | 1406910 | AVN89451.1 | K9N7C7 |
| AATEHSNIVRGW | 1 | 0 | 0 | 1406916 | AVP78042.1 | Q3LZX1 |
| AATEKSNVIRGW | 1 | 0 | 0 | 1406917 | ATO98181.1 | Q3LZX1 |
| AATGVTSSMTDN | 1 | 0 | 0 | 1406921 | AUM60024.1 | K9N5Q8 |
| AATNGEMQKESD | 1 | 0 | 0 | 1406932 | APO40578.1 | P0C6X7 |
| AATSANLFPPWS | 1 | 0 | 0 | 1406936 | CAA83661.1 | P36334 |
| AATSASLFPPWS | 1 | 0 | 0 | 1406938 | CAA83660.1 | P36334 |
| AATTCVTGPTTE | 1 | 0 | 0 | 1406941 | ATO98145.1 | Q3LZX1 |

| | | | | | | |
|---|---|---|---|---|---|---|
| AATVRLQAGSAT | 1 | 0 | 0 | 1406947 | ACV88184.1 | P0C6X7 |
| AAVAAAMATPQP | 1 | 0 | 0 | 1406951 | AVV62544.1 | K9N4V7 |
| AAVDEAFPLNEA | 1 | 0 | 0 | 1406955 | AHI48527.1 | K9N638 |
| AAVDLSMDKLCR | 1 | 0 | 0 | 1406956 | AVN89451.1 | K9N7C7 |
| AAVDLSMDRLRR | 1 | 0 | 0 | 1406958 | ALA49384.1 | K9N7C7 |
| AAVDSLCAKAMT | 1 | 0 | 0 | 1406959 | AWK59942.1 | H9EJ42 |
| AAVHEWKMVVCD | 1 | 0 | 0 | 1406968 | AXN92258.1 | K9N7C7 |
| AAVHKWKMVVCD | 1 | 0 | 0 | 1406970 | AXN92247.1 | K9N7C7 |
| AAVHRWKMVVCD | 1 | 0 | 0 | 1406971 | AVN89451.1 | K9N7C7 |
| AAVIAGDRWFLN | 1 | 0 | 0 | 1406972 | 3M3S_A | P0C6U8 |
| AAVINGERWFLN | 1 | 0 | 0 | 1406974 | APO40578.1 | P0C6X7 |
| AAVINIPDLPIC | 1 | 0 | 0 | 1406975 | AGT51520.1 | P30215 |
| AAVLEWDIKQCP | 1 | 0 | 0 | 1406981 | AUM60013.1 | K9N7C7 |
| AAVLNSLCICFV | 1 | 0 | 0 | 1406983 | AXN92247.1 | K9N7C7 |
| AAVLNSLCICLV | 1 | 0 | 0 | 1406984 | ASU91085.1 | K9N7C7 |
| AAVNLALKNLGF | 1 | 0 | 0 | 1406985 | ABK63972.1 | H9EJ69 |
| AAVNSVPWSKIF | 1 | 0 | 0 | 1406987 | ARO76381.1 | P0C6U8 |
| AAVRSVTVEYNI | 1 | 0 | 0 | 1406992 | AXN92247.1 | K9N7C7 |
| AAVSAMMRISYF | 1 | 0 | 0 | 1406994 | ALA49424.1 | K9N7A1 |
| AAVSAMMWISYF | 1 | 0 | 0 | 1406995 | ASY99815.1 | K9N7A1 |
| AAVSAMMWLSYF | 1 | 0 | 0 | 1406996 | AVN89358.1 | K9N7A1 |
| AAVSKWKMLMCD | 1 | 0 | 0 | 1406997 | AUM60013.1 | K9N7C7 |
| AAVSRINWVTGG | 1 | 0 | 0 | 1406998 | AFR58690.1 | P59596 |
| AAVTILDGISEH | 1 | 0 | 0 | 1407000 | ARO76381.1 | P0C6U8 |
| AAVYRINWATGG | 1 | 0 | 0 | 1407004 | APO40582.1 | P59596 |
| AAYAEVHAEPKG | 1 | 0 | 0 | 1407011 | P0C6U5.1 | A0A140H1G9 |
| AAYDIEVTKGSF | 1 | 0 | 0 | 1407014 | AUM60013.1 | K9N7C7 |
| AAYDIGVTVGPY | 1 | 0 | 0 | 1407015 | AVV62535.1 | K9N7C7 |
| AAYENFNSQDTL | 1 | 0 | 0 | 1407017 | ATO98229.1 | P0C6W2 |
| AAYENFNSQEIL | 1 | 0 | 0 | 1407018 | ARO76381.1 | P0C6U8 |
| AAYGLCNSEVPP | 1 | 0 | 0 | 1407024 | AUM60014.1 | K9N5Q8 |
| AAYHANSPEPAL | 1 | 0 | 0 | 1407026 | AIX10763.1 | P36334 |
| AAYHANSSDPAL | 1 | 0 | 0 | 1407027 | AIL49515.1 | P36334 |
| AAYHANSSEPAL | 1 | 0 | 0 | 1407028 | AEN19366.1 | P36334 |
| AAYHANSSEPTL | 1 | 0 | 0 | 1407029 | AIL49500.1 | P36334 |
| AAYINIPDLPIC | 1 | 0 | 0 | 1407031 | AEN19365.1 | P30215 |
| AAYINIPDLPVC | 1 | 0 | 0 | 1407032 | ARB07432.1 | P30215 |
| AAYNGKPQGAFH | 1 | 0 | 0 | 1407041 | AEN19363.1 | P0C6X6 |
| AAYNGPPQGAFH | 1 | 0 | 0 | 1407042 | ABD75599.1 | A0A140H1G9 |
| AAYNNCTGGAEI | 1 | 0 | 0 | 1407043 | AVQ05264.1 | P36334 |
| AAYREAAACHLA | 1 | 0 | 0 | 1407046 | AXN92247.1 | K9N7C7 |
| AAYTSSSLGSIA | 1 | 0 | 0 | 1407056 | AOR17480.1 | K9N5Q8 |
| AAYVQEESITIK | 1 | 0 | 0 | 1407062 | APO40578.1 | P0C6X7 |
| ACADGTKHTYQL | 1 | 0 | 0 | 1407069 | ARO76387.1 | P59635 |
| ACADGTRHTYQL | 1 | 0 | 0 | 1407070 | ATO98237.1 | Q3LZX7 |
| ACAHAAVDSLCA | 1 | 0 | 0 | 1407072 | AWK59942.1 | H9EJ42 |
| ACAQYYNGIMVV | 1 | 0 | 0 | 1407076 | BAL45639.1 | A0A0P0K6L9 |
| ACAVDVNVCHSV | 1 | 0 | 0 | 1407077 | ARU07575.1 | A0A140H1G9 |
| ACCHLAKAINDF | 1 | 0 | 0 | 1407078 | AVP78030.1 | P0C6W2 |
| ACCYVGVQTVED | 1 | 0 | 0 | 1407089 | AXN92247.1 | K9N7C7 |
| ACDCGACFSAND | 1 | 0 | 0 | 1407090 | AVV62524.1 | K9N7C7 |
| ACDCGAVFSGND | 1 | 0 | 0 | 1407091 | AVV62535.1 | K9N7C7 |
| ACDGLALAYRAN | 1 | 0 | 0 | 1407092 | ALA49659.1 | K9N7C7 |
| ACDGLARDYRAN | 1 | 0 | 0 | 1407093 | AVV62535.1 | K9N7C7 |
| ACDGLASNYREN | 1 | 0 | 0 | 1407095 | AVV62524.1 | K9N7C7 |
| ACDYDRLSNEVT | 1 | 0 | 0 | 1407100 | AMQ49021.1 | K9N7A1 |
| ACDYLYNPLIVD | 1 | 0 | 0 | 1407101 | AGT51488.1 | P0C6X6 |
| ACECGAVYTAND | 1 | 0 | 0 | 1407103 | AUM60013.1 | K9N7C7 |
| ACEHISSTMFQY | 1 | 0 | 0 | 1407106 | ASU90747.1 | K9N5Q8 |
| ACEHISSTMSQH | 1 | 0 | 0 | 1407107 | ASU90241.1 | K9N5Q8 |
| ACEHITTMLHQF | 1 | 0 | 0 | 1407108 | AVV62537.1 | K9N5Q8 |
| ACEHITTMMSQF | 1 | 0 | 0 | 1407109 | AVV62526.1 | K9N5Q8 |
| ACENSFMVNQLA | 1 | 0 | 0 | 1407114 | AVN89312.1 | K9N638 |
| ACEVLKPVSEEV | 1 | 0 | 0 | 1407118 | AVP78041.1 | P0C6W2 |
| ACEYAEEGICFN | 1 | 0 | 0 | 1407119 | ARU07575.1 | A0A140H1G9 |
| ACFSANDVEIQS | 1 | 0 | 0 | 1407128 | AVV62524.1 | K9N7C7 |
| ACFYVGVQSIEE | 1 | 0 | 0 | 1407136 | AUM60013.1 | K9N7C7 |
| ACGCGTSYTAND | 1 | 0 | 0 | 1407138 | QBM11736.1 | K9N638 |
| ACGNCTFGAPDD | 1 | 0 | 0 | 1407139 | AID50427.1 | K9N7C7 |
| ACIIATIFLLWF | 1 | 0 | 0 | 1407157 | AEN19363.1 | P0C6X6 |
| ACIKIAQADTAG | 1 | 0 | 0 | 1407158 | ASU89922.1 | K9N796 |
| ACINGHAVPTLF | 1 | 0 | 0 | 1407165 | QBF80553.1 | K9N7C7 |
| ACIRSAKIDTSG | 1 | 0 | 0 | 1407167 | AVV62527.1 | K9N796 |
| ACKFAVCGDGFV | 1 | 0 | 0 | 1407178 | ARU07575.1 | A0A140H1G9 |
| ACKGFYSNASSD | 1 | 0 | 0 | 1407179 | AIW52757.1 | A0A0P0K6L9 |
| ACKGFYSNSSSD | 1 | 0 | 0 | 1407180 | AIW52747.1 | A0A0P0K6L9 |
| ACKSQLGEYGSF | 1 | 0 | 0 | 1407184 | AHF54658.1 | P36334 |
| ACKSQLVEYGSF | 1 | 0 | 0 | 1407185 | AVQ05264.1 | P36334 |
| ACLKGGLILKFD | 1 | 0 | 0 | 1407195 | AXN92226.1 | K9N7C7 |
| ACNANRFNVAIT | 1 | 0 | 0 | 1407207 | NP_073549.1 | A0A0P0LKV0 |
| ACNFELCDNPFI | 1 | 0 | 0 | 1407209 | ABD73001.1 | P59594 |
| ACNGHKIEDFSI | 1 | 0 | 0 | 1407210 | AIG96659.1 | A0A140H1G9 |
| ACNGHKIEDLSI | 1 | 0 | 0 | 1407211 | AFK23449.1 | A0A140H1G9 |
| ACNGISVTQIMF | 1 | 0 | 0 | 1407212 | P0C6U5.1 | A0A140H1G9 |
| ACNGISVTQTMF | 1 | 0 | 0 | 1407213 | ARU07575.1 | A0A140H1G9 |
| ACNKVSSRSAIE | 1 | 0 | 0 | 1407214 | CAA83660.1 | P36334 |
| ACNTVKDILSKE | 1 | 0 | 0 | 1407218 | AUM60013.1 | K9N7C7 |
| ACNVNRFNVAIT | 1 | 0 | 0 | 1407220 | AWK59942.1 | H9EJ42 |
| ACPDGVRHTFQL | 1 | 0 | 0 | 1407223 | AVP78047.1 | Q3LZX7 |
| ACQNLDCNCLGF | 1 | 0 | 0 | 1407239 | AWW13564.1 | Q80872 |
| ACQNPKIGPDHS | 1 | 0 | 0 | 1407240 | APO40578.1 | P0C6X7 |

| | | | | | | |
|---|---|---|---|---|---|---|
| ACQYTMCEFPHT | 1 | 0 | 0 | 1407242 | ARB07438.1 | A0A140H1H1 |
| ACQYTMCEYPYT | 1 | 0 | 0 | 1407244 | AMM42415.1 | A0A140H1H1 |
| ACRAYLDSCTTQ | 1 | 0 | 0 | 1407246 | AHI48794.1 | K9N638 |
| ACRAYLNSRTTQ | 1 | 0 | 0 | 1407249 | ALW82708.1 | K9N638 |
| ACRFIAFGLQDC | 1 | 0 | 0 | 1407250 | AVA26872.1 | H9EJ42 |
| ACRFVAFGLQDC | 1 | 0 | 0 | 1407251 | AWK59942.1 | H9EJ42 |
| ACSCIKDALSYY | 1 | 0 | 0 | 1407260 | AVV62535.1 | K9N7C7 |
| ACSHITTFMPQF | 1 | 0 | 0 | 1407264 | AUM60014.1 | K9N5Q8 |
| ACSYVGIQTIEE | 1 | 0 | 0 | 1407272 | AVV62524.1 | K9N7C7 |
| ACSYVGIQTLDE | 1 | 0 | 0 | 1407273 | AVV62535.1 | K9N7C7 |
| ACTNGHALPTLF | 1 | 0 | 0 | 1407281 | ALL26408.1 | K9N638 |
| ACTNGHAVPTLF | 1 | 0 | 0 | 1407282 | AXN92247.1 | K9N7C7 |
| ACTPESKPLADD | 1 | 0 | 0 | 1407283 | ABK63972.1 | H9EJ69 |
| ACTRLEGLGGNN | 1 | 0 | 0 | 1407286 | NP_073549.1 | A0A0P0LKV0 |
| ACTSFTCQCGGV | 1 | 0 | 0 | 1407288 | AVV62535.1 | K9N7C7 |
| ACTYTMCEYPHT | 1 | 0 | 0 | 1407291 | ABA60802.1 | A0A140H1H1 |
| ACVCGDYAACKS | 1 | 0 | 0 | 1407292 | ABD43177.1 | P36334 |
| ACVKNYEFKFYD | 1 | 0 | 0 | 1407297 | AGW27834.1 | A0A140H1G9 |
| ACVVAESVVKTL | 1 | 0 | 0 | 1407306 | ARO76381.1 | P0C6U8 |
| ACVVSDTVVKTL | 1 | 0 | 0 | 1407309 | APO40578.1 | P0C6X7 |
| ACVYYSQVLCKS | 1 | 0 | 0 | 1407314 | AXN92247.1 | K9N7C7 |
| ACYNGAPSGVYQ | 1 | 0 | 0 | 1407324 | APO40578.1 | P0C6X7 |
| ACYNGRPIGTFT | 1 | 0 | 0 | 1407325 | ASU45838.1 | K9N7C7 |
| ACYNGRPTGTFT | 1 | 0 | 0 | 1407326 | AXN92247.1 | K9N7C7 |
| ACYNGSPTGTFT | 1 | 0 | 0 | 1407328 | AUM60013.1 | K9N7C7 |
| ADAALSIKENVD | 1 | 0 | 0 | 1407330 | AIV41889.1 | Q80872 |
| ADAEALNNLLQQ | 1 | 0 | 0 | 1407331 | CAA83661.1 | P36334 |
| ADAEGYINFKEK | 1 | 0 | 0 | 1407332 | AVV62524.1 | K9N7C7 |
| ADAEIVQPATSK | 1 | 0 | 0 | 1407333 | AUM60013.1 | K9N7C7 |
| ADAERMAMYTGS | 1 | 0 | 0 | 1407334 | AIW52843.1 | A0A0P0G321 |
| ADAETSEQVVAK | 1 | 0 | 0 | 1407335 | AVV62535.1 | K9N7C7 |
| ADAMSAKEQRAP | 1 | 0 | 0 | 1407344 | AUM60031.1 | K9N4V7 |
| ADAMSAKEQRVP | 1 | 0 | 0 | 1407345 | AUM60021.1 | K9N4V7 |
| ADAPEPQRGRQG | 1 | 0 | 0 | 1407346 | APT69853.1 | A0A0N7J131 |
| ADAQSFLNRVCG | 1 | 0 | 0 | 1407347 | QYC64967.1 | P0DTD1 |
| ADASEPQRGRQS | 1 | 0 | 0 | 1407349 | ALK28785.1 | A0A0N7J131 |
| ADASETQRGRHG | 1 | 0 | 0 | 1407350 | AIW52713.1 | A0A0N7J131 |
| ADASETQRGRQG | 1 | 0 | 0 | 1407351 | AIW52715.1 | A0A0N7J131 |
| ADAYARYLAMFN | 1 | 0 | 0 | 1407358 | AVV62524.1 | K9N7C7 |
| ADAYLADTVWYV | 1 | 0 | 0 | 1407360 | AOL02455.1 | Q04854 |
| ADCQTTNGLNTS | 1 | 0 | 0 | 1407364 | ABB90509.1 | A0A0P0K6L9 |
| ADCTSCSNGRWL | 1 | 0 | 0 | 1407365 | AUM60013.1 | K9N7C7 |
| ADCVACSKSARL | 1 | 0 | 0 | 1407366 | AWK59942.1 | H9EJ42 |
| ADDEGFITLKNN | 1 | 0 | 0 | 1407368 | AXN92247.1 | K9N7C7 |
| ADDFVIASDTTL | 1 | 0 | 0 | 1407369 | AVL25554.1 | H9EJ42 |
| ADDKCFANKYRS | 1 | 0 | 0 | 1407370 | AUM60013.1 | K9N7C7 |
| ADDKPIVLFTDS | 1 | 0 | 0 | 1407371 | NP_073549.1 | A0A0P0LKV0 |
| ADDLNQLSGYKK | 1 | 0 | 0 | 1407372 | AVP78041.1 | P0C6W2 |
| ADDRCVLHCANF | 1 | 0 | 0 | 1407377 | AVV62524.1 | K9N7C7 |
| ADDSGNPVVEYL | 1 | 0 | 0 | 1407378 | QGV13494.1 | K9N7C7 |
| ADDSGTPVVECL | 1 | 0 | 0 | 1407379 | AVN89451.1 | K9N7C7 |
| ADDSGTPVVEYL | 1 | 0 | 0 | 1407380 | AHI48798.1 | K9N7C7 |
| ADEIKALKELYG | 1 | 0 | 0 | 1407385 | AVN89363.1 | K9N7C7 |
| ADELQALPQRQK | 1 | 0 | 0 | 1407386 | AVP78038.1 | Q3LZX4 |
| ADEPDSRAPSRS | 1 | 0 | 0 | 1407387 | AST12968.1 | A0A0N7J131 |
| ADESEPQRGRQG | 1 | 0 | 0 | 1407389 | ABB90491.1 | A0A0N7J131 |
| ADETCEHEYGTE | 1 | 0 | 0 | 1407390 | ARO76381.1 | P0C6U8 |
| ADETQALPQRQKKQQTVTLLPAADL | 1 | 0 | 0 | 1407393 | P0DTC9.1 | P0DTC9 |
| ADEVQPLPQRQK | 1 | 0 | 0 | 1407394 | APO40586.1 | P59595 |
| ADFACKFAVCGD | 1 | 0 | 0 | 1407396 | ARU07575.1 | A0A140H1G9 |
| ADFDSWFSQRGG | 1 | 0 | 0 | 1407398 | AVP78041.1 | P0C6W2 |
| ADFHLSGCDEYI | 1 | 0 | 0 | 1407400 | AEN19365.1 | P30215 |
| ADFIKHVRTNGG | 1 | 0 | 0 | 1407401 | ATQ39389.1 | K9N7C7 |
| ADFKSFTLIFGG | 1 | 0 | 0 | 1407402 | AXN73435.1 | K9N7C7 |
| ADFLAGSSDVLV | 1 | 0 | 0 | 1407403 | NP_073549.1 | A0A0P0LKV0 |
| ADFNAIATCDWT | 1 | 0 | 0 | 1407404 | APO40578.1 | P0C6X7 |
| ADFTPVSGTTLH | 1 | 0 | 0 | 1407406 | AVV62528.1 | K9N4V0 |
| ADFVQIENCDVT | 1 | 0 | 0 | 1407407 | BAL45640.1 | A0A0P0K6L9 |
| ADFVQIENCNVT | 1 | 0 | 0 | 1407408 | AIW52757.1 | A0A0P0K6L9 |
| ADFYLSGCDEYI | 1 | 0 | 0 | 1407410 | AEN19357.1 | P30215 |
| ADGAEEEVTVDT | 1 | 0 | 0 | 1407411 | ATQ39389.1 | K9N7C7 |
| ADGATACEMVRA | 1 | 0 | 0 | 1407412 | AVV62535.1 | K9N7C7 |
| ADGDEAWGSTMT | 1 | 0 | 0 | 1407413 | AUM60013.1 | K9N7C7 |
| ADGDQRQLLPRW | 1 | 0 | 0 | 1407414 | AJQ24083.1 | P33469 |
| ADGETACNTVKD | 1 | 0 | 0 | 1407416 | AUM60013.1 | K9N7C7 |
| ADGFMPFLLDDL | 1 | 0 | 0 | 1407417 | AEN19363.1 | P0C6X6 |
| ADGIIHPQGRTY | 1 | 0 | 0 | 1407418 | AGV08524.1 | K9N5Q8 |
| ADGIIYPTGKSF | 1 | 0 | 0 | 1407420 | AVV62526.1 | K9N5Q8 |
| ADGIIYPTGKSY | 1 | 0 | 0 | 1407421 | AVV62537.1 | K9N5Q8 |
| ADGINQMTGFKK | 1 | 0 | 0 | 1407423 | APO40578.1 | P0C6X7 |
| ADGIQEAKPSGS | 1 | 0 | 0 | 1407424 | 4KQZ_A | K9N5Q8 |
| ADGKPVISEDLW | 1 | 0 | 0 | 1407425 | AGT21366.1 | A0A0P0LKV0 |
| ADGLKITEELQH | 1 | 0 | 0 | 1407427 | APO40578.1 | P0C6X7 |
| ADGNLRQLLPRW | 1 | 0 | 0 | 1407428 | QRK03816.1 | P33469 |
| ADGQQKHLLPRW | 1 | 0 | 0 | 1407430 | ABD75597.1 | Q19U29 |
| ADGQQKPLLPRW | 1 | 0 | 0 | 1407431 | ABA60796.1 | Q19U29 |
| ADGSIIAVQPRN | 1 | 0 | 0 | 1407433 | AIW52757.1 | A0A0P0K6L9 |
| ADGSLIPVRLRN | 1 | 0 | 0 | 1407434 | AIW52843.1 | A0A0P0G321 |
| ADGSLQPYCYTD | 1 | 0 | 0 | 1407435 | ARU07583.1 | P0C6X6 |
| ADGSPQPYCYTD | 1 | 0 | 0 | 1407436 | AEN19363.1 | P0C6X6 |
| ADGSPQPYCYTE | 1 | 0 | 0 | 1407437 | AGT51787.1 | P0C6X6 |

| | | | | | | |
|---|---|---|---|---|---|---|
| ADGSPQSYCYTD | 1 | 0 | 0 | 1407438 | AGT51488.1 | P0C6X6 |
| ADGSQRQLLPRW | 1 | 0 | 0 | 1407439 | AIX10728.1 | P33469 |
| ADGTKHTYQLRA | 1 | 0 | 0 | 1407440 | ARO76387.1 | P59635 |
| ADGTRHTYQLRA | 1 | 0 | 0 | 1407441 | ATO98237.1 | Q3LZX7 |
| ADGVECDFSPLL | 1 | 0 | 0 | 1407442 | ATG84679.1 | K9N5Q8 |
| ADGVQCYTPHSQ | 1 | 0 | 0 | 1407444 | AEN19363.1 | P0C6X6 |
| ADHIASLVLAKL | 1 | 0 | 0 | 1407445 | AIL49401.1 | P33469 |
| ADIASRVDDEGF | 1 | 0 | 0 | 1407446 | ATQ39389.1 | K9N7C7 |
| ADICYKFTLWDY | 1 | 0 | 0 | 1407447 | AUM60013.1 | K9N7C7 |
| ADIDYRNPSIDE | 1 | 0 | 0 | 1407449 | AJQ24074.1 | P33469 |
| ADIGANHTGITG | 1 | 0 | 0 | 1407450 | ARO76381.1 | P0C6U8 |
| ADIGDRDPRNDE | 1 | 0 | 0 | 1407451 | AJQ24068.1 | P33469 |
| ADIKTQATQPLD | 1 | 0 | 0 | 1407454 | AVV62526.1 | K9N5Q8 |
| ADILDRDPSSDE | 1 | 0 | 0 | 1407457 | QBP84708.1 | P33469 |
| ADILVVDEVSMC | 1 | 0 | 0 | 1407458 | AXN92247.1 | K9N7C7 |
| ADINGELHEDSK | 1 | 0 | 0 | 1407459 | APO40578.1 | P0C6X7 |
| ADINGKLYQDSQ | 1 | 0 | 0 | 1407461 | ARO76381.1 | P0C6U8 |
| ADINIHDLPICV | 1 | 0 | 0 | 1407462 | AGT51530.1 | P30215 |
| ADINIPDLPISV | 1 | 0 | 0 | 1407464 | AIL49333.1 | P30215 |
| ADINIPHLPICV | 1 | 0 | 0 | 1407465 | AXX83356.1 | P30215 |
| ADINNPDLPICV | 1 | 0 | 0 | 1407466 | P30215.1 | P30215 |
| ADINTPDVPICV | 1 | 0 | 0 | 1407467 | AAA73152.1 | P30215 |
| ADITLFGDRVTV | 1 | 0 | 0 | 1407469 | AGV08523.1 | K9N7C7 |
| ADITLLGDCVTV | 1 | 0 | 0 | 1407470 | ALL26405.1 | K9N7C7 |
| ADIVDRDPSNDE | 1 | 0 | 0 | 1407471 | AJQ24082.1 | P33469 |
| ADIVDRDQSSEE | 1 | 0 | 0 | 1407473 | ANZ78879.1 | P33469 |
| ADIVEEAMVVKP | 1 | 0 | 0 | 1407475 | APO40578.1 | P0C6X7 |
| ADIVHRDPSNDE | 1 | 0 | 0 | 1407476 | AJQ24067.1 | P33469 |
| ADIVVFDEVSMA | 1 | 0 | 0 | 1407479 | APO40578.1 | P0C6X7 |
| ADKANFLNAVVF | 1 | 0 | 0 | 1407480 | AXX83300.1 | P0C6X6 |
| ADKEETNILSEL | 1 | 0 | 0 | 1407482 | AVV62524.1 | K9N7C7 |
| ADKEKLISLADV | 1 | 0 | 0 | 1407483 | AUM60013.1 | K9N7C7 |
| ADKEKLISLTDV | 1 | 0 | 0 | 1407484 | AUM60023.1 | K9N7C7 |
| ADKFPILHDIGN | 1 | 0 | 0 | 1407485 | APO40578.1 | P0C6X7 |
| ADKICIVDNVYM | 1 | 0 | 0 | 1407488 | AEN19363.1 | P0C6X6 |
| ADKLCCAVVSKS | 1 | 0 | 0 | 1407489 | AIV41888.1 | P0C6X6 |
| ADKLCHAVVSKS | 1 | 0 | 0 | 1407490 | AEN19363.1 | P0C6X6 |
| ADKLCHAVVYKS | 1 | 0 | 0 | 1407491 | AIV41786.1 | P0C6X6 |
| ADKLVEVGLVGV | 1 | 0 | 0 | 1407492 | AEN19363.1 | P0C6X6 |
| ADKNKQDSTDDG | 1 | 0 | 0 | 1407493 | AVV62533.1 | K9N4V7 |
| ADKPSQLKRPRW | 1 | 0 | 0 | 1407496 | ADK37677.1 | H9EJ69 |
| ADKTLQEMKALL | 1 | 0 | 0 | 1407497 | QEG03790.1 | P0C6X6 |
| ADKTLQEMKALY | 1 | 0 | 0 | 1407498 | AGT51518.1 | P0C6X6 |
| ADLDEFSKQLQQ | 1 | 0 | 0 | 1407503 | AVP78038.1 | Q3LZX4 |
| ADLHSSQVCGLF | 1 | 0 | 0 | 1407506 | AWK59942.1 | H9EJ42 |
| ADLPAYMSVAKV | 1 | 0 | 0 | 1407510 | AWW13547.1 | Q01455 |
| ADLPAYMTVAKV | 1 | 0 | 0 | 1407511 | BAM16435.1 | Q01455 |
| ADLPCTLHFSLF | 1 | 0 | 0 | 1407512 | ARU07575.1 | A0A140H1G9 |
| ADLQHEDSDDDG | 1 | 0 | 0 | 1407514 | ATQ39389.1 | K9N7C7 |
| ADLSGAICELEL | 1 | 0 | 0 | 1407515 | ARU07575.1 | A0A140H1G9 |
| ADLYELMGKFAY | 1 | 0 | 0 | 1407517 | AVV62524.1 | K9N7C7 |
| ADLYELMGRFAW | 1 | 0 | 0 | 1407518 | AUM60013.1 | K9N7C7 |
| ADLYHYQECARG | 1 | 0 | 0 | 1407519 | APO40584.1 | P59635 |
| ADLYKLMGYFAW | 1 | 0 | 0 | 1407522 | AVP78041.1 | P0C6W2 |
| ADMRNSSSEFFN | 1 | 0 | 0 | 1407524 | AVV62537.1 | K9N5Q8 |
| ADNGAITVEELK | 1 | 0 | 0 | 1407525 | BAE93405.1 | P59596 |
| ADNGTITVEKLK | 1 | 0 | 0 | 1407528 | AAS48455.1 | P59596 |
| ADNKFALACFST | 1 | 0 | 0 | 1407529 | AVP78036.1 | Q3LZX7 |
| ADNKFALTCFST | 1 | 0 | 0 | 1407530 | AVP78047.1 | Q3LZX7 |
| ADNKFALTCIST | 1 | 0 | 0 | 1407531 | ARO76387.1 | P59635 |
| ADNKFALTCTST | 1 | 0 | 0 | 1407532 | ATO98237.1 | Q3LZX7 |
| ADNKFAVTCTTT | 1 | 0 | 0 | 1407533 | APO40584.1 | P59635 |
| ADNLKNVLVWLE | 1 | 0 | 0 | 1407535 | APO40578.1 | P0C6X7 |
| ADNLNADETKAL | 1 | 0 | 0 | 1407536 | ATQ39389.1 | K9N7C7 |
| ADNLSAAEKEVL | 1 | 0 | 0 | 1407537 | AVV62524.1 | K9N7C7 |
| ADNLSASEKEVL | 1 | 0 | 0 | 1407538 | AUM60013.1 | K9N7C7 |
| ADNLTADEIKAL | 1 | 0 | 0 | 1407539 | AVN89363.1 | K9N7C7 |
| ADNLTDYSVSSC | 1 | 0 | 0 | 1407540 | BBA20983.1 | A0A140H1H1 |
| ADNNCYLSNVLL | 1 | 0 | 0 | 1407542 | APO40578.1 | P0C6X7 |
| ADNNDITNTNMS | 1 | 0 | 0 | 1407544 | ATG84708.1 | K9N4V7 |
| ADNNDNNNQQSR | 1 | 0 | 0 | 1407545 | AUM60021.1 | K9N4V7 |
| ADNNIEKPKPDG | 1 | 0 | 0 | 1407547 | AVV62526.1 | K9N5Q8 |
| ADNNQNGGRNGA | 1 | 0 | 0 | 1407548 | ARO76389.1 | P59595 |
| ADNNSYLSSVLL | 1 | 0 | 0 | 1407549 | 4M0W_A | P0C6U8 |
| ADNPNHFINFPL | 1 | 0 | 0 | 1407550 | ARU07566.1 | Q80872 |
| ADNSIFRSFHFT | 1 | 0 | 0 | 1407551 | AMK59676.1 | P30215 |
| ADNSTASAVQTC | 1 | 0 | 0 | 1407552 | CAA83661.1 | P36334 |
| ADNSYTSFATYH | 1 | 0 | 0 | 1407554 | AID55096.1 | K9N5Q8 |
| ADPGYMQGYDDC | 1 | 0 | 0 | 1407556 | AXN92228.1 | K9N5Q8 |
| ADPGYMQGYDEC | 1 | 0 | 0 | 1407557 | AUM60014.1 | K9N5Q8 |
| ADPKCKYAQNLL | 1 | 0 | 0 | 1407559 | ALA49857.1 | K9N7C7 |
| ADPKGKYSQSLL | 1 | 0 | 0 | 1407561 | AUM60013.1 | K9N7C7 |
| ADPKSFTLIFGG | 1 | 0 | 0 | 1407562 | AQZ41321.1 | K9N7C7 |
| ADPNHFTLGKLY | 1 | 0 | 0 | 1407564 | AUM60024.1 | K9N5Q8 |
| ADPNITNLCPFG | 1 | 0 | 0 | 1407565 | 2GHV_E | P59594 |
| ADPNRFILGKLY | 1 | 0 | 0 | 1407566 | AUM60014.1 | K9N5Q8 |
| ADPNYDALLVSM | 1 | 0 | 0 | 1407567 | AUM60013.1 | K9N7C7 |
| ADPSVVNSCVKS | 1 | 0 | 0 | 1407568 | BBA20986.1 | A0A140H1H1 |
| ADPTFLHRFYSL | 1 | 0 | 0 | 1407569 | AHI48582.1 | K9N7C7 |
| ADPVHSLRVCVE | 1 | 0 | 0 | 1407571 | AVP78030.1 | P0C6W2 |
| ADQESYGGASVC | 1 | 0 | 0 | 1407574 | AUM60013.1 | K9N7C7 |

| | | | | | | |
|---|---|---|---|---|---|---|
| ADQLADPNYDAL | 1 | 0 | 0 | 1407576 | AUM60013.1 | K9N7C7 |
| ADQLSDPNYDAL | 1 | 0 | 0 | 1407577 | AXN92247.1 | K9N7C7 |
| ADQLTPAWRIYA | 1 | 0 | 0 | 1407578 | AVP78031.1 | Q3LZX1 |
| ADQLTPAWRVYS | 1 | 0 | 0 | 1407580 | ATO98181.1 | Q3LZX1 |
| ADQNGYITLKEK | 1 | 0 | 0 | 1407582 | AUM60013.1 | K9N7C7 |
| ADQNKQESSDDV | 1 | 0 | 0 | 1407583 | AVV62544.1 | K9N4V7 |
| ADQSDQFRNFQT | 1 | 0 | 0 | 1407585 | AWW13570.1 | P33469 |
| ADQSDQFRNVQI | 1 | 0 | 0 | 1407586 | ARB07416.1 | P33469 |
| ADQSDQFRSVQT | 1 | 0 | 0 | 1407588 | AIL49389.1 | P33469 |
| ADQSDQVRNVQT | 1 | 0 | 0 | 1407590 | P33469.1 | P33469 |
| ADQSRKWTGHCT | 1 | 0 | 0 | 1407593 | AIV41775.1 | Q80872 |
| ADQTMTQMYKQA | 1 | 0 | 0 | 1407594 | AVP78041.1 | P0C6W2 |
| ADRGQESLARSA | 1 | 0 | 0 | 1407595 | ATQ39389.1 | K9N7C7 |
| ADRNAWAAFYVY | 1 | 0 | 0 | 1407598 | AUM60014.1 | K9N5Q8 |
| ADRRAFVDYTGV | 1 | 0 | 0 | 1407599 | AEN19363.1 | P0C6X6 |
| ADRRVFVDYTGV | 1 | 0 | 0 | 1407600 | QBP84701.1 | P0C6X6 |
| ADRSVVNSCVKS | 1 | 0 | 0 | 1407601 | ABD75545.1 | A0A140H1H1 |
| ADRTLQEMKALF | 1 | 0 | 0 | 1407602 | AIV41774.1 | P0C6X6 |
| ADRVAWAAFYIY | 1 | 0 | 0 | 1407603 | AVV62526.1 | K9N5Q8 |
| ADSCLQGNKCNI | 1 | 0 | 0 | 1407605 | AXX83303.1 | P36334 |
| ADSFTCNIIDAA | 1 | 0 | 0 | 1407606 | AIV41909.1 | P36334 |
| ADSFTCNNIDVA | 1 | 0 | 0 | 1407608 | ABD43176.1 | P36334 |
| ADSGDSSESGAK | 1 | 0 | 0 | 1407611 | QEG03790.1 | P0C6X6 |
| ADSGMPTTFLFS | 1 | 0 | 0 | 1407612 | BBA20983.1 | A0A140H1H1 |
| ADSKSFTLIFGG | 1 | 0 | 0 | 1407617 | AXN92247.1 | K9N7C7 |
| ADSLSPSHYWSP | 1 | 0 | 0 | 1407619 | ARB07436.1 | A0A140H1G9 |
| ADSNGFIRFPDV | 1 | 0 | 0 | 1407620 | ABD75543.1 | A0A140H1G9 |
| ADSPSKDTFELR | 1 | 0 | 0 | 1407623 | AMM42397.1 | Q19U29 |
| ADSPVKDVFELH | 1 | 0 | 0 | 1407624 | ABD75589.1 | Q19U29 |
| ADSPVKDVFELR | 1 | 0 | 0 | 1407625 | ARU07581.1 | Q19U29 |
| ADSTLIGDCVTV | 1 | 0 | 0 | 1407628 | QYB08949.1 | P0DTD1 |
| ADSTLIGHCATV | 1 | 0 | 0 | 1407629 | AVP78030.1 | P0C6W2 |
| ADSVLSAVSAGL | 1 | 0 | 0 | 1407630 | AIV41810.1 | P0C6X6 |
| ADSVMSAVSAGL | 1 | 0 | 0 | 1407631 | AEN19363.1 | P0C6X6 |
| ADSVMSAVSSGL | 1 | 0 | 0 | 1407632 | AXX83372.1 | P0C6X6 |
| ADSYFSYMMPML | 1 | 0 | 0 | 1407633 | ABD75559.1 | A0A140H1G9 |
| ADSYQKYVEAYN | 1 | 0 | 0 | 1407634 | QED88024.1 | H9EJ42 |
| ADTAGLFRIDVP | 1 | 0 | 0 | 1407637 | ALW82670.1 | K9N796 |
| ADTAGLFTNFRI | 1 | 0 | 0 | 1407638 | ALA49936.1 | K9N796 |
| ADTAGLYTNFPI | 1 | 0 | 0 | 1407639 | AZK15901.1 | K9N796 |
| ADTAGLYTNSRI | 1 | 0 | 0 | 1407641 | AGV08439.1 | K9N796 |
| ADTEQTQSAIAK | 1 | 0 | 0 | 1407644 | ATQ39389.1 | K9N7C7 |
| ADTEQTQSVIVK | 1 | 0 | 0 | 1407645 | AVN89451.1 | K9N7C7 |
| ADTEQTQSVNVK | 1 | 0 | 0 | 1407646 | ATG84700.1 | K9N638 |
| ADTEQTQSVTVK | 1 | 0 | 0 | 1407647 | AXN92247.1 | K9N7C7 |
| ADTGINNATFFL | 1 | 0 | 0 | 1407650 | APO40580.1 | P59632 |
| ADTLQEIPVVSD | 1 | 0 | 0 | 1407652 | AQZ41284.1 | K9N7C7 |
| ADTQHISNDFTP | 1 | 0 | 0 | 1407653 | ARU07581.1 | Q19U29 |
| ADTSIPSDVSAR | 1 | 0 | 0 | 1407654 | AMM42397.1 | Q19U29 |
| ADTVMSAVSAGL | 1 | 0 | 0 | 1407658 | AQN78757.1 | P0C6X6 |
| ADTVWYVGQIIF | 1 | 0 | 0 | 1407660 | AOL02455.1 | Q04854 |
| ADTYACWHHSVG | 1 | 0 | 0 | 1407661 | APO40578.1 | P0C6X7 |
| ADVAARADDEGF | 1 | 0 | 0 | 1407662 | AXN92247.1 | K9N7C7 |
| ADVAARADYEGF | 1 | 0 | 0 | 1407663 | ANI69867.1 | K9N638 |
| ADVATDVQSRAD | 1 | 0 | 0 | 1407664 | AUM60013.1 | K9N7C7 |
| ADVEAAEQVVAK | 1 | 0 | 0 | 1407666 | AVV62524.1 | K9N7C7 |
| ADVEADIASRVD | 1 | 0 | 0 | 1407667 | ATQ39389.1 | K9N7C7 |
| ADVEADVAARAD | 1 | 0 | 0 | 1407668 | AXN92247.1 | K9N7C7 |
| ADVGMPTTFLFS | 1 | 0 | 0 | 1407671 | BBA20986.1 | A0A140H1H1 |
| ADVLVNVSQTAI | 1 | 0 | 0 | 1407672 | AIW52757.1 | A0A0P0K6L9 |
| ADVMRYNFNFSS | 1 | 0 | 0 | 1407674 | AVA26873.1 | A0A0P0G321 |
| ADVMRYNLNFSA | 1 | 0 | 0 | 1407675 | AIW52841.1 | A0A0P0G321 |
| ADVMRYNLNFSS | 1 | 0 | 0 | 1407676 | AIW52843.1 | A0A0P0G321 |
| ADVNIPADIGDR | 1 | 0 | 0 | 1407677 | AJQ24068.1 | P33469 |
| ADVNIPADIVDR | 1 | 0 | 0 | 1407678 | AIX10718.1 | P33469 |
| ADVNTPADIDYR | 1 | 0 | 0 | 1407679 | AJQ24074.1 | P33469 |
| ADVNTPADILDR | 1 | 0 | 0 | 1407680 | QBP84708.1 | P33469 |
| ADVNTPADIVHR | 1 | 0 | 0 | 1407682 | AJQ24067.1 | P33469 |
| ADVNTSADIVDR | 1 | 0 | 0 | 1407683 | ARK08674.1 | P33469 |
| ADVPDAFQPVYR | 1 | 0 | 0 | 1407684 | AGT51379.1 | H9EJ42 |
| ADVPDAFQSVYH | 1 | 0 | 0 | 1407685 | ARK08626.1 | H9EJ42 |
| ADVTKIKPHSTH | 1 | 0 | 0 | 1407690 | APO40578.1 | P0C6X7 |
| ADVYARYLAMFN | 1 | 0 | 0 | 1407694 | AVV62535.1 | K9N7C7 |
| ADVYRRIPNLPD | 1 | 0 | 0 | 1407695 | CAA83660.1 | P36334 |
| ADVYRRKLNLPN | 1 | 0 | 0 | 1407696 | AAR01015.1 | P36334 |
| ADVYRRKPDLPN | 1 | 0 | 0 | 1407697 | AEN19366.1 | P36334 |
| ADVYRRKPDPPN | 1 | 0 | 0 | 1407698 | AHA80201.1 | P36334 |
| ADVYRRRPDLPN | 1 | 0 | 0 | 1407700 | AHA80225.1 | P36334 |
| ADWIELFPHNDR | 1 | 0 | 0 | 1407701 | QEO75983.1 | A0A0P0LKV0 |
| ADWKPGHSMPSL | 1 | 0 | 0 | 1407703 | AVN89289.1 | K9N7C7 |
| ADWKPGQAMPSL | 1 | 0 | 0 | 1407704 | ATQ39389.1 | K9N7C7 |
| ADWKPGYAMPSL | 1 | 0 | 0 | 1407705 | ALA49934.1 | K9N7C7 |
| ADYAFIADKEKL | 1 | 0 | 0 | 1407708 | AUM60013.1 | K9N7C7 |
| ADYAVLYNSSSS | 1 | 0 | 0 | 1407709 | APO40579.1 | P59594 |
| ADYDIAVTAGPF | 1 | 0 | 0 | 1407710 | AQZ41321.1 | K9N7C7 |
| ADYDIVVTAGPF | 1 | 0 | 0 | 1407711 | ALA49901.1 | K9N7C7 |
| ADYDVAVTVGPF | 1 | 0 | 0 | 1407713 | AXN92258.1 | K9N7C7 |
| ADYDVVVTAGPF | 1 | 0 | 0 | 1407714 | AVN89363.1 | K9N7C7 |
| ADYEGFITLKNN | 1 | 0 | 0 | 1407715 | ANI69867.1 | K9N638 |
| ADYHNHSGIISQ | 1 | 0 | 0 | 1407717 | APO40578.1 | P0C6X7 |
| ADYILPLVLSSL | 1 | 0 | 0 | 1407720 | NP_073549.1 | A0A0P0LKV0 |

| | | | | | | |
|---|---|---|---|---|---|---|
| ADYLAGSSDVLV | 1 | 0 | 0 | 1407724 | ATU80780.1 | A0A0P0LKV0 |
| ADYNAKDEISVA | 1 | 0 | 0 | 1407726 | APT69850.1 | A0A0P0LKV0 |
| ADYRCACYAYLA | 1 | 0 | 0 | 1407730 | NP_073549.1 | A0A0P0LKV0 |
| ADYSVLYNSTSF | 1 | 0 | 0 | 1407733 | ATO98231.1 | Q3LZX1 |
| ADYTAFYNSTSF | 1 | 0 | 0 | 1407734 | ATO98145.1 | Q3LZX1 |
| ADYTPVIGTSTH | 1 | 0 | 0 | 1407736 | AVV62539.1 | K9N4V0 |
| ADYTQCAVIAFA | 1 | 0 | 0 | 1407737 | AVV62524.1 | K9N7C7 |
| ADYTVFYNSTSF | 1 | 0 | 0 | 1407740 | AVP78031.1 | Q3LZX1 |
| ADYTVLYNSTSF | 1 | 0 | 0 | 1407741 | ATO98181.1 | Q3LZX1 |
| AEAAVKPLLAPH | 1 | 0 | 0 | 1407744 | ARO76392.1 | Q7TLC7 |
| AEAEWIELFPHN | 1 | 0 | 0 | 1407746 | AST12962.1 | A0A0P0LKV0 |
| AEAFEKFVCLFA | 1 | 0 | 0 | 1407747 | AVV62524.1 | K9N7C7 |
| AEAHNNLLQQLS | 1 | 0 | 0 | 1407749 | ARB66064.1 | P36334 |
| AEALNNLLQQLS | 1 | 0 | 0 | 1407753 | CAA79896.1 | P36334 |
| AEANAVQQEFSV | 1 | 0 | 0 | 1407754 | AVV62535.1 | K9N7C7 |
| AEAQIIAIIKDW | 1 | 0 | 0 | 1407756 | AHI48545.1 | K9N7A1 |
| AECEEEEIADET | 1 | 0 | 0 | 1407760 | ARO76381.1 | P0C6U8 |
| AECKAALGLTVS | 1 | 0 | 0 | 1407762 | AGT51337.1 | H9EJ42 |
| AECKATLGLTVS | 1 | 0 | 0 | 1407763 | AWK59942.1 | H9EJ42 |
| AECTIFKDSSGK | 1 | 0 | 0 | 1407767 | AVP78041.1 | P0C6W2 |
| AECTIFKNAQGK | 1 | 0 | 0 | 1407768 | APO40578.1 | P0C6X7 |
| AECTMFRDAEGR | 1 | 0 | 0 | 1407769 | ATQ39389.1 | K9N7C7 |
| AEDCMSDFMSEI | 1 | 0 | 0 | 1407770 | P36334.1 | P36334 |
| AEDGITKLADVE | 1 | 0 | 0 | 1407772 | AKN58609.1 | K9N638 |
| AEDIFCPCWSAL | 1 | 0 | 0 | 1407773 | AST12962.1 | A0A0P0LKV0 |
| AEDIFGPCWYAL | 1 | 0 | 0 | 1407775 | AGT21344.1 | A0A0P0LKV0 |
| AEDISLLRKMDE | 1 | 0 | 0 | 1407777 | AHA80151.1 | P33469 |
| AEDVTEFVQEEA | 1 | 0 | 0 | 1407780 | AVN89451.1 | K9N7C7 |
| AEDVTEFVQEEV | 1 | 0 | 0 | 1407781 | AXN92247.1 | K9N7C7 |
| AEDVTESVQEKA | 1 | 0 | 0 | 1407783 | AMQ49025.1 | K9N7C7 |
| AEEAHGGEVDIQ | 1 | 0 | 0 | 1407785 | AUM60018.1 | K9N7D2 |
| AEEEVTVDTTVE | 1 | 0 | 0 | 1407788 | ATQ39389.1 | K9N7C7 |
| AEEFDDSTGDEA | 1 | 0 | 0 | 1407789 | AVV62524.1 | K9N7C7 |
| AEEKKIKASIEE | 1 | 0 | 0 | 1407790 | AVP78041.1 | P0C6W2 |
| AEEKLDNPNSSE | 1 | 0 | 0 | 1407791 | AOL02450.1 | P0C6X6 |
| AEELASPMKSDE | 1 | 0 | 0 | 1407792 | ARU07575.1 | A0A140H1G9 |
| AEELHGGEVDIQ | 1 | 0 | 0 | 1407793 | AUM60028.1 | K9N7D2 |
| AEEPAKDIPEKV | 1 | 0 | 0 | 1407794 | ATQ39389.1 | K9N7C7 |
| AEEPDSRAPSRS | 1 | 0 | 0 | 1407795 | AIW52721.1 | A0A0N7J131 |
| AEEPDSRVPSRS | 1 | 0 | 0 | 1407796 | APT69887.1 | A0A0N7J131 |
| AEEPVVGIVYSS | 1 | 0 | 0 | 1407797 | AVA26874.1 | H9EJ24 |
| AEETRKIMPICM | 1 | 0 | 0 | 1407798 | APO40578.1 | P0C6X7 |
| AEFDENGTITDA | 1 | 0 | 0 | 1407804 | APO40579.1 | P59594 |
| AEFDTYDPTYKN | 1 | 0 | 0 | 1407806 | AVN89451.1 | K9N7C7 |
| AEFSLNNHHSTV | 1 | 0 | 0 | 1407810 | AVV62541.1 | K9N7D2 |
| AEFSLWFKNHTS | 1 | 0 | 0 | 1407811 | BBA20986.1 | A0A140H1H1 |
| AEFSTHDPIYKN | 1 | 0 | 0 | 1407812 | AVV62524.1 | K9N7C7 |
| AEFVHYDPMYKN | 1 | 0 | 0 | 1407814 | AUM60013.1 | K9N7C7 |
| AEGAHATLDSIG | 1 | 0 | 0 | 1407815 | AEN19363.1 | P0C6X6 |
| AEGDAFWSSTMI | 1 | 0 | 0 | 1407817 | AVN89333.1 | K9N7C7 |
| AEGDASWSSTMI | 1 | 0 | 0 | 1407818 | AXN92247.1 | K9N7C7 |
| AEGDASWYSTMI | 1 | 0 | 0 | 1407819 | AHI48789.1 | K9N638 |
| AEGDGGVLGDGN | 1 | 0 | 0 | 1407820 | AWK59942.1 | H9EJ42 |
| AEGEETDLTVTS | 1 | 0 | 0 | 1407821 | ATP16754.1 | P0C6X6 |
| AEGFECDFSPLL | 1 | 0 | 0 | 1407822 | AQZ41282.1 | K9N5Q8 |
| AEGGASWSSTMI | 1 | 0 | 0 | 1407823 | AKN24802.1 | K9N7C7 |
| AEGISLLKKMDE | 1 | 0 | 0 | 1407824 | AHA80174.1 | P33469 |
| AEGKMVPYCYDP | 1 | 0 | 0 | 1407825 | AVV62524.1 | K9N7C7 |
| AEGQGVPIAPGV | 1 | 0 | 0 | 1407826 | AEN19370.1 | P33469 |
| AEGRITPYCHDP | 1 | 0 | 0 | 1407827 | AHL18091.1 | K9N7C7 |
| AEGRMVPYCYDP | 1 | 0 | 0 | 1407830 | AUM60013.1 | K9N7C7 |
| AEGSHATLDSIG | 1 | 0 | 0 | 1407831 | AGT51518.1 | P0C6X6 |
| AEGSRGGSQTAS | 1 | 0 | 0 | 1407833 | APO40586.1 | P59595 |
| AEGTKQIIHSHP | 1 | 0 | 0 | 1407834 | ATQ39396.1 | K9N643 |
| AEGTPVCVNGLM | 1 | 0 | 0 | 1407836 | APO40578.1 | P0C6X7 |
| AEGVECDFSLLL | 1 | 0 | 0 | 1407837 | ALJ54500.1 | K9N5Q8 |
| AEGVECDFSPLL | 1 | 0 | 0 | 1407838 | AXN92228.1 | K9N5Q8 |
| AEGVIYPTGRTY | 1 | 0 | 0 | 1407839 | AUM60014.1 | K9N5Q8 |
| AEGVPFYLNVQY | 1 | 0 | 0 | 1407840 | AIL49550.1 | P36334 |
| AEGYINFKEKLY | 1 | 0 | 0 | 1407841 | AVV62524.1 | K9N7C7 |
| AEHLSPWFADGA | 1 | 0 | 0 | 1407844 | AVV62535.1 | K9N7C7 |
| AEHVNASYECDI | 1 | 0 | 0 | 1407850 | AVP78031.1 | Q3LZX1 |
| AEIHRDCDFNKP | 1 | 0 | 0 | 1407852 | AVV62535.1 | K9N7C7 |
| AEIKDLICVQSY | 1 | 0 | 0 | 1407853 | AGT51590.1 | P36334 |
| AEILIIIMRTFR | 1 | 0 | 0 | 1407855 | ATO98186.1 | Q3LZX8 |
| AEISKINTTLLD | 1 | 0 | 0 | 1407861 | AUM60014.1 | K9N5Q8 |
| AEIVDTVSTLVY | 1 | 0 | 0 | 1407863 | APO40578.1 | P0C6X7 |
| AEKEVLRELYGT | 1 | 0 | 0 | 1407866 | AVV62524.1 | K9N7C7 |
| AEKLASPMKSDE | 1 | 0 | 0 | 1407868 | AZS52616.1 | A0A140H1G9 |
| AEKLNAFLVHDN | 1 | 0 | 0 | 1407869 | 3EJG_A | A0A0P0LKV0 |
| AEKPNLNSLCLL | 1 | 0 | 0 | 1407870 | AET99209.1 | Q0ZJ80 |
| AEKPTENACSSL | 1 | 0 | 0 | 1407871 | AVP78041.1 | P0C6W2 |
| AELAPPAGAFFF | 1 | 0 | 0 | 1407873 | ANZ78874.1 | P33469 |
| AELAPSASAFMG | 1 | 0 | 0 | 1407874 | AUM60021.1 | K9N4V7 |
| AELAPSASAFMS | 1 | 0 | 0 | 1407875 | AVV62544.1 | K9N4V7 |
| AELAPSSSAFIG | 1 | 0 | 0 | 1407876 | AVV62533.1 | K9N4V7 |
| AELAPTAGAFFF | 1 | 0 | 0 | 1407877 | AEN19370.1 | P33469 |
| AELAPTASAFMG | 1 | 0 | 0 | 1407878 | AXN92267.1 | K9N4V7 |
| AELCCFARMVWR | 1 | 0 | 0 | 1407880 | ANI69888.1 | K9N638 |
| AELCCSACMVWR | 1 | 0 | 0 | 1407881 | QBF80553.1 | K9N7C7 |
| AELCCSARMIWR | 1 | 0 | 0 | 1407882 | AZK15899.1 | K9N638 |

| | | | | | | |
|---|---|---|---|---|---|---|
| AELCCSARMVWR | 1 | 0 | 0 | 1407883 | AXN92247.1 | K9N7C7 |
| AELDDFSKQLQD | 1 | 0 | 0 | 1407884 | APO40586.1 | P59595 |
| AELDGIQYGRSG | 1 | 0 | 0 | 1407885 | AVP78041.1 | P0C6W2 |
| AELDGVQFGRSG | 1 | 0 | 0 | 1407886 | ARO76381.1 | P0C6U8 |
| AELEAAQKSYQE | 1 | 0 | 0 | 1407887 | AVV62524.1 | K9N7C7 |
| AELEAARKAYQE | 1 | 0 | 0 | 1407888 | AMQ49036.1 | K9N638 |
| AELEPPCRFVTD | 1 | 0 | 0 | 1407890 | AEA10817.1 | P0C6X7 |
| AELETAQKSYQE | 1 | 0 | 0 | 1407891 | AUM60013.1 | K9N7C7 |
| AELGYIADISSF | 1 | 0 | 0 | 1407894 | ARB07423.1 | A0A0P0LKV0 |
| AELIPNHAALFF | 1 | 0 | 0 | 1407896 | AMM42619.1 | H9EJ69 |
| AELNGVMVDKVG | 1 | 0 | 0 | 1407905 | QBP84746.1 | P0C6X6 |
| AELNQLNEPLIT | 1 | 0 | 0 | 1407906 | APO40578.1 | P0C6X7 |
| AELNYTVQKLQS | 1 | 0 | 0 | 1407907 | BAL45640.1 | A0A0P0K6L9 |
| AELSTINYKLQS | 1 | 0 | 0 | 1407910 | AUM60013.1 | K9N7C7 |
| AELSTKNYKLQS | 1 | 0 | 0 | 1407911 | AVV62535.1 | K9N7C7 |
| AELSTNNYKLQS | 1 | 0 | 0 | 1407912 | AVV62524.1 | K9N7C7 |
| AELSVSARMVWR | 1 | 0 | 0 | 1407914 | AVV62524.1 | K9N7C7 |
| AELTGTQSVSVD | 1 | 0 | 0 | 1407915 | AVN89314.1 | K9N796 |
| AELVPSTAAMLS | 1 | 0 | 0 | 1407917 | ABB90484.1 | A0A0N7J131 |
| AELVPTASAFMG | 1 | 0 | 0 | 1407918 | AXG21672.1 | K9N4V7 |
| AEMNRLQEAIKV | 1 | 0 | 0 | 1407921 | AGT51521.1 | P36334 |
| AENDTITVDQLK | 1 | 0 | 0 | 1407924 | APO40582.1 | P59596 |
| AENGTISVEELK | 1 | 0 | 0 | 1407925 | ATO98197.1 | Q3LZX9 |
| AENKDGFYETIV | 1 | 0 | 0 | 1407926 | AIV41955.1 | Q80872 |
| AENPKEEVKPFF | 1 | 0 | 0 | 1407927 | AVP78041.1 | P0C6W2 |
| AENPNPNSLCLL | 1 | 0 | 0 | 1407928 | AET99208.1 | Q0ZJ80 |
| AENPNQNSLCLL | 1 | 0 | 0 | 1407929 | Q0ZME2.1 | Q0ZJ80 |
| AENSIAYANNSI | 1 | 0 | 0 | 1407930 | AVP78031.1 | Q3LZX1 |
| AENVTESVQEEA | 1 | 0 | 0 | 1407932 | QBF80465.1 | K9N7C7 |
| AEPLVQDRAVEP | 1 | 0 | 0 | 1407934 | Q4VID0.1 | Q4VID0 |
| AEPLVQDRVVEP | 1 | 0 | 0 | 1407935 | AVR40343.1 | Q4VID0 |
| AEPVRDEFSIET | 1 | 0 | 0 | 1407936 | AAA45463.1 | A0A0N7J131 |
| AEQAMTSMYKQA | 1 | 0 | 0 | 1407937 | AVV62524.1 | K9N7C7 |
| AEQCIRGAIGDV | 1 | 0 | 0 | 1407938 | ARB07424.1 | A0A0P0ILY9 |
| AERDCYIFKRTF | 1 | 0 | 0 | 1407943 | AZS52616.1 | A0A140H1G9 |
| AERDCYIFKRTV | 1 | 0 | 0 | 1407944 | ARB07436.1 | A0A140H1G9 |
| AERDCYTLKKTV | 1 | 0 | 0 | 1407946 | ABD75615.1 | A0A140H1G9 |
| AERHRQLIEHST | 1 | 0 | 0 | 1407947 | AUM60013.1 | K9N7C7 |
| AESFKKVAAVPY | 1 | 0 | 0 | 1407949 | AUM60013.1 | K9N7C7 |
| AESHIDSDVTKP | 1 | 0 | 0 | 1407950 | APO40578.1 | P0C6X7 |
| AESKPRFVRKAE | 1 | 0 | 0 | 1407954 | AGT21359.1 | A0A0P0LKV0 |
| AESVVKTLQPVS | 1 | 0 | 0 | 1407957 | ARO76381.1 | P0C6U8 |
| AETGTSDLDRCT | 1 | 0 | 0 | 1407959 | 6NB6_A | P59594 |
| AETGTVDVGPDS | 1 | 0 | 0 | 1407960 | 6NB3_A | K9N5Q8 |
| AETLKANEETFK | 1 | 0 | 0 | 1407963 | APO40578.1 | P0C6X7 |
| AETLRATEEASR | 1 | 0 | 0 | 1407968 | AIL23989.1 | K9N7C7 |
| AETPALQVSLTQ | 1 | 0 | 0 | 1407969 | ATQ39389.1 | K9N7C7 |
| AETQKATEEAFKQSYA | 1 | 0 | 0 | 1407970 | YP_009555238.1 | P0C6X6 |
| AETSEQVVAKPK | 1 | 0 | 0 | 1407972 | AVV62535.1 | K9N7C7 |
| AETTDFCTVALA | 1 | 0 | 0 | 1407973 | NP_073551.1 | A0A0P0K6L9 |
| AETTDFFTVALA | 1 | 0 | 0 | 1407974 | CAA71056.1 | A0A0P0K6L9 |
| AETTDFLTVALA | 1 | 0 | 0 | 1407975 | CAA71147.1 | A0A0P0K6L9 |
| AETTVCTVALAS | 1 | 0 | 0 | 1407976 | CAA71146.1 | A0A0P0K6L9 |
| AEVASDSDSLPS | 1 | 0 | 0 | 1407980 | AVV62541.1 | K9N7D2 |
| AEVDWKFYDAQP | 1 | 0 | 0 | 1407982 | AVP78041.1 | P0C6W2 |
| AEVNPDLSAYYV | 1 | 0 | 0 | 1407985 | AVV62535.1 | K9N7C7 |
| AEVQIDRLITGG | 1 | 0 | 0 | 1407986 | 1WYY_A | P59594 |
| AEVQQSAEEFDD | 1 | 0 | 0 | 1407988 | AVV62524.1 | K9N7C7 |
| AEVSRQVIQEVK | 1 | 0 | 0 | 1407990 | ABM64804.1 | A0A0P0KWE8 |
| AEVVKLPSRYCC | 1 | 0 | 0 | 1407991 | ARO76392.1 | Q7TLC7 |
| AEVVKQMYGVNH | 1 | 0 | 0 | 1407992 | 6FV1_A | H9EJ42 |
| AEVVVNSANGHM | 1 | 0 | 0 | 1407994 | AIV41936.1 | P0C6X6 |
| AEWFGITQNAQG | 1 | 0 | 0 | 1407995 | AUM60014.1 | K9N5Q8 |
| AEWFGITQNTQG | 1 | 0 | 0 | 1407996 | AVV62537.1 | K9N5Q8 |
| AEWKCGYAMPQI | 1 | 0 | 0 | 1408002 | AWK59942.1 | H9EJ42 |
| AEWKCGYSMPGI | 1 | 0 | 0 | 1408003 | NP_073549.1 | A0A0P0LKV0 |
| AEYHNESGLKTV | 1 | 0 | 0 | 1408007 | AVP78041.1 | P0C6W2 |
| AEYIFGPCWSAL | 1 | 0 | 0 | 1408008 | APT69850.1 | A0A0P0LKV0 |
| AEYREYMEAYNL | 1 | 0 | 0 | 1408012 | AVV62535.1 | K9N7C7 |
| AEYREYVEAYNL | 1 | 0 | 0 | 1408013 | AVV62524.1 | K9N7C7 |
| AEYSTYDPIYKN | 1 | 0 | 0 | 1408014 | AVV62535.1 | K9N7C7 |
| AEYVTEFVQEEA | 1 | 0 | 0 | 1408016 | AVN89322.1 | K9N7C7 |
| AFAAIGDLKCPL | 1 | 0 | 0 | 1408017 | AIL49486.1 | P36334 |
| AFAAIGDLNCTL | 1 | 0 | 0 | 1408018 | BBO58773.1 | P36334 |
| AFAATIMNSLCL | 1 | 0 | 0 | 1408019 | AVV62524.1 | K9N7C7 |
| AFACADGTKHTY | 1 | 0 | 0 | 1408020 | ARO76387.1 | P59635 |
| AFACADGTRHTY | 1 | 0 | 0 | 1408021 | ATO98237.1 | Q3LZX7 |
| AFACPDGVRHTF | 1 | 0 | 0 | 1408023 | AVP78047.1 | Q3LZX7 |
| AFADCTSCSNGR | 1 | 0 | 0 | 1408024 | AUM60013.1 | K9N7C7 |
| AFADYTPCAVIA | 1 | 0 | 0 | 1408025 | ARQ84755.1 | K9N7C7 |
| AFADYTQCAVIA | 1 | 0 | 0 | 1408026 | AXN92247.1 | K9N7C7 |
| AFAECTGCGNGR | 1 | 0 | 0 | 1408027 | AVV62535.1 | K9N7C7 |
| AFAEELHGGEVD | 1 | 0 | 0 | 1408029 | AUM60028.1 | K9N7D2 |
| AFAKVQEAVNAN | 1 | 0 | 0 | 1408031 | AUM60014.1 | K9N5Q8 |
| AFALHSGYYDAN | 1 | 0 | 0 | 1408032 | AWO68966.1 | A0A0P0G321 |
| AFALHTGYYDAN | 1 | 0 | 0 | 1408033 | AVL25587.1 | A0A0P0G321 |
| AFALHTGYYDVN | 1 | 0 | 0 | 1408034 | AIW52843.1 | A0A0P0G321 |
| AFANWLAPTNAY | 1 | 0 | 0 | 1408036 | QBF80585.1 | K9N7C7 |
| AFASDSVQCYTP | 1 | 0 | 0 | 1408037 | ARU07575.1 | A0A140H1G9 |
| AFASEAARVVRS | 1 | 0 | 0 | 1408038 | AVP78041.1 | P0C6W2 |
| AFAVDPAAAYLD | 1 | 0 | 0 | 1408042 | AST12962.1 | A0A0P0LKV0 |

| | | | | | | |
|---|---|---|---|---|---|---|
| AFAVEPAKAYKD | 1 | 0 | 0 | 1408044 | APO40578.1 | P0C6X7 |
| AFAVIGDLNCPL | 1 | 0 | 0 | 1408047 | AEN19366.1 | P36334 |
| AFAVVGDLNCPL | 1 | 0 | 0 | 1408049 | QBP84704.1 | P36334 |
| AFAYADKPNHFI | 1 | 0 | 0 | 1408051 | AGT51410.1 | Q80872 |
| AFAYESAVVNGI | 1 | 0 | 0 | 1408052 | NP_073549.1 | A0A0P0LKV0 |
| AFCDEMVLKGVV | 1 | 0 | 0 | 1408055 | AIW52776.1 | A0A0P0LKV0 |
| AFCDYADKLCCA | 1 | 0 | 0 | 1408056 | AIV41888.1 | P0C6X6 |
| AFCDYADKLCHA | 1 | 0 | 0 | 1408057 | AEN19363.1 | P0C6X6 |
| AFCDYTDKLCHA | 1 | 0 | 0 | 1408059 | AGT51737.1 | P0C6X6 |
| AFCGDQVHEVAA | 1 | 0 | 0 | 1408060 | ALT66879.1 | K9N7C7 |
| AFCKPGTLVLNN | 1 | 0 | 0 | 1408061 | AGT51421.1 | P30215 |
| AFCKSGSLVLNN | 1 | 0 | 0 | 1408063 | AEN19365.1 | P30215 |
| AFCVYKLQPLTF | 1 | 0 | 0 | 1408068 | AKN24812.1 | K9N5Q8 |
| AFDCGDYAACKS | 1 | 0 | 0 | 1408070 | ARB66062.1 | P36334 |
| AFDFSSDVFKIG | 1 | 0 | 0 | 1408073 | QDH43737.1 | H9EJ42 |
| AFDGLRGGVRKA | 1 | 0 | 0 | 1408074 | AUM60013.1 | K9N7C7 |
| AFDGNMYGAFQD | 1 | 0 | 0 | 1408075 | AVV62535.1 | K9N7C7 |
| AFDGTMYGAFMD | 1 | 0 | 0 | 1408076 | AXN92247.1 | K9N7C7 |
| AFDIQQWGYVGS | 1 | 0 | 0 | 1408078 | AWK59942.1 | H9EJ42 |
| AFDLSKELKRPI | 1 | 0 | 0 | 1408088 | QBQ01836.1 | P0C6X6 |
| AFDQDGIIFNAV | 1 | 0 | 0 | 1408089 | BBA84088.1 | P36334 |
| AFDVPDGFYSTS | 1 | 0 | 0 | 1408092 | AFR79257.1 | A0A0P0K6L9 |
| AFEAFKRSNNGV | 1 | 0 | 0 | 1408095 | AEN19363.1 | P0C6X6 |
| AFEFAEGQGVPI | 1 | 0 | 0 | 1408097 | AIV41850.1 | P33469 |
| AFENHSYIYHSA | 1 | 0 | 0 | 1408107 | AVV62524.1 | K9N7C7 |
| AFETPAVQFSTA | 1 | 0 | 0 | 1408109 | AVV62535.1 | K9N7C7 |
| AFFFGSKFELAK | 1 | 0 | 0 | 1408113 | AMM42445.1 | P33469 |
| AFFFGSKLELAK | 1 | 0 | 0 | 1408115 | AEN19370.1 | P33469 |
| AFGAILSNLGVP | 1 | 0 | 0 | 1408130 | APO40578.1 | P0C6X7 |
| AFGEYNHVVAAN | 1 | 0 | 0 | 1408131 | P0C6W6.1 | P0C6W2 |
| AFGGDQIHEVAA | 1 | 0 | 0 | 1408135 | ALA49340.1 | K9N7C7 |
| AFGGDQVHEVVA | 1 | 0 | 0 | 1408137 | ALA49384.1 | K9N7C7 |
| AFGIPPSEAKGY | 1 | 0 | 0 | 1408139 | ABA60794.1 | Q19U29 |
| AFGLQDCVTGVN | 1 | 0 | 0 | 1408141 | AGT51337.1 | H9EJ42 |
| AFGNAGVCYDIF | 1 | 0 | 0 | 1408145 | NP_073549.1 | A0A0P0LKV0 |
| AFGNTGVCYDFD | 1 | 0 | 0 | 1408146 | AWK59942.1 | H9EJ42 |
| AFGRRGPGALQG | 1 | 0 | 0 | 1408149 | AVV62544.1 | K9N4V7 |
| AFGSPFITNGMS | 1 | 0 | 0 | 1408152 | AWK59942.1 | H9EJ42 |
| AFHANSSEPALI | 1 | 0 | 0 | 1408157 | AGT51481.1 | P36334 |
| AFHKPAYDVSAF | 1 | 0 | 0 | 1408160 | AHI48661.1 | K9N7C7 |
| AFHKVQDAVNNN | 1 | 0 | 0 | 1408161 | AHI48803.1 | K9N5Q8 |
| AFHRIESTDPIV | 1 | 0 | 0 | 1408168 | AVN89337.1 | K9N7D2 |
| AFHTPAYDGSAF | 1 | 0 | 0 | 1408172 | AHI48736.1 | K9N7C7 |
| AFHVNSSEPALL | 1 | 0 | 0 | 1408175 | AIX10749.1 | P36334 |
| AFIADKEKLISL | 1 | 0 | 0 | 1408177 | AUM60013.1 | K9N7C7 |
| AFIAFNHPIQVD | 1 | 0 | 0 | 1408178 | ASU90142.1 | K9N5Q8 |
| AFIDTVKGLDYK | 1 | 0 | 0 | 1408180 | ARO76381.1 | P0C6U8 |
| AFIECGSCGNDS | 1 | 0 | 0 | 1408181 | AXN92247.1 | K9N7C7 |
| AFIECGSCGNES | 1 | 0 | 0 | 1408182 | AVN89333.1 | K9N7C7 |
| AFIECGSCGNNS | 1 | 0 | 0 | 1408183 | ALA49340.1 | K9N7C7 |
| AFIETVKNLDYS | 1 | 0 | 0 | 1408184 | APO40578.1 | P0C6X7 |
| AFIGVLDITDAA | 1 | 0 | 0 | 1408186 | AFR79249.1 | A0A0P0ILY9 |
| AFIKHKGGDSHE | 1 | 0 | 0 | 1408188 | AUM60013.1 | K9N7C7 |
| AFIKHKGGDSTE | 1 | 0 | 0 | 1408189 | AVV62524.1 | K9N7C7 |
| AFIKTAQADTAG | 1 | 0 | 0 | 1408190 | AXN92261.1 | K9N796 |
| AFISVFDITDAA | 1 | 0 | 0 | 1408196 | APT69868.1 | A0A0P0ILY9 |
| AFKAVCSVIDLK | 1 | 0 | 0 | 1408203 | AXN92247.1 | K9N7C7 |
| AFKCNRFGSNDI | 1 | 0 | 0 | 1408207 | AXX83326.1 | P30215 |
| AFKQSYASATIQ | 1 | 0 | 0 | 1408216 | AEN19363.1 | P0C6X6 |
| AFKRQIRIACRK | 1 | 0 | 0 | 1408218 | ASU91217.1 | K9N7C7 |
| AFLMAEDGITKL | 1 | 0 | 0 | 1408247 | AKN58609.1 | K9N638 |
| AFLNGSCGSVGF | 1 | 0 | 0 | 1408249 | 3F9E_A | P0C6U8 |
| AFLPAQDTWGTS | 1 | 0 | 0 | 1408251 | ATO98205.1 | Q3LZX1 |
| AFLQKLEDNPLF | 1 | 0 | 0 | 1408257 | AEN19363.1 | P0C6X6 |
| AFLQKLEDNSLF | 1 | 0 | 0 | 1408258 | AIV41864.1 | P0C6X6 |
| AFLTAARLCVQC | 1 | 0 | 0 | 1408260 | AVV62531.1 | K9N5R3 |
| AFLTLLFYYINK | 1 | 0 | 0 | 1408264 | AXN92247.1 | K9N7C7 |
| AFLTQYITRMTA | 1 | 0 | 0 | 1408265 | AVV62529.1 | K9N643 |
| AFLVVLIILWFV | 1 | 0 | 0 | 1408267 | AZK15904.1 | K9N7D2 |
| AFMGMSQFKLTH | 1 | 0 | 0 | 1408271 | AXN92267.1 | K9N4V7 |
| AFMGMSQFKVTH | 1 | 0 | 0 | 1408272 | AUM60021.1 | K9N4V7 |
| AFMKHKGGDLTD | 1 | 0 | 0 | 1408273 | AHI48527.1 | K9N638 |
| AFMPVQSRKFIV | 1 | 0 | 0 | 1408283 | AEN19363.1 | P0C6X6 |
| AFNHPIQIDQLN | 1 | 0 | 0 | 1408294 | ASU91208.1 | K9N5Q8 |
| AFNHPIQVDQFN | 1 | 0 | 0 | 1408296 | ASU90362.1 | K9N5Q8 |
| AFNHPIQVGQLN | 1 | 0 | 0 | 1408298 | ALJ54452.1 | K9N5Q8 |
| AFNKLLGYDPSK | 1 | 0 | 0 | 1408300 | AVV62535.1 | K9N7C7 |
| AFNNALDAIQEG | 1 | 0 | 0 | 1408301 | CAA79896.1 | P36334 |
| AFNNALGAIQEG | 1 | 0 | 0 | 1408302 | AVI15011.1 | P36334 |
| AFNNALHAIQQG | 1 | 0 | 0 | 1408303 | AEN19366.1 | P36334 |
| AFNNALHSIQQG | 1 | 0 | 0 | 1408304 | AIX10759.1 | P36334 |
| AFNNALNAIQQG | 1 | 0 | 0 | 1408305 | AGT51491.1 | P36334 |
| AFNNALYAIQQG | 1 | 0 | 0 | 1408307 | AIX10750.1 | P36334 |
| AFNPETNAIISL | 1 | 0 | 0 | 1408311 | ARE29965.1 | U3M6Q8 |
| AFNQDGIIFNAE | 1 | 0 | 0 | 1408312 | P36334.1 | P36334 |
| AFNQDGVIFNAV | 1 | 0 | 0 | 1408314 | CAA83660.1 | P36334 |
| AFNQLTADLQHK | 1 | 0 | 0 | 1408317 | ARU07575.1 | A0A140H1G9 |
| AFNSYGNMFQTQ | 1 | 0 | 0 | 1408320 | APO40579.1 | P59594 |
| AFNVVKHGRFNG | 1 | 0 | 0 | 1408333 | AVV62524.1 | K9N7C7 |
| AFNWLLLAGTLH | 1 | 0 | 0 | 1408337 | AXN92247.1 | K9N7C7 |
| AFPFETFSACRA | 1 | 0 | 0 | 1408342 | ATG84733.1 | K9N638 |

| | | | | | | |
|---|---|---|---|---|---|---|
| AFPLDEAEDFTE | 1 | 0 | 0 | 1408346 | AIL23988.1 | K9N7C7 |
| AFPLDEAENVTE | 1 | 0 | 0 | 1408348 | QBF80465.1 | K9N7C7 |
| AFPLDEAEYVTE | 1 | 0 | 0 | 1408349 | AVN89322.1 | K9N7C7 |
| AFPLNEAEDVTE | 1 | 0 | 0 | 1408352 | AHI48527.1 | K9N638 |
| AFPSAKVEGCMV | 1 | 0 | 0 | 1408354 | 2PWX_A | P0C6X7 |
| AFPSDAAHVVRS | 1 | 0 | 0 | 1408355 | AVP78030.1 | P0C6W2 |
| AFPSGKVECCMV | 1 | 0 | 0 | 1408356 | APO40578.1 | P0C6X7 |
| AFQDRQIHQAQL | 1 | 0 | 0 | 1408366 | AVV62524.1 | K9N7C7 |
| AFQDRQIYQSQL | 1 | 0 | 0 | 1408367 | AUM60013.1 | K9N7C7 |
| AFQKVQDAVNNI | 1 | 0 | 0 | 1408370 | QFS19596.1 | K9N5Q8 |
| AFQNHSYIYHSA | 1 | 0 | 0 | 1408373 | AVV62535.1 | K9N7C7 |
| AFQPVYRTFTQA | 1 | 0 | 0 | 1408374 | AGT51379.1 | H9EJ42 |
| AFQSVYRTFTQA | 1 | 0 | 0 | 1408381 | AWK59942.1 | H9EJ42 |
| AFRDLKPLPFFY | 1 | 0 | 0 | 1408385 | AXN92247.1 | K9N7C7 |
| AFRELTRELNGG | 1 | 0 | 0 | 1408391 | ARO76381.1 | P0C6U8 |
| AFRNACGTNVPL | 1 | 0 | 0 | 1408394 | AXN92247.1 | K9N7C7 |
| AFRNVVFELCDF | 1 | 0 | 0 | 1408395 | AVV62529.1 | K9N643 |
| AFSHSANDPLLG | 1 | 0 | 0 | 1408403 | AVV62524.1 | K9N7C7 |
| AFSLSLFRPIQL | 1 | 0 | 0 | 1408414 | AVV62541.1 | K9N7D2 |
| AFSMSLFRPIQL | 1 | 0 | 0 | 1408415 | AVV62530.1 | K9N7D2 |
| AFSNALDAIQEG | 1 | 0 | 0 | 1408416 | CAA83660.1 | P36334 |
| AFSPAQDTWGTS | 1 | 0 | 0 | 1408419 | 5X58_A | P59594 |
| AFTAPHGHVMVE | 1 | 0 | 0 | 1408425 | APO40578.1 | P0C6X7 |
| AFTREMQQQPLL | 1 | 0 | 0 | 1408438 | AIW52723.1 | A0A0N7J131 |
| AFTRYVDNNFCG | 1 | 0 | 0 | 1408439 | ATO98229.1 | P0C6W2 |
| AFVAQQLVRSEA | 1 | 0 | 0 | 1408445 | AUM60014.1 | K9N5Q8 |
| AFVAQQLVRSET | 1 | 0 | 0 | 1408447 | AVV62526.1 | K9N5Q8 |
| AFVCGDCAACKS | 1 | 0 | 0 | 1408448 | CAA83661.1 | P36334 |
| AFVCGDFAACKS | 1 | 0 | 0 | 1408449 | AAX85669.1 | P36334 |
| AFVCGDYAACKL | 1 | 0 | 0 | 1408450 | AEN19366.1 | P36334 |
| AFVCGDYAACKS | 1 | 0 | 0 | 1408451 | CAA79896.1 | P36334 |
| AFVECGSCGNDS | 1 | 0 | 0 | 1408455 | AXN73479.1 | K9N7C7 |
| AFVECTGCGNGR | 1 | 0 | 0 | 1408456 | AVV62524.1 | K9N7C7 |
| AFVKCNCGSENW | 1 | 0 | 0 | 1408460 | AGT51379.1 | H9EJ42 |
| AFVKCNCGSESW | 1 | 0 | 0 | 1408461 | AWK59942.1 | H9EJ42 |
| AFVNSVFNICQA | 1 | 0 | 0 | 1408469 | ADO24692.1 | P0C6X6 |
| AFVPQPTGVFTN | 1 | 0 | 0 | 1408470 | AGT51481.1 | P36334 |
| AFVRGDYAACKL | 1 | 0 | 0 | 1408471 | ANZ78843.1 | P36334 |
| AFVVTGGAQTST | 1 | 0 | 0 | 1408480 | ARO76382.1 | P59594 |
| AFWVFSYCRKLG | 1 | 0 | 0 | 1408487 | AEN19363.1 | P0C6X6 |
| AFYGVALKDCGF | 1 | 0 | 0 | 1408492 | AEN19363.1 | P0C6X6 |
| AFYILPSIISNEKQEI | 1 | 0 | 0 | 1408495 | P0DTC1.1 | P0DTD1 |
| AFYPACTNGHAV | 1 | 0 | 0 | 1408502 | AXN92247.1 | K9N7C7 |
| AFYRDLYSSPTT | 1 | 0 | 0 | 1408505 | AXN92247.1 | K9N7C7 |
| AFYVLPSETPNA | 1 | 0 | 0 | 1408509 | P0C6W6.1 | P0C6W2 |
| AGAGWTAGLSSF | 1 | 0 | 0 | 1408522 | AVV62526.1 | K9N5Q8 |
| AGAGWTAGLSSS | 1 | 0 | 0 | 1408523 | AOR17480.1 | K9N5Q8 |
| AGALNAATNGEM | 1 | 0 | 0 | 1408533 | APO40578.1 | P0C6X7 |
| AGALWDIAVINT | 1 | 0 | 0 | 1408537 | AVV62524.1 | K9N7C7 |
| AGAMLSENFTSY | 1 | 0 | 0 | 1408540 | ADK37701.1 | A0A0P0K6L9 |
| AGAPFYLNVQYR | 1 | 0 | 0 | 1408541 | ARB66070.1 | P36334 |
| AGASRAIEKVNE | 1 | 0 | 0 | 1408544 | BBA20986.1 | A0A140H1H1 |
| AGAWNIGSNKTL | 1 | 0 | 0 | 1408546 | APO40578.1 | P0C6X7 |
| AGAYSSHLQSYA | 1 | 0 | 0 | 1408547 | AVV62524.1 | K9N7C7 |
| AGAYSSPLQSYA | 1 | 0 | 0 | 1408548 | AVV62535.1 | K9N7C7 |
| AGCEQSYFTKKD | 1 | 0 | 0 | 1408550 | AEN19363.1 | P0C6X6 |
| AGCFVNDLLKTD | 1 | 0 | 0 | 1408552 | ANZ78855.1 | P0C6X6 |
| AGCLIGAEHVNA | 1 | 0 | 0 | 1408557 | AVP78031.1 | Q3LZX1 |
| AGCQTTNGMNTS | 1 | 0 | 0 | 1408561 | AIW52756.1 | A0A0P0K6L9 |
| AGCQTTNGTNTI | 1 | 0 | 0 | 1408562 | AIW52741.1 | A0A0P0K6L9 |
| AGDAVHSLTWLW | 1 | 0 | 0 | 1408565 | ARA15428.1 | A0A0P0ILY9 |
| AGDGISRPKLHE | 1 | 0 | 0 | 1408567 | ARO76383.1 | P59632 |
| AGDGISTPKLKE | 1 | 0 | 0 | 1408568 | ATO98194.1 | Q3LZX0 |
| AGDGVTIPIKTQ | 1 | 0 | 0 | 1408569 | APO40580.1 | P59632 |
| AGDQAAKLSPWF | 1 | 0 | 0 | 1408573 | AVV62524.1 | K9N7C7 |
| AGDSSWHIYLKS | 1 | 0 | 0 | 1408579 | AWO68966.1 | A0A0P0G321 |
| AGEAANFCALIL | 1 | 0 | 0 | 1408582 | AVP78041.1 | P0C6W2 |
| AGEEKLSSRMYC | 1 | 0 | 0 | 1408589 | ATO98229.1 | P0C6W2 |
| AGEENLSSRMYC | 1 | 0 | 0 | 1408591 | ARO76381.1 | P0C6U8 |
| AGEEVFAPKLYC | 1 | 0 | 0 | 1408592 | AEN19363.1 | P0C6X6 |
| AGEPCVASEQEE | 1 | 0 | 0 | 1408594 | AVI15051.1 | P0C6X6 |
| AGFDSWFSQRGG | 1 | 0 | 0 | 1408601 | P0C6W6.1 | P0C6W2 |
| AGFDTWFSQRGG | 1 | 0 | 0 | 1408602 | ARO76381.1 | P0C6U8 |
| AGFGVDEEKCGV | 1 | 0 | 0 | 1408604 | BBA20983.1 | A0A140H1H1 |
| AGFKRTYNFVWY | 1 | 0 | 0 | 1408606 | ARK08626.1 | H9EJ42 |
| AGFNIWVPHSFD | 1 | 0 | 0 | 1408611 | AWK59942.1 | H9EJ42 |
| AGFNTEFASNSS | 1 | 0 | 0 | 1408612 | ANI69888.1 | K9N638 |
| AGGHCYGADLKS | 1 | 0 | 0 | 1408625 | APO40578.1 | P0C6X7 |
| AGGIADAINKAS | 1 | 0 | 0 | 1408630 | AVV62535.1 | K9N7C7 |
| AGGTCSSISQLG | 1 | 0 | 0 | 1408637 | APO40579.1 | P59594 |
| AGGTTEMLAKAL | 1 | 0 | 0 | 1408638 | AVP78041.1 | P0C6W2 |
| AGGTTEMLARAL | 1 | 0 | 0 | 1408639 | ARO76381.1 | P0C6U8 |
| AGGVYTNSCLVS | 1 | 0 | 0 | 1408644 | QBM11736.1 | K9N638 |
| AGHADNNIEKPK | 1 | 0 | 0 | 1408646 | AVV62526.1 | K9N5Q8 |
| AGHDDFSTEIIG | 1 | 0 | 0 | 1408652 | APO40578.1 | P0C6X7 |
| AGHYDGRIGELP | 1 | 0 | 0 | 1408662 | ARB07436.1 | A0A140H1G9 |
| AGHYTGQAGEMP | 1 | 0 | 0 | 1408663 | AQN78669.1 | P0C6X6 |
| AGIAGARIPGVP | 1 | 0 | 0 | 1408664 | AVV62535.1 | K9N7C7 |
| AGICASYHTVSS | 1 | 0 | 0 | 1408670 | ATO98231.1 | Q3LZX1 |
| AGIFGAKPLDSL | 1 | 0 | 0 | 1408677 | ARO76381.1 | P0C6U8 |
| AGIFGVDPGLSL | 1 | 0 | 0 | 1408679 | APO40578.1 | P0C6X7 |

| | | | | | | |
|---|---|---|---|---|---|---|
| AGIFGVDPVHSL | 1 | 0 | 0 | 1408681 | AVP78041.1 | P0C6W2 |
| AGIGNTFICEEV | 1 | 0 | 0 | 1408686 | AVV62524.1 | K9N7C7 |
| AGIRDLICVQSY | 1 | 0 | 0 | 1408698 | CAA83661.1 | P36334 |
| AGKASYALSEQL | 1 | 0 | 0 | 1408702 | AVP78030.1 | P0C6W2 |
| AGKFDSFKGECP | 1 | 0 | 0 | 1408704 | APO40578.1 | P0C6X7 |
| AGKNKAAPKVVT | 1 | 0 | 0 | 1408706 | AVV62533.1 | K9N4V7 |
| AGKSCAPLSDQI | 1 | 0 | 0 | 1408707 | APO40578.1 | P0C6X7 |
| AGLDQTSFNVNS | 1 | 0 | 0 | 1408711 | AVV62535.1 | K9N7C7 |
| AGLFRIDVPSAE | 1 | 0 | 0 | 1408716 | ALW82670.1 | K9N796 |
| AGLFRIFLRKAY | 1 | 0 | 0 | 1408718 | AHY04433.1 | K9N638 |
| AGLKLPSGIMFN | 1 | 0 | 0 | 1408728 | NP_073549.1 | A0A0P0LKV0 |
| AGLNGDFNLTLL | 1 | 0 | 0 | 1408732 | AVV62537.1 | K9N5Q8 |
| AGLNGDFNLTML | 1 | 0 | 0 | 1408733 | AVV62526.1 | K9N5Q8 |
| AGLSSSAAIPFA | 1 | 0 | 0 | 1408741 | AOR17480.1 | K9N5Q8 |
| AGLVTDSVIVMA | 1 | 0 | 0 | 1408747 | APO40578.1 | P0C6X7 |
| AGLWLDDTVYCP | 1 | 0 | 0 | 1408750 | 3FZD_A | P0C6X7 |
| AGLYTNFPIDFP | 1 | 0 | 0 | 1408751 | AZK15901.1 | K9N796 |
| AGLYTNFRIDIP | 1 | 0 | 0 | 1408753 | AXP07346.1 | K9N796 |
| AGLYTNSRIDVP | 1 | 0 | 0 | 1408755 | AGV08439.1 | K9N796 |
| AGMHFGQCDFRS | 1 | 0 | 0 | 1408760 | AVV62543.1 | K9N7A1 |
| AGMHFGQCDYAQ | 1 | 0 | 0 | 1408761 | AUM60030.1 | K9N7A1 |
| AGNGGDTALALL | 1 | 0 | 0 | 1408768 | AVP78038.1 | Q3LZX4 |
| AGNIFVVNKDTY | 1 | 0 | 0 | 1408771 | AVV62524.1 | K9N7C7 |
| AGNKFGLYKNSA | 1 | 0 | 0 | 1408772 | QED88010.1 | H9EJ42 |
| AGNYRRPAIVQD | 1 | 0 | 0 | 1408788 | AVV62543.1 | K9N7A1 |
| AGPDLEGKFYGP | 1 | 0 | 0 | 1408791 | ACV88184.1 | P0C6X7 |
| AGPFMDDAINVG | 1 | 0 | 0 | 1408792 | AVN89322.1 | K9N7C7 |
| AGPFMGNAINVG | 1 | 0 | 0 | 1408794 | ATQ39389.1 | K9N7C7 |
| AGPSVETSDSQV | 1 | 0 | 0 | 1408800 | AGT51617.1 | P0C6X6 |
| AGPYVEGAINVG | 1 | 0 | 0 | 1408802 | AVV62524.1 | K9N7C7 |
| AGQDISLLDVAY | 1 | 0 | 0 | 1408805 | APO40578.1 | P0C6X7 |
| AGQEQTNCNISS | 1 | 0 | 0 | 1408806 | ASU45728.1 | K9N7C7 |
| AGQEQTNCNTSA | 1 | 0 | 0 | 1408807 | AXN73479.1 | K9N7C7 |
| AGQEQTNCNTSS | 1 | 0 | 0 | 1408808 | AXN92247.1 | K9N7C7 |
| AGQPCVASEQDE | 1 | 0 | 0 | 1408815 | AIV41846.1 | P0C6X6 |
| AGQPCVASEQEE | 1 | 0 | 0 | 1408816 | AEN19363.1 | P0C6X6 |
| AGQPCVVSEQEE | 1 | 0 | 0 | 1408817 | AVQ05261.1 | P0C6X6 |
| AGQPYVASEQEE | 1 | 0 | 0 | 1408818 | AGT51707.1 | P0C6X6 |
| AGQQFVKETTDM | 1 | 0 | 0 | 1408819 | AVQ05261.1 | P0C6X6 |
| AGQQFVKETTNM | 1 | 0 | 0 | 1408820 | AEN19363.1 | P0C6X6 |
| AGQSCVVSEQEE | 1 | 0 | 0 | 1408821 | ATN39862.1 | P0C6X6 |
| AGQSIQALQTAA | 1 | 0 | 0 | 1408822 | APO40578.1 | P0C6X7 |
| AGQSTYERHPLS | 1 | 0 | 0 | 1408823 | APO40578.1 | P0C6X7 |
| AGQVHVLNHVDP | 1 | 0 | 0 | 1408824 | P0C5B5.1 | Q0ZJ80 |
| AGQVQTFYPRLQ | 1 | 0 | 0 | 1408825 | AIL23945.1 | K9N7C7 |
| AGRPCVASEQEE | 1 | 0 | 0 | 1408827 | QRK03919.1 | P0C6X6 |
| AGRQTTNGMNTS | 1 | 0 | 0 | 1408828 | AKA09607.1 | A0A0P0K6L9 |
| AGSAAAPFTKPF | 1 | 0 | 0 | 1408835 | 2VRI_A | H9EJ42 |
| AGSALQIPFAMQ | 1 | 0 | 0 | 1408836 | Q3I5J5.1 | Q3LZX1 |
| AGSATEVPANST | 1 | 0 | 0 | 1408837 | ACV88184.1 | P0C6X7 |
| AGSDKGITPGTS | 1 | 0 | 0 | 1408839 | AXN92247.1 | K9N7C7 |
| AGSDKGVAPGSA | 1 | 0 | 0 | 1408840 | APO40578.1 | P0C6X7 |
| AGSLELTTMRYF | 1 | 0 | 0 | 1408852 | AWK59942.1 | H9EJ42 |
| AGSNDIWMQNKG | 1 | 0 | 0 | 1408853 | AGT51729.1 | P30215 |
| AGSNISAVIYSG | 1 | 0 | 0 | 1408855 | AXN92247.1 | K9N7C7 |
| AGSNTEFAANSS | 1 | 0 | 0 | 1408856 | AVV62524.1 | K9N7C7 |
| AGSNTEFAFNSS | 1 | 0 | 0 | 1408857 | QAT98908.1 | K9N7C7 |
| AGSNTELAANSS | 1 | 0 | 0 | 1408859 | AVV62535.1 | K9N7C7 |
| AGSQTTNGTNTN | 1 | 0 | 0 | 1408861 | AIW52735.1 | A0A0P0K6L9 |
| AGSTFISDEVAR | 1 | 0 | 0 | 1408867 | P0C6W6.1 | P0C6W2 |
| AGSVGFNIDYDC | 1 | 0 | 0 | 1408870 | 1Z1J_A | P0C6U8 |
| AGSVGYTKEGSV | 1 | 0 | 0 | 1408871 | 4WMD_A | K9N7C7 |
| AGSVYTNSCLVA | 1 | 0 | 0 | 1408873 | AVV62524.1 | K9N7C7 |
| AGSVYTNSCLVS | 1 | 0 | 0 | 1408874 | AXN92247.1 | K9N7C7 |
| AGTAVINLRSDQ | 1 | 0 | 0 | 1408879 | AQN78733.1 | P0C6X6 |
| AGTAVINLRTDQ | 1 | 0 | 0 | 1408880 | AIX09796.1 | P0C6X6 |
| AGTDIEGNFYGP | 1 | 0 | 0 | 1408881 | APO40578.1 | P0C6X7 |
| AGTILVDNDLHP | 1 | 0 | 0 | 1408891 | QDH43722.1 | P0C6X6 |
| AGTKLRSLNLGE | 1 | 0 | 0 | 1408895 | ATO98179.1 | P0C6W2 |
| AGTKYRHCDLDT | 1 | 0 | 0 | 1408896 | BBA20986.1 | A0A140H1H1 |
| AGTKYRHCDVDS | 1 | 0 | 0 | 1408897 | ARB07438.1 | A0A140H1H1 |
| AGTKYRHCDVDT | 1 | 0 | 0 | 1408898 | AGT17758.1 | A0A140H1H1 |
| AGTNHLTCDNLC | 1 | 0 | 0 | 1408901 | ARE30008.1 | P36334 |
| AGTNHLTCHNLC | 1 | 0 | 0 | 1408902 | BBO58900.1 | P36334 |
| AGTNSLTCDNLC | 1 | 0 | 0 | 1408903 | AWW13566.1 | P36334 |
| AGTNSLTCGNLC | 1 | 0 | 0 | 1408904 | BBA20976.1 | P36334 |
| AGTNYLNCDNLC | 1 | 0 | 0 | 1408905 | AIL49520.1 | P36334 |
| AGTNYLTCHNAA | 1 | 0 | 0 | 1408907 | CAA83660.1 | P36334 |
| AGTNYLTCHNAV | 1 | 0 | 0 | 1408908 | CAA83661.1 | P36334 |
| AGTNYLTCHNLC | 1 | 0 | 0 | 1408909 | AVQ05264.1 | P36334 |
| AGVAGARVPGVP | 1 | 0 | 0 | 1408913 | AVV62524.1 | K9N7C7 |
| AGVATQGARSMY | 1 | 0 | 0 | 1408916 | AUM60013.1 | K9N7C7 |
| AGVCLGAGQFMG | 1 | 0 | 0 | 1408917 | AIV41774.1 | P0C6X6 |
| AGVESDIDTGDI | 1 | 0 | 0 | 1408922 | AVV62535.1 | K9N7C7 |
| AGVESDIETGDI | 1 | 0 | 0 | 1408923 | AVV62524.1 | K9N7C7 |
| AGVFTDHDVVYA | 1 | 0 | 0 | 1408925 | CAA83661.1 | P36334 |
| AGVFTNHDVVYA | 1 | 0 | 0 | 1408926 | ATP16757.1 | P36334 |
| AGVGVIRISNDT | 1 | 0 | 0 | 1408928 | AIW52741.1 | A0A0P0K6L9 |
| AGVGWTTGLSSF | 1 | 0 | 0 | 1408931 | AOR17479.1 | K9N5Q8 |
| AGVIAQGARGTY | 1 | 0 | 0 | 1408932 | AXN92247.1 | K9N7C7 |
| AGVISQYNYKQS | 1 | 0 | 0 | 1408935 | AVV62537.1 | K9N5Q8 |

| | | | | | | |
|---|---|---|---|---|---|---|
| AGVKLRSLNLGE | 1 | 0 | 0 | 1408936 | ATO98217.1 | P0C6W2 |
| AGVKYFCGMTLK | 1 | 0 | 0 | 1408937 | AST12962.1 | A0A0P0LKV0 |
| AGVLTNHDVVYA | 1 | 0 | 0 | 1408939 | CAA79896.1 | P36334 |
| AGVSILSTVTGR | 1 | 0 | 0 | 1408946 | AIL49624.1 | P0C6X6 |
| AGVSVLSTMTGR | 1 | 0 | 0 | 1408947 | AJS10811.1 | P0C6X6 |
| AGVTAQGARGTY | 1 | 0 | 0 | 1408949 | AHI48793.1 | K9N7C7 |
| AGVVANGVTAKG | 1 | 0 | 0 | 1408950 | ABB90495.1 | A0A0N7J131 |
| AGVVAQFNYKQS | 1 | 0 | 0 | 1408951 | AVV62526.1 | K9N5Q8 |
| AGVVAQGARNMY | 1 | 0 | 0 | 1408952 | AVV62524.1 | K9N7C7 |
| AGVVAQGARNRY | 1 | 0 | 0 | 1408953 | AVV62535.1 | K9N7C7 |
| AGVVTGNVLKHC | 1 | 0 | 0 | 1408955 | AUM60013.1 | K9N7C7 |
| AGVVTSNVLKHC | 1 | 0 | 0 | 1408957 | AUM60023.1 | K9N7C7 |
| AGWTAGLSSSAA | 1 | 0 | 0 | 1408965 | AOR17480.1 | K9N5Q8 |
| AGYGGFGCKDSF | 1 | 0 | 0 | 1408970 | AEN19363.1 | P0C6X6 |
| AGYKVLPPLYDV | 1 | 0 | 0 | 1408976 | AUM60014.1 | K9N5Q8 |
| AGYNTNATLNSF | 1 | 0 | 0 | 1408977 | AVV62537.1 | K9N5Q8 |
| AGYTLDEARTAL | 1 | 0 | 0 | 1408983 | ARO76381.1 | P0C6U8 |
| AHADPNHFTLGK | 1 | 0 | 0 | 1408991 | AUM60024.1 | K9N5Q8 |
| AHADPNRFILGK | 1 | 0 | 0 | 1408992 | AUM60014.1 | K9N5Q8 |
| AHASKNACGTNV | 1 | 0 | 0 | 1409002 | AVV62524.1 | K9N7C7 |
| AHATLDSIGTNF | 1 | 0 | 0 | 1409006 | AEN19363.1 | P0C6X6 |
| AHATRDNIGTNF | 1 | 0 | 0 | 1409007 | ARB07436.1 | A0A140H1G9 |
| AHAYGSMKPHVR | 1 | 0 | 0 | 1409011 | AVV62524.1 | K9N7C7 |
| AHDNILSVKFSV | 1 | 0 | 0 | 1409015 | AUM60013.1 | K9N7C7 |
| AHELHHHMRVPI | 1 | 0 | 0 | 1409017 | AVV62529.1 | K9N643 |
| AHGGEVDIQPQV | 1 | 0 | 0 | 1409023 | AUM60018.1 | K9N7D2 |
| AHIPSGTVLGNF | 1 | 0 | 0 | 1409031 | AIW52757.1 | A0A0P0K6L9 |
| AHKHDINLTNEG | 1 | 0 | 0 | 1409040 | AUM60013.1 | K9N7C7 |
| AHKHDIQLTNES | 1 | 0 | 0 | 1409043 | AVN89451.1 | K9N7C7 |
| AHKHDIQLTTEG | 1 | 0 | 0 | 1409044 | AVV62535.1 | K9N7C7 |
| AHLNEIAKNLNE | 1 | 0 | 0 | 1409050 | APO40579.1 | P59594 |
| AHNSRSTSRTSS | 1 | 0 | 0 | 1409058 | AIL49429.1 | P33469 |
| AHPAEPLVQDRV | 1 | 0 | 0 | 1409061 | AVR40343.1 | Q4VID0 |
| AHPVEPLVQDRA | 1 | 0 | 0 | 1409065 | AHF54653.1 | Q4VID0 |
| AHPVEPLVQDRV | 1 | 0 | 0 | 1409066 | ARE29989.1 | Q4VID0 |
| AHSSIKQGTQIY | 1 | 0 | 0 | 1409078 | AEN19363.1 | P0C6X6 |
| AHTAGVCIKYYA | 1 | 0 | 0 | 1409081 | AWK59942.1 | H9EJ42 |
| AHTGRNYPKCDC | 1 | 0 | 0 | 1409082 | AJS10812.1 | P0C6X6 |
| AHTPSGTVLGNF | 1 | 0 | 0 | 1409084 | AMM42538.1 | A0A0P0K6L9 |
| AHVMFKHNDFSS | 1 | 0 | 0 | 1409092 | AVP78041.1 | P0C6W2 |
| AHVNATAAYGLC | 1 | 0 | 0 | 1409093 | AUM60014.1 | K9N5Q8 |
| AIAFCKSGNLVL | 1 | 0 | 0 | 1409106 | AIL49349.1 | P30215 |
| AIAKIVANWLSV | 1 | 0 | 0 | 1409110 | ARU07575.1 | A0A140H1G9 |
| AIAKPKRLRKKR | 1 | 0 | 0 | 1409111 | ATQ39389.1 | K9N7C7 |
| AIANIIYCNSVI | 1 | 0 | 0 | 1409115 | AIW52757.1 | A0A0P0K6L9 |
| AIASKFSSLPSY | 1 | 0 | 0 | 1409120 | AVP78041.1 | P0C6W2 |
| AIASNCYSSLIV | 1 | 0 | 0 | 1409122 | ANI69889.1 | K9N5Q8 |
| AICAAFDFSSDV | 1 | 0 | 0 | 1409129 | QDH43737.1 | H9EJ42 |
| AIDCAQNPLSEL | 1 | 0 | 0 | 1409155 | ATO98193.1 | Q3LZX1 |
| AIDCGHDDLAQL | 1 | 0 | 0 | 1409157 | AVV62526.1 | K9N5Q8 |
| AIDCGYDDLSQL | 1 | 0 | 0 | 1409158 | AUM60014.1 | K9N5Q8 |
| AIDCSDGNYNRN | 1 | 0 | 0 | 1409159 | ASU91010.1 | K9N5Q8 |
| AIDFITWRRQEL | 1 | 0 | 0 | 1409160 | QBP84714.1 | Q04853 |
| AIDVEHLELKET | 1 | 0 | 0 | 1409166 | AWK59942.1 | H9EJ42 |
| AIDVVQYEADRR | 1 | 0 | 0 | 1409168 | AEN19363.1 | P0C6X6 |
| AIEDLLFNKITI | 1 | 0 | 0 | 1409175 | AUM60014.1 | K9N5Q8 |
| AIEDLLFSKVKL | 1 | 0 | 0 | 1409176 | CAA83660.1 | P36334 |
| AIEDLLFSKVRL | 1 | 0 | 0 | 1409177 | CAA83661.1 | P36334 |
| AIEEKLSPCKEI | 1 | 0 | 0 | 1409180 | AGT51707.1 | P0C6X6 |
| AIENALNSCKDH | 1 | 0 | 0 | 1409184 | ARB07436.1 | A0A140H1G9 |
| AIENMWLSYKVG | 1 | 0 | 0 | 1409186 | ARU07575.1 | A0A140H1G9 |
| AIETAAYREAAA | 1 | 0 | 0 | 1409188 | ALW82690.1 | K9N7C7 |
| AIFAYSPQLTLV | 1 | 0 | 0 | 1409192 | AXN92247.1 | K9N7C7 |
| AIFNDFHANRPG | 1 | 0 | 0 | 1409199 | AVV62535.1 | K9N7C7 |
| AIFNDFHASRPG | 1 | 0 | 0 | 1409200 | AUM60013.1 | K9N7C7 |
| AIFNDYHANRPG | 1 | 0 | 0 | 1409202 | AVV62524.1 | K9N7C7 |
| AIFTGCVGSWNK | 1 | 0 | 0 | 1409207 | AUM60023.1 | K9N7C7 |
| AIGATAALPTQF | 1 | 0 | 0 | 1409210 | ARO76382.1 | P59594 |
| AIGDLKCPLDSR | 1 | 0 | 0 | 1409211 | AIL49486.1 | P36334 |
| AIGDLNCTLDPR | 1 | 0 | 0 | 1409212 | BBO58773.1 | P36334 |
| AIGDVGLCMYRL | 1 | 0 | 0 | 1409213 | ARB07424.1 | A0A0P0ILY9 |
| AIGKIQESLTST | 1 | 0 | 0 | 1409216 | AVP78031.1 | Q3LZX1 |
| AIGSSLQEILSR | 1 | 0 | 0 | 1409223 | CAA83660.1 | P36334 |
| AIGSTFVNTSYS | 1 | 0 | 0 | 1409224 | AKA09659.1 | P36334 |
| AIGTANLSIPSN | 1 | 0 | 0 | 1409225 | AAK32190.1 | A0A0P0K6L9 |
| AIINDKVLTKID | 1 | 0 | 0 | 1409249 | QBP84719.1 | P0C6X6 |
| AIINDKVVTKID | 1 | 0 | 0 | 1409251 | AEN19363.1 | P0C6X6 |
| AIIRDGDVVKQP | 1 | 0 | 0 | 1409252 | ALA49648.1 | K9N7C7 |
| AIIRDSDFVKQP | 1 | 0 | 0 | 1409253 | AVN89451.1 | K9N7C7 |
| AIIRDSEAIKQP | 1 | 0 | 0 | 1409255 | AVV62524.1 | K9N7C7 |
| AIKDDVLYSEFP | 1 | 0 | 0 | 1409271 | AKA09643.1 | P36334 |
| AIKDFGGFNFSQ | 1 | 0 | 0 | 1409272 | ATO98181.1 | Q3LZX1 |
| AIKITEHSWSAD | 1 | 0 | 0 | 1409276 | APO40578.1 | P0C6X7 |
| AIKLAKCYKDAV | 1 | 0 | 0 | 1409281 | AVV62524.1 | K9N7C7 |
| AIKLDSKNPNYK | 1 | 0 | 0 | 1409285 | AVV62544.1 | K9N4V7 |
| AIKLVPTTTGAL | 1 | 0 | 0 | 1409287 | AGT51358.1 | H9EJ42 |
| AIKNLTPIKLNF | 1 | 0 | 0 | 1409290 | AGT51393.1 | H9EJ42 |
| AIKSDHCGGNPC | 1 | 0 | 0 | 1409298 | AVI15011.1 | P36334 |
| AIKSDYCGGNPC | 1 | 0 | 0 | 1409299 | CAA83660.1 | P36334 |
| AIKVTEYSWNKY | 1 | 0 | 0 | 1409301 | AFD98819.1 | H9EJ42 |
| AILCATCPFAKG | 1 | 0 | 0 | 1409304 | QBF80531.1 | K9N7C7 |

| | | | | | | |
|---|---|---|---|---|---|---|
| AILNNCNWFIQS | 1 | 0 | 0 | 1409316 | AEN19363.1 | P0C6X6 |
| AILNSFKEYFNL | 1 | 0 | 0 | 1409318 | ALJ54448.1 | K9N5Q8 |
| AILNTVKFCDHI | 1 | 0 | 0 | 1409320 | AVV61898.1 | K9N7C7 |
| AILNTVTFCDHM | 1 | 0 | 0 | 1409321 | AGV08514.1 | K9N7C7 |
| AILSGTTTNVDK | 1 | 0 | 0 | 1409328 | AXN92247.1 | K9N7C7 |
| AILSGTTTNVEK | 1 | 0 | 0 | 1409329 | AVN89333.1 | K9N7C7 |
| AILSNLGVPSYC | 1 | 0 | 0 | 1409330 | APO40578.1 | P0C6X7 |
| AILTAFLPAQDT | 1 | 0 | 0 | 1409332 | ATO98205.1 | Q3LZX1 |
| AILTAFSPAQDT | 1 | 0 | 0 | 1409334 | 5X58_A | P59594 |
| AIMSMAECKATL | 1 | 0 | 0 | 1409355 | QED88025.1 | H9EJ42 |
| AINDFSNSGSDV | 1 | 0 | 0 | 1409363 | AVP78030.1 | P0C6W2 |
| AINDLNTTIPRI | 1 | 0 | 0 | 1409365 | ACV87140.1 | A0A140H1H1 |
| AINDVDTGVPST | 1 | 0 | 0 | 1409366 | CAA83661.1 | P36334 |
| AINFNQPLVVDT | 1 | 0 | 0 | 1409367 | AUM60024.1 | K9N5Q8 |
| AINKASNDAVQL | 1 | 0 | 0 | 1409370 | AVV62524.1 | K9N7C7 |
| AINLPIGCMMNV | 1 | 0 | 0 | 1409372 | ABD75623.1 | A0A140H1G9 |
| AINLPTGCMMNV | 1 | 0 | 0 | 1409373 | ARB07436.1 | A0A140H1G9 |
| AINNASNGAVQI | 1 | 0 | 0 | 1409374 | AUM60013.1 | K9N7C7 |
| AINNIVASFSSV | 1 | 0 | 0 | 1409375 | AIW52843.1 | A0A0P0G321 |
| AINSEEIYNKLT | 1 | 0 | 0 | 1409380 | AVV62535.1 | K9N7C7 |
| AINSVPWDTIAN | 1 | 0 | 0 | 1409381 | AVP78041.1 | P0C6W2 |
| AINSVPWSKICD | 1 | 0 | 0 | 1409382 | APO40578.1 | P0C6X7 |
| AINSVPWSKIFA | 1 | 0 | 0 | 1409383 | ATO98143.1 | P0C6W2 |
| AINVGKNGLQFA | 1 | 0 | 0 | 1409387 | AUM60013.1 | K9N7C7 |
| AINVGKQGLEFA | 1 | 0 | 0 | 1409388 | AVV62524.1 | K9N7C7 |
| AINVGKQGLEYA | 1 | 0 | 0 | 1409389 | AVV62535.1 | K9N7C7 |
| AIPDYNYYKYNL | 1 | 0 | 0 | 1409392 | ACV87150.1 | A0A140H1G9 |
| AIPNGRKVDLHL | 1 | 0 | 0 | 1409396 | AIL49485.1 | P36334 |
| AIPNGRRVDLQL | 1 | 0 | 0 | 1409398 | AGT51600.1 | P36334 |
| AIPNRRKFDLQL | 1 | 0 | 0 | 1409399 | BBO58726.1 | P36334 |
| AIPNRRKVDLQL | 1 | 0 | 0 | 1409400 | AEN19366.1 | P36334 |
| AIPNRRRDDLQL | 1 | 0 | 0 | 1409401 | BBA20986.1 | A0A140H1H1 |
| AIPNRRRVDLQL | 1 | 0 | 0 | 1409402 | ANZ78849.1 | P36334 |
| AIPNSRKVDLQL | 1 | 0 | 0 | 1409403 | AIX10751.1 | P36334 |
| AIPNSRRSDLQF | 1 | 0 | 0 | 1409404 | AMM42415.1 | A0A140H1H1 |
| AIPSMFPPGTVL | 1 | 0 | 0 | 1409408 | AJQ24068.1 | P33469 |
| AIQDGVTRDILS | 1 | 0 | 0 | 1409422 | ARO76381.1 | P0C6U8 |
| AIQDGVTRDIVS | 1 | 0 | 0 | 1409424 | P0C6W6.1 | P0C6W2 |
| AIQEVIDDMLWG | 1 | 0 | 0 | 1409426 | ARK08668.1 | Q80872 |
| AIQGFACGCGAS | 1 | 0 | 0 | 1409427 | AXN92247.1 | K9N7C7 |
| AIQLAKCFQECV | 1 | 0 | 0 | 1409429 | AUM60013.1 | K9N7C7 |
| AIQNCAWDYHIT | 1 | 0 | 0 | 1409430 | AST12962.1 | A0A0P0LKV0 |
| AIQNCAWDYHVT | 1 | 0 | 0 | 1409431 | NP_073549.1 | A0A0P0LKV0 |
| AIQQGFDATNFA | 1 | 0 | 0 | 1409432 | QBP84722.1 | P36334 |
| AIQQGSDATNSA | 1 | 0 | 0 | 1409433 | ARB66064.1 | P36334 |
| AIQQLHQGFKGK | 1 | 0 | 0 | 1409434 | AUM60023.1 | K9N7C7 |
| AIQQLYTGFQGK | 1 | 0 | 0 | 1409437 | AXN92247.1 | K9N7C7 |
| AIQQQFNVALPG | 1 | 0 | 0 | 1409438 | AUM60013.1 | K9N7C7 |
| AIQVAKCYGESV | 1 | 0 | 0 | 1409443 | AXN92247.1 | K9N7C7 |
| AIRCVPQSEVDW | 1 | 0 | 0 | 1409446 | AVP78030.1 | P0C6W2 |
| AIRGSCEKNLQL | 1 | 0 | 0 | 1409449 | AVA26874.1 | H9EJ24 |
| AIRISNDTFLNG | 1 | 0 | 0 | 1409451 | ADK37703.1 | A0A0P0K6L9 |
| AIRNFDEQNCDV | 1 | 0 | 0 | 1409453 | AGT51330.1 | H9EJ42 |
| AISDLGSLIDSN | 1 | 0 | 0 | 1409467 | AVV62524.1 | K9N7C7 |
| AISFADNNDITN | 1 | 0 | 0 | 1409469 | ANI69896.1 | K9N4V7 |
| AISKIQDSLTTT | 1 | 0 | 0 | 1409475 | APO40579.1 | P59594 |
| AISNELFLTVPT | 1 | 0 | 0 | 1409481 | AGT51520.1 | P30215 |
| AISNEQHYKSWV | 1 | 0 | 0 | 1409482 | ATO98193.1 | Q3LZX1 |
| AISNEWYIGAII | 1 | 0 | 0 | 1409483 | NP_073549.1 | A0A0P0LKV0 |
| AISPKIDNETLE | 1 | 0 | 0 | 1409486 | AMM42407.1 | A0A140H1H1 |
| AISPKTDKETLE | 1 | 0 | 0 | 1409487 | AZS52618.1 | A0A140H1H1 |
| AISPNTDNETLQ | 1 | 0 | 0 | 1409489 | ABA60801.1 | A0A140H1H1 |
| AISSNTDNETLE | 1 | 0 | 0 | 1409493 | 5I08_A | A0A140H1H1 |
| AISSSISDILQR | 1 | 0 | 0 | 1409495 | AVV62526.1 | K9N5Q8 |
| AISSTTDNETLE | 1 | 0 | 0 | 1409497 | ARB07438.1 | A0A140H1H1 |
| AISVKNRARTVA | 1 | 0 | 0 | 1409501 | BAI44145.1 | P0C6X6 |
| AISVNCCHDGIF | 1 | 0 | 0 | 1409503 | AGT51675.1 | Q4VID0 |
| AITDYSYYAYNL | 1 | 0 | 0 | 1409511 | AUM60013.1 | K9N7C7 |
| AITEARQHLIDG | 1 | 0 | 0 | 1409512 | APO40578.1 | P0C6X7 |
| AITIKPEANINQ | 1 | 0 | 0 | 1409515 | ABD75615.1 | A0A140H1G9 |
| AITPYVCNASGC | 1 | 0 | 0 | 1409520 | AWK59942.1 | H9EJ42 |
| AITQTSPALQTV | 1 | 0 | 0 | 1409522 | AIW52728.1 | A0A0P0K6L9 |
| AITTENRFTNFE | 1 | 0 | 0 | 1409527 | AIL49500.1 | P36334 |
| AITVEELKQLLE | 1 | 0 | 0 | 1409531 | BAE93405.1 | P59596 |
| AIVAYTMSLGAE | 1 | 0 | 0 | 1409534 | AVP78031.1 | Q3LZX1 |
| AIVESCGNYRVT | 1 | 0 | 0 | 1409538 | ARO76381.1 | P0C6U8 |
| AIVGAMLSENFT | 1 | 0 | 0 | 1409549 | ABB90509.1 | A0A0P0K6L9 |
| AIVIDNDLNEFV | 1 | 0 | 0 | 1409552 | ASY99818.1 | K9N7C7 |
| AIVKETNLICKA | 1 | 0 | 0 | 1409557 | AEN19363.1 | P0C6X6 |
| AIVLPQGGFVAD | 1 | 0 | 0 | 1409561 | ARU07575.1 | A0A140H1G9 |
| AIVLQLPQGTSL | 1 | 0 | 0 | 1409562 | APO40586.1 | P59595 |
| AIVLVNGVHHPT | 1 | 0 | 0 | 1409565 | QBM11740.1 | K9N7D2 |
| AIVLVNGVHYPT | 1 | 0 | 0 | 1409566 | AXG21669.1 | K9N7D2 |
| AIVLVNGVRYPT | 1 | 0 | 0 | 1409567 | AXN73419.1 | K9N7D2 |
| AIVQATESKVLL | 1 | 0 | 0 | 1409575 | ABE97136.1 | H9EJ42 |
| AIVTGLLPTHWI | 1 | 0 | 0 | 1409581 | AKA09625.1 | A0A0P0G321 |
| AIVTQFAPGTQL | 1 | 0 | 0 | 1409583 | AVV62533.1 | K9N4V7 |
| AIVYGDFSHGQL | 1 | 0 | 0 | 1409587 | 2RHB_A | P0C6X7 |
| AIWEAPHNDCTP | 1 | 0 | 0 | 1409590 | AVV62526.1 | K9N5Q8 |
| AIYHRYKAGNYR | 1 | 0 | 0 | 1409606 | ASY99815.1 | K9N7A1 |
| AIYNGVTRDILS | 1 | 0 | 0 | 1409611 | APO40578.1 | P0C6X7 |

| | | | | | | |
|---|---|---|---|---|---|---|
| AIYPIDLASQII | 1 | 0 | 0 | 1409612 | ASY99815.1 | K9N7A1 |
| AIYPIDLSSQII | 1 | 0 | 0 | 1409613 | ALA49908.1 | K9N7A1 |
| AIYPIHLASQII | 1 | 0 | 0 | 1409614 | QEJ82232.1 | K9N7A1 |
| AKAELCCFARMV | 1 | 0 | 0 | 1409622 | ANI69888.1 | K9N638 |
| AKAELCCSACMV | 1 | 0 | 0 | 1409623 | QBF80553.1 | K9N7C7 |
| AKAKNTKVIKDD | 1 | 0 | 0 | 1409631 | AKA09644.1 | P36334 |
| AKAQEGCGIISR | 1 | 0 | 0 | 1409643 | Q3I5J5.1 | Q3LZX1 |
| AKCYGEPVLVNA | 1 | 0 | 0 | 1409652 | AVN89451.1 | K9N7C7 |
| AKDDLYVKRYER | 1 | 0 | 0 | 1409655 | ATN39862.1 | P0C6X6 |
| AKDDYGTDIDGV | 1 | 0 | 0 | 1409656 | AJQ24074.1 | P33469 |
| AKDNTVLQALQS | 1 | 0 | 0 | 1409667 | AEN19363.1 | P0C6X6 |
| AKDQYGSDINGV | 1 | 0 | 0 | 1409668 | AIX10729.1 | P33469 |
| AKDQYGTDFDGV | 1 | 0 | 0 | 1409669 | AGT51416.1 | P33469 |
| AKDQYGTDIDGI | 1 | 0 | 0 | 1409670 | AIL49434.1 | P33469 |
| AKDQYGTDINGV | 1 | 0 | 0 | 1409672 | AEN19370.1 | P33469 |
| AKDYGFTMFVCI | 1 | 0 | 0 | 1409675 | AUM60013.1 | K9N7C7 |
| AKDYGFTVFVCT | 1 | 0 | 0 | 1409676 | AXN92247.1 | K9N7C7 |
| AKEGLNSLADVL | 1 | 0 | 0 | 1409682 | AVV62524.1 | K9N7C7 |
| AKEIRQKILNKP | 1 | 0 | 0 | 1409686 | BBM61433.1 | P33469 |
| AKELVDGYGNTM | 1 | 0 | 0 | 1409687 | AUM60027.1 | K9N643 |
| AKERCTHWCTYL | 1 | 0 | 0 | 1409691 | NP_073549.1 | A0A0P0LKV0 |
| AKFERQHMDSPD | 1 | 0 | 0 | 1409700 | 3R24_A | P0C6X7 |
| AKFGFTPLNKQS | 1 | 0 | 0 | 1409702 | NP_073549.1 | A0A0P0LKV0 |
| AKFKNSVTSINS | 1 | 0 | 0 | 1409706 | AST12962.1 | A0A0P0LKV0 |
| AKFWEESFYREL | 1 | 0 | 0 | 1409717 | AVV62524.1 | K9N7C7 |
| AKGFPLLAELIP | 1 | 0 | 0 | 1409724 | AMM42619.1 | H9EJ69 |
| AKGFPQFAELIP | 1 | 0 | 0 | 1409725 | AGT51335.1 | H9EJ69 |
| AKGLLQVGDSVL | 1 | 0 | 0 | 1409729 | ALA49901.1 | K9N7C7 |
| AKGPLQVGDAVL | 1 | 0 | 0 | 1409731 | AVV62524.1 | K9N7C7 |
| AKGVSLDNVLST | 1 | 0 | 0 | 1409733 | APO40578.1 | P0C6X7 |
| AKHGDFSGVASQ | 1 | 0 | 0 | 1409742 | AWK59934.1 | U3M6Q8 |
| AKIDTSGIHNRV | 1 | 0 | 0 | 1409755 | AVV62527.1 | K9N796 |
| AKILAPSRSSET | 1 | 0 | 0 | 1409757 | AIW52721.1 | A0A0N7J131 |
| AKINQALHGANL | 1 | 0 | 0 | 1409765 | AVV62526.1 | K9N5Q8 |
| AKIVGGAPIWFN | 1 | 0 | 0 | 1409769 | AVN89451.1 | K9N7C7 |
| AKKGAWNIGEQS | 1 | 0 | 0 | 1409786 | AVP78030.1 | P0C6W2 |
| AKKNLDEARLSG | 1 | 0 | 0 | 1409792 | AIV41942.1 | P0C6X6 |
| AKKNNIPFKLTC | 1 | 0 | 0 | 1409794 | APO40578.1 | P0C6X7 |
| AKKNNLPFRLTC | 1 | 0 | 0 | 1409797 | AVP78041.1 | P0C6W2 |
| AKLGGDASKPHQ | 1 | 0 | 0 | 1409813 | AJQ24068.1 | P33469 |
| AKLGKESKPQQV | 1 | 0 | 0 | 1409819 | ARU07581.1 | Q19U29 |
| AKLHNLTDSQRA | 1 | 0 | 0 | 1409821 | AUM60027.1 | K9N643 |
| AKMIERCYTAEQ | 1 | 0 | 0 | 1409840 | NP_073549.1 | A0A0P0LKV0 |
| AKNILHVVGPDA | 1 | 0 | 0 | 1409850 | AXN92247.1 | K9N7C7 |
| AKNKMRHKRIST | 1 | 0 | 0 | 1409852 | AID66496.1 | K9N4V7 |
| AKNSGSVNQQQI | 1 | 0 | 0 | 1409862 | ARU07575.1 | A0A140H1G9 |
| AKPKRLRKKRNV | 1 | 0 | 0 | 1409874 | ATQ39389.1 | K9N7C7 |
| AKPNVLIALKMV | 1 | 0 | 0 | 1409881 | ASY99815.1 | K9N7A1 |
| AKPNVVIALKMV | 1 | 0 | 0 | 1409882 | AIL23940.1 | K9N7A1 |
| AKPVSEYHVTLA | 1 | 0 | 0 | 1409894 | AVV62529.1 | K9N643 |
| AKPVTEYHITIA | 1 | 0 | 0 | 1409896 | AVV62540.1 | K9N643 |
| AKQDTGHYFYRS | 1 | 0 | 0 | 1409904 | AVP78042.1 | Q3LZX1 |
| AKQDVGNYFYRS | 1 | 0 | 0 | 1409905 | AVP78031.1 | Q3LZX1 |
| AKQEILGTVSWN | 1 | 0 | 0 | 1409910 | AVP78030.1 | P0C6W2 |
| AKQNVSLLSKCY | 1 | 0 | 0 | 1409913 | ASU91141.1 | K9N7C7 |
| AKRALRSHPDFK | 1 | 0 | 0 | 1409917 | ASU90074.1 | K9N7C7 |
| AKRSTEKVLERA | 1 | 0 | 0 | 1409932 | AUM60013.1 | K9N7C7 |
| AKRVVRSHPDFK | 1 | 0 | 0 | 1409937 | ASU90569.1 | K9N7C7 |
| AKSIQQDLDVIL | 1 | 0 | 0 | 1409947 | ALA49934.1 | K9N7C7 |
| AKSIVSTIGEAI | 1 | 0 | 0 | 1409950 | AUM60023.1 | K9N7C7 |
| AKSIVSTIGESI | 1 | 0 | 0 | 1409951 | AUM60013.1 | K9N7C7 |
| AKTELCCSARMV | 1 | 0 | 0 | 1409977 | AXN73469.1 | K9N7C7 |
| AKTSVDCTMYIC | 1 | 0 | 0 | 1409986 | AVP78031.1 | Q3LZX1 |
| AKVANWKMFMCD | 1 | 0 | 0 | 1409993 | AVV62524.1 | K9N7C7 |
| AKVEGCMVQVTC | 1 | 0 | 0 | 1409997 | 2PWX_A | P0C6X7 |
| AKVEGKDGFVTV | 1 | 0 | 0 | 1409998 | AUM60013.1 | K9N7C7 |
| AKVGDKYYPVVV | 1 | 0 | 0 | 1410001 | AEN19363.1 | P0C6X6 |
| AKVHNLSGNPDE | 1 | 0 | 0 | 1410005 | AHA80155.1 | P33469 |
| AKVKNTKAIKDD | 1 | 0 | 0 | 1410008 | AKA09643.1 | P36334 |
| AKVKNTKVFIDG | 1 | 0 | 0 | 1410010 | QDH43719.1 | P36334 |
| AKVKNTKVFKHG | 1 | 0 | 0 | 1410012 | AIL49520.1 | P36334 |
| AKVKNTKVIKDD | 1 | 0 | 0 | 1410013 | ATP16757.1 | P36334 |
| AKVKNTKVIKDG | 1 | 0 | 0 | 1410014 | CAA79896.1 | P36334 |
| AKVKNTKVVGKD | 1 | 0 | 0 | 1410018 | AWW13511.1 | P36334 |
| AKVRAVIYDSKE | 1 | 0 | 0 | 1410024 | AVV62524.1 | K9N7C7 |
| AKVRVDCYDKFKINDTTCKY | 1 | 0 | 0 | 1410027 | ABD75503.1 | A0A140H1G9 |
| AKVRVECYDKFKINDTTRKYV | 1 | 0 | 0 | 1410028 | AQN78733.1 | P0C6X6 |
| AKVSPGLCISGD | 1 | 0 | 0 | 1410033 | AGT51551.1 | P36334 |
| AKVSTGLCIAGD | 1 | 0 | 0 | 1410034 | AMK59677.1 | P36334 |
| AKWKMLMCDKVR | 1 | 0 | 0 | 1410038 | AVV62535.1 | K9N7C7 |
| AKWVFDSSKEFV | 1 | 0 | 0 | 1410040 | AIV41984.1 | P0C6X6 |
| AKYIPLVGTPLH | 1 | 0 | 0 | 1410043 | ANI69891.1 | K9N4V0 |
| AKYTPLVGTSLH | 1 | 0 | 0 | 1410048 | AXN92262.1 | K9N4V0 |
| ALAAKEGLNSLA | 1 | 0 | 0 | 1410054 | AVV62524.1 | K9N7C7 |
| ALADLYHYQECA | 1 | 0 | 0 | 1410056 | APO40584.1 | P59635 |
| ALADYTQCAIIA | 1 | 0 | 0 | 1410057 | AUM60013.1 | K9N7C7 |
| ALADYTQCAVIA | 1 | 0 | 0 | 1410058 | AVV62524.1 | K9N7C7 |
| ALAEHDFFTWKD | 1 | 0 | 0 | 1410061 | AIW52827.1 | H9EJ42 |
| ALAFAWVLFHVL | 1 | 0 | 0 | 1410062 | AEN19363.1 | P0C6X6 |
| ALAKQYAAQRIA | 1 | 0 | 0 | 1410072 | AMQ48995.1 | K9N4V0 |
| ALAMAYGDCTFG | 1 | 0 | 0 | 1410085 | AVV62535.1 | K9N7C7 |

| | | | | | | |
|---|---|---|---|---|---|---|
| ALANQFTEFVST | 1 | 0 | 0 | 1410089 | ATG84865.1 | K9N638 |
| ALASDIFATPSV | 1 | 0 | 0 | 1410094 | ASY99818.1 | K9N7C7 |
| ALASGCYSSLTV | 1 | 0 | 0 | 1410096 | AUM60014.1 | K9N5Q8 |
| ALAVNACLIPFL | 1 | 0 | 0 | 1410101 | AUM60018.1 | K9N7D2 |
| ALAVNACLIPLL | 1 | 0 | 0 | 1410102 | ALB08293.1 | K9N7D2 |
| ALAVNVCIIPLL | 1 | 0 | 0 | 1410103 | AHI48676.1 | K9N7D2 |
| ALAVNVCLIPLM | 1 | 0 | 0 | 1410105 | AVV62541.1 | K9N7D2 |
| ALAVRLTANVGR | 1 | 0 | 0 | 1410106 | ARU07575.1 | A0A140H1G9 |
| ALAYRANSFDVP | 1 | 0 | 0 | 1410109 | ALA49659.1 | K9N7C7 |
| ALCFNKPKQFVP | 1 | 0 | 0 | 1410121 | ABD75512.1 | A0A140H1H0 |
| ALDAESHIDSDV | 1 | 0 | 0 | 1410137 | APO40578.1 | P0C6X7 |
| ALDAYVSQQLSD | 1 | 0 | 0 | 1410139 | CAA83661.1 | P36334 |
| ALDDATRNYIIT | 1 | 0 | 0 | 1410141 | AHL18094.1 | K9N643 |
| ALDDDARTFIQS | 1 | 0 | 0 | 1410142 | AGT51409.1 | P0C6X6 |
| ALDDDARTFVQS | 1 | 0 | 0 | 1410143 | AEN19363.1 | P0C6X6 |
| ALDDVTLNYIIT | 1 | 0 | 0 | 1410144 | ALA49575.1 | K9N643 |
| ALDDVTRNYIIT | 1 | 0 | 0 | 1410145 | AXN92231.1 | K9N643 |
| ALDLSLDYINVT | 1 | 0 | 0 | 1410148 | ARA15421.1 | P36334 |
| ALDQDVFIQKYD | 1 | 0 | 0 | 1410154 | ARU07564.1 | P0C6X6 |
| ALDQDVFIQKYG | 1 | 0 | 0 | 1410155 | AEN19363.1 | P0C6X6 |
| ALDSGDASPQVL | 1 | 0 | 0 | 1410156 | AUM60013.1 | K9N7C7 |
| ALDSLGAAIHYL | 1 | 0 | 0 | 1410157 | AVQ05261.1 | P0C6X6 |
| ALDTILQEANKS | 1 | 0 | 0 | 1410159 | ATQ39389.1 | K9N7C7 |
| ALDTTIPFGPSS | 1 | 0 | 0 | 1410160 | AIW52835.1 | A0A0P0G321 |
| ALDWELYVNNAY | 1 | 0 | 0 | 1410164 | AIL49596.1 | P0C6X6 |
| ALEDEFTPFDVV | 1 | 0 | 0 | 1410171 | 3M3V_A | P0C6X7 |
| ALEKFMSTSNAI | 1 | 0 | 0 | 1410175 | AVV62524.1 | K9N7C7 |
| ALESKVLLERFL | 1 | 0 | 0 | 1410194 | QED88024.1 | H9EJ42 |
| ALFASLNVKPYG | 1 | 0 | 0 | 1410204 | AWK59948.1 | H9EJ42 |
| ALFCATCPFAKG | 1 | 0 | 0 | 1410206 | AUM60013.1 | K9N7C7 |
| ALFDSLEFTELS | 1 | 0 | 0 | 1410208 | AVN89451.1 | K9N7C7 |
| ALFVVEQKWCVI | 1 | 0 | 0 | 1410239 | ABF82439.1 | H9EJ42 |
| ALGAIQEGFDAT | 1 | 0 | 0 | 1410242 | AVI15011.1 | P36334 |
| ALGGSVAVKITE | 1 | 0 | 0 | 1410250 | P0C6W6.1 | P0C6W2 |
| ALGKLQDVINQN | 1 | 0 | 0 | 1410251 | APO40579.1 | P59594 |
| ALGLFTLHFVSA | 1 | 0 | 0 | 1410255 | ARA15430.1 | A0A0P0KWE8 |
| ALHGANLMQDES | 1 | 0 | 0 | 1410267 | AVV62537.1 | K9N5Q8 |
| ALHGANLRQDES | 1 | 0 | 0 | 1410270 | AVV62526.1 | K9N5Q8 |
| ALHLVDPQIQVT | 1 | 0 | 0 | 1410275 | AVP78050.1 | Q3LZX3 |
| ALHNRWQLALYK | 1 | 0 | 0 | 1410279 | ARO76383.1 | P59632 |
| ALHSGYYDANQY | 1 | 0 | 0 | 1410283 | AWO68966.1 | A0A0P0G321 |
| ALHSIQQGFDAT | 1 | 0 | 0 | 1410284 | AIX10759.1 | P36334 |
| ALHSKCLTSIEE | 1 | 0 | 0 | 1410285 | ATP16764.1 | P0C6X6 |
| ALHTGYYDVNQY | 1 | 0 | 0 | 1410287 | AIW52843.1 | A0A0P0G321 |
| ALIGANGNKIVD | 1 | 0 | 0 | 1410296 | AVN89322.1 | K9N7C7 |
| ALIMACGNCTFG | 1 | 0 | 0 | 1410301 | AID50427.1 | K9N7C7 |
| ALIPDYAKILVD | 1 | 0 | 0 | 1410307 | AMN88692.1 | A0A140H1G9 |
| ALISCNGNKIED | 1 | 0 | 0 | 1410312 | AIL49553.1 | P0C6X6 |
| ALISGTATAGYT | 1 | 0 | 0 | 1410314 | APO40579.1 | P59594 |
| ALKELYGPADPT | 1 | 0 | 0 | 1410336 | AHI48582.1 | K9N7C7 |
| ALKELYGPIDPT | 1 | 0 | 0 | 1410337 | ATG84722.1 | K9N7C7 |
| ALKELYGPLDPT | 1 | 0 | 0 | 1410338 | AXN92247.1 | K9N7C7 |
| ALKELYGSVDPT | 1 | 0 | 0 | 1410340 | AHC74096.1 | K9N7C7 |
| ALKKVPIDEYIT | 1 | 0 | 0 | 1410350 | AER30328.1 | P0C6X7 |
| ALKMVQTHISHY | 1 | 0 | 0 | 1410362 | AVV62524.1 | K9N7C7 |
| ALKNAFQTFRWS | 1 | 0 | 0 | 1410364 | ARU07575.1 | A0A140H1G9 |
| ALKNEKRQDHVS | 1 | 0 | 0 | 1410365 | AVV62524.1 | K9N7C7 |
| ALKNLGFDNQSK | 1 | 0 | 0 | 1410366 | AIW52805.1 | H9EJ69 |
| ALKSLGFEKPQE | 1 | 0 | 0 | 1410374 | ARB07429.1 | A0A0N7J131 |
| ALLHIADCQTTN | 1 | 0 | 0 | 1410405 | ABB90509.1 | A0A0P0K6L9 |
| ALLHIAGSQTTN | 1 | 0 | 0 | 1410407 | AIW52735.1 | A0A0P0K6L9 |
| ALLLLDRLNQLESKMSG | 1 | 0 | 0 | 1410414 | P0DTC9.1 | P0DTC9 |
| ALLQNGMNGRTI | 1 | 0 | 0 | 1410424 | AEA10817.1 | P0C6X7 |
| ALLRKTLVLNAS | 1 | 0 | 0 | 1410426 | AXN92231.1 | K9N643 |
| ALLSDHLDLKWA | 1 | 0 | 0 | 1410427 | APO40578.1 | P0C6X7 |
| ALLTKSSEYKGS | 1 | 0 | 0 | 1410438 | AVP78041.1 | P0C6W2 |
| ALLVPKIFKARA | 1 | 0 | 0 | 1410443 | AMM42360.1 | A0A140H1G9 |
| ALMQLFFVHFST | 1 | 0 | 0 | 1410448 | APO40578.1 | P0C6X7 |
| ALMSCNGNKIGD | 1 | 0 | 0 | 1410449 | AJE26218.1 | P0C6X6 |
| ALMSCNGNKSDD | 1 | 0 | 0 | 1410450 | AKL87224.1 | P0C6X6 |
| ALNAATNGEMQK | 1 | 0 | 0 | 1410452 | APO40578.1 | P0C6X7 |
| ALNAIQQGFDAT | 1 | 0 | 0 | 1410454 | AGT51491.1 | P36334 |
| ALNCYWPLKDYG | 1 | 0 | 0 | 1410456 | 3D0H_E | P59594 |
| ALNDFSNLGADV | 1 | 0 | 0 | 1410460 | BAE93400.1 | P0C6U8 |
| ALNGSCGSVGFN | 1 | 0 | 0 | 1410472 | 3F9G_A | P0C6X7 |
| ALNKATNNAMQL | 1 | 0 | 0 | 1410482 | AVP78041.1 | P0C6W2 |
| ALNKIHDVVNQQ | 1 | 0 | 0 | 1410484 | ABB90526.1 | A0A0P0K6L9 |
| ALNNVYFGLSIV | 1 | 0 | 0 | 1410488 | QDH43729.1 | Q01455 |
| ALNNWFLLTNTS | 1 | 0 | 0 | 1410489 | ADK37704.1 | A0A0P0K6L9 |
| ALNSCKDHPVVG | 1 | 0 | 0 | 1410490 | ARB07436.1 | A0A140H1G9 |
| ALNVCFDVRDNG | 1 | 0 | 0 | 1410503 | AVV62524.1 | K9N7C7 |
| ALPDTHSTLTPR | 1 | 0 | 0 | 1410507 | AID55099.1 | K9N5Q8 |
| ALPDTPITLTPR | 1 | 0 | 0 | 1410508 | ANI69900.1 | K9N5Q8 |
| ALPDTPSPLTPR | 1 | 0 | 0 | 1410509 | AJG44080.1 | K9N5Q8 |
| ALPDTPSTFTPR | 1 | 0 | 0 | 1410510 | AVK87439.1 | K9N5Q8 |
| ALPDTPSTLAPR | 1 | 0 | 0 | 1410511 | ALJ54472.1 | K9N5Q8 |
| ALPDTPSTLKPR | 1 | 0 | 0 | 1410512 | ALJ54490.1 | K9N5Q8 |
| ALPDTPSTLTLR | 1 | 0 | 0 | 1410513 | ANI69889.1 | K9N5Q8 |
| ALPDTPSTLTPA | 1 | 0 | 0 | 1410514 | 5W9H_A | K9N5Q8 |
| ALPDTPSTLTPC | 1 | 0 | 0 | 1410515 | AKN24812.1 | K9N5Q8 |
| ALPFEDLNKCRA | 1 | 0 | 0 | 1410518 | AVV62524.1 | K9N7C7 |

| | | | | | | |
|---|---|---|---|---|---|---|
| ALPFEDLSKCRA | 1 | 0 | 0 | 1410519 | AVV62535.1 | K9N7C7 |
| ALPGVFCGVDAM | 1 | 0 | 0 | 1410523 | P0C6W6.1 | P0C6W2 |
| ALPGVFCGVDSM | 1 | 0 | 0 | 1410524 | ARO76381.1 | P0C6U8 |
| ALPNMIRMISAM | 1 | 0 | 0 | 1410529 | AVP77959.1 | P0C6W2 |
| ALPNMIRTISAM | 1 | 0 | 0 | 1410530 | ADK37697.1 | H9EJ42 |
| ALPNRRKVDLQL | 1 | 0 | 0 | 1410532 | AIL49495.1 | P36334 |
| ALPQSKDSNFLK | 1 | 0 | 0 | 1410535 | ALT66879.1 | K9N7C7 |
| ALPQSKESNFLN | 1 | 0 | 0 | 1410537 | AQZ41321.1 | K9N7C7 |
| ALPVYYQHTDIN | 1 | 0 | 0 | 1410542 | AWO68966.1 | A0A0P0G321 |
| ALPYGANKEGIV | 1 | 0 | 0 | 1410543 | APO40586.1 | P59595 |
| ALQGNFGDLQLN | 1 | 0 | 0 | 1410555 | AVV62544.1 | K9N4V7 |
| ALQKAVNVAKNA | 1 | 0 | 0 | 1410560 | AUM60013.1 | K9N7C7 |
| ALQQIDIKFNAP | 1 | 0 | 0 | 1410569 | APO40578.1 | P0C6X7 |
| ALQQIEVKFNAP | 1 | 0 | 0 | 1410570 | P0C6W6.1 | P0C6W2 |
| ALQTAAMTIFSD | 1 | 0 | 0 | 1410574 | APO40578.1 | P0C6X7 |
| ALQTDVLQEDQK | 1 | 0 | 0 | 1410575 | BAL45639.1 | A0A0P0K6L9 |
| ALQTDVLQENQR | 1 | 0 | 0 | 1410577 | AIW52747.1 | A0A0P0K6L9 |
| ALRAGSFDKVYD | 1 | 0 | 0 | 1410583 | AIY60536.1 | K9N7C7 |
| ALRGGKIVNNWL | 1 | 0 | 0 | 1410593 | AVP78041.1 | P0C6W2 |
| ALRHAFMKHKGG | 1 | 0 | 0 | 1410594 | AHL18091.1 | K9N7C7 |
| ALRHFDQNIEVL | 1 | 0 | 0 | 1410598 | AVN89363.1 | K9N7C7 |
| ALRHFDQNNCEV | 1 | 0 | 0 | 1410599 | AVV62524.1 | K9N7C7 |
| ALRHFDRNDCML | 1 | 0 | 0 | 1410601 | AEN19363.1 | P0C6X6 |
| ALRNAINADVYA | 1 | 0 | 0 | 1410614 | AVV62535.1 | K9N7C7 |
| ALRNDTSVEDKL | 1 | 0 | 0 | 1410615 | AUM60014.1 | K9N5Q8 |
| ALRNSINADAYA | 1 | 0 | 0 | 1410616 | AVV62524.1 | K9N7C7 |
| ALRNSISTDVYA | 1 | 0 | 0 | 1410617 | AUM60013.1 | K9N7C7 |
| ALRTAGVVTGNV | 1 | 0 | 0 | 1410623 | AUM60013.1 | K9N7C7 |
| ALSEAREHLKSG | 1 | 0 | 0 | 1410629 | P0C6W6.1 | P0C6W2 |
| ALSEQLDFIDTK | 1 | 0 | 0 | 1410631 | AVP78030.1 | P0C6W2 |
| ALSGIAIEQDKN | 1 | 0 | 0 | 1410636 | AVP78031.1 | Q3LZX1 |
| ALSGIAVEQDRN | 1 | 0 | 0 | 1410637 | ATO98231.1 | Q3LZX1 |
| ALSNGFSQVKSD | 1 | 0 | 0 | 1410659 | AEN19363.1 | P0C6X6 |
| ALTGIAIEQDKN | 1 | 0 | 0 | 1410679 | ATO98181.1 | Q3LZX1 |
| ALTGTNQLCAKI | 1 | 0 | 0 | 1410681 | AWK59942.1 | H9EJ42 |
| ALTGVALEQDKN | 1 | 0 | 0 | 1410682 | ATO98145.1 | Q3LZX1 |
| ALTKRNVMPTIT | 1 | 0 | 0 | 1410684 | AUM60013.1 | K9N7C7 |
| ALTNNVAFQTVKPGNF | 1 | 0 | 0 | 1410687 | YP_009724389.1 | P0DTD1 |
| ALTPALLATNNV | 1 | 0 | 0 | 1410689 | APO40578.1 | P0C6X7 |
| ALTQHGKENLTF | 1 | 0 | 0 | 1410692 | AVP78038.1 | Q3LZX4 |
| ALTQSKDTNFLN | 1 | 0 | 0 | 1410695 | AUM60023.1 | K9N7C7 |
| ALTVEEFVDVVK | 1 | 0 | 0 | 1410698 | AVV62524.1 | K9N7C7 |
| ALVDQRTICFSV | 1 | 0 | 0 | 1410708 | AIW52827.1 | H9EJ42 |
| ALVFIILCFTIK | 1 | 0 | 0 | 1410709 | ATO98237.1 | Q3LZX7 |
| ALVFMIVQFMPF | 1 | 0 | 0 | 1410711 | AMM42568.1 | H9EJ42 |
| ALVKSLNLNCVS | 1 | 0 | 0 | 1410719 | AVQ05261.1 | P0C6X6 |
| ALVNSQCDLTGR | 1 | 0 | 0 | 1410725 | AVP78042.1 | Q3LZX1 |
| ALVNSQCVNLTG | 1 | 0 | 0 | 1410726 | AVP78031.1 | Q3LZX1 |
| ALVQCNCGTKSW | 1 | 0 | 0 | 1410727 | AFR79256.1 | A0A0P0ILY9 |
| ALVYDNKLKAKN | 1 | 0 | 0 | 1410739 | ARU07574.1 | A0A140H1G9 |
| ALWDIAVINNVD | 1 | 0 | 0 | 1410744 | AVV62535.1 | K9N7C7 |
| ALYAIQQGFDAT | 1 | 0 | 0 | 1410750 | AIX10750.1 | P36334 |
| ALYTPWFYPLLA | 1 | 0 | 0 | 1410777 | AIV41960.1 | P0C6X6 |
| AMAASQAWQPGV | 1 | 0 | 0 | 1410784 | 3R24_A | P0C6X7 |
| AMAEKLNAFLVH | 1 | 0 | 0 | 1410789 | 3EJG_A | A0A0P0LKV0 |
| AMANQFTEFIGT | 1 | 0 | 0 | 1410793 | AVV62524.1 | K9N7C7 |
| AMANQFTEFVGT | 1 | 0 | 0 | 1410794 | AUM60013.1 | K9N7C7 |
| AMANWLAPNNMY | 1 | 0 | 0 | 1410795 | AVV62524.1 | K9N7C7 |
| AMASQKPQRAPR | 1 | 0 | 0 | 1410796 | AVV62533.1 | K9N4V7 |
| AMATPQPQRAPK | 1 | 0 | 0 | 1410798 | AVV62544.1 | K9N4V7 |
| AMAYGDCTFGVP | 1 | 0 | 0 | 1410800 | AVV62535.1 | K9N7C7 |
| AMCNWCLISQDS | 1 | 0 | 0 | 1410807 | AXN92247.1 | K9N7C7 |
| AMCNWCLVSQDS | 1 | 0 | 0 | 1410808 | AHL18091.1 | K9N7C7 |
| AMDEFIERYKLD | 1 | 0 | 0 | 1410809 | AVP78041.1 | P0C6W2 |
| AMDTAAYREAAA | 1 | 0 | 0 | 1410816 | AVV62524.1 | K9N7C7 |
| AMETFTNNNGSD | 1 | 0 | 0 | 1410825 | AQN78669.1 | P0C6X6 |
| AMFPPWSAAAGV | 1 | 0 | 0 | 1410833 | BBA20986.1 | A0A140H1H1 |
| AMFQPRLPVLSD | 1 | 0 | 0 | 1410834 | ARU07594.1 | A0A0P0G321 |
| AMGCNTLLVQPA | 1 | 0 | 0 | 1410837 | AVV62542.1 | K9N5R3 |
| AMGDVGLCMYRL | 1 | 0 | 0 | 1410838 | NP_073549.1 | A0A0P0LKV0 |
| AMGLPNNTASWF | 1 | 0 | 0 | 1410842 | 1SSK_A | P59595 |
| AMGQGQNNADPK | 1 | 0 | 0 | 1410843 | ARO76391.1 | P59636 |
| AMILGSKHVGCC | 1 | 0 | 0 | 1410850 | AAV28291.1 | A0A0P0LKV0 |
| AMIMPKTCFGPI | 1 | 0 | 0 | 1410855 | AVN89289.1 | K9N7C7 |
| AMINTAGLVTDS | 1 | 0 | 0 | 1410856 | APO40578.1 | P0C6X7 |
| AMLDDATGVMAL | 1 | 0 | 0 | 1410865 | AVV62524.1 | K9N7C7 |
| AMLFKKEPLLYV | 1 | 0 | 0 | 1410870 | ALA49373.1 | K9N7C7 |
| AMLNGKPVLWVN | 1 | 0 | 0 | 1410875 | AUM60013.1 | K9N7C7 |
| AMLNGKPVLWVT | 1 | 0 | 0 | 1410876 | AVV62535.1 | K9N7C7 |
| AMLTGFTTTNEA | 1 | 0 | 0 | 1410880 | QDP16206.1 | K9N5Q8 |
| AMLVNGTHTPLG | 1 | 0 | 0 | 1410882 | AVV62541.1 | K9N7D2 |
| AMMFTSDLVTNN | 1 | 0 | 0 | 1410884 | AVP78041.1 | P0C6W2 |
| AMMRISYFVQSI | 1 | 0 | 0 | 1410886 | ALA49424.1 | K9N7A1 |
| AMMWLSYFVQSI | 1 | 0 | 0 | 1410888 | AVN89358.1 | K9N7A1 |
| AMNGEDAFSILS | 1 | 0 | 0 | 1410893 | APT69882.1 | A0A0P0ILY9 |
| AMNSHPLVITPL | 1 | 0 | 0 | 1410896 | AVV62535.1 | K9N7C7 |
| AMNSYPLVITPL | 1 | 0 | 0 | 1410897 | AVV62524.1 | K9N7C7 |
| AMNVIGQHIELP | 1 | 0 | 0 | 1410898 | ARE29978.1 | H9EJ42 |
| AMNVIGQHIKLP | 1 | 0 | 0 | 1410899 | AWK59942.1 | H9EJ42 |
| AMPIMANLANLN | 1 | 0 | 0 | 1410900 | CAA71146.1 | A0A0P0K6L9 |
| AMPIMANLVNSK | 1 | 0 | 0 | 1410902 | AOG74783.1 | A0A0P0K6L9 |

| | | | | | | |
|---|---|---|---|---|---|---|
| AMPIVANLAYIN | 1 | 0 | 0 | 1410904 | ABB90509.1 | A0A0P0K6L9 |
| AMPIVANLAYLN | 1 | 0 | 0 | 1410905 | ABB90510.1 | A0A0P0K6L9 |
| AMPNIIRIVSSL | 1 | 0 | 0 | 1410907 | AKL87224.1 | P0C6X6 |
| AMPTLFKVQNVN | 1 | 0 | 0 | 1410915 | AVV62524.1 | K9N7C7 |
| AMQLNLISNITT | 1 | 0 | 0 | 1410924 | AVP78041.1 | P0C6W2 |
| AMQTGFTTTNVA | 1 | 0 | 0 | 1410929 | AUM60014.1 | K9N5Q8 |
| AMQVESDKYITT | 1 | 0 | 0 | 1410933 | AVP78030.1 | P0C6W2 |
| AMRPNFTIKGSF | 1 | 0 | 0 | 1410940 | AVP78041.1 | P0C6W2 |
| AMRPNYTIKGSF | 1 | 0 | 0 | 1410945 | AEA10817.1 | P0C6X7 |
| AMSAKEQRVPKG | 1 | 0 | 0 | 1410949 | AUM60021.1 | K9N4V7 |
| AMSGQILLNCAL | 1 | 0 | 0 | 1410951 | NP_073549.1 | A0A0P0LKV0 |
| AMTKRNVFPTIT | 1 | 0 | 0 | 1410960 | AVV62524.1 | K9N7C7 |
| AMTYTSDLLTNS | 1 | 0 | 0 | 1410968 | ARO76381.1 | P0C6U8 |
| AMVAKGVVGVLT | 1 | 0 | 0 | 1410969 | AIW52827.1 | H9EJ42 |
| AMVHDLNNVDPK | 1 | 0 | 0 | 1410970 | AVP78050.1 | Q3LZX3 |
| AMVLGSKHVNCC | 1 | 0 | 0 | 1410973 | AIW52827.1 | H9EJ42 |
| AMVVKPTVIVNA | 1 | 0 | 0 | 1410976 | APO40578.1 | P0C6X7 |
| ANAEAHNNLLQQ | 1 | 0 | 0 | 1410985 | ARB66064.1 | P36334 |
| ANAFNNALHSIQ | 1 | 0 | 0 | 1410987 | AIX10759.1 | P36334 |
| ANAFRNVVFELC | 1 | 0 | 0 | 1410988 | AVV62529.1 | K9N643 |
| ANAFSKPFYIPE | 1 | 0 | 0 | 1410989 | ATQ39394.1 | K9N796 |
| ANAFSKPLYVPE | 1 | 0 | 0 | 1410990 | AVV62527.1 | K9N796 |
| ANAFSNALDAIQ | 1 | 0 | 0 | 1410991 | CAA83660.1 | P36334 |
| ANAKIVTLPGSD | 1 | 0 | 0 | 1410997 | ATQ39390.1 | K9N5Q8 |
| ANALPECNADIV | 1 | 0 | 0 | 1411001 | ABE97136.1 | H9EJ42 |
| ANANSKPLIGYI | 1 | 0 | 0 | 1411003 | AUM60025.1 | K9N796 |
| ANANSKPLVGYI | 1 | 0 | 0 | 1411004 | AUM60015.1 | K9N796 |
| ANAQALNSLLQQ | 1 | 0 | 0 | 1411005 | BBA20986.1 | A0A140H1H1 |
| ANARNGCIPLSV | 1 | 0 | 0 | 1411006 | AUM60013.1 | K9N7C7 |
| ANARYVSPDVQF | 1 | 0 | 0 | 1411007 | AVV62526.1 | K9N5Q8 |
| ANATGSVIISTV | 1 | 0 | 0 | 1411011 | AVV62537.1 | K9N5Q8 |
| ANCLATPSGGVS | 1 | 0 | 0 | 1411017 | AVV62529.1 | K9N643 |
| ANCVNCADDRCV | 1 | 0 | 0 | 1411019 | AVV62524.1 | K9N7C7 |
| ANCVNCSDDRCV | 1 | 0 | 0 | 1411020 | AUM60013.1 | K9N7C7 |
| ANCVNCTDDRCV | 1 | 0 | 0 | 1411021 | AXN92247.1 | K9N7C7 |
| ANDDLYVKRYER | 1 | 0 | 0 | 1411022 | AEN19363.1 | P0C6X6 |
| ANDEKDTLRLFA | 1 | 0 | 0 | 1411023 | QED88024.1 | H9EJ42 |
| ANDFTTALRRPI | 1 | 0 | 0 | 1411024 | AKI29254.1 | K9N638 |
| ANDHYGTDIDGV | 1 | 0 | 0 | 1411026 | AJQ24067.1 | P33469 |
| ANDKFYRLSNEL | 1 | 0 | 0 | 1411027 | ALJ99930.1 | A0A0P0LKV0 |
| ANDLKDTLRLFA | 1 | 0 | 0 | 1411029 | QED88010.1 | H9EJ42 |
| ANDLTASLRRLV | 1 | 0 | 0 | 1411030 | AUM60013.1 | K9N7C7 |
| ANDLTTTLRRLV | 1 | 0 | 0 | 1411032 | AVV62524.1 | K9N7C7 |
| ANDLTTTLRRPI | 1 | 0 | 0 | 1411033 | AXN92247.1 | K9N7C7 |
| ANDNILSVKFTA | 1 | 0 | 0 | 1411034 | AVN89312.1 | K9N638 |
| ANDPVGFILRNT | 1 | 0 | 0 | 1411036 | ARO76381.1 | P0C6U8 |
| ANDTKIASQLGM | 1 | 0 | 0 | 1411039 | AWF93655.1 | K9N5Q8 |
| ANDTKIVSQLGN | 1 | 0 | 0 | 1411041 | AVN89313.1 | K9N5Q8 |
| ANDVDLQSSGLL | 1 | 0 | 0 | 1411042 | AUM60023.1 | K9N7C7 |
| ANDVDLQSSGLV | 1 | 0 | 0 | 1411043 | AUM60013.1 | K9N7C7 |
| ANDVEIQSSGVV | 1 | 0 | 0 | 1411044 | AVV62524.1 | K9N7C7 |
| ANDVEVQSSGII | 1 | 0 | 0 | 1411045 | AHY22544.1 | K9N7C7 |
| ANDWKPGLAMPS | 1 | 0 | 0 | 1411048 | AVV62535.1 | K9N7C7 |
| ANDWKPGLAMPT | 1 | 0 | 0 | 1411049 | AVV62524.1 | K9N7C7 |
| ANECAQILSEIV | 1 | 0 | 0 | 1411052 | AIL49554.1 | P0C6X6 |
| ANECAQVLSEIV | 1 | 0 | 0 | 1411053 | QXY02687.1 | P0DTD1 |
| ANECAQVLSETV | 1 | 0 | 0 | 1411055 | ADO24634.1 | A0A140H1G9 |
| ANECQHPNMTVH | 1 | 0 | 0 | 1411056 | AVV62540.1 | K9N643 |
| ANEETFKLSYGI | 1 | 0 | 0 | 1411058 | APO40578.1 | P0C6X7 |
| ANEKFADGINQM | 1 | 0 | 0 | 1411059 | APO40578.1 | P0C6X7 |
| ANELTTALRRPI | 1 | 0 | 0 | 1411062 | QBM11736.1 | K9N638 |
| ANFFDVPTFCAN | 1 | 0 | 0 | 1411071 | AMQ49014.1 | K9N7C7 |
| ANFILHDVNNGL | 1 | 0 | 0 | 1411074 | AEN19366.1 | P36334 |
| ANFILHDVNSGT | 1 | 0 | 0 | 1411076 | CAA83660.1 | P36334 |
| ANFNETNGPLCV | 1 | 0 | 0 | 1411081 | CAA71147.1 | A0A0P0K6L9 |
| ANFNILFSMILP | 1 | 0 | 0 | 1411082 | ABD75551.1 | A0A140H1G9 |
| ANFQHDNVLPET | 1 | 0 | 0 | 1411085 | AWO68987.1 | A0A0P0G321 |
| ANFRDLALKGSK | 1 | 0 | 0 | 1411090 | ARB07438.1 | A0A140H1H1 |
| ANFTDLALKGST | 1 | 0 | 0 | 1411093 | AZS52618.1 | A0A140H1H1 |
| ANGACIWNAAAY | 1 | 0 | 0 | 1411097 | AVV62524.1 | K9N7C7 |
| ANGACIWNCSAY | 1 | 0 | 0 | 1411098 | AUM60013.1 | K9N7C7 |
| ANGACVWNAAAY | 1 | 0 | 0 | 1411099 | AVV62535.1 | K9N7C7 |
| ANGFFYIDVGKH | 1 | 0 | 0 | 1411105 | AWO68966.1 | A0A0P0G321 |
| ANGFTAMNGEDA | 1 | 0 | 0 | 1411109 | NP_073549.1 | A0A0P0LKV0 |
| ANGFTVMNGEDA | 1 | 0 | 0 | 1411110 | QEO75983.1 | A0A0P0LKV0 |
| ANGFVLRIGAAA | 1 | 0 | 0 | 1411111 | AHI48723.1 | K9N5Q8 |
| ANGLFYIDVGNH | 1 | 0 | 0 | 1411122 | ACY56731.1 | A0A0P0G321 |
| ANGLHAPHNSWE | 1 | 0 | 0 | 1411124 | AVV62535.1 | K9N7C7 |
| ANGLHAPRNSWE | 1 | 0 | 0 | 1411125 | AVV62524.1 | K9N7C7 |
| ANGRLVGTTLQG | 1 | 0 | 0 | 1411129 | AJG44057.1 | K9N638 |
| ANGSLDGTTLQG | 1 | 0 | 0 | 1411131 | AXG21642.1 | K9N638 |
| ANGSLVGTTLQG | 1 | 0 | 0 | 1411134 | AXN92247.1 | K9N7C7 |
| ANGTHTGSSFDG | 1 | 0 | 0 | 1411137 | AVV62524.1 | K9N7C7 |
| ANHVANFQKAGF | 1 | 0 | 0 | 1411146 | AVV62524.1 | K9N7C7 |
| ANHYSPCVSIVP | 1 | 0 | 0 | 1411149 | AID55087.1 | K9N5Q8 |
| ANICHSNILSLN | 1 | 0 | 0 | 1411150 | ABD75616.1 | A0A140H1H0 |
| ANICNSNILSLN | 1 | 0 | 0 | 1411151 | AGT17757.1 | A0A140H1H0 |
| ANIGSNHTGVVG | 1 | 0 | 0 | 1411158 | APO40578.1 | P0C6X7 |
| ANIKSDILSLNN | 1 | 0 | 0 | 1411164 | AZS52617.1 | A0A140H1H0 |
| ANINQDSYGGAS | 1 | 0 | 0 | 1411166 | ABD75615.1 | A0A140H1G9 |
| ANISMLQLGAPD | 1 | 0 | 0 | 1411171 | AKA09625.1 | A0A0P0G321 |

| | | | | | | |
|---|---|---|---|---|---|---|
| ANISMLQLGVPD | 1 | 0 | 0 | 1411172 | AKA09626.1 | A0A0P0G321 |
| ANISTLQLGVPD | 1 | 0 | 0 | 1411173 | AKA09623.1 | A0A0P0G321 |
| ANIVFPQGGYVA | 1 | 0 | 0 | 1411174 | AGT51419.1 | P0C6X6 |
| ANIVHEPTTPIS | 1 | 0 | 0 | 1411175 | ALA49956.1 | K9N7C7 |
| ANIVYEPSTPIS | 1 | 0 | 0 | 1411177 | AUM60013.1 | K9N7C7 |
| ANIVYEPSTPVS | 1 | 0 | 0 | 1411178 | AVV62524.1 | K9N7C7 |
| ANIYLKHGGGVA | 1 | 0 | 0 | 1411180 | AVP78041.1 | P0C6W2 |
| ANKEQSLVLTST | 1 | 0 | 0 | 1411189 | AVV62524.1 | K9N7C7 |
| ANKFFQFNQWYE | 1 | 0 | 0 | 1411192 | ABD75567.1 | A0A140H1G9 |
| ANKFNQAPGAMQ | 1 | 0 | 0 | 1411195 | AVV62526.1 | K9N5Q8 |
| ANKHADFDSWFS | 1 | 0 | 0 | 1411197 | AVP78041.1 | P0C6W2 |
| ANKHAGFDSWFS | 1 | 0 | 0 | 1411200 | P0C6W6.1 | P0C6W2 |
| ANKHAGFDTWFS | 1 | 0 | 0 | 1411201 | ARO76381.1 | P0C6U8 |
| ANKIVGGAHTWF | 1 | 0 | 0 | 1411203 | ALA49340.1 | K9N7C7 |
| ANKTKGIVAHPL | 1 | 0 | 0 | 1411212 | AVV62524.1 | K9N7C7 |
| ANLANLNSHNIG | 1 | 0 | 0 | 1411218 | CAA71146.1 | A0A0P0K6L9 |
| ANLANLNSHTIG | 1 | 0 | 0 | 1411219 | AGT21338.1 | A0A0P0K6L9 |
| ANLAYINSYTIG | 1 | 0 | 0 | 1411221 | ABB90509.1 | A0A0P0K6L9 |
| ANLAYSNSYTIG | 1 | 0 | 0 | 1411223 | AAQ90006.1 | A0A0P0K6L9 |
| ANLFLEQSQRSL | 1 | 0 | 0 | 1411229 | AVV62535.1 | K9N7C7 |
| ANLFPPWSAAAG | 1 | 0 | 0 | 1411232 | CAA83661.1 | P36334 |
| ANLFSDIKTHKS | 1 | 0 | 0 | 1411234 | AVV62537.1 | K9N5Q8 |
| ANLFVEQSQRSL | 1 | 0 | 0 | 1411235 | AVV62524.1 | K9N7C7 |
| ANLGERVRQALL | 1 | 0 | 0 | 1411237 | P0C6W6.1 | P0C6W2 |
| ANLILHDVNSGL | 1 | 0 | 0 | 1411239 | AVQ05264.1 | P36334 |
| ANLKLPSHDVTV | 1 | 0 | 0 | 1411241 | AVA26874.1 | H9EJ24 |
| ANLMQDESVANL | 1 | 0 | 0 | 1411246 | AVV62537.1 | K9N5Q8 |
| ANLNFVALIPDY | 1 | 0 | 0 | 1411247 | P0C6U5.1 | A0A140H1G9 |
| ANLNILFSMVLP | 1 | 0 | 0 | 1411248 | AIL49625.1 | P0C6X6 |
| ANLRQDDFVRNL | 1 | 0 | 0 | 1411252 | AMW90843.1 | K9N5Q8 |
| ANLRQDDSVRDL | 1 | 0 | 0 | 1411253 | QBF80611.1 | K9N5Q8 |
| ANLRQDESVSNL | 1 | 0 | 0 | 1411255 | AVV62526.1 | K9N5Q8 |
| ANLSIPSNWTIS | 1 | 0 | 0 | 1411258 | AAY43188.1 | A0A0P0G321 |
| ANLSMLQLSVPD | 1 | 0 | 0 | 1411261 | BAM16475.1 | A0A0P0G321 |
| ANLTISKPSSYS | 1 | 0 | 0 | 1411262 | AVV62526.1 | K9N5Q8 |
| ANLTITKPSSYS | 1 | 0 | 0 | 1411263 | AVV62537.1 | K9N5Q8 |
| ANLVNHKSHNIG | 1 | 0 | 0 | 1411264 | AIW52757.1 | A0A0P0K6L9 |
| ANLVNLNSHNIG | 1 | 0 | 0 | 1411265 | AIW52756.1 | A0A0P0K6L9 |
| ANLVNYKSHNIG | 1 | 0 | 0 | 1411267 | QOP39313.1 | A0A0P0K6L9 |
| ANMGIDLDEWSV | 1 | 0 | 0 | 1411273 | ARO76381.1 | P0C6U8 |
| ANMRNANGACIW | 1 | 0 | 0 | 1411275 | AVV62524.1 | K9N7C7 |
| ANMRNANGACVW | 1 | 0 | 0 | 1411276 | AVV62535.1 | K9N7C7 |
| ANMSMAECKAAL | 1 | 0 | 0 | 1411280 | AGT51337.1 | H9EJ42 |
| ANNGHKQIVKSI | 1 | 0 | 0 | 1411284 | AUM60027.1 | K9N643 |
| ANNINRFNVAVT | 1 | 0 | 0 | 1411286 | AUM60013.1 | K9N7C7 |
| ANNLMNGVTLST | 1 | 0 | 0 | 1411288 | AIL49500.1 | P36334 |
| ANNLNRFNVAVT | 1 | 0 | 0 | 1411289 | AVV62524.1 | K9N7C7 |
| ANNSIAIPTNFS | 1 | 0 | 0 | 1411292 | AVP78031.1 | Q3LZX1 |
| ANNTIGIVSQPL | 1 | 0 | 0 | 1411294 | AUM60013.1 | K9N7C7 |
| ANNYSKLFDEGH | 1 | 0 | 0 | 1411297 | AFR79256.1 | A0A0P0ILY9 |
| ANNYSKLSDEGY | 1 | 0 | 0 | 1411299 | AST12962.1 | A0A0P0LKV0 |
| ANPAAVDGKCLT | 1 | 0 | 0 | 1411301 | AMK59674.1 | P0C6X6 |
| ANPAAVDSKCLA | 1 | 0 | 0 | 1411302 | AGT51687.1 | P0C6X6 |
| ANPAAVDSKCLT | 1 | 0 | 0 | 1411303 | AEN19363.1 | P0C6X6 |
| ANPLWHKAAFIS | 1 | 0 | 0 | 1411308 | AXX83330.1 | P0C6X6 |
| ANPNSVLTTAVY | 1 | 0 | 0 | 1411310 | QRK03774.1 | A0A0P0ILY9 |
| ANPSTPPYTFTT | 1 | 0 | 0 | 1411312 | AUM60013.1 | K9N7C7 |
| ANPSTPVYTFTT | 1 | 0 | 0 | 1411313 | QBF80585.1 | K9N7C7 |
| ANPTCRIFATAP | 1 | 0 | 0 | 1411314 | AVV62526.1 | K9N5Q8 |
| ANPTNCITPVNG | 1 | 0 | 0 | 1411317 | ALJ76278.1 | K9N5Q8 |
| ANPTVIVNAANI | 1 | 0 | 0 | 1411318 | ATO98229.1 | P0C6W2 |
| ANPVKEYEADAE | 1 | 0 | 0 | 1411319 | AVV62535.1 | K9N7C7 |
| ANQERYVKITGL | 1 | 0 | 0 | 1411322 | AXG21641.1 | K9N7C7 |
| ANQFNNAISKIQ | 1 | 0 | 0 | 1411325 | APO40579.1 | P59594 |
| ANQFTEFIGTQA | 1 | 0 | 0 | 1411327 | AVV62524.1 | K9N7C7 |
| ANQFTEFVGTQA | 1 | 0 | 0 | 1411328 | AUM60013.1 | K9N7C7 |
| ANQFTEFVSTQS | 1 | 0 | 0 | 1411330 | ATG84865.1 | K9N638 |
| ANQNTSVYSANG | 1 | 0 | 0 | 1411332 | BAM16475.1 | A0A0P0G321 |
| ANQSTSSYPANG | 1 | 0 | 0 | 1411333 | AWO68991.1 | A0A0P0G321 |
| ANQSTSTYLANG | 1 | 0 | 0 | 1411334 | AMM42642.1 | A0A0P0G321 |
| ANQSTSTYPANG | 1 | 0 | 0 | 1411335 | AWO68966.1 | A0A0P0G321 |
| ANQSTSTYSANG | 1 | 0 | 0 | 1411336 | AWO68998.1 | A0A0P0G321 |
| ANQSTSVYSAKG | 1 | 0 | 0 | 1411337 | AKA09625.1 | A0A0P0G321 |
| ANQYSPCVPIVP | 1 | 0 | 0 | 1411339 | QAT98898.1 | K9N5Q8 |
| ANQYSPCVSIAP | 1 | 0 | 0 | 1411340 | QEJ82215.1 | K9N5Q8 |
| ANQYSPCVSILP | 1 | 0 | 0 | 1411341 | AKL59401.1 | K9N5Q8 |
| ANQYSPCVSTVP | 1 | 0 | 0 | 1411343 | ANC28634.1 | K9N5Q8 |
| ANRCNEVSATVL | 1 | 0 | 0 | 1411349 | AVQ05261.1 | P0C6X6 |
| ANRPGVYCGSDY | 1 | 0 | 0 | 1411352 | AVV62524.1 | K9N7C7 |
| ANRSALCQWCLI | 1 | 0 | 0 | 1411354 | AUM60013.1 | K9N7C7 |
| ANRSAMCNWCLV | 1 | 0 | 0 | 1411356 | AHL18091.1 | K9N7C7 |
| ANRYNEVPATVL | 1 | 0 | 0 | 1411358 | ATN39877.2 | P0C6X6 |
| ANSEALNNLLQQ | 1 | 0 | 0 | 1411364 | AWW13566.1 | P36334 |
| ANSFDAPTFCAN | 1 | 0 | 0 | 1411365 | AHI48604.1 | K9N7C7 |
| ANSFDISNFCAN | 1 | 0 | 0 | 1411366 | AVV62535.1 | K9N7C7 |
| ANSFDISSYCAN | 1 | 0 | 0 | 1411367 | AUM60013.1 | K9N7C7 |
| ANSGDYYHKVEA | 1 | 0 | 0 | 1411370 | AGT51421.1 | P30215 |
| ANSGDYYYKVEA | 1 | 0 | 0 | 1411371 | P30215.1 | P30215 |
| ANSGGYYYKVEA | 1 | 0 | 0 | 1411372 | AGT51669.1 | P30215 |
| ANSGNRAPTSGV | 1 | 0 | 0 | 1411373 | AIX10723.1 | P33469 |
| ANSGNRTPTPGV | 1 | 0 | 0 | 1411376 | AIX10722.1 | P33469 |

| | | | | | | |
|---|---|---|---|---|---|---|
| ANSIDVPTFCAN | 1 | 0 | 0 | 1411379 | AQZ41320.1 | K9N7C7 |
| ANSKPLIGYIPE | 1 | 0 | 0 | 1411385 | AUM60025.1 | K9N796 |
| ANSKPLVGYIPE | 1 | 0 | 0 | 1411386 | AUM60015.1 | K9N796 |
| ANSLDNLKSGVI | 1 | 0 | 0 | 1411387 | YP_003767.1 | A0A0P0G321 |
| ANSLHATATIPL | 1 | 0 | 0 | 1411388 | APO40580.1 | P59632 |
| ANSNLRTFCVDK | 1 | 0 | 0 | 1411390 | AVV62524.1 | K9N7C7 |
| ANSNLRTFSVDK | 1 | 0 | 0 | 1411391 | AUM60013.1 | K9N7C7 |
| ANSNSVCYTPTN | 1 | 0 | 0 | 1411392 | AUM60013.1 | K9N7C7 |
| ANSPEPALLFRN | 1 | 0 | 0 | 1411393 | AIX10763.1 | P36334 |
| ANSSDPALLFRN | 1 | 0 | 0 | 1411396 | AIL49515.1 | P36334 |
| ANSSEPTLLFRN | 1 | 0 | 0 | 1411397 | AIL49500.1 | P36334 |
| ANSTGNLIISSS | 1 | 0 | 0 | 1411399 | AVV62526.1 | K9N5Q8 |
| ANSTGTVIIRPS | 1 | 0 | 0 | 1411400 | QBF80608.1 | K9N5Q8 |
| ANSTPSQNAGYW | 1 | 0 | 0 | 1411403 | AQZ41279.1 | K9N4V7 |
| ANSVFNIFQAVS | 1 | 0 | 0 | 1411407 | AVP78019.1 | P0C6W2 |
| ANSVFNILQVTT | 1 | 0 | 0 | 1411409 | AVV62524.1 | K9N7C7 |
| ANSVLNICQAVT | 1 | 0 | 0 | 1411410 | AVP77956.1 | P0C6W2 |
| ANTDIILGVCVN | 1 | 0 | 0 | 1411412 | CAA79896.1 | P36334 |
| ANTDIKLDVCVN | 1 | 0 | 0 | 1411413 | BBO58740.1 | P36334 |
| ANTEIELGVCVN | 1 | 0 | 0 | 1411415 | AEN19366.1 | P36334 |
| ANTLTIIIPDKQ | 1 | 0 | 0 | 1411422 | ARU07575.1 | A0A140H1G9 |
| ANTTTAPLKLFA | 1 | 0 | 0 | 1411427 | AGV08437.1 | K9N7C7 |
| ANTVIWDFNREA | 1 | 0 | 0 | 1411429 | APO40578.1 | P0C6X7 |
| ANTVIWDHKREA | 1 | 0 | 0 | 1411430 | AVP78041.1 | P0C6W2 |
| ANVCALISCNGN | 1 | 0 | 0 | 1411434 | AIL49553.1 | P0C6X6 |
| ANVCSPMACNGH | 1 | 0 | 0 | 1411436 | ACR82761.1 | A0A140H1G9 |
| ANVISFGDYNLS | 1 | 0 | 0 | 1411441 | AFD64760.1 | A0A0P0G321 |
| ANVNWADDRAAR | 1 | 0 | 0 | 1411446 | AVL25604.1 | H9EJ69 |
| ANVRSLTACNGH | 1 | 0 | 0 | 1411450 | ADO24641.1 | A0A140H1G9 |
| ANVSALIGANGN | 1 | 0 | 0 | 1411451 | AVN89322.1 | K9N7C7 |
| ANVSVSRFDPST | 1 | 0 | 0 | 1411452 | AIV41831.1 | P36334 |
| ANVSVSRFNPSI | 1 | 0 | 0 | 1411453 | AIL49507.1 | P36334 |
| ANVSVSRFNPST | 1 | 0 | 0 | 1411454 | CAA79896.1 | P36334 |
| ANVSYVYNGSAQ | 1 | 0 | 0 | 1411455 | QBP84739.1 | P30215 |
| ANVTQAFGRRGP | 1 | 0 | 0 | 1411456 | 2GIB_A | P59595 |
| ANVTSFGDYNLS | 1 | 0 | 0 | 1411457 | AIW52843.1 | A0A0P0G321 |
| ANVYRRKPDLPN | 1 | 0 | 0 | 1411460 | ABD43177.1 | P36334 |
| ANWLAPNNMYMR | 1 | 0 | 0 | 1411465 | AVV62524.1 | K9N7C7 |
| ANWLAPTNAYTR | 1 | 0 | 0 | 1411467 | ANI69888.1 | K9N638 |
| ANWLTPTNMYMR | 1 | 0 | 0 | 1411470 | AUM60013.1 | K9N7C7 |
| ANYFQDVKQFAN | 1 | 0 | 0 | 1411473 | AID55096.1 | K9N5Q8 |
| ANYNTYDVVLSP | 1 | 0 | 0 | 1411489 | APO40578.1 | P0C6X7 |
| ANYQKVGVQKYS | 1 | 0 | 0 | 1411491 | APO40578.1 | P0C6X7 |
| APAAPRAVSFAD | 1 | 0 | 0 | 1411496 | AQZ41292.1 | K9N4V7 |
| APAICHEGKAHF | 1 | 0 | 0 | 1411501 | AVP78031.1 | Q3LZX1 |
| APAICHNGKAYF | 1 | 0 | 0 | 1411503 | APO40579.1 | P59594 |
| APALPKTEVIDT | 1 | 0 | 0 | 1411504 | AVV62535.1 | K9N7C7 |
| APALQDAYYRAR | 1 | 0 | 0 | 1411505 | APO40578.1 | P0C6X7 |
| APANLRVVGHSM | 1 | 0 | 0 | 1411508 | AVV62524.1 | K9N7C7 |
| APANLTISKPSS | 1 | 0 | 0 | 1411509 | AVV62526.1 | K9N5Q8 |
| APANLTITKPSS | 1 | 0 | 0 | 1411510 | AVV62537.1 | K9N5Q8 |
| APAQTGVIADYN | 1 | 0 | 0 | 1411512 | APO40579.1 | P59594 |
| APCAVSFADNND | 1 | 0 | 0 | 1411520 | AKK52599.1 | K9N4V7 |
| APCIKVAAMQSK | 1 | 0 | 0 | 1411522 | ASY99852.1 | K9N7C7 |
| APDDASRLLHTV | 1 | 0 | 0 | 1411527 | AXN92247.1 | K9N7C7 |
| APDDATRLLHVV | 1 | 0 | 0 | 1411528 | AUM60013.1 | K9N7C7 |
| APDFSLDYINVT | 1 | 0 | 0 | 1411530 | BBA20976.1 | P36334 |
| APDLSFDYINVT | 1 | 0 | 0 | 1411535 | AIX10753.1 | P36334 |
| APDLSLDYINAT | 1 | 0 | 0 | 1411536 | QEG03776.1 | P36334 |
| APDNPSAIVTGL | 1 | 0 | 0 | 1411538 | AKA09625.1 | A0A0P0G321 |
| APDVMLNISTPN | 1 | 0 | 0 | 1411541 | CAA83660.1 | P36334 |
| APEDDDFLEGSD | 1 | 0 | 0 | 1411546 | AIV41846.1 | P0C6X6 |
| APEPQRGRQGRI | 1 | 0 | 0 | 1411555 | APT69853.1 | A0A0N7J131 |
| APESKSFTMTFG | 1 | 0 | 0 | 1411556 | AVV62524.1 | K9N7C7 |
| APEYDDFLEESD | 1 | 0 | 0 | 1411558 | AGT51707.1 | P0C6X6 |
| APFIYLNNSIPN | 1 | 0 | 0 | 1411560 | BBA20986.1 | A0A140H1H1 |
| APGETGVIADYN | 1 | 0 | 0 | 1411577 | ATO98181.1 | Q3LZX1 |
| APGVPATQAKGY | 1 | 0 | 0 | 1411590 | ATN39868.1 | P33469 |
| APHDFFIFDVDK | 1 | 0 | 0 | 1411592 | AXN92247.1 | K9N7C7 |
| APHGPFDALWLS | 1 | 0 | 0 | 1411596 | AWK59942.1 | H9EJ42 |
| APHISLAMLDIQ | 1 | 0 | 0 | 1411600 | NP_150075.1 | Q80872 |
| APHNDCTPINTN | 1 | 0 | 0 | 1411602 | AVV62526.1 | K9N5Q8 |
| APHNSWEALALN | 1 | 0 | 0 | 1411603 | AVV62535.1 | K9N7C7 |
| APIDGYFVLNET | 1 | 0 | 0 | 1411608 | AVV62526.1 | K9N5Q8 |
| APIDGYFVLNNT | 1 | 0 | 0 | 1411609 | AVV62537.1 | K9N5Q8 |
| APIDIDAFPVRP | 1 | 0 | 0 | 1411611 | AWK59942.1 | H9EJ42 |
| APIELDNKFTPL | 1 | 0 | 0 | 1411612 | AUM60013.1 | K9N7C7 |
| APKNFCPCKLIS | 1 | 0 | 0 | 1411638 | AIL49492.1 | P36334 |
| APKNFCPCKLNS | 1 | 0 | 0 | 1411640 | AVQ05264.1 | P36334 |
| APKNFCPCKSND | 1 | 0 | 0 | 1411641 | AIV41891.1 | P36334 |
| APKNFCPCKSNG | 1 | 0 | 0 | 1411642 | AEN19358.1 | P36334 |
| APKNFCPCSSCF | 1 | 0 | 0 | 1411644 | QEG03814.1 | P36334 |
| APKNFCPCSSCH | 1 | 0 | 0 | 1411645 | ARB66068.1 | P36334 |
| APKNFCPCSSCP | 1 | 0 | 0 | 1411646 | AWW13566.1 | P36334 |
| APKNFCPCSSCS | 1 | 0 | 0 | 1411647 | AVR40344.1 | P36334 |
| APKQGYFIKHND | 1 | 0 | 0 | 1411651 | BBA20983.1 | A0A140H1H1 |
| APKSRVQQNKIR | 1 | 0 | 0 | 1411655 | AGT51784.1 | P33469 |
| APKVITKKDAAA | 1 | 0 | 0 | 1411659 | AVV62544.1 | K9N4V7 |
| APKVVTKKDAAA | 1 | 0 | 0 | 1411660 | AVV62533.1 | K9N4V7 |
| APLSDQIDFLQS | 1 | 0 | 0 | 1411675 | APO40578.1 | P0C6X7 |
| APLTNCVKMLTP | 1 | 0 | 0 | 1411676 | AXN92247.1 | K9N7C7 |

| | | | | | | |
|---|---|---|---|---|---|---|
| APLVGTPVCING | 1 | 0 | 0 | 1411678 | QYC64967.1 | P0DTD1 |
| APNAKEEILGAV | 1 | 0 | 0 | 1411681 | BAE93400.1 | P0C6U8 |
| APNGFSQDGDYF | 1 | 0 | 0 | 1411687 | AVV62537.1 | K9N5Q8 |
| APNGLSEDGQVF | 1 | 0 | 0 | 1411689 | AVV62526.1 | K9N5Q8 |
| APNGLYFMHVGY | 1 | 0 | 0 | 1411691 | AXN92228.1 | K9N5Q8 |
| APNGTYNCTDAV | 1 | 0 | 0 | 1411692 | ADK37704.1 | A0A0P0K6L9 |
| APNLTFNSHINA | 1 | 0 | 0 | 1411693 | BBA20986.1 | A0A140H1H1 |
| APNLTLNLHTIN | 1 | 0 | 0 | 1411695 | BBA20983.1 | A0A140H1H1 |
| APNNMYMRTTHT | 1 | 0 | 0 | 1411697 | AVV62524.1 | K9N7C7 |
| APNNSVSWYTGL | 1 | 0 | 0 | 1411698 | AVV62544.1 | K9N4V7 |
| APNNTVSWYTGL | 1 | 0 | 0 | 1411699 | AXN92267.1 | K9N4V7 |
| APNTSVSWYTGL | 1 | 0 | 0 | 1411701 | AVV62533.1 | K9N4V7 |
| APNTTVSWYTGL | 1 | 0 | 0 | 1411702 | AUM60021.1 | K9N4V7 |
| APPAQYELKHGT | 1 | 0 | 0 | 1411711 | AVP78041.1 | P0C6W2 |
| APPGDNFKHLIP | 1 | 0 | 0 | 1411712 | ARB07436.1 | A0A140H1G9 |
| APPGEQFTHIVP | 1 | 0 | 0 | 1411715 | NP_073549.1 | A0A0P0LKV0 |
| APPIADFTPVSG | 1 | 0 | 0 | 1411716 | AVV62528.1 | K9N4V0 |
| APPSECRLQNGD | 1 | 0 | 0 | 1411718 | APO40578.1 | P0C6X7 |
| APPTGVYELHGY | 1 | 0 | 0 | 1411720 | AIL49485.1 | P36334 |
| APQANFGDYYYK | 1 | 0 | 0 | 1411726 | AGT51599.1 | P30215 |
| APQANSGGYYYK | 1 | 0 | 0 | 1411729 | AGT51669.1 | P30215 |
| APQATYQNISTN | 1 | 0 | 0 | 1411731 | ALM26400.1 | K9N5Q8 |
| APQVTYQNISIN | 1 | 0 | 0 | 1411738 | ATG84888.1 | K9N5Q8 |
| APRAISFADNND | 1 | 0 | 0 | 1411740 | ANI69896.1 | K9N4V7 |
| APRAVSFAYNND | 1 | 0 | 0 | 1411744 | AVN89381.1 | K9N4V7 |
| APRAVTFGDNND | 1 | 0 | 0 | 1411745 | AVV62533.1 | K9N4V7 |
| APRGSITQRSHA | 1 | 0 | 0 | 1411747 | AVV62533.1 | K9N4V7 |
| APRGSITQRSRP | 1 | 0 | 0 | 1411748 | AUM60031.1 | K9N4V7 |
| APRNSWEALALN | 1 | 0 | 0 | 1411755 | AVV62524.1 | K9N7C7 |
| APRSVRSVPGEM | 1 | 0 | 0 | 1411758 | ALJ54472.1 | K9N5Q8 |
| APSASAFMSMSQ | 1 | 0 | 0 | 1411766 | AVV62544.1 | K9N4V7 |
| APSRSSETKEQK | 1 | 0 | 0 | 1411774 | AIW52721.1 | A0A0N7J131 |
| APSTVWEDGDYY | 1 | 0 | 0 | 1411778 | QEJ82215.1 | K9N5Q8 |
| APSWVTALYNFT | 1 | 0 | 0 | 1411779 | AUM60013.1 | K9N7C7 |
| APTDTCFANKHA | 1 | 0 | 0 | 1411788 | AVP78041.1 | P0C6W2 |
| APTIANQERYVK | 1 | 0 | 0 | 1411793 | AXG21663.1 | K9N7C7 |
| APTLIPQENYSS | 1 | 0 | 0 | 1411798 | AWW13571.1 | P0C6X6 |
| APTNAYTRTTHT | 1 | 0 | 0 | 1411805 | ANI69888.1 | K9N638 |
| APTNFCPCKLDG | 1 | 0 | 0 | 1411806 | CAA83660.1 | P36334 |
| APTNVYMRTTHT | 1 | 0 | 0 | 1411808 | ANC28699.1 | K9N638 |
| APTRSVTVEYNI | 1 | 0 | 0 | 1411809 | AVV62535.1 | K9N7C7 |
| APTSGVIPDMAD | 1 | 0 | 0 | 1411810 | AMK59681.1 | P33469 |
| APTTCGYLPNNA | 1 | 0 | 0 | 1411812 | APO40578.1 | P0C6X7 |
| APVDGYFILNQT | 1 | 0 | 0 | 1411818 | AUM60014.1 | K9N5Q8 |
| APVEDIAQVVIA | 1 | 0 | 0 | 1411819 | AVN89451.1 | K9N7C7 |
| APVELENKFTPL | 1 | 0 | 0 | 1411822 | AVV62524.1 | K9N7C7 |
| APVPVKEYEADV | 1 | 0 | 0 | 1411830 | AVV62524.1 | K9N7C7 |
| APVSPAFHRIES | 1 | 0 | 0 | 1411835 | AKM76243.1 | K9N7D2 |
| APYIVGDVVHNG | 1 | 0 | 0 | 1411848 | APO40578.1 | P0C6X7 |
| APYMVGDVINSG | 1 | 0 | 0 | 1411850 | ARO76381.1 | P0C6U8 |
| APYVMLNTSSPN | 1 | 0 | 0 | 1411857 | AIL49492.1 | P36334 |
| APYVMLNTSTSN | 1 | 0 | 0 | 1411858 | AMK59677.1 | P36334 |
| AQADTAGLFRID | 1 | 0 | 0 | 1411866 | ALW82670.1 | K9N796 |
| AQADTAGLFTNF | 1 | 0 | 0 | 1411867 | ALA49936.1 | K9N796 |
| AQADTAGLYTNS | 1 | 0 | 0 | 1411869 | AGV08439.1 | K9N796 |
| AQAFETPAVQFS | 1 | 0 | 0 | 1411873 | AVV62535.1 | K9N7C7 |
| AQAFGLRGPGDL | 1 | 0 | 0 | 1411874 | AXG21672.1 | K9N4V7 |
| AQALAKLASELS | 1 | 0 | 0 | 1411879 | AVK87450.1 | K9N5Q8 |
| AQALTEVVYSNG | 1 | 0 | 0 | 1411883 | ADO24700.1 | A0A0P0LKV0 |
| AQANGFTVMNGE | 1 | 0 | 0 | 1411886 | QEO75983.1 | A0A0P0LKV0 |
| AQAQIDRLINGR | 1 | 0 | 0 | 1411887 | CAA83660.1 | P36334 |
| AQATSFCKSGNL | 1 | 0 | 0 | 1411891 | AIL49332.1 | P30215 |
| AQAVRRYSEAAS | 1 | 0 | 0 | 1411894 | NP_073549.1 | A0A0P0LKV0 |
| AQCDCLCTPDPI | 1 | 0 | 0 | 1411896 | CAA83660.1 | P36334 |
| AQCTRDPVGFCL | 1 | 0 | 0 | 1411903 | AVV62524.1 | K9N7C7 |
| AQDGCGHISTKP | 1 | 0 | 0 | 1411905 | ATO98193.1 | Q3LZX1 |
| AQDGHAAITDYS | 1 | 0 | 0 | 1411906 | AUM60013.1 | K9N7C7 |
| AQDGNAAISDYDYYRYN | 1 | 0 | 0 | 1411908 | P0DTD1.1 | P0DTD1 |
| AQDTWGTSAAAY | 1 | 0 | 0 | 1411911 | ATO98205.1 | Q3LZX1 |
| AQEAYEQAVVNG | 1 | 0 | 0 | 1411913 | AVP78041.1 | P0C6W2 |
| AQEGCGILSNKS | 1 | 0 | 0 | 1411916 | APO40579.1 | P59594 |
| AQEGICITTNGS | 1 | 0 | 0 | 1411917 | ALA49758.1 | K9N7C7 |
| AQEGVCITTNGL | 1 | 0 | 0 | 1411918 | AVK87438.1 | K9N638 |
| AQEGVCITTNGS | 1 | 0 | 0 | 1411919 | AXN92247.1 | K9N7C7 |
| AQFNYKQSFANP | 1 | 0 | 0 | 1411927 | AVV62526.1 | K9N5Q8 |
| AQFNYRRDGQPC | 1 | 0 | 0 | 1411928 | QAT98908.1 | K9N7C7 |
| AQGAHLFSSRYV | 1 | 0 | 0 | 1411930 | ALJ54512.1 | K9N5Q8 |
| AQGFQACRFIAF | 1 | 0 | 0 | 1411935 | AVA26872.1 | H9EJ42 |
| AQGFQACRFVAF | 1 | 0 | 0 | 1411936 | AWK59942.1 | H9EJ42 |
| AQGKPVPYCYDT | 1 | 0 | 0 | 1411937 | APO40578.1 | P0C6X7 |
| AQGMETLACESQ | 1 | 0 | 0 | 1411941 | APO40578.1 | P0C6X7 |
| AQGTGHGHKVCE | 1 | 0 | 0 | 1411946 | ARO76381.1 | P0C6U8 |
| AQHFSDFGTSFV | 1 | 0 | 0 | 1411961 | AEN19363.1 | P0C6X6 |
| AQHQPYVVDDPC | 1 | 0 | 0 | 1411962 | AVP78037.1 | Q3LZX5 |
| AQILSEIVMCGG | 1 | 0 | 0 | 1411970 | AIL49554.1 | P0C6X6 |
| AQIQNKTVDVVV | 1 | 0 | 0 | 1411971 | AVV62535.1 | K9N7C7 |
| AQKLETDGICPE | 1 | 0 | 0 | 1411979 | AEN19363.1 | P0C6X6 |
| AQKLKTDGICPE | 1 | 0 | 0 | 1411982 | AIX09796.1 | P0C6X6 |
| AQKSYQEALASG | 1 | 0 | 0 | 1411984 | AVV62524.1 | K9N7C7 |
| AQKSYQEALDSG | 1 | 0 | 0 | 1411985 | AUM60013.1 | K9N7C7 |
| AQKSYQEAVDSG | 1 | 0 | 0 | 1411986 | AVV62535.1 | K9N7C7 |

| | | | | | | |
|---|---|---|---|---|---|---|
| AQLASDKVNECV | 1 | 0 | 0 | 1411987 | AVV62526.1 | K9N5Q8 |
| AQLGCVFLNGVD | 1 | 0 | 0 | 1411989 | AVV62524.1 | K9N7C7 |
| AQLNAKLGFNVD | 1 | 0 | 0 | 1411991 | ABD75519.1 | A0A140H1G9 |
| AQLPAPRTLLTR | 1 | 0 | 0 | 1411994 | QWK59326.1 | P0DTD1 |
| AQLTDKYCTINI | 1 | 0 | 0 | 1411997 | AVV62524.1 | K9N7C7 |
| AQLTDKYCTTNI | 1 | 0 | 0 | 1411998 | AVV62535.1 | K9N7C7 |
| AQNANPMVIVNA | 1 | 0 | 0 | 1412009 | P0C6W6.1 | P0C6W2 |
| AQNANPTVIVNA | 1 | 0 | 0 | 1412010 | ATO98229.1 | P0C6W2 |
| AQNPLSELKCTI | 1 | 0 | 0 | 1412015 | ATO98193.1 | Q3LZX1 |
| AQNTVLNACTPE | 1 | 0 | 0 | 1412017 | ABK63972.1 | H9EJ69 |
| AQNTVLNASISE | 1 | 0 | 0 | 1412019 | AMM42596.1 | H9EJ69 |
| AQNTVLNDSIPE | 1 | 0 | 0 | 1412020 | AMM42614.1 | H9EJ69 |
| AQPCSDKAYKIA | 1 | 0 | 0 | 1412021 | QRU91444.1 | P0DTD1 |
| AQPEGCVLTNTG | 1 | 0 | 0 | 1412023 | NP_073549.1 | A0A0P0LKV0 |
| AQPKQTSTSQQP | 1 | 0 | 0 | 1412027 | ATP16771.1 | P33469 |
| AQPLTRKVQQLF | 1 | 0 | 0 | 1412035 | AXN92247.1 | K9N7C7 |
| AQPNQTATSQQP | 1 | 0 | 0 | 1412036 | AIX10721.1 | P33469 |
| AQQEATHIAEDV | 1 | 0 | 0 | 1412048 | AST12962.1 | A0A0P0LKV0 |
| AQQEVQLKVIKC | 1 | 0 | 0 | 1412050 | AUM60013.1 | K9N7C7 |
| AQQEVTHIAEDV | 1 | 0 | 0 | 1412051 | APT69889.1 | A0A0P0LKV0 |
| AQQKVNECVESQ | 1 | 0 | 0 | 1412054 | ABB90528.1 | A0A0P0K6L9 |
| AQQLVRSEAAAR | 1 | 0 | 0 | 1412057 | AUM60014.1 | K9N5Q8 |
| AQQLVRSESAAI | 1 | 0 | 0 | 1412058 | AVN89365.1 | K9N5Q8 |
| AQQLVRSETAAR | 1 | 0 | 0 | 1412060 | AVV62526.1 | K9N5Q8 |
| AQRISWLLHKDG | 1 | 0 | 0 | 1412063 | AMQ49017.1 | K9N4V0 |
| AQRTRTRPSVQP | 1 | 0 | 0 | 1412065 | QEJ82222.1 | K9N4V7 |
| AQRVFNNYMPYV | 1 | 0 | 0 | 1412066 | AEA10817.1 | P0C6X7 |
| AQSELAKGVSLD | 1 | 0 | 0 | 1412069 | APO40578.1 | P0C6X7 |
| AQSFLNRVCGVS | 1 | 0 | 0 | 1412070 | QYC64967.1 | P0DTD1 |
| AQSIFYRMNGIG | 1 | 0 | 0 | 1412072 | AVV62537.1 | K9N5Q8 |
| AQSSETKEQKHE | 1 | 0 | 0 | 1412080 | APT69860.1 | A0A0N7J131 |
| AQTGSSKCGCSV | 1 | 0 | 0 | 1412086 | 2H85_A | P0C6X7 |
| AQTGVIADYNYK | 1 | 0 | 0 | 1412089 | APO40579.1 | P59594 |
| AQTSTAIYTNLR | 1 | 0 | 0 | 1412094 | ATO98145.1 | Q3LZX1 |
| AQVAKSHSVSLI | 1 | 0 | 0 | 1412099 | APO40578.1 | P0C6X7 |
| AQVISYLSERCV | 1 | 0 | 0 | 1412101 | AUM60013.1 | K9N7C7 |
| AQVLSEIVMCGG | 1 | 0 | 0 | 1412105 | QXY02687.1 | P0DTD1 |
| AQVLSETVMCGG | 1 | 0 | 0 | 1412107 | ADO24634.1 | A0A140H1G9 |
| AQVLTEVVHCTG | 1 | 0 | 0 | 1412111 | AAV28291.1 | A0A0P0LKV0 |
| AQVQLEHYSSCV | 1 | 0 | 0 | 1412114 | NP_073549.1 | A0A0P0LKV0 |
| AQVVQLPIQDYI | 1 | 0 | 0 | 1412119 | AEN19363.1 | P0C6X6 |
| AQVVQLPVKDYI | 1 | 0 | 0 | 1412120 | ABD75583.1 | A0A140H1G9 |
| AQVVSHLSEKCM | 1 | 0 | 0 | 1412121 | AVV62535.1 | K9N7C7 |
| AQVVSYLSEKCM | 1 | 0 | 0 | 1412122 | AVV62524.1 | K9N7C7 |
| AQYNYKQDFSNP | 1 | 0 | 0 | 1412130 | AUM60014.1 | K9N5Q8 |
| ARADAEGYINFK | 1 | 0 | 0 | 1412138 | AVV62524.1 | K9N7C7 |
| ARADDEGFITLK | 1 | 0 | 0 | 1412139 | AXN92247.1 | K9N7C7 |
| ARADYEGFITLK | 1 | 0 | 0 | 1412140 | ANI69867.1 | K9N638 |
| ARAGEAANFCAL | 1 | 0 | 0 | 1412147 | AVP78041.1 | P0C6W2 |
| ARALQICVALCD | 1 | 0 | 0 | 1412162 | QED88024.1 | H9EJ42 |
| ARAMLDYAKDHN | 1 | 0 | 0 | 1412164 | AVA26872.1 | H9EJ42 |
| ARANCLATPSGG | 1 | 0 | 0 | 1412165 | AVV62529.1 | K9N643 |
| ARAQSSETKEQK | 1 | 0 | 0 | 1412166 | APT69860.1 | A0A0N7J131 |
| ARAYKHLNNYDC | 1 | 0 | 0 | 1412171 | ARB07605.1 | A0A140H1G9 |
| ARDALDTILQEA | 1 | 0 | 0 | 1412176 | ATQ39389.1 | K9N7C7 |
| ARDGIRRGDNFH | 1 | 0 | 0 | 1412180 | ANI69888.1 | K9N638 |
| ARDLICSQYVAG | 1 | 0 | 0 | 1412186 | APT69978.1 | K9N5Q8 |
| ARDLSLDYINVT | 1 | 0 | 0 | 1412187 | AIX10750.1 | P36334 |
| ARDPIIQEAIPT | 1 | 0 | 0 | 1412191 | AMM42389.1 | Q19U29 |
| ARDPTIQEAIPT | 1 | 0 | 0 | 1412192 | AMM42397.1 | Q19U29 |
| ARDYRANSFDIS | 1 | 0 | 0 | 1412193 | AVV62535.1 | K9N7C7 |
| AREANFGDYYYK | 1 | 0 | 0 | 1412194 | AAA73152.1 | P30215 |
| AREHEFCCSHGD | 1 | 0 | 0 | 1412195 | ABU39953.1 | A0A140H1G9 |
| ARELICAQYVAG | 1 | 0 | 0 | 1412199 | ARA67740.1 | K9N5Q8 |
| ARESIFCSNTYG | 1 | 0 | 0 | 1412200 | AEN19363.1 | P0C6X6 |
| ARFHDFTPLANA | 1 | 0 | 0 | 1412203 | AVV62529.1 | K9N643 |
| ARGPLQVGESAL | 1 | 0 | 0 | 1412220 | ATQ39389.1 | K9N7C7 |
| ARGWVVGTISST | 1 | 0 | 0 | 1412224 | AEN19363.1 | P0C6X6 |
| ARHDFFIFDVDK | 1 | 0 | 0 | 1412225 | AVV62535.1 | K9N7C7 |
| ARHSDNMTAVAC | 1 | 0 | 0 | 1412230 | AQN78751.1 | P30215 |
| ARKHDTCCSQSD | 1 | 0 | 0 | 1412239 | ANZ78857.1 | P0C6X6 |
| ARKHETCCSQID | 1 | 0 | 0 | 1412240 | QBP84719.1 | P0C6X6 |
| ARKHETCCSQRD | 1 | 0 | 0 | 1412241 | AKL87224.1 | P0C6X6 |
| ARKHETCCSQSD | 1 | 0 | 0 | 1412243 | AEN19363.1 | P0C6X6 |
| ARKHGTCCSTSD | 1 | 0 | 0 | 1412245 | AVV62524.1 | K9N7C7 |
| ARKHNTCCTTRD | 1 | 0 | 0 | 1412248 | AUM60013.1 | K9N7C7 |
| ARKRSYSPTKKL | 1 | 0 | 0 | 1412258 | ANF29275.1 | K9N643 |
| ARKSCSYDSDVD | 1 | 0 | 0 | 1412262 | AIV41810.1 | P0C6X6 |
| ARLELVKLGAEK | 1 | 0 | 0 | 1412273 | AWW13564.1 | Q80872 |
| ARLNCGLILKFD | 1 | 0 | 0 | 1412286 | AVN89312.1 | K9N638 |
| ARLNYVALQTDV | 1 | 0 | 0 | 1412287 | AIW52843.1 | A0A0P0G321 |
| ARLVKGDKGDAE | 1 | 0 | 0 | 1412298 | AST12962.1 | A0A0P0LKV0 |
| ARLVQLCAGEYG | 1 | 0 | 0 | 1412299 | ATO98227.1 | Q3LZX5 |
| ARMTYTCQCGSE | 1 | 0 | 0 | 1412308 | ATQ39389.1 | K9N7C7 |
| ARNHGTCCTTRD | 1 | 0 | 0 | 1412313 | QDP16215.1 | K9N7C7 |
| ARNLNESLIDLQ | 1 | 0 | 0 | 1412316 | AVP78031.1 | Q3LZX1 |
| ARNLQAGGDTAL | 1 | 0 | 0 | 1412317 | APO40586.1 | P59595 |
| ARNPVDLPPSHA | 1 | 0 | 0 | 1412320 | QEG03783.1 | A0A0P0LKV0 |
| ARQEYENAVANS | 1 | 0 | 0 | 1412328 | AGT21359.1 | A0A0P0LKV0 |
| ARQFDNMTAVAC | 1 | 0 | 0 | 1412329 | AXX83326.1 | P30215 |
| ARQHLIDGTCGI | 1 | 0 | 0 | 1412333 | APO40578.1 | P0C6X7 |

| | | | | | | |
|---|---|---|---|---|---|---|
| ARQIAWQLHRQG | 1 | 0 | 0 | 1412335 | AVV62528.1 | K9N4V0 |
| ARRHETCCSQSD | 1 | 0 | 0 | 1412340 | AJE26227.1 | P0C6X6 |
| ARRNIGFTLEYW | 1 | 0 | 0 | 1412342 | AMM42485.1 | P36334 |
| ARRYETCCSQSD | 1 | 0 | 0 | 1412347 | AJE26237.1 | P0C6X6 |
| ARSHRGRAALIV | 1 | 0 | 0 | 1412353 | APD51508.1 | A0A0P0KWE8 |
| ARSMLGLITQTV | 1 | 0 | 0 | 1412357 | ASU90899.1 | K9N7C7 |
| ARSMYRAALNSE | 1 | 0 | 0 | 1412360 | AUM60013.1 | K9N7C7 |
| ARSQSSETKEQK | 1 | 0 | 0 | 1412361 | AIW52723.1 | A0A0N7J131 |
| ARSVASQSIIAYTMSLGAENSVA | 1 | 0 | 0 | 1412364 | P0DTC2.1 | P0DTC2 |
| ARSVVSRIGDSI | 1 | 0 | 0 | 1412367 | ATG84876.1 | K9N638 |
| ARTFVQSNVDVL | 1 | 0 | 0 | 1412375 | AEN19363.1 | P0C6X6 |
| ARTFWAWNPEVN | 1 | 0 | 0 | 1412378 | AST12967.1 | A0A0P0KXN3 |
| ARTTSSEFQLAA | 1 | 0 | 0 | 1412389 | AUM60024.1 | K9N5Q8 |
| ARTTSSEFQLAS | 1 | 0 | 0 | 1412390 | AUM60014.1 | K9N5Q8 |
| ARTVYDDATRRV | 1 | 0 | 0 | 1412395 | APO40578.1 | P0C6X7 |
| ARTVYDDSARRV | 1 | 0 | 0 | 1412397 | AVP78041.1 | P0C6W2 |
| ARVRTINLGEVL | 1 | 0 | 0 | 1412406 | APO40578.1 | P0C6X7 |
| ARWSASIDTGNC | 1 | 0 | 0 | 1412413 | QRK03782.1 | A0A0P0K6L9 |
| ARYVSPDVQFEQ | 1 | 0 | 0 | 1412422 | AVV62526.1 | K9N5Q8 |
| ARYVSQDVKFEN | 1 | 0 | 0 | 1412423 | AVV62537.1 | K9N5Q8 |
| ASADIGANHTGI | 1 | 0 | 0 | 1412425 | ARO76381.1 | P0C6U8 |
| ASADWKPGQAMP | 1 | 0 | 0 | 1412427 | ATQ39389.1 | K9N7C7 |
| ASADWKPGYAMP | 1 | 0 | 0 | 1412428 | ALA49934.1 | K9N7C7 |
| ASAFRDLKPLPF | 1 | 0 | 0 | 1412434 | AUM60013.1 | K9N7C7 |
| ASAPATGWLLYQ | 1 | 0 | 0 | 1412444 | ARU07575.1 | A0A140H1G9 |
| ASAPVEPSTVDV | 1 | 0 | 0 | 1412445 | QBF80531.1 | K9N7C7 |
| ASARDLICSQYV | 1 | 0 | 0 | 1412449 | APT69978.1 | K9N5Q8 |
| ASARELICAQYV | 1 | 0 | 0 | 1412450 | ARA67740.1 | K9N5Q8 |
| ASARNLICAQYV | 1 | 0 | 0 | 1412452 | QFS19619.1 | K9N5Q8 |
| ASATDFWTVAFA | 1 | 0 | 0 | 1412454 | AIW52843.1 | A0A0P0G321 |
| ASAVAQAFQSVA | 1 | 0 | 0 | 1412457 | AEN19363.1 | P0C6X6 |
| ASAVAQAFQSVV | 1 | 0 | 0 | 1412458 | AVQ05261.1 | P0C6X6 |
| ASAVKLQNNEIR | 1 | 0 | 0 | 1412459 | AVV62524.1 | K9N7C7 |
| ASAVQTCDLTVG | 1 | 0 | 0 | 1412460 | CAA83661.1 | P36334 |
| ASAYGLCDAANP | 1 | 0 | 0 | 1412462 | ANF29261.1 | K9N5Q8 |
| ASAYRANFFDVP | 1 | 0 | 0 | 1412463 | AMQ49014.1 | K9N7C7 |
| ASAYRANSFDAP | 1 | 0 | 0 | 1412464 | AHI48604.1 | K9N7C7 |
| ASAYRANSIDVP | 1 | 0 | 0 | 1412466 | AQZ41320.1 | K9N7C7 |
| ASCELYHYQECV | 1 | 0 | 0 | 1412469 | ATO98237.1 | Q3LZX7 |
| ASCFFTCNSNAN | 1 | 0 | 0 | 1412470 | AIW52843.1 | A0A0P0G321 |
| ASCFSTCNSNAN | 1 | 0 | 0 | 1412472 | AVL25555.1 | A0A0P0G321 |
| ASDAIVGTTLQG | 1 | 0 | 0 | 1412480 | AVV62535.1 | K9N7C7 |
| ASDFIDSVRDPQ | 1 | 0 | 0 | 1412482 | AVP78031.1 | Q3LZX1 |
| ASDGKPGGDAIS | 1 | 0 | 0 | 1412483 | AUM60013.1 | K9N7C7 |
| ASDGSPSGDAIS | 1 | 0 | 0 | 1412485 | AVV62535.1 | K9N7C7 |
| ASDGVPSNDNIS | 1 | 0 | 0 | 1412486 | AVV62524.1 | K9N7C7 |
| ASDGYFRLMANP | 1 | 0 | 0 | 1412487 | QEO75983.1 | A0A0P0LKV0 |
| ASDIFATPSVLQ | 1 | 0 | 0 | 1412488 | ASY99818.1 | K9N7C7 |
| ASDISLSVACQL | 1 | 0 | 0 | 1412490 | ARU07576.1 | A0A140H1H0 |
| ASDKVNECVKSQ | 1 | 0 | 0 | 1412491 | AVV62526.1 | K9N5Q8 |
| ASDLLDNASVLQ | 1 | 0 | 0 | 1412494 | AVV62535.1 | K9N7C7 |
| ASDLLDNVSVLQ | 1 | 0 | 0 | 1412495 | AVV62524.1 | K9N7C7 |
| ASDLSTLAVTAI | 1 | 0 | 0 | 1412496 | AWK59942.1 | H9EJ42 |
| ASDRGQENLARS | 1 | 0 | 0 | 1412500 | AVV62524.1 | K9N7C7 |
| ASDSDSLPSVPF | 1 | 0 | 0 | 1412502 | AVV62541.1 | K9N7D2 |
| ASDSVQCYTPHI | 1 | 0 | 0 | 1412505 | ARU07575.1 | A0A140H1G9 |
| ASDTSLSDACQL | 1 | 0 | 0 | 1412506 | AGT17757.1 | A0A140H1H0 |
| ASDTYACWHHSI | 1 | 0 | 0 | 1412508 | QHD43415.1 | P0DTD1 |
| ASDVMFEDAQGL | 1 | 0 | 0 | 1412509 | ANI69888.1 | K9N638 |
| ASDWKSGYSMPV | 1 | 0 | 0 | 1412512 | AGT51587.1 | P0C6X6 |
| ASDYRANSFDIS | 1 | 0 | 0 | 1412513 | AUM60013.1 | K9N7C7 |
| ASEAARVVRSIF | 1 | 0 | 0 | 1412514 | AVP78041.1 | P0C6W2 |
| ASEIYCANIHRF | 1 | 0 | 0 | 1412522 | AXN92231.1 | K9N643 |
| ASEIYCANIHSF | 1 | 0 | 0 | 1412523 | QDI73613.1 | K9N643 |
| ASEIYCANLHRF | 1 | 0 | 0 | 1412524 | QDP16209.1 | K9N643 |
| ASEMYCANIHRF | 1 | 0 | 0 | 1412526 | ATQ39396.1 | K9N643 |
| ASEPQRGRQSRI | 1 | 0 | 0 | 1412528 | ALK28785.1 | A0A0N7J131 |
| ASETQRGRHGRI | 1 | 0 | 0 | 1412531 | AIW52713.1 | A0A0N7J131 |
| ASETQRGRQGRI | 1 | 0 | 0 | 1412532 | AIW52715.1 | A0A0N7J131 |
| ASEYRQYLDAYN | 1 | 0 | 0 | 1412533 | APO40578.1 | P0C6X7 |
| ASFDNFRFVCDN | 1 | 0 | 0 | 1412539 | AVP78041.1 | P0C6W2 |
| ASFGNRSGNGIL | 1 | 0 | 0 | 1412542 | QBP84763.1 | P33469 |
| ASFIKLFSLFRH | 1 | 0 | 0 | 1412547 | AGT51787.1 | P0C6X6 |
| ASFNEWAMANQF | 1 | 0 | 0 | 1412550 | AVV62535.1 | K9N7C7 |
| ASFSSVNDAITQ | 1 | 0 | 0 | 1412559 | AIW52843.1 | A0A0P0G321 |
| ASGDCPSVDMQP | 1 | 0 | 0 | 1412570 | ATQ39390.1 | K9N5Q8 |
| ASGNLLLDKRTS | 1 | 0 | 0 | 1412580 | APO40578.1 | P0C6X7 |
| ASGPTVAMTEQL | 1 | 0 | 0 | 1412582 | AKN24830.1 | K9N5Q8 |
| ASGTHTGSAFDG | 1 | 0 | 0 | 1412591 | AVN89451.1 | K9N7C7 |
| ASGYGGFGCKDS | 1 | 0 | 0 | 1412596 | AIV41888.1 | P0C6X6 |
| ASGYVAASAVNV | 1 | 0 | 0 | 1412599 | AHI48543.1 | K9N7D2 |
| ASHLAKALQTYS | 1 | 0 | 0 | 1412603 | AVV62524.1 | K9N7C7 |
| ASHYVVTDIKQV | 1 | 0 | 0 | 1412609 | ARU07575.1 | A0A140H1G9 |
| ASIAFNHPIQID | 1 | 0 | 0 | 1412610 | ASU91208.1 | K9N5Q8 |
| ASIAFNHPIQVD | 1 | 0 | 0 | 1412611 | AXN92228.1 | K9N5Q8 |
| ASIEEVCDDYAK | 1 | 0 | 0 | 1412615 | AGT51687.1 | P0C6X6 |
| ASIGDIIQRLDF | 1 | 0 | 0 | 1412617 | AXN92260.1 | K9N5Q8 |
| ASIGDIIQRLDG | 1 | 0 | 0 | 1412618 | AHI48733.1 | K9N5Q8 |
| ASIGDIIQRLDP | 1 | 0 | 0 | 1412619 | 5W9H_A | K9N5Q8 |
| ASIGDIIRRLDV | 1 | 0 | 0 | 1412621 | ALB08246.1 | K9N5Q8 |
| ASINFNQPLVVD | 1 | 0 | 0 | 1412630 | AUM60014.1 | K9N5Q8 |

| | | | | | | |
|---|---|---|---|---|---|---|
| ASIPLCILPYTA | 1 | 0 | 0 | 1412635 | ALD51903.1 | K9N7C7 |
| ASIPLCIVPYTV | 1 | 0 | 0 | 1412637 | AXN92247.1 | K9N7C7 |
| ASIQADVNTPAD | 1 | 0 | 0 | 1412638 | QBQ01843.1 | P33469 |
| ASISAFVETVKG | 1 | 0 | 0 | 1412641 | AVP78030.1 | P0C6W2 |
| ASISESKPLADD | 1 | 0 | 0 | 1412642 | AVL25595.1 | H9EJ69 |
| ASIVDIIQRLDV | 1 | 0 | 0 | 1412647 | AGV08438.1 | K9N5Q8 |
| ASKADGIIYPTG | 1 | 0 | 0 | 1412649 | AVV62537.1 | K9N5Q8 |
| ASKCVCSVIDLL | 1 | 0 | 0 | 1412650 | APO40578.1 | P0C6X7 |
| ASKMFDAYVDTF | 1 | 0 | 0 | 1412661 | APO40578.1 | P0C6X7 |
| ASKNACGTNVPL | 1 | 0 | 0 | 1412662 | AVV62524.1 | K9N7C7 |
| ASKVIALHKRWQ | 1 | 0 | 0 | 1412666 | ATO98194.1 | Q3LZX0 |
| ASKVIALHNRWQ | 1 | 0 | 0 | 1412667 | ARO76383.1 | P59632 |
| ASKVIPFNSLWQ | 1 | 0 | 0 | 1412668 | APO40580.1 | P59632 |
| ASLFKPVQLAPV | 1 | 0 | 0 | 1412678 | AKM76243.1 | K9N7D2 |
| ASLFPLWTAAAG | 1 | 0 | 0 | 1412680 | AAR01015.1 | P36334 |
| ASLFPPWSAAAG | 1 | 0 | 0 | 1412681 | CAA83660.1 | P36334 |
| ASLFPPWTAAEG | 1 | 0 | 0 | 1412683 | AIL49550.1 | P36334 |
| ASLGERVRQAML | 1 | 0 | 0 | 1412688 | APO40578.1 | P0C6X7 |
| ASLINTLNNLNE | 1 | 0 | 0 | 1412693 | AVP78041.1 | P0C6W2 |
| ASLNAFKQYFNL | 1 | 0 | 0 | 1412695 | AVV62526.1 | K9N5Q8 |
| ASLNGVTLIGES | 1 | 0 | 0 | 1412697 | APO40578.1 | P0C6X7 |
| ASLNSLKEYFNL | 1 | 0 | 0 | 1412699 | AXN92228.1 | K9N5Q8 |
| ASLNSLTYFNKP | 1 | 0 | 0 | 1412700 | AIV41786.1 | P0C6X6 |
| ASLNSLTYFNRS | 1 | 0 | 0 | 1412702 | AGT51737.1 | P0C6X6 |
| ASLNVKPYGYPN | 1 | 0 | 0 | 1412703 | AWK59948.1 | H9EJ42 |
| ASLPVYDTIKYY | 1 | 0 | 0 | 1412709 | AVV62526.1 | K9N5Q8 |
| ASLQEIISRLDA | 1 | 0 | 0 | 1412712 | ANZ78843.1 | P36334 |
| ASLRQIFDVPPI | 1 | 0 | 0 | 1412722 | AKJ80136.2 | K9N7C7 |
| ASLRQIYDLAPI | 1 | 0 | 0 | 1412723 | AUM60013.1 | K9N7C7 |
| ASLRQIYDPAPV | 1 | 0 | 0 | 1412724 | AVV62535.1 | K9N7C7 |
| ASMEVKTIKVFT | 1 | 0 | 0 | 1412734 | 5TL6_B | P0C6X7 |
| ASMLNGTPPPVY | 1 | 0 | 0 | 1412735 | AVV62526.1 | K9N5Q8 |
| ASMRNANGACIW | 1 | 0 | 0 | 1412737 | AUM60013.1 | K9N7C7 |
| ASMYENYTILQA | 1 | 0 | 0 | 1412742 | QED88024.1 | H9EJ42 |
| ASNDAIMTRCLA | 1 | 0 | 0 | 1412745 | AXN92247.1 | K9N7C7 |
| ASNDAVQLESDA | 1 | 0 | 0 | 1412746 | AVV62524.1 | K9N7C7 |
| ASNDWKPGLAMP | 1 | 0 | 0 | 1412747 | AUM60013.1 | K9N7C7 |
| ASNFPSVYAWER | 1 | 0 | 0 | 1412748 | APO40579.1 | P59594 |
| ASNKLRVVIPDL | 1 | 0 | 0 | 1412756 | AVV62535.1 | K9N7C7 |
| ASNPTNCIAPVN | 1 | 0 | 0 | 1412760 | AHY22565.1 | K9N5Q8 |
| ASNQADVNIPAD | 1 | 0 | 0 | 1412761 | AIX10718.1 | P33469 |
| ASNQADVNTSAD | 1 | 0 | 0 | 1412763 | ARK08674.1 | P33469 |
| ASNSKLWQYCST | 1 | 0 | 0 | 1412765 | AGT51737.1 | P0C6X6 |
| ASNSRLWQYCSI | 1 | 0 | 0 | 1412766 | ARB07436.1 | A0A140H1G9 |
| ASNSRSTSRTSS | 1 | 0 | 0 | 1412767 | AXX83347.1 | P33469 |
| ASNSSVLSLVNF | 1 | 0 | 0 | 1412768 | AXN92247.1 | K9N7C7 |
| ASNVPSIVSEKI | 1 | 0 | 0 | 1412769 | AVA26872.1 | H9EJ42 |
| ASNVSPIVSEKI | 1 | 0 | 0 | 1412770 | QEG03746.1 | H9EJ42 |
| ASNYRENSFDIS | 1 | 0 | 0 | 1412771 | AVV62524.1 | K9N7C7 |
| ASPAAPCAVSFA | 1 | 0 | 0 | 1412772 | AKK52599.1 | K9N4V7 |
| ASPAAPRAISFA | 1 | 0 | 0 | 1412773 | ANI69896.1 | K9N4V7 |
| ASPAAPRAVFFA | 1 | 0 | 0 | 1412774 | ASY99807.1 | K9N4V7 |
| ASPAAPRAVSFA | 1 | 0 | 0 | 1412775 | AXN92267.1 | K9N4V7 |
| ASPASTVHAIAT | 1 | 0 | 0 | 1412777 | ACQ82726.1 | P59632 |
| ASPASTVHTTAT | 1 | 0 | 0 | 1412780 | ARO76383.1 | P59632 |
| ASPGPSGIGAVG | 1 | 0 | 0 | 1412781 | AVV62533.1 | K9N4V7 |
| ASPRECNDMHLS | 1 | 0 | 0 | 1412787 | ARO76381.1 | P0C6U8 |
| ASPSECNSMHLS | 1 | 0 | 0 | 1412789 | APO40578.1 | P0C6X7 |
| ASQEGVLSEREKLLHLI | 1 | 0 | 0 | 1412797 | QED88043.1 | U3M6Q8 |
| ASQGLVASIKNF | 1 | 0 | 0 | 1412798 | QYC64967.1 | P0DTD1 |
| ASQIISGTVAAV | 1 | 0 | 0 | 1412804 | AVN89380.1 | K9N7A1 |
| ASQLGNCVEYHH | 1 | 0 | 0 | 1412807 | 4KQZ_A | K9N5Q8 |
| ASQLGNCVEYSL | 1 | 0 | 0 | 1412808 | AXN92228.1 | K9N5Q8 |
| ASQLTKQLLDIL | 1 | 0 | 0 | 1412809 | AVV62524.1 | K9N7C7 |
| ASQVIVNNYEKS | 1 | 0 | 0 | 1412815 | AIL49570.1 | P0C6X6 |
| ASRAIEKVNECV | 1 | 0 | 0 | 1412816 | BBA20986.1 | A0A140H1H1 |
| ASRDPVPLHIIA | 1 | 0 | 0 | 1412817 | AVV62530.1 | K9N7D2 |
| ASRDSVPLHIIA | 1 | 0 | 0 | 1412818 | AVV62541.1 | K9N7D2 |
| ASRDVNVTFFPD | 1 | 0 | 0 | 1412819 | APO40578.1 | P0C6X7 |
| ASRFPNVYAWER | 1 | 0 | 0 | 1412823 | ATO98181.1 | Q3LZX1 |
| ASRNSSRSNSRG | 1 | 0 | 0 | 1412833 | AUM60021.1 | K9N4V7 |
| ASRNSSRSSSQG | 1 | 0 | 0 | 1412834 | AXN92267.1 | K9N4V7 |
| ASRPGVYCGHDY | 1 | 0 | 0 | 1412836 | AUM60013.1 | K9N7C7 |
| ASSAAPRAVSFA | 1 | 0 | 0 | 1412847 | AHL18098.1 | K9N4V7 |
| ASSAGPLQTTVG | 1 | 0 | 0 | 1412848 | AVV62526.1 | K9N5Q8 |
| ASSAGSRSKANS | 1 | 0 | 0 | 1412849 | AJQ24081.1 | P33469 |
| ASSAINRPQLAF | 1 | 0 | 0 | 1412851 | AVV62524.1 | K9N7C7 |
| ASSASRGSSRSS | 1 | 0 | 0 | 1412853 | AVV62533.1 | K9N4V7 |
| ASSAVKLQNNEI | 1 | 0 | 0 | 1412855 | AUM60013.1 | K9N7C7 |
| ASSDAIMTRCLA | 1 | 0 | 0 | 1412858 | QXR89047.1 | P0DTD1 |
| ASSDVIHYNINF | 1 | 0 | 0 | 1412859 | BAL45639.1 | A0A0P0K6L9 |
| ASSDVIRYNINF | 1 | 0 | 0 | 1412860 | AIW52757.1 | A0A0P0K6L9 |
| ASSEFAVLYQDV | 1 | 0 | 0 | 1412861 | BAE93401.1 | P59594 |
| ASSEVAVLYQGV | 1 | 0 | 0 | 1412864 | ACQ82725.1 | P59594 |
| ASSGNRAPTSGV | 1 | 0 | 0 | 1412867 | ATP16771.1 | P33469 |
| ASSGNRASTSGV | 1 | 0 | 0 | 1412868 | AIL49428.1 | P33469 |
| ASSGNRPGNGIL | 1 | 0 | 0 | 1412869 | AMM42456.1 | P33469 |
| ASSGNRSSNGIL | 1 | 0 | 0 | 1412871 | AMM42463.1 | P33469 |
| ASSGNRSVNGIL | 1 | 0 | 0 | 1412872 | AGT51505.1 | P33469 |
| ASSGNRTPTSGV | 1 | 0 | 0 | 1412873 | AJQ24068.1 | P33469 |
| ASSGYRSGNGIL | 1 | 0 | 0 | 1412875 | AIV41850.1 | P33469 |

| | | | | | | |
|---|---|---|---|---|---|---|
| ASSNKTGNNVIS | 1 | 0 | 0 | 1412884 | AUM60014.1 | K9N5Q8 |
| ASSNQSPQGNLG | 1 | 0 | 0 | 1412885 | AIX09796.1 | P0C6X6 |
| ASSQADVNTPAD | 1 | 0 | 0 | 1412889 | ANZ78879.1 | P33469 |
| ASSQSAAKILAP | 1 | 0 | 0 | 1412890 | AIW52721.1 | A0A0N7J131 |
| ASSQSAAKILAR | 1 | 0 | 0 | 1412891 | AIW52722.1 | A0A0N7J131 |
| ASSQSSAKSLAR | 1 | 0 | 0 | 1412894 | ABB90490.1 | A0A0N7J131 |
| ASSQSSAKSLSR | 1 | 0 | 0 | 1412895 | ABB90484.1 | A0A0N7J131 |
| ASSSEAFLIGCN | 1 | 0 | 0 | 1412903 | AVP78041.1 | P0C6W2 |
| ASSSEAFLIGVN | 1 | 0 | 0 | 1412904 | P0C6W6.1 | P0C6W2 |
| ASSSGPISPTNL | 1 | 0 | 0 | 1412905 | AOL02458.1 | Q4VID0 |
| ASSSLRTFVVDK | 1 | 0 | 0 | 1412906 | AXN92247.1 | K9N7C7 |
| ASSTFVEEEHVN | 1 | 0 | 0 | 1412907 | NP_073549.1 | A0A0P0LKV0 |
| ASSTGSRSRANS | 1 | 0 | 0 | 1412908 | AIX10719.1 | P33469 |
| ASTDFHQLECIV | 1 | 0 | 0 | 1412913 | ARU07593.1 | H9EJ42 |
| ASTDFHQLEFVR | 1 | 0 | 0 | 1412914 | AWK59942.1 | H9EJ42 |
| ASTDFHQLEYDS | 1 | 0 | 0 | 1412915 | AGT51337.1 | H9EJ42 |
| ASTDSIVFTYIP | 1 | 0 | 0 | 1412916 | QDI73614.1 | K9N7D2 |
| ASTEELPDEFVV | 1 | 0 | 0 | 1412918 | AAT76156.1 | Q3LZX3 |
| ASTFLFSCIITA | 1 | 0 | 0 | 1412924 | AEN19363.1 | P0C6X6 |
| ASTGVIDISAGM | 1 | 0 | 0 | 1412928 | AWK59942.1 | H9EJ42 |
| ASTPESKPLADD | 1 | 0 | 0 | 1412933 | AIW52805.1 | H9EJ69 |
| ASTPVEPPTVDV | 1 | 0 | 0 | 1412934 | AXN92247.1 | K9N7C7 |
| ASTPVEPSTVDV | 1 | 0 | 0 | 1412935 | AXN92258.1 | K9N7C7 |
| ASTPVESPTVDV | 1 | 0 | 0 | 1412936 | ATG84853.1 | K9N7C7 |
| ASTQNEELEQTI | 1 | 0 | 0 | 1412938 | APO40578.1 | P0C6X7 |
| ASTSAFIDTVKG | 1 | 0 | 0 | 1412942 | ARO76381.1 | P0C6U8 |
| ASTSGVTPDMAD | 1 | 0 | 0 | 1412945 | AIL49428.1 | P33469 |
| ASTSLDWFPRPH | 1 | 0 | 0 | 1412946 | AUM60016.1 | K9N4V0 |
| ASTSLDWFPRPY | 1 | 0 | 0 | 1412947 | AUM60026.1 | K9N4V0 |
| ASTSQKSIVAYT | 1 | 0 | 0 | 1412948 | 5WRG_A | P59594 |
| ASTVHAIATIPL | 1 | 0 | 0 | 1412949 | ACQ82726.1 | P59632 |
| ASTVHTTATIPL | 1 | 0 | 0 | 1412952 | ARO76383.1 | P59632 |
| ASTWFNALRDFT | 1 | 0 | 0 | 1412957 | AHI48593.1 | K9N638 |
| ASVDARLVPCAN | 1 | 0 | 0 | 1412963 | AOL02450.1 | P0C6X6 |
| ASVDARLVPCAS | 1 | 0 | 0 | 1412964 | AEN19363.1 | P0C6X6 |
| ASVFKDGCDVGF | 1 | 0 | 0 | 1412966 | ABE97136.1 | H9EJ42 |
| ASVGSVPGEMRL | 1 | 0 | 0 | 1412968 | 5W9H_A | K9N5Q8 |
| ASVIAELNQLNE | 1 | 0 | 0 | 1412969 | APO40578.1 | P0C6X7 |
| ASVIAGGLIAIL | 1 | 0 | 0 | 1412970 | APO40578.1 | P0C6X7 |
| ASVIQDYENVCF | 1 | 0 | 0 | 1412971 | AIV41852.1 | P0C6X6 |
| ASVKIYQSSPII | 1 | 0 | 0 | 1412973 | ARA67741.1 | K9N5Q8 |
| ASVKLPKGVGSA | 1 | 0 | 0 | 1412974 | AEN19363.1 | P0C6X6 |
| ASVKNSQSSPII | 1 | 0 | 0 | 1412976 | QBF80598.1 | K9N5Q8 |
| ASVKSYQSSPII | 1 | 0 | 0 | 1412979 | AVN89387.1 | K9N5Q8 |
| ASVLGAAAKSVV | 1 | 0 | 0 | 1412980 | AVV62535.1 | K9N7C7 |
| ASVLKDGCDVGF | 1 | 0 | 0 | 1412981 | AWK59942.1 | H9EJ42 |
| ASVLQATLSEFS | 1 | 0 | 0 | 1412982 | AVV62535.1 | K9N7C7 |
| ASVNPANMRNAN | 1 | 0 | 0 | 1412985 | AVV62524.1 | K9N7C7 |
| ASVNPASMRNAN | 1 | 0 | 0 | 1412986 | AUM60013.1 | K9N7C7 |
| ASVPANVSIPKP | 1 | 0 | 0 | 1412990 | AUM60014.1 | K9N5Q8 |
| ASVRDLICAQYV | 1 | 0 | 0 | 1412991 | ALJ54502.1 | K9N5Q8 |
| ASVVDIKKEIAH | 1 | 0 | 0 | 1413004 | APO40579.1 | P59594 |
| ASVVEVHKAVVV | 1 | 0 | 0 | 1413006 | AVV62535.1 | K9N7C7 |
| ASYADVLVNVSQ | 1 | 0 | 0 | 1413021 | AIW52757.1 | A0A0P0K6L9 |
| ASYDINLNKFNR | 1 | 0 | 0 | 1413024 | AMQ48992.1 | K9N7C7 |
| ASYDTNLNKFNG | 1 | 0 | 0 | 1413025 | AVN89385.1 | K9N7C7 |
| ASYECDIPIGAG | 1 | 0 | 0 | 1413026 | AVP78031.1 | Q3LZX1 |
| ASYGESLEGVFW | 1 | 0 | 0 | 1413031 | ARU07581.1 | Q19U29 |
| ASYHTASILRST | 1 | 0 | 0 | 1413034 | AVP78031.1 | Q3LZX1 |
| ASYLQQGSTGEI | 1 | 0 | 0 | 1413042 | AUM60014.1 | K9N5Q8 |
| ASYSDQQPSVLF | 1 | 0 | 0 | 1413046 | AVV62530.1 | K9N7D2 |
| ATAALPTQFLPL | 1 | 0 | 0 | 1413049 | ARO76382.1 | P59594 |
| ATAAYGLCNSEV | 1 | 0 | 0 | 1413050 | AUM60014.1 | K9N5Q8 |
| ATAEPVRDEFSI | 1 | 0 | 0 | 1413053 | AAA45463.1 | A0A0N7J131 |
| ATAGWTFGAGSA | 1 | 0 | 0 | 1413058 | Q3I5J5.1 | Q3LZX1 |
| ATAGYTFGAGAA | 1 | 0 | 0 | 1413060 | APO40579.1 | P59594 |
| ATAHSSIKQGTL | 1 | 0 | 0 | 1413062 | AIV41996.1 | P0C6X6 |
| ATAKVSPGLCIA | 1 | 0 | 0 | 1413065 | AQN78776.1 | P36334 |
| ATALNKIHDVVN | 1 | 0 | 0 | 1413068 | ABB90526.1 | A0A0P0K6L9 |
| ATAPANLTISKP | 1 | 0 | 0 | 1413070 | AVV62526.1 | K9N5Q8 |
| ATAPANLTITKP | 1 | 0 | 0 | 1413071 | AVV62537.1 | K9N5Q8 |
| ATAQEAYEQAVV | 1 | 0 | 0 | 1413074 | AVP78041.1 | P0C6W2 |
| ATATIPIQATLP | 1 | 0 | 0 | 1413076 | AVP78043.1 | Q3LZX0 |
| ATATIPLQASLP | 1 | 0 | 0 | 1413077 | ATO98194.1 | Q3LZX0 |
| ATCELCGTQNAV | 1 | 0 | 0 | 1413086 | APO40578.1 | P0C6X7 |
| ATCEQCGTENLV | 1 | 0 | 0 | 1413088 | P0C6W6.1 | P0C6W2 |
| ATCFSTSADTYA | 1 | 0 | 0 | 1413090 | APO40578.1 | P0C6X7 |
| ATCPFAKSDGCS | 1 | 0 | 0 | 1413095 | AVV62524.1 | K9N7C7 |
| ATCPFVKGDSCS | 1 | 0 | 0 | 1413096 | AKN24784.1 | K9N7C7 |
| ATCVNCCHDGIF | 1 | 0 | 0 | 1413097 | QRK03817.1 | Q4VID0 |
| ATCYNSVSNEYC | 1 | 0 | 0 | 1413098 | AWK59942.1 | H9EJ42 |
| ATDAPSTFGTRN | 1 | 0 | 0 | 1413100 | AXN92267.1 | K9N4V7 |
| ATDCSDGNYNCN | 1 | 0 | 0 | 1413101 | AHL31360.1 | K9N5Q8 |
| ATDCSDGNYNRY | 1 | 0 | 0 | 1413103 | ALJ54486.1 | K9N5Q8 |
| ATDPSEAFEKFI | 1 | 0 | 0 | 1413110 | ANI69899.1 | K9N7C7 |
| ATDRFCRLGNEL | 1 | 0 | 0 | 1413111 | ABO28724.1 | H9EJ42 |
| ATDVQSRADQNG | 1 | 0 | 0 | 1413113 | AUM60013.1 | K9N7C7 |
| ATDYREYMEAYN | 1 | 0 | 0 | 1413114 | ASU91085.1 | K9N7C7 |
| ATEAKGYWYRHN | 1 | 0 | 0 | 1413117 | AEN19370.1 | P33469 |
| ATEATFNYLLDL | 1 | 0 | 0 | 1413119 | AZK15899.1 | K9N638 |
| ATEEAFKQSYAS | 1 | 0 | 0 | 1413121 | AEN19363.1 | P0C6X6 |

| | | | | | | |
|---|---|---|---|---|---|---|
| ATEFADKLVEVG | 1 | 0 | 0 | 1413123 | AEN19363.1 | P0C6X6 |
| ATEFANKLVEVG | 1 | 0 | 0 | 1413124 | AXX83348.1 | P0C6X6 |
| ATEGALNTPKEH | 1 | 0 | 0 | 1413126 | APO40586.1 | P59595 |
| ATEHSNIIRGWI | 1 | 0 | 0 | 1413127 | AVP78031.1 | Q3LZX1 |
| ATEHSNIVRGWI | 1 | 0 | 0 | 1413128 | AVP78042.1 | Q3LZX1 |
| ATEKSNVIRGWI | 1 | 0 | 0 | 1413129 | ATO98181.1 | Q3LZX1 |
| ATEQLLYAIQQL | 1 | 0 | 0 | 1413132 | AUM60023.1 | K9N7C7 |
| ATEQLLYVIQQL | 1 | 0 | 0 | 1413133 | AUM60013.1 | K9N7C7 |
| ATESKGYWYRHN | 1 | 0 | 0 | 1413135 | AIX09811.1 | P33469 |
| ATEYRDYMEAYN | 1 | 0 | 0 | 1413140 | AUM60013.1 | K9N7C7 |
| ATFAELEAARKA | 1 | 0 | 0 | 1413143 | AMQ49036.1 | K9N638 |
| ATFNFLLDLTGL | 1 | 0 | 0 | 1413157 | AGV08389.1 | K9N638 |
| ATFNYLLDLAGL | 1 | 0 | 0 | 1413158 | AZK15899.1 | K9N638 |
| ATGDLAVCLGIS | 1 | 0 | 0 | 1413171 | QEG03763.1 | P0C6X6 |
| ATGDLAVCLGVG | 1 | 0 | 0 | 1413172 | AIV41816.1 | P0C6X6 |
| ATGGIAIAMACL | 1 | 0 | 0 | 1413176 | APO40582.1 | P59596 |
| ATGITPQKLFVA | 1 | 0 | 0 | 1413178 | ATG84701.1 | K9N5Q8 |
| ATGLFAERDCYI | 1 | 0 | 0 | 1413179 | ARB07436.1 | A0A140H1G9 |
| ATGSTPQKLFVA | 1 | 0 | 0 | 1413186 | AZU90731.1 | K9N5Q8 |
| ATGTNVFQTQAG | 1 | 0 | 0 | 1413188 | AVP78031.1 | Q3LZX1 |
| ATGTTLQKLFVA | 1 | 0 | 0 | 1413189 | AID55095.1 | K9N5Q8 |
| ATGTTPRKLFVA | 1 | 0 | 0 | 1413191 | AJG44069.1 | K9N5Q8 |
| ATGTYKCPQTKS | 1 | 0 | 0 | 1413193 | CAA79896.1 | P36334 |
| ATGVTSSMTDNL | 1 | 0 | 0 | 1413196 | AUM60024.1 | K9N5Q8 |
| ATHGIAAVNSVP | 1 | 0 | 0 | 1413200 | ARO76381.1 | P0C6U8 |
| ATHGLTAINSVP | 1 | 0 | 0 | 1413203 | APO40578.1 | P0C6X7 |
| ATHHDKFTDGIC | 1 | 0 | 0 | 1413205 | APO40578.1 | P0C6X7 |
| ATHIAEDVVDQV | 1 | 0 | 0 | 1413207 | AST12962.1 | A0A0P0LKV0 |
| ATHVCFKDCYNI | 1 | 0 | 0 | 1413211 | AWK59942.1 | H9EJ42 |
| ATHVCFKDCYNV | 1 | 0 | 0 | 1413212 | P0C6U6.1 | H9EJ42 |
| ATHVCFKDCYSI | 1 | 0 | 0 | 1413213 | AGT51337.1 | H9EJ42 |
| ATIEELPDEFVV | 1 | 0 | 0 | 1413218 | APO40587.1 | P59636 |
| ATIKEIVGERQL | 1 | 0 | 0 | 1413224 | AXN92247.1 | K9N7C7 |
| ATILDYCDGNLP | 1 | 0 | 0 | 1413230 | APO40578.1 | P0C6X7 |
| ATINFIFDLAQL | 1 | 0 | 0 | 1413234 | AVV62535.1 | K9N7C7 |
| ATINYTQPEVLS | 1 | 0 | 0 | 1413236 | AVV62526.1 | K9N5Q8 |
| ATINYTQPQLLS | 1 | 0 | 0 | 1413237 | AVV62537.1 | K9N5Q8 |
| ATINYVIDLANL | 1 | 0 | 0 | 1413238 | AUM60023.1 | K9N7C7 |
| ATIPIQATLPFG | 1 | 0 | 0 | 1413241 | QWS71388.1 | P0DTC3 |
| ATIPLQASLPFG | 1 | 0 | 0 | 1413242 | ATO98219.1 | Q3LZX0 |
| ATKFTLWNYLFE | 1 | 0 | 0 | 1413260 | AXN92247.1 | K9N7C7 |
| ATKGEGDGGIIS | 1 | 0 | 0 | 1413263 | AST12962.1 | A0A0P0LKV0 |
| ATKIQNLLYCDS | 1 | 0 | 0 | 1413266 | AIW52843.1 | A0A0P0G321 |
| ATKRTDNPILDF | 1 | 0 | 0 | 1413276 | AVP78031.1 | Q3LZX1 |
| ATKSFNMVQAFG | 1 | 0 | 0 | 1413277 | AVV62533.1 | K9N4V7 |
| ATKSFNVVQAFG | 1 | 0 | 0 | 1413278 | AUM60021.1 | K9N4V7 |
| ATKTVGGAPNWL | 1 | 0 | 0 | 1413280 | AVV62524.1 | K9N7C7 |
| ATKVQQQQDDLF | 1 | 0 | 0 | 1413281 | APO40584.1 | P59635 |
| ATKVVPFNNGFV | 1 | 0 | 0 | 1413282 | AVV62537.1 | K9N5Q8 |
| ATKVYFDNPTLP | 1 | 0 | 0 | 1413283 | ATO98193.1 | Q3LZX1 |
| ATLDKCTKYNIY | 1 | 0 | 0 | 1413289 | CAA71147.1 | A0A0P0K6L9 |
| ATLINNISYVVT | 1 | 0 | 0 | 1413294 | AWK59942.1 | H9EJ42 |
| ATLMAPTIVNQE | 1 | 0 | 0 | 1413298 | AVN89451.1 | K9N7C7 |
| ATLNSFKEYFDL | 1 | 0 | 0 | 1413300 | AVV62537.1 | K9N5Q8 |
| ATLPDIAEDVVD | 1 | 0 | 0 | 1413301 | NP_073549.1 | A0A0P0LKV0 |
| ATLPVYDGIKYY | 1 | 0 | 0 | 1413306 | AVV62537.1 | K9N5Q8 |
| ATLPVYDTIKYY | 1 | 0 | 0 | 1413307 | AXN92228.1 | K9N5Q8 |
| ATLQEILSRLDA | 1 | 0 | 0 | 1413311 | ARB66071.1 | P36334 |
| ATLTASTIVNQE | 1 | 0 | 0 | 1413321 | ALA50055.1 | K9N7C7 |
| ATNCSDGNYNRN | 1 | 0 | 0 | 1413331 | ALA49671.1 | K9N5Q8 |
| ATNEWYYTGSSY | 1 | 0 | 0 | 1413333 | AUM60014.1 | K9N5Q8 |
| ATNHDRYCAVHQ | 1 | 0 | 0 | 1413341 | AVV62524.1 | K9N7C7 |
| ATNHDRYCSVHQ | 1 | 0 | 0 | 1413342 | AXN92247.1 | K9N7C7 |
| ATNILNSCFAIV | 1 | 0 | 0 | 1413343 | AVV62524.1 | K9N7C7 |
| ATNKATYKPNIW | 1 | 0 | 0 | 1413345 | AVP78041.1 | P0C6W2 |
| ATNNAMQLNLIS | 1 | 0 | 0 | 1413348 | AVP78041.1 | P0C6W2 |
| ATNNAMQVESDK | 1 | 0 | 0 | 1413350 | AVP78030.1 | P0C6W2 |
| ATNNVFTLKGGA | 1 | 0 | 0 | 1413355 | APO40578.1 | P0C6X7 |
| ATPAAPRAVTFG | 1 | 0 | 0 | 1413363 | AVV62533.1 | K9N4V7 |
| ATPFDVVRQCSG | 1 | 0 | 0 | 1413364 | 3EA9_A | P0C6U8 |
| ATPHKMVLSVSP | 1 | 0 | 0 | 1413366 | AXN92247.1 | K9N7C7 |
| ATPQPQRAPKGS | 1 | 0 | 0 | 1413370 | AVV62544.1 | K9N4V7 |
| ATPSDSIRATAT | 1 | 0 | 0 | 1413372 | AVP78043.1 | Q3LZX0 |
| ATPSTTIVCDRV | 1 | 0 | 0 | 1413374 | AWK59934.1 | U3M6Q8 |
| ATPSVLQATLSE | 1 | 0 | 0 | 1413375 | ASY99818.1 | K9N7C7 |
| ATPTNSVRTTAT | 1 | 0 | 0 | 1413376 | AVP78032.1 | Q3LZX0 |
| ATPVQPRAVSFA | 1 | 0 | 0 | 1413377 | AUM60021.1 | K9N4V7 |
| ATQAKSGIITKV | 1 | 0 | 0 | 1413379 | AIV41824.1 | P30215 |
| ATQAMEKVNECV | 1 | 0 | 0 | 1413380 | AIL49512.1 | P36334 |
| ATQDGVNKLQGL | 1 | 0 | 0 | 1413381 | AXX83321.1 | P36334 |
| ATQGFQACRFVA | 1 | 0 | 0 | 1413383 | AFD98756.1 | H9EJ42 |
| ATQPLDAGLNGD | 1 | 0 | 0 | 1413386 | AVV62526.1 | K9N5Q8 |
| ATQYLVQQESTF | 1 | 0 | 0 | 1413390 | APO40578.1 | P0C6X7 |
| ATRDTVPLHIIA | 1 | 0 | 0 | 1413394 | AUM60018.1 | K9N7D2 |
| ATRFPSVYAWER | 1 | 0 | 0 | 1413397 | AVP78031.1 | Q3LZX1 |
| ATRGVTCVIGTT | 1 | 0 | 0 | 1413400 | ALA49384.1 | K9N7C7 |
| ATRLCVQCAVGF | 1 | 0 | 0 | 1413401 | AUM60019.1 | K9N5R3 |
| ATRLCVQCITGF | 1 | 0 | 0 | 1413402 | ALR69646.1 | K9N5R3 |
| ATRLCVQCMTGF | 1 | 0 | 0 | 1413404 | ASU90554.1 | K9N5R3 |
| ATRLCVQCMTGL | 1 | 0 | 0 | 1413405 | ALA49390.1 | K9N5R3 |
| ATRLMSIPQLTL | 1 | 0 | 0 | 1413406 | AVR40345.1 | Q4VID0 |

| | | | | | | |
|---|---|---|---|---|---|---|
| ATRNNCWLRSVL | 1 | 0 | 0 | 1413410 | AEN19363.1 | P0C6X6 |
| ATRNYIITMPCF | 1 | 0 | 0 | 1413411 | AHL18094.1 | K9N643 |
| ATRSVTVEYNIH | 1 | 0 | 0 | 1413417 | AVV62524.1 | K9N7C7 |
| ATRTFTATTVVG | 1 | 0 | 0 | 1413421 | AXN92247.1 | K9N7C7 |
| ATRVQVGQLPKY | 1 | 0 | 0 | 1413425 | AWK59934.1 | U3M6Q8 |
| ATRYCRVGECRD | 1 | 0 | 0 | 1413427 | AWK59942.1 | H9EJ42 |
| ATSANCCHDGIF | 1 | 0 | 0 | 1413430 | ATP16762.1 | Q4VID0 |
| ATSANLFPPWSA | 1 | 0 | 0 | 1413431 | CAA83661.1 | P36334 |
| ATSASLFPLWTA | 1 | 0 | 0 | 1413432 | AAR01015.1 | P36334 |
| ATSASLFPPWSA | 1 | 0 | 0 | 1413433 | CAA83660.1 | P36334 |
| ATSCKEADMRNS | 1 | 0 | 0 | 1413435 | AVV62537.1 | K9N5Q8 |
| ATSCQLYYNLPS | 1 | 0 | 0 | 1413437 | BBO58721.1 | P36334 |
| ATSCQLYYSLPA | 1 | 0 | 0 | 1413438 | ARB66070.1 | P36334 |
| ATSDLGSLIDCN | 1 | 0 | 0 | 1413439 | AUM60013.1 | K9N7C7 |
| ATSDLGSLIDSN | 1 | 0 | 0 | 1413440 | AVV62535.1 | K9N7C7 |
| ATSDRTDVRDYK | 1 | 0 | 0 | 1413441 | QED88010.1 | H9EJ42 |
| ATSFCKSGNLVL | 1 | 0 | 0 | 1413444 | AIL49332.1 | P30215 |
| ATSLSFNNLLGF | 1 | 0 | 0 | 1413455 | ANI69888.1 | K9N638 |
| ATSNVTQCFGPR | 1 | 0 | 0 | 1413456 | APT69866.1 | A0A0N7J131 |
| ATSQQPSGGNVI | 1 | 0 | 0 | 1413458 | QEG03818.1 | P33469 |
| ATSQQPSGRNVV | 1 | 0 | 0 | 1413460 | ARU07591.1 | P33469 |
| ATSSYDALLHRN | 1 | 0 | 0 | 1413464 | AGT51379.1 | H9EJ42 |
| ATSVNCCHDGIF | 1 | 0 | 0 | 1413471 | AVR40343.1 | Q4VID0 |
| ATSVNCFHDGIF | 1 | 0 | 0 | 1413472 | AHF54653.1 | Q4VID0 |
| ATSYGMCFSSIT | 1 | 0 | 0 | 1413473 | AIV41909.1 | P36334 |
| ATTANVSALIGA | 1 | 0 | 0 | 1413475 | AVN89322.1 | K9N7C7 |
| ATTAYANSVLNI | 1 | 0 | 0 | 1413483 | AVP77956.1 | P0C6W2 |
| ATTCVTGPTTEN | 1 | 0 | 0 | 1413485 | ATO98145.1 | Q3LZX1 |
| ATTSQDSYGGAS | 1 | 0 | 0 | 1413496 | AEN19363.1 | P0C6X6 |
| ATTVDVNVLNDT | 1 | 0 | 0 | 1413499 | AUM60013.1 | K9N7C7 |
| ATTVELPDEFVA | 1 | 0 | 0 | 1413501 | AVP78050.1 | Q3LZX3 |
| ATTVELPDEFVV | 1 | 0 | 0 | 1413502 | AVP78039.1 | Q3LZX3 |
| ATTYCRVGECVD | 1 | 0 | 0 | 1413506 | APT69868.1 | A0A0P0ILY9 |
| ATVCGPKLSTGL | 1 | 0 | 0 | 1413512 | ATO98193.1 | Q3LZX1 |
| ATVCGPKLSTQL | 1 | 0 | 0 | 1413513 | AVP78031.1 | Q3LZX1 |
| ATVCNSLCLCFV | 1 | 0 | 0 | 1413514 | AUM60013.1 | K9N7C7 |
| ATVDHSAFAYES | 1 | 0 | 0 | 1413515 | NP_073549.1 | A0A0P0LKV0 |
| ATVDVQIQNNTV | 1 | 0 | 0 | 1413516 | AVV62524.1 | K9N7C7 |
| ATVDVTNGLGTY | 1 | 0 | 0 | 1413517 | AVQ05264.1 | P36334 |
| ATVEPVRDEVSI | 1 | 0 | 0 | 1413518 | AIW52723.1 | A0A0N7J131 |
| ATVEPVRDEVST | 1 | 0 | 0 | 1413519 | ABB90487.1 | A0A0N7J131 |
| ATVIAGSVYTNS | 1 | 0 | 0 | 1413524 | AVV62524.1 | K9N7C7 |
| ATVKWADAPEPQ | 1 | 0 | 0 | 1413528 | APT69853.1 | A0A0N7J131 |
| ATVKWADESEPQ | 1 | 0 | 0 | 1413530 | ABB90491.1 | A0A0N7J131 |
| ATVLQNNELMPA | 1 | 0 | 0 | 1413532 | AEN19363.1 | P0C6X6 |
| ATVLQNNELMPV | 1 | 0 | 0 | 1413533 | AXX83324.1 | P0C6X6 |
| ATVLSGSVYTNS | 1 | 0 | 0 | 1413534 | AUM60013.1 | K9N7C7 |
| ATVMNSLCLCFV | 1 | 0 | 0 | 1413535 | AVV62535.1 | K9N7C7 |
| ATVNLRLVDCAP | 1 | 0 | 0 | 1413537 | AWK59942.1 | H9EJ42 |
| ATVNLRLVDCVP | 1 | 0 | 0 | 1413538 | AFD98812.1 | H9EJ42 |
| ATVTANLSIPSN | 1 | 0 | 0 | 1413555 | AAK32189.1 | A0A0P0K6L9 |
| ATVYRRIPDLPD | 1 | 0 | 0 | 1413561 | ABD75593.1 | A0A140H1H1 |
| ATVYTANKWDLI | 1 | 0 | 0 | 1413562 | APO40578.1 | P0C6X7 |
| ATYDNFRIVCAN | 1 | 0 | 0 | 1413566 | APO40578.1 | P0C6X7 |
| ATYEGNSPFHPL | 1 | 0 | 0 | 1413568 | ABI96964.1 | P59635 |
| ATYHTPAIDCSD | 1 | 0 | 0 | 1413570 | ASU91010.1 | K9N5Q8 |
| ATYHTPATNCSD | 1 | 0 | 0 | 1413572 | ALA49671.1 | K9N5Q8 |
| ATYHTPSTDCSD | 1 | 0 | 0 | 1413573 | ALA49803.1 | K9N5Q8 |
| ATYIECTFISRF | 1 | 0 | 0 | 1413575 | AUM60018.1 | K9N7D2 |
| ATYQNISTNLPP | 1 | 0 | 0 | 1413582 | ALM26400.1 | K9N5Q8 |
| ATYYTPATDCSD | 1 | 0 | 0 | 1413586 | ALA49352.1 | K9N5Q8 |
| AVAAAMASQKPQ | 1 | 0 | 0 | 1413587 | AVV62533.1 | K9N4V7 |
| AVAAAMATPQPQ | 1 | 0 | 0 | 1413588 | AVV62544.1 | K9N4V7 |
| AVADAMSAKEQR | 1 | 0 | 0 | 1413591 | AUM60021.1 | K9N4V7 |
| AVADSYFSYMMP | 1 | 0 | 0 | 1413592 | ABD75559.1 | A0A140H1G9 |
| AVAKAPPGEQFK | 1 | 0 | 0 | 1413595 | AEN19363.1 | P0C6X6 |
| AVALNTLVKQLS | 1 | 0 | 0 | 1413600 | APO40579.1 | P59594 |
| AVANGSHFKSWV | 1 | 0 | 0 | 1413603 | APO40579.1 | P59594 |
| AVANSTGTVIIS | 1 | 0 | 0 | 1413604 | AGV08520.1 | K9N5Q8 |
| AVANWLTPTNMY | 1 | 0 | 0 | 1413606 | AUM60013.1 | K9N7C7 |
| AVAYADNPNHFI | 1 | 0 | 0 | 1413621 | ARU07566.1 | Q80872 |
| AVAYSSPLQSYA | 1 | 0 | 0 | 1413622 | ALD51902.1 | K9N7C7 |
| AVAYVDKPNHFI | 1 | 0 | 0 | 1413624 | AXX83373.1 | Q80872 |
| AVCITTCNLGGA | 1 | 0 | 0 | 1413630 | AHI48776.1 | K9N7C7 |
| AVCLGISDSAVT | 1 | 0 | 0 | 1413632 | QXL74882.1 | P0C6X6 |
| AVCNGCVGHSEN | 1 | 0 | 0 | 1413635 | QEG03785.1 | A0A0P0K6L9 |
| AVCRHHASEYRQ | 1 | 0 | 0 | 1413639 | APO40578.1 | P0C6X7 |
| AVCRKHAAEYRE | 1 | 0 | 0 | 1413640 | AVV62524.1 | K9N7C7 |
| AVCRKHATDYRE | 1 | 0 | 0 | 1413641 | ASU91085.1 | K9N7C7 |
| AVCRKHATEYRD | 1 | 0 | 0 | 1413642 | AUM60013.1 | K9N7C7 |
| AVCSKHAEDYCK | 1 | 0 | 0 | 1413648 | ARB07436.1 | A0A140H1G9 |
| AVCSKHAEEYCN | 1 | 0 | 0 | 1413649 | ARU07574.1 | A0A140H1G9 |
| AVCSVIDLKLDD | 1 | 0 | 0 | 1413653 | AXN92247.1 | K9N7C7 |
| AVCVTACNLGGA | 1 | 0 | 0 | 1413655 | AVV62524.1 | K9N7C7 |
| AVDALCEKALKYLP | 1 | 0 | 0 | 1413658 | P0DTD1.1 | P0DTD1 |
| AVDCGHDDLSQL | 1 | 0 | 0 | 1413662 | AVV62537.1 | K9N5Q8 |
| AVDCKSDFMSEI | 1 | 0 | 0 | 1413663 | CAA83660.1 | P36334 |
| AVDCMSDFMSGI | 1 | 0 | 0 | 1413665 | AIL49491.1 | P36334 |
| AVDCSQDPLAEL | 1 | 0 | 0 | 1413666 | ATO98145.1 | Q3LZX1 |
| AVDEAFPLNEAE | 1 | 0 | 0 | 1413673 | AHI48527.1 | K9N638 |
| AVDGAKTEPTGH | 1 | 0 | 0 | 1413676 | AAA45463.1 | A0A0N7J131 |

| | | | | | | |
|---|---|---|---|---|---|---|
| AVDINKLCDEML | 1 | 0 | 0 | 1413678 | APO40578.1 | P0C6X7 |
| AVDIVSIRSSNQ | 1 | 0 | 0 | 1413682 | AKQ21067.1 | K9N643 |
| AVDLNKLCEEML | 1 | 0 | 0 | 1413683 | AVP78041.1 | P0C6W2 |
| AVDLSMDKLCRV | 1 | 0 | 0 | 1413684 | AVN89451.1 | K9N7C7 |
| AVDLSMDRLRRV | 1 | 0 | 0 | 1413687 | ALA49384.1 | K9N7C7 |
| AVDSLCAKAVTS | 1 | 0 | 0 | 1413691 | APT69884.1 | A0A0P0LKV0 |
| AVDVVSIRTSNQ | 1 | 0 | 0 | 1413694 | AHI48842.1 | K9N643 |
| AVEENKIPSVEV | 1 | 0 | 0 | 1413699 | AVP78030.1 | P0C6W2 |
| AVEKLRPVFAWV | 1 | 0 | 0 | 1413701 | ARO76381.1 | P0C6U8 |
| AVELTKDKLQRI | 1 | 0 | 0 | 1413708 | AUM60013.1 | K9N7C7 |
| AVEPILAIEPPP | 1 | 0 | 0 | 1413714 | Q4VID0.1 | Q4VID0 |
| AVEQDRNTRDVF | 1 | 0 | 0 | 1413716 | APO40579.1 | P59594 |
| AVEQRITVEVFE | 1 | 0 | 0 | 1413719 | AVV62535.1 | K9N7C7 |
| AVESCMVQVTCG | 1 | 0 | 0 | 1413720 | AVV62524.1 | K9N7C7 |
| AVESGGYIPPNF | 1 | 0 | 0 | 1413721 | AIW52740.1 | A0A0P0K6L9 |
| AVESGGYIPSNF | 1 | 0 | 0 | 1413722 | AIW52757.1 | A0A0P0K6L9 |
| AVESNVMDVNDC | 1 | 0 | 0 | 1413723 | AFD98812.1 | H9EJ42 |
| AVFDTPHTDCTS | 1 | 0 | 0 | 1413729 | AVV62537.1 | K9N5Q8 |
| AVFISPYNSQNT | 1 | 0 | 0 | 1413735 | AVP78041.1 | P0C6W2 |
| AVFQSASKVIAL | 1 | 0 | 0 | 1413740 | ATO98194.1 | Q3LZX0 |
| AVFTGCVGTWDK | 1 | 0 | 0 | 1413745 | AVV62535.1 | K9N7C7 |
| AVGDAGAAAFST | 1 | 0 | 0 | 1413752 | AVV62544.1 | K9N4V7 |
| AVGDIFVLTSHS | 1 | 0 | 0 | 1413753 | AVV62524.1 | K9N7C7 |
| AVGFNTLLVQPA | 1 | 0 | 0 | 1413756 | AUM60019.1 | K9N5R3 |
| AVGHPAPTHYLR | 1 | 0 | 0 | 1413759 | AVV62540.1 | K9N643 |
| AVGHTEFYDSQP | 1 | 0 | 0 | 1413760 | AUM60025.1 | K9N796 |
| AVGNICYTPSTL | 1 | 0 | 0 | 1413771 | APO40578.1 | P0C6X7 |
| AVGQMEDVNLND | 1 | 0 | 0 | 1413774 | AVV62544.1 | K9N4V7 |
| AVHEWKMVVCDK | 1 | 0 | 0 | 1413788 | AXN92258.1 | K9N7C7 |
| AVHSDGMFVASS | 1 | 0 | 0 | 1413796 | AWK59942.1 | H9EJ42 |
| AVIAGVAGARIP | 1 | 0 | 0 | 1413803 | AXN92247.1 | K9N7C7 |
| AVIAGVAGARVP | 1 | 0 | 0 | 1413804 | AVV62524.1 | K9N7C7 |
| AVIDEHVFYKKD | 1 | 0 | 0 | 1413809 | NP_073549.1 | A0A0P0LKV0 |
| AVIGDFNCTNSF | 1 | 0 | 0 | 1413811 | BBA20986.1 | A0A140H1H1 |
| AVIGDLKCPLDS | 1 | 0 | 0 | 1413812 | AVQ05264.1 | P36334 |
| AVIGDLKCPLDT | 1 | 0 | 0 | 1413813 | AKA09663.1 | P36334 |
| AVIGDLKCPLES | 1 | 0 | 0 | 1413814 | QEG03803.1 | P36334 |
| AVIGDLKCTSDT | 1 | 0 | 0 | 1413815 | CAA79896.1 | P36334 |
| AVIGDLKCTTVA | 1 | 0 | 0 | 1413816 | CAA83661.1 | P36334 |
| AVIGDLKCTTVS | 1 | 0 | 0 | 1413817 | CAA83660.1 | P36334 |
| AVIGDLNCPLDP | 1 | 0 | 0 | 1413818 | AWW13511.1 | P36334 |
| AVIGDLNCPLDS | 1 | 0 | 0 | 1413819 | AIL49507.1 | P36334 |
| AVIGDLNCPLDT | 1 | 0 | 0 | 1413820 | AEN19366.1 | P36334 |
| AVIGDLNCTLDP | 1 | 0 | 0 | 1413821 | AWW13566.1 | P36334 |
| AVIGDLNCTLHP | 1 | 0 | 0 | 1413822 | BBA84086.1 | P36334 |
| AVILLGRFFHTA | 1 | 0 | 0 | 1413828 | ARB07393.1 | A0A0P0KWE8 |
| AVINGERWFLNR | 1 | 0 | 0 | 1413832 | APO40578.1 | P0C6X7 |
| AVINIPDLPICV | 1 | 0 | 0 | 1413833 | AGT51520.1 | P30215 |
| AVKDGIVWVHEE | 1 | 0 | 0 | 1413854 | AVV62533.1 | K9N4V7 |
| AVKHGAKPVSNC | 1 | 0 | 0 | 1413860 | AWK59942.1 | H9EJ42 |
| AVKLQNNEIKPT | 1 | 0 | 0 | 1413865 | AUM60013.1 | K9N7C7 |
| AVKLQNNEIRPT | 1 | 0 | 0 | 1413866 | AVV62524.1 | K9N7C7 |
| AVKMFDAYVNIF | 1 | 0 | 0 | 1413868 | AVP78041.1 | P0C6W2 |
| AVKPLLAPHHVV | 1 | 0 | 0 | 1413873 | ARO76392.1 | Q7TLC7 |
| AVKSDYCGGNLC | 1 | 0 | 0 | 1413875 | QQY99180.1 | P36334 |
| AVKTGVATEQLL | 1 | 0 | 0 | 1413882 | AUM60013.1 | K9N7C7 |
| AVLATVCNSLCL | 1 | 0 | 0 | 1413887 | AUM60013.1 | K9N7C7 |
| AVLDTLLASSSF | 1 | 0 | 0 | 1413894 | ANI69888.1 | K9N638 |
| AVLEWDIKQCPA | 1 | 0 | 0 | 1413895 | AUM60013.1 | K9N7C7 |
| AVLFCTPIKKAF | 1 | 0 | 0 | 1413896 | APT69868.1 | A0A0P0ILY9 |
| AVLFSTVIIKKL | 1 | 0 | 0 | 1413900 | AWK59978.1 | H9EJ42 |
| AVLIDNDLNEYV | 1 | 0 | 0 | 1413902 | AVV62524.1 | K9N7C7 |
| AVLNSLCICFVA | 1 | 0 | 0 | 1413916 | AXN92247.1 | K9N7C7 |
| AVLVNGTHTPLG | 1 | 0 | 0 | 1413933 | AVV62530.1 | K9N7D2 |
| AVLYNSSSSFST | 1 | 0 | 0 | 1413936 | APO40579.1 | P59594 |
| AVMFEDAQGLKK | 1 | 0 | 0 | 1413945 | AXG21642.1 | K9N638 |
| AVMSLKDGQIND | 1 | 0 | 0 | 1413950 | AVP78041.1 | P0C6W2 |
| AVMSLKDNQVND | 1 | 0 | 0 | 1413951 | APO40578.1 | P0C6X7 |
| AVNANAQALSKL | 1 | 0 | 0 | 1413963 | AVV62537.1 | K9N5Q8 |
| AVNDQKLYDQVV | 1 | 0 | 0 | 1413966 | ARO76381.1 | P0C6U8 |
| AVNESRYGLHNL | 1 | 0 | 0 | 1413967 | ARU07594.1 | A0A0P0G321 |
| AVNFNVTNAATI | 1 | 0 | 0 | 1413969 | ARU07601.1 | A0A0P0K6L9 |
| AVNFNVTTASAT | 1 | 0 | 0 | 1413970 | AIW52843.1 | A0A0P0G321 |
| AVNLALKNLGFD | 1 | 0 | 0 | 1413978 | ABK63972.1 | H9EJ69 |
| AVNLNICEPVLT | 1 | 0 | 0 | 1413980 | ARU07574.1 | A0A140H1G9 |
| AVNNIAQALSKL | 1 | 0 | 0 | 1413984 | QFS19596.1 | K9N5Q8 |
| AVNNNAHALSKL | 1 | 0 | 0 | 1413985 | ATG84679.1 | K9N5Q8 |
| AVNNNAQALAKL | 1 | 0 | 0 | 1413986 | AVK87450.1 | K9N5Q8 |
| AVNNSKPYFIKN | 1 | 0 | 0 | 1413988 | AVV62530.1 | K9N7D2 |
| AVNNSKPYFLKN | 1 | 0 | 0 | 1413989 | AVV62541.1 | K9N7D2 |
| AVNPTNCIAPVN | 1 | 0 | 0 | 1413990 | ATG84723.1 | K9N5Q8 |
| AVNQSLIKSKVS | 1 | 0 | 0 | 1413991 | ARB07393.1 | A0A0P0KWE8 |
| AVNSEDIYNTLT | 1 | 0 | 0 | 1413995 | AUM60013.1 | K9N7C7 |
| AVNSEEIYNKLT | 1 | 0 | 0 | 1413996 | AVV62524.1 | K9N7C7 |
| AVNSTKALAKQY | 1 | 0 | 0 | 1413999 | AMQ48995.1 | K9N4V0 |
| AVNTGVFVAPPI | 1 | 0 | 0 | 1414002 | AVV62528.1 | K9N4V0 |
| AVNYTKAPDVML | 1 | 0 | 0 | 1414013 | CAA83660.1 | P36334 |
| AVPANSLHATAT | 1 | 0 | 0 | 1414016 | APO40580.1 | P59632 |
| AVPDNSSTIVTG | 1 | 0 | 0 | 1414018 | AKA09630.1 | A0A0P0G321 |
| AVPGGCNFPLEA | 1 | 0 | 0 | 1414020 | AWO68993.1 | A0A0P0G321 |
| AVPGSCNFPLEA | 1 | 0 | 0 | 1414021 | AIW52843.1 | A0A0P0G321 |

| | | | | | | |
|---|---|---|---|---|---|---|
| AVPPTSPTARTT | 1 | 0 | 0 | 1414025 | AUM60024.1 | K9N5Q8 |
| AVPSTTRSSSQL | 1 | 0 | 0 | 1414028 | AVV62537.1 | K9N5Q8 |
| AVPTGYVDTEHS | 1 | 0 | 0 | 1414029 | AVP78030.1 | P0C6W2 |
| AVPTGYVDTSAN | 1 | 0 | 0 | 1414032 | APO40578.1 | P0C6X7 |
| AVQGFACDCGAC | 1 | 0 | 0 | 1414040 | AVV62524.1 | K9N7C7 |
| AVQGFACDCGAV | 1 | 0 | 0 | 1414041 | AVV62535.1 | K9N7C7 |
| AVQGFACECGAV | 1 | 0 | 0 | 1414042 | AUM60013.1 | K9N7C7 |
| AVQLESDAYLDA | 1 | 0 | 0 | 1414046 | AVV62524.1 | K9N7C7 |
| AVQMGSNNVCTY | 1 | 0 | 0 | 1414047 | AGT51323.1 | H9EJ42 |
| AVQNTVFNASIS | 1 | 0 | 0 | 1414048 | AVL25595.1 | H9EJ69 |
| AVQQESDEYILA | 1 | 0 | 0 | 1414053 | ATQ39389.1 | K9N7C7 |
| AVQQLNNGFQGK | 1 | 0 | 0 | 1414054 | AVV62524.1 | K9N7C7 |
| AVQTCDLTVGSG | 1 | 0 | 0 | 1414057 | CAA83661.1 | P36334 |
| AVRKFLDGLDLP | 1 | 0 | 0 | 1414063 | QED88038.1 | H9EJ42 |
| AVRLQPSMALQL | 1 | 0 | 0 | 1414064 | AVV62540.1 | K9N643 |
| AVRNKISACFNF | 1 | 0 | 0 | 1414066 | AGT51787.1 | P0C6X6 |
| AVRNKTFACFNF | 1 | 0 | 0 | 1414068 | AGT51617.1 | P0C6X6 |
| AVRNKTSACFNF | 1 | 0 | 0 | 1414069 | AEN19363.1 | P0C6X6 |
| AVRTYSEAAAQG | 1 | 0 | 0 | 1414080 | AWK59942.1 | H9EJ42 |
| AVSAGLEFTDES | 1 | 0 | 0 | 1414083 | QDH43722.1 | P0C6X6 |
| AVSANVCALISC | 1 | 0 | 0 | 1414087 | AIL49553.1 | P0C6X6 |
| AVSANVSALMSC | 1 | 0 | 0 | 1414088 | AJE26220.1 | P0C6X6 |
| AVSDNVRALMSC | 1 | 0 | 0 | 1414091 | ADO24673.1 | P0C6X6 |
| AVSFADNNDNNN | 1 | 0 | 0 | 1414093 | AUM60021.1 | K9N4V7 |
| AVSFADNNDTTN | 1 | 0 | 0 | 1414094 | ALW82676.1 | K9N4V7 |
| AVSFAYNNDITN | 1 | 0 | 0 | 1414096 | AVN89381.1 | K9N4V7 |
| AVSFDYLIREAE | 1 | 0 | 0 | 1414098 | AXN73523.1 | K9N7C7 |
| AVSFDYLIREAR | 1 | 0 | 0 | 1414100 | QDP16204.1 | K9N7C7 |
| AVSKLVESASDF | 1 | 0 | 0 | 1414111 | Q14EA9.1 | Q0ZJ84 |
| AVSKVVKVPIDL | 1 | 0 | 0 | 1414114 | ANI69887.1 | K9N7C7 |
| AVSPDFKAFNVF | 1 | 0 | 0 | 1414117 | AVV62535.1 | K9N7C7 |
| AVSRINWVTGGI | 1 | 0 | 0 | 1414118 | AFR58690.1 | P59596 |
| AVSRLVESASEF | 1 | 0 | 0 | 1414119 | ADN03340.1 | Q0ZJ84 |
| AVSSNINMLLSV | 1 | 0 | 0 | 1414124 | AGH06535.1 | H9EJ42 |
| AVSSNINRILGV | 1 | 0 | 0 | 1414125 | AUT76985.1 | A0A0P0LKV0 |
| AVSSNINRLFSV | 1 | 0 | 0 | 1414126 | AFD98777.1 | H9EJ42 |
| AVSSNINRVLSV | 1 | 0 | 0 | 1414127 | ABQ16583.1 | H9EJ42 |
| AVSWNLREMLAH | 1 | 0 | 0 | 1414131 | BAE93400.1 | P0C6U8 |
| AVTANVCSPMAC | 1 | 0 | 0 | 1414136 | ACR82761.1 | A0A140H1G9 |
| AVTANVRSLTAC | 1 | 0 | 0 | 1414138 | ADO24641.1 | A0A140H1G9 |
| AVTFGDNNDNNS | 1 | 0 | 0 | 1414144 | AVV62533.1 | K9N4V7 |
| AVTFGDNNDNNT | 1 | 0 | 0 | 1414145 | AVV62544.1 | K9N4V7 |
| AVTKTSEQNNFT | 1 | 0 | 0 | 1414150 | AVP78041.1 | P0C6W2 |
| AVVATVMNSLCL | 1 | 0 | 0 | 1414160 | AVV62535.1 | K9N7C7 |
| AVVCASVFKDGC | 1 | 0 | 0 | 1414161 | ABE97136.1 | H9EJ42 |
| AVVDGKQIDDYR | 1 | 0 | 0 | 1414166 | AIV41978.1 | P0C6X6 |
| AVVDSYYSYLMP | 1 | 0 | 0 | 1414167 | AVV62524.1 | K9N7C7 |
| AVVEAPTTCGYL | 1 | 0 | 0 | 1414168 | APO40578.1 | P0C6X7 |
| AVVGDLNCPLDP | 1 | 0 | 0 | 1414173 | QBP84704.1 | P36334 |
| AVVGTPVCINGL | 1 | 0 | 0 | 1414175 | P0C6W6.1 | P0C6W2 |
| AVVGTPVCVNGL | 1 | 0 | 0 | 1414176 | ATO98229.1 | P0C6W2 |
| AVVIREPVHLLS | 1 | 0 | 0 | 1414182 | ARB07436.1 | A0A140H1G9 |
| AVVKIYCPACHN | 1 | 0 | 0 | 1414183 | AVP78041.1 | P0C6W2 |
| AVVKLYCPACHN | 1 | 0 | 0 | 1414184 | AVP78030.1 | P0C6W2 |
| AVVKMACPACQN | 1 | 0 | 0 | 1414185 | APO40578.1 | P0C6X7 |
| AVVNADAEALNN | 1 | 0 | 0 | 1414190 | CAA83661.1 | P36334 |
| AVVNANAEAHNN | 1 | 0 | 0 | 1414191 | ARB66064.1 | P36334 |
| AVVNDKVFTKSG | 1 | 0 | 0 | 1414193 | ANF29215.1 | K9N7C7 |
| AVVNGDSEVVLK | 1 | 0 | 0 | 1414195 | AVP78041.1 | P0C6W2 |
| AVVNIRETKCAP | 1 | 0 | 0 | 1414198 | AVV62524.1 | K9N7C7 |
| AVVNSQDVYKSL | 1 | 0 | 0 | 1414199 | ANF29227.1 | K9N638 |
| AVVYFGGCVYWS | 1 | 0 | 0 | 1414210 | QBP84710.1 | P0C6X6 |
| AVYTANDVDLQS | 1 | 0 | 0 | 1414238 | AUM60013.1 | K9N7C7 |
| AWAAFYIYKLHQ | 1 | 0 | 0 | 1414246 | AVV62526.1 | K9N5Q8 |
| AWAALVDEEFPL | 1 | 0 | 0 | 1414251 | AVV62524.1 | K9N7C7 |
| AWAHCVKCHNDI | 1 | 0 | 0 | 1414253 | AUM60013.1 | K9N7C7 |
| AWATLVDEEFPL | 1 | 0 | 0 | 1414255 | AVV62535.1 | K9N7C7 |
| AWCVKCHNDILT | 1 | 0 | 0 | 1414260 | AVV62524.1 | K9N7C7 |
| AWDFMLKTLYKD | 1 | 0 | 0 | 1414261 | QCZ25053.1 | K9N7C7 |
| AWDKVSKIANLF | 1 | 0 | 0 | 1414263 | AVV62524.1 | K9N7C7 |
| AWDYHITKVLAE | 1 | 0 | 0 | 1414265 | AST12962.1 | A0A0P0LKV0 |
| AWEDGDYYRKQL | 1 | 0 | 0 | 1414267 | AGV08584.1 | K9N5Q8 |
| AWFVQNDVCSTE | 1 | 0 | 0 | 1414289 | ARU07575.1 | A0A140H1G9 |
| AWGSTMTFSVHP | 1 | 0 | 0 | 1414291 | AUM60013.1 | K9N7C7 |
| AWLLHKDGGIPD | 1 | 0 | 0 | 1414304 | AXN92262.1 | K9N4V0 |
| AWLLHKDGGVPD | 1 | 0 | 0 | 1414305 | QBF80567.1 | K9N4V0 |
| AWLNDKAVPSPL | 1 | 0 | 0 | 1414308 | AIV41909.1 | P36334 |
| AWLNDKLVPSPL | 1 | 0 | 0 | 1414309 | ATN39872.1 | P36334 |
| AWLNGKSVPSPL | 1 | 0 | 0 | 1414311 | AIL49492.1 | P36334 |
| AWLSYYVPSVEY | 1 | 0 | 0 | 1414313 | AEN19363.1 | P0C6X6 |
| AWLTKRKPLDYK | 1 | 0 | 0 | 1414314 | NP_073549.1 | A0A0P0LKV0 |
| AWNIGEQSSILS | 1 | 0 | 0 | 1414322 | AVP78030.1 | P0C6W2 |
| AWNIGSNKTLLT | 1 | 0 | 0 | 1414326 | APO40578.1 | P0C6X7 |
| AWNLREMLAYAE | 1 | 0 | 0 | 1414329 | APO40578.1 | P0C6X7 |
| AWNPEVNAITVT | 1 | 0 | 0 | 1414330 | AST12967.1 | A0A0P0KXN3 |
| AWNTAKQDQGQY | 1 | 0 | 0 | 1414332 | ATO98193.1 | Q3LZX1 |
| AWNTAKQDTGHY | 1 | 0 | 0 | 1414333 | AVP78042.1 | Q3LZX1 |
| AWNTAKQDVGNY | 1 | 0 | 0 | 1414334 | AVP78031.1 | Q3LZX1 |
| AWNTAKQDVGSY | 1 | 0 | 0 | 1414335 | ATO98145.1 | Q3LZX1 |
| AWNTNSVDSKSG | 1 | 0 | 0 | 1414338 | APO40579.1 | P59594 |
| AWSGLCVDGRNG | 1 | 0 | 0 | 1414355 | AIW52756.1 | A0A0P0K6L9 |

| | | | | | | |
|---|---|---|---|---|---|---|
| AWVESRLSDGVE | 1 | 0 | 0 | 1414365 | ARO76381.1 | P0C6U8 |
| AYADNPNHFINF | 1 | 0 | 0 | 1414382 | ARU07566.1 | Q80872 |
| AYADRTLQEMKA | 1 | 0 | 0 | 1414383 | AIV41774.1 | P0C6X6 |
| AYAEETRKIMPI | 1 | 0 | 0 | 1414384 | APO40578.1 | P0C6X7 |
| AYALIRGYRGVK | 1 | 0 | 0 | 1414392 | AEN19363.1 | P0C6X6 |
| AYANNSIAIPTN | 1 | 0 | 0 | 1414399 | AVP78031.1 | Q3LZX1 |
| AYANSVFNIFQA | 1 | 0 | 0 | 1414402 | AVP78019.1 | P0C6W2 |
| AYANSVFNILQV | 1 | 0 | 0 | 1414404 | AVV62524.1 | K9N7C7 |
| AYANYNTYDVVL | 1 | 0 | 0 | 1414406 | APO40578.1 | P0C6X7 |
| AYDGSAFRDLKP | 1 | 0 | 0 | 1414424 | AHI48736.1 | K9N7C7 |
| AYDQVHEEPKGK | 1 | 0 | 0 | 1414433 | ARK08632.1 | P0C6X6 |
| AYEASAFRDLKP | 1 | 0 | 0 | 1414435 | AUM60013.1 | K9N7C7 |
| AYENFNQHEILL | 1 | 0 | 0 | 1414440 | AVP78041.1 | P0C6W2 |
| AYENFNSQDTLL | 1 | 0 | 0 | 1414443 | ATO98229.1 | P0C6W2 |
| AYENFNSQEILL | 1 | 0 | 0 | 1414444 | ARO76381.1 | P0C6U8 |
| AYEQAVVNGDSE | 1 | 0 | 0 | 1414446 | AVP78041.1 | P0C6W2 |
| AYESSAFRDLKP | 1 | 0 | 0 | 1414449 | AUM60023.1 | K9N7C7 |
| AYFNGAIIRDGD | 1 | 0 | 0 | 1414459 | ALA49648.1 | K9N7C7 |
| AYFNGAIIRDSE | 1 | 0 | 0 | 1414460 | AVV62524.1 | K9N7C7 |
| AYFNMIYMPASW | 1 | 0 | 0 | 1414463 | APO40578.1 | P0C6X7 |
| AYFSDSKFGTTE | 1 | 0 | 0 | 1414470 | AHI48527.1 | K9N638 |
| AYFTDNGVVNKF | 1 | 0 | 0 | 1414477 | AHA80218.1 | P36334 |
| AYFTDTGFVTKF | 1 | 0 | 0 | 1414478 | AEN19366.1 | P36334 |
| AYGIPLSEAKGY | 1 | 0 | 0 | 1414491 | AMM42386.1 | Q19U29 |
| AYGIPPSEAKGY | 1 | 0 | 0 | 1414492 | AMM42397.1 | Q19U29 |
| AYGLCDASNPTN | 1 | 0 | 0 | 1414494 | AHY22565.1 | K9N5Q8 |
| AYGLCDLANPTN | 1 | 0 | 0 | 1414496 | AVN89387.1 | K9N5Q8 |
| AYGLCDVANPTN | 1 | 0 | 0 | 1414498 | AKM76239.1 | K9N5Q8 |
| AYGLCNKGSPQQ | 1 | 0 | 0 | 1414499 | AVV62526.1 | K9N5Q8 |
| AYGNCTFGDPDD | 1 | 0 | 0 | 1414503 | AVN89333.1 | K9N7C7 |
| AYGNCTFGSPDD | 1 | 0 | 0 | 1414504 | AHI48794.1 | K9N638 |
| AYHANSPEPALL | 1 | 0 | 0 | 1414509 | AIX10763.1 | P36334 |
| AYHANSSDPALL | 1 | 0 | 0 | 1414510 | AIL49515.1 | P36334 |
| AYHANSSEPTLL | 1 | 0 | 0 | 1414511 | AIL49500.1 | P36334 |
| AYHNLVGYYSDD | 1 | 0 | 0 | 1414513 | ALA49649.1 | K9N5Q8 |
| AYHTVIDVKQVG | 1 | 0 | 0 | 1414514 | AIV41894.1 | P0C6X6 |
| AYIAPQAKSGII | 1 | 0 | 0 | 1414517 | ATN39864.2 | P30215 |
| AYIAPQANFGDY | 1 | 0 | 0 | 1414519 | AGT51599.1 | P30215 |
| AYIAPQANSGGY | 1 | 0 | 0 | 1414520 | AGT51669.1 | P30215 |
| AYIAPQANSGII | 1 | 0 | 0 | 1414521 | AEN19365.1 | P30215 |
| AYIAREANFGDY | 1 | 0 | 0 | 1414523 | AAA73152.1 | P30215 |
| AYIDFVNSGGTP | 1 | 0 | 0 | 1414527 | AUM60023.1 | K9N7C7 |
| AYINILDLPICV | 1 | 0 | 0 | 1414538 | ARK08634.1 | P30215 |
| AYINIPDLPICV | 1 | 0 | 0 | 1414539 | AEN19365.1 | P30215 |
| AYINIPDLPVCV | 1 | 0 | 0 | 1414540 | ARB07432.1 | P30215 |
| AYITFSGSFDNG | 1 | 0 | 0 | 1414546 | ABE97129.1 | H9EJ42 |
| AYITGGVVQMTS | 1 | 0 | 0 | 1414548 | AVP78041.1 | P0C6W2 |
| AYKHLNKYDCCL | 1 | 0 | 0 | 1414556 | ARU07575.1 | A0A140H1G9 |
| AYKHLNNYDCCL | 1 | 0 | 0 | 1414557 | ARB07605.1 | A0A140H1G9 |
| AYLAGTVWYVGQ | 1 | 0 | 0 | 1414571 | AGT51533.1 | Q04854 |
| AYLDAKGPLQVG | 1 | 0 | 0 | 1414574 | AVV62524.1 | K9N7C7 |
| AYLDLRTTQQLT | 1 | 0 | 0 | 1414575 | AHI48765.1 | K9N638 |
| AYLDSCTTQQLT | 1 | 0 | 0 | 1414576 | AHI48794.1 | K9N638 |
| AYLDSRATQQLT | 1 | 0 | 0 | 1414577 | ANF29194.1 | K9N7C7 |
| AYLDSRTTQQLT | 1 | 0 | 0 | 1414578 | AXN92247.1 | K9N7C7 |
| AYLKLSDALKRQ | 1 | 0 | 0 | 1414587 | AVV62535.1 | K9N7C7 |
| AYLNSRTTQQLT | 1 | 0 | 0 | 1414595 | ALW82708.1 | K9N638 |
| AYLQSSNYRIDT | 1 | 0 | 0 | 1414597 | ARB66070.1 | P36334 |
| AYMDFVNSGGTP | 1 | 0 | 0 | 1414605 | AUM60013.1 | K9N7C7 |
| AYNCTYDRVEQS | 1 | 0 | 0 | 1414620 | ATO98120.1 | Q3LZX1 |
| AYNGKPQGAFHV | 1 | 0 | 0 | 1414623 | AEN19363.1 | P0C6X6 |
| AYNGPPQGAFHV | 1 | 0 | 0 | 1414624 | ABD75599.1 | A0A140H1G9 |
| AYNGYVTSSSKS | 1 | 0 | 0 | 1414627 | APO40578.1 | P0C6X7 |
| AYNKDHGCGGFG | 1 | 0 | 0 | 1414629 | AEN19363.1 | P0C6X6 |
| AYNLPTMCDIKQ | 1 | 0 | 0 | 1414634 | AUM60013.1 | K9N7C7 |
| AYNNCTGGAEIK | 1 | 0 | 0 | 1414636 | AGT51590.1 | P36334 |
| AYNNCTGGAGIR | 1 | 0 | 0 | 1414638 | CAA83661.1 | P36334 |
| AYNNDITNTNLS | 1 | 0 | 0 | 1414639 | AVN89381.1 | K9N4V7 |
| AYNNTAISVQTC | 1 | 0 | 0 | 1414640 | AIL49538.1 | P36334 |
| AYNSILVENGIP | 1 | 0 | 0 | 1414644 | ARU07593.1 | H9EJ42 |
| AYNSTAISVKTC | 1 | 0 | 0 | 1414647 | AIL49550.1 | P36334 |
| AYNTFTQAGFNI | 1 | 0 | 0 | 1414648 | AWK59942.1 | H9EJ42 |
| AYNVINKGHFDG | 1 | 0 | 0 | 1414650 | AVP78041.1 | P0C6W2 |
| AYNVVKQGHFTG | 1 | 0 | 0 | 1414652 | AUM60023.1 | K9N7C7 |
| AYQGLFPKQGDQ | 1 | 0 | 0 | 1414664 | AUM60014.1 | K9N5Q8 |
| AYQLYMIDPAGV | 1 | 0 | 0 | 1414666 | AEN19363.1 | P0C6X6 |
| AYQNLLGYYSDD | 1 | 0 | 0 | 1414669 | QEU56412.1 | K9N5Q8 |
| AYQPYDHAIGAT | 1 | 0 | 0 | 1414671 | ATO98145.1 | Q3LZX1 |
| AYQQQDGMMNIS | 1 | 0 | 0 | 1414672 | AIX10729.1 | P33469 |
| AYQQQDGMMNMS | 1 | 0 | 0 | 1414673 | AEN19362.1 | P33469 |
| AYQQQDGMNMSP | 1 | 0 | 0 | 1414674 | AIV41850.1 | P33469 |
| AYQQQDGMVNMS | 1 | 0 | 0 | 1414675 | AEN19370.1 | P33469 |
| AYRANFFDVPTF | 1 | 0 | 0 | 1414678 | AMQ49014.1 | K9N7C7 |
| AYRANSFDAPTF | 1 | 0 | 0 | 1414679 | AHI48604.1 | K9N7C7 |
| AYRANSIDVPTF | 1 | 0 | 0 | 1414681 | AQZ41320.1 | K9N7C7 |
| AYRPVPGTSTHP | 1 | 0 | 0 | 1414690 | AUM60016.1 | K9N4V0 |
| AYSGDIPHYVQP | 1 | 0 | 0 | 1414696 | AUM60024.1 | K9N5Q8 |
| AYSKYYTIGTLY | 1 | 0 | 0 | 1414698 | 6ATK_E | A0A0P0K6L9 |
| AYSNNIIAIPTN | 1 | 0 | 0 | 1414703 | APO40579.1 | P59594 |
| AYSPIDIPRGLP | 1 | 0 | 0 | 1414708 | ATO98193.1 | Q3LZX1 |
| AYSPQLTLVFLE | 1 | 0 | 0 | 1414710 | ALL26408.1 | K9N638 |

| | | | | | | |
|---|---|---|---|---|---|---|
| AYSSPILPFGHG | 1 | 0 | 0 | 1414719 | ATO98145.1 | Q3LZX1 |
| AYSSSANGILVG | 1 | 0 | 0 | 1414722 | AVN89312.1 | K9N638 |
| AYSSSANGRLVG | 1 | 0 | 0 | 1414723 | AJG44057.1 | K9N638 |
| AYSSSANGSLDG | 1 | 0 | 0 | 1414724 | AXG21642.1 | K9N638 |
| AYSSSANGSLVC | 1 | 0 | 0 | 1414725 | AVN89333.1 | K9N7C7 |
| AYSSSASDAIVG | 1 | 0 | 0 | 1414727 | AVV62535.1 | K9N7C7 |
| AYSSSASEGLVG | 1 | 0 | 0 | 1414728 | AUM60013.1 | K9N7C7 |
| AYTFTTVKPSAA | 1 | 0 | 0 | 1414740 | AZK15899.1 | K9N638 |
| AYTQGRSCSDFE | 1 | 0 | 0 | 1414749 | ATQ39389.1 | K9N7C7 |
| AYVANPTEDLCN | 1 | 0 | 0 | 1414761 | AGT51737.1 | P0C6X6 |
| AYVDKPNHFINF | 1 | 0 | 0 | 1414762 | AXX83373.1 | Q80872 |
| AYVTGGLVQQVE | 1 | 0 | 0 | 1414787 | APO40578.1 | P0C6X7 |
| AYVTGGLVQQVS | 1 | 0 | 0 | 1414788 | ARO76381.1 | P0C6U8 |
| AYWFLNQFDINA | 1 | 0 | 0 | 1414794 | ARU07575.1 | A0A140H1G9 |
| AYWVPRASANIG | 1 | 0 | 0 | 1414799 | AVP78041.1 | P0C6W2 |
| AYYADSGMPTTF | 1 | 0 | 0 | 1414800 | BBA20983.1 | A0A140H1H1 |
| AYYADVGMPTTF | 1 | 0 | 0 | 1414801 | BBA20986.1 | A0A140H1H1 |
| AYYNDWQNLLYD | 1 | 0 | 0 | 1414814 | AZS52618.1 | A0A140H1H1 |
| AYYNNWQNLLYD | 1 | 0 | 0 | 1414819 | BBA20986.1 | A0A140H1H1 |
| AYYSQSILYRVY | 1 | 0 | 0 | 1414824 | AVV62524.1 | K9N7C7 |
| AYYVKDGKYFTS | 1 | 0 | 0 | 1414829 | AVV62524.1 | K9N7C7 |
| CAACVCGDYAAC | 1 | 0 | 0 | 1414831 | ABD43177.1 | P36334 |
| CAAFDCGDYAAC | 1 | 0 | 0 | 1414832 | ARB66062.1 | P36334 |
| CAAGTTQTACTDDN | 1 | 0 | 0 | 1414840 | P0DTC1.1 | P0DTD1 |
| CADALHMYVKSS | 1 | 0 | 0 | 1414853 | APO40578.1 | P0C6X7 |
| CAGYTLDEARTA | 1 | 0 | 0 | 1414890 | ARO76381.1 | P0C6U8 |
| CAIVQASESKVF | 1 | 0 | 0 | 1414906 | ARU07593.1 | H9EJ42 |
| CAIVQATESKVL | 1 | 0 | 0 | 1414908 | ABE97136.1 | H9EJ42 |
| CAKILPFSDRPL | 1 | 0 | 0 | 1414914 | AWK59942.1 | H9EJ42 |
| CAKSPPGEQFRR | 1 | 0 | 0 | 1414924 | AWK59942.1 | H9EJ42 |
| CALISCNGNKIE | 1 | 0 | 0 | 1414937 | AIL49553.1 | P0C6X6 |
| CALMSCNGNKIG | 1 | 0 | 0 | 1414939 | AJE26218.1 | P0C6X6 |
| CALMSCNGNKSD | 1 | 0 | 0 | 1414940 | AKL87224.1 | P0C6X6 |
| CALPDTPSTFTP | 1 | 0 | 0 | 1414944 | AVK87439.1 | K9N5Q8 |
| CALPDTPSTLAP | 1 | 0 | 0 | 1414945 | ALJ54472.1 | K9N5Q8 |
| CALPDTPSTLTL | 1 | 0 | 0 | 1414947 | ANI69889.1 | K9N5Q8 |
| CALPFEDLNKCR | 1 | 0 | 0 | 1414949 | AVV62524.1 | K9N7C7 |
| CALTPALLATNN | 1 | 0 | 0 | 1414955 | APO40578.1 | P0C6X7 |
| CAMGIVNSSMVV | 1 | 0 | 0 | 1414958 | AUM60014.1 | K9N5Q8 |
| CAMPIMANLANL | 1 | 0 | 0 | 1414960 | CAA71146.1 | A0A0P0K6L9 |
| CAMRPNYTIKGS | 1 | 0 | 0 | 1414973 | AEA10817.1 | P0C6X7 |
| CANEKFADGINQ | 1 | 0 | 0 | 1414979 | APO40578.1 | P0C6X7 |
| CANFNVLFAMTM | 1 | 0 | 0 | 1414983 | AXN92247.1 | K9N7C7 |
| CANFTILFSMVL | 1 | 0 | 0 | 1414985 | AIL49570.1 | P0C6X6 |
| CANQSTSTYLAN | 1 | 0 | 0 | 1414996 | AMM42642.1 | A0A0P0G321 |
| CANQSTSVYSAK | 1 | 0 | 0 | 1414999 | AKA09625.1 | A0A0P0G321 |
| CARKSCSYDSDV | 1 | 0 | 0 | 1415029 | AIV41810.1 | P0C6X6 |
| CARKSFSIDSDV | 1 | 0 | 0 | 1415030 | AJC98124.1 | P0C6X6 |
| CARNPVDLPPSH | 1 | 0 | 0 | 1415033 | QEG03783.1 | A0A0P0LKV0 |
| CASVFKDGCDVG | 1 | 0 | 0 | 1415040 | ABE97136.1 | H9EJ42 |
| CATCPFAKSDGC | 1 | 0 | 0 | 1415052 | AVV62524.1 | K9N7C7 |
| CATVYTANKWDL | 1 | 0 | 0 | 1415063 | APO40578.1 | P0C6X7 |
| CAVHQGAHVASG | 1 | 0 | 0 | 1415067 | AVV62524.1 | K9N7C7 |
| CAVPPTSSTART | 1 | 0 | 0 | 1415081 | AUM60014.1 | K9N5Q8 |
| CAVSFADNNDIT | 1 | 0 | 0 | 1415084 | AKK52599.1 | K9N4V7 |
| CAVYPIDLASQI | 1 | 0 | 0 | 1415087 | AVV62532.1 | K9N7A1 |
| CAVYPIDLATQI | 1 | 0 | 0 | 1415088 | AVV62543.1 | K9N7A1 |
| CAYTQGRSCSDF | 1 | 0 | 0 | 1415099 | ATQ39389.1 | K9N7C7 |
| CCCTGCGPSCFK | 1 | 0 | 0 | 1415105 | AXX83351.1 | P36334 |
| CCCTGCGTSCFK | 1 | 0 | 0 | 1415107 | CAA79896.1 | P36334 |
| CCDASYFDNKLW | 1 | 0 | 0 | 1415112 | AVV62524.1 | K9N7C7 |
| CCDDNYFSKKDW | 1 | 0 | 0 | 1415117 | APO40578.1 | P0C6X7 |
| CCDDYTGHQELV | 1 | 0 | 0 | 1415118 | CAA83660.1 | P36334 |
| CCDDYTGYQEFV | 1 | 0 | 0 | 1415119 | AIV41909.1 | P36334 |
| CCDDYTGYQESV | 1 | 0 | 0 | 1415121 | AIL49487.1 | P36334 |
| CCDEYGGHNDFV | 1 | 0 | 0 | 1415123 | BBA20986.1 | A0A140H1H1 |
| CCDGDYFNKKDW | 1 | 0 | 0 | 1415124 | AVP78041.1 | P0C6W2 |
| CCDKYDDLEPLK | 1 | 0 | 0 | 1415125 | AVV62537.1 | K9N5Q8 |
| CCDRYEEYDLQP | 1 | 0 | 0 | 1415131 | AHI48550.1 | K9N5Q8 |
| CCEASYFDNKLW | 1 | 0 | 0 | 1415133 | AUM60013.1 | K9N7C7 |
| CCEDYTGYQELV | 1 | 0 | 0 | 1415134 | AIL49528.1 | P36334 |
| CCENQPYQIEDP | 1 | 0 | 0 | 1415136 | Q3I5I8.1 | Q3LZX5 |
| CCGAGDQAAKLS | 1 | 0 | 0 | 1415141 | AVV62524.1 | K9N7C7 |
| CCGTPAKKLCVV | 1 | 0 | 0 | 1415146 | AWK59942.1 | H9EJ42 |
| CCHLAKAINDFS | 1 | 0 | 0 | 1415151 | AVP78030.1 | P0C6W2 |
| CCKCCYDHVIAT | 1 | 0 | 0 | 1415156 | QWA55870.1 | P0DTD1 |
| CCLDVVRKDLEE | 1 | 0 | 0 | 1415164 | ARK08633.1 | Q80872 |
| CCLDVVRKELEE | 1 | 0 | 0 | 1415165 | AWW13564.1 | Q80872 |
| CCLEVVRKELEE | 1 | 0 | 0 | 1415167 | NP_150075.1 | Q80872 |
| CCMTNCCSCFKG | 1 | 0 | 0 | 1415178 | APO40579.1 | P59594 |
| CCRLVEKNVVRA | 1 | 0 | 0 | 1415204 | AUM60013.1 | K9N7C7 |
| CCRVVERNIVRA | 1 | 0 | 0 | 1415206 | AVV62524.1 | K9N7C7 |
| CCRVVERNVVRA | 1 | 0 | 0 | 1415207 | AVV62535.1 | K9N7C7 |
| CCSACMVWREWC | 1 | 0 | 0 | 1415208 | QBF80553.1 | K9N7C7 |
| CCSQIDRFYRLA | 1 | 0 | 0 | 1415219 | QBP84719.1 | P0C6X6 |
| CCTAAIDLAHTK | 1 | 0 | 0 | 1415225 | AVV62535.1 | K9N7C7 |
| CCTAAIDLAYNK | 1 | 0 | 0 | 1415226 | AVV62524.1 | K9N7C7 |
| CCTEQDLHKDLT | 1 | 0 | 0 | 1415232 | AUM60027.1 | K9N643 |
| CCVVEASTDFHQ | 1 | 0 | 0 | 1415252 | AGT51386.1 | H9EJ42 |
| CCYDHVIATSHK | 1 | 0 | 0 | 1415253 | QWA55870.1 | P0DTD1 |
| CDAKFKNSVTSI | 1 | 0 | 0 | 1415270 | AST12962.1 | A0A0P0LKV0 |

| | | | | | | |
|---|---|---|---|---|---|---|
| CDCGACFSANDV | 1 | 0 | 0 | 1415281 | AVV62524.1 | K9N7C7 |
| CDCGAVFSGNDV | 1 | 0 | 0 | 1415282 | AVV62535.1 | K9N7C7 |
| CDCGIKQESRVG | 1 | 0 | 0 | 1415284 | ARU07575.1 | A0A140H1G9 |
| CDCGKKFDEQVG | 1 | 0 | 0 | 1415286 | AWK59942.1 | H9EJ42 |
| CDCGSTKLPYYEFEKVHVQ | 1 | 0 | 0 | 1415288 | Q6Q1S2.1 | A0A0P0G321 |
| CDCLCTPDPITS | 1 | 0 | 0 | 1415289 | CAA83660.1 | P36334 |
| CDDNYFNKKDWY | 1 | 0 | 0 | 1415292 | AVP78030.1 | P0C6W2 |
| CDDYTGHQELVI | 1 | 0 | 0 | 1415296 | CAA83660.1 | P36334 |
| CDDYTGYQESVI | 1 | 0 | 0 | 1415299 | AIL49487.1 | P36334 |
| CDEYEEEYEEQV | 1 | 0 | 0 | 1415313 | AVV62535.1 | K9N7C7 |
| CDEYGGHNDFVI | 1 | 0 | 0 | 1415315 | BBA20986.1 | A0A140H1H1 |
| CDFKGWIPGNMI | 1 | 0 | 0 | 1415321 | AEN19363.1 | P0C6X6 |
| CDFTKLLSGTPP | 1 | 0 | 0 | 1415332 | AVV62537.1 | K9N5Q8 |
| CDGGKYYTQRII | 1 | 0 | 0 | 1415338 | AEN19363.1 | P0C6X6 |
| CDGISPDAIARG | 1 | 0 | 0 | 1415342 | AVV62537.1 | K9N5Q8 |
| CDGLARDYRANS | 1 | 0 | 0 | 1415345 | AVV62535.1 | K9N7C7 |
| CDGLASAYRANF | 1 | 0 | 0 | 1415346 | AMQ49014.1 | K9N7C7 |
| CDGLASNYRENS | 1 | 0 | 0 | 1415348 | AVV62524.1 | K9N7C7 |
| CDGNLPCGICLS | 1 | 0 | 0 | 1415349 | APO40578.1 | P0C6X7 |
| CDHMVKDGLVGV | 1 | 0 | 0 | 1415361 | AVN89451.1 | K9N7C7 |
| CDIDKWLNNFSV | 1 | 0 | 0 | 1415366 | AMM42415.1 | A0A140H1H1 |
| CDIDNWLNKVSV | 1 | 0 | 0 | 1415367 | AMM42407.1 | A0A140H1H1 |
| CDINKPLQVFVY | 1 | 0 | 0 | 1415373 | AWK59942.1 | H9EJ42 |
| CDIPIGAGICAK | 1 | 0 | 0 | 1415375 | APO40579.1 | P59594 |
| CDIPVGAGICAS | 1 | 0 | 0 | 1415377 | ACQ82725.1 | P59594 |
| CDISYKGKVFFQ | 1 | 0 | 0 | 1415379 | AWW13571.1 | P0C6X6 |
| CDKVRSLKLSDC | 1 | 0 | 0 | 1415390 | AVV62535.1 | K9N7C7 |
| CDKYDDLEPLKI | 1 | 0 | 0 | 1415392 | AVV62537.1 | K9N5Q8 |
| CDLALKLKGLDA | 1 | 0 | 0 | 1415395 | AEN19363.1 | P0C6X6 |
| CDLHNYGENAVL | 1 | 0 | 0 | 1415399 | APO40578.1 | P0C6X7 |
| CDLMQARAFVNM | 1 | 0 | 0 | 1415403 | AVV62524.1 | K9N7C7 |
| CDLMQARAFVNQ | 1 | 0 | 0 | 1415404 | AVV62535.1 | K9N7C7 |
| CDLQNYGDSAIL | 1 | 0 | 0 | 1415405 | AVP78030.1 | P0C6W2 |
| CDLRMGSGFCID | 1 | 0 | 0 | 1415408 | BBA20986.1 | A0A140H1H1 |
| CDLYSIHDVGFK | 1 | 0 | 0 | 1415413 | AAX85666.1 | P0C6X6 |
| CDLYSMQDVGFK | 1 | 0 | 0 | 1415414 | AAR01012.1 | P0C6X6 |
| CDMCKSTVVEVK | 1 | 0 | 0 | 1415415 | QED88039.1 | H9EJ42 |
| CDNDRCIIHCAN | 1 | 0 | 0 | 1415417 | ARU07574.1 | A0A140H1G9 |
| CDNLCTLDPIAL | 1 | 0 | 0 | 1415423 | BBO58891.1 | P36334 |
| CDNPAFVVTGGA | 1 | 0 | 0 | 1415428 | ARO76382.1 | P59594 |
| CDNPFIAVSKPM | 1 | 0 | 0 | 1415431 | ABD73001.1 | P59594 |
| CDNYDDYDFEPQ | 1 | 0 | 0 | 1415433 | AUM60014.1 | K9N5Q8 |
| CDQLAFDVPDGF | 1 | 0 | 0 | 1415435 | AFR79257.1 | A0A0P0K6L9 |
| CDRAMPNIIRIV | 1 | 0 | 0 | 1415444 | AKL87224.1 | P0C6X6 |
| CDRTAIQSFDSS | 1 | 0 | 0 | 1415453 | AST12962.1 | A0A0P0LKV0 |
| CDRYEEYDLQPH | 1 | 0 | 0 | 1415459 | AHI48550.1 | K9N5Q8 |
| CDSLNAKWIFDS | 1 | 0 | 0 | 1415464 | AIV41816.1 | P0C6X6 |
| CDSLNAKWVFDS | 1 | 0 | 0 | 1415465 | AIV41984.1 | P0C6X6 |
| CDSLNFKLGFDS | 1 | 0 | 0 | 1415466 | AAX85666.1 | P0C6X6 |
| CDSLRQAALPQS | 1 | 0 | 0 | 1415469 | AXN92247.1 | K9N7C7 |
| CDSLRQVALPQS | 1 | 0 | 0 | 1415471 | AXN92258.1 | K9N7C7 |
| CDSLRQVALTQS | 1 | 0 | 0 | 1415472 | AUM60023.1 | K9N7C7 |
| CDVCPNRATCYN | 1 | 0 | 0 | 1415492 | ARB07436.1 | A0A140H1G9 |
| CDVDTTTLYVNN | 1 | 0 | 0 | 1415494 | AGT17758.1 | A0A140H1H1 |
| CDYAQLPNEITV | 1 | 0 | 0 | 1415513 | AUM60030.1 | K9N7A1 |
| CDYLADYDVAVT | 1 | 0 | 0 | 1415517 | ASU90174.1 | K9N638 |
| CDYTGSLADGLE | 1 | 0 | 0 | 1415522 | AEN19363.1 | P0C6X6 |
| CDYTGSLADGLG | 1 | 0 | 0 | 1415523 | AIV42008.1 | P0C6X6 |
| CECGAVYTANDV | 1 | 0 | 0 | 1415528 | AUM60013.1 | K9N7C7 |
| CEDFYASMYENS | 1 | 0 | 0 | 1415530 | AIW52827.1 | H9EJ42 |
| CEDFYASMYENY | 1 | 0 | 0 | 1415531 | QED88024.1 | H9EJ42 |
| CEEADEGICFNL | 1 | 0 | 0 | 1415536 | ATP16754.1 | P0C6X6 |
| CEESSAKSASVYYSQ | 1 | 0 | 0 | 1415548 | P0DTC1.1 | P0DTD1 |
| CEEVANDFTTAL | 1 | 0 | 0 | 1415549 | AKI29254.1 | K9N638 |
| CEHITTMMSQFS | 1 | 0 | 0 | 1415576 | AVV62526.1 | K9N5Q8 |
| CEHLQFGLQDGF | 1 | 0 | 0 | 1415577 | AIW52843.1 | A0A0P0G321 |
| CEIATKMFDSFV | 1 | 0 | 0 | 1415584 | AUM60013.1 | K9N7C7 |
| CEIVGGQIVTRA | 1 | 0 | 0 | 1415587 | AVP78041.1 | P0C6W2 |
| CELCGTQNAVVE | 1 | 0 | 0 | 1415599 | APO40578.1 | P0C6X7 |
| CELELICDCGIK | 1 | 0 | 0 | 1415600 | ARU07575.1 | A0A140H1G9 |
| CELHGNGSMIED | 1 | 0 | 0 | 1415601 | ALW82645.1 | K9N7C7 |
| CELVAELDGVQF | 1 | 0 | 0 | 1415602 | ARO76381.1 | P0C6U8 |
| CELVNYKQSIPM | 1 | 0 | 0 | 1415603 | QAT98908.1 | K9N7C7 |
| CELYVNNAYRIF | 1 | 0 | 0 | 1415605 | QDH43722.1 | P0C6X6 |
| CEMVRAILKKKE | 1 | 0 | 0 | 1415607 | AVV62535.1 | K9N7C7 |
| CENQPYQIEDPC | 1 | 0 | 0 | 1415613 | Q3I5I8.1 | Q3LZX5 |
| CEPSTQYEYGTE | 1 | 0 | 0 | 1415618 | AVP78030.1 | P0C6W2 |
| CEPVLTYSRLIS | 1 | 0 | 0 | 1415619 | ARU07574.1 | A0A140H1G9 |
| CEQAYQLYDASN | 1 | 0 | 0 | 1415620 | AIV42008.1 | P0C6X6 |
| CEQFYQLYDASN | 1 | 0 | 0 | 1415625 | AWW13571.1 | P0C6X6 |
| CEQSYQLYDACN | 1 | 0 | 0 | 1415628 | QBP84746.1 | P0C6X6 |
| CEQTPCGFKGGI | 1 | 0 | 0 | 1415630 | AIV42002.1 | P0C6X6 |
| CERLTGGNYSDP | 1 | 0 | 0 | 1415632 | AUM60013.1 | K9N7C7 |
| CESIYEFDTYKA | 1 | 0 | 0 | 1415642 | ARU07575.1 | A0A140H1G9 |
| CESTTVLDHTDW | 1 | 0 | 0 | 1415647 | 5GNB_A | A0A140H1H1 |
| CESTTVLDRTDW | 1 | 0 | 0 | 1415648 | BBA20983.1 | A0A140H1H1 |
| CETSNVPPIVSE | 1 | 0 | 0 | 1415651 | ACA49954.1 | H9EJ42 |
| CEVEYGTEDDYN | 1 | 0 | 0 | 1415652 | APO40578.1 | P0C6X7 |
| CEYPQTICHPNL | 1 | 0 | 0 | 1415673 | CAA79896.1 | P36334 |
| CFALRNFDERDC | 1 | 0 | 0 | 1415679 | AIW52766.1 | A0A0P0LKV0 |
| CFANKHADFDSW | 1 | 0 | 0 | 1415682 | AVP78041.1 | P0C6W2 |

| | | | | | | |
|---|---|---|---|---|---|---|
| CFCPRDFNHNMG | 1 | 0 | 0 | 1415698 | AFD98754.1 | H9EJ69 |
| CFDIRDNGSLEK | 1 | 0 | 0 | 1415704 | AXN92247.1 | K9N7C7 |
| CFDIRDNGSLER | 1 | 0 | 0 | 1415705 | AUM60013.1 | K9N7C7 |
| CFDIRDNVSLEK | 1 | 0 | 0 | 1415706 | ANF29193.1 | K9N7C7 |
| CFDNSIQGSYER | 1 | 0 | 0 | 1415711 | AWK59942.1 | H9EJ42 |
| CFEATEATFNYL | 1 | 0 | 0 | 1415721 | AZK15899.1 | K9N638 |
| CFEATTATINYL | 1 | 0 | 0 | 1415724 | AUM60013.1 | K9N7C7 |
| CFEIDGNVMHAN | 1 | 0 | 0 | 1415726 | ARB07436.1 | A0A140H1G9 |
| CFFTCNSNANIS | 1 | 0 | 0 | 1415732 | AKA09626.1 | A0A0P0G321 |
| CFFTCNSNANLS | 1 | 0 | 0 | 1415733 | AIW52843.1 | A0A0P0G321 |
| CFGKNNGIGTCP | 1 | 0 | 0 | 1415741 | QEG03814.1 | P36334 |
| CFGPRDFNHNMG | 1 | 0 | 0 | 1415750 | AIW52805.1 | H9EJ69 |
| CFINSTINTTHV | 1 | 0 | 0 | 1415762 | AIW52843.1 | A0A0P0G321 |
| CFINSTINTTPV | 1 | 0 | 0 | 1415763 | AMM42635.1 | A0A0P0G321 |
| CFKGMCSCGSCC | 1 | 0 | 0 | 1415778 | APO40579.1 | P59594 |
| CFKKDHVVLKIT | 1 | 0 | 0 | 1415790 | QED88038.1 | H9EJ42 |
| CFLPTFNMQTVS | 1 | 0 | 0 | 1415811 | AVV62524.1 | K9N7C7 |
| CFNADGDEAWGS | 1 | 0 | 0 | 1415814 | AUM60013.1 | K9N7C7 |
| CFNAEGDAFWSS | 1 | 0 | 0 | 1415815 | AVN89333.1 | K9N7C7 |
| CFNKRKYFTPVQ | 1 | 0 | 0 | 1415829 | AQN78743.1 | P30215 |
| CFNRDGDEQWGS | 1 | 0 | 0 | 1415833 | AVV62524.1 | K9N7C7 |
| CFNRDGDVQWSS | 1 | 0 | 0 | 1415834 | AVV62535.1 | K9N7C7 |
| CFRIYPQQMFVT | 1 | 0 | 0 | 1415846 | ALA49355.1 | K9N643 |
| CFRTYPQQMFVT | 1 | 0 | 0 | 1415850 | AXN92231.1 | K9N643 |
| CFSANDVEIQSS | 1 | 0 | 0 | 1415851 | AVV62524.1 | K9N7C7 |
| CFSFVKMLFMLS | 1 | 0 | 0 | 1415856 | ALT66869.1 | K9N7C7 |
| CFSTCNSNANIS | 1 | 0 | 0 | 1415872 | AKA09625.1 | A0A0P0G321 |
| CFSTCNSNASIS | 1 | 0 | 0 | 1415873 | AWO68966.1 | A0A0P0G321 |
| CFSTRNSNASIS | 1 | 0 | 0 | 1415875 | AKA09629.1 | A0A0P0G321 |
| CFSTVAVPGGCN | 1 | 0 | 0 | 1415878 | AWO68993.1 | A0A0P0G321 |
| CFSTVEVPGSCN | 1 | 0 | 0 | 1415879 | AWO68966.1 | A0A0P0G321 |
| CFVDDIVKTDGT | 1 | 0 | 0 | 1415895 | AEA10817.1 | P0C6X7 |
| CFVDDLLKTDSI | 1 | 0 | 0 | 1415896 | ABD75559.1 | A0A140H1G9 |
| CFVLSLVCFLGL | 1 | 0 | 0 | 1415912 | AXX83324.1 | P0C6X6 |
| CFVNDLLKTDSV | 1 | 0 | 0 | 1415915 | ANZ78855.1 | P0C6X6 |
| CFVNFNVDWIFF | 1 | 0 | 0 | 1415916 | AWK59934.1 | U3M6Q8 |
| CFVNTTIDNETT | 1 | 0 | 0 | 1415918 | AIW52752.1 | A0A0P0K6L9 |
| CGAERHRQLIEH | 1 | 0 | 0 | 1415941 | AUM60013.1 | K9N7C7 |
| CGAGDQAAKLSP | 1 | 0 | 0 | 1415943 | AVV62524.1 | K9N7C7 |
| CGAVYTANDVDL | 1 | 0 | 0 | 1415947 | AUM60013.1 | K9N7C7 |
| CGCGTSYTANDV | 1 | 0 | 0 | 1415951 | QBM11736.1 | K9N638 |
| CGDCAACKSQLV | 1 | 0 | 0 | 1415954 | CAA83661.1 | P36334 |
| CGDNSAKFWEEA | 1 | 0 | 0 | 1415964 | AXN92247.1 | K9N7C7 |
| CGDQVHEVAAVR | 1 | 0 | 0 | 1415966 | ALT66879.1 | K9N7C7 |
| CGESSWQTGDFL | 1 | 0 | 0 | 1415978 | APO40578.1 | P0C6X7 |
| CGFCCKFDEDDS | 1 | 0 | 0 | 1415980 | AVP78042.1 | Q3LZX1 |
| CGFDGKPMLPKN | 1 | 0 | 0 | 1415983 | AFD98826.1 | H9EJ42 |
| CGFDGKPVLPKN | 1 | 0 | 0 | 1415984 | AWK59942.1 | H9EJ42 |
| CGFVMQFGYIDC | 1 | 0 | 0 | 1415997 | AEN19363.1 | P0C6X6 |
| CGFVMQYGYVDC | 1 | 0 | 0 | 1415998 | ARU07575.1 | A0A140H1G9 |
| CGGCCDDYTGHQ | 1 | 0 | 0 | 1416000 | CAA83660.1 | P36334 |
| CGGCCEDYTGYQ | 1 | 0 | 0 | 1416002 | AIL49528.1 | P36334 |
| CGGCYYVTPGGT | 1 | 0 | 0 | 1416005 | ACR82746.1 | A0A140H1G9 |
| CGGERHQQLVEH | 1 | 0 | 0 | 1416006 | AVN89322.1 | K9N7C7 |
| CGGFCDDYTGYQ | 1 | 0 | 0 | 1416009 | AQN78672.1 | P36334 |
| CGGILAVIDEHV | 1 | 0 | 0 | 1416010 | NP_073549.1 | A0A0P0LKV0 |
| CGGSLYVKPSGT | 1 | 0 | 0 | 1416023 | AVP78041.1 | P0C6W2 |
| CGGVRHRQIVEH | 1 | 0 | 0 | 1416024 | AVV62524.1 | K9N7C7 |
| CGHDDLAQLRCS | 1 | 0 | 0 | 1416026 | AVV62526.1 | K9N5Q8 |
| CGHVYEVSDLIA | 1 | 0 | 0 | 1416032 | AIV41990.1 | P0C6X6 |
| CGHYHTNSYSQN | 1 | 0 | 0 | 1416033 | AST12962.1 | A0A0P0LKV0 |
| CGHYTHITAKET | 1 | 0 | 0 | 1416036 | AEA10817.1 | P0C6X7 |
| CGIKDVVLQGLR | 1 | 0 | 0 | 1416049 | AMQ48992.1 | K9N7C7 |
| CGIKQESRVGVD | 1 | 0 | 0 | 1416050 | ARU07575.1 | A0A140H1G9 |
| CGIVELEKGVLP | 1 | 0 | 0 | 1416053 | P0C6W6.1 | P0C6W2 |
| CGKDGNPIRDYA | 1 | 0 | 0 | 1416055 | AVV62524.1 | K9N7C7 |
| CGLILKFDSGTV | 1 | 0 | 0 | 1416063 | AVN89312.1 | K9N638 |
| CGNCTFGAPDDA | 1 | 0 | 0 | 1416071 | AID50427.1 | K9N7C7 |
| CGNDSWLTGHAI | 1 | 0 | 0 | 1416072 | AHI48785.1 | K9N638 |
| CGNESWLTGNAI | 1 | 0 | 0 | 1416074 | AVN89333.1 | K9N7C7 |
| CGNLCTLDPITL | 1 | 0 | 0 | 1416080 | BBA20976.1 | P36334 |
| CGNYRVTKGKPV | 1 | 0 | 0 | 1416088 | ARO76381.1 | P0C6U8 |
| CGPKKSTELVKN | 1 | 0 | 0 | 1416091 | APO40579.1 | P59594 |
| CGQKSTTLIGVE | 1 | 0 | 0 | 1416099 | APO40578.1 | P0C6X7 |
| CGREATQYLVQQ | 1 | 0 | 0 | 1416106 | ARO76381.1 | P0C6U8 |
| CGSACFSKCHNC | 1 | 0 | 0 | 1416108 | BBA20983.1 | A0A140H1H1 |
| CGSDGKPIADYA | 1 | 0 | 0 | 1416117 | AUM60013.1 | K9N7C7 |
| CGSENWSVGAWD | 1 | 0 | 0 | 1416120 | AGT51379.1 | H9EJ42 |
| CGSVGYTKDGSV | 1 | 0 | 0 | 1416133 | ATG84865.1 | K9N638 |
| CGSVGYTKEGNV | 1 | 0 | 0 | 1416134 | AVV62535.1 | K9N7C7 |
| CGTGALVSKLSP | 1 | 0 | 0 | 1416141 | AUM60023.1 | K9N7C7 |
| CGTQLNHSSCSC | 1 | 0 | 0 | 1416157 | ARB07438.1 | A0A140H1H1 |
| CGTQLNHTSCSC | 1 | 0 | 0 | 1416158 | BBA20986.1 | A0A140H1H1 |
| CGTSYTANDVEV | 1 | 0 | 0 | 1416168 | QBM11736.1 | K9N638 |
| CGVDSMNLIANI | 1 | 0 | 0 | 1416179 | ARO76381.1 | P0C6U8 |
| CGVHGNGSMIED | 1 | 0 | 0 | 1416181 | AGV08556.1 | K9N7C7 |
| CGVLDGSYNASC | 1 | 0 | 0 | 1416187 | BBA20983.1 | A0A140H1H1 |
| CGVLDGSYNVSC | 1 | 0 | 0 | 1416188 | 5GNB_A | A0A140H1H1 |
| CGYDDLSQLRCS | 1 | 0 | 0 | 1416196 | AUM60014.1 | K9N5Q8 |
| CHDPTVLPGAFA | 1 | 0 | 0 | 1416223 | AXN92247.1 | K9N7C7 |
| CHGKNNGIGTCP | 1 | 0 | 0 | 1416227 | ARB66068.1 | P36334 |

| | | | | | | |
|---|---|---|---|---|---|---|
| CHLAKAINDFSN | 1 | 0 | 0 | 1416233 | AVP78030.1 | P0C6W2 |
| CHNAAQCDCLCT | 1 | 0 | 0 | 1416239 | CAA83660.1 | P36334 |
| CHNDILTANDPS | 1 | 0 | 0 | 1416245 | AUM60013.1 | K9N7C7 |
| CHNDILTATDPA | 1 | 0 | 0 | 1416246 | AVV62524.1 | K9N7C7 |
| CHNGKAYFPREG | 1 | 0 | 0 | 1416248 | APO40579.1 | P59594 |
| CHNLCNPDPITF | 1 | 0 | 0 | 1416251 | AVQ05264.1 | P36334 |
| CHPKLGNHFKEL | 1 | 0 | 0 | 1416256 | AEN19366.1 | P36334 |
| CHPKLGNHFKES | 1 | 0 | 0 | 1416257 | AIL49486.1 | P36334 |
| CHPNLGNHRKEL | 1 | 0 | 0 | 1416259 | CAA79896.1 | P36334 |
| CHPNLGNRRVEL | 1 | 0 | 0 | 1416261 | CAA83660.1 | P36334 |
| CHQVSPSALASG | 1 | 0 | 0 | 1416263 | AUM60014.1 | K9N5Q8 |
| CHSNILSLNNPI | 1 | 0 | 0 | 1416267 | ABD75616.1 | A0A140H1H0 |
| CHSQTSLRCGTC | 1 | 0 | 0 | 1416269 | AXN92247.1 | K9N7C7 |
| CHTGTAFNGDFY | 1 | 0 | 0 | 1416272 | AIV41912.1 | P0C6X6 |
| CHTGTDFNGDFY | 1 | 0 | 0 | 1416273 | AEN19363.1 | P0C6X6 |
| CHTGTDFTGNFY | 1 | 0 | 0 | 1416274 | ARU07575.1 | A0A140H1G9 |
| CHYLVLPSGNYL | 1 | 0 | 0 | 1416279 | AEN19365.1 | P30215 |
| CIADYTVFYNST | 1 | 0 | 0 | 1416283 | AVP78031.1 | Q3LZX1 |
| CIAGNRGIAPKS | 1 | 0 | 0 | 1416285 | AEN19366.1 | P36334 |
| CIALQYSKPHFI | 1 | 0 | 0 | 1416288 | NP_073549.1 | A0A0P0LKV0 |
| CIAPVNGYFIKT | 1 | 0 | 0 | 1416292 | AXN92228.1 | K9N5Q8 |
| CICLVASIPLCI | 1 | 0 | 0 | 1416300 | ASU91085.1 | K9N7C7 |
| CICTVVQRCASN | 1 | 0 | 0 | 1416303 | NP_849176.1 | Q7TFA0 |
| CIDCSARHINAH | 1 | 0 | 0 | 1416306 | AVP78041.1 | P0C6W2 |
| CIDYVTIAGTKL | 1 | 0 | 0 | 1416323 | ARO76381.1 | P0C6U8 |
| CIERLKLFAAET | 1 | 0 | 0 | 1416326 | AGT51787.1 | P0C6X6 |
| CIESTDSIVFTY | 1 | 0 | 0 | 1416327 | ANI69893.1 | K9N7D2 |
| CIFNGKSFSNTK | 1 | 0 | 0 | 1416336 | ARK08650.1 | P30215 |
| CIGKGSLTASSF | 1 | 0 | 0 | 1416341 | AUM60027.1 | K9N643 |
| CIGRGNIGFTLE | 1 | 0 | 0 | 1416343 | AHA80218.1 | P36334 |
| CIGRRNIGFTLE | 1 | 0 | 0 | 1416345 | AHA80213.1 | P36334 |
| CIHKECSIQECC | 1 | 0 | 0 | 1416348 | Q3I5I8.1 | Q3LZX5 |
| CIKDADYNAKDE | 1 | 0 | 0 | 1416364 | AST12962.1 | A0A0P0LKV0 |
| CIKDLLARAGKS | 1 | 0 | 0 | 1416368 | P0C6W6.1 | P0C6W2 |
| CIKIAQADTAGL | 1 | 0 | 0 | 1416371 | ASU89922.1 | K9N796 |
| CIKPVPEMKILQ | 1 | 0 | 0 | 1416374 | APO40578.1 | P0C6X7 |
| CILEPRPGNHCP | 1 | 0 | 0 | 1416393 | QBF80608.1 | K9N5Q8 |
| CILEPRSGNYCP | 1 | 0 | 0 | 1416395 | AVN89387.1 | K9N5Q8 |
| CILSVNSSNCNN | 1 | 0 | 0 | 1416409 | AIW52766.1 | A0A0P0LKV0 |
| CINCDTFCAGST | 1 | 0 | 0 | 1416416 | AVP78041.1 | P0C6W2 |
| CIPLSVIPLCAS | 1 | 0 | 0 | 1416425 | AXN92247.1 | K9N7C7 |
| CIPLSVVPLCAS | 1 | 0 | 0 | 1416426 | AUM60013.1 | K9N7C7 |
| CIPTVGYSAALA | 1 | 0 | 0 | 1416427 | AVV62530.1 | K9N7D2 |
| CIPYNSITDTIV | 1 | 0 | 0 | 1416428 | ATO98146.1 | Q3LZX0 |
| CIRGAIGDVGLC | 1 | 0 | 0 | 1416437 | ARB07424.1 | A0A0P0ILY9 |
| CIRRAKDGFSPE | 1 | 0 | 0 | 1416440 | QEG03767.1 | P36334 |
| CIRRPKDGFSPE | 1 | 0 | 0 | 1416442 | QEG03814.1 | P36334 |
| CIRSAKIDTSGI | 1 | 0 | 0 | 1416444 | AVV62527.1 | K9N796 |
| CISRLKDGFSLE | 1 | 0 | 0 | 1416454 | AVR40342.1 | P36334 |
| CITIIKLTKLCF | 1 | 0 | 0 | 1416470 | AGW80935.1 | A0A0P0KH29 |
| CITPNVCFVKGD | 1 | 0 | 0 | 1416473 | AEN19363.1 | P0C6X6 |
| CITSREVVVTNLNKSA | 1 | 0 | 0 | 1416476 | ALK43115.1 | A0A0P0LKV0 |
| CIVFDVKGVEEL | 1 | 0 | 0 | 1416485 | AOL02451.1 | Q80872 |
| CIVNLSFTEQLG | 1 | 0 | 0 | 1416493 | ACY56724.1 | A0A0P0G321 |
| CIWNAAVYMKLS | 1 | 0 | 0 | 1416504 | QFQ59585.1 | K9N7C7 |
| CIWNCSAYMQLS | 1 | 0 | 0 | 1416505 | AUM60013.1 | K9N7C7 |
| CKAGQVQTFYPR | 1 | 0 | 0 | 1416511 | AIL23945.1 | K9N7C7 |
| CKAISSNTDNET | 1 | 0 | 0 | 1416513 | Q14EB0.1 | A0A140H1H1 |
| CKCGNLEQNHIL | 1 | 0 | 0 | 1416525 | ARU07575.1 | A0A140H1G9 |
| CKCGVKQEQCTG | 1 | 0 | 0 | 1416526 | ARE30011.1 | P0C6X6 |
| CKCGVKQEQRIG | 1 | 0 | 0 | 1416527 | AVQ05261.1 | P0C6X6 |
| CKDGKVQTFYPR | 1 | 0 | 0 | 1416530 | ATQ39389.1 | K9N7C7 |
| CKEPMYAISNEQ | 1 | 0 | 0 | 1416545 | ATO98193.1 | Q3LZX1 |
| CKFAVCGDGFVP | 1 | 0 | 0 | 1416548 | ARU07575.1 | A0A140H1G9 |
| CKFDEDHSEPVS | 1 | 0 | 0 | 1416550 | APO40579.1 | P59594 |
| CKFGNTTFDFLS | 1 | 0 | 0 | 1416555 | AWO69005.1 | A0A0P0G321 |
| CKFGTNTTFDFL | 1 | 0 | 0 | 1416556 | AVL25589.1 | A0A0P0G321 |
| CKFKGKFVQIPA | 1 | 0 | 0 | 1416557 | AVV62524.1 | K9N7C7 |
| CKFSSNTTFDFL | 1 | 0 | 0 | 1416564 | ACY56730.1 | A0A0P0G321 |
| CKGFSSNVSSDV | 1 | 0 | 0 | 1416571 | ABB90509.1 | A0A0P0K6L9 |
| CKGFYSNASSDV | 1 | 0 | 0 | 1416572 | AIW52757.1 | A0A0P0K6L9 |
| CKGFYSNSSSDV | 1 | 0 | 0 | 1416573 | AIW52747.1 | A0A0P0K6L9 |
| CKHTTAQVVSHL | 1 | 0 | 0 | 1416590 | AVV62535.1 | K9N7C7 |
| CKLDFLDCTGQA | 1 | 0 | 0 | 1416604 | AVV62539.1 | K9N4V0 |
| CKLNGSCPDKNN | 1 | 0 | 0 | 1416613 | AIX10751.1 | P36334 |
| CKLNGSCPGKNN | 1 | 0 | 0 | 1416614 | AEN19366.1 | P36334 |
| CKMCLFRYQEFN | 1 | 0 | 0 | 1416627 | QED88024.1 | H9EJ42 |
| CKNCGQKSTTLI | 1 | 0 | 0 | 1416630 | APO40578.1 | P0C6X7 |
| CKPDQQFHNFSY | 1 | 0 | 0 | 1416634 | AVV62540.1 | K9N643 |
| CKPHQVNLSLNG | 1 | 0 | 0 | 1416637 | AWO69005.1 | A0A0P0G321 |
| CKQYICNGFQKC | 1 | 0 | 0 | 1416640 | AKM76239.1 | K9N5Q8 |
| CKSALYVLPSET | 1 | 0 | 0 | 1416652 | ATO98143.1 | P0C6W2 |
| CKSGSKFKNLVL | 1 | 0 | 0 | 1416658 | ARU07609.1 | P30215 |
| CKSGSLVLNNPA | 1 | 0 | 0 | 1416660 | AEN19365.1 | P30215 |
| CKSGSLVLNNPV | 1 | 0 | 0 | 1416662 | AVQ05263.1 | P30215 |
| CKSHGGKDSLTM | 1 | 0 | 0 | 1416664 | ABB90496.1 | A0A0N7J131 |
| CKSIVKETALIT | 1 | 0 | 0 | 1416670 | ARU07575.1 | A0A140H1G9 |
| CKSKGSFRNESW | 1 | 0 | 0 | 1416671 | AMN88694.1 | A0A140H1H1 |
| CKSKNPLIYDAS | 1 | 0 | 0 | 1416672 | APO40580.1 | P59632 |
| CKSNDSCPGKNN | 1 | 0 | 0 | 1416677 | AIV41891.1 | P36334 |
| CKSNGSCPGKNN | 1 | 0 | 0 | 1416678 | AEN19358.1 | P36334 |

| | | | | | | |
|---|---|---|---|---|---|---|
| CKSNSSLCVGSG | 1 | 0 | 0 | 1416679 | ARB66062.1 | P36334 |
| CKSQLGEYGSFC | 1 | 0 | 0 | 1416680 | AHF54658.1 | P36334 |
| CKSQNPLLYDAN | 1 | 0 | 0 | 1416683 | AAP30031.1 | P59632 |
| CKSRNPLLYDAN | 1 | 0 | 0 | 1416684 | ARO76383.1 | P59632 |
| CKSTTGIPEHNF | 1 | 0 | 0 | 1416685 | AVV62524.1 | K9N7C7 |
| CKSTTGIPEHNL | 1 | 0 | 0 | 1416686 | AUM60013.1 | K9N7C7 |
| CKSVFYILPSIL | 1 | 0 | 0 | 1416687 | AVP78041.1 | P0C6W2 |
| CKTAVTTGDVKI | 1 | 0 | 0 | 1416691 | NP_073549.1 | A0A0P0LKV0 |
| CKTQFIAPPTGV | 1 | 0 | 0 | 1416699 | AHA80218.1 | P36334 |
| CKVTDILYPGQK | 1 | 0 | 0 | 1416713 | AVV62524.1 | K9N7C7 |
| CKVTDVLFSGQK | 1 | 0 | 0 | 1416716 | AQZ41321.1 | K9N7C7 |
| CKVTDVLYPQQK | 1 | 0 | 0 | 1416717 | AUM60013.1 | K9N7C7 |
| CKYTDIDVSPAL | 1 | 0 | 0 | 1416729 | AVV62524.1 | K9N7C7 |
| CLAAEIHRDCDF | 1 | 0 | 0 | 1416733 | AVV62535.1 | K9N7C7 |
| CLAAETHRDCDL | 1 | 0 | 0 | 1416735 | AVV62524.1 | K9N7C7 |
| CLARSACVYYSQ | 1 | 0 | 0 | 1416746 | AZK15899.1 | K9N638 |
| CLASIAFNHPIQ | 1 | 0 | 0 | 1416748 | AVN89313.1 | K9N5Q8 |
| CLASIEEVCDDY | 1 | 0 | 0 | 1416749 | AGT51687.1 | P0C6X6 |
| CLATPSGGVSTN | 1 | 0 | 0 | 1416750 | AVV62529.1 | K9N643 |
| CLDVVRKELEEG | 1 | 0 | 0 | 1416773 | AWW13564.1 | Q80872 |
| CLEAGINYVKSP | 1 | 0 | 0 | 1416776 | ARO76381.1 | P0C6U8 |
| CLEASFNYLKSP | 1 | 0 | 0 | 1416777 | QYC64967.1 | P0DTD1 |
| CLENHVMVDCRR | 1 | 0 | 0 | 1416778 | ARK08667.1 | P0C6X6 |
| CLEVQVDPNAFI | 1 | 0 | 0 | 1416781 | AUM60014.1 | K9N5Q8 |
| CLFKKNFTYNVS | 1 | 0 | 0 | 1416786 | BBA20983.1 | A0A140H1H1 |
| CLFKKNFTYNVY | 1 | 0 | 0 | 1416787 | ARB07438.1 | A0A140H1H1 |
| CLFKSNFNQYYY | 1 | 0 | 0 | 1416788 | AZS52617.1 | A0A140H1H0 |
| CLFSYQEFNDLD | 1 | 0 | 0 | 1416796 | AWK59942.1 | H9EJ42 |
| CLFSYQEFNDLV | 1 | 0 | 0 | 1416797 | AMM42571.1 | H9EJ42 |
| CLGDISARDLIC | 1 | 0 | 0 | 1416815 | AVP78042.1 | Q3LZX1 |
| CLGGLDSLDSYP | 1 | 0 | 0 | 1416823 | ARO76381.1 | P0C6U8 |
| CLGSECNKVSSR | 1 | 0 | 0 | 1416835 | CAA83661.1 | P36334 |
| CLGSECSKASSR | 1 | 0 | 0 | 1416836 | CAA79896.1 | P36334 |
| CLGSLIYSTAAL | 1 | 0 | 0 | 1416839 | AVP78041.1 | P0C6W2 |
| CLGSLIYVTAAL | 1 | 0 | 0 | 1416841 | P0C6W6.1 | P0C6W2 |
| CLGVGDSAVTYS | 1 | 0 | 0 | 1416842 | AIV41816.1 | P0C6X6 |
| CLILATVSHNLT | 1 | 0 | 0 | 1416859 | AJG44102.1 | K9N5Q8 |
| CLKGVCSCGSCC | 1 | 0 | 0 | 1416881 | ARO76382.1 | P59594 |
| CLKSIAATRGVP | 1 | 0 | 0 | 1416890 | AEN19363.1 | P0C6X6 |
| CLLANKDQSLVL | 1 | 0 | 0 | 1416893 | AVV62535.1 | K9N7C7 |
| CLLSGHNLANKC | 1 | 0 | 0 | 1416920 | ARO76381.1 | P0C6U8 |
| CLMFYNFVLSAV | 1 | 0 | 0 | 1416932 | AVV62530.1 | K9N7D2 |
| CLMPTYTVHKSD | 1 | 0 | 0 | 1416938 | AEN19363.1 | P0C6X6 |
| CLPDPISTYSPN | 1 | 0 | 0 | 1416953 | BBA20986.1 | A0A140H1H1 |
| CLQAGKQTELAV | 1 | 0 | 0 | 1416959 | AAW33912.1 | H9EJ42 |
| CLQGDKCNIFSN | 1 | 0 | 0 | 1416962 | BBA84101.1 | P36334 |
| CLSDLDSLDFYP | 1 | 0 | 0 | 1416980 | APO40578.1 | P0C6X7 |
| CLSGRKIYEVAR | 1 | 0 | 0 | 1416985 | AWW13571.1 | P0C6X6 |
| CLSNNPCNVCQY | 1 | 0 | 0 | 1416992 | ANI69899.1 | K9N7C7 |
| CLSNVSTSHHCI | 1 | 0 | 0 | 1416994 | QED88040.1 | A0A0P0G321 |
| CLSSMNIAANSN | 1 | 0 | 0 | 1416996 | AGT51337.1 | H9EJ42 |
| CLSSNMNEPDYS | 1 | 0 | 0 | 1416998 | ARB07436.1 | A0A140H1G9 |
| CLSTGCCGCCNC | 1 | 0 | 0 | 1416999 | AIW52843.1 | A0A0P0G321 |
| CLTFVVKHKVLF | 1 | 0 | 0 | 1417007 | NP_073549.1 | A0A0P0LKV0 |
| CLTSSMRGCCDC | 1 | 0 | 0 | 1417016 | AIW52843.1 | A0A0P0G321 |
| CLTYANIVHEPT | 1 | 0 | 0 | 1417017 | ALA49956.1 | K9N7C7 |
| CLTYANIVYEPS | 1 | 0 | 0 | 1417018 | AVV62524.1 | K9N7C7 |
| CLVASDGKPGGD | 1 | 0 | 0 | 1417020 | AUM60013.1 | K9N7C7 |
| CLVSSNGQPGGD | 1 | 0 | 0 | 1417032 | QAT98908.1 | K9N7C7 |
| CLWFLRRFYNHA | 1 | 0 | 0 | 1417034 | AVV62524.1 | K9N7C7 |
| CLYCRAHIEHPD | 1 | 0 | 0 | 1417052 | AXN92247.1 | K9N7C7 |
| CLYKRNFTYDVN | 1 | 0 | 0 | 1417058 | CAA79896.1 | P36334 |
| CLYKRNFTYDVS | 1 | 0 | 0 | 1417059 | AKA09662.1 | P36334 |
| CLYLTSSGFDKS | 1 | 0 | 0 | 1417061 | AUM60024.1 | K9N5Q8 |
| CLYRNRDVDQEF | 1 | 0 | 0 | 1417066 | AVP77994.1 | P0C6W2 |
| CMHKECSIQECC | 1 | 0 | 0 | 1417090 | ATO98227.1 | Q3LZX5 |
| CMKQGPASARDL | 1 | 0 | 0 | 1417095 | QEJ82215.1 | K9N5Q8 |
| CMKQGPPSARDL | 1 | 0 | 0 | 1417096 | AVV62537.1 | K9N5Q8 |
| CMLRAFIKTAQA | 1 | 0 | 0 | 1417103 | AXN92261.1 | K9N796 |
| CMNQGPPSARDL | 1 | 0 | 0 | 1417111 | AVV62526.1 | K9N5Q8 |
| CMQQGPASAREL | 1 | 0 | 0 | 1417114 | ARA67740.1 | K9N5Q8 |
| CMQQGPASARNL | 1 | 0 | 0 | 1417115 | QFS19619.1 | K9N5Q8 |
| CMQQGPQSARDL | 1 | 0 | 0 | 1417117 | AUM60014.1 | K9N5Q8 |
| CMRAGSKVINHN | 1 | 0 | 0 | 1417119 | QDH43737.1 | H9EJ42 |
| CMVGPDCAYFNG | 1 | 0 | 0 | 1417133 | AUM60023.1 | K9N7C7 |
| CMYTDLNFIDKL | 1 | 0 | 0 | 1417139 | ARK08667.1 | P0C6X6 |
| CNAASVNPASMR | 1 | 0 | 0 | 1417141 | AUM60013.1 | K9N7C7 |
| CNAISPKTDNET | 1 | 0 | 0 | 1417147 | BBA20986.1 | A0A140H1H1 |
| CNAVEQQITVEV | 1 | 0 | 0 | 1417161 | AVV62524.1 | K9N7C7 |
| CNCDSLRQAALP | 1 | 0 | 0 | 1417165 | AXN92247.1 | K9N7C7 |
| CNCGSENWSVGA | 1 | 0 | 0 | 1417169 | AGT51379.1 | H9EJ42 |
| CNCLCTPDPITS | 1 | 0 | 0 | 1417172 | CAA83661.1 | P36334 |
| CNCLTSSMRGCC | 1 | 0 | 0 | 1417174 | AIW52843.1 | A0A0P0G321 |
| CNCTFGAPDDAS | 1 | 0 | 0 | 1417175 | ANI69888.1 | K9N638 |
| CNESIACQFCLA | 1 | 0 | 0 | 1417185 | AXX83294.1 | P0C6X6 |
| CNFELCDNPFIA | 1 | 0 | 0 | 1417190 | ABD73001.1 | P59594 |
| CNFSNLDKLKFK | 1 | 0 | 0 | 1417197 | AVV62524.1 | K9N7C7 |
| CNFYFCQDPMLA | 1 | 0 | 0 | 1417199 | APO40579.1 | P59594 |
| CNGCVGHSENVF | 1 | 0 | 0 | 1417201 | AIW52757.1 | A0A0P0K6L9 |
| CNGFEKCEQLLR | 1 | 0 | 0 | 1417205 | AVV62526.1 | K9N5Q8 |
| CNGISPDAIARG | 1 | 0 | 0 | 1417212 | AVV62526.1 | K9N5Q8 |

| | | | | | | |
|---|---|---|---|---|---|---|
| CNGNKIEDFSIR | 1 | 0 | 0 | 1417215 | AIV41978.1 | P0C6X6 |
| CNGNKIEDLSIR | 1 | 0 | 0 | 1417216 | AEN19363.1 | P0C6X6 |
| CNGNKIEDLSMR | 1 | 0 | 0 | 1417217 | AJE26223.1 | P0C6X6 |
| CNGNKIGDLSIR | 1 | 0 | 0 | 1417218 | AJE26218.1 | P0C6X6 |
| CNGNKSDDLSIS | 1 | 0 | 0 | 1417219 | AKL87224.1 | P0C6X6 |
| CNGSIACQFCLA | 1 | 0 | 0 | 1417225 | AEN19363.1 | P0C6X6 |
| CNGTDIDYCVRA | 1 | 0 | 0 | 1417228 | AIW52776.1 | A0A0P0LKV0 |
| CNGTDIDYCVRV | 1 | 0 | 0 | 1417229 | ARX11394.1 | A0A0P0LKV0 |
| CNGYQHHSVADV | 1 | 0 | 0 | 1417235 | AIW52843.1 | A0A0P0G321 |
| CNGYQHNSFADV | 1 | 0 | 0 | 1417236 | ALJ53431.1 | A0A0P0G321 |
| CNGYQHNSVVDV | 1 | 0 | 0 | 1417238 | YP_003767.1 | A0A0P0G321 |
| CNHCGESSWQTG | 1 | 0 | 0 | 1417242 | APO40578.1 | P0C6X7 |
| CNHIGNIKPHNY | 1 | 0 | 0 | 1417243 | AIL49382.1 | P30215 |
| CNHINNLKVKNY | 1 | 0 | 0 | 1417247 | ABD75544.1 | A0A140H1H0 |
| CNIEAWLNDKAV | 1 | 0 | 0 | 1417254 | AIV41909.1 | P36334 |
| CNIEAWLNDKLV | 1 | 0 | 0 | 1417255 | ATN39872.1 | P36334 |
| CNIEAWLNGKSV | 1 | 0 | 0 | 1417257 | AIL49492.1 | P36334 |
| CNIESWLNDKSV | 1 | 0 | 0 | 1417258 | AIX10749.1 | P36334 |
| CNIGAWLNDKSV | 1 | 0 | 0 | 1417264 | AHA80222.1 | P36334 |
| CNNIDVAKIYGM | 1 | 0 | 0 | 1417300 | ABD43176.1 | P36334 |
| CNNLVQTYLKSD | 1 | 0 | 0 | 1417304 | AIV42002.1 | P0C6X6 |
| CNNSDYASKGYI | 1 | 0 | 0 | 1417306 | AIX10745.1 | P0C6X6 |
| CNPDPIRFTGPY | 1 | 0 | 0 | 1417309 | AGT51422.1 | P36334 |
| CNRAGSNDIWMQ | 1 | 0 | 0 | 1417319 | AGT51729.1 | P30215 |
| CNRCCDKYDDYD | 1 | 0 | 0 | 1417321 | AUM60024.1 | K9N5Q8 |
| CNRCCDNYDDYD | 1 | 0 | 0 | 1417323 | AUM60014.1 | K9N5Q8 |
| CNRRAGAYSSPL | 1 | 0 | 0 | 1417328 | AUM60013.1 | K9N7C7 |
| CNSFFKDLTANM | 1 | 0 | 0 | 1417335 | ARK08626.1 | H9EJ42 |
| CNSFFKELTAIM | 1 | 0 | 0 | 1417336 | QED88024.1 | H9EJ42 |
| CNSKVKNGFTLE | 1 | 0 | 0 | 1417343 | CAA79896.1 | P36334 |
| CNSMHLSTYMNC | 1 | 0 | 0 | 1417345 | APO40578.1 | P0C6X7 |
| CNSNANISMLQL | 1 | 0 | 0 | 1417346 | AKA09626.1 | A0A0P0G321 |
| CNSNANISTLQL | 1 | 0 | 0 | 1417347 | AKA09623.1 | A0A0P0G321 |
| CNSVKDTLTYYA | 1 | 0 | 0 | 1417357 | AXN92247.1 | K9N7C7 |
| CNTDTPPRCIAP | 1 | 0 | 0 | 1417359 | AVV62537.1 | K9N5Q8 |
| CNTGSCTVNKHV | 1 | 0 | 0 | 1417361 | AGT51697.1 | P0C6X6 |
| CNTVKDILSKEG | 1 | 0 | 0 | 1417371 | AUM60013.1 | K9N7C7 |
| CNVIADFKSFTL | 1 | 0 | 0 | 1417382 | AXN73435.1 | K9N7C7 |
| CNVIADPKSFTL | 1 | 0 | 0 | 1417383 | AQZ41321.1 | K9N7C7 |
| CNVPRYPNNAIV | 1 | 0 | 0 | 1417391 | AVV62524.1 | K9N7C7 |
| CNWFIQSDKCSI | 1 | 0 | 0 | 1417407 | AWW13571.1 | P0C6X6 |
| CNWFIQSDKCSV | 1 | 0 | 0 | 1417408 | AEN19363.1 | P0C6X6 |
| CNWFVQSDKCSV | 1 | 0 | 0 | 1417410 | AGT51787.1 | P0C6X6 |
| CNYVFNNSLTRQ | 1 | 0 | 0 | 1417422 | CAA79896.1 | P36334 |
| CPADNSYTSFAT | 1 | 0 | 0 | 1417430 | AID55096.1 | K9N5Q8 |
| CPADQLSDPNYD | 1 | 0 | 0 | 1417431 | AXN92247.1 | K9N7C7 |
| CPAEIVDTVSTL | 1 | 0 | 0 | 1417433 | APO40578.1 | P0C6X7 |
| CPAGNSHTSFAT | 1 | 0 | 0 | 1417435 | AHX71946.1 | K9N5Q8 |
| CPAGNSYTSFAT | 1 | 0 | 0 | 1417437 | AXN92228.1 | K9N5Q8 |
| CPAGTNSLTCGN | 1 | 0 | 0 | 1417444 | BBA20976.1 | P36334 |
| CPAGTNYLNCDN | 1 | 0 | 0 | 1417445 | AIL49520.1 | P36334 |
| CPCADPSVVNSC | 1 | 0 | 0 | 1417451 | BBA20986.1 | A0A140H1H1 |
| CPCADRSVVNSC | 1 | 0 | 0 | 1417452 | ABD75545.1 | A0A140H1H1 |
| CPCSSCFGKNNG | 1 | 0 | 0 | 1417466 | QEG03814.1 | P36334 |
| CPCSSCHGKNNG | 1 | 0 | 0 | 1417467 | ARB66068.1 | P36334 |
| CPCSSCPGKNNG | 1 | 0 | 0 | 1417468 | AWW13566.1 | P36334 |
| CPCSSCSCPGKN | 1 | 0 | 0 | 1417470 | AVR40344.1 | P36334 |
| CPCSSCSGKNNG | 1 | 0 | 0 | 1417471 | ANZ78838.1 | P36334 |
| CPDKNNGIGICP | 1 | 0 | 0 | 1417475 | AIX10751.1 | P36334 |
| CPDKNNGIGTCP | 1 | 0 | 0 | 1417476 | AIX10750.1 | P36334 |
| CPEWMDEFEADP | 1 | 0 | 0 | 1417481 | ATG84853.1 | K9N7C7 |
| CPEWMDYFEADP | 1 | 0 | 0 | 1417482 | ASY99841.1 | K9N638 |
| CPFAKGDSCSSS | 1 | 0 | 0 | 1417484 | AUM60013.1 | K9N7C7 |
| CPFAKSDGCSCS | 1 | 0 | 0 | 1417485 | AVV62524.1 | K9N7C7 |
| CPFVKGDSCSSN | 1 | 0 | 0 | 1417496 | AKN24784.1 | K9N7C7 |
| CPGKNNGIGTCP | 1 | 0 | 0 | 1417497 | AEN19366.1 | P36334 |
| CPGKNNGIGTRP | 1 | 0 | 0 | 1417498 | ARB66066.1 | P36334 |
| CPGKNNGISTCP | 1 | 0 | 0 | 1417499 | BBA84094.1 | P36334 |
| CPGKNNGLGTCP | 1 | 0 | 0 | 1417500 | AIL49503.1 | P36334 |
| CPGNSNYNTFAV | 1 | 0 | 0 | 1417502 | AVV62537.1 | K9N5Q8 |
| CPKFEFANDTKI | 1 | 0 | 0 | 1417516 | AVN89387.1 | K9N5Q8 |
| CPLDPKLKGSFN | 1 | 0 | 0 | 1417526 | AWW13511.1 | P36334 |
| CPLDPRLKGSFN | 1 | 0 | 0 | 1417527 | BAM16463.1 | P36334 |
| CPLDSRTGSFNN | 1 | 0 | 0 | 1417529 | AIX10756.1 | P36334 |
| CPMMDLGNSSTI | 1 | 0 | 0 | 1417542 | AVV62526.1 | K9N5Q8 |
| CPMQALRNDTSV | 1 | 0 | 0 | 1417544 | AUM60014.1 | K9N5Q8 |
| CPNDNAYTSYGV | 1 | 0 | 0 | 1417545 | AUM60014.1 | K9N5Q8 |
| CPNDNAYVSHAI | 1 | 0 | 0 | 1417546 | AVV62526.1 | K9N5Q8 |
| CPNFVFPLNSII | 1 | 0 | 0 | 1417547 | QYC64967.1 | P0DTD1 |
| CPPGMGIPYCTS | 1 | 0 | 0 | 1417550 | AIW52776.1 | A0A0P0LKV0 |
| CPQSKSLVGIGE | 1 | 0 | 0 | 1417555 | AWW13566.1 | P36334 |
| CPRHVICPADQL | 1 | 0 | 0 | 1417561 | QGV13494.1 | K9N7C7 |
| CPRHVMCPADQL | 1 | 0 | 0 | 1417566 | AXN92247.1 | K9N7C7 |
| CPRHVMVPADQL | 1 | 0 | 0 | 1417567 | AUM60013.1 | K9N7C7 |
| CPSATYEGNSPF | 1 | 0 | 0 | 1417568 | ABI96964.1 | P59635 |
| CPSKNNGIGTCP | 1 | 0 | 0 | 1417572 | QDH43719.1 | P36334 |
| CPTAADINIHDL | 1 | 0 | 0 | 1417575 | AGT51530.1 | P30215 |
| CPTAADINIPHL | 1 | 0 | 0 | 1417577 | AXX83356.1 | P30215 |
| CPTAADINNPDL | 1 | 0 | 0 | 1417578 | P30215.1 | P30215 |
| CPTAADINTPDV | 1 | 0 | 0 | 1417579 | AAA73152.1 | P30215 |
| CPTGTKYRHCDL | 1 | 0 | 0 | 1417583 | AIG96691.1 | A0A140H1H1 |

| | | | | | | |
|---|---|---|---|---|---|---|
| CPWLLRKFYQHV | 1 | 0 | 0 | 1417594 | AMQ49058.1 | K9N638 |
| CQAVSANVCALI | 1 | 0 | 0 | 1417597 | AIL49553.1 | P0C6X6 |
| CQAVTANVCSPM | 1 | 0 | 0 | 1417599 | ACR82761.1 | A0A140H1G9 |
| CQAVTANVRSLT | 1 | 0 | 0 | 1417601 | ADO24641.1 | A0A140H1G9 |
| CQDPMLAVANGS | 1 | 0 | 0 | 1417614 | APO40579.1 | P59594 |
| CQFKDKNLQGLW | 1 | 0 | 0 | 1417618 | AGT51787.1 | P0C6X6 |
| CQFNMCEYPHTI | 1 | 0 | 0 | 1417619 | ANZ78837.1 | P36334 |
| CQITSVVGTKAL | 1 | 0 | 0 | 1417626 | ARU07575.1 | A0A140H1G9 |
| CQKAVSAFITTC | 1 | 0 | 0 | 1417629 | AUM60013.1 | K9N7C7 |
| CQLANYKQSIPM | 1 | 0 | 0 | 1417630 | AVV62524.1 | K9N7C7 |
| CQLYYNLPSANV | 1 | 0 | 0 | 1417637 | BBO58721.1 | P36334 |
| CQNLDCNCLGFY | 1 | 0 | 0 | 1417644 | AWW13564.1 | Q80872 |
| CQPVSEYHITLA | 1 | 0 | 0 | 1417666 | AHE78102.1 | K9N643 |
| CQTADTAHANNL | 1 | 0 | 0 | 1417674 | AVV62524.1 | K9N7C7 |
| CQTTNGMNTSQS | 1 | 0 | 0 | 1417681 | ABB90520.1 | A0A0P0K6L9 |
| CQTTNGTNTIHS | 1 | 0 | 0 | 1417682 | AIW52741.1 | A0A0P0K6L9 |
| CQTTNGTNTSNS | 1 | 0 | 0 | 1417684 | AST12964.1 | A0A0P0K6L9 |
| CQVHGNGSMIED | 1 | 0 | 0 | 1417687 | AGV08533.1 | K9N7C7 |
| CQYNMCDYPHTI | 1 | 0 | 0 | 1417691 | ANZ78839.1 | P36334 |
| CQYNMCEHPHTI | 1 | 0 | 0 | 1417692 | BAM16463.1 | P36334 |
| CQYNMCEHPHTS | 1 | 0 | 0 | 1417693 | AWW13511.1 | P36334 |
| CQYNMCEYPHTS | 1 | 0 | 0 | 1417695 | AKA09651.1 | P36334 |
| CQYNMCEYPHTT | 1 | 0 | 0 | 1417696 | BBA84088.1 | P36334 |
| CQYNMCEYPQTI | 1 | 0 | 0 | 1417697 | CAA79896.1 | P36334 |
| CQYNMCEYPYTI | 1 | 0 | 0 | 1417698 | BBA20979.1 | P36334 |
| CQYTMCEFPHTV | 1 | 0 | 0 | 1417699 | ARB07438.1 | A0A140H1H1 |
| CQYTMCEYPHTV | 1 | 0 | 0 | 1417701 | BBA20986.1 | A0A140H1H1 |
| CQYTMCEYPNTI | 1 | 0 | 0 | 1417702 | CAA83661.1 | P36334 |
| CQYWIGYGCNCD | 1 | 0 | 0 | 1417704 | AXN92247.1 | K9N7C7 |
| CQYWKGYGCNCD | 1 | 0 | 0 | 1417705 | AVV62535.1 | K9N7C7 |
| CRAYLDSCTTQQ | 1 | 0 | 0 | 1417711 | AHI48794.1 | K9N638 |
| CRAYLTSRAAQQ | 1 | 0 | 0 | 1417715 | AVV62535.1 | K9N7C7 |
| CRFQELDEEGNL | 1 | 0 | 0 | 1417743 | APO40578.1 | P0C6X7 |
| CRFVEKDDQGHN | 1 | 0 | 0 | 1417749 | AVV62524.1 | K9N7C7 |
| CRGDQACYERFP | 1 | 0 | 0 | 1417754 | AUM60013.1 | K9N7C7 |
| CRGFAPNGLSED | 1 | 0 | 0 | 1417755 | AVV62526.1 | K9N5Q8 |
| CRHNWNCVDCDT | 1 | 0 | 0 | 1417764 | AVV62535.1 | K9N7C7 |
| CRILASVPANVS | 1 | 0 | 0 | 1417772 | AUM60014.1 | K9N5Q8 |
| CRIPNGFPFNNW | 1 | 0 | 0 | 1417773 | AKA09611.1 | A0A0P0G321 |
| CRKCNLAFRLTS | 1 | 0 | 0 | 1417776 | AGV08522.1 | K9N7C7 |
| CRKHAAEYREYM | 1 | 0 | 0 | 1417782 | AVV62535.1 | K9N7C7 |
| CRKHAAEYREYV | 1 | 0 | 0 | 1417783 | AVV62524.1 | K9N7C7 |
| CRKIGVNVCNDS | 1 | 0 | 0 | 1417787 | ARB07436.1 | A0A140H1G9 |
| CRKLGSSVRSDG | 1 | 0 | 0 | 1417789 | ATP16754.1 | P0C6X6 |
| CRKLGTAVRSDG | 1 | 0 | 0 | 1417790 | AQN78773.1 | P0C6X6 |
| CRLQNGDFLCAN | 1 | 0 | 0 | 1417802 | APO40578.1 | P0C6X7 |
| CRPDARFHDFTP | 1 | 0 | 0 | 1417811 | AVV62529.1 | K9N643 |
| CRRLLKQECCVQ | 1 | 0 | 0 | 1417827 | AEN19363.1 | P0C6X6 |
| CRSGHLIDEVQN | 1 | 0 | 0 | 1417832 | AUM60027.1 | K9N643 |
| CRSKNPVLYDAN | 1 | 0 | 0 | 1417834 | AVP78043.1 | Q3LZX0 |
| CRVGECRDSHKG | 1 | 0 | 0 | 1417842 | AVA26872.1 | H9EJ42 |
| CSAFVCGDCAAC | 1 | 0 | 0 | 1417855 | CAA83661.1 | P36334 |
| CSCDQLREPMMQ | 1 | 0 | 0 | 1417868 | P0C6W6.1 | P0C6W2 |
| CSCGFCCKFDED | 1 | 0 | 0 | 1417871 | AVP78042.1 | Q3LZX1 |
| CSCIKDALSYYS | 1 | 0 | 0 | 1417874 | AVV62535.1 | K9N7C7 |
| CSCPGKNNGIGT | 1 | 0 | 0 | 1417881 | AVR40344.1 | P36334 |
| CSCSCKHTTAQV | 1 | 0 | 0 | 1417882 | AVV62524.1 | K9N7C7 |
| CSCSPHAFLGWS | 1 | 0 | 0 | 1417884 | ABD75617.1 | A0A140H1H1 |
| CSCVSNDTTVQS | 1 | 0 | 0 | 1417888 | AXX83324.1 | P0C6X6 |
| CSDCYDDMCVIH | 1 | 0 | 0 | 1417890 | AIW52827.1 | H9EJ42 |
| CSDFEPMTDMEK | 1 | 0 | 0 | 1417894 | AVV62524.1 | K9N7C7 |
| CSDFIPLSDMEK | 1 | 0 | 0 | 1417896 | ANI69887.1 | K9N7C7 |
| CSDFYADKILYQ | 1 | 0 | 0 | 1417900 | ARU07575.1 | A0A140H1G9 |
| CSDGNYNCNASL | 1 | 0 | 0 | 1417901 | AHL31360.1 | K9N5Q8 |
| CSDGNYNRNAIL | 1 | 0 | 0 | 1417902 | ALJ54448.1 | K9N5Q8 |
| CSDGNYNRYASL | 1 | 0 | 0 | 1417904 | ALJ54486.1 | K9N5Q8 |
| CSEVEASGLEEG | 1 | 0 | 0 | 1417915 | AQZ41341.1 | K9N7C7 |
| CSFTNLDKLKFK | 1 | 0 | 0 | 1417923 | AUM60013.1 | K9N7C7 |
| CSFYPPDEEEDC | 1 | 0 | 0 | 1417931 | P0C6W6.1 | P0C6W2 |
| CSGKNNGIGTCP | 1 | 0 | 0 | 1417935 | ANZ78838.1 | P36334 |
| CSGLAIKSDYCG | 1 | 0 | 0 | 1417936 | CAA83660.1 | P36334 |
| CSHITTFMPQFS | 1 | 0 | 0 | 1417946 | AUM60014.1 | K9N5Q8 |
| CSIEDFNVWALS | 1 | 0 | 0 | 1417951 | AWW13571.1 | P0C6X6 |
| CSIEDFNVWAMT | 1 | 0 | 0 | 1417952 | ARB07436.1 | A0A140H1G9 |
| CSIGIGVYYPGA | 1 | 0 | 0 | 1417954 | NP_073549.1 | A0A0P0LKV0 |
| CSIQECCENQPF | 1 | 0 | 0 | 1417957 | ATO98239.1 | Q3LZX5 |
| CSIQECCENQPY | 1 | 0 | 0 | 1417958 | Q3I5I8.1 | Q3LZX5 |
| CSKEEDGFAFYP | 1 | 0 | 0 | 1417966 | QAT98908.1 | K9N7C7 |
| CSKIINGLHPTQ | 1 | 0 | 0 | 1417971 | ATO98167.1 | P0C6W2 |
| CSNDLLYSNTDV | 1 | 0 | 0 | 1418000 | AZS52618.1 | A0A140H1H1 |
| CSNDLLYSNTEI | 1 | 0 | 0 | 1418001 | 5I08_A | A0A140H1H1 |
| CSNGRWLTGNAV | 1 | 0 | 0 | 1418004 | AUM60013.1 | K9N7C7 |
| CSNSSSGALDTT | 1 | 0 | 0 | 1418007 | AIW52835.1 | A0A0P0G321 |
| CSNSYSGYHPAH | 1 | 0 | 0 | 1418009 | AGT51577.1 | P0C6X6 |
| CSPMACNGHKIE | 1 | 0 | 0 | 1418016 | ACR82761.1 | A0A140H1G9 |
| CSPNTYEGNSPY | 1 | 0 | 0 | 1418017 | APO40584.1 | P59635 |
| CSQHTHYIKDGD | 1 | 0 | 0 | 1418021 | AHI48773.1 | K9N7C7 |
| CSQISPAAIASN | 1 | 0 | 0 | 1418026 | AXN92228.1 | K9N5Q8 |
| CSQISPAAISSN | 1 | 0 | 0 | 1418027 | AHL18090.1 | K9N5Q8 |
| CSQLDEEQPMEI | 1 | 0 | 0 | 1418035 | AVP78035.1 | Q3LZX8 |
| CSRRAGAYSSHL | 1 | 0 | 0 | 1418064 | AVV62524.1 | K9N7C7 |

| | | | | | | |
|---|---|---|---|---|---|---|
| CSSCFGKNNGIG | 1 | 0 | 0 | 1418066 | QEG03814.1 | P36334 |
| CSSCHGKNNGIG | 1 | 0 | 0 | 1418067 | ARB66068.1 | P36334 |
| CSSCPGKNNGIG | 1 | 0 | 0 | 1418068 | AWW13566.1 | P36334 |
| CSSCPGKNNGIS | 1 | 0 | 0 | 1418069 | BBA84094.1 | P36334 |
| CSSCPGKNNGLG | 1 | 0 | 0 | 1418070 | AIL49503.1 | P36334 |
| CSSCPSKNNGIG | 1 | 0 | 0 | 1418071 | QDH43719.1 | P36334 |
| CSSCSCPGKNNG | 1 | 0 | 0 | 1418072 | AVR40344.1 | P36334 |
| CSSCSGKNNGIG | 1 | 0 | 0 | 1418073 | ANZ78838.1 | P36334 |
| CSSCSGKNNGVG | 1 | 0 | 0 | 1418074 | QEG03757.1 | P36334 |
| CSSQGIVAVNST | 1 | 0 | 0 | 1418085 | AUM60018.1 | K9N7D2 |
| CSSSFLSEIQCK | 1 | 0 | 0 | 1418089 | BBA20986.1 | A0A140H1H1 |
| CSSYTVETGTQL | 1 | 0 | 0 | 1418097 | APO40578.1 | P0C6X7 |
| CSTDLQKANIEI | 1 | 0 | 0 | 1418098 | BBO58790.1 | P36334 |
| CSTDLQKANTEI | 1 | 0 | 0 | 1418100 | AEN19366.1 | P36334 |
| CSTLHNEILATS | 1 | 0 | 0 | 1418109 | AEN19363.1 | P0C6X6 |
| CSTNLFKDCSNS | 1 | 0 | 0 | 1418113 | AGT51577.1 | P0C6X6 |
| CSYDYDFVEHHD | 1 | 0 | 0 | 1418150 | Q3I5I8.1 | Q3LZX5 |
| CSYESFDVESGV | 1 | 0 | 0 | 1418153 | AXN92228.1 | K9N5Q8 |
| CSYESFDVGSGV | 1 | 0 | 0 | 1418154 | AVV62537.1 | K9N5Q8 |
| CSYVGIQTLDEL | 1 | 0 | 0 | 1418160 | AVV62535.1 | K9N7C7 |
| CSYYEDFLEYHD | 1 | 0 | 0 | 1418164 | AVP78037.1 | Q3LZX5 |
| CTAAIDLAYNKL | 1 | 0 | 0 | 1418166 | AVV62524.1 | K9N7C7 |
| CTANDKFYRLSN | 1 | 0 | 0 | 1418168 | ALJ99930.1 | A0A0P0LKV0 |
| CTATDRFCRLGN | 1 | 0 | 0 | 1418169 | ABO28724.1 | H9EJ42 |
| CTAVGVKQQRFV | 1 | 0 | 0 | 1418172 | AVV62526.1 | K9N5Q8 |
| CTCYDNSIQGSY | 1 | 0 | 0 | 1418186 | NP_073549.1 | A0A0P0LKV0 |
| CTDVPTAIRADQ | 1 | 0 | 0 | 1418195 | ATO98181.1 | Q3LZX1 |
| CTDVPTMIHVEQ | 1 | 0 | 0 | 1418196 | APO40579.1 | P59594 |
| CTDYILLAGARV | 1 | 0 | 0 | 1418204 | APO40578.1 | P0C6X7 |
| CTEQDLHKDLTP | 1 | 0 | 0 | 1418207 | AUM60027.1 | K9N643 |
| CTFGAPDDATRL | 1 | 0 | 0 | 1418219 | AUM60013.1 | K9N7C7 |
| CTFGSPDDASRL | 1 | 0 | 0 | 1418221 | AHI48794.1 | K9N638 |
| CTFGVPDDAARL | 1 | 0 | 0 | 1418222 | AVV62535.1 | K9N7C7 |
| CTFQEAAANIFV | 1 | 0 | 0 | 1418229 | AVV62535.1 | K9N7C7 |
| CTFQEAAGNIFV | 1 | 0 | 0 | 1418230 | AVV62524.1 | K9N7C7 |
| CTFQEAASNIFL | 1 | 0 | 0 | 1418231 | AUM60013.1 | K9N7C7 |
| CTGGIDICSQDK | 1 | 0 | 0 | 1418246 | AEN19363.1 | P0C6X6 |
| CTGLDAVMHFGT | 1 | 0 | 0 | 1418247 | ARE30011.1 | P0C6X6 |
| CTGQAVNTSKAL | 1 | 0 | 0 | 1418249 | AVV62539.1 | K9N4V0 |
| CTGSWNKVNQIA | 1 | 0 | 0 | 1418255 | QBF80541.1 | K9N7C7 |
| CTGSWNKVTQIA | 1 | 0 | 0 | 1418256 | AXN92247.1 | K9N7C7 |
| CTHWCTYLFSLI | 1 | 0 | 0 | 1418262 | NP_073549.1 | A0A0P0LKV0 |
| CTIANASSSEGF | 1 | 0 | 0 | 1418264 | AGV08523.1 | K9N7C7 |
| CTIKNFNVSKGI | 1 | 0 | 0 | 1418274 | ATO98181.1 | Q3LZX1 |
| CTKHTMLVKMDG | 1 | 0 | 0 | 1418284 | AIL49625.1 | P0C6X6 |
| CTKKFLTVDGVQ | 1 | 0 | 0 | 1418285 | AXN92247.1 | K9N7C7 |
| CTKYNIYDVAGV | 1 | 0 | 0 | 1418293 | AIW52741.1 | A0A0P0K6L9 |
| CTKYNIYDVVGV | 1 | 0 | 0 | 1418295 | BAL45638.1 | A0A0P0K6L9 |
| CTLDPIALKAPD | 1 | 0 | 0 | 1418300 | BBO58891.1 | P36334 |
| CTLDPISFKAPD | 1 | 0 | 0 | 1418301 | ARE30008.1 | P36334 |
| CTLDPITLKTPD | 1 | 0 | 0 | 1418302 | AXX83339.1 | P36334 |
| CTLDSITFKAPG | 1 | 0 | 0 | 1418305 | AIV41891.1 | P36334 |
| CTLFRDAEGKMV | 1 | 0 | 0 | 1418306 | AVV62524.1 | K9N7C7 |
| CTLRDTLEYYAQ | 1 | 0 | 0 | 1418320 | AUM60013.1 | K9N7C7 |
| CTLSEQLDFIES | 1 | 0 | 0 | 1418322 | ARO76381.1 | P0C6U8 |
| CTMFMDAEGRMT | 1 | 0 | 0 | 1418327 | AKJ80136.2 | K9N7C7 |
| CTMFRDAEGKMV | 1 | 0 | 0 | 1418328 | AVV62535.1 | K9N7C7 |
| CTMFRDAEGRIT | 1 | 0 | 0 | 1418329 | AHL18091.1 | K9N7C7 |
| CTMFRDAEGRMT | 1 | 0 | 0 | 1418330 | AXN92247.1 | K9N7C7 |
| CTMFRDAEGRMV | 1 | 0 | 0 | 1418331 | AUM60013.1 | K9N7C7 |
| CTMFTMADGSLQ | 1 | 0 | 0 | 1418332 | ARU07583.1 | P0C6X6 |
| CTMFTTADGSPQ | 1 | 0 | 0 | 1418334 | AGT51707.1 | P0C6X6 |
| CTMTDIAEKPTE | 1 | 0 | 0 | 1418335 | AVP78041.1 | P0C6W2 |
| CTMYICGDSQEC | 1 | 0 | 0 | 1418339 | ATO98181.1 | Q3LZX1 |
| CTNGHALPTLFR | 1 | 0 | 0 | 1418344 | ALL26408.1 | K9N638 |
| CTNGKVVPTLFK | 1 | 0 | 0 | 1418347 | AVV62524.1 | K9N7C7 |
| CTPEDMLNPNYE | 1 | 0 | 0 | 1418359 | APO40578.1 | P0C6X7 |
| CTPESKPLADDD | 1 | 0 | 0 | 1418360 | ABK63972.1 | H9EJ69 |
| CTPINTNASLNA | 1 | 0 | 0 | 1418362 | AVV62526.1 | K9N5Q8 |
| CTRAAASAVKLQ | 1 | 0 | 0 | 1418378 | AVV62524.1 | K9N7C7 |
| CTRAASSAVKLQ | 1 | 0 | 0 | 1418379 | AUM60013.1 | K9N7C7 |
| CTRKLNRLMCDI | 1 | 0 | 0 | 1418387 | NP_073549.1 | A0A0P0LKV0 |
| CTRTPLNLPPTH | 1 | 0 | 0 | 1418391 | AWK59942.1 | H9EJ42 |
| CTSAGYNTNATL | 1 | 0 | 0 | 1418394 | AVV62537.1 | K9N5Q8 |
| CTSPGPSGIGAV | 1 | 0 | 0 | 1418402 | ALA49909.1 | K9N4V7 |
| CTSYYAYMMPVM | 1 | 0 | 0 | 1418408 | AIW52776.1 | A0A0P0LKV0 |
| CTSYYSYMMPIM | 1 | 0 | 0 | 1418409 | AIW52813.1 | H9EJ42 |
| CTSYYSYMMPVM | 1 | 0 | 0 | 1418410 | AIW52827.1 | H9EJ42 |
| CTTAVMTYSNFG | 1 | 0 | 0 | 1418411 | AIW52843.1 | A0A0P0G321 |
| CTTCGHVYEVGD | 1 | 0 | 0 | 1418412 | AEN19363.1 | P0C6X6 |
| CTTIVNGMKKSF | 1 | 0 | 0 | 1418417 | APO40578.1 | P0C6X7 |
| CTYDRVEKSFQL | 1 | 0 | 0 | 1418447 | Q3I5J5.1 | Q3LZX1 |
| CTYDRVEQSFQL | 1 | 0 | 0 | 1418448 | ATO98120.1 | Q3LZX1 |
| CTYISRFSKISL | 1 | 0 | 0 | 1418455 | AVK87480.1 | K9N7D2 |
| CTYTMCEYPHTI | 1 | 0 | 0 | 1418466 | ABA60802.1 | A0A140H1H1 |
| CTYVDTLRDLPL | 1 | 0 | 0 | 1418467 | ATO98145.1 | Q3LZX1 |
| CVADQSRKWTGH | 1 | 0 | 0 | 1418472 | AIV41775.1 | Q80872 |
| CVADYAVLYNSS | 1 | 0 | 0 | 1418473 | APO40579.1 | P59594 |
| CVAETGTSDLDR | 1 | 0 | 0 | 1418478 | 6NB6_A | P59594 |
| CVALCDAMVAKG | 1 | 0 | 0 | 1418482 | AIW52827.1 | H9EJ42 |
| CVCFDNSIQGSY | 1 | 0 | 0 | 1418489 | AWK59942.1 | H9EJ42 |

| | | | | | | |
|---|---|---|---|---|---|---|
| CVDCDAAGVGNT | 1 | 0 | 0 | 1418496 | AVN89451.1 | K9N7C7 |
| CVDCDIAGVGNT | 1 | 0 | 0 | 1418497 | AXN92247.1 | K9N7C7 |
| CVDCDTAGIGNT | 1 | 0 | 0 | 1418498 | AVV62524.1 | K9N7C7 |
| CVDCDTAGIGST | 1 | 0 | 0 | 1418499 | AUM60013.1 | K9N7C7 |
| CVDCDTAGVGNT | 1 | 0 | 0 | 1418500 | AHI48798.1 | K9N7C7 |
| CVDGFGLNKIQP | 1 | 0 | 0 | 1418505 | AST12962.1 | A0A0P0LKV0 |
| CVDGIYGYVLRQ | 1 | 0 | 0 | 1418508 | AIW52843.1 | A0A0P0G321 |
| CVDGRNGYVLRQ | 1 | 0 | 0 | 1418509 | AIW52756.1 | A0A0P0K6L9 |
| CVDHRPVCSFPL | 1 | 0 | 0 | 1418510 | AXN92247.1 | K9N7C7 |
| CVDLSVASTYFL | 1 | 0 | 0 | 1418517 | AXN92247.1 | K9N7C7 |
| CVDYSKNGGSGG | 1 | 0 | 0 | 1418528 | 6NZK_A | P36334 |
| CVDYSKNRRSRK | 1 | 0 | 0 | 1418531 | AIL49542.1 | P36334 |
| CVESQSKRYGFG | 1 | 0 | 0 | 1418540 | ABB90528.1 | A0A0P0K6L9 |
| CVFAYVGCYNKC | 1 | 0 | 0 | 1418555 | AVP78041.1 | P0C6W2 |
| CVFENKTTLPTN | 1 | 0 | 0 | 1418558 | AVV62524.1 | K9N7C7 |
| CVFFNGVDISDT | 1 | 0 | 0 | 1418562 | AQZ41321.1 | K9N7C7 |
| CVFNADNLTDYS | 1 | 0 | 0 | 1418564 | BBA20983.1 | A0A140H1H1 |
| CVFSVVNTTVTV | 1 | 0 | 0 | 1418565 | AIW52831.1 | A0A0P0G321 |
| CVGHSENVFVVE | 1 | 0 | 0 | 1418575 | AGT21353.1 | A0A0P0K6L9 |
| CVGNGPGIDAGY | 1 | 0 | 0 | 1418577 | CAA83660.1 | P36334 |
| CVGSGPGKNNGM | 1 | 0 | 0 | 1418581 | AIL49487.1 | P36334 |
| CVIAWNTAKQDT | 1 | 0 | 0 | 1418593 | AVP78042.1 | Q3LZX1 |
| CVIAWNTAKQDV | 1 | 0 | 0 | 1418594 | AVP78031.1 | Q3LZX1 |
| CVIGTTKFYGGW | 1 | 0 | 0 | 1418599 | AXN92247.1 | K9N7C7 |
| CVKCHNDILTAT | 1 | 0 | 0 | 1418608 | AVV62524.1 | K9N7C7 |
| CVKSFIDVINKA | 1 | 0 | 0 | 1418616 | ARO76381.1 | P0C6U8 |
| CVLAWNTNSVDS | 1 | 0 | 0 | 1418629 | APO40579.1 | P59594 |
| CVLEPRNGTRCP | 1 | 0 | 0 | 1418633 | AVV62526.1 | K9N5Q8 |
| CVLGLVNSSLSV | 1 | 0 | 0 | 1418635 | ALL26396.1 | K9N5Q8 |
| CVMLVYCCLGYF | 1 | 0 | 0 | 1418648 | APO40578.1 | P0C6X7 |
| CVMPTYTVHKSD | 1 | 0 | 0 | 1418649 | ATN39877.2 | P0C6X6 |
| CVNFNFNGLKGT | 1 | 0 | 0 | 1418666 | AVP78031.1 | Q3LZX1 |
| CVNLNIPANVPY | 1 | 0 | 0 | 1418674 | AUM60013.1 | K9N7C7 |
| CVNLTGRTPLNP | 1 | 0 | 0 | 1418677 | AVP78031.1 | Q3LZX1 |
| CVPLSAVPLCAS | 1 | 0 | 0 | 1418689 | AVV62524.1 | K9N7C7 |
| CVPLSVVPLCAS | 1 | 0 | 0 | 1418690 | AVV62535.1 | K9N7C7 |
| CVPQAEVDWKFY | 1 | 0 | 0 | 1418692 | AVP78041.1 | P0C6W2 |
| CVPQSEVDWKFY | 1 | 0 | 0 | 1418694 | AVP78030.1 | P0C6W2 |
| CVPTVGYSAALA | 1 | 0 | 0 | 1418695 | AUM60018.1 | K9N7D2 |
| CVQTVRTHVYIA | 1 | 0 | 0 | 1418709 | ARO76381.1 | P0C6U8 |
| CVRAFDGYNKDA | 1 | 0 | 0 | 1418714 | AST12962.1 | A0A0P0LKV0 |
| CVRPCVSVPVSV | 1 | 0 | 0 | 1418724 | AUM60014.1 | K9N5Q8 |
| CVRVAKNKVYFV | 1 | 0 | 0 | 1418730 | APO40578.1 | P0C6X7 |
| CVSDFMSEIKCK | 1 | 0 | 0 | 1418732 | AHA80206.1 | P36334 |
| CVSIVPPTVWED | 1 | 0 | 0 | 1418737 | QBF80598.1 | K9N5Q8 |
| CVSIVPSTAWED | 1 | 0 | 0 | 1418738 | AGV08584.1 | K9N5Q8 |
| CVSKVVNLNVDF | 1 | 0 | 0 | 1418742 | AOL02450.1 | P0C6X6 |
| CVSNDTTVQSKD | 1 | 0 | 0 | 1418744 | AXX83324.1 | P0C6X6 |
| CVSVPLSVVYDK | 1 | 0 | 0 | 1418748 | AVV62526.1 | K9N5Q8 |
| CVTCPFAKGDSC | 1 | 0 | 0 | 1418752 | ATG84865.1 | K9N638 |
| CVTGPTTENKLN | 1 | 0 | 0 | 1418756 | ATO98145.1 | Q3LZX1 |
| CVTRFFLAPIAV | 1 | 0 | 0 | 1418760 | AUM60017.1 | K9N643 |
| CVTSNVPPIISE | 1 | 0 | 0 | 1418763 | AMM42546.1 | H9EJ42 |
| CVTSNVPPIVSE | 1 | 0 | 0 | 1418764 | AWK59942.1 | H9EJ42 |
| CVTSNVSPIVNE | 1 | 0 | 0 | 1418768 | AGT51337.1 | H9EJ42 |
| CVTSNVSPIVSE | 1 | 0 | 0 | 1418769 | ARU07593.1 | H9EJ42 |
| CVTTKAKSIQQD | 1 | 0 | 0 | 1418770 | ALA49934.1 | K9N7C7 |
| CVVDQSRKWIGH | 1 | 0 | 0 | 1418783 | ATP16755.1 | Q80872 |
| CVVDQSRKWISH | 1 | 0 | 0 | 1418784 | AIV41836.1 | Q80872 |
| CVVDYSVLYNST | 1 | 0 | 0 | 1418787 | ABD72995.1 | P59594 |
| CVVKPDGKSFIL | 1 | 0 | 0 | 1418792 | AUM60023.1 | K9N7C7 |
| CVVNAYNNTAIS | 1 | 0 | 0 | 1418797 | AIL49538.1 | P36334 |
| CVYELNGYTVQP | 1 | 0 | 0 | 1418818 | AQN78656.1 | P36334 |
| CVYLFCEHPAVG | 1 | 0 | 0 | 1418820 | NP_073553.1 | A0A0P0KWE8 |
| CVYTVAMCFRHF | 1 | 0 | 0 | 1418823 | AXN92247.1 | K9N7C7 |
| CVYYSQVLSKSI | 1 | 0 | 0 | 1418830 | ATQ39389.1 | K9N7C7 |
| CWETGKVKPPLNKNYVFTG | 1 | 0 | 0 | 1418834 | ABD75503.1 | A0A140H1G9 |
| CWHTNCFDYCIP | 1 | 0 | 0 | 1418842 | Q3I5J4.1 | Q3LZX0 |
| CWLYKTWDFIVF | 1 | 0 | 0 | 1418863 | ARA15430.1 | A0A0P0KWE8 |
| CWYVERKDVPYT | 1 | 0 | 0 | 1418877 | AUM60013.1 | K9N7C7 |
| CYALRHFDRNDC | 1 | 0 | 0 | 1418883 | AEN19363.1 | P0C6X6 |
| CYCFNADGDEAW | 1 | 0 | 0 | 1418885 | AUM60013.1 | K9N7C7 |
| CYCFNRDGDEQW | 1 | 0 | 0 | 1418889 | AVV62524.1 | K9N7C7 |
| CYDIFGVTTPEK | 1 | 0 | 0 | 1418900 | NP_073549.1 | A0A0P0LKV0 |
| CYDPTVLPGAHA | 1 | 0 | 0 | 1418910 | AVV62524.1 | K9N7C7 |
| CYDPTVLPGAHP | 1 | 0 | 0 | 1418911 | AUM60013.1 | K9N7C7 |
| CYELLKDCDAVA | 1 | 0 | 0 | 1418920 | AUM60013.1 | K9N7C7 |
| CYFNLCKEPMYA | 1 | 0 | 0 | 1418939 | ATO98193.1 | Q3LZX1 |
| CYGADLKSPDLG | 1 | 0 | 0 | 1418951 | APO40578.1 | P0C6X7 |
| CYGEPVLVNAAN | 1 | 0 | 0 | 1418952 | AVN89451.1 | K9N7C7 |
| CYGFNEPLNVVS | 1 | 0 | 0 | 1418954 | ARU07576.1 | A0A140H1H0 |
| CYLFIGYVISCY | 1 | 0 | 0 | 1418987 | AIV41774.1 | P0C6X6 |
| CYLNVVIMMLDL | 1 | 0 | 0 | 1418989 | AVV62524.1 | K9N7C7 |
| CYMHQMELANGT | 1 | 0 | 0 | 1418995 | AXN92247.1 | K9N7C7 |
| CYNCRPAVVNIT | 1 | 0 | 0 | 1418999 | AIW52756.1 | A0A0P0K6L9 |
| CYNGAPSGVYQC | 1 | 0 | 0 | 1419003 | APO40578.1 | P0C6X7 |
| CYNSEYAAKGYV | 1 | 0 | 0 | 1419015 | AVV62535.1 | K9N7C7 |
| CYRCPREIVNTV | 1 | 0 | 0 | 1419024 | AVV62524.1 | K9N7C7 |
| CYRKSNVDESFV | 1 | 0 | 0 | 1419027 | AIG96641.1 | A0A0P0LKV0 |
| CYRNSNVDEAFV | 1 | 0 | 0 | 1419029 | ALJ99927.1 | A0A0P0LKV0 |
| CYRNSNVGESFV | 1 | 0 | 0 | 1419030 | ADO24705.1 | A0A0P0LKV0 |

| | | | | | | |
|---|---|---|---|---|---|---|
| CYSAKDTLTELV | 1 | 0 | 0 | 1419035 | AVV62535.1 | K9N7C7 |
| CYSFSAYSGDVP | 1 | 0 | 0 | 1419037 | AUM60014.1 | K9N5Q8 |
| CYSSLILDYFSY | 1 | 0 | 0 | 1419047 | AXN92228.1 | K9N5Q8 |
| CYTAEQCIRGAI | 1 | 0 | 0 | 1419052 | ARB07424.1 | A0A0P0ILY9 |
| CYTAEQCIRGAM | 1 | 0 | 0 | 1419053 | NP_073549.1 | A0A0P0LKV0 |
| CYTARDALDTIL | 1 | 0 | 0 | 1419054 | ATQ39389.1 | K9N7C7 |
| CYTDGLMQNAFL | 1 | 0 | 0 | 1419055 | AIV41774.1 | P0C6X6 |
| CYTEGLMQNASL | 1 | 0 | 0 | 1419058 | AGT51787.1 | P0C6X6 |
| CYVCKPHQVNLS | 1 | 0 | 0 | 1419068 | AWO69005.1 | A0A0P0G321 |
| CYVGVQTLEDLH | 1 | 0 | 0 | 1419073 | ATQ39389.1 | K9N7C7 |
| CYVGVQTVEDLC | 1 | 0 | 0 | 1419074 | AKN24802.1 | K9N7C7 |
| CYWPLKDYGFYT | 1 | 0 | 0 | 1419081 | 3D0H_E | P59594 |
| CYYVKPGGTGSG | 1 | 0 | 0 | 1419090 | ADO24677.1 | P0C6X6 |
| CYYVTPGGTSSG | 1 | 0 | 0 | 1419092 | ACR82746.1 | A0A140H1G9 |
| DAAEELASPMKS | 1 | 0 | 0 | 1419095 | ARU07575.1 | A0A140H1G9 |
| DAAKIYGMGFSS | 1 | 0 | 0 | 1419102 | AIL49487.1 | P36334 |
| DAANHSIPDLQK | 1 | 0 | 0 | 1419108 | ARO76381.1 | P0C6U8 |
| DAANPTNCITPV | 1 | 0 | 0 | 1419110 | ALJ76278.1 | K9N5Q8 |
| DAARQVAWELHR | 1 | 0 | 0 | 1419119 | AUM60016.1 | K9N4V0 |
| DAARQVAWQLHR | 1 | 0 | 0 | 1419120 | AVV62539.1 | K9N4V0 |
| DACNVKKVTDVT | 1 | 0 | 0 | 1419132 | QBP84746.1 | P0C6X6 |
| DACQLPYCYFRN | 1 | 0 | 0 | 1419133 | AGT17757.1 | A0A140H1H0 |
| DADSTLIGHCAT | 1 | 0 | 0 | 1419146 | AVP78030.1 | P0C6W2 |
| DADYKMCTKGLS | 1 | 0 | 0 | 1419148 | APT69849.1 | A0A0P0K6L9 |
| DAEGKMVPYCYD | 1 | 0 | 0 | 1419156 | AVV62524.1 | K9N7C7 |
| DAEIVQPATSKS | 1 | 0 | 0 | 1419161 | AUM60013.1 | K9N7C7 |
| DAFNQLTADLQH | 1 | 0 | 0 | 1419178 | ARU07575.1 | A0A140H1G9 |
| DAFQSVYHTFTQ | 1 | 0 | 0 | 1419183 | ARK08626.1 | H9EJ42 |
| DAGCGVKYYAGL | 1 | 0 | 0 | 1419188 | AGT51379.1 | H9EJ42 |
| DAGEEKLSSRMY | 1 | 0 | 0 | 1419189 | ATO98229.1 | P0C6W2 |
| DAGEENLSSRMY | 1 | 0 | 0 | 1419191 | ARO76381.1 | P0C6U8 |
| DAGLNGDFNLTM | 1 | 0 | 0 | 1419199 | AVV62526.1 | K9N5Q8 |
| DAGNHRSAFALH | 1 | 0 | 0 | 1419200 | AKA09621.1 | A0A0P0G321 |
| DAGVVSCLYKRN | 1 | 0 | 0 | 1419202 | AKA09652.1 | P36334 |
| DAINVGGTGLQY | 1 | 0 | 0 | 1419226 | AVN89322.1 | K9N7C7 |
| DAIQVAKCYGEP | 1 | 0 | 0 | 1419230 | AVN89451.1 | K9N7C7 |
| DAISAAFNNLLG | 1 | 0 | 0 | 1419232 | AUM60013.1 | K9N7C7 |
| DAISQAFNKLLG | 1 | 0 | 0 | 1419233 | AVV62535.1 | K9N7C7 |
| DAITQTSPALQT | 1 | 0 | 0 | 1419235 | AIW52728.1 | A0A0P0K6L9 |
| DAITRTAEAIHT | 1 | 0 | 0 | 1419236 | QED88026.1 | A0A0P0G321 |
| DAIVIDNDLNEF | 1 | 0 | 0 | 1419238 | ASY99818.1 | K9N7C7 |
| DAKETKEINIIK | 1 | 0 | 0 | 1419241 | AGT51737.1 | P0C6X6 |
| DAKFKNSVTSIN | 1 | 0 | 0 | 1419243 | AST12962.1 | A0A0P0LKV0 |
| DAKGFPQFAELI | 1 | 0 | 0 | 1419246 | AGT51335.1 | H9EJ69 |
| DALDTILQEANK | 1 | 0 | 0 | 1419261 | ATQ39389.1 | K9N7C7 |
| DALKLGDTQGMD | 1 | 0 | 0 | 1419269 | AVP78041.1 | P0C6W2 |
| DALQSREAVLVT | 1 | 0 | 0 | 1419281 | AEN19363.1 | P0C6X6 |
| DALRNSAMLESA | 1 | 0 | 0 | 1419284 | AIW52757.1 | A0A0P0K6L9 |
| DAMLDDATGVMA | 1 | 0 | 0 | 1419298 | AVV62524.1 | K9N7C7 |
| DAMSAKEQRVPK | 1 | 0 | 0 | 1419306 | AUM60021.1 | K9N4V7 |
| DANIPDYPKLFI | 1 | 0 | 0 | 1419311 | AVV62535.1 | K9N7C7 |
| DANYKKCTKGLS | 1 | 0 | 0 | 1419321 | CAA71147.1 | A0A0P0K6L9 |
| DAPCYCFNAEGG | 1 | 0 | 0 | 1419325 | AKN24802.1 | K9N7C7 |
| DAPSTFGTRNPN | 1 | 0 | 0 | 1419327 | AXN92267.1 | K9N4V7 |
| DAPYMVGDVINS | 1 | 0 | 0 | 1419333 | ARO76381.1 | P0C6U8 |
| DAQPLRKNVQQL | 1 | 0 | 0 | 1419344 | AVV62524.1 | K9N7C7 |
| DAQSFLNRVCGV | 1 | 0 | 0 | 1419345 | QYC64967.1 | P0DTD1 |
| DAQTGSSKCGCS | 1 | 0 | 0 | 1419346 | 2H85_A | P0C6X7 |
| DAQVVQLPIQDY | 1 | 0 | 0 | 1419350 | AEN19363.1 | P0C6X6 |
| DARAKQDVSLLG | 1 | 0 | 0 | 1419352 | AKN24802.1 | K9N7C7 |
| DARAKQNVSLLS | 1 | 0 | 0 | 1419354 | ASU91141.1 | K9N7C7 |
| DARFHDFTPLAN | 1 | 0 | 0 | 1419356 | AVV62529.1 | K9N643 |
| DARTAHHGHVMV | 1 | 0 | 0 | 1419358 | AVP78041.1 | P0C6W2 |
| DASAPATGWLLY | 1 | 0 | 0 | 1419361 | ARU07575.1 | A0A140H1G9 |
| DASAQCFKILYK | 1 | 0 | 0 | 1419362 | AUM60023.1 | K9N7C7 |
| DASEPQRGRQSR | 1 | 0 | 0 | 1419368 | ALK28785.1 | A0A0N7J131 |
| DASGDCPSVDMQ | 1 | 0 | 0 | 1419372 | ATQ39390.1 | K9N5Q8 |
| DASHYVVTDIKQ | 1 | 0 | 0 | 1419375 | ARU07575.1 | A0A140H1G9 |
| DASKPHQVTKHT | 1 | 0 | 0 | 1419377 | AJQ24068.1 | P33469 |
| DASNVKKVIDVT | 1 | 0 | 0 | 1419379 | AGT51687.1 | P0C6X6 |
| DASNVKNVTDVT | 1 | 0 | 0 | 1419381 | AVQ05261.1 | P0C6X6 |
| DATGVMALYKEI | 1 | 0 | 0 | 1419407 | AVV62524.1 | K9N7C7 |
| DATKPKQVTKHT | 1 | 0 | 0 | 1419409 | AGT51505.1 | P33469 |
| DATNEWYYTGSS | 1 | 0 | 0 | 1419412 | AUM60014.1 | K9N5Q8 |
| DATTAYANSVLN | 1 | 0 | 0 | 1419432 | AVP77956.1 | P0C6W2 |
| DAVARHDFFTFD | 1 | 0 | 0 | 1419439 | AVV62524.1 | K9N7C7 |
| DAVARHDFFVFD | 1 | 0 | 0 | 1419440 | AUM60013.1 | K9N7C7 |
| DAVFKPQPAGVL | 1 | 0 | 0 | 1419447 | QEG03794.1 | P36334 |
| DAVFNATRFPSV | 1 | 0 | 0 | 1419448 | ATO98145.1 | Q3LZX1 |
| DAVLIDNDLNEY | 1 | 0 | 0 | 1419452 | AVV62524.1 | K9N7C7 |
| DAVLVNAANTHL | 1 | 0 | 0 | 1419456 | AVV62524.1 | K9N7C7 |
| DAVNANAQALSK | 1 | 0 | 0 | 1419458 | AVV62537.1 | K9N5Q8 |
| DAVNNNAHALSK | 1 | 0 | 0 | 1419461 | ATG84679.1 | K9N5Q8 |
| DAVNPTNCIAPV | 1 | 0 | 0 | 1419463 | ATG84723.1 | K9N5Q8 |
| DAVQYAHKHDIQ | 1 | 0 | 0 | 1419466 | AVV62524.1 | K9N7C7 |
| DAWEILKFLIRG | 1 | 0 | 0 | 1419479 | ARO76381.1 | P0C6U8 |
| DAYDLVHDEPKG | 1 | 0 | 0 | 1419489 | AVI15051.1 | P0C6X6 |
| DAYKLFLSGDDC | 1 | 0 | 0 | 1419495 | ARU07575.1 | A0A140H1G9 |
| DAYKSFPVKEKK | 1 | 0 | 0 | 1419497 | AVV62533.1 | K9N4V7 |
| DAYKTFPKKENK | 1 | 0 | 0 | 1419498 | AHL18098.1 | K9N4V7 |
| DAYKTFPVKEKK | 1 | 0 | 0 | 1419500 | AVV62544.1 | K9N4V7 |

| | | | | | | |
|---|---|---|---|---|---|---|
| DAYLDAKGPLQV | 1 | 0 | 0 | 1419503 | AVV62524.1 | K9N7C7 |
| DAYVSQQLSDST | 1 | 0 | 0 | 1419518 | CAA83661.1 | P36334 |
| DCAACVCGDYAA | 1 | 0 | 0 | 1419522 | ABD43177.1 | P36334 |
| DCAAFVCGDFAA | 1 | 0 | 0 | 1419524 | AAX85669.1 | P36334 |
| DCATVYTANKWD | 1 | 0 | 0 | 1419534 | APO40578.1 | P0C6X7 |
| DCDAVARHDFFT | 1 | 0 | 0 | 1419542 | AVV62524.1 | K9N7C7 |
| DCDFNKPLIVWP | 1 | 0 | 0 | 1419545 | QBF80519.1 | K9N7C7 |
| DCDIDKWLNNFS | 1 | 0 | 0 | 1419549 | AMM42415.1 | A0A140H1H1 |
| DCDIDNWLNNVS | 1 | 0 | 0 | 1419551 | BBA20986.1 | A0A140H1H1 |
| DCEQTPCGFKGG | 1 | 0 | 0 | 1419567 | AIV42002.1 | P0C6X6 |
| DCFANKHAGFDS | 1 | 0 | 0 | 1419570 | P0C6W6.1 | P0C6W2 |
| DCGAVFSGNDVE | 1 | 0 | 0 | 1419582 | AVV62535.1 | K9N7C7 |
| DCGHDDLAQLRC | 1 | 0 | 0 | 1419589 | AVV62526.1 | K9N5Q8 |
| DCGIKQESRVGV | 1 | 0 | 0 | 1419591 | ARU07575.1 | A0A140H1G9 |
| DCGKKFDEQVGC | 1 | 0 | 0 | 1419593 | AWK59942.1 | H9EJ42 |
| DCGVNALDGDIY | 1 | 0 | 0 | 1419596 | AWK59942.1 | H9EJ42 |
| DCGVNALDGDTY | 1 | 0 | 0 | 1419597 | AFD98777.1 | H9EJ42 |
| DCINGQYDKNAS | 1 | 0 | 0 | 1419607 | AUM60014.1 | K9N5Q8 |
| DCKQYICNGFQK | 1 | 0 | 0 | 1419614 | AKM76239.1 | K9N5Q8 |
| DCKQYVCNGFKK | 1 | 0 | 0 | 1419618 | AUM60014.1 | K9N5Q8 |
| DCLFMYTFNITE | 1 | 0 | 0 | 1419624 | AUM60014.1 | K9N5Q8 |
| DCMHQGPASARD | 1 | 0 | 0 | 1419635 | ATG84866.1 | K9N5Q8 |
| DCMKQGPASARD | 1 | 0 | 0 | 1419636 | QEJ82215.1 | K9N5Q8 |
| DCMKQGPPSARD | 1 | 0 | 0 | 1419637 | AVV62537.1 | K9N5Q8 |
| DCMNQGPPSARD | 1 | 0 | 0 | 1419640 | AVV62526.1 | K9N5Q8 |
| DCMQQGPASARE | 1 | 0 | 0 | 1419642 | ARA67740.1 | K9N5Q8 |
| DCMQQGPASARN | 1 | 0 | 0 | 1419643 | QFS19619.1 | K9N5Q8 |
| DCMSDFMSGIKC | 1 | 0 | 0 | 1419647 | AIL49491.1 | P36334 |
| DCNAASVNPASM | 1 | 0 | 0 | 1419648 | AUM60013.1 | K9N7C7 |
| DCNHIGNINPHN | 1 | 0 | 0 | 1419658 | AEN19365.1 | P30215 |
| DCNHINNLKVKN | 1 | 0 | 0 | 1419659 | ABD75544.1 | A0A140H1H0 |
| DCNHVVNTNPRN | 1 | 0 | 0 | 1419661 | AAA73152.1 | P30215 |
| DCNIEAWLNDKS | 1 | 0 | 0 | 1419662 | CAA83660.1 | P36334 |
| DCNLPLGQSLCA | 1 | 0 | 0 | 1419664 | ALA49451.1 | K9N5Q8 |
| DCNVVRYSLDGN | 1 | 0 | 0 | 1419669 | AXN92247.1 | K9N7C7 |
| DCRIPNGFPFNN | 1 | 0 | 0 | 1419684 | AKA09611.1 | A0A0P0G321 |
| DCSAFVCGDCAA | 1 | 0 | 0 | 1419687 | CAA83661.1 | P36334 |
| DCSAFVCGDYAA | 1 | 0 | 0 | 1419688 | CAA83660.1 | P36334 |
| DCSDGNYNCNAS | 1 | 0 | 0 | 1419693 | AHL31360.1 | K9N5Q8 |
| DCSDGNYNRNAI | 1 | 0 | 0 | 1419694 | ALJ54448.1 | K9N5Q8 |
| DCSDGNYNRYAS | 1 | 0 | 0 | 1419696 | ALJ54486.1 | K9N5Q8 |
| DCSKVITGLHPT | 1 | 0 | 0 | 1419702 | QYC64967.1 | P0DTD1 |
| DCSNSYSGYHPA | 1 | 0 | 0 | 1419704 | AGT51577.1 | P0C6X6 |
| DCSSSFLSEIQC | 1 | 0 | 0 | 1419712 | BBA20986.1 | A0A140H1H1 |
| DCTGQAVNTSKA | 1 | 0 | 0 | 1419717 | AVV62539.1 | K9N4V0 |
| DCVGLSVASTYF | 1 | 0 | 0 | 1419738 | ATQ39389.1 | K9N7C7 |
| DCVPVVCPKGKI | 1 | 0 | 0 | 1419744 | AFD98812.1 | H9EJ42 |
| DCWTATDRFYRL | 1 | 0 | 0 | 1419756 | ABQ16583.1 | H9EJ42 |
| DCYIFKRTFAKA | 1 | 0 | 0 | 1419761 | AZS52616.1 | A0A140H1G9 |
| DCYTLKKTVAKA | 1 | 0 | 0 | 1419768 | ABD75615.1 | A0A140H1G9 |
| DDAINVGGTGLQ | 1 | 0 | 0 | 1419777 | AVN89322.1 | K9N7C7 |
| DDAQGMETLACE | 1 | 0 | 0 | 1419778 | APO40578.1 | P0C6X7 |
| DDATRNYIITMP | 1 | 0 | 0 | 1419788 | AHL18094.1 | K9N643 |
| DDATRRVWTFMN | 1 | 0 | 0 | 1419789 | APO40578.1 | P0C6X7 |
| DDATSNVTQCFG | 1 | 0 | 0 | 1419791 | APT69866.1 | A0A0N7J131 |
| DDCMHQGPASAR | 1 | 0 | 0 | 1419799 | ATG84866.1 | K9N5Q8 |
| DDCMKQGPASAR | 1 | 0 | 0 | 1419800 | QEJ82215.1 | K9N5Q8 |
| DDCMKQGPPSAR | 1 | 0 | 0 | 1419801 | AVV62537.1 | K9N5Q8 |
| DDCMNQGPPSAR | 1 | 0 | 0 | 1419802 | AVV62526.1 | K9N5Q8 |
| DDCMQQGPASVR | 1 | 0 | 0 | 1419804 | ALJ54502.1 | K9N5Q8 |
| DDCMQQGPSSAR | 1 | 0 | 0 | 1419805 | AHL18090.1 | K9N5Q8 |
| DDDDSQPRADKP | 1 | 0 | 0 | 1419816 | 5EPW_A | H9EJ69 |
| DDEDVVTGDNDQ | 1 | 0 | 0 | 1419837 | AZS52616.1 | A0A140H1G9 |
| DDEGFITLKNNL | 1 | 0 | 0 | 1419841 | AXN92247.1 | K9N7C7 |
| DDEWAAAVDEVF | 1 | 0 | 0 | 1419845 | ASU90174.1 | K9N638 |
| DDEWTAAVDEAF | 1 | 0 | 0 | 1419846 | AVN89451.1 | K9N7C7 |
| DDEWVAAVDEAF | 1 | 0 | 0 | 1419847 | ASU45728.1 | K9N7C7 |
| DDFFKNVSTSIP | 1 | 0 | 0 | 1419853 | AMW90852.1 | K9N5Q8 |
| DDFNEHVTSVNL | 1 | 0 | 0 | 1419859 | AXG21666.1 | K9N796 |
| DDFQGYLQKHFS | 1 | 0 | 0 | 1419862 | AIG96641.1 | A0A0P0LKV0 |
| DDFSKQLQDSMN | 1 | 0 | 0 | 1419864 | APO40586.1 | P59595 |
| DDFSTEIIGYKA | 1 | 0 | 0 | 1419867 | APO40578.1 | P0C6X7 |
| DDFTEHVTSVNL | 1 | 0 | 0 | 1419868 | AJG44103.1 | K9N796 |
| DDGSLAFDEAVA | 1 | 0 | 0 | 1419892 | AVV62533.1 | K9N4V7 |
| DDHIPSGTVLGN | 1 | 0 | 0 | 1419904 | AIW52748.1 | A0A0P0K6L9 |
| DDIDTINCESIY | 1 | 0 | 0 | 1419911 | ARU07575.1 | A0A140H1G9 |
| DDIFAESKPRFV | 1 | 0 | 0 | 1419914 | AGT21359.1 | A0A0P0LKV0 |
| DDINFAPVLGCL | 1 | 0 | 0 | 1419923 | QXL74886.1 | P36334 |
| DDISESVTSVNL | 1 | 0 | 0 | 1419926 | ATQ39394.1 | K9N796 |
| DDKASVNWADDR | 1 | 0 | 0 | 1419934 | 5N4K_A | H9EJ69 |
| DDKYKTIDELCV | 1 | 0 | 0 | 1419942 | ALA50066.1 | K9N7C7 |
| DDKYKTSDELCV | 1 | 0 | 0 | 1419943 | AXN92247.1 | K9N7C7 |
| DDLHQLYVDSKY | 1 | 0 | 0 | 1419948 | AUM60013.1 | K9N7C7 |
| DDLNQLSGYKKP | 1 | 0 | 0 | 1419952 | AVP78041.1 | P0C6W2 |
| DDLQLGSSGFLQ | 1 | 0 | 0 | 1419956 | BBA20986.1 | A0A140H1H1 |
| DDMHWGDGFLIK | 1 | 0 | 0 | 1419968 | AGT51559.1 | Q80872 |
| DDMLWGDGFQIK | 1 | 0 | 0 | 1419974 | ARK08668.1 | Q80872 |
| DDNKFDGLKVDD | 1 | 0 | 0 | 1419982 | ATP16754.1 | P0C6X6 |
| DDNKFDVLKLDD | 1 | 0 | 0 | 1419983 | AIV42008.1 | P0C6X6 |
| DDNQALLVPKIF | 1 | 0 | 0 | 1419985 | AMM42360.1 | A0A140H1G9 |
| DDNQTLLVPQIF | 1 | 0 | 0 | 1419987 | AMM42357.1 | A0A140H1G9 |

| | | | | | | |
|---|---|---|---|---|---|---|
| DDNQTLLVSKIF | 1 | 0 | 0 | 1419988 | AMM42372.1 | A0A140H1G9 |
| DDNTDSIVIDGV | 1 | 0 | 0 | 1419989 | AWK59930.1 | H9EJ42 |
| DDNVLPDTYVAL | 1 | 0 | 0 | 1419991 | AWO68966.1 | A0A0P0G321 |
| DDQGHHLDSHFV | 1 | 0 | 0 | 1420005 | AUM60013.1 | K9N7C7 |
| DDQGHNLDSYFV | 1 | 0 | 0 | 1420007 | AVV62524.1 | K9N7C7 |
| DDQIGYYRRASR | 1 | 0 | 0 | 1420010 | APO40586.1 | P59595 |
| DDRGVITNAVDC | 1 | 0 | 0 | 1420022 | ADN03339.1 | A0A140H1H1 |
| DDSGDSSEGGAK | 1 | 0 | 0 | 1420034 | AVQ05261.1 | P0C6X6 |
| DDSGNPVVEYLS | 1 | 0 | 0 | 1420037 | QGV13494.1 | K9N7C7 |
| DDSQPRADKPSQ | 1 | 0 | 0 | 1420043 | 5EPW_A | H9EJ69 |
| DDSQYFFNKDTG | 1 | 0 | 0 | 1420045 | AGT51759.1 | P30215 |
| DDSQYYLNKDTG | 1 | 0 | 0 | 1420047 | AGT51480.1 | P30215 |
| DDSTGDEAWAAL | 1 | 0 | 0 | 1420050 | AVV62524.1 | K9N7C7 |
| DDSVREEVDIIE | 1 | 0 | 0 | 1420051 | ARU07593.1 | H9EJ42 |
| DDTLRTEAFDYY | 1 | 0 | 0 | 1420060 | APO40578.1 | P0C6X7 |
| DDVDDGGDSSES | 1 | 0 | 0 | 1420073 | AGT51787.1 | P0C6X6 |
| DDVDDIDPINCE | 1 | 0 | 0 | 1420074 | ABD75591.1 | A0A140H1G9 |
| DDVDDIESIYEF | 1 | 0 | 0 | 1420075 | ABD75535.1 | A0A140H1G9 |
| DDVIFSRADSLS | 1 | 0 | 0 | 1420082 | ARB07436.1 | A0A140H1G9 |
| DDVITSLCKKSF | 1 | 0 | 0 | 1420083 | AWK59942.1 | H9EJ42 |
| DDVIVHDVEPTH | 1 | 0 | 0 | 1420084 | AWK59942.1 | H9EJ42 |
| DDVNFSPVLGCL | 1 | 0 | 0 | 1420089 | CAA83661.1 | P36334 |
| DDVQRWLAPYGA | 1 | 0 | 0 | 1420097 | AVV62540.1 | K9N643 |
| DDVRQIAPAQTG | 1 | 0 | 0 | 1420098 | APO40579.1 | P59594 |
| DDYDFSVEDVYA | 1 | 0 | 0 | 1420113 | ARB07605.1 | A0A140H1G9 |
| DDYGTDIDGVYW | 1 | 0 | 0 | 1420117 | AJQ24074.1 | P33469 |
| DDYKAFISVFDI | 1 | 0 | 0 | 1420123 | APT69868.1 | A0A0P0ILY9 |
| DDYNGTPLEFGA | 1 | 0 | 0 | 1420127 | APO40578.1 | P0C6X7 |
| DDYQIGGYTEKW | 1 | 0 | 0 | 1420130 | AVP78043.1 | Q3LZX0 |
| DDYRITSITSDK | 1 | 0 | 0 | 1420132 | AIV41978.1 | P0C6X6 |
| DEAFPLDEAENV | 1 | 0 | 0 | 1420148 | QBF80465.1 | K9N7C7 |
| DEAIKRVKGWVG | 1 | 0 | 0 | 1420159 | ABD75519.1 | A0A140H1G9 |
| DEAIPTKFPPGT | 1 | 0 | 0 | 1420161 | AJQ24067.1 | P33469 |
| DEAMASRMYCTF | 1 | 0 | 0 | 1420162 | ARU07575.1 | A0A140H1G9 |
| DEARFSGTANQQ | 1 | 0 | 0 | 1420167 | AGT51787.1 | P0C6X6 |
| DEATPFDVVRQC | 1 | 0 | 0 | 1420170 | 3EA9_A | P0C6U8 |
| DECMQQGPQSAR | 1 | 0 | 0 | 1420175 | AUM60014.1 | K9N5Q8 |
| DECQLPLGQSLC | 1 | 0 | 0 | 1420176 | AVV62526.1 | K9N5Q8 |
| DEDVVTGDNDDC | 1 | 0 | 0 | 1420191 | ARB07436.1 | A0A140H1G9 |
| DEDVVTGDNDQI | 1 | 0 | 0 | 1420193 | AZS52616.1 | A0A140H1G9 |
| DEEDLYDVSKPV | 1 | 0 | 0 | 1420197 | QED88039.1 | H9EJ42 |
| DEEFPLDDEESV | 1 | 0 | 0 | 1420205 | AVV62535.1 | K9N7C7 |
| DEEGDNVQITYT | 1 | 0 | 0 | 1420207 | AMM42619.1 | H9EJ69 |
| DEFIQRYKLESY | 1 | 0 | 0 | 1420222 | APO40578.1 | P0C6X7 |
| DEFTPFDVVAQC | 1 | 0 | 0 | 1420228 | 3M3T_A | P0C6X7 |
| DEGSELTLKHFL | 1 | 0 | 0 | 1420244 | QED88038.1 | H9EJ42 |
| DEHEFKFWDESF | 1 | 0 | 0 | 1420249 | APT69863.1 | A0A0P0LKV0 |
| DEIKALKELYGP | 1 | 0 | 0 | 1420255 | AVN89363.1 | K9N7C7 |
| DEIPYKSFFDSG | 1 | 0 | 0 | 1420258 | AMD02834.1 | K9N638 |
| DELCVNLNIPAN | 1 | 0 | 0 | 1420269 | AUM60013.1 | K9N7C7 |
| DELDDFFKNVST | 1 | 0 | 0 | 1420270 | AMW90852.1 | K9N5Q8 |
| DELDEFFKNFST | 1 | 0 | 0 | 1420271 | AKM76239.1 | K9N5Q8 |
| DELFAMTKRNVF | 1 | 0 | 0 | 1420273 | AVV62524.1 | K9N7C7 |
| DELKINSACRKV | 1 | 0 | 0 | 1420282 | P0C6W6.1 | P0C6W2 |
| DEMLNNRATLQA | 1 | 0 | 0 | 1420291 | APO40578.1 | P0C6X7 |
| DEMNRLQEAIKV | 1 | 0 | 0 | 1420292 | CAA83660.1 | P36334 |
| DENAMFDKAVAD | 1 | 0 | 0 | 1420294 | AUM60021.1 | K9N4V7 |
| DENGVRTLSTYD | 1 | 0 | 0 | 1420297 | AVP78031.1 | Q3LZX1 |
| DEPDSRAPSRSQ | 1 | 0 | 0 | 1420299 | AST12968.1 | A0A0N7J131 |
| DESAAKSASVYY | 1 | 0 | 0 | 1420315 | P0C6W6.1 | P0C6W2 |
| DESCNNLVQTYL | 1 | 0 | 0 | 1420318 | AIV42002.1 | P0C6X6 |
| DESFVDDFQGYL | 1 | 0 | 0 | 1420323 | AIG96641.1 | A0A0P0LKV0 |
| DESGEPVYFLSY | 1 | 0 | 0 | 1420329 | AVV62544.1 | K9N4V7 |
| DESVANLFSDIK | 1 | 0 | 0 | 1420332 | AVV62537.1 | K9N5Q8 |
| DESVSNLFADIK | 1 | 0 | 0 | 1420334 | AVV62526.1 | K9N5Q8 |
| DETAEWFGITQN | 1 | 0 | 0 | 1420335 | AUM60014.1 | K9N5Q8 |
| DETKALKELYGS | 1 | 0 | 0 | 1420340 | AHC74096.1 | K9N7C7 |
| DEVARDLSVQFK | 1 | 0 | 0 | 1420344 | APO40578.1 | P0C6X7 |
| DEVGDNVQITHT | 1 | 0 | 0 | 1420346 | AMM42628.1 | H9EJ69 |
| DEWVAAVDEAFP | 1 | 0 | 0 | 1420373 | ASU45728.1 | K9N7C7 |
| DEYLTQRGPLQV | 1 | 0 | 0 | 1420382 | AUM60013.1 | K9N7C7 |
| DFDLEDFIDNPC | 1 | 0 | 0 | 1420415 | AVV62524.1 | K9N7C7 |
| DFEEDDVEGEET | 1 | 0 | 0 | 1420427 | AIV41978.1 | P0C6X6 |
| DFFIFDVDKFKT | 1 | 0 | 0 | 1420436 | AVN89322.1 | K9N7C7 |
| DFGDFIQVAPGA | 1 | 0 | 0 | 1420448 | APO40578.1 | P0C6X7 |
| DFGYVCDGSFCK | 1 | 0 | 0 | 1420462 | ARB07436.1 | A0A140H1G9 |
| DFGYVCDGSFYK | 1 | 0 | 0 | 1420463 | ARU07575.1 | A0A140H1G9 |
| DFHANRPGVYCG | 1 | 0 | 0 | 1420464 | AVV62535.1 | K9N7C7 |
| DFHSVEIEVGEV | 1 | 0 | 0 | 1420472 | AKT07951.1 | H9EJ42 |
| DFIDMEDSLFIA | 1 | 0 | 0 | 1420483 | ARB07436.1 | A0A140H1G9 |
| DFIDSVRDPQTL | 1 | 0 | 0 | 1420486 | AVP78031.1 | Q3LZX1 |
| DFIKHVQLEGSY | 1 | 0 | 0 | 1420494 | AUM60013.1 | K9N7C7 |
| DFIPIDQYMCGK | 1 | 0 | 0 | 1420506 | AVV62535.1 | K9N7C7 |
| DFIQVAPGAGIP | 1 | 0 | 0 | 1420513 | APO40578.1 | P0C6X7 |
| DFKDFQFMLWGN | 1 | 0 | 0 | 1420521 | AIV41912.1 | P0C6X6 |
| DFKEELDQWLKN | 1 | 0 | 0 | 1420524 | ARB07414.1 | P36334 |
| DFKEELGQWFKN | 1 | 0 | 0 | 1420526 | AIX10763.1 | P36334 |
| DFKFLHNLQADI | 1 | 0 | 0 | 1420528 | AVN89451.1 | K9N7C7 |
| DFKFSDGQGVPL | 1 | 0 | 0 | 1420529 | ABA60796.1 | Q19U29 |
| DFKGELDQWFKN | 1 | 0 | 0 | 1420530 | QDH43719.1 | P36334 |
| DFKVEYAYDQVH | 1 | 0 | 0 | 1420546 | APU51933.1 | P0C6X6 |

| | | | | | | |
|---|---|---|---|---|---|---|
| DFLDCTGQAVNT | 1 | 0 | 0 | 1420554 | AVV62539.1 | K9N4V0 |
| DFLEASDVEEDD | 1 | 0 | 0 | 1420555 | ATN39862.1 | P0C6X6 |
| DFLNIDMGVFIN | 1 | 0 | 0 | 1420574 | AGT51365.1 | H9EJ42 |
| DFLPIDSTVKNY | 1 | 0 | 0 | 1420576 | APO40578.1 | P0C6X7 |
| DFLSFDSDVFVK | 1 | 0 | 0 | 1420580 | AHI48614.1 | K9N7C7 |
| DFLSLDSDVFIK | 1 | 0 | 0 | 1420581 | AVV62535.1 | K9N7C7 |
| DFLSNSSSSFDC | 1 | 0 | 0 | 1420584 | AIW52843.1 | A0A0P0G321 |
| DFLSSDSDVFIK | 1 | 0 | 0 | 1420585 | ANI69887.1 | K9N7C7 |
| DFMLKTLYKDVE | 1 | 0 | 0 | 1420596 | AVV62524.1 | K9N7C7 |
| DFNCTNSFINDY | 1 | 0 | 0 | 1420612 | ARB07438.1 | A0A140H1H1 |
| DFNEDVCVCFDN | 1 | 0 | 0 | 1420614 | AWK59942.1 | H9EJ42 |
| DFNHNMGDSDLV | 1 | 0 | 0 | 1420617 | AIW52805.1 | H9EJ69 |
| DFNKPTSKTCTY | 1 | 0 | 0 | 1420621 | AUM60013.1 | K9N7C7 |
| DFNPNVPLDYQA | 1 | 0 | 0 | 1420626 | ATO98145.1 | Q3LZX1 |
| DFNPNVPLEYQA | 1 | 0 | 0 | 1420627 | AVP78031.1 | Q3LZX1 |
| DFNREAPAYIST | 1 | 0 | 0 | 1420628 | APO40578.1 | P0C6X7 |
| DFNVNVIGTRPE | 1 | 0 | 0 | 1420633 | AST12962.1 | A0A0P0LKV0 |
| DFNVNVMGTRPE | 1 | 0 | 0 | 1420634 | AGT21366.1 | A0A0P0LKV0 |
| DFPFAESTGTQS | 1 | 0 | 0 | 1420637 | QBF80599.1 | K9N796 |
| DFQENWNTKHGS | 1 | 0 | 0 | 1420642 | AVP78041.1 | P0C6W2 |
| DFQGYLQKHFSM | 1 | 0 | 0 | 1420649 | AIG96641.1 | A0A0P0LKV0 |
| DFQNKMAATGVI | 1 | 0 | 0 | 1420655 | AUM60014.1 | K9N5Q8 |
| DFSNLGADVLYQ | 1 | 0 | 0 | 1420679 | BAE93400.1 | P0C6U8 |
| DFSSDVFKIGDV | 1 | 0 | 0 | 1420690 | QDH43737.1 | H9EJ42 |
| DFTEERLCLFNR | 1 | 0 | 0 | 1420710 | APO40578.1 | P0C6X7 |
| DFTIGNMEEFIQ | 1 | 0 | 0 | 1420719 | ARB66065.1 | P36334 |
| DFTIGNMGEFIQ | 1 | 0 | 0 | 1420720 | ARB66062.1 | P36334 |
| DFTKLLSGTPPQ | 1 | 0 | 0 | 1420722 | AVV62537.1 | K9N5Q8 |
| DFTVLFDRLSLT | 1 | 0 | 0 | 1420732 | AEN19363.1 | P0C6X6 |
| DFVDFESVIQDY | 1 | 0 | 0 | 1420739 | APU51933.1 | P0C6X6 |
| DFVDLASVIQDY | 1 | 0 | 0 | 1420740 | AIV41852.1 | P0C6X6 |
| DFYTTEPEFVKV | 1 | 0 | 0 | 1420798 | QBQ01836.1 | P0C6X6 |
| DGADVTKIKPHS | 1 | 0 | 0 | 1420801 | APO40578.1 | P0C6X7 |
| DGAKTEPTGHGA | 1 | 0 | 0 | 1420806 | AAA45463.1 | A0A0N7J131 |
| DGALLTKMSEYK | 1 | 0 | 0 | 1420808 | APO40578.1 | P0C6X7 |
| DGARLFSSDLST | 1 | 0 | 0 | 1420811 | P0C6U6.1 | H9EJ42 |
| DGARLSASDLST | 1 | 0 | 0 | 1420812 | AAS58186.1 | H9EJ42 |
| DGCGTLLGAFYC | 1 | 0 | 0 | 1420821 | AHN10826.1 | K9N5Q8 |
| DGDVQWSSTMIF | 1 | 0 | 0 | 1420855 | AVV62535.1 | K9N7C7 |
| DGDVVKQPVKFY | 1 | 0 | 0 | 1420857 | ALA49648.1 | K9N7C7 |
| DGEIISLEKLKT | 1 | 0 | 0 | 1420864 | APO40578.1 | P0C6X7 |
| DGETITFDNLKT | 1 | 0 | 0 | 1420869 | AVP78030.1 | P0C6W2 |
| DGFGLNKIQPWT | 1 | 0 | 0 | 1420885 | AST12962.1 | A0A0P0LKV0 |
| DGFLSVYNAYSP | 1 | 0 | 0 | 1420890 | ATO98193.1 | Q3LZX1 |
| DGFLSVYQTYTA | 1 | 0 | 0 | 1420891 | ATO98181.1 | Q3LZX1 |
| DGFNEWAIANGY | 1 | 0 | 0 | 1420896 | AGT51358.1 | H9EJ42 |
| DGFNHQFVNASS | 1 | 0 | 0 | 1420900 | AVA26872.1 | H9EJ42 |
| DGFNLEYWVTPL | 1 | 0 | 0 | 1420902 | ARK08651.1 | P36334 |
| DGFSFVDDVNGM | 1 | 0 | 0 | 1420906 | AVV62535.1 | K9N7C7 |
| DGFSFVNDDTGV | 1 | 0 | 0 | 1420907 | AVV62524.1 | K9N7C7 |
| DGFTEHVTSVNL | 1 | 0 | 0 | 1420911 | ASU89978.1 | K9N796 |
| DGFYSTSPIQPA | 1 | 0 | 0 | 1420922 | ADK37704.1 | A0A0P0K6L9 |
| DGFYSTSSIQPV | 1 | 0 | 0 | 1420926 | AMM42536.1 | A0A0P0K6L9 |
| DGFYTQGRNLSD | 1 | 0 | 0 | 1420927 | AWK59942.1 | H9EJ42 |
| DGGIPDGCSLYL | 1 | 0 | 0 | 1420932 | AXN92262.1 | K9N4V0 |
| DGGKYYTHRIIK | 1 | 0 | 0 | 1420935 | AGT51448.1 | P0C6X6 |
| DGGRSGARPKQR | 1 | 0 | 0 | 1420937 | Q3I5I7.1 | Q3LZX4 |
| DGHVGLLDQAWR | 1 | 0 | 0 | 1420947 | AVZ61116.1 | P0C6X6 |
| DGIASDYRANSF | 1 | 0 | 0 | 1420949 | AUM60013.1 | K9N7C7 |
| DGICSENHVMVD | 1 | 0 | 0 | 1420952 | AIV41942.1 | P0C6X6 |
| DGIFKTEVDPNL | 1 | 0 | 0 | 1420957 | ATQ39389.1 | K9N7C7 |
| DGIFTIWEQDRM | 1 | 0 | 0 | 1420960 | ATP16762.1 | Q4VID0 |
| DGIIYPTGKSFS | 1 | 0 | 0 | 1420970 | AVV62526.1 | K9N5Q8 |
| DGINQMTGFKKP | 1 | 0 | 0 | 1420976 | APO40578.1 | P0C6X7 |
| DGINVKDVFVES | 1 | 0 | 0 | 1420977 | QEG03747.1 | H9EJ42 |
| DGIRRGDNFHSV | 1 | 0 | 0 | 1420982 | ANI69888.1 | K9N638 |
| DGISEHSLRLVD | 1 | 0 | 0 | 1420984 | ARO76381.1 | P0C6U8 |
| DGKCLTSIEEVC | 1 | 0 | 0 | 1421004 | AMK59674.1 | P0C6X6 |
| DGKHVDHYDGFY | 1 | 0 | 0 | 1421006 | AST12962.1 | A0A0P0LKV0 |
| DGKMGLFFNHTL | 1 | 0 | 0 | 1421009 | ANF29217.1 | K9N5Q8 |
| DGKMGRFFYHTL | 1 | 0 | 0 | 1421012 | AID55079.1 | K9N5Q8 |
| DGKPCTPPAFNC | 1 | 0 | 0 | 1421017 | ATO98231.1 | Q3LZX1 |
| DGKPMLPKNMWE | 1 | 0 | 0 | 1421024 | AFD98826.1 | H9EJ42 |
| DGKPVISEDLWQ | 1 | 0 | 0 | 1421025 | AGT21366.1 | A0A0P0LKV0 |
| DGKPVLPKNMWE | 1 | 0 | 0 | 1421026 | AWK59942.1 | H9EJ42 |
| DGKQIDDHRITS | 1 | 0 | 0 | 1421027 | AEN19363.1 | P0C6X6 |
| DGKYFTSAPPVT | 1 | 0 | 0 | 1421038 | AVK87459.1 | K9N7C7 |
| DGLEQDAQIDRL | 1 | 0 | 0 | 1421054 | AHI48733.1 | K9N5Q8 |
| DGLKITEELQHE | 1 | 0 | 0 | 1421061 | APO40578.1 | P0C6X7 |
| DGLMQNAFLYSS | 1 | 0 | 0 | 1421065 | AIV41774.1 | P0C6X6 |
| DGLMQNASLYSS | 1 | 0 | 0 | 1421066 | AEN19363.1 | P0C6X6 |
| DGLRGGVRKAKD | 1 | 0 | 0 | 1421069 | AUM60013.1 | K9N7C7 |
| DGLTEHVTSVNL | 1 | 0 | 0 | 1421073 | AXN92229.1 | K9N796 |
| DGMMNMSPKPQL | 1 | 0 | 0 | 1421085 | AMM42447.1 | P33469 |
| DGMNMSPKPQRQ | 1 | 0 | 0 | 1421087 | AIV41850.1 | P33469 |
| DGNAAIPDYNYY | 1 | 0 | 0 | 1421089 | ACV87150.1 | A0A140H1G9 |
| DGNMYGAFQDRQ | 1 | 0 | 0 | 1421101 | AVV62524.1 | K9N7C7 |
| DGNPIRDYAALA | 1 | 0 | 0 | 1421102 | AVV62524.1 | K9N7C7 |
| DGNYNCNASLNS | 1 | 0 | 0 | 1421107 | AHL31360.1 | K9N5Q8 |
| DGNYNRNAILNS | 1 | 0 | 0 | 1421108 | ALJ54448.1 | K9N5Q8 |
| DGNYYCVRPCVS | 1 | 0 | 0 | 1421114 | AVV62526.1 | K9N5Q8 |

| | | | | | | |
|---|---|---|---|---|---|---|
| DGPNEHVTSVNL | 1 | 0 | 0 | 1421120 | AGV08391.1 | K9N796 |
| DGPSVDTSDSQV | 1 | 0 | 0 | 1421121 | AIV41888.1 | P0C6X6 |
| DGPTEHVNLFDV | 1 | 0 | 0 | 1421124 | ANI69890.1 | K9N796 |
| DGQGVPIAFGIP | 1 | 0 | 0 | 1421132 | ABA60794.1 | Q19U29 |
| DGQGVPIAYGIP | 1 | 0 | 0 | 1421133 | AMM42397.1 | Q19U29 |
| DGQGVPLAFGVP | 1 | 0 | 0 | 1421134 | ABA60796.1 | Q19U29 |
| DGQNGEAPVSIV | 1 | 0 | 0 | 1421137 | AVP78030.1 | P0C6W2 |
| DGQTTFYPMCTN | 1 | 0 | 0 | 1421150 | AVV62524.1 | K9N7C7 |
| DGQVQTFYPRFQ | 1 | 0 | 0 | 1421155 | ALA49945.1 | K9N7C7 |
| DGRDNGAFEAFK | 1 | 0 | 0 | 1421156 | AEN19363.1 | P0C6X6 |
| DGRIPNDFSFNN | 1 | 0 | 0 | 1421160 | ABE97137.1 | A0A0P0G321 |
| DGRPVHVKEITR | 1 | 0 | 0 | 1421168 | AGT51393.1 | H9EJ42 |
| DGRTPNGFSFNN | 1 | 0 | 0 | 1421172 | AKA09627.1 | A0A0P0G321 |
| DGSAFRDLKPLP | 1 | 0 | 0 | 1421179 | AHI48736.1 | K9N7C7 |
| DGSGVGVIRISN | 1 | 0 | 0 | 1421184 | BAL45639.1 | A0A0P0K6L9 |
| DGSPQPYCYTDG | 1 | 0 | 0 | 1421193 | AEN19363.1 | P0C6X6 |
| DGSPQSYCYTDG | 1 | 0 | 0 | 1421195 | AGT51488.1 | P0C6X6 |
| DGTMYGAFMDKQ | 1 | 0 | 0 | 1421220 | AXN92247.1 | K9N7C7 |
| DGTTQPKLKEDY | 1 | 0 | 0 | 1421231 | Q3I5J4.1 | Q3LZX0 |
| DGVDCYNWDSAE | 1 | 0 | 0 | 1421237 | QED88024.1 | H9EJ42 |
| DGVEFLKDAWEI | 1 | 0 | 0 | 1421243 | ARO76381.1 | P0C6U8 |
| DGVKRGENLQSV | 1 | 0 | 0 | 1421251 | AUM60013.1 | K9N7C7 |
| DGVNFNIDDINF | 1 | 0 | 0 | 1421258 | ARB66060.1 | P36334 |
| DGVNFNVDDVNF | 1 | 0 | 0 | 1421259 | CAA83661.1 | P36334 |
| DGVNFRTCVVNN | 1 | 0 | 0 | 1421260 | AVV62524.1 | K9N7C7 |
| DGVNKLQGLLEI | 1 | 0 | 0 | 1421263 | AVQ05264.1 | P36334 |
| DGVNKLQGLLEV | 1 | 0 | 0 | 1421264 | AEN19366.1 | P36334 |
| DGVNVTKHRCDI | 1 | 0 | 0 | 1421266 | QBP84746.1 | P0C6X6 |
| DGVQCYTPHSQI | 1 | 0 | 0 | 1421272 | AEN19363.1 | P0C6X6 |
| DGVTRDIVSTDD | 1 | 0 | 0 | 1421287 | P0C6W6.1 | P0C6W2 |
| DGVYTNFKLIGH | 1 | 0 | 0 | 1421300 | AEN19363.1 | P0C6X6 |
| DGVYWVASNQAD | 1 | 0 | 0 | 1421303 | AIV41850.1 | P33469 |
| DGYAFEHIVYGD | 1 | 0 | 0 | 1421308 | AVP78041.1 | P0C6W2 |
| DGYGNTMYVNMA | 1 | 0 | 0 | 1421316 | AUM60027.1 | K9N643 |
| DGYIRRAIDCGH | 1 | 0 | 0 | 1421320 | AVV62526.1 | K9N5Q8 |
| DHAIGATAALPT | 1 | 0 | 0 | 1421340 | ARO76382.1 | P59594 |
| DHAYSTMRLHNF | 1 | 0 | 0 | 1421345 | AWK59942.1 | H9EJ42 |
| DHDVYYDESRAT | 1 | 0 | 0 | 1421351 | AVV62527.1 | K9N796 |
| DHEISVQKKINR | 1 | 0 | 0 | 1421356 | AWK59942.1 | H9EJ42 |
| DHFGENEEIVIN | 1 | 0 | 0 | 1421361 | AST12962.1 | A0A0P0LKV0 |
| DHFKELWHLDTG | 1 | 0 | 0 | 1421362 | AKA09663.1 | P36334 |
| DHFNYNVSVLQM | 1 | 0 | 0 | 1421363 | AWK59942.1 | H9EJ42 |
| DHIGTRNPNNNV | 1 | 0 | 0 | 1421380 | ATO98117.1 | Q3LZX4 |
| DHNDMLYSPPTI | 1 | 0 | 0 | 1421396 | AWK59942.1 | H9EJ42 |
| DHPHLGKFLEEL | 1 | 0 | 0 | 1421399 | AIW52723.1 | A0A0N7J131 |
| DHPNPKGFCDFK | 1 | 0 | 0 | 1421401 | APO40578.1 | P0C6X7 |
| DHQDLKWSRFPK | 1 | 0 | 0 | 1421405 | AVP78041.1 | P0C6W2 |
| DHSGSNDNIALL | 1 | 0 | 0 | 1421414 | Q3I5J2.1 | Q3LZX9 |
| DHSQFYFNKDTG | 1 | 0 | 0 | 1421415 | QEG03793.1 | P30215 |
| DHSQYYFNKDTG | 1 | 0 | 0 | 1421416 | AEN19365.1 | P30215 |
| DHSQYYLNKDTG | 1 | 0 | 0 | 1421417 | AIX09798.1 | P30215 |
| DHSVADYHNHSG | 1 | 0 | 0 | 1421419 | APO40578.1 | P0C6X7 |
| DHTSLVWKHIRD | 1 | 0 | 0 | 1421422 | AWK59942.1 | H9EJ42 |
| DHVSLTVSCCGA | 1 | 0 | 0 | 1421437 | AVV62524.1 | K9N7C7 |
| DHYDGFYTQGRN | 1 | 0 | 0 | 1421443 | AWK59942.1 | H9EJ42 |
| DHYIVLGGETIT | 1 | 0 | 0 | 1421445 | ARU07595.1 | H9EJ24 |
| DHYVVLGGETIT | 1 | 0 | 0 | 1421446 | AVA26874.1 | H9EJ24 |
| DIAANTVIWDHK | 1 | 0 | 0 | 1421448 | AVP78041.1 | P0C6W2 |
| DIAKKPTEHACS | 1 | 0 | 0 | 1421458 | APO40578.1 | P0C6X7 |
| DICAQLNAKLGF | 1 | 0 | 0 | 1421470 | ABD75519.1 | A0A140H1G9 |
| DIDDFDIDTFID | 1 | 0 | 0 | 1421486 | AUM60013.1 | K9N7C7 |
| DIDKWLNNFSVP | 1 | 0 | 0 | 1421500 | AMM42415.1 | A0A140H1H1 |
| DIDTGDIVDAVQ | 1 | 0 | 0 | 1421506 | AVV62535.1 | K9N7C7 |
| DIEGNFYGPFVD | 1 | 0 | 0 | 1421519 | APO40578.1 | P0C6X7 |
| DIESIYEFDTYK | 1 | 0 | 0 | 1421524 | ABD75535.1 | A0A140H1G9 |
| DIETGDIVDAVQ | 1 | 0 | 0 | 1421525 | AVV62524.1 | K9N7C7 |
| DIFAESKPRFVR | 1 | 0 | 0 | 1421533 | AGT21359.1 | A0A0P0LKV0 |
| DIFATPSVLQAT | 1 | 0 | 0 | 1421535 | ASY99818.1 | K9N7C7 |
| DIFRSDVLHLTQ | 1 | 0 | 0 | 1421547 | ATO98181.1 | Q3LZX1 |
| DIGDRDPRNDEA | 1 | 0 | 0 | 1421551 | AJQ24068.1 | P33469 |
| DIGIDTTGVLLK | 1 | 0 | 0 | 1421557 | ABD75582.1 | Q0ZJ80 |
| DIGNPKGIRCAV | 1 | 0 | 0 | 1421564 | NP_073549.1 | A0A0P0LKV0 |
| DIGNPKGVRCAV | 1 | 0 | 0 | 1421566 | AWK59942.1 | H9EJ42 |
| DIGNYTVSCSPF | 1 | 0 | 0 | 1421569 | AVP78037.1 | Q3LZX5 |
| DIIEQPFGEVEH | 1 | 0 | 0 | 1421580 | AVA26872.1 | H9EJ42 |
| DIILKPASDGLK | 1 | 0 | 0 | 1421585 | AVP78030.1 | P0C6W2 |
| DIIQADQQVDRL | 1 | 0 | 0 | 1421586 | AIW52757.1 | A0A0P0K6L9 |
| DIIQRLDGLEQD | 1 | 0 | 0 | 1421588 | AHI48733.1 | K9N5Q8 |
| DIIQRLDPPEQD | 1 | 0 | 0 | 1421589 | 5W9H_A | K9N5Q8 |
| DIKDAPALPKTE | 1 | 0 | 0 | 1421596 | AVV62535.1 | K9N7C7 |
| DIKKEIAHLNEI | 1 | 0 | 0 | 1421602 | APO40579.1 | P59594 |
| DILKRLDAVEQE | 1 | 0 | 0 | 1421619 | AVV62537.1 | K9N5Q8 |
| DILQHANKSVGI | 1 | 0 | 0 | 1421624 | AHC74096.1 | K9N7C7 |
| DILSTDDCFANK | 1 | 0 | 0 | 1421634 | ARO76381.1 | P0C6U8 |
| DILYPGQKYTSD | 1 | 0 | 0 | 1421641 | AVV62524.1 | K9N7C7 |
| DINGELHEDSKS | 1 | 0 | 0 | 1421657 | APO40578.1 | P0C6X7 |
| DINGKLYQDSQN | 1 | 0 | 0 | 1421659 | ARO76381.1 | P0C6U8 |
| DINHGDNGFTSI | 1 | 0 | 0 | 1421662 | AGT17757.1 | A0A140H1H0 |
| DINHGDSGFISI | 1 | 0 | 0 | 1421663 | ARU07576.1 | A0A140H1H0 |
| DINNGPHEFCTK | 1 | 0 | 0 | 1421674 | AIL49625.1 | P0C6X6 |
| DINTPDVPICVY | 1 | 0 | 0 | 1421681 | AAA73152.1 | P30215 |

| | | | | | | |
|---|---|---|---|---|---|---|
| DIPHYVQPGQYT | 1 | 0 | 0 | 1421689 | AUM60024.1 | K9N5Q8 |
| DIPTTENLYFQG | 1 | 0 | 0 | 1421696 | 3EJG_A | A0A0P0LKV0 |
| DIPVGAGICASY | 1 | 0 | 0 | 1421697 | ACQ82725.1 | P59594 |
| DIPYNVSLFKFN | 1 | 0 | 0 | 1421698 | ABD75496.1 | A0A140H1H0 |
| DIQLLKNAYENF | 1 | 0 | 0 | 1421702 | AVP78041.1 | P0C6W2 |
| DIQLTNESYNNY | 1 | 0 | 0 | 1421704 | AVN89451.1 | K9N7C7 |
| DIQPQVLFTCVP | 1 | 0 | 0 | 1421707 | AUM60018.1 | K9N7D2 |
| DIQQWGYVGSLS | 1 | 0 | 0 | 1421710 | AWK59942.1 | H9EJ42 |
| DIRDNGSLEKFM | 1 | 0 | 0 | 1421714 | AXN92247.1 | K9N7C7 |
| DIRDNGSLERFM | 1 | 0 | 0 | 1421715 | AUM60013.1 | K9N7C7 |
| DIRDNVSLEKFM | 1 | 0 | 0 | 1421716 | ANF29193.1 | K9N7C7 |
| DISDLSNRLLDA | 1 | 0 | 0 | 1421729 | APO40578.1 | P0C6X7 |
| DISFERLRDYLA | 1 | 0 | 0 | 1421732 | AUM60013.1 | K9N7C7 |
| DISFIDYTTTVT | 1 | 0 | 0 | 1421733 | QED88024.1 | H9EJ42 |
| DISLFGDCVTVR | 1 | 0 | 0 | 1421740 | AVV62524.1 | K9N7C7 |
| DISNFCANRSAL | 1 | 0 | 0 | 1421747 | AVV62535.1 | K9N7C7 |
| DISNVLYNSAGG | 1 | 0 | 0 | 1421748 | APO40579.1 | P59594 |
| DISNVPFSPNGK | 1 | 0 | 0 | 1421750 | ABD72972.1 | P59594 |
| DISYLKRARGSS | 1 | 0 | 0 | 1421763 | QDH43737.1 | H9EJ42 |
| DITFLKKDVPYI | 1 | 0 | 0 | 1421773 | AVP78041.1 | P0C6W2 |
| DITNINLSRGRG | 1 | 0 | 0 | 1421779 | ASJ26617.1 | K9N4V7 |
| DITNTNLSRDRG | 1 | 0 | 0 | 1421780 | AVN89370.1 | K9N4V7 |
| DITNTNMSRGRG | 1 | 0 | 0 | 1421783 | ATG84708.1 | K9N4V7 |
| DITSAGKFDSFK | 1 | 0 | 0 | 1421786 | APO40578.1 | P0C6X7 |
| DIVCKVENKAEA | 1 | 0 | 0 | 1421788 | AST12962.1 | A0A0P0LKV0 |
| DIVDRDPSNDEA | 1 | 0 | 0 | 1421792 | AJQ24082.1 | P33469 |
| DIVDVPPQVVEL | 1 | 0 | 0 | 1421796 | AVN89352.1 | K9N7C7 |
| DIVEEAMVVKPT | 1 | 0 | 0 | 1421799 | APO40578.1 | P0C6X7 |
| DIVEVPPQVVEL | 1 | 0 | 0 | 1421801 | AVN89451.1 | K9N7C7 |
| DIVKRLATGLFQ | 1 | 0 | 0 | 1421809 | AVV62535.1 | K9N7C7 |
| DIVNGVAEGTPV | 1 | 0 | 0 | 1421812 | APO40578.1 | P0C6X7 |
| DIVRGQIQVTSD | 1 | 0 | 0 | 1421815 | ARO76381.1 | P0C6U8 |
| DIWRPEKKFPPY | 1 | 0 | 0 | 1421835 | AIV41989.1 | Q04853 |
| DIWRPEKKYLRY | 1 | 0 | 0 | 1421836 | YP_009555242.1 | Q04853 |
| DKANFLNAVVFY | 1 | 0 | 0 | 1421863 | AXX83300.1 | P0C6X6 |
| DKAVPSPLNWER | 1 | 0 | 0 | 1421867 | AIV41909.1 | P36334 |
| DKDTGPPPISTD | 1 | 0 | 0 | 1421892 | CAA79896.1 | P36334 |
| DKEEDLQQAQQE | 1 | 0 | 0 | 1421894 | APT69868.1 | A0A0P0ILY9 |
| DKEKLISLTDVA | 1 | 0 | 0 | 1421898 | AUM60023.1 | K9N7C7 |
| DKETLEYWVTPL | 1 | 0 | 0 | 1421901 | AZS52618.1 | A0A140H1H1 |
| DKFAIPNRRRVD | 1 | 0 | 0 | 1421911 | ANZ78849.1 | P36334 |
| DKGALEKFMSTS | 1 | 0 | 0 | 1421932 | AVV62535.1 | K9N7C7 |
| DKGCVVDQSRKW | 1 | 0 | 0 | 1421934 | ATP16755.1 | Q80872 |
| DKGDAEDTLNKL | 1 | 0 | 0 | 1421935 | AST12962.1 | A0A0P0LKV0 |
| DKGVAPGSAVLK | 1 | 0 | 0 | 1421941 | APO40578.1 | P0C6X7 |
| DKIEEIESKQKK | 1 | 0 | 0 | 1421949 | 5X58_A | P59594 |
| DKIFTKSGFNDI | 1 | 0 | 0 | 1421950 | AVN89289.1 | K9N7C7 |
| DKIHHHHHHNST | 1 | 0 | 0 | 1421956 | 3R24_B | P0C6X7 |
| DKIHHHHHHNTA | 1 | 0 | 0 | 1421957 | 2OFZ_A | P59595 |
| DKIIEFQLDYSI | 1 | 0 | 0 | 1421958 | AGT21366.1 | A0A0P0LKV0 |
| DKIRHLLKGVTL | 1 | 0 | 0 | 1421963 | AHI48785.1 | K9N638 |
| DKISDTSGFAVY | 1 | 0 | 0 | 1421966 | BAM16435.1 | Q01455 |
| DKIVYHTSGNVI | 1 | 0 | 0 | 1421970 | APO40578.1 | P0C6X7 |
| DKKKKADEVQPL | 1 | 0 | 0 | 1421974 | APO40586.1 | P59595 |
| DKLCHAVVSKSK | 1 | 0 | 0 | 1421982 | AEN19363.1 | P0C6X6 |
| DKLEKLIFTARD | 1 | 0 | 0 | 1421986 | AGV08389.1 | K9N638 |
| DKLGVCIDYTLY | 1 | 0 | 0 | 1421995 | AUM60014.1 | K9N5Q8 |
| DKLGVCVEYNLY | 1 | 0 | 0 | 1421996 | AVV62537.1 | K9N5Q8 |
| DKLIATAKDGLK | 1 | 0 | 0 | 1421997 | AVV62524.1 | K9N7C7 |
| DKLISTARDGVK | 1 | 0 | 0 | 1421998 | AUM60013.1 | K9N7C7 |
| DKLPGNCIGKGS | 1 | 0 | 0 | 1422004 | AUM60027.1 | K9N643 |
| DKLQFASLEVPR | 1 | 0 | 0 | 1422005 | APO40578.1 | P0C6X7 |
| DKLVPSPLNWER | 1 | 0 | 0 | 1422018 | ATN39872.1 | P36334 |
| DKNASLNSFKEY | 1 | 0 | 0 | 1422020 | AUM60014.1 | K9N5Q8 |
| DKNICLLDQAWR | 1 | 0 | 0 | 1422023 | ARU07575.1 | A0A140H1G9 |
| DKNKLIGYWNIQ | 1 | 0 | 0 | 1422024 | APT69880.1 | A0A0N7J131 |
| DKNKLIGYWNVQ | 1 | 0 | 0 | 1422025 | AIW52723.1 | A0A0N7J131 |
| DKNKQDSTDDGS | 1 | 0 | 0 | 1422026 | AVV62533.1 | K9N4V7 |
| DKNLQDLWLLYK | 1 | 0 | 0 | 1422028 | AGT51568.1 | P0C6X6 |
| DKNNGIGICPAG | 1 | 0 | 0 | 1422032 | AIX10751.1 | P36334 |
| DKNNGIGTCPAG | 1 | 0 | 0 | 1422033 | AIX10750.1 | P36334 |
| DKPNHFINFPLS | 1 | 0 | 0 | 1422038 | AQN78734.1 | Q80872 |
| DKQVHQVQLIDK | 1 | 0 | 0 | 1422049 | ATG84700.1 | K9N638 |
| DKSAIPNGRKVD | 1 | 0 | 0 | 1422064 | BBA84104.1 | P36334 |
| DKSELTNGVYYR | 1 | 0 | 0 | 1422068 | AEN19363.1 | P0C6X6 |
| DKSKIFGSCFNS | 1 | 0 | 0 | 1422072 | BBA20986.1 | A0A140H1H1 |
| DKSWPRPIDPAA | 1 | 0 | 0 | 1422086 | AVV62526.1 | K9N5Q8 |
| DKTGDFMSTIII | 1 | 0 | 0 | 1422091 | AVN89363.1 | K9N7C7 |
| DKTLQEMKALFL | 1 | 0 | 0 | 1422096 | ARB07430.1 | P0C6X6 |
| DKTWPRPIDVSK | 1 | 0 | 0 | 1422105 | AXN92228.1 | K9N5Q8 |
| DKVAGFAKFLKT | 1 | 0 | 0 | 1422107 | APO40578.1 | P0C6X7 |
| DKVAKIANLFLE | 1 | 0 | 0 | 1422108 | AVV62535.1 | K9N7C7 |
| DKVFNASRFPNV | 1 | 0 | 0 | 1422116 | ATO98181.1 | Q3LZX1 |
| DKVGDTDCVFYF | 1 | 0 | 0 | 1422121 | AEN19363.1 | P0C6X6 |
| DKVGNTDCVFYF | 1 | 0 | 0 | 1422123 | AGT51498.1 | P0C6X6 |
| DKVIIADPGYMQ | 1 | 0 | 0 | 1422124 | AXP07345.1 | K9N5Q8 |
| DKVKTPHIVRQR | 1 | 0 | 0 | 1422126 | AXN92247.1 | K9N7C7 |
| DKVMTFYPRLQA | 1 | 0 | 0 | 1422132 | AVQ05261.1 | P0C6X6 |
| DKVPQAVETRVQ | 1 | 0 | 0 | 1422136 | AVI15051.1 | P0C6X6 |
| DKVPQSVETKVQ | 1 | 0 | 0 | 1422137 | AIV41864.1 | P0C6X6 |
| DKVRSLKLSDCN | 1 | 0 | 0 | 1422141 | AVV62535.1 | K9N7C7 |

| | | | | | | |
|---|---|---|---|---|---|---|
| DKVSKIANLFVE | 1 | 0 | 0 | 1422143 | AVV62524.1 | K9N7C7 |
| DKVYCPRHVICL | 1 | 0 | 0 | 1422148 | ARB07436.1 | A0A140H1G9 |
| DKWLNNFSVPSP | 1 | 0 | 0 | 1422155 | AMM42415.1 | A0A140H1H1 |
| DKYDDLEPLKIH | 1 | 0 | 0 | 1422161 | AVV62537.1 | K9N5Q8 |
| DKYDDYDLEPQK | 1 | 0 | 0 | 1422162 | AUM60024.1 | K9N5Q8 |
| DKYKATGDLAVC | 1 | 0 | 0 | 1422171 | AEN19363.1 | P0C6X6 |
| DKYKTIDELCVN | 1 | 0 | 0 | 1422172 | ALA50066.1 | K9N7C7 |
| DKYMCGFDGKPM | 1 | 0 | 0 | 1422174 | AFD98826.1 | H9EJ42 |
| DLAGLFRIFLRK | 1 | 0 | 0 | 1422184 | AHY04433.1 | K9N638 |
| DLAHTKLQSVDW | 1 | 0 | 0 | 1422187 | AVV62535.1 | K9N7C7 |
| DLAQLFKVFLRQ | 1 | 0 | 0 | 1422200 | AVV62524.1 | K9N7C7 |
| DLCTHSFRKKQV | 1 | 0 | 0 | 1422235 | AYV99777.1 | P59633 |
| DLDCTYISLKPG | 1 | 0 | 0 | 1422239 | AGT17757.1 | A0A140H1H0 |
| DLDDATRSYIGS | 1 | 0 | 0 | 1422240 | AVV62540.1 | K9N643 |
| DLDEFSKQLQQS | 1 | 0 | 0 | 1422244 | AVP78038.1 | Q3LZX4 |
| DLDVMLPGEFSK | 1 | 0 | 0 | 1422257 | AKK52590.1 | K9N7C7 |
| DLEADPKGKYAQ | 1 | 0 | 0 | 1422258 | ATQ39389.1 | K9N7C7 |
| DLEAGKNKAAPK | 1 | 0 | 0 | 1422260 | AVV62533.1 | K9N4V7 |
| DLERDFIDMEDS | 1 | 0 | 0 | 1422274 | ARB07436.1 | A0A140H1G9 |
| DLGAVSKVVKVP | 1 | 0 | 0 | 1422307 | ANI69887.1 | K9N7C7 |
| DLGKQYVYSAGH | 1 | 0 | 0 | 1422315 | AVV62526.1 | K9N5Q8 |
| DLGNQILDSYSV | 1 | 0 | 0 | 1422318 | AEN19363.1 | P0C6X6 |
| DLGNSSTITNQL | 1 | 0 | 0 | 1422319 | AVV62526.1 | K9N5Q8 |
| DLGNSTTITDKL | 1 | 0 | 0 | 1422320 | AVV62537.1 | K9N5Q8 |
| DLGSLIDCNAAS | 1 | 0 | 0 | 1422321 | AXN92247.1 | K9N7C7 |
| DLGSLIDSNAAS | 1 | 0 | 0 | 1422322 | AVV62524.1 | K9N7C7 |
| DLGTENLYFQSM | 1 | 0 | 0 | 1422324 | 5X29_A | P59637 |
| DLHKDLTPLANI | 1 | 0 | 0 | 1422333 | AUM60027.1 | K9N643 |
| DLHQLYVDSKYQ | 1 | 0 | 0 | 1422338 | AUM60013.1 | K9N7C7 |
| DLHTLYVDSKYQ | 1 | 0 | 0 | 1422341 | ALA49340.1 | K9N7C7 |
| DLIAQSSGVLPV | 1 | 0 | 0 | 1422342 | AEN19363.1 | P0C6X6 |
| DLICVQSYNGIK | 1 | 0 | 0 | 1422348 | CAA83660.1 | P36334 |
| DLISRCQITAVE | 1 | 0 | 0 | 1422363 | QBP84728.1 | P0C6X6 |
| DLKCPLDTRYKG | 1 | 0 | 0 | 1422368 | AKA09647.1 | P36334 |
| DLKCPLDTSLKG | 1 | 0 | 0 | 1422369 | AIX10753.1 | P36334 |
| DLKCPLDTSYKG | 1 | 0 | 0 | 1422372 | AKA09658.1 | P36334 |
| DLKELGNYTYYK | 1 | 0 | 0 | 1422379 | AHL18090.1 | K9N5Q8 |
| DLKFVKWEYEGG | 1 | 0 | 0 | 1422386 | AWK59942.1 | H9EJ42 |
| DLKGKFVQIPTT | 1 | 0 | 0 | 1422387 | 2XYV_B | P0C6U8 |
| DLKWSRFPKSDG | 1 | 0 | 0 | 1422402 | AVP78041.1 | P0C6W2 |
| DLLANMGIDLDE | 1 | 0 | 0 | 1422404 | ARO76381.1 | P0C6U8 |
| DLLARAGKASCT | 1 | 0 | 0 | 1422406 | QYC64967.1 | P0DTD1 |
| DLLCVADDKPIV | 1 | 0 | 0 | 1422411 | NP_073549.1 | A0A0P0LKV0 |
| DLLCVANDKPIV | 1 | 0 | 0 | 1422412 | APT69868.1 | A0A0P0ILY9 |
| DLLDKGMQLLHT | 1 | 0 | 0 | 1422414 | ASU91196.1 | K9N638 |
| DLLFNKVTIADP | 1 | 0 | 0 | 1422423 | AVV62526.1 | K9N5Q8 |
| DLLHAHPVEPLV | 1 | 0 | 0 | 1422427 | AHF54653.1 | Q4VID0 |
| DLLLGDFVEIIK | 1 | 0 | 0 | 1422441 | 2H85_A | P0C6X7 |
| DLLNRLQSLESG | 1 | 0 | 0 | 1422447 | ASU90457.1 | K9N4V7 |
| DLLQILNRQCVN | 1 | 0 | 0 | 1422450 | APO40578.1 | P0C6X7 |
| DLMAQSSGVLPV | 1 | 0 | 0 | 1422478 | YP_009555247.1 | P0C6X6 |
| DLMQARAFVNMV | 1 | 0 | 0 | 1422483 | AVV62524.1 | K9N7C7 |
| DLMQARAFVNQV | 1 | 0 | 0 | 1422484 | AVV62535.1 | K9N7C7 |
| DLNCHYLVLPFG | 1 | 0 | 0 | 1422488 | AGT51619.1 | P30215 |
| DLNCPLDSRLKG | 1 | 0 | 0 | 1422491 | AIL49507.1 | P36334 |
| DLNCPLDTRLKG | 1 | 0 | 0 | 1422492 | AEN19366.1 | P36334 |
| DLNKLCEEMLDN | 1 | 0 | 0 | 1422504 | AVP78041.1 | P0C6W2 |
| DLNKPLIEWPLM | 1 | 0 | 0 | 1422505 | AUM60013.1 | K9N7C7 |
| DLNKPLIEWPLT | 1 | 0 | 0 | 1422506 | AVV62524.1 | K9N7C7 |
| DLNNVDPKAYPI | 1 | 0 | 0 | 1422507 | AVP78050.1 | Q3LZX3 |
| DLNPALCDSGKI | 1 | 0 | 0 | 1422508 | AEN19365.1 | P30215 |
| DLNWLNRVETYI | 1 | 0 | 0 | 1422518 | AGT21353.1 | A0A0P0K6L9 |
| DLPENDDDAWFK | 1 | 0 | 0 | 1422522 | AWW13564.1 | Q80872 |
| DLPICVYDRLPV | 1 | 0 | 0 | 1422528 | AIL49324.1 | P30215 |
| DLPLSLTEVDGG | 1 | 0 | 0 | 1422534 | ATO98145.1 | Q3LZX1 |
| DLPNCNIEAWLN | 1 | 0 | 0 | 1422535 | AEN19366.1 | P36334 |
| DLPNCNIGAWLN | 1 | 0 | 0 | 1422536 | AHA80222.1 | P36334 |
| DLQELGKYEHYI | 1 | 0 | 0 | 1422547 | AVP78042.1 | Q3LZX1 |
| DLQELGKYEQYV | 1 | 0 | 0 | 1422550 | APO40579.1 | P59594 |
| DLQFNKLGTEDP | 1 | 0 | 0 | 1422552 | ASY99807.1 | K9N4V7 |
| DLQGNFGDLKFN | 1 | 0 | 0 | 1422553 | AWM99589.1 | K9N4V7 |
| DLQGNFGDLQFN | 1 | 0 | 0 | 1422554 | ASY99807.1 | K9N4V7 |
| DLQGNFGDLQLT | 1 | 0 | 0 | 1422556 | AVN89320.1 | K9N4V7 |
| DLQGNFGDMQFN | 1 | 0 | 0 | 1422557 | AUM60021.1 | K9N4V7 |
| DLQKANIEIELG | 1 | 0 | 0 | 1422561 | BBO58790.1 | P36334 |
| DLQKANTDIILG | 1 | 0 | 0 | 1422562 | CAA79896.1 | P36334 |
| DLQKANTDIKLD | 1 | 0 | 0 | 1422563 | BBO58740.1 | P36334 |
| DLQKANTDIKLG | 1 | 0 | 0 | 1422564 | AVQ05264.1 | P36334 |
| DLQKSNTDIILG | 1 | 0 | 0 | 1422566 | CAA83660.1 | P36334 |
| DLQLGNLAYLQS | 1 | 0 | 0 | 1422567 | ARB66070.1 | P36334 |
| DLQLGNLGNLQS | 1 | 0 | 0 | 1422568 | AIL49519.1 | P36334 |
| DLQLNKLGTDDP | 1 | 0 | 0 | 1422571 | ATG84774.1 | K9N4V7 |
| DLQLTKLGIEDP | 1 | 0 | 0 | 1422573 | QBM11743.1 | K9N4V7 |
| DLQLTKLGTEDP | 1 | 0 | 0 | 1422574 | AVN89320.1 | K9N4V7 |
| DLQNCSFIYSFN | 1 | 0 | 0 | 1422575 | AVV62537.1 | K9N5Q8 |
| DLQNYGDSAILP | 1 | 0 | 0 | 1422576 | AVP78030.1 | P0C6W2 |
| DLQQAQQEATHI | 1 | 0 | 0 | 1422580 | APT69868.1 | A0A0P0ILY9 |
| DLQVEMNRLQDA | 1 | 0 | 0 | 1422587 | QEG03776.1 | P36334 |
| DLREYVFKNKDG | 1 | 0 | 0 | 1422591 | ATO98193.1 | Q3LZX1 |
| DLSAYYVKDGKY | 1 | 0 | 0 | 1422600 | AVV62524.1 | K9N7C7 |
| DLSDEMKVLNAV | 1 | 0 | 0 | 1422601 | AVV62526.1 | K9N5Q8 |

| | | | | | | |
|---|---|---|---|---|---|---|
| DLSSDENGVRTL | 1 | 0 | 0 | 1422634 | AVP78031.1 | Q3LZX1 |
| DLSVFYVKDGKY | 1 | 0 | 0 | 1422641 | ALA49857.1 | K9N7C7 |
| DLSVSSAGPIFQ | 1 | 0 | 0 | 1422647 | ASU90747.1 | K9N5Q8 |
| DLTAFYVKDGKY | 1 | 0 | 0 | 1422661 | AUM60013.1 | K9N7C7 |
| DLTGAICDFEIA | 1 | 0 | 0 | 1422667 | AEN19363.1 | P0C6X6 |
| DLTKCRTYLTSR | 1 | 0 | 0 | 1422675 | AUM60013.1 | K9N7C7 |
| DLTSDENGVRTL | 1 | 0 | 0 | 1422688 | ATO98120.1 | Q3LZX1 |
| DLTVGSGYCVDC | 1 | 0 | 0 | 1422695 | ABD43176.1 | P36334 |
| DLTYEMLSLQQV | 1 | 0 | 0 | 1422698 | AXN92228.1 | K9N5Q8 |
| DLVHDEPKGKFS | 1 | 0 | 0 | 1422705 | AVI15051.1 | P0C6X6 |
| DLVKRDSEADSP | 1 | 0 | 0 | 1422709 | ABD75589.1 | Q19U29 |
| DLYDKLQFASLE | 1 | 0 | 0 | 1422738 | APO40578.1 | P0C6X7 |
| DLYDKLQFMSLE | 1 | 0 | 0 | 1422739 | AVP78030.1 | P0C6W2 |
| DLYHYQECARGT | 1 | 0 | 0 | 1422751 | APO40584.1 | P59635 |
| DLYSAPTTLQAV | 1 | 0 | 0 | 1422757 | AUM60013.1 | K9N7C7 |
| DLYSMQDVGFKN | 1 | 0 | 0 | 1422760 | AAR01012.1 | P0C6X6 |
| DMADQIASLVLA | 1 | 0 | 0 | 1422771 | AEN19370.1 | P33469 |
| DMDDNLFIAKYG | 1 | 0 | 0 | 1422782 | ARU07574.1 | A0A140H1G9 |
| DMDSGGSSFSEL | 1 | 0 | 0 | 1422784 | AHX71940.1 | K9N7C7 |
| DMDYVPLKSAVC | 1 | 0 | 0 | 1422788 | AXG21663.1 | K9N7C7 |
| DMEKDFLSLDSD | 1 | 0 | 0 | 1422796 | AVV62535.1 | K9N7C7 |
| DMHWGDGFQIKF | 1 | 0 | 0 | 1422812 | AWW13564.1 | Q80872 |
| DMICSTTAQKLE | 1 | 0 | 0 | 1422816 | AEN19363.1 | P0C6X6 |
| DMLRPLIKDVDN | 1 | 0 | 0 | 1422829 | AHF54667.1 | P0C6X6 |
| DMNDFVSDADST | 1 | 0 | 0 | 1422835 | APO40578.1 | P0C6X7 |
| DMQFNKLGTEDP | 1 | 0 | 0 | 1422839 | AUM60021.1 | K9N4V7 |
| DMQNKMAATGVT | 1 | 0 | 0 | 1422840 | AUM60024.1 | K9N5Q8 |
| DMRNSSSEFFNK | 1 | 0 | 0 | 1422843 | AVV62537.1 | K9N5Q8 |
| DMSSVDNILNMA | 1 | 0 | 0 | 1422858 | AST12962.1 | A0A0P0LKV0 |
| DMYDPITKNVKG | 1 | 0 | 0 | 1422871 | AVV62535.1 | K9N7C7 |
| DMYDPVTKNING | 1 | 0 | 0 | 1422878 | AVV62524.1 | K9N7C7 |
| DMYVYSAGYATG | 1 | 0 | 0 | 1422884 | ANC28634.1 | K9N5Q8 |
| DMYVYSSGHATG | 1 | 0 | 0 | 1422885 | AVN89334.1 | K9N5Q8 |
| DNASPASTVHAI | 1 | 0 | 0 | 1422895 | ACQ82726.1 | P59632 |
| DNAYYTQQPIDL | 1 | 0 | 0 | 1422904 | APO40578.1 | P0C6X7 |
| DNCQLPLGQSLC | 1 | 0 | 0 | 1422906 | AVV62537.1 | K9N5Q8 |
| DNCYRKSNVDES | 1 | 0 | 0 | 1422908 | AIG96641.1 | A0A0P0LKV0 |
| DNCYRNSNVDEA | 1 | 0 | 0 | 1422910 | ALJ99927.1 | A0A0P0LKV0 |
| DNCYRNSNVGES | 1 | 0 | 0 | 1422911 | ADO24705.1 | A0A0P0LKV0 |
| DNCYRNSNVNKS | 1 | 0 | 0 | 1422912 | AJE26205.1 | A0A0P0LKV0 |
| DNCYRSTSVEES | 1 | 0 | 0 | 1422914 | AIG96682.1 | H9EJ42 |
| DNDDDNDDQIVV | 1 | 0 | 0 | 1422916 | ARU07575.1 | A0A140H1G9 |
| DNDDEHVVTGDN | 1 | 0 | 0 | 1422921 | ABD75543.1 | A0A140H1G9 |
| DNDLNEYVSDAD | 1 | 0 | 0 | 1422930 | AVV62524.1 | K9N7C7 |
| DNDVFIKKYGLE | 1 | 0 | 0 | 1422936 | QBM11746.1 | K9N7C7 |
| DNEALTVSTDVL | 1 | 0 | 0 | 1422939 | AVV62524.1 | K9N7C7 |
| DNETLEYWVTPL | 1 | 0 | 0 | 1422943 | BBA20986.1 | A0A140H1H1 |
| DNFCAFCTPKKV | 1 | 0 | 0 | 1422948 | AZS52616.1 | A0A140H1G9 |
| DNFDNNQNGDRT | 1 | 0 | 0 | 1422949 | APO40586.1 | P59595 |
| DNFNQWHEAKFG | 1 | 0 | 0 | 1422957 | AWK59942.1 | H9EJ42 |
| DNFPQGFSVLKP | 1 | 0 | 0 | 1422958 | APO40579.1 | P59594 |
| DNFTSIWPYYSF | 1 | 0 | 0 | 1422963 | ARU07576.1 | A0A140H1H0 |
| DNGAITVEELKQ | 1 | 0 | 0 | 1422966 | BAE93405.1 | P59596 |
| DNGALEKFMSTS | 1 | 0 | 0 | 1422969 | AVV62524.1 | K9N7C7 |
| DNGNYYCVRPCV | 1 | 0 | 0 | 1422979 | AUM60014.1 | K9N5Q8 |
| DNGPQNQSSAPR | 1 | 0 | 0 | 1422982 | AVP78049.1 | Q3LZX4 |
| DNGPQQNQRSAP | 1 | 0 | 0 | 1422984 | ARO76389.1 | P59595 |
| DNGSLEKFMSTP | 1 | 0 | 0 | 1422987 | AXN92247.1 | K9N7C7 |
| DNHITTYPGQGL | 1 | 0 | 0 | 1422999 | AVP78041.1 | P0C6W2 |
| DNIFSVQEDGRI | 1 | 0 | 0 | 1423002 | ABE97137.1 | A0A0P0G321 |
| DNIFSVQQDCRI | 1 | 0 | 0 | 1423003 | AKA09611.1 | A0A0P0G321 |
| DNIFSVQQDGRT | 1 | 0 | 0 | 1423005 | AKA09627.1 | A0A0P0G321 |
| DNINSTLVDLNW | 1 | 0 | 0 | 1423020 | AGT21353.1 | A0A0P0K6L9 |
| DNISVASPKSRV | 1 | 0 | 0 | 1423025 | ATP16771.1 | P33469 |
| DNISVATPKSRV | 1 | 0 | 0 | 1423026 | AIX10720.1 | P33469 |
| DNKFALTCFSTQ | 1 | 0 | 0 | 1423032 | AVP78047.1 | Q3LZX7 |
| DNKFALTCTSTH | 1 | 0 | 0 | 1423034 | ATO98237.1 | Q3LZX7 |
| DNKFAVTCTTTK | 1 | 0 | 0 | 1423035 | APO40584.1 | P59635 |
| DNKFTPLKVEEV | 1 | 0 | 0 | 1423037 | AUM60013.1 | K9N7C7 |
| DNKIMTFYPKMQ | 1 | 0 | 0 | 1423038 | ARU07574.1 | A0A140H1G9 |
| DNKLKAHKEKSA | 1 | 0 | 0 | 1423041 | P0C6W6.1 | P0C6W2 |
| DNKLKAHKEKSS | 1 | 0 | 0 | 1423042 | APO40578.1 | P0C6X7 |
| DNLCTLDPITLK | 1 | 0 | 0 | 1423057 | AWW13566.1 | P36334 |
| DNLEPVSGLYEI | 1 | 0 | 0 | 1423060 | AIL49538.1 | P36334 |
| DNLIGYYSDDGN | 1 | 0 | 0 | 1423061 | AVV62537.1 | K9N5Q8 |
| DNLRANNTKGVL | 1 | 0 | 0 | 1423073 | APO40578.1 | P0C6X7 |
| DNLSASEKEVLL | 1 | 0 | 0 | 1423077 | AUM60013.1 | K9N7C7 |
| DNLTADEIKALK | 1 | 0 | 0 | 1423080 | AVN89363.1 | K9N7C7 |
| DNLTDYSVSSCA | 1 | 0 | 0 | 1423081 | BBA20983.1 | A0A140H1H1 |
| DNNDITNTNMSR | 1 | 0 | 0 | 1423093 | ATG84708.1 | K9N4V7 |
| DNNDNNNQQSRG | 1 | 0 | 0 | 1423094 | AUM60021.1 | K9N4V7 |
| DNNDNNSNNQQS | 1 | 0 | 0 | 1423095 | AVV62533.1 | K9N4V7 |
| DNNDNNTNQQSR | 1 | 0 | 0 | 1423096 | AVV62544.1 | K9N4V7 |
| DNNDTTNTNLSR | 1 | 0 | 0 | 1423097 | ALW82676.1 | K9N4V7 |
| DNNGEDVVTGDN | 1 | 0 | 0 | 1423100 | ABD75631.1 | A0A140H1G9 |
| DNNNQQSRGRGR | 1 | 0 | 0 | 1423103 | AUM60021.1 | K9N4V7 |
| DNNQDGGRSGAR | 1 | 0 | 0 | 1423104 | Q3I5I7.1 | Q3LZX4 |
| DNNSNNQQSRGR | 1 | 0 | 0 | 1423108 | AVV62533.1 | K9N4V7 |
| DNNSTLQSVASS | 1 | 0 | 0 | 1423109 | AGT51379.1 | H9EJ42 |
| DNNTDSIVIDGV | 1 | 0 | 0 | 1423111 | AWK59942.1 | H9EJ42 |
| DNNTNQQSRGRG | 1 | 0 | 0 | 1423112 | AVV62544.1 | K9N4V7 |

| | | | | | | |
|---|---|---|---|---|---|---|
| DNNVQRCTRKLN | 1 | 0 | 0 | 1423113 | NP_073549.1 | A0A0P0LKV0 |
| DNPAFVVTGGAQ | 1 | 0 | 0 | 1423114 | ARO76382.1 | P59594 |
| DNPCYCFNADGD | 1 | 0 | 0 | 1423116 | AUM60013.1 | K9N7C7 |
| DNPNHFINFPLT | 1 | 0 | 0 | 1423125 | ARU07566.1 | Q80872 |
| DNPNIPFKDGLY | 1 | 0 | 0 | 1423126 | APO40579.1 | P59594 |
| DNPNSSEVDMIC | 1 | 0 | 0 | 1423127 | AOL02450.1 | P0C6X6 |
| DNPSAIVTGLLP | 1 | 0 | 0 | 1423130 | AKA09625.1 | A0A0P0G321 |
| DNPTLPFGDGIY | 1 | 0 | 0 | 1423133 | ATO98193.1 | Q3LZX1 |
| DNPVIPFKDGIY | 1 | 0 | 0 | 1423134 | 5X58_A | P59594 |
| DNQHIETPITIN | 1 | 0 | 0 | 1423140 | AVV62526.1 | K9N5Q8 |
| DNSANTKVLKNK | 1 | 0 | 0 | 1423162 | ATQ39389.1 | K9N7C7 |
| DNSHIETPIVIN | 1 | 0 | 0 | 1423165 | AVV62537.1 | K9N5Q8 |
| DNSTSSVVQTCD | 1 | 0 | 0 | 1423179 | CAA83660.1 | P36334 |
| DNTDSIVIDGVT | 1 | 0 | 0 | 1423185 | AWK59930.1 | H9EJ42 |
| DNTKFADDLNQL | 1 | 0 | 0 | 1423190 | AVP78030.1 | P0C6W2 |
| DNTNQYATLFGS | 1 | 0 | 0 | 1423194 | ATQ39390.1 | K9N5Q8 |
| DNTTQSAVLFNN | 1 | 0 | 0 | 1423205 | APO40579.1 | P59594 |
| DNVADFNAIATC | 1 | 0 | 0 | 1423207 | APO40578.1 | P0C6X7 |
| DNVFIKNADIVE | 1 | 0 | 0 | 1423214 | AVP78041.1 | P0C6W2 |
| DNVIMLNKHIDA | 1 | 0 | 0 | 1423218 | ARO76389.1 | P59595 |
| DNVSSVWPFYPY | 1 | 0 | 0 | 1423233 | AGT51679.1 | P30215 |
| DNVYCYNTGLIE | 1 | 0 | 0 | 1423239 | AFV53147.1 | H9EJ42 |
| DNVYMAKSGDKY | 1 | 0 | 0 | 1423243 | AXX83372.1 | P0C6X6 |
| DNVYMAKVGDKY | 1 | 0 | 0 | 1423244 | AEN19363.1 | P0C6X6 |
| DNWLNKVSVPSP | 1 | 0 | 0 | 1423245 | AMM42407.1 | A0A140H1H1 |
| DNYDDYDFEPQK | 1 | 0 | 0 | 1423246 | AUM60014.1 | K9N5Q8 |
| DNYFSKKDWYDF | 1 | 0 | 0 | 1423248 | APO40578.1 | P0C6X7 |
| DPAGVCLGAGQF | 1 | 0 | 0 | 1423259 | AEN19363.1 | P0C6X6 |
| DPALETGEYTFL | 1 | 0 | 0 | 1423263 | AVV62524.1 | K9N7C7 |
| DPAQLPTPRTLL | 1 | 0 | 0 | 1423271 | ATG84864.1 | K9N7C7 |
| DPDDASRLLHTV | 1 | 0 | 0 | 1423278 | AVN89333.1 | K9N7C7 |
| DPFMVNQLAYSS | 1 | 0 | 0 | 1423290 | AUM60013.1 | K9N7C7 |
| DPIALKAPDTYK | 1 | 0 | 0 | 1423297 | BBO58891.1 | P36334 |
| DPINCESIYEFD | 1 | 0 | 0 | 1423304 | ABD75591.1 | A0A140H1G9 |
| DPIRSCLENTVC | 1 | 0 | 0 | 1423305 | CAA49377.1 | A0A0P0LKV0 |
| DPISFKAPDTYK | 1 | 0 | 0 | 1423306 | ARE30008.1 | P36334 |
| DPITFAGPYKCP | 1 | 0 | 0 | 1423308 | ATP16757.1 | P36334 |
| DPITFKAPDTYK | 1 | 0 | 0 | 1423309 | ARB07414.1 | P36334 |
| DPITKNVKGNNE | 1 | 0 | 0 | 1423310 | AVV62535.1 | K9N7C7 |
| DPITKTEKDTNE | 1 | 0 | 0 | 1423311 | AUM60013.1 | K9N7C7 |
| DPITLKTPDTYK | 1 | 0 | 0 | 1423313 | AXX83339.1 | P36334 |
| DPKLKGSFNDRD | 1 | 0 | 0 | 1423330 | AIL49495.1 | P36334 |
| DPKVSFEFLQRD | 1 | 0 | 0 | 1423345 | AVV62524.1 | K9N7C7 |
| DPLFSNIQPFIV | 1 | 0 | 0 | 1423351 | AST12962.1 | A0A0P0LKV0 |
| DPLNLSADTQHI | 1 | 0 | 0 | 1423358 | ADN03343.1 | Q19U29 |
| DPMPNNGWTAFS | 1 | 0 | 0 | 1423371 | AVP78031.1 | Q3LZX1 |
| DPMYKNGAMLNG | 1 | 0 | 0 | 1423373 | AUM60013.1 | K9N7C7 |
| DPNAFIYEWPMP | 1 | 0 | 0 | 1423374 | AUM60014.1 | K9N5Q8 |
| DPNGNLYGFRDY | 1 | 0 | 0 | 1423376 | ARB66062.1 | P36334 |
| DPNITNLCPFGE | 1 | 0 | 0 | 1423378 | 2GHV_E | P59594 |
| DPNKTNVVPPAL | 1 | 0 | 0 | 1423380 | ARO76391.1 | P59636 |
| DPNLSGFYVKDG | 1 | 0 | 0 | 1423381 | ATQ39389.1 | K9N7C7 |
| DPNMEGAYTSSL | 1 | 0 | 0 | 1423385 | AVV62537.1 | K9N5Q8 |
| DPPNCNIEAWLN | 1 | 0 | 0 | 1423398 | AHA80201.1 | P36334 |
| DPPQADQQVDRL | 1 | 0 | 0 | 1423400 | 6U7H_A | A0A0P0K6L9 |
| DPQIQLTTTRME | 1 | 0 | 0 | 1423410 | Q3I5I6.1 | Q3LZX3 |
| DPQIQVTIAKTE | 1 | 0 | 0 | 1423412 | AVP78050.1 | Q3LZX3 |
| DPQKAYIDFVNS | 1 | 0 | 0 | 1423413 | AUM60023.1 | K9N7C7 |
| DPQKAYMDFVNS | 1 | 0 | 0 | 1423415 | AUM60013.1 | K9N7C7 |
| DPQQLPAPRVLI | 1 | 0 | 0 | 1423416 | NP_073549.1 | A0A0P0LKV0 |
| DPQTLEILDISP | 1 | 0 | 0 | 1423418 | Q3I5J5.1 | Q3LZX1 |
| DPRLKGSFNDRD | 1 | 0 | 0 | 1423422 | AIL49520.1 | P36334 |
| DPRLRGSFNNRD | 1 | 0 | 0 | 1423424 | BBA84088.1 | P36334 |
| DPRNDEAIPSMF | 1 | 0 | 0 | 1423425 | AJQ24068.1 | P33469 |
| DPSGLSPAYAPT | 1 | 0 | 0 | 1423434 | AVV62524.1 | K9N7C7 |
| DPTVLPGAHAYE | 1 | 0 | 0 | 1423457 | AVV62535.1 | K9N7C7 |
| DPTVLPGAHAYG | 1 | 0 | 0 | 1423458 | AVV62524.1 | K9N7C7 |
| DPVGFCLTNTPC | 1 | 0 | 0 | 1423465 | AUM60013.1 | K9N7C7 |
| DPVGFVLRNTVC | 1 | 0 | 0 | 1423469 | APO40578.1 | P0C6X7 |
| DPVLMGWDYPKC | 1 | 0 | 0 | 1423473 | AJS10811.1 | P0C6X6 |
| DPVTKNINGSNE | 1 | 0 | 0 | 1423477 | AVV62524.1 | K9N7C7 |
| DPYLSGYYHNNK | 1 | 0 | 0 | 1423482 | AVP78031.1 | Q3LZX1 |
| DQACYERFPLCS | 1 | 0 | 0 | 1423484 | AUM60013.1 | K9N7C7 |
| DQFRSVQTRGRR | 1 | 0 | 0 | 1423524 | AIL49389.1 | P33469 |
| DQGHILDSYFVV | 1 | 0 | 0 | 1423526 | AVV62535.1 | K9N7C7 |
| DQGLKKFYKEVR | 1 | 0 | 0 | 1423528 | AGV08522.1 | K9N7C7 |
| DQGQYYYRSSRK | 1 | 0 | 0 | 1423530 | ATO98193.1 | Q3LZX1 |
| DQKLYDQVVMDY | 1 | 0 | 0 | 1423551 | ARO76381.1 | P0C6U8 |
| DQLNSSYKLSIP | 1 | 0 | 0 | 1423556 | ATW75477.1 | K9N5Q8 |
| DQLNSTYFKLSI | 1 | 0 | 0 | 1423557 | ANI69889.1 | K9N5Q8 |
| DQLREPMMQSAD | 1 | 0 | 0 | 1423560 | P0C6W6.1 | P0C6W2 |
| DQLREPMMQSSD | 1 | 0 | 0 | 1423561 | APO40578.1 | P0C6X7 |
| DQMSELPKEQRV | 1 | 0 | 0 | 1423573 | ASJ26617.1 | K9N4V7 |
| DQMSEPPNEQRV | 1 | 0 | 0 | 1423577 | AVN89381.1 | K9N4V7 |
| DQNKQESSDDVD | 1 | 0 | 0 | 1423580 | AVV62544.1 | K9N4V7 |
| DQPYHANCVNCA | 1 | 0 | 0 | 1423587 | AVV62535.1 | K9N7C7 |
| DQQAQQEATHIA | 1 | 0 | 0 | 1423588 | APT69896.1 | A0A0P0LKV0 |
| DQQPSVLFTCIP | 1 | 0 | 0 | 1423591 | AVV62530.1 | K9N7D2 |
| DQSDQVRNVQTR | 1 | 0 | 0 | 1423604 | P33469.1 | P33469 |
| DQSFKDELEEFF | 1 | 0 | 0 | 1423609 | AVV62526.1 | K9N5Q8 |
| DQSRKWISHCTI | 1 | 0 | 0 | 1423614 | AIV41836.1 | Q80872 |

| | | | | | | |
|---|---|---|---|---|---|---|
| DQSRKWTGHCTI | 1 | 0 | 0 | 1423615 | AIV41775.1 | Q80872 |
| DQTMTQMYKQAR | 1 | 0 | 0 | 1423628 | AVP78041.1 | P0C6W2 |
| DQTSFNVSSLAY | 1 | 0 | 0 | 1423631 | AVV62524.1 | K9N7C7 |
| DQVVMDYLDSLK | 1 | 0 | 0 | 1423656 | ARO76381.1 | P0C6U8 |
| DQWFKNQTLVAP | 1 | 0 | 0 | 1423658 | CAA83661.1 | P36334 |
| DQWFKNQTSVAR | 1 | 0 | 0 | 1423661 | AIX10750.1 | P36334 |
| DQWLKNQTSVAP | 1 | 0 | 0 | 1423662 | ARB07414.1 | P36334 |
| DQYGTDINGVYW | 1 | 0 | 0 | 1423672 | AEN19370.1 | P33469 |
| DQYLDSKGPLQV | 1 | 0 | 0 | 1423673 | AVV62535.1 | K9N7C7 |
| DQYMCGSDGKPI | 1 | 0 | 0 | 1423676 | AUM60013.1 | K9N7C7 |
| DRAAFVCGDYAA | 1 | 0 | 0 | 1423678 | AIL49498.1 | P36334 |
| DRAMRLALRLVS | 1 | 0 | 0 | 1423689 | APO40587.1 | P59636 |
| DRDLYDKLQFAS | 1 | 0 | 0 | 1423703 | APO40578.1 | P0C6X7 |
| DRDLYDKLQFMS | 1 | 0 | 0 | 1423704 | AVP78030.1 | P0C6W2 |
| DRDPRNDEAIPS | 1 | 0 | 0 | 1423706 | AJQ24068.1 | P33469 |
| DRDPSNDEAIPT | 1 | 0 | 0 | 1423707 | AJQ24082.1 | P33469 |
| DRDSAVGHTEFY | 1 | 0 | 0 | 1423710 | AUM60025.1 | K9N796 |
| DRDSTSTHDGPT | 1 | 0 | 0 | 1423712 | QDI73611.1 | K9N796 |
| DRDTGPPSISTD | 1 | 0 | 0 | 1423713 | AIL49520.1 | P36334 |
| DRESTSTHDGFN | 1 | 0 | 0 | 1423724 | AHI48551.1 | K9N796 |
| DRESTSTHDGLN | 1 | 0 | 0 | 1423727 | ANF29196.1 | K9N796 |
| DRESTSTHDGPN | 1 | 0 | 0 | 1423730 | AHE78098.1 | K9N796 |
| DRESTSTHDGPT | 1 | 0 | 0 | 1423731 | AHI48573.1 | K9N796 |
| DRESTSTHDVPT | 1 | 0 | 0 | 1423733 | AMO03402.1 | K9N796 |
| DRFKVNETTSQY | 1 | 0 | 0 | 1423735 | ALW82700.1 | K9N7C7 |
| DRGQESLARSAC | 1 | 0 | 0 | 1423748 | AXN92247.1 | K9N7C7 |
| DRGRNPKPRAAP | 1 | 0 | 0 | 1423749 | AVN89370.1 | K9N4V7 |
| DRILDYKVLDNG | 1 | 0 | 0 | 1423757 | AVV62524.1 | K9N7C7 |
| DRIRFCLMLNSY | 1 | 0 | 0 | 1423763 | AXG21669.1 | K9N7D2 |
| DRKINTGNGTKA | 1 | 0 | 0 | 1423778 | AVV62533.1 | K9N4V7 |
| DRKINTGNGTKQ | 1 | 0 | 0 | 1423780 | AUM60021.1 | K9N4V7 |
| DRLDIIQADQQV | 1 | 0 | 0 | 1423785 | AIW52757.1 | A0A0P0K6L9 |
| DRLDKLIATAKD | 1 | 0 | 0 | 1423786 | AVV62524.1 | K9N7C7 |
| DRLNEVARNLNE | 1 | 0 | 0 | 1423796 | AVP78031.1 | Q3LZX1 |
| DRLNQLENKISG | 1 | 0 | 0 | 1423797 | AVP78049.1 | Q3LZX4 |
| DRLNQLENKVSG | 1 | 0 | 0 | 1423798 | AVP78038.1 | Q3LZX4 |
| DRMLKTSMAPIL | 1 | 0 | 0 | 1423808 | AVR40343.1 | Q4VID0 |
| DRNQHSKLDEEV | 1 | 0 | 0 | 1423817 | APO40583.1 | P59634 |
| DRQIHQAQLTDK | 1 | 0 | 0 | 1423825 | AVV62524.1 | K9N7C7 |
| DRQIYQSQLTDK | 1 | 0 | 0 | 1423826 | AUM60013.1 | K9N7C7 |
| DRSEKSYEQQTP | 1 | 0 | 0 | 1423835 | APO40578.1 | P0C6X7 |
| DRSHYYVDSVTV | 1 | 0 | 0 | 1423837 | AXN92247.1 | K9N7C7 |
| DRTAIQSFDSSY | 1 | 0 | 0 | 1423842 | AST12962.1 | A0A0P0LKV0 |
| DRTELPQLVNAN | 1 | 0 | 0 | 1423848 | ATG84679.1 | K9N5Q8 |
| DRVEKSFQLDTA | 1 | 0 | 0 | 1423858 | Q3I5J5.1 | Q3LZX1 |
| DRVLPNMIRMIS | 1 | 0 | 0 | 1423861 | QED88038.1 | H9EJ42 |
| DRVVEPILATEP | 1 | 0 | 0 | 1423867 | AHF54652.1 | Q4VID0 |
| DRWSASINTGNC | 1 | 0 | 0 | 1423880 | AIW52757.1 | A0A0P0K6L9 |
| DRYCAVHQGAHV | 1 | 0 | 0 | 1423881 | AVV62524.1 | K9N7C7 |
| DRYEEYDLQPHK | 1 | 0 | 0 | 1423885 | AHI48550.1 | K9N5Q8 |
| DRYEEYEIEPHK | 1 | 0 | 0 | 1423886 | AVV62526.1 | K9N5Q8 |
| DRYHNSQDDIMK | 1 | 0 | 0 | 1423888 | AAA45463.1 | A0A0N7J131 |
| DRYVYSAGHATG | 1 | 0 | 0 | 1423891 | ALA49803.1 | K9N5Q8 |
| DSADNNQNGGRN | 1 | 0 | 0 | 1423894 | ARO76389.1 | P59595 |
| DSAFVPQPTGVF | 1 | 0 | 0 | 1423898 | AGT51481.1 | P36334 |
| DSAILPKGIMMN | 1 | 0 | 0 | 1423900 | AVP78030.1 | P0C6W2 |
| DSAKSACIEVDI | 1 | 0 | 0 | 1423901 | AVN89387.1 | K9N5Q8 |
| DSAKSVCIEVDI | 1 | 0 | 0 | 1423902 | QAT98898.1 | K9N5Q8 |
| DSANPTNCIAPV | 1 | 0 | 0 | 1423903 | AVN89334.1 | K9N5Q8 |
| DSATKVYFDNPT | 1 | 0 | 0 | 1423905 | ATO98193.1 | Q3LZX1 |
| DSAVGHTEFYDS | 1 | 0 | 0 | 1423910 | AUM60025.1 | K9N796 |
| DSCISNNRCNIF | 1 | 0 | 0 | 1423913 | BBA20986.1 | A0A140H1H1 |
| DSCPGKNNGIGT | 1 | 0 | 0 | 1423921 | AIV41891.1 | P36334 |
| DSDFVKQPVKFY | 1 | 0 | 0 | 1423931 | AVN89451.1 | K9N7C7 |
| DSDMNDFVSDAD | 1 | 0 | 0 | 1423935 | APO40578.1 | P0C6X7 |
| DSDQPWKVIPRN | 1 | 0 | 0 | 1423936 | AST12968.1 | A0A0N7J131 |
| DSDSLPSVPFTC | 1 | 0 | 0 | 1423937 | AVV62541.1 | K9N7D2 |
| DSDTPSDPLNNF | 1 | 0 | 0 | 1423939 | AUM60013.1 | K9N7C7 |
| DSDVTKPLIKWD | 1 | 0 | 0 | 1423946 | APO40578.1 | P0C6X7 |
| DSDVVKQPVKFY | 1 | 0 | 0 | 1423947 | AXN92247.1 | K9N7C7 |
| DSDVVKQPVNFY | 1 | 0 | 0 | 1423948 | AXN73479.1 | K9N7C7 |
| DSEAIKQPVKFY | 1 | 0 | 0 | 1423949 | AVV62524.1 | K9N7C7 |
| DSEATKQPVKFY | 1 | 0 | 0 | 1423950 | AVV62535.1 | K9N7C7 |
| DSESSVDPLQSF | 1 | 0 | 0 | 1423959 | AVV62535.1 | K9N7C7 |
| DSFRSTSRQQSR | 1 | 0 | 0 | 1423978 | ABE97134.1 | H9EJ69 |
| DSFSCNNLDKSK | 1 | 0 | 0 | 1423980 | BBA20986.1 | A0A140H1H1 |
| DSFTCNNIDVAK | 1 | 0 | 0 | 1423982 | ABD43176.1 | P36334 |
| DSFYTQGRSCSD | 1 | 0 | 0 | 1423994 | AVV62524.1 | K9N7C7 |
| DSGSLTKASDWK | 1 | 0 | 0 | 1424019 | AUM60013.1 | K9N7C7 |
| DSGSLTKSSEWK | 1 | 0 | 0 | 1424020 | AVV62524.1 | K9N7C7 |
| DSIKSACIEVDI | 1 | 0 | 0 | 1424050 | QBF80543.1 | K9N5Q8 |
| DSKCLTSIEEVC | 1 | 0 | 0 | 1424069 | AEN19363.1 | P0C6X6 |
| DSKNPNYKKWLE | 1 | 0 | 0 | 1424081 | AVV62544.1 | K9N4V7 |
| DSKNPNYQKWME | 1 | 0 | 0 | 1424082 | AUM60021.1 | K9N4V7 |
| DSKPPLPPDEWF | 1 | 0 | 0 | 1424083 | ANF29266.1 | K9N5R3 |
| DSKPQQVTKQNA | 1 | 0 | 0 | 1424087 | AMM42397.1 | Q19U29 |
| DSKSGNNFYYRL | 1 | 0 | 0 | 1424091 | APO40579.1 | P59594 |
| DSLCKTIQSALS | 1 | 0 | 0 | 1424100 | QEO75984.1 | A0A0P0ILY9 |
| DSLECSNLLLQY | 1 | 0 | 0 | 1424106 | Q3I5J5.1 | Q3LZX1 |
| DSLGASSNQSPQ | 1 | 0 | 0 | 1424112 | AIX09796.1 | P0C6X6 |
| DSLIYKFDSGSL | 1 | 0 | 0 | 1424114 | AVV62524.1 | K9N7C7 |

| | | | | | | |
|---|---|---|---|---|---|---|
| DSLIYKYDSGSL | 1 | 0 | 0 | 1424115 | AVV62535.1 | K9N7C7 |
| DSLKPTVETPKE | 1 | 0 | 0 | 1424118 | ARO76381.1 | P0C6U8 |
| DSLKPVGGLYEI | 1 | 0 | 0 | 1424119 | AIL49500.1 | P36334 |
| DSLKTGVSVPCV | 1 | 0 | 0 | 1424121 | ARO76381.1 | P0C6U8 |
| DSLLKLEDFIPM | 1 | 0 | 0 | 1424123 | AVP78041.1 | P0C6W2 |
| DSLNAKWIFDSS | 1 | 0 | 0 | 1424125 | AIV41816.1 | P0C6X6 |
| DSLNAKWVFDSS | 1 | 0 | 0 | 1424126 | AIV41984.1 | P0C6X6 |
| DSLNSKLGFDSS | 1 | 0 | 0 | 1424131 | AEN19363.1 | P0C6X6 |
| DSLQFAELSTNN | 1 | 0 | 0 | 1424136 | AVV62524.1 | K9N7C7 |
| DSLRQATLPQSK | 1 | 0 | 0 | 1424140 | AKI29254.1 | K9N638 |
| DSLRQVSLTQSK | 1 | 0 | 0 | 1424143 | AUM60013.1 | K9N7C7 |
| DSLRVTFIDGLS | 1 | 0 | 0 | 1424145 | AEN19363.1 | P0C6X6 |
| DSLTHYPALKMV | 1 | 0 | 0 | 1424150 | ANI69867.1 | K9N638 |
| DSMNGASDSTQA | 1 | 0 | 0 | 1424154 | APO40586.1 | P59595 |
| DSMNLIANIFTP | 1 | 0 | 0 | 1424155 | ARO76381.1 | P0C6U8 |
| DSNAASVNPANM | 1 | 0 | 0 | 1424158 | AVV62524.1 | K9N7C7 |
| DSNAFSLANVIS | 1 | 0 | 0 | 1424159 | AFD64760.1 | A0A0P0G321 |
| DSNAFSLANVTS | 1 | 0 | 0 | 1424160 | AIW52843.1 | A0A0P0G321 |
| DSNCCVVEASTD | 1 | 0 | 0 | 1424161 | AGT51386.1 | H9EJ42 |
| DSNFLYKYYSLK | 1 | 0 | 0 | 1424168 | AVV62524.1 | K9N7C7 |
| DSNFRTEVDPDL | 1 | 0 | 0 | 1424169 | ASU90302.1 | K9N7C7 |
| DSNGFIRFPDVI | 1 | 0 | 0 | 1424171 | ABD75543.1 | A0A140H1G9 |
| DSNGNLYGLRDY | 1 | 0 | 0 | 1424173 | ABD43179.1 | P36334 |
| DSNIDAYKNFPK | 1 | 0 | 0 | 1424175 | AUM60021.1 | K9N4V7 |
| DSNVKEEPTTQK | 1 | 0 | 0 | 1424183 | ARO76381.1 | P0C6U8 |
| DSNWAFVAFSFL | 1 | 0 | 0 | 1424185 | AST12967.1 | A0A0P0KXN3 |
| DSPCFSQQGVFR | 1 | 0 | 0 | 1424190 | AAA73152.1 | P30215 |
| DSPDLGTDDDDK | 1 | 0 | 0 | 1424192 | 3R24_A | P0C6X7 |
| DSPVKDVFELHY | 1 | 0 | 0 | 1424198 | ABD75589.1 | Q19U29 |
| DSQDGQETAVTK | 1 | 0 | 0 | 1424201 | AVP78041.1 | P0C6W2 |
| DSQFEEDVEMSD | 1 | 0 | 0 | 1424203 | AGT51787.1 | P0C6X6 |
| DSQNMLRGDDMS | 1 | 0 | 0 | 1424206 | AFR58685.1 | P0C6X7 |
| DSQNMLRGEDVS | 1 | 0 | 0 | 1424209 | ARO76381.1 | P0C6U8 |
| DSQPRADKPSQL | 1 | 0 | 0 | 1424210 | 5EPW_A | H9EJ69 |
| DSQPSTSFYDIA | 1 | 0 | 0 | 1424211 | AUM60025.1 | K9N796 |
| DSQRAYIGSTAY | 1 | 0 | 0 | 1424213 | AUM60027.1 | K9N643 |
| DSQYFFNKDTGV | 1 | 0 | 0 | 1424216 | AGT51759.1 | P30215 |
| DSRLKGSFNNRD | 1 | 0 | 0 | 1424224 | AIL49507.1 | P36334 |
| DSRSDCNHIGNI | 1 | 0 | 0 | 1424225 | AEN19365.1 | P30215 |
| DSRTGSFNNRDT | 1 | 0 | 0 | 1424229 | ABD43177.1 | P36334 |
| DSRTGSLNNVDT | 1 | 0 | 0 | 1424230 | AMK59677.1 | P36334 |
| DSRWNNARHSDN | 1 | 0 | 0 | 1424234 | AQN78751.1 | P30215 |
| DSRWNNARLSDN | 1 | 0 | 0 | 1424235 | AAX85668.1 | P30215 |
| DSSDHGQESLAR | 1 | 0 | 0 | 1424240 | AHI48549.1 | K9N7C7 |
| DSSDNSQNGERN | 1 | 0 | 0 | 1424242 | AVP78049.1 | Q3LZX4 |
| DSSEGDAKETKE | 1 | 0 | 0 | 1424246 | AGT51737.1 | P0C6X6 |
| DSSEPLCLFKKN | 1 | 0 | 0 | 1424248 | BBA20986.1 | A0A140H1H1 |
| DSSFEKLHCEHL | 1 | 0 | 0 | 1424251 | AWO68974.1 | A0A0P0G321 |
| DSSGKPVPYCYD | 1 | 0 | 0 | 1424255 | AVP78041.1 | P0C6W2 |
| DSSKSVTKKYTY | 1 | 0 | 0 | 1424264 | AVV61898.1 | K9N7C7 |
| DSTDNNQDGGRS | 1 | 0 | 0 | 1424279 | Q3I5I7.1 | Q3LZX4 |
| DSTLIGDCVTVH | 1 | 0 | 0 | 1424298 | QSV83056.1 | P0DTD1 |
| DSTQDGVNKLQG | 1 | 0 | 0 | 1424304 | QBQ01839.1 | P36334 |
| DSTRHTYQLRAR | 1 | 0 | 0 | 1424305 | APO40584.1 | P59635 |
| DSTYGPLQTPVS | 1 | 0 | 0 | 1424314 | ALX27228.1 | K9N5Q8 |
| DSTYGPLQTSVG | 1 | 0 | 0 | 1424315 | ASU91284.1 | K9N5Q8 |
| DSVFKPQPAGVL | 1 | 0 | 0 | 1424327 | ARB66062.1 | P36334 |
| DSVISAVSAGIE | 1 | 0 | 0 | 1424334 | AGT51787.1 | P0C6X6 |
| DSVIVNDDTSES | 1 | 0 | 0 | 1424337 | AWW13571.1 | P0C6X6 |
| DSVIVNNDTSET | 1 | 0 | 0 | 1424338 | AOL02450.1 | P0C6X6 |
| DSVKSACIKVDI | 1 | 0 | 0 | 1424344 | QCQ29075.1 | K9N5Q8 |
| DSVQYAHKHDIN | 1 | 0 | 0 | 1424353 | AUM60013.1 | K9N7C7 |
| DSVRDPQTLDIL | 1 | 0 | 0 | 1424358 | ATO98145.1 | Q3LZX1 |
| DSVRDPQTLQIL | 1 | 0 | 0 | 1424360 | ATO98193.1 | Q3LZX1 |
| DSVRNLFESVKS | 1 | 0 | 0 | 1424363 | QBM11737.1 | K9N5Q8 |
| DSVSAIGTANLS | 1 | 0 | 0 | 1424366 | AAK32190.1 | A0A0P0K6L9 |
| DSVTVKETVAQF | 1 | 0 | 0 | 1424374 | QAT98908.1 | K9N7C7 |
| DSVTVKNGTLYM | 1 | 0 | 0 | 1424377 | APO40578.1 | P0C6X7 |
| DSYLGCVFNADN | 1 | 0 | 0 | 1424394 | BBA20983.1 | A0A140H1H1 |
| DTALALLLLDRL | 1 | 0 | 0 | 1424417 | AVP78038.1 | Q3LZX4 |
| DTAPKTGNFKDL | 1 | 0 | 0 | 1424420 | ATO98181.1 | Q3LZX1 |
| DTCFGVSKPNAI | 1 | 0 | 0 | 1424430 | AWK59942.1 | H9EJ42 |
| DTCVRSMIKTIV | 1 | 0 | 0 | 1424436 | AVV62530.1 | K9N7D2 |
| DTCVSNNLCNIF | 1 | 0 | 0 | 1424438 | ABD75609.1 | A0A140H1H1 |
| DTCVSNNRCNIF | 1 | 0 | 0 | 1424439 | BBA20983.1 | A0A140H1H1 |
| DTENSTEFTRVN | 1 | 0 | 0 | 1424451 | AVP78041.1 | P0C6W2 |
| DTEQTQSAIAKP | 1 | 0 | 0 | 1424452 | ATQ39389.1 | K9N7C7 |
| DTEQTQSVIVKP | 1 | 0 | 0 | 1424453 | AVN89451.1 | K9N7C7 |
| DTEQTQSVNVKL | 1 | 0 | 0 | 1424454 | ANF29194.1 | K9N7C7 |
| DTEQTQSVNVKP | 1 | 0 | 0 | 1424455 | ATG84700.1 | K9N638 |
| DTEWGNMMTCIS | 1 | 0 | 0 | 1424460 | AVV62524.1 | K9N7C7 |
| DTEWGNMMTGIS | 1 | 0 | 0 | 1424461 | AVV62535.1 | K9N7C7 |
| DTFIDNPCYCFN | 1 | 0 | 0 | 1424467 | AUM60013.1 | K9N7C7 |
| DTFLIRFSEVRQ | 1 | 0 | 0 | 1424469 | AVP78031.1 | Q3LZX1 |
| DTFNQLTADLQH | 1 | 0 | 0 | 1424474 | ARB07436.1 | A0A140H1G9 |
| DTGHPSISTDTV | 1 | 0 | 0 | 1424491 | AIX10759.1 | P36334 |
| DTGLPSISTDTV | 1 | 0 | 0 | 1424496 | BBA84092.1 | P36334 |
| DTGNCPFSFGKV | 1 | 0 | 0 | 1424497 | ALK28781.1 | A0A0P0K6L9 |
| DTGRPSISTDTV | 1 | 0 | 0 | 1424504 | BBO58746.1 | P36334 |
| DTGSFYGFSNVV | 1 | 0 | 0 | 1424505 | ABD75600.1 | A0A140H1H0 |
| DTGVIYGFNSTE | 1 | 0 | 0 | 1424511 | AIL49343.1 | P30215 |

| | | | | | | |
|---|---|---|---|---|---|---|
| DTGVPSTSTDIV | 1 | 0 | 0 | 1424513 | CAA83661.1 | P36334 |
| DTGVQVVDYLTR | 1 | 0 | 0 | 1424514 | AVV62524.1 | K9N7C7 |
| DTIDVLLKQPMA | 1 | 0 | 0 | 1424523 | AGT51735.1 | Q4VID0 |
| DTIPQQSQNGFS | 1 | 0 | 0 | 1424536 | AVV62535.1 | K9N7C7 |
| DTITVDQLKHLL | 1 | 0 | 0 | 1424539 | APO40582.1 | P59596 |
| DTIVVTAGDGIS | 1 | 0 | 0 | 1424543 | ATO98194.1 | Q3LZX0 |
| DTKCLADTVMSA | 1 | 0 | 0 | 1424549 | AQN78757.1 | P0C6X6 |
| DTKIASQLGMGG | 1 | 0 | 0 | 1424554 | AWF93655.1 | K9N5Q8 |
| DTKIVSQLGNCV | 1 | 0 | 0 | 1424556 | AVN89313.1 | K9N5Q8 |
| DTLDDGPSVDTS | 1 | 0 | 0 | 1424560 | AIV41888.1 | P0C6X6 |
| DTLDKLSDYCTF | 1 | 0 | 0 | 1424563 | AXN92247.1 | K9N7C7 |
| DTLFASSSLRTF | 1 | 0 | 0 | 1424566 | ATG84909.1 | K9N7C7 |
| DTLLTSEEVVIK | 1 | 0 | 0 | 1424575 | APO40578.1 | P0C6X7 |
| DTLNDGPSVETS | 1 | 0 | 0 | 1424577 | AGT51737.1 | P0C6X6 |
| DTLNKLSKYLVN | 1 | 0 | 0 | 1424578 | AST12962.1 | A0A0P0LKV0 |
| DTLQETSVVSDT | 1 | 0 | 0 | 1424584 | ASU45706.1 | K9N7C7 |
| DTLRDLPLSLTE | 1 | 0 | 0 | 1424587 | ATO98145.1 | Q3LZX1 |
| DTLTELVVAANK | 1 | 0 | 0 | 1424593 | AVV62535.1 | K9N7C7 |
| DTNGDPVYFLRY | 1 | 0 | 0 | 1424606 | AUM60021.1 | K9N4V7 |
| DTNHGDAGFTSI | 1 | 0 | 0 | 1424607 | AQN78743.1 | P30215 |
| DTNLNKFNGASL | 1 | 0 | 0 | 1424610 | AVN89385.1 | K9N7C7 |
| DTNSACPKLEFA | 1 | 0 | 0 | 1424613 | AXN92260.1 | K9N5Q8 |
| DTNSVCPKFEFA | 1 | 0 | 0 | 1424614 | AVN89387.1 | K9N5Q8 |
| DTNSVCPKLEHH | 1 | 0 | 0 | 1424618 | 4ZS6_A | K9N5Q8 |
| DTNSVCPKLHHH | 1 | 0 | 0 | 1424619 | 5GMQ_A | K9N5Q8 |
| DTNSVCPMMDLG | 1 | 0 | 0 | 1424620 | AVV62526.1 | K9N5Q8 |
| DTNSVCPMQALR | 1 | 0 | 0 | 1424621 | AUM60014.1 | K9N5Q8 |
| DTNSVSPKLEFA | 1 | 0 | 0 | 1424622 | AKN24803.1 | K9N5Q8 |
| DTPSDPLNNFKH | 1 | 0 | 0 | 1424636 | AUM60013.1 | K9N7C7 |
| DTPSPLTPRSVR | 1 | 0 | 0 | 1424637 | AJG44080.1 | K9N5Q8 |
| DTPSSANPVKEY | 1 | 0 | 0 | 1424638 | AVV62535.1 | K9N7C7 |
| DTPSTLTPCSVR | 1 | 0 | 0 | 1424644 | AKN24812.1 | K9N5Q8 |
| DTPTEHVTSVNL | 1 | 0 | 0 | 1424648 | AJG44059.1 | K9N796 |
| DTQVLLRRRDAS | 1 | 0 | 0 | 1424654 | AVV62526.1 | K9N5Q8 |
| DTRVLSSLNLPG | 1 | 0 | 0 | 1424659 | APO40578.1 | P0C6X7 |
| DTSDFTDSVRDP | 1 | 0 | 0 | 1424664 | ATO98181.1 | Q3LZX1 |
| DTSGFAVYVKSK | 1 | 0 | 0 | 1424670 | BAM16435.1 | Q01455 |
| DTSGIHNRVIAI | 1 | 0 | 0 | 1424671 | AVV62527.1 | K9N796 |
| DTSLSDACQLPY | 1 | 0 | 0 | 1424681 | AGT17757.1 | A0A140H1H0 |
| DTSRKGSFNNRD | 1 | 0 | 0 | 1424691 | AIX10751.1 | P36334 |
| DTSYINNKDTGP | 1 | 0 | 0 | 1424701 | AKA09646.1 | P36334 |
| DTSYKGTFNNKD | 1 | 0 | 0 | 1424702 | AKA09658.1 | P36334 |
| DTTDVLLKQLMA | 1 | 0 | 0 | 1424708 | AGT51695.1 | Q4VID0 |
| DTTGVLLKQLMV | 1 | 0 | 0 | 1424717 | ABD75582.1 | Q0ZJ80 |
| DTTLYVNNWCRC | 1 | 0 | 0 | 1424725 | Q14EB0.1 | A0A140H1H1 |
| DTTNTNLSRGRG | 1 | 0 | 0 | 1424726 | ALW82676.1 | K9N4V7 |
| DTVDVTNGLGTY | 1 | 0 | 0 | 1424738 | CAA79896.1 | P36334 |
| DTVEEAITEARQ | 1 | 0 | 0 | 1424739 | APO40578.1 | P0C6X7 |
| DTVEQEAQIDRL | 1 | 0 | 0 | 1424741 | AVV62526.1 | K9N5Q8 |
| DTVIEVQGYKSVNI | 1 | 0 | 0 | 1424755 | P0DTC1.1 | P0DTD1 |
| DTVNRDSFNLYG | 1 | 0 | 0 | 1424762 | ADN03337.1 | A0A140H1G9 |
| DTVVKTLQPISD | 1 | 0 | 0 | 1424768 | APO40578.1 | P0C6X7 |
| DTVVQLNYSRGD | 1 | 0 | 0 | 1424772 | AUM60023.1 | K9N7C7 |
| DTVYCPRAVICT | 1 | 0 | 0 | 1424776 | 2Q6G_A | P0C6X7 |
| DTYAALRNSLTN | 1 | 0 | 0 | 1424784 | AXN92247.1 | K9N7C7 |
| DTYDPTYKNGAM | 1 | 0 | 0 | 1424789 | AVN89451.1 | K9N7C7 |
| DTYNLWNTFTKL | 1 | 0 | 0 | 1424794 | AVP78041.1 | P0C6W2 |
| DTYVALRNSINA | 1 | 0 | 0 | 1424801 | AVV62524.1 | K9N7C7 |
| DVAARADYEGFI | 1 | 0 | 0 | 1424805 | ANI69867.1 | K9N638 |
| DVAFNRYLSLYN | 1 | 0 | 0 | 1424808 | AEN19363.1 | P0C6X6 |
| DVAIQEVIDDML | 1 | 0 | 0 | 1424812 | ARK08668.1 | Q80872 |
| DVALDSLGAAIH | 1 | 0 | 0 | 1424814 | AVQ05261.1 | P0C6X6 |
| DVANNAVYDGAR | 1 | 0 | 0 | 1424816 | AFD98763.1 | H9EJ42 |
| DVANPTNCIAPV | 1 | 0 | 0 | 1424818 | AKM76239.1 | K9N5Q8 |
| DVATDVQSRADQ | 1 | 0 | 0 | 1424820 | AUM60013.1 | K9N7C7 |
| DVAYANYNTYDV | 1 | 0 | 0 | 1424824 | APO40578.1 | P0C6X7 |
| DVDDIDPINCES | 1 | 0 | 0 | 1424841 | ABD75591.1 | A0A140H1G9 |
| DVDDYILANECI | 1 | 0 | 0 | 1424847 | AGT51787.1 | P0C6X6 |
| DVDHEFVGEFYA | 1 | 0 | 0 | 1424856 | P0C6W6.1 | P0C6W2 |
| DVDKFKTPHIVR | 1 | 0 | 0 | 1424858 | AVN89322.1 | K9N7C7 |
| DVDNIDFKSLLG | 1 | 0 | 0 | 1424864 | BBA20986.1 | A0A140H1H1 |
| DVDNPAHTGRNY | 1 | 0 | 0 | 1424865 | AJS10812.1 | P0C6X6 |
| DVDQEFVDEFYA | 1 | 0 | 0 | 1424868 | AVP77995.1 | P0C6W2 |
| DVDTGVPSTSTD | 1 | 0 | 0 | 1424875 | CAA83661.1 | P36334 |
| DVDTGVYSVSSF | 1 | 0 | 0 | 1424876 | AVV62526.1 | K9N5Q8 |
| DVDTHRYCLSLK | 1 | 0 | 0 | 1424877 | ANZ78856.1 | P0C6X6 |
| DVDTKDVVECLK | 1 | 0 | 0 | 1424881 | APO40578.1 | P0C6X7 |
| DVDTNCLADSVM | 1 | 0 | 0 | 1424882 | AIV41996.1 | P0C6X6 |
| DVDTTTLYVNNW | 1 | 0 | 0 | 1424884 | AGT17758.1 | A0A140H1H1 |
| DVEEDDAEGEET | 1 | 0 | 0 | 1424900 | ATP16754.1 | P0C6X6 |
| DVETPNLMGWDY | 1 | 0 | 0 | 1424920 | ASB51350.1 | P0C6X7 |
| DVFDLIYQLFKG | 1 | 0 | 0 | 1424927 | AEN19363.1 | P0C6X6 |
| DVFDLSYQLFKG | 1 | 0 | 0 | 1424929 | AIV41918.1 | P0C6X6 |
| DVFHETSDDEWT | 1 | 0 | 0 | 1424933 | AVN89352.1 | K9N7C7 |
| DVFTDGKLNCTF | 1 | 0 | 0 | 1424942 | AVV62524.1 | K9N7C7 |
| DVGLDSAKSACI | 1 | 0 | 0 | 1424963 | AHL18090.1 | K9N5Q8 |
| DVGNHRGAFALH | 1 | 0 | 0 | 1424965 | AKA09616.1 | A0A0P0G321 |
| DVGNHRSAFALH | 1 | 0 | 0 | 1424966 | AIW52843.1 | A0A0P0G321 |
| DVGNRRSAFALH | 1 | 0 | 0 | 1424967 | AMM42641.1 | A0A0P0G321 |
| DVGNYFYRSHRS | 1 | 0 | 0 | 1424968 | AVP78031.1 | Q3LZX1 |
| DVHDTVLTSEEV | 1 | 0 | 0 | 1424983 | ARO76381.1 | P0C6U8 |

| | | | | | | |
|---|---|---|---|---|---|---|
| DVHSTYGTQAYS | 1 | 0 | 0 | 1424984 | ATQ39396.1 | K9N643 |
| DVIKEGALTAVV | 1 | 0 | 0 | 1425001 | AVP78041.1 | P0C6W2 |
| DVINQNAVALNT | 1 | 0 | 0 | 1425004 | APO40579.1 | P59594 |
| DVIVHDVEPTHK | 1 | 0 | 0 | 1425018 | AWK59942.1 | H9EJ42 |
| DVKFENLTNQLP | 1 | 0 | 0 | 1425028 | AVV62537.1 | K9N5Q8 |
| DVKGLEELHYDL | 1 | 0 | 0 | 1425033 | AIV41836.1 | Q80872 |
| DVLFSGQKYSSD | 1 | 0 | 0 | 1425064 | AQZ41321.1 | K9N7C7 |
| DVLLKQPMATCV | 1 | 0 | 0 | 1425081 | QRK03817.1 | Q4VID0 |
| DVLNKNQKLIAN | 1 | 0 | 0 | 1425084 | BBA20986.1 | A0A140H1H1 |
| DVLPEGWHVVNK | 1 | 0 | 0 | 1425086 | AIV41876.1 | P0C6X6 |
| DVLTDVSARADA | 1 | 0 | 0 | 1425098 | AVV62524.1 | K9N7C7 |
| DVLYQPPQTSITSAVL | 1 | 0 | 0 | 1425105 | P0DTC1.1 | P0DTD1 |
| DVMFEDDQGLKK | 1 | 0 | 0 | 1425112 | AGV08522.1 | K9N7C7 |
| DVMLEDATGLKW | 1 | 0 | 0 | 1425115 | AUM60013.1 | K9N7C7 |
| DVMRYNFNFSSN | 1 | 0 | 0 | 1425120 | AVA26873.1 | A0A0P0G321 |
| DVNCNGYQHHSF | 1 | 0 | 0 | 1425125 | ABE97137.1 | A0A0P0G321 |
| DVNCNGYQHHSI | 1 | 0 | 0 | 1425126 | AVL25583.1 | A0A0P0G321 |
| DVNCNGYQHNSF | 1 | 0 | 0 | 1425128 | ALJ53431.1 | A0A0P0G321 |
| DVNDSHSMPVVD | 1 | 0 | 0 | 1425144 | AGT51409.1 | P0C6X6 |
| DVNFSPVLGCLG | 1 | 0 | 0 | 1425146 | CAA83661.1 | P36334 |
| DVNGMKVVDYLT | 1 | 0 | 0 | 1425147 | AVV62535.1 | K9N7C7 |
| DVNKTLQEFAKN | 1 | 0 | 0 | 1425152 | AWK59979.1 | A0A0P0G321 |
| DVNMEAAYTSSL | 1 | 0 | 0 | 1425155 | AXN92228.1 | K9N5Q8 |
| DVNMQLMGVVMQ | 1 | 0 | 0 | 1425160 | AQZ41284.1 | K9N7C7 |
| DVNSVLNICFDI | 1 | 0 | 0 | 1425168 | QBF80475.1 | K9N7C7 |
| DVNTLLSNQLRL | 1 | 0 | 0 | 1425172 | AWK59942.1 | H9EJ42 |
| DVNTPADIDYRN | 1 | 0 | 0 | 1425173 | AJQ24074.1 | P33469 |
| DVNVTFFPDLNG | 1 | 0 | 0 | 1425177 | APO40578.1 | P0C6X7 |
| DVPFLICSNTPA | 1 | 0 | 0 | 1425187 | AEN19363.1 | P0C6X6 |
| DVPHYVLPGQYT | 1 | 0 | 0 | 1425190 | AUM60014.1 | K9N5Q8 |
| DVPSISGTQVYS | 1 | 0 | 0 | 1425198 | ATG84770.1 | K9N643 |
| DVPSTYGTQVYS | 1 | 0 | 0 | 1425202 | AIA22866.1 | K9N643 |
| DVPTAIRADQLT | 1 | 0 | 0 | 1425203 | ATO98181.1 | Q3LZX1 |
| DVPTEHVTSVNL | 1 | 0 | 0 | 1425204 | AMO03402.1 | K9N796 |
| DVPTTLAWLNNQ | 1 | 0 | 0 | 1425209 | AQZ41321.1 | K9N7C7 |
| DVPTTLAWVNNH | 1 | 0 | 0 | 1425210 | AHI48582.1 | K9N7C7 |
| DVPTVIRADQLT | 1 | 0 | 0 | 1425212 | ATO98145.1 | Q3LZX1 |
| DVQFEQLDNKLP | 1 | 0 | 0 | 1425220 | AVV62526.1 | K9N5Q8 |
| DVQQAQQEATHI | 1 | 0 | 0 | 1425228 | AST12962.1 | A0A0P0LKV0 |
| DVQQAQQEVTHI | 1 | 0 | 0 | 1425229 | APT69889.1 | A0A0P0LKV0 |
| DVQQWGYIGNLA | 1 | 0 | 0 | 1425231 | AVV62524.1 | K9N7C7 |
| DVQQWGYVGNLA | 1 | 0 | 0 | 1425232 | AXN92247.1 | K9N7C7 |
| DVQSRADQNGYI | 1 | 0 | 0 | 1425234 | AUM60013.1 | K9N7C7 |
| DVQVSAQQEVQL | 1 | 0 | 0 | 1425235 | AUM60013.1 | K9N7C7 |
| DVRDNGALEKFM | 1 | 0 | 0 | 1425241 | AVV62524.1 | K9N7C7 |
| DVRETMAHLLQH | 1 | 0 | 0 | 1425243 | ARO76381.1 | P0C6U8 |
| DVRQIAPAQTGV | 1 | 0 | 0 | 1425248 | APO40579.1 | P59594 |
| DVRSGHMLDDVQ | 1 | 0 | 0 | 1425250 | AVV62540.1 | K9N643 |
| DVSARDPTIQEA | 1 | 0 | 0 | 1425255 | AMM42397.1 | Q19U29 |
| DVSDTIPEQKQN | 1 | 0 | 0 | 1425261 | AVV62524.1 | K9N7C7 |
| DVSDTIPQQSQN | 1 | 0 | 0 | 1425262 | AVV62535.1 | K9N7C7 |
| DVSENNEHLTWP | 1 | 0 | 0 | 1425265 | AVV62524.1 | K9N7C7 |
| DVSFLAHIQWFA | 1 | 0 | 0 | 1425267 | APO40578.1 | P0C6X7 |
| DVSFLERDAPYM | 1 | 0 | 0 | 1425270 | ARO76381.1 | P0C6U8 |
| DVSGVGVIRVSN | 1 | 0 | 0 | 1425277 | CAA71056.1 | A0A0P0K6L9 |
| DVSHETSDDDWT | 1 | 0 | 0 | 1425280 | AVN89374.1 | K9N7C7 |
| DVSKADGIIHPQ | 1 | 0 | 0 | 1425288 | AGV08524.1 | K9N5Q8 |
| DVSKPNAIDVEH | 1 | 0 | 0 | 1425290 | AGT51337.1 | H9EJ42 |
| DVSNSGPLQTAV | 1 | 0 | 0 | 1425295 | AVV62537.1 | K9N5Q8 |
| DVTGKLSDCLYL | 1 | 0 | 0 | 1425319 | AEN19363.1 | P0C6X6 |
| DVTLEKLRDYLA | 1 | 0 | 0 | 1425328 | ATQ39389.1 | K9N7C7 |
| DVTQLAEEPAKD | 1 | 0 | 0 | 1425333 | ATQ39389.1 | K9N7C7 |
| DVVIAILSGTTT | 1 | 0 | 0 | 1425361 | AVN89333.1 | K9N7C7 |
| DVVLSPLLSAGI | 1 | 0 | 0 | 1425375 | APO40578.1 | P0C6X7 |
| DVVNQQGNSLNY | 1 | 0 | 0 | 1425380 | AAK32190.1 | A0A0P0K6L9 |
| DVVNQQGNSLSH | 1 | 0 | 0 | 1425381 | ARA15429.1 | A0A0P0K6L9 |
| DVVQYEADRRVF | 1 | 0 | 0 | 1425384 | QBP84701.1 | P0C6X6 |
| DVVSIRSSNRGN | 1 | 0 | 0 | 1425396 | ATG84704.1 | K9N643 |
| DVVSIRTSNQGN | 1 | 0 | 0 | 1425397 | AHI48842.1 | K9N643 |
| DVVSSEAVEDTI | 1 | 0 | 0 | 1425398 | AZS52616.1 | A0A140H1G9 |
| DVVTGDNDDDND | 1 | 0 | 0 | 1425400 | ARU07575.1 | A0A140H1G9 |
| DVVTGDNNDEDV | 1 | 0 | 0 | 1425402 | P0C6U5.1 | A0A140H1G9 |
| DVVTGDNNGEDV | 1 | 0 | 0 | 1425405 | ABD75631.1 | A0A140H1G9 |
| DVVYAQHCFKVP | 1 | 0 | 0 | 1425412 | AGT51590.1 | P36334 |
| DVYARYLAMFNK | 1 | 0 | 0 | 1425422 | AVV62535.1 | K9N7C7 |
| DVYDMWLTGCKY | 1 | 0 | 0 | 1425425 | AEN19363.1 | P0C6X6 |
| DVYDNFPQGFSV | 1 | 0 | 0 | 1425426 | APO40579.1 | P59594 |
| DVYEADAEIVQP | 1 | 0 | 0 | 1425427 | AUM60013.1 | K9N7C7 |
| DVYNLTMPSQVQ | 1 | 0 | 0 | 1425434 | AAX85666.1 | P0C6X6 |
| DVYRRIPNLPDC | 1 | 0 | 0 | 1425436 | CAA83660.1 | P36334 |
| DVYRRKPDPPNC | 1 | 0 | 0 | 1425439 | AHA80201.1 | P36334 |
| DVYRRRPDLPNC | 1 | 0 | 0 | 1425441 | AHA80225.1 | P36334 |
| DWDSFCKTIQSA | 1 | 0 | 0 | 1425445 | NP_073549.1 | A0A0P0LKV0 |
| DWDSFGKTIQSA | 1 | 0 | 0 | 1425446 | AST12962.1 | A0A0P0LKV0 |
| DWKCKVTDVLYP | 1 | 0 | 0 | 1425471 | AUM60013.1 | K9N7C7 |
| DWKPGLAMPSLF | 1 | 0 | 0 | 1425475 | AVV62535.1 | K9N7C7 |
| DWKPGLAMPTLF | 1 | 0 | 0 | 1425476 | AVV62524.1 | K9N7C7 |
| DWLDVQEESEPL | 1 | 0 | 0 | 1425487 | APO40578.1 | P0C6X7 |
| DWLWFALELLFA | 1 | 0 | 0 | 1425496 | AVV62535.1 | K9N7C7 |
| DWNSFCKTIQSA | 1 | 0 | 0 | 1425505 | APT69882.1 | A0A0P0ILY9 |
| DWNTKHGSGLKR | 1 | 0 | 0 | 1425506 | APO40578.1 | P0C6X7 |

| | | | | | | |
|---|---|---|---|---|---|---|
| DWQNLLYDSNGN | 1 | 0 | 0 | 1425509 | AZS52618.1 | A0A140H1H1 |
| DWRSCNYAVSSA | 1 | 0 | 0 | 1425511 | ALD51903.1 | K9N7C7 |
| DWRSYNYAVSSA | 1 | 0 | 0 | 1425512 | AXN92247.1 | K9N7C7 |
| DWSYSGQNTELG | 1 | 0 | 0 | 1425525 | APO40578.1 | P0C6X7 |
| DWYDFVENPDVL | 1 | 0 | 0 | 1425542 | AVP78030.1 | P0C6W2 |
| DYAAKGYVAGIQ | 1 | 0 | 0 | 1425547 | AVV62524.1 | K9N7C7 |
| DYADKLCHAVVS | 1 | 0 | 0 | 1425550 | AEN19363.1 | P0C6X6 |
| DYADKLCHAVVY | 1 | 0 | 0 | 1425551 | AIV41786.1 | P0C6X6 |
| DYAFIADKEKLI | 1 | 0 | 0 | 1425554 | AUM60013.1 | K9N7C7 |
| DYAKDNTVLQAL | 1 | 0 | 0 | 1425557 | AEN19363.1 | P0C6X6 |
| DYANKGYIANIS | 1 | 0 | 0 | 1425562 | ABD75519.1 | A0A140H1G9 |
| DYASKGYIVNIS | 1 | 0 | 0 | 1425565 | AIL49553.1 | P0C6X6 |
| DYASLAAKEGIN | 1 | 0 | 0 | 1425566 | AVV62535.1 | K9N7C7 |
| DYATKGYVAGIQ | 1 | 0 | 0 | 1425568 | AUM60013.1 | K9N7C7 |
| DYCDGNLPCGIC | 1 | 0 | 0 | 1425575 | APO40578.1 | P0C6X7 |
| DYCGGNLCTCQP | 1 | 0 | 0 | 1425576 | QQY99180.1 | P36334 |
| DYCGGNPCICQP | 1 | 0 | 0 | 1425577 | BBO58900.1 | P36334 |
| DYCGGNPCTCQP | 1 | 0 | 0 | 1425578 | AVQ05264.1 | P36334 |
| DYCGGNSCTCQP | 1 | 0 | 0 | 1425580 | AGT51690.1 | P36334 |
| DYCGNNSCTCQP | 1 | 0 | 0 | 1425582 | AEN19366.1 | P36334 |
| DYCRGNFCTCQP | 1 | 0 | 0 | 1425587 | AGT51600.1 | P36334 |
| DYCRGNSCICQP | 1 | 0 | 0 | 1425588 | AIX10751.1 | P36334 |
| DYCRGNSCTCQP | 1 | 0 | 0 | 1425589 | AIX10750.1 | P36334 |
| DYCVRAFDGYNK | 1 | 0 | 0 | 1425591 | AST12962.1 | A0A0P0LKV0 |
| DYDIAVTAGPFM | 1 | 0 | 0 | 1425597 | AQZ41321.1 | K9N7C7 |
| DYDIPTTENLYF | 1 | 0 | 0 | 1425598 | 3EJG_A | A0A0P0LKV0 |
| DYDLEPQKIHIH | 1 | 0 | 0 | 1425600 | AUM60024.1 | K9N5Q8 |
| DYDVAVTVGPFM | 1 | 0 | 0 | 1425603 | AXN92258.1 | K9N7C7 |
| DYDVVVTAGPFM | 1 | 0 | 0 | 1425604 | AVN89363.1 | K9N7C7 |
| DYEAERPFTSYT | 1 | 0 | 0 | 1425606 | AWK59942.1 | H9EJ42 |
| DYENDWNTKHGS | 1 | 0 | 0 | 1425612 | APO40578.1 | P0C6X7 |
| DYERCNIYGSAT | 1 | 0 | 0 | 1425617 | AVV62524.1 | K9N7C7 |
| DYERSNIYGTAT | 1 | 0 | 0 | 1425618 | AXN92247.1 | K9N7C7 |
| DYFAYPLSMKSY | 1 | 0 | 0 | 1425619 | AVV62537.1 | K9N5Q8 |
| DYFEMKNWFDPI | 1 | 0 | 0 | 1425624 | AIW52776.1 | A0A0P0LKV0 |
| DYFSYPFSMKSD | 1 | 0 | 0 | 1425632 | AVN89313.1 | K9N5Q8 |
| DYFSYPLSMKSD | 1 | 0 | 0 | 1425633 | AXN92228.1 | K9N5Q8 |
| DYFTRQLSQLEG | 1 | 0 | 0 | 1425635 | AVV62537.1 | K9N5Q8 |
| DYHANRPGVYCG | 1 | 0 | 0 | 1425652 | AVV62524.1 | K9N7C7 |
| DYIKQNGPLRVG | 1 | 0 | 0 | 1425670 | ARO76381.1 | P0C6U8 |
| DYKNVKAWSGIC | 1 | 0 | 0 | 1425701 | AIW52843.1 | A0A0P0G321 |
| DYKVLDNGVIRD | 1 | 0 | 0 | 1425714 | AVV62524.1 | K9N7C7 |
| DYLAAYDIEVTK | 1 | 0 | 0 | 1425719 | AUM60013.1 | K9N7C7 |
| DYLADYDVAVTV | 1 | 0 | 0 | 1425722 | AXN92258.1 | K9N7C7 |
| DYLAVYDIEVTK | 1 | 0 | 0 | 1425725 | AUM60023.1 | K9N7C7 |
| DYLTRDDMHQLY | 1 | 0 | 0 | 1425736 | AVV62535.1 | K9N7C7 |
| DYLTREDMHQLY | 1 | 0 | 0 | 1425738 | AVV62524.1 | K9N7C7 |
| DYLWYVVDDIYY | 1 | 0 | 0 | 1425740 | AWK59942.1 | H9EJ42 |
| DYMDIVKRLATG | 1 | 0 | 0 | 1425744 | AVV62535.1 | K9N7C7 |
| DYMTHGQMEGLC | 1 | 0 | 0 | 1425751 | AST12962.1 | A0A0P0LKV0 |
| DYNAILQSDNPQ | 1 | 0 | 0 | 1425752 | QDH43737.1 | H9EJ42 |
| DYNGTPLEFGAS | 1 | 0 | 0 | 1425756 | APO40578.1 | P0C6X7 |
| DYNKTIPRISED | 1 | 0 | 0 | 1425757 | ARB07438.1 | A0A140H1H1 |
| DYPIISNEVSIN | 1 | 0 | 0 | 1425769 | ABD75551.1 | A0A140H1G9 |
| DYPKLFITREEA | 1 | 0 | 0 | 1425776 | AVV62535.1 | K9N7C7 |
| DYQIGGYSEDWH | 1 | 0 | 0 | 1425782 | ATO98182.1 | Q3LZX0 |
| DYRITSITSDKF | 1 | 0 | 0 | 1425792 | AIV41978.1 | P0C6X6 |
| DYSIEMQLFKVG | 1 | 0 | 0 | 1425798 | AWK59978.1 | H9EJ42 |
| DYTGSLADGLGA | 1 | 0 | 0 | 1425827 | AIV42008.1 | P0C6X6 |
| DYTGYQEFVIKT | 1 | 0 | 0 | 1425829 | AIV41909.1 | P36334 |
| DYTGYQESVIKT | 1 | 0 | 0 | 1425831 | AIL49487.1 | P36334 |
| DYTKEGSVINFC | 1 | 0 | 0 | 1425833 | AKN24802.1 | K9N7C7 |
| DYTLANTTTAPL | 1 | 0 | 0 | 1425836 | AGV08437.1 | K9N7C7 |
| DYTQYYFNKDTG | 1 | 0 | 0 | 1425846 | QEG03802.1 | P30215 |
| DYTWTPYPELME | 1 | 0 | 0 | 1425851 | AUM60013.1 | K9N7C7 |
| DYVKPFLNHTQT | 1 | 0 | 0 | 1425865 | APO40578.1 | P0C6X7 |
| DYVSLLNEVWQK | 1 | 0 | 0 | 1425878 | AUM60016.1 | K9N4V0 |
| DYVSLLNQIWQK | 1 | 0 | 0 | 1425879 | AXN92262.1 | K9N4V0 |
| DYVTIAGTKLRS | 1 | 0 | 0 | 1425881 | ARO76381.1 | P0C6U8 |
| DYYNMLVNCFKW | 1 | 0 | 0 | 1425894 | ATN39862.1 | P0C6X6 |
| EAAAQGFQACRF | 1 | 0 | 0 | 1425905 | AWK59942.1 | H9EJ42 |
| EAAGNIFVVNKD | 1 | 0 | 0 | 1425913 | AVV62524.1 | K9N7C7 |
| EAALPQSKDSNF | 1 | 0 | 0 | 1425917 | AVV62535.1 | K9N7C7 |
| EAALPYGANKEG | 1 | 0 | 0 | 1425918 | APO40586.1 | P59595 |
| EAARKAYQEAMD | 1 | 0 | 0 | 1425927 | AMQ49036.1 | K9N638 |
| EAASNGFRACRF | 1 | 0 | 0 | 1425931 | NP_073549.1 | A0A0P0LKV0 |
| EAATQGFQACRF | 1 | 0 | 0 | 1425933 | AFD98756.1 | H9EJ42 |
| EAAYAEVHAEPK | 1 | 0 | 0 | 1425937 | P0C6U5.1 | A0A140H1G9 |
| EAAYTSSSLGSI | 1 | 0 | 0 | 1425939 | AOR17480.1 | K9N5Q8 |
| EACYERGCNPKG | 1 | 0 | 0 | 1425946 | AAM95936.1 | P0C6U7 |
| EADMRNSSSEFF | 1 | 0 | 0 | 1425956 | AVV62537.1 | K9N5Q8 |
| EADPKCKYAQNL | 1 | 0 | 0 | 1425957 | ALA49857.1 | K9N7C7 |
| EADTEQTQSVNV | 1 | 0 | 0 | 1425966 | ATG84700.1 | K9N638 |
| EADWIELFPHND | 1 | 0 | 0 | 1425971 | QEO75983.1 | A0A0P0LKV0 |
| EAFDYYHTLDDS | 1 | 0 | 0 | 1425990 | APO40578.1 | P0C6X7 |
| EAFHKVQDAVNN | 1 | 0 | 0 | 1425997 | AHI48803.1 | K9N5Q8 |
| EAFKRFNNGVYI | 1 | 0 | 0 | 1425998 | AGT51558.1 | P0C6X6 |
| EAFPLDEAENVT | 1 | 0 | 0 | 1426005 | QBF80465.1 | K9N7C7 |
| EAFTKPVFISAL | 1 | 0 | 0 | 1426012 | NP_073549.1 | A0A0P0LKV0 |
| EAHNNLLQQLSN | 1 | 0 | 0 | 1426031 | ARB66064.1 | P36334 |
| EAIKHVRAWIGF | 1 | 0 | 0 | 1426041 | APO40578.1 | P0C6X7 |

| | | | | | | |
|---|---|---|---|---|---|---|
| EAIKRVKGWVGF | 1 | 0 | 0 | 1426044 | ABD75519.1 | A0A140H1G9 |
| EAKCWVEPDLKK | 1 | 0 | 0 | 1426058 | AVV62524.1 | K9N7C7 |
| EAKFKNSVTSIN | 1 | 0 | 0 | 1426061 | ARB07424.1 | A0A0P0ILY9 |
| EAKGYWYKLNRR | 1 | 0 | 0 | 1426063 | AJQ24067.1 | P33469 |
| EAKGYWYRHNRR | 1 | 0 | 0 | 1426066 | AEN19370.1 | P33469 |
| EAKTASLFKTTV | 1 | 0 | 0 | 1426077 | ARU07594.1 | A0A0P0G321 |
| EAKTASLFQTTV | 1 | 0 | 0 | 1426078 | AIW52843.1 | A0A0P0G321 |
| EALESGKQKTAA | 1 | 0 | 0 | 1426087 | AUM60021.1 | K9N4V7 |
| EALQFTTLTLDK | 1 | 0 | 0 | 1426095 | AEN19363.1 | P0C6X6 |
| EALQFTTLTVDK | 1 | 0 | 0 | 1426096 | AXX83324.1 | P0C6X6 |
| EANLPFRAVKDG | 1 | 0 | 0 | 1426123 | AVV62533.1 | K9N4V7 |
| EANLPFRAVKEG | 1 | 0 | 0 | 1426124 | AVV62544.1 | K9N4V7 |
| EAPAPELQVPVT | 1 | 0 | 0 | 1426133 | APO40578.1 | P0C6X7 |
| EAPHNDCTPINT | 1 | 0 | 0 | 1426137 | AVV62526.1 | K9N5Q8 |
| EAPNAKEEILGA | 1 | 0 | 0 | 1426144 | BAE93400.1 | P0C6U8 |
| EAPRDGQAYVRK | 1 | 0 | 0 | 1426146 | 6CRW_A | P59594 |
| EAQAFETPAVQF | 1 | 0 | 0 | 1426153 | AVV62535.1 | K9N7C7 |
| EAQAQIDRLING | 1 | 0 | 0 | 1426154 | CAA83660.1 | P36334 |
| EAQLVEVPVEDI | 1 | 0 | 0 | 1426164 | ATG84788.1 | K9N7C7 |
| EAQNANPMVIVN | 1 | 0 | 0 | 1426165 | P0C6W6.1 | P0C6W2 |
| EAQNANPTVIVN | 1 | 0 | 0 | 1426166 | ATO98229.1 | P0C6W2 |
| EAQPVEVHVEDI | 1 | 0 | 0 | 1426172 | ANF29161.1 | K9N7C7 |
| EARFSGTANQQQ | 1 | 0 | 0 | 1426182 | AGT51787.1 | P0C6X6 |
| EARLSGSANQQQ | 1 | 0 | 0 | 1426184 | AIV41942.1 | P0C6X6 |
| EASAFRDLKPLP | 1 | 0 | 0 | 1426188 | AUM60013.1 | K9N7C7 |
| EASLNSLTYFNK | 1 | 0 | 0 | 1426194 | ARU07575.1 | A0A140H1G9 |
| EASTDFHQLECI | 1 | 0 | 0 | 1426200 | ARU07593.1 | H9EJ42 |
| EASTVVCGGKRT | 1 | 0 | 0 | 1426204 | AXN92247.1 | K9N7C7 |
| EASTVVCGTKRT | 1 | 0 | 0 | 1426205 | AVV62535.1 | K9N7C7 |
| EATEATFNYLLD | 1 | 0 | 0 | 1426211 | AZK15899.1 | K9N638 |
| EAVDSGDASPQV | 1 | 0 | 0 | 1426228 | AVV62535.1 | K9N7C7 |
| EAVKHGAKPISN | 1 | 0 | 0 | 1426232 | AFD98833.1 | H9EJ42 |
| EAVNANAQALSK | 1 | 0 | 0 | 1426244 | AUM60014.1 | K9N5Q8 |
| EAVSRLVESASE | 1 | 0 | 0 | 1426247 | ADN03340.1 | Q0ZJ84 |
| EAWLEFRAGRPA | 1 | 0 | 0 | 1426252 | AVQ05261.1 | P0C6X6 |
| EAWLNDKAVPSP | 1 | 0 | 0 | 1426254 | AIV41909.1 | P36334 |
| EAWLNGKSVPSP | 1 | 0 | 0 | 1426257 | AIL49492.1 | P36334 |
| EAWSGLCVDGRN | 1 | 0 | 0 | 1426259 | AIW52756.1 | A0A0P0K6L9 |
| EAWSGLCVDGTN | 1 | 0 | 0 | 1426260 | CAA71056.1 | A0A0P0K6L9 |
| EAWYEFRAGRPH | 1 | 0 | 0 | 1426261 | ARU07575.1 | A0A140H1G9 |
| EAYADRTLQEMK | 1 | 0 | 0 | 1426263 | AIV41774.1 | P0C6X6 |
| EAYEQAVVNGDS | 1 | 0 | 0 | 1426265 | AVP78041.1 | P0C6W2 |
| ECCEASYFDNKL | 1 | 0 | 0 | 1426283 | AUM60013.1 | K9N7C7 |
| ECDFTKLLSGTP | 1 | 0 | 0 | 1426302 | AVV62537.1 | K9N5Q8 |
| ECFDKFKVNSAL | 1 | 0 | 0 | 1426312 | APO40578.1 | P0C6X7 |
| ECGAVYTANDVD | 1 | 0 | 0 | 1426318 | AUM60013.1 | K9N7C7 |
| ECMQQGPQSARD | 1 | 0 | 0 | 1426347 | AUM60014.1 | K9N5Q8 |
| ECNLSLGQSLCA | 1 | 0 | 0 | 1426351 | ATQ39390.1 | K9N5Q8 |
| ECNQMCLSTLMK | 1 | 0 | 0 | 1426354 | AVP78041.1 | P0C6W2 |
| ECPNSIYICSQG | 1 | 0 | 0 | 1426362 | ARU07575.1 | A0A140H1G9 |
| ECQHPNMTVHHI | 1 | 0 | 0 | 1426363 | AVV62540.1 | K9N643 |
| ECQLPLGQSLCA | 1 | 0 | 0 | 1426364 | AVV62526.1 | K9N5Q8 |
| ECRDSRKGVCFG | 1 | 0 | 0 | 1426366 | AWK59942.1 | H9EJ42 |
| ECTERLKLFAEE | 1 | 0 | 0 | 1426376 | AIV41984.1 | P0C6X6 |
| ECTIFKDSSGKP | 1 | 0 | 0 | 1426383 | AVP78041.1 | P0C6W2 |
| ECTIFKNAQGKP | 1 | 0 | 0 | 1426384 | APO40578.1 | P0C6X7 |
| ECTMFKDAEGRM | 1 | 0 | 0 | 1426386 | AVN89322.1 | K9N7C7 |
| ECTMFRDAEGKM | 1 | 0 | 0 | 1426388 | AVV62535.1 | K9N7C7 |
| ECTMFRDAEGRI | 1 | 0 | 0 | 1426389 | AHL18091.1 | K9N7C7 |
| ECTMFRDAEGRM | 1 | 0 | 0 | 1426390 | AXN92247.1 | K9N7C7 |
| ECTTIVNGMKKS | 1 | 0 | 0 | 1426393 | APO40578.1 | P0C6X7 |
| ECTYIGRFSKIS | 1 | 0 | 0 | 1426396 | ATG84705.1 | K9N7D2 |
| ECTYITRFSKTQ | 1 | 0 | 0 | 1426399 | AVV62541.1 | K9N7D2 |
| EDACFKFNYQFP | 1 | 0 | 0 | 1426425 | AQN78665.1 | Q04853 |
| EDACFKFNYQLP | 1 | 0 | 0 | 1426426 | AXX83304.1 | Q04853 |
| EDALRNSAMLES | 1 | 0 | 0 | 1426436 | AIW52757.1 | A0A0P0K6L9 |
| EDAYDLVHDEPK | 1 | 0 | 0 | 1426450 | AVI15051.1 | P0C6X6 |
| EDCKLPLGRSLC | 1 | 0 | 0 | 1426455 | ATG84888.1 | K9N5Q8 |
| EDCNLPLGQSLC | 1 | 0 | 0 | 1426457 | ALA49451.1 | K9N5Q8 |
| EDDCEEGQFEPS | 1 | 0 | 0 | 1426460 | AVP78041.1 | P0C6W2 |
| EDDDFLEENDVE | 1 | 0 | 0 | 1426462 | QBQ01836.1 | P0C6X6 |
| EDDGTPVYFLRY | 1 | 0 | 0 | 1426464 | AVV62533.1 | K9N4V7 |
| EDDQGLKKFYKE | 1 | 0 | 0 | 1426467 | AGV08522.1 | K9N7C7 |
| EDDYNGTPLEFG | 1 | 0 | 0 | 1426475 | APO40578.1 | P0C6X7 |
| EDDYQIGGYTEK | 1 | 0 | 0 | 1426478 | AVP78043.1 | Q3LZX0 |
| EDENAEWFGITQ | 1 | 0 | 0 | 1426492 | AVV62537.1 | K9N5Q8 |
| EDENAMFDKAVA | 1 | 0 | 0 | 1426493 | AUM60021.1 | K9N4V7 |
| EDFDLEDFIDNP | 1 | 0 | 0 | 1426497 | AVV62524.1 | K9N7C7 |
| EDFIKHIRTNGG | 1 | 0 | 0 | 1426503 | ASU90174.1 | K9N638 |
| EDFIKHVRTNGD | 1 | 0 | 0 | 1426506 | QDI73607.1 | K9N7C7 |
| EDFLPIDSTVKN | 1 | 0 | 0 | 1426516 | APO40578.1 | P0C6X7 |
| EDFNLEDFIDSP | 1 | 0 | 0 | 1426517 | AVV62535.1 | K9N7C7 |
| EDFQENWNTKHG | 1 | 0 | 0 | 1426518 | AVP78041.1 | P0C6W2 |
| EDGDTVYYPMCT | 1 | 0 | 0 | 1426526 | AUM60023.1 | K9N7C7 |
| EDGELNSLDGDC | 1 | 0 | 0 | 1426530 | AUM60013.1 | K9N7C7 |
| EDGFAFYPACIN | 1 | 0 | 0 | 1426532 | QBF80553.1 | K9N7C7 |
| EDGINVKDVFVE | 1 | 0 | 0 | 1426535 | QEG03747.1 | H9EJ42 |
| EDGQTTFYPMCT | 1 | 0 | 0 | 1426539 | AVV62524.1 | K9N7C7 |
| EDGQTVFYPMCT | 1 | 0 | 0 | 1426540 | AVV62535.1 | K9N7C7 |
| EDGQVFTRQLSD | 1 | 0 | 0 | 1426541 | AVV62526.1 | K9N5Q8 |
| EDGRIPNDFSFN | 1 | 0 | 0 | 1426542 | ABE97137.1 | A0A0P0G321 |

| | | | | | | |
|---|---|---|---|---|---|---|
| EDIMEVDISQPT | 1 | 0 | 0 | 1426555 | AUM60017.1 | K9N643 |
| EDIQLLKNAYEN | 1 | 0 | 0 | 1426558 | AVP78041.1 | P0C6W2 |
| EDISLLRKMDEP | 1 | 0 | 0 | 1426562 | AHA80151.1 | P33469 |
| EDKFWCEDFYAS | 1 | 0 | 0 | 1426566 | AIW52827.1 | H9EJ42 |
| EDKVCRLIPTVA | 1 | 0 | 0 | 1426573 | APO40587.1 | P59636 |
| EDLEAGKNKAAP | 1 | 0 | 0 | 1426585 | AVV62533.1 | K9N4V7 |
| EDLHTLYVDSKY | 1 | 0 | 0 | 1426589 | ALA49340.1 | K9N7C7 |
| EDLLAAYVQEES | 1 | 0 | 0 | 1426593 | APO40578.1 | P0C6X7 |
| EDLLFNKVTIAD | 1 | 0 | 0 | 1426597 | QWB73837.1 | P0DTC2 |
| EDLLYNKVTLSD | 1 | 0 | 0 | 1426605 | APO40579.1 | P59594 |
| EDLMEVDHSGPT | 1 | 0 | 0 | 1426611 | AVV62529.1 | K9N643 |
| EDLQQAQQEATH | 1 | 0 | 0 | 1426614 | APT69868.1 | A0A0P0ILY9 |
| EDLSIRALQKHL | 1 | 0 | 0 | 1426618 | AXX83366.1 | P0C6X6 |
| EDLYDVSKPVMI | 1 | 0 | 0 | 1426633 | QED88038.1 | H9EJ42 |
| EDMHQLYVNPKH | 1 | 0 | 0 | 1426638 | AVV62524.1 | K9N7C7 |
| EDNPLFLFDEAG | 1 | 0 | 0 | 1426642 | AEN19363.1 | P0C6X6 |
| EDPPWPQIAELV | 1 | 0 | 0 | 1426646 | QDP16213.1 | K9N4V7 |
| EDPRWPQIAELV | 1 | 0 | 0 | 1426649 | AXG21672.1 | K9N4V7 |
| EDPVVDWHYYDA | 1 | 0 | 0 | 1426650 | AUM60023.1 | K9N7C7 |
| EDQSIAYSNNII | 1 | 0 | 0 | 1426655 | APO40579.1 | P59594 |
| EDRILEFKVLDN | 1 | 0 | 0 | 1426659 | AUM60013.1 | K9N7C7 |
| EDSKSLLVSEDV | 1 | 0 | 0 | 1426663 | APO40578.1 | P0C6X7 |
| EDSTGDEAWATL | 1 | 0 | 0 | 1426668 | AVV62535.1 | K9N7C7 |
| EDSVFKPQPAGV | 1 | 0 | 0 | 1426670 | ARB66062.1 | P36334 |
| EDSVFVPQPTGV | 1 | 0 | 0 | 1426673 | AEN19366.1 | P36334 |
| EDSVIINDDTSE | 1 | 0 | 0 | 1426674 | AXX83318.1 | P0C6X6 |
| EDSVIVNDDTGE | 1 | 0 | 0 | 1426675 | AXX83366.1 | P0C6X6 |
| EDSVIVNDDTSD | 1 | 0 | 0 | 1426676 | AIV41780.1 | P0C6X6 |
| EDTQVVIADTLQ | 1 | 0 | 0 | 1426683 | AVN89312.1 | K9N638 |
| EDVDYVPLKSAV | 1 | 0 | 0 | 1426689 | AMQ49035.1 | K9N7C7 |
| EDVKLNPSEEFI | 1 | 0 | 0 | 1426697 | ALW82730.1 | K9N7C7 |
| EDVQQAQQEATH | 1 | 0 | 0 | 1426703 | AST12962.1 | A0A0P0LKV0 |
| EDVQQAQQEVTH | 1 | 0 | 0 | 1426704 | APT69889.1 | A0A0P0LKV0 |
| EDVSFLERDAPY | 1 | 0 | 0 | 1426705 | ARO76381.1 | P0C6U8 |
| EDVSFLKNDAPY | 1 | 0 | 0 | 1426706 | APO40578.1 | P0C6X7 |
| EDVTEFVQEEVQ | 1 | 0 | 0 | 1426707 | AXN92247.1 | K9N7C7 |
| EDWHYAKFGFTP | 1 | 0 | 0 | 1426721 | NP_073549.1 | A0A0P0LKV0 |
| EDWLDVQEESEP | 1 | 0 | 0 | 1426725 | APO40578.1 | P0C6X7 |
| EDWLESDSQDGQ | 1 | 0 | 0 | 1426727 | AVP78041.1 | P0C6W2 |
| EDYAFEHVVYGD | 1 | 0 | 0 | 1426730 | AUM60013.1 | K9N7C7 |
| EDYCKYLESYNV | 1 | 0 | 0 | 1426731 | ARB07436.1 | A0A140H1G9 |
| EDYQIGGYSEDW | 1 | 0 | 0 | 1426738 | ATO98182.1 | Q3LZX0 |
| EDYTGYQELVIK | 1 | 0 | 0 | 1426739 | AIL49528.1 | P36334 |
| EEAARCMRSLRA | 1 | 0 | 0 | 1426742 | 2W2G_A | P0C6U8 |
| EEADGAEEEVTV | 1 | 0 | 0 | 1426747 | ATQ39389.1 | K9N7C7 |
| EEAFYRDLYSSP | 1 | 0 | 0 | 1426749 | AXN92247.1 | K9N7C7 |
| EEAHGGEVDIQP | 1 | 0 | 0 | 1426751 | AUM60018.1 | K9N7D2 |
| EEDENAMFDKAV | 1 | 0 | 0 | 1426782 | AUM60021.1 | K9N4V7 |
| EEDLTKGPYEFC | 1 | 0 | 0 | 1426788 | AVL25535.1 | H9EJ42 |
| EEEIVPTLELCN | 1 | 0 | 0 | 1426826 | AVV62535.1 | K9N7C7 |
| EEEQEEDWLESD | 1 | 0 | 0 | 1426830 | AVP78041.1 | P0C6W2 |
| EEFACVVSDTVV | 1 | 0 | 0 | 1426835 | APO40578.1 | P0C6X7 |
| EEFIKHVRTNGG | 1 | 0 | 0 | 1426842 | ALW82730.1 | K9N7C7 |
| EEFPLEEEVQEE | 1 | 0 | 0 | 1426848 | AVV62524.1 | K9N7C7 |
| EEFVAASDITLK | 1 | 0 | 0 | 1426851 | NP_073549.1 | A0A0P0LKV0 |
| EEGESECISETS | 1 | 0 | 0 | 1426860 | AXN92247.1 | K9N7C7 |
| EEGQCEPSTQYE | 1 | 0 | 0 | 1426864 | AVP78030.1 | P0C6W2 |
| EEIADETCEHEY | 1 | 0 | 0 | 1426876 | ARO76381.1 | P0C6U8 |
| EEIVINGRTYVC | 1 | 0 | 0 | 1426889 | AST12962.1 | A0A0P0LKV0 |
| EEIYNKLTEVKV | 1 | 0 | 0 | 1426891 | AVV62524.1 | K9N7C7 |
| EEKKIKASIEEV | 1 | 0 | 0 | 1426898 | AVP78041.1 | P0C6W2 |
| EELASPMKSDEG | 1 | 0 | 0 | 1426908 | ARU07575.1 | A0A140H1G9 |
| EELDQWFKNQIS | 1 | 0 | 0 | 1426910 | ANZ78835.1 | P36334 |
| EELDQWFKNQTL | 1 | 0 | 0 | 1426911 | CAA83661.1 | P36334 |
| EELEQTIQQKEL | 1 | 0 | 0 | 1426915 | APO40578.1 | P0C6X7 |
| EELKRLLEQWNL | 1 | 0 | 0 | 1426928 | ATO98197.1 | Q3LZX9 |
| EELQHEDLLAAY | 1 | 0 | 0 | 1426939 | APO40578.1 | P0C6X7 |
| EEMLKRSSTIHN | 1 | 0 | 0 | 1426950 | ASU90393.1 | K9N7C7 |
| EENISRGTILFK | 1 | 0 | 0 | 1426956 | CAA71146.1 | A0A0P0K6L9 |
| EENLSSRMYCSF | 1 | 0 | 0 | 1426964 | ARO76381.1 | P0C6U8 |
| EENVIQCFCPRD | 1 | 0 | 0 | 1426965 | AFD98754.1 | H9EJ69 |
| EEPIVEQPVEEF | 1 | 0 | 0 | 1426974 | AVV62535.1 | K9N7C7 |
| EEPTTQKPVDVK | 1 | 0 | 0 | 1426977 | ARO76381.1 | P0C6U8 |
| EESITIKKPNEL | 1 | 0 | 0 | 1426995 | APO40578.1 | P0C6X7 |
| EESKLALKKCKS | 1 | 0 | 0 | 1426996 | APO40578.1 | P0C6X7 |
| EESLMDVPSTYG | 1 | 0 | 0 | 1426998 | AIA22866.1 | K9N643 |
| EESLTDVPSISG | 1 | 0 | 0 | 1426999 | ATG84770.1 | K9N643 |
| EESNDQEMTTPP | 1 | 0 | 0 | 1427000 | ATQ39393.1 | K9N4V7 |
| EESQPVEVPVED | 1 | 0 | 0 | 1427002 | AHI48792.1 | K9N638 |
| EESTDQMSEPPM | 1 | 0 | 0 | 1427008 | AXP07352.1 | K9N4V7 |
| EETIYNLLKDCP | 1 | 0 | 0 | 1427024 | QYC64967.1 | P0DTD1 |
| EETRKLMPVCMD | 1 | 0 | 0 | 1427038 | P0C6W6.1 | P0C6W2 |
| EEVIVQQAAELH | 1 | 0 | 0 | 1427056 | AUM60013.1 | K9N7C7 |
| EEVKPFFTENKP | 1 | 0 | 0 | 1427057 | AVP78041.1 | P0C6W2 |
| EEVQEEKTSISV | 1 | 0 | 0 | 1427062 | AVV62524.1 | K9N7C7 |
| EEYEIEPHKIHV | 1 | 0 | 0 | 1427090 | AVV62526.1 | K9N5Q8 |
| EFACVVADAVIKTLQPVS | 1 | 0 | 0 | 1427096 | P0DTC1.1 | P0DTD1 |
| EFACVVSDTVVK | 1 | 0 | 0 | 1427099 | APO40578.1 | P0C6X7 |
| EFADKLVEVGLV | 1 | 0 | 0 | 1427100 | AEN19363.1 | P0C6X6 |
| EFAKNLPKYVKP | 1 | 0 | 0 | 1427103 | AWK59979.1 | A0A0P0G321 |
| EFANDTKIVSQL | 1 | 0 | 0 | 1427106 | AVN89313.1 | K9N5Q8 |

| | | | | | | |
|---|---|---|---|---|---|---|
| EFANHVANFQKS | 1 | 0 | 0 | 1427107 | AKN24802.1 | K9N7C7 |
| EFASRTVSAASV | 1 | 0 | 0 | 1427114 | AVV62535.1 | K9N7C7 |
| EFAVLYQDVNCT | 1 | 0 | 0 | 1427118 | BAE93401.1 | P59594 |
| EFAVNSTKALAK | 1 | 0 | 0 | 1427119 | ASU90276.1 | K9N4V0 |
| EFCSQHTHYIKD | 1 | 0 | 0 | 1427125 | AHI48773.1 | K9N7C7 |
| EFCTKHTMLVKM | 1 | 0 | 0 | 1427129 | AIL49625.1 | P0C6X6 |
| EFDENGTITDAV | 1 | 0 | 0 | 1427132 | APO40579.1 | P59594 |
| EFDTYDPTYKNG | 1 | 0 | 0 | 1427138 | AVN89451.1 | K9N7C7 |
| EFDVETNEIVDS | 1 | 0 | 0 | 1427140 | AIL23988.1 | K9N7C7 |
| EFEFAEGQGVPI | 1 | 0 | 0 | 1427147 | AEN19370.1 | P33469 |
| EFEFVEGQGPPI | 1 | 0 | 0 | 1427148 | P33469.1 | P33469 |
| EFFKNFSTSIPN | 1 | 0 | 0 | 1427152 | AKM76239.1 | K9N5Q8 |
| EFFKNVSSQGPN | 1 | 0 | 0 | 1427153 | AVV62526.1 | K9N5Q8 |
| EFGASTEMQVEE | 1 | 0 | 0 | 1427159 | ATO98229.1 | P0C6W2 |
| EFGASTETLQVE | 1 | 0 | 0 | 1427160 | ARO76381.1 | P0C6U8 |
| EFIEQPTSVECD | 1 | 0 | 0 | 1427173 | AVV62526.1 | K9N5Q8 |
| EFIERYKLDGYA | 1 | 0 | 0 | 1427174 | AVP78041.1 | P0C6W2 |
| EFIFEKVEYGSD | 1 | 0 | 0 | 1427176 | AWK59942.1 | H9EJ42 |
| EFIGTQALDMLA | 1 | 0 | 0 | 1427177 | AVV62524.1 | K9N7C7 |
| EFIKHVRTNGGY | 1 | 0 | 0 | 1427179 | ALW82730.1 | K9N7C7 |
| EFINFSDSFYTQ | 1 | 0 | 0 | 1427180 | AVV62524.1 | K9N7C7 |
| EFKFWDESFYAS | 1 | 0 | 0 | 1427184 | APT69863.1 | A0A0P0LKV0 |
| EFKVLDNGIIRD | 1 | 0 | 0 | 1427189 | AUM60013.1 | K9N7C7 |
| EFLATFIVCKIV | 1 | 0 | 0 | 1427191 | AWK59942.1 | H9EJ42 |
| EFLKRGEKIVYH | 1 | 0 | 0 | 1427202 | P0C6W6.1 | P0C6W2 |
| EFMLWCKDGKVQ | 1 | 0 | 0 | 1427213 | ATQ39389.1 | K9N7C7 |
| EFMLWCKNGQVQ | 1 | 0 | 0 | 1427215 | AVV62535.1 | K9N7C7 |
| EFNDLAHTSLVW | 1 | 0 | 0 | 1427216 | AVA26872.1 | H9EJ42 |
| EFPAITIGSNFV | 1 | 0 | 0 | 1427228 | BBO58761.1 | P36334 |
| EFRDYFDNNTDS | 1 | 0 | 0 | 1427244 | AWK59942.1 | H9EJ42 |
| EFRDYFNDNTDS | 1 | 0 | 0 | 1427245 | AVA26872.1 | H9EJ42 |
| EFSCDGISPDAI | 1 | 0 | 0 | 1427260 | AVV62537.1 | K9N5Q8 |
| EFSCHQVSPSAL | 1 | 0 | 0 | 1427261 | AUM60014.1 | K9N5Q8 |
| EFSKKQLGLLQS | 1 | 0 | 0 | 1427268 | AXG21653.1 | K9N7C7 |
| EFSKQLQQSMSG | 1 | 0 | 0 | 1427269 | AVP78038.1 | Q3LZX4 |
| EFSTHDPIYKNG | 1 | 0 | 0 | 1427277 | AVV62524.1 | K9N7C7 |
| EFSVVLPGEMAK | 1 | 0 | 0 | 1427278 | AVV62535.1 | K9N7C7 |
| EFSVVLPGEVAK | 1 | 0 | 0 | 1427279 | AVV62524.1 | K9N7C7 |
| EFTIGNMEEFIQ | 1 | 0 | 0 | 1427286 | CAA79896.1 | P36334 |
| EFTPEDVNMQLM | 1 | 0 | 0 | 1427291 | AQZ41284.1 | K9N7C7 |
| EFTTGFVYFNGN | 1 | 0 | 0 | 1427298 | APT69890.1 | A0A0P0K6L9 |
| EFVDVVKDEVSD | 1 | 0 | 0 | 1427304 | AVV62535.1 | K9N7C7 |
| EFVEYKITQWPT | 1 | 0 | 0 | 1427311 | AVQ05261.1 | P0C6X6 |
| EFVFEKVDYGSD | 1 | 0 | 0 | 1427313 | NP_073549.1 | A0A0P0LKV0 |
| EFVGEFYAYLRK | 1 | 0 | 0 | 1427320 | P0C6W6.1 | P0C6W2 |
| EFVHYDPMYKNG | 1 | 0 | 0 | 1427322 | AUM60013.1 | K9N7C7 |
| EFVIARTGHFYI | 1 | 0 | 0 | 1427323 | AIW52745.1 | A0A0P0K6L9 |
| EFVQEEAQPVEA | 1 | 0 | 0 | 1427328 | AVN89451.1 | K9N7C7 |
| EFVQEEVQPVEV | 1 | 0 | 0 | 1427330 | AXN92247.1 | K9N7C7 |
| EFVREEVDIIEQ | 1 | 0 | 0 | 1427331 | AWK59942.1 | H9EJ42 |
| EFVSDADITLLG | 1 | 0 | 0 | 1427332 | ALL26405.1 | K9N7C7 |
| EFVVNDVVIAIL | 1 | 0 | 0 | 1427337 | AVN89333.1 | K9N7C7 |
| EFYDDRILEFKV | 1 | 0 | 0 | 1427342 | ATQ39389.1 | K9N7C7 |
| EFYGFKDAVTFA | 1 | 0 | 0 | 1427346 | NP_073549.1 | A0A0P0LKV0 |
| EFYNDIKPPVLD | 1 | 0 | 0 | 1427349 | AOL02455.1 | Q04854 |
| EFYTTEPEFVKV | 1 | 0 | 0 | 1427351 | AEN19363.1 | P0C6X6 |
| EGAINVGKNGLQ | 1 | 0 | 0 | 1427355 | AUM60013.1 | K9N7C7 |
| EGAINVGKQGLE | 1 | 0 | 0 | 1427356 | AVV62524.1 | K9N7C7 |
| EGALNTPKEHIG | 1 | 0 | 0 | 1427359 | APO40586.1 | P59595 |
| EGALTAVVIPTK | 1 | 0 | 0 | 1427360 | AVP78041.1 | P0C6W2 |
| EGCGILSNKSNP | 1 | 0 | 0 | 1427365 | APO40579.1 | P59594 |
| EGCHATRDAVGT | 1 | 0 | 0 | 1427366 | AEA10817.1 | P0C6X7 |
| EGDASWYSTMIF | 1 | 0 | 0 | 1427376 | AHI48789.1 | K9N638 |
| EGDLPENDDDAW | 1 | 0 | 0 | 1427384 | AWW13564.1 | Q80872 |
| EGDWDSFGKTIQ | 1 | 0 | 0 | 1427394 | AST12962.1 | A0A0P0LKV0 |
| EGDYYRKQLSPL | 1 | 0 | 0 | 1427398 | ALJ76277.1 | K9N5Q8 |
| EGEDVIFSQFDS | 1 | 0 | 0 | 1427401 | AVQ05261.1 | P0C6X6 |
| EGFGGKNILGYA | 1 | 0 | 0 | 1427413 | 5NH0_A | H9EJ42 |
| EGFGGKNILGYS | 1 | 0 | 0 | 1427414 | AWK59942.1 | H9EJ42 |
| EGFMGRIRSLYP | 1 | 0 | 0 | 1427421 | ARO76381.1 | P0C6U8 |
| EGGAKETKEINI | 1 | 0 | 0 | 1427430 | AVQ05261.1 | P0C6X6 |
| EGGASWSSTMIF | 1 | 0 | 0 | 1427431 | AKN24802.1 | K9N7C7 |
| EGHIIMEEMLKE | 1 | 0 | 0 | 1427453 | AUM60013.1 | K9N7C7 |
| EGHIIMEEMLRG | 1 | 0 | 0 | 1427456 | AVN89322.1 | K9N7C7 |
| EGIEQVPPHSFT | 1 | 0 | 0 | 1427466 | AVV62535.1 | K9N7C7 |
| EGIKVTQELGHE | 1 | 0 | 0 | 1427470 | ARO76381.1 | P0C6U8 |
| EGILSEREKLLH | 1 | 0 | 0 | 1427471 | AGT51369.1 | U3M6Q8 |
| EGIVWVHEDGAT | 1 | 0 | 0 | 1427479 | AVV62544.1 | K9N4V7 |
| EGKDGFVSVELQ | 1 | 0 | 0 | 1427482 | AXN92247.1 | K9N7C7 |
| EGKDGFVTVELQ | 1 | 0 | 0 | 1427483 | AUM60013.1 | K9N7C7 |
| EGKVFQSTPIAL | 1 | 0 | 0 | 1427494 | AVP78039.1 | Q3LZX3 |
| EGLASDLLDNAS | 1 | 0 | 0 | 1427495 | AVV62535.1 | K9N7C7 |
| EGLASDLLDNVS | 1 | 0 | 0 | 1427496 | AVV62524.1 | K9N7C7 |
| EGLDCKIQTAPH | 1 | 0 | 0 | 1427499 | ARK08668.1 | Q80872 |
| EGLLLIPDLLHA | 1 | 0 | 0 | 1427506 | AHF54653.1 | Q4VID0 |
| EGLLLIPDLLRA | 1 | 0 | 0 | 1427507 | AVR40343.1 | Q4VID0 |
| EGLVRIVRTRSM | 1 | 0 | 0 | 1427516 | AVQ05261.1 | P0C6X6 |
| EGMLYPTILGSL | 1 | 0 | 0 | 1427519 | ARE29989.1 | Q4VID0 |
| EGMLYPTTLGSL | 1 | 0 | 0 | 1427520 | AGT51745.1 | Q4VID0 |
| EGNSPYHPLADN | 1 | 0 | 0 | 1427532 | APO40584.1 | P59635 |
| EGNVINFCYMHQ | 1 | 0 | 0 | 1427534 | AVV62535.1 | K9N7C7 |

| | | | | | | |
|---|---|---|---|---|---|---|
| EGQFEPSTQYEY | 1 | 0 | 0 | 1427545 | AVP78041.1 | P0C6W2 |
| EGQLQSLSRDYI | 1 | 0 | 0 | 1427550 | ABE97136.1 | H9EJ42 |
| EGQQIIFYEGVN | 1 | 0 | 0 | 1427552 | AEN19365.1 | P30215 |
| EGRITPYCHDPT | 1 | 0 | 0 | 1427554 | AHL18091.1 | K9N7C7 |
| EGRMVPYCYDPT | 1 | 0 | 0 | 1427556 | AUM60013.1 | K9N7C7 |
| EGSFPCSVCLGG | 1 | 0 | 0 | 1427562 | ARO76381.1 | P0C6U8 |
| EGSGKSAPNSRS | 1 | 0 | 0 | 1427564 | AJQ24087.1 | P33469 |
| EGSGKYAPNSRS | 1 | 0 | 0 | 1427565 | AJQ24067.1 | P33469 |
| EGSGRPAPNSRS | 1 | 0 | 0 | 1427566 | AIL49470.1 | P33469 |
| EGSYHNWHLVKG | 1 | 0 | 0 | 1427584 | AVV62524.1 | K9N7C7 |
| EGTGGNIQSSSR | 1 | 0 | 0 | 1427588 | ANI69874.1 | K9N4V7 |
| EGVCITTNGLWA | 1 | 0 | 0 | 1427594 | AVK87438.1 | K9N638 |
| EGVCLGAGQFLG | 1 | 0 | 0 | 1427596 | AWW13571.1 | P0C6X6 |
| EGVDCKIQTAPH | 1 | 0 | 0 | 1427597 | AWW13564.1 | Q80872 |
| EGVFVMNGTHWF | 1 | 0 | 0 | 1427602 | APO40579.1 | P59594 |
| EGVFVSNGTSWF | 1 | 0 | 0 | 1427604 | ATO98181.1 | Q3LZX1 |
| EGVGARVSAFLQ | 1 | 0 | 0 | 1427608 | AXX83306.1 | P0C6X6 |
| EGVLTAVVIPTK | 1 | 0 | 0 | 1427612 | QYC64967.1 | P0DTD1 |
| EGVMYSEFPAIA | 1 | 0 | 0 | 1427613 | AKA09659.1 | P36334 |
| EGVNFNPHHRFK | 1 | 0 | 0 | 1427615 | AGT17757.1 | A0A140H1H0 |
| EGVSSFMNVTLN | 1 | 0 | 0 | 1427623 | AIW52757.1 | A0A0P0K6L9 |
| EGWHVVNKFYQI | 1 | 0 | 0 | 1427627 | AIV41876.1 | P0C6X6 |
| EHALSIKQPFSF | 1 | 0 | 0 | 1427638 | AGT51337.1 | H9EJ42 |
| EHCQNYSGSMLR | 1 | 0 | 0 | 1427646 | ASU45697.1 | K9N796 |
| EHCSGLAIKSDH | 1 | 0 | 0 | 1427650 | AVI15011.1 | P36334 |
| EHCSGLAIKSDY | 1 | 0 | 0 | 1427651 | CAA83660.1 | P36334 |
| EHDEWFGITQDT | 1 | 0 | 0 | 1427654 | AVV62526.1 | K9N5Q8 |
| EHDYQIGGYTEKWESGVKDCVVL | 1 | 0 | 0 | 1427659 | P0DTC3.1 | P0DTC3 |
| EHEIAWFTERSE | 1 | 0 | 0 | 1427668 | ARO76381.1 | P0C6U8 |
| EHGATDAPSTFG | 1 | 0 | 0 | 1427683 | AXN92267.1 | K9N4V7 |
| EHLSPWFADGAT | 1 | 0 | 0 | 1427712 | AVV62535.1 | K9N7C7 |
| EHPDVSGVCKYK | 1 | 0 | 0 | 1427722 | AXN92247.1 | K9N7C7 |
| EHPHTICHPKLG | 1 | 0 | 0 | 1427723 | AKA09654.1 | P36334 |
| EHPHTICHPNLG | 1 | 0 | 0 | 1427724 | BAM16463.1 | P36334 |
| EHRRGPGKCYNC | 1 | 0 | 0 | 1427732 | AOG74783.1 | A0A0P0K6L9 |
| EHSLRLVDAMTY | 1 | 0 | 0 | 1427737 | ARO76381.1 | P0C6U8 |
| EHSTEFTRVNAK | 1 | 0 | 0 | 1427743 | AVP78030.1 | P0C6W2 |
| EHSWSADLYELM | 1 | 0 | 0 | 1427747 | AVV62524.1 | K9N7C7 |
| EHVLSIKQPFSF | 1 | 0 | 0 | 1427763 | ARU07593.1 | H9EJ42 |
| EHVTSGNLFDVG | 1 | 0 | 0 | 1427773 | AGV08439.1 | K9N796 |
| EHVTSVNLFDIG | 1 | 0 | 0 | 1427775 | AID50419.1 | K9N796 |
| EHVTSVNLLDVG | 1 | 0 | 0 | 1427777 | AHI48551.1 | K9N796 |
| EHVVTGDNDDEH | 1 | 0 | 0 | 1427778 | ABD75543.1 | A0A140H1G9 |
| EHYRALPGVFCG | 1 | 0 | 0 | 1427783 | P0C6W6.1 | P0C6W2 |
| EIADETCEHEYG | 1 | 0 | 0 | 1427790 | ARO76381.1 | P0C6U8 |
| EIAPVYEADTEQ | 1 | 0 | 0 | 1427796 | ALA49362.1 | K9N7C7 |
| EIAQYNYKQDFS | 1 | 0 | 0 | 1427798 | AUM60014.1 | K9N5Q8 |
| EIAWFTERSEKS | 1 | 0 | 0 | 1427805 | ARO76381.1 | P0C6U8 |
| EIDITVRDAKFL | 1 | 0 | 0 | 1427811 | APO40578.1 | P0C6X7 |
| EIDRLNEVARNL | 1 | 0 | 0 | 1427818 | AVP78031.1 | Q3LZX1 |
| EIEGVGAKVSAF | 1 | 0 | 0 | 1427821 | AGT51707.1 | P0C6X6 |
| EIEKPRWKRQPN | 1 | 0 | 0 | 1427822 | AAA45463.1 | A0A0N7J131 |
| EIGTYEMYVKWP | 1 | 0 | 0 | 1427842 | BBA20986.1 | A0A140H1H1 |
| EIIGYKAIYNGV | 1 | 0 | 0 | 1427851 | APO40578.1 | P0C6X7 |
| EIIKSSDVSENN | 1 | 0 | 0 | 1427856 | AVV62524.1 | K9N7C7 |
| EIKALKELYGPV | 1 | 0 | 0 | 1427865 | AVN89363.1 | K9N7C7 |
| EIKCKTQFIAPP | 1 | 0 | 0 | 1427868 | AHA80218.1 | P36334 |
| EIKPGKMKVKAT | 1 | 0 | 0 | 1427881 | 2J98_B | A0A0P0LKV0 |
| EIKPSGLKTMIV | 1 | 0 | 0 | 1427882 | AQZ41321.1 | K9N7C7 |
| EIKPTGLKTMVV | 1 | 0 | 0 | 1427884 | AUM60013.1 | K9N7C7 |
| EILLAPLLSAGI | 1 | 0 | 0 | 1427906 | AVP78041.1 | P0C6W2 |
| EINRLQEAIKVL | 1 | 0 | 0 | 1427931 | AIL49487.1 | P36334 |
| EIPSLMTVNVMF | 1 | 0 | 0 | 1427938 | ABD75639.1 | A0A140H1G9 |
| EIPVVSDTVEVP | 1 | 0 | 0 | 1427940 | AQZ41284.1 | K9N7C7 |
| EIQIPSDFTIGN | 1 | 0 | 0 | 1427946 | ARB66062.1 | P36334 |
| EIQSSGVINSSA | 1 | 0 | 0 | 1427954 | AVV62535.1 | K9N7C7 |
| EIRDLICVQSYN | 1 | 0 | 0 | 1427958 | CAA83660.1 | P36334 |
| EIRPTGLKTMIV | 1 | 0 | 0 | 1427962 | AVV62535.1 | K9N7C7 |
| EIRQKILNKPRQ | 1 | 0 | 0 | 1427964 | BBM61433.1 | P33469 |
| EISCDCGKKFDE | 1 | 0 | 0 | 1427969 | QED88024.1 | H9EJ42 |
| EITACQYTMCEF | 1 | 0 | 0 | 1427988 | ARB07438.1 | A0A140H1H1 |
| EITKENVETLTW | 1 | 0 | 0 | 1427990 | AWK59942.1 | H9EJ42 |
| EIVGERELILVW | 1 | 0 | 0 | 1428001 | AVV62524.1 | K9N7C7 |
| EIVGERQLLLLW | 1 | 0 | 0 | 1428002 | AZU90729.1 | K9N7C7 |
| EIVTGDNDDEDV | 1 | 0 | 0 | 1428019 | ABD75503.1 | A0A140H1G9 |
| EIYNKLTEVKVP | 1 | 0 | 0 | 1428028 | AVV62524.1 | K9N7C7 |
| EKCGVLDGSYNA | 1 | 0 | 0 | 1428043 | BBA20983.1 | A0A140H1H1 |
| EKCSSYTVETGT | 1 | 0 | 0 | 1428051 | APO40578.1 | P0C6X7 |
| EKDDQGHNLDSY | 1 | 0 | 0 | 1428057 | AVV62524.1 | K9N7C7 |
| EKDFLSFDNDVF | 1 | 0 | 0 | 1428061 | QBM11746.1 | K9N7C7 |
| EKDFLSFDSAVF | 1 | 0 | 0 | 1428062 | AIZ74426.1 | K9N7C7 |
| EKDFLSLDSDVF | 1 | 0 | 0 | 1428064 | AVV62535.1 | K9N7C7 |
| EKDFLSSDSDVF | 1 | 0 | 0 | 1428067 | ANI69887.1 | K9N7C7 |
| EKDKSVARKLER | 1 | 0 | 0 | 1428072 | AUM60013.1 | K9N7C7 |
| EKEGSYHNWHLV | 1 | 0 | 0 | 1428076 | AVV62524.1 | K9N7C7 |
| EKEVLSELYGTP | 1 | 0 | 0 | 1428080 | AVV62535.1 | K9N7C7 |
| EKIVYHTLESPV | 1 | 0 | 0 | 1428112 | P0C6W6.1 | P0C6W2 |
| EKKFPPYINGFN | 1 | 0 | 0 | 1428115 | AIV41989.1 | Q04853 |
| EKKIKASIEEVT | 1 | 0 | 0 | 1428116 | AVP78041.1 | P0C6W2 |
| EKKQKASEKQGG | 1 | 0 | 0 | 1428126 | AUM60021.1 | K9N4V7 |
| EKLASPMKSDEG | 1 | 0 | 0 | 1428130 | AZS52616.1 | A0A140H1G9 |

| | | | | | | |
|---|---|---|---|---|---|---|
| EKLDNPSGSEVD | 1 | 0 | 0 | 1428133 | AGT51717.1 | P0C6X6 |
| EKLHCEHLQFGL | 1 | 0 | 0 | 1428139 | AWO68974.1 | A0A0P0G321 |
| EKLISLADVATD | 1 | 0 | 0 | 1428148 | AUM60013.1 | K9N7C7 |
| EKLISLTDVATD | 1 | 0 | 0 | 1428149 | AUM60023.1 | K9N7C7 |
| EKLISTARDGLK | 1 | 0 | 0 | 1428151 | AVV62535.1 | K9N7C7 |
| EKLKQLLEQWNL | 1 | 0 | 0 | 1428157 | AAS48455.1 | P59596 |
| EKLKTILALREV | 1 | 0 | 0 | 1428159 | APO40578.1 | P0C6X7 |
| EKLNAFLVHGNV | 1 | 0 | 0 | 1428165 | AST12962.1 | A0A0P0LKV0 |
| EKLSPCKEIEGV | 1 | 0 | 0 | 1428176 | AGT51707.1 | P0C6X6 |
| EKLYKDLTGYML | 1 | 0 | 0 | 1428191 | QAT98908.1 | K9N7C7 |
| EKNYYELTKSCG | 1 | 0 | 0 | 1428200 | AZS52616.1 | A0A140H1G9 |
| EKPNLNSLCLLN | 1 | 0 | 0 | 1428202 | AET99209.1 | Q0ZJ80 |
| EKPQEKDKKSAK | 1 | 0 | 0 | 1428203 | ARB07429.1 | A0A0N7J131 |
| EKQNGLSLYLAD | 1 | 0 | 0 | 1428213 | AUM60013.1 | K9N7C7 |
| EKRKTEGFMGRI | 1 | 0 | 0 | 1428218 | APO40578.1 | P0C6X7 |
| EKSAGYPFNQFG | 1 | 0 | 0 | 1428222 | AIL49570.1 | P0C6X6 |
| EKSFQLDTAPKT | 1 | 0 | 0 | 1428224 | Q3I5J5.1 | Q3LZX1 |
| EKSYEQQTPFDI | 1 | 0 | 0 | 1428239 | APO40578.1 | P0C6X7 |
| EKSYERQSPFEI | 1 | 0 | 0 | 1428240 | ARO76381.1 | P0C6U8 |
| EKTHGPLSLPVL | 1 | 0 | 0 | 1428243 | AVP78030.1 | P0C6W2 |
| EKYFKHWDQAYH | 1 | 0 | 0 | 1428264 | AVV62524.1 | K9N7C7 |
| EKYFKYWDQTYY | 1 | 0 | 0 | 1428266 | ATG84699.1 | K9N7C7 |
| ELAKGVSLDNVL | 1 | 0 | 0 | 1428274 | APO40578.1 | P0C6X7 |
| ELANGTHTGSAF | 1 | 0 | 0 | 1428281 | AXN92247.1 | K9N7C7 |
| ELANGTHTGSSF | 1 | 0 | 0 | 1428282 | AVV62524.1 | K9N7C7 |
| ELAPPAGAFFFG | 1 | 0 | 0 | 1428284 | ANZ78874.1 | P33469 |
| ELAPSASAFMSM | 1 | 0 | 0 | 1428286 | AVV62544.1 | K9N4V7 |
| ELAPTAGAFFFG | 1 | 0 | 0 | 1428288 | AEN19370.1 | P33469 |
| ELAPTAGAFFFR | 1 | 0 | 0 | 1428289 | AHA80162.1 | P33469 |
| ELAQALTEVVYS | 1 | 0 | 0 | 1428291 | ADO24700.1 | A0A0P0LKV0 |
| ELASPMKSDEGG | 1 | 0 | 0 | 1428298 | ARU07575.1 | A0A140H1G9 |
| ELCNAVEQRITV | 1 | 0 | 0 | 1428311 | AVV62535.1 | K9N7C7 |
| ELCVDEVGSKSP | 1 | 0 | 0 | 1428314 | AVP78037.1 | Q3LZX5 |
| ELDDEELMELDY | 1 | 0 | 0 | 1428315 | AAP30035.1 | P59634 |
| ELDDFSKQLQDS | 1 | 0 | 0 | 1428319 | APO40586.1 | P59595 |
| ELDEEGNLLDSY | 1 | 0 | 0 | 1428321 | APO40578.1 | P0C6X7 |
| ELDNKFTPLKVE | 1 | 0 | 0 | 1428330 | AUM60013.1 | K9N7C7 |
| ELEAAQKSYQEA | 1 | 0 | 0 | 1428340 | AVV62524.1 | K9N7C7 |
| ELEAARKAYQEA | 1 | 0 | 0 | 1428341 | AMQ49036.1 | K9N638 |
| ELEKRCYELVKD | 1 | 0 | 0 | 1428358 | AVV62524.1 | K9N7C7 |
| ELELICDCGIKQ | 1 | 0 | 0 | 1428359 | ARU07575.1 | A0A140H1G9 |
| ELEQTIQQKELE | 1 | 0 | 0 | 1428367 | APO40578.1 | P0C6X7 |
| ELESFKQELDKY | 1 | 0 | 0 | 1428368 | APO40579.1 | P59594 |
| ELFDSLNLSADT | 1 | 0 | 0 | 1428375 | ARU07581.1 | Q19U29 |
| ELFTKRSIRHHP | 1 | 0 | 0 | 1428389 | ARU07574.1 | A0A140H1G9 |
| ELGDVRETMAHL | 1 | 0 | 0 | 1428396 | ARO76381.1 | P0C6U8 |
| ELGKYEHYIKWP | 1 | 0 | 0 | 1428404 | AVP78042.1 | Q3LZX1 |
| ELGKYEQYIKGS | 1 | 0 | 0 | 1428406 | 6NB6_A | P59594 |
| ELGKYEQYIKIK | 1 | 0 | 0 | 1428407 | 5X58_A | P59594 |
| ELGKYEQYVKWP | 1 | 0 | 0 | 1428408 | APO40579.1 | P59594 |
| ELGNYSYYQKWP | 1 | 0 | 0 | 1428413 | AVV62537.1 | K9N5Q8 |
| ELGNYTYYNKEF | 1 | 0 | 0 | 1428416 | 5X59_A | K9N5Q8 |
| ELGNYTYYNKGS | 1 | 0 | 0 | 1428417 | 6NB3_A | K9N5Q8 |
| ELGVVMNFDVDT | 1 | 0 | 0 | 1428431 | ACV87155.1 | A0A140H1G9 |
| ELGYIADISSFK | 1 | 0 | 0 | 1428433 | ARB07423.1 | A0A0P0LKV0 |
| ELHGNGSMIEDI | 1 | 0 | 0 | 1428443 | ALW82645.1 | K9N7C7 |
| ELHGYTVQPIAD | 1 | 0 | 0 | 1428444 | AIL49485.1 | P36334 |
| ELICDCGIKQES | 1 | 0 | 0 | 1428456 | ARU07575.1 | A0A140H1G9 |
| ELIPNHAALFFD | 1 | 0 | 0 | 1428460 | AMM42619.1 | H9EJ69 |
| ELKCTTKSLTVE | 1 | 0 | 0 | 1428475 | APO40579.1 | P59594 |
| ELKERDQYCALT | 1 | 0 | 0 | 1428478 | APO40578.1 | P0C6X7 |
| ELKETVFVEPKN | 1 | 0 | 0 | 1428479 | ARU07593.1 | H9EJ42 |
| ELKRELNGGAYT | 1 | 0 | 0 | 1428492 | AVP78041.1 | P0C6W2 |
| ELLEQNIDAYKS | 1 | 0 | 0 | 1428507 | AVV62533.1 | K9N4V7 |
| ELLNAPATVSGL | 1 | 0 | 0 | 1428532 | 2GHV_E | P59594 |
| ELLQLRTQTASL | 1 | 0 | 0 | 1428535 | QED88040.1 | A0A0P0G321 |
| ELLSELLKDKYT | 1 | 0 | 0 | 1428539 | ABE73410.1 | H9EJ42 |
| ELLSFKREKSLS | 1 | 0 | 0 | 1428541 | AGT51407.1 | Q4VID0 |
| ELLSFKRERSFS | 1 | 0 | 0 | 1428543 | QBP84718.1 | Q4VID0 |
| ELLTKAIDVDWV | 1 | 0 | 0 | 1428546 | NP_073549.1 | A0A0P0LKV0 |
| ELLVQDLCLNKL | 1 | 0 | 0 | 1428550 | ATG84700.1 | K9N638 |
| ELNAFTREMQQQ | 1 | 0 | 0 | 1428563 | AIW52723.1 | A0A0N7J131 |
| ELNGVMVDKVGD | 1 | 0 | 0 | 1428572 | QBP84746.1 | P0C6X6 |
| ELNGYPVQPIAD | 1 | 0 | 0 | 1428573 | AMM42466.1 | P36334 |
| ELNGYTVQPIAD | 1 | 0 | 0 | 1428574 | CAA79896.1 | P36334 |
| ELPDEFVVVTEK | 1 | 0 | 0 | 1428586 | APO40587.1 | P59636 |
| ELPFEPQTIQPE | 1 | 0 | 0 | 1428589 | QGV13494.1 | K9N7C7 |
| ELPYEPRTIQPE | 1 | 0 | 0 | 1428609 | ANI69888.1 | K9N638 |
| ELQALPQRQKKQ | 1 | 0 | 0 | 1428610 | AVP78038.1 | Q3LZX4 |
| ELQHEDLLAAYV | 1 | 0 | 0 | 1428615 | APO40578.1 | P0C6X7 |
| ELQVPVTLEQKV | 1 | 0 | 0 | 1428621 | APO40578.1 | P0C6X7 |
| ELRRGPGRCYNC | 1 | 0 | 0 | 1428627 | AIW52757.1 | A0A0P0K6L9 |
| ELRRGPGTCYNC | 1 | 0 | 0 | 1428628 | BAL45638.1 | A0A0P0K6L9 |
| ELSFTNYKLQSQ | 1 | 0 | 0 | 1428637 | AXN92247.1 | K9N7C7 |
| ELSNTFGAISSS | 1 | 0 | 0 | 1428649 | AVV62526.1 | K9N5Q8 |
| ELTAEDISLLRK | 1 | 0 | 0 | 1428668 | AHA80151.1 | P33469 |
| ELTASMSMAECK | 1 | 0 | 0 | 1428672 | AFD98819.1 | H9EJ42 |
| ELTKDKLQRIDW | 1 | 0 | 0 | 1428677 | AUM60013.1 | K9N7C7 |
| ELVDGYGNTMYV | 1 | 0 | 0 | 1428695 | AUM60027.1 | K9N643 |
| ELVDSVQYAHKH | 1 | 0 | 0 | 1428696 | AUM60013.1 | K9N7C7 |
| ELVVAANKTKGI | 1 | 0 | 0 | 1428716 | AVV62535.1 | K9N7C7 |

| | | | | | | |
|---|---|---|---|---|---|---|
| ELVYSYATHHDK | 1 | 0 | 0 | 1428718 | APO40578.1 | P0C6X7 |
| ELYAKRAIRSHP | 1 | 0 | 0 | 1428729 | AUM60013.1 | K9N7C7 |
| ELYDAPTTLQAV | 1 | 0 | 0 | 1428733 | AVV62535.1 | K9N7C7 |
| ELYEAPTTLQAV | 1 | 0 | 0 | 1428734 | AVV62524.1 | K9N7C7 |
| ELYGPLDPTFLH | 1 | 0 | 0 | 1428740 | AXN92247.1 | K9N7C7 |
| ELYGTPDSNFLY | 1 | 0 | 0 | 1428745 | AVV62524.1 | K9N7C7 |
| ELYLSWEVGKPR | 1 | 0 | 0 | 1428750 | AVP78041.1 | P0C6W2 |
| EMAKKQIGLLEP | 1 | 0 | 0 | 1428759 | AVV62535.1 | K9N7C7 |
| EMCILHCSNFNT | 1 | 0 | 0 | 1428765 | AIW52776.1 | A0A0P0LKV0 |
| EMCLASIAFNHP | 1 | 0 | 0 | 1428766 | AVN89313.1 | K9N5Q8 |
| EMEEYPMDVHST | 1 | 0 | 0 | 1428769 | ATQ39396.1 | K9N643 |
| EMFKPEVEELNP | 1 | 0 | 0 | 1428775 | AGT51436.1 | Q4VID0 |
| EMFKPGVEELNP | 1 | 0 | 0 | 1428776 | AOL02458.1 | Q4VID0 |
| EMKALFPIWSQE | 1 | 0 | 0 | 1428783 | AVQ05261.1 | P0C6X6 |
| EMKALLPTWSQE | 1 | 0 | 0 | 1428785 | QEG03790.1 | P0C6X6 |
| EMKSEKQVEQKV | 1 | 0 | 0 | 1428792 | AVP78030.1 | P0C6W2 |
| EMKVLNAVVKQL | 1 | 0 | 0 | 1428795 | AVV62526.1 | K9N5Q8 |
| EMKVLNEVVKQL | 1 | 0 | 0 | 1428796 | AVV62537.1 | K9N5Q8 |
| EMLKGSSTIHNY | 1 | 0 | 0 | 1428807 | AXN92247.1 | K9N7C7 |
| EMLKLGTSDSQF | 1 | 0 | 0 | 1428810 | AGT51764.1 | P33469 |
| EMLKRSSTIHNY | 1 | 0 | 0 | 1428811 | ASU90393.1 | K9N7C7 |
| EMNRLQEAIKVL | 1 | 0 | 0 | 1428820 | CAA79896.1 | P36334 |
| EMPCAIINDKVL | 1 | 0 | 0 | 1428821 | QBP84719.1 | P0C6X6 |
| EMPKFFYAPNGT | 1 | 0 | 0 | 1428824 | ADK37704.1 | A0A0P0K6L9 |
| EMPKFFYASNGT | 1 | 0 | 0 | 1428825 | AIW52757.1 | A0A0P0K6L9 |
| EMQQHPLLNPGA | 1 | 0 | 0 | 1428829 | AGT21342.1 | A0A0N7J131 |
| EMQQQPLLNPSA | 1 | 0 | 0 | 1428831 | AIW52723.1 | A0A0N7J131 |
| EMQSQPSHAVQN | 1 | 0 | 0 | 1428833 | AVL25602.1 | H9EJ69 |
| EMQSQPSHVAQN | 1 | 0 | 0 | 1428834 | AVA26877.1 | H9EJ69 |
| EMRLASIAFNHP | 1 | 0 | 0 | 1428841 | AXN92228.1 | K9N5Q8 |
| EMSCDCGKKFDE | 1 | 0 | 0 | 1428843 | AWK59942.1 | H9EJ42 |
| EMSDFVDFESVI | 1 | 0 | 0 | 1428844 | APU51933.1 | P0C6X6 |
| EMSDFVDLESVI | 1 | 0 | 0 | 1428846 | AEN19363.1 | P0C6X6 |
| EMSPFEMPFEEL | 1 | 0 | 0 | 1428847 | AST12962.1 | A0A0P0LKV0 |
| ENAFMPFAMGII | 1 | 0 | 0 | 1428868 | AVP78041.1 | P0C6W2 |
| ENALNSCKDHPV | 1 | 0 | 0 | 1428869 | ARB07436.1 | A0A140H1G9 |
| ENAMFDKAVADA | 1 | 0 | 0 | 1428873 | AUM60021.1 | K9N4V7 |
| ENASPASTVHTT | 1 | 0 | 0 | 1428876 | ARO76383.1 | P59632 |
| ENAVLPKGIMMN | 1 | 0 | 0 | 1428880 | APO40578.1 | P0C6X7 |
| ENAYEVVKAMLF | 1 | 0 | 0 | 1428881 | ALA49373.1 | K9N7C7 |
| ENAYLKWAKVEG | 1 | 0 | 0 | 1428882 | AUM60013.1 | K9N7C7 |
| ENAYLKYAKVEG | 1 | 0 | 0 | 1428883 | AVV62524.1 | K9N7C7 |
| ENDDDAWFKPSY | 1 | 0 | 0 | 1428889 | AQN78686.1 | Q80872 |
| ENDNISVAAPKS | 1 | 0 | 0 | 1428891 | AIX10718.1 | P33469 |
| ENDTITVDQLKH | 1 | 0 | 0 | 1428899 | APO40582.1 | P59596 |
| ENDWNTKHGSGL | 1 | 0 | 0 | 1428901 | APO40578.1 | P0C6X7 |
| ENEEIVINGHTY | 1 | 0 | 0 | 1428903 | ARB07390.1 | A0A0P0LKV0 |
| ENEEIVINGRTY | 1 | 0 | 0 | 1428904 | AST12962.1 | A0A0P0LKV0 |
| ENEIARIKKIKL | 1 | 0 | 0 | 1428905 | 5X58_A | P59594 |
| ENFIVYNNSGSN | 1 | 0 | 0 | 1428909 | AWK59942.1 | H9EJ42 |
| ENFNQHEILLAP | 1 | 0 | 0 | 1428911 | AVP78041.1 | P0C6W2 |
| ENFNSQDTLLAP | 1 | 0 | 0 | 1428914 | ATO98229.1 | P0C6W2 |
| ENFNSVTRLMCN | 1 | 0 | 0 | 1428916 | AXN92247.1 | K9N7C7 |
| ENGTISVEELKR | 1 | 0 | 0 | 1428924 | ATO98197.1 | Q3LZX9 |
| ENGTITDAIDCA | 1 | 0 | 0 | 1428925 | ATO98181.1 | Q3LZX1 |
| ENGTMSNAVDCS | 1 | 0 | 0 | 1428928 | ATO98145.1 | Q3LZX1 |
| ENGVRTLSTYDF | 1 | 0 | 0 | 1428929 | AVP78031.1 | Q3LZX1 |
| ENIAFNVFKQGH | 1 | 0 | 0 | 1428934 | ANI69909.1 | K9N7C7 |
| ENIAYNVVKQGH | 1 | 0 | 0 | 1428936 | AUM60023.1 | K9N7C7 |
| ENIDGDAMHANY | 1 | 0 | 0 | 1428937 | AVN89322.1 | K9N7C7 |
| ENIVYNVVKQGH | 1 | 0 | 0 | 1428941 | AUM60013.1 | K9N7C7 |
| ENIYNLIKDCPA | 1 | 0 | 0 | 1428942 | APO40578.1 | P0C6X7 |
| ENKFTPLSVASA | 1 | 0 | 0 | 1428948 | QBF80531.1 | K9N7C7 |
| ENKIPSVEVPPV | 1 | 0 | 0 | 1428953 | AVP78030.1 | P0C6W2 |
| ENKISGKGQQQQ | 1 | 0 | 0 | 1428954 | AVP78049.1 | Q3LZX4 |
| ENKLNISSGSRG | 1 | 0 | 0 | 1428955 | ATO98145.1 | Q3LZX1 |
| ENKLNSCCRIVE | 1 | 0 | 0 | 1428956 | ALA49351.1 | K9N7C7 |
| ENKLWFDFIENP | 1 | 0 | 0 | 1428957 | AXN92226.1 | K9N7C7 |
| ENKLWFDFVENP | 1 | 0 | 0 | 1428958 | AXN92247.1 | K9N7C7 |
| ENKTEAEWIELF | 1 | 0 | 0 | 1428962 | NP_073549.1 | A0A0P0LKV0 |
| ENLFVRSKEEDG | 1 | 0 | 0 | 1428971 | ANF29194.1 | K9N7C7 |
| ENLIHGGGVARA | 1 | 0 | 0 | 1428972 | AWK59942.1 | H9EJ42 |
| ENLIKRMKQIED | 1 | 0 | 0 | 1428973 | 5SZS_A | A0A0P0G321 |
| ENLLHGGGVARA | 1 | 0 | 0 | 1428976 | P0C6U6.1 | H9EJ42 |
| ENLMHGGGVARA | 1 | 0 | 0 | 1428978 | AVA26872.1 | H9EJ42 |
| ENLNILFDGRDT | 1 | 0 | 0 | 1428981 | ARU07574.1 | A0A140H1G9 |
| ENLPWPLVLEFA | 1 | 0 | 0 | 1428982 | ALD51903.1 | K9N7C7 |
| ENLSSEDLKAVR | 1 | 0 | 0 | 1428986 | AIV41978.1 | P0C6X6 |
| ENLTFPRGQGVP | 1 | 0 | 0 | 1428989 | AVP78038.1 | Q3LZX4 |
| ENLTNQLPPPLL | 1 | 0 | 0 | 1428992 | AVV62537.1 | K9N5Q8 |
| ENLYFQGGGGSG | 1 | 0 | 0 | 1428998 | 6NB6_A | P59594 |
| ENLYFQSMETGT | 1 | 0 | 0 | 1429000 | 5X29_A | P59637 |
| ENNCSQLDEEQP | 1 | 0 | 0 | 1429003 | AVP78035.1 | Q3LZX8 |
| ENNRVIISSDVL | 1 | 0 | 0 | 1429007 | AVP78041.1 | P0C6W2 |
| ENNRVVVSSDVL | 1 | 0 | 0 | 1429010 | ATO98179.1 | P0C6W2 |
| ENPKEEVKPFFT | 1 | 0 | 0 | 1429030 | AVP78041.1 | P0C6W2 |
| ENPNQNSLCLLN | 1 | 0 | 0 | 1429032 | Q0ZME2.1 | Q0ZJ80 |
| ENQINGYTLAAT | 1 | 0 | 0 | 1429042 | ANZ78836.1 | P36334 |
| ENQKQIANQFNN | 1 | 0 | 0 | 1429049 | APO40579.1 | P59594 |
| ENQPYQIEDPCP | 1 | 0 | 0 | 1429051 | Q3I5I8.1 | Q3LZX5 |
| ENSESAVLFVND | 1 | 0 | 0 | 1429055 | APT69882.1 | A0A0P0ILY9 |

| | | | | | | |
|---|---|---|---|---|---|---|
| ENSFMVNQLAYS | 1 | 0 | 0 | 1429057 | AVN89312.1 | K9N638 |
| ENTIPHYSWFSG | 1 | 0 | 0 | 1429068 | ABD75589.1 | Q19U29 |
| ENVGGNFDNDIN | 1 | 0 | 0 | 1429083 | AVV62535.1 | K9N7C7 |
| ENVIQGFGPRDF | 1 | 0 | 0 | 1429086 | QED88044.1 | H9EJ69 |
| ENYAFEHVVYGD | 1 | 0 | 0 | 1429095 | AXN92247.1 | K9N7C7 |
| ENYELEKHCYEL | 1 | 0 | 0 | 1429097 | AVV62535.1 | K9N7C7 |
| ENYELEKRCYEL | 1 | 0 | 0 | 1429098 | AVV62524.1 | K9N7C7 |
| EPCEVEYGTEDD | 1 | 0 | 0 | 1429111 | APO40578.1 | P0C6X7 |
| EPCNLWHYGAGI | 1 | 0 | 0 | 1429113 | QEG03729.1 | H9EJ42 |
| EPCPSGTYEGNS | 1 | 0 | 0 | 1429117 | ATO98237.1 | Q3LZX7 |
| EPFNVSFVNDSV | 1 | 0 | 0 | 1429145 | BBA20986.1 | A0A140H1H1 |
| EPFTVNSINDSL | 1 | 0 | 0 | 1429146 | AIL49542.1 | P36334 |
| EPFTVNSKNDNL | 1 | 0 | 0 | 1429147 | AIL49538.1 | P36334 |
| EPFTVNSVSDSL | 1 | 0 | 0 | 1429149 | ARB66068.1 | P36334 |
| EPIISQSSKLLN | 1 | 0 | 0 | 1429159 | AFO70496.1 | H9EJ42 |
| EPITLANARYVS | 1 | 0 | 0 | 1429168 | AVV62526.1 | K9N5Q8 |
| EPITMANARYVS | 1 | 0 | 0 | 1429169 | AVV62537.1 | K9N5Q8 |
| EPITSFNTKYVA | 1 | 0 | 0 | 1429170 | AVN89387.1 | K9N5Q8 |
| EPITYLNTKYVA | 1 | 0 | 0 | 1429173 | AGV08438.1 | K9N5Q8 |
| EPIVQHNTEPKV | 1 | 0 | 0 | 1429175 | APO40578.1 | P0C6X7 |
| EPKDGGHFFVSD | 1 | 0 | 0 | 1429177 | AFD98791.1 | H9EJ42 |
| EPKDGGHFFVSG | 1 | 0 | 0 | 1429178 | ARE29978.1 | H9EJ42 |
| EPKDGGQFFVSD | 1 | 0 | 0 | 1429179 | AWK59942.1 | H9EJ42 |
| EPKDGGQFFVSG | 1 | 0 | 0 | 1429180 | ARK08626.1 | H9EJ42 |
| EPMYAISNEQHY | 1 | 0 | 0 | 1429217 | ATO98193.1 | Q3LZX1 |
| EPPKEHRVQGTQ | 1 | 0 | 0 | 1429224 | YP_007188586.1 | K9N4V7 |
| EPPKEQRVQGII | 1 | 0 | 0 | 1429226 | AHI48535.1 | K9N4V7 |
| EPPKEQRVQSSI | 1 | 0 | 0 | 1429229 | ANI69896.1 | K9N4V7 |
| EPPKVAAKICIV | 1 | 0 | 0 | 1429232 | YP_009555247.1 | P0C6X6 |
| EPPLYVPIRLAG | 1 | 0 | 0 | 1429234 | AVN89312.1 | K9N638 |
| EPPNEQRVQGSI | 1 | 0 | 0 | 1429236 | AVN89381.1 | K9N4V7 |
| EPQRGRQSRIPY | 1 | 0 | 0 | 1429246 | ALK28785.1 | A0A0N7J131 |
| EPQTIKPEVKEV | 1 | 0 | 0 | 1429249 | AIZ74432.1 | K9N7C7 |
| EPQTIQPEVKEV | 1 | 0 | 0 | 1429251 | AXN92247.1 | K9N7C7 |
| EPRNGSYCPGNS | 1 | 0 | 0 | 1429256 | AVV62537.1 | K9N5Q8 |
| EPRNGTRCPNDN | 1 | 0 | 0 | 1429257 | AVV62526.1 | K9N5Q8 |
| EPRPGNHCPAGN | 1 | 0 | 0 | 1429258 | QBF80608.1 | K9N5Q8 |
| EPRSGNHCPADN | 1 | 0 | 0 | 1429259 | AID55096.1 | K9N5Q8 |
| EPSITYSPATIL | 1 | 0 | 0 | 1429266 | ATQ39389.1 | K9N7C7 |
| EPSTPISSALIA | 1 | 0 | 0 | 1429268 | AUM60013.1 | K9N7C7 |
| EPSTPVSSALIA | 1 | 0 | 0 | 1429269 | AVV62524.1 | K9N7C7 |
| EPTISQPSKLLN | 1 | 0 | 0 | 1429275 | ABE97136.1 | H9EJ42 |
| EPTTQKPVDVKP | 1 | 0 | 0 | 1429282 | ARO76381.1 | P0C6U8 |
| EPVHVYADDSII | 1 | 0 | 0 | 1429287 | AEN19363.1 | P0C6X6 |
| EPVHVYANDSII | 1 | 0 | 0 | 1429288 | QDH43722.1 | P0C6X6 |
| EPVLVNAANTHL | 1 | 0 | 0 | 1429292 | AVN89451.1 | K9N7C7 |
| EPVQGRKMLMGI | 1 | 0 | 0 | 1429297 | AVV62524.1 | K9N7C7 |
| EPVQGRKMLMGL | 1 | 0 | 0 | 1429298 | AUM60013.1 | K9N7C7 |
| EPVSGLYEIQIP | 1 | 0 | 0 | 1429302 | AIL49538.1 | P36334 |
| EPVSISTGSLSA | 1 | 0 | 0 | 1429303 | AVN89324.1 | K9N5Q8 |
| EPVSISTGSPSA | 1 | 0 | 0 | 1429304 | QFS19619.1 | K9N5Q8 |
| EPVSISTGSRSA | 1 | 0 | 0 | 1429305 | AXN92228.1 | K9N5Q8 |
| EQAEGFECDFSP | 1 | 0 | 0 | 1429312 | AQZ41282.1 | K9N5Q8 |
| EQAEGVECDFSL | 1 | 0 | 0 | 1429313 | ALJ54500.1 | K9N5Q8 |
| EQCIRGAIGDVG | 1 | 0 | 0 | 1429322 | ARB07424.1 | A0A0P0ILY9 |
| EQCTGLDSVMHF | 1 | 0 | 0 | 1429325 | AIV42002.1 | P0C6X6 |
| EQDRMLKTSMAP | 1 | 0 | 0 | 1429333 | QBQ01844.1 | Q4VID0 |
| EQDRMLKTSTAP | 1 | 0 | 0 | 1429334 | ATP16762.1 | Q4VID0 |
| EQDRNTRDVFAQ | 1 | 0 | 0 | 1429337 | APO40579.1 | P59594 |
| EQGSGYIPEAPR | 1 | 0 | 0 | 1429361 | 6CRW_A | P59594 |
| EQISAFTKPSSV | 1 | 0 | 0 | 1429373 | ABE97141.1 | H9EJ69 |
| EQKCCMCNRRAA | 1 | 0 | 0 | 1429377 | AXN92247.1 | K9N7C7 |
| EQKCCMCNRRAG | 1 | 0 | 0 | 1429378 | AUM60013.1 | K9N7C7 |
| EQKCCMCNRRAV | 1 | 0 | 0 | 1429379 | ALD51902.1 | K9N7C7 |
| EQKCCVCNRRAA | 1 | 0 | 0 | 1429381 | AHY22543.1 | K9N7C7 |
| EQKHEIEKPRWK | 1 | 0 | 0 | 1429382 | AAA45463.1 | A0A0N7J131 |
| EQLDFIESKRGV | 1 | 0 | 0 | 1429395 | ARO76381.1 | P0C6U8 |
| EQLDTTQLQVAN | 1 | 0 | 0 | 1429397 | AHF54662.1 | P36334 |
| EQLGAPLGITIS | 1 | 0 | 0 | 1429402 | AIW52843.1 | A0A0P0G321 |
| EQLLREYGQFCA | 1 | 0 | 0 | 1429408 | AVV62526.1 | K9N5Q8 |
| EQLLYAVQQLNN | 1 | 0 | 0 | 1429415 | AVV62524.1 | K9N7C7 |
| EQNHILGVNNSW | 1 | 0 | 0 | 1429423 | ARU07575.1 | A0A140H1G9 |
| EQPFGEVEYSLP | 1 | 0 | 0 | 1429436 | AWK59942.1 | H9EJ42 |
| EQPKIEDSNVKE | 1 | 0 | 0 | 1429440 | ARO76381.1 | P0C6U8 |
| EQPNSVECDFTK | 1 | 0 | 0 | 1429441 | AVV62537.1 | K9N5Q8 |
| EQPTVKEIISKP | 1 | 0 | 0 | 1429446 | AIV41888.1 | P0C6X6 |
| EQPTVKEIISMH | 1 | 0 | 0 | 1429447 | APU51933.1 | P0C6X6 |
| EQQNVLAIESIH | 1 | 0 | 0 | 1429460 | AWK59942.1 | H9EJ42 |
| EQRQQAEEKKIK | 1 | 0 | 0 | 1429467 | AVP78041.1 | P0C6W2 |
| EQRVPKGSITQR | 1 | 0 | 0 | 1429470 | AUM60021.1 | K9N4V7 |
| EQRVQGSITQHT | 1 | 0 | 0 | 1429475 | AKN24774.1 | K9N4V7 |
| EQRVQGSITQRA | 1 | 0 | 0 | 1429476 | AVN89381.1 | K9N4V7 |
| EQRVQSSIIQRT | 1 | 0 | 0 | 1429481 | QCZ25056.1 | K9N4V7 |
| EQRVRGSITQRT | 1 | 0 | 0 | 1429483 | ALA49392.1 | K9N4V7 |
| EQSFGEVEHSLP | 1 | 0 | 0 | 1429486 | ABE73411.1 | H9EJ42 |
| EQSFYENRFAAI | 1 | 0 | 0 | 1429488 | AVA26874.1 | H9EJ24 |
| EQSQRSLNFVGE | 1 | 0 | 0 | 1429495 | AVV62524.1 | K9N7C7 |
| EQTNMHSHDVIV | 1 | 0 | 0 | 1429508 | ASU90888.1 | K9N7C7 |
| EQTNMHSPDLIV | 1 | 0 | 0 | 1429510 | AUM60013.1 | K9N7C7 |
| EQTQHSLNFVGE | 1 | 0 | 0 | 1429516 | AXN92247.1 | K9N7C7 |
| EQTQSAIAKPKR | 1 | 0 | 0 | 1429520 | ATQ39389.1 | K9N7C7 |

| | | | | | | |
|---|---|---|---|---|---|---|
| EQTQSVNVKLKR | 1 | 0 | 0 | 1429522 | ANF29194.1 | K9N7C7 |
| EQTQSVNVKPKR | 1 | 0 | 0 | 1429523 | ATG84700.1 | K9N638 |
| EQTQSVTVKPKR | 1 | 0 | 0 | 1429524 | AXN92247.1 | K9N7C7 |
| EQVIPPTSDYLI | 1 | 0 | 0 | 1429536 | YP_173240.1 | Q0ZJ83 |
| EQYSCDFKGWIP | 1 | 0 | 0 | 1429562 | QBP84701.1 | P0C6X6 |
| ERAAVLEWDIKQ | 1 | 0 | 0 | 1429567 | AUM60013.1 | K9N7C7 |
| ERARTVGGVPLL | 1 | 0 | 0 | 1429571 | ADK37692.1 | H9EJ42 |
| ERCNVIADFKSF | 1 | 0 | 0 | 1429580 | AXN73435.1 | K9N7C7 |
| ERCVVRVCYGST | 1 | 0 | 0 | 1429585 | AWK59942.1 | H9EJ42 |
| ERCYTAEQCIRG | 1 | 0 | 0 | 1429586 | NP_073549.1 | A0A0P0LKV0 |
| ERDFIDMEDSLF | 1 | 0 | 0 | 1429591 | ARB07436.1 | A0A140H1G9 |
| ERDISNVLYNSA | 1 | 0 | 0 | 1429593 | APO40579.1 | P59594 |
| ERDISNVPFSPN | 1 | 0 | 0 | 1429595 | ABD72972.1 | P59594 |
| ERDLSSDENGVR | 1 | 0 | 0 | 1429598 | AVP78031.1 | Q3LZX1 |
| ERDLTSDENGVR | 1 | 0 | 0 | 1429600 | ATO98120.1 | Q3LZX1 |
| ERFMNTPNAILI | 1 | 0 | 0 | 1429607 | AUM60013.1 | K9N7C7 |
| ERGLLHSSQLPL | 1 | 0 | 0 | 1429616 | AEN19363.1 | P0C6X6 |
| ERGLLHSSRLPL | 1 | 0 | 0 | 1429617 | AGT51419.1 | P0C6X6 |
| ERHPLSYFVNLD | 1 | 0 | 0 | 1429625 | APO40578.1 | P0C6X7 |
| ERHRQLIEHSTP | 1 | 0 | 0 | 1429627 | AUM60013.1 | K9N7C7 |
| ERILYTFYGKNA | 1 | 0 | 0 | 1429635 | AVV62535.1 | K9N7C7 |
| ERINVVPGVSTN | 1 | 0 | 0 | 1429636 | AKT07951.1 | H9EJ42 |
| ERKDVPYSKQSI | 1 | 0 | 0 | 1429639 | ATQ39389.1 | K9N7C7 |
| ERKDVPYTKQTI | 1 | 0 | 0 | 1429640 | AUM60013.1 | K9N7C7 |
| ERKSMYDGDRFV | 1 | 0 | 0 | 1429645 | AMM42508.1 | A0A0P0ILY9 |
| ERKTFSNCNFNM | 1 | 0 | 0 | 1429646 | CAA79896.1 | P36334 |
| ERLINTISPEKL | 1 | 0 | 0 | 1429653 | AWK59942.1 | H9EJ42 |
| ERLKLFAAETQK | 1 | 0 | 0 | 1429656 | AEN19363.1 | P0C6X6 |
| ERLRDYLAAYDI | 1 | 0 | 0 | 1429658 | AUM60013.1 | K9N7C7 |
| ERLRDYLAVYDI | 1 | 0 | 0 | 1429659 | AUM60023.1 | K9N7C7 |
| ERLRISDCVADY | 1 | 0 | 0 | 1429660 | APO40579.1 | P59594 |
| ERLTGGNYSDPF | 1 | 0 | 0 | 1429663 | AUM60013.1 | K9N7C7 |
| ERNGARPKQRRP | 1 | 0 | 0 | 1429672 | AVP78038.1 | Q3LZX4 |
| ERNIVRAALLAG | 1 | 0 | 0 | 1429673 | AVV62524.1 | K9N7C7 |
| ERNVVRASLLAG | 1 | 0 | 0 | 1429680 | QBF80596.1 | K9N7C7 |
| ERQSPFEIKSAK | 1 | 0 | 0 | 1429687 | ARO76381.1 | P0C6U8 |
| ERSEKSYERQSP | 1 | 0 | 0 | 1429697 | ARO76381.1 | P0C6U8 |
| ERSEKSYESQTP | 1 | 0 | 0 | 1429699 | AVP78041.1 | P0C6W2 |
| ERSFSLQKDKVC | 1 | 0 | 0 | 1429701 | QBP84718.1 | Q4VID0 |
| ERSNIYGTATIG | 1 | 0 | 0 | 1429705 | AXN92247.1 | K9N7C7 |
| ERSVSLAIDAYP | 1 | 0 | 0 | 1429708 | ASU90624.1 | K9N7C7 |
| ERVDVYDNFPQG | 1 | 0 | 0 | 1429715 | APO40579.1 | P59594 |
| ERVEWDIEYPYI | 1 | 0 | 0 | 1429717 | AUM60013.1 | K9N7C7 |
| ERVKISNCVADY | 1 | 0 | 0 | 1429721 | ATO98145.1 | Q3LZX1 |
| ERVNVVPGVPTN | 1 | 0 | 0 | 1429725 | AFD98763.1 | H9EJ42 |
| ERVRKAILNTVK | 1 | 0 | 0 | 1429727 | ANI69887.1 | K9N7C7 |
| ERVRQAILNTVT | 1 | 0 | 0 | 1429728 | AGV08514.1 | K9N7C7 |
| ERVSLWNYGKAI | 1 | 0 | 0 | 1429732 | ARB07436.1 | A0A140H1G9 |
| ERVYKHFNNYDC | 1 | 0 | 0 | 1429738 | AEN19363.1 | P0C6X6 |
| ERVYNHFNNYDC | 1 | 0 | 0 | 1429739 | AIV41936.1 | P0C6X6 |
| ERYVKMFWVVPT | 1 | 0 | 0 | 1429750 | AXG21663.1 | K9N7C7 |
| ESAAQADLQHED | 1 | 0 | 0 | 1429754 | ATQ39389.1 | K9N7C7 |
| ESASAVNMQQIY | 1 | 0 | 0 | 1429765 | AIV41846.1 | P0C6X6 |
| ESATVRNYFITE | 1 | 0 | 0 | 1429768 | AUM60013.1 | K9N7C7 |
| ESCLQGDRCNIF | 1 | 0 | 0 | 1429777 | AMQ23491.1 | P36334 |
| ESCVRVAKNKVY | 1 | 0 | 0 | 1429780 | APO40578.1 | P0C6X7 |
| ESDDYIKQNGPL | 1 | 0 | 0 | 1429785 | ARO76381.1 | P0C6U8 |
| ESDKEEDLQQAQ | 1 | 0 | 0 | 1429792 | APT69868.1 | A0A0P0ILY9 |
| ESDKEEDVQQAQ | 1 | 0 | 0 | 1429793 | AST12962.1 | A0A0P0LKV0 |
| ESDQWFKNQTSV | 1 | 0 | 0 | 1429797 | BBA20973.1 | P36334 |
| ESDSQDGQETAV | 1 | 0 | 0 | 1429798 | AVP78041.1 | P0C6W2 |
| ESDTYITKNGPL | 1 | 0 | 0 | 1429799 | APO40578.1 | P0C6X7 |
| ESEPLCDQFIDY | 1 | 0 | 0 | 1429808 | APO40578.1 | P0C6X7 |
| ESEPQRGRQGRI | 1 | 0 | 0 | 1429809 | ABB90491.1 | A0A0N7J131 |
| ESFDVDTGVYSV | 1 | 0 | 0 | 1429814 | AVV62526.1 | K9N5Q8 |
| ESFDVESGVYSV | 1 | 0 | 0 | 1429815 | AXN92228.1 | K9N5Q8 |
| ESFENPTYIYHS | 1 | 0 | 0 | 1429817 | AVN89322.1 | K9N7C7 |
| ESFIDDYYGYLM | 1 | 0 | 0 | 1429821 | ADO24650.1 | H9EJ42 |
| ESFKPRSQMEVD | 1 | 0 | 0 | 1429827 | APO40578.1 | P0C6X7 |
| ESFKQELDKYFK | 1 | 0 | 0 | 1429828 | APO40579.1 | P59594 |
| ESFLGRYMSAFN | 1 | 0 | 0 | 1429829 | BAE93400.1 | P0C6U8 |
| ESFVDDFQGYLQ | 1 | 0 | 0 | 1429832 | AIG96641.1 | A0A0P0LKV0 |
| ESFYKNMYLSSA | 1 | 0 | 0 | 1429835 | ANZ78864.1 | P0C6X6 |
| ESFYRELYDAPT | 1 | 0 | 0 | 1429837 | AVV62535.1 | K9N7C7 |
| ESFYRELYEAPT | 1 | 0 | 0 | 1429838 | AVV62524.1 | K9N7C7 |
| ESGFVVRIGAAA | 1 | 0 | 0 | 1429849 | AVV62526.1 | K9N5Q8 |
| ESGGYIPSNFAF | 1 | 0 | 0 | 1429852 | AIW52757.1 | A0A0P0K6L9 |
| ESGGYIPSNFSF | 1 | 0 | 0 | 1429854 | ARU07601.1 | A0A0P0K6L9 |
| ESGKLKQSQPKV | 1 | 0 | 0 | 1429855 | ATG84807.1 | K9N4V7 |
| ESGKQKTAAPKV | 1 | 0 | 0 | 1429856 | AUM60021.1 | K9N4V7 |
| ESGKTKAAPKVI | 1 | 0 | 0 | 1429857 | AVV62544.1 | K9N4V7 |
| ESGLKTVLRKGG | 1 | 0 | 0 | 1429861 | AVP78041.1 | P0C6W2 |
| ESIPIVWGAKDF | 1 | 0 | 0 | 1429883 | AFD98777.1 | H9EJ42 |
| ESIPLADDDSAI | 1 | 0 | 0 | 1429885 | AMM42614.1 | H9EJ69 |
| ESIVAYTMSIGE | 1 | 0 | 0 | 1429888 | APO40579.1 | P59594 |
| ESIVSLPVYHKH | 1 | 0 | 0 | 1429890 | ARA15429.1 | A0A0P0K6L9 |
| ESIYEFDTYKAL | 1 | 0 | 0 | 1429892 | ARU07575.1 | A0A140H1G9 |
| ESKPQQVTKQNA | 1 | 0 | 0 | 1429907 | ARU07581.1 | Q19U29 |
| ESKQKKIENEIA | 1 | 0 | 0 | 1429913 | 5X58_A | P59594 |
| ESKVFLERFLPK | 1 | 0 | 0 | 1429921 | ARU07593.1 | H9EJ42 |
| ESKYVKSLSIVD | 1 | 0 | 0 | 1429927 | AWW13571.1 | P0C6X6 |

| | | | | | | |
|---|---|---|---|---|---|---|
| ESLIFPSGTYYC | 1 | 0 | 0 | 1429939 | AXN92247.1 | K9N7C7 |
| ESLNDNLLEILQ | 1 | 0 | 0 | 1429946 | AVP78041.1 | P0C6W2 |
| ESLRPDIRYVLM | 1 | 0 | 0 | 1429948 | AVP78041.1 | P0C6W2 |
| ESNMFIKFPEVV | 1 | 0 | 0 | 1429971 | AVV62524.1 | K9N7C7 |
| ESNVVDVNDCFK | 1 | 0 | 0 | 1429974 | AGT51379.1 | H9EJ42 |
| ESPLLKQKDIGI | 1 | 0 | 0 | 1429981 | AGW27859.1 | Q0ZJ80 |
| ESQGNVVTPVME | 1 | 0 | 0 | 1429986 | ARB07605.1 | A0A140H1G9 |
| ESQLQKGVSFLK | 1 | 0 | 0 | 1429990 | APO40578.1 | P0C6X7 |
| ESRLSDGVEFLK | 1 | 0 | 0 | 1429999 | ARO76381.1 | P0C6U8 |
| ESSDDVDPFDAA | 1 | 0 | 0 | 1430006 | AVV62544.1 | K9N4V7 |
| ESSEVLEDTLDA | 1 | 0 | 0 | 1430007 | AGT51617.1 | P0C6X6 |
| ESSGSLATRLMS | 1 | 0 | 0 | 1430012 | ATP16762.1 | Q4VID0 |
| ESTDQMFEPPKE | 1 | 0 | 0 | 1430021 | QCZ25056.1 | K9N4V7 |
| ESTDQMSELPKE | 1 | 0 | 0 | 1430022 | ASJ26617.1 | K9N4V7 |
| ESTDSTVLTYIP | 1 | 0 | 0 | 1430029 | AVN89378.1 | K9N7D2 |
| ESTFVMMSAPPS | 1 | 0 | 0 | 1430032 | APO40578.1 | P0C6X7 |
| ESTGSLAARLMS | 1 | 0 | 0 | 1430033 | QRK03827.1 | Q4VID0 |
| ESTGTQSVSVDR | 1 | 0 | 0 | 1430037 | AXN92229.1 | K9N796 |
| ESTLRITRTLST | 1 | 0 | 0 | 1430041 | AXN92247.1 | K9N7C7 |
| ESTPVELPTVDV | 1 | 0 | 0 | 1430045 | AVN89312.1 | K9N638 |
| ESTPVEPSTVDV | 1 | 0 | 0 | 1430047 | AIY60577.1 | K9N638 |
| ESTRSYLGSSAF | 1 | 0 | 0 | 1430053 | AVV62529.1 | K9N643 |
| ESTSTHDGFNEH | 1 | 0 | 0 | 1430057 | AHI48551.1 | K9N796 |
| ESTSTHDGFTEH | 1 | 0 | 0 | 1430058 | ASU89978.1 | K9N796 |
| ESTSTHDGPNEH | 1 | 0 | 0 | 1430063 | AHE78098.1 | K9N796 |
| ESTSTHDTPTEH | 1 | 0 | 0 | 1430066 | AJG44059.1 | K9N796 |
| ESTSVEPPTVDV | 1 | 0 | 0 | 1430068 | AJG44112.1 | K9N638 |
| ESTTDPLLSFKH | 1 | 0 | 0 | 1430069 | AVV62524.1 | K9N7C7 |
| ESVANLFSDIKT | 1 | 0 | 0 | 1430071 | AVV62537.1 | K9N5Q8 |
| ESVKSSQSSPII | 1 | 0 | 0 | 1430080 | QBM11737.1 | K9N5Q8 |
| ESVLVNAVNTHL | 1 | 0 | 0 | 1430083 | AHI48549.1 | K9N7C7 |
| ESVSNLFADIKT | 1 | 0 | 0 | 1430089 | AVV62526.1 | K9N5Q8 |
| ESVTSVNLFDVG | 1 | 0 | 0 | 1430090 | ATQ39394.1 | K9N796 |
| ESVVDLQQAVVN | 1 | 0 | 0 | 1430092 | AVV62524.1 | K9N7C7 |
| ESVYDFDTYKAL | 1 | 0 | 0 | 1430094 | ARB07436.1 | A0A140H1G9 |
| ESWLNDKSVPSP | 1 | 0 | 0 | 1430100 | AIX10749.1 | P36334 |
| ESYIDLKKLGNY | 1 | 0 | 0 | 1430105 | AVV62526.1 | K9N5Q8 |
| ESYVDVGLDSVK | 1 | 0 | 0 | 1430109 | AXN73437.1 | K9N5Q8 |
| ESYVDVGPDSAK | 1 | 0 | 0 | 1430110 | AVN89387.1 | K9N5Q8 |
| ESYVDVGPDSLK | 1 | 0 | 0 | 1430113 | QBF80608.1 | K9N5Q8 |
| ETAAAKFERQHM | 1 | 0 | 0 | 1430117 | 3R24_A | P0C6X7 |
| ETAHHSVHVLQK | 1 | 0 | 0 | 1430121 | AVP78030.1 | P0C6W2 |
| ETALDLSKELKR | 1 | 0 | 0 | 1430125 | AXX83306.1 | P0C6X6 |
| ETAVTKTSEQNN | 1 | 0 | 0 | 1430135 | AVP78041.1 | P0C6W2 |
| ETDGICLENHVM | 1 | 0 | 0 | 1430145 | ARK08667.1 | P0C6X6 |
| ETFSACRAYLDS | 1 | 0 | 0 | 1430162 | ATG84733.1 | K9N638 |
| ETFTNNNGSDVL | 1 | 0 | 0 | 1430163 | AQN78669.1 | P0C6X6 |
| ETFTVCADGFMP | 1 | 0 | 0 | 1430165 | AEN19363.1 | P0C6X6 |
| ETGDIVDAVQYA | 1 | 0 | 0 | 1430174 | AVV62524.1 | K9N7C7 |
| ETGLLLPLKAPK | 1 | 0 | 0 | 1430177 | AVP78041.1 | P0C6W2 |
| ETGTFTFKIRKN | 1 | 0 | 0 | 1430182 | AUM60013.1 | K9N7C7 |
| ETGTSDLDRCTT | 1 | 0 | 0 | 1430189 | 6NB6_A | P59594 |
| ETIMKVLTENLN | 1 | 0 | 0 | 1430199 | AIL49396.1 | P33469 |
| ETIPTQLTPLMF | 1 | 0 | 0 | 1430202 | AUM60013.1 | K9N7C7 |
| ETITFDNLKTLL | 1 | 0 | 0 | 1430207 | AVP78030.1 | P0C6W2 |
| ETIVLTAGDGVT | 1 | 0 | 0 | 1430213 | APO40580.1 | P59632 |
| ETIYNLLKDCPA | 1 | 0 | 0 | 1430220 | QYC64967.1 | P0DTD1 |
| ETKCAPELTKTE | 1 | 0 | 0 | 1430231 | AVV62524.1 | K9N7C7 |
| ETKCTLKSLTVQ | 1 | 0 | 0 | 1430234 | AVP78031.1 | Q3LZX1 |
| ETLACESQTLVS | 1 | 0 | 0 | 1430245 | APO40578.1 | P0C6X7 |
| ETLKANEETFKL | 1 | 0 | 0 | 1430250 | APO40578.1 | P0C6X7 |
| ETLNEDLLEILH | 1 | 0 | 0 | 1430258 | ARO76381.1 | P0C6U8 |
| ETLSACRAYLNS | 1 | 0 | 0 | 1430272 | ALW82708.1 | K9N638 |
| ETLTDMEKDFLS | 1 | 0 | 0 | 1430275 | AUM60013.1 | K9N7C7 |
| ETMAHLLQHANL | 1 | 0 | 0 | 1430285 | ARO76381.1 | P0C6U8 |
| ETMFDVYVDTFL | 1 | 0 | 0 | 1430286 | AEN19363.1 | P0C6X6 |
| ETMNDDLLQILN | 1 | 0 | 0 | 1430289 | APO40578.1 | P0C6X7 |
| ETNCLLNVPIGG | 1 | 0 | 0 | 1430297 | ASY99815.1 | K9N7A1 |
| ETNCLLNVPVGG | 1 | 0 | 0 | 1430299 | AVV62543.1 | K9N7A1 |
| ETNTAAFNAVCS | 1 | 0 | 0 | 1430318 | ANI69909.1 | K9N7C7 |
| ETPNAKEEILGT | 1 | 0 | 0 | 1430331 | P0C6W6.1 | P0C6W2 |
| ETPNLMGWDYPK | 1 | 0 | 0 | 1430332 | ASB51350.1 | P0C6X7 |
| ETPTGYRKVLVR | 1 | 0 | 0 | 1430333 | APO40578.1 | P0C6X7 |
| ETPTIQKDVIEC | 1 | 0 | 0 | 1430334 | APO40578.1 | P0C6X7 |
| ETPVDSFTNGAV | 1 | 0 | 0 | 1430339 | P0C6W6.1 | P0C6W2 |
| ETPVVSDTVEVP | 1 | 0 | 0 | 1430347 | AXN92247.1 | K9N7C7 |
| ETRKLMPVCMDV | 1 | 0 | 0 | 1430360 | P0C6W6.1 | P0C6W2 |
| ETSNSFDALKLG | 1 | 0 | 0 | 1430381 | AVP78041.1 | P0C6W2 |
| ETSNSFEVLAVE | 1 | 0 | 0 | 1430383 | ACV88184.1 | P0C6X7 |
| ETSTEQVDVSHE | 1 | 0 | 0 | 1430391 | AXN92247.1 | K9N7C7 |
| ETSWQTGDFVRA | 1 | 0 | 0 | 1430396 | AVP78041.1 | P0C6W2 |
| ETSYTSDIKPVT | 1 | 0 | 0 | 1430397 | APO40578.1 | P0C6X7 |
| ETTDNNCWVNAT | 1 | 0 | 0 | 1430405 | AGT51365.1 | H9EJ42 |
| ETVFVEPKDGGH | 1 | 0 | 0 | 1430424 | ARE29978.1 | H9EJ42 |
| ETVFVEPKNGGQ | 1 | 0 | 0 | 1430426 | ARU07593.1 | H9EJ42 |
| ETVHTNVYLVVF | 1 | 0 | 0 | 1430427 | AVP78041.1 | P0C6W2 |
| ETVRLHLYNATR | 1 | 0 | 0 | 1430436 | AWO68966.1 | A0A0P0G321 |
| ETVSLAGSYKDW | 1 | 0 | 0 | 1430443 | AVP78041.1 | P0C6W2 |
| ETYIKSGGYIPE | 1 | 0 | 0 | 1430457 | 6U7H_A | A0A0P0K6L9 |
| EVAAVRSVTVEY | 1 | 0 | 0 | 1430463 | AXN92247.1 | K9N7C7 |
| EVAKKQIGLLEP | 1 | 0 | 0 | 1430469 | AVV62524.1 | K9N7C7 |

| | | | | | | |
|---|---|---|---|---|---|---|
| EVALPQSKDSNF | 1 | 0 | 0 | 1430472 | AVV62524.1 | K9N7C7 |
| EVARDLSVQFKR | 1 | 0 | 0 | 1430480 | APO40578.1 | P0C6X7 |
| EVARNLNESLID | 1 | 0 | 0 | 1430481 | AVP78031.1 | Q3LZX1 |
| EVASYSDQQPSV | 1 | 0 | 0 | 1430486 | AVV62530.1 | K9N7D2 |
| EVDFLELAMDEF | 1 | 0 | 0 | 1430500 | APO40578.1 | P0C6X7 |
| EVDHSGPTHYLR | 1 | 0 | 0 | 1430502 | AVV62529.1 | K9N643 |
| EVDWKFYDAQPC | 1 | 0 | 0 | 1430517 | AVP78041.1 | P0C6W2 |
| EVDWVAFYGFEK | 1 | 0 | 0 | 1430518 | AWK59942.1 | H9EJ42 |
| EVEAGEVKPFAV | 1 | 0 | 0 | 1430519 | ARU07593.1 | H9EJ42 |
| EVEELNPSKLLL | 1 | 0 | 0 | 1430527 | AGT51436.1 | Q4VID0 |
| EVEEQAEVQQSA | 1 | 0 | 0 | 1430528 | AVV62524.1 | K9N7C7 |
| EVEPEPEPTLEE | 1 | 0 | 0 | 1430538 | ARO76381.1 | P0C6U8 |
| EVFDALQDDPKG | 1 | 0 | 0 | 1430548 | AVV62524.1 | K9N7C7 |
| EVFDDEWAAAVD | 1 | 0 | 0 | 1430549 | AQZ41284.1 | K9N7C7 |
| EVFEAQAFETPA | 1 | 0 | 0 | 1430550 | AVV62535.1 | K9N7C7 |
| EVFFNNTVYQKL | 1 | 0 | 0 | 1430552 | AVV62524.1 | K9N7C7 |
| EVFVAFVYYVAR | 1 | 0 | 0 | 1430560 | ARK08640.1 | A0A0P0LKV0 |
| EVGDNVQITHTY | 1 | 0 | 0 | 1430566 | AMM42628.1 | H9EJ69 |
| EVGEVKPFAVYK | 1 | 0 | 0 | 1430568 | AKT07951.1 | H9EJ42 |
| EVHGNGSMIEDM | 1 | 0 | 0 | 1430583 | AXG21663.1 | K9N7C7 |
| EVHGNGSMIEDV | 1 | 0 | 0 | 1430584 | AMQ49035.1 | K9N7C7 |
| EVHIQQTFFDKT | 1 | 0 | 0 | 1430587 | ANI69922.1 | K9N5Q8 |
| EVHKAVVVTKDI | 1 | 0 | 0 | 1430588 | AVV62535.1 | K9N7C7 |
| EVHQELYSPLFL | 1 | 0 | 0 | 1430589 | ATO98237.1 | Q3LZX7 |
| EVKFSVIELATG | 1 | 0 | 0 | 1430620 | ABQ16581.1 | H9EJ42 |
| EVKPFFTENKPS | 1 | 0 | 0 | 1430627 | AVP78041.1 | P0C6W2 |
| EVKTASLFQTTV | 1 | 0 | 0 | 1430633 | AGT51366.1 | A0A0P0G321 |
| EVKTFETPAGQF | 1 | 0 | 0 | 1430634 | AVV62524.1 | K9N7C7 |
| EVKVSALKTAGV | 1 | 0 | 0 | 1430638 | AXN92247.1 | K9N7C7 |
| EVLEDTLDAGPS | 1 | 0 | 0 | 1430643 | AGT51617.1 | P0C6X6 |
| EVLEDTLNDGPS | 1 | 0 | 0 | 1430645 | AGT51737.1 | P0C6X6 |
| EVLLDVCNTKEF | 1 | 0 | 0 | 1430656 | ARX11394.1 | A0A0P0LKV0 |
| EVLLKDALQSRE | 1 | 0 | 0 | 1430658 | AXX83318.1 | P0C6X6 |
| EVLRELYGTPDS | 1 | 0 | 0 | 1430667 | AVV62524.1 | K9N7C7 |
| EVLSELYGTPDP | 1 | 0 | 0 | 1430671 | AVV62535.1 | K9N7C7 |
| EVLVTVDGINFR | 1 | 0 | 0 | 1430680 | AVV62535.1 | K9N7C7 |
| EVLVTVDGVNFR | 1 | 0 | 0 | 1430681 | AXN92247.1 | K9N7C7 |
| EVNAPYYNSWQN | 1 | 0 | 0 | 1430685 | CAA83660.1 | P36334 |
| EVNPDLSAYYVK | 1 | 0 | 0 | 1430692 | AVV62535.1 | K9N7C7 |
| EVPGSCNFPLEA | 1 | 0 | 0 | 1430700 | AWO68966.1 | A0A0P0G321 |
| EVPPQLVELPSE | 1 | 0 | 0 | 1430704 | AVN89312.1 | K9N638 |
| EVPPQVVELPYE | 1 | 0 | 0 | 1430708 | ANI69888.1 | K9N638 |
| EVPPRCIAPVDG | 1 | 0 | 0 | 1430713 | AUM60014.1 | K9N5Q8 |
| EVPQLVNSNQYS | 1 | 0 | 0 | 1430717 | AHY22555.1 | K9N5Q8 |
| EVQPVEVPVEDI | 1 | 0 | 0 | 1430742 | AXN92247.1 | K9N7C7 |
| EVQVDPNAFIYE | 1 | 0 | 0 | 1430749 | AUM60014.1 | K9N5Q8 |
| EVRQVAPGETGV | 1 | 0 | 0 | 1430758 | ATO98181.1 | Q3LZX1 |
| EVRSKGEFIEQP | 1 | 0 | 0 | 1430761 | AVV62526.1 | K9N5Q8 |
| EVSALRTAGVVT | 1 | 0 | 0 | 1430764 | AUM60013.1 | K9N7C7 |
| EVSDDEWVAAVD | 1 | 0 | 0 | 1430768 | ASU45728.1 | K9N7C7 |
| EVSDLIAQSSGV | 1 | 0 | 0 | 1430769 | AIV41990.1 | P0C6X6 |
| EVSMATNYDLSV | 1 | 0 | 0 | 1430778 | APO40578.1 | P0C6X7 |
| EVSRQVIQEVKD | 1 | 0 | 0 | 1430780 | ABM64804.1 | A0A0P0KWE8 |
| EVSVKMVDAYVD | 1 | 0 | 0 | 1430791 | BAE93400.1 | P0C6U8 |
| EVSVQKKINRMA | 1 | 0 | 0 | 1430792 | ARE29978.1 | H9EJ42 |
| EVTKGPFVEGAI | 1 | 0 | 0 | 1430803 | AUM60023.1 | K9N7C7 |
| EVTQKILNNPRR | 1 | 0 | 0 | 1430812 | AJQ24083.1 | P33469 |
| EVTVDTTVESAA | 1 | 0 | 0 | 1430821 | ATQ39389.1 | K9N7C7 |
| EVVAVRSVTVEY | 1 | 0 | 0 | 1430823 | ALA49384.1 | K9N7C7 |
| EVVENPTVQKEV | 1 | 0 | 0 | 1430829 | ARO76381.1 | P0C6U8 |
| EVVETPTIQKDV | 1 | 0 | 0 | 1430830 | APO40578.1 | P0C6X7 |
| EVVGDIILKPAI | 1 | 0 | 0 | 1430833 | AVP78041.1 | P0C6W2 |
| EVVGDIILKPAS | 1 | 0 | 0 | 1430835 | AVP78030.1 | P0C6W2 |
| EVVKAMLFKKEP | 1 | 0 | 0 | 1430844 | ALA49373.1 | K9N7C7 |
| EVVLKKLKKSLNVAK | 1 | 0 | 0 | 1430849 | P0DTC1.1 | P0DTD1 |
| EVVRFPNITNLC | 1 | 0 | 0 | 1430858 | ATO98231.1 | Q3LZX1 |
| EVVSAYGLCDSA | 1 | 0 | 0 | 1430864 | AVN89334.1 | K9N5Q8 |
| EVVSAYGLCDVA | 1 | 0 | 0 | 1430865 | AKM76239.1 | K9N5Q8 |
| EVWQKYTNSAST | 1 | 0 | 0 | 1430873 | AUM60016.1 | K9N4V0 |
| EVYLPYPNPSRI | 1 | 0 | 0 | 1430876 | AIL49620.1 | P0C6X6 |
| EWAMANQFTEFV | 1 | 0 | 0 | 1430889 | AUM60013.1 | K9N7C7 |
| EWDIEYPYISHE | 1 | 0 | 0 | 1430898 | AUM60013.1 | K9N7C7 |
| EWFGITQDTQGV | 1 | 0 | 0 | 1430900 | AVV62526.1 | K9N5Q8 |
| EWFGITQNAQGV | 1 | 0 | 0 | 1430901 | AUM60014.1 | K9N5Q8 |
| EWFGITQNTQGV | 1 | 0 | 0 | 1430902 | AVV62537.1 | K9N5Q8 |
| EWFGITQTAQGA | 1 | 0 | 0 | 1430903 | ALJ54512.1 | K9N5Q8 |
| EWFGITQTAQGV | 1 | 0 | 0 | 1430904 | AXN92228.1 | K9N5Q8 |
| EWKCKVTDILYP | 1 | 0 | 0 | 1430922 | AVV62524.1 | K9N7C7 |
| EWKMVVCDKVRS | 1 | 0 | 0 | 1430925 | AXN92258.1 | K9N7C7 |
| EWPLFDQQAQQE | 1 | 0 | 0 | 1430945 | APT69896.1 | A0A0P0LKV0 |
| EWPLMEYDFTDY | 1 | 0 | 0 | 1430947 | AUM60013.1 | K9N7C7 |
| EWVAAVDEAFPL | 1 | 0 | 0 | 1430958 | ASU45728.1 | K9N7C7 |
| EYAYDQVHDEPK | 1 | 0 | 0 | 1430972 | APU51933.1 | P0C6X6 |
| EYDLEPHKVHVH | 1 | 0 | 0 | 1430981 | AXN92228.1 | K9N5Q8 |
| EYDLQPHKVHVH | 1 | 0 | 0 | 1430982 | AHI48550.1 | K9N5Q8 |
| EYDNISVAVPKS | 1 | 0 | 0 | 1430983 | AXX83299.1 | P33469 |
| EYGGHNDFVIKA | 1 | 0 | 0 | 1431001 | BBA20986.1 | A0A140H1H1 |
| EYGQFCSKINQA | 1 | 0 | 0 | 1431004 | AXN92228.1 | K9N5Q8 |
| EYGTEDDYNGTP | 1 | 0 | 0 | 1431010 | APO40578.1 | P0C6X7 |
| EYHITIALLICK | 1 | 0 | 0 | 1431017 | AVV62540.1 | K9N643 |
| EYIRYGIKPITI | 1 | 0 | 0 | 1431029 | ARU07575.1 | A0A140H1G9 |

| | | | | | | |
|---|---|---|---|---|---|---|
| EYKGSITDVFYK | 1 | 0 | 0 | 1431038 | AVP78041.1 | P0C6W2 |
| EYLLREVKTRAL | 1 | 0 | 0 | 1431050 | AUM60013.1 | K9N7C7 |
| EYLQITSIPIVV | 1 | 0 | 0 | 1431051 | APF29071.1 | A0A0P0G321 |
| EYLTKDDLHQLY | 1 | 0 | 0 | 1431055 | AUM60013.1 | K9N7C7 |
| EYLTQRGPLQVG | 1 | 0 | 0 | 1431056 | AUM60013.1 | K9N7C7 |
| EYNPPTDKLAMI | 1 | 0 | 0 | 1431074 | NP_150075.1 | Q80872 |
| EYPFLKQKDIGI | 1 | 0 | 0 | 1431076 | ABD75542.1 | Q0ZJ80 |
| EYPMDVHSTYGT | 1 | 0 | 0 | 1431088 | ATQ39396.1 | K9N643 |
| EYPNTICHPNLG | 1 | 0 | 0 | 1431089 | CAA83661.1 | P36334 |
| EYPYISHENKLN | 1 | 0 | 0 | 1431093 | ALA49351.1 | K9N7C7 |
| EYPYISHETKLN | 1 | 0 | 0 | 1431094 | ASU45783.1 | K9N7C7 |
| EYREYMEAYNFV | 1 | 0 | 0 | 1431101 | AHY22563.1 | K9N7C7 |
| EYREYVEAYNLV | 1 | 0 | 0 | 1431102 | AVV62524.1 | K9N7C7 |
| EYSKRSAIEDLL | 1 | 0 | 0 | 1431113 | AVV62526.1 | K9N5Q8 |
| EYTYTDEVIYGM | 1 | 0 | 0 | 1431134 | AEN19363.1 | P0C6X6 |
| EYTYTDEVIYSM | 1 | 0 | 0 | 1431135 | AIV41834.1 | P0C6X6 |
| EYVFKNKDGFLS | 1 | 0 | 0 | 1431139 | ATO98193.1 | Q3LZX1 |
| EYVSYFDGVKTV | 1 | 0 | 0 | 1431144 | NP_073549.1 | A0A0P0LKV0 |
| EYVTEFVQEEAQ | 1 | 0 | 0 | 1431145 | AVN89322.1 | K9N7C7 |
| EYYEFLNKHFNM | 1 | 0 | 0 | 1431161 | AIL49554.1 | P0C6X6 |
| FAAIGDLKCPLD | 1 | 0 | 0 | 1431176 | AIL49486.1 | P36334 |
| FAAIGDLNCTLD | 1 | 0 | 0 | 1431177 | BBO58773.1 | P36334 |
| FACADGTRHTYQ | 1 | 0 | 0 | 1431192 | ATO98237.1 | Q3LZX7 |
| FACDCGACFSAN | 1 | 0 | 0 | 1431193 | AVV62524.1 | K9N7C7 |
| FACDCGAVFSGN | 1 | 0 | 0 | 1431194 | AVV62535.1 | K9N7C7 |
| FACGCGTSYTAN | 1 | 0 | 0 | 1431198 | QBM11736.1 | K9N638 |
| FACKFAVCGDGF | 1 | 0 | 0 | 1431199 | ARU07575.1 | A0A140H1G9 |
| FACTTCGHVYEV | 1 | 0 | 0 | 1431204 | AEN19363.1 | P0C6X6 |
| FADCTSCSNGRW | 1 | 0 | 0 | 1431210 | AUM60013.1 | K9N7C7 |
| FADDLNQLSGYK | 1 | 0 | 0 | 1431211 | AVP78041.1 | P0C6W2 |
| FADGINQMTGFK | 1 | 0 | 0 | 1431217 | APO40578.1 | P0C6X7 |
| FADIKTQATQPL | 1 | 0 | 0 | 1431220 | AVV62526.1 | K9N5Q8 |
| FADINGKLYQDS | 1 | 0 | 0 | 1431222 | ARO76381.1 | P0C6U8 |
| FADNNDTTNTNL | 1 | 0 | 0 | 1431226 | ALW82676.1 | K9N4V7 |
| FADVVKGQVSDL | 1 | 0 | 0 | 1431232 | ALA49659.1 | K9N7C7 |
| FAECTGCGNGRW | 1 | 0 | 0 | 1431235 | AVV62535.1 | K9N7C7 |
| FAELEAARKAYQ | 1 | 0 | 0 | 1431242 | AMQ49036.1 | K9N638 |
| FAELIPNQAALF | 1 | 0 | 0 | 1431244 | AGT51335.1 | H9EJ69 |
| FAFACADGTKHT | 1 | 0 | 0 | 1431254 | ARO76387.1 | P59635 |
| FAFACPDGVRHT | 1 | 0 | 0 | 1431257 | AVP78047.1 | Q3LZX7 |
| FAFNSSVLSLVN | 1 | 0 | 0 | 1431264 | QAT98908.1 | K9N7C7 |
| FAGFVGQAVNSS | 1 | 0 | 0 | 1431274 | AUM60016.1 | K9N4V0 |
| FAGYTESAVNFT | 1 | 0 | 0 | 1431281 | AVN89336.1 | K9N4V0 |
| FAINDLNTTIPR | 1 | 0 | 0 | 1431287 | ACV87140.1 | A0A140H1H1 |
| FAIPNGRKVDLH | 1 | 0 | 0 | 1431290 | AIL49485.1 | P36334 |
| FAIPNRRKFDLQ | 1 | 0 | 0 | 1431293 | BBO58726.1 | P36334 |
| FAIPNRRKVDLQ | 1 | 0 | 0 | 1431294 | AEN19366.1 | P36334 |
| FAIPNRRRDDLQ | 1 | 0 | 0 | 1431295 | BBA20986.1 | A0A140H1H1 |
| FAIPNRRRVDLQ | 1 | 0 | 0 | 1431296 | ANZ78849.1 | P36334 |
| FAKVKNTKVFGK | 1 | 0 | 0 | 1431316 | AWW13566.1 | P36334 |
| FAKVKNTKVIKD | 1 | 0 | 0 | 1431319 | CAA79896.1 | P36334 |
| FAKVKNTKVVGK | 1 | 0 | 0 | 1431322 | AWW13511.1 | P36334 |
| FALTCFSTQFAF | 1 | 0 | 0 | 1431345 | AVP78047.1 | Q3LZX7 |
| FAMMFVKHKHAF | 1 | 0 | 0 | 1431359 | AVP78041.1 | P0C6W2 |
| FANFILHDVNNG | 1 | 0 | 0 | 1431371 | AEN19366.1 | P36334 |
| FANGFVVRIGAV | 1 | 0 | 0 | 1431375 | AGV08520.1 | K9N5Q8 |
| FANHVANFQKSG | 1 | 0 | 0 | 1431376 | AKN24802.1 | K9N7C7 |
| FANPAALHSKCL | 1 | 0 | 0 | 1431388 | ATP16764.1 | P0C6X6 |
| FANPAAVDSKCL | 1 | 0 | 0 | 1431389 | AEN19363.1 | P0C6X6 |
| FANSNSVCYTPT | 1 | 0 | 0 | 1431393 | AUM60013.1 | K9N7C7 |
| FANSPNVCYTPR | 1 | 0 | 0 | 1431394 | AVV62535.1 | K9N7C7 |
| FANVSYVYNGSA | 1 | 0 | 0 | 1431397 | QBP84739.1 | P30215 |
| FANWLAPTNAYM | 1 | 0 | 0 | 1431398 | QBF80585.1 | K9N7C7 |
| FAQSEKEESFSN | 1 | 0 | 0 | 1431421 | ATG84802.1 | K9N4V0 |
| FAQTKSIYKTPN | 1 | 0 | 0 | 1431428 | APO40579.1 | P59594 |
| FASDSVQCYTPH | 1 | 0 | 0 | 1431443 | ARU07575.1 | A0A140H1G9 |
| FASLEVPRKSVA | 1 | 0 | 0 | 1431453 | APO40578.1 | P0C6X7 |
| FASLPVYETVKY | 1 | 0 | 0 | 1431456 | AUM60014.1 | K9N5Q8 |
| FASMLNGTPPPV | 1 | 0 | 0 | 1431457 | AVV62526.1 | K9N5Q8 |
| FASSCKSHKPSS | 1 | 0 | 0 | 1431464 | AFR45556.1 | A0A140H1H1 |
| FASVKSYQSSPI | 1 | 0 | 0 | 1431473 | AVN89387.1 | K9N5Q8 |
| FATAPANLTISK | 1 | 0 | 0 | 1431475 | AVV62526.1 | K9N5Q8 |
| FATKVQQQQDDL | 1 | 0 | 0 | 1431485 | APO40584.1 | P59635 |
| FATLPVYDGIKY | 1 | 0 | 0 | 1431488 | AVV62537.1 | K9N5Q8 |
| FATTIPNTAFGP | 1 | 0 | 0 | 1431499 | AIW52767.1 | A0A0P0LKV0 |
| FATYHTPSTDCS | 1 | 0 | 0 | 1431506 | ALA49803.1 | K9N5Q8 |
| FAVEPAKAYKDY | 1 | 0 | 0 | 1431511 | APO40578.1 | P0C6X7 |
| FAVFDTPHTDCT | 1 | 0 | 0 | 1431513 | AVV62537.1 | K9N5Q8 |
| FAVIGDLKCPLD | 1 | 0 | 0 | 1431517 | AVQ05264.1 | P36334 |
| FAVIGDLKCPLE | 1 | 0 | 0 | 1431518 | QEG03803.1 | P36334 |
| FAVIGDLKCTTV | 1 | 0 | 0 | 1431520 | CAA83660.1 | P36334 |
| FAVIGDLNCPLD | 1 | 0 | 0 | 1431521 | AEN19366.1 | P36334 |
| FAVIGDLNCTLD | 1 | 0 | 0 | 1431522 | AWW13566.1 | P36334 |
| FAVLYQDVNCTD | 1 | 0 | 0 | 1431526 | BAE93401.1 | P59594 |
| FAVVGDLNCPLD | 1 | 0 | 0 | 1431535 | QBP84704.1 | P36334 |
| FAVYHRYRIGNY | 1 | 0 | 0 | 1431536 | APO40582.1 | P59596 |
| FAVYKNVKFYLG | 1 | 0 | 0 | 1431537 | AWK59942.1 | H9EJ42 |
| FAWWTVFCTIAN | 1 | 0 | 0 | 1431550 | AGV08523.1 | K9N7C7 |
| FAYADKPNHFIN | 1 | 0 | 0 | 1431551 | AGT51410.1 | Q80872 |
| FAYNNDITNTNL | 1 | 0 | 0 | 1431557 | AVN89381.1 | K9N4V7 |
| FAYPLSMKSYMQ | 1 | 0 | 0 | 1431558 | AVV62537.1 | K9N5Q8 |

| | | | | | | |
|---|---|---|---|---|---|---|
| FAYVKPFLGQAA | 1 | 0 | 0 | 1431570 | ATO98143.1 | P0C6W2 |
| FCAFAVEPAKAY | 1 | 0 | 0 | 1431578 | APO40578.1 | P0C6X7 |
| FCAFCTPKKVFI | 1 | 0 | 0 | 1431580 | AZS52616.1 | A0A140H1G9 |
| FCALCVISVAML | 1 | 0 | 0 | 1431585 | AGT51757.1 | P0C6X6 |
| FCATCPFAKGDS | 1 | 0 | 0 | 1431594 | AUM60013.1 | K9N7C7 |
| FCATCPFAKSDG | 1 | 0 | 0 | 1431595 | AVV62524.1 | K9N7C7 |
| FCCFLVTKFRRM | 1 | 0 | 0 | 1431599 | NP_073549.1 | A0A0P0LKV0 |
| FCCKFDEDDSEP | 1 | 0 | 0 | 1431600 | AVP78042.1 | Q3LZX1 |
| FCCRHNWNCVDC | 1 | 0 | 0 | 1431602 | AVV62535.1 | K9N7C7 |
| FCDHMVKDGLVG | 1 | 0 | 0 | 1431611 | AVN89451.1 | K9N7C7 |
| FCDYADKLCHAV | 1 | 0 | 0 | 1431619 | AEN19363.1 | P0C6X6 |
| FCDYTDKLCHAV | 1 | 0 | 0 | 1431621 | AGT51737.1 | P0C6X6 |
| FCGDQVHEVAAV | 1 | 0 | 0 | 1431630 | ALT66879.1 | K9N7C7 |
| FCGRGYHLMSFP | 1 | 0 | 0 | 1431642 | ATO98181.1 | Q3LZX1 |
| FCGVDALDLVFN | 1 | 0 | 0 | 1431645 | APO40578.1 | P0C6X7 |
| FCGVDSMNLIAN | 1 | 0 | 0 | 1431648 | ARO76381.1 | P0C6U8 |
| FCHRHNWNCVDC | 1 | 0 | 0 | 1431651 | AVN89333.1 | K9N7C7 |
| FCIDYALPSSRR | 1 | 0 | 0 | 1431654 | BBA20986.1 | A0A140H1H1 |
| FCKIGKEQKCCV | 1 | 0 | 0 | 1431660 | AHY22543.1 | K9N7C7 |
| FCKIGREQKCCM | 1 | 0 | 0 | 1431661 | ALA50033.1 | K9N7C7 |
| FCKSGSKFKNLV | 1 | 0 | 0 | 1431666 | ARU07609.1 | P30215 |
| FCLEASFNYLKS | 1 | 0 | 0 | 1431672 | QYC64967.1 | P0DTD1 |
| FCPCADRSVVNS | 1 | 0 | 0 | 1431701 | ABD75545.1 | A0A140H1H1 |
| FCPCKSNGSCPG | 1 | 0 | 0 | 1431712 | AEN19358.1 | P36334 |
| FCPCSSCFGKNN | 1 | 0 | 0 | 1431714 | QEG03814.1 | P36334 |
| FCPCSSCHGKNN | 1 | 0 | 0 | 1431715 | ARB66068.1 | P36334 |
| FCPCSSCPGKNN | 1 | 0 | 0 | 1431716 | AWW13566.1 | P36334 |
| FCPCSSCPSKNN | 1 | 0 | 0 | 1431717 | QDH43719.1 | P36334 |
| FCPCSSCSGKNN | 1 | 0 | 0 | 1431719 | ANZ78838.1 | P36334 |
| FCQDPMLAVANG | 1 | 0 | 0 | 1431723 | APO40579.1 | P59594 |
| FCRLYIEESDLI | 1 | 0 | 0 | 1431728 | ARU07575.1 | A0A140H1G9 |
| FCTKHTMLVKMD | 1 | 0 | 0 | 1431749 | AIL49625.1 | P0C6X6 |
| FCVDKALTVEEF | 1 | 0 | 0 | 1431761 | AVV62524.1 | K9N7C7 |
| FCVDKSLTVEEF | 1 | 0 | 0 | 1431762 | AVV62535.1 | K9N7C7 |
| FCVNCDSFGPGN | 1 | 0 | 0 | 1431769 | AWK59942.1 | H9EJ42 |
| FCVNSKANAIQQ | 1 | 0 | 0 | 1431770 | AUM60013.1 | K9N7C7 |
| FDAAVAAAMATP | 1 | 0 | 0 | 1431786 | AVV62544.1 | K9N4V7 |
| FDALKLGDTQGM | 1 | 0 | 0 | 1431792 | AVP78041.1 | P0C6W2 |
| FDAQPLTRKVQQ | 1 | 0 | 0 | 1431796 | AXN92247.1 | K9N7C7 |
| FDCGVNALDGDT | 1 | 0 | 0 | 1431813 | AFD98777.1 | H9EJ42 |
| FDCYNLWQTFTE | 1 | 0 | 0 | 1431824 | NP_073549.1 | A0A0P0LKV0 |
| FDDNFYAFCTPK | 1 | 0 | 0 | 1431832 | ABD75647.1 | A0A140H1G9 |
| FDDNTDSIVIDG | 1 | 0 | 0 | 1431833 | AWK59930.1 | H9EJ42 |
| FDDSTGDEAWAA | 1 | 0 | 0 | 1431838 | AVV62524.1 | K9N7C7 |
| FDEAVAAAMASQ | 1 | 0 | 0 | 1431843 | AVV62533.1 | K9N4V7 |
| FDEHGVITKAVD | 1 | 0 | 0 | 1431848 | AZS52618.1 | A0A140H1H1 |
| FDENGDKLDQFF | 1 | 0 | 0 | 1431851 | AIX10734.1 | P0C6X6 |
| FDENGTITDAVD | 1 | 0 | 0 | 1431852 | APO40579.1 | P59594 |
| FDFIIGHGMSYS | 1 | 0 | 0 | 1431860 | AIV41888.1 | P0C6X6 |
| FDFLSNVSTSHH | 1 | 0 | 0 | 1431864 | AWO68966.1 | A0A0P0G321 |
| FDFSSDVFKIGD | 1 | 0 | 0 | 1431867 | QDH43737.1 | H9EJ42 |
| FDFYTTEPEFVK | 1 | 0 | 0 | 1431869 | QBQ01836.1 | P0C6X6 |
| FDGDVGEKPVSI | 1 | 0 | 0 | 1431871 | APO40578.1 | P0C6X7 |
| FDGKPMLPKNMW | 1 | 0 | 0 | 1431874 | AFD98826.1 | H9EJ42 |
| FDGKPVLPKNMW | 1 | 0 | 0 | 1431875 | AWK59942.1 | H9EJ42 |
| FDGKSKCDESAA | 1 | 0 | 0 | 1431876 | P0C6W6.1 | P0C6W2 |
| FDGNMYGAFQDR | 1 | 0 | 0 | 1431881 | AVV62524.1 | K9N7C7 |
| FDGRVEGQANLF | 1 | 0 | 0 | 1431888 | AVP78041.1 | P0C6W2 |
| FDGTMYGAFMDK | 1 | 0 | 0 | 1431891 | AXN92247.1 | K9N7C7 |
| FDHSQYYFNKDT | 1 | 0 | 0 | 1431901 | AIV41896.1 | P30215 |
| FDIRDNGSLEKF | 1 | 0 | 0 | 1431913 | AXN92247.1 | K9N7C7 |
| FDIRDNGSLERF | 1 | 0 | 0 | 1431914 | AUM60013.1 | K9N7C7 |
| FDIRDNVSLEKF | 1 | 0 | 0 | 1431915 | ANF29193.1 | K9N7C7 |
| FDITDAAVKAAE | 1 | 0 | 0 | 1431920 | APT69868.1 | A0A0P0ILY9 |
| FDKAVADAMSAK | 1 | 0 | 0 | 1431930 | AUM60021.1 | K9N4V7 |
| FDKFKVNSALEQ | 1 | 0 | 0 | 1431931 | APO40578.1 | P0C6X7 |
| FDKSELTNGVYY | 1 | 0 | 0 | 1431945 | AEN19363.1 | P0C6X6 |
| FDKSEPLCLFKK | 1 | 0 | 0 | 1431946 | BBA20983.1 | A0A140H1H1 |
| FDKSWPRPIDPA | 1 | 0 | 0 | 1431948 | AVV62526.1 | K9N5Q8 |
| FDKSYQTNRDMQ | 1 | 0 | 0 | 1431949 | AUM60024.1 | K9N5Q8 |
| FDKTWPRPIDVF | 1 | 0 | 0 | 1431950 | AHI48583.1 | K9N5Q8 |
| FDKTWPRPIDVS | 1 | 0 | 0 | 1431951 | AXN92228.1 | K9N5Q8 |
| FDKWFVNDGRVG | 1 | 0 | 0 | 1431955 | AST12962.1 | A0A0P0LKV0 |
| FDLEDFIDNPCY | 1 | 0 | 0 | 1431961 | AVV62524.1 | K9N7C7 |
| FDLNGLDALFFY | 1 | 0 | 0 | 1431976 | ARB07605.1 | A0A140H1G9 |
| FDLNGLDAVFFY | 1 | 0 | 0 | 1431977 | ARU07575.1 | A0A140H1G9 |
| FDLQLGNLGYLQ | 1 | 0 | 0 | 1431978 | BBO58726.1 | P36334 |
| FDLQNCSFIYSF | 1 | 0 | 0 | 1431979 | AVV62537.1 | K9N5Q8 |
| FDLSKELKRPIQ | 1 | 0 | 0 | 1431980 | QBQ01836.1 | P0C6X6 |
| FDLTPFNLTYFN | 1 | 0 | 0 | 1431981 | ARE29967.1 | A0A0P0G321 |
| FDLTPFNLTYLN | 1 | 0 | 0 | 1431982 | AIW52843.1 | A0A0P0G321 |
| FDLVQYDFTDYKLEL | 1 | 0 | 0 | 1431984 | ABD75503.1 | A0A140H1G9 |
| FDNLSSEDLTAV | 1 | 0 | 0 | 1432012 | AIV41792.1 | P0C6X6 |
| FDNLVGYYSDDG | 1 | 0 | 0 | 1432014 | AVV62526.1 | K9N5Q8 |
| FDNMTAVACQPP | 1 | 0 | 0 | 1432015 | AXX83326.1 | P30215 |
| FDNNQNGDRTGA | 1 | 0 | 0 | 1432016 | APO40586.1 | P59595 |
| FDNNTDSIVIDG | 1 | 0 | 0 | 1432017 | AWK59942.1 | H9EJ42 |
| FDNPILDFGDGV | 1 | 0 | 0 | 1432018 | ATO98181.1 | Q3LZX1 |
| FDNPNIPFKDGL | 1 | 0 | 0 | 1432019 | APO40579.1 | P59594 |
| FDNPPTNVVSHV | 1 | 0 | 0 | 1432020 | AGT51679.1 | P30215 |
| FDNPVIPFKDGI | 1 | 0 | 0 | 1432022 | 5X58_A | P59594 |

| | | | | | | |
|---|---|---|---|---|---|---|
| FDNQSKLPSSSG | 1 | 0 | 0 | 1432024 | AIW52795.1 | H9EJ69 |
| FDNQSKPPSSSG | 1 | 0 | 0 | 1432025 | AGT51335.1 | H9EJ69 |
| FDNQSKSLSSSG | 1 | 0 | 0 | 1432026 | ARU07598.1 | H9EJ69 |
| FDNSIQGSYERF | 1 | 0 | 0 | 1432028 | AWK59942.1 | H9EJ42 |
| FDNSYQTNRDFQ | 1 | 0 | 0 | 1432029 | AUM60014.1 | K9N5Q8 |
| FDPAPVELENKF | 1 | 0 | 0 | 1432031 | AVV62524.1 | K9N7C7 |
| FDPITLKAPDTY | 1 | 0 | 0 | 1432033 | AVR40342.1 | P36334 |
| FDQQAQQEATHI | 1 | 0 | 0 | 1432038 | APT69896.1 | A0A0P0LKV0 |
| FDQQQLLAYYNF | 1 | 0 | 0 | 1432040 | ARU07575.1 | A0A140H1G9 |
| FDQRCRCSRCFY | 1 | 0 | 0 | 1432042 | AUM60018.1 | K9N7D2 |
| FDQRCSCSRCFY | 1 | 0 | 0 | 1432043 | AXG21669.1 | K9N7D2 |
| FDRNDCMLLCDI | 1 | 0 | 0 | 1432051 | AEN19363.1 | P0C6X6 |
| FDSCISNNRCNI | 1 | 0 | 0 | 1432056 | BBA20986.1 | A0A140H1H1 |
| FDSELNKFNRAS | 1 | 0 | 0 | 1432059 | AUM60013.1 | K9N7C7 |
| FDSEVSTDEEGD | 1 | 0 | 0 | 1432060 | AMM42619.1 | H9EJ69 |
| FDSGTVTKISDW | 1 | 0 | 0 | 1432070 | AVN89441.1 | K9N7C7 |
| FDSNAFSLANVI | 1 | 0 | 0 | 1432078 | AFD64760.1 | A0A0P0G321 |
| FDSNAFSLANVT | 1 | 0 | 0 | 1432079 | AIW52843.1 | A0A0P0G321 |
| FDTNMFIKFPEV | 1 | 0 | 0 | 1432096 | AUM60013.1 | K9N7C7 |
| FDTPHTDCTSAG | 1 | 0 | 0 | 1432097 | AVV62537.1 | K9N5Q8 |
| FDTRVHSEFNLP | 1 | 0 | 0 | 1432099 | AXN92247.1 | K9N7C7 |
| FDTRVLSNLNLPGCD | 1 | 0 | 0 | 1432101 | P0DTD1.1 | P0DTD1 |
| FDTVEVPPQVVK | 1 | 0 | 0 | 1432103 | AMQ48992.1 | K9N7C7 |
| FDVDKFKTPHIV | 1 | 0 | 0 | 1432113 | AVN89322.1 | K9N7C7 |
| FDVDKVKTPHIV | 1 | 0 | 0 | 1432117 | AXN92247.1 | K9N7C7 |
| FDVEGAHASKNA | 1 | 0 | 0 | 1432121 | AVV62524.1 | K9N7C7 |
| FDVKGVEELHDD | 1 | 0 | 0 | 1432130 | AOL02451.1 | Q80872 |
| FDVVFGHGAGSV | 1 | 0 | 0 | 1432142 | AWK59942.1 | H9EJ42 |
| FDWIKISADNKV | 1 | 0 | 0 | 1432152 | AGT51707.1 | P0C6X6 |
| FDYCIPYNSITD | 1 | 0 | 0 | 1432153 | Q3I5J4.1 | Q3LZX0 |
| FDYLIREAETRV | 1 | 0 | 0 | 1432157 | AXN73523.1 | K9N7C7 |
| FDYVYNPFMIDVQQWG | 1 | 0 | 0 | 1432163 | P0DTD1.1 | P0DTD1 |
| FDYYHTLDDSFL | 1 | 0 | 0 | 1432164 | APO40578.1 | P0C6X7 |
| FDYYRYNRPTIL | 1 | 0 | 0 | 1432165 | AIW52776.1 | A0A0P0LKV0 |
| FEAHKDFFKDGL | 1 | 0 | 0 | 1432170 | ARK08657.1 | P0C6X6 |
| FEAIPSGSVVEQ | 1 | 0 | 0 | 1432173 | ALA49649.1 | K9N5Q8 |
| FEAQAFETPAVQ | 1 | 0 | 0 | 1432181 | AVV62535.1 | K9N7C7 |
| FEAQPRGEFVEQ | 1 | 0 | 0 | 1432182 | AUM60014.1 | K9N5Q8 |
| FEDAQGLKEFYK | 1 | 0 | 0 | 1432196 | AMQ49036.1 | K9N638 |
| FEDDQGLKKFYK | 1 | 0 | 0 | 1432199 | AGV08522.1 | K9N7C7 |
| FEDDVITSLCKK | 1 | 0 | 0 | 1432200 | AWK59942.1 | H9EJ42 |
| FEDEKLVDVCEK | 1 | 0 | 0 | 1432204 | NP_073549.1 | A0A0P0LKV0 |
| FEDLNKCRAYMT | 1 | 0 | 0 | 1432206 | AVV62524.1 | K9N7C7 |
| FEDLSKCRAYLT | 1 | 0 | 0 | 1432207 | AVV62535.1 | K9N7C7 |
| FEDLTKCRTYLT | 1 | 0 | 0 | 1432208 | AUM60013.1 | K9N7C7 |
| FEFEDDVITSLC | 1 | 0 | 0 | 1432228 | AWK59942.1 | H9EJ42 |
| FEFVEGQGPPIA | 1 | 0 | 0 | 1432234 | P33469.1 | P33469 |
| FEFYTTEPEFVK | 1 | 0 | 0 | 1432236 | AEN19363.1 | P0C6X6 |
| FEGAQGLKKFYK | 1 | 0 | 0 | 1432237 | ASU90140.1 | K9N7C7 |
| FEISCDCGKKFD | 1 | 0 | 0 | 1432268 | QED88024.1 | H9EJ42 |
| FEKCEQLLREYG | 1 | 0 | 0 | 1432271 | AVV62526.1 | K9N5Q8 |
| FEKLHCEHLQFG | 1 | 0 | 0 | 1432274 | AWO68974.1 | A0A0P0G321 |
| FEKPQEKDKKSA | 1 | 0 | 0 | 1432278 | ARB07429.1 | A0A0N7J131 |
| FELAKVQNLSGN | 1 | 0 | 0 | 1432283 | AMM42445.1 | P33469 |
| FELLNAPATVSG | 1 | 0 | 0 | 1432300 | 2GHV_E | P59594 |
| FELTCLRYFAKL | 1 | 0 | 0 | 1432310 | ARU07574.1 | A0A140H1G9 |
| FEMKNWFDPIEN | 1 | 0 | 0 | 1432321 | AIW52776.1 | A0A0P0LKV0 |
| FEMLKLGTNDPQ | 1 | 0 | 0 | 1432322 | AMM42397.1 | Q19U29 |
| FEMNRLQEAIKV | 1 | 0 | 0 | 1432323 | AIL49493.1 | P36334 |
| FENLTNNLPPPL | 1 | 0 | 0 | 1432333 | AUM60014.1 | K9N5Q8 |
| FENLTNQLPPPL | 1 | 0 | 0 | 1432334 | AVV62537.1 | K9N5Q8 |
| FEPFTVNSINDS | 1 | 0 | 0 | 1432341 | AIL49542.1 | P36334 |
| FEPFTVNSKNDN | 1 | 0 | 0 | 1432342 | AIL49538.1 | P36334 |
| FEPFTVNSVSDS | 1 | 0 | 0 | 1432344 | ARB66068.1 | P36334 |
| FEPPKEQRVQSS | 1 | 0 | 0 | 1432348 | QCZ25056.1 | K9N4V7 |
| FEPRIPTIADFV | 1 | 0 | 0 | 1432350 | AIW52757.1 | A0A0P0K6L9 |
| FEQQCRCSRCFY | 1 | 0 | 0 | 1432359 | AVV62530.1 | K9N7D2 |
| FERDLSSDENGV | 1 | 0 | 0 | 1432367 | AVP78031.1 | Q3LZX1 |
| FERDLTSDENGV | 1 | 0 | 0 | 1432368 | ATO98120.1 | Q3LZX1 |
| FERNLRRGTILF | 1 | 0 | 0 | 1432372 | ABB90515.1 | A0A0P0K6L9 |
| FERVYDIGNPKG | 1 | 0 | 0 | 1432375 | AVV62524.1 | K9N7C7 |
| FESNMFIKFPEV | 1 | 0 | 0 | 1432383 | AVV62524.1 | K9N7C7 |
| FESTINKSVANL | 1 | 0 | 0 | 1432384 | AGT51388.1 | H9EJ24 |
| FESVKSSQSSPI | 1 | 0 | 0 | 1432388 | QBM11737.1 | K9N5Q8 |
| FETIPTQLTPLM | 1 | 0 | 0 | 1432400 | AUM60013.1 | K9N7C7 |
| FETLTDMEKDFL | 1 | 0 | 0 | 1432405 | AUM60013.1 | K9N7C7 |
| FETPAGQFCTES | 1 | 0 | 0 | 1432406 | AVV62524.1 | K9N7C7 |
| FEVKTFETPAGQ | 1 | 0 | 0 | 1432411 | AVV62524.1 | K9N7C7 |
| FEVPGVSIVKAK | 1 | 0 | 0 | 1432418 | AUM60013.1 | K9N7C7 |
| FEYNPPTDKLAM | 1 | 0 | 0 | 1432429 | NP_150075.1 | Q80872 |
| FFADNNDITNTN | 1 | 0 | 0 | 1432437 | ASY99807.1 | K9N4V7 |
| FFDKTWPRPIDV | 1 | 0 | 0 | 1432459 | AXN92228.1 | K9N5Q8 |
| FFEIKHADYTLA | 1 | 0 | 0 | 1432468 | AHF51980.1 | H9EJ42 |
| FFFLTHINTTLL | 1 | 0 | 0 | 1432479 | ANI69922.1 | K9N5Q8 |
| FFFPYDPALETG | 1 | 0 | 0 | 1432481 | AVV62535.1 | K9N7C7 |
| FFFPYEEGLETG | 1 | 0 | 0 | 1432482 | AUM60013.1 | K9N7C7 |
| FFFTQDGNAAIP | 1 | 0 | 0 | 1432484 | ACV87150.1 | A0A140H1G9 |
| FFFVYENAFMPF | 1 | 0 | 0 | 1432487 | AVP78041.1 | P0C6W2 |
| FFGSKLELVKRE | 1 | 0 | 0 | 1432499 | ARU07581.1 | Q19U29 |
| FFICCCTGCGPS | 1 | 0 | 0 | 1432508 | AXX83351.1 | P36334 |
| FFILCCTGCGTN | 1 | 0 | 0 | 1432519 | AXN92228.1 | K9N5Q8 |

| | | | | | | |
|---|---|---|---|---|---|---|
| FFKNVSSQGPNF | 1 | 0 | 0 | 1432539 | AVV62526.1 | K9N5Q8 |
| FFKNVSTSISNF | 1 | 0 | 0 | 1432542 | QBF80598.1 | K9N5Q8 |
| FFKTFTTSTALA | 1 | 0 | 0 | 1432545 | AEN19363.1 | P0C6X6 |
| FFLADNLSAAEK | 1 | 0 | 0 | 1432546 | AVV62524.1 | K9N7C7 |
| FFNLEPITLANA | 1 | 0 | 0 | 1432577 | AVV62526.1 | K9N5Q8 |
| FFNPEPITMANA | 1 | 0 | 0 | 1432580 | AVV62537.1 | K9N5Q8 |
| FFPQFTSDQAIT | 1 | 0 | 0 | 1432585 | ABD75596.1 | Q0ZJ82 |
| FFPYDIEFVTGK | 1 | 0 | 0 | 1432587 | ALA50044.1 | K9N7C7 |
| FFPYDPALETGD | 1 | 0 | 0 | 1432589 | AVV62535.1 | K9N7C7 |
| FFPYDPALETGE | 1 | 0 | 0 | 1432590 | AVV62524.1 | K9N7C7 |
| FFQFGDFVMNNI | 1 | 0 | 0 | 1432592 | AWK59942.1 | H9EJ42 |
| FFQPVTYFQLTT | 1 | 0 | 0 | 1432596 | AUM60013.1 | K9N7C7 |
| FFSNTKYYDHSQ | 1 | 0 | 0 | 1432607 | AIL49348.1 | P30215 |
| FFTCNSNANISM | 1 | 0 | 0 | 1432619 | AKA09626.1 | A0A0P0G321 |
| FFTPYFMYDIVS | 1 | 0 | 0 | 1432635 | AGW80947.1 | A0A0P0ILY9 |
| FFTYLCNFIKNN | 1 | 0 | 0 | 1432649 | AUM60013.1 | K9N7C7 |
| FFVNDFTCSQIS | 1 | 0 | 0 | 1432661 | ANI69889.1 | K9N5Q8 |
| FFVSRVFANSPN | 1 | 0 | 0 | 1432666 | AVV62535.1 | K9N7C7 |
| FFYGDVVSHICK | 1 | 0 | 0 | 1432688 | AEN19363.1 | P0C6X6 |
| FFYIDVGNHRGA | 1 | 0 | 0 | 1432694 | AKA09616.1 | A0A0P0G321 |
| FFYYSTTPCGVH | 1 | 0 | 0 | 1432710 | AGV08556.1 | K9N7C7 |
| FGACDYDRLSNE | 1 | 0 | 0 | 1432713 | AMQ49021.1 | K9N7A1 |
| FGAKPLDSLKVC | 1 | 0 | 0 | 1432732 | ARO76381.1 | P0C6U8 |
| FGAPDDATRLLH | 1 | 0 | 0 | 1432738 | AUM60013.1 | K9N7C7 |
| FGCENPDCIACS | 1 | 0 | 0 | 1432751 | NP_073549.1 | A0A0P0LKV0 |
| FGCKDSFTLYGQ | 1 | 0 | 0 | 1432755 | AEN19363.1 | P0C6X6 |
| FGDFIQTTPGFG | 1 | 0 | 0 | 1432763 | ABD75607.1 | A0A140H1G9 |
| FGDGIYFAATEK | 1 | 0 | 0 | 1432773 | ATO98193.1 | Q3LZX1 |
| FGDGVYFAATEK | 1 | 0 | 0 | 1432774 | ATO98181.1 | Q3LZX1 |
| FGDLQFNKLGTE | 1 | 0 | 0 | 1432776 | ASY99807.1 | K9N4V7 |
| FGDMQFNKLGTE | 1 | 0 | 0 | 1432780 | AUM60021.1 | K9N4V7 |
| FGDNNDNNTNQQ | 1 | 0 | 0 | 1432782 | AVV62544.1 | K9N4V7 |
| FGDPDDASRLLH | 1 | 0 | 0 | 1432784 | AVN89333.1 | K9N7C7 |
| FGDQELIRLGTD | 1 | 0 | 0 | 1432787 | APO40586.1 | P59595 |
| FGDVKPVTEYHL | 1 | 0 | 0 | 1432800 | AUM60027.1 | K9N643 |
| FGDVNKPVCAKS | 1 | 0 | 0 | 1432801 | QED88038.1 | H9EJ42 |
| FGEDTVLEVQGY | 1 | 0 | 0 | 1432807 | ATO98229.1 | P0C6W2 |
| FGEVEYSLPIKQ | 1 | 0 | 0 | 1432820 | AWK59942.1 | H9EJ42 |
| FGFIEDAVFKPQ | 1 | 0 | 0 | 1432831 | QEG03794.1 | P36334 |
| FGFIENSVFKPQ | 1 | 0 | 0 | 1432836 | AVQ05264.1 | P36334 |
| FGGPSDSSDNSK | 1 | 0 | 0 | 1432865 | AVP78038.1 | Q3LZX4 |
| FGGPTDSADNNQ | 1 | 0 | 0 | 1432869 | ARO76389.1 | P59595 |
| FGGQPVSEYHIT | 1 | 0 | 0 | 1432871 | AKN24752.1 | K9N643 |
| FGGSCYVCKPRQ | 1 | 0 | 0 | 1432873 | AVL25589.1 | A0A0P0G321 |
| FGINLQSGKTTS | 1 | 0 | 0 | 1432894 | ARB07390.1 | A0A0P0LKV0 |
| FGIPPSEAKGYW | 1 | 0 | 0 | 1432895 | ABA60794.1 | Q19U29 |
| FGITFQYGTDTN | 1 | 0 | 0 | 1432896 | AID55088.1 | K9N5Q8 |
| FGITQNAQGVHL | 1 | 0 | 0 | 1432899 | AUM60014.1 | K9N5Q8 |
| FGKDASDFIDSV | 1 | 0 | 0 | 1432906 | AVP78031.1 | Q3LZX1 |
| FGKDGVMYSEFP | 1 | 0 | 0 | 1432907 | AWW13566.1 | P36334 |
| FGKNNGIGTCPA | 1 | 0 | 0 | 1432910 | QEG03814.1 | P36334 |
| FGKRGGGNVTYT | 1 | 0 | 0 | 1432913 | NP_073549.1 | A0A0P0LKV0 |
| FGKRGPSHNFGG | 1 | 0 | 0 | 1432919 | AJQ24068.1 | P33469 |
| FGKSIIYTGDWE | 1 | 0 | 0 | 1432921 | AWK59942.1 | H9EJ42 |
| FGLHTVQSGDLN | 1 | 0 | 0 | 1432929 | AVV62535.1 | K9N7C7 |
| FGLQDCVTGVND | 1 | 0 | 0 | 1432938 | AGT51337.1 | H9EJ42 |
| FGLSYYSNSMDC | 1 | 0 | 0 | 1432946 | AXX83324.1 | P0C6X6 |
| FGNYTISCEPLE | 1 | 0 | 0 | 1432976 | ATO98227.1 | Q3LZX5 |
| FGPFKFCSETVN | 1 | 0 | 0 | 1432983 | ARB07424.1 | A0A0P0ILY9 |
| FGPTYMDGADVT | 1 | 0 | 0 | 1432994 | APO40578.1 | P0C6X7 |
| FGQCDFRSLPNE | 1 | 0 | 0 | 1432995 | AVV62543.1 | K9N7A1 |
| FGQCDYAQLPNE | 1 | 0 | 0 | 1432996 | AUM60030.1 | K9N7A1 |
| FGRDTSDFTDSV | 1 | 0 | 0 | 1433000 | ATO98181.1 | Q3LZX1 |
| FGRRGPGDLQGN | 1 | 0 | 0 | 1433006 | AVV62533.1 | K9N4V7 |
| FGSAGVVANGVT | 1 | 0 | 0 | 1433009 | ABB90495.1 | A0A0N7J131 |
| FGSDKIRMGQES | 1 | 0 | 0 | 1433015 | APO40579.1 | P59594 |
| FGSKLELAKVQN | 1 | 0 | 0 | 1433020 | AEN19370.1 | P33469 |
| FGSKLELVKRES | 1 | 0 | 0 | 1433023 | ARU07581.1 | Q19U29 |
| FGSNDIWMQNKG | 1 | 0 | 0 | 1433032 | AXX83326.1 | P30215 |
| FGSPDDASRLLH | 1 | 0 | 0 | 1433033 | AHI48794.1 | K9N638 |
| FGSSGFLQSSNY | 1 | 0 | 0 | 1433039 | AMM42415.1 | A0A140H1H1 |
| FGSTMDNTTQSA | 1 | 0 | 0 | 1433042 | ATO98193.1 | Q3LZX1 |
| FGTDHRFILAIT | 1 | 0 | 0 | 1433052 | ARU07599.1 | A0A0P0LKV0 |
| FGTRNPNNDAAI | 1 | 0 | 0 | 1433056 | AVV62533.1 | K9N4V7 |
| FGTRNPNNDLAI | 1 | 0 | 0 | 1433057 | AKN24774.1 | K9N4V7 |
| FGTRNPNNDPAI | 1 | 0 | 0 | 1433058 | AUM60021.1 | K9N4V7 |
| FGTTLDNTSQSL | 1 | 0 | 0 | 1433063 | AVP78031.1 | Q3LZX1 |
| FGTTLDNTTQSA | 1 | 0 | 0 | 1433064 | ATO98145.1 | Q3LZX1 |
| FGVPLSEAKGYW | 1 | 0 | 0 | 1433097 | ABA60796.1 | Q19U29 |
| FGVSDNFDNNQN | 1 | 0 | 0 | 1433103 | APO40586.1 | P59595 |
| FGVSKPNAIDVE | 1 | 0 | 0 | 1433104 | AWK59942.1 | H9EJ42 |
| FGVTDLEDLCFK | 1 | 0 | 0 | 1433106 | Q14EA9.1 | Q0ZJ84 |
| FGVYDYLVSTQE | 1 | 0 | 0 | 1433114 | APO40578.1 | P0C6X7 |
| FGWVKISADNKV | 1 | 0 | 0 | 1433123 | QBQ01836.1 | P0C6X6 |
| FGYVCDGSFCKA | 1 | 0 | 0 | 1433131 | ARB07436.1 | A0A140H1G9 |
| FGYVCDGSFYKA | 1 | 0 | 0 | 1433132 | ARU07575.1 | A0A140H1G9 |
| FGYVQQGVFNKA | 1 | 0 | 0 | 1433134 | ATN39862.1 | P0C6X6 |
| FHANSSEPALIF | 1 | 0 | 0 | 1433137 | AGT51481.1 | P36334 |
| FHASRPGVYCGH | 1 | 0 | 0 | 1433139 | AUM60013.1 | K9N7C7 |
| FHEHVGGNFDND | 1 | 0 | 0 | 1433145 | AVV62524.1 | K9N7C7 |
| FHENVGGNFDND | 1 | 0 | 0 | 1433146 | AVV62535.1 | K9N7C7 |

| | | | | | | |
|---|---|---|---|---|---|---|
| FHFRKACQNLDC | 1 | 0 | 0 | 1433149 | AWW13564.1 | Q80872 |
| FHFRKSCQNLDC | 1 | 0 | 0 | 1433150 | AGT51688.1 | Q80872 |
| FHKVFNATRFPS | 1 | 0 | 0 | 1433172 | AVP78031.1 | Q3LZX1 |
| FHLDGEIISLEK | 1 | 0 | 0 | 1433173 | APO40578.1 | P0C6X7 |
| FHLDGETITFDN | 1 | 0 | 0 | 1433174 | AVP78030.1 | P0C6W2 |
| FHNYTGEGQQII | 1 | 0 | 0 | 1433188 | AXX83296.1 | P30215 |
| FHRIESTDSTVL | 1 | 0 | 0 | 1433207 | AVN89378.1 | K9N7D2 |
| FHSSQLPLDVYD | 1 | 0 | 0 | 1433212 | AJC98124.1 | P0C6X6 |
| FHSVEIEVGEVK | 1 | 0 | 0 | 1433214 | AKT07951.1 | H9EJ42 |
| FHTPAYDVSAFR | 1 | 0 | 0 | 1433230 | AXN92247.1 | K9N7C7 |
| FHTVLLPTQYKD | 1 | 0 | 0 | 1433233 | AIW52753.1 | A0A0P0K6L9 |
| FHYERDNTSYPE | 1 | 0 | 0 | 1433245 | AHI48797.1 | K9N638 |
| FIADKEKLISLA | 1 | 0 | 0 | 1433248 | AUM60013.1 | K9N7C7 |
| FIAFNHPIQVDQ | 1 | 0 | 0 | 1433251 | ASU90142.1 | K9N5Q8 |
| FIAPPTGVYELN | 1 | 0 | 0 | 1433266 | AHA80218.1 | P36334 |
| FIAVSKPMGTQT | 1 | 0 | 0 | 1433270 | ABD73001.1 | P59594 |
| FICCCTGCGPSC | 1 | 0 | 0 | 1433272 | AXX83351.1 | P36334 |
| FICCCTGCGTSC | 1 | 0 | 0 | 1433273 | CAA79896.1 | P36334 |
| FICSQISPAAIA | 1 | 0 | 0 | 1433282 | AVN89334.1 | K9N5Q8 |
| FIDDYYGYLMKR | 1 | 0 | 0 | 1433286 | ADO24650.1 | H9EJ42 |
| FIDGVMYSEFPA | 1 | 0 | 0 | 1433290 | QDH43719.1 | P36334 |
| FIDVGNHRSAFA | 1 | 0 | 0 | 1433307 | QEG03748.1 | A0A0P0G321 |
| FIECGSCGNNSW | 1 | 0 | 0 | 1433314 | ALA49340.1 | K9N7C7 |
| FIEDAVFKPQPA | 1 | 0 | 0 | 1433315 | QEG03794.1 | P36334 |
| FIEDLLFDKVKL | 1 | 0 | 0 | 1433316 | ARU07577.1 | A0A140H1H1 |
| FIEDSVFKPQPA | 1 | 0 | 0 | 1433321 | ARB66062.1 | P36334 |
| FIEDSVFVPQPT | 1 | 0 | 0 | 1433323 | AEN19366.1 | P36334 |
| FIEFTDSFYTQG | 1 | 0 | 0 | 1433324 | AUM60013.1 | K9N7C7 |
| FIENSVFKPQPA | 1 | 0 | 0 | 1433327 | AVQ05264.1 | P36334 |
| FIEQPNSVECDF | 1 | 0 | 0 | 1433329 | AVV62537.1 | K9N5Q8 |
| FIEQPTSVECDF | 1 | 0 | 0 | 1433330 | AVV62526.1 | K9N5Q8 |
| FIFDVDKFKTPH | 1 | 0 | 0 | 1433340 | AVN89322.1 | K9N7C7 |
| FIFDVDKVKTPH | 1 | 0 | 0 | 1433341 | AXN92247.1 | K9N7C7 |
| FIGDNVGHYVHV | 1 | 0 | 0 | 1433358 | AEN19363.1 | P0C6X6 |
| FIGTQALDMLTV | 1 | 0 | 0 | 1433368 | AVV62535.1 | K9N7C7 |
| FIKDGDDGYFLP | 1 | 0 | 0 | 1433396 | AVV62524.1 | K9N7C7 |
| FIKHKGGDSHEF | 1 | 0 | 0 | 1433405 | AUM60013.1 | K9N7C7 |
| FIKKYKLEDYAF | 1 | 0 | 0 | 1433418 | AUM60013.1 | K9N7C7 |
| FIKKYKLENFAF | 1 | 0 | 0 | 1433419 | AVV62524.1 | K9N7C7 |
| FIKTAQADTAGL | 1 | 0 | 0 | 1433439 | AXN92261.1 | K9N796 |
| FIKTNNIRIVDE | 1 | 0 | 0 | 1433440 | ASU90175.1 | K9N5Q8 |
| FIKTNNTMIVDE | 1 | 0 | 0 | 1433441 | ATG84866.1 | K9N5Q8 |
| FIKTNNTSIVDE | 1 | 0 | 0 | 1433445 | ANF29261.1 | K9N5Q8 |
| FIKTNNTTIVDE | 1 | 0 | 0 | 1433446 | ALA49957.1 | K9N5Q8 |
| FIKYNWPEPIDM | 1 | 0 | 0 | 1433451 | ATQ39390.1 | K9N5Q8 |
| FILHDVNNGLTC | 1 | 0 | 0 | 1433457 | AEN19366.1 | P36334 |
| FILHDVNSGTTC | 1 | 0 | 0 | 1433459 | CAA83660.1 | P36334 |
| FILLFGWVKISA | 1 | 0 | 0 | 1433462 | QBQ01836.1 | P0C6X6 |
| FILPLVYFSDSK | 1 | 0 | 0 | 1433466 | ANI69921.1 | K9N7C7 |
| FILVSCIIGSLG | 1 | 0 | 0 | 1433471 | AEN19365.1 | P30215 |
| FIMVLGSKHVNC | 1 | 0 | 0 | 1433481 | ABQ16582.1 | H9EJ42 |
| FINGISAKVKNT | 1 | 0 | 0 | 1433495 | AKA09642.1 | P36334 |
| FINSTINTTPVS | 1 | 0 | 0 | 1433500 | AMM42635.1 | A0A0P0G321 |
| FIPIDQYMCGKD | 1 | 0 | 0 | 1433505 | AVV62535.1 | K9N7C7 |
| FIQSIRLFRRTG | 1 | 0 | 0 | 1433519 | AVV62543.1 | K9N7A1 |
| FIRANAKIVTLP | 1 | 0 | 0 | 1433523 | ATQ39390.1 | K9N5Q8 |
| FISDEVARDLSV | 1 | 0 | 0 | 1433541 | APO40578.1 | P0C6X7 |
| FISPYNSQNAVS | 1 | 0 | 0 | 1433559 | 5WWP_B | K9N7C7 |
| FISPYNSQNTVA | 1 | 0 | 0 | 1433561 | AVP78041.1 | P0C6W2 |
| FITDAQTGSSKCVCSVID | 1 | 0 | 0 | 1433574 | P0DTD1.1 | P0DTD1 |
| FITETNTAAFNA | 1 | 0 | 0 | 1433580 | ANI69909.1 | K9N7C7 |
| FITLEEAVRQVR | 1 | 0 | 0 | 1433591 | AHL18091.1 | K9N7C7 |
| FITMHSINHDIF | 1 | 0 | 0 | 1433595 | ARU07575.1 | A0A140H1G9 |
| FITNGMSLLDII | 1 | 0 | 0 | 1433598 | AWK59942.1 | H9EJ42 |
| FITQRNFYSPQV | 1 | 0 | 0 | 1433603 | APO40579.1 | P59594 |
| FITREEAIRQVR | 1 | 0 | 0 | 1433606 | QWP96910.1 | P0DTD1 |
| FITTCFDATTAT | 1 | 0 | 0 | 1433607 | AUM60023.1 | K9N7C7 |
| FITVEAAFDLSK | 1 | 0 | 0 | 1433610 | QBQ01836.1 | P0C6X6 |
| FITVETAIELSK | 1 | 0 | 0 | 1433611 | ABD75535.1 | A0A140H1G9 |
| FITVETALDLSK | 1 | 0 | 0 | 1433612 | AXX83306.1 | P0C6X6 |
| FIVVLQYGHYKF | 1 | 0 | 0 | 1433651 | QDH43733.1 | U3M6Q8 |
| FIVYNNSGSNVT | 1 | 0 | 0 | 1433656 | ARK08626.1 | H9EJ42 |
| FIYFITMSSKGQ | 1 | 0 | 0 | 1433659 | AWK59942.1 | H9EJ42 |
| FIYVFSLLHFTN | 1 | 0 | 0 | 1433667 | ARU07575.1 | A0A140H1G9 |
| FKADGIIYPQGR | 1 | 0 | 0 | 1433671 | AHI48583.1 | K9N5Q8 |
| FKAPKNFCPCKL | 1 | 0 | 0 | 1433678 | CAA79896.1 | P36334 |
| FKASALFKEFAS | 1 | 0 | 0 | 1433687 | AVV62524.1 | K9N7C7 |
| FKATGTYKCPQT | 1 | 0 | 0 | 1433688 | CAA79896.1 | P36334 |
| FKCSSNGSNDVW | 1 | 0 | 0 | 1433698 | AGT17757.1 | A0A140H1H0 |
| FKDAEGRMTPYC | 1 | 0 | 0 | 1433703 | AVN89322.1 | K9N7C7 |
| FKDCSKVITGLH | 1 | 0 | 0 | 1433711 | QYC64967.1 | P0DTD1 |
| FKDCSRDPSGLP | 1 | 0 | 0 | 1433713 | AVV62535.1 | K9N7C7 |
| FKDCSRDPSGLS | 1 | 0 | 0 | 1433714 | AVV62524.1 | K9N7C7 |
| FKDFQFMLWGND | 1 | 0 | 0 | 1433723 | AIV41912.1 | P0C6X6 |
| FKDGIYFAATEH | 1 | 0 | 0 | 1433726 | AVP78031.1 | Q3LZX1 |
| FKDGLYFAATER | 1 | 0 | 0 | 1433728 | APO40579.1 | P59594 |
| FKDGVMHSEFPA | 1 | 0 | 0 | 1433729 | AKA09651.1 | P36334 |
| FKDGVMYSEFPA | 1 | 0 | 0 | 1433730 | AEN19366.1 | P36334 |
| FKDGVMYSEFPT | 1 | 0 | 0 | 1433731 | AIL49548.1 | P36334 |
| FKDLTANMSMAE | 1 | 0 | 0 | 1433737 | ARK08626.1 | H9EJ42 |
| FKDSSGKPVPYC | 1 | 0 | 0 | 1433741 | AVP78041.1 | P0C6W2 |

| | | | | | | |
|---|---|---|---|---|---|---|
| FKEELDQWLKNQ | 1 | 0 | 0 | 1433743 | ARB07414.1 | P36334 |
| FKEESDQWFKNQ | 1 | 0 | 0 | 1433745 | BBA20973.1 | P36334 |
| FKEFASRTVSAA | 1 | 0 | 0 | 1433749 | AVV62535.1 | K9N7C7 |
| FKELWHLDTGAV | 1 | 0 | 0 | 1433758 | ANZ78840.1 | P36334 |
| FKELWHMDTGVV | 1 | 0 | 0 | 1433759 | AVQ05264.1 | P36334 |
| FKENVDFINSMQ | 1 | 0 | 0 | 1433760 | AXX83325.1 | Q80872 |
| FKEVSAAYYNDW | 1 | 0 | 0 | 1433767 | AZS52618.1 | A0A140H1H1 |
| FKFCSETVNGYA | 1 | 0 | 0 | 1433774 | ARB07424.1 | A0A0P0ILY9 |
| FKFDDYDFSVEA | 1 | 0 | 0 | 1433775 | P0C6U5.1 | A0A140H1G9 |
| FKFNYQLPKVGY | 1 | 0 | 0 | 1433784 | AXX83304.1 | Q04853 |
| FKFPDGQGVPIA | 1 | 0 | 0 | 1433785 | AMM42397.1 | Q19U29 |
| FKFRDLAIENMW | 1 | 0 | 0 | 1433786 | ABD75647.1 | A0A140H1G9 |
| FKGCETFGKPVI | 1 | 0 | 0 | 1433797 | ARU07575.1 | A0A140H1G9 |
| FKGGIPGNMIDG | 1 | 0 | 0 | 1433800 | AIV42002.1 | P0C6X6 |
| FKGKFVQIPAQC | 1 | 0 | 0 | 1433801 | AVV62524.1 | K9N7C7 |
| FKHGVMYSEFPA | 1 | 0 | 0 | 1433811 | AIL49520.1 | P36334 |
| FKHKVCTDSVTL | 1 | 0 | 0 | 1433812 | AVV62535.1 | K9N7C7 |
| FKHLREFVFKTS | 1 | 0 | 0 | 1433819 | ATO98145.1 | Q3LZX1 |
| FKHWSFDYHPNC | 1 | 0 | 0 | 1433828 | AIW52827.1 | H9EJ42 |
| FKIGRRRICLSG | 1 | 0 | 0 | 1433835 | AEN19363.1 | P0C6X6 |
| FKILYKGSVTHD | 1 | 0 | 0 | 1433839 | AUM60023.1 | K9N7C7 |
| FKKCGGFCDDYT | 1 | 0 | 0 | 1433849 | AQN78672.1 | P36334 |
| FKKNFTYNVSTD | 1 | 0 | 0 | 1433863 | BBA20983.1 | A0A140H1H1 |
| FKKNFTYNVYAD | 1 | 0 | 0 | 1433864 | ARB07438.1 | A0A140H1H1 |
| FKLDAIVPGYPK | 1 | 0 | 0 | 1433876 | QBM11746.1 | K9N7C7 |
| FKLGFDSSKEFV | 1 | 0 | 0 | 1433885 | AAX85666.1 | P0C6X6 |
| FKLRSGVGRCYN | 1 | 0 | 0 | 1433907 | CAA71146.1 | A0A0P0K6L9 |
| FKLSDNNCYLNA | 1 | 0 | 0 | 1433908 | AVN89451.1 | K9N7C7 |
| FKLTHQSNEDDG | 1 | 0 | 0 | 1433917 | AVV62533.1 | K9N4V7 |
| FKMFVLWLLWPS | 1 | 0 | 0 | 1433922 | ASY99815.1 | K9N7A1 |
| FKNADLKDGYFV | 1 | 0 | 0 | 1433928 | AIW52827.1 | H9EJ42 |
| FKNFSTSIPNFG | 1 | 0 | 0 | 1433931 | AKM76239.1 | K9N5Q8 |
| FKNLQYDVLFYC | 1 | 0 | 0 | 1433945 | QEG59359.1 | A0A0P0G321 |
| FKNQHCNGSIAC | 1 | 0 | 0 | 1433948 | AWW13571.1 | P0C6X6 |
| FKNQTLVAPDLS | 1 | 0 | 0 | 1433950 | CAA83661.1 | P36334 |
| FKNQTSVAPDFS | 1 | 0 | 0 | 1433951 | BBA20976.1 | P36334 |
| FKNSVTSINSAI | 1 | 0 | 0 | 1433960 | AST12962.1 | A0A0P0LKV0 |
| FKNVSTSIPNFF | 1 | 0 | 0 | 1433966 | ANI69922.1 | K9N5Q8 |
| FKNVSTSISNFG | 1 | 0 | 0 | 1433969 | QBF80598.1 | K9N5Q8 |
| FKPGGTTSGEAI | 1 | 0 | 0 | 1433972 | ADK37693.1 | H9EJ42 |
| FKPQPAGVFTDH | 1 | 0 | 0 | 1433985 | CAA83661.1 | P36334 |
| FKPQPAGVFTNH | 1 | 0 | 0 | 1433986 | ATP16757.1 | P36334 |
| FKPQPVGVFTHH | 1 | 0 | 0 | 1433989 | CAA83660.1 | P36334 |
| FKPQSGGGKCYN | 1 | 0 | 0 | 1433992 | ABB90509.1 | A0A0P0K6L9 |
| FKPQSGGGRCYN | 1 | 0 | 0 | 1433993 | ABB90508.1 | A0A0P0K6L9 |
| FKPVHLVPVSPA | 1 | 0 | 0 | 1434000 | AVN89337.1 | K9N7D2 |
| FKPVHLVPVSPV | 1 | 0 | 0 | 1434001 | AVN89455.1 | K9N7D2 |
| FKPVQLAPVSPA | 1 | 0 | 0 | 1434003 | AKM76243.1 | K9N7D2 |
| FKQGHFIGVEGE | 1 | 0 | 0 | 1434011 | ANI69909.1 | K9N7C7 |
| FKRDSDADSPSK | 1 | 0 | 0 | 1434015 | AMM42394.1 | Q19U29 |
| FKSIVNTRNATV | 1 | 0 | 0 | 1434041 | ADK37693.1 | H9EJ42 |
| FKTADGQQKHLL | 1 | 0 | 0 | 1434053 | ABD75597.1 | Q19U29 |
| FKTADGQQKPLL | 1 | 0 | 0 | 1434054 | ABA60796.1 | Q19U29 |
| FKTAVFVYTCVV | 1 | 0 | 0 | 1434056 | AWK59942.1 | H9EJ42 |
| FKTEGLCVDIPS | 1 | 0 | 0 | 1434058 | APO40578.1 | P0C6X7 |
| FKTEVDPNLSGF | 1 | 0 | 0 | 1434059 | ATQ39389.1 | K9N7C7 |
| FKVEDAYDQVHE | 1 | 0 | 0 | 1434071 | ARK08632.1 | P0C6X6 |
| FKVEDSVIINDD | 1 | 0 | 0 | 1434072 | AXX83318.1 | P0C6X6 |
| FKVEDSVIVNDA | 1 | 0 | 0 | 1434073 | AIV41846.1 | P0C6X6 |
| FKVEDSVIVNND | 1 | 0 | 0 | 1434075 | AOL02450.1 | P0C6X6 |
| FKVLDNGIIRDI | 1 | 0 | 0 | 1434084 | AUM60013.1 | K9N7C7 |
| FKVNETTSQYLF | 1 | 0 | 0 | 1434086 | ALW82700.1 | K9N7C7 |
| FLAFVVFLLAIL | 1 | 0 | 0 | 1434127 | AAS48454.1 | P59637 |
| FLANANSKPLVG | 1 | 0 | 0 | 1434136 | AUM60015.1 | K9N796 |
| FLDCTGQAVNTS | 1 | 0 | 0 | 1434173 | AVV62539.1 | K9N4V0 |
| FLDGNFRTEVDP | 1 | 0 | 0 | 1434177 | AVN89363.1 | K9N7C7 |
| FLDLQAEMNRLQ | 1 | 0 | 0 | 1434183 | AGT51521.1 | P36334 |
| FLDLQDEMNRLQ | 1 | 0 | 0 | 1434184 | CAA83660.1 | P36334 |
| FLDTCFDVSKPN | 1 | 0 | 0 | 1434192 | AGT51337.1 | H9EJ42 |
| FLEENDVEEDDV | 1 | 0 | 0 | 1434197 | QBQ01836.1 | P0C6X6 |
| FLEMKSGKQVEQ | 1 | 0 | 0 | 1434209 | AVP78041.1 | P0C6W2 |
| FLEQDAQIDRLI | 1 | 0 | 0 | 1434211 | AXN92260.1 | K9N5Q8 |
| FLGLWCLMPTYT | 1 | 0 | 0 | 1434251 | AXX83324.1 | P0C6X6 |
| FLIAGPKGPAIR | 1 | 0 | 0 | 1434293 | ASU45717.1 | K9N7C7 |
| FLIAGPKGPEIR | 1 | 0 | 0 | 1434294 | AXN92247.1 | K9N7C7 |
| FLIHNFVIFFTW | 1 | 0 | 0 | 1434308 | NP_073549.1 | A0A0P0LKV0 |
| FLISGPKGPEIR | 1 | 0 | 0 | 1434319 | AVV62524.1 | K9N7C7 |
| FLKKAYIYTSTG | 1 | 0 | 0 | 1434339 | AUM60013.1 | K9N7C7 |
| FLKLVDDHALIV | 1 | 0 | 0 | 1434343 | AOG74785.1 | A0A0P0KH29 |
| FLKNDAPYIVGD | 1 | 0 | 0 | 1434344 | APO40578.1 | P0C6X7 |
| FLKNFDQQCHCS | 1 | 0 | 0 | 1434345 | AVV62541.1 | K9N7D2 |
| FLKPLYVPEHCQ | 1 | 0 | 0 | 1434346 | ATY74382.1 | K9N796 |
| FLLCCTGCGTNC | 1 | 0 | 0 | 1434364 | AVV62526.1 | K9N5Q8 |
| FLLDFSVDGYIR | 1 | 0 | 0 | 1434367 | AXN92228.1 | K9N5Q8 |
| FLLNGYYPTSGS | 1 | 0 | 0 | 1434394 | CAA79896.1 | P36334 |
| FLMAEDGITKLA | 1 | 0 | 0 | 1434425 | AKN58609.1 | K9N638 |
| FLMLFVFQVYPI | 1 | 0 | 0 | 1434431 | ARU07575.1 | A0A140H1G9 |
| FLNATNCTYNRV | 1 | 0 | 0 | 1434440 | APO40579.1 | P59594 |
| FLNGADISDTIP | 1 | 0 | 0 | 1434443 | AUM60013.1 | K9N7C7 |
| FLNGSAGSVGFN | 1 | 0 | 0 | 1434445 | 1Z1J_A | P0C6U8 |
| FLNKVVSTTTNIVTRC | 1 | 0 | 0 | 1434453 | P0DTC1.1 | P0DTD1 |

| | | | | | | |
|---|---|---|---|---|---|---|
| FLNSENLYFQSG | 1 | 0 | 0 | 1434462 | 6U7H_A | A0A0P0K6L9 |
| FLPAQDTWGTSA | 1 | 0 | 0 | 1434463 | ATO98205.1 | Q3LZX1 |
| FLPESAAYYVGN | 1 | 0 | 0 | 1434464 | ATO98181.1 | Q3LZX1 |
| FLPEVAAYFVGN | 1 | 0 | 0 | 1434465 | ATO98193.1 | Q3LZX1 |
| FLPFDSNLTQYF | 1 | 0 | 0 | 1434468 | ATO98181.1 | Q3LZX1 |
| FLPIDSTVKNYF | 1 | 0 | 0 | 1434481 | APO40578.1 | P0C6X7 |
| FLPTFIGKIAQW | 1 | 0 | 0 | 1434494 | AXX83336.1 | P0C6X6 |
| FLRYINGFNVSE | 1 | 0 | 0 | 1434535 | AIV41839.1 | Q04853 |
| FLSDDRTEVPQL | 1 | 0 | 0 | 1434541 | 6C6Z_A | K9N5Q8 |
| FLSDFINAIFAK | 1 | 0 | 0 | 1434542 | QBP84704.1 | P36334 |
| FLSKHSDFGLDG | 1 | 0 | 0 | 1434554 | AWK59942.1 | H9EJ42 |
| FLSNASSSFDCI | 1 | 0 | 0 | 1434560 | AVL25587.1 | A0A0P0G321 |
| FLSNTKYYDHSQ | 1 | 0 | 0 | 1434565 | AEN19365.1 | P30215 |
| FLSNTKYYDYSQ | 1 | 0 | 0 | 1434566 | AGT51729.1 | P30215 |
| FLSNVSTSHHCI | 1 | 0 | 0 | 1434567 | AWO68966.1 | A0A0P0G321 |
| FLTLFLLPTALC | 1 | 0 | 0 | 1434589 | AVV62524.1 | K9N7C7 |
| FLTSLTSDYTVT | 1 | 0 | 0 | 1434605 | AUM60013.1 | K9N7C7 |
| FLTYIIFSYAPH | 1 | 0 | 0 | 1434618 | AUM60013.1 | K9N7C7 |
| FLVKNGIVLPFE | 1 | 0 | 0 | 1434630 | AUM60013.1 | K9N7C7 |
| FLVKNPSWSNAV | 1 | 0 | 0 | 1434631 | ABE97136.1 | H9EJ42 |
| FLVVDDKYKVNE | 1 | 0 | 0 | 1434655 | ABD75575.1 | A0A140H1G9 |
| FMDGENAYEMVK | 1 | 0 | 0 | 1434695 | AVN89352.1 | K9N7C7 |
| FMIRSCYSGHVS | 1 | 0 | 0 | 1434728 | AIL49489.1 | P36334 |
| FMKHKGGDLTDF | 1 | 0 | 0 | 1434736 | AHI48527.1 | K9N638 |
| FMKQYGDCLGGI | 1 | 0 | 0 | 1434741 | APO40579.1 | P59594 |
| FMLGSSVGNFSN | 1 | 0 | 0 | 1434750 | AVV62537.1 | K9N5Q8 |
| FMLKYNENGTIT | 1 | 0 | 0 | 1434759 | AVP78031.1 | Q3LZX1 |
| FMLNFNENGTIT | 1 | 0 | 0 | 1434760 | ATO98193.1 | Q3LZX1 |
| FMLNISGNGGLF | 1 | 0 | 0 | 1434761 | AVP78031.1 | Q3LZX1 |
| FMLSFNENGTIT | 1 | 0 | 0 | 1434766 | ATO98181.1 | Q3LZX1 |
| FMLSTLHWSFRL | 1 | 0 | 0 | 1434768 | AEN19363.1 | P0C6X6 |
| FMNSCSVNFTKA | 1 | 0 | 0 | 1434783 | BBA20986.1 | A0A140H1H1 |
| FMPQFSRMTQSA | 1 | 0 | 0 | 1434792 | AUM60014.1 | K9N5Q8 |
| FMPVQSRKFIVP | 1 | 0 | 0 | 1434794 | AEN19363.1 | P0C6X6 |
| FMPYFMYDVVSS | 1 | 0 | 0 | 1434799 | AGT21366.1 | A0A0P0LKV0 |
| FMSLEVPRRNVA | 1 | 0 | 0 | 1434827 | AVP78030.1 | P0C6W2 |
| FMVDNGTRLHDA | 1 | 0 | 0 | 1434840 | P30215.1 | P30215 |
| FMVDPTTVLNDP | 1 | 0 | 0 | 1434841 | AWK59942.1 | H9EJ42 |
| FMVDPTTVLNGP | 1 | 0 | 0 | 1434842 | ABE97136.1 | H9EJ42 |
| FMVGNVTRLHDA | 1 | 0 | 0 | 1434844 | AWW13510.1 | P30215 |
| FMYAYISNKADL | 1 | 0 | 0 | 1434850 | AWK59942.1 | H9EJ42 |
| FNADGDEAWGST | 1 | 0 | 0 | 1434862 | AUM60013.1 | K9N7C7 |
| FNADNLTDYSVS | 1 | 0 | 0 | 1434863 | BBA20983.1 | A0A140H1H1 |
| FNAEDCMSDFMS | 1 | 0 | 0 | 1434864 | P36334.1 | P36334 |
| FNAEGDAFWSST | 1 | 0 | 0 | 1434865 | AVN89333.1 | K9N7C7 |
| FNAEGDASWSST | 1 | 0 | 0 | 1434866 | AXN92247.1 | K9N7C7 |
| FNAFVKCNCGSE | 1 | 0 | 0 | 1434869 | AWK59942.1 | H9EJ42 |
| FNASISESKPLA | 1 | 0 | 0 | 1434876 | AVL25595.1 | H9EJ69 |
| FNAVDCKSDFMS | 1 | 0 | 0 | 1434885 | CAA83660.1 | P36334 |
| FNAVDCVSDFMS | 1 | 0 | 0 | 1434887 | AHA80206.1 | P36334 |
| FNCDTFCTGSTF | 1 | 0 | 0 | 1434890 | BAE93400.1 | P0C6U8 |
| FNCHSFKPGNTF | 1 | 0 | 0 | 1434894 | ARU07575.1 | A0A140H1G9 |
| FNCTNFAINDKN | 1 | 0 | 0 | 1434900 | BBA20983.1 | A0A140H1H1 |
| FNCTYDRVEKSF | 1 | 0 | 0 | 1434902 | Q3I5J5.1 | Q3LZX1 |
| FNDLSQLYCSYE | 1 | 0 | 0 | 1434914 | ANI69889.1 | K9N5Q8 |
| FNDYHANRPGVY | 1 | 0 | 0 | 1434920 | AVV62524.1 | K9N7C7 |
| FNEHVTSVNLFD | 1 | 0 | 0 | 1434922 | AXG21666.1 | K9N796 |
| FNEHVTSVNLFS | 1 | 0 | 0 | 1434923 | QBM11745.1 | K9N796 |
| FNEKTHGPLSLP | 1 | 0 | 0 | 1434926 | AVP78030.1 | P0C6W2 |
| FNEWAMANQFTE | 1 | 0 | 0 | 1434939 | AVV62524.1 | K9N7C7 |
| FNFCKSDILSLN | 1 | 0 | 0 | 1434941 | AMN88685.1 | A0A140H1H0 |
| FNFCYGFNEPLN | 1 | 0 | 0 | 1434943 | AGT17757.1 | A0A140H1H0 |
| FNFDQKELLDYY | 1 | 0 | 0 | 1434945 | ATN39862.1 | P0C6X6 |
| FNFSQILPDSSK | 1 | 0 | 0 | 1434960 | ARO76382.1 | P59594 |
| FNGAIIRDSEAI | 1 | 0 | 0 | 1434969 | AVV62524.1 | K9N7C7 |
| FNGATIRGSDVK | 1 | 0 | 0 | 1434973 | AUM60013.1 | K9N7C7 |
| FNGLTGTGVLTS | 1 | 0 | 0 | 1434987 | ATO98231.1 | Q3LZX1 |
| FNGSNDVWIFNK | 1 | 0 | 0 | 1434991 | AZS52617.1 | A0A140H1H0 |
| FNGTGRGACKGF | 1 | 0 | 0 | 1434993 | AIW52757.1 | A0A0P0K6L9 |
| FNGTGRGVCKGF | 1 | 0 | 0 | 1434996 | AIW52750.1 | A0A0P0K6L9 |
| FNHNMGDSDLVQ | 1 | 0 | 0 | 1435003 | AIW52805.1 | H9EJ69 |
| FNHPIQIDQLNS | 1 | 0 | 0 | 1435005 | ASU91208.1 | K9N5Q8 |
| FNHPIQVDQFNS | 1 | 0 | 0 | 1435007 | ASU90362.1 | K9N5Q8 |
| FNHPIQVGQLNS | 1 | 0 | 0 | 1435009 | ALJ54452.1 | K9N5Q8 |
| FNILQATTANVS | 1 | 0 | 0 | 1435032 | AXN92247.1 | K9N7C7 |
| FNITEDENAEWF | 1 | 0 | 0 | 1435036 | AVV62537.1 | K9N5Q8 |
| FNKLLGYDPSKP | 1 | 0 | 0 | 1435054 | AVV62535.1 | K9N7C7 |
| FNKPLIEWPLIE | 1 | 0 | 0 | 1435058 | QBF80541.1 | K9N7C7 |
| FNKQWPMPINAS | 1 | 0 | 0 | 1435063 | AVV62537.1 | K9N5Q8 |
| FNKSKQFVPVQV | 1 | 0 | 0 | 1435066 | AGT17757.1 | A0A140H1H0 |
| FNKWQWQEAWYE | 1 | 0 | 0 | 1435072 | ARU07575.1 | A0A140H1G9 |
| FNLEDFIDSPCY | 1 | 0 | 0 | 1435080 | AVV62535.1 | K9N7C7 |
| FNMAQAFGLRGP | 1 | 0 | 0 | 1435113 | AXG21672.1 | K9N4V7 |
| FNMQTVSFHEDR | 1 | 0 | 0 | 1435120 | AVV62524.1 | K9N7C7 |
| FNNALDAIQEGF | 1 | 0 | 0 | 1435132 | CAA79896.1 | P36334 |
| FNNALGAIQEGF | 1 | 0 | 0 | 1435133 | AVI15011.1 | P36334 |
| FNNALHAIQQGS | 1 | 0 | 0 | 1435135 | ARB66064.1 | P36334 |
| FNNALHSIQQGF | 1 | 0 | 0 | 1435136 | AIX10759.1 | P36334 |
| FNNALNAIQQGF | 1 | 0 | 0 | 1435137 | AGT51491.1 | P36334 |
| FNNGFVVRIGAA | 1 | 0 | 0 | 1435144 | AVV62537.1 | K9N5Q8 |
| FNNGVYISTTKV | 1 | 0 | 0 | 1435146 | AGT51558.1 | P0C6X6 |

| | | | | | | |
|---|---|---|---|---|---|---|
| FNNKNTGPPFIS | 1 | 0 | 0 | 1435148 | AKA09659.1 | P36334 |
| FNNLLGFDFSKP | 1 | 0 | 0 | 1435149 | AVN89352.1 | K9N7C7 |
| FNNLLGFDSSKS | 1 | 0 | 0 | 1435151 | AVV61898.1 | K9N7C7 |
| FNNMLGYNPSKP | 1 | 0 | 0 | 1435153 | AVV62524.1 | K9N7C7 |
| FNNNTDSIIIDG | 1 | 0 | 0 | 1435154 | AGT51351.1 | H9EJ42 |
| FNNRDTGHPSIS | 1 | 0 | 0 | 1435157 | AWW13566.1 | P36334 |
| FNNRDTGPLSIS | 1 | 0 | 0 | 1435159 | ANZ78836.1 | P36334 |
| FNNRDTGPPSIS | 1 | 0 | 0 | 1435160 | AEN19366.1 | P36334 |
| FNNWFLLTNDST | 1 | 0 | 0 | 1435167 | ARE29967.1 | A0A0P0G321 |
| FNNYDCVITTLI | 1 | 0 | 0 | 1435170 | QBP84737.1 | P0C6X6 |
| FNPEPITMANAR | 1 | 0 | 0 | 1435176 | AVV62537.1 | K9N5Q8 |
| FNPHHRFKCSSN | 1 | 0 | 0 | 1435184 | AGT17757.1 | A0A140H1H0 |
| FNPNVPLDYQAT | 1 | 0 | 0 | 1435185 | ATO98145.1 | Q3LZX1 |
| FNPNVPLEYQAT | 1 | 0 | 0 | 1435186 | AVP78031.1 | Q3LZX1 |
| FNQALEAMQTGF | 1 | 0 | 0 | 1435198 | QBF80499.1 | K9N5Q8 |
| FNQAPGAMQTGF | 1 | 0 | 0 | 1435200 | AVV62526.1 | K9N5Q8 |
| FNQDFYDFVVAK | 1 | 0 | 0 | 1435201 | AUM60013.1 | K9N7C7 |
| FNQGGIIFNAVD | 1 | 0 | 0 | 1435205 | ABD43183.1 | P36334 |
| FNQVLSSDVMFE | 1 | 0 | 0 | 1435212 | AXN92247.1 | K9N7C7 |
| FNQYYYNMATGF | 1 | 0 | 0 | 1435217 | AZS52617.1 | A0A140H1H0 |
| FNRCCDRYEEYD | 1 | 0 | 0 | 1435222 | ALA49473.1 | K9N5Q8 |
| FNRDGDEQWGST | 1 | 0 | 0 | 1435223 | AVV62524.1 | K9N7C7 |
| FNRSSFVDDNKF | 1 | 0 | 0 | 1435236 | AGT51657.1 | P0C6X6 |
| FNSCQNAVSSFI | 1 | 0 | 0 | 1435242 | AVV62524.1 | K9N7C7 |
| FNSNLTGYSSRN | 1 | 0 | 0 | 1435250 | APO40579.1 | P59594 |
| FNSYGNMFQTQA | 1 | 0 | 0 | 1435262 | APO40579.1 | P59594 |
| FNTEFASNSSVL | 1 | 0 | 0 | 1435263 | ANI69888.1 | K9N638 |
| FNTLFSTTIPNT | 1 | 0 | 0 | 1435266 | AIW52776.1 | A0A0P0LKV0 |
| FNVAITRAKVGI | 1 | 0 | 0 | 1435280 | APO40578.1 | P0C6X7 |
| FNVDWVFFGFSI | 1 | 0 | 0 | 1435286 | ARE29965.1 | U3M6Q8 |
| FNVKKLQKQLYD | 1 | 0 | 0 | 1435301 | ALJ99896.1 | A0A0P0LKV0 |
| FNVNSLAYYEPV | 1 | 0 | 0 | 1435307 | AVV62535.1 | K9N7C7 |
| FNVSKGIYQTSN | 1 | 0 | 0 | 1435316 | ATO98181.1 | Q3LZX1 |
| FNVTTAETTDFL | 1 | 0 | 0 | 1435327 | CAA71147.1 | A0A0P0K6L9 |
| FNVVGPRTGKHE | 1 | 0 | 0 | 1435330 | AWK59942.1 | H9EJ42 |
| FNVVPYYSWFSG | 1 | 0 | 0 | 1435334 | 4LM9_A | P33469 |
| FNVWNQVVTYPS | 1 | 0 | 0 | 1435337 | AVV62524.1 | K9N7C7 |
| FNYFKKVDGVVQ | 1 | 0 | 0 | 1435349 | APO40578.1 | P0C6X7 |
| FPCAGRRVTFKE | 1 | 0 | 0 | 1435379 | AEN19363.1 | P0C6X6 |
| FPDGQGVPIAYG | 1 | 0 | 0 | 1435385 | AMM42397.1 | Q19U29 |
| FPEWIDDLEADP | 1 | 0 | 0 | 1435396 | ATQ39389.1 | K9N7C7 |
| FPFAESTGTQSV | 1 | 0 | 0 | 1435397 | QBF80599.1 | K9N796 |
| FPGPSGIGAVGG | 1 | 0 | 0 | 1435403 | ANI69896.1 | K9N4V7 |
| FPIDFPSAESTG | 1 | 0 | 0 | 1435407 | AZK15901.1 | K9N796 |
| FPIVMNDKNICL | 1 | 0 | 0 | 1435413 | ARU07575.1 | A0A140H1G9 |
| FPKHGDLGEQYI | 1 | 0 | 0 | 1435417 | AVV62537.1 | K9N5Q8 |
| FPKSDGTGTIYA | 1 | 0 | 0 | 1435422 | AEA10817.1 | P0C6X7 |
| FPKSGANFRDLA | 1 | 0 | 0 | 1435424 | BBA20986.1 | A0A140H1H1 |
| FPKSGANFTDLA | 1 | 0 | 0 | 1435425 | AZS52618.1 | A0A140H1H1 |
| FPLKLAGTAVINLRAD | 1 | 0 | 0 | 1435437 | YP_009555238.1 | P0C6X6 |
| FPLTFPPGQGVP | 1 | 0 | 0 | 1435456 | AVV62544.1 | K9N4V7 |
| FPNITNLCPFGQ | 1 | 0 | 0 | 1435464 | APO40579.1 | P59594 |
| FPNITNVCPFHK | 1 | 0 | 0 | 1435466 | AVP78031.1 | Q3LZX1 |
| FPNSNYVLQDFD | 1 | 0 | 0 | 1435469 | AFV53147.1 | H9EJ42 |
| FPPGQGVPLNSN | 1 | 0 | 0 | 1435474 | AHI48825.1 | K9N4V7 |
| FPPGTILPQGYY | 1 | 0 | 0 | 1435475 | AOL02457.1 | P33469 |
| FPPGTVLPQGYY | 1 | 0 | 0 | 1435477 | AEN19370.1 | P33469 |
| FPPSRNWDVRSG | 1 | 0 | 0 | 1435481 | AVV62540.1 | K9N643 |
| FPPSRNWHVSSG | 1 | 0 | 0 | 1435482 | AVV62529.1 | K9N643 |
| FPPWTAAAGAPF | 1 | 0 | 0 | 1435488 | ARB66070.1 | P36334 |
| FPPWTAAAGVPL | 1 | 0 | 0 | 1435490 | ARB66061.1 | P36334 |
| FPPWTAAEGVPF | 1 | 0 | 0 | 1435491 | AIL49550.1 | P36334 |
| FPQFAELIPNQA | 1 | 0 | 0 | 1435494 | AGT51335.1 | H9EJ69 |
| FPQFTSDQAITF | 1 | 0 | 0 | 1435495 | ABD75596.1 | Q0ZJ82 |
| FPREGVFVMNGT | 1 | 0 | 0 | 1435506 | APO40579.1 | P59594 |
| FPSAKVEGCMVQ | 1 | 0 | 0 | 1435515 | 2PWX_A | P0C6X7 |
| FPSGQGVPLNAN | 1 | 0 | 0 | 1435520 | AZK15907.1 | K9N4V7 |
| FPTTVTCENLES | 1 | 0 | 0 | 1435537 | NP_073549.1 | A0A0P0LKV0 |
| FPTWSQELPFDV | 1 | 0 | 0 | 1435541 | AEN19355.1 | P0C6X6 |
| FPVAKSVKQLFY | 1 | 0 | 0 | 1435543 | ARU07574.1 | A0A140H1G9 |
| FPYDIEFVTGKQ | 1 | 0 | 0 | 1435553 | ALA50044.1 | K9N7C7 |
| FPYDMDFGVSSF | 1 | 0 | 0 | 1435556 | AHY22554.1 | K9N638 |
| FPYDMDSGGSSF | 1 | 0 | 0 | 1435557 | AHX71940.1 | K9N7C7 |
| FPYDPALETGDF | 1 | 0 | 0 | 1435561 | AVV62535.1 | K9N7C7 |
| FPYDPALETGEY | 1 | 0 | 0 | 1435562 | AVV62524.1 | K9N7C7 |
| FPYQGDHGDMYV | 1 | 0 | 0 | 1435568 | AXN92228.1 | K9N5Q8 |
| FPYQGDHGDRYV | 1 | 0 | 0 | 1435569 | ALA49803.1 | K9N5Q8 |
| FQASADWKPGHA | 1 | 0 | 0 | 1435575 | ALA49945.1 | K9N7C7 |
| FQAVSSNINCIL | 1 | 0 | 0 | 1435576 | AIW52766.1 | A0A0P0LKV0 |
| FQDELDDFFKNV | 1 | 0 | 0 | 1435584 | AMW90852.1 | K9N5Q8 |
| FQDELDEFFKNF | 1 | 0 | 0 | 1435585 | AKM76239.1 | K9N5Q8 |
| FQDRQIHQAQLT | 1 | 0 | 0 | 1435587 | AVV62524.1 | K9N7C7 |
| FQFATFPVYDTI | 1 | 0 | 0 | 1435606 | ALA49363.1 | K9N5Q8 |
| FQFMLWCNDEKV | 1 | 0 | 0 | 1435618 | AEN19363.1 | P0C6X6 |
| FQFMLWGNDEKV | 1 | 0 | 0 | 1435621 | AIV41912.1 | P0C6X6 |
| FQGAMAEKLNAF | 1 | 0 | 0 | 1435624 | 3EJG_A | A0A0P0LKV0 |
| FQGGGGSGYIPE | 1 | 0 | 0 | 1435626 | 6NB6_A | P59594 |
| FQGIETAVGHYV | 1 | 0 | 0 | 1435628 | AXN92247.1 | K9N7C7 |
| FQGPGHHHHHHH | 1 | 0 | 0 | 1435634 | 6CRV_A | P59594 |
| FQGRQLMGSCSF | 1 | 0 | 0 | 1435637 | AIV41816.1 | P0C6X6 |
| FQGYLQKHFSMM | 1 | 0 | 0 | 1435639 | AIG96641.1 | A0A0P0LKV0 |

| | | | | | | |
|---|---|---|---|---|---|---|
| FQHKLKKACCKT | 1 | 0 | 0 | 1435642 | AEN19363.1 | P0C6X6 |
| FQILDEGLDCKI | 1 | 0 | 0 | 1435647 | ARK08668.1 | Q80872 |
| FQKAGQSTYERH | 1 | 0 | 0 | 1435653 | APO40578.1 | P0C6X7 |
| FQKGKEFEFAEG | 1 | 0 | 0 | 1435660 | AEN19370.1 | P33469 |
| FQLDTAPQTGNF | 1 | 0 | 0 | 1435675 | ATO98193.1 | Q3LZX1 |
| FQLTFESTINKS | 1 | 0 | 0 | 1435686 | AGT51388.1 | H9EJ24 |
| FQLTLESTINKS | 1 | 0 | 0 | 1435687 | AVA26874.1 | H9EJ24 |
| FQLTPIALKMTK | 1 | 0 | 0 | 1435688 | AVP78050.1 | Q3LZX3 |
| FQMVKEYPLLTG | 1 | 0 | 0 | 1435693 | AET99208.1 | Q0ZJ80 |
| FQNASKVIPFNS | 1 | 0 | 0 | 1435694 | APO40580.1 | P59632 |
| FQNCTAVGLRQQ | 1 | 0 | 0 | 1435696 | AUM60014.1 | K9N5Q8 |
| FQNCTAVGVPQQ | 1 | 0 | 0 | 1435698 | AVN89313.1 | K9N5Q8 |
| FQNCTAVGVRQQ | 1 | 0 | 0 | 1435699 | AXN92228.1 | K9N5Q8 |
| FQNLTNNLPPPL | 1 | 0 | 0 | 1435704 | AUM60024.1 | K9N5Q8 |
| FQQFGKDASDFI | 1 | 0 | 0 | 1435714 | AVP78031.1 | Q3LZX1 |
| FQQFGRDTSDFT | 1 | 0 | 0 | 1435716 | ATO98181.1 | Q3LZX1 |
| FQSFQQFGKDAS | 1 | 0 | 0 | 1435727 | AVP78031.1 | Q3LZX1 |
| FQTIPTQLTPIM | 1 | 0 | 0 | 1435741 | AVV62524.1 | K9N7C7 |
| FQTIPTQLTPLM | 1 | 0 | 0 | 1435742 | AVV62535.1 | K9N7C7 |
| FQTVRSGNFNQD | 1 | 0 | 0 | 1435752 | AHI48750.1 | K9N7C7 |
| FRDAEGKMVPYC | 1 | 0 | 0 | 1435773 | AVV62524.1 | K9N7C7 |
| FRDLALKGSKYL | 1 | 0 | 0 | 1435780 | ACV87130.1 | A0A140H1H1 |
| FRDYFDDNTDSI | 1 | 0 | 0 | 1435789 | AWK59930.1 | H9EJ42 |
| FREELDQWFKNQ | 1 | 0 | 0 | 1435797 | AVQ05264.1 | P36334 |
| FRHSDSIVKPDM | 1 | 0 | 0 | 1435828 | ARU07581.1 | Q19U29 |
| FRIDFPFAESTG | 1 | 0 | 0 | 1435836 | QBF80599.1 | K9N796 |
| FRIDFPSAELTG | 1 | 0 | 0 | 1435837 | QBF80533.1 | K9N796 |
| FRIDFPSAESTG | 1 | 0 | 0 | 1435838 | AXN92229.1 | K9N796 |
| FRKACQNLDCNC | 1 | 0 | 0 | 1435852 | AWW13564.1 | Q80872 |
| FRKMAFPSAKVE | 1 | 0 | 0 | 1435853 | 2PWX_A | P0C6X7 |
| FRKSCQNLDCNC | 1 | 0 | 0 | 1435856 | AGT51688.1 | Q80872 |
| FRKVQDAVNNNA | 1 | 0 | 0 | 1435858 | AXN92228.1 | K9N5Q8 |
| FRPFNSLLCGDI | 1 | 0 | 0 | 1435904 | ABW71808.1 | H9EJ42 |
| FRPIQLVPAFPE | 1 | 0 | 0 | 1435906 | AVV62541.1 | K9N7D2 |
| FRRAFGEYNHVV | 1 | 0 | 0 | 1435914 | P0C6W6.1 | P0C6W2 |
| FRRRSVCSTGRC | 1 | 0 | 0 | 1435922 | AIV42008.1 | P0C6X6 |
| FRSDVLHLTQDY | 1 | 0 | 0 | 1435930 | ATO98181.1 | Q3LZX1 |
| FRSLPNEVTVAK | 1 | 0 | 0 | 1435936 | AVV62543.1 | K9N7A1 |
| FRTFPQQMFVTP | 1 | 0 | 0 | 1435948 | AVN89294.1 | K9N643 |
| FRTGVDPDLSAF | 1 | 0 | 0 | 1435950 | ASU90030.1 | K9N7C7 |
| FRTRKGKRVDLP | 1 | 0 | 0 | 1435952 | ALK28797.1 | A0A0N7J131 |
| FRTRKGKRVDLS | 1 | 0 | 0 | 1435953 | AIW52723.1 | A0A0N7J131 |
| FRVTSRVMFQPR | 1 | 0 | 0 | 1435977 | AIW52843.1 | A0A0P0G321 |
| FRYDNVSSVWPL | 1 | 0 | 0 | 1435987 | P30215.1 | P30215 |
| FSAAQAMEKVNE | 1 | 0 | 0 | 1436000 | CAA79896.1 | P36334 |
| FSAFYPIDLASQ | 1 | 0 | 0 | 1436003 | AJG44108.1 | K9N7A1 |
| FSAGSRSRASSG | 1 | 0 | 0 | 1436004 | AMK59681.1 | P33469 |
| FSANDVEIQSSG | 1 | 0 | 0 | 1436009 | AVV62524.1 | K9N7C7 |
| FSASLFKPVQLA | 1 | 0 | 0 | 1436013 | AKM76243.1 | K9N7D2 |
| FSAYSGDVPHYV | 1 | 0 | 0 | 1436027 | AUM60014.1 | K9N5Q8 |
| FSDDVIVHDVEP | 1 | 0 | 0 | 1436039 | AWK59942.1 | H9EJ42 |
| FSDDVIVYDVEP | 1 | 0 | 0 | 1436040 | AFD98798.1 | H9EJ42 |
| FSDGKMGRFFNY | 1 | 0 | 0 | 1436044 | AVN89387.1 | K9N5Q8 |
| FSDIKTHKSQPL | 1 | 0 | 0 | 1436047 | AVV62537.1 | K9N5Q8 |
| FSDPTTFGSSFT | 1 | 0 | 0 | 1436054 | AVN89383.1 | K9N643 |
| FSDSKFGTTEVK | 1 | 0 | 0 | 1436062 | AHI48527.1 | K9N638 |
| FSFFDCGVNALA | 1 | 0 | 0 | 1436082 | ARU07593.1 | H9EJ42 |
| FSFGTVNNFVKF | 1 | 0 | 0 | 1436089 | BAL45640.1 | A0A0P0K6L9 |
| FSFIMFVGHLIS | 1 | 0 | 0 | 1436093 | AMK59674.1 | P0C6X6 |
| FSFISDSNGVPV | 1 | 0 | 0 | 1436095 | AUM60013.1 | K9N7C7 |
| FSFLNLKLRAPM | 1 | 0 | 0 | 1436096 | AVK87459.1 | K9N7C7 |
| FSFVDDVNGMKV | 1 | 0 | 0 | 1436105 | AVV62535.1 | K9N7C7 |
| FSGITQFQKGKE | 1 | 0 | 0 | 1436120 | AEN19370.1 | P33469 |
| FSGNDVEIQSSG | 1 | 0 | 0 | 1436122 | AVV62535.1 | K9N7C7 |
| FSGNVDLEGLAS | 1 | 0 | 0 | 1436125 | AVV62524.1 | K9N7C7 |
| FSHQQLGGLHLM | 1 | 0 | 0 | 1436144 | APO40578.1 | P0C6X7 |
| FSHSANDPLLGT | 1 | 0 | 0 | 1436145 | AVV62524.1 | K9N7C7 |
| FSIETDIIDEVN | 1 | 0 | 0 | 1436149 | AAA45463.1 | A0A0N7J131 |
| FSIKANIKSDIL | 1 | 0 | 0 | 1436158 | AZS52617.1 | A0A140H1H0 |
| FSKKQLGLLQST | 1 | 0 | 0 | 1436178 | AXG21653.1 | K9N7C7 |
| FSKLLYGLKMSI | 1 | 0 | 0 | 1436184 | QDH43733.1 | U3M6Q8 |
| FSKPFYVPEHCQ | 1 | 0 | 0 | 1436188 | AVN89384.1 | K9N796 |
| FSKPIYVPEHCQ | 1 | 0 | 0 | 1436189 | QAT98905.1 | K9N796 |
| FSKQLQDSMNGA | 1 | 0 | 0 | 1436192 | APO40586.1 | P59595 |
| FSKVQDAVNANA | 1 | 0 | 0 | 1436198 | AVV62537.1 | K9N5Q8 |
| FSKYVTVQGPPG | 1 | 0 | 0 | 1436203 | AVV62524.1 | K9N7C7 |
| FSLDVSEKSGNFKHLREFVF | 1 | 0 | 0 | 1436214 | 6NB6_A | P59594 |
| FSLELQDIEEPC | 1 | 0 | 0 | 1436216 | Q3I5I9.1 | Q3LZX6 |
| FSLGDVEAVNFN | 1 | 0 | 0 | 1436228 | AST12964.1 | A0A0P0K6L9 |
| FSLGNVEAVNFN | 1 | 0 | 0 | 1436230 | AIW52757.1 | A0A0P0K6L9 |
| FSLHPVECDEEC | 1 | 0 | 0 | 1436231 | AXN92247.1 | K9N7C7 |
| FSLHPVECDEGC | 1 | 0 | 0 | 1436232 | ALA50033.1 | K9N7C7 |
| FSLSVLVTASSK | 1 | 0 | 0 | 1436267 | ANI69890.1 | K9N796 |
| FSNALDAIQEGF | 1 | 0 | 0 | 1436298 | CAA83660.1 | P36334 |
| FSNFILHDVNNG | 1 | 0 | 0 | 1436303 | BBA84101.1 | P36334 |
| FSNLDKLKFKEV | 1 | 0 | 0 | 1436308 | AVV62524.1 | K9N7C7 |
| FSNMVETVVGQL | 1 | 0 | 0 | 1436312 | AVN89363.1 | K9N7C7 |
| FSPESVFNVFGD | 1 | 0 | 0 | 1436335 | AFD98763.1 | H9EJ42 |
| FSPGQGVPLNAN | 1 | 0 | 0 | 1436339 | AHI48535.1 | K9N4V7 |
| FSPGTILPQGYY | 1 | 0 | 0 | 1436340 | AMM42397.1 | Q19U29 |
| FSPLLSGTPPQV | 1 | 0 | 0 | 1436350 | AXN92228.1 | K9N5Q8 |

| | | | | | | |
|---|---|---|---|---|---|---|
| FSPVLGCLGSDC | 1 | 0 | 0 | 1436355 | AHF54664.1 | P36334 |
| FSQDGDYFTRQL | 1 | 0 | 0 | 1436357 | AVV62537.1 | K9N5Q8 |
| FSQFDSLGASSN | 1 | 0 | 0 | 1436359 | AIX09796.1 | P0C6X6 |
| FSQILPDPSKPT | 1 | 0 | 0 | 1436365 | ATO98181.1 | Q3LZX1 |
| FSQILPDSSKPT | 1 | 0 | 0 | 1436366 | ARO76382.1 | P59594 |
| FSQILPSPLKPT | 1 | 0 | 0 | 1436367 | 5XJK_A | P59594 |
| FSQQGVFRYDNV | 1 | 0 | 0 | 1436370 | P30215.1 | P30215 |
| FSQSDSLGVSSN | 1 | 0 | 0 | 1436377 | QBP84719.1 | P0C6X6 |
| FSQVDLTEAICD | 1 | 0 | 0 | 1436379 | AAX85666.1 | P0C6X6 |
| FSQVKSDLVIDA | 1 | 0 | 0 | 1436381 | AEN19363.1 | P0C6X6 |
| FSRMTQSALRMR | 1 | 0 | 0 | 1436388 | AUM60014.1 | K9N5Q8 |
| FSRTLETAQYSV | 1 | 0 | 0 | 1436394 | AVP78041.1 | P0C6W2 |
| FSRTLYQPVYKI | 1 | 0 | 0 | 1436395 | AAV67329.1 | H9EJA2 |
| FSRTVDTAGQSI | 1 | 0 | 0 | 1436398 | APO40578.1 | P0C6X7 |
| FSSDVFKIGDVK | 1 | 0 | 0 | 1436405 | QDH43737.1 | H9EJ42 |
| FSSITIDKSAIP | 1 | 0 | 0 | 1436409 | BBA84104.1 | P36334 |
| FSSSRRGVYYND | 1 | 0 | 0 | 1436422 | ATO98193.1 | Q3LZX1 |
| FSSTFNVPMEKLKT | 1 | 0 | 0 | 1436426 | P0DTC1.1 | P0DTD1 |
| FSTCNSNANISM | 1 | 0 | 0 | 1436436 | AKA09625.1 | A0A0P0G321 |
| FSTCNSNANIST | 1 | 0 | 0 | 1436437 | AKA09623.1 | A0A0P0G321 |
| FSTCNSNANLSM | 1 | 0 | 0 | 1436438 | AVL25555.1 | A0A0P0G321 |
| FSTINALPEISA | 1 | 0 | 0 | 1436458 | ALW82729.1 | K9N7C7 |
| FSTMSQYSRSTR | 1 | 0 | 0 | 1436464 | AHZ64057.1 | K9N5Q8 |
| FSTRNSNASISM | 1 | 0 | 0 | 1436468 | AKA09629.1 | A0A0P0G321 |
| FSTTIPNTAFGP | 1 | 0 | 0 | 1436471 | AIW52776.1 | A0A0P0LKV0 |
| FSVEDVYAEVHA | 1 | 0 | 0 | 1436496 | ARB07605.1 | A0A140H1G9 |
| FSVHPVECDEYE | 1 | 0 | 0 | 1436500 | AVV62535.1 | K9N7C7 |
| FSVNDFICSQIS | 1 | 0 | 0 | 1436512 | AVN89334.1 | K9N5Q8 |
| FSVNDFTCRQIS | 1 | 0 | 0 | 1436513 | ATG84888.1 | K9N5Q8 |
| FSVNSDFCPCAD | 1 | 0 | 0 | 1436516 | BBA20986.1 | A0A140H1H1 |
| FSVPSPLNWERK | 1 | 0 | 0 | 1436520 | AMM42415.1 | A0A140H1H1 |
| FSVQEDGRIPND | 1 | 0 | 0 | 1436521 | ABE97137.1 | A0A0P0G321 |
| FSVQQDGRTPNG | 1 | 0 | 0 | 1436529 | AKA09627.1 | A0A0P0G321 |
| FSVTQEYIQTTI | 1 | 0 | 0 | 1436537 | AUM60014.1 | K9N5Q8 |
| FSVVLPGEVAKK | 1 | 0 | 0 | 1436540 | AVV62524.1 | K9N7C7 |
| FSWNAELYNLMG | 1 | 0 | 0 | 1436545 | AWW13571.1 | P0C6X6 |
| FSWNLILTVFIV | 1 | 0 | 0 | 1436547 | AWK59934.1 | U3M6Q8 |
| FSYPLSMKSDLG | 1 | 0 | 0 | 1436564 | AHX00711.1 | K9N5Q8 |
| FSYQEFNDLAHT | 1 | 0 | 0 | 1436567 | AVA26872.1 | H9EJ42 |
| FSYQEFNDLVHT | 1 | 0 | 0 | 1436569 | AMM42571.1 | H9EJ42 |
| FTALTQHGKETL | 1 | 0 | 0 | 1436580 | APO40586.1 | P59595 |
| FTARDGVRRGDN | 1 | 0 | 0 | 1436589 | AGV08389.1 | K9N638 |
| FTCASEYTGNYQCGHY | 1 | 0 | 0 | 1436594 | P0DTC1.1 | P0DTD1 |
| FTCIPTVGYSAA | 1 | 0 | 0 | 1436598 | AVV62530.1 | K9N7D2 |
| FTCISTVGYSAA | 1 | 0 | 0 | 1436599 | AVV62541.1 | K9N7D2 |
| FTCNNIDVAKIY | 1 | 0 | 0 | 1436602 | ABD43176.1 | P36334 |
| FTCNSNANISML | 1 | 0 | 0 | 1436603 | AKA09626.1 | A0A0P0G321 |
| FTCRQISPAAIA | 1 | 0 | 0 | 1436607 | ATG84888.1 | K9N5Q8 |
| FTCSQISPAAIS | 1 | 0 | 0 | 1436610 | AHL18090.1 | K9N5Q8 |
| FTCSQISPPAIA | 1 | 0 | 0 | 1436612 | AHI48528.1 | K9N5Q8 |
| FTCVPTVGYSAA | 1 | 0 | 0 | 1436614 | AUM60018.1 | K9N7D2 |
| FTDFHNYTGEGQ | 1 | 0 | 0 | 1436618 | AXX83296.1 | P30215 |
| FTDSVRDPQTLE | 1 | 0 | 0 | 1436638 | ATO98181.1 | Q3LZX1 |
| FTECAYTQGRSC | 1 | 0 | 0 | 1436649 | ATQ39389.1 | K9N7C7 |
| FTEDLLCVANDK | 1 | 0 | 0 | 1436650 | APT69868.1 | A0A0P0ILY9 |
| FTEFVGTQSIDM | 1 | 0 | 0 | 1436657 | ATG84733.1 | K9N638 |
| FTEHVTSVNLFD | 1 | 0 | 0 | 1436659 | ASU89978.1 | K9N796 |
| FTENKPSVEQRQ | 1 | 0 | 0 | 1436664 | AVP78041.1 | P0C6W2 |
| FTEQLGAPLGIT | 1 | 0 | 0 | 1436665 | AIW52843.1 | A0A0P0G321 |
| FTEQSVFKPQPA | 1 | 0 | 0 | 1436669 | CAA83661.1 | P36334 |
| FTEQSVFKPQPV | 1 | 0 | 0 | 1436670 | CAA83660.1 | P36334 |
| FTETITGFDLN | 1 | 0 | 0 | 1436675 | ATP16766.1 | P30215 |
| FTFKIRKNGRGG | 1 | 0 | 0 | 1436682 | AUM60013.1 | K9N7C7 |
| FTFLLRKNGRGG | 1 | 0 | 0 | 1436685 | AVV62535.1 | K9N7C7 |
| FTGCVGTWDKVA | 1 | 0 | 0 | 1436693 | AVV62535.1 | K9N7C7 |
| FTGDLSYTIKFS | 1 | 0 | 0 | 1436700 | QQY99387.1 | H9EJ42 |
| FTGNFPKSGANF | 1 | 0 | 0 | 1436702 | ABA60802.1 | A0A140H1H1 |
| FTGNLQSNHDLY | 1 | 0 | 0 | 1436705 | APO40578.1 | P0C6X7 |
| FTGTYKCPQTKS | 1 | 0 | 0 | 1436713 | P36334.1 | P36334 |
| FTGYFPKAGANF | 1 | 0 | 0 | 1436715 | BBA20983.1 | A0A140H1H1 |
| FTHHDVVYAQHC | 1 | 0 | 0 | 1436719 | CAA83660.1 | P36334 |
| FTHIPLDGLVRA | 1 | 0 | 0 | 1436720 | AUM60013.1 | K9N7C7 |
| FTIAGHEEFIQT | 1 | 0 | 0 | 1436727 | BBA20986.1 | A0A140H1H1 |
| FTIGNIEEFIQT | 1 | 0 | 0 | 1436730 | BBA20979.1 | P36334 |
| FTIGNMEEFIQT | 1 | 0 | 0 | 1436731 | CAA79896.1 | P36334 |
| FTIGNMGEFIQT | 1 | 0 | 0 | 1436732 | ARB66062.1 | P36334 |
| FTIGNMVEFIQT | 1 | 0 | 0 | 1436733 | P36334.1 | P36334 |
| FTILEQDRMPKT | 1 | 0 | 0 | 1436738 | AHF54653.1 | Q4VID0 |
| FTIPALQQAFIK | 1 | 0 | 0 | 1436745 | AVV62524.1 | K9N7C7 |
| FTIWEPDRMLKT | 1 | 0 | 0 | 1436750 | AVR40343.1 | Q4VID0 |
| FTIWEQDRMLKT | 1 | 0 | 0 | 1436751 | ATP16762.1 | Q4VID0 |
| FTKFFYVLGLFA | 1 | 0 | 0 | 1436756 | APO40578.1 | P0C6X7 |
| FTKLLSGTPPQV | 1 | 0 | 0 | 1436761 | AVV62537.1 | K9N5Q8 |
| FTKLQSLENVAY | 1 | 0 | 0 | 1436763 | AVP78041.1 | P0C6W2 |
| FTKPSSVKEMQS | 1 | 0 | 0 | 1436768 | ABE97141.1 | H9EJ69 |
| FTKVQGLENIAF | 1 | 0 | 0 | 1436776 | AXN92258.1 | K9N7C7 |
| FTKVQGLENIAY | 1 | 0 | 0 | 1436777 | AUM60023.1 | K9N7C7 |
| FTKVQGLENIVF | 1 | 0 | 0 | 1436778 | AXN92247.1 | K9N7C7 |
| FTLANVSSFGEY | 1 | 0 | 0 | 1436781 | ALK28787.1 | A0A0P0K6L9 |
| FTLCCTGCGTNC | 1 | 0 | 0 | 1436782 | ATG84866.1 | K9N5Q8 |
| FTLQAAFGNTGV | 1 | 0 | 0 | 1436820 | AWK59942.1 | H9EJ42 |

| | | | | | | |
|---|---|---|---|---|---|---|
| FTLRSITAQPVK | 1 | 0 | 0 | 1436825 | NP_828852.2 | P59632 |
| FTLWDYERSNIY | 1 | 0 | 0 | 1436832 | AUM60013.1 | K9N7C7 |
| FTLYGQTVVHFG | 1 | 0 | 0 | 1436839 | AIV41888.1 | P0C6X6 |
| FTMADGSLQPYC | 1 | 0 | 0 | 1436842 | ARU07583.1 | P0C6X6 |
| FTNFEPFTVNSK | 1 | 0 | 0 | 1436856 | AIL49538.1 | P36334 |
| FTNRFVPVGDSF | 1 | 0 | 0 | 1436871 | ATN39877.2 | P0C6X6 |
| FTPEDVNMQLMG | 1 | 0 | 0 | 1436879 | AQZ41284.1 | K9N7C7 |
| FTPFDVVAQCSG | 1 | 0 | 0 | 1436880 | 3M3T_A | P0C6X7 |
| FTPIDQYMCGKD | 1 | 0 | 0 | 1436883 | AVV62524.1 | K9N7C7 |
| FTPIDQYMCGSD | 1 | 0 | 0 | 1436884 | AUM60013.1 | K9N7C7 |
| FTPKKPNKPLSQ | 1 | 0 | 0 | 1436885 | ABE73425.1 | H9EJ69 |
| FTPLKVEEVQLQ | 1 | 0 | 0 | 1436889 | AUM60013.1 | K9N7C7 |
| FTPLSVEPTPIE | 1 | 0 | 0 | 1436892 | ATQ39389.1 | K9N7C7 |
| FTPRLFATKVQQ | 1 | 0 | 0 | 1436897 | APO40584.1 | P59635 |
| FTPRSVRSVPGE | 1 | 0 | 0 | 1436898 | AVK87439.1 | K9N5Q8 |
| FTPVNLNRGLPT | 1 | 0 | 0 | 1436900 | AVP78031.1 | Q3LZX1 |
| FTQDGNAAIPDY | 1 | 0 | 0 | 1436912 | ACV87150.1 | A0A140H1G9 |
| FTQSRNLQEFKP | 1 | 0 | 0 | 1436918 | QYC64967.1 | P0DTD1 |
| FTQWYHEHIGGV | 1 | 0 | 0 | 1436921 | AVN89333.1 | K9N7C7 |
| FTREMQQQPLLN | 1 | 0 | 0 | 1436927 | AIW52723.1 | A0A0N7J131 |
| FTRNYIITMPCF | 1 | 0 | 0 | 1436935 | AIY60561.1 | K9N643 |
| FTRQEEVQELYS | 1 | 0 | 0 | 1436938 | QVM71556.1 | P0DTC7 |
| FTRQLSDYEGGG | 1 | 0 | 0 | 1436939 | AVV62526.1 | K9N5Q8 |
| FTRQLSQLEGGG | 1 | 0 | 0 | 1436940 | AVV62537.1 | K9N5Q8 |
| FTSACTRLEGLG | 1 | 0 | 0 | 1436947 | NP_073549.1 | A0A0P0LKV0 |
| FTSDQAITFLKE | 1 | 0 | 0 | 1436955 | ABD75596.1 | Q0ZJ82 |
| FTSEPPITYSPA | 1 | 0 | 0 | 1436958 | QBF80553.1 | K9N7C7 |
| FTSEPPVTYSTA | 1 | 0 | 0 | 1436960 | AXN92247.1 | K9N7C7 |
| FTSYGFSNVVEM | 1 | 0 | 0 | 1436975 | AIW52757.1 | A0A0P0K6L9 |
| FTTADGSPQPYC | 1 | 0 | 0 | 1436977 | AGT51707.1 | P0C6X6 |
| FTTAQEGCGILS | 1 | 0 | 0 | 1436981 | APO40579.1 | P59594 |
| FTTQFVDNVKLA | 1 | 0 | 0 | 1436996 | AIW52747.1 | A0A0P0K6L9 |
| FTTTNVAFAKVQ | 1 | 0 | 0 | 1437007 | AUM60014.1 | K9N5Q8 |
| FTVALASYADVL | 1 | 0 | 0 | 1437013 | CAA71056.1 | A0A0P0K6L9 |
| FTVCADGFMPFL | 1 | 0 | 0 | 1437014 | AEN19363.1 | P0C6X6 |
| FTVDGVQYYCYT | 1 | 0 | 0 | 1437015 | AKI29254.1 | K9N638 |
| FTVDPKKAYMDF | 1 | 0 | 0 | 1437016 | AVV62524.1 | K9N7C7 |
| FTVDPQKAYMDF | 1 | 0 | 0 | 1437018 | AUM60013.1 | K9N7C7 |
| FTVLAAYNGPPQ | 1 | 0 | 0 | 1437028 | ABD75599.1 | A0A140H1G9 |
| FTVMNGEDAFSI | 1 | 0 | 0 | 1437030 | QEO75983.1 | A0A0P0LKV0 |
| FTVNSVNDSLEH | 1 | 0 | 0 | 1437033 | CAA83661.1 | P36334 |
| FTVNSVNDSLKP | 1 | 0 | 0 | 1437034 | AIL49500.1 | P36334 |
| FTVNSVSDSLEP | 1 | 0 | 0 | 1437035 | ARB66068.1 | P36334 |
| FTVNSVVQREGI | 1 | 0 | 0 | 1437037 | AVV62535.1 | K9N7C7 |
| FTVSTQPQENTI | 1 | 0 | 0 | 1437039 | ABD75589.1 | Q19U29 |
| FTVSTQPQGNII | 1 | 0 | 0 | 1437040 | ABA60796.1 | Q19U29 |
| FTYICYMIRDKL | 1 | 0 | 0 | 1437056 | AQN78757.1 | P0C6X6 |
| FVAAYNNCTGGA | 1 | 0 | 0 | 1437071 | AVQ05264.1 | P36334 |
| FVAFNVFQGIET | 1 | 0 | 0 | 1437080 | AXN92247.1 | K9N7C7 |
| FVAGNCDVVIGI | 1 | 0 | 0 | 1437085 | ATO98181.1 | Q3LZX1 |
| FVANYFQDVKQF | 1 | 0 | 0 | 1437090 | AID55096.1 | K9N5Q8 |
| FVAQQLVRSEAA | 1 | 0 | 0 | 1437095 | AUM60014.1 | K9N5Q8 |
| FVCASEYTGNYQ | 1 | 0 | 0 | 1437106 | AVP78041.1 | P0C6W2 |
| FVCIDNSANTKF | 1 | 0 | 0 | 1437116 | AUM60013.1 | K9N7C7 |
| FVCQDSTRHTYQ | 1 | 0 | 0 | 1437121 | APO40584.1 | P59635 |
| FVDDFQGYLQKH | 1 | 0 | 0 | 1437132 | AIG96641.1 | A0A0P0LKV0 |
| FVDDIVKTDGTL | 1 | 0 | 0 | 1437134 | AEA10817.1 | P0C6X7 |
| FVDGVPFVVSYG | 1 | 0 | 0 | 1437143 | ASU90459.1 | K9N7C7 |
| FVDHFGENEEIV | 1 | 0 | 0 | 1437144 | AST12962.1 | A0A0P0LKV0 |
| FVDIQTAQAAGT | 1 | 0 | 0 | 1437146 | 3ATW_A | P0C6X7 |
| FVDYLMTTIPKA | 1 | 0 | 0 | 1437167 | ARB07605.1 | A0A140H1G9 |
| FVEFMLWCKDGQ | 1 | 0 | 0 | 1437177 | AVV62524.1 | K9N7C7 |
| FVEGAINVGKNG | 1 | 0 | 0 | 1437178 | AUM60013.1 | K9N7C7 |
| FVEKDDQGHILD | 1 | 0 | 0 | 1437183 | AVV62535.1 | K9N7C7 |
| FVEKWESGVKDY | 1 | 0 | 0 | 1437186 | APO40580.1 | P59632 |
| FVELDDQGHHLD | 1 | 0 | 0 | 1437187 | AXN92247.1 | K9N7C7 |
| FVEPKDGGHFFV | 1 | 0 | 0 | 1437195 | ARE29978.1 | H9EJ42 |
| FVEPKNGGQFFV | 1 | 0 | 0 | 1437197 | ARU07593.1 | H9EJ42 |
| FVEQPQGVECDF | 1 | 0 | 0 | 1437199 | AUM60014.1 | K9N5Q8 |
| FVEVDDQGHHLD | 1 | 0 | 0 | 1437205 | AUM60013.1 | K9N7C7 |
| FVFDVDKVKTPH | 1 | 0 | 0 | 1437215 | AUM60013.1 | K9N7C7 |
| FVGGFRVLRTTD | 1 | 0 | 0 | 1437246 | AWK59942.1 | H9EJ42 |
| FVGTQALDMLAV | 1 | 0 | 0 | 1437257 | AUM60013.1 | K9N7C7 |
| FVIDTPTGPQIK | 1 | 0 | 0 | 1437275 | NP_073549.1 | A0A0P0LKV0 |
| FVKCNCGSENWS | 1 | 0 | 0 | 1437293 | AGT51379.1 | H9EJ42 |
| FVKDGFSFVNDD | 1 | 0 | 0 | 1437296 | AVV62524.1 | K9N7C7 |
| FVKDNVSFVVDD | 1 | 0 | 0 | 1437299 | AXN92247.1 | K9N7C7 |
| FVKGDFSFISDS | 1 | 0 | 0 | 1437304 | AUM60013.1 | K9N7C7 |
| FVKNIPCYASAV | 1 | 0 | 0 | 1437321 | QBP84719.1 | P0C6X6 |
| FVKNLNNLHRGQ | 1 | 0 | 0 | 1437323 | AXN92247.1 | K9N7C7 |
| FVKPNRTSVASF | 1 | 0 | 0 | 1437326 | AVV62535.1 | K9N7C7 |
| FVKQPVKFYLYK | 1 | 0 | 0 | 1437331 | AVN89451.1 | K9N7C7 |
| FVKRRADFACKF | 1 | 0 | 0 | 1437332 | ARU07575.1 | A0A140H1G9 |
| FVKWEYEGGCNT | 1 | 0 | 0 | 1437340 | AWK59942.1 | H9EJ42 |
| FVLKNGCALPFE | 1 | 0 | 0 | 1437359 | AVV62524.1 | K9N7C7 |
| FVLLFFICCCTG | 1 | 0 | 0 | 1437362 | AIX10760.1 | P36334 |
| FVLLRFYIVVTA | 1 | 0 | 0 | 1437364 | ARU07575.1 | A0A140H1G9 |
| FVLNNTTTRDVV | 1 | 0 | 0 | 1437367 | AVV62537.1 | K9N5Q8 |
| FVLPSDDTLRTE | 1 | 0 | 0 | 1437373 | APO40578.1 | P0C6X7 |
| FVLSDIVNGVAE | 1 | 0 | 0 | 1437378 | APO40578.1 | P0C6X7 |
| FVLTSHTVMPLT | 1 | 0 | 0 | 1437384 | ATO98130.1 | P0C6W2 |

| | | | | | | |
|---|---|---|---|---|---|---|
| FVMMSALPAEYK | 1 | 0 | 0 | 1437399 | ARO76381.1 | P0C6U8 |
| FVNAGGVPLTNC | 1 | 0 | 0 | 1437415 | AHI48749.1 | K9N638 |
| FVNDDTGVQVVD | 1 | 0 | 0 | 1437416 | AVV62524.1 | K9N7C7 |
| FVNEYYEFLFFH | 1 | 0 | 0 | 1437422 | AIG96649.1 | A0A140H1G9 |
| FVNFNVDWIFFG | 1 | 0 | 0 | 1437423 | AWK59934.1 | U3M6Q8 |
| FVNIDNSANTKT | 1 | 0 | 0 | 1437427 | AVV62524.1 | K9N7C7 |
| FVNIGVWISPPV | 1 | 0 | 0 | 1437429 | AVV62539.1 | K9N4V0 |
| FVNMVKVPYVCL | 1 | 0 | 0 | 1437435 | AVV62524.1 | K9N7C7 |
| FVNQVKVPYVCL | 1 | 0 | 0 | 1437436 | AVV62535.1 | K9N7C7 |
| FVNSGGTPLTNC | 1 | 0 | 0 | 1437439 | AUM60013.1 | K9N7C7 |
| FVNTTIGNETTF | 1 | 0 | 0 | 1437442 | ABB90514.1 | A0A0P0K6L9 |
| FVNVPYVYNGSV | 1 | 0 | 0 | 1437450 | AGT51729.1 | P30215 |
| FVNVSYFYNGSA | 1 | 0 | 0 | 1437452 | ARK08634.1 | P30215 |
| FVNVSYVYNGSA | 1 | 0 | 0 | 1437453 | AEN19365.1 | P30215 |
| FVPATRDTVPLH | 1 | 0 | 0 | 1437458 | AUM60018.1 | K9N7D2 |
| FVPQPTGVFTNH | 1 | 0 | 0 | 1437465 | AEN19366.1 | P36334 |
| FVPRAKSIVSTI | 1 | 0 | 0 | 1437468 | AUM60013.1 | K9N7C7 |
| FVPVQVVDSRWN | 1 | 0 | 0 | 1437471 | ARU07576.1 | A0A140H1H0 |
| FVQDPSPIDIDA | 1 | 0 | 0 | 1437473 | AFD98763.1 | H9EJ42 |
| FVQEQIGSFIVN | 1 | 0 | 0 | 1437476 | AVV62531.1 | K9N5R3 |
| FVQERVGLFIVN | 1 | 0 | 0 | 1437481 | AGV08472.1 | K9N5R3 |
| FVQIENCDVTFV | 1 | 0 | 0 | 1437482 | BAL45640.1 | A0A0P0K6L9 |
| FVQIPTTCANDP | 1 | 0 | 0 | 1437486 | 2XYV_B | P0C6U8 |
| FVQPAIYVYNTG | 1 | 0 | 0 | 1437487 | AVV62531.1 | K9N5R3 |
| FVQQQIGSFIVN | 1 | 0 | 0 | 1437489 | AUM60019.1 | K9N5R3 |
| FVQTCDLTVGSG | 1 | 0 | 0 | 1437494 | AAB27260.2 | P36334 |
| FVQWFHENVGGN | 1 | 0 | 0 | 1437500 | AVV62535.1 | K9N7C7 |
| FVRCEEDGQTVF | 1 | 0 | 0 | 1437507 | AVV62535.1 | K9N7C7 |
| FVRKAEDIFCPC | 1 | 0 | 0 | 1437519 | AST12962.1 | A0A0P0LKV0 |
| FVRKASGLFDAI | 1 | 0 | 0 | 1437521 | AWK59942.1 | H9EJ42 |
| FVRPIIEDYHTL | 1 | 0 | 0 | 1437523 | AJC98130.1 | Q01455 |
| FVRSVEDGDTVY | 1 | 0 | 0 | 1437528 | AUM60023.1 | K9N7C7 |
| FVRVSTCSSQGI | 1 | 0 | 0 | 1437530 | AUM60018.1 | K9N7D2 |
| FVSAYNNCTGGA | 1 | 0 | 0 | 1437536 | AIL49488.1 | P36334 |
| FVSDADITLLGD | 1 | 0 | 0 | 1437538 | ALL26405.1 | K9N7C7 |
| FVSKDVKFENLT | 1 | 0 | 0 | 1437557 | AUM60014.1 | K9N5Q8 |
| FVSTFNVPMEKL | 1 | 0 | 0 | 1437576 | APO40578.1 | P0C6X7 |
| FVTQRNFYEPKI | 1 | 0 | 0 | 1437606 | AVP78031.1 | Q3LZX1 |
| FVVAFLATIKLC | 1 | 0 | 0 | 1437618 | AGT17765.1 | Q0ZJ83 |
| FVVAYPFCIAPY | 1 | 0 | 0 | 1437619 | AUM60013.1 | K9N7C7 |
| FVVESGGYIPSN | 1 | 0 | 0 | 1437626 | AGT21353.1 | A0A0P0K6L9 |
| FVVKRHTLSNYQ | 1 | 0 | 0 | 1437637 | APO40578.1 | P0C6X7 |
| FVVNDKKLFHQL | 1 | 0 | 0 | 1437644 | APO40578.1 | P0C6X7 |
| FVVQIPTNFTFG | 1 | 0 | 0 | 1437649 | AVV62526.1 | K9N5Q8 |
| FVVQTEGCVSTN | 1 | 0 | 0 | 1437650 | AWK59942.1 | H9EJ42 |
| FVVRIGAAANAT | 1 | 0 | 0 | 1437653 | AVK87439.1 | K9N5Q8 |
| FVYDAYHNLVGY | 1 | 0 | 0 | 1437684 | ALA49649.1 | K9N5Q8 |
| FVYDGFDNLIGY | 1 | 0 | 0 | 1437686 | AVV62537.1 | K9N5Q8 |
| FVYDGFDNLVGY | 1 | 0 | 0 | 1437687 | AVV62526.1 | K9N5Q8 |
| FVYENAFMPFAM | 1 | 0 | 0 | 1437693 | AVP78041.1 | P0C6W2 |
| FVYFNGTGRGAC | 1 | 0 | 0 | 1437696 | AIW52757.1 | A0A0P0K6L9 |
| FVYTDTEVCKVK | 1 | 0 | 0 | 1437701 | NP_073549.1 | A0A0P0LKV0 |
| FVYVIKMIILWL | 1 | 0 | 0 | 1437707 | AWW13547.1 | Q01455 |
| FVYVRKDGKHVD | 1 | 0 | 0 | 1437711 | AST12962.1 | A0A0P0LKV0 |
| FVYYVARLMKGD | 1 | 0 | 0 | 1437712 | NP_073549.1 | A0A0P0LKV0 |
| FWAWNPEVNAIT | 1 | 0 | 0 | 1437717 | AST12967.1 | A0A0P0KXN3 |
| FWPKQETSTQVE | 1 | 0 | 0 | 1437748 | AUM60013.1 | K9N7C7 |
| FWRYDFDITESK | 1 | 0 | 0 | 1437757 | AWK59942.1 | H9EJ42 |
| FWSSTMIFSLHP | 1 | 0 | 0 | 1437759 | AVN89333.1 | K9N7C7 |
| FWVASNQADVNI | 1 | 0 | 0 | 1437763 | AIX10718.1 | P33469 |
| FWVASNQADVNT | 1 | 0 | 0 | 1437764 | ATP16771.1 | P33469 |
| FYAPEPITSFNT | 1 | 0 | 0 | 1437783 | AVN89387.1 | K9N5Q8 |
| FYAYFPDTGFVT | 1 | 0 | 0 | 1437795 | AMM42466.1 | P36334 |
| FYCIIEPRNGSR | 1 | 0 | 0 | 1437806 | AUM60014.1 | K9N5Q8 |
| FYCSNSSSSVLD | 1 | 0 | 0 | 1437811 | AVL25581.1 | A0A0P0G321 |
| FYCTNTTLVSGD | 1 | 0 | 0 | 1437812 | BAL45640.1 | A0A0P0K6L9 |
| FYDRDGQRIYDD | 1 | 0 | 0 | 1437822 | AEN19363.1 | P0C6X6 |
| FYEFYNDIKPPV | 1 | 0 | 0 | 1437835 | AOL02455.1 | Q04854 |
| FYEGVNFSPHHR | 1 | 0 | 0 | 1437838 | ABD75512.1 | A0A140H1H0 |
| FYENRFAAIYGG | 1 | 0 | 0 | 1437842 | AVA26874.1 | H9EJ24 |
| FYEPEPITSLNT | 1 | 0 | 0 | 1437843 | ALM26400.1 | K9N5Q8 |
| FYFAVRKEGEDV | 1 | 0 | 0 | 1437850 | AVQ05261.1 | P0C6X6 |
| FYFCQDPMLAVA | 1 | 0 | 0 | 1437852 | APO40579.1 | P59594 |
| FYFKPGGTTSGE | 1 | 0 | 0 | 1437861 | ADK37693.1 | H9EJ42 |
| FYGGWDDMLRPL | 1 | 0 | 0 | 1437877 | AHF54667.1 | P0C6X6 |
| FYGKDAFENHSY | 1 | 0 | 0 | 1437880 | AVV62524.1 | K9N7C7 |
| FYGKESFENPTY | 1 | 0 | 0 | 1437881 | AVN89322.1 | K9N7C7 |
| FYGKESLENPTY | 1 | 0 | 0 | 1437883 | AXN92247.1 | K9N7C7 |
| FYGKNAFQNHSY | 1 | 0 | 0 | 1437886 | AVV62535.1 | K9N7C7 |
| FYGPFVDIQTAQ | 1 | 0 | 0 | 1437888 | 3ATW_A | P0C6X7 |
| FYGVALKDCGFV | 1 | 0 | 0 | 1437892 | AEN19363.1 | P0C6X6 |
| FYIDVGNHRGAF | 1 | 0 | 0 | 1437901 | AKA09616.1 | A0A0P0G321 |
| FYIEDMASRFAD | 1 | 0 | 0 | 1437904 | ATG84699.1 | K9N7C7 |
| FYILPSILPNEK | 1 | 0 | 0 | 1437913 | AVP78041.1 | P0C6W2 |
| FYINGYRYFSLG | 1 | 0 | 0 | 1437917 | AIW52757.1 | A0A0P0K6L9 |
| FYIYKLHQLTYL | 1 | 0 | 0 | 1437925 | AVV62526.1 | K9N5Q8 |
| FYKETSYTSDIK | 1 | 0 | 0 | 1437932 | APO40578.1 | P0C6X7 |
| FYLFDDAGEENL | 1 | 0 | 0 | 1437947 | ARO76381.1 | P0C6U8 |
| FYLGDISHLINC | 1 | 0 | 0 | 1437948 | QEG03730.1 | H9EJ42 |
| FYLGDISHLVNC | 1 | 0 | 0 | 1437949 | AWK59942.1 | H9EJ42 |
| FYLLGCDEYIVP | 1 | 0 | 0 | 1437954 | QBP84758.1 | P30215 |

| | | | | | | |
|---|---|---|---|---|---|---|
| FYLTMYIIPVLC | 1 | 0 | 0 | 1437964 | ARU07575.1 | A0A140H1G9 |
| FYNDIKPPVLDV | 1 | 0 | 0 | 1437976 | AOL02455.1 | Q04854 |
| FYNLWSTFTKLQ | 1 | 0 | 0 | 1437986 | QBP84746.1 | P0C6X6 |
| FYPACTNGHALP | 1 | 0 | 0 | 1437997 | ALL26408.1 | K9N638 |
| FYPACTNGHAVP | 1 | 0 | 0 | 1437998 | AXN92247.1 | K9N7C7 |
| FYPNVPIEYQAT | 1 | 0 | 0 | 1438010 | ATO98120.1 | Q3LZX1 |
| FYPPDEEEDCEE | 1 | 0 | 0 | 1438013 | P0C6W6.1 | P0C6W2 |
| FYPPEDEGEDDC | 1 | 0 | 0 | 1438016 | AVP78041.1 | P0C6W2 |
| FYQLYDASNVKK | 1 | 0 | 0 | 1438034 | AWW13571.1 | P0C6X6 |
| FYRELYEAPTTL | 1 | 0 | 0 | 1438042 | AVV62524.1 | K9N7C7 |
| FYRLANGCAQVL | 1 | 0 | 0 | 1438050 | AJE26242.1 | P0C6X6 |
| FYRMNGIGITQQ | 1 | 0 | 0 | 1438055 | AVV62537.1 | K9N5Q8 |
| FYRPANECAQVL | 1 | 0 | 0 | 1438056 | AIL49562.1 | P0C6X6 |
| FYSPEPITSLNT | 1 | 0 | 0 | 1438080 | AHI48672.1 | K9N5Q8 |
| FYSPQIITTDNT | 1 | 0 | 0 | 1438081 | ATO98181.1 | Q3LZX1 |
| FYSPQVITTDNT | 1 | 0 | 0 | 1438083 | APO40579.1 | P59594 |
| FYSSKQSVASVI | 1 | 0 | 0 | 1438085 | APO40578.1 | P0C6X7 |
| FYSTSPIQPAEL | 1 | 0 | 0 | 1438086 | ADK37704.1 | A0A0P0K6L9 |
| FYSTSSIQPVEL | 1 | 0 | 0 | 1438089 | AMM42536.1 | A0A0P0K6L9 |
| FYVGVQSIEELH | 1 | 0 | 0 | 1438110 | AUM60013.1 | K9N7C7 |
| FYVKDGKYFISE | 1 | 0 | 0 | 1438114 | ALA49340.1 | K9N7C7 |
| FYVKDGKYFTSA | 1 | 0 | 0 | 1438115 | AVK87459.1 | K9N7C7 |
| FYVKDGKYFTSP | 1 | 0 | 0 | 1438117 | AUM60013.1 | K9N7C7 |
| FYVLPNDDTLRA | 1 | 0 | 0 | 1438122 | AVP78030.1 | P0C6W2 |
| FYVYANGGKGFC | 1 | 0 | 0 | 1438137 | AVP78041.1 | P0C6W2 |
| FYYLGTGPHAND | 1 | 0 | 0 | 1438154 | AJQ24067.1 | P33469 |
| FYYLGTGPYANA | 1 | 0 | 0 | 1438162 | ARU07581.1 | Q19U29 |
| FYYRLFRHGKIK | 1 | 0 | 0 | 1438165 | APO40579.1 | P59594 |
| GAAALEDEATPF | 1 | 0 | 0 | 1438177 | 3EA9_A | P0C6U8 |
| GAAANSTGNLII | 1 | 0 | 0 | 1438181 | AVV62526.1 | K9N5Q8 |
| GACIWNAAVYMK | 1 | 0 | 0 | 1438200 | QFQ59585.1 | K9N7C7 |
| GACKGFYSNSSS | 1 | 0 | 0 | 1438202 | AIW52747.1 | A0A0P0K6L9 |
| GADLKSPDLGDE | 1 | 0 | 0 | 1438212 | APO40578.1 | P0C6X7 |
| GAEIKDLICVQS | 1 | 0 | 0 | 1438232 | AGT51590.1 | P36334 |
| GAEIRDLICVQS | 1 | 0 | 0 | 1438233 | CAA79896.1 | P36334 |
| GAEKEDGFYETI | 1 | 0 | 0 | 1438235 | AMK59675.1 | Q80872 |
| GAENKDGFYETI | 1 | 0 | 0 | 1438239 | AIV41955.1 | Q80872 |
| GAENSIAYANNS | 1 | 0 | 0 | 1438240 | AVP78031.1 | Q3LZX1 |
| GAFLNGSCGSVG | 1 | 0 | 0 | 1438260 | 3F9E_A | P0C6U8 |
| GAFTRYVDNNFC | 1 | 0 | 0 | 1438266 | ATO98229.1 | P0C6W2 |
| GAGCFVNDLLKT | 1 | 0 | 0 | 1438271 | ANZ78855.1 | P0C6X6 |
| GAGDQAAKLSPW | 1 | 0 | 0 | 1438277 | AVV62524.1 | K9N7C7 |
| GAGGHCYGADLK | 1 | 0 | 0 | 1438280 | APO40578.1 | P0C6X7 |
| GAGIPIVDSYYS | 1 | 0 | 0 | 1438290 | APO40578.1 | P0C6X7 |
| GAGIRDLICVQS | 1 | 0 | 0 | 1438291 | CAA83661.1 | P36334 |
| GAGLKLPSGIMF | 1 | 0 | 0 | 1438292 | NP_073549.1 | A0A0P0LKV0 |
| GAGSDKGVAPGS | 1 | 0 | 0 | 1438301 | APO40578.1 | P0C6X7 |
| GAGWTAGLSSSA | 1 | 0 | 0 | 1438307 | AOR17480.1 | K9N5Q8 |
| GAHAFRNACGTN | 1 | 0 | 0 | 1438309 | AXN92247.1 | K9N7C7 |
| GAHAYETMKPHV | 1 | 0 | 0 | 1438312 | AVV62535.1 | K9N7C7 |
| GAIGDVGLCMYR | 1 | 0 | 0 | 1438325 | ARB07424.1 | A0A0P0ILY9 |
| GAIGSSLQEILS | 1 | 0 | 0 | 1438326 | CAA83660.1 | P36334 |
| GAIIRDGDVVKQ | 1 | 0 | 0 | 1438327 | ALA49648.1 | K9N7C7 |
| GAIIRDSEAIKQ | 1 | 0 | 0 | 1438330 | AVV62524.1 | K9N7C7 |
| GAILSNLGVPSY | 1 | 0 | 0 | 1438338 | APO40578.1 | P0C6X7 |
| GAILSTMTGRMF | 1 | 0 | 0 | 1438339 | AJS10812.1 | P0C6X6 |
| GAINVGKNGLQF | 1 | 0 | 0 | 1438341 | AUM60013.1 | K9N7C7 |
| GAINVGKQGLEF | 1 | 0 | 0 | 1438342 | AVV62524.1 | K9N7C7 |
| GAINVGKQGLEY | 1 | 0 | 0 | 1438343 | AVV62535.1 | K9N7C7 |
| GAIRISNDTFLN | 1 | 0 | 0 | 1438347 | ADK37703.1 | A0A0P0K6L9 |
| GAIRSDSTLSGF | 1 | 0 | 0 | 1438349 | AHA80176.1 | P33469 |
| GAISSSLQEILS | 1 | 0 | 0 | 1438357 | AVQ05264.1 | P36334 |
| GAITVEELKQLL | 1 | 0 | 0 | 1438363 | BAE93405.1 | P59596 |
| GAKFGRWSASIN | 1 | 0 | 0 | 1438375 | AGT21338.1 | A0A0P0K6L9 |
| GALAEHDFFTWK | 1 | 0 | 0 | 1438399 | AIW52827.1 | H9EJ42 |
| GALDTTIPFGPS | 1 | 0 | 0 | 1438403 | AIW52835.1 | A0A0P0G321 |
| GALNAATNGEMQ | 1 | 0 | 0 | 1438418 | APO40578.1 | P0C6X7 |
| GALNTPKEHIGT | 1 | 0 | 0 | 1438423 | APO40586.1 | P59595 |
| GALQGNFGDLQL | 1 | 0 | 0 | 1438425 | AVV62544.1 | K9N4V7 |
| GALWDIAVINNV | 1 | 0 | 0 | 1438432 | AVV62535.1 | K9N7C7 |
| GAMQTGFTTTNL | 1 | 0 | 0 | 1438451 | AVV62526.1 | K9N5Q8 |
| GANHTGITGDNV | 1 | 0 | 0 | 1438456 | ARO76381.1 | P0C6U8 |
| GANLMQDESVAN | 1 | 0 | 0 | 1438461 | AVV62537.1 | K9N5Q8 |
| GANLRQDDFVRN | 1 | 0 | 0 | 1438462 | AMW90843.1 | K9N5Q8 |
| GANLRQDDSVRD | 1 | 0 | 0 | 1438463 | QBF80611.1 | K9N5Q8 |
| GANLRQDESVSN | 1 | 0 | 0 | 1438465 | AVV62526.1 | K9N5Q8 |
| GAPDDASRLLHT | 1 | 0 | 0 | 1438473 | AXN92247.1 | K9N7C7 |
| GAPDDATRLLHV | 1 | 0 | 0 | 1438474 | AUM60013.1 | K9N7C7 |
| GAPDNPSAIVTG | 1 | 0 | 0 | 1438475 | AKA09625.1 | A0A0P0G321 |
| GAPIKSVTFGED | 1 | 0 | 0 | 1438480 | ARO76381.1 | P0C6U8 |
| GAPLGITISGET | 1 | 0 | 0 | 1438482 | AIW52843.1 | A0A0P0G321 |
| GAPNWLMSLYSF | 1 | 0 | 0 | 1438484 | AVV62524.1 | K9N7C7 |
| GAPNWLMSLYSL | 1 | 0 | 0 | 1438485 | AVV62535.1 | K9N7C7 |
| GAPTKGVTFGED | 1 | 0 | 0 | 1438493 | P0C6W6.1 | P0C6W2 |
| GAPVKKVAFCGD | 1 | 0 | 0 | 1438499 | ALT66879.1 | K9N7C7 |
| GAQGLKKFYKEV | 1 | 0 | 0 | 1438503 | ASU90140.1 | K9N7C7 |
| GAQSDHDVYYDE | 1 | 0 | 0 | 1438505 | AVV62527.1 | K9N796 |
| GAQTSTAIYTNL | 1 | 0 | 0 | 1438506 | ATO98145.1 | Q3LZX1 |
| GARLFASDLSTL | 1 | 0 | 0 | 1438515 | AWK59942.1 | H9EJ42 |
| GARLFSSDLSTL | 1 | 0 | 0 | 1438516 | P0C6U6.1 | H9EJ42 |
| GARLSASDLSTL | 1 | 0 | 0 | 1438517 | AAS58186.1 | H9EJ42 |

| | | | | | | |
|---|---|---|---|---|---|---|
| GARNMYRAALKN | 1 | 0 | 0 | 1438518 | AVV62524.1 | K9N7C7 |
| GARNRYRAALNN | 1 | 0 | 0 | 1438519 | AVV62535.1 | K9N7C7 |
| GARPKHRRPQGP | 1 | 0 | 0 | 1438520 | APO40586.1 | P59595 |
| GARRVWTLMNVL | 1 | 0 | 0 | 1438523 | AVP78030.1 | P0C6W2 |
| GASLRQIFDVAP | 1 | 0 | 0 | 1438536 | AVN89385.1 | K9N7C7 |
| GASRAIEKVNEC | 1 | 0 | 0 | 1438541 | BBA20986.1 | A0A140H1H1 |
| GASSNKTGNNVI | 1 | 0 | 0 | 1438542 | AUM60014.1 | K9N5Q8 |
| GASSNQSPQGNL | 1 | 0 | 0 | 1438543 | AIX09796.1 | P0C6X6 |
| GASTQNEELEQT | 1 | 0 | 0 | 1438547 | APO40578.1 | P0C6X7 |
| GATAALPTQFLP | 1 | 0 | 0 | 1438554 | ARO76382.1 | P59594 |
| GATIRGSDVKQP | 1 | 0 | 0 | 1438558 | AUM60013.1 | K9N7C7 |
| GATIRLQTGKQT | 1 | 0 | 0 | 1438559 | AAW33902.1 | H9EJ42 |
| GATSFSSSSQEE | 1 | 0 | 0 | 1438563 | AVP78041.1 | P0C6W2 |
| GAVANSTGTVII | 1 | 0 | 0 | 1438567 | AGV08520.1 | K9N5Q8 |
| GAVCRHHASEYR | 1 | 0 | 0 | 1438569 | APO40578.1 | P0C6X7 |
| GAVDINKLCDEM | 1 | 0 | 0 | 1438580 | APO40578.1 | P0C6X7 |
| GAVGDAGAAAFS | 1 | 0 | 0 | 1438589 | AVV62544.1 | K9N4V7 |
| GAVQIESDEYLT | 1 | 0 | 0 | 1438602 | AUM60013.1 | K9N7C7 |
| GAVSCLYKRNFT | 1 | 0 | 0 | 1438607 | ANZ78840.1 | P36334 |
| GAVVGTPVCING | 1 | 0 | 0 | 1438616 | P0C6W6.1 | P0C6W2 |
| GAVVGTPVCVNG | 1 | 0 | 0 | 1438617 | ATO98229.1 | P0C6W2 |
| GAVYTANDVDLQ | 1 | 0 | 0 | 1438619 | AUM60013.1 | K9N7C7 |
| GAWLNDKSVPSP | 1 | 0 | 0 | 1438621 | AHA80222.1 | P36334 |
| GAWNIGEQSSIL | 1 | 0 | 0 | 1438623 | AVP78030.1 | P0C6W2 |
| GAYNKDHGCGGF | 1 | 0 | 0 | 1438633 | AEN19363.1 | P0C6X6 |
| GAYQPYDHAIGA | 1 | 0 | 0 | 1438636 | ATO98145.1 | Q3LZX1 |
| GAYSSHLQSYAC | 1 | 0 | 0 | 1438638 | AVV62524.1 | K9N7C7 |
| GCAGYTLDEART | 1 | 0 | 0 | 1438648 | ARO76381.1 | P0C6U8 |
| GCALPFEDLNKC | 1 | 0 | 0 | 1438651 | AVV62524.1 | K9N7C7 |
| GCAMPIMANLVN | 1 | 0 | 0 | 1438655 | AIW52757.1 | A0A0P0K6L9 |
| GCCDASYFDNKL | 1 | 0 | 0 | 1438661 | AVV62524.1 | K9N7C7 |
| GCCDDYTGYQES | 1 | 0 | 0 | 1438666 | AIL49487.1 | P36334 |
| GCCDVTYFENKL | 1 | 0 | 0 | 1438668 | AXN92247.1 | K9N7C7 |
| GCCNCLTSSMRG | 1 | 0 | 0 | 1438673 | AIW52843.1 | A0A0P0G321 |
| GCCNTDYFEMKN | 1 | 0 | 0 | 1438674 | AIW52776.1 | A0A0P0LKV0 |
| GCCSTDYFEMKN | 1 | 0 | 0 | 1438677 | NP_073549.1 | A0A0P0LKV0 |
| GCDLMQARAFVN | 1 | 0 | 0 | 1438682 | AVV62524.1 | K9N7C7 |
| GCDVTDVTQLYLGG | 1 | 0 | 0 | 1438686 | P0DTD1.1 | P0DTD1 |
| GCEQSYFTKKDW | 1 | 0 | 0 | 1438694 | AEN19363.1 | P0C6X6 |
| GCETSNVPPIVS | 1 | 0 | 0 | 1438696 | ACA49954.1 | H9EJ42 |
| GCFVNDLLKTDS | 1 | 0 | 0 | 1438706 | ANZ78855.1 | P0C6X6 |
| GCGNGRWLTGNA | 1 | 0 | 0 | 1438715 | AVV62524.1 | K9N7C7 |
| GCHTGTAFNGDF | 1 | 0 | 0 | 1438742 | AIV41912.1 | P0C6X6 |
| GCILPSECTMFK | 1 | 0 | 0 | 1438748 | AVN89322.1 | K9N7C7 |
| GCKAIVKETNFI | 1 | 0 | 0 | 1438755 | AXX83336.1 | P0C6X6 |
| GCKYVVRTANAL | 1 | 0 | 0 | 1438767 | AEN19363.1 | P0C6X6 |
| GCLFCYKRNCSV | 1 | 0 | 0 | 1438770 | ARU07575.1 | A0A140H1G9 |
| GCLGSDCNRVSS | 1 | 0 | 0 | 1438776 | AMQ23491.1 | P36334 |
| GCLGSECNKVSS | 1 | 0 | 0 | 1438777 | CAA83661.1 | P36334 |
| GCLIGAEHVNAS | 1 | 0 | 0 | 1438784 | AVP78031.1 | Q3LZX1 |
| GCMLVLTVTLQN | 1 | 0 | 0 | 1438796 | AEN19363.1 | P0C6X6 |
| GCNNADCVACSK | 1 | 0 | 0 | 1438805 | AWK59942.1 | H9EJ42 |
| GCQTTNGLNTSH | 1 | 0 | 0 | 1438816 | ABB90513.1 | A0A0P0K6L9 |
| GCQTTNGTNTIH | 1 | 0 | 0 | 1438820 | AIW52741.1 | A0A0P0K6L9 |
| GCQTTNGTNTSN | 1 | 0 | 0 | 1438822 | AST12964.1 | A0A0P0K6L9 |
| GCRWWLCSTRVN | 1 | 0 | 0 | 1438824 | 5NH0_C | H9EJ42 |
| GCRYFTLGNVEA | 1 | 0 | 0 | 1438826 | AKA09609.1 | A0A0P0K6L9 |
| GCSVGNFSDGKM | 1 | 0 | 0 | 1438841 | QEU56412.1 | K9N5Q8 |
| GCSVIDLLLGDF | 1 | 0 | 0 | 1438842 | 2H85_A | P0C6X7 |
| GCTGSWNKVTQI | 1 | 0 | 0 | 1438848 | AXN92247.1 | K9N7C7 |
| GCVADQSRKWTG | 1 | 0 | 0 | 1438855 | AIV41775.1 | Q80872 |
| GCVASNVSPIVS | 1 | 0 | 0 | 1438858 | QEG03746.1 | H9EJ42 |
| GCVGHSENVFVV | 1 | 0 | 0 | 1438867 | AGT21353.1 | A0A0P0K6L9 |
| GCVGTWDKVAKI | 1 | 0 | 0 | 1438870 | AVV62535.1 | K9N7C7 |
| GCVIAWNTAKQD | 1 | 0 | 0 | 1438873 | AVP78031.1 | Q3LZX1 |
| GCVIAWNTAQQD | 1 | 0 | 0 | 1438874 | ATO98108.1 | Q3LZX1 |
| GCVLAWNTNSVD | 1 | 0 | 0 | 1438877 | APO40579.1 | P59594 |
| GCVLNAVNLTSY | 1 | 0 | 0 | 1438884 | BBA20986.1 | A0A140H1H1 |
| GCVSTNFGDVNK | 1 | 0 | 0 | 1438894 | QED88038.1 | H9EJ42 |
| GCVTSNVPPIIS | 1 | 0 | 0 | 1438896 | AMM42546.1 | H9EJ42 |
| GCVTSNVPPIVS | 1 | 0 | 0 | 1438897 | AWK59942.1 | H9EJ42 |
| GCVTSNVPSLVS | 1 | 0 | 0 | 1438898 | AFD98756.1 | H9EJ42 |
| GCVTSNVPTIVS | 1 | 0 | 0 | 1438899 | P0C6U6.1 | H9EJ42 |
| GCVTSNVSPIVN | 1 | 0 | 0 | 1438900 | AGT51337.1 | H9EJ42 |
| GCVTSNVSPIVS | 1 | 0 | 0 | 1438901 | ARU07593.1 | H9EJ42 |
| GCVVDQSRKWIG | 1 | 0 | 0 | 1438902 | ATP16755.1 | Q80872 |
| GCVVNADNSTAS | 1 | 0 | 0 | 1438904 | CAA83661.1 | P36334 |
| GCVVNAYNNTAI | 1 | 0 | 0 | 1438905 | AIL49538.1 | P36334 |
| GCVVNAYNSTAI | 1 | 0 | 0 | 1438906 | CAA79896.1 | P36334 |
| GCVVNAYNSTSI | 1 | 0 | 0 | 1438907 | ARK08679.1 | P36334 |
| GCWRHSVACDYL | 1 | 0 | 0 | 1438911 | AGT51488.1 | P0C6X6 |
| GDAFWSSTMIFS | 1 | 0 | 0 | 1438927 | AVN89333.1 | K9N7C7 |
| GDAGAAAFSTLF | 1 | 0 | 0 | 1438928 | AVV62544.1 | K9N4V7 |
| GDAGFTSILIGL | 1 | 0 | 0 | 1438929 | QBP84721.1 | P30215 |
| GDAHEGIFWVAS | 1 | 0 | 0 | 1438932 | AMM42397.1 | Q19U29 |
| GDAHIPSGTSFG | 1 | 0 | 0 | 1438934 | AIG59196.1 | A0A0P0K6L9 |
| GDAHIPSGTVLG | 1 | 0 | 0 | 1438935 | AIW52757.1 | A0A0P0K6L9 |
| GDAHTPSGTVLG | 1 | 0 | 0 | 1438936 | AMM42538.1 | A0A0P0K6L9 |
| GDAMHANYIFWR | 1 | 0 | 0 | 1438945 | AVN89322.1 | K9N7C7 |
| GDASKPHQVTKH | 1 | 0 | 0 | 1438947 | AJQ24068.1 | P33469 |
| GDASPQVLKALQ | 1 | 0 | 0 | 1438948 | AVV62524.1 | K9N7C7 |

| | | | | | | |
|---|---|---|---|---|---|---|
| GDASSLLYLNLL | 1 | 0 | 0 | 1438949 | AUM60021.1 | K9N4V7 |
| GDASTAFANSVF | 1 | 0 | 0 | 1438950 | AKL87224.1 | P0C6X6 |
| GDASWSSTMILS | 1 | 0 | 0 | 1438953 | ALW82719.1 | K9N7C7 |
| GDASWYSTMIFS | 1 | 0 | 0 | 1438954 | AHI48789.1 | K9N638 |
| GDATTAYANSVL | 1 | 0 | 0 | 1438957 | AVP77956.1 | P0C6W2 |
| GDAVFTGCVGTW | 1 | 0 | 0 | 1438958 | AVV62535.1 | K9N7C7 |
| GDCATVVYTANKW | 1 | 0 | 0 | 1438964 | APO40578.1 | P0C6X7 |
| GDCKGFSSNVSS | 1 | 0 | 0 | 1438968 | ABB90509.1 | A0A0P0K6L9 |
| GDCKGFYSNASS | 1 | 0 | 0 | 1438969 | AIW52756.1 | A0A0P0K6L9 |
| GDCLGGINARDL | 1 | 0 | 0 | 1438972 | APO40579.1 | P59594 |
| GDCQGFYSNASS | 1 | 0 | 0 | 1438976 | BAL45639.1 | A0A0P0K6L9 |
| GDCTFGVPDDAA | 1 | 0 | 0 | 1438978 | AVV62535.1 | K9N7C7 |
| GDCTFGVPDDAS | 1 | 0 | 0 | 1438979 | AVV62524.1 | K9N7C7 |
| GDCVTVHTANKW | 1 | 0 | 0 | 1438980 | QYB08949.1 | P0DTD1 |
| GDCVVEASTDFH | 1 | 0 | 0 | 1438983 | AWK59942.1 | H9EJ42 |
| GDDDDSQPRADK | 1 | 0 | 0 | 1438986 | 5EPW_A | H9EJ69 |
| GDDVDDIDTINC | 1 | 0 | 0 | 1438993 | ARU07575.1 | A0A140H1G9 |
| GDDVRQIAPAQT | 1 | 0 | 0 | 1438996 | APO40579.1 | P59594 |
| GDEQWGSTMIFS | 1 | 0 | 0 | 1439010 | AVV62524.1 | K9N7C7 |
| GDEVYLPYPNPS | 1 | 0 | 0 | 1439012 | AIL49620.1 | P0C6X6 |
| GDFAACKSQLVE | 1 | 0 | 0 | 1439013 | AAX85669.1 | P36334 |
| GDFIPIDQYMCG | 1 | 0 | 0 | 1439017 | AVV62535.1 | K9N7C7 |
| GDFIQTTPGSGVPV | 1 | 0 | 0 | 1439020 | P0DTD1.1 | P0DTD1 |
| GDFIQVAPGAGI | 1 | 0 | 0 | 1439021 | APO40578.1 | P0C6X7 |
| GDFMQGPFIAGN | 1 | 0 | 0 | 1439029 | AFD98763.1 | H9EJ42 |
| GDFNCTNSFIND | 1 | 0 | 0 | 1439034 | ARB07438.1 | A0A140H1H1 |
| GDFNCTNSFINH | 1 | 0 | 0 | 1439035 | BBA20986.1 | A0A140H1H1 |
| GDFNLTLLQVPE | 1 | 0 | 0 | 1439037 | AUM60014.1 | K9N5Q8 |
| GDFNLTLLQVPQ | 1 | 0 | 0 | 1439038 | AVV62537.1 | K9N5Q8 |
| GDFNLTMLQVPK | 1 | 0 | 0 | 1439039 | AVV62526.1 | K9N5Q8 |
| GDFQLNEEVAII | 1 | 0 | 0 | 1439041 | P0C6W6.1 | P0C6W2 |
| GDFQLTEEVAII | 1 | 0 | 0 | 1439042 | ATO98229.1 | P0C6W2 |
| GDFSHSQLGGLH | 1 | 0 | 0 | 1439046 | QYC64967.1 | P0DTD1 |
| GDFTPIDQYMCG | 1 | 0 | 0 | 1439049 | AVV62524.1 | K9N7C7 |
| GDFTSTIITSCQ | 1 | 0 | 0 | 1439050 | AJO62148.1 | K9N7C7 |
| GDGISRPKLHED | 1 | 0 | 0 | 1439064 | ARO76383.1 | P59632 |
| GDGIYFAATEKS | 1 | 0 | 0 | 1439069 | ATO98193.1 | Q3LZX1 |
| GDGLIDLLINVL | 1 | 0 | 0 | 1439074 | ARU07593.1 | H9EJ42 |
| GDGLIDLLVNVL | 1 | 0 | 0 | 1439075 | AWK59942.1 | H9EJ42 |
| GDGVYFAATEKS | 1 | 0 | 0 | 1439084 | ATO98181.1 | Q3LZX1 |
| GDHFKELWHLDT | 1 | 0 | 0 | 1439085 | AKA09663.1 | P36334 |
| GDHGDMYVYSSG | 1 | 0 | 0 | 1439088 | AVN89334.1 | K9N5Q8 |
| GDHGDRYVYSAG | 1 | 0 | 0 | 1439089 | ALA49803.1 | K9N5Q8 |
| GDIARELGNVVK | 1 | 0 | 0 | 1439093 | AWK59942.1 | H9EJ42 |
| GDIGIAPKQGYF | 1 | 0 | 0 | 1439095 | AZS52618.1 | A0A140H1H1 |
| GDIILKPASDGL | 1 | 0 | 0 | 1439098 | AVP78030.1 | P0C6W2 |
| GDIIQRLDGLEQ | 1 | 0 | 0 | 1439100 | AHI48733.1 | K9N5Q8 |
| GDIIQRLDPPEQ | 1 | 0 | 0 | 1439101 | 5W9H_A | K9N5Q8 |
| GDINVIAASGSL | 1 | 0 | 0 | 1439105 | ANI69921.1 | K9N7C7 |
| GDINVVAASGSL | 1 | 0 | 0 | 1439106 | AXN92247.1 | K9N7C7 |
| GDIPHYVQPGQY | 1 | 0 | 0 | 1439107 | AUM60024.1 | K9N5Q8 |
| GDIVDAVQYAHK | 1 | 0 | 0 | 1439117 | AVV62524.1 | K9N7C7 |
| GDIVSHVCKCGH | 1 | 0 | 0 | 1439120 | ARB07605.1 | A0A140H1G9 |
| GDIVTHLKNWNF | 1 | 0 | 0 | 1439121 | AST12967.1 | A0A0P0KXN3 |
| GDKIVYHTSGNV | 1 | 0 | 0 | 1439129 | APO40578.1 | P0C6X7 |
| GDLAVCLGVGDS | 1 | 0 | 0 | 1439141 | AIV41816.1 | P0C6X6 |
| GDLAVQMGSNNV | 1 | 0 | 0 | 1439143 | AGT51323.1 | H9EJ42 |
| GDLKCPLDSRTG | 1 | 0 | 0 | 1439149 | AVQ05264.1 | P36334 |
| GDLKCPLDTRYK | 1 | 0 | 0 | 1439150 | AKA09647.1 | P36334 |
| GDLKCPLDTSLK | 1 | 0 | 0 | 1439151 | AIX10753.1 | P36334 |
| GDLKCPLDTSYK | 1 | 0 | 0 | 1439154 | AKA09658.1 | P36334 |
| GDLKCPLESRTG | 1 | 0 | 0 | 1439155 | QEG03803.1 | P36334 |
| GDLKCTTVAIND | 1 | 0 | 0 | 1439158 | CAA83661.1 | P36334 |
| GDLNCPLDPKLK | 1 | 0 | 0 | 1439162 | AWW13511.1 | P36334 |
| GDLNCPLDPRLK | 1 | 0 | 0 | 1439163 | BAM16463.1 | P36334 |
| GDLNCPLDSRLK | 1 | 0 | 0 | 1439164 | AIL49507.1 | P36334 |
| GDLNCPLDTRLK | 1 | 0 | 0 | 1439165 | AEN19366.1 | P36334 |
| GDLNCTLDPRLK | 1 | 0 | 0 | 1439166 | AWW13566.1 | P36334 |
| GDLNCTLDPRLR | 1 | 0 | 0 | 1439167 | BBA84088.1 | P36334 |
| GDLNCTLHPRLK | 1 | 0 | 0 | 1439168 | BBA84086.1 | P36334 |
| GDLNVVAVSGSL | 1 | 0 | 0 | 1439169 | AVV62535.1 | K9N7C7 |
| GDLPENDDDAWF | 1 | 0 | 0 | 1439170 | AWW13564.1 | Q80872 |
| GDLQFNKLGTED | 1 | 0 | 0 | 1439171 | ASY99807.1 | K9N4V7 |
| GDLQGNFGDLKF | 1 | 0 | 0 | 1439172 | AWM99589.1 | K9N4V7 |
| GDLQGNFGDLQF | 1 | 0 | 0 | 1439173 | ASY99807.1 | K9N4V7 |
| GDLQGNFGDMQF | 1 | 0 | 0 | 1439175 | AUM60021.1 | K9N4V7 |
| GDLQLTKLGIED | 1 | 0 | 0 | 1439176 | QBM11743.1 | K9N4V7 |
| GDLQVVADTGSL | 1 | 0 | 0 | 1439178 | AUM60013.1 | K9N7C7 |
| GDLVSKLSPWFA | 1 | 0 | 0 | 1439184 | AUM60013.1 | K9N7C7 |
| GDMQFNKLGTED | 1 | 0 | 0 | 1439189 | AUM60021.1 | K9N4V7 |
| GDNDDEDNKDEE | 1 | 0 | 0 | 1439194 | AGW27834.1 | A0A140H1G9 |
| GDNDDEEIVTGD | 1 | 0 | 0 | 1439197 | P0C6U4.1 | A0A140H1G9 |
| GDNGTITVEELK | 1 | 0 | 0 | 1439203 | AVP78034.1 | Q3LZX9 |
| GDNNDNNSNNQQ | 1 | 0 | 0 | 1439209 | AVV62533.1 | K9N4V7 |
| GDNNDNNTNQQS | 1 | 0 | 0 | 1439210 | AVV62544.1 | K9N4V7 |
| GDNNGEDVVTGD | 1 | 0 | 0 | 1439211 | ABD75631.1 | A0A140H1G9 |
| GDNQHIETPITI | 1 | 0 | 0 | 1439212 | AVV62526.1 | K9N5Q8 |
| GDNSWRYDFDIT | 1 | 0 | 0 | 1439216 | QED88018.1 | H9EJ42 |
| GDNVGHYVHVKC | 1 | 0 | 0 | 1439219 | AEN19363.1 | P0C6X6 |
| GDNVQITHTYKM | 1 | 0 | 0 | 1439220 | AMM42628.1 | H9EJ69 |
| GDNYLRCFLLYF | 1 | 0 | 0 | 1439223 | NP_073549.1 | A0A0P0LKV0 |

| | | | | | | |
|---|---|---|---|---|---|---|
| GDPAQLPTPRTL | 1 | 0 | 0 | 1439227 | ATG84864.1 | K9N7C7 |
| GDPMPNNGWTAF | 1 | 0 | 0 | 1439229 | AVP78031.1 | Q3LZX1 |
| GDPMSNNGWTAF | 1 | 0 | 0 | 1439230 | AVP78042.1 | Q3LZX1 |
| GDPVYFLRYSGA | 1 | 0 | 0 | 1439232 | AUM60021.1 | K9N4V7 |
| GDQAAKLSPWFM | 1 | 0 | 0 | 1439233 | AVV62524.1 | K9N7C7 |
| GDQACYERFPLC | 1 | 0 | 0 | 1439234 | AUM60013.1 | K9N7C7 |
| GDQGQMYLYSAA | 1 | 0 | 0 | 1439240 | AUM60014.1 | K9N5Q8 |
| GDQIHEVAAVRS | 1 | 0 | 0 | 1439241 | ALA49340.1 | K9N7C7 |
| GDRDPRNDEAIP | 1 | 0 | 0 | 1439248 | AJQ24068.1 | P33469 |
| GDSAILPKGIMM | 1 | 0 | 0 | 1439267 | AVP78030.1 | P0C6W2 |
| GDSCSSSCKHTM | 1 | 0 | 0 | 1439272 | AUM60013.1 | K9N7C7 |
| GDSGELLSELLK | 1 | 0 | 0 | 1439279 | ABE73410.1 | H9EJ42 |
| GDSIFKSYHFTR | 1 | 0 | 0 | 1439289 | AGT17757.1 | A0A140H1H0 |
| GDSIFTGCTGSW | 1 | 0 | 0 | 1439290 | AXN92247.1 | K9N7C7 |
| GDSLECSNLLLQ | 1 | 0 | 0 | 1439292 | Q3I5J5.1 | Q3LZX1 |
| GDSQECSNLLLQ | 1 | 0 | 0 | 1439293 | ATO98181.1 | Q3LZX1 |
| GDSVFTGCTGSW | 1 | 0 | 0 | 1439313 | AQZ41321.1 | K9N7C7 |
| GDTDCVFYFAVR | 1 | 0 | 0 | 1439318 | AEN19363.1 | P0C6X6 |
| GDTQGMDNLACE | 1 | 0 | 0 | 1439320 | AVP78041.1 | P0C6W2 |
| GDTVGHYVHVKC | 1 | 0 | 0 | 1439324 | AIV41930.1 | P0C6X6 |
| GDTVYYPMCTNG | 1 | 0 | 0 | 1439326 | AUM60023.1 | K9N7C7 |
| GDTYLLFTNSIL | 1 | 0 | 0 | 1439327 | ARU07593.1 | H9EJ42 |
| GDVDSGPPSSAT | 1 | 0 | 0 | 1439331 | AVV62537.1 | K9N5Q8 |
| GDVDTVVNGVEF | 1 | 0 | 0 | 1439333 | AGT21350.1 | A0A0P0LKV0 |
| GDVGEKPVSIIN | 1 | 0 | 0 | 1439336 | APO40578.1 | P0C6X7 |
| GDVGLCMYRLLK | 1 | 0 | 0 | 1439337 | NP_073549.1 | A0A0P0LKV0 |
| GDVLLAEFSTHD | 1 | 0 | 0 | 1439347 | AVV62524.1 | K9N7C7 |
| GDVNKPVCAKSP | 1 | 0 | 0 | 1439350 | QED88038.1 | H9EJ42 |
| GDVNVVAASGSL | 1 | 0 | 0 | 1439351 | ASU45805.1 | K9N7C7 |
| GDVPHYVLPGQY | 1 | 0 | 0 | 1439352 | AUM60014.1 | K9N5Q8 |
| GDVQWSSTMIFS | 1 | 0 | 0 | 1439353 | AVV62535.1 | K9N7C7 |
| GDVVSHICKCGE | 1 | 0 | 0 | 1439368 | AEN19363.1 | P0C6X6 |
| GDWDSFGKTIQS | 1 | 0 | 0 | 1439371 | AST12962.1 | A0A0P0LKV0 |
| GDWNSFCKTIQS | 1 | 0 | 0 | 1439375 | APT69882.1 | A0A0P0ILY9 |
| GDWTGFKSSCCN | 1 | 0 | 0 | 1439376 | NP_073549.1 | A0A0P0LKV0 |
| GDYTLANTTTAP | 1 | 0 | 0 | 1439391 | AGV08437.1 | K9N7C7 |
| GDYVIISDKLFV | 1 | 0 | 0 | 1439393 | AUM60013.1 | K9N7C7 |
| GECLGDISARDL | 1 | 0 | 0 | 1439416 | ATO98145.1 | Q3LZX1 |
| GEDAFSILAVKT | 1 | 0 | 0 | 1439425 | AST12962.1 | A0A0P0LKV0 |
| GEDIQLLKNAYE | 1 | 0 | 0 | 1439432 | AVP78041.1 | P0C6W2 |
| GEDVDDIESIYD | 1 | 0 | 0 | 1439444 | ADN03337.1 | A0A140H1G9 |
| GEDVVTGDNNDE | 1 | 0 | 0 | 1439447 | ABD75631.1 | A0A140H1G9 |
| GEEKVHEVAPTR | 1 | 0 | 0 | 1439451 | AVV62535.1 | K9N7C7 |
| GEEKVHQVAATR | 1 | 0 | 0 | 1439452 | AVV62524.1 | K9N7C7 |
| GEENLSSRMYCS | 1 | 0 | 0 | 1439454 | ARO76381.1 | P0C6U8 |
| GEFASHVANFQK | 1 | 0 | 0 | 1439458 | AXN73435.1 | K9N7C7 |
| GEFIFEKVEYGS | 1 | 0 | 0 | 1439461 | AWK59942.1 | H9EJ42 |
| GEFIQTSSPKVT | 1 | 0 | 0 | 1439462 | ARB66062.1 | P36334 |
| GEFVFEKVDYGS | 1 | 0 | 0 | 1439469 | NP_073549.1 | A0A0P0LKV0 |
| GEGDQIIFYEGV | 1 | 0 | 0 | 1439478 | ARU07576.1 | A0A140H1H0 |
| GEHCSGLAIKSD | 1 | 0 | 0 | 1439485 | CAA83660.1 | P36334 |
| GEHFKHLVPLMH | 1 | 0 | 0 | 1439487 | ALW82707.1 | K9N7C7 |
| GEIAQYNYKQDF | 1 | 0 | 0 | 1439489 | AUM60014.1 | K9N5Q8 |
| GEKGAYNKDHGC | 1 | 0 | 0 | 1439497 | AEN19363.1 | P0C6X6 |
| GELDQWFKNQTS | 1 | 0 | 0 | 1439504 | QDH43719.1 | P36334 |
| GELLVDNVKYNK | 1 | 0 | 0 | 1439517 | AUM60013.1 | K9N7C7 |
| GELLVQDLRFNK | 1 | 0 | 0 | 1439519 | AKK52620.1 | K9N7C7 |
| GELNSLDGDCSD | 1 | 0 | 0 | 1439522 | AUM60013.1 | K9N7C7 |
| GELNVEFWFAMR | 1 | 0 | 0 | 1439523 | ARB07436.1 | A0A140H1G9 |
| GELYHYQECIKG | 1 | 0 | 0 | 1439526 | AVP78047.1 | Q3LZX7 |
| GEMQKESDTYIT | 1 | 0 | 0 | 1439533 | APO40578.1 | P0C6X7 |
| GENAVLPKGIMM | 1 | 0 | 0 | 1439537 | APO40578.1 | P0C6X7 |
| GENDNISVAAPK | 1 | 0 | 0 | 1439540 | AIX10718.1 | P33469 |
| GENDNISVATPK | 1 | 0 | 0 | 1439542 | AIX10720.1 | P33469 |
| GENDNISVAVSK | 1 | 0 | 0 | 1439544 | AIX10717.1 | P33469 |
| GENDNTSVAVPK | 1 | 0 | 0 | 1439545 | AIX09811.1 | P33469 |
| GENTVVEVQGYK | 1 | 0 | 0 | 1439552 | APO40578.1 | P0C6X7 |
| GEQSSILSPLYA | 1 | 0 | 0 | 1439566 | AVP78030.1 | P0C6W2 |
| GERELILVWEAG | 1 | 0 | 0 | 1439568 | AVV62524.1 | K9N7C7 |
| GERHQQLVEHTT | 1 | 0 | 0 | 1439570 | AVN89322.1 | K9N7C7 |
| GERNGARPKQRR | 1 | 0 | 0 | 1439574 | AVP78038.1 | Q3LZX4 |
| GERQLLLLWEAG | 1 | 0 | 0 | 1439575 | AZU90729.1 | K9N7C7 |
| GERWFLNRFTTT | 1 | 0 | 0 | 1439585 | APO40578.1 | P0C6X7 |
| GESKSQSRNSSS | 1 | 0 | 0 | 1439596 | AMM42526.1 | A0A0N7J131 |
| GESSWQTGDFLK | 1 | 0 | 0 | 1439600 | APO40578.1 | P0C6X7 |
| GETGVIADYNYK | 1 | 0 | 0 | 1439606 | ATO98181.1 | Q3LZX1 |
| GETKALKELYGP | 1 | 0 | 0 | 1439609 | QBF80609.1 | K9N7C7 |
| GETVRLHLYNAT | 1 | 0 | 0 | 1439620 | AWO68966.1 | A0A0P0G321 |
| GEVITFDNLKTI | 1 | 0 | 0 | 1439630 | AVP78041.1 | P0C6W2 |
| GEVPVAYRKVLL | 1 | 0 | 0 | 1439640 | AVP78041.1 | P0C6W2 |
| GEWNAVYKAFGS | 1 | 0 | 0 | 1439642 | AGT51379.1 | H9EJ42 |
| GEWNAVYRAFGS | 1 | 0 | 0 | 1439643 | AWK59942.1 | H9EJ42 |
| GEYDNISVAVPK | 1 | 0 | 0 | 1439647 | AXX83299.1 | P33469 |
| GEYGSFCDNINA | 1 | 0 | 0 | 1439649 | AHF54658.1 | P36334 |
| GEYNHVVFANAL | 1 | 0 | 0 | 1439651 | APO40578.1 | P0C6X7 |
| GEYSICRGFAPN | 1 | 0 | 0 | 1439654 | AVV62526.1 | K9N5Q8 |
| GFACDCGACFSA | 1 | 0 | 0 | 1439660 | AVV62524.1 | K9N7C7 |
| GFACDCGAVFSG | 1 | 0 | 0 | 1439661 | AVV62535.1 | K9N7C7 |
| GFACGCGTSYTA | 1 | 0 | 0 | 1439664 | QBM11736.1 | K9N638 |
| GFAVYHRYRIGN | 1 | 0 | 0 | 1439676 | APO40582.1 | P59596 |
| GFCDLKGKFVQI | 1 | 0 | 0 | 1439683 | 2XYV_B | P0C6U8 |

| | | | | | | |
|---|---|---|---|---|---|---|
| GFCEEADEGICF | 1 | 0 | 0 | 1439685 | AWW13571.1 | P0C6X6 |
| GFCLSNNPCNVC | 1 | 0 | 0 | 1439697 | ANI69899.1 | K9N7C7 |
| GFCVKHQWNCFI | 1 | 0 | 0 | 1439703 | ABD75543.1 | A0A140H1G9 |
| GFDKPQEKDKSQ | 1 | 0 | 0 | 1439709 | ABB90496.1 | A0A0N7J131 |
| GFDKPQEKKKSA | 1 | 0 | 0 | 1439710 | AMM42528.1 | A0A0N7J131 |
| GFDKSYQTNRDM | 1 | 0 | 0 | 1439711 | AUM60024.1 | K9N5Q8 |
| GFDKWYVNDGRV | 1 | 0 | 0 | 1439712 | AWK59942.1 | H9EJ42 |
| GFDLNCYYLVLP | 1 | 0 | 0 | 1439714 | P30215.1 | P30215 |
| GFDNQSKLPSSS | 1 | 0 | 0 | 1439719 | AIW52795.1 | H9EJ69 |
| GFDNQSKPPSSS | 1 | 0 | 0 | 1439720 | AGT51335.1 | H9EJ69 |
| GFDSSKSVTKKY | 1 | 0 | 0 | 1439726 | AVV61898.1 | K9N7C7 |
| GFECDFSPLLSG | 1 | 0 | 0 | 1439733 | AQZ41282.1 | K9N5Q8 |
| GFEIDPSPNTGS | 1 | 0 | 0 | 1439735 | APO40579.1 | P59594 |
| GFEKAALFASLN | 1 | 0 | 0 | 1439737 | AWK59948.1 | H9EJ42 |
| GFEKPQEKDKKS | 1 | 0 | 0 | 1439739 | ARB07429.1 | A0A0N7J131 |
| GFENPPTNVVSH | 1 | 0 | 0 | 1439741 | AAA73152.1 | P30215 |
| GFETIMKVLEEN | 1 | 0 | 0 | 1439742 | ABD75589.1 | Q19U29 |
| GFFFPYDPALET | 1 | 0 | 0 | 1439748 | AVV62535.1 | K9N7C7 |
| GFFYIDVGNHRG | 1 | 0 | 0 | 1439763 | AKA09616.1 | A0A0P0G321 |
| GFGGKNILGYAS | 1 | 0 | 0 | 1439777 | 5NH0_A | H9EJ42 |
| GFGGKNILGYSS | 1 | 0 | 0 | 1439778 | AWK59942.1 | H9EJ42 |
| GFGGNFNLKLLE | 1 | 0 | 0 | 1439780 | AOR17482.1 | K9N5Q8 |
| GFGPRDFNHNMG | 1 | 0 | 0 | 1439786 | QED88044.1 | H9EJ69 |
| GFHNPPTNVVSH | 1 | 0 | 0 | 1439796 | AVQ05263.1 | P30215 |
| GFIEAVNFNVTP | 1 | 0 | 0 | 1439813 | QEG59359.1 | A0A0P0G321 |
| GFIEAVNFNVTT | 1 | 0 | 0 | 1439814 | AIW52843.1 | A0A0P0G321 |
| GFIEDAVFKPQP | 1 | 0 | 0 | 1439815 | QEG03794.1 | P36334 |
| GFIEDSAFVPQP | 1 | 0 | 0 | 1439816 | AGT51481.1 | P36334 |
| GFIEDSVFKPQP | 1 | 0 | 0 | 1439817 | ARB66062.1 | P36334 |
| GFIEDSVFVPQP | 1 | 0 | 0 | 1439819 | AEN19366.1 | P36334 |
| GFIENSVFKPQP | 1 | 0 | 0 | 1439820 | AVQ05264.1 | P36334 |
| GFIKQYGDCLGG | 1 | 0 | 0 | 1439828 | AVP78031.1 | Q3LZX1 |
| GFITLKNNLYRL | 1 | 0 | 0 | 1439835 | AXN92247.1 | K9N7C7 |
| GFKDVTKGTIYS | 1 | 0 | 0 | 1439839 | CAA71056.1 | A0A0P0K6L9 |
| GFKDVTNGTIYS | 1 | 0 | 0 | 1439840 | AIW52757.1 | A0A0P0K6L9 |
| GFKEELDQWFKN | 1 | 0 | 0 | 1439841 | AIL49523.1 | P36334 |
| GFKGKQILGNTM | 1 | 0 | 0 | 1439845 | AUM60013.1 | K9N7C7 |
| GFKLDAIVPGYP | 1 | 0 | 0 | 1439851 | QBM11746.1 | K9N7C7 |
| GFKLDANIPDYP | 1 | 0 | 0 | 1439852 | AVV62535.1 | K9N7C7 |
| GFKLDANIPGYP | 1 | 0 | 0 | 1439853 | AVV62524.1 | K9N7C7 |
| GFKLDASIPGYP | 1 | 0 | 0 | 1439855 | AUM60013.1 | K9N7C7 |
| GFKNQHCNGSIA | 1 | 0 | 0 | 1439861 | AWW13571.1 | P0C6X6 |
| GFKNQYCNESIA | 1 | 0 | 0 | 1439862 | AXX83294.1 | P0C6X6 |
| GFKNVSTGNIFI | 1 | 0 | 0 | 1439864 | AIW52843.1 | A0A0P0G321 |
| GFLDKISDTSGF | 1 | 0 | 0 | 1439877 | BAM16435.1 | Q01455 |
| GFLETVCSVAHT | 1 | 0 | 0 | 1439879 | AWK59942.1 | H9EJ42 |
| GFLIGCNYLGFI | 1 | 0 | 0 | 1439891 | AUM60013.1 | K9N7C7 |
| GFLPTFVGKIAQ | 1 | 0 | 0 | 1439903 | AEN19363.1 | P0C6X6 |
| GFMGATICLQAG | 1 | 0 | 0 | 1439918 | AAW33912.1 | H9EJ42 |
| GFMKQYGDCLGG | 1 | 0 | 0 | 1439924 | APO40579.1 | P59594 |
| GFNDICMFENKT | 1 | 0 | 0 | 1439932 | AVN89289.1 | K9N7C7 |
| GFNDLSQLYCSY | 1 | 0 | 0 | 1439934 | ANI69889.1 | K9N5Q8 |
| GFNFSQILPDPK | 1 | 0 | 0 | 1439940 | APO40579.1 | P59594 |
| GFNFSQILPDSS | 1 | 0 | 0 | 1439943 | ARO76382.1 | P59594 |
| GFNNYVPSYIKP | 1 | 0 | 0 | 1439953 | AUM60013.1 | K9N7C7 |
| GFNTEFASNSSV | 1 | 0 | 0 | 1439956 | ANI69888.1 | K9N638 |
| GFPSQAASVIRA | 1 | 0 | 0 | 1439973 | ARO76381.1 | P0C6U8 |
| GFQGKQILGSSM | 1 | 0 | 0 | 1439979 | AVV62524.1 | K9N7C7 |
| GFQGKQILGSTM | 1 | 0 | 0 | 1439980 | AXN92247.1 | K9N7C7 |
| GFQIKFENPHIL | 1 | 0 | 0 | 1439985 | AWW13564.1 | Q80872 |
| GFSFVDDVNGMK | 1 | 0 | 0 | 1440009 | AVV62535.1 | K9N7C7 |
| GFSFVNDDTGVQ | 1 | 0 | 0 | 1440010 | AVV62524.1 | K9N7C7 |
| GFSNVVEMPKFF | 1 | 0 | 0 | 1440020 | AIW52757.1 | A0A0P0K6L9 |
| GFSQVKSDLVID | 1 | 0 | 0 | 1440024 | AEN19363.1 | P0C6X6 |
| GFSSDVLSDVIR | 1 | 0 | 0 | 1440026 | CAA71146.1 | A0A0P0K6L9 |
| GFSTGVNFVVQP | 1 | 0 | 0 | 1440030 | AXN92247.1 | K9N7C7 |
| GFSVLRPILKLP | 1 | 0 | 0 | 1440038 | ATO98181.1 | Q3LZX1 |
| GFTEQSVFKPQP | 1 | 0 | 0 | 1440044 | CAA83660.1 | P36334 |
| GFTPTVGVGYQP | 1 | 0 | 0 | 1440060 | APO40579.1 | P59594 |
| GFTVMNGEDAFS | 1 | 0 | 0 | 1440074 | QEO75983.1 | A0A0P0LKV0 |
| GFVLNYKGLQFQ | 1 | 0 | 0 | 1440085 | AWW13564.1 | Q80872 |
| GFVLRNTVCTVC | 1 | 0 | 0 | 1440089 | APO40578.1 | P0C6X7 |
| GFVLSIPTKALC | 1 | 0 | 0 | 1440090 | AGT17757.1 | A0A140H1H0 |
| GFVTVELQPPCK | 1 | 0 | 0 | 1440103 | AUM60013.1 | K9N7C7 |
| GFVVRIGASSNK | 1 | 0 | 0 | 1440110 | AUM60014.1 | K9N5Q8 |
| GFVYDNHVKIDC | 1 | 0 | 0 | 1440116 | ARU07575.1 | A0A140H1G9 |
| GFYNPLTNVVSH | 1 | 0 | 0 | 1440128 | AGT51799.1 | P30215 |
| GFYRLYSGDNSW | 1 | 0 | 0 | 1440134 | QED88018.1 | H9EJ42 |
| GFYSNSSSDVIR | 1 | 0 | 0 | 1440141 | AIW52747.1 | A0A0P0K6L9 |
| GFYSTSPIQPVE | 1 | 0 | 0 | 1440143 | AIW52757.1 | A0A0P0K6L9 |
| GFYVKPGGTTSG | 1 | 0 | 0 | 1440152 | AVP78019.1 | P0C6W2 |
| GGAEIRDLICVS | 1 | 0 | 0 | 1440156 | AIL49490.1 | P36334 |
| GGAFTRYVDNNF | 1 | 0 | 0 | 1440157 | ATO98229.1 | P0C6W2 |
| GGAPIWFNALRD | 1 | 0 | 0 | 1440168 | AVN89451.1 | K9N7C7 |
| GGAPNWLMSLYS | 1 | 0 | 0 | 1440169 | AVV62524.1 | K9N7C7 |
| GGCAMPIMANLV | 1 | 0 | 0 | 1440198 | AIW52757.1 | A0A0P0K6L9 |
| GGCCEDYTGYQE | 1 | 0 | 0 | 1440203 | AIL49528.1 | P36334 |
| GGCNFPLEATWH | 1 | 0 | 0 | 1440207 | AWO68993.1 | A0A0P0G321 |
| GGDASKPHQVTK | 1 | 0 | 0 | 1440222 | AJQ24068.1 | P33469 |
| GGDATSLSFNNL | 1 | 0 | 0 | 1440223 | ANI69888.1 | K9N638 |
| GGDFIPIDQYMC | 1 | 0 | 0 | 1440224 | AVV62535.1 | K9N7C7 |

| | | | | | | |
|---|---|---|---|---|---|---|
| GGDFTPIDQYMC | 1 | 0 | 0 | 1440225 | AVV62524.1 | K9N7C7 |
| GGDNVYCYNTGL | 1 | 0 | 0 | 1440236 | AFV53147.1 | H9EJ42 |
| GGDQIHEVAAVR | 1 | 0 | 0 | 1440238 | ALA49340.1 | K9N7C7 |
| GGDQVHEVVAVR | 1 | 0 | 0 | 1440240 | ALA49384.1 | K9N7C7 |
| GGDVLLAEFVHY | 1 | 0 | 0 | 1440253 | AUM60013.1 | K9N7C7 |
| GGEEDENAMFDK | 1 | 0 | 0 | 1440255 | AUM60021.1 | K9N4V7 |
| GGEVDIQPQVLF | 1 | 0 | 0 | 1440264 | AUM60018.1 | K9N7D2 |
| GGFNFSQILPDS | 1 | 0 | 0 | 1440277 | ARO76382.1 | P59594 |
| GGGIAAAINNAS | 1 | 0 | 0 | 1440296 | AUM60013.1 | K9N7C7 |
| GGGKCFNCAPAV | 1 | 0 | 0 | 1440299 | AAQ90006.1 | A0A0P0K6L9 |
| GGGSGYIPEAPR | 1 | 0 | 0 | 1440307 | 6NB6_A | P59594 |
| GGGVAGAFNKAT | 1 | 0 | 0 | 1440310 | QOJ87595.1 | P0DTD1 |
| GGGWLVASGPTV | 1 | 0 | 0 | 1440314 | AKN24830.1 | K9N5Q8 |
| GGHCYGADLKSP | 1 | 0 | 0 | 1440319 | APO40578.1 | P0C6X7 |
| GGIADAINKASN | 1 | 0 | 0 | 1440329 | AVV62535.1 | K9N7C7 |
| GGISARDLICAQ | 1 | 0 | 0 | 1440344 | AVP78031.1 | Q3LZX1 |
| GGITQRTRTRPS | 1 | 0 | 0 | 1440347 | ANF29213.1 | K9N4V7 |
| GGKIVNNWLKQL | 1 | 0 | 0 | 1440355 | AVP78041.1 | P0C6W2 |
| GGKNILGYASLC | 1 | 0 | 0 | 1440360 | 5NH0_A | H9EJ42 |
| GGKNILGYSSLC | 1 | 0 | 0 | 1440361 | AWK59942.1 | H9EJ42 |
| GGKVEVLVPSNK | 1 | 0 | 0 | 1440371 | AVV62535.1 | K9N7C7 |
| GGLDSLDSYPAL | 1 | 0 | 0 | 1440378 | ARO76381.1 | P0C6U8 |
| GGLFQLTFESTI | 1 | 0 | 0 | 1440383 | AGT51388.1 | H9EJ24 |
| GGNCAGNASGND | 1 | 0 | 0 | 1440411 | ARB07436.1 | A0A140H1G9 |
| GGNEALSISTIF | 1 | 0 | 0 | 1440416 | AIV41792.1 | P0C6X6 |
| GGNFNLKLLEPV | 1 | 0 | 0 | 1440418 | AOR17482.1 | K9N5Q8 |
| GGNFTPIDQYMC | 1 | 0 | 0 | 1440419 | AUM60013.1 | K9N7C7 |
| GGNGTHIFSLVN | 1 | 0 | 0 | 1440420 | ABB90528.1 | A0A0P0K6L9 |
| GGNHFRHPLDED | 1 | 0 | 0 | 1440421 | AVV62524.1 | K9N7C7 |
| GGNHFRHPLEED | 1 | 0 | 0 | 1440422 | AVV62535.1 | K9N7C7 |
| GGNLCTCQPQAF | 1 | 0 | 0 | 1440424 | QQY99180.1 | P36334 |
| GGNMFQFATFPV | 1 | 0 | 0 | 1440427 | ALA49363.1 | K9N5Q8 |
| GGNMFRFASLPV | 1 | 0 | 0 | 1440429 | AVV62526.1 | K9N5Q8 |
| GGNSCTCQPQAF | 1 | 0 | 0 | 1440435 | AGT51690.1 | P36334 |
| GGNVIPHYSWFS | 1 | 0 | 0 | 1440438 | QEG03818.1 | P33469 |
| GGNVVPHYSWFS | 1 | 0 | 0 | 1440440 | AWW13570.1 | P33469 |
| GGNYSDPFMVNQ | 1 | 0 | 0 | 1440443 | AUM60013.1 | K9N7C7 |
| GGPSDSSDNSQN | 1 | 0 | 0 | 1440446 | AVP78049.1 | Q3LZX4 |
| GGPTDSTDNNQD | 1 | 0 | 0 | 1440450 | Q3I5I7.1 | Q3LZX4 |
| GGRGGVPDLVVE | 1 | 0 | 0 | 1440472 | 5YL9_B | A0A0P0K6L9 |
| GGRPLPVYHKHM | 1 | 0 | 0 | 1440477 | 6U7G_C | A0A0P0K6L9 |
| GGSDVDLGDISG | 1 | 0 | 0 | 1440492 | 1WYY_A | P59594 |
| GGSGYIPEAPRD | 1 | 0 | 0 | 1440496 | 6NB6_A | P59594 |
| GGSLYVKPSGTS | 1 | 0 | 0 | 1440500 | AVP78041.1 | P0C6W2 |
| GGSQTASRSSSR | 1 | 0 | 0 | 1440503 | APO40586.1 | P59595 |
| GGSYKNDKACPI | 1 | 0 | 0 | 1440510 | APO40578.1 | P0C6X7 |
| GGTFYAYFTDNG | 1 | 0 | 0 | 1440521 | AHA80218.1 | P36334 |
| GGTYSTPFMVNQ | 1 | 0 | 0 | 1440545 | AVV62535.1 | K9N7C7 |
| GGVAGALNAATN | 1 | 0 | 0 | 1440547 | APO40578.1 | P0C6X7 |
| GGVTRDIAPTDT | 1 | 0 | 0 | 1440567 | AVP78041.1 | P0C6W2 |
| GGVVTRYVDNNF | 1 | 0 | 0 | 1440574 | ARO76381.1 | P0C6U8 |
| GGWGNMLKNLMA | 1 | 0 | 0 | 1440583 | ADK37686.1 | A0A0P0LKV0 |
| GGWNNMLRTLID | 1 | 0 | 0 | 1440590 | AIW52827.1 | H9EJ42 |
| GGYIPEAPRDGQ | 1 | 0 | 0 | 1440597 | 6U7H_A | A0A0P0K6L9 |
| GGYIPPNFAFNN | 1 | 0 | 0 | 1440598 | AIW52740.1 | A0A0P0K6L9 |
| GGYIPSNFAFNN | 1 | 0 | 0 | 1440600 | AIW52757.1 | A0A0P0K6L9 |
| GGYNSWHLVEGG | 1 | 0 | 0 | 1440604 | QDP16215.1 | K9N7C7 |
| GGYSEDWHSGVK | 1 | 0 | 0 | 1440606 | ATO98182.1 | Q3LZX0 |
| GGYYYKVEADFY | 1 | 0 | 0 | 1440610 | AGT51669.1 | P30215 |
| GHAAITDYSYYA | 1 | 0 | 0 | 1440611 | AUM60013.1 | K9N7C7 |
| GHADNNIEKPKP | 1 | 0 | 0 | 1440612 | AVV62526.1 | K9N5Q8 |
| GHATGTTPKKLF | 1 | 0 | 0 | 1440621 | ALA49649.1 | K9N5Q8 |
| GHCTIAQLTDAA | 1 | 0 | 0 | 1440630 | AWW13564.1 | Q80872 |
| GHCTIVQLTDAA | 1 | 0 | 0 | 1440632 | AIV41895.1 | Q80872 |
| GHCYGADLKSPD | 1 | 0 | 0 | 1440633 | APO40578.1 | P0C6X7 |
| GHDICAQLNAKL | 1 | 0 | 0 | 1440637 | ABD75519.1 | A0A140H1G9 |
| GHDICAQLNDKL | 1 | 0 | 0 | 1440638 | ARU07575.1 | A0A140H1G9 |
| GHEEFIQTSSPK | 1 | 0 | 0 | 1440643 | BBA20986.1 | A0A140H1H1 |
| GHFDGDVGEKPV | 1 | 0 | 0 | 1440645 | APO40578.1 | P0C6X7 |
| GHFLAKNILHVV | 1 | 0 | 0 | 1440657 | AHI48765.1 | K9N638 |
| GHHLDSHFVVKR | 1 | 0 | 0 | 1440679 | AUM60013.1 | K9N7C7 |
| GHIIMEEMLRGS | 1 | 0 | 0 | 1440691 | AVN89322.1 | K9N7C7 |
| GHIVCDSLNSKL | 1 | 0 | 0 | 1440695 | QBP84746.1 | P0C6X6 |
| GHKNGQGEYDNI | 1 | 0 | 0 | 1440704 | AXX83299.1 | P33469 |
| GHKTGQGENDNI | 1 | 0 | 0 | 1440708 | AHA80176.1 | P33469 |
| GHLIDEVQNWLS | 1 | 0 | 0 | 1440715 | AUM60027.1 | K9N643 |
| GHLKMAGMHFGA | 1 | 0 | 0 | 1440717 | ASY99815.1 | K9N7A1 |
| GHMAHDGGVAKA | 1 | 0 | 0 | 1440729 | AMK59674.1 | P0C6X6 |
| GHMVHGGGVAKA | 1 | 0 | 0 | 1440736 | AEN19363.1 | P0C6X6 |
| GHPAPTHYLRVV | 1 | 0 | 0 | 1440748 | AVV62540.1 | K9N643 |
| GHRVPIHYEMFG | 1 | 0 | 0 | 1440760 | ATO98227.1 | Q3LZX5 |
| GHSEIVFDVESG | 1 | 0 | 0 | 1440763 | AST12964.1 | A0A0P0K6L9 |
| GHSENVFAVESG | 1 | 0 | 0 | 1440764 | AIW52757.1 | A0A0P0K6L9 |
| GHTEFYDSQPST | 1 | 0 | 0 | 1440782 | AUM60025.1 | K9N796 |
| GHTLKSGCKLIN | 1 | 0 | 0 | 1440785 | AWK59942.1 | H9EJ42 |
| GHTVCDSLNAKW | 1 | 0 | 0 | 1440789 | AIV41984.1 | P0C6X6 |
| GHTVCDSLNSKL | 1 | 0 | 0 | 1440791 | AEN19363.1 | P0C6X6 |
| GHVYEVSDLIAQ | 1 | 0 | 0 | 1440806 | AIV41990.1 | P0C6X6 |
| GIAAVNSVPWSK | 1 | 0 | 0 | 1440840 | ARO76381.1 | P0C6U8 |
| GIACVKDFDFKF | 1 | 0 | 0 | 1440841 | AEN19363.1 | P0C6X6 |
| GIADAINKASNG | 1 | 0 | 0 | 1440843 | AVV62535.1 | K9N7C7 |

| | | | | | | |
|---|---|---|---|---|---|---|
| GIAIEQDKNTQE | 1 | 0 | 0 | 1440848 | AVP78031.1 | Q3LZX1 |
| GIAISVKPESNA | 1 | 0 | 0 | 1440849 | AVV62524.1 | K9N7C7 |
| GIAISVKPESTA | 1 | 0 | 0 | 1440850 | AXN92247.1 | K9N7C7 |
| GIAISVKPESTV | 1 | 0 | 0 | 1440851 | QBF80597.1 | K9N638 |
| GIALGGLTSAAS | 1 | 0 | 0 | 1440853 | AIW52757.1 | A0A0P0K6L9 |
| GIALTGTNQLCA | 1 | 0 | 0 | 1440854 | QED88010.1 | H9EJ42 |
| GIAVEQDRNTRD | 1 | 0 | 0 | 1440860 | APO40579.1 | P59594 |
| GICFEYTSPTDK | 1 | 0 | 0 | 1440876 | AGT51509.1 | Q80872 |
| GICPAGTNYLTC | 1 | 0 | 0 | 1440884 | AIX10751.1 | P36334 |
| GICSENHVMVDC | 1 | 0 | 0 | 1440886 | AIV41942.1 | P0C6X6 |
| GIDAGYKNSGIG | 1 | 0 | 0 | 1440890 | CAA83660.1 | P36334 |
| GIEDPRWPQIAE | 1 | 0 | 0 | 1440905 | QBM11743.1 | K9N4V7 |
| GIERVPPHSFTL | 1 | 0 | 0 | 1440914 | AVV62524.1 | K9N7C7 |
| GIFAKAKNTKVI | 1 | 0 | 0 | 1440917 | AKA09644.1 | P36334 |
| GIFAKVKNTKVT | 1 | 0 | 0 | 1440922 | AMQ23491.1 | P36334 |
| GIFAKVKNTKVV | 1 | 0 | 0 | 1440923 | AWW13511.1 | P36334 |
| GIFDGALSRFNR | 1 | 0 | 0 | 1440927 | AVV62535.1 | K9N7C7 |
| GIFGAKPLDSLK | 1 | 0 | 0 | 1440933 | ARO76381.1 | P0C6U8 |
| GIFGVDPGLSLE | 1 | 0 | 0 | 1440937 | APO40578.1 | P0C6X7 |
| GIFGVDPKVSFE | 1 | 0 | 0 | 1440938 | AVV62524.1 | K9N7C7 |
| GIFKEVSAVYYN | 1 | 0 | 0 | 1440942 | BBA20983.1 | A0A140H1H1 |
| GIFKTEVDPNLS | 1 | 0 | 0 | 1440943 | ATQ39389.1 | K9N7C7 |
| GIFVASSSYDAL | 1 | 0 | 0 | 1440949 | AGT51372.1 | H9EJ42 |
| GIFVEVNAPYYN | 1 | 0 | 0 | 1440950 | CAA83660.1 | P36334 |
| GIFVEVNATYCN | 1 | 0 | 0 | 1440951 | ABU39950.1 | P36334 |
| GIGAVGDAGAAA | 1 | 0 | 0 | 1440955 | AVV62544.1 | K9N4V7 |
| GIGEHCSGLAIK | 1 | 0 | 0 | 1440960 | CAA83660.1 | P36334 |
| GIGGIPCIKVAS | 1 | 0 | 0 | 1440970 | AUM60013.1 | K9N7C7 |
| GIGICPAGTNYL | 1 | 0 | 0 | 1440976 | AIX10751.1 | P36334 |
| GIGLCAFLTLLF | 1 | 0 | 0 | 1440980 | AXN92247.1 | K9N7C7 |
| GIGLHLKVNCCR | 1 | 0 | 0 | 1440983 | AEN19363.1 | P0C6X6 |
| GIGNTFICEEVA | 1 | 0 | 0 | 1440985 | AVV62524.1 | K9N7C7 |
| GIGSTFICEEVA | 1 | 0 | 0 | 1440986 | AUM60013.1 | K9N7C7 |
| GIGTCPAGTNSL | 1 | 0 | 0 | 1440988 | AWW13566.1 | P36334 |
| GIGYSLADLPAY | 1 | 0 | 0 | 1440999 | AWW13547.1 | Q01455 |
| GIHNRVIAIPTS | 1 | 0 | 0 | 1441002 | AVV62527.1 | K9N796 |
| GIIAAISGMMWV | 1 | 0 | 0 | 1441004 | AVV62543.1 | K9N7A1 |
| GIIFNAVDCKSD | 1 | 0 | 0 | 1441009 | AHA80218.1 | P36334 |
| GIITGNVVKQCC | 1 | 0 | 0 | 1441026 | AVV62535.1 | K9N7C7 |
| GIIYPTGKSYSN | 1 | 0 | 0 | 1441038 | AVV62537.1 | K9N5Q8 |
| GIKDVVLQGLRA | 1 | 0 | 0 | 1441044 | AMQ48992.1 | K9N7C7 |
| GIKLGIGYSLAD | 1 | 0 | 0 | 1441048 | AWW13547.1 | Q01455 |
| GIKLGIGYSWAD | 1 | 0 | 0 | 1441049 | AAQ93015.1 | Q01455 |
| GIKLQSKRTRFF | 1 | 0 | 0 | 1441054 | AIV41996.1 | P0C6X6 |
| GIKLQSKRTRLF | 1 | 0 | 0 | 1441055 | AEN19363.1 | P0C6X6 |
| GIKPFTIPIDLL | 1 | 0 | 0 | 1441057 | AMM42367.1 | A0A140H1G9 |
| GIKQESRVGVDA | 1 | 0 | 0 | 1441059 | ARU07575.1 | A0A140H1G9 |
| GIKVLPPLLSDN | 1 | 0 | 0 | 1441061 | CAA83661.1 | P36334 |
| GIKYYSRVKSVR | 1 | 0 | 0 | 1441066 | AST12962.1 | A0A0P0LKV0 |
| GILGQGIFVEVN | 1 | 0 | 0 | 1441077 | CAA79896.1 | P36334 |
| GILKKTSWVDQS | 1 | 0 | 0 | 1441081 | AMM42397.1 | Q19U29 |
| GILLCLALLFVV | 1 | 0 | 0 | 1441085 | ARU07576.1 | A0A140H1H0 |
| GILPSPGMPALL | 1 | 0 | 0 | 1441094 | 6NB6_A | P59594 |
| GILSENAYLKYA | 1 | 0 | 0 | 1441095 | AVV62524.1 | K9N7C7 |
| GILTTVAAFHQE | 1 | 0 | 0 | 1441106 | AVP78037.1 | Q3LZX5 |
| GIMVLPGAADAE | 1 | 0 | 0 | 1441113 | APT69876.1 | A0A0P0K6L9 |
| GINEKTHVQLSL | 1 | 0 | 0 | 1441122 | P0C6W6.1 | P0C6W2 |
| GINFRTCILNNQ | 1 | 0 | 0 | 1441123 | AUM60013.1 | K9N7C7 |
| GINFRTSVVNNQ | 1 | 0 | 0 | 1441124 | AVV62535.1 | K9N7C7 |
| GINTTGVLLKQL | 1 | 0 | 0 | 1441136 | AET99208.1 | Q0ZJ80 |
| GINYLGDFMQGP | 1 | 0 | 0 | 1441139 | AFD98763.1 | H9EJ42 |
| GIPFEYTSPTDK | 1 | 0 | 0 | 1441148 | AXX83349.1 | Q80872 |
| GIPGNMIDGFAC | 1 | 0 | 0 | 1441150 | AIV42002.1 | P0C6X6 |
| GIPLSEAKGYWY | 1 | 0 | 0 | 1441156 | AMM42386.1 | Q19U29 |
| GIPPSEAKGYWY | 1 | 0 | 0 | 1441157 | AMM42397.1 | Q19U29 |
| GIPVVEYLSKED | 1 | 0 | 0 | 1441160 | ATQ39389.1 | K9N7C7 |
| GIQEAKPSGSVV | 1 | 0 | 0 | 1441163 | 4KQZ_A | K9N5Q8 |
| GIRFEYISPTDK | 1 | 0 | 0 | 1441179 | AGT51578.1 | Q80872 |
| GIRKAKDYGFTV | 1 | 0 | 0 | 1441183 | AVV62524.1 | K9N7C7 |
| GISGRGVFQNCT | 1 | 0 | 0 | 1441198 | AXN92260.1 | K9N5Q8 |
| GISLLKKMDEPF | 1 | 0 | 0 | 1441202 | AHA80174.1 | P33469 |
| GISPDAIARGCY | 1 | 0 | 0 | 1441203 | AVV62526.1 | K9N5Q8 |
| GISRPKLHEDYQ | 1 | 0 | 0 | 1441206 | ARO76383.1 | P59632 |
| GISRQGIFVEVN | 1 | 0 | 0 | 1441207 | BBA20982.1 | P36334 |
| GISSACDGLALA | 1 | 0 | 0 | 1441208 | ALA49659.1 | K9N7C7 |
| GITFQYGTDTNS | 1 | 0 | 0 | 1441220 | AID55088.1 | K9N5Q8 |
| GITIQKGLVDYR | 1 | 0 | 0 | 1441226 | APO40578.1 | P0C6X7 |
| GITQFQKGKAFE | 1 | 0 | 0 | 1441234 | AIV41850.1 | P33469 |
| GITQFQKGKEFK | 1 | 0 | 0 | 1441236 | ARA15425.1 | P33469 |
| GITQFQKGKEFQ | 1 | 0 | 0 | 1441237 | QBP84717.1 | P33469 |
| GITQQVLSDNQK | 1 | 0 | 0 | 1441241 | ALA49671.1 | K9N5Q8 |
| GITQQVLSKNQK | 1 | 0 | 0 | 1441243 | AVN89376.1 | K9N5Q8 |
| GIVAAVSAMMRI | 1 | 0 | 0 | 1441254 | ALA49424.1 | K9N7A1 |
| GIVEDTINDDED | 1 | 0 | 0 | 1441263 | AGW27861.1 | A0A140H1G9 |
| GIVELEKGVLPQ | 1 | 0 | 0 | 1441264 | P0C6W6.1 | P0C6W2 |
| GIVKMVSPTSKI | 1 | 0 | 0 | 1441270 | ARU07575.1 | A0A140H1G9 |
| GIVLPFEDLTKC | 1 | 0 | 0 | 1441272 | AUM60013.1 | K9N7C7 |
| GIVNSSMVVDEC | 1 | 0 | 0 | 1441274 | AUM60014.1 | K9N5Q8 |
| GIVTGNVVKQCC | 1 | 0 | 0 | 1441282 | AVV62524.1 | K9N7C7 |
| GIYGYVLRQSNL | 1 | 0 | 0 | 1441292 | AFD98771.1 | A0A0P0G321 |
| GIYKPLLENIDY | 1 | 0 | 0 | 1441293 | AIV41840.1 | P0C6X6 |

| | | | | | | |
|---|---|---|---|---|---|---|
| GIYQTSNFRVSP | 1 | 0 | 0 | 1441297 | ATO98181.1 | Q3LZX1 |
| GKAFEFAEGQGV | 1 | 0 | 0 | 1441301 | AIV41850.1 | P33469 |
| GKASSRSAIEDL | 1 | 0 | 0 | 1441310 | BBA20982.1 | P36334 |
| GKASYALSEQLD | 1 | 0 | 0 | 1441311 | AVP78030.1 | P0C6W2 |
| GKCYNCRPAVVN | 1 | 0 | 0 | 1441324 | ABB90509.1 | A0A0P0K6L9 |
| GKDAFENHSYIY | 1 | 0 | 0 | 1441326 | AVV62524.1 | K9N7C7 |
| GKDASDFIDSVR | 1 | 0 | 0 | 1441328 | AVP78031.1 | Q3LZX1 |
| GKDGNPIRDYAS | 1 | 0 | 0 | 1441337 | AVV62535.1 | K9N7C7 |
| GKDGVMYSEFPA | 1 | 0 | 0 | 1441338 | AWW13566.1 | P36334 |
| GKDLNANMSLAD | 1 | 0 | 0 | 1441339 | AST12962.1 | A0A0P0LKV0 |
| GKEFEFAEGQGV | 1 | 0 | 0 | 1441344 | AEN19370.1 | P33469 |
| GKEFKFAEGQGV | 1 | 0 | 0 | 1441347 | ARA15425.1 | P33469 |
| GKEFQFAEGQGV | 1 | 0 | 0 | 1441348 | QBP84717.1 | P33469 |
| GKEQKCCVCNRR | 1 | 0 | 0 | 1441351 | AHY22543.1 | K9N7C7 |
| GKERARTVGGVP | 1 | 0 | 0 | 1441354 | ADK37692.1 | H9EJ42 |
| GKESFEQHSYIY | 1 | 0 | 0 | 1441356 | AUM60023.1 | K9N7C7 |
| GKESLENPTYIY | 1 | 0 | 0 | 1441358 | AXN92247.1 | K9N7C7 |
| GKEVEFVEGQGV | 1 | 0 | 0 | 1441361 | AJQ24087.1 | P33469 |
| GKFPLSFPPGQG | 1 | 0 | 0 | 1441381 | AVV62533.1 | K9N4V7 |
| GKFPLTFPPGQG | 1 | 0 | 0 | 1441382 | AVV62544.1 | K9N4V7 |
| GKGSKDEQIGYW | 1 | 0 | 0 | 1441399 | AIW52795.1 | H9EJ69 |
| GKIAHWIKNTFS | 1 | 0 | 0 | 1441408 | AIV41978.1 | P0C6X6 |
| GKIKPYERDISN | 1 | 0 | 0 | 1441411 | APO40579.1 | P59594 |
| GKIVNNWLKQLI | 1 | 0 | 0 | 1441418 | AVP78041.1 | P0C6W2 |
| GKLADCLQAYGH | 1 | 0 | 0 | 1441434 | ARU07575.1 | A0A140H1G9 |
| GKLFVRCDEDGQ | 1 | 0 | 0 | 1441436 | AVV62524.1 | K9N7C7 |
| GKLFVSDYATKV | 1 | 0 | 0 | 1441438 | AVV62537.1 | K9N5Q8 |
| GKLKQKPMKAEG | 1 | 0 | 0 | 1441444 | AWK59942.1 | H9EJ42 |
| GKLLIRDNEALS | 1 | 0 | 0 | 1441451 | AVV62535.1 | K9N7C7 |
| GKLLIRDNEALT | 1 | 0 | 0 | 1441452 | AVV62524.1 | K9N7C7 |
| GKLLIRDNEGLS | 1 | 0 | 0 | 1441453 | AUM60013.1 | K9N7C7 |
| GKLLIRDNGTLS | 1 | 0 | 0 | 1441454 | AKN24802.1 | K9N7C7 |
| GKLLVRDTNKEV | 1 | 0 | 0 | 1441457 | AVI15051.1 | P0C6X6 |
| GKLYQDSQNMLR | 1 | 0 | 0 | 1441474 | ARO76381.1 | P0C6U8 |
| GKMGLFFNHTLV | 1 | 0 | 0 | 1441478 | ANF29217.1 | K9N5Q8 |
| GKMVPYCYDPTV | 1 | 0 | 0 | 1441488 | AVV62524.1 | K9N7C7 |
| GKNILGYASLCD | 1 | 0 | 0 | 1441492 | 5NH0_A | H9EJ42 |
| GKNNGIGTCPVG | 1 | 0 | 0 | 1441497 | AIL49486.1 | P36334 |
| GKNNGIGTRPAG | 1 | 0 | 0 | 1441498 | ARB66066.1 | P36334 |
| GKNNGLGTCPAG | 1 | 0 | 0 | 1441500 | AIL49503.1 | P36334 |
| GKNNGVGTCPAG | 1 | 0 | 0 | 1441502 | QEG03757.1 | P36334 |
| GKPGNVGGNDDL | 1 | 0 | 0 | 1441510 | AIV41912.1 | P0C6X6 |
| GKPLEFGATSFS | 1 | 0 | 0 | 1441523 | AVP78041.1 | P0C6W2 |
| GKPNAIDVEHLE | 1 | 0 | 0 | 1441525 | ARU07593.1 | H9EJ42 |
| GKPVISEDLWQF | 1 | 0 | 0 | 1441532 | AGT21366.1 | A0A0P0LKV0 |
| GKQGLEFAAITA | 1 | 0 | 0 | 1441542 | AVV62524.1 | K9N7C7 |
| GKQGLEYAAITA | 1 | 0 | 0 | 1441543 | AVV62535.1 | K9N7C7 |
| GKQILGNTMLED | 1 | 0 | 0 | 1441547 | AUM60013.1 | K9N7C7 |
| GKQNILLRKYGR | 1 | 0 | 0 | 1441553 | AXN92247.1 | K9N7C7 |
| GKQSSSRASFGN | 1 | 0 | 0 | 1441554 | QBP84763.1 | P33469 |
| GKQTEFVSNSHL | 1 | 0 | 0 | 1441556 | NP_073549.1 | A0A0P0LKV0 |
| GKRGGGSVTYTD | 1 | 0 | 0 | 1441562 | ARB07390.1 | A0A0P0LKV0 |
| GKRGHHKNFGGG | 1 | 0 | 0 | 1441563 | AJQ24090.1 | P33469 |
| GKRGPNHKFGGG | 1 | 0 | 0 | 1441566 | AJQ24079.1 | P33469 |
| GKSAPNSRSTSR | 1 | 0 | 0 | 1441577 | AJQ24087.1 | P33469 |
| GKSFSNISLSYQ | 1 | 0 | 0 | 1441582 | AVV62526.1 | K9N5Q8 |
| GKSKHYDKYFGV | 1 | 0 | 0 | 1441593 | AQN78665.1 | Q04853 |
| GKTLVLNAKELV | 1 | 0 | 0 | 1441611 | AUM60027.1 | K9N643 |
| GKVISTWFKLML | 1 | 0 | 0 | 1441627 | ARO76381.1 | P0C6U8 |
| GKVKQSQPKVIT | 1 | 0 | 0 | 1441629 | AXN92267.1 | K9N4V7 |
| GKVNNFVKFGSV | 1 | 0 | 0 | 1441631 | AIW52757.1 | A0A0P0K6L9 |
| GKVPLIFPPGQG | 1 | 0 | 0 | 1441632 | ASU90622.1 | K9N4V7 |
| GKVSTLVKQVLD | 1 | 0 | 0 | 1441638 | AXN92247.1 | K9N7C7 |
| GKVVPTLFKLKG | 1 | 0 | 0 | 1441639 | AVV62524.1 | K9N7C7 |
| GKYDINHGDNGF | 1 | 0 | 0 | 1441644 | AGT17757.1 | A0A140H1H0 |
| GKYEQYIKIKRM | 1 | 0 | 0 | 1441649 | 5X58_A | P59594 |
| GLAAINSVPWDT | 1 | 0 | 0 | 1441673 | AVP78041.1 | P0C6W2 |
| GLAIKSDHCGGN | 1 | 0 | 0 | 1441683 | AVI15011.1 | P36334 |
| GLAIKSDYCGGN | 1 | 0 | 0 | 1441684 | CAA83660.1 | P36334 |
| GLAMPSLFKVQN | 1 | 0 | 0 | 1441694 | AVV62535.1 | K9N7C7 |
| GLAMPTLFKVQN | 1 | 0 | 0 | 1441695 | AVV62524.1 | K9N7C7 |
| GLASAYRANSID | 1 | 0 | 0 | 1441700 | AQZ41320.1 | K9N7C7 |
| GLASDLLDNVSV | 1 | 0 | 0 | 1441702 | AVV62524.1 | K9N7C7 |
| GLASNYRENSFD | 1 | 0 | 0 | 1441703 | AVV62524.1 | K9N7C7 |
| GLAVKSDYCRGN | 1 | 0 | 0 | 1441707 | AIX10751.1 | P36334 |
| GLCDASNPTNCI | 1 | 0 | 0 | 1441712 | AHY22565.1 | K9N5Q8 |
| GLCDVANPTNCI | 1 | 0 | 0 | 1441716 | AKM76239.1 | K9N5Q8 |
| GLCNKGSPQQCI | 1 | 0 | 0 | 1441727 | AVV62526.1 | K9N5Q8 |
| GLDAMFFYGDVV | 1 | 0 | 0 | 1441737 | AEN19363.1 | P0C6X6 |
| GLDARTQAKQSY | 1 | 0 | 0 | 1441738 | AIV41936.1 | P0C6X6 |
| GLDLIQAGSVVR | 1 | 0 | 0 | 1441742 | AXN92247.1 | K9N7C7 |
| GLDLMQAGTIVR | 1 | 0 | 0 | 1441744 | ATQ39389.1 | K9N7C7 |
| GLDLTPVIDDVD | 1 | 0 | 0 | 1441747 | AWK59942.1 | H9EJ42 |
| GLDQTSFNVNSL | 1 | 0 | 0 | 1441748 | AVV62535.1 | K9N7C7 |
| GLDYKTFKAIVE | 1 | 0 | 0 | 1441756 | ARO76381.1 | P0C6U8 |
| GLEAYADRTLQE | 1 | 0 | 0 | 1441767 | AIV41774.1 | P0C6X6 |
| GLENHAFEHVVY | 1 | 0 | 0 | 1441777 | AMQ49013.1 | K9N7C7 |
| GLENIAFNVVKH | 1 | 0 | 0 | 1441780 | AVV62524.1 | K9N7C7 |
| GLENIAYNVVKQ | 1 | 0 | 0 | 1441782 | AUM60023.1 | K9N7C7 |
| GLENIVFNVVKQ | 1 | 0 | 0 | 1441783 | AXN92247.1 | K9N7C7 |
| GLENYAFEHVVY | 1 | 0 | 0 | 1441785 | AXN92247.1 | K9N7C7 |

| | | | | | | |
|---|---|---|---|---|---|---|
| GLEYAAITAPFL | 1 | 0 | 0 | 1441790 | AVV62535.1 | K9N7C7 |
| GLFHSSQLPLDV | 1 | 0 | 0 | 1441798 | AJC98124.1 | P0C6X6 |
| GLFKDCSKIING | 1 | 0 | 0 | 1441802 | ATO98167.1 | P0C6W2 |
| GLFKDCSREISG | 1 | 0 | 0 | 1441806 | AUM60013.1 | K9N7C7 |
| GLFPKHGDLGEQ | 1 | 0 | 0 | 1441813 | AVV62537.1 | K9N5Q8 |
| GLFPYQGDHGDR | 1 | 0 | 0 | 1441818 | ALA49803.1 | K9N5Q8 |
| GLFRIDVPSAES | 1 | 0 | 0 | 1441821 | ALW82670.1 | K9N796 |
| GLFRIFFRNAYV | 1 | 0 | 0 | 1441822 | ALA49901.1 | K9N7C7 |
| GLFRILLRNAYV | 1 | 0 | 0 | 1441825 | ALA50044.1 | K9N7C7 |
| GLFRRRSVGNTG | 1 | 0 | 0 | 1441828 | AZP73837.1 | P0C6X6 |
| GLFYIDVGNHRS | 1 | 0 | 0 | 1441836 | ACY56731.1 | A0A0P0G321 |
| GLGLIQAGSVVR | 1 | 0 | 0 | 1441847 | ASU45706.1 | K9N7C7 |
| GLGTVDADYKMC | 1 | 0 | 0 | 1441850 | APT69849.1 | A0A0P0K6L9 |
| GLGTVDANYKKC | 1 | 0 | 0 | 1441851 | CAA71147.1 | A0A0P0K6L9 |
| GLHAPHNSWEAL | 1 | 0 | 0 | 1441856 | AVV62535.1 | K9N7C7 |
| GLHAPRNSWEAM | 1 | 0 | 0 | 1441858 | AUM60013.1 | K9N7C7 |
| GLHPTQSPTYLS | 1 | 0 | 0 | 1441872 | APO40578.1 | P0C6X7 |
| GLHTVQSGDLNV | 1 | 0 | 0 | 1441874 | AVV62535.1 | K9N7C7 |
| GLHYPTGSCLIA | 1 | 0 | 0 | 1441875 | ALD51908.1 | K9N7D2 |
| GLIASKGPKQAS | 1 | 0 | 0 | 1441881 | APO40578.1 | P0C6X7 |
| GLIDLIVNVLSI | 1 | 0 | 0 | 1441882 | QED88011.1 | H9EJ42 |
| GLILKFDSGTVT | 1 | 0 | 0 | 1441894 | AVN89451.1 | K9N7C7 |
| GLINSSLFVEDC | 1 | 0 | 0 | 1441896 | ALK80261.1 | K9N5Q8 |
| GLKGGIRKAKDY | 1 | 0 | 0 | 1441906 | AVV62524.1 | K9N7C7 |
| GLKNGQGENDNI | 1 | 0 | 0 | 1441919 | ATP16771.1 | P33469 |
| GLLDLAWRVPCA | 1 | 0 | 0 | 1441940 | AIV41888.1 | P0C6X6 |
| GLLDQAWRFPCA | 1 | 0 | 0 | 1441941 | AEN19363.1 | P0C6X6 |
| GLLEPVKHSSTV | 1 | 0 | 0 | 1441946 | AVV62535.1 | K9N7C7 |
| GLLEPVNHSATV | 1 | 0 | 0 | 1441947 | AVV62524.1 | K9N7C7 |
| GLLHSSQLPLDI | 1 | 0 | 0 | 1441957 | AIV41960.1 | P0C6X6 |
| GLLLSSIYHLYV | 1 | 0 | 0 | 1441968 | AVV62524.1 | K9N7C7 |
| GLLNVQKVAQPK | 1 | 0 | 0 | 1441971 | APT69882.1 | A0A0P0ILY9 |
| GLLQPTANSTTV | 1 | 0 | 0 | 1441977 | AHI48527.1 | K9N638 |
| GLLRPTDNSTTV | 1 | 0 | 0 | 1441985 | ANI69823.1 | K9N638 |
| GLLSENAYLKWA | 1 | 0 | 0 | 1441986 | AUM60013.1 | K9N7C7 |
| GLLTSVYCIHKE | 1 | 0 | 0 | 1441988 | Q3I5I8.1 | Q3LZX5 |
| GLLYDSPCFSQQ | 1 | 0 | 0 | 1441992 | AAA73152.1 | P30215 |
| GLLYNSPCFSQQ | 1 | 0 | 0 | 1441995 | P30215.1 | P30215 |
| GLLYNSSCFSHQ | 1 | 0 | 0 | 1441996 | QBQ01838.1 | P30215 |
| GLLYNSSCFSQQ | 1 | 0 | 0 | 1441997 | AEN19365.1 | P30215 |
| GLMLLELKERDQ | 1 | 0 | 0 | 1442005 | APO40578.1 | P0C6X7 |
| GLMQNAFLYSSL | 1 | 0 | 0 | 1442006 | AIV41774.1 | P0C6X6 |
| GLNFTETITTGF | 1 | 0 | 0 | 1442015 | ATP16766.1 | P30215 |
| GLNGDFNLTLLQ | 1 | 0 | 0 | 1442016 | AVV62537.1 | K9N5Q8 |
| GLNGDFNLTMLQ | 1 | 0 | 0 | 1442017 | AVV62526.1 | K9N5Q8 |
| GLNNLNRGMVLE | 1 | 0 | 0 | 1442027 | 3EE7_A | P0C6X7 |
| GLNTLLVQPALY | 1 | 0 | 0 | 1442032 | ALA49390.1 | K9N5R3 |
| GLPGTVIRTLNG | 1 | 0 | 0 | 1442038 | APO40578.1 | P0C6X7 |
| GLPLEFGASTEM | 1 | 0 | 0 | 1442042 | ATO98229.1 | P0C6W2 |
| GLPLEFGASTET | 1 | 0 | 0 | 1442043 | ARO76381.1 | P0C6U8 |
| GLPNFKKVETLN | 1 | 0 | 0 | 1442047 | AET99208.1 | Q0ZJ80 |
| GLPNNIASWFTA | 1 | 0 | 0 | 1442049 | AAP49024.1 | P59595 |
| GLPNTCFGPLVR | 1 | 0 | 0 | 1442051 | AIV41996.1 | P0C6X6 |
| GLQDGFYSANFQ | 1 | 0 | 0 | 1442064 | AWO68987.1 | A0A0P0G321 |
| GLQFQILDEGLD | 1 | 0 | 0 | 1442066 | ARK08668.1 | Q80872 |
| GLQKCEQLLREY | 1 | 0 | 0 | 1442068 | QFS19588.1 | K9N5Q8 |
| GLQLFTTWLPDL | 1 | 0 | 0 | 1442069 | ARU07575.1 | A0A140H1G9 |
| GLQMAFIISVQY | 1 | 0 | 0 | 1442070 | AVV62526.1 | K9N5Q8 |
| GLQYAAITAPYV | 1 | 0 | 0 | 1442072 | AXN92247.1 | K9N7C7 |
| GLRDYITNRTFM | 1 | 0 | 0 | 1442075 | ABD43179.1 | P36334 |
| GLRGSGDLQGNF | 1 | 0 | 0 | 1442082 | AUM60021.1 | K9N4V7 |
| GLSEDGQVFTRQ | 1 | 0 | 0 | 1442096 | AVV62526.1 | K9N5Q8 |
| GLSGIGAVGGDL | 1 | 0 | 0 | 1442097 | ALA49601.1 | K9N4V7 |
| GLSKPFVKGDFS | 1 | 0 | 0 | 1442106 | AUM60013.1 | K9N7C7 |
| GLSSFAAIPFAQ | 1 | 0 | 0 | 1442109 | AXN92228.1 | K9N5Q8 |
| GLSSFASIPFAQ | 1 | 0 | 0 | 1442110 | AVV62526.1 | K9N5Q8 |
| GLSSSAAIPFAQ | 1 | 0 | 0 | 1442111 | AOR17480.1 | K9N5Q8 |
| GLTCSTDLQKAN | 1 | 0 | 0 | 1442116 | CAA79896.1 | P36334 |
| GLTCSTDSQKAN | 1 | 0 | 0 | 1442117 | BBO58736.1 | P36334 |
| GLTCVKDYEFKF | 1 | 0 | 0 | 1442118 | ABD75503.1 | A0A140H1G9 |
| GLTFQTVRSGNF | 1 | 0 | 0 | 1442127 | AHI48750.1 | K9N7C7 |
| GLTFSYDGLKGG | 1 | 0 | 0 | 1442128 | AVV62524.1 | K9N7C7 |
| GLTNLNEITKPL | 1 | 0 | 0 | 1442135 | ARU07582.1 | Q0ZJ80 |
| GLVGTTLQGKAI | 1 | 0 | 0 | 1442165 | AUM60013.1 | K9N7C7 |
| GLVNSSLLVEDC | 1 | 0 | 0 | 1442176 | ATW75477.1 | K9N5Q8 |
| GLVNSSLSVEDC | 1 | 0 | 0 | 1442177 | ALL26396.1 | K9N5Q8 |
| GLVNSSMVVDNC | 1 | 0 | 0 | 1442178 | AVV62537.1 | K9N5Q8 |
| GLVNTSMVVDEC | 1 | 0 | 0 | 1442179 | AVV62526.1 | K9N5Q8 |
| GLVNVQKFAQPK | 1 | 0 | 0 | 1442180 | ARB07390.1 | A0A0P0LKV0 |
| GLVNVQKVAQPK | 1 | 0 | 0 | 1442181 | AST12962.1 | A0A0P0LKV0 |
| GLVPRGSAMGLP | 1 | 0 | 0 | 1442183 | 1SSK_A | P59595 |
| GLVRMYNLLACP | 1 | 0 | 0 | 1442195 | AMQ49058.1 | K9N638 |
| GLWSPTITHTNC | 1 | 0 | 0 | 1442214 | ARU07575.1 | A0A140H1G9 |
| GLYEIKIPTNFT | 1 | 0 | 0 | 1442217 | BBA20983.1 | A0A140H1H1 |
| GLYEIQIPSDFT | 1 | 0 | 0 | 1442218 | ARB66062.1 | P36334 |
| GLYKNMCTGSPP | 1 | 0 | 0 | 1442231 | AHI48748.1 | K9N7C7 |
| GLYKNTCVGSDNVT | 1 | 0 | 0 | 1442233 | P0DTD1.1 | P0DTD1 |
| GLYTNRVVSPPN | 1 | 0 | 0 | 1442252 | AUM60015.1 | K9N796 |
| GLYVPIATRNNC | 1 | 0 | 0 | 1442254 | AXX83378.1 | P0C6X6 |
| GMAITIKPEANI | 1 | 0 | 0 | 1442262 | ABD75615.1 | A0A140H1G9 |
| GMDNLACEVLKP | 1 | 0 | 0 | 1442274 | AVP78041.1 | P0C6W2 |

| | | | | | | |
|---|---|---|---|---|---|---|
| GMEVTPTGTWLT | 1 | 0 | 0 | 1442287 | APO40586.1 | P59595 |
| GMGTCPAGTNYL | 1 | 0 | 0 | 1442298 | AIL49487.1 | P36334 |
| GMHFGQCDYAQL | 1 | 0 | 0 | 1442301 | AUM60030.1 | K9N7A1 |
| GMKETAAAKFER | 1 | 0 | 0 | 1442312 | 3R24_A | P0C6X7 |
| GMKYVKWEHDSG | 1 | 0 | 0 | 1442316 | NP_073549.1 | A0A0P0LKV0 |
| GMLNGMPVSSIK | 1 | 0 | 0 | 1442318 | AWK59942.1 | H9EJ42 |
| GMLYPITLGSLE | 1 | 0 | 0 | 1442321 | AGT51685.1 | Q4VID0 |
| GMLYPTTLGSLE | 1 | 0 | 0 | 1442323 | AGT51745.1 | Q4VID0 |
| GMMNISPKPQRQ | 1 | 0 | 0 | 1442324 | AIX10729.1 | P33469 |
| GMMNMSPKPQLQ | 1 | 0 | 0 | 1442325 | AMM42447.1 | P33469 |
| GMNGRTILGSALLED | 1 | 0 | 0 | 1442331 | P0DTC1.1 | P0DTD1 |
| GMNMSPKPQRQR | 1 | 0 | 0 | 1442333 | AIV41850.1 | P33469 |
| GMNTSHSVCNGC | 1 | 0 | 0 | 1442334 | AIW52756.1 | A0A0P0K6L9 |
| GMNTSQSVCNGC | 1 | 0 | 0 | 1442335 | ABB90520.1 | A0A0P0K6L9 |
| GMNTSYPVCNGR | 1 | 0 | 0 | 1442336 | ADK37701.1 | A0A0P0K6L9 |
| GMNTSYSVCNGC | 1 | 0 | 0 | 1442337 | ADK37704.1 | A0A0P0K6L9 |
| GMPVSSIKSDKG | 1 | 0 | 0 | 1442346 | AGT51393.1 | H9EJ42 |
| GMQTVEDLRARM | 1 | 0 | 0 | 1442350 | AVN89451.1 | K9N7C7 |
| GMSFYCEDHKPQ | 1 | 0 | 0 | 1442354 | AGT51438.1 | P0C6X6 |
| GMSYSMTTFEIA | 1 | 0 | 0 | 1442368 | AIV41888.1 | P0C6X6 |
| GMSYYCKMHKPP | 1 | 0 | 0 | 1442371 | APO40578.1 | P0C6X7 |
| GMTNCLASECFM | 1 | 0 | 0 | 1442376 | AIW52776.1 | A0A0P0LKV0 |
| GMVNMSPKPQRQ | 1 | 0 | 0 | 1442381 | AEN19370.1 | P33469 |
| GNAAIPDYNYYK | 1 | 0 | 0 | 1442390 | ACV87150.1 | A0A140H1G9 |
| GNAINVGGRGLQ | 1 | 0 | 0 | 1442394 | ATQ39389.1 | K9N7C7 |
| GNALYNNECGRA | 1 | 0 | 0 | 1442397 | AGT21359.1 | A0A0P0LKV0 |
| GNASGNDALARF | 1 | 0 | 0 | 1442401 | ARB07436.1 | A0A140H1G9 |
| GNAVQGFACDCG | 1 | 0 | 0 | 1442405 | AVV62524.1 | K9N7C7 |
| GNCAKRFRRVLV | 1 | 0 | 0 | 1442409 | AWK59942.1 | H9EJ42 |
| GNCIGKGSLTAS | 1 | 0 | 0 | 1442414 | AUM60027.1 | K9N643 |
| GNCTFGSPDDAS | 1 | 0 | 0 | 1442421 | AHI48794.1 | K9N638 |
| GNCVEYHHHHHH | 1 | 0 | 0 | 1442422 | 4KQZ_A | K9N5Q8 |
| GNDDLSISTIFT | 1 | 0 | 0 | 1442431 | AIV41912.1 | P0C6X6 |
| GNDEKVMTFYPR | 1 | 0 | 0 | 1442432 | AIV41912.1 | P0C6X6 |
| GNDLSLPVMFSE | 1 | 0 | 0 | 1442433 | AST12962.1 | A0A0P0LKV0 |
| GNEALSISTIFT | 1 | 0 | 0 | 1442440 | AIV41792.1 | P0C6X6 |
| GNESWLTGNAIQ | 1 | 0 | 0 | 1442442 | AVN89333.1 | K9N7C7 |
| GNETTFAFVGAL | 1 | 0 | 0 | 1442443 | ABB90514.1 | A0A0P0K6L9 |
| GNEWAQVLSEIV | 1 | 0 | 0 | 1442446 | AKL87224.1 | P0C6X6 |
| GNFDNDINCPVT | 1 | 0 | 0 | 1442451 | AVV62524.1 | K9N7C7 |
| GNFGDLQLTKLG | 1 | 0 | 0 | 1442455 | AVN89320.1 | K9N4V7 |
| GNFNQDFYDFIL | 1 | 0 | 0 | 1442470 | AGT51787.1 | P0C6X6 |
| GNFPKSGANFRD | 1 | 0 | 0 | 1442471 | ABA60802.1 | A0A140H1H1 |
| GNFRTGVDPDLS | 1 | 0 | 0 | 1442473 | ASU90030.1 | K9N7C7 |
| GNFTARANCLAT | 1 | 0 | 0 | 1442481 | AVV62529.1 | K9N643 |
| GNFTPIDQYMCG | 1 | 0 | 0 | 1442482 | AUM60013.1 | K9N7C7 |
| GNGGDTALALLL | 1 | 0 | 0 | 1442490 | AVP78038.1 | Q3LZX4 |
| GNGILKWSDQSD | 1 | 0 | 0 | 1442493 | ARK08639.1 | P33469 |
| GNGNHIISLVQN | 1 | 0 | 0 | 1442495 | CAA79896.1 | P36334 |
| GNGNHILSLVQN | 1 | 0 | 0 | 1442496 | BBA20983.1 | A0A140H1H1 |
| GNGPGIDAGYKN | 1 | 0 | 0 | 1442497 | CAA83660.1 | P36334 |
| GNGSMIEDMDYV | 1 | 0 | 0 | 1442502 | AXG21663.1 | K9N7C7 |
| GNGSMIEDVDYV | 1 | 0 | 0 | 1442503 | AMQ49035.1 | K9N7C7 |
| GNGSMLEDIDYV | 1 | 0 | 0 | 1442504 | AVV62524.1 | K9N7C7 |
| GNGTKQLAPRWY | 1 | 0 | 0 | 1442508 | AUM60021.1 | K9N4V7 |
| GNHCPAGNSHTS | 1 | 0 | 0 | 1442513 | AHX71946.1 | K9N5Q8 |
| GNHCPAGNSYTS | 1 | 0 | 0 | 1442514 | AXN92228.1 | K9N5Q8 |
| GNHFKELWHLDT | 1 | 0 | 0 | 1442517 | AEN19366.1 | P36334 |
| GNHFKESWHMDT | 1 | 0 | 0 | 1442520 | AIL49486.1 | P36334 |
| GNHFRHPLDEDE | 1 | 0 | 0 | 1442521 | AVV62524.1 | K9N7C7 |
| GNHRSAFALHTG | 1 | 0 | 0 | 1442528 | AIW52843.1 | A0A0P0G321 |
| GNIIPHYSWFSG | 1 | 0 | 0 | 1442540 | ABA60796.1 | Q19U29 |
| GNIKPHNYSYMD | 1 | 0 | 0 | 1442542 | AIL49382.1 | P30215 |
| GNITKTNVQPTG | 1 | 0 | 0 | 1442546 | NP_073549.1 | A0A0P0LKV0 |
| GNKGAGGHSYGADLKSFDL | 1 | 0 | 0 | 1442554 | P0DTC1.1 | P0DTD1 |
| GNKIGDLSIRAL | 1 | 0 | 0 | 1442558 | AJE26218.1 | P0C6X6 |
| GNLGGNCAGNAS | 1 | 0 | 0 | 1442577 | ARB07436.1 | A0A140H1G9 |
| GNLGNLQSFNYR | 1 | 0 | 0 | 1442579 | AIL49492.1 | P36334 |
| GNLGNLQSSNYR | 1 | 0 | 0 | 1442580 | AIL49519.1 | P36334 |
| GNLKHTTMMLSF | 1 | 0 | 0 | 1442589 | ATO98145.1 | Q3LZX1 |
| GNLKMAGMHFGQ | 1 | 0 | 0 | 1442590 | AUM60030.1 | K9N7A1 |
| GNLQSSNYRIDT | 1 | 0 | 0 | 1442601 | AIL49519.1 | P36334 |
| GNMEEFIQTSSP | 1 | 0 | 0 | 1442615 | CAA79896.1 | P36334 |
| GNMFQFATFPVY | 1 | 0 | 0 | 1442616 | ALA49363.1 | K9N5Q8 |
| GNMFQTQAGCLV | 1 | 0 | 0 | 1442618 | APO40579.1 | P59594 |
| GNMFRFASLPVY | 1 | 0 | 0 | 1442619 | AVV62526.1 | K9N5Q8 |
| GNMGEFIQTSSP | 1 | 0 | 0 | 1442620 | ARB66062.1 | P36334 |
| GNMLTGIAAHPP | 1 | 0 | 0 | 1442625 | AVN89289.1 | K9N7C7 |
| GNMLTGIAARPP | 1 | 0 | 0 | 1442626 | AXN92247.1 | K9N7C7 |
| GNMMTCISARPP | 1 | 0 | 0 | 1442628 | AVV62524.1 | K9N7C7 |
| GNMMTGISARPP | 1 | 0 | 0 | 1442629 | AVV62535.1 | K9N7C7 |
| GNNEAKALFFTY | 1 | 0 | 0 | 1442635 | AVV62535.1 | K9N7C7 |
| GNNSCTCQPLAF | 1 | 0 | 0 | 1442637 | BBO58769.1 | P36334 |
| GNNSCTCQPQAF | 1 | 0 | 0 | 1442638 | AEN19366.1 | P36334 |
| GNNSWLTGNAIQ | 1 | 0 | 0 | 1442639 | ALA49340.1 | K9N7C7 |
| GNNVVVMSTCAV | 1 | 0 | 0 | 1442642 | CAA83660.1 | P36334 |
| GNNVYCYNTELM | 1 | 0 | 0 | 1442644 | AST12962.1 | A0A0P0LKV0 |
| GNPDEPHKDVYE | 1 | 0 | 0 | 1442648 | AIX10716.1 | P33469 |
| GNPKGIPIFDDP | 1 | 0 | 0 | 1442657 | AVN89322.1 | K9N7C7 |
| GNPNNHKKIVQS | 1 | 0 | 0 | 1442663 | AVV62529.1 | K9N643 |
| GNPVVEYLSKED | 1 | 0 | 0 | 1442667 | QGV13494.1 | K9N7C7 |

| | | | | | | |
|---|---|---|---|---|---|---|
| GNPVYFLRYSGA | 1 | 0 | 0 | 1442668 | AXN92267.1 | K9N4V7 |
| GNRNPTSGVTPD | 1 | 0 | 0 | 1442680 | AJQ24067.1 | P33469 |
| GNRPGNGILKWA | 1 | 0 | 0 | 1442681 | AMM42456.1 | P33469 |
| GNRRSAFALHSG | 1 | 0 | 0 | 1442682 | AMM42641.1 | A0A0P0G321 |
| GNRRVELWHWDT | 1 | 0 | 0 | 1442683 | CAA83660.1 | P36334 |
| GNRSGNGILKWS | 1 | 0 | 0 | 1442686 | ARK08639.1 | P33469 |
| GNRTPTPGVTPD | 1 | 0 | 0 | 1442689 | AIX10722.1 | P33469 |
| GNSLNHLTFQLR | 1 | 0 | 0 | 1442706 | ABB90520.1 | A0A0P0K6L9 |
| GNSLNYLTSQLR | 1 | 0 | 0 | 1442708 | AAK32190.1 | A0A0P0K6L9 |
| GNSNYNTFAVFD | 1 | 0 | 0 | 1442711 | AVV62537.1 | K9N5Q8 |
| GNSPYHPLADNK | 1 | 0 | 0 | 1442717 | APO40584.1 | P59635 |
| GNSSRGSSPGPA | 1 | 0 | 0 | 1442724 | AUM60021.1 | K9N4V7 |
| GNSSRNASPGPS | 1 | 0 | 0 | 1442726 | AVV62533.1 | K9N4V7 |
| GNSSTITNQLGV | 1 | 0 | 0 | 1442727 | AVV62526.1 | K9N5Q8 |
| GNSTRGTFPGPS | 1 | 0 | 0 | 1442731 | ANI69896.1 | K9N4V7 |
| GNSTRGTSPCPS | 1 | 0 | 0 | 1442732 | QCI31476.1 | K9N4V7 |
| GNSTRGTSPVPS | 1 | 0 | 0 | 1442735 | AIY60565.1 | K9N4V7 |
| GNSTRSTSPGPS | 1 | 0 | 0 | 1442737 | AVN89381.1 | K9N4V7 |
| GNSTTITDKLGV | 1 | 0 | 0 | 1442738 | AVV62537.1 | K9N5Q8 |
| GNSYTSFAIYHT | 1 | 0 | 0 | 1442739 | ATG84866.1 | K9N5Q8 |
| GNTMYVNMAVPH | 1 | 0 | 0 | 1442758 | AUM60027.1 | K9N643 |
| GNTSFACLSNVS | 1 | 0 | 0 | 1442760 | QED88040.1 | A0A0P0G321 |
| GNTSFDFLSNVS | 1 | 0 | 0 | 1442761 | AWO68966.1 | A0A0P0G321 |
| GNVDLDNLASEL | 1 | 0 | 0 | 1442772 | AUM60013.1 | K9N7C7 |
| GNVEAVNFNVTN | 1 | 0 | 0 | 1442774 | AIW52757.1 | A0A0P0K6L9 |
| GNVGGNDALSTS | 1 | 0 | 0 | 1442779 | AWW13571.1 | P0C6X6 |
| GNVGGNEALSIS | 1 | 0 | 0 | 1442780 | AIV41792.1 | P0C6X6 |
| GNVILKPAADGL | 1 | 0 | 0 | 1442783 | APO40578.1 | P0C6X7 |
| GNVILKSSDEGI | 1 | 0 | 0 | 1442786 | ARO76381.1 | P0C6U8 |
| GNVLDQAIAMWA | 1 | 0 | 0 | 1442790 | APO40578.1 | P0C6X7 |
| GNVVQQCCTAAV | 1 | 0 | 0 | 1442808 | ALA49384.1 | K9N7C7 |
| GNYFTFKQANNN | 1 | 0 | 0 | 1442816 | AIV41846.1 | P0C6X6 |
| GNYNCNASLNSF | 1 | 0 | 0 | 1442824 | AHL31360.1 | K9N5Q8 |
| GNYRVTKGKPVQ | 1 | 0 | 0 | 1442847 | ARO76381.1 | P0C6U8 |
| GNYTISCEPLEI | 1 | 0 | 0 | 1442852 | ATO98227.1 | Q3LZX5 |
| GPAGVESDIETG | 1 | 0 | 0 | 1442873 | AVV62524.1 | K9N7C7 |
| GPASARELICAQ | 1 | 0 | 0 | 1442881 | ARA67740.1 | K9N5Q8 |
| GPASARNLICAQ | 1 | 0 | 0 | 1442882 | QFS19619.1 | K9N5Q8 |
| GPASVRDLICAQ | 1 | 0 | 0 | 1442883 | ALJ54502.1 | K9N5Q8 |
| GPCWSALVSALK | 1 | 0 | 0 | 1442887 | ARX11394.1 | A0A0P0LKV0 |
| GPDATNNTCLEV | 1 | 0 | 0 | 1442893 | AUM60014.1 | K9N5Q8 |
| GPDDQIGFYRRA | 1 | 0 | 0 | 1442895 | AAT76155.1 | Q3LZX4 |
| GPDHSVADYHNH | 1 | 0 | 0 | 1442900 | APO40578.1 | P0C6X7 |
| GPDLEGKFYGPF | 1 | 0 | 0 | 1442904 | ACV88184.1 | P0C6X7 |
| GPDQICNTPTQC | 1 | 0 | 0 | 1442908 | ARB07411.1 | P0C6X6 |
| GPDVFLHKCYRY | 1 | 0 | 0 | 1442921 | QED88024.1 | H9EJ42 |
| GPDYAYFNGATI | 1 | 0 | 0 | 1442924 | AUM60013.1 | K9N7C7 |
| GPEAALPYGANK | 1 | 0 | 0 | 1442926 | APO40586.1 | P59595 |
| GPELYHYQECVR | 1 | 0 | 0 | 1442934 | 1YO4_A | P59635 |
| GPFMGNAINVGG | 1 | 0 | 0 | 1442947 | ATQ39389.1 | K9N7C7 |
| GPFVDIQTAQAA | 1 | 0 | 0 | 1442948 | 3ATW_A | P0C6X7 |
| GPFVEGAINVGK | 1 | 0 | 0 | 1442950 | AUM60023.1 | K9N7C7 |
| GPGALQGNFGDL | 1 | 0 | 0 | 1442951 | AVV62544.1 | K9N4V7 |
| GPGESKPQSRNP | 1 | 0 | 0 | 1442953 | AAA45463.1 | A0A0N7J131 |
| GPGIDAGYKNSG | 1 | 0 | 0 | 1442955 | CAA83660.1 | P36334 |
| GPGKNNGIGTCP | 1 | 0 | 0 | 1442958 | CAA79896.1 | P36334 |
| GPGKNNGMGTCP | 1 | 0 | 0 | 1442959 | AIL49487.1 | P36334 |
| GPHAKDQYGSDI | 1 | 0 | 0 | 1442964 | AIX10729.1 | P33469 |
| GPHANDHYGTDI | 1 | 0 | 0 | 1442967 | AJQ24067.1 | P33469 |
| GPHKAAKFRERV | 1 | 0 | 0 | 1442974 | BAP75906.1 | A0A0N7J131 |
| GPHKDAKFRERV | 1 | 0 | 0 | 1442975 | AIW52723.1 | A0A0N7J131 |
| GPHKDLKFRQRS | 1 | 0 | 0 | 1442978 | AIW52795.1 | H9EJ69 |
| GPIEHVTSVNLF | 1 | 0 | 0 | 1442984 | AVN89292.1 | K9N796 |
| GPIQTPVGCAMG | 1 | 0 | 0 | 1442990 | AUM60014.1 | K9N5Q8 |
| GPISPISLEMFK | 1 | 0 | 0 | 1442991 | AGT51805.1 | Q4VID0 |
| GPISPTNLEMFK | 1 | 0 | 0 | 1442992 | AOL02458.1 | Q4VID0 |
| GPISPTSLEMFK | 1 | 0 | 0 | 1442994 | ARE29989.1 | Q4VID0 |
| GPIYLDGADVTK | 1 | 0 | 0 | 1443000 | AVP78030.1 | P0C6W2 |
| GPKKSTELVKNK | 1 | 0 | 0 | 1443003 | APO40579.1 | P59594 |
| GPKLSTQLVKNQ | 1 | 0 | 0 | 1443007 | AVP78031.1 | Q3LZX1 |
| GPLGSSGLKKMA | 1 | 0 | 0 | 1443020 | 5GWY_A | H9EJ42 |
| GPLQNFGNSEML | 1 | 0 | 0 | 1443028 | AMM42394.1 | Q19U29 |
| GPLQVGDAVLLK | 1 | 0 | 0 | 1443035 | AVV62524.1 | K9N7C7 |
| GPLQVGDSVLLQ | 1 | 0 | 0 | 1443036 | AXN92247.1 | K9N7C7 |
| GPMQDVNQDDDR | 1 | 0 | 0 | 1443046 | AVV62533.1 | K9N4V7 |
| GPNEHVTSVNLL | 1 | 0 | 0 | 1443049 | AGV08391.1 | K9N796 |
| GPNFAEISKINT | 1 | 0 | 0 | 1443050 | AUM60014.1 | K9N5Q8 |
| GPNFQEISKINT | 1 | 0 | 0 | 1443051 | AVV62526.1 | K9N5Q8 |
| GPNKNAGQDISL | 1 | 0 | 0 | 1443052 | APO40578.1 | P0C6X7 |
| GPNKYCYCGNFA | 1 | 0 | 0 | 1443053 | QED88017.1 | H9EJ42 |
| GPNTRSLSRSNS | 1 | 0 | 0 | 1443060 | AGW27840.1 | Q19U29 |
| GPPFISTDTVDV | 1 | 0 | 0 | 1443064 | AKA09663.1 | P36334 |
| GPPIAPGVPATE | 1 | 0 | 0 | 1443067 | P33469.1 | P33469 |
| GPPNNTASWFTA | 1 | 0 | 0 | 1443068 | APO40586.1 | P59595 |
| GPPQGAFHVTMR | 1 | 0 | 0 | 1443070 | ABD75599.1 | A0A140H1G9 |
| GPPSISTATVDV | 1 | 0 | 0 | 1443074 | AVQ05264.1 | P36334 |
| GPPSISTDTVDV | 1 | 0 | 0 | 1443075 | AEN19366.1 | P36334 |
| GPQNQRSAPRIT | 1 | 0 | 0 | 1443078 | AVP78038.1 | Q3LZX4 |
| GPQNQSSAPRIT | 1 | 0 | 0 | 1443079 | AVP78049.1 | Q3LZX4 |
| GPSGIGTVGGDL | 1 | 0 | 0 | 1443106 | ATG84708.1 | K9N4V7 |
| GPSSARDLICAQ | 1 | 0 | 0 | 1443110 | AHL18090.1 | K9N5Q8 |

| | | | | | | |
|---|---|---|---|---|---|---|
| GPSVETSDSQFE | 1 | 0 | 0 | 1443112 | AGT51787.1 | P0C6X6 |
| GPTDSADNNQNG | 1 | 0 | 0 | 1443114 | ARO76389.1 | P59595 |
| GPVFTGDLFYTI | 1 | 0 | 0 | 1443134 | ABE97136.1 | H9EJ42 |
| GPYAKDQYGTDI | 1 | 0 | 0 | 1443137 | AIL49482.1 | P33469 |
| GPYANASYGQSL | 1 | 0 | 0 | 1443139 | ADN03343.1 | Q19U29 |
| GPYEFCSQHTMQ | 1 | 0 | 0 | 1443142 | AVL25535.1 | H9EJ42 |
| GPYKDAQVVQLP | 1 | 0 | 0 | 1443149 | AEN19363.1 | P0C6X6 |
| GPYVEGAINVGK | 1 | 0 | 0 | 1443151 | AVV62524.1 | K9N7C7 |
| GQAFFYSGGFYR | 1 | 0 | 0 | 1443158 | AWK59942.1 | H9EJ42 |
| GQAMESVGFVYD | 1 | 0 | 0 | 1443162 | ARU07575.1 | A0A140H1G9 |
| GQANLFRNARNG | 1 | 0 | 0 | 1443164 | AVP78041.1 | P0C6W2 |
| GQAVPTLFRLKG | 1 | 0 | 0 | 1443168 | AUM60013.1 | K9N7C7 |
| GQAYVRKDGEWV | 1 | 0 | 0 | 1443170 | 6CRW_A | P59594 |
| GQCDFRSLPNEV | 1 | 0 | 0 | 1443171 | AVV62543.1 | K9N7A1 |
| GQCDYAQLPNEI | 1 | 0 | 0 | 1443172 | AUM60030.1 | K9N7A1 |
| GQCTFGAPDDAT | 1 | 0 | 0 | 1443176 | AUM60013.1 | K9N7C7 |
| GQECLARSACVY | 1 | 0 | 0 | 1443183 | AZK15899.1 | K9N638 |
| GQEENDNISVAV | 1 | 0 | 0 | 1443185 | AGT51525.1 | P33469 |
| GQFCTESVVDLQ | 1 | 0 | 0 | 1443197 | AVV62524.1 | K9N7C7 |
| GQFVGWVIPLAF | 1 | 0 | 0 | 1443205 | AEN19363.1 | P0C6X6 |
| GQFYINGFKYFD | 1 | 0 | 0 | 1443206 | AIW52843.1 | A0A0P0G321 |
| GQGENDNTSVAV | 1 | 0 | 0 | 1443211 | AIX09811.1 | P33469 |
| GQGEYDNISVAV | 1 | 0 | 0 | 1443212 | AXX83299.1 | P33469 |
| GQGHLGYTLEES | 1 | 0 | 0 | 1443213 | APO40578.1 | P0C6X7 |
| GQGIFFEVNATY | 1 | 0 | 0 | 1443214 | AAX84795.1 | P36334 |
| GQGQNNADPKVY | 1 | 0 | 0 | 1443220 | ARO76391.1 | P59636 |
| GQGVPIAPGVPS | 1 | 0 | 0 | 1443228 | AJQ24074.1 | P33469 |
| GQGVPIAYGIPP | 1 | 0 | 0 | 1443230 | AMM42397.1 | Q19U29 |
| GQGVPLAFGVPL | 1 | 0 | 0 | 1443235 | ABA60796.1 | Q19U29 |
| GQIQVTSDNIKE | 1 | 0 | 0 | 1443246 | ARO76381.1 | P0C6U8 |
| GQIVTRAKEVKE | 1 | 0 | 0 | 1443249 | AVP78041.1 | P0C6W2 |
| GQKAIVAYTMSL | 1 | 0 | 0 | 1443250 | AVP78042.1 | Q3LZX1 |
| GQKNGQGENDNI | 1 | 0 | 0 | 1443253 | BBM61433.1 | P33469 |
| GQKSTTLIGVEA | 1 | 0 | 0 | 1443255 | APO40578.1 | P0C6X7 |
| GQKTYERHSLSH | 1 | 0 | 0 | 1443260 | AVP78041.1 | P0C6W2 |
| GQLKNFEAPLKC | 1 | 0 | 0 | 1443272 | NP_073549.1 | A0A0P0LKV0 |
| GQMEDVNLNDSR | 1 | 0 | 0 | 1443281 | AVV62544.1 | K9N4V7 |
| GQNNADPKVYPI | 1 | 0 | 0 | 1443285 | ARO76391.1 | P59636 |
| GQNTELGVEFLK | 1 | 0 | 0 | 1443287 | APO40578.1 | P0C6X7 |
| GQPCVVSEQEES | 1 | 0 | 0 | 1443290 | AVQ05261.1 | P0C6X6 |
| GQQFGPTYMDGA | 1 | 0 | 0 | 1443301 | APO40578.1 | P0C6X7 |
| GQQFVKETTDMV | 1 | 0 | 0 | 1443302 | AVQ05261.1 | P0C6X6 |
| GQQKHLLPRWYF | 1 | 0 | 0 | 1443307 | ABD75597.1 | Q19U29 |
| GQQKQLLPRWYF | 1 | 0 | 0 | 1443309 | ARU07581.1 | Q19U29 |
| GQRIYDDVNASL | 1 | 0 | 0 | 1443315 | AEN19363.1 | P0C6X6 |
| GQSIQALQTAAM | 1 | 0 | 0 | 1443322 | APO40578.1 | P0C6X7 |
| GQSLCALPDTHS | 1 | 0 | 0 | 1443327 | AID55099.1 | K9N5Q8 |
| GQSLCALPDTPI | 1 | 0 | 0 | 1443328 | ANI69900.1 | K9N5Q8 |
| GQSLEGVFWVAN | 1 | 0 | 0 | 1443334 | ADN03343.1 | Q19U29 |
| GQSTELGIEFLK | 1 | 0 | 0 | 1443338 | AVP78041.1 | P0C6W2 |
| GQVCYERFSLCN | 1 | 0 | 0 | 1443351 | AVV62524.1 | K9N7C7 |
| GQVFNASNFPSV | 1 | 0 | 0 | 1443356 | APO40579.1 | P59594 |
| GQVFTRQLSDYE | 1 | 0 | 0 | 1443357 | AVV62526.1 | K9N5Q8 |
| GQVHVLNHVDPI | 1 | 0 | 0 | 1443358 | P0C5B5.1 | Q0ZJ80 |
| GQWFKNQTSVAP | 1 | 0 | 0 | 1443362 | AIX10763.1 | P36334 |
| GRCFNCRPAVVN | 1 | 0 | 0 | 1443374 | AGT21338.1 | A0A0P0K6L9 |
| GRCIVFDVKGVE | 1 | 0 | 0 | 1443375 | AOL02451.1 | Q80872 |
| GRCPTAADINIH | 1 | 0 | 0 | 1443379 | AGT51530.1 | P30215 |
| GRCPTAADINNP | 1 | 0 | 0 | 1443381 | P30215.1 | P30215 |
| GRCPTAADINTP | 1 | 0 | 0 | 1443382 | AAA73152.1 | P30215 |
| GRCYNCRPAVIN | 1 | 0 | 0 | 1443389 | AIW52757.1 | A0A0P0K6L9 |
| GRCYNCRPAVVN | 1 | 0 | 0 | 1443390 | AIW52756.1 | A0A0P0K6L9 |
| GRDFKFPDGQGV | 1 | 0 | 0 | 1443394 | AMM42397.1 | Q19U29 |
| GRDTSDFTDSVR | 1 | 0 | 0 | 1443402 | ATO98181.1 | Q3LZX1 |
| GREATQYLVQQE | 1 | 0 | 0 | 1443408 | ARO76381.1 | P0C6U8 |
| GRELNCDVCPNR | 1 | 0 | 0 | 1443410 | ARB07436.1 | A0A140H1G9 |
| GRELNCNVCANR | 1 | 0 | 0 | 1443411 | ABD75615.1 | A0A140H1G9 |
| GRENLYFQGGGG | 1 | 0 | 0 | 1443412 | 6NB6_A | P59594 |
| GRESLIFPLGTY | 1 | 0 | 0 | 1443415 | AHY22564.1 | K9N638 |
| GRGGNHFRHPLD | 1 | 0 | 0 | 1443444 | AVV62524.1 | K9N7C7 |
| GRGGNHFRHPLE | 1 | 0 | 0 | 1443445 | AVV62535.1 | K9N7C7 |
| GRGPGKCYNCRP | 1 | 0 | 0 | 1443458 | ARA15429.1 | A0A0P0K6L9 |
| GRITPYCHDPTV | 1 | 0 | 0 | 1443487 | AHL18091.1 | K9N7C7 |
| GRIVNYTVCDDC | 1 | 0 | 0 | 1443490 | AIW52843.1 | A0A0P0G321 |
| GRKSLIFPSGTY | 1 | 0 | 0 | 1443503 | AVV61898.1 | K9N7C7 |
| GRLAALNVFVSH | 1 | 0 | 0 | 1443509 | AIW52757.1 | A0A0P0K6L9 |
| GRLIQFDNPNIP | 1 | 0 | 0 | 1443512 | APO40579.1 | P59594 |
| GRLNCSFQDAAS | 1 | 0 | 0 | 1443518 | AVN89333.1 | K9N7C7 |
| GRLNCTFQEAAS | 1 | 0 | 0 | 1443519 | AUM60013.1 | K9N7C7 |
| GRLVGTTLQGKP | 1 | 0 | 0 | 1443526 | AJG44057.1 | K9N638 |
| GRMTPYCHDPTV | 1 | 0 | 0 | 1443533 | AXN92247.1 | K9N7C7 |
| GRMVPYCYDPTV | 1 | 0 | 0 | 1443534 | AUM60013.1 | K9N7C7 |
| GRPQGAFHVTMR | 1 | 0 | 0 | 1443552 | ARU07575.1 | A0A140H1G9 |
| GRPVHVKEITRE | 1 | 0 | 0 | 1443557 | AGT51393.1 | H9EJ42 |
| GRQFYEFYNDIK | 1 | 0 | 0 | 1443559 | AOL02455.1 | Q04854 |
| GRQLYKSYSEHV | 1 | 0 | 0 | 1443564 | BBA20987.1 | Q0ZJ83 |
| GRQTTNGMNTSH | 1 | 0 | 0 | 1443566 | AKA09607.1 | A0A0P0K6L9 |
| GRRAQPKQTSTS | 1 | 0 | 0 | 1443568 | ATP16771.1 | P33469 |
| GRRAQPKQTVTS | 1 | 0 | 0 | 1443569 | AIL49418.1 | P33469 |
| GRSAYVKFQESK | 1 | 0 | 0 | 1443584 | AVV62531.1 | K9N5R3 |
| GRSVYVKFQESK | 1 | 0 | 0 | 1443601 | AVV62542.1 | K9N5R3 |

| | | | | | | |
|---|---|---|---|---|---|---|
| GRTEVPQLVNAN | 1 | 0 | 0 | 1443602 | ANC28678.1 | K9N5Q8 |
| GRTGELPCAIIN | 1 | 0 | 0 | 1443603 | AGW27861.1 | A0A140H1G9 |
| GRTPKPRPAPNN | 1 | 0 | 0 | 1443613 | AVV62544.1 | K9N4V7 |
| GRTPNGFSFNNW | 1 | 0 | 0 | 1443615 | AKA09627.1 | A0A0P0G321 |
| GRTPTSGVTPDM | 1 | 0 | 0 | 1443616 | AXX83311.1 | P33469 |
| GRTVVRPLVEDS | 1 | 0 | 0 | 1443619 | ASY99815.1 | K9N7A1 |
| GRVDDDYICGDG | 1 | 0 | 0 | 1443625 | QED88011.1 | H9EJ42 |
| GRVEGQANLFRN | 1 | 0 | 0 | 1443628 | AVP78041.1 | P0C6W2 |
| GRVGNGYVCGTG | 1 | 0 | 0 | 1443632 | AST12962.1 | A0A0P0LKV0 |
| GRVIYGNVSRHN | 1 | 0 | 0 | 1443634 | AIW52827.1 | H9EJ42 |
| GRVLRPYHANSS | 1 | 0 | 0 | 1443638 | AIV41873.1 | P36334 |
| GRVVNYTVCDDC | 1 | 0 | 0 | 1443649 | AIW52841.1 | A0A0P0G321 |
| GRWSTSINTGNC | 1 | 0 | 0 | 1443652 | AAQ90004.1 | A0A0P0K6L9 |
| GRYMSALNHTKKWKYPQV | 1 | 0 | 0 | 1443660 | P0DTC1.1 | P0DTD1 |
| GRYMSALNHTKR | 1 | 0 | 0 | 1443661 | APO40578.1 | P0C6X7 |
| GSAFDGNMYGAF | 1 | 0 | 0 | 1443665 | AVV62535.1 | K9N7C7 |
| GSAFRDLKPLPF | 1 | 0 | 0 | 1443667 | AHI48736.1 | K9N7C7 |
| GSAGNATEVPAN | 1 | 0 | 0 | 1443668 | 5C8S_A | P0C6U8 |
| GSAIASEFSSLP | 1 | 0 | 0 | 1443675 | 2AHM_E | P0C6U8 |
| GSALNGSCGSVG | 1 | 0 | 0 | 1443678 | 3F9G_A | P0C6X7 |
| GSALQIPFAMQM | 1 | 0 | 0 | 1443680 | Q3I5J5.1 | Q3LZX1 |
| GSAMGLPNNTAS | 1 | 0 | 0 | 1443681 | 1SSK_A | P59595 |
| GSANIFIGDNVG | 1 | 0 | 0 | 1443683 | AEN19363.1 | P0C6X6 |
| GSANIFIGDTVG | 1 | 0 | 0 | 1443684 | AIV41930.1 | P0C6X6 |
| GSANQQQLKQLE | 1 | 0 | 0 | 1443685 | AEN19363.1 | P0C6X6 |
| GSAQVTSFCKSG | 1 | 0 | 0 | 1443690 | QBP84703.1 | P30215 |
| GSAVLWANGVKA | 1 | 0 | 0 | 1443698 | AMM42513.1 | A0A0N7J131 |
| GSCCKFDEDHSE | 1 | 0 | 0 | 1443701 | APO40579.1 | P59594 |
| GSCEYAQEGVCI | 1 | 0 | 0 | 1443706 | AXN92247.1 | K9N7C7 |
| GSCFNSITVDKF | 1 | 0 | 0 | 1443707 | BBA20986.1 | A0A140H1H1 |
| GSCGSVGYTKDG | 1 | 0 | 0 | 1443711 | ATG84865.1 | K9N638 |
| GSCLLSGHNLAN | 1 | 0 | 0 | 1443724 | ARO76381.1 | P0C6U8 |
| GSCNFPLEATWH | 1 | 0 | 0 | 1443729 | AIW52843.1 | A0A0P0G321 |
| GSCPDKNNGIGI | 1 | 0 | 0 | 1443730 | AIX10751.1 | P36334 |
| GSCPDKNNGIGT | 1 | 0 | 0 | 1443731 | AIX10750.1 | P36334 |
| GSCPGKNNGIGT | 1 | 0 | 0 | 1443732 | AEN19366.1 | P36334 |
| GSDGKPIADYAF | 1 | 0 | 0 | 1443745 | AUM60013.1 | K9N7C7 |
| GSDINCNIPTQC | 1 | 0 | 0 | 1443746 | P0C6U5.1 | A0A140H1G9 |
| GSDKIRMGQESI | 1 | 0 | 0 | 1443753 | APO40579.1 | P59594 |
| GSDLLYQLPNCS | 1 | 0 | 0 | 1443755 | AGV08515.1 | K9N7C7 |
| GSDNVADFNAIA | 1 | 0 | 0 | 1443757 | APO40578.1 | P0C6X7 |
| GSDVDLGDISGI | 1 | 0 | 0 | 1443763 | 1WYY_A | P59594 |
| GSDYMDIVKRLA | 1 | 0 | 0 | 1443771 | AVV62535.1 | K9N7C7 |
| GSENWSVGAWDG | 1 | 0 | 0 | 1443779 | AGT51379.1 | H9EJ42 |
| GSERDNINAILT | 1 | 0 | 0 | 1443780 | AHF54662.1 | P36334 |
| GSESLNDNLLEI | 1 | 0 | 0 | 1443782 | AVP78041.1 | P0C6W2 |
| GSFERVYDIGNP | 1 | 0 | 0 | 1443797 | AVV62524.1 | K9N7C7 |
| GSFKTADGNQRQ | 1 | 0 | 0 | 1443802 | P33469.1 | P33469 |
| GSFNHKVIGGLH | 1 | 0 | 0 | 1443810 | ARU07574.1 | A0A140H1G9 |
| GSFNNIDTGPPS | 1 | 0 | 0 | 1443811 | AIX10756.1 | P36334 |
| GSFNNKDTGPPF | 1 | 0 | 0 | 1443812 | AKA09663.1 | P36334 |
| GSFNNKNTGPPF | 1 | 0 | 0 | 1443813 | AKA09659.1 | P36334 |
| GSFNNRDTGHPS | 1 | 0 | 0 | 1443814 | AWW13566.1 | P36334 |
| GSFNNRDTGLPS | 1 | 0 | 0 | 1443816 | BBA84092.1 | P36334 |
| GSFNNRDTGPLS | 1 | 0 | 0 | 1443817 | ANZ78836.1 | P36334 |
| GSFNNRDTGPPS | 1 | 0 | 0 | 1443818 | AEN19366.1 | P36334 |
| GSFNNRDTGRPS | 1 | 0 | 0 | 1443819 | BBO58746.1 | P36334 |
| GSFNNRDTGSPP | 1 | 0 | 0 | 1443820 | QBP84713.1 | P36334 |
| GSFNNRDTGSPS | 1 | 0 | 0 | 1443821 | AVR40342.1 | P36334 |
| GSFVEGAINVGK | 1 | 0 | 0 | 1443829 | AUM60013.1 | K9N7C7 |
| GSFYKATVNQVC | 1 | 0 | 0 | 1443833 | ARU07575.1 | A0A140H1G9 |
| GSGDATTAFANS | 1 | 0 | 0 | 1443839 | ADO24677.1 | P0C6X6 |
| GSGFCIDYALPS | 1 | 0 | 0 | 1443842 | BBA20986.1 | A0A140H1H1 |
| GSGKSAPNSRST | 1 | 0 | 0 | 1443852 | AJQ24087.1 | P33469 |
| GSGRENLYFQGG | 1 | 0 | 0 | 1443864 | 6NB6_A | P59594 |
| GSGRPAPNSRST | 1 | 0 | 0 | 1443866 | AIL49470.1 | P33469 |
| GSGSMLEDIDYV | 1 | 0 | 0 | 1443873 | AUM60013.1 | K9N7C7 |
| GSGYIPEAPRDG | 1 | 0 | 0 | 1443889 | 6CRW_A | P59594 |
| GSGYYYPEPIIE | 1 | 0 | 0 | 1443891 | AIX09799.1 | P36334 |
| GSGYYYPEPITG | 1 | 0 | 0 | 1443894 | CAA83660.1 | P36334 |
| GSHMDPLSNFEH | 1 | 0 | 0 | 1443896 | 5HIH_A | K9N638 |
| GSHMVLPSEAPN | 1 | 0 | 0 | 1443899 | 2JZF_A | P0C6U8 |
| GSIAGAGWTAGL | 1 | 0 | 0 | 1443902 | AVV62526.1 | K9N5Q8 |
| GSILVDNDLNPF | 1 | 0 | 0 | 1443916 | ARU07574.1 | A0A140H1G9 |
| GSIPNFGSLTQI | 1 | 0 | 0 | 1443918 | 4NJL_A | K9N5Q8 |
| GSITQRSRPRAA | 1 | 0 | 0 | 1443931 | AVV62544.1 | K9N4V7 |
| GSITQRTRSRPS | 1 | 0 | 0 | 1443933 | ASY99807.1 | K9N4V7 |
| GSIVDARIEPCA | 1 | 0 | 0 | 1443939 | AVV62524.1 | K9N7C7 |
| GSIVNARIEPCA | 1 | 0 | 0 | 1443940 | AUM60013.1 | K9N7C7 |
| GSKDEQIGYWNV | 1 | 0 | 0 | 1443944 | AIW52795.1 | H9EJ69 |
| GSKHVTCCTAND | 1 | 0 | 0 | 1443954 | ALJ99930.1 | A0A0P0LKV0 |
| GSKLELFKRDSD | 1 | 0 | 0 | 1443963 | AMM42394.1 | Q19U29 |
| GSKLKNLVLNNP | 1 | 0 | 0 | 1443964 | AWW13558.1 | P30215 |
| GSKSPIQYIDIGNYTVSCLPF | 1 | 0 | 0 | 1443968 | P0DTC8.1 | P0DTC8 |
| GSLAFDEAVAAA | 1 | 0 | 0 | 1443976 | AVV62533.1 | K9N4V7 |
| GSLASRLMSIPR | 1 | 0 | 0 | 1443977 | QBQ01844.1 | Q4VID0 |
| GSLATRLMSIPR | 1 | 0 | 0 | 1443979 | AVR40343.1 | Q4VID0 |
| GSLCVGSGPGKN | 1 | 0 | 0 | 1443982 | ATP16757.1 | P36334 |
| GSLEKFMSTPNA | 1 | 0 | 0 | 1443984 | AXN92247.1 | K9N7C7 |
| GSLERFMNTPNA | 1 | 0 | 0 | 1443988 | AUM60013.1 | K9N7C7 |
| GSLEWSHPQFEK | 1 | 0 | 0 | 1443989 | 5X58_A | P59594 |

| | | | | | | |
|---|---|---|---|---|---|---|
| GSLGDSVKLLSE | 1 | 0 | 0 | 1443993 | ABE73416.1 | H9EJ42 |
| GSLGFHNPPTNV | 1 | 0 | 0 | 1443994 | AVQ05263.1 | P30215 |
| GSLNNIDTGPPS | 1 | 0 | 0 | 1444007 | AVQ05264.1 | P36334 |
| GSLNNVDTGPPS | 1 | 0 | 0 | 1444008 | AMK59677.1 | P36334 |
| GSLTHINTTLLD | 1 | 0 | 0 | 1444019 | ALJ54471.1 | K9N5Q8 |
| GSLTKKVKMLFK | 1 | 0 | 0 | 1444023 | AVV62524.1 | K9N7C7 |
| GSLVTRLMSIPR | 1 | 0 | 0 | 1444033 | AHF54653.1 | Q4VID0 |
| GSMIEDVDYVPL | 1 | 0 | 0 | 1444044 | AMQ49035.1 | K9N7C7 |
| GSMLTSIAARPP | 1 | 0 | 0 | 1444051 | AUM60013.1 | K9N7C7 |
| GSNDVWIFNKVR | 1 | 0 | 0 | 1444056 | AGT17757.1 | A0A140H1H0 |
| GSNEPGNVGGND | 1 | 0 | 0 | 1444058 | AVI15051.1 | P0C6X6 |
| GSPLSLNMVKKL | 1 | 0 | 0 | 1444086 | APO40587.1 | P59636 |
| GSPQSYCYTDGL | 1 | 0 | 0 | 1444090 | AGT51488.1 | P0C6X6 |
| GSQLELAKVQNL | 1 | 0 | 0 | 1444100 | AHA80186.1 | P33469 |
| GSQTASRSSSRS | 1 | 0 | 0 | 1444106 | APO40586.1 | P59595 |
| GSQTTNGTNTNH | 1 | 0 | 0 | 1444107 | AIW52735.1 | A0A0P0K6L9 |
| GSQTVLRCGDCL | 1 | 0 | 0 | 1444108 | AWK59942.1 | H9EJ42 |
| GSRSGNSSRNAS | 1 | 0 | 0 | 1444122 | AVV62533.1 | K9N4V7 |
| GSRSGNSTRGTF | 1 | 0 | 0 | 1444124 | ANI69896.1 | K9N4V7 |
| GSRSGNSTRSTS | 1 | 0 | 0 | 1444125 | AVN89381.1 | K9N4V7 |
| GSRSRANSGNRA | 1 | 0 | 0 | 1444130 | AIX10723.1 | P33469 |
| GSSFFNLEPITL | 1 | 0 | 0 | 1444147 | AVV62526.1 | K9N5Q8 |
| GSSFSELLFDCV | 1 | 0 | 0 | 1444150 | AHX71940.1 | K9N7C7 |
| GSSFYAPEPITY | 1 | 0 | 0 | 1444153 | AGV08438.1 | K9N5Q8 |
| GSSLQEILSRLD | 1 | 0 | 0 | 1444170 | CAA83660.1 | P36334 |
| GSSMVCKMCLFS | 1 | 0 | 0 | 1444173 | AGT51337.1 | H9EJ42 |
| GSSSRSFIEDLL | 1 | 0 | 0 | 1444185 | ARU07577.1 | A0A140H1H1 |
| GSSVGNFSNGVS | 1 | 0 | 0 | 1444188 | AVV62537.1 | K9N5Q8 |
| GSSVGNYYNGYP | 1 | 0 | 0 | 1444190 | AVV62526.1 | K9N5Q8 |
| GSTADNLKNVLV | 1 | 0 | 0 | 1444195 | APO40578.1 | P0C6X7 |
| GSTLDNTTQSAV | 1 | 0 | 0 | 1444205 | APO40579.1 | P59594 |
| GSTLVDGVSRRY | 1 | 0 | 0 | 1444208 | QED88026.1 | A0A0P0G321 |
| GSTMTFSVHPVV | 1 | 0 | 0 | 1444215 | AUM60013.1 | K9N7C7 |
| GSVAYESLRPDI | 1 | 0 | 0 | 1444231 | AVP78041.1 | P0C6W2 |
| GSVCFSLKYIPG | 1 | 0 | 0 | 1444234 | APT69876.1 | A0A0P0K6L9 |
| GSVDPTFLHRFY | 1 | 0 | 0 | 1444236 | AHC74096.1 | K9N7C7 |
| GSVELSLDRGAQ | 1 | 0 | 0 | 1444238 | AVV62527.1 | K9N796 |
| GSVVEQADGVEC | 1 | 0 | 0 | 1444267 | ATG84679.1 | K9N5Q8 |
| GSWNKVTQIANM | 1 | 0 | 0 | 1444286 | AXN92247.1 | K9N7C7 |
| GSWWSFNPETNC | 1 | 0 | 0 | 1444289 | ASY99815.1 | K9N7A1 |
| GSYCPGNSNYNT | 1 | 0 | 0 | 1444291 | AVV62537.1 | K9N5Q8 |
| GSYHNWHLVKGE | 1 | 0 | 0 | 1444294 | AVV62524.1 | K9N7C7 |
| GSYNASCLCSPD | 1 | 0 | 0 | 1444304 | AFR45556.1 | A0A140H1H1 |
| GSYRDWSYSGQN | 1 | 0 | 0 | 1444310 | APO40578.1 | P0C6X7 |
| GTANLSIPSNWT | 1 | 0 | 0 | 1444318 | AAK32190.1 | A0A0P0K6L9 |
| GTANQQQLKQLE | 1 | 0 | 0 | 1444319 | AGT51787.1 | P0C6X6 |
| GTATAGYTFGAG | 1 | 0 | 0 | 1444322 | APO40579.1 | P59594 |
| GTAVMSLKDNQV | 1 | 0 | 0 | 1444328 | APO40578.1 | P0C6X7 |
| GTCAKRFKRWLG | 1 | 0 | 0 | 1444332 | NP_073549.1 | A0A0P0LKV0 |
| GTCCTTSYKFYR | 1 | 0 | 0 | 1444335 | AWU59319.1 | K9N7C7 |
| GTCGIVELEKGV | 1 | 0 | 0 | 1444345 | P0C6W6.1 | P0C6W2 |
| GTCYNCRPAVVN | 1 | 0 | 0 | 1444360 | BAL45638.1 | A0A0P0K6L9 |
| GTDDDDKAMAAS | 1 | 0 | 0 | 1444361 | 3R24_A | P0C6X7 |
| GTDDPRWPQIAE | 1 | 0 | 0 | 1444362 | ATG84774.1 | K9N4V7 |
| GTDIEGNFYGPF | 1 | 0 | 0 | 1444369 | APO40578.1 | P0C6X7 |
| GTDKSSTLWFKP | 1 | 0 | 0 | 1444372 | AKA09652.1 | P36334 |
| GTDLLSTLWFKP | 1 | 0 | 0 | 1444376 | AWW13566.1 | P36334 |
| GTDPIEDFQENW | 1 | 0 | 0 | 1444378 | AVP78041.1 | P0C6W2 |
| GTDPIEDYENDW | 1 | 0 | 0 | 1444379 | APO40578.1 | P0C6X7 |
| GTDYKNWPQIAQ | 1 | 0 | 0 | 1444397 | APO40586.1 | P59595 |
| GTEDDYNGTPLE | 1 | 0 | 0 | 1444402 | APO40578.1 | P0C6X7 |
| GTEDPPWPQIAE | 1 | 0 | 0 | 1444405 | QDP16213.1 | K9N4V7 |
| GTENLVCEGPTT | 1 | 0 | 0 | 1444411 | ARO76381.1 | P0C6U8 |
| GTENLYFQSMET | 1 | 0 | 0 | 1444413 | 5X29_A | P59637 |
| GTFDNDIACPLT | 1 | 0 | 0 | 1444421 | AUM60013.1 | K9N7C7 |
| GTFEEMALTTFM | 1 | 0 | 0 | 1444422 | AEN19363.1 | P0C6X6 |
| GTFNNKDTGPPF | 1 | 0 | 0 | 1444426 | AKA09658.1 | P36334 |
| GTFPGPSGIGAV | 1 | 0 | 0 | 1444427 | ANI69896.1 | K9N4V7 |
| GTFTPKKPNKPL | 1 | 0 | 0 | 1444430 | ABE73425.1 | H9EJ69 |
| GTGALVSKLSPW | 1 | 0 | 0 | 1444449 | AUM60023.1 | K9N7C7 |
| GTGDLAEHLSPW | 1 | 0 | 0 | 1444450 | AVV62535.1 | K9N7C7 |
| GTGGNIQSSSRA | 1 | 0 | 0 | 1444453 | ANI69874.1 | K9N4V7 |
| GTGIAISVKPES | 1 | 0 | 0 | 1444456 | AXN92247.1 | K9N7C7 |
| GTGLQYAAITAP | 1 | 0 | 0 | 1444458 | AXN92247.1 | K9N7C7 |
| GTGPHAKDDYGT | 1 | 0 | 0 | 1444465 | AJQ24074.1 | P33469 |
| GTGPHKDLKFRQ | 1 | 0 | 0 | 1444471 | AIW52805.1 | H9EJ69 |
| GTGPHSKDQYGT | 1 | 0 | 0 | 1444473 | AIV41994.1 | P33469 |
| GTGPYANASYGQ | 1 | 0 | 0 | 1444477 | ADN03343.1 | Q19U29 |
| GTGRGDCKGFYS | 1 | 0 | 0 | 1444482 | AIW52756.1 | A0A0P0K6L9 |
| GTGRGDCQGFYS | 1 | 0 | 0 | 1444483 | BAL45639.1 | A0A0P0K6L9 |
| GTGSGDATTAFA | 1 | 0 | 0 | 1444486 | ADO24677.1 | P0C6X6 |
| GTGTDTIDVLLK | 1 | 0 | 0 | 1444487 | AGT51735.1 | Q4VID0 |
| GTGTIYAELEPP | 1 | 0 | 0 | 1444490 | AEA10817.1 | P0C6X7 |
| GTGVFVQDPSPI | 1 | 0 | 0 | 1444495 | AFD98763.1 | H9EJ42 |
| GTGVLTSSTKKF | 1 | 0 | 0 | 1444499 | APO40579.1 | P59594 |
| GTHTGSAFDGNM | 1 | 0 | 0 | 1444516 | AVV62535.1 | K9N7C7 |
| GTHTGSSFDGNM | 1 | 0 | 0 | 1444518 | AVV62524.1 | K9N7C7 |
| GTILQGKPIGMF | 1 | 0 | 0 | 1444526 | AIZ48759.1 | K9N638 |
| GTILRTINGDFL | 1 | 0 | 0 | 1444527 | AVP78041.1 | P0C6W2 |
| GTISVEELKRLL | 1 | 0 | 0 | 1444533 | ATO98197.1 | Q3LZX9 |
| GTITDAIDCAQN | 1 | 0 | 0 | 1444534 | ATO98181.1 | Q3LZX1 |

| | | | | | | |
|---|---|---|---|---|---|---|
| GTIYAELEPPCR | 1 | 0 | 0 | 1444546 | AEA10817.1 | P0C6X7 |
| GTKDVVIEGLKA | 1 | 0 | 0 | 1444550 | AVV62524.1 | K9N7C7 |
| GTKQIIHSHPIL | 1 | 0 | 0 | 1444558 | ATQ39396.1 | K9N643 |
| GTLLVDSDMNDF | 1 | 0 | 0 | 1444584 | APO40578.1 | P0C6X7 |
| GTLMVERSVSLA | 1 | 0 | 0 | 1444589 | ASU90624.1 | K9N7C7 |
| GTLVGVGNTVPM | 1 | 0 | 0 | 1444606 | AVV62526.1 | K9N5Q8 |
| GTLYMYFQKAGQ | 1 | 0 | 0 | 1444608 | APO40578.1 | P0C6X7 |
| GTMSNAVDCSQD | 1 | 0 | 0 | 1444611 | ATO98145.1 | Q3LZX1 |
| GTNCMGKLKRNR | 1 | 0 | 0 | 1444621 | AHL18090.1 | K9N5Q8 |
| GTNDICVFENKT | 1 | 0 | 0 | 1444622 | ATQ39389.1 | K9N7C7 |
| GTNGNKIVDKEI | 1 | 0 | 0 | 1444627 | AUM60013.1 | K9N7C7 |
| GTNGNKIVDKEV | 1 | 0 | 0 | 1444628 | AVV62524.1 | K9N7C7 |
| GTNSGVAIYHRY | 1 | 0 | 0 | 1444640 | ASY99815.1 | K9N7A1 |
| GTNTSNSVCNGC | 1 | 0 | 0 | 1444648 | AST12964.1 | A0A0P0K6L9 |
| GTNTTFDFLSNS | 1 | 0 | 0 | 1444650 | AVL25589.1 | A0A0P0G321 |
| GTNVFQTQAGCL | 1 | 0 | 0 | 1444651 | AVP78031.1 | Q3LZX1 |
| GTNYLTCHNAVQ | 1 | 0 | 0 | 1444656 | CAA83661.1 | P36334 |
| GTPDSNFLYKYY | 1 | 0 | 0 | 1444661 | AVV62524.1 | K9N7C7 |
| GTPKPSRNHSPA | 1 | 0 | 0 | 1444664 | ABB90490.1 | A0A0N7J131 |
| GTPKPSRNQSPV | 1 | 0 | 0 | 1444667 | AMM42518.1 | A0A0N7J131 |
| GTPLEFGASTQN | 1 | 0 | 0 | 1444668 | APO40578.1 | P0C6X7 |
| GTPLHPVLWNCQ | 1 | 0 | 0 | 1444669 | ANI69891.1 | K9N4V0 |
| GTPNEKLVTTST | 1 | 0 | 0 | 1444671 | AXN92247.1 | K9N7C7 |
| GTPNNPKQIVQS | 1 | 0 | 0 | 1444672 | AVV62540.1 | K9N643 |
| GTPPQVYNFNRL | 1 | 0 | 0 | 1444676 | AVV62537.1 | K9N5Q8 |
| GTQAYSLRVRKR | 1 | 0 | 0 | 1444684 | ATQ39396.1 | K9N643 |
| GTQLNHTSCSCS | 1 | 0 | 0 | 1444691 | BBA20986.1 | A0A140H1H1 |
| GTQLPKNFHIEG | 1 | 0 | 0 | 1444692 | AVV62533.1 | K9N4V7 |
| GTQNAVVEAPTT | 1 | 0 | 0 | 1444693 | APO40578.1 | P0C6X7 |
| GTQSVSVDRELT | 1 | 0 | 0 | 1444700 | AVN89314.1 | K9N796 |
| GTQYVYSPSNHK | 1 | 0 | 0 | 1444717 | ATQ39390.1 | K9N5Q8 |
| GTRCPNDNAYVS | 1 | 0 | 0 | 1444718 | AVV62526.1 | K9N5Q8 |
| GTRNPNNDAAIV | 1 | 0 | 0 | 1444724 | AVV62533.1 | K9N4V7 |
| GTRNPNNDLAIV | 1 | 0 | 0 | 1444725 | AKN24774.1 | K9N4V7 |
| GTRNPNNDPAIV | 1 | 0 | 0 | 1444726 | AUM60021.1 | K9N4V7 |
| GTSDPQFPILAE | 1 | 0 | 0 | 1444742 | AEN19370.1 | P33469 |
| GTSDSQFPILAE | 1 | 0 | 0 | 1444743 | AGT51764.1 | P33469 |
| GTSLHPVLWNCQ | 1 | 0 | 0 | 1444750 | AXN92262.1 | K9N4V0 |
| GTSLPKGFYAEG | 1 | 0 | 0 | 1444753 | APO40586.1 | P59595 |
| GTSPCPSGIGAV | 1 | 0 | 0 | 1444755 | QCI31476.1 | K9N4V7 |
| GTSPGLSGIGAV | 1 | 0 | 0 | 1444756 | ALA49601.1 | K9N4V7 |
| GTSPGPSGIGTV | 1 | 0 | 0 | 1444758 | ATG84708.1 | K9N4V7 |
| GTSSGDASTAFA | 1 | 0 | 0 | 1444761 | AKL87224.1 | P0C6X6 |
| GTSVLRQWLPTD | 1 | 0 | 0 | 1444772 | AXN92247.1 | K9N7C7 |
| GTSVTEDVKFAA | 1 | 0 | 0 | 1444774 | AWK59948.1 | H9EJ42 |
| GTSYTANDVEVQ | 1 | 0 | 0 | 1444778 | QBM11736.1 | K9N638 |
| GTTCSNDLLYSN | 1 | 0 | 0 | 1444780 | BBA20986.1 | A0A140H1H1 |
| GTTEVKVSALKT | 1 | 0 | 0 | 1444787 | AXN92247.1 | K9N7C7 |
| GTTKFYGGWGFM | 1 | 0 | 0 | 1444792 | ASU45728.1 | K9N7C7 |
| GTTLDNTSQSLL | 1 | 0 | 0 | 1444794 | AVP78031.1 | Q3LZX1 |
| GTTLDNTTQSAL | 1 | 0 | 0 | 1444795 | ATO98145.1 | Q3LZX1 |
| GTTLHPVQWQCK | 1 | 0 | 0 | 1444798 | AVV62528.1 | K9N4V0 |
| GTTLQGKPVGYF | 1 | 0 | 0 | 1444803 | AVV62524.1 | K9N7C7 |
| GTTVILKEPCPS | 1 | 0 | 0 | 1444821 | 1XAK_A | P59635 |
| GTTVLLKEPCSS | 1 | 0 | 0 | 1444823 | AVP78047.1 | Q3LZX7 |
| GTTVLLSEPCSP | 1 | 0 | 0 | 1444824 | APO40584.1 | P59635 |
| GTVDADYKMCTK | 1 | 0 | 0 | 1444830 | APT69849.1 | A0A0P0K6L9 |
| GTVIINPSTSAT | 1 | 0 | 0 | 1444838 | AJD81440.1 | K9N5Q8 |
| GTVIISPFTSAT | 1 | 0 | 0 | 1444840 | AHI48702.1 | K9N5Q8 |
| GTVIRTLNGDFL | 1 | 0 | 0 | 1444844 | APO40578.1 | P0C6X7 |
| GTVLPRGYYIEG | 1 | 0 | 0 | 1444847 | AJQ24068.1 | P33469 |
| GTVNNFVKFGSV | 1 | 0 | 0 | 1444852 | BAL45640.1 | A0A0P0K6L9 |
| GTVSLKQGEIKD | 1 | 0 | 0 | 1444854 | AVP78043.1 | Q3LZX0 |
| GTVWYVGQIIFI | 1 | 0 | 0 | 1444861 | AGT51533.1 | Q04854 |
| GTWDKVAKIANL | 1 | 0 | 0 | 1444863 | AVV62535.1 | K9N7C7 |
| GTYHCVTTKAKS | 1 | 0 | 0 | 1444875 | ALA49758.1 | K9N7C7 |
| GTYSTPFMVNQL | 1 | 0 | 0 | 1444884 | AVV62535.1 | K9N7C7 |
| GTYTWTTVLSMA | 1 | 0 | 0 | 1444886 | AEN19363.1 | P0C6X6 |
| GVALEQDKNTQE | 1 | 0 | 0 | 1444908 | ATO98108.1 | Q3LZX1 |
| GVATSILYSCFT | 1 | 0 | 0 | 1444927 | AVV62535.1 | K9N7C7 |
| GVCKGFYSNASS | 1 | 0 | 0 | 1444941 | AIW52750.1 | A0A0P0K6L9 |
| GVCKYTDIDVNT | 1 | 0 | 0 | 1444947 | AUM60013.1 | K9N7C7 |
| GVCMYTDLNFID | 1 | 0 | 0 | 1444956 | ARK08667.1 | P0C6X6 |
| GVCVDYKLYGIT | 1 | 0 | 0 | 1444962 | AVV62526.1 | K9N5Q8 |
| GVDAKGFPLLAE | 1 | 0 | 0 | 1444968 | AMM42619.1 | H9EJ69 |
| GVDAKGFPQFAE | 1 | 0 | 0 | 1444969 | AGT51335.1 | H9EJ69 |
| GVDCYNWDSAEL | 1 | 0 | 0 | 1444977 | QED88024.1 | H9EJ42 |
| GVDLGTENLYFQ | 1 | 0 | 0 | 1444985 | 5X29_A | P59637 |
| GVDSMNLIANIF | 1 | 0 | 0 | 1444992 | ARO76381.1 | P0C6U8 |
| GVDYTKKQITVN | 1 | 0 | 0 | 1445002 | AUM60013.1 | K9N7C7 |
| GVDYTKKSLNVD | 1 | 0 | 0 | 1445003 | ATQ39389.1 | K9N7C7 |
| GVEDTPPHYFTL | 1 | 0 | 0 | 1445009 | ATQ39389.1 | K9N7C7 |
| GVEIEDFDLEDF | 1 | 0 | 0 | 1445020 | AVV62524.1 | K9N7C7 |
| GVENTPPHFFTL | 1 | 0 | 0 | 1445024 | ATG84821.1 | K9N638 |
| GVENTPPHYFTL | 1 | 0 | 0 | 1445025 | AXN92247.1 | K9N7C7 |
| GVESDIETGDIV | 1 | 0 | 0 | 1445027 | AVV62524.1 | K9N7C7 |
| GVFLGVVGVTMH | 1 | 0 | 0 | 1445040 | AWK59942.1 | H9EJ42 |
| GVFQNCTAVGVP | 1 | 0 | 0 | 1445047 | AVN89313.1 | K9N5Q8 |
| GVFRVTSRAMFQ | 1 | 0 | 0 | 1445049 | ARU07594.1 | A0A0P0G321 |
| GVFYFKPGGTTS | 1 | 0 | 0 | 1445076 | AWK59978.1 | H9EJ42 |
| GVGAVSGDASSL | 1 | 0 | 0 | 1445082 | AUM60021.1 | K9N4V7 |

| | | | | | | |
|---|---|---|---|---|---|---|
| GVGITQQVLSKN | 1 | 0 | 0 | 1445089 | AVN89376.1 | K9N5Q8 |
| GVGNTVPMTEGL | 1 | 0 | 0 | 1445092 | AVV62526.1 | K9N5Q8 |
| GVGSANIFIGDN | 1 | 0 | 0 | 1445097 | AEN19363.1 | P0C6X6 |
| GVGWTTGLSSFA | 1 | 0 | 0 | 1445107 | AOR17479.1 | K9N5Q8 |
| GVHGNGSMIEDI | 1 | 0 | 0 | 1445115 | AGV08556.1 | K9N7C7 |
| GVHKGAHVASSD | 1 | 0 | 0 | 1445117 | AXX83294.1 | P0C6X6 |
| GVHMNIAKYMQL | 1 | 0 | 0 | 1445121 | AXN92247.1 | K9N7C7 |
| GVIAQGARGTYR | 1 | 0 | 0 | 1445130 | AXN92247.1 | K9N7C7 |
| GVIFNAVDCKSD | 1 | 0 | 0 | 1445133 | CAA83660.1 | P36334 |
| GVIKNVNYVRDW | 1 | 0 | 0 | 1445136 | AST12962.1 | A0A0P0LKV0 |
| GVILTRSFKYNI | 1 | 0 | 0 | 1445137 | AWK59942.1 | H9EJ42 |
| GVINGFTAFANT | 1 | 0 | 0 | 1445140 | AQN78665.1 | Q04853 |
| GVINSSAMFCAT | 1 | 0 | 0 | 1445142 | AVV62535.1 | K9N7C7 |
| GVIPDMADQIAS | 1 | 0 | 0 | 1445143 | AMK59681.1 | P33469 |
| GVISNARNGCIP | 1 | 0 | 0 | 1445146 | AVN89322.1 | K9N7C7 |
| GVISSMTDNLQM | 1 | 0 | 0 | 1445148 | AUM60014.1 | K9N5Q8 |
| GVIYGFNSTETI | 1 | 0 | 0 | 1445151 | AIL49343.1 | P30215 |
| GVIYGLNFTETI | 1 | 0 | 0 | 1445152 | ATP16766.1 | P30215 |
| GVIYGLNSTETI | 1 | 0 | 0 | 1445153 | AEN19365.1 | P30215 |
| GVKDYVTLIGLF | 1 | 0 | 0 | 1445162 | APO40580.1 | P59632 |
| GVKKTFKVEDSV | 1 | 0 | 0 | 1445171 | AQN78677.1 | P0C6X6 |
| GVKLRSLNLGEV | 1 | 0 | 0 | 1445173 | ATO98217.1 | P0C6W2 |
| GVKPAVSFDYLI | 1 | 0 | 0 | 1445174 | AXN92247.1 | K9N7C7 |
| GVKPAVSFDYLT | 1 | 0 | 0 | 1445175 | ANI69910.1 | K9N7C7 |
| GVKQEQCTGLDS | 1 | 0 | 0 | 1445181 | AIV42002.1 | P0C6X6 |
| GVKQSPELLDPL | 1 | 0 | 0 | 1445188 | ADN03343.1 | Q19U29 |
| GVKQSPESFDSL | 1 | 0 | 0 | 1445189 | AMM42397.1 | Q19U29 |
| GVKSGFKFCADA | 1 | 0 | 0 | 1445191 | APO40578.1 | P0C6X7 |
| GVLDGSYNASCL | 1 | 0 | 0 | 1445198 | BBA20983.1 | A0A140H1H1 |
| GVLDGYNVSCLC | 1 | 0 | 0 | 1445200 | AGT17769.1 | A0A140H1H1 |
| GVLDTTIPFGHS | 1 | 0 | 0 | 1445202 | AWO68990.1 | A0A0P0G321 |
| GVLEVSVCQYNM | 1 | 0 | 0 | 1445206 | AMM42485.1 | P36334 |
| GVLKADDFVIAS | 1 | 0 | 0 | 1445214 | AVL25554.1 | H9EJ42 |
| GVLLVDGHKIAT | 1 | 0 | 0 | 1445219 | AWK59934.1 | U3M6Q8 |
| GVLQQRFVYDAY | 1 | 0 | 0 | 1445225 | ALJ54472.1 | K9N5Q8 |
| GVLVPHAGETPI | 1 | 0 | 0 | 1445241 | ARO76381.1 | P0C6U8 |
| GVLVPHTYETPT | 1 | 0 | 0 | 1445242 | APO40578.1 | P0C6X7 |
| GVLVPHVGEVPV | 1 | 0 | 0 | 1445246 | AVP78041.1 | P0C6W2 |
| GVMHSEFPAITI | 1 | 0 | 0 | 1445252 | AKA09651.1 | P36334 |
| GVMYSEFPAIAI | 1 | 0 | 0 | 1445260 | AKA09659.1 | P36334 |
| GVMYSEFPAITI | 1 | 0 | 0 | 1445261 | CAA79896.1 | P36334 |
| GVMYSEFPAIVI | 1 | 0 | 0 | 1445262 | AIL49490.1 | P36334 |
| GVMYSEFPTITI | 1 | 0 | 0 | 1445263 | AIL49548.1 | P36334 |
| GVNDAITQTSPA | 1 | 0 | 0 | 1445265 | AIW52728.1 | A0A0P0K6L9 |
| GVNDDDYVIALT | 1 | 0 | 0 | 1445266 | AGT51337.1 | H9EJ42 |
| GVNDICIFENKT | 1 | 0 | 0 | 1445267 | ANF29215.1 | K9N7C7 |
| GVNFNVDDINFA | 1 | 0 | 0 | 1445275 | QEG03814.1 | P36334 |
| GVNFNVDDVNFS | 1 | 0 | 0 | 1445276 | CAA83661.1 | P36334 |
| GVNFRTCVVNNQ | 1 | 0 | 0 | 1445277 | AVV62524.1 | K9N7C7 |
| GVNKLQGLLEIS | 1 | 0 | 0 | 1445284 | AVQ05264.1 | P36334 |
| GVNLVTVPTGYV | 1 | 0 | 0 | 1445287 | AVP78041.1 | P0C6W2 |
| GVNSSNCNNLNV | 1 | 0 | 0 | 1445288 | AUT76985.1 | A0A0P0LKV0 |
| GVNVFQTQAGCL | 1 | 0 | 0 | 1445291 | ATO98145.1 | Q3LZX1 |
| GVNVTKHRCDIN | 1 | 0 | 0 | 1445295 | QBP84746.1 | P0C6X6 |
| GVNYLGDFIQGP | 1 | 0 | 0 | 1445296 | AVL25588.1 | H9EJ42 |
| GVPATEAKGYWY | 1 | 0 | 0 | 1445301 | AEN19370.1 | P33469 |
| GVPATQAKGYWY | 1 | 0 | 0 | 1445303 | ATN39868.1 | P33469 |
| GVPDNFSTIVTG | 1 | 0 | 0 | 1445310 | AWO68998.1 | A0A0P0G321 |
| GVPIAPGVPSTE | 1 | 0 | 0 | 1445318 | AJQ24074.1 | P33469 |
| GVPIAYGIPLSE | 1 | 0 | 0 | 1445319 | AMM42386.1 | Q19U29 |
| GVPINTNSSKDD | 1 | 0 | 0 | 1445324 | AVP78038.1 | Q3LZX4 |
| GVPLGITISGET | 1 | 0 | 0 | 1445329 | AWO68966.1 | A0A0P0G321 |
| GVPLLSTMTTRQ | 1 | 0 | 0 | 1445330 | ADK37692.1 | H9EJ42 |
| GVPLNANSTPSQ | 1 | 0 | 0 | 1445332 | AQZ41279.1 | K9N4V7 |
| GVPLSEAKGYWY | 1 | 0 | 0 | 1445334 | ABA60796.1 | Q19U29 |
| GVPLTNCVKMLT | 1 | 0 | 0 | 1445335 | AHI48749.1 | K9N638 |
| GVPSNDNISLPF | 1 | 0 | 0 | 1445341 | AVV62524.1 | K9N7C7 |
| GVPSTEAKGYWY | 1 | 0 | 0 | 1445342 | AJQ24074.1 | P33469 |
| GVQCYTPHSQIS | 1 | 0 | 0 | 1445350 | AEN19363.1 | P0C6X6 |
| GVQIVEDLRARM | 1 | 0 | 0 | 1445354 | ASY99852.1 | K9N7C7 |
| GVQKYSTLQGPP | 1 | 0 | 0 | 1445355 | APO40578.1 | P0C6X7 |
| GVQSIEELHQPM | 1 | 0 | 0 | 1445357 | AUM60013.1 | K9N7C7 |
| GVQTIEDLRARM | 1 | 0 | 0 | 1445358 | AJG44112.1 | K9N638 |
| GVQTLEDLHARM | 1 | 0 | 0 | 1445359 | ATQ39389.1 | K9N7C7 |
| GVQTVEDLCARM | 1 | 0 | 0 | 1445360 | AKN24802.1 | K9N7C7 |
| GVQVHNLPEYMA | 1 | 0 | 0 | 1445366 | APT69865.1 | A0A0P0KXN3 |
| GVQYYCYSAKDT | 1 | 0 | 0 | 1445368 | AVV62524.1 | K9N7C7 |
| GVRGASVDARLV | 1 | 0 | 0 | 1445377 | AIV41780.1 | P0C6X6 |
| GVRGSGHYQTNL | 1 | 0 | 0 | 1445378 | AFD98812.1 | H9EJ42 |
| GVRIENSLKALF | 1 | 0 | 0 | 1445381 | AWK59942.1 | H9EJ42 |
| GVRTLSTYDFNP | 1 | 0 | 0 | 1445394 | AVP78031.1 | Q3LZX1 |
| GVRYPTGSCLIA | 1 | 0 | 0 | 1445398 | AXN73419.1 | K9N7D2 |
| GVSCIWSVDAFN | 1 | 0 | 0 | 1445402 | AEN19363.1 | P0C6X6 |
| GVSDNFDNNQNG | 1 | 0 | 0 | 1445403 | APO40586.1 | P59595 |
| GVSERVNVVPGV | 1 | 0 | 0 | 1445408 | AFD98763.1 | H9EJ42 |
| GVSFSTFEEAAL | 1 | 0 | 0 | 1445412 | AVP78041.1 | P0C6W2 |
| GVSFVVQPVGVV | 1 | 0 | 0 | 1445413 | ALA49417.1 | K9N7C7 |
| GVSIEQLLSAIQ | 1 | 0 | 0 | 1445421 | AVV62535.1 | K9N7C7 |
| GVSILSTMTGRV | 1 | 0 | 0 | 1445425 | AIL49573.1 | P0C6X6 |
| GVSILSTVTGRM | 1 | 0 | 0 | 1445426 | AIL49624.1 | P0C6X6 |
| GVSLDNVLSTFI | 1 | 0 | 0 | 1445430 | APO40578.1 | P0C6X7 |

| | | | | | | |
|---|---|---|---|---|---|---|
| GVSNLSLKQKLL | 1 | 0 | 0 | 1445432 | QAT98908.1 | K9N7C7 |
| GVSSAGPISQFN | 1 | 0 | 0 | 1445438 | AHX00711.1 | K9N5Q8 |
| GVSSFMNVTLNK | 1 | 0 | 0 | 1445440 | AIW52757.1 | A0A0P0K6L9 |
| GVTAKGYPQFAE | 1 | 0 | 0 | 1445452 | ABB90495.1 | A0A0N7J131 |
| GVTHEYIQTTIQ | 1 | 0 | 0 | 1445464 | ATG84679.1 | K9N5Q8 |
| GVTIDKLCDYLA | 1 | 0 | 0 | 1445465 | ASU90174.1 | K9N638 |
| GVTIDKLRDYLA | 1 | 0 | 0 | 1445466 | AXN92247.1 | K9N7C7 |
| GVTIPIKTQDYQ | 1 | 0 | 0 | 1445468 | APO40580.1 | P59632 |
| GVTLKIKVSQTN | 1 | 0 | 0 | 1445474 | NP_073549.1 | A0A0P0LKV0 |
| GVTPDMADQIAS | 1 | 0 | 0 | 1445480 | AEN19370.1 | P33469 |
| GVTRDILSTDDC | 1 | 0 | 0 | 1445488 | ARO76381.1 | P0C6U8 |
| GVTRDMADQIAS | 1 | 0 | 0 | 1445491 | AIL49470.1 | P33469 |
| GVTSSMTDNLQM | 1 | 0 | 0 | 1445495 | AUM60024.1 | K9N5Q8 |
| GVTTSILSNCFT | 1 | 0 | 0 | 1445498 | AUM60013.1 | K9N7C7 |
| GVTVAEEPDSRV | 1 | 0 | 0 | 1445500 | APT69887.1 | A0A0N7J131 |
| GVTVGPYVEGAI | 1 | 0 | 0 | 1445501 | AVV62535.1 | K9N7C7 |
| GVVAQGARNRYR | 1 | 0 | 0 | 1445509 | AVV62535.1 | K9N7C7 |
| GVVCNNSDYASK | 1 | 0 | 0 | 1445510 | AIX10745.1 | P0C6X6 |
| GVVHNQDVNLHN | 1 | 0 | 0 | 1445529 | AVP78030.1 | P0C6W2 |
| GVVMNFDVDTHR | 1 | 0 | 0 | 1445532 | ACV87155.1 | A0A140H1G9 |
| GVVNPIYDEPTP | 1 | 0 | 0 | 1445539 | APO40580.1 | P59632 |
| GVVNPVMDPIYD | 1 | 0 | 0 | 1445540 | ARO76383.1 | P59632 |
| GVVQQLPETYFTQSRN | 1 | 0 | 0 | 1445547 | P0DTD1.1 | P0DTD1 |
| GVVTCNVVKQCC | 1 | 0 | 0 | 1445556 | AVN89451.1 | K9N7C7 |
| GVVTGNVLKHCC | 1 | 0 | 0 | 1445557 | AUM60013.1 | K9N7C7 |
| GVVTGNVVQQCC | 1 | 0 | 0 | 1445559 | ALA49384.1 | K9N7C7 |
| GVVYVNATGSDV | 1 | 0 | 0 | 1445575 | ABU88826.1 | A0A0P0G321 |
| GVWDTPTLDCIN | 1 | 0 | 0 | 1445576 | AUM60014.1 | K9N5Q8 |
| GVWEINACTYTM | 1 | 0 | 0 | 1445577 | ABA60802.1 | A0A140H1H1 |
| GVWISPPVADYT | 1 | 0 | 0 | 1445578 | AVV62539.1 | K9N4V0 |
| GVYDTNHGDAGF | 1 | 0 | 0 | 1445588 | AQN78743.1 | P30215 |
| GVYDYKVSTQEF | 1 | 0 | 0 | 1445589 | AUM60013.1 | K9N7C7 |
| GVYELHGYTVQP | 1 | 0 | 0 | 1445591 | AIL49485.1 | P36334 |
| GVYELNGYPVQP | 1 | 0 | 0 | 1445592 | AMM42466.1 | P36334 |
| GVYFSTIDKSNV | 1 | 0 | 0 | 1445595 | ATO98145.1 | Q3LZX1 |
| GVYHKLGERLRQ | 1 | 0 | 0 | 1445598 | QBF80563.1 | K9N7C7 |
| GVYSVSSFEAQP | 1 | 0 | 0 | 1445616 | AUM60014.1 | K9N5Q8 |
| GVYSVSSLEAKP | 1 | 0 | 0 | 1445619 | AIA22864.1 | K9N5Q8 |
| GVYYRATTTYKL | 1 | 0 | 0 | 1445636 | AEN19363.1 | P0C6X6 |
| GWALYVRAKHGD | 1 | 0 | 0 | 1445640 | AFO70499.1 | U3M6Q8 |
| GWGNMLKNLMAD | 1 | 0 | 0 | 1445649 | ADK37686.1 | A0A0P0LKV0 |
| GWQNLLYDSNGN | 1 | 0 | 0 | 1445681 | ABU39950.1 | P36334 |
| GWTAGLSSFASI | 1 | 0 | 0 | 1445693 | AVV62526.1 | K9N5Q8 |
| GYCCLDVVRKEL | 1 | 0 | 0 | 1445722 | AWW13564.1 | Q80872 |
| GYCVDYSKIRRS | 1 | 0 | 0 | 1445730 | QBP84759.1 | P36334 |
| GYDDCMHQGPAS | 1 | 0 | 0 | 1445740 | ATG84866.1 | K9N5Q8 |
| GYDDCMNQGPPS | 1 | 0 | 0 | 1445742 | AVV62526.1 | K9N5Q8 |
| GYDDCMQQGPSS | 1 | 0 | 0 | 1445744 | AHL18090.1 | K9N5Q8 |
| GYDECMQQGPQS | 1 | 0 | 0 | 1445746 | AUM60014.1 | K9N5Q8 |
| GYDFNKPTSKTC | 1 | 0 | 0 | 1445748 | AUM60013.1 | K9N7C7 |
| GYFFPYDPALET | 1 | 0 | 0 | 1445756 | AVV62524.1 | K9N7C7 |
| GYFIKQNDSWMF | 1 | 0 | 0 | 1445759 | BBA20986.1 | A0A140H1H1 |
| GYFLPFNSNLTG | 1 | 0 | 0 | 1445765 | APO40579.1 | P59594 |
| GYFPKAGANFRD | 1 | 0 | 0 | 1445768 | BBA20983.1 | A0A140H1H1 |
| GYGCNCDSLRQV | 1 | 0 | 0 | 1445783 | AXN92258.1 | K9N7C7 |
| GYGKSTFNKDIY | 1 | 0 | 0 | 1445789 | AAY32939.1 | H9EJ42 |
| GYGNTMYVNMAV | 1 | 0 | 0 | 1445790 | AUM60027.1 | K9N643 |
| GYHPAHAPSFLV | 1 | 0 | 0 | 1445799 | ABD75575.1 | A0A140H1G9 |
| GYHYKELGLVMN | 1 | 0 | 0 | 1445804 | AXN92247.1 | K9N7C7 |
| GYHYNELGIEMN | 1 | 0 | 0 | 1445806 | AIL49563.1 | P0C6X6 |
| GYIRRAVDCGHD | 1 | 0 | 0 | 1445837 | AVV62537.1 | K9N5Q8 |
| GYIRTAIDCGFN | 1 | 0 | 0 | 1445839 | AXN92260.1 | K9N5Q8 |
| GYKIGCNCAGRI | 1 | 0 | 0 | 1445850 | ARU07575.1 | A0A140H1G9 |
| GYKVLPPLYDPN | 1 | 0 | 0 | 1445859 | AVV62526.1 | K9N5Q8 |
| GYLDIHPSLCNN | 1 | 0 | 0 | 1445862 | AGT17757.1 | A0A140H1H0 |
| GYLEAVNFNVTN | 1 | 0 | 0 | 1445864 | AMM42536.1 | A0A0P0K6L9 |
| GYLNSTNVTTAT | 1 | 0 | 0 | 1445876 | AVP78030.1 | P0C6W2 |
| GYLPNNAVVKMA | 1 | 0 | 0 | 1445878 | APO40578.1 | P0C6X7 |
| GYMQGYDDCMKQ | 1 | 0 | 0 | 1445897 | AVV62537.1 | K9N5Q8 |
| GYMQGYDECMQQ | 1 | 0 | 0 | 1445900 | AUM60014.1 | K9N5Q8 |
| GYNKDASFIGKN | 1 | 0 | 0 | 1445902 | AST12962.1 | A0A0P0LKV0 |
| GYNMTLIAADLP | 1 | 0 | 0 | 1445903 | ARU07575.1 | A0A140H1G9 |
| GYNTNATLNSFK | 1 | 0 | 0 | 1445911 | AVV62537.1 | K9N5Q8 |
| GYPLECIKNFLA | 1 | 0 | 0 | 1445923 | AVP78041.1 | P0C6W2 |
| GYPNDFVGGFRV | 1 | 0 | 0 | 1445924 | AWK59942.1 | H9EJ42 |
| GYPNDIVGGFRV | 1 | 0 | 0 | 1445925 | AWK59978.1 | H9EJ42 |
| GYPNMFITRDEA | 1 | 0 | 0 | 1445926 | APO40578.1 | P0C6X7 |
| GYQELSQFSHLD | 1 | 0 | 0 | 1445933 | NP_073549.1 | A0A0P0LKV0 |
| GYQESVIKTSHD | 1 | 0 | 0 | 1445935 | AIL49487.1 | P36334 |
| GYQHHSIADVMR | 1 | 0 | 0 | 1445937 | AVL25583.1 | A0A0P0G321 |
| GYQHHSVADVMR | 1 | 0 | 0 | 1445938 | AIW52843.1 | A0A0P0G321 |
| GYQHNSFADVMR | 1 | 0 | 0 | 1445939 | ALJ53431.1 | A0A0P0G321 |
| GYRLKDCVMYAI | 1 | 0 | 0 | 1445954 | APO40578.1 | P0C6X7 |
| GYRYFTLGNVEA | 1 | 0 | 0 | 1445960 | CAA71056.1 | A0A0P0K6L9 |
| GYSEDWHSGVKD | 1 | 0 | 0 | 1445964 | ATO98182.1 | Q3LZX0 |
| GYSEPPKVADKI | 1 | 0 | 0 | 1445966 | AEN19363.1 | P0C6X6 |
| GYSLADLPAYMS | 1 | 0 | 0 | 1445969 | AWW13547.1 | Q01455 |
| GYSNFVSSDLDC | 1 | 0 | 0 | 1445974 | AGT17757.1 | A0A140H1H0 |
| GYTDNIFSVQED | 1 | 0 | 0 | 1445982 | ABE97137.1 | A0A0P0G321 |
| GYTDNIFSVQKD | 1 | 0 | 0 | 1445983 | AMM42643.1 | A0A0P0G321 |
| GYTKEGNVINFC | 1 | 0 | 0 | 1445993 | AVV62535.1 | K9N7C7 |

| | | | | | | |
|---|---|---|---|---|---|---|
| GYTLDEARTALK | 1 | 0 | 0 | 1446000 | ARO76381.1 | P0C6U8 |
| GYTLEESKLALK | 1 | 0 | 0 | 1446005 | APO40578.1 | P0C6X7 |
| GYTVQPVADVYR | 1 | 0 | 0 | 1446016 | AVR40344.1 | P36334 |
| GYVAGIQNFKET | 1 | 0 | 0 | 1446022 | AUM60013.1 | K9N7C7 |
| GYVDTENSTEFT | 1 | 0 | 0 | 1446031 | AVP78041.1 | P0C6W2 |
| GYVDTSANTEFT | 1 | 0 | 0 | 1446033 | APO40578.1 | P0C6X7 |
| GYVPTNHVNVTA | 1 | 0 | 0 | 1446043 | AVV62537.1 | K9N5Q8 |
| GYVQQGVFNKAT | 1 | 0 | 0 | 1446046 | ATN39862.1 | P0C6X6 |
| GYVRRAIDCGYD | 1 | 0 | 0 | 1446047 | AUM60014.1 | K9N5Q8 |
| GYVTSSSKSSEE | 1 | 0 | 0 | 1446055 | APO40578.1 | P0C6X7 |
| GYYIEGSGKYAP | 1 | 0 | 0 | 1446079 | AJQ24067.1 | P33469 |
| GYYIEGSGRSAS | 1 | 0 | 0 | 1446083 | AXX83347.1 | P33469 |
| GYYIEGSGRSVP | 1 | 0 | 0 | 1446084 | AIL49415.1 | P33469 |
| GYYKKDNSYFTE | 1 | 0 | 0 | 1446089 | AVP78041.1 | P0C6W2 |
| GYYPSNHIEVAS | 1 | 0 | 0 | 1446091 | ANF29261.1 | K9N5Q8 |
| GYYVEGSGRSAS | 1 | 0 | 0 | 1446104 | ARU07581.1 | Q19U29 |
| GYYYPEPIIENN | 1 | 0 | 0 | 1446109 | AIX09799.1 | P36334 |
| GYYYPEPITGNN | 1 | 0 | 0 | 1446112 | CAA83660.1 | P36334 |
| HAAEYREYMEAY | 1 | 0 | 0 | 1446114 | AVV62535.1 | K9N7C7 |
| HAAITDYNYYSY | 1 | 0 | 0 | 1446116 | AVV62524.1 | K9N7C7 |
| HACLKGGLILKF | 1 | 0 | 0 | 1446124 | AXN92226.1 | K9N7C7 |
| HACTSFTCQCGG | 1 | 0 | 0 | 1446126 | AVV62535.1 | K9N7C7 |
| HADPNRFILGKL | 1 | 0 | 0 | 1446133 | AUM60014.1 | K9N5Q8 |
| HAENVTGLFKDC | 1 | 0 | 0 | 1446143 | 5NFY_A | P0C6X7 |
| HAFHTPAYEASA | 1 | 0 | 0 | 1446151 | AUM60013.1 | K9N7C7 |
| HAFKCTTSGSND | 1 | 0 | 0 | 1446157 | AAA73152.1 | P30215 |
| HAFMKHKGGDLT | 1 | 0 | 0 | 1446160 | AHI48527.1 | K9N638 |
| HAGPDLEGKFYG | 1 | 0 | 0 | 1446174 | ACV88184.1 | P0C6X7 |
| HAHPVEPLVQDR | 1 | 0 | 0 | 1446182 | AHF54653.1 | Q4VID0 |
| HAIATIPLQASL | 1 | 0 | 0 | 1446184 | ACQ82726.1 | P59632 |
| HAIGATAALPTQ | 1 | 0 | 0 | 1446186 | ARO76382.1 | P59594 |
| HAIQQGFDATNF | 1 | 0 | 0 | 1446190 | QBP84722.1 | P36334 |
| HAIQQGSDATNS | 1 | 0 | 0 | 1446192 | ARB66064.1 | P36334 |
| HAKDQYGTDFDG | 1 | 0 | 0 | 1446197 | AGT51416.1 | P33469 |
| HAKDQYGTDING | 1 | 0 | 0 | 1446199 | AEN19370.1 | P33469 |
| HALDDATRNYII | 1 | 0 | 0 | 1446203 | AHL18094.1 | K9N643 |
| HALDDFTRNYII | 1 | 0 | 0 | 1446204 | AIY60561.1 | K9N643 |
| HALDDVTLNYII | 1 | 0 | 0 | 1446205 | ALA49575.1 | K9N643 |
| HALDDVTRNYII | 1 | 0 | 0 | 1446206 | AXN92231.1 | K9N643 |
| HANCVNCADDRC | 1 | 0 | 0 | 1446219 | AVV62524.1 | K9N7C7 |
| HANCVNCSDDRC | 1 | 0 | 0 | 1446220 | AUM60013.1 | K9N7C7 |
| HANSSEPAMLFR | 1 | 0 | 0 | 1446236 | AIV41873.1 | P36334 |
| HANSYQKYVEAY | 1 | 0 | 0 | 1446238 | QEG03729.1 | H9EJ42 |
| HANYIFWRNSTP | 1 | 0 | 0 | 1446241 | AXN92247.1 | K9N7C7 |
| HASEYRQYLDAY | 1 | 0 | 0 | 1446261 | APO40578.1 | P0C6X7 |
| HATGTTPKKLFV | 1 | 0 | 0 | 1446276 | ALA49649.1 | K9N5Q8 |
| HATRLTITELLQ | 1 | 0 | 0 | 1446287 | AIW52763.1 | A0A0P0LKV0 |
| HATTVDVNVLND | 1 | 0 | 0 | 1446289 | AUM60013.1 | K9N7C7 |
| HAVVSKSKELLN | 1 | 0 | 0 | 1446297 | QBP84755.1 | P0C6U7 |
| HAWQPGVAMPNL | 1 | 0 | 0 | 1446302 | AVP78041.1 | P0C6W2 |
| HAYPLVYHENEE | 1 | 0 | 0 | 1446306 | ANZ78855.1 | P0C6X6 |
| HCAFAVDPAAAY | 1 | 0 | 0 | 1446308 | AST12962.1 | A0A0P0LKV0 |
| HCATVHTANKWD | 1 | 0 | 0 | 1446316 | AVP78030.1 | P0C6W2 |
| HCEHLQFGLQDG | 1 | 0 | 0 | 1446323 | AWO68974.1 | A0A0P0G321 |
| HCGSSSRSFIED | 1 | 0 | 0 | 1446339 | ARU07577.1 | A0A140H1H1 |
| HCPADNSYTSFA | 1 | 0 | 0 | 1446352 | AID55096.1 | K9N5Q8 |
| HCPAGNFYTSFA | 1 | 0 | 0 | 1446353 | ALJ54446.1 | K9N5Q8 |
| HCPAGNSHTSFA | 1 | 0 | 0 | 1446354 | AHX71946.1 | K9N5Q8 |
| HCPAGNSYTSFA | 1 | 0 | 0 | 1446355 | AXN92228.1 | K9N5Q8 |
| HCSFAVDPAAAY | 1 | 0 | 0 | 1446372 | NP_073549.1 | A0A0P0LKV0 |
| HCSFAVDPAVAY | 1 | 0 | 0 | 1446373 | AGT21359.1 | A0A0P0LKV0 |
| HCSGLAIKSDYC | 1 | 0 | 0 | 1446374 | CAA83660.1 | P36334 |
| HCSNFNTLFSTT | 1 | 0 | 0 | 1446378 | AIW52776.1 | A0A0P0LKV0 |
| HCTIAQLADAAL | 1 | 0 | 0 | 1446383 | AIV41889.1 | Q80872 |
| HCTIAQLTDAAL | 1 | 0 | 0 | 1446384 | AWW13564.1 | Q80872 |
| HCVKCHNDILTA | 1 | 0 | 0 | 1446390 | AUM60013.1 | K9N7C7 |
| HCVSTQANAVQQ | 1 | 0 | 0 | 1446394 | AVV62524.1 | K9N7C7 |
| HCYGADLKSPDL | 1 | 0 | 0 | 1446402 | APO40578.1 | P0C6X7 |
| HDDFSTEIIGYK | 1 | 0 | 0 | 1446407 | APO40578.1 | P0C6X7 |
| HDFFIFDVDKFK | 1 | 0 | 0 | 1446418 | AVN89322.1 | K9N7C7 |
| HDFKEELDQWFK | 1 | 0 | 0 | 1446427 | CAA83660.1 | P36334 |
| HDGDWDSFCKTI | 1 | 0 | 0 | 1446429 | AGT21366.1 | A0A0P0LKV0 |
| HDGLNEHVTSVN | 1 | 0 | 0 | 1446436 | ANF29196.1 | K9N796 |
| HDGPIEHVTSVN | 1 | 0 | 0 | 1446438 | AVN89292.1 | K9N796 |
| HDGPNEHVTSVN | 1 | 0 | 0 | 1446440 | AGV08391.1 | K9N796 |
| HDGPTEHVNLFD | 1 | 0 | 0 | 1446441 | ANI69890.1 | K9N796 |
| HDIGNPKGVRCA | 1 | 0 | 0 | 1446454 | AWK59942.1 | H9EJ42 |
| HDINNGPHEFCT | 1 | 0 | 0 | 1446457 | AIL49625.1 | P0C6X6 |
| HDIQLTNENYNN | 1 | 0 | 0 | 1446460 | QGV13494.1 | K9N7C7 |
| HDIQLTNESYNN | 1 | 0 | 0 | 1446461 | AVN89451.1 | K9N7C7 |
| HDIQLTTEGYNN | 1 | 0 | 0 | 1446462 | AVV62535.1 | K9N7C7 |
| HDIQLTVEGYNN | 1 | 0 | 0 | 1446463 | AVV62524.1 | K9N7C7 |
| HDITLDDDARTF | 1 | 0 | 0 | 1446465 | AQN78773.1 | P0C6X6 |
| HDLDDATRSYIG | 1 | 0 | 0 | 1446468 | AVV62540.1 | K9N643 |
| HDLNNVDPKAYP | 1 | 0 | 0 | 1446472 | AVP78050.1 | Q3LZX3 |
| HDNVLPETYVAL | 1 | 0 | 0 | 1446480 | AWO68987.1 | A0A0P0G321 |
| HDPIYKNGAMFH | 1 | 0 | 0 | 1446481 | AVV62524.1 | K9N7C7 |
| HDPTVLPGAFAY | 1 | 0 | 0 | 1446483 | AXN92247.1 | K9N7C7 |
| HDSQNMLRGDDM | 1 | 0 | 0 | 1446492 | AFR58685.1 | P0C6X7 |
| HDTSHKNNFGVI | 1 | 0 | 0 | 1446496 | Q14EA9.1 | Q0ZJ84 |
| HDVIVLRDNLKP | 1 | 0 | 0 | 1446504 | AFD98800.1 | H9EJ24 |

| | | | | | | |
|---|---|---|---|---|---|---|
| HDVNNGLTCSTD | 1 | 0 | 0 | 1446506 | AEN19366.1 | P36334 |
| HDVPTEHVTSVN | 1 | 0 | 0 | 1446509 | AMO03402.1 | K9N796 |
| HDVVSSFEGYDF | 1 | 0 | 0 | 1446515 | AST12962.1 | A0A0P0LKV0 |
| HDYCIPYNSITE | 1 | 0 | 0 | 1446522 | APO40580.1 | P59632 |
| HDYCIPYNSVTD | 1 | 0 | 0 | 1446523 | ARO76383.1 | P59632 |
| HEEGATDAPSTF | 1 | 0 | 0 | 1446542 | AVV62533.1 | K9N4V7 |
| HEENIYNLIKDC | 1 | 0 | 0 | 1446543 | APO40578.1 | P0C6X7 |
| HEETIYNLLKDC | 1 | 0 | 0 | 1446548 | QYC64967.1 | P0DTD1 |
| HEFAFLRKTLVL | 1 | 0 | 0 | 1446552 | ALA50059.1 | K9N643 |
| HEGDWDSFGKTI | 1 | 0 | 0 | 1446567 | AST12962.1 | A0A0P0LKV0 |
| HEGDWNSFCKTI | 1 | 0 | 0 | 1446569 | APT69882.1 | A0A0P0ILY9 |
| HEGFGGKNILGY | 1 | 0 | 0 | 1446570 | AWK59942.1 | H9EJ42 |
| HEHIGGVYDNSI | 1 | 0 | 0 | 1446586 | AVN89333.1 | K9N7C7 |
| HEHVGGNFDNDI | 1 | 0 | 0 | 1446588 | AVV62524.1 | K9N7C7 |
| HEIAWFTERSEK | 1 | 0 | 0 | 1446593 | ARO76381.1 | P0C6U8 |
| HEIEKPRWKRQP | 1 | 0 | 0 | 1446596 | AAA45463.1 | A0A0N7J131 |
| HEISDDEWAAAV | 1 | 0 | 0 | 1446598 | AHI48784.1 | K9N7C7 |
| HEKKLNSCCRLV | 1 | 0 | 0 | 1446604 | AUM60013.1 | K9N7C7 |
| HEMSPFEMPFEE | 1 | 0 | 0 | 1446610 | AST12962.1 | A0A0P0LKV0 |
| HENKLNSCCRIV | 1 | 0 | 0 | 1446611 | ALA49351.1 | K9N7C7 |
| HEPTTPISSALI | 1 | 0 | 0 | 1446613 | ALA49956.1 | K9N7C7 |
| HETCCSQRTRFY | 1 | 0 | 0 | 1446619 | AAD32993.1 | P0C6X6 |
| HETSDDDWTAAV | 1 | 0 | 0 | 1446623 | AVN89374.1 | K9N7C7 |
| HETSDDEWTAAV | 1 | 0 | 0 | 1446625 | AVN89451.1 | K9N7C7 |
| HEVFDDEWAAAV | 1 | 0 | 0 | 1446630 | AQZ41284.1 | K9N7C7 |
| HEWKMVVCDKVR | 1 | 0 | 0 | 1446642 | AXN92258.1 | K9N7C7 |
| HEYGATDAPSTF | 1 | 0 | 0 | 1446643 | QFQ59594.1 | K9N4V7 |
| HFCPCKLNGSCV | 1 | 0 | 0 | 1446659 | AIX10748.1 | P36334 |
| HFDCNDCSVLCE | 1 | 0 | 0 | 1446660 | ARB07436.1 | A0A140H1G9 |
| HFDGQSGEAPVS | 1 | 0 | 0 | 1446669 | AVP78041.1 | P0C6W2 |
| HFFFTQDGNAAI | 1 | 0 | 0 | 1446682 | QWY52192.1 | P0DTD1 |
| HFGAGSDKGVAPGTA | 1 | 0 | 0 | 1446694 | P0DTD1.1 | P0DTD1 |
| HFGENEEIVING | 1 | 0 | 0 | 1446696 | AST12962.1 | A0A0P0LKV0 |
| HFGQCDFRSLPN | 1 | 0 | 0 | 1446699 | AVV62543.1 | K9N7A1 |
| HFGQCDYAQLPN | 1 | 0 | 0 | 1446700 | AUM60030.1 | K9N7A1 |
| HFISQGLDAAWN | 1 | 0 | 0 | 1446716 | NP_073549.1 | A0A0P0LKV0 |
| HFKELWHLDAGV | 1 | 0 | 0 | 1446718 | AKA09652.1 | P36334 |
| HFKELWHLDTGA | 1 | 0 | 0 | 1446719 | ANZ78840.1 | P36334 |
| HFKESWHMDTGV | 1 | 0 | 0 | 1446723 | AIL49486.1 | P36334 |
| HFMVGNVTRLHD | 1 | 0 | 0 | 1446736 | AWW13510.1 | P30215 |
| HFNQKLPNGVNV | 1 | 0 | 0 | 1446742 | AMM42518.1 | A0A0N7J131 |
| HFNYVPTKYVTA | 1 | 0 | 0 | 1446747 | AEN19366.1 | P36334 |
| HFRHPLDEDEII | 1 | 0 | 0 | 1446754 | AVV62524.1 | K9N7C7 |
| HFRKACQNLDCN | 1 | 0 | 0 | 1446756 | AWW13564.1 | Q80872 |
| HFRKSCQNLDCN | 1 | 0 | 0 | 1446757 | AGT51688.1 | Q80872 |
| HFSLFDDNFYAF | 1 | 0 | 0 | 1446760 | ABD75647.1 | A0A140H1G9 |
| HFSMMILSDEGG | 1 | 0 | 0 | 1446765 | ARO76377.1 | P0C6X7 |
| HFSYKPTSFKTV | 1 | 0 | 0 | 1446770 | BBA20986.1 | A0A140H1H1 |
| HFTNDKTIFYTT | 1 | 0 | 0 | 1446780 | ARU07575.1 | A0A140H1G9 |
| HFTTQFVDNVKL | 1 | 0 | 0 | 1446786 | AIW52747.1 | A0A0P0K6L9 |
| HFTTQFVGNVKF | 1 | 0 | 0 | 1446789 | AIW52729.1 | A0A0P0K6L9 |
| HFTTQFVGVKFD | 1 | 0 | 0 | 1446791 | AIW52757.1 | A0A0P0K6L9 |
| HFYYLGTGPHKA | 1 | 0 | 0 | 1446818 | BAP75906.1 | A0A0N7J131 |
| HGAIKLDDKDPN | 1 | 0 | 0 | 1446823 | APO40586.1 | P59595 |
| HGAKPISNCIKM | 1 | 0 | 0 | 1446825 | AFD98833.1 | H9EJ42 |
| HGANLKQDESVA | 1 | 0 | 0 | 1446827 | AUM60014.1 | K9N5Q8 |
| HGCDLMQARAFV | 1 | 0 | 0 | 1446836 | AVV62524.1 | K9N7C7 |
| HGDFSGVASQEG | 1 | 0 | 0 | 1446843 | AWK59934.1 | U3M6Q8 |
| HGFEIDPSPNTG | 1 | 0 | 0 | 1446853 | APO40579.1 | P59594 |
| HGGGVAGAFNKA | 1 | 0 | 0 | 1446863 | QOJ87595.1 | P0DTD1 |
| HGIAAVNSVPWS | 1 | 0 | 0 | 1446872 | ARO76381.1 | P0C6U8 |
| HGKENLTFPRGQ | 1 | 0 | 0 | 1446878 | AVP78038.1 | Q3LZX4 |
| HGKETLTFPRGQ | 1 | 0 | 0 | 1446879 | APO40586.1 | P59595 |
| HGKIKPYERDIS | 1 | 0 | 0 | 1446882 | APO40579.1 | P59594 |
| HGKLLIRDNDTL | 1 | 0 | 0 | 1446883 | AHL31379.1 | K9N7C7 |
| HGKNNGIGTCPA | 1 | 0 | 0 | 1446885 | ARB66068.1 | P36334 |
| HGLAAINSVPWD | 1 | 0 | 0 | 1446893 | AVP78041.1 | P0C6W2 |
| HGLDLMQASQIV | 1 | 0 | 0 | 1446898 | AUM60013.1 | K9N7C7 |
| HGLGLIQAGSVV | 1 | 0 | 0 | 1446899 | ASU45706.1 | K9N7C7 |
| HGLLHSSQLPLD | 1 | 0 | 0 | 1446901 | AIV41780.1 | P0C6X6 |
| HGMSFSMTTFEI | 1 | 0 | 0 | 1446907 | AEN19363.1 | P0C6X6 |
| HGNGSMIEDMDY | 1 | 0 | 0 | 1446913 | AXG21663.1 | K9N7C7 |
| HGNGSMIEDVDY | 1 | 0 | 0 | 1446914 | AMQ49035.1 | K9N7C7 |
| HGNGSMLEDIDY | 1 | 0 | 0 | 1446915 | AVV62524.1 | K9N7C7 |
| HGNVAFYQGDVD | 1 | 0 | 0 | 1446919 | ARB07424.1 | A0A0P0ILY9 |
| HGTCERSEAGIC | 1 | 0 | 0 | 1446940 | P0C6W6.1 | P0C6W2 |
| HGTKKDAAAAKN | 1 | 0 | 0 | 1446947 | 6G13_A | K9N4V7 |
| HGVITKAVDCSS | 1 | 0 | 0 | 1446954 | AZS52618.1 | A0A140H1H1 |
| HGVITNAVDCSS | 1 | 0 | 0 | 1446955 | BBA20986.1 | A0A140H1H1 |
| HGYTIHNSGLPL | 1 | 0 | 0 | 1446969 | AVV62529.1 | K9N643 |
| HHAENVTGLFKD | 1 | 0 | 0 | 1446971 | 5NFY_A | P0C6X7 |
| HHASEYRQYLDA | 1 | 0 | 0 | 1446977 | APO40578.1 | P0C6X7 |
| HHCIVNSSFTEQ | 1 | 0 | 0 | 1446981 | AWO68966.1 | A0A0P0G321 |
| HHHAGNATEVPA | 1 | 0 | 0 | 1446990 | 5NFY_M | P0C6U8 |
| HHHAWSHPQFEK | 1 | 0 | 0 | 1446992 | 6NUR_A | P0C6X7 |
| HHHEEPVNQFTG | 1 | 0 | 0 | 1446993 | 2FAV_A | P0C6U8 |
| HHHHAGNATEVP | 1 | 0 | 0 | 1446995 | 5NFY_M | P0C6U8 |
| HHHHEEPVNQFT | 1 | 0 | 0 | 1446998 | 2FAV_A | P0C6U8 |
| HHHHHAENVTGL | 1 | 0 | 0 | 1446999 | 5NFY_A | P0C6X7 |
| HHHHHAGNATEV | 1 | 0 | 0 | 1447000 | 5NFY_M | P0C6U8 |
| HHHHHHAGNATE | 1 | 0 | 0 | 1447005 | 5NFY_M | P0C6U8 |

| | | | | | | |
|---|---|---|---|---|---|---|
| HHHHHHEEPVNQ | 1 | 0 | 0 | 1447008 | 2FAV_A | P0C6U8 |
| HHHHHHMPVNQF | 1 | 0 | 0 | 1447012 | 2ACF_A | P0C6U8 |
| HHHMPVNQFTGY | 1 | 0 | 0 | 1447041 | 2ACF_A | P0C6U8 |
| HHHSAWSHPQFE | 1 | 0 | 0 | 1447047 | 6CRV_A | P59594 |
| HHHSGDDDDKAS | 1 | 0 | 0 | 1447048 | 5N4K_A | H9EJ69 |
| HHLDSHFVVKRH | 1 | 0 | 0 | 1447055 | AUM60013.1 | K9N7C7 |
| HHLHSVYNWLKP | 1 | 0 | 0 | 1447062 | ATY74384.1 | K9N643 |
| HHMPVNQFTGYL | 1 | 0 | 0 | 1447066 | 2ACF_A | P0C6U8 |
| HHMRVPINMALT | 1 | 0 | 0 | 1447067 | AVV62529.1 | K9N643 |
| HHPMSDYDIPTT | 1 | 0 | 0 | 1447072 | 3EJG_A | A0A0P0LKV0 |
| HHRFKCSSNGSN | 1 | 0 | 0 | 1447076 | AGT17757.1 | A0A140H1H0 |
| HHSDLEVTSDSC | 1 | 0 | 0 | 1447079 | APO40578.1 | P0C6X7 |
| HHVINGCKDTAC | 1 | 0 | 0 | 1447094 | AVN89363.1 | K9N7C7 |
| HIADCQTTNGLN | 1 | 0 | 0 | 1447104 | ABB90509.1 | A0A0P0K6L9 |
| HIAGCQTTNGTN | 1 | 0 | 0 | 1447108 | AIW52757.1 | A0A0P0K6L9 |
| HIAYRPVPGTST | 1 | 0 | 0 | 1447115 | AUM60016.1 | K9N4V0 |
| HIDPFPKRVIDL | 1 | 0 | 0 | 1447123 | CAA46114.1 | A0A0P0KH29 |
| HIDSSEPLCLFK | 1 | 0 | 0 | 1447125 | BBA20986.1 | A0A140H1H1 |
| HIEVASAYGLCD | 1 | 0 | 0 | 1447133 | ANF29261.1 | K9N5Q8 |
| HIGNINPHNYSY | 1 | 0 | 0 | 1447147 | AEN19365.1 | P30215 |
| HIGPPANLRVVG | 1 | 0 | 0 | 1447148 | QBF80508.1 | K9N7C7 |
| HIGTRNPNNNVA | 1 | 0 | 0 | 1447152 | ATO98117.1 | Q3LZX4 |
| HIIPGSLGLPNF | 1 | 0 | 0 | 1447163 | AET99208.1 | Q0ZJ80 |
| HINAHVAKSHNI | 1 | 0 | 0 | 1447184 | AVP78041.1 | P0C6W2 |
| HINNLKIKNYGY | 1 | 0 | 0 | 1447192 | AGT17757.1 | A0A140H1H0 |
| HINNLKVKNYGY | 1 | 0 | 0 | 1447193 | ABD75544.1 | A0A140H1H0 |
| HISSTMSQHSRS | 1 | 0 | 0 | 1447221 | ASU90241.1 | K9N5Q8 |
| HISTKPQPLMDK | 1 | 0 | 0 | 1447224 | ATO98193.1 | Q3LZX1 |
| HITAKETLYRID | 1 | 0 | 0 | 1447226 | AEA10817.1 | P0C6X7 |
| HITDPLFSNMQP | 1 | 0 | 0 | 1447229 | NP_073549.1 | A0A0P0LKV0 |
| HITKVLAEKFDY | 1 | 0 | 0 | 1447232 | AST12962.1 | A0A0P0LKV0 |
| HITTYPGQGLNG | 1 | 0 | 0 | 1447241 | AVP78041.1 | P0C6W2 |
| HIVCDSLNSKLG | 1 | 0 | 0 | 1447243 | QBP84746.1 | P0C6X6 |
| HIVGPDARAKQD | 1 | 0 | 0 | 1447244 | AVK87459.1 | K9N7C7 |
| HIVNINPHNYSY | 1 | 0 | 0 | 1447248 | AWW13558.1 | P30215 |
| HIVSRETLYKID | 1 | 0 | 0 | 1447251 | APO40578.1 | P0C6X7 |
| HIVYGNFNQKII | 1 | 0 | 0 | 1447255 | AEN19363.1 | P0C6X6 |
| HKAYVDVLCNSF | 1 | 0 | 0 | 1447268 | AWK59942.1 | H9EJ42 |
| HKCGNLHQRELL | 1 | 0 | 0 | 1447270 | AEN19363.1 | P0C6X6 |
| HKDFFKDGLCMF | 1 | 0 | 0 | 1447275 | ARK08657.1 | P0C6X6 |
| HKDITECVTIVL | 1 | 0 | 0 | 1447278 | ALA49659.1 | K9N7C7 |
| HKEILSHIIPGS | 1 | 0 | 0 | 1447287 | AET99209.1 | Q0ZJ80 |
| HKELKFRQRSDG | 1 | 0 | 0 | 1447290 | AMM42628.1 | H9EJ69 |
| HKGAAWPIVRRR | 1 | 0 | 0 | 1447292 | AXN92247.1 | K9N7C7 |
| HKHDINLTNEGF | 1 | 0 | 0 | 1447302 | AUM60013.1 | K9N7C7 |
| HKHFYLTMYIIP | 1 | 0 | 0 | 1447308 | ARU07575.1 | A0A140H1G9 |
| HKLGERVRQAVL | 1 | 0 | 0 | 1447333 | AUM60013.1 | K9N7C7 |
| HKMQTYKLVNMK | 1 | 0 | 0 | 1447342 | AGT51330.1 | H9EJ42 |
| HKNGQGEYDNIS | 1 | 0 | 0 | 1447348 | AXX83299.1 | P33469 |
| HKNVQGENDNIS | 1 | 0 | 0 | 1447353 | AIX10722.1 | P33469 |
| HKPSSASCPIGT | 1 | 0 | 0 | 1447363 | AFR45556.1 | A0A140H1H1 |
| HKQCSVQQCFGK | 1 | 0 | 0 | 1447364 | AJQ24068.1 | P33469 |
| HKSDFQLPVYAS | 1 | 0 | 0 | 1447378 | AEN19363.1 | P0C6X6 |
| HKSNTNDAFISN | 1 | 0 | 0 | 1447383 | ATQ39390.1 | K9N5Q8 |
| HLAKALQTFSET | 1 | 0 | 0 | 1447410 | AUM60013.1 | K9N7C7 |
| HLDGEIISLEKL | 1 | 0 | 0 | 1447422 | APO40578.1 | P0C6X7 |
| HLDGEVLPLDKL | 1 | 0 | 0 | 1447427 | P0C6W6.1 | P0C6W2 |
| HLDILGPLSAQT | 1 | 0 | 0 | 1447429 | AVP78041.1 | P0C6W2 |
| HLGKFLEELNAF | 1 | 0 | 0 | 1447454 | AIW52723.1 | A0A0N7J131 |
| HLGSPLSLNMAR | 1 | 0 | 0 | 1447456 | AVP78050.1 | Q3LZX3 |
| HLHSVHNWLNGG | 1 | 0 | 0 | 1447467 | AVN89460.1 | K9N643 |
| HLIDEVQNWLSA | 1 | 0 | 0 | 1447472 | AUM60027.1 | K9N643 |
| HLKMAGMHFGAC | 1 | 0 | 0 | 1447487 | ASY99815.1 | K9N7A1 |
| HLNEIAKNLNES | 1 | 0 | 0 | 1447532 | APO40579.1 | P59594 |
| HLSEKCMIAPDG | 1 | 0 | 0 | 1447570 | AVV62535.1 | K9N7C7 |
| HLSNDTNGDPVY | 1 | 0 | 0 | 1447581 | AUM60021.1 | K9N4V7 |
| HLSPWFADGATA | 1 | 0 | 0 | 1447582 | AVV62535.1 | K9N7C7 |
| HLVSVTEFTLNN | 1 | 0 | 0 | 1447631 | AVV62530.1 | K9N7D2 |
| HMIDDVHSWLQP | 1 | 0 | 0 | 1447665 | AVV62529.1 | K9N643 |
| HMLDDVQRWLAP | 1 | 0 | 0 | 1447669 | AVV62540.1 | K9N643 |
| HMPLSNFEHKVI | 1 | 0 | 0 | 1447678 | 5DUS_A | K9N638 |
| HMPMHPMVNPSK | 1 | 0 | 0 | 1447679 | AET99209.1 | Q0ZJ80 |
| HMPVNQFTGYLK | 1 | 0 | 0 | 1447682 | 2ACF_A | P0C6U8 |
| HMRVPINMALTH | 1 | 0 | 0 | 1447683 | AVV62529.1 | K9N643 |
| HMSVAKLHNLTD | 1 | 0 | 0 | 1447688 | AUM60027.1 | K9N643 |
| HMVKDGLVGVLT | 1 | 0 | 0 | 1447690 | AVN89451.1 | K9N7C7 |
| HMVLPSEAPNAK | 1 | 0 | 0 | 1447692 | 2JZF_A | P0C6U8 |
| HMYCSFYPPEDE | 1 | 0 | 0 | 1447699 | AVP78041.1 | P0C6W2 |
| HNAAQCDCLCTP | 1 | 0 | 0 | 1447701 | CAA83660.1 | P36334 |
| HNAVQCNCLCTP | 1 | 0 | 0 | 1447703 | CAA83661.1 | P36334 |
| HNDCTPINTNAS | 1 | 0 | 0 | 1447706 | AVV62526.1 | K9N5Q8 |
| HNDFSSEIIGYK | 1 | 0 | 0 | 1447708 | AVP78041.1 | P0C6W2 |
| HNDILTATDPAE | 1 | 0 | 0 | 1447711 | AVV62524.1 | K9N7C7 |
| HNESGLKTVLRK | 1 | 0 | 0 | 1447715 | AVP78041.1 | P0C6W2 |
| HNFIIYDASDRG | 1 | 0 | 0 | 1447718 | AVV62524.1 | K9N7C7 |
| HNFLVQAGNVQL | 1 | 0 | 0 | 1447719 | AVP78041.1 | P0C6W2 |
| HNGVLDITACQY | 1 | 0 | 0 | 1447731 | AGW27872.1 | A0A140H1H1 |
| HNHSGIISQLRK | 1 | 0 | 0 | 1447735 | APO40578.1 | P0C6X7 |
| HNIGTLYVSWSD | 1 | 0 | 0 | 1447740 | 6U7G_C | A0A0P0K6L9 |
| HNKCAYWVPRAS | 1 | 0 | 0 | 1447741 | QHD43415.1 | P0DTD1 |
| HNLCTLDPITFK | 1 | 0 | 0 | 1447753 | AIL49502.1 | P36334 |

| | | | | | | |
|---|---|---|---|---|---|---|
| HNLIIYDSSDRG | 1 | 0 | 0 | 1447757 | AUM60013.1 | K9N7C7 |
| HNLPEYMAVAVP | 1 | 0 | 0 | 1447759 | APT69865.1 | A0A0P0KXN3 |
| HNLSGNPDEPQK | 1 | 0 | 0 | 1447762 | AHA80155.1 | P33469 |
| HNMGDSDLVQNG | 1 | 0 | 0 | 1447769 | AIW52805.1 | H9EJ69 |
| HNNKTWSIREFA | 1 | 0 | 0 | 1447774 | AVP78031.1 | Q3LZX1 |
| HNRRSFKTADGN | 1 | 0 | 0 | 1447783 | AEN19370.1 | P33469 |
| HNRRSFKTADGS | 1 | 0 | 0 | 1447784 | AIX10728.1 | P33469 |
| HNRVIAIPTSTG | 1 | 0 | 0 | 1447785 | AVV62527.1 | K9N796 |
| HNRWQLALYKGI | 1 | 0 | 0 | 1447786 | ARO76383.1 | P59632 |
| HNTASWFTALTQ | 1 | 0 | 0 | 1447800 | 2OFZ_A | P59595 |
| HNVCYTPRSEIP | 1 | 0 | 0 | 1447806 | AVV62524.1 | K9N7C7 |
| HNWLNGGQPVSE | 1 | 0 | 0 | 1447814 | AVN89361.1 | K9N643 |
| HNYSYMDLNSAL | 1 | 0 | 0 | 1447833 | AIL49334.1 | P30215 |
| HPAEPLVQDRVV | 1 | 0 | 0 | 1447837 | AVR40343.1 | Q4VID0 |
| HPDCVDCHDEMC | 1 | 0 | 0 | 1447846 | AIW52776.1 | A0A0P0LKV0 |
| HPHLGKFLEELN | 1 | 0 | 0 | 1447859 | AIW52723.1 | A0A0N7J131 |
| HPHTICHPNLGN | 1 | 0 | 0 | 1447861 | BAM16463.1 | P36334 |
| HPIQIDQLNSSY | 1 | 0 | 0 | 1447866 | ASU91208.1 | K9N5Q8 |
| HPIQVDQFNSSY | 1 | 0 | 0 | 1447868 | ASU90362.1 | K9N5Q8 |
| HPIQVDQLNSTY | 1 | 0 | 0 | 1447870 | ANI69889.1 | K9N5Q8 |
| HPIQVGQLNSSY | 1 | 0 | 0 | 1447871 | ALJ54452.1 | K9N5Q8 |
| HPKLGNHFKESW | 1 | 0 | 0 | 1447876 | AIL49486.1 | P36334 |
| HPLLNPGALEFN | 1 | 0 | 0 | 1447884 | AGT21342.1 | A0A0N7J131 |
| HPMSDYDIPTTE | 1 | 0 | 0 | 1447891 | 3EJG_A | A0A0P0LKV0 |
| HPNCSNCLDDRC | 1 | 0 | 0 | 1447896 | APO40578.1 | P0C6X7 |
| HPNLGNHFKELW | 1 | 0 | 0 | 1447898 | AWW13566.1 | P36334 |
| HPPPGEQFKHLV | 1 | 0 | 0 | 1447907 | AVN89289.1 | K9N7C7 |
| HPSISTDTVDVT | 1 | 0 | 0 | 1447917 | AIX10759.1 | P36334 |
| HPVECDEGCSEV | 1 | 0 | 0 | 1447926 | ALA50033.1 | K9N7C7 |
| HPVEPLVQDRAV | 1 | 0 | 0 | 1447930 | AHF54653.1 | Q4VID0 |
| HPVEPLVQDRVV | 1 | 0 | 0 | 1447931 | ARE29989.1 | Q4VID0 |
| HPVLMGWDYPKC | 1 | 0 | 0 | 1447932 | ACR82747.1 | A0A140H1G9 |
| HQEGMLYPITLG | 1 | 0 | 0 | 1447957 | AGT51685.1 | Q4VID0 |
| HQFVNASSATDA | 1 | 0 | 0 | 1447965 | AVA26872.1 | H9EJ42 |
| HQGVFRYDNISS | 1 | 0 | 0 | 1447971 | QBQ01838.1 | P30215 |
| HQLECIVDDSVR | 1 | 0 | 0 | 1447982 | ARU07593.1 | H9EJ42 |
| HQMELASGTHTG | 1 | 0 | 0 | 1447996 | AVN89451.1 | K9N7C7 |
| HQTYNRGRKPQP | 1 | 0 | 0 | 1448017 | AMM42397.1 | Q19U29 |
| HQVQLIDKYCSV | 1 | 0 | 0 | 1448021 | ATG84700.1 | K9N638 |
| HQVQLTDKYCSV | 1 | 0 | 0 | 1448022 | AXN92247.1 | K9N7C7 |
| HQWNCIDCDSYK | 1 | 0 | 0 | 1448027 | AEN19363.1 | P0C6X6 |
| HRCDINYKGKVF | 1 | 0 | 0 | 1448029 | QBP84746.1 | P0C6X6 |
| HRDCDLNKPLIE | 1 | 0 | 0 | 1448032 | AVV62524.1 | K9N7C7 |
| HRFKCFFNGSND | 1 | 0 | 0 | 1448038 | AZS52617.1 | A0A140H1H0 |
| HRFYSLKAAVHK | 1 | 0 | 0 | 1448045 | AXN92247.1 | K9N7C7 |
| HRGDPMPNNGWT | 1 | 0 | 0 | 1448050 | AVP78031.1 | Q3LZX1 |
| HRGDPMSNNGWT | 1 | 0 | 0 | 1448051 | AVP78042.1 | Q3LZX1 |
| HRIESIDSIVFT | 1 | 0 | 0 | 1448062 | AXP07349.1 | K9N7D2 |
| HRIIKAQFKTFE | 1 | 0 | 0 | 1448069 | AGT51448.1 | P0C6X6 |
| HRITSITSDKFD | 1 | 0 | 0 | 1448070 | AEN19363.1 | P0C6X6 |
| HRITSVTSDKFD | 1 | 0 | 0 | 1448071 | AXX83348.1 | P0C6X6 |
| HRNLGNHFKELW | 1 | 0 | 0 | 1448079 | BBO58769.1 | P36334 |
| HRSAIEDLLFNK | 1 | 0 | 0 | 1448095 | AVV62537.1 | K9N5Q8 |
| HRSFTQWYHEHI | 1 | 0 | 0 | 1448096 | AVN89333.1 | K9N7C7 |
| HRVPIHYEMFGN | 1 | 0 | 0 | 1448106 | ATO98227.1 | Q3LZX5 |
| HRWKMVVCDKVR | 1 | 0 | 0 | 1448112 | AVN89451.1 | K9N7C7 |
| HRYYSIKAAVSK | 1 | 0 | 0 | 1448122 | AUM60013.1 | K9N7C7 |
| HSAFADCTSCSN | 1 | 0 | 0 | 1448123 | AUM60013.1 | K9N7C7 |
| HSAFAECTGCGN | 1 | 0 | 0 | 1448124 | AVV62535.1 | K9N7C7 |
| HSAFVECGSCGN | 1 | 0 | 0 | 1448127 | AXN73479.1 | K9N7C7 |
| HSANDPLLGTTL | 1 | 0 | 0 | 1448130 | AVV62524.1 | K9N7C7 |
| HSDFGLDGLIDS | 1 | 0 | 0 | 1448141 | AWK59942.1 | H9EJ42 |
| HSDRFYRLRNEL | 1 | 0 | 0 | 1448152 | AAV28291.1 | A0A0P0LKV0 |
| HSDVDNIDFKSL | 1 | 0 | 0 | 1448157 | BBA20986.1 | A0A140H1H1 |
| HSEFNLPGCNGG | 1 | 0 | 0 | 1448159 | AUM60013.1 | K9N7C7 |
| HSEIVFDVESGG | 1 | 0 | 0 | 1448160 | AST12964.1 | A0A0P0K6L9 |
| HSFRTLKSGEGF | 1 | 0 | 0 | 1448170 | NP_073549.1 | A0A0P0LKV0 |
| HSFTLGGKVDLL | 1 | 0 | 0 | 1448174 | AUM60013.1 | K9N7C7 |
| HSGDDDDSQPRA | 1 | 0 | 0 | 1448178 | 5EPW_A | H9EJ69 |
| HSGSNDNIALLV | 1 | 0 | 0 | 1448189 | Q3I5J2.1 | Q3LZX9 |
| HSGYYDANQYYI | 1 | 0 | 0 | 1448193 | AWO68966.1 | A0A0P0G321 |
| HSIADVMRYNLN | 1 | 0 | 0 | 1448198 | AVL25583.1 | A0A0P0G321 |
| HSIPDLQKAAVT | 1 | 0 | 0 | 1448204 | ARO76381.1 | P0C6U8 |
| HSIQQGFDATNS | 1 | 0 | 0 | 1448207 | AIX10759.1 | P36334 |
| HSIRSFQSDRKA | 1 | 0 | 0 | 1448209 | ANF29195.1 | K9N5Q8 |
| HSIRSIKSDRKA | 1 | 0 | 0 | 1448210 | AJG44080.1 | K9N5Q8 |
| HSIRSIQSGRKA | 1 | 0 | 0 | 1448212 | QBF80543.1 | K9N5Q8 |
| HSKCLTSIEEVC | 1 | 0 | 0 | 1448213 | ATP16764.1 | P0C6X6 |
| HSKEQYGTDIDG | 1 | 0 | 0 | 1448215 | ANZ78874.1 | P33469 |
| HSKLDEEVPMEI | 1 | 0 | 0 | 1448219 | APO40583.1 | P59634 |
| HSKMSDVKCTSV | 1 | 0 | 0 | 1448221 | 1YSY_A | P0C6X7 |
| HSLNFVGEFVVN | 1 | 0 | 0 | 1448231 | AXN92247.1 | K9N7C7 |
| HSMPVVDGKQID | 1 | 0 | 0 | 1448249 | AGT51409.1 | P0C6X6 |
| HSPPTKQRYVKR | 1 | 0 | 0 | 1448266 | AUM60027.1 | K9N643 |
| HSQFYFNKDTGV | 1 | 0 | 0 | 1448269 | QEG03793.1 | P30215 |
| HSQTSLRCGTCI | 1 | 0 | 0 | 1448272 | AXN92247.1 | K9N7C7 |
| HSSIKEGTQIYK | 1 | 0 | 0 | 1448289 | AGT51787.1 | P0C6X6 |
| HSSIKQGTQIYK | 1 | 0 | 0 | 1448292 | AEN19363.1 | P0C6X6 |
| HSSLFAQSEEEE | 1 | 0 | 0 | 1448295 | AVN89326.1 | K9N4V0 |
| HSSQLPLDIYDL | 1 | 0 | 0 | 1448307 | AIV41960.1 | P0C6X6 |
| HSSQLPLDVYNL | 1 | 0 | 0 | 1448309 | AAX85666.1 | P0C6X6 |

| | | | | | | |
|---|---|---|---|---|---|---|
| HSTRSIQSDRKA | 1 | 0 | 0 | 1448329 | ATG84899.1 | K9N5Q8 |
| HSTVFVPASRDP | 1 | 0 | 0 | 1448330 | AVV62530.1 | K9N7D2 |
| HSVADVMRYNFN | 1 | 0 | 0 | 1448334 | AVA26873.1 | A0A0P0G321 |
| HSVADYHNHSGI | 1 | 0 | 0 | 1448336 | APO40578.1 | P0C6X7 |
| HSVCKGCVGHSE | 1 | 0 | 0 | 1448344 | AIW52752.1 | A0A0P0K6L9 |
| HSVCNGCVGHSE | 1 | 0 | 0 | 1448345 | AIW52757.1 | A0A0P0K6L9 |
| HSVHNRLKPYGG | 1 | 0 | 0 | 1448353 | ALW82723.1 | K9N643 |
| HSVHNWLKPFGG | 1 | 0 | 0 | 1448354 | AKN24752.1 | K9N643 |
| HSVHNWLNGGQP | 1 | 0 | 0 | 1448357 | AVN89361.1 | K9N643 |
| HSVSLIWNVKDY | 1 | 0 | 0 | 1448365 | APO40578.1 | P0C6X7 |
| HSWSADLYELMG | 1 | 0 | 0 | 1448376 | AVV62524.1 | K9N7C7 |
| HSYIYHSAFADC | 1 | 0 | 0 | 1448384 | AUM60013.1 | K9N7C7 |
| HTASTLRSVGQK | 1 | 0 | 0 | 1448396 | ATO98181.1 | Q3LZX1 |
| HTDCTSAGYNTN | 1 | 0 | 0 | 1448400 | AVV62537.1 | K9N5Q8 |
| HTDSFSCNNFDE | 1 | 0 | 0 | 1448407 | AET99191.1 | A0A140H1H1 |
| HTGFQGKQILGS | 1 | 0 | 0 | 1448430 | ATQ39389.1 | K9N7C7 |
| HTGRNYPKCDCA | 1 | 0 | 0 | 1448433 | AJS10812.1 | P0C6X6 |
| HTGSLLKACIET | 1 | 0 | 0 | 1448437 | AUM60025.1 | K9N796 |
| HTGSSFDGNMYG | 1 | 0 | 0 | 1448438 | AVV62524.1 | K9N7C7 |
| HTHTHDYCIPYN | 1 | 0 | 0 | 1448448 | APO40580.1 | P59632 |
| HTICKSIGSSRN | 1 | 0 | 0 | 1448456 | ADN03339.1 | A0A140H1H1 |
| HTIKGSFLNGSA | 1 | 0 | 0 | 1448465 | 1Z1J_A | P0C6U8 |
| HTILNKPRPKRS | 1 | 0 | 0 | 1448467 | AJQ24068.1 | P33469 |
| HTKKWKIPQVGG | 1 | 0 | 0 | 1448482 | BAE93400.1 | P0C6U8 |
| HTKSAAGYGGFG | 1 | 0 | 0 | 1448487 | AEN19363.1 | P0C6X6 |
| HTLDDSFLGRYM | 1 | 0 | 0 | 1448494 | APO40578.1 | P0C6X7 |
| HTLDDVTRNYII | 1 | 0 | 0 | 1448495 | ASU90464.1 | K9N643 |
| HTLSNYQHEENI | 1 | 0 | 0 | 1448510 | APO40578.1 | P0C6X7 |
| HTLSVHDGYTNE | 1 | 0 | 0 | 1448511 | ARO76381.1 | P0C6U8 |
| HTLTKHTEVRAS | 1 | 0 | 0 | 1448513 | APT69856.1 | A0A0P0K6L9 |
| HTLTKYTEVRAS | 1 | 0 | 0 | 1448514 | AIW52757.1 | A0A0P0K6L9 |
| HTLYIKDGDYGY | 1 | 0 | 0 | 1448519 | ALW82667.1 | K9N7C7 |
| HTMENYELEKRC | 1 | 0 | 0 | 1448525 | AVV62524.1 | K9N7C7 |
| HTNPFTRTVFEN | 1 | 0 | 0 | 1448542 | ARU07574.1 | A0A140H1G9 |
| HTPAIDCSDGNY | 1 | 0 | 0 | 1448550 | ASU91010.1 | K9N5Q8 |
| HTPATNCSDGNY | 1 | 0 | 0 | 1448552 | ALA49671.1 | K9N5Q8 |
| HTPAYDGSAFRD | 1 | 0 | 0 | 1448553 | AHI48736.1 | K9N7C7 |
| HTPSTDCSDGNY | 1 | 0 | 0 | 1448562 | ALA49803.1 | K9N5Q8 |
| HTRYSLADSNGF | 1 | 0 | 0 | 1448572 | ABD75543.1 | A0A140H1G9 |
| HTSCSCSPDAFL | 1 | 0 | 0 | 1448575 | BBA20986.1 | A0A140H1H1 |
| HTSFATYHTPAT | 1 | 0 | 0 | 1448576 | AHX71946.1 | K9N5Q8 |
| HTSGNVIEFHLD | 1 | 0 | 0 | 1448577 | APO40578.1 | P0C6X7 |
| HTSIAPNLTFNS | 1 | 0 | 0 | 1448578 | BBA20986.1 | A0A140H1H1 |
| HTSPDIDLGDIS | 1 | 0 | 0 | 1448582 | AVP78031.1 | Q3LZX1 |
| HTTAQVVSYLSE | 1 | 0 | 0 | 1448587 | AVV62524.1 | K9N7C7 |
| HTTATIPLQASL | 1 | 0 | 0 | 1448588 | ARO76383.1 | P59632 |
| HTTLGGLHLFIG | 1 | 0 | 0 | 1448594 | QAT98908.1 | K9N7C7 |
| HTVCDSLNAKWI | 1 | 0 | 0 | 1448600 | AIV41816.1 | P0C6X6 |
| HTVCDSLNAKWV | 1 | 0 | 0 | 1448601 | AIV41984.1 | P0C6X6 |
| HTVCDSLNFKLG | 1 | 0 | 0 | 1448602 | AAX85666.1 | P0C6X6 |
| HTVIDVKQVGCS | 1 | 0 | 0 | 1448608 | AIV41894.1 | P0C6X6 |
| HTVTDVKQVGCF | 1 | 0 | 0 | 1448624 | AGT51518.1 | P0C6X6 |
| HTVTDVKQVGCS | 1 | 0 | 0 | 1448625 | AEN19363.1 | P0C6X6 |
| HTVYPDYVDVNK | 1 | 0 | 0 | 1448629 | AWK59979.1 | A0A0P0G321 |
| HVAQNTVLNACT | 1 | 0 | 0 | 1448645 | ABK63972.1 | H9EJ69 |
| HVCFKDCYNIWE | 1 | 0 | 0 | 1448653 | AWK59942.1 | H9EJ42 |
| HVDHYDGFYTQG | 1 | 0 | 0 | 1448661 | AST12962.1 | A0A0P0LKV0 |
| HVDVFTDGKLNC | 1 | 0 | 0 | 1448671 | AVV62524.1 | K9N7C7 |
| HVGLLDQAWRFP | 1 | 0 | 0 | 1448700 | AEN19363.1 | P0C6X6 |
| HVGYQPTAHVNA | 1 | 0 | 0 | 1448704 | AUM60014.1 | K9N5Q8 |
| HVGYYPSNHIDV | 1 | 0 | 0 | 1448707 | ALX27232.1 | K9N5Q8 |
| HVGYYPSNHIEV | 1 | 0 | 0 | 1448708 | AXN92228.1 | K9N5Q8 |
| HVHVGGVYDNSI | 1 | 0 | 0 | 1448710 | ASU90174.1 | K9N638 |
| HVICPADQLSDP | 1 | 0 | 0 | 1448711 | QGV13494.1 | K9N7C7 |
| HVIWDYVRESIF | 1 | 0 | 0 | 1448720 | AXX83324.1 | P0C6X6 |
| HVKEITRENVET | 1 | 0 | 0 | 1448725 | AGT51393.1 | H9EJ42 |
| HVMCPADQLNDP | 1 | 0 | 0 | 1448746 | AVV62524.1 | K9N7C7 |
| HVMFKHNDFSSE | 1 | 0 | 0 | 1448748 | AVP78041.1 | P0C6W2 |
| HVNASYECDIPI | 1 | 0 | 0 | 1448756 | AVP78031.1 | Q3LZX1 |
| HVNQSTNKTTFR | 1 | 0 | 0 | 1448771 | APO40578.1 | P0C6X7 |
| HVNSSEPALLFR | 1 | 0 | 0 | 1448773 | AIX10749.1 | P36334 |
| HVQIHTIDGTSG | 1 | 0 | 0 | 1448778 | AVP78043.1 | Q3LZX0 |
| HVRNNGGYNSWH | 1 | 0 | 0 | 1448788 | AXN92247.1 | K9N7C7 |
| HVSSGHMIDDVH | 1 | 0 | 0 | 1448806 | AVV62529.1 | K9N643 |
| HVTCCTANDKFY | 1 | 0 | 0 | 1448814 | ALJ99930.1 | A0A0P0LKV0 |
| HVTSVNLLDVGY | 1 | 0 | 0 | 1448824 | AHI48551.1 | K9N796 |
| HVTYIPSQEKNF | 1 | 0 | 0 | 1448827 | AVP78031.1 | Q3LZX1 |
| HVTYVPSQEKNF | 1 | 0 | 0 | 1448831 | ATO98181.1 | Q3LZX1 |
| HVTYVPSQQQNF | 1 | 0 | 0 | 1448833 | APO40579.1 | P59594 |
| HVVPLENDADKA | 1 | 0 | 0 | 1448858 | AIV41996.1 | P0C6X6 |
| HVYDFKVEYAYD | 1 | 0 | 0 | 1448871 | APU51933.1 | P0C6X6 |
| HVYEVGDLIAQS | 1 | 0 | 0 | 1448872 | AEN19363.1 | P0C6X6 |
| HVYEVSDLIAQS | 1 | 0 | 0 | 1448874 | AIV41990.1 | P0C6X6 |
| HVYLLYVNFELR | 1 | 0 | 0 | 1448877 | ADK37703.1 | A0A0P0K6L9 |
| HWDQPYHANCVN | 1 | 0 | 0 | 1448888 | AVV62535.1 | K9N7C7 |
| HWFVTQRNFYEPQIITT | 1 | 0 | 0 | 1448897 | P0DTC2.1 | P0DTC2 |
| HWICANQSTSTY | 1 | 0 | 0 | 1448905 | AWO68966.1 | A0A0P0G321 |
| HWLCANQSTSVY | 1 | 0 | 0 | 1448908 | ABE73442.1 | A0A0P0G321 |
| HWSMPYHPNPVD | 1 | 0 | 0 | 1448937 | ANZ78865.1 | P0C6X6 |
| HYCSTLHNEILA | 1 | 0 | 0 | 1448943 | AEN19363.1 | P0C6X6 |
| HYDLVNYIRDKG | 1 | 0 | 0 | 1448951 | AIV41836.1 | Q80872 |

| | | | | | | |
|---|---|---|---|---|---|---|
| HYERDNTSYPEW | 1 | 0 | 0 | 1448957 | AHI48797.1 | K9N638 |
| HYERNNTSCPEW | 1 | 0 | 0 | 1448958 | AVN89451.1 | K9N7C7 |
| HYFDAQPLTRKV | 1 | 0 | 0 | 1448960 | AXN92247.1 | K9N7C7 |
| HYIKDGDDGYFL | 1 | 0 | 0 | 1448971 | AHI48773.1 | K9N7C7 |
| HYKELGIVMNMD | 1 | 0 | 0 | 1448977 | AEN19363.1 | P0C6X6 |
| HYKELGVVMNFD | 1 | 0 | 0 | 1448979 | ACV87155.1 | A0A140H1G9 |
| HYKYSRFLYGLK | 1 | 0 | 0 | 1448985 | AFQ55887.1 | U3M6Q8 |
| HYMYSRFLYGLK | 1 | 0 | 0 | 1448999 | AWK59934.1 | U3M6Q8 |
| HYTPVFKKGAKL | 1 | 0 | 0 | 1449043 | APO40578.1 | P0C6X7 |
| HYTPVHYERDNT | 1 | 0 | 0 | 1449044 | AXN92247.1 | K9N7C7 |
| HYTPVRYERDNT | 1 | 0 | 0 | 1449045 | ALA50044.1 | K9N7C7 |
| HYVHARLKCGLI | 1 | 0 | 0 | 1449050 | AVN89290.1 | K9N7C7 |
| HYVNRLVTQFKL | 1 | 0 | 0 | 1449065 | ARU07575.1 | A0A140H1G9 |
| HYVRVTFSDPNI | 1 | 0 | 0 | 1449071 | AVN89373.1 | K9N643 |
| HYVRVTFSDPNL | 1 | 0 | 0 | 1449072 | ALA49564.1 | K9N643 |
| HYVRVTFSDPNT | 1 | 0 | 0 | 1449073 | ASU90321.1 | K9N643 |
| HYVVYDAANNAV | 1 | 0 | 0 | 1449077 | AWK59942.1 | H9EJ42 |
| IAAAINKASNDA | 1 | 0 | 0 | 1449103 | AVV62524.1 | K9N7C7 |
| IAAAINNASNGA | 1 | 0 | 0 | 1449104 | AUM60013.1 | K9N7C7 |
| IAADLPCTLHFS | 1 | 0 | 0 | 1449106 | ARU07575.1 | A0A140H1G9 |
| IAAHPPPGEQFK | 1 | 0 | 0 | 1449109 | AVN89289.1 | K9N7C7 |
| IAANTVIWDFNR | 1 | 0 | 0 | 1449116 | APO40578.1 | P0C6X7 |
| IAARPPPGEHFK | 1 | 0 | 0 | 1449121 | ALW82707.1 | K9N7C7 |
| IAAVITREVGFI | 1 | 0 | 0 | 1449127 | APO40578.1 | P0C6X7 |
| IAAVNSVPWSKI | 1 | 0 | 0 | 1449129 | ARO76381.1 | P0C6U8 |
| IACVKDYEFKFY | 1 | 0 | 0 | 1449152 | ARB07436.1 | A0A140H1G9 |
| IADAINKASNGA | 1 | 0 | 0 | 1449153 | AVV62535.1 | K9N7C7 |
| IADETCEHEYGT | 1 | 0 | 0 | 1449156 | ARO76381.1 | P0C6U8 |
| IADPGYMQGYDD | 1 | 0 | 0 | 1449165 | AXN92228.1 | K9N5Q8 |
| IADPGYMQGYDE | 1 | 0 | 0 | 1449166 | AUM60014.1 | K9N5Q8 |
| IADTLQEIPVVS | 1 | 0 | 0 | 1449173 | AQZ41284.1 | K9N7C7 |
| IADTLQETSVVS | 1 | 0 | 0 | 1449177 | ASU45706.1 | K9N7C7 |
| IADVMRYNLNFS | 1 | 0 | 0 | 1449179 | AVL25583.1 | A0A0P0G321 |
| IADVYRRKLNLP | 1 | 0 | 0 | 1449181 | AAR01015.1 | P36334 |
| IADVYRRKPDLP | 1 | 0 | 0 | 1449182 | AEN19366.1 | P36334 |
| IADVYRRKPNLP | 1 | 0 | 0 | 1449184 | CAA79896.1 | P36334 |
| IADVYRRRPDLP | 1 | 0 | 0 | 1449185 | AHA80225.1 | P36334 |
| IAFANWLAPTNA | 1 | 0 | 0 | 1449202 | QBF80585.1 | K9N7C7 |
| IAFCKSGNLVLN | 1 | 0 | 0 | 1449203 | AIL49349.1 | P30215 |
| IAFELYAKRSVR | 1 | 0 | 0 | 1449209 | AHY22543.1 | K9N7C7 |
| IAFGIPPSEAKG | 1 | 0 | 0 | 1449211 | ABA60794.1 | Q19U29 |
| IAFGLQDCVTGI | 1 | 0 | 0 | 1449212 | AVA26872.1 | H9EJ42 |
| IAFGVPLSEAKG | 1 | 0 | 0 | 1449213 | ABD75541.1 | Q19U29 |
| IAFNHPIQIDQL | 1 | 0 | 0 | 1449218 | ASU91208.1 | K9N5Q8 |
| IAFNHPIQVDHI | 1 | 0 | 0 | 1449219 | AMQ49004.1 | K9N5Q8 |
| IAFNHPIQVDQF | 1 | 0 | 0 | 1449220 | ASU90362.1 | K9N5Q8 |
| IAFSHSANDPLL | 1 | 0 | 0 | 1449226 | AVV62524.1 | K9N7C7 |
| IAGHEEFIQTSS | 1 | 0 | 0 | 1449241 | BBA20986.1 | A0A140H1H1 |
| IAGPKGPEIRYL | 1 | 0 | 0 | 1449256 | AXN92247.1 | K9N7C7 |
| IAGQAFFYSDGF | 1 | 0 | 0 | 1449257 | AVA26872.1 | H9EJ42 |
| IAGQVHVLNHVD | 1 | 0 | 0 | 1449259 | P0C5B5.1 | Q0ZJ80 |
| IAHWIKNTFSLV | 1 | 0 | 0 | 1449274 | AIV41978.1 | P0C6X6 |
| IAIEDLLFDKVT | 1 | 0 | 0 | 1449277 | AHL18090.1 | K9N5Q8 |
| IAIEQDKNTQEV | 1 | 0 | 0 | 1449278 | AVP78031.1 | Q3LZX1 |
| IAILSGTTTNVD | 1 | 0 | 0 | 1449287 | AVV62524.1 | K9N7C7 |
| IAISDLGSLIDS | 1 | 0 | 0 | 1449296 | AVV62524.1 | K9N7C7 |
| IAISVKPESNAD | 1 | 0 | 0 | 1449297 | AVV62524.1 | K9N7C7 |
| IAIVKDWNFAWS | 1 | 0 | 0 | 1449301 | AUM60030.1 | K9N7A1 |
| IAKPKRLRKKRN | 1 | 0 | 0 | 1449321 | ATQ39389.1 | K9N7C7 |
| IAKTESAMVHDL | 1 | 0 | 0 | 1449323 | AVP78050.1 | Q3LZX3 |
| IALAMAYGDCTF | 1 | 0 | 0 | 1449327 | AVV62535.1 | K9N7C7 |
| IALDDDARTFIQ | 1 | 0 | 0 | 1449330 | AGT51409.1 | P0C6X6 |
| IALDDDARTFVQ | 1 | 0 | 0 | 1449331 | AEN19363.1 | P0C6X6 |
| IALKMTKLATTV | 1 | 0 | 0 | 1449352 | AVP78050.1 | Q3LZX3 |
| IAMANWLAPNNM | 1 | 0 | 0 | 1449367 | AVV62524.1 | K9N7C7 |
| IANAFNNALDAI | 1 | 0 | 0 | 1449372 | CAA79896.1 | P36334 |
| IANAFNNALHAI | 1 | 0 | 0 | 1449373 | AEN19366.1 | P36334 |
| IANAFSNALDAI | 1 | 0 | 0 | 1449375 | CAA83660.1 | P36334 |
| IANKFNQAPGAM | 1 | 0 | 0 | 1449385 | AVV62526.1 | K9N5Q8 |
| IANLFVEQSQRS | 1 | 0 | 0 | 1449388 | AVV62524.1 | K9N7C7 |
| IANQERYVKISG | 1 | 0 | 0 | 1449394 | AXG21652.1 | K9N7C7 |
| IANQERYVKITG | 1 | 0 | 0 | 1449395 | AXG21641.1 | K9N7C7 |
| IANQERYVKMFW | 1 | 0 | 0 | 1449396 | AXG21663.1 | K9N7C7 |
| IANQFNNAISKI | 1 | 0 | 0 | 1449398 | APO40579.1 | P59594 |
| IANVYRRKPDLP | 1 | 0 | 0 | 1449405 | ABD43177.1 | P36334 |
| IAPDGKSFTMTF | 1 | 0 | 0 | 1449409 | AVV62535.1 | K9N7C7 |
| IAPNLTLNLHTI | 1 | 0 | 0 | 1449424 | BBA20983.1 | A0A140H1H1 |
| IAPQANFGDYYY | 1 | 0 | 0 | 1449430 | AGT51599.1 | P30215 |
| IAPQANSGDYYH | 1 | 0 | 0 | 1449431 | AGT51421.1 | P30215 |
| IAPQANSGDYYY | 1 | 0 | 0 | 1449432 | P30215.1 | P30215 |
| IAPQANSGGYYY | 1 | 0 | 0 | 1449433 | AGT51669.1 | P30215 |
| IAPQANSGIITK | 1 | 0 | 0 | 1449434 | AEN19365.1 | P30215 |
| IAPSTVWEDGDY | 1 | 0 | 0 | 1449436 | QEJ82215.1 | K9N5Q8 |
| IAQRTRTRPSVQ | 1 | 0 | 0 | 1449453 | QEJ82222.1 | K9N4V7 |
| IAQVDVVNFNLT | 1 | 0 | 0 | 1449457 | YP_009725255.1 | A0A663DJA2 |
| IAQYNYKQDFSN | 1 | 0 | 0 | 1449461 | AUM60014.1 | K9N5Q8 |
| IASNCYSSLILD | 1 | 0 | 0 | 1449484 | AXN92228.1 | K9N5Q8 |
| IASQLGNCVEYH | 1 | 0 | 0 | 1449488 | 4KQZ_A | K9N5Q8 |
| IASQLGNCVEYS | 1 | 0 | 0 | 1449489 | AXN92228.1 | K9N5Q8 |
| IATAKDGLKRGE | 1 | 0 | 0 | 1449499 | AVV62524.1 | K9N7C7 |
| IATPHKMVLSVS | 1 | 0 | 0 | 1449511 | AXN92247.1 | K9N7C7 |

| | | | | | | |
|---|---|---|---|---|---|---|
| IATRNNCWLRSV | 1 | 0 | 0 | 1449513 | AXX83378.1 | P0C6X6 |
| IATSHKMVLSVS | 1 | 0 | 0 | 1449516 | ATG84699.1 | K9N7C7 |
| IAVANWLAPTNV | 1 | 0 | 0 | 1449523 | ANC28699.1 | K9N638 |
| IAVANWLTPTNM | 1 | 0 | 0 | 1449524 | AUM60013.1 | K9N7C7 |
| IAVDIQQTFFDK | 1 | 0 | 0 | 1449528 | ALJ54490.1 | K9N5Q8 |
| IAVEQDRNTRDV | 1 | 0 | 0 | 1449533 | APO40579.1 | P59594 |
| IAVNDQKLYDQV | 1 | 0 | 0 | 1449546 | ARO76381.1 | P0C6U8 |
| IAWNTAKQDTGH | 1 | 0 | 0 | 1449568 | AVP78042.1 | Q3LZX1 |
| IAWNTAKQDVGN | 1 | 0 | 0 | 1449569 | AVP78031.1 | Q3LZX1 |
| IAYAILYFFATR | 1 | 0 | 0 | 1449575 | NP_073549.1 | A0A0P0LKV0 |
| IAYANNSIAIPT | 1 | 0 | 0 | 1449576 | AVP78031.1 | Q3LZX1 |
| IAYFNMIYMPAS | 1 | 0 | 0 | 1449578 | APO40578.1 | P0C6X7 |
| IAYGIPLSEAKG | 1 | 0 | 0 | 1449580 | AMM42386.1 | Q19U29 |
| ICDCGIKQESRV | 1 | 0 | 0 | 1449630 | ARU07575.1 | A0A140H1G9 |
| ICEKLAIGGSIA | 1 | 0 | 0 | 1449643 | NP_073549.1 | A0A0P0LKV0 |
| ICFDIRDNGSLE | 1 | 0 | 0 | 1449650 | AXN92247.1 | K9N7C7 |
| ICFEYTSPTDKL | 1 | 0 | 0 | 1449654 | AGT51509.1 | Q80872 |
| ICFIGLWCLMPT | 1 | 0 | 0 | 1449655 | AGT51737.1 | P0C6X6 |
| ICGGCCDDYTGH | 1 | 0 | 0 | 1449674 | CAA83660.1 | P36334 |
| ICHPKLGNHFKE | 1 | 0 | 0 | 1449685 | AEN19366.1 | P36334 |
| ICHPNLGNRRVE | 1 | 0 | 0 | 1449689 | CAA83660.1 | P36334 |
| ICIKDADYNAKD | 1 | 0 | 0 | 1449697 | AST12962.1 | A0A0P0LKV0 |
| ICKFGNTTFDFL | 1 | 0 | 0 | 1449711 | AWO69005.1 | A0A0P0G321 |
| ICKFSTNTTFDF | 1 | 0 | 0 | 1449718 | ACV87106.1 | A0A0P0G321 |
| ICKNFAPNGFSQ | 1 | 0 | 0 | 1449720 | AVV62537.1 | K9N5Q8 |
| ICKPDQQFHNFS | 1 | 0 | 0 | 1449721 | AVV62540.1 | K9N643 |
| ICKTTTGIPEYN | 1 | 0 | 0 | 1449724 | ATG84821.1 | K9N638 |
| ICLSGRKIYEIE | 1 | 0 | 0 | 1449739 | AGT51737.1 | P0C6X6 |
| ICNSNILSLNNP | 1 | 0 | 0 | 1449758 | AGT17757.1 | A0A140H1H0 |
| ICPAGTKYRHCD | 1 | 0 | 0 | 1449763 | BBA20986.1 | A0A140H1H1 |
| ICPTGTKYRHCD | 1 | 0 | 0 | 1449766 | AIG96691.1 | A0A140H1H1 |
| ICQAVSDNVRAL | 1 | 0 | 0 | 1449767 | ADO24673.1 | P0C6X6 |
| ICQAVTANVRSL | 1 | 0 | 0 | 1449770 | ADO24641.1 | A0A140H1G9 |
| ICSQISPAAIAS | 1 | 0 | 0 | 1449785 | AVN89334.1 | K9N5Q8 |
| ICSQYVAGYKVL | 1 | 0 | 0 | 1449786 | APT69978.1 | K9N5Q8 |
| ICSVAYVANPTE | 1 | 0 | 0 | 1449790 | AEN19363.1 | P0C6X6 |
| ICVALCDAMVAK | 1 | 0 | 0 | 1449799 | QED88024.1 | H9EJ42 |
| IDAAKIYGMGFS | 1 | 0 | 0 | 1449833 | AIL49487.1 | P36334 |
| IDAGNHRSAFAL | 1 | 0 | 0 | 1449842 | AKA09621.1 | A0A0P0G321 |
| IDAYKTFPKKEN | 1 | 0 | 0 | 1449859 | AHL18098.1 | K9N4V7 |
| IDAYKTFPVKEK | 1 | 0 | 0 | 1449861 | AVV62544.1 | K9N4V7 |
| IDCAACVCGDYA | 1 | 0 | 0 | 1449867 | ABD43177.1 | P36334 |
| IDCAAFVRGDYA | 1 | 0 | 0 | 1449871 | ANZ78843.1 | P36334 |
| IDCEQDLCDFKG | 1 | 0 | 0 | 1449873 | AXX83372.1 | P0C6X6 |
| IDCEQTPCGFKG | 1 | 0 | 0 | 1449875 | AIV42002.1 | P0C6X6 |
| IDCGFNDLSQLY | 1 | 0 | 0 | 1449878 | ANI69889.1 | K9N5Q8 |
| IDCGHDDLAQLR | 1 | 0 | 0 | 1449879 | AVV62526.1 | K9N5Q8 |
| IDCNAASVNPAS | 1 | 0 | 0 | 1449881 | AUM60013.1 | K9N7C7 |
| IDCSAFVCGDCA | 1 | 0 | 0 | 1449887 | CAA83661.1 | P36334 |
| IDCSAFVCGDYA | 1 | 0 | 0 | 1449888 | CAA83660.1 | P36334 |
| IDCSDGNYNRNA | 1 | 0 | 0 | 1449891 | ASU91010.1 | K9N5Q8 |
| IDDCYGYLRKHF | 1 | 0 | 0 | 1449893 | ADO24649.1 | H9EJ42 |
| IDDGKIWNGVIK | 1 | 0 | 0 | 1449899 | QEO75984.1 | A0A0P0ILY9 |
| IDEIPYKSFSDS | 1 | 0 | 0 | 1449918 | AXN92247.1 | K9N7C7 |
| IDEVQNWLSAFG | 1 | 0 | 0 | 1449922 | AUM60027.1 | K9N643 |
| IDEYITTYPGQG | 1 | 0 | 0 | 1449925 | AER30328.1 | P0C6X7 |
| IDFITWRRQELN | 1 | 0 | 0 | 1449927 | QBP84714.1 | Q04853 |
| IDFPFAESTGTQ | 1 | 0 | 0 | 1449931 | QBF80599.1 | K9N796 |
| IDFPSAELTGTQ | 1 | 0 | 0 | 1449932 | QBF80533.1 | K9N796 |
| IDFQDELDDFFK | 1 | 0 | 0 | 1449934 | AMW90852.1 | K9N5Q8 |
| IDFVNSGGTPLT | 1 | 0 | 0 | 1449936 | AUM60023.1 | K9N7C7 |
| IDGVYWVASSQA | 1 | 0 | 0 | 1449976 | ANZ78879.1 | P33469 |
| IDHPNPKGFCDF | 1 | 0 | 0 | 1449984 | APO40578.1 | P0C6X7 |
| IDINPALNVCFD | 1 | 0 | 0 | 1449995 | AVV62535.1 | K9N7C7 |
| IDKFAIPNRRKF | 1 | 0 | 0 | 1450012 | BBO58726.1 | P36334 |
| IDKLRDYLADYD | 1 | 0 | 0 | 1450020 | AXN92247.1 | K9N7C7 |
| IDKSAIPNGRKV | 1 | 0 | 0 | 1450021 | BBA84104.1 | P36334 |
| IDKVLNEKCSNY | 1 | 0 | 0 | 1450025 | AVP78030.1 | P0C6W2 |
| IDKWLNNFSVPS | 1 | 0 | 0 | 1450027 | AMM42415.1 | A0A140H1H1 |
| IDLGDISGINAS | 1 | 0 | 0 | 1450043 | AVP78031.1 | Q3LZX1 |
| IDLQELGKYEHY | 1 | 0 | 0 | 1450058 | AVP78042.1 | Q3LZX1 |
| IDLQELGKYEQG | 1 | 0 | 0 | 1450059 | 6CRW_A | P59594 |
| IDLTECIYTQSR | 1 | 0 | 0 | 1450067 | AZK15898.1 | K9N7C7 |
| IDLTMIEFMLWC | 1 | 0 | 0 | 1450070 | AXN92247.1 | K9N7C7 |
| IDMRSSFIRVST | 1 | 0 | 0 | 1450083 | AVV62541.1 | K9N7D2 |
| IDNMYMGKCGDK | 1 | 0 | 0 | 1450095 | ARU07575.1 | A0A140H1G9 |
| IDPAGVCLGAGQ | 1 | 0 | 0 | 1450102 | AEN19363.1 | P0C6X6 |
| IDQYMCGSDGKP | 1 | 0 | 0 | 1450119 | AUM60013.1 | K9N7C7 |
| IDSSEPLCLFKK | 1 | 0 | 0 | 1450133 | BBA20986.1 | A0A140H1H1 |
| IDSTQDGVNKLQ | 1 | 0 | 0 | 1450135 | QBQ01839.1 | P36334 |
| IDSTVLTYIPAS | 1 | 0 | 0 | 1450137 | AVN89455.1 | K9N7D2 |
| IDSVRDPQTLEI | 1 | 0 | 0 | 1450138 | AVP78031.1 | Q3LZX1 |
| IDTFNQLTADLQ | 1 | 0 | 0 | 1450143 | ARB07436.1 | A0A140H1G9 |
| IDTINCESIYEF | 1 | 0 | 0 | 1450150 | ARU07575.1 | A0A140H1G9 |
| IDTTLLDLTYEM | 1 | 0 | 0 | 1450157 | ASU91284.1 | K9N5Q8 |
| IDTVDVTNGLGT | 1 | 0 | 0 | 1450158 | AWW13566.1 | P36334 |
| IDVGNHRSAFAL | 1 | 0 | 0 | 1450172 | AIW52843.1 | A0A0P0G321 |
| IDVGNRRSAFAL | 1 | 0 | 0 | 1450173 | AMM42641.1 | A0A0P0G321 |
| IDVINKALEMCI | 1 | 0 | 0 | 1450174 | ARO76381.1 | P0C6U8 |
| IDVNPALNICFD | 1 | 0 | 0 | 1450176 | ANI69887.1 | K9N7C7 |
| IDVNTALNICFD | 1 | 0 | 0 | 1450179 | AUM60013.1 | K9N7C7 |

| | | | | | | |
|---|---|---|---|---|---|---|
| IDVPFAELTGTQ | 1 | 0 | 0 | 1450180 | QAT98905.1 | K9N796 |
| IDVTGKLSDCLY | 1 | 0 | 0 | 1450189 | AGT51687.1 | P0C6X6 |
| IDVVSAYGLCDA | 1 | 0 | 0 | 1450194 | ALX27232.1 | K9N5Q8 |
| IDYRNPSIDEAF | 1 | 0 | 0 | 1450211 | AJQ24074.1 | P33469 |
| IDYVTIAGTKLR | 1 | 0 | 0 | 1450218 | ARO76381.1 | P0C6U8 |
| IEAAAQGFQACR | 1 | 0 | 0 | 1450219 | QED88024.1 | H9EJ42 |
| IEAENPNQNSLC | 1 | 0 | 0 | 1450223 | Q0ZME2.1 | Q0ZJ80 |
| IEASTVVCGAKR | 1 | 0 | 0 | 1450229 | AVV62524.1 | K9N7C7 |
| IECGSCGNESWL | 1 | 0 | 0 | 1450238 | AVN89333.1 | K9N7C7 |
| IECGSCGNNSWL | 1 | 0 | 0 | 1450239 | ALA49340.1 | K9N7C7 |
| IECTFISRFEKV | 1 | 0 | 0 | 1450243 | AUM60018.1 | K9N7D2 |
| IEDAETPALQVS | 1 | 0 | 0 | 1450250 | ATQ39389.1 | K9N7C7 |
| IEDFQENWNTKH | 1 | 0 | 0 | 1450254 | AVP78041.1 | P0C6W2 |
| IEDILFSKIVTS | 1 | 0 | 0 | 1450259 | AGW80932.1 | A0A0P0K6L9 |
| IEDKIEEIESKQ | 1 | 0 | 0 | 1450260 | 5X58_A | P59594 |
| IEDLLFNKVTIA | 1 | 0 | 0 | 1450262 | AVV62526.1 | K9N5Q8 |
| IEDLLFSKVRLS | 1 | 0 | 0 | 1450266 | CAA83661.1 | P36334 |
| IEDLRARMTYVC | 1 | 0 | 0 | 1450269 | AJG44112.1 | K9N638 |
| IEDLSIRAFQKR | 1 | 0 | 0 | 1450270 | ARB07411.1 | P0C6X6 |
| IEDLSIRALQKR | 1 | 0 | 0 | 1450272 | AEN19363.1 | P0C6X6 |
| IEDPCPIHYYSD | 1 | 0 | 0 | 1450279 | Q3I5I8.1 | Q3LZX5 |
| IEDSNVKEEPTT | 1 | 0 | 0 | 1450282 | ARO76381.1 | P0C6U8 |
| IEDSVFVPQPTD | 1 | 0 | 0 | 1450285 | ARB66060.1 | P36334 |
| IEDVEDVISSEV | 1 | 0 | 0 | 1450288 | ARU07575.1 | A0A140H1G9 |
| IEEFADVVKGQV | 1 | 0 | 0 | 1450295 | ALA49659.1 | K9N7C7 |
| IEEFIQTSSPKV | 1 | 0 | 0 | 1450297 | BBA20979.1 | P36334 |
| IEFANDTKIASQ | 1 | 0 | 0 | 1450312 | ATG84701.1 | K9N5Q8 |
| IEFTDSFYTQGR | 1 | 0 | 0 | 1450319 | AUM60013.1 | K9N7C7 |
| IEGLKACSYVGI | 1 | 0 | 0 | 1450325 | AVV62524.1 | K9N7C7 |
| IEGNFYGPFVDR | 1 | 0 | 0 | 1450326 | APO40578.1 | P0C6X7 |
| IEGSGRPAPNSR | 1 | 0 | 0 | 1450330 | AIL49470.1 | P33469 |
| IEGSGRSASNSR | 1 | 0 | 0 | 1450333 | AXX83347.1 | P33469 |
| IEGSGRSVPNSR | 1 | 0 | 0 | 1450334 | AIL49415.1 | P33469 |
| IEGTGGNIQSSS | 1 | 0 | 0 | 1450337 | ANI69874.1 | K9N4V7 |
| IEHVTSVNLFDV | 1 | 0 | 0 | 1450344 | AVN89292.1 | K9N796 |
| IEKDFLSFDSDV | 1 | 0 | 0 | 1450349 | AHI48581.1 | K9N7C7 |
| IEKPKPDGVSKI | 1 | 0 | 0 | 1450353 | AVV62526.1 | K9N5Q8 |
| IELCVDEVGSKS | 1 | 0 | 0 | 1450359 | AVP78037.1 | Q3LZX5 |
| IELDNKFTPLKV | 1 | 0 | 0 | 1450360 | AUM60013.1 | K9N7C7 |
| IELGSGAHVGSD | 1 | 0 | 0 | 1450364 | AWK59942.1 | H9EJ42 |
| IELGVCVNYDLY | 1 | 0 | 0 | 1450366 | AEN19366.1 | P36334 |
| IELQPPCKFLIS | 1 | 0 | 0 | 1450371 | AVV62524.1 | K9N7C7 |
| IELTAEDISLLR | 1 | 0 | 0 | 1450373 | AHA80151.1 | P33469 |
| IENASPASTVHT | 1 | 0 | 0 | 1450382 | ARO76383.1 | P59632 |
| IENSLNALFSCD | 1 | 0 | 0 | 1450393 | QED88017.1 | H9EJ42 |
| IERGLLHSSRLP | 1 | 0 | 0 | 1450422 | AGT51419.1 | P0C6X6 |
| IESIYEFDTYKA | 1 | 0 | 0 | 1450435 | ABD75535.1 | A0A140H1G9 |
| IESSSKLWTQCV | 1 | 0 | 0 | 1450441 | APO40578.1 | P0C6X7 |
| IESWLNDKSVPS | 1 | 0 | 0 | 1450443 | AIX10749.1 | P36334 |
| IETAKFDTIGLY | 1 | 0 | 0 | 1450445 | AUM60025.1 | K9N796 |
| IETAVGHYFHAR | 1 | 0 | 0 | 1450446 | ALA49351.1 | K9N7C7 |
| IETAVGHYVHAC | 1 | 0 | 0 | 1450447 | AXN92226.1 | K9N7C7 |
| IETVDVKHEMSP | 1 | 0 | 0 | 1450460 | AST12962.1 | A0A0P0LKV0 |
| IETVSLAGSYKD | 1 | 0 | 0 | 1450463 | AVP78041.1 | P0C6W2 |
| IETVSLAGSYRD | 1 | 0 | 0 | 1450464 | APO40578.1 | P0C6X7 |
| IEVGEVKPFAVY | 1 | 0 | 0 | 1450468 | AKT07951.1 | H9EJ42 |
| IEVKFNAPALQE | 1 | 0 | 0 | 1450470 | P0C6W6.1 | P0C6W2 |
| IEVNELLDTTQL | 1 | 0 | 0 | 1450472 | AIX10755.1 | P36334 |
| IEVVSAYGLCDA | 1 | 0 | 0 | 1450481 | AXN92228.1 | K9N5Q8 |
| IEVVSAYGLCDS | 1 | 0 | 0 | 1450483 | AVN89334.1 | K9N5Q8 |
| IEWPLMEYDFTD | 1 | 0 | 0 | 1450485 | AUM60013.1 | K9N7C7 |
| IEWSGRSANTSR | 1 | 0 | 0 | 1450487 | AJQ24074.1 | P33469 |
| IEYPYISHENKL | 1 | 0 | 0 | 1450492 | ALA49351.1 | K9N7C7 |
| IEYPYISHETKL | 1 | 0 | 0 | 1450493 | ASU45783.1 | K9N7C7 |
| IFAEEPVVGIVY | 1 | 0 | 0 | 1450499 | AVA26874.1 | H9EJ24 |
| IFAKAKNTKVIK | 1 | 0 | 0 | 1450502 | AKA09644.1 | P36334 |
| IFAKVKNNKGFK | 1 | 0 | 0 | 1450503 | AIL49539.1 | P36334 |
| IFAKVKNTKAIK | 1 | 0 | 0 | 1450504 | AKA09643.1 | P36334 |
| IFAKVKNTKVVG | 1 | 0 | 0 | 1450507 | AWW13511.1 | P36334 |
| IFANFILHDVNN | 1 | 0 | 0 | 1450511 | AEN19366.1 | P36334 |
| IFANFILHDVNS | 1 | 0 | 0 | 1450512 | CAA79896.1 | P36334 |
| IFASVPEKFLNA | 1 | 0 | 0 | 1450520 | ARB07424.1 | A0A0P0ILY9 |
| IFATAPANLTIS | 1 | 0 | 0 | 1450521 | AVV62526.1 | K9N5Q8 |
| IFATPSVLQATL | 1 | 0 | 0 | 1450523 | ASY99818.1 | K9N7C7 |
| IFCATCPFAKSD | 1 | 0 | 0 | 1450531 | AVV62524.1 | K9N7C7 |
| IFDSSKEFVEYK | 1 | 0 | 0 | 1450551 | AIV41816.1 | P0C6X6 |
| IFDVDKFKTPHI | 1 | 0 | 0 | 1450557 | AVN89322.1 | K9N7C7 |
| IFDVDKVKTPHI | 1 | 0 | 0 | 1450558 | AXN92247.1 | K9N7C7 |
| IFERVDYSDAVT | 1 | 0 | 0 | 1450568 | AUM60013.1 | K9N7C7 |
| IFGSTLDNTTQS | 1 | 0 | 0 | 1450604 | APO40579.1 | P59594 |
| IFGTTLDNTTQS | 1 | 0 | 0 | 1450608 | ATO98145.1 | Q3LZX1 |
| IFGVDPGLSLES | 1 | 0 | 0 | 1450611 | APO40578.1 | P0C6X7 |
| IFICLFVALLLT | 1 | 0 | 0 | 1450622 | AGW80947.1 | A0A0P0ILY9 |
| IFIGDNVGHYVH | 1 | 0 | 0 | 1450624 | AEN19363.1 | P0C6X6 |
| IFIGDTVGHYVH | 1 | 0 | 0 | 1450625 | AIV41930.1 | P0C6X6 |
| IFKDDGNFVVLE | 1 | 0 | 0 | 1450643 | AMK59674.1 | P0C6X6 |
| IFKDSSGKPVPY | 1 | 0 | 0 | 1450644 | AVP78041.1 | P0C6W2 |
| IFKPGVEELNPS | 1 | 0 | 0 | 1450650 | APU51922.1 | Q4VID0 |
| IFKTEVDPNLSG | 1 | 0 | 0 | 1450654 | ATQ39389.1 | K9N7C7 |
| IFLAWIMLLQFA | 1 | 0 | 0 | 1450655 | Q3I5J2.1 | Q3LZX9 |
| IFNAVDCKSDFM | 1 | 0 | 0 | 1450685 | CAA83660.1 | P36334 |

| | | | | | | |
|---|---|---|---|---|---|---|
| IFNDFHANRPGV | 1 | 0 | 0 | 1450692 | AVV62535.1 | K9N7C7 |
| IFNDFHASRPGV | 1 | 0 | 0 | 1450693 | AUM60013.1 | K9N7C7 |
| IFNDYHANRPGV | 1 | 0 | 0 | 1450695 | AVV62524.1 | K9N7C7 |
| IFNGINSGTTCS | 1 | 0 | 0 | 1450698 | BBA20986.1 | A0A140H1H1 |
| IFNGKFFSNTKY | 1 | 0 | 0 | 1450699 | AIL49348.1 | P30215 |
| IFNIFQGVSSNI | 1 | 0 | 0 | 1450703 | AIG96678.1 | H9EJ42 |
| IFPPGQGVPLNA | 1 | 0 | 0 | 1450714 | ASU90622.1 | K9N4V7 |
| IFPSGTYYCVTT | 1 | 0 | 0 | 1450717 | AXN92247.1 | K9N7C7 |
| IFQAVSSNINCI | 1 | 0 | 0 | 1450720 | AIW52766.1 | A0A0P0LKV0 |
| IFQGVSSNINRL | 1 | 0 | 0 | 1450726 | AIG96678.1 | H9EJ42 |
| IFRSDVLHLTQD | 1 | 0 | 0 | 1450736 | ATO98181.1 | Q3LZX1 |
| IFRSDVSVLVTG | 1 | 0 | 0 | 1450737 | ATO98145.1 | Q3LZX1 |
| IFSAIYPIDLAS | 1 | 0 | 0 | 1450741 | ASY99815.1 | K9N7A1 |
| IFSAVYPIDLAS | 1 | 0 | 0 | 1450744 | AHI48833.1 | K9N7A1 |
| IFSDISDLSNRL | 1 | 0 | 0 | 1450747 | APO40578.1 | P0C6X7 |
| IFSIVNSAPDGL | 1 | 0 | 0 | 1450751 | AIW52843.1 | A0A0P0G321 |
| IFSLHPVECDEE | 1 | 0 | 0 | 1450753 | AXN92247.1 | K9N7C7 |
| IFSLHPVECDEG | 1 | 0 | 0 | 1450754 | ALA50033.1 | K9N7C7 |
| IFSLVNAAPEGL | 1 | 0 | 0 | 1450757 | AIW52757.1 | A0A0P0K6L9 |
| IFSQSDSLGVSS | 1 | 0 | 0 | 1450766 | QBP84719.1 | P0C6X6 |
| IFSSTFNVPMEK | 1 | 0 | 0 | 1450773 | AVP78041.1 | P0C6W2 |
| IFSTANALPECN | 1 | 0 | 0 | 1450775 | ABE97136.1 | H9EJ42 |
| IFSVHPVECDEY | 1 | 0 | 0 | 1450784 | AVV62535.1 | K9N7C7 |
| IFTARDGVRRGD | 1 | 0 | 0 | 1450793 | AGV08389.1 | K9N638 |
| IFTGCVGAWDKV | 1 | 0 | 0 | 1450796 | AVV62524.1 | K9N7C7 |
| IFTHIPLDGLVR | 1 | 0 | 0 | 1450798 | AVV62524.1 | K9N7C7 |
| IFTKSGFNDICM | 1 | 0 | 0 | 1450804 | AVN89289.1 | K9N7C7 |
| IFTKSGVNDICV | 1 | 0 | 0 | 1450806 | AUM60013.1 | K9N7C7 |
| IFTPLLKPVGTL | 1 | 0 | 0 | 1450816 | APO40578.1 | P0C6X7 |
| IFTVNSVVQREG | 1 | 0 | 0 | 1450827 | AVV62535.1 | K9N7C7 |
| IFVDGVPFVVSY | 1 | 0 | 0 | 1450834 | ASU90459.1 | K9N7C7 |
| IFVEVNAPYYNS | 1 | 0 | 0 | 1450837 | CAA83660.1 | P36334 |
| IFYRMNGIGITQ | 1 | 0 | 0 | 1450872 | AVV62537.1 | K9N5Q8 |
| IFYTTEIASKFT | 1 | 0 | 0 | 1450873 | ARU07575.1 | A0A140H1G9 |
| IGAAANATGSVI | 1 | 0 | 0 | 1450874 | AVV62537.1 | K9N5Q8 |
| IGAAANATGTVI | 1 | 0 | 0 | 1450875 | AVK87439.1 | K9N5Q8 |
| IGAEHVNASYEC | 1 | 0 | 0 | 1450883 | AVP78031.1 | Q3LZX1 |
| IGANGNKIVDKE | 1 | 0 | 0 | 1450893 | AVN89322.1 | K9N7C7 |
| IGANHTGITGDN | 1 | 0 | 0 | 1450894 | ARO76381.1 | P0C6U8 |
| IGAVGDAGAAAF | 1 | 0 | 0 | 1450905 | AVV62544.1 | K9N4V7 |
| IGAWLNDKSVPS | 1 | 0 | 0 | 1450911 | AHA80222.1 | P36334 |
| IGDCVTVHTANK | 1 | 0 | 0 | 1450922 | QYB08949.1 | P0DTD1 |
| IGDDVDDIESIY | 1 | 0 | 0 | 1450924 | ABD75535.1 | A0A140H1G9 |
| IGDFMSTIITSC | 1 | 0 | 0 | 1450929 | AXG21664.1 | K9N638 |
| IGDIIQRLDFLE | 1 | 0 | 0 | 1450931 | AXN92260.1 | K9N5Q8 |
| IGDIIQRLDGLE | 1 | 0 | 0 | 1450932 | AHI48733.1 | K9N5Q8 |
| IGDIIQRLDPPE | 1 | 0 | 0 | 1450933 | 5W9H_A | K9N5Q8 |
| IGDLKCPLDTSY | 1 | 0 | 0 | 1450941 | AKA09658.1 | P36334 |
| IGDLKCTTVAIN | 1 | 0 | 0 | 1450945 | CAA83661.1 | P36334 |
| IGDLNCPLDPKL | 1 | 0 | 0 | 1450947 | AWW13511.1 | P36334 |
| IGDLNCPLDPRL | 1 | 0 | 0 | 1450948 | BAM16463.1 | P36334 |
| IGDLNCPLDSRL | 1 | 0 | 0 | 1450949 | AIL49507.1 | P36334 |
| IGDLNCPLDTRL | 1 | 0 | 0 | 1450950 | AEN19366.1 | P36334 |
| IGDLNCTLDPRL | 1 | 0 | 0 | 1450951 | AWW13566.1 | P36334 |
| IGDLNCTLHPRL | 1 | 0 | 0 | 1450952 | BBA84086.1 | P36334 |
| IGDLSIRALQKR | 1 | 0 | 0 | 1450953 | AJE26218.1 | P0C6X6 |
| IGDRDPRNDEAI | 1 | 0 | 0 | 1450957 | AJQ24068.1 | P33469 |
| IGDTVGHYVHVK | 1 | 0 | 0 | 1450963 | AIV41930.1 | P0C6X6 |
| IGEASSPIAYLV | 1 | 0 | 0 | 1450969 | AXN92247.1 | K9N7C7 |
| IGEDQSIAYSNN | 1 | 0 | 0 | 1450971 | APO40579.1 | P59594 |
| IGELPCAIMNDK | 1 | 0 | 0 | 1450975 | ARB07436.1 | A0A140H1G9 |
| IGFDVEGAHAFR | 1 | 0 | 0 | 1450983 | AXN92247.1 | K9N7C7 |
| IGFTLEYWATPL | 1 | 0 | 0 | 1451001 | AHA80225.1 | P36334 |
| IGGFVEKWESGV | 1 | 0 | 0 | 1451016 | APO40580.1 | P59632 |
| IGGGDLLYLDLL | 1 | 0 | 0 | 1451017 | AVN89359.1 | K9N4V7 |
| IGGTIVVRPLVE | 1 | 0 | 0 | 1451026 | ATY74387.1 | K9N7A1 |
| IGHGMSYSMTTF | 1 | 0 | 0 | 1451043 | AIV41888.1 | P0C6X6 |
| IGHTVCDSLNFK | 1 | 0 | 0 | 1451050 | AAX85666.1 | P0C6X6 |
| IGHTVCDSLNSK | 1 | 0 | 0 | 1451051 | AEN19363.1 | P0C6X6 |
| IGICPAGTNYLT | 1 | 0 | 0 | 1451054 | AIX10751.1 | P36334 |
| IGIDFQDELDEF | 1 | 0 | 0 | 1451055 | AGV08467.1 | K9N5Q8 |
| IGIDTTGVLLKQ | 1 | 0 | 0 | 1451058 | ABD75582.1 | Q0ZJ80 |
| IGINYLGKSCFE | 1 | 0 | 0 | 1451066 | ARB07436.1 | A0A140H1G9 |
| IGKGSKDEQIGY | 1 | 0 | 0 | 1451077 | AIW52795.1 | H9EJ69 |
| IGKGSLTASSFD | 1 | 0 | 0 | 1451078 | AUM60027.1 | K9N643 |
| IGKIAQWIKNTF | 1 | 0 | 0 | 1451079 | AXX83336.1 | P0C6X6 |
| IGKIQESLTSTA | 1 | 0 | 0 | 1451082 | AVP78031.1 | Q3LZX1 |
| IGLLEPVKHSST | 1 | 0 | 0 | 1451108 | AVV62535.1 | K9N7C7 |
| IGLVNSSMVVDN | 1 | 0 | 0 | 1451117 | AVV62537.1 | K9N5Q8 |
| IGLYKKQQEGHI | 1 | 0 | 0 | 1451122 | AXN92247.1 | K9N7C7 |
| IGLYTNRVVSPP | 1 | 0 | 0 | 1451128 | AUM60025.1 | K9N796 |
| IGNIEEFIQTSS | 1 | 0 | 0 | 1451142 | BBA20979.1 | P36334 |
| IGNYTVSCSPFT | 1 | 0 | 0 | 1451166 | AVP78037.1 | Q3LZX5 |
| IGPDHSVADYHN | 1 | 0 | 0 | 1451167 | APO40578.1 | P0C6X7 |
| IGPHEFCSQHTM | 1 | 0 | 0 | 1451174 | AIW52776.1 | A0A0P0LKV0 |
| IGPIKYCYCGNF | 1 | 0 | 0 | 1451175 | AWK59942.1 | H9EJ42 |
| IGPNKYCYCGNF | 1 | 0 | 0 | 1451177 | QED88017.1 | H9EJ42 |
| IGQDSLNRYSSL | 1 | 0 | 0 | 1451181 | AUM60013.1 | K9N7C7 |
| IGREQKCCMCNR | 1 | 0 | 0 | 1451191 | ALA50033.1 | K9N7C7 |
| IGRGNIGFTLEY | 1 | 0 | 0 | 1451193 | AHA80218.1 | P36334 |
| IGRRNIGFTLEY | 1 | 0 | 0 | 1451196 | AHA80213.1 | P36334 |

| | | | | | | |
|---|---|---|---|---|---|---|
| IGSLGFHNPPTN | 1 | 0 | 0 | 1451206 | AVQ05263.1 | P30215 |
| IGSNFVNTSYSV | 1 | 0 | 0 | 1451210 | BBO58761.1 | P36334 |
| IGSNKTLLTPLC | 1 | 0 | 0 | 1451212 | APO40578.1 | P0C6X7 |
| IGSSLQEILSRL | 1 | 0 | 0 | 1451214 | CAA83660.1 | P36334 |
| IGSTAYIQQCLT | 1 | 0 | 0 | 1451217 | AUM60027.1 | K9N643 |
| IGSTAYIQQCVT | 1 | 0 | 0 | 1451218 | AUM60017.1 | K9N643 |
| IGTANLSIPSNW | 1 | 0 | 0 | 1451222 | AAK32190.1 | A0A0P0K6L9 |
| IGTCPAGTNHLT | 1 | 0 | 0 | 1451223 | ARE30008.1 | P36334 |
| IGTCPAGTNYLN | 1 | 0 | 0 | 1451225 | AIL49520.1 | P36334 |
| IGTCPVGTNYLT | 1 | 0 | 0 | 1451227 | AIL49486.1 | P36334 |
| IGTSTHPVQWQC | 1 | 0 | 0 | 1451254 | AVV62539.1 | K9N4V0 |
| IGTTKFCGGWNN | 1 | 0 | 0 | 1451257 | ADK37693.1 | H9EJ42 |
| IGTTKFYGGWGN | 1 | 0 | 0 | 1451259 | ADK37686.1 | A0A0P0LKV0 |
| IGTYEMYVKWPW | 1 | 0 | 0 | 1451262 | BBA20983.1 | A0A140H1H1 |
| IGVCTMTDIAEK | 1 | 0 | 0 | 1451269 | AVP78041.1 | P0C6W2 |
| IGVFVQKYGLED | 1 | 0 | 0 | 1451274 | AGT21344.1 | A0A0P0LKV0 |
| IGVNYLGKPKEI | 1 | 0 | 0 | 1451279 | APO40578.1 | P0C6X7 |
| IGVYHKLGERIR | 1 | 0 | 0 | 1451292 | AVV62524.1 | K9N7C7 |
| IGVYHKLGERVR | 1 | 0 | 0 | 1451294 | AXN92247.1 | K9N7C7 |
| IGYIPEHCQSHT | 1 | 0 | 0 | 1451303 | AUM60025.1 | K9N796 |
| IGYSLADLPAYM | 1 | 0 | 0 | 1451308 | AWW13547.1 | Q01455 |
| IHAVLDALLANS | 1 | 0 | 0 | 1451330 | AUM60013.1 | K9N7C7 |
| IHDVVNQQGNSL | 1 | 0 | 0 | 1451346 | ABB90526.1 | A0A0P0K6L9 |
| IHIVSFVVNAPN | 1 | 0 | 0 | 1451366 | ALJ54495.1 | K9N5Q8 |
| IHRDCDFNKPLI | 1 | 0 | 0 | 1451388 | AVV62535.1 | K9N7C7 |
| IHRGDPMSNNGW | 1 | 0 | 0 | 1451393 | AVP78042.1 | Q3LZX1 |
| IHSVCNGCVGHS | 1 | 0 | 0 | 1451405 | AIW52741.1 | A0A0P0K6L9 |
| IHSVYLLMFLLT | 1 | 0 | 0 | 1451407 | ALJ54471.1 | K9N5Q8 |
| IHTKPLFIRNFD | 1 | 0 | 0 | 1451413 | AXG21669.1 | K9N7D2 |
| IHVEQVSSDWRV | 1 | 0 | 0 | 1451416 | APO40579.1 | P59594 |
| IHYNINFEENLR | 1 | 0 | 0 | 1451423 | BAL45639.1 | A0A0P0K6L9 |
| IIANENAINKGG | 1 | 0 | 0 | 1451443 | AGT21359.1 | A0A0P0LKV0 |
| IIANRCNEVSAT | 1 | 0 | 0 | 1451445 | AVQ05261.1 | P0C6X6 |
| IIDAATSYGMCF | 1 | 0 | 0 | 1451461 | AIV41909.1 | P36334 |
| IIEAENPNPNSL | 1 | 0 | 0 | 1451469 | AET99208.1 | Q0ZJ80 |
| IIEAENPNQNSL | 1 | 0 | 0 | 1451470 | Q0ZME2.1 | Q0ZJ80 |
| IIEQPFGEVEHS | 1 | 0 | 0 | 1451477 | AVA26872.1 | H9EJ42 |
| IIEQPFGEVEYS | 1 | 0 | 0 | 1451478 | AWK59942.1 | H9EJ42 |
| IIEQSFGEVEHS | 1 | 0 | 0 | 1451479 | ABE73411.1 | H9EJ42 |
| IIFNAEDCMSDF | 1 | 0 | 0 | 1451495 | P36334.1 | P36334 |
| IIFNAVDCKSDF | 1 | 0 | 0 | 1451496 | AHA80218.1 | P36334 |
| IIGDELKINSAC | 1 | 0 | 0 | 1451513 | P0C6W6.1 | P0C6W2 |
| IIGVPYPVSGIR | 1 | 0 | 0 | 1451529 | AVL25553.1 | A0A0P0G321 |
| IIHDIGNPKGVR | 1 | 0 | 0 | 1451537 | ABE97136.1 | H9EJ42 |
| IIHSHPILHDPG | 1 | 0 | 0 | 1451541 | ATQ39396.1 | K9N643 |
| IIIDKVDFVNGF | 1 | 0 | 0 | 1451543 | AGT51379.1 | H9EJ42 |
| IIISCQTAVSKF | 1 | 0 | 0 | 1451562 | AVN89363.1 | K9N7C7 |
| IIKPNALLCATC | 1 | 0 | 0 | 1451582 | AHY22544.1 | K9N7C7 |
| IIKSSDVSENNE | 1 | 0 | 0 | 1451587 | AVV62524.1 | K9N7C7 |
| IILKPASDGLKI | 1 | 0 | 0 | 1451613 | AVP78030.1 | P0C6W2 |
| IINARVKAKHIV | 1 | 0 | 0 | 1451656 | AUM60013.1 | K9N7C7 |
| IINDKVLTKIDK | 1 | 0 | 0 | 1451659 | QBP84719.1 | P0C6X6 |
| IINNAVYTKMDG | 1 | 0 | 0 | 1451670 | APO40578.1 | P0C6X7 |
| IIPGFGGNFNLK | 1 | 0 | 0 | 1451682 | AOR17482.1 | K9N5Q8 |
| IIPHSTRSIQSD | 1 | 0 | 0 | 1451690 | ATG84899.1 | K9N5Q8 |
| IIPLGIDLDEWS | 1 | 0 | 0 | 1451692 | AVP78030.1 | P0C6W2 |
| IIQADQQVDRLI | 1 | 0 | 0 | 1451698 | AIW52757.1 | A0A0P0K6L9 |
| IIRDGDVVKQPV | 1 | 0 | 0 | 1451713 | ALA49648.1 | K9N7C7 |
| IISEKISVMDKL | 1 | 0 | 0 | 1451750 | AMM42546.1 | H9EJ42 |
| IISPSTRATIRK | 1 | 0 | 0 | 1451780 | AKL59401.1 | K9N5Q8 |
| IISQLRKGGRVL | 1 | 0 | 0 | 1451783 | APO40578.1 | P0C6X7 |
| IISTQDGVNKLQ | 1 | 0 | 0 | 1451789 | BBA20973.1 | P36334 |
| IISTVQKTIKKI | 1 | 0 | 0 | 1451790 | AVV62537.1 | K9N5Q8 |
| IITGNVVKQCCT | 1 | 0 | 0 | 1451797 | AVV62535.1 | K9N7C7 |
| IITMPCFRTFPQ | 1 | 0 | 0 | 1451807 | AVN89294.1 | K9N643 |
| IITSCQTAVSKL | 1 | 0 | 0 | 1451817 | AXN92247.1 | K9N7C7 |
| IITSTDAGCGVK | 1 | 0 | 0 | 1451818 | AGT51379.1 | H9EJ42 |
| IITTDNTFVAGN | 1 | 0 | 0 | 1451821 | ATO98181.1 | Q3LZX1 |
| IIVFLCAVLMYL | 1 | 0 | 0 | 1451828 | AXN92247.1 | K9N7C7 |
| IIVPDKSVYDQI | 1 | 0 | 0 | 1451836 | AEN19363.1 | P0C6X6 |
| IIYDASDRGQEN | 1 | 0 | 0 | 1451847 | AVV62524.1 | K9N7C7 |
| IIYDSADRGQES | 1 | 0 | 0 | 1451848 | ATQ39389.1 | K9N7C7 |
| IIYPTGKSYSNI | 1 | 0 | 0 | 1451856 | AVV62537.1 | K9N5Q8 |
| IKANIKSDILSL | 1 | 0 | 0 | 1451872 | AZS52617.1 | A0A140H1H0 |
| IKCNYVFNNTLS | 1 | 0 | 0 | 1451888 | CAA83661.1 | P36334 |
| IKDATPSDSIRA | 1 | 0 | 0 | 1451901 | AVP78043.1 | Q3LZX0 |
| IKDGDYGYFLPY | 1 | 0 | 0 | 1451911 | ALW82667.1 | K9N7C7 |
| IKDGVMYSEFPA | 1 | 0 | 0 | 1451912 | CAA79896.1 | P36334 |
| IKDLLARAGKSC | 1 | 0 | 0 | 1451916 | APO40578.1 | P0C6X7 |
| IKDWNFVWSLIF | 1 | 0 | 0 | 1451934 | AGV08590.1 | K9N7A1 |
| IKEGALTAVVIP | 1 | 0 | 0 | 1451937 | AVP78041.1 | P0C6W2 |
| IKENVDFINSKQ | 1 | 0 | 0 | 1451950 | ARB07431.1 | Q80872 |
| IKEPVHVYANDS | 1 | 0 | 0 | 1451952 | QDH43722.1 | P0C6X6 |
| IKFADDLNQLSG | 1 | 0 | 0 | 1451959 | AVP78041.1 | P0C6W2 |
| IKFETDAYKLFL | 1 | 0 | 0 | 1451968 | ARU07575.1 | A0A140H1G9 |
| IKFPEVVFESTL | 1 | 0 | 0 | 1451974 | AXN92247.1 | K9N7C7 |
| IKGATPTNSVRT | 1 | 0 | 0 | 1451980 | AVP78032.1 | Q3LZX0 |
| IKGCCESTKLPY | 1 | 0 | 0 | 1451982 | AWH62679.1 | A0A0P0K6L9 |
| IKGTTVLLKEPC | 1 | 0 | 0 | 1451995 | AVP78047.1 | Q3LZX7 |
| IKHEGDWNSFCK | 1 | 0 | 0 | 1452003 | APT69882.1 | A0A0P0ILY9 |
| IKHGVMYSEFPA | 1 | 0 | 0 | 1452004 | CAA83661.1 | P36334 |

| | | | | | | |
|---|---|---|---|---|---|---|
| IKHVEKEGSYHN | 1 | 0 | 0 | 1452012 | AVV62524.1 | K9N7C7 |
| IKHVRNNGGYNS | 1 | 0 | 0 | 1452015 | AXN92247.1 | K9N7C7 |
| IKHVRTNGDYNS | 1 | 0 | 0 | 1452016 | QDI73607.1 | K9N7C7 |
| IKIAQADTAGLY | 1 | 0 | 0 | 1452018 | ASU89922.1 | K9N796 |
| IKITEHSWSADL | 1 | 0 | 0 | 1452028 | APO40578.1 | P0C6X7 |
| IKITEYSWSADL | 1 | 0 | 0 | 1452031 | AVV62535.1 | K9N7C7 |
| IKKAMFTKVVVG | 1 | 0 | 0 | 1452035 | AWK59942.1 | H9EJ42 |
| IKKGVMYSEFPA | 1 | 0 | 0 | 1452037 | CAA83660.1 | P36334 |
| IKKLTPIKLNFG | 1 | 0 | 0 | 1452042 | AWK59978.1 | H9EJ42 |
| IKKPSELSRVLG | 1 | 0 | 0 | 1452047 | AVP78041.1 | P0C6W2 |
| IKKVSFGPEEVH | 1 | 0 | 0 | 1452048 | AUM60013.1 | K9N7C7 |
| IKKYPCIVGPDY | 1 | 0 | 0 | 1452055 | AXN92258.1 | K9N7C7 |
| IKLDDKDPNFKDQVIL | 1 | 0 | 0 | 1452063 | YP_009724397.2 | P0DTC9 |
| IKLDSKNPNYKK | 1 | 0 | 0 | 1452066 | AVV62544.1 | K9N4V7 |
| IKLKRAFGDYTS | 1 | 0 | 0 | 1452082 | AEN19363.1 | P0C6X6 |
| IKLQSKRTRLFK | 1 | 0 | 0 | 1452092 | AEN19363.1 | P0C6X6 |
| IKNLSKPPTENN | 1 | 0 | 0 | 1452122 | AVP78035.1 | Q3LZX8 |
| IKNTVDTIPKEK | 1 | 0 | 0 | 1452130 | AGT21366.1 | A0A0P0LKV0 |
| IKNYPCMVGPDC | 1 | 0 | 0 | 1452135 | AUM60023.1 | K9N7C7 |
| IKPEVKEVAPVY | 1 | 0 | 0 | 1452139 | AIZ74432.1 | K9N7C7 |
| IKPNAILCATCP | 1 | 0 | 0 | 1452149 | QBF80531.1 | K9N7C7 |
| IKPNALLCATCP | 1 | 0 | 0 | 1452151 | AXN92247.1 | K9N7C7 |
| IKPNARLCATCP | 1 | 0 | 0 | 1452153 | AHI48793.1 | K9N7C7 |
| IKPQQVTKHTAK | 1 | 0 | 0 | 1452156 | AOL02457.1 | P33469 |
| IKPYERDISNVL | 1 | 0 | 0 | 1452169 | APO40579.1 | P59594 |
| IKQGTLIYKVLD | 1 | 0 | 0 | 1452172 | AIV41996.1 | P0C6X6 |
| IKRMKQIEDKIE | 1 | 0 | 0 | 1452192 | 5X58_A | P59594 |
| IKRSDARTAHHG | 1 | 0 | 0 | 1452199 | AVP78041.1 | P0C6W2 |
| IKRVKGWVGFDV | 1 | 0 | 0 | 1452202 | ABD75519.1 | A0A140H1G9 |
| IKSDHCGGNPCT | 1 | 0 | 0 | 1452206 | AVI15011.1 | P36334 |
| IKSGGYIPEAPR | 1 | 0 | 0 | 1452211 | 6U7H_A | A0A0P0K6L9 |
| IKSGRLLLRNNG | 1 | 0 | 0 | 1452212 | AWK59942.1 | H9EJ42 |
| IKSQDLGVVSKV | 1 | 0 | 0 | 1452219 | AVV62535.1 | K9N7C7 |
| IKSQDLSVVSKVVKV | 1 | 0 | 0 | 1452222 | P0DTD1.1 | P0DTD1 |
| IKSTQDGDNKLQ | 1 | 0 | 0 | 1452225 | AIX10748.1 | P36334 |
| IKTAQADTAGLF | 1 | 0 | 0 | 1452233 | ALW82670.1 | K9N796 |
| IKTHKSQPLNAG | 1 | 0 | 0 | 1452238 | AVV62537.1 | K9N5Q8 |
| IKTNNIRIVDEW | 1 | 0 | 0 | 1452243 | ASU90175.1 | K9N5Q8 |
| IKTNNTMIVDEW | 1 | 0 | 0 | 1452244 | ATG84866.1 | K9N5Q8 |
| IKTNNTRVVDEW | 1 | 0 | 0 | 1452247 | ALA49803.1 | K9N5Q8 |
| IKTNNTSIVDEW | 1 | 0 | 0 | 1452248 | ANF29261.1 | K9N5Q8 |
| IKTQATQPLDAG | 1 | 0 | 0 | 1452250 | AVV62526.1 | K9N5Q8 |
| IKTSKYGLGFKW | 1 | 0 | 0 | 1452252 | ARU07575.1 | A0A140H1G9 |
| IKVLPPLLSDNQ | 1 | 0 | 0 | 1452266 | CAA83661.1 | P36334 |
| IKVQEGIVDYGV | 1 | 0 | 0 | 1452269 | ATO98229.1 | P0C6W2 |
| IKVSQTNMHTPR | 1 | 0 | 0 | 1452272 | NP_073549.1 | A0A0P0LKV0 |
| IKVTQELGHEDL | 1 | 0 | 0 | 1452278 | ARO76381.1 | P0C6U8 |
| IKWLFVLLFDWI | 1 | 0 | 0 | 1452287 | AGT51707.1 | P0C6X6 |
| ILALREVKTIKV | 1 | 0 | 0 | 1452315 | APO40578.1 | P0C6X7 |
| ILARSQSSETKE | 1 | 0 | 0 | 1452322 | AIW52723.1 | A0A0N7J131 |
| ILCATCPFAKGD | 1 | 0 | 0 | 1452336 | QBF80531.1 | K9N7C7 |
| ILDYCDGNLPCG | 1 | 0 | 0 | 1452378 | APO40578.1 | P0C6X7 |
| ILEFKVLDNGII | 1 | 0 | 0 | 1452384 | AUM60013.1 | K9N7C7 |
| ILEQDRMPKTRM | 1 | 0 | 0 | 1452392 | AHF54653.1 | Q4VID0 |
| ILEQDRMPKTSM | 1 | 0 | 0 | 1452393 | AHF54652.1 | Q4VID0 |
| ILFANPAAVDSK | 1 | 0 | 0 | 1452396 | AIV42008.1 | P0C6X6 |
| ILFTGYFPKAGA | 1 | 0 | 0 | 1452417 | BBA20983.1 | A0A140H1H1 |
| ILGGCIVLDVKG | 1 | 0 | 0 | 1452427 | AIV41836.1 | Q80872 |
| ILHDVNNGLTCS | 1 | 0 | 0 | 1452452 | AEN19366.1 | P36334 |
| ILHDVNSGTTCS | 1 | 0 | 0 | 1452454 | CAA83660.1 | P36334 |
| ILHHPGFLNPSS | 1 | 0 | 0 | 1452456 | AXN92231.1 | K9N643 |
| ILHLGSPLSLNM | 1 | 0 | 0 | 1452460 | AVP78050.1 | Q3LZX3 |
| ILHRECVNINIV | 1 | 0 | 0 | 1452462 | ARO76381.1 | P0C6U8 |
| ILIIIMRTFRIA | 1 | 0 | 0 | 1452472 | ATO98186.1 | Q3LZX8 |
| ILIQSTQWSLFF | 1 | 0 | 0 | 1452478 | APO40578.1 | P0C6X7 |
| ILKPAADGLKIT | 1 | 0 | 0 | 1452499 | APO40578.1 | P0C6X7 |
| ILKPAIDGLKIT | 1 | 0 | 0 | 1452500 | AVP78041.1 | P0C6W2 |
| ILKSQDLGAVSK | 1 | 0 | 0 | 1452509 | ANI69887.1 | K9N7C7 |
| ILKWSDQSDQFR | 1 | 0 | 0 | 1452514 | ARK08639.1 | P33469 |
| ILLAGARVRTIN | 1 | 0 | 0 | 1452516 | APO40578.1 | P0C6X7 |
| ILLEAYNAFLNT | 1 | 0 | 0 | 1452525 | AST12962.1 | A0A0P0LKV0 |
| ILLKQPMATSVN | 1 | 0 | 0 | 1452545 | AHF54651.1 | Q4VID0 |
| ILNGCTWWLKGE | 1 | 0 | 0 | 1452595 | NP_073549.1 | A0A0P0LKV0 |
| ILNGINSGTTCS | 1 | 0 | 0 | 1452596 | BBA20983.1 | A0A140H1H1 |
| ILNKPRQKRSPN | 1 | 0 | 0 | 1452601 | AEN19370.1 | P33469 |
| ILNKPRQKSSPN | 1 | 0 | 0 | 1452602 | AJQ24079.1 | P33469 |
| ILNQTTRDATNE | 1 | 0 | 0 | 1452612 | AUM60014.1 | K9N5Q8 |
| ILNRQCVNINIV | 1 | 0 | 0 | 1452614 | APO40578.1 | P0C6X7 |
| ILPFSDRPLNLR | 1 | 0 | 0 | 1452635 | AWK59942.1 | H9EJ42 |
| ILPLAYFSDSKL | 1 | 0 | 0 | 1452637 | AXN92247.1 | K9N7C7 |
| ILPLVSCFSTRN | 1 | 0 | 0 | 1452642 | AKA09629.1 | A0A0P0G321 |
| ILPQGYYIEGSG | 1 | 0 | 0 | 1452648 | AOL02457.1 | P33469 |
| ILQDKTGDFMST | 1 | 0 | 0 | 1452662 | YP_009047214.1 | K9N638 |
| ILQEVKDNDGKN | 1 | 0 | 0 | 1452664 | QEO75983.1 | A0A0P0LKV0 |
| ILQGKPIGMFFP | 1 | 0 | 0 | 1452667 | AIZ48759.1 | K9N638 |
| ILQVTTANVSAL | 1 | 0 | 0 | 1452679 | AVV62524.1 | K9N7C7 |
| ILRSTGQKAIVA | 1 | 0 | 0 | 1452701 | AVP78042.1 | Q3LZX1 |
| ILSCNFNMSSLM | 1 | 0 | 0 | 1452710 | AIV41909.1 | P36334 |
| ILSKAMQLLHTK | 1 | 0 | 0 | 1452734 | AVV62524.1 | K9N7C7 |
| ILSKTDVSTPVD | 1 | 0 | 0 | 1452736 | AUM60013.1 | K9N7C7 |
| ILSNKSNPALTQ | 1 | 0 | 0 | 1452745 | APO40579.1 | P59594 |

| | | | | | | |
|---|---|---|---|---|---|---|
| ILSNLGVPSYCD | 1 | 0 | 0 | 1452746 | APO40578.1 | P0C6X7 |
| ILSRLDPPEAEV | 1 | 0 | 0 | 1452753 | 6CRW_A | P59594 |
| ILSRLDVLEAQV | 1 | 0 | 0 | 1452754 | ABD75553.1 | A0A140H1H1 |
| ILSVKFSVTKIV | 1 | 0 | 0 | 1452760 | AUM60013.1 | K9N7C7 |
| ILSWELGKVKPP | 1 | 0 | 0 | 1452766 | AGT51677.1 | P0C6X6 |
| ILTAFSPAQDTW | 1 | 0 | 0 | 1452770 | 5X58_A | P59594 |
| ILVGTTLQGKPI | 1 | 0 | 0 | 1452813 | AVN89312.1 | K9N638 |
| ILVTYKCCDDNY | 1 | 0 | 0 | 1452827 | AVP78030.1 | P0C6W2 |
| ILYKGSVTHDAS | 1 | 0 | 0 | 1452851 | AUM60023.1 | K9N7C7 |
| ILYPGQKYTSDC | 1 | 0 | 0 | 1452854 | AVV62524.1 | K9N7C7 |
| ILYTFYGKDAFE | 1 | 0 | 0 | 1452860 | AVV62524.1 | K9N7C7 |
| IMACGNCTFGAP | 1 | 0 | 0 | 1452868 | AID50427.1 | K9N7C7 |
| IMANLANLNSHT | 1 | 0 | 0 | 1452871 | AGT21338.1 | A0A0P0K6L9 |
| IMAYGDCTFGVP | 1 | 0 | 0 | 1452881 | AVV62524.1 | K9N7C7 |
| IMAYGQCTFGAP | 1 | 0 | 0 | 1452885 | AUM60013.1 | K9N7C7 |
| IMFQPRLPVLSD | 1 | 0 | 0 | 1452908 | AKT07952.1 | A0A0P0G321 |
| IMKHDIKVIAKA | 1 | 0 | 0 | 1452933 | AWK59942.1 | H9EJ42 |
| IMLNKHIDAYKT | 1 | 0 | 0 | 1452950 | ARO76389.1 | P59595 |
| IMPTHVLAGHDD | 1 | 0 | 0 | 1452964 | APO40578.1 | P0C6X7 |
| IMPVHTLSVHDG | 1 | 0 | 0 | 1452966 | ARO76381.1 | P0C6U8 |
| IMRTFRIAIWNL | 1 | 0 | 0 | 1452983 | ATO98186.1 | Q3LZX8 |
| IMSMAECKATLG | 1 | 0 | 0 | 1452989 | QED88024.1 | H9EJ42 |
| INADAYARYLAM | 1 | 0 | 0 | 1453017 | AVV62524.1 | K9N7C7 |
| INADDKCFANKY | 1 | 0 | 0 | 1453018 | AUM60013.1 | K9N7C7 |
| INASVVDIKKEI | 1 | 0 | 0 | 1453040 | APO40579.1 | P59594 |
| INCILSVNSSNC | 1 | 0 | 0 | 1453049 | AIW52766.1 | A0A0P0LKV0 |
| INCQAPPVGSLI | 1 | 0 | 0 | 1453053 | Q3I5I8.1 | Q3LZX5 |
| INCQNPPVGSLI | 1 | 0 | 0 | 1453056 | ATO98227.1 | Q3LZX5 |
| INDKVLTKIDKE | 1 | 0 | 0 | 1453069 | QBP84719.1 | P0C6X6 |
| INDYNKTIPRIS | 1 | 0 | 0 | 1453080 | ARB07438.1 | A0A140H1H1 |
| INECVKSQSNRY | 1 | 0 | 0 | 1453082 | AIW52841.1 | A0A0P0G321 |
| INFRTCILNNQS | 1 | 0 | 0 | 1453104 | AUM60013.1 | K9N7C7 |
| INFRTSVVNNQT | 1 | 0 | 0 | 1453105 | AVV62535.1 | K9N7C7 |
| INFVKFGSVCFS | 1 | 0 | 0 | 1453110 | APT69876.1 | A0A0P0K6L9 |
| INGERWFLNRFT | 1 | 0 | 0 | 1453118 | APO40578.1 | P0C6X7 |
| INGIFAKAKNTK | 1 | 0 | 0 | 1453128 | AKA09644.1 | P36334 |
| INGKLYQDSQNM | 1 | 0 | 0 | 1453135 | ARO76381.1 | P0C6U8 |
| INGLHPTQSPTY | 1 | 0 | 0 | 1453136 | APO40578.1 | P0C6X7 |
| INGQYDKNASLN | 1 | 0 | 0 | 1453141 | AUM60014.1 | K9N5Q8 |
| INGYTLAATSAS | 1 | 0 | 0 | 1453152 | ANZ78836.1 | P36334 |
| INKGHFDGQNGE | 1 | 0 | 0 | 1453205 | AVP78030.1 | P0C6W2 |
| INKVKRALADYT | 1 | 0 | 0 | 1453215 | AVV62524.1 | K9N7C7 |
| INLCDDPEKAQG | 1 | 0 | 0 | 1453218 | AWK59942.1 | H9EJ42 |
| INLKDIGTYEMY | 1 | 0 | 0 | 1453226 | BBA20983.1 | A0A140H1H1 |
| INLKDIGTYEYY | 1 | 0 | 0 | 1453227 | CAA79896.1 | P36334 |
| INLKFNKWQWQE | 1 | 0 | 0 | 1453228 | ARU07575.1 | A0A140H1G9 |
| INLPPPLLGNST | 1 | 0 | 0 | 1453232 | ATG84888.1 | K9N5Q8 |
| INLTNEGFNNYV | 1 | 0 | 0 | 1453238 | AUM60013.1 | K9N7C7 |
| INNFNPSSWNRR | 1 | 0 | 0 | 1453251 | BBA20986.1 | A0A140H1H1 |
| INNGPHEFCTKH | 1 | 0 | 0 | 1453253 | AIL49625.1 | P0C6X6 |
| INNKDTGPPFIS | 1 | 0 | 0 | 1453255 | AKA09646.1 | P36334 |
| INNVDNEIIKSS | 1 | 0 | 0 | 1453264 | AVV62535.1 | K9N7C7 |
| INNVDNEIVKSS | 1 | 0 | 0 | 1453265 | AXN92247.1 | K9N7C7 |
| INPNALLCATCP | 1 | 0 | 0 | 1453275 | ASU90746.1 | K9N7C7 |
| INQALHGANLMQ | 1 | 0 | 0 | 1453282 | AVV62537.1 | K9N5Q8 |
| INQMTGFKKPAS | 1 | 0 | 0 | 1453286 | APO40578.1 | P0C6X7 |
| INRFNVAVTRAQ | 1 | 0 | 0 | 1453294 | AUM60013.1 | K9N7C7 |
| INRGIYKPLLEN | 1 | 0 | 0 | 1453295 | AIV41840.1 | P0C6X6 |
| INRLFSVPSDSC | 1 | 0 | 0 | 1453300 | AFD98777.1 | H9EJ42 |
| INRMAEQAAAAM | 1 | 0 | 0 | 1453305 | NP_073549.1 | A0A0P0LKV0 |
| INRVLSVPSDSC | 1 | 0 | 0 | 1453312 | ABQ16583.1 | H9EJ42 |
| INSILNEVNDLL | 1 | 0 | 0 | 1453319 | BBA20986.1 | A0A140H1H1 |
| INSNDKCFANKY | 1 | 0 | 0 | 1453323 | AVV62524.1 | K9N7C7 |
| INSQDGDNKLQG | 1 | 0 | 0 | 1453324 | AKA09654.1 | P36334 |
| INSQDGVNKLQG | 1 | 0 | 0 | 1453325 | QDH43726.1 | P36334 |
| INSSAMFCATCP | 1 | 0 | 0 | 1453326 | AVV62535.1 | K9N7C7 |
| INSTINTTPVST | 1 | 0 | 0 | 1453329 | AMM42635.1 | A0A0P0G321 |
| INSTQDGVNKLQ | 1 | 0 | 0 | 1453333 | AEN19358.1 | P36334 |
| INSVIQKQDIDD | 1 | 0 | 0 | 1453337 | AUM60013.1 | K9N7C7 |
| INSVPWSKICDY | 1 | 0 | 0 | 1453340 | APO40578.1 | P0C6X7 |
| INSVVQKDGVED | 1 | 0 | 0 | 1453343 | ATQ39389.1 | K9N7C7 |
| INTGNGIKQLAP | 1 | 0 | 0 | 1453351 | AXN92267.1 | K9N4V7 |
| INTGNGTKPLAP | 1 | 0 | 0 | 1453353 | AVV62544.1 | K9N4V7 |
| INTGNGTKQLAP | 1 | 0 | 0 | 1453354 | AUM60021.1 | K9N4V7 |
| INTLNNLNETLV | 1 | 0 | 0 | 1453356 | AVP78041.1 | P0C6W2 |
| INTNSSKDDQIG | 1 | 0 | 0 | 1453360 | AVP78038.1 | Q3LZX4 |
| INTTPVSTFVGV | 1 | 0 | 0 | 1453371 | AMM42635.1 | A0A0P0G321 |
| INVGKNGLQFAA | 1 | 0 | 0 | 1453378 | AUM60013.1 | K9N7C7 |
| INVGKQGLEFAA | 1 | 0 | 0 | 1453379 | AVV62524.1 | K9N7C7 |
| INVGKQGLEYAA | 1 | 0 | 0 | 1453380 | AVV62535.1 | K9N7C7 |
| INVKDVFVESSK | 1 | 0 | 0 | 1453384 | QEG03747.1 | H9EJ42 |
| INVLSIFSSSFS | 1 | 0 | 0 | 1453385 | ARU07593.1 | H9EJ42 |
| INVTINNYNPSF | 1 | 0 | 0 | 1453390 | ABD75609.1 | A0A140H1H1 |
| INVVAASGSLTR | 1 | 0 | 0 | 1453393 | AXN92247.1 | K9N7C7 |
| INYTQPQLLSPL | 1 | 0 | 0 | 1453412 | AVV62537.1 | K9N5Q8 |
| IPAHCVRDPVGF | 1 | 0 | 0 | 1453419 | ALA49417.1 | K9N7C7 |
| IPAKARVECYDR | 1 | 0 | 0 | 1453420 | AXN92247.1 | K9N7C7 |
| IPASGYVAALAV | 1 | 0 | 0 | 1453430 | AXG21669.1 | K9N7D2 |
| IPDFNVWNQVVT | 1 | 0 | 0 | 1453447 | AVV62524.1 | K9N7C7 |
| IPDLPICVYDSL | 1 | 0 | 0 | 1453461 | AXX83314.1 | P30215 |
| IPDLQKAAVTIL | 1 | 0 | 0 | 1453464 | ARO76381.1 | P0C6U8 |

| | | | | | | |
|---|---|---|---|---|---|---|
| IPDMADQIASLV | 1 | 0 | 0 | 1453468 | AMK59681.1 | P33469 |
| IPDPKFVDKYYA | 1 | 0 | 0 | 1453469 | AUM60013.1 | K9N7C7 |
| IPDYNYYKYNLP | 1 | 0 | 0 | 1453474 | ACV87150.1 | A0A140H1G9 |
| IPEAPRDGQAYV | 1 | 0 | 0 | 1453477 | 6CRW_A | P59594 |
| IPEQKQNGFSFF | 1 | 0 | 0 | 1453484 | AVV62524.1 | K9N7C7 |
| IPFGTVLGNFYC | 1 | 0 | 0 | 1453496 | CAA71146.1 | A0A0P0K6L9 |
| IPFNWTTSVQVE | 1 | 0 | 0 | 1453503 | AOG74783.1 | A0A0P0K6L9 |
| IPGGCAMPIMAN | 1 | 0 | 0 | 1453510 | AIW52757.1 | A0A0P0K6L9 |
| IPGSLGLPNFKK | 1 | 0 | 0 | 1453516 | AET99208.1 | Q0ZJ80 |
| IPHYSWFSGITQ | 1 | 0 | 0 | 1453532 | ARU07581.1 | Q19U29 |
| IPHYVQPGQYTP | 1 | 0 | 0 | 1453533 | AUM60024.1 | K9N5Q8 |
| IPIDLLCLRDDN | 1 | 0 | 0 | 1453535 | ARU07575.1 | A0A140H1G9 |
| IPIDQYMCGKDG | 1 | 0 | 0 | 1453537 | AVV62535.1 | K9N7C7 |
| IPIFDDPVVDWH | 1 | 0 | 0 | 1453538 | AVN89322.1 | K9N7C7 |
| IPIVDDLVVDWH | 1 | 0 | 0 | 1453549 | ASU91162.1 | K9N7C7 |
| IPKPTAKYTPFA | 1 | 0 | 0 | 1453565 | ARQ84748.1 | K9N4V0 |
| IPLADDDSAIIE | 1 | 0 | 0 | 1453567 | AMM42614.1 | H9EJ69 |
| IPLGIDLDEWSM | 1 | 0 | 0 | 1453578 | AVP78030.1 | P0C6W2 |
| IPLLVDSEQPWK | 1 | 0 | 0 | 1453582 | ABB90504.1 | A0A0N7J131 |
| IPLSDMEKDFLS | 1 | 0 | 0 | 1453593 | ANI69887.1 | K9N7C7 |
| IPMPRGVHMNVA | 1 | 0 | 0 | 1453608 | AVV62535.1 | K9N7C7 |
| IPNFGSLTQINI | 1 | 0 | 0 | 1453616 | QBF80554.1 | K9N5Q8 |
| IPNLPDFKEELG | 1 | 0 | 0 | 1453629 | AIX10763.1 | P36334 |
| IPNLPDFKEESD | 1 | 0 | 0 | 1453630 | BBA20973.1 | P36334 |
| IPNLPDFKGELD | 1 | 0 | 0 | 1453631 | QDH43719.1 | P36334 |
| IPNLSDFEAEFS | 1 | 0 | 0 | 1453634 | BBA20986.1 | A0A140H1H1 |
| IPNLSDFEAELS | 1 | 0 | 0 | 1453635 | 5I08_A | A0A140H1H1 |
| IPNRRKVDLQLG | 1 | 0 | 0 | 1453638 | AEN19366.1 | P36334 |
| IPNRRRVDLQLG | 1 | 0 | 0 | 1453640 | ANZ78849.1 | P36334 |
| IPQQSQNGFSFF | 1 | 0 | 0 | 1453661 | AVV62535.1 | K9N7C7 |
| IPQVGGLTSIKW | 1 | 0 | 0 | 1453663 | BAE93400.1 | P0C6U8 |
| IPRISEDGVDVS | 1 | 0 | 0 | 1453665 | AZS52618.1 | A0A140H1H1 |
| IPRYASAVAQAF | 1 | 0 | 0 | 1453679 | AEN19363.1 | P0C6X6 |
| IPSDVSARDPII | 1 | 0 | 0 | 1453684 | AMM42389.1 | Q19U29 |
| IPSDVSARDPTI | 1 | 0 | 0 | 1453685 | AMM42397.1 | Q19U29 |
| IPSEFTIGNIEE | 1 | 0 | 0 | 1453686 | BBA20979.1 | P36334 |
| IPSEFTIGNMEE | 1 | 0 | 0 | 1453687 | CAA79896.1 | P36334 |
| IPSNWTTSVQVE | 1 | 0 | 0 | 1453702 | AIW52843.1 | A0A0P0G321 |
| IPSQEKNFTTAP | 1 | 0 | 0 | 1453704 | AVP78031.1 | Q3LZX1 |
| IPTIEDWLLVQG | 1 | 0 | 0 | 1453712 | AHI48842.1 | K9N643 |
| IPTIKDWLLVQG | 1 | 0 | 0 | 1453713 | AXN92231.1 | K9N643 |
| IPTKFPPGTVLP | 1 | 0 | 0 | 1453718 | AJQ24067.1 | P33469 |
| IPTMTHMNLKYA | 1 | 0 | 0 | 1453721 | AZK15898.1 | K9N7C7 |
| IPTNFTFSVTQE | 1 | 0 | 0 | 1453727 | AUM60014.1 | K9N5Q8 |
| IPTQCYYNNVSS | 1 | 0 | 0 | 1453731 | P0C6U5.1 | A0A140H1G9 |
| IPTSTGSVELSL | 1 | 0 | 0 | 1453735 | AVV62527.1 | K9N796 |
| IPYNSITDTIVL | 1 | 0 | 0 | 1453764 | Q3I5J4.1 | Q3LZX0 |
| IQADVNTPADIV | 1 | 0 | 0 | 1453778 | QBQ01843.1 | P33469 |
| IQAGSIVRRVNV | 1 | 0 | 0 | 1453779 | AXN92258.1 | K9N7C7 |
| IQDYIQSVNFVA | 1 | 0 | 0 | 1453812 | AGT51787.1 | P0C6X6 |
| IQEAIPTRFSPG | 1 | 0 | 0 | 1453813 | AMM42397.1 | Q19U29 |
| IQEVIDDMHWGE | 1 | 0 | 0 | 1453832 | NP_150075.1 | Q80872 |
| IQFDNPNIPFKD | 1 | 0 | 0 | 1453837 | APO40579.1 | P59594 |
| IQGFGPRDFNHN | 1 | 0 | 0 | 1453847 | QED88044.1 | H9EJ69 |
| IQIDQLNSSYFK | 1 | 0 | 0 | 1453858 | ASU91208.1 | K9N5Q8 |
| IQIPSDFTIGNM | 1 | 0 | 0 | 1453861 | ARB66062.1 | P36334 |
| IQIPSKFTIGNM | 1 | 0 | 0 | 1453864 | AIL49542.1 | P36334 |
| IQKQDIDDVPAH | 1 | 0 | 0 | 1453877 | AUM60013.1 | K9N7C7 |
| IQLLKNAYENFN | 1 | 0 | 0 | 1453889 | AVP78041.1 | P0C6W2 |
| IQLQLTKLLDYS | 1 | 0 | 0 | 1453896 | AWK59978.1 | H9EJ42 |
| IQLTITDPKVYP | 1 | 0 | 0 | 1453906 | 2CME_E | P59636 |
| IQLTTEGYNNYV | 1 | 0 | 0 | 1453912 | AVV62535.1 | K9N7C7 |
| IQLTVEGYNNYV | 1 | 0 | 0 | 1453915 | AVV62524.1 | K9N7C7 |
| IQNQTLPRLHCT | 1 | 0 | 0 | 1453928 | ARU07574.1 | A0A140H1G9 |
| IQNQTLSRLHCT | 1 | 0 | 0 | 1453929 | ARB07436.1 | A0A140H1G9 |
| IQPVELPVSIVS | 1 | 0 | 0 | 1453954 | AIW52757.1 | A0A0P0K6L9 |
| IQQDLDVILPGE | 1 | 0 | 0 | 1453959 | ALA49934.1 | K9N7C7 |
| IQQGFDATNFAL | 1 | 0 | 0 | 1453960 | QBP84722.1 | P36334 |
| IQQGFDATNSAL | 1 | 0 | 0 | 1453961 | AEN19366.1 | P36334 |
| IQQGGIPIANCV | 1 | 0 | 0 | 1453962 | AIV41966.1 | P0C6X6 |
| IQQGSDATNSAL | 1 | 0 | 0 | 1453964 | ARB66064.1 | P36334 |
| IQQIELKFNPPA | 1 | 0 | 0 | 1453965 | AVP78030.1 | P0C6W2 |
| IQQQFNVALPGE | 1 | 0 | 0 | 1453975 | AUM60013.1 | K9N7C7 |
| IQQWGYVGSLSQ | 1 | 0 | 0 | 1453982 | AWK59942.1 | H9EJ42 |
| IQQWGYVGSLST | 1 | 0 | 0 | 1453983 | NP_073549.1 | A0A0P0LKV0 |
| IQRLDGLEQDAQ | 1 | 0 | 0 | 1453991 | AHI48733.1 | K9N5Q8 |
| IQSFDSSYLNES | 1 | 0 | 0 | 1454006 | AST12963.1 | A0A0P0LKV0 |
| IQSKLTDLKCTN | 1 | 0 | 0 | 1454012 | AGT51337.1 | H9EJ42 |
| IQSNVDVVPEGW | 1 | 0 | 0 | 1454013 | AGT51409.1 | P0C6X6 |
| IQSSGVVKPNAI | 1 | 0 | 0 | 1454017 | AVV62524.1 | K9N7C7 |
| IQSVELPVSIVA | 1 | 0 | 0 | 1454023 | AAQ90008.1 | A0A0P0K6L9 |
| IQTAQAAGTDTT | 1 | 0 | 0 | 1454028 | 3ATW_A | P0C6X7 |
| IQTLDELHACTS | 1 | 0 | 0 | 1454034 | AVV62535.1 | K9N7C7 |
| IQTTIQKITVDC | 1 | 0 | 0 | 1454037 | AUM60014.1 | K9N5Q8 |
| IQTTPGFGVAVA | 1 | 0 | 0 | 1454039 | ABD75607.1 | A0A140H1G9 |
| IQTVFDCAVETC | 1 | 0 | 0 | 1454042 | NP_073549.1 | A0A0P0LKV0 |
| IQVDQFNSSYFK | 1 | 0 | 0 | 1454051 | ASU90362.1 | K9N5Q8 |
| IQVDQLNSSYKL | 1 | 0 | 0 | 1454053 | ATW75477.1 | K9N5Q8 |
| IQVDQLNSTYFK | 1 | 0 | 0 | 1454054 | ANI69889.1 | K9N5Q8 |
| IQVTIAKTESAM | 1 | 0 | 0 | 1454057 | AVP78050.1 | Q3LZX3 |
| IQVVNEGLLHFV | 1 | 0 | 0 | 1454060 | APO40578.1 | P0C6X7 |

| | | | | | | |
|---|---|---|---|---|---|---|
| IQVVNSGPDQIC | 1 | 0 | 0 | 1454061 | ARB07411.1 | P0C6X6 |
| IQVVNSGSDINC | 1 | 0 | 0 | 1454063 | P0C6U5.1 | A0A140H1G9 |
| IRANAKIVTLPG | 1 | 0 | 0 | 1454086 | ATQ39390.1 | K9N5Q8 |
| IRASMPTTIAKN | 1 | 0 | 0 | 1454089 | AVP78041.1 | P0C6W2 |
| IRDNGSLEKFMS | 1 | 0 | 0 | 1454113 | AXN92247.1 | K9N7C7 |
| IRDNGSLERFMN | 1 | 0 | 0 | 1454114 | AUM60013.1 | K9N7C7 |
| IRDNVSLEKFMS | 1 | 0 | 0 | 1454115 | ANF29193.1 | K9N7C7 |
| IRDSEATKQPVK | 1 | 0 | 0 | 1454123 | AVV62535.1 | K9N7C7 |
| IRDYAALAAKEG | 1 | 0 | 0 | 1454126 | AVV62524.1 | K9N7C7 |
| IRDYASLAAKEG | 1 | 0 | 0 | 1454127 | AVV62535.1 | K9N7C7 |
| IREIVMNASPYD | 1 | 0 | 0 | 1454137 | AGT51737.1 | P0C6X6 |
| IRFEYNPPTDKL | 1 | 0 | 0 | 1454159 | NP_150075.1 | Q80872 |
| IRGAIGDVGLCM | 1 | 0 | 0 | 1454169 | ARB07424.1 | A0A0P0ILY9 |
| IRGNPNNHKKIV | 1 | 0 | 0 | 1454183 | AVV62529.1 | K9N643 |
| IRGSDVKQPVKF | 1 | 0 | 0 | 1454187 | AUM60013.1 | K9N7C7 |
| IRHKILTKPRQK | 1 | 0 | 0 | 1454199 | ARU07581.1 | Q19U29 |
| IRIVDEWSYTGS | 1 | 0 | 0 | 1454205 | ASU90175.1 | K9N5Q8 |
| IRKNGRGGNLFR | 1 | 0 | 0 | 1454218 | AUM60013.1 | K9N7C7 |
| IRLTITELLQFV | 1 | 0 | 0 | 1454236 | AIW52813.1 | H9EJ42 |
| IRMEESLMDVPS | 1 | 0 | 0 | 1454238 | AIZ74435.1 | K9N643 |
| IRNESWHIDSSE | 1 | 0 | 0 | 1454244 | BBA20986.1 | A0A140H1H1 |
| IRNLQKRLYFNV | 1 | 0 | 0 | 1454250 | ABD75527.1 | A0A140H1G9 |
| IRPIDFFSLTAS | 1 | 0 | 0 | 1454258 | ARU07575.1 | A0A140H1G9 |
| IRQKILNKPRQK | 1 | 0 | 0 | 1454270 | BBM61433.1 | P33469 |
| IRRAIDCGHDDL | 1 | 0 | 0 | 1454278 | AVV62526.1 | K9N5Q8 |
| IRRLDVLEQDAQ | 1 | 0 | 0 | 1454286 | ALB08246.1 | K9N5Q8 |
| IRRPKDGFNLEY | 1 | 0 | 0 | 1454290 | ARK08651.1 | P36334 |
| IRSDSTLSGFET | 1 | 0 | 0 | 1454306 | AHA80176.1 | P33469 |
| IRSHPDFKLLHN | 1 | 0 | 0 | 1454308 | AUM60013.1 | K9N7C7 |
| IRSKAADRVAWA | 1 | 0 | 0 | 1454314 | AVV62526.1 | K9N5Q8 |
| IRSYEKLANSIS | 1 | 0 | 0 | 1454326 | AST12962.1 | A0A0P0LKV0 |
| IRTRTLVAINSE | 1 | 0 | 0 | 1454343 | AVV62535.1 | K9N7C7 |
| IRYGIKPITIPI | 1 | 0 | 0 | 1454359 | ARU07575.1 | A0A140H1G9 |
| ISAKVKNTKVIK | 1 | 0 | 0 | 1454387 | AKA09642.1 | P36334 |
| ISAMQTMLFGMI | 1 | 0 | 0 | 1454392 | AVN89333.1 | K9N7C7 |
| ISAQLAKDKVNE | 1 | 0 | 0 | 1454395 | AVN89365.1 | K9N5Q8 |
| ISARPPPGEQFK | 1 | 0 | 0 | 1454397 | AVV62524.1 | K9N7C7 |
| ISASYRFVTFEPFNVS | 1 | 0 | 0 | 1454404 | YP_173238.1 | A0A140H1H1 |
| ISAYGCSTIAQA | 1 | 0 | 0 | 1454411 | AST12962.1 | A0A0P0LKV0 |
| ISCDCGKKFDEQ | 1 | 0 | 0 | 1454412 | QED88024.1 | H9EJ42 |
| ISCIWFIDAFNQ | 1 | 0 | 0 | 1454415 | ARU07575.1 | A0A140H1G9 |
| ISCNGNKIEDLS | 1 | 0 | 0 | 1454417 | AIL49553.1 | P0C6X6 |
| ISDFVDVGPDAT | 1 | 0 | 0 | 1454433 | AUM60014.1 | K9N5Q8 |
| ISDGNQRQLLPL | 1 | 0 | 0 | 1454434 | AJQ24067.1 | P33469 |
| ISDILKRLDAVE | 1 | 0 | 0 | 1454436 | AVV62537.1 | K9N5Q8 |
| ISDILQRLDTVE | 1 | 0 | 0 | 1454437 | AVV62526.1 | K9N5Q8 |
| ISDKNVVFMNSC | 1 | 0 | 0 | 1454439 | BBA20986.1 | A0A140H1H1 |
| ISDLLTPMGIDL | 1 | 0 | 0 | 1454441 | APO40578.1 | P0C6X7 |
| ISDLSNRLLDAM | 1 | 0 | 0 | 1454442 | APO40578.1 | P0C6X7 |
| ISDMYDGRIKFC | 1 | 0 | 0 | 1454443 | AWK59942.1 | H9EJ42 |
| ISDMYDPITKNV | 1 | 0 | 0 | 1454444 | AVV62535.1 | K9N7C7 |
| ISDMYDPITKTE | 1 | 0 | 0 | 1454445 | AUM60013.1 | K9N7C7 |
| ISDMYDPKTKHV | 1 | 0 | 0 | 1454446 | ATO98229.1 | P0C6W2 |
| ISDMYDPVTKNI | 1 | 0 | 0 | 1454451 | AVV62524.1 | K9N7C7 |
| ISDRKIKKYPCI | 1 | 0 | 0 | 1454453 | AXN92258.1 | K9N7C7 |
| ISDSNGVPVVEY | 1 | 0 | 0 | 1454458 | AUM60013.1 | K9N7C7 |
| ISDTSGFAVYVK | 1 | 0 | 0 | 1454460 | AWW13547.1 | Q01455 |
| ISEDLWQFVDHF | 1 | 0 | 0 | 1454464 | AGT21366.1 | A0A0P0LKV0 |
| ISEGNALYNNEG | 1 | 0 | 0 | 1454467 | AST12962.1 | A0A0P0LKV0 |
| ISEHSLRLVDAM | 1 | 0 | 0 | 1454469 | ARO76381.1 | P0C6U8 |
| ISEWPLFDQQAQ | 1 | 0 | 0 | 1454484 | APT69896.1 | A0A0P0LKV0 |
| ISFADNNDITNT | 1 | 0 | 0 | 1454490 | ANI69896.1 | K9N4V7 |
| ISFCCRHNWNCV | 1 | 0 | 0 | 1454492 | ASU90975.1 | K9N7C7 |
| ISFCKSGSLVLN | 1 | 0 | 0 | 1454494 | QBP84748.1 | P30215 |
| ISFERLRDYLAA | 1 | 0 | 0 | 1454497 | AUM60013.1 | K9N7C7 |
| ISFSDDVIVHDV | 1 | 0 | 0 | 1454508 | AWK59942.1 | H9EJ42 |
| ISGDRGIAPKSG | 1 | 0 | 0 | 1454512 | AGT51551.1 | P36334 |
| ISGFLSRVIRRE | 1 | 0 | 0 | 1454515 | QED88025.1 | H9EJ42 |
| ISGIVSAVSAMM | 1 | 0 | 0 | 1454524 | ANF29190.1 | K9N7A1 |
| ISGKELARTVGG | 1 | 0 | 0 | 1454526 | AIW52767.1 | A0A0P0LKV0 |
| ISGKGQQQQGQT | 1 | 0 | 0 | 1454527 | AVP78049.1 | Q3LZX4 |
| ISGNGGLFNTLR | 1 | 0 | 0 | 1454533 | AVP78031.1 | Q3LZX1 |
| ISIMKHDIKVIA | 1 | 0 | 0 | 1454559 | AVA26872.1 | H9EJ42 |
| ISINLPPPLLGN | 1 | 0 | 0 | 1454561 | ATG84888.1 | K9N5Q8 |
| ISIVNKALELCT | 1 | 0 | 0 | 1454569 | APO40578.1 | P0C6X7 |
| ISKCSRLTGDNQ | 1 | 0 | 0 | 1454573 | AVV62526.1 | K9N5Q8 |
| ISLADVATDVQS | 1 | 0 | 0 | 1454590 | AUM60013.1 | K9N7C7 |
| ISLICFILLWAL | 1 | 0 | 0 | 1454605 | ARU07575.1 | A0A140H1G9 |
| ISLNAPVTLKIC | 1 | 0 | 0 | 1454618 | ARU07594.1 | A0A0P0G321 |
| ISLNASVTLRIC | 1 | 0 | 0 | 1454619 | AMM42653.1 | A0A0P0G321 |
| ISLPTAFAAIGD | 1 | 0 | 0 | 1454624 | AIL49486.1 | P36334 |
| ISLPTAFAVVGD | 1 | 0 | 0 | 1454626 | QBP84704.1 | P36334 |
| ISLPTAFSVIGD | 1 | 0 | 0 | 1454627 | BAM16455.1 | P36334 |
| ISLQKGVLPQLE | 1 | 0 | 0 | 1454628 | APO40578.1 | P0C6X7 |
| ISLTDVATDVQS | 1 | 0 | 0 | 1454635 | AUM60023.1 | K9N7C7 |
| ISMLQLGAPDNP | 1 | 0 | 0 | 1454644 | AKA09625.1 | A0A0P0G321 |
| ISMQGAVDINKL | 1 | 0 | 0 | 1454651 | ARO76381.1 | P0C6U8 |
| ISMSLVLVIVVK | 1 | 0 | 0 | 1454653 | AXN92247.1 | K9N7C7 |
| ISMTNHSFSVQK | 1 | 0 | 0 | 1454654 | AXN92247.1 | K9N7C7 |
| ISNCVADYTAFY | 1 | 0 | 0 | 1454662 | ATO98145.1 | Q3LZX1 |
| ISNDTVDVTNGL | 1 | 0 | 0 | 1454667 | ANZ78836.1 | P36334 |

| | | | | | | |
|---|---|---|---|---|---|---|
| ISNVPFSPNGKP | 1 | 0 | 0 | 1454686 | ABD72972.1 | P59594 |
| ISPAAISSNCYS | 1 | 0 | 0 | 1454691 | AHL18090.1 | K9N5Q8 |
| ISPDAIARGCYS | 1 | 0 | 0 | 1454696 | AVV62526.1 | K9N5Q8 |
| ISPKIDNETLEY | 1 | 0 | 0 | 1454703 | AMM42407.1 | A0A140H1H1 |
| ISPKTDKETLEY | 1 | 0 | 0 | 1454706 | AZS52618.1 | A0A140H1H1 |
| ISPNTDNETLQY | 1 | 0 | 0 | 1454708 | ABA60801.1 | A0A140H1H1 |
| ISPSTSAIIRKI | 1 | 0 | 0 | 1454715 | AHL31378.1 | K9N5Q8 |
| ISPSTSATIQKI | 1 | 0 | 0 | 1454716 | ALW82742.1 | K9N5Q8 |
| ISPTAIASNCYS | 1 | 0 | 0 | 1454719 | AWF93654.1 | K9N5Q8 |
| ISPTNLEMFKPG | 1 | 0 | 0 | 1454720 | AOL02458.1 | Q4VID0 |
| ISPTSLEIFKPG | 1 | 0 | 0 | 1454721 | APU51922.1 | Q4VID0 |
| ISPTSLEMFKPE | 1 | 0 | 0 | 1454722 | AGT51436.1 | Q4VID0 |
| ISPTSLEMFKSG | 1 | 0 | 0 | 1454724 | ARE29989.1 | Q4VID0 |
| ISPYNSQNTVAS | 1 | 0 | 0 | 1454728 | AVP78041.1 | P0C6W2 |
| ISPYNSQNYVAA | 1 | 0 | 0 | 1454729 | NP_073549.1 | A0A0P0LKV0 |
| ISPYNSQNYVAKRVLGVQ | 1 | 0 | 0 | 1454730 | ABD75543.1 | A0A140H1G9 |
| ISQDSITHYSAL | 1 | 0 | 0 | 1454734 | ASU91185.1 | K9N7C7 |
| ISQFNYKQSSSN | 1 | 0 | 0 | 1454737 | AXN92228.1 | K9N5Q8 |
| ISQGLDAAWNKF | 1 | 0 | 0 | 1454739 | NP_073549.1 | A0A0P0LKV0 |
| ISQLGCYEPLKS | 1 | 0 | 0 | 1454743 | APO40579.1 | P59594 |
| ISQWPISNDSNC | 1 | 0 | 0 | 1454750 | AGT51386.1 | H9EJ42 |
| ISRCQITAVEGT | 1 | 0 | 0 | 1454755 | QBP84728.1 | P0C6X6 |
| ISRMGFKLDANI | 1 | 0 | 0 | 1454767 | AVV62524.1 | K9N7C7 |
| ISRRNIGFTLEY | 1 | 0 | 0 | 1454776 | AMK59677.1 | P36334 |
| ISRVELHTVYPD | 1 | 0 | 0 | 1454781 | AWK59979.1 | A0A0P0G321 |
| ISSCDGIASDYR | 1 | 0 | 0 | 1454786 | AUM60023.1 | K9N7C7 |
| ISSGIFSVPSDV | 1 | 0 | 0 | 1454792 | ARK08632.1 | P0C6X6 |
| ISSGSRGVYYPD | 1 | 0 | 0 | 1454793 | ATO98145.1 | Q3LZX1 |
| ISSKVGNSIFRS | 1 | 0 | 0 | 1454797 | AIL49309.1 | P30215 |
| ISSLAYYEPVQG | 1 | 0 | 0 | 1454798 | ASU45728.1 | K9N7C7 |
| ISSMTDNLQMAF | 1 | 0 | 0 | 1454801 | AUM60014.1 | K9N5Q8 |
| ISSNCYSSLILD | 1 | 0 | 0 | 1454802 | AHL18090.1 | K9N5Q8 |
| ISSNGPLKVGTG | 1 | 0 | 0 | 1454804 | ARU07593.1 | H9EJ42 |
| ISTDIVDVTNGL | 1 | 0 | 0 | 1454835 | CAA83660.1 | P36334 |
| ISTDVLVNTYKR | 1 | 0 | 0 | 1454836 | AUM60013.1 | K9N7C7 |
| ISTGVCVNYDLY | 1 | 0 | 0 | 1454845 | 5I08_A | A0A140H1H1 |
| ISTNLPPPLLSN | 1 | 0 | 0 | 1454859 | ALL26396.1 | K9N5Q8 |
| ISTNYSKDVYFT | 1 | 0 | 0 | 1454860 | AGT17757.1 | A0A140H1H0 |
| ISTPKKPNKPLS | 1 | 0 | 0 | 1454861 | AMM42629.1 | H9EJ69 |
| ISTPNLPDFKEE | 1 | 0 | 0 | 1454864 | CAA83661.1 | P36334 |
| ISTQDGVNKLQG | 1 | 0 | 0 | 1454865 | BBA20973.1 | P36334 |
| ISTRENKSAELN | 1 | 0 | 0 | 1454867 | BAL45640.1 | A0A0P0K6L9 |
| ISTSKVKSLSMI | 1 | 0 | 0 | 1454869 | AIV41780.1 | P0C6X6 |
| ISTTLIQLQLTK | 1 | 0 | 0 | 1454873 | AWK59942.1 | H9EJ42 |
| ISVAPIKNTVDT | 1 | 0 | 0 | 1454883 | APT69850.1 | A0A0P0LKV0 |
| ISVDRDSAVGHT | 1 | 0 | 0 | 1454891 | AUM60025.1 | K9N796 |
| ISVEELKRLLEQ | 1 | 0 | 0 | 1454893 | ATO98197.1 | Q3LZX9 |
| ISVPAHMPMPPM | 1 | 0 | 0 | 1454906 | ARU07582.1 | Q0ZJ80 |
| ISVVATSGSLTK | 1 | 0 | 0 | 1454918 | AVV62524.1 | K9N7C7 |
| ISWLLHKDGGIP | 1 | 0 | 0 | 1454927 | AMQ49017.1 | K9N4V0 |
| ISYEEQDAIFLL | 1 | 0 | 0 | 1454932 | AIW52776.1 | A0A0P0LKV0 |
| ISYLNEQGVLVQ | 1 | 0 | 0 | 1454937 | AVA26878.1 | H9EJ42 |
| ISYLNRARGSSAARLEPC | 1 | 0 | 0 | 1454938 | QED88024.1 | H9EJ42 |
| ISYNSTLQSGLK | 1 | 0 | 0 | 1454940 | AWK59942.1 | H9EJ42 |
| ITACQYTMCEFP | 1 | 0 | 0 | 1454945 | ARB07438.1 | A0A140H1H1 |
| ITAGDGISRPKL | 1 | 0 | 0 | 1454950 | ARO76383.1 | P59632 |
| ITANGGISFCCR | 1 | 0 | 0 | 1454955 | ASU90975.1 | K9N7C7 |
| ITANGGISFCRR | 1 | 0 | 0 | 1454957 | AXN92247.1 | K9N7C7 |
| ITARETLYRIDG | 1 | 0 | 0 | 1454968 | ARO76381.1 | P0C6U8 |
| ITCEEAVRQVRS | 1 | 0 | 0 | 1454972 | AQZ41330.1 | K9N7C7 |
| ITCGDGTTNPIS | 1 | 0 | 0 | 1454973 | AVP78043.1 | Q3LZX0 |
| ITDPLFSNIQPF | 1 | 0 | 0 | 1454994 | AST12962.1 | A0A0P0LKV0 |
| ITEDEILEWFGI | 1 | 0 | 0 | 1455005 | AXN92228.1 | K9N5Q8 |
| ITEDENAEWFGI | 1 | 0 | 0 | 1455006 | AVV62537.1 | K9N5Q8 |
| ITEDETAEWFGI | 1 | 0 | 0 | 1455007 | AUM60014.1 | K9N5Q8 |
| ITEDGINVKDVF | 1 | 0 | 0 | 1455008 | QEG03747.1 | H9EJ42 |
| ITEDGINVKDVV | 1 | 0 | 0 | 1455009 | AWK59942.1 | H9EJ42 |
| ITEELQHEDLLA | 1 | 0 | 0 | 1455010 | APO40578.1 | P0C6X7 |
| ITEGPTTCGYLP | 1 | 0 | 0 | 1455014 | ATO98179.1 | P0C6W2 |
| ITEGSVKGLIAS | 1 | 0 | 0 | 1455015 | APO40578.1 | P0C6X7 |
| ITEHSWSADLYK | 1 | 0 | 0 | 1455021 | APO40578.1 | P0C6X7 |
| ITENNVVVTSTC | 1 | 0 | 0 | 1455023 | ARB66070.1 | P36334 |
| ITETNTAAFKAV | 1 | 0 | 0 | 1455027 | AXN92247.1 | K9N7C7 |
| ITFDNLKTILSL | 1 | 0 | 0 | 1455038 | AVP78041.1 | P0C6W2 |
| ITGLYPTITVPG | 1 | 0 | 0 | 1455081 | AXN73435.1 | K9N7C7 |
| ITHALSADGVQC | 1 | 0 | 0 | 1455097 | AEN19363.1 | P0C6X6 |
| ITIADPGYMQGY | 1 | 0 | 0 | 1455106 | AUM60014.1 | K9N5Q8 |
| ITIDKFAIPNRR | 1 | 0 | 0 | 1455111 | AEN19366.1 | P36334 |
| ITIDKFAIPNSR | 1 | 0 | 0 | 1455112 | AIX10751.1 | P36334 |
| ITIDKFALPNRR | 1 | 0 | 0 | 1455113 | AIL49495.1 | P36334 |
| ITIDKSAIPNGR | 1 | 0 | 0 | 1455114 | BBA84104.1 | P36334 |
| ITIEAALDLSKE | 1 | 0 | 0 | 1455115 | AXX83372.1 | P0C6X6 |
| ITIGSNFVNTSY | 1 | 0 | 0 | 1455116 | BBO58761.1 | P36334 |
| ITIKKPNELSIV | 1 | 0 | 0 | 1455121 | APO40578.1 | P0C6X7 |
| ITINPSSPARLE | 1 | 0 | 0 | 1455127 | AWW13564.1 | Q80872 |
| ITKDTYVALRNS | 1 | 0 | 0 | 1455138 | AUM60013.1 | K9N7C7 |
| ITKKHAAAAKNK | 1 | 0 | 0 | 1455145 | ANI69874.1 | K9N4V7 |
| ITKLADVEADIA | 1 | 0 | 0 | 1455146 | ATQ39389.1 | K9N7C7 |
| ITKLEADFYLSG | 1 | 0 | 0 | 1455148 | QQY99179.1 | P30215 |
| ITKPLKYSYIDK | 1 | 0 | 0 | 1455157 | ASY99811.1 | K9N5Q8 |
| ITKTEKDTNEAK | 1 | 0 | 0 | 1455162 | AUM60013.1 | K9N7C7 |

| | | | | | | |
|---|---|---|---|---|---|---|
| ITKVEADFYLSG | 1 | 0 | 0 | 1455165 | AEN19357.1 | P30215 |
| ITLANFNETNGP | 1 | 0 | 0 | 1455171 | CAA71147.1 | A0A0P0K6L9 |
| ITLDDDARTFVQ | 1 | 0 | 0 | 1455177 | AQN78773.1 | P0C6X6 |
| ITLEEAVRQVRS | 1 | 0 | 0 | 1455178 | AHL18091.1 | K9N7C7 |
| ITLFGDCVTVRV | 1 | 0 | 0 | 1455179 | AXN92247.1 | K9N7C7 |
| ITLKKRWQLALSKG | 1 | 0 | 0 | 1455189 | P0DTC3.1 | P0DTC3 |
| ITLKTPDTYKCP | 1 | 0 | 0 | 1455191 | AXX83339.1 | P36334 |
| ITMANTRFVSKD | 1 | 0 | 0 | 1455210 | AUM60014.1 | K9N5Q8 |
| ITMPCFITYPQQ | 1 | 0 | 0 | 1455212 | QDI73613.1 | K9N643 |
| ITNINLSRGRGR | 1 | 0 | 0 | 1455228 | ASJ26617.1 | K9N4V7 |
| ITNLDIEKLRQY | 1 | 0 | 0 | 1455233 | AVV62535.1 | K9N7C7 |
| ITNQLGVCVDYK | 1 | 0 | 0 | 1455234 | AVV62526.1 | K9N5Q8 |
| ITNVGEPTISQP | 1 | 0 | 0 | 1455243 | ABE97136.1 | H9EJ42 |
| ITPAHVMFKHND | 1 | 0 | 0 | 1455244 | AVP78041.1 | P0C6W2 |
| ITPGTNASSEFA | 1 | 0 | 0 | 1455248 | BAE93401.1 | P59594 |
| ITPGTNTSKAVA | 1 | 0 | 0 | 1455250 | APO40579.1 | P59594 |
| ITPGTNTSLEVA | 1 | 0 | 0 | 1455251 | AVP78031.1 | Q3LZX1 |
| ITPYCHDPTVLP | 1 | 0 | 0 | 1455259 | AHL18091.1 | K9N7C7 |
| ITQEYIQTTIQK | 1 | 0 | 0 | 1455262 | AVV62526.1 | K9N5Q8 |
| ITQFQKGKEFQF | 1 | 0 | 0 | 1455267 | QBP84717.1 | P33469 |
| ITQHTRTRPSVQ | 1 | 0 | 0 | 1455269 | AKN24774.1 | K9N4V7 |
| ITQPGSGVAVVD | 1 | 0 | 0 | 1455275 | AVV62524.1 | K9N7C7 |
| ITQQVLSDNQKL | 1 | 0 | 0 | 1455276 | ALA49671.1 | K9N5Q8 |
| ITQQVLSENQKQ | 1 | 0 | 0 | 1455278 | AVV62526.1 | K9N5Q8 |
| ITQQVLSKNQKL | 1 | 0 | 0 | 1455279 | AVN89376.1 | K9N5Q8 |
| ITQRNFYSPQII | 1 | 0 | 0 | 1455283 | ATO98181.1 | Q3LZX1 |
| ITQRNFYSPQTI | 1 | 0 | 0 | 1455284 | ARO76382.1 | P59594 |
| ITQRNFYSPQVI | 1 | 0 | 0 | 1455285 | APO40579.1 | P59594 |
| ITQTSPALQTVA | 1 | 0 | 0 | 1455295 | AIW52728.1 | A0A0P0K6L9 |
| ITQTSQALQTVS | 1 | 0 | 0 | 1455296 | ABB90519.1 | A0A0P0K6L9 |
| ITQWPTATGDVV | 1 | 0 | 0 | 1455300 | AVQ05261.1 | P0C6X6 |
| ITRDEAIKHVRA | 1 | 0 | 0 | 1455307 | APO40578.1 | P0C6X7 |
| ITRQSSKIENAS | 1 | 0 | 0 | 1455323 | ARO76383.1 | P59632 |
| ITSAGQPCVASE | 1 | 0 | 0 | 1455331 | AEN19363.1 | P0C6X6 |
| ITSFKVVMSMFS | 1 | 0 | 0 | 1455342 | ATO98193.1 | Q3LZX1 |
| ITSFNTKYVAPQ | 1 | 0 | 0 | 1455343 | AVN89387.1 | K9N5Q8 |
| ITSVTSDKFDFI | 1 | 0 | 0 | 1455368 | AXX83348.1 | P0C6X6 |
| ITTCFDATTATI | 1 | 0 | 0 | 1455372 | AUM60023.1 | K9N7C7 |
| ITTFMPQFSRMT | 1 | 0 | 0 | 1455381 | AUM60014.1 | K9N5Q8 |
| ITTKISLKGGKV | 1 | 0 | 0 | 1455392 | ARO76381.1 | P0C6U8 |
| ITTNGPLIVGGS | 1 | 0 | 0 | 1455398 | AVP78041.1 | P0C6W2 |
| ITVDCKQYVCNG | 1 | 0 | 0 | 1455412 | AUM60014.1 | K9N5Q8 |
| ITVDKFAIPNRR | 1 | 0 | 0 | 1455413 | BBA20986.1 | A0A140H1H1 |
| ITVDQLKHLLEQ | 1 | 0 | 0 | 1455414 | APO40582.1 | P59596 |
| ITVDVLKDMPVL | 1 | 0 | 0 | 1455415 | ARU07575.1 | A0A140H1G9 |
| ITVEAAFDLSKE | 1 | 0 | 0 | 1455416 | QBQ01836.1 | P0C6X6 |
| ITVEAALDLSRE | 1 | 0 | 0 | 1455417 | AOL02450.1 | P0C6X6 |
| ITVEKLKQLLEQ | 1 | 0 | 0 | 1455420 | AAS48455.1 | P59596 |
| ITVETAIELSKE | 1 | 0 | 0 | 1455421 | ABD75535.1 | A0A140H1G9 |
| ITVETALDLSKE | 1 | 0 | 0 | 1455422 | AXX83306.1 | P0C6X6 |
| ITVEVFEAQAFE | 1 | 0 | 0 | 1455423 | AVV62535.1 | K9N7C7 |
| ITVPGEFASHVA | 1 | 0 | 0 | 1455436 | AXN73435.1 | K9N7C7 |
| ITVQYGTDTISV | 1 | 0 | 0 | 1455437 | AGV08438.1 | K9N5Q8 |
| ITVQYGTDTNSA | 1 | 0 | 0 | 1455438 | AXN92260.1 | K9N5Q8 |
| ITYSPATILAGS | 1 | 0 | 0 | 1455456 | QBF80553.1 | K9N7C7 |
| IVAAVSAMMRIS | 1 | 0 | 0 | 1455465 | ALA49424.1 | K9N7A1 |
| IVAICLLVTIVL | 1 | 0 | 0 | 1455471 | AMK59679.1 | Q04854 |
| IVAMTEQLQMGF | 1 | 0 | 0 | 1455480 | AMQ49004.1 | K9N5Q8 |
| IVANWAYSKYYT | 1 | 0 | 0 | 1455481 | CAA71056.1 | A0A0P0K6L9 |
| IVAYTMSIGEDQ | 1 | 0 | 0 | 1455493 | APO40579.1 | P59594 |
| IVCRLDTRVHSE | 1 | 0 | 0 | 1455513 | AKN24802.1 | K9N7C7 |
| IVDARIEPCASG | 1 | 0 | 0 | 1455516 | AVV62524.1 | K9N7C7 |
| IVDIIQRLDVLE | 1 | 0 | 0 | 1455525 | AGV08438.1 | K9N5Q8 |
| IVDMSMTYGQQF | 1 | 0 | 0 | 1455535 | AVP78041.1 | P0C6W2 |
| IVDNVYMAKSGD | 1 | 0 | 0 | 1455537 | AXX83372.1 | P0C6X6 |
| IVDNVYMAKVGD | 1 | 0 | 0 | 1455538 | AEN19363.1 | P0C6X6 |
| IVDRDQSSEEAI | 1 | 0 | 0 | 1455543 | ANZ78879.1 | P33469 |
| IVDTLQETPVVS | 1 | 0 | 0 | 1455549 | AVK87459.1 | K9N7C7 |
| IVDVTNGLGTYY | 1 | 0 | 0 | 1455554 | CAA83660.1 | P36334 |
| IVEDTINDDEDV | 1 | 0 | 0 | 1455560 | AGW27861.1 | A0A140H1G9 |
| IVEEAMVVKPTV | 1 | 0 | 0 | 1455562 | APO40578.1 | P0C6X7 |
| IVFNVVKQGHFI | 1 | 0 | 0 | 1455599 | AXN92247.1 | K9N7C7 |
| IVGCNNADCVAC | 1 | 0 | 0 | 1455613 | AWK59942.1 | H9EJ42 |
| IVGDVIKEGALT | 1 | 0 | 0 | 1455619 | AVP78041.1 | P0C6W2 |
| IVGDVVHNGDIT | 1 | 0 | 0 | 1455621 | APO40578.1 | P0C6X7 |
| IVGERQLLLLWE | 1 | 0 | 0 | 1455625 | AZU90729.1 | K9N7C7 |
| IVGGQIVTRAKE | 1 | 0 | 0 | 1455637 | AVP78041.1 | P0C6W2 |
| IVHSHPILHHPG | 1 | 0 | 0 | 1455663 | ASU90849.1 | K9N643 |
| IVHSYPILHHSG | 1 | 0 | 0 | 1455667 | ASU90607.1 | K9N643 |
| IVIDNDLNEFVS | 1 | 0 | 0 | 1455672 | ASY99818.1 | K9N7C7 |
| IVIRDIVCKVEN | 1 | 0 | 0 | 1455687 | NP_073549.1 | A0A0P0LKV0 |
| IVKEAQNANPTV | 1 | 0 | 0 | 1455705 | ATO98229.1 | P0C6W2 |
| IVKLGAEKEDGF | 1 | 0 | 0 | 1455715 | AMK59675.1 | Q80872 |
| IVKMVSPTSKIE | 1 | 0 | 0 | 1455718 | ARU07575.1 | A0A140H1G9 |
| IVLPQGGYVADF | 1 | 0 | 0 | 1455760 | AEN19363.1 | P0C6X6 |
| IVLTAGDGVTIP | 1 | 0 | 0 | 1455771 | APO40580.1 | P59632 |
| IVLVCYLFIGYI | 1 | 0 | 0 | 1455776 | AVI15051.1 | P0C6X6 |
| IVLYVDFKPQSG | 1 | 0 | 0 | 1455784 | CAA71056.1 | A0A0P0K6L9 |
| IVMNASPYDLEV | 1 | 0 | 0 | 1455797 | AGT51737.1 | P0C6X6 |
| IVNGYAKSNFVK | 1 | 0 | 0 | 1455827 | APT69868.1 | A0A0P0ILY9 |
| IVNNYEKSAGYP | 1 | 0 | 0 | 1455843 | AIL49570.1 | P0C6X6 |

| | | | | | | |
|---|---|---|---|---|---|---|
| IVNQEKYVKITG | 1 | 0 | 0 | 1455845 | AVV62524.1 | K9N7C7 |
| IVNSSFTEQLGV | 1 | 0 | 0 | 1455849 | AWO68966.1 | A0A0P0G321 |
| IVPPCPSILKVI | 1 | 0 | 0 | 1455876 | NP_073549.1 | A0A0P0LKV0 |
| IVPSTAWEDGDY | 1 | 0 | 0 | 1455878 | AGV08584.1 | K9N5Q8 |
| IVPSTVWEEGDY | 1 | 0 | 0 | 1455882 | ALJ76277.1 | K9N5Q8 |
| IVQASESKVFLE | 1 | 0 | 0 | 1455890 | ARU07593.1 | H9EJ42 |
| IVQDEELALLRA | 1 | 0 | 0 | 1455894 | AVV62543.1 | K9N7A1 |
| IVQHNTEPKVPE | 1 | 0 | 0 | 1455897 | APO40578.1 | P0C6X7 |
| IVQPATSKSKRL | 1 | 0 | 0 | 1455907 | AUM60013.1 | K9N7C7 |
| IVRIVRTRSMTY | 1 | 0 | 0 | 1455929 | ARU07575.1 | A0A140H1G9 |
| IVRRDAAFATHV | 1 | 0 | 0 | 1455934 | ABE97136.1 | H9EJ42 |
| IVSIRSSNQGNK | 1 | 0 | 0 | 1455958 | AKQ21067.1 | K9N643 |
| IVSLPVYHKHTF | 1 | 0 | 0 | 1455961 | AIW52757.1 | A0A0P0K6L9 |
| IVTAGLDQTSFN | 1 | 0 | 0 | 1455984 | AVV62535.1 | K9N7C7 |
| IVTGLFKDCSRD | 1 | 0 | 0 | 1455991 | AVV62524.1 | K9N7C7 |
| IVTGNVVKQCCT | 1 | 0 | 0 | 1455997 | AVV62524.1 | K9N7C7 |
| IVTLPGSDASGD | 1 | 0 | 0 | 1456004 | ATQ39390.1 | K9N5Q8 |
| IVVDCATYVCNG | 1 | 0 | 0 | 1456015 | AIW52843.1 | A0A0P0G321 |
| IVVGDYVVISGK | 1 | 0 | 0 | 1456018 | AVV62535.1 | K9N7C7 |
| IVVQPHNGVLEI | 1 | 0 | 0 | 1456036 | BBA20983.1 | A0A140H1H1 |
| IVWGAKDFNTLS | 1 | 0 | 0 | 1456050 | AFD98777.1 | H9EJ42 |
| IVWPLTEYDFTD | 1 | 0 | 0 | 1456054 | QBF80519.1 | K9N7C7 |
| IVYEPSTPISSA | 1 | 0 | 0 | 1456065 | AUM60013.1 | K9N7C7 |
| IVYEPSTPVSSA | 1 | 0 | 0 | 1456066 | AVV62524.1 | K9N7C7 |
| IVYGDFSHQQLG | 1 | 0 | 0 | 1456072 | APO40578.1 | P0C6X7 |
| IVYHTLESPIEF | 1 | 0 | 0 | 1456077 | ARO76381.1 | P0C6U8 |
| IVYHTSGNVIEF | 1 | 0 | 0 | 1456079 | APO40578.1 | P0C6X7 |
| IVYVGDPAQLPT | 1 | 0 | 0 | 1456083 | ATG84864.1 | K9N7C7 |
| IWDFNREAPAYI | 1 | 0 | 0 | 1456088 | APO40578.1 | P0C6X7 |
| IWDYKRDAPAHI | 1 | 0 | 0 | 1456092 | QYC64967.1 | P0DTD1 |
| IWEAPHNDCTPI | 1 | 0 | 0 | 1456096 | AVV62526.1 | K9N5Q8 |
| IWHIAYIVAYFL | 1 | 0 | 0 | 1456107 | AWK59942.1 | H9EJ42 |
| IWLSHEKASLSS | 1 | 0 | 0 | 1456123 | AIV41990.1 | P0C6X6 |
| IWNKRFGFIEDS | 1 | 0 | 0 | 1456132 | AIL49507.1 | P36334 |
| IWSVDAFNQFSS | 1 | 0 | 0 | 1456154 | AEN19363.1 | P0C6X6 |
| IYCRAHVAHPTM | 1 | 0 | 0 | 1456185 | NP_073549.1 | A0A0P0LKV0 |
| IYDDNTNQYATL | 1 | 0 | 0 | 1456189 | ATQ39390.1 | K9N5Q8 |
| IYDEEDLYDVSK | 1 | 0 | 0 | 1456190 | QED88039.1 | H9EJ42 |
| IYDEPTPTTSVP | 1 | 0 | 0 | 1456194 | APO40580.1 | P59632 |
| IYDKKTKTHATL | 1 | 0 | 0 | 1456201 | AHL18090.1 | K9N5Q8 |
| IYDLAPIELDNK | 1 | 0 | 0 | 1456204 | AUM60013.1 | K9N7C7 |
| IYDRLDIIQADQ | 1 | 0 | 0 | 1456207 | AIW52757.1 | A0A0P0K6L9 |
| IYDSSDRGQECL | 1 | 0 | 0 | 1456214 | AZK15899.1 | K9N638 |
| IYDSSDRGQENL | 1 | 0 | 0 | 1456215 | AUM60013.1 | K9N7C7 |
| IYENTTYECDIP | 1 | 0 | 0 | 1456225 | APO40579.1 | P59594 |
| IYEWPMPIETSK | 1 | 0 | 0 | 1456231 | AUM60014.1 | K9N5Q8 |
| IYFLVRLLQDKT | 1 | 0 | 0 | 1456240 | AVN89451.1 | K9N7C7 |
| IYGAYQPYDHAI | 1 | 0 | 0 | 1456246 | ATO98145.1 | Q3LZX1 |
| IYGFNSTETITT | 1 | 0 | 0 | 1456247 | AIL49343.1 | P30215 |
| IYGLNSTETIST | 1 | 0 | 0 | 1456252 | AGT51729.1 | P30215 |
| IYGLNSTETITT | 1 | 0 | 0 | 1456253 | AEN19365.1 | P30215 |
| IYHSAFADCTSC | 1 | 0 | 0 | 1456272 | AUM60013.1 | K9N7C7 |
| IYKFDSGSLTKS | 1 | 0 | 0 | 1456289 | AVV62524.1 | K9N7C7 |
| IYKNGAMLNGKP | 1 | 0 | 0 | 1456298 | AVV62535.1 | K9N7C7 |
| IYKTPNIKDFGG | 1 | 0 | 0 | 1456302 | APO40579.1 | P59594 |
| IYKYDSGSLTKS | 1 | 0 | 0 | 1456306 | AVV62535.1 | K9N7C7 |
| IYLINKFLKANP | 1 | 0 | 0 | 1456311 | AEN19363.1 | P0C6X6 |
| IYLLFTHIPLDG | 1 | 0 | 0 | 1456317 | AVV62535.1 | K9N7C7 |
| IYLTNKISLNAS | 1 | 0 | 0 | 1456323 | AMM42653.1 | A0A0P0G321 |
| IYLTNNISLNAP | 1 | 0 | 0 | 1456324 | QEG59356.1 | A0A0P0G321 |
| IYNDKVAGFAKF | 1 | 0 | 0 | 1456334 | APO40578.1 | P0C6X7 |
| IYNGVTRDILST | 1 | 0 | 0 | 1456339 | APO40578.1 | P0C6X7 |
| IYNKLTEVKVPQ | 1 | 0 | 0 | 1456343 | AVV62524.1 | K9N7C7 |
| IYNLFNVGHYDG | 1 | 0 | 0 | 1456344 | ABD75583.1 | A0A140H1G9 |
| IYNLIKDCPAVA | 1 | 0 | 0 | 1456345 | APO40578.1 | P0C6X7 |
| IYPQGRTYSNIT | 1 | 0 | 0 | 1456370 | AXN92228.1 | K9N5Q8 |
| IYQLDENAQKFF | 1 | 0 | 0 | 1456377 | AMM42491.1 | A0A0P0ILY9 |
| IYRSDTLVLSQG | 1 | 0 | 0 | 1456387 | AVP78031.1 | Q3LZX1 |
| IYSKFTPVNLNR | 1 | 0 | 0 | 1456395 | AVP78031.1 | Q3LZX1 |
| IYVPEHCQNYSG | 1 | 0 | 0 | 1456428 | QAT98905.1 | K9N796 |
| IYVYNTGRSAYV | 1 | 0 | 0 | 1456434 | AVV62531.1 | K9N5R3 |
| KAAPKVVTKKDA | 1 | 0 | 0 | 1456457 | AVV62533.1 | K9N4V7 |
| KAASGSDSESTT | 1 | 0 | 0 | 1456460 | AVV62524.1 | K9N7C7 |
| KAAYENFNSQDT | 1 | 0 | 0 | 1456471 | ATO98229.1 | P0C6W2 |
| KACCYVGMQTVE | 1 | 0 | 0 | 1456474 | AVN89451.1 | K9N7C7 |
| KACKSIKDTLAY | 1 | 0 | 0 | 1456484 | AVV62524.1 | K9N7C7 |
| KACQNLDCNCLG | 1 | 0 | 0 | 1456490 | AWW13564.1 | Q80872 |
| KACSCIKDALSY | 1 | 0 | 0 | 1456491 | AVV62535.1 | K9N7C7 |
| KADDFVIASDTT | 1 | 0 | 0 | 1456495 | AVL25554.1 | H9EJ42 |
| KADELQALPQRQ | 1 | 0 | 0 | 1456497 | AVP78038.1 | Q3LZX4 |
| KADEVQPLPQRQ | 1 | 0 | 0 | 1456499 | APO40586.1 | P59595 |
| KADGIIYPTGKS | 1 | 0 | 0 | 1456502 | AVV62537.1 | K9N5Q8 |
| KADLVLDALASM | 1 | 0 | 0 | 1456506 | ARU07575.1 | A0A140H1G9 |
| KADQNKQESSDD | 1 | 0 | 0 | 1456508 | AVV62544.1 | K9N4V7 |
| KAEDIFGPCWSA | 1 | 0 | 0 | 1456511 | NP_073549.1 | A0A0P0LKV0 |
| KAEFDRESSVQK | 1 | 0 | 0 | 1456513 | NP_073549.1 | A0A0P0LKV0 |
| KAEGVIYPTGRT | 1 | 0 | 0 | 1456515 | AUM60014.1 | K9N5Q8 |
| KAFEFAEGQGVP | 1 | 0 | 0 | 1456527 | AIV41850.1 | P33469 |
| KAFGSPFITNGI | 1 | 0 | 0 | 1456528 | AGT51379.1 | H9EJ42 |
| KAGFSKYVTVQG | 1 | 0 | 0 | 1456543 | AVV62524.1 | K9N7C7 |
| KAGHYTGQAGEM | 1 | 0 | 0 | 1456548 | AQN78669.1 | P0C6X6 |

| | | | | | | |
|---|---|---|---|---|---|---|
| KAGNYRRPPVVL | 1 | 0 | 0 | 1456552 | AUM60030.1 | K9N7A1 |
| KAIDGGVTRDIA | 1 | 0 | 0 | 1456568 | AVP78041.1 | P0C6W2 |
| KAIGFFFPYEEG | 1 | 0 | 0 | 1456570 | AUM60013.1 | K9N7C7 |
| KAILNTVKFCDH | 1 | 0 | 0 | 1456578 | ANI69887.1 | K9N7C7 |
| KAINDFSNSGSD | 1 | 0 | 0 | 1456583 | AVP78030.1 | P0C6W2 |
| KAIQDGVTRDIL | 1 | 0 | 0 | 1456587 | ARO76381.1 | P0C6U8 |
| KAIQDGVTRDIV | 1 | 0 | 0 | 1456589 | P0C6W6.1 | P0C6W2 |
| KAIRCVPQAEVD | 1 | 0 | 0 | 1456590 | AVP78041.1 | P0C6W2 |
| KAIRCVPQSEVD | 1 | 0 | 0 | 1456591 | AVP78030.1 | P0C6W2 |
| KAIVAYTMSLGA | 1 | 0 | 0 | 1456594 | AVP78031.1 | Q3LZX1 |
| KAIVESCGNYRV | 1 | 0 | 0 | 1456595 | ARO76381.1 | P0C6U8 |
| KAIYNGVTRDIL | 1 | 0 | 0 | 1456597 | APO40578.1 | P0C6X7 |
| KAKRSTEKVLER | 1 | 0 | 0 | 1456610 | AUM60013.1 | K9N7C7 |
| KALCFNKPKQFV | 1 | 0 | 0 | 1456626 | ABD75512.1 | A0A140H1H0 |
| KALKELYGPIDP | 1 | 0 | 0 | 1456646 | ATG84722.1 | K9N7C7 |
| KALNDFSNLGAD | 1 | 0 | 0 | 1456658 | BAE93400.1 | P0C6U8 |
| KALSKLASELSN | 1 | 0 | 0 | 1456668 | AVV62526.1 | K9N5Q8 |
| KAMETFTNNNGS | 1 | 0 | 0 | 1456677 | AQN78669.1 | P0C6X6 |
| KANAIQQQFNVA | 1 | 0 | 0 | 1456693 | AUM60013.1 | K9N7C7 |
| KANEETFKLSYG | 1 | 0 | 0 | 1456694 | APO40578.1 | P0C6X7 |
| KANSGNRTPTSG | 1 | 0 | 0 | 1456704 | AJQ24081.1 | P33469 |
| KANTDIKLGVCV | 1 | 0 | 0 | 1456708 | AVQ05264.1 | P36334 |
| KAPKEESTDQIS | 1 | 0 | 0 | 1456718 | AQZ41303.1 | K9N4V7 |
| KAQEGCGIISRK | 1 | 0 | 0 | 1456739 | ATO98181.1 | Q3LZX1 |
| KARSHRGRAALI | 1 | 0 | 0 | 1456759 | APD51508.1 | A0A0P0KWE8 |
| KASDWKCKVTDV | 1 | 0 | 0 | 1456765 | AUM60013.1 | K9N7C7 |
| KATGDLAVCLGV | 1 | 0 | 0 | 1456789 | AIV41816.1 | P0C6X6 |
| KAVARKLERMAE | 1 | 0 | 0 | 1456809 | AVV62524.1 | K9N7C7 |
| KAVDCSSSFLSE | 1 | 0 | 0 | 1456811 | AZS52618.1 | A0A140H1H1 |
| KAVPSPLNWERK | 1 | 0 | 0 | 1456818 | AIV41909.1 | P36334 |
| KAVRNKISVCFN | 1 | 0 | 0 | 1456820 | AAR01012.1 | P0C6X6 |
| KAWAHCVKCHND | 1 | 0 | 0 | 1456830 | AUM60013.1 | K9N7C7 |
| KAYMDFVNAGGA | 1 | 0 | 0 | 1456848 | AVV62535.1 | K9N7C7 |
| KCCDDNYFNKKD | 1 | 0 | 0 | 1456864 | AVP78030.1 | P0C6W2 |
| KCDLQNYGDSAI | 1 | 0 | 0 | 1456883 | AVP78030.1 | P0C6W2 |
| KCDQLSFDVPDG | 1 | 0 | 0 | 1456886 | AMM42536.1 | A0A0P0K6L9 |
| KCEQFYQLYDAS | 1 | 0 | 0 | 1456901 | AWW13571.1 | P0C6X6 |
| KCFFNGSNDVWI | 1 | 0 | 0 | 1456912 | AZS52617.1 | A0A140H1H0 |
| KCFPNGSNDVWL | 1 | 0 | 0 | 1456917 | ARU07576.1 | A0A140H1H0 |
| KCGGCCDDYTGH | 1 | 0 | 0 | 1456922 | AIL49486.1 | P36334 |
| KCGVKQEQCTGL | 1 | 0 | 0 | 1456932 | ARE30011.1 | P0C6X6 |
| KCGVKQEQRTGV | 1 | 0 | 0 | 1456935 | AVI15051.1 | P0C6X6 |
| KCGVLDGSYNAS | 1 | 0 | 0 | 1456936 | BBA20983.1 | A0A140H1H1 |
| KCHNDILAATDP | 1 | 0 | 0 | 1456941 | AXN92247.1 | K9N7C7 |
| KCKVTDVIFPGQ | 1 | 0 | 0 | 1456969 | AXN92226.1 | K9N7C7 |
| KCKVTDVLFSGQ | 1 | 0 | 0 | 1456971 | AQZ41321.1 | K9N7C7 |
| KCLNVAKSEFDR | 1 | 0 | 0 | 1456980 | APO40578.1 | P0C6X7 |
| KCNLPFRLTTSK | 1 | 0 | 0 | 1457001 | AUM60013.1 | K9N7C7 |
| KCNRAGSNDIWM | 1 | 0 | 0 | 1457002 | AGT51729.1 | P30215 |
| KCNRCCDKYDDL | 1 | 0 | 0 | 1457003 | AVV62537.1 | K9N5Q8 |
| KCNRCCDKYDDY | 1 | 0 | 0 | 1457004 | AUM60024.1 | K9N5Q8 |
| KCNYVFNNTLSR | 1 | 0 | 0 | 1457011 | CAA83661.1 | P36334 |
| KCPLDSRTGSFN | 1 | 0 | 0 | 1457015 | AIX10756.1 | P36334 |
| KCPLDTSRKGSF | 1 | 0 | 0 | 1457019 | AKA09663.1 | P36334 |
| KCPQSKSIVGIG | 1 | 0 | 0 | 1457023 | ARB66065.1 | P36334 |
| KCSRFLSDDRTE | 1 | 0 | 0 | 1457038 | 6C6Z_A | K9N5Q8 |
| KCSRLTGDNSHI | 1 | 0 | 0 | 1457042 | AVV62537.1 | K9N5Q8 |
| KCSSNGSNDVWI | 1 | 0 | 0 | 1457043 | AGT17757.1 | A0A140H1H0 |
| KCSVKSHHHHHH | 1 | 0 | 0 | 1457051 | 5X4S_A | P59594 |
| KCTIKNFNVSKG | 1 | 0 | 0 | 1457055 | ATO98181.1 | Q3LZX1 |
| KCTKYNIYDVAG | 1 | 0 | 0 | 1457058 | AIW52741.1 | A0A0P0K6L9 |
| KCTKYNIYDVVG | 1 | 0 | 0 | 1457060 | BAL45638.1 | A0A0P0K6L9 |
| KCTLKSLSVQKG | 1 | 0 | 0 | 1457064 | AVP78042.1 | Q3LZX1 |
| KCTSDTSYINDV | 1 | 0 | 0 | 1457069 | ATP16757.1 | P36334 |
| KCTTKSLTVEKG | 1 | 0 | 0 | 1457071 | APO40579.1 | P59594 |
| KCVALCDAMVAK | 1 | 0 | 0 | 1457076 | AIW52827.1 | H9EJ42 |
| KCVRSREETGLL | 1 | 0 | 0 | 1457091 | AVP78041.1 | P0C6W2 |
| KDAIKPQQVTKH | 1 | 0 | 0 | 1457124 | AOL02457.1 | P33469 |
| KDALQSREAVLV | 1 | 0 | 0 | 1457126 | AXX83318.1 | P0C6X6 |
| KDAQVVQLPIQD | 1 | 0 | 0 | 1457133 | AEN19363.1 | P0C6X6 |
| KDASDFIDSVRD | 1 | 0 | 0 | 1457134 | AVP78031.1 | Q3LZX1 |
| KDAVLVNAANTH | 1 | 0 | 0 | 1457142 | AVV62524.1 | K9N7C7 |
| KDCSKIINGLHP | 1 | 0 | 0 | 1457151 | ATO98167.1 | P0C6W2 |
| KDCSKVITGLHP | 1 | 0 | 0 | 1457155 | QYC64967.1 | P0DTD1 |
| KDCSNSYSGYHP | 1 | 0 | 0 | 1457156 | AGT51577.1 | P0C6X6 |
| KDCSREISGLSP | 1 | 0 | 0 | 1457159 | AUM60013.1 | K9N7C7 |
| KDCVMYAITAIL | 1 | 0 | 0 | 1457163 | APO40578.1 | P0C6X7 |
| KDDLHQLYVDSK | 1 | 0 | 0 | 1457172 | AUM60013.1 | K9N7C7 |
| KDDQGHILDSYF | 1 | 0 | 0 | 1457174 | AVV62535.1 | K9N7C7 |
| KDDQGHNLDSYF | 1 | 0 | 0 | 1457175 | AVV62524.1 | K9N7C7 |
| KDDQIGYYRRAS | 1 | 0 | 0 | 1457176 | APO40586.1 | P59595 |
| KDDQIGYYRRAT | 1 | 0 | 0 | 1457177 | AVP78038.1 | Q3LZX4 |
| KDDYGTDIDGVY | 1 | 0 | 0 | 1457180 | AJQ24074.1 | P33469 |
| KDEAIKRVKGWV | 1 | 0 | 0 | 1457181 | ABD75519.1 | A0A140H1G9 |
| KDEISVTPIKNT | 1 | 0 | 0 | 1457185 | AST12962.1 | A0A0P0LKV0 |
| KDFGYVCDGSFY | 1 | 0 | 0 | 1457193 | ARU07575.1 | A0A140H1G9 |
| KDFKPQQVTKQN | 1 | 0 | 0 | 1457194 | ABD75549.1 | Q19U29 |
| KDFLSFDNDVFI | 1 | 0 | 0 | 1457197 | QBM11746.1 | K9N7C7 |
| KDFLSLDSDVFI | 1 | 0 | 0 | 1457200 | AVV62535.1 | K9N7C7 |
| KDFLSMDSDVFI | 1 | 0 | 0 | 1457201 | AVV62524.1 | K9N7C7 |
| KDFQFMLWCNDD | 1 | 0 | 0 | 1457207 | AVQ05261.1 | P0C6X6 |

| | | | | | | |
|---|---|---|---|---|---|---|
| KDFQFMLWCNDE | 1 | 0 | 0 | 1457208 | AEN19363.1 | P0C6X6 |
| KDFQFMLWCNDH | 1 | 0 | 0 | 1457209 | ABD75631.1 | A0A140H1G9 |
| KDFQFMLWGNDE | 1 | 0 | 0 | 1457211 | AIV41912.1 | P0C6X6 |
| KDGDYYRKQLSP | 1 | 0 | 0 | 1457219 | ATW75480.1 | K9N5Q8 |
| KDGFNLEYWVTP | 1 | 0 | 0 | 1457224 | ARK08651.1 | P36334 |
| KDGFSFVDDVNG | 1 | 0 | 0 | 1457226 | AVV62535.1 | K9N7C7 |
| KDGFSFVNDDTG | 1 | 0 | 0 | 1457227 | AVV62524.1 | K9N7C7 |
| KDGFSPEYWVTP | 1 | 0 | 0 | 1457229 | QEG03814.1 | P36334 |
| KDGFVSVELQPP | 1 | 0 | 0 | 1457231 | AXN92247.1 | K9N7C7 |
| KDGFYETIVSYW | 1 | 0 | 0 | 1457234 | AIV41848.1 | Q80872 |
| KDGGHFFVSGDY | 1 | 0 | 0 | 1457236 | ARE29978.1 | H9EJ42 |
| KDGITKLADVEA | 1 | 0 | 0 | 1457243 | AXN92247.1 | K9N7C7 |
| KDGIVWVHEEGA | 1 | 0 | 0 | 1457245 | AVV62533.1 | K9N4V7 |
| KDGIVWVHEHGA | 1 | 0 | 0 | 1457246 | AXN92267.1 | K9N4V7 |
| KDGIVWVHEYGA | 1 | 0 | 0 | 1457247 | QFQ59594.1 | K9N4V7 |
| KDGKVQTFYPRL | 1 | 0 | 0 | 1457252 | ATQ39389.1 | K9N7C7 |
| KDGKYFTSEPPI | 1 | 0 | 0 | 1457255 | QBF80553.1 | K9N7C7 |
| KDGNPIRDYAAL | 1 | 0 | 0 | 1457264 | AVV62524.1 | K9N7C7 |
| KDGNPIRDYASL | 1 | 0 | 0 | 1457265 | AVV62535.1 | K9N7C7 |
| KDGQINDMILSL | 1 | 0 | 0 | 1457266 | AVP78041.1 | P0C6W2 |
| KDGQVQTFYPRF | 1 | 0 | 0 | 1457267 | ALA49945.1 | K9N7C7 |
| KDGVEDTPPHYF | 1 | 0 | 0 | 1457273 | ATQ39389.1 | K9N7C7 |
| KDGVENTPPHFF | 1 | 0 | 0 | 1457274 | ATG84821.1 | K9N638 |
| KDGVENTPPHYF | 1 | 0 | 0 | 1457275 | AXN92247.1 | K9N7C7 |
| KDGVMHSEFPAI | 1 | 0 | 0 | 1457276 | AKA09651.1 | P36334 |
| KDGVMYSEFPAI | 1 | 0 | 0 | 1457277 | CAA79896.1 | P36334 |
| KDGVMYSEFPTI | 1 | 0 | 0 | 1457278 | AIL49548.1 | P36334 |
| KDGVNFNIDDIN | 1 | 0 | 0 | 1457279 | ARB66060.1 | P36334 |
| KDGVNFNVDDVN | 1 | 0 | 0 | 1457281 | CAA83661.1 | P36334 |
| KDIDITFLKKDV | 1 | 0 | 0 | 1457293 | AVP78041.1 | P0C6W2 |
| KDIGTYEMYVKW | 1 | 0 | 0 | 1457299 | BBA20983.1 | A0A140H1H1 |
| KDIKDAPALPKT | 1 | 0 | 0 | 1457301 | AVV62535.1 | K9N7C7 |
| KDIPEKVSLVSV | 1 | 0 | 0 | 1457308 | ATQ39389.1 | K9N7C7 |
| KDIPGGCAMPIM | 1 | 0 | 0 | 1457309 | AIW52757.1 | A0A0P0K6L9 |
| KDKKKKADEVQP | 1 | 0 | 0 | 1457323 | APO40586.1 | P59595 |
| KDKLQRIDWKST | 1 | 0 | 0 | 1457327 | AUM60013.1 | K9N7C7 |
| KDKNKLIGYWNI | 1 | 0 | 0 | 1457328 | APT69880.1 | A0A0N7J131 |
| KDLKFRQRSDGV | 1 | 0 | 0 | 1457344 | AIW52795.1 | H9EJ69 |
| KDLLARAGKASY | 1 | 0 | 0 | 1457346 | AVP78030.1 | P0C6W2 |
| KDLNANMSLADC | 1 | 0 | 0 | 1457349 | AST12962.1 | A0A0P0LKV0 |
| KDLREYVFKNKD | 1 | 0 | 0 | 1457353 | ATO98193.1 | Q3LZX1 |
| KDMQFDLYVNVY | 1 | 0 | 0 | 1457363 | AXN92247.1 | K9N7C7 |
| KDNAYYTQQPID | 1 | 0 | 0 | 1457369 | APO40578.1 | P0C6X7 |
| KDNQVNDMIYSL | 1 | 0 | 0 | 1457373 | APO40578.1 | P0C6X7 |
| KDNVIFVADDSG | 1 | 0 | 0 | 1457376 | AVN89451.1 | K9N7C7 |
| KDQSLVLTSNDE | 1 | 0 | 0 | 1457392 | AVV62535.1 | K9N7C7 |
| KDQTPIVWHAKD | 1 | 0 | 0 | 1457394 | NP_073549.1 | A0A0P0LKV0 |
| KDQYGSDINGVY | 1 | 0 | 0 | 1457396 | AIX10729.1 | P33469 |
| KDQYGTDIDGIY | 1 | 0 | 0 | 1457398 | AIL49434.1 | P33469 |
| KDQYGTDINGVY | 1 | 0 | 0 | 1457401 | AEN19370.1 | P33469 |
| KDSFTLYDQTVV | 1 | 0 | 0 | 1457406 | AIV41846.1 | P0C6X6 |
| KDSLIYKYDSGS | 1 | 0 | 0 | 1457412 | AVV62535.1 | K9N7C7 |
| KDSNFLNGSGVL | 1 | 0 | 0 | 1457415 | AVV62536.1 | K9N638 |
| KDSSGKPVPYCY | 1 | 0 | 0 | 1457417 | AVP78041.1 | P0C6W2 |
| KDTGPPPISTDT | 1 | 0 | 0 | 1457425 | CAA79896.1 | P36334 |
| KDTGPPSISTAT | 1 | 0 | 0 | 1457426 | ACV60203.1 | P36334 |
| KDTGVIYGFNST | 1 | 0 | 0 | 1457427 | AIL49343.1 | P30215 |
| KDTGVIYGLNFT | 1 | 0 | 0 | 1457428 | ATP16766.1 | P30215 |
| KDTNFLNGFGVR | 1 | 0 | 0 | 1457441 | ATP16763.1 | P0C6X6 |
| KDTNFLNGVRGA | 1 | 0 | 0 | 1457442 | AIV41780.1 | P0C6X6 |
| KDTNFLNRVRGA | 1 | 0 | 0 | 1457443 | AEN19363.1 | P0C6X6 |
| KDTNFLNRVRGS | 1 | 0 | 0 | 1457444 | AUM60013.1 | K9N7C7 |
| KDTVVQLNYTRD | 1 | 0 | 0 | 1457450 | AVV62535.1 | K9N7C7 |
| KDVKFENLTNNL | 1 | 0 | 0 | 1457467 | AUM60014.1 | K9N5Q8 |
| KDVNTHATRLTI | 1 | 0 | 0 | 1457469 | AIW52763.1 | A0A0P0LKV0 |
| KDVPYIVGDVIK | 1 | 0 | 0 | 1457472 | AVP78041.1 | P0C6W2 |
| KDVTFLEGDVHD | 1 | 0 | 0 | 1457476 | ARO76381.1 | P0C6U8 |
| KDWSYSGQSTEL | 1 | 0 | 0 | 1457496 | AVP78041.1 | P0C6W2 |
| KDYGFTMFVCID | 1 | 0 | 0 | 1457502 | AUM60013.1 | K9N7C7 |
| KEADMRNSSSEF | 1 | 0 | 0 | 1457516 | AVV62537.1 | K9N5Q8 |
| KEAQNANPMVIV | 1 | 0 | 0 | 1457518 | ARO76381.1 | P0C6U8 |
| KECVKSFIDVIN | 1 | 0 | 0 | 1457528 | ARO76381.1 | P0C6U8 |
| KEDDVDDSGDSS | 1 | 0 | 0 | 1457529 | AMM42440.1 | P0C6X6 |
| KEDGDVVLAEFD | 1 | 0 | 0 | 1457532 | AHI48794.1 | K9N638 |
| KEDLHTLYVDSK | 1 | 0 | 0 | 1457535 | ALA49340.1 | K9N7C7 |
| KEEDVQQAQQEA | 1 | 0 | 0 | 1457542 | AST12962.1 | A0A0P0LKV0 |
| KEEGDVLLAEFA | 1 | 0 | 0 | 1457545 | ATQ39389.1 | K9N7C7 |
| KEELDQWFKNQI | 1 | 0 | 0 | 1457549 | ANZ78835.1 | P36334 |
| KEELTDQMSEPP | 1 | 0 | 0 | 1457553 | AGV08397.1 | K9N4V7 |
| KEEPTTQKPVDV | 1 | 0 | 0 | 1457554 | ARO76381.1 | P0C6U8 |
| KEEQPKIEDSNV | 1 | 0 | 0 | 1457555 | ARO76381.1 | P0C6U8 |
| KEESTDQISEPP | 1 | 0 | 0 | 1457558 | AQZ41303.1 | K9N4V7 |
| KEESTDQMFEPP | 1 | 0 | 0 | 1457559 | QCZ25056.1 | K9N4V7 |
| KEESTDQVSEPP | 1 | 0 | 0 | 1457562 | AHI48535.1 | K9N4V7 |
| KEEVKPFFTENK | 1 | 0 | 0 | 1457565 | AVP78041.1 | P0C6W2 |
| KEFEFAEGQGVP | 1 | 0 | 0 | 1457573 | AEN19370.1 | P33469 |
| KEFQFAEGQGVP | 1 | 0 | 0 | 1457581 | QBP84717.1 | P33469 |
| KEFVEYKITEWP | 1 | 0 | 0 | 1457587 | AEN19363.1 | P0C6X6 |
| KEGALTAVVIPT | 1 | 0 | 0 | 1457592 | AVP78041.1 | P0C6W2 |
| KEGIEFLRDGWE | 1 | 0 | 0 | 1457600 | AVP78030.1 | P0C6W2 |
| KEGIIWVHEDGA | 1 | 0 | 0 | 1457602 | ATQ39393.1 | K9N4V7 |

| | | | | | | |
|---|---|---|---|---|---|---|
| KEGIVWVHEDGA | 1 | 0 | 0 | 1457607 | AVV62544.1 | K9N4V7 |
| KEGVMYSEFPAI | 1 | 0 | 0 | 1457621 | AMK59677.1 | P36334 |
| KEHRVQGTQRTR | 1 | 0 | 0 | 1457626 | YP_007188586.1 | K9N4V7 |
| KEIAHLNEIAKN | 1 | 0 | 0 | 1457627 | APO40579.1 | P59594 |
| KEIAPVYEADTE | 1 | 0 | 0 | 1457628 | ALA49362.1 | K9N7C7 |
| KEIDRLNEVARN | 1 | 0 | 0 | 1457631 | AVP78031.1 | Q3LZX1 |
| KEIGTYEMYVKW | 1 | 0 | 0 | 1457633 | BBA20986.1 | A0A140H1H1 |
| KEIISIPNIIKV | 1 | 0 | 0 | 1457638 | AIX09804.1 | P0C6X6 |
| KEIKDMQFELYV | 1 | 0 | 0 | 1457642 | AUM60013.1 | K9N7C7 |
| KEITRENVETLT | 1 | 0 | 0 | 1457656 | AGT51393.1 | H9EJ42 |
| KEKILFLPLNLV | 1 | 0 | 0 | 1457664 | AVV62535.1 | K9N7C7 |
| KEKLISLADVAT | 1 | 0 | 0 | 1457670 | AUM60013.1 | K9N7C7 |
| KELEQPIEEEDW | 1 | 0 | 0 | 1457686 | APO40578.1 | P0C6X7 |
| KELGLVMNMDVS | 1 | 0 | 0 | 1457688 | AXN92247.1 | K9N7C7 |
| KELGNYTYYNKE | 1 | 0 | 0 | 1457692 | 5X59_A | K9N5Q8 |
| KELGNYTYYNKG | 1 | 0 | 0 | 1457693 | 6NB3_A | K9N5Q8 |
| KELGVVMNFDVD | 1 | 0 | 0 | 1457695 | ACV87155.1 | A0A140H1G9 |
| KELKFRQRSDGV | 1 | 0 | 0 | 1457696 | AMM42628.1 | H9EJ69 |
| KELLDYYNMLVN | 1 | 0 | 0 | 1457700 | ATN39862.1 | P0C6X6 |
| KELVDGYGNTMY | 1 | 0 | 0 | 1457712 | AUM60027.1 | K9N643 |
| KELWHLDAGVVS | 1 | 0 | 0 | 1457713 | AKA09652.1 | P36334 |
| KELWHLDTGAVS | 1 | 0 | 0 | 1457714 | ANZ78840.1 | P36334 |
| KELWHLDTGVVS | 1 | 0 | 0 | 1457716 | CAA79896.1 | P36334 |
| KELWHSDTGVVS | 1 | 0 | 0 | 1457718 | BAM16460.1 | P36334 |
| KELYGPADPTFL | 1 | 0 | 0 | 1457721 | AHI48582.1 | K9N7C7 |
| KELYGSVDPTFL | 1 | 0 | 0 | 1457726 | AHC74096.1 | K9N7C7 |
| KENIDGDAMHAN | 1 | 0 | 0 | 1457737 | AVN89322.1 | K9N7C7 |
| KEPCSSGTYEGNSPFHP | 1 | 0 | 0 | 1457748 | P0DTC7.1 | P0DTC7 |
| KEPVHVYANDSI | 1 | 0 | 0 | 1457757 | QDH43722.1 | P0C6X6 |
| KEQRVQGGITQR | 1 | 0 | 0 | 1457779 | ANF29213.1 | K9N4V7 |
| KEQRVQGSTTQR | 1 | 0 | 0 | 1457784 | AQZ41292.1 | K9N4V7 |
| KEQRVQSSIIQR | 1 | 0 | 0 | 1457785 | QCZ25056.1 | K9N4V7 |
| KEQRVQSSITQR | 1 | 0 | 0 | 1457786 | ANI69896.1 | K9N4V7 |
| KEQYGTDIDGVY | 1 | 0 | 0 | 1457793 | ANZ78874.1 | P33469 |
| KERILYTFYGKD | 1 | 0 | 0 | 1457798 | AVV62524.1 | K9N7C7 |
| KESFEQHSYIYH | 1 | 0 | 0 | 1457808 | AUM60023.1 | K9N7C7 |
| KESIPIVWGAKD | 1 | 0 | 0 | 1457811 | AFD98777.1 | H9EJ42 |
| KESWHMDTGVVP | 1 | 0 | 0 | 1457825 | AIL49486.1 | P36334 |
| KETSYTSDIKPV | 1 | 0 | 0 | 1457839 | APO40578.1 | P0C6X7 |
| KETSYTTTIKPL | 1 | 0 | 0 | 1457840 | 5E6J_A | P0C6X7 |
| KETVFVEPKDGG | 1 | 0 | 0 | 1457846 | AWK59942.1 | H9EJ42 |
| KETVFVEPKNGG | 1 | 0 | 0 | 1457847 | ARU07593.1 | H9EJ42 |
| KEVKESVQTFFK | 1 | 0 | 0 | 1457863 | AVP78041.1 | P0C6W2 |
| KEVLRELYGTPD | 1 | 0 | 0 | 1457867 | AVV62524.1 | K9N7C7 |
| KEVLVLTGCCDN | 1 | 0 | 0 | 1457869 | AIW52827.1 | H9EJ42 |
| KEVRQKILNKPR | 1 | 0 | 0 | 1457872 | AEN19370.1 | P33469 |
| KEVSAAYYNDWQ | 1 | 0 | 0 | 1457873 | AZS52618.1 | A0A140H1H1 |
| KEVSAVYYNSWQ | 1 | 0 | 0 | 1457875 | BBA20983.1 | A0A140H1H1 |
| KEVTQKILNNPR | 1 | 0 | 0 | 1457878 | AJQ24083.1 | P33469 |
| KEWLKVQGFTLY | 1 | 0 | 0 | 1457881 | AUM60027.1 | K9N643 |
| KEYFDLQNCSFI | 1 | 0 | 0 | 1457887 | AVV62537.1 | K9N5Q8 |
| KFADDLNQLSGY | 1 | 0 | 0 | 1457892 | AVP78041.1 | P0C6W2 |
| KFADGINQMTGF | 1 | 0 | 0 | 1457895 | APO40578.1 | P0C6X7 |
| KFAEGQGVPIAP | 1 | 0 | 0 | 1457896 | ARA15425.1 | P33469 |
| KFAIPNRRKFDL | 1 | 0 | 0 | 1457900 | BBO58726.1 | P36334 |
| KFAVESNVMDVN | 1 | 0 | 0 | 1457911 | AFD98812.1 | H9EJ42 |
| KFCDHMVKDGLV | 1 | 0 | 0 | 1457920 | AVN89451.1 | K9N7C7 |
| KFCLEASFNYLK | 1 | 0 | 0 | 1457926 | QYC64967.1 | P0DTD1 |
| KFCSETVNGYAK | 1 | 0 | 0 | 1457928 | ARB07424.1 | A0A0P0ILY9 |
| KFDDYDFSVEDV | 1 | 0 | 0 | 1457931 | ARB07605.1 | A0A140H1G9 |
| KFDLQLGNLGYL | 1 | 0 | 0 | 1457940 | BBO58726.1 | P36334 |
| KFEFANDTKIAS | 1 | 0 | 0 | 1457954 | AVN89387.1 | K9N5Q8 |
| KFEHGRGPGKCY | 1 | 0 | 0 | 1457955 | ARA15429.1 | A0A0P0K6L9 |
| KFEHQRGPGKCY | 1 | 0 | 0 | 1457956 | AST12964.1 | A0A0P0K6L9 |
| KFELAKVQNLSG | 1 | 0 | 0 | 1457957 | AMM42445.1 | P33469 |
| KFENLTNQLPPP | 1 | 0 | 0 | 1457959 | AVV62537.1 | K9N5Q8 |
| KFFEIKHADYTL | 1 | 0 | 0 | 1457968 | AHF51980.1 | H9EJ42 |
| KFGFIEDSVFVP | 1 | 0 | 0 | 1457987 | ARB66070.1 | P36334 |
| KFGFTPLNKQRC | 1 | 0 | 0 | 1457988 | APT69896.1 | A0A0P0LKV0 |
| KFGIITKVEADF | 1 | 0 | 0 | 1457990 | AIV41992.1 | P30215 |
| KFGNTSFACLSN | 1 | 0 | 0 | 1458000 | QED88040.1 | A0A0P0G321 |
| KFGNTSFDFLSN | 1 | 0 | 0 | 1458001 | AWO68966.1 | A0A0P0G321 |
| KFGNTTFDFLSN | 1 | 0 | 0 | 1458002 | AWO69005.1 | A0A0P0G321 |
| KFGSDKIRMGQE | 1 | 0 | 0 | 1458004 | APO40579.1 | P59594 |
| KFGSVCFSLKDI | 1 | 0 | 0 | 1458005 | AIW52757.1 | A0A0P0K6L9 |
| KFIGTFESAAAG | 1 | 0 | 0 | 1458011 | AWK59942.1 | H9EJ42 |
| KFKTPHIVRQRL | 1 | 0 | 0 | 1458041 | AVN89322.1 | K9N7C7 |
| KFKVNSALEQYV | 1 | 0 | 0 | 1458043 | APO40578.1 | P0C6X7 |
| KFLADHFNYNVS | 1 | 0 | 0 | 1458045 | AWK59942.1 | H9EJ42 |
| KFLDGLDLTPVI | 1 | 0 | 0 | 1458048 | AWK59942.1 | H9EJ42 |
| KFLSNTKYFDHS | 1 | 0 | 0 | 1458072 | AIV41896.1 | P30215 |
| KFMSTSNAILIS | 1 | 0 | 0 | 1458088 | AVV62524.1 | K9N7C7 |
| KFNKWQWQEAWY | 1 | 0 | 0 | 1458098 | ARU07575.1 | A0A140H1G9 |
| KFNQALGAMLTG | 1 | 0 | 0 | 1458102 | QDP16206.1 | K9N5Q8 |
| KFNYQLPKVGYC | 1 | 0 | 0 | 1458113 | AXX83304.1 | Q04853 |
| KFPPPSFYMPLL | 1 | 0 | 0 | 1458124 | AIW52805.1 | H9EJ69 |
| KFQDSKPPLPPH | 1 | 0 | 0 | 1458134 | ANI69894.1 | K9N5R3 |
| KFQNLTNNLPPP | 1 | 0 | 0 | 1458140 | AUM60024.1 | K9N5Q8 |
| KFRNTTFDFLSN | 1 | 0 | 0 | 1458146 | AKA09625.1 | A0A0P0G321 |
| KFTIGNMEEFIQ | 1 | 0 | 0 | 1458182 | AIL49542.1 | P36334 |
| KFTPLKVEEVQL | 1 | 0 | 0 | 1458187 | AUM60013.1 | K9N7C7 |

| | | | | | | |
|---|---|---|---|---|---|---|
| KFTPLSVASTLV | 1 | 0 | 0 | 1458189 | ATG84722.1 | K9N7C7 |
| KFTPLSVEDLVP | 1 | 0 | 0 | 1458191 | AVV62524.1 | K9N7C7 |
| KFTPLSVESTPV | 1 | 0 | 0 | 1458193 | AHI48796.1 | K9N7C7 |
| KFTPLSVESTSV | 1 | 0 | 0 | 1458194 | AJG44112.1 | K9N638 |
| KFTVSTQPQENT | 1 | 0 | 0 | 1458197 | ABD75589.1 | Q19U29 |
| KFTVSTQPQGNS | 1 | 0 | 0 | 1458200 | AMM42386.1 | Q19U29 |
| KFVGTFESAAAG | 1 | 0 | 0 | 1458212 | NP_073549.1 | A0A0P0LKV0 |
| KFVVRHLGVTTS | 1 | 0 | 0 | 1458230 | AUM60013.1 | K9N7C7 |
| KFVYMHQLELST | 1 | 0 | 0 | 1458232 | AEN19363.1 | P0C6X6 |
| KFYGGWDDMLRP | 1 | 0 | 0 | 1458242 | AHF54667.1 | P0C6X6 |
| KFYNHAINGCKD | 1 | 0 | 0 | 1458256 | AVV62535.1 | K9N7C7 |
| KGAFLNGSCGSV | 1 | 0 | 0 | 1458270 | 3F9E_A | P0C6U8 |
| KGALEKFMSTSN | 1 | 0 | 0 | 1458280 | AVV62535.1 | K9N7C7 |
| KGCCSCGFCCKF | 1 | 0 | 0 | 1458292 | AVP78042.1 | Q3LZX1 |
| KGCPAVAVHDFF | 1 | 0 | 0 | 1458298 | AVP78030.1 | P0C6W2 |
| KGCVVDQSRKWI | 1 | 0 | 0 | 1458303 | ATP16755.1 | Q80872 |
| KGDFSFISDSNG | 1 | 0 | 0 | 1458308 | AUM60013.1 | K9N7C7 |
| KGDKLFVEHYNE | 1 | 0 | 0 | 1458313 | ARX11394.1 | A0A0P0LKV0 |
| KGECCICHKMQT | 1 | 0 | 0 | 1458325 | AWK59942.1 | H9EJ42 |
| KGEFIEQPTSVE | 1 | 0 | 0 | 1458329 | AVV62526.1 | K9N5Q8 |
| KGEGDGGIISEG | 1 | 0 | 0 | 1458330 | AST12962.1 | A0A0P0LKV0 |
| KGFCDFKGKYVQ | 1 | 0 | 0 | 1458336 | APO40578.1 | P0C6X7 |
| KGFKKGVMYSEF | 1 | 0 | 0 | 1458344 | AIL49539.1 | P36334 |
| KGFPQFAELIPN | 1 | 0 | 0 | 1458346 | AGT51335.1 | H9EJ69 |
| KGFSSDVLSDVI | 1 | 0 | 0 | 1458351 | CAA71146.1 | A0A0P0K6L9 |
| KGGAPTKVTFGDDTVIEV | 1 | 0 | 0 | 1458359 | P0DTC1.1 | P0DTD1 |
| KGGIRKAKDYGF | 1 | 0 | 0 | 1458372 | AVV62524.1 | K9N7C7 |
| KGGLILRFDSGT | 1 | 0 | 0 | 1458379 | AGV08389.1 | K9N638 |
| KGIFANCFSFVK | 1 | 0 | 0 | 1458402 | ATQ39389.1 | K9N7C7 |
| KGILTDCFSFVK | 1 | 0 | 0 | 1458411 | AXN92247.1 | K9N7C7 |
| KGIPIVDDLVVD | 1 | 0 | 0 | 1458416 | ASU91162.1 | K9N7C7 |
| KGIPIVEDPVVD | 1 | 0 | 0 | 1458418 | AUM60023.1 | K9N7C7 |
| KGIVAHPLGYIS | 1 | 0 | 0 | 1458423 | AVV62524.1 | K9N7C7 |
| KGKEFQFAEGQG | 1 | 0 | 0 | 1458435 | QBP84717.1 | P33469 |
| KGKFVQIPTQCT | 1 | 0 | 0 | 1458440 | AUM60013.1 | K9N7C7 |
| KGKQILGNTMLE | 1 | 0 | 0 | 1458451 | AUM60013.1 | K9N7C7 |
| KGKVFFQFENLS | 1 | 0 | 0 | 1458456 | AIV41978.1 | P0C6X6 |
| KGKYVQIPTTCV | 1 | 0 | 0 | 1458460 | AVP78030.1 | P0C6W2 |
| KGLEELHDDLVN | 1 | 0 | 0 | 1458466 | ATP16755.1 | Q80872 |
| KGLEELHYDLVN | 1 | 0 | 0 | 1458467 | AIV41836.1 | Q80872 |
| KGLIASKGPKQA | 1 | 0 | 0 | 1458470 | APO40578.1 | P0C6X7 |
| KGLQFQILDEGL | 1 | 0 | 0 | 1458476 | ARK08668.1 | Q80872 |
| KGLSKPFVKDNV | 1 | 0 | 0 | 1458483 | ATQ39389.1 | K9N7C7 |
| KGLSKPFVKGDF | 1 | 0 | 0 | 1458484 | AUM60013.1 | K9N7C7 |
| KGLTFSYDGLKG | 1 | 0 | 0 | 1458487 | AVV62535.1 | K9N7C7 |
| KGLWSPTITHTN | 1 | 0 | 0 | 1458491 | ARU07575.1 | A0A140H1G9 |
| KGPKQASLNGVT | 1 | 0 | 0 | 1458509 | APO40578.1 | P0C6X7 |
| KGPYEFCSQHTM | 1 | 0 | 0 | 1458514 | AVL25535.1 | H9EJ42 |
| KGQVSDLLVKLL | 1 | 0 | 0 | 1458520 | ALA49659.1 | K9N7C7 |
| KGSALNGSCGSV | 1 | 0 | 0 | 1458528 | 3F9G_A | P0C6X7 |
| KGSFNDRDTGPP | 1 | 0 | 0 | 1458532 | AIL49520.1 | P36334 |
| KGSFNNKDTGPP | 1 | 0 | 0 | 1458533 | AKA09663.1 | P36334 |
| KGSFNNRDTGPL | 1 | 0 | 0 | 1458537 | ANZ78836.1 | P36334 |
| KGSFVEGAINVG | 1 | 0 | 0 | 1458541 | AUM60013.1 | K9N7C7 |
| KGSITQRSRPRA | 1 | 0 | 0 | 1458548 | AVV62544.1 | K9N4V7 |
| KGSITQRSRPVK | 1 | 0 | 0 | 1458549 | AUM60021.1 | K9N4V7 |
| KGSKDEQIGYWN | 1 | 0 | 0 | 1458551 | AIW52795.1 | H9EJ69 |
| KGSLTASSFDGV | 1 | 0 | 0 | 1458553 | AUM60027.1 | K9N643 |
| KGSNLEEELLLM | 1 | 0 | 0 | 1458556 | AGT51707.1 | P0C6X6 |
| KGSPQQCIAPID | 1 | 0 | 0 | 1458557 | AVV62526.1 | K9N5Q8 |
| KGTDKLSTLWFK | 1 | 0 | 0 | 1458564 | AEN19366.1 | P36334 |
| KGTGVFVQDPSP | 1 | 0 | 0 | 1458570 | AFD98763.1 | H9EJ42 |
| KGTGVLTDSSKR | 1 | 0 | 0 | 1458571 | AVP78031.1 | Q3LZX1 |
| KGTVCWIMASTF | 1 | 0 | 0 | 1458582 | AEN19363.1 | P0C6X6 |
| KGVDCTKKFLTV | 1 | 0 | 0 | 1458592 | AXN92247.1 | K9N7C7 |
| KGVDYTKKFLIV | 1 | 0 | 0 | 1458594 | QBF80541.1 | K9N7C7 |
| KGVDYTKKQITV | 1 | 0 | 0 | 1458596 | AUM60013.1 | K9N7C7 |
| KGVDYTKKSLNV | 1 | 0 | 0 | 1458597 | ATQ39389.1 | K9N7C7 |
| KGVDYTKRQVIA | 1 | 0 | 0 | 1458598 | AVV62524.1 | K9N7C7 |
| KGVEKLVSWYIY | 1 | 0 | 0 | 1458601 | AGT51337.1 | H9EJ42 |
| KGVMYSEFPAIT | 1 | 0 | 0 | 1458612 | CAA83660.1 | P36334 |
| KGVQIPCMCGKQ | 1 | 0 | 0 | 1458613 | AVP78041.1 | P0C6W2 |
| KGVRCAVTDAKW | 1 | 0 | 0 | 1458615 | AWK59942.1 | H9EJ42 |
| KGVREGGLNKVK | 1 | 0 | 0 | 1458616 | AST12962.1 | A0A0P0LKV0 |
| KGVRKTVKIEDA | 1 | 0 | 0 | 1458618 | ARU07575.1 | A0A140H1G9 |
| KGVTFGENTVVE | 1 | 0 | 0 | 1458624 | APO40578.1 | P0C6X7 |
| KGVTIDKLRDYL | 1 | 0 | 0 | 1458625 | AXN92247.1 | K9N7C7 |
| KGVVGVLTLDNQ | 1 | 0 | 0 | 1458627 | AIW52827.1 | H9EJ42 |
| KGWVGFDVEGAH | 1 | 0 | 0 | 1458630 | ABD75519.1 | A0A140H1G9 |
| KGYWYPSNRGSF | 1 | 0 | 0 | 1458646 | AJQ24074.1 | P33469 |
| KHDIQLTTEGYN | 1 | 0 | 0 | 1458698 | AVV62535.1 | K9N7C7 |
| KHDIQLTVEGYN | 1 | 0 | 0 | 1458699 | AVV62524.1 | K9N7C7 |
| KHEGDWDSFGKT | 1 | 0 | 0 | 1458705 | AST12962.1 | A0A0P0LKV0 |
| KHEMSPFEMPFE | 1 | 0 | 0 | 1458712 | AST12962.1 | A0A0P0LKV0 |
| KHGGGVAGAFNK | 1 | 0 | 0 | 1458741 | QOJ87595.1 | P0DTD1 |
| KHGGGVAGALNA | 1 | 0 | 0 | 1458742 | APO40578.1 | P0C6X7 |
| KHILHVVGPDAR | 1 | 0 | 0 | 1458761 | AUM60013.1 | K9N7C7 |
| KHKFLFLQVFLL | 1 | 0 | 0 | 1458774 | AWK59942.1 | H9EJ42 |
| KHKGGDSIDFIA | 1 | 0 | 0 | 1458777 | AKK52590.1 | K9N7C7 |
| KHLIPLMSKGQK | 1 | 0 | 0 | 1458790 | ARU07574.1 | A0A140H1G9 |
| KHLLEQWNLVIG | 1 | 0 | 0 | 1458794 | APO40582.1 | P59596 |

| | | | | | | |
|---|---|---|---|---|---|---|
| KHLLPRWYFYYL | 1 | 0 | 0 | 1458795 | ABD75597.1 | Q19U29 |
| KHLNKYDCCLST | 1 | 0 | 0 | 1458797 | ARU07575.1 | A0A140H1G9 |
| KHLVPLIHKGAA | 1 | 0 | 0 | 1458804 | AHL18091.1 | K9N7C7 |
| KHLVPLMHKGAA | 1 | 0 | 0 | 1458805 | AXN92247.1 | K9N7C7 |
| KHNDFSSEIIGY | 1 | 0 | 0 | 1458812 | AVP78041.1 | P0C6W2 |
| KHPCMVGPDYAY | 1 | 0 | 0 | 1458817 | AHL18091.1 | K9N7C7 |
| KHQIIVLRDNFL | 1 | 0 | 0 | 1458823 | AVV62535.1 | K9N7C7 |
| KHQWNCFNCHSF | 1 | 0 | 0 | 1458826 | ARU07575.1 | A0A140H1G9 |
| KHRCDINYKGKV | 1 | 0 | 0 | 1458829 | QBP84746.1 | P0C6X6 |
| KHSDFGLGDLVDSYFEND | 1 | 0 | 0 | 1458838 | ALK43115.1 | A0A0P0LKV0 |
| KHTYQLRARSVS | 1 | 0 | 0 | 1458864 | ARO76387.1 | P59635 |
| KHVAYGCSKSGC | 1 | 0 | 0 | 1458865 | AEN19363.1 | P0C6X6 |
| KHVDHYDGFYTQ | 1 | 0 | 0 | 1458868 | AST12962.1 | A0A0P0LKV0 |
| KHVEKEGSYHNW | 1 | 0 | 0 | 1458870 | AVV62524.1 | K9N7C7 |
| KHVMHDNDSKEG | 1 | 0 | 0 | 1458877 | APO40578.1 | P0C6X7 |
| KHVNCCNATDRF | 1 | 0 | 0 | 1458878 | AKL87221.1 | H9EJ42 |
| KHVRAWIGFDVE | 1 | 0 | 0 | 1458887 | APO40578.1 | P0C6X7 |
| KHVRTNGDYNSW | 1 | 0 | 0 | 1458890 | QDI73607.1 | K9N7C7 |
| KHVTVFTDGRLN | 1 | 0 | 0 | 1458895 | AUM60013.1 | K9N7C7 |
| KHWDQQYHANCV | 1 | 0 | 0 | 1458900 | AUM60013.1 | K9N7C7 |
| KHWSMPYHPNPV | 1 | 0 | 0 | 1458903 | ANZ78865.1 | P0C6X6 |
| KIAQADTAGLYT | 1 | 0 | 0 | 1458921 | ASU89922.1 | K9N796 |
| KICDYVKPFLNH | 1 | 0 | 0 | 1458928 | APO40578.1 | P0C6X7 |
| KICGGCCDDYTG | 1 | 0 | 0 | 1458929 | CAA83660.1 | P36334 |
| KICKFGNTSFDF | 1 | 0 | 0 | 1458933 | AWO68991.1 | A0A0P0G321 |
| KICKFGTNTTFD | 1 | 0 | 0 | 1458935 | AVL25589.1 | A0A0P0G321 |
| KIDNETLEYWVT | 1 | 0 | 0 | 1458948 | AMM42407.1 | A0A140H1H1 |
| KIDTSGIHNRVI | 1 | 0 | 0 | 1458949 | AVV62527.1 | K9N796 |
| KIEDLSIRAFQK | 1 | 0 | 0 | 1458953 | ARB07411.1 | P0C6X6 |
| KIEDLSMRALQK | 1 | 0 | 0 | 1458956 | AJE26223.1 | P0C6X6 |
| KIEDSNVKEEPT | 1 | 0 | 0 | 1458957 | ARO76381.1 | P0C6U8 |
| KIENASPASTVH | 1 | 0 | 0 | 1458965 | ATO98194.1 | Q3LZX0 |
| KIEYKPDLSQYY | 1 | 0 | 0 | 1458968 | AEN19363.1 | P0C6X6 |
| KIFAYVKPFLGQ | 1 | 0 | 0 | 1458969 | ATO98143.1 | P0C6W2 |
| KIFIAACAVDVN | 1 | 0 | 0 | 1458971 | ARB07605.1 | A0A140H1G9 |
| KIFISNYANETE | 1 | 0 | 0 | 1458972 | AVV62526.1 | K9N5Q8 |
| KIFKARAIECYG | 1 | 0 | 0 | 1458973 | AMM42360.1 | A0A140H1G9 |
| KIFKDDGNFVVL | 1 | 0 | 0 | 1458975 | AMK59674.1 | P0C6X6 |
| KIFPTGFVLSIP | 1 | 0 | 0 | 1458977 | ABD75632.1 | A0A140H1H0 |
| KIGDLSIRALQK | 1 | 0 | 0 | 1458991 | AJE26218.1 | P0C6X6 |
| KIGPDHSVADYH | 1 | 0 | 0 | 1459003 | APO40578.1 | P0C6X7 |
| KIGTTEVKVSAL | 1 | 0 | 0 | 1459011 | ALD51903.1 | K9N7C7 |
| KIGVNVCNDSTF | 1 | 0 | 0 | 1459012 | ARB07436.1 | A0A140H1G9 |
| KIHDVVNQQGNS | 1 | 0 | 0 | 1459013 | ABB90526.1 | A0A0P0K6L9 |
| KIHHHHHHNTAS | 1 | 0 | 0 | 1459015 | 2OFZ_A | P59595 |
| KIINGLHPTQSP | 1 | 0 | 0 | 1459031 | APO40578.1 | P0C6X7 |
| KIISKHLDRNQH | 1 | 0 | 0 | 1459033 | APO40583.1 | P59634 |
| KIITTDNTFVSG | 1 | 0 | 0 | 1459036 | AVP78031.1 | Q3LZX1 |
| KIKHDGDWDSFC | 1 | 0 | 0 | 1459042 | AGT21366.1 | A0A0P0LKV0 |
| KIKHEGDWDSLC | 1 | 0 | 0 | 1459045 | QEO75984.1 | A0A0P0ILY9 |
| KIKNYGYLDIHP | 1 | 0 | 0 | 1459053 | AGT17757.1 | A0A140H1H0 |
| KIKPHSTHEGKT | 1 | 0 | 0 | 1459057 | APO40578.1 | P0C6X7 |
| KIKPYERDISNV | 1 | 0 | 0 | 1459059 | APO40579.1 | P59594 |
| KIKVSMPTVIAK | 1 | 0 | 0 | 1459061 | APO40578.1 | P0C6X7 |
| KILARAQSSETK | 1 | 0 | 0 | 1459066 | APT69860.1 | A0A0N7J131 |
| KILARSQSSETK | 1 | 0 | 0 | 1459067 | AIW52722.1 | A0A0N7J131 |
| KILGNVFCDGID | 1 | 0 | 0 | 1459073 | ARU07575.1 | A0A140H1G9 |
| KILPFSDRPLNL | 1 | 0 | 0 | 1459085 | AWK59942.1 | H9EJ42 |
| KILQNLGVDIAA | 1 | 0 | 0 | 1459086 | APO40578.1 | P0C6X7 |
| KILSMIRGPPRA | 1 | 0 | 0 | 1459089 | AGT51409.1 | P0C6X6 |
| KILTLTPRNKWN | 1 | 0 | 0 | 1459092 | ASU90779.1 | K9N638 |
| KILYKGSVTHDA | 1 | 0 | 0 | 1459096 | AUM60023.1 | K9N7C7 |
| KINQALHGANLK | 1 | 0 | 0 | 1459111 | AUM60014.1 | K9N5Q8 |
| KINRALHGANLR | 1 | 0 | 0 | 1459113 | AID55087.1 | K9N5Q8 |
| KINSACRKVQHM | 1 | 0 | 0 | 1459115 | P0C6W6.1 | P0C6W2 |
| KINTGNGIKQLA | 1 | 0 | 0 | 1459116 | AXN92267.1 | K9N4V7 |
| KINTTLLDLSTE | 1 | 0 | 0 | 1459122 | AVV62537.1 | K9N5Q8 |
| KIPANIVLSQGG | 1 | 0 | 0 | 1459124 | AJC98133.1 | P0C6X6 |
| KIQAVVNADAEA | 1 | 0 | 0 | 1459131 | CAA83661.1 | P36334 |
| KIQESLTSTASA | 1 | 0 | 0 | 1459139 | AVP78031.1 | Q3LZX1 |
| KIQTAPHISLAM | 1 | 0 | 0 | 1459147 | NP_150075.1 | Q80872 |
| KIQTAPHISLTM | 1 | 0 | 0 | 1459148 | AWW13564.1 | Q80872 |
| KIQVVNSGPDQI | 1 | 0 | 0 | 1459150 | ARB07411.1 | P0C6X6 |
| KIQVVNSGSDIN | 1 | 0 | 0 | 1459151 | P0C6U5.1 | A0A140H1G9 |
| KIRELTAEDISL | 1 | 0 | 0 | 1459154 | AGT51784.1 | P33469 |
| KIRMGQESIVAY | 1 | 0 | 0 | 1459157 | APO40579.1 | P59594 |
| KISDCIADYTVF | 1 | 0 | 0 | 1459165 | AVP78031.1 | Q3LZX1 |
| KISGKGQQQQGQ | 1 | 0 | 0 | 1459170 | AVP78049.1 | Q3LZX4 |
| KISLKGGKVIST | 1 | 0 | 0 | 1459175 | ARO76381.1 | P0C6U8 |
| KISSKAGNSIFR | 1 | 0 | 0 | 1459183 | AEN19365.1 | P30215 |
| KISWAGAKIRAV | 1 | 0 | 0 | 1459188 | AUM60023.1 | K9N7C7 |
| KITEDGINVKDV | 1 | 0 | 0 | 1459192 | AWK59942.1 | H9EJ42 |
| KITEELQHEDLL | 1 | 0 | 0 | 1459193 | APO40578.1 | P0C6X7 |
| KITEYSWSADLY | 1 | 0 | 0 | 1459198 | AVV62535.1 | K9N7C7 |
| KITIADPGYMQG | 1 | 0 | 0 | 1459200 | AUM60014.1 | K9N5Q8 |
| KITQWPTATGDV | 1 | 0 | 0 | 1459205 | AVQ05261.1 | P0C6X6 |
| KIVARAMLKCVA | 1 | 0 | 0 | 1459208 | AIW52827.1 | H9EJ42 |
| KIVGGAHTWFNA | 1 | 0 | 0 | 1459218 | ALA49340.1 | K9N7C7 |
| KIVGGAPSWVTA | 1 | 0 | 0 | 1459220 | AUM60013.1 | K9N7C7 |
| KIVQWIKNTFSL | 1 | 0 | 0 | 1459235 | ARK08667.1 | P0C6X6 |
| KIVTLPGSDASG | 1 | 0 | 0 | 1459241 | ATQ39390.1 | K9N5Q8 |

| | | | | | | |
|---|---|---|---|---|---|---|
| KIVYHTSGNVIE | 1 | 0 | 0 | 1459247 | APO40578.1 | P0C6X7 |
| KIYEIERGLLHS | 1 | 0 | 0 | 1459252 | AGT51737.1 | P0C6X6 |
| KIYSKFTPVNLN | 1 | 0 | 0 | 1459266 | AVP78031.1 | Q3LZX1 |
| KKAGGTTEMLAK | 1 | 0 | 0 | 1459274 | AVP78041.1 | P0C6W2 |
| KKCGCCCDDYTG | 1 | 0 | 0 | 1459284 | AIX09807.1 | P36334 |
| KKDKKKKADELQ | 1 | 0 | 0 | 1459303 | AVP78038.1 | Q3LZX4 |
| KKDNAYYTQQPI | 1 | 0 | 0 | 1459311 | APO40578.1 | P0C6X7 |
| KKDVPYIVGDVI | 1 | 0 | 0 | 1459313 | AVP78041.1 | P0C6W2 |
| KKEAAAAKNKMR | 1 | 0 | 0 | 1459315 | AXG21672.1 | K9N4V7 |
| KKEFSGQCFKIL | 1 | 0 | 0 | 1459316 | AVN89322.1 | K9N7C7 |
| KKEIAHLNEIAK | 1 | 0 | 0 | 1459317 | APO40579.1 | P59594 |
| KKENKQKAPKEE | 1 | 0 | 0 | 1459325 | AHL18098.1 | K9N4V7 |
| KKEPLLYVPIRL | 1 | 0 | 0 | 1459326 | AXN92247.1 | K9N7C7 |
| KKFLRYINGFNV | 1 | 0 | 0 | 1459340 | AIV41839.1 | Q04853 |
| KKGAKLLHKPIL | 1 | 0 | 0 | 1459350 | AID16715.1 | P0C6W2 |
| KKGAWNIGEQSS | 1 | 0 | 0 | 1459353 | AVP78030.1 | P0C6W2 |
| KKGVMYSEFPAI | 1 | 0 | 0 | 1459357 | CAA83660.1 | P36334 |
| KKHPCMVGPDYA | 1 | 0 | 0 | 1459361 | AHL18091.1 | K9N7C7 |
| KKHVEVFTDGRL | 1 | 0 | 0 | 1459365 | AVN89333.1 | K9N7C7 |
| KKIYPAFMLGSS | 1 | 0 | 0 | 1459385 | AVV62526.1 | K9N5Q8 |
| KKKADEVQPLPQ | 1 | 0 | 0 | 1459389 | APO40586.1 | P59595 |
| KKKKADELQALP | 1 | 0 | 0 | 1459391 | AVP78038.1 | Q3LZX4 |
| KKKKADEVQPLP | 1 | 0 | 0 | 1459393 | APO40586.1 | P59595 |
| KKLFVANYSQDV | 1 | 0 | 0 | 1459411 | ALA49649.1 | K9N5Q8 |
| KKLKKCLNVAKS | 1 | 0 | 0 | 1459418 | APO40578.1 | P0C6X7 |
| KKLSPSREDKVC | 1 | 0 | 0 | 1459432 | APO40587.1 | P59636 |
| KKLVGGNFTPID | 1 | 0 | 0 | 1459437 | AUM60013.1 | K9N7C7 |
| KKMDEPFTEDTS | 1 | 0 | 0 | 1459442 | AEN19370.1 | P33469 |
| KKNFTYNVSTDF | 1 | 0 | 0 | 1459448 | AGW27863.1 | A0A140H1H1 |
| KKNFTYNVSTDG | 1 | 0 | 0 | 1459449 | ABA60800.1 | A0A140H1H1 |
| KKNFTYNVSTDW | 1 | 0 | 0 | 1459450 | BBA20983.1 | A0A140H1H1 |
| KKNLDEARFSGT | 1 | 0 | 0 | 1459454 | AGT51787.1 | P0C6X6 |
| KKNYSELDDEEL | 1 | 0 | 0 | 1459462 | AAP30035.1 | P59634 |
| KKPFKVEDSVII | 1 | 0 | 0 | 1459469 | AXX83318.1 | P0C6X6 |
| KKQIGLLEPVKH | 1 | 0 | 0 | 1459487 | AVV62535.1 | K9N7C7 |
| KKQIGLLEPVNH | 1 | 0 | 0 | 1459488 | AVV62524.1 | K9N7C7 |
| KKQITVNGVQYY | 1 | 0 | 0 | 1459489 | AUM60013.1 | K9N7C7 |
| KKRAVKAASGSD | 1 | 0 | 0 | 1459508 | AVV62524.1 | K9N7C7 |
| KKRNVDLLSNFE | 1 | 0 | 0 | 1459510 | AKN58609.1 | K9N638 |
| KKRVEVFTDGKL | 1 | 0 | 0 | 1459513 | AQZ41341.1 | K9N7C7 |
| KKRWQLALSKGV | 1 | 0 | 0 | 1459515 | AVP78043.1 | Q3LZX0 |
| KKSFEQHSYIYH | 1 | 0 | 0 | 1459519 | AUM60013.1 | K9N7C7 |
| KKSFGKSIIYTS | 1 | 0 | 0 | 1459521 | AWK59972.1 | H9EJ42 |
| KKSTELVKNKCV | 1 | 0 | 0 | 1459530 | APO40579.1 | P59594 |
| KKVAAIPYKACK | 1 | 0 | 0 | 1459548 | AVV62524.1 | K9N7C7 |
| KKVAFCGDQVHE | 1 | 0 | 0 | 1459552 | ALT66879.1 | K9N7C7 |
| KKVPIDEYITTY | 1 | 0 | 0 | 1459570 | AER30328.1 | P0C6X7 |
| KKWKIPQVGGLT | 1 | 0 | 0 | 1459586 | BAE93400.1 | P0C6U8 |
| KKYGLEKYAFEH | 1 | 0 | 0 | 1459589 | AVN89352.1 | K9N7C7 |
| KKYKLEDYAFEH | 1 | 0 | 0 | 1459592 | AUM60013.1 | K9N7C7 |
| KKYPCIVGPDYA | 1 | 0 | 0 | 1459598 | AXN92258.1 | K9N7C7 |
| KKYSELDDEEPM | 1 | 0 | 0 | 1459600 | Q3I5J1.1 | Q3LZX8 |
| KLADVEADIASR | 1 | 0 | 0 | 1459611 | ATQ39389.1 | K9N7C7 |
| KLAGVSCIWSVD | 1 | 0 | 0 | 1459614 | AGT51737.1 | P0C6X6 |
| KLAKCYKDSVLV | 1 | 0 | 0 | 1459616 | AVV62535.1 | K9N7C7 |
| KLARWSASIDTG | 1 | 0 | 0 | 1459625 | QRK03775.1 | A0A0P0K6L9 |
| KLASPMKSDEGG | 1 | 0 | 0 | 1459628 | AZS52616.1 | A0A140H1G9 |
| KLASTEELPDEF | 1 | 0 | 0 | 1459629 | AAT76156.1 | Q3LZX3 |
| KLATIEELPDEF | 1 | 0 | 0 | 1459630 | APO40587.1 | P59636 |
| KLCDEMLNNRAT | 1 | 0 | 0 | 1459634 | APO40578.1 | P0C6X7 |
| KLCHAVVSKSKE | 1 | 0 | 0 | 1459642 | AEN19363.1 | P0C6X6 |
| KLCHAVVYKSKE | 1 | 0 | 0 | 1459643 | AIV41786.1 | P0C6X6 |
| KLCMQLCGFCNF | 1 | 0 | 0 | 1459650 | AGT17765.1 | Q0ZJ83 |
| KLDANIPDYPKL | 1 | 0 | 0 | 1459653 | AVV62535.1 | K9N7C7 |
| KLDANIPGYPKL | 1 | 0 | 0 | 1459654 | AVV62524.1 | K9N7C7 |
| KLDGSLCVGNGP | 1 | 0 | 0 | 1459668 | CAA83660.1 | P36334 |
| KLDGYYKKDNSY | 1 | 0 | 0 | 1459676 | AVP78041.1 | P0C6W2 |
| KLDNPNSSEVDM | 1 | 0 | 0 | 1459686 | AOL02450.1 | P0C6X6 |
| KLDNPSGSEVDM | 1 | 0 | 0 | 1459687 | AGT51717.1 | P0C6X6 |
| KLDNPSSSEVDM | 1 | 0 | 0 | 1459688 | AEN19363.1 | P0C6X6 |
| KLDSKNPNYQKW | 1 | 0 | 0 | 1459694 | AUM60021.1 | K9N4V7 |
| KLEDIMEVDISQ | 1 | 0 | 0 | 1459703 | AUM60017.1 | K9N643 |
| KLETDGICLENH | 1 | 0 | 0 | 1459727 | ARK08667.1 | P0C6X6 |
| KLETDRICPENH | 1 | 0 | 0 | 1459730 | AXX83372.1 | P0C6X6 |
| KLFAAESQKATE | 1 | 0 | 0 | 1459732 | AIV41834.1 | P0C6X6 |
| KLFAAETLKANE | 1 | 0 | 0 | 1459733 | APO40578.1 | P0C6X7 |
| KLFCAFITKRIV | 1 | 0 | 0 | 1459737 | AEN19363.1 | P0C6X6 |
| KLFVRCEEDGQT | 1 | 0 | 0 | 1459769 | AVV62535.1 | K9N7C7 |
| KLFVSDYATKVV | 1 | 0 | 0 | 1459775 | AVV62537.1 | K9N5Q8 |
| KLGAEKEDGFYE | 1 | 0 | 0 | 1459777 | AMK59675.1 | Q80872 |
| KLGAENKDGFYE | 1 | 0 | 0 | 1459779 | AIV41955.1 | Q80872 |
| KLGDTQGMDNLA | 1 | 0 | 0 | 1459785 | AVP78041.1 | P0C6W2 |
| KLGERVRKAILN | 1 | 0 | 0 | 1459789 | ANI69887.1 | K9N7C7 |
| KLGNHFKESWHM | 1 | 0 | 0 | 1459805 | AIL49486.1 | P36334 |
| KLGRELNCNVCA | 1 | 0 | 0 | 1459809 | ABD75615.1 | A0A140H1G9 |
| KLGTSDSQFPIL | 1 | 0 | 0 | 1459824 | AGT51764.1 | P33469 |
| KLHALDDATRNY | 1 | 0 | 0 | 1459828 | AHL18094.1 | K9N643 |
| KLHCEHLQFGLQ | 1 | 0 | 0 | 1459831 | AWO68974.1 | A0A0P0G321 |
| KLHDLDDATRSY | 1 | 0 | 0 | 1459834 | AVV62540.1 | K9N643 |
| KLHNWNCINCDT | 1 | 0 | 0 | 1459839 | AVP78041.1 | P0C6W2 |
| KLIASAFNNALH | 1 | 0 | 0 | 1459849 | AIL49544.1 | P36334 |

| | | | | | | |
|---|---|---|---|---|---|---|
| KLIFEFEDDVIT | 1 | 0 | 0 | 1459854 | AWK59942.1 | H9EJ42 |
| KLIQLCFTCHYF | 1 | 0 | 0 | 1459880 | AAV67329.1 | H9EJA2 |
| KLISLADVATDV | 1 | 0 | 0 | 1459881 | AUM60013.1 | K9N7C7 |
| KLISLTDVATDV | 1 | 0 | 0 | 1459882 | AUM60023.1 | K9N7C7 |
| KLKCNRCCDNYD | 1 | 0 | 0 | 1459896 | AUM60014.1 | K9N5Q8 |
| KLKCSDFYADKI | 1 | 0 | 0 | 1459898 | ARU07575.1 | A0A140H1G9 |
| KLKDCVMYASAV | 1 | 0 | 0 | 1459899 | AVP78041.1 | P0C6W2 |
| KLKDGVNFNIDD | 1 | 0 | 0 | 1459900 | ARB66060.1 | P36334 |
| KLKKSLNVAKSEFDRD | 1 | 0 | 0 | 1459917 | P0DTC1.1 | P0DTD1 |
| KLKLTYNKQMAN | 1 | 0 | 0 | 1459920 | AEN19363.1 | P0C6X6 |
| KLLDICFEATEA | 1 | 0 | 0 | 1459945 | ASU45805.1 | K9N7C7 |
| KLLDVYNGFLET | 1 | 0 | 0 | 1459948 | AWK59942.1 | H9EJ42 |
| KLLRGMPIPDFD | 1 | 0 | 0 | 1459994 | AXN92247.1 | K9N7C7 |
| KLLRGVELEDFN | 1 | 0 | 0 | 1459997 | AVV62535.1 | K9N7C7 |
| KLLSKKDASAQC | 1 | 0 | 0 | 1459999 | AUM60013.1 | K9N7C7 |
| KLMFKASALFKE | 1 | 0 | 0 | 1460016 | AUM60013.1 | K9N7C7 |
| KLNCTFQEAAAN | 1 | 0 | 0 | 1460032 | AVV62535.1 | K9N7C7 |
| KLNGSCPGKNNG | 1 | 0 | 0 | 1460044 | AEN19366.1 | P36334 |
| KLNISSGSRGVY | 1 | 0 | 0 | 1460050 | ATO98145.1 | Q3LZX1 |
| KLNLPNCNIEAW | 1 | 0 | 0 | 1460052 | AAR01015.1 | P36334 |
| KLNPSEDFIKHI | 1 | 0 | 0 | 1460055 | ASU90174.1 | K9N638 |
| KLNPSEEFIKHV | 1 | 0 | 0 | 1460057 | ALW82730.1 | K9N7C7 |
| KLNSSLCVGSGP | 1 | 0 | 0 | 1460063 | AVQ05264.1 | P36334 |
| KLPNGVTVADEP | 1 | 0 | 0 | 1460090 | AST12968.1 | A0A0N7J131 |
| KLPPHDVIVLRD | 1 | 0 | 0 | 1460092 | AFD98800.1 | H9EJ24 |
| KLPPPLLSNQTD | 1 | 0 | 0 | 1460094 | AVV62526.1 | K9N5Q8 |
| KLPSEPRTIQPE | 1 | 0 | 0 | 1460101 | AMQ49036.1 | K9N638 |
| KLPSSSGTSTPK | 1 | 0 | 0 | 1460102 | AIW52795.1 | H9EJ69 |
| KLQAGSNTEFAA | 1 | 0 | 0 | 1460106 | AVV62524.1 | K9N7C7 |
| KLQDVINQNAVA | 1 | 0 | 0 | 1460109 | APO40579.1 | P59594 |
| KLQGVLEVSVCQ | 1 | 0 | 0 | 1460122 | AMM42485.1 | P36334 |
| KLQNNEIKPTGL | 1 | 0 | 0 | 1460126 | AUM60013.1 | K9N7C7 |
| KLQNNEIMPGKM | 1 | 0 | 0 | 1460128 | NP_073549.1 | A0A0P0LKV0 |
| KLQNNEIRPTGL | 1 | 0 | 0 | 1460129 | AVV62524.1 | K9N7C7 |
| KLQSGVGRCFNC | 1 | 0 | 0 | 1460137 | AGT21338.1 | A0A0P0K6L9 |
| KLQSKRTRLFKS | 1 | 0 | 0 | 1460140 | AGT51787.1 | P0C6X6 |
| KLQSSHAWQPGV | 1 | 0 | 0 | 1460143 | AVP78041.1 | P0C6W2 |
| KLRANDNILSVR | 1 | 0 | 0 | 1460150 | AXN92247.1 | K9N7C7 |
| KLRDYLADYDIA | 1 | 0 | 0 | 1460153 | AQZ41321.1 | K9N7C7 |
| KLRDYLADYDVV | 1 | 0 | 0 | 1460156 | AVN89363.1 | K9N7C7 |
| KLRQYLAAYDIG | 1 | 0 | 0 | 1460166 | AVV62535.1 | K9N7C7 |
| KLRSETLLPITQ | 1 | 0 | 0 | 1460173 | APO40578.1 | P0C6X7 |
| KLSDALKRQIRI | 1 | 0 | 0 | 1460193 | AXN92247.1 | K9N7C7 |
| KLSDCLYFKNLK | 1 | 0 | 0 | 1460195 | AGT51737.1 | P0C6X6 |
| KLSDNNCYFNAV | 1 | 0 | 0 | 1460198 | ASU90086.1 | K9N638 |
| KLSDVGKGIYQT | 1 | 0 | 0 | 1460200 | ATO98145.1 | Q3LZX1 |
| KLSGYTIRWRFN | 1 | 0 | 0 | 1460208 | AVV62528.1 | K9N4V0 |
| KLSHHSDLEVTS | 1 | 0 | 0 | 1460210 | APO40578.1 | P0C6X7 |
| KLSPWFMDGQNA | 1 | 0 | 0 | 1460225 | AVV62524.1 | K9N7C7 |
| KLSSLSMIKGPQ | 1 | 0 | 0 | 1460228 | ARB07436.1 | A0A140H1G9 |
| KLTPIKLNFGML | 1 | 0 | 0 | 1460270 | AWK59978.1 | H9EJ42 |
| KLTYNKQMANVS | 1 | 0 | 0 | 1460273 | AEN19363.1 | P0C6X6 |
| KLVFLAIDFITW | 1 | 0 | 0 | 1460279 | QBP84714.1 | Q04853 |
| KLVGGDFTPIDQ | 1 | 0 | 0 | 1460284 | AVV62524.1 | K9N7C7 |
| KLVGGNFTPIDQ | 1 | 0 | 0 | 1460285 | AUM60013.1 | K9N7C7 |
| KLVGHTVCDSLN | 1 | 0 | 0 | 1460287 | AGT51757.1 | P0C6X6 |
| KLVKDVDHVQIH | 1 | 0 | 0 | 1460292 | Q3I5J4.1 | Q3LZX0 |
| KLVTTSTNPEFV | 1 | 0 | 0 | 1460312 | AUM60013.1 | K9N7C7 |
| KLWFDFIENPSV | 1 | 0 | 0 | 1460317 | AXN92226.1 | K9N7C7 |
| KLWFDFVENPGI | 1 | 0 | 0 | 1460318 | AVV62524.1 | K9N7C7 |
| KLWTQCVQLHND | 1 | 0 | 0 | 1460324 | APO40578.1 | P0C6X7 |
| KLYRLVWYVERK | 1 | 0 | 0 | 1460342 | ATQ39389.1 | K9N7C7 |
| KLYVSNYSHEVN | 1 | 0 | 0 | 1460345 | AUM60014.1 | K9N5Q8 |
| KMADQTMTQMYK | 1 | 0 | 0 | 1460350 | AVP78041.1 | P0C6W2 |
| KMCLFRYQEFND | 1 | 0 | 0 | 1460360 | QED88024.1 | H9EJ42 |
| KMCLFSYQEFND | 1 | 0 | 0 | 1460361 | AWK59942.1 | H9EJ42 |
| KMDEPFTEDTSD | 1 | 0 | 0 | 1460365 | ANZ78880.1 | P33469 |
| KMDEPFTEHTSE | 1 | 0 | 0 | 1460367 | AHA80186.1 | P33469 |
| KMDEPFTVDTSE | 1 | 0 | 0 | 1460368 | AIX10719.1 | P33469 |
| KMDGDEVYLPYP | 1 | 0 | 0 | 1460370 | AIL49620.1 | P0C6X6 |
| KMFDSFVNSFVL | 1 | 0 | 0 | 1460381 | ANI69899.1 | K9N7C7 |
| KMGVLKADDFVI | 1 | 0 | 0 | 1460397 | AVL25554.1 | H9EJ42 |
| KMLMGILSENAY | 1 | 0 | 0 | 1460436 | AVV62524.1 | K9N7C7 |
| KMLMGLLSENAY | 1 | 0 | 0 | 1460437 | AUM60013.1 | K9N7C7 |
| KMLTPKTGTGIA | 1 | 0 | 0 | 1460440 | AXN92247.1 | K9N7C7 |
| KMQTYKLVSMKG | 1 | 0 | 0 | 1460456 | AWK59942.1 | H9EJ42 |
| KMVDAYVDTFSA | 1 | 0 | 0 | 1460477 | BAE93400.1 | P0C6U8 |
| KMVSPTSKIEPC | 1 | 0 | 0 | 1460492 | ARU07575.1 | A0A140H1G9 |
| KMVVCDKVRSLK | 1 | 0 | 0 | 1460498 | AXN92247.1 | K9N7C7 |
| KNAFQTFRWSIF | 1 | 0 | 0 | 1460506 | ARU07575.1 | A0A140H1G9 |
| KNAQGKPVPYCY | 1 | 0 | 0 | 1460508 | APO40578.1 | P0C6X7 |
| KNASLNSFKEYF | 1 | 0 | 0 | 1460509 | AUM60014.1 | K9N5Q8 |
| KNCGQKSTTLIG | 1 | 0 | 0 | 1460513 | APO40578.1 | P0C6X7 |
| KNFCPCSSCPSK | 1 | 0 | 0 | 1460534 | QDH43719.1 | P36334 |
| KNFEQQCRCSRC | 1 | 0 | 0 | 1460540 | AVV62530.1 | K9N7D2 |
| KNFHIEGTGGNI | 1 | 0 | 0 | 1460541 | ANI69874.1 | K9N4V7 |
| KNFIAACAVDVN | 1 | 0 | 0 | 1460543 | ARU07575.1 | A0A140H1G9 |
| KNFLARAGKASC | 1 | 0 | 0 | 1460548 | AVP78041.1 | P0C6W2 |
| KNFSTSIPNFGS | 1 | 0 | 0 | 1460551 | AKM76239.1 | K9N5Q8 |
| KNFTYNVSTDFL | 1 | 0 | 0 | 1460555 | AGW27863.1 | A0A140H1H1 |
| KNGCALPFEDLN | 1 | 0 | 0 | 1460564 | AVV62524.1 | K9N7C7 |

| | | | | | | |
|---|---|---|---|---|---|---|
| KNGCALPFEDLS | 1 | 0 | 0 | 1460565 | AVV62535.1 | K9N7C7 |
| KNGFQGRQLMGS | 1 | 0 | 0 | 1460572 | AIV41816.1 | P0C6X6 |
| KNGLQFAAITAP | 1 | 0 | 0 | 1460579 | AUM60013.1 | K9N7C7 |
| KNGQEENDNISV | 1 | 0 | 0 | 1460586 | AGT51525.1 | P33469 |
| KNGQGENDNTSV | 1 | 0 | 0 | 1460589 | AIX09811.1 | P33469 |
| KNGQGEYDNISV | 1 | 0 | 0 | 1460590 | AXX83299.1 | P33469 |
| KNGRGGNHFRHP | 1 | 0 | 0 | 1460592 | AVV62524.1 | K9N7C7 |
| KNIDLFVKRRAD | 1 | 0 | 0 | 1460606 | ARU07575.1 | A0A140H1G9 |
| KNIPCYASAVAQ | 1 | 0 | 0 | 1460614 | QBP84719.1 | P0C6X6 |
| KNIPRYASAVAQ | 1 | 0 | 0 | 1460615 | AEN19363.1 | P0C6X6 |
| KNKAAPKVVTKK | 1 | 0 | 0 | 1460617 | AVV62533.1 | K9N4V7 |
| KNKDGFLSVYNA | 1 | 0 | 0 | 1460620 | ATO98193.1 | Q3LZX1 |
| KNKMRHKRTSTK | 1 | 0 | 0 | 1460630 | AXN92267.1 | K9N4V7 |
| KNKQDSTDDGSL | 1 | 0 | 0 | 1460632 | AVV62533.1 | K9N4V7 |
| KNLDEARFSGTA | 1 | 0 | 0 | 1460637 | AGT51787.1 | P0C6X6 |
| KNLGFDNQSKPP | 1 | 0 | 0 | 1460643 | AGT51335.1 | H9EJ69 |
| KNLNEGVLEYFY | 1 | 0 | 0 | 1460653 | QED88017.1 | H9EJ42 |
| KNLNTLRRGAVP | 1 | 0 | 0 | 1460661 | AAW33908.1 | H9EJ42 |
| KNLPKYVKPNFD | 1 | 0 | 0 | 1460663 | AWK59979.1 | A0A0P0G321 |
| KNLSKPPTENNC | 1 | 0 | 0 | 1460672 | AVP78035.1 | Q3LZX8 |
| KNNFGVITSFTS | 1 | 0 | 0 | 1460689 | Q14EA9.1 | Q0ZJ84 |
| KNNGIGTCPVGT | 1 | 0 | 0 | 1460692 | AIL49486.1 | P36334 |
| KNNTSLPTNIAV | 1 | 0 | 0 | 1460707 | ARB07436.1 | A0A140H1G9 |
| KNQISVAPDLSL | 1 | 0 | 0 | 1460724 | ANZ78835.1 | P36334 |
| KNQKLIANAFNK | 1 | 0 | 0 | 1460725 | BBA20986.1 | A0A140H1H1 |
| KNQTLVAPDLSL | 1 | 0 | 0 | 1460728 | CAA83661.1 | P36334 |
| KNSGIGTCPAGT | 1 | 0 | 0 | 1460757 | CAA83660.1 | P36334 |
| KNSGSVNQQQIK | 1 | 0 | 0 | 1460758 | ARU07575.1 | A0A140H1G9 |
| KNSLSDVAFNRH | 1 | 0 | 0 | 1460761 | ARU07583.1 | P0C6X6 |
| KNSQSSPIIPGF | 1 | 0 | 0 | 1460763 | QBF80598.1 | K9N5Q8 |
| KNTKAIKDDVLY | 1 | 0 | 0 | 1460780 | AKA09643.1 | P36334 |
| KNTVDTIPKEEF | 1 | 0 | 0 | 1460801 | AST12962.1 | A0A0P0LKV0 |
| KNTYISQLGCVF | 1 | 0 | 0 | 1460805 | AVN89333.1 | K9N7C7 |
| KNVDWSITYPII | 1 | 0 | 0 | 1460808 | AGT21359.1 | A0A0P0LKV0 |
| KNVKAWSGICVD | 1 | 0 | 0 | 1460812 | AIW52843.1 | A0A0P0G321 |
| KNVLVWLESQLQ | 1 | 0 | 0 | 1460816 | APO40578.1 | P0C6X7 |
| KNVSFLGKCLKM | 1 | 0 | 0 | 1460824 | AIW52827.1 | H9EJ42 |
| KNVSSQGPNFQE | 1 | 0 | 0 | 1460827 | AVV62526.1 | K9N5Q8 |
| KNVSTSIPNFSS | 1 | 0 | 0 | 1460832 | ANF29272.1 | K9N5Q8 |
| KNVTDVTGNLSD | 1 | 0 | 0 | 1460833 | AVQ05261.1 | P0C6X6 |
| KNYEFKFYDAFP | 1 | 0 | 0 | 1460843 | AGW27834.1 | A0A140H1G9 |
| KNYPCMVGPDCA | 1 | 0 | 0 | 1460848 | AUM60023.1 | K9N7C7 |
| KNYVFTGYHFTK | 1 | 0 | 0 | 1460853 | AEN19363.1 | P0C6X6 |
| KPAFNVVKAVRN | 1 | 0 | 0 | 1460859 | AIV41792.1 | P0C6X6 |
| KPAIDGLKITEE | 1 | 0 | 0 | 1460860 | AVP78041.1 | P0C6W2 |
| KPAMADEIASLV | 1 | 0 | 0 | 1460864 | AMM42397.1 | Q19U29 |
| KPAVGPMVDVNS | 1 | 0 | 0 | 1460871 | AUM60021.1 | K9N4V7 |
| KPDSIAISDLGS | 1 | 0 | 0 | 1460890 | AVV62524.1 | K9N7C7 |
| KPEANINQDSYG | 1 | 0 | 0 | 1460893 | ABD75615.1 | A0A140H1G9 |
| KPEDLMEVDHSG | 1 | 0 | 0 | 1460894 | AVV62529.1 | K9N643 |
| KPEYRKVFYALL | 1 | 0 | 0 | 1460903 | AIW52776.1 | A0A0P0LKV0 |
| KPFFTENKPSVE | 1 | 0 | 0 | 1460909 | AVP78041.1 | P0C6W2 |
| KPFLSDFNNGIF | 1 | 0 | 0 | 1460920 | BBA20983.1 | A0A140H1H1 |
| KPFSQPRADKPS | 1 | 0 | 0 | 1460922 | AVA26877.1 | H9EJ69 |
| KPFVKDGFSFVD | 1 | 0 | 0 | 1460925 | AVV62535.1 | K9N7C7 |
| KPFVKDGFSFVN | 1 | 0 | 0 | 1460926 | AVV62524.1 | K9N7C7 |
| KPFYIPEHCQNY | 1 | 0 | 0 | 1460931 | ATQ39394.1 | K9N796 |
| KPFYVPEHCQNY | 1 | 0 | 0 | 1460932 | AVN89384.1 | K9N796 |
| KPGASFSVLACY | 1 | 0 | 0 | 1460934 | AVV62524.1 | K9N7C7 |
| KPGGDAISAAFN | 1 | 0 | 0 | 1460936 | AUM60013.1 | K9N7C7 |
| KPGHSMPSLFKV | 1 | 0 | 0 | 1460945 | AVN89289.1 | K9N7C7 |
| KPGNTFITIEAA | 1 | 0 | 0 | 1460951 | AXX83372.1 | P0C6X6 |
| KPGNVGGNDDLS | 1 | 0 | 0 | 1460953 | AIV41912.1 | P0C6X6 |
| KPGNVGGNEALS | 1 | 0 | 0 | 1460954 | AIV41792.1 | P0C6X6 |
| KPHQVNLSLNGN | 1 | 0 | 0 | 1460967 | AWO69005.1 | A0A0P0G321 |
| KPHVRYDMFDTN | 1 | 0 | 0 | 1460976 | AUM60013.1 | K9N7C7 |
| KPIADYAFIADK | 1 | 0 | 0 | 1460978 | AUM60013.1 | K9N7C7 |
| KPICSSVYLGVR | 1 | 0 | 0 | 1460981 | AFD98812.1 | H9EJ42 |
| KPIGMFFPYDIE | 1 | 0 | 0 | 1460986 | AXN92247.1 | K9N7C7 |
| KPIYLKGSGFDF | 1 | 0 | 0 | 1461012 | ARE30011.1 | P0C6X6 |
| KPIYVPEHCQNY | 1 | 0 | 0 | 1461014 | QAT98905.1 | K9N796 |
| KPKPVISQLDEN | 1 | 0 | 0 | 1461023 | AST12962.1 | A0A0P0LKV0 |
| KPLEPKFSTALP | 1 | 0 | 0 | 1461038 | AFD98810.1 | H9EJ69 |
| KPLLVDDNKFDV | 1 | 0 | 0 | 1461052 | AIV41786.1 | P0C6X6 |
| KPLSAICPAGTK | 1 | 0 | 0 | 1461060 | BBA20986.1 | A0A140H1H1 |
| KPLSQPRSDKPS | 1 | 0 | 0 | 1461063 | AVL25604.1 | H9EJ69 |
| KPLVDFNVNVMG | 1 | 0 | 0 | 1461071 | AGT21366.1 | A0A0P0LKV0 |
| KPNFDLTPFNLT | 1 | 0 | 0 | 1461094 | AIW52843.1 | A0A0P0G321 |
| KPNKPLSQPRSD | 1 | 0 | 0 | 1461102 | AVL25604.1 | H9EJ69 |
| KPNLPNCNIESW | 1 | 0 | 0 | 1461105 | AIX10749.1 | P36334 |
| KPNRTSVASFNE | 1 | 0 | 0 | 1461107 | AVV62535.1 | K9N7C7 |
| KPPSAICPAGTK | 1 | 0 | 0 | 1461120 | 5I08_A | A0A140H1H1 |
| KPQEKDKSQRKR | 1 | 0 | 0 | 1461125 | ABB90496.1 | A0A0N7J131 |
| KPQPAGVFTDHD | 1 | 0 | 0 | 1461129 | CAA83661.1 | P36334 |
| KPQPAGVFTNHD | 1 | 0 | 0 | 1461130 | ATP16757.1 | P36334 |
| KPQPAGVLTNHD | 1 | 0 | 0 | 1461132 | AVQ05264.1 | P36334 |
| KPQPKFTVSTQP | 1 | 0 | 0 | 1461133 | AMM42397.1 | Q19U29 |
| KPQPVGVFTHHD | 1 | 0 | 0 | 1461136 | CAA83660.1 | P36334 |
| KPQRAPRGSITQ | 1 | 0 | 0 | 1461139 | AVV62533.1 | K9N4V7 |
| KPRAAPNNTVSW | 1 | 0 | 0 | 1461156 | AXN92267.1 | K9N4V7 |
| KPRPAPNNSVSW | 1 | 0 | 0 | 1461161 | AVV62544.1 | K9N4V7 |

| | | | | | | |
|---|---|---|---|---|---|---|
| KPRPAPNTTVSW | 1 | 0 | 0 | 1461163 | AUM60021.1 | K9N4V7 |
| KPRQKRSPNTQR | 1 | 0 | 0 | 1461167 | AJQ24090.1 | P33469 |
| KPRSQMEVDFLE | 1 | 0 | 0 | 1461179 | APO40578.1 | P0C6X7 |
| KPRTFMLKYNEN | 1 | 0 | 0 | 1461181 | AVP78031.1 | Q3LZX1 |
| KPSDKHATLFGS | 1 | 0 | 0 | 1461191 | AUM60014.1 | K9N5Q8 |
| KPSELSRVLGLK | 1 | 0 | 0 | 1461193 | AVP78041.1 | P0C6W2 |
| KPSGMDTALLRA | 1 | 0 | 0 | 1461199 | YP_173241.1 | Q0ZJ82 |
| KPSGSFVEQAEG | 1 | 0 | 0 | 1461200 | AID55084.1 | K9N5Q8 |
| KPSIIHDIGNPK | 1 | 0 | 0 | 1461203 | ABE97136.1 | H9EJ42 |
| KPSSASCPIGTN | 1 | 0 | 0 | 1461212 | AFR45556.1 | A0A140H1H1 |
| KPSSVKEMQSQS | 1 | 0 | 0 | 1461216 | ABE97141.1 | H9EJ69 |
| KPTAKYTPLVGT | 1 | 0 | 0 | 1461230 | AXN92262.1 | K9N4V0 |
| KPTAKYTPLVRT | 1 | 0 | 0 | 1461231 | QBF80613.1 | K9N4V0 |
| KPTDRSHYYVDS | 1 | 0 | 0 | 1461233 | AVV62524.1 | K9N7C7 |
| KPTDRSHYYVNS | 1 | 0 | 0 | 1461234 | AVV62535.1 | K9N7C7 |
| KPTPAPVYIWTA | 1 | 0 | 0 | 1461245 | AWW13562.1 | Q01455 |
| KPTVETPKEEQP | 1 | 0 | 0 | 1461252 | ARO76381.1 | P0C6U8 |
| KPVDHYDGFYTQ | 1 | 0 | 0 | 1461261 | NP_073549.1 | A0A0P0LKV0 |
| KPVFFNAFVKCN | 1 | 0 | 0 | 1461268 | AWK59942.1 | H9EJ42 |
| KPVGFFFPYDPA | 1 | 0 | 0 | 1461273 | AVV62535.1 | K9N7C7 |
| KPVGYFFPYDPA | 1 | 0 | 0 | 1461276 | AVV62524.1 | K9N7C7 |
| KPVISQLDENAQ | 1 | 0 | 0 | 1461281 | AST12962.1 | A0A0P0LKV0 |
| KPVIWLSHEQAS | 1 | 0 | 0 | 1461285 | AVI15051.1 | P0C6X6 |
| KPVMISQWPISD | 1 | 0 | 0 | 1461292 | AWK59942.1 | H9EJ42 |
| KPVQGAWNIGQQ | 1 | 0 | 0 | 1461302 | ARO76381.1 | P0C6U8 |
| KPVSEYHVTLAL | 1 | 0 | 0 | 1461310 | AVV62529.1 | K9N643 |
| KPVYRVNTAIPT | 1 | 0 | 0 | 1461324 | AXN92231.1 | K9N643 |
| KPYERDISNVLY | 1 | 0 | 0 | 1461329 | APO40579.1 | P59594 |
| KQADLSGAICEL | 1 | 0 | 0 | 1461339 | ARU07575.1 | A0A140H1G9 |
| KQANDLKDTLRL | 1 | 0 | 0 | 1461340 | QED88010.1 | H9EJ42 |
| KQCCTAAVDLSM | 1 | 0 | 0 | 1461352 | AXN92247.1 | K9N7C7 |
| KQDQGQYYYRSS | 1 | 0 | 0 | 1461368 | ATO98193.1 | Q3LZX1 |
| KQDVGNYFYRSH | 1 | 0 | 0 | 1461371 | AVP78031.1 | Q3LZX1 |
| KQDVSLLGKCYK | 1 | 0 | 0 | 1461373 | AKN24802.1 | K9N7C7 |
| KQEQRIGLDAVM | 1 | 0 | 0 | 1461385 | AVQ05261.1 | P0C6X6 |
| KQFVPVQVVDSR | 1 | 0 | 0 | 1461395 | ARU07576.1 | A0A140H1H0 |
| KQGDLGKQYVYS | 1 | 0 | 0 | 1461399 | AVV62526.1 | K9N5Q8 |
| KQGEIKGATPTN | 1 | 0 | 0 | 1461402 | AVP78032.1 | Q3LZX0 |
| KQGHFIGVDGEL | 1 | 0 | 0 | 1461404 | AVN89289.1 | K9N7C7 |
| KQGLEFAAITAP | 1 | 0 | 0 | 1461409 | AVV62524.1 | K9N7C7 |
| KQGPASARDLIC | 1 | 0 | 0 | 1461412 | QEJ82215.1 | K9N5Q8 |
| KQGPPSARDLIC | 1 | 0 | 0 | 1461413 | AVV62537.1 | K9N5Q8 |
| KQGTLIYKVLDT | 1 | 0 | 0 | 1461416 | AIV41996.1 | P0C6X6 |
| KQGTQIYKVLDT | 1 | 0 | 0 | 1461418 | AEN19363.1 | P0C6X6 |
| KQIANKFNQAPG | 1 | 0 | 0 | 1461425 | AVV62526.1 | K9N5Q8 |
| KQIANQFNNAIS | 1 | 0 | 0 | 1461427 | APO40579.1 | P59594 |
| KQIGLLEPVKHS | 1 | 0 | 0 | 1461430 | AVV62535.1 | K9N7C7 |
| KQIGLLEPVNHS | 1 | 0 | 0 | 1461431 | AVV62524.1 | K9N7C7 |
| KQIIHSHPILHD | 1 | 0 | 0 | 1461432 | ATQ39396.1 | K9N643 |
| KQILGNTMLEDE | 1 | 0 | 0 | 1461433 | AUM60013.1 | K9N7C7 |
| KQILGSSMLEDE | 1 | 0 | 0 | 1461434 | AVV62524.1 | K9N7C7 |
| KQILGSTMLEDE | 1 | 0 | 0 | 1461435 | AXN92247.1 | K9N7C7 |
| KQKASEKQGGEE | 1 | 0 | 0 | 1461456 | AUM60021.1 | K9N4V7 |
| KQLEAKTASLFK | 1 | 0 | 0 | 1461476 | ARU07594.1 | A0A0P0G321 |
| KQLEVKTASLFQ | 1 | 0 | 0 | 1461479 | AGT51366.1 | A0A0P0G321 |
| KQLGLVWNKDVN | 1 | 0 | 0 | 1461487 | AIW52827.1 | H9EJ42 |
| KQLLINRGVYKP | 1 | 0 | 0 | 1461494 | AEN19363.1 | P0C6X6 |
| KQLNESYIDLKE | 1 | 0 | 0 | 1461504 | AVV62537.1 | K9N5Q8 |
| KQLNESYIDLKK | 1 | 0 | 0 | 1461505 | AVV62526.1 | K9N5Q8 |
| KQLSPLEGGGRL | 1 | 0 | 0 | 1461512 | AXG21654.1 | K9N5Q8 |
| KQNDSWMFTGSS | 1 | 0 | 0 | 1461525 | BBA20986.1 | A0A140H1H1 |
| KQNGCSLYLADN | 1 | 0 | 0 | 1461526 | ATQ39389.1 | K9N7C7 |
| KQNGFSFFLADN | 1 | 0 | 0 | 1461527 | AVV62524.1 | K9N7C7 |
| KQNGLSLYLADN | 1 | 0 | 0 | 1461529 | AUM60013.1 | K9N7C7 |
| KQPMATSANCCH | 1 | 0 | 0 | 1461536 | AGT51735.1 | Q4VID0 |
| KQPMATSVNCCH | 1 | 0 | 0 | 1461537 | APU51922.1 | Q4VID0 |
| KQPMATSVNCFH | 1 | 0 | 0 | 1461538 | AHF54653.1 | Q4VID0 |
| KQPTVTLLPAAE | 1 | 0 | 0 | 1461541 | APO40586.1 | P59595 |
| KQPTVTPLPAAD | 1 | 0 | 0 | 1461542 | AAS48456.1 | P59595 |
| KQQRFVYDGFDN | 1 | 0 | 0 | 1461549 | AVV62526.1 | K9N5Q8 |
| KQQYSQLFVDTL | 1 | 0 | 0 | 1461551 | AEN19363.1 | P0C6X6 |
| KQTATSQQPSRG | 1 | 0 | 0 | 1461583 | ANZ78871.1 | P33469 |
| KQTIFTVNSVVQ | 1 | 0 | 0 | 1461587 | AVV62524.1 | K9N7C7 |
| KQTSTSQQPLGG | 1 | 0 | 0 | 1461589 | AMK59681.1 | P33469 |
| KQTVTSQQPSGG | 1 | 0 | 0 | 1461592 | AIL49418.1 | P33469 |
| KQVGCYMRLFYD | 1 | 0 | 0 | 1461599 | AVZ61116.1 | P0C6X6 |
| KQVLDLLDKGMQ | 1 | 0 | 0 | 1461604 | ASU91196.1 | K9N638 |
| KQWPMPINASKA | 1 | 0 | 0 | 1461613 | AVV62537.1 | K9N5Q8 |
| KQYLLAFNQDGV | 1 | 0 | 0 | 1461626 | CAA83660.1 | P36334 |
| KQYVCNGFEKCE | 1 | 0 | 0 | 1461630 | AVV62526.1 | K9N5Q8 |
| KRARGSSAARLE | 1 | 0 | 0 | 1461646 | QDH43737.1 | H9EJ42 |
| KRDAAAAKNKMR | 1 | 0 | 0 | 1461659 | ANF29235.1 | K9N4V7 |
| KRESEADSPVKD | 1 | 0 | 0 | 1461671 | ARU07581.1 | Q19U29 |
| KRFIHKNSTWSK | 1 | 0 | 0 | 1461679 | NP_073549.1 | A0A0P0LKV0 |
| KRFQSFQQFGRD | 1 | 0 | 0 | 1461685 | ATO98181.1 | Q3LZX1 |
| KRFRRVLVKLLD | 1 | 0 | 0 | 1461687 | AWK59942.1 | H9EJ42 |
| KRGFLDKISDTS | 1 | 0 | 0 | 1461700 | BAM16435.1 | Q01455 |
| KRHTLSNYQHEE | 1 | 0 | 0 | 1461718 | APO40578.1 | P0C6X7 |
| KRKTEGFMGRIR | 1 | 0 | 0 | 1461732 | APO40578.1 | P0C6X7 |
| KRLDAVEQEAQI | 1 | 0 | 0 | 1461739 | AVV62537.1 | K9N5Q8 |
| KRLKNGFQGRQL | 1 | 0 | 0 | 1461744 | AIV41816.1 | P0C6X6 |

| | | | | | | |
|---|---|---|---|---|---|---|
| KRLRKRRDAKVS | 1 | 0 | 0 | 1461750 | AUM60013.1 | K9N7C7 |
| KRLYPNVYRTDY | 1 | 0 | 0 | 1461759 | ACR82756.1 | A0A140H1G9 |
| KRMKQIEDKIEE | 1 | 0 | 0 | 1461766 | 5X58_A | P59594 |
| KRNFTYDVNAIY | 1 | 0 | 0 | 1461768 | BBO58839.1 | P36334 |
| KRNFTYDVNANY | 1 | 0 | 0 | 1461769 | AKA09658.1 | P36334 |
| KRNFTYDVSATY | 1 | 0 | 0 | 1461771 | AKA09662.1 | P36334 |
| KRPRWKRVPTRE | 1 | 0 | 0 | 1461786 | ADK37677.1 | H9EJ69 |
| KRQIRIACRKFN | 1 | 0 | 0 | 1461792 | ASU90547.1 | K9N7C7 |
| KRQNYGTNSGVA | 1 | 0 | 0 | 1461794 | AUM60030.1 | K9N7A1 |
| KRQPNDDATSNV | 1 | 0 | 0 | 1461795 | APT69866.1 | A0A0N7J131 |
| KRRFSLLRPEDL | 1 | 0 | 0 | 1461808 | AXN92231.1 | K9N643 |
| KRSVRSHPDFKL | 1 | 0 | 0 | 1461850 | AHY22543.1 | K9N7C7 |
| KRTDNPILDFKD | 1 | 0 | 0 | 1461856 | AVP78031.1 | Q3LZX1 |
| KRTHSPTKKLRY | 1 | 0 | 0 | 1461864 | AMD02849.1 | K9N643 |
| KRTRLFKSTVCW | 1 | 0 | 0 | 1461873 | AGT51787.1 | P0C6X6 |
| KRVDLPPKLHFY | 1 | 0 | 0 | 1461887 | ALK28797.1 | A0A0N7J131 |
| KRVPTIEENVIQ | 1 | 0 | 0 | 1461902 | ADK37676.1 | H9EJ69 |
| KSAAGYGGFGCK | 1 | 0 | 0 | 1461925 | AEN19363.1 | P0C6X6 |
| KSAGYPFNQFGK | 1 | 0 | 0 | 1461940 | AIL49570.1 | P0C6X6 |
| KSCAPLSDQIDF | 1 | 0 | 0 | 1461970 | APO40578.1 | P0C6X7 |
| KSCFEIDGNVMH | 1 | 0 | 0 | 1461972 | ARB07436.1 | A0A140H1G9 |
| KSCQNLDCNCLG | 1 | 0 | 0 | 1461975 | AGT51688.1 | Q80872 |
| KSDYCGGNPCIC | 1 | 0 | 0 | 1462002 | BBO58900.1 | P36334 |
| KSDYCRGNSCIC | 1 | 0 | 0 | 1462004 | AIX10751.1 | P36334 |
| KSEFDHEISVQK | 1 | 0 | 0 | 1462008 | AWK59942.1 | H9EJ42 |
| KSFFPQFTSDQA | 1 | 0 | 0 | 1462021 | ABD75500.1 | Q0ZJ82 |
| KSFFSQFTSDQA | 1 | 0 | 0 | 1462022 | ABD75588.1 | Q0ZJ82 |
| KSFIDVINKALE | 1 | 0 | 0 | 1462028 | ARO76381.1 | P0C6U8 |
| KSFQLDTAPKTG | 1 | 0 | 0 | 1462047 | Q3I5J5.1 | Q3LZX1 |
| KSGANFRDLSLK | 1 | 0 | 0 | 1462061 | AMN88694.1 | A0A140H1H1 |
| KSGCKLINAKPP | 1 | 0 | 0 | 1462063 | AWK59942.1 | H9EJ42 |
| KSGGYIPEAPRD | 1 | 0 | 0 | 1462069 | 6U7H_A | A0A0P0K6L9 |
| KSGSKFKNLVLN | 1 | 0 | 0 | 1462081 | ARU07609.1 | P30215 |
| KSGTCGIVELEK | 1 | 0 | 0 | 1462089 | P0C6W6.1 | P0C6W2 |
| KSGTITKVEADF | 1 | 0 | 0 | 1462090 | AWW13510.1 | P30215 |
| KSGTLVLNNPAY | 1 | 0 | 0 | 1462091 | AGT51729.1 | P30215 |
| KSGVNDVCVFEN | 1 | 0 | 0 | 1462097 | AVV62524.1 | K9N7C7 |
| KSIELTAEDISL | 1 | 0 | 0 | 1462116 | AEN19370.1 | P33469 |
| KSIYNVDLCDCG | 1 | 0 | 0 | 1462140 | ARU07575.1 | A0A140H1G9 |
| KSKELLDVALDS | 1 | 0 | 0 | 1462145 | AVQ05261.1 | P0C6X6 |
| KSKGSFRNESWH | 1 | 0 | 0 | 1462148 | AMN88694.1 | A0A140H1H1 |
| KSKVSAEVSRQV | 1 | 0 | 0 | 1462166 | ARB07393.1 | A0A0P0KWE8 |
| KSLGFDKPQEKK | 1 | 0 | 0 | 1462175 | AMM42528.1 | A0A0N7J131 |
| KSLNVDGVQYYC | 1 | 0 | 0 | 1462185 | ATQ39389.1 | K9N7C7 |
| KSLSRSQSSETK | 1 | 0 | 0 | 1462194 | ABB90484.1 | A0A0N7J131 |
| KSLSVDEFVDVV | 1 | 0 | 0 | 1462196 | AUM60013.1 | K9N7C7 |
| KSLTIVDIPLSL | 1 | 0 | 0 | 1462200 | AVN89312.1 | K9N638 |
| KSLTTVSIPQGL | 1 | 0 | 0 | 1462202 | ATQ39389.1 | K9N7C7 |
| KSNDSCPGKNNG | 1 | 0 | 0 | 1462214 | AIV41891.1 | P36334 |
| KSNTNDAFISNY | 1 | 0 | 0 | 1462229 | ATQ39390.1 | K9N5Q8 |
| KSNVDESFVDDF | 1 | 0 | 0 | 1462230 | AIG96641.1 | A0A0P0LKV0 |
| KSQSNRYGFCGN | 1 | 0 | 0 | 1462259 | AIW52843.1 | A0A0P0G321 |
| KSRNPLLYDANY | 1 | 0 | 0 | 1462271 | ARO76383.1 | P59632 |
| KSRVQQNKIREL | 1 | 0 | 0 | 1462275 | AGT51784.1 | P33469 |
| KSSCCNLISNKL | 1 | 0 | 0 | 1462279 | AST12962.1 | A0A0P0LKV0 |
| KSSDEGIKVTQE | 1 | 0 | 0 | 1462281 | ARO76381.1 | P0C6U8 |
| KSSEYKGSITDV | 1 | 0 | 0 | 1462287 | AVP78041.1 | P0C6W2 |
| KSSIDAFKLNIR | 1 | 0 | 0 | 1462291 | ARO76381.1 | P0C6U8 |
| KSSQCFKIFYKG | 1 | 0 | 0 | 1462296 | APO40578.1 | P0C6X7 |
| KSTELVKNKCVN | 1 | 0 | 0 | 1462303 | APO40579.1 | P59594 |
| KSVDVAIQEVID | 1 | 0 | 0 | 1462330 | AWW13564.1 | Q80872 |
| KSVGKFCIEASF | 1 | 0 | 0 | 1462337 | AVP78041.1 | P0C6W2 |
| KSVVSKIGDSIF | 1 | 0 | 0 | 1462366 | ATG84700.1 | K9N638 |
| KSVVSKIGNSIF | 1 | 0 | 0 | 1462367 | ATQ39389.1 | K9N7C7 |
| KSVYDQIVDNVY | 1 | 0 | 0 | 1462370 | AIV42002.1 | P0C6X6 |
| KSWPRPIDPAAA | 1 | 0 | 0 | 1462375 | AVV62526.1 | K9N5Q8 |
| KSWYDPVENEDI | 1 | 0 | 0 | 1462380 | AWK59948.1 | H9EJ42 |
| KSYEQQTPFDIT | 1 | 0 | 0 | 1462387 | APO40578.1 | P0C6X7 |
| KSYERQSPFEIK | 1 | 0 | 0 | 1462388 | ARO76381.1 | P0C6U8 |
| KSYGFTPTVGVG | 1 | 0 | 0 | 1462391 | APO40579.1 | P59594 |
| KSYQEALDSGDA | 1 | 0 | 0 | 1462397 | AUM60013.1 | K9N7C7 |
| KSYSEHVIPSTL | 1 | 0 | 0 | 1462400 | BBA20987.1 | Q0ZJ83 |
| KTADGDQRQLLP | 1 | 0 | 0 | 1462410 | AJQ24083.1 | P33469 |
| KTADGQQKPLLP | 1 | 0 | 0 | 1462413 | ABA60796.1 | Q19U29 |
| KTAGVVTCNVVK | 1 | 0 | 0 | 1462417 | AVN89451.1 | K9N7C7 |
| KTAQADTAGLFT | 1 | 0 | 0 | 1462422 | ALA49936.1 | K9N796 |
| KTAQAYTAGLYT | 1 | 0 | 0 | 1462425 | AJG44059.1 | K9N796 |
| KTEADWIELFPH | 1 | 0 | 0 | 1462456 | QEO75983.1 | A0A0P0LKV0 |
| KTEKDTNEAKAL | 1 | 0 | 0 | 1462466 | AUM60013.1 | K9N7C7 |
| KTEPTGYGVRRK | 1 | 0 | 0 | 1462470 | AIW52723.1 | A0A0N7J131 |
| KTESAMVHDLNN | 1 | 0 | 0 | 1462471 | AVP78050.1 | Q3LZX3 |
| KTEVDPNLSGFY | 1 | 0 | 0 | 1462472 | ATQ39389.1 | K9N7C7 |
| KTFSNCNFNMSS | 1 | 0 | 0 | 1462493 | CAA79896.1 | P36334 |
| KTGNFKDLREYV | 1 | 0 | 0 | 1462507 | ATO98181.1 | Q3LZX1 |
| KTGQGENDNISV | 1 | 0 | 0 | 1462508 | AHA80176.1 | P33469 |
| KTGVATEQLLYA | 1 | 0 | 0 | 1462516 | AUM60023.1 | K9N7C7 |
| KTGVLQPLEQPT | 1 | 0 | 0 | 1462518 | AVP78041.1 | P0C6W2 |
| KTIATHGIAAVN | 1 | 0 | 0 | 1462532 | ARO76381.1 | P0C6U8 |
| KTIDELCVNLNL | 1 | 0 | 0 | 1462538 | ALA50066.1 | K9N7C7 |
| KTIEDALRNSAM | 1 | 0 | 0 | 1462540 | AIW52757.1 | A0A0P0K6L9 |
| KTIWAFNPETNA | 1 | 0 | 0 | 1462556 | ARE29965.1 | U3M6Q8 |

| | | | | | | |
|---|---|---|---|---|---|---|
| KTKAAPKVITKK | 1 | 0 | 0 | 1462557 | AVV62544.1 | K9N4V7 |
| KTKGIVAHPLGY | 1 | 0 | 0 | 1462559 | AVV62524.1 | K9N7C7 |
| KTLATHGLAAIN | 1 | 0 | 0 | 1462569 | AVP78041.1 | P0C6W2 |
| KTLFDSLKFFEI | 1 | 0 | 0 | 1462572 | AWK59942.1 | H9EJ42 |
| KTLLTPLCGFSS | 1 | 0 | 0 | 1462578 | APO40578.1 | P0C6X7 |
| KTLQEMKALFLT | 1 | 0 | 0 | 1462584 | ARB07430.1 | P0C6X6 |
| KTLQPISDLLTP | 1 | 0 | 0 | 1462590 | APO40578.1 | P0C6X7 |
| KTLRSLEGKVFQ | 1 | 0 | 0 | 1462596 | AVP78050.1 | Q3LZX3 |
| KTMVVTAGLDQT | 1 | 0 | 0 | 1462615 | AVV62524.1 | K9N7C7 |
| KTNNIRIVDEWS | 1 | 0 | 0 | 1462618 | ASU90175.1 | K9N5Q8 |
| KTNNTSIVDEWS | 1 | 0 | 0 | 1462623 | ANF29261.1 | K9N5Q8 |
| KTNVVHPALHLV | 1 | 0 | 0 | 1462626 | AVP78039.1 | Q3LZX3 |
| KTPDTYKCPQSK | 1 | 0 | 0 | 1462629 | AXX83339.1 | P36334 |
| KTPEEHFIETVS | 1 | 0 | 0 | 1462631 | AVP78041.1 | P0C6W2 |
| KTPEEHFVETVS | 1 | 0 | 0 | 1462632 | ARO76381.1 | P0C6U8 |
| KTPTLKDFGGFN | 1 | 0 | 0 | 1462640 | ATO98231.1 | Q3LZX1 |
| KTQDYQIGGFVE | 1 | 0 | 0 | 1462644 | APO40580.1 | P59632 |
| KTQFIAPPTGVY | 1 | 0 | 0 | 1462645 | AHA80218.1 | P36334 |
| KTRALVAVNSED | 1 | 0 | 0 | 1462656 | AUM60013.1 | K9N7C7 |
| KTRVLAVVNSQD | 1 | 0 | 0 | 1462663 | ANF29227.1 | K9N638 |
| KTRVVHPALHLV | 1 | 0 | 0 | 1462665 | AVP78050.1 | Q3LZX3 |
| KTSDELCVNLNI | 1 | 0 | 0 | 1462668 | AUM60013.1 | K9N7C7 |
| KTSDGFLHIYGA | 1 | 0 | 0 | 1462670 | ATO98145.1 | Q3LZX1 |
| KTSEQNNFTGYL | 1 | 0 | 0 | 1462677 | AVP78041.1 | P0C6W2 |
| KTSMAPILTEST | 1 | 0 | 0 | 1462683 | AVR40343.1 | Q4VID0 |
| KTSQADTAGLYT | 1 | 0 | 0 | 1462685 | AXN92229.1 | K9N796 |
| KTSTAPILMEST | 1 | 0 | 0 | 1462688 | AGT51765.1 | Q4VID0 |
| KTSTAPILTESS | 1 | 0 | 0 | 1462689 | ATP16762.1 | Q4VID0 |
| KTSTAPILTEST | 1 | 0 | 0 | 1462690 | ARE29989.1 | Q4VID0 |
| KTSWVDQFERSH | 1 | 0 | 0 | 1462698 | AZS52622.1 | Q19U29 |
| KTSWVDQSERSH | 1 | 0 | 0 | 1462699 | AMM42397.1 | Q19U29 |
| KTVKYFECPNSI | 1 | 0 | 0 | 1462720 | ARU07575.1 | A0A140H1G9 |
| KTVQADTAGLYT | 1 | 0 | 0 | 1462731 | AHC74089.1 | K9N796 |
| KTVQFCDVMRDA | 1 | 0 | 0 | 1462734 | APO40578.1 | P0C6X7 |
| KTVREFVIARTG | 1 | 0 | 0 | 1462735 | AIW52745.1 | A0A0P0K6L9 |
| KTVYSDVENPHL | 1 | 0 | 0 | 1462737 | AID16714.1 | P0C6W2 |
| KTVYSDVESPHL | 1 | 0 | 0 | 1462738 | AVP78041.1 | P0C6W2 |
| KTVYSDVETPNL | 1 | 0 | 0 | 1462740 | ASB51350.1 | P0C6X7 |
| KTYAELGYIADI | 1 | 0 | 0 | 1462746 | AIW52776.1 | A0A0P0LKV0 |
| KTYLDFIQQGGI | 1 | 0 | 0 | 1462754 | AIV41966.1 | P0C6X6 |
| KVAKIANLFLEQ | 1 | 0 | 0 | 1462771 | AVV62535.1 | K9N7C7 |
| KVANLFLTQAQR | 1 | 0 | 0 | 1462774 | AUM60013.1 | K9N7C7 |
| KVAQPKIEQKPL | 1 | 0 | 0 | 1462776 | AST12962.1 | A0A0P0LKV0 |
| KVDIQQTFFDKT | 1 | 0 | 0 | 1462809 | QCQ29075.1 | K9N5Q8 |
| KVDLLVPRNKWN | 1 | 0 | 0 | 1462811 | AUM60013.1 | K9N7C7 |
| KVEDAYDQVHEE | 1 | 0 | 0 | 1462825 | ARK08632.1 | P0C6X6 |
| KVEDSVIVNDAT | 1 | 0 | 0 | 1462827 | AIV41846.1 | P0C6X6 |
| KVEDSVIVNNDT | 1 | 0 | 0 | 1462828 | AOL02450.1 | P0C6X6 |
| KVELYNGECVYL | 1 | 0 | 0 | 1462834 | NP_073553.1 | A0A0P0KWE8 |
| KVESLMSTILSS | 1 | 0 | 0 | 1462840 | AVV62535.1 | K9N7C7 |
| KVETANPKTPKY | 1 | 0 | 0 | 1462841 | APO40578.1 | P0C6X7 |
| KVEVNALRTTGI | 1 | 0 | 0 | 1462845 | AVV62524.1 | K9N7C7 |
| KVFFQFENLSSE | 1 | 0 | 0 | 1462851 | AIV41978.1 | P0C6X6 |
| KVFGKDGVMYSE | 1 | 0 | 0 | 1462852 | AWW13566.1 | P36334 |
| KVFGYVQQGVFN | 1 | 0 | 0 | 1462856 | ATN39862.1 | P0C6X6 |
| KVFKEGVMYSEF | 1 | 0 | 0 | 1462861 | AMK59677.1 | P36334 |
| KVFNASRFPNVY | 1 | 0 | 0 | 1462866 | ATO98181.1 | Q3LZX1 |
| KVHEVAPTRSVT | 1 | 0 | 0 | 1462905 | AVV62535.1 | K9N7C7 |
| KVHNLSGNPDEP | 1 | 0 | 0 | 1462909 | AHA80155.1 | P33469 |
| KVHQVAATRSVT | 1 | 0 | 0 | 1462910 | AVV62524.1 | K9N7C7 |
| KVIDGGKIWNGV | 1 | 0 | 0 | 1462916 | NP_073549.1 | A0A0P0LKV0 |
| KVIKDGVMYSEF | 1 | 0 | 0 | 1462921 | CAA79896.1 | P36334 |
| KVIKHGVMYSEF | 1 | 0 | 0 | 1462923 | CAA83661.1 | P36334 |
| KVINHNVLIKES | 1 | 0 | 0 | 1462927 | QDH43737.1 | H9EJ42 |
| KVITKKDAAAAK | 1 | 0 | 0 | 1462939 | AXN92267.1 | K9N4V7 |
| KVITKRDAAAAK | 1 | 0 | 0 | 1462942 | ANF29235.1 | K9N4V7 |
| KVKDFVSGLLNV | 1 | 0 | 0 | 1462945 | APT69882.1 | A0A0P0ILY9 |
| KVKNTKAIKDDV | 1 | 0 | 0 | 1462952 | AKA09643.1 | P36334 |
| KVKNTKVFKDGV | 1 | 0 | 0 | 1462957 | AEN19366.1 | P36334 |
| KVKNTKVIKDDV | 1 | 0 | 0 | 1462960 | ATP16757.1 | P36334 |
| KVKNTKVIKDGV | 1 | 0 | 0 | 1462961 | CAA79896.1 | P36334 |
| KVKNTKVIKHGV | 1 | 0 | 0 | 1462963 | CAA83661.1 | P36334 |
| KVKPTVVVNAANVY | 1 | 0 | 0 | 1462970 | P0DTC1.1 | P0DTD1 |
| KVLPPILSETQI | 1 | 0 | 0 | 1463009 | BBA20986.1 | A0A140H1H1 |
| KVLPPLLSDNQI | 1 | 0 | 0 | 1463010 | CAA83661.1 | P36334 |
| KVLPPLYDPNME | 1 | 0 | 0 | 1463014 | AVV62526.1 | K9N5Q8 |
| KVLSENLNAYQQ | 1 | 0 | 0 | 1463021 | AEN19370.1 | P33469 |
| KVLTENLNAYQQ | 1 | 0 | 0 | 1463023 | AIL49396.1 | P33469 |
| KVNGLWSPTITH | 1 | 0 | 0 | 1463041 | ARB07605.1 | A0A140H1G9 |
| KVPAVVSVSSPD | 1 | 0 | 0 | 1463056 | APO40578.1 | P0C6X7 |
| KVQNLFGNPDEP | 1 | 0 | 0 | 1463102 | AHA80176.1 | P33469 |
| KVQQLFYTEDMA | 1 | 0 | 0 | 1463109 | AXN92247.1 | K9N7C7 |
| KVQQQQDDLFTP | 1 | 0 | 0 | 1463110 | APO40584.1 | P59635 |
| KVQTFYPRLQAS | 1 | 0 | 0 | 1463112 | ATQ39389.1 | K9N7C7 |
| KVRNLSGNPDEP | 1 | 0 | 0 | 1463116 | AHA80174.1 | P33469 |
| KVSAGSDSDTPS | 1 | 0 | 0 | 1463120 | AUM60013.1 | K9N7C7 |
| KVSCVCGRDATQ | 1 | 0 | 0 | 1463122 | APO40578.1 | P0C6X7 |
| KVSELEDACFKF | 1 | 0 | 0 | 1463123 | AQN78665.1 | Q04853 |
| KVSKIANLFVEQ | 1 | 0 | 0 | 1463131 | AVV62524.1 | K9N7C7 |
| KVSKVANLFLTQ | 1 | 0 | 0 | 1463132 | AUM60013.1 | K9N7C7 |
| KVSMPTVIAKNG | 1 | 0 | 0 | 1463137 | APO40578.1 | P0C6X7 |

| | | | | | | |
|---|---|---|---|---|---|---|
| KVSPDLCTHSFR | 1 | 0 | 0 | 1463138 | AYV99777.1 | P59633 |
| KVSQLTKQLLDI | 1 | 0 | 0 | 1463142 | AVV62535.1 | K9N7C7 |
| KVSVPSPLNWER | 1 | 0 | 0 | 1463152 | AMM42407.1 | A0A140H1H1 |
| KVTDILYPGQKY | 1 | 0 | 0 | 1463159 | AVV62524.1 | K9N7C7 |
| KVTDVLFSGQKY | 1 | 0 | 0 | 1463162 | AQZ41321.1 | K9N7C7 |
| KVTDVLYPQQKY | 1 | 0 | 0 | 1463163 | AUM60013.1 | K9N7C7 |
| KVTIDCAAFDCG | 1 | 0 | 0 | 1463170 | ARB66062.1 | P36334 |
| KVVGKDGVMYSE | 1 | 0 | 0 | 1463208 | AWW13511.1 | P36334 |
| KVVKGTHHWMLL | 1 | 0 | 0 | 1463209 | APO40578.1 | P0C6X7 |
| KVVMSMFSRTTS | 1 | 0 | 0 | 1463220 | ATO98193.1 | Q3LZX1 |
| KVVTTLKPTNQA | 1 | 0 | 0 | 1463229 | ATO98145.1 | Q3LZX1 |
| KVVVGPTTEVKS | 1 | 0 | 0 | 1463230 | AWK59942.1 | H9EJ42 |
| KVYFDNPTLPFG | 1 | 0 | 0 | 1463236 | ATO98193.1 | Q3LZX1 |
| KWADASETQRGR | 1 | 0 | 0 | 1463249 | AIW52715.1 | A0A0N7J131 |
| KWADESEPQRGR | 1 | 0 | 0 | 1463250 | ABB90491.1 | A0A0N7J131 |
| KWADNNCYLATA | 1 | 0 | 0 | 1463251 | AVP78030.1 | P0C6W2 |
| KWADNNSYLSSV | 1 | 0 | 0 | 1463254 | 4M0W_A | P0C6U8 |
| KWADQSDQFRSV | 1 | 0 | 0 | 1463257 | AIL49389.1 | P33469 |
| KWADQSDQVRNV | 1 | 0 | 0 | 1463258 | P33469.1 | P33469 |
| KWESGVKDYVTL | 1 | 0 | 0 | 1463275 | APO40580.1 | P59632 |
| KWEYEGGCNTIE | 1 | 0 | 0 | 1463276 | AWK59942.1 | H9EJ42 |
| KWIFDSSKEFVE | 1 | 0 | 0 | 1463280 | AIV41816.1 | P0C6X6 |
| KWLELLEANIDA | 1 | 0 | 0 | 1463294 | AVV62544.1 | K9N4V7 |
| KWNGVSDLSLKQ | 1 | 0 | 0 | 1463309 | ASU90779.1 | K9N638 |
| KWSGISDLSLKQ | 1 | 0 | 0 | 1463323 | ALA49758.1 | K9N7C7 |
| KWSGVSDLSSNK | 1 | 0 | 0 | 1463325 | QGV13494.1 | K9N7C7 |
| KWTDVDDYILAN | 1 | 0 | 0 | 1463329 | AEN19363.1 | P0C6X6 |
| KYAPNSRSTSRT | 1 | 0 | 0 | 1463354 | AJQ24067.1 | P33469 |
| KYDDYDLEPQKI | 1 | 0 | 0 | 1463371 | AUM60024.1 | K9N5Q8 |
| KYDSGSLTKSSE | 1 | 0 | 0 | 1463381 | AVV62535.1 | K9N7C7 |
| KYFDLGFIEAVN | 1 | 0 | 0 | 1463397 | AIW52843.1 | A0A0P0G321 |
| KYFISEPPVTYS | 1 | 0 | 0 | 1463404 | ALA49340.1 | K9N7C7 |
| KYFNTVTKLMCC | 1 | 0 | 0 | 1463413 | AEN19363.1 | P0C6X6 |
| KYFTSPPPITFS | 1 | 0 | 0 | 1463427 | AUM60013.1 | K9N7C7 |
| KYKLENFAFEHV | 1 | 0 | 0 | 1463462 | AVV62524.1 | K9N7C7 |
| KYMCGFDGKPML | 1 | 0 | 0 | 1463489 | AFD98826.1 | H9EJ42 |
| KYNENGTITDAVDCALDPLS | 1 | 0 | 0 | 1463497 | P0DTC2.1 | P0DTC2 |
| KYNIYDVAGVGV | 1 | 0 | 0 | 1463501 | AIW52741.1 | A0A0P0K6L9 |
| KYPQVGGLTSIK | 1 | 0 | 0 | 1463517 | APO40578.1 | P0C6X7 |
| KYQSFVQWFHEN | 1 | 0 | 0 | 1463522 | AVV62535.1 | K9N7C7 |
| KYRHCDLDTTLY | 1 | 0 | 0 | 1463527 | 5I08_A | A0A140H1H1 |
| KYRHCDVDTTTL | 1 | 0 | 0 | 1463530 | AGT17758.1 | A0A140H1H1 |
| KYSELDDEEPME | 1 | 0 | 0 | 1463537 | Q3I5J1.1 | Q3LZX8 |
| KYSTFMLNFNEN | 1 | 0 | 0 | 1463553 | ATO98193.1 | Q3LZX1 |
| KYTDFNEDVCVC | 1 | 0 | 0 | 1463564 | AWK59942.1 | H9EJ42 |
| KYTDIDINPALN | 1 | 0 | 0 | 1463565 | AVV62535.1 | K9N7C7 |
| KYTDIDVSPALN | 1 | 0 | 0 | 1463570 | AVV62524.1 | K9N7C7 |
| KYTDLNFIENLN | 1 | 0 | 0 | 1463572 | ARB07436.1 | A0A140H1G9 |
| KYTGVSGCLTDC | 1 | 0 | 0 | 1463578 | ARU07575.1 | A0A140H1G9 |
| KYTKIFKDDGNF | 1 | 0 | 0 | 1463579 | AMK59674.1 | P0C6X6 |
| KYTKIIKDDGNC | 1 | 0 | 0 | 1463580 | ARB07436.1 | A0A140H1G9 |
| KYVIVATPSTTI | 1 | 0 | 0 | 1463617 | AWK59934.1 | U3M6Q8 |
| KYVKWEHDSGVV | 1 | 0 | 0 | 1463623 | NP_073549.1 | A0A0P0LKV0 |
| KYYDDSQYFFNK | 1 | 0 | 0 | 1463646 | AGT51759.1 | P30215 |
| KYYDDSQYYLNK | 1 | 0 | 0 | 1463648 | AGT51480.1 | P30215 |
| KYYTVIPRSIRS | 1 | 0 | 0 | 1463670 | AVV62526.1 | K9N5Q8 |
| LAAINSVPWDTI | 1 | 0 | 0 | 1463692 | AVP78041.1 | P0C6W2 |
| LAAKEGINSLAD | 1 | 0 | 0 | 1463693 | AVV62535.1 | K9N7C7 |
| LAALESGKTKAA | 1 | 0 | 0 | 1463695 | AVV62544.1 | K9N4V7 |
| LAATSANLFPPW | 1 | 0 | 0 | 1463708 | CAA83661.1 | P36334 |
| LAAYDIEVTKGS | 1 | 0 | 0 | 1463721 | AUM60013.1 | K9N7C7 |
| LAAYDIGVTVGP | 1 | 0 | 0 | 1463722 | AVV62535.1 | K9N7C7 |
| LAAYNGPPQGAF | 1 | 0 | 0 | 1463723 | ABD75599.1 | A0A140H1G9 |
| LACEVLKPVSEE | 1 | 0 | 0 | 1463732 | AVP78041.1 | P0C6W2 |
| LACYNGRPTGTF | 1 | 0 | 0 | 1463756 | AXN92247.1 | K9N7C7 |
| LADGLEAYADRT | 1 | 0 | 0 | 1463766 | AIV41774.1 | P0C6X6 |
| LADKFPVLHDIGNPKAIKCVP | 1 | 0 | 0 | 1463772 | P0DTD1.1 | P0DTD1 |
| LADLYHYQECAR | 1 | 0 | 0 | 1463777 | APO40584.1 | P59635 |
| LADNLTADEIKA | 1 | 0 | 0 | 1463787 | AVN89363.1 | K9N7C7 |
| LADNLTADETKA | 1 | 0 | 0 | 1463788 | AXN92247.1 | K9N7C7 |
| LADSNGFIRFPD | 1 | 0 | 0 | 1463791 | ABD75543.1 | A0A140H1G9 |
| LADSVMSAVSSG | 1 | 0 | 0 | 1463793 | AXX83372.1 | P0C6X6 |
| LADTVMSAVSAG | 1 | 0 | 0 | 1463797 | AQN78757.1 | P0C6X6 |
| LADVATDVQSRA | 1 | 0 | 0 | 1463799 | AUM60013.1 | K9N7C7 |
| LADVEADVAARA | 1 | 0 | 0 | 1463801 | AXN92247.1 | K9N7C7 |
| LADYDIAVTAGP | 1 | 0 | 0 | 1463803 | AQZ41321.1 | K9N7C7 |
| LADYTQCAVIAV | 1 | 0 | 0 | 1463810 | AVV62535.1 | K9N7C7 |
| LAEFDENGTITD | 1 | 0 | 0 | 1463817 | APO40579.1 | P59594 |
| LAEFDTYDPTYK | 1 | 0 | 0 | 1463819 | AVN89451.1 | K9N7C7 |
| LAEFSLNNHHST | 1 | 0 | 0 | 1463820 | AVV62541.1 | K9N7D2 |
| LAEFVHYDPMYK | 1 | 0 | 0 | 1463821 | AUM60013.1 | K9N7C7 |
| LAELAPPAGAFF | 1 | 0 | 0 | 1463823 | ANZ78874.1 | P33469 |
| LAELIPNHAALF | 1 | 0 | 0 | 1463824 | AMM42619.1 | H9EJ69 |
| LAESFKKVAAVP | 1 | 0 | 0 | 1463832 | AUM60013.1 | K9N7C7 |
| LAFDEAVAAAMA | 1 | 0 | 0 | 1463839 | AVV62533.1 | K9N4V7 |
| LAFGVPLSEAKG | 1 | 0 | 0 | 1463844 | ABA60796.1 | Q19U29 |
| LAFMPVQSRRFI | 1 | 0 | 0 | 1463851 | AEN19355.1 | P0C6X6 |
| LAFSKVQDAVNA | 1 | 0 | 0 | 1463855 | AVV62537.1 | K9N5Q8 |
| LAFSKVQEAVNA | 1 | 0 | 0 | 1463856 | AVV62526.1 | K9N5Q8 |
| LAGHDDFSTEII | 1 | 0 | 0 | 1463866 | APO40578.1 | P0C6X7 |
| LAGHTGHLPGPR | 1 | 0 | 0 | 1463868 | AVN89290.1 | K9N7C7 |

| | | | | | | |
|---|---|---|---|---|---|---|
| LAGLGESFKKVA | 1 | 0 | 0 | 1463877 | AVV62535.1 | K9N7C7 |
| LAIDAYPLVHHE | 1 | 0 | 0 | 1463903 | ARB07436.1 | A0A140H1G9 |
| LAIKSDHCGGNP | 1 | 0 | 0 | 1463912 | AVI15011.1 | P36334 |
| LAIKSDYCGGNP | 1 | 0 | 0 | 1463913 | CAA83660.1 | P36334 |
| LAILTALRLAAY | 1 | 0 | 0 | 1463917 | 5X29_A | P59637 |
| LAKAELCCSARM | 1 | 0 | 0 | 1463933 | AXN92247.1 | K9N7C7 |
| LAKAINDFSNSG | 1 | 0 | 0 | 1463934 | AVP78030.1 | P0C6W2 |
| LAKALNDFSNLG | 1 | 0 | 0 | 1463936 | BAE93400.1 | P0C6U8 |
| LAKDDLYVKRYE | 1 | 0 | 0 | 1463943 | ATN39862.1 | P0C6X6 |
| LAKLGKDFKPQQ | 1 | 0 | 0 | 1463960 | ABD75549.1 | Q19U29 |
| LAKLGRDATKPQ | 1 | 0 | 0 | 1463963 | AGT51674.1 | P33469 |
| LAKNCLHVVGPN | 1 | 0 | 0 | 1463965 | AVP78041.1 | P0C6W2 |
| LAKQDAAQRISW | 1 | 0 | 0 | 1463972 | AMQ49017.1 | K9N4V0 |
| LAKQDAARQIAW | 1 | 0 | 0 | 1463973 | AVV62528.1 | K9N4V0 |
| LAKQDAARQVAW | 1 | 0 | 0 | 1463974 | AVV62539.1 | K9N4V0 |
| LAKTELCCSARM | 1 | 0 | 0 | 1463982 | AXN73469.1 | K9N7C7 |
| LAKVHNLSGNPD | 1 | 0 | 0 | 1463985 | AHA80155.1 | P33469 |
| LAKVQNLFGNPD | 1 | 0 | 0 | 1463986 | AHA80176.1 | P33469 |
| LAKVRNLSGNPD | 1 | 0 | 0 | 1463988 | AHA80174.1 | P33469 |
| LALLNLTDEDLA | 1 | 0 | 0 | 1464022 | AXN92231.1 | K9N643 |
| LALLSDLQDLKWARF | 1 | 0 | 0 | 1464027 | P0DTC1.1 | P0DTD1 |
| LALQQIDIKFNA | 1 | 0 | 0 | 1464031 | APO40578.1 | P0C6X7 |
| LAMAYGDCTFGV | 1 | 0 | 0 | 1464049 | AVV62535.1 | K9N7C7 |
| LANAKGFIRFPE | 1 | 0 | 0 | 1464066 | AEN19363.1 | P0C6X6 |
| LANANSKPLIGY | 1 | 0 | 0 | 1464067 | AUM60025.1 | K9N796 |
| LANANSKPLVGY | 1 | 0 | 0 | 1464068 | AUM60015.1 | K9N796 |
| LANARYVSPDVQ | 1 | 0 | 0 | 1464069 | AVV62526.1 | K9N5Q8 |
| LANDDLYVKRYE | 1 | 0 | 0 | 1464070 | AEN19363.1 | P0C6X6 |
| LANFNETNGPLC | 1 | 0 | 0 | 1464079 | CAA71147.1 | A0A0P0K6L9 |
| LANGTHTGSAFD | 1 | 0 | 0 | 1464082 | AXN92247.1 | K9N7C7 |
| LANLNSHNIGSL | 1 | 0 | 0 | 1464092 | CAA71146.1 | A0A0P0K6L9 |
| LANLNSHTIGSL | 1 | 0 | 0 | 1464093 | AGT21338.1 | A0A0P0K6L9 |
| LANMGIDLDEWS | 1 | 0 | 0 | 1464094 | ARO76381.1 | P0C6U8 |
| LANPTNCIAPVN | 1 | 0 | 0 | 1464096 | AVN89387.1 | K9N5Q8 |
| LANQFTEFVSTQ | 1 | 0 | 0 | 1464098 | ATG84865.1 | K9N638 |
| LANSNLRTFCVD | 1 | 0 | 0 | 1464100 | AVV62524.1 | K9N7C7 |
| LANTTTAPLKLF | 1 | 0 | 0 | 1464102 | AGV08437.1 | K9N7C7 |
| LANVISFGDYNL | 1 | 0 | 0 | 1464103 | AFD64760.1 | A0A0P0G321 |
| LAPQAKSGIITK | 1 | 0 | 0 | 1464121 | AIL49375.1 | P30215 |
| LAPRSVRSVPGE | 1 | 0 | 0 | 1464122 | ALJ54472.1 | K9N5Q8 |
| LAPSASAFMSMS | 1 | 0 | 0 | 1464127 | AVV62544.1 | K9N4V7 |
| LARAGKSCAPLS | 1 | 0 | 0 | 1464157 | APO40578.1 | P0C6X7 |
| LARALKKVPIDE | 1 | 0 | 0 | 1464161 | ARO76381.1 | P0C6U8 |
| LARDYRANSFDI | 1 | 0 | 0 | 1464167 | AVV62535.1 | K9N7C7 |
| LARKHDTCCSQS | 1 | 0 | 0 | 1464172 | ANZ78857.1 | P0C6X6 |
| LARKHSTCCNLS | 1 | 0 | 0 | 1464176 | P0C6W6.1 | P0C6W2 |
| LARRYETCCSQS | 1 | 0 | 0 | 1464178 | AJE26237.1 | P0C6X6 |
| LASCFFTCNSNA | 1 | 0 | 0 | 1464187 | AIW52843.1 | A0A0P0G321 |
| LASCFSTCNSNA | 1 | 0 | 0 | 1464189 | AVL25555.1 | A0A0P0G321 |
| LASDKVNECVKS | 1 | 0 | 0 | 1464193 | AVV62526.1 | K9N5Q8 |
| LASDLLDNASVL | 1 | 0 | 0 | 1464194 | AVV62535.1 | K9N7C7 |
| LASFAELEAAQK | 1 | 0 | 0 | 1464200 | AVV62524.1 | K9N7C7 |
| LASIAFNHPIQI | 1 | 0 | 0 | 1464209 | ASU91208.1 | K9N5Q8 |
| LASIAFNHPIQV | 1 | 0 | 0 | 1464210 | AXN92228.1 | K9N5Q8 |
| LASNYRENSFDI | 1 | 0 | 0 | 1464216 | AVV62524.1 | K9N7C7 |
| LASPMKSDEGGL | 1 | 0 | 0 | 1464217 | ARU07575.1 | A0A140H1G9 |
| LASRFADGLCLF | 1 | 0 | 0 | 1464224 | AUM60013.1 | K9N7C7 |
| LASSSLRTFVVD | 1 | 0 | 0 | 1464228 | AXN92247.1 | K9N7C7 |
| LASVIQDYENVC | 1 | 0 | 0 | 1464231 | AIV41852.1 | P0C6X6 |
| LASYLQQGSTGE | 1 | 0 | 0 | 1464234 | AUM60014.1 | K9N5Q8 |
| LATFAELEAARK | 1 | 0 | 0 | 1464241 | AMQ49036.1 | K9N638 |
| LATIEELPDEFV | 1 | 0 | 0 | 1464252 | APO40587.1 | P59636 |
| LATINYTQPQLL | 1 | 0 | 0 | 1464256 | AVV62537.1 | K9N5Q8 |
| LAVIGDLKCTSD | 1 | 0 | 0 | 1464291 | 6NZK_A | P36334 |
| LAVIGDLKCTTV | 1 | 0 | 0 | 1464292 | CAA83661.1 | P36334 |
| LAVNTGVFVAPP | 1 | 0 | 0 | 1464301 | AVV62528.1 | K9N4V0 |
| LAVSGQAFCDYT | 1 | 0 | 0 | 1464312 | AGT51737.1 | P0C6X6 |
| LAVTAIVVVSGC | 1 | 0 | 0 | 1464315 | ACA49954.1 | H9EJ42 |
| LAVVNSQDVYKS | 1 | 0 | 0 | 1464316 | ANF29227.1 | K9N638 |
| LAVYDIEVTKGP | 1 | 0 | 0 | 1464319 | AUM60023.1 | K9N7C7 |
| LAWLYAAVINGE | 1 | 0 | 0 | 1464326 | APO40578.1 | P0C6X7 |
| LAYAEETRKIMP | 1 | 0 | 0 | 1464335 | APO40578.1 | P0C6X7 |
| LAYFSDSKFGTT | 1 | 0 | 0 | 1464338 | AHI48527.1 | K9N638 |
| LAYNKLQSVDWK | 1 | 0 | 0 | 1464351 | AVV62524.1 | K9N7C7 |
| LAYSSSANGSLD | 1 | 0 | 0 | 1464361 | AXG21642.1 | K9N638 |
| LCAKILPFSDRP | 1 | 0 | 0 | 1464385 | AWK59942.1 | H9EJ42 |
| LCALPDTPSTLA | 1 | 0 | 0 | 1464391 | ALJ54472.1 | K9N5Q8 |
| LCALPDTPSTLK | 1 | 0 | 0 | 1464392 | ALJ54490.1 | K9N5Q8 |
| LCATCPFVKGDS | 1 | 0 | 0 | 1464400 | AKN24784.1 | K9N7C7 |
| LCAVPPTSSTAR | 1 | 0 | 0 | 1464407 | AUM60014.1 | K9N5Q8 |
| LCCKCCYDHVMA | 1 | 0 | 0 | 1464417 | AEN19363.1 | P0C6X6 |
| LCCMTNCCSCFK | 1 | 0 | 0 | 1464419 | APO40579.1 | P59594 |
| LCDDPEKAQGML | 1 | 0 | 0 | 1464436 | AWK59942.1 | H9EJ42 |
| LCDEMLNNRATL | 1 | 0 | 0 | 1464439 | APO40578.1 | P0C6X7 |
| LCDLANPTNCIA | 1 | 0 | 0 | 1464447 | AVN89387.1 | K9N5Q8 |
| LCDNPFIAVSKP | 1 | 0 | 0 | 1464450 | ABD73001.1 | P59594 |
| LCDVANPTNCIA | 1 | 0 | 0 | 1464458 | AKM76239.1 | K9N5Q8 |
| LCEKALKYLPIDKC | 1 | 0 | 0 | 1464469 | P0DTD1.1 | P0DTD1 |
| LCERLTGGNYSD | 1 | 0 | 0 | 1464471 | AUM60013.1 | K9N7C7 |
| LCGDIVSGYGKS | 1 | 0 | 0 | 1464503 | AAY32939.1 | H9EJ42 |
| LCGDNSAKFWEE | 1 | 0 | 0 | 1464504 | AXN92247.1 | K9N7C7 |

| | | | | | | |
|---|---|---|---|---|---|---|
| LCIFNGKSFSNT | 1 | 0 | 0 | 1464528 | ARK08650.1 | P30215 |
| LCIFNGKSLSNT | 1 | 0 | 0 | 1464529 | ATN39871.1 | P30215 |
| LCKEPMYAISNE | 1 | 0 | 0 | 1464540 | ATO98193.1 | Q3LZX1 |
| LCLEAGINYVKS | 1 | 0 | 0 | 1464564 | ARO76381.1 | P0C6U8 |
| LCLRDDNQALLV | 1 | 0 | 0 | 1464585 | AMM42360.1 | A0A140H1G9 |
| LCNPDPIRFTGP | 1 | 0 | 0 | 1464603 | AGT51422.1 | P36334 |
| LCNPDPITFAGP | 1 | 0 | 0 | 1464604 | ATP16757.1 | P36334 |
| LCNPDPITFTGP | 1 | 0 | 0 | 1464605 | BBA84099.1 | P36334 |
| LCNPDPITFTGS | 1 | 0 | 0 | 1464606 | AVQ05264.1 | P36334 |
| LCNTDPITFTGP | 1 | 0 | 0 | 1464612 | BBO58736.1 | P36334 |
| LCPFDAVFNATR | 1 | 0 | 0 | 1464614 | ATO98145.1 | Q3LZX1 |
| LCPPKNSFEALM | 1 | 0 | 0 | 1464619 | ATN39862.1 | P0C6X6 |
| LCRVTLGDFTIM | 1 | 0 | 0 | 1464626 | ARU07575.1 | A0A140H1G9 |
| LCTLDPIALKAP | 1 | 0 | 0 | 1464646 | BBO58891.1 | P36334 |
| LCTPDPITFTGT | 1 | 0 | 0 | 1464657 | P36334.1 | P36334 |
| LCVADDKPIVLF | 1 | 0 | 0 | 1464665 | NP_073549.1 | A0A0P0LKV0 |
| LCVDIPSIPKDM | 1 | 0 | 0 | 1464671 | APO40578.1 | P0C6X7 |
| LCVGNGPGIDAG | 1 | 0 | 0 | 1464678 | CAA83660.1 | P36334 |
| LCVGSGPGKNNG | 1 | 0 | 0 | 1464679 | AVQ05264.1 | P36334 |
| LCVQCAMGCNTL | 1 | 0 | 0 | 1464693 | AVV62542.1 | K9N5R3 |
| LCVQCAVGFNTL | 1 | 0 | 0 | 1464694 | AUM60019.1 | K9N5R3 |
| LCVQCGMGFNTL | 1 | 0 | 0 | 1464695 | AVV62531.1 | K9N5R3 |
| LCVTKGILTGCF | 1 | 0 | 0 | 1464702 | AVN89322.1 | K9N7C7 |
| LCWKCKSKNPLI | 1 | 0 | 0 | 1464707 | APO40580.1 | P59632 |
| LCYALRHFDCND | 1 | 0 | 0 | 1464713 | ARB07436.1 | A0A140H1G9 |
| LCYDIGNPKAIA | 1 | 0 | 0 | 1464715 | AGT51787.1 | P0C6X6 |
| LDANIPDYPKLF | 1 | 0 | 0 | 1464737 | AVV62535.1 | K9N7C7 |
| LDANIPGYPKLF | 1 | 0 | 0 | 1464738 | AVV62524.1 | K9N7C7 |
| LDCTGQAVNTSK | 1 | 0 | 0 | 1464758 | AVV62539.1 | K9N4V0 |
| LDDATRNYIITM | 1 | 0 | 0 | 1464766 | AHL18094.1 | K9N643 |
| LDDATRSYIGST | 1 | 0 | 0 | 1464767 | AVV62540.1 | K9N643 |
| LDDEELMELDYP | 1 | 0 | 0 | 1464771 | AAP30035.1 | P59634 |
| LDDEESVEEEIV | 1 | 0 | 0 | 1464774 | AVV62535.1 | K9N7C7 |
| LDDFSKQLQDSM | 1 | 0 | 0 | 1464780 | APO40586.1 | P59595 |
| LDDSIEMQLFKV | 1 | 0 | 0 | 1464794 | AWK59942.1 | H9EJ42 |
| LDEAEDVTEFVQ | 1 | 0 | 0 | 1464806 | AXN92247.1 | K9N7C7 |
| LDEARTALKKCK | 1 | 0 | 0 | 1464811 | ARO76381.1 | P0C6U8 |
| LDEEGNLLDSYF | 1 | 0 | 0 | 1464813 | APO40578.1 | P0C6X7 |
| LDEFFKNFSTSI | 1 | 0 | 0 | 1464815 | AKM76239.1 | K9N5Q8 |
| LDEGVDVKIQTA | 1 | 0 | 0 | 1464821 | AIV41842.1 | Q80872 |
| LDELHACTSFTC | 1 | 0 | 0 | 1464825 | AVV62535.1 | K9N7C7 |
| LDEQESKFWDEN | 1 | 0 | 0 | 1464829 | AGT21344.1 | A0A0P0LKV0 |
| LDEQESKFWDES | 1 | 0 | 0 | 1464830 | AIW52776.1 | A0A0P0LKV0 |
| LDFKEELEEFFK | 1 | 0 | 0 | 1464845 | AVV62537.1 | K9N5Q8 |
| LDFLDCTGQAVN | 1 | 0 | 0 | 1464848 | AVV62539.1 | K9N4V0 |
| LDFVNAGGVPLT | 1 | 0 | 0 | 1464855 | AHI48749.1 | K9N638 |
| LDFYPALETVQV | 1 | 0 | 0 | 1464857 | APO40578.1 | P0C6X7 |
| LDGETITFDNLK | 1 | 0 | 0 | 1464863 | AVP78030.1 | P0C6W2 |
| LDGIFKTEVDPN | 1 | 0 | 0 | 1464871 | ATQ39389.1 | K9N7C7 |
| LDGIQYGRSGET | 1 | 0 | 0 | 1464872 | AVP78041.1 | P0C6W2 |
| LDGISKAEVDPD | 1 | 0 | 0 | 1464877 | AVV62524.1 | K9N7C7 |
| LDGLEQDAQIDR | 1 | 0 | 0 | 1464880 | AHI48733.1 | K9N5Q8 |
| LDGNFRTGVDPD | 1 | 0 | 0 | 1464885 | ASU90030.1 | K9N7C7 |
| LDGSLCVGNGPG | 1 | 0 | 0 | 1464886 | CAA83660.1 | P36334 |
| LDGSYNASCLCS | 1 | 0 | 0 | 1464887 | BBA20983.1 | A0A140H1H1 |
| LDGYYKKDNSYF | 1 | 0 | 0 | 1464901 | AVP78041.1 | P0C6W2 |
| LDHTSLVWKHIR | 1 | 0 | 0 | 1464907 | AWK59942.1 | H9EJ42 |
| LDICFEATEATF | 1 | 0 | 0 | 1464908 | ASU45805.1 | K9N7C7 |
| LDIEKLRQYLAA | 1 | 0 | 0 | 1464910 | AVV62535.1 | K9N7C7 |
| LDILDITPCSFG | 1 | 0 | 0 | 1464919 | ATO98145.1 | Q3LZX1 |
| LDILGPLSAQTG | 1 | 0 | 0 | 1464920 | AVP78041.1 | P0C6W2 |
| LDIVDTSLSGFK | 1 | 0 | 0 | 1464936 | AVP78041.1 | P0C6W2 |
| LDIYDLTMPSQV | 1 | 0 | 0 | 1464937 | AIV41960.1 | P0C6X6 |
| LDKIQNQTLPRL | 1 | 0 | 0 | 1464943 | ARU07574.1 | A0A140H1G9 |
| LDKLIATAKDGL | 1 | 0 | 0 | 1464944 | AVV62524.1 | K9N7C7 |
| LDLLKDIKLVIP | 1 | 0 | 0 | 1464978 | AVV61898.1 | K9N7C7 |
| LDLQAEMNRLQE | 1 | 0 | 0 | 1464986 | AGT51521.1 | P36334 |
| LDLQDEMNRLQE | 1 | 0 | 0 | 1464988 | CAA83660.1 | P36334 |
| LDLQVEINRLQE | 1 | 0 | 0 | 1464990 | AIL49487.1 | P36334 |
| LDLQVEMNRLQD | 1 | 0 | 0 | 1464991 | QEG03776.1 | P36334 |
| LDLRTSCFSVAA | 1 | 0 | 0 | 1464993 | AXN92247.1 | K9N7C7 |
| LDLSDEMKVLNA | 1 | 0 | 0 | 1464995 | AVV62526.1 | K9N5Q8 |
| LDLSTEMKVLNE | 1 | 0 | 0 | 1465001 | AVV62537.1 | K9N5Q8 |
| LDLTPVIDDVDV | 1 | 0 | 0 | 1465006 | AWK59942.1 | H9EJ42 |
| LDLVKRDSEADS | 1 | 0 | 0 | 1465012 | ABD75589.1 | Q19U29 |
| LDNKFTPLKVEE | 1 | 0 | 0 | 1465038 | AUM60013.1 | K9N7C7 |
| LDNKLPPPLLSN | 1 | 0 | 0 | 1465039 | AVV62526.1 | K9N5Q8 |
| LDNLRANNTKGV | 1 | 0 | 0 | 1465044 | APO40578.1 | P0C6X7 |
| LDNTTQSALLVN | 1 | 0 | 0 | 1465054 | ATO98145.1 | Q3LZX1 |
| LDNTTQSAVLFN | 1 | 0 | 0 | 1465055 | APO40579.1 | P59594 |
| LDPISFKAPDTY | 1 | 0 | 0 | 1465067 | ARE30008.1 | P36334 |
| LDPITFKAPDTY | 1 | 0 | 0 | 1465068 | ARB07414.1 | P36334 |
| LDPKLKGSFNDR | 1 | 0 | 0 | 1465073 | AIL49495.1 | P36334 |
| LDPRLKGSFNDR | 1 | 0 | 0 | 1465084 | AIL49520.1 | P36334 |
| LDQPPTASVSTS | 1 | 0 | 0 | 1465100 | AIV41798.1 | P0C6X6 |
| LDQTSFNVNSLA | 1 | 0 | 0 | 1465105 | AVV62535.1 | K9N7C7 |
| LDQWLKNQTSVA | 1 | 0 | 0 | 1465109 | ARB07414.1 | P36334 |
| LDRLNQLENKIS | 1 | 0 | 0 | 1465114 | AVP78049.1 | Q3LZX4 |
| LDRLNQLENKVS | 1 | 0 | 0 | 1465115 | AVP78038.1 | Q3LZX4 |
| LDRNQHSKLDEE | 1 | 0 | 0 | 1465119 | APO40583.1 | P59634 |
| LDSNFRTEVDPD | 1 | 0 | 0 | 1465158 | ASU90302.1 | K9N7C7 |

| | | | | | | |
|---|---|---|---|---|---|---|
| LDSNIDAYKNFP | 1 | 0 | 0 | 1465159 | AUM60021.1 | K9N4V7 |
| LDSRTGSFNNID | 1 | 0 | 0 | 1465162 | AIX10756.1 | P36334 |
| LDSTYGPLQTPV | 1 | 0 | 0 | 1465164 | AXG21654.1 | K9N5Q8 |
| LDTAPQTGNFKD | 1 | 0 | 0 | 1465172 | ATO98193.1 | Q3LZX1 |
| LDTCFEAIEATF | 1 | 0 | 0 | 1465173 | AXN92226.1 | K9N7C7 |
| LDTCFEATEATL | 1 | 0 | 0 | 1465176 | ATQ39389.1 | K9N7C7 |
| LDTCFGVSKPNA | 1 | 0 | 0 | 1465178 | AWK59942.1 | H9EJ42 |
| LDTCFSVSKPNA | 1 | 0 | 0 | 1465179 | AVA26872.1 | H9EJ42 |
| LDTLLASSSFRT | 1 | 0 | 0 | 1465189 | ANI69888.1 | K9N638 |
| LDTRYKGTFNNK | 1 | 0 | 0 | 1465194 | AKA09647.1 | P36334 |
| LDTTLYVNNWCR | 1 | 0 | 0 | 1465203 | Q14EB0.1 | A0A140H1H1 |
| LDTTQLQVANNL | 1 | 0 | 0 | 1465204 | AIL49500.1 | P36334 |
| LDVAYANYNTYD | 1 | 0 | 0 | 1465209 | APO40578.1 | P0C6X7 |
| LDVCNTKEFKVF | 1 | 0 | 0 | 1465211 | ARX11394.1 | A0A0P0LKV0 |
| LDVKGLEELHDD | 1 | 0 | 0 | 1465218 | ATP16755.1 | Q80872 |
| LDVKGLEELHYD | 1 | 0 | 0 | 1465219 | AIV41836.1 | Q80872 |
| LDVLEQSGGRGG | 1 | 0 | 0 | 1465224 | 4NJL_A | K9N5Q8 |
| LDVMLPGEFSKK | 1 | 0 | 0 | 1465228 | AKK52590.1 | K9N7C7 |
| LDVQEESEPLCD | 1 | 0 | 0 | 1465231 | APO40578.1 | P0C6X7 |
| LDVVRKDLEEGD | 1 | 0 | 0 | 1465237 | ARK08633.1 | Q80872 |
| LDVVRKELEEGD | 1 | 0 | 0 | 1465238 | AWW13564.1 | Q80872 |
| LDVYNLTMPSQV | 1 | 0 | 0 | 1465242 | AAX85666.1 | P0C6X6 |
| LDYCDGNLPCGI | 1 | 0 | 0 | 1465252 | APO40578.1 | P0C6X7 |
| LDYPKCDRAMPN | 1 | 0 | 0 | 1465268 | ARO76377.1 | P0C6X7 |
| LDYPKCDRAMPS | 1 | 0 | 0 | 1465269 | AKL87217.1 | A0A0P0LKV0 |
| LEAAQKSYQEAL | 1 | 0 | 0 | 1465280 | AVV62524.1 | K9N7C7 |
| LEAAQKSYQEAV | 1 | 0 | 0 | 1465281 | AVV62535.1 | K9N7C7 |
| LEACYERGCNPK | 1 | 0 | 0 | 1465282 | AAM95936.1 | P0C6U7 |
| LEALESGKQKTA | 1 | 0 | 0 | 1465299 | AUM60021.1 | K9N4V7 |
| LEAMQTGFTTTN | 1 | 0 | 0 | 1465302 | QBF80499.1 | K9N5Q8 |
| LEANIDAYKTFP | 1 | 0 | 0 | 1465304 | AVV62544.1 | K9N4V7 |
| LEANSKAWAWCV | 1 | 0 | 0 | 1465307 | AVV62524.1 | K9N7C7 |
| LEASFNYLKSPNFS | 1 | 0 | 0 | 1465316 | P0DTC1.1 | P0DTD1 |
| LEAYADRTLQEM | 1 | 0 | 0 | 1465327 | AIV41774.1 | P0C6X6 |
| LECTRASTSAVK | 1 | 0 | 0 | 1465335 | AXN92247.1 | K9N7C7 |
| LECVGFTTASVY | 1 | 0 | 0 | 1465336 | AUM60013.1 | K9N7C7 |
| LEDACFKFNYQF | 1 | 0 | 0 | 1465339 | AQN78665.1 | Q04853 |
| LEDEFTPFDVVA | 1 | 0 | 0 | 1465344 | 3M3T_A | P0C6X7 |
| LEDFIDNPCYCF | 1 | 0 | 0 | 1465346 | AVV62524.1 | K9N7C7 |
| LEDFLPIDSTVK | 1 | 0 | 0 | 1465352 | APO40578.1 | P0C6X7 |
| LEDFNLEDFIDS | 1 | 0 | 0 | 1465353 | AVV62535.1 | K9N7C7 |
| LEDIDYVPLKSA | 1 | 0 | 0 | 1465354 | AVV62524.1 | K9N7C7 |
| LEDLEAGKNKAA | 1 | 0 | 0 | 1465360 | AVV62533.1 | K9N4V7 |
| LEEAARCMRSLR | 1 | 0 | 0 | 1465373 | 2W2G_A | P0C6U8 |
| LEEAVRQVRSWI | 1 | 0 | 0 | 1465381 | AHL18091.1 | K9N7C7 |
| LEEDDIVYFPYE | 1 | 0 | 0 | 1465383 | AVV62535.1 | K9N7C7 |
| LEELHDDLVNYI | 1 | 0 | 0 | 1465390 | ATP16755.1 | Q80872 |
| LEESGAVLFCTP | 1 | 0 | 0 | 1465398 | NP_073549.1 | A0A0P0LKV0 |
| LEETNMHSPDLV | 1 | 0 | 0 | 1465405 | AVV62524.1 | K9N7C7 |
| LEFGASTEMQVE | 1 | 0 | 0 | 1465411 | ATO98229.1 | P0C6W2 |
| LEFKVLDNGIIR | 1 | 0 | 0 | 1465416 | AUM60013.1 | K9N7C7 |
| LEFRAGRPARFV | 1 | 0 | 0 | 1465417 | AVQ05261.1 | P0C6X6 |
| LEFVREEVDIIE | 1 | 0 | 0 | 1465422 | AWK59942.1 | H9EJ42 |
| LEGGGILVGVGS | 1 | 0 | 0 | 1465427 | AVV62537.1 | K9N5Q8 |
| LEGGGWLVASGP | 1 | 0 | 0 | 1465429 | AKN24830.1 | K9N5Q8 |
| LEGLASDLLDNA | 1 | 0 | 0 | 1465438 | AVV62535.1 | K9N7C7 |
| LEGLASDLLDNV | 1 | 0 | 0 | 1465439 | AVV62524.1 | K9N7C7 |
| LEISVCQYNMCE | 1 | 0 | 0 | 1465466 | AVQ05264.1 | P36334 |
| LEIVKLGAEKED | 1 | 0 | 0 | 1465469 | AMK59675.1 | Q80872 |
| LEKFMSTPNAVF | 1 | 0 | 0 | 1465479 | ATG84897.1 | K9N7C7 |
| LEKFMSTSNAIL | 1 | 0 | 0 | 1465480 | AVV62524.1 | K9N7C7 |
| LEKLKTILALRE | 1 | 0 | 0 | 1465489 | APO40578.1 | P0C6X7 |
| LEKLQCEHLQFG | 1 | 0 | 0 | 1465491 | AWO68978.1 | A0A0P0G321 |
| LEKRCYELLKDC | 1 | 0 | 0 | 1465495 | AUM60013.1 | K9N7C7 |
| LEKRCYELVKDC | 1 | 0 | 0 | 1465496 | AVV62524.1 | K9N7C7 |
| LELAKVRNLSGN | 1 | 0 | 0 | 1465502 | AHA80174.1 | P33469 |
| LELDPPCKFSIQ | 1 | 0 | 0 | 1465507 | ARU07575.1 | A0A140H1G9 |
| LELFVGYALYTA | 1 | 0 | 0 | 1465513 | AUM60013.1 | K9N7C7 |
| LELGDTQGMDNL | 1 | 0 | 0 | 1465514 | AVP78030.1 | P0C6W2 |
| LELICDCGIKQE | 1 | 0 | 0 | 1465517 | ARU07575.1 | A0A140H1G9 |
| LELKERDQYCAL | 1 | 0 | 0 | 1465520 | APO40578.1 | P0C6X7 |
| LELLSFKRERSL | 1 | 0 | 0 | 1465530 | ARE29989.1 | Q4VID0 |
| LELLSKAMQLLH | 1 | 0 | 0 | 1465531 | AUM60013.1 | K9N7C7 |
| LELTTMRYFVRI | 1 | 0 | 0 | 1465537 | AFD98819.1 | H9EJ42 |
| LELVTIRGTPNN | 1 | 0 | 0 | 1465541 | AVV62540.1 | K9N643 |
| LEMADTSLSGYR | 1 | 0 | 0 | 1465542 | APO40578.1 | P0C6X7 |
| LEMFKPEVEELN | 1 | 0 | 0 | 1465548 | AGT51436.1 | Q4VID0 |
| LENGRLIIRENN | 1 | 0 | 0 | 1465554 | ATO98143.1 | P0C6W2 |
| LENHVMVDCRRL | 1 | 0 | 0 | 1465556 | ARK08667.1 | P0C6X6 |
| LENIAFNVFKQG | 1 | 0 | 0 | 1465557 | ANI69909.1 | K9N7C7 |
| LENIAFNVVKHG | 1 | 0 | 0 | 1465558 | AVV62524.1 | K9N7C7 |
| LENIVYNVVKQG | 1 | 0 | 0 | 1465561 | AUM60013.1 | K9N7C7 |
| LENKALKPFVCF | 1 | 0 | 0 | 1465562 | NP_073553.1 | A0A0P0KWE8 |
| LENKISGKGQQQ | 1 | 0 | 0 | 1465564 | AVP78049.1 | Q3LZX4 |
| LENPTYIYHSAF | 1 | 0 | 0 | 1465567 | AXN92247.1 | K9N7C7 |
| LENVIYNLFNVG | 1 | 0 | 0 | 1465571 | ABD75583.1 | A0A140H1G9 |
| LEPCNLWHYGAG | 1 | 0 | 0 | 1465576 | QEG03729.1 | H9EJ42 |
| LEPCQLANYKQS | 1 | 0 | 0 | 1465578 | AVV62524.1 | K9N7C7 |
| LEPQTHATTVDV | 1 | 0 | 0 | 1465591 | AUM60013.1 | K9N7C7 |
| LEPRSGNHCPAD | 1 | 0 | 0 | 1465595 | AID55096.1 | K9N5Q8 |
| LEPRSGNYCPAG | 1 | 0 | 0 | 1465597 | AVN89387.1 | K9N5Q8 |

| | | | | | | |
|---|---|---|---|---|---|---|
| LEPVKHSSTVDA | 1 | 0 | 0 | 1465599 | AVV62535.1 | K9N7C7 |
| LEQDKNTQEVFA | 1 | 0 | 0 | 1465605 | ATO98108.1 | Q3LZX1 |
| LEQDRMPKTRMA | 1 | 0 | 0 | 1465607 | AHF54653.1 | Q4VID0 |
| LEQNIDAYKSFP | 1 | 0 | 0 | 1465615 | AVV62533.1 | K9N4V7 |
| LEQPTNEAVEAP | 1 | 0 | 0 | 1465618 | AVP78041.1 | P0C6W2 |
| LEQTIQQKELEQ | 1 | 0 | 0 | 1465625 | APO40578.1 | P0C6X7 |
| LEQTNMHSHDVI | 1 | 0 | 0 | 1465627 | ASU90888.1 | K9N7C7 |
| LEQTNMHSPDLI | 1 | 0 | 0 | 1465629 | AUM60013.1 | K9N7C7 |
| LEQTQHSLNFVR | 1 | 0 | 0 | 1465633 | ASU90790.1 | K9N638 |
| LERAAVLEWDIK | 1 | 0 | 0 | 1465642 | AUM60013.1 | K9N7C7 |
| LERFMNTPNAIL | 1 | 0 | 0 | 1465649 | AUM60013.1 | K9N7C7 |
| LERMAEQAMTSM | 1 | 0 | 0 | 1465652 | AVV62524.1 | K9N7C7 |
| LERVYKHFNNYD | 1 | 0 | 0 | 1465655 | AEN19363.1 | P0C6X6 |
| LERVYNHFNNYD | 1 | 0 | 0 | 1465657 | AIV41936.1 | P0C6X6 |
| LESFKPRSQMEV | 1 | 0 | 0 | 1465666 | APO40578.1 | P0C6X7 |
| LESFKQELDKYF | 1 | 0 | 0 | 1465667 | APO40579.1 | P59594 |
| LESKVLLERFLP | 1 | 0 | 0 | 1465676 | QED88024.1 | H9EJ42 |
| LESKVSGKTQQQ | 1 | 0 | 0 | 1465678 | APO40586.1 | P59595 |
| LESTQLSTDTGA | 1 | 0 | 0 | 1465689 | Q3I5J4.1 | Q3LZX0 |
| LESYAFEHIVYG | 1 | 0 | 0 | 1465695 | APO40578.1 | P0C6X7 |
| LETAHHSVHVLQ | 1 | 0 | 0 | 1465696 | AVP78030.1 | P0C6W2 |
| LETAQKSYQEAL | 1 | 0 | 0 | 1465697 | AUM60013.1 | K9N7C7 |
| LETDGICLENHV | 1 | 0 | 0 | 1465700 | ARK08667.1 | P0C6X6 |
| LETPVGCVLGLV | 1 | 0 | 0 | 1465718 | ASU90230.1 | K9N5Q8 |
| LETVCSVAHTAG | 1 | 0 | 0 | 1465720 | AWK59942.1 | H9EJ42 |
| LETVCSVAYTAG | 1 | 0 | 0 | 1465721 | ARU07593.1 | H9EJ42 |
| LETVCSVVHTAG | 1 | 0 | 0 | 1465722 | AGT51379.1 | H9EJ42 |
| LETVCSVVRTAG | 1 | 0 | 0 | 1465723 | AFD98763.1 | H9EJ42 |
| LETVCSVVYTAG | 1 | 0 | 0 | 1465724 | AVA26872.1 | H9EJ42 |
| LEVKTASLFQTT | 1 | 0 | 0 | 1465736 | AGT51366.1 | A0A0P0G321 |
| LEVLFQGPGHHH | 1 | 0 | 0 | 1465739 | 6CRV_A | P59594 |
| LEVLLKDALQSR | 1 | 0 | 0 | 1465742 | AXX83318.1 | P0C6X6 |
| LEVLTFEVPGVS | 1 | 0 | 0 | 1465745 | AUM60013.1 | K9N7C7 |
| LEVQGYKNVRIT | 1 | 0 | 0 | 1465751 | ATO98229.1 | P0C6W2 |
| LEVSICQYNMCE | 1 | 0 | 0 | 1465753 | BBO58751.1 | P36334 |
| LEVSVCQFNMCE | 1 | 0 | 0 | 1465754 | ANZ78837.1 | P36334 |
| LEWDIKQCPAKS | 1 | 0 | 0 | 1465766 | AUM60013.1 | K9N7C7 |
| LEWLAEVVKLPS | 1 | 0 | 0 | 1465768 | ARO76392.1 | Q7TLC7 |
| LEYLLREVKTRA | 1 | 0 | 0 | 1465779 | AUM60013.1 | K9N7C7 |
| LEYWVTPLTPRQ | 1 | 0 | 0 | 1465786 | AWW13566.1 | P36334 |
| LFAAETLKANEE | 1 | 0 | 0 | 1465793 | APO40578.1 | P0C6X7 |
| LFAAETLRATEE | 1 | 0 | 0 | 1465795 | QWY57050.1 | P0DTD1 |
| LFADIKTQATQP | 1 | 0 | 0 | 1465796 | AVV62526.1 | K9N5Q8 |
| LFADINGKLYQD | 1 | 0 | 0 | 1465798 | ARO76381.1 | P0C6U8 |
| LFANPAAVDSKC | 1 | 0 | 0 | 1465814 | AEN19363.1 | P0C6X6 |
| LFAQSEEQEPFS | 1 | 0 | 0 | 1465818 | AHX00713.1 | K9N4V0 |
| LFAQSEEQESFS | 1 | 0 | 0 | 1465819 | ANF29197.1 | K9N4V0 |
| LFAQSEKEESFS | 1 | 0 | 0 | 1465820 | ATG84802.1 | K9N4V0 |
| LFAQSQEEESFS | 1 | 0 | 0 | 1465821 | AXN92262.1 | K9N4V0 |
| LFASVKNSQSSP | 1 | 0 | 0 | 1465827 | QBF80598.1 | K9N5Q8 |
| LFASVKSYQSSP | 1 | 0 | 0 | 1465830 | AVN89387.1 | K9N5Q8 |
| LFCATCPFAKGD | 1 | 0 | 0 | 1465841 | AUM60013.1 | K9N7C7 |
| LFCNIMKFSDRP | 1 | 0 | 0 | 1465845 | NP_073549.1 | A0A0P0LKV0 |
| LFCSSTYGVCKY | 1 | 0 | 0 | 1465846 | ARU07574.1 | A0A140H1G9 |
| LFDCVGLSVAST | 1 | 0 | 0 | 1465856 | ATQ39389.1 | K9N7C7 |
| LFDEAGEEVFAP | 1 | 0 | 0 | 1465862 | AEN19363.1 | P0C6X6 |
| LFDGRVEGQANL | 1 | 0 | 0 | 1465868 | AVP78041.1 | P0C6W2 |
| LFDQQAQQEATH | 1 | 0 | 0 | 1465878 | APT69896.1 | A0A0P0LKV0 |
| LFDSELNKFNRA | 1 | 0 | 0 | 1465880 | AUM60013.1 | K9N7C7 |
| LFDSLNLSADTQ | 1 | 0 | 0 | 1465884 | ARU07581.1 | Q19U29 |
| LFDSLQFAELST | 1 | 0 | 0 | 1465885 | AVV62524.1 | K9N7C7 |
| LFDWIKISADNK | 1 | 0 | 0 | 1465888 | AGT51707.1 | P0C6X6 |
| LFETIPTQLTPL | 1 | 0 | 0 | 1465903 | AUM60013.1 | K9N7C7 |
| LFFDSEVSTDEE | 1 | 0 | 0 | 1465908 | AMM42619.1 | H9EJ69 |
| LFFDSEVSTEEV | 1 | 0 | 0 | 1465909 | ABK63972.1 | H9EJ69 |
| LFFICCCTGCGP | 1 | 0 | 0 | 1465914 | AXX83351.1 | P36334 |
| LFFNDTAWYIGQ | 1 | 0 | 0 | 1465923 | AGT17765.1 | Q0ZJ83 |
| LFGCENPDCIAC | 1 | 0 | 0 | 1465941 | NP_073549.1 | A0A0P0LKV0 |
| LFGNCVTVRVGQ | 1 | 0 | 0 | 1465949 | AUM60013.1 | K9N7C7 |
| LFGNPDEPQKDV | 1 | 0 | 0 | 1465950 | AHA80176.1 | P33469 |
| LFGSVACEHIFS | 1 | 0 | 0 | 1465951 | AHZ64057.1 | K9N5Q8 |
| LFGWIKISADNK | 1 | 0 | 0 | 1465957 | AEN19363.1 | P0C6X6 |
| LFGWIKISADNN | 1 | 0 | 0 | 1465958 | AGT51508.1 | P0C6X6 |
| LFIVDVYDMWLT | 1 | 0 | 0 | 1465988 | AMM42427.1 | P0C6U7 |
| LFIYTNTIWINP | 1 | 0 | 0 | 1465993 | AWK59942.1 | H9EJ42 |
| LFKAAPPQIYNF | 1 | 0 | 0 | 1465994 | AUM60014.1 | K9N5Q8 |
| LFKDCARNPVDL | 1 | 0 | 0 | 1465999 | QEG03783.1 | A0A0P0LKV0 |
| LFKDCSNSYSGY | 1 | 0 | 0 | 1466004 | AGT51577.1 | P0C6X6 |
| LFKDCSREISGL | 1 | 0 | 0 | 1466005 | AUM60013.1 | K9N7C7 |
| LFKKNFTYNVSA | 1 | 0 | 0 | 1466018 | BBA20986.1 | A0A140H1H1 |
| LFKPLTARGRVV | 1 | 0 | 0 | 1466026 | ARE29970.1 | H9EJ24 |
| LFKRDSDADSPS | 1 | 0 | 0 | 1466036 | AMM42394.1 | Q19U29 |
| LFLFDEAGEEVF | 1 | 0 | 0 | 1466056 | AEN19363.1 | P0C6X6 |
| LFLTLIVLASCE | 1 | 0 | 0 | 1466094 | ARO76387.1 | P59635 |
| LFLTLTNGSDLD | 1 | 0 | 0 | 1466095 | ABD72988.1 | P59594 |
| LFMLPLVYFSDS | 1 | 0 | 0 | 1466116 | ATQ39389.1 | K9N7C7 |
| LFMRIFTLGTVT | 1 | 0 | 0 | 1466125 | AVP78032.1 | Q3LZX0 |
| LFNKYFKHWSFD | 1 | 0 | 0 | 1466138 | AIW52827.1 | H9EJ42 |
| LFNVTKLCFISG | 1 | 0 | 0 | 1466147 | AFD98835.1 | H9EJ24 |
| LFNVTKLCFVLG | 1 | 0 | 0 | 1466148 | AGT51332.1 | H9EJ24 |
| LFNVYLGMALSQ | 1 | 0 | 0 | 1466150 | AHA80218.1 | P36334 |

| | | | | | | |
|---|---|---|---|---|---|---|
| LFPKQGDLGKQY | 1 | 0 | 0 | 1466159 | AVV62526.1 | K9N5Q8 |
| LFPPWSAAAGVP | 1 | 0 | 0 | 1466167 | CAA83660.1 | P36334 |
| LFPPWTAAAGAP | 1 | 0 | 0 | 1466168 | ARB66070.1 | P36334 |
| LFPTWSQELPFD | 1 | 0 | 0 | 1466175 | AEN19355.1 | P0C6X6 |
| LFPYQGDHGDRY | 1 | 0 | 0 | 1466178 | ALA49803.1 | K9N5Q8 |
| LFQGPGHHHHHH | 1 | 0 | 0 | 1466182 | 6CRV_A | P59594 |
| LFQLCTFTKGAG | 1 | 0 | 0 | 1466187 | APO40578.1 | P0C6X7 |
| LFQPVTYFQLTT | 1 | 0 | 0 | 1466192 | AVV62535.1 | K9N7C7 |
| LFQTIPTQLTPL | 1 | 0 | 0 | 1466194 | AVV62535.1 | K9N7C7 |
| LFRDAEGKMVPY | 1 | 0 | 0 | 1466199 | AVV62524.1 | K9N7C7 |
| LFRELNGGVYTR | 1 | 0 | 0 | 1466201 | APO40578.1 | P0C6X7 |
| LFRIDVPSAEST | 1 | 0 | 0 | 1466206 | ALW82670.1 | K9N796 |
| LFRLKGGAPIKK | 1 | 0 | 0 | 1466214 | AUM60013.1 | K9N7C7 |
| LFRRLKKSNLLI | 1 | 0 | 0 | 1466222 | ARB07605.1 | A0A140H1G9 |
| LFSCDINKLLQV | 1 | 0 | 0 | 1466232 | ARU07593.1 | H9EJ42 |
| LFSDRPLNLRGW | 1 | 0 | 0 | 1466234 | AVA26872.1 | H9EJ42 |
| LFSSDLSTLAVT | 1 | 0 | 0 | 1466255 | AAY32939.1 | H9EJ42 |
| LFSTINALPEIS | 1 | 0 | 0 | 1466257 | ALW82729.1 | K9N7C7 |
| LFSTVAIKNLTP | 1 | 0 | 0 | 1466260 | AGT51393.1 | H9EJ42 |
| LFSTVFPPTCFG | 1 | 0 | 0 | 1466261 | APO40578.1 | P0C6X7 |
| LFSTVIIKKLTP | 1 | 0 | 0 | 1466263 | AWK59978.1 | H9EJ42 |
| LFSVPSDSCNNV | 1 | 0 | 0 | 1466270 | AFD98777.1 | H9EJ42 |
| LFSYQEFNDLVH | 1 | 0 | 0 | 1466274 | AMM42571.1 | H9EJ42 |
| LFTGYFPKAGAN | 1 | 0 | 0 | 1466280 | BBA20983.1 | A0A140H1H1 |
| LFTRQEEVQELY | 1 | 0 | 0 | 1466304 | QTS27337.1 | P0DTC7 |
| LFVCSNYAACHD | 1 | 0 | 0 | 1466313 | BBA20983.1 | A0A140H1H1 |
| LFVEDCKLPLGR | 1 | 0 | 0 | 1466319 | ATG84888.1 | K9N5Q8 |
| LFVEDCNLPLGQ | 1 | 0 | 0 | 1466320 | ALA49451.1 | K9N5Q8 |
| LFVIEVVDKYFD | 1 | 0 | 0 | 1466328 | AVP78041.1 | P0C6W2 |
| LFVNDKIIEFQL | 1 | 0 | 0 | 1466335 | AGT21366.1 | A0A0P0LKV0 |
| LFVRCEEDGQTV | 1 | 0 | 0 | 1466341 | AVV62535.1 | K9N7C7 |
| LFVRSKEEDGFA | 1 | 0 | 0 | 1466343 | AXN92247.1 | K9N7C7 |
| LFVSDYATKVVP | 1 | 0 | 0 | 1466348 | AVV62537.1 | K9N5Q8 |
| LFWNCNVPKYPN | 1 | 0 | 0 | 1466367 | AXN92247.1 | K9N7C7 |
| LFWNCNVPRYPN | 1 | 0 | 0 | 1466368 | AVV62524.1 | K9N7C7 |
| LFYCSNSSSSVL | 1 | 0 | 0 | 1466375 | AVL25581.1 | A0A0P0G321 |
| LFYTEDLASRFA | 1 | 0 | 0 | 1466391 | AUM60013.1 | K9N7C7 |
| LFYVNANGGSKF | 1 | 0 | 0 | 1466397 | AST12962.1 | A0A0P0LKV0 |
| LGAAALEDEATP | 1 | 0 | 0 | 1466400 | 3EA9_A | P0C6U8 |
| LGADVLYQPPQT | 1 | 0 | 0 | 1466406 | BAE93400.1 | P0C6U8 |
| LGAENSIAYANN | 1 | 0 | 0 | 1466410 | AVP78031.1 | Q3LZX1 |
| LGAMLTGFTTTN | 1 | 0 | 0 | 1466420 | QDP16206.1 | K9N5Q8 |
| LGAPDNPSAIVT | 1 | 0 | 0 | 1466422 | AKA09625.1 | A0A0P0G321 |
| LGASSNQSPQGN | 1 | 0 | 0 | 1466427 | AIX09796.1 | P0C6X6 |
| LGAVSKVVKVPI | 1 | 0 | 0 | 1466428 | ANI69887.1 | K9N7C7 |
| LGAYADKTLQEM | 1 | 0 | 0 | 1466430 | AIV42008.1 | P0C6X6 |
| LGCDVTDVTQLY | 1 | 0 | 0 | 1466434 | AVP78041.1 | P0C6W2 |
| LGCLGSECSKAS | 1 | 0 | 0 | 1466445 | CAA79896.1 | P36334 |
| LGCLGSQCGSSS | 1 | 0 | 0 | 1466447 | BBA20986.1 | A0A140H1H1 |
| LGCSIEMEEYPM | 1 | 0 | 0 | 1466449 | ATQ39396.1 | K9N643 |
| LGCVVNADNSTS | 1 | 0 | 0 | 1466455 | CAA83660.1 | P36334 |
| LGCVVNAYNNTA | 1 | 0 | 0 | 1466456 | AIL49538.1 | P36334 |
| LGDAIQVAKCYG | 1 | 0 | 0 | 1466463 | AXN92247.1 | K9N7C7 |
| LGDISARDLICA | 1 | 0 | 0 | 1466478 | AVP78042.1 | Q3LZX1 |
| LGDSVKLLSELL | 1 | 0 | 0 | 1466484 | ABE73416.1 | H9EJ42 |
| LGESFKKVATIP | 1 | 0 | 0 | 1466502 | AXN92247.1 | K9N7C7 |
| LGEYGSFCDNIN | 1 | 0 | 0 | 1466506 | AHF54658.1 | P36334 |
| LGFDKPQEKDKS | 1 | 0 | 0 | 1466513 | ABB90496.1 | A0A0N7J131 |
| LGFDKPQEKKKS | 1 | 0 | 0 | 1466514 | AMM42528.1 | A0A0N7J131 |
| LGFDNPPTNVVS | 1 | 0 | 0 | 1466515 | AGT51679.1 | P30215 |
| LGFDNQSKLPSS | 1 | 0 | 0 | 1466516 | AIW52795.1 | H9EJ69 |
| LGFDNQSKPPSS | 1 | 0 | 0 | 1466517 | AGT51335.1 | H9EJ69 |
| LGFDNQSKSLSS | 1 | 0 | 0 | 1466518 | ARU07598.1 | H9EJ69 |
| LGFDSSKPVAKK | 1 | 0 | 0 | 1466521 | ANI69899.1 | K9N7C7 |
| LGFDSSKPVIKK | 1 | 0 | 0 | 1466522 | AHI48784.1 | K9N7C7 |
| LGFEKPQEKDKK | 1 | 0 | 0 | 1466525 | ARB07429.1 | A0A0N7J131 |
| LGFENPPTNVVS | 1 | 0 | 0 | 1466526 | AAA73152.1 | P30215 |
| LGFHNPPTNVVS | 1 | 0 | 0 | 1466528 | AVQ05263.1 | P30215 |
| LGFYNPLTNVVS | 1 | 0 | 0 | 1466552 | AGT51799.1 | P30215 |
| LGGDASKPHQVT | 1 | 0 | 0 | 1466560 | AJQ24068.1 | P33469 |
| LGGLDSLDSYPA | 1 | 0 | 0 | 1466570 | ARO76381.1 | P0C6U8 |
| LGGMSYFCVDHR | 1 | 0 | 0 | 1466580 | AXN92247.1 | K9N7C7 |
| LGIGDSAVTYSR | 1 | 0 | 0 | 1466602 | AEN19363.1 | P0C6X6 |
| LGIGYSLADLPA | 1 | 0 | 0 | 1466613 | AWW13547.1 | Q01455 |
| LGINEEKCGTQL | 1 | 0 | 0 | 1466623 | BBA20986.1 | A0A140H1H1 |
| LGINEKTHVQLS | 1 | 0 | 0 | 1466624 | P0C6W6.1 | P0C6W2 |
| LGISDSAVTYSR | 1 | 0 | 0 | 1466629 | QEG03763.1 | P0C6X6 |
| LGISSACDGLAS | 1 | 0 | 0 | 1466630 | AXN92247.1 | K9N7C7 |
| LGITSSCDGIAS | 1 | 0 | 0 | 1466631 | AUM60013.1 | K9N7C7 |
| LGKCYKAMNAYP | 1 | 0 | 0 | 1466633 | AKN24802.1 | K9N7C7 |
| LGKDAIKPQQVT | 1 | 0 | 0 | 1466634 | AOL02457.1 | P33469 |
| LGKLQDVINQNA | 1 | 0 | 0 | 1466643 | APO40579.1 | P59594 |
| LGKLYVSNYSHE | 1 | 0 | 0 | 1466647 | AUM60014.1 | K9N5Q8 |
| LGKPKEIIDGYI | 1 | 0 | 0 | 1466649 | APO40578.1 | P0C6X7 |
| LGKVKPPLNKNY | 1 | 0 | 0 | 1466653 | AGT51677.1 | P0C6X6 |
| LGKYEQGSGYIP | 1 | 0 | 0 | 1466656 | 6CRW_A | P59594 |
| LGLLQPTDNSTT | 1 | 0 | 0 | 1466677 | AXN92247.1 | K9N7C7 |
| LGLSIEDRKALL | 1 | 0 | 0 | 1466690 | APO40587.1 | P59636 |
| LGLVNSSLFVED | 1 | 0 | 0 | 1466695 | AXN92228.1 | K9N5Q8 |
| LGLVNSSLLVED | 1 | 0 | 0 | 1466696 | ATW75477.1 | K9N5Q8 |
| LGLYVSLTIALA | 1 | 0 | 0 | 1466704 | AUM60013.1 | K9N7C7 |

| | | | | | | |
|---|---|---|---|---|---|---|
| LGNCVEYHHHHH | 1 | 0 | 0 | 1466720 | 4KQZ_A | K9N5Q8 |
| LGNHFKELWHSD | 1 | 0 | 0 | 1466729 | BAM16460.1 | P36334 |
| LGNHRKELWHLD | 1 | 0 | 0 | 1466732 | CAA79896.1 | P36334 |
| LGNLGNLQSSNY | 1 | 0 | 0 | 1466736 | AIL49519.1 | P36334 |
| LGNSIGIDFQDE | 1 | 0 | 0 | 1466744 | AGV08467.1 | K9N5Q8 |
| LGNSSTITNQLG | 1 | 0 | 0 | 1466745 | AVV62526.1 | K9N5Q8 |
| LGNSTTITDKLG | 1 | 0 | 0 | 1466748 | AVV62537.1 | K9N5Q8 |
| LGNTMLEDEFTP | 1 | 0 | 0 | 1466749 | AUM60013.1 | K9N7C7 |
| LGNYTYYNKGSG | 1 | 0 | 0 | 1466756 | 6NB3_A | K9N5Q8 |
| LGQAATTTSNCT | 1 | 0 | 0 | 1466766 | ARO76381.1 | P0C6U8 |
| LGRELNCDVCPN | 1 | 0 | 0 | 1466782 | ARB07436.1 | A0A140H1G9 |
| LGRELNCNVCAN | 1 | 0 | 0 | 1466783 | ABD75615.1 | A0A140H1G9 |
| LGSAGNATEVPA | 1 | 0 | 0 | 1466798 | 5C8S_A | P0C6U8 |
| LGSCSFEDELTP | 1 | 0 | 0 | 1466804 | AIV41834.1 | P0C6X6 |
| LGSGAHVGSDFT | 1 | 0 | 0 | 1466809 | AWK59942.1 | H9EJ42 |
| LGSGCGKASSRS | 1 | 0 | 0 | 1466810 | BBA20982.1 | P36334 |
| LGSIAGASWTAG | 1 | 0 | 0 | 1466813 | AUM60014.1 | K9N5Q8 |
| LGSNEPGNVGGN | 1 | 0 | 0 | 1466838 | AVI15051.1 | P0C6X6 |
| LGSNLSLSMARR | 1 | 0 | 0 | 1466840 | 2CME_A | P59636 |
| LGTDPIEDFQEN | 1 | 0 | 0 | 1466867 | AVP78041.1 | P0C6W2 |
| LGTDPIEDYEND | 1 | 0 | 0 | 1466868 | APO40578.1 | P0C6X7 |
| LGTEDPRCPQIA | 1 | 0 | 0 | 1466874 | AVV61907.1 | K9N4V7 |
| LGTGPEAALPYG | 1 | 0 | 0 | 1466878 | APO40586.1 | P59595 |
| LGTGPHAKDDYG | 1 | 0 | 0 | 1466881 | AJQ24074.1 | P33469 |
| LGTGPHSKDQYG | 1 | 0 | 0 | 1466888 | AIV41994.1 | P33469 |
| LGTGPYASSSYG | 1 | 0 | 0 | 1466892 | AMM42397.1 | Q19U29 |
| LGTHCLHVVGPN | 1 | 0 | 0 | 1466898 | APO40578.1 | P0C6X7 |
| LGTSDPQFPIFA | 1 | 0 | 0 | 1466907 | AAQ93017.1 | P33469 |
| LGTSDSQFPILA | 1 | 0 | 0 | 1466909 | AGT51764.1 | P33469 |
| LGTVDADYKMCT | 1 | 0 | 0 | 1466918 | APT69849.1 | A0A0P0K6L9 |
| LGTVDANYKKCT | 1 | 0 | 0 | 1466919 | CAA71147.1 | A0A0P0K6L9 |
| LGVGDSAVTYSR | 1 | 0 | 0 | 1466944 | AIV41816.1 | P0C6X6 |
| LGVNSSNCNNLN | 1 | 0 | 0 | 1466960 | AUT76985.1 | A0A0P0LKV0 |
| LGVRGSGHYQTN | 1 | 0 | 0 | 1466966 | AFD98812.1 | H9EJ42 |
| LGVSDVWHKQLL | 1 | 0 | 0 | 1466968 | AEN19363.1 | P0C6X6 |
| LGWSADSCLQGN | 1 | 0 | 0 | 1466987 | AXX83303.1 | P36334 |
| LGWSVDSCLQGD | 1 | 0 | 0 | 1466989 | CAA83660.1 | P36334 |
| LGWSVESCLQGD | 1 | 0 | 0 | 1466990 | AMQ23491.1 | P36334 |
| LGWSYDTCVSNN | 1 | 0 | 0 | 1466991 | BBA20983.1 | A0A140H1H1 |
| LGYDFNKPTSKT | 1 | 0 | 0 | 1466995 | AUM60013.1 | K9N7C7 |
| LGYISHGCDLMQ | 1 | 0 | 0 | 1467000 | AVV62524.1 | K9N7C7 |
| LHAIQQGFDATN | 1 | 0 | 0 | 1467015 | AEN19366.1 | P36334 |
| LHALDDATRNYI | 1 | 0 | 0 | 1467017 | AHL18094.1 | K9N643 |
| LHALDDVTRNYI | 1 | 0 | 0 | 1467020 | AXN92231.1 | K9N643 |
| LHARMTYTCQCG | 1 | 0 | 0 | 1467027 | ATQ39389.1 | K9N7C7 |
| LHCANFNVLFAM | 1 | 0 | 0 | 1467029 | AXN92247.1 | K9N7C7 |
| LHCTTNLFKDCS | 1 | 0 | 0 | 1467038 | ARU07574.1 | A0A140H1G9 |
| LHCVRNVFDNFN | 1 | 0 | 0 | 1467039 | AWK59942.1 | H9EJ42 |
| LHDLDDATRSYI | 1 | 0 | 0 | 1467048 | AVV62540.1 | K9N643 |
| LHDVNNGLTCST | 1 | 0 | 0 | 1467049 | AEN19366.1 | P36334 |
| LHDVNSGLTCST | 1 | 0 | 0 | 1467050 | CAA79896.1 | P36334 |
| LHFSLFDDNFCA | 1 | 0 | 0 | 1467066 | ARU07575.1 | A0A140H1G9 |
| LHFSTYIECTYI | 1 | 0 | 0 | 1467068 | AKM76243.1 | K9N7D2 |
| LHFTNDKTIFYT | 1 | 0 | 0 | 1467069 | ARU07575.1 | A0A140H1G9 |
| LHGANLMQDESV | 1 | 0 | 0 | 1467073 | AVV62537.1 | K9N5Q8 |
| LHGANLRQDDFV | 1 | 0 | 0 | 1467074 | AMW90843.1 | K9N5Q8 |
| LHGNGSMIEDID | 1 | 0 | 0 | 1467080 | ALW82645.1 | K9N7C7 |
| LHIAGCQTTNGT | 1 | 0 | 0 | 1467092 | AIW52757.1 | A0A0P0K6L9 |
| LHIAVCQTTNGT | 1 | 0 | 0 | 1467095 | AIW52747.1 | A0A0P0K6L9 |
| LHLEANGRAWAF | 1 | 0 | 0 | 1467115 | AYO86698.1 | K9N638 |
| LHLTQDYFLPFD | 1 | 0 | 0 | 1467141 | ATO98181.1 | Q3LZX1 |
| LHLVDPQIQLTT | 1 | 0 | 0 | 1467144 | Q3I5I6.1 | Q3LZX3 |
| LHNVKFDVGGFI | 1 | 0 | 0 | 1467167 | AUM60027.1 | K9N643 |
| LHNYGENAVLPK | 1 | 0 | 0 | 1467170 | APO40578.1 | P0C6X7 |
| LHPRLKGSFNNR | 1 | 0 | 0 | 1467176 | BBA84086.1 | P36334 |
| LHPVQWQCKLDF | 1 | 0 | 0 | 1467184 | AVV62528.1 | K9N4V0 |
| LHQESQLLKLRD | 1 | 0 | 0 | 1467187 | AGT51675.1 | Q4VID0 |
| LHQFSRTTQASL | 1 | 0 | 0 | 1467189 | AVV62537.1 | K9N5Q8 |
| LHQLLGSTADNL | 1 | 0 | 0 | 1467193 | APO40578.1 | P0C6X7 |
| LHRFKPVYRVNT | 1 | 0 | 0 | 1467201 | QDP16209.1 | K9N643 |
| LHSSQLPLDVYN | 1 | 0 | 0 | 1467220 | AAX85666.1 | P0C6X6 |
| LHTGFQGKQILG | 1 | 0 | 0 | 1467235 | ATQ39389.1 | K9N7C7 |
| LHTGYYDANQYY | 1 | 0 | 0 | 1467236 | AVL25587.1 | A0A0P0G321 |
| LHTLDDVTRNYI | 1 | 0 | 0 | 1467247 | ASU90464.1 | K9N643 |
| LHTLSIHDGYTN | 1 | 0 | 0 | 1467249 | AAP30028.1 | P0C6X7 |
| LHTVLLPTDYKN | 1 | 0 | 0 | 1467260 | AIW52843.1 | A0A0P0G321 |
| LHVTYIPSQEKN | 1 | 0 | 0 | 1467273 | AVP78031.1 | Q3LZX1 |
| LHVTYVPSQEKN | 1 | 0 | 0 | 1467277 | ATO98181.1 | Q3LZX1 |
| LHVTYVPSQQQN | 1 | 0 | 0 | 1467279 | APO40579.1 | P59594 |
| LIAGPKGPEIRY | 1 | 0 | 0 | 1467307 | AXN92247.1 | K9N7C7 |
| LIANAFNNALDA | 1 | 0 | 0 | 1467322 | CAA79896.1 | P36334 |
| LIANAFNNALNA | 1 | 0 | 0 | 1467326 | AGT51491.1 | P36334 |
| LIANKFNQALEA | 1 | 0 | 0 | 1467330 | QBF80499.1 | K9N5Q8 |
| LIAQSRGFYRKC | 1 | 0 | 0 | 1467333 | APO40578.1 | P0C6X7 |
| LIAVANWLTPTN | 1 | 0 | 0 | 1467340 | AUM60013.1 | K9N7C7 |
| LICDCGIKQESR | 1 | 0 | 0 | 1467347 | ARU07575.1 | A0A140H1G9 |
| LICKPDQQFHNF | 1 | 0 | 0 | 1467350 | AVV62540.1 | K9N643 |
| LICSNTPLSKDL | 1 | 0 | 0 | 1467354 | ARU07575.1 | A0A140H1G9 |
| LIDCNAASVNQT | 1 | 0 | 0 | 1467366 | AVN89451.1 | K9N7C7 |
| LIDEVQNWLSAF | 1 | 0 | 0 | 1467368 | AUM60027.1 | K9N643 |
| LIDLANLFYIFL | 1 | 0 | 0 | 1467372 | AUM60013.1 | K9N7C7 |

| | | | | | | |
|---|---|---|---|---|---|---|
| LIDLQELGKYEH | 1 | 0 | 0 | 1467376 | AVP78042.1 | Q3LZX1 |
| LIDSNAASVNPA | 1 | 0 | 0 | 1467383 | AVV62524.1 | K9N7C7 |
| LIEHVTSVNLFD | 1 | 0 | 0 | 1467390 | QBF80533.1 | K9N796 |
| LIELCVDEAGSKSPIQYID | 1 | 0 | 0 | 1467392 | P0DTC8.1 | P0DTC8 |
| LIELENKFTPLS | 1 | 0 | 0 | 1467394 | ASU91261.1 | K9N638 |
| LIEYDFTDYKVQ | 1 | 0 | 0 | 1467398 | QBF80541.1 | K9N7C7 |
| LIFEFEDDVITS | 1 | 0 | 0 | 1467402 | AWK59942.1 | H9EJ42 |
| LIFPNSNYVLQD | 1 | 0 | 0 | 1467414 | AFV53147.1 | H9EJ42 |
| LIFPPGQGVPLN | 1 | 0 | 0 | 1467416 | ASU90622.1 | K9N4V7 |
| LIFRNIKCNYVF | 1 | 0 | 0 | 1467420 | AGT51481.1 | P36334 |
| LIGAEHVNASYE | 1 | 0 | 0 | 1467435 | AVP78031.1 | Q3LZX1 |
| LIGANGNKIVDK | 1 | 0 | 0 | 1467438 | AVN89322.1 | K9N7C7 |
| LIGCNYLGFIKD | 1 | 0 | 0 | 1467442 | AUM60013.1 | K9N7C7 |
| LIGHCATVHTAN | 1 | 0 | 0 | 1467451 | AVP78030.1 | P0C6W2 |
| LIGYIPEHCQSH | 1 | 0 | 0 | 1467475 | AUM60025.1 | K9N796 |
| LIGYTAATTCVT | 1 | 0 | 0 | 1467476 | ATO98145.1 | Q3LZX1 |
| LIGYYSDDGNYY | 1 | 0 | 0 | 1467480 | AVV62537.1 | K9N5Q8 |
| LIIKNLSKPPTE | 1 | 0 | 0 | 1467502 | AVP78035.1 | Q3LZX8 |
| LIIPLGIDLDEW | 1 | 0 | 0 | 1467509 | AVP78030.1 | P0C6W2 |
| LIKDCPAVAVHD | 1 | 0 | 0 | 1467522 | APO40578.1 | P0C6X7 |
| LIKDVDNPAHTG | 1 | 0 | 0 | 1467524 | AJS10812.1 | P0C6X6 |
| LIKSKVSAEVSR | 1 | 0 | 0 | 1467543 | ARB07393.1 | A0A0P0KWE8 |
| LILDYFSYPFSM | 1 | 0 | 0 | 1467556 | AVN89313.1 | K9N5Q8 |
| LIMAYGNCTFGD | 1 | 0 | 0 | 1467600 | AVN89333.1 | K9N7C7 |
| LIMRCWLCWKCK | 1 | 0 | 0 | 1467607 | ARO76383.1 | P59632 |
| LIMVLLVLLIYK | 1 | 0 | 0 | 1467609 | AFR45573.1 | P36334 |
| LINNISYVVTQN | 1 | 0 | 0 | 1467617 | AWK59942.1 | H9EJ42 |
| LINRGVYKPLLE | 1 | 0 | 0 | 1467620 | AEN19363.1 | P0C6X6 |
| LINSSLFVEDCK | 1 | 0 | 0 | 1467621 | ALK80261.1 | K9N5Q8 |
| LINTISPEKLKN | 1 | 0 | 0 | 1467624 | AWK59942.1 | H9EJ42 |
| LINTLNNLNETL | 1 | 0 | 0 | 1467626 | AVP78041.1 | P0C6W2 |
| LIPDLLHAHPVE | 1 | 0 | 0 | 1467628 | AHF54653.1 | Q4VID0 |
| LIPDLLRAHPVE | 1 | 0 | 0 | 1467630 | ARE29989.1 | Q4VID0 |
| LIPDLLYAHPAE | 1 | 0 | 0 | 1467631 | AHF54648.1 | Q4VID0 |
| LIPNHAALFFDS | 1 | 0 | 0 | 1467644 | AMM42619.1 | H9EJ69 |
| LIQQWIPFMMSR | 1 | 0 | 0 | 1467673 | AAP30032.1 | P59633 |
| LIRDNEALTVST | 1 | 0 | 0 | 1467685 | AVV62524.1 | K9N7C7 |
| LIRGYRGIKPLL | 1 | 0 | 0 | 1467697 | AXX83336.1 | P0C6X6 |
| LIRQLQPINYFD | 1 | 0 | 0 | 1467708 | AVQ05264.1 | P36334 |
| LISCNGNKIEDL | 1 | 0 | 0 | 1467712 | AIL49553.1 | P0C6X6 |
| LISGPKGPEIRY | 1 | 0 | 0 | 1467718 | AVV62524.1 | K9N7C7 |
| LISGSLGDSGEL | 1 | 0 | 0 | 1467719 | ABE73410.1 | H9EJ42 |
| LISLADVATDVQ | 1 | 0 | 0 | 1467726 | AUM60013.1 | K9N7C7 |
| LISLTDVATDVQ | 1 | 0 | 0 | 1467733 | AUM60023.1 | K9N7C7 |
| LISQFRLSKMGV | 1 | 0 | 0 | 1467744 | AWK59942.1 | H9EJ42 |
| LISQVRLSKMGI | 1 | 0 | 0 | 1467745 | NP_073549.1 | A0A0P0LKV0 |
| LISSGIFSVPSD | 1 | 0 | 0 | 1467748 | ARK08632.1 | P0C6X6 |
| LITLLFANAFSK | 1 | 0 | 0 | 1467769 | AVV62527.1 | K9N796 |
| LITNLDIEKLRQ | 1 | 0 | 0 | 1467772 | AVV62535.1 | K9N7C7 |
| LIVNVLSIFSSS | 1 | 0 | 0 | 1467814 | QED88011.1 | H9EJ42 |
| LKAAVHEWKMVV | 1 | 0 | 0 | 1467843 | AXN92258.1 | K9N7C7 |
| LKAAYENFNSQE | 1 | 0 | 0 | 1467850 | ARO76381.1 | P0C6U8 |
| LKAHKEKSSQCF | 1 | 0 | 0 | 1467862 | APO40578.1 | P0C6X7 |
| LKAIMLKAQEGL | 1 | 0 | 0 | 1467863 | ARU07582.1 | Q0ZJ80 |
| LKANEETFKLSY | 1 | 0 | 0 | 1467871 | APO40578.1 | P0C6X7 |
| LKAPKDVTFLEG | 1 | 0 | 0 | 1467877 | ARO76381.1 | P0C6U8 |
| LKCPLDTSRKGS | 1 | 0 | 0 | 1467903 | AKA09663.1 | P36334 |
| LKCPLDTSYKGT | 1 | 0 | 0 | 1467905 | AKA09658.1 | P36334 |
| LKCTTVAINDVD | 1 | 0 | 0 | 1467917 | CAA83661.1 | P36334 |
| LKDALQSREAVL | 1 | 0 | 0 | 1467923 | AXX83318.1 | P0C6X6 |
| LKDGQINDMILS | 1 | 0 | 0 | 1467934 | AVP78041.1 | P0C6W2 |
| LKDGVNFNIDDI | 1 | 0 | 0 | 1467936 | ARB66060.1 | P36334 |
| LKDGVNFNVDDV | 1 | 0 | 0 | 1467937 | CAA83661.1 | P36334 |
| LKDKILYTFYGK | 1 | 0 | 0 | 1467947 | AUM60013.1 | K9N7C7 |
| LKDNQVNDMIYS | 1 | 0 | 0 | 1467952 | APO40578.1 | P0C6X7 |
| LKDNVLSFMLRL | 1 | 0 | 0 | 1467953 | AHI48786.1 | K9N7C7 |
| LKEDDVDDSGDS | 1 | 0 | 0 | 1467964 | AMM42440.1 | P0C6X6 |
| LKELYGPADPTF | 1 | 0 | 0 | 1467979 | AHI48582.1 | K9N7C7 |
| LKELYGPIDPTF | 1 | 0 | 0 | 1467980 | ATG84722.1 | K9N7C7 |
| LKELYGPLDPTF | 1 | 0 | 0 | 1467981 | AXN92247.1 | K9N7C7 |
| LKETVFVEPKNG | 1 | 0 | 0 | 1467994 | ARU07593.1 | H9EJ42 |
| LKFFEIKHADYT | 1 | 0 | 0 | 1468004 | AHF51980.1 | H9EJ42 |
| LKGGKIVSTWFK | 1 | 0 | 0 | 1468040 | P0C6W6.1 | P0C6W2 |
| LKGGLILRFDSG | 1 | 0 | 0 | 1468044 | AGV08389.1 | K9N638 |
| LKGLDAMFFYGD | 1 | 0 | 0 | 1468049 | AEN19363.1 | P0C6X6 |
| LKGSFNNRDTGH | 1 | 0 | 0 | 1468052 | AWW13566.1 | P36334 |
| LKGSFNNRDTGL | 1 | 0 | 0 | 1468053 | AWW13511.1 | P36334 |
| LKGSFNNRDTGP | 1 | 0 | 0 | 1468054 | AEN19366.1 | P36334 |
| LKICKFGNTSFA | 1 | 0 | 0 | 1468096 | QED88040.1 | A0A0P0G321 |
| LKICKFGNTTFD | 1 | 0 | 0 | 1468098 | AWO69005.1 | A0A0P0G321 |
| LKICKFSSNTTF | 1 | 0 | 0 | 1468100 | ACY56730.1 | A0A0P0G321 |
| LKKEPLLYVPIR | 1 | 0 | 0 | 1468132 | AXN92247.1 | K9N7C7 |
| LKKLVGGNFTPI | 1 | 0 | 0 | 1468145 | AUM60013.1 | K9N7C7 |
| LKKMDEPFTEDT | 1 | 0 | 0 | 1468147 | AEN19370.1 | P33469 |
| LKKPRWKRVPTR | 1 | 0 | 0 | 1468152 | AIW52805.1 | H9EJ69 |
| LKKTSWVDQSER | 1 | 0 | 0 | 1468159 | AMM42397.1 | Q19U29 |
| LKLEDIMEVDIS | 1 | 0 | 0 | 1468169 | AUM60017.1 | K9N643 |
| LKLLEPVSISTG | 1 | 0 | 0 | 1468185 | AOR17482.1 | K9N5Q8 |
| LKLLGIGGDRCI | 1 | 0 | 0 | 1468187 | AKT07951.1 | H9EJ42 |
| LKLPPHDVIVLR | 1 | 0 | 0 | 1468194 | AFD98800.1 | H9EJ24 |
| LKLRSETLLPIT | 1 | 0 | 0 | 1468202 | APO40578.1 | P0C6X7 |

| | | | | | | |
|---|---|---|---|---|---|---|
| LKLTHQSNDESG | 1 | 0 | 0 | 1468219 | AVV62544.1 | K9N4V7 |
| LKLTLTSNGLLG | 1 | 0 | 0 | 1468220 | AWK59942.1 | H9EJ42 |
| LKLTYNKQKANV | 1 | 0 | 0 | 1468221 | AXX83324.1 | P0C6X6 |
| LKLVDDHALIVN | 1 | 0 | 0 | 1468222 | AOG74785.1 | A0A0P0KH29 |
| LKMAGMHFGACD | 1 | 0 | 0 | 1468227 | ASY99815.1 | K9N7A1 |
| LKMAGMHFGQCD | 1 | 0 | 0 | 1468228 | AUM60030.1 | K9N7A1 |
| LKMIQTHLSHYV | 1 | 0 | 0 | 1468231 | AUM60013.1 | K9N7C7 |
| LKMVQTHISHYV | 1 | 0 | 0 | 1468240 | AVV62524.1 | K9N7C7 |
| LKMVQTHLSHYV | 1 | 0 | 0 | 1468241 | AXN92247.1 | K9N7C7 |
| LKNFDQQCHCSR | 1 | 0 | 0 | 1468248 | AVV62541.1 | K9N7D2 |
| LKNGELETLETP | 1 | 0 | 0 | 1468252 | AAP33695.1 | P0C6X7 |
| LKPFERDLSSDE | 1 | 0 | 0 | 1468278 | AVP78031.1 | Q3LZX1 |
| LKPFERDLTSDE | 1 | 0 | 0 | 1468279 | ATO98120.1 | Q3LZX1 |
| LKPLPFLYYSTT | 1 | 0 | 0 | 1468290 | ANI69887.1 | K9N7C7 |
| LKPLYVPEHCQN | 1 | 0 | 0 | 1468293 | ATY74382.1 | K9N796 |
| LKPPFLSDFING | 1 | 0 | 0 | 1468294 | AKA09633.1 | P36334 |
| LKPTFKSKGLNV | 1 | 0 | 0 | 1468303 | AWK59942.1 | H9EJ42 |
| LKQADLSGAICE | 1 | 0 | 0 | 1468318 | ARU07575.1 | A0A140H1G9 |
| LKQPMATSANCC | 1 | 0 | 0 | 1468339 | AGT51735.1 | Q4VID0 |
| LKQPMATSVNCC | 1 | 0 | 0 | 1468340 | APU51922.1 | Q4VID0 |
| LKQPMATSVNCF | 1 | 0 | 0 | 1468341 | AHF54653.1 | Q4VID0 |
| LKQRDSTYGPLQ | 1 | 0 | 0 | 1468343 | AVN89365.1 | K9N5Q8 |
| LKRGEKIVYHTL | 1 | 0 | 0 | 1468356 | P0C6W6.1 | P0C6W2 |
| LKRLLEQWNLVI | 1 | 0 | 0 | 1468363 | ATO98197.1 | Q3LZX9 |
| LKRQIRIACHKC | 1 | 0 | 0 | 1468367 | ALA49758.1 | K9N7C7 |
| LKRRDSTYGPLE | 1 | 0 | 0 | 1468372 | ASU90230.1 | K9N5Q8 |
| LKSFIPLVDMRS | 1 | 0 | 0 | 1468394 | AVV62530.1 | K9N7D2 |
| LKSLKLNLTQQG | 1 | 0 | 0 | 1468404 | NP_073549.1 | A0A0P0LKV0 |
| LKSNRCCDRYEE | 1 | 0 | 0 | 1468412 | AKN24812.1 | K9N5Q8 |
| LKTGVLQPLEQP | 1 | 0 | 0 | 1468435 | AVP78041.1 | P0C6W2 |
| LKTLATHGLAAV | 1 | 0 | 0 | 1468448 | QYC64967.1 | P0DTD1 |
| LKTLKLTLTSNG | 1 | 0 | 0 | 1468449 | AWK59942.1 | H9EJ42 |
| LKTMVVTAGLDQ | 1 | 0 | 0 | 1468460 | AVV62524.1 | K9N7C7 |
| LKTPDTYKCPQS | 1 | 0 | 0 | 1468463 | AXX83339.1 | P36334 |
| LKTSTAPILMES | 1 | 0 | 0 | 1468466 | AGT51765.1 | Q4VID0 |
| LKTVLRKGGRTI | 1 | 0 | 0 | 1468469 | AVP78041.1 | P0C6W2 |
| LKVNCCRFQRVG | 1 | 0 | 0 | 1468498 | ATN39877.2 | P0C6X6 |
| LKVPATVSVSSPDA | 1 | 0 | 0 | 1468500 | P0DTC1.1 | P0DTD1 |
| LKWADQSDQFRS | 1 | 0 | 0 | 1468505 | AIL49389.1 | P33469 |
| LKWADQSDQVRN | 1 | 0 | 0 | 1468506 | P33469.1 | P33469 |
| LKWSDQSDQFRN | 1 | 0 | 0 | 1468515 | ARK08639.1 | P33469 |
| LLADKFPILHDI | 1 | 0 | 0 | 1468564 | APO40578.1 | P0C6X7 |
| LLAGGLEATFLF | 1 | 0 | 0 | 1468579 | AVP78032.1 | Q3LZX0 |
| LLANAFSKPFYI | 1 | 0 | 0 | 1468604 | ATQ39394.1 | K9N796 |
| LLANANSKPLIG | 1 | 0 | 0 | 1468605 | AUM60025.1 | K9N796 |
| LLANMGIDLDEW | 1 | 0 | 0 | 1468611 | ARO76381.1 | P0C6U8 |
| LLARAGKSCAPL | 1 | 0 | 0 | 1468617 | APO40578.1 | P0C6X7 |
| LLAVFQNASKVI | 1 | 0 | 0 | 1468630 | APO40580.1 | P59632 |
| LLCATCPFVKGD | 1 | 0 | 0 | 1468639 | AKN24784.1 | K9N7C7 |
| LLCCTEQDLHKD | 1 | 0 | 0 | 1468644 | AUM60027.1 | K9N643 |
| LLCEPIKLVNSE | 1 | 0 | 0 | 1468650 | AWK59942.1 | H9EJ42 |
| LLCVANDKPIVL | 1 | 0 | 0 | 1468662 | APT69868.1 | A0A0P0ILY9 |
| LLCVTCPFAKGD | 1 | 0 | 0 | 1468667 | ATG84865.1 | K9N638 |
| LLDEGVDCKIQT | 1 | 0 | 0 | 1468673 | NP_150075.1 | Q80872 |
| LLDLAGLFRILL | 1 | 0 | 0 | 1468693 | ALA50044.1 | K9N7C7 |
| LLDVCNTKEFKV | 1 | 0 | 0 | 1468727 | ARX11394.1 | A0A0P0LKV0 |
| LLEAYNAFLNTV | 1 | 0 | 0 | 1468732 | AST12962.1 | A0A0P0LKV0 |
| LLEKCDLHNYGE | 1 | 0 | 0 | 1468746 | APO40578.1 | P0C6X7 |
| LLELKERDQYCA | 1 | 0 | 0 | 1468752 | APO40578.1 | P0C6X7 |
| LLENGRLIIREN | 1 | 0 | 0 | 1468754 | ATO98143.1 | P0C6W2 |
| LLEPVKHSSTVD | 1 | 0 | 0 | 1468758 | AVV62535.1 | K9N7C7 |
| LLEYDFTEHKEN | 1 | 0 | 0 | 1468780 | AGT51365.1 | H9EJ42 |
| LLFANAFSKPLY | 1 | 0 | 0 | 1468784 | AVV62527.1 | K9N796 |
| LLFDWIKISADN | 1 | 0 | 0 | 1468789 | AGT51707.1 | P0C6X6 |
| LLGFDSSKPVAK | 1 | 0 | 0 | 1468855 | ANI69899.1 | K9N7C7 |
| LLGFDSSKPVIK | 1 | 0 | 0 | 1468856 | AHI48784.1 | K9N7C7 |
| LLGNSIGIDFQD | 1 | 0 | 0 | 1468875 | AGV08467.1 | K9N5Q8 |
| LLGSIAGAGWTA | 1 | 0 | 0 | 1468886 | AVV62537.1 | K9N5Q8 |
| LLGSTADNLKNV | 1 | 0 | 0 | 1468891 | APO40578.1 | P0C6X7 |
| LLGVSDVWHKQL | 1 | 0 | 0 | 1468897 | AEN19363.1 | P0C6X6 |
| LLHFTNDKTIFY | 1 | 0 | 0 | 1468905 | ARU07575.1 | A0A140H1G9 |
| LLHIAGRQTTNG | 1 | 0 | 0 | 1468910 | AKA09607.1 | A0A0P0K6L9 |
| LLHNLQADICYN | 1 | 0 | 0 | 1468923 | QGV13494.1 | K9N7C7 |
| LLHSKVKVVPNLYVVV | 1 | 0 | 0 | 1468930 | ABD75503.1 | A0A140H1G9 |
| LLHTKVSWAGSK | 1 | 0 | 0 | 1468937 | AVN89451.1 | K9N7C7 |
| LLICVHDPLPVI | 1 | 0 | 0 | 1468951 | AGT51729.1 | P30215 |
| LLIKRFVSLAIH | 1 | 0 | 0 | 1468970 | ANZ78855.1 | P0C6X6 |
| LLIKRMKQIEDK | 1 | 0 | 0 | 1468971 | 6NZK_A | P36334 |
| LLIPDLLHAHPV | 1 | 0 | 0 | 1468981 | AHF54653.1 | Q4VID0 |
| LLIPDLLRAHPA | 1 | 0 | 0 | 1468982 | AVR40343.1 | Q4VID0 |
| LLKDALQSREAV | 1 | 0 | 0 | 1469025 | AXX83318.1 | P0C6X6 |
| LLKDIKLVIPAL | 1 | 0 | 0 | 1469029 | AVV61898.1 | K9N7C7 |
| LLKGHNLAKNIL | 1 | 0 | 0 | 1469039 | AVV62524.1 | K9N7C7 |
| LLKKMDEPFTED | 1 | 0 | 0 | 1469051 | AEN19370.1 | P33469 |
| LLKKMDEPFTEH | 1 | 0 | 0 | 1469052 | AHA80186.1 | P33469 |
| LLKKMDEPFTVD | 1 | 0 | 0 | 1469053 | AIX10719.1 | P33469 |
| LLKNAYENFNQH | 1 | 0 | 0 | 1469059 | AVP78041.1 | P0C6W2 |
| LLKQPMAISVNC | 1 | 0 | 0 | 1469070 | AGT51675.1 | Q4VID0 |
| LLKQPTATSVNC | 1 | 0 | 0 | 1469074 | ARE29989.1 | Q4VID0 |
| LLLANANSKPLI | 1 | 0 | 0 | 1469097 | AUM60025.1 | K9N796 |
| LLLDCVGFTAAS | 1 | 0 | 0 | 1469100 | AVV62524.1 | K9N7C7 |

| | | | | | | |
|---|---|---|---|---|---|---|
| LLLECITVDVLK | 1 | 0 | 0 | 1469113 | ARU07575.1 | A0A140H1G9 |
| LLLEYGQFCSKI | 1 | 0 | 0 | 1469116 | QAT98898.1 | K9N5Q8 |
| LLLGDFVEIIKS | 1 | 0 | 0 | 1469123 | 2H85_A | P0C6X7 |
| LLLILQKLWPNR | 1 | 0 | 0 | 1469136 | AIZ74435.1 | K9N643 |
| LLLIPDLLHAHP | 1 | 0 | 0 | 1469137 | AHF54653.1 | Q4VID0 |
| LLLIVDQVHVLN | 1 | 0 | 0 | 1469140 | ARU07582.1 | Q0ZJ80 |
| LLLLANANSKPL | 1 | 0 | 0 | 1469147 | AUM60025.1 | K9N796 |
| LLLLDRLNQLEN | 1 | 0 | 0 | 1469148 | AVP78038.1 | Q3LZX4 |
| LLLMLGTTMVLL | 1 | 0 | 0 | 1469163 | ATQ39389.1 | K9N7C7 |
| LLLPLKAPKEII | 1 | 0 | 0 | 1469175 | AVP78041.1 | P0C6W2 |
| LLLSRLWFKPPF | 1 | 0 | 0 | 1469194 | CAA83660.1 | P36334 |
| LLMFLLTPIKGD | 1 | 0 | 0 | 1469227 | AVV62537.1 | K9N5Q8 |
| LLMPLKAPKDVT | 1 | 0 | 0 | 1469239 | ARO76381.1 | P0C6U8 |
| LLNHKKILSHII | 1 | 0 | 0 | 1469269 | ABD75590.1 | Q0ZJ80 |
| LLNKLVGGDFTP | 1 | 0 | 0 | 1469277 | AVV62524.1 | K9N7C7 |
| LLNNQTDLDFKE | 1 | 0 | 0 | 1469281 | AVV62537.1 | K9N5Q8 |
| LLNRYFRFTFGV | 1 | 0 | 0 | 1469293 | APO40578.1 | P0C6X7 |
| LLNVPIGGTTVV | 1 | 0 | 0 | 1469303 | AUM60030.1 | K9N7A1 |
| LLNVPLRGTILT | 1 | 0 | 0 | 1469307 | Q3I5J2.1 | Q3LZX9 |
| LLPAAELDDFSK | 1 | 0 | 0 | 1469316 | APO40586.1 | P59595 |
| LLPLKAPKEIIF | 1 | 0 | 0 | 1469328 | AVP78041.1 | P0C6W2 |
| LLPLLFVTMAFV | 1 | 0 | 0 | 1469330 | ATG84886.1 | K9N7C7 |
| LLPTHWLCANQS | 1 | 0 | 0 | 1469355 | ABE73442.1 | A0A0P0G321 |
| LLQDKAGDFMST | 1 | 0 | 0 | 1469362 | AQZ41321.1 | K9N7C7 |
| LLQDKIGDFMST | 1 | 0 | 0 | 1469363 | AXG21664.1 | K9N638 |
| LLQDKTGDFMSI | 1 | 0 | 0 | 1469364 | AMO03400.1 | K9N7C7 |
| LLQGKTGDFMST | 1 | 0 | 0 | 1469370 | AHI48793.1 | K9N7C7 |
| LLQPTANSTTVS | 1 | 0 | 0 | 1469385 | AHI48527.1 | K9N638 |
| LLQVGDSVLLQG | 1 | 0 | 0 | 1469397 | ALA49901.1 | K9N7C7 |
| LLRGMPIPDFDL | 1 | 0 | 0 | 1469411 | AXN92247.1 | K9N7C7 |
| LLRGVEIEDFDL | 1 | 0 | 0 | 1469413 | AVV62524.1 | K9N7C7 |
| LLRKMDEPFTED | 1 | 0 | 0 | 1469420 | AHA80151.1 | P33469 |
| LLRLVHTDSFSC | 1 | 0 | 0 | 1469428 | AET99191.1 | A0A140H1H1 |
| LLRNTSQKSIVA | 1 | 0 | 0 | 1469436 | AAT76147.1 | Q3LZX1 |
| LLRTSIMKRDIK | 1 | 0 | 0 | 1469449 | ABV45235.1 | H9EJ42 |
| LLSENQFSGYTL | 1 | 0 | 0 | 1469470 | AIX10748.1 | P36334 |
| LLSESPLLKQKD | 1 | 0 | 0 | 1469474 | AGW27859.1 | Q0ZJ80 |
| LLSFKREKSLSL | 1 | 0 | 0 | 1469479 | AGT51407.1 | Q4VID0 |
| LLSGDRTEVPQL | 1 | 0 | 0 | 1469482 | ANF29261.1 | K9N5Q8 |
| LLSKCYKAMNSY | 1 | 0 | 0 | 1469502 | AVV62524.1 | K9N7C7 |
| LLSKLWFKPPFL | 1 | 0 | 0 | 1469506 | AWW13511.1 | P36334 |
| LLSNQTDQSFKD | 1 | 0 | 0 | 1469531 | AVV62526.1 | K9N5Q8 |
| LLSSLTLTVKFA | 1 | 0 | 0 | 1469540 | AFD98812.1 | H9EJ42 |
| LLSVLISMQGAV | 1 | 0 | 0 | 1469551 | ARO76381.1 | P0C6U8 |
| LLTHCAFAVDPA | 1 | 0 | 0 | 1469573 | AST12962.1 | A0A0P0LKV0 |
| LLTHCSFAVDPA | 1 | 0 | 0 | 1469574 | NP_073549.1 | A0A0P0LKV0 |
| LLTHIKGDVDSG | 1 | 0 | 0 | 1469575 | AVV62526.1 | K9N5Q8 |
| LLTIHRGDPMSN | 1 | 0 | 0 | 1469578 | AVP78042.1 | Q3LZX1 |
| LLTNTSSVVDSV | 1 | 0 | 0 | 1469596 | AAQ90005.1 | A0A0P0K6L9 |
| LLTPIKGDVDSG | 1 | 0 | 0 | 1469597 | AVV62537.1 | K9N5Q8 |
| LLVDSEQPWKVV | 1 | 0 | 0 | 1469633 | AIW52723.1 | A0A0N7J131 |
| LLVNNGSAITIQ | 1 | 0 | 0 | 1469670 | ATO98145.1 | Q3LZX1 |
| LLVQDLCLNKLL | 1 | 0 | 0 | 1469675 | ATG84700.1 | K9N638 |
| LLVQDLRLNKLL | 1 | 0 | 0 | 1469677 | AXN92247.1 | K9N7C7 |
| LLVQNQSFNKLL | 1 | 0 | 0 | 1469679 | ATQ39389.1 | K9N7C7 |
| LLVSSDKAPHRV | 1 | 0 | 0 | 1469686 | AVL25594.1 | H9EJ69 |
| LLYAHPAEPLVQ | 1 | 0 | 0 | 1469714 | AHF54648.1 | Q4VID0 |
| LLYCDSSFEKLH | 1 | 0 | 0 | 1469723 | AWO68974.1 | A0A0P0G321 |
| LLYDPNGNLYGF | 1 | 0 | 0 | 1469728 | ARB66062.1 | P36334 |
| LLYDSNGNLYGL | 1 | 0 | 0 | 1469730 | ABD43179.1 | P36334 |
| LLYDSPCFSQQG | 1 | 0 | 0 | 1469731 | AAA73152.1 | P30215 |
| LLYFMVDNGTRL | 1 | 0 | 0 | 1469734 | P30215.1 | P30215 |
| LLYGLKMLVLWL | 1 | 0 | 0 | 1469736 | AAA45461.1 | A0A0P0KXN3 |
| LLYNSPCFSQQG | 1 | 0 | 0 | 1469751 | P30215.1 | P30215 |
| LLYNTPCFSQQG | 1 | 0 | 0 | 1469755 | AGT51579.1 | P30215 |
| LLYTFYGKESFE | 1 | 0 | 0 | 1469766 | AVN89322.1 | K9N7C7 |
| LMACNGHKIEDF | 1 | 0 | 0 | 1469785 | AIG96659.1 | A0A140H1G9 |
| LMAQSSGVLPVN | 1 | 0 | 0 | 1469794 | YP_009555247.1 | P0C6X6 |
| LMATSVNCCHDG | 1 | 0 | 0 | 1469796 | AGT51695.1 | Q4VID0 |
| LMFKASALFKEF | 1 | 0 | 0 | 1469832 | AUM60013.1 | K9N7C7 |
| LMFLLTPIKGDV | 1 | 0 | 0 | 1469834 | AVV62537.1 | K9N5Q8 |
| LMGCVAETGTSD | 1 | 0 | 0 | 1469840 | 6NB6_A | P59594 |
| LMKPRQKRTPNK | 1 | 0 | 0 | 1469865 | AMM42397.1 | Q19U29 |
| LMLDKQGQLLDI | 1 | 0 | 0 | 1469873 | ARK08626.1 | H9EJ42 |
| LMLLELKDKEQY | 1 | 0 | 0 | 1469883 | ARO76381.1 | P0C6U8 |
| LMLLELKERDQY | 1 | 0 | 0 | 1469884 | APO40578.1 | P0C6X7 |
| LMMYAADPAMHI | 1 | 0 | 0 | 1469893 | AXN92247.1 | K9N7C7 |
| LMPLKAPKDVTF | 1 | 0 | 0 | 1469907 | ARO76381.1 | P0C6U8 |
| LMPTYTVHKSDF | 1 | 0 | 0 | 1469912 | AEN19363.1 | P0C6X6 |
| LMPTYTVHKSGF | 1 | 0 | 0 | 1469913 | AIV41822.1 | P0C6X6 |
| LMQASQIVKMVT | 1 | 0 | 0 | 1469922 | AUM60013.1 | K9N7C7 |
| LMQDESVANLFS | 1 | 0 | 0 | 1469923 | AVV62537.1 | K9N5Q8 |
| LMSCNGNKIGDL | 1 | 0 | 0 | 1469931 | AJE26218.1 | P0C6X6 |
| LMSCNGNKSDDL | 1 | 0 | 0 | 1469932 | AKL87224.1 | P0C6X6 |
| LMTFSGNVDLDN | 1 | 0 | 0 | 1469958 | AUM60013.1 | K9N7C7 |
| LMTFSGNVDLEG | 1 | 0 | 0 | 1469959 | AVV62524.1 | K9N7C7 |
| LMTIDALRDIRF | 1 | 0 | 0 | 1469960 | AUM60013.1 | K9N7C7 |
| LMVNKSSCYQDG | 1 | 0 | 0 | 1469967 | AET99208.1 | Q0ZJ80 |
| LMYLCLPTFSMA | 1 | 0 | 0 | 1469983 | AVN89451.1 | K9N7C7 |
| LNAATNGEMQKE | 1 | 0 | 0 | 1469989 | APO40578.1 | P0C6X7 |
| LNADETKALKEL | 1 | 0 | 0 | 1469992 | ATQ39389.1 | K9N7C7 |

| | | | | | | |
|---|---|---|---|---|---|---|
| LNAGLNGDFNLT | 1 | 0 | 0 | 1470002 | AVV62537.1 | K9N5Q8 |
| LNAPATVCGPKK | 1 | 0 | 0 | 1470014 | APO40579.1 | P59594 |
| LNASTPESKPLA | 1 | 0 | 0 | 1470021 | AIW52805.1 | H9EJ69 |
| LNAVVKQLNESY | 1 | 0 | 0 | 1470026 | AVV62526.1 | K9N5Q8 |
| LNAYQQQDGMNM | 1 | 0 | 0 | 1470028 | AIV41850.1 | P33469 |
| LNAYQQQDGMVN | 1 | 0 | 0 | 1470029 | AEN19370.1 | P33469 |
| LNCPLDPRLKGS | 1 | 0 | 0 | 1470048 | BAM16463.1 | P36334 |
| LNCPLDSRLKGS | 1 | 0 | 0 | 1470049 | AIL49507.1 | P36334 |
| LNCTFQEAAGNI | 1 | 0 | 0 | 1470053 | AVV62524.1 | K9N7C7 |
| LNCTFQEAASNI | 1 | 0 | 0 | 1470054 | AUM60013.1 | K9N7C7 |
| LNCTLHPRLKGS | 1 | 0 | 0 | 1470056 | BBA84086.1 | P36334 |
| LNDKLGFNVDLP | 1 | 0 | 0 | 1470076 | ARU07575.1 | A0A140H1G9 |
| LNDKLVPSPLNW | 1 | 0 | 0 | 1470077 | ATN39872.1 | P36334 |
| LNDPNYDGLLIS | 1 | 0 | 0 | 1470087 | AVV62524.1 | K9N7C7 |
| LNEHVTSVNLFD | 1 | 0 | 0 | 1470110 | ANF29196.1 | K9N796 |
| LNEPLITMPLGY | 1 | 0 | 0 | 1470121 | APO40578.1 | P0C6X7 |
| LNEVNDLLDITQ | 1 | 0 | 0 | 1470130 | BBA20986.1 | A0A140H1H1 |
| LNEVVKQLNESY | 1 | 0 | 0 | 1470131 | AVV62537.1 | K9N5Q8 |
| LNFNENGTITDA | 1 | 0 | 0 | 1470147 | ATO98193.1 | Q3LZX1 |
| LNFSGALAEHDF | 1 | 0 | 0 | 1470151 | AIW52827.1 | H9EJ42 |
| LNFVGEFVINDV | 1 | 0 | 0 | 1470155 | ATQ39389.1 | K9N7C7 |
| LNGCTWWLKGEK | 1 | 0 | 0 | 1470165 | NP_073549.1 | A0A0P0LKV0 |
| LNGIAYITFSGS | 1 | 0 | 0 | 1470181 | ABE97129.1 | H9EJ42 |
| LNGMPVSSIKGD | 1 | 0 | 0 | 1470197 | AWK59942.1 | H9EJ42 |
| LNGMPVSSIKSD | 1 | 0 | 0 | 1470198 | AGT51393.1 | H9EJ42 |
| LNGSAGSVGFNI | 1 | 0 | 0 | 1470208 | 1Z1J_A | P0C6U8 |
| LNGSCPDKNNGI | 1 | 0 | 0 | 1470211 | AIX10751.1 | P36334 |
| LNGSCPGKNNGI | 1 | 0 | 0 | 1470212 | AEN19366.1 | P36334 |
| LNGTVSTVFVPA | 1 | 0 | 0 | 1470216 | ANI69893.1 | K9N7D2 |
| LNGVIREKLAIG | 1 | 0 | 0 | 1470220 | AWK59942.1 | H9EJ42 |
| LNGVMVDKVGDT | 1 | 0 | 0 | 1470221 | QBP84746.1 | P0C6X6 |
| LNGYPVQPIADV | 1 | 0 | 0 | 1470225 | AMM42466.1 | P36334 |
| LNHTQTVVTNCV | 1 | 0 | 0 | 1470246 | APO40578.1 | P0C6X7 |
| LNILQATTANVS | 1 | 0 | 0 | 1470263 | AXG21663.1 | K9N7C7 |
| LNKHFNMMILSD | 1 | 0 | 0 | 1470291 | AIL49554.1 | P0C6X6 |
| LNKIHDVVNQQG | 1 | 0 | 0 | 1470295 | ABB90526.1 | A0A0P0K6L9 |
| LNKLGTEDPPWP | 1 | 0 | 0 | 1470301 | QDP16213.1 | K9N4V7 |
| LNKLVGGDFIPI | 1 | 0 | 0 | 1470306 | AVV62535.1 | K9N7C7 |
| LNKVSVPSPLNW | 1 | 0 | 0 | 1470321 | AMM42407.1 | A0A140H1H1 |
| LNLGEIFIAQSK | 1 | 0 | 0 | 1470328 | ATO98143.1 | P0C6W2 |
| LNLGETFVTHSK | 1 | 0 | 0 | 1470329 | AVP78041.1 | P0C6W2 |
| LNLQFEMNRLQE | 1 | 0 | 0 | 1470342 | AIL49493.1 | P36334 |
| LNLSADTQHISN | 1 | 0 | 0 | 1470347 | ARU07581.1 | Q19U29 |
| LNLSDRLAALES | 1 | 0 | 0 | 1470349 | AVV62544.1 | K9N4V7 |
| LNNDYYRSVPGT | 1 | 0 | 0 | 1470371 | AGW27852.1 | A0A140H1G9 |
| LNNFKHKVLTDS | 1 | 0 | 0 | 1470375 | AUM60013.1 | K9N7C7 |
| LNNGQGENDNIS | 1 | 0 | 0 | 1470378 | AGT51734.1 | P33469 |
| LNNKNIYRSQLG | 1 | 0 | 0 | 1470382 | ASU90570.1 | K9N638 |
| LNNKSTYRSQLG | 1 | 0 | 0 | 1470386 | AZK15899.1 | K9N638 |
| LNNPAYIAPQAN | 1 | 0 | 0 | 1470394 | AEN19365.1 | P30215 |
| LNNPAYIAPRAN | 1 | 0 | 0 | 1470395 | AIL49334.1 | P30215 |
| LNNPAYIAREAN | 1 | 0 | 0 | 1470396 | AAA73152.1 | P30215 |
| LNNPAYIASQAK | 1 | 0 | 0 | 1470397 | AIV41992.1 | P30215 |
| LNNSIPNLSDFE | 1 | 0 | 0 | 1470409 | BBA20986.1 | A0A140H1H1 |
| LNNVSVPSPFNW | 1 | 0 | 0 | 1470414 | ABU39973.1 | A0A140H1H1 |
| LNPGALEFNPSQ | 1 | 0 | 0 | 1470420 | AGT21342.1 | A0A0N7J131 |
| LNPSEEFIKHVR | 1 | 0 | 0 | 1470426 | ALW82730.1 | K9N7C7 |
| LNQIWQKYLNSS | 1 | 0 | 0 | 1470435 | AXN92262.1 | K9N4V0 |
| LNQLNEPLITMP | 1 | 0 | 0 | 1470440 | APO40578.1 | P0C6X7 |
| LNQSYINLKYIG | 1 | 0 | 0 | 1470446 | AGT51412.1 | P36334 |
| LNQVWQHHINLA | 1 | 0 | 0 | 1470448 | AVV62528.1 | K9N4V0 |
| LNRALTGVALEQ | 1 | 0 | 0 | 1470454 | ATO98145.1 | Q3LZX1 |
| LNRRSFRISDGN | 1 | 0 | 0 | 1470471 | AJQ24067.1 | P33469 |
| LNSENLYFQSGS | 1 | 0 | 0 | 1470489 | 6U7H_A | A0A0P0K6L9 |
| LNSFKEYFDLQN | 1 | 0 | 0 | 1470490 | AVV62537.1 | K9N5Q8 |
| LNSNSTPAQNAG | 1 | 0 | 0 | 1470512 | AHI48825.1 | K9N4V7 |
| LNSSQGIPDLLV | 1 | 0 | 0 | 1470525 | APO40581.1 | P59637 |
| LNSSSCVGSGPG | 1 | 0 | 0 | 1470526 | AIX10756.1 | P36334 |
| LNSTNVTTATYC | 1 | 0 | 0 | 1470532 | AVP78030.1 | P0C6W2 |
| LNSYIPLFDMRY | 1 | 0 | 0 | 1470536 | AMQ49019.1 | K9N7D2 |
| LNTDHSGSNDNI | 1 | 0 | 0 | 1470542 | Q3I5J2.1 | Q3LZX9 |
| LNTNLHSDVDNI | 1 | 0 | 0 | 1470559 | BBA20986.1 | A0A140H1H1 |
| LNTPIPNLPDFR | 1 | 0 | 0 | 1470560 | AIL49487.1 | P36334 |
| LNTSIPNLHDFK | 1 | 0 | 0 | 1470565 | AAB27260.2 | P36334 |
| LNTSYSVCNGCV | 1 | 0 | 0 | 1470570 | NP_073551.1 | A0A0P0K6L9 |
| LNTTFLLNGYYP | 1 | 0 | 0 | 1470571 | CAA79896.1 | P36334 |
| LNTTIPRISEYV | 1 | 0 | 0 | 1470573 | ACV87140.1 | A0A140H1H1 |
| LNVCFDVRDKGA | 1 | 0 | 0 | 1470586 | AVV62535.1 | K9N7C7 |
| LNVCFDVRDNGA | 1 | 0 | 0 | 1470587 | AVV62524.1 | K9N7C7 |
| LNVKPYGYPNDF | 1 | 0 | 0 | 1470593 | AWK59948.1 | H9EJ42 |
| LNVLFDGRVDGQ | 1 | 0 | 0 | 1470596 | APO40578.1 | P0C6X7 |
| LNVQKVAQPKIE | 1 | 0 | 0 | 1470609 | APT69882.1 | A0A0P0ILY9 |
| LNVTINNFNPSS | 1 | 0 | 0 | 1470613 | BBA20986.1 | A0A140H1H1 |
| LNVVGLDARTQA | 1 | 0 | 0 | 1470619 | AIV41936.1 | P0C6X6 |
| LNWLNRVETYIK | 1 | 0 | 0 | 1470626 | AGT21353.1 | A0A0P0K6L9 |
| LNYAGALWDITV | 1 | 0 | 0 | 1470629 | AXN92247.1 | K9N7C7 |
| LPAADLDEFSKQ | 1 | 0 | 0 | 1470640 | AVP78038.1 | Q3LZX4 |
| LPAAELDDFSKQ | 1 | 0 | 0 | 1470642 | APO40586.1 | P59595 |
| LPAEYKLRQGTF | 1 | 0 | 0 | 1470645 | ARO76381.1 | P0C6U8 |
| LPAQDTWGTSAA | 1 | 0 | 0 | 1470652 | ATO98205.1 | Q3LZX1 |
| LPCGICLSDLDS | 1 | 0 | 0 | 1470660 | APO40578.1 | P0C6X7 |

| | | | | | | |
|---|---|---|---|---|---|---|
| LPDAAEKLASPM | 1 | 0 | 0 | 1470668 | AZS52616.1 | A0A140H1G9 |
| LPDCDIDNWLNN | 1 | 0 | 0 | 1470671 | BBA20986.1 | A0A140H1H1 |
| LPDFKEELDQWL | 1 | 0 | 0 | 1470679 | ARB07414.1 | P36334 |
| LPDFKEELGQWF | 1 | 0 | 0 | 1470680 | AIX10763.1 | P36334 |
| LPDFKEESDQWF | 1 | 0 | 0 | 1470681 | BBA20973.1 | P36334 |
| LPDIAEDVVDQV | 1 | 0 | 0 | 1470691 | NP_073549.1 | A0A0P0LKV0 |
| LPDTPSTLKPRS | 1 | 0 | 0 | 1470706 | ALJ54490.1 | K9N5Q8 |
| LPELVIAANKTK | 1 | 0 | 0 | 1470722 | AVV62524.1 | K9N7C7 |
| LPENDDDAWFKP | 1 | 0 | 0 | 1470725 | AQN78686.1 | Q80872 |
| LPEYMAVAVPST | 1 | 0 | 0 | 1470735 | APT69865.1 | A0A0P0KXN3 |
| LPEYMTVAVPST | 1 | 0 | 0 | 1470736 | AST12967.1 | A0A0P0KXN3 |
| LPFDSNLTQYFS | 1 | 0 | 0 | 1470740 | ATO98181.1 | Q3LZX1 |
| LPFEDLNKCRAY | 1 | 0 | 0 | 1470742 | AVV62524.1 | K9N7C7 |
| LPFEDLSKCRAY | 1 | 0 | 0 | 1470743 | AVV62535.1 | K9N7C7 |
| LPFEDLTKCRTY | 1 | 0 | 0 | 1470744 | AUM60013.1 | K9N7C7 |
| LPFFYYSGSPCE | 1 | 0 | 0 | 1470749 | AER30333.1 | P0C6X7 |
| LPFNNMLGYNPS | 1 | 0 | 0 | 1470770 | AVV62524.1 | K9N7C7 |
| LPFVQEQIGTFI | 1 | 0 | 0 | 1470783 | AVV62542.1 | K9N5R3 |
| LPGAHAYETMKP | 1 | 0 | 0 | 1470793 | AVV62535.1 | K9N7C7 |
| LPGAHPYSAMKP | 1 | 0 | 0 | 1470795 | AUM60013.1 | K9N7C7 |
| LPGELSKKQLGL | 1 | 0 | 0 | 1470799 | AUM60013.1 | K9N7C7 |
| LPGTVIRTLNGD | 1 | 0 | 0 | 1470817 | APO40578.1 | P0C6X7 |
| LPICVYHPLPVI | 1 | 0 | 0 | 1470834 | QEG03802.1 | P30215 |
| LPIQDYIQSVNF | 1 | 0 | 0 | 1470847 | AEN19363.1 | P0C6X6 |
| LPKEEGDVLLAE | 1 | 0 | 0 | 1470857 | ATQ39389.1 | K9N7C7 |
| LPKGVGSANIFK | 1 | 0 | 0 | 1470868 | AVI15051.1 | P0C6X6 |
| LPLAYFSDSKFG | 1 | 0 | 0 | 1470883 | AHI48527.1 | K9N638 |
| LPLDIYDLTMPS | 1 | 0 | 0 | 1470888 | AIV41960.1 | P0C6X6 |
| LPLDKIQNQTLP | 1 | 0 | 0 | 1470889 | ARU07574.1 | A0A140H1G9 |
| LPLDVYNLTMPS | 1 | 0 | 0 | 1470892 | AAX85666.1 | P0C6X6 |
| LPLEFGASTEMQ | 1 | 0 | 0 | 1470895 | ATO98229.1 | P0C6W2 |
| LPLSDIEKDFLS | 1 | 0 | 0 | 1470924 | AHI48581.1 | K9N7C7 |
| LPLSLTEVDGGF | 1 | 0 | 0 | 1470927 | ATO98145.1 | Q3LZX1 |
| LPLVSCFSTRNS | 1 | 0 | 0 | 1470933 | AKA09629.1 | A0A0P0G321 |
| LPMSDMEKDFLS | 1 | 0 | 0 | 1470942 | QBF80541.1 | K9N7C7 |
| LPNCSITSGVLQ | 1 | 0 | 0 | 1470950 | AGV08515.1 | K9N7C7 |
| LPNRRKVDLQLG | 1 | 0 | 0 | 1470969 | AIL49495.1 | P36334 |
| LPPAYAPTYVSV | 1 | 0 | 0 | 1470976 | AVV62535.1 | K9N7C7 |
| LPPLLSDNQISG | 1 | 0 | 0 | 1470989 | CAA83661.1 | P36334 |
| LPPLLSENQFSG | 1 | 0 | 0 | 1470990 | AIX10748.1 | P36334 |
| LPPLLSENQING | 1 | 0 | 0 | 1470991 | ANZ78836.1 | P36334 |
| LPPLLTDEMIAA | 1 | 0 | 0 | 1470995 | AVP78031.1 | Q3LZX1 |
| LPPLYDPNMEAA | 1 | 0 | 0 | 1470998 | AVV62526.1 | K9N5Q8 |
| LPPLYDPNMEGA | 1 | 0 | 0 | 1470999 | AVV62537.1 | K9N5Q8 |
| LPPLYDVNMEAA | 1 | 0 | 0 | 1471000 | AUM60014.1 | K9N5Q8 |
| LPPPLLGNSIGI | 1 | 0 | 0 | 1471001 | AGV08467.1 | K9N5Q8 |
| LPPPLLNNQTDL | 1 | 0 | 0 | 1471003 | AVV62537.1 | K9N5Q8 |
| LPPPLLSNQTDQ | 1 | 0 | 0 | 1471004 | AVV62526.1 | K9N5Q8 |
| LPPPLLSNQTDV | 1 | 0 | 0 | 1471005 | AUM60014.1 | K9N5Q8 |
| LPPPLLSNSTGI | 1 | 0 | 0 | 1471006 | ALL26396.1 | K9N5Q8 |
| LPQGGFVADFAY | 1 | 0 | 0 | 1471014 | ARU07575.1 | A0A140H1G9 |
| LPQGYYIEGSGK | 1 | 0 | 0 | 1471018 | AJQ24087.1 | P33469 |
| LPQLVNANQYSP | 1 | 0 | 0 | 1471024 | ATG84679.1 | K9N5Q8 |
| LPQSKDSNFLNG | 1 | 0 | 0 | 1471036 | AVV62536.1 | K9N638 |
| LPQSKESNFLNE | 1 | 0 | 0 | 1471038 | AQZ41321.1 | K9N7C7 |
| LPQSKESNFLNR | 1 | 0 | 0 | 1471039 | AQZ41330.1 | K9N7C7 |
| LPSANVSVSRFN | 1 | 0 | 0 | 1471054 | BBO58721.1 | P36334 |
| LPSGNYLAISNE | 1 | 0 | 0 | 1471066 | AEN19365.1 | P30215 |
| LPSGNYLSISNE | 1 | 0 | 0 | 1471067 | AVQ05263.1 | P30215 |
| LPSPGMPALLSL | 1 | 0 | 0 | 1471079 | 6NB6_A | P59594 |
| LPSSKSSGMDTA | 1 | 0 | 0 | 1471082 | AZS52621.1 | Q0ZJ82 |
| LPSVSLNSKEEL | 1 | 0 | 0 | 1471090 | APO40578.1 | P0C6X7 |
| LPTAFAAIGDLK | 1 | 0 | 0 | 1471093 | AIL49486.1 | P36334 |
| LPTAFAAIGDLN | 1 | 0 | 0 | 1471094 | BBO58773.1 | P36334 |
| LPTAFAVVGDLN | 1 | 0 | 0 | 1471097 | QBP84704.1 | P36334 |
| LPTDYKNVKAWS | 1 | 0 | 0 | 1471101 | AIW52843.1 | A0A0P0G321 |
| LPTHWICANQGT | 1 | 0 | 0 | 1471121 | AKA09620.1 | A0A0P0G321 |
| LPTHWICANQNT | 1 | 0 | 0 | 1471122 | BAM16475.1 | A0A0P0G321 |
| LPTHWICANQST | 1 | 0 | 0 | 1471123 | AIW52843.1 | A0A0P0G321 |
| LPTHWLCANQST | 1 | 0 | 0 | 1471124 | ABE73442.1 | A0A0P0G321 |
| LPTMCDIKQMLF | 1 | 0 | 0 | 1471128 | AXN92247.1 | K9N7C7 |
| LPTNAVMKMPCP | 1 | 0 | 0 | 1471131 | ARO76381.1 | P0C6U8 |
| LPTWSQELPFDV | 1 | 0 | 0 | 1471140 | QQY99294.1 | P0C6X6 |
| LPVAVVNDKVFT | 1 | 0 | 0 | 1471146 | ANF29215.1 | K9N7C7 |
| LPVGFSVLKPIL | 1 | 0 | 0 | 1471150 | ATO98193.1 | Q3LZX1 |
| LPVKDYIQTVNV | 1 | 0 | 0 | 1471155 | ABD75583.1 | A0A140H1G9 |
| LPVMISEWPLFE | 1 | 0 | 0 | 1471164 | ARX11394.1 | A0A0P0LKV0 |
| LPVYASYKVLDN | 1 | 0 | 0 | 1471174 | AEN19363.1 | P0C6X6 |
| LPVYTSYKVLDN | 1 | 0 | 0 | 1471180 | ARK08667.1 | P0C6X6 |
| LPVYYQHTDINF | 1 | 0 | 0 | 1471182 | AWO68966.1 | A0A0P0G321 |
| LPYPNPSRILGA | 1 | 0 | 0 | 1471193 | QVU04458.1 | P0DTD1 |
| LQAAFGNTGVCY | 1 | 0 | 0 | 1471199 | AWK59942.1 | H9EJ42 |
| LQAANDWKPGLA | 1 | 0 | 0 | 1471200 | AVV62524.1 | K9N7C7 |
| LQALESGKVKQS | 1 | 0 | 0 | 1471216 | AXN92267.1 | K9N4V7 |
| LQALPQRQKKQQ | 1 | 0 | 0 | 1471217 | AVP78038.1 | Q3LZX4 |
| LQASNDWKPGLA | 1 | 0 | 0 | 1471224 | AUM60013.1 | K9N7C7 |
| LQDFDVVFGHGA | 1 | 0 | 0 | 1471245 | AWK59942.1 | H9EJ42 |
| LQDGFYSANFQH | 1 | 0 | 0 | 1471248 | AWO68987.1 | A0A0P0G321 |
| LQDKIGDFMSTI | 1 | 0 | 0 | 1471251 | AXG21664.1 | K9N638 |
| LQDKTGDFMSII | 1 | 0 | 0 | 1471253 | AMO03400.1 | K9N7C7 |
| LQDKTGDFTSTI | 1 | 0 | 0 | 1471255 | AJO62148.1 | K9N7C7 |

| | | | | | | |
|---|---|---|---|---|---|---|
| LQDSMNGASDST | 1 | 0 | 0 | 1471263 | APO40586.1 | P59595 |
| LQDVQVSAQQEV | 1 | 0 | 0 | 1471267 | AUM60013.1 | K9N7C7 |
| LQEIISRLDALE | 1 | 0 | 0 | 1471277 | ANZ78843.1 | P36334 |
| LQELGKYEQGSG | 1 | 0 | 0 | 1471282 | 6CRW_A | P59594 |
| LQENQRILAASF | 1 | 0 | 0 | 1471298 | AIW52747.1 | A0A0P0K6L9 |
| LQFQLLDEGVDC | 1 | 0 | 0 | 1471330 | NP_150075.1 | Q80872 |
| LQFTTLTLDKVP | 1 | 0 | 0 | 1471333 | AEN19363.1 | P0C6X6 |
| LQGDKCNIFANL | 1 | 0 | 0 | 1471337 | AVQ05264.1 | P36334 |
| LQGDRCNIFANF | 1 | 0 | 0 | 1471340 | CAA83660.1 | P36334 |
| LQGHFLAKNILH | 1 | 0 | 0 | 1471341 | AHI48765.1 | K9N638 |
| LQGKAIGFFFPY | 1 | 0 | 0 | 1471343 | AUM60013.1 | K9N7C7 |
| LQGLKACCYVGM | 1 | 0 | 0 | 1471351 | AVN89451.1 | K9N7C7 |
| LQGLLEVSICQY | 1 | 0 | 0 | 1471354 | BBO58751.1 | P36334 |
| LQGLLEVSVCQI | 1 | 0 | 0 | 1471356 | AIG59210.1 | P36334 |
| LQGNFGDLQFNK | 1 | 0 | 0 | 1471360 | ASY99807.1 | K9N4V7 |
| LQGNFGDLQLTK | 1 | 0 | 0 | 1471362 | AVN89320.1 | K9N4V7 |
| LQGNFGDMQFNK | 1 | 0 | 0 | 1471363 | AUM60021.1 | K9N4V7 |
| LQGTIIFVQQKP | 1 | 0 | 0 | 1471366 | NP_073549.1 | A0A0P0LKV0 |
| LQHAFMKHKGGG | 1 | 0 | 0 | 1471369 | AIY60587.1 | K9N638 |
| LQHAGGIADAIN | 1 | 0 | 0 | 1471370 | AVV62535.1 | K9N7C7 |
| LQHEDLLAAYVQ | 1 | 0 | 0 | 1471373 | APO40578.1 | P0C6X7 |
| LQILNRQCVNIN | 1 | 0 | 0 | 1471380 | APO40578.1 | P0C6X7 |
| LQKHLYSHVYRS | 1 | 0 | 0 | 1471402 | AXX83366.1 | P0C6X6 |
| LQKLFVANYSQD | 1 | 0 | 0 | 1471403 | AID55095.1 | K9N5Q8 |
| LQLATINYTQPE | 1 | 0 | 0 | 1471412 | AVV62526.1 | K9N5Q8 |
| LQLGAPDNPSAI | 1 | 0 | 0 | 1471419 | AKA09625.1 | A0A0P0G321 |
| LQLGFSTGVSFV | 1 | 0 | 0 | 1471422 | ALA49417.1 | K9N7C7 |
| LQLKESQINELV | 1 | 0 | 0 | 1471427 | AXN92247.1 | K9N7C7 |
| LQLNKLGTDDPR | 1 | 0 | 0 | 1471436 | ATG84774.1 | K9N4V7 |
| LQLNKLGTEDPP | 1 | 0 | 0 | 1471437 | QDP16213.1 | K9N4V7 |
| LQLVNANQYSPC | 1 | 0 | 0 | 1471449 | QBM11748.1 | K9N5Q8 |
| LQMDVQGLVNVL | 1 | 0 | 0 | 1471452 | AWK59942.1 | H9EJ42 |
| LQNFGNFEMLKL | 1 | 0 | 0 | 1471459 | AMM42397.1 | Q19U29 |
| LQNGDFLCANEY | 1 | 0 | 0 | 1471461 | APO40578.1 | P0C6X7 |
| LQNNEIKPTGLK | 1 | 0 | 0 | 1471464 | AUM60013.1 | K9N7C7 |
| LQNNEIMPGKMK | 1 | 0 | 0 | 1471465 | NP_073549.1 | A0A0P0LKV0 |
| LQNYGDSAILPK | 1 | 0 | 0 | 1471473 | AVP78030.1 | P0C6W2 |
| LQPLEQPTNEAV | 1 | 0 | 0 | 1471484 | AVP78041.1 | P0C6W2 |
| LQPNAYYKYDVK | 1 | 0 | 0 | 1471488 | AWK59942.1 | H9EJ42 |
| LQPNTEVYTDVC | 1 | 0 | 0 | 1471489 | AGW27863.1 | A0A140H1H1 |
| LQQAQQEATHIA | 1 | 0 | 0 | 1471506 | APT69868.1 | A0A0P0ILY9 |
| LQQAVVNIRETK | 1 | 0 | 0 | 1471507 | AVV62524.1 | K9N7C7 |
| LQQGSTGEIAQY | 1 | 0 | 0 | 1471513 | AUM60014.1 | K9N5Q8 |
| LQQIDIKFNAPA | 1 | 0 | 0 | 1471516 | APO40578.1 | P0C6X7 |
| LQQIEVKFNAPA | 1 | 0 | 0 | 1471518 | P0C6W6.1 | P0C6W2 |
| LQQLHLEANSKA | 1 | 0 | 0 | 1471521 | AVV62524.1 | K9N7C7 |
| LQQLRIESSSKL | 1 | 0 | 0 | 1471523 | APO40578.1 | P0C6X7 |
| LQQLTLVRCKGL | 1 | 0 | 0 | 1471527 | AVV62535.1 | K9N7C7 |
| LQQMFVTPLAVD | 1 | 0 | 0 | 1471528 | ATG84858.1 | K9N643 |
| LQSGVGRCFNCR | 1 | 0 | 0 | 1471565 | AGT21338.1 | A0A0P0K6L9 |
| LQSKRTRLFKST | 1 | 0 | 0 | 1471570 | AGT51787.1 | P0C6X6 |
| LQSKTKRFIKET | 1 | 0 | 0 | 1471571 | ARU07575.1 | A0A140H1G9 |
| LQSYACWTHSCG | 1 | 0 | 0 | 1471597 | AXN92247.1 | K9N7C7 |
| LQTDVLQEDQKI | 1 | 0 | 0 | 1471601 | BAL45639.1 | A0A0P0K6L9 |
| LQTFSETGSDVL | 1 | 0 | 0 | 1471604 | AUM60013.1 | K9N7C7 |
| LQTGKQTELAVN | 1 | 0 | 0 | 1471605 | AAW33902.1 | H9EJ42 |
| LQTHVGCVLGLV | 1 | 0 | 0 | 1471606 | ATG84701.1 | K9N5Q8 |
| LQTIINGMHKSF | 1 | 0 | 0 | 1471607 | AWK59942.1 | H9EJ42 |
| LQTYSETGSDLL | 1 | 0 | 0 | 1471624 | AXN92247.1 | K9N7C7 |
| LQVANNLMNGVT | 1 | 0 | 0 | 1471628 | AIL49500.1 | P36334 |
| LQVTTANVSALM | 1 | 0 | 0 | 1471647 | AVV62524.1 | K9N7C7 |
| LQVVADTGSLTK | 1 | 0 | 0 | 1471648 | AUM60013.1 | K9N7C7 |
| LRAANNGHKQIV | 1 | 0 | 0 | 1471672 | AUM60017.1 | K9N643 |
| LRACCYVGVQTV | 1 | 0 | 0 | 1471674 | AMQ48992.1 | K9N7C7 |
| LRACIKIAQADT | 1 | 0 | 0 | 1471676 | ASU89922.1 | K9N796 |
| LRACIKTVQADT | 1 | 0 | 0 | 1471681 | AHC74089.1 | K9N796 |
| LRAFDIYNANVA | 1 | 0 | 0 | 1471688 | AGT51737.1 | P0C6X6 |
| LRAFIKTAQADT | 1 | 0 | 0 | 1471690 | AXN92261.1 | K9N796 |
| LRAVSFADNNDI | 1 | 0 | 0 | 1471712 | QDP16202.1 | K9N4V7 |
| LRCSYESFDVDT | 1 | 0 | 0 | 1471724 | AVV62526.1 | K9N5Q8 |
| LRDDNQALLVPK | 1 | 0 | 0 | 1471729 | AMM42360.1 | A0A140H1G9 |
| LRDFMLKGYVLA | 1 | 0 | 0 | 1471733 | AVN89312.1 | K9N638 |
| LRDTLEYYAQSI | 1 | 0 | 0 | 1471745 | AUM60013.1 | K9N7C7 |
| LREAALPQSKDS | 1 | 0 | 0 | 1471753 | AVV62535.1 | K9N7C7 |
| LRELYGTPDSNF | 1 | 0 | 0 | 1471769 | AVV62524.1 | K9N7C7 |
| LREPMMQSSDAS | 1 | 0 | 0 | 1471776 | APO40578.1 | P0C6X7 |
| LRGDDMSFLEKD | 1 | 0 | 0 | 1471796 | AFR58685.1 | P0C6X7 |
| LRGVDIDDFDID | 1 | 0 | 0 | 1471825 | AUM60013.1 | K9N7C7 |
| LRHDITLDDDAR | 1 | 0 | 0 | 1471833 | AQN78773.1 | P0C6X6 |
| LRHFDEGNCDVL | 1 | 0 | 0 | 1471837 | APO40578.1 | P0C6X7 |
| LRHFDRNDCMLL | 1 | 0 | 0 | 1471839 | AEN19363.1 | P0C6X6 |
| LRHSSLFAQSKE | 1 | 0 | 0 | 1471849 | ARQ84748.1 | K9N4V0 |
| LRIFNVVGPRKG | 1 | 0 | 0 | 1471860 | NP_073549.1 | A0A0P0LKV0 |
| LRIKVSQSNVHT | 1 | 0 | 0 | 1471866 | AWK59942.1 | H9EJ42 |
| LRISDCVADYAV | 1 | 0 | 0 | 1471868 | APO40579.1 | P59594 |
| LRITSACDGLAS | 1 | 0 | 0 | 1471873 | AVV62524.1 | K9N7C7 |
| LRKGGRTIAYGG | 1 | 0 | 0 | 1471884 | AVP78041.1 | P0C6W2 |
| LRKMDEPFTEDT | 1 | 0 | 0 | 1471894 | AHA80151.1 | P33469 |
| LRKNVQQLFYTE | 1 | 0 | 0 | 1471903 | AVV62524.1 | K9N7C7 |
| LRKTLVLNASEM | 1 | 0 | 0 | 1471909 | ATQ39396.1 | K9N643 |
| LRKVPTDNHITT | 1 | 0 | 0 | 1471910 | AVP78041.1 | P0C6W2 |

| | | | | | | |
|---|---|---|---|---|---|---|
| LRKVPTDNYITTYP | 1 | 0 | 0 | 1471912 | P0DTC1.1 | P0DTD1 |
| LRKYGRGGYHYT | 1 | 0 | 0 | 1471914 | AXN92247.1 | K9N7C7 |
| LRLGSNLSLSMA | 1 | 0 | 0 | 1471923 | 2CME_A | P59636 |
| LRLHTVESGDVN | 1 | 0 | 0 | 1471928 | ASU45805.1 | K9N7C7 |
| LRLLLMLCTTMV | 1 | 0 | 0 | 1471936 | AXN92247.1 | K9N7C7 |
| LRNDTSVEDKLG | 1 | 0 | 0 | 1471973 | AUM60014.1 | K9N5Q8 |
| LRNKGVDCTKKF | 1 | 0 | 0 | 1471979 | AXN92247.1 | K9N7C7 |
| LRNLQADICYKF | 1 | 0 | 0 | 1471983 | ATG84842.1 | K9N7C7 |
| LRNNGVDYTKKF | 1 | 0 | 0 | 1471987 | AVN89441.1 | K9N7C7 |
| LRNSGRFGGFSN | 1 | 0 | 0 | 1471988 | AVL25576.1 | H9EJ42 |
| LRNSINADAYAR | 1 | 0 | 0 | 1471989 | AVV62524.1 | K9N7C7 |
| LRNVTFELHEFA | 1 | 0 | 0 | 1472003 | ANF29275.1 | K9N643 |
| LRPILKLPFGIN | 1 | 0 | 0 | 1472018 | ATO98181.1 | Q3LZX1 |
| LRPLIKDVDNPV | 1 | 0 | 0 | 1472020 | AHF54667.1 | P0C6X6 |
| LRPPKNSFEALV | 1 | 0 | 0 | 1472022 | APU51939.1 | P0C6X6 |
| LRPPRNSFEAIL | 1 | 0 | 0 | 1472023 | ARU07575.1 | A0A140H1G9 |
| LRQIFDPAPVEL | 1 | 0 | 0 | 1472041 | AVV62524.1 | K9N7C7 |
| LRQIYDPAPVEL | 1 | 0 | 0 | 1472047 | AVV62535.1 | K9N7C7 |
| LRQTYVYTTTGF | 1 | 0 | 0 | 1472054 | AVV62524.1 | K9N7C7 |
| LRRLVKPTDRSH | 1 | 0 | 0 | 1472084 | AVV62524.1 | K9N7C7 |
| LRSETLLPITQY | 1 | 0 | 0 | 1472097 | APO40578.1 | P0C6X7 |
| LRSGHHLHSVHN | 1 | 0 | 0 | 1472099 | AXN92231.1 | K9N643 |
| LRSHDTSHKNNF | 1 | 0 | 0 | 1472106 | Q14EA9.1 | Q0ZJ84 |
| LRSINHDLFSFI | 1 | 0 | 0 | 1472108 | AEN19363.1 | P0C6X6 |
| LRSLVTNMDIEK | 1 | 0 | 0 | 1472117 | AVV62524.1 | K9N7C7 |
| LRSTRVNVDGFN | 1 | 0 | 0 | 1472125 | AWK59942.1 | H9EJ42 |
| LRTDQINDMVYS | 1 | 0 | 0 | 1472136 | AIX09796.1 | P0C6X6 |
| LRTEAFDYYHTL | 1 | 0 | 0 | 1472137 | APO40578.1 | P0C6X7 |
| LRTSIMKRDIKV | 1 | 0 | 0 | 1472156 | ABV45235.1 | H9EJ42 |
| LRVSSNQSPQGN | 1 | 0 | 0 | 1472187 | AVI15051.1 | P0C6X6 |
| LRVVFPPSRNWD | 1 | 0 | 0 | 1472193 | AVV62540.1 | K9N643 |
| LRVVIPDFNVWN | 1 | 0 | 0 | 1472198 | AVV62524.1 | K9N7C7 |
| LRYMNANGLCPP | 1 | 0 | 0 | 1472211 | ATN39862.1 | P0C6X6 |
| LSAFGDVKPVTE | 1 | 0 | 0 | 1472228 | AUM60027.1 | K9N643 |
| LSAGIFGADPVH | 1 | 0 | 0 | 1472233 | AVP78030.1 | P0C6W2 |
| LSAIQQLNNGFQ | 1 | 0 | 0 | 1472244 | AVV62535.1 | K9N7C7 |
| LSAMISGSKHVT | 1 | 0 | 0 | 1472246 | AIG96641.1 | A0A0P0LKV0 |
| LSAMLVNGTHTP | 1 | 0 | 0 | 1472247 | AVV62541.1 | K9N7D2 |
| LSARSAIEDLLF | 1 | 0 | 0 | 1472253 | AVN89324.1 | K9N5Q8 |
| LSAVLVNGTHTP | 1 | 0 | 0 | 1472255 | AVV62530.1 | K9N7D2 |
| LSCARKSCSIDS | 1 | 0 | 0 | 1472258 | AEN19363.1 | P0C6X6 |
| LSCNFNMSSLMS | 1 | 0 | 0 | 1472268 | AIV41909.1 | P36334 |
| LSDEMKVLNAVV | 1 | 0 | 0 | 1472291 | AVV62526.1 | K9N5Q8 |
| LSDFHILGRTLV | 1 | 0 | 0 | 1472292 | AVV62540.1 | K9N643 |
| LSDIEKDFLSFD | 1 | 0 | 0 | 1472303 | AHI48581.1 | K9N7C7 |
| LSDIVNGVAEGT | 1 | 0 | 0 | 1472306 | APO40578.1 | P0C6X7 |
| LSDMEKDFLSSD | 1 | 0 | 0 | 1472313 | ANI69887.1 | K9N7C7 |
| LSDNQKLIANKF | 1 | 0 | 0 | 1472318 | ALA49671.1 | K9N5Q8 |
| LSDTLDKLSDYC | 1 | 0 | 0 | 1472333 | AXN92247.1 | K9N7C7 |
| LSDVAGLKYTKI | 1 | 0 | 0 | 1472338 | ARB07411.1 | P0C6X6 |
| LSEEFRVLNEVV | 1 | 0 | 0 | 1472354 | AUM60014.1 | K9N5Q8 |
| LSEIQCKTQSFA | 1 | 0 | 0 | 1472360 | BBA20986.1 | A0A140H1H1 |
| LSEKCMIAPESK | 1 | 0 | 0 | 1472363 | AVV62524.1 | K9N7C7 |
| LSELYGTPDPMF | 1 | 0 | 0 | 1472372 | AVV62535.1 | K9N7C7 |
| LSENAYLKYAKV | 1 | 0 | 0 | 1472375 | AVV62524.1 | K9N7C7 |
| LSENLNAYQQQD | 1 | 0 | 0 | 1472376 | AEN19370.1 | P33469 |
| LSENQKQIANKF | 1 | 0 | 0 | 1472380 | AVV62526.1 | K9N5Q8 |
| LSETKCTLKSLT | 1 | 0 | 0 | 1472395 | AVP78031.1 | Q3LZX1 |
| LSFCAFAVDAAKAY | 1 | 0 | 0 | 1472407 | P0DTC1.1 | P0DTD1 |
| LSFCAFAVEPAK | 1 | 0 | 0 | 1472409 | APO40578.1 | P0C6X7 |
| LSFDNDVFIKKY | 1 | 0 | 0 | 1472411 | QBM11746.1 | K9N7C7 |
| LSFINICFEATT | 1 | 0 | 0 | 1472422 | AVV62535.1 | K9N7C7 |
| LSFKENVDFINS | 1 | 0 | 0 | 1472424 | AXX83325.1 | Q80872 |
| LSFKREKSLSLQ | 1 | 0 | 0 | 1472428 | AGT51407.1 | Q4VID0 |
| LSFNENGTITDA | 1 | 0 | 0 | 1472437 | ATO98181.1 | Q3LZX1 |
| LSFNENGTMSNA | 1 | 0 | 0 | 1472438 | ATO98145.1 | Q3LZX1 |
| LSFTEQLGAPLG | 1 | 0 | 0 | 1472448 | ACY56724.1 | A0A0P0G321 |
| LSFTNYKLQSQI | 1 | 0 | 0 | 1472449 | AXN92247.1 | K9N7C7 |
| LSGDISVVATSG | 1 | 0 | 0 | 1472455 | AVV62524.1 | K9N7C7 |
| LSGDRTEVPQLV | 1 | 0 | 0 | 1472457 | ANF29261.1 | K9N5Q8 |
| LSGEWNAVYKAF | 1 | 0 | 0 | 1472459 | AGT51379.1 | H9EJ42 |
| LSGIAIEQDKNT | 1 | 0 | 0 | 1472474 | AVP78031.1 | Q3LZX1 |
| LSGIAVEQDRNT | 1 | 0 | 0 | 1472475 | ATO98231.1 | Q3LZX1 |
| LSGIGAVGGDLL | 1 | 0 | 0 | 1472476 | ALA49601.1 | K9N4V7 |
| LSGNLDEPQKDV | 1 | 0 | 0 | 1472483 | AOL02457.1 | P33469 |
| LSGRKIYEVERG | 1 | 0 | 0 | 1472493 | AEN19363.1 | P0C6X6 |
| LSGRKIYEVVRG | 1 | 0 | 0 | 1472494 | AWW13508.1 | P0C6X6 |
| LSGTPNEKLVTT | 1 | 0 | 0 | 1472499 | AXN92247.1 | K9N7C7 |
| LSGTPPQVYNFK | 1 | 0 | 0 | 1472501 | AXN92228.1 | K9N5Q8 |
| LSGVAYTAFSGP | 1 | 0 | 0 | 1472507 | NP_073549.1 | A0A0P0LKV0 |
| LSHEKASLNSLT | 1 | 0 | 0 | 1472516 | AEN19363.1 | P0C6X6 |
| LSHEKASLSSLT | 1 | 0 | 0 | 1472517 | AIV41990.1 | P0C6X6 |
| LSIDDGRLWNLF | 1 | 0 | 0 | 1472549 | AWK59942.1 | H9EJ42 |
| LSIDEFADVVKE | 1 | 0 | 0 | 1472550 | AZK15899.1 | K9N638 |
| LSIEEFADVVKG | 1 | 0 | 0 | 1472553 | ALA49659.1 | K9N7C7 |
| LSIEEFAVVVKE | 1 | 0 | 0 | 1472554 | AHI48778.1 | K9N7C7 |
| LSILWFVDRIRF | 1 | 0 | 0 | 1472573 | AVV62541.1 | K9N7D2 |
| LSINTSCRVLQR | 1 | 0 | 0 | 1472574 | AEN19363.1 | P0C6X6 |
| LSIPSNWTISVQ | 1 | 0 | 0 | 1472575 | AAY43188.1 | A0A0P0G321 |
| LSISNDTVDVTN | 1 | 0 | 0 | 1472591 | ANZ78836.1 | P36334 |
| LSIVLGLKTIAT | 1 | 0 | 0 | 1472599 | APO40578.1 | P0C6X7 |

| | | | | | | |
|---|---|---|---|---|---|---|
| LSKFSLKLKGTP | 1 | 0 | 0 | 1472619 | AUM60013.1 | K9N7C7 |
| LSKKQLGLLEPQ | 1 | 0 | 0 | 1472629 | AUM60013.1 | K9N7C7 |
| LSKPFVKDNVSF | 1 | 0 | 0 | 1472641 | ATQ39389.1 | K9N7C7 |
| LSLDRGAQSDHD | 1 | 0 | 0 | 1472658 | AVV62527.1 | K9N796 |
| LSLFAQSEEEEP | 1 | 0 | 0 | 1472663 | AVN89336.1 | K9N4V0 |
| LSLFLLMFLLTH | 1 | 0 | 0 | 1472666 | AVV62526.1 | K9N5Q8 |
| LSLHPVECDEEC | 1 | 0 | 0 | 1472677 | ALW82719.1 | K9N7C7 |
| LSLLSFTTAQEG | 1 | 0 | 0 | 1472690 | APO40579.1 | P59594 |
| LSLSMVRRNLDS | 1 | 0 | 0 | 1472721 | Q3I5I6.1 | Q3LZX3 |
| LSLYLADNLSAS | 1 | 0 | 0 | 1472738 | AUM60013.1 | K9N7C7 |
| LSMKSDLSVSSS | 1 | 0 | 0 | 1472754 | ALA50067.1 | K9N5Q8 |
| LSMVRRNLDSLE | 1 | 0 | 0 | 1472764 | Q3I5I6.1 | Q3LZX3 |
| LSNGFSQVKSDL | 1 | 0 | 0 | 1472777 | AEN19363.1 | P0C6X6 |
| LSNKSNPALTQY | 1 | 0 | 0 | 1472780 | APO40579.1 | P59594 |
| LSNRLLDAMINT | 1 | 0 | 0 | 1472798 | APO40578.1 | P0C6X7 |
| LSNTKYYDHSQY | 1 | 0 | 0 | 1472806 | AEN19365.1 | P30215 |
| LSPLEGGGRLVA | 1 | 0 | 0 | 1472824 | AXG21654.1 | K9N5Q8 |
| LSPSHYWSPQGN | 1 | 0 | 0 | 1472835 | ARB07436.1 | A0A140H1G9 |
| LSPVDEFIKHVE | 1 | 0 | 0 | 1472839 | AVV62535.1 | K9N7C7 |
| LSPWFADGATAC | 1 | 0 | 0 | 1472840 | AVV62535.1 | K9N7C7 |
| LSPWFMDGQNAY | 1 | 0 | 0 | 1472844 | AVV62524.1 | K9N7C7 |
| LSQPRSDKPSQL | 1 | 0 | 0 | 1472867 | AVL25604.1 | H9EJ69 |
| LSQPRVDKPSQL | 1 | 0 | 0 | 1472868 | QED88037.1 | H9EJ69 |
| LSRDYISSNGPL | 1 | 0 | 0 | 1472875 | ABE97136.1 | H9EJ42 |
| LSRLDVLEAQVQ | 1 | 0 | 0 | 1472884 | ABD75553.1 | A0A140H1H1 |
| LSRLEALESGKQ | 1 | 0 | 0 | 1472885 | AUM60021.1 | K9N4V7 |
| LSRLIIFFSPES | 1 | 0 | 0 | 1472887 | AGT51393.1 | H9EJ42 |
| LSRSQSSETKEQ | 1 | 0 | 0 | 1472899 | ABB90484.1 | A0A0N7J131 |
| LSSACTMFTTAD | 1 | 0 | 0 | 1472906 | AGT51707.1 | P0C6X6 |
| LSSDENGVRTLS | 1 | 0 | 0 | 1472909 | AVP78031.1 | Q3LZX1 |
| LSSDSDVFIKKY | 1 | 0 | 0 | 1472911 | ANI69887.1 | K9N7C7 |
| LSSDVMFEDAQG | 1 | 0 | 0 | 1472912 | AXN92247.1 | K9N7C7 |
| LSSDVMFEDDQG | 1 | 0 | 0 | 1472914 | AGV08522.1 | K9N7C7 |
| LSSDVMFEGAQG | 1 | 0 | 0 | 1472915 | ASU90140.1 | K9N7C7 |
| LSSDVMLEDATG | 1 | 0 | 0 | 1472917 | AUM60013.1 | K9N7C7 |
| LSSFFGLHTVQS | 1 | 0 | 0 | 1472922 | AVV62535.1 | K9N7C7 |
| LSSLNLPGCDGG | 1 | 0 | 0 | 1472929 | APO40578.1 | P0C6X7 |
| LSSYSLFDMSKFPLKL | 1 | 0 | 0 | 1472955 | YP_009724389.1 | P0DTD1 |
| LSTCDGQKFTDESFYKNMY | 1 | 0 | 0 | 1472958 | AQN78733.1 | P0C6X6 |
| LSTDDCFANKHA | 1 | 0 | 0 | 1472959 | ARO76381.1 | P0C6U8 |
| LSTDTGAENATF | 1 | 0 | 0 | 1472962 | Q3I5J4.1 | Q3LZX0 |
| LSTEQEDAWSVA | 1 | 0 | 0 | 1472965 | AUM60013.1 | K9N7C7 |
| LSTFEPRSDLER | 1 | 0 | 0 | 1472966 | ARB07436.1 | A0A140H1G9 |
| LSTGCCGCCNCL | 1 | 0 | 0 | 1472972 | AIW52843.1 | A0A0P0G321 |
| LSTNGRWVLNND | 1 | 0 | 0 | 1472989 | APO40578.1 | P0C6X7 |
| LSVEDCKLPLGQ | 1 | 0 | 0 | 1473017 | ALL26396.1 | K9N5Q8 |
| LSVEDLVPSADK | 1 | 0 | 0 | 1473018 | AVV62524.1 | K9N7C7 |
| LSVHDGYTNEII | 1 | 0 | 0 | 1473026 | ARO76381.1 | P0C6U8 |
| LSVLVTASSKPL | 1 | 0 | 0 | 1473041 | ANI69890.1 | K9N796 |
| LSVNSSNCNNLN | 1 | 0 | 0 | 1473047 | ATU79932.1 | A0A0P0LKV0 |
| LSVQQAQQEATH | 1 | 0 | 0 | 1473053 | ARB07424.1 | A0A0P0ILY9 |
| LSYDSLKTGVSV | 1 | 0 | 0 | 1473088 | ARO76381.1 | P0C6U8 |
| LSYRSFIEDLLY | 1 | 0 | 0 | 1473107 | APO40579.1 | P59594 |
| LSYSGAIKLDSK | 1 | 0 | 0 | 1473108 | AVV62544.1 | K9N4V7 |
| LSYYRLGASQRV | 1 | 0 | 0 | 1473111 | APO40582.1 | P59596 |
| LTAATIVNQERY | 1 | 0 | 0 | 1473118 | AGV08597.1 | K9N7C7 |
| LTACNGHKIEDL | 1 | 0 | 0 | 1473120 | ADO24641.1 | A0A140H1G9 |
| LTADEIKALKEL | 1 | 0 | 0 | 1473121 | AVN89363.1 | K9N7C7 |
| LTAEDISLLRKM | 1 | 0 | 0 | 1473124 | AHA80151.1 | P33469 |
| LTAFSPAQDTWG | 1 | 0 | 0 | 1473130 | 5X58_A | P59594 |
| LTAGETKALKEL | 1 | 0 | 0 | 1473133 | QBF80609.1 | K9N7C7 |
| LTALDAYVSQQL | 1 | 0 | 0 | 1473136 | CAA83661.1 | P36334 |
| LTASMSMAECKA | 1 | 0 | 0 | 1473150 | AFD98819.1 | H9EJ42 |
| LTASTIVNQERY | 1 | 0 | 0 | 1473151 | ALA50055.1 | K9N7C7 |
| LTCHNAAQCDCL | 1 | 0 | 0 | 1473165 | CAA83660.1 | P36334 |
| LTCHNLCNTDPI | 1 | 0 | 0 | 1473168 | BBO58740.1 | P36334 |
| LTCIRRPKDGFN | 1 | 0 | 0 | 1473176 | ARK08651.1 | P36334 |
| LTCKAISSNTDN | 1 | 0 | 0 | 1473187 | Q14EB0.1 | A0A140H1H1 |
| LTCLCPVPGLKS | 1 | 0 | 0 | 1473190 | ABU88845.1 | A0A0P0G321 |
| LTCNAISPKTDK | 1 | 0 | 0 | 1473197 | AZS52618.1 | A0A140H1H1 |
| LTCSTDLQKANI | 1 | 0 | 0 | 1473205 | BBO58790.1 | P36334 |
| LTCSTDLQKANT | 1 | 0 | 0 | 1473206 | CAA79896.1 | P36334 |
| LTCVKDYEFKFY | 1 | 0 | 0 | 1473210 | ABD75503.1 | A0A140H1G9 |
| LTDESCNNLVQT | 1 | 0 | 0 | 1473220 | AIV42002.1 | P0C6X6 |
| LTDHDVVYAQHC | 1 | 0 | 0 | 1473222 | QEG03776.1 | P36334 |
| LTDQMSEPPKEQ | 1 | 0 | 0 | 1473237 | AGV08397.1 | K9N4V7 |
| LTDSSKRFQSFQ | 1 | 0 | 0 | 1473240 | AVP78031.1 | Q3LZX1 |
| LTDVATDVQSRA | 1 | 0 | 0 | 1473241 | AUM60023.1 | K9N7C7 |
| LTDYNAILQSDN | 1 | 0 | 0 | 1473247 | QDH43737.1 | H9EJ42 |
| LTEGQLQSLSRD | 1 | 0 | 0 | 1473258 | ABE97136.1 | H9EJ42 |
| LTEHVTSVNFFD | 1 | 0 | 0 | 1473259 | AXN73427.1 | K9N796 |
| LTEHVTSVNLFD | 1 | 0 | 0 | 1473260 | AXN92229.1 | K9N796 |
| LTESSGSLATRL | 1 | 0 | 0 | 1473269 | ATP16762.1 | Q4VID0 |
| LTFDSNAFSLAN | 1 | 0 | 0 | 1473287 | AIW52843.1 | A0A0P0G321 |
| LTFSYDGLKGGI | 1 | 0 | 0 | 1473313 | AVV62524.1 | K9N7C7 |
| LTGHAIQGFACG | 1 | 0 | 0 | 1473344 | AHI48785.1 | K9N638 |
| LTGIAIEQDKNT | 1 | 0 | 0 | 1473347 | ATO98181.1 | Q3LZX1 |
| LTGNAIQGFACG | 1 | 0 | 0 | 1473352 | AXN92247.1 | K9N7C7 |
| LTGRTPLNPNYT | 1 | 0 | 0 | 1473355 | AVP78031.1 | Q3LZX1 |
| LTGTQSVSVDRE | 1 | 0 | 0 | 1473362 | AVN89314.1 | K9N796 |
| LTGVALEQDKNT | 1 | 0 | 0 | 1473363 | ATO98145.1 | Q3LZX1 |

| | | | | | | |
|---|---|---|---|---|---|---|
| LTHCAFAVDPAA | 1 | 0 | 0 | 1473367 | AST12962.1 | A0A0P0LKV0 |
| LTHCSFAVDPAA | 1 | 0 | 0 | 1473368 | NP_073549.1 | A0A0P0LKV0 |
| LTHCSFAVDPAV | 1 | 0 | 0 | 1473369 | AGT21359.1 | A0A0P0LKV0 |
| LTHFQGFVLNYK | 1 | 0 | 0 | 1473370 | AWW13564.1 | Q80872 |
| LTIHRGDPMSNN | 1 | 0 | 0 | 1473388 | AVP78042.1 | Q3LZX1 |
| LTIQQIELKFNP | 1 | 0 | 0 | 1473399 | AVP78030.1 | P0C6W2 |
| LTITADPKVYPI | 1 | 0 | 0 | 1473401 | 2CME_A | P59636 |
| LTITDPKVYPII | 1 | 0 | 0 | 1473402 | 2CME_E | P59636 |
| LTITSAGQPCVA | 1 | 0 | 0 | 1473408 | AEN19363.1 | P0C6X6 |
| LTKCRTYLTSRT | 1 | 0 | 0 | 1473420 | AUM60013.1 | K9N7C7 |
| LTKDDLHQLYVD | 1 | 0 | 0 | 1473422 | AUM60013.1 | K9N7C7 |
| LTKDQTPIVWHA | 1 | 0 | 0 | 1473424 | NP_073549.1 | A0A0P0LKV0 |
| LTKKKYSELDDE | 1 | 0 | 0 | 1473434 | Q3I5J1.1 | Q3LZX8 |
| LTKLGTEDPRWP | 1 | 0 | 0 | 1473441 | AVN89320.1 | K9N4V7 |
| LTKLSVKCYFNY | 1 | 0 | 0 | 1473450 | AIW52841.1 | A0A0P0G321 |
| LTKNLLLYADIN | 1 | 0 | 0 | 1473454 | APO40578.1 | P0C6X7 |
| LTKQLLEVLSKA | 1 | 0 | 0 | 1473459 | AUM60023.1 | K9N7C7 |
| LTLLQVPQVSTS | 1 | 0 | 0 | 1473499 | AVV62537.1 | K9N5Q8 |
| LTLQQIELKFNP | 1 | 0 | 0 | 1473503 | AVP78041.1 | P0C6W2 |
| LTLTPRNKWSGV | 1 | 0 | 0 | 1473517 | AXN92247.1 | K9N7C7 |
| LTLTVKFAVESN | 1 | 0 | 0 | 1473522 | AFD98812.1 | H9EJ42 |
| LTMLAEFDENGT | 1 | 0 | 0 | 1473545 | APO40579.1 | P59594 |
| LTMVEFMLWCKD | 1 | 0 | 0 | 1473551 | ATQ39389.1 | K9N7C7 |
| LTMVEFMLWCKN | 1 | 0 | 0 | 1473552 | AVV62535.1 | K9N7C7 |
| LTNEGFNNYVPS | 1 | 0 | 0 | 1473564 | AUM60013.1 | K9N7C7 |
| LTNGSDLDRCTT | 1 | 0 | 0 | 1473566 | ABD72988.1 | P59594 |
| LTNIFGTVYEKLKPVLD | 1 | 0 | 0 | 1473569 | P0DTC1.1 | P0DTD1 |
| LTNKISLNAPVT | 1 | 0 | 0 | 1473572 | ARU07594.1 | A0A0P0G321 |
| LTNLNEITKPLI | 1 | 0 | 0 | 1473574 | ARU07582.1 | Q0ZJ80 |
| LTNNISLNAPVT | 1 | 0 | 0 | 1473578 | QEG59356.1 | A0A0P0G321 |
| LTNQLPPPLLNN | 1 | 0 | 0 | 1473583 | AVV62537.1 | K9N5Q8 |
| LTPALLATNNVF | 1 | 0 | 0 | 1473594 | APO40578.1 | P0C6X7 |
| LTPASVGSVPGE | 1 | 0 | 0 | 1473595 | 5W9H_A | K9N5Q8 |
| LTPAWRVYSTGV | 1 | 0 | 0 | 1473599 | ATO98145.1 | Q3LZX1 |
| LTPIKGDVDSGP | 1 | 0 | 0 | 1473609 | AVV62537.1 | K9N5Q8 |
| LTPKTGTGIAIS | 1 | 0 | 0 | 1473612 | AXN92247.1 | K9N7C7 |
| LTPLANILHNVK | 1 | 0 | 0 | 1473613 | AUM60027.1 | K9N643 |
| LTPLCGFSSQAA | 1 | 0 | 0 | 1473615 | APO40578.1 | P0C6X7 |
| LTPRSVSSVPGE | 1 | 0 | 0 | 1473633 | 5X59_A | K9N5Q8 |
| LTPTESYVDVGL | 1 | 0 | 0 | 1473637 | AHL18090.1 | K9N5Q8 |
| LTQDHLDILGPL | 1 | 0 | 0 | 1473646 | AVP78041.1 | P0C6W2 |
| LTQDYFLPFDSN | 1 | 0 | 0 | 1473651 | ATO98181.1 | Q3LZX1 |
| LTQDYFLPFDTN | 1 | 0 | 0 | 1473652 | ATO98193.1 | Q3LZX1 |
| LTQRGPLQVGDA | 1 | 0 | 0 | 1473671 | AUM60013.1 | K9N7C7 |
| LTRDDMHQLYVN | 1 | 0 | 0 | 1473679 | AVV62535.1 | K9N7C7 |
| LTREDMHQLYVN | 1 | 0 | 0 | 1473680 | AVV62524.1 | K9N7C7 |
| LTRGSDLDRCTT | 1 | 0 | 0 | 1473684 | ABD72995.1 | P59594 |
| LTRIEASTVVCG | 1 | 0 | 0 | 1473686 | AVV62524.1 | K9N7C7 |
| LTRVEASTVVCG | 1 | 0 | 0 | 1473698 | AXN92247.1 | K9N7C7 |
| LTSCFFTCNSNA | 1 | 0 | 0 | 1473704 | BAM16477.1 | A0A0P0G321 |
| LTSDENGVRTLS | 1 | 0 | 0 | 1473705 | ATO98120.1 | Q3LZX1 |
| LTSDYMITVFVT | 1 | 0 | 0 | 1473706 | ANI69888.1 | K9N638 |
| LTSEEVVIKTGT | 1 | 0 | 0 | 1473712 | APO40578.1 | P0C6X7 |
| LTSEISILENKS | 1 | 0 | 0 | 1473715 | AIW52757.1 | A0A0P0K6L9 |
| LTSEQIVAIIKD | 1 | 0 | 0 | 1473719 | AVV62532.1 | K9N7A1 |
| LTSGVTPDMADQ | 1 | 0 | 0 | 1473723 | AAX85673.1 | P33469 |
| LTSLLILIQSTQ | 1 | 0 | 0 | 1473734 | APO40578.1 | P0C6X7 |
| LTSLTSDYTVTV | 1 | 0 | 0 | 1473741 | AUM60013.1 | K9N7C7 |
| LTSRAAQQLTIE | 1 | 0 | 0 | 1473750 | AVV62535.1 | K9N7C7 |
| LTSSGFDNSYQT | 1 | 0 | 0 | 1473755 | AUM60014.1 | K9N5Q8 |
| LTSSMRGCCDCG | 1 | 0 | 0 | 1473756 | AIW52843.1 | A0A0P0G321 |
| LTSSTKKFQPFQ | 1 | 0 | 0 | 1473761 | APO40579.1 | P59594 |
| LTSTHDGPTEHV | 1 | 0 | 0 | 1473764 | AVN89314.1 | K9N796 |
| LTTAQSELAKGV | 1 | 0 | 0 | 1473771 | APO40578.1 | P0C6X7 |
| LTTCAFSVDPAT | 1 | 0 | 0 | 1473773 | QED88017.1 | H9EJ42 |
| LTTEGYNNYVPS | 1 | 0 | 0 | 1473775 | AVV62535.1 | K9N7C7 |
| LTTGYFLPFNSN | 1 | 0 | 0 | 1473784 | APO40579.1 | P59594 |
| LTTLNAFVAQQL | 1 | 0 | 0 | 1473789 | AXN92228.1 | K9N5Q8 |
| LTVDAANPSTPP | 1 | 0 | 0 | 1473815 | AUM60013.1 | K9N7C7 |
| LTVDGVQYYCYT | 1 | 0 | 0 | 1473817 | AXN92247.1 | K9N7C7 |
| LTVDSSNPNTPA | 1 | 0 | 0 | 1473819 | AVV62524.1 | K9N7C7 |
| LTVDYFAYPLSM | 1 | 0 | 0 | 1473823 | AVV62537.1 | K9N5Q8 |
| LTVEGYNNYVPS | 1 | 0 | 0 | 1473826 | AVV62524.1 | K9N7C7 |
| LTVKFAVESNVM | 1 | 0 | 0 | 1473836 | AFD98812.1 | H9EJ42 |
| LTVMSYQMQGCM | 1 | 0 | 0 | 1473848 | AVQ05261.1 | P0C6X6 |
| LTVMSYQMRGCM | 1 | 0 | 0 | 1473849 | AEN19363.1 | P0C6X6 |
| LTVPFCGSGNLV | 1 | 0 | 0 | 1473851 | ASU45728.1 | K9N7C7 |
| LTVQKGIYQTSN | 1 | 0 | 0 | 1473856 | AVP78031.1 | Q3LZX1 |
| LTVTLQNSCTPK | 1 | 0 | 0 | 1473863 | AQN78773.1 | P0C6X6 |
| LTVTSAGQPCVV | 1 | 0 | 0 | 1473866 | AVQ05261.1 | P0C6X6 |
| LTVTSAGQPYVA | 1 | 0 | 0 | 1473867 | AGT51707.1 | P0C6X6 |
| LTVTSDGQPYVA | 1 | 0 | 0 | 1473868 | AGT51657.1 | P0C6X6 |
| LTWPLGLECTRA | 1 | 0 | 0 | 1473878 | AHI48747.1 | K9N638 |
| LTWSQELPFDVI | 1 | 0 | 0 | 1473882 | ARB07430.1 | P0C6X6 |
| LTYANIVHEPTT | 1 | 0 | 0 | 1473885 | ALA49956.1 | K9N7C7 |
| LTYEMLSLQQVV | 1 | 0 | 0 | 1473888 | AXN92228.1 | K9N5Q8 |
| LTYLNLSSELKQ | 1 | 0 | 0 | 1473904 | AIW52843.1 | A0A0P0G321 |
| LTYNKGENMTPR | 1 | 0 | 0 | 1473907 | BAE93400.1 | P0C6U8 |
| LTYQNISTNLPP | 1 | 0 | 0 | 1473910 | AVN89324.1 | K9N5Q8 |
| LTYYLHSVLYRV | 1 | 0 | 0 | 1473916 | ANI69823.1 | K9N638 |
| LVAELDGIQYGR | 1 | 0 | 0 | 1473922 | AVP78041.1 | P0C6W2 |

| | | | | | | |
|---|---|---|---|---|---|---|
| LVAELDGVQFGR | 1 | 0 | 0 | 1473923 | ARO76381.1 | P0C6U8 |
| LVAKCYGESVLV | 1 | 0 | 0 | 1473932 | QBF80520.1 | K9N638 |
| LVAQEFVSYDSS | 1 | 0 | 0 | 1473947 | APU51933.1 | P0C6X6 |
| LVAQNTVLNDSI | 1 | 0 | 0 | 1473948 | AMM42614.1 | H9EJ69 |
| LVASGPTVAMTE | 1 | 0 | 0 | 1473951 | AKN24830.1 | K9N5Q8 |
| LVASGSIVAMTE | 1 | 0 | 0 | 1473952 | AMQ49004.1 | K9N5Q8 |
| LVAVNNSKPYFI | 1 | 0 | 0 | 1473964 | AVV62530.1 | K9N7D2 |
| LVAVNSEDIYNT | 1 | 0 | 0 | 1473966 | AUM60013.1 | K9N7C7 |
| LVAYDIAVTAGP | 1 | 0 | 0 | 1473975 | AVV62524.1 | K9N7C7 |
| LVCITIIKLTKL | 1 | 0 | 0 | 1473982 | AGW80935.1 | A0A0P0KH29 |
| LVCTTLQGKPIG | 1 | 0 | 0 | 1473992 | AVN89333.1 | K9N7C7 |
| LVCYIVMPLHTL | 1 | 0 | 0 | 1473995 | AAP30028.1 | P0C6X7 |
| LVDDNKFDGLKV | 1 | 0 | 0 | 1474004 | ATP16754.1 | P0C6X6 |
| LVDEEFPLDDEE | 1 | 0 | 0 | 1474005 | AVV62535.1 | K9N7C7 |
| LVDEEFPLEEEV | 1 | 0 | 0 | 1474006 | AVV62524.1 | K9N7C7 |
| LVDGYGNTMYVN | 1 | 0 | 0 | 1474012 | AUM60027.1 | K9N643 |
| LVDSDMNDFVSD | 1 | 0 | 0 | 1474032 | APO40578.1 | P0C6X7 |
| LVDSEQPCLVIP | 1 | 0 | 0 | 1474034 | AMM42513.1 | A0A0N7J131 |
| LVDSEQPWKVIP | 1 | 0 | 0 | 1474035 | AIW52721.1 | A0A0N7J131 |
| LVDSEQSWKVIP | 1 | 0 | 0 | 1474037 | AAA45463.1 | A0A0N7J131 |
| LVDSVQYAHKHD | 1 | 0 | 0 | 1474040 | AUM60013.1 | K9N7C7 |
| LVEAYNSILVEN | 1 | 0 | 0 | 1474044 | ARU07593.1 | H9EJ42 |
| LVEDSTSVTAVV | 1 | 0 | 0 | 1474046 | ASY99815.1 | K9N7A1 |
| LVEKNVVRAALL | 1 | 0 | 0 | 1474055 | AUM60013.1 | K9N7C7 |
| LVESASDFIAWR | 1 | 0 | 0 | 1474064 | Q14EA9.1 | Q0ZJ84 |
| LVEVPVEDIAQV | 1 | 0 | 0 | 1474068 | ATG84788.1 | K9N7C7 |
| LVEYGQFCAKIN | 1 | 0 | 0 | 1474070 | AUM60014.1 | K9N5Q8 |
| LVFLILCFTIKR | 1 | 0 | 0 | 1474082 | ARO76387.1 | P59635 |
| LVFVTTLTSDYT | 1 | 0 | 0 | 1474114 | AXN92247.1 | K9N7C7 |
| LVGCLGPHCGSS | 1 | 0 | 0 | 1474117 | BBA20983.1 | A0A140H1H1 |
| LVGFKDVTNGTI | 1 | 0 | 0 | 1474118 | CAA71146.1 | A0A0P0K6L9 |
| LVGGNFTPIDQY | 1 | 0 | 0 | 1474122 | AUM60013.1 | K9N7C7 |
| LVGHDICAQLNA | 1 | 0 | 0 | 1474123 | ABD75519.1 | A0A140H1G9 |
| LVGHDICAQLND | 1 | 0 | 0 | 1474124 | ARU07575.1 | A0A140H1G9 |
| LVGIGGTPCIKV | 1 | 0 | 0 | 1474126 | AVV62524.1 | K9N7C7 |
| LVGQLEQTNMHS | 1 | 0 | 0 | 1474130 | ASU90174.1 | K9N638 |
| LVGTTLQGKAIG | 1 | 0 | 0 | 1474142 | AUM60013.1 | K9N7C7 |
| LVGYHSDNGNYY | 1 | 0 | 0 | 1474148 | AUM60014.1 | K9N5Q8 |
| LVGYIPEHCQSH | 1 | 0 | 0 | 1474149 | AUM60015.1 | K9N796 |
| LVHTDSFSCNNF | 1 | 0 | 0 | 1474158 | AET99191.1 | A0A140H1H1 |
| LVILLIFGNMYL | 1 | 0 | 0 | 1474183 | AWK59942.1 | H9EJ42 |
| LVISDMYDPVTK | 1 | 0 | 0 | 1474194 | AVV62524.1 | K9N7C7 |
| LVKDCDAVARHD | 1 | 0 | 0 | 1474202 | AVV62524.1 | K9N7C7 |
| LVKDVDHVQIHT | 1 | 0 | 0 | 1474207 | Q3I5J4.1 | Q3LZX0 |
| LVKFSATQAMEK | 1 | 0 | 0 | 1474212 | AIL49512.1 | P36334 |
| LVKGCPAVAVHD | 1 | 0 | 0 | 1474213 | AVP78030.1 | P0C6W2 |
| LVKIQAVVNADA | 1 | 0 | 0 | 1474218 | CAA83661.1 | P36334 |
| LVKTGHYIGQAG | 1 | 0 | 0 | 1474247 | ARU07606.1 | P0C6X6 |
| LVLAKLGGDASK | 1 | 0 | 0 | 1474255 | AJQ24068.1 | P33469 |
| LVLAKLGRDATK | 1 | 0 | 0 | 1474258 | AGT51674.1 | P33469 |
| LVLECTRAAASA | 1 | 0 | 0 | 1474268 | AVV62524.1 | K9N7C7 |
| LVLECTRAASSA | 1 | 0 | 0 | 1474269 | AUM60013.1 | K9N7C7 |
| LVLGDAIQLAKC | 1 | 0 | 0 | 1474272 | AUM60013.1 | K9N7C7 |
| LVLGINEKTHVQ | 1 | 0 | 0 | 1474274 | P0C6W6.1 | P0C6W2 |
| LVLGSKHVNCCT | 1 | 0 | 0 | 1474275 | ABQ16584.1 | H9EJ42 |
| LVLNAKELVDGY | 1 | 0 | 0 | 1474293 | AUM60027.1 | K9N643 |
| LVLNANECQHPN | 1 | 0 | 0 | 1474294 | AVV62540.1 | K9N643 |
| LVLNASEMYCAN | 1 | 0 | 0 | 1474297 | ATQ39396.1 | K9N643 |
| LVLNNPAYIATQ | 1 | 0 | 0 | 1474304 | AIV41824.1 | P30215 |
| LVLNNPAYLAPQ | 1 | 0 | 0 | 1474305 | AIL49375.1 | P30215 |
| LVLNNPSYIAPQ | 1 | 0 | 0 | 1474306 | AIX09798.1 | P30215 |
| LVLNNPVYIAPQ | 1 | 0 | 0 | 1474307 | AVQ05263.1 | P30215 |
| LVLPSGNYLSIS | 1 | 0 | 0 | 1474313 | AVQ05263.1 | P30215 |
| LVLTSNDERLSP | 1 | 0 | 0 | 1474324 | AVV62535.1 | K9N7C7 |
| LVNANHYSPCVS | 1 | 0 | 0 | 1474351 | AID55087.1 | K9N5Q8 |
| LVNANQYSPCVP | 1 | 0 | 0 | 1474352 | QAT98898.1 | K9N5Q8 |
| LVNGLHYPTGSC | 1 | 0 | 0 | 1474363 | ALD51908.1 | K9N7D2 |
| LVNGTHTPLGSC | 1 | 0 | 0 | 1474364 | AVV62530.1 | K9N7D2 |
| LVNGVHHPTGSC | 1 | 0 | 0 | 1474366 | QBM11740.1 | K9N7D2 |
| LVNGVHYPTGGC | 1 | 0 | 0 | 1474367 | ALA49906.1 | K9N7D2 |
| LVNGVHYPTGIC | 1 | 0 | 0 | 1474368 | ALW82735.1 | K9N7D2 |
| LVNGVHYPTGSC | 1 | 0 | 0 | 1474369 | AXG21669.1 | K9N7D2 |
| LVNPTDKSHYYV | 1 | 0 | 0 | 1474378 | ATQ39389.1 | K9N7C7 |
| LVNSNQYSPCVS | 1 | 0 | 0 | 1474379 | AHY22555.1 | K9N5Q8 |
| LVNTSMVVDECQ | 1 | 0 | 0 | 1474387 | AVV62526.1 | K9N5Q8 |
| LVNWYIYVRKNG | 1 | 0 | 0 | 1474392 | AWK59942.1 | H9EJ42 |
| LVNYKQSIPMPR | 1 | 0 | 0 | 1474395 | QAT98908.1 | K9N7C7 |
| LVPAFAEELHGG | 1 | 0 | 0 | 1474398 | AUM60028.1 | K9N7D2 |
| LVPAFPETEVAS | 1 | 0 | 0 | 1474400 | AVV62530.1 | K9N7D2 |
| LVPHAGETPIAY | 1 | 0 | 0 | 1474407 | ARO76381.1 | P0C6U8 |
| LVPHVGEVPVAY | 1 | 0 | 0 | 1474414 | AVP78041.1 | P0C6W2 |
| LVPIGKGSKDEQ | 1 | 0 | 0 | 1474416 | AIW52795.1 | H9EJ69 |
| LVPRNKWNGKSS | 1 | 0 | 0 | 1474438 | AUM60013.1 | K9N7C7 |
| LVPSNKWTGKKN | 1 | 0 | 0 | 1474440 | AVV62535.1 | K9N7C7 |
| LVPTQPLPNATY | 1 | 0 | 0 | 1474445 | APO40578.1 | P0C6X7 |
| LVPTTTGGLGRF | 1 | 0 | 0 | 1474447 | QED88038.1 | H9EJ42 |
| LVPTTTGGLVRF | 1 | 0 | 0 | 1474448 | AGT51379.1 | H9EJ42 |
| LVQCTCGTKSWS | 1 | 0 | 0 | 1474464 | NP_073549.1 | A0A0P0LKV0 |
| LVQDHVDILGPL | 1 | 0 | 0 | 1474465 | APO40578.1 | P0C6X7 |
| LVQNQSFNKLLH | 1 | 0 | 0 | 1474481 | ATQ39389.1 | K9N7C7 |
| LVQPAVYMYNTG | 1 | 0 | 0 | 1474485 | AUM60019.1 | K9N5R3 |

| | | | | | | |
|---|---|---|---|---|---|---|
| LVQQESTFVMMS | 1 | 0 | 0 | 1474490 | APO40578.1 | P0C6X7 |
| LVQQVEKWLHQL | 1 | 0 | 0 | 1474493 | APO40578.1 | P0C6X7 |
| LVQTYLKSDNIV | 1 | 0 | 0 | 1474498 | AIV42002.1 | P0C6X6 |
| LVRLLQDKTGDF | 1 | 0 | 0 | 1474520 | AXN92247.1 | K9N7C7 |
| LVRLLQGKTGDF | 1 | 0 | 0 | 1474522 | AHI48793.1 | K9N7C7 |
| LVSCFSTRNSNA | 1 | 0 | 0 | 1474539 | AKA09629.1 | A0A0P0G321 |
| LVSGDAHTPSGT | 1 | 0 | 0 | 1474552 | AMM42538.1 | A0A0P0K6L9 |
| LVSKIFKARAIE | 1 | 0 | 0 | 1474562 | AMM42372.1 | A0A140H1G9 |
| LVSSDKAPHRVI | 1 | 0 | 0 | 1474578 | AVL25594.1 | H9EJ69 |
| LVSSNGQPGGDA | 1 | 0 | 0 | 1474583 | QAT98908.1 | K9N7C7 |
| LVSSTFVPLGSC | 1 | 0 | 0 | 1474586 | AUM60018.1 | K9N7D2 |
| LVTAQDGCGHIS | 1 | 0 | 0 | 1474601 | ATO98193.1 | Q3LZX1 |
| LVTQFKLLGTTL | 1 | 0 | 0 | 1474631 | ARU07575.1 | A0A140H1G9 |
| LVTTEVKVSALK | 1 | 0 | 0 | 1474637 | AHI48796.1 | K9N7C7 |
| LVTTSMAPDFVA | 1 | 0 | 0 | 1474642 | QBF80609.1 | K9N7C7 |
| LVTTSTAPDFVA | 1 | 0 | 0 | 1474643 | AXN92247.1 | K9N7C7 |
| LVTTSTVPDFVA | 1 | 0 | 0 | 1474646 | ATG84754.1 | K9N7C7 |
| LVTVDGINFRTC | 1 | 0 | 0 | 1474647 | AUM60013.1 | K9N7C7 |
| LVTYNCCDGDYF | 1 | 0 | 0 | 1474656 | AVP78041.1 | P0C6W2 |
| LVVAANKTKGII | 1 | 0 | 0 | 1474658 | AVV62535.1 | K9N7C7 |
| LVVDDKYKVNEN | 1 | 0 | 0 | 1474659 | ABD75575.1 | A0A140H1G9 |
| LVVDTLNSSGFV | 1 | 0 | 0 | 1474660 | AUM60014.1 | K9N5Q8 |
| LVVDWHYFDAQP | 1 | 0 | 0 | 1474661 | ASU91162.1 | K9N7C7 |
| LVVGDYVIISGK | 1 | 0 | 0 | 1474666 | AVV62524.1 | K9N7C7 |
| LVWLESQLQKGV | 1 | 0 | 0 | 1474692 | APO40578.1 | P0C6X7 |
| LWAVGHPAPTHY | 1 | 0 | 0 | 1474735 | AVV62540.1 | K9N643 |
| LWCKDGKVQTFY | 1 | 0 | 0 | 1474738 | ATQ39389.1 | K9N7C7 |
| LWDYEAERPFTS | 1 | 0 | 0 | 1474748 | AWK59942.1 | H9EJ42 |
| LWFDFVENPGII | 1 | 0 | 0 | 1474764 | AVV62524.1 | K9N7C7 |
| LWNKFVTGDVGP | 1 | 0 | 0 | 1474824 | AWK59942.1 | H9EJ42 |
| LWNTFTKLQSLE | 1 | 0 | 0 | 1474828 | AVP78041.1 | P0C6W2 |
| LWNYLFETIPTQ | 1 | 0 | 0 | 1474832 | AXN92247.1 | K9N7C7 |
| LWPLVLALSIFD | 1 | 0 | 0 | 1474836 | AWK59934.1 | U3M6Q8 |
| LWQTFTDVNLQG | 1 | 0 | 0 | 1474848 | AST12962.1 | A0A0P0LKV0 |
| LWSVSGLRFTTG | 1 | 0 | 0 | 1474864 | NP_073551.1 | A0A0P0K6L9 |
| LWSVSGSRFTTG | 1 | 0 | 0 | 1474869 | AIW52754.1 | A0A0P0K6L9 |
| LWVKKASYDTNL | 1 | 0 | 0 | 1474876 | AXN92226.1 | K9N7C7 |
| LWVNKASYDINL | 1 | 0 | 0 | 1474880 | AMQ48992.1 | K9N7C7 |
| LWYKGSNLEEEI | 1 | 0 | 0 | 1474890 | AEN19363.1 | P0C6X6 |
| LWYVVDDIYYPA | 1 | 0 | 0 | 1474898 | AWK59942.1 | H9EJ42 |
| LYADINGELHED | 1 | 0 | 0 | 1474907 | APO40578.1 | P0C6X7 |
| LYAIQQGFDATN | 1 | 0 | 0 | 1474913 | AIX10750.1 | P36334 |
| LYAKRALRSHPD | 1 | 0 | 0 | 1474920 | ASU90074.1 | K9N7C7 |
| LYAVQQLNNGFQ | 1 | 0 | 0 | 1474928 | AVV62524.1 | K9N7C7 |
| LYCGVHKGAHVA | 1 | 0 | 0 | 1474933 | AXX83294.1 | P0C6X6 |
| LYDACNVKKVTD | 1 | 0 | 0 | 1474946 | QBP84746.1 | P0C6X6 |
| LYDASNVKNVTD | 1 | 0 | 0 | 1474951 | AVQ05261.1 | P0C6X6 |
| LYDFNGNIIGFK | 1 | 0 | 0 | 1474953 | ADN03339.1 | A0A140H1H1 |
| LYDKLQFASLEV | 1 | 0 | 0 | 1474956 | APO40578.1 | P0C6X7 |
| LYDNCYRNSNVG | 1 | 0 | 0 | 1474962 | ADO24705.1 | A0A0P0LKV0 |
| LYDNCYRNSNVN | 1 | 0 | 0 | 1474963 | AJE26205.1 | A0A0P0LKV0 |
| LYDPNGNLYGFR | 1 | 0 | 0 | 1474969 | ARB66062.1 | P36334 |
| LYDPNMEAAYTS | 1 | 0 | 0 | 1474970 | AVV62526.1 | K9N5Q8 |
| LYDPNMEGAYTS | 1 | 0 | 0 | 1474971 | AVV62537.1 | K9N5Q8 |
| LYDQVVMDYLDS | 1 | 0 | 0 | 1474972 | ARO76381.1 | P0C6U8 |
| LYDSNGNLYGLR | 1 | 0 | 0 | 1474975 | ABD43179.1 | P36334 |
| LYDSPCFSQQGV | 1 | 0 | 0 | 1474976 | AAA73152.1 | P30215 |
| LYDVNMEAAYTS | 1 | 0 | 0 | 1474980 | AUM60014.1 | K9N5Q8 |
| LYEAPTTLQAVG | 1 | 0 | 0 | 1474985 | AVV62524.1 | K9N7C7 |
| LYENAFLPFAMG | 1 | 0 | 0 | 1474994 | QHD43415.1 | P0DTD1 |
| LYFAATERSNVI | 1 | 0 | 0 | 1475001 | APO40579.1 | P59594 |
| LYFQGGGGSGYI | 1 | 0 | 0 | 1475026 | 6NB6_A | P59594 |
| LYFQSGSHHHHH | 1 | 0 | 0 | 1475027 | 6U7H_A | A0A0P0K6L9 |
| LYFQSMETGTLI | 1 | 0 | 0 | 1475028 | 5X29_A | P59637 |
| LYGISGQGIFVE | 1 | 0 | 0 | 1475048 | AEN19366.1 | P36334 |
| LYGQTVFYFGGC | 1 | 0 | 0 | 1475066 | ATN39862.1 | P0C6X6 |
| LYHDSQNMLRGD | 1 | 0 | 0 | 1475080 | AFR58685.1 | P0C6X7 |
| LYHYQECARGTT | 1 | 0 | 0 | 1475084 | APO40584.1 | P59635 |
| LYIPKPTAKYTP | 1 | 0 | 0 | 1475103 | AXN92262.1 | K9N4V0 |
| LYIYALIYILQI | 1 | 0 | 0 | 1475110 | Q3I5J4.1 | Q3LZX0 |
| LYKDLTGYMLDS | 1 | 0 | 0 | 1475115 | QAT98908.1 | K9N7C7 |
| LYKGNVTHDASS | 1 | 0 | 0 | 1475123 | AXN92247.1 | K9N7C7 |
| LYKNMCTGSPPI | 1 | 0 | 0 | 1475138 | AHI48748.1 | K9N7C7 |
| LYKRNFTYDVNA | 1 | 0 | 0 | 1475145 | CAA79896.1 | P36334 |
| LYKRNFTYDVSA | 1 | 0 | 0 | 1475146 | AKA09662.1 | P36334 |
| LYLADNLNADET | 1 | 0 | 0 | 1475155 | ATQ39389.1 | K9N7C7 |
| LYLADNLTAGET | 1 | 0 | 0 | 1475160 | QBF80609.1 | K9N7C7 |
| LYLGGMRYFCVD | 1 | 0 | 0 | 1475170 | AHI48744.1 | K9N7C7 |
| LYLGVMSYFCVD | 1 | 0 | 0 | 1475179 | AHI48776.1 | K9N7C7 |
| LYLKNLKQTLKS | 1 | 0 | 0 | 1475181 | AEN19355.1 | P0C6X6 |
| LYLTSSGFDKSY | 1 | 0 | 0 | 1475203 | AUM60024.1 | K9N5Q8 |
| LYMIDPAGVCLG | 1 | 0 | 0 | 1475215 | AEN19363.1 | P0C6X6 |
| LYNAPCFSQQGV | 1 | 0 | 0 | 1475219 | AGT51729.1 | P30215 |
| LYNCVLDYYIVK | 1 | 0 | 0 | 1475221 | AWK59942.1 | H9EJ42 |
| LYNSPCFSQQGV | 1 | 0 | 0 | 1475245 | P30215.1 | P30215 |
| LYNSSCFSHQGV | 1 | 0 | 0 | 1475248 | QBQ01838.1 | P30215 |
| LYNSSCFSQQGV | 1 | 0 | 0 | 1475249 | AEN19365.1 | P30215 |
| LYNTEGGKTFMY | 1 | 0 | 0 | 1475254 | AWK59942.1 | H9EJ42 |
| LYNTPCFSQQGV | 1 | 0 | 0 | 1475257 | AGT51579.1 | P30215 |
| LYNVTRDKLEKL | 1 | 0 | 0 | 1475260 | AXN92247.1 | K9N7C7 |
| LYPNVYRTDYVD | 1 | 0 | 0 | 1475268 | ACR82756.1 | A0A140H1G9 |

| | | | | | | |
|---|---|---|---|---|---|---|
| LYPQQKYTSQCN | 1 | 0 | 0 | 1475269 | AUM60013.1 | K9N7C7 |
| LYPVASPRECND | 1 | 0 | 0 | 1475274 | ARO76381.1 | P0C6U8 |
| LYQDVNCTDVPT | 1 | 0 | 0 | 1475279 | AVP78031.1 | Q3LZX1 |
| LYSAAHADPNHF | 1 | 0 | 0 | 1475314 | AUM60024.1 | K9N5Q8 |
| LYSAAHADPNRF | 1 | 0 | 0 | 1475315 | AUM60014.1 | K9N5Q8 |
| LYSFDKAIDGFG | 1 | 0 | 0 | 1475321 | AWK59942.1 | H9EJ42 |
| LYSIQDVGFKNQ | 1 | 0 | 0 | 1475327 | AEN19363.1 | P0C6X6 |
| LYSPLLVDSDQP | 1 | 0 | 0 | 1475335 | AST12968.1 | A0A0N7J131 |
| LYSPLLVDSEQP | 1 | 0 | 0 | 1475336 | AIW52723.1 | A0A0N7J131 |
| LYSSRKGDLYGG | 1 | 0 | 0 | 1475341 | AVV62526.1 | K9N5Q8 |
| LYSTKPITTSNP | 1 | 0 | 0 | 1475344 | APO40578.1 | P0C6X7 |
| LYVLPSVSLNSK | 1 | 0 | 0 | 1475406 | APO40578.1 | P0C6X7 |
| LYVNFEHQRGPG | 1 | 0 | 0 | 1475407 | APT69876.1 | A0A0P0K6L9 |
| LYVNVYRSTNPD | 1 | 0 | 0 | 1475419 | ATU80202.1 | K9N7C7 |
| LYVPEHCQSYTG | 1 | 0 | 0 | 1475421 | AVV62527.1 | K9N796 |
| LYVPIRLAGHTR | 1 | 0 | 0 | 1475425 | AXN92247.1 | K9N7C7 |
| LYVSNYSHEVNL | 1 | 0 | 0 | 1475432 | AUM60014.1 | K9N5Q8 |
| MAATGVISSMTD | 1 | 0 | 0 | 1475465 | AUM60014.1 | K9N5Q8 |
| MACNGHKIEDFS | 1 | 0 | 0 | 1475477 | AIG96659.1 | A0A140H1G9 |
| MACPACQNPKIG | 1 | 0 | 0 | 1475479 | APO40578.1 | P0C6X7 |
| MADGSPQSYCYT | 1 | 0 | 0 | 1475490 | AGT51488.1 | P0C6X6 |
| MADNGTITVEELKQLLEQWNLVIG | 1 | 0 | 0 | 1475496 | AFR58690.1 | P59596 |
| MADPNITNLCPF | 1 | 0 | 0 | 1475499 | 2GHV_E | P59594 |
| MADQTMTQMYKQ | 1 | 0 | 0 | 1475502 | AVP78041.1 | P0C6W2 |
| MAENVTGLFKDC | 1 | 0 | 0 | 1475513 | 5C8T_B | P0C6X7 |
| MAEQAMTSMYKQ | 1 | 0 | 0 | 1475514 | AVV62524.1 | K9N7C7 |
| MAFAYADKPNHF | 1 | 0 | 0 | 1475516 | AGT51410.1 | Q80872 |
| MAFPSAKVEGCM | 1 | 0 | 0 | 1475521 | 2PWX_A | P0C6X7 |
| MAFSASLFKPVH | 1 | 0 | 0 | 1475525 | AVN89455.1 | K9N7D2 |
| MAGNGGDAALALLLLDR | 1 | 0 | 0 | 1475537 | P0DTC9.1 | P0DTC9 |
| MAGRSALEDLLF | 1 | 0 | 0 | 1475539 | AIW52841.1 | A0A0P0G321 |
| MAILTATRLCVQ | 1 | 0 | 0 | 1475548 | AUM60019.1 | K9N5R3 |
| MAITIKPEANIN | 1 | 0 | 0 | 1475551 | ABD75615.1 | A0A140H1G9 |
| MAKTSVDCTMYI | 1 | 0 | 0 | 1475563 | AVP78031.1 | Q3LZX1 |
| MALKGTDKSSTL | 1 | 0 | 0 | 1475575 | AKA09652.1 | P36334 |
| MALNFKLIGVGG | 1 | 0 | 0 | 1475581 | AQZ41320.1 | K9N7C7 |
| MALSIFSAFYPI | 1 | 0 | 0 | 1475587 | AJG44108.1 | K9N7A1 |
| MALSQYYVMPLT | 1 | 0 | 0 | 1475591 | AHA80218.1 | P36334 |
| MALSRYYVMPLT | 1 | 0 | 0 | 1475592 | AKA09640.1 | P36334 |
| MANLANLNSHNI | 1 | 0 | 0 | 1475600 | CAA71146.1 | A0A0P0K6L9 |
| MANQFTEFVGTQ | 1 | 0 | 0 | 1475605 | AUM60013.1 | K9N7C7 |
| MAPILTESSGSL | 1 | 0 | 0 | 1475611 | AHF54652.1 | Q4VID0 |
| MAPPAAPRAVSF | 1 | 0 | 0 | 1475617 | AQZ41303.1 | K9N4V7 |
| MAQAFGLRGPGD | 1 | 0 | 0 | 1475622 | AXG21672.1 | K9N4V7 |
| MAQSSGVLPVNP | 1 | 0 | 0 | 1475623 | YP_009555247.1 | P0C6X6 |
| MAQVISYLSERC | 1 | 0 | 0 | 1475624 | AUM60013.1 | K9N7C7 |
| MASPAAPRAVSF | 1 | 0 | 0 | 1475648 | AXN92267.1 | K9N4V7 |
| MASSAAPRAVSF | 1 | 0 | 0 | 1475651 | AHL18098.1 | K9N4V7 |
| MASSNGPISPTS | 1 | 0 | 0 | 1475652 | AGT51605.1 | Q4VID0 |
| MATPAAPRAVTF | 1 | 0 | 0 | 1475667 | AVV62533.1 | K9N4V7 |
| MATPQPQRAPKG | 1 | 0 | 0 | 1475668 | AVV62544.1 | K9N4V7 |
| MATSANCCHDGI | 1 | 0 | 0 | 1475670 | ATP16762.1 | Q4VID0 |
| MATSVNCCHDGI | 1 | 0 | 0 | 1475671 | AVR40343.1 | Q4VID0 |
| MATVKWADESEP | 1 | 0 | 0 | 1475676 | ABB90491.1 | A0A0N7J131 |
| MAVAYADNPNHF | 1 | 0 | 0 | 1475681 | ARU07566.1 | Q80872 |
| MAVAYVDKPNHF | 1 | 0 | 0 | 1475682 | AXX83373.1 | Q80872 |
| MAVNLAKFLTYL | 1 | 0 | 0 | 1475684 | AVV62535.1 | K9N7C7 |
| MAVVDGKQIDDY | 1 | 0 | 0 | 1475690 | AIV41978.1 | P0C6X6 |
| MAYGDCTFGVPD | 1 | 0 | 0 | 1475696 | AVV62524.1 | K9N7C7 |
| MAYGNCTFGDPD | 1 | 0 | 0 | 1475698 | AVN89333.1 | K9N7C7 |
| MAYGNCTFGSPD | 1 | 0 | 0 | 1475699 | AHI48794.1 | K9N638 |
| MCDYPHTICHPN | 1 | 0 | 0 | 1475723 | ANZ78839.1 | P36334 |
| MCEHPHTICHPK | 1 | 0 | 0 | 1475725 | AKA09654.1 | P36334 |
| MCEHPHTICHPN | 1 | 0 | 0 | 1475726 | BAM16463.1 | P36334 |
| MCEHPHTSCHPN | 1 | 0 | 0 | 1475728 | AWW13511.1 | P36334 |
| MCEYPHTICHPN | 1 | 0 | 0 | 1475730 | AWW13566.1 | P36334 |
| MCEYPHTTCHPN | 1 | 0 | 0 | 1475736 | BBA84088.1 | P36334 |
| MCEYPNTICHPN | 1 | 0 | 0 | 1475738 | CAA83661.1 | P36334 |
| MCEYPQTICHPN | 1 | 0 | 0 | 1475740 | CAA79896.1 | P36334 |
| MCEYPYTICHPN | 1 | 0 | 0 | 1475741 | BBA20979.1 | P36334 |
| MCGFDGKPMLPK | 1 | 0 | 0 | 1475749 | AFD98826.1 | H9EJ42 |
| MCGSDGKPIADY | 1 | 0 | 0 | 1475757 | AUM60013.1 | K9N7C7 |
| MCIDQVTIAGVK | 1 | 0 | 0 | 1475771 | ATO98217.1 | P0C6W2 |
| MCLASIAFNHPI | 1 | 0 | 0 | 1475777 | AVN89313.1 | K9N5Q8 |
| MCPADQLNDPNY | 1 | 0 | 0 | 1475796 | AVV62524.1 | K9N7C7 |
| MCPADQLSDPNY | 1 | 0 | 0 | 1475797 | AXN92247.1 | K9N7C7 |
| MCSRRAGAYSSH | 1 | 0 | 0 | 1475807 | AVV62524.1 | K9N7C7 |
| MCTNGQVVPTLF | 1 | 0 | 0 | 1475818 | AVV62535.1 | K9N7C7 |
| MCVVSSINDLSF | 1 | 0 | 0 | 1475825 | AVA26872.1 | H9EJ42 |
| MDAEGRMTPYCH | 1 | 0 | 0 | 1475832 | AKJ80136.2 | K9N7C7 |
| MDDAINVGGTGL | 1 | 0 | 0 | 1475837 | AVN89322.1 | K9N7C7 |
| MDEFIQRYKLES | 1 | 0 | 0 | 1475846 | APO40578.1 | P0C6X7 |
| MDFCEGSFPCSV | 1 | 0 | 0 | 1475856 | P0C6W6.1 | P0C6W2 |
| MDFVNAGGAPLT | 1 | 0 | 0 | 1475857 | AVV62535.1 | K9N7C7 |
| MDFVNAGGVPLT | 1 | 0 | 0 | 1475858 | AVV62524.1 | K9N7C7 |
| MDFVNSGGTPLT | 1 | 0 | 0 | 1475859 | AUM60013.1 | K9N7C7 |
| MDFVYALRHFDQ | 1 | 0 | 0 | 1475860 | ATG84732.1 | K9N7C7 |
| MDIEKLRQYLVA | 1 | 0 | 0 | 1475878 | AVV62524.1 | K9N7C7 |
| MDIVKRLATGLF | 1 | 0 | 0 | 1475881 | AVV62535.1 | K9N7C7 |
| MDIWRPEKKFPP | 1 | 0 | 0 | 1475884 | AIV41989.1 | Q04853 |
| MDKQVHQVQLID | 1 | 0 | 0 | 1475893 | ATG84700.1 | K9N638 |

| | | | | | | |
|---|---|---|---|---|---|---|
| MDNKDRAMRLAL | 1 | 0 | 0 | 1475917 | APO40587.1 | P59636 |
| MDNTTQSAIIVN | 1 | 0 | 0 | 1475925 | ATO98193.1 | Q3LZX1 |
| MDPKTNVVHPAL | 1 | 0 | 0 | 1475929 | AVP78039.1 | Q3LZX3 |
| MDPLSNFEHKVI | 1 | 0 | 0 | 1475932 | 5HIH_A | K9N638 |
| MDPNKTNVVPPA | 1 | 0 | 0 | 1475933 | ARO76391.1 | P59636 |
| MDSGGSSFSELL | 1 | 0 | 0 | 1475944 | AHX71940.1 | K9N7C7 |
| MDSPDLGTDDDD | 1 | 0 | 0 | 1475948 | 3R24_A | P0C6X7 |
| MDTAAYREAAAS | 1 | 0 | 0 | 1475950 | AVV62524.1 | K9N7C7 |
| MDVPSTYGTQVY | 1 | 0 | 0 | 1475971 | AIA22866.1 | K9N643 |
| MDVQGLVNVLAC | 1 | 0 | 0 | 1475972 | AGT51372.1 | H9EJ42 |
| MDVSVHRHRLSL | 1 | 0 | 0 | 1475977 | AVN89342.1 | K9N7C7 |
| MDYKCSCTSGRL | 1 | 0 | 0 | 1475980 | NP_073549.1 | A0A0P0LKV0 |
| MDYLDSLKPTVE | 1 | 0 | 0 | 1475983 | ARO76381.1 | P0C6U8 |
| MDYVSLLNQVWQ | 1 | 0 | 0 | 1475988 | AVV62539.1 | K9N4V0 |
| MEAKPSGSVVEQ | 1 | 0 | 0 | 1475996 | 5GMQ_A | K9N5Q8 |
| MECDFLNMDMGV | 1 | 0 | 0 | 1476001 | AVL25566.1 | H9EJ42 |
| MEDAMGQGQNNA | 1 | 0 | 0 | 1476002 | ARO76391.1 | P59636 |
| MEEDFLNMDMGV | 1 | 0 | 0 | 1476007 | 4RS4_A | A0A0P0LKV0 |
| MEEFIQTNSPKV | 1 | 0 | 0 | 1476008 | AIL49500.1 | P36334 |
| MEESLTDVPSTS | 1 | 0 | 0 | 1476019 | AXN92231.1 | K9N643 |
| MEESPTDVPSTS | 1 | 0 | 0 | 1476021 | ASU90409.1 | K9N643 |
| MEEYPMDVHSTY | 1 | 0 | 0 | 1476022 | ATQ39396.1 | K9N643 |
| MEKDFLSLDSDV | 1 | 0 | 0 | 1476030 | AVV62535.1 | K9N7C7 |
| MEKDFLSMDSDV | 1 | 0 | 0 | 1476031 | AVV62524.1 | K9N7C7 |
| MEKVNERVKSQS | 1 | 0 | 0 | 1476037 | ARB66064.1 | P36334 |
| MELANGTHTGSS | 1 | 0 | 0 | 1476040 | AVV62524.1 | K9N7C7 |
| MELLDSNIDAYK | 1 | 0 | 0 | 1476042 | AUM60021.1 | K9N4V7 |
| MENFELEKHCYD | 1 | 0 | 0 | 1476050 | ALW82707.1 | K9N7C7 |
| MESTGSLATRLM | 1 | 0 | 0 | 1476071 | AGT51765.1 | Q4VID0 |
| METAAYREAAAC | 1 | 0 | 0 | 1476073 | AXN92247.1 | K9N7C7 |
| METFTNNNGSDV | 1 | 0 | 0 | 1476078 | AQN78669.1 | P0C6X6 |
| MEVDHSGPTHYL | 1 | 0 | 0 | 1476085 | AVV62529.1 | K9N643 |
| MEYDFLNIDMGV | 1 | 0 | 0 | 1476095 | AGT51365.1 | H9EJ42 |
| MFADHLIDLSDC | 1 | 0 | 0 | 1476099 | AEN19363.1 | P0C6X6 |
| MFADHLTDLSDC | 1 | 0 | 0 | 1476100 | AXX83336.1 | P0C6X6 |
| MFCATCPFAKSD | 1 | 0 | 0 | 1476102 | AVV62535.1 | K9N7C7 |
| MFDAYVDTFVST | 1 | 0 | 0 | 1476106 | APO40578.1 | P0C6X7 |
| MFDKAVADAMSA | 1 | 0 | 0 | 1476110 | AUM60021.1 | K9N4V7 |
| MFDTNMFIKFPE | 1 | 0 | 0 | 1476112 | AUM60013.1 | K9N7C7 |
| MFDVDKKILNAL | 1 | 0 | 0 | 1476113 | AIV41972.1 | P0C6X6 |
| MFDVDKKSLNVL | 1 | 0 | 0 | 1476115 | AIV41834.1 | P0C6X6 |
| MFEDAQGLKEFY | 1 | 0 | 0 | 1476118 | AMQ49036.1 | K9N638 |
| MFEDAQSLKKFY | 1 | 0 | 0 | 1476120 | ALA50066.1 | K9N7C7 |
| MFEDDQGLKKFY | 1 | 0 | 0 | 1476121 | AGV08522.1 | K9N7C7 |
| MFEPPKEQRVQS | 1 | 0 | 0 | 1476127 | QCZ25056.1 | K9N4V7 |
| MFFPYDIEFVTG | 1 | 0 | 0 | 1476134 | ALA50044.1 | K9N7C7 |
| MFGHIVPIWMTC | 1 | 0 | 0 | 1476139 | ALA49351.1 | K9N7C7 |
| MFHQKCLKSIAA | 1 | 0 | 0 | 1476146 | AEN19363.1 | P0C6X6 |
| MFITRDEAIKHV | 1 | 0 | 0 | 1476155 | APO40578.1 | P0C6X7 |
| MFIVLYVDFKPQ | 1 | 0 | 0 | 1476160 | 6ATK_E | A0A0P0K6L9 |
| MFKDAEGRMTPY | 1 | 0 | 0 | 1476162 | AVN89322.1 | K9N7C7 |
| MFKPEVEELNPS | 1 | 0 | 0 | 1476164 | AGT51436.1 | Q4VID0 |
| MFKPGVEELNPS | 1 | 0 | 0 | 1476165 | AOL02458.1 | Q4VID0 |
| MFLLPRFVLVSC | 1 | 0 | 0 | 1476189 | AAA73152.1 | P30215 |
| MFLLTPIKGDVD | 1 | 0 | 0 | 1476192 | AVV62537.1 | K9N5Q8 |
| MFLLTPTESYVD | 1 | 0 | 0 | 1476193 | AXN92228.1 | K9N5Q8 |
| MFMADAYLADTV | 1 | 0 | 0 | 1476199 | AOL02455.1 | Q04854 |
| MFMADAYLAGTV | 1 | 0 | 0 | 1476200 | AGT51533.1 | Q04854 |
| MFMDAEGRMTPY | 1 | 0 | 0 | 1476202 | AKJ80136.2 | K9N7C7 |
| MFPPGTVLPRGY | 1 | 0 | 0 | 1476212 | AJQ24068.1 | P33469 |
| MFQPRLPVLSDF | 1 | 0 | 0 | 1476218 | AIW52843.1 | A0A0P0G321 |
| MFRDAEGKMVPY | 1 | 0 | 0 | 1476221 | AVV62535.1 | K9N7C7 |
| MFRDAEGRITPY | 1 | 0 | 0 | 1476222 | AHL18091.1 | K9N7C7 |
| MFRDAEGRMTPY | 1 | 0 | 0 | 1476223 | AXN92247.1 | K9N7C7 |
| MFRDAEGRMVPY | 1 | 0 | 0 | 1476224 | AUM60013.1 | K9N7C7 |
| MFRFASLPVYDT | 1 | 0 | 0 | 1476225 | AVV62526.1 | K9N5Q8 |
| MFSAMILGSKHV | 1 | 0 | 0 | 1476228 | AAV28291.1 | A0A0P0LKV0 |
| MFTGSGYYYPEP | 1 | 0 | 0 | 1476245 | CAA83660.1 | P36334 |
| MFTGSRYYYPEP | 1 | 0 | 0 | 1476246 | CAA83661.1 | P36334 |
| MFTPLIQPIGAL | 1 | 0 | 0 | 1476253 | AVP78041.1 | P0C6W2 |
| MFVATSSYDALL | 1 | 0 | 0 | 1476259 | AGT51379.1 | H9EJ42 |
| MFVCIDNSANTK | 1 | 0 | 0 | 1476260 | AUM60013.1 | K9N7C7 |
| MGATICLQAGKQ | 1 | 0 | 0 | 1476290 | AAW33912.1 | H9EJ42 |
| MGDCVKFVYMHQ | 1 | 0 | 0 | 1476296 | AEN19363.1 | P0C6X6 |
| MGEFIQTSSPKV | 1 | 0 | 0 | 1476299 | ARB66062.1 | P36334 |
| MGFKLDANIPDY | 1 | 0 | 0 | 1476304 | AVV62535.1 | K9N7C7 |
| MGFKLDANIPGY | 1 | 0 | 0 | 1476305 | AVV62524.1 | K9N7C7 |
| MGFRFDISIPGS | 1 | 0 | 0 | 1476313 | AWK59942.1 | H9EJ42 |
| MGIDLDEWSVST | 1 | 0 | 0 | 1476321 | APO40578.1 | P0C6X7 |
| MGILPSPGMPAL | 1 | 0 | 0 | 1476326 | 6NB6_A | P59594 |
| MGILSENAYLKY | 1 | 0 | 0 | 1476327 | AVV62524.1 | K9N7C7 |
| MGLLSENAYLKW | 1 | 0 | 0 | 1476353 | AUM60013.1 | K9N7C7 |
| MGLPNNTASWFT | 1 | 0 | 0 | 1476356 | 1SSK_A | P59595 |
| MGLTICLASECF | 1 | 0 | 0 | 1476357 | QED88024.1 | H9EJ42 |
| MGLTNCLASECF | 1 | 0 | 0 | 1476358 | AIW52827.1 | H9EJ42 |
| MGMSQFKVTHLS | 1 | 0 | 0 | 1476361 | AUM60021.1 | K9N4V7 |
| MGNAINVGGRGL | 1 | 0 | 0 | 1476363 | ATQ39389.1 | K9N7C7 |
| MGQESIVAYTMS | 1 | 0 | 0 | 1476365 | APO40579.1 | P59594 |
| MGQGQNNADPKV | 1 | 0 | 0 | 1476366 | ARO76391.1 | P59636 |
| MGQLEETNMHSP | 1 | 0 | 0 | 1476368 | AUM60023.1 | K9N7C7 |
| MGRFFNYTLVLL | 1 | 0 | 0 | 1476372 | AVN89387.1 | K9N5Q8 |

| | | | | | | |
|---|---|---|---|---|---|---|
| MGRIRSVYPVAT | 1 | 0 | 0 | 1476377 | P0C6W6.1 | P0C6W2 |
| MGSNNVCTYEHV | 1 | 0 | 0 | 1476387 | AGT51323.1 | H9EJ42 |
| MGSSHHHHHHSS | 1 | 0 | 0 | 1476388 | 1SSK_A | P59595 |
| MGTKKSAAEASK | 1 | 0 | 0 | 1476390 | 2CJR_A | P59595 |
| MGTLSYDDLKRG | 1 | 0 | 0 | 1476392 | APO40578.1 | P0C6X7 |
| MGTLSYDSLKTG | 1 | 0 | 0 | 1476394 | ARO76381.1 | P0C6U8 |
| MGTLSYEQFKKGVQIPC | 1 | 0 | 0 | 1476397 | P0DTC1.1 | P0DTD1 |
| MGTNGNKIVDKE | 1 | 0 | 0 | 1476399 | AVV62524.1 | K9N7C7 |
| MGVLKADDFVIA | 1 | 0 | 0 | 1476406 | AVL25554.1 | H9EJ42 |
| MGVVMQSGVKRV | 1 | 0 | 0 | 1476409 | AVV62524.1 | K9N7C7 |
| MHDNDSKEGFFT | 1 | 0 | 0 | 1476426 | APO40578.1 | P0C6X7 |
| MHFGAGSDKGIA | 1 | 0 | 0 | 1476429 | AVV62524.1 | K9N7C7 |
| MHGSVLRIKVSQ | 1 | 0 | 0 | 1476437 | AWK59942.1 | H9EJ42 |
| MHHHHHHSSGVD | 1 | 0 | 0 | 1476443 | 5X29_A | P59637 |
| MHKIIKVFYELD | 1 | 0 | 0 | 1476449 | APU51933.1 | P0C6X6 |
| MHKPPISFPLCA | 1 | 0 | 0 | 1476450 | APO40578.1 | P0C6X7 |
| MHSEFPAITIGS | 1 | 0 | 0 | 1476463 | AKA09651.1 | P36334 |
| MHSINHDIFSLM | 1 | 0 | 0 | 1476465 | ARU07575.1 | A0A140H1G9 |
| MHSPDIVVGDYV | 1 | 0 | 0 | 1476468 | AVV62535.1 | K9N7C7 |
| MHSPDLVVGDYV | 1 | 0 | 0 | 1476470 | AVV62524.1 | K9N7C7 |
| MHSPDVIVGDYV | 1 | 0 | 0 | 1476471 | AXN92247.1 | K9N7C7 |
| MHVGYYPSNHID | 1 | 0 | 0 | 1476476 | ALX27232.1 | K9N5Q8 |
| MHVGYYPSNHIE | 1 | 0 | 0 | 1476477 | AXN92228.1 | K9N5Q8 |
| MHVGYYPSNQIE | 1 | 0 | 0 | 1476478 | QBF80554.1 | K9N5Q8 |
| MIAPESKSFTMT | 1 | 0 | 0 | 1476495 | AVV62524.1 | K9N7C7 |
| MICSTTAQKLKT | 1 | 0 | 0 | 1476498 | AIX09796.1 | P0C6X6 |
| MIDPEGVCLGAG | 1 | 0 | 0 | 1476503 | AWW13571.1 | P0C6X6 |
| MIDVQQWGFTGNLQ | 1 | 0 | 0 | 1476506 | P0DTD1.1 | P0DTD1 |
| MIEDMDYVPLKS | 1 | 0 | 0 | 1476509 | AXG21663.1 | K9N7C7 |
| MIEDVDYVPLKS | 1 | 0 | 0 | 1476510 | AMQ49035.1 | K9N7C7 |
| MIFSVHPVECDE | 1 | 0 | 0 | 1476523 | AVV62524.1 | K9N7C7 |
| MIKPNAILCATC | 1 | 0 | 0 | 1476539 | QBF80531.1 | K9N7C7 |
| MIKPNALLCATC | 1 | 0 | 0 | 1476541 | AXN92247.1 | K9N7C7 |
| MIKPNARLCATC | 1 | 0 | 0 | 1476543 | AHI48793.1 | K9N7C7 |
| MILKNQDLGVVS | 1 | 0 | 0 | 1476549 | AHI48724.1 | K9N7C7 |
| MILKSQDLGAVS | 1 | 0 | 0 | 1476550 | ANI69887.1 | K9N7C7 |
| MILPTFNMQSVS | 1 | 0 | 0 | 1476558 | AVV62535.1 | K9N7C7 |
| MILSDDGVVCNN | 1 | 0 | 0 | 1476562 | AIX10745.1 | P0C6X6 |
| MILSLHPVECDE | 1 | 0 | 0 | 1476563 | ALW82719.1 | K9N7C7 |
| MINPNALLCATC | 1 | 0 | 0 | 1476570 | ASU90746.1 | K9N7C7 |
| MIQTHLSHYVLN | 1 | 0 | 0 | 1476580 | AUM60013.1 | K9N7C7 |
| MIRMLSAMISGS | 1 | 0 | 0 | 1476584 | AIG96641.1 | A0A0P0LKV0 |
| MIRSCYSGHVSA | 1 | 0 | 0 | 1476585 | AIL49489.1 | P36334 |
| MISTVGVFLGYK | 1 | 0 | 0 | 1476602 | NP_073549.1 | A0A0P0LKV0 |
| MIVDEWSYTGSS | 1 | 0 | 0 | 1476607 | ATG84866.1 | K9N5Q8 |
| MKALFPIWSQEL | 1 | 0 | 0 | 1476621 | AVQ05261.1 | P0C6X6 |
| MKALYPTWSQEL | 1 | 0 | 0 | 1476625 | AGT51518.1 | P0C6X6 |
| MKGTGVFVQDPS | 1 | 0 | 0 | 1476648 | AFD98763.1 | H9EJ42 |
| MKHKGGDSIDFI | 1 | 0 | 0 | 1476654 | AKK52590.1 | K9N7C7 |
| MKQGPPSARDLI | 1 | 0 | 0 | 1476694 | AVV62537.1 | K9N5Q8 |
| MKQIEDKIEEIE | 1 | 0 | 0 | 1476695 | 5X58_A | P59594 |
| MKQYGDCLGGIN | 1 | 0 | 0 | 1476696 | APO40579.1 | P59594 |
| MKQYGECLGDIS | 1 | 0 | 0 | 1476699 | ATO98145.1 | Q3LZX1 |
| MKSDLSVSFAGP | 1 | 0 | 0 | 1476708 | AHX71943.1 | K9N5Q8 |
| MKVLTENLNAYQ | 1 | 0 | 0 | 1476728 | AIL49396.1 | P33469 |
| MLAEFDENGTIT | 1 | 0 | 0 | 1476737 | APO40579.1 | P59594 |
| MLAKKELSGQCF | 1 | 0 | 0 | 1476743 | AGV08405.1 | K9N7C7 |
| MLAVKTGVATEQ | 1 | 0 | 0 | 1476749 | AUM60013.1 | K9N7C7 |
| MLDDVQRWLAPY | 1 | 0 | 0 | 1476760 | AVV62540.1 | K9N643 |
| MLDLLRDIRFTI | 1 | 0 | 0 | 1476766 | AVV62524.1 | K9N7C7 |
| MLEDIDYVPLKS | 1 | 0 | 0 | 1476774 | AVV62524.1 | K9N7C7 |
| MLFKASALFKEF | 1 | 0 | 0 | 1476782 | AVV62524.1 | K9N7C7 |
| MLGCSVGNFSDG | 1 | 0 | 0 | 1476796 | QEU56412.1 | K9N5Q8 |
| MLGSSVGNFSNG | 1 | 0 | 0 | 1476809 | AVV62537.1 | K9N5Q8 |
| MLGSSVGNYYNG | 1 | 0 | 0 | 1476811 | AVV62526.1 | K9N5Q8 |
| MLGYNPSKPISK | 1 | 0 | 0 | 1476816 | AVV62524.1 | K9N7C7 |
| MLKDSSTIHNYF | 1 | 0 | 0 | 1476838 | AQZ41305.1 | K9N7C7 |
| MLKHFFFAQDGN | 1 | 0 | 0 | 1476843 | QAT98908.1 | K9N7C7 |
| MLKRRDSTYSPL | 1 | 0 | 0 | 1476852 | QEU56411.1 | K9N5Q8 |
| MLKTVYSDVENP | 1 | 0 | 0 | 1476858 | AID16714.1 | P0C6W2 |
| MLLALLAFFLSK | 1 | 0 | 0 | 1476865 | AWK59942.1 | H9EJ42 |
| MLLELKDKEQYC | 1 | 0 | 0 | 1476873 | ARO76381.1 | P0C6U8 |
| MLLELKERDQYC | 1 | 0 | 0 | 1476874 | APO40578.1 | P0C6X7 |
| MLLSVPSDSCNN | 1 | 0 | 0 | 1476886 | AGH06535.1 | H9EJ42 |
| MLMGLLSENAYL | 1 | 0 | 0 | 1476894 | AUM60013.1 | K9N7C7 |
| MLNGMPVSSIKS | 1 | 0 | 0 | 1476899 | AGT51393.1 | H9EJ42 |
| MLNISGNGGLFN | 1 | 0 | 0 | 1476901 | AVP78031.1 | Q3LZX1 |
| MLNISTPNLHDF | 1 | 0 | 0 | 1476902 | CAA83660.1 | P36334 |
| MLNKHIDAYKTF | 1 | 0 | 0 | 1476903 | ARO76389.1 | P59595 |
| MLNTPIPNLPDF | 1 | 0 | 0 | 1476910 | AIL49487.1 | P36334 |
| MLNTSSPNLPDF | 1 | 0 | 0 | 1476914 | AIL49492.1 | P36334 |
| MLNTSTPNLPDF | 1 | 0 | 0 | 1476915 | AVQ05264.1 | P36334 |
| MLPFVQEQIGSF | 1 | 0 | 0 | 1476918 | AVV62531.1 | K9N5R3 |
| MLPFVQQQIGSF | 1 | 0 | 0 | 1476922 | AUM60019.1 | K9N5R3 |
| MLPGEFSKKQLG | 1 | 0 | 0 | 1476923 | AKK52590.1 | K9N7C7 |
| MLPKNMWEFRDY | 1 | 0 | 0 | 1476924 | AFD98826.1 | H9EJ42 |
| MLPSCYNFLKEQ | 1 | 0 | 0 | 1476929 | ARO76392.1 | Q7TLC7 |
| MLQIQLTGILDG | 1 | 0 | 0 | 1476930 | NP_073549.1 | A0A0P0LKV0 |
| MLRACIKIAQAD | 1 | 0 | 0 | 1476941 | ASU89922.1 | K9N796 |
| MLRACIKTVQAD | 1 | 0 | 0 | 1476946 | AHC74089.1 | K9N796 |
| MLRGDDMSFLEK | 1 | 0 | 0 | 1476949 | AFR58685.1 | P0C6X7 |

| | | | | | | |
|---|---|---|---|---|---|---|
| MLRPLIKDVDNP | 1 | 0 | 0 | 1476960 | AHF54667.1 | P0C6X6 |
| MLSFNENGTMSN | 1 | 0 | 0 | 1476971 | ATO98145.1 | Q3LZX1 |
| MLTFDSNAFSLA | 1 | 0 | 0 | 1476984 | AIW52843.1 | A0A0P0G321 |
| MLTGIAARPPPG | 1 | 0 | 0 | 1476986 | AXN92247.1 | K9N7C7 |
| MLTKALRSVPTD | 1 | 0 | 0 | 1476988 | APO40578.1 | P0C6X7 |
| MLVKMDGDEVYL | 1 | 0 | 0 | 1477002 | AIL49620.1 | P0C6X6 |
| MLVNCFKWQVVV | 1 | 0 | 0 | 1477012 | AEN19363.1 | P0C6X6 |
| MLWCKDGKVQTF | 1 | 0 | 0 | 1477023 | ATQ39389.1 | K9N7C7 |
| MLWGDGFQIKFE | 1 | 0 | 0 | 1477025 | ARK08668.1 | Q80872 |
| MLYPTTLGSLEL | 1 | 0 | 0 | 1477029 | AGT51745.1 | Q4VID0 |
| MMDLGNSTTITD | 1 | 0 | 0 | 1477036 | AVV62537.1 | K9N5Q8 |
| MMNISPKPQRQR | 1 | 0 | 0 | 1477056 | AIX10729.1 | P33469 |
| MMQSADASTFLN | 1 | 0 | 0 | 1477065 | P0C6W6.1 | P0C6W2 |
| MMSALPAEYKLR | 1 | 0 | 0 | 1477069 | ARO76381.1 | P0C6U8 |
| MMTGISARPPPG | 1 | 0 | 0 | 1477078 | AVV62535.1 | K9N7C7 |
| MMWVSYFIQSIR | 1 | 0 | 0 | 1477083 | AVV62543.1 | K9N7A1 |
| MMYAADPAMHIA | 1 | 0 | 0 | 1477085 | AXN92247.1 | K9N7C7 |
| MNATLDKCTKYN | 1 | 0 | 0 | 1477090 | CAA71147.1 | A0A0P0K6L9 |
| MNAYPFVVTPLV | 1 | 0 | 0 | 1477091 | AHY22534.1 | K9N7C7 |
| MNCNHCGESSWQ | 1 | 0 | 0 | 1477093 | APO40578.1 | P0C6X7 |
| MNDDLLQILNRQ | 1 | 0 | 0 | 1477096 | APO40578.1 | P0C6X7 |
| MNDKNICLLDHA | 1 | 0 | 0 | 1477098 | P0C6U5.1 | A0A140H1G9 |
| MNGEDAFSILSA | 1 | 0 | 0 | 1477108 | APT69882.1 | A0A0P0ILY9 |
| MNIAKSEFDHEV | 1 | 0 | 0 | 1477114 | ARE29978.1 | H9EJ42 |
| MNIAKYMQLCQY | 1 | 0 | 0 | 1477115 | AXN92247.1 | K9N7C7 |
| MNKSFFPQFTSD | 1 | 0 | 0 | 1477118 | ABD75500.1 | Q0ZJ82 |
| MNKSFFSQFTSD | 1 | 0 | 0 | 1477119 | ABD75588.1 | Q0ZJ82 |
| MNMDVDTHRYCL | 1 | 0 | 0 | 1477127 | ANZ78856.1 | P0C6X6 |
| MNMDVSLHRHRL | 1 | 0 | 0 | 1477128 | AXN92247.1 | K9N7C7 |
| MNTSHSVCNGCV | 1 | 0 | 0 | 1477153 | AIW52756.1 | A0A0P0K6L9 |
| MNTSYSVCNGCV | 1 | 0 | 0 | 1477156 | ADK37704.1 | A0A0P0K6L9 |
| MNVIGQHIELPQ | 1 | 0 | 0 | 1477159 | ARE29978.1 | H9EJ42 |
| MNVIGQHIKLPQ | 1 | 0 | 0 | 1477160 | AWK59942.1 | H9EJ42 |
| MNVTLNKCTKYN | 1 | 0 | 0 | 1477164 | AIW52757.1 | A0A0P0K6L9 |
| MPALLSLVSLLS | 1 | 0 | 0 | 1477170 | 6NB6_A | P59594 |
| MPIMANLVNLNS | 1 | 0 | 0 | 1477210 | AIW52756.1 | A0A0P0K6L9 |
| MPKFFYASNGTY | 1 | 0 | 0 | 1477229 | AIW52757.1 | A0A0P0K6L9 |
| MPLKAPKDVTFL | 1 | 0 | 0 | 1477239 | ARO76381.1 | P0C6U8 |
| MPLTCIGRGNIG | 1 | 0 | 0 | 1477253 | AHA80218.1 | P36334 |
| MPLTRIARRNIG | 1 | 0 | 0 | 1477272 | AHA80201.1 | P36334 |
| MPMLTMCHALDC | 1 | 0 | 0 | 1477277 | AEN19363.1 | P0C6X6 |
| MPMLTMCHVLDC | 1 | 0 | 0 | 1477279 | ARU07574.1 | A0A140H1G9 |
| MPNNGWTAFSAA | 1 | 0 | 0 | 1477290 | AVP78031.1 | Q3LZX1 |
| MPQFSRMTQSAL | 1 | 0 | 0 | 1477293 | AUM60014.1 | K9N5Q8 |
| MPQGVHMNVAKY | 1 | 0 | 0 | 1477294 | AVV62524.1 | K9N7C7 |
| MPTHVLAGHDDF | 1 | 0 | 0 | 1477305 | APO40578.1 | P0C6X7 |
| MPTITQMNLKYA | 1 | 0 | 0 | 1477306 | AUM60013.1 | K9N7C7 |
| MPTYTVHKSDFQ | 1 | 0 | 0 | 1477314 | AEN19363.1 | P0C6X6 |
| MPVSSIKSDKGV | 1 | 0 | 0 | 1477335 | AGT51393.1 | H9EJ42 |
| MPVVDGKQIDDH | 1 | 0 | 0 | 1477336 | AGT51409.1 | P0C6X6 |
| MPYFMYDVVSSF | 1 | 0 | 0 | 1477340 | AGT21366.1 | A0A0P0LKV0 |
| MPYHPNPVDCQD | 1 | 0 | 0 | 1477341 | ANZ78865.1 | P0C6X6 |
| MQCFGPRDFNHN | 1 | 0 | 0 | 1477357 | AMM42619.1 | H9EJ69 |
| MQFGYIDCEQDL | 1 | 0 | 0 | 1477364 | AXX83372.1 | P0C6X6 |
| MQFNYKITINPS | 1 | 0 | 0 | 1477369 | AWW13564.1 | Q80872 |
| MQGAVDINKLCD | 1 | 0 | 0 | 1477370 | APO40578.1 | P0C6X7 |
| MQGAVDLNKLCE | 1 | 0 | 0 | 1477373 | AVP78041.1 | P0C6W2 |
| MQGPFIAGNTVH | 1 | 0 | 0 | 1477375 | AFD98763.1 | H9EJ42 |
| MQGYDDCMNQGP | 1 | 0 | 0 | 1477380 | AVV62526.1 | K9N5Q8 |
| MQGYDECMQQGP | 1 | 0 | 0 | 1477382 | AUM60014.1 | K9N5Q8 |
| MQIVDKDGNYYL | 1 | 0 | 0 | 1477391 | QED88010.1 | H9EJ42 |
| MQLLHTKVSWAG | 1 | 0 | 0 | 1477410 | AXN92247.1 | K9N7C7 |
| MQLNLISNITTN | 1 | 0 | 0 | 1477412 | AVP78041.1 | P0C6W2 |
| MQNKGLFYTQVY | 1 | 0 | 0 | 1477424 | AEN19365.1 | P30215 |
| MQPDKLPGNCIG | 1 | 0 | 0 | 1477427 | AUM60027.1 | K9N643 |
| MQQGPASARDLI | 1 | 0 | 0 | 1477433 | AXN92228.1 | K9N5Q8 |
| MQQGPQSARDLI | 1 | 0 | 0 | 1477437 | AUM60014.1 | K9N5Q8 |
| MQQHPLLNPGAL | 1 | 0 | 0 | 1477439 | AGT21342.1 | A0A0N7J131 |
| MQQQPLLNPSAL | 1 | 0 | 0 | 1477442 | AIW52723.1 | A0A0N7J131 |
| MQSGVKRVTYGT | 1 | 0 | 0 | 1477451 | AVV62524.1 | K9N7C7 |
| MQSQSSHAVQNT | 1 | 0 | 0 | 1477455 | AIW52795.1 | H9EJ69 |
| MQSSDASTFLNR | 1 | 0 | 0 | 1477459 | APO40578.1 | P0C6X7 |
| MQTGFTTTNVAF | 1 | 0 | 0 | 1477464 | AUM60014.1 | K9N5Q8 |
| MQTVEDLRARMT | 1 | 0 | 0 | 1477467 | AVN89451.1 | K9N7C7 |
| MQTYKLVNMKGT | 1 | 0 | 0 | 1477469 | AGT51330.1 | H9EJ42 |
| MRDVSNSGPLQT | 1 | 0 | 0 | 1477490 | AVV62537.1 | K9N5Q8 |
| MRHKRVATKSFN | 1 | 0 | 0 | 1477502 | AVV62533.1 | K9N4V7 |
| MRISYFVQSIRL | 1 | 0 | 0 | 1477513 | ALA49424.1 | K9N7A1 |
| MRLFYNRDGQRT | 1 | 0 | 0 | 1477525 | AEN19355.1 | P0C6X6 |
| MRNANGACIWNC | 1 | 0 | 0 | 1477535 | AUM60013.1 | K9N7C7 |
| MRPNYTIKGSFL | 1 | 0 | 0 | 1477543 | AEA10817.1 | P0C6X7 |
| MRSTGPIQTPVG | 1 | 0 | 0 | 1477559 | AUM60014.1 | K9N5Q8 |
| MRYNLNFSSNSV | 1 | 0 | 0 | 1477583 | AIW52843.1 | A0A0P0G321 |
| MSCDCGKKFDEQ | 1 | 0 | 0 | 1477604 | AWK59942.1 | H9EJ42 |
| MSCNGNKIEDFS | 1 | 0 | 0 | 1477607 | AIV41978.1 | P0C6X6 |
| MSCNGNKIGDLS | 1 | 0 | 0 | 1477608 | AJE26218.1 | P0C6X6 |
| MSCNGNKSDDLS | 1 | 0 | 0 | 1477609 | AKL87224.1 | P0C6X6 |
| MSDFMSEIKCET | 1 | 0 | 0 | 1477610 | AIV41831.1 | P36334 |
| MSDNGPQNQRSA | 1 | 0 | 0 | 1477620 | AVP78038.1 | Q3LZX4 |
| MSDNGPQNQSSA | 1 | 0 | 0 | 1477621 | AVP78049.1 | Q3LZX4 |
| MSDNGPQPNQRS | 1 | 0 | 0 | 1477622 | ATO98228.1 | Q3LZX4 |

| | | | | | | |
|---|---|---|---|---|---|---|
| MSDNGPQQNQRS | 1 | 0 | 0 | 1477623 | ARO76389.1 | P59595 |
| MSDYDIPTTENL | 1 | 0 | 0 | 1477628 | 3EJG_A | A0A0P0LKV0 |
| MSEIKCKTLSIA | 1 | 0 | 0 | 1477640 | CAA83660.1 | P36334 |
| MSELPKEQRVQG | 1 | 0 | 0 | 1477642 | ASJ26617.1 | K9N4V7 |
| MSEPPKEHRVQG | 1 | 0 | 0 | 1477643 | YP_007188586.1 | K9N4V7 |
| MSEPPNEQRVQG | 1 | 0 | 0 | 1477648 | AVN89381.1 | K9N4V7 |
| MSISKLHALDDV | 1 | 0 | 0 | 1477693 | AXN92231.1 | K9N643 |
| MSLEACYERGCN | 1 | 0 | 0 | 1477702 | AAM95936.1 | P0C6U7 |
| MSLKDGQINDMI | 1 | 0 | 0 | 1477714 | AVP78041.1 | P0C6W2 |
| MSLKDNQVNDMI | 1 | 0 | 0 | 1477715 | APO40578.1 | P0C6X7 |
| MSMAECKAALGL | 1 | 0 | 0 | 1477729 | AGT51337.1 | H9EJ42 |
| MSMFSRTTSNFL | 1 | 0 | 0 | 1477731 | ATO98193.1 | Q3LZX1 |
| MSMTYGQQFGPI | 1 | 0 | 0 | 1477735 | AVP78030.1 | P0C6W2 |
| MSNDNCTGDIVT | 1 | 0 | 0 | 1477738 | AST12967.1 | A0A0P0KXN3 |
| MSNDTLTQQQII | 1 | 0 | 0 | 1477739 | AUM60030.1 | K9N7A1 |
| MSNITQLTEAQI | 1 | 0 | 0 | 1477740 | AKM76245.1 | K9N7A1 |
| MSNLGMPSYCTV | 1 | 0 | 0 | 1477742 | AVP78041.1 | P0C6W2 |
| MSNMTHLTEAQI | 1 | 0 | 0 | 1477744 | AZK15906.1 | K9N7A1 |
| MSNNGWTAFSAA | 1 | 0 | 0 | 1477750 | AVP78042.1 | Q3LZX1 |
| MSPFEMPFEELN | 1 | 0 | 0 | 1477756 | AST12962.1 | A0A0P0LKV0 |
| MSPKPQRQRGLN | 1 | 0 | 0 | 1477761 | AGT51734.1 | P33469 |
| MSPVDEFIKHVE | 1 | 0 | 0 | 1477764 | AVV62524.1 | K9N7C7 |
| MSQFKLTHQSNE | 1 | 0 | 0 | 1477766 | AVV62533.1 | K9N4V7 |
| MSQFSQEVSDMC | 1 | 0 | 0 | 1477769 | ARU07575.1 | A0A140H1G9 |
| MSQYSRSTRSML | 1 | 0 | 0 | 1477775 | AXN92228.1 | K9N5Q8 |
| MSQYSRSTRSTL | 1 | 0 | 0 | 1477776 | AHL18090.1 | K9N5Q8 |
| MSSVDNILNMAR | 1 | 0 | 0 | 1477794 | AST12962.1 | A0A0P0LKV0 |
| MSTPNAIFISDR | 1 | 0 | 0 | 1477803 | AXN92247.1 | K9N7C7 |
| MSTPNAVFISDR | 1 | 0 | 0 | 1477805 | ATG84897.1 | K9N7C7 |
| MSWLEMADTSLS | 1 | 0 | 0 | 1477814 | APO40578.1 | P0C6X7 |
| MSYYCENHKPHY | 1 | 0 | 0 | 1477827 | ARU07574.1 | A0A140H1G9 |
| MTAVNESRYGLH | 1 | 0 | 0 | 1477838 | ARU07594.1 | A0A0P0G321 |
| MTDCLAAEIHRD | 1 | 0 | 0 | 1477847 | AVV62535.1 | K9N7C7 |
| MTDIAEKPTENA | 1 | 0 | 0 | 1477848 | AVP78041.1 | P0C6W2 |
| MTDIAKKPTEHA | 1 | 0 | 0 | 1477849 | APO40578.1 | P0C6X7 |
| MTDMEKDFLSLD | 1 | 0 | 0 | 1477853 | AVV62535.1 | K9N7C7 |
| MTDMEKDFLSMD | 1 | 0 | 0 | 1477854 | AVV62524.1 | K9N7C7 |
| MTDNGQQGPRNA | 1 | 0 | 0 | 1477855 | APO40586.1 | P59595 |
| MTDNLQMAFVIS | 1 | 0 | 0 | 1477857 | AUM60014.1 | K9N5Q8 |
| MTEQLQMGFGIT | 1 | 0 | 0 | 1477860 | AXN92228.1 | K9N5Q8 |
| MTFGGVVYAYLG | 1 | 0 | 0 | 1477861 | AVV62524.1 | K9N7C7 |
| MTFSGNVDLEGL | 1 | 0 | 0 | 1477866 | AVV62524.1 | K9N7C7 |
| MTFSVHPVVCDE | 1 | 0 | 0 | 1477867 | AUM60013.1 | K9N7C7 |
| MTFYPKMQATND | 1 | 0 | 0 | 1477869 | ARU07574.1 | A0A140H1G9 |
| MTGRVFHQKCLK | 1 | 0 | 0 | 1477878 | AIL49573.1 | P0C6X6 |
| MTHMDDFSIKSI | 1 | 0 | 0 | 1477882 | ARU07575.1 | A0A140H1G9 |
| MTIDALRDIRFT | 1 | 0 | 0 | 1477884 | AUM60013.1 | K9N7C7 |
| MTKLASTEELPD | 1 | 0 | 0 | 1477891 | AAT76156.1 | Q3LZX3 |
| MTKLATIEELPD | 1 | 0 | 0 | 1477892 | APO40587.1 | P59636 |
| MTKRNVIPTMTQ | 1 | 0 | 0 | 1477898 | AXN92247.1 | K9N7C7 |
| MTLDLLKDIKLV | 1 | 0 | 0 | 1477902 | AVV61898.1 | K9N7C7 |
| MTLFSPTTSSFN | 1 | 0 | 0 | 1477904 | APO40579.1 | P59594 |
| MTLNGLWLDNTV | 1 | 0 | 0 | 1477907 | AXN92247.1 | K9N7C7 |
| MTLNGLWLDNYV | 1 | 0 | 0 | 1477908 | AVV62524.1 | K9N7C7 |
| MTNIVDAFTGVN | 1 | 0 | 0 | 1477921 | AIW52757.1 | A0A0P0K6L9 |
| MTPYCHDPTVLP | 1 | 0 | 0 | 1477925 | AXN92247.1 | K9N7C7 |
| MTSQALFDSLEF | 1 | 0 | 0 | 1477950 | AVN89451.1 | K9N7C7 |
| MTTSTTNFGEEF | 1 | 0 | 0 | 1477959 | ABD75599.1 | A0A140H1G9 |
| MTYVCQCGGERH | 1 | 0 | 0 | 1477980 | AXN92247.1 | K9N7C7 |
| MVCKMCLFRYQE | 1 | 0 | 0 | 1477985 | QEG03730.1 | H9EJ42 |
| MVDCRRLLKQDC | 1 | 0 | 0 | 1477988 | AZP73837.1 | P0C6X6 |
| MVDPTTVLNDPV | 1 | 0 | 0 | 1477993 | AWK59942.1 | H9EJ42 |
| MVDPTTVLNGPV | 1 | 0 | 0 | 1477994 | ABE97136.1 | H9EJ42 |
| MVDVFFTDTAWY | 1 | 0 | 0 | 1477998 | BBA20987.1 | Q0ZJ83 |
| MVERSVSLAIDA | 1 | 0 | 0 | 1478008 | ASU90624.1 | K9N7C7 |
| MVETLVGQLEQT | 1 | 0 | 0 | 1478009 | ASU90174.1 | K9N638 |
| MVETPNGPQVKY | 1 | 0 | 0 | 1478010 | AWK59942.1 | H9EJ42 |
| MVGDVINSGDIT | 1 | 0 | 0 | 1478015 | ARO76381.1 | P0C6U8 |
| MVGPDCAYFNGA | 1 | 0 | 0 | 1478020 | AUM60023.1 | K9N7C7 |
| MVHDLNNVDPKA | 1 | 0 | 0 | 1478022 | AVP78050.1 | Q3LZX3 |
| MVKEYPLLTGYP | 1 | 0 | 0 | 1478031 | AET99208.1 | Q0ZJ80 |
| MVKRQDYGTNSG | 1 | 0 | 0 | 1478033 | AVV62543.1 | K9N7A1 |
| MVLESLAATVRL | 1 | 0 | 0 | 1478039 | 3EE7_A | P0C6X7 |
| MVLPGVADAERM | 1 | 0 | 0 | 1478049 | AIW52843.1 | A0A0P0G321 |
| MVLTDVDVPFTA | 1 | 0 | 0 | 1478054 | AGT51637.1 | P0C6X6 |
| MVNKSSCYQDGI | 1 | 0 | 0 | 1478064 | AET99208.1 | Q0ZJ80 |
| MVNMSPKPQRQR | 1 | 0 | 0 | 1478066 | AEN19370.1 | P33469 |
| MVNPSKGSSGLLITKLT | 1 | 0 | 0 | 1478068 | AGW27877.1 | Q0ZJ80 |
| MVNQLAYSSSAS | 1 | 0 | 0 | 1478072 | AVV62535.1 | K9N7C7 |
| MVPADQLADPNY | 1 | 0 | 0 | 1478074 | AUM60013.1 | K9N7C7 |
| MVPVEFYEDRIL | 1 | 0 | 0 | 1478077 | AXN92247.1 | K9N7C7 |
| MVPYCYDPTVLP | 1 | 0 | 0 | 1478081 | AVV62524.1 | K9N7C7 |
| MVQAFGRRGPGD | 1 | 0 | 0 | 1478085 | AVV62533.1 | K9N4V7 |
| MVQTHISHYVLN | 1 | 0 | 0 | 1478090 | AVV62524.1 | K9N7C7 |
| MVQTHLSHYVFN | 1 | 0 | 0 | 1478091 | ALT66812.1 | K9N7C7 |
| MVVDNCQLPLGQ | 1 | 0 | 0 | 1478118 | AVV62537.1 | K9N5Q8 |
| MVVSAGQEQTNC | 1 | 0 | 0 | 1478126 | AXN92247.1 | K9N7C7 |
| MWCLRCLYSTKP | 1 | 0 | 0 | 1478137 | APO40578.1 | P0C6X7 |
| MWVSYFVQSIRL | 1 | 0 | 0 | 1478173 | AVV62532.1 | K9N7A1 |
| MYAISNEQHYKS | 1 | 0 | 0 | 1478178 | ATO98193.1 | Q3LZX1 |
| MYDPVTKNINGS | 1 | 0 | 0 | 1478203 | AVV62524.1 | K9N7C7 |

| | | | | | | |
|---|---|---|---|---|---|---|
| MYFQKAGQSTYE | 1 | 0 | 0 | 1478207 | APO40578.1 | P0C6X7 |
| MYGAFMDKQVHQ | 1 | 0 | 0 | 1478212 | AXN92247.1 | K9N7C7 |
| MYGAFQDRQIYQ | 1 | 0 | 0 | 1478214 | AUM60013.1 | K9N7C7 |
| MYGVNLQSGKVI | 1 | 0 | 0 | 1478219 | AWK59930.1 | H9EJ42 |
| MYKGLPWNIVRV | 1 | 0 | 0 | 1478240 | APO40578.1 | P0C6X7 |
| MYKNGAMLNGKP | 1 | 0 | 0 | 1478243 | AUM60013.1 | K9N7C7 |
| MYKTPTLKDFGG | 1 | 0 | 0 | 1478248 | ATO98231.1 | Q3LZX1 |
| MYKVVGFIRHIV | 1 | 0 | 0 | 1478251 | ARU07575.1 | A0A140H1G9 |
| MYLYSAAHADPN | 1 | 0 | 0 | 1478261 | AUM60014.1 | K9N5Q8 |
| MYMGKCGDKFFP | 1 | 0 | 0 | 1478262 | ARU07575.1 | A0A140H1G9 |
| MYMGTLSYDSLK | 1 | 0 | 0 | 1478265 | ARO76381.1 | P0C6U8 |
| MYMGTLSYEQLK | 1 | 0 | 0 | 1478268 | AVP78041.1 | P0C6W2 |
| MYTILPTFNTRT | 1 | 0 | 0 | 1478297 | AUM60013.1 | K9N7C7 |
| MYTYNITEDEIL | 1 | 0 | 0 | 1478302 | AXN92228.1 | K9N5Q8 |
| MYVNMAVPHIKE | 1 | 0 | 0 | 1478307 | AUM60027.1 | K9N643 |
| MYVYSAGHATGS | 1 | 0 | 0 | 1478315 | AZU90731.1 | K9N5Q8 |
| NAAAYMKLSDAF | 1 | 0 | 0 | 1478320 | ASU91217.1 | K9N7C7 |
| NAAIPDYNYYKY | 1 | 0 | 0 | 1478322 | ACV87150.1 | A0A140H1G9 |
| NAAQCDCLCTPD | 1 | 0 | 0 | 1478332 | CAA83660.1 | P36334 |
| NAASVNPASMRN | 1 | 0 | 0 | 1478335 | AUM60013.1 | K9N7C7 |
| NAASVNQTVLRN | 1 | 0 | 0 | 1478338 | AVN89451.1 | K9N7C7 |
| NAATKRTDNPIL | 1 | 0 | 0 | 1478340 | AVP78031.1 | Q3LZX1 |
| NACTPESKPLAD | 1 | 0 | 0 | 1478350 | ABK63972.1 | H9EJ69 |
| NADAEALNNLLQ | 1 | 0 | 0 | 1478353 | CAA83661.1 | P36334 |
| NADYLYFHFYQE | 1 | 0 | 0 | 1478372 | CAA79896.1 | P36334 |
| NAEAHNNLLQQL | 1 | 0 | 0 | 1478374 | ARB66064.1 | P36334 |
| NAFYGVALKDCG | 1 | 0 | 0 | 1478410 | AEN19363.1 | P0C6X6 |
| NAGYWRRQYRKI | 1 | 0 | 0 | 1478426 | AVN89320.1 | K9N4V7 |
| NAHRYDALLSDL | 1 | 0 | 0 | 1478428 | QED88018.1 | H9EJ42 |
| NAISPKTDNETL | 1 | 0 | 0 | 1478455 | BBA20986.1 | A0A140H1H1 |
| NAKEEILGAVSW | 1 | 0 | 0 | 1478465 | BAE93400.1 | P0C6U8 |
| NAKEIRHKILMK | 1 | 0 | 0 | 1478467 | AMM42397.1 | Q19U29 |
| NAKIVTLPGSDA | 1 | 0 | 0 | 1478471 | ATQ39390.1 | K9N5Q8 |
| NAKLGFNVDLPF | 1 | 0 | 0 | 1478473 | ABD75519.1 | A0A140H1G9 |
| NAKWIFDSSKEF | 1 | 0 | 0 | 1478477 | AIV41816.1 | P0C6X6 |
| NALFCATCPFAK | 1 | 0 | 0 | 1478484 | ASU91323.1 | K9N7C7 |
| NALHAIQQGSDA | 1 | 0 | 0 | 1478488 | ARB66064.1 | P36334 |
| NALLCVTCPFAK | 1 | 0 | 0 | 1478493 | ATG84865.1 | K9N638 |
| NALMQGVTLSSN | 1 | 0 | 0 | 1478499 | BBA20986.1 | A0A140H1H1 |
| NANAKALSKLAS | 1 | 0 | 0 | 1478517 | AVV62526.1 | K9N5Q8 |
| NANGACVWNAAA | 1 | 0 | 0 | 1478524 | AVV62535.1 | K9N7C7 |
| NANSEALNNLLQ | 1 | 0 | 0 | 1478542 | AWW13566.1 | P36334 |
| NAPATVCGPKKS | 1 | 0 | 0 | 1478550 | APO40579.1 | P59594 |
| NAPRITFGVSDN | 1 | 0 | 0 | 1478560 | APO40586.1 | P59595 |
| NAPYYNSWQNLL | 1 | 0 | 0 | 1478566 | CAA83660.1 | P36334 |
| NAQGKPVPYCYD | 1 | 0 | 0 | 1478570 | APO40578.1 | P0C6X7 |
| NAQVAKSHSVSL | 1 | 0 | 0 | 1478578 | APO40578.1 | P0C6X7 |
| NARQFDNMTAVA | 1 | 0 | 0 | 1478591 | AXX83326.1 | P30215 |
| NASFDNFRFVCD | 1 | 0 | 0 | 1478604 | AVP78041.1 | P0C6W2 |
| NASGNDALARFT | 1 | 0 | 0 | 1478605 | ARB07436.1 | A0A140H1G9 |
| NASISESKPLAD | 1 | 0 | 0 | 1478608 | AVL25595.1 | H9EJ69 |
| NASKADGIIYPT | 1 | 0 | 0 | 1478610 | AVV62537.1 | K9N5Q8 |
| NASRFPNVYAWE | 1 | 0 | 0 | 1478623 | ATO98181.1 | Q3LZX1 |
| NASVVDIKKEIA | 1 | 0 | 0 | 1478640 | APO40579.1 | P59594 |
| NASYECDIPIGA | 1 | 0 | 0 | 1478643 | AVP78031.1 | Q3LZX1 |
| NATFLDLQVEMN | 1 | 0 | 0 | 1478654 | QEG03776.1 | P36334 |
| NATFLDLYYEMN | 1 | 0 | 0 | 1478655 | BBA20983.1 | A0A140H1H1 |
| NATNCTYNRVHG | 1 | 0 | 0 | 1478663 | APO40579.1 | P59594 |
| NAVALNTLVKQL | 1 | 0 | 0 | 1478680 | APO40579.1 | P59594 |
| NAVCSVIDLKLD | 1 | 0 | 0 | 1478685 | ANI69909.1 | K9N7C7 |
| NAVDCMSDFMSG | 1 | 0 | 0 | 1478688 | AIL49491.1 | P36334 |
| NAVEQRITVEVF | 1 | 0 | 0 | 1478692 | AVV62535.1 | K9N7C7 |
| NAVLTIANNYSK | 1 | 0 | 0 | 1478700 | AST12962.1 | A0A0P0LKV0 |
| NAVMQNNELMPH | 1 | 0 | 0 | 1478702 | ARU07575.1 | A0A140H1G9 |
| NAVPANSLHATA | 1 | 0 | 0 | 1478703 | APO40580.1 | P59632 |
| NAVQGFACDCGA | 1 | 0 | 0 | 1478706 | AVV62524.1 | K9N7C7 |
| NAVVKQLNESYI | 1 | 0 | 0 | 1478720 | AVV62526.1 | K9N5Q8 |
| NAVYDGVRLFAS | 1 | 0 | 0 | 1478725 | AMM42570.1 | H9EJ42 |
| NAVYTKMDGVDV | 1 | 0 | 0 | 1478727 | APO40578.1 | P0C6X7 |
| NAYNNTAISVQT | 1 | 0 | 0 | 1478744 | AIL49538.1 | P36334 |
| NAYNSTSISVQT | 1 | 0 | 0 | 1478749 | ARK08679.1 | P36334 |
| NAYQQQDGMMNI | 1 | 0 | 0 | 1478752 | AIX10729.1 | P33469 |
| NAYSPIDIPRGL | 1 | 0 | 0 | 1478758 | ATO98193.1 | Q3LZX1 |
| NAYYTQQPIDLV | 1 | 0 | 0 | 1478769 | APO40578.1 | P0C6X7 |
| NCAGNASGNDAL | 1 | 0 | 0 | 1478771 | ARB07436.1 | A0A140H1G9 |
| NCCDDNYFSKKD | 1 | 0 | 0 | 1478783 | APO40578.1 | P0C6X7 |
| NCCDGDYFNKKD | 1 | 0 | 0 | 1478785 | AVP78041.1 | P0C6W2 |
| NCESIYEFDTYK | 1 | 0 | 0 | 1478819 | ARU07575.1 | A0A140H1G9 |
| NCFICHSFKPGN | 1 | 0 | 0 | 1478824 | ABD75543.1 | A0A140H1G9 |
| NCGQKSTTLIGV | 1 | 0 | 0 | 1478838 | APO40578.1 | P0C6X7 |
| NCIGKGSLTASS | 1 | 0 | 0 | 1478846 | AUM60027.1 | K9N643 |
| NCLASECFMKSD | 1 | 0 | 0 | 1478859 | AIW52776.1 | A0A0P0LKV0 |
| NCNGYQHNSFAD | 1 | 0 | 0 | 1478903 | ALJ53431.1 | A0A0P0G321 |
| NCNIEAWLNGKS | 1 | 0 | 0 | 1478910 | AIL49492.1 | P36334 |
| NCNIESWLNDKS | 1 | 0 | 0 | 1478911 | AIX10749.1 | P36334 |
| NCNWFVQSDKCS | 1 | 0 | 0 | 1478929 | AGT51787.1 | P0C6X6 |
| NCPLDPRLKGSF | 1 | 0 | 0 | 1478936 | BAM16463.1 | P36334 |
| NCQLPLGQSLCA | 1 | 0 | 0 | 1478945 | AVV62537.1 | K9N5Q8 |
| NCSDCYDDMCVI | 1 | 0 | 0 | 1478954 | AIW52827.1 | H9EJ42 |
| NCSDGNYNRNAS | 1 | 0 | 0 | 1478956 | ALA49671.1 | K9N5Q8 |
| NCTAVGVKQQRF | 1 | 0 | 0 | 1478968 | AVV62526.1 | K9N5Q8 |

| | | | | | | |
|---|---|---|---|---|---|---|
| NCTDVPTMIHVE | 1 | 0 | 0 | 1478977 | APO40579.1 | P59594 |
| NCTDVPTTIHAD | 1 | 0 | 0 | 1478979 | AVP78031.1 | Q3LZX1 |
| NCTFGDPDDASR | 1 | 0 | 0 | 1478991 | AVN89333.1 | K9N7C7 |
| NCTFGSPDDASR | 1 | 0 | 0 | 1478992 | AHI48794.1 | K9N638 |
| NCTFMYIYNITE | 1 | 0 | 0 | 1478993 | AID55097.1 | K9N5Q8 |
| NCTFMYTFNITE | 1 | 0 | 0 | 1478994 | AQZ41296.1 | K9N5Q8 |
| NCTFQEAAANIF | 1 | 0 | 0 | 1478996 | AVV62535.1 | K9N7C7 |
| NCTYVDTLRDLP | 1 | 0 | 0 | 1479018 | ATO98145.1 | Q3LZX1 |
| NCVDCDIAGVGN | 1 | 0 | 0 | 1479022 | AXN92247.1 | K9N7C7 |
| NCVDCDTAGIGN | 1 | 0 | 0 | 1479023 | AVV62524.1 | K9N7C7 |
| NCVDCDTAGVGN | 1 | 0 | 0 | 1479025 | AHI48798.1 | K9N7C7 |
| NCVLKLKVDTAN | 1 | 0 | 0 | 1479032 | AVP78041.1 | P0C6W2 |
| NCYRNSNVDEAF | 1 | 0 | 0 | 1479057 | ALJ99927.1 | A0A0P0LKV0 |
| NCYRNSNVGESF | 1 | 0 | 0 | 1479058 | ADO24705.1 | A0A0P0LKV0 |
| NDADKANFLNAV | 1 | 0 | 0 | 1479074 | AXX83300.1 | P0C6X6 |
| NDAITRTAEAIH | 1 | 0 | 0 | 1479081 | QED88026.1 | A0A0P0G321 |
| NDDDYGIALTGT | 1 | 0 | 0 | 1479097 | QED88010.1 | H9EJ42 |
| NDDLYVKRYERG | 1 | 0 | 0 | 1479108 | AEN19363.1 | P0C6X6 |
| NDEESVTGDNDD | 1 | 0 | 0 | 1479120 | P0C6U5.1 | A0A140H1G9 |
| NDFSSEIIGYKA | 1 | 0 | 0 | 1479135 | AVP78041.1 | P0C6W2 |
| NDGRVDDDYICG | 1 | 0 | 0 | 1479148 | QED88011.1 | H9EJ42 |
| NDILSRLDPPEA | 1 | 0 | 0 | 1479161 | 6CRW_A | P59594 |
| NDKDTGPPSIST | 1 | 0 | 0 | 1479182 | ACV60203.1 | P36334 |
| NDKLGFNVDLPF | 1 | 0 | 0 | 1479189 | ARU07575.1 | A0A140H1G9 |
| NDKNICLLDHAW | 1 | 0 | 0 | 1479191 | P0C6U5.1 | A0A140H1G9 |
| NDKNTTVPRISE | 1 | 0 | 0 | 1479193 | BBA20983.1 | A0A140H1H1 |
| NDLLDITQLQVA | 1 | 0 | 0 | 1479211 | BBA20986.1 | A0A140H1H1 |
| NDLLQPNTEVYT | 1 | 0 | 0 | 1479214 | AGW27863.1 | A0A140H1H1 |
| NDLNTTIPRISE | 1 | 0 | 0 | 1479219 | ACV87140.1 | A0A140H1H1 |
| NDNKIMTFYPKM | 1 | 0 | 0 | 1479245 | ARU07574.1 | A0A140H1G9 |
| NDNNNQQSRGRG | 1 | 0 | 0 | 1479248 | AUM60021.1 | K9N4V7 |
| NDNNSNNQQSRG | 1 | 0 | 0 | 1479249 | AVV62533.1 | K9N4V7 |
| NDNNTNQQSRGR | 1 | 0 | 0 | 1479250 | AVV62544.1 | K9N4V7 |
| NDNVVYLFDEAG | 1 | 0 | 0 | 1479256 | ARB07605.1 | A0A140H1G9 |
| NDPAIVTQFAPG | 1 | 0 | 0 | 1479257 | AUM60021.1 | K9N4V7 |
| NDPCNVCQYWKG | 1 | 0 | 0 | 1479258 | AVV62535.1 | K9N7C7 |
| NDPIRSCLENTV | 1 | 0 | 0 | 1479260 | CAA49377.1 | A0A0P0LKV0 |
| NDQEMTTPPPKE | 1 | 0 | 0 | 1479269 | ATQ39393.1 | K9N4V7 |
| NDSCPGKNNGIG | 1 | 0 | 0 | 1479278 | AIV41891.1 | P36334 |
| NDTITVDQLKHL | 1 | 0 | 0 | 1479297 | APO40582.1 | P59596 |
| NDTKIASQLGMG | 1 | 0 | 0 | 1479298 | AWF93655.1 | K9N5Q8 |
| NDTKIVSQLGNC | 1 | 0 | 0 | 1479300 | AVN89313.1 | K9N5Q8 |
| NDTNGDPVYFLR | 1 | 0 | 0 | 1479302 | AUM60021.1 | K9N4V7 |
| NDTSETKYVKSL | 1 | 0 | 0 | 1479303 | AOL02450.1 | P0C6X6 |
| NDTTNTNLSRGR | 1 | 0 | 0 | 1479305 | ALW82676.1 | K9N4V7 |
| NDTVDVTNGLGT | 1 | 0 | 0 | 1479308 | ANZ78836.1 | P36334 |
| NDVSSMLTFDSN | 1 | 0 | 0 | 1479333 | AIW52843.1 | A0A0P0G321 |
| NDWKPGLAMPSL | 1 | 0 | 0 | 1479341 | AVV62535.1 | K9N7C7 |
| NDWNTKHGSGLK | 1 | 0 | 0 | 1479344 | APO40578.1 | P0C6X7 |
| NDWQNLLYDSNG | 1 | 0 | 0 | 1479345 | AZS52618.1 | A0A140H1H1 |
| NDYNKTIPRISE | 1 | 0 | 0 | 1479350 | ARB07438.1 | A0A140H1H1 |
| NEAFHKVQDAVN | 1 | 0 | 0 | 1479357 | AHI48803.1 | K9N5Q8 |
| NECTERLKLFAE | 1 | 0 | 0 | 1479375 | AIV41984.1 | P0C6X6 |
| NECVESQSKRYG | 1 | 0 | 0 | 1479376 | ABB90528.1 | A0A0P0K6L9 |
| NEDLAVCLNICE | 1 | 0 | 0 | 1479388 | ARB07436.1 | A0A140H1G9 |
| NEEKCGTQLNHT | 1 | 0 | 0 | 1479396 | BBA20986.1 | A0A140H1H1 |
| NEELEQTIQQKE | 1 | 0 | 0 | 1479397 | APO40578.1 | P0C6X7 |
| NEFINFSDSFYT | 1 | 0 | 0 | 1479405 | AVV62524.1 | K9N7C7 |
| NEGLLHFVRSFI | 1 | 0 | 0 | 1479416 | APO40578.1 | P0C6X7 |
| NEHVTSVNLLDV | 1 | 0 | 0 | 1479422 | AHI48551.1 | K9N796 |
| NEIRPTGLKTMV | 1 | 0 | 0 | 1479438 | AVV62524.1 | K9N7C7 |
| NEKLVTTSTAPD | 1 | 0 | 0 | 1479457 | AXN92247.1 | K9N7C7 |
| NEKLVTTSTEPD | 1 | 0 | 0 | 1479458 | ATG84711.1 | K9N7C7 |
| NEKRPDHVSLTV | 1 | 0 | 0 | 1479463 | AVV62535.1 | K9N7C7 |
| NELMPVKLKIQV | 1 | 0 | 0 | 1479475 | AXX83324.1 | P0C6X6 |
| NENGTITDAIDC | 1 | 0 | 0 | 1479487 | ATO98181.1 | Q3LZX1 |
| NENGTMSNAVDC | 1 | 0 | 0 | 1479489 | ATO98145.1 | Q3LZX1 |
| NENLTWPLGLEC | 1 | 0 | 0 | 1479492 | AHI48747.1 | K9N638 |
| NEQHYKSWVYQN | 1 | 0 | 0 | 1479503 | ATO98193.1 | Q3LZX1 |
| NESIACQFCLAG | 1 | 0 | 0 | 1479511 | AXX83294.1 | P0C6X6 |
| NESWFEDDYKAF | 1 | 0 | 0 | 1479519 | AST12962.1 | A0A0P0LKV0 |
| NESWLTGNAIQG | 1 | 0 | 0 | 1479523 | AVN89333.1 | K9N7C7 |
| NETNTRDVDQQW | 1 | 0 | 0 | 1479536 | AVV62526.1 | K9N5Q8 |
| NEVNDLLDITQL | 1 | 0 | 0 | 1479546 | BBA20986.1 | A0A140H1H1 |
| NEVWQKYTNSAS | 1 | 0 | 0 | 1479551 | AUM60016.1 | K9N4V0 |
| NEWAIANGYTSV | 1 | 0 | 0 | 1479552 | AGT51358.1 | H9EJ42 |
| NEWAMANQFTEF | 1 | 0 | 0 | 1479554 | AVV62524.1 | K9N7C7 |
| NEWAQANGFTVM | 1 | 0 | 0 | 1479556 | QEO75983.1 | A0A0P0LKV0 |
| NFAWSIIFLLIT | 1 | 0 | 0 | 1479580 | AVV62543.1 | K9N7A1 |
| NFCPCKSNGSCP | 1 | 0 | 0 | 1479602 | AEN19358.1 | P36334 |
| NFCPCSSCFGKN | 1 | 0 | 0 | 1479604 | QEG03814.1 | P36334 |
| NFCPCSSCHGKN | 1 | 0 | 0 | 1479605 | ARB66068.1 | P36334 |
| NFCPCSSCPGKN | 1 | 0 | 0 | 1479606 | AWW13566.1 | P36334 |
| NFCPCSSCPSKN | 1 | 0 | 0 | 1479607 | QDH43719.1 | P36334 |
| NFCPCSSCSGKN | 1 | 0 | 0 | 1479609 | ANZ78838.1 | P36334 |
| NFCYMHQMELAS | 1 | 0 | 0 | 1479613 | AVN89451.1 | K9N7C7 |
| NFDNNQNGDRTG | 1 | 0 | 0 | 1479622 | APO40586.1 | P59595 |
| NFDQKELLDYYN | 1 | 0 | 0 | 1479624 | ATN39862.1 | P0C6X6 |
| NFEHKDITECVT | 1 | 0 | 0 | 1479639 | ALA49659.1 | K9N7C7 |
| NFELCDNPFIAV | 1 | 0 | 0 | 1479649 | ABD73001.1 | P59594 |
| NFEPFTVNSIND | 1 | 0 | 0 | 1479655 | AIL49542.1 | P36334 |

| | | | | | | |
|---|---|---|---|---|---|---|
| NFEPFTVNSKND | 1 | 0 | 0 | 1479656 | AIL49538.1 | P36334 |
| NFEPFTVNSVSD | 1 | 0 | 0 | 1479658 | ARB66068.1 | P36334 |
| NFFFLTHINTTL | 1 | 0 | 0 | 1479663 | ANI69922.1 | K9N5Q8 |
| NFGDLQLTKLGI | 1 | 0 | 0 | 1479680 | QBM11743.1 | K9N4V7 |
| NFGDVNKPVCAK | 1 | 0 | 0 | 1479690 | QED88038.1 | H9EJ42 |
| NFIDKLNVLFDG | 1 | 0 | 0 | 1479712 | ARK08667.1 | P0C6X6 |
| NFIIYDASDRGQ | 1 | 0 | 0 | 1479714 | AVV62524.1 | K9N7C7 |
| NFILHDVNNGLT | 1 | 0 | 0 | 1479723 | AEN19366.1 | P36334 |
| NFILHDVNSGTT | 1 | 0 | 0 | 1479725 | CAA83660.1 | P36334 |
| NFINNNLALGGS | 1 | 0 | 0 | 1479727 | AVV62524.1 | K9N7C7 |
| NFKDLREYVFKN | 1 | 0 | 0 | 1479737 | ATO98181.1 | Q3LZX1 |
| NFKRERSLSLQK | 1 | 0 | 0 | 1479759 | AGT51735.1 | Q4VID0 |
| NFLGYHPAHAPS | 1 | 0 | 0 | 1479769 | ARB07436.1 | A0A140H1G9 |
| NFLSDILRVHTV | 1 | 0 | 0 | 1479787 | AUM60013.1 | K9N7C7 |
| NFLSSFFGLHTV | 1 | 0 | 0 | 1479789 | AVV62535.1 | K9N7C7 |
| NFNENGTITDAI | 1 | 0 | 0 | 1479797 | ATO98193.1 | Q3LZX1 |
| NFNETNGPLCVD | 1 | 0 | 0 | 1479798 | CAA71147.1 | A0A0P0K6L9 |
| NFNFNGLKGTGV | 1 | 0 | 0 | 1479799 | AVP78031.1 | Q3LZX1 |
| NFNFSSNSVDNL | 1 | 0 | 0 | 1479802 | AVA26873.1 | A0A0P0G321 |
| NFNMSCLMSFIQ | 1 | 0 | 0 | 1479811 | AHA80204.1 | P36334 |
| NFNPSSWNRRYG | 1 | 0 | 0 | 1479817 | BBA20986.1 | A0A140H1H1 |
| NFNQYYYNMATG | 1 | 0 | 0 | 1479824 | AZS52617.1 | A0A140H1H0 |
| NFNRLVFTNCNY | 1 | 0 | 0 | 1479825 | AVV62526.1 | K9N5Q8 |
| NFNTLFSTTIPN | 1 | 0 | 0 | 1479830 | AIW52776.1 | A0A0P0LKV0 |
| NFNVDDINFAPV | 1 | 0 | 0 | 1479831 | QEG03814.1 | P36334 |
| NFNVTNAETTDF | 1 | 0 | 0 | 1479842 | AAQ90006.1 | A0A0P0K6L9 |
| NFPKKEKKQKAS | 1 | 0 | 0 | 1479845 | AUM60021.1 | K9N4V7 |
| NFRDLALKGSTY | 1 | 0 | 0 | 1479870 | BBA20986.1 | A0A140H1H1 |
| NFRIDIPSAELT | 1 | 0 | 0 | 1479877 | AXP07346.1 | K9N796 |
| NFRTEVDPDLSA | 1 | 0 | 0 | 1479886 | AXN92247.1 | K9N7C7 |
| NFRVSPSTEVVR | 1 | 0 | 0 | 1479897 | APO40579.1 | P59594 |
| NFSDGKMGLFFN | 1 | 0 | 0 | 1479905 | ANF29217.1 | K9N5Q8 |
| NFSGALAEHDFF | 1 | 0 | 0 | 1479915 | AIW52827.1 | H9EJ42 |
| NFSNLDKLKFKE | 1 | 0 | 0 | 1479927 | AVV62524.1 | K9N7C7 |
| NFSQILPDPSKPSKRS | 1 | 0 | 0 | 1479936 | P0DTC2.1 | P0DTC2 |
| NFSQILPDSSKP | 1 | 0 | 0 | 1479938 | ARO76382.1 | P59594 |
| NFSSNSVDNLKS | 1 | 0 | 0 | 1479943 | AIW52843.1 | A0A0P0G321 |
| NFSSVLGCLGSG | 1 | 0 | 0 | 1479946 | BBA20982.1 | P36334 |
| NFTKAPFIYLNN | 1 | 0 | 0 | 1479971 | BBA20986.1 | A0A140H1H1 |
| NFVFPLNSTIKT | 1 | 0 | 0 | 1480005 | AVP78041.1 | P0C6W2 |
| NFVGEFVINDVV | 1 | 0 | 0 | 1480006 | ATQ39389.1 | K9N7C7 |
| NFVKQGHFIGVE | 1 | 0 | 0 | 1480009 | AVK87459.1 | K9N7C7 |
| NFYFCQDPMLAV | 1 | 0 | 0 | 1480032 | APO40579.1 | P59594 |
| NFYQHVINGCKD | 1 | 0 | 0 | 1480037 | AZK15899.1 | K9N638 |
| NFYSPQIITTDN | 1 | 0 | 0 | 1480038 | ATO98181.1 | Q3LZX1 |
| NFYSPQVITTDN | 1 | 0 | 0 | 1480040 | APO40579.1 | P59594 |
| NFYYVSNGSNSC | 1 | 0 | 0 | 1480045 | AWO69003.1 | A0A0P0G321 |
| NGAITVEELKQL | 1 | 0 | 0 | 1480059 | BAE93405.1 | P59596 |
| NGALEKFMSTSN | 1 | 0 | 0 | 1480063 | AVV62524.1 | K9N7C7 |
| NGAVQLESDQYL | 1 | 0 | 0 | 1480081 | AVV62535.1 | K9N7C7 |
| NGCNWFVKPSRT | 1 | 0 | 0 | 1480092 | AUM60013.1 | K9N7C7 |
| NGCVGHSENVFA | 1 | 0 | 0 | 1480102 | AIW52757.1 | A0A0P0K6L9 |
| NGCVPLSAVPLC | 1 | 0 | 0 | 1480106 | AVV62524.1 | K9N7C7 |
| NGFFYIDVGKRR | 1 | 0 | 0 | 1480142 | AKA09629.1 | A0A0P0G321 |
| NGFICEKLAIGG | 1 | 0 | 0 | 1480146 | NP_073549.1 | A0A0P0LKV0 |
| NGFQGRQLLGSC | 1 | 0 | 0 | 1480159 | AIV41834.1 | P0C6X6 |
| NGFQKCEQLLLE | 1 | 0 | 0 | 1480161 | QAT98898.1 | K9N5Q8 |
| NGFTAMNGEDAF | 1 | 0 | 0 | 1480168 | NP_073549.1 | A0A0P0LKV0 |
| NGFVYFNGTGRG | 1 | 0 | 0 | 1480178 | AAQ90005.1 | A0A0P0K6L9 |
| NGGDTALALLLL | 1 | 0 | 0 | 1480187 | AVP78038.1 | Q3LZX4 |
| NGGTSFCCRHNW | 1 | 0 | 0 | 1480203 | AVV62535.1 | K9N7C7 |
| NGGVVTRYVDNN | 1 | 0 | 0 | 1480205 | ARO76381.1 | P0C6U8 |
| NGHKIEDFSIRN | 1 | 0 | 0 | 1480211 | AIG96659.1 | A0A140H1G9 |
| NGIAYITFSGSF | 1 | 0 | 0 | 1480218 | ABE97129.1 | H9EJ42 |
| NGIFAKVKNTKA | 1 | 0 | 0 | 1480222 | AKA09643.1 | P36334 |
| NGIGTCPAGTNY | 1 | 0 | 0 | 1480231 | CAA79896.1 | P36334 |
| NGIGTRPAGTNS | 1 | 0 | 0 | 1480233 | ARB66066.1 | P36334 |
| NGIQYGRSGETL | 1 | 0 | 0 | 1480252 | APO40578.1 | P0C6X7 |
| NGISTCPAGTNS | 1 | 0 | 0 | 1480258 | BBA84094.1 | P36334 |
| NGKIVYAILSDV | 1 | 0 | 0 | 1480266 | AEN19363.1 | P0C6X6 |
| NGKPCTPPALNC | 1 | 0 | 0 | 1480269 | ABD72972.1 | P59594 |
| NGKSSLSLKDKI | 1 | 0 | 0 | 1480277 | AUM60013.1 | K9N7C7 |
| NGKSVPSPLNWE | 1 | 0 | 0 | 1480278 | AIL49492.1 | P36334 |
| NGLHPTQSPTYL | 1 | 0 | 0 | 1480299 | APO40578.1 | P0C6X7 |
| NGLTCSTDLQKA | 1 | 0 | 0 | 1480318 | AEN19366.1 | P36334 |
| NGMGTCPAGTNY | 1 | 0 | 0 | 1480337 | AIL49487.1 | P36334 |
| NGMKVVDYLTRD | 1 | 0 | 0 | 1480340 | AVV62535.1 | K9N7C7 |
| NGMNGRTILGAA | 1 | 0 | 0 | 1480341 | 3M3V_A | P0C6X7 |
| NGMNTSQSVCNG | 1 | 0 | 0 | 1480345 | ABB90520.1 | A0A0P0K6L9 |
| NGMPVSSIKSDK | 1 | 0 | 0 | 1480348 | AGT51393.1 | H9EJ42 |
| NGNFDCALSRYN | 1 | 0 | 0 | 1480351 | AVV62524.1 | K9N7C7 |
| NGNKIGDLSIRA | 1 | 0 | 0 | 1480359 | AJE26218.1 | P0C6X6 |
| NGNVSTVFVPAM | 1 | 0 | 0 | 1480371 | AKN24834.1 | K9N7D2 |
| NGPGIDAGYKNS | 1 | 0 | 0 | 1480379 | CAA83660.1 | P36334 |
| NGPHEFCSQHTM | 1 | 0 | 0 | 1480380 | QUH45707.1 | P0DTD1 |
| NGPHEFCTKHTM | 1 | 0 | 0 | 1480381 | AIL49625.1 | P0C6X6 |
| NGPQNQSSAPRI | 1 | 0 | 0 | 1480393 | AVP78049.1 | Q3LZX4 |
| NGSAQAISFCKS | 1 | 0 | 0 | 1480441 | QBP84748.1 | P30215 |
| NGSAQATALCKS | 1 | 0 | 0 | 1480443 | AGT51769.1 | P30215 |
| NGSAQATSFCKS | 1 | 0 | 0 | 1480444 | AWW13558.1 | P30215 |
| NGSAQVTSFCKS | 1 | 0 | 0 | 1480446 | QBP84703.1 | P30215 |

| | | | | | | |
|---|---|---|---|---|---|---|
| NGSCPDKNNGIG | 1 | 0 | 0 | 1480449 | AIX10751.1 | P36334 |
| NGSCPGKNNGIG | 1 | 0 | 0 | 1480450 | AEN19366.1 | P36334 |
| NGSDVLDQPPTA | 1 | 0 | 0 | 1480452 | AIV41798.1 | P0C6X6 |
| NGSMIEDVDYVP | 1 | 0 | 0 | 1480463 | AMQ49035.1 | K9N7C7 |
| NGSNEAKALFFT | 1 | 0 | 0 | 1480465 | AVV62524.1 | K9N7C7 |
| NGSYCPGNSNYN | 1 | 0 | 0 | 1480481 | AVV62537.1 | K9N5Q8 |
| NGTHIVINVCNF | 1 | 0 | 0 | 1480493 | APO40579.1 | P59594 |
| NGTHTGSSFDGN | 1 | 0 | 0 | 1480496 | AVV62524.1 | K9N7C7 |
| NGTISVEELKRL | 1 | 0 | 0 | 1480501 | ATO98197.1 | Q3LZX9 |
| NGTITDAIDCAQ | 1 | 0 | 0 | 1480502 | ATO98181.1 | Q3LZX1 |
| NGTITVEKLKQL | 1 | 0 | 0 | 1480507 | AAS48455.1 | P59596 |
| NGTNTSHSVCKG | 1 | 0 | 0 | 1480518 | AIW52752.1 | A0A0P0K6L9 |
| NGTNTSHSVCNG | 1 | 0 | 0 | 1480519 | AIW52757.1 | A0A0P0K6L9 |
| NGTPLEFGASTQ | 1 | 0 | 0 | 1480522 | APO40578.1 | P0C6X7 |
| NGVDAKGFPLLA | 1 | 0 | 0 | 1480531 | AMM42619.1 | H9EJ69 |
| NGVDAKGFPQFA | 1 | 0 | 0 | 1480532 | AGT51335.1 | H9EJ69 |
| NGVDVSDTIPQQ | 1 | 0 | 0 | 1480536 | AVV62535.1 | K9N7C7 |
| NGVDYTKKFLTV | 1 | 0 | 0 | 1480537 | AVN89441.1 | K9N7C7 |
| NGVEGELPVAVV | 1 | 0 | 0 | 1480539 | AVV62524.1 | K9N7C7 |
| NGVGITQQVLSK | 1 | 0 | 0 | 1480546 | AVN89376.1 | K9N5Q8 |
| NGVIRDINSNDK | 1 | 0 | 0 | 1480556 | AVV62524.1 | K9N7C7 |
| NGVPVVEYLTKD | 1 | 0 | 0 | 1480567 | AUM60013.1 | K9N7C7 |
| NGVRTLSTYDFN | 1 | 0 | 0 | 1480571 | AVP78031.1 | Q3LZX1 |
| NGVSDLSLKQKL | 1 | 0 | 0 | 1480575 | ASU90779.1 | K9N638 |
| NGVYWVASNQAD | 1 | 0 | 0 | 1480590 | AEN19370.1 | P33469 |
| NGWQNLLYDSNG | 1 | 0 | 0 | 1480592 | ABU39950.1 | P36334 |
| NGYFIKTNNTMI | 1 | 0 | 0 | 1480598 | ATG84866.1 | K9N5Q8 |
| NGYFIKTNNTRV | 1 | 0 | 0 | 1480600 | ALA49803.1 | K9N5Q8 |
| NGYLEAVNFNVT | 1 | 0 | 0 | 1480606 | AMM42536.1 | A0A0P0K6L9 |
| NGYPGRYFNHTL | 1 | 0 | 0 | 1480612 | AVV62526.1 | K9N5Q8 |
| NGYPVQPIADVY | 1 | 0 | 0 | 1480616 | AMM42466.1 | P36334 |
| NGYQHNSFADVM | 1 | 0 | 0 | 1480620 | ALJ53431.1 | A0A0P0G321 |
| NGYQHNSVVDVM | 1 | 0 | 0 | 1480622 | YP_003767.1 | A0A0P0G321 |
| NGYVTSSSKSSE | 1 | 0 | 0 | 1480633 | APO40578.1 | P0C6X7 |
| NGYYPTSGSTYL | 1 | 0 | 0 | 1480635 | BBA20982.1 | P36334 |
| NHFKELWHMDTG | 1 | 0 | 0 | 1480676 | AVQ05264.1 | P36334 |
| NHFKESWHMDTG | 1 | 0 | 0 | 1480679 | AIL49486.1 | P36334 |
| NHFRHPLEEDDI | 1 | 0 | 0 | 1480682 | AVV62535.1 | K9N7C7 |
| NHIEVVSAYGLC | 1 | 0 | 0 | 1480698 | AXN92228.1 | K9N5Q8 |
| NHINNLKVKNYG | 1 | 0 | 0 | 1480707 | ABD75544.1 | A0A140H1H0 |
| NHLHSVHNWLKP | 1 | 0 | 0 | 1480716 | AWU59324.1 | K9N643 |
| NHNFLVQAGNVQ | 1 | 0 | 0 | 1480731 | AVP78041.1 | P0C6W2 |
| NHNMGDSDLVQN | 1 | 0 | 0 | 1480732 | AIW52805.1 | H9EJ69 |
| NHPIQIDQLNSS | 1 | 0 | 0 | 1480737 | ASU91208.1 | K9N5Q8 |
| NHPIQVDQFNSS | 1 | 0 | 0 | 1480739 | ASU90362.1 | K9N5Q8 |
| NHPIQVDQLNST | 1 | 0 | 0 | 1480741 | ANI69889.1 | K9N5Q8 |
| NHQFVTASSATD | 1 | 0 | 0 | 1480749 | AWK59942.1 | H9EJ42 |
| NHSGSNDIWMQN | 1 | 0 | 0 | 1480763 | AQN78687.1 | P30215 |
| NHTGITGDNVET | 1 | 0 | 0 | 1480786 | ARO76381.1 | P0C6U8 |
| NHTIKGAFLNGS | 1 | 0 | 0 | 1480790 | 3F9E_A | P0C6U8 |
| NHVMVDCRRLLK | 1 | 0 | 0 | 1480816 | AEN19363.1 | P0C6X6 |
| NICFEATTATIN | 1 | 0 | 0 | 1480842 | AVV62535.1 | K9N7C7 |
| NICQAVSDNVRA | 1 | 0 | 0 | 1480848 | ADO24673.1 | P0C6X6 |
| NICQAVTANVRS | 1 | 0 | 0 | 1480850 | ADO24641.1 | A0A140H1G9 |
| NIDAAKIYGMGF | 1 | 0 | 0 | 1480855 | AIL49487.1 | P36334 |
| NIDMGVFINKYG | 1 | 0 | 0 | 1480869 | AGT51365.1 | H9EJ42 |
| NIDTGSFYGFSN | 1 | 0 | 0 | 1480872 | ABD75600.1 | A0A140H1H0 |
| NIDVAKIYGMGF | 1 | 0 | 0 | 1480873 | ABD43176.1 | P36334 |
| NIEEFIQTSSPK | 1 | 0 | 0 | 1480883 | BBA20979.1 | P36334 |
| NIEKPKPDGVSK | 1 | 0 | 0 | 1480885 | AVV62526.1 | K9N5Q8 |
| NIESWLNDKSVP | 1 | 0 | 0 | 1480886 | AIX10749.1 | P36334 |
| NIFQAVSSNINM | 1 | 0 | 0 | 1480901 | AGH06535.1 | H9EJ42 |
| NIFSSTFNVPME | 1 | 0 | 0 | 1480906 | AVP78041.1 | P0C6W2 |
| NIFSVQEDGRIP | 1 | 0 | 0 | 1480907 | ABE97137.1 | A0A0P0G321 |
| NIFSVQQDGRTP | 1 | 0 | 0 | 1480910 | AKA09627.1 | A0A0P0G321 |
| NIFTPLLKPVGT | 1 | 0 | 0 | 1480911 | APO40578.1 | P0C6X7 |
| NIGSNKTLLTPL | 1 | 0 | 0 | 1480932 | APO40578.1 | P0C6X7 |
| NIHAVLDTLLAS | 1 | 0 | 0 | 1480938 | AXN92247.1 | K9N7C7 |
| NIHSSRIAGRSA | 1 | 0 | 0 | 1480945 | AIW52843.1 | A0A0P0G321 |
| NIIACLWFLRRF | 1 | 0 | 0 | 1480949 | AUM60013.1 | K9N7C7 |
| NIIDAATSYGMC | 1 | 0 | 0 | 1480951 | AIV41909.1 | P36334 |
| NIKFADDLNQLS | 1 | 0 | 0 | 1480970 | AVP78041.1 | P0C6W2 |
| NILACYDGCAQG | 1 | 0 | 0 | 1480981 | NP_073549.1 | A0A0P0LKV0 |
| NILGYSSLCDEF | 1 | 0 | 0 | 1480989 | AWK59942.1 | H9EJ42 |
| NINAILTEVKEQ | 1 | 0 | 0 | 1481012 | AHF54662.1 | P36334 |
| NINCILSVNSSN | 1 | 0 | 0 | 1481014 | AIW52766.1 | A0A0P0LKV0 |
| NINFERNLRRGT | 1 | 0 | 0 | 1481022 | ABB90515.1 | A0A0P0K6L9 |
| NINGSNEAKALF | 1 | 0 | 0 | 1481023 | AVV62524.1 | K9N7C7 |
| NINRFNVAITRT | 1 | 0 | 0 | 1481037 | QBM11735.1 | K9N7C7 |
| NINRILGVNSSN | 1 | 0 | 0 | 1481038 | AUT76985.1 | A0A0P0LKV0 |
| NIPADIGDRDPR | 1 | 0 | 0 | 1481047 | AJQ24068.1 | P33469 |
| NIPADIVDRDPS | 1 | 0 | 0 | 1481048 | AIX10718.1 | P33469 |
| NIPCYASAVAQA | 1 | 0 | 0 | 1481050 | QBP84719.1 | P0C6X6 |
| NIPDLPICVYDR | 1 | 0 | 0 | 1481055 | AIL49324.1 | P30215 |
| NIPDLPISVYDP | 1 | 0 | 0 | 1481058 | AIL49333.1 | P30215 |
| NIPYSRVISRMG | 1 | 0 | 0 | 1481070 | AXN92258.1 | K9N7C7 |
| NIRETKCAPELT | 1 | 0 | 0 | 1481078 | AVV62524.1 | K9N7C7 |
| NIRIVDEWSYTG | 1 | 0 | 0 | 1481079 | ASU90175.1 | K9N5Q8 |
| NISLPFNNMLGY | 1 | 0 | 0 | 1481097 | AVV62524.1 | K9N7C7 |
| NISMLQLGVPDN | 1 | 0 | 0 | 1481103 | AKA09626.1 | A0A0P0G321 |
| NISRGTILFKTS | 1 | 0 | 0 | 1481106 | CAA71146.1 | A0A0P0K6L9 |

| | | | | | | |
|---|---|---|---|---|---|---|
| NISTLQLGVPDN | 1 | 0 | 0 | 1481114 | AKA09623.1 | A0A0P0G321 |
| NISTPNLHDFKE | 1 | 0 | 0 | 1481117 | CAA83660.1 | P36334 |
| NISTPNLPDFKE | 1 | 0 | 0 | 1481118 | CAA83661.1 | P36334 |
| NISYVVTQNLLF | 1 | 0 | 0 | 1481125 | AVA26872.1 | H9EJ42 |
| NITEDETAEWFG | 1 | 0 | 0 | 1481129 | AUM60014.1 | K9N5Q8 |
| NIVGDFQLTEEV | 1 | 0 | 0 | 1481162 | ATO98229.1 | P0C6W2 |
| NIVYEPSTPISS | 1 | 0 | 0 | 1481179 | AUM60013.1 | K9N7C7 |
| NIYGTATIGVCK | 1 | 0 | 0 | 1481193 | AXN92247.1 | K9N7C7 |
| NKALEMCIDYVT | 1 | 0 | 0 | 1481212 | ARO76381.1 | P0C6U8 |
| NKATNNAMQLNL | 1 | 0 | 0 | 1481218 | AVP78041.1 | P0C6W2 |
| NKCSRFLSDDRT | 1 | 0 | 0 | 1481229 | 6C6Z_A | K9N5Q8 |
| NKCTKYNIYDGS | 1 | 0 | 0 | 1481232 | BAL45639.1 | A0A0P0K6L9 |
| NKCTKYNIYDVS | 1 | 0 | 0 | 1481234 | AIW52757.1 | A0A0P0K6L9 |
| NKCTKYNIYDVV | 1 | 0 | 0 | 1481235 | BAL45638.1 | A0A0P0K6L9 |
| NKCTKYNIYGVS | 1 | 0 | 0 | 1481236 | APT69862.1 | A0A0P0K6L9 |
| NKDQEAILMSED | 1 | 0 | 0 | 1481255 | ALA49351.1 | K9N7C7 |
| NKEVFVGDSLVN | 1 | 0 | 0 | 1481275 | AEN19363.1 | P0C6X6 |
| NKFCNVQQCFGK | 1 | 0 | 0 | 1481281 | AMM42397.1 | Q19U29 |
| NKFGAISSSLQE | 1 | 0 | 0 | 1481285 | BBA20983.1 | A0A140H1H1 |
| NKFNQALEAMQT | 1 | 0 | 0 | 1481292 | QBF80499.1 | K9N5Q8 |
| NKFTPLIVEETT | 1 | 0 | 0 | 1481299 | AVV62535.1 | K9N7C7 |
| NKFTPLSVASAP | 1 | 0 | 0 | 1481300 | QBF80531.1 | K9N7C7 |
| NKGAGGHCYGAD | 1 | 0 | 0 | 1481310 | APO40578.1 | P0C6X7 |
| NKGEDIQLLKSAYENF | 1 | 0 | 0 | 1481315 | P0DTC1.1 | P0DTD1 |
| NKGHFDGDVGEK | 1 | 0 | 0 | 1481319 | APO40578.1 | P0C6X7 |
| NKGSPQQCIAPI | 1 | 0 | 0 | 1481332 | AVV62526.1 | K9N5Q8 |
| NKGVDCTKKFLT | 1 | 0 | 0 | 1481333 | AXN92247.1 | K9N7C7 |
| NKGVDYTKKFLT | 1 | 0 | 0 | 1481336 | AXN92258.1 | K9N7C7 |
| NKHAFHTPAYDG | 1 | 0 | 0 | 1481345 | AHI48736.1 | K9N7C7 |
| NKHRSFTQWYHE | 1 | 0 | 0 | 1481359 | AXN92247.1 | K9N7C7 |
| NKIEDLSMRALQ | 1 | 0 | 0 | 1481364 | AJE26223.1 | P0C6X6 |
| NKIGDLSIRALQ | 1 | 0 | 0 | 1481365 | AJE26218.1 | P0C6X6 |
| NKIHDVVNQQGN | 1 | 0 | 0 | 1481366 | ABB90526.1 | A0A0P0K6L9 |
| NKIPANIVLPQG | 1 | 0 | 0 | 1481375 | AEN19363.1 | P0C6X6 |
| NKISGKGQQQQG | 1 | 0 | 0 | 1481383 | AVP78049.1 | Q3LZX4 |
| NKISLNAPVTLK | 1 | 0 | 0 | 1481384 | ARU07594.1 | A0A0P0G321 |
| NKITIADPGYMQ | 1 | 0 | 0 | 1481387 | AUM60014.1 | K9N5Q8 |
| NKKFGFIEDSVF | 1 | 0 | 0 | 1481397 | ARB66070.1 | P36334 |
| NKLKAHKEKSSQ | 1 | 0 | 0 | 1481416 | APO40578.1 | P0C6X7 |
| NKLTEVKVPQGL | 1 | 0 | 0 | 1481435 | AVV62524.1 | K9N7C7 |
| NKLWFDFIENPS | 1 | 0 | 0 | 1481440 | AXN92226.1 | K9N7C7 |
| NKNAGQDISLLD | 1 | 0 | 0 | 1481451 | APO40578.1 | P0C6X7 |
| NKNTGPPFISTD | 1 | 0 | 0 | 1481459 | AKA09659.1 | P36334 |
| NKNTYRSQLGCV | 1 | 0 | 0 | 1481461 | AXN92247.1 | K9N7C7 |
| NKPLSQPRVDKP | 1 | 0 | 0 | 1481477 | QED88037.1 | H9EJ69 |
| NKPSVEQRQQAE | 1 | 0 | 0 | 1481483 | AVP78041.1 | P0C6W2 |
| NKRFGFIEDSAF | 1 | 0 | 0 | 1481506 | AGT51481.1 | P36334 |
| NKSFFPQFTSDQ | 1 | 0 | 0 | 1481511 | ABD75500.1 | Q0ZJ82 |
| NKSFFSQFTSDQ | 1 | 0 | 0 | 1481512 | ABD75588.1 | Q0ZJ82 |
| NKSMKCKSHGGK | 1 | 0 | 0 | 1481521 | ABB90496.1 | A0A0N7J131 |
| NKSNPALTQYFS | 1 | 0 | 0 | 1481522 | APO40579.1 | P59594 |
| NKTEADWIELFP | 1 | 0 | 0 | 1481536 | QEO75983.1 | A0A0P0LKV0 |
| NKTLQEFAKNLP | 1 | 0 | 0 | 1481547 | AWK59979.1 | A0A0P0G321 |
| NKVSVPSPLNWE | 1 | 0 | 0 | 1481568 | AMM42407.1 | A0A140H1H1 |
| NKVTLSDPGFMK | 1 | 0 | 0 | 1481573 | APO40579.1 | P59594 |
| NKWQWQEAWYEF | 1 | 0 | 0 | 1481585 | ARU07575.1 | A0A140H1G9 |
| NKWSGVSDLSLK | 1 | 0 | 0 | 1481588 | AXN92247.1 | K9N7C7 |
| NKWSGVSDLSSN | 1 | 0 | 0 | 1481589 | QGV13494.1 | K9N7C7 |
| NKWSGVSNLSLK | 1 | 0 | 0 | 1481590 | QAT98908.1 | K9N7C7 |
| NKYLYELIQKFA | 1 | 0 | 0 | 1481603 | AGT51393.1 | H9EJ42 |
| NLAKFLTYIIFS | 1 | 0 | 0 | 1481622 | AUM60013.1 | K9N7C7 |
| NLAKNILHVVGP | 1 | 0 | 0 | 1481629 | AVV62524.1 | K9N7C7 |
| NLANLNSHNIGT | 1 | 0 | 0 | 1481638 | 6U7G_C | A0A0P0K6L9 |
| NLANLNSHTIGT | 1 | 0 | 0 | 1481640 | 6U7F_C | A0A0P0K6L9 |
| NLAYSNSYTIGS | 1 | 0 | 0 | 1481647 | AAQ90006.1 | A0A0P0K6L9 |
| NLCDDPEKAQGM | 1 | 0 | 0 | 1481648 | AWK59942.1 | H9EJ42 |
| NLCNPDPIRFTG | 1 | 0 | 0 | 1481655 | AGT51422.1 | P36334 |
| NLCVDGFGLNKI | 1 | 0 | 0 | 1481673 | AST12962.1 | A0A0P0LKV0 |
| NLDAYVNSNQNT | 1 | 0 | 0 | 1481675 | ARU07581.1 | Q19U29 |
| NLDEARFSGTAN | 1 | 0 | 0 | 1481679 | AGT51787.1 | P0C6X6 |
| NLDEARLSGSAN | 1 | 0 | 0 | 1481680 | AIV41942.1 | P0C6X6 |
| NLDEPQKDVYEL | 1 | 0 | 0 | 1481681 | AOL02457.1 | P33469 |
| NLEAQVQIDRLI | 1 | 0 | 0 | 1481701 | BBA20986.1 | A0A140H1H1 |
| NLEDFIDSPCYC | 1 | 0 | 0 | 1481703 | AVV62535.1 | K9N7C7 |
| NLERCELVNYKQ | 1 | 0 | 0 | 1481716 | QAT98908.1 | K9N7C7 |
| NLFADIKTQATQ | 1 | 0 | 0 | 1481723 | AVV62526.1 | K9N5Q8 |
| NLFASVKNSQSS | 1 | 0 | 0 | 1481725 | QBF80598.1 | K9N5Q8 |
| NLFGNPDEPQKD | 1 | 0 | 0 | 1481734 | AHA80176.1 | P33469 |
| NLFIAKYGLEDY | 1 | 0 | 0 | 1481736 | ARU07574.1 | A0A140H1G9 |
| NLFKDCSNSYSG | 1 | 0 | 0 | 1481738 | AGT51577.1 | P0C6X6 |
| NLGFDNQSKLPS | 1 | 0 | 0 | 1481764 | AIW52795.1 | H9EJ69 |
| NLGGAVCRKHAA | 1 | 0 | 0 | 1481770 | AVV62524.1 | K9N7C7 |
| NLGGNCAGNASG | 1 | 0 | 0 | 1481774 | ARB07436.1 | A0A140H1G9 |
| NLGNHRKELWHL | 1 | 0 | 0 | 1481782 | CAA79896.1 | P36334 |
| NLGNLQSFNYRI | 1 | 0 | 0 | 1481784 | AIL49492.1 | P36334 |
| NLGSNEPGNVGG | 1 | 0 | 0 | 1481789 | AVI15051.1 | P0C6X6 |
| NLHPDSATLVKD | 1 | 0 | 0 | 1481799 | AVP78041.1 | P0C6W2 |
| NLICAQYVAGYK | 1 | 0 | 0 | 1481809 | QFS19619.1 | K9N5Q8 |
| NLIGYYSDDGNY | 1 | 0 | 0 | 1481812 | AVV62537.1 | K9N5Q8 |
| NLIISSSTQKTI | 1 | 0 | 0 | 1481815 | AVV62526.1 | K9N5Q8 |
| NLIKRMKQIEDK | 1 | 0 | 0 | 1481818 | 5SZS_A | A0A0P0G321 |

| | | | | | | |
|---|---|---|---|---|---|---|
| NLILHDVNSGLT | 1 | 0 | 0 | 1481819 | AVQ05264.1 | P36334 |
| NLISNITTNGPL | 1 | 0 | 0 | 1481822 | AVP78041.1 | P0C6W2 |
| NLLFTEQLGAPL | 1 | 0 | 0 | 1481869 | AIW52843.1 | A0A0P0G321 |
| NLLGFDSSKPVI | 1 | 0 | 0 | 1481871 | AHI48784.1 | K9N7C7 |
| NLLGYDFNKPTS | 1 | 0 | 0 | 1481874 | AUM60013.1 | K9N7C7 |
| NLLSRLEALESG | 1 | 0 | 0 | 1481896 | AUM60021.1 | K9N4V7 |
| NLLYCDSSLEKL | 1 | 0 | 0 | 1481901 | AWO68978.1 | A0A0P0G321 |
| NLLYDFNGNIIG | 1 | 0 | 0 | 1481902 | ADN03339.1 | A0A140H1H1 |
| NLLYDPNGNLYG | 1 | 0 | 0 | 1481903 | ARB66062.1 | P36334 |
| NLMCIDMKGKMY | 1 | 0 | 0 | 1481910 | AZS52621.1 | Q0ZJ82 |
| NLMQDESVANLF | 1 | 0 | 0 | 1481916 | AVV62537.1 | K9N5Q8 |
| NLNAYQQQDGMN | 1 | 0 | 0 | 1481923 | AIV41850.1 | P33469 |
| NLNFEENISRGT | 1 | 0 | 0 | 1481930 | CAA71146.1 | A0A0P0K6L9 |
| NLNRGLPTGLSV | 1 | 0 | 0 | 1481941 | AVP78031.1 | Q3LZX1 |
| NLNSSEGVSDLL | 1 | 0 | 0 | 1481947 | AAS48454.1 | P59637 |
| NLNTLRRGAALG | 1 | 0 | 0 | 1481949 | AAW33939.1 | H9EJ42 |
| NLPDFKEELGQW | 1 | 0 | 0 | 1481959 | AIX10763.1 | P36334 |
| NLPEYMAVAVPS | 1 | 0 | 0 | 1481964 | APT69865.1 | A0A0P0KXN3 |
| NLPFRAVKDGIV | 1 | 0 | 0 | 1481967 | AVV62533.1 | K9N4V7 |
| NLPPPLLGNSTR | 1 | 0 | 0 | 1481982 | ALJ54517.1 | K9N5Q8 |
| NLPPPLLSNQTD | 1 | 0 | 0 | 1481983 | AUM60014.1 | K9N5Q8 |
| NLPPPLLSNSTG | 1 | 0 | 0 | 1481984 | ALL26396.1 | K9N5Q8 |
| NLQFEMNRLQEA | 1 | 0 | 0 | 1482000 | AIL49493.1 | P36334 |
| NLRANNTKGVLP | 1 | 0 | 0 | 1482018 | APO40578.1 | P0C6X7 |
| NLSAAEKEVLRE | 1 | 0 | 0 | 1482050 | AVV62524.1 | K9N7C7 |
| NLSAPTLIPQEN | 1 | 0 | 0 | 1482052 | AWW13571.1 | P0C6X6 |
| NLSASEKEVLLQ | 1 | 0 | 0 | 1482054 | AUM60013.1 | K9N7C7 |
| NLSGNLDEPQKD | 1 | 0 | 0 | 1482061 | AOL02457.1 | P33469 |
| NLSGNPDEPHKD | 1 | 0 | 0 | 1482062 | AIX10716.1 | P33469 |
| NLSLSMARRNLD | 1 | 0 | 0 | 1482074 | 2CME_A | P59636 |
| NLSMLQLSVPDN | 1 | 0 | 0 | 1482077 | BAM16475.1 | A0A0P0G321 |
| NLSSVVPSLPTS | 1 | 0 | 0 | 1482090 | ABB90513.1 | A0A0P0K6L9 |
| NLTADEIKALKE | 1 | 0 | 0 | 1482097 | AVN89363.1 | K9N7C7 |
| NLTKLLSLFMVN | 1 | 0 | 0 | 1482114 | AVV62526.1 | K9N5Q8 |
| NLTNQLPPPLLN | 1 | 0 | 0 | 1482130 | AVV62537.1 | K9N5Q8 |
| NLTYLNFSFELL | 1 | 0 | 0 | 1482144 | QED88040.1 | A0A0P0G321 |
| NLVGFKDVTNGT | 1 | 0 | 0 | 1482152 | CAA71146.1 | A0A0P0K6L9 |
| NLVGSAVEKLRP | 1 | 0 | 0 | 1482153 | ARO76381.1 | P0C6U8 |
| NLVGYHSDNGNY | 1 | 0 | 0 | 1482154 | AUM60014.1 | K9N5Q8 |
| NLVNSKSHNIGS | 1 | 0 | 0 | 1482174 | AOG74783.1 | A0A0P0K6L9 |
| NLVNYKSHNIGS | 1 | 0 | 0 | 1482175 | QRK03789.1 | A0A0P0K6L9 |
| NLVPAFAEELHG | 1 | 0 | 0 | 1482177 | AUM60028.1 | K9N7D2 |
| NLVPINKKDKNK | 1 | 0 | 0 | 1482180 | AIW52723.1 | A0A0N7J131 |
| NLVQMAPISAMVRMY | 1 | 0 | 0 | 1482186 | P0DTC1.1 | P0DTD1 |
| NLYFQGGGGSGY | 1 | 0 | 0 | 1482206 | 6NB6_A | P59594 |
| NLYGFRDYIINR | 1 | 0 | 0 | 1482209 | P36334.1 | P36334 |
| NLYSFDKAIDGF | 1 | 0 | 0 | 1482222 | AWK59942.1 | H9EJ42 |
| NMALKGTDKSST | 1 | 0 | 0 | 1482227 | AKA09652.1 | P36334 |
| NMALKGTDLLSK | 1 | 0 | 0 | 1482228 | AWW13511.1 | P36334 |
| NMAVPHIKEWLK | 1 | 0 | 0 | 1482240 | AUM60027.1 | K9N643 |
| NMCEHPHTICHP | 1 | 0 | 0 | 1482245 | BAM16463.1 | P36334 |
| NMCEHPHTSCHP | 1 | 0 | 0 | 1482247 | AWW13511.1 | P36334 |
| NMCEYPHTICHS | 1 | 0 | 0 | 1482249 | AKA09663.1 | P36334 |
| NMCEYPHTTCHP | 1 | 0 | 0 | 1482251 | BBA84088.1 | P36334 |
| NMCEYPQTICHP | 1 | 0 | 0 | 1482252 | CAA79896.1 | P36334 |
| NMCEYPYTICHP | 1 | 0 | 0 | 1482253 | BBA20979.1 | P36334 |
| NMDTGFAYGYSN | 1 | 0 | 0 | 1482268 | AGT17757.1 | A0A140H1H0 |
| NMEAAYTSSSLG | 1 | 0 | 0 | 1482274 | AOR17480.1 | K9N5Q8 |
| NMEEFIQTNSPK | 1 | 0 | 0 | 1482276 | AIL49500.1 | P36334 |
| NMEGAYTSSLLG | 1 | 0 | 0 | 1482278 | AVV62537.1 | K9N5Q8 |
| NMFTPLIQPIGA | 1 | 0 | 0 | 1482291 | AVP78041.1 | P0C6W2 |
| NMHSPDIVVGDY | 1 | 0 | 0 | 1482302 | AVV62535.1 | K9N7C7 |
| NMHSPDVVGDYV | 1 | 0 | 0 | 1482304 | AKK52620.1 | K9N7C7 |
| NMKAEVDPDLTA | 1 | 0 | 0 | 1482312 | AUM60013.1 | K9N7C7 |
| NMLGYNPSKPIS | 1 | 0 | 0 | 1482316 | AVV62524.1 | K9N7C7 |
| NMQLFEALQFTT | 1 | 0 | 0 | 1482347 | AEN19363.1 | P0C6X6 |
| NMQSVSFHEDRI | 1 | 0 | 0 | 1482352 | AVV62535.1 | K9N7C7 |
| NMQTVSFHEDRI | 1 | 0 | 0 | 1482353 | AVV62524.1 | K9N7C7 |
| NMRNANGACVWN | 1 | 0 | 0 | 1482355 | AVV62535.1 | K9N7C7 |
| NMSPKPQRQRGQ | 1 | 0 | 0 | 1482368 | BBM61433.1 | P33469 |
| NMSSLMSFIQAD | 1 | 0 | 0 | 1482371 | CAA79896.1 | P36334 |
| NMWEFRDYFNDN | 1 | 0 | 0 | 1482400 | AVA26872.1 | H9EJ42 |
| NMYMGKCGDKFF | 1 | 0 | 0 | 1482412 | ARU07575.1 | A0A140H1G9 |
| NNADLSPISVDR | 1 | 0 | 0 | 1482420 | AUM60015.1 | K9N796 |
| NNAHALSKLASE | 1 | 0 | 0 | 1482423 | ATG84679.1 | K9N5Q8 |
| NNAISKIQDSLT | 1 | 0 | 0 | 1482424 | APO40579.1 | P59594 |
| NNALDAIQEGFD | 1 | 0 | 0 | 1482425 | CAA79896.1 | P36334 |
| NNALHAIQQGFD | 1 | 0 | 0 | 1482426 | AEN19366.1 | P36334 |
| NNALHSIQQGFD | 1 | 0 | 0 | 1482428 | AIX10759.1 | P36334 |
| NNALNAIQQGFD | 1 | 0 | 0 | 1482429 | AGT51491.1 | P36334 |
| NNALYAIQQGFD | 1 | 0 | 0 | 1482430 | AIX10750.1 | P36334 |
| NNAQALAKLASE | 1 | 0 | 0 | 1482434 | AVK87450.1 | K9N5Q8 |
| NNARQSDNKTAV | 1 | 0 | 0 | 1482439 | AIL49307.1 | P30215 |
| NNATNVIIKVCN | 1 | 0 | 0 | 1482443 | AVP78031.1 | Q3LZX1 |
| NNAVVKMACPAC | 1 | 0 | 0 | 1482444 | APO40578.1 | P0C6X7 |
| NNCTTAVMTYSN | 1 | 0 | 0 | 1482467 | AIW52843.1 | A0A0P0G321 |
| NNDNNNQQSRGR | 1 | 0 | 0 | 1482487 | AUM60021.1 | K9N4V7 |
| NNDNNSNNQQSR | 1 | 0 | 0 | 1482488 | AVV62533.1 | K9N4V7 |
| NNDNNTNQQSRG | 1 | 0 | 0 | 1482489 | AVV62544.1 | K9N4V7 |
| NNDTTNTNLSRG | 1 | 0 | 0 | 1482493 | ALW82676.1 | K9N4V7 |
| NNDYCCFKHALG | 1 | 0 | 0 | 1482495 | ARB07390.1 | A0A0P0LKV0 |

| | | | | | | |
|---|---|---|---|---|---|---|
| NNECGRAFMYAY | 1 | 0 | 0 | 1482499 | AGT21359.1 | A0A0P0LKV0 |
| NNEFIDLTECIY | 1 | 0 | 0 | 1482500 | AZK15898.1 | K9N7C7 |
| NNEHYRALPGVF | 1 | 0 | 0 | 1482506 | P0C6W6.1 | P0C6W2 |
| NNEIKPTGLKTM | 1 | 0 | 0 | 1482510 | AUM60013.1 | K9N7C7 |
| NNEIRPTGLKTM | 1 | 0 | 0 | 1482512 | AVV62524.1 | K9N7C7 |
| NNEKRPDHVSLT | 1 | 0 | 0 | 1482513 | AVV62535.1 | K9N7C7 |
| NNELMPVKLKIQ | 1 | 0 | 0 | 1482514 | AXX83324.1 | P0C6X6 |
| NNEQGTPLTPIL | 1 | 0 | 0 | 1482517 | NP_073549.1 | A0A0P0LKV0 |
| NNGHKQIVKSIP | 1 | 0 | 0 | 1482547 | AUM60027.1 | K9N643 |
| NNGIGICPAGTN | 1 | 0 | 0 | 1482548 | AIX10751.1 | P36334 |
| NNGIGTCPAGTN | 1 | 0 | 0 | 1482549 | CAA79896.1 | P36334 |
| NNGIGTCPVGTN | 1 | 0 | 0 | 1482550 | AIL49486.1 | P36334 |
| NNGISTCPAGTN | 1 | 0 | 0 | 1482552 | BBA84094.1 | P36334 |
| NNGLGTCPAGTN | 1 | 0 | 0 | 1482555 | AIL49503.1 | P36334 |
| NNGLTCSTDLQK | 1 | 0 | 0 | 1482556 | AEN19366.1 | P36334 |
| NNGMGTCPAGTN | 1 | 0 | 0 | 1482557 | AIL49487.1 | P36334 |
| NNGQGENDNISV | 1 | 0 | 0 | 1482560 | AGT51734.1 | P33469 |
| NNGRFGGFSNHL | 1 | 0 | 0 | 1482561 | AWK59942.1 | H9EJ42 |
| NNGTHIVINVCN | 1 | 0 | 0 | 1482563 | APO40579.1 | P59594 |
| NNGVGTCPAGTN | 1 | 0 | 0 | 1482566 | QEG03757.1 | P36334 |
| NNIASWFTALTQ | 1 | 0 | 0 | 1482573 | AAP49024.1 | P59595 |
| NNIDAAKIYGMG | 1 | 0 | 0 | 1482575 | AIL49487.1 | P36334 |
| NNIDVAKIYGMG | 1 | 0 | 0 | 1482577 | ABD43176.1 | P36334 |
| NNIEKPKPDGVS | 1 | 0 | 0 | 1482578 | AVV62526.1 | K9N5Q8 |
| NNIVASFSSVND | 1 | 0 | 0 | 1482590 | AIW52843.1 | A0A0P0G321 |
| NNKDTGPPFIST | 1 | 0 | 0 | 1482594 | AKA09663.1 | P36334 |
| NNKNTYRSQLGC | 1 | 0 | 0 | 1482604 | AXN92247.1 | K9N7C7 |
| NNKSTYRSQLGC | 1 | 0 | 0 | 1482610 | AZK15899.1 | K9N638 |
| NNKTWSIREFAV | 1 | 0 | 0 | 1482613 | AVP78031.1 | Q3LZX1 |
| NNLDKSAGYPFN | 1 | 0 | 0 | 1482619 | QWK72597.1 | P0DTD1 |
| NNLKIKNYGYLD | 1 | 0 | 0 | 1482627 | AGT17757.1 | A0A140H1H0 |
| NNLMNGVTLSTK | 1 | 0 | 0 | 1482637 | AIL49500.1 | P36334 |
| NNNGSDVLDQPP | 1 | 0 | 0 | 1482659 | AIV41798.1 | P0C6X6 |
| NNNLALGGSVAI | 1 | 0 | 0 | 1482662 | AXN92247.1 | K9N7C7 |
| NNNTDSIIIDGV | 1 | 0 | 0 | 1482664 | AGT51351.1 | H9EJ42 |
| NNNTDSIVIDGV | 1 | 0 | 0 | 1482665 | AKT07951.1 | H9EJ42 |
| NNPAYIAPQANF | 1 | 0 | 0 | 1482669 | AGT51599.1 | P30215 |
| NNPAYIAPQANS | 1 | 0 | 0 | 1482670 | AEN19365.1 | P30215 |
| NNPAYIASQAKF | 1 | 0 | 0 | 1482673 | AIV41992.1 | P30215 |
| NNPAYIATQAKS | 1 | 0 | 0 | 1482674 | AIV41824.1 | P30215 |
| NNPKQIVQSIPI | 1 | 0 | 0 | 1482680 | AVV62540.1 | K9N643 |
| NNPQCAIVQATE | 1 | 0 | 0 | 1482683 | ABE97136.1 | H9EJ42 |
| NNQEDFDLISRC | 1 | 0 | 0 | 1482689 | QBP84728.1 | P0C6X6 |
| NNRDTGHPSISI | 1 | 0 | 0 | 1482702 | AWW13566.1 | P36334 |
| NNRDTGLPSIST | 1 | 0 | 0 | 1482705 | BBA84092.1 | P36334 |
| NNRDTGPLSISN | 1 | 0 | 0 | 1482706 | ANZ78836.1 | P36334 |
| NNRDTGRPSIST | 1 | 0 | 0 | 1482709 | BBO58746.1 | P36334 |
| NNSCTCQPQAFL | 1 | 0 | 0 | 1482719 | AEN19366.1 | P36334 |
| NNSCTCQPQAFV | 1 | 0 | 0 | 1482720 | BBA84088.1 | P36334 |
| NNSIPNLSDFEA | 1 | 0 | 0 | 1482726 | BBA20986.1 | A0A140H1H1 |
| NNSKPYFIKNFE | 1 | 0 | 0 | 1482728 | AVV62530.1 | K9N7D2 |
| NNSLIRQLQPIN | 1 | 0 | 0 | 1482731 | AVQ05264.1 | P36334 |
| NNSLTRQLRPIN | 1 | 0 | 0 | 1482734 | AIL49543.1 | P36334 |
| NNSTHIIIRVCN | 1 | 0 | 0 | 1482740 | ATO98181.1 | Q3LZX1 |
| NNSWLTGNAIQG | 1 | 0 | 0 | 1482746 | ALA49340.1 | K9N7C7 |
| NNSYTIVVQPHN | 1 | 0 | 0 | 1482750 | BBA20983.1 | A0A140H1H1 |
| NNTAISVQTCDL | 1 | 0 | 0 | 1482751 | AIL49538.1 | P36334 |
| NNTDSIVIDGVT | 1 | 0 | 0 | 1482755 | AWK59942.1 | H9EJ42 |
| NNTLVSGDDHIP | 1 | 0 | 0 | 1482769 | AIW52748.1 | A0A0P0K6L9 |
| NNTRVVDEWSYT | 1 | 0 | 0 | 1482774 | ALA49803.1 | K9N5Q8 |
| NNTSCPEWMDDF | 1 | 0 | 0 | 1482776 | AVN89451.1 | K9N7C7 |
| NNTTIVDEWSYT | 1 | 0 | 0 | 1482779 | ALA49957.1 | K9N5Q8 |
| NNTTTRDVVDQW | 1 | 0 | 0 | 1482780 | AVV62537.1 | K9N5Q8 |
| NNTVSWYTGLTQ | 1 | 0 | 0 | 1482782 | AXN92267.1 | K9N4V7 |
| NNTWMHTGSGYY | 1 | 0 | 0 | 1482789 | ARB66068.1 | P36334 |
| NNVDNEIIKSSD | 1 | 0 | 0 | 1482794 | AVV62535.1 | K9N7C7 |
| NNVDPKAYPIIL | 1 | 0 | 0 | 1482796 | AVP78050.1 | Q3LZX3 |
| NNVFMSEAKCWL | 1 | 0 | 0 | 1482797 | AVN89342.1 | K9N7C7 |
| NNVSVPSPFNWE | 1 | 0 | 0 | 1482809 | ABU39973.1 | A0A140H1H1 |
| NNWFLLTTGFVY | 1 | 0 | 0 | 1482825 | AIG59204.1 | A0A0P0K6L9 |
| NNWLKQLIKVTL | 1 | 0 | 0 | 1482826 | AVP78041.1 | P0C6W2 |
| NNWQNLLYDSNG | 1 | 0 | 0 | 1482827 | BBA20986.1 | A0A140H1H1 |
| NNYEKSAGYPFN | 1 | 0 | 0 | 1482830 | AIL49570.1 | P0C6X6 |
| NNYVPSYIKPES | 1 | 0 | 0 | 1482843 | AUM60013.1 | K9N7C7 |
| NPANMRNANGAC | 1 | 0 | 0 | 1482857 | AVV62524.1 | K9N7C7 |
| NPAYIAPQANFG | 1 | 0 | 0 | 1482865 | AGT51599.1 | P30215 |
| NPAYIAPRANSG | 1 | 0 | 0 | 1482867 | AIL49334.1 | P30215 |
| NPAYIATQAKSG | 1 | 0 | 0 | 1482870 | AIV41824.1 | P30215 |
| NPCTCRPQAFLG | 1 | 0 | 0 | 1482877 | CAA79896.1 | P36334 |
| NPCYCFNRDGDE | 1 | 0 | 0 | 1482879 | AVV62524.1 | K9N7C7 |
| NPDEPHKDVYEL | 1 | 0 | 0 | 1482883 | AIX10716.1 | P33469 |
| NPDPITFAGPYK | 1 | 0 | 0 | 1482892 | ATP16757.1 | P36334 |
| NPDPKFVDRYYA | 1 | 0 | 0 | 1482896 | ATU80202.1 | K9N7C7 |
| NPEPITMANARY | 1 | 0 | 0 | 1482902 | AVV62537.1 | K9N5Q8 |
| NPHHRFKCSSNG | 1 | 0 | 0 | 1482930 | AGT17757.1 | A0A140H1H0 |
| NPHNYSYMDLNS | 1 | 0 | 0 | 1482939 | AIL49334.1 | P30215 |
| NPILDFKDGIYF | 1 | 0 | 0 | 1482945 | AVP78031.1 | Q3LZX1 |
| NPIRDYAALAAK | 1 | 0 | 0 | 1482948 | AVV62524.1 | K9N7C7 |
| NPITFYLDGEVI | 1 | 0 | 0 | 1482951 | AVP78041.1 | P0C6W2 |
| NPITKNIGEYNV | 1 | 0 | 0 | 1482952 | ATN39862.1 | P0C6X6 |
| NPIYDEPTPTTS | 1 | 0 | 0 | 1482953 | APO40580.1 | P59632 |

| | | | | | | |
|---|---|---|---|---|---|---|
| NPKEEVKPFFTE | 1 | 0 | 0 | 1482958 | AVP78041.1 | P0C6W2 |
| NPKGIPIFDDPV | 1 | 0 | 0 | 1482962 | AVN89322.1 | K9N7C7 |
| NPKIGPDHSVAD | 1 | 0 | 0 | 1482970 | APO40578.1 | P0C6X7 |
| NPKLDGYYKKDN | 1 | 0 | 0 | 1482971 | APO40578.1 | P0C6X7 |
| NPKPRAAPNNTV | 1 | 0 | 0 | 1482972 | AXN92267.1 | K9N4V7 |
| NPNIPFKDGLYF | 1 | 0 | 0 | 1482995 | APO40579.1 | P59594 |
| NPNNDPAIVTQF | 1 | 0 | 0 | 1482998 | AUM60021.1 | K9N4V7 |
| NPNNHKKIVQSI | 1 | 0 | 0 | 1483001 | AVV62529.1 | K9N643 |
| NPNSSEVDMICS | 1 | 0 | 0 | 1483008 | AOL02450.1 | P0C6X6 |
| NPNVPLDYQATR | 1 | 0 | 0 | 1483010 | ATO98145.1 | Q3LZX1 |
| NPNVPLEYQATR | 1 | 0 | 0 | 1483011 | AVP78031.1 | Q3LZX1 |
| NPPDQLVVYQQA | 1 | 0 | 0 | 1483019 | AIW52757.1 | A0A0P0K6L9 |
| NPPTDKLAMIMG | 1 | 0 | 0 | 1483020 | NP_150075.1 | Q80872 |
| NPPTNVVSHLNG | 1 | 0 | 0 | 1483021 | AAA73152.1 | P30215 |
| NPPTNVVSHVNG | 1 | 0 | 0 | 1483022 | AEN19365.1 | P30215 |
| NPSEDFIKHVRN | 1 | 0 | 0 | 1483035 | AXN92247.1 | K9N7C7 |
| NPSEEFIKHVRT | 1 | 0 | 0 | 1483037 | ALW82730.1 | K9N7C7 |
| NPSSPARLEIVK | 1 | 0 | 0 | 1483052 | ATP16755.1 | Q80872 |
| NPSTWNKKFGFI | 1 | 0 | 0 | 1483058 | ARB66070.1 | P36334 |
| NPTEKLVTTSTN | 1 | 0 | 0 | 1483075 | AUM60013.1 | K9N7C7 |
| NPTSKVESCVVS | 1 | 0 | 0 | 1483085 | AIV41876.1 | P0C6X6 |
| NPVDCQDDRCII | 1 | 0 | 0 | 1483092 | ANZ78865.1 | P0C6X6 |
| NPVDLPPSHATT | 1 | 0 | 0 | 1483093 | QEG03783.1 | A0A0P0LKV0 |
| NPVMDPIYDEPT | 1 | 0 | 0 | 1483101 | ARO76383.1 | P59632 |
| NQADVNIPADIV | 1 | 0 | 0 | 1483115 | AIX10718.1 | P33469 |
| NQALHGANLMQD | 1 | 0 | 0 | 1483120 | AVV62537.1 | K9N5Q8 |
| NQALHGANLRQD | 1 | 0 | 0 | 1483121 | AXN92228.1 | K9N5Q8 |
| NQAPGAMQTGFT | 1 | 0 | 0 | 1483123 | AVV62526.1 | K9N5Q8 |
| NQDFYDFVVAKG | 1 | 0 | 0 | 1483130 | AUM60013.1 | K9N7C7 |
| NQDTCVRSMIKT | 1 | 0 | 0 | 1483137 | AVV62530.1 | K9N7D2 |
| NQDTCVRSMVKT | 1 | 0 | 0 | 1483138 | AVV62541.1 | K9N7D2 |
| NQFTEFVSTQSV | 1 | 0 | 0 | 1483169 | ATG84865.1 | K9N638 |
| NQGNKQIVHSCP | 1 | 0 | 0 | 1483174 | ASU90211.1 | K9N643 |
| NQGPPSARDLIC | 1 | 0 | 0 | 1483177 | AVV62526.1 | K9N5Q8 |
| NQHCNGSIACQF | 1 | 0 | 0 | 1483180 | AWW13571.1 | P0C6X6 |
| NQISVAPDLSLD | 1 | 0 | 0 | 1483194 | ANZ78835.1 | P36334 |
| NQIVLRNSNGAC | 1 | 0 | 0 | 1483196 | AXN92247.1 | K9N7C7 |
| NQKLIANAFSNA | 1 | 0 | 0 | 1483204 | CAA83660.1 | P36334 |
| NQLAYSSSANGI | 1 | 0 | 0 | 1483215 | AVN89312.1 | K9N638 |
| NQLESKMSGKGQQQQGQTVTKKSA | 1 | 0 | 0 | 1483226 | P0DTC9.1 | P0DTC9 |
| NQLNEPLITMPL | 1 | 0 | 0 | 1483230 | APO40578.1 | P0C6X7 |
| NQLPPPLLNNQT | 1 | 0 | 0 | 1483231 | AVV62537.1 | K9N5Q8 |
| NQSKPPSSSGTS | 1 | 0 | 0 | 1483279 | AGT51335.1 | H9EJ69 |
| NQSLIKSKVSAE | 1 | 0 | 0 | 1483284 | ARB07393.1 | A0A0P0KWE8 |
| NQSPASSQSAAT | 1 | 0 | 0 | 1483289 | AIW52723.1 | A0A0N7J131 |
| NQSPASSQSVAK | 1 | 0 | 0 | 1483290 | ABB90499.1 | A0A0N7J131 |
| NQSSAPRITFGG | 1 | 0 | 0 | 1483293 | AVP78049.1 | Q3LZX4 |
| NQSTNKTTFRPN | 1 | 0 | 0 | 1483296 | APO40578.1 | P0C6X7 |
| NQSTSSYPANGF | 1 | 0 | 0 | 1483297 | AWO68991.1 | A0A0P0G321 |
| NQSTSTYPANGF | 1 | 0 | 0 | 1483299 | AWO68966.1 | A0A0P0G321 |
| NQTDLDFKEELE | 1 | 0 | 0 | 1483307 | AVV62537.1 | K9N5Q8 |
| NQTSVAPDFSLD | 1 | 0 | 0 | 1483320 | BBA20976.1 | P36334 |
| NQTSVAPDLALD | 1 | 0 | 0 | 1483321 | ANZ78847.1 | P36334 |
| NQTSVAPDLSLG | 1 | 0 | 0 | 1483324 | AIL49488.1 | P36334 |
| NQTSVAPHLSLD | 1 | 0 | 0 | 1483325 | AQN78744.1 | P36334 |
| NQVLSSDVMFEG | 1 | 0 | 0 | 1483343 | ASU90140.1 | K9N7C7 |
| NQVLSSDVMFEN | 1 | 0 | 0 | 1483344 | AXN92226.1 | K9N7C7 |
| NQVNDMIYSLLE | 1 | 0 | 0 | 1483346 | APO40578.1 | P0C6X7 |
| NQYCNGSIACQF | 1 | 0 | 0 | 1483354 | AEN19363.1 | P0C6X6 |
| NQYYYNMATGFA | 1 | 0 | 0 | 1483367 | AZS52617.1 | A0A140H1H0 |
| NRAGSNDIWMQN | 1 | 0 | 0 | 1483369 | AGT51729.1 | P30215 |
| NRALTGIAIEQD | 1 | 0 | 0 | 1483374 | ATO98181.1 | Q3LZX1 |
| NRARTVAGVSVL | 1 | 0 | 0 | 1483382 | AJS10811.1 | P0C6X6 |
| NRASLRQIFDPA | 1 | 0 | 0 | 1483383 | AVV62524.1 | K9N7C7 |
| NRASLRQIFDVP | 1 | 0 | 0 | 1483385 | AKJ80136.2 | K9N7C7 |
| NRASLRQIYDLA | 1 | 0 | 0 | 1483386 | AUM60013.1 | K9N7C7 |
| NRASLRQIYDPA | 1 | 0 | 0 | 1483387 | AVV62535.1 | K9N7C7 |
| NRATCYNSRTGY | 1 | 0 | 0 | 1483388 | ARU07574.1 | A0A140H1G9 |
| NRCCDKYDDLEP | 1 | 0 | 0 | 1483392 | AVV62537.1 | K9N5Q8 |
| NRCCDKYDDYDL | 1 | 0 | 0 | 1483393 | AUM60024.1 | K9N5Q8 |
| NRCNEVSATVLQ | 1 | 0 | 0 | 1483398 | AVQ05261.1 | P0C6X6 |
| NRCPFDKVFNAS | 1 | 0 | 0 | 1483401 | ATO98181.1 | Q3LZX1 |
| NRDGDVQWSSTM | 1 | 0 | 0 | 1483404 | AVV62535.1 | K9N7C7 |
| NRDTGRPSISTD | 1 | 0 | 0 | 1483417 | BBO58746.1 | P36334 |
| NRDVDQEFVDEF | 1 | 0 | 0 | 1483422 | AVP77994.1 | P0C6W2 |
| NRDVDTDFVNEFYAYL | 1 | 0 | 0 | 1483424 | P0DTD1.1 | P0DTD1 |
| NREAPAYISTIG | 1 | 0 | 0 | 1483426 | APO40578.1 | P0C6X7 |
| NRFGSNDIWMQN | 1 | 0 | 0 | 1483437 | AXX83326.1 | P30215 |
| NRFNVAITRAKV | 1 | 0 | 0 | 1483442 | APO40578.1 | P0C6X7 |
| NRFTTTLNDFNLVAMK | 1 | 0 | 0 | 1483448 | P0DTC1.1 | P0DTD1 |
| NRILGVNSSNCN | 1 | 0 | 0 | 1483467 | AUT76985.1 | A0A0P0LKV0 |
| NRLLDAMINTAG | 1 | 0 | 0 | 1483479 | APO40578.1 | P0C6X7 |
| NRLSRRAITTGY | 1 | 0 | 0 | 1483490 | QBP84713.1 | P36334 |
| NRMAEQAAAAMY | 1 | 0 | 0 | 1483494 | NP_073549.1 | A0A0P0LKV0 |
| NRQCVNINIVGD | 1 | 0 | 0 | 1483516 | APO40578.1 | P0C6X7 |
| NRRKVDLQLGNL | 1 | 0 | 0 | 1483523 | AEN19366.1 | P36334 |
| NRRSFKTADGSQ | 1 | 0 | 0 | 1483531 | AIX10728.1 | P33469 |
| NRRSFRISDGNQ | 1 | 0 | 0 | 1483532 | AJQ24067.1 | P33469 |
| NRRSRRAITTEN | 1 | 0 | 0 | 1483535 | AIL49500.1 | P36334 |
| NRRYGFNNFILS | 1 | 0 | 0 | 1483540 | ARU07577.1 | A0A140H1H1 |
| NRSSFVDDNKFD | 1 | 0 | 0 | 1483552 | AGT51657.1 | P0C6X6 |

| | | | | | | |
|---|---|---|---|---|---|---|
| NRTLTSGVTPDM | 1 | 0 | 0 | 1483561 | AAX85673.1 | P33469 |
| NRTPTSGVTRDM | 1 | 0 | 0 | 1483564 | AIL49470.1 | P33469 |
| NRVHGFEIDPSP | 1 | 0 | 0 | 1483575 | APO40579.1 | P59594 |
| NRVIAIPTSTGS | 1 | 0 | 0 | 1483576 | AVV62527.1 | K9N796 |
| NRVLSVPSDSCN | 1 | 0 | 0 | 1483581 | ABQ16583.1 | H9EJ42 |
| NRVVSPPNNADL | 1 | 0 | 0 | 1483588 | AUM60015.1 | K9N796 |
| NRYGFCGNGTHI | 1 | 0 | 0 | 1483597 | AIW52843.1 | A0A0P0G321 |
| NSACPKLEFAND | 1 | 0 | 0 | 1483605 | AXN92260.1 | K9N5Q8 |
| NSANTKVLKNKG | 1 | 0 | 0 | 1483624 | ATQ39389.1 | K9N7C7 |
| NSAPVPVKEYEA | 1 | 0 | 0 | 1483628 | AVV62524.1 | K9N7C7 |
| NSCCRLVEKNVV | 1 | 0 | 0 | 1483639 | AUM60013.1 | K9N7C7 |
| NSCLVASDGSPS | 1 | 0 | 0 | 1483648 | AVV62535.1 | K9N7C7 |
| NSCLVSSDGQPD | 1 | 0 | 0 | 1483651 | AVN89451.1 | K9N7C7 |
| NSCLVSSNGQPG | 1 | 0 | 0 | 1483653 | QAT98908.1 | K9N7C7 |
| NSCQNAVSSFIN | 1 | 0 | 0 | 1483654 | AVV62524.1 | K9N7C7 |
| NSCVKSKPPSAI | 1 | 0 | 0 | 1483659 | ABD75545.1 | A0A140H1H1 |
| NSDFCPCADPSV | 1 | 0 | 0 | 1483660 | BBA20986.1 | A0A140H1H1 |
| NSDYANKGYIAN | 1 | 0 | 0 | 1483669 | ABD75519.1 | A0A140H1G9 |
| NSEMLKLGTSDP | 1 | 0 | 0 | 1483677 | AZS52622.1 | Q19U29 |
| NSENLYFQSGSH | 1 | 0 | 0 | 1483678 | 6U7H_A | A0A0P0K6L9 |
| NSEVPPRCIAPV | 1 | 0 | 0 | 1483682 | AUM60014.1 | K9N5Q8 |
| NSEYAAKGYVAG | 1 | 0 | 0 | 1483683 | AVV62535.1 | K9N7C7 |
| NSFDVLKSEDAQGMDNL | 1 | 0 | 0 | 1483693 | P0DTC1.1 | P0DTD1 |
| NSFKEYFDLQNC | 1 | 0 | 0 | 1483703 | AVV62537.1 | K9N5Q8 |
| NSFKEYFNLRDC | 1 | 0 | 0 | 1483704 | AUM60014.1 | K9N5Q8 |
| NSFKEYFNLRNS | 1 | 0 | 0 | 1483706 | AGV08520.1 | K9N5Q8 |
| NSGGTPLTNCVK | 1 | 0 | 0 | 1483721 | AUM60013.1 | K9N7C7 |
| NSGIGTCPAGTN | 1 | 0 | 0 | 1483723 | CAA83660.1 | P36334 |
| NSGLTCSTDLQK | 1 | 0 | 0 | 1483728 | CAA79896.1 | P36334 |
| NSGLTCSTDSQK | 1 | 0 | 0 | 1483729 | BBO58736.1 | P36334 |
| NSGNRNPTSGVT | 1 | 0 | 0 | 1483732 | AJQ24067.1 | P33469 |
| NSGNRTSTSGVT | 1 | 0 | 0 | 1483736 | ARB07416.1 | P33469 |
| NSGTTCSTDLQK | 1 | 0 | 0 | 1483751 | CAA83660.1 | P36334 |
| NSHNIGSLYVSW | 1 | 0 | 0 | 1483760 | AIW52756.1 | A0A0P0K6L9 |
| NSIAYANNSIAI | 1 | 0 | 0 | 1483769 | AVP78031.1 | Q3LZX1 |
| NSISTDVYARYL | 1 | 0 | 0 | 1483792 | AUM60013.1 | K9N7C7 |
| NSITDTIVLTSG | 1 | 0 | 0 | 1483794 | Q3I5J4.1 | Q3LZX0 |
| NSKPLVGYIPEH | 1 | 0 | 0 | 1483817 | AUM60015.1 | K9N796 |
| NSKPYFIKNFEQ | 1 | 0 | 0 | 1483818 | AVV62530.1 | K9N7D2 |
| NSKQFNYKITIN | 1 | 0 | 0 | 1483820 | ARB07431.1 | Q80872 |
| NSKRKDGFTLEY | 1 | 0 | 0 | 1483821 | AKA09664.1 | P36334 |
| NSKRKGGFTLEY | 1 | 0 | 0 | 1483822 | AKA09645.1 | P36334 |
| NSKRKNGFTLEY | 1 | 0 | 0 | 1483823 | ATP16757.1 | P36334 |
| NSLADVLTDVSA | 1 | 0 | 0 | 1483827 | AVV62524.1 | K9N7C7 |
| NSNANISMLQLG | 1 | 0 | 0 | 1483879 | AKA09626.1 | A0A0P0G321 |
| NSNANISTLQLG | 1 | 0 | 0 | 1483880 | AKA09623.1 | A0A0P0G321 |
| NSNASISMLQLA | 1 | 0 | 0 | 1483881 | AKA09630.1 | A0A0P0G321 |
| NSNASISMLQLG | 1 | 0 | 0 | 1483882 | AWO68966.1 | A0A0P0G321 |
| NSNVGESFVDDF | 1 | 0 | 0 | 1483901 | ADO24705.1 | A0A0P0LKV0 |
| NSPKVTIDCAAF | 1 | 0 | 0 | 1483912 | AIL49500.1 | P36334 |
| NSPYHPLADNKF | 1 | 0 | 0 | 1483921 | APO40584.1 | P59635 |
| NSQLRQNFQAIS | 1 | 0 | 0 | 1483937 | BAL45637.1 | A0A0P0K6L9 |
| NSQNTVASKILG | 1 | 0 | 0 | 1483943 | AVP78041.1 | P0C6W2 |
| NSRIDVPSAEST | 1 | 0 | 0 | 1483958 | AGV08439.1 | K9N796 |
| NSRKVDLQLGNL | 1 | 0 | 0 | 1483962 | AIX10751.1 | P36334 |
| NSSDNGISAIVT | 1 | 0 | 0 | 1483985 | AVL25577.1 | A0A0P0G321 |
| NSSFINLKEIGT | 1 | 0 | 0 | 1483994 | BBA20986.1 | A0A140H1H1 |
| NSSFTEQLGVPL | 1 | 0 | 0 | 1483995 | AWO68966.1 | A0A0P0G321 |
| NSSKDDQIGYYR | 1 | 0 | 0 | 1484000 | AVP78038.1 | Q3LZX4 |
| NSSLFVEDCNLP | 1 | 0 | 0 | 1484003 | ALA49451.1 | K9N5Q8 |
| NSSLLVEDCKLP | 1 | 0 | 0 | 1484004 | ATW75477.1 | K9N5Q8 |
| NSSLSVEDCKLP | 1 | 0 | 0 | 1484005 | ALL26396.1 | K9N5Q8 |
| NSSRGSSPGPAG | 1 | 0 | 0 | 1484018 | AUM60021.1 | K9N4V7 |
| NSSRGSSPGPSG | 1 | 0 | 0 | 1484019 | AVV62544.1 | K9N4V7 |
| NSSRSNSRGSRS | 1 | 0 | 0 | 1484022 | AUM60021.1 | K9N4V7 |
| NSSSDRNHNSQD | 1 | 0 | 0 | 1484025 | AMM42526.1 | A0A0N7J131 |
| NSSSEFFNKQWP | 1 | 0 | 0 | 1484027 | AVV62537.1 | K9N5Q8 |
| NSSSGALDTTIP | 1 | 0 | 0 | 1484028 | AIW52835.1 | A0A0P0G321 |
| NSSSSFDCIVNL | 1 | 0 | 0 | 1484030 | AIW52843.1 | A0A0P0G321 |
| NSSVPLSEVYVH | 1 | 0 | 0 | 1484035 | AWK59934.1 | U3M6Q8 |
| NSTASAVQTCDL | 1 | 0 | 0 | 1484044 | CAA83661.1 | P36334 |
| NSTDTVNPDSFN | 1 | 0 | 0 | 1484046 | ARB07436.1 | A0A140H1G9 |
| NSTKPCFIRNFD | 1 | 0 | 0 | 1484065 | AUM60018.1 | K9N7D2 |
| NSTQDGYNKLQG | 1 | 0 | 0 | 1484078 | CAA79896.1 | P36334 |
| NSTRGTFPGPSG | 1 | 0 | 0 | 1484082 | ANI69896.1 | K9N4V7 |
| NSTTITDKLGVC | 1 | 0 | 0 | 1484094 | AVV62537.1 | K9N5Q8 |
| NSTYASQGLVAS | 1 | 0 | 0 | 1484100 | QYC64967.1 | P0DTD1 |
| NSVADVMRYNLN | 1 | 0 | 0 | 1484103 | AIW52841.1 | A0A0P0G321 |
| NSVCNGCVGHSE | 1 | 0 | 0 | 1484106 | AST12964.1 | A0A0P0K6L9 |
| NSVCPKFEFAND | 1 | 0 | 0 | 1484107 | AVN89387.1 | K9N5Q8 |
| NSVCPMMDLGNS | 1 | 0 | 0 | 1484111 | AVV62526.1 | K9N5Q8 |
| NSVCPMQALRND | 1 | 0 | 0 | 1484112 | AUM60014.1 | K9N5Q8 |
| NSVFKPQPAGVL | 1 | 0 | 0 | 1484119 | AVQ05264.1 | P36334 |
| NSVIVMAYVTGG | 1 | 0 | 0 | 1484129 | ARO76381.1 | P0C6U8 |
| NSVPVNKKDKNK | 1 | 0 | 0 | 1484147 | AMM42513.1 | A0A0N7J131 |
| NSVPWSKICDYV | 1 | 0 | 0 | 1484150 | APO40578.1 | P0C6X7 |
| NSVPWSKIFAYV | 1 | 0 | 0 | 1484151 | ATO98143.1 | P0C6W2 |
| NSVSDSLEPVGG | 1 | 0 | 0 | 1484155 | ARB66068.1 | P36334 |
| NSVSPKLEFAND | 1 | 0 | 0 | 1484157 | AKN24803.1 | K9N5Q8 |
| NSWLTGNAIQGF | 1 | 0 | 0 | 1484180 | ALA49340.1 | K9N7C7 |
| NSWQNLLYDSSG | 1 | 0 | 0 | 1484184 | ARK08661.1 | P36334 |

| | | | | | | |
|---|---|---|---|---|---|---|
| NSWRYDFDITES | 1 | 0 | 0 | 1484185 | QED88017.1 | H9EJ42 |
| NSYTSFATYYTP | 1 | 0 | 0 | 1484206 | ALA49352.1 | K9N5Q8 |
| NTAKQDQGQYYY | 1 | 0 | 0 | 1484216 | ATO98193.1 | Q3LZX1 |
| NTAKQDTGHYFY | 1 | 0 | 0 | 1484217 | AVP78042.1 | Q3LZX1 |
| NTAKQDVGNYFY | 1 | 0 | 0 | 1484218 | AVP78031.1 | Q3LZX1 |
| NTAKQDVGSYFY | 1 | 0 | 0 | 1484219 | ATO98145.1 | Q3LZX1 |
| NTCDGTTFTYAS | 1 | 0 | 0 | 1484228 | QHD43415.1 | P0DTD1 |
| NTCLEVQVDPNA | 1 | 0 | 0 | 1484232 | AUM60014.1 | K9N5Q8 |
| NTCVGSDNVTDFNAIATCDW | 1 | 0 | 0 | 1484242 | P0DTD1.1 | P0DTD1 |
| NTDNETLQYGVT | 1 | 0 | 0 | 1484254 | ABA60801.1 | A0A140H1H1 |
| NTDTPPRCIAPI | 1 | 0 | 0 | 1484258 | AVV62537.1 | K9N5Q8 |
| NTEIELGVCVNY | 1 | 0 | 0 | 1484267 | AEN19366.1 | P36334 |
| NTEPKVPEEAPA | 1 | 0 | 0 | 1484269 | APO40578.1 | P0C6X7 |
| NTETMNDDLLQI | 1 | 0 | 0 | 1484270 | APO40578.1 | P0C6X7 |
| NTFAVFDTPHTD | 1 | 0 | 0 | 1484272 | AVV62537.1 | K9N5Q8 |
| NTFICEEVANDF | 1 | 0 | 0 | 1484279 | AKI29254.1 | K9N638 |
| NTFTKLQSLENV | 1 | 0 | 0 | 1484290 | AVP78041.1 | P0C6W2 |
| NTGNGTKALAPR | 1 | 0 | 0 | 1484304 | AVV62533.1 | K9N4V7 |
| NTGPPFISTDTV | 1 | 0 | 0 | 1484309 | AKA09659.1 | P36334 |
| NTGRCNVNKHVA | 1 | 0 | 0 | 1484310 | QBP84746.1 | P0C6X6 |
| NTGVFVAPPIAD | 1 | 0 | 0 | 1484322 | AVV62528.1 | K9N4V0 |
| NTIHSVCNGCVG | 1 | 0 | 0 | 1484336 | AIW52741.1 | A0A0P0K6L9 |
| NTKLYVDNTLYS | 1 | 0 | 0 | 1484358 | ACV87129.1 | A0A140H1H1 |
| NTKVFKDGVMYS | 1 | 0 | 0 | 1484364 | AEN19366.1 | P36334 |
| NTKVIKDDVMYS | 1 | 0 | 0 | 1484367 | ACV60189.1 | P36334 |
| NTKVIKDGVMYS | 1 | 0 | 0 | 1484368 | CAA79896.1 | P36334 |
| NTLNNLNETLVT | 1 | 0 | 0 | 1484404 | AVP78041.1 | P0C6W2 |
| NTLTIIIPDKQV | 1 | 0 | 0 | 1484415 | ARU07575.1 | A0A140H1G9 |
| NTMIVDEWSYTG | 1 | 0 | 0 | 1484425 | ATG84866.1 | K9N5Q8 |
| NTMLEDEFTPED | 1 | 0 | 0 | 1484426 | AUM60013.1 | K9N7C7 |
| NTNANLSMLQLS | 1 | 0 | 0 | 1484428 | BAM16475.1 | A0A0P0G321 |
| NTNATLNSFKEY | 1 | 0 | 0 | 1484431 | AVV62537.1 | K9N5Q8 |
| NTNSGPDDQIGF | 1 | 0 | 0 | 1484448 | AAT76155.1 | Q3LZX4 |
| NTNSSKDDQIGY | 1 | 0 | 0 | 1484450 | AVP78038.1 | Q3LZX4 |
| NTNSVDSKSGNN | 1 | 0 | 0 | 1484453 | APO40579.1 | P59594 |
| NTPADIDYRNPS | 1 | 0 | 0 | 1484455 | AJQ24074.1 | P33469 |
| NTPADILDRDPS | 1 | 0 | 0 | 1484456 | QBP84708.1 | P33469 |
| NTPADIVDRDQS | 1 | 0 | 0 | 1484457 | ANZ78879.1 | P33469 |
| NTPADIVHRDPS | 1 | 0 | 0 | 1484458 | AJQ24067.1 | P33469 |
| NTPCNVCQYWIG | 1 | 0 | 0 | 1484461 | AXN92247.1 | K9N7C7 |
| NTPCNVCQYWVG | 1 | 0 | 0 | 1484462 | AUM60013.1 | K9N7C7 |
| NTPDVPICVYDP | 1 | 0 | 0 | 1484463 | AAA73152.1 | P30215 |
| NTPIPNLPDFRE | 1 | 0 | 0 | 1484464 | AIL49487.1 | P36334 |
| NTPKEHIGTRNP | 1 | 0 | 0 | 1484466 | APO40586.1 | P59595 |
| NTPNAILISDRK | 1 | 0 | 0 | 1484468 | AUM60013.1 | K9N7C7 |
| NTPVSVKLPKGV | 1 | 0 | 0 | 1484471 | AOL02450.1 | P0C6X6 |
| NTQEVFAQVKQM | 1 | 0 | 0 | 1484475 | ATO98181.1 | Q3LZX1 |
| NTQTVDSSQGSE | 1 | 0 | 0 | 1484478 | AVV62524.1 | K9N7C7 |
| NTRDVFAQTKSI | 1 | 0 | 0 | 1484481 | APO40579.1 | P59594 |
| NTRFVSKDVKFE | 1 | 0 | 0 | 1484484 | AUM60014.1 | K9N5Q8 |
| NTSADIVDRDPS | 1 | 0 | 0 | 1484494 | ARK08674.1 | P33469 |
| NTSCPEWMDEFE | 1 | 0 | 0 | 1484498 | ATG84853.1 | K9N7C7 |
| NTSFPEWIDDLE | 1 | 0 | 0 | 1484505 | ATQ39389.1 | K9N7C7 |
| NTSGPEWMDDFE | 1 | 0 | 0 | 1484506 | AWM99580.1 | K9N7C7 |
| NTSHAVCNGCVG | 1 | 0 | 0 | 1484507 | QEG03785.1 | A0A0P0K6L9 |
| NTSHSVCNGCVG | 1 | 0 | 0 | 1484509 | AIW52757.1 | A0A0P0K6L9 |
| NTSIPNLPDFKG | 1 | 0 | 0 | 1484513 | QDH43719.1 | P36334 |
| NTSIPNLPDFRE | 1 | 0 | 0 | 1484514 | AIX10756.1 | P36334 |
| NTSIPNLPGFKE | 1 | 0 | 0 | 1484515 | AIL49523.1 | P36334 |
| NTSKAVAVLYQD | 1 | 0 | 0 | 1484519 | APO40579.1 | P59594 |
| NTSSPNLPDFRE | 1 | 0 | 0 | 1484533 | AIL49492.1 | P36334 |
| NTSTPNLPDFKE | 1 | 0 | 0 | 1484536 | BBA20982.1 | P36334 |
| NTSTPNLPDFRE | 1 | 0 | 0 | 1484537 | AVQ05264.1 | P36334 |
| NTSTSNLPDFRE | 1 | 0 | 0 | 1484538 | AMK59677.1 | P36334 |
| NTSYSVCNGCVG | 1 | 0 | 0 | 1484545 | NP_073551.1 | A0A0P0K6L9 |
| NTTLLDLSEEFR | 1 | 0 | 0 | 1484562 | AUM60014.1 | K9N5Q8 |
| NTTQLTEAQIIA | 1 | 0 | 0 | 1484571 | ALT66808.1 | K9N7A1 |
| NTTSFLNGYYPT | 1 | 0 | 0 | 1484576 | AKA09662.1 | P36334 |
| NTTVPRISEYVV | 1 | 0 | 0 | 1484579 | BBA20983.1 | A0A140H1H1 |
| NTTVTVNVTTHN | 1 | 0 | 0 | 1484581 | AIW52831.1 | A0A0P0G321 |
| NTVDCQDDRCFI | 1 | 0 | 0 | 1484587 | AIL49573.1 | P0C6X6 |
| NTVKFCDHIVKA | 1 | 0 | 0 | 1484599 | AVV61898.1 | K9N7C7 |
| NTVLNASISESK | 1 | 0 | 0 | 1484610 | AMM42596.1 | H9EJ69 |
| NTVLNASTPESK | 1 | 0 | 0 | 1484611 | AIW52805.1 | H9EJ69 |
| NTYEGNSPYHPL | 1 | 0 | 0 | 1484635 | APO40584.1 | P59635 |
| NVADFNAIATCD | 1 | 0 | 0 | 1484645 | APO40578.1 | P0C6X7 |
| NVAITRAKVGILCI | 1 | 0 | 0 | 1484658 | P0DTD1.1 | P0DTD1 |
| NVCFDVRDKGAL | 1 | 0 | 0 | 1484677 | AVV62535.1 | K9N7C7 |
| NVCGIKDVFIQG | 1 | 0 | 0 | 1484683 | ATQ39389.1 | K9N7C7 |
| NVCNFYFCQDPM | 1 | 0 | 0 | 1484691 | APO40579.1 | P59594 |
| NVDDINFAPVLG | 1 | 0 | 0 | 1484703 | QXL74886.1 | P36334 |
| NVDKIRQLLKDV | 1 | 0 | 0 | 1484714 | ATQ39389.1 | K9N7C7 |
| NVFVSHTLTKYT | 1 | 0 | 0 | 1484775 | AIW52757.1 | A0A0P0K6L9 |
| NVGEPIISQPSKLLNG | 1 | 0 | 0 | 1484780 | QED88024.1 | H9EJ42 |
| NVGKNGLQFAAI | 1 | 0 | 0 | 1484788 | AUM60013.1 | K9N7C7 |
| NVGKQGLEYAAI | 1 | 0 | 0 | 1484790 | AVV62535.1 | K9N7C7 |
| NVIADSKSFTLI | 1 | 0 | 0 | 1484802 | AXN92247.1 | K9N7C7 |
| NVIEFHLDGEII | 1 | 0 | 0 | 1484804 | APO40578.1 | P0C6X7 |
| NVIRGWIFGSTL | 1 | 0 | 0 | 1484829 | APO40579.1 | P59594 |
| NVIYTTEIASKL | 1 | 0 | 0 | 1484837 | AGT51508.1 | P0C6X6 |
| NVKAWSGICVDD | 1 | 0 | 0 | 1484838 | ARE29979.1 | A0A0P0G321 |

| | | | | | | |
|---|---|---|---|---|---|---|
| NVKEEPTTQKPV | 1 | 0 | 0 | 1484842 | ARO76381.1 | P0C6U8 |
| NVKLPRWSASID | 1 | 0 | 0 | 1484854 | QNT54756.1 | A0A0P0K6L9 |
| NVKNVTDVTGNL | 1 | 0 | 0 | 1484856 | AVQ05261.1 | P0C6X6 |
| NVLDQAIAMWAL | 1 | 0 | 0 | 1484868 | APO40578.1 | P0C6X7 |
| NVLKHCCIAAVE | 1 | 0 | 0 | 1484877 | AUM60013.1 | K9N7C7 |
| NVLSSMLRLHTV | 1 | 0 | 0 | 1484886 | AXN92247.1 | K9N7C7 |
| NVMDVNDCFKKD | 1 | 0 | 0 | 1484900 | QED88039.1 | H9EJ42 |
| NVMFDLDSTFDD | 1 | 0 | 0 | 1484902 | ABD75639.1 | A0A140H1G9 |
| NVMQCFGPRDFN | 1 | 0 | 0 | 1484905 | AMM42619.1 | H9EJ69 |
| NVNNTWMFTGSG | 1 | 0 | 0 | 1484911 | CAA83660.1 | P36334 |
| NVPMEKLKTFLT | 1 | 0 | 0 | 1484945 | APO40578.1 | P0C6X7 |
| NVPYSRVISRMG | 1 | 0 | 0 | 1484956 | AXN92247.1 | K9N7C7 |
| NVPYVYNGSVQA | 1 | 0 | 0 | 1484960 | AGT51729.1 | P30215 |
| NVQKVAQPKIEQ | 1 | 0 | 0 | 1484969 | AST12962.1 | A0A0P0LKV0 |
| NVSALIGANGNK | 1 | 0 | 0 | 1484995 | AVN89322.1 | K9N7C7 |
| NVSFVVDDSGTP | 1 | 0 | 0 | 1485004 | AXN92247.1 | K9N7C7 |
| NVSHVYNGSAQA | 1 | 0 | 0 | 1485005 | QQY99179.1 | P30215 |
| NVSLEKFMSTPN | 1 | 0 | 0 | 1485011 | ANF29193.1 | K9N7C7 |
| NVSSDVIRYNIN | 1 | 0 | 0 | 1485026 | ABB90509.1 | A0A0P0K6L9 |
| NVSSQGPNFQEI | 1 | 0 | 0 | 1485029 | AVV62526.1 | K9N5Q8 |
| NVSSVWPHYPYG | 1 | 0 | 0 | 1485031 | AGT51769.1 | P30215 |
| NVSTDGLYFHFY | 1 | 0 | 0 | 1485036 | ABA60800.1 | A0A140H1H1 |
| NVSTSIPNFGSL | 1 | 0 | 0 | 1485041 | AXN92228.1 | K9N5Q8 |
| NVSTSIPNFSSL | 1 | 0 | 0 | 1485042 | ANF29272.1 | K9N5Q8 |
| NVSTSISNFGSL | 1 | 0 | 0 | 1485043 | QBF80598.1 | K9N5Q8 |
| NVSTVFVPVTRD | 1 | 0 | 0 | 1485046 | AHI48554.1 | K9N7D2 |
| NVSVPSPFNWER | 1 | 0 | 0 | 1485050 | ABU39973.1 | A0A140H1H1 |
| NVSYDSVSAIGT | 1 | 0 | 0 | 1485055 | AAK32190.1 | A0A0P0K6L9 |
| NVSYDSVSATVT | 1 | 0 | 0 | 1485056 | AAK32189.1 | A0A0P0K6L9 |
| NVTAAYGLCNTD | 1 | 0 | 0 | 1485061 | AVV62537.1 | K9N5Q8 |
| NVTFFPDLNGDV | 1 | 0 | 0 | 1485067 | APO40578.1 | P0C6X7 |
| NVTGSNDSKALF | 1 | 0 | 0 | 1485076 | AZU90729.1 | K9N7C7 |
| NVTINNYNPSFW | 1 | 0 | 0 | 1485082 | ABD75609.1 | A0A140H1H1 |
| NVTKHKCDISYK | 1 | 0 | 0 | 1485085 | AWW13571.1 | P0C6X6 |
| NVTQAFGRRGPG | 1 | 0 | 0 | 1485095 | AVV62544.1 | K9N4V7 |
| NVTQCFGPRDLD | 1 | 0 | 0 | 1485096 | AIW52723.1 | A0A0N7J131 |
| NVTRDKLEKLIF | 1 | 0 | 0 | 1485097 | AGV08389.1 | K9N638 |
| NVTREKLEKLIS | 1 | 0 | 0 | 1485102 | ALD51903.1 | K9N7C7 |
| NVTSFGDYNLSS | 1 | 0 | 0 | 1485103 | AIW52843.1 | A0A0P0G321 |
| NVTTAETTDFLT | 1 | 0 | 0 | 1485105 | CAA71147.1 | A0A0P0K6L9 |
| NVTTASATDFWT | 1 | 0 | 0 | 1485106 | AIW52843.1 | A0A0P0G321 |
| NVVDVNDCFKND | 1 | 0 | 0 | 1485124 | AGT51379.1 | H9EJ42 |
| NVVEMPKFFYAS | 1 | 0 | 0 | 1485127 | AIW52757.1 | A0A0P0K6L9 |
| NVVKAVRNKISA | 1 | 0 | 0 | 1485141 | AGT51787.1 | P0C6X6 |
| NVVKHGRFNGVE | 1 | 0 | 0 | 1485144 | AVV62524.1 | K9N7C7 |
| NVVPGDVIITST | 1 | 0 | 0 | 1485158 | AWK59942.1 | H9EJ42 |
| NVVPGVSTNVYL | 1 | 0 | 0 | 1485160 | AKT07951.1 | H9EJ42 |
| NVVTTKIALRGG | 1 | 0 | 0 | 1485181 | AVP78041.1 | P0C6W2 |
| NVYAWERTKISD | 1 | 0 | 0 | 1485199 | ATO98181.1 | Q3LZX1 |
| NVYMAKVGDKYY | 1 | 0 | 0 | 1485218 | AEN19363.1 | P0C6X6 |
| NVYMRTTHTDIG | 1 | 0 | 0 | 1485220 | ANC28699.1 | K9N638 |
| NVYRSSIPDPKF | 1 | 0 | 0 | 1485223 | AUM60013.1 | K9N7C7 |
| NVYVDSFVVKGD | 1 | 0 | 0 | 1485226 | ABD72969.1 | P59594 |
| NWFIQSDKCSVE | 1 | 0 | 0 | 1485248 | AEN19363.1 | P0C6X6 |
| NWLAPTNAYTRT | 1 | 0 | 0 | 1485262 | ANI69888.1 | K9N638 |
| NWQNLLYDSNGN | 1 | 0 | 0 | 1485309 | BBA20986.1 | A0A140H1H1 |
| NYCQPIVGYCIV | 1 | 0 | 0 | 1485333 | ADN03340.1 | Q0ZJ84 |
| NYDDYDFEPQKI | 1 | 0 | 0 | 1485338 | AUM60014.1 | K9N5Q8 |
| NYEKSAGYPFNQ | 1 | 0 | 0 | 1485348 | AIL49570.1 | P0C6X6 |
| NYELEKRCYELL | 1 | 0 | 0 | 1485351 | AUM60013.1 | K9N7C7 |
| NYEPLVQDHVDI | 1 | 0 | 0 | 1485357 | APO40578.1 | P0C6X7 |
| NYFITDAKTGAS | 1 | 0 | 0 | 1485360 | APO40578.1 | P0C6X7 |
| NYFTFKQANNNC | 1 | 0 | 0 | 1485371 | AIV41846.1 | P0C6X6 |
| NYKGKVFFQFEN | 1 | 0 | 0 | 1485406 | AIV41978.1 | P0C6X6 |
| NYKQDFSNPTCR | 1 | 0 | 0 | 1485419 | AUM60014.1 | K9N5Q8 |
| NYKQSIPMPQGV | 1 | 0 | 0 | 1485422 | AVV62524.1 | K9N7C7 |
| NYLGKPKEIIDG | 1 | 0 | 0 | 1485444 | APO40578.1 | P0C6X7 |
| NYLTCHNAAQCD | 1 | 0 | 0 | 1485462 | CAA83660.1 | P36334 |
| NYLVVYKHTFRG | 1 | 0 | 0 | 1485468 | AEN19363.1 | P0C6X6 |
| NYNCNASLNSFK | 1 | 0 | 0 | 1485477 | AHL31360.1 | K9N5Q8 |
| NYNLTKLLSLFH | 1 | 0 | 0 | 1485480 | AUM60014.1 | K9N5Q8 |
| NYPCMVGPDCAY | 1 | 0 | 0 | 1485497 | AUM60023.1 | K9N7C7 |
| NYQKVGVQKYST | 1 | 0 | 0 | 1485510 | APO40578.1 | P0C6X7 |
| NYQKWMELLDSN | 1 | 0 | 0 | 1485511 | AUM60021.1 | K9N4V7 |
| NYQLPKVGYCRV | 1 | 0 | 0 | 1485512 | AXX83304.1 | Q04853 |
| NYSGSMLRACIK | 1 | 0 | 0 | 1485547 | ASU45697.1 | K9N796 |
| NYSKDVYFTLSG | 1 | 0 | 0 | 1485551 | AGT17757.1 | A0A140H1H0 |
| NYSKLFDEGHTV | 1 | 0 | 0 | 1485554 | AFR79256.1 | A0A0P0ILY9 |
| NYSKLSDEGYTV | 1 | 0 | 0 | 1485556 | AST12962.1 | A0A0P0LKV0 |
| NYSYMDLNPALC | 1 | 0 | 0 | 1485561 | AEN19365.1 | P30215 |
| NYTVQKLQSLID | 1 | 0 | 0 | 1485587 | BAL45640.1 | A0A0P0K6L9 |
| NYVPSYVKPDSI | 1 | 0 | 0 | 1485606 | AVV62524.1 | K9N7C7 |
| NYYCVRPCVSVP | 1 | 0 | 0 | 1485624 | AVV62526.1 | K9N5Q8 |
| PAAADGIIYPTG | 1 | 0 | 0 | 1485634 | AVV62526.1 | K9N5Q8 |
| PAADLDEFSKQL | 1 | 0 | 0 | 1485637 | AVP78038.1 | Q3LZX4 |
| PAAELDDFSKQL | 1 | 0 | 0 | 1485639 | APO40586.1 | P59595 |
| PAAISSNCYSSL | 1 | 0 | 0 | 1485641 | AHL18090.1 | K9N5Q8 |
| PAANVSVSRFDP | 1 | 0 | 0 | 1485646 | AIV41831.1 | P36334 |
| PAANVSVSRFNP | 1 | 0 | 0 | 1485647 | CAA79896.1 | P36334 |
| PAAPRAISFADN | 1 | 0 | 0 | 1485649 | ANI69896.1 | K9N4V7 |
| PAAPRAVTFGDN | 1 | 0 | 0 | 1485653 | AVV62533.1 | K9N4V7 |

| | | | | | | |
|---|---|---|---|---|---|---|
| PACIEVDIQQTF | 1 | 0 | 0 | 1485657 | ASU91284.1 | K9N5Q8 |
| PACINGHAVPTL | 1 | 0 | 0 | 1485658 | QBF80553.1 | K9N7C7 |
| PACQNPKIGPDH | 1 | 0 | 0 | 1485661 | APO40578.1 | P0C6X7 |
| PACTNGHALPTL | 1 | 0 | 0 | 1485663 | ALL26408.1 | K9N638 |
| PACTNGHAVPTL | 1 | 0 | 0 | 1485664 | AXN92247.1 | K9N7C7 |
| PADIDYRNPSID | 1 | 0 | 0 | 1485665 | AJQ24074.1 | P33469 |
| PADIGDRDPRND | 1 | 0 | 0 | 1485666 | AJQ24068.1 | P33469 |
| PADILDRDPSSD | 1 | 0 | 0 | 1485667 | QBP84708.1 | P33469 |
| PADIVDRDPSND | 1 | 0 | 0 | 1485668 | AJQ24082.1 | P33469 |
| PADIVDRDPSSD | 1 | 0 | 0 | 1485669 | AEN19370.1 | P33469 |
| PADIVDRDQSSE | 1 | 0 | 0 | 1485670 | ANZ78879.1 | P33469 |
| PADIVHRDPSND | 1 | 0 | 0 | 1485671 | AJQ24067.1 | P33469 |
| PADNSYTSFATY | 1 | 0 | 0 | 1485672 | AID55096.1 | K9N5Q8 |
| PADQLADPNYDA | 1 | 0 | 0 | 1485674 | AUM60013.1 | K9N7C7 |
| PADQLNDPNYDG | 1 | 0 | 0 | 1485675 | AVV62524.1 | K9N7C7 |
| PADQLSDPNYDA | 1 | 0 | 0 | 1485676 | AXN92247.1 | K9N7C7 |
| PADVNTPADIDY | 1 | 0 | 0 | 1485678 | AJQ24074.1 | P33469 |
| PADYILPLVLSS | 1 | 0 | 0 | 1485679 | NP_073549.1 | A0A0P0LKV0 |
| PAEAFEKFVCLF | 1 | 0 | 0 | 1485680 | AVV62524.1 | K9N7C7 |
| PAECTMFRDAEG | 1 | 0 | 0 | 1485682 | ATQ39389.1 | K9N7C7 |
| PAEIVDTVSTLV | 1 | 0 | 0 | 1485685 | APO40578.1 | P0C6X7 |
| PAELPVSIVSLP | 1 | 0 | 0 | 1485686 | ADK37704.1 | A0A0P0K6L9 |
| PAELTGTQSVSV | 1 | 0 | 0 | 1485687 | ANI69890.1 | K9N796 |
| PAEPLVQDRAVE | 1 | 0 | 0 | 1485688 | Q4VID0.1 | Q4VID0 |
| PAEPLVQDRVVE | 1 | 0 | 0 | 1485689 | AVR40343.1 | Q4VID0 |
| PAEYKLRQGTFL | 1 | 0 | 0 | 1485691 | ARO76381.1 | P0C6U8 |
| PAFAEEAHGGEV | 1 | 0 | 0 | 1485692 | AUM60018.1 | K9N7D2 |
| PAFHRIESPDSI | 1 | 0 | 0 | 1485702 | ASU91080.1 | K9N7D2 |
| PAFHRIESTDPI | 1 | 0 | 0 | 1485703 | AVN89337.1 | K9N7D2 |
| PAFVVTGGAQTS | 1 | 0 | 0 | 1485717 | ARO76382.1 | P59594 |
| PAGNSHTSFATY | 1 | 0 | 0 | 1485725 | AHX71946.1 | K9N5Q8 |
| PAGTILVDNDLH | 1 | 0 | 0 | 1485730 | QDH43722.1 | P0C6X6 |
| PAGTKYRHCDLD | 1 | 0 | 0 | 1485732 | BBA20986.1 | A0A140H1H1 |
| PAGTKYRHCDVD | 1 | 0 | 0 | 1485733 | ARB07438.1 | A0A140H1H1 |
| PAGTNHLTCDNL | 1 | 0 | 0 | 1485734 | ARE30008.1 | P36334 |
| PAGTNHLTCHNL | 1 | 0 | 0 | 1485735 | BBO58900.1 | P36334 |
| PAGTNSLTCDNL | 1 | 0 | 0 | 1485736 | AWW13566.1 | P36334 |
| PAGTNSLTCGNL | 1 | 0 | 0 | 1485737 | BBA20976.1 | P36334 |
| PAGTNYLNCDNL | 1 | 0 | 0 | 1485738 | AIL49520.1 | P36334 |
| PAGTNYLTCHNA | 1 | 0 | 0 | 1485740 | CAA83660.1 | P36334 |
| PAGVCLGAGQFM | 1 | 0 | 0 | 1485743 | AIV41774.1 | P0C6X6 |
| PAGVCLGAGQFV | 1 | 0 | 0 | 1485744 | AEN19363.1 | P0C6X6 |
| PAHMPMHPMVNP | 1 | 0 | 0 | 1485756 | AET99209.1 | Q0ZJ80 |
| PAHMPMPPMVNP | 1 | 0 | 0 | 1485757 | ARU07582.1 | Q0ZJ80 |
| PAHVMFKHNDFS | 1 | 0 | 0 | 1485762 | AVP78041.1 | P0C6W2 |
| PAICHEGKAHFP | 1 | 0 | 0 | 1485767 | AVP78031.1 | Q3LZX1 |
| PAIDCSDGNYNR | 1 | 0 | 0 | 1485770 | ASU91010.1 | K9N5Q8 |
| PAIDGLKITEEV | 1 | 0 | 0 | 1485771 | AVP78041.1 | P0C6W2 |
| PAIKDFGGFNFS | 1 | 0 | 0 | 1485772 | ATO98181.1 | Q3LZX1 |
| PAITIGSNFVNT | 1 | 0 | 0 | 1485773 | BBO58761.1 | P36334 |
| PAKDIPEKVSLV | 1 | 0 | 0 | 1485781 | ATQ39389.1 | K9N7C7 |
| PAKSQTSDVYEA | 1 | 0 | 0 | 1485783 | AUM60013.1 | K9N7C7 |
| PALHLVDPQIQV | 1 | 0 | 0 | 1485794 | AVP78050.1 | Q3LZX3 |
| PALLFRNIKCNY | 1 | 0 | 0 | 1485802 | AEN19366.1 | P36334 |
| PALNCYWPLKDY | 1 | 0 | 0 | 1485808 | 3D0H_E | P59594 |
| PALQTVATALNK | 1 | 0 | 0 | 1485820 | AIW52728.1 | A0A0P0K6L9 |
| PAMRDSVPLHII | 1 | 0 | 0 | 1485835 | AKN24834.1 | K9N7D2 |
| PANECAQVLSEI | 1 | 0 | 0 | 1485838 | AIL49562.1 | P0C6X6 |
| PANGHMAHDGGV | 1 | 0 | 0 | 1485841 | AMK59674.1 | P0C6X6 |
| PANGHMVHGGGV | 1 | 0 | 0 | 1485844 | AEN19363.1 | P0C6X6 |
| PANLTISKPSSY | 1 | 0 | 0 | 1485848 | AVV62526.1 | K9N5Q8 |
| PANLTITKPSSY | 1 | 0 | 0 | 1485849 | AVV62537.1 | K9N5Q8 |
| PANMRNANGACI | 1 | 0 | 0 | 1485850 | AVV62524.1 | K9N7C7 |
| PANMRNANGACV | 1 | 0 | 0 | 1485851 | AVV62535.1 | K9N7C7 |
| PANMRVMHFGAG | 1 | 0 | 0 | 1485853 | AVV62524.1 | K9N7C7 |
| PANSLHATATIP | 1 | 0 | 0 | 1485858 | APO40580.1 | P59632 |
| PAPIDIDAFPVR | 1 | 0 | 0 | 1485864 | AWK59942.1 | H9EJ42 |
| PAPNNSVSWYTG | 1 | 0 | 0 | 1485869 | AVV62544.1 | K9N4V7 |
| PAPTHYLRVVFP | 1 | 0 | 0 | 1485875 | AVV62540.1 | K9N643 |
| PAQCIRDPVGFC | 1 | 0 | 0 | 1485879 | AHY22544.1 | K9N7C7 |
| PAQCTRDPVGFC | 1 | 0 | 0 | 1485880 | AVV62524.1 | K9N7C7 |
| PAQDTWGTSAAA | 1 | 0 | 0 | 1485884 | ATO98205.1 | Q3LZX1 |
| PAQTGVIADYNY | 1 | 0 | 0 | 1485890 | APO40579.1 | P59594 |
| PAQYELKHGTFV | 1 | 0 | 0 | 1485892 | AVP78041.1 | P0C6W2 |
| PARFVIDTPTGP | 1 | 0 | 0 | 1485895 | 2J98_B | A0A0P0LKV0 |
| PARMAGNGGDTA | 1 | 0 | 0 | 1485899 | AVP78038.1 | Q3LZX4 |
| PARNLQAGGDTA | 1 | 0 | 0 | 1485904 | APO40586.1 | P59595 |
| PASARDLICSQY | 1 | 0 | 0 | 1485907 | APT69978.1 | K9N5Q8 |
| PASARELICAQY | 1 | 0 | 0 | 1485908 | ARA67740.1 | K9N5Q8 |
| PASARNLICAQY | 1 | 0 | 0 | 1485909 | QFS19619.1 | K9N5Q8 |
| PASCNGVLPVAF | 1 | 0 | 0 | 1485910 | AWK59942.1 | H9EJ42 |
| PASDGLKITKEV | 1 | 0 | 0 | 1485911 | AVP78030.1 | P0C6W2 |
| PASGYVAASAVN | 1 | 0 | 0 | 1485913 | AHI48543.1 | K9N7D2 |
| PASKQCFKVFFK | 1 | 0 | 0 | 1485915 | AWK59942.1 | H9EJ42 |
| PASMRNANGACI | 1 | 0 | 0 | 1485916 | AUM60013.1 | K9N7C7 |
| PASQVIVNNYDKSAGY | 1 | 0 | 0 | 1485918 | ABD75503.1 | A0A140H1G9 |
| PASQVIVNNYEK | 1 | 0 | 0 | 1485919 | AIL49570.1 | P0C6X6 |
| PASRDSVPLHII | 1 | 0 | 0 | 1485921 | AVV62541.1 | K9N7D2 |
| PASRDVNVTFFP | 1 | 0 | 0 | 1485922 | APO40578.1 | P0C6X7 |
| PASSQSAAKILA | 1 | 0 | 0 | 1485925 | AIW52722.1 | A0A0N7J131 |
| PASSQSAATILA | 1 | 0 | 0 | 1485926 | AIW52723.1 | A0A0N7J131 |

| | | | | | | |
|---|---|---|---|---|---|---|
| PASSQSSAKILA | 1 | 0 | 0 | 1485927 | ABB90486.1 | A0A0N7J131 |
| PASTVHTTATIP | 1 | 0 | 0 | 1485937 | ARO76383.1 | P59632 |
| PASVGSVPGEMR | 1 | 0 | 0 | 1485938 | 5W9H_A | K9N5Q8 |
| PASVKLPKGVGS | 1 | 0 | 0 | 1485939 | AEN19363.1 | P0C6X6 |
| PASVRDLICAQY | 1 | 0 | 0 | 1485941 | ALJ54502.1 | K9N5Q8 |
| PATAEPVRDEFS | 1 | 0 | 0 | 1485944 | AAA45463.1 | A0A0N7J131 |
| PATAEPVRDEVS | 1 | 0 | 0 | 1485945 | NP_073556.1 | A0A0N7J131 |
| PATDCSDGNYNC | 1 | 0 | 0 | 1485947 | AHL31360.1 | K9N5Q8 |
| PATNCSDGNYNR | 1 | 0 | 0 | 1485957 | ALA49671.1 | K9N5Q8 |
| PATRDTVPLHII | 1 | 0 | 0 | 1485960 | AUM60018.1 | K9N7D2 |
| PATSKSKRLRKR | 1 | 0 | 0 | 1485963 | AUM60013.1 | K9N7C7 |
| PAVNVSVSRFNP | 1 | 0 | 0 | 1485980 | AIV41873.1 | P36334 |
| PAVQFSTASVVE | 1 | 0 | 0 | 1485982 | AVV62535.1 | K9N7C7 |
| PAVSFDYLTREA | 1 | 0 | 0 | 1485984 | ANI69910.1 | K9N7C7 |
| PAVYMYNTGRSV | 1 | 0 | 0 | 1485986 | AUM60019.1 | K9N5R3 |
| PAYDGSAFRDLK | 1 | 0 | 0 | 1485994 | AHI48736.1 | K9N7C7 |
| PAYEASAFRDLK | 1 | 0 | 0 | 1485995 | AUM60013.1 | K9N7C7 |
| PAYESSAFRDLK | 1 | 0 | 0 | 1485996 | AUM60023.1 | K9N7C7 |
| PAYIAPQANSGD | 1 | 0 | 0 | 1486000 | P30215.1 | P30215 |
| PAYIAPQANSGG | 1 | 0 | 0 | 1486001 | AGT51669.1 | P30215 |
| PAYIAPQANSGI | 1 | 0 | 0 | 1486002 | AEN19365.1 | P30215 |
| PCADPSVVNSCA | 1 | 0 | 0 | 1486015 | ARB07438.1 | A0A140H1H1 |
| PCADPSVVNSCV | 1 | 0 | 0 | 1486016 | BBA20986.1 | A0A140H1H1 |
| PCADRSVVNSCV | 1 | 0 | 0 | 1486017 | ABD75545.1 | A0A140H1H1 |
| PCAIINDKVLTK | 1 | 0 | 0 | 1486021 | QBP84719.1 | P0C6X6 |
| PCAVSFADNNDI | 1 | 0 | 0 | 1486027 | AKK52599.1 | K9N4V7 |
| PCCGTGDLVSKL | 1 | 0 | 0 | 1486030 | AUM60013.1 | K9N7C7 |
| PCCTSYYSYMMP | 1 | 0 | 0 | 1486031 | AIW52827.1 | H9EJ42 |
| PCELHGNGSMIE | 1 | 0 | 0 | 1486032 | ALW82645.1 | K9N7C7 |
| PCEVEYGTEDDY | 1 | 0 | 0 | 1486034 | APO40578.1 | P0C6X7 |
| PCFITYPQQMFV | 1 | 0 | 0 | 1486038 | QDI73613.1 | K9N643 |
| PCFRIYPQQMFV | 1 | 0 | 0 | 1486040 | ALA49355.1 | K9N643 |
| PCFRTFPQQMFV | 1 | 0 | 0 | 1486041 | AVN89294.1 | K9N643 |
| PCGFKGGIPGNM | 1 | 0 | 0 | 1486046 | AIV42002.1 | P0C6X6 |
| PCGICLSDLDSL | 1 | 0 | 0 | 1486047 | APO40578.1 | P0C6X7 |
| PCGVHGNGSMIE | 1 | 0 | 0 | 1486050 | AGV08556.1 | K9N7C7 |
| PCGYSEPPKVAD | 1 | 0 | 0 | 1486052 | AEN19363.1 | P0C6X6 |
| PCKEIEGVGAKV | 1 | 0 | 0 | 1486058 | AGT51707.1 | P0C6X6 |
| PCKELEGVGARV | 1 | 0 | 0 | 1486060 | AXX83306.1 | P0C6X6 |
| PCKLDGSLCVGN | 1 | 0 | 0 | 1486063 | CAA83660.1 | P36334 |
| PCKLNSSLCVGS | 1 | 0 | 0 | 1486068 | AVQ05264.1 | P36334 |
| PCKLNSSSCVGS | 1 | 0 | 0 | 1486069 | AIX10756.1 | P36334 |
| PCKSNDSCPGKN | 1 | 0 | 0 | 1486070 | AIV41891.1 | P36334 |
| PCKSNGSCPGKN | 1 | 0 | 0 | 1486071 | AEN19358.1 | P36334 |
| PCKSNSSLCVGS | 1 | 0 | 0 | 1486072 | ARB66062.1 | P36334 |
| PCMCGKQATQYL | 1 | 0 | 0 | 1486077 | AVP78041.1 | P0C6W2 |
| PCMVGPDCAYFN | 1 | 0 | 0 | 1486078 | AUM60023.1 | K9N7C7 |
| PCNAQKQQIYFF | 1 | 0 | 0 | 1486081 | ABD75519.1 | A0A140H1G9 |
| PCNGTDIDYCVR | 1 | 0 | 0 | 1486083 | AIW52776.1 | A0A0P0LKV0 |
| PCNPPDQLVVYQ | 1 | 0 | 0 | 1486086 | AIW52757.1 | A0A0P0K6L9 |
| PCPSATYEGNSP | 1 | 0 | 0 | 1486096 | ABI96964.1 | P59635 |
| PCPSGIGAVGGD | 1 | 0 | 0 | 1486097 | QCI31476.1 | K9N4V7 |
| PCQFKDKNLQGL | 1 | 0 | 0 | 1486103 | AGT51787.1 | P0C6X6 |
| PCQLANYKQSIP | 1 | 0 | 0 | 1486104 | AVV62524.1 | K9N7C7 |
| PCSDKAYKIAEL | 1 | 0 | 0 | 1486107 | QVO99997.1 | P0DTD1 |
| PCSPNTYEGNSP | 1 | 0 | 0 | 1486111 | APO40584.1 | P59635 |
| PCSSCFGKNNGI | 1 | 0 | 0 | 1486112 | QEG03814.1 | P36334 |
| PCSSCHGKNNGI | 1 | 0 | 0 | 1486113 | ARB66068.1 | P36334 |
| PCSSCPGKNNGI | 1 | 0 | 0 | 1486114 | AWW13566.1 | P36334 |
| PCSSCPGKNNGL | 1 | 0 | 0 | 1486115 | AIL49503.1 | P36334 |
| PCSSCPSKNNGI | 1 | 0 | 0 | 1486116 | QDH43719.1 | P36334 |
| PCSSCSCPGKNN | 1 | 0 | 0 | 1486117 | AVR40344.1 | P36334 |
| PCSSCSGKNNGI | 1 | 0 | 0 | 1486118 | ANZ78838.1 | P36334 |
| PCSSCSGKNNGV | 1 | 0 | 0 | 1486119 | QEG03757.1 | P36334 |
| PCSVCLGGLDSL | 1 | 0 | 0 | 1486121 | ARO76381.1 | P0C6U8 |
| PCSVRSVPGEMR | 1 | 0 | 0 | 1486123 | AKN24812.1 | K9N5Q8 |
| PCYCFNADGDEA | 1 | 0 | 0 | 1486162 | AUM60013.1 | K9N7C7 |
| PCYCFNAEGGAS | 1 | 0 | 0 | 1486165 | AKN24802.1 | K9N7C7 |
| PDAAEKLASPMK | 1 | 0 | 0 | 1486174 | AZS52616.1 | A0A140H1G9 |
| PDAFQSVYHTFT | 1 | 0 | 0 | 1486176 | ARK08626.1 | H9EJ42 |
| PDAIARGCYSSL | 1 | 0 | 0 | 1486178 | AVV62526.1 | K9N5Q8 |
| PDARAKQDFSLL | 1 | 0 | 0 | 1486180 | ATG84722.1 | K9N7C7 |
| PDARFHDFTPLA | 1 | 0 | 0 | 1486183 | AVV62529.1 | K9N643 |
| PDATNNTCLEVQ | 1 | 0 | 0 | 1486185 | AUM60014.1 | K9N5Q8 |
| PDAVTAYNGYVT | 1 | 0 | 0 | 1486187 | APO40578.1 | P0C6X7 |
| PDCDIDNWLNKV | 1 | 0 | 0 | 1486191 | AMM42407.1 | A0A140H1H1 |
| PDCDIDNWLNNV | 1 | 0 | 0 | 1486192 | BBA20986.1 | A0A140H1H1 |
| PDCNIEAWLNDK | 1 | 0 | 0 | 1486195 | CAA83660.1 | P36334 |
| PDDAARLLHILL | 1 | 0 | 0 | 1486196 | AVV62535.1 | K9N7C7 |
| PDDASRLIHTLL | 1 | 0 | 0 | 1486197 | AVV62524.1 | K9N7C7 |
| PDDQIGFYRRAT | 1 | 0 | 0 | 1486208 | AAT76155.1 | Q3LZX4 |
| PDDSSTIVTGLL | 1 | 0 | 0 | 1486210 | AKA09622.1 | A0A0P0G321 |
| PDEEEDDDAECE | 1 | 0 | 0 | 1486217 | ARO76381.1 | P0C6U8 |
| PDEGGDVLLAEF | 1 | 0 | 0 | 1486219 | AUM60013.1 | K9N7C7 |
| PDEKQNGCSLYL | 1 | 0 | 0 | 1486221 | ATQ39389.1 | K9N7C7 |
| PDEKQNGHSLYL | 1 | 0 | 0 | 1486222 | AXN92247.1 | K9N7C7 |
| PDEKQNGLSLYL | 1 | 0 | 0 | 1486223 | AUM60013.1 | K9N7C7 |
| PDEPHKDVYELR | 1 | 0 | 0 | 1486224 | AIX10716.1 | P33469 |
| PDFKEELDQWLK | 1 | 0 | 0 | 1486232 | ARB07414.1 | P36334 |
| PDFKEESDQWFK | 1 | 0 | 0 | 1486233 | BBA20973.1 | P36334 |
| PDFKGELDQWFK | 1 | 0 | 0 | 1486235 | QDH43719.1 | P36334 |

| | | | | | | |
|---|---|---|---|---|---|---|
| PDFKLLRNLQAD | 1 | 0 | 0 | 1486237 | ATG84842.1 | K9N7C7 |
| PDFNVWNQVVTY | 1 | 0 | 0 | 1486238 | AVV62524.1 | K9N7C7 |
| PDFREELDQWFK | 1 | 0 | 0 | 1486239 | AVQ05264.1 | P36334 |
| PDGFYSTSPIQP | 1 | 0 | 0 | 1486256 | AIW52757.1 | A0A0P0K6L9 |
| PDGFYSTSPTQP | 1 | 0 | 0 | 1486258 | AIW52741.1 | A0A0P0K6L9 |
| PDGFYSTSSIQP | 1 | 0 | 0 | 1486259 | AMM42536.1 | A0A0P0K6L9 |
| PDGKPCTPPAFN | 1 | 0 | 0 | 1486260 | ATO98231.1 | Q3LZX1 |
| PDGKSFTMTFGG | 1 | 0 | 0 | 1486265 | AVV62535.1 | K9N7C7 |
| PDGQGVPIAYGI | 1 | 0 | 0 | 1486267 | AMM42397.1 | Q19U29 |
| PDGYPLECIKNF | 1 | 0 | 0 | 1486275 | AVP78041.1 | P0C6W2 |
| PDHSVADYHNHS | 1 | 0 | 0 | 1486277 | APO40578.1 | P0C6X7 |
| PDHTCNTPTQCY | 1 | 0 | 0 | 1486278 | AIV41780.1 | P0C6X6 |
| PDHVSLTVPCCG | 1 | 0 | 0 | 1486279 | AVV62535.1 | K9N7C7 |
| PDHVSRTVPCCG | 1 | 0 | 0 | 1486280 | AUM60013.1 | K9N7C7 |
| PDIDLGDISGIN | 1 | 0 | 0 | 1486282 | AVP78031.1 | Q3LZX1 |
| PDIVDVPPQVVE | 1 | 0 | 0 | 1486289 | AVN89352.1 | K9N7C7 |
| PDKNNGIGICPA | 1 | 0 | 0 | 1486293 | AIX10751.1 | P36334 |
| PDKNNGIGTCPA | 1 | 0 | 0 | 1486294 | AIX10750.1 | P36334 |
| PDKSVYDQIVDN | 1 | 0 | 0 | 1486296 | AEN19363.1 | P0C6X6 |
| PDLALDYINVTF | 1 | 0 | 0 | 1486299 | ANZ78847.1 | P36334 |
| PDLEGKFYGPFV | 1 | 0 | 0 | 1486302 | ACV88184.1 | P0C6X7 |
| PDLESYSKIVCD | 1 | 0 | 0 | 1486303 | AWK59942.1 | H9EJ42 |
| PDLGTDDDDKAM | 1 | 0 | 0 | 1486306 | 3R24_A | P0C6X7 |
| PDLKKGPHEFCS | 1 | 0 | 0 | 1486307 | AVV62524.1 | K9N7C7 |
| PDLLRAHPAEPL | 1 | 0 | 0 | 1486310 | AVR40343.1 | Q4VID0 |
| PDLLRAHPVEPL | 1 | 0 | 0 | 1486311 | ARE29989.1 | Q4VID0 |
| PDLLYAHPAEPL | 1 | 0 | 0 | 1486312 | AHF54648.1 | Q4VID0 |
| PDLPICVHDPLP | 1 | 0 | 0 | 1486315 | AIV41818.1 | P30215 |
| PDLPICVYDPLP | 1 | 0 | 0 | 1486317 | AEN19365.1 | P30215 |
| PDLPICVYDRLP | 1 | 0 | 0 | 1486318 | AIL49324.1 | P30215 |
| PDLPICVYDSDP | 1 | 0 | 0 | 1486319 | 5N11_A | P30215 |
| PDLPICVYDSLP | 1 | 0 | 0 | 1486320 | AXX83314.1 | P30215 |
| PDLPICVYHPLP | 1 | 0 | 0 | 1486321 | QEG03802.1 | P30215 |
| PDLPISVYDPLP | 1 | 0 | 0 | 1486322 | AIL49333.1 | P30215 |
| PDLPNCNIGAWL | 1 | 0 | 0 | 1486324 | AHA80222.1 | P36334 |
| PDLPVCVYDPLP | 1 | 0 | 0 | 1486325 | ARB07432.1 | P30215 |
| PDLQKAAVTILD | 1 | 0 | 0 | 1486326 | ARO76381.1 | P0C6U8 |
| PDLSAYYVKDGK | 1 | 0 | 0 | 1486330 | AVV62524.1 | K9N7C7 |
| PDLSFDYINVTF | 1 | 0 | 0 | 1486331 | AIX10753.1 | P36334 |
| PDLSLDYINATF | 1 | 0 | 0 | 1486332 | QEG03776.1 | P36334 |
| PDLTAFYVKDGK | 1 | 0 | 0 | 1486338 | AUM60013.1 | K9N7C7 |
| PDMADEIANLVL | 1 | 0 | 0 | 1486348 | ARU07581.1 | Q19U29 |
| PDMADHIASLVL | 1 | 0 | 0 | 1486349 | AIL49401.1 | P33469 |
| PDMADQIASLVL | 1 | 0 | 0 | 1486350 | AEN19370.1 | P33469 |
| PDPITFAGPYKC | 1 | 0 | 0 | 1486358 | ATP16757.1 | P36334 |
| PDPITSKSTGPY | 1 | 0 | 0 | 1486363 | CAA83660.1 | P36334 |
| PDPKFVDRYYAF | 1 | 0 | 0 | 1486365 | ATU80202.1 | K9N7C7 |
| PDPPNCNIEAWL | 1 | 0 | 0 | 1486372 | AHA80201.1 | P36334 |
| PDPSKPTKRSFI | 1 | 0 | 0 | 1486375 | ATO98181.1 | Q3LZX1 |
| PDQICNTPTQCY | 1 | 0 | 0 | 1486377 | ARB07411.1 | P0C6X6 |
| PDQLNDLVYSLI | 1 | 0 | 0 | 1486378 | ARU07574.1 | A0A140H1G9 |
| PDQLVVYQQAIV | 1 | 0 | 0 | 1486379 | ABB90509.1 | A0A0P0K6L9 |
| PDQLVVYQQAVA | 1 | 0 | 0 | 1486380 | ADK37701.1 | A0A0P0K6L9 |
| PDQLVVYQQAVV | 1 | 0 | 0 | 1486381 | AIW52757.1 | A0A0P0K6L9 |
| PDQQFHNFSYLA | 1 | 0 | 0 | 1486382 | AVV62540.1 | K9N643 |
| PDRMLKTSMAPI | 1 | 0 | 0 | 1486385 | AVR40343.1 | Q4VID0 |
| PDSAKSACIEVD | 1 | 0 | 0 | 1486386 | AVN89387.1 | K9N5Q8 |
| PDSAKSVCIEVD | 1 | 0 | 0 | 1486387 | QAT98898.1 | K9N5Q8 |
| PDSATLVKDIDI | 1 | 0 | 0 | 1486388 | AVP78041.1 | P0C6W2 |
| PDSATLVKDIDT | 1 | 0 | 0 | 1486389 | AVP78030.1 | P0C6W2 |
| PDSIAISDLGSL | 1 | 0 | 0 | 1486393 | AVV62524.1 | K9N7C7 |
| PDSNFLYKYYSL | 1 | 0 | 0 | 1486398 | AVV62524.1 | K9N7C7 |
| PDSSKPTKRSFI | 1 | 0 | 0 | 1486401 | ARO76382.1 | P59594 |
| PDSVKPACIEVD | 1 | 0 | 0 | 1486402 | ASU91284.1 | K9N5Q8 |
| PDSVKSACIAVD | 1 | 0 | 0 | 1486403 | ALJ54490.1 | K9N5Q8 |
| PDSVKSACIEVH | 1 | 0 | 0 | 1486405 | ANI69922.1 | K9N5Q8 |
| PDSVKSACIKVD | 1 | 0 | 0 | 1486406 | QCQ29075.1 | K9N5Q8 |
| PDSVSTSDLGSL | 1 | 0 | 0 | 1486408 | AXN92247.1 | K9N7C7 |
| PDTHSTLTPRSV | 1 | 0 | 0 | 1486410 | AID55099.1 | K9N5Q8 |
| PDTIYRSDTLVL | 1 | 0 | 0 | 1486411 | AVP78031.1 | Q3LZX1 |
| PDTPITLTPRSV | 1 | 0 | 0 | 1486412 | ANI69900.1 | K9N5Q8 |
| PDTPSPLTPRSV | 1 | 0 | 0 | 1486413 | AJG44080.1 | K9N5Q8 |
| PDTPSTFTPRSV | 1 | 0 | 0 | 1486414 | AVK87439.1 | K9N5Q8 |
| PDTPSTLAPRSV | 1 | 0 | 0 | 1486415 | ALJ54472.1 | K9N5Q8 |
| PDTPSTLKPRSV | 1 | 0 | 0 | 1486416 | ALJ54490.1 | K9N5Q8 |
| PDTPSTLTLRSV | 1 | 0 | 0 | 1486417 | ANI69889.1 | K9N5Q8 |
| PDTPSTLTPASV | 1 | 0 | 0 | 1486418 | 5W9H_A | K9N5Q8 |
| PDTPSTLTPCSV | 1 | 0 | 0 | 1486419 | AKN24812.1 | K9N5Q8 |
| PDTVGVPPQVVE | 1 | 0 | 0 | 1486426 | AHL18091.1 | K9N7C7 |
| PDTYKCPQTKSL | 1 | 0 | 0 | 1486428 | ARE30008.1 | P36334 |
| PDTYVALPIYYQ | 1 | 0 | 0 | 1486429 | AWO68991.1 | A0A0P0G321 |
| PDTYVALPVYYQ | 1 | 0 | 0 | 1486430 | AWO68966.1 | A0A0P0G321 |
| PDVFLHKCYRYP | 1 | 0 | 0 | 1486434 | QED88024.1 | H9EJ42 |
| PDVLRVYANLGE | 1 | 0 | 0 | 1486436 | AVP78030.1 | P0C6W2 |
| PDVMLNISTPNL | 1 | 0 | 0 | 1486437 | CAA83660.1 | P36334 |
| PDVPICVYDPLP | 1 | 0 | 0 | 1486438 | AAA73152.1 | P30215 |
| PDVQFEQLDNKL | 1 | 0 | 0 | 1486441 | AVV62526.1 | K9N5Q8 |
| PDVSGVCKFKGK | 1 | 0 | 0 | 1486442 | AVV62524.1 | K9N7C7 |
| PDVSGVCKFKGR | 1 | 0 | 0 | 1486443 | AVV62535.1 | K9N7C7 |
| PDYGTYKNTCEG | 1 | 0 | 0 | 1486449 | APO40578.1 | P0C6X7 |
| PDYNYYKYNLPT | 1 | 0 | 0 | 1486451 | ACV87150.1 | A0A140H1G9 |

| | | | | | | |
|---|---|---|---|---|---|---|
| PDYSTYKNTCDG | 1 | 0 | 0 | 1486455 | QXU68417.1 | P0DTD1 |
| PEAALPYGANKE | 1 | 0 | 0 | 1486460 | APO40586.1 | P59595 |
| PEAEVQIDRLIT | 1 | 0 | 0 | 1486462 | 6CRW_A | P59594 |
| PEANINQDSYGG | 1 | 0 | 0 | 1486467 | ABD75615.1 | A0A140H1G9 |
| PEANLPFRAVKD | 1 | 0 | 0 | 1486468 | AVV62533.1 | K9N4V7 |
| PEANLPFRAVKE | 1 | 0 | 0 | 1486469 | AVV62544.1 | K9N4V7 |
| PEAPRDGQAYVR | 1 | 0 | 0 | 1486472 | 6CRW_A | P59594 |
| PEDDDFLEASDV | 1 | 0 | 0 | 1486476 | ATN39862.1 | P0C6X6 |
| PEDDDFLEENDV | 1 | 0 | 0 | 1486477 | QBQ01836.1 | P0C6X6 |
| PEDDDFLEGSDV | 1 | 0 | 0 | 1486480 | AIV41846.1 | P0C6X6 |
| PEDEGEDDCEEG | 1 | 0 | 0 | 1486481 | AVP78041.1 | P0C6W2 |
| PEDFSVIVQPTH | 1 | 0 | 0 | 1486482 | AKN24752.1 | K9N643 |
| PEDLMEVDHSGP | 1 | 0 | 0 | 1486485 | AVV62529.1 | K9N643 |
| PEDLSVIIQPTH | 1 | 0 | 0 | 1486486 | ALA49344.1 | K9N643 |
| PEDLSVIQPTHY | 1 | 0 | 0 | 1486487 | QBF80458.1 | K9N643 |
| PEDLSVMVQPTH | 1 | 0 | 0 | 1486488 | QBF80535.1 | K9N643 |
| PEDLSVVTQPTH | 1 | 0 | 0 | 1486489 | ATQ39396.1 | K9N643 |
| PEDVNMQLMGVV | 1 | 0 | 0 | 1486493 | AQZ41284.1 | K9N7C7 |
| PEEAPAPELQVP | 1 | 0 | 0 | 1486494 | APO40578.1 | P0C6X7 |
| PEEFANHVANFQ | 1 | 0 | 0 | 1486496 | AVV62524.1 | K9N7C7 |
| PEEHFIETVSLA | 1 | 0 | 0 | 1486499 | AVP78041.1 | P0C6W2 |
| PEEHFVETVSLA | 1 | 0 | 0 | 1486500 | ARO76381.1 | P0C6U8 |
| PEELSVIVQPTH | 1 | 0 | 0 | 1486503 | ASU90849.1 | K9N643 |
| PEEVHEVTPVRS | 1 | 0 | 0 | 1486506 | AUM60013.1 | K9N7C7 |
| PEFSIVCRFDTR | 1 | 0 | 0 | 1486509 | AWK59942.1 | H9EJ42 |
| PEGCVLTNTGSV | 1 | 0 | 0 | 1486513 | NP_073549.1 | A0A0P0LKV0 |
| PEGLVFFHTVLL | 1 | 0 | 0 | 1486514 | AIW52753.1 | A0A0P0K6L9 |
| PEGVCLGAGQFL | 1 | 0 | 0 | 1486515 | AWW13571.1 | P0C6X6 |
| PEGVCLGAGQFV | 1 | 0 | 0 | 1486516 | AQN78773.1 | P0C6X6 |
| PEGWHVVNKFYQ | 1 | 0 | 0 | 1486517 | AIV41876.1 | P0C6X6 |
| PEHCQNYSGCML | 1 | 0 | 0 | 1486519 | AXN92229.1 | K9N796 |
| PEHCQNYSGFML | 1 | 0 | 0 | 1486520 | AVN89325.1 | K9N796 |
| PEHCQNYSGSML | 1 | 0 | 0 | 1486522 | ASU45697.1 | K9N796 |
| PEHCQSHTGSLL | 1 | 0 | 0 | 1486523 | AUM60025.1 | K9N796 |
| PEHCQSYTGKLL | 1 | 0 | 0 | 1486524 | AVV62527.1 | K9N796 |
| PEHNFIIYDASD | 1 | 0 | 0 | 1486526 | AVV62524.1 | K9N7C7 |
| PEHNLIIYDSSD | 1 | 0 | 0 | 1486527 | AUM60013.1 | K9N7C7 |
| PEKAQGMLLALL | 1 | 0 | 0 | 1486536 | AWK59942.1 | H9EJ42 |
| PEKAQSMLLALL | 1 | 0 | 0 | 1486537 | P0C6U6.1 | H9EJ42 |
| PEKCIFTSACTR | 1 | 0 | 0 | 1486538 | NP_073549.1 | A0A0P0LKV0 |
| PEKFPTTVTCEN | 1 | 0 | 0 | 1486540 | NP_073549.1 | A0A0P0LKV0 |
| PEKKFLRYINGF | 1 | 0 | 0 | 1486541 | AQN78665.1 | Q04853 |
| PEKKFPPYINGF | 1 | 0 | 0 | 1486542 | AIV41989.1 | Q04853 |
| PELESFKQELDK | 1 | 0 | 0 | 1486551 | APO40579.1 | P59594 |
| PELLDPLNLSAD | 1 | 0 | 0 | 1486556 | ADN03343.1 | Q19U29 |
| PELMEDLEADPK | 1 | 0 | 0 | 1486557 | AUM60013.1 | K9N7C7 |
| PELQVPVTLEQK | 1 | 0 | 0 | 1486558 | APO40578.1 | P0C6X7 |
| PELVIAANKTKG | 1 | 0 | 0 | 1486559 | AVV62524.1 | K9N7C7 |
| PELYHYQECVRG | 1 | 0 | 0 | 1486560 | 1YO4_A | P59635 |
| PEMKILQNLGVD | 1 | 0 | 0 | 1486561 | APO40578.1 | P0C6X7 |
| PENDDDAWFKLS | 1 | 0 | 0 | 1486563 | AWW13564.1 | Q80872 |
| PENDDDAWFKPS | 1 | 0 | 0 | 1486564 | AQN78686.1 | Q80872 |
| PEPALLFRNIKC | 1 | 0 | 0 | 1486566 | AIX10763.1 | P36334 |
| PEPEPTLEEPVN | 1 | 0 | 0 | 1486568 | ARO76381.1 | P0C6U8 |
| PEPIIENNVVVM | 1 | 0 | 0 | 1486571 | AIX09799.1 | P36334 |
| PEPINENNVVVM | 1 | 0 | 0 | 1486572 | AGT51800.1 | P36334 |
| PEPITENNVVVT | 1 | 0 | 0 | 1486575 | ARB66070.1 | P36334 |
| PEPITGNNVVVM | 1 | 0 | 0 | 1486576 | CAA83660.1 | P36334 |
| PEPITMANARYV | 1 | 0 | 0 | 1486577 | AVV62537.1 | K9N5Q8 |
| PEPITMANTRFV | 1 | 0 | 0 | 1486578 | AUM60014.1 | K9N5Q8 |
| PEPITSFNTKYV | 1 | 0 | 0 | 1486579 | AVN89387.1 | K9N5Q8 |
| PEPITSLNIKYV | 1 | 0 | 0 | 1486580 | AHI48737.1 | K9N5Q8 |
| PEPQRGRQGRIP | 1 | 0 | 0 | 1486585 | APT69853.1 | A0A0N7J131 |
| PEPTLEEPVNQF | 1 | 0 | 0 | 1486586 | ARO76381.1 | P0C6U8 |
| PEQDAQIDRLIN | 1 | 0 | 0 | 1486589 | 5W9H_A | K9N5Q8 |
| PEQKQNGFSFFL | 1 | 0 | 0 | 1486591 | AVV62524.1 | K9N7C7 |
| PESAAYYVGNLK | 1 | 0 | 0 | 1486597 | ATO98181.1 | Q3LZX1 |
| PESFDSLNLSAD | 1 | 0 | 0 | 1486598 | AMM42397.1 | Q19U29 |
| PESIPLADDDSA | 1 | 0 | 0 | 1486599 | AMM42614.1 | H9EJ69 |
| PESKSFTMTFGG | 1 | 0 | 0 | 1486601 | AVV62524.1 | K9N7C7 |
| PESNADQESYGG | 1 | 0 | 0 | 1486602 | AUM60013.1 | K9N7C7 |
| PESNADQETYGG | 1 | 0 | 0 | 1486603 | AVV62524.1 | K9N7C7 |
| PETEVASYSDQQ | 1 | 0 | 0 | 1486605 | AVV62530.1 | K9N7D2 |
| PETNAIISLQVY | 1 | 0 | 0 | 1486607 | ARE29965.1 | U3M6Q8 |
| PETNCLLNVPIG | 1 | 0 | 0 | 1486609 | ASY99815.1 | K9N7A1 |
| PETNCLLNVPVG | 1 | 0 | 0 | 1486611 | AVV62543.1 | K9N7A1 |
| PETYFTQSRNLE | 1 | 0 | 0 | 1486618 | APO40578.1 | P0C6X7 |
| PEVEELNPSKLL | 1 | 0 | 0 | 1486624 | AGT51436.1 | Q4VID0 |
| PEVKEIAPVYEA | 1 | 0 | 0 | 1486627 | ALA49362.1 | K9N7C7 |
| PEVKEVAPVYEA | 1 | 0 | 0 | 1486628 | AXN92247.1 | K9N7C7 |
| PEVLSPINSSGF | 1 | 0 | 0 | 1486634 | AVV62526.1 | K9N5Q8 |
| PEVTTSQYSHRS | 1 | 0 | 0 | 1486635 | AUM60024.1 | K9N5Q8 |
| PEVTTSQYSYRS | 1 | 0 | 0 | 1486636 | AUM60014.1 | K9N5Q8 |
| PEWIDDLEADPK | 1 | 0 | 0 | 1486641 | ATQ39389.1 | K9N7C7 |
| PEWMDEFEADPK | 1 | 0 | 0 | 1486643 | ATG84853.1 | K9N7C7 |
| PEWMDYFEADPK | 1 | 0 | 0 | 1486644 | ASY99841.1 | K9N638 |
| PEYDDFLEESDV | 1 | 0 | 0 | 1486648 | AGT51707.1 | P0C6X6 |
| PEYMAVAVPSTT | 1 | 0 | 0 | 1486650 | APT69865.1 | A0A0P0KXN3 |
| PEYMTVAVPSTT | 1 | 0 | 0 | 1486651 | AST12967.1 | A0A0P0KXN3 |
| PEYNFIIYDSSD | 1 | 0 | 0 | 1486653 | AXN92247.1 | K9N7C7 |
| PFAELTGTQSVS | 1 | 0 | 0 | 1486656 | QAT98905.1 | K9N796 |

| | | | | | | |
|---|---|---|---|---|---|---|
| PFAGYTESAVNS | 1 | 0 | 0 | 1486657 | ARQ84748.1 | K9N4V0 |
| PFAKGDSCSSSC | 1 | 0 | 0 | 1486660 | AUM60013.1 | K9N7C7 |
| PFAKSDGCSCSC | 1 | 0 | 0 | 1486661 | AVV62524.1 | K9N7C7 |
| PFAVYKNVKFYL | 1 | 0 | 0 | 1486671 | AWK59942.1 | H9EJ42 |
| PFCGSGNLVEKL | 1 | 0 | 0 | 1486672 | ASU45728.1 | K9N7C7 |
| PFDAAVAAAMAT | 1 | 0 | 0 | 1486674 | AVV62544.1 | K9N4V7 |
| PFDAVFNATRFP | 1 | 0 | 0 | 1486677 | ATO98145.1 | Q3LZX1 |
| PFDITSAGKFDS | 1 | 0 | 0 | 1486680 | APO40578.1 | P0C6X7 |
| PFDSNLTQYFSL | 1 | 0 | 0 | 1486682 | ATO98181.1 | Q3LZX1 |
| PFDVVAQCSGVT | 1 | 0 | 0 | 1486685 | 3M3T_A | P0C6X7 |
| PFDVVRQCSGVV | 1 | 0 | 0 | 1486687 | 2ALV_A | P0C6X7 |
| PFEDLNKCRAYM | 1 | 0 | 0 | 1486688 | AVV62524.1 | K9N7C7 |
| PFEDLSKCRAYL | 1 | 0 | 0 | 1486689 | AVV62535.1 | K9N7C7 |
| PFEDLTKCRTYL | 1 | 0 | 0 | 1486690 | AUM60013.1 | K9N7C7 |
| PFEEVEHVLSIK | 1 | 0 | 0 | 1486691 | ARU07593.1 | H9EJ42 |
| PFEEVEHVLSIR | 1 | 0 | 0 | 1486692 | ARE29978.1 | H9EJ42 |
| PFEPQTIQPEVK | 1 | 0 | 0 | 1486699 | ALW82701.1 | K9N7C7 |
| PFERDLSSDENG | 1 | 0 | 0 | 1486702 | AVP78031.1 | Q3LZX1 |
| PFERDLTSDENG | 1 | 0 | 0 | 1486703 | ATO98120.1 | Q3LZX1 |
| PFEVLCLDDAQG | 1 | 0 | 0 | 1486708 | APO40578.1 | P0C6X7 |
| PFEYTSPTDKLA | 1 | 0 | 0 | 1486709 | AXX83349.1 | Q80872 |
| PFFTENKPSVEQ | 1 | 0 | 0 | 1486713 | AVP78041.1 | P0C6W2 |
| PFFVDVQQWGYV | 1 | 0 | 0 | 1486715 | AXN92247.1 | K9N7C7 |
| PFFYYSDTSCVY | 1 | 0 | 0 | 1486720 | QBP84728.1 | P0C6X6 |
| PFGEVEHALSIK | 1 | 0 | 0 | 1486727 | AGT51337.1 | H9EJ42 |
| PFGEVEHSLPIK | 1 | 0 | 0 | 1486728 | AVA26872.1 | H9EJ42 |
| PFGEVEHSLTIK | 1 | 0 | 0 | 1486729 | AFD98763.1 | H9EJ42 |
| PFGEVEYSLPIK | 1 | 0 | 0 | 1486730 | AWK59942.1 | H9EJ42 |
| PFGGQPVSEYHI | 1 | 0 | 0 | 1486735 | AKN24752.1 | K9N643 |
| PFGHGVYFSTID | 1 | 0 | 0 | 1486737 | ATO98145.1 | Q3LZX1 |
| PFGQVFNASNFP | 1 | 0 | 0 | 1486743 | APO40579.1 | P59594 |
| PFGVVDTEWGNM | 1 | 0 | 0 | 1486746 | AXG21652.1 | K9N7C7 |
| PFHYERDNTSYP | 1 | 0 | 0 | 1486756 | AHI48797.1 | K9N638 |
| PFHYERNNTSCP | 1 | 0 | 0 | 1486757 | AVN89451.1 | K9N7C7 |
| PFIAVSKPMGTQ | 1 | 0 | 0 | 1486759 | ABD73001.1 | P59594 |
| PFKDGLYFAATE | 1 | 0 | 0 | 1486768 | APO40579.1 | P59594 |
| PFKFCSETVNGY | 1 | 0 | 0 | 1486769 | ARB07424.1 | A0A0P0ILY9 |
| PFLGQAATTTSN | 1 | 0 | 0 | 1486781 | ARO76381.1 | P0C6U8 |
| PFLSDFINGISA | 1 | 0 | 0 | 1486796 | AKA09642.1 | P36334 |
| PFLYLLALIYFL | 1 | 0 | 0 | 1486802 | APO40580.1 | P59632 |
| PFMDDAINVGGT | 1 | 0 | 0 | 1486805 | AVN89322.1 | K9N7C7 |
| PFMGNAINVGGR | 1 | 0 | 0 | 1486807 | ATQ39389.1 | K9N7C7 |
| PFNNMLGYNPSK | 1 | 0 | 0 | 1486823 | AVV62524.1 | K9N7C7 |
| PFSDRPLNLRGW | 1 | 0 | 0 | 1486842 | AWK59942.1 | H9EJ42 |
| PFSPNGKPCTPP | 1 | 0 | 0 | 1486848 | ABD72972.1 | P59594 |
| PFTVNSINDSLE | 1 | 0 | 0 | 1486864 | AIL49542.1 | P36334 |
| PFTVNSKNDNLE | 1 | 0 | 0 | 1486865 | AIL49538.1 | P36334 |
| PFTVNSVNDSLK | 1 | 0 | 0 | 1486866 | AIL49500.1 | P36334 |
| PFTVNSVSDSLE | 1 | 0 | 0 | 1486867 | ARB66068.1 | P36334 |
| PFVDIQTAQAAG | 1 | 0 | 0 | 1486869 | 3ATW_A | P0C6X7 |
| PFVEGAINVGKN | 1 | 0 | 0 | 1486871 | AUM60023.1 | K9N7C7 |
| PFVKDGFSFVDD | 1 | 0 | 0 | 1486874 | AVV62535.1 | K9N7C7 |
| PFVKDGFSFVND | 1 | 0 | 0 | 1486875 | AVV62524.1 | K9N7C7 |
| PFVKGDFSFISD | 1 | 0 | 0 | 1486879 | AUM60013.1 | K9N7C7 |
| PFVKGDSCSSNC | 1 | 0 | 0 | 1486880 | AKN24784.1 | K9N7C7 |
| PFVQEQIGSFIV | 1 | 0 | 0 | 1486883 | AVV62531.1 | K9N5R3 |
| PFVSDSVATYFG | 1 | 0 | 0 | 1486888 | AEN19363.1 | P0C6X6 |
| PFVSFIYFITMS | 1 | 0 | 0 | 1486889 | AWK59942.1 | H9EJ42 |
| PFYLNVQYRING | 1 | 0 | 0 | 1486905 | CAA79896.1 | P36334 |
| PFYVPEHCQNYS | 1 | 0 | 0 | 1486910 | AVN89384.1 | K9N796 |
| PFYYERDNTSCP | 1 | 0 | 0 | 1486911 | AVN89322.1 | K9N7C7 |
| PGAADAERMAMY | 1 | 0 | 0 | 1486912 | APT69876.1 | A0A0P0K6L9 |
| PGAFAYSQMRPH | 1 | 0 | 0 | 1486915 | AXN92247.1 | K9N7C7 |
| PGAGIPIVDSYY | 1 | 0 | 0 | 1486918 | APO40578.1 | P0C6X7 |
| PGAHAYETMKPH | 1 | 0 | 0 | 1486919 | AVV62535.1 | K9N7C7 |
| PGAHAYGSMKPH | 1 | 0 | 0 | 1486920 | AVV62524.1 | K9N7C7 |
| PGAHPYSAMKPH | 1 | 0 | 0 | 1486921 | AUM60013.1 | K9N7C7 |
| PGALEFNPSQTS | 1 | 0 | 0 | 1486922 | AGT21342.1 | A0A0N7J131 |
| PGALQGNFGDLQ | 1 | 0 | 0 | 1486923 | AVV62544.1 | K9N4V7 |
| PGAMQTGFTTTN | 1 | 0 | 0 | 1486924 | AVV62526.1 | K9N5Q8 |
| PGCGGILSVIDD | 1 | 0 | 0 | 1486928 | AFR79256.1 | A0A0P0ILY9 |
| PGDLQGNFGDLK | 1 | 0 | 0 | 1486931 | AWM99589.1 | K9N4V7 |
| PGDVIITSTDAG | 1 | 0 | 0 | 1486935 | AGT51379.1 | H9EJ42 |
| PGDVIITSTGAG | 1 | 0 | 0 | 1486936 | AWK59942.1 | H9EJ42 |
| PGEFASHVANFQ | 1 | 0 | 0 | 1486938 | AXN73435.1 | K9N7C7 |
| PGEHFKHLVPLM | 1 | 0 | 0 | 1486940 | ALW82707.1 | K9N7C7 |
| PGEMAKKQIGLL | 1 | 0 | 0 | 1486942 | AVV62535.1 | K9N7C7 |
| PGEMRLASIAFN | 1 | 0 | 0 | 1486945 | AXN92228.1 | K9N5Q8 |
| PGEQFKHLIPLM | 1 | 0 | 0 | 1486946 | QXH28115.1 | P0DTD1 |
| PGEQFKHLIPLV | 1 | 0 | 0 | 1486947 | QBP84719.1 | P0C6X6 |
| PGEQFKHLVPLI | 1 | 0 | 0 | 1486948 | AHL18091.1 | K9N7C7 |
| PGESKPQSRNPS | 1 | 0 | 0 | 1486950 | AAA45463.1 | A0A0N7J131 |
| PGETGVIADYNY | 1 | 0 | 0 | 1486951 | ATO98181.1 | Q3LZX1 |
| PGEVAKKQIGLL | 1 | 0 | 0 | 1486952 | AVV62524.1 | K9N7C7 |
| PGEYSICKNFAP | 1 | 0 | 0 | 1486953 | AVV62537.1 | K9N5Q8 |
| PGEYSICRGFAP | 1 | 0 | 0 | 1486954 | AVV62526.1 | K9N5Q8 |
| PGFGGNFNLKLL | 1 | 0 | 0 | 1486958 | AOR17482.1 | K9N5Q8 |
| PGFGVAVADSYF | 1 | 0 | 0 | 1486959 | ABD75559.1 | A0A140H1G9 |
| PGFKEELDQWFK | 1 | 0 | 0 | 1486961 | AIL49523.1 | P36334 |
| PGFLTPFMILLV | 1 | 0 | 0 | 1486962 | NP_073549.1 | A0A0P0LKV0 |
| PGFMKQYGDCLG | 1 | 0 | 0 | 1486963 | APO40579.1 | P59594 |

| | | | | | | |
|---|---|---|---|---|---|---|
| PGFNEKTHGPLS | 1 | 0 | 0 | 1486964 | AVP78030.1 | P0C6W2 |
| PGGCAMPIMANL | 1 | 0 | 0 | 1486969 | AIW52757.1 | A0A0P0K6L9 |
| PGGCAMPIVANL | 1 | 0 | 0 | 1486970 | ABB90509.1 | A0A0P0K6L9 |
| PGGCNFPLEATW | 1 | 0 | 0 | 1486972 | AWO68993.1 | A0A0P0G321 |
| PGGDAISAAFNN | 1 | 0 | 0 | 1486973 | AUM60013.1 | K9N7C7 |
| PGGDATSLSFNN | 1 | 0 | 0 | 1486975 | ANI69888.1 | K9N638 |
| PGGIDICSQDKV | 1 | 0 | 0 | 1486978 | AVI15051.1 | P0C6X6 |
| PGGTGSGDATTA | 1 | 0 | 0 | 1486984 | ADO24677.1 | P0C6X6 |
| PGGTSSGDASTA | 1 | 0 | 0 | 1486985 | AKL87224.1 | P0C6X6 |
| PGGTTSGDATTA | 1 | 0 | 0 | 1486990 | AVP78019.1 | P0C6W2 |
| PGHHHHHHHHSA | 1 | 0 | 0 | 1486993 | 6CRV_A | P59594 |
| PGIDAGYKNSGI | 1 | 0 | 0 | 1486996 | CAA83660.1 | P36334 |
| PGKCYNCRPAVI | 1 | 0 | 0 | 1486999 | AIW52747.1 | A0A0P0K6L9 |
| PGKCYNCRPSVI | 1 | 0 | 0 | 1487000 | AWH62679.1 | A0A0P0K6L9 |
| PGKNNGIGTCPA | 1 | 0 | 0 | 1487004 | CAA79896.1 | P36334 |
| PGKNNGIGTCPV | 1 | 0 | 0 | 1487005 | AIL49486.1 | P36334 |
| PGKNNGIGTRPA | 1 | 0 | 0 | 1487006 | ARB66066.1 | P36334 |
| PGKNNGISTCPA | 1 | 0 | 0 | 1487007 | BBA84094.1 | P36334 |
| PGKNNGLGTCPA | 1 | 0 | 0 | 1487008 | AIL49503.1 | P36334 |
| PGKNNGMGTCPA | 1 | 0 | 0 | 1487009 | AIL49487.1 | P36334 |
| PGLCIAGNRGIA | 1 | 0 | 0 | 1487015 | AEN19366.1 | P36334 |
| PGLCISGDRGIA | 1 | 0 | 0 | 1487017 | AGT51551.1 | P36334 |
| PGLCLSGDIGIA | 1 | 0 | 0 | 1487019 | AZS52618.1 | A0A140H1H1 |
| PGLCLSGDRGIA | 1 | 0 | 0 | 1487020 | BBA20986.1 | A0A140H1H1 |
| PGLGINEEKCGT | 1 | 0 | 0 | 1487022 | BBA20986.1 | A0A140H1H1 |
| PGLPGTVIRTLN | 1 | 0 | 0 | 1487028 | APO40578.1 | P0C6X7 |
| PGLSGIGAVGGD | 1 | 0 | 0 | 1487031 | ALA49601.1 | K9N4V7 |
| PGLSLESCVRVA | 1 | 0 | 0 | 1487032 | APO40578.1 | P0C6X7 |
| PGMIFVLTSHNV | 1 | 0 | 0 | 1487036 | AWK59942.1 | H9EJ42 |
| PGMKYVKWEHDS | 1 | 0 | 0 | 1487037 | NP_073549.1 | A0A0P0LKV0 |
| PGNCIGKGSLTA | 1 | 0 | 0 | 1487040 | AUM60027.1 | K9N643 |
| PGNGILKWADQS | 1 | 0 | 0 | 1487042 | AMM42456.1 | P33469 |
| PGNHCPAGNSYT | 1 | 0 | 0 | 1487043 | QBF80608.1 | K9N5Q8 |
| PGNMIDGFACTS | 1 | 0 | 0 | 1487044 | AIV41774.1 | P0C6X6 |
| PGNVGGNDALST | 1 | 0 | 0 | 1487053 | AWW13571.1 | P0C6X6 |
| PGNVGGNDDLSI | 1 | 0 | 0 | 1487054 | AIV41912.1 | P0C6X6 |
| PGNVGGNEALSI | 1 | 0 | 0 | 1487055 | AIV41792.1 | P0C6X6 |
| PGNVVPGDVIIT | 1 | 0 | 0 | 1487056 | AWK59942.1 | H9EJ42 |
| PGPRVFLVEWLT | 1 | 0 | 0 | 1487058 | AVV62524.1 | K9N7C7 |
| PGPSGIGAVGDA | 1 | 0 | 0 | 1487062 | AVV62544.1 | K9N4V7 |
| PGPSGIGGGDLL | 1 | 0 | 0 | 1487064 | AVN89359.1 | K9N4V7 |
| PGPSGIGTVGGD | 1 | 0 | 0 | 1487065 | ATG84708.1 | K9N4V7 |
| PGQAMPSLFKVQ | 1 | 0 | 0 | 1487067 | ATQ39389.1 | K9N7C7 |
| PGQGHLGYTLEE | 1 | 0 | 0 | 1487070 | APO40578.1 | P0C6X7 |
| PGQGVPLNSNST | 1 | 0 | 0 | 1487072 | AHI48825.1 | K9N4V7 |
| PGQKYSSDCNVV | 1 | 0 | 0 | 1487073 | AXN92247.1 | K9N7C7 |
| PGQKYTSDCNVV | 1 | 0 | 0 | 1487074 | AVV62524.1 | K9N7C7 |
| PGQYTPCLYLTS | 1 | 0 | 0 | 1487080 | AUM60014.1 | K9N5Q8 |
| PGRCYNCRPAVI | 1 | 0 | 0 | 1487081 | AIW52757.1 | A0A0P0K6L9 |
| PGRCYNCRPAVV | 1 | 0 | 0 | 1487082 | AIW52756.1 | A0A0P0K6L9 |
| PGRTSSGDATTA | 1 | 0 | 0 | 1487083 | AFR45548.1 | P0C6X6 |
| PGSAGVISQYNY | 1 | 0 | 0 | 1487085 | AVV62537.1 | K9N5Q8 |
| PGSAVLKQWLPV | 1 | 0 | 0 | 1487088 | APO40578.1 | P0C6X7 |
| PGSCNFPLEATW | 1 | 0 | 0 | 1487091 | AIW52843.1 | A0A0P0G321 |
| PGSDASGDCPSV | 1 | 0 | 0 | 1487092 | ATQ39390.1 | K9N5Q8 |
| PGSGYIPEAPRD | 1 | 0 | 0 | 1487094 | 5X59_A | K9N5Q8 |
| PGTCYNCRPAVV | 1 | 0 | 0 | 1487108 | BAL45638.1 | A0A0P0K6L9 |
| PGTGPHKDLKFR | 1 | 0 | 0 | 1487111 | ABK63972.1 | H9EJ69 |
| PGTNTSKAVAVL | 1 | 0 | 0 | 1487119 | APO40579.1 | P59594 |
| PGTNTSSEVAVL | 1 | 0 | 0 | 1487121 | AVP78042.1 | Q3LZX1 |
| PGTQLPKNFHIE | 1 | 0 | 0 | 1487122 | AVV62533.1 | K9N4V7 |
| PGTVIRTLNGDF | 1 | 0 | 0 | 1487127 | APO40578.1 | P0C6X7 |
| PGVADAERMAMY | 1 | 0 | 0 | 1487134 | AIW52843.1 | A0A0P0G321 |
| PGVEELNPSKLL | 1 | 0 | 0 | 1487137 | APU51922.1 | Q4VID0 |
| PGVPATEAKGYW | 1 | 0 | 0 | 1487144 | AEN19370.1 | P33469 |
| PGVPATESKGYW | 1 | 0 | 0 | 1487145 | AIX09811.1 | P33469 |
| PGVPSTEAKGYW | 1 | 0 | 0 | 1487147 | AJQ24074.1 | P33469 |
| PGVSTNVYLVGK | 1 | 0 | 0 | 1487152 | AKT07951.1 | H9EJ42 |
| PGVTPDMADQIA | 1 | 0 | 0 | 1487153 | AIX10722.1 | P33469 |
| PGVYCGSDYMDI | 1 | 0 | 0 | 1487156 | AVV62535.1 | K9N7C7 |
| PGYMQGYDDCMH | 1 | 0 | 0 | 1487163 | ATG84866.1 | K9N5Q8 |
| PGYMQGYDDCMK | 1 | 0 | 0 | 1487164 | AVV62537.1 | K9N5Q8 |
| PGYMQGYDDCMN | 1 | 0 | 0 | 1487165 | AVV62526.1 | K9N5Q8 |
| PGYMQGYDECMQ | 1 | 0 | 0 | 1487167 | AUM60014.1 | K9N5Q8 |
| PHAGETPIAYRT | 1 | 0 | 0 | 1487172 | ARO76381.1 | P0C6U8 |
| PHAKDQYGSDIN | 1 | 0 | 0 | 1487174 | AIX10729.1 | P33469 |
| PHAKDQYGTDFD | 1 | 0 | 0 | 1487175 | AGT51416.1 | P33469 |
| PHAKDQYGTDIN | 1 | 0 | 0 | 1487177 | AEN19370.1 | P33469 |
| PHDFFIFDVDKV | 1 | 0 | 0 | 1487184 | AXN92247.1 | K9N7C7 |
| PHDVIVLRDNLK | 1 | 0 | 0 | 1487186 | AFD98800.1 | H9EJ24 |
| PHEADTEQTQSV | 1 | 0 | 0 | 1487188 | QBF80585.1 | K9N7C7 |
| PHEFCSQHTHYI | 1 | 0 | 0 | 1487189 | AHI48773.1 | K9N7C7 |
| PHEFCTKHTMLV | 1 | 0 | 0 | 1487194 | AIL49625.1 | P0C6X6 |
| PHFNQKLPNGVN | 1 | 0 | 0 | 1487198 | AMM42518.1 | A0A0N7J131 |
| PHKDLKFRQRSD | 1 | 0 | 0 | 1487224 | AIW52795.1 | H9EJ69 |
| PHKDVYELRYNG | 1 | 0 | 0 | 1487225 | AIX10716.1 | P33469 |
| PHNDCTPINTNA | 1 | 0 | 0 | 1487237 | AVV62526.1 | K9N5Q8 |
| PHNGVLDITACQ | 1 | 0 | 0 | 1487240 | AGW27872.1 | A0A140H1H1 |
| PHNGVWEINACT | 1 | 0 | 0 | 1487242 | ABA60802.1 | A0A140H1H1 |
| PHNYFYMDLNPA | 1 | 0 | 0 | 1487248 | QBQ01838.1 | P30215 |
| PHNYSYMDLNPA | 1 | 0 | 0 | 1487249 | AEN19365.1 | P30215 |

| | | | | | | |
|---|---|---|---|---|---|---|
| PHNYSYMDLNSA | 1 | 0 | 0 | 1487251 | AIL49334.1 | P30215 |
| PHQVNLSLNGNT | 1 | 0 | 0 | 1487255 | AWO69005.1 | A0A0P0G321 |
| PHSFTLGGKVEV | 1 | 0 | 0 | 1487262 | AVV62535.1 | K9N7C7 |
| PHSIRSIKSDRK | 1 | 0 | 0 | 1487266 | AJG44080.1 | K9N5Q8 |
| PHSIRSIQSDRK | 1 | 0 | 0 | 1487267 | AXN92228.1 | K9N5Q8 |
| PHSIRSIQSGRK | 1 | 0 | 0 | 1487268 | QBF80543.1 | K9N5Q8 |
| PHSKDQYGTDIN | 1 | 0 | 0 | 1487269 | AIV41994.1 | P33469 |
| PHSKEQYGTDID | 1 | 0 | 0 | 1487270 | ANZ78874.1 | P33469 |
| PHTDCTSAGYNT | 1 | 0 | 0 | 1487274 | AVV62537.1 | K9N5Q8 |
| PHTRYSLADSNG | 1 | 0 | 0 | 1487282 | ABD75543.1 | A0A140H1G9 |
| PHTSCHPNLGNH | 1 | 0 | 0 | 1487284 | AWW13511.1 | P36334 |
| PHTTNLDNKLQG | 1 | 0 | 0 | 1487287 | CAA83660.1 | P36334 |
| PHTVLLVPRGSG | 1 | 0 | 0 | 1487289 | 6NUR_A | P0C6X7 |
| PHVRYDMFDTNM | 1 | 0 | 0 | 1487303 | AUM60013.1 | K9N7C7 |
| PHYVQPGQYTPC | 1 | 0 | 0 | 1487315 | AUM60024.1 | K9N5Q8 |
| PIADFTPVSGTT | 1 | 0 | 0 | 1487317 | AVV62528.1 | K9N4V0 |
| PIADVYRRKPDL | 1 | 0 | 0 | 1487321 | AEN19366.1 | P36334 |
| PIADVYRRRPDL | 1 | 0 | 0 | 1487324 | AHA80225.1 | P36334 |
| PIAFGIPPSEAK | 1 | 0 | 0 | 1487326 | ABA60794.1 | Q19U29 |
| PIALKAPDTYKC | 1 | 0 | 0 | 1487334 | BBO58891.1 | P36334 |
| PIAPGVPSTEAK | 1 | 0 | 0 | 1487339 | AJQ24074.1 | P33469 |
| PIAYGIPLSEAK | 1 | 0 | 0 | 1487344 | AMM42386.1 | Q19U29 |
| PIAYGIPPSEAK | 1 | 0 | 0 | 1487345 | AMM42397.1 | Q19U29 |
| PIDAIIIDNDLN | 1 | 0 | 0 | 1487363 | ATG84699.1 | K9N7C7 |
| PIDEIPYKSFFD | 1 | 0 | 0 | 1487365 | AMD02834.1 | K9N638 |
| PIDEIPYKSFSD | 1 | 0 | 0 | 1487366 | AXN92247.1 | K9N7C7 |
| PIDEYITTYPGQ | 1 | 0 | 0 | 1487367 | AER30328.1 | P0C6X7 |
| PIDFPSAESTGT | 1 | 0 | 0 | 1487369 | AZK15901.1 | K9N796 |
| PIDGYFVLNETN | 1 | 0 | 0 | 1487370 | AVV62526.1 | K9N5Q8 |
| PIDGYFVLNNTT | 1 | 0 | 0 | 1487371 | AVV62537.1 | K9N5Q8 |
| PIDIDAFPVRPI | 1 | 0 | 0 | 1487373 | AWK59942.1 | H9EJ42 |
| PIDIPRGLPVGF | 1 | 0 | 0 | 1487374 | ATO98193.1 | Q3LZX1 |
| PIDKCTRIIPAR | 1 | 0 | 0 | 1487377 | APO40578.1 | P0C6X7 |
| PIDLASQIISGT | 1 | 0 | 0 | 1487378 | AVN89380.1 | K9N7A1 |
| PIDLATQIISGI | 1 | 0 | 0 | 1487379 | AVV62543.1 | K9N7A1 |
| PIDLLCLRDDNQ | 1 | 0 | 0 | 1487381 | ARU07575.1 | A0A140H1G9 |
| PIDLSSQIISGI | 1 | 0 | 0 | 1487382 | ALA49908.1 | K9N7A1 |
| PIDPAAADGIIY | 1 | 0 | 0 | 1487388 | AVV62526.1 | K9N5Q8 |
| PIDQYMCGKDGN | 1 | 0 | 0 | 1487390 | AVV62524.1 | K9N7C7 |
| PIDQYMCGSDGK | 1 | 0 | 0 | 1487391 | AUM60013.1 | K9N7C7 |
| PIDSTVKNYFIT | 1 | 0 | 0 | 1487393 | APO40578.1 | P0C6X7 |
| PIDVFKADGIIY | 1 | 0 | 0 | 1487395 | AHI48583.1 | K9N5Q8 |
| PIEDAETPALQV | 1 | 0 | 0 | 1487398 | ATQ39389.1 | K9N7C7 |
| PIEDFQENWNTK | 1 | 0 | 0 | 1487399 | AVP78041.1 | P0C6W2 |
| PIEDYENDWNTK | 1 | 0 | 0 | 1487400 | APO40578.1 | P0C6X7 |
| PIEEEDWLDVQE | 1 | 0 | 0 | 1487402 | APO40578.1 | P0C6X7 |
| PIEFHLDGEVLP | 1 | 0 | 0 | 1487403 | ARO76381.1 | P0C6U8 |
| PIEHVTSVNLFD | 1 | 0 | 0 | 1487404 | AVN89292.1 | K9N796 |
| PIETSKAEGVIY | 1 | 0 | 0 | 1487407 | AUM60014.1 | K9N5Q8 |
| PIFDDPVVDWHY | 1 | 0 | 0 | 1487413 | AVN89322.1 | K9N7C7 |
| PIGKGSKDEQIG | 1 | 0 | 0 | 1487439 | AIW52795.1 | H9EJ69 |
| PIGQSLCAVPPT | 1 | 0 | 0 | 1487443 | AUM60014.1 | K9N5Q8 |
| PIGTKYRHCDLD | 1 | 0 | 0 | 1487444 | AZS52618.1 | A0A140H1H1 |
| PIGTNDPIRSCL | 1 | 0 | 0 | 1487446 | CAA49377.1 | A0A0P0LKV0 |
| PIHYEMFGNYTI | 1 | 0 | 0 | 1487454 | ATO98227.1 | Q3LZX5 |
| PIIEAENPNPNS | 1 | 0 | 0 | 1487458 | AET99208.1 | Q0ZJ80 |
| PIIEAENPNQNS | 1 | 0 | 0 | 1487459 | Q0ZME2.1 | Q0ZJ80 |
| PIIGDELKINSA | 1 | 0 | 0 | 1487463 | P0C6W6.1 | P0C6W2 |
| PIIQEAIPTRFS | 1 | 0 | 0 | 1487476 | AMM42389.1 | Q19U29 |
| PIISEKISVMDK | 1 | 0 | 0 | 1487477 | AMM42546.1 | H9EJ42 |
| PIKGDVDSGPPS | 1 | 0 | 0 | 1487483 | AVV62537.1 | K9N5Q8 |
| PIKKAFPYGTCL | 1 | 0 | 0 | 1487485 | APT69868.1 | A0A0P0ILY9 |
| PIKKVSFGPEEV | 1 | 0 | 0 | 1487486 | AUM60013.1 | K9N7C7 |
| PIKNTVDTIPKE | 1 | 0 | 0 | 1487488 | AST12962.1 | A0A0P0LKV0 |
| PIKTQDYQIGGF | 1 | 0 | 0 | 1487494 | APO40580.1 | P59632 |
| PILAELAPPAGA | 1 | 0 | 0 | 1487497 | ANZ78874.1 | P33469 |
| PILAELAPTAGA | 1 | 0 | 0 | 1487498 | AEN19370.1 | P33469 |
| PILDFKDGIYFA | 1 | 0 | 0 | 1487500 | AVP78031.1 | Q3LZX1 |
| PILHHPGFLNPS | 1 | 0 | 0 | 1487503 | AXN92231.1 | K9N643 |
| PILKLPFGINIT | 1 | 0 | 0 | 1487506 | ATO98181.1 | Q3LZX1 |
| PILKTFIDAARG | 1 | 0 | 0 | 1487510 | AVV62524.1 | K9N7C7 |
| PILLVDSNLVTS | 1 | 0 | 0 | 1487515 | AVV62524.1 | K9N7C7 |
| PILMESTGSLAT | 1 | 0 | 0 | 1487516 | AGT51765.1 | Q4VID0 |
| PILWVKKASYDT | 1 | 0 | 0 | 1487530 | AXN92226.1 | K9N7C7 |
| PIMANLANLNSH | 1 | 0 | 0 | 1487534 | CAA71146.1 | A0A0P0K6L9 |
| PIMANLVNHKSH | 1 | 0 | 0 | 1487535 | AIW52757.1 | A0A0P0K6L9 |
| PIMANLVNLNSH | 1 | 0 | 0 | 1487536 | AIW52756.1 | A0A0P0K6L9 |
| PIMANLVNSKSH | 1 | 0 | 0 | 1487537 | AOG74783.1 | A0A0P0K6L9 |
| PINADLSPISVD | 1 | 0 | 0 | 1487540 | AUM60025.1 | K9N796 |
| PINASKADGIIY | 1 | 0 | 0 | 1487541 | AVV62537.1 | K9N5Q8 |
| PINKKDKNKLIG | 1 | 0 | 0 | 1487543 | AIW52723.1 | A0A0N7J131 |
| PINKKEKNKLIG | 1 | 0 | 0 | 1487544 | AIW52704.1 | A0A0N7J131 |
| PINSSGFVVQIP | 1 | 0 | 0 | 1487549 | AVV62526.1 | K9N5Q8 |
| PINTNSGKDDQI | 1 | 0 | 0 | 1487552 | Q3I5I7.1 | Q3LZX4 |
| PINTNSSKDDQI | 1 | 0 | 0 | 1487554 | AVP78038.1 | Q3LZX4 |
| PINYFWPKQETS | 1 | 0 | 0 | 1487557 | AUM60013.1 | K9N7C7 |
| PIPNLPDFREEL | 1 | 0 | 0 | 1487562 | AIL49487.1 | P36334 |
| PIQDYIQSVNFL | 1 | 0 | 0 | 1487568 | AEN19363.1 | P0C6X6 |
| PIQIDQLNSSYF | 1 | 0 | 0 | 1487570 | ASU91208.1 | K9N5Q8 |
| PIQLVPAFPEAE | 1 | 0 | 0 | 1487573 | AVV62541.1 | K9N7D2 |
| PIQLVPAFPETE | 1 | 0 | 0 | 1487574 | AVV62530.1 | K9N7D2 |

| | | | | | | |
|---|---|---|---|---|---|---|
| PIQPAELPVSIV | 1 | 0 | 0 | 1487575 | ADK37704.1 | A0A0P0K6L9 |
| PIQPVELPESIV | 1 | 0 | 0 | 1487578 | ARA15429.1 | A0A0P0K6L9 |
| PIQPVELPMSIV | 1 | 0 | 0 | 1487579 | AST12964.1 | A0A0P0K6L9 |
| PIQPVELPVSIV | 1 | 0 | 0 | 1487580 | AIW52757.1 | A0A0P0K6L9 |
| PIQPVELPVSMC | 1 | 0 | 0 | 1487581 | ADK37703.1 | A0A0P0K6L9 |
| PIQSAELPVSIV | 1 | 0 | 0 | 1487582 | AGT21353.1 | A0A0P0K6L9 |
| PIQTPVGCAMGI | 1 | 0 | 0 | 1487583 | AUM60014.1 | K9N5Q8 |
| PIQVDHIFAWQH | 1 | 0 | 0 | 1487584 | AMQ49004.1 | K9N5Q8 |
| PIQVDQFNSSYF | 1 | 0 | 0 | 1487585 | ASU90362.1 | K9N5Q8 |
| PIQVDQLNSSYK | 1 | 0 | 0 | 1487587 | ATW75477.1 | K9N5Q8 |
| PIQVDQLNSTYF | 1 | 0 | 0 | 1487588 | ANI69889.1 | K9N5Q8 |
| PIQVGQLNSSYF | 1 | 0 | 0 | 1487589 | ALJ54452.1 | K9N5Q8 |
| PIRDYAALAAKE | 1 | 0 | 0 | 1487591 | AVV62524.1 | K9N7C7 |
| PISAYAFLMAED | 1 | 0 | 0 | 1487601 | AKN58609.1 | K9N638 |
| PISDLLTPMGID | 1 | 0 | 0 | 1487605 | APO40578.1 | P0C6X7 |
| PISEDDYQIGGY | 1 | 0 | 0 | 1487606 | AVP78043.1 | Q3LZX0 |
| PISLEMFKPGVE | 1 | 0 | 0 | 1487612 | AGT51805.1 | Q4VID0 |
| PISNCIKMLSNG | 1 | 0 | 0 | 1487613 | AFD98833.1 | H9EJ42 |
| PISNDSNCCVVE | 1 | 0 | 0 | 1487614 | AGT51386.1 | H9EJ42 |
| PISPISLEMFKP | 1 | 0 | 0 | 1487616 | AGT51805.1 | Q4VID0 |
| PISTYSPNTCPQ | 1 | 0 | 0 | 1487624 | BBA20986.1 | A0A140H1H1 |
| PISVDRDSAVGH | 1 | 0 | 0 | 1487625 | AUM60025.1 | K9N796 |
| PITENNVVVTST | 1 | 0 | 0 | 1487631 | ARB66070.1 | P36334 |
| PITFAGPYKCPQ | 1 | 0 | 0 | 1487632 | ATP16757.1 | P36334 |
| PITFKAPDTYKC | 1 | 0 | 0 | 1487633 | ARB07414.1 | P36334 |
| PITFSPATVLSG | 1 | 0 | 0 | 1487634 | AUM60013.1 | K9N7C7 |
| PITGNNVVVMST | 1 | 0 | 0 | 1487639 | CAA83660.1 | P36334 |
| PITINPGEYSIC | 1 | 0 | 0 | 1487641 | AVV62526.1 | K9N5Q8 |
| PITKNVKGNNEA | 1 | 0 | 0 | 1487644 | AVV62535.1 | K9N7C7 |
| PITKTEKDTNEA | 1 | 0 | 0 | 1487645 | AUM60013.1 | K9N7C7 |
| PITLANARYVSP | 1 | 0 | 0 | 1487647 | AVV62526.1 | K9N5Q8 |
| PITLGSLELLSF | 1 | 0 | 0 | 1487648 | AGT51685.1 | Q4VID0 |
| PITLKTPDTYKC | 1 | 0 | 0 | 1487650 | AXX83339.1 | P36334 |
| PITLTPRSVRSV | 1 | 0 | 0 | 1487652 | ANI69900.1 | K9N5Q8 |
| PITSFNTKYVAP | 1 | 0 | 0 | 1487657 | AVN89387.1 | K9N5Q8 |
| PIVDDPVVDWHY | 1 | 0 | 0 | 1487671 | AXN92258.1 | K9N7C7 |
| PIVEDPVVDWHY | 1 | 0 | 0 | 1487673 | AUM60023.1 | K9N7C7 |
| PIVEQPVEEFED | 1 | 0 | 0 | 1487676 | AVV62535.1 | K9N7C7 |
| PIVNEKISVMDK | 1 | 0 | 0 | 1487683 | AGT51337.1 | H9EJ42 |
| PIVQHNTEPKVP | 1 | 0 | 0 | 1487688 | APO40578.1 | P0C6X7 |
| PIVRKIFVDGVP | 1 | 0 | 0 | 1487690 | QXN12055.1 | P0DTD1 |
| PIVSEKISVMDK | 1 | 0 | 0 | 1487692 | AWK59942.1 | H9EJ42 |
| PIVWGAKDFNTL | 1 | 0 | 0 | 1487700 | AFD98777.1 | H9EJ42 |
| PIVYETDANKLI | 1 | 0 | 0 | 1487703 | AQN78717.1 | P0C6X6 |
| PIYDEPTPTTSV | 1 | 0 | 0 | 1487712 | APO40580.1 | P59632 |
| PIYLKGSGFDFS | 1 | 0 | 0 | 1487719 | ARE30011.1 | P0C6X6 |
| PKAGANFRDLSL | 1 | 0 | 0 | 1487723 | BBA20983.1 | A0A140H1H1 |
| PKAIACVKDFDF | 1 | 0 | 0 | 1487724 | AGT51787.1 | P0C6X6 |
| PKAIRCVPQAEV | 1 | 0 | 0 | 1487727 | AVP78041.1 | P0C6W2 |
| PKAIRCVPQSEV | 1 | 0 | 0 | 1487728 | AVP78030.1 | P0C6W2 |
| PKCDRAMPNIIR | 1 | 0 | 0 | 1487733 | AKL87224.1 | P0C6X6 |
| PKDGFNLEYWVT | 1 | 0 | 0 | 1487741 | ARK08651.1 | P36334 |
| PKDGFSPEYWVT | 1 | 0 | 0 | 1487743 | QEG03814.1 | P36334 |
| PKDGGHFFVSDD | 1 | 0 | 0 | 1487744 | AFD98791.1 | H9EJ42 |
| PKDGGHFFVSGD | 1 | 0 | 0 | 1487745 | ARE29978.1 | H9EJ42 |
| PKDGGQFFVSDD | 1 | 0 | 0 | 1487746 | AWK59942.1 | H9EJ42 |
| PKDGGQFFVSGD | 1 | 0 | 0 | 1487747 | ARK08626.1 | H9EJ42 |
| PKEDGDVVLAEF | 1 | 0 | 0 | 1487755 | AHI48794.1 | K9N638 |
| PKEEGDVLLAEF | 1 | 0 | 0 | 1487756 | ATQ39389.1 | K9N7C7 |
| PKEELTDQMSEP | 1 | 0 | 0 | 1487757 | AGV08397.1 | K9N4V7 |
| PKEEQPKIEDSN | 1 | 0 | 0 | 1487758 | ARO76381.1 | P0C6U8 |
| PKEESNDQEMTT | 1 | 0 | 0 | 1487759 | ATQ39393.1 | K9N4V7 |
| PKEESTDQISEP | 1 | 0 | 0 | 1487760 | AQZ41303.1 | K9N4V7 |
| PKEESTDQMFEP | 1 | 0 | 0 | 1487761 | QCZ25056.1 | K9N4V7 |
| PKEESTDQMSEL | 1 | 0 | 0 | 1487762 | ASJ26617.1 | K9N4V7 |
| PKEEVKPFFTEN | 1 | 0 | 0 | 1487764 | AVP78041.1 | P0C6W2 |
| PKEHIGTRNPNN | 1 | 0 | 0 | 1487765 | APO40586.1 | P59595 |
| PKEHRVQGTQRT | 1 | 0 | 0 | 1487766 | YP_007188586.1 | K9N4V7 |
| PKEIIDGYIMHA | 1 | 0 | 0 | 1487767 | APO40578.1 | P0C6X7 |
| PKEITVATSRML | 1 | 0 | 0 | 1487770 | AYV99793.1 | P59596 |
| PKEQRVQGGITQ | 1 | 0 | 0 | 1487775 | ANF29213.1 | K9N4V7 |
| PKEQRVQGIITQ | 1 | 0 | 0 | 1487776 | AHI48535.1 | K9N4V7 |
| PKEQRVQGSIAQ | 1 | 0 | 0 | 1487777 | QEJ82222.1 | K9N4V7 |
| PKEQRVQGSITH | 1 | 0 | 0 | 1487778 | AXN73444.1 | K9N4V7 |
| PKEQRVQGSTTQ | 1 | 0 | 0 | 1487780 | AQZ41292.1 | K9N4V7 |
| PKFEFANDTKIA | 1 | 0 | 0 | 1487786 | AVN89387.1 | K9N5Q8 |
| PKFFYAPNGTYN | 1 | 0 | 0 | 1487787 | ADK37704.1 | A0A0P0K6L9 |
| PKFTVSTQPQEN | 1 | 0 | 0 | 1487794 | ABD75589.1 | Q19U29 |
| PKGFCDFKGKYV | 1 | 0 | 0 | 1487799 | APO40578.1 | P0C6X7 |
| PKGIPIFDDPVV | 1 | 0 | 0 | 1487804 | AVN89322.1 | K9N7C7 |
| PKGIPIVEDPVV | 1 | 0 | 0 | 1487805 | AUM60023.1 | K9N7C7 |
| PKGKYSQSLLKK | 1 | 0 | 0 | 1487807 | AUM60013.1 | K9N7C7 |
| PKHGDLGEQYIY | 1 | 0 | 0 | 1487820 | AVV62537.1 | K9N5Q8 |
| PKIDNETLEYWV | 1 | 0 | 0 | 1487824 | AMM42407.1 | A0A140H1H1 |
| PKIEDSNVKEEP | 1 | 0 | 0 | 1487825 | ARO76381.1 | P0C6U8 |
| PKIEFANDTKIA | 1 | 0 | 0 | 1487827 | ATG84701.1 | K9N5Q8 |
| PKIFKARAIECY | 1 | 0 | 0 | 1487828 | AMM42360.1 | A0A140H1G9 |
| PKIFKARAIEFY | 1 | 0 | 0 | 1487829 | ARU07575.1 | A0A140H1G9 |
| PKIGPDHSVADY | 1 | 0 | 0 | 1487833 | APO40578.1 | P0C6X7 |
| PKIITTDNTFVS | 1 | 0 | 0 | 1487835 | AVP78031.1 | Q3LZX1 |
| PKKAYMDFVNAG | 1 | 0 | 0 | 1487839 | AVV62524.1 | K9N7C7 |

| | | | | | | |
|---|---|---|---|---|---|---|
| PKKDKKKKADEL | 1 | 0 | 0 | 1487840 | AVP78038.1 | Q3LZX4 |
| PKKDKKKKADEV | 1 | 0 | 0 | 1487842 | APO40586.1 | P59595 |
| PKKIFIAACAVD | 1 | 0 | 0 | 1487845 | ARB07605.1 | A0A140H1G9 |
| PKLEFANDTKIV | 1 | 0 | 0 | 1487858 | AVN89313.1 | K9N5Q8 |
| PKLFIRQEEVQQ | 1 | 0 | 0 | 1487862 | 1XAK_A | P59635 |
| PKLFTRQEEVQE | 1 | 0 | 0 | 1487865 | AVP78047.1 | Q3LZX7 |
| PKLGNHFKESWH | 1 | 0 | 0 | 1487867 | AIL49486.1 | P36334 |
| PKLHEDYQIGGY | 1 | 0 | 0 | 1487870 | ARO76383.1 | P59632 |
| PKLKGSFNNRDT | 1 | 0 | 0 | 1487874 | AWW13511.1 | P36334 |
| PKLQSSHAWQPG | 1 | 0 | 0 | 1487876 | AVP78041.1 | P0C6W2 |
| PKLSTGLVKNQC | 1 | 0 | 0 | 1487880 | ATO98193.1 | Q3LZX1 |
| PKLYCAFTAPEY | 1 | 0 | 0 | 1487883 | AGT51707.1 | P0C6X6 |
| PKNFCPCSSCFG | 1 | 0 | 0 | 1487897 | QEG03814.1 | P36334 |
| PKNFCPCSSCHG | 1 | 0 | 0 | 1487898 | ARB66068.1 | P36334 |
| PKNFCPCSSCPG | 1 | 0 | 0 | 1487899 | AWW13566.1 | P36334 |
| PKNFCPCSSCPS | 1 | 0 | 0 | 1487900 | QDH43719.1 | P36334 |
| PKNFCPCSSCSC | 1 | 0 | 0 | 1487901 | AVR40344.1 | P36334 |
| PKNFCPCSSCSG | 1 | 0 | 0 | 1487902 | ANZ78838.1 | P36334 |
| PKNFHIEGTGGN | 1 | 0 | 0 | 1487903 | AXN92267.1 | K9N4V7 |
| PKNGGQFFVSGD | 1 | 0 | 0 | 1487905 | ARU07593.1 | H9EJ42 |
| PKNGQGENDNIS | 1 | 0 | 0 | 1487906 | QEG03805.1 | P33469 |
| PKNMWEFRDYFD | 1 | 0 | 0 | 1487907 | AWK59942.1 | H9EJ42 |
| PKPQRQRGHRNG | 1 | 0 | 0 | 1487918 | ANZ78875.1 | P33469 |
| PKPSRNQSPVSS | 1 | 0 | 0 | 1487931 | AMM42518.1 | A0A0N7J131 |
| PKPVEGVSSFMN | 1 | 0 | 0 | 1487937 | AIW52757.1 | A0A0P0K6L9 |
| PKPVIYQLDENA | 1 | 0 | 0 | 1487940 | AMM42491.1 | A0A0P0ILY9 |
| PKQETSTQVEVS | 1 | 0 | 0 | 1487946 | AUM60013.1 | K9N7C7 |
| PKQFVPVQVVDS | 1 | 0 | 0 | 1487947 | ABD75512.1 | A0A140H1H0 |
| PKQGDLGKQYVY | 1 | 0 | 0 | 1487948 | AVV62526.1 | K9N5Q8 |
| PKQGDQGQMYLY | 1 | 0 | 0 | 1487949 | AUM60014.1 | K9N5Q8 |
| PKQGYFIKHNDH | 1 | 0 | 0 | 1487950 | BBA20983.1 | A0A140H1H1 |
| PKQGYFIKQNDS | 1 | 0 | 0 | 1487951 | BBA20986.1 | A0A140H1H1 |
| PKQTSTSQQPLG | 1 | 0 | 0 | 1487958 | AMK59681.1 | P33469 |
| PKQTSTSQQPSG | 1 | 0 | 0 | 1487959 | ATP16771.1 | P33469 |
| PKQTSTSQQPSR | 1 | 0 | 0 | 1487960 | AGT51784.1 | P33469 |
| PKQTVTSQQPSG | 1 | 0 | 0 | 1487961 | AIL49418.1 | P33469 |
| PKRLRIKRNVDP | 1 | 0 | 0 | 1487962 | AGN72639.1 | K9N7C7 |
| PKRSRHKQCSVQ | 1 | 0 | 0 | 1487965 | AJQ24068.1 | P33469 |
| PKRTGVILTRSL | 1 | 0 | 0 | 1487967 | AMM42558.1 | H9EJ42 |
| PKSDGTGTIYAE | 1 | 0 | 0 | 1487968 | AEA10817.1 | P0C6X7 |
| PKSRSTSRTSGL | 1 | 0 | 0 | 1487972 | AJQ24068.1 | P33469 |
| PKSRVQQNKSIE | 1 | 0 | 0 | 1487974 | AEN19370.1 | P33469 |
| PKTDNETLEYWV | 1 | 0 | 0 | 1487979 | BBA20986.1 | A0A140H1H1 |
| PKTEVIDTPSSA | 1 | 0 | 0 | 1487981 | AVV62535.1 | K9N7C7 |
| PKTGTGIAISVK | 1 | 0 | 0 | 1487985 | AXN92247.1 | K9N7C7 |
| PKTKHVMHDNDS | 1 | 0 | 0 | 1487986 | APO40578.1 | P0C6X7 |
| PKTKHVTKENDS | 1 | 0 | 0 | 1487987 | ATO98229.1 | P0C6W2 |
| PKTRVVHPALHL | 1 | 0 | 0 | 1487994 | AVP78050.1 | Q3LZX3 |
| PKTSMAPILTES | 1 | 0 | 0 | 1487998 | AHF54652.1 | Q4VID0 |
| PKVADKICIVDN | 1 | 0 | 0 | 1488004 | AEN19363.1 | P0C6X6 |
| PKVHFYYPGTGP | 1 | 0 | 0 | 1488008 | ABK63972.1 | H9EJ69 |
| PKVITKRDAAAA | 1 | 0 | 0 | 1488012 | ANF29235.1 | K9N4V7 |
| PKVPEEAPAPEL | 1 | 0 | 0 | 1488014 | APO40578.1 | P0C6X7 |
| PKVTIDCAAFDC | 1 | 0 | 0 | 1488018 | ARB66062.1 | P36334 |
| PLADNKFALACF | 1 | 0 | 0 | 1488033 | AVP78036.1 | Q3LZX7 |
| PLADNKFALTCI | 1 | 0 | 0 | 1488035 | ARO76387.1 | P59635 |
| PLADNKFAVTCT | 1 | 0 | 0 | 1488037 | APO40584.1 | P59635 |
| PLAELKCTLKLS | 1 | 0 | 0 | 1488041 | ATO98145.1 | Q3LZX1 |
| PLAFGVPLSEAK | 1 | 0 | 0 | 1488042 | ABA60796.1 | Q19U29 |
| PLAFMPVQSRRF | 1 | 0 | 0 | 1488045 | AEN19355.1 | P0C6X6 |
| PLASCFFTCNSN | 1 | 0 | 0 | 1488053 | AIW52843.1 | A0A0P0G321 |
| PLASCFFTCNTN | 1 | 0 | 0 | 1488054 | BAM16475.1 | A0A0P0G321 |
| PLASCFSTCNSN | 1 | 0 | 0 | 1488055 | AVL25555.1 | A0A0P0G321 |
| PLAYFSDSKIGT | 1 | 0 | 0 | 1488061 | ALD51903.1 | K9N7C7 |
| PLCDQFIDYHQL | 1 | 0 | 0 | 1488066 | APO40578.1 | P0C6X7 |
| PLCGFPSQAASV | 1 | 0 | 0 | 1488069 | ARO76381.1 | P0C6U8 |
| PLCLFKSNFNQY | 1 | 0 | 0 | 1488077 | AZS52617.1 | A0A140H1H0 |
| PLDAGLNGDFNL | 1 | 0 | 0 | 1488083 | AVV62526.1 | K9N5Q8 |
| PLDEAEDFTESV | 1 | 0 | 0 | 1488086 | AIL23988.1 | K9N7C7 |
| PLDEAEDVTEFV | 1 | 0 | 0 | 1488087 | AXN92247.1 | K9N7C7 |
| PLDEAENVTESV | 1 | 0 | 0 | 1488088 | QBF80465.1 | K9N7C7 |
| PLDEAEYVTEFV | 1 | 0 | 0 | 1488089 | AVN89322.1 | K9N7C7 |
| PLDEDEIIYFPY | 1 | 0 | 0 | 1488090 | AVV62524.1 | K9N7C7 |
| PLDGLVRAYNII | 1 | 0 | 0 | 1488093 | AUM60013.1 | K9N7C7 |
| PLDGLVRIYNVL | 1 | 0 | 0 | 1488094 | AVV62524.1 | K9N7C7 |
| PLDIYDLTMPSQ | 1 | 0 | 0 | 1488095 | AIV41960.1 | P0C6X6 |
| PLDKIQNQTLSR | 1 | 0 | 0 | 1488096 | ARB07436.1 | A0A140H1G9 |
| PLDPKLKGSFND | 1 | 0 | 0 | 1488099 | AIL49495.1 | P36334 |
| PLDPRLKGSFND | 1 | 0 | 0 | 1488100 | AIL49520.1 | P36334 |
| PLDSLKVCVQTV | 1 | 0 | 0 | 1488102 | ARO76381.1 | P0C6U8 |
| PLDSRLKGSFNN | 1 | 0 | 0 | 1488103 | AIL49507.1 | P36334 |
| PLDSRTGSFNNI | 1 | 0 | 0 | 1488104 | AIX10756.1 | P36334 |
| PLDSRTGSFNNR | 1 | 0 | 0 | 1488105 | ABD43177.1 | P36334 |
| PLDTRLKGSFNN | 1 | 0 | 0 | 1488106 | AEN19366.1 | P36334 |
| PLDTRYKGTFNN | 1 | 0 | 0 | 1488107 | AKA09647.1 | P36334 |
| PLDTSLKGSFNN | 1 | 0 | 0 | 1488108 | AIX10753.1 | P36334 |
| PLDTSRKGSFNN | 1 | 0 | 0 | 1488109 | AKA09663.1 | P36334 |
| PLDTSYINNKDT | 1 | 0 | 0 | 1488110 | AKA09646.1 | P36334 |
| PLDTSYKGTFNN | 1 | 0 | 0 | 1488111 | AKA09658.1 | P36334 |
| PLDVYNLTMPSQ | 1 | 0 | 0 | 1488113 | AAX85666.1 | P0C6X6 |
| PLEEDDIVYFPY | 1 | 0 | 0 | 1488120 | AVV62535.1 | K9N7C7 |

| | | | | | | |
|---|---|---|---|---|---|---|
| PLEEEVQEEKTS | 1 | 0 | 0 | 1488121 | AVV62524.1 | K9N7C7 |
| PLEFGASAETVR | 1 | 0 | 0 | 1488122 | AEA10817.1 | P0C6X7 |
| PLEFGASTEMQV | 1 | 0 | 0 | 1488124 | ATO98229.1 | P0C6W2 |
| PLEFGASTETLQ | 1 | 0 | 0 | 1488125 | ARO76381.1 | P0C6U8 |
| PLEFGASTQNEE | 1 | 0 | 0 | 1488126 | APO40578.1 | P0C6X7 |
| PLEFGATSFSSS | 1 | 0 | 0 | 1488128 | AVP78041.1 | P0C6W2 |
| PLEGGGRLVASG | 1 | 0 | 0 | 1488129 | AXG21654.1 | K9N5Q8 |
| PLEGGGWLVVSG | 1 | 0 | 0 | 1488131 | ALJ54495.1 | K9N5Q8 |
| PLEINCQAPPVG | 1 | 0 | 0 | 1488132 | Q3I5I8.1 | Q3LZX5 |
| PLENDADKANFL | 1 | 0 | 0 | 1488133 | AIV41996.1 | P0C6X6 |
| PLEQPTNEAVEA | 1 | 0 | 0 | 1488134 | AVP78041.1 | P0C6W2 |
| PLESRTGSLNNI | 1 | 0 | 0 | 1488136 | QEG03803.1 | P36334 |
| PLETPVGCVLGL | 1 | 0 | 0 | 1488137 | ASU90230.1 | K9N5Q8 |
| PLFALISIQILT | 1 | 0 | 0 | 1488140 | AEN19363.1 | P0C6X6 |
| PLFDQQAQQEAT | 1 | 0 | 0 | 1488143 | APT69896.1 | A0A0P0LKV0 |
| PLFESDKEEDVQ | 1 | 0 | 0 | 1488145 | AST12962.1 | A0A0P0LKV0 |
| PLFIRNFDQRCS | 1 | 0 | 0 | 1488146 | AXG21669.1 | K9N7D2 |
| PLGIKDPILYVL | 1 | 0 | 0 | 1488155 | ARU07575.1 | A0A140H1G9 |
| PLGQSLCAVPAT | 1 | 0 | 0 | 1488164 | AVV62526.1 | K9N5Q8 |
| PLGQSLCAVPST | 1 | 0 | 0 | 1488165 | AVV62537.1 | K9N5Q8 |
| PLGRSLCALPDT | 1 | 0 | 0 | 1488167 | ATG84888.1 | K9N5Q8 |
| PLGSAGNATEVP | 1 | 0 | 0 | 1488168 | 5C8S_A | P0C6U8 |
| PLGSAIASEFSS | 1 | 0 | 0 | 1488169 | 2AHM_E | P0C6U8 |
| PLGSMYKQARSE | 1 | 0 | 0 | 1488172 | 5F22_B | P0C6U8 |
| PLGSSKMSDVKC | 1 | 0 | 0 | 1488175 | 2AHM_A | P0C6U8 |
| PLGVYDYKVSTQ | 1 | 0 | 0 | 1488179 | AUM60013.1 | K9N7C7 |
| PLHLSVSKLHDL | 1 | 0 | 0 | 1488194 | AVV62540.1 | K9N643 |
| PLHMSISKLHAL | 1 | 0 | 0 | 1488195 | AXN92231.1 | K9N643 |
| PLHPYGRCPTAA | 1 | 0 | 0 | 1488200 | AQN78751.1 | P30215 |
| PLHTLSIHDGYT | 1 | 0 | 0 | 1488203 | AAP30028.1 | P0C6X7 |
| PLIKDVDNPVLM | 1 | 0 | 0 | 1488217 | AHF54667.1 | P0C6X6 |
| PLKLEDFLPIDS | 1 | 0 | 0 | 1488236 | APO40578.1 | P0C6X7 |
| PLKPEDLMEVDH | 1 | 0 | 0 | 1488240 | AVV62529.1 | K9N643 |
| PLKPLWKLPLGL | 1 | 0 | 0 | 1488242 | ATO98145.1 | Q3LZX1 |
| PLKVEEVQLQDV | 1 | 0 | 0 | 1488253 | AUM60013.1 | K9N7C7 |
| PLLAELIPNHAA | 1 | 0 | 0 | 1488261 | AMM42619.1 | H9EJ69 |
| PLLGNSTGIDFQ | 1 | 0 | 0 | 1488275 | AXN92228.1 | K9N5Q8 |
| PLLKPVGTLDIS | 1 | 0 | 0 | 1488281 | APO40578.1 | P0C6X7 |
| PLLNNQTDLDFK | 1 | 0 | 0 | 1488289 | AVV62537.1 | K9N5Q8 |
| PLLNPGALEFNP | 1 | 0 | 0 | 1488290 | AGT21342.1 | A0A0N7J131 |
| PLLSDNQISGYT | 1 | 0 | 0 | 1488299 | CAA83661.1 | P36334 |
| PLLSENQFSGYT | 1 | 0 | 0 | 1488300 | AIX10748.1 | P36334 |
| PLLSENQINGYT | 1 | 0 | 0 | 1488301 | ANZ78836.1 | P36334 |
| PLLSGTPPQVYN | 1 | 0 | 0 | 1488305 | AXN92228.1 | K9N5Q8 |
| PLLSNQTDQSFK | 1 | 0 | 0 | 1488306 | AVV62526.1 | K9N5Q8 |
| PLLSNSTGIDFQ | 1 | 0 | 0 | 1488307 | ALL26396.1 | K9N5Q8 |
| PLLVDDNKFDVL | 1 | 0 | 0 | 1488316 | AEN19363.1 | P0C6X6 |
| PLLVDSDQPWKV | 1 | 0 | 0 | 1488317 | AST12968.1 | A0A0N7J131 |
| PLLVDSEQPCLV | 1 | 0 | 0 | 1488318 | AMM42513.1 | A0A0N7J131 |
| PLLVDSEQPWKV | 1 | 0 | 0 | 1488319 | AIW52723.1 | A0A0N7J131 |
| PLLVDSEQSWKV | 1 | 0 | 0 | 1488320 | AAA45463.1 | A0A0N7J131 |
| PLLVLFGMYLDS | 1 | 0 | 0 | 1488322 | AVA26874.1 | H9EJ24 |
| PLLVSSDKAPHR | 1 | 0 | 0 | 1488323 | AVL25594.1 | H9EJ69 |
| PLMEYDFTDYKV | 1 | 0 | 0 | 1488336 | AUM60013.1 | K9N7C7 |
| PLMKRGHRWDVV | 1 | 0 | 0 | 1488339 | AGT51787.1 | P0C6X6 |
| PLMSKGQKWDVV | 1 | 0 | 0 | 1488347 | ARB07436.1 | A0A140H1G9 |
| PLNAGLNGDFNL | 1 | 0 | 0 | 1488351 | AVV62537.1 | K9N5Q8 |
| PLNANSTPSQNA | 1 | 0 | 0 | 1488353 | AQZ41279.1 | K9N4V7 |
| PLNLPPTHAHTF | 1 | 0 | 0 | 1488362 | AWK59942.1 | H9EJ42 |
| PLNLPPTQAHTF | 1 | 0 | 0 | 1488363 | QED88038.1 | H9EJ42 |
| PLNLVGYIKHLP | 1 | 0 | 0 | 1488366 | AVV62535.1 | K9N7C7 |
| PLNSNSTPAQNA | 1 | 0 | 0 | 1488375 | AHI48825.1 | K9N4V7 |
| PLNWERKTFSNC | 1 | 0 | 0 | 1488379 | CAA79896.1 | P36334 |
| PLPAADMDDFSR | 1 | 0 | 0 | 1488381 | AAS48456.1 | P59595 |
| PLQPELESFKQE | 1 | 0 | 0 | 1488402 | APO40579.1 | P59594 |
| PLQSFKHKVCTD | 1 | 0 | 0 | 1488403 | AVV62535.1 | K9N7C7 |
| PLQSYACWAHSC | 1 | 0 | 0 | 1488407 | AVV62535.1 | K9N7C7 |
| PLSAICPTGTKY | 1 | 0 | 0 | 1488441 | AIG96691.1 | A0A140H1H1 |
| PLSDIEKDFLSF | 1 | 0 | 0 | 1488447 | AHI48581.1 | K9N7C7 |
| PLSDMEKDFLSS | 1 | 0 | 0 | 1488450 | ANI69887.1 | K9N7C7 |
| PLSDQIDFLQSK | 1 | 0 | 0 | 1488451 | APO40578.1 | P0C6X7 |
| PLSEVYVHLRNW | 1 | 0 | 0 | 1488459 | AWK59934.1 | U3M6Q8 |
| PLSEVYVQLRNW | 1 | 0 | 0 | 1488460 | QED88043.1 | U3M6Q8 |
| PLSFPPGQGVPL | 1 | 0 | 0 | 1488461 | AVV62533.1 | K9N4V7 |
| PLSHFQGFVLNY | 1 | 0 | 0 | 1488462 | AQN78734.1 | Q80872 |
| PLSISNDTVDVT | 1 | 0 | 0 | 1488464 | ANZ78836.1 | P36334 |
| PLSLPVLQVRDV | 1 | 0 | 0 | 1488468 | AVP78030.1 | P0C6W2 |
| PLSLTEVDGGFK | 1 | 0 | 0 | 1488469 | ATO98145.1 | Q3LZX1 |
| PLSMKSDLGVSS | 1 | 0 | 0 | 1488470 | AHX00711.1 | K9N5Q8 |
| PLSMKSDLSVSF | 1 | 0 | 0 | 1488471 | AHX71943.1 | K9N5Q8 |
| PLSMKSDLSVSS | 1 | 0 | 0 | 1488472 | AXN92228.1 | K9N5Q8 |
| PLSMKSDLSVTS | 1 | 0 | 0 | 1488473 | ATW75479.1 | K9N5Q8 |
| PLSMKSYMDPGS | 1 | 0 | 0 | 1488474 | AVV62526.1 | K9N5Q8 |
| PLSMKSYMQPGS | 1 | 0 | 0 | 1488475 | AVV62537.1 | K9N5Q8 |
| PLSNFEHKDITE | 1 | 0 | 0 | 1488476 | ALA49659.1 | K9N7C7 |
| PLSQPRVDKPSQ | 1 | 0 | 0 | 1488484 | QED88037.1 | H9EJ69 |
| PLSVASAPVEPS | 1 | 0 | 0 | 1488485 | QBF80531.1 | K9N7C7 |
| PLSVASTLVEPP | 1 | 0 | 0 | 1488486 | ATG84722.1 | K9N7C7 |
| PLSVASTPVEPP | 1 | 0 | 0 | 1488487 | AXN92247.1 | K9N7C7 |
| PLSVASTPVEPS | 1 | 0 | 0 | 1488488 | AXN92258.1 | K9N7C7 |
| PLSVASTPVESP | 1 | 0 | 0 | 1488489 | ATG84853.1 | K9N7C7 |

| | | | | | | |
|---|---|---|---|---|---|---|
| PLSVEPTPIEDA | 1 | 0 | 0 | 1488491 | ATQ39389.1 | K9N7C7 |
| PLSVESTPVELP | 1 | 0 | 0 | 1488492 | AVN89312.1 | K9N638 |
| PLSVESTPVEPP | 1 | 0 | 0 | 1488493 | AHI48796.1 | K9N7C7 |
| PLSVESTPVEPS | 1 | 0 | 0 | 1488494 | AIY60577.1 | K9N638 |
| PLSVVYDKTTNS | 1 | 0 | 0 | 1488500 | AVV62537.1 | K9N5Q8 |
| PLSVVYDKTTNT | 1 | 0 | 0 | 1488501 | AVV62526.1 | K9N5Q8 |
| PLTAPTLVPQEH | 1 | 0 | 0 | 1488503 | ATO98130.1 | P0C6W2 |
| PLTCISRLKDGF | 1 | 0 | 0 | 1488516 | AVR40342.1 | P36334 |
| PLTCKAISSNTD | 1 | 0 | 0 | 1488520 | Q14EB0.1 | A0A140H1H1 |
| PLTCNAISPKTD | 1 | 0 | 0 | 1488522 | BBA20986.1 | A0A140H1H1 |
| PLTCNAISPNTD | 1 | 0 | 0 | 1488523 | ABA60801.1 | A0A140H1H1 |
| PLTCNSAMTLEY | 1 | 0 | 0 | 1488524 | CAA83660.1 | P36334 |
| PLTCNSKRKDGF | 1 | 0 | 0 | 1488525 | AKA09664.1 | P36334 |
| PLTCNSKRKGGF | 1 | 0 | 0 | 1488526 | AKA09645.1 | P36334 |
| PLTCNSKVKNGF | 1 | 0 | 0 | 1488528 | CAA79896.1 | P36334 |
| PLTFPPGQGVPL | 1 | 0 | 0 | 1488533 | AXN92267.1 | K9N4V7 |
| PLTFPSGQGVPL | 1 | 0 | 0 | 1488534 | AZK15907.1 | K9N4V7 |
| PLTFSPGQGVPL | 1 | 0 | 0 | 1488535 | AHI48535.1 | K9N4V7 |
| PLTKKKYSELDD | 1 | 0 | 0 | 1488543 | Q3I5J1.1 | Q3LZX8 |
| PLTMLAEFDENG | 1 | 0 | 0 | 1488547 | APO40579.1 | P59594 |
| PLTNVVSHVNGD | 1 | 0 | 0 | 1488549 | AGT51799.1 | P30215 |
| PLTQDHLDILGP | 1 | 0 | 0 | 1488555 | AVP78041.1 | P0C6W2 |
| PLTYLPATVIAG | 1 | 0 | 0 | 1488574 | AVV62535.1 | K9N7C7 |
| PLVDFNVNVMGT | 1 | 0 | 0 | 1488576 | AGT21366.1 | A0A0P0LKV0 |
| PLVEDSTSVTAV | 1 | 0 | 0 | 1488579 | ASY99815.1 | K9N7A1 |
| PLVELPVSINIT | 1 | 0 | 0 | 1488580 | AVP78031.1 | Q3LZX1 |
| PLVGTPVCINGL | 1 | 0 | 0 | 1488584 | QYC64967.1 | P0DTD1 |
| PLVGYIPEHCQS | 1 | 0 | 0 | 1488588 | AUM60015.1 | K9N796 |
| PLVLECTRAAAS | 1 | 0 | 0 | 1488595 | AVV62524.1 | K9N7C7 |
| PLVQDHVDILGP | 1 | 0 | 0 | 1488599 | APO40578.1 | P0C6X7 |
| PLVSCFSTCNSN | 1 | 0 | 0 | 1488604 | AWO68966.1 | A0A0P0G321 |
| PLVYFSDSKPST | 1 | 0 | 0 | 1488611 | ATQ39389.1 | K9N7C7 |
| PLWFCLLYIAVV | 1 | 0 | 0 | 1488613 | AEN19363.1 | P0C6X6 |
| PLYAFASEAARV | 1 | 0 | 0 | 1488619 | AVP78041.1 | P0C6W2 |
| PLYAFPSDAAHV | 1 | 0 | 0 | 1488620 | AVP78030.1 | P0C6W2 |
| PLYDPNMEAAYT | 1 | 0 | 0 | 1488622 | AVV62526.1 | K9N5Q8 |
| PLYDPNMEGAYT | 1 | 0 | 0 | 1488623 | AVV62537.1 | K9N5Q8 |
| PLYDVNMEAAYT | 1 | 0 | 0 | 1488624 | AUM60014.1 | K9N5Q8 |
| PMACNGHKIEDL | 1 | 0 | 0 | 1488635 | ACR82761.1 | A0A140H1G9 |
| PMAISVNCCHDG | 1 | 0 | 0 | 1488637 | AGT51675.1 | Q4VID0 |
| PMATSANCCHDG | 1 | 0 | 0 | 1488640 | AGT51735.1 | Q4VID0 |
| PMATSVNCCHDG | 1 | 0 | 0 | 1488641 | APU51922.1 | Q4VID0 |
| PMATSVNCFHDG | 1 | 0 | 0 | 1488642 | AHF54653.1 | Q4VID0 |
| PMCTNGKVVPTL | 1 | 0 | 0 | 1488645 | AVV62524.1 | K9N7C7 |
| PMCTNGQAVPTL | 1 | 0 | 0 | 1488646 | AUM60013.1 | K9N7C7 |
| PMCTNGQVVPTL | 1 | 0 | 0 | 1488647 | AVV62535.1 | K9N7C7 |
| PMDVHSTYGTQA | 1 | 0 | 0 | 1488649 | ATQ39396.1 | K9N643 |
| PMEQRVQGSITQ | 1 | 0 | 0 | 1488654 | AXP07352.1 | K9N4V7 |
| PMERVSLWNYGK | 1 | 0 | 0 | 1488655 | AEN19363.1 | P0C6X6 |
| PMEYVSYFDGVK | 1 | 0 | 0 | 1488656 | AGT21344.1 | A0A0P0LKV0 |
| PMGSLQPLATLY | 1 | 0 | 0 | 1488661 | 6NZK_A | P36334 |
| PMGVYDFKVSTQ | 1 | 0 | 0 | 1488664 | AXN92247.1 | K9N7C7 |
| PMKAEGDGGVLG | 1 | 0 | 0 | 1488667 | AWK59942.1 | H9EJ42 |
| PMLPKNMWEFRD | 1 | 0 | 0 | 1488672 | AFD98826.1 | H9EJ42 |
| PMMDLGNSSTIT | 1 | 0 | 0 | 1488678 | AVV62526.1 | K9N5Q8 |
| PMMDLGNSTTIT | 1 | 0 | 0 | 1488679 | AVV62537.1 | K9N5Q8 |
| PMMQSADASTFL | 1 | 0 | 0 | 1488680 | P0C6W6.1 | P0C6W2 |
| PMMQSSDASTFL | 1 | 0 | 0 | 1488681 | APO40578.1 | P0C6X7 |
| PMPINASKADGI | 1 | 0 | 0 | 1488686 | AVV62537.1 | K9N5Q8 |
| PMPNNGWTAFSA | 1 | 0 | 0 | 1488687 | AVP78031.1 | Q3LZX1 |
| PMPPMVNPSKES | 1 | 0 | 0 | 1488688 | ARU07582.1 | Q0ZJ80 |
| PMPQGVHMNVAK | 1 | 0 | 0 | 1488690 | AVV62524.1 | K9N7C7 |
| PMQALRNDTSVE | 1 | 0 | 0 | 1488694 | AUM60014.1 | K9N5Q8 |
| PMQDVNQDDDRQ | 1 | 0 | 0 | 1488695 | AVV62533.1 | K9N4V7 |
| PMSDMEKDFLSF | 1 | 0 | 0 | 1488696 | QBF80541.1 | K9N7C7 |
| PMSDYDIPTTEN | 1 | 0 | 0 | 1488698 | 3EJG_A | A0A0P0LKV0 |
| PMSNNGWTAFSA | 1 | 0 | 0 | 1488700 | AVP78042.1 | Q3LZX1 |
| PMTDMEKDFLSL | 1 | 0 | 0 | 1488701 | AVV62535.1 | K9N7C7 |
| PMTDMEKDFLSM | 1 | 0 | 0 | 1488702 | AVV62524.1 | K9N7C7 |
| PMTDTLQMGFII | 1 | 0 | 0 | 1488703 | AVV62537.1 | K9N5Q8 |
| PMTEGLQMAFII | 1 | 0 | 0 | 1488704 | AVV62526.1 | K9N5Q8 |
| PMVNPSKESSGL | 1 | 0 | 0 | 1488708 | AET99203.1 | Q0ZJ80 |
| PNAILISDRKIK | 1 | 0 | 0 | 1488720 | AUM60013.1 | K9N7C7 |
| PNARLCATCPFA | 1 | 0 | 0 | 1488727 | AHI48793.1 | K9N7C7 |
| PNCNIEAWLNGK | 1 | 0 | 0 | 1488737 | AIL49492.1 | P36334 |
| PNCNIESWLNDK | 1 | 0 | 0 | 1488738 | AIX10749.1 | P36334 |
| PNCNIGAWLNDK | 1 | 0 | 0 | 1488739 | AHA80222.1 | P36334 |
| PNCSNCLDDRCI | 1 | 0 | 0 | 1488740 | APO40578.1 | P0C6X7 |
| PNDDATSNVTQC | 1 | 0 | 0 | 1488746 | APT69866.1 | A0A0N7J131 |
| PNDFSFNNWFLL | 1 | 0 | 0 | 1488750 | ABE97137.1 | A0A0P0G321 |
| PNDFVGGFRVLG | 1 | 0 | 0 | 1488752 | AVA26872.1 | H9EJ42 |
| PNDIVGGFRVLR | 1 | 0 | 0 | 1488754 | AWK59978.1 | H9EJ42 |
| PNDNAYVSHAIW | 1 | 0 | 0 | 1488757 | AVV62526.1 | K9N5Q8 |
| PNECNQMCLSIL | 1 | 0 | 0 | 1488758 | AVP78030.1 | P0C6W2 |
| PNEHVTSVNLLD | 1 | 0 | 0 | 1488761 | AGV08391.1 | K9N796 |
| PNEITVAKPSVL | 1 | 0 | 0 | 1488763 | AUM60030.1 | K9N7A1 |
| PNEKLVTTFTAP | 1 | 0 | 0 | 1488764 | AMQ48992.1 | K9N7C7 |
| PNEKLVTTSMAP | 1 | 0 | 0 | 1488765 | QBF80609.1 | K9N7C7 |
| PNEKLVTTSTAP | 1 | 0 | 0 | 1488766 | AXN92247.1 | K9N7C7 |
| PNEKLVTTSTVP | 1 | 0 | 0 | 1488767 | ATG84754.1 | K9N7C7 |
| PNEKQEILGTVS | 1 | 0 | 0 | 1488768 | AVP78041.1 | P0C6W2 |

| | | | | | | |
|---|---|---|---|---|---|---|
| PNEQRVQGSITQ | 1 | 0 | 0 | 1488773 | AVN89381.1 | K9N4V7 |
| PNESGDVLLAEY | 1 | 0 | 0 | 1488774 | AVV62535.1 | K9N7C7 |
| PNEVTVAKPNVV | 1 | 0 | 0 | 1488776 | AIL23940.1 | K9N7A1 |
| PNFAEISKINTT | 1 | 0 | 0 | 1488777 | AUM60014.1 | K9N5Q8 |
| PNFDLTPFNLTY | 1 | 0 | 0 | 1488779 | AIW52843.1 | A0A0P0G321 |
| PNFGSLTHINTT | 1 | 0 | 0 | 1488782 | ALJ54471.1 | K9N5Q8 |
| PNFKKVETLNFQ | 1 | 0 | 0 | 1488787 | AET99208.1 | Q0ZJ80 |
| PNFQEISKINTT | 1 | 0 | 0 | 1488788 | AVV62526.1 | K9N5Q8 |
| PNFVFPLNSIIK | 1 | 0 | 0 | 1488792 | QYC64967.1 | P0DTD1 |
| PNGFSQDGDYFT | 1 | 0 | 0 | 1488799 | AVV62537.1 | K9N5Q8 |
| PNGLSEDGQVFT | 1 | 0 | 0 | 1488803 | AVV62526.1 | K9N5Q8 |
| PNGRRVDLQLGN | 1 | 0 | 0 | 1488810 | AGT51600.1 | P36334 |
| PNGTYNCTDAVL | 1 | 0 | 0 | 1488812 | ADK37704.1 | A0A0P0K6L9 |
| PNGVNVAEEPDS | 1 | 0 | 0 | 1488813 | AMM42518.1 | A0A0N7J131 |
| PNHAALFFDSEV | 1 | 0 | 0 | 1488818 | AMM42619.1 | H9EJ69 |
| PNHFINFPLSHF | 1 | 0 | 0 | 1488819 | AQN78734.1 | Q80872 |
| PNHTIKGAFLNG | 1 | 0 | 0 | 1488823 | 3F9E_A | P0C6U8 |
| PNHTIKGSALNG | 1 | 0 | 0 | 1488824 | 3F9G_A | P0C6X7 |
| PNIIKVFYELDN | 1 | 0 | 0 | 1488826 | AIX09804.1 | P0C6X6 |
| PNIIRIVSSLVL | 1 | 0 | 0 | 1488827 | AKL87224.1 | P0C6X6 |
| PNIKDFGGFNFS | 1 | 0 | 0 | 1488828 | APO40579.1 | P59594 |
| PNITNLCPFDAV | 1 | 0 | 0 | 1488832 | ATO98145.1 | Q3LZX1 |
| PNITNLCPFGQV | 1 | 0 | 0 | 1488834 | APO40579.1 | P59594 |
| PNITNRCPFDKV | 1 | 0 | 0 | 1488835 | ATO98181.1 | Q3LZX1 |
| PNKFCNVQQCFG | 1 | 0 | 0 | 1488840 | AMM42397.1 | Q19U29 |
| PNKNAGQDISLL | 1 | 0 | 0 | 1488842 | APO40578.1 | P0C6X7 |
| PNKYCYCGNFAT | 1 | 0 | 0 | 1488849 | QED88017.1 | H9EJ42 |
| PNLGNHRTELWH | 1 | 0 | 0 | 1488853 | ATP16757.1 | P36334 |
| PNLGNRRVELWH | 1 | 0 | 0 | 1488854 | CAA83660.1 | P36334 |
| PNLHDFKEELDQ | 1 | 0 | 0 | 1488855 | CAA83660.1 | P36334 |
| PNLMGWDYPKCD | 1 | 0 | 0 | 1488856 | ASB51350.1 | P0C6X7 |
| PNLPDFKEELGQ | 1 | 0 | 0 | 1488864 | AIX10763.1 | P36334 |
| PNLPDFKEESDQ | 1 | 0 | 0 | 1488865 | BBA20973.1 | P36334 |
| PNLPDFKGELDQ | 1 | 0 | 0 | 1488866 | QDH43719.1 | P36334 |
| PNLPDFREELDQ | 1 | 0 | 0 | 1488867 | AVQ05264.1 | P36334 |
| PNLPGFKEELDQ | 1 | 0 | 0 | 1488868 | AIL49523.1 | P36334 |
| PNLVPAFAEELH | 1 | 0 | 0 | 1488877 | AUM60028.1 | K9N7D2 |
| PNMEGAYTSSLL | 1 | 0 | 0 | 1488886 | AVV62537.1 | K9N5Q8 |
| PNMFITRDEAIK | 1 | 0 | 0 | 1488887 | APO40578.1 | P0C6X7 |
| PNMGVPCCTSYY | 1 | 0 | 0 | 1488891 | AIW52827.1 | H9EJ42 |
| PNMHVVPLENDA | 1 | 0 | 0 | 1488892 | AIV41996.1 | P0C6X6 |
| PNMMVTNNTFTL | 1 | 0 | 0 | 1488897 | AVP78041.1 | P0C6W2 |
| PNMTVHHIKEWL | 1 | 0 | 0 | 1488898 | AVV62540.1 | K9N643 |
| PNMWCLRSGHHL | 1 | 0 | 0 | 1488900 | AXG21646.1 | K9N643 |
| PNMWYLRSGYHL | 1 | 0 | 0 | 1488903 | ANF29275.1 | K9N643 |
| PNNADLSPISVD | 1 | 0 | 0 | 1488904 | AUM60015.1 | K9N796 |
| PNNAVVKMACPA | 1 | 0 | 0 | 1488906 | APO40578.1 | P0C6X7 |
| PNNDAAIVTQFA | 1 | 0 | 0 | 1488907 | AVV62533.1 | K9N4V7 |
| PNNDPAIVTQFA | 1 | 0 | 0 | 1488909 | AUM60021.1 | K9N4V7 |
| PNNGWTAFSAAY | 1 | 0 | 0 | 1488913 | AVP78031.1 | Q3LZX1 |
| PNNMYMRTTHTD | 1 | 0 | 0 | 1488917 | AVV62524.1 | K9N7C7 |
| PNNPKQIVQSIP | 1 | 0 | 0 | 1488922 | AVV62540.1 | K9N643 |
| PNNSVSWYTGLT | 1 | 0 | 0 | 1488924 | AVV62544.1 | K9N4V7 |
| PNQAALFFDSEN | 1 | 0 | 0 | 1488931 | ADK37678.1 | H9EJ69 |
| PNQNSLCLLNHK | 1 | 0 | 0 | 1488936 | Q0ZME2.1 | Q0ZJ80 |
| PNQTATSQQPSG | 1 | 0 | 0 | 1488939 | AIX10721.1 | P33469 |
| PNRRKVDLQLGN | 1 | 0 | 0 | 1488944 | AEN19366.1 | P36334 |
| PNRRRVDLQLGN | 1 | 0 | 0 | 1488946 | ANZ78849.1 | P36334 |
| PNSIYICSQGKD | 1 | 0 | 0 | 1488951 | ARU07575.1 | A0A140H1G9 |
| PNSNYVLQDFDV | 1 | 0 | 0 | 1488953 | AFV53147.1 | H9EJ42 |
| PNSRRSDLQLAS | 1 | 0 | 0 | 1488956 | ALJ77702.1 | A0A140H1H1 |
| PNSSEVDMICST | 1 | 0 | 0 | 1488959 | AOL02450.1 | P0C6X6 |
| PNSVECDFTKLL | 1 | 0 | 0 | 1488960 | AVV62537.1 | K9N5Q8 |
| PNTDNETLQYGV | 1 | 0 | 0 | 1488963 | ABA60801.1 | A0A140H1H1 |
| PNTRSLSRSNSN | 1 | 0 | 0 | 1488973 | AGW27840.1 | Q19U29 |
| PNTVDCQDDRCF | 1 | 0 | 0 | 1488976 | AIL49573.1 | P0C6X6 |
| PNTWCIRCLWST | 1 | 0 | 0 | 1488978 | QYC64967.1 | P0DTD1 |
| PNTWYLRSGHHL | 1 | 0 | 0 | 1488980 | ASU90321.1 | K9N643 |
| PNTYEGNSPYHP | 1 | 0 | 0 | 1488981 | APO40584.1 | P59635 |
| PNVCYTPRSEIP | 1 | 0 | 0 | 1488984 | AVV62535.1 | K9N7C7 |
| PNVPIEYQATRV | 1 | 0 | 0 | 1488988 | ATO98120.1 | Q3LZX1 |
| PNVPLDYQATRV | 1 | 0 | 0 | 1488989 | ATO98145.1 | Q3LZX1 |
| PNWLMSLYSFTL | 1 | 0 | 0 | 1488998 | AVV62524.1 | K9N7C7 |
| PNYKKWLELLEA | 1 | 0 | 0 | 1489002 | AVV62544.1 | K9N4V7 |
| PNYQKWMELLDS | 1 | 0 | 0 | 1489004 | AUM60021.1 | K9N4V7 |
| PNYRQWHCRSGH | 1 | 0 | 0 | 1489005 | AUM60017.1 | K9N643 |
| PNYTNSSQRGVY | 1 | 0 | 0 | 1489007 | AVP78031.1 | Q3LZX1 |
| PPAAPRAVSFAD | 1 | 0 | 0 | 1489011 | AQZ41303.1 | K9N4V7 |
| PPADVNTPADID | 1 | 0 | 0 | 1489012 | AJQ24074.1 | P33469 |
| PPAIASNCYSSL | 1 | 0 | 0 | 1489018 | AHI48528.1 | K9N5Q8 |
| PPALNCYWPLKD | 1 | 0 | 0 | 1489022 | 3D0H_E | P59594 |
| PPANLRVVGHAM | 1 | 0 | 0 | 1489027 | QBF80508.1 | K9N7C7 |
| PPARFVIDTPTG | 1 | 0 | 0 | 1489033 | 2J98_B | A0A0P0LKV0 |
| PPCKFLISGPKG | 1 | 0 | 0 | 1489036 | AVV62524.1 | K9N7C7 |
| PPDEEEDCEEYE | 1 | 0 | 0 | 1489044 | P0C6W6.1 | P0C6W2 |
| PPDQLVVYQQAI | 1 | 0 | 0 | 1489047 | ABB90509.1 | A0A0P0K6L9 |
| PPDQLVVYQQAV | 1 | 0 | 0 | 1489048 | AIW52757.1 | A0A0P0K6L9 |
| PPFLSDFINGIF | 1 | 0 | 0 | 1489054 | CAA79896.1 | P36334 |
| PPGEHFKHLVPL | 1 | 0 | 0 | 1489057 | ALW82707.1 | K9N7C7 |
| PPGEQFKHLIPL | 1 | 0 | 0 | 1489058 | QXH28115.1 | P0DTD1 |
| PPGMGIPYCTSY | 1 | 0 | 0 | 1489060 | AIW52776.1 | A0A0P0LKV0 |

| | | | | | | |
|---|---|---|---|---|---|---|
| PPGQGVPLNSNS | 1 | 0 | 0 | 1489062 | AHI48825.1 | K9N4V7 |
| PPHDVIVLRDNL | 1 | 0 | 0 | 1489066 | AFD98800.1 | H9EJ24 |
| PPHSFTLGGKVD | 1 | 0 | 0 | 1489071 | AVV62524.1 | K9N7C7 |
| PPIADFTPVSGT | 1 | 0 | 0 | 1489075 | AVV62528.1 | K9N4V0 |
| PPIAPGVPATEA | 1 | 0 | 0 | 1489076 | P33469.1 | P33469 |
| PPIELENKFTPL | 1 | 0 | 0 | 1489078 | AKJ80136.2 | K9N7C7 |
| PPILSETQISGY | 1 | 0 | 0 | 1489082 | BBA20986.1 | A0A140H1H1 |
| PPINADLSPISV | 1 | 0 | 0 | 1489083 | AUM60025.1 | K9N796 |
| PPITFSPATVLS | 1 | 0 | 0 | 1489086 | AUM60013.1 | K9N7C7 |
| PPKEQRVQGIIT | 1 | 0 | 0 | 1489093 | AHI48535.1 | K9N4V7 |
| PPKEQRVQSSIT | 1 | 0 | 0 | 1489096 | ANI69896.1 | K9N4V7 |
| PPLLGNSIGIDF | 1 | 0 | 0 | 1489110 | AGV08467.1 | K9N5Q8 |
| PPLLGNSTRIDF | 1 | 0 | 0 | 1489111 | ALJ54517.1 | K9N5Q8 |
| PPLLNNQTDLDF | 1 | 0 | 0 | 1489112 | AVV62537.1 | K9N5Q8 |
| PPLLSDNQISGY | 1 | 0 | 0 | 1489113 | CAA83661.1 | P36334 |
| PPLLSENQFSGY | 1 | 0 | 0 | 1489114 | AIX10748.1 | P36334 |
| PPLLSENQINGY | 1 | 0 | 0 | 1489115 | ANZ78836.1 | P36334 |
| PPLLSNQTDQSF | 1 | 0 | 0 | 1489116 | AVV62526.1 | K9N5Q8 |
| PPLLSNQTDVDF | 1 | 0 | 0 | 1489117 | AUM60014.1 | K9N5Q8 |
| PPLLSNSTGIDF | 1 | 0 | 0 | 1489118 | ALL26396.1 | K9N5Q8 |
| PPLLSVNQISGY | 1 | 0 | 0 | 1489120 | CAA83660.1 | P36334 |
| PPLYDPNMEAAY | 1 | 0 | 0 | 1489131 | AVV62526.1 | K9N5Q8 |
| PPLYDPNMEGAY | 1 | 0 | 0 | 1489132 | AVV62537.1 | K9N5Q8 |
| PPLYDVNMEAAY | 1 | 0 | 0 | 1489133 | AUM60014.1 | K9N5Q8 |
| PPLYVPIRLAGH | 1 | 0 | 0 | 1489134 | AVN89312.1 | K9N638 |
| PPMEQRVQGSIT | 1 | 0 | 0 | 1489135 | AXP07352.1 | K9N4V7 |
| PPMVNPSKESSG | 1 | 0 | 0 | 1489138 | ARU07582.1 | Q0ZJ80 |
| PPNCNIEAWLND | 1 | 0 | 0 | 1489140 | AHA80201.1 | P36334 |
| PPNEQRVQGSIT | 1 | 0 | 0 | 1489141 | AVN89381.1 | K9N4V7 |
| PPNLVPAFAEEL | 1 | 0 | 0 | 1489143 | AUM60028.1 | K9N7D2 |
| PPNNADLSPISV | 1 | 0 | 0 | 1489144 | AUM60015.1 | K9N796 |
| PPNNTASWFTAL | 1 | 0 | 0 | 1489145 | APO40586.1 | P59595 |
| PPPGEHFKHLVP | 1 | 0 | 0 | 1489150 | ALW82707.1 | K9N7C7 |
| PPPITFSPATVL | 1 | 0 | 0 | 1489153 | AUM60013.1 | K9N7C7 |
| PPPLLGNSIGID | 1 | 0 | 0 | 1489155 | AGV08467.1 | K9N5Q8 |
| PPPLLGNSTRID | 1 | 0 | 0 | 1489157 | ALJ54517.1 | K9N5Q8 |
| PPPLLNNQTDLD | 1 | 0 | 0 | 1489158 | AVV62537.1 | K9N5Q8 |
| PPPLLSNQTDQS | 1 | 0 | 0 | 1489159 | AVV62526.1 | K9N5Q8 |
| PPPLLSNQTDVD | 1 | 0 | 0 | 1489160 | AUM60014.1 | K9N5Q8 |
| PPPLLSNSTGID | 1 | 0 | 0 | 1489161 | ALL26396.1 | K9N5Q8 |
| PPPSFYMPLLVS | 1 | 0 | 0 | 1489163 | AIW52805.1 | H9EJ69 |
| PPPVYNFNRLVF | 1 | 0 | 0 | 1489164 | AVV62526.1 | K9N5Q8 |
| PPQADQQVDRLI | 1 | 0 | 0 | 1489165 | 6U7H_A | A0A0P0K6L9 |
| PPQLVELPSEPQ | 1 | 0 | 0 | 1489168 | AVN89312.1 | K9N638 |
| PPQTSITSAILQ | 1 | 0 | 0 | 1489170 | AVP78041.1 | P0C6W2 |
| PPQVMKLPSEPQ | 1 | 0 | 0 | 1489172 | ALA50044.1 | K9N7C7 |
| PPQVVELPSEPQ | 1 | 0 | 0 | 1489174 | AVN89451.1 | K9N7C7 |
| PPQVVELPYEPR | 1 | 0 | 0 | 1489175 | ANI69888.1 | K9N638 |
| PPQVVKLPFEPQ | 1 | 0 | 0 | 1489176 | ALW82701.1 | K9N7C7 |
| PPQVVKLPSEPQ | 1 | 0 | 0 | 1489178 | AXN92247.1 | K9N7C7 |
| PPQVVKLPSEPR | 1 | 0 | 0 | 1489179 | AMQ49036.1 | K9N638 |
| PPQVVKLQFEPQ | 1 | 0 | 0 | 1489180 | QBF80520.1 | K9N638 |
| PPQVYNFNRLVF | 1 | 0 | 0 | 1489183 | AVV62537.1 | K9N5Q8 |
| PPRAELNGVMVD | 1 | 0 | 0 | 1489185 | QBP84746.1 | P0C6X6 |
| PPRCIAPIDGYF | 1 | 0 | 0 | 1489186 | AVV62537.1 | K9N5Q8 |
| PPRCIAPVDGYF | 1 | 0 | 0 | 1489187 | AUM60014.1 | K9N5Q8 |
| PPSAICPAGTKY | 1 | 0 | 0 | 1489188 | 5I08_A | A0A140H1H1 |
| PPSARDLICAQY | 1 | 0 | 0 | 1489189 | AVV62526.1 | K9N5Q8 |
| PPSASCPIGTKY | 1 | 0 | 0 | 1489190 | AZS52618.1 | A0A140H1H1 |
| PPSECRLQNGDF | 1 | 0 | 0 | 1489194 | APO40578.1 | P0C6X7 |
| PPSSATSCKEAD | 1 | 0 | 0 | 1489199 | AVV62537.1 | K9N5Q8 |
| PPSSSGTSTPKK | 1 | 0 | 0 | 1489200 | AGT51335.1 | H9EJ69 |
| PPTCVYELNGYT | 1 | 0 | 0 | 1489202 | AQN78656.1 | P36334 |
| PPTDKLAMIMGY | 1 | 0 | 0 | 1489203 | NP_150075.1 | Q80872 |
| PPTENNCSQLDE | 1 | 0 | 0 | 1489204 | AVP78035.1 | Q3LZX8 |
| PPTGVYELHGYT | 1 | 0 | 0 | 1489206 | AIL49485.1 | P36334 |
| PPTGVYELNGYP | 1 | 0 | 0 | 1489207 | AMM42466.1 | P36334 |
| PPTGVYELNGYT | 1 | 0 | 0 | 1489208 | CAA79896.1 | P36334 |
| PPTKQRYAKRRF | 1 | 0 | 0 | 1489211 | AUM60017.1 | K9N643 |
| PPTNVVSHLNGD | 1 | 0 | 0 | 1489221 | AAA73152.1 | P30215 |
| PPTNVVSHVNGD | 1 | 0 | 0 | 1489222 | AEN19365.1 | P30215 |
| PPTSPTARTTSS | 1 | 0 | 0 | 1489225 | AUM60024.1 | K9N5Q8 |
| PPTSSTARTTSS | 1 | 0 | 0 | 1489226 | AUM60014.1 | K9N5Q8 |
| PPVADYTPVIGT | 1 | 0 | 0 | 1489230 | AVV62539.1 | K9N4V0 |
| PPVIDDGDGVKP | 1 | 0 | 0 | 1489232 | QED88038.1 | H9EJ42 |
| PPVIDDVDVVKP | 1 | 0 | 0 | 1489233 | QED88031.1 | H9EJ42 |
| PPVLQVESTPVV | 1 | 0 | 0 | 1489235 | AVP78030.1 | P0C6W2 |
| PPVTYLPATILA | 1 | 0 | 0 | 1489236 | ANF29227.1 | K9N638 |
| PPVTYSTATILA | 1 | 0 | 0 | 1489239 | AXN92247.1 | K9N7C7 |
| PPWPQIAELVPT | 1 | 0 | 0 | 1489243 | QDP16213.1 | K9N4V7 |
| PPWSAAAGVPFS | 1 | 0 | 0 | 1489245 | BBA20986.1 | A0A140H1H1 |
| PPYINGFNVSEL | 1 | 0 | 0 | 1489252 | AIV41989.1 | Q04853 |
| PQADQQVDRLIT | 1 | 0 | 0 | 1489255 | 6U7H_A | A0A0P0K6L9 |
| PQAEVDWKFYDA | 1 | 0 | 0 | 1489257 | AVP78041.1 | P0C6W2 |
| PQAKSGIITKLE | 1 | 0 | 0 | 1489264 | QEG03802.1 | P30215 |
| PQAKSLVGIGEH | 1 | 0 | 0 | 1489267 | BBO58715.1 | P36334 |
| PQANSGDYYHKV | 1 | 0 | 0 | 1489269 | AGT51421.1 | P30215 |
| PQANSGDYYYKV | 1 | 0 | 0 | 1489270 | P30215.1 | P30215 |
| PQANSGIITKVE | 1 | 0 | 0 | 1489272 | AEN19365.1 | P30215 |
| PQATYQNISTNL | 1 | 0 | 0 | 1489273 | ALM26400.1 | K9N5Q8 |
| PQAVETKVQCST | 1 | 0 | 0 | 1489274 | AEN19363.1 | P0C6X6 |

| | | | | | | |
|---|---|---|---|---|---|---|
| PQAVETRVQCST | 1 | 0 | 0 | 1489275 | AVI15051.1 | P0C6X6 |
| PQCAIVQALESK | 1 | 0 | 0 | 1489276 | QED88024.1 | H9EJ42 |
| PQCAIVQASESK | 1 | 0 | 0 | 1489277 | AWK59942.1 | H9EJ42 |
| PQCAIVQATESK | 1 | 0 | 0 | 1489278 | ABE97136.1 | H9EJ42 |
| PQENTIPHYSWF | 1 | 0 | 0 | 1489285 | ABD75589.1 | Q19U29 |
| PQFAELIPNQAA | 1 | 0 | 0 | 1489288 | AGT51335.1 | H9EJ69 |
| PQFPILAELAPP | 1 | 0 | 0 | 1489292 | ANZ78874.1 | P33469 |
| PQFTSDQAITFL | 1 | 0 | 0 | 1489295 | ABD75596.1 | Q0ZJ82 |
| PQGAFHVTMRSS | 1 | 0 | 0 | 1489299 | AEN19363.1 | P0C6X6 |
| PQGGYVADFAYW | 1 | 0 | 0 | 1489302 | AEN19363.1 | P0C6X6 |
| PQGLPNNIASWF | 1 | 0 | 0 | 1489304 | AAP49024.1 | P59595 |
| PQGLTFAFDGLR | 1 | 0 | 0 | 1489306 | AUM60013.1 | K9N7C7 |
| PQGLTFSYDGLK | 1 | 0 | 0 | 1489307 | AVV62524.1 | K9N7C7 |
| PQGNSIPHYSWF | 1 | 0 | 0 | 1489312 | AMM42386.1 | Q19U29 |
| PQGPPNNTASWF | 1 | 0 | 0 | 1489315 | APO40586.1 | P59595 |
| PQGRTYSNITIT | 1 | 0 | 0 | 1489316 | AXN92228.1 | K9N5Q8 |
| PQGVECDFSKLF | 1 | 0 | 0 | 1489318 | AUM60014.1 | K9N5Q8 |
| PQGVHMNVAKYM | 1 | 0 | 0 | 1489319 | AVV62524.1 | K9N7C7 |
| PQGYYIEGSGKS | 1 | 0 | 0 | 1489320 | AJQ24087.1 | P33469 |
| PQGYYIEGSGKY | 1 | 0 | 0 | 1489321 | AJQ24067.1 | P33469 |
| PQGYYIEGSGRP | 1 | 0 | 0 | 1489322 | AIL49470.1 | P33469 |
| PQGYYVEGSGRS | 1 | 0 | 0 | 1489325 | AIX10731.1 | P33469 |
| PQIAELAPSASA | 1 | 0 | 0 | 1489326 | AVV62544.1 | K9N4V7 |
| PQIAELAPSSSA | 1 | 0 | 0 | 1489327 | AVV62533.1 | K9N4V7 |
| PQIAELVPTASA | 1 | 0 | 0 | 1489329 | AXG21672.1 | K9N4V7 |
| PQIITTDNTFVA | 1 | 0 | 0 | 1489334 | ATO98181.1 | Q3LZX1 |
| PQIQLTITDPKV | 1 | 0 | 0 | 1489339 | 2CME_E | P59636 |
| PQIQLTITRADP | 1 | 0 | 0 | 1489340 | 2CME_B | P59636 |
| PQIQLTTTRMED | 1 | 0 | 0 | 1489342 | Q3I5I6.1 | Q3LZX3 |
| PQKAYIDFVNSG | 1 | 0 | 0 | 1489346 | AUM60023.1 | K9N7C7 |
| PQKAYMDFVNSG | 1 | 0 | 0 | 1489348 | AUM60013.1 | K9N7C7 |
| PQLEQPYVFLKR | 1 | 0 | 0 | 1489359 | APO40578.1 | P0C6X7 |
| PQLLSPLNSSGF | 1 | 0 | 0 | 1489362 | AVV62537.1 | K9N5Q8 |
| PQLSFPLCSAGN | 1 | 0 | 0 | 1489365 | AWK59942.1 | H9EJ42 |
| PQLTYQNISTNL | 1 | 0 | 0 | 1489371 | AVN89324.1 | K9N5Q8 |
| PQLVELPSEPQT | 1 | 0 | 0 | 1489372 | AVN89312.1 | K9N638 |
| PQLVNANHYSPC | 1 | 0 | 0 | 1489373 | AID55087.1 | K9N5Q8 |
| PQLVNSNQYSPC | 1 | 0 | 0 | 1489375 | AHY22555.1 | K9N5Q8 |
| PQNAVVKLYCPA | 1 | 0 | 0 | 1489378 | AVP78030.1 | P0C6W2 |
| PQNQSSAPRITF | 1 | 0 | 0 | 1489381 | AVP78049.1 | Q3LZX4 |
| PQPAGVFTDHDV | 1 | 0 | 0 | 1489382 | CAA83661.1 | P36334 |
| PQPAGVFTNHDV | 1 | 0 | 0 | 1489383 | ATP16757.1 | P36334 |
| PQPAGVLTNHDV | 1 | 0 | 0 | 1489384 | AVQ05264.1 | P36334 |
| PQPKFTVSTQPQ | 1 | 0 | 0 | 1489385 | AMM42397.1 | Q19U29 |
| PQPLMDKFSSSR | 1 | 0 | 0 | 1489387 | ATO98193.1 | Q3LZX1 |
| PQPQRAPKGSIT | 1 | 0 | 0 | 1489389 | AVV62544.1 | K9N4V7 |
| PQPTDVFTNHSV | 1 | 0 | 0 | 1489390 | ARB66060.1 | P36334 |
| PQPTGVFTNHSV | 1 | 0 | 0 | 1489391 | AEN19366.1 | P36334 |
| PQPVEGVSSFMN | 1 | 0 | 0 | 1489392 | AIW52753.1 | A0A0P0K6L9 |
| PQPVGVFTHHDV | 1 | 0 | 0 | 1489393 | CAA83660.1 | P36334 |
| PQPYCYTDGLMQ | 1 | 0 | 0 | 1489394 | AEN19363.1 | P0C6X6 |
| PQPYCYTEGLMQ | 1 | 0 | 0 | 1489395 | AGT51787.1 | P0C6X6 |
| PQQCIAPIDGYF | 1 | 0 | 0 | 1489396 | AVV62526.1 | K9N5Q8 |
| PQQKYTSQCNVV | 1 | 0 | 0 | 1489397 | AUM60013.1 | K9N7C7 |
| PQQMFVTPLAVD | 1 | 0 | 0 | 1489398 | AXN92231.1 | K9N643 |
| PQQMFVTPLSVD | 1 | 0 | 0 | 1489399 | ALA49399.1 | K9N643 |
| PQQNQRSAPRIT | 1 | 0 | 0 | 1489402 | ARO76389.1 | P59595 |
| PQQRFVYDAYQN | 1 | 0 | 0 | 1489403 | AVN89313.1 | K9N5Q8 |
| PQQSQNGFSFFL | 1 | 0 | 0 | 1489404 | AVV62535.1 | K9N7C7 |
| PQQVTKQNAKEI | 1 | 0 | 0 | 1489406 | ARU07581.1 | Q19U29 |
| PQRKRGVKQSPE | 1 | 0 | 0 | 1489414 | ARU07581.1 | Q19U29 |
| PQRQKKQHTVTL | 1 | 0 | 0 | 1489418 | ARO76389.1 | P59595 |
| PQRQRGHKNGQE | 1 | 0 | 0 | 1489423 | AGT51525.1 | P33469 |
| PQRQRGHKNGQV | 1 | 0 | 0 | 1489425 | AIL49482.1 | P33469 |
| PQRQRGHKNVQG | 1 | 0 | 0 | 1489426 | AIX10722.1 | P33469 |
| PQRQRGHKYGQG | 1 | 0 | 0 | 1489428 | AIX10721.1 | P33469 |
| PQRQRGHRNGQG | 1 | 0 | 0 | 1489429 | ANZ78875.1 | P33469 |
| PQRQRGQKNGQG | 1 | 0 | 0 | 1489432 | BBM61433.1 | P33469 |
| PQSARDLICAQY | 1 | 0 | 0 | 1489434 | AUM60014.1 | K9N5Q8 |
| PQSEVDWKFYDA | 1 | 0 | 0 | 1489435 | AVP78030.1 | P0C6W2 |
| PQSGGGKCFNCA | 1 | 0 | 0 | 1489436 | AAQ90006.1 | A0A0P0K6L9 |
| PQSGGGKCYNCR | 1 | 0 | 0 | 1489438 | ABB90509.1 | A0A0P0K6L9 |
| PQSGGGRCYNCR | 1 | 0 | 0 | 1489439 | ABB90508.1 | A0A0P0K6L9 |
| PQSKDSNFLNGS | 1 | 0 | 0 | 1489442 | AVV62536.1 | K9N638 |
| PQSKESNFLNES | 1 | 0 | 0 | 1489444 | AQZ41321.1 | K9N7C7 |
| PQSMLVFPEVKM | 1 | 0 | 0 | 1489448 | AVV62524.1 | K9N7C7 |
| PQSVETKVQCST | 1 | 0 | 0 | 1489452 | AIV41864.1 | P0C6X6 |
| PQSYCYTDGLMQ | 1 | 0 | 0 | 1489453 | AGT51488.1 | P0C6X6 |
| PQTGNFKDLREY | 1 | 0 | 0 | 1489454 | ATO98193.1 | Q3LZX1 |
| PQTHATTVDVNV | 1 | 0 | 0 | 1489455 | AUM60013.1 | K9N7C7 |
| PQTICHPKLGNH | 1 | 0 | 0 | 1489457 | AMK59677.1 | P36334 |
| PQTICHPNLGNH | 1 | 0 | 0 | 1489458 | CAA79896.1 | P36334 |
| PQTIKPEVKEVA | 1 | 0 | 0 | 1489459 | AIZ74432.1 | K9N7C7 |
| PQTIQPEVKEIA | 1 | 0 | 0 | 1489460 | ALA49362.1 | K9N7C7 |
| PQTITTDNTFVA | 1 | 0 | 0 | 1489462 | ARO76382.1 | P59594 |
| PQTKYLVGIGEH | 1 | 0 | 0 | 1489464 | CAA83660.1 | P36334 |
| PQTLDILDITPC | 1 | 0 | 0 | 1489465 | ATO98145.1 | Q3LZX1 |
| PQTSITSAILQS | 1 | 0 | 0 | 1489472 | AVP78041.1 | P0C6W2 |
| PQVITTDNTFES | 1 | 0 | 0 | 1489478 | APO40579.1 | P59594 |
| PQVLFTCVPTVG | 1 | 0 | 0 | 1489479 | AUM60018.1 | K9N7D2 |
| PQVMKLPSEPQT | 1 | 0 | 0 | 1489481 | ALA50044.1 | K9N7C7 |

| | | | | | | |
|---|---|---|---|---|---|---|
| PQVVELLSEPQT | 1 | 0 | 0 | 1489489 | AVN89374.1 | K9N7C7 |
| PQVVELPSEPQT | 1 | 0 | 0 | 1489490 | AVN89451.1 | K9N7C7 |
| PQVVELPYEPRT | 1 | 0 | 0 | 1489491 | ANI69888.1 | K9N638 |
| PQVVKLPFEPQT | 1 | 0 | 0 | 1489492 | ALW82701.1 | K9N7C7 |
| PQVVKLPSEPQT | 1 | 0 | 0 | 1489494 | AXN92247.1 | K9N7C7 |
| PQVVKLPSEPRT | 1 | 0 | 0 | 1489495 | AMQ49036.1 | K9N638 |
| PQVVKLQFEPQT | 1 | 0 | 0 | 1489496 | QBF80520.1 | K9N638 |
| PQVYNFKRLVFT | 1 | 0 | 0 | 1489499 | AXN92228.1 | K9N5Q8 |
| PRAAANIGSNHT | 1 | 0 | 0 | 1489501 | APO40578.1 | P0C6X7 |
| PRAAPNNTVSWY | 1 | 0 | 0 | 1489502 | AXN92267.1 | K9N4V7 |
| PRAELNGVMVDK | 1 | 0 | 0 | 1489507 | QBP84746.1 | P0C6X6 |
| PRAGKHEHSLLV | 1 | 0 | 0 | 1489508 | ARU07593.1 | H9EJ42 |
| PRAISFADNNDI | 1 | 0 | 0 | 1489509 | ANI69896.1 | K9N4V7 |
| PRANSGIITKVE | 1 | 0 | 0 | 1489517 | AIL49334.1 | P30215 |
| PRASADIGANHT | 1 | 0 | 0 | 1489519 | ARO76381.1 | P0C6U8 |
| PRAVSFADNNDN | 1 | 0 | 0 | 1489526 | AUM60021.1 | K9N4V7 |
| PRAVSFADNNDT | 1 | 0 | 0 | 1489527 | ALW82676.1 | K9N4V7 |
| PRAVTFGDNNDN | 1 | 0 | 0 | 1489529 | AVV62533.1 | K9N4V7 |
| PRCPQIAELVPT | 1 | 0 | 0 | 1489535 | AVV61907.1 | K9N4V7 |
| PRDFNHNMGDSD | 1 | 0 | 0 | 1489536 | AIW52805.1 | H9EJ69 |
| PRDGQAYVRKDG | 1 | 0 | 0 | 1489537 | 6CRW_A | P59594 |
| PRDLDHNFGSAV | 1 | 0 | 0 | 1489538 | AMM42513.1 | A0A0N7J131 |
| PRECNDMHLSTL | 1 | 0 | 0 | 1489541 | ARO76381.1 | P0C6U8 |
| PREGVFVMNGTH | 1 | 0 | 0 | 1489544 | APO40579.1 | P59594 |
| PREGVFVSNGTS | 1 | 0 | 0 | 1489546 | ATO98181.1 | Q3LZX1 |
| PRGEFVEQPQGV | 1 | 0 | 0 | 1489553 | AUM60014.1 | K9N5Q8 |
| PRGIACVKDFDF | 1 | 0 | 0 | 1489555 | ARE30011.1 | P0C6X6 |
| PRGSLEWSHPQF | 1 | 0 | 0 | 1489566 | 5X58_A | P59594 |
| PRGVGSANIFIG | 1 | 0 | 0 | 1489568 | ARU07606.1 | P0C6X6 |
| PRHVMCPADQLN | 1 | 0 | 0 | 1489578 | AVV62524.1 | K9N7C7 |
| PRHVMCPADQLS | 1 | 0 | 0 | 1489579 | AXN92247.1 | K9N7C7 |
| PRHVMVPADQLA | 1 | 0 | 0 | 1489580 | AUM60013.1 | K9N7C7 |
| PRITFGGPSDSS | 1 | 0 | 0 | 1489589 | AVP78038.1 | Q3LZX4 |
| PRITFGGPTDSA | 1 | 0 | 0 | 1489591 | ARO76389.1 | P59595 |
| PRIVQMFADHLT | 1 | 0 | 0 | 1489595 | AXX83336.1 | P0C6X6 |
| PRLKGSFNNRDT | 1 | 0 | 0 | 1489605 | AWW13566.1 | P36334 |
| PRLPVLSDFVQI | 1 | 0 | 0 | 1489606 | AIW52843.1 | A0A0P0G321 |
| PRLQAANDWKPG | 1 | 0 | 0 | 1489607 | AVV62524.1 | K9N7C7 |
| PRNDEAIPSMFP | 1 | 0 | 0 | 1489616 | AJQ24068.1 | P33469 |
| PRNGSRCPNDNA | 1 | 0 | 0 | 1489617 | AUM60014.1 | K9N5Q8 |
| PRNGSYCPGNSN | 1 | 0 | 0 | 1489618 | AVV62537.1 | K9N5Q8 |
| PRNGTRCPNDNA | 1 | 0 | 0 | 1489619 | AVV62526.1 | K9N5Q8 |
| PRNKWNGVSDLS | 1 | 0 | 0 | 1489621 | ASU90779.1 | K9N638 |
| PRNKWSGISDLS | 1 | 0 | 0 | 1489623 | ALA49758.1 | K9N7C7 |
| PRNKWSGVSDLS | 1 | 0 | 0 | 1489624 | AXN92247.1 | K9N7C7 |
| PRNLVPIGKGSK | 1 | 0 | 0 | 1489629 | AIW52795.1 | H9EJ69 |
| PRNLVPINKKDK | 1 | 0 | 0 | 1489630 | AIW52723.1 | A0A0N7J131 |
| PRNSWEALALNF | 1 | 0 | 0 | 1489635 | AVV62524.1 | K9N7C7 |
| PRNVLYDSVSAI | 1 | 0 | 0 | 1489639 | BAL45640.1 | A0A0P0K6L9 |
| PRNVSYDSVSAI | 1 | 0 | 0 | 1489640 | AIW52757.1 | A0A0P0K6L9 |
| PRNVSYDSVSAT | 1 | 0 | 0 | 1489641 | AAK32189.1 | A0A0P0K6L9 |
| PRNYSYMDLNPA | 1 | 0 | 0 | 1489642 | AAA73152.1 | P30215 |
| PRPAGVLTNHDV | 1 | 0 | 0 | 1489643 | CAA79896.1 | P36334 |
| PRPAPNNSVSWY | 1 | 0 | 0 | 1489644 | AVV62544.1 | K9N4V7 |
| PRPGNHCPAGNS | 1 | 0 | 0 | 1489648 | QBF80608.1 | K9N5Q8 |
| PRPIDPAAADGI | 1 | 0 | 0 | 1489650 | AVV62526.1 | K9N5Q8 |
| PRPIDVFKADGI | 1 | 0 | 0 | 1489651 | AHI48583.1 | K9N5Q8 |
| PRQKRTPNKHCN | 1 | 0 | 0 | 1489664 | ARU07581.1 | Q19U29 |
| PRQYSLAFNQDG | 1 | 0 | 0 | 1489668 | AHA80190.1 | P36334 |
| PRRKTNPHKHCN | 1 | 0 | 0 | 1489670 | AJQ24083.1 | P33469 |
| PRSDMECDFLNM | 1 | 0 | 0 | 1489674 | AVL25566.1 | H9EJ42 |
| PRSDMEYDFLNI | 1 | 0 | 0 | 1489675 | AGT51365.1 | H9EJ42 |
| PRSEIPYKSFSD | 1 | 0 | 0 | 1489676 | AVV62524.1 | K9N7C7 |
| PRSGNHCPADNS | 1 | 0 | 0 | 1489678 | AID55096.1 | K9N5Q8 |
| PRSGNHCPAGNF | 1 | 0 | 0 | 1489679 | ALJ54446.1 | K9N5Q8 |
| PRSGNHCPAGNS | 1 | 0 | 0 | 1489680 | AXN92228.1 | K9N5Q8 |
| PRSGNHCPAGNY | 1 | 0 | 0 | 1489681 | ATG84899.1 | K9N5Q8 |
| PRSGNYCPAGNS | 1 | 0 | 0 | 1489683 | AVN89387.1 | K9N5Q8 |
| PRSIRSYHADRN | 1 | 0 | 0 | 1489687 | AUM60014.1 | K9N5Q8 |
| PRSQMEVDFLEL | 1 | 0 | 0 | 1489692 | APO40578.1 | P0C6X7 |
| PRSVRSVPGEMC | 1 | 0 | 0 | 1489694 | AVN89313.1 | K9N5Q8 |
| PRTESMCPGNSN | 1 | 0 | 0 | 1489698 | ATQ39390.1 | K9N5Q8 |
| PRTIDSTQDGVN | 1 | 0 | 0 | 1489702 | QBQ01839.1 | P36334 |
| PRTINATQDGVN | 1 | 0 | 0 | 1489704 | AXX83321.1 | P36334 |
| PRTINSQDGDNK | 1 | 0 | 0 | 1489706 | AKA09654.1 | P36334 |
| PRTINSQDGVNK | 1 | 0 | 0 | 1489707 | QDH43726.1 | P36334 |
| PRTINSTQDGVN | 1 | 0 | 0 | 1489709 | AEN19358.1 | P36334 |
| PRTINSTQDGYN | 1 | 0 | 0 | 1489710 | CAA79896.1 | P36334 |
| PRTIQPEVKEVA | 1 | 0 | 0 | 1489713 | ANI69888.1 | K9N638 |
| PRTTQDGDNKLQ | 1 | 0 | 0 | 1489718 | AKA09658.1 | P36334 |
| PRVEKRKTEGFM | 1 | 0 | 0 | 1489723 | APO40578.1 | P0C6X7 |
| PRWKRQPNDDAT | 1 | 0 | 0 | 1489734 | APT69866.1 | A0A0N7J131 |
| PRWPQIAELAPS | 1 | 0 | 0 | 1489738 | AVV62533.1 | K9N4V7 |
| PRWPQIAELVPT | 1 | 0 | 0 | 1489740 | AXG21672.1 | K9N4V7 |
| PSADKEETNILS | 1 | 0 | 0 | 1489752 | AVV62524.1 | K9N7C7 |
| PSAELTGTQSVS | 1 | 0 | 0 | 1489753 | AVN89314.1 | K9N796 |
| PSAKVEGCMVQV | 1 | 0 | 0 | 1489757 | 2PWX_A | P0C6X7 |
| PSALASGCYSSL | 1 | 0 | 0 | 1489758 | AUM60014.1 | K9N5Q8 |
| PSANVSVSRFNP | 1 | 0 | 0 | 1489760 | BBO58721.1 | P36334 |
| PSARDLICAQYV | 1 | 0 | 0 | 1489762 | AVV62526.1 | K9N5Q8 |
| PSASAFMGMSQF | 1 | 0 | 0 | 1489766 | AUM60021.1 | K9N4V7 |

| | | | | | | |
|---|---|---|---|---|---|---|
| PSASAFMSMSQL | 1 | 0 | 0 | 1489767 | AVV62544.1 | K9N4V7 |
| PSASCPIGTKYR | 1 | 0 | 0 | 1489768 | AZS52618.1 | A0A140H1H1 |
| PSASVVEQAEGV | 1 | 0 | 0 | 1489770 | ATG84767.1 | K9N5Q8 |
| PSATYEGNSPFH | 1 | 0 | 0 | 1489771 | ABI96964.1 | P59635 |
| PSCFKKCGGCCD | 1 | 0 | 0 | 1489775 | AXX83351.1 | P36334 |
| PSCYNFLKEQHC | 1 | 0 | 0 | 1489777 | ARO76392.1 | Q7TLC7 |
| PSDDRTEVPQLV | 1 | 0 | 0 | 1489780 | ALJ54518.1 | K9N5Q8 |
| PSDDTLRTEAFD | 1 | 0 | 0 | 1489782 | APO40578.1 | P0C6X7 |
| PSDFTIGNMEEF | 1 | 0 | 0 | 1489785 | ARB66065.1 | P36334 |
| PSDFTIGNMGEF | 1 | 0 | 0 | 1489786 | ARB66062.1 | P36334 |
| PSDKHATLFGSV | 1 | 0 | 0 | 1489788 | AUM60014.1 | K9N5Q8 |
| PSDLNTSELLTK | 1 | 0 | 0 | 1489789 | NP_073549.1 | A0A0P0LKV0 |
| PSDPLNNFKHKV | 1 | 0 | 0 | 1489790 | AUM60013.1 | K9N7C7 |
| PSDSCNNVNVRD | 1 | 0 | 0 | 1489791 | AIW52827.1 | H9EJ42 |
| PSDSIRATATIP | 1 | 0 | 0 | 1489792 | AVP78043.1 | Q3LZX0 |
| PSDSSDNSKNGE | 1 | 0 | 0 | 1489793 | AVP78038.1 | Q3LZX4 |
| PSDSSDNSQNGE | 1 | 0 | 0 | 1489794 | AVP78049.1 | Q3LZX4 |
| PSDVSARDPIIQ | 1 | 0 | 0 | 1489797 | AMM42389.1 | Q19U29 |
| PSDVSARDPTIQ | 1 | 0 | 0 | 1489798 | AMM42397.1 | Q19U29 |
| PSEAKGYWYKHN | 1 | 0 | 0 | 1489802 | AMM42397.1 | Q19U29 |
| PSEAKGYWYRHN | 1 | 0 | 0 | 1489803 | ABD75581.1 | Q19U29 |
| PSECNSMHLSTY | 1 | 0 | 0 | 1489806 | APO40578.1 | P0C6X7 |
| PSECRLQNGDFL | 1 | 0 | 0 | 1489807 | APO40578.1 | P0C6X7 |
| PSECTLFRDAEG | 1 | 0 | 0 | 1489808 | AVV62524.1 | K9N7C7 |
| PSECTMFMDAEG | 1 | 0 | 0 | 1489809 | AKJ80136.2 | K9N7C7 |
| PSECTMFRDAEG | 1 | 0 | 0 | 1489810 | AXN92247.1 | K9N7C7 |
| PSEDFIKHIRTN | 1 | 0 | 0 | 1489811 | ASU90174.1 | K9N638 |
| PSEDFIKHVRNN | 1 | 0 | 0 | 1489812 | AXN92247.1 | K9N7C7 |
| PSEEFIKHVRTN | 1 | 0 | 0 | 1489814 | ALW82730.1 | K9N7C7 |
| PSEFTIGNIEEF | 1 | 0 | 0 | 1489816 | BBA20979.1 | P36334 |
| PSEFTIGNMEEF | 1 | 0 | 0 | 1489817 | CAA79896.1 | P36334 |
| PSEFTIGNMVEF | 1 | 0 | 0 | 1489818 | P36334.1 | P36334 |
| PSEPQTIQPEVK | 1 | 0 | 0 | 1489821 | AXN92247.1 | K9N7C7 |
| PSEPRTIQPEVK | 1 | 0 | 0 | 1489822 | AMQ49036.1 | K9N638 |
| PSETPNAKEEIL | 1 | 0 | 0 | 1489823 | P0C6W6.1 | P0C6W2 |
| PSFLVVDDKYKV | 1 | 0 | 0 | 1489829 | ABD75575.1 | A0A140H1G9 |
| PSFSSSRRKRRS | 1 | 0 | 0 | 1489830 | AGW27863.1 | A0A140H1H1 |
| PSGDAISQAFNK | 1 | 0 | 0 | 1489836 | AVV62535.1 | K9N7C7 |
| PSGDVEACMVQV | 1 | 0 | 0 | 1489838 | AXN92247.1 | K9N7C7 |
| PSGGNVIPHYSW | 1 | 0 | 0 | 1489841 | QEG03818.1 | P33469 |
| PSGGNVVPHYSW | 1 | 0 | 0 | 1489842 | AWW13570.1 | P33469 |
| PSGIGAVGDAGA | 1 | 0 | 0 | 1489845 | AVV62544.1 | K9N4V7 |
| PSGIGAVGGDAS | 1 | 0 | 0 | 1489846 | AVV62533.1 | K9N4V7 |
| PSGIGGGDLLYL | 1 | 0 | 0 | 1489848 | AVN89359.1 | K9N4V7 |
| PSGIGTVGGDLL | 1 | 0 | 0 | 1489849 | ATG84708.1 | K9N4V7 |
| PSGKVECCMVQV | 1 | 0 | 0 | 1489850 | APO40578.1 | P0C6X7 |
| PSGLKTMIVSAG | 1 | 0 | 0 | 1489852 | AQZ41321.1 | K9N7C7 |
| PSGLPPAYAPTY | 1 | 0 | 0 | 1489854 | AVV62535.1 | K9N7C7 |
| PSGLSPAYAPTY | 1 | 0 | 0 | 1489855 | AVV62524.1 | K9N7C7 |
| PSGNLIKFDIDS | 1 | 0 | 0 | 1489856 | AST12962.1 | A0A0P0LKV0 |
| PSGNYLAISNEL | 1 | 0 | 0 | 1489858 | AEN19365.1 | P30215 |
| PSGNYLSISNEL | 1 | 0 | 0 | 1489860 | AVQ05263.1 | P30215 |
| PSGQGVPLNANS | 1 | 0 | 0 | 1489862 | AZK15907.1 | K9N4V7 |
| PSGSFVEQAEGV | 1 | 0 | 0 | 1489864 | AID55084.1 | K9N5Q8 |
| PSGSILVDNDLN | 1 | 0 | 0 | 1489865 | ARU07574.1 | A0A140H1G9 |
| PSGSVVEQADGV | 1 | 0 | 0 | 1489866 | ATG84679.1 | K9N5Q8 |
| PSGSVVEQAEGF | 1 | 0 | 0 | 1489867 | AQZ41282.1 | K9N5Q8 |
| PSGSVVEQAEGV | 1 | 0 | 0 | 1489868 | AXN92228.1 | K9N5Q8 |
| PSGTSSGDATTA | 1 | 0 | 0 | 1489869 | AVP78041.1 | P0C6W2 |
| PSHAVQNTVLNA | 1 | 0 | 0 | 1489879 | AVL25602.1 | H9EJ69 |
| PSHDVTVLRDNL | 1 | 0 | 0 | 1489883 | AVA26874.1 | H9EJ24 |
| PSHVAQNTVLNA | 1 | 0 | 0 | 1489885 | AVA26877.1 | H9EJ69 |
| PSIDEAFPTRFP | 1 | 0 | 0 | 1489887 | AJQ24074.1 | P33469 |
| PSIIHDIGNPKG | 1 | 0 | 0 | 1489888 | ABE97136.1 | H9EJ42 |
| PSIISNAKQEIL | 1 | 0 | 0 | 1489889 | AVP78030.1 | P0C6W2 |
| PSILKVIDDGKI | 1 | 0 | 0 | 1489891 | QEO75984.1 | A0A0P0ILY9 |
| PSILPNEKQEIL | 1 | 0 | 0 | 1489892 | AVP78041.1 | P0C6W2 |
| PSIVEFNRLSTC | 1 | 0 | 0 | 1489902 | AUM60013.1 | K9N7C7 |
| PSKDTFELRYSG | 1 | 0 | 0 | 1489907 | AMM42397.1 | Q19U29 |
| PSKLLNGIAYIT | 1 | 0 | 0 | 1489919 | ABE97129.1 | H9EJ42 |
| PSKNNGIGTCPA | 1 | 0 | 0 | 1489921 | QDH43719.1 | P36334 |
| PSKPTKRSFIED | 1 | 0 | 0 | 1489924 | ATO98181.1 | Q3LZX1 |
| PSLVDDNKFDGL | 1 | 0 | 0 | 1489944 | ATP16754.1 | P0C6X6 |
| PSNDEAIPTKFP | 1 | 0 | 0 | 1489952 | AJQ24067.1 | P33469 |
| PSNDEAIPTRFP | 1 | 0 | 0 | 1489953 | AJQ24082.1 | P33469 |
| PSNDNISLPFNN | 1 | 0 | 0 | 1489954 | AVV62524.1 | K9N7C7 |
| PSNFSFNNWFLL | 1 | 0 | 0 | 1489957 | ARU07601.1 | A0A0P0K6L9 |
| PSNGTNILPVAF | 1 | 0 | 0 | 1489958 | NP_073549.1 | A0A0P0LKV0 |
| PSNHIDVVSAYG | 1 | 0 | 0 | 1489959 | ALX27232.1 | K9N5Q8 |
| PSNHIEVASAYG | 1 | 0 | 0 | 1489960 | ANF29261.1 | K9N5Q8 |
| PSNHKSNTNDAF | 1 | 0 | 0 | 1489961 | ATQ39390.1 | K9N5Q8 |
| PSNKWTGKKTLT | 1 | 0 | 0 | 1489963 | AVV62524.1 | K9N7C7 |
| PSNRGSFSSATG | 1 | 0 | 0 | 1489965 | AJQ24074.1 | P33469 |
| PSNWTTSVQVEY | 1 | 0 | 0 | 1489967 | AIW52843.1 | A0A0P0G321 |
| PSPFNWERRIFS | 1 | 0 | 0 | 1489968 | ABU39973.1 | A0A140H1H1 |
| PSPGMPALLSLV | 1 | 0 | 0 | 1489969 | 6NB6_A | P59594 |
| PSPIDIDAFPVR | 1 | 0 | 0 | 1489970 | AFD98763.1 | H9EJ42 |
| PSQEKNFTTAPA | 1 | 0 | 0 | 1489982 | AVP78031.1 | Q3LZX1 |
| PSQLKKPRWKRV | 1 | 0 | 0 | 1489985 | AIW52805.1 | H9EJ69 |
| PSQNAGYWRRQD | 1 | 0 | 0 | 1489987 | AQZ41279.1 | K9N4V7 |
| PSQNFGNSEMLK | 1 | 0 | 0 | 1489989 | ABD75597.1 | Q19U29 |

| | | | | | | |
|---|---|---|---|---|---|---|
| PSQQQNFTTAPA | 1 | 0 | 0 | 1489990 | APO40579.1 | P59594 |
| PSQTSPATVEPV | 1 | 0 | 0 | 1489992 | AIW52723.1 | A0A0N7J131 |
| PSREDKVCRLIP | 1 | 0 | 0 | 1489994 | APO40587.1 | P59636 |
| PSRNQSPASSQS | 1 | 0 | 0 | 1489999 | AIW52723.1 | A0A0N7J131 |
| PSRNQSPSSFQS | 1 | 0 | 0 | 1490001 | ABB90505.1 | A0A0N7J131 |
| PSRNQSPSSSQS | 1 | 0 | 0 | 1490002 | BAP75906.1 | A0A0N7J131 |
| PSRNQSPVSSQS | 1 | 0 | 0 | 1490003 | AMM42518.1 | A0A0N7J131 |
| PSRNWDVRSGHM | 1 | 0 | 0 | 1490004 | AVV62540.1 | K9N643 |
| PSRSQSRSQSRS | 1 | 0 | 0 | 1490010 | AIW52723.1 | A0A0N7J131 |
| PSSANPVKEYEA | 1 | 0 | 0 | 1490012 | AVV62535.1 | K9N7C7 |
| PSSARDLICAQY | 1 | 0 | 0 | 1490013 | AHL18090.1 | K9N5Q8 |
| PSSASCPIGTNY | 1 | 0 | 0 | 1490014 | AFR45556.1 | A0A140H1H1 |
| PSSDEAISTRFP | 1 | 0 | 0 | 1490017 | AIL49389.1 | P33469 |
| PSSDRYHNSQDD | 1 | 0 | 0 | 1490019 | AAA45463.1 | A0A0N7J131 |
| PSSIKEMQSQLS | 1 | 0 | 0 | 1490022 | AMM42613.1 | H9EJ69 |
| PSSIKEMQSQPS | 1 | 0 | 0 | 1490023 | AVL25602.1 | H9EJ69 |
| PSSKSSGMDTAL | 1 | 0 | 0 | 1490026 | AZS52621.1 | Q0ZJ82 |
| PSSPARLEIVKL | 1 | 0 | 0 | 1490032 | ATP16755.1 | Q80872 |
| PSSPARLELVKL | 1 | 0 | 0 | 1490033 | AWW13564.1 | Q80872 |
| PSSSAFIGMSQF | 1 | 0 | 0 | 1490037 | AVV62533.1 | K9N4V7 |
| PSSSGISTPKKP | 1 | 0 | 0 | 1490038 | AMM42629.1 | H9EJ69 |
| PSSSRGSSPARN | 1 | 0 | 0 | 1490042 | APO40586.1 | P59595 |
| PSSVKEMQSQSS | 1 | 0 | 0 | 1490043 | ABE97141.1 | H9EJ69 |
| PSTAAMLSDSHI | 1 | 0 | 0 | 1490050 | ABB90484.1 | A0A0N7J131 |
| PSTAWEDGDYYR | 1 | 0 | 0 | 1490051 | AGV08584.1 | K9N5Q8 |
| PSTDCSDGNYNR | 1 | 0 | 0 | 1490052 | ALA49803.1 | K9N5Q8 |
| PSTEAKGYWYPS | 1 | 0 | 0 | 1490054 | AJQ24074.1 | P33469 |
| PSTEVVRFPNIT | 1 | 0 | 0 | 1490055 | APO40579.1 | P59594 |
| PSTGVYELNGYT | 1 | 0 | 0 | 1490058 | CAA83660.1 | P36334 |
| PSTNTVIKKIYP | 1 | 0 | 0 | 1490066 | AUM60014.1 | K9N5Q8 |
| PSTQKVSGMDTA | 1 | 0 | 0 | 1490072 | AQN78779.1 | Q01455 |
| PSTSFYDIAVSQ | 1 | 0 | 0 | 1490080 | AUM60025.1 | K9N796 |
| PSTSGTQVYSHK | 1 | 0 | 0 | 1490083 | AVN89383.1 | K9N643 |
| PSTVDVVALQQE | 1 | 0 | 0 | 1490091 | AXN92258.1 | K9N7C7 |
| PSTVWEDGNYYR | 1 | 0 | 0 | 1490093 | ALJ54517.1 | K9N5Q8 |
| PSTVWKDGDYYR | 1 | 0 | 0 | 1490095 | ATW75480.1 | K9N5Q8 |
| PSTWNKKFGFIE | 1 | 0 | 0 | 1490096 | ARB66070.1 | P36334 |
| PSTWNRRFGFTE | 1 | 0 | 0 | 1490098 | CAA83660.1 | P36334 |
| PSVDMQPDYFIK | 1 | 0 | 0 | 1490103 | ATQ39390.1 | K9N5Q8 |
| PSVEQRQQAEEK | 1 | 0 | 0 | 1490105 | AVP78041.1 | P0C6W2 |
| PSVEVPPVLQVE | 1 | 0 | 0 | 1490106 | AVP78030.1 | P0C6W2 |
| PSVGPKQASLNGVTL | 1 | 0 | 0 | 1490109 | P0DTD1.1 | P0DTD1 |
| PSVLQATLSEFS | 1 | 0 | 0 | 1490113 | AXN92247.1 | K9N7C7 |
| PSVQPGPMIDVN | 1 | 0 | 0 | 1490115 | AXN92267.1 | K9N4V7 |
| PSVSLNSKEELL | 1 | 0 | 0 | 1490117 | APO40578.1 | P0C6X7 |
| PSVSNLALAFCS | 1 | 0 | 0 | 1490118 | AVV62524.1 | K9N7C7 |
| PSVVEFNRLATC | 1 | 0 | 0 | 1490119 | AVV62524.1 | K9N7C7 |
| PSYIKPESVATS | 1 | 0 | 0 | 1490138 | AUM60013.1 | K9N7C7 |
| PSYVKPDSIAIS | 1 | 0 | 0 | 1490144 | AVV62524.1 | K9N7C7 |
| PSYVKPDSIATS | 1 | 0 | 0 | 1490145 | AVV62535.1 | K9N7C7 |
| PTAADINIHDLP | 1 | 0 | 0 | 1490147 | AGT51530.1 | P30215 |
| PTAADINIPDLL | 1 | 0 | 0 | 1490148 | AGT51729.1 | P30215 |
| PTAADINIPHLP | 1 | 0 | 0 | 1490150 | AXX83356.1 | P30215 |
| PTAADINNPDLP | 1 | 0 | 0 | 1490151 | P30215.1 | P30215 |
| PTAAYINIPDLP | 1 | 0 | 0 | 1490153 | AEN19365.1 | P30215 |
| PTAFAAIGDLKC | 1 | 0 | 0 | 1490154 | AIL49486.1 | P36334 |
| PTAFAAIGDLNC | 1 | 0 | 0 | 1490155 | BBO58773.1 | P36334 |
| PTAFAVVGDLNC | 1 | 0 | 0 | 1490158 | QBP84704.1 | P36334 |
| PTAFSVIGDLKC | 1 | 0 | 0 | 1490159 | BAM16455.1 | P36334 |
| PTAGSTYRNMAL | 1 | 0 | 0 | 1490165 | AWW13566.1 | P36334 |
| PTAHVNATAAYG | 1 | 0 | 0 | 1490166 | AUM60014.1 | K9N5Q8 |
| PTAIASNCYSSL | 1 | 0 | 0 | 1490167 | AWF93654.1 | K9N5Q8 |
| PTAIRADQLTPA | 1 | 0 | 0 | 1490168 | ATO98181.1 | Q3LZX1 |
| PTAKYTPFAGYT | 1 | 0 | 0 | 1490171 | ARQ84748.1 | K9N4V0 |
| PTAKYTPLVGTS | 1 | 0 | 0 | 1490173 | AXN92262.1 | K9N4V0 |
| PTARTTSSEFQL | 1 | 0 | 0 | 1490179 | AUM60024.1 | K9N5Q8 |
| PTASAFMGMSQF | 1 | 0 | 0 | 1490180 | AXN92267.1 | K9N4V7 |
| PTATGDVVLAKD | 1 | 0 | 0 | 1490181 | ATN39862.1 | P0C6X6 |
| PTATSVNCCHDG | 1 | 0 | 0 | 1490183 | ARE29989.1 | Q4VID0 |
| PTDAIVIDNDLN | 1 | 0 | 0 | 1490192 | ASY99818.1 | K9N7C7 |
| PTDIYLTTYPGQ | 1 | 0 | 0 | 1490193 | APO40578.1 | P0C6X7 |
| PTDKLAMIMGYC | 1 | 0 | 0 | 1490194 | AWW13564.1 | Q80872 |
| PTDNHITTYPGQ | 1 | 0 | 0 | 1490196 | AVP78041.1 | P0C6W2 |
| PTDQSSYTVDSV | 1 | 0 | 0 | 1490200 | APO40578.1 | P0C6X7 |
| PTDSADNNQNGG | 1 | 0 | 0 | 1490202 | ARO76389.1 | P59595 |
| PTDSTDNNQDGG | 1 | 0 | 0 | 1490203 | Q3I5I7.1 | Q3LZX4 |
| PTDTCFANKHAD | 1 | 0 | 0 | 1490207 | AVP78041.1 | P0C6W2 |
| PTDVAYHTVIDV | 1 | 0 | 0 | 1490208 | AIV41894.1 | P0C6X6 |
| PTDVFTNHSVVY | 1 | 0 | 0 | 1490209 | ARB66060.1 | P36334 |
| PTDVPSTSGTQV | 1 | 0 | 0 | 1490210 | ASU90409.1 | K9N643 |
| PTDYKNVKAWSG | 1 | 0 | 0 | 1490211 | AIW52843.1 | A0A0P0G321 |
| PTECIYTQSRSC | 1 | 0 | 0 | 1490212 | AXN92247.1 | K9N7C7 |
| PTEDLCNGSVVI | 1 | 0 | 0 | 1490214 | AGT51737.1 | P0C6X6 |
| PTEHACSSLNVL | 1 | 0 | 0 | 1490216 | APO40578.1 | P0C6X7 |
| PTEHVNLFDVGY | 1 | 0 | 0 | 1490217 | ANI69890.1 | K9N796 |
| PTEHVTSGNLFD | 1 | 0 | 0 | 1490218 | AGV08439.1 | K9N796 |
| PTEKLVTTSTNP | 1 | 0 | 0 | 1490220 | AUM60013.1 | K9N7C7 |
| PTENACSSLTVL | 1 | 0 | 0 | 1490221 | AVP78041.1 | P0C6W2 |
| PTENNCSQLDEE | 1 | 0 | 0 | 1490222 | AVP78035.1 | Q3LZX8 |
| PTEQTVSTAVSP | 1 | 0 | 0 | 1490225 | AVV62535.1 | K9N7C7 |
| PTEQVVSTSSDP | 1 | 0 | 0 | 1490226 | AVV62524.1 | K9N7C7 |

| | | | | | | |
|---|---|---|---|---|---|---|
| PTESYVDVGSDS | 1 | 0 | 0 | 1490230 | AMW90853.1 | K9N5Q8 |
| PTGKSYSNISLT | 1 | 0 | 0 | 1490258 | AVV62537.1 | K9N5Q8 |
| PTGLKTMVVSAG | 1 | 0 | 0 | 1490260 | AUM60013.1 | K9N7C7 |
| PTGLKTMVVTAG | 1 | 0 | 0 | 1490261 | AVV62524.1 | K9N7C7 |
| PTGLSVLQPLVE | 1 | 0 | 0 | 1490262 | AVP78031.1 | Q3LZX1 |
| PTGPAYVMIDKV | 1 | 0 | 0 | 1490263 | NP_073549.1 | A0A0P0LKV0 |
| PTGTKYRHCDLD | 1 | 0 | 0 | 1490270 | AIG96691.1 | A0A140H1H1 |
| PTGTYHCVSTQA | 1 | 0 | 0 | 1490274 | AVV62524.1 | K9N7C7 |
| PTGVCLGAGQFV | 1 | 0 | 0 | 1490276 | AVQ05261.1 | P0C6X6 |
| PTGVHAGPDLEG | 1 | 0 | 0 | 1490278 | ACV88184.1 | P0C6X7 |
| PTGVYELNGYPV | 1 | 0 | 0 | 1490282 | AMM42466.1 | P36334 |
| PTGYVDTEHSTE | 1 | 0 | 0 | 1490287 | AVP78030.1 | P0C6W2 |
| PTGYVDTENSTE | 1 | 0 | 0 | 1490289 | AVP78041.1 | P0C6W2 |
| PTGYVDTSANTE | 1 | 0 | 0 | 1490291 | APO40578.1 | P0C6X7 |
| PTHEVIRFPNIT | 1 | 0 | 0 | 1490293 | ATO98193.1 | Q3LZX1 |
| PTHWICANQGTS | 1 | 0 | 0 | 1490299 | AKA09620.1 | A0A0P0G321 |
| PTHWICANQNTS | 1 | 0 | 0 | 1490300 | BAM16475.1 | A0A0P0G321 |
| PTIADFVQIENC | 1 | 0 | 0 | 1490310 | AIW52757.1 | A0A0P0K6L9 |
| PTIEDWLLVQGF | 1 | 0 | 0 | 1490314 | AHI48842.1 | K9N643 |
| PTIEENVIQCFG | 1 | 0 | 0 | 1490315 | ADK37676.1 | H9EJ69 |
| PTIQKDVIECDV | 1 | 0 | 0 | 1490320 | APO40578.1 | P0C6X7 |
| PTISQPSKLLNG | 1 | 0 | 0 | 1490325 | ABE97136.1 | H9EJ42 |
| PTITVPEEFANH | 1 | 0 | 0 | 1490332 | AVV62524.1 | K9N7C7 |
| PTKLNDLCFSSV | 1 | 0 | 0 | 1490343 | APO40579.1 | P59594 |
| PTKQRYAKRRFS | 1 | 0 | 0 | 1490345 | AUM60017.1 | K9N643 |
| PTKVLRYGYHVL | 1 | 0 | 0 | 1490349 | AEN19363.1 | P0C6X6 |
| PTLDCINGQYDK | 1 | 0 | 0 | 1490352 | AUM60014.1 | K9N5Q8 |
| PTLEEPVNQFTG | 1 | 0 | 0 | 1490353 | ARO76381.1 | P0C6U8 |
| PTLELCNAVEQR | 1 | 0 | 0 | 1490354 | AVV62535.1 | K9N7C7 |
| PTLIPQENYSSI | 1 | 0 | 0 | 1490359 | AWW13571.1 | P0C6X6 |
| PTLKDFGGFNFS | 1 | 0 | 0 | 1490360 | ATO98231.1 | Q3LZX1 |
| PTLTESSGSLVT | 1 | 0 | 0 | 1490377 | AHF54653.1 | Q4VID0 |
| PTMADFVQIENC | 1 | 0 | 0 | 1490382 | CAA71056.1 | A0A0P0K6L9 |
| PTMCDIKQMLFC | 1 | 0 | 0 | 1490383 | AXN92247.1 | K9N7C7 |
| PTMIHVEQVSSD | 1 | 0 | 0 | 1490386 | APO40579.1 | P59594 |
| PTMTHMNLKYAI | 1 | 0 | 0 | 1490387 | AZK15898.1 | K9N7C7 |
| PTNAVMKMPCPA | 1 | 0 | 0 | 1490388 | ARO76381.1 | P0C6U8 |
| PTNAYTRTTHTD | 1 | 0 | 0 | 1490391 | ANI69888.1 | K9N638 |
| PTNCITPVNGYF | 1 | 0 | 0 | 1490393 | ALJ76278.1 | K9N5Q8 |
| PTNEAVEAPLIG | 1 | 0 | 0 | 1490394 | AVP78041.1 | P0C6W2 |
| PTNEIPYKSFSD | 1 | 0 | 0 | 1490395 | AUM60013.1 | K9N7C7 |
| PTNFCPCKLDGS | 1 | 0 | 0 | 1490396 | CAA83660.1 | P36334 |
| PTNFSISVTTEV | 1 | 0 | 0 | 1490402 | AVP78031.1 | Q3LZX1 |
| PTNHVNVTAAYG | 1 | 0 | 0 | 1490408 | AVV62537.1 | K9N5Q8 |
| PTNLEMFKPGVE | 1 | 0 | 0 | 1490410 | AOL02458.1 | Q4VID0 |
| PTNMYMRTTHTD | 1 | 0 | 0 | 1490412 | AUM60013.1 | K9N7C7 |
| PTNSVRTTATIP | 1 | 0 | 0 | 1490414 | AVP78032.1 | Q3LZX0 |
| PTNVVSHVNGDW | 1 | 0 | 0 | 1490415 | AEN19365.1 | P30215 |
| PTNVYMRTTHTD | 1 | 0 | 0 | 1490417 | ANC28699.1 | K9N638 |
| PTPAPVYIWTAD | 1 | 0 | 0 | 1490418 | AWW13562.1 | Q01455 |
| PTPIEDAETPAL | 1 | 0 | 0 | 1490420 | ATQ39389.1 | K9N7C7 |
| PTQAPTHLSVDI | 1 | 0 | 0 | 1490423 | AEA10817.1 | P0C6X7 |
| PTQCTRDPVGFC | 1 | 0 | 0 | 1490425 | AUM60013.1 | K9N7C7 |
| PTQEVVRFPNIT | 1 | 0 | 0 | 1490427 | ATO98108.1 | Q3LZX1 |
| PTQPLPNATYDN | 1 | 0 | 0 | 1490437 | APO40578.1 | P0C6X7 |
| PTQSPTYLSVDT | 1 | 0 | 0 | 1490439 | APO40578.1 | P0C6X7 |
| PTQSVVRFPNIT | 1 | 0 | 0 | 1490440 | AVP78031.1 | Q3LZX1 |
| PTREENVMQCFG | 1 | 0 | 0 | 1490444 | AMM42619.1 | H9EJ69 |
| PTSEEVVENPTV | 1 | 0 | 0 | 1490453 | ARO76381.1 | P0C6U8 |
| PTSGSTYLNMAL | 1 | 0 | 0 | 1490456 | BBA20982.1 | P36334 |
| PTSGSTYRNMAL | 1 | 0 | 0 | 1490457 | CAA79896.1 | P36334 |
| PTSGVIPDMADQ | 1 | 0 | 0 | 1490458 | AMK59681.1 | P33469 |
| PTSGVTPDMADE | 1 | 0 | 0 | 1490459 | AMM42457.1 | P33469 |
| PTSGVTPDMADH | 1 | 0 | 0 | 1490460 | AIL49401.1 | P33469 |
| PTSKIEPCIVSV | 1 | 0 | 0 | 1490464 | ARU07575.1 | A0A140H1G9 |
| PTSKIEPCLVSV | 1 | 0 | 0 | 1490465 | P0C6U5.1 | A0A140H1G9 |
| PTSLEIFKPGVE | 1 | 0 | 0 | 1490468 | APU51922.1 | Q4VID0 |
| PTSPTARTTSSE | 1 | 0 | 0 | 1490469 | AUM60024.1 | K9N5Q8 |
| PTSSSTYRNMAL | 1 | 0 | 0 | 1490471 | AIL49538.1 | P36334 |
| PTTCANDPVGFI | 1 | 0 | 0 | 1490476 | ARO76381.1 | P0C6U8 |
| PTTCGYLPNNAV | 1 | 0 | 0 | 1490479 | APO40578.1 | P0C6X7 |
| PTTCVNDPVGFT | 1 | 0 | 0 | 1490481 | AVP78030.1 | P0C6W2 |
| PTTCVNDPVGFV | 1 | 0 | 0 | 1490482 | APO40578.1 | P0C6X7 |
| PTTDCSDGNYNR | 1 | 0 | 0 | 1490483 | ALA49473.1 | K9N5Q8 |
| PTTENKLNISSG | 1 | 0 | 0 | 1490484 | ATO98145.1 | Q3LZX1 |
| PTTENLYFQGAM | 1 | 0 | 0 | 1490485 | 3EJG_A | A0A0P0LKV0 |
| PTTFHLDGETIT | 1 | 0 | 0 | 1490487 | AVP78030.1 | P0C6W2 |
| PTTIHADQLTPA | 1 | 0 | 0 | 1490496 | AVP78031.1 | Q3LZX1 |
| PTTPISSALIAF | 1 | 0 | 0 | 1490506 | QBF80585.1 | K9N7C7 |
| PTTSSFNADASV | 1 | 0 | 0 | 1490510 | APO40579.1 | P59594 |
| PTTVLNGPVFTG | 1 | 0 | 0 | 1490515 | ABE97136.1 | H9EJ42 |
| PTVAMTEQLQMG | 1 | 0 | 0 | 1490518 | AKN24830.1 | K9N5Q8 |
| PTVDVVALQQEM | 1 | 0 | 0 | 1490519 | AXN92247.1 | K9N7C7 |
| PTVDVVRFPNIT | 1 | 0 | 0 | 1490520 | ATO98145.1 | Q3LZX1 |
| PTVETPKEEQPK | 1 | 0 | 0 | 1490521 | ARO76381.1 | P0C6U8 |
| PTVIRADQLTPA | 1 | 0 | 0 | 1490525 | ATO98145.1 | Q3LZX1 |
| PTVKEIISIPNI | 1 | 0 | 0 | 1490530 | AIX09804.1 | P0C6X6 |
| PTVKEIISKPKI | 1 | 0 | 0 | 1490531 | AIV41888.1 | P0C6X6 |
| PTVLPGAHAYET | 1 | 0 | 0 | 1490536 | AVV62535.1 | K9N7C7 |
| PTVLPGAHAYGS | 1 | 0 | 0 | 1490537 | AVV62524.1 | K9N7C7 |
| PTVLPGAHPYSA | 1 | 0 | 0 | 1490538 | AUM60013.1 | K9N7C7 |

| | | | | | | |
|---|---|---|---|---|---|---|
| PTVNEIASTPKT | 1 | 0 | 0 | 1490539 | AVI15051.1 | P0C6X6 |
| PTVQKEVIECDV | 1 | 0 | 0 | 1490540 | ARO76381.1 | P0C6U8 |
| PTVTLLPAAELD | 1 | 0 | 0 | 1490544 | APO40586.1 | P59595 |
| PTVTPLPAADMD | 1 | 0 | 0 | 1490545 | AAS48456.1 | P59595 |
| PTVWEDGDYYRK | 1 | 0 | 0 | 1490548 | QBF80598.1 | K9N5Q8 |
| PTYKNGAMYKGK | 1 | 0 | 0 | 1490562 | AVN89451.1 | K9N7C7 |
| PTYMDGADVTKI | 1 | 0 | 0 | 1490567 | APO40578.1 | P0C6X7 |
| PVADYTPVIGTS | 1 | 0 | 0 | 1490572 | AVV62539.1 | K9N4V0 |
| PVASPRECNDMH | 1 | 0 | 0 | 1490582 | ARO76381.1 | P0C6U8 |
| PVCNGRVGHSGN | 1 | 0 | 0 | 1490593 | ADK37701.1 | A0A0P0K6L9 |
| PVCSFPLCANGL | 1 | 0 | 0 | 1490594 | AXN92247.1 | K9N7C7 |
| PVDAVLIDNDLN | 1 | 0 | 0 | 1490601 | AVV62524.1 | K9N7C7 |
| PVDCQDDRCIIH | 1 | 0 | 0 | 1490602 | ANZ78865.1 | P0C6X6 |
| PVDDFIKHVQLE | 1 | 0 | 0 | 1490603 | AUM60013.1 | K9N7C7 |
| PVDGYFILNQTT | 1 | 0 | 0 | 1490606 | AUM60014.1 | K9N5Q8 |
| PVDHGVESNDSV | 1 | 0 | 0 | 1490607 | ATQ39397.1 | K9N7D2 |
| PVDHYDGFYTQG | 1 | 0 | 0 | 1490608 | NP_073549.1 | A0A0P0LKV0 |
| PVDLKHFFFTQD | 1 | 0 | 0 | 1490610 | QEG03736.1 | P0C6X6 |
| PVDLPPKVYFYY | 1 | 0 | 0 | 1490611 | AMM42619.1 | H9EJ69 |
| PVDSFTNGAVVG | 1 | 0 | 0 | 1490617 | P0C6W6.1 | P0C6W2 |
| PVEAPVEDIAQV | 1 | 0 | 0 | 1490623 | AVN89451.1 | K9N7C7 |
| PVECDEGCSEVE | 1 | 0 | 0 | 1490624 | ALA50033.1 | K9N7C7 |
| PVECDEQEEEVE | 1 | 0 | 0 | 1490625 | AVV62524.1 | K9N7C7 |
| PVEDTQVVIADT | 1 | 0 | 0 | 1490629 | AVN89312.1 | K9N638 |
| PVEEFEDSTGDE | 1 | 0 | 0 | 1490630 | AVV62535.1 | K9N7C7 |
| PVEFHLDGEVLP | 1 | 0 | 0 | 1490631 | ATO98229.1 | P0C6W2 |
| PVEGVSSFMNAT | 1 | 0 | 0 | 1490634 | CAA71147.1 | A0A0P0K6L9 |
| PVEGVSSFMNVT | 1 | 0 | 0 | 1490635 | AIW52757.1 | A0A0P0K6L9 |
| PVELENKFTPLS | 1 | 0 | 0 | 1490637 | AVV62524.1 | K9N7C7 |
| PVELPTVDVVAL | 1 | 0 | 0 | 1490638 | AVN89312.1 | K9N638 |
| PVELPVSMCRYL | 1 | 0 | 0 | 1490639 | ADK37703.1 | A0A0P0K6L9 |
| PVEPLVQDRAVE | 1 | 0 | 0 | 1490641 | AHF54653.1 | Q4VID0 |
| PVEPLVQDRVVE | 1 | 0 | 0 | 1490642 | ARE29989.1 | Q4VID0 |
| PVEPPTVDVVAL | 1 | 0 | 0 | 1490643 | AXN92247.1 | K9N7C7 |
| PVEPSTVDVVAL | 1 | 0 | 0 | 1490644 | AXN92258.1 | K9N7C7 |
| PVESPTVDVVAL | 1 | 0 | 0 | 1490645 | ATG84853.1 | K9N7C7 |
| PVETSNSFDALE | 1 | 0 | 0 | 1490646 | AVP78030.1 | P0C6W2 |
| PVETSNSFDALK | 1 | 0 | 0 | 1490647 | AVP78041.1 | P0C6W2 |
| PVETSNSFEVLA | 1 | 0 | 0 | 1490649 | ACV88184.1 | P0C6X7 |
| PVEVHVEDIVQV | 1 | 0 | 0 | 1490651 | ANF29161.1 | K9N7C7 |
| PVEVPVEDIAQV | 1 | 0 | 0 | 1490652 | AXN92247.1 | K9N7C7 |
| PVEVPVEDIVQV | 1 | 0 | 0 | 1490653 | AVK87459.1 | K9N7C7 |
| PVEVPVEDTQVV | 1 | 0 | 0 | 1490654 | AVN89312.1 | K9N638 |
| PVEYVSYFDGVK | 1 | 0 | 0 | 1490656 | NP_073549.1 | A0A0P0LKV0 |
| PVFKKGAKLLHK | 1 | 0 | 0 | 1490665 | APO40578.1 | P0C6X7 |
| PVFPAFHRIEST | 1 | 0 | 0 | 1490667 | AGV08496.1 | K9N7D2 |
| PVGCAMGIVNSS | 1 | 0 | 0 | 1490675 | AUM60014.1 | K9N5Q8 |
| PVGCVLGLINSS | 1 | 0 | 0 | 1490676 | ALK80261.1 | K9N5Q8 |
| PVGCVLGLVNSS | 1 | 0 | 0 | 1490677 | AXN92228.1 | K9N5Q8 |
| PVGDSFGKSLGN | 1 | 0 | 0 | 1490678 | ATN39877.2 | P0C6X6 |
| PVGESFGKSLGN | 1 | 0 | 0 | 1490679 | AEN19363.1 | P0C6X6 |
| PVGFCLSNNPCN | 1 | 0 | 0 | 1490680 | ANI69899.1 | K9N7C7 |
| PVGFCLTNDPCN | 1 | 0 | 0 | 1490683 | AVV62535.1 | K9N7C7 |
| PVGFFFPYDPAL | 1 | 0 | 0 | 1490685 | AVV62535.1 | K9N7C7 |
| PVGGLYEIQIPS | 1 | 0 | 0 | 1490691 | CAA79896.1 | P36334 |
| PVGGTTVVRPLV | 1 | 0 | 0 | 1490692 | AVV62543.1 | K9N7A1 |
| PVGIRDPVSYVL | 1 | 0 | 0 | 1490694 | ATN39862.1 | P0C6X6 |
| PVGTLDISASVI | 1 | 0 | 0 | 1490697 | APO40578.1 | P0C6X7 |
| PVGTNYLTCHNL | 1 | 0 | 0 | 1490699 | AIL49486.1 | P36334 |
| PVGYFFPYDPAL | 1 | 0 | 0 | 1490703 | AVV62524.1 | K9N7C7 |
| PVHMPVHPMVMP | 1 | 0 | 0 | 1490709 | AET99208.1 | Q0ZJ80 |
| PVHVYADDSIIL | 1 | 0 | 0 | 1490718 | AEN19363.1 | P0C6X6 |
| PVHYERDNTSCP | 1 | 0 | 0 | 1490721 | AXN92247.1 | K9N7C7 |
| PVHYERDNTSGP | 1 | 0 | 0 | 1490722 | AWM99580.1 | K9N7C7 |
| PVIDDGDGVKPF | 1 | 0 | 0 | 1490723 | QED88039.1 | H9EJ42 |
| PVIGTSTHPVQW | 1 | 0 | 0 | 1490725 | AVV62539.1 | K9N4V0 |
| PVISQLDENAQK | 1 | 0 | 0 | 1490735 | AST12962.1 | A0A0P0LKV0 |
| PVIWLSHEQASL | 1 | 0 | 0 | 1490739 | AVI15051.1 | P0C6X6 |
| PVIYQLDENAQK | 1 | 0 | 0 | 1490740 | AMM42491.1 | A0A0P0ILY9 |
| PVKDVFELRYSG | 1 | 0 | 0 | 1490742 | ARU07581.1 | Q19U29 |
| PVKDYIQTVNVI | 1 | 0 | 0 | 1490743 | ABD75583.1 | A0A140H1G9 |
| PVKKVAFGGDQI | 1 | 0 | 0 | 1490758 | ALA49340.1 | K9N7C7 |
| PVKPAVGPMVDV | 1 | 0 | 0 | 1490762 | AUM60021.1 | K9N4V7 |
| PVLQVCDVLVRG | 1 | 0 | 0 | 1490781 | AVP78041.1 | P0C6W2 |
| PVLQVESTPVVT | 1 | 0 | 0 | 1490782 | AVP78030.1 | P0C6W2 |
| PVLVNAANTHLK | 1 | 0 | 0 | 1490789 | AVN89451.1 | K9N7C7 |
| PVLWVTNGNFDC | 1 | 0 | 0 | 1490793 | AVV62524.1 | K9N7C7 |
| PVLYKYLNSPME | 1 | 0 | 0 | 1490795 | AEN19363.1 | P0C6X6 |
| PVMAAPTGITLT | 1 | 0 | 0 | 1490796 | ABD34826.1 | U3M6Q8 |
| PVMAAPTGVTLT | 1 | 0 | 0 | 1490797 | AWK59934.1 | U3M6Q8 |
| PVMDPIYDEPTT | 1 | 0 | 0 | 1490798 | ARO76383.1 | P59632 |
| PVMESQISTKEV | 1 | 0 | 0 | 1490800 | ARB07605.1 | A0A140H1G9 |
| PVMISEWPLFDQ | 1 | 0 | 0 | 1490802 | APT69896.1 | A0A0P0LKV0 |
| PVMISQWPISDD | 1 | 0 | 0 | 1490806 | AWK59942.1 | H9EJ42 |
| PVMSEDLWQFVD | 1 | 0 | 0 | 1490808 | NP_073549.1 | A0A0P0LKV0 |
| PVNGYFIKTNNI | 1 | 0 | 0 | 1490812 | ASU90175.1 | K9N5Q8 |
| PVNLNRGLPTGL | 1 | 0 | 0 | 1490815 | AVP78031.1 | Q3LZX1 |
| PVNPTDQSSYTV | 1 | 0 | 0 | 1490817 | APO40578.1 | P0C6X7 |
| PVNPVLHTKSAA | 1 | 0 | 0 | 1490818 | AEN19363.1 | P0C6X6 |
| PVPEMKILQNLG | 1 | 0 | 0 | 1490826 | APO40578.1 | P0C6X7 |
| PVPGTSTHPVQW | 1 | 0 | 0 | 1490829 | AUM60016.1 | K9N4V0 |

| | | | | | | |
|---|---|---|---|---|---|---|
| PVPVKEYEADVE | 1 | 0 | 0 | 1490834 | AVV62524.1 | K9N7C7 |
| PVQGAWNIGQQK | 1 | 0 | 0 | 1490837 | ARO76381.1 | P0C6U8 |
| PVQLAPVSPAFH | 1 | 0 | 0 | 1490841 | AKM76243.1 | K9N7D2 |
| PVQSRKFIVPWV | 1 | 0 | 0 | 1490849 | AEN19363.1 | P0C6X6 |
| PVRDEVSTETDI | 1 | 0 | 0 | 1490860 | ABB90487.1 | A0A0N7J131 |
| PVRLRNSSDNGI | 1 | 0 | 0 | 1490863 | AIW52843.1 | A0A0P0G321 |
| PVRNIWIPILKS | 1 | 0 | 0 | 1490864 | AXX83306.1 | P0C6X6 |
| PVRYERDNTSCP | 1 | 0 | 0 | 1490868 | ALA50044.1 | K9N7C7 |
| PVSDLLANMGID | 1 | 0 | 0 | 1490871 | ARO76381.1 | P0C6U8 |
| PVSGIREFSNLV | 1 | 0 | 0 | 1490880 | AIW52843.1 | A0A0P0G321 |
| PVSGLKSSTGFV | 1 | 0 | 0 | 1490881 | QEG59359.1 | A0A0P0G321 |
| PVSGLYEIQIPS | 1 | 0 | 0 | 1490882 | AIL49538.1 | P36334 |
| PVSGTTLHPVQW | 1 | 0 | 0 | 1490883 | AVV62528.1 | K9N4V0 |
| PVSNCIKMLSNG | 1 | 0 | 0 | 1490902 | AWK59942.1 | H9EJ42 |
| PVSPAFHCIEST | 1 | 0 | 0 | 1490903 | ANI69893.1 | K9N7D2 |
| PVSPAFHRIESP | 1 | 0 | 0 | 1490906 | ASU91080.1 | K9N7D2 |
| PVSPVDHGVESN | 1 | 0 | 0 | 1490909 | ATQ39397.1 | K9N7D2 |
| PVSPVFHRIESI | 1 | 0 | 0 | 1490910 | AVN89455.1 | K9N7D2 |
| PVSSIKGDKGVE | 1 | 0 | 0 | 1490913 | AWK59942.1 | H9EJ42 |
| PVSVIYDDNTNQ | 1 | 0 | 0 | 1490920 | ATQ39390.1 | K9N5Q8 |
| PVSVIYDKPSDK | 1 | 0 | 0 | 1490923 | AUM60014.1 | K9N5Q8 |
| PVTKNINGSNEA | 1 | 0 | 0 | 1490935 | AVV62524.1 | K9N7C7 |
| PVTLEQKVKPIV | 1 | 0 | 0 | 1490938 | APO40578.1 | P0C6X7 |
| PVTRDSVPLHII | 1 | 0 | 0 | 1490941 | AHI48554.1 | K9N7D2 |
| PVVCDEDGELNS | 1 | 0 | 0 | 1490951 | AUM60013.1 | K9N7C7 |
| PVVDGKQIDDHR | 1 | 0 | 0 | 1490953 | AGT51409.1 | P0C6X6 |
| PVVDWHYYDAQP | 1 | 0 | 0 | 1490956 | AVV62524.1 | K9N7C7 |
| PVVMEIPSLMTV | 1 | 0 | 0 | 1490968 | ARU07575.1 | A0A140H1G9 |
| PVVPDIVDVPPQ | 1 | 0 | 0 | 1490971 | AVN89352.1 | K9N7C7 |
| PVVPDTVGVPPQ | 1 | 0 | 0 | 1490974 | AHL18091.1 | K9N7C7 |
| PVVSDTVDVPPQ | 1 | 0 | 0 | 1490976 | ASU91261.1 | K9N638 |
| PVVSDTVEVPPH | 1 | 0 | 0 | 1490977 | QBF80585.1 | K9N7C7 |
| PVVSDTVEVPPQ | 1 | 0 | 0 | 1490978 | AXN92247.1 | K9N7C7 |
| PVVSDTVEVSPQ | 1 | 0 | 0 | 1490979 | ASU45805.1 | K9N7C7 |
| PVVSDTVEVTPQ | 1 | 0 | 0 | 1490980 | ALA50033.1 | K9N7C7 |
| PVVTETSEQNNF | 1 | 0 | 0 | 1490982 | AVP78030.1 | P0C6W2 |
| PVYFLSYSGAIK | 1 | 0 | 0 | 1490992 | AVV62544.1 | K9N4V7 |
| PVYFLWYSGAIK | 1 | 0 | 0 | 1490993 | AQZ41280.1 | K9N4V7 |
| PVYIAPQANSGI | 1 | 0 | 0 | 1490995 | AVQ05263.1 | P30215 |
| PWDTIANYVKPF | 1 | 0 | 0 | 1491006 | AVP78030.1 | P0C6W2 |
| PWLLLSGNPTEK | 1 | 0 | 0 | 1491023 | AUM60013.1 | K9N7C7 |
| PWPQIAELVPTA | 1 | 0 | 0 | 1491033 | QDP16213.1 | K9N4V7 |
| PWSAAAGVPFSL | 1 | 0 | 0 | 1491035 | BBA20986.1 | A0A140H1H1 |
| PWSKICDYVKPF | 1 | 0 | 0 | 1491036 | APO40578.1 | P0C6X7 |
| PWTAAEGVPFYL | 1 | 0 | 0 | 1491042 | AIL49550.1 | P36334 |
| PWYVWLLIGFAG | 1 | 0 | 0 | 1491058 | CAA83660.1 | P36334 |
| PYAKDQYGTDID | 1 | 0 | 0 | 1491062 | AIL49482.1 | P33469 |
| PYANASYGQSLE | 1 | 0 | 0 | 1491065 | ADN03343.1 | Q19U29 |
| PYCHDPTVLPGA | 1 | 0 | 0 | 1491069 | AXN92247.1 | K9N7C7 |
| PYCYTEGLMQNA | 1 | 0 | 0 | 1491074 | AGT51787.1 | P0C6X6 |
| PYDHAIGATAAL | 1 | 0 | 0 | 1491076 | ATO98145.1 | Q3LZX1 |
| PYDIEFVTGKQN | 1 | 0 | 0 | 1491077 | ALA50044.1 | K9N7C7 |
| PYDIELVTGKQN | 1 | 0 | 0 | 1491078 | AXN92247.1 | K9N7C7 |
| PYDMDFGVSSFS | 1 | 0 | 0 | 1491081 | AHY22554.1 | K9N638 |
| PYDMDSGGSSFS | 1 | 0 | 0 | 1491082 | AHX71940.1 | K9N7C7 |
| PYDMDSGVSSIS | 1 | 0 | 0 | 1491084 | AHI48797.1 | K9N638 |
| PYDMSSDVSSFS | 1 | 0 | 0 | 1491085 | AUM60013.1 | K9N7C7 |
| PYDPALETGDFT | 1 | 0 | 0 | 1491086 | AVV62535.1 | K9N7C7 |
| PYEEGLETGTFT | 1 | 0 | 0 | 1491089 | AUM60013.1 | K9N7C7 |
| PYEFCSQHTMQI | 1 | 0 | 0 | 1491090 | AVL25535.1 | H9EJ42 |
| PYEISDDVSAFS | 1 | 0 | 0 | 1491091 | AVV62524.1 | K9N7C7 |
| PYEPRTIQPEVK | 1 | 0 | 0 | 1491093 | ANI69888.1 | K9N638 |
| PYERDISNVLYN | 1 | 0 | 0 | 1491094 | APO40579.1 | P59594 |
| PYEVFDALQDDP | 1 | 0 | 0 | 1491095 | AVV62524.1 | K9N7C7 |
| PYEVSDALQDDP | 1 | 0 | 0 | 1491096 | AVV62535.1 | K9N7C7 |
| PYFMHDVVSSFE | 1 | 0 | 0 | 1491102 | AST12962.1 | A0A0P0LKV0 |
| PYFMYDVVSSFE | 1 | 0 | 0 | 1491104 | AGT21366.1 | A0A0P0LKV0 |
| PYGAKPVSEYHV | 1 | 0 | 0 | 1491105 | AVV62529.1 | K9N643 |
| PYGAKPVTEYHI | 1 | 0 | 0 | 1491106 | AVV62540.1 | K9N643 |
| PYGCQPVSEYHI | 1 | 0 | 0 | 1491111 | AHE78102.1 | K9N643 |
| PYGRCPTAAVIN | 1 | 0 | 0 | 1491118 | AGT51520.1 | P30215 |
| PYGYPNDIVGGF | 1 | 0 | 0 | 1491122 | AWK59978.1 | H9EJ42 |
| PYHANCVNCADD | 1 | 0 | 0 | 1491128 | AVV62535.1 | K9N7C7 |
| PYHANSSEPAML | 1 | 0 | 0 | 1491129 | AIV41873.1 | P36334 |
| PYHPLADNKFAV | 1 | 0 | 0 | 1491131 | APO40584.1 | P59635 |
| PYISHENKLNSC | 1 | 0 | 0 | 1491141 | ALA49351.1 | K9N7C7 |
| PYISHETKLNSC | 1 | 0 | 0 | 1491142 | ASU45783.1 | K9N7C7 |
| PYIVGDVIKEGA | 1 | 0 | 0 | 1491143 | AVP78041.1 | P0C6W2 |
| PYKACSCIKDAL | 1 | 0 | 0 | 1491149 | AVV62535.1 | K9N7C7 |
| PYKDAQVVQLPI | 1 | 0 | 0 | 1491156 | AEN19363.1 | P0C6X6 |
| PYLYNGSAQATA | 1 | 0 | 0 | 1491174 | AVQ05263.1 | P30215 |
| PYMVGDVINSGD | 1 | 0 | 0 | 1491175 | ARO76381.1 | P0C6U8 |
| PYNLRVIHFGAG | 1 | 0 | 0 | 1491177 | QUH73894.1 | P0DTD1 |
| PYNSQNAVARSM | 1 | 0 | 0 | 1491181 | AXN92247.1 | K9N7C7 |
| PYNSQNTVASKI | 1 | 0 | 0 | 1491184 | AVP78041.1 | P0C6W2 |
| PYNVSLSNFNSC | 1 | 0 | 0 | 1491191 | ADN03338.1 | A0A140H1H0 |
| PYPELMEDLEAD | 1 | 0 | 0 | 1491193 | AUM60013.1 | K9N7C7 |
| PYPNPSRILGAG | 1 | 0 | 0 | 1491197 | QVU04458.1 | P0DTD1 |
| PYPVSGIREFSN | 1 | 0 | 0 | 1491198 | AIW52843.1 | A0A0P0G321 |
| PYQGDHGDIYVY | 1 | 0 | 0 | 1491199 | ALJ54488.1 | K9N5Q8 |
| PYQGDHGDMYVY | 1 | 0 | 0 | 1491200 | AXN92228.1 | K9N5Q8 |

| | | | | | | |
|---|---|---|---|---|---|---|
| PYQGDHGDRYVY | 1 | 0 | 0 | 1491201 | ALA49803.1 | K9N5Q8 |
| PYQIEDPCPIHY | 1 | 0 | 0 | 1491203 | Q3I5I8.1 | Q3LZX5 |
| PYQRNNTSFPEW | 1 | 0 | 0 | 1491204 | ATQ39389.1 | K9N7C7 |
| PYSAMKPHVRYD | 1 | 0 | 0 | 1491214 | AUM60013.1 | K9N7C7 |
| PYTALYYYATFY | 1 | 0 | 0 | 1491222 | AXN92258.1 | K9N7C7 |
| PYTKLFQKGECY | 1 | 0 | 0 | 1491229 | QED88024.1 | H9EJ42 |
| PYVASEQEESSD | 1 | 0 | 0 | 1491234 | AGT51657.1 | P0C6X6 |
| PYVASEQEESSE | 1 | 0 | 0 | 1491235 | AGT51707.1 | P0C6X6 |
| PYVCNAPGCDVS | 1 | 0 | 0 | 1491242 | AVV62524.1 | K9N7C7 |
| PYVCNAPGCGAS | 1 | 0 | 0 | 1491244 | AVN89289.1 | K9N7C7 |
| PYVCNTPGCDVN | 1 | 0 | 0 | 1491245 | QEG03773.1 | P0C6X6 |
| PYVEGAINVGKQ | 1 | 0 | 0 | 1491246 | AVV62524.1 | K9N7C7 |
| PYVFIKRSDAQG | 1 | 0 | 0 | 1491249 | P0C6W6.1 | P0C6W2 |
| PYVMLNTPIPNL | 1 | 0 | 0 | 1491256 | AIL49487.1 | P36334 |
| PYVMLNTSSPNL | 1 | 0 | 0 | 1491258 | AIL49492.1 | P36334 |
| PYVMLNTSTSNL | 1 | 0 | 0 | 1491260 | AMK59677.1 | P36334 |
| PYVYNGSAHATA | 1 | 0 | 0 | 1491264 | AIV41890.1 | P30215 |
| PYVYNGSAQAIA | 1 | 0 | 0 | 1491265 | AIL49349.1 | P30215 |
| PYVYNGSAQSTA | 1 | 0 | 0 | 1491267 | P30215.1 | P30215 |
| QAAGTDITITLN | 1 | 0 | 0 | 1491282 | APO40578.1 | P0C6X7 |
| QAAKLSPWFMDG | 1 | 0 | 0 | 1491289 | AVV62524.1 | K9N7C7 |
| QADSFTCNNIDV | 1 | 0 | 0 | 1491312 | ABD43176.1 | P36334 |
| QADTSIPSDVSA | 1 | 0 | 0 | 1491318 | AMM42397.1 | Q19U29 |
| QADVNIPADIGD | 1 | 0 | 0 | 1491320 | AJQ24068.1 | P33469 |
| QADVNIPADIVD | 1 | 0 | 0 | 1491321 | AIX10718.1 | P33469 |
| QAEMNRLQEAIK | 1 | 0 | 0 | 1491328 | AGT51521.1 | P36334 |
| QAEVDWKFYDAQ | 1 | 0 | 0 | 1491333 | AVP78041.1 | P0C6W2 |
| QAFCDYADKLCC | 1 | 0 | 0 | 1491335 | AIV41888.1 | P0C6X6 |
| QAFCDYADKLCH | 1 | 0 | 0 | 1491336 | AEN19363.1 | P0C6X6 |
| QAFCDYTDKLCH | 1 | 0 | 0 | 1491338 | AGT51737.1 | P0C6X6 |
| QAFETPAVQFST | 1 | 0 | 0 | 1491339 | AVV62535.1 | K9N7C7 |
| QAFGLRGSGDLQ | 1 | 0 | 0 | 1491345 | AUM60021.1 | K9N4V7 |
| QAFGRRGPGALQ | 1 | 0 | 0 | 1491347 | AVV62544.1 | K9N4V7 |
| QAFIKHKGGDST | 1 | 0 | 0 | 1491350 | AVV62524.1 | K9N7C7 |
| QAFNKLLGYDPS | 1 | 0 | 0 | 1491354 | AVV62535.1 | K9N7C7 |
| QAFQAVYIVGNL | 1 | 0 | 0 | 1491355 | ATO98145.1 | Q3LZX1 |
| QAGNVQLRVVGH | 1 | 0 | 0 | 1491377 | AVP78041.1 | P0C6W2 |
| QAGSIVRRVNVP | 1 | 0 | 0 | 1491379 | AXN92258.1 | K9N7C7 |
| QAILNTVKFCDH | 1 | 0 | 0 | 1491393 | AXN92247.1 | K9N7C7 |
| QAIYDRLDPPQA | 1 | 0 | 0 | 1491403 | 6U7H_A | A0A0P0K6L9 |
| QAKFGIITKVEA | 1 | 0 | 0 | 1491407 | AIV41992.1 | P30215 |
| QAKSGIITKLEA | 1 | 0 | 0 | 1491409 | QEG03802.1 | P30215 |
| QAKSGIITKVEA | 1 | 0 | 0 | 1491410 | ATN39864.2 | P30215 |
| QALAKLASELSN | 1 | 0 | 0 | 1491411 | AVK87450.1 | K9N5Q8 |
| QALFDSLEFTEL | 1 | 0 | 0 | 1491418 | AVN89451.1 | K9N7C7 |
| QALHGANLKQDE | 1 | 0 | 0 | 1491423 | AUM60014.1 | K9N5Q8 |
| QALHGANLRQDE | 1 | 0 | 0 | 1491426 | AVV62526.1 | K9N5Q8 |
| QALNSILDNAVR | 1 | 0 | 0 | 1491431 | ATN39862.1 | P0C6X6 |
| QALQAGLNGDFN | 1 | 0 | 0 | 1491437 | AUM60014.1 | K9N5Q8 |
| QALQTVSTALNK | 1 | 0 | 0 | 1491440 | ABB90519.1 | A0A0P0K6L9 |
| QALRNDTSVEDK | 1 | 0 | 0 | 1491441 | AUM60014.1 | K9N5Q8 |
| QAMESVGFVYDN | 1 | 0 | 0 | 1491449 | ARU07575.1 | A0A140H1G9 |
| QANLFRNARNGV | 1 | 0 | 0 | 1491459 | AVP78041.1 | P0C6W2 |
| QANLNFVALIPD | 1 | 0 | 0 | 1491460 | P0C6U5.1 | A0A140H1G9 |
| QANSGGYYYKVE | 1 | 0 | 0 | 1491464 | AGT51669.1 | P30215 |
| QANSGIITKVEA | 1 | 0 | 0 | 1491465 | AEN19365.1 | P30215 |
| QAPGAMQTGFTT | 1 | 0 | 0 | 1491466 | AVV62526.1 | K9N5Q8 |
| QAPTHLSVDTKF | 1 | 0 | 0 | 1491471 | P0C6W6.1 | P0C6W2 |
| QARAFVNQVKVP | 1 | 0 | 0 | 1491483 | AVV62535.1 | K9N7C7 |
| QASADWKPGQAM | 1 | 0 | 0 | 1491489 | ATQ39389.1 | K9N7C7 |
| QASADWKPGYAM | 1 | 0 | 0 | 1491490 | ALA49934.1 | K9N7C7 |
| QASLPFGWLIVG | 1 | 0 | 0 | 1491495 | QTS17443.1 | P0DTC3 |
| QASNDWKPGLAM | 1 | 0 | 0 | 1491500 | AUM60013.1 | K9N7C7 |
| QATLPQSKDSNF | 1 | 0 | 0 | 1491514 | AKI29254.1 | K9N638 |
| QATSFCKSGSKF | 1 | 0 | 0 | 1491520 | ARU07609.1 | P30215 |
| QAVGVCVLCNSQ | 1 | 0 | 0 | 1491531 | AEA10817.1 | P0C6X7 |
| QAVRLQPSMALQ | 1 | 0 | 0 | 1491538 | AVV62540.1 | K9N643 |
| QAVRRYSEAASN | 1 | 0 | 0 | 1491539 | NP_073549.1 | A0A0P0LKV0 |
| QAVSDNVRALMS | 1 | 0 | 0 | 1491541 | ADO24673.1 | P0C6X6 |
| QAVSSNINMLLS | 1 | 0 | 0 | 1491542 | AGH06535.1 | H9EJ42 |
| QAVSSNINRILG | 1 | 0 | 0 | 1491543 | AUT76985.1 | A0A0P0LKV0 |
| QAVVNANAEAHN | 1 | 0 | 0 | 1491552 | ARB66064.1 | P36334 |
| QAVVNIRETKCA | 1 | 0 | 0 | 1491554 | AVV62524.1 | K9N7C7 |
| QAYHANCVNCAD | 1 | 0 | 0 | 1491564 | AVV62524.1 | K9N7C7 |
| QAYSLRVRKRRH | 1 | 0 | 0 | 1491569 | ATQ39396.1 | K9N643 |
| QCAIVQALESKV | 1 | 0 | 0 | 1491573 | QED88024.1 | H9EJ42 |
| QCAIVQATESKV | 1 | 0 | 0 | 1491575 | ABE97136.1 | H9EJ42 |
| QCCTAAIDLAHT | 1 | 0 | 0 | 1491589 | AVV62535.1 | K9N7C7 |
| QCDCLCTPDPIT | 1 | 0 | 0 | 1491592 | CAA83660.1 | P36334 |
| QCDYAQLPNEIT | 1 | 0 | 0 | 1491596 | AUM60030.1 | K9N7A1 |
| QCGAERHRQLIE | 1 | 0 | 0 | 1491613 | AUM60013.1 | K9N7C7 |
| QCGGVRHRQIVE | 1 | 0 | 0 | 1491617 | AVV62524.1 | K9N7C7 |
| QCGTENLVCEGP | 1 | 0 | 0 | 1491626 | ARO76381.1 | P0C6U8 |
| QCIRGAIGDVGL | 1 | 0 | 0 | 1491633 | ARB07424.1 | A0A0P0ILY9 |
| QCKCGNLEQNHI | 1 | 0 | 0 | 1491639 | ARU07575.1 | A0A140H1G9 |
| QCKFFYLYNITD | 1 | 0 | 0 | 1491640 | AVV62526.1 | K9N5Q8 |
| QCNCGTKSWSVG | 1 | 0 | 0 | 1491650 | AFR79256.1 | A0A0P0ILY9 |
| QCSTNLFKDCSN | 1 | 0 | 0 | 1491660 | AGT51577.1 | P0C6X6 |
| QCTCGTKSWSVG | 1 | 0 | 0 | 1491662 | NP_073549.1 | A0A0P0LKV0 |
| QCYSFSAYSGDV | 1 | 0 | 0 | 1491685 | AUM60014.1 | K9N5Q8 |
| QDAARQIAWQLH | 1 | 0 | 0 | 1491690 | AVV62528.1 | K9N4V0 |

| | | | | | | |
|---|---|---|---|---|---|---|
| QDAARQVAWELH | 1 | 0 | 0 | 1491691 | AUM60016.1 | K9N4V0 |
| QDAARQVAWQLH | 1 | 0 | 0 | 1491692 | AVV62539.1 | K9N4V0 |
| QDAVNANAQALS | 1 | 0 | 0 | 1491702 | AVV62537.1 | K9N5Q8 |
| QDAVNNNAHALS | 1 | 0 | 0 | 1491703 | ATG84679.1 | K9N5Q8 |
| QDAVNNNAQALA | 1 | 0 | 0 | 1491704 | AVK87450.1 | K9N5Q8 |
| QDCRIPNGFPFN | 1 | 0 | 0 | 1491711 | AKA09611.1 | A0A0P0G321 |
| QDDFVRNLFASV | 1 | 0 | 0 | 1491712 | AMW90843.1 | K9N5Q8 |
| QDDRCIIHCANF | 1 | 0 | 0 | 1491718 | AEN19363.1 | P0C6X6 |
| QDDRCIIHCANL | 1 | 0 | 0 | 1491719 | AIL49625.1 | P0C6X6 |
| QDELDDFFKNVS | 1 | 0 | 0 | 1491722 | AMW90852.1 | K9N5Q8 |
| QDELDEFFKNFS | 1 | 0 | 0 | 1491723 | AKM76239.1 | K9N5Q8 |
| QDEMNRLQEAIK | 1 | 0 | 0 | 1491725 | CAA83660.1 | P36334 |
| QDFDVVFGHGAG | 1 | 0 | 0 | 1491730 | AWK59942.1 | H9EJ42 |
| QDFDVVFGHVAG | 1 | 0 | 0 | 1491731 | AWK59936.1 | H9EJ42 |
| QDFYDFVVAKGF | 1 | 0 | 0 | 1491737 | AUM60013.1 | K9N7C7 |
| QDGDYFTRQLSQ | 1 | 0 | 0 | 1491741 | AVV62537.1 | K9N5Q8 |
| QDGMMNISPKPQ | 1 | 0 | 0 | 1491753 | AIX10729.1 | P33469 |
| QDGNAAIPDYNY | 1 | 0 | 0 | 1491757 | ACV87150.1 | A0A140H1G9 |
| QDGQETAVTKTS | 1 | 0 | 0 | 1491760 | AVP78041.1 | P0C6W2 |
| QDGRTPNGFSFN | 1 | 0 | 0 | 1491764 | AKA09627.1 | A0A0P0G321 |
| QDGVIFNAVDCK | 1 | 0 | 0 | 1491766 | CAA83660.1 | P36334 |
| QDGYNKLQGLLE | 1 | 0 | 0 | 1491773 | CAA79896.1 | P36334 |
| QDHLDILGPLSA | 1 | 0 | 0 | 1491775 | AVP78041.1 | P0C6W2 |
| QDKSLGNFTARA | 1 | 0 | 0 | 1491790 | AVV62529.1 | K9N643 |
| QDKTGDFISTII | 1 | 0 | 0 | 1491791 | AXN92258.1 | K9N7C7 |
| QDKVFGYVQQGV | 1 | 0 | 0 | 1491798 | ATN39862.1 | P0C6X6 |
| QDLCDFKGWIPG | 1 | 0 | 0 | 1491799 | AXX83372.1 | P0C6X6 |
| QDLDVILPGEFS | 1 | 0 | 0 | 1491802 | AXN92247.1 | K9N7C7 |
| QDLDVMLPGEFS | 1 | 0 | 0 | 1491803 | AKK52590.1 | K9N7C7 |
| QDLKWSRFPKSD | 1 | 0 | 0 | 1491815 | AVP78041.1 | P0C6W2 |
| QDLTKYTMMDLC | 1 | 0 | 0 | 1491824 | AIW52776.1 | A0A0P0LKV0 |
| QDPLAELKCTLK | 1 | 0 | 0 | 1491835 | ATO98145.1 | Q3LZX1 |
| QDPMLAVANGSH | 1 | 0 | 0 | 1491837 | APO40579.1 | P59594 |
| QDPSPIDIDAFP | 1 | 0 | 0 | 1491838 | AFD98763.1 | H9EJ42 |
| QDQTTCNVNTLA | 1 | 0 | 0 | 1491844 | AUM60013.1 | K9N7C7 |
| QDRKINTGNGTK | 1 | 0 | 0 | 1491849 | AVV62533.1 | K9N4V7 |
| QDRMLKTSTAPI | 1 | 0 | 0 | 1491852 | ATP16762.1 | Q4VID0 |
| QDRNTRDVFAQT | 1 | 0 | 0 | 1491856 | APO40579.1 | P59594 |
| QDRQIHQAQLTD | 1 | 0 | 0 | 1491859 | AVV62524.1 | K9N7C7 |
| QDRQIYQSQLTD | 1 | 0 | 0 | 1491860 | AUM60013.1 | K9N7C7 |
| QDRVVEPILVIE | 1 | 0 | 0 | 1491863 | AGT51745.1 | Q4VID0 |
| QDRVVEPVLAIE | 1 | 0 | 0 | 1491864 | ATP16772.1 | Q4VID0 |
| QDSKPPLPPHEW | 1 | 0 | 0 | 1491870 | ANI69894.1 | K9N5R3 |
| QDSKPPLPSDEW | 1 | 0 | 0 | 1491871 | ALX27237.1 | K9N5R3 |
| QDSLLKLEDFIP | 1 | 0 | 0 | 1491872 | AVP78041.1 | P0C6W2 |
| QDSQNMLRGEDV | 1 | 0 | 0 | 1491880 | ARO76381.1 | P0C6U8 |
| QDSTVLQALQSE | 1 | 0 | 0 | 1491883 | ARU07575.1 | A0A140H1G9 |
| QDTGHYFYRSHR | 1 | 0 | 0 | 1491890 | AVP78042.1 | Q3LZX1 |
| QDTQGVHLYSSR | 1 | 0 | 0 | 1491892 | AVV62526.1 | K9N5Q8 |
| QDVFIQKYDLED | 1 | 0 | 0 | 1491894 | ARU07564.1 | P0C6X6 |
| QDVFIQKYGLEY | 1 | 0 | 0 | 1491896 | QBQ01836.1 | P0C6X6 |
| QDVGFKNQHCNG | 1 | 0 | 0 | 1491897 | AWW13571.1 | P0C6X6 |
| QDVKQFANGFVL | 1 | 0 | 0 | 1491907 | AHI48723.1 | K9N5Q8 |
| QDVNCTDVPTAI | 1 | 0 | 0 | 1491913 | ATO98181.1 | Q3LZX1 |
| QDVNCTDVPTMI | 1 | 0 | 0 | 1491914 | APO40579.1 | P59594 |
| QDVNCTDVPTTI | 1 | 0 | 0 | 1491916 | AVP78031.1 | Q3LZX1 |
| QDVNCTDVPTVI | 1 | 0 | 0 | 1491917 | ATO98145.1 | Q3LZX1 |
| QDVQVSAQQEVQ | 1 | 0 | 0 | 1491926 | AUM60013.1 | K9N7C7 |
| QDVSLLGKCYKA | 1 | 0 | 0 | 1491927 | AKN24802.1 | K9N7C7 |
| QDYFLPFDTNLT | 1 | 0 | 0 | 1491940 | ATO98193.1 | Q3LZX1 |
| QDYIQSVNFLAW | 1 | 0 | 0 | 1491942 | AEN19363.1 | P0C6X6 |
| QDYQIGGFVEKW | 1 | 0 | 0 | 1491945 | APO40580.1 | P59632 |
| QEAMDSGDTSPQ | 1 | 0 | 0 | 1491954 | AXN92247.1 | K9N7C7 |
| QEAQIDRLINGR | 1 | 0 | 0 | 1491956 | AVV62526.1 | K9N5Q8 |
| QEAVNANAQALS | 1 | 0 | 0 | 1491960 | AUM60014.1 | K9N5Q8 |
| QEAWLEFRAGRP | 1 | 0 | 0 | 1491962 | AVQ05261.1 | P0C6X6 |
| QEDFDLISRCQI | 1 | 0 | 0 | 1491984 | QBP84728.1 | P0C6X6 |
| QEDQKILAASFN | 1 | 0 | 0 | 1491987 | BAL45639.1 | A0A0P0K6L9 |
| QEEDWLESDSQD | 1 | 0 | 0 | 1491993 | AVP78041.1 | P0C6W2 |
| QEESITIKKPNE | 1 | 0 | 0 | 1492002 | APO40578.1 | P0C6X7 |
| QEFNDLAHTSLV | 1 | 0 | 0 | 1492013 | AVA26872.1 | H9EJ42 |
| QEFSVVLPGEMA | 1 | 0 | 0 | 1492019 | AVV62535.1 | K9N7C7 |
| QEGCGIISRKPQ | 1 | 0 | 0 | 1492024 | ATO98181.1 | Q3LZX1 |
| QEGCGILSNKSN | 1 | 0 | 0 | 1492025 | APO40579.1 | P59594 |
| QEGMLYPITLGS | 1 | 0 | 0 | 1492038 | AGT51685.1 | Q4VID0 |
| QEIISRLDALEA | 1 | 0 | 0 | 1492047 | ANZ78843.1 | P36334 |
| QEILLAPLLSAG | 1 | 0 | 0 | 1492049 | ARO76381.1 | P0C6U8 |
| QEILSRLDVLEA | 1 | 0 | 0 | 1492050 | ABD75553.1 | A0A140H1H1 |
| QEISKINTTLLD | 1 | 0 | 0 | 1492053 | AVV62526.1 | K9N5Q8 |
| QELGKYEHYIKW | 1 | 0 | 0 | 1492071 | AVP78042.1 | Q3LZX1 |
| QELPFDVIVAWH | 1 | 0 | 0 | 1492085 | AEN19355.1 | P0C6X6 |
| QEMKALYPTWSQ | 1 | 0 | 0 | 1492101 | AGT51518.1 | P0C6X6 |
| QEMTTPPPKEQR | 1 | 0 | 0 | 1492105 | ATQ39393.1 | K9N4V7 |
| QENQRILAASFN | 1 | 0 | 0 | 1492109 | AIW52747.1 | A0A0P0K6L9 |
| QENWNTKHGSGV | 1 | 0 | 0 | 1492111 | AVP78041.1 | P0C6W2 |
| QEQDELFAMTKR | 1 | 0 | 0 | 1492121 | AXN92247.1 | K9N7C7 |
| QEQRTGLDAVMY | 1 | 0 | 0 | 1492127 | ARK08647.1 | P0C6X6 |
| QEQTNCNTSALA | 1 | 0 | 0 | 1492129 | AXN73479.1 | K9N7C7 |
| QESIVAYTMSIG | 1 | 0 | 0 | 1492149 | APO40579.1 | P59594 |
| QESKFWDENFYA | 1 | 0 | 0 | 1492150 | AGT21344.1 | A0A0P0LKV0 |
| QESQLLKLRGTG | 1 | 0 | 0 | 1492159 | ARE29989.1 | Q4VID0 |

| | | | | | | |
|---|---|---|---|---|---|---|
| QESSDDVDPFDA | 1 | 0 | 0 | 1492161 | AVV62544.1 | K9N4V7 |
| QETPVVFDTVEV | 1 | 0 | 0 | 1492171 | AMQ48992.1 | K9N7C7 |
| QEVFAQVKQMYK | 1 | 0 | 0 | 1492174 | ATO98181.1 | Q3LZX1 |
| QEYIQTTIQKIT | 1 | 0 | 0 | 1492189 | AUM60014.1 | K9N5Q8 |
| QFAELSTNNYKL | 1 | 0 | 0 | 1492194 | AVV62524.1 | K9N7C7 |
| QFAFACPDGVRH | 1 | 0 | 0 | 1492197 | AVP78047.1 | Q3LZX7 |
| QFANGFVLRIGA | 1 | 0 | 0 | 1492200 | AHI48723.1 | K9N5Q8 |
| QFAPGTQLPKNF | 1 | 0 | 0 | 1492202 | AVV62533.1 | K9N4V7 |
| QFATFPVYDTIK | 1 | 0 | 0 | 1492205 | ALA49363.1 | K9N5Q8 |
| QFCDVMRDAGIV | 1 | 0 | 0 | 1492213 | APO40578.1 | P0C6X7 |
| QFCIDNCGEPWF | 1 | 0 | 0 | 1492214 | AWK59942.1 | H9EJ42 |
| QFCLAGFDMLDN | 1 | 0 | 0 | 1492215 | AEN19363.1 | P0C6X6 |
| QFDQWYESTFGS | 1 | 0 | 0 | 1492228 | ARU07575.1 | A0A140H1G9 |
| QFEMNRLQEAIK | 1 | 0 | 0 | 1492237 | AIL49493.1 | P36334 |
| QFGKDASDFIDS | 1 | 0 | 0 | 1492253 | AVP78031.1 | Q3LZX1 |
| QFGLQDGFYSANFLDDNVL | 1 | 0 | 0 | 1492255 | Q6Q1S2.1 | A0A0P0G321 |
| QFGPTYMDGADV | 1 | 0 | 0 | 1492258 | APO40578.1 | P0C6X7 |
| QFGYIDCEQDLC | 1 | 0 | 0 | 1492264 | AXX83372.1 | P0C6X6 |
| QFHNFSYLAHML | 1 | 0 | 0 | 1492269 | AVV62540.1 | K9N643 |
| QFISTFGSFLGF | 1 | 0 | 0 | 1492277 | AWK59942.1 | H9EJ42 |
| QFKDKNLQDLWL | 1 | 0 | 0 | 1492278 | AGT51568.1 | P0C6X6 |
| QFKDKNLQGLWV | 1 | 0 | 0 | 1492280 | AGT51787.1 | P0C6X6 |
| QFKLTHQSNEDD | 1 | 0 | 0 | 1492290 | AVV62533.1 | K9N4V7 |
| QFKRPVNPTDQS | 1 | 0 | 0 | 1492292 | APO40578.1 | P0C6X7 |
| QFKTTGDLAVQM | 1 | 0 | 0 | 1492296 | AGT51323.1 | H9EJ42 |
| QFMLWGNDEKVM | 1 | 0 | 0 | 1492314 | AIV41912.1 | P0C6X6 |
| QFNKLGTEDPRW | 1 | 0 | 0 | 1492321 | ASY99807.1 | K9N4V7 |
| QFNNAISKIQDS | 1 | 0 | 0 | 1492323 | APO40579.1 | P59594 |
| QFNSAIGKIQES | 1 | 0 | 0 | 1492326 | AVP78031.1 | Q3LZX1 |
| QFNVALPGELSK | 1 | 0 | 0 | 1492329 | AUM60013.1 | K9N7C7 |
| QFNYFKKVDGVV | 1 | 0 | 0 | 1492332 | APO40578.1 | P0C6X7 |
| QFNYKQSSSNPT | 1 | 0 | 0 | 1492336 | AXN92228.1 | K9N5Q8 |
| QFQILDEGLDCK | 1 | 0 | 0 | 1492352 | ARK08668.1 | Q80872 |
| QFQKGKAFEFAE | 1 | 0 | 0 | 1492356 | AIV41850.1 | P33469 |
| QFQKGKEFEFAE | 1 | 0 | 0 | 1492357 | AEN19370.1 | P33469 |
| QFQKGKEFQFAE | 1 | 0 | 0 | 1492360 | QBP84717.1 | P33469 |
| QFTEFVGTQSID | 1 | 0 | 0 | 1492381 | ATG84733.1 | K9N638 |
| QFTTLTLDKVPQ | 1 | 0 | 0 | 1492391 | AEN19363.1 | P0C6X6 |
| QFTTLTVDKVPQ | 1 | 0 | 0 | 1492392 | AXX83324.1 | P0C6X6 |
| QFVKETTDMVKS | 1 | 0 | 0 | 1492401 | AVQ05261.1 | P0C6X6 |
| QFVKETTNMVKS | 1 | 0 | 0 | 1492402 | AEN19363.1 | P0C6X6 |
| QFVSAVNQSLIK | 1 | 0 | 0 | 1492406 | ARB07393.1 | A0A0P0KWE8 |
| QFYEFYNDIKPP | 1 | 0 | 0 | 1492410 | AOL02455.1 | Q04854 |
| QGAHLFSSRYVD | 1 | 0 | 0 | 1492423 | ALJ54512.1 | K9N5Q8 |
| QGAWNIGQQKSI | 1 | 0 | 0 | 1492436 | ARO76381.1 | P0C6U8 |
| QGDHGDMYVYSS | 1 | 0 | 0 | 1492445 | AVN89334.1 | K9N5Q8 |
| QGDLGKQYVYSA | 1 | 0 | 0 | 1492451 | AVV62526.1 | K9N5Q8 |
| QGDRCNIFANFI | 1 | 0 | 0 | 1492453 | CAA83660.1 | P36334 |
| QGDVDTVVNGVE | 1 | 0 | 0 | 1492454 | AGT21350.1 | A0A0P0LKV0 |
| QGEIKGATPTNS | 1 | 0 | 0 | 1492457 | AVP78032.1 | Q3LZX0 |
| QGENDNISVAAP | 1 | 0 | 0 | 1492458 | AIX10718.1 | P33469 |
| QGENDNTSVAVP | 1 | 0 | 0 | 1492463 | AIX09811.1 | P33469 |
| QGFACDCGACFS | 1 | 0 | 0 | 1492466 | AVV62524.1 | K9N7C7 |
| QGFACDCGAVFS | 1 | 0 | 0 | 1492467 | AVV62535.1 | K9N7C7 |
| QGFDATNSALVK | 1 | 0 | 0 | 1492472 | AEN19366.1 | P36334 |
| QGFGPRDFNHNM | 1 | 0 | 0 | 1492474 | QED88044.1 | H9EJ69 |
| QGFVVVNGKVST | 1 | 0 | 0 | 1492487 | AXN92247.1 | K9N7C7 |
| QGGITQRTRTRP | 1 | 0 | 0 | 1492493 | ANF29213.1 | K9N4V7 |
| QGHFIGVDGELP | 1 | 0 | 0 | 1492495 | AVN89289.1 | K9N7C7 |
| QGHFTGVDGELP | 1 | 0 | 0 | 1492498 | AUM60013.1 | K9N7C7 |
| QGHLGYTLEESK | 1 | 0 | 0 | 1492503 | APO40578.1 | P0C6X7 |
| QGIETAVGHYFH | 1 | 0 | 0 | 1492508 | ALA49351.1 | K9N7C7 |
| QGIETAVGHYVH | 1 | 0 | 0 | 1492509 | AXN92247.1 | K9N7C7 |
| QGIFVEVNAPYY | 1 | 0 | 0 | 1492512 | CAA83660.1 | P36334 |
| QGKCWFLENKAL | 1 | 0 | 0 | 1492525 | NP_073553.1 | A0A0P0KWE8 |
| QGKFKKVVKGTH | 1 | 0 | 0 | 1492528 | APO40578.1 | P0C6X7 |
| QGKLLIRDNEAL | 1 | 0 | 0 | 1492531 | AVV62524.1 | K9N7C7 |
| QGKPIGMFFPYD | 1 | 0 | 0 | 1492536 | AXN92247.1 | K9N7C7 |
| QGKPLEFGATSF | 1 | 0 | 0 | 1492538 | AVP78041.1 | P0C6W2 |
| QGKPVPYCYDTN | 1 | 0 | 0 | 1492541 | APO40578.1 | P0C6X7 |
| QGKTGDFMSTII | 1 | 0 | 0 | 1492544 | AHI48793.1 | K9N7C7 |
| QGLKEFYKEVRA | 1 | 0 | 0 | 1492562 | AMQ49036.1 | K9N638 |
| QGLLEVSVCQYN | 1 | 0 | 0 | 1492570 | CAA79896.1 | P36334 |
| QGLLPPKNSIDAFKLNIK | 1 | 0 | 0 | 1492573 | P0DTC1.1 | P0DTD1 |
| QGLPLEFGASAE | 1 | 0 | 0 | 1492576 | AEA10817.1 | P0C6X7 |
| QGMDNLACEVLK | 1 | 0 | 0 | 1492591 | AVP78041.1 | P0C6W2 |
| QGMETLACESQT | 1 | 0 | 0 | 1492592 | APO40578.1 | P0C6X7 |
| QGNFGDLQFNKL | 1 | 0 | 0 | 1492596 | ASY99807.1 | K9N4V7 |
| QGNFGDLQLTKL | 1 | 0 | 0 | 1492598 | AVN89320.1 | K9N4V7 |
| QGNKQIVHSCPI | 1 | 0 | 0 | 1492606 | ASU90211.1 | K9N643 |
| QGNSLNYLTSQL | 1 | 0 | 0 | 1492614 | AAK32190.1 | A0A0P0K6L9 |
| QGPASARDLICS | 1 | 0 | 0 | 1492621 | APT69978.1 | K9N5Q8 |
| QGPASARELICA | 1 | 0 | 0 | 1492622 | ARA67740.1 | K9N5Q8 |
| QGPASVRDLICA | 1 | 0 | 0 | 1492624 | ALJ54502.1 | K9N5Q8 |
| QGPNFAEISKIN | 1 | 0 | 0 | 1492630 | AUM60014.1 | K9N5Q8 |
| QGPPIAPGVPAT | 1 | 0 | 0 | 1492635 | P33469.1 | P33469 |
| QGPPNNTASWFT | 1 | 0 | 0 | 1492636 | APO40586.1 | P59595 |
| QGPQSARDLICA | 1 | 0 | 0 | 1492637 | AUM60014.1 | K9N5Q8 |
| QGPSSARDLICA | 1 | 0 | 0 | 1492639 | AHL18090.1 | K9N5Q8 |
| QGQNNADPKVYP | 1 | 0 | 0 | 1492641 | ARO76391.1 | P59636 |
| QGRSCSDFEPMT | 1 | 0 | 0 | 1492658 | AVV62524.1 | K9N7C7 |

| | | | | | | |
|---|---|---|---|---|---|---|
| QGRSCSDFETLT | 1 | 0 | 0 | 1492659 | AUM60013.1 | K9N7C7 |
| QGSDATNSALVK | 1 | 0 | 0 | 1492661 | ARB66064.1 | P36334 |
| QGSGYIPEAPRD | 1 | 0 | 0 | 1492668 | 6CRW_A | P59594 |
| QGSIAQRTRTRP | 1 | 0 | 0 | 1492669 | QEJ82222.1 | K9N4V7 |
| QGSITQRARTRP | 1 | 0 | 0 | 1492672 | AVN89381.1 | K9N4V7 |
| QGSRSRNSTRGT | 1 | 0 | 0 | 1492680 | AHY22571.1 | K9N4V7 |
| QGTPLTPILSCG | 1 | 0 | 0 | 1492693 | NP_073549.1 | A0A0P0LKV0 |
| QGTSLPKGFYAE | 1 | 0 | 0 | 1492695 | APO40586.1 | P59595 |
| QGVFNKATVAQI | 1 | 0 | 0 | 1492701 | ATN39862.1 | P0C6X6 |
| QGVFRYDSVSSV | 1 | 0 | 0 | 1492704 | AGT51619.1 | P30215 |
| QGVFRYNNVSSV | 1 | 0 | 0 | 1492705 | AAA73152.1 | P30215 |
| QGVPIAFGIPPS | 1 | 0 | 0 | 1492714 | ABA60794.1 | Q19U29 |
| QGVPIAPGVPST | 1 | 0 | 0 | 1492717 | AJQ24074.1 | P33469 |
| QGVPINTNSSKD | 1 | 0 | 0 | 1492722 | AVP78038.1 | Q3LZX4 |
| QGYDDCMHQGPA | 1 | 0 | 0 | 1492731 | ATG84866.1 | K9N5Q8 |
| QGYDDCMKQGPA | 1 | 0 | 0 | 1492732 | QEJ82215.1 | K9N5Q8 |
| QGYDDCMNQGPP | 1 | 0 | 0 | 1492734 | AVV62526.1 | K9N5Q8 |
| QGYDDCMQQGPS | 1 | 0 | 0 | 1492736 | AHL18090.1 | K9N5Q8 |
| QGYDECMQQGPQ | 1 | 0 | 0 | 1492737 | AUM60014.1 | K9N5Q8 |
| QGYFLPFYSNVS | 1 | 0 | 0 | 1492740 | AVP78031.1 | Q3LZX1 |
| QGYYIEGSGKSA | 1 | 0 | 0 | 1492745 | AJQ24087.1 | P33469 |
| QGYYIEGSGKYA | 1 | 0 | 0 | 1492746 | AJQ24067.1 | P33469 |
| QGYYIEGSGRSV | 1 | 0 | 0 | 1492749 | AIL49415.1 | P33469 |
| QGYYVEGSGRSA | 1 | 0 | 0 | 1492751 | AIX10731.1 | P33469 |
| QHAGGIAAAINK | 1 | 0 | 0 | 1492755 | AVV62524.1 | K9N7C7 |
| QHCFKAPKNFCP | 1 | 0 | 0 | 1492762 | CAA79896.1 | P36334 |
| QHCFKAPKTFCP | 1 | 0 | 0 | 1492763 | BBO58793.1 | P36334 |
| QHCNGSIACQFC | 1 | 0 | 0 | 1492767 | AWW13571.1 | P0C6X6 |
| QHEDLLAAYVQE | 1 | 0 | 0 | 1492774 | APO40578.1 | P0C6X7 |
| QHEENIYNLIKD | 1 | 0 | 0 | 1492776 | APO40578.1 | P0C6X7 |
| QHEENIYNLVKE | 1 | 0 | 0 | 1492777 | AVP78041.1 | P0C6W2 |
| QHEFCCSHGDRF | 1 | 0 | 0 | 1492782 | QCU72857.1 | A0A140H1G9 |
| QHKCGNLHQREL | 1 | 0 | 0 | 1492809 | AEN19363.1 | P0C6X6 |
| QHMVVKSVLLAD | 1 | 0 | 0 | 1492819 | APO40578.1 | P0C6X7 |
| QHPLLNPGALEF | 1 | 0 | 0 | 1492825 | AGT21342.1 | A0A0N7J131 |
| QHPNMTVHHIKE | 1 | 0 | 0 | 1492827 | AVV62540.1 | K9N643 |
| QHTDINFTATAS | 1 | 0 | 0 | 1492842 | AIW52843.1 | A0A0P0G321 |
| QHTMLVKMDGDE | 1 | 0 | 0 | 1492847 | AIL49620.1 | P0C6X6 |
| QHVIKGCKDTAC | 1 | 0 | 0 | 1492854 | AHI48777.1 | K9N7C7 |
| QIANMFLEQTQH | 1 | 0 | 0 | 1492868 | AXN92247.1 | K9N7C7 |
| QIANQFNNAISK | 1 | 0 | 0 | 1492870 | APO40579.1 | P59594 |
| QIDDYRITSITS | 1 | 0 | 0 | 1492886 | AIV41978.1 | P0C6X6 |
| QIDQLNSSYFKL | 1 | 0 | 0 | 1492892 | ASU91208.1 | K9N5Q8 |
| QIDRFYRLANEC | 1 | 0 | 0 | 1492893 | QBP84719.1 | P0C6X6 |
| QIDRLITGGRGG | 1 | 0 | 0 | 1492897 | 1WYY_A | P59594 |
| QIEDKIEEIESK | 1 | 0 | 0 | 1492900 | 5X58_A | P59594 |
| QIELGSGAHVGS | 1 | 0 | 0 | 1492902 | AWK59942.1 | H9EJ42 |
| QIESDEYLTQRG | 1 | 0 | 0 | 1492906 | AUM60013.1 | K9N7C7 |
| QIEVKFNAPALQ | 1 | 0 | 0 | 1492907 | P0C6W6.1 | P0C6W2 |
| QIFASVPEKFLN | 1 | 0 | 0 | 1492908 | ARB07424.1 | A0A0P0ILY9 |
| QIFDPAPVELEN | 1 | 0 | 0 | 1492909 | AVV62524.1 | K9N7C7 |
| QIGLLEPVNHSA | 1 | 0 | 0 | 1492926 | AVV62524.1 | K9N7C7 |
| QIIAIIKDWSFA | 1 | 0 | 0 | 1492944 | ALX27238.1 | K9N7A1 |
| QIIHSHPILHDP | 1 | 0 | 0 | 1492953 | ATQ39396.1 | K9N643 |
| QIIKSQDLGVVS | 1 | 0 | 0 | 1492954 | AVV62535.1 | K9N7C7 |
| QIILNLTSEIST | 1 | 0 | 0 | 1492955 | ABB90514.1 | A0A0P0K6L9 |
| QIKFENPHILGT | 1 | 0 | 0 | 1492969 | AWW13564.1 | Q80872 |
| QILDEGLDCKIQ | 1 | 0 | 0 | 1492974 | ARK08668.1 | Q80872 |
| QILDITPCSFGG | 1 | 0 | 0 | 1492976 | ATO98193.1 | Q3LZX1 |
| QILIDNINSTLV | 1 | 0 | 0 | 1492980 | ARB07425.1 | A0A0P0K6L9 |
| QILLNCALGALA | 1 | 0 | 0 | 1492982 | AFR79256.1 | A0A0P0ILY9 |
| QILPDPSKPTKR | 1 | 0 | 0 | 1492988 | ATO98181.1 | Q3LZX1 |
| QILPDSSKPTKR | 1 | 0 | 0 | 1492989 | ARO76382.1 | P59594 |
| QILPSPLKPTKR | 1 | 0 | 0 | 1492991 | 5XJK_A | P59594 |
| QIMGVVMQSSVR | 1 | 0 | 0 | 1492999 | AUM60013.1 | K9N7C7 |
| QINGYTLAATSA | 1 | 0 | 0 | 1493006 | ANZ78836.1 | P36334 |
| QIPAHCVRDPVG | 1 | 0 | 0 | 1493012 | ALA49417.1 | K9N7C7 |
| QIPAQCTRDPVG | 1 | 0 | 0 | 1493013 | AVV62524.1 | K9N7C7 |
| QIPCMCGKQATQ | 1 | 0 | 0 | 1493015 | AVP78041.1 | P0C6W2 |
| QIPSDFTIGNMG | 1 | 0 | 0 | 1493021 | ARB66062.1 | P36334 |
| QIPSEFTIGNMV | 1 | 0 | 0 | 1493024 | P36334.1 | P36334 |
| QIPSKFTIGNME | 1 | 0 | 0 | 1493025 | AIL49542.1 | P36334 |
| QIQNNTVDVVVG | 1 | 0 | 0 | 1493045 | AVV62524.1 | K9N7C7 |
| QISPTAIASNCY | 1 | 0 | 0 | 1493069 | AWF93654.1 | K9N5Q8 |
| QIVDKDGNYYLP | 1 | 0 | 0 | 1493091 | QED88010.1 | H9EJ42 |
| QIVDMSMTYGQQ | 1 | 0 | 0 | 1493093 | AVP78041.1 | P0C6W2 |
| QIVFFVSRVFAN | 1 | 0 | 0 | 1493096 | AVV62535.1 | K9N7C7 |
| QIVHSHPILHHP | 1 | 0 | 0 | 1493099 | ASU90849.1 | K9N643 |
| QIVKSIPIHHDT | 1 | 0 | 0 | 1493108 | AUM60027.1 | K9N643 |
| QIVNMFLEQTQH | 1 | 0 | 0 | 1493110 | ALA49362.1 | K9N7C7 |
| QIYDPAPVELEN | 1 | 0 | 0 | 1493119 | AVV62535.1 | K9N7C7 |
| QKANTEIELGVC | 1 | 0 | 0 | 1493144 | AEN19366.1 | P36334 |
| QKASTQREAEAA | 1 | 0 | 0 | 1493153 | ARO76392.1 | Q7TLC7 |
| QKAVNVAKNAYE | 1 | 0 | 0 | 1493155 | AUM60013.1 | K9N7C7 |
| QKAYMDFVNSGG | 1 | 0 | 0 | 1493159 | AUM60013.1 | K9N7C7 |
| QKDIGIDTTGVL | 1 | 0 | 0 | 1493177 | ABD75582.1 | Q0ZJ80 |
| QKEIDRLNEVAR | 1 | 0 | 0 | 1493184 | AVP78031.1 | Q3LZX1 |
| QKESDDYIKQNG | 1 | 0 | 0 | 1493189 | ARO76381.1 | P0C6U8 |
| QKFKTICFSTVA | 1 | 0 | 0 | 1493201 | AIW52843.1 | A0A0P0G321 |
| QKGILCVMTSQA | 1 | 0 | 0 | 1493213 | AXN92258.1 | K9N7C7 |
| QKGIYQTSNFRV | 1 | 0 | 0 | 1493215 | AVP78031.1 | Q3LZX1 |

| | | | | | | |
|---|---|---|---|---|---|---|
| QKGKAFEFAEGQ | 1 | 0 | 0 | 1493216 | AIV41850.1 | P33469 |
| QKGKEFEFAEGQ | 1 | 0 | 0 | 1493217 | AEN19370.1 | P33469 |
| QKGRDFKFPDGQ | 1 | 0 | 0 | 1493222 | AMM42397.1 | Q19U29 |
| QKHLKSIVNTRN | 1 | 0 | 0 | 1493234 | AIW52827.1 | H9EJ42 |
| QKHVGAPANLRV | 1 | 0 | 0 | 1493237 | AVV62524.1 | K9N7C7 |
| QKITTIQGPPGS | 1 | 0 | 0 | 1493246 | AWK59942.1 | H9EJ42 |
| QKKQPTVTPLPA | 1 | 0 | 0 | 1493253 | AAS48456.1 | P59595 |
| QKLALGGSVAVK | 1 | 0 | 0 | 1493259 | P0C6W6.1 | P0C6W2 |
| QKLEDNSLFLFD | 1 | 0 | 0 | 1493261 | AIV41864.1 | P0C6X6 |
| QKLIANAFNNAL | 1 | 0 | 0 | 1493269 | CAA79896.1 | P36334 |
| QKLQSLIDNINS | 1 | 0 | 0 | 1493285 | BAL45640.1 | A0A0P0K6L9 |
| QKLYDQVVMDYL | 1 | 0 | 0 | 1493289 | ARO76381.1 | P0C6U8 |
| QKNGQGENDNIS | 1 | 0 | 0 | 1493291 | BBM61433.1 | P33469 |
| QKQIANKFNQAP | 1 | 0 | 0 | 1493305 | AVV62526.1 | K9N5Q8 |
| QKQIANQFNNAI | 1 | 0 | 0 | 1493307 | APO40579.1 | P59594 |
| QKRSPNTQRTVQ | 1 | 0 | 0 | 1493326 | AJQ24090.1 | P33469 |
| QKRTPNKFCNVQ | 1 | 0 | 0 | 1493330 | AMM42397.1 | Q19U29 |
| QKSGYSKYVTVQ | 1 | 0 | 0 | 1493332 | AXN92247.1 | K9N7C7 |
| QKVAENPKEEVK | 1 | 0 | 0 | 1493352 | AVP78041.1 | P0C6W2 |
| QKWMELLDSNID | 1 | 0 | 0 | 1493369 | AUM60021.1 | K9N4V7 |
| QKYDLEDYAFEH | 1 | 0 | 0 | 1493371 | ARU07564.1 | P0C6X6 |
| QKYLNSSYTTCL | 1 | 0 | 0 | 1493377 | AXN92262.1 | K9N4V0 |
| QLAAINFNQPLV | 1 | 0 | 0 | 1493384 | AUM60024.1 | K9N5Q8 |
| QLADAALSIKEN | 1 | 0 | 0 | 1493385 | AIV41889.1 | Q80872 |
| QLAEEPAKDIPE | 1 | 0 | 0 | 1493388 | ATQ39389.1 | K9N7C7 |
| QLAFDVPDGFYS | 1 | 0 | 0 | 1493390 | AFR79257.1 | A0A0P0K6L9 |
| QLAGIKLQSKHT | 1 | 0 | 0 | 1493392 | AIV41894.1 | P0C6X6 |
| QLAGVKLQSKTK | 1 | 0 | 0 | 1493394 | ARU07575.1 | A0A140H1G9 |
| QLAVPDNSSTIV | 1 | 0 | 0 | 1493412 | AKA09630.1 | A0A0P0G321 |
| QLCQYLNTCTLA | 1 | 0 | 0 | 1493431 | AXN92247.1 | K9N7C7 |
| QLDENAQKFFDF | 1 | 0 | 0 | 1493439 | AST12962.1 | A0A0P0LKV0 |
| QLDFIESKRGVY | 1 | 0 | 0 | 1493442 | ARO76381.1 | P0C6U8 |
| QLDTAPKTGNFK | 1 | 0 | 0 | 1493445 | ATO98181.1 | Q3LZX1 |
| QLDTTQLQVANS | 1 | 0 | 0 | 1493447 | AHF54662.1 | P36334 |
| QLEAKTASLFKT | 1 | 0 | 0 | 1493450 | ARU07594.1 | A0A0P0G321 |
| QLECIVDDSREE | 1 | 0 | 0 | 1493452 | ABE73415.1 | H9EJ42 |
| QLEFVREEVDII | 1 | 0 | 0 | 1493457 | AWK59942.1 | H9EJ42 |
| QLELAKVQNLSG | 1 | 0 | 0 | 1493465 | AHA80186.1 | P33469 |
| QLESDAYLDAKG | 1 | 0 | 0 | 1493478 | AVV62524.1 | K9N7C7 |
| QLESDQYLDSKG | 1 | 0 | 0 | 1493479 | AVV62535.1 | K9N7C7 |
| QLESKVSGKTQQ | 1 | 0 | 0 | 1493482 | APO40586.1 | P59595 |
| QLEVKTASLFQT | 1 | 0 | 0 | 1493488 | AGT51366.1 | A0A0P0G321 |
| QLFYTEDLASRF | 1 | 0 | 0 | 1493504 | AUM60013.1 | K9N7C7 |
| QLGAPDNPSAIV | 1 | 0 | 0 | 1493508 | AKA09625.1 | A0A0P0G321 |
| QLGCYEPLKSYG | 1 | 0 | 0 | 1493512 | APO40579.1 | P59594 |
| QLGNLGNLQSFN | 1 | 0 | 0 | 1493530 | AIL49492.1 | P36334 |
| QLGVPLGITISG | 1 | 0 | 0 | 1493538 | AWO68966.1 | A0A0P0G321 |
| QLHTGFQGKQIL | 1 | 0 | 0 | 1493548 | ATQ39389.1 | K9N7C7 |
| QLIDKYCSVNVV | 1 | 0 | 0 | 1493552 | ATG84700.1 | K9N638 |
| QLKKGVQIPCMC | 1 | 0 | 0 | 1493567 | AVP78041.1 | P0C6W2 |
| QLKLTHQSNDES | 1 | 0 | 0 | 1493571 | AVV62544.1 | K9N4V7 |
| QLLDEGVDCKIQ | 1 | 0 | 0 | 1493580 | NP_150075.1 | Q80872 |
| QLLFVVEVVDKYFDCYDGGC | 1 | 0 | 0 | 1493592 | P0DTD1.1 | P0DTD1 |
| QLLGSTADNLKN | 1 | 0 | 0 | 1493597 | APO40578.1 | P0C6X7 |
| QLLHTKISWAGA | 1 | 0 | 0 | 1493598 | AUM60013.1 | K9N7C7 |
| QLLHTKISWAGS | 1 | 0 | 0 | 1493599 | AVN89363.1 | K9N7C7 |
| QLLHTKVSWAGF | 1 | 0 | 0 | 1493601 | AHI48793.1 | K9N7C7 |
| QLLIQDYIQSVN | 1 | 0 | 0 | 1493606 | AAR01012.1 | P0C6X6 |
| QLLKDVTLEKLR | 1 | 0 | 0 | 1493610 | ATQ39389.1 | K9N7C7 |
| QLLLEYGQFCSK | 1 | 0 | 0 | 1493615 | QAT98898.1 | K9N5Q8 |
| QLLMPLKAPKEI | 1 | 0 | 0 | 1493626 | ATO98130.1 | P0C6W2 |
| QLLYAIQQLHTG | 1 | 0 | 0 | 1493641 | ATQ39389.1 | K9N7C7 |
| QLLYAVQQLNNG | 1 | 0 | 0 | 1493643 | AVV62524.1 | K9N7C7 |
| QLMATSVNCCHD | 1 | 0 | 0 | 1493645 | AGT51695.1 | Q4VID0 |
| QLNHSSCFCSPN | 1 | 0 | 0 | 1493659 | AIG96691.1 | A0A140H1H1 |
| QLNHTSCSCSPD | 1 | 0 | 0 | 1493662 | BBA20986.1 | A0A140H1H1 |
| QLNLISNITTNG | 1 | 0 | 0 | 1493667 | AVP78041.1 | P0C6W2 |
| QLNNGFQGKQIL | 1 | 0 | 0 | 1493670 | AVV62524.1 | K9N7C7 |
| QLNSSYKLSIPT | 1 | 0 | 0 | 1493677 | ATW75477.1 | K9N5Q8 |
| QLPIQDYIQSVN | 1 | 0 | 0 | 1493690 | AEN19363.1 | P0C6X6 |
| QLPLDVYNLTMP | 1 | 0 | 0 | 1493696 | AAX85666.1 | P0C6X6 |
| QLPNEITVAKPS | 1 | 0 | 0 | 1493699 | AUM60030.1 | K9N7A1 |
| QLPPPLLNNQTD | 1 | 0 | 0 | 1493702 | AVV62537.1 | K9N5Q8 |
| QLPVKDYIQTVN | 1 | 0 | 0 | 1493705 | ABD75583.1 | A0A140H1G9 |
| QLPVYTSYKVLD | 1 | 0 | 0 | 1493707 | ARK08667.1 | P0C6X6 |
| QLPYCYFRNSSG | 1 | 0 | 0 | 1493709 | AGT17757.1 | A0A140H1H0 |
| QLQDSMNGASDS | 1 | 0 | 0 | 1493711 | APO40586.1 | P59595 |
| QLQDVQVSAQQE | 1 | 0 | 0 | 1493712 | AUM60013.1 | K9N7C7 |
| QLQLATINYTQP | 1 | 0 | 0 | 1493715 | AVV62537.1 | K9N5Q8 |
| QLQPINYSFDSY | 1 | 0 | 0 | 1493722 | AEN19366.1 | P36334 |
| QLQSAEWKCGYA | 1 | 0 | 0 | 1493728 | AWK59942.1 | H9EJ42 |
| QLQVANALMQGV | 1 | 0 | 0 | 1493732 | BBA20986.1 | A0A140H1H1 |
| QLQVANSLMNGV | 1 | 0 | 0 | 1493734 | CAA79896.1 | P36334 |
| QLRAFDIYNANV | 1 | 0 | 0 | 1493735 | AGT51737.1 | P0C6X6 |
| QLSDAMKRQIRI | 1 | 0 | 0 | 1493758 | AUM60013.1 | K9N7C7 |
| QLSDPNYDALLI | 1 | 0 | 0 | 1493760 | AXN92247.1 | K9N7C7 |
| QLSNRFGAISAT | 1 | 0 | 0 | 1493788 | ARB66071.1 | P36334 |
| QLSPLEGGGRLV | 1 | 0 | 0 | 1493789 | AXG21654.1 | K9N5Q8 |
| QLTDKYCSVNVA | 1 | 0 | 0 | 1493804 | AHI48798.1 | K9N7C7 |
| QLTITDPKVYPI | 1 | 0 | 0 | 1493821 | 2CME_E | P59636 |
| QLTKQLLDILSK | 1 | 0 | 0 | 1493828 | AVV62524.1 | K9N7C7 |

| | | | | | | |
|---|---|---|---|---|---|---|
| QLTPAWRVYSTG | 1 | 0 | 0 | 1493838 | ATO98181.1 | Q3LZX1 |
| QLTVEGYNNYVP | 1 | 0 | 0 | 1493850 | AVV62524.1 | K9N7C7 |
| QLTVSAVVILIQ | 1 | 0 | 0 | 1493852 | AVV62540.1 | K9N643 |
| QLTYQNISTNLP | 1 | 0 | 0 | 1493854 | AVN89324.1 | K9N5Q8 |
| QLVEHTTPWLLL | 1 | 0 | 0 | 1493858 | AVN89451.1 | K9N7C7 |
| QLVNANHYSPCV | 1 | 0 | 0 | 1493864 | AID55087.1 | K9N5Q8 |
| QLVNANQYSPCV | 1 | 0 | 0 | 1493865 | AXN92228.1 | K9N5Q8 |
| QLVNSNQYSPCV | 1 | 0 | 0 | 1493866 | AHY22555.1 | K9N5Q8 |
| QLVPAFPEAEVA | 1 | 0 | 0 | 1493867 | AVV62541.1 | K9N7D2 |
| QLVPAFPETEVA | 1 | 0 | 0 | 1493868 | AVV62530.1 | K9N7D2 |
| QLVSLAEFSLNN | 1 | 0 | 0 | 1493881 | AVV62541.1 | K9N7D2 |
| QLVTYPSLNYAG | 1 | 0 | 0 | 1493885 | AHL18091.1 | K9N7C7 |
| QLYCSYESFDVG | 1 | 0 | 0 | 1493891 | AVV62537.1 | K9N5Q8 |
| QLYDASNVKNVT | 1 | 0 | 0 | 1493894 | AVQ05261.1 | P0C6X6 |
| QLYDNCYRKSNV | 1 | 0 | 0 | 1493895 | AIG96641.1 | A0A0P0LKV0 |
| QLYYNLPAVNVS | 1 | 0 | 0 | 1493913 | AIV41873.1 | P36334 |
| QMDIQGFVNIFI | 1 | 0 | 0 | 1493926 | NP_073549.1 | A0A0P0LKV0 |
| QMEVDFLELAMD | 1 | 0 | 0 | 1493936 | APO40578.1 | P0C6X7 |
| QMFVTPLSVDVV | 1 | 0 | 0 | 1493942 | ALA49399.1 | K9N643 |
| QMGSNNVCTYEH | 1 | 0 | 0 | 1493948 | AGT51323.1 | H9EJ42 |
| QMSELPKEQRVQ | 1 | 0 | 0 | 1493963 | ASJ26617.1 | K9N4V7 |
| QMTGFKKPASRD | 1 | 0 | 0 | 1493969 | APO40578.1 | P0C6X7 |
| QMYKTPTLKDFG | 1 | 0 | 0 | 1493982 | ATO98231.1 | Q3LZX1 |
| QNAGYWRRQYRK | 1 | 0 | 0 | 1493987 | AVN89320.1 | K9N4V7 |
| QNAVARSMLGLN | 1 | 0 | 0 | 1494000 | AVV62524.1 | K9N7C7 |
| QNAVPANSLHAT | 1 | 0 | 0 | 1494003 | APO40580.1 | P59632 |
| QNAVVKIYCPAC | 1 | 0 | 0 | 1494008 | AVP78041.1 | P0C6W2 |
| QNAVVKLYCPAC | 1 | 0 | 0 | 1494009 | AVP78030.1 | P0C6W2 |
| QNCTAVGVLQQR | 1 | 0 | 0 | 1494020 | ALJ54472.1 | K9N5Q8 |
| QNCTAVGVPQQR | 1 | 0 | 0 | 1494021 | AVN89313.1 | K9N5Q8 |
| QNEELEQTIQQK | 1 | 0 | 0 | 1494029 | APO40578.1 | P0C6X7 |
| QNFGNFEMLKLG | 1 | 0 | 0 | 1494033 | AMM42397.1 | Q19U29 |
| QNFGNSEMLKLG | 1 | 0 | 0 | 1494034 | AMM42394.1 | Q19U29 |
| QNFTTAPAICHN | 1 | 0 | 0 | 1494037 | APO40579.1 | P59594 |
| QNGCSLYLADNL | 1 | 0 | 0 | 1494039 | ATQ39389.1 | K9N7C7 |
| QNGDRTGARPKH | 1 | 0 | 0 | 1494042 | APO40586.1 | P59595 |
| QNGEAPVSIVNN | 1 | 0 | 0 | 1494043 | AVP78030.1 | P0C6W2 |
| QNGMNGRTILGA | 1 | 0 | 0 | 1494050 | 3M3V_A | P0C6X7 |
| QNHSYIYHSAFA | 1 | 0 | 0 | 1494060 | AVV62535.1 | K9N7C7 |
| QNIDAYKSFPVK | 1 | 0 | 0 | 1494062 | AVV62533.1 | K9N4V7 |
| QNILLRKYGRGG | 1 | 0 | 0 | 1494065 | AXN92247.1 | K9N7C7 |
| QNISINLPPPLL | 1 | 0 | 0 | 1494067 | ATG84888.1 | K9N5Q8 |
| QNKQESSDDVDP | 1 | 0 | 0 | 1494074 | AVV62544.1 | K9N4V7 |
| QNKSIELTAEGI | 1 | 0 | 0 | 1494076 | AHA80174.1 | P33469 |
| QNKSRELTSEDI | 1 | 0 | 0 | 1494079 | AGT51674.1 | P33469 |
| QNKTVDVVVGQL | 1 | 0 | 0 | 1494080 | AVV62535.1 | K9N7C7 |
| QNLDCNCLGFYE | 1 | 0 | 0 | 1494084 | AWW13564.1 | Q80872 |
| QNLLYCDSSFEK | 1 | 0 | 0 | 1494094 | AWO68987.1 | A0A0P0G321 |
| QNLLYDPNGNLY | 1 | 0 | 0 | 1494096 | ARB66062.1 | P36334 |
| QNLTNNLPPPLL | 1 | 0 | 0 | 1494101 | AUM60024.1 | K9N5Q8 |
| QNNADPKVYPII | 1 | 0 | 0 | 1494108 | ARO76391.1 | P59636 |
| QNNDDHGNPVYF | 1 | 0 | 0 | 1494111 | AXN92267.1 | K9N4V7 |
| QNNEIMPGKLKQ | 1 | 0 | 0 | 1494112 | AWK59942.1 | H9EJ42 |
| QNNEIMPGKMKV | 1 | 0 | 0 | 1494113 | NP_073549.1 | A0A0P0LKV0 |
| QNNELMPVKLKI | 1 | 0 | 0 | 1494114 | AXX83324.1 | P0C6X6 |
| QNNFTGYLKLTD | 1 | 0 | 0 | 1494116 | AVP78041.1 | P0C6W2 |
| QNQKLIASAFNN | 1 | 0 | 0 | 1494131 | AIL49544.1 | P36334 |
| QNTSVYSANGFF | 1 | 0 | 0 | 1494154 | BAM16475.1 | A0A0P0G321 |
| QNTVLNDSIPES | 1 | 0 | 0 | 1494161 | AMM42614.1 | H9EJ69 |
| QNVNLEPCQLAN | 1 | 0 | 0 | 1494168 | AVV62524.1 | K9N7C7 |
| QNVNLERCELVN | 1 | 0 | 0 | 1494170 | QAT98908.1 | K9N7C7 |
| QNWLSAFGDVKP | 1 | 0 | 0 | 1494173 | AUM60027.1 | K9N643 |
| QNYGDSAILPKG | 1 | 0 | 0 | 1494179 | AVP78030.1 | P0C6W2 |
| QPAGVFTDHDVV | 1 | 0 | 0 | 1494192 | CAA83661.1 | P36334 |
| QPALYLYNTGRL | 1 | 0 | 0 | 1494197 | QBM11741.1 | K9N5R3 |
| QPALYLYNTGRS | 1 | 0 | 0 | 1494198 | ASU90334.1 | K9N5R3 |
| QPCLVIPRNSVP | 1 | 0 | 0 | 1494200 | AMM42513.1 | A0A0N7J131 |
| QPCSDKAYKIAE | 1 | 0 | 0 | 1494201 | QVO99997.1 | P0DTD1 |
| QPCVASEQDESS | 1 | 0 | 0 | 1494203 | AIV41846.1 | P0C6X6 |
| QPEDYKSVDVAI | 1 | 0 | 0 | 1494213 | AWW13564.1 | Q80872 |
| QPELESFKQELD | 1 | 0 | 0 | 1494218 | APO40579.1 | P59594 |
| QPEVLSPINSSG | 1 | 0 | 0 | 1494222 | AVV62526.1 | K9N5Q8 |
| QPFEEVEHVLSI | 1 | 0 | 0 | 1494226 | ARU07593.1 | H9EJ42 |
| QPFGEVEHSLPI | 1 | 0 | 0 | 1494228 | AVA26872.1 | H9EJ42 |
| QPFGEVEYSLPI | 1 | 0 | 0 | 1494230 | AWK59942.1 | H9EJ42 |
| QPGGDATSLSFN | 1 | 0 | 0 | 1494242 | ANI69888.1 | K9N638 |
| QPIADVYRRKPD | 1 | 0 | 0 | 1494256 | AEN19366.1 | P36334 |
| QPIADVYRRRPD | 1 | 0 | 0 | 1494258 | AHA80225.1 | P36334 |
| QPIANVYRRKPD | 1 | 0 | 0 | 1494259 | ABD43177.1 | P36334 |
| QPIEEEDWLDVQ | 1 | 0 | 0 | 1494263 | APO40578.1 | P0C6X7 |
| QPISDLLTPMGI | 1 | 0 | 0 | 1494270 | APO40578.1 | P0C6X7 |
| QPKIEQKPLSLI | 1 | 0 | 0 | 1494276 | AST12962.1 | A0A0P0LKV0 |
| QPKQTSTSQQPS | 1 | 0 | 0 | 1494283 | ATP16771.1 | P33469 |
| QPKQTVTSQQPS | 1 | 0 | 0 | 1494284 | AIL49418.1 | P33469 |
| QPKVITKKEAAA | 1 | 0 | 0 | 1494286 | AXG21672.1 | K9N4V7 |
| QPKVITKRDAAA | 1 | 0 | 0 | 1494288 | ANF29235.1 | K9N4V7 |
| QPLDAGLNGDFN | 1 | 0 | 0 | 1494291 | AVV62526.1 | K9N5Q8 |
| QPLGGNVVPYYS | 1 | 0 | 0 | 1494294 | AMK59681.1 | P33469 |
| QPLNAGLNGDFN | 1 | 0 | 0 | 1494303 | AVV62537.1 | K9N5Q8 |
| QPLPNATYDNFR | 1 | 0 | 0 | 1494306 | APO40578.1 | P0C6X7 |
| QPLRLTCLWPVS | 1 | 0 | 0 | 1494313 | QED88026.1 | A0A0P0G321 |

| | | | | | | |
|---|---|---|---|---|---|---|
| QPLVVDTLNSSG | 1 | 0 | 0 | 1494317 | AUM60014.1 | K9N5Q8 |
| QPMAISVNCCHD | 1 | 0 | 0 | 1494318 | AGT51675.1 | Q4VID0 |
| QPMATSANCCHD | 1 | 0 | 0 | 1494320 | AGT51735.1 | Q4VID0 |
| QPMLTTCQCGAE | 1 | 0 | 0 | 1494323 | AUM60013.1 | K9N7C7 |
| QPNDDATSNVTQ | 1 | 0 | 0 | 1494326 | APT69866.1 | A0A0N7J131 |
| QPNLALYKEGNY | 1 | 0 | 0 | 1494328 | AIW52757.1 | A0A0P0K6L9 |
| QPNQTATSQQPS | 1 | 0 | 0 | 1494331 | AIX10721.1 | P33469 |
| QPNTEVFTDVCV | 1 | 0 | 0 | 1494333 | BBA20983.1 | A0A140H1H1 |
| QPPQTSITSAIL | 1 | 0 | 0 | 1494337 | AVP78041.1 | P0C6W2 |
| QPQAFVGWSADS | 1 | 0 | 0 | 1494343 | BBA84088.1 | P36334 |
| QPQLLSPLNSSG | 1 | 0 | 0 | 1494352 | AVV62537.1 | K9N5Q8 |
| QPQRAPKGSITQ | 1 | 0 | 0 | 1494353 | AVV62544.1 | K9N4V7 |
| QPQVVTRKSASE | 1 | 0 | 0 | 1494355 | APO40586.1 | P59595 |
| QPRLPVLSDFVQ | 1 | 0 | 0 | 1494360 | AIW52843.1 | A0A0P0G321 |
| QPRNVLYDSVSA | 1 | 0 | 0 | 1494361 | BAL45640.1 | A0A0P0K6L9 |
| QPRNVSYDSVSA | 1 | 0 | 0 | 1494362 | AIW52757.1 | A0A0P0K6L9 |
| QPRSDKPSQLKK | 1 | 0 | 0 | 1494363 | AVL25604.1 | H9EJ69 |
| QPRTIDSTQDGV | 1 | 0 | 0 | 1494365 | QBQ01839.1 | P36334 |
| QPRTIISTQDGV | 1 | 0 | 0 | 1494366 | BBA20973.1 | P36334 |
| QPRTINATQDGV | 1 | 0 | 0 | 1494368 | AXX83321.1 | P36334 |
| QPRTINSTQDGV | 1 | 0 | 0 | 1494374 | AEN19358.1 | P36334 |
| QPRTINSTQDGY | 1 | 0 | 0 | 1494375 | CAA79896.1 | P36334 |
| QPSGGNVIPHYS | 1 | 0 | 0 | 1494383 | QEG03818.1 | P33469 |
| QPSGGNVVPHYS | 1 | 0 | 0 | 1494384 | AWW13570.1 | P33469 |
| QPSHAVQNTVLN | 1 | 0 | 0 | 1494387 | AVL25602.1 | H9EJ69 |
| QPSKLLNGIAYT | 1 | 0 | 0 | 1494391 | AWK59942.1 | H9EJ42 |
| QPSNQRDAWAAF | 1 | 0 | 0 | 1494394 | ATQ39390.1 | K9N5Q8 |
| QPSRGNVVPHYS | 1 | 0 | 0 | 1494395 | ANZ78871.1 | P33469 |
| QPSRGNVVPYYS | 1 | 0 | 0 | 1494396 | AGT51784.1 | P33469 |
| QPSTSFYDIAVS | 1 | 0 | 0 | 1494397 | AUM60025.1 | K9N796 |
| QPTAHVNATAAY | 1 | 0 | 0 | 1494400 | AUM60014.1 | K9N5Q8 |
| QPTANSTTVSVT | 1 | 0 | 0 | 1494401 | AHI48527.1 | K9N638 |
| QPTGPAYVMIDK | 1 | 0 | 0 | 1494412 | NP_073549.1 | A0A0P0LKV0 |
| QPTQSIVRFPNI | 1 | 0 | 0 | 1494421 | AVP78042.1 | Q3LZX1 |
| QPTRNSTTVSVT | 1 | 0 | 0 | 1494423 | AVN89322.1 | K9N7C7 |
| QPTVKEIISIPK | 1 | 0 | 0 | 1494428 | AIV42008.1 | P0C6X6 |
| QPTVKEIISIPN | 1 | 0 | 0 | 1494429 | AIX09804.1 | P0C6X6 |
| QPVDLPPKVYFY | 1 | 0 | 0 | 1494437 | AMM42619.1 | H9EJ69 |
| QPVEVHVEDIVQ | 1 | 0 | 0 | 1494448 | ANF29161.1 | K9N7C7 |
| QPVEVPVEDTQV | 1 | 0 | 0 | 1494451 | AVN89312.1 | K9N638 |
| QPVSDLLANMGI | 1 | 0 | 0 | 1494466 | ARO76381.1 | P0C6U8 |
| QPVYRTFTQAIC | 1 | 0 | 0 | 1494473 | AGT51379.1 | H9EJ42 |
| QPYCYTEGLMQN | 1 | 0 | 0 | 1494479 | AGT51787.1 | P0C6X6 |
| QPYPNASFDNFR | 1 | 0 | 0 | 1494484 | AVP78041.1 | P0C6W2 |
| QPYQIEDPCPIH | 1 | 0 | 0 | 1494485 | Q3I5I8.1 | Q3LZX5 |
| QPYVFIKRSDAQ | 1 | 0 | 0 | 1494491 | P0C6W6.1 | P0C6W2 |
| QQAFIKHKGGDS | 1 | 0 | 0 | 1494498 | AVV62524.1 | K9N7C7 |
| QQAGAGIKYFCG | 1 | 0 | 0 | 1494499 | NP_073549.1 | A0A0P0LKV0 |
| QQAQQEATHIAE | 1 | 0 | 0 | 1494504 | AST12962.1 | A0A0P0LKV0 |
| QQAVVNIRETKC | 1 | 0 | 0 | 1494508 | AVV62524.1 | K9N7C7 |
| QQCFGKRGPNHK | 1 | 0 | 0 | 1494512 | AJQ24079.1 | P33469 |
| QQDCRIPNGFPF | 1 | 0 | 0 | 1494523 | AKA09611.1 | A0A0P0G321 |
| QQDGMNMSPKPQ | 1 | 0 | 0 | 1494529 | AIV41850.1 | P33469 |
| QQFGKDASDFID | 1 | 0 | 0 | 1494553 | AVP78031.1 | Q3LZX1 |
| QQFGRDTSDFTD | 1 | 0 | 0 | 1494558 | ATO98181.1 | Q3LZX1 |
| QQFNVALPGELS | 1 | 0 | 0 | 1494562 | AUM60013.1 | K9N7C7 |
| QQFVKETTNMVK | 1 | 0 | 0 | 1494564 | AEN19363.1 | P0C6X6 |
| QQGFDATNFALV | 1 | 0 | 0 | 1494568 | QBP84722.1 | P36334 |
| QQGGIPIANCVK | 1 | 0 | 0 | 1494569 | AIV41966.1 | P0C6X6 |
| QQGLLGTCAKRF | 1 | 0 | 0 | 1494572 | NP_073549.1 | A0A0P0LKV0 |
| QQGNLLNHLNSQ | 1 | 0 | 0 | 1494573 | BAL45637.1 | A0A0P0K6L9 |
| QQGNSLNHLTFQ | 1 | 0 | 0 | 1494574 | ABB90520.1 | A0A0P0K6L9 |
| QQGNSLNYLTSQ | 1 | 0 | 0 | 1494576 | AAK32190.1 | A0A0P0K6L9 |
| QQGNSLSHLTSQ | 1 | 0 | 0 | 1494577 | ARA15429.1 | A0A0P0K6L9 |
| QQGPASARDLIC | 1 | 0 | 0 | 1494578 | AXN92228.1 | K9N5Q8 |
| QQGPASVRDLIC | 1 | 0 | 0 | 1494581 | ALJ54502.1 | K9N5Q8 |
| QQGPQSARDLIC | 1 | 0 | 0 | 1494582 | AUM60014.1 | K9N5Q8 |
| QQGSFNKATVAQ | 1 | 0 | 0 | 1494588 | AVZ61116.1 | P0C6X6 |
| QQGVFNKATVAQ | 1 | 0 | 0 | 1494591 | ATN39862.1 | P0C6X6 |
| QQGVFRYDSVSS | 1 | 0 | 0 | 1494594 | AGT51619.1 | P30215 |
| QQGVFRYNNVSS | 1 | 0 | 0 | 1494595 | AAA73152.1 | P30215 |
| QQHFVYDAYQNL | 1 | 0 | 0 | 1494596 | AKN11074.1 | K9N5Q8 |
| QQHPLLNPGALE | 1 | 0 | 0 | 1494597 | AGT21342.1 | A0A0N7J131 |
| QQIDIKFNAPAL | 1 | 0 | 0 | 1494600 | APO40578.1 | P0C6X7 |
| QQIEVKFNAPAL | 1 | 0 | 0 | 1494602 | P0C6W6.1 | P0C6W2 |
| QQKELEQPIEEE | 1 | 0 | 0 | 1494611 | APO40578.1 | P0C6X7 |
| QQKQLLPRWYFY | 1 | 0 | 0 | 1494619 | ARU07581.1 | Q19U29 |
| QQLAGIKLQSKH | 1 | 0 | 0 | 1494626 | AIV41894.1 | P0C6X6 |
| QQLHLEANGRAW | 1 | 0 | 0 | 1494637 | AYO86698.1 | K9N638 |
| QQLHTGFQGKQI | 1 | 0 | 0 | 1494640 | ATQ39389.1 | K9N7C7 |
| QQLNNGFQGKQI | 1 | 0 | 0 | 1494645 | AVV62524.1 | K9N7C7 |
| QQLSNRFGAIGS | 1 | 0 | 0 | 1494654 | CAA83660.1 | P36334 |
| QQLVEHTTPWLL | 1 | 0 | 0 | 1494659 | AVN89322.1 | K9N7C7 |
| QQMFVTPLSVDV | 1 | 0 | 0 | 1494668 | ALA49399.1 | K9N643 |
| QQNKSIELTAEG | 1 | 0 | 0 | 1494674 | AHA80174.1 | P33469 |
| QQNQRSAPRITF | 1 | 0 | 0 | 1494678 | ARO76389.1 | P59595 |
| QQPLGGNVVPYY | 1 | 0 | 0 | 1494681 | AMK59681.1 | P33469 |
| QQPSGGNVIPHY | 1 | 0 | 0 | 1494684 | QEG03818.1 | P33469 |
| QQPSRGNVVPHY | 1 | 0 | 0 | 1494688 | ANZ78871.1 | P33469 |
| QQQDDLFTPVVL | 1 | 0 | 0 | 1494692 | APO40584.1 | P59635 |
| QQQDGMMNISPK | 1 | 0 | 0 | 1494693 | AIX10729.1 | P33469 |

| | | | | | | |
|---|---|---|---|---|---|---|
| QQQDGMMNMSPK | 1 | 0 | 0 | 1494694 | AEN19362.1 | P33469 |
| QQQFNVALPGEL | 1 | 0 | 0 | 1494697 | AUM60013.1 | K9N7C7 |
| QQQNFTTAPAIC | 1 | 0 | 0 | 1494704 | APO40579.1 | P59594 |
| QQQQPQVVTRKS | 1 | 0 | 0 | 1494709 | APO40586.1 | P59595 |
| QQRFVYDSFDNL | 1 | 0 | 0 | 1494714 | AUM60014.1 | K9N5Q8 |
| QQSQNGFSFFLA | 1 | 0 | 0 | 1494721 | AVV62535.1 | K9N7C7 |
| QQTPFDITSAGK | 1 | 0 | 0 | 1494728 | APO40578.1 | P0C6X7 |
| QQVLSENQKQIA | 1 | 0 | 0 | 1494740 | AVV62526.1 | K9N5Q8 |
| QQVSQWLSNLVG | 1 | 0 | 0 | 1494743 | ARO76381.1 | P0C6U8 |
| QQVTKHTAKEVT | 1 | 0 | 0 | 1494745 | AJQ24083.1 | P33469 |
| QQWGYVGSLSQN | 1 | 0 | 0 | 1494750 | AWK59942.1 | H9EJ42 |
| QRGLNNGQGEND | 1 | 0 | 0 | 1494805 | AGT51734.1 | P33469 |
| QRILAASFNKAM | 1 | 0 | 0 | 1494820 | AIW52747.1 | A0A0P0K6L9 |
| QRKLEKMADQTM | 1 | 0 | 0 | 1494830 | AVP78041.1 | P0C6W2 |
| QRKYKGITIQKG | 1 | 0 | 0 | 1494839 | APO40578.1 | P0C6X7 |
| QRMLLEKCDLHN | 1 | 0 | 0 | 1494861 | APO40578.1 | P0C6X7 |
| QRNNTSFPEWID | 1 | 0 | 0 | 1494876 | ATQ39389.1 | K9N7C7 |
| QRQKKQPTVTPL | 1 | 0 | 0 | 1494883 | AAS48456.1 | P59595 |
| QRQRGHKNGQEE | 1 | 0 | 0 | 1494891 | AGT51525.1 | P33469 |
| QRQRGHRNGQGE | 1 | 0 | 0 | 1494897 | ANZ78875.1 | P33469 |
| QRQRGLKNGQGE | 1 | 0 | 0 | 1494898 | ATP16771.1 | P33469 |
| QRQRGLNNGQGE | 1 | 0 | 0 | 1494899 | AGT51734.1 | P33469 |
| QRVFNNYMPYVL | 1 | 0 | 0 | 1494933 | ATO98229.1 | P0C6W2 |
| QRVPKGSITQRS | 1 | 0 | 0 | 1494941 | AUM60021.1 | K9N4V7 |
| QRVQGGITQRTR | 1 | 0 | 0 | 1494942 | ANF29213.1 | K9N4V7 |
| QRVQGIITQRTR | 1 | 0 | 0 | 1494943 | AHI48535.1 | K9N4V7 |
| QRVQGSITQHTR | 1 | 0 | 0 | 1494946 | AKN24774.1 | K9N4V7 |
| QRYKLESYAFEH | 1 | 0 | 0 | 1494959 | APO40578.1 | P0C6X7 |
| QSAIAKPKRLRK | 1 | 0 | 0 | 1494974 | ATQ39389.1 | K9N7C7 |
| QSALLVNNGSAI | 1 | 0 | 0 | 1494976 | ATO98145.1 | Q3LZX1 |
| QSALRMRSTGPI | 1 | 0 | 0 | 1494977 | AUM60014.1 | K9N5Q8 |
| QSAVKRTIKGTH | 1 | 0 | 0 | 1494987 | AVP78041.1 | P0C6W2 |
| QSAVLFNNGTHI | 1 | 0 | 0 | 1494988 | APO40579.1 | P59594 |
| QSCPIVVGVSEI | 1 | 0 | 0 | 1494991 | NP_073549.1 | A0A0P0LKV0 |
| QSDHDVYYDESR | 1 | 0 | 0 | 1494996 | AVV62527.1 | K9N796 |
| QSDQVRNVQTRG | 1 | 0 | 0 | 1495006 | P33469.1 | P33469 |
| QSDRFYRLANEC | 1 | 0 | 0 | 1495007 | AEN19363.1 | P0C6X6 |
| QSDRFYRPANEC | 1 | 0 | 0 | 1495009 | AIL49562.1 | P0C6X6 |
| QSDSLGVSSNQS | 1 | 0 | 0 | 1495013 | QBP84719.1 | P0C6X6 |
| QSEEVIVQQAAE | 1 | 0 | 0 | 1495014 | AUM60013.1 | K9N7C7 |
| QSELAKGVSLDN | 1 | 0 | 0 | 1495016 | APO40578.1 | P0C6X7 |
| QSERYGFCGNGT | 1 | 0 | 0 | 1495021 | ABB90523.1 | A0A0P0K6L9 |
| QSEVDWKFYDAQ | 1 | 0 | 0 | 1495023 | AVP78030.1 | P0C6W2 |
| QSFKDELEEFFK | 1 | 0 | 0 | 1495030 | AVV62526.1 | K9N5Q8 |
| QSFKETLYYQNN | 1 | 0 | 0 | 1495031 | AVV62524.1 | K9N7C7 |
| QSFQQFGKDASD | 1 | 0 | 0 | 1495038 | AVP78031.1 | Q3LZX1 |
| QSFQQFGRDTSD | 1 | 0 | 0 | 1495039 | ATO98181.1 | Q3LZX1 |
| QSFVQWFHEHVG | 1 | 0 | 0 | 1495042 | AVV62524.1 | K9N7C7 |
| QSFVQWFHENVG | 1 | 0 | 0 | 1495043 | AVV62535.1 | K9N7C7 |
| QSGGGKCFNCAP | 1 | 0 | 0 | 1495050 | AAQ90006.1 | A0A0P0K6L9 |
| QSGVRKVTYGTV | 1 | 0 | 0 | 1495065 | AQZ41320.1 | K9N7C7 |
| QSIPMPRGVHMN | 1 | 0 | 0 | 1495082 | AXN92247.1 | K9N7C7 |
| QSIQALQTAAMT | 1 | 0 | 0 | 1495084 | APO40578.1 | P0C6X7 |
| QSKDSNFLKRVR | 1 | 0 | 0 | 1495088 | ALT66879.1 | K9N7C7 |
| QSKDTNFLNAFG | 1 | 0 | 0 | 1495092 | ATN39869.1 | P0C6X6 |
| QSKDTNFLNGFG | 1 | 0 | 0 | 1495093 | ATP16763.1 | P0C6X6 |
| QSKESNFLNRVR | 1 | 0 | 0 | 1495096 | AQZ41330.1 | K9N7C7 |
| QSKGLYRHCIRG | 1 | 0 | 0 | 1495097 | ARO76381.1 | P0C6U8 |
| QSKLPSSSGTST | 1 | 0 | 0 | 1495100 | AIW52795.1 | H9EJ69 |
| QSKRGVYCCREH | 1 | 0 | 0 | 1495106 | APO40578.1 | P0C6X7 |
| QSKRSGFCGQGI | 1 | 0 | 0 | 1495108 | ALJ54495.1 | K9N5Q8 |
| QSKRTRFFKGTV | 1 | 0 | 0 | 1495110 | AIV41996.1 | P0C6X6 |
| QSKRTRLFKGTV | 1 | 0 | 0 | 1495112 | AEN19363.1 | P0C6X6 |
| QSKRVDFCGRGY | 1 | 0 | 0 | 1495115 | ATO98181.1 | Q3LZX1 |
| QSKRYGFGGNGT | 1 | 0 | 0 | 1495117 | ABB90528.1 | A0A0P0K6L9 |
| QSKSPSSSGIST | 1 | 0 | 0 | 1495121 | AMM42629.1 | H9EJ69 |
| QSKTKRFIKETI | 1 | 0 | 0 | 1495124 | ARU07575.1 | A0A140H1G9 |
| QSLCALPDTHST | 1 | 0 | 0 | 1495128 | AID55099.1 | K9N5Q8 |
| QSLCALPDTPSP | 1 | 0 | 0 | 1495130 | AJG44080.1 | K9N5Q8 |
| QSLENVIYNLFN | 1 | 0 | 0 | 1495141 | ABD75583.1 | A0A140H1G9 |
| QSLENVVYNLVN | 1 | 0 | 0 | 1495142 | ABD75615.1 | A0A140H1G9 |
| QSLIDNINSTLV | 1 | 0 | 0 | 1495148 | BAL45640.1 | A0A0P0K6L9 |
| QSLVLTSNDERL | 1 | 0 | 0 | 1495169 | AVV62535.1 | K9N7C7 |
| QSLVLTSTDERM | 1 | 0 | 0 | 1495170 | AVV62524.1 | K9N7C7 |
| QSNEDDGTPVYF | 1 | 0 | 0 | 1495181 | AVV62533.1 | K9N4V7 |
| QSNRYGFCGNGT | 1 | 0 | 0 | 1495187 | AIW52843.1 | A0A0P0G321 |
| QSNVDVLPEGWR | 1 | 0 | 0 | 1495188 | AEN19363.1 | P0C6X6 |
| QSNVEVLPEGWR | 1 | 0 | 0 | 1495190 | QEG03790.1 | P0C6X6 |
| QSPASSQSAAKI | 1 | 0 | 0 | 1495192 | AIW52722.1 | A0A0N7J131 |
| QSPASSQSAATI | 1 | 0 | 0 | 1495193 | AIW52723.1 | A0A0N7J131 |
| QSPASSQSSAKI | 1 | 0 | 0 | 1495194 | ABB90486.1 | A0A0N7J131 |
| QSPASSQSSAKS | 1 | 0 | 0 | 1495195 | ABB90484.1 | A0A0N7J131 |
| QSPASSQSVAKI | 1 | 0 | 0 | 1495196 | ABB90499.1 | A0A0N7J131 |
| QSPELFDSLNLS | 1 | 0 | 0 | 1495198 | ARU07581.1 | Q19U29 |
| QSPFEIKSAKKF | 1 | 0 | 0 | 1495199 | ARO76381.1 | P0C6U8 |
| QSPQGNLGSNEP | 1 | 0 | 0 | 1495200 | AVI15051.1 | P0C6X6 |
| QSPTYLSVDTKF | 1 | 0 | 0 | 1495205 | APO40578.1 | P0C6X7 |
| QSPVSSQSAAKI | 1 | 0 | 0 | 1495206 | AMM42518.1 | A0A0N7J131 |
| QSQPKVITKRDA | 1 | 0 | 0 | 1495214 | ANF29235.1 | K9N4V7 |
| QSQSSHAVQNTV | 1 | 0 | 0 | 1495218 | AIW52795.1 | H9EJ69 |
| QSQSSHVVQNTV | 1 | 0 | 0 | 1495220 | AIW52805.1 | H9EJ69 |

| | | | | | | |
|---|---|---|---|---|---|---|
| QSQSSLVAQNTV | 1 | 0 | 0 | 1495221 | AMM42614.1 | H9EJ69 |
| QSRADQNGYITL | 1 | 0 | 0 | 1495222 | AUM60013.1 | K9N7C7 |
| QSRGRGRNPKPR | 1 | 0 | 0 | 1495232 | AVV62533.1 | K9N4V7 |
| QSRNPSSDRYHN | 1 | 0 | 0 | 1495244 | AAA45463.1 | A0A0N7J131 |
| QSRRFIVPWVMY | 1 | 0 | 0 | 1495247 | AEN19355.1 | P0C6X6 |
| QSRSCSDFIPLS | 1 | 0 | 0 | 1495248 | ANI69887.1 | K9N7C7 |
| QSRSQSRSRGES | 1 | 0 | 0 | 1495255 | AIW52723.1 | A0A0N7J131 |
| QSRVISSFTCRT | 1 | 0 | 0 | 1495259 | AEN19363.1 | P0C6X6 |
| QSSAPRITFGGP | 1 | 0 | 0 | 1495263 | AVP78049.1 | Q3LZX4 |
| QSSETKEQKHEI | 1 | 0 | 0 | 1495266 | AAA45463.1 | A0A0N7J131 |
| QSSGMIKPNALF | 1 | 0 | 0 | 1495274 | ASU91323.1 | K9N7C7 |
| QSSGMINPNALL | 1 | 0 | 0 | 1495277 | ASU90746.1 | K9N7C7 |
| QSSGVINSSAMF | 1 | 0 | 0 | 1495278 | AVV62535.1 | K9N7C7 |
| QSSHAVQNTVFN | 1 | 0 | 0 | 1495281 | AVL25595.1 | H9EJ69 |
| QSSHAWQPGVAM | 1 | 0 | 0 | 1495283 | AVP78041.1 | P0C6W2 |
| QSSITQRTRTRP | 1 | 0 | 0 | 1495289 | ANI69896.1 | K9N4V7 |
| QSSLVAQNTVLN | 1 | 0 | 0 | 1495293 | AMM42614.1 | H9EJ69 |
| QSSNYRIDTTAT | 1 | 0 | 0 | 1495295 | AEN19366.1 | P36334 |
| QSSPIIPGFGGN | 1 | 0 | 0 | 1495297 | AOR17482.1 | K9N5Q8 |
| QSSSDLVAAVNL | 1 | 0 | 0 | 1495300 | ABK63972.1 | H9EJ69 |
| QSTALCKSGSLV | 1 | 0 | 0 | 1495313 | P30215.1 | P30215 |
| QSVNVKPKRLRK | 1 | 0 | 0 | 1495355 | ATG84700.1 | K9N638 |
| QSVSVDRDSTST | 1 | 0 | 0 | 1495361 | QDI73611.1 | K9N796 |
| QSVTVKPKRLRK | 1 | 0 | 0 | 1495368 | AXN92247.1 | K9N7C7 |
| QTAQAAGTDITI | 1 | 0 | 0 | 1495403 | APO40578.1 | P0C6X7 |
| QTAQGAHLFSSR | 1 | 0 | 0 | 1495405 | ALJ54512.1 | K9N5Q8 |
| QTASRSSSRSRG | 1 | 0 | 0 | 1495407 | APO40586.1 | P59595 |
| QTDVDFKDELEE | 1 | 0 | 0 | 1495424 | AUM60014.1 | K9N5Q8 |
| QTFQAISSSIQA | 1 | 0 | 0 | 1495436 | AGW80677.1 | A0A0P0K6L9 |
| QTFYPRLQAAND | 1 | 0 | 0 | 1495441 | AVV62524.1 | K9N7C7 |
| QTGDFLKATCEL | 1 | 0 | 0 | 1495443 | APO40578.1 | P0C6X7 |
| QTGNFKDLREYV | 1 | 0 | 0 | 1495455 | ATO98193.1 | Q3LZX1 |
| QTGSSKCGCSVI | 1 | 0 | 0 | 1495456 | 2H85_A | P0C6X7 |
| QTIEDLRARMTY | 1 | 0 | 0 | 1495477 | AJG44112.1 | K9N638 |
| QTIEELHVRSSF | 1 | 0 | 0 | 1495478 | AVV62524.1 | K9N7C7 |
| QTIINGMHKSFY | 1 | 0 | 0 | 1495483 | AWK59942.1 | H9EJ42 |
| QTIKPEVKEVAP | 1 | 0 | 0 | 1495484 | AIZ74432.1 | K9N7C7 |
| QTIQPEVKEIAP | 1 | 0 | 0 | 1495490 | ALA49362.1 | K9N7C7 |
| QTIQQKELEQPI | 1 | 0 | 0 | 1495492 | APO40578.1 | P0C6X7 |
| QTITTDNTFVAG | 1 | 0 | 0 | 1495493 | ARO76382.1 | P59594 |
| QTKSIYKTPNIK | 1 | 0 | 0 | 1495496 | APO40579.1 | P59594 |
| QTKYLVGIGEHC | 1 | 0 | 0 | 1495498 | CAA83660.1 | P36334 |
| QTLDELHACTSF | 1 | 0 | 0 | 1495500 | AVV62535.1 | K9N7C7 |
| QTLDILDITPCS | 1 | 0 | 0 | 1495501 | ATO98145.1 | Q3LZX1 |
| QTLEDLHARMTY | 1 | 0 | 0 | 1495502 | ATQ39389.1 | K9N7C7 |
| QTLEILDISPCS | 1 | 0 | 0 | 1495503 | Q3I5J5.1 | Q3LZX1 |
| QTMTQMYKQARS | 1 | 0 | 0 | 1495523 | AVP78041.1 | P0C6W2 |
| QTNLYSFDKAID | 1 | 0 | 0 | 1495528 | AWK59942.1 | H9EJ42 |
| QTNSPKVTIDCA | 1 | 0 | 0 | 1495541 | AIL49500.1 | P36334 |
| QTPFDITSAGKF | 1 | 0 | 0 | 1495545 | APO40578.1 | P0C6X7 |
| QTQHSLNFVGEF | 1 | 0 | 0 | 1495559 | AXN92247.1 | K9N7C7 |
| QTQSAIAKPKRL | 1 | 0 | 0 | 1495563 | ATQ39389.1 | K9N7C7 |
| QTQSVIVKPKRL | 1 | 0 | 0 | 1495564 | AVN89451.1 | K9N7C7 |
| QTQSVTVKPKRL | 1 | 0 | 0 | 1495567 | AXN92247.1 | K9N7C7 |
| QTSDVYEADAEI | 1 | 0 | 0 | 1495574 | AUM60013.1 | K9N7C7 |
| QTSNFRVQPTQS | 1 | 0 | 0 | 1495587 | AVP78031.1 | Q3LZX1 |
| QTSNFRVSPSTE | 1 | 0 | 0 | 1495589 | APO40579.1 | P59594 |
| QTSNFRVSPTHE | 1 | 0 | 0 | 1495590 | ATO98193.1 | Q3LZX1 |
| QTSTSQQPSGGN | 1 | 0 | 0 | 1495607 | ATP16771.1 | P33469 |
| QTSVAPDLSLGY | 1 | 0 | 0 | 1495611 | AIL49488.1 | P36334 |
| QTTNGMNTSHSV | 1 | 0 | 0 | 1495626 | AIW52756.1 | A0A0P0K6L9 |
| QTTNGTNTIHSV | 1 | 0 | 0 | 1495627 | AIW52741.1 | A0A0P0K6L9 |
| QTVATALNKIHD | 1 | 0 | 0 | 1495641 | ABB90526.1 | A0A0P0K6L9 |
| QTVAWNLREMLA | 1 | 0 | 0 | 1495643 | APO40578.1 | P0C6X7 |
| QTVDSSQGSEYP | 1 | 0 | 0 | 1495647 | AVV62524.1 | K9N7C7 |
| QTVEDLCARMTY | 1 | 0 | 0 | 1495649 | AKN24802.1 | K9N7C7 |
| QTVEDLRVRMTY | 1 | 0 | 0 | 1495651 | QDP16215.1 | K9N7C7 |
| QTVRSGNFNQDF | 1 | 0 | 0 | 1495663 | AHI48750.1 | K9N7C7 |
| QTVSFHEDRILD | 1 | 0 | 0 | 1495667 | AVV62524.1 | K9N7C7 |
| QTVSTALNKIQD | 1 | 0 | 0 | 1495668 | ABB90519.1 | A0A0P0K6L9 |
| QTVSTAVSPDFK | 1 | 0 | 0 | 1495669 | AVV62535.1 | K9N7C7 |
| QTVTKKSAAEASKKPRQK | 1 | 0 | 0 | 1495671 | P0DTC9.1 | P0DTC9 |
| QTVTSQQPSGGN | 1 | 0 | 0 | 1495673 | AIL49418.1 | P33469 |
| QTYLKSDNIVAA | 1 | 0 | 0 | 1495683 | AIV42002.1 | P0C6X6 |
| QTYSETGSDVLY | 1 | 0 | 0 | 1495686 | AVV62535.1 | K9N7C7 |
| QTYYQPIQQAPT | 1 | 0 | 0 | 1495692 | AST12967.1 | A0A0P0KXN3 |
| QVALPQSKDSNF | 1 | 0 | 0 | 1495700 | AXN92258.1 | K9N7C7 |
| QVALTQSKDTNF | 1 | 0 | 0 | 1495701 | AUM60023.1 | K9N7C7 |
| QVANNLMNGVTL | 1 | 0 | 0 | 1495703 | AIL49500.1 | P36334 |
| QVAPGETGVIAD | 1 | 0 | 0 | 1495706 | ATO98181.1 | Q3LZX1 |
| QVCAFLNKLNEN | 1 | 0 | 0 | 1495719 | ABD75607.1 | A0A140H1G9 |
| QVDLTEAICDFE | 1 | 0 | 0 | 1495732 | AAX85666.1 | P0C6X6 |
| QVDQLNSSYKLS | 1 | 0 | 0 | 1495737 | ATW75477.1 | K9N5Q8 |
| QVDQLNSTYFKL | 1 | 0 | 0 | 1495738 | ANI69889.1 | K9N5Q8 |
| QVDSFTCNNIDA | 1 | 0 | 0 | 1495740 | AMK59677.1 | P36334 |
| QVDVSHEVFDDE | 1 | 0 | 0 | 1495746 | AQZ41284.1 | K9N7C7 |
| QVEEDVEMSDFV | 1 | 0 | 0 | 1495748 | AEN19363.1 | P0C6X6 |
| QVEINRLQEAIK | 1 | 0 | 0 | 1495750 | AIL49487.1 | P36334 |
| QVEMNRLQEAIK | 1 | 0 | 0 | 1495753 | CAA79896.1 | P36334 |
| QVEQKVAENPKE | 1 | 0 | 0 | 1495756 | AVP78041.1 | P0C6W2 |
| QVESDDYIATNGPLK | 1 | 0 | 0 | 1495759 | P0DTC1.1 | P0DTD1 |

| | | | | | | |
|---|---|---|---|---|---|---|
| QVEYLQITSKPI | 1 | 0 | 0 | 1495764 | APT69883.1 | A0A0P0K6L9 |
| QVGDAVLLKGHG | 1 | 0 | 0 | 1495781 | AUM60013.1 | K9N7C7 |
| QVGDSVLLQGHF | 1 | 0 | 0 | 1495783 | AHI48765.1 | K9N638 |
| QVGDSVLLQGHS | 1 | 0 | 0 | 1495784 | AXN92247.1 | K9N7C7 |
| QVHGNGSMIEDI | 1 | 0 | 0 | 1495799 | AGV08533.1 | K9N7C7 |
| QVHNLPEYMAVA | 1 | 0 | 0 | 1495801 | APT69865.1 | A0A0P0KXN3 |
| QVIQDVKDGTVF | 1 | 0 | 0 | 1495814 | CAA46113.1 | A0A0P0KWE8 |
| QVIVMKDNFLSD | 1 | 0 | 0 | 1495822 | AUM60013.1 | K9N7C7 |
| QVKNASVYFSQL | 1 | 0 | 0 | 1495825 | NP_073549.1 | A0A0P0LKV0 |
| QVKQMYKTPAIK | 1 | 0 | 0 | 1495827 | ATO98181.1 | Q3LZX1 |
| QVLSSDVMFEGA | 1 | 0 | 0 | 1495860 | ASU90140.1 | K9N7C7 |
| QVLSSDVMFENA | 1 | 0 | 0 | 1495861 | AXN92226.1 | K9N7C7 |
| QVMKLPSEPQTI | 1 | 0 | 0 | 1495868 | ALA50044.1 | K9N7C7 |
| QVNLSLNGNTSV | 1 | 0 | 0 | 1495876 | AWO69005.1 | A0A0P0G321 |
| QVPIGTNDPIRS | 1 | 0 | 0 | 1495881 | CAA49377.1 | A0A0P0LKV0 |
| QVPTNFSFGITQ | 1 | 0 | 0 | 1495885 | AVV62537.1 | K9N5Q8 |
| QVQLIDKYCSVN | 1 | 0 | 0 | 1495892 | ATG84700.1 | K9N638 |
| QVQLTDKYCSVN | 1 | 0 | 0 | 1495893 | AXN92247.1 | K9N7C7 |
| QVRAFLNKLNDN | 1 | 0 | 0 | 1495896 | ARB07605.1 | A0A140H1G9 |
| QVSEPPKEQRVQ | 1 | 0 | 0 | 1495906 | AHI48535.1 | K9N4V7 |
| QVSLTQSKDTNF | 1 | 0 | 0 | 1495908 | AUM60013.1 | K9N7C7 |
| QVSTSQYSHRSA | 1 | 0 | 0 | 1495913 | AVV62537.1 | K9N5Q8 |
| QVTIDYIEISFM | 1 | 0 | 0 | 1495925 | APO40578.1 | P0C6X7 |
| QVTTANVSALMG | 1 | 0 | 0 | 1495934 | AVV62524.1 | K9N7C7 |
| QVTYQNISFNLP | 1 | 0 | 0 | 1495936 | ANI69889.1 | K9N5Q8 |
| QVVDSRWNNARL | 1 | 0 | 0 | 1495946 | AAX85668.1 | P30215 |
| QVVDYLTREDMH | 1 | 0 | 0 | 1495950 | AVV62524.1 | K9N7C7 |
| QVVELLSEPQTI | 1 | 0 | 0 | 1495951 | AVN89374.1 | K9N7C7 |
| QVVIADTLQETS | 1 | 0 | 0 | 1495956 | ASU45706.1 | K9N7C7 |
| QVVKLPSEPRTI | 1 | 0 | 0 | 1495962 | AMQ49036.1 | K9N638 |
| QVVMDYLDSLKP | 1 | 0 | 0 | 1495965 | P0C6W6.1 | P0C6W2 |
| QVVNSGPDHTCN | 1 | 0 | 0 | 1495969 | AIV41780.1 | P0C6X6 |
| QVVNSGSDINCN | 1 | 0 | 0 | 1495972 | P0C6U5.1 | A0A140H1G9 |
| QVVQLPVKDYIQ | 1 | 0 | 0 | 1495978 | ABD75583.1 | A0A140H1G9 |
| QVVSDIDYVPLKSATC | 1 | 0 | 0 | 1495980 | YP_009724389.1 | P0DTD1 |
| QVVTRKSASEAS | 1 | 0 | 0 | 1495987 | APO40586.1 | P59595 |
| QWDLIISDMYNP | 1 | 0 | 0 | 1496019 | ATN39862.1 | P0C6X6 |
| QWFHENVGGNFD | 1 | 0 | 0 | 1496023 | AVV62535.1 | K9N7C7 |
| QWFKNQISVAPD | 1 | 0 | 0 | 1496024 | ANZ78835.1 | P36334 |
| QWFKNQTSVAPV | 1 | 0 | 0 | 1496030 | AIV41873.1 | P36334 |
| QWFKNQTSVARD | 1 | 0 | 0 | 1496031 | AIX10750.1 | P36334 |
| QWGSTMIFSVHP | 1 | 0 | 0 | 1496034 | AVV62524.1 | K9N7C7 |
| QWLKNQTSVAPD | 1 | 0 | 0 | 1496042 | ARB07414.1 | P36334 |
| QWNCIDCDSYKP | 1 | 0 | 0 | 1496058 | AEN19363.1 | P0C6X6 |
| QWYDFGDFIQTT | 1 | 0 | 0 | 1496082 | ABD75607.1 | A0A140H1G9 |
| QYAHKHDIQLTN | 1 | 0 | 0 | 1496095 | AVN89451.1 | K9N7C7 |
| QYAHKHDIQLTT | 1 | 0 | 0 | 1496096 | AVV62535.1 | K9N7C7 |
| QYAHKHDIQLTV | 1 | 0 | 0 | 1496097 | AVV62524.1 | K9N7C7 |
| QYCFKAPKNFCP | 1 | 0 | 0 | 1496101 | ABD43182.1 | P36334 |
| QYCNGSIACQFC | 1 | 0 | 0 | 1496103 | AEN19363.1 | P0C6X6 |
| QYCSTLHNEILA | 1 | 0 | 0 | 1496107 | AGT51737.1 | P0C6X6 |
| QYDKNASLNSFK | 1 | 0 | 0 | 1496109 | AUM60014.1 | K9N5Q8 |
| QYELKHGTFVCA | 1 | 0 | 0 | 1496115 | AVP78041.1 | P0C6W2 |
| QYGCDYSGKLAA | 1 | 0 | 0 | 1496124 | ABD75583.1 | A0A140H1G9 |
| QYGECLGDISAR | 1 | 0 | 0 | 1496133 | ATO98145.1 | Q3LZX1 |
| QYGTDTISVCPK | 1 | 0 | 0 | 1496151 | AGV08438.1 | K9N5Q8 |
| QYIKGSGRENLY | 1 | 0 | 0 | 1496167 | 6NB6_A | P59594 |
| QYIYSVGHSDSN | 1 | 0 | 0 | 1496174 | AVV62537.1 | K9N5Q8 |
| QYLLKFDDRGVI | 1 | 0 | 0 | 1496188 | ADN03339.1 | A0A140H1H1 |
| QYMCGKDGNPIR | 1 | 0 | 0 | 1496199 | AVV62524.1 | K9N7C7 |
| QYMCGSDGKPIA | 1 | 0 | 0 | 1496200 | AUM60013.1 | K9N7C7 |
| QYNMCEYPHTSC | 1 | 0 | 0 | 1496208 | AKA09651.1 | P36334 |
| QYNMCEYPQTIC | 1 | 0 | 0 | 1496210 | CAA79896.1 | P36334 |
| QYNYKQSFANPT | 1 | 0 | 0 | 1496216 | AVV62537.1 | K9N5Q8 |
| QYRKINTGNGIK | 1 | 0 | 0 | 1496221 | AVN89320.1 | K9N4V7 |
| QYSCDFKGWIPG | 1 | 0 | 0 | 1496222 | QBP84701.1 | P0C6X6 |
| QYSHRSAIEDLL | 1 | 0 | 0 | 1496226 | AVV62537.1 | K9N5Q8 |
| QYSLAFNQDGII | 1 | 0 | 0 | 1496228 | AHA80190.1 | P36334 |
| QYSPCVSIAPST | 1 | 0 | 0 | 1496232 | QEJ82215.1 | K9N5Q8 |
| QYSPCVSTVPST | 1 | 0 | 0 | 1496236 | ANC28634.1 | K9N5Q8 |
| QYTMCEYPNTIC | 1 | 0 | 0 | 1496250 | CAA83661.1 | P36334 |
| QYTPCLYLTSSG | 1 | 0 | 0 | 1496252 | AUM60014.1 | K9N5Q8 |
| QYVCNGFKKCEQ | 1 | 0 | 0 | 1496261 | AUM60014.1 | K9N5Q8 |
| QYWKGYGCSCDS | 1 | 0 | 0 | 1496272 | AVV62524.1 | K9N7C7 |
| QYYCDGGKYYTQ | 1 | 0 | 0 | 1496276 | AEN19363.1 | P0C6X6 |
| QYYCYSAKDTLP | 1 | 0 | 0 | 1496278 | AVV62524.1 | K9N7C7 |
| QYYFYSAKDDIK | 1 | 0 | 0 | 1496283 | AUM60013.1 | K9N7C7 |
| QYYNGIMVVPGV | 1 | 0 | 0 | 1496290 | BAL45639.1 | A0A0P0K6L9 |
| QYYYNMATGFAY | 1 | 0 | 0 | 1496294 | AZS52617.1 | A0A140H1H0 |
| QYYYNMDTGFAY | 1 | 0 | 0 | 1496295 | AGT17757.1 | A0A140H1H0 |
| RAAFEYLKPMPF | 1 | 0 | 0 | 1496303 | AGT51737.1 | P0C6X6 |
| RAALNNEKRPDH | 1 | 0 | 0 | 1496317 | AVV62535.1 | K9N7C7 |
| RAALNSEKRPDH | 1 | 0 | 0 | 1496319 | AUM60013.1 | K9N7C7 |
| RAANNGHKQIVK | 1 | 0 | 0 | 1496321 | AUM60017.1 | K9N643 |
| RAAPNNTVSWYT | 1 | 0 | 0 | 1496322 | AXN92267.1 | K9N4V7 |
| RAAVGPMQDVNQ | 1 | 0 | 0 | 1496326 | AVV62533.1 | K9N4V7 |
| RAAVLEWDIKQC | 1 | 0 | 0 | 1496329 | AUM60013.1 | K9N7C7 |
| RACIKTAQAGLY | 1 | 0 | 0 | 1496337 | ALT66847.1 | K9N796 |
| RACIKTVQADTA | 1 | 0 | 0 | 1496340 | AHC74089.1 | K9N796 |
| RADQNGYITLKE | 1 | 0 | 0 | 1496353 | AUM60013.1 | K9N7C7 |
| RAELNGVMVDKV | 1 | 0 | 0 | 1496359 | QBP84746.1 | P0C6X6 |

| | | | | | | |
|---|---|---|---|---|---|---|
| RAFADYTPCAVI | 1 | 0 | 0 | 1496361 | ARQ84755.1 | K9N7C7 |
| RAFVNQVKVPYV | 1 | 0 | 0 | 1496382 | AVV62535.1 | K9N7C7 |
| RAGAWNIGSNKT | 1 | 0 | 0 | 1496385 | APO40578.1 | P0C6X7 |
| RAGEAANFCALI | 1 | 0 | 0 | 1496390 | AVP78041.1 | P0C6W2 |
| RAGKHEHSLLVE | 1 | 0 | 0 | 1496394 | ARU07593.1 | H9EJ42 |
| RAHPAEPLVQDR | 1 | 0 | 0 | 1496402 | AVR40343.1 | Q4VID0 |
| RAHPVEPLVQDR | 1 | 0 | 0 | 1496403 | ARE29989.1 | Q4VID0 |
| RAILKKKEKILF | 1 | 0 | 0 | 1496415 | AVV62535.1 | K9N7C7 |
| RAITTENRFTNF | 1 | 0 | 0 | 1496423 | AIL49500.1 | P36334 |
| RAITTGYRFTNF | 1 | 0 | 0 | 1496425 | AEN19366.1 | P36334 |
| RAKSIVSTIGEA | 1 | 0 | 0 | 1496439 | AUM60023.1 | K9N7C7 |
| RAKSIVSTIGES | 1 | 0 | 0 | 1496440 | AUM60013.1 | K9N7C7 |
| RAKSVVSKIGNS | 1 | 0 | 0 | 1496442 | ATQ39389.1 | K9N7C7 |
| RALADYTQCAII | 1 | 0 | 0 | 1496446 | AUM60013.1 | K9N7C7 |
| RALADYTQCAVI | 1 | 0 | 0 | 1496447 | AVV62524.1 | K9N7C7 |
| RALMSCNGNKIE | 1 | 0 | 0 | 1496456 | ADO24673.1 | P0C6X6 |
| RALSGIAIEQDK | 1 | 0 | 0 | 1496473 | AVP78031.1 | Q3LZX1 |
| RALSGIAVEQDR | 1 | 0 | 0 | 1496474 | ATO98231.1 | Q3LZX1 |
| RALTGIAIEQDK | 1 | 0 | 0 | 1496477 | ATO98181.1 | Q3LZX1 |
| RALTGIAVEQDKNTQEVFAQV | 1 | 0 | 0 | 1496479 | P0DTC2.1 | P0DTC2 |
| RALTGVALEQDK | 1 | 0 | 0 | 1496480 | ATO98145.1 | Q3LZX1 |
| RALVAVNSEDIY | 1 | 0 | 0 | 1496482 | AUM60013.1 | K9N7C7 |
| RAMFQPRLPVLS | 1 | 0 | 0 | 1496486 | ARU07594.1 | A0A0P0G321 |
| RAMLDYAKDHND | 1 | 0 | 0 | 1496487 | AVA26872.1 | H9EJ42 |
| RANFFDVPTFCA | 1 | 0 | 0 | 1496499 | AMQ49014.1 | K9N7C7 |
| RANSGNRTPTPG | 1 | 0 | 0 | 1496509 | AIX10722.1 | P33469 |
| RAPRGSITQRSR | 1 | 0 | 0 | 1496517 | AUM60031.1 | K9N4V7 |
| RAQPKQTVTSQQ | 1 | 0 | 0 | 1496527 | AIL49418.1 | P33469 |
| RARTRPSVQPGP | 1 | 0 | 0 | 1496541 | AVN89381.1 | K9N4V7 |
| RASADIGANHTG | 1 | 0 | 0 | 1496547 | ARO76381.1 | P0C6U8 |
| RASDTSLSDACQ | 1 | 0 | 0 | 1496552 | AGT17757.1 | A0A140H1H0 |
| RASLRQIFDPAP | 1 | 0 | 0 | 1496558 | AVV62524.1 | K9N7C7 |
| RASLRQIYDPAP | 1 | 0 | 0 | 1496563 | AVV62535.1 | K9N7C7 |
| RASSGNRPGNGI | 1 | 0 | 0 | 1496578 | AMM42456.1 | P33469 |
| RASSGNRSVNGI | 1 | 0 | 0 | 1496580 | AGT51505.1 | P33469 |
| RATTVLITDLIQ | 1 | 0 | 0 | 1496606 | AVV62527.1 | K9N796 |
| RAVEPILAIEPL | 1 | 0 | 0 | 1496611 | AHF54653.1 | Q4VID0 |
| RAVICTAEDMLN | 1 | 0 | 0 | 1496614 | 2Q6G_A | P0C6X7 |
| RAVIYDSKESLV | 1 | 0 | 0 | 1496615 | AVV62524.1 | K9N7C7 |
| RAVIYNSRESLV | 1 | 0 | 0 | 1496617 | AVV62535.1 | K9N7C7 |
| RAVRSHPDFKFL | 1 | 0 | 0 | 1496623 | AVN89451.1 | K9N7C7 |
| RAVSFAYNNDIT | 1 | 0 | 0 | 1496627 | AVN89381.1 | K9N4V7 |
| RAYIEAAAQGFQ | 1 | 0 | 0 | 1496631 | QED88024.1 | H9EJ42 |
| RAYIGSTAYIQQ | 1 | 0 | 0 | 1496632 | AUM60027.1 | K9N643 |
| RAYKHLNNYDCC | 1 | 0 | 0 | 1496634 | ARB07605.1 | A0A140H1G9 |
| RAYLDLRTTQQL | 1 | 0 | 0 | 1496635 | AHI48765.1 | K9N638 |
| RAYLDSCTTQQL | 1 | 0 | 0 | 1496636 | AHI48794.1 | K9N638 |
| RAYSEAAAQGFQ | 1 | 0 | 0 | 1496646 | AVA26872.1 | H9EJ42 |
| RAYSEAATQGFQ | 1 | 0 | 0 | 1496647 | AFD98756.1 | H9EJ42 |
| RCDINYKGKVFF | 1 | 0 | 0 | 1496670 | QBP84746.1 | P0C6X6 |
| RCDQLAFDVPDG | 1 | 0 | 0 | 1496672 | AFR79257.1 | A0A0P0K6L9 |
| RCDQLSFDVPDG | 1 | 0 | 0 | 1496673 | AIW52757.1 | A0A0P0K6L9 |
| RCIVFDVKGVEE | 1 | 0 | 0 | 1496701 | AOL02451.1 | Q80872 |
| RCIVLDVKGLEE | 1 | 0 | 0 | 1496702 | ATP16755.1 | Q80872 |
| RCNVIADFKSFT | 1 | 0 | 0 | 1496727 | AXN73435.1 | K9N7C7 |
| RCNVIADPKSFT | 1 | 0 | 0 | 1496728 | AQZ41321.1 | K9N7C7 |
| RCPNDNAYTSYG | 1 | 0 | 0 | 1496737 | AUM60014.1 | K9N5Q8 |
| RCPREIVNTVSA | 1 | 0 | 0 | 1496740 | AVV62524.1 | K9N7C7 |
| RCPTAADINIHD | 1 | 0 | 0 | 1496741 | AGT51530.1 | P30215 |
| RCPTAADINIPH | 1 | 0 | 0 | 1496743 | AXX83356.1 | P30215 |
| RCPTAADINNPD | 1 | 0 | 0 | 1496744 | P30215.1 | P30215 |
| RCPTAADINTPD | 1 | 0 | 0 | 1496745 | AAA73152.1 | P30215 |
| RCPTAAYINILD | 1 | 0 | 0 | 1496747 | ARK08634.1 | P30215 |
| RCSRCFYVHSAI | 1 | 0 | 0 | 1496758 | AUM60028.1 | K9N7D2 |
| RCVVKPDGKSFI | 1 | 0 | 0 | 1496779 | AUM60023.1 | K9N7C7 |
| RCYTAEQCIRGA | 1 | 0 | 0 | 1496795 | NP_073549.1 | A0A0P0LKV0 |
| RDAPYMVGDVIN | 1 | 0 | 0 | 1496809 | ARO76381.1 | P0C6U8 |
| RDATKPQQVTKH | 1 | 0 | 0 | 1496812 | AGT51674.1 | P33469 |
| RDCDLNKPLIEW | 1 | 0 | 0 | 1496821 | AVV62524.1 | K9N7C7 |
| RDDLQLGSSGFL | 1 | 0 | 0 | 1496827 | BBA20986.1 | A0A140H1H1 |
| RDDNQALLVPKI | 1 | 0 | 0 | 1496829 | AMM42360.1 | A0A140H1G9 |
| RDDNQTLLVPQI | 1 | 0 | 0 | 1496831 | AMM42357.1 | A0A140H1G9 |
| RDEFSIETDIID | 1 | 0 | 0 | 1496834 | AAA45463.1 | A0A0N7J131 |
| RDFKFPDGQGVP | 1 | 0 | 0 | 1496840 | AMM42397.1 | Q19U29 |
| RDFNHNMGDSDL | 1 | 0 | 0 | 1496843 | AIW52805.1 | H9EJ69 |
| RDGDVVKQPVKF | 1 | 0 | 0 | 1496855 | ALA49648.1 | K9N7C7 |
| RDGQAYVRKDGE | 1 | 0 | 0 | 1496861 | 6CRW_A | P59594 |
| RDGQPCYEQFPL | 1 | 0 | 0 | 1496862 | AVN89451.1 | K9N7C7 |
| RDGQPCYERFPL | 1 | 0 | 0 | 1496863 | AVN89312.1 | K9N638 |
| RDGQPFYEPFPL | 1 | 0 | 0 | 1496864 | ALD51903.1 | K9N7C7 |
| RDGQPFYERFPL | 1 | 0 | 0 | 1496865 | AXN92247.1 | K9N7C7 |
| RDGQRIYDDVNA | 1 | 0 | 0 | 1496866 | AEN19363.1 | P0C6X6 |
| RDGQVCYERFSL | 1 | 0 | 0 | 1496868 | AVV62535.1 | K9N7C7 |
| RDGVRRGDNFHN | 1 | 0 | 0 | 1496876 | AVN89451.1 | K9N7C7 |
| RDINADDKCFAN | 1 | 0 | 0 | 1496893 | AUM60013.1 | K9N7C7 |
| RDINSNDKCFAN | 1 | 0 | 0 | 1496895 | AVV62524.1 | K9N7C7 |
| RDLQRRLYDNCY | 1 | 0 | 0 | 1496940 | AIW52827.1 | H9EJ42 |
| RDLSSDENGVRT | 1 | 0 | 0 | 1496944 | AVP78031.1 | Q3LZX1 |
| RDLTSDENGVRT | 1 | 0 | 0 | 1496947 | ATO98120.1 | Q3LZX1 |
| RDLYDKLQFMSL | 1 | 0 | 0 | 1496949 | AVP78030.1 | P0C6W2 |
| RDMADQIASLVL | 1 | 0 | 0 | 1496952 | AIL49470.1 | P33469 |

| | | | | | | |
|---|---|---|---|---|---|---|
| RDMQFDLYVNVY | 1 | 0 | 0 | 1496953 | AVV62524.1 | K9N7C7 |
| RDNEALSVSTDV | 1 | 0 | 0 | 1496957 | AVV62535.1 | K9N7C7 |
| RDNEALTVSTDV | 1 | 0 | 0 | 1496958 | AVV62524.1 | K9N7C7 |
| RDNFLSSFFGLH | 1 | 0 | 0 | 1496960 | AVV62535.1 | K9N7C7 |
| RDNGSLEKFMST | 1 | 0 | 0 | 1496963 | AXN92247.1 | K9N7C7 |
| RDNIGTNFPLQI | 1 | 0 | 0 | 1496965 | ARB07436.1 | A0A140H1G9 |
| RDPILISLQPFV | 1 | 0 | 0 | 1496975 | AWK59942.1 | H9EJ42 |
| RDPQTLDILDIT | 1 | 0 | 0 | 1496979 | ATO98145.1 | Q3LZX1 |
| RDPQTLEILDIS | 1 | 0 | 0 | 1496980 | Q3I5J5.1 | Q3LZX1 |
| RDPRNDEAIPSM | 1 | 0 | 0 | 1496983 | AJQ24068.1 | P33469 |
| RDPSGLPPAYAP | 1 | 0 | 0 | 1496985 | AVV62535.1 | K9N7C7 |
| RDPSGLSPAYAP | 1 | 0 | 0 | 1496986 | AVV62524.1 | K9N7C7 |
| RDPSNDEAIPTK | 1 | 0 | 0 | 1496987 | AJQ24067.1 | P33469 |
| RDPSNDEAIPTR | 1 | 0 | 0 | 1496988 | AJQ24082.1 | P33469 |
| RDPTIQEAIPTR | 1 | 0 | 0 | 1496989 | AMM42397.1 | Q19U29 |
| RDPVGFCLSNTP | 1 | 0 | 0 | 1496991 | AXN92247.1 | K9N7C7 |
| RDPVPLHIIAPS | 1 | 0 | 0 | 1496994 | AVV62530.1 | K9N7D2 |
| RDSAVGHTEFYD | 1 | 0 | 0 | 1497003 | AUM60025.1 | K9N796 |
| RDSDFVKQPVKF | 1 | 0 | 0 | 1497004 | AVN89451.1 | K9N7C7 |
| RDSDVVKQPVNF | 1 | 0 | 0 | 1497006 | AXN73479.1 | K9N7C7 |
| RDSEAIKQPVKF | 1 | 0 | 0 | 1497007 | AVV62524.1 | K9N7C7 |
| RDSSRSTSRQHS | 1 | 0 | 0 | 1497017 | AWK59971.1 | H9EJ69 |
| RDSTSTHDGPTE | 1 | 0 | 0 | 1497019 | QDI73611.1 | K9N796 |
| RDSTYSPLQTPV | 1 | 0 | 0 | 1497024 | QEU56411.1 | K9N5Q8 |
| RDTGHPSISTDT | 1 | 0 | 0 | 1497026 | AIX10759.1 | P36334 |
| RDTQVLLRRRDA | 1 | 0 | 0 | 1497039 | AVV62526.1 | K9N5Q8 |
| RDVNVTFFPDLN | 1 | 0 | 0 | 1497055 | APO40578.1 | P0C6X7 |
| RDWSYSGQNTEL | 1 | 0 | 0 | 1497062 | APO40578.1 | P0C6X7 |
| RDYFDDNTDSIV | 1 | 0 | 0 | 1497065 | AWK59930.1 | H9EJ42 |
| RDYFDNNTDSIV | 1 | 0 | 0 | 1497066 | AWK59942.1 | H9EJ42 |
| RDYLADYDIAVT | 1 | 0 | 0 | 1497080 | AQZ41321.1 | K9N7C7 |
| RDYLADYDIVVT | 1 | 0 | 0 | 1497081 | ALA49901.1 | K9N7C7 |
| RDYLADYDVVVT | 1 | 0 | 0 | 1497083 | AVN89363.1 | K9N7C7 |
| REAACCHLAKAI | 1 | 0 | 0 | 1497089 | AVP78030.1 | P0C6W2 |
| RECNDMHLSTLM | 1 | 0 | 0 | 1497102 | ARO76381.1 | P0C6U8 |
| REELDQWFKNQT | 1 | 0 | 0 | 1497109 | AVQ05264.1 | P36334 |
| REENVMQCFGPR | 1 | 0 | 0 | 1497113 | AMM42619.1 | H9EJ69 |
| REEVDIIEQPFG | 1 | 0 | 0 | 1497117 | AWK59942.1 | H9EJ42 |
| REGGLNKVKYAT | 1 | 0 | 0 | 1497131 | AST12962.1 | A0A0P0LKV0 |
| REGIEQVPPHSF | 1 | 0 | 0 | 1497132 | AVV62535.1 | K9N7C7 |
| REHEFCCSHGDR | 1 | 0 | 0 | 1497144 | ABU39953.1 | A0A140H1G9 |
| REHLKSGTCGIV | 1 | 0 | 0 | 1497150 | P0C6W6.1 | P0C6W2 |
| REIKPAFNVVKA | 1 | 0 | 0 | 1497155 | AIV41792.1 | P0C6X6 |
| REIVMNASPYDL | 1 | 0 | 0 | 1497159 | AGT51737.1 | P0C6X6 |
| REIVMNASPYHL | 1 | 0 | 0 | 1497160 | AEN19363.1 | P0C6X6 |
| REIVMNASPYYL | 1 | 0 | 0 | 1497161 | AWW13571.1 | P0C6X6 |
| REKLEKLISTAR | 1 | 0 | 0 | 1497164 | ALD51903.1 | K9N7C7 |
| RELNCDVCPNRA | 1 | 0 | 0 | 1497176 | ARB07436.1 | A0A140H1G9 |
| RELTRELNGGVV | 1 | 0 | 0 | 1497186 | ARO76381.1 | P0C6U8 |
| RELTSTHDGPTE | 1 | 0 | 0 | 1497187 | AVN89314.1 | K9N796 |
| RELYDAPTTLQA | 1 | 0 | 0 | 1497188 | AVV62535.1 | K9N7C7 |
| REMQQQPLLNPS | 1 | 0 | 0 | 1497200 | AIW52723.1 | A0A0N7J131 |
| RENKSAELNYTV | 1 | 0 | 0 | 1497201 | BAL45640.1 | A0A0P0K6L9 |
| REPMMQSADAST | 1 | 0 | 0 | 1497211 | P0C6W6.1 | P0C6W2 |
| REPMMQSSDAST | 1 | 0 | 0 | 1497212 | APO40578.1 | P0C6X7 |
| REPVHLLSADAI | 1 | 0 | 0 | 1497213 | ARU07575.1 | A0A140H1G9 |
| RERGLNKVKYAT | 1 | 0 | 0 | 1497220 | NP_073549.1 | A0A0P0LKV0 |
| RESTSTHDDFTE | 1 | 0 | 0 | 1497238 | AJG44103.1 | K9N796 |
| RESTSTHDGFNE | 1 | 0 | 0 | 1497239 | AHI48551.1 | K9N796 |
| RESTSTHDVPTE | 1 | 0 | 0 | 1497248 | AMO03402.1 | K9N796 |
| REVALPQSKDSN | 1 | 0 | 0 | 1497260 | AVV62524.1 | K9N7C7 |
| REWCNVCGIKDV | 1 | 0 | 0 | 1497277 | AXN92247.1 | K9N7C7 |
| REWCNVCGTKDV | 1 | 0 | 0 | 1497278 | AVV62524.1 | K9N7C7 |
| REWCNVCGTRDV | 1 | 0 | 0 | 1497279 | AUM60013.1 | K9N7C7 |
| RFCLLLKSFIPL | 1 | 0 | 0 | 1497302 | AVV62541.1 | K9N7D2 |
| RFDENGDKLDQF | 1 | 0 | 0 | 1497308 | AIX10734.1 | P0C6X6 |
| RFEYISPTDKLA | 1 | 0 | 0 | 1497326 | AGT51578.1 | Q80872 |
| RFEYNPPTDKLA | 1 | 0 | 0 | 1497327 | NP_150075.1 | Q80872 |
| RFGAISASLQEI | 1 | 0 | 0 | 1497344 | CAA79896.1 | P36334 |
| RFGFIEDAVFKP | 1 | 0 | 0 | 1497347 | QEG03794.1 | P36334 |
| RFGFIEDSAFVP | 1 | 0 | 0 | 1497348 | AGT51481.1 | P36334 |
| RFGFIEDSVFVP | 1 | 0 | 0 | 1497350 | AEN19366.1 | P36334 |
| RFGPFEFCSEIV | 1 | 0 | 0 | 1497355 | APT69868.1 | A0A0P0ILY9 |
| RFKCSSNGSNDV | 1 | 0 | 0 | 1497375 | AGT17757.1 | A0A140H1H0 |
| RFLSDDRTEVPQ | 1 | 0 | 0 | 1497390 | 6C6Z_A | K9N5Q8 |
| RFMNTPNAILIS | 1 | 0 | 0 | 1497395 | AUM60013.1 | K9N7C7 |
| RFMVDPTTVLNG | 1 | 0 | 0 | 1497398 | ABE97136.1 | H9EJ42 |
| RFPNITNRCPFD | 1 | 0 | 0 | 1497423 | ATO98181.1 | Q3LZX1 |
| RFPPGTVLPQGY | 1 | 0 | 0 | 1497428 | AEN19370.1 | P33469 |
| RFQRFDENGDKL | 1 | 0 | 0 | 1497436 | AIX10734.1 | P0C6X6 |
| RFQRVGENGDKL | 1 | 0 | 0 | 1497438 | ATN39877.2 | P0C6X6 |
| RFSKVSLVSVAD | 1 | 0 | 0 | 1497461 | ATQ39397.1 | K9N7D2 |
| RFSPLKLEDIME | 1 | 0 | 0 | 1497468 | AUM60027.1 | K9N643 |
| RFTGPYKCPQTK | 1 | 0 | 0 | 1497477 | AGT51422.1 | P36334 |
| RFVEKDDQGHIL | 1 | 0 | 0 | 1497488 | AVV62535.1 | K9N7C7 |
| RFVELDDQGHHL | 1 | 0 | 0 | 1497490 | AXN92247.1 | K9N7C7 |
| RFVSKDVKFENL | 1 | 0 | 0 | 1497503 | AUM60014.1 | K9N5Q8 |
| RFVSKDVKFQNL | 1 | 0 | 0 | 1497504 | AUM60024.1 | K9N5Q8 |
| RFVYDGFDNLIG | 1 | 0 | 0 | 1497514 | AVV62537.1 | K9N5Q8 |
| RFVYDSFDNLVG | 1 | 0 | 0 | 1497516 | AUM60014.1 | K9N5Q8 |
| RFYRLANECAQI | 1 | 0 | 0 | 1497528 | AIL49554.1 | P0C6X6 |

| | | | | | | |
|---|---|---|---|---|---|---|
| RGAFALHTGYYD | 1 | 0 | 0 | 1497545 | AKA09616.1 | A0A0P0G321 |
| RGAIGDVGLCMY | 1 | 0 | 0 | 1497547 | ARB07424.1 | A0A0P0ILY9 |
| RGAVSCGHYHTN | 1 | 0 | 0 | 1497556 | AST12962.1 | A0A0P0LKV0 |
| RGDDMSFLEKDA | 1 | 0 | 0 | 1497573 | AFR58685.1 | P0C6X7 |
| RGDKIVYHTSGN | 1 | 0 | 0 | 1497578 | APO40578.1 | P0C6X7 |
| RGDPMPNNGWTA | 1 | 0 | 0 | 1497583 | AVP78031.1 | Q3LZX1 |
| RGDPMSNNGWTA | 1 | 0 | 0 | 1497584 | AVP78042.1 | Q3LZX1 |
| RGEFVEQPQGVE | 1 | 0 | 0 | 1497592 | AUM60014.1 | K9N5Q8 |
| RGFCKTHNWNCF | 1 | 0 | 0 | 1497602 | BAE93400.1 | P0C6U8 |
| RGFFDEGSELTL | 1 | 0 | 0 | 1497604 | AIW52827.1 | H9EJ42 |
| RGGGSVTYTDQY | 1 | 0 | 0 | 1497623 | ARB07390.1 | A0A0P0LKV0 |
| RGGSDVDLGDIS | 1 | 0 | 0 | 1497630 | 1WYY_A | P59594 |
| RGGSYKNDKACP | 1 | 0 | 0 | 1497634 | APO40578.1 | P0C6X7 |
| RGGYHYTPVRYE | 1 | 0 | 0 | 1497645 | ALA50044.1 | K9N7C7 |
| RGIACVKDFDFK | 1 | 0 | 0 | 1497662 | ARE30011.1 | P0C6X6 |
| RGLFHSSQLPLD | 1 | 0 | 0 | 1497684 | AJC98124.1 | P0C6X6 |
| RGLKNGQGENDN | 1 | 0 | 0 | 1497685 | ATP16771.1 | P33469 |
| RGNPNNHKKIVQ | 1 | 0 | 0 | 1497704 | AVV62529.1 | K9N643 |
| RGPAGVESDVDT | 1 | 0 | 0 | 1497719 | AUM60013.1 | K9N7C7 |
| RGPGALQGNFGD | 1 | 0 | 0 | 1497723 | AVV62544.1 | K9N4V7 |
| RGPGKCYNCRPA | 1 | 0 | 0 | 1497726 | AIW52747.1 | A0A0P0K6L9 |
| RGPGKCYNCRPS | 1 | 0 | 0 | 1497727 | AWH62679.1 | A0A0P0K6L9 |
| RGPGRCYNCRPA | 1 | 0 | 0 | 1497728 | AIW52757.1 | A0A0P0K6L9 |
| RGPKNGQGENDN | 1 | 0 | 0 | 1497730 | QEG03805.1 | P33469 |
| RGPLQNFGNFEM | 1 | 0 | 0 | 1497732 | AMM42397.1 | Q19U29 |
| RGPRSGNSSRGS | 1 | 0 | 0 | 1497742 | AVV62544.1 | K9N4V7 |
| RGRESKSQSRNP | 1 | 0 | 0 | 1497755 | ABB90484.1 | A0A0N7J131 |
| RGRKTQPKFTVS | 1 | 0 | 0 | 1497765 | ARU07581.1 | Q19U29 |
| RGSDVKQPVKFY | 1 | 0 | 0 | 1497782 | AUM60013.1 | K9N7C7 |
| RGSFNNRDTGPP | 1 | 0 | 0 | 1497785 | BBA84088.1 | P36334 |
| RGSGHYQTNLYS | 1 | 0 | 0 | 1497791 | AFD98812.1 | H9EJ42 |
| RGSLEWSHPQFE | 1 | 0 | 0 | 1497800 | 5X58_A | P59594 |
| RGSRSGNSSRGS | 1 | 0 | 0 | 1497804 | AUM60021.1 | K9N4V7 |
| RGTAVMSLKDGQ | 1 | 0 | 0 | 1497814 | AVP78041.1 | P0C6W2 |
| RGTAVMSLKDNQ | 1 | 0 | 0 | 1497815 | APO40578.1 | P0C6X7 |
| RGTFPGPSGIGA | 1 | 0 | 0 | 1497818 | ANI69896.1 | K9N4V7 |
| RGTGTDTIDVLL | 1 | 0 | 0 | 1497820 | AGT51735.1 | Q4VID0 |
| RGTGTDTTDILL | 1 | 0 | 0 | 1497821 | AHF54651.1 | Q4VID0 |
| RGTILFKTSYGA | 1 | 0 | 0 | 1497825 | AIW52757.1 | A0A0P0K6L9 |
| RGTTTYKLNVGDYF | 1 | 0 | 0 | 1497847 | P0DTD1.1 | P0DTD1 |
| RGVEIEDFDLED | 1 | 0 | 0 | 1497856 | AVV62524.1 | K9N7C7 |
| RGVKQSPELFDS | 1 | 0 | 0 | 1497867 | ARU07581.1 | Q19U29 |
| RGVKQSPELLDP | 1 | 0 | 0 | 1497868 | ADN03343.1 | Q19U29 |
| RGVKQSPESFDS | 1 | 0 | 0 | 1497869 | AMM42397.1 | Q19U29 |
| RHCDLDTTLYVK | 1 | 0 | 0 | 1497901 | 5I08_A | A0A140H1H1 |
| RHFFWVLAYFSK | 1 | 0 | 0 | 1497922 | AXN92247.1 | K9N7C7 |
| RHKRTSTKSFNM | 1 | 0 | 0 | 1497947 | AXN92267.1 | K9N4V7 |
| RHPLEEDDIVYF | 1 | 0 | 0 | 1497972 | AVV62535.1 | K9N7C7 |
| RHSFRTLKSGEG | 1 | 0 | 0 | 1497988 | NP_073549.1 | A0A0P0LKV0 |
| RHSSLFAQSKEE | 1 | 0 | 0 | 1498000 | ARQ84748.1 | K9N4V0 |
| RHSSLSAQSEEE | 1 | 0 | 0 | 1498004 | AKN24751.1 | K9N4V0 |
| RHTYQLRARAFT | 1 | 0 | 0 | 1498016 | APO40584.1 | P59635 |
| RHVICPADQLSD | 1 | 0 | 0 | 1498019 | QGV13494.1 | K9N7C7 |
| RHVICTPEDMLN | 1 | 0 | 0 | 1498021 | APO40578.1 | P0C6X7 |
| RHVMCPADQLND | 1 | 0 | 0 | 1498024 | AVV62524.1 | K9N7C7 |
| RHVMCPADQLSD | 1 | 0 | 0 | 1498025 | AXN92247.1 | K9N7C7 |
| RHVMVPADQLAD | 1 | 0 | 0 | 1498026 | AUM60013.1 | K9N7C7 |
| RIAIEDLLFDKV | 1 | 0 | 0 | 1498040 | AHL18090.1 | K9N5Q8 |
| RIASCKWTDVDD | 1 | 0 | 0 | 1498045 | AEN19363.1 | P0C6X6 |
| RIDFQDELDEFF | 1 | 0 | 0 | 1498063 | ALJ54517.1 | K9N5Q8 |
| RIDLPSTDSTGT | 1 | 0 | 0 | 1498072 | ATQ39394.1 | K9N796 |
| RIDTTATSCQLY | 1 | 0 | 0 | 1498074 | CAA79896.1 | P36334 |
| RIDVPPAELTGT | 1 | 0 | 0 | 1498076 | ANI69890.1 | K9N796 |
| RIESSSKLWTQC | 1 | 0 | 0 | 1498088 | APO40578.1 | P0C6X7 |
| RIGELPCAIMND | 1 | 0 | 0 | 1498114 | ARB07436.1 | A0A140H1G9 |
| RIGPIKYCYCGN | 1 | 0 | 0 | 1498125 | AFD98819.1 | H9EJ42 |
| RIKFCDGENVSK | 1 | 0 | 0 | 1498138 | AWK59942.1 | H9EJ42 |
| RILGAGCFVNDL | 1 | 0 | 0 | 1498151 | ANZ78855.1 | P0C6X6 |
| RILGVNSSNCNN | 1 | 0 | 0 | 1498152 | AUT76985.1 | A0A0P0LKV0 |
| RINWATGGIAIA | 1 | 0 | 0 | 1498172 | APO40582.1 | P59596 |
| RIPDLPDCDIDK | 1 | 0 | 0 | 1498175 | AET99191.1 | A0A140H1H1 |
| RIPNDFSFNNWF | 1 | 0 | 0 | 1498180 | ABE97137.1 | A0A0P0G321 |
| RIPNLPDCDIDN | 1 | 0 | 0 | 1498183 | BBA20986.1 | A0A140H1H1 |
| RIPTIADFVQIE | 1 | 0 | 0 | 1498185 | AIW52757.1 | A0A0P0K6L9 |
| RIQSRNPSSDRN | 1 | 0 | 0 | 1498190 | QEO75989.1 | A0A0N7J131 |
| RIRTRPSVQPGP | 1 | 0 | 0 | 1498208 | AKI29262.1 | K9N4V7 |
| RISDGNQRQLLP | 1 | 0 | 0 | 1498211 | AJQ24067.1 | P33469 |
| RISEYVVDVSHG | 1 | 0 | 0 | 1498215 | ARU07577.1 | A0A140H1H1 |
| RITFGGPTDSAD | 1 | 0 | 0 | 1498229 | ARO76389.1 | P59595 |
| RITPYCHDPTVL | 1 | 0 | 0 | 1498234 | AHL18091.1 | K9N7C7 |
| RITRTLSTQYCR | 1 | 0 | 0 | 1498235 | AXN92247.1 | K9N7C7 |
| RIVCANEKFADG | 1 | 0 | 0 | 1498244 | APO40578.1 | P0C6X7 |
| RIVHCTKLNVPF | 1 | 0 | 0 | 1498249 | ARU07575.1 | A0A140H1G9 |
| RIYPQQMFVTPL | 1 | 0 | 0 | 1498273 | ALA49355.1 | K9N643 |
| RKACQNLDCNCL | 1 | 0 | 0 | 1498275 | AWW13564.1 | Q80872 |
| RKAEDIFCPCWS | 1 | 0 | 0 | 1498276 | AST12962.1 | A0A0P0LKV0 |
| RKARNRSHSPTK | 1 | 0 | 0 | 1498294 | ATG84704.1 | K9N643 |
| RKASGLFDAIWDAFVAAI | 1 | 0 | 0 | 1498296 | QED88024.1 | H9EJ42 |
| RKATKRSHSPTK | 1 | 0 | 0 | 1498297 | AHC74084.1 | K9N643 |
| RKCNLPFRLTTS | 1 | 0 | 0 | 1498307 | AUM60013.1 | K9N7C7 |
| RKCVRSREETGL | 1 | 0 | 0 | 1498310 | AVP78041.1 | P0C6W2 |

| | | | | | | |
|---|---|---|---|---|---|---|
| RKDFTPVQVVDS | 1 | 0 | 0 | 1498311 | AEN19365.1 | P30215 |
| RKEIFVGDSLVN | 1 | 0 | 0 | 1498322 | ARU07574.1 | A0A140H1G9 |
| RKELWHLDTGVV | 1 | 0 | 0 | 1498325 | CAA79896.1 | P36334 |
| RKFIKFGMTFVS | 1 | 0 | 0 | 1498330 | QBP84710.1 | P0C6X6 |
| RKFYPAFMLGSS | 1 | 0 | 0 | 1498337 | ALA49660.1 | K9N5Q8 |
| RKGKRVDLSPKL | 1 | 0 | 0 | 1498352 | AIW52723.1 | A0A0N7J131 |
| RKGSFNNKDTGP | 1 | 0 | 0 | 1498354 | AKA09663.1 | P36334 |
| RKHAAEYREYME | 1 | 0 | 0 | 1498358 | AVV62535.1 | K9N7C7 |
| RKHATDYREYME | 1 | 0 | 0 | 1498361 | ASU91085.1 | K9N7C7 |
| RKINTGNGIKQL | 1 | 0 | 0 | 1498386 | AXN92267.1 | K9N4V7 |
| RKINTGNGTKPL | 1 | 0 | 0 | 1498388 | AVV62544.1 | K9N4V7 |
| RKINTGNGTKQL | 1 | 0 | 0 | 1498389 | AUM60021.1 | K9N4V7 |
| RKLLGLTNLNEI | 1 | 0 | 0 | 1498421 | ARU07582.1 | Q0ZJ80 |
| RKLNLPNCNIEA | 1 | 0 | 0 | 1498427 | AAR01015.1 | P36334 |
| RKMAFPSAKVEG | 1 | 0 | 0 | 1498430 | 2PWX_A | P0C6X7 |
| RKMDEPFTEDTS | 1 | 0 | 0 | 1498435 | AHA80151.1 | P33469 |
| RKRGVKQSPESF | 1 | 0 | 0 | 1498476 | AMM42397.1 | Q19U29 |
| RKRTHSPTKKLR | 1 | 0 | 0 | 1498489 | AMD02849.1 | K9N643 |
| RKSASEASKKPR | 1 | 0 | 0 | 1498495 | APO40586.1 | P59595 |
| RKSCQNLDCNCL | 1 | 0 | 0 | 1498497 | AGT51688.1 | Q80872 |
| RKSCSYDSDVDT | 1 | 0 | 0 | 1498499 | AIV41810.1 | P0C6X6 |
| RKSNVDESFVDD | 1 | 0 | 0 | 1498508 | AIG96641.1 | A0A0P0LKV0 |
| RKTKLKPFERDL | 1 | 0 | 0 | 1498515 | ATO98181.1 | Q3LZX1 |
| RKTLVLNASEIY | 1 | 0 | 0 | 1498518 | AXN92231.1 | K9N643 |
| RKTLVLNASEMY | 1 | 0 | 0 | 1498519 | ATQ39396.1 | K9N643 |
| RKTNPHKHCNVQ | 1 | 0 | 0 | 1498520 | AJQ24083.1 | P33469 |
| RKTQPKFTVSTQ | 1 | 0 | 0 | 1498522 | ARU07581.1 | Q19U29 |
| RKVQDAVNNNAH | 1 | 0 | 0 | 1498540 | ATG84679.1 | K9N5Q8 |
| RKVQQLFYIEDM | 1 | 0 | 0 | 1498545 | ATG84699.1 | K9N7C7 |
| RLANELAQVLTE | 1 | 0 | 0 | 1498584 | AFK23434.1 | A0A0P0LKV0 |
| RLASIAFNHPIQ | 1 | 0 | 0 | 1498592 | AXN92228.1 | K9N5Q8 |
| RLCATCPFAKGD | 1 | 0 | 0 | 1498596 | AHI48793.1 | K9N7C7 |
| RLCVQCAMGCNT | 1 | 0 | 0 | 1498605 | AVV62542.1 | K9N5R3 |
| RLDKLIATAKDG | 1 | 0 | 0 | 1498622 | AVV62524.1 | K9N7C7 |
| RLDMSSVDNILN | 1 | 0 | 0 | 1498624 | AST12962.1 | A0A0P0LKV0 |
| RLDSTYGPLQTP | 1 | 0 | 0 | 1498630 | AXG21654.1 | K9N5Q8 |
| RLDVLEAQVQID | 1 | 0 | 0 | 1498633 | ABD75553.1 | A0A140H1H1 |
| RLEDLEAGKNKA | 1 | 0 | 0 | 1498637 | AVV62533.1 | K9N4V7 |
| RLEESGAVLFCT | 1 | 0 | 0 | 1498638 | NP_073549.1 | A0A0P0LKV0 |
| RLEIVKLGAEKE | 1 | 0 | 0 | 1498641 | AMK59675.1 | Q80872 |
| RLEIVKLGAENK | 1 | 0 | 0 | 1498643 | AIV41955.1 | Q80872 |
| RLELVKLGAEKK | 1 | 0 | 0 | 1498646 | AWW13564.1 | Q80872 |
| RLEPCNGTDIDY | 1 | 0 | 0 | 1498648 | AIW52776.1 | A0A0P0LKV0 |
| RLFASDLSTLAV | 1 | 0 | 0 | 1498653 | AWK59942.1 | H9EJ42 |
| RLFATKVQQQQD | 1 | 0 | 0 | 1498654 | APO40584.1 | P59635 |
| RLFRRTGSWWSF | 1 | 0 | 0 | 1498668 | AVV62543.1 | K9N7A1 |
| RLFSVPSDSCNN | 1 | 0 | 0 | 1498669 | AFD98777.1 | H9EJ42 |
| RLGASQRVGTDS | 1 | 0 | 0 | 1498674 | APO40582.1 | P59596 |
| RLGNEWAQVLSE | 1 | 0 | 0 | 1498678 | AKL87224.1 | P0C6X6 |
| RLGSQLSLSMVR | 1 | 0 | 0 | 1498681 | Q3I5I6.1 | Q3LZX3 |
| RLHCTTNLFKDC | 1 | 0 | 0 | 1498685 | ARU07574.1 | A0A140H1G9 |
| RLHTVESGDVNV | 1 | 0 | 0 | 1498691 | ASU45805.1 | K9N7C7 |
| RLIFPNYRQWHC | 1 | 0 | 0 | 1498697 | AUM60017.1 | K9N643 |
| RLIKDVDDPVLM | 1 | 0 | 0 | 1498704 | AJS10811.1 | P0C6X6 |
| RLIMRCWLCWKC | 1 | 0 | 0 | 1498708 | ARO76383.1 | P59632 |
| RLINGRLTALDA | 1 | 0 | 0 | 1498709 | CAA83661.1 | P36334 |
| RLIPTVAKMTKL | 1 | 0 | 0 | 1498711 | APO40587.1 | P59636 |
| RLIQFDNPNIPF | 1 | 0 | 0 | 1498712 | APO40579.1 | P59594 |
| RLKGSFNDRDTG | 1 | 0 | 0 | 1498732 | AIL49520.1 | P36334 |
| RLKGSFNNRDTG | 1 | 0 | 0 | 1498733 | AEN19366.1 | P36334 |
| RLKKSNLLIQEF | 1 | 0 | 0 | 1498734 | ARB07605.1 | A0A140H1G9 |
| RLKKSNLVIQEF | 1 | 0 | 0 | 1498735 | ARB07436.1 | A0A140H1G9 |
| RLLEDKVEALMS | 1 | 0 | 0 | 1498751 | AVV62524.1 | K9N7C7 |
| RLLEDKVESLMS | 1 | 0 | 0 | 1498752 | AVV62535.1 | K9N7C7 |
| RLLLRNSGRFGG | 1 | 0 | 0 | 1498771 | AVL25576.1 | H9EJ42 |
| RLLQDKAGDFMS | 1 | 0 | 0 | 1498776 | AQZ41321.1 | K9N7C7 |
| RLLQDKIGDFMS | 1 | 0 | 0 | 1498777 | AXG21664.1 | K9N638 |
| RLLQDKTGDFTS | 1 | 0 | 0 | 1498780 | AJO62148.1 | K9N7C7 |
| RLLSDGRTEVPQ | 1 | 0 | 0 | 1498784 | ANC28678.1 | K9N5Q8 |
| RLNQLENKVSGK | 1 | 0 | 0 | 1498810 | AVP78038.1 | Q3LZX4 |
| RLPSSKSSGMDT | 1 | 0 | 0 | 1498819 | AZS52621.1 | Q0ZJ82 |
| RLQAGFNTEFAS | 1 | 0 | 0 | 1498825 | ANI69888.1 | K9N638 |
| RLQASADWKPGY | 1 | 0 | 0 | 1498833 | ALA49934.1 | K9N7C7 |
| RLQASNDWKPGL | 1 | 0 | 0 | 1498834 | AUM60013.1 | K9N7C7 |
| RLQEAIKVLNQS | 1 | 0 | 0 | 1498836 | CAA79896.1 | P36334 |
| RLQNGDFLCANE | 1 | 0 | 0 | 1498837 | APO40578.1 | P0C6X7 |
| RLQTGKQTELAV | 1 | 0 | 0 | 1498843 | AAW33902.1 | H9EJ42 |
| RLSDVGFVEAYN | 1 | 0 | 0 | 1498870 | CAA83661.1 | P36334 |
| RLSPVDEFIKHV | 1 | 0 | 0 | 1498882 | AVV62535.1 | K9N7C7 |
| RLSVCLLMFLLT | 1 | 0 | 0 | 1498890 | AVV62537.1 | K9N5Q8 |
| RLTGDNQHIETP | 1 | 0 | 0 | 1498901 | AVV62526.1 | K9N5Q8 |
| RLTTLNAFVAQQ | 1 | 0 | 0 | 1498918 | AXN92228.1 | K9N5Q8 |
| RLTTSKLRANDN | 1 | 0 | 0 | 1498922 | AXN92247.1 | K9N7C7 |
| RLVASGSTVAMT | 1 | 0 | 0 | 1498925 | AXG21654.1 | K9N5Q8 |
| RLVEYGSFCDNI | 1 | 0 | 0 | 1498932 | ARB66066.1 | P36334 |
| RLVHTDSFSCNN | 1 | 0 | 0 | 1498937 | AET99191.1 | A0A140H1H1 |
| RLYKRNFTYDVN | 1 | 0 | 0 | 1498972 | AHA80190.1 | P36334 |
| RLYSPSLSNLAL | 1 | 0 | 0 | 1498986 | AVV62535.1 | K9N7C7 |
| RMAGNGGDTALA | 1 | 0 | 0 | 1498998 | AVP78038.1 | Q3LZX4 |
| RMEDAMGQGQNN | 1 | 0 | 0 | 1499016 | ARO76391.1 | P59636 |
| RMGFKLDAIVPG | 1 | 0 | 0 | 1499022 | QBM11746.1 | K9N7C7 |

| | | | | | | |
|---|---|---|---|---|---|---|
| RMGFKLDASIPG | 1 | 0 | 0 | 1499026 | AUM60013.1 | K9N7C7 |
| RMGFKLDATIPG | 1 | 0 | 0 | 1499027 | AVN89352.1 | K9N7C7 |
| RMLLEKCDLHNY | 1 | 0 | 0 | 1499041 | APO40578.1 | P0C6X7 |
| RMQIGSVFLNGA | 1 | 0 | 0 | 1499053 | AUM60013.1 | K9N7C7 |
| RMTAKPIALELV | 1 | 0 | 0 | 1499060 | AVV62529.1 | K9N643 |
| RMTPYCHDPTVL | 1 | 0 | 0 | 1499061 | AXN92247.1 | K9N7C7 |
| RMVWREWCNVCG | 1 | 0 | 0 | 1499066 | AXN92247.1 | K9N7C7 |
| RMYCSFYPPDDE | 1 | 0 | 0 | 1499067 | ATO98229.1 | P0C6W2 |
| RNANGACIWNAA | 1 | 0 | 0 | 1499080 | AVV62524.1 | K9N7C7 |
| RNANGACVWNAA | 1 | 0 | 0 | 1499082 | AVV62535.1 | K9N7C7 |
| RNASPGPSGIGA | 1 | 0 | 0 | 1499088 | AVV62533.1 | K9N4V7 |
| RNAWAAFYVYKL | 1 | 0 | 0 | 1499089 | AUM60014.1 | K9N5Q8 |
| RNAYVYTSQGFV | 1 | 0 | 0 | 1499090 | AXN92247.1 | K9N7C7 |
| RNDEAIPSMFPP | 1 | 0 | 0 | 1499096 | AJQ24068.1 | P33469 |
| RNFTYDVNANYL | 1 | 0 | 0 | 1499113 | AKA09658.1 | P36334 |
| RNFTYDVNATYL | 1 | 0 | 0 | 1499114 | AEN19366.1 | P36334 |
| RNFYEPKIITTD | 1 | 0 | 0 | 1499116 | AVP78031.1 | Q3LZX1 |
| RNFYSPQIITTD | 1 | 0 | 0 | 1499119 | ATO98181.1 | Q3LZX1 |
| RNFYSPQVITTD | 1 | 0 | 0 | 1499121 | APO40579.1 | P59594 |
| RNGQGENDNISV | 1 | 0 | 0 | 1499126 | ANZ78875.1 | P33469 |
| RNGSRCPNDNAY | 1 | 0 | 0 | 1499127 | AUM60014.1 | K9N5Q8 |
| RNGSYCPGNSNY | 1 | 0 | 0 | 1499128 | AVV62537.1 | K9N5Q8 |
| RNGTRCPNDNAY | 1 | 0 | 0 | 1499129 | AVV62526.1 | K9N5Q8 |
| RNKWNGKSSLSL | 1 | 0 | 0 | 1499159 | AUM60013.1 | K9N7C7 |
| RNKWSGVSDLSL | 1 | 0 | 0 | 1499162 | AXN92247.1 | K9N7C7 |
| RNKWSGVSDLSS | 1 | 0 | 0 | 1499163 | QGV13494.1 | K9N7C7 |
| RNKWSGVSNLSL | 1 | 0 | 0 | 1499164 | QAT98908.1 | K9N7C7 |
| RNLESFKPRSQM | 1 | 0 | 0 | 1499166 | APO40578.1 | P0C6X7 |
| RNLQAGGDTALA | 1 | 0 | 0 | 1499177 | APO40586.1 | P59595 |
| RNLQKRLYPNVY | 1 | 0 | 0 | 1499180 | ACR82756.1 | A0A140H1G9 |
| RNLSGNPDEPQK | 1 | 0 | 0 | 1499185 | AHA80174.1 | P33469 |
| RNLTDRVVFVLW | 1 | 0 | 0 | 1499187 | APO40578.1 | P0C6X7 |
| RNMALKGTDKSS | 1 | 0 | 0 | 1499196 | AKA09652.1 | P36334 |
| RNNGCFGGFSNH | 1 | 0 | 0 | 1499202 | AGT51393.1 | H9EJ42 |
| RNNSRDSFRSTS | 1 | 0 | 0 | 1499206 | ABE97134.1 | H9EJ69 |
| RNNTSCPEWMDD | 1 | 0 | 0 | 1499208 | AVN89451.1 | K9N7C7 |
| RNPNNDAAIVTQ | 1 | 0 | 0 | 1499219 | AVV62533.1 | K9N4V7 |
| RNPNNDLAIVTQ | 1 | 0 | 0 | 1499220 | AKN24774.1 | K9N4V7 |
| RNPNNDPAIVTQ | 1 | 0 | 0 | 1499221 | AUM60021.1 | K9N4V7 |
| RNPSIDEAFPTR | 1 | 0 | 0 | 1499227 | AJQ24074.1 | P33469 |
| RNPVDLPPSHAT | 1 | 0 | 0 | 1499231 | QEG03783.1 | A0A0P0LKV0 |
| RNQHSKLDEEVP | 1 | 0 | 0 | 1499235 | APO40583.1 | P59634 |
| RNQSPASSQSAA | 1 | 0 | 0 | 1499236 | AIW52723.1 | A0A0N7J131 |
| RNQSPASSQSVA | 1 | 0 | 0 | 1499237 | ABB90499.1 | A0A0N7J131 |
| RNQSPASSQTSA | 1 | 0 | 0 | 1499238 | NP_073556.1 | A0A0N7J131 |
| RNQSPSSSQSAA | 1 | 0 | 0 | 1499240 | BAP75906.1 | A0A0N7J131 |
| RNQSPVSSQSAA | 1 | 0 | 0 | 1499241 | AMM42518.1 | A0A0N7J131 |
| RNSLDPFCFDVN | 1 | 0 | 0 | 1499259 | AWK59942.1 | H9EJ42 |
| RNSNGACIWNAA | 1 | 0 | 0 | 1499262 | AXN92247.1 | K9N7C7 |
| RNSNGVCIWNAA | 1 | 0 | 0 | 1499263 | ATG84677.1 | K9N7C7 |
| RNSSDNGISAIV | 1 | 0 | 0 | 1499269 | AVL25577.1 | A0A0P0G321 |
| RNSSSEFFNKQW | 1 | 0 | 0 | 1499275 | AVV62537.1 | K9N5Q8 |
| RNSTTVSVTVSS | 1 | 0 | 0 | 1499282 | AVN89322.1 | K9N7C7 |
| RNSTVWNGGAYS | 1 | 0 | 0 | 1499284 | AEN19363.1 | P0C6X6 |
| RNSVTGRLIQFD | 1 | 0 | 0 | 1499288 | APO40579.1 | P59594 |
| RNSWEAMALNFT | 1 | 0 | 0 | 1499291 | ALA49417.1 | K9N7C7 |
| RNVDGHFKIYSK | 1 | 0 | 0 | 1499302 | AVP78031.1 | Q3LZX1 |
| RNVSYDSVSAIV | 1 | 0 | 0 | 1499322 | AIW52757.1 | A0A0P0K6L9 |
| RNYIITMPCFRT | 1 | 0 | 0 | 1499338 | AXN92231.1 | K9N643 |
| RPAGTNSLTCDN | 1 | 0 | 0 | 1499342 | ARB66066.1 | P36334 |
| RPAIVQDEELAL | 1 | 0 | 0 | 1499344 | AVV62543.1 | K9N7A1 |
| RPAPNNSVSWYT | 1 | 0 | 0 | 1499346 | AVV62544.1 | K9N4V7 |
| RPAPNTSVSWYT | 1 | 0 | 0 | 1499348 | AVV62533.1 | K9N4V7 |
| RPAPNTTVSWYT | 1 | 0 | 0 | 1499349 | AUM60021.1 | K9N4V7 |
| RPAVINITLANF | 1 | 0 | 0 | 1499351 | AIW52757.1 | A0A0P0K6L9 |
| RPDARFHDFTPL | 1 | 0 | 0 | 1499357 | AVV62529.1 | K9N643 |
| RPDLPNCNIEAW | 1 | 0 | 0 | 1499361 | AHA80225.1 | P36334 |
| RPEDLSVIQPTH | 1 | 0 | 0 | 1499367 | QBF80458.1 | K9N643 |
| RPEDLVQSTHYV | 1 | 0 | 0 | 1499371 | AVN89383.1 | K9N643 |
| RPGNGILKWADQ | 1 | 0 | 0 | 1499385 | AMM42456.1 | P33469 |
| RPGVYCGHDYMD | 1 | 0 | 0 | 1499387 | AUM60013.1 | K9N7C7 |
| RPGVYCGSDYFD | 1 | 0 | 0 | 1499389 | AVV62524.1 | K9N7C7 |
| RPGVYCGSDYMD | 1 | 0 | 0 | 1499390 | AVV62535.1 | K9N7C7 |
| RPIDPAAADGII | 1 | 0 | 0 | 1499397 | AVV62526.1 | K9N5Q8 |
| RPIDVFKADGII | 1 | 0 | 0 | 1499398 | AHI48583.1 | K9N5Q8 |
| RPIQLVPAFPET | 1 | 0 | 0 | 1499413 | AVV62530.1 | K9N7D2 |
| RPIVYETDANKL | 1 | 0 | 0 | 1499416 | AQN78717.1 | P0C6X6 |
| RPLIKDVDNPVL | 1 | 0 | 0 | 1499424 | AHF54667.1 | P0C6X6 |
| RPLLVDDNKFDV | 1 | 0 | 0 | 1499427 | AEN19363.1 | P0C6X6 |
| RPLNLVPAFAEE | 1 | 0 | 0 | 1499431 | AUM60018.1 | K9N7D2 |
| RPNFTIKGSFLN | 1 | 0 | 0 | 1499435 | AVP78041.1 | P0C6W2 |
| RPNYTIKGSFLN | 1 | 0 | 0 | 1499441 | AEA10817.1 | P0C6X7 |
| RPPKNSFEALIL | 1 | 0 | 0 | 1499442 | AGT51747.1 | P0C6X6 |
| RPPPGEHFKHLV | 1 | 0 | 0 | 1499447 | ALW82707.1 | K9N7C7 |
| RPQGLPNNIASW | 1 | 0 | 0 | 1499457 | AAP49024.1 | P59595 |
| RPQGPPNNTASW | 1 | 0 | 0 | 1499459 | APO40586.1 | P59595 |
| RPSLVDDNKFDG | 1 | 0 | 0 | 1499470 | ATP16754.1 | P0C6X6 |
| RPSVQPGPMIDV | 1 | 0 | 0 | 1499475 | AXN92267.1 | K9N4V7 |
| RPSYKYSLITRE | 1 | 0 | 0 | 1499476 | Q14EA9.1 | Q0ZJ84 |
| RPTDNSTTVSVT | 1 | 0 | 0 | 1499478 | ANI69823.1 | K9N638 |
| RPVIWLSHEKAS | 1 | 0 | 0 | 1499491 | AQN78773.1 | P0C6X6 |

| | | | | | | |
|---|---|---|---|---|---|---|
| RPVNPTDQSSYT | 1 | 0 | 0 | 1499494 | APO40578.1 | P0C6X7 |
| RPVPGTSTHPVQ | 1 | 0 | 0 | 1499495 | AUM60016.1 | K9N4V0 |
| RPYHANSSEPAM | 1 | 0 | 0 | 1499499 | AIV41873.1 | P36334 |
| RQAALPQSKESN | 1 | 0 | 0 | 1499502 | AQZ41321.1 | K9N7C7 |
| RQDYGTNSGVAI | 1 | 0 | 0 | 1499527 | AVV62543.1 | K9N7A1 |
| RQEEVQQELYSR | 1 | 0 | 0 | 1499532 | 1YO4_A | P59635 |
| RQFDNMTAVACQ | 1 | 0 | 0 | 1499533 | AXX83326.1 | P30215 |
| RQGIFVEVNATY | 1 | 0 | 0 | 1499537 | BBA20982.1 | P36334 |
| RQGTFLCANEYT | 1 | 0 | 0 | 1499547 | ARO76381.1 | P0C6U8 |
| RQIAPAQTGVIA | 1 | 0 | 0 | 1499557 | APO40579.1 | P59594 |
| RQIAWQLHRQGK | 1 | 0 | 0 | 1499560 | AVV62528.1 | K9N4V0 |
| RQIFDPAPVELE | 1 | 0 | 0 | 1499561 | AVV62524.1 | K9N7C7 |
| RQIFDVPPIELE | 1 | 0 | 0 | 1499565 | AKJ80136.2 | K9N7C7 |
| RQIHQAQLTDKY | 1 | 0 | 0 | 1499566 | AVV62524.1 | K9N7C7 |
| RQIYDLAPIELD | 1 | 0 | 0 | 1499574 | AUM60013.1 | K9N7C7 |
| RQKKQHTVTLLP | 1 | 0 | 0 | 1499579 | ARO76389.1 | P59595 |
| RQKRTPNKFCNV | 1 | 0 | 0 | 1499589 | AMM42397.1 | Q19U29 |
| RQLMGSCSFEDE | 1 | 0 | 0 | 1499604 | AIV41816.1 | P0C6X6 |
| RQLRPINYSFDS | 1 | 0 | 0 | 1499610 | AIL49543.1 | P36334 |
| RQPFSFSFRDEL | 1 | 0 | 0 | 1499622 | ARE29978.1 | H9EJ42 |
| RQQAEEKKIKAS | 1 | 0 | 0 | 1499626 | AVP78041.1 | P0C6W2 |
| RQQEGHIIMEEM | 1 | 0 | 0 | 1499627 | 5YVD_A | K9N7C7 |
| RQQTSNLVAQEF | 1 | 0 | 0 | 1499635 | APU51933.1 | P0C6X6 |
| RQRGLKNGQGEN | 1 | 0 | 0 | 1499646 | ATP16771.1 | P33469 |
| RQRGPKNGQGEN | 1 | 0 | 0 | 1499648 | QEG03805.1 | P33469 |
| RQRLTEYTMMDL | 1 | 0 | 0 | 1499652 | AXN92247.1 | K9N7C7 |
| RQSSKIENASPA | 1 | 0 | 0 | 1499661 | ARO76383.1 | P59632 |
| RQVAPGETGVIA | 1 | 0 | 0 | 1499674 | ATO98181.1 | Q3LZX1 |
| RQVIQEVKDGTV | 1 | 0 | 0 | 1499680 | ABM64804.1 | A0A0P0KWE8 |
| RQVSLTQSKDTN | 1 | 0 | 0 | 1499685 | AUM60013.1 | K9N7C7 |
| RRAGAYSSPLQS | 1 | 0 | 0 | 1499717 | AVV62535.1 | K9N7C7 |
| RRAITTENRFTN | 1 | 0 | 0 | 1499721 | AIL49500.1 | P36334 |
| RRAQPKQTVTSQ | 1 | 0 | 0 | 1499729 | AIL49418.1 | P33469 |
| RRAQPNQTATSQ | 1 | 0 | 0 | 1499730 | AIX10721.1 | P33469 |
| RRATCFSTSADT | 1 | 0 | 0 | 1499737 | APO40578.1 | P0C6X7 |
| RRAVDCGHDDLS | 1 | 0 | 0 | 1499743 | AVV62537.1 | K9N5Q8 |
| RRDGQPCYEQFP | 1 | 0 | 0 | 1499750 | AVN89451.1 | K9N7C7 |
| RRDGQPFYEPFP | 1 | 0 | 0 | 1499752 | ALD51903.1 | K9N7C7 |
| RRFLVKLLDVYN | 1 | 0 | 0 | 1499767 | ABE97136.1 | H9EJ42 |
| RRFSPLKPEDLM | 1 | 0 | 0 | 1499775 | AVV62529.1 | K9N643 |
| RRGPGDLQGNFG | 1 | 0 | 0 | 1499791 | AVV62533.1 | K9N4V7 |
| RRGQPVDLPPKV | 1 | 0 | 0 | 1499795 | AMM42619.1 | H9EJ69 |
| RRGVYYNDDIFR | 1 | 0 | 0 | 1499801 | ATO98181.1 | Q3LZX1 |
| RRIPDLPDCDID | 1 | 0 | 0 | 1499813 | AET99191.1 | A0A140H1H1 |
| RRKPDPPNCNIE | 1 | 0 | 0 | 1499822 | AHA80201.1 | P36334 |
| RRLKKSNLVIQE | 1 | 0 | 0 | 1499842 | ARB07436.1 | A0A140H1G9 |
| RRLSKERFTHWF | 1 | 0 | 0 | 1499849 | QED88010.1 | H9EJ42 |
| RRLVNPTDKSHY | 1 | 0 | 0 | 1499853 | ATQ39389.1 | K9N7C7 |
| RRQYLLNFDEHG | 1 | 0 | 0 | 1499879 | BBA20986.1 | A0A140H1H1 |
| RRRPDLPNCNIE | 1 | 0 | 0 | 1499886 | AHA80225.1 | P36334 |
| RRSICRTLVLYL | 1 | 0 | 0 | 1499905 | ATQ39397.1 | K9N7D2 |
| RRSIDCGFNDLS | 1 | 0 | 0 | 1499906 | AHZ90568.1 | K9N5Q8 |
| RRTSPDPKFVDK | 1 | 0 | 0 | 1499926 | AVV62524.1 | K9N7C7 |
| RRYSEAASNGFR | 1 | 0 | 0 | 1499955 | NP_073549.1 | A0A0P0LKV0 |
| RSAIEDILFSKI | 1 | 0 | 0 | 1499966 | AGW80932.1 | A0A0P0K6L9 |
| RSAPRITFGGPS | 1 | 0 | 0 | 1499983 | AVP78038.1 | Q3LZX4 |
| RSCNYAVSSAFW | 1 | 0 | 0 | 1499995 | ALD51903.1 | K9N7C7 |
| RSCSDFEPMTDM | 1 | 0 | 0 | 1499997 | AVV62524.1 | K9N7C7 |
| RSCSDFLPLSDI | 1 | 0 | 0 | 1500000 | AHI48581.1 | K9N7C7 |
| RSDCNHIGNIKP | 1 | 0 | 0 | 1500014 | AIL49382.1 | P30215 |
| RSDCNHIGNINP | 1 | 0 | 0 | 1500015 | AEN19365.1 | P30215 |
| RSDCNHIVNINP | 1 | 0 | 0 | 1500018 | AWW13558.1 | P30215 |
| RSDLERDFIDME | 1 | 0 | 0 | 1500025 | ARB07436.1 | A0A140H1G9 |
| RSDTLVLSQGYF | 1 | 0 | 0 | 1500033 | AVP78031.1 | Q3LZX1 |
| RSDVLHLTQDYF | 1 | 0 | 0 | 1500035 | ATO98181.1 | Q3LZX1 |
| RSETLLPITQYN | 1 | 0 | 0 | 1500058 | APO40578.1 | P0C6X7 |
| RSFKTADGDQRQ | 1 | 0 | 0 | 1500069 | AJQ24083.1 | P33469 |
| RSFKTADGQQKH | 1 | 0 | 0 | 1500072 | ABD75597.1 | Q19U29 |
| RSGILKKTSWVD | 1 | 0 | 0 | 1500100 | AMM42397.1 | Q19U29 |
| RSGNSTRCTSPG | 1 | 0 | 0 | 1500114 | ALA49909.1 | K9N4V7 |
| RSGNSTRSTSPG | 1 | 0 | 0 | 1500119 | AVN89381.1 | K9N4V7 |
| RSGVGRCYNCRP | 1 | 0 | 0 | 1500126 | CAA71146.1 | A0A0P0K6L9 |
| RSHDTSHKNNFG | 1 | 0 | 0 | 1500130 | Q14EA9.1 | Q0ZJ84 |
| RSHFIRVSTVSS | 1 | 0 | 0 | 1500132 | AXG21669.1 | K9N7D2 |
| RSHRSTKLKPFE | 1 | 0 | 0 | 1500142 | AVP78031.1 | Q3LZX1 |
| RSHYYVDSVTVK | 1 | 0 | 0 | 1500145 | AXN92247.1 | K9N7C7 |
| RSIDCGFNDLSQ | 1 | 0 | 0 | 1500149 | AHZ90568.1 | K9N5Q8 |
| RSIFSRTLETAH | 1 | 0 | 0 | 1500151 | AVP78030.1 | P0C6W2 |
| RSIFSRTVDTAG | 1 | 0 | 0 | 1500153 | APO40578.1 | P0C6X7 |
| RSIKSDRKAWAA | 1 | 0 | 0 | 1500159 | AJG44080.1 | K9N5Q8 |
| RSIQSGRKAWAA | 1 | 0 | 0 | 1500167 | QBF80543.1 | K9N5Q8 |
| RSITTGYRFTNF | 1 | 0 | 0 | 1500173 | ABD43183.1 | P36334 |
| RSKEEDGFAFYP | 1 | 0 | 0 | 1500178 | AXN92247.1 | K9N7C7 |
| RSKNPVLYDANY | 1 | 0 | 0 | 1500185 | AVP78043.1 | Q3LZX0 |
| RSLESPLKLEDF | 1 | 0 | 0 | 1500194 | APO40578.1 | P0C6X7 |
| RSLYPVASPREC | 1 | 0 | 0 | 1500225 | ARO76381.1 | P0C6U8 |
| RSMLGLITQTVD | 1 | 0 | 0 | 1500231 | ASU90899.1 | K9N7C7 |
| RSMLGLNTQTVD | 1 | 0 | 0 | 1500232 | AVV62524.1 | K9N7C7 |
| RSMLKQRDSTYG | 1 | 0 | 0 | 1500234 | AVN89365.1 | K9N5Q8 |
| RSNIYGTATIGV | 1 | 0 | 0 | 1500250 | AXN92247.1 | K9N7C7 |
| RSNSNFRHSASI | 1 | 0 | 0 | 1500252 | AJQ24072.1 | Q19U29 |

| | | | | | | |
|---|---|---|---|---|---|---|
| RSPNKQCTVQQC | 1 | 0 | 0 | 1500256 | AEN19370.1 | P33469 |
| RSPPITADIELA | 1 | 0 | 0 | 1500258 | ASY99815.1 | K9N7A1 |
| RSQDSLLKLEDF | 1 | 0 | 0 | 1500260 | AVP78041.1 | P0C6W2 |
| RSQSRGPGESKP | 1 | 0 | 0 | 1500269 | AAA45463.1 | A0A0N7J131 |
| RSRASSGNRAPT | 1 | 0 | 0 | 1500287 | ATP16771.1 | P33469 |
| RSREETGLLLPL | 1 | 0 | 0 | 1500292 | AVP78041.1 | P0C6W2 |
| RSRGNSRNSTPS | 1 | 0 | 0 | 1500297 | APO40586.1 | P59595 |
| RSRRAITTENRF | 1 | 0 | 0 | 1500305 | AIL49500.1 | P36334 |
| RSRRSITTGYRF | 1 | 0 | 0 | 1500309 | ABD43183.1 | P36334 |
| RSSFVDDNKFDV | 1 | 0 | 0 | 1500319 | AGT51657.1 | P0C6X6 |
| RSSRIAIEDLLF | 1 | 0 | 0 | 1500333 | AHL18090.1 | K9N5Q8 |
| RSSSQGSRSRNS | 1 | 0 | 0 | 1500337 | AHY22571.1 | K9N4V7 |
| RSSTRNNSRDSF | 1 | 0 | 0 | 1500346 | ABE97134.1 | H9EJ69 |
| RSTEKVLERAAV | 1 | 0 | 0 | 1500351 | AUM60013.1 | K9N7C7 |
| RSVAYVANPTED | 1 | 0 | 0 | 1500381 | AEN19355.1 | P0C6X6 |
| RSVDVAIQEVID | 1 | 0 | 0 | 1500387 | NP_150075.1 | Q80872 |
| RSVEDGDTVYYP | 1 | 0 | 0 | 1500388 | AUM60023.1 | K9N7C7 |
| RSVEDGETVYYP | 1 | 0 | 0 | 1500389 | AUM60013.1 | K9N7C7 |
| RSVKVPAVVSVS | 1 | 0 | 0 | 1500395 | APO40578.1 | P0C6X7 |
| RSVRSHPDFKLL | 1 | 0 | 0 | 1500411 | AHY22543.1 | K9N7C7 |
| RSVRSVPGEMCL | 1 | 0 | 0 | 1500412 | AVN89313.1 | K9N5Q8 |
| RSVRSVPGQMRL | 1 | 0 | 0 | 1500414 | AHL18090.1 | K9N5Q8 |
| RSVSSVPGEMRL | 1 | 0 | 0 | 1500418 | 5X59_A | K9N5Q8 |
| RSVTVEYNIHAV | 1 | 0 | 0 | 1500419 | AXN92247.1 | K9N7C7 |
| RSVYPVATPQEC | 1 | 0 | 0 | 1500427 | P0C6W6.1 | P0C6W2 |
| RSYERLINTISP | 1 | 0 | 0 | 1500430 | AWK59942.1 | H9EJ42 |
| RTAEAIHTVTIA | 1 | 0 | 0 | 1500451 | QED88026.1 | A0A0P0G321 |
| RTAGVVTSNVLK | 1 | 0 | 0 | 1500454 | AUM60023.1 | K9N7C7 |
| RTAIDCGFNDLS | 1 | 0 | 0 | 1500456 | AXN92260.1 | K9N5Q8 |
| RTDQINDMVYSL | 1 | 0 | 0 | 1500477 | AIX09796.1 | P0C6X6 |
| RTDVRDYKQAND | 1 | 0 | 0 | 1500479 | QED88010.1 | H9EJ42 |
| RTESMCPGNSNY | 1 | 0 | 0 | 1500493 | ATQ39390.1 | K9N5Q8 |
| RTEVDPDLSAFY | 1 | 0 | 0 | 1500494 | AXN92247.1 | K9N7C7 |
| RTEVDPDLSVFY | 1 | 0 | 0 | 1500495 | ALA49857.1 | K9N7C7 |
| RTEVLQLVNANQ | 1 | 0 | 0 | 1500496 | QBM11748.1 | K9N5Q8 |
| RTFAKAPPGDNF | 1 | 0 | 0 | 1500500 | AZS52616.1 | A0A140H1G9 |
| RTFMLKYNENGT | 1 | 0 | 0 | 1500509 | AVP78031.1 | Q3LZX1 |
| RTFPQQMFVTPL | 1 | 0 | 0 | 1500510 | AVN89294.1 | K9N643 |
| RTFVVDKSLSID | 1 | 0 | 0 | 1500519 | AZK15899.1 | K9N638 |
| RTGARPKHRRPQ | 1 | 0 | 0 | 1500527 | APO40586.1 | P59595 |
| RTGSFNNIDTGP | 1 | 0 | 0 | 1500537 | AIX10756.1 | P36334 |
| RTGVDPDLSAFY | 1 | 0 | 0 | 1500544 | ASU90030.1 | K9N7C7 |
| RTIDSTQDGVNK | 1 | 0 | 0 | 1500554 | QBQ01839.1 | P36334 |
| RTIKSTQDGDNK | 1 | 0 | 0 | 1500558 | AIX10748.1 | P36334 |
| RTILGAAALEDE | 1 | 0 | 0 | 1500561 | 3M3V_A | P0C6X7 |
| RTIQPEVKEVAP | 1 | 0 | 0 | 1500574 | ANI69888.1 | K9N638 |
| RTKGVDYTKKQI | 1 | 0 | 0 | 1500582 | AUM60013.1 | K9N7C7 |
| RTLETAHHSVHV | 1 | 0 | 0 | 1500593 | AVP78030.1 | P0C6W2 |
| RTLFDALKFFEI | 1 | 0 | 0 | 1500596 | NP_073549.1 | A0A0P0LKV0 |
| RTLFDALKFFEV | 1 | 0 | 0 | 1500597 | AST12962.1 | A0A0P0LKV0 |
| RTLIDGVENPML | 1 | 0 | 0 | 1500598 | AIW52827.1 | H9EJ42 |
| RTLQEMKALFPT | 1 | 0 | 0 | 1500605 | AIV41774.1 | P0C6X6 |
| RTLSTYDFNPNV | 1 | 0 | 0 | 1500607 | AVP78031.1 | Q3LZX1 |
| RTLVLNANECQH | 1 | 0 | 0 | 1500616 | AVV62540.1 | K9N643 |
| RTLYQPVYKIFL | 1 | 0 | 0 | 1500618 | AAV67329.1 | H9EJA2 |
| RTPLNLPPTHAH | 1 | 0 | 0 | 1500635 | AWK59942.1 | H9EJ42 |
| RTPLNPNYTNSS | 1 | 0 | 0 | 1500637 | AVP78031.1 | Q3LZX1 |
| RTPTPGVTPDMA | 1 | 0 | 0 | 1500641 | AIX10722.1 | P33469 |
| RTPTSGVTRDMA | 1 | 0 | 0 | 1500643 | AIL49470.1 | P33469 |
| RTRKGKRVDLSP | 1 | 0 | 0 | 1500657 | AIW52723.1 | A0A0N7J131 |
| RTSIASFNEWAM | 1 | 0 | 0 | 1500680 | AUM60013.1 | K9N7C7 |
| RTSLHPVLWNCQ | 1 | 0 | 0 | 1500683 | QBF80613.1 | K9N4V0 |
| RTSNQGNKQIVH | 1 | 0 | 0 | 1500684 | AHI48842.1 | K9N643 |
| RTSPDPKFVDKY | 1 | 0 | 0 | 1500685 | AVV62524.1 | K9N7C7 |
| RTSSRASSTGSR | 1 | 0 | 0 | 1500691 | AIX10719.1 | P33469 |
| RTSSRTSSAGSR | 1 | 0 | 0 | 1500692 | ARK08665.1 | P33469 |
| RTSTSGVTPDMA | 1 | 0 | 0 | 1500693 | ARB07416.1 | P33469 |
| RTSVASFNEWAM | 1 | 0 | 0 | 1500694 | AVV62535.1 | K9N7C7 |
| RTSVVSFNEWAM | 1 | 0 | 0 | 1500699 | AVV62524.1 | K9N7C7 |
| RTTATIPIQATL | 1 | 0 | 0 | 1500701 | AVP78032.1 | Q3LZX0 |
| RTTGIITGNVVK | 1 | 0 | 0 | 1500704 | AVV62535.1 | K9N7C7 |
| RTTSNFLPEVAA | 1 | 0 | 0 | 1500716 | ATO98193.1 | Q3LZX1 |
| RTVGGVPLLSTM | 1 | 0 | 0 | 1500726 | ADK37692.1 | H9EJ42 |
| RTVSFYEDRILE | 1 | 0 | 0 | 1500734 | AUM60013.1 | K9N7C7 |
| RTVVRPLVEDST | 1 | 0 | 0 | 1500741 | ASY99815.1 | K9N7A1 |
| RTYSEAAAQGFQ | 1 | 0 | 0 | 1500757 | AWK59942.1 | H9EJ42 |
| RVATKSFNVVQA | 1 | 0 | 0 | 1500770 | AUM60021.1 | K9N4V7 |
| RVCGVSAARLTPCGT | 1 | 0 | 0 | 1500775 | P0DTD1.1 | P0DTD1 |
| RVCIKTAQADTA | 1 | 0 | 0 | 1500776 | AMD02845.1 | K9N796 |
| RVCYFNLCKEPM | 1 | 0 | 0 | 1500781 | ATO98193.1 | Q3LZX1 |
| RVDLNTTILFTG | 1 | 0 | 0 | 1500796 | ABA60800.1 | A0A140H1H1 |
| RVDLQLGNLGYL | 1 | 0 | 0 | 1500799 | AGT51600.1 | P36334 |
| RVDYSDAVTYKS | 1 | 0 | 0 | 1500808 | AUM60013.1 | K9N7C7 |
| RVEVFTDGKLNC | 1 | 0 | 0 | 1500838 | AQZ41341.1 | K9N7C7 |
| RVFPYDMSSDVS | 1 | 0 | 0 | 1500851 | AUM60013.1 | K9N7C7 |
| RVFTDYMPYIFT | 1 | 0 | 0 | 1500854 | APO40578.1 | P0C6X7 |
| RVHGFEIDPSPN | 1 | 0 | 0 | 1500874 | APO40579.1 | P59594 |
| RVIAIPTSTGSV | 1 | 0 | 0 | 1500878 | AVV62527.1 | K9N796 |
| RVIFPPSRNWHV | 1 | 0 | 0 | 1500880 | AVV62529.1 | K9N643 |
| RVISSFTCRTDM | 1 | 0 | 0 | 1500893 | AEN19363.1 | P0C6X6 |
| RVIYGNVSRHNL | 1 | 0 | 0 | 1500894 | AIW52827.1 | H9EJ42 |

| | | | | | | |
|---|---|---|---|---|---|---|
| RVKGWVGFDVEG | 1 | 0 | 0 | 1500901 | ABD75519.1 | A0A140H1G9 |
| RVKLHTVIPDYV | 1 | 0 | 0 | 1500905 | QEG03731.1 | A0A0P0G321 |
| RVKNLNSSEGVS | 1 | 0 | 0 | 1500907 | AAS48454.1 | P59637 |
| RVKNLNSSQGIP | 1 | 0 | 0 | 1500908 | APO40581.1 | P59637 |
| RVKSVNGRVVIT | 1 | 0 | 0 | 1500914 | QED88038.1 | H9EJ42 |
| RVKSVSPGDSSW | 1 | 0 | 0 | 1500917 | AVL25553.1 | A0A0P0G321 |
| RVLDQSENNCWI | 1 | 0 | 0 | 1500922 | ABE73416.1 | H9EJ42 |
| RVLSSLNLPGCD | 1 | 0 | 0 | 1500944 | APO40578.1 | P0C6X7 |
| RVLSTFEPRSDL | 1 | 0 | 0 | 1500945 | ARB07436.1 | A0A140H1G9 |
| RVLTTGYFLPFN | 1 | 0 | 0 | 1500947 | APO40579.1 | P59594 |
| RVMITKGVMEPI | 1 | 0 | 0 | 1500954 | AGT51393.1 | H9EJ42 |
| RVNTAIPTIKDW | 1 | 0 | 0 | 1500967 | AXN92231.1 | K9N643 |
| RVPIHYEMFGNY | 1 | 0 | 0 | 1500975 | ATO98227.1 | Q3LZX5 |
| RVPSRSQSRSQS | 1 | 0 | 0 | 1500982 | APT69887.1 | A0A0N7J131 |
| RVPTIEENVIQC | 1 | 0 | 0 | 1500983 | ADK37676.1 | H9EJ69 |
| RVPYQRNNTSFP | 1 | 0 | 0 | 1500988 | ATQ39389.1 | K9N7C7 |
| RVQGGITQRTRT | 1 | 0 | 0 | 1500990 | ANF29213.1 | K9N4V7 |
| RVQGSITQHTRT | 1 | 0 | 0 | 1500994 | AKN24774.1 | K9N4V7 |
| RVQGSITQRIRT | 1 | 0 | 0 | 1500996 | AKI29262.1 | K9N4V7 |
| RVQGSITQRTRS | 1 | 0 | 0 | 1500998 | ASY99807.1 | K9N4V7 |
| RVQQNKIRELTA | 1 | 0 | 0 | 1501007 | AGT51784.1 | P33469 |
| RVQSSIIQRTRT | 1 | 0 | 0 | 1501015 | QCZ25056.1 | K9N4V7 |
| RVRNPKPRAAPN | 1 | 0 | 0 | 1501028 | AXG21672.1 | K9N4V7 |
| RVRTINLGEVLI | 1 | 0 | 0 | 1501034 | APO40578.1 | P0C6X7 |
| RVSNLTKQLLEL | 1 | 0 | 0 | 1501048 | AUM60013.1 | K9N7C7 |
| RVSNLTKQLLEV | 1 | 0 | 0 | 1501049 | AUM60023.1 | K9N7C7 |
| RVSSNQSPQGNL | 1 | 0 | 0 | 1501055 | AVI15051.1 | P0C6X6 |
| RVTFSDPTTFGS | 1 | 0 | 0 | 1501075 | AVN89383.1 | K9N643 |
| RVTLGDFTIMSG | 1 | 0 | 0 | 1501081 | ARU07575.1 | A0A140H1G9 |
| RVTYGTFHWLFA | 1 | 0 | 0 | 1501090 | AVV62524.1 | K9N7C7 |
| RVVDEWSYTGSS | 1 | 0 | 0 | 1501093 | ALA49803.1 | K9N5Q8 |
| RVVEPILAIEPL | 1 | 0 | 0 | 1501094 | AVR40343.1 | Q4VID0 |
| RVVEPILATEPL | 1 | 0 | 0 | 1501096 | AHF54652.1 | Q4VID0 |
| RVVERNVVRAAL | 1 | 0 | 0 | 1501100 | AVV62535.1 | K9N7C7 |
| RVVFPPSRNWDV | 1 | 0 | 0 | 1501103 | AVV62540.1 | K9N643 |
| RVVIPDFTVWNQ | 1 | 0 | 0 | 1501109 | AXN92247.1 | K9N7C7 |
| RVVITNVGEPTI | 1 | 0 | 0 | 1501112 | ABE97136.1 | H9EJ42 |
| RVVSPPINADLS | 1 | 0 | 0 | 1501123 | AUM60025.1 | K9N796 |
| RVVSPPNNADLS | 1 | 0 | 0 | 1501124 | AUM60015.1 | K9N796 |
| RVYLAYIKKLYN | 1 | 0 | 0 | 1501145 | AEN19363.1 | P0C6X6 |
| RVYNHFNNYDCV | 1 | 0 | 0 | 1501157 | AIV41936.1 | P0C6X6 |
| RVYSTGVNVFQT | 1 | 0 | 0 | 1501161 | ATO98145.1 | Q3LZX1 |
| RWKRQPNDDATS | 1 | 0 | 0 | 1501171 | APT69866.1 | A0A0N7J131 |
| RWNNARHSDNMT | 1 | 0 | 0 | 1501183 | AQN78751.1 | P30215 |
| RWNNARQSDNKT | 1 | 0 | 0 | 1501186 | AIL49307.1 | P30215 |
| RWSTSINTGNCP | 1 | 0 | 0 | 1501203 | AAQ90004.1 | A0A0P0K6L9 |
| RYAKRRFSPLKL | 1 | 0 | 0 | 1501217 | AUM60017.1 | K9N643 |
| RYASAVAQAFQS | 1 | 0 | 0 | 1501218 | AEN19363.1 | P0C6X6 |
| RYDALLSDLSFN | 1 | 0 | 0 | 1501229 | QED88018.1 | H9EJ42 |
| RYDMFESNMFIK | 1 | 0 | 0 | 1501233 | AVV62524.1 | K9N7C7 |
| RYDNISSVWPLH | 1 | 0 | 0 | 1501234 | AQN78751.1 | P30215 |
| RYDNVSSVWPFY | 1 | 0 | 0 | 1501236 | AGT51679.1 | P30215 |
| RYFTLGNVEAVN | 1 | 0 | 0 | 1501259 | CAA71056.1 | A0A0P0K6L9 |
| RYGIKPFTIPID | 1 | 0 | 0 | 1501269 | AMM42367.1 | A0A140H1G9 |
| RYIYNRVKSVSA | 1 | 0 | 0 | 1501280 | AWO68966.1 | A0A0P0G321 |
| RYIYNRVKSVSP | 1 | 0 | 0 | 1501281 | AVL25553.1 | A0A0P0G321 |
| RYMRSVKVPAVV | 1 | 0 | 0 | 1501309 | APO40578.1 | P0C6X7 |
| RYMSAFNHTKKW | 1 | 0 | 0 | 1501310 | BAE93400.1 | P0C6U8 |
| RYNINFEENQRR | 1 | 0 | 0 | 1501323 | ACO44679.1 | A0A0P0K6L9 |
| RYQEFNDLDHTS | 1 | 0 | 0 | 1501343 | QED88024.1 | H9EJ42 |
| RYRAALNNEKRP | 1 | 0 | 0 | 1501345 | AVV62535.1 | K9N7C7 |
| RYSEAASNGFRA | 1 | 0 | 0 | 1501349 | NP_073549.1 | A0A0P0LKV0 |
| RYSLADSNGFIR | 1 | 0 | 0 | 1501354 | ABD75543.1 | A0A140H1G9 |
| RYVSPDVQFEQL | 1 | 0 | 0 | 1501381 | AVV62526.1 | K9N5Q8 |
| RYYSIKAAVSKW | 1 | 0 | 0 | 1501391 | AUM60013.1 | K9N7C7 |
| SAAEKEVLSELY | 1 | 0 | 0 | 1501399 | AVV62535.1 | K9N7C7 |
| SAAFHQNASSLA | 1 | 0 | 0 | 1501401 | BBA20983.1 | A0A140H1H1 |
| SAAFHQSASSLA | 1 | 0 | 0 | 1501402 | AGT17777.1 | A0A140H1H1 |
| SAAGYGGFGCKD | 1 | 0 | 0 | 1501405 | AEN19363.1 | P0C6X6 |
| SAAKKNNIPFKL | 1 | 0 | 0 | 1501415 | APO40578.1 | P0C6X7 |
| SAALAVNACLIP | 1 | 0 | 0 | 1501422 | AUM60018.1 | K9N7D2 |
| SAAQADLQHEDS | 1 | 0 | 0 | 1501430 | ATQ39389.1 | K9N7C7 |
| SAARNGVVDSDV | 1 | 0 | 0 | 1501437 | APO40578.1 | P0C6X7 |
| SAAYHANSPEPA | 1 | 0 | 0 | 1501446 | AIX10763.1 | P36334 |
| SAAYHANSSDPA | 1 | 0 | 0 | 1501447 | AIL49515.1 | P36334 |
| SACPKLEFANDT | 1 | 0 | 0 | 1501467 | AXN92260.1 | K9N5Q8 |
| SACTRLEGLGGN | 1 | 0 | 0 | 1501480 | NP_073549.1 | A0A0P0LKV0 |
| SADFIKHVRTNG | 1 | 0 | 0 | 1501490 | ATQ39389.1 | K9N7C7 |
| SADGVQCYTPHS | 1 | 0 | 0 | 1501492 | AEN19363.1 | P0C6X6 |
| SADIGANHTGIT | 1 | 0 | 0 | 1501493 | ARO76381.1 | P0C6U8 |
| SADIVDRDPSSD | 1 | 0 | 0 | 1501495 | ARK08674.1 | P33469 |
| SADLYELMGRFA | 1 | 0 | 0 | 1501498 | AUM60013.1 | K9N7C7 |
| SADWKPGHSMPS | 1 | 0 | 0 | 1501511 | AVN89289.1 | K9N7C7 |
| SAEGRKYIVKTS | 1 | 0 | 0 | 1501517 | NP_073549.1 | A0A0P0LKV0 |
| SAEWKCGYAMPQ | 1 | 0 | 0 | 1501526 | AWK59942.1 | H9EJ42 |
| SAEWKCGYSMPG | 1 | 0 | 0 | 1501527 | NP_073549.1 | A0A0P0LKV0 |
| SAFDGNMYGAFQ | 1 | 0 | 0 | 1501532 | AVV62535.1 | K9N7C7 |
| SAFGAILSNLGV | 1 | 0 | 0 | 1501535 | APO40578.1 | P0C6X7 |
| SAFGDVKPVTEY | 1 | 0 | 0 | 1501536 | AUM60027.1 | K9N643 |
| SAFIDTVKGLDY | 1 | 0 | 0 | 1501538 | ARO76381.1 | P0C6U8 |
| SAFIECGSCGND | 1 | 0 | 0 | 1501539 | AXN92247.1 | K9N7C7 |

| | | | | | | |
|---|---|---|---|---|---|---|
| SAFIECGSCGNN | 1 | 0 | 0 | 1501541 | ALA49340.1 | K9N7C7 |
| SAFIETVKNLDY | 1 | 0 | 0 | 1501542 | APO40578.1 | P0C6X7 |
| SAFMSMSQLKLT | 1 | 0 | 0 | 1501555 | AVV62544.1 | K9N4V7 |
| SAFVECGSCGND | 1 | 0 | 0 | 1501573 | AXN73479.1 | K9N7C7 |
| SAGDSIFKSFHF | 1 | 0 | 0 | 1501586 | AMN88685.1 | A0A140H1H0 |
| SAGGTCSSISQL | 1 | 0 | 0 | 1501593 | APO40579.1 | P59594 |
| SAGHADNNIEKP | 1 | 0 | 0 | 1501595 | AVV62526.1 | K9N5Q8 |
| SAGHATGITPQK | 1 | 0 | 0 | 1501596 | ATG84701.1 | K9N5Q8 |
| SAGHATGTTLQK | 1 | 0 | 0 | 1501598 | AID55095.1 | K9N5Q8 |
| SAGIFGVDPGLS | 1 | 0 | 0 | 1501610 | APO40578.1 | P0C6X7 |
| SAGKFDSFKGEC | 1 | 0 | 0 | 1501617 | APO40578.1 | P0C6X7 |
| SAGNATEVPANS | 1 | 0 | 0 | 1501620 | 5C8S_A | P0C6U8 |
| SAGQDQTTCNVN | 1 | 0 | 0 | 1501625 | AUM60013.1 | K9N7C7 |
| SAGQEQTNCNTS | 1 | 0 | 0 | 1501627 | AXN92247.1 | K9N7C7 |
| SAGSVGFNIDYD | 1 | 0 | 0 | 1501639 | 1Z1J_A | P0C6U8 |
| SAGYNTNATLNS | 1 | 0 | 0 | 1501647 | AVV62537.1 | K9N5Q8 |
| SAGYPFNQFGKA | 1 | 0 | 0 | 1501648 | AIL49570.1 | P0C6X6 |
| SAIAKPKRLRKK | 1 | 0 | 0 | 1501654 | ATQ39389.1 | K9N7C7 |
| SAIEDLLFSKVR | 1 | 0 | 0 | 1501660 | CAA83661.1 | P36334 |
| SAILPKGIMMNV | 1 | 0 | 0 | 1501666 | AVP78030.1 | P0C6W2 |
| SAIPNGRKVDLQ | 1 | 0 | 0 | 1501671 | BBA84104.1 | P36334 |
| SAIQQLNNGFQG | 1 | 0 | 0 | 1501672 | AVV62535.1 | K9N7C7 |
| SAIYPIDLSSQI | 1 | 0 | 0 | 1501679 | ALA49908.1 | K9N7A1 |
| SAKDDIKSVVAA | 1 | 0 | 0 | 1501681 | AUM60013.1 | K9N7C7 |
| SAKFWEESFYRE | 1 | 0 | 0 | 1501689 | AVV62524.1 | K9N7C7 |
| SAKGFFYIDAGN | 1 | 0 | 0 | 1501690 | AKA09621.1 | A0A0P0G321 |
| SAKIDTSGIHNR | 1 | 0 | 0 | 1501692 | AVV62527.1 | K9N796 |
| SAKVKNTKVIKD | 1 | 0 | 0 | 1501707 | AKA09642.1 | P36334 |
| SALEQYVFCTVN | 1 | 0 | 0 | 1501716 | APO40578.1 | P0C6X7 |
| SALFCATCPFAK | 1 | 0 | 0 | 1501717 | AUM60013.1 | K9N7C7 |
| SALFVVKQKWCV | 1 | 0 | 0 | 1501722 | ARK08626.1 | H9EJ42 |
| SALIAVANWLTP | 1 | 0 | 0 | 1501729 | AUM60013.1 | K9N7C7 |
| SALIGANGNKIV | 1 | 0 | 0 | 1501730 | AVN89322.1 | K9N7C7 |
| SALKMVQTHISH | 1 | 0 | 0 | 1501732 | AVV62524.1 | K9N7C7 |
| SALLVNNGSAIT | 1 | 0 | 0 | 1501741 | ATO98145.1 | Q3LZX1 |
| SALMGTNGNKIV | 1 | 0 | 0 | 1501742 | AVV62524.1 | K9N7C7 |
| SALMSCNGNKIE | 1 | 0 | 0 | 1501743 | AJE26220.1 | P0C6X6 |
| SALNGSCGSVGF | 1 | 0 | 0 | 1501744 | 3F9G_A | P0C6X7 |
| SAMFCATCPFAK | 1 | 0 | 0 | 1501767 | AVV62535.1 | K9N7C7 |
| SAMGLPNNTASW | 1 | 0 | 0 | 1501768 | 1SSK_A | P59595 |
| SAMNVIGQHIEL | 1 | 0 | 0 | 1501780 | ARE29978.1 | H9EJ42 |
| SAMNVIGQHIKL | 1 | 0 | 0 | 1501781 | AWK59942.1 | H9EJ42 |
| SAMVHDLNNVDP | 1 | 0 | 0 | 1501786 | AVP78050.1 | Q3LZX3 |
| SANDPLLGTTLQ | 1 | 0 | 0 | 1501791 | AVV62524.1 | K9N7C7 |
| SANDVEIQSSGV | 1 | 0 | 0 | 1501792 | AVV62524.1 | K9N7C7 |
| SANFLDDNVLPD | 1 | 0 | 0 | 1501793 | AWO68966.1 | A0A0P0G321 |
| SANFQHDNVLPE | 1 | 0 | 0 | 1501795 | AWO68987.1 | A0A0P0G321 |
| SANIFIGDNVGH | 1 | 0 | 0 | 1501806 | AEN19363.1 | P0C6X6 |
| SANLFPPWSAAA | 1 | 0 | 0 | 1501811 | CAA83661.1 | P36334 |
| SANPVKEYEADA | 1 | 0 | 0 | 1501816 | AVV62535.1 | K9N7C7 |
| SANTKTLRNKGV | 1 | 0 | 0 | 1501821 | AVV62524.1 | K9N7C7 |
| SANVCALISCNG | 1 | 0 | 0 | 1501826 | AIL49553.1 | P0C6X6 |
| SANVSALMSCNG | 1 | 0 | 0 | 1501828 | AJE26220.1 | P0C6X6 |
| SAPPAQYELKHG | 1 | 0 | 0 | 1501841 | AVP78041.1 | P0C6W2 |
| SAPPVTYSPATI | 1 | 0 | 0 | 1501844 | AVK87459.1 | K9N7C7 |
| SAPTLIPQENYS | 1 | 0 | 0 | 1501850 | AWW13571.1 | P0C6X6 |
| SAQAIAFCKSGS | 1 | 0 | 0 | 1501857 | AIL49345.1 | P30215 |
| SAQSEEEESFSN | 1 | 0 | 0 | 1501869 | AKN24751.1 | K9N4V0 |
| SAQYIFSTANAL | 1 | 0 | 0 | 1501874 | ABE97136.1 | H9EJ42 |
| SARDPIIQEAIP | 1 | 0 | 0 | 1501880 | AMM42389.1 | Q19U29 |
| SARDPTIQEAIP | 1 | 0 | 0 | 1501881 | AMM42397.1 | Q19U29 |
| SARLVQLCAGEY | 1 | 0 | 0 | 1501890 | ATO98227.1 | Q3LZX5 |
| SARNLICAQYVA | 1 | 0 | 0 | 1501894 | QFS19619.1 | K9N5Q8 |
| SARPPPGEQFKH | 1 | 0 | 0 | 1501895 | AVV62524.1 | K9N7C7 |
| SASAVNMQQIYL | 1 | 0 | 0 | 1501904 | AIV41846.1 | P0C6X6 |
| SASDFIAWRAEA | 1 | 0 | 0 | 1501908 | Q14EA9.1 | Q0ZJ84 |
| SASEGLVGTTLQ | 1 | 0 | 0 | 1501914 | AUM60013.1 | K9N7C7 |
| SASEKEVLLQLY | 1 | 0 | 0 | 1501915 | AUM60013.1 | K9N7C7 |
| SASGYGGFGCKD | 1 | 0 | 0 | 1501922 | AIV41888.1 | P0C6X6 |
| SASIGDIIRRLD | 1 | 0 | 0 | 1501924 | ALB08246.1 | K9N5Q8 |
| SASISAFVETVK | 1 | 0 | 0 | 1501926 | AVP78030.1 | P0C6W2 |
| SASIVKPDMADE | 1 | 0 | 0 | 1501930 | AJQ24072.1 | Q19U29 |
| SASKVIALHNRW | 1 | 0 | 0 | 1501933 | ARO76383.1 | P59632 |
| SASLFKPVHLVP | 1 | 0 | 0 | 1501934 | AVN89455.1 | K9N7D2 |
| SASLFKPVHLVS | 1 | 0 | 0 | 1501935 | AHL18095.1 | K9N7D2 |
| SASLFKPVQLAP | 1 | 0 | 0 | 1501936 | AKM76243.1 | K9N7D2 |
| SASNSRSTSRTS | 1 | 0 | 0 | 1501943 | AXX83347.1 | P33469 |
| SASSGYLQLATI | 1 | 0 | 0 | 1501947 | AVV62526.1 | K9N5Q8 |
| SASTSAFIDTVK | 1 | 0 | 0 | 1501952 | ARO76381.1 | P0C6U8 |
| SASTSLDWFPRP | 1 | 0 | 0 | 1501955 | AUM60016.1 | K9N4V0 |
| SASYFCKIGREQ | 1 | 0 | 0 | 1501960 | ALA50033.1 | K9N7C7 |
| SATEFADKLVEV | 1 | 0 | 0 | 1501967 | AEN19363.1 | P0C6X6 |
| SATGLHRQLVTL | 1 | 0 | 0 | 1501972 | AJQ24074.1 | P33469 |
| SATIRKFYPAFM | 1 | 0 | 0 | 1501976 | ALA49660.1 | K9N5Q8 |
| SATKTVGGAPNW | 1 | 0 | 0 | 1501981 | AVV62524.1 | K9N7C7 |
| SATKVYFDNPTL | 1 | 0 | 0 | 1501982 | ATO98193.1 | Q3LZX1 |
| SATLVKDIDITF | 1 | 0 | 0 | 1501985 | AVP78041.1 | P0C6W2 |
| SATLVKDIDTTF | 1 | 0 | 0 | 1501986 | AVP78030.1 | P0C6W2 |
| SATNSALAKIQS | 1 | 0 | 0 | 1501989 | BBA20983.1 | A0A140H1H1 |
| SATQPYFDSYLG | 1 | 0 | 0 | 1501991 | AGW27863.1 | A0A140H1H1 |
| SATVTANLSIPS | 1 | 0 | 0 | 1501996 | AAK32189.1 | A0A0P0K6L9 |

| | | | | | | |
|---|---|---|---|---|---|---|
| SATYEGNSPFHP | 1 | 0 | 0 | 1501997 | ABI96964.1 | P59635 |
| SAVANAHRYDVL | 1 | 0 | 0 | 1502002 | AWK59942.1 | H9EJ42 |
| SAVEKLRPVFAW | 1 | 0 | 0 | 1502007 | ARO76381.1 | P0C6U8 |
| SAVGNICYTPST | 1 | 0 | 0 | 1502013 | APO40578.1 | P0C6X7 |
| SAVGVRQQRFVY | 1 | 0 | 0 | 1502014 | ASU90791.1 | K9N5Q8 |
| SAVLWANGVKAK | 1 | 0 | 0 | 1502025 | AMM42513.1 | A0A0N7J131 |
| SAVNFTKALAKQ | 1 | 0 | 0 | 1502029 | AVN89336.1 | K9N4V0 |
| SAVSFDYLIREA | 1 | 0 | 0 | 1502041 | ANI69899.1 | K9N7C7 |
| SAYGCSTIAQAV | 1 | 0 | 0 | 1502052 | AST12962.1 | A0A0P0LKV0 |
| SAYGLCDLANPT | 1 | 0 | 0 | 1502056 | AVN89387.1 | K9N5Q8 |
| SAYIKLNCFYDP | 1 | 0 | 0 | 1502059 | AGT17757.1 | A0A140H1H0 |
| SAYRANSFDAPT | 1 | 0 | 0 | 1502066 | AHI48604.1 | K9N7C7 |
| SAYSGDIPHYVQ | 1 | 0 | 0 | 1502070 | AUM60024.1 | K9N5Q8 |
| SAYVKFQESKPP | 1 | 0 | 0 | 1502073 | AVV62531.1 | K9N5R3 |
| SAYYVKDGKYFT | 1 | 0 | 0 | 1502076 | AVV62524.1 | K9N7C7 |
| SCAHEYGTEEDY | 1 | 0 | 0 | 1502078 | ATO98229.1 | P0C6W2 |
| SCAKSKPLSAIC | 1 | 0 | 0 | 1502079 | ARB07438.1 | A0A140H1H1 |
| SCAPLSDQIDFL | 1 | 0 | 0 | 1502083 | APO40578.1 | P0C6X7 |
| SCARKSFSIDSD | 1 | 0 | 0 | 1502087 | AJC98124.1 | P0C6X6 |
| SCCKFDEDHSEP | 1 | 0 | 0 | 1502091 | APO40579.1 | P59594 |
| SCDCGKKFDEQV | 1 | 0 | 0 | 1502105 | AWK59942.1 | H9EJ42 |
| SCDFKGWIPGNM | 1 | 0 | 0 | 1502106 | AEN19363.1 | P0C6X6 |
| SCDGIASDYRAN | 1 | 0 | 0 | 1502107 | AUM60013.1 | K9N7C7 |
| SCDINKPLQVFV | 1 | 0 | 0 | 1502111 | AWK59942.1 | H9EJ42 |
| SCDLRMGSGFCI | 1 | 0 | 0 | 1502112 | BBA20986.1 | A0A140H1H1 |
| SCDQLREPMLQS | 1 | 0 | 0 | 1502114 | QYC64967.1 | P0DTD1 |
| SCDQLREPMMQS | 1 | 0 | 0 | 1502115 | P0C6W6.1 | P0C6W2 |
| SCELYHYQECVR | 1 | 0 | 0 | 1502120 | ATO98237.1 | Q3LZX7 |
| SCESTTVLDRTD | 1 | 0 | 0 | 1502124 | BBA20983.1 | A0A140H1H1 |
| SCFEIDGNVMHA | 1 | 0 | 0 | 1502132 | ARB07436.1 | A0A140H1G9 |
| SCFFTCNSNANI | 1 | 0 | 0 | 1502133 | AKA09626.1 | A0A0P0G321 |
| SCFFTCNSNANL | 1 | 0 | 0 | 1502134 | AIW52843.1 | A0A0P0G321 |
| SCFGKNNGIGTC | 1 | 0 | 0 | 1502136 | QEG03814.1 | P36334 |
| SCFIERVEWDIE | 1 | 0 | 0 | 1502139 | AUM60013.1 | K9N7C7 |
| SCFSTCNSNANL | 1 | 0 | 0 | 1502151 | AVL25555.1 | A0A0P0G321 |
| SCFSTCNSNASI | 1 | 0 | 0 | 1502152 | AWO68966.1 | A0A0P0G321 |
| SCGNESWLTGNA | 1 | 0 | 0 | 1502168 | AVN89333.1 | K9N7C7 |
| SCGNNSWLTGNA | 1 | 0 | 0 | 1502171 | ALA49340.1 | K9N7C7 |
| SCGYDYVYSPFF | 1 | 0 | 0 | 1502185 | AGV08437.1 | K9N7C7 |
| SCGYHYKELGLV | 1 | 0 | 0 | 1502186 | AXN92247.1 | K9N7C7 |
| SCHGKNNGIGTC | 1 | 0 | 0 | 1502187 | ARB66068.1 | P36334 |
| SCHQVSPSALAS | 1 | 0 | 0 | 1502191 | AUM60014.1 | K9N5Q8 |
| SCIWSVDAFNQF | 1 | 0 | 0 | 1502214 | AEN19363.1 | P0C6X6 |
| SCIYWSPRPGLW | 1 | 0 | 0 | 1502216 | ARU07575.1 | A0A140H1G9 |
| SCKHTTAQVVSH | 1 | 0 | 0 | 1502222 | AVV62535.1 | K9N7C7 |
| SCLKGVCSCGSC | 1 | 0 | 0 | 1502241 | ARO76382.1 | P59594 |
| SCLVASDGVPSN | 1 | 0 | 0 | 1502260 | AVV62524.1 | K9N7C7 |
| SCLVSSNGQPGG | 1 | 0 | 0 | 1502264 | QAT98908.1 | K9N7C7 |
| SCNFNMSSLMSF | 1 | 0 | 0 | 1502268 | AIV41909.1 | P36334 |
| SCNFPLEATWHY | 1 | 0 | 0 | 1502269 | AIW52843.1 | A0A0P0G321 |
| SCNGNKIEDFSI | 1 | 0 | 0 | 1502271 | AIV41978.1 | P0C6X6 |
| SCNGNKIEDLSI | 1 | 0 | 0 | 1502272 | AEN19363.1 | P0C6X6 |
| SCNGNKIEDLSM | 1 | 0 | 0 | 1502273 | AJE26223.1 | P0C6X6 |
| SCNGNKIGDLSI | 1 | 0 | 0 | 1502274 | AJE26218.1 | P0C6X6 |
| SCNGNKSDDLSI | 1 | 0 | 0 | 1502275 | AKL87224.1 | P0C6X6 |
| SCPDKNNGIGIC | 1 | 0 | 0 | 1502284 | AIX10751.1 | P36334 |
| SCPDKNNGIGTC | 1 | 0 | 0 | 1502285 | AIX10750.1 | P36334 |
| SCPEWMDEFEAD | 1 | 0 | 0 | 1502287 | ATG84853.1 | K9N7C7 |
| SCPGKNNGIGTC | 1 | 0 | 0 | 1502289 | AEN19366.1 | P36334 |
| SCPGKNNGIGTR | 1 | 0 | 0 | 1502290 | ARB66066.1 | P36334 |
| SCPGKNNGISTC | 1 | 0 | 0 | 1502291 | BBA84094.1 | P36334 |
| SCPGKNNGLGTC | 1 | 0 | 0 | 1502292 | AIL49503.1 | P36334 |
| SCQLYYSLPLLN | 1 | 0 | 0 | 1502307 | BBA20986.1 | A0A140H1H1 |
| SCQNLDCNCLGF | 1 | 0 | 0 | 1502310 | AGT51688.1 | Q80872 |
| SCRLLQRVMLKA | 1 | 0 | 0 | 1502317 | ARU07574.1 | A0A140H1G9 |
| SCSCKHTTAQVV | 1 | 0 | 0 | 1502321 | AVV62524.1 | K9N7C7 |
| SCSCPGKNNGIG | 1 | 0 | 0 | 1502322 | AVR40344.1 | P36334 |
| SCSCVGSGVAVQ | 1 | 0 | 0 | 1502323 | ARB07436.1 | A0A140H1G9 |
| SCSDFIPLSDME | 1 | 0 | 0 | 1502327 | ANI69887.1 | K9N7C7 |
| SCSDFLPMSDME | 1 | 0 | 0 | 1502330 | QBF80541.1 | K9N7C7 |
| SCSGKNNGIGTC | 1 | 0 | 0 | 1502332 | ANZ78838.1 | P36334 |
| SCSGKNNGVGTC | 1 | 0 | 0 | 1502333 | QEG03757.1 | P36334 |
| SCSIDSDVDTKC | 1 | 0 | 0 | 1502334 | AEN19363.1 | P0C6X6 |
| SCSNGRWLTGNA | 1 | 0 | 0 | 1502337 | AUM60013.1 | K9N7C7 |
| SCSSSCKHTMAQ | 1 | 0 | 0 | 1502347 | AUM60013.1 | K9N7C7 |
| SCTCQPQAFLGW | 1 | 0 | 0 | 1502349 | AEN19366.1 | P36334 |
| SCTCQPQAFVGW | 1 | 0 | 0 | 1502350 | BBA84088.1 | P36334 |
| SCTCRPQAFLGW | 1 | 0 | 0 | 1502351 | P36334.1 | P36334 |
| SCTTQQLTIEVL | 1 | 0 | 0 | 1502361 | AHI48794.1 | K9N638 |
| SCVCGRDATQYL | 1 | 0 | 0 | 1502362 | APO40578.1 | P0C6X7 |
| SCVVCHSQTSLR | 1 | 0 | 0 | 1502382 | AXN92247.1 | K9N7C7 |
| SCVYAICYFYAT | 1 | 0 | 0 | 1502383 | AEN19363.1 | P0C6X6 |
| SCYNFLKEQHCQ | 1 | 0 | 0 | 1502387 | ARO76392.1 | Q7TLC7 |
| SCYVCKPHQVNL | 1 | 0 | 0 | 1502395 | AWO69005.1 | A0A0P0G321 |
| SDALQDDPKGKY | 1 | 0 | 0 | 1502414 | AVV62535.1 | K9N7C7 |
| SDAMLDDVTGFM | 1 | 0 | 0 | 1502420 | AVV62535.1 | K9N7C7 |
| SDAQGTGHGHKV | 1 | 0 | 0 | 1502421 | ARO76381.1 | P0C6U8 |
| SDASTFLNRVCG | 1 | 0 | 0 | 1502427 | APO40578.1 | P0C6X7 |
| SDCNCYVNVVVM | 1 | 0 | 0 | 1502436 | AVV62535.1 | K9N7C7 |
| SDCNHIGNIKPH | 1 | 0 | 0 | 1502437 | AIL49382.1 | P30215 |
| SDCNHIGNINPH | 1 | 0 | 0 | 1502438 | AEN19365.1 | P30215 |

| | | | | | | |
|---|---|---|---|---|---|---|
| SDCNHIVNINPH | 1 | 0 | 0 | 1502440 | AWW13558.1 | P30215 |
| SDCNHVVNTNPR | 1 | 0 | 0 | 1502441 | AAA73152.1 | P30215 |
| SDCNRVSSRSAI | 1 | 0 | 0 | 1502443 | AVI15011.1 | P36334 |
| SDCNVVRYFLDG | 1 | 0 | 0 | 1502444 | AVN89363.1 | K9N7C7 |
| SDCNVVRYSLDG | 1 | 0 | 0 | 1502445 | AXN92247.1 | K9N7C7 |
| SDDEWVAAVDEA | 1 | 0 | 0 | 1502462 | ASU45728.1 | K9N7C7 |
| SDDVDPFDAAVA | 1 | 0 | 0 | 1502478 | AVV62544.1 | K9N4V7 |
| SDDVIVHDVEPT | 1 | 0 | 0 | 1502479 | AWK59942.1 | H9EJ42 |
| SDDVIVYDVEPT | 1 | 0 | 0 | 1502480 | AFD98798.1 | H9EJ42 |
| SDDYIKQNGPLR | 1 | 0 | 0 | 1502485 | ARO76381.1 | P0C6U8 |
| SDEAISTRFPPG | 1 | 0 | 0 | 1502487 | AIL49389.1 | P33469 |
| SDENGVRTLSTY | 1 | 0 | 0 | 1502497 | AVP78031.1 | Q3LZX1 |
| SDEYILAKGLLQ | 1 | 0 | 0 | 1502501 | ALA49901.1 | K9N7C7 |
| SDEYLTQRGPLQ | 1 | 0 | 0 | 1502503 | AUM60013.1 | K9N7C7 |
| SDFINAIFAKVK | 1 | 0 | 0 | 1502518 | QBP84704.1 | P36334 |
| SDFLPLSDIEKD | 1 | 0 | 0 | 1502526 | AHI48581.1 | K9N7C7 |
| SDFQLPVYASYK | 1 | 0 | 0 | 1502533 | AEN19363.1 | P0C6X6 |
| SDFQLPVYTSYK | 1 | 0 | 0 | 1502534 | ARK08667.1 | P0C6X6 |
| SDFVDFESVIQD | 1 | 0 | 0 | 1502543 | APU51933.1 | P0C6X6 |
| SDGCSCSCKHTT | 1 | 0 | 0 | 1502552 | AVV62524.1 | K9N7C7 |
| SDGFLHIYGAYQ | 1 | 0 | 0 | 1502558 | ATO98145.1 | Q3LZX1 |
| SDGMFVATSSYD | 1 | 0 | 0 | 1502570 | AGT51379.1 | H9EJ42 |
| SDGNYNCNASLN | 1 | 0 | 0 | 1502572 | AHL31360.1 | K9N5Q8 |
| SDGNYNRNAILN | 1 | 0 | 0 | 1502573 | ALJ54448.1 | K9N5Q8 |
| SDGNYNRYASLN | 1 | 0 | 0 | 1502575 | ALJ54486.1 | K9N5Q8 |
| SDGQGVPIAFGI | 1 | 0 | 0 | 1502576 | ABA60794.1 | Q19U29 |
| SDGQGVPLAFGV | 1 | 0 | 0 | 1502577 | ABA60796.1 | Q19U29 |
| SDGQPYVASEQE | 1 | 0 | 0 | 1502580 | AGT51657.1 | P0C6X6 |
| SDGSPSGDAISQ | 1 | 0 | 0 | 1502583 | AVV62535.1 | K9N7C7 |
| SDGVEFLKDAWE | 1 | 0 | 0 | 1502587 | ARO76381.1 | P0C6U8 |
| SDHDVYYDESRA | 1 | 0 | 0 | 1502593 | AVV62527.1 | K9N796 |
| SDHQDLKWSRFP | 1 | 0 | 0 | 1502599 | AVP78041.1 | P0C6W2 |
| SDIDYVPLKSST | 1 | 0 | 0 | 1502602 | APO40578.1 | P0C6X7 |
| SDIFATPSVLQA | 1 | 0 | 0 | 1502604 | ASY99818.1 | K9N7C7 |
| SDIKPVTYKLDG | 1 | 0 | 0 | 1502607 | APO40578.1 | P0C6X7 |
| SDILKRLDAVEQ | 1 | 0 | 0 | 1502609 | AVV62537.1 | K9N5Q8 |
| SDILQRLDTVEQ | 1 | 0 | 0 | 1502610 | AVV62526.1 | K9N5Q8 |
| SDINCNIPTQCY | 1 | 0 | 0 | 1502611 | P0C6U5.1 | A0A140H1G9 |
| SDINGVYWVASN | 1 | 0 | 0 | 1502612 | AIX10729.1 | P33469 |
| SDISDLSNRLLD | 1 | 0 | 0 | 1502614 | APO40578.1 | P0C6X7 |
| SDIVNGVAEGTP | 1 | 0 | 0 | 1502617 | APO40578.1 | P0C6X7 |
| SDIYFVKNIPCY | 1 | 0 | 0 | 1502618 | QBP84719.1 | P0C6X6 |
| SDKCSIEDFNVW | 1 | 0 | 0 | 1502627 | AWW13571.1 | P0C6X6 |
| SDKIHHHHHHNT | 1 | 0 | 0 | 1502638 | 2OFZ_A | P59595 |
| SDKIRMGQESIV | 1 | 0 | 0 | 1502639 | APO40579.1 | P59594 |
| SDKNVVFMNSCS | 1 | 0 | 0 | 1502641 | BBA20986.1 | A0A140H1H1 |
| SDKVNECVKSQS | 1 | 0 | 0 | 1502646 | AVV62526.1 | K9N5Q8 |
| SDLEEGESECIS | 1 | 0 | 0 | 1502653 | AXN92247.1 | K9N7C7 |
| SDLEESESECIS | 1 | 0 | 0 | 1502654 | AXN92258.1 | K9N7C7 |
| SDLQFGSSGFLQ | 1 | 0 | 0 | 1502675 | AMM42415.1 | A0A140H1H1 |
| SDLVIDALASMT | 1 | 0 | 0 | 1502687 | AEN19363.1 | P0C6X6 |
| SDMNDFVSDADS | 1 | 0 | 0 | 1502695 | APO40578.1 | P0C6X7 |
| SDMVETVVGQLE | 1 | 0 | 0 | 1502696 | ASU45706.1 | K9N7C7 |
| SDMYDPKTKHVT | 1 | 0 | 0 | 1502700 | ATO98229.1 | P0C6W2 |
| SDMYDPVTKNIN | 1 | 0 | 0 | 1502705 | AVV62524.1 | K9N7C7 |
| SDNFDNNQNGDR | 1 | 0 | 0 | 1502707 | APO40586.1 | P59595 |
| SDNGPQNQRSAP | 1 | 0 | 0 | 1502710 | AVP78038.1 | Q3LZX4 |
| SDNGPQPNQRSA | 1 | 0 | 0 | 1502712 | ATO98228.1 | Q3LZX4 |
| SDNIKECVKSFI | 1 | 0 | 0 | 1502717 | ARO76381.1 | P0C6U8 |
| SDNKTAVACQPP | 1 | 0 | 0 | 1502719 | AIL49307.1 | P30215 |
| SDNNCYFNAVIM | 1 | 0 | 0 | 1502722 | ASU90086.1 | K9N638 |
| SDNPQCAIVQAS | 1 | 0 | 0 | 1502725 | QDH43737.1 | H9EJ42 |
| SDNQISGYTLAA | 1 | 0 | 0 | 1502726 | CAA83661.1 | P36334 |
| SDNQKLIANKFN | 1 | 0 | 0 | 1502727 | ALA49671.1 | K9N5Q8 |
| SDPFMVNQLAYS | 1 | 0 | 0 | 1502737 | AUM60013.1 | K9N7C7 |
| SDPGFMKQYGDC | 1 | 0 | 0 | 1502738 | APO40579.1 | P59594 |
| SDPLNNFKHKVL | 1 | 0 | 0 | 1502741 | AUM60013.1 | K9N7C7 |
| SDPNLWYLRSGH | 1 | 0 | 0 | 1502743 | ALA49564.1 | K9N643 |
| SDPNMCLLSGHH | 1 | 0 | 0 | 1502744 | AVN89460.1 | K9N643 |
| SDPNMWCLRSGH | 1 | 0 | 0 | 1502745 | AXG21646.1 | K9N643 |
| SDPNMWYLRSGH | 1 | 0 | 0 | 1502746 | AXN92231.1 | K9N643 |
| SDPNMWYLRSGN | 1 | 0 | 0 | 1502747 | AWU59324.1 | K9N643 |
| SDPNTWYLRSGH | 1 | 0 | 0 | 1502749 | ASU90321.1 | K9N643 |
| SDPNYDALLISM | 1 | 0 | 0 | 1502750 | AXN92247.1 | K9N7C7 |
| SDQQPSVLFTCI | 1 | 0 | 0 | 1502763 | AVV62530.1 | K9N7D2 |
| SDQSDQFRNVQT | 1 | 0 | 0 | 1502764 | ARK08639.1 | P33469 |
| SDRDLYDKLQFM | 1 | 0 | 0 | 1502773 | AVP78030.1 | P0C6W2 |
| SDRFYRLANGCA | 1 | 0 | 0 | 1502779 | AJE26242.1 | P0C6X6 |
| SDRGQECLARSA | 1 | 0 | 0 | 1502781 | AZK15899.1 | K9N638 |
| SDRKIKKHPCMV | 1 | 0 | 0 | 1502787 | AHL18091.1 | K9N7C7 |
| SDRLAALESGKT | 1 | 0 | 0 | 1502791 | AVV62544.1 | K9N4V7 |
| SDRYHNSQDDIM | 1 | 0 | 0 | 1502798 | AAA45463.1 | A0A0N7J131 |
| SDSDSLPSVPFT | 1 | 0 | 0 | 1502802 | AVV62541.1 | K9N7D2 |
| SDSFYTQGRSCS | 1 | 0 | 0 | 1502808 | AVV62524.1 | K9N7C7 |
| SDSLGVSSNQSP | 1 | 0 | 0 | 1502823 | QBP84719.1 | P0C6X6 |
| SDSLPSVPFTCI | 1 | 0 | 0 | 1502824 | AVV62541.1 | K9N7D2 |
| SDSNCYMNVVLM | 1 | 0 | 0 | 1502827 | AUM60013.1 | K9N7C7 |
| SDSNGVPVVEYL | 1 | 0 | 0 | 1502828 | AUM60013.1 | K9N7C7 |
| SDSNMWYLRSGH | 1 | 0 | 0 | 1502829 | ALA49894.1 | K9N643 |
| SDSNYDYLGKLF | 1 | 0 | 0 | 1502831 | AVV62537.1 | K9N5Q8 |
| SDSQDGQETAVT | 1 | 0 | 0 | 1502834 | AVP78041.1 | P0C6W2 |

| | | | | | | |
|---|---|---|---|---|---|---|
| SDSQFEEDVEMS | 1 | 0 | 0 | 1502835 | AGT51787.1 | P0C6X6 |
| SDSQFPILAELA | 1 | 0 | 0 | 1502836 | AGT51764.1 | P33469 |
| SDSVKSACIEVD | 1 | 0 | 0 | 1502844 | AMW90853.1 | K9N5Q8 |
| SDTIPDEKQNGL | 1 | 0 | 0 | 1502850 | AUM60013.1 | K9N7C7 |
| SDTIPQQSQNGF | 1 | 0 | 0 | 1502852 | AVV62535.1 | K9N7C7 |
| SDTPCVYMDGMD | 1 | 0 | 0 | 1502859 | AEN19363.1 | P0C6X6 |
| SDTSLSDACQLP | 1 | 0 | 0 | 1502863 | AGT17757.1 | A0A140H1H0 |
| SDTSYINDKDTG | 1 | 0 | 0 | 1502864 | CAA79896.1 | P36334 |
| SDTVEVPPHEAD | 1 | 0 | 0 | 1502870 | QBF80585.1 | K9N7C7 |
| SDTVEVSPQVVK | 1 | 0 | 0 | 1502875 | ASU45805.1 | K9N7C7 |
| SDVDLGDISGIN | 1 | 0 | 0 | 1502883 | 1WYY_A | P59594 |
| SDVFVKKYGLEN | 1 | 0 | 0 | 1502897 | AHI48614.1 | K9N7C7 |
| SDVGFVSAYNNC | 1 | 0 | 0 | 1502902 | AIL49488.1 | P36334 |
| SDVGKGIYQTSN | 1 | 0 | 0 | 1502903 | ATO98145.1 | Q3LZX1 |
| SDVIHYNINFEE | 1 | 0 | 0 | 1502904 | BAL45639.1 | A0A0P0K6L9 |
| SDVMFEDAQSLK | 1 | 0 | 0 | 1502919 | ALA50066.1 | K9N7C7 |
| SDVMFEGAQGLK | 1 | 0 | 0 | 1502921 | ASU90140.1 | K9N7C7 |
| SDVNCNGYQHHS | 1 | 0 | 0 | 1502924 | AIW52843.1 | A0A0P0G321 |
| SDVSARDPIIQE | 1 | 0 | 0 | 1502928 | AMM42389.1 | Q19U29 |
| SDVYEADAEIVQ | 1 | 0 | 0 | 1502944 | AUM60013.1 | K9N7C7 |
| SDYANKGYIANI | 1 | 0 | 0 | 1502957 | ABD75519.1 | A0A140H1G9 |
| SDYATKVVPFNN | 1 | 0 | 0 | 1502959 | AVV62537.1 | K9N5Q8 |
| SDYCGGNPCICQ | 1 | 0 | 0 | 1502961 | BBO58900.1 | P36334 |
| SDYCGGNPCTCR | 1 | 0 | 0 | 1502963 | CAA79896.1 | P36334 |
| SDYCGGNSCTCQ | 1 | 0 | 0 | 1502964 | AGT51690.1 | P36334 |
| SDYDIPTTENLY | 1 | 0 | 0 | 1502970 | 3EJG_A | A0A0P0LKV0 |
| SDYFDIVRRLAL | 1 | 0 | 0 | 1502973 | AVV62524.1 | K9N7C7 |
| SEAAAQGFQACR | 1 | 0 | 0 | 1502985 | AWK59942.1 | H9EJ42 |
| SEAAARSSQLAS | 1 | 0 | 0 | 1502986 | AUM60014.1 | K9N5Q8 |
| SEAASNGFRACR | 1 | 0 | 0 | 1502988 | NP_073549.1 | A0A0P0LKV0 |
| SEAFEKFISLFA | 1 | 0 | 0 | 1502992 | ANI69899.1 | K9N7C7 |
| SEAFEKFVSLFA | 1 | 0 | 0 | 1502994 | AXN92247.1 | K9N7C7 |
| SEAIKPIIEAEN | 1 | 0 | 0 | 1503005 | P0C5B5.1 | Q0ZJ80 |
| SEAKCWVEPDLK | 1 | 0 | 0 | 1503009 | AVV62524.1 | K9N7C7 |
| SECRLQNGDFLC | 1 | 0 | 0 | 1503028 | APO40578.1 | P0C6X7 |
| SECTMFKDAEGR | 1 | 0 | 0 | 1503031 | AVN89322.1 | K9N7C7 |
| SECTMFRDAEGK | 1 | 0 | 0 | 1503033 | AVV62535.1 | K9N7C7 |
| SEDIYNTLTEVK | 1 | 0 | 0 | 1503044 | AUM60013.1 | K9N7C7 |
| SEEAIPTRFPPG | 1 | 0 | 0 | 1503058 | ANZ78879.1 | P33469 |
| SEEPQGNVVTSV | 1 | 0 | 0 | 1503065 | ARB07436.1 | A0A140H1G9 |
| SEEVVENPTIQKDVLECN | 1 | 0 | 0 | 1503077 | P0DTC1.1 | P0DTD1 |
| SEEVVENPTVQK | 1 | 0 | 0 | 1503078 | ARO76381.1 | P0C6U8 |
| SEFNLPGCNGGS | 1 | 0 | 0 | 1503091 | AUM60013.1 | K9N7C7 |
| SEFPAITIGSNF | 1 | 0 | 0 | 1503093 | BBO58761.1 | P36334 |
| SEFQLASINFNQ | 1 | 0 | 0 | 1503097 | AUM60014.1 | K9N5Q8 |
| SEFTIGNMEEFI | 1 | 0 | 0 | 1503103 | CAA79896.1 | P36334 |
| SEGLNVLWNKFV | 1 | 0 | 0 | 1503111 | ARU07593.1 | H9EJ42 |
| SEGLVGTTLQGK | 1 | 0 | 0 | 1503112 | AUM60013.1 | K9N7C7 |
| SEGNALYNNECG | 1 | 0 | 0 | 1503113 | AGT21359.1 | A0A0P0LKV0 |
| SEGNSIIGVPYP | 1 | 0 | 0 | 1503115 | AVL25553.1 | A0A0P0G321 |
| SEKLFVRSREED | 1 | 0 | 0 | 1503160 | ATG84722.1 | K9N7C7 |
| SEKQVEQKVAEN | 1 | 0 | 0 | 1503163 | AVP78030.1 | P0C6W2 |
| SEKSYERQSPFE | 1 | 0 | 0 | 1503169 | ARO76381.1 | P0C6U8 |
| SELAKGVSLDNV | 1 | 0 | 0 | 1503174 | APO40578.1 | P0C6X7 |
| SELPKEQRVQGS | 1 | 0 | 0 | 1503189 | ASJ26617.1 | K9N4V7 |
| SELYGTPDPMFL | 1 | 0 | 0 | 1503202 | AVV62535.1 | K9N7C7 |
| SEMLKLGTSDPQ | 1 | 0 | 0 | 1503205 | AZS52622.1 | Q19U29 |
| SENAYLKYAKVE | 1 | 0 | 0 | 1503211 | AVV62524.1 | K9N7C7 |
| SENLFVRSKEED | 1 | 0 | 0 | 1503215 | ANF29194.1 | K9N7C7 |
| SENQKQIANKFN | 1 | 0 | 0 | 1503224 | AVV62526.1 | K9N5Q8 |
| SEPALLSRNIKC | 1 | 0 | 0 | 1503232 | CAA79896.1 | P36334 |
| SEPIVQHNTEPK | 1 | 0 | 0 | 1503236 | APO40578.1 | P0C6X7 |
| SEPPKEQRVQGI | 1 | 0 | 0 | 1503242 | AHI48535.1 | K9N4V7 |
| SEPQRGRQSRIP | 1 | 0 | 0 | 1503255 | ALK28785.1 | A0A0N7J131 |
| SEQLDFIESKRG | 1 | 0 | 0 | 1503269 | ARO76381.1 | P0C6U8 |
| SEQNNFTGYLKL | 1 | 0 | 0 | 1503273 | AVP78041.1 | P0C6W2 |
| SEQPWKVVPRNL | 1 | 0 | 0 | 1503276 | AIW52723.1 | A0A0N7J131 |
| SEQSWKVIPRNL | 1 | 0 | 0 | 1503279 | AAA45463.1 | A0A0N7J131 |
| SEQVIPPTSDYL | 1 | 0 | 0 | 1503280 | YP_173240.1 | Q0ZJ83 |
| SERCNVIADFKS | 1 | 0 | 0 | 1503281 | AXN73435.1 | K9N7C7 |
| SERDNINAILTE | 1 | 0 | 0 | 1503285 | AHF54662.1 | P36334 |
| SERINVVPGVST | 1 | 0 | 0 | 1503287 | AKT07951.1 | H9EJ42 |
| SERVNVVPGVPT | 1 | 0 | 0 | 1503291 | AFD98763.1 | H9EJ42 |
| SESAAISAQLAK | 1 | 0 | 0 | 1503294 | AVN89365.1 | K9N5Q8 |
| SESAALSAQLAK | 1 | 0 | 0 | 1503295 | AXN92228.1 | K9N5Q8 |
| SESGVKETKEIN | 1 | 0 | 0 | 1503302 | ATN39877.2 | P0C6X6 |
| SESLNDNLLEIL | 1 | 0 | 0 | 1503309 | AVP78041.1 | P0C6W2 |
| SETKCTLKSLSV | 1 | 0 | 0 | 1503320 | AVP78042.1 | Q3LZX1 |
| SETKEQKHEIEK | 1 | 0 | 0 | 1503322 | AAA45463.1 | A0A0N7J131 |
| SETPNAKEEILG | 1 | 0 | 0 | 1503328 | P0C6W6.1 | P0C6W2 |
| SETQRGRQGRIP | 1 | 0 | 0 | 1503331 | AIW52715.1 | A0A0N7J131 |
| SEVDWKFYDAQP | 1 | 0 | 0 | 1503340 | AVP78030.1 | P0C6W2 |
| SEVEPEPEPTLE | 1 | 0 | 0 | 1503342 | ARO76381.1 | P0C6U8 |
| SEVPPRCIAPVD | 1 | 0 | 0 | 1503349 | AUM60014.1 | K9N5Q8 |
| SEVSTDEVGDNV | 1 | 0 | 0 | 1503353 | AIW52805.1 | H9EJ69 |
| SEYKGSITDVFY | 1 | 0 | 0 | 1503371 | AVP78041.1 | P0C6W2 |
| SEYLVGDSIVTL | 1 | 0 | 0 | 1503373 | AVA26872.1 | H9EJ42 |
| SEYPYVIFCQTA | 1 | 0 | 0 | 1503377 | AVV62524.1 | K9N7C7 |
| SEYRQYLDAYNM | 1 | 0 | 0 | 1503380 | APO40578.1 | P0C6X7 |
| SEYSKRSAIEDL | 1 | 0 | 0 | 1503381 | AVV62526.1 | K9N5Q8 |
| SFADNNDITNIN | 1 | 0 | 0 | 1503388 | ASJ26617.1 | K9N4V7 |

| | | | | | | |
|---|---|---|---|---|---|---|
| SFADNNDNNNQQ | 1 | 0 | 0 | 1503390 | AUM60021.1 | K9N4V7 |
| SFADNNDTTNTN | 1 | 0 | 0 | 1503391 | ALW82676.1 | K9N4V7 |
| SFADVMRYNLNF | 1 | 0 | 0 | 1503393 | ALJ53431.1 | A0A0P0G321 |
| SFAELETAQKSY | 1 | 0 | 0 | 1503395 | AUM60013.1 | K9N7C7 |
| SFAGPISQFNYK | 1 | 0 | 0 | 1503396 | AHX71943.1 | K9N5Q8 |
| SFAGYTESAVNF | 1 | 0 | 0 | 1503398 | AVN89336.1 | K9N4V0 |
| SFAIYHTPATDC | 1 | 0 | 0 | 1503400 | ATG84866.1 | K9N5Q8 |
| SFASIPFAQSIF | 1 | 0 | 0 | 1503407 | AVV62526.1 | K9N5Q8 |
| SFATYHTPAIDC | 1 | 0 | 0 | 1503409 | ASU91010.1 | K9N5Q8 |
| SFAYNNDITNTN | 1 | 0 | 0 | 1503417 | AVN89381.1 | K9N4V7 |
| SFCAFAVEPAKA | 1 | 0 | 0 | 1503420 | APO40578.1 | P0C6X7 |
| SFCKSGNLVLNN | 1 | 0 | 0 | 1503425 | AIL49332.1 | P30215 |
| SFCKTIQSALSV | 1 | 0 | 0 | 1503428 | NP_073549.1 | A0A0P0LKV0 |
| SFDALELGDTQG | 1 | 0 | 0 | 1503437 | AVP78030.1 | P0C6W2 |
| SFDALKLGDTQG | 1 | 0 | 0 | 1503438 | AVP78041.1 | P0C6W2 |
| SFDGNMYGAFQD | 1 | 0 | 0 | 1503446 | AVV62524.1 | K9N7C7 |
| SFDNLVGYHSDN | 1 | 0 | 0 | 1503462 | AUM60014.1 | K9N5Q8 |
| SFDSSYLNESGA | 1 | 0 | 0 | 1503469 | AST12963.1 | A0A0P0LKV0 |
| SFDVDTGVYSVS | 1 | 0 | 0 | 1503471 | AVV62526.1 | K9N5Q8 |
| SFDVESGVYSVS | 1 | 0 | 0 | 1503472 | AXN92228.1 | K9N5Q8 |
| SFDYLTREAKTR | 1 | 0 | 0 | 1503483 | ANI69910.1 | K9N7C7 |
| SFEAHKDFFKDG | 1 | 0 | 0 | 1503484 | ARK08657.1 | P0C6X6 |
| SFEKLHCEHLQF | 1 | 0 | 0 | 1503496 | AWO68974.1 | A0A0P0G321 |
| SFELLNAPATVS | 1 | 0 | 0 | 1503501 | 2GHV_E | P59594 |
| SFERLRDYLAVY | 1 | 0 | 0 | 1503509 | AUM60023.1 | K9N7C7 |
| SFEVLEDTLDDG | 1 | 0 | 0 | 1503512 | AVQ05261.1 | P0C6X6 |
| SFFDCGVNALAG | 1 | 0 | 0 | 1503516 | ARU07593.1 | H9EJ42 |
| SFFDCGVNALDG | 1 | 0 | 0 | 1503517 | AWK59942.1 | H9EJ42 |
| SFFLADNLSAAE | 1 | 0 | 0 | 1503520 | AVV62524.1 | K9N7C7 |
| SFFNPEPITMAN | 1 | 0 | 0 | 1503523 | AVV62537.1 | K9N5Q8 |
| SFFPQFTSDQAI | 1 | 0 | 0 | 1503524 | ABD75596.1 | Q0ZJ82 |
| SFFSWYGPCHTH | 1 | 0 | 0 | 1503528 | AMQ49019.1 | K9N7D2 |
| SFGITQEYIQTT | 1 | 0 | 0 | 1503535 | AVV62537.1 | K9N5Q8 |
| SFGKSIIYTGDW | 1 | 0 | 0 | 1503536 | AWK59942.1 | H9EJ42 |
| SFGNRSGNGILK | 1 | 0 | 0 | 1503542 | QBP84763.1 | P33469 |
| SFHFTRFYNYTG | 1 | 0 | 0 | 1503558 | ARU07576.1 | A0A140H1H0 |
| SFIDDYYGYLMK | 1 | 0 | 0 | 1503564 | ADO24650.1 | H9EJ42 |
| SFIDVINKALEM | 1 | 0 | 0 | 1503566 | ARO76381.1 | P0C6U8 |
| SFIEDLLFDKVK | 1 | 0 | 0 | 1503567 | ARU07577.1 | A0A140H1H1 |
| SFINICFEATTA | 1 | 0 | 0 | 1503585 | AVV62535.1 | K9N7C7 |
| SFINVCFEATAA | 1 | 0 | 0 | 1503587 | AVV62524.1 | K9N7C7 |
| SFISIVNKALEL | 1 | 0 | 0 | 1503595 | APO40578.1 | P0C6X7 |
| SFISQPFYFDSY | 1 | 0 | 0 | 1503597 | BBA20986.1 | A0A140H1H1 |
| SFKAPDTYKCPQ | 1 | 0 | 0 | 1503604 | ARE30008.1 | P36334 |
| SFKDELEEFFKN | 1 | 0 | 0 | 1503605 | AVV62526.1 | K9N5Q8 |
| SFKENVDFINSM | 1 | 0 | 0 | 1503608 | AXX83325.1 | Q80872 |
| SFKEYFDLQNCS | 1 | 0 | 0 | 1503609 | AVV62537.1 | K9N5Q8 |
| SFKEYFNLRDCL | 1 | 0 | 0 | 1503610 | AUM60014.1 | K9N5Q8 |
| SFKEYFNLRNST | 1 | 0 | 0 | 1503611 | AGV08520.1 | K9N5Q8 |
| SFKHKVFTDSVT | 1 | 0 | 0 | 1503615 | AVV62524.1 | K9N7C7 |
| SFKKVAAIPYKA | 1 | 0 | 0 | 1503621 | AVV62524.1 | K9N7C7 |
| SFKPRSQMEVDF | 1 | 0 | 0 | 1503626 | APO40578.1 | P0C6X7 |
| SFKTADGDQRQL | 1 | 0 | 0 | 1503636 | AJQ24083.1 | P33469 |
| SFKTADGQQKPL | 1 | 0 | 0 | 1503638 | ABA60796.1 | Q19U29 |
| SFLEMKSGKQVE | 1 | 0 | 0 | 1503654 | AVP78041.1 | P0C6W2 |
| SFLNGYYPTSGS | 1 | 0 | 0 | 1503670 | AKA09662.1 | P36334 |
| SFLPKEEGDVLL | 1 | 0 | 0 | 1503677 | ATQ39389.1 | K9N7C7 |
| SFLQSGIVKMVS | 1 | 0 | 0 | 1503681 | ARU07575.1 | A0A140H1G9 |
| SFNENGTMSNAV | 1 | 0 | 0 | 1503705 | ATO98145.1 | Q3LZX1 |
| SFNFDQKELLDY | 1 | 0 | 0 | 1503710 | ATN39862.1 | P0C6X6 |
| SFNIGRSIVYET | 1 | 0 | 0 | 1503713 | AIV41804.1 | P0C6X6 |
| SFNKAIDGFGVF | 1 | 0 | 0 | 1503716 | ARU07593.1 | H9EJ42 |
| SFNNIDTGPPSI | 1 | 0 | 0 | 1503726 | AIX10756.1 | P36334 |
| SFNNKNTGPPFI | 1 | 0 | 0 | 1503727 | AKA09659.1 | P36334 |
| SFNNRDTGPLSI | 1 | 0 | 0 | 1503729 | ANZ78836.1 | P36334 |
| SFNNRDTGRPSI | 1 | 0 | 0 | 1503730 | BBO58746.1 | P36334 |
| SFNVNSLAYYEP | 1 | 0 | 0 | 1503740 | AVV62535.1 | K9N7C7 |
| SFPPGQGVPLNA | 1 | 0 | 0 | 1503753 | AVV62533.1 | K9N4V7 |
| SFQQFGKDASDF | 1 | 0 | 0 | 1503762 | AVP78031.1 | Q3LZX1 |
| SFRTLKSGEGFN | 1 | 0 | 0 | 1503776 | NP_073549.1 | A0A0P0LKV0 |
| SFSASISAFVET | 1 | 0 | 0 | 1503777 | AVP78030.1 | P0C6W2 |
| SFSAYSGDIPHY | 1 | 0 | 0 | 1503781 | AUM60024.1 | K9N5Q8 |
| SFSIKANICNSN | 1 | 0 | 0 | 1503796 | AGT17757.1 | A0A140H1H0 |
| SFSNTKYYDHSQ | 1 | 0 | 0 | 1503802 | ARK08650.1 | P30215 |
| SFSSVNDAITQT | 1 | 0 | 0 | 1503808 | AIW52843.1 | A0A0P0G321 |
| SFSVQKHIGPPA | 1 | 0 | 0 | 1503816 | QBF80508.1 | K9N7C7 |
| SFTCNNIDVAKI | 1 | 0 | 0 | 1503821 | ABD43176.1 | P36334 |
| SFTEQLGVPLVI | 1 | 0 | 0 | 1503825 | AMM42637.1 | A0A0P0G321 |
| SFTNLDKLKFKE | 1 | 0 | 0 | 1503846 | AUM60013.1 | K9N7C7 |
| SFTQWYHEHIGG | 1 | 0 | 0 | 1503849 | AVN89333.1 | K9N7C7 |
| SFTSYGNTVSEA | 1 | 0 | 0 | 1503853 | Q14EA9.1 | Q0ZJ84 |
| SFVCQDSTRHTY | 1 | 0 | 0 | 1503860 | APO40584.1 | P59635 |
| SFVDDFQGYLQK | 1 | 0 | 0 | 1503861 | AIG96641.1 | A0A0P0LKV0 |
| SFVDDNKFDVLK | 1 | 0 | 0 | 1503863 | AGT51657.1 | P0C6X6 |
| SFVDDVNGMKVV | 1 | 0 | 0 | 1503864 | AVV62535.1 | K9N7C7 |
| SFVEGAINVGKN | 1 | 0 | 0 | 1503867 | AUM60013.1 | K9N7C7 |
| SFVMMSALPAEY | 1 | 0 | 0 | 1503884 | ARO76381.1 | P0C6U8 |
| SFVNDDTGVQVV | 1 | 0 | 0 | 1503891 | AVV62524.1 | K9N7C7 |
| SFVSKIIHFFKT | 1 | 0 | 0 | 1503901 | QBQ01836.1 | P0C6X6 |
| SFVSKIVHFFET | 1 | 0 | 0 | 1503902 | ATN39862.1 | P0C6X6 |
| SFYAPDEDEEEP | 1 | 0 | 0 | 1503918 | APO40578.1 | P0C6X7 |

| | | | | | | |
|---|---|---|---|---|---|---|
| SFYCEDHKPQYS | 1 | 0 | 0 | 1503923 | AGT51438.1 | P0C6X6 |
| SFYEDRILEFKV | 1 | 0 | 0 | 1503924 | AUM60013.1 | K9N7C7 |
| SFYGFSNVVYPD | 1 | 0 | 0 | 1503929 | ABD75600.1 | A0A140H1H0 |
| SFYPPEDEGEDD | 1 | 0 | 0 | 1503940 | AVP78041.1 | P0C6W2 |
| SFYRDLYSAPTT | 1 | 0 | 0 | 1503942 | AUM60013.1 | K9N7C7 |
| SFYRELYDAPTT | 1 | 0 | 0 | 1503943 | AVV62535.1 | K9N7C7 |
| SFYVYANGGKGF | 1 | 0 | 0 | 1503948 | AVP78041.1 | P0C6W2 |
| SGAFRELTRELN | 1 | 0 | 0 | 1503956 | ARO76381.1 | P0C6U8 |
| SGALAEHDFFTW | 1 | 0 | 0 | 1503964 | AIW52827.1 | H9EJ42 |
| SGCETSNVPPIV | 1 | 0 | 0 | 1503978 | ACA49954.1 | H9EJ42 |
| SGCMLRVCIKTA | 1 | 0 | 0 | 1503992 | AMD02845.1 | K9N796 |
| SGDAHIPSGTSF | 1 | 0 | 0 | 1503996 | AIG59196.1 | A0A0P0K6L9 |
| SGDASTAFANSV | 1 | 0 | 0 | 1504000 | AKL87224.1 | P0C6X6 |
| SGDCPSVDMQPD | 1 | 0 | 0 | 1504003 | ATQ39390.1 | K9N5Q8 |
| SGDCVVEASTDF | 1 | 0 | 0 | 1504004 | AWK59942.1 | H9EJ42 |
| SGDKYYPVVVDD | 1 | 0 | 0 | 1504017 | AXX83372.1 | P0C6X6 |
| SGDNGTITVEEL | 1 | 0 | 0 | 1504021 | AVP78034.1 | Q3LZX9 |
| SGDRTEVPQLVN | 1 | 0 | 0 | 1504025 | ANF29261.1 | K9N5Q8 |
| SGDSSEGGAKET | 1 | 0 | 0 | 1504026 | AVQ05261.1 | P0C6X6 |
| SGDVNVVAASGS | 1 | 0 | 0 | 1504034 | ASU45805.1 | K9N7C7 |
| SGDVPHYVLPGQ | 1 | 0 | 0 | 1504035 | AUM60014.1 | K9N5Q8 |
| SGDYTLANTTTA | 1 | 0 | 0 | 1504039 | AGV08437.1 | K9N7C7 |
| SGEIATKMFDSF | 1 | 0 | 0 | 1504044 | QBM11736.1 | K9N638 |
| SGETLGVLVPHT | 1 | 0 | 0 | 1504051 | APO40578.1 | P0C6X7 |
| SGFAVYVKSKVG | 1 | 0 | 0 | 1504062 | BAM16435.1 | Q01455 |
| SGFCGQGIHIVS | 1 | 0 | 0 | 1504063 | ALJ54495.1 | K9N5Q8 |
| SGGFYRFMVDST | 1 | 0 | 0 | 1504099 | AGT51337.1 | H9EJ42 |
| SGGGKCFNCAPA | 1 | 0 | 0 | 1504101 | AAQ90006.1 | A0A0P0K6L9 |
| SGGGKCYNCRPA | 1 | 0 | 0 | 1504103 | ABB90509.1 | A0A0P0K6L9 |
| SGGNVVPHYSWF | 1 | 0 | 0 | 1504108 | AWW13570.1 | P33469 |
| SGGYIPEAPRDG | 1 | 0 | 0 | 1504121 | 6U7H_A | A0A0P0K6L9 |
| SGHLIDEVQNWL | 1 | 0 | 0 | 1504134 | AUM60027.1 | K9N643 |
| SGHMLDDVQRWL | 1 | 0 | 0 | 1504136 | AVV62540.1 | K9N643 |
| SGHNLAKNCLHV | 1 | 0 | 0 | 1504140 | AVP78041.1 | P0C6W2 |
| SGIAIEQDKNTQ | 1 | 0 | 0 | 1504147 | AVP78031.1 | Q3LZX1 |
| SGIAVEQDRNTR | 1 | 0 | 0 | 1504150 | ATO98231.1 | Q3LZX1 |
| SGIGAVGDAGAA | 1 | 0 | 0 | 1504154 | AVV62544.1 | K9N4V7 |
| SGIGGGDLLYLD | 1 | 0 | 0 | 1504157 | AVN89359.1 | K9N4V7 |
| SGIGTCPAGTNY | 1 | 0 | 0 | 1504158 | CAA83660.1 | P36334 |
| SGISDLSLKQKL | 1 | 0 | 0 | 1504176 | ALA49758.1 | K9N7C7 |
| SGITQQVLSENQ | 1 | 0 | 0 | 1504183 | 4NJL_A | K9N5Q8 |
| SGKDDQIGYYRR | 1 | 0 | 0 | 1504192 | Q3I5I7.1 | Q3LZX4 |
| SGKISSKADNSI | 1 | 0 | 0 | 1504195 | AMK59676.1 | P30215 |
| SGKNNGIGTCPA | 1 | 0 | 0 | 1504202 | ANZ78838.1 | P36334 |
| SGKNNGVGTCPA | 1 | 0 | 0 | 1504203 | QEG03757.1 | P36334 |
| SGKTKAAPKVIT | 1 | 0 | 0 | 1504210 | AVV62544.1 | K9N4V7 |
| SGKYAPNSRSTS | 1 | 0 | 0 | 1504219 | AJQ24067.1 | P33469 |
| SGLAIKSDYCGG | 1 | 0 | 0 | 1504220 | CAA83660.1 | P36334 |
| SGLCVDGRNGYV | 1 | 0 | 0 | 1504226 | AIW52756.1 | A0A0P0K6L9 |
| SGLEEGESECIS | 1 | 0 | 0 | 1504231 | AQZ41341.1 | K9N7C7 |
| SGLGTVDADYKK | 1 | 0 | 0 | 1504233 | AIW52757.1 | A0A0P0K6L9 |
| SGLGTVDANYKK | 1 | 0 | 0 | 1504234 | CAA71147.1 | A0A0P0K6L9 |
| SGLKTMIVSAGQ | 1 | 0 | 0 | 1504241 | AQZ41321.1 | K9N7C7 |
| SGLLNVQKVAQP | 1 | 0 | 0 | 1504245 | APT69882.1 | A0A0P0ILY9 |
| SGLTCSTDLQKA | 1 | 0 | 0 | 1504265 | CAA79896.1 | P36334 |
| SGLTCSTDSQKA | 1 | 0 | 0 | 1504266 | BBO58736.1 | P36334 |
| SGLVPRGSAMGL | 1 | 0 | 0 | 1504272 | 1SSK_A | P59595 |
| SGLVPRGSGMKE | 1 | 0 | 0 | 1504273 | 3R24_A | P0C6X7 |
| SGLYPTITVPEE | 1 | 0 | 0 | 1504275 | AXG21652.1 | K9N7C7 |
| SGMIRYYDVMAN | 1 | 0 | 0 | 1504283 | AIV41888.1 | P0C6X6 |
| SGMKETAAAKFE | 1 | 0 | 0 | 1504284 | 3R24_A | P0C6X7 |
| SGNDALARFTIF | 1 | 0 | 0 | 1504292 | ARB07436.1 | A0A140H1G9 |
| SGNGGLFNTLRE | 1 | 0 | 0 | 1504296 | AVP78031.1 | Q3LZX1 |
| SGNHCPADNSYT | 1 | 0 | 0 | 1504299 | AID55096.1 | K9N5Q8 |
| SGNHCPAGNFYT | 1 | 0 | 0 | 1504300 | ALJ54446.1 | K9N5Q8 |
| SGNHCPAGNSYT | 1 | 0 | 0 | 1504302 | AXN92228.1 | K9N5Q8 |
| SGNHCPAGNYYT | 1 | 0 | 0 | 1504303 | ATG84899.1 | K9N5Q8 |
| SGNHLHSVHNWL | 1 | 0 | 0 | 1504304 | AWU59324.1 | K9N643 |
| SGNLDEPQKDVY | 1 | 0 | 0 | 1504305 | AOL02457.1 | P33469 |
| SGNLLLDKRTSC | 1 | 0 | 0 | 1504309 | APO40578.1 | P0C6X7 |
| SGNLVGFKDVTN | 1 | 0 | 0 | 1504312 | CAA71146.1 | A0A0P0K6L9 |
| SGNLYGFRDYIT | 1 | 0 | 0 | 1504313 | ARK08661.1 | P36334 |
| SGNPDEPHKDVY | 1 | 0 | 0 | 1504316 | AIX10716.1 | P33469 |
| SGNPVVEYLSKE | 1 | 0 | 0 | 1504319 | QGV13494.1 | K9N7C7 |
| SGNRAPTSGVIP | 1 | 0 | 0 | 1504320 | AMK59681.1 | P33469 |
| SGNRNPTSGVTP | 1 | 0 | 0 | 1504323 | AJQ24067.1 | P33469 |
| SGNRPGNGILKW | 1 | 0 | 0 | 1504324 | AMM42456.1 | P33469 |
| SGNRSGILKKTS | 1 | 0 | 0 | 1504325 | ARU07581.1 | Q19U29 |
| SGNSSRNASPGP | 1 | 0 | 0 | 1504335 | AVV62533.1 | K9N4V7 |
| SGNTVVLTFTTK | 1 | 0 | 0 | 1504345 | AIW52712.1 | A0A0N7J131 |
| SGNVIEFHLDGE | 1 | 0 | 0 | 1504349 | APO40578.1 | P0C6X7 |
| SGNYCPAGNSYT | 1 | 0 | 0 | 1504350 | AVN89387.1 | K9N5Q8 |
| SGNYLSISNELL | 1 | 0 | 0 | 1504352 | AVQ05263.1 | P30215 |
| SGPDQICNTPTQ | 1 | 0 | 0 | 1504356 | ARB07411.1 | P0C6X6 |
| SGPDQTCNTPTQ | 1 | 0 | 0 | 1504357 | AEN19363.1 | P0C6X6 |
| SGPEWMDDFEAD | 1 | 0 | 0 | 1504358 | AWM99580.1 | K9N7C7 |
| SGPGKNNGIGTC | 1 | 0 | 0 | 1504359 | CAA79896.1 | P36334 |
| SGPGKNNGMGTC | 1 | 0 | 0 | 1504360 | AIL49487.1 | P36334 |
| SGPISPTSLEIF | 1 | 0 | 0 | 1504363 | APU51922.1 | Q4VID0 |
| SGPISQFNYKQS | 1 | 0 | 0 | 1504365 | ALA50067.1 | K9N5Q8 |
| SGPLQTAVGCVI | 1 | 0 | 0 | 1504367 | AVV62537.1 | K9N5Q8 |

| | | | | | | |
|---|---|---|---|---|---|---|
| SGQAFCDYTDKL | 1 | 0 | 0 | 1504376 | AGT51737.1 | P0C6X6 |
| SGQGIFFEVNAT | 1 | 0 | 0 | 1504378 | AAX84795.1 | P36334 |
| SGQGVPLNANST | 1 | 0 | 0 | 1504379 | AZK15907.1 | K9N4V7 |
| SGRENLYFQGGG | 1 | 0 | 0 | 1504387 | 6NB6_A | P59594 |
| SGRKIYEIERGL | 1 | 0 | 0 | 1504396 | AGT51737.1 | P0C6X6 |
| SGRKIYEVERGL | 1 | 0 | 0 | 1504399 | AEN19363.1 | P0C6X6 |
| SGRSANTSRSTL | 1 | 0 | 0 | 1504409 | AJQ24074.1 | P33469 |
| SGRTPTSGVTPD | 1 | 0 | 0 | 1504413 | AXX83311.1 | P33469 |
| SGSDINCNIPTQ | 1 | 0 | 0 | 1504424 | P0C6U5.1 | A0A140H1G9 |
| SGSIRFDSTLPG | 1 | 0 | 0 | 1504434 | ARU07581.1 | Q19U29 |
| SGSLATRLMSIP | 1 | 0 | 0 | 1504438 | ATP16762.1 | Q4VID0 |
| SGSPCESHGKQV | 1 | 0 | 0 | 1504458 | AER30333.1 | P0C6X7 |
| SGTCGIVELEKG | 1 | 0 | 0 | 1504480 | P0C6W6.1 | P0C6W2 |
| SGTHTGSAFDGT | 1 | 0 | 0 | 1504488 | AVN89451.1 | K9N7C7 |
| SGTNDICVFENK | 1 | 0 | 0 | 1504491 | ATQ39389.1 | K9N7C7 |
| SGTPPQVYNFNR | 1 | 0 | 0 | 1504496 | AVV62537.1 | K9N5Q8 |
| SGTTCSTDLQKS | 1 | 0 | 0 | 1504511 | CAA83660.1 | P36334 |
| SGTTTNVEKIRQ | 1 | 0 | 0 | 1504516 | AVN89333.1 | K9N7C7 |
| SGTYHCVTTKAK | 1 | 0 | 0 | 1504526 | ALA49758.1 | K9N7C7 |
| SGVCKFKGKFVQ | 1 | 0 | 0 | 1504535 | AVV62524.1 | K9N7C7 |
| SGVGRCYNCRPA | 1 | 0 | 0 | 1504546 | CAA71146.1 | A0A0P0K6L9 |
| SGVIPDMADQIA | 1 | 0 | 0 | 1504550 | AMK59681.1 | P33469 |
| SGVKETKEINII | 1 | 0 | 0 | 1504555 | ATN39877.2 | P0C6X6 |
| SGVKKPFKVEDS | 1 | 0 | 0 | 1504556 | AEN19363.1 | P0C6X6 |
| SGVKKTFKVEDS | 1 | 0 | 0 | 1504557 | AQN78677.1 | P0C6X6 |
| SGVLPVNPVLHT | 1 | 0 | 0 | 1504562 | AEN19363.1 | P0C6X6 |
| SGVNDICVFENK | 1 | 0 | 0 | 1504569 | AUM60013.1 | K9N7C7 |
| SGVNDVCVFENK | 1 | 0 | 0 | 1504570 | AVV62524.1 | K9N7C7 |
| SGVSDLSLKQKL | 1 | 0 | 0 | 1504575 | AXN92247.1 | K9N7C7 |
| SGVSNLSLKQKL | 1 | 0 | 0 | 1504576 | QAT98908.1 | K9N7C7 |
| SGVTPDMADEIA | 1 | 0 | 0 | 1504584 | AMM42457.1 | P33469 |
| SGVTPDMADHIA | 1 | 0 | 0 | 1504585 | AIL49401.1 | P33469 |
| SGVTRDMADQIA | 1 | 0 | 0 | 1504588 | AIL49470.1 | P33469 |
| SGVVGCKSIVKE | 1 | 0 | 0 | 1504591 | ARU07575.1 | A0A140H1G9 |
| SGVVNPIYDEPT | 1 | 0 | 0 | 1504595 | APO40580.1 | P59632 |
| SGVVNPVMDPIY | 1 | 0 | 0 | 1504596 | ARO76383.1 | P59632 |
| SGYCVDYSKNGG | 1 | 0 | 0 | 1504621 | 6NZK_A | P36334 |
| SGYGKSTFNKDI | 1 | 0 | 0 | 1504627 | AAY32939.1 | H9EJ42 |
| SGYIPEAPRDGQ | 1 | 0 | 0 | 1504630 | 6CRW_A | P59594 |
| SGYYYPEPIIEN | 1 | 0 | 0 | 1504648 | AIX09799.1 | P36334 |
| SHATTYLSLSDR | 1 | 0 | 0 | 1504657 | NP_073549.1 | A0A0P0LKV0 |
| SHAVQNTVFNAS | 1 | 0 | 0 | 1504658 | AVL25595.1 | H9EJ69 |
| SHAVQNTVLNAS | 1 | 0 | 0 | 1504659 | AIW52795.1 | H9EJ69 |
| SHAWCRNQGRFC | 1 | 0 | 0 | 1504662 | AQN78665.1 | Q04853 |
| SHAWQPGVAMPN | 1 | 0 | 0 | 1504663 | AVP78041.1 | P0C6W2 |
| SHDVSDDEWAAA | 1 | 0 | 0 | 1504671 | AXG21664.1 | K9N638 |
| SHEISDDEWAAA | 1 | 0 | 0 | 1504675 | AHI48784.1 | K9N7C7 |
| SHENKLNSCCRI | 1 | 0 | 0 | 1504678 | ALA49351.1 | K9N7C7 |
| SHETSDDDWTAA | 1 | 0 | 0 | 1504680 | AVN89374.1 | K9N7C7 |
| SHEVSDDEWAAA | 1 | 0 | 0 | 1504685 | AXN92247.1 | K9N7C7 |
| SHFTTQFVDNVK | 1 | 0 | 0 | 1504704 | AIW52747.1 | A0A0P0K6L9 |
| SHFTTQFVGNVK | 1 | 0 | 0 | 1504706 | AIW52740.1 | A0A0P0K6L9 |
| SHFTTQFVSNVK | 1 | 0 | 0 | 1504708 | AIW52741.1 | A0A0P0K6L9 |
| SHGCDLMQARAF | 1 | 0 | 0 | 1504714 | AVV62524.1 | K9N7C7 |
| SHGLDLMQASQI | 1 | 0 | 0 | 1504723 | AUM60013.1 | K9N7C7 |
| SHHSDLEVTSDS | 1 | 0 | 0 | 1504738 | APO40578.1 | P0C6X7 |
| SHKNNFGVITSF | 1 | 0 | 0 | 1504755 | Q14EA9.1 | Q0ZJ84 |
| SHLLLLAGGLEA | 1 | 0 | 0 | 1504772 | AVP78032.1 | Q3LZX0 |
| SHMPLSNFEHKV | 1 | 0 | 0 | 1504793 | 5DUS_A | K9N638 |
| SHSMAVVDGKQI | 1 | 0 | 0 | 1504829 | AEN19363.1 | P0C6X6 |
| SHSVVYSRYCFS | 1 | 0 | 0 | 1504838 | BBA20983.1 | A0A140H1H1 |
| SHTIGTLYVSWS | 1 | 0 | 0 | 1504841 | 6U7F_C | A0A0P0K6L9 |
| SHTLTKHTEVRA | 1 | 0 | 0 | 1504842 | APT69856.1 | A0A0P0K6L9 |
| SHVAQNTVLNAC | 1 | 0 | 0 | 1504850 | ABK63972.1 | H9EJ69 |
| SHYVLNIDWLWF | 1 | 0 | 0 | 1504872 | AXN92247.1 | K9N7C7 |
| SHYVVTDIKQVG | 1 | 0 | 0 | 1504874 | ARU07575.1 | A0A140H1G9 |
| SIAFNHPIQIDQ | 1 | 0 | 0 | 1504892 | ASU91208.1 | K9N5Q8 |
| SIAFNHPIQVDH | 1 | 0 | 0 | 1504893 | AMQ49004.1 | K9N5Q8 |
| SIAFNHPIQVGQ | 1 | 0 | 0 | 1504895 | ALJ54452.1 | K9N5Q8 |
| SIAGVGWTTGLS | 1 | 0 | 0 | 1504901 | AOR17479.1 | K9N5Q8 |
| SIAPSTVWEDGD | 1 | 0 | 0 | 1504913 | QEJ82215.1 | K9N5Q8 |
| SIASFNEWAMAN | 1 | 0 | 0 | 1504915 | AUM60013.1 | K9N7C7 |
| SIDEFADVVKEQ | 1 | 0 | 0 | 1504938 | AZK15899.1 | K9N638 |
| SIDSTVLTYIPA | 1 | 0 | 0 | 1504946 | AVN89455.1 | K9N7D2 |
| SIDTGNCPFSFG | 1 | 0 | 0 | 1504947 | ALK28781.1 | A0A0P0K6L9 |
| SIEDFNVWALSN | 1 | 0 | 0 | 1504952 | AWW13571.1 | P0C6X6 |
| SIEEFADVVKGQ | 1 | 0 | 0 | 1504955 | ALA49659.1 | K9N7C7 |
| SIEEVTTTLEET | 1 | 0 | 0 | 1504958 | AVP78041.1 | P0C6W2 |
| SIEGHIIMEEML | 1 | 0 | 0 | 1504959 | AUM60013.1 | K9N7C7 |
| SIFNIFQGVSSN | 1 | 0 | 0 | 1504980 | AIG96678.1 | H9EJ42 |
| SIFPHWNSDQAI | 1 | 0 | 0 | 1504981 | AZS52621.1 | Q0ZJ82 |
| SIFSRTLETAHH | 1 | 0 | 0 | 1504990 | AVP78030.1 | P0C6W2 |
| SIFSRTLETAQY | 1 | 0 | 0 | 1504992 | AVP78041.1 | P0C6W2 |
| SIGDIIQRLDFL | 1 | 0 | 0 | 1505010 | AXN92260.1 | K9N5Q8 |
| SIGDIIQRLDGL | 1 | 0 | 0 | 1505011 | AHI48733.1 | K9N5Q8 |
| SIGDIIQRLDPP | 1 | 0 | 0 | 1505012 | 5W9H_A | K9N5Q8 |
| SIGDIIRRLDVL | 1 | 0 | 0 | 1505014 | ALB08246.1 | K9N5Q8 |
| SIINNAVYTKMD | 1 | 0 | 0 | 1505055 | APO40578.1 | P0C6X7 |
| SIIPHSILSIQS | 1 | 0 | 0 | 1505059 | AHI48727.1 | K9N5Q8 |
| SIKEMQSQPSHA | 1 | 0 | 0 | 1505087 | AVL25602.1 | H9EJ69 |
| SIKSDKGVEKLV | 1 | 0 | 0 | 1505105 | AGT51393.1 | H9EJ42 |

| | | | | | | |
|---|---|---|---|---|---|---|
| SIKSDRKAWAAF | 1 | 0 | 0 | 1505106 | AJG44080.1 | K9N5Q8 |
| SIKVFTTVDNIN | 1 | 0 | 0 | 1505108 | AVP78041.1 | P0C6W2 |
| SILLSTLWFKPP | 1 | 0 | 0 | 1505121 | AKA09643.1 | P36334 |
| SILMKCDHCGET | 1 | 0 | 0 | 1505125 | AVP78030.1 | P0C6W2 |
| SILNEVNDLLDI | 1 | 0 | 0 | 1505127 | BBA20986.1 | A0A140H1H1 |
| SILSGLLYNSPC | 1 | 0 | 0 | 1505135 | P30215.1 | P30215 |
| SILSIQSDRKAW | 1 | 0 | 0 | 1505136 | AHI48727.1 | K9N5Q8 |
| SILSNCFTIMKM | 1 | 0 | 0 | 1505137 | AUM60013.1 | K9N7C7 |
| SILSPLYAFPSD | 1 | 0 | 0 | 1505139 | AVP78030.1 | P0C6W2 |
| SILVENGIPLMP | 1 | 0 | 0 | 1505144 | ARU07593.1 | H9EJ42 |
| SILVKYGCCDAS | 1 | 0 | 0 | 1505145 | AVV62524.1 | K9N7C7 |
| SILWCLWPLVLA | 1 | 0 | 0 | 1505146 | QDH43733.1 | U3M6Q8 |
| SINDIDTGAPSI | 1 | 0 | 0 | 1505160 | CAA83660.1 | P36334 |
| SINDSLEPVGGL | 1 | 0 | 0 | 1505162 | AIL49542.1 | P36334 |
| SINTTFDFLSNA | 1 | 0 | 0 | 1505178 | AKA09614.1 | A0A0P0G321 |
| SIPDLQKAAVTI | 1 | 0 | 0 | 1505181 | ARO76381.1 | P0C6U8 |
| SIPIVWGAKDFN | 1 | 0 | 0 | 1505196 | AFD98777.1 | H9EJ42 |
| SIPLADDDSAII | 1 | 0 | 0 | 1505201 | AMM42614.1 | H9EJ69 |
| SIPLCIVPYTAL | 1 | 0 | 0 | 1505203 | AXN92258.1 | K9N7C7 |
| SIPNFGSLTHIN | 1 | 0 | 0 | 1505210 | ALJ54471.1 | K9N5Q8 |
| SIPNFGSLTQID | 1 | 0 | 0 | 1505211 | ASU91284.1 | K9N5Q8 |
| SIPNLPDFKEES | 1 | 0 | 0 | 1505216 | BBA20973.1 | P36334 |
| SIPNLPDFKGEL | 1 | 0 | 0 | 1505217 | QDH43719.1 | P36334 |
| SIPNLPDFREEL | 1 | 0 | 0 | 1505218 | AIX10756.1 | P36334 |
| SIPNLPGFKEEL | 1 | 0 | 0 | 1505219 | AIL49523.1 | P36334 |
| SIPNLSDFEAEF | 1 | 0 | 0 | 1505220 | BBA20986.1 | A0A140H1H1 |
| SIQADVNTPADI | 1 | 0 | 0 | 1505235 | QBQ01843.1 | P33469 |
| SIQALQTAAMTI | 1 | 0 | 0 | 1505236 | APO40578.1 | P0C6X7 |
| SIQECCENQPFQ | 1 | 0 | 0 | 1505239 | ATO98239.1 | Q3LZX5 |
| SIQECCENQPYQ | 1 | 0 | 0 | 1505240 | Q3I5I8.1 | Q3LZX5 |
| SIQNGFTATNSA | 1 | 0 | 0 | 1505247 | BBA20986.1 | A0A140H1H1 |
| SIQQGFDATNSA | 1 | 0 | 0 | 1505250 | AIX10759.1 | P36334 |
| SIQSGRKAWAAF | 1 | 0 | 0 | 1505253 | QBF80543.1 | K9N5Q8 |
| SIRNESWHIDSS | 1 | 0 | 0 | 1505274 | BBA20986.1 | A0A140H1H1 |
| SIRNLQKRLYPN | 1 | 0 | 0 | 1505276 | ACR82756.1 | A0A140H1G9 |
| SIRSIKSDRKAW | 1 | 0 | 0 | 1505282 | AJG44080.1 | K9N5Q8 |
| SIRSIQSDRKAW | 1 | 0 | 0 | 1505283 | AXN92228.1 | K9N5Q8 |
| SIRSKYSERQAW | 1 | 0 | 0 | 1505286 | AVV62537.1 | K9N5Q8 |
| SIRWADNNCYLA | 1 | 0 | 0 | 1505291 | AVP78041.1 | P0C6W2 |
| SISALQKRLYSH | 1 | 0 | 0 | 1505295 | AKL87224.1 | P0C6X6 |
| SISDILQRLDTV | 1 | 0 | 0 | 1505301 | AVV62526.1 | K9N5Q8 |
| SISKLHALDDAT | 1 | 0 | 0 | 1505305 | AHL18094.1 | K9N643 |
| SISNDTVDVTNG | 1 | 0 | 0 | 1505313 | ANZ78836.1 | P36334 |
| SISNELLLTVPT | 1 | 0 | 0 | 1505314 | AVQ05263.1 | P30215 |
| SISNFGSLTQIN | 1 | 0 | 0 | 1505315 | QBF80598.1 | K9N5Q8 |
| SISTDIVDVTNG | 1 | 0 | 0 | 1505319 | CAA83660.1 | P36334 |
| SISTDVLVNTYK | 1 | 0 | 0 | 1505321 | AUM60013.1 | K9N7C7 |
| SISTGSLSARSA | 1 | 0 | 0 | 1505323 | AVN89324.1 | K9N5Q8 |
| SITFCALCVISV | 1 | 0 | 0 | 1505339 | AGT51757.1 | P0C6X6 |
| SITIDKSAIPNG | 1 | 0 | 0 | 1505352 | BBA84104.1 | P36334 |
| SITQRTRSRPSV | 1 | 0 | 0 | 1505364 | ASY99807.1 | K9N4V7 |
| SITRQPSKIENA | 1 | 0 | 0 | 1505367 | ATO98194.1 | Q3LZX0 |
| SITSGMLQSGLV | 1 | 0 | 0 | 1505372 | AVV62535.1 | K9N7C7 |
| SITTGYRFTNFE | 1 | 0 | 0 | 1505375 | ABD43183.1 | P36334 |
| SITVDKFAIPNR | 1 | 0 | 0 | 1505377 | BBA20986.1 | A0A140H1H1 |
| SIVAYTMSIGED | 1 | 0 | 0 | 1505386 | APO40579.1 | P59594 |
| SIVDEWSYTGSS | 1 | 0 | 0 | 1505392 | ANF29261.1 | K9N5Q8 |
| SIVDIIQRLDVL | 1 | 0 | 0 | 1505393 | AGV08438.1 | K9N5Q8 |
| SIVDVYDMWLTG | 1 | 0 | 0 | 1505394 | AEN19363.1 | P0C6X6 |
| SIVITCGDGTTN | 1 | 0 | 0 | 1505402 | AVP78043.1 | Q3LZX0 |
| SIVKPDMADEIA | 1 | 0 | 0 | 1505406 | ARU07581.1 | Q19U29 |
| SIVLGLKTIATH | 1 | 0 | 0 | 1505408 | APO40578.1 | P0C6X7 |
| SIVPSTAWEDGD | 1 | 0 | 0 | 1505414 | AGV08584.1 | K9N5Q8 |
| SIVPSTLWEDGD | 1 | 0 | 0 | 1505415 | ALJ54501.1 | K9N5Q8 |
| SIVPSTVWEEGD | 1 | 0 | 0 | 1505418 | ALJ76277.1 | K9N5Q8 |
| SIVSEKISVMDK | 1 | 0 | 0 | 1505428 | AVA26872.1 | H9EJ42 |
| SIYDFDTYKALL | 1 | 0 | 0 | 1505445 | ARB07605.1 | A0A140H1G9 |
| SIYICSQGKDFG | 1 | 0 | 0 | 1505450 | ARU07575.1 | A0A140H1G9 |
| SIYLSTLWYKPP | 1 | 0 | 0 | 1505455 | 5I08_A | A0A140H1H1 |
| SKAADRVAWAAF | 1 | 0 | 0 | 1505458 | AVV62526.1 | K9N5Q8 |
| SKADGIIYPTGK | 1 | 0 | 0 | 1505461 | AVV62537.1 | K9N5Q8 |
| SKAEGVIYPTGR | 1 | 0 | 0 | 1505463 | AUM60014.1 | K9N5Q8 |
| SKAGNSIFRSFH | 1 | 0 | 0 | 1505466 | AEN19365.1 | P30215 |
| SKAKVSAEVSRQ | 1 | 0 | 0 | 1505468 | ALK28782.1 | A0A0P0KWE8 |
| SKANAIQQQFNV | 1 | 0 | 0 | 1505474 | AUM60013.1 | K9N7C7 |
| SKANSGNRTPTS | 1 | 0 | 0 | 1505475 | AJQ24081.1 | P33469 |
| SKCLTSIEEVCD | 1 | 0 | 0 | 1505491 | AEN19363.1 | P0C6X6 |
| SKDQYGTDINGV | 1 | 0 | 0 | 1505512 | AIV41994.1 | P33469 |
| SKDSNFLKRVRG | 1 | 0 | 0 | 1505513 | ALT66879.1 | K9N7C7 |
| SKDVKFQNLTNN | 1 | 0 | 0 | 1505521 | AUM60024.1 | K9N5Q8 |
| SKESNFLNESGV | 1 | 0 | 0 | 1505541 | AQZ41321.1 | K9N7C7 |
| SKESNFLNRVRG | 1 | 0 | 0 | 1505542 | AQZ41330.1 | K9N7C7 |
| SKETAAMVKSKG | 1 | 0 | 0 | 1505543 | ARU07575.1 | A0A140H1G9 |
| SKGIYQTSNFRV | 1 | 0 | 0 | 1505570 | ATO98181.1 | Q3LZX1 |
| SKGLVAVNNSKP | 1 | 0 | 0 | 1505572 | AVV62541.1 | K9N7D2 |
| SKGSFRNESWHF | 1 | 0 | 0 | 1505583 | AMN88694.1 | A0A140H1H1 |
| SKGTLEPKYFNT | 1 | 0 | 0 | 1505588 | AEN19363.1 | P0C6X6 |
| SKGVTFGENTVV | 1 | 0 | 0 | 1505592 | APO40578.1 | P0C6X7 |
| SKHSDFGLDGLI | 1 | 0 | 0 | 1505604 | AWK59942.1 | H9EJ42 |
| SKIANLFVEQSQ | 1 | 0 | 0 | 1505620 | AVV62524.1 | K9N7C7 |
| SKIENASPASTV | 1 | 0 | 0 | 1505623 | ATO98194.1 | Q3LZX0 |

| | | | | | | |
|---|---|---|---|---|---|---|
| SKIGNSIFTGCT | 1 | 0 | 0 | 1505631 | ATQ39389.1 | K9N7C7 |
| SKIGTTEVKVSA | 1 | 0 | 0 | 1505632 | ALD51903.1 | K9N7C7 |
| SKIINGLHPTQS | 1 | 0 | 0 | 1505636 | APO40578.1 | P0C6X7 |
| SKIQDSLTTTSA | 1 | 0 | 0 | 1505648 | APO40579.1 | P59594 |
| SKIVCDGSVYYA | 1 | 0 | 0 | 1505653 | ABE97136.1 | H9EJ42 |
| SKIVTSGLGTVD | 1 | 0 | 0 | 1505660 | AGW80932.1 | A0A0P0K6L9 |
| SKKHVDVFTDGK | 1 | 0 | 0 | 1505671 | AVV62524.1 | K9N7C7 |
| SKKLLNKLVGGD | 1 | 0 | 0 | 1505674 | AVV62524.1 | K9N7C7 |
| SKKYTYSKLPDE | 1 | 0 | 0 | 1505686 | AVV62524.1 | K9N7C7 |
| SKKYTYSLLPNE | 1 | 0 | 0 | 1505687 | AVV62535.1 | K9N7C7 |
| SKLELAKVHNLS | 1 | 0 | 0 | 1505693 | AHA80155.1 | P33469 |
| SKLELAKVRNLS | 1 | 0 | 0 | 1505696 | AHA80174.1 | P33469 |
| SKLELVKRESEA | 1 | 0 | 0 | 1505698 | ARU07581.1 | Q19U29 |
| SKLHALDDATRN | 1 | 0 | 0 | 1505705 | AHL18094.1 | K9N643 |
| SKLHALDDVTLN | 1 | 0 | 0 | 1505707 | ALA49575.1 | K9N643 |
| SKLHDLDDATRS | 1 | 0 | 0 | 1505710 | AVV62540.1 | K9N643 |
| SKLKNLVLNNPA | 1 | 0 | 0 | 1505717 | AWW13558.1 | P30215 |
| SKLKPFERDLSS | 1 | 0 | 0 | 1505718 | ATO98145.1 | Q3LZX1 |
| SKLNNFQKFKTI | 1 | 0 | 0 | 1505727 | AIW52843.1 | A0A0P0G321 |
| SKLPSSSGTSTP | 1 | 0 | 0 | 1505729 | AIW52795.1 | H9EJ69 |
| SKLRAHDNILSV | 1 | 0 | 0 | 1505731 | AUM60013.1 | K9N7C7 |
| SKLRANDNILSV | 1 | 0 | 0 | 1505732 | AXN92247.1 | K9N7C7 |
| SKLVTSGLGTVD | 1 | 0 | 0 | 1505743 | AIW52757.1 | A0A0P0K6L9 |
| SKNDNLEPVSGL | 1 | 0 | 0 | 1505761 | AIL49538.1 | P36334 |
| SKNGGSGGAITT | 1 | 0 | 0 | 1505764 | 6NZK_A | P36334 |
| SKNPVLYDANYF | 1 | 0 | 0 | 1505772 | AVP78043.1 | Q3LZX0 |
| SKPFVKGDFSFI | 1 | 0 | 0 | 1505784 | AUM60013.1 | K9N7C7 |
| SKPFYIPEHCQN | 1 | 0 | 0 | 1505785 | ATQ39394.1 | K9N796 |
| SKPFYVPEHCQN | 1 | 0 | 0 | 1505786 | AVN89384.1 | K9N796 |
| SKPIYVPEHCQN | 1 | 0 | 0 | 1505791 | QAT98905.1 | K9N796 |
| SKPLYVPEHCQS | 1 | 0 | 0 | 1505800 | AVV62527.1 | K9N796 |
| SKPTKRSFIEDL | 1 | 0 | 0 | 1505815 | ATO98181.1 | Q3LZX1 |
| SKQLQDSMNGAS | 1 | 0 | 0 | 1505832 | APO40586.1 | P59595 |
| SKQSVASVIAEL | 1 | 0 | 0 | 1505837 | APO40578.1 | P0C6X7 |
| SKRFQSFQQFGK | 1 | 0 | 0 | 1505841 | AVP78031.1 | Q3LZX1 |
| SKRFQSFQQFGR | 1 | 0 | 0 | 1505842 | ATO98181.1 | Q3LZX1 |
| SKRLRKRRDAKV | 1 | 0 | 0 | 1505848 | AUM60013.1 | K9N7C7 |
| SKRQYLLKFDDR | 1 | 0 | 0 | 1505849 | ADN03339.1 | A0A140H1H1 |
| SKRSAIEDLLFN | 1 | 0 | 0 | 1505851 | AVV62526.1 | K9N5Q8 |
| SKSHNIGSLYVS | 1 | 0 | 0 | 1505873 | AOG74783.1 | A0A0P0K6L9 |
| SKSKELLNVALD | 1 | 0 | 0 | 1505877 | QBP84755.1 | P0C6U7 |
| SKSQSRNPSSDR | 1 | 0 | 0 | 1505889 | AIW52723.1 | A0A0N7J131 |
| SKSSEEHFIETV | 1 | 0 | 0 | 1505894 | APO40578.1 | P0C6X7 |
| SKSSGMDTALLR | 1 | 0 | 0 | 1505895 | AZS52621.1 | Q0ZJ82 |
| SKTDVSTPVDDF | 1 | 0 | 0 | 1505901 | AUM60013.1 | K9N7C7 |
| SKTPEEHFIETV | 1 | 0 | 0 | 1505906 | AVP78041.1 | P0C6W2 |
| SKTRRSRRAITT | 1 | 0 | 0 | 1505912 | AVQ05264.1 | P36334 |
| SKVEPCVVSVTY | 1 | 0 | 0 | 1505922 | AEN19363.1 | P0C6X6 |
| SKVKNGFTLEYW | 1 | 0 | 0 | 1505936 | CAA79896.1 | P36334 |
| SKVQDAVNANAQ | 1 | 0 | 0 | 1505943 | AVV62537.1 | K9N5Q8 |
| SKVRLSDVGFVE | 1 | 0 | 0 | 1505948 | CAA83661.1 | P36334 |
| SKVSGKTQQQQP | 1 | 0 | 0 | 1505950 | APO40586.1 | P59595 |
| SKVVKFPIDLTM | 1 | 0 | 0 | 1505952 | ATG84897.1 | K9N7C7 |
| SKVVKVPIDLTI | 1 | 0 | 0 | 1505955 | ANI69887.1 | K9N7C7 |
| SKVVKVPIDLTM | 1 | 0 | 0 | 1505956 | AXN92247.1 | K9N7C7 |
| SKVVLSGEWNAV | 1 | 0 | 0 | 1505961 | AWK59942.1 | H9EJ42 |
| SKYQVIVLKDNV | 1 | 0 | 0 | 1505972 | ALA49340.1 | K9N7C7 |
| SKYTMLDLCYAL | 1 | 0 | 0 | 1505976 | ARU07574.1 | A0A140H1G9 |
| SKYVTVQGPPGT | 1 | 0 | 0 | 1505977 | AXN92247.1 | K9N7C7 |
| SLACEHISSTMS | 1 | 0 | 0 | 1505986 | QDI73610.1 | K9N5Q8 |
| SLADGLEAYADR | 1 | 0 | 0 | 1505988 | AIV41774.1 | P0C6X6 |
| SLADGLGAYADK | 1 | 0 | 0 | 1505989 | AIV42008.1 | P0C6X6 |
| SLADLPAYMSVA | 1 | 0 | 0 | 1505990 | AWW13547.1 | Q01455 |
| SLADSNGFIRFP | 1 | 0 | 0 | 1505992 | ABD75543.1 | A0A140H1G9 |
| SLADVATDVQSR | 1 | 0 | 0 | 1505993 | AUM60013.1 | K9N7C7 |
| SLAEFSLNNHHS | 1 | 0 | 0 | 1505996 | AVV62541.1 | K9N7D2 |
| SLAIDAYPLVHH | 1 | 0 | 0 | 1506006 | ARB07436.1 | A0A140H1G9 |
| SLANVISFGDYN | 1 | 0 | 0 | 1506021 | AFD64760.1 | A0A0P0G321 |
| SLATRLMSIPQL | 1 | 0 | 0 | 1506026 | AVR40345.1 | Q4VID0 |
| SLAYYEPVQGRK | 1 | 0 | 0 | 1506031 | AXN92247.1 | K9N7C7 |
| SLCKTIQSALSV | 1 | 0 | 0 | 1506050 | QEO75984.1 | A0A0P0ILY9 |
| SLCLCFVMTCPL | 1 | 0 | 0 | 1506052 | AVV62524.1 | K9N7C7 |
| SLCLLNHKETLS | 1 | 0 | 0 | 1506055 | AET99208.1 | Q0ZJ80 |
| SLCNFSNLDKLK | 1 | 0 | 0 | 1506057 | AVV62524.1 | K9N7C7 |
| SLCVGSGPGKNN | 1 | 0 | 0 | 1506064 | AVQ05264.1 | P36334 |
| SLDFYPALETVQ | 1 | 0 | 0 | 1506067 | APO40578.1 | P0C6X7 |
| SLDGISKAEVDP | 1 | 0 | 0 | 1506070 | AVV62524.1 | K9N7C7 |
| SLDGNFRTGVDP | 1 | 0 | 0 | 1506072 | ASU90030.1 | K9N7C7 |
| SLDGTTLQGKPI | 1 | 0 | 0 | 1506074 | AXG21642.1 | K9N638 |
| SLDSNFRTEVDP | 1 | 0 | 0 | 1506082 | ASU90302.1 | K9N7C7 |
| SLDYINVTFLDL | 1 | 0 | 0 | 1506090 | CAA79896.1 | P36334 |
| SLEACYLRGCNP | 1 | 0 | 0 | 1506096 | QBQ01836.1 | P0C6X6 |
| SLEGKVFQLTPI | 1 | 0 | 0 | 1506103 | AVP78050.1 | Q3LZX3 |
| SLEGKVFQSTPI | 1 | 0 | 0 | 1506104 | AVP78039.1 | Q3LZX3 |
| SLEKLQCEHLQF | 1 | 0 | 0 | 1506112 | AWO68978.1 | A0A0P0G321 |
| SLELLNFKRERS | 1 | 0 | 0 | 1506113 | AGT51735.1 | Q4VID0 |
| SLEMFKPGVEEL | 1 | 0 | 0 | 1506123 | AGT51805.1 | Q4VID0 |
| SLENPTYIYHSA | 1 | 0 | 0 | 1506125 | AXN92247.1 | K9N7C7 |
| SLESPLKLEDFL | 1 | 0 | 0 | 1506136 | APO40578.1 | P0C6X7 |
| SLFAQSDEEESF | 1 | 0 | 0 | 1506153 | ANF29241.1 | K9N4V0 |
| SLFAQSEEQESF | 1 | 0 | 0 | 1506160 | ANF29197.1 | K9N4V0 |

| | | | | | | |
|---|---|---|---|---|---|---|
| SLFKPVQLVPVF | 1 | 0 | 0 | 1506188 | AGV08496.1 | K9N7D2 |
| SLFLFDEAGEEV | 1 | 0 | 0 | 1506192 | AIV41864.1 | P0C6X6 |
| SLFLLMFLLTHI | 1 | 0 | 0 | 1506193 | AVV62526.1 | K9N5Q8 |
| SLFPLWTAAAGV | 1 | 0 | 0 | 1506198 | AAR01015.1 | P36334 |
| SLFSNIKTSSTQ | 1 | 0 | 0 | 1506209 | AUM60014.1 | K9N5Q8 |
| SLFSVNDFTCRQ | 1 | 0 | 0 | 1506211 | ATG84888.1 | K9N5Q8 |
| SLGASSNQSPQG | 1 | 0 | 0 | 1506226 | AIX09796.1 | P0C6X6 |
| SLGDSVKLLSEL | 1 | 0 | 0 | 1506232 | ABE73416.1 | H9EJ42 |
| SLGFDKPQEKKK | 1 | 0 | 0 | 1506235 | AMM42528.1 | A0A0N7J131 |
| SLGFDNPPTNVV | 1 | 0 | 0 | 1506236 | AGT51679.1 | P30215 |
| SLGNFTARANCL | 1 | 0 | 0 | 1506253 | AVV62529.1 | K9N643 |
| SLHKLTYLLDFS | 1 | 0 | 0 | 1506264 | AVV62537.1 | K9N5Q8 |
| SLHPVECDEECS | 1 | 0 | 0 | 1506266 | AXN92247.1 | K9N7C7 |
| SLHSVLWNCQLS | 1 | 0 | 0 | 1506272 | AVN89336.1 | K9N4V0 |
| SLIPVRLRNSSD | 1 | 0 | 0 | 1506293 | AIW52843.1 | A0A0P0G321 |
| SLKDGQINDMIL | 1 | 0 | 0 | 1506317 | AVP78041.1 | P0C6W2 |
| SLKDNQVNDMIY | 1 | 0 | 0 | 1506319 | APO40578.1 | P0C6X7 |
| SLKFFEIKHADY | 1 | 0 | 0 | 1506327 | AHF51980.1 | H9EJ42 |
| SLKGGKLNNTWF | 1 | 0 | 0 | 1506331 | APO40578.1 | P0C6X7 |
| SLKQGEIKDATP | 1 | 0 | 0 | 1506347 | AVP78043.1 | Q3LZX0 |
| SLKYNLKASQAV | 1 | 0 | 0 | 1506358 | CAA49377.1 | A0A0P0LKV0 |
| SLLATLDDPYVEDSVA | 1 | 0 | 0 | 1506363 | YP_173242.1 | Q19U29 |
| SLLGCLGSQCGS | 1 | 0 | 0 | 1506372 | BBA20986.1 | A0A140H1H1 |
| SLLGSIAGASWT | 1 | 0 | 0 | 1506377 | AUM60014.1 | K9N5Q8 |
| SLLIVNNATNVI | 1 | 0 | 0 | 1506381 | AVP78031.1 | Q3LZX1 |
| SLLKLEDFIPMD | 1 | 0 | 0 | 1506389 | AVP78041.1 | P0C6W2 |
| SLLNEVWQKYTN | 1 | 0 | 0 | 1506403 | AUM60016.1 | K9N4V0 |
| SLLNQVWQHHIN | 1 | 0 | 0 | 1506408 | AVV62528.1 | K9N4V0 |
| SLLNQVWQHYTN | 1 | 0 | 0 | 1506409 | AVV62539.1 | K9N4V0 |
| SLLSVLLMGCVA | 1 | 0 | 0 | 1506434 | 6NB6_A | P59594 |
| SLLTKCYKAMNS | 1 | 0 | 0 | 1506436 | AVV62535.1 | K9N7C7 |
| SLLVEDCKLPLG | 1 | 0 | 0 | 1506439 | ATW75477.1 | K9N5Q8 |
| SLLVTASSKPLY | 1 | 0 | 0 | 1506451 | YP_009047205.1 | K9N796 |
| SLMNSLFRMPLG | 1 | 0 | 0 | 1506459 | AEN19363.1 | P0C6X6 |
| SLNDNLLEILQK | 1 | 0 | 0 | 1506468 | AVP78041.1 | P0C6W2 |
| SLNFKLGFDSSK | 1 | 0 | 0 | 1506472 | AAX85666.1 | P0C6X6 |
| SLNFVGEFVIND | 1 | 0 | 0 | 1506474 | ATQ39389.1 | K9N7C7 |
| SLNFVGEFVVND | 1 | 0 | 0 | 1506476 | AXN92247.1 | K9N7C7 |
| SLNGTVSTVFVP | 1 | 0 | 0 | 1506478 | ANI69893.1 | K9N7D2 |
| SLNMVKKLSPSR | 1 | 0 | 0 | 1506490 | APO40587.1 | P59636 |
| SLNNHHSTVFVP | 1 | 0 | 0 | 1506491 | AVV62541.1 | K9N7D2 |
| SLNNPIFISTNY | 1 | 0 | 0 | 1506493 | AGT17757.1 | A0A140H1H0 |
| SLNSKEELLQTV | 1 | 0 | 0 | 1506495 | APO40578.1 | P0C6X7 |
| SLPFNNMLGYNP | 1 | 0 | 0 | 1506513 | AVV62524.1 | K9N7C7 |
| SLPIKQPFSFSF | 1 | 0 | 0 | 1506516 | AWK59942.1 | H9EJ42 |
| SLPNEVTVAKPN | 1 | 0 | 0 | 1506524 | AVV62543.1 | K9N7A1 |
| SLPNMGVPCCTS | 1 | 0 | 0 | 1506525 | AIW52827.1 | H9EJ42 |
| SLPTAFAAIGDL | 1 | 0 | 0 | 1506530 | AIL49486.1 | P36334 |
| SLPTSGSRVAGR | 1 | 0 | 0 | 1506533 | CAA71056.1 | A0A0P0K6L9 |
| SLPVMISEWPLS | 1 | 0 | 0 | 1506542 | NP_073549.1 | A0A0P0LKV0 |
| SLPVYDTIKYYT | 1 | 0 | 0 | 1506543 | AVV62526.1 | K9N5Q8 |
| SLPVYHRHTFIV | 1 | 0 | 0 | 1506546 | AMM42532.1 | A0A0P0K6L9 |
| SLQDMRQKQSVW | 1 | 0 | 0 | 1506551 | ARU07575.1 | A0A140H1G9 |
| SLQEILSRLDVL | 1 | 0 | 0 | 1506554 | ABD75553.1 | A0A140H1H1 |
| SLQKGVLPQLEQ | 1 | 0 | 0 | 1506561 | APO40578.1 | P0C6X7 |
| SLREAALPQSKD | 1 | 0 | 0 | 1506580 | AVV62535.1 | K9N7C7 |
| SLREVALPQSKD | 1 | 0 | 0 | 1506581 | AVV62524.1 | K9N7C7 |
| SLRLSDSNCYMN | 1 | 0 | 0 | 1506594 | AUM60013.1 | K9N7C7 |
| SLRQIFDPAPVE | 1 | 0 | 0 | 1506604 | AVV62524.1 | K9N7C7 |
| SLRVCVETVHTN | 1 | 0 | 0 | 1506620 | AVP78041.1 | P0C6W2 |
| SLRVTMLKAQEG | 1 | 0 | 0 | 1506626 | QBP84745.1 | Q4VID0 |
| SLRYAWIWCAAY | 1 | 0 | 0 | 1506627 | NP_073549.1 | A0A0P0LKV0 |
| SLSHLTSQLRQN | 1 | 0 | 0 | 1506645 | ARA15429.1 | A0A0P0K6L9 |
| SLSLLVTASSKP | 1 | 0 | 0 | 1506660 | YP_009047205.1 | K9N796 |
| SLSMVRRNLDSL | 1 | 0 | 0 | 1506665 | Q3I5I6.1 | Q3LZX3 |
| SLSNFALALCSG | 1 | 0 | 0 | 1506666 | AXN92247.1 | K9N7C7 |
| SLSNTKYYDHSQ | 1 | 0 | 0 | 1506670 | ATN39871.1 | P30215 |
| SLSRSQSSETKE | 1 | 0 | 0 | 1506677 | ABB90484.1 | A0A0N7J131 |
| SLSSSGTSTPKK | 1 | 0 | 0 | 1506681 | ARU07598.1 | H9EJ69 |
| SLSVESTPVEPP | 1 | 0 | 0 | 1506688 | AGV08515.1 | K9N7C7 |
| SLTASSIFGAIL | 1 | 0 | 0 | 1506695 | ARU07575.1 | A0A140H1G9 |
| SLTDVATDVQSR | 1 | 0 | 0 | 1506700 | AUM60023.1 | K9N7C7 |
| SLTDVRSTSGTQ | 1 | 0 | 0 | 1506703 | QBF80601.1 | K9N643 |
| SLTHINTTLLDL | 1 | 0 | 0 | 1506712 | ALJ54471.1 | K9N5Q8 |
| SLTQIDTTLLDL | 1 | 0 | 0 | 1506728 | ASU91284.1 | K9N5Q8 |
| SLTQSKDTNFLN | 1 | 0 | 0 | 1506732 | AUM60013.1 | K9N7C7 |
| SLTTNNAATKRT | 1 | 0 | 0 | 1506742 | AVP78031.1 | Q3LZX1 |
| SLTTTSAALGKL | 1 | 0 | 0 | 1506744 | APO40579.1 | P59594 |
| SLTTVSIPQGLT | 1 | 0 | 0 | 1506746 | ATQ39389.1 | K9N7C7 |
| SLTWLWLLCGLV | 1 | 0 | 0 | 1506760 | NP_073549.1 | A0A0P0LKV0 |
| SLVAQNTVLNDS | 1 | 0 | 0 | 1506767 | AMM42614.1 | H9EJ69 |
| SLVCTTLQGKPI | 1 | 0 | 0 | 1506771 | AVN89333.1 | K9N7C7 |
| SLVEFQVTIAEL | 1 | 0 | 0 | 1506772 | APO40583.1 | P59634 |
| SLVGSIAGAGWT | 1 | 0 | 0 | 1506779 | AVV62526.1 | K9N5Q8 |
| SLVLAKLGKDFK | 1 | 0 | 0 | 1506794 | ABD75549.1 | Q19U29 |
| SLVLAKLGKDSK | 1 | 0 | 0 | 1506795 | AMM42397.1 | Q19U29 |
| SLVLNNPAYIAS | 1 | 0 | 0 | 1506803 | AIV41992.1 | P30215 |
| SLVLNNPSYIAP | 1 | 0 | 0 | 1506805 | AIX09798.1 | P30215 |
| SLVSLLSVLLMG | 1 | 0 | 0 | 1506824 | 6NB6_A | P59594 |
| SLVSVQELVQNT | 1 | 0 | 0 | 1506828 | ATQ39389.1 | K9N7C7 |
| SLVTFVGDSSEI | 1 | 0 | 0 | 1506830 | ALB08245.1 | K9N7C7 |

| | | | | | | |
|---|---|---|---|---|---|---|
| SLVTRLMSIPRL | 1 | 0 | 0 | 1506837 | AHF54653.1 | Q4VID0 |
| SLWNYGKAINLP | 1 | 0 | 0 | 1506850 | ARB07436.1 | A0A140H1G9 |
| SLWNYGKPVTLP | 1 | 0 | 0 | 1506852 | AEN19363.1 | P0C6X6 |
| SLWYKGSNLEEE | 1 | 0 | 0 | 1506857 | AEN19363.1 | P0C6X6 |
| SLYLADNLNADE | 1 | 0 | 0 | 1506866 | ATQ39389.1 | K9N7C7 |
| SLYLADNLTAGE | 1 | 0 | 0 | 1506870 | QBF80609.1 | K9N7C7 |
| SLYNVTREKLDK | 1 | 0 | 0 | 1506885 | AUM60013.1 | K9N7C7 |
| SLYPVASPRECN | 1 | 0 | 0 | 1506887 | ARO76381.1 | P0C6U8 |
| SLYVKPGGTSSGDATTAYA | 1 | 0 | 0 | 1506893 | P0DTD1.1 | P0DTD1 |
| SLYVKPSGTSSG | 1 | 0 | 0 | 1506894 | AVP78041.1 | P0C6W2 |
| SMAFLTAARLCV | 1 | 0 | 0 | 1506905 | AVV62542.1 | K9N5R3 |
| SMALSIFSAFYP | 1 | 0 | 0 | 1506910 | AJG44108.1 | K9N7A1 |
| SMALSIFSAIYP | 1 | 0 | 0 | 1506911 | ASY99815.1 | K9N7A1 |
| SMALSIFSAVYP | 1 | 0 | 0 | 1506912 | AHI48833.1 | K9N7A1 |
| SMAVAKVIAKWV | 1 | 0 | 0 | 1506923 | AEN19363.1 | P0C6X6 |
| SMCPGNSNYVSY | 1 | 0 | 0 | 1506926 | ATQ39390.1 | K9N5Q8 |
| SMCRYLFITTHV | 1 | 0 | 0 | 1506927 | ADK37703.1 | A0A0P0K6L9 |
| SMCYRCPREIVN | 1 | 0 | 0 | 1506931 | AVV62524.1 | K9N7C7 |
| SMDVLSQNQKLI | 1 | 0 | 0 | 1506943 | AEN19366.1 | P36334 |
| SMEVKTIKVFTT | 1 | 0 | 0 | 1506946 | 5TL6_B | P0C6X7 |
| SMFDVDKKILNA | 1 | 0 | 0 | 1506948 | AIV41972.1 | P0C6X6 |
| SMFSRTTSNFLP | 1 | 0 | 0 | 1506956 | ATO98193.1 | Q3LZX1 |
| SMGLPNTCFGPL | 1 | 0 | 0 | 1506963 | AIV41996.1 | P0C6X6 |
| SMHLSTYMNCNH | 1 | 0 | 0 | 1506965 | APO40578.1 | P0C6X7 |
| SMIEDMDYVPLK | 1 | 0 | 0 | 1506967 | AXG21663.1 | K9N7C7 |
| SMKSDLSVSFAG | 1 | 0 | 0 | 1506981 | AHX71943.1 | K9N5Q8 |
| SMKSDLSVSSSG | 1 | 0 | 0 | 1506983 | ALA50067.1 | K9N5Q8 |
| SMLGLITQTVDS | 1 | 0 | 0 | 1506989 | ASU90899.1 | K9N7C7 |
| SMLGLNTQTVDS | 1 | 0 | 0 | 1506990 | AVV62524.1 | K9N7C7 |
| SMLKQRDSTYGP | 1 | 0 | 0 | 1506992 | AVN89365.1 | K9N5Q8 |
| SMLTFDSNAFSL | 1 | 0 | 0 | 1507005 | AIW52843.1 | A0A0P0G321 |
| SMLTSIAARPPP | 1 | 0 | 0 | 1507007 | AUM60013.1 | K9N7C7 |
| SMMGFKMNYQVNGYPNM | 1 | 0 | 0 | 1507010 | P0DTD1.1 | P0DTD1 |
| SMNIAANSNEWA | 1 | 0 | 0 | 1507014 | AGT51337.1 | H9EJ42 |
| SMPVVDGKQIDD | 1 | 0 | 0 | 1507028 | AGT51409.1 | P0C6X6 |
| SMPYHPNPVDCQ | 1 | 0 | 0 | 1507029 | ANZ78865.1 | P0C6X6 |
| SMQDVGFKNQYC | 1 | 0 | 0 | 1507031 | AAR01012.1 | P0C6X6 |
| SMRNANGACIWN | 1 | 0 | 0 | 1507046 | AUM60013.1 | K9N7C7 |
| SMTDNLQMAFVI | 1 | 0 | 0 | 1507064 | AUM60014.1 | K9N5Q8 |
| SMTLNGLWLDNY | 1 | 0 | 0 | 1507072 | AVV62524.1 | K9N7C7 |
| SMTTSTTNFGEE | 1 | 0 | 0 | 1507079 | ABD75599.1 | A0A140H1G9 |
| SMVCKMCLFRYQ | 1 | 0 | 0 | 1507086 | QEG03730.1 | H9EJ42 |
| SMVPVEFYEDRI | 1 | 0 | 0 | 1507092 | AXN92247.1 | K9N7C7 |
| SMVVDECQLPLG | 1 | 0 | 0 | 1507095 | AVV62526.1 | K9N5Q8 |
| SMVVDNCQLPLG | 1 | 0 | 0 | 1507097 | AVV62537.1 | K9N5Q8 |
| SMYKQARSEDKR | 1 | 0 | 0 | 1507107 | 5F22_B | P0C6U8 |
| SNAFSLANVISF | 1 | 0 | 0 | 1507115 | AFD64760.1 | A0A0P0G321 |
| SNAHRCDVLLSD | 1 | 0 | 0 | 1507117 | NP_073549.1 | A0A0P0LKV0 |
| SNAILISDRKIK | 1 | 0 | 0 | 1507118 | AVV62524.1 | K9N7C7 |
| SNANISMLQLGA | 1 | 0 | 0 | 1507124 | AKA09625.1 | A0A0P0G321 |
| SNANISMLQLGV | 1 | 0 | 0 | 1507125 | AKA09626.1 | A0A0P0G321 |
| SNASISMLQLGV | 1 | 0 | 0 | 1507131 | AWO68966.1 | A0A0P0G321 |
| SNASSDVIHYNI | 1 | 0 | 0 | 1507132 | BAL45639.1 | A0A0P0K6L9 |
| SNATILDYCDGN | 1 | 0 | 0 | 1507135 | APO40578.1 | P0C6X7 |
| SNCNFNLSTLLR | 1 | 0 | 0 | 1507146 | AET99191.1 | A0A140H1H1 |
| SNCNFNMSCLMS | 1 | 0 | 0 | 1507147 | AHA80204.1 | P36334 |
| SNCQKAVSAFIT | 1 | 0 | 0 | 1507151 | AUM60013.1 | K9N7C7 |
| SNDAVQLESDAY | 1 | 0 | 0 | 1507162 | AVV62524.1 | K9N7C7 |
| SNDEAIPTKFPP | 1 | 0 | 0 | 1507163 | AJQ24067.1 | P33469 |
| SNDLLQPNTEVY | 1 | 0 | 0 | 1507170 | AGW27863.1 | A0A140H1H1 |
| SNDNISLPFNNM | 1 | 0 | 0 | 1507175 | AVV62524.1 | K9N7C7 |
| SNDQEMTTPPPK | 1 | 0 | 0 | 1507176 | ATQ39393.1 | K9N4V7 |
| SNDSCPGKNNGI | 1 | 0 | 0 | 1507177 | AIV41891.1 | P36334 |
| SNDSKALFFTYL | 1 | 0 | 0 | 1507178 | AZU90729.1 | K9N7C7 |
| SNDTLTQQQIIA | 1 | 0 | 0 | 1507181 | AUM60030.1 | K9N7A1 |
| SNDWKPGLAMPS | 1 | 0 | 0 | 1507187 | AUM60013.1 | K9N7C7 |
| SNENLTWPLGLE | 1 | 0 | 0 | 1507196 | AHI48747.1 | K9N638 |
| SNENLTWPLVLE | 1 | 0 | 0 | 1507197 | AXN92247.1 | K9N7C7 |
| SNEPGNVGGNDA | 1 | 0 | 0 | 1507198 | AVI15051.1 | P0C6X6 |
| SNESWFEDDYKA | 1 | 0 | 0 | 1507202 | AST12962.1 | A0A0P0LKV0 |
| SNEVTVAKPNVL | 1 | 0 | 0 | 1507205 | AMQ49021.1 | K9N7A1 |
| SNFILHDVNNGL | 1 | 0 | 0 | 1507224 | BBA84101.1 | P36334 |
| SNFNQYYYNMAT | 1 | 0 | 0 | 1507229 | AZS52617.1 | A0A140H1H0 |
| SNFNSCKSDILS | 1 | 0 | 0 | 1507230 | ADN03338.1 | A0A140H1H0 |
| SNGILKWADQSD | 1 | 0 | 0 | 1507264 | AMM42463.1 | P33469 |
| SNGKPGNVGGNE | 1 | 0 | 0 | 1507266 | AIV41792.1 | P0C6X6 |
| SNGPLKVGTGVM | 1 | 0 | 0 | 1507273 | ARU07593.1 | H9EJ42 |
| SNGSCPGKNNGI | 1 | 0 | 0 | 1507276 | AEN19358.1 | P36334 |
| SNGVFYFKPGGT | 1 | 0 | 0 | 1507285 | AWK59978.1 | H9EJ42 |
| SNGVIIMCADVP | 1 | 0 | 0 | 1507286 | AWK59942.1 | H9EJ42 |
| SNHIEVASAYGL | 1 | 0 | 0 | 1507300 | ANF29261.1 | K9N5Q8 |
| SNHQEGMLYPTI | 1 | 0 | 0 | 1507305 | ARE29989.1 | Q4VID0 |
| SNHQEGMLYPTT | 1 | 0 | 0 | 1507306 | AGT51745.1 | Q4VID0 |
| SNINRVLSVPSD | 1 | 0 | 0 | 1507325 | ABQ16583.1 | H9EJ42 |
| SNITTNGPLIVG | 1 | 0 | 0 | 1507336 | AVP78041.1 | P0C6W2 |
| SNKLRVVIPDFN | 1 | 0 | 0 | 1507340 | AVV62524.1 | K9N7C7 |
| SNLDKLKFKEVC | 1 | 0 | 0 | 1507357 | AVV62524.1 | K9N7C7 |
| SNLFADIKTQAT | 1 | 0 | 0 | 1507359 | AVV62526.1 | K9N5Q8 |
| SNLSLSMARRNL | 1 | 0 | 0 | 1507381 | 2CME_A | P59636 |
| SNLTKQLLELLS | 1 | 0 | 0 | 1507383 | AUM60013.1 | K9N7C7 |
| SNLTKQLLEVLS | 1 | 0 | 0 | 1507384 | AUM60023.1 | K9N7C7 |

| | | | | | | |
|---|---|---|---|---|---|---|
| SNMTQLAEAQII | 1 | 0 | 0 | 1507398 | AHI48545.1 | K9N7A1 |
| SNMTQLTEAQII | 1 | 0 | 0 | 1507401 | ASY99815.1 | K9N7A1 |
| SNNGWTAFSAAY | 1 | 0 | 0 | 1507410 | AVP78042.1 | Q3LZX1 |
| SNNKTGRYMNYT | 1 | 0 | 0 | 1507414 | AUM60014.1 | K9N5Q8 |
| SNNPQCAIVQAL | 1 | 0 | 0 | 1507420 | QED88024.1 | H9EJ42 |
| SNNQEDFDLISR | 1 | 0 | 0 | 1507424 | QBP84728.1 | P0C6X6 |
| SNPAAVDTKCLA | 1 | 0 | 0 | 1507430 | P0C6U5.1 | A0A140H1G9 |
| SNPNTPAYTFTT | 1 | 0 | 0 | 1507436 | AVV62524.1 | K9N7C7 |
| SNPTNCIAPVNG | 1 | 0 | 0 | 1507442 | AHY22565.1 | K9N5Q8 |
| SNQADVNIPADI | 1 | 0 | 0 | 1507445 | AIX10718.1 | P33469 |
| SNQGNKQIVHSC | 1 | 0 | 0 | 1507447 | ASU90211.1 | K9N643 |
| SNQGNKQIVHSH | 1 | 0 | 0 | 1507448 | ASU90849.1 | K9N643 |
| SNSCTTAVMTYS | 1 | 0 | 0 | 1507474 | AWO69003.1 | A0A0P0G321 |
| SNSFDALKLGDT | 1 | 0 | 0 | 1507477 | AVP78041.1 | P0C6W2 |
| SNSNFRHSASIV | 1 | 0 | 0 | 1507492 | AJQ24072.1 | Q19U29 |
| SNSSLCVGSGPG | 1 | 0 | 0 | 1507499 | ARB66062.1 | P36334 |
| SNSSVPLSEVYV | 1 | 0 | 0 | 1507507 | AWK59934.1 | U3M6Q8 |
| SNSTPAQNAGYW | 1 | 0 | 0 | 1507511 | AHI48825.1 | K9N4V7 |
| SNSVCNGCVGHS | 1 | 0 | 0 | 1507512 | AST12964.1 | A0A0P0K6L9 |
| SNTDNETLEYWV | 1 | 0 | 0 | 1507520 | 5I08_A | A0A140H1H1 |
| SNTTFDFLSNAS | 1 | 0 | 0 | 1507536 | ACY56730.1 | A0A0P0G321 |
| SNVCALMSCNGN | 1 | 0 | 0 | 1507546 | AIV41816.1 | P0C6X6 |
| SNVDVLPEGWHV | 1 | 0 | 0 | 1507548 | AIV41876.1 | P0C6X6 |
| SNVDVLPEGWRV | 1 | 0 | 0 | 1507549 | AEN19363.1 | P0C6X6 |
| SNVEKLRSLIQD | 1 | 0 | 0 | 1507551 | AUM60013.1 | K9N7C7 |
| SNVEVLPEGWRV | 1 | 0 | 0 | 1507553 | QEG03790.1 | P0C6X6 |
| SNVKEEPTTQKP | 1 | 0 | 0 | 1507559 | ARO76381.1 | P0C6U8 |
| SNVMDVNDCFKK | 1 | 0 | 0 | 1507566 | QED88039.1 | H9EJ42 |
| SNVMDVNDCFKN | 1 | 0 | 0 | 1507567 | AWK59942.1 | H9EJ42 |
| SNVSPIVNEKIS | 1 | 0 | 0 | 1507577 | AGT51337.1 | H9EJ42 |
| SNVSWYYSLTTN | 1 | 0 | 0 | 1507580 | AVP78031.1 | Q3LZX1 |
| SNVTQIKNACVY | 1 | 0 | 0 | 1507583 | ARK08626.1 | H9EJ42 |
| SNVVDVNDCFKN | 1 | 0 | 0 | 1507586 | AGT51379.1 | H9EJ42 |
| SNVYVDSFVVKG | 1 | 0 | 0 | 1507594 | ABD72969.1 | P59594 |
| SNYNTFAVFDTP | 1 | 0 | 0 | 1507610 | AVV62537.1 | K9N5Q8 |
| SNYQHEENIYNL | 1 | 0 | 0 | 1507611 | AVP78041.1 | P0C6W2 |
| SPAAISSNCYSS | 1 | 0 | 0 | 1507623 | AHL18090.1 | K9N5Q8 |
| SPAAPRAISFAD | 1 | 0 | 0 | 1507626 | ANI69896.1 | K9N4V7 |
| SPAQDTWGTSAA | 1 | 0 | 0 | 1507642 | 5X58_A | P59594 |
| SPARMAGNGGDT | 1 | 0 | 0 | 1507645 | AVP78038.1 | Q3LZX4 |
| SPASSQSAATIL | 1 | 0 | 0 | 1507651 | AIW52723.1 | A0A0N7J131 |
| SPASSQSSAKSL | 1 | 0 | 0 | 1507652 | ABB90484.1 | A0A0N7J131 |
| SPATAEPVRDEF | 1 | 0 | 0 | 1507660 | AAA45463.1 | A0A0N7J131 |
| SPCFSQQGVFRY | 1 | 0 | 0 | 1507671 | P30215.1 | P30215 |
| SPCKELEGVGAR | 1 | 0 | 0 | 1507675 | AXX83306.1 | P0C6X6 |
| SPCPSGIGAVGG | 1 | 0 | 0 | 1507676 | QCI31476.1 | K9N4V7 |
| SPCVSILPSTVW | 1 | 0 | 0 | 1507681 | AKL59401.1 | K9N5Q8 |
| SPDAVTAYNGYV | 1 | 0 | 0 | 1507691 | APO40578.1 | P0C6X7 |
| SPDFKAFNVFIG | 1 | 0 | 0 | 1507695 | AVV62535.1 | K9N7C7 |
| SPDGKPCTPPAF | 1 | 0 | 0 | 1507697 | ATO98231.1 | Q3LZX1 |
| SPDIDLGDISGI | 1 | 0 | 0 | 1507700 | AVP78031.1 | Q3LZX1 |
| SPDVQFEQLDNK | 1 | 0 | 0 | 1507712 | AVV62526.1 | K9N5Q8 |
| SPELLDPLNLSA | 1 | 0 | 0 | 1507716 | ADN03343.1 | Q19U29 |
| SPEPALLFRNIK | 1 | 0 | 0 | 1507717 | AIX10763.1 | P36334 |
| SPGLSGIGAVGG | 1 | 0 | 0 | 1507744 | ALA49601.1 | K9N4V7 |
| SPGPSGIGAVGD | 1 | 0 | 0 | 1507747 | AVV62544.1 | K9N4V7 |
| SPGPSGIGGGDL | 1 | 0 | 0 | 1507749 | AVN89359.1 | K9N4V7 |
| SPGPSGIGTVGG | 1 | 0 | 0 | 1507750 | ATG84708.1 | K9N4V7 |
| SPGQGVPLNANS | 1 | 0 | 0 | 1507752 | AHI48535.1 | K9N4V7 |
| SPGSGYIPEAPR | 1 | 0 | 0 | 1507753 | 5X59_A | K9N5Q8 |
| SPIDIPRGLPVG | 1 | 0 | 0 | 1507771 | ATO98193.1 | Q3LZX1 |
| SPKIDNETLEYW | 1 | 0 | 0 | 1507794 | AMM42407.1 | A0A140H1H1 |
| SPKPQRKRGVKQ | 1 | 0 | 0 | 1507803 | ARU07581.1 | Q19U29 |
| SPKPQRQRGHKT | 1 | 0 | 0 | 1507805 | AHA80176.1 | P33469 |
| SPKPQRQRGLKN | 1 | 0 | 0 | 1507808 | ATP16771.1 | P33469 |
| SPKSGYFVNVNN | 1 | 0 | 0 | 1507812 | AIL49520.1 | P36334 |
| SPKTDKETLEYW | 1 | 0 | 0 | 1507814 | AZS52618.1 | A0A140H1H1 |
| SPKVTIDCAACV | 1 | 0 | 0 | 1507816 | ABD43177.1 | P36334 |
| SPKVTIDCAAFV | 1 | 0 | 0 | 1507818 | CAA79896.1 | P36334 |
| SPKVTIDCVAFV | 1 | 0 | 0 | 1507821 | AIL49523.1 | P36334 |
| SPKVTIDRAAFV | 1 | 0 | 0 | 1507822 | AIL49498.1 | P36334 |
| SPLEGGGRLVAS | 1 | 0 | 0 | 1507823 | AXG21654.1 | K9N5Q8 |
| SPLKPTKRSFIE | 1 | 0 | 0 | 1507832 | 5XJK_A | P59594 |
| SPLLKQKDIGID | 1 | 0 | 0 | 1507834 | AGW27859.1 | Q0ZJ80 |
| SPLLVDSEQPWK | 1 | 0 | 0 | 1507839 | AIW52723.1 | A0A0N7J131 |
| SPLLVDSEQSWK | 1 | 0 | 0 | 1507840 | AAA45463.1 | A0A0N7J131 |
| SPLNWERKTILS | 1 | 0 | 0 | 1507844 | AIV41909.1 | P36334 |
| SPLQSYACWAHS | 1 | 0 | 0 | 1507845 | AVV62535.1 | K9N7C7 |
| SPLYAFASEAAR | 1 | 0 | 0 | 1507852 | AVP78041.1 | P0C6W2 |
| SPMACNGHKIED | 1 | 0 | 0 | 1507854 | ACR82761.1 | A0A140H1G9 |
| SPNECNQMCLSI | 1 | 0 | 0 | 1507860 | AVP78030.1 | P0C6W2 |
| SPNGKPCTPPAL | 1 | 0 | 0 | 1507864 | ABD72972.1 | P59594 |
| SPNLPDFREELD | 1 | 0 | 0 | 1507867 | AIL49492.1 | P36334 |
| SPNTDNETLQYG | 1 | 0 | 0 | 1507869 | ABA60801.1 | A0A140H1H1 |
| SPNTQRTVQHWF | 1 | 0 | 0 | 1507871 | AJQ24090.1 | P33469 |
| SPNTYEGNSPYH | 1 | 0 | 0 | 1507872 | APO40584.1 | P59635 |
| SPNVCYTPRSEI | 1 | 0 | 0 | 1507873 | AVV62535.1 | K9N7C7 |
| SPPADVNTPADI | 1 | 0 | 0 | 1507874 | AJQ24074.1 | P33469 |
| SPPAIASNCYSS | 1 | 0 | 0 | 1507875 | AHI48528.1 | K9N5Q8 |
| SPPGEQFKHLVP | 1 | 0 | 0 | 1507876 | QED88038.1 | H9EJ42 |
| SPPISTDTVDVT | 1 | 0 | 0 | 1507881 | QBP84713.1 | P36334 |

| | | | | | | |
|---|---|---|---|---|---|---|
| SPPNNADLSPIS | 1 | 0 | 0 | 1507885 | AUM60015.1 | K9N796 |
| SPPTKQRYAKRR | 1 | 0 | 0 | 1507888 | AUM60017.1 | K9N643 |
| SPPVADYTPVIG | 1 | 0 | 0 | 1507890 | AVV62539.1 | K9N4V0 |
| SPQQCIAPIDGY | 1 | 0 | 0 | 1507899 | AVV62526.1 | K9N5Q8 |
| SPQSMLVFPEVK | 1 | 0 | 0 | 1507901 | AVV62524.1 | K9N7C7 |
| SPQVITTDNTFE | 1 | 0 | 0 | 1507904 | APO40579.1 | P59594 |
| SPSECNSMHLST | 1 | 0 | 0 | 1507919 | APO40578.1 | P0C6X7 |
| SPSGDAISQAFN | 1 | 0 | 0 | 1507921 | AVV62535.1 | K9N7C7 |
| SPSHYWSPQGNL | 1 | 0 | 0 | 1507924 | ARB07436.1 | A0A140H1G9 |
| SPSKDTFELRYS | 1 | 0 | 0 | 1507928 | AMM42397.1 | Q19U29 |
| SPSLSNLALALC | 1 | 0 | 0 | 1507930 | AVV62535.1 | K9N7C7 |
| SPSNHKSNTNDA | 1 | 0 | 0 | 1507931 | ATQ39390.1 | K9N5Q8 |
| SPSSFQSAAKIL | 1 | 0 | 0 | 1507933 | ABB90505.1 | A0A0N7J131 |
| SPSSSGTFTPKK | 1 | 0 | 0 | 1507935 | ABE73425.1 | H9EJ69 |
| SPSTNTVIKKIY | 1 | 0 | 0 | 1507939 | AUM60014.1 | K9N5Q8 |
| SPTNLEMFKPGV | 1 | 0 | 0 | 1507955 | AOL02458.1 | Q4VID0 |
| SPTSKIEPCLVS | 1 | 0 | 0 | 1507959 | P0C6U5.1 | A0A140H1G9 |
| SPTSLEIFKPGV | 1 | 0 | 0 | 1507960 | APU51922.1 | Q4VID0 |
| SPTSLEMFKPEV | 1 | 0 | 0 | 1507961 | AGT51436.1 | Q4VID0 |
| SPTTSSFNADAS | 1 | 0 | 0 | 1507964 | APO40579.1 | P59594 |
| SPVDEFIKHVEK | 1 | 0 | 0 | 1507970 | AVV62524.1 | K9N7C7 |
| SPVFHRIESIDS | 1 | 0 | 0 | 1507973 | AVN89455.1 | K9N7D2 |
| SPVFHRIESTDS | 1 | 0 | 0 | 1507974 | AVN89422.1 | K9N7D2 |
| SPVLGCLGSDCN | 1 | 0 | 0 | 1507977 | AHF54664.1 | P36334 |
| SPWFADGETACN | 1 | 0 | 0 | 1507984 | AUM60013.1 | K9N7C7 |
| SPYRFVTFEPFN | 1 | 0 | 0 | 1507997 | BBA20986.1 | A0A140H1H1 |
| SPYVCNAPGCGA | 1 | 0 | 0 | 1508000 | AVN89289.1 | K9N7C7 |
| SQAKFGIITKVE | 1 | 0 | 0 | 1508008 | AIV41992.1 | P30215 |
| SQALQTVSTALN | 1 | 0 | 0 | 1508012 | ABB90519.1 | A0A0P0K6L9 |
| SQDGDYFTRQLS | 1 | 0 | 0 | 1508031 | AVV62537.1 | K9N5Q8 |
| SQDSITHYSALK | 1 | 0 | 0 | 1508045 | ASU91185.1 | K9N7C7 |
| SQDSLLKLEDFI | 1 | 0 | 0 | 1508047 | AVP78041.1 | P0C6W2 |
| SQDTLLAPLLSA | 1 | 0 | 0 | 1508051 | ATO98229.1 | P0C6W2 |
| SQDVYKSLTTVS | 1 | 0 | 0 | 1508057 | ATQ39389.1 | K9N7C7 |
| SQECSNLLLQYG | 1 | 0 | 0 | 1508059 | ATO98181.1 | Q3LZX1 |
| SQEEEQEEDWLE | 1 | 0 | 0 | 1508060 | AVP78041.1 | P0C6W2 |
| SQEGILSEREKL | 1 | 0 | 0 | 1508061 | AGT51369.1 | U3M6Q8 |
| SQEKNFTTAPAI | 1 | 0 | 0 | 1508065 | AVP78031.1 | Q3LZX1 |
| SQFDSLGASSNQ | 1 | 0 | 0 | 1508070 | AIX09796.1 | P0C6X6 |
| SQFVNLDNLRAN | 1 | 0 | 0 | 1508087 | ARO76381.1 | P0C6U8 |
| SQGFVVVNGKFS | 1 | 0 | 0 | 1508090 | QAT98908.1 | K9N7C7 |
| SQGKLLIRDNEG | 1 | 0 | 0 | 1508096 | AUM60013.1 | K9N7C7 |
| SQGLVASIKNFKSV | 1 | 0 | 0 | 1508101 | P0DTD1.1 | P0DTD1 |
| SQGSRSGNSTRC | 1 | 0 | 0 | 1508106 | ALA49909.1 | K9N4V7 |
| SQHTHYIKDGDD | 1 | 0 | 0 | 1508111 | AHI48773.1 | K9N7C7 |
| SQHTLFIKDGDD | 1 | 0 | 0 | 1508112 | AVV62524.1 | K9N7C7 |
| SQHTMQIVDEKG | 1 | 0 | 0 | 1508115 | AST12962.1 | A0A0P0LKV0 |
| SQIISGIVAAVS | 1 | 0 | 0 | 1508120 | ASY99815.1 | K9N7A1 |
| SQILPDPKKLSY | 1 | 0 | 0 | 1508123 | APO40579.1 | P59594 |
| SQILPDPSKPTK | 1 | 0 | 0 | 1508127 | ATO98181.1 | Q3LZX1 |
| SQILPDSSKPTK | 1 | 0 | 0 | 1508128 | ARO76382.1 | P59594 |
| SQILPSPLKPTK | 1 | 0 | 0 | 1508129 | 5XJK_A | P59594 |
| SQISPAAISSNC | 1 | 0 | 0 | 1508136 | AHL18090.1 | K9N5Q8 |
| SQISPTAIASNC | 1 | 0 | 0 | 1508139 | AWF93654.1 | K9N5Q8 |
| SQKAIVAYTMSL | 1 | 0 | 0 | 1508144 | AVP78031.1 | Q3LZX1 |
| SQLAKAMDTFTN | 1 | 0 | 0 | 1508151 | AEN19363.1 | P0C6X6 |
| SQLGCVFFNGAD | 1 | 0 | 0 | 1508161 | AXN92247.1 | K9N7C7 |
| SQLGEYGSFCDN | 1 | 0 | 0 | 1508164 | AHF54658.1 | P36334 |
| SQLGNCVEYHHH | 1 | 0 | 0 | 1508166 | 4KQZ_A | K9N5Q8 |
| SQLPLDIYDLTM | 1 | 0 | 0 | 1508179 | AIV41960.1 | P0C6X6 |
| SQLPLDVYNLTM | 1 | 0 | 0 | 1508181 | AAX85666.1 | P0C6X6 |
| SQLQKGVSFLKE | 1 | 0 | 0 | 1508182 | APO40578.1 | P0C6X7 |
| SQLQLATINYTQ | 1 | 0 | 0 | 1508183 | AVV62537.1 | K9N5Q8 |
| SQLRKGGRVLPF | 1 | 0 | 0 | 1508187 | APO40578.1 | P0C6X7 |
| SQLTDKYCTVNV | 1 | 0 | 0 | 1508193 | AUM60013.1 | K9N7C7 |
| SQLTKQLLDILS | 1 | 0 | 0 | 1508194 | AVV62524.1 | K9N7C7 |
| SQMEVDFLELAM | 1 | 0 | 0 | 1508201 | APO40578.1 | P0C6X7 |
| SQNMLRGDDMSF | 1 | 0 | 0 | 1508214 | AFR58685.1 | P0C6X7 |
| SQNMLRGEDVSF | 1 | 0 | 0 | 1508217 | ARO76381.1 | P0C6U8 |
| SQNVKQFANGFV | 1 | 0 | 0 | 1508227 | AIY60528.1 | K9N5Q8 |
| SQPKVITKRDAA | 1 | 0 | 0 | 1508233 | ANF29235.1 | K9N4V7 |
| SQPLNAGLNGDF | 1 | 0 | 0 | 1508235 | AVV62537.1 | K9N5Q8 |
| SQPRVDKPSQLK | 1 | 0 | 0 | 1508238 | QED88037.1 | H9EJ69 |
| SQPSHAVQNTVL | 1 | 0 | 0 | 1508239 | AVL25602.1 | H9EJ69 |
| SQQGVFRYDNIS | 1 | 0 | 0 | 1508246 | AEN19365.1 | P30215 |
| SQQGVFRYDSVS | 1 | 0 | 0 | 1508248 | AGT51619.1 | P30215 |
| SQQPLGGNVVPY | 1 | 0 | 0 | 1508252 | AMK59681.1 | P33469 |
| SQQPSGGNVVPH | 1 | 0 | 0 | 1508254 | AWW13570.1 | P33469 |
| SQQPSGRNVVPH | 1 | 0 | 0 | 1508256 | ARU07591.1 | P33469 |
| SQQPSRGNVVPH | 1 | 0 | 0 | 1508257 | ANZ78871.1 | P33469 |
| SQQPSRGNVVPY | 1 | 0 | 0 | 1508258 | AGT51784.1 | P33469 |
| SQQQNFTTAPAI | 1 | 0 | 0 | 1508260 | APO40579.1 | P59594 |
| SQRDRFYRLGNE | 1 | 0 | 0 | 1508264 | AKL87224.1 | P0C6X6 |
| SQRGGSYKNDKA | 1 | 0 | 0 | 1508265 | APO40578.1 | P0C6X7 |
| SQRGVYYPDTIY | 1 | 0 | 0 | 1508270 | AVP78031.1 | Q3LZX1 |
| SQSAAKILAPSR | 1 | 0 | 0 | 1508278 | AIW52721.1 | A0A0N7J131 |
| SQSAAKILARSQ | 1 | 0 | 0 | 1508280 | AIW52722.1 | A0A0N7J131 |
| SQSDRFYRPANE | 1 | 0 | 0 | 1508284 | AIL49562.1 | P0C6X6 |
| SQSDSLGVSSNQ | 1 | 0 | 0 | 1508285 | QBP84719.1 | P0C6X6 |
| SQSERYGFCGNG | 1 | 0 | 0 | 1508286 | ABB90523.1 | A0A0P0K6L9 |
| SQSILYRVFPYE | 1 | 0 | 0 | 1508288 | AVV62535.1 | K9N7C7 |

| | | | | | | |
|---|---|---|---|---|---|---|
| SQSNRYGFCGNG | 1 | 0 | 0 | 1508295 | AIW52843.1 | A0A0P0G321 |
| SQSRGPGESKPQ | 1 | 0 | 0 | 1508298 | AAA45463.1 | A0A0N7J131 |
| SQSRGRGESKSQ | 1 | 0 | 0 | 1508301 | ARB07429.1 | A0A0N7J131 |
| SQSRNPSSDRNH | 1 | 0 | 0 | 1508303 | AIW52723.1 | A0A0N7J131 |
| SQSRNSSSDRNH | 1 | 0 | 0 | 1508304 | AMM42526.1 | A0A0N7J131 |
| SQSRSQSRGRES | 1 | 0 | 0 | 1508306 | ABB90484.1 | A0A0N7J131 |
| SQSRSQSRIQSR | 1 | 0 | 0 | 1508308 | QEO75989.1 | A0A0N7J131 |
| SQSSAKSLARSQ | 1 | 0 | 0 | 1508314 | ABB90490.1 | A0A0N7J131 |
| SQSSAKSPARSQ | 1 | 0 | 0 | 1508316 | ABB90485.1 | A0A0N7J131 |
| SQSSHAVQNTVF | 1 | 0 | 0 | 1508318 | AVL25595.1 | H9EJ69 |
| SQSSHVVQNTVL | 1 | 0 | 0 | 1508322 | AIW52805.1 | H9EJ69 |
| SQSVAKILARSQ | 1 | 0 | 0 | 1508329 | ABB90499.1 | A0A0N7J131 |
| SQSVCNGCVGHS | 1 | 0 | 0 | 1508330 | ABB90520.1 | A0A0P0K6L9 |
| SQTSIANIIYCN | 1 | 0 | 0 | 1508344 | CAA71056.1 | A0A0P0K6L9 |
| SQVIVNNYDKSAGYPF | 1 | 0 | 0 | 1508356 | YP_009555238.1 | P0C6X6 |
| SQVIVNNYEKSA | 1 | 0 | 0 | 1508357 | AIL49570.1 | P0C6X6 |
| SQVKADLVLDAL | 1 | 0 | 0 | 1508358 | ARU07575.1 | A0A140H1G9 |
| SQWLSNLVGSAV | 1 | 0 | 0 | 1508366 | ARO76381.1 | P0C6U8 |
| SQWPISDDSDGC | 1 | 0 | 0 | 1508368 | AGT51337.1 | H9EJ42 |
| SQWPISDDSGDC | 1 | 0 | 0 | 1508369 | AWK59942.1 | H9EJ42 |
| SQWPISNDSNCC | 1 | 0 | 0 | 1508370 | AGT51386.1 | H9EJ42 |
| SQYSRSTRSILK | 1 | 0 | 0 | 1508377 | ATG84723.1 | K9N5Q8 |
| SQYSRSTRSMLK | 1 | 0 | 0 | 1508379 | AXN92228.1 | K9N5Q8 |
| SRADQNGYITLK | 1 | 0 | 0 | 1508388 | AUM60013.1 | K9N7C7 |
| SRALKKVPIDEY | 1 | 0 | 0 | 1508392 | AER30328.1 | P0C6X7 |
| SRANSGNRAPTS | 1 | 0 | 0 | 1508397 | AIX10723.1 | P33469 |
| SRANSGNRNPTS | 1 | 0 | 0 | 1508398 | AJQ24067.1 | P33469 |
| SRANSGNRTSTS | 1 | 0 | 0 | 1508402 | ARB07416.1 | P33469 |
| SRASSAGSRSKA | 1 | 0 | 0 | 1508406 | AJQ24081.1 | P33469 |
| SRASSGNRSSNG | 1 | 0 | 0 | 1508412 | AMM42463.1 | P33469 |
| SRAWAFCVKCHN | 1 | 0 | 0 | 1508421 | AXN92247.1 | K9N7C7 |
| SRDSVPLHIIAP | 1 | 0 | 0 | 1508444 | AVV62541.1 | K9N7D2 |
| SRDTQVLLRRRD | 1 | 0 | 0 | 1508445 | AVV62526.1 | K9N5Q8 |
| SREISGLSPAYA | 1 | 0 | 0 | 1508457 | AUM60013.1 | K9N7C7 |
| SRETFGLSPAYA | 1 | 0 | 0 | 1508462 | QGV13494.1 | K9N7C7 |
| SRFNPSTWNKKF | 1 | 0 | 0 | 1508473 | ARB66070.1 | P36334 |
| SRGESKSQSRNP | 1 | 0 | 0 | 1508486 | AIW52723.1 | A0A0N7J131 |
| SRGNSRNSTPSS | 1 | 0 | 0 | 1508494 | APO40586.1 | P59595 |
| SRGNVVPHYSWF | 1 | 0 | 0 | 1508495 | ANZ78871.1 | P33469 |
| SRGPGESKPQSR | 1 | 0 | 0 | 1508497 | AAA45463.1 | A0A0N7J131 |
| SRGQFIEQPNSV | 1 | 0 | 0 | 1508499 | AVV62537.1 | K9N5Q8 |
| SRGRGRNPKPRP | 1 | 0 | 0 | 1508502 | AVV62533.1 | K9N4V7 |
| SRGRGRTPKPRP | 1 | 0 | 0 | 1508503 | AVV62544.1 | K9N4V7 |
| SRGSSPARNLQA | 1 | 0 | 0 | 1508506 | APO40586.1 | P59595 |
| SRGSSPGPAGVG | 1 | 0 | 0 | 1508507 | AUM60021.1 | K9N4V7 |
| SRGVYYPDDIFR | 1 | 0 | 0 | 1508512 | ATO98145.1 | Q3LZX1 |
| SRHKQCSVQQCF | 1 | 0 | 0 | 1508513 | AJQ24068.1 | P33469 |
| SRILGAGCFVND | 1 | 0 | 0 | 1508527 | ANZ78855.1 | P0C6X6 |
| SRIRASMPTTIA | 1 | 0 | 0 | 1508536 | AVP78041.1 | P0C6W2 |
| SRKAMKRSHSPT | 1 | 0 | 0 | 1508539 | ANF29242.1 | K9N643 |
| SRKARNRSHSPT | 1 | 0 | 0 | 1508546 | ATG84704.1 | K9N643 |
| SRKFIVPWVMYL | 1 | 0 | 0 | 1508548 | AEN19363.1 | P0C6X6 |
| SRKLLGLTNLNE | 1 | 0 | 0 | 1508556 | ARU07582.1 | Q0ZJ80 |
| SRLPLDVYDLTM | 1 | 0 | 0 | 1508597 | AGT51419.1 | P0C6X6 |
| SRLRSDDRTEVP | 1 | 0 | 0 | 1508600 | ALW82691.1 | K9N5Q8 |
| SRLVESASEFIV | 1 | 0 | 0 | 1508606 | ADN03340.1 | Q0ZJ84 |
| SRMGFKLDAIVP | 1 | 0 | 0 | 1508616 | QBM11746.1 | K9N7C7 |
| SRNASPGPSGIG | 1 | 0 | 0 | 1508627 | AVV62533.1 | K9N4V7 |
| SRNQSPASSQSV | 1 | 0 | 0 | 1508638 | ABB90499.1 | A0A0N7J131 |
| SRNQSPSSSQSA | 1 | 0 | 0 | 1508640 | BAP75906.1 | A0A0N7J131 |
| SRNQSPVSSQSA | 1 | 0 | 0 | 1508641 | AMM42518.1 | A0A0N7J131 |
| SRNSSRSNSRGS | 1 | 0 | 0 | 1508643 | AUM60021.1 | K9N4V7 |
| SRNSTPSSSRGS | 1 | 0 | 0 | 1508648 | APO40586.1 | P59595 |
| SRNSTRGTSPGP | 1 | 0 | 0 | 1508649 | AHY22571.1 | K9N4V7 |
| SRQLQPINYFDS | 1 | 0 | 0 | 1508667 | CAA83660.1 | P36334 |
| SRQVIQEVKDGT | 1 | 0 | 0 | 1508671 | ABM64804.1 | A0A0P0KWE8 |
| SRRAITTENRFT | 1 | 0 | 0 | 1508677 | AIL49500.1 | P36334 |
| SRRFIVPWVMYL | 1 | 0 | 0 | 1508682 | AEN19355.1 | P0C6X6 |
| SRRGVYYNDDIF | 1 | 0 | 0 | 1508684 | ATO98181.1 | Q3LZX1 |
| SRRSDLQFGSSG | 1 | 0 | 0 | 1508691 | AMM42415.1 | A0A140H1H1 |
| SRSCSDFIPLSD | 1 | 0 | 0 | 1508697 | ANI69887.1 | K9N7C7 |
| SRSCSDFLPLSD | 1 | 0 | 0 | 1508698 | AXN92247.1 | K9N7C7 |
| SRSDCNHIGNIN | 1 | 0 | 0 | 1508701 | AEN19365.1 | P30215 |
| SRSFFEDLLFDKVKLS | 1 | 0 | 0 | 1508706 | AXT92537.1 | A0A140H1H1 |
| SRSGNSTRCTSP | 1 | 0 | 0 | 1508710 | ALA49909.1 | K9N4V7 |
| SRSNSNFRHSAS | 1 | 0 | 0 | 1508721 | AJQ24072.1 | Q19U29 |
| SRSQSRIQSRNP | 1 | 0 | 0 | 1508726 | QEO75989.1 | A0A0N7J131 |
| SRSQSRSQSRGP | 1 | 0 | 0 | 1508727 | AAA45463.1 | A0A0N7J131 |
| SRSQSSETKEQK | 1 | 0 | 0 | 1508731 | ABB90484.1 | A0A0N7J131 |
| SRSRANSGNRNP | 1 | 0 | 0 | 1508734 | AJQ24067.1 | P33469 |
| SRSRANSGNRTL | 1 | 0 | 0 | 1508735 | AAX85673.1 | P33469 |
| SRSRASSGNRAP | 1 | 0 | 0 | 1508738 | ATP16771.1 | P33469 |
| SRSRASSGNRTP | 1 | 0 | 0 | 1508739 | AJQ24068.1 | P33469 |
| SRSRNSTRGTSP | 1 | 0 | 0 | 1508744 | AHY22571.1 | K9N4V7 |
| SRSRVSSGNRAP | 1 | 0 | 0 | 1508745 | AIL49426.1 | P33469 |
| SRSSSQGSRSRN | 1 | 0 | 0 | 1508749 | AHY22571.1 | K9N4V7 |
| SRSTRSILKRRD | 1 | 0 | 0 | 1508756 | ATG84723.1 | K9N5Q8 |
| SRTGHFYINGYR | 1 | 0 | 0 | 1508773 | AIW52757.1 | A0A0P0K6L9 |
| SRTLSYYRLGAS | 1 | 0 | 0 | 1508784 | APO40582.1 | P59596 |
| SRTPKYTFGVVK | 1 | 0 | 0 | 1508785 | AEN19363.1 | P0C6X6 |
| SRTSSRASSTGS | 1 | 0 | 0 | 1508790 | AIX10719.1 | P33469 |

| | | | | | | |
|---|---|---|---|---|---|---|
| SRVESKSQSRNP | 1 | 0 | 0 | 1508804 | AST12968.1 | A0A0N7J131 |
| SRVGRSVNSQNC | 1 | 0 | 0 | 1508810 | CAA33521.1 | A0A0P0KXN3 |
| SRVIRRERCDMT | 1 | 0 | 0 | 1508812 | AWK59942.1 | H9EJ42 |
| SRVSSGNRAPTS | 1 | 0 | 0 | 1508832 | AIL49426.1 | P33469 |
| SRYVDLYGGNMF | 1 | 0 | 0 | 1508851 | AXN92228.1 | K9N5Q8 |
| SSAFLTQYITRM | 1 | 0 | 0 | 1508864 | AVV62529.1 | K9N643 |
| SSAGPLQTTVGC | 1 | 0 | 0 | 1508871 | AVV62526.1 | K9N5Q8 |
| SSALIAMANWLA | 1 | 0 | 0 | 1508882 | AVV62524.1 | K9N7C7 |
| SSAMFCATCPFA | 1 | 0 | 0 | 1508885 | AVV62535.1 | K9N7C7 |
| SSANGSLDGTTL | 1 | 0 | 0 | 1508887 | AXG21642.1 | K9N638 |
| SSANGSLVCTTL | 1 | 0 | 0 | 1508888 | AVN89333.1 | K9N7C7 |
| SSANGSLVGTIL | 1 | 0 | 0 | 1508889 | AIZ48759.1 | K9N638 |
| SSAPRITFGGPS | 1 | 0 | 0 | 1508893 | AVP78049.1 | Q3LZX4 |
| SSASEGLVGTTL | 1 | 0 | 0 | 1508897 | AUM60013.1 | K9N7C7 |
| SSASRGSSRSSS | 1 | 0 | 0 | 1508898 | AVV62533.1 | K9N4V7 |
| SSATQPYFDSYL | 1 | 0 | 0 | 1508903 | AGW27863.1 | A0A140H1H1 |
| SSCFGKNNGIGT | 1 | 0 | 0 | 1508916 | QEG03814.1 | P36334 |
| SSCKSHKPSSAS | 1 | 0 | 0 | 1508924 | AFR45556.1 | A0A140H1H1 |
| SSCPGKNNGIGT | 1 | 0 | 0 | 1508925 | AWW13566.1 | P36334 |
| SSCPGKNNGIST | 1 | 0 | 0 | 1508926 | BBA84094.1 | P36334 |
| SSCPGKNNGLGT | 1 | 0 | 0 | 1508927 | AIL49503.1 | P36334 |
| SSCSCPGKNNGI | 1 | 0 | 0 | 1508932 | AVR40344.1 | P36334 |
| SSCSGKNNGIGT | 1 | 0 | 0 | 1508933 | ANZ78838.1 | P36334 |
| SSCSGKNNGVGT | 1 | 0 | 0 | 1508934 | QEG03757.1 | P36334 |
| SSCVGSGPGKNN | 1 | 0 | 0 | 1508937 | AIX10756.1 | P36334 |
| SSDAMLDDATGV | 1 | 0 | 0 | 1508941 | AVV62524.1 | K9N7C7 |
| SSDDVDPFDAAV | 1 | 0 | 0 | 1508947 | AVV62544.1 | K9N4V7 |
| SSDENGVRTLST | 1 | 0 | 0 | 1508949 | AVP78031.1 | Q3LZX1 |
| SSDKAPYRVIPR | 1 | 0 | 0 | 1508959 | AIW52805.1 | H9EJ69 |
| SSDMVETVVGQL | 1 | 0 | 0 | 1508963 | ASU45706.1 | K9N7C7 |
| SSDRYHNSQDDI | 1 | 0 | 0 | 1508972 | AAA45463.1 | A0A0N7J131 |
| SSDSDVFIKKYG | 1 | 0 | 0 | 1508973 | ANI69887.1 | K9N7C7 |
| SSDVFKIGDVKF | 1 | 0 | 0 | 1508975 | QDH43737.1 | H9EJ42 |
| SSDVIHYNINFE | 1 | 0 | 0 | 1508976 | BAL45639.1 | A0A0P0K6L9 |
| SSDVMLEDATGL | 1 | 0 | 0 | 1508984 | AUM60013.1 | K9N7C7 |
| SSDWRVYAFNSY | 1 | 0 | 0 | 1508988 | APO40579.1 | P59594 |
| SSEPALIFRNIK | 1 | 0 | 0 | 1509012 | AGT51481.1 | P36334 |
| SSEPTLLFRNIK | 1 | 0 | 0 | 1509017 | AIL49500.1 | P36334 |
| SSEYKGSITDVF | 1 | 0 | 0 | 1509030 | AVP78041.1 | P0C6W2 |
| SSFDGNMYGAFQ | 1 | 0 | 0 | 1509037 | AVV62524.1 | K9N7C7 |
| SSFGEYNLSSVV | 1 | 0 | 0 | 1509056 | ABB90513.1 | A0A0P0K6L9 |
| SSFGFDQQQLLA | 1 | 0 | 0 | 1509057 | ARU07575.1 | A0A140H1G9 |
| SSFIELLFDCVG | 1 | 0 | 0 | 1509059 | ATQ39389.1 | K9N7C7 |
| SSFKATLYYQNG | 1 | 0 | 0 | 1509062 | ARB07423.1 | A0A0P0LKV0 |
| SSFLGRYMSALN | 1 | 0 | 0 | 1509066 | AVP78041.1 | P0C6W2 |
| SSFMNATLDKCT | 1 | 0 | 0 | 1509069 | CAA71147.1 | A0A0P0K6L9 |
| SSFMNVTLNKCT | 1 | 0 | 0 | 1509071 | AIW52757.1 | A0A0P0K6L9 |
| SSFNFDQKELLD | 1 | 0 | 0 | 1509074 | ATN39862.1 | P0C6X6 |
| SSFQSAAKILAR | 1 | 0 | 0 | 1509075 | ABB90505.1 | A0A0N7J131 |
| SSFSEILFDCVD | 1 | 0 | 0 | 1509077 | AZL96751.1 | K9N638 |
| SSFSELLFDCVD | 1 | 0 | 0 | 1509078 | AXN92247.1 | K9N7C7 |
| SSFTEQLGVPLG | 1 | 0 | 0 | 1509087 | AWO68966.1 | A0A0P0G321 |
| SSFTEQLGVPLV | 1 | 0 | 0 | 1509088 | AMM42637.1 | A0A0P0G321 |
| SSFYAPEPITSF | 1 | 0 | 0 | 1509093 | AVN89387.1 | K9N5Q8 |
| SSFYSPEPITSL | 1 | 0 | 0 | 1509098 | AHI48672.1 | K9N5Q8 |
| SSGFDKSYQTNR | 1 | 0 | 0 | 1509103 | AUM60024.1 | K9N5Q8 |
| SSGFDNSYQTNR | 1 | 0 | 0 | 1509104 | AUM60014.1 | K9N5Q8 |
| SSGFRKMAFPSG | 1 | 0 | 0 | 1509108 | 3M3V_A | P0C6X7 |
| SSGIFSVPSDVS | 1 | 0 | 0 | 1509114 | ARK08632.1 | P0C6X6 |
| SSGKPVPYCYDT | 1 | 0 | 0 | 1509117 | AVP78041.1 | P0C6W2 |
| SSGLVKPSALFC | 1 | 0 | 0 | 1509121 | AUM60013.1 | K9N7C7 |
| SSGLVPRGSAMG | 1 | 0 | 0 | 1509122 | 1SSK_A | P59595 |
| SSGMINPNALLC | 1 | 0 | 0 | 1509128 | ASU90746.1 | K9N7C7 |
| SSGNRPGNGILK | 1 | 0 | 0 | 1509132 | AMM42456.1 | P33469 |
| SSGNRSVNGILK | 1 | 0 | 0 | 1509134 | AGT51505.1 | P33469 |
| SSGNYVGKYDIN | 1 | 0 | 0 | 1509136 | AGT17757.1 | A0A140H1H0 |
| SSGPISPTNLEM | 1 | 0 | 0 | 1509138 | AOL02458.1 | Q4VID0 |
| SSGPISPTSLEM | 1 | 0 | 0 | 1509140 | ARE29989.1 | Q4VID0 |
| SSGPISQFNYKQ | 1 | 0 | 0 | 1509141 | ALA50067.1 | K9N5Q8 |
| SSGSLATRLMSI | 1 | 0 | 0 | 1509143 | ATP16762.1 | Q4VID0 |
| SSGVDLGTENLY | 1 | 0 | 0 | 1509153 | 5X29_A | P59637 |
| SSGVINSSAMFC | 1 | 0 | 0 | 1509154 | AVV62535.1 | K9N7C7 |
| SSGYLQLATINY | 1 | 0 | 0 | 1509162 | AVV62526.1 | K9N5Q8 |
| SSGYRSGNGILK | 1 | 0 | 0 | 1509164 | AIV41850.1 | P33469 |
| SSHAVQNTVFNA | 1 | 0 | 0 | 1509165 | AVL25595.1 | H9EJ69 |
| SSHAWQPGVAMP | 1 | 0 | 0 | 1509167 | AVP78041.1 | P0C6W2 |
| SSIIQRTRTRPS | 1 | 0 | 0 | 1509193 | QCZ25056.1 | K9N4V7 |
| SSIKSDKGVEKL | 1 | 0 | 0 | 1509202 | AGT51393.1 | H9EJ42 |
| SSINDLSFDKNL | 1 | 0 | 0 | 1509204 | AVA26872.1 | H9EJ42 |
| SSIPDPKFVDKY | 1 | 0 | 0 | 1509205 | AUM60013.1 | K9N7C7 |
| SSIQAIYDRLDI | 1 | 0 | 0 | 1509206 | AIW52757.1 | A0A0P0K6L9 |
| SSISDILKRLDA | 1 | 0 | 0 | 1509212 | AVV62537.1 | K9N5Q8 |
| SSISDILQRLDT | 1 | 0 | 0 | 1509213 | AVV62526.1 | K9N5Q8 |
| SSIVRQLFKPLT | 1 | 0 | 0 | 1509223 | ATO98186.1 | Q3LZX8 |
| SSKADNSIFRSF | 1 | 0 | 0 | 1509228 | AMK59676.1 | P30215 |
| SSKAGNSIFRSF | 1 | 0 | 0 | 1509229 | AEN19365.1 | P30215 |
| SSKALAKQDAAR | 1 | 0 | 0 | 1509230 | AUM60016.1 | K9N4V0 |
| SSKEFVEYKITQ | 1 | 0 | 0 | 1509236 | AVQ05261.1 | P0C6X6 |
| SSKIENASPAST | 1 | 0 | 0 | 1509239 | ARO76383.1 | P59632 |
| SSKMSDVKCTSV | 1 | 0 | 0 | 1509243 | 2AHM_A | P0C6U8 |
| SSKPTKRSFIED | 1 | 0 | 0 | 1509246 | ARO76382.1 | P59594 |

| | | | | | | |
|---|---|---|---|---|---|---|
| SSKPTPAPVYIW | 1 | 0 | 0 | 1509247 | AWW13562.1 | Q01455 |
| SSKPVAKKYTYS | 1 | 0 | 0 | 1509248 | ANI69899.1 | K9N7C7 |
| SSKPVIKKYTYS | 1 | 0 | 0 | 1509249 | AHI48784.1 | K9N7C7 |
| SSKQSVASVIAE | 1 | 0 | 0 | 1509251 | APO40578.1 | P0C6X7 |
| SSKRFQSFQQFG | 1 | 0 | 0 | 1509253 | AVP78031.1 | Q3LZX1 |
| SSKTPEEHFVET | 1 | 0 | 0 | 1509258 | ARO76381.1 | P0C6U8 |
| SSLCVGSGPGKN | 1 | 0 | 0 | 1509268 | AVQ05264.1 | P36334 |
| SSLEKLQCEHLQ | 1 | 0 | 0 | 1509272 | AWO68978.1 | A0A0P0G321 |
| SSLFAQSEEDES | 1 | 0 | 0 | 1509274 | AKK52584.1 | K9N4V0 |
| SSLFAQSEEEEL | 1 | 0 | 0 | 1509275 | ANF29274.1 | K9N4V0 |
| SSLFAQSEEQES | 1 | 0 | 0 | 1509280 | ANF29197.1 | K9N4V0 |
| SSLFAQSKEEEL | 1 | 0 | 0 | 1509282 | ALT66804.1 | K9N4V0 |
| SSLFAQSKEEES | 1 | 0 | 0 | 1509283 | ARQ84748.1 | K9N4V0 |
| SSLILDYFSYPF | 1 | 0 | 0 | 1509291 | AVN89313.1 | K9N5Q8 |
| SSLIREIVMNAS | 1 | 0 | 0 | 1509293 | AEN19363.1 | P0C6X6 |
| SSLIVDYFSYPL | 1 | 0 | 0 | 1509295 | AVV62526.1 | K9N5Q8 |
| SSLKMIQTHLSH | 1 | 0 | 0 | 1509297 | AUM60013.1 | K9N7C7 |
| SSLNLPGCDGGS | 1 | 0 | 0 | 1509307 | APO40578.1 | P0C6X7 |
| SSLQEILSRLDV | 1 | 0 | 0 | 1509312 | ABD75553.1 | A0A140H1H1 |
| SSLQQVVKALNE | 1 | 0 | 0 | 1509313 | ATG84756.1 | K9N5Q8 |
| SSLSAQSEEEES | 1 | 0 | 0 | 1509317 | AKN24751.1 | K9N4V0 |
| SSLTSEQIVAII | 1 | 0 | 0 | 1509324 | AVV62532.1 | K9N7A1 |
| SSLVLAREHEFC | 1 | 0 | 0 | 1509334 | ABU39953.1 | A0A140H1G9 |
| SSMLRLHTVESG | 1 | 0 | 0 | 1509345 | AXN92247.1 | K9N7C7 |
| SSMLTFDGNAFS | 1 | 0 | 0 | 1509346 | AVL25589.1 | A0A0P0G321 |
| SSMNIAANSNEW | 1 | 0 | 0 | 1509348 | AGT51337.1 | H9EJ42 |
| SSMRGCCDCGST | 1 | 0 | 0 | 1509350 | AIW52843.1 | A0A0P0G321 |
| SSMTDNLQMAFV | 1 | 0 | 0 | 1509352 | AUM60014.1 | K9N5Q8 |
| SSMVVDNCQLPL | 1 | 0 | 0 | 1509356 | AVV62537.1 | K9N5Q8 |
| SSNATILDYCDG | 1 | 0 | 0 | 1509358 | APO40578.1 | P0C6X7 |
| SSNGILKWADQS | 1 | 0 | 0 | 1509363 | AMM42463.1 | P33469 |
| SSNGPISPTSLE | 1 | 0 | 0 | 1509364 | AGT51605.1 | Q4VID0 |
| SSNGPLKVGTGV | 1 | 0 | 0 | 1509366 | ARU07593.1 | H9EJ42 |
| SSNGQPGGDAIS | 1 | 0 | 0 | 1509367 | QAT98908.1 | K9N7C7 |
| SSNGSNDVWIFN | 1 | 0 | 0 | 1509368 | AGT17757.1 | A0A140H1H0 |
| SSNINRILGVNS | 1 | 0 | 0 | 1509373 | AUT76985.1 | A0A0P0LKV0 |
| SSNINRVLSVPS | 1 | 0 | 0 | 1509374 | ABQ16583.1 | H9EJ42 |
| SSNMVETLVGQL | 1 | 0 | 0 | 1509378 | ASU90174.1 | K9N638 |
| SSNPNTPAYTFT | 1 | 0 | 0 | 1509380 | AVV62524.1 | K9N7C7 |
| SSNRGNKQIVHS | 1 | 0 | 0 | 1509385 | ATG84704.1 | K9N643 |
| SSNTDNETLEYW | 1 | 0 | 0 | 1509387 | 5I08_A | A0A140H1H1 |
| SSNVANYQKVGV | 1 | 0 | 0 | 1509392 | APO40578.1 | P0C6X7 |
| SSPIAYLVFLTS | 1 | 0 | 0 | 1509407 | AUM60013.1 | K9N7C7 |
| SSPIIPGFGGNF | 1 | 0 | 0 | 1509411 | AOR17482.1 | K9N5Q8 |
| SSPKVTIDCAAC | 1 | 0 | 0 | 1509413 | ABD43177.1 | P36334 |
| SSPKVTIDCVAF | 1 | 0 | 0 | 1509416 | AIL49523.1 | P36334 |
| SSPNKQCTVQQC | 1 | 0 | 0 | 1509421 | AJQ24079.1 | P33469 |
| SSPNLPDFREEL | 1 | 0 | 0 | 1509422 | AIL49492.1 | P36334 |
| SSPQIIKQLKKA | 1 | 0 | 0 | 1509423 | NP_073549.1 | A0A0P0LKV0 |
| SSPTTLQAVGSC | 1 | 0 | 0 | 1509424 | AXN92247.1 | K9N7C7 |
| SSPYRFVTFEPF | 1 | 0 | 0 | 1509425 | BBA20986.1 | A0A140H1H1 |
| SSQGIVAVNSTK | 1 | 0 | 0 | 1509431 | AUM60018.1 | K9N7D2 |
| SSQGPNFAEISK | 1 | 0 | 0 | 1509432 | AUM60014.1 | K9N5Q8 |
| SSQGPNFQEISK | 1 | 0 | 0 | 1509433 | AVV62526.1 | K9N5Q8 |
| SSQGSRSRNSTR | 1 | 0 | 0 | 1509436 | AHY22571.1 | K9N4V7 |
| SSQLPLDIYDLT | 1 | 0 | 0 | 1509439 | AIV41960.1 | P0C6X6 |
| SSQLPLDVYNLT | 1 | 0 | 0 | 1509440 | AAX85666.1 | P0C6X6 |
| SSQSSAKILARS | 1 | 0 | 0 | 1509450 | ABB90486.1 | A0A0N7J131 |
| SSRASSGNRPGN | 1 | 0 | 0 | 1509461 | AMM42456.1 | P33469 |
| SSRASSGNRSSN | 1 | 0 | 0 | 1509462 | AMM42463.1 | P33469 |
| SSRASSGNRSVN | 1 | 0 | 0 | 1509463 | AGT51505.1 | P33469 |
| SSRGSSPARNLQ | 1 | 0 | 0 | 1509470 | APO40586.1 | P59595 |
| SSRKGDLYGGNM | 1 | 0 | 0 | 1509476 | AVV62526.1 | K9N5Q8 |
| SSRRGFYYFDDT | 1 | 0 | 0 | 1509488 | APO40579.1 | P59594 |
| SSRRKRRGISSP | 1 | 0 | 0 | 1509490 | BBA20986.1 | A0A140H1H1 |
| SSRRLAQQKVNE | 1 | 0 | 0 | 1509493 | AIW52843.1 | A0A0P0G321 |
| SSRVERSNAVLT | 1 | 0 | 0 | 1509505 | AST12962.1 | A0A0P0LKV0 |
| SSSAAIPFAQSI | 1 | 0 | 0 | 1509508 | AOR17480.1 | K9N5Q8 |
| SSSAFIGMSQFK | 1 | 0 | 0 | 1509509 | AVV62533.1 | K9N4V7 |
| SSSANGRLVGTT | 1 | 0 | 0 | 1509514 | AJG44057.1 | K9N638 |
| SSSANGSLDGTT | 1 | 0 | 0 | 1509515 | AXG21642.1 | K9N638 |
| SSSANGSLVCTT | 1 | 0 | 0 | 1509516 | AVN89333.1 | K9N7C7 |
| SSSANGSLVGTI | 1 | 0 | 0 | 1509517 | AIZ48759.1 | K9N638 |
| SSSCVGSGPGKN | 1 | 0 | 0 | 1509526 | AIX10756.1 | P36334 |
| SSSEAFLIGVNY | 1 | 0 | 0 | 1509534 | P0C6W6.1 | P0C6W2 |
| SSSEFFNKQWPM | 1 | 0 | 0 | 1509535 | AVV62537.1 | K9N5Q8 |
| SSSFSTFKCYGV | 1 | 0 | 0 | 1509541 | APO40579.1 | P59594 |
| SSSFSVVAMSGH | 1 | 0 | 0 | 1509543 | AWK59942.1 | H9EJ42 |
| SSSGPISPISLE | 1 | 0 | 0 | 1509547 | AGT51805.1 | Q4VID0 |
| SSSGPISPTNLE | 1 | 0 | 0 | 1509548 | AOL02458.1 | Q4VID0 |
| SSSGPISPTSLE | 1 | 0 | 0 | 1509549 | ARE29989.1 | Q4VID0 |
| SSSGPISQFNYK | 1 | 0 | 0 | 1509550 | ALA50067.1 | K9N5Q8 |
| SSSKSSEEHFIE | 1 | 0 | 0 | 1509558 | APO40578.1 | P0C6X7 |
| SSSKTPEEHFVE | 1 | 0 | 0 | 1509560 | ARO76381.1 | P0C6U8 |
| SSSLRTFVVDKS | 1 | 0 | 0 | 1509566 | AXN92247.1 | K9N7C7 |
| SSSNPTCLILAT | 1 | 0 | 0 | 1509571 | AXN92228.1 | K9N5Q8 |
| SSSPALVDQRTI | 1 | 0 | 0 | 1509573 | ARE29984.1 | H9EJ42 |
| SSSRASFGNRSG | 1 | 0 | 0 | 1509579 | QBP84763.1 | P33469 |
| SSSRASSGNRPG | 1 | 0 | 0 | 1509583 | AMM42456.1 | P33469 |
| SSSSFDCIVNLS | 1 | 0 | 0 | 1509594 | ACY56724.1 | A0A0P0G321 |
| SSSSFSTFKCYG | 1 | 0 | 0 | 1509595 | APO40579.1 | P59594 |

| | | | | | | |
|---|---|---|---|---|---|---|
| SSSTYRNMALKG | 1 | 0 | 0 | 1509600 | AIL49538.1 | P36334 |
| SSSVLDTTIPFG | 1 | 0 | 0 | 1509602 | AVL25581.1 | A0A0P0G321 |
| SSSYGDAHEGIF | 1 | 0 | 0 | 1509605 | AMM42397.1 | Q19U29 |
| SSTKKFQPFQQF | 1 | 0 | 0 | 1509621 | APO40579.1 | P59594 |
| SSTMIFSLHPVE | 1 | 0 | 0 | 1509625 | AXN92247.1 | K9N7C7 |
| SSTQKTIKKIYP | 1 | 0 | 0 | 1509632 | AVV62526.1 | K9N5Q8 |
| SSTVDAQIQNKT | 1 | 0 | 0 | 1509637 | AVV62535.1 | K9N7C7 |
| SSVACEHITTMM | 1 | 0 | 0 | 1509646 | AVV62526.1 | K9N5Q8 |
| SSVAGVVAQGAR | 1 | 0 | 0 | 1509648 | AVV62524.1 | K9N7C7 |
| SSVDNILNMARN | 1 | 0 | 0 | 1509649 | AST12962.1 | A0A0P0LKV0 |
| SSVGNFSNGVSG | 1 | 0 | 0 | 1509655 | AVV62537.1 | K9N5Q8 |
| SSVGNYYNGYPG | 1 | 0 | 0 | 1509657 | AVV62526.1 | K9N5Q8 |
| SSVLDTTIPFGP | 1 | 0 | 0 | 1509664 | AVL25581.1 | A0A0P0G321 |
| SSVNDAITRTAE | 1 | 0 | 0 | 1509674 | QED88026.1 | A0A0P0G321 |
| SSVPGEMRLASI | 1 | 0 | 0 | 1509675 | 5X59_A | K9N5Q8 |
| SSVPLSEVYVHL | 1 | 0 | 0 | 1509677 | AWK59934.1 | U3M6Q8 |
| SSVRSDGTFEEM | 1 | 0 | 0 | 1509680 | AVQ05261.1 | P0C6X6 |
| SSVSNFSYGKMG | 1 | 0 | 0 | 1509682 | QGV13484.1 | K9N5Q8 |
| SSVVPSLPTSGS | 1 | 0 | 0 | 1509687 | ABB90513.1 | A0A0P0K6L9 |
| SSVYSVPLLFQT | 1 | 0 | 0 | 1509698 | AZS52616.1 | A0A140H1G9 |
| SSWQTGDFLKAT | 1 | 0 | 0 | 1509704 | APO40578.1 | P0C6X7 |
| SSWVLNNDYYRS | 1 | 0 | 0 | 1509705 | ARU07575.1 | A0A140H1G9 |
| SSWVYQSAFNCT | 1 | 0 | 0 | 1509706 | ATO98181.1 | Q3LZX1 |
| SSYKLDLTFLGL | 1 | 0 | 0 | 1509722 | ARO76381.1 | P0C6U8 |
| SSYTVDSVTVKN | 1 | 0 | 0 | 1509737 | APO40578.1 | P0C6X7 |
| SSYTVELGTDVN | 1 | 0 | 0 | 1509738 | AVP78041.1 | P0C6W2 |
| STADNLKNVLVW | 1 | 0 | 0 | 1509746 | APO40578.1 | P0C6X7 |
| STAIYTNLRNCT | 1 | 0 | 0 | 1509755 | ATO98145.1 | Q3LZX1 |
| STALCKSGSLVL | 1 | 0 | 0 | 1509758 | P30215.1 | P30215 |
| STANALPECNAD | 1 | 0 | 0 | 1509764 | ABE97136.1 | H9EJ42 |
| STAPILTESTGS | 1 | 0 | 0 | 1509767 | ARE29989.1 | Q4VID0 |
| STARDGIRRGDN | 1 | 0 | 0 | 1509768 | ANI69888.1 | K9N638 |
| STARDGLKRGEN | 1 | 0 | 0 | 1509769 | AVV62535.1 | K9N7C7 |
| STARTTSSEFQL | 1 | 0 | 0 | 1509772 | AUM60014.1 | K9N5Q8 |
| STASAVQTCDLT | 1 | 0 | 0 | 1509774 | CAA83661.1 | P36334 |
| STAVSPDFKAFN | 1 | 0 | 0 | 1509780 | AVV62535.1 | K9N7C7 |
| STAYIQQCVTRF | 1 | 0 | 0 | 1509787 | AUM60017.1 | K9N643 |
| STCNSNANISML | 1 | 0 | 0 | 1509800 | AKA09625.1 | A0A0P0G321 |
| STCNSNANISTL | 1 | 0 | 0 | 1509801 | AKA09623.1 | A0A0P0G321 |
| STCNSNANLSML | 1 | 0 | 0 | 1509802 | AVL25555.1 | A0A0P0G321 |
| STCSSKGLVAVN | 1 | 0 | 0 | 1509805 | AVV62541.1 | K9N7D2 |
| STCSSQGIVAVN | 1 | 0 | 0 | 1509806 | AUM60018.1 | K9N7D2 |
| STDCSDGNYNRN | 1 | 0 | 0 | 1509810 | ALA49803.1 | K9N5Q8 |
| STDDGSLAFDEA | 1 | 0 | 0 | 1509812 | AVV62533.1 | K9N4V7 |
| STDEEGDNVQIT | 1 | 0 | 0 | 1509813 | AMM42619.1 | H9EJ69 |
| STDFHQLECIVD | 1 | 0 | 0 | 1509816 | ARU07593.1 | H9EJ42 |
| STDFHQLEYDSV | 1 | 0 | 0 | 1509817 | AGT51337.1 | H9EJ42 |
| STDGLYFHFYQE | 1 | 0 | 0 | 1509823 | ABA60800.1 | A0A140H1H1 |
| STDLQKANTDII | 1 | 0 | 0 | 1509827 | CAA79896.1 | P36334 |
| STDLQKANTDIK | 1 | 0 | 0 | 1509828 | AVQ05264.1 | P36334 |
| STDLQKSNTDII | 1 | 0 | 0 | 1509830 | CAA83660.1 | P36334 |
| STDNNQDGGRSG | 1 | 0 | 0 | 1509831 | Q3I5I7.1 | Q3LZX4 |
| STDQMSELPKEQ | 1 | 0 | 0 | 1509838 | ASJ26617.1 | K9N4V7 |
| STDQMSEPPKEH | 1 | 0 | 0 | 1509839 | YP_007188586.1 | K9N4V7 |
| STDQMSEPPMEQ | 1 | 0 | 0 | 1509841 | AXP07352.1 | K9N4V7 |
| STDSTVFTYIPA | 1 | 0 | 0 | 1509848 | AVN89422.1 | K9N7D2 |
| STDTVNPDSFNL | 1 | 0 | 0 | 1509855 | ARB07436.1 | A0A140H1G9 |
| STDVLVNTYKRK | 1 | 0 | 0 | 1509858 | AUM60013.1 | K9N7C7 |
| STEEVGDNVQIT | 1 | 0 | 0 | 1509867 | ABK63972.1 | H9EJ69 |
| STEQVDVFHETS | 1 | 0 | 0 | 1509885 | AVN89352.1 | K9N7C7 |
| STEVASKMFDAY | 1 | 0 | 0 | 1509893 | APO40578.1 | P0C6X7 |
| STEVVRFPNITN | 1 | 0 | 0 | 1509897 | APO40579.1 | P59594 |
| STFDDILGKVCS | 1 | 0 | 0 | 1509898 | ARU07575.1 | A0A140H1G9 |
| STFEPRSDLERD | 1 | 0 | 0 | 1509900 | ARB07436.1 | A0A140H1G9 |
| STFGSFYYHNSM | 1 | 0 | 0 | 1509906 | ARB07436.1 | A0A140H1G9 |
| STFGTRNPNNDA | 1 | 0 | 0 | 1509908 | AVV62533.1 | K9N4V7 |
| STFGTRNPNNDL | 1 | 0 | 0 | 1509909 | AKN24774.1 | K9N4V7 |
| STFGTRNPNNDP | 1 | 0 | 0 | 1509910 | AUM60021.1 | K9N4V7 |
| STFISAARNGVV | 1 | 0 | 0 | 1509912 | APO40578.1 | P0C6X7 |
| STFTPRSVRSVP | 1 | 0 | 0 | 1509933 | AVK87439.1 | K9N5Q8 |
| STFVGVLPPTVR | 1 | 0 | 0 | 1509936 | AIW52841.1 | A0A0P0G321 |
| STGEIAQYNYKQ | 1 | 0 | 0 | 1509952 | AUM60014.1 | K9N5Q8 |
| STGFDFNCHYLV | 1 | 0 | 0 | 1509953 | AGT51729.1 | P30215 |
| STGLVKNQCVNF | 1 | 0 | 0 | 1509962 | ATO98193.1 | Q3LZX1 |
| STGNIFIVTPCN | 1 | 0 | 0 | 1509964 | AIW52843.1 | A0A0P0G321 |
| STGNLIISSSTQ | 1 | 0 | 0 | 1509965 | AVV62526.1 | K9N5Q8 |
| STGRCTVNKHVA | 1 | 0 | 0 | 1509979 | AIV42008.1 | P0C6X6 |
| STGSLASRLMSI | 1 | 0 | 0 | 1509981 | QBQ01844.1 | Q4VID0 |
| STGSLATRLMSI | 1 | 0 | 0 | 1509982 | AVR40343.1 | Q4VID0 |
| STGSRSRANSGN | 1 | 0 | 0 | 1509987 | AIX10719.1 | P33469 |
| STGSYFCVNSKA | 1 | 0 | 0 | 1509991 | AUM60013.1 | K9N7C7 |
| STGSYYCVNSKA | 1 | 0 | 0 | 1509992 | AUM60023.1 | K9N7C7 |
| STGTQSVSVDRE | 1 | 0 | 0 | 1509997 | AXN92229.1 | K9N796 |
| STGTVIIRPSTS | 1 | 0 | 0 | 1509999 | QBF80608.1 | K9N5Q8 |
| STHDDFNEHVTS | 1 | 0 | 0 | 1510014 | AXG21666.1 | K9N796 |
| STHDDFTEHVTS | 1 | 0 | 0 | 1510015 | AJG44103.1 | K9N796 |
| STHDGFNEHVTS | 1 | 0 | 0 | 1510017 | AHI48551.1 | K9N796 |
| STHDGLIEHVTS | 1 | 0 | 0 | 1510019 | QBF80533.1 | K9N796 |
| STHDGPTEHVNL | 1 | 0 | 0 | 1510025 | ANI69890.1 | K9N796 |
| STHDGPTEYVTI | 1 | 0 | 0 | 1510027 | AVN89362.1 | K9N796 |
| STHDTPTEHVTS | 1 | 0 | 0 | 1510029 | AJG44059.1 | K9N796 |

| | | | | | | |
|---|---|---|---|---|---|---|
| STIDKSNVVRGW | 1 | 0 | 0 | 1510040 | ATO98145.1 | Q3LZX1 |
| STIFNSCQNAVS | 1 | 0 | 0 | 1510042 | AVV62524.1 | K9N7C7 |
| STIGDSIFTGCT | 1 | 0 | 0 | 1510045 | QAT98908.1 | K9N7C7 |
| STIGEAIFTGCV | 1 | 0 | 0 | 1510046 | AUM60023.1 | K9N7C7 |
| STINALPEISAD | 1 | 0 | 0 | 1510062 | ALW82729.1 | K9N7C7 |
| STISVPAHMPMH | 1 | 0 | 0 | 1510067 | AET99209.1 | Q0ZJ80 |
| STKLPYYDVKKI | 1 | 0 | 0 | 1510084 | BAL45640.1 | A0A0P0K6L9 |
| STKPCFIRNFDQ | 1 | 0 | 0 | 1510088 | AUM60018.1 | K9N7D2 |
| STLLGSIAGVGW | 1 | 0 | 0 | 1510121 | AWU59321.1 | K9N5Q8 |
| STLPGFETIMKV | 1 | 0 | 0 | 1510127 | ARU07581.1 | Q19U29 |
| STLRDTLEYYAQ | 1 | 0 | 0 | 1510134 | AUM60023.1 | K9N7C7 |
| STLSGFETIMKV | 1 | 0 | 0 | 1510148 | AEN19370.1 | P33469 |
| STLTPASVGSVP | 1 | 0 | 0 | 1510151 | 5W9H_A | K9N5Q8 |
| STLVDLNWLNRV | 1 | 0 | 0 | 1510157 | AGT21353.1 | A0A0P0K6L9 |
| STMDNTTQSAII | 1 | 0 | 0 | 1510170 | ATO98193.1 | Q3LZX1 |
| STMEEDFLNMDI | 1 | 0 | 0 | 1510171 | NP_073549.1 | A0A0P0LKV0 |
| STMTGRVFHQKC | 1 | 0 | 0 | 1510184 | AIL49573.1 | P0C6X6 |
| STMTNRQYHQKM | 1 | 0 | 0 | 1510186 | AXN92247.1 | K9N7C7 |
| STNKTTFRPNMW | 1 | 0 | 0 | 1510196 | APO40578.1 | P0C6X7 |
| STNLFKDCSNSY | 1 | 0 | 0 | 1510200 | AGT51577.1 | P0C6X6 |
| STNLPPPLLSNS | 1 | 0 | 0 | 1510202 | ALL26396.1 | K9N5Q8 |
| STNPDPKFVDRY | 1 | 0 | 0 | 1510207 | ATU80202.1 | K9N7C7 |
| STNTTFDFLSNA | 1 | 0 | 0 | 1510213 | ACV87106.1 | A0A0P0G321 |
| STNVTTATYCTG | 1 | 0 | 0 | 1510215 | AVP78030.1 | P0C6W2 |
| STPNAIFISDRK | 1 | 0 | 0 | 1510235 | AXN92247.1 | K9N7C7 |
| STPNLPDFKEEL | 1 | 0 | 0 | 1510239 | BBA20982.1 | P36334 |
| STPNLPDFREEL | 1 | 0 | 0 | 1510240 | AVQ05264.1 | P36334 |
| STPPYTFTTVKP | 1 | 0 | 0 | 1510241 | AUM60013.1 | K9N7C7 |
| STPSQNAGYWRR | 1 | 0 | 0 | 1510242 | AQZ41279.1 | K9N4V7 |
| STPSTVYSPATV | 1 | 0 | 0 | 1510244 | AVV62524.1 | K9N7C7 |
| STPVELPTVDVV | 1 | 0 | 0 | 1510245 | AVN89312.1 | K9N638 |
| STPWLLLSGNPT | 1 | 0 | 0 | 1510251 | AUM60013.1 | K9N7C7 |
| STQANAVQQEFS | 1 | 0 | 0 | 1510252 | AVV62524.1 | K9N7C7 |
| STQDGVNKLQGL | 1 | 0 | 0 | 1510254 | AEN19358.1 | P36334 |
| STQDGVNKLQGV | 1 | 0 | 0 | 1510255 | AMM42485.1 | P36334 |
| STQDGYNKLQGL | 1 | 0 | 0 | 1510256 | CAA79896.1 | P36334 |
| STQISADTGINN | 1 | 0 | 0 | 1510262 | APO40580.1 | P59632 |
| STQLVKNQCVNF | 1 | 0 | 0 | 1510273 | AVP78031.1 | Q3LZX1 |
| STQNEELEQTIQ | 1 | 0 | 0 | 1510274 | APO40578.1 | P0C6X7 |
| STQPQGNIIPHY | 1 | 0 | 0 | 1510278 | ABA60796.1 | Q19U29 |
| STQREAEAAVKP | 1 | 0 | 0 | 1510282 | ARO76392.1 | Q7TLC7 |
| STQSSPIIPGFG | 1 | 0 | 0 | 1510284 | AOR17484.1 | K9N5Q8 |
| STRATIRKIYPA | 1 | 0 | 0 | 1510291 | AKL59401.1 | K9N5Q8 |
| STRCTSPGPSGI | 1 | 0 | 0 | 1510292 | ALA49909.1 | K9N4V7 |
| STRGSSPGPSGI | 1 | 0 | 0 | 1510294 | ALA49392.1 | K9N4V7 |
| STRGTFPGPSGI | 1 | 0 | 0 | 1510295 | ANI69896.1 | K9N4V7 |
| STRGTSPCPSGI | 1 | 0 | 0 | 1510296 | QCI31476.1 | K9N4V7 |
| STRGTSPGLSGI | 1 | 0 | 0 | 1510297 | ALA49601.1 | K9N4V7 |
| STRGTSPVPSGI | 1 | 0 | 0 | 1510299 | AIY60565.1 | K9N4V7 |
| STRHTYQLRARA | 1 | 0 | 0 | 1510301 | APO40584.1 | P59635 |
| STRNNSRDSFRS | 1 | 0 | 0 | 1510304 | ABE97134.1 | H9EJ69 |
| STRSMLKRLDST | 1 | 0 | 0 | 1510312 | AXG21654.1 | K9N5Q8 |
| STRSTSPGPSGI | 1 | 0 | 0 | 1510315 | AVN89381.1 | K9N4V7 |
| STSFSTFKCYGV | 1 | 0 | 0 | 1510331 | AVP78031.1 | Q3LZX1 |
| STSGNLVGFKDV | 1 | 0 | 0 | 1510336 | CAA71146.1 | A0A0P0K6L9 |
| STSGTQVYPRKA | 1 | 0 | 0 | 1510339 | AKN24752.1 | K9N643 |
| STSGVTPDMADQ | 1 | 0 | 0 | 1510343 | ARB07416.1 | P33469 |
| STSINTGNCPFS | 1 | 0 | 0 | 1510346 | AAQ90004.1 | A0A0P0K6L9 |
| STSISNFGSLTQ | 1 | 0 | 0 | 1510351 | QBF80598.1 | K9N5Q8 |
| STSISVQTCDLT | 1 | 0 | 0 | 1510352 | ARK08679.1 | P36334 |
| STSNAILISDRK | 1 | 0 | 0 | 1510359 | AVV62524.1 | K9N7C7 |
| STSNLPDFREEL | 1 | 0 | 0 | 1510360 | AMK59677.1 | P36334 |
| STSPGPSGIGAV | 1 | 0 | 0 | 1510362 | AVN89381.1 | K9N4V7 |
| STSPGPSGIGGG | 1 | 0 | 0 | 1510363 | AVN89359.1 | K9N4V7 |
| STSQQPLGGNVV | 1 | 0 | 0 | 1510371 | AMK59681.1 | P33469 |
| STSRTSSKASSA | 1 | 0 | 0 | 1510376 | AJQ24067.1 | P33469 |
| STSSDPGFVAFN | 1 | 0 | 0 | 1510380 | AVV62524.1 | K9N7C7 |
| STSTHDDFNEHV | 1 | 0 | 0 | 1510386 | AXG21666.1 | K9N796 |
| STSTHDGLIEHV | 1 | 0 | 0 | 1510390 | QBF80533.1 | K9N796 |
| STSTHDGLNEHV | 1 | 0 | 0 | 1510391 | ANF29196.1 | K9N796 |
| STSTHDGPIEHV | 1 | 0 | 0 | 1510393 | AVN89292.1 | K9N796 |
| STSTHDGPNEHV | 1 | 0 | 0 | 1510394 | AHE78098.1 | K9N796 |
| STSTYPANGFFY | 1 | 0 | 0 | 1510401 | AWO68966.1 | A0A0P0G321 |
| STSVEESFIDDY | 1 | 0 | 0 | 1510403 | AIG96682.1 | H9EJ42 |
| STTAQKLETDRI | 1 | 0 | 0 | 1510411 | AXX83372.1 | P0C6X6 |
| STTDNETLEYWV | 1 | 0 | 0 | 1510413 | ARB07438.1 | A0A140H1H1 |
| STTITDKLGVCV | 1 | 0 | 0 | 1510422 | AVV62537.1 | K9N5Q8 |
| STTNYVGVYDTN | 1 | 0 | 0 | 1510440 | AQN78743.1 | P30215 |
| STTPCELHGNGS | 1 | 0 | 0 | 1510441 | ALW82645.1 | K9N7C7 |
| STTPCQVHGNGS | 1 | 0 | 0 | 1510444 | AGV08533.1 | K9N7C7 |
| STTRSSSQLQLA | 1 | 0 | 0 | 1510446 | AVV62537.1 | K9N5Q8 |
| STVAMTEQLQMG | 1 | 0 | 0 | 1510457 | AXN92228.1 | K9N5Q8 |
| STVDAQIQNKTV | 1 | 0 | 0 | 1510461 | AVV62535.1 | K9N7C7 |
| STVDVVALQQEM | 1 | 0 | 0 | 1510463 | AXN92258.1 | K9N7C7 |
| STVNALPETSAD | 1 | 0 | 0 | 1510485 | AUM60013.1 | K9N7C7 |
| STVPSTVWEDGD | 1 | 0 | 0 | 1510487 | ANC28634.1 | K9N5Q8 |
| STVRLQAGNATE | 1 | 0 | 0 | 1510492 | QWO62522.1 | P0DTC1 |
| STVSALVYNNKM | 1 | 0 | 0 | 1510495 | AGV08405.1 | K9N7C7 |
| STVTGRMFHQKC | 1 | 0 | 0 | 1510501 | AIL49624.1 | P0C6X6 |
| STVWEDGNYYRK | 1 | 0 | 0 | 1510509 | ALJ54517.1 | K9N5Q8 |
| STVYSPATVIAG | 1 | 0 | 0 | 1510514 | AVV62524.1 | K9N7C7 |

| | | | | | | |
|---|---|---|---|---|---|---|
| STWNKKFGFIED | 1 | 0 | 0 | 1510519 | ARB66070.1 | P36334 |
| STYDFNPNVPLD | 1 | 0 | 0 | 1510526 | ATO98145.1 | Q3LZX1 |
| STYDFNPNVPLE | 1 | 0 | 0 | 1510527 | AVP78031.1 | Q3LZX1 |
| STYDFYPNVPIE | 1 | 0 | 0 | 1510528 | ATO98120.1 | Q3LZX1 |
| STYDPIYKNGAM | 1 | 0 | 0 | 1510530 | AVV62535.1 | K9N7C7 |
| STYFLVRILQDK | 1 | 0 | 0 | 1510534 | YP_009047214.1 | K9N638 |
| STYRNMALKGTD | 1 | 0 | 0 | 1510553 | AEN19366.1 | P36334 |
| SVAAPKSRVQQN | 1 | 0 | 0 | 1510567 | AIX10718.1 | P33469 |
| SVACEHIFSTMS | 1 | 0 | 0 | 1510569 | AHZ64057.1 | K9N5Q8 |
| SVACEHISSTIS | 1 | 0 | 0 | 1510570 | AJG44102.1 | K9N5Q8 |
| SVACEHITTMMS | 1 | 0 | 0 | 1510574 | AVV62526.1 | K9N5Q8 |
| SVACSHITTFMP | 1 | 0 | 0 | 1510577 | AUM60014.1 | K9N5Q8 |
| SVADVMRYNLNF | 1 | 0 | 0 | 1510580 | AIW52843.1 | A0A0P0G321 |
| SVAKLHNLTDSQ | 1 | 0 | 0 | 1510599 | AUM60027.1 | K9N643 |
| SVANLFSDIKTH | 1 | 0 | 0 | 1510603 | AVV62537.1 | K9N5Q8 |
| SVASFNEWAMAN | 1 | 0 | 0 | 1510620 | AVV62535.1 | K9N7C7 |
| SVASLFSNIKTS | 1 | 0 | 0 | 1510621 | AUM60014.1 | K9N5Q8 |
| SVASPKSRVQQN | 1 | 0 | 0 | 1510622 | ATP16771.1 | P33469 |
| SVASTPVEPSTV | 1 | 0 | 0 | 1510624 | AXN92258.1 | K9N7C7 |
| SVASTYFIIRLL | 1 | 0 | 0 | 1510626 | ATQ39389.1 | K9N7C7 |
| SVATSDLGSLID | 1 | 0 | 0 | 1510634 | AUM60013.1 | K9N7C7 |
| SVAVKSGALHLY | 1 | 0 | 0 | 1510640 | ARO76381.1 | P0C6U8 |
| SVAYTAGVCIKY | 1 | 0 | 0 | 1510647 | ARU07593.1 | H9EJ42 |
| SVAYVANPTEDL | 1 | 0 | 0 | 1510648 | AEN19363.1 | P0C6X6 |
| SVCLGGLDSLDS | 1 | 0 | 0 | 1510655 | ARO76381.1 | P0C6U8 |
| SVCLSGLDSLDS | 1 | 0 | 0 | 1510659 | P0C6W6.1 | P0C6W2 |
| SVCNGCVGHSEN | 1 | 0 | 0 | 1510662 | AIW52757.1 | A0A0P0K6L9 |
| SVCNGCVGHSKN | 1 | 0 | 0 | 1510663 | AIW52741.1 | A0A0P0K6L9 |
| SVCNTGSCTVNK | 1 | 0 | 0 | 1510668 | AGT51697.1 | P0C6X6 |
| SVCPKFEFANDT | 1 | 0 | 0 | 1510669 | AVN89387.1 | K9N5Q8 |
| SVCPKLHHHHHH | 1 | 0 | 0 | 1510674 | 5GMQ_A | K9N5Q8 |
| SVCPMMDLGNST | 1 | 0 | 0 | 1510676 | AVV62537.1 | K9N5Q8 |
| SVCQYNMCDYPH | 1 | 0 | 0 | 1510679 | ANZ78839.1 | P36334 |
| SVCQYNMCEHPH | 1 | 0 | 0 | 1510680 | AWW13511.1 | P36334 |
| SVCQYNMCEYPH | 1 | 0 | 0 | 1510681 | AEN19366.1 | P36334 |
| SVCQYNMCEYPQ | 1 | 0 | 0 | 1510682 | CAA79896.1 | P36334 |
| SVCVRTSHFSIR | 1 | 0 | 0 | 1510689 | AIW52843.1 | A0A0P0G321 |
| SVDARLVPCASG | 1 | 0 | 0 | 1510693 | AEN19363.1 | P0C6X6 |
| SVDDKYKTIDEL | 1 | 0 | 0 | 1510696 | ALA50066.1 | K9N7C7 |
| SVDDKYKVNEDL | 1 | 0 | 0 | 1510698 | ARB07436.1 | A0A140H1G9 |
| SVDEFVDVVKDE | 1 | 0 | 0 | 1510699 | AUM60013.1 | K9N7C7 |
| SVDGYIRRAIDC | 1 | 0 | 0 | 1510702 | AXN92228.1 | K9N5Q8 |
| SVDGYIRRAVDC | 1 | 0 | 0 | 1510703 | AVV62537.1 | K9N5Q8 |
| SVDGYIRRSIDC | 1 | 0 | 0 | 1510704 | AHZ90568.1 | K9N5Q8 |
| SVDISYLKRARG | 1 | 0 | 0 | 1510708 | QDH43737.1 | H9EJ42 |
| SVDLESTSTHDG | 1 | 0 | 0 | 1510711 | YP_009047205.1 | K9N796 |
| SVDRDSAVGHTE | 1 | 0 | 0 | 1510718 | AUM60025.1 | K9N796 |
| SVDRESTSTHDD | 1 | 0 | 0 | 1510723 | AXG21666.1 | K9N796 |
| SVDRESTSTHDG | 1 | 0 | 0 | 1510724 | AXN92229.1 | K9N796 |
| SVDVKDTVVQFN | 1 | 0 | 0 | 1510734 | AVV62524.1 | K9N7C7 |
| SVEAAYAEVHAE | 1 | 0 | 0 | 1510738 | P0C6U5.1 | A0A140H1G9 |
| SVEEEIVPTLEL | 1 | 0 | 0 | 1510751 | AVV62535.1 | K9N7C7 |
| SVEELKRLLEQW | 1 | 0 | 0 | 1510752 | ATO98197.1 | Q3LZX9 |
| SVEGHIIMEEML | 1 | 0 | 0 | 1510757 | AVV62524.1 | K9N7C7 |
| SVELSLDRGAQS | 1 | 0 | 0 | 1510762 | AVV62527.1 | K9N796 |
| SVEQLEPCVQSR | 1 | 0 | 0 | 1510767 | AST12962.1 | A0A0P0LKV0 |
| SVEVPPVLQVES | 1 | 0 | 0 | 1510777 | AVP78030.1 | P0C6W2 |
| SVFDITDAAVKA | 1 | 0 | 0 | 1510782 | APT69868.1 | A0A0P0ILY9 |
| SVFKPQPAGVFT | 1 | 0 | 0 | 1510788 | ATP16757.1 | P36334 |
| SVFKPQPAGVLT | 1 | 0 | 0 | 1510789 | AVQ05264.1 | P36334 |
| SVFPIFVGRIVM | 1 | 0 | 0 | 1510802 | ARU07575.1 | A0A140H1G9 |
| SVGDSTEIATKM | 1 | 0 | 0 | 1510825 | AVV62524.1 | K9N7C7 |
| SVGEHSSPIAYL | 1 | 0 | 0 | 1510827 | AVV62524.1 | K9N7C7 |
| SVGESFGKSLGN | 1 | 0 | 0 | 1510828 | QEG03773.1 | P0C6X6 |
| SVGKFCLEASFN | 1 | 0 | 0 | 1510841 | QYC64967.1 | P0DTD1 |
| SVGNFSNGVSGR | 1 | 0 | 0 | 1510844 | AVV62537.1 | K9N5Q8 |
| SVGNFSNNKTGR | 1 | 0 | 0 | 1510845 | AUM60014.1 | K9N5Q8 |
| SVGNYYNGYPGR | 1 | 0 | 0 | 1510849 | AVV62526.1 | K9N5Q8 |
| SVGQKSIVAYTM | 1 | 0 | 0 | 1510851 | ATO98181.1 | Q3LZX1 |
| SVGSVPGEMRLA | 1 | 0 | 0 | 1510852 | 5W9H_A | K9N5Q8 |
| SVGYTKEGNVIN | 1 | 0 | 0 | 1510855 | AVV62535.1 | K9N7C7 |
| SVGYTKGGSVIN | 1 | 0 | 0 | 1510858 | AHI48747.1 | K9N638 |
| SVHDGYTNEIIG | 1 | 0 | 0 | 1510859 | ARO76381.1 | P0C6U8 |
| SVHPVVCDEDGE | 1 | 0 | 0 | 1510867 | AUM60013.1 | K9N7C7 |
| SVIAELNQLNEP | 1 | 0 | 0 | 1510872 | APO40578.1 | P0C6X7 |
| SVIDLRLDDFVE | 1 | 0 | 0 | 1510878 | AVV62524.1 | K9N7C7 |
| SVIDLRLDDFVQ | 1 | 0 | 0 | 1510880 | AVV62535.1 | K9N7C7 |
| SVIGDLKCPLDS | 1 | 0 | 0 | 1510881 | BAM16455.1 | P36334 |
| SVIQDYENVCFD | 1 | 0 | 0 | 1510904 | QBQ01836.1 | P0C6X6 |
| SVIQNYENVCFE | 1 | 0 | 0 | 1510907 | AIV41930.1 | P0C6X6 |
| SVITPGTNTSSE | 1 | 0 | 0 | 1510918 | AVP78042.1 | Q3LZX1 |
| SVIVNDATSDTK | 1 | 0 | 0 | 1510924 | AIV41846.1 | P0C6X6 |
| SVIVNDDTSDTK | 1 | 0 | 0 | 1510926 | AIV41780.1 | P0C6X6 |
| SVIYDDNTNQYA | 1 | 0 | 0 | 1510931 | ATQ39390.1 | K9N5Q8 |
| SVKGLIASKGPK | 1 | 0 | 0 | 1510944 | APO40578.1 | P0C6X7 |
| SVKLPKGVGSAN | 1 | 0 | 0 | 1510952 | AEN19363.1 | P0C6X6 |
| SVKMVDAYVDTF | 1 | 0 | 0 | 1510955 | BAE93400.1 | P0C6U8 |
| SVKPESNADQES | 1 | 0 | 0 | 1510959 | AUM60013.1 | K9N7C7 |
| SVKTCDLTVGSG | 1 | 0 | 0 | 1510977 | AIL49550.1 | P36334 |
| SVLAVMQKLPCQ | 1 | 0 | 0 | 1510985 | AEN19363.1 | P0C6X6 |
| SVLDLSKFECKH | 1 | 0 | 0 | 1510990 | AWK59942.1 | H9EJ42 |

| | | | | | | |
|---|---|---|---|---|---|---|
| SVLFTCIPTVGY | 1 | 0 | 0 | 1510994 | AVV62530.1 | K9N7D2 |
| SVLGAAAKSVVR | 1 | 0 | 0 | 1510995 | AVV62535.1 | K9N7C7 |
| SVLGCLGSECSK | 1 | 0 | 0 | 1510996 | AVQ05264.1 | P36334 |
| SVLLALQQIEVK | 1 | 0 | 0 | 1511008 | P0C6W6.1 | P0C6W2 |
| SVLNDILSRLDP | 1 | 0 | 0 | 1511021 | 6CRW_A | P59594 |
| SVLQMDVRGLVN | 1 | 0 | 0 | 1511031 | QED88024.1 | H9EJ42 |
| SVLQQLHLEANG | 1 | 0 | 0 | 1511033 | AYO86698.1 | K9N638 |
| SVLRQWLPTDAI | 1 | 0 | 0 | 1511039 | AXN92247.1 | K9N7C7 |
| SVLTTAVYKPLV | 1 | 0 | 0 | 1511046 | AST12962.1 | A0A0P0LKV0 |
| SVLVNAANTHLQ | 1 | 0 | 0 | 1511048 | AVV62535.1 | K9N7C7 |
| SVMHFGTLSRED | 1 | 0 | 0 | 1511062 | AIV42002.1 | P0C6X6 |
| SVMLCGDNSAKF | 1 | 0 | 0 | 1511064 | AXN92247.1 | K9N7C7 |
| SVMQMDIQGFVN | 1 | 0 | 0 | 1511067 | NP_073549.1 | A0A0P0LKV0 |
| SVNDSLEHVGGL | 1 | 0 | 0 | 1511079 | CAA83661.1 | P36334 |
| SVNDSLKPVGGL | 1 | 0 | 0 | 1511080 | AIL49500.1 | P36334 |
| SVNETSQTGWAL | 1 | 0 | 0 | 1511081 | AFO70499.1 | U3M6Q8 |
| SVNLNICEPVLT | 1 | 0 | 0 | 1511094 | AGW27870.1 | A0A140H1G9 |
| SVNPANMRNANG | 1 | 0 | 0 | 1511096 | AVV62524.1 | K9N7C7 |
| SVNPASMRNANG | 1 | 0 | 0 | 1511097 | AUM60013.1 | K9N7C7 |
| SVNQISGYTLAA | 1 | 0 | 0 | 1511100 | CAA83660.1 | P36334 |
| SVPNMHVVPLEN | 1 | 0 | 0 | 1511135 | AIV41996.1 | P0C6X6 |
| SVPNSRSTSRTS | 1 | 0 | 0 | 1511137 | AIL49415.1 | P33469 |
| SVPSPFNWERRI | 1 | 0 | 0 | 1511139 | ABU39973.1 | A0A140H1H1 |
| SVPSPLNWERKI | 1 | 0 | 0 | 1511140 | AMM42415.1 | A0A140H1H1 |
| SVQFKRPVNPTD | 1 | 0 | 0 | 1511163 | APO40578.1 | P0C6X7 |
| SVQNAVPANSLH | 1 | 0 | 0 | 1511172 | APO40580.1 | P59632 |
| SVQQAQQEATHI | 1 | 0 | 0 | 1511174 | ARB07424.1 | A0A0P0ILY9 |
| SVQQDCRIPNGF | 1 | 0 | 0 | 1511176 | AKA09611.1 | A0A0P0G321 |
| SVQQDGRTPNGF | 1 | 0 | 0 | 1511180 | AKA09627.1 | A0A0P0G321 |
| SVQQDLDVICPG | 1 | 0 | 0 | 1511181 | AZL96750.1 | K9N638 |
| SVRDPKTFEILD | 1 | 0 | 0 | 1511196 | APO40579.1 | P59594 |
| SVRDPQTLDILD | 1 | 0 | 0 | 1511199 | ATO98145.1 | Q3LZX1 |
| SVRDPQTLQILD | 1 | 0 | 0 | 1511201 | ATO98193.1 | Q3LZX1 |
| SVRNLFESVKSS | 1 | 0 | 0 | 1511214 | QBM11737.1 | K9N5Q8 |
| SVRSDGTFEEMA | 1 | 0 | 0 | 1511216 | AEN19363.1 | P0C6X6 |
| SVRSHPDFKLLH | 1 | 0 | 0 | 1511217 | AHY22543.1 | K9N7C7 |
| SVSARMIWREWC | 1 | 0 | 0 | 1511231 | AUM60013.1 | K9N7C7 |
| SVSDSLEPVGGL | 1 | 0 | 0 | 1511235 | ARB66068.1 | P36334 |
| SVSFDYLIREIE | 1 | 0 | 0 | 1511238 | ATQ39389.1 | K9N7C7 |
| SVSFHEDRILDY | 1 | 0 | 0 | 1511239 | AVV62535.1 | K9N7C7 |
| SVSKLHDLDDAT | 1 | 0 | 0 | 1511248 | AVV62540.1 | K9N643 |
| SVSKPNAIDVEH | 1 | 0 | 0 | 1511249 | AVA26872.1 | H9EJ42 |
| SVSLAIDAYPLT | 1 | 0 | 0 | 1511250 | ASU90624.1 | K9N7C7 |
| SVSNFSYGKMGR | 1 | 0 | 0 | 1511256 | QGV13484.1 | K9N5Q8 |
| SVSNLALAFCSF | 1 | 0 | 0 | 1511257 | AVV62524.1 | K9N7C7 |
| SVSNLFADIKTQ | 1 | 0 | 0 | 1511258 | AVV62526.1 | K9N5Q8 |
| SVSPKLEFANDT | 1 | 0 | 0 | 1511260 | AKN24803.1 | K9N5Q8 |
| SVSSCALRIGSG | 1 | 0 | 0 | 1511271 | ARU07577.1 | A0A140H1H1 |
| SVSSCDLRMGSG | 1 | 0 | 0 | 1511273 | BBA20986.1 | A0A140H1H1 |
| SVSSVPGEMRLA | 1 | 0 | 0 | 1511284 | 5X59_A | K9N5Q8 |
| SVSSVWPLYPYG | 1 | 0 | 0 | 1511285 | AGT51619.1 | P30215 |
| SVSTDVLVNTYK | 1 | 0 | 0 | 1511287 | AVV62535.1 | K9N7C7 |
| SVTAVVTNGNLK | 1 | 0 | 0 | 1511304 | AUM60030.1 | K9N7A1 |
| SVTDECRLAMLG | 1 | 0 | 0 | 1511306 | NP_073549.1 | A0A0P0LKV0 |
| SVTDFSLGGKSS | 1 | 0 | 0 | 1511307 | AUM60018.1 | K9N7D2 |
| SVTDFSLNGTVS | 1 | 0 | 0 | 1511309 | ANI69893.1 | K9N7D2 |
| SVTEFTLNNHHS | 1 | 0 | 0 | 1511314 | AVV62530.1 | K9N7D2 |
| SVTGRLIQFDNP | 1 | 0 | 0 | 1511318 | APO40579.1 | P59594 |
| SVTLKICKFGIN | 1 | 0 | 0 | 1511326 | AIW52843.1 | A0A0P0G321 |
| SVTPIKNTVDTI | 1 | 0 | 0 | 1511339 | AST12962.1 | A0A0P0LKV0 |
| SVTSAGPISQFN | 1 | 0 | 0 | 1511344 | ATW75479.1 | K9N5Q8 |
| SVTSNYSGVVTA | 1 | 0 | 0 | 1511346 | APO40578.1 | P0C6X7 |
| SVTSVNLFDVGY | 1 | 0 | 0 | 1511351 | ATQ39394.1 | K9N796 |
| SVTVKNGTLYMY | 1 | 0 | 0 | 1511362 | APO40578.1 | P0C6X7 |
| SVTVKRKRLRKK | 1 | 0 | 0 | 1511364 | AXN92258.1 | K9N7C7 |
| SVTYTDQYLCGA | 1 | 0 | 0 | 1511366 | ARB07390.1 | A0A0P0LKV0 |
| SVVDIKKEIAHL | 1 | 0 | 0 | 1511373 | APO40579.1 | P59594 |
| SVVEVHKAVVVT | 1 | 0 | 0 | 1511382 | AVV62535.1 | K9N7C7 |
| SVVHTAGVCIKY | 1 | 0 | 0 | 1511391 | AGT51379.1 | H9EJ42 |
| SVVILSKTDVST | 1 | 0 | 0 | 1511394 | AUM60013.1 | K9N7C7 |
| SVVIQPRTINST | 1 | 0 | 0 | 1511395 | BBO58758.1 | P36334 |
| SVVLPGEMAKKQ | 1 | 0 | 0 | 1511401 | AVV62535.1 | K9N7C7 |
| SVVMVGDYVAPV | 1 | 0 | 0 | 1511403 | AST12962.1 | A0A0P0LKV0 |
| SVVNTTVTVNVT | 1 | 0 | 0 | 1511410 | AIW52831.1 | A0A0P0G321 |
| SVVQKDGVEDTP | 1 | 0 | 0 | 1511416 | ATQ39389.1 | K9N7C7 |
| SVVQREGIEQVP | 1 | 0 | 0 | 1511419 | AVV62535.1 | K9N7C7 |
| SVVSDTVEVPPQ | 1 | 0 | 0 | 1511430 | ASU45706.1 | K9N7C7 |
| SVVSFNEWAMAN | 1 | 0 | 0 | 1511433 | AVV62524.1 | K9N7C7 |
| SVVVQPRTIDST | 1 | 0 | 0 | 1511442 | QBQ01839.1 | P36334 |
| SVVVQPRTIIST | 1 | 0 | 0 | 1511443 | BBA20973.1 | P36334 |
| SVVVQPRTINSQ | 1 | 0 | 0 | 1511448 | AKA09654.1 | P36334 |
| SVVVQPRTTQDG | 1 | 0 | 0 | 1511451 | AKA09658.1 | P36334 |
| SVYADYFVVKGD | 1 | 0 | 0 | 1511469 | APO40579.1 | P59594 |
| SVYDIACCMRAS | 1 | 0 | 0 | 1511479 | AWK59942.1 | H9EJ42 |
| SVYDQIVDNIYV | 1 | 0 | 0 | 1511481 | AEN19363.1 | P0C6X6 |
| SVYGFSNVVYPD | 1 | 0 | 0 | 1511485 | ARU07576.1 | A0A140H1H0 |
| SVYNAYSPIDIP | 1 | 0 | 0 | 1511493 | ATO98193.1 | Q3LZX1 |
| SVYSVPLLFQTN | 1 | 0 | 0 | 1511507 | AZS52616.1 | A0A140H1G9 |
| SVYSVPLVFQNN | 1 | 0 | 0 | 1511508 | ARU07574.1 | A0A140H1G9 |
| SVYTNSCLVSSN | 1 | 0 | 0 | 1511511 | QAT98908.1 | K9N7C7 |
| SVYTVESGTEVS | 1 | 0 | 0 | 1511512 | ARO76381.1 | P0C6U8 |

| | | | | | | |
|---|---|---|---|---|---|---|
| SWADLPAYMTVA | 1 | 0 | 0 | 1511520 | Q01455.1 | Q01455 |
| SWAGSKINAVIY | 1 | 0 | 0 | 1511527 | ATQ39389.1 | K9N7C7 |
| SWAIFNDYHANR | 1 | 0 | 0 | 1511534 | AVV62524.1 | K9N7C7 |
| SWEIGKVKPPLNKNYVFTG | 1 | 0 | 0 | 1511542 | AQN78733.1 | P0C6X6 |
| SWFEDDYKAFIG | 1 | 0 | 0 | 1511544 | AFR79249.1 | A0A0P0ILY9 |
| SWHMDTGVVPCL | 1 | 0 | 0 | 1511566 | AIL49486.1 | P36334 |
| SWLEMADTSLSG | 1 | 0 | 0 | 1511570 | APO40578.1 | P0C6X7 |
| SWLLHKDGGIPD | 1 | 0 | 0 | 1511571 | AMQ49017.1 | K9N4V0 |
| SWLMWLVINVVQ | 1 | 0 | 0 | 1511574 | APO40578.1 | P0C6X7 |
| SWLNDKSVPSPL | 1 | 0 | 0 | 1511575 | AIX10749.1 | P36334 |
| SWNKVTQIANMF | 1 | 0 | 0 | 1511587 | AXN92247.1 | K9N7C7 |
| SWNKYLYELIQK | 1 | 0 | 0 | 1511589 | AGT51393.1 | H9EJ42 |
| SWNQAYSSPILP | 1 | 0 | 0 | 1511593 | ATO98145.1 | Q3LZX1 |
| SWNRRYGFGSFN | 1 | 0 | 0 | 1511594 | BBA20986.1 | A0A140H1H1 |
| SWPRPIDPAAAD | 1 | 0 | 0 | 1511598 | AVV62526.1 | K9N5Q8 |
| SWQNLLYDFNGN | 1 | 0 | 0 | 1511599 | ADN03339.1 | A0A140H1H1 |
| SWQNLLYDPNGN | 1 | 0 | 0 | 1511600 | ARB66062.1 | P36334 |
| SWQNLLYDSSGN | 1 | 0 | 0 | 1511602 | ARK08661.1 | P36334 |
| SWSADLYELMGK | 1 | 0 | 0 | 1511609 | AVV62524.1 | K9N7C7 |
| SWSDGDVITGVP | 1 | 0 | 0 | 1511613 | AIW52757.1 | A0A0P0K6L9 |
| SWSSTMIFSLHP | 1 | 0 | 0 | 1511618 | AXN92247.1 | K9N7C7 |
| SWSSTMILSLHP | 1 | 0 | 0 | 1511620 | ALW82719.1 | K9N7C7 |
| SWVDQFERSHQT | 1 | 0 | 0 | 1511627 | AZS52622.1 | Q19U29 |
| SWVDQSERSHQT | 1 | 0 | 0 | 1511628 | AMM42397.1 | Q19U29 |
| SWWSFNPETNNL | 1 | 0 | 0 | 1511643 | AZS52621.1 | Q0ZJ82 |
| SWYDPVENEDIH | 1 | 0 | 0 | 1511645 | AWK59948.1 | H9EJ42 |
| SYAFEHIVYGDF | 1 | 0 | 0 | 1511660 | APO40578.1 | P0C6X7 |
| SYAIHHDKFTDG | 1 | 0 | 0 | 1511662 | 5NFY_A | P0C6X7 |
| SYALYTAWFYPL | 1 | 0 | 0 | 1511666 | AEN19363.1 | P0C6X6 |
| SYALYTPWFYPL | 1 | 0 | 0 | 1511667 | AIV41960.1 | P0C6X6 |
| SYAPHFALVLPE | 1 | 0 | 0 | 1511670 | AUM60013.1 | K9N7C7 |
| SYCPGNSNYNTF | 1 | 0 | 0 | 1511680 | AVV62537.1 | K9N5Q8 |
| SYCRKLGSSVRS | 1 | 0 | 0 | 1511682 | ATP16754.1 | P0C6X6 |
| SYDINLNKFNRA | 1 | 0 | 0 | 1511697 | AMQ48992.1 | K9N7C7 |
| SYDISHSNNFGV | 1 | 0 | 0 | 1511698 | ADN03340.1 | Q0ZJ84 |
| SYDTNLNKFNGA | 1 | 0 | 0 | 1511708 | AVN89385.1 | K9N7C7 |
| SYERQSPFEIKS | 1 | 0 | 0 | 1511726 | ARO76381.1 | P0C6U8 |
| SYESFDVDTGVY | 1 | 0 | 0 | 1511728 | AVV62526.1 | K9N5Q8 |
| SYESQTPFEIKL | 1 | 0 | 0 | 1511730 | AVP78041.1 | P0C6W2 |
| SYFDKSWPRPID | 1 | 0 | 0 | 1511736 | AVV62526.1 | K9N5Q8 |
| SYFNPEPITMAN | 1 | 0 | 0 | 1511744 | AUM60014.1 | K9N5Q8 |
| SYFRKSCQNLDC | 1 | 0 | 0 | 1511745 | NP_150075.1 | Q80872 |
| SYFVQSIRLFMR | 1 | 0 | 0 | 1511755 | ASY99815.1 | K9N7A1 |
| SYFYNGSAQATS | 1 | 0 | 0 | 1511761 | ARK08634.1 | P30215 |
| SYGFSNVVELPK | 1 | 0 | 0 | 1511772 | CAA71056.1 | A0A0P0K6L9 |
| SYGFSNVVEMPK | 1 | 0 | 0 | 1511773 | AIW52757.1 | A0A0P0K6L9 |
| SYGGVSVITPGT | 1 | 0 | 0 | 1511775 | APO40579.1 | P59594 |
| SYGNMFQTQAGC | 1 | 0 | 0 | 1511782 | APO40579.1 | P59594 |
| SYIDLKELGNYS | 1 | 0 | 0 | 1511808 | AVV62537.1 | K9N5Q8 |
| SYINDVDTGVPP | 1 | 0 | 0 | 1511816 | ATP16757.1 | P36334 |
| SYINNKDTGPPF | 1 | 0 | 0 | 1511820 | AKA09646.1 | P36334 |
| SYIPLFDMRYSF | 1 | 0 | 0 | 1511822 | AMQ49019.1 | K9N7D2 |
| SYIYHSAFADCT | 1 | 0 | 0 | 1511828 | AUM60013.1 | K9N7C7 |
| SYKGTFNNKDTG | 1 | 0 | 0 | 1511837 | AKA09658.1 | P36334 |
| SYKIFSTGFVLS | 1 | 0 | 0 | 1511840 | AGT17757.1 | A0A140H1H0 |
| SYLGCVLNAVNL | 1 | 0 | 0 | 1511864 | BBA20986.1 | A0A140H1H1 |
| SYLGCVVNAYNS | 1 | 0 | 0 | 1511866 | CAA79896.1 | P36334 |
| SYMMPVMGMTNC | 1 | 0 | 0 | 1511900 | NP_073549.1 | A0A0P0LKV0 |
| SYMQPGSAGVIS | 1 | 0 | 0 | 1511901 | AVV62537.1 | K9N5Q8 |
| SYNASCLCSTDA | 1 | 0 | 0 | 1511904 | BBA20983.1 | A0A140H1H1 |
| SYNVSCLCSTDA | 1 | 0 | 0 | 1511915 | 5GNB_A | A0A140H1H1 |
| SYPLSMKSDLGV | 1 | 0 | 0 | 1511932 | AHX00711.1 | K9N5Q8 |
| SYPLSMKSDLSV | 1 | 0 | 0 | 1511933 | AXN92228.1 | K9N5Q8 |
| SYQEALDSGDAS | 1 | 0 | 0 | 1511939 | AUM60013.1 | K9N7C7 |
| SYQEFNDLDHTS | 1 | 0 | 0 | 1511942 | AWK59942.1 | H9EJ42 |
| SYQLYDASNVKN | 1 | 0 | 0 | 1511951 | AVQ05261.1 | P0C6X6 |
| SYQTNRDFQNKM | 1 | 0 | 0 | 1511955 | AUM60014.1 | K9N5Q8 |
| SYQTNRDMQNKM | 1 | 0 | 0 | 1511956 | AUM60024.1 | K9N5Q8 |
| SYRNDKSCPVVA | 1 | 0 | 0 | 1511965 | ARO76381.1 | P0C6U8 |
| SYSQNLCVDGFG | 1 | 0 | 0 | 1511989 | AST12962.1 | A0A0P0LKV0 |
| SYSVVVQPRTID | 1 | 0 | 0 | 1511995 | QBQ01839.1 | P36334 |
| SYTGSSFYEPEP | 1 | 0 | 0 | 1512004 | ALM26400.1 | K9N5Q8 |
| SYTSDIKPVTYK | 1 | 0 | 0 | 1512014 | APO40578.1 | P0C6X7 |
| SYTTTIKPLEHH | 1 | 0 | 0 | 1512018 | 5E6J_A | P0C6X7 |
| SYTTTILEHHHH | 1 | 0 | 0 | 1512022 | 5Y3E_A | P0C6U8 |
| SYVGCYNKCAYW | 1 | 0 | 0 | 1512041 | APO40578.1 | P0C6X7 |
| SYVGIQTIEELH | 1 | 0 | 0 | 1512042 | AVV62524.1 | K9N7C7 |
| SYVLLERVYKHF | 1 | 0 | 0 | 1512053 | AEN19363.1 | P0C6X6 |
| SYVLLERVYNHF | 1 | 0 | 0 | 1512055 | AIV41936.1 | P0C6X6 |
| SYVPTKYVTAKV | 1 | 0 | 0 | 1512060 | CAA79896.1 | P36334 |
| SYVVDSVAVKSG | 1 | 0 | 0 | 1512063 | ARO76381.1 | P0C6U8 |
| SYVYNGSAQVTS | 1 | 0 | 0 | 1512067 | QBP84703.1 | P30215 |
| SYYAYMMPVMGM | 1 | 0 | 0 | 1512069 | AIW52776.1 | A0A0P0LKV0 |
| SYYEDFLEYHDI | 1 | 0 | 0 | 1512077 | AVP78037.1 | Q3LZX5 |
| SYYLIQSGIFFT | 1 | 0 | 0 | 1512083 | ARU07575.1 | A0A140H1G9 |
| SYYRLGASQRVG | 1 | 0 | 0 | 1512085 | APO40582.1 | P59596 |
| TAAAKFERQHMD | 1 | 0 | 0 | 1512096 | 3R24_A | P0C6X7 |
| TAADINIHDLPI | 1 | 0 | 0 | 1512097 | AGT51530.1 | P30215 |
| TAADINIPHLPI | 1 | 0 | 0 | 1512100 | AXX83356.1 | P30215 |
| TAADINNPDLPI | 1 | 0 | 0 | 1512101 | P30215.1 | P30215 |
| TAADINTPDVPI | 1 | 0 | 0 | 1512102 | AAA73152.1 | P30215 |

| | | | | | | |
|---|---|---|---|---|---|---|
| TAAIDLAYNKLQ | 1 | 0 | 0 | 1512112 | AVV62524.1 | K9N7C7 |
| TAARLCVQCAMG | 1 | 0 | 0 | 1512125 | AVV62542.1 | K9N5R3 |
| TAATIVNQERYV | 1 | 0 | 0 | 1512128 | AGV08597.1 | K9N7C7 |
| TAATTCVTGPTT | 1 | 0 | 0 | 1512130 | ATO98145.1 | Q3LZX1 |
| TAAVDLSMDKLC | 1 | 0 | 0 | 1512133 | AVN89451.1 | K9N7C7 |
| TAAVINIPDLPI | 1 | 0 | 0 | 1512136 | AGT51520.1 | P30215 |
| TAAYGLCNTDTP | 1 | 0 | 0 | 1512141 | AVV62537.1 | K9N5Q8 |
| TAAYINIPDLPI | 1 | 0 | 0 | 1512143 | AEN19365.1 | P30215 |
| TAAYREAAASHL | 1 | 0 | 0 | 1512146 | AVV62524.1 | K9N7C7 |
| TACNGHKIEDLS | 1 | 0 | 0 | 1512151 | ADO24641.1 | A0A140H1G9 |
| TACNGISVTQIM | 1 | 0 | 0 | 1512152 | P0C6U5.1 | A0A140H1G9 |
| TACQYTMCEYPY | 1 | 0 | 0 | 1512158 | AMM42415.1 | A0A140H1H1 |
| TADEIKALKELY | 1 | 0 | 0 | 1512161 | AVN89363.1 | K9N7C7 |
| TADGSPQPYCYT | 1 | 0 | 0 | 1512169 | AGT51707.1 | P0C6X6 |
| TADNLKNVLVWL | 1 | 0 | 0 | 1512174 | APO40578.1 | P0C6X7 |
| TAEPVRDEFSIE | 1 | 0 | 0 | 1512184 | AAA45463.1 | A0A0N7J131 |
| TAEQCIRGAIGD | 1 | 0 | 0 | 1512186 | ARB07424.1 | A0A0P0ILY9 |
| TAETTDFLTVAL | 1 | 0 | 0 | 1512190 | CAA71147.1 | A0A0P0K6L9 |
| TAEWFGITQNAQ | 1 | 0 | 0 | 1512191 | AUM60014.1 | K9N5Q8 |
| TAFAAIGDLKCP | 1 | 0 | 0 | 1512192 | AIL49486.1 | P36334 |
| TAFAVIGDLKCP | 1 | 0 | 0 | 1512193 | AVQ05264.1 | P36334 |
| TAFAVIGDLNCT | 1 | 0 | 0 | 1512196 | AWW13566.1 | P36334 |
| TAFSKPLYVPEH | 1 | 0 | 0 | 1512215 | AXN92229.1 | K9N796 |
| TAFSPAQDTWGT | 1 | 0 | 0 | 1512217 | 5X58_A | P59594 |
| TAFSVIGDLKCP | 1 | 0 | 0 | 1512219 | BAM16455.1 | P36334 |
| TAGDGISRPKLH | 1 | 0 | 0 | 1512228 | ARO76383.1 | P59632 |
| TAGLFTNFRIDV | 1 | 0 | 0 | 1512238 | ALA49936.1 | K9N796 |
| TAGLSSFASIPF | 1 | 0 | 0 | 1512241 | AVV62526.1 | K9N5Q8 |
| TAGLYTNSRIDV | 1 | 0 | 0 | 1512248 | AGV08439.1 | K9N796 |
| TAGVVTCNVVKQ | 1 | 0 | 0 | 1512256 | AVN89451.1 | K9N7C7 |
| TAGVVTGNVLKH | 1 | 0 | 0 | 1512257 | AUM60013.1 | K9N7C7 |
| TAGWTFGAGSAL | 1 | 0 | 0 | 1512262 | Q3I5J5.1 | Q3LZX1 |
| TAHVNATAAYGL | 1 | 0 | 0 | 1512274 | AUM60014.1 | K9N5Q8 |
| TAIDCGFNDLSQ | 1 | 0 | 0 | 1512280 | AXN92260.1 | K9N5Q8 |
| TAIPTIKDWLLV | 1 | 0 | 0 | 1512291 | AXN92231.1 | K9N643 |
| TAIQTVFDCAVE | 1 | 0 | 0 | 1512292 | NP_073549.1 | A0A0P0LKV0 |
| TAIRADQLTPAW | 1 | 0 | 0 | 1512293 | ATO98181.1 | Q3LZX1 |
| TAKEVRQKILNK | 1 | 0 | 0 | 1512304 | AEN19370.1 | P33469 |
| TAKEVTQKILNN | 1 | 0 | 0 | 1512305 | AJQ24083.1 | P33469 |
| TAKQDVGSYFYR | 1 | 0 | 0 | 1512316 | ATO98145.1 | Q3LZX1 |
| TAKYIPLVGTPL | 1 | 0 | 0 | 1512320 | ANI69891.1 | K9N4V0 |
| TALDAYVSQQLS | 1 | 0 | 0 | 1512331 | CAA83661.1 | P36334 |
| TALLTLQQIELK | 1 | 0 | 0 | 1512342 | AVP78041.1 | P0C6W2 |
| TALNICFDIRDN | 1 | 0 | 0 | 1512345 | AUM60013.1 | K9N7C7 |
| TALRRPINATDR | 1 | 0 | 0 | 1512357 | AXN92258.1 | K9N7C7 |
| TALTQHGKEDLKFPR | 1 | 0 | 0 | 1512360 | P0DTC9.1 | P0DTC9 |
| TANDPSEAFEKF | 1 | 0 | 0 | 1512374 | AUM60013.1 | K9N7C7 |
| TANVCSPMACNG | 1 | 0 | 0 | 1512396 | ACR82761.1 | A0A140H1G9 |
| TAPANLTISKPS | 1 | 0 | 0 | 1512404 | AVV62526.1 | K9N5Q8 |
| TAPANLTITKPS | 1 | 0 | 0 | 1512405 | AVV62537.1 | K9N5Q8 |
| TAPEDDDFLEEN | 1 | 0 | 0 | 1512411 | QBQ01836.1 | P0C6X6 |
| TAPEYDDFLEES | 1 | 0 | 0 | 1512412 | AGT51707.1 | P0C6X6 |
| TAPFLVLAGLGE | 1 | 0 | 0 | 1512414 | AVV62535.1 | K9N7C7 |
| TAPKTGNFKDLR | 1 | 0 | 0 | 1512423 | ATO98181.1 | Q3LZX1 |
| TAPQTGNFKDLR | 1 | 0 | 0 | 1512425 | ATO98193.1 | Q3LZX1 |
| TAQAAGTDITIT | 1 | 0 | 0 | 1512433 | APO40578.1 | P0C6X7 |
| TAQADTAGLFTN | 1 | 0 | 0 | 1512436 | ALA49936.1 | K9N796 |
| TAQEAYEQAVVN | 1 | 0 | 0 | 1512439 | AVP78041.1 | P0C6W2 |
| TAQELLLALLAF | 1 | 0 | 0 | 1512441 | NP_073549.1 | A0A0P0LKV0 |
| TAQNSVRVLQKA | 1 | 0 | 0 | 1512450 | QYC64967.1 | P0DTD1 |
| TARDALDTILQE | 1 | 0 | 0 | 1512461 | ATQ39389.1 | K9N7C7 |
| TARDGLKRGENL | 1 | 0 | 0 | 1512463 | AVV62535.1 | K9N7C7 |
| TARTTSSEFQLA | 1 | 0 | 0 | 1512474 | AUM60014.1 | K9N5Q8 |
| TARYFDCYEGGC | 1 | 0 | 0 | 1512480 | ALJ99935.1 | A0A0P0LKV0 |
| TASLFQTTVKLQ | 1 | 0 | 0 | 1512492 | AVA26873.1 | A0A0P0G321 |
| TASLRRLVKPTD | 1 | 0 | 0 | 1512493 | AUM60013.1 | K9N7C7 |
| TASMSMAECKAT | 1 | 0 | 0 | 1512494 | AFD98819.1 | H9EJ42 |
| TASRSSSRSRGN | 1 | 0 | 0 | 1512495 | APO40586.1 | P59595 |
| TASTLRGVGQKS | 1 | 0 | 0 | 1512501 | ATO98145.1 | Q3LZX1 |
| TASVVEVHKAVV | 1 | 0 | 0 | 1512504 | AVV62535.1 | K9N7C7 |
| TATDPAEAFEKF | 1 | 0 | 0 | 1512512 | AVV62524.1 | K9N7C7 |
| TATINYLIDLAN | 1 | 0 | 0 | 1512520 | AUM60013.1 | K9N7C7 |
| TATINYVIDLAN | 1 | 0 | 0 | 1512521 | AUM60023.1 | K9N7C7 |
| TATIPIQATLPF | 1 | 0 | 0 | 1512523 | QWS71388.1 | P0DTC3 |
| TATSVNCCHDGI | 1 | 0 | 0 | 1512539 | ARE29989.1 | Q4VID0 |
| TATTVVGSCIKG | 1 | 0 | 0 | 1512542 | ATQ39389.1 | K9N7C7 |
| TATTVVGSCIKS | 1 | 0 | 0 | 1512543 | AXN92247.1 | K9N7C7 |
| TAVGVPQQRFVY | 1 | 0 | 0 | 1512563 | AVN89313.1 | K9N5Q8 |
| TAVHSDGMFVAT | 1 | 0 | 0 | 1512569 | AGT51379.1 | H9EJ42 |
| TAVMSLKDGQIN | 1 | 0 | 0 | 1512579 | AVP78041.1 | P0C6W2 |
| TAVNESRYGLHN | 1 | 0 | 0 | 1512584 | ARU07594.1 | A0A0P0G321 |
| TAVRSDGTFEEM | 1 | 0 | 0 | 1512588 | AQN78773.1 | P0C6X6 |
| TAVRSSFNFDQK | 1 | 0 | 0 | 1512589 | AIV41792.1 | P0C6X6 |
| TAVSPDFKAFNV | 1 | 0 | 0 | 1512594 | AVV62535.1 | K9N7C7 |
| TAVTKTSEQNNF | 1 | 0 | 0 | 1512595 | AVP78041.1 | P0C6W2 |
| TAVTTGDVKIMA | 1 | 0 | 0 | 1512597 | NP_073549.1 | A0A0P0LKV0 |
| TAVVTNGYLKIA | 1 | 0 | 0 | 1512602 | AVV62543.1 | K9N7A1 |
| TAVVTNGYLKMA | 1 | 0 | 0 | 1512603 | AVN89380.1 | K9N7A1 |
| TAVYKPLVDFNV | 1 | 0 | 0 | 1512606 | AST12962.1 | A0A0P0LKV0 |
| TAWYVGQIFFLV | 1 | 0 | 0 | 1512614 | AGT17760.1 | Q0ZJ83 |
| TCACEIVGGQIVTCA | 1 | 0 | 0 | 1512629 | P0DTC1.1 | P0DTD1 |

| | | | | | | |
|---|---|---|---|---|---|---|
| TCAKRFKRWLCI | 1 | 0 | 0 | 1512632 | AFR79249.1 | A0A0P0ILY9 |
| TCATTRQVISVI | 1 | 0 | 0 | 1512639 | APO40578.1 | P0C6X7 |
| TCAVNYTKAPDV | 1 | 0 | 0 | 1512643 | CAA83660.1 | P36334 |
| TCDGTTFTYASA | 1 | 0 | 0 | 1512660 | QYC64967.1 | P0DTD1 |
| TCDNLCTLDPIA | 1 | 0 | 0 | 1512664 | BBO58891.1 | P36334 |
| TCDNLCTLDSIT | 1 | 0 | 0 | 1512666 | AIV41891.1 | P36334 |
| TCEGNTFTYASA | 1 | 0 | 0 | 1512676 | APO40578.1 | P0C6X7 |
| TCELCGTQNAVV | 1 | 0 | 0 | 1512680 | APO40578.1 | P0C6X7 |
| TCFANKHADFDS | 1 | 0 | 0 | 1512688 | AVP78041.1 | P0C6W2 |
| TCFDATTATINY | 1 | 0 | 0 | 1512692 | AUM60023.1 | K9N7C7 |
| TCFDVSKPNAID | 1 | 0 | 0 | 1512693 | AGT51337.1 | H9EJ42 |
| TCFGVGKPNAID | 1 | 0 | 0 | 1512702 | ARU07593.1 | H9EJ42 |
| TCFGVSKPNAID | 1 | 0 | 0 | 1512703 | AWK59942.1 | H9EJ42 |
| TCFRCPKEIVDT | 1 | 0 | 0 | 1512707 | QBQ01836.1 | P0C6X6 |
| TCFSTSADTYAC | 1 | 0 | 0 | 1512710 | APO40578.1 | P0C6X7 |
| TCFSVSKPNAID | 1 | 0 | 0 | 1512713 | AVA26872.1 | H9EJ42 |
| TCGIVELEKGVL | 1 | 0 | 0 | 1512720 | P0C6W6.1 | P0C6W2 |
| TCGTTTLAGLWL | 1 | 0 | 0 | 1512727 | 3FZD_A | P0C6X7 |
| TCGYLPNNAVVK | 1 | 0 | 0 | 1512730 | APO40578.1 | P0C6X7 |
| TCHNAAQCDCLC | 1 | 0 | 0 | 1512736 | CAA83660.1 | P36334 |
| TCHNLCNTDPIT | 1 | 0 | 0 | 1512740 | BBO58740.1 | P36334 |
| TCIPTVGYSAAL | 1 | 0 | 0 | 1512752 | AVV62530.1 | K9N7D2 |
| TCIVLDVKGVEE | 1 | 0 | 0 | 1512769 | AWW13564.1 | Q80872 |
| TCLEVQVDPNAF | 1 | 0 | 0 | 1512775 | AUM60014.1 | K9N5Q8 |
| TCLILPSKSAGG | 1 | 0 | 0 | 1512776 | APO40578.1 | P0C6X7 |
| TCNAISPKIDNE | 1 | 0 | 0 | 1512791 | AMM42407.1 | A0A140H1H1 |
| TCNAISPKTDKE | 1 | 0 | 0 | 1512792 | AZS52618.1 | A0A140H1H1 |
| TCNNIDVAKIYG | 1 | 0 | 0 | 1512799 | ABD43176.1 | P36334 |
| TCNSKVKNGFTL | 1 | 0 | 0 | 1512806 | CAA79896.1 | P36334 |
| TCNSNANISMLQ | 1 | 0 | 0 | 1512807 | AKA09626.1 | A0A0P0G321 |
| TCNSNANISTLQ | 1 | 0 | 0 | 1512808 | AKA09623.1 | A0A0P0G321 |
| TCNVVKQCCTAA | 1 | 0 | 0 | 1512814 | AVN89451.1 | K9N7C7 |
| TCPAGTNYLNCD | 1 | 0 | 0 | 1512819 | AIL49520.1 | P36334 |
| TCPFAKSDGCSC | 1 | 0 | 0 | 1512823 | AVV62524.1 | K9N7C7 |
| TCRFVELDDQGH | 1 | 0 | 0 | 1512842 | AXN92247.1 | K9N7C7 |
| TCRQISPAAIAS | 1 | 0 | 0 | 1512848 | ATG84888.1 | K9N5Q8 |
| TCSQISPAAIVS | 1 | 0 | 0 | 1512860 | AHI48733.1 | K9N5Q8 |
| TCSTDLQKANIE | 1 | 0 | 0 | 1512862 | BBO58790.1 | P36334 |
| TCTIDSNTTQDT | 1 | 0 | 0 | 1512871 | NP_073549.1 | A0A0P0LKV0 |
| TCTKVEVNALRT | 1 | 0 | 0 | 1512872 | AVV62524.1 | K9N7C7 |
| TCTSTHFAFACA | 1 | 0 | 0 | 1512876 | ATO98237.1 | Q3LZX7 |
| TCTTTKFSFVCQ | 1 | 0 | 0 | 1512877 | APO40584.1 | P59635 |
| TCVGSDNVADFN | 1 | 0 | 0 | 1512879 | APO40578.1 | P0C6X7 |
| TCVPTVGYSAAL | 1 | 0 | 0 | 1512887 | AUM60018.1 | K9N7D2 |
| TDAALSFKENVD | 1 | 0 | 0 | 1512911 | AXX83325.1 | Q80872 |
| TDAFLGWSYDTC | 1 | 0 | 0 | 1512912 | BBA20983.1 | A0A140H1H1 |
| TDAIIAVELLLS | 1 | 0 | 0 | 1512916 | AVA26872.1 | H9EJ42 |
| TDAPSTFGTRNP | 1 | 0 | 0 | 1512921 | AXN92267.1 | K9N4V7 |
| TDAQTGSSKCGC | 1 | 0 | 0 | 1512922 | 2H85_A | P0C6X7 |
| TDAYKLFLSGDD | 1 | 0 | 0 | 1512932 | ARU07575.1 | A0A140H1G9 |
| TDCLAAEIHRDC | 1 | 0 | 0 | 1512936 | AVV62535.1 | K9N7C7 |
| TDCSDGNYNCNA | 1 | 0 | 0 | 1512939 | AHL31360.1 | K9N5Q8 |
| TDCSDGNYNRYA | 1 | 0 | 0 | 1512941 | ALJ54486.1 | K9N5Q8 |
| TDDEHDEWFGIT | 1 | 0 | 0 | 1512947 | AVV62526.1 | K9N5Q8 |
| TDDNALAYYNTS | 1 | 0 | 0 | 1512952 | APO40578.1 | P0C6X7 |
| TDDPRWPQIAEL | 1 | 0 | 0 | 1512954 | ATG84774.1 | K9N4V7 |
| TDESFYKNMYLS | 1 | 0 | 0 | 1512966 | ANZ78864.1 | P0C6X6 |
| TDFHNYTGEGQQ | 1 | 0 | 0 | 1512976 | AXX83296.1 | P30215 |
| TDFNGNFYGPYR | 1 | 0 | 0 | 1512992 | P0C6U5.1 | A0A140H1G9 |
| TDFSLGGKSSTV | 1 | 0 | 0 | 1512993 | AUM60018.1 | K9N7D2 |
| TDGICSENHVMV | 1 | 0 | 0 | 1513005 | AIV41942.1 | P0C6X6 |
| TDHDVVYAQHCF | 1 | 0 | 0 | 1513020 | CAA83661.1 | P36334 |
| TDHSGSNDNIAL | 1 | 0 | 0 | 1513024 | Q3I5J2.1 | Q3LZX9 |
| TDIAEKPTENAC | 1 | 0 | 0 | 1513026 | AVP78041.1 | P0C6W2 |
| TDIDVNPALNIC | 1 | 0 | 0 | 1513036 | ANI69887.1 | K9N7C7 |
| TDIDVNTALNIC | 1 | 0 | 0 | 1513040 | AUM60013.1 | K9N7C7 |
| TDIEGNFYGPFV | 1 | 0 | 0 | 1513044 | APO40578.1 | P0C6X7 |
| TDILLKQPMATS | 1 | 0 | 0 | 1513051 | AHF54651.1 | Q4VID0 |
| TDILYPGQKYTS | 1 | 0 | 0 | 1513052 | AVV62524.1 | K9N7C7 |
| TDINFTATASFG | 1 | 0 | 0 | 1513053 | AIW52843.1 | A0A0P0G321 |
| TDINGVYWVASN | 1 | 0 | 0 | 1513054 | AEN19370.1 | P33469 |
| TDIVDVTNGLGT | 1 | 0 | 0 | 1513059 | CAA83660.1 | P36334 |
| TDIYLTTYPGQG | 1 | 0 | 0 | 1513062 | APO40578.1 | P0C6X7 |
| TDKETLEYWVTP | 1 | 0 | 0 | 1513063 | AZS52618.1 | A0A140H1H1 |
| TDKLCHAVVSKS | 1 | 0 | 0 | 1513065 | AGT51737.1 | P0C6X6 |
| TDKSSTLWFKPP | 1 | 0 | 0 | 1513071 | AKA09652.1 | P36334 |
| TDLLSTLWFKPP | 1 | 0 | 0 | 1513092 | AWW13566.1 | P36334 |
| TDLQKANIEIEL | 1 | 0 | 0 | 1513096 | BBO58790.1 | P36334 |
| TDLQKANTDIIL | 1 | 0 | 0 | 1513097 | CAA79896.1 | P36334 |
| TDLQKANTDIKL | 1 | 0 | 0 | 1513098 | AVQ05264.1 | P36334 |
| TDLQKANTEIEL | 1 | 0 | 0 | 1513099 | AEN19366.1 | P36334 |
| TDLQKSNTDIIL | 1 | 0 | 0 | 1513100 | CAA83660.1 | P36334 |
| TDLTVTSDGQPY | 1 | 0 | 0 | 1513108 | AGT51657.1 | P0C6X6 |
| TDNETLEYWVTP | 1 | 0 | 0 | 1513116 | BBA20986.1 | A0A140H1H1 |
| TDNETLQYGVTP | 1 | 0 | 0 | 1513117 | ABA60801.1 | A0A140H1H1 |
| TDNHITTYPGQG | 1 | 0 | 0 | 1513122 | AVP78041.1 | P0C6W2 |
| TDNIFSVQEDGR | 1 | 0 | 0 | 1513123 | ABE97137.1 | A0A0P0G321 |
| TDNIFSVQKDGR | 1 | 0 | 0 | 1513124 | AMM42643.1 | A0A0P0G321 |
| TDNNQDGGRSGA | 1 | 0 | 0 | 1513131 | Q3I5I7.1 | Q3LZX4 |
| TDNTFVAGNCDV | 1 | 0 | 0 | 1513139 | ATO98181.1 | Q3LZX1 |
| TDNTFVSGKCDV | 1 | 0 | 0 | 1513140 | AAT76147.1 | Q3LZX1 |

| | | | | | | |
|---|---|---|---|---|---|---|
| TDNVFIKNADIV | 1 | 0 | 0 | 1513145 | AVP78041.1 | P0C6W2 |
| TDPIEDYENDWN | 1 | 0 | 0 | 1513151 | APO40578.1 | P0C6X7 |
| TDPIVFTYIPAS | 1 | 0 | 0 | 1513153 | AVN89337.1 | K9N7D2 |
| TDQINDMVYSLL | 1 | 0 | 0 | 1513162 | AIX09796.1 | P0C6X6 |
| TDQSSYTVDSVT | 1 | 0 | 0 | 1513173 | APO40578.1 | P0C6X7 |
| TDSADNNQNGGR | 1 | 0 | 0 | 1513185 | ARO76389.1 | P59595 |
| TDSGFAVYHRYR | 1 | 0 | 0 | 1513190 | APO40582.1 | P59596 |
| TDSIVIDGVTYQ | 1 | 0 | 0 | 1513195 | AWK59942.1 | H9EJ42 |
| TDSQKANTDIKL | 1 | 0 | 0 | 1513198 | BBO58736.1 | P36334 |
| TDSQRAYIGSTA | 1 | 0 | 0 | 1513199 | AUM60027.1 | K9N643 |
| TDSTDNNQDGGR | 1 | 0 | 0 | 1513202 | Q3I5I7.1 | Q3LZX4 |
| TDSTGTESVTVD | 1 | 0 | 0 | 1513205 | ATQ39394.1 | K9N796 |
| TDSTVFTYIPAS | 1 | 0 | 0 | 1513206 | AVN89422.1 | K9N7D2 |
| TDSTVLTYIPAS | 1 | 0 | 0 | 1513208 | AVN89378.1 | K9N7D2 |
| TDSVRDPQTLDI | 1 | 0 | 0 | 1513215 | ATO98145.1 | Q3LZX1 |
| TDSVRDPQTLEI | 1 | 0 | 0 | 1513216 | ATO98181.1 | Q3LZX1 |
| TDTIVVTAGDGI | 1 | 0 | 0 | 1513234 | ATO98194.1 | Q3LZX0 |
| TDTNSVCPKFEF | 1 | 0 | 0 | 1513238 | AVN89387.1 | K9N5Q8 |
| TDTNSVCPKIEF | 1 | 0 | 0 | 1513239 | ATG84701.1 | K9N5Q8 |
| TDTNSVCPKLEG | 1 | 0 | 0 | 1513241 | 6C6Z_A | K9N5Q8 |
| TDTNSVCPKLEH | 1 | 0 | 0 | 1513242 | 4ZS6_A | K9N5Q8 |
| TDTNSVCPMQAL | 1 | 0 | 0 | 1513245 | AUM60014.1 | K9N5Q8 |
| TDTNSVSPKLEF | 1 | 0 | 0 | 1513246 | AKN24803.1 | K9N5Q8 |
| TDTVDVTNGLGT | 1 | 0 | 0 | 1513258 | CAA79896.1 | P36334 |
| TDTVNRDSFNLY | 1 | 0 | 0 | 1513263 | ADN03337.1 | A0A140H1G9 |
| TDVATDVQSRAD | 1 | 0 | 0 | 1513264 | AUM60023.1 | K9N7C7 |
| TDVKQVGCFMRL | 1 | 0 | 0 | 1513280 | AGT51518.1 | P0C6X6 |
| TDVKQVGCSMRL | 1 | 0 | 0 | 1513281 | AEN19363.1 | P0C6X6 |
| TDVLFSGQKYSS | 1 | 0 | 0 | 1513283 | AQZ41321.1 | K9N7C7 |
| TDVLLKQPMATC | 1 | 0 | 0 | 1513286 | QRK03817.1 | Q4VID0 |
| TDVLLKQPTATS | 1 | 0 | 0 | 1513288 | ARE29989.1 | Q4VID0 |
| TDVLQEDQKILA | 1 | 0 | 0 | 1513289 | BAL45639.1 | A0A0P0K6L9 |
| TDVPLKQPMATS | 1 | 0 | 0 | 1513298 | AHF54653.1 | Q4VID0 |
| TDVPSISGTQVY | 1 | 0 | 0 | 1513299 | ATG84770.1 | K9N643 |
| TDVPTAIRADQL | 1 | 0 | 0 | 1513301 | ATO98181.1 | Q3LZX1 |
| TDVPTMIHVEQV | 1 | 0 | 0 | 1513302 | APO40579.1 | P59594 |
| TDVPTTIHADQL | 1 | 0 | 0 | 1513304 | AVP78031.1 | Q3LZX1 |
| TDVQSRADQNGY | 1 | 0 | 0 | 1513306 | AUM60013.1 | K9N7C7 |
| TDYKNWPQIAQF | 1 | 0 | 0 | 1513330 | APO40586.1 | P59595 |
| TDYNAILQSDNP | 1 | 0 | 0 | 1513335 | QDH43737.1 | H9EJ42 |
| TEADWIELFPHN | 1 | 0 | 0 | 1513342 | QEO75983.1 | A0A0P0LKV0 |
| TEDDYNGTPLEF | 1 | 0 | 0 | 1513366 | APO40578.1 | P0C6X7 |
| TEDETAEWFGIT | 1 | 0 | 0 | 1513371 | AUM60014.1 | K9N5Q8 |
| TEDPPWPQIAEL | 1 | 0 | 0 | 1513380 | QDP16213.1 | K9N4V7 |
| TEFAVNSTKALA | 1 | 0 | 0 | 1513407 | ASU90276.1 | K9N4V0 |
| TEFIGTQALDML | 1 | 0 | 0 | 1513410 | AVV62524.1 | K9N7C7 |
| TEFVGTQSIDML | 1 | 0 | 0 | 1513418 | ATG84733.1 | K9N638 |
| TEGGKTFMYAYI | 1 | 0 | 0 | 1513431 | AWK59942.1 | H9EJ42 |
| TEGQLQSLSRDY | 1 | 0 | 0 | 1513438 | ABE97136.1 | H9EJ42 |
| TEGSVKGLIASK | 1 | 0 | 0 | 1513439 | APO40578.1 | P0C6X7 |
| TEIELGVCVNYD | 1 | 0 | 0 | 1513457 | AEN19366.1 | P36334 |
| TEKDTNEAKALF | 1 | 0 | 0 | 1513468 | AUM60013.1 | K9N7C7 |
| TELAVNSGLLTA | 1 | 0 | 0 | 1513484 | AWK59942.1 | H9EJ42 |
| TELPQLVNANQY | 1 | 0 | 0 | 1513496 | ATG84679.1 | K9N5Q8 |
| TENKLNISSGSR | 1 | 0 | 0 | 1513511 | ATO98145.1 | Q3LZX1 |
| TENLNAYQQQDG | 1 | 0 | 0 | 1513515 | AIL49396.1 | P33469 |
| TENLYFQSMETG | 1 | 0 | 0 | 1513520 | 5X29_A | P59637 |
| TENRFTNFEPFT | 1 | 0 | 0 | 1513527 | AIL49500.1 | P36334 |
| TEPKVPEEAPAP | 1 | 0 | 0 | 1513532 | APO40578.1 | P0C6X7 |
| TEQLGAPLGITI | 1 | 0 | 0 | 1513537 | AIW52843.1 | A0A0P0G321 |
| TEQTQSAIAKPK | 1 | 0 | 0 | 1513551 | ATQ39389.1 | K9N7C7 |
| TEQTQSVNVKLK | 1 | 0 | 0 | 1513552 | ANF29194.1 | K9N7C7 |
| TEQTQSVNVKPK | 1 | 0 | 0 | 1513553 | ATG84700.1 | K9N638 |
| TERLKLFAEETQ | 1 | 0 | 0 | 1513564 | AIV41984.1 | P0C6X6 |
| TESNKKFLPFQQFGRDIAD | 1 | 0 | 0 | 1513583 | P0DTC2.1 | P0DTC2 |
| TESSGSLATRLM | 1 | 0 | 0 | 1513585 | ATP16762.1 | Q4VID0 |
| TESSGSLVTRLM | 1 | 0 | 0 | 1513586 | AHF54653.1 | Q4VID0 |
| TESYVDVGPDSF | 1 | 0 | 0 | 1513597 | AQZ41296.1 | K9N5Q8 |
| TETITTGFDFNC | 1 | 0 | 0 | 1513605 | AGT51679.1 | P30215 |
| TETITTGFDLNC | 1 | 0 | 0 | 1513606 | AEN19365.1 | P30215 |
| TETIVLTAGDGV | 1 | 0 | 0 | 1513608 | APO40580.1 | P59632 |
| TETTSAFVGALP | 1 | 0 | 0 | 1513614 | AAK32191.1 | A0A0P0K6L9 |
| TEVDGGFKHLRE | 1 | 0 | 0 | 1513620 | ATO98145.1 | Q3LZX1 |
| TEVKEQLDTTQL | 1 | 0 | 0 | 1513626 | AHF54662.1 | P36334 |
| TEVKIPQGLTFA | 1 | 0 | 0 | 1513628 | AUM60013.1 | K9N7C7 |
| TEVKVSALKTAG | 1 | 0 | 0 | 1513631 | AXN92247.1 | K9N7C7 |
| TEVLPVSMTKTS | 1 | 0 | 0 | 1513632 | APO40579.1 | P59594 |
| TEVPQLVNANHY | 1 | 0 | 0 | 1513642 | AID55087.1 | K9N5Q8 |
| TEVSVKMVDAYV | 1 | 0 | 0 | 1513652 | BAE93400.1 | P0C6U8 |
| TEVVGNVILKPA | 1 | 0 | 0 | 1513656 | AID16715.1 | P0C6W2 |
| TEWGNMMTGISA | 1 | 0 | 0 | 1513666 | AVV62535.1 | K9N7C7 |
| TEYREYMEAYNF | 1 | 0 | 0 | 1513674 | AHY22563.1 | K9N7C7 |
| TEYYEFLNKHFN | 1 | 0 | 0 | 1513684 | AIL49554.1 | P0C6X6 |
| TFAELEAARKAY | 1 | 0 | 0 | 1513687 | AMQ49036.1 | K9N638 |
| TFCSGSTFISDE | 1 | 0 | 0 | 1513711 | APO40578.1 | P0C6X7 |
| TFCVDKALTVEE | 1 | 0 | 0 | 1513713 | AVV62524.1 | K9N7C7 |
| TFDDILGKVCSE | 1 | 0 | 0 | 1513718 | ARU07575.1 | A0A140H1G9 |
| TFDFYNLWSTFT | 1 | 0 | 0 | 1513724 | QBP84746.1 | P0C6X6 |
| TFDNDIACPLTV | 1 | 0 | 0 | 1513730 | AUM60013.1 | K9N7C7 |
| TFDNPVIPFKDG | 1 | 0 | 0 | 1513732 | 5X58_A | P59594 |
| TFDPITLKAPDT | 1 | 0 | 0 | 1513733 | AVR40342.1 | P36334 |

| | | | | | | |
|---|---|---|---|---|---|---|
| TFEPRSDLERDF | 1 | 0 | 0 | 1513751 | ARB07436.1 | A0A140H1G9 |
| TFERGSVQVDNG | 1 | 0 | 0 | 1513752 | AFR79255.1 | A0A0P0LKV0 |
| TFETPAGQFCTE | 1 | 0 | 0 | 1513756 | AVV62524.1 | K9N7C7 |
| TFGAPDDATRLL | 1 | 0 | 0 | 1513783 | AUM60013.1 | K9N7C7 |
| TFGEEKVHQVAA | 1 | 0 | 0 | 1513794 | AVV62524.1 | K9N7C7 |
| TFGGPSDSSDNS | 1 | 0 | 0 | 1513798 | AVP78038.1 | Q3LZX4 |
| TFGITQEYIQTT | 1 | 0 | 0 | 1513803 | AVV62526.1 | K9N5Q8 |
| TFGTRNPNNDAA | 1 | 0 | 0 | 1513816 | AVV62533.1 | K9N4V7 |
| TFGTRNPNNDLA | 1 | 0 | 0 | 1513817 | AKN24774.1 | K9N4V7 |
| TFGTRNPNNDPA | 1 | 0 | 0 | 1513818 | AUM60021.1 | K9N4V7 |
| TFGVSDNFDNNQ | 1 | 0 | 0 | 1513822 | APO40586.1 | P59595 |
| TFGVVKPGDTFT | 1 | 0 | 0 | 1513824 | ATN39877.2 | P0C6X6 |
| TFGVYDYLVSTQ | 1 | 0 | 0 | 1513826 | APO40578.1 | P0C6X7 |
| TFIQSNVDVVPE | 1 | 0 | 0 | 1513837 | AGT51409.1 | P0C6X6 |
| TFIVLYVDFKPQ | 1 | 0 | 0 | 1513851 | CAA71056.1 | A0A0P0K6L9 |
| TFKAPDTYKCPQ | 1 | 0 | 0 | 1513857 | ARB07414.1 | P36334 |
| TFKATGTYKCPQ | 1 | 0 | 0 | 1513859 | CAA79896.1 | P36334 |
| TFKIVQWQEAWF | 1 | 0 | 0 | 1513868 | ARK08632.1 | P0C6X6 |
| TFKQTNNNCFVN | 1 | 0 | 0 | 1513877 | AGT51787.1 | P0C6X6 |
| TFLDLQAEMNRL | 1 | 0 | 0 | 1513889 | AGT51521.1 | P36334 |
| TFLDLQDEMNRL | 1 | 0 | 0 | 1513890 | CAA83660.1 | P36334 |
| TFLEGDVHDTVL | 1 | 0 | 0 | 1513895 | ARO76381.1 | P0C6U8 |
| TFLSCARKSFSI | 1 | 0 | 0 | 1513926 | AJC98124.1 | P0C6X6 |
| TFLSMFDVDKKI | 1 | 0 | 0 | 1513929 | AIV41972.1 | P0C6X6 |
| TFMLKYNENGTI | 1 | 0 | 0 | 1513942 | AVP78031.1 | Q3LZX1 |
| TFNITEDEILEW | 1 | 0 | 0 | 1513956 | AQZ41296.1 | K9N5Q8 |
| TFNMQTVSFHED | 1 | 0 | 0 | 1513962 | AVV62524.1 | K9N7C7 |
| TFNQLTADLQHK | 1 | 0 | 0 | 1513964 | ARB07436.1 | A0A140H1G9 |
| TFNRGRKTQPKF | 1 | 0 | 0 | 1513965 | ARU07581.1 | Q19U29 |
| TFPGPSGIGAVG | 1 | 0 | 0 | 1513975 | ANI69896.1 | K9N4V7 |
| TFPQQMFVTPLA | 1 | 0 | 0 | 1513980 | AVN89294.1 | K9N643 |
| TFPSGQGVPLNA | 1 | 0 | 0 | 1513982 | AZK15907.1 | K9N4V7 |
| TFQGKFKKVVKG | 1 | 0 | 0 | 1513991 | APO40578.1 | P0C6X7 |
| TFQTVRPGNFNQ | 1 | 0 | 0 | 1513997 | AXN92247.1 | K9N7C7 |
| TFQTVRSGNFNQ | 1 | 0 | 0 | 1513998 | AHI48750.1 | K9N7C7 |
| TFSDPNMCLLSG | 1 | 0 | 0 | 1514016 | AVN89460.1 | K9N643 |
| TFSDPTTFGSSF | 1 | 0 | 0 | 1514020 | AVN89383.1 | K9N643 |
| TFSETGSDVLYQ | 1 | 0 | 0 | 1514023 | AUM60013.1 | K9N7C7 |
| TFSGNVDLDNLA | 1 | 0 | 0 | 1514026 | AUM60013.1 | K9N7C7 |
| TFSGNVDLEGLA | 1 | 0 | 0 | 1514027 | AVV62524.1 | K9N7C7 |
| TFSPGQGVPLNA | 1 | 0 | 0 | 1514037 | AHI48535.1 | K9N4V7 |
| TFSVHPVVCDED | 1 | 0 | 0 | 1514040 | AUM60013.1 | K9N7C7 |
| TFSVTQEYIQTT | 1 | 0 | 0 | 1514046 | AUM60014.1 | K9N5Q8 |
| TFTATTVVGSCI | 1 | 0 | 0 | 1514050 | AXN92247.1 | K9N7C7 |
| TFTDVNLQGLEN | 1 | 0 | 0 | 1514053 | AST12962.1 | A0A0P0LKV0 |
| TFTEVNLQGLEN | 1 | 0 | 0 | 1514054 | NP_073549.1 | A0A0P0LKV0 |
| TFTFAGFVGQAV | 1 | 0 | 0 | 1514055 | AUM60016.1 | K9N4V0 |
| TFTGSYKCPQTK | 1 | 0 | 0 | 1514060 | AVQ05264.1 | P36334 |
| TFTKLQSLENVA | 1 | 0 | 0 | 1514063 | AVP78041.1 | P0C6W2 |
| TFTQAGFNIWVP | 1 | 0 | 0 | 1514071 | AWK59942.1 | H9EJ42 |
| TFTRSTNSRIKA | 1 | 0 | 0 | 1514075 | QYC64967.1 | P0DTD1 |
| TFTTKVTVPKDH | 1 | 0 | 0 | 1514077 | AIW52712.1 | A0A0N7J131 |
| TFTVDPKKAYMD | 1 | 0 | 0 | 1514086 | AVV62524.1 | K9N7C7 |
| TFVCASEYTGNY | 1 | 0 | 0 | 1514093 | AVP78041.1 | P0C6W2 |
| TFVMMSAPPSEC | 1 | 0 | 0 | 1514107 | APO40578.1 | P0C6X7 |
| TFVNISRVELHT | 1 | 0 | 0 | 1514110 | AIW52843.1 | A0A0P0G321 |
| TFVQSNVEVLPE | 1 | 0 | 0 | 1514120 | QEG03790.1 | P0C6X6 |
| TFVSTFNVPMEK | 1 | 0 | 0 | 1514125 | APO40578.1 | P0C6X7 |
| TFVVDKSLSIDE | 1 | 0 | 0 | 1514129 | AZK15899.1 | K9N638 |
| TFYAYFTDTGVV | 1 | 0 | 0 | 1514144 | CAA79896.1 | P36334 |
| TFYGKDAFENHS | 1 | 0 | 0 | 1514149 | AVV62524.1 | K9N7C7 |
| TFYGKESFENPT | 1 | 0 | 0 | 1514150 | AVN89322.1 | K9N7C7 |
| TFYGKESFEQHS | 1 | 0 | 0 | 1514151 | AUM60023.1 | K9N7C7 |
| TFYGKESLENPT | 1 | 0 | 0 | 1514152 | AXN92247.1 | K9N7C7 |
| TFYGKNAFQNHS | 1 | 0 | 0 | 1514155 | AVV62535.1 | K9N7C7 |
| TFYIAANGGTSF | 1 | 0 | 0 | 1514158 | AVV62535.1 | K9N7C7 |
| TFYLDGEVITFD | 1 | 0 | 0 | 1514163 | AVP78041.1 | P0C6W2 |
| TGCFSFVKMLFM | 1 | 0 | 0 | 1514198 | AXN92258.1 | K9N7C7 |
| TGCGNGRWLTGN | 1 | 0 | 0 | 1514199 | AVV62524.1 | K9N7C7 |
| TGCTGSWNKVNQ | 1 | 0 | 0 | 1514209 | QBF80541.1 | K9N7C7 |
| TGCVGAWDKVSK | 1 | 0 | 0 | 1514211 | AVV62524.1 | K9N7C7 |
| TGCVGTWDKVAK | 1 | 0 | 0 | 1514213 | AVV62535.1 | K9N7C7 |
| TGCVIAWNTAQQ | 1 | 0 | 0 | 1514215 | ATO98108.1 | Q3LZX1 |
| TGDCATVYLEDK | 1 | 0 | 0 | 1514218 | AWK59942.1 | H9EJ42 |
| TGDLQPLEQPTSEAVEAPLV | 1 | 0 | 0 | 1514240 | P0DTC1.1 | P0DTD1 |
| TGDVKIMAKAPQ | 1 | 0 | 0 | 1514257 | NP_073549.1 | A0A0P0LKV0 |
| TGEGDQIIFYEG | 1 | 0 | 0 | 1514262 | ARU07576.1 | A0A140H1H0 |
| TGEIAQYNYKQD | 1 | 0 | 0 | 1514264 | AUM60014.1 | K9N5Q8 |
| TGFCSKHQWNCI | 1 | 0 | 0 | 1514268 | AEN19363.1 | P0C6X6 |
| TGFHTINHTFDN | 1 | 0 | 0 | 1514275 | 5X58_A | P59594 |
| TGFINHLPGPRV | 1 | 0 | 0 | 1514279 | AVV62524.1 | K9N7C7 |
| TGFTTTNEAFRK | 1 | 0 | 0 | 1514291 | AXN92228.1 | K9N5Q8 |
| TGGAQTSTAIYT | 1 | 0 | 0 | 1514303 | ARO76382.1 | P59594 |
| TGGFMSTIITSC | 1 | 0 | 0 | 1514307 | AXN92236.1 | K9N7C7 |
| TGGIDICSQDKV | 1 | 0 | 0 | 1514311 | AEN19363.1 | P0C6X6 |
| TGGLVQQVEKWL | 1 | 0 | 0 | 1514315 | APO40578.1 | P0C6X7 |
| TGGNIQSSSRAS | 1 | 0 | 0 | 1514317 | ANI69874.1 | K9N4V7 |
| TGGTYSAPFMVN | 1 | 0 | 0 | 1514323 | AVV62524.1 | K9N7C7 |
| TGHFYINGYRYF | 1 | 0 | 0 | 1514331 | AIW52757.1 | A0A0P0K6L9 |
| TGHYIGQAGEMP | 1 | 0 | 0 | 1514341 | ARU07606.1 | P0C6X6 |
| TGIAIEQDKNTQ | 1 | 0 | 0 | 1514342 | ATO98181.1 | Q3LZX1 |

| | | | | | | |
|---|---|---|---|---|---|---|
| TGIDFQDELDDF | 1 | 0 | 0 | 1514348 | AMW90852.1 | K9N5Q8 |
| TGKLLQACIRSA | 1 | 0 | 0 | 1514368 | AVV62527.1 | K9N796 |
| TGKLSDCLYFKN | 1 | 0 | 0 | 1514369 | AGT51737.1 | P0C6X6 |
| TGKSFSNISLSY | 1 | 0 | 0 | 1514373 | AVV62526.1 | K9N5Q8 |
| TGLCIAGDRGIA | 1 | 0 | 0 | 1514380 | AMK59677.1 | P36334 |
| TGLFKDCSREIS | 1 | 0 | 0 | 1514386 | AUM60013.1 | K9N7C7 |
| TGLTFQTVRSGN | 1 | 0 | 0 | 1514414 | AHI48750.1 | K9N7C7 |
| TGLVKNQCVNFN | 1 | 0 | 0 | 1514420 | ATO98193.1 | Q3LZX1 |
| TGMAITIKPEAN | 1 | 0 | 0 | 1514423 | ABD75615.1 | A0A140H1G9 |
| TGNAVQGFACDC | 1 | 0 | 0 | 1514426 | AVV62524.1 | K9N7C7 |
| TGNGTKALAPRW | 1 | 0 | 0 | 1514435 | AVV62533.1 | K9N4V7 |
| TGNGTKPLAPRW | 1 | 0 | 0 | 1514436 | AVV62544.1 | K9N4V7 |
| TGNLIISSSTQK | 1 | 0 | 0 | 1514443 | AVV62526.1 | K9N5Q8 |
| TGNLQSNHDLYCQVH | 1 | 0 | 0 | 1514445 | P0DTD1.1 | P0DTD1 |
| TGNVLKHCCIAA | 1 | 0 | 0 | 1514452 | AUM60013.1 | K9N7C7 |
| TGNVVQQCCTAA | 1 | 0 | 0 | 1514455 | ALA49384.1 | K9N7C7 |
| TGPHANDHYGTD | 1 | 0 | 0 | 1514476 | AJQ24067.1 | P33469 |
| TGPHKDAKFRER | 1 | 0 | 0 | 1514478 | AIW52723.1 | A0A0N7J131 |
| TGPYAKDQYGTD | 1 | 0 | 0 | 1514489 | AIL49482.1 | P33469 |
| TGPYANASYGQS | 1 | 0 | 0 | 1514491 | ADN03343.1 | Q19U29 |
| TGPYASSSYGEA | 1 | 0 | 0 | 1514493 | AMM42394.1 | Q19U29 |
| TGPYKCPQSKSL | 1 | 0 | 0 | 1514496 | BBA84107.1 | P36334 |
| TGQGIFKEVSAV | 1 | 0 | 0 | 1514502 | BBA20983.1 | A0A140H1H1 |
| TGRCTVNKHVAY | 1 | 0 | 0 | 1514508 | AEN19363.1 | P0C6X6 |
| TGRLIQFDNPNI | 1 | 0 | 0 | 1514516 | APO40579.1 | P59594 |
| TGSAFDGNMYGA | 1 | 0 | 0 | 1514528 | AVV62535.1 | K9N7C7 |
| TGSAFDGTMYGA | 1 | 0 | 0 | 1514529 | AXN92247.1 | K9N7C7 |
| TGSFNNIDTGPP | 1 | 0 | 0 | 1514540 | AIX10756.1 | P36334 |
| TGSLAARLMSIP | 1 | 0 | 0 | 1514550 | QRK03817.1 | Q4VID0 |
| TGSLADGLGAYA | 1 | 0 | 0 | 1514551 | AIV42008.1 | P0C6X6 |
| TGSLASRLMSIP | 1 | 0 | 0 | 1514552 | QBQ01844.1 | Q4VID0 |
| TGSNDSKALFFT | 1 | 0 | 0 | 1514560 | AZU90729.1 | K9N7C7 |
| TGSPSVVEFNRL | 1 | 0 | 0 | 1514567 | AVV62524.1 | K9N7C7 |
| TGSRYYYPEPIT | 1 | 0 | 0 | 1514573 | CAA83661.1 | P36334 |
| TGSSFDGNMYGA | 1 | 0 | 0 | 1514574 | AVV62524.1 | K9N7C7 |
| TGSSFFNLEPIT | 1 | 0 | 0 | 1514575 | AVV62526.1 | K9N5Q8 |
| TGSSFYEPEPIT | 1 | 0 | 0 | 1514578 | ALM26400.1 | K9N5Q8 |
| TGSSFYSPEPIT | 1 | 0 | 0 | 1514579 | AHI48672.1 | K9N5Q8 |
| TGSTPQKLFVAN | 1 | 0 | 0 | 1514587 | AZU90731.1 | K9N5Q8 |
| TGSWNKVTQIAN | 1 | 0 | 0 | 1514596 | AXN92247.1 | K9N7C7 |
| TGSWWSFNPETN | 1 | 0 | 0 | 1514598 | ASY99815.1 | K9N7A1 |
| TGTAFNGDFYGP | 1 | 0 | 0 | 1514603 | AIV41912.1 | P0C6X6 |
| TGTDFNGNFYGP | 1 | 0 | 0 | 1514606 | P0C6U5.1 | A0A140H1G9 |
| TGTDTTDVLLKQ | 1 | 0 | 0 | 1514610 | ARE29989.1 | Q4VID0 |
| TGTESVTVDRGS | 1 | 0 | 0 | 1514612 | ATQ39394.1 | K9N796 |
| TGTGVLTSSTKK | 1 | 0 | 0 | 1514623 | APO40579.1 | P59594 |
| TGTIYAELEPPC | 1 | 0 | 0 | 1514624 | AEA10817.1 | P0C6X7 |
| TGTLIVNSVLPF | 1 | 0 | 0 | 1514632 | AAS48454.1 | P59637 |
| TGTNVFQTQAGC | 1 | 0 | 0 | 1514639 | AVP78031.1 | Q3LZX1 |
| TGTQSVSVDREL | 1 | 0 | 0 | 1514645 | AVN89314.1 | K9N796 |
| TGTVIIRPSTSA | 1 | 0 | 0 | 1514657 | QBF80608.1 | K9N5Q8 |
| TGTVIISPFTSA | 1 | 0 | 0 | 1514658 | AHI48702.1 | K9N5Q8 |
| TGTYKCPQTKSL | 1 | 0 | 0 | 1514662 | CAA79896.1 | P36334 |
| TGVAIEQLLYAI | 1 | 0 | 0 | 1514665 | AXN92247.1 | K9N7C7 |
| TGVALEQDKNTQ | 1 | 0 | 0 | 1514666 | ATO98145.1 | Q3LZX1 |
| TGVATEQLLYAI | 1 | 0 | 0 | 1514667 | AUM60023.1 | K9N7C7 |
| TGVFVAPPIADF | 1 | 0 | 0 | 1514678 | AVV62528.1 | K9N4V0 |
| TGVISSMTDNLQ | 1 | 0 | 0 | 1514688 | AUM60014.1 | K9N5Q8 |
| TGVIYGFNSTET | 1 | 0 | 0 | 1514689 | AIL49343.1 | P30215 |
| TGVIYGLNFTET | 1 | 0 | 0 | 1514690 | ATP16766.1 | P30215 |
| TGVLTSSTKKFQ | 1 | 0 | 0 | 1514700 | APO40579.1 | P59594 |
| TGVNDDDYVIAL | 1 | 0 | 0 | 1514706 | AGT51337.1 | H9EJ42 |
| TGVNFVVQPVGV | 1 | 0 | 0 | 1514708 | AXN92247.1 | K9N7C7 |
| TGVSIEILLAAI | 1 | 0 | 0 | 1514720 | ABD75639.1 | A0A140H1G9 |
| TGVSVPCVCGRE | 1 | 0 | 0 | 1514725 | ARO76381.1 | P0C6U8 |
| TGVTSSMTDNLQ | 1 | 0 | 0 | 1514727 | AUM60024.1 | K9N5Q8 |
| TGVVYVNATGSD | 1 | 0 | 0 | 1514730 | ABU88826.1 | A0A0P0G321 |
| TGWLLYQLLNGF | 1 | 0 | 0 | 1514739 | P0C6U5.1 | A0A140H1G9 |
| TGYFPKAGANFR | 1 | 0 | 0 | 1514743 | BBA20983.1 | A0A140H1H1 |
| TGYSSRNSVTGR | 1 | 0 | 0 | 1514761 | APO40579.1 | P59594 |
| TGYSWADLPAYM | 1 | 0 | 0 | 1514762 | Q01455.1 | Q01455 |
| TGYTLADLPVYV | 1 | 0 | 0 | 1514763 | AZS52621.1 | Q0ZJ82 |
| TGYVDTEHSTEF | 1 | 0 | 0 | 1514765 | AVP78030.1 | P0C6W2 |
| THATTVDVNVLN | 1 | 0 | 0 | 1514781 | AUM60013.1 | K9N7C7 |
| THCAFAVDPAAA | 1 | 0 | 0 | 1514783 | AST12962.1 | A0A0P0LKV0 |
| THCSFAVDPAAA | 1 | 0 | 0 | 1514785 | NP_073549.1 | A0A0P0LKV0 |
| THCSFAVDPAVA | 1 | 0 | 0 | 1514786 | AGT21359.1 | A0A0P0LKV0 |
| THDGPNEHVTSV | 1 | 0 | 0 | 1514798 | AGV08391.1 | K9N796 |
| THDPIYKNGAMF | 1 | 0 | 0 | 1514803 | AVV62524.1 | K9N7C7 |
| THFAFACADGTK | 1 | 0 | 0 | 1514812 | ARO76387.1 | P59635 |
| THGIAAVNSVPW | 1 | 0 | 0 | 1514822 | ARO76381.1 | P0C6U8 |
| THGLNLEEAARYMRS | 1 | 0 | 0 | 1514827 | P0DTC1.1 | P0DTD1 |
| THHDKFTDGICL | 1 | 0 | 0 | 1514831 | APO40578.1 | P0C6X7 |
| THIAEDVVDQVE | 1 | 0 | 0 | 1514838 | AST12962.1 | A0A0P0LKV0 |
| THLSHYVFNIDW | 1 | 0 | 0 | 1514866 | ALT66812.1 | K9N7C7 |
| THLSNDTNGDPV | 1 | 0 | 0 | 1514869 | AUM60021.1 | K9N4V7 |
| THMNLKYAISAK | 1 | 0 | 0 | 1514875 | AZK15898.1 | K9N7C7 |
| THNGRVVNYTVC | 1 | 0 | 0 | 1514876 | AIW52841.1 | A0A0P0G321 |
| THPVQWQCKLDF | 1 | 0 | 0 | 1514883 | AVV62539.1 | K9N4V0 |
| THQNNDDHGNPV | 1 | 0 | 0 | 1514885 | AXN92267.1 | K9N4V7 |
| THQSNDESGEPV | 1 | 0 | 0 | 1514886 | AVV62544.1 | K9N4V7 |

| | | | | | | |
|---|---|---|---|---|---|---|
| THSKGLYRKCVR | 1 | 0 | 0 | 1514899 | AVP78041.1 | P0C6W2 |
| THTGSSFDGNMY | 1 | 0 | 0 | 1514911 | AVV62524.1 | K9N7C7 |
| THWICANQGTSV | 1 | 0 | 0 | 1514935 | AKA09620.1 | A0A0P0G321 |
| THYIKDGDDGYF | 1 | 0 | 0 | 1514940 | AHI48773.1 | K9N7C7 |
| THYVRVTFSDSN | 1 | 0 | 0 | 1514954 | ALA49894.1 | K9N643 |
| TIADFVQIENCD | 1 | 0 | 0 | 1514955 | BAL45640.1 | A0A0P0K6L9 |
| TIADFVQIENCN | 1 | 0 | 0 | 1514956 | AIW52757.1 | A0A0P0K6L9 |
| TIANYAKPFLNKVVSTTT | 1 | 0 | 0 | 1514983 | P0DTC1.1 | P0DTD1 |
| TIATYCTGSIPC | 1 | 0 | 0 | 1514994 | AVP78041.1 | P0C6W2 |
| TICDLYSIHDVG | 1 | 0 | 0 | 1515002 | AAX85666.1 | P0C6X6 |
| TICDLYSIQDVG | 1 | 0 | 0 | 1515003 | AEN19363.1 | P0C6X6 |
| TICDLYSMQDVG | 1 | 0 | 0 | 1515004 | AAR01012.1 | P0C6X6 |
| TICHPNLGNHRK | 1 | 0 | 0 | 1515012 | CAA79896.1 | P36334 |
| TIDCAAFDCGDY | 1 | 0 | 0 | 1515030 | ARB66062.1 | P36334 |
| TIDCSAFVCGDC | 1 | 0 | 0 | 1515034 | CAA83661.1 | P36334 |
| TIDELCVNLNLP | 1 | 0 | 0 | 1515038 | ALA50066.1 | K9N7C7 |
| TIDGIVEDTIND | 1 | 0 | 0 | 1515039 | AGW27861.1 | A0A140H1G9 |
| TIDKFAIPNRRK | 1 | 0 | 0 | 1515048 | AEN19366.1 | P36334 |
| TIDKLCDYLADY | 1 | 0 | 0 | 1515052 | ASU90174.1 | K9N638 |
| TIDKLRDYLADY | 1 | 0 | 0 | 1515053 | AXN92247.1 | K9N7C7 |
| TIDSTQDGVNKL | 1 | 0 | 0 | 1515060 | QBQ01839.1 | P36334 |
| TIEDALRNSAML | 1 | 0 | 0 | 1515067 | AIW52757.1 | A0A0P0K6L9 |
| TIEDLRARMTYV | 1 | 0 | 0 | 1515068 | AJG44112.1 | K9N638 |
| TIEDWLLVQGFS | 1 | 0 | 0 | 1515069 | AHI48842.1 | K9N643 |
| TIEENVIQCFGP | 1 | 0 | 0 | 1515072 | ADK37676.1 | H9EJ69 |
| TIFKDSSGKPVP | 1 | 0 | 0 | 1515081 | AVP78041.1 | P0C6W2 |
| TIFYTTEIASKF | 1 | 0 | 0 | 1515096 | ARU07575.1 | A0A140H1G9 |
| TIGDSIFTGCTG | 1 | 0 | 0 | 1515097 | QAT98908.1 | K9N7C7 |
| TIGESIFTGCVG | 1 | 0 | 0 | 1515099 | AUM60013.1 | K9N7C7 |
| TIGNIEEFIQTS | 1 | 0 | 0 | 1515107 | BBA20979.1 | P36334 |
| TIGNMEEFIQTN | 1 | 0 | 0 | 1515108 | AIL49500.1 | P36334 |
| TIGNMEEFIQTS | 1 | 0 | 0 | 1515109 | CAA79896.1 | P36334 |
| TIGNMGEFIQTS | 1 | 0 | 0 | 1515110 | ARB66062.1 | P36334 |
| TIGTVTLKQGEIKDAT | 1 | 0 | 0 | 1515118 | YP_009724391.1 | P0DTC3 |
| TIHRGDPMSNNG | 1 | 0 | 0 | 1515128 | AVP78042.1 | Q3LZX1 |
| TIHSVCNGCVGH | 1 | 0 | 0 | 1515130 | AIW52741.1 | A0A0P0K6L9 |
| TIIIPDKQVFDK | 1 | 0 | 0 | 1515133 | ARU07575.1 | A0A140H1G9 |
| TIILTTFNCVYA | 1 | 0 | 0 | 1515138 | AWW13547.1 | Q01455 |
| TIISTQDGVNKL | 1 | 0 | 0 | 1515143 | BBA20973.1 | P36334 |
| TIKKPNELSRVL | 1 | 0 | 0 | 1515164 | AVP78030.1 | P0C6W2 |
| TIKKPSELSRVL | 1 | 0 | 0 | 1515165 | AVP78041.1 | P0C6W2 |
| TIKPEANINQDS | 1 | 0 | 0 | 1515169 | ABD75615.1 | A0A140H1G9 |
| TIKPLEHHHHHH | 1 | 0 | 0 | 1515170 | 5E6J_A | P0C6X7 |
| TIKPVSLEHHHH | 1 | 0 | 0 | 1515171 | 4M0W_A | P0C6U8 |
| TIKVFTTVDNVN | 1 | 0 | 0 | 1515180 | APO40578.1 | P0C6X7 |
| TILDYCDGNLPC | 1 | 0 | 0 | 1515197 | APO40578.1 | P0C6X7 |
| TILEQDRMPKTR | 1 | 0 | 0 | 1515200 | AHF54653.1 | Q4VID0 |
| TILEQDRMPKTS | 1 | 0 | 0 | 1515201 | AHF54652.1 | Q4VID0 |
| TILFITSYGAVV | 1 | 0 | 0 | 1515203 | AII82118.1 | A0A0P0K6L9 |
| TILNKPRPKRSR | 1 | 0 | 0 | 1515219 | AJQ24068.1 | P33469 |
| TILPCYSGRVSA | 1 | 0 | 0 | 1515225 | BBA20986.1 | A0A140H1H1 |
| TILPTFNTRTVS | 1 | 0 | 0 | 1515226 | AUM60013.1 | K9N7C7 |
| TILRTINGDFLH | 1 | 0 | 0 | 1515230 | AVP78041.1 | P0C6W2 |
| TINCESIYEFDT | 1 | 0 | 0 | 1515250 | ARU07575.1 | A0A140H1G9 |
| TINNFNPSSWNR | 1 | 0 | 0 | 1515265 | BBA20986.1 | A0A140H1H1 |
| TINPSSPARLEI | 1 | 0 | 0 | 1515269 | ATP16755.1 | Q80872 |
| TINSQDGVNKLQ | 1 | 0 | 0 | 1515272 | QDH43726.1 | P36334 |
| TINSTQDGVNKL | 1 | 0 | 0 | 1515273 | AEN19358.1 | P36334 |
| TINSTQDGYNKL | 1 | 0 | 0 | 1515274 | CAA79896.1 | P36334 |
| TINSVIQKQDID | 1 | 0 | 0 | 1515275 | AUM60013.1 | K9N7C7 |
| TINYTQPEVLSP | 1 | 0 | 0 | 1515280 | AVV62526.1 | K9N5Q8 |
| TIPDEKQNGHSL | 1 | 0 | 0 | 1515287 | AXN92247.1 | K9N7C7 |
| TIPDEKQNGLSL | 1 | 0 | 0 | 1515288 | AUM60013.1 | K9N7C7 |
| TIPEQKQNGFSF | 1 | 0 | 0 | 1515289 | AVV62524.1 | K9N7C7 |
| TIPQQSQNGFSF | 1 | 0 | 0 | 1515304 | AVV62535.1 | K9N7C7 |
| TIPTQLTPIMFV | 1 | 0 | 0 | 1515310 | AVV62524.1 | K9N7C7 |
| TIQTVFDCAVET | 1 | 0 | 0 | 1515338 | AST12962.1 | A0A0P0LKV0 |
| TIREVIDGNAMH | 1 | 0 | 0 | 1515341 | AVV62524.1 | K9N7C7 |
| TISTGFDFNCHY | 1 | 0 | 0 | 1515370 | AGT51729.1 | P30215 |
| TISVEELKRLLE | 1 | 0 | 0 | 1515373 | ATO98197.1 | Q3LZX9 |
| TISVPAHMPMPP | 1 | 0 | 0 | 1515375 | ARU07582.1 | Q0ZJ80 |
| TISVPVHMPVHP | 1 | 0 | 0 | 1515376 | AET99208.1 | Q0ZJ80 |
| TITDAIDCAQNP | 1 | 0 | 0 | 1515380 | ATO98181.1 | Q3LZX1 |
| TITDPKVYPIIL | 1 | 0 | 0 | 1515385 | 2CME_E | P59636 |
| TITIGSTFVNTS | 1 | 0 | 0 | 1515389 | AIL49548.1 | P36334 |
| TITSAGRPCVAS | 1 | 0 | 0 | 1515408 | QRK03919.1 | P0C6X6 |
| TITTDNTFVAGN | 1 | 0 | 0 | 1515411 | ARO76382.1 | P59594 |
| TITTGFDFNCHY | 1 | 0 | 0 | 1515412 | AGT51679.1 | P30215 |
| TITVDQLKHLLE | 1 | 0 | 0 | 1515415 | APO40582.1 | P59596 |
| TITVEKLKQLLE | 1 | 0 | 0 | 1515418 | AAS48455.1 | P59596 |
| TITVPEEFANHV | 1 | 0 | 0 | 1515420 | AVV62524.1 | K9N7C7 |
| TIVVQPHNGVWE | 1 | 0 | 0 | 1515472 | ABA60802.1 | A0A140H1H1 |
| TIWEPDRMLKTS | 1 | 0 | 0 | 1515480 | AVR40343.1 | Q4VID0 |
| TIWEQDRMLKTS | 1 | 0 | 0 | 1515481 | ATP16762.1 | Q4VID0 |
| TIYAELEPPCRF | 1 | 0 | 0 | 1515486 | AEA10817.1 | P0C6X7 |
| TKAIKDDVLYSE | 1 | 0 | 0 | 1515511 | AKA09643.1 | P36334 |
| TKALKELYGPID | 1 | 0 | 0 | 1515528 | ATG84722.1 | K9N7C7 |
| TKALKELYGSVD | 1 | 0 | 0 | 1515531 | AHC74096.1 | K9N7C7 |
| TKCTLKSLSVQK | 1 | 0 | 0 | 1515551 | AVP78042.1 | Q3LZX1 |
| TKDVVIEGLKAC | 1 | 0 | 0 | 1515564 | AVV62524.1 | K9N7C7 |
| TKEVGHTDLMAA | 1 | 0 | 0 | 1515574 | AVP78030.1 | P0C6W2 |

| | | | | | | |
|---|---|---|---|---|---|---|
| TKFADDLNQLSG | 1 | 0 | 0 | 1515578 | AVP78030.1 | P0C6W2 |
| TKFQGLENIAFN | 1 | 0 | 0 | 1515592 | AXG21663.1 | K9N7C7 |
| TKFVGANFGRWS | 1 | 0 | 0 | 1515598 | ABB90515.1 | A0A0P0K6L9 |
| TKGKPVQGAWNI | 1 | 0 | 0 | 1515620 | ARO76381.1 | P0C6U8 |
| TKGPFVEGAINV | 1 | 0 | 0 | 1515622 | AUM60023.1 | K9N7C7 |
| TKGPYEFCSQHT | 1 | 0 | 0 | 1515624 | AVL25535.1 | H9EJ42 |
| TKGVMEPIDYNV | 1 | 0 | 0 | 1515631 | AGT51393.1 | H9EJ42 |
| TKGVMEPVDYNV | 1 | 0 | 0 | 1515632 | AWK59942.1 | H9EJ42 |
| TKGVTFGEDTVL | 1 | 0 | 0 | 1515633 | ATO98229.1 | P0C6W2 |
| TKGYVAGIQNFK | 1 | 0 | 0 | 1515634 | AUM60013.1 | K9N7C7 |
| TKHTAREVRHTI | 1 | 0 | 0 | 1515650 | AJQ24068.1 | P33469 |
| TKIFKDDGNFVV | 1 | 0 | 0 | 1515661 | AMK59674.1 | P0C6X6 |
| TKISLKGGKVIS | 1 | 0 | 0 | 1515676 | ARO76381.1 | P0C6U8 |
| TKKFQPFQQFGR | 1 | 0 | 0 | 1515692 | APO40579.1 | P59594 |
| TKKKYSELDDEE | 1 | 0 | 0 | 1515694 | Q3I5J1.1 | Q3LZX8 |
| TKKSLNVDGVQY | 1 | 0 | 0 | 1515704 | ATQ39389.1 | K9N7C7 |
| TKLASTEELPDE | 1 | 0 | 0 | 1515716 | AAT76156.1 | Q3LZX3 |
| TKLATTVELPDE | 1 | 0 | 0 | 1515719 | AVP78050.1 | Q3LZX3 |
| TKLKDGVNFNID | 1 | 0 | 0 | 1515735 | ARB66060.1 | P36334 |
| TKLLDYSIEMQL | 1 | 0 | 0 | 1515739 | AWK59978.1 | H9EJ42 |
| TKLLFNVYLGMA | 1 | 0 | 0 | 1515740 | AXX83339.1 | P36334 |
| TKLLRGVELEDF | 1 | 0 | 0 | 1515742 | AVV62535.1 | K9N7C7 |
| TKLMCCIGPDIF | 1 | 0 | 0 | 1515746 | QEG03773.1 | P0C6X6 |
| TKLPYYDVKKIH | 1 | 0 | 0 | 1515756 | BAL45640.1 | A0A0P0K6L9 |
| TKLQSLENVAYN | 1 | 0 | 0 | 1515758 | AVP78041.1 | P0C6W2 |
| TKNVTGSNDSKA | 1 | 0 | 0 | 1515790 | AZU90729.1 | K9N7C7 |
| TKPFVRAAFEHL | 1 | 0 | 0 | 1515799 | ATN39862.1 | P0C6X6 |
| TKPITTSNPFEV | 1 | 0 | 0 | 1515802 | APO40578.1 | P0C6X7 |
| TKPQPLMDKFSS | 1 | 0 | 0 | 1515811 | ATO98193.1 | Q3LZX1 |
| TKPVETSNSFDA | 1 | 0 | 0 | 1515818 | AVP78041.1 | P0C6W2 |
| TKRTDNPILDFK | 1 | 0 | 0 | 1515849 | AVP78031.1 | Q3LZX1 |
| TKSAAGYGGFGC | 1 | 0 | 0 | 1515852 | AEN19363.1 | P0C6X6 |
| TKSCGVVAEHDF | 1 | 0 | 0 | 1515854 | ARU07574.1 | A0A140H1G9 |
| TKSGTNDICVFE | 1 | 0 | 0 | 1515860 | ATQ39389.1 | K9N7C7 |
| TKSGVNDICIFE | 1 | 0 | 0 | 1515861 | ANF29215.1 | K9N7C7 |
| TKSIYKTPNIKD | 1 | 0 | 0 | 1515865 | APO40579.1 | P59594 |
| TKSSEYKGSITD | 1 | 0 | 0 | 1515872 | AVP78041.1 | P0C6W2 |
| TKSTTSRIRASM | 1 | 0 | 0 | 1515875 | AVP78041.1 | P0C6W2 |
| TKSWSVGDWTGF | 1 | 0 | 0 | 1515878 | NP_073549.1 | A0A0P0LKV0 |
| TKTAVKSGTAIL | 1 | 0 | 0 | 1515880 | ARU07575.1 | A0A140H1G9 |
| TKTEKDTNEAKA | 1 | 0 | 0 | 1515882 | AUM60013.1 | K9N7C7 |
| TKTHATLFGSVA | 1 | 0 | 0 | 1515884 | AXN92228.1 | K9N5Q8 |
| TKVFKDGVMYSE | 1 | 0 | 0 | 1515907 | AEN19366.1 | P36334 |
| TKVFKEGVMYSE | 1 | 0 | 0 | 1515908 | AMK59677.1 | P36334 |
| TKVIKDGVMYSE | 1 | 0 | 0 | 1515911 | CAA79896.1 | P36334 |
| TKVLRYGYHVLY | 1 | 0 | 0 | 1515920 | AQN78693.1 | P0C6X6 |
| TKVQQQQDDLFT | 1 | 0 | 0 | 1515926 | APO40584.1 | P59635 |
| TKVTVPKDHPHL | 1 | 0 | 0 | 1515934 | AIW52712.1 | A0A0N7J131 |
| TKYNIYDGSGVG | 1 | 0 | 0 | 1515945 | BAL45639.1 | A0A0P0K6L9 |
| TKYNIYDVAGVG | 1 | 0 | 0 | 1515946 | AIW52741.1 | A0A0P0K6L9 |
| TKYVAPHVTYQN | 1 | 0 | 0 | 1515959 | AID55097.1 | K9N5Q8 |
| TKYVAPQATYQN | 1 | 0 | 0 | 1515960 | ALM26400.1 | K9N5Q8 |
| TKYYDHSQYYLN | 1 | 0 | 0 | 1515971 | AIX09798.1 | P30215 |
| TLACFVLAVVYR | 1 | 0 | 0 | 1515981 | ATO98209.1 | Q3LZX9 |
| TLALLNLTDEDL | 1 | 0 | 0 | 1515993 | AXN92231.1 | K9N643 |
| TLANFNETNGPL | 1 | 0 | 0 | 1515997 | CAA71147.1 | A0A0P0K6L9 |
| TLANTTTAPLKL | 1 | 0 | 0 | 1515999 | AGV08437.1 | K9N7C7 |
| TLAVIGDFNCTN | 1 | 0 | 0 | 1516008 | BBA20983.1 | A0A140H1H1 |
| TLDCINGQYDKN | 1 | 0 | 0 | 1516033 | AUM60014.1 | K9N5Q8 |
| TLDDILQHANKS | 1 | 0 | 0 | 1516036 | AHC74096.1 | K9N7C7 |
| TLDDSFLGRYMS | 1 | 0 | 0 | 1516039 | APO40578.1 | P0C6X7 |
| TLDDVTRNYIIT | 1 | 0 | 0 | 1516040 | ASU90464.1 | K9N643 |
| TLDELHACTSFT | 1 | 0 | 0 | 1516042 | AVV62535.1 | K9N7C7 |
| TLDILDITPCSF | 1 | 0 | 0 | 1516045 | ATO98145.1 | Q3LZX1 |
| TLDKLSDYCTFV | 1 | 0 | 0 | 1516049 | AXN92247.1 | K9N7C7 |
| TLEDLHARMTYT | 1 | 0 | 0 | 1516067 | ATQ39389.1 | K9N7C7 |
| TLEEAVRQVRSW | 1 | 0 | 0 | 1516072 | AHL18091.1 | K9N7C7 |
| TLEQKVKPIVEE | 1 | 0 | 0 | 1516088 | APO40578.1 | P0C6X7 |
| TLEQYSTCDMCK | 1 | 0 | 0 | 1516090 | QED88039.1 | H9EJ42 |
| TLESGKVKQSQP | 1 | 0 | 0 | 1516093 | ARQ84742.1 | K9N4V7 |
| TLFRDAEGKMVP | 1 | 0 | 0 | 1516142 | AVV62524.1 | K9N7C7 |
| TLGSITRQPSKI | 1 | 0 | 0 | 1516167 | ATO98194.1 | Q3LZX0 |
| TLGSLELLNFKR | 1 | 0 | 0 | 1516169 | AGT51735.1 | Q4VID0 |
| TLGTVSLKQGEI | 1 | 0 | 0 | 1516170 | AVP78043.1 | Q3LZX0 |
| TLHPVQWQCKLD | 1 | 0 | 0 | 1516185 | AVV62528.1 | K9N4V0 |
| TLIPQENYSSIR | 1 | 0 | 0 | 1516214 | AWW13571.1 | P0C6X6 |
| TLIVFTSCELYH | 1 | 0 | 0 | 1516220 | ABI96964.1 | P59635 |
| TLKANEETFKLS | 1 | 0 | 0 | 1516227 | APO40578.1 | P0C6X7 |
| TLKHFFFAQDGH | 1 | 0 | 0 | 1516242 | AVV62524.1 | K9N7C7 |
| TLKHFFFAQDGN | 1 | 0 | 0 | 1516243 | AXN92247.1 | K9N7C7 |
| TLKIKVSQTNMY | 1 | 0 | 0 | 1516255 | APT69868.1 | A0A0P0ILY9 |
| TLKQGEIKGATP | 1 | 0 | 0 | 1516267 | AVP78032.1 | Q3LZX0 |
| TLLAKAELSVSA | 1 | 0 | 0 | 1516280 | AVV62524.1 | K9N7C7 |
| TLLCCTEQDLHK | 1 | 0 | 0 | 1516283 | AUM60027.1 | K9N643 |
| TLLDLQVEMNRL | 1 | 0 | 0 | 1516291 | AVQ05264.1 | P36334 |
| TLLDNQEDKFWC | 1 | 0 | 0 | 1516297 | AIW52827.1 | H9EJ42 |
| TLLPAADLDDFSKQLQQSMSSADS | 1 | 0 | 0 | 1516334 | P0DTC9.1 | P0DTC9 |
| TLLPAAELDDFS | 1 | 0 | 0 | 1516338 | APO40586.1 | P59595 |
| TLLQVPQVSTSQ | 1 | 0 | 0 | 1516347 | AVV62537.1 | K9N5Q8 |
| TLLVDSDMNDFV | 1 | 0 | 0 | 1516365 | APO40578.1 | P0C6X7 |
| TLLVPQIFKARA | 1 | 0 | 0 | 1516369 | AMM42357.1 | A0A140H1G9 |

| | | | | | | |
|---|---|---|---|---|---|---|
| TLLYSFEAHKDF | 1 | 0 | 0 | 1516374 | ARK08657.1 | P0C6X6 |
| TLNEDLLEILHR | 1 | 0 | 0 | 1516395 | ARO76381.1 | P0C6U8 |
| TLNSFKEYFDLQ | 1 | 0 | 0 | 1516416 | AVV62537.1 | K9N5Q8 |
| TLPDIAEDVVDQ | 1 | 0 | 0 | 1516424 | NP_073549.1 | A0A0P0LKV0 |
| TLPQSKDSNFLN | 1 | 0 | 0 | 1516439 | AKI29254.1 | K9N638 |
| TLQAAFGNTGVC | 1 | 0 | 0 | 1516446 | AWK59942.1 | H9EJ42 |
| TLQCVMLVYCCL | 1 | 0 | 0 | 1516452 | APO40578.1 | P0C6X7 |
| TLQLGVPDNSST | 1 | 0 | 0 | 1516473 | AKA09623.1 | A0A0P0G321 |
| TLQPISDLLTPM | 1 | 0 | 0 | 1516478 | APO40578.1 | P0C6X7 |
| TLQPVSDLLANM | 1 | 0 | 0 | 1516481 | ARO76381.1 | P0C6U8 |
| TLQSGLKKMAQP | 1 | 0 | 0 | 1516488 | AWK59942.1 | H9EJ42 |
| TLQYFFSQTSLF | 1 | 0 | 0 | 1516495 | AVV62524.1 | K9N7C7 |
| TLRATEEASRQS | 1 | 0 | 0 | 1516498 | AIL23989.1 | K9N7C7 |
| TLRVTKTLNTQY | 1 | 0 | 0 | 1516542 | AVV62535.1 | K9N7C7 |
| TLSHRFYRLANE | 1 | 0 | 0 | 1516560 | APO40578.1 | P0C6X7 |
| TLSLLDQYLLAF | 1 | 0 | 0 | 1516567 | AAB27260.2 | P36334 |
| TLSTYDFYPNVP | 1 | 0 | 0 | 1516578 | ATO98120.1 | Q3LZX1 |
| TLSVDFNGVLHK | 1 | 0 | 0 | 1516581 | AWK59942.1 | H9EJ42 |
| TLSVHDGYTNEI | 1 | 0 | 0 | 1516582 | ARO76381.1 | P0C6U8 |
| TLTAPTIANQER | 1 | 0 | 0 | 1516593 | AXG21663.1 | K9N7C7 |
| TLTDMEKDFLSL | 1 | 0 | 0 | 1516596 | AUM60013.1 | K9N7C7 |
| TLTEVKIPQGLT | 1 | 0 | 0 | 1516599 | AUM60013.1 | K9N7C7 |
| TLTLDKVPQAVE | 1 | 0 | 0 | 1516613 | AEN19363.1 | P0C6X6 |
| TLTLKERILYTF | 1 | 0 | 0 | 1516614 | AVV62524.1 | K9N7C7 |
| TLTNGSDLDRCT | 1 | 0 | 0 | 1516625 | ABD72988.1 | P59594 |
| TLTPCSVRSVPG | 1 | 0 | 0 | 1516627 | AKN24812.1 | K9N5Q8 |
| TLTRGSDLDRCT | 1 | 0 | 0 | 1516637 | ABD72995.1 | P59594 |
| TLTTPTIVNQER | 1 | 0 | 0 | 1516645 | ALA49912.1 | K9N7C7 |
| TLTTQYCRFGSC | 1 | 0 | 0 | 1516646 | AUM60013.1 | K9N7C7 |
| TLVAPDLSLDYI | 1 | 0 | 0 | 1516656 | CAA83661.1 | P36334 |
| TLVATAEAELAKNVSL | 1 | 0 | 0 | 1516658 | P0DTC1.1 | P0DTD1 |
| TLVCCDAIKLAK | 1 | 0 | 0 | 1516660 | AVV62524.1 | K9N7C7 |
| TLVDLNWLNRVE | 1 | 0 | 0 | 1516664 | AGT21353.1 | A0A0P0K6L9 |
| TLVGQLEQTNMH | 1 | 0 | 0 | 1516671 | ASU90174.1 | K9N638 |
| TLVGVGNTVPMT | 1 | 0 | 0 | 1516672 | AVV62526.1 | K9N5Q8 |
| TLVLNAHELHHH | 1 | 0 | 0 | 1516686 | AVV62529.1 | K9N643 |
| TLVLNANECQHP | 1 | 0 | 0 | 1516687 | AVV62540.1 | K9N643 |
| TLVSGDDHIPSG | 1 | 0 | 0 | 1516710 | AIW52748.1 | A0A0P0K6L9 |
| TLWNYLFETIPT | 1 | 0 | 0 | 1516725 | AXN92247.1 | K9N7C7 |
| TLWYQKPFLSDF | 1 | 0 | 0 | 1516729 | BBA20983.1 | A0A140H1H1 |
| TLYFNNEPAFIM | 1 | 0 | 0 | 1516732 | AXG21664.1 | K9N638 |
| TLYGQAVVYFGG | 1 | 0 | 0 | 1516738 | QBP84710.1 | P0C6X6 |
| TLYKIDGALLTK | 1 | 0 | 0 | 1516744 | APO40578.1 | P0C6X7 |
| TLYVKNWCRCSC | 1 | 0 | 0 | 1516755 | BBA20986.1 | A0A140H1H1 |
| TMADFVQIENCN | 1 | 0 | 0 | 1516761 | CAA71056.1 | A0A0P0K6L9 |
| TMCDIKQMLFCM | 1 | 0 | 0 | 1516771 | AXN92247.1 | K9N7C7 |
| TMCEYPHTICKS | 1 | 0 | 0 | 1516776 | BBA20983.1 | A0A140H1H1 |
| TMCEYPNTICHP | 1 | 0 | 0 | 1516778 | CAA83661.1 | P36334 |
| TMDGFCQYKGKW | 1 | 0 | 0 | 1516788 | NP_073549.1 | A0A0P0LKV0 |
| TMENFELEKHCY | 1 | 0 | 0 | 1516794 | ALW82707.1 | K9N7C7 |
| TMFDVYVDTFLS | 1 | 0 | 0 | 1516798 | AEN19363.1 | P0C6X6 |
| TMFMDAEGRMTP | 1 | 0 | 0 | 1516804 | AKJ80136.2 | K9N7C7 |
| TMFRDAEGKMVP | 1 | 0 | 0 | 1516810 | AVV62535.1 | K9N7C7 |
| TMFRDAEGRITP | 1 | 0 | 0 | 1516811 | AHL18091.1 | K9N7C7 |
| TMFTMADGSPQS | 1 | 0 | 0 | 1516815 | AGT51488.1 | P0C6X6 |
| TMFTTADGSPQP | 1 | 0 | 0 | 1516816 | AGT51707.1 | P0C6X6 |
| TMFVCIDNSANT | 1 | 0 | 0 | 1516817 | AUM60013.1 | K9N7C7 |
| TMHSINHDIFSL | 1 | 0 | 0 | 1516827 | ARU07575.1 | A0A140H1G9 |
| TMIFSVHPVECD | 1 | 0 | 0 | 1516838 | AVV62524.1 | K9N7C7 |
| TMILSLHPVECD | 1 | 0 | 0 | 1516840 | ALW82719.1 | K9N7C7 |
| TMIVDEWSYTGS | 1 | 0 | 0 | 1516847 | ATG84866.1 | K9N5Q8 |
| TMLAEFDENGTI | 1 | 0 | 0 | 1516853 | APO40579.1 | P59594 |
| TMNDDLLQILNR | 1 | 0 | 0 | 1516868 | APO40578.1 | P0C6X7 |
| TMQIVDKDGNYY | 1 | 0 | 0 | 1516883 | QED88010.1 | H9EJ42 |
| TMRYFVKIGPNK | 1 | 0 | 0 | 1516890 | QED88017.1 | H9EJ42 |
| TMSQHSRSTRSM | 1 | 0 | 0 | 1516901 | ASU90241.1 | K9N5Q8 |
| TMTDIAEKPTEN | 1 | 0 | 0 | 1516909 | AVP78041.1 | P0C6W2 |
| TMTFSVHPVVCD | 1 | 0 | 0 | 1516912 | AUM60013.1 | K9N7C7 |
| TMTGRVFHQKCL | 1 | 0 | 0 | 1516913 | AIL49573.1 | P0C6X6 |
| TMYGAFMDKQVH | 1 | 0 | 0 | 1516937 | AXN92247.1 | K9N7C7 |
| TNAETTDFFTVA | 1 | 0 | 0 | 1516949 | AAQ90006.1 | A0A0P0K6L9 |
| TNANLSMLQLSV | 1 | 0 | 0 | 1516956 | BAM16475.1 | A0A0P0G321 |
| TNCLAAETHRDC | 1 | 0 | 0 | 1516977 | AVV62524.1 | K9N7C7 |
| TNCSDGNYNRNA | 1 | 0 | 0 | 1516994 | ALA49671.1 | K9N5Q8 |
| TNDICVFENKTT | 1 | 0 | 0 | 1517004 | ATQ39389.1 | K9N7C7 |
| TNDPCNVCQYWK | 1 | 0 | 0 | 1517009 | AVV62535.1 | K9N7C7 |
| TNDPIRSCLENT | 1 | 0 | 0 | 1517010 | CAA49377.1 | A0A0P0LKV0 |
| TNFAINDKNTTV | 1 | 0 | 0 | 1517033 | BBA20983.1 | A0A140H1H1 |
| TNFAINDLNTTI | 1 | 0 | 0 | 1517034 | ACV87140.1 | A0A140H1H1 |
| TNFEPFTVNSIN | 1 | 0 | 0 | 1517038 | AIL49542.1 | P36334 |
| TNFEPFTVNSVS | 1 | 0 | 0 | 1517041 | ARB66068.1 | P36334 |
| TNFGDVIKPVCA | 1 | 0 | 0 | 1517043 | AWK59942.1 | H9EJ42 |
| TNFGDVNKPVCA | 1 | 0 | 0 | 1517044 | QED88038.1 | H9EJ42 |
| TNFGEDFLGWIV | 1 | 0 | 0 | 1517047 | ARU07575.1 | A0A140H1G9 |
| TNFPIDFPSAES | 1 | 0 | 0 | 1517056 | AZK15901.1 | K9N796 |
| TNFRIDIPSAEL | 1 | 0 | 0 | 1517062 | AXP07346.1 | K9N796 |
| TNFRIDVPFAEL | 1 | 0 | 0 | 1517063 | QAT98905.1 | K9N796 |
| TNFRIDVPPAEL | 1 | 0 | 0 | 1517064 | ANI69890.1 | K9N796 |
| TNFRIDVPSAEL | 1 | 0 | 0 | 1517065 | AVN89314.1 | K9N796 |
| TNFSFGVTQEYI | 1 | 0 | 0 | 1517069 | AXN92228.1 | K9N5Q8 |
| TNGDYNSWHLVE | 1 | 0 | 0 | 1517090 | QDI73607.1 | K9N7C7 |

| | | | | | | |
|---|---|---|---|---|---|---|
| TNGHAVPTLFKL | 1 | 0 | 0 | 1517096 | AHL18091.1 | K9N7C7 |
| TNGKVVPTLFKL | 1 | 0 | 0 | 1517099 | AVV62524.1 | K9N7C7 |
| TNGMNTSHSVCN | 1 | 0 | 0 | 1517104 | AIW52756.1 | A0A0P0K6L9 |
| TNGMNTSQSVCN | 1 | 0 | 0 | 1517105 | ABB90520.1 | A0A0P0K6L9 |
| TNGMNTSYPVCN | 1 | 0 | 0 | 1517106 | ADK37701.1 | A0A0P0K6L9 |
| TNGMNTSYSVCN | 1 | 0 | 0 | 1517107 | ADK37704.1 | A0A0P0K6L9 |
| TNGNFDCALSRY | 1 | 0 | 0 | 1517109 | AVV62524.1 | K9N7C7 |
| TNGPLCVDTSHF | 1 | 0 | 0 | 1517112 | CAA71147.1 | A0A0P0K6L9 |
| TNGPLIVGGGCV | 1 | 0 | 0 | 1517113 | AVP78030.1 | P0C6W2 |
| TNGQAVPTLFRL | 1 | 0 | 0 | 1517116 | AUM60013.1 | K9N7C7 |
| TNGSDLDRCTTF | 1 | 0 | 0 | 1517119 | ABD72988.1 | P59594 |
| TNGSWAIFNDFH | 1 | 0 | 0 | 1517123 | AVV62535.1 | K9N7C7 |
| TNGTNTSHAVCN | 1 | 0 | 0 | 1517130 | QEG03785.1 | A0A0P0K6L9 |
| TNGTNTSHSVCK | 1 | 0 | 0 | 1517131 | AIW52752.1 | A0A0P0K6L9 |
| TNHSVVYAQHCF | 1 | 0 | 0 | 1517156 | AEN19366.1 | P36334 |
| TNHVNVTAAYGL | 1 | 0 | 0 | 1517159 | AVV62537.1 | K9N5Q8 |
| TNILLNVPLRGS | 1 | 0 | 0 | 1517165 | APO40582.1 | P59596 |
| TNKISLNASVTL | 1 | 0 | 0 | 1517185 | AMM42653.1 | A0A0P0G321 |
| TNLDIEKLRQYL | 1 | 0 | 0 | 1517198 | AVV62535.1 | K9N7C7 |
| TNLPPPLLGNSI | 1 | 0 | 0 | 1517218 | AGV08467.1 | K9N5Q8 |
| TNMDIEKLRQYL | 1 | 0 | 0 | 1517235 | AVV62524.1 | K9N7C7 |
| TNMHSPDIVVGD | 1 | 0 | 0 | 1517243 | AVV62535.1 | K9N7C7 |
| TNMSRGRGRNPK | 1 | 0 | 0 | 1517250 | ATG84708.1 | K9N4V7 |
| TNNAMQVESDKY | 1 | 0 | 0 | 1517256 | AVP78030.1 | P0C6W2 |
| TNNNGSDVLDQP | 1 | 0 | 0 | 1517264 | AIV41798.1 | P0C6X6 |
| TNNTLVSGDDHI | 1 | 0 | 0 | 1517269 | AIW52748.1 | A0A0P0K6L9 |
| TNNTRIVDDWSY | 1 | 0 | 0 | 1517271 | ALJ54496.1 | K9N5Q8 |
| TNNTSIVDEWSY | 1 | 0 | 0 | 1517274 | ANF29261.1 | K9N5Q8 |
| TNNVFTLKGGAP | 1 | 0 | 0 | 1517280 | APO40578.1 | P0C6X7 |
| TNPDPKFVDRYY | 1 | 0 | 0 | 1517283 | ATU80202.1 | K9N7C7 |
| TNPHKHCNVQRC | 1 | 0 | 0 | 1517287 | AJQ24083.1 | P33469 |
| TNPRNYSYMDLN | 1 | 0 | 0 | 1517290 | AAA73152.1 | P30215 |
| TNQLPPPLLNNQ | 1 | 0 | 0 | 1517297 | AVV62537.1 | K9N5Q8 |
| TNRCPFDKVFNA | 1 | 0 | 0 | 1517302 | ATO98181.1 | Q3LZX1 |
| TNRFVSVGESFG | 1 | 0 | 0 | 1517307 | QEG03779.1 | P0C6X6 |
| TNRVVSPPINAD | 1 | 0 | 0 | 1517316 | AUM60025.1 | K9N796 |
| TNRVVSPPNNAD | 1 | 0 | 0 | 1517317 | AUM60015.1 | K9N796 |
| TNSASTSLDWFP | 1 | 0 | 0 | 1517319 | AUM60016.1 | K9N4V0 |
| TNSCLVASDGKP | 1 | 0 | 0 | 1517320 | AUM60013.1 | K9N7C7 |
| TNSCLVASDGSP | 1 | 0 | 0 | 1517321 | AVV62535.1 | K9N7C7 |
| TNSCLVASDGVP | 1 | 0 | 0 | 1517322 | AVV62524.1 | K9N7C7 |
| TNSCLVSSNGQP | 1 | 0 | 0 | 1517325 | QAT98908.1 | K9N7C7 |
| TNSGPDDQIGFY | 1 | 0 | 0 | 1517331 | AAT76155.1 | Q3LZX4 |
| TNSGVAIYHRYK | 1 | 0 | 0 | 1517333 | ASY99815.1 | K9N7A1 |
| TNSSKDDQIGYY | 1 | 0 | 0 | 1517347 | AVP78038.1 | Q3LZX4 |
| TNSSQRGVYYPD | 1 | 0 | 0 | 1517349 | AVP78031.1 | Q3LZX1 |
| TNSVCPMMDLGN | 1 | 0 | 0 | 1517356 | AVV62526.1 | K9N5Q8 |
| TNSVDSKSGNNF | 1 | 0 | 0 | 1517358 | APO40579.1 | P59594 |
| TNSYSQNLCVDG | 1 | 0 | 0 | 1517363 | AST12962.1 | A0A0P0LKV0 |
| TNTAAFNAVCSV | 1 | 0 | 0 | 1517364 | ANI69909.1 | K9N7C7 |
| TNTIHSVCNGCV | 1 | 0 | 0 | 1517368 | AIW52741.1 | A0A0P0K6L9 |
| TNTNLSRDRGRN | 1 | 0 | 0 | 1517371 | AVN89370.1 | K9N4V7 |
| TNTRDVDQQWYY | 1 | 0 | 0 | 1517376 | AVV62526.1 | K9N5Q8 |
| TNTSHAVCNGCV | 1 | 0 | 0 | 1517377 | QEG03785.1 | A0A0P0K6L9 |
| TNTSHSVCKGCV | 1 | 0 | 0 | 1517378 | AIW52752.1 | A0A0P0K6L9 |
| TNTSNSVCNGCV | 1 | 0 | 0 | 1517383 | AST12964.1 | A0A0P0K6L9 |
| TNTTFDFLSNAS | 1 | 0 | 0 | 1517387 | AKA09616.1 | A0A0P0G321 |
| TNVDKIRHLLKG | 1 | 0 | 0 | 1517393 | AHI48785.1 | K9N638 |
| TNVEKIRQLLKG | 1 | 0 | 0 | 1517397 | AVN89333.1 | K9N7C7 |
| TNVFKDGVMYSE | 1 | 0 | 0 | 1517398 | BBA84088.1 | P36334 |
| TNVGEPIISQSS | 1 | 0 | 0 | 1517401 | AFO70496.1 | H9EJ42 |
| TNVTIATYCTGS | 1 | 0 | 0 | 1517409 | AVP78041.1 | P0C6W2 |
| TNVTTATYCTGS | 1 | 0 | 0 | 1517411 | AVP78030.1 | P0C6W2 |
| TNVYMRTTHTDI | 1 | 0 | 0 | 1517422 | ANC28699.1 | K9N638 |
| TNYDLSIINARV | 1 | 0 | 0 | 1517425 | AUM60013.1 | K9N7C7 |
| TNYLTCHNAAQC | 1 | 0 | 0 | 1517432 | CAA83660.1 | P36334 |
| TPADIDYRNPSI | 1 | 0 | 0 | 1517445 | AJQ24074.1 | P33469 |
| TPADILDRDPSS | 1 | 0 | 0 | 1517446 | QBP84708.1 | P33469 |
| TPADIVDRDPSN | 1 | 0 | 0 | 1517447 | AJQ24082.1 | P33469 |
| TPADIVDRDPSS | 1 | 0 | 0 | 1517448 | AEN19370.1 | P33469 |
| TPADIVDRDQSS | 1 | 0 | 0 | 1517449 | ANZ78879.1 | P33469 |
| TPADIVHRDPSN | 1 | 0 | 0 | 1517450 | AJQ24067.1 | P33469 |
| TPAFDKSAFTHL | 1 | 0 | 0 | 1517452 | AVP78041.1 | P0C6W2 |
| TPAIDCSDGNYN | 1 | 0 | 0 | 1517458 | ASU91010.1 | K9N5Q8 |
| TPAIKDFGGFNF | 1 | 0 | 0 | 1517459 | ATO98181.1 | Q3LZX1 |
| TPALLATNNVFT | 1 | 0 | 0 | 1517461 | APO40578.1 | P0C6X7 |
| TPASVGSVPGEM | 1 | 0 | 0 | 1517467 | 5W9H_A | K9N5Q8 |
| TPATNCSDGNYN | 1 | 0 | 0 | 1517471 | ALA49671.1 | K9N5Q8 |
| TPAYDGSAFRDL | 1 | 0 | 0 | 1517476 | AHI48736.1 | K9N7C7 |
| TPAYTFTTVKPS | 1 | 0 | 0 | 1517480 | AZK15899.1 | K9N638 |
| TPCELHGNGSMI | 1 | 0 | 0 | 1517482 | ALW82645.1 | K9N7C7 |
| TPCGFKGGIPGN | 1 | 0 | 0 | 1517487 | AIV42002.1 | P0C6X6 |
| TPCNPPDQLVVY | 1 | 0 | 0 | 1517497 | AIW52757.1 | A0A0P0K6L9 |
| TPDMADHIASLV | 1 | 0 | 0 | 1517506 | AIL49401.1 | P33469 |
| TPDMADQIASLV | 1 | 0 | 0 | 1517507 | AEN19370.1 | P33469 |
| TPDPITFKATGT | 1 | 0 | 0 | 1517508 | CAA79896.1 | P36334 |
| TPEDMLNPNYED | 1 | 0 | 0 | 1517518 | APO40578.1 | P0C6X7 |
| TPEDVNMQLMGV | 1 | 0 | 0 | 1517520 | AQZ41284.1 | K9N7C7 |
| TPEEHFVETVSL | 1 | 0 | 0 | 1517523 | ARO76381.1 | P0C6U8 |
| TPESKPLADDDS | 1 | 0 | 0 | 1517526 | AIW52805.1 | H9EJ69 |
| TPFGSCIYWSPR | 1 | 0 | 0 | 1517535 | ARU07575.1 | A0A140H1G9 |

| | | | | | | |
|---|---|---|---|---|---|---|
| TPFHYERDNTSY | 1 | 0 | 0 | 1517537 | AHI48797.1 | K9N638 |
| TPFMVNQLAYSS | 1 | 0 | 0 | 1517540 | AVV62535.1 | K9N7C7 |
| TPFYYERDNTSC | 1 | 0 | 0 | 1517546 | AVN89322.1 | K9N7C7 |
| TPGFGVAVADSY | 1 | 0 | 0 | 1517550 | ABD75607.1 | A0A140H1G9 |
| TPGKQSSSRASF | 1 | 0 | 0 | 1517553 | QBP84763.1 | P33469 |
| TPGTNASSEFAV | 1 | 0 | 0 | 1517558 | BAE93401.1 | P59594 |
| TPGTNTSKAVAV | 1 | 0 | 0 | 1517560 | APO40579.1 | P59594 |
| TPIDEIPYKSFF | 1 | 0 | 0 | 1517581 | AMD02834.1 | K9N638 |
| TPIDQYMCGSDG | 1 | 0 | 0 | 1517583 | AUM60013.1 | K9N7C7 |
| TPINTNASLNAF | 1 | 0 | 0 | 1517595 | AVV62526.1 | K9N5Q8 |
| TPKEHIGTRNPN | 1 | 0 | 0 | 1517611 | APO40586.1 | P59595 |
| TPKKPNKPFSQP | 1 | 0 | 0 | 1517619 | AVA26877.1 | H9EJ69 |
| TPKPSRNQSPVS | 1 | 0 | 0 | 1517626 | AMM42518.1 | A0A0N7J131 |
| TPLCGFPSQAAS | 1 | 0 | 0 | 1517637 | ARO76381.1 | P0C6U8 |
| TPLEFGASTQNE | 1 | 0 | 0 | 1517639 | APO40578.1 | P0C6X7 |
| TPLNLPPTQAHT | 1 | 0 | 0 | 1517657 | QED88038.1 | H9EJ42 |
| TPLNPNYTNSSQ | 1 | 0 | 0 | 1517658 | AVP78031.1 | Q3LZX1 |
| TPLPAADMDDFS | 1 | 0 | 0 | 1517659 | AAS48456.1 | P59595 |
| TPLSKRQYLLKF | 1 | 0 | 0 | 1517660 | BBA20983.1 | A0A140H1H1 |
| TPLSVASAPVEP | 1 | 0 | 0 | 1517662 | QBF80531.1 | K9N7C7 |
| TPLSVASTLVEP | 1 | 0 | 0 | 1517663 | ATG84722.1 | K9N7C7 |
| TPLSVEDLVPSA | 1 | 0 | 0 | 1517667 | AVV62524.1 | K9N7C7 |
| TPLSVESTPVEL | 1 | 0 | 0 | 1517669 | AVN89312.1 | K9N638 |
| TPLVGTSLHPVL | 1 | 0 | 0 | 1517681 | AXN92262.1 | K9N4V0 |
| TPLVPFWITIAY | 1 | 0 | 0 | 1517683 | QYC64967.1 | P0DTD1 |
| TPMTDTLQMGFI | 1 | 0 | 0 | 1517693 | AVV62537.1 | K9N5Q8 |
| TPNAILISDRKI | 1 | 0 | 0 | 1517695 | AUM60013.1 | K9N7C7 |
| TPNAVFISDRKI | 1 | 0 | 0 | 1517697 | ATG84897.1 | K9N7C7 |
| TPNEKLVTTSMA | 1 | 0 | 0 | 1517699 | QBF80609.1 | K9N7C7 |
| TPNIKDFGGFNF | 1 | 0 | 0 | 1517703 | APO40579.1 | P59594 |
| TPNKFCNVQQCF | 1 | 0 | 0 | 1517704 | AMM42397.1 | Q19U29 |
| TPNLPDFKEELD | 1 | 0 | 0 | 1517708 | BBA20982.1 | P36334 |
| TPPALNCYWPLK | 1 | 0 | 0 | 1517716 | 3D0H_E | P59594 |
| TPPRCIAPIDGY | 1 | 0 | 0 | 1517734 | AVV62537.1 | K9N5Q8 |
| TPQPQRAPKGSI | 1 | 0 | 0 | 1517739 | AVV62544.1 | K9N4V7 |
| TPRNKWNGVSDL | 1 | 0 | 0 | 1517747 | ASU90779.1 | K9N638 |
| TPRNKWSGISDL | 1 | 0 | 0 | 1517749 | ALA49758.1 | K9N7C7 |
| TPRQYLLAFNQD | 1 | 0 | 0 | 1517753 | AWW13566.1 | P36334 |
| TPRQYLLSFNQD | 1 | 0 | 0 | 1517754 | AQN78656.1 | P36334 |
| TPRQYSLAFNQD | 1 | 0 | 0 | 1517755 | AHA80190.1 | P36334 |
| TPRSEIPYKSFS | 1 | 0 | 0 | 1517758 | AVV62524.1 | K9N7C7 |
| TPSDPLNNFKHK | 1 | 0 | 0 | 1517764 | AUM60013.1 | K9N7C7 |
| TPSPLTPRSVRS | 1 | 0 | 0 | 1517774 | AJG44080.1 | K9N5Q8 |
| TPSSANPVKEYE | 1 | 0 | 0 | 1517776 | AVV62535.1 | K9N7C7 |
| TPSTDCSDGNYN | 1 | 0 | 0 | 1517779 | ALA49803.1 | K9N5Q8 |
| TPSTLTLRSVRS | 1 | 0 | 0 | 1517784 | ANI69889.1 | K9N5Q8 |
| TPTEHVTSVNLF | 1 | 0 | 0 | 1517790 | AJG44059.1 | K9N796 |
| TPTEQVVSTSSD | 1 | 0 | 0 | 1517791 | AVV62524.1 | K9N7C7 |
| TPTESYVDVGLD | 1 | 0 | 0 | 1517792 | AHL18090.1 | K9N5Q8 |
| TPTESYVDVGSD | 1 | 0 | 0 | 1517794 | AMW90853.1 | K9N5Q8 |
| TPTNMYMRTTHT | 1 | 0 | 0 | 1517804 | AUM60013.1 | K9N7C7 |
| TPTPGVTPDMAD | 1 | 0 | 0 | 1517806 | AIX10722.1 | P33469 |
| TPTTDCSDGNYN | 1 | 0 | 0 | 1517813 | ALA49473.1 | K9N5Q8 |
| TPVDSFTNGAVV | 1 | 0 | 0 | 1517821 | P0C6W6.1 | P0C6W2 |
| TPVHYERDNTSC | 1 | 0 | 0 | 1517828 | AXN92247.1 | K9N7C7 |
| TPVHYERDNTSG | 1 | 0 | 0 | 1517829 | AWM99580.1 | K9N7C7 |
| TPVIDDVDVVKP | 1 | 0 | 0 | 1517830 | AWK59942.1 | H9EJ42 |
| TPVNLNRGLPTG | 1 | 0 | 0 | 1517834 | AVP78031.1 | Q3LZX1 |
| TPVSGTTLHPVQ | 1 | 0 | 0 | 1517839 | AVV62528.1 | K9N4V0 |
| TPVSTFVGVLPP | 1 | 0 | 0 | 1517841 | AMM42635.1 | A0A0P0G321 |
| TPVSVKLPKGVG | 1 | 0 | 0 | 1517842 | AOL02450.1 | P0C6X6 |
| TPVVSDTVEVSP | 1 | 0 | 0 | 1517850 | ASU45805.1 | K9N7C7 |
| TPVVTETSEQNN | 1 | 0 | 0 | 1517851 | AVP78030.1 | P0C6W2 |
| TPWLLLSGKPTE | 1 | 0 | 0 | 1517856 | AVV62535.1 | K9N7C7 |
| TPYCHDPTVLPG | 1 | 0 | 0 | 1517861 | AXN92247.1 | K9N7C7 |
| TQALQAGLNGDF | 1 | 0 | 0 | 1517883 | AUM60014.1 | K9N5Q8 |
| TQANAVQQEFSV | 1 | 0 | 0 | 1517885 | AVV62524.1 | K9N7C7 |
| TQASLRMRDVSN | 1 | 0 | 0 | 1517889 | AVV62537.1 | K9N5Q8 |
| TQATQPLDAGLN | 1 | 0 | 0 | 1517890 | AVV62526.1 | K9N5Q8 |
| TQCFGPRDLDHN | 1 | 0 | 0 | 1517897 | AIW52723.1 | A0A0N7J131 |
| TQDGNAAIPDYN | 1 | 0 | 0 | 1517903 | ACV87150.1 | A0A140H1G9 |
| TQDHLDILGPLS | 1 | 0 | 0 | 1517908 | AVP78041.1 | P0C6W2 |
| TQDTQGVHLYSS | 1 | 0 | 0 | 1517913 | AVV62526.1 | K9N5Q8 |
| TQDYFLPFDTNL | 1 | 0 | 0 | 1517916 | ATO98193.1 | Q3LZX1 |
| TQDYQIGGFVEK | 1 | 0 | 0 | 1517917 | APO40580.1 | P59632 |
| TQEVFAQVKQMY | 1 | 0 | 0 | 1517928 | ATO98181.1 | Q3LZX1 |
| TQFAFACPDGVR | 1 | 0 | 0 | 1517934 | AVP78047.1 | Q3LZX7 |
| TQFAPGTQLPKN | 1 | 0 | 0 | 1517936 | AVV62533.1 | K9N4V7 |
| TQFNYFKKVDGV | 1 | 0 | 0 | 1517947 | APO40578.1 | P0C6X7 |
| TQFQGFVLNYKG | 1 | 0 | 0 | 1517949 | ATP16755.1 | Q80872 |
| TQFQKGKEFEFA | 1 | 0 | 0 | 1517951 | AEN19370.1 | P33469 |
| TQFQKGKEFKFA | 1 | 0 | 0 | 1517953 | ARA15425.1 | P33469 |
| TQFQKGKEFQFA | 1 | 0 | 0 | 1517954 | QBP84717.1 | P33469 |
| TQFVGNVKFDRW | 1 | 0 | 0 | 1517962 | AIW52729.1 | A0A0P0K6L9 |
| TQFVGNVKLDRW | 1 | 0 | 0 | 1517963 | AIW52740.1 | A0A0P0K6L9 |
| TQFVGVKFDRWS | 1 | 0 | 0 | 1517964 | AIW52757.1 | A0A0P0K6L9 |
| TQFVSNVKLDRW | 1 | 0 | 0 | 1517965 | AIW52741.1 | A0A0P0K6L9 |
| TQGRSCSDFEPM | 1 | 0 | 0 | 1517975 | AVV62524.1 | K9N7C7 |
| TQHGKVPLSFPP | 1 | 0 | 0 | 1517985 | AUM60021.1 | K9N4V7 |
| TQHGKVPLTFPS | 1 | 0 | 0 | 1517986 | AZK15907.1 | K9N4V7 |
| TQKPVDVKPKIK | 1 | 0 | 0 | 1518015 | ARO76381.1 | P0C6U8 |

| | | | | | | |
|---|---|---|---|---|---|---|
| TQLAEEPAKDIP | 1 | 0 | 0 | 1518020 | ATQ39389.1 | K9N7C7 |
| TQLFPLLFVTMA | 1 | 0 | 0 | 1518026 | AXN92247.1 | K9N7C7 |
| TQLLFVIEVVDK | 1 | 0 | 0 | 1518032 | AVP78041.1 | P0C6W2 |
| TQLPKNFHIEGT | 1 | 0 | 0 | 1518047 | AVV62533.1 | K9N4V7 |
| TQLVSLAEFSLN | 1 | 0 | 0 | 1518064 | AVV62541.1 | K9N7D2 |
| TQNEELEQTIQQ | 1 | 0 | 0 | 1518072 | APO40578.1 | P0C6X7 |
| TQNLFFMLLYAI | 1 | 0 | 0 | 1518074 | AWK59942.1 | H9EJ42 |
| TQPGSGVAVVDS | 1 | 0 | 0 | 1518086 | AVV62524.1 | K9N7C7 |
| TQPKLKEDYQIG | 1 | 0 | 0 | 1518088 | Q3I5J4.1 | Q3LZX0 |
| TQPLPNATYDNF | 1 | 0 | 0 | 1518091 | APO40578.1 | P0C6X7 |
| TQPQGNSIPHYS | 1 | 0 | 0 | 1518095 | AMM42386.1 | Q19U29 |
| TQQQQPQVVTRK | 1 | 0 | 0 | 1518111 | APO40586.1 | P59595 |
| TQQVLSENQKQI | 1 | 0 | 0 | 1518116 | AVV62526.1 | K9N5Q8 |
| TQRIIKAQFKTF | 1 | 0 | 0 | 1518124 | AEN19363.1 | P0C6X6 |
| TQRNFYSPQIIT | 1 | 0 | 0 | 1518130 | ATO98181.1 | Q3LZX1 |
| TQRSRPRAAVGQ | 1 | 0 | 0 | 1518136 | AVV62544.1 | K9N4V7 |
| TQSAIAKPKRLR | 1 | 0 | 0 | 1518141 | ATQ39389.1 | K9N7C7 |
| TQSALLVNNGSA | 1 | 0 | 0 | 1518143 | ATO98145.1 | Q3LZX1 |
| TQSAVLFNNGTH | 1 | 0 | 0 | 1518145 | APO40579.1 | P59594 |
| TQSFAPNTGVYD | 1 | 0 | 0 | 1518147 | BBA20986.1 | A0A140H1H1 |
| TQSIVRFPNITN | 1 | 0 | 0 | 1518150 | AVP78042.1 | Q3LZX1 |
| TQSRNLQEFKPR | 1 | 0 | 0 | 1518158 | QYC64967.1 | P0DTD1 |
| TQSSPIIPGFGG | 1 | 0 | 0 | 1518162 | AOR17484.1 | K9N5Q8 |
| TQSVNVKLKRLR | 1 | 0 | 0 | 1518166 | ANF29194.1 | K9N7C7 |
| TQSVTVKPKRLR | 1 | 0 | 0 | 1518173 | AXN92247.1 | K9N7C7 |
| TQTAQGAHLFSS | 1 | 0 | 0 | 1518177 | ALJ54512.1 | K9N5Q8 |
| TQTSQALQTVST | 1 | 0 | 0 | 1518186 | ABB90519.1 | A0A0P0K6L9 |
| TQVSTDTGVEHV | 1 | 0 | 0 | 1518198 | AVP78032.1 | Q3LZX0 |
| TQVYSRKARNRS | 1 | 0 | 0 | 1518215 | ATG84704.1 | K9N643 |
| TQVYSRKATKRS | 1 | 0 | 0 | 1518216 | AHC74084.1 | K9N643 |
| TQYITRMTAKPI | 1 | 0 | 0 | 1518229 | AVV62529.1 | K9N643 |
| TQYVYSPSNHKS | 1 | 0 | 0 | 1518236 | ATQ39390.1 | K9N5Q8 |
| TQYYFNKDTGVI | 1 | 0 | 0 | 1518237 | QEG03802.1 | P30215 |
| TRAASSAVKLQN | 1 | 0 | 0 | 1518239 | AUM60013.1 | K9N7C7 |
| TRCTSPGPSGIG | 1 | 0 | 0 | 1518267 | ALA49909.1 | K9N4V7 |
| TRDEAIKHVRAW | 1 | 0 | 0 | 1518271 | APO40578.1 | P0C6X7 |
| TRDGQVCYERFS | 1 | 0 | 0 | 1518274 | AVV62535.1 | K9N7C7 |
| TRDIAPTDTCFA | 1 | 0 | 0 | 1518275 | AVP78041.1 | P0C6W2 |
| TRDIVSTDDCFA | 1 | 0 | 0 | 1518280 | P0C6W6.1 | P0C6W2 |
| TRDMADQIASLV | 1 | 0 | 0 | 1518284 | AIL49470.1 | P33469 |
| TRDTVPLHIIAP | 1 | 0 | 0 | 1518292 | AUM60018.1 | K9N7D2 |
| TRDVFAQTKSIY | 1 | 0 | 0 | 1518294 | APO40579.1 | P59594 |
| TRDVVDQWYYTG | 1 | 0 | 0 | 1518296 | AVV62537.1 | K9N5Q8 |
| TREENVIQCFCP | 1 | 0 | 0 | 1518301 | AFD98754.1 | H9EJ69 |
| TREFGVTDLEDL | 1 | 0 | 0 | 1518304 | Q14EA9.1 | Q0ZJ84 |
| TREKLDKLISTA | 1 | 0 | 0 | 1518306 | AUM60013.1 | K9N7C7 |
| TREKLEKLISTA | 1 | 0 | 0 | 1518307 | ALD51903.1 | K9N7C7 |
| TRELKRELNGGA | 1 | 0 | 0 | 1518308 | AVP78041.1 | P0C6W2 |
| TRENKSAELNYT | 1 | 0 | 0 | 1518314 | BAL45640.1 | A0A0P0K6L9 |
| TRENVETLTWPL | 1 | 0 | 0 | 1518315 | AGT51393.1 | H9EJ42 |
| TRFITKSMISAV | 1 | 0 | 0 | 1518326 | ARU07575.1 | A0A140H1G9 |
| TRFPSVYAWERV | 1 | 0 | 0 | 1518331 | ATO98145.1 | Q3LZX1 |
| TRFVSKDVKFEN | 1 | 0 | 0 | 1518338 | AUM60014.1 | K9N5Q8 |
| TRGTFPGPSGIG | 1 | 0 | 0 | 1518355 | ANI69896.1 | K9N4V7 |
| TRGTSPGLSGIG | 1 | 0 | 0 | 1518357 | ALA49601.1 | K9N4V7 |
| TRHPGSPLSLNM | 1 | 0 | 0 | 1518361 | APO40587.1 | P59636 |
| TRILAKPTALEL | 1 | 0 | 0 | 1518370 | AVV62540.1 | K9N643 |
| TRKSASEASKKP | 1 | 0 | 0 | 1518380 | APO40586.1 | P59595 |
| TRLKGSFNNRDT | 1 | 0 | 0 | 1518395 | AEN19366.1 | P36334 |
| TRNNSRDSFRST | 1 | 0 | 0 | 1518415 | ABE97134.1 | H9EJ69 |
| TRNPNNDAAIVT | 1 | 0 | 0 | 1518420 | AVV62533.1 | K9N4V7 |
| TRNPNNDPAIVT | 1 | 0 | 0 | 1518422 | AUM60021.1 | K9N4V7 |
| TRPAGTNSLTCD | 1 | 0 | 0 | 1518431 | ARB66066.1 | P36334 |
| TRSKLKRRDSTY | 1 | 0 | 0 | 1518467 | ALJ54455.1 | K9N5Q8 |
| TRSLKYNIRSAL | 1 | 0 | 0 | 1518469 | AMM42558.1 | H9EJ42 |
| TRTAEAIHTVTI | 1 | 0 | 0 | 1518491 | QED88026.1 | A0A0P0G321 |
| TRTFTATTVVGS | 1 | 0 | 0 | 1518494 | AXN92247.1 | K9N7C7 |
| TRTLVAVNSEEI | 1 | 0 | 0 | 1518504 | AVV62524.1 | K9N7C7 |
| TRTTHTDIGVYI | 1 | 0 | 0 | 1518510 | ANI69888.1 | K9N638 |
| TRYSLADSNGFI | 1 | 0 | 0 | 1518529 | ABD75543.1 | A0A140H1G9 |
| TRYSLANSNGFI | 1 | 0 | 0 | 1518530 | ARU07575.1 | A0A140H1G9 |
| TSAAAIPFSLAL | 1 | 0 | 0 | 1518535 | AIW52843.1 | A0A0P0G321 |
| TSADIVDRDPSS | 1 | 0 | 0 | 1518543 | ARK08674.1 | P33469 |
| TSAFIDTVKGLD | 1 | 0 | 0 | 1518546 | ARO76381.1 | P0C6U8 |
| TSAGQPCVVSEQ | 1 | 0 | 0 | 1518554 | AVQ05261.1 | P0C6X6 |
| TSAGQPYVASEQ | 1 | 0 | 0 | 1518555 | AGT51707.1 | P0C6X6 |
| TSAGRPCVASEQ | 1 | 0 | 0 | 1518557 | QRK03919.1 | P0C6X6 |
| TSANLFPPWSAA | 1 | 0 | 0 | 1518568 | CAA83661.1 | P36334 |
| TSASLFPLWTAA | 1 | 0 | 0 | 1518574 | AAR01015.1 | P36334 |
| TSASLFPPWSAA | 1 | 0 | 0 | 1518575 | CAA83660.1 | P36334 |
| TSATSCQLYYNL | 1 | 0 | 0 | 1518581 | AIL49487.1 | P36334 |
| TSCELYHYQECV | 1 | 0 | 0 | 1518588 | ABI96964.1 | P59635 |
| TSCFFTCNSNAN | 1 | 0 | 0 | 1518589 | BAM16477.1 | A0A0P0G321 |
| TSCKEADMRNSS | 1 | 0 | 0 | 1518600 | AVV62537.1 | K9N5Q8 |
| TSCQIAVSKFLD | 1 | 0 | 0 | 1518606 | AHL18091.1 | K9N7C7 |
| TSCQTAVSKFLD | 1 | 0 | 0 | 1518613 | AVN89451.1 | K9N7C7 |
| TSCSNGRWLTGN | 1 | 0 | 0 | 1518620 | AUM60013.1 | K9N7C7 |
| TSDDDWTAAVDE | 1 | 0 | 0 | 1518624 | AVN89374.1 | K9N7C7 |
| TSDDEWAAAVDE | 1 | 0 | 0 | 1518625 | AVN89312.1 | K9N638 |
| TSDFTDSVRDPQ | 1 | 0 | 0 | 1518631 | ATO98181.1 | Q3LZX1 |
| TSDIKPVTYKLD | 1 | 0 | 0 | 1518634 | APO40578.1 | P0C6X7 |

| | | | | | | |
|---|---|---|---|---|---|---|
| TSDNNCWVNAVC | 1 | 0 | 0 | 1518643 | NP_073549.1 | A0A0P0LKV0 |
| TSDPQFPILAEL | 1 | 0 | 0 | 1518644 | AEN19370.1 | P33469 |
| TSDTSYINDKDT | 1 | 0 | 0 | 1518655 | CAA79896.1 | P36334 |
| TSDVYEADAEIV | 1 | 0 | 0 | 1518656 | AUM60013.1 | K9N7C7 |
| TSEPIVQHNTEP | 1 | 0 | 0 | 1518686 | APO40578.1 | P0C6X7 |
| TSEPPVTYLPAT | 1 | 0 | 0 | 1518688 | ANF29227.1 | K9N638 |
| TSEQIVAIIKDW | 1 | 0 | 0 | 1518691 | AVV62532.1 | K9N7A1 |
| TSEQIVAIVKDW | 1 | 0 | 0 | 1518692 | AVV62543.1 | K9N7A1 |
| TSEQNNFTGYLK | 1 | 0 | 0 | 1518693 | AVP78041.1 | P0C6W2 |
| TSFACLSNVSTS | 1 | 0 | 0 | 1518698 | QED88040.1 | A0A0P0G321 |
| TSFAIYHTPATD | 1 | 0 | 0 | 1518699 | ATG84866.1 | K9N5Q8 |
| TSFATYHTPATN | 1 | 0 | 0 | 1518701 | ALA49671.1 | K9N5Q8 |
| TSFATYHTPSTD | 1 | 0 | 0 | 1518702 | ALA49803.1 | K9N5Q8 |
| TSFATYHTPTTD | 1 | 0 | 0 | 1518703 | ALA49473.1 | K9N5Q8 |
| TSFATYYTPATD | 1 | 0 | 0 | 1518704 | ALA49352.1 | K9N5Q8 |
| TSFDFLSNVSTS | 1 | 0 | 0 | 1518708 | AWO68966.1 | A0A0P0G321 |
| TSGDGTTQPKLK | 1 | 0 | 0 | 1518729 | Q3I5J4.1 | Q3LZX0 |
| TSGFAVYVKSKV | 1 | 0 | 0 | 1518733 | BAM16435.1 | Q01455 |
| TSGNLVGFKDVT | 1 | 0 | 0 | 1518747 | CAA71146.1 | A0A0P0K6L9 |
| TSGNVIEFHLDG | 1 | 0 | 0 | 1518748 | APO40578.1 | P0C6X7 |
| TSGSTYLNMALK | 1 | 0 | 0 | 1518754 | BBA20982.1 | P36334 |
| TSGSTYRNMALK | 1 | 0 | 0 | 1518755 | CAA79896.1 | P36334 |
| TSGVTRDMADQI | 1 | 0 | 0 | 1518766 | AIL49470.1 | P33469 |
| TSHFTTQFVGVK | 1 | 0 | 0 | 1518777 | AIW52757.1 | A0A0P0K6L9 |
| TSHTVMPLTAPT | 1 | 0 | 0 | 1518790 | ATO98130.1 | P0C6W2 |
| TSIASFNEWAMA | 1 | 0 | 0 | 1518795 | AUM60013.1 | K9N7C7 |
| TSIEEVCDDYAK | 1 | 0 | 0 | 1518796 | AEN19363.1 | P0C6X6 |
| TSIKWADNNSYL | 1 | 0 | 0 | 1518802 | 4M0W_A | P0C6U8 |
| TSINTGNCPFSF | 1 | 0 | 0 | 1518813 | AAQ90004.1 | A0A0P0K6L9 |
| TSIPNLHDFKEE | 1 | 0 | 0 | 1518820 | AAB27260.2 | P36334 |
| TSIPNLPDFKGE | 1 | 0 | 0 | 1518822 | QDH43719.1 | P36334 |
| TSIRWADNNCYL | 1 | 0 | 0 | 1518827 | AVP78041.1 | P0C6W2 |
| TSKALAKQDAAR | 1 | 0 | 0 | 1518840 | AVV62539.1 | K9N4V0 |
| TSKLRAHDNILS | 1 | 0 | 0 | 1518853 | AUM60013.1 | K9N7C7 |
| TSKVKSLSMIRG | 1 | 0 | 0 | 1518861 | AIV41780.1 | P0C6X6 |
| TSLDWFPRPHIA | 1 | 0 | 0 | 1518864 | AUM60016.1 | K9N4V0 |
| TSLNTKYVAPHV | 1 | 0 | 0 | 1518887 | AID55097.1 | K9N5Q8 |
| TSLRRLVNPTDK | 1 | 0 | 0 | 1518895 | ATQ39389.1 | K9N7C7 |
| TSLSDACQLPYC | 1 | 0 | 0 | 1518896 | AGT17757.1 | A0A140H1H0 |
| TSLSFNNLLGFD | 1 | 0 | 0 | 1518898 | ANI69888.1 | K9N638 |
| TSLTSDYTITVF | 1 | 0 | 0 | 1518904 | AVV62524.1 | K9N7C7 |
| TSLVMGIGLCGF | 1 | 0 | 0 | 1518909 | AUM60013.1 | K9N7C7 |
| TSNCTKRLVQRV | 1 | 0 | 0 | 1518926 | ARO76381.1 | P0C6U8 |
| TSNDERLSPVDE | 1 | 0 | 0 | 1518927 | AVV62535.1 | K9N7C7 |
| TSNFRVSPTHEV | 1 | 0 | 0 | 1518934 | ATO98193.1 | Q3LZX1 |
| TSNGLLGNCAKR | 1 | 0 | 0 | 1518938 | AWK59942.1 | H9EJ42 |
| TSNLPDFREELD | 1 | 0 | 0 | 1518940 | AMK59677.1 | P36334 |
| TSNSFDALKLGD | 1 | 0 | 0 | 1518948 | AVP78041.1 | P0C6W2 |
| TSNVPPIVSEKI | 1 | 0 | 0 | 1518957 | AWK59942.1 | H9EJ42 |
| TSNVPSIVSEKI | 1 | 0 | 0 | 1518958 | AMM42558.1 | H9EJ42 |
| TSNVPSLVSEKI | 1 | 0 | 0 | 1518959 | AFD98756.1 | H9EJ42 |
| TSNVPTIVSEKI | 1 | 0 | 0 | 1518960 | P0C6U6.1 | H9EJ42 |
| TSNVSPIVNEKI | 1 | 0 | 0 | 1518961 | AGT51337.1 | H9EJ42 |
| TSPATVEPVRDE | 1 | 0 | 0 | 1518970 | AIW52723.1 | A0A0N7J131 |
| TSPCPSGIGAVG | 1 | 0 | 0 | 1518971 | QCI31476.1 | K9N4V7 |
| TSPGLSGIGAVG | 1 | 0 | 0 | 1518976 | ALA49601.1 | K9N4V7 |
| TSPGPSGIGTVG | 1 | 0 | 0 | 1518979 | ATG84708.1 | K9N4V7 |
| TSPIQPVELPAS | 1 | 0 | 0 | 1518981 | APT69862.1 | A0A0P0K6L9 |
| TSPPADVNTPAD | 1 | 0 | 0 | 1518987 | AJQ24074.1 | P33469 |
| TSQGFVVVNGKF | 1 | 0 | 0 | 1519003 | QAT98908.1 | K9N7C7 |
| TSQGFVVVNGKV | 1 | 0 | 0 | 1519004 | AXN92247.1 | K9N7C7 |
| TSQKAIVAYTMS | 1 | 0 | 0 | 1519005 | AVP78031.1 | Q3LZX1 |
| TSQQPLGGNVVP | 1 | 0 | 0 | 1519015 | AMK59681.1 | P33469 |
| TSQQPSGGNVIP | 1 | 0 | 0 | 1519016 | QEG03818.1 | P33469 |
| TSQSVCNGCVGH | 1 | 0 | 0 | 1519020 | ABB90520.1 | A0A0P0K6L9 |
| TSRKGSFNNKDT | 1 | 0 | 0 | 1519037 | AKA09663.1 | P36334 |
| TSRMLSYYKLGA | 1 | 0 | 0 | 1519042 | AYV99793.1 | P59596 |
| TSRTSGLASSAG | 1 | 0 | 0 | 1519055 | AJQ24068.1 | P33469 |
| TSSCDGIASDYR | 1 | 0 | 0 | 1519064 | AUM60013.1 | K9N7C7 |
| TSSGDASTAFAN | 1 | 0 | 0 | 1519073 | AKL87224.1 | P0C6X6 |
| TSSGFDKSYQTN | 1 | 0 | 0 | 1519075 | AUM60024.1 | K9N5Q8 |
| TSSIQPVELPVS | 1 | 0 | 0 | 1519078 | AMM42536.1 | A0A0P0K6L9 |
| TSSLAYYEPVQG | 1 | 0 | 0 | 1519082 | AXN92247.1 | K9N7C7 |
| TSSMRGCCDCGS | 1 | 0 | 0 | 1519086 | AIW52843.1 | A0A0P0G321 |
| TSSMTDNLQMAF | 1 | 0 | 0 | 1519088 | AUM60024.1 | K9N5Q8 |
| TSSPKVTIDCVA | 1 | 0 | 0 | 1519092 | AIL49523.1 | P36334 |
| TSSPKVTIDRAA | 1 | 0 | 0 | 1519093 | AIL49498.1 | P36334 |
| TSSRTSSAGSRS | 1 | 0 | 0 | 1519099 | ARK08665.1 | P33469 |
| TSSSLGSIAGAG | 1 | 0 | 0 | 1519106 | AOR17480.1 | K9N5Q8 |
| TSSSTYRNMALK | 1 | 0 | 0 | 1519107 | AIL49538.1 | P36334 |
| TSSTARTTSSEF | 1 | 0 | 0 | 1519108 | AUM60014.1 | K9N5Q8 |
| TSSTKKFQPFQQ | 1 | 0 | 0 | 1519111 | APO40579.1 | P59594 |
| TSTASALGKLQD | 1 | 0 | 0 | 1519126 | AVP78031.1 | Q3LZX1 |
| TSTDRFYRLGNE | 1 | 0 | 0 | 1519131 | AVL25540.1 | H9EJ42 |
| TSTEQVDVSHDV | 1 | 0 | 0 | 1519135 | AXG21664.1 | K9N638 |
| TSTGNYNYKYRC | 1 | 0 | 0 | 1519146 | BAE93401.1 | P59594 |
| TSTGNYNYKYRF | 1 | 0 | 0 | 1519147 | 3SCI_E | P59594 |
| TSTGNYNYKYRS | 1 | 0 | 0 | 1519148 | ATO98231.1 | Q3LZX1 |
| TSTHDGFNEHVT | 1 | 0 | 0 | 1519153 | AHI48551.1 | K9N796 |
| TSTHDGLIEHVT | 1 | 0 | 0 | 1519155 | QBF80533.1 | K9N796 |
| TSTHDGLNEHVT | 1 | 0 | 0 | 1519156 | ANF29196.1 | K9N796 |

| | | | | | | |
|---|---|---|---|---|---|---|
| TSTHDGLTEHVT | 1 | 0 | 0 | 1519157 | AXN92229.1 | K9N796 |
| TSTHDGPNEHVT | 1 | 0 | 0 | 1519159 | AHE78098.1 | K9N796 |
| TSTHDGPTEHVN | 1 | 0 | 0 | 1519160 | ANI69890.1 | K9N796 |
| TSTHPVQWECTF | 1 | 0 | 0 | 1519166 | AUM60016.1 | K9N4V0 |
| TSTNPEFVAFNV | 1 | 0 | 0 | 1519178 | AUM60013.1 | K9N7C7 |
| TSTPIVVDCATY | 1 | 0 | 0 | 1519179 | AIW52843.1 | A0A0P0G321 |
| TSTPIVVDCSTY | 1 | 0 | 0 | 1519180 | AIW52757.1 | A0A0P0K6L9 |
| TSTPNLPDFKEE | 1 | 0 | 0 | 1519182 | BBA20982.1 | P36334 |
| TSTPSTVYSPAT | 1 | 0 | 0 | 1519184 | AVV62524.1 | K9N7C7 |
| TSTSNLPDFREE | 1 | 0 | 0 | 1519191 | AMK59677.1 | P36334 |
| TSTTNFGEEFLG | 1 | 0 | 0 | 1519200 | ABD75599.1 | A0A140H1G9 |
| TSVAPDLSFDYI | 1 | 0 | 0 | 1519209 | AIX10753.1 | P36334 |
| TSVASFNEWAMA | 1 | 0 | 0 | 1519213 | AVV62535.1 | K9N7C7 |
| TSVECDFASMLN | 1 | 0 | 0 | 1519218 | AVV62526.1 | K9N5Q8 |
| TSVEESFIDDYY | 1 | 0 | 0 | 1519221 | AIW52827.1 | H9EJ42 |
| TSVLRQWLPTDA | 1 | 0 | 0 | 1519229 | AXN92247.1 | K9N7C7 |
| TSVTEDVKFAAS | 1 | 0 | 0 | 1519246 | AWK59948.1 | H9EJ42 |
| TSWVDQSERSHQ | 1 | 0 | 0 | 1519265 | AMM42397.1 | Q19U29 |
| TSYECDIPVGAG | 1 | 0 | 0 | 1519268 | ACQ82725.1 | P59594 |
| TSYGFSNVVEMP | 1 | 0 | 0 | 1519272 | AIW52757.1 | A0A0P0K6L9 |
| TSYGVWDTPTLD | 1 | 0 | 0 | 1519276 | AUM60014.1 | K9N5Q8 |
| TSYINDKDTGPP | 1 | 0 | 0 | 1519277 | CAA79896.1 | P36334 |
| TSYINDVDTGVP | 1 | 0 | 0 | 1519278 | ATP16757.1 | P36334 |
| TSYINNKDTGPP | 1 | 0 | 0 | 1519279 | AKA09646.1 | P36334 |
| TSYKGTFNNKDT | 1 | 0 | 0 | 1519281 | AKA09658.1 | P36334 |
| TSYSVCNGCVGH | 1 | 0 | 0 | 1519288 | ADK37704.1 | A0A0P0K6L9 |
| TSYSVVIQPRTI | 1 | 0 | 0 | 1519291 | BBO58758.1 | P36334 |
| TSYTANDVEVQS | 1 | 0 | 0 | 1519293 | QBM11736.1 | K9N638 |
| TSYTSDIKPVTY | 1 | 0 | 0 | 1519295 | APO40578.1 | P0C6X7 |
| TSYTTTILEHHH | 1 | 0 | 0 | 1519299 | 5Y3E_A | P0C6U8 |
| TTAETTDFLTVA | 1 | 0 | 0 | 1519310 | CAA71147.1 | A0A0P0K6L9 |
| TTAPLKLFAAET | 1 | 0 | 0 | 1519321 | AGV08437.1 | K9N7C7 |
| TTAQKLETDRIC | 1 | 0 | 0 | 1519323 | AXX83372.1 | P0C6X6 |
| TTATIPIQATLP | 1 | 0 | 0 | 1519331 | AVP78032.1 | Q3LZX0 |
| TTATIPLQASLP | 1 | 0 | 0 | 1519332 | ARO76383.1 | P59632 |
| TTAVYKPLVDFN | 1 | 0 | 0 | 1519340 | AST12962.1 | A0A0P0LKV0 |
| TTCAFSVDPATT | 1 | 0 | 0 | 1519344 | QED88017.1 | H9EJ42 |
| TTCANDPVGFIL | 1 | 0 | 0 | 1519345 | ARO76381.1 | P0C6U8 |
| TTCFEATTATIN | 1 | 0 | 0 | 1519350 | AUM60013.1 | K9N7C7 |
| TTCGYLPNNAVV | 1 | 0 | 0 | 1519355 | APO40578.1 | P0C6X7 |
| TTCGYLPTNAVM | 1 | 0 | 0 | 1519357 | ARO76381.1 | P0C6U8 |
| TTCLYIPKPTAK | 1 | 0 | 0 | 1519362 | AXN92262.1 | K9N4V0 |
| TTCQCGAERHRQ | 1 | 0 | 0 | 1519365 | AUM60013.1 | K9N7C7 |
| TTCSTDLQKSNT | 1 | 0 | 0 | 1519367 | CAA83660.1 | P36334 |
| TTCVNDPVGFTL | 1 | 0 | 0 | 1519369 | AVP78030.1 | P0C6W2 |
| TTCVTGPTTENK | 1 | 0 | 0 | 1519371 | ATO98145.1 | Q3LZX1 |
| TTDCSDGNYNRN | 1 | 0 | 0 | 1519373 | ALA49473.1 | K9N5Q8 |
| TTDILLKQPMAT | 1 | 0 | 0 | 1519377 | AHF54651.1 | Q4VID0 |
| TTDNTFVAGNCD | 1 | 0 | 0 | 1519383 | ATO98181.1 | Q3LZX1 |
| TTDVLLKQPMAI | 1 | 0 | 0 | 1519391 | AGT51675.1 | Q4VID0 |
| TTDVLLKQPMAT | 1 | 0 | 0 | 1519392 | APU51922.1 | Q4VID0 |
| TTDVPLKQPMAT | 1 | 0 | 0 | 1519393 | AHF54653.1 | Q4VID0 |
| TTEGYNNYVPSY | 1 | 0 | 0 | 1519396 | AVV62535.1 | K9N7C7 |
| TTENKLNISSGS | 1 | 0 | 0 | 1519403 | ATO98145.1 | Q3LZX1 |
| TTENLYFQGAMA | 1 | 0 | 0 | 1519404 | 3EJG_A | A0A0P0LKV0 |
| TTENRFTNFEPF | 1 | 0 | 0 | 1519405 | AIL49500.1 | P36334 |
| TTEVLPVSMTKT | 1 | 0 | 0 | 1519411 | APO40579.1 | P59594 |
| TTFDFLSNSSSS | 1 | 0 | 0 | 1519420 | AIW52843.1 | A0A0P0G321 |
| TTFLLNGYYPTS | 1 | 0 | 0 | 1519429 | CAA79896.1 | P36334 |
| TTFMPQFSRMTQ | 1 | 0 | 0 | 1519434 | AUM60014.1 | K9N5Q8 |
| TTFRPNMWCLRC | 1 | 0 | 0 | 1519435 | APO40578.1 | P0C6X7 |
| TTFTAPDFVAFN | 1 | 0 | 0 | 1519437 | AMQ48992.1 | K9N7C7 |
| TTFYPMCTNGKV | 1 | 0 | 0 | 1519439 | AVV62524.1 | K9N7C7 |
| TTGIITGNVVKQ | 1 | 0 | 0 | 1519451 | AVV62535.1 | K9N7C7 |
| TTGIPEHNFIIY | 1 | 0 | 0 | 1519453 | AVV62524.1 | K9N7C7 |
| TTGIVTGNVVKQ | 1 | 0 | 0 | 1519456 | AVV62524.1 | K9N7C7 |
| TTGLLLSSIYHL | 1 | 0 | 0 | 1519457 | AVV62524.1 | K9N7C7 |
| TTIHADQLTPAW | 1 | 0 | 0 | 1519481 | AVP78031.1 | Q3LZX1 |
| TTIQSVDISYLK | 1 | 0 | 0 | 1519500 | QDH43737.1 | H9EJ42 |
| TTIQSVDISYLN | 1 | 0 | 0 | 1519501 | AWK59942.1 | H9EJ42 |
| TTIQTVFDCAVE | 1 | 0 | 0 | 1519503 | AST12962.1 | A0A0P0LKV0 |
| TTITDKLGVCVE | 1 | 0 | 0 | 1519506 | AVV62537.1 | K9N5Q8 |
| TTKISLKGGKLN | 1 | 0 | 0 | 1519532 | APO40578.1 | P0C6X7 |
| TTKISLKGGKVI | 1 | 0 | 0 | 1519533 | ARO76381.1 | P0C6U8 |
| TTKVTVPKDHPH | 1 | 0 | 0 | 1519542 | AIW52712.1 | A0A0N7J131 |
| TTLAGLWLDDTV | 1 | 0 | 0 | 1519546 | 3FZD_A | P0C6X7 |
| TTLAVIGDFNCT | 1 | 0 | 0 | 1519547 | BBA20983.1 | A0A140H1H1 |
| TTLDNTSQSLLI | 1 | 0 | 0 | 1519554 | AVP78031.1 | Q3LZX1 |
| TTLDNTTQSALL | 1 | 0 | 0 | 1519555 | ATO98145.1 | Q3LZX1 |
| TTLGGLHLLISR | 1 | 0 | 0 | 1519566 | QED88024.1 | H9EJ42 |
| TTLLDLSDEMKV | 1 | 0 | 0 | 1519572 | AVV62526.1 | K9N5Q8 |
| TTLLDLSEEFRV | 1 | 0 | 0 | 1519573 | AUM60014.1 | K9N5Q8 |
| TTLLDLSTEMKV | 1 | 0 | 0 | 1519575 | AVV62537.1 | K9N5Q8 |
| TTLLDLTYEMLS | 1 | 0 | 0 | 1519576 | AXN92228.1 | K9N5Q8 |
| TTLLDLTYEMSS | 1 | 0 | 0 | 1519577 | ATG84756.1 | K9N5Q8 |
| TTLNGLWLDDIV | 1 | 0 | 0 | 1519580 | AID16715.1 | P0C6W2 |
| TTLPTFNTRTVS | 1 | 0 | 0 | 1519587 | AUM60023.1 | K9N7C7 |
| TTLQSLENVVYN | 1 | 0 | 0 | 1519595 | ABD75615.1 | A0A140H1G9 |
| TTLSTITAYLLV | 1 | 0 | 0 | 1519599 | AVA26874.1 | H9EJ24 |
| TTLTSDYTITVF | 1 | 0 | 0 | 1519603 | AXN92247.1 | K9N7C7 |
| TTLTSEQIVAIV | 1 | 0 | 0 | 1519605 | AVV62543.1 | K9N7A1 |

| | | | | | | |
|---|---|---|---|---|---|---|
| TTMMLSFNENGT | 1 | 0 | 0 | 1519615 | ATO98145.1 | Q3LZX1 |
| TTMMSQFSRDTQ | 1 | 0 | 0 | 1519616 | AVV62526.1 | K9N5Q8 |
| TTMSVIPPIDTK | 1 | 0 | 0 | 1519619 | AVV62535.1 | K9N7C7 |
| TTNEAFHKVQDA | 1 | 0 | 0 | 1519624 | AHI48803.1 | K9N5Q8 |
| TTNEAFRKVQDA | 1 | 0 | 0 | 1519626 | AXN92228.1 | K9N5Q8 |
| TTNFGEDFLGWI | 1 | 0 | 0 | 1519628 | ARU07575.1 | A0A140H1G9 |
| TTNGPLIVGGGC | 1 | 0 | 0 | 1519632 | AVP78030.1 | P0C6W2 |
| TTNGSWAIFNDF | 1 | 0 | 0 | 1519634 | AVV62535.1 | K9N7C7 |
| TTNGSWAIFNDY | 1 | 0 | 0 | 1519636 | AVV62524.1 | K9N7C7 |
| TTNLAFSKVQDA | 1 | 0 | 0 | 1519642 | AVV62537.1 | K9N5Q8 |
| TTNLDNKLQGLL | 1 | 0 | 0 | 1519644 | CAA83660.1 | P36334 |
| TTNPISEDDYQI | 1 | 0 | 0 | 1519648 | AVP78043.1 | Q3LZX0 |
| TTNVAFAKVQEA | 1 | 0 | 0 | 1519655 | AUM60014.1 | K9N5Q8 |
| TTPCELHGNGSM | 1 | 0 | 0 | 1519665 | ALW82645.1 | K9N7C7 |
| TTPCGVHGNGSM | 1 | 0 | 0 | 1519668 | AGV08556.1 | K9N7C7 |
| TTPCQVHGNGSM | 1 | 0 | 0 | 1519669 | AGV08533.1 | K9N7C7 |
| TTPLGMSLEACY | 1 | 0 | 0 | 1519677 | AEN19363.1 | P0C6X6 |
| TTPLGMSLGACY | 1 | 0 | 0 | 1519678 | QBP84728.1 | P0C6X6 |
| TTPVSTFVGVLP | 1 | 0 | 0 | 1519682 | AMM42635.1 | A0A0P0G321 |
| TTQASLRMRDVS | 1 | 0 | 0 | 1519685 | AVV62537.1 | K9N5Q8 |
| TTQDGDNKLQGL | 1 | 0 | 0 | 1519686 | AKA09658.1 | P36334 |
| TTQFVDNVKLAR | 1 | 0 | 0 | 1519690 | AIW52747.1 | A0A0P0K6L9 |
| TTQFVGAKFDRW | 1 | 0 | 0 | 1519691 | CAA71146.1 | A0A0P0K6L9 |
| TTQLQVANNLMN | 1 | 0 | 0 | 1519698 | AIL49500.1 | P36334 |
| TTQPKLKEDYQI | 1 | 0 | 0 | 1519700 | Q3I5J4.1 | Q3LZX0 |
| TTQSAIIVNNST | 1 | 0 | 0 | 1519705 | ATO98193.1 | Q3LZX1 |
| TTQSALLVNNGS | 1 | 0 | 0 | 1519706 | ATO98145.1 | Q3LZX1 |
| TTQSAVLFNNGT | 1 | 0 | 0 | 1519707 | APO40579.1 | P59594 |
| TTRDATNEWYYT | 1 | 0 | 0 | 1519711 | AUM60014.1 | K9N5Q8 |
| TTRDVVDQWYYT | 1 | 0 | 0 | 1519713 | AVV62537.1 | K9N5Q8 |
| TTRILAKPTALE | 1 | 0 | 0 | 1519715 | AVV62540.1 | K9N643 |
| TTSAALGKLQDV | 1 | 0 | 0 | 1519729 | APO40579.1 | P59594 |
| TTSGSNDIWMQN | 1 | 0 | 0 | 1519736 | AAA73152.1 | P30215 |
| TTSMAPDFVAFN | 1 | 0 | 0 | 1519745 | QBF80609.1 | K9N7C7 |
| TTSNFLPEVAAY | 1 | 0 | 0 | 1519749 | ATO98193.1 | Q3LZX1 |
| TTSQYSHRSAIE | 1 | 0 | 0 | 1519753 | AUM60024.1 | K9N5Q8 |
| TTSSFNADASVY | 1 | 0 | 0 | 1519758 | APO40579.1 | P59594 |
| TTSTNPEFVAFN | 1 | 0 | 0 | 1519761 | AUM60013.1 | K9N7C7 |
| TTTAPLKLFAAE | 1 | 0 | 0 | 1519766 | AGV08437.1 | K9N7C7 |
| TTTGIPEYNFII | 1 | 0 | 0 | 1519770 | AXN92247.1 | K9N7C7 |
| TTTIKPLEHHHH | 1 | 0 | 0 | 1519772 | 5E6J_A | P0C6X7 |
| TTTIKPVSLEHH | 1 | 0 | 0 | 1519773 | 4M0W_A | P0C6U8 |
| TTTLNGLWLDDI | 1 | 0 | 0 | 1519784 | AID16715.1 | P0C6W2 |
| TTTNEAFHKVQD | 1 | 0 | 0 | 1519793 | AHI48803.1 | K9N5Q8 |
| TTTNEAFRKVQD | 1 | 0 | 0 | 1519795 | AXN92228.1 | K9N5Q8 |
| TTTNLAFSKVQE | 1 | 0 | 0 | 1519798 | AVV62526.1 | K9N5Q8 |
| TTTNVAFAKVQE | 1 | 0 | 0 | 1519800 | AUM60014.1 | K9N5Q8 |
| TTTNVDKIRHLL | 1 | 0 | 0 | 1519801 | AHI48785.1 | K9N638 |
| TTTSAALGKLQD | 1 | 0 | 0 | 1519807 | APO40579.1 | P59594 |
| TTTVTSFHGYDF | 1 | 0 | 0 | 1519811 | AWK59942.1 | H9EJ42 |
| TTVAINDVDTGV | 1 | 0 | 0 | 1519815 | CAA83661.1 | P36334 |
| TTVASLINTLNN | 1 | 0 | 0 | 1519817 | AVP78041.1 | P0C6W2 |
| TTVDNINLHTQI | 1 | 0 | 0 | 1519820 | AVP78041.1 | P0C6W2 |
| TTVDNTNLHTQI | 1 | 0 | 0 | 1519822 | ARO76381.1 | P0C6U8 |
| TTVDNVNLHTQV | 1 | 0 | 0 | 1519824 | APO40578.1 | P0C6X7 |
| TTVDVNVLNDTV | 1 | 0 | 0 | 1519825 | AUM60013.1 | K9N7C7 |
| TTVDVSVLNDTV | 1 | 0 | 0 | 1519826 | AUM60023.1 | K9N7C7 |
| TTVESAAQADLQ | 1 | 0 | 0 | 1519831 | ATQ39389.1 | K9N7C7 |
| TTVLDRTDWCRC | 1 | 0 | 0 | 1519835 | BBA20983.1 | A0A140H1H1 |
| TTVLLKEPCSSG | 1 | 0 | 0 | 1519840 | AVP78047.1 | Q3LZX7 |
| TTVLNDPVFTGE | 1 | 0 | 0 | 1519843 | AGT51337.1 | H9EJ42 |
| TTVLNGPVFTGD | 1 | 0 | 0 | 1519844 | ABE97136.1 | H9EJ42 |
| TTVRSDGTFEEM | 1 | 0 | 0 | 1519848 | ATN39877.2 | P0C6X6 |
| TTVSINDIDTGA | 1 | 0 | 0 | 1519849 | CAA83660.1 | P36334 |
| TTVTVNVTTHNG | 1 | 0 | 0 | 1519857 | AIW52831.1 | A0A0P0G321 |
| TTYECDIPIGAG | 1 | 0 | 0 | 1519866 | APO40579.1 | P59594 |
| TTYPGQGHLGYT | 1 | 0 | 0 | 1519871 | APO40578.1 | P0C6X7 |
| TVAEEPDSRVPS | 1 | 0 | 0 | 1519881 | APT69887.1 | A0A0N7J131 |
| TVAKAPPGEKFK | 1 | 0 | 0 | 1519889 | ARU07574.1 | A0A140H1G9 |
| TVAQFNYRRDGQ | 1 | 0 | 0 | 1519902 | QAT98908.1 | K9N7C7 |
| TVASLINTLNNL | 1 | 0 | 0 | 1519906 | AVP78041.1 | P0C6W2 |
| TVATALNKIHDV | 1 | 0 | 0 | 1519907 | ABB90526.1 | A0A0P0K6L9 |
| TVATSRMLSYYK | 1 | 0 | 0 | 1519908 | AYV99793.1 | P59596 |
| TVAVPGGCNFPL | 1 | 0 | 0 | 1519911 | AWO68993.1 | A0A0P0G321 |
| TVCDSLNSKLGF | 1 | 0 | 0 | 1519922 | AEN19363.1 | P0C6X6 |
| TVCGPKLSTQLV | 1 | 0 | 0 | 1519929 | AVP78031.1 | Q3LZX1 |
| TVDANYKKCTKG | 1 | 0 | 0 | 1519948 | CAA71147.1 | A0A0P0K6L9 |
| TVDAQIQNKTVD | 1 | 0 | 0 | 1519949 | AVV62535.1 | K9N7C7 |
| TVDGINFRTCIL | 1 | 0 | 0 | 1519956 | AUM60013.1 | K9N7C7 |
| TVDNINLHTQIV | 1 | 0 | 0 | 1519965 | AVP78041.1 | P0C6W2 |
| TVDNTNLHTQIV | 1 | 0 | 0 | 1519967 | ARO76381.1 | P0C6U8 |
| TVDPKKAYMDFV | 1 | 0 | 0 | 1519970 | AVV62524.1 | K9N7C7 |
| TVDQLKHLLEQW | 1 | 0 | 0 | 1519975 | APO40582.1 | P59596 |
| TVDTTVESAAQA | 1 | 0 | 0 | 1519982 | ATQ39389.1 | K9N7C7 |
| TVDVQIQNNTVD | 1 | 0 | 0 | 1519988 | AVV62524.1 | K9N7C7 |
| TVDYFAYPLSMK | 1 | 0 | 0 | 1519992 | AVV62537.1 | K9N5Q8 |
| TVEAAFDLSKEL | 1 | 0 | 0 | 1519993 | QBQ01836.1 | P0C6X6 |
| TVEGYNNYVPSY | 1 | 0 | 0 | 1520004 | AVV62524.1 | K9N7C7 |
| TVELGTDVNEFA | 1 | 0 | 0 | 1520009 | AVP78041.1 | P0C6W2 |
| TVELQPPCKFLI | 1 | 0 | 0 | 1520013 | AUM60013.1 | K9N7C7 |
| TVEPVRDEVSTE | 1 | 0 | 0 | 1520016 | ABB90487.1 | A0A0N7J131 |

| | | | | | | |
|---|---|---|---|---|---|---|
| TVEQEAQIDRLI | 1 | 0 | 0 | 1520017 | AVV62526.1 | K9N5Q8 |
| TVESAAQADLQH | 1 | 0 | 0 | 1520018 | ATQ39389.1 | K9N7C7 |
| TVETALDLSKEL | 1 | 0 | 0 | 1520022 | AXX83306.1 | P0C6X6 |
| TVETPKEEQPKI | 1 | 0 | 0 | 1520024 | ARO76381.1 | P0C6U8 |
| TVEVSPQVVKLP | 1 | 0 | 0 | 1520036 | ASU45805.1 | K9N7C7 |
| TVFANTVEDAVN | 1 | 0 | 0 | 1520045 | AGT51403.1 | Q04853 |
| TVFSNMVETVVG | 1 | 0 | 0 | 1520059 | AVN89363.1 | K9N7C7 |
| TVFTDGRLNCTF | 1 | 0 | 0 | 1520060 | AUM60013.1 | K9N7C7 |
| TVFVCTDNSANT | 1 | 0 | 0 | 1520064 | AXN92247.1 | K9N7C7 |
| TVFVEPKDGGHF | 1 | 0 | 0 | 1520065 | ARE29978.1 | H9EJ42 |
| TVFVEPKDGGQF | 1 | 0 | 0 | 1520066 | AWK59942.1 | H9EJ42 |
| TVFVEPKNGGQF | 1 | 0 | 0 | 1520067 | ARU07593.1 | H9EJ42 |
| TVFVNIDNSANT | 1 | 0 | 0 | 1520068 | AVV62524.1 | K9N7C7 |
| TVFVPATRDTVP | 1 | 0 | 0 | 1520073 | AUM60018.1 | K9N7D2 |
| TVFVTVNLAKFL | 1 | 0 | 0 | 1520075 | AUM60013.1 | K9N7C7 |
| TVGGVPLLSTMT | 1 | 0 | 0 | 1520088 | ADK37692.1 | H9EJ42 |
| TVGPFMDNAINV | 1 | 0 | 0 | 1520091 | AXN92258.1 | K9N7C7 |
| TVGPYVEGAINV | 1 | 0 | 0 | 1520092 | AVV62535.1 | K9N7C7 |
| TVGQQDGSEDNQTT | 1 | 0 | 0 | 1520094 | P0DTC1.1 | P0DTD1 |
| TVGSGYCLDYSK | 1 | 0 | 0 | 1520095 | ARB66063.1 | P36334 |
| TVGVFLGYKETD | 1 | 0 | 0 | 1520101 | AMM42494.1 | A0A0P0ILY9 |
| TVHKSGFQLPVY | 1 | 0 | 0 | 1520115 | AIV41822.1 | P0C6X6 |
| TVIAKNGVKSGF | 1 | 0 | 0 | 1520123 | APO40578.1 | P0C6X7 |
| TVIISPSTRATI | 1 | 0 | 0 | 1520139 | AKL59401.1 | K9N5Q8 |
| TVILKEPCPSGT | 1 | 0 | 0 | 1520141 | 1XAK_A | P59635 |
| TVINNVDNEIIK | 1 | 0 | 0 | 1520144 | ATG84853.1 | K9N7C7 |
| TVINNVDNEIVR | 1 | 0 | 0 | 1520146 | AUM60013.1 | K9N7C7 |
| TVIPRSIRSYHA | 1 | 0 | 0 | 1520150 | AUM60014.1 | K9N5Q8 |
| TVIRADQLTPAW | 1 | 0 | 0 | 1520152 | ATO98145.1 | Q3LZX1 |
| TVIRTLNGDFLH | 1 | 0 | 0 | 1520154 | APO40578.1 | P0C6X7 |
| TVIWDFNREAPA | 1 | 0 | 0 | 1520159 | APO40578.1 | P0C6X7 |
| TVKATLRNKEKL | 1 | 0 | 0 | 1520164 | AVV62524.1 | K9N7C7 |
| TVKEIISIPNII | 1 | 0 | 0 | 1520167 | AIX09804.1 | P0C6X6 |
| TVKFCDHIVKAG | 1 | 0 | 0 | 1520174 | AVV61898.1 | K9N7C7 |
| TVKFCDQMVSAG | 1 | 0 | 0 | 1520178 | AVV62524.1 | K9N7C7 |
| TVKRKRLRKKRN | 1 | 0 | 0 | 1520194 | AXN92258.1 | K9N7C7 |
| TVKWADASETQR | 1 | 0 | 0 | 1520202 | AIW52715.1 | A0A0N7J131 |
| TVKWADESEPQR | 1 | 0 | 0 | 1520203 | ABB90491.1 | A0A0N7J131 |
| TVKYFECTGGID | 1 | 0 | 0 | 1520206 | AEN19363.1 | P0C6X6 |
| TVLAAYNGPPQG | 1 | 0 | 0 | 1520208 | ABD75599.1 | A0A140H1G9 |
| TVLEVQGYKNVR | 1 | 0 | 0 | 1520216 | ATO98229.1 | P0C6W2 |
| TVLLSEPCSPNT | 1 | 0 | 0 | 1520231 | APO40584.1 | P59635 |
| TVLNASIPESKPLADDDSA | 1 | 0 | 0 | 1520236 | AMM42589.1 | H9EJ69 |
| TVLNASTPESKP | 1 | 0 | 0 | 1520238 | AIW52805.1 | H9EJ69 |
| TVLNDPVFTGEL | 1 | 0 | 0 | 1520240 | AGT51337.1 | H9EJ42 |
| TVLNDPVFTGNL | 1 | 0 | 0 | 1520241 | AWK59948.1 | H9EJ42 |
| TVLPGAFAYSQM | 1 | 0 | 0 | 1520244 | AXN92247.1 | K9N7C7 |
| TVLPGAHAYETM | 1 | 0 | 0 | 1520245 | AVV62535.1 | K9N7C7 |
| TVLPGAHAYGSM | 1 | 0 | 0 | 1520246 | AVV62524.1 | K9N7C7 |
| TVLPGAHPYSAM | 1 | 0 | 0 | 1520247 | AUM60013.1 | K9N7C7 |
| TVLQNNELMPVK | 1 | 0 | 0 | 1520256 | AXX83324.1 | P0C6X6 |
| TVLRNSNGACIW | 1 | 0 | 0 | 1520260 | AVN89451.1 | K9N7C7 |
| TVLSVAPEVDWV | 1 | 0 | 0 | 1520268 | AWK59942.1 | H9EJ42 |
| TVLSYYYVMPLT | 1 | 0 | 0 | 1520269 | CAA83661.1 | P36334 |
| TVMPLTAPTLVP | 1 | 0 | 0 | 1520285 | ATO98130.1 | P0C6W2 |
| TVNLAKVCTYAV | 1 | 0 | 0 | 1520296 | AGV08596.1 | K9N7C7 |
| TVNNFVKFGSVC | 1 | 0 | 0 | 1520299 | BAL45640.1 | A0A0P0K6L9 |
| TVNSINDSLEPV | 1 | 0 | 0 | 1520300 | AIL49542.1 | P36334 |
| TVNSVNDSLEHV | 1 | 0 | 0 | 1520302 | CAA83661.1 | P36334 |
| TVNSVNDSLKPV | 1 | 0 | 0 | 1520303 | AIL49500.1 | P36334 |
| TVNVTTLNGRLV | 1 | 0 | 0 | 1520309 | AMM42642.1 | A0A0P0G321 |
| TVPEEFANHVAN | 1 | 0 | 0 | 1520315 | AVV62524.1 | K9N7C7 |
| TVPSTVWEDGDY | 1 | 0 | 0 | 1520332 | ANC28634.1 | K9N5Q8 |
| TVPTGYVDTENS | 1 | 0 | 0 | 1520333 | AVP78041.1 | P0C6W2 |
| TVPYVCLLANKE | 1 | 0 | 0 | 1520336 | AUM60013.1 | K9N7C7 |
| TVQADTAGLYTN | 1 | 0 | 0 | 1520337 | AHC74089.1 | K9N796 |
| TVQFCDAMRNAG | 1 | 0 | 0 | 1520342 | ATO98179.1 | P0C6W2 |
| TVQKGIYQTSNF | 1 | 0 | 0 | 1520347 | AVP78031.1 | Q3LZX1 |
| TVQKLQSLIDNI | 1 | 0 | 0 | 1520350 | BAL45640.1 | A0A0P0K6L9 |
| TVQPIANVYRRK | 1 | 0 | 0 | 1520355 | ABD43177.1 | P36334 |
| TVQSKDTNFLNA | 1 | 0 | 0 | 1520360 | ATN39869.1 | P0C6X6 |
| TVQSKMSDIKCT | 1 | 0 | 0 | 1520361 | APO40578.1 | P0C6X7 |
| TVREYIRYGIKP | 1 | 0 | 0 | 1520373 | ARU07575.1 | A0A140H1G9 |
| TVRSGNFNQDFY | 1 | 0 | 0 | 1520380 | AHI48750.1 | K9N7C7 |
| TVRVTKTLNTQY | 1 | 0 | 0 | 1520390 | AVV62524.1 | K9N7C7 |
| TVSINDIDTGAP | 1 | 0 | 0 | 1520403 | CAA83660.1 | P36334 |
| TVSLKQGEIKDA | 1 | 0 | 0 | 1520412 | AVP78043.1 | Q3LZX0 |
| TVSLLRNTSQKS | 1 | 0 | 0 | 1520414 | AAT76147.1 | Q3LZX1 |
| TVSSHGMVPVIH | 1 | 0 | 0 | 1520423 | AXG21669.1 | K9N7D2 |
| TVSTALNKIQDV | 1 | 0 | 0 | 1520430 | ABB90519.1 | A0A0P0K6L9 |
| TVSTAVSPDFKA | 1 | 0 | 0 | 1520431 | AVV62535.1 | K9N7C7 |
| TVTANLSIPSNW | 1 | 0 | 0 | 1520446 | AAK32189.1 | A0A0P0K6L9 |
| TVTDVKQVGCFM | 1 | 0 | 0 | 1520449 | AGT51518.1 | P0C6X6 |
| TVTKTSDWKCKV | 1 | 0 | 0 | 1520457 | AVN89451.1 | K9N7C7 |
| TVTSDGQPYVAS | 1 | 0 | 0 | 1520471 | AGT51657.1 | P0C6X6 |
| TVTYLNDPSSKT | 1 | 0 | 0 | 1520479 | NP_073549.1 | A0A0P0LKV0 |
| TVVEEPDSRAPS | 1 | 0 | 0 | 1520487 | AIW52723.1 | A0A0N7J131 |
| TVVFTVKIGGLT | 1 | 0 | 0 | 1520489 | AGT21344.1 | A0A0P0LKV0 |
| TVVIGTTKFCGG | 1 | 0 | 0 | 1520496 | ADK37693.1 | H9EJ42 |
| TVVLNNKNTYRS | 1 | 0 | 0 | 1520502 | AXN92247.1 | K9N7C7 |
| TVVLNNKSTYRS | 1 | 0 | 0 | 1520503 | AZK15899.1 | K9N638 |

| | | | | | | |
|---|---|---|---|---|---|---|
| TVVQLNYTRDGQ | 1 | 0 | 0 | 1520517 | AVV62535.1 | K9N7C7 |
| TVVVPGSCNFPL | 1 | 0 | 0 | 1520525 | AFD98750.1 | A0A0P0G321 |
| TVWKDGDYYRKQ | 1 | 0 | 0 | 1520534 | ATW75480.1 | K9N5Q8 |
| TVYDDATRRVWT | 1 | 0 | 0 | 1520542 | APO40578.1 | P0C6X7 |
| TVYDDSARRVWT | 1 | 0 | 0 | 1520544 | AVP78041.1 | P0C6W2 |
| TVYDPLQPELES | 1 | 0 | 0 | 1520546 | APO40579.1 | P59594 |
| TVYPDYVDVNKT | 1 | 0 | 0 | 1520553 | AWK59979.1 | A0A0P0G321 |
| TVYQVASPSECN | 1 | 0 | 0 | 1520556 | APO40578.1 | P0C6X7 |
| TVYREGYLNSTN | 1 | 0 | 0 | 1520557 | AVP78041.1 | P0C6W2 |
| TVYRRIPDLPDC | 1 | 0 | 0 | 1520558 | ABD75593.1 | A0A140H1H1 |
| TVYRRIPNLPDC | 1 | 0 | 0 | 1520559 | BBA20986.1 | A0A140H1H1 |
| TVYSDVESPHLM | 1 | 0 | 0 | 1520561 | AVP78041.1 | P0C6W2 |
| TVYSDVETPNLM | 1 | 0 | 0 | 1520563 | ASB51350.1 | P0C6X7 |
| TVYYPMCTNGQA | 1 | 0 | 0 | 1520576 | AUM60013.1 | K9N7C7 |
| TWLPELFMLNTL | 1 | 0 | 0 | 1520606 | AGT51548.1 | P0C6X6 |
| TWNKKFGFIEDS | 1 | 0 | 0 | 1520616 | ARB66070.1 | P36334 |
| TWNKRFGFIEDA | 1 | 0 | 0 | 1520617 | QEG03794.1 | P36334 |
| TWPLGLECTRAS | 1 | 0 | 0 | 1520621 | AHI48747.1 | K9N638 |
| TWPLVLECTRAA | 1 | 0 | 0 | 1520623 | AVV62524.1 | K9N7C7 |
| TWPLVLECTRAS | 1 | 0 | 0 | 1520624 | AXN92247.1 | K9N7C7 |
| TWPRPIDVFKAD | 1 | 0 | 0 | 1520625 | AHI48583.1 | K9N5Q8 |
| TWTPYPELMEDL | 1 | 0 | 0 | 1520639 | AUM60013.1 | K9N7C7 |
| TWWLKGDKLFVE | 1 | 0 | 0 | 1520644 | ARX11394.1 | A0A0P0LKV0 |
| TWWLKGDKLSVE | 1 | 0 | 0 | 1520645 | AST12962.1 | A0A0P0LKV0 |
| TYANIVHEPTTP | 1 | 0 | 0 | 1520662 | ALA49956.1 | K9N7C7 |
| TYANIVYEPSTP | 1 | 0 | 0 | 1520663 | AVV62524.1 | K9N7C7 |
| TYANIVYEPTTP | 1 | 0 | 0 | 1520664 | AXN92247.1 | K9N7C7 |
| TYDFNPNVPLEY | 1 | 0 | 0 | 1520675 | AVP78031.1 | Q3LZX1 |
| TYDPTYKNGAMY | 1 | 0 | 0 | 1520681 | AVN89451.1 | K9N7C7 |
| TYECDIPIGAGI | 1 | 0 | 0 | 1520691 | APO40579.1 | P59594 |
| TYEGNSPYHPLA | 1 | 0 | 0 | 1520694 | APO40584.1 | P59635 |
| TYERHPLSYFVN | 1 | 0 | 0 | 1520700 | APO40578.1 | P0C6X7 |
| TYFDNPILDFGD | 1 | 0 | 0 | 1520707 | ATO98181.1 | Q3LZX1 |
| TYFKVNGLWSPI | 1 | 0 | 0 | 1520714 | ADN03337.1 | A0A140H1G9 |
| TYFLVRILQDKT | 1 | 0 | 0 | 1520716 | YP_009047214.1 | K9N638 |
| TYFNKPLLVDDN | 1 | 0 | 0 | 1520721 | AIV41786.1 | P0C6X6 |
| TYFNLSSELKQL | 1 | 0 | 0 | 1520723 | ARE29967.1 | A0A0P0G321 |
| TYFQLTTSLVMG | 1 | 0 | 0 | 1520729 | AVV62524.1 | K9N7C7 |
| TYGQQFGPTYMD | 1 | 0 | 0 | 1520746 | APO40578.1 | P0C6X7 |
| TYGSGYIPEAPR | 1 | 0 | 0 | 1520747 | 5W9H_A | K9N5Q8 |
| TYGVCMYTDLNF | 1 | 0 | 0 | 1520756 | ARK08667.1 | P0C6X6 |
| TYINGFICEKLA | 1 | 0 | 0 | 1520776 | NP_073549.1 | A0A0P0LKV0 |
| TYIPASGYVAAS | 1 | 0 | 0 | 1520779 | AHI48543.1 | K9N7D2 |
| TYIPSQEKNFTT | 1 | 0 | 0 | 1520780 | AVP78031.1 | Q3LZX1 |
| TYKCPQSKSLVG | 1 | 0 | 0 | 1520794 | AWW13566.1 | P36334 |
| TYKLAMIMGYCC | 1 | 0 | 0 | 1520797 | AXX83319.1 | Q80872 |
| TYKLDGVTYTEL | 1 | 0 | 0 | 1520798 | APO40578.1 | P0C6X7 |
| TYKLVSMKGTGV | 1 | 0 | 0 | 1520801 | AWK59942.1 | H9EJ42 |
| TYLDFIQQGGIP | 1 | 0 | 0 | 1520819 | AIV41966.1 | P0C6X6 |
| TYLPATILAGSV | 1 | 0 | 0 | 1520837 | ANF29227.1 | K9N638 |
| TYLSVDTKFKTE | 1 | 0 | 0 | 1520845 | APO40578.1 | P0C6X7 |
| TYLTSRTSQQLT | 1 | 0 | 0 | 1520846 | AUM60013.1 | K9N7C7 |
| TYNCCDGDYFNK | 1 | 0 | 0 | 1520856 | AVP78041.1 | P0C6W2 |
| TYNKGENMTPRD | 1 | 0 | 0 | 1520863 | BAE93400.1 | P0C6U8 |
| TYNRVHGFEIDP | 1 | 0 | 0 | 1520870 | APO40579.1 | P59594 |
| TYPQQMFVTPLS | 1 | 0 | 0 | 1520882 | ALA49399.1 | K9N643 |
| TYQLRARAFTPR | 1 | 0 | 0 | 1520889 | APO40584.1 | P59635 |
| TYQNISFNLPPP | 1 | 0 | 0 | 1520891 | ANI69889.1 | K9N5Q8 |
| TYQNISINLPPP | 1 | 0 | 0 | 1520892 | ATG84888.1 | K9N5Q8 |
| TYRNMALKGTDL | 1 | 0 | 0 | 1520899 | AWW13566.1 | P36334 |
| TYSEAAAQGFQA | 1 | 0 | 0 | 1520904 | AWK59942.1 | H9EJ42 |
| TYSKLPDESGDV | 1 | 0 | 0 | 1520913 | AVV62524.1 | K9N7C7 |
| TYSNITLAYQGL | 1 | 0 | 0 | 1520920 | AUM60014.1 | K9N5Q8 |
| TYSPATILAGGV | 1 | 0 | 0 | 1520922 | QBM11736.1 | K9N638 |
| TYSVGEHSSPIA | 1 | 0 | 0 | 1520932 | AVV62524.1 | K9N7C7 |
| TYTELNPKLDGY | 1 | 0 | 0 | 1520939 | APO40578.1 | P0C6X7 |
| TYTSDLLTNSVI | 1 | 0 | 0 | 1520944 | ARO76381.1 | P0C6U8 |
| TYTWTTVLSMAV | 1 | 0 | 0 | 1520949 | AEN19363.1 | P0C6X6 |
| TYVCNGNPRCKN | 1 | 0 | 0 | 1520956 | AIW52843.1 | A0A0P0G321 |
| TYVCQCGGERHQ | 1 | 0 | 0 | 1520958 | AVN89322.1 | K9N7C7 |
| TYVCSLIAVAYT | 1 | 0 | 0 | 1520961 | AWK59942.1 | H9EJ42 |
| TYVPSQEKNFTT | 1 | 0 | 0 | 1520971 | ATO98181.1 | Q3LZX1 |
| TYVPSQQQNFTT | 1 | 0 | 0 | 1520973 | APO40579.1 | P59594 |
| TYVSVDDKYKTI | 1 | 0 | 0 | 1520974 | ALA50066.1 | K9N7C7 |
| TYVYTTSGFVVV | 1 | 0 | 0 | 1520983 | AVV62535.1 | K9N7C7 |
| TYVYTTTGFVVV | 1 | 0 | 0 | 1520984 | AVV62524.1 | K9N7C7 |
| TYYAHSVLYRVF | 1 | 0 | 0 | 1520986 | AXN92247.1 | K9N7C7 |
| TYYCVTTKAKSI | 1 | 0 | 0 | 1520988 | ALA49934.1 | K9N7C7 |
| TYYQPIQQAPTG | 1 | 0 | 0 | 1521003 | AST12967.1 | A0A0P0KXN3 |
| VAAAMASQKPQR | 1 | 0 | 0 | 1521008 | AVV62533.1 | K9N4V7 |
| VAAGQQFVKETT | 1 | 0 | 0 | 1521016 | AEN19363.1 | P0C6X6 |
| VAAIPYKACKSI | 1 | 0 | 0 | 1521020 | AVV62524.1 | K9N7C7 |
| VAALSTGLTNQV | 1 | 0 | 0 | 1521030 | AIW52827.1 | H9EJ42 |
| VAANVGRWSASI | 1 | 0 | 0 | 1521041 | ABB90509.1 | A0A0P0K6L9 |
| VAARADYEGFIT | 1 | 0 | 0 | 1521044 | ANI69867.1 | K9N638 |
| VAASDITLKCCT | 1 | 0 | 0 | 1521046 | NP_073549.1 | A0A0P0LKV0 |
| VAATRSVTVEYN | 1 | 0 | 0 | 1521049 | AVV62524.1 | K9N7C7 |
| VAAVDEAFPLDE | 1 | 0 | 0 | 1521050 | ASU45728.1 | K9N7C7 |
| VAAYNNCTGGAE | 1 | 0 | 0 | 1521066 | AVQ05264.1 | P36334 |
| VACEHITTMMSQ | 1 | 0 | 0 | 1521073 | AVV62526.1 | K9N5Q8 |
| VACSHITTFMPQ | 1 | 0 | 0 | 1521079 | AUM60014.1 | K9N5Q8 |

| | | | | | | |
|---|---|---|---|---|---|---|
| VADAMSAKEQRA | 1 | 0 | 0 | 1521082 | AUM60031.1 | K9N4V7 |
| VADDSGTPVVEC | 1 | 0 | 0 | 1521087 | AVN89451.1 | K9N7C7 |
| VADDSGTPVVEY | 1 | 0 | 0 | 1521088 | AHI48798.1 | K9N7C7 |
| VADFNAIATCDW | 1 | 0 | 0 | 1521090 | APO40578.1 | P0C6X7 |
| VADQSRKWTGHC | 1 | 0 | 0 | 1521093 | AIV41775.1 | Q80872 |
| VADSYFSYMMPM | 1 | 0 | 0 | 1521094 | ABD75559.1 | A0A140H1G9 |
| VADVMRYNFNFS | 1 | 0 | 0 | 1521097 | AVA26873.1 | A0A0P0G321 |
| VADVMRYNLNFS | 1 | 0 | 0 | 1521098 | AIW52843.1 | A0A0P0G321 |
| VAFCGDQVHEVA | 1 | 0 | 0 | 1521131 | ALT66879.1 | K9N7C7 |
| VAFGGDQIHEVA | 1 | 0 | 0 | 1521135 | ALA49340.1 | K9N7C7 |
| VAFGGDQVHEVV | 1 | 0 | 0 | 1521137 | ALA49384.1 | K9N7C7 |
| VAFNVFIGSETS | 1 | 0 | 0 | 1521148 | AUM60013.1 | K9N7C7 |
| VAGGLEATFLFL | 1 | 0 | 0 | 1521169 | AVP78043.1 | Q3LZX0 |
| VAHPLGYISHGC | 1 | 0 | 0 | 1521191 | AVV62524.1 | K9N7C7 |
| VAICLTYANIVH | 1 | 0 | 0 | 1521195 | ALA49956.1 | K9N7C7 |
| VAIIKDWNFAWS | 1 | 0 | 0 | 1521202 | AVV62532.1 | K9N7A1 |
| VAINDVDTGVPS | 1 | 0 | 0 | 1521217 | CAA83661.1 | P36334 |
| VAIQEVIDDMLW | 1 | 0 | 0 | 1521220 | ARK08668.1 | Q80872 |
| VAITRAQKGILC | 1 | 0 | 0 | 1521225 | AXN92247.1 | K9N7C7 |
| VAKAIAVAAGQQ | 1 | 0 | 0 | 1521235 | AEN19363.1 | P0C6X6 |
| VAKAPPGEQFKH | 1 | 0 | 0 | 1521236 | AEN19363.1 | P0C6X6 |
| VAKILARSQSSE | 1 | 0 | 0 | 1521248 | ABB90499.1 | A0A0N7J131 |
| VAKIYGMGFSSI | 1 | 0 | 0 | 1521249 | ABD43176.1 | P36334 |
| VAKKQIGLLEPV | 1 | 0 | 0 | 1521251 | AVV62524.1 | K9N7C7 |
| VAKLHNLTDSQR | 1 | 0 | 0 | 1521257 | AUM60027.1 | K9N643 |
| VAKNAYEKDKSV | 1 | 0 | 0 | 1521261 | AUM60013.1 | K9N7C7 |
| VAKPNVLIALKM | 1 | 0 | 0 | 1521266 | ASY99815.1 | K9N7A1 |
| VALADLYHYQEC | 1 | 0 | 0 | 1521282 | APO40584.1 | P59635 |
| VALEQDKNTQEV | 1 | 0 | 0 | 1521291 | ATO98108.1 | Q3LZX1 |
| VALPGELSKKQL | 1 | 0 | 0 | 1521301 | AUM60013.1 | K9N7C7 |
| VALPQSKDSNFL | 1 | 0 | 0 | 1521303 | AXN92258.1 | K9N7C7 |
| VALQTDVLQEDQ | 1 | 0 | 0 | 1521306 | BAL45639.1 | A0A0P0K6L9 |
| VALQTDVLQENQ | 1 | 0 | 0 | 1521307 | AIW52843.1 | A0A0P0G321 |
| VALRNSINADAY | 1 | 0 | 0 | 1521309 | AVV62524.1 | K9N7C7 |
| VALTQSKDTNFL | 1 | 0 | 0 | 1521312 | AUM60023.1 | K9N7C7 |
| VAMKYNYEPLVQ | 1 | 0 | 0 | 1521320 | APO40578.1 | P0C6X7 |
| VANDFTTALRRP | 1 | 0 | 0 | 1521328 | AKI29254.1 | K9N638 |
| VANGVTAKGYPQ | 1 | 0 | 0 | 1521340 | ABB90495.1 | A0A0N7J131 |
| VANLFSDIKTHK | 1 | 0 | 0 | 1521345 | AVV62537.1 | K9N5Q8 |
| VANLKLPPHDVI | 1 | 0 | 0 | 1521346 | AFD98800.1 | H9EJ24 |
| VANNLMNGVTLS | 1 | 0 | 0 | 1521349 | AIL49500.1 | P36334 |
| VANPTEDLCNGS | 1 | 0 | 0 | 1521355 | AGT51737.1 | P0C6X6 |
| VANPTNCIAPVN | 1 | 0 | 0 | 1521356 | AKM76239.1 | K9N5Q8 |
| VANSTGTVIISP | 1 | 0 | 0 | 1521358 | AGV08520.1 | K9N5Q8 |
| VANWLAPTNAYT | 1 | 0 | 0 | 1521361 | ANI69888.1 | K9N638 |
| VANWLAPTNVYM | 1 | 0 | 0 | 1521362 | ANC28699.1 | K9N638 |
| VANYFQDVKQFA | 1 | 0 | 0 | 1521364 | AID55096.1 | K9N5Q8 |
| VANYSQNVKQFA | 1 | 0 | 0 | 1521368 | AIY60528.1 | K9N5Q8 |
| VAPDLSFDYINV | 1 | 0 | 0 | 1521373 | AIX10753.1 | P36334 |
| VAPGETGVIADY | 1 | 0 | 0 | 1521378 | ATO98181.1 | Q3LZX1 |
| VAPGQTGVIADY | 1 | 0 | 0 | 1521380 | AVP78031.1 | Q3LZX1 |
| VAPHLSLDYINV | 1 | 0 | 0 | 1521386 | AQN78744.1 | P36334 |
| VAPLLSLDYINV | 1 | 0 | 0 | 1521390 | AQN78680.1 | P36334 |
| VAPPIADFTPVS | 1 | 0 | 0 | 1521391 | AVV62528.1 | K9N4V0 |
| VAPQATYQNIST | 1 | 0 | 0 | 1521392 | ALM26400.1 | K9N5Q8 |
| VAPTRSVTVEYN | 1 | 0 | 0 | 1521398 | AVV62535.1 | K9N7C7 |
| VAQFNYKQSFAN | 1 | 0 | 0 | 1521407 | AVV62526.1 | K9N5Q8 |
| VAQNTVLNACTP | 1 | 0 | 0 | 1521414 | ABK63972.1 | H9EJ69 |
| VAQNTVLNDSIP | 1 | 0 | 0 | 1521417 | AMM42614.1 | H9EJ69 |
| VAQQLVRSEAAA | 1 | 0 | 0 | 1521421 | AUM60014.1 | K9N5Q8 |
| VAQQLVRSETAA | 1 | 0 | 0 | 1521423 | AVV62526.1 | K9N5Q8 |
| VARDLSVQFKRP | 1 | 0 | 0 | 1521431 | APO40578.1 | P0C6X7 |
| VARNLNESLIDL | 1 | 0 | 0 | 1521439 | AVP78031.1 | Q3LZX1 |
| VASDGSPSGDAI | 1 | 0 | 0 | 1521447 | AVV62535.1 | K9N7C7 |
| VASDLTTSLRRL | 1 | 0 | 0 | 1521449 | ATQ39389.1 | K9N7C7 |
| VASDSDSLPSVP | 1 | 0 | 0 | 1521452 | AVV62541.1 | K9N7D2 |
| VASIPLCILPYT | 1 | 0 | 0 | 1521468 | ALD51903.1 | K9N7C7 |
| VASIQADVNTPA | 1 | 0 | 0 | 1521470 | QBQ01843.1 | P33469 |
| VASNDAIMTRCL | 1 | 0 | 0 | 1521477 | AXN92247.1 | K9N7C7 |
| VASNQADVNIPA | 1 | 0 | 0 | 1521478 | AIX10718.1 | P33469 |
| VASPKSRVQQNK | 1 | 0 | 0 | 1521480 | ATP16771.1 | P33469 |
| VASTLVEPPTVD | 1 | 0 | 0 | 1521490 | ATG84722.1 | K9N7C7 |
| VASVIAELNQLN | 1 | 0 | 0 | 1521497 | APO40578.1 | P0C6X7 |
| VATALNKIHDVV | 1 | 0 | 0 | 1521501 | ABB90526.1 | A0A0P0K6L9 |
| VATCYNSVNNDY | 1 | 0 | 0 | 1521502 | ARB07390.1 | A0A0P0LKV0 |
| VATDVQSRADQN | 1 | 0 | 0 | 1521503 | AUM60013.1 | K9N7C7 |
| VATKSYNVTQAF | 1 | 0 | 0 | 1521514 | AVV62544.1 | K9N4V7 |
| VATLMAPTIVNQ | 1 | 0 | 0 | 1521515 | AVN89451.1 | K9N7C7 |
| VATLTAPTIANQ | 1 | 0 | 0 | 1521518 | AXG21663.1 | K9N7C7 |
| VATSRTLSYYRL | 1 | 0 | 0 | 1521531 | APO40582.1 | P59596 |
| VATSSYDALLHR | 1 | 0 | 0 | 1521532 | AGT51379.1 | H9EJ42 |
| VATVFLFWFNFL | 1 | 0 | 0 | 1521533 | ARU07575.1 | A0A140H1G9 |
| VAVADSYFSYMM | 1 | 0 | 0 | 1521541 | ABD75559.1 | A0A140H1G9 |
| VAVKSGALHLYF | 1 | 0 | 0 | 1521556 | ARO76381.1 | P0C6U8 |
| VAVNNSKPYFIK | 1 | 0 | 0 | 1521563 | AVV62530.1 | K9N7D2 |
| VAVNNSKPYFLK | 1 | 0 | 0 | 1521564 | AVV62541.1 | K9N7D2 |
| VAVSKSRVQQNK | 1 | 0 | 0 | 1521581 | AIX10717.1 | P33469 |
| VAVTAGPFMGNA | 1 | 0 | 0 | 1521584 | ATQ39389.1 | K9N7C7 |
| VAWNLREMLAYA | 1 | 0 | 0 | 1521594 | APO40578.1 | P0C6X7 |
| VAWVKKQIVFFV | 1 | 0 | 0 | 1521596 | AVV62524.1 | K9N7C7 |
| VAYADNPNHFIN | 1 | 0 | 0 | 1521598 | ARU07566.1 | Q80872 |

| | | | | | | |
|---|---|---|---|---|---|---|
| VAYESLRPDIRY | 1 | 0 | 0 | 1521608 | AVP78041.1 | P0C6W2 |
| VAYHTVIDVKQV | 1 | 0 | 0 | 1521615 | AIV41894.1 | P0C6X6 |
| VAYQLYMIDPAG | 1 | 0 | 0 | 1521621 | AEN19363.1 | P0C6X6 |
| VAYVANPTEDLC | 1 | 0 | 0 | 1521636 | AEN19363.1 | P0C6X6 |
| VCAFLNKLNENV | 1 | 0 | 0 | 1521641 | ABD75607.1 | A0A140H1G9 |
| VCANEKFADGIN | 1 | 0 | 0 | 1521650 | APO40578.1 | P0C6X7 |
| VCAWLTKRKPLD | 1 | 0 | 0 | 1521655 | NP_073549.1 | A0A0P0LKV0 |
| VCDKVCSLKLSD | 1 | 0 | 0 | 1521666 | ALA49384.1 | K9N7C7 |
| VCDSLNAKWIFD | 1 | 0 | 0 | 1521673 | AIV41816.1 | P0C6X6 |
| VCDSLNAKWVFD | 1 | 0 | 0 | 1521674 | AIV41984.1 | P0C6X6 |
| VCDSLNFKLGFD | 1 | 0 | 0 | 1521675 | AAX85666.1 | P0C6X6 |
| VCEDVTQLAEEP | 1 | 0 | 0 | 1521677 | ATQ39389.1 | K9N7C7 |
| VCFDNSIQGSYE | 1 | 0 | 0 | 1521684 | AWK59942.1 | H9EJ42 |
| VCFDVRDKGALE | 1 | 0 | 0 | 1521685 | AVV62535.1 | K9N7C7 |
| VCFVDVDFHSVK | 1 | 0 | 0 | 1521702 | AWK59942.1 | H9EJ42 |
| VCGDYAACKSQL | 1 | 0 | 0 | 1521712 | CAA79896.1 | P36334 |
| VCGREATQYLVQ | 1 | 0 | 0 | 1521726 | ARO76381.1 | P0C6U8 |
| VCKFKGRFVQIP | 1 | 0 | 0 | 1521755 | AVV62535.1 | K9N7C7 |
| VCKGCVGHSENV | 1 | 0 | 0 | 1521756 | AIW52752.1 | A0A0P0K6L9 |
| VCKNCGQKSTTL | 1 | 0 | 0 | 1521762 | APO40578.1 | P0C6X7 |
| VCKVCGCWLNHG | 1 | 0 | 0 | 1521773 | NP_073549.1 | A0A0P0LKV0 |
| VCKVENKAEAEW | 1 | 0 | 0 | 1521774 | AST12962.1 | A0A0P0LKV0 |
| VCKYKGKFVQIP | 1 | 0 | 0 | 1521779 | AXN92247.1 | K9N7C7 |
| VCKYTDIDINPA | 1 | 0 | 0 | 1521781 | AVV62535.1 | K9N7C7 |
| VCKYTDIDVNTA | 1 | 0 | 0 | 1521784 | AUM60013.1 | K9N7C7 |
| VCLGGLDSLDSY | 1 | 0 | 0 | 1521794 | ARO76381.1 | P0C6U8 |
| VCLGISDSAVTY | 1 | 0 | 0 | 1521795 | QXL74882.1 | P0C6X6 |
| VCLLANKEQSLV | 1 | 0 | 0 | 1521805 | AVV62524.1 | K9N7C7 |
| VCLLANKEQSVV | 1 | 0 | 0 | 1521806 | AUM60013.1 | K9N7C7 |
| VCLSGLDSLDSY | 1 | 0 | 0 | 1521810 | P0C6W6.1 | P0C6W2 |
| VCMAFLTATRLC | 1 | 0 | 0 | 1521817 | ASU90554.1 | K9N5R3 |
| VCMETKAIVSTI | 1 | 0 | 0 | 1521820 | AVP78041.1 | P0C6W2 |
| VCNFNLCKEPMY | 1 | 0 | 0 | 1521827 | ATO98181.1 | Q3LZX1 |
| VCNFYFCQDPML | 1 | 0 | 0 | 1521828 | APO40579.1 | P59594 |
| VCNGCVGHSEIV | 1 | 0 | 0 | 1521829 | AST12964.1 | A0A0P0K6L9 |
| VCNGCVGHSKNV | 1 | 0 | 0 | 1521831 | AIW52741.1 | A0A0P0K6L9 |
| VCNGFEKCEQLL | 1 | 0 | 0 | 1521833 | AVV62526.1 | K9N5Q8 |
| VCNGFQKCEQLL | 1 | 0 | 0 | 1521836 | AXN92228.1 | K9N5Q8 |
| VCNGLQKCEQLL | 1 | 0 | 0 | 1521837 | QFS19588.1 | K9N5Q8 |
| VCNGNPRCKNLL | 1 | 0 | 0 | 1521838 | AIW52843.1 | A0A0P0G321 |
| VCNGRVGHSGNV | 1 | 0 | 0 | 1521840 | ADK37701.1 | A0A0P0K6L9 |
| VCNNSDYASKGY | 1 | 0 | 0 | 1521845 | AIX10745.1 | P0C6X6 |
| VCPKLEGSLEVL | 1 | 0 | 0 | 1521859 | 6C6Z_A | K9N5Q8 |
| VCPMQALRNDTS | 1 | 0 | 0 | 1521864 | AUM60014.1 | K9N5Q8 |
| VCQDSTRHTYQL | 1 | 0 | 0 | 1521869 | APO40584.1 | P59635 |
| VCQTTNGTNTSH | 1 | 0 | 0 | 1521871 | AIW52747.1 | A0A0P0K6L9 |
| VCQYNMCEHPHT | 1 | 0 | 0 | 1521873 | AWW13511.1 | P36334 |
| VCQYNMCEYPQT | 1 | 0 | 0 | 1521875 | CAA79896.1 | P36334 |
| VCRKHATDYREY | 1 | 0 | 0 | 1521890 | ASU91085.1 | K9N7C7 |
| VCRKHATEYREY | 1 | 0 | 0 | 1521892 | AXN92247.1 | K9N7C7 |
| VCSLKLSDNNCY | 1 | 0 | 0 | 1521905 | ALA49384.1 | K9N7C7 |
| VCSSVKDTLTYY | 1 | 0 | 0 | 1521911 | ATG84733.1 | K9N638 |
| VCVEYNLYGVSG | 1 | 0 | 0 | 1521940 | AVV62537.1 | K9N5Q8 |
| VCVFENKTTLPT | 1 | 0 | 0 | 1521942 | AVV62524.1 | K9N7C7 |
| VCVYDPLPVILL | 1 | 0 | 0 | 1521952 | ARB07432.1 | P30215 |
| VCYNSDYATKGY | 1 | 0 | 0 | 1521972 | AUM60013.1 | K9N7C7 |
| VDAKGFPLLAEL | 1 | 0 | 0 | 1521985 | AMM42619.1 | H9EJ69 |
| VDAKGFPQFAEL | 1 | 0 | 0 | 1521986 | AGT51335.1 | H9EJ69 |
| VDALDLVFNIFT | 1 | 0 | 0 | 1521990 | APO40578.1 | P0C6X7 |
| VDAMTYTSDLLT | 1 | 0 | 0 | 1521993 | ARO76381.1 | P0C6U8 |
| VDANYKKCTKGL | 1 | 0 | 0 | 1521998 | CAA71147.1 | A0A0P0K6L9 |
| VDARIEPCASGL | 1 | 0 | 0 | 1521999 | AVV62524.1 | K9N7C7 |
| VDAVQYAHKHDI | 1 | 0 | 0 | 1522002 | AVV62524.1 | K9N7C7 |
| VDAYVDTFSATF | 1 | 0 | 0 | 1522004 | BAE93400.1 | P0C6U8 |
| VDCATYVCNGNP | 1 | 0 | 0 | 1522006 | AIW52843.1 | A0A0P0G321 |
| VDCDAAGVGNTF | 1 | 0 | 0 | 1522007 | AVN89451.1 | K9N7C7 |
| VDCDIAGVGNTF | 1 | 0 | 0 | 1522008 | AXN92247.1 | K9N7C7 |
| VDCDTAGIGNTF | 1 | 0 | 0 | 1522009 | AVV62524.1 | K9N7C7 |
| VDCDTAGIGSTF | 1 | 0 | 0 | 1522010 | AUM60013.1 | K9N7C7 |
| VDCDTAGVGNTF | 1 | 0 | 0 | 1522011 | AHI48798.1 | K9N7C7 |
| VDCGHDDLSQLY | 1 | 0 | 0 | 1522013 | AVV62537.1 | K9N5Q8 |
| VDCKIQTAPHIS | 1 | 0 | 0 | 1522014 | AWW13564.1 | Q80872 |
| VDCKQYICNGFQ | 1 | 0 | 0 | 1522015 | AKM76239.1 | K9N5Q8 |
| VDCKQYVCNGFE | 1 | 0 | 0 | 1522017 | AVV62526.1 | K9N5Q8 |
| VDCKQYVCNGFH | 1 | 0 | 0 | 1522018 | ASU45807.1 | K9N5Q8 |
| VDCKQYVCNGLQ | 1 | 0 | 0 | 1522021 | QFS19588.1 | K9N5Q8 |
| VDCKSDFMSEIK | 1 | 0 | 0 | 1522022 | CAA83660.1 | P36334 |
| VDCMSDFMSGIK | 1 | 0 | 0 | 1522024 | AIL49491.1 | P36334 |
| VDCQDDRCFIHC | 1 | 0 | 0 | 1522026 | AIL49573.1 | P0C6X6 |
| VDCRRLLKQDCC | 1 | 0 | 0 | 1522027 | AZP73837.1 | P0C6X6 |
| VDCSQDPLAELK | 1 | 0 | 0 | 1522031 | ATO98145.1 | Q3LZX1 |
| VDCTMYICGDSI | 1 | 0 | 0 | 1522036 | AVP78031.1 | Q3LZX1 |
| VDCTMYICGDSL | 1 | 0 | 0 | 1522037 | Q3I5J5.1 | Q3LZX1 |
| VDCTMYICGDSQ | 1 | 0 | 0 | 1522038 | ATO98181.1 | Q3LZX1 |
| VDCVSDFMSEIK | 1 | 0 | 0 | 1522041 | AHA80206.1 | P36334 |
| VDCWTATDRFYR | 1 | 0 | 0 | 1522042 | ABQ16583.1 | H9EJ42 |
| VDDDYICGDGLI | 1 | 0 | 0 | 1522044 | QED88011.1 | H9EJ42 |
| VDDFQGYLQKHF | 1 | 0 | 0 | 1522046 | AIG96641.1 | A0A0P0LKV0 |
| VDDFYGYLQRHF | 1 | 0 | 0 | 1522047 | ATU79905.1 | A0A0P0LKV0 |
| VDDGYICGDGLI | 1 | 0 | 0 | 1522050 | AWK59942.1 | H9EJ42 |
| VDDHALIVNVLL | 1 | 0 | 0 | 1522052 | AFR79252.1 | A0A0P0KH29 |

| | | | | | | |
|---|---|---|---|---|---|---|
| VDDHVGLLDLAW | 1 | 0 | 0 | 1522054 | AIV41888.1 | P0C6X6 |
| VDDHVRLLDQAW | 1 | 0 | 0 | 1522056 | AGT51797.1 | P0C6X6 |
| VDDIDPINCESI | 1 | 0 | 0 | 1522057 | ABD75591.1 | A0A140H1G9 |
| VDDIDTINCESI | 1 | 0 | 0 | 1522058 | ARU07575.1 | A0A140H1G9 |
| VDDIESIYEFDT | 1 | 0 | 0 | 1522060 | ABD75535.1 | A0A140H1G9 |
| VDDIYGYVLRQP | 1 | 0 | 0 | 1522063 | ARE29979.1 | A0A0P0G321 |
| VDDKYKTIDELC | 1 | 0 | 0 | 1522064 | ALA50066.1 | K9N7C7 |
| VDDKYKTSDELC | 1 | 0 | 0 | 1522065 | AXN92247.1 | K9N7C7 |
| VDDLLKTDSILL | 1 | 0 | 0 | 1522067 | ABD75559.1 | A0A140H1G9 |
| VDDLVVDWHYFD | 1 | 0 | 0 | 1522068 | ASU91162.1 | K9N7C7 |
| VDDPCPIHFYSR | 1 | 0 | 0 | 1522070 | AVP78037.1 | Q3LZX5 |
| VDDPVLMGWDYP | 1 | 0 | 0 | 1522071 | AJS10811.1 | P0C6X6 |
| VDDPVVDWHYFD | 1 | 0 | 0 | 1522072 | AXN92247.1 | K9N7C7 |
| VDDPVVDWHYYD | 1 | 0 | 0 | 1522073 | AVV62524.1 | K9N7C7 |
| VDDQGHHLDSHF | 1 | 0 | 0 | 1522074 | AUM60013.1 | K9N7C7 |
| VDDQGHHLDSYF | 1 | 0 | 0 | 1522075 | AUM60023.1 | K9N7C7 |
| VDDSGDSSEGGA | 1 | 0 | 0 | 1522076 | AVQ05261.1 | P0C6X6 |
| VDDSGDSSESGA | 1 | 0 | 0 | 1522077 | AEN19363.1 | P0C6X6 |
| VDDSGTPVVEYL | 1 | 0 | 0 | 1522078 | AXN92247.1 | K9N7C7 |
| VDDSIILRQYNL | 1 | 0 | 0 | 1522079 | ARA15418.1 | P0C6X6 |
| VDDVNFSPVLGC | 1 | 0 | 0 | 1522082 | CAA83661.1 | P36334 |
| VDDVNGMKVVDY | 1 | 0 | 0 | 1522083 | AVV62535.1 | K9N7C7 |
| VDDWSYTGSSFY | 1 | 0 | 0 | 1522084 | ALJ54496.1 | K9N5Q8 |
| VDEAFPLDEAEN | 1 | 0 | 0 | 1522086 | QBF80465.1 | K9N7C7 |
| VDEAFPLNEAED | 1 | 0 | 0 | 1522089 | AHI48527.1 | K9N638 |
| VDECQLPLGQSL | 1 | 0 | 0 | 1522090 | AVV62526.1 | K9N5Q8 |
| VDECTLPIGQSL | 1 | 0 | 0 | 1522091 | AUM60014.1 | K9N5Q8 |
| VDEEFPLDDEES | 1 | 0 | 0 | 1522093 | AVV62535.1 | K9N7C7 |
| VDEEFPLEEEVQ | 1 | 0 | 0 | 1522094 | AVV62524.1 | K9N7C7 |
| VDESFVDDFQGY | 1 | 0 | 0 | 1522101 | AIG96641.1 | A0A0P0LKV0 |
| VDFCGRGYHLMS | 1 | 0 | 0 | 1522109 | ATO98181.1 | Q3LZX1 |
| VDFESVIQDYEN | 1 | 0 | 0 | 1522110 | APU51933.1 | P0C6X6 |
| VDFKDFQFMLWG | 1 | 0 | 0 | 1522117 | AIV41912.1 | P0C6X6 |
| VDFMEYHDVRVV | 1 | 0 | 0 | 1522124 | ATO98227.1 | Q3LZX5 |
| VDFNVNVIGTRP | 1 | 0 | 0 | 1522125 | AST12962.1 | A0A0P0LKV0 |
| VDFNVNVMGTRP | 1 | 0 | 0 | 1522126 | AGT21366.1 | A0A0P0LKV0 |
| VDFTECAYTQGR | 1 | 0 | 0 | 1522130 | ATQ39389.1 | K9N7C7 |
| VDGAKTEPTGHG | 1 | 0 | 0 | 1522132 | AAA45463.1 | A0A0N7J131 |
| VDGCTSSTCMMC | 1 | 0 | 0 | 1522137 | AVP78041.1 | P0C6W2 |
| VDGDVVIREPVH | 1 | 0 | 0 | 1522139 | ARU07575.1 | A0A140H1G9 |
| VDGFGLNKIQPW | 1 | 0 | 0 | 1522141 | AST12962.1 | A0A0P0LKV0 |
| VDGFNEWAIANG | 1 | 0 | 0 | 1522144 | AGT51358.1 | H9EJ42 |
| VDGGFKHLREFV | 1 | 0 | 0 | 1522145 | ATO98145.1 | Q3LZX1 |
| VDGHVGLLDQAW | 1 | 0 | 0 | 1522148 | AVZ61116.1 | P0C6X6 |
| VDGINFRTCILN | 1 | 0 | 0 | 1522153 | AUM60013.1 | K9N7C7 |
| VDGKPISAYAFL | 1 | 0 | 0 | 1522154 | AXN92247.1 | K9N7C7 |
| VDGKQIDDHRIT | 1 | 0 | 0 | 1522156 | AEN19363.1 | P0C6X6 |
| VDGQVDLFRNAR | 1 | 0 | 0 | 1522160 | APO40578.1 | P0C6X7 |
| VDGVNFRTCVVN | 1 | 0 | 0 | 1522163 | AVV62524.1 | K9N7C7 |
| VDGVPFVVSYGY | 1 | 0 | 0 | 1522168 | ASU90459.1 | K9N7C7 |
| VDGVSRRYQPLR | 1 | 0 | 0 | 1522171 | QED88026.1 | A0A0P0G321 |
| VDGYGNTMYVNM | 1 | 0 | 0 | 1522174 | AUM60027.1 | K9N643 |
| VDGYIRRAIDCG | 1 | 0 | 0 | 1522175 | AXN92228.1 | K9N5Q8 |
| VDGYIRRAVDCG | 1 | 0 | 0 | 1522176 | AVV62537.1 | K9N5Q8 |
| VDGYIRRSIDCG | 1 | 0 | 0 | 1522177 | AHZ90568.1 | K9N5Q8 |
| VDGYIRTAIDCG | 1 | 0 | 0 | 1522178 | AXN92260.1 | K9N5Q8 |
| VDGYVRRAIDCG | 1 | 0 | 0 | 1522179 | AUM60014.1 | K9N5Q8 |
| VDHIFAWQHSRC | 1 | 0 | 0 | 1522185 | AMQ49004.1 | K9N5Q8 |
| VDHKPHLSFPLC | 1 | 0 | 0 | 1522186 | NP_073549.1 | A0A0P0LKV0 |
| VDHRPVCSFPLC | 1 | 0 | 0 | 1522187 | AXN92247.1 | K9N7C7 |
| VDHSAFAYESAV | 1 | 0 | 0 | 1522188 | NP_073549.1 | A0A0P0LKV0 |
| VDHSGPTHYLRV | 1 | 0 | 0 | 1522189 | AVV62529.1 | K9N643 |
| VDHVQIHTIDGS | 1 | 0 | 0 | 1522190 | Q3I5J4.1 | Q3LZX0 |
| VDIIEQPFGEVE | 1 | 0 | 0 | 1522196 | AWK59942.1 | H9EJ42 |
| VDIIEQPFGEVK | 1 | 0 | 0 | 1522197 | QEG03747.1 | H9EJ42 |
| VDIIEQSFGEVE | 1 | 0 | 0 | 1522198 | ABE73411.1 | H9EJ42 |
| VDIIQRLDVLEQ | 1 | 0 | 0 | 1522199 | AGV08438.1 | K9N5Q8 |
| VDINKLCDEMLN | 1 | 0 | 0 | 1522206 | APO40578.1 | P0C6X7 |
| VDIPSIPKDMTY | 1 | 0 | 0 | 1522213 | APO40578.1 | P0C6X7 |
| VDIQPQVLFTCV | 1 | 0 | 0 | 1522214 | AUM60018.1 | K9N7D2 |
| VDIQQWGYIGSL | 1 | 0 | 0 | 1522216 | AEN19363.1 | P0C6X6 |
| VDIQTAQAAGTD | 1 | 0 | 0 | 1522217 | 3ATW_A | P0C6X7 |
| VDKDGNYYLPYP | 1 | 0 | 0 | 1522226 | QED88010.1 | H9EJ42 |
| VDKEIKDMQFEL | 1 | 0 | 0 | 1522227 | AUM60013.1 | K9N7C7 |
| VDKEVRDMQFDL | 1 | 0 | 0 | 1522230 | AVV62524.1 | K9N7C7 |
| VDKFAIPNRRRD | 1 | 0 | 0 | 1522231 | BBA20986.1 | A0A140H1H1 |
| VDKFKTPHIVRQ | 1 | 0 | 0 | 1522232 | AVN89322.1 | K9N7C7 |
| VDKIRHLLKGVT | 1 | 0 | 0 | 1522234 | AHI48785.1 | K9N638 |
| VDKPNHFINFPL | 1 | 0 | 0 | 1522238 | AXX83373.1 | Q80872 |
| VDKQVILVSNNKEDFDIIQ | 1 | 0 | 0 | 1522241 | ABD75503.1 | A0A140H1G9 |
| VDKVLNEKCSSY | 1 | 0 | 0 | 1522247 | APO40578.1 | P0C6X7 |
| VDLDALASDIFA | 1 | 0 | 0 | 1522253 | ASY99818.1 | K9N7C7 |
| VDLGTENLYFQS | 1 | 0 | 0 | 1522262 | 5X29_A | P59637 |
| VDLLSNFEHKVI | 1 | 0 | 0 | 1522268 | AKN58609.1 | K9N638 |
| VDLNTTILFTGY | 1 | 0 | 0 | 1522272 | ABA60800.1 | A0A140H1H1 |
| VDLNWLNRVETY | 1 | 0 | 0 | 1522273 | AGT21353.1 | A0A0P0K6L9 |
| VDLPFVEYKVTV | 1 | 0 | 0 | 1522274 | ARU07575.1 | A0A140H1G9 |
| VDLPPKLHFYYL | 1 | 0 | 0 | 1522275 | ALK28797.1 | A0A0N7J131 |
| VDLPPKVHFYYP | 1 | 0 | 0 | 1522276 | ABK63972.1 | H9EJ69 |
| VDLPPSHATTYL | 1 | 0 | 0 | 1522278 | QEG03783.1 | A0A0P0LKV0 |
| VDLQLGNLAYLQ | 1 | 0 | 0 | 1522280 | ARB66070.1 | P36334 |

| | | | | | | |
|---|---|---|---|---|---|---|
| VDLQQAVVNIRE | 1 | 0 | 0 | 1522282 | AVV62524.1 | K9N7C7 |
| VDLQSSGLLKPS | 1 | 0 | 0 | 1522283 | AUM60023.1 | K9N7C7 |
| VDLQSSGLVKPS | 1 | 0 | 0 | 1522284 | AUM60013.1 | K9N7C7 |
| VDLSMDKLRRAD | 1 | 0 | 0 | 1522286 | ATG84688.1 | K9N7C7 |
| VDLTEAICDFEI | 1 | 0 | 0 | 1522293 | AAX85666.1 | P0C6X6 |
| VDLTGAICDFEI | 1 | 0 | 0 | 1522294 | AEN19363.1 | P0C6X6 |
| VDLVISDMYDPV | 1 | 0 | 0 | 1522297 | AVV62524.1 | K9N7C7 |
| VDMQPDYFIKYN | 1 | 0 | 0 | 1522303 | ATQ39390.1 | K9N5Q8 |
| VDMYPEFSIVCR | 1 | 0 | 0 | 1522306 | AWK59942.1 | H9EJ42 |
| VDNCQLPLGQSL | 1 | 0 | 0 | 1522307 | AVV62537.1 | K9N5Q8 |
| VDNDLHPFVSDS | 1 | 0 | 0 | 1522308 | QDH43722.1 | P0C6X6 |
| VDNDLNPFVSDS | 1 | 0 | 0 | 1522309 | ARU07574.1 | A0A140H1G9 |
| VDNEIVRSSDVV | 1 | 0 | 0 | 1522311 | AUM60013.1 | K9N7C7 |
| VDNIYVTYAGNV | 1 | 0 | 0 | 1522315 | AEN19363.1 | P0C6X6 |
| VDNPAHTGRNYP | 1 | 0 | 0 | 1522318 | AJS10812.1 | P0C6X6 |
| VDNTLYSEFSTI | 1 | 0 | 0 | 1522322 | ACV87129.1 | A0A140H1H1 |
| VDNTNLHTQIVD | 1 | 0 | 0 | 1522323 | ARO76381.1 | P0C6U8 |
| VDNVKLPRWSAS | 1 | 0 | 0 | 1522325 | QNT54756.1 | A0A0P0K6L9 |
| VDNVNLHTQVVD | 1 | 0 | 0 | 1522327 | APO40578.1 | P0C6X7 |
| VDNVYMAKVGDK | 1 | 0 | 0 | 1522330 | AEN19363.1 | P0C6X6 |
| VDPDLSAYYVKD | 1 | 0 | 0 | 1522334 | AVV62524.1 | K9N7C7 |
| VDPFDAAVAAAM | 1 | 0 | 0 | 1522335 | AVV62544.1 | K9N4V7 |
| VDPLSNFEHKDI | 1 | 0 | 0 | 1522343 | ALA49659.1 | K9N7C7 |
| VDPNAFIYEWPM | 1 | 0 | 0 | 1522345 | AUM60014.1 | K9N5Q8 |
| VDPNLSGFYVKD | 1 | 0 | 0 | 1522346 | ATQ39389.1 | K9N7C7 |
| VDPQIQLTITAD | 1 | 0 | 0 | 1522348 | 2CME_A | P59636 |
| VDPQIQLTITDP | 1 | 0 | 0 | 1522349 | 2CME_E | P59636 |
| VDPQIQLTITRA | 1 | 0 | 0 | 1522350 | 2CME_B | P59636 |
| VDPQIQLTITRM | 1 | 0 | 0 | 1522351 | ARO76391.1 | P59636 |
| VDPQIQLTTTRM | 1 | 0 | 0 | 1522352 | Q3I5I6.1 | Q3LZX3 |
| VDPQKAYIDFVN | 1 | 0 | 0 | 1522355 | AUM60023.1 | K9N7C7 |
| VDPQKAYMDFVN | 1 | 0 | 0 | 1522357 | AUM60013.1 | K9N7C7 |
| VDPTTVLNGPVF | 1 | 0 | 0 | 1522359 | ABE97136.1 | H9EJ42 |
| VDQFERSHQTYN | 1 | 0 | 0 | 1522362 | AZS52622.1 | Q19U29 |
| VDQFNSSYFKLS | 1 | 0 | 0 | 1522363 | ASU90362.1 | K9N5Q8 |
| VDQLKHLLEQWN | 1 | 0 | 0 | 1522364 | APO40582.1 | P59596 |
| VDQLNSSYKLSI | 1 | 0 | 0 | 1522365 | ATW75477.1 | K9N5Q8 |
| VDQLNSTYFKLS | 1 | 0 | 0 | 1522366 | ANI69889.1 | K9N5Q8 |
| VDQQWYYTGSSF | 1 | 0 | 0 | 1522367 | AVV62526.1 | K9N5Q8 |
| VDQSERSHQTYN | 1 | 0 | 0 | 1522369 | AMM42397.1 | Q19U29 |
| VDQSRKWIGHCT | 1 | 0 | 0 | 1522370 | ATP16755.1 | Q80872 |
| VDQSRKWISHCT | 1 | 0 | 0 | 1522371 | AIV41836.1 | Q80872 |
| VDQWYYTGSSFF | 1 | 0 | 0 | 1522373 | AVV62537.1 | K9N5Q8 |
| VDRDPSNDEAIP | 1 | 0 | 0 | 1522375 | AJQ24082.1 | P33469 |
| VDRDSAVGHTEF | 1 | 0 | 0 | 1522379 | AUM60025.1 | K9N796 |
| VDRELTSTHDGP | 1 | 0 | 0 | 1522380 | AVN89314.1 | K9N796 |
| VDRESTSIHDGP | 1 | 0 | 0 | 1522382 | AVN89384.1 | K9N796 |
| VDRIRFCLMLGS | 1 | 0 | 0 | 1522392 | ATQ39397.1 | K9N7D2 |
| VDRIRFCLMLNS | 1 | 0 | 0 | 1522393 | AXG21669.1 | K9N7D2 |
| VDSCLQGDRCNI | 1 | 0 | 0 | 1522400 | CAA83660.1 | P36334 |
| VDSDQPWKVIPR | 1 | 0 | 0 | 1522404 | AST12968.1 | A0A0N7J131 |
| VDSEQPCLVIPR | 1 | 0 | 0 | 1522407 | AMM42513.1 | A0A0N7J131 |
| VDSEQPWKVVPR | 1 | 0 | 0 | 1522409 | AIW52723.1 | A0A0N7J131 |
| VDSEQSWKVIPR | 1 | 0 | 0 | 1522410 | AAA45463.1 | A0A0N7J131 |
| VDSFTCNNIDAA | 1 | 0 | 0 | 1522413 | AMK59677.1 | P36334 |
| VDSFTNGAVVGT | 1 | 0 | 0 | 1522417 | P0C6W6.1 | P0C6W2 |
| VDSFVVKGDDVR | 1 | 0 | 0 | 1522418 | ABD72969.1 | P59594 |
| VDSGDASPQVLK | 1 | 0 | 0 | 1522419 | AVV62535.1 | K9N7C7 |
| VDSGPPSSATSC | 1 | 0 | 0 | 1522420 | AVV62537.1 | K9N5Q8 |
| VDSIVQRCYELS | 1 | 0 | 0 | 1522423 | ABE73414.1 | H9EJ42 |
| VDSKCLTSIEGV | 1 | 0 | 0 | 1522425 | QEG03737.1 | P0C6X6 |
| VDSKSGNNFYYR | 1 | 0 | 0 | 1522426 | APO40579.1 | P59594 |
| VDSKYQVIVLKD | 1 | 0 | 0 | 1522427 | ALA49340.1 | K9N7C7 |
| VDSKYQVIVMKD | 1 | 0 | 0 | 1522428 | AUM60013.1 | K9N7C7 |
| VDSMNLIANIFT | 1 | 0 | 0 | 1522429 | ARO76381.1 | P0C6U8 |
| VDSNENLTWPLG | 1 | 0 | 0 | 1522430 | AHI48747.1 | K9N638 |
| VDSRWNNARHSD | 1 | 0 | 0 | 1522433 | AQN78751.1 | P30215 |
| VDSRWNNARLSD | 1 | 0 | 0 | 1522434 | AAX85668.1 | P30215 |
| VDSRWNNARQFD | 1 | 0 | 0 | 1522435 | AXX83326.1 | P30215 |
| VDSRWNNLRASD | 1 | 0 | 0 | 1522438 | AGT17757.1 | A0A140H1H0 |
| VDSTPVEPPTVD | 1 | 0 | 0 | 1522443 | QGV13494.1 | K9N7C7 |
| VDSVAVKSGALH | 1 | 0 | 0 | 1522445 | ARO76381.1 | P0C6U8 |
| VDSVSEEPQGNV | 1 | 0 | 0 | 1522448 | ARB07436.1 | A0A140H1G9 |
| VDSVTVKETVAQ | 1 | 0 | 0 | 1522449 | QAT98908.1 | K9N7C7 |
| VDSVTVKETVVQ | 1 | 0 | 0 | 1522450 | AXN92247.1 | K9N7C7 |
| VDSVTVKNGTLY | 1 | 0 | 0 | 1522452 | APO40578.1 | P0C6X7 |
| VDSVVRSFQPLL | 1 | 0 | 0 | 1522453 | AAQ90005.1 | A0A0P0K6L9 |
| VDSVVVKDTVVQ | 1 | 0 | 0 | 1522454 | AUM60013.1 | K9N7C7 |
| VDSYYSYLMPAL | 1 | 0 | 0 | 1522456 | QBF80541.1 | K9N7C7 |
| VDTAGQSIQALQ | 1 | 0 | 0 | 1522457 | APO40578.1 | P0C6X7 |
| VDTEHSTEFTRV | 1 | 0 | 0 | 1522463 | AVP78030.1 | P0C6W2 |
| VDTENSTEFTRV | 1 | 0 | 0 | 1522466 | AVP78041.1 | P0C6W2 |
| VDTEWGNMMTCI | 1 | 0 | 0 | 1522468 | AVV62524.1 | K9N7C7 |
| VDTEWGNMMTGI | 1 | 0 | 0 | 1522469 | AVV62535.1 | K9N7C7 |
| VDTEWGSMLTSI | 1 | 0 | 0 | 1522470 | AUM60013.1 | K9N7C7 |
| VDTFVSTFNVPM | 1 | 0 | 0 | 1522475 | APO40578.1 | P0C6X7 |
| VDTGPPSISTAT | 1 | 0 | 0 | 1522476 | AMK59677.1 | P36334 |
| VDTGVPPISTDT | 1 | 0 | 0 | 1522477 | ATP16757.1 | P36334 |
| VDTGVPSTSTDI | 1 | 0 | 0 | 1522478 | CAA83661.1 | P36334 |
| VDTHRYCLSLKD | 1 | 0 | 0 | 1522480 | ANZ78856.1 | P0C6X6 |
| VDTHRYRLSLKD | 1 | 0 | 0 | 1522481 | AEN19363.1 | P0C6X6 |

| | | | | | | |
|---|---|---|---|---|---|---|
| VDTKDVVECLKL | 1 | 0 | 0 | 1522484 | APO40578.1 | P0C6X7 |
| VDTLQETPVVSD | 1 | 0 | 0 | 1522486 | AVK87459.1 | K9N7C7 |
| VDTLRDLPLSLT | 1 | 0 | 0 | 1522487 | ATO98145.1 | Q3LZX1 |
| VDTNCLADSVMS | 1 | 0 | 0 | 1522489 | AIV41996.1 | P0C6X6 |
| VDTSANTEFTRV | 1 | 0 | 0 | 1522491 | APO40578.1 | P0C6X7 |
| VDTSHFTTQFFG | 1 | 0 | 0 | 1522493 | ABB90529.1 | A0A0P0K6L9 |
| VDTSHFTTQFVS | 1 | 0 | 0 | 1522494 | AIW52741.1 | A0A0P0K6L9 |
| VDTSQPTHYLRL | 1 | 0 | 0 | 1522499 | AUM60027.1 | K9N643 |
| VDTSYECDIPVG | 1 | 0 | 0 | 1522502 | ACQ82725.1 | P59594 |
| VDTTVESAAQAD | 1 | 0 | 0 | 1522507 | ATQ39389.1 | K9N7C7 |
| VDTVSTLVYDNK | 1 | 0 | 0 | 1522512 | APO40578.1 | P0C6X7 |
| VDTVVNGVEFDF | 1 | 0 | 0 | 1522513 | AGT21350.1 | A0A0P0LKV0 |
| VDVANPSTPVYT | 1 | 0 | 0 | 1522514 | QBF80585.1 | K9N7C7 |
| VDVDFHSVEVEA | 1 | 0 | 0 | 1522518 | ARU07593.1 | H9EJ42 |
| VDVDFHSVKPFA | 1 | 0 | 0 | 1522519 | AWK59942.1 | H9EJ42 |
| VDVFHETSDDEW | 1 | 0 | 0 | 1522525 | AVN89352.1 | K9N7C7 |
| VDVFTDGKLNCT | 1 | 0 | 0 | 1522526 | AVV62524.1 | K9N7C7 |
| VDVGPDSAKSVC | 1 | 0 | 0 | 1522531 | QAT98898.1 | K9N5Q8 |
| VDVGPDSFKSAC | 1 | 0 | 0 | 1522532 | AQZ41296.1 | K9N5Q8 |
| VDVGPDSIKSAC | 1 | 0 | 0 | 1522533 | QBF80543.1 | K9N5Q8 |
| VDVGPDSLKSAC | 1 | 0 | 0 | 1522534 | QBF80608.1 | K9N5Q8 |
| VDVGPDSVKPAC | 1 | 0 | 0 | 1522535 | ASU91284.1 | K9N5Q8 |
| VDVLPEGWHVVN | 1 | 0 | 0 | 1522545 | AIV41876.1 | P0C6X6 |
| VDVLPEGWRVVN | 1 | 0 | 0 | 1522546 | AEN19363.1 | P0C6X6 |
| VDVLVPSNKWTG | 1 | 0 | 0 | 1522547 | AVV62524.1 | K9N7C7 |
| VDVNDCFKNDNV | 1 | 0 | 0 | 1522548 | AGT51379.1 | H9EJ42 |
| VDVNKTLQEFAK | 1 | 0 | 0 | 1522549 | AWK59979.1 | A0A0P0G321 |
| VDVNVLNDTVEV | 1 | 0 | 0 | 1522551 | AUM60013.1 | K9N7C7 |
| VDVPPQVVELPS | 1 | 0 | 0 | 1522554 | AVN89352.1 | K9N7C7 |
| VDVPPQVVKLPS | 1 | 0 | 0 | 1522555 | ASU91261.1 | K9N638 |
| VDVQIQNNTVDV | 1 | 0 | 0 | 1522556 | AVV62524.1 | K9N7C7 |
| VDVQQWGYIGNL | 1 | 0 | 0 | 1522557 | AVV62524.1 | K9N7C7 |
| VDVQQWGYVGNL | 1 | 0 | 0 | 1522558 | AXN92247.1 | K9N7C7 |
| VDVQQWGYVSNL | 1 | 0 | 0 | 1522559 | ATG84677.1 | K9N7C7 |
| VDVSDTIPEQKQ | 1 | 0 | 0 | 1522560 | AVV62524.1 | K9N7C7 |
| VDVSHDVSDDEW | 1 | 0 | 0 | 1522562 | AXG21664.1 | K9N638 |
| VDVSHETSDDDW | 1 | 0 | 0 | 1522564 | AVN89374.1 | K9N7C7 |
| VDVSHEVFDDEW | 1 | 0 | 0 | 1522566 | AQZ41284.1 | K9N7C7 |
| VDVSHGLGTYYI | 1 | 0 | 0 | 1522568 | ARU07577.1 | A0A140H1H1 |
| VDVTNGLGTYYI | 1 | 0 | 0 | 1522570 | AIX10756.1 | P36334 |
| VDVVRFPNITNL | 1 | 0 | 0 | 1522575 | ATO98145.1 | Q3LZX1 |
| VDVVVGQLEETN | 1 | 0 | 0 | 1522579 | AVV62524.1 | K9N7C7 |
| VDVWDMYLTGCD | 1 | 0 | 0 | 1522580 | ARU07575.1 | A0A140H1G9 |
| VDVYDMWLTGCK | 1 | 0 | 0 | 1522581 | AEN19363.1 | P0C6X6 |
| VDVYDNFPQGFS | 1 | 0 | 0 | 1522582 | APO40579.1 | P59594 |
| VDWHYYDAQPLR | 1 | 0 | 0 | 1522587 | AVV62524.1 | K9N7C7 |
| VDWKASLRLLLI | 1 | 0 | 0 | 1522589 | AVV62524.1 | K9N7C7 |
| VDWKFYDAQPCS | 1 | 0 | 0 | 1522592 | AVP78041.1 | P0C6W2 |
| VDWRSCNYAVSS | 1 | 0 | 0 | 1522598 | ALD51903.1 | K9N7C7 |
| VDYFSYPLSMKS | 1 | 0 | 0 | 1522615 | AVV62526.1 | K9N5Q8 |
| VDYLTREDMHQL | 1 | 0 | 0 | 1522621 | AVV62524.1 | K9N7C7 |
| VDYNVVTQRMCA | 1 | 0 | 0 | 1522623 | AWK59942.1 | H9EJ42 |
| VDYSKNGGSGGA | 1 | 0 | 0 | 1522628 | 6NZK_A | P36334 |
| VDYSTKRRSRRA | 1 | 0 | 0 | 1522636 | CAA83660.1 | P36334 |
| VDYTKEGSVINF | 1 | 0 | 0 | 1522640 | AKN24802.1 | K9N7C7 |
| VDYTKKQITVNG | 1 | 0 | 0 | 1522644 | AUM60013.1 | K9N7C7 |
| VDYTKKSLNVDG | 1 | 0 | 0 | 1522646 | ATQ39389.1 | K9N7C7 |
| VEAAEQVVAKPK | 1 | 0 | 0 | 1522650 | AVV62524.1 | K9N7C7 |
| VEAAFDLSKELK | 1 | 0 | 0 | 1522651 | QBQ01836.1 | P0C6X6 |
| VEACIVQVTCGS | 1 | 0 | 0 | 1522652 | AGV08515.1 | K9N7C7 |
| VEACMVQVTCGG | 1 | 0 | 0 | 1522653 | ALB08298.1 | K9N7C7 |
| VEACVVQVTCGS | 1 | 0 | 0 | 1522656 | ATG84853.1 | K9N7C7 |
| VEADFHLSGCDE | 1 | 0 | 0 | 1522657 | AEN19365.1 | P30215 |
| VEADVAARADYE | 1 | 0 | 0 | 1522662 | ANI69867.1 | K9N638 |
| VEAFEYYHTTDS | 1 | 0 | 0 | 1522665 | AVP78041.1 | P0C6W2 |
| VEAGEVKPFAVY | 1 | 0 | 0 | 1522666 | ARU07593.1 | H9EJ42 |
| VEAHFYLSGCDE | 1 | 0 | 0 | 1522667 | AIV42010.1 | P30215 |
| VEAPTTCGYLPN | 1 | 0 | 0 | 1522673 | APO40578.1 | P0C6X7 |
| VEASDLEEGESE | 1 | 0 | 0 | 1522675 | AXN92247.1 | K9N7C7 |
| VEASTDFHQLEC | 1 | 0 | 0 | 1522678 | ARU07593.1 | H9EJ42 |
| VEASTDFHQLEF | 1 | 0 | 0 | 1522679 | AWK59942.1 | H9EJ42 |
| VEASTDFHQLEY | 1 | 0 | 0 | 1522680 | AGT51337.1 | H9EJ42 |
| VEASTVVCGGKR | 1 | 0 | 0 | 1522681 | AXN92247.1 | K9N7C7 |
| VEATGLFADRDG | 1 | 0 | 0 | 1522682 | AEN19363.1 | P0C6X6 |
| VEATGLFAERDC | 1 | 0 | 0 | 1522683 | ARU07574.1 | A0A140H1G9 |
| VEAVNFNVTNAE | 1 | 0 | 0 | 1522685 | CAA71146.1 | A0A0P0K6L9 |
| VEAWSGLCVDGI | 1 | 0 | 0 | 1522687 | AIW52757.1 | A0A0P0K6L9 |
| VEAWSGLCVDGR | 1 | 0 | 0 | 1522688 | AIW52756.1 | A0A0P0K6L9 |
| VEAWSGLCVDGT | 1 | 0 | 0 | 1522689 | CAA71056.1 | A0A0P0K6L9 |
| VEAYNLVSASGF | 1 | 0 | 0 | 1522690 | AVV62524.1 | K9N7C7 |
| VEAYNNCTGGAG | 1 | 0 | 0 | 1522692 | CAA83661.1 | P36334 |
| VEAYNSILVENG | 1 | 0 | 0 | 1522695 | ARU07593.1 | H9EJ42 |
| VECCMVQVTCGT | 1 | 0 | 0 | 1522697 | APO40578.1 | P0C6X7 |
| VECDEECSEVEA | 1 | 0 | 0 | 1522700 | AXN92247.1 | K9N7C7 |
| VECDFASMLNGT | 1 | 0 | 0 | 1522704 | AVV62526.1 | K9N5Q8 |
| VECDFSPLLFGT | 1 | 0 | 0 | 1522707 | AVN89365.1 | K9N5Q8 |
| VECFDKFKVNSA | 1 | 0 | 0 | 1522711 | APO40578.1 | P0C6X7 |
| VECGSCGNDSWL | 1 | 0 | 0 | 1522713 | AXN73479.1 | K9N7C7 |
| VECLKLSHHSDL | 1 | 0 | 0 | 1522714 | APO40578.1 | P0C6X7 |
| VECLSKEDLHTL | 1 | 0 | 0 | 1522716 | AVN89451.1 | K9N7C7 |
| VECTTIVNGMKK | 1 | 0 | 0 | 1522717 | APO40578.1 | P0C6X7 |

| | | | | | | |
|---|---|---|---|---|---|---|
| VEDAYDLVHDEP | 1 | 0 | 0 | 1522723 | AVI15051.1 | P0C6X6 |
| VEDCKLPLGRSL | 1 | 0 | 0 | 1522727 | ATG84888.1 | K9N5Q8 |
| VEDCNLPLGQSL | 1 | 0 | 0 | 1522728 | ALA49451.1 | K9N5Q8 |
| VEDGDTVYYPMC | 1 | 0 | 0 | 1522730 | AUM60023.1 | K9N7C7 |
| VEDLCARMTYVC | 1 | 0 | 0 | 1522736 | AKN24802.1 | K9N7C7 |
| VEDLVPSADKEE | 1 | 0 | 0 | 1522739 | AVV62524.1 | K9N7C7 |
| VEDSTSVTAVVT | 1 | 0 | 0 | 1522741 | ASY99815.1 | K9N7A1 |
| VEDSVIVNDATS | 1 | 0 | 0 | 1522743 | AIV41846.1 | P0C6X6 |
| VEDSVIVNDDTG | 1 | 0 | 0 | 1522744 | AXX83366.1 | P0C6X6 |
| VEDSVIVNNDTS | 1 | 0 | 0 | 1522745 | AOL02450.1 | P0C6X6 |
| VEEAITEARQHL | 1 | 0 | 0 | 1522752 | APO40578.1 | P0C6X7 |
| VEEAMVVKPTVI | 1 | 0 | 0 | 1522758 | APO40578.1 | P0C6X7 |
| VEECNLSLGQSL | 1 | 0 | 0 | 1522760 | ATQ39390.1 | K9N5Q8 |
| VEEDLTKGPYEF | 1 | 0 | 0 | 1522761 | AVL25535.1 | H9EJ42 |
| VEEEIVPTLELC | 1 | 0 | 0 | 1522765 | AVV62535.1 | K9N7C7 |
| VEEFEDSTGDEA | 1 | 0 | 0 | 1522766 | AVV62535.1 | K9N7C7 |
| VEENKIPSVEVP | 1 | 0 | 0 | 1522772 | AVP78030.1 | P0C6W2 |
| VEEQAEVQQSAE | 1 | 0 | 0 | 1522774 | AVV62524.1 | K9N7C7 |
| VEESFIDDCYGY | 1 | 0 | 0 | 1522775 | ADO24649.1 | H9EJ42 |
| VEESFIDDYYGY | 1 | 0 | 0 | 1522776 | AIW52827.1 | H9EJ42 |
| VEETTMSVIPPI | 1 | 0 | 0 | 1522778 | AVV62535.1 | K9N7C7 |
| VEEVQLQDVQVS | 1 | 0 | 0 | 1522781 | AUM60013.1 | K9N7C7 |
| VEFDFIVNAANE | 1 | 0 | 0 | 1522783 | ALK43115.1 | A0A0P0LKV0 |
| VEFHLDGEVLPL | 1 | 0 | 0 | 1522785 | ATO98229.1 | P0C6W2 |
| VEFMLWCKDGKV | 1 | 0 | 0 | 1522793 | ATQ39389.1 | K9N7C7 |
| VEFMLWCKDGQV | 1 | 0 | 0 | 1522794 | AVV62524.1 | K9N7C7 |
| VEFMLWCKNGQV | 1 | 0 | 0 | 1522795 | AVV62535.1 | K9N7C7 |
| VEFNRLSTCDWS | 1 | 0 | 0 | 1522797 | AUM60013.1 | K9N7C7 |
| VEFVEGQGVPIA | 1 | 0 | 0 | 1522799 | AJQ24087.1 | P33469 |
| VEGAHAFRNACG | 1 | 0 | 0 | 1522803 | AXN92247.1 | K9N7C7 |
| VEGAHASKNACG | 1 | 0 | 0 | 1522804 | AVV62524.1 | K9N7C7 |
| VEGAHASRNACG | 1 | 0 | 0 | 1522805 | AXN92258.1 | K9N7C7 |
| VEGAINVGKNGL | 1 | 0 | 0 | 1522806 | AUM60013.1 | K9N7C7 |
| VEGAINVGKQGL | 1 | 0 | 0 | 1522807 | AVV62524.1 | K9N7C7 |
| VEGCHATRDAVG | 1 | 0 | 0 | 1522808 | AEA10817.1 | P0C6X7 |
| VEGCHATREAVG | 1 | 0 | 0 | 1522809 | QYC64967.1 | P0DTD1 |
| VEGELLVQDLRF | 1 | 0 | 0 | 1522813 | AKK52620.1 | K9N7C7 |
| VEGELLVQDLRL | 1 | 0 | 0 | 1522814 | AXN92247.1 | K9N7C7 |
| VEGELLVQNQSF | 1 | 0 | 0 | 1522815 | ATQ39389.1 | K9N7C7 |
| VEGGLLVQDLRL | 1 | 0 | 0 | 1522818 | QDP16215.1 | K9N7C7 |
| VEGHIIMEEMLK | 1 | 0 | 0 | 1522819 | AVV62524.1 | K9N7C7 |
| VEGKDGFVSVEL | 1 | 0 | 0 | 1522820 | AXN92247.1 | K9N7C7 |
| VEGKDGFVTVEL | 1 | 0 | 0 | 1522821 | AUM60013.1 | K9N7C7 |
| VEGNFSFFDCGV | 1 | 0 | 0 | 1522822 | AWK59942.1 | H9EJ42 |
| VEGQANLFRNAR | 1 | 0 | 0 | 1522823 | AVP78041.1 | P0C6W2 |
| VEGQDGFVTIEL | 1 | 0 | 0 | 1522825 | AVV62524.1 | K9N7C7 |
| VEGQGPPIAPGV | 1 | 0 | 0 | 1522826 | P33469.1 | P33469 |
| VEGSGRSAPNSR | 1 | 0 | 0 | 1522829 | AIX10731.1 | P33469 |
| VEGVSSFMNATL | 1 | 0 | 0 | 1522832 | CAA71147.1 | A0A0P0K6L9 |
| VEGVSSFMNVTL | 1 | 0 | 0 | 1522833 | AIW52757.1 | A0A0P0K6L9 |
| VEGVVWVAVDGA | 1 | 0 | 0 | 1522834 | AIW52723.1 | A0A0N7J131 |
| VEGYNNYVPSYV | 1 | 0 | 0 | 1522835 | AVV62524.1 | K9N7C7 |
| VEHALSIKQPFS | 1 | 0 | 0 | 1522836 | AGT51337.1 | H9EJ42 |
| VEHGLLHSSQLP | 1 | 0 | 0 | 1522837 | AIV41780.1 | P0C6X6 |
| VEHSLPIKQPFS | 1 | 0 | 0 | 1522841 | AVA26872.1 | H9EJ42 |
| VEHSLTIKQPFS | 1 | 0 | 0 | 1522842 | AFD98763.1 | H9EJ42 |
| VEIEVGEVKPFA | 1 | 0 | 0 | 1522847 | AKT07951.1 | H9EJ42 |
| VEINRLQEAIKV | 1 | 0 | 0 | 1522854 | AIL49487.1 | P36334 |
| VEIQSSGVINSS | 1 | 0 | 0 | 1522855 | AVV62535.1 | K9N7C7 |
| VEIQSSGVVKPN | 1 | 0 | 0 | 1522856 | AVV62524.1 | K9N7C7 |
| VEKCVVRVCYGN | 1 | 0 | 0 | 1522858 | NP_073549.1 | A0A0P0LKV0 |
| VEKDDQGHNLDS | 1 | 0 | 0 | 1522860 | AVV62524.1 | K9N7C7 |
| VEKEGSYHNWHL | 1 | 0 | 0 | 1522861 | AVV62524.1 | K9N7C7 |
| VEKEGSYHNWYL | 1 | 0 | 0 | 1522862 | AVV62535.1 | K9N7C7 |
| VEKFNFVTDWLK | 1 | 0 | 0 | 1522863 | AWK59942.1 | H9EJ42 |
| VEKFTFVTDWLK | 1 | 0 | 0 | 1522864 | AFV53147.1 | H9EJ42 |
| VEKICIIDNMYM | 1 | 0 | 0 | 1522867 | ARU07575.1 | A0A140H1G9 |
| VEKLKQLLEQWN | 1 | 0 | 0 | 1522871 | AAS48455.1 | P59596 |
| VEKLRPVFAWVE | 1 | 0 | 0 | 1522874 | ARO76381.1 | P0C6U8 |
| VEKLVSWYIYVR | 1 | 0 | 0 | 1522878 | AGT51337.1 | H9EJ42 |
| VEKNVVRAALLA | 1 | 0 | 0 | 1522879 | AUM60013.1 | K9N7C7 |
| VEKSFQLDTAPK | 1 | 0 | 0 | 1522881 | Q3I5J5.1 | Q3LZX1 |
| VEKWESGVKDYV | 1 | 0 | 0 | 1522882 | APO40580.1 | P59632 |
| VEKWLHQLLGST | 1 | 0 | 0 | 1522883 | APO40578.1 | P0C6X7 |
| VELENKFTPLSV | 1 | 0 | 0 | 1522886 | AVV62524.1 | K9N7C7 |
| VELFAKRNVRHH | 1 | 0 | 0 | 1522887 | AGT51498.1 | P0C6X6 |
| VELGSKHVDCWT | 1 | 0 | 0 | 1522892 | ABQ16583.1 | H9EJ42 |
| VELGTDVNEFAC | 1 | 0 | 0 | 1522894 | AVP78041.1 | P0C6W2 |
| VELHTVIPDYVD | 1 | 0 | 0 | 1522896 | AIW52843.1 | A0A0P0G321 |
| VELHTVYPDYVD | 1 | 0 | 0 | 1522897 | AWK59979.1 | A0A0P0G321 |
| VELLSEPQTIQP | 1 | 0 | 0 | 1522902 | AVN89374.1 | K9N7C7 |
| VELSLDRGAQSD | 1 | 0 | 0 | 1522913 | AVV62527.1 | K9N796 |
| VELVAELDGIQY | 1 | 0 | 0 | 1522916 | AVP78041.1 | P0C6W2 |
| VELYELMGKFAW | 1 | 0 | 0 | 1522924 | AXN92247.1 | K9N7C7 |
| VELYNGECVYLF | 1 | 0 | 0 | 1522926 | NP_073553.1 | A0A0P0KWE8 |
| VEMNRLQDAIKV | 1 | 0 | 0 | 1522928 | QEG03776.1 | P36334 |
| VEMNRLQEAIKV | 1 | 0 | 0 | 1522929 | CAA79896.1 | P36334 |
| VEMPKFFYAPNG | 1 | 0 | 0 | 1522930 | ADK37704.1 | A0A0P0K6L9 |
| VENDNISVAVPK | 1 | 0 | 0 | 1522934 | AIL49482.1 | P33469 |
| VENGIPLMPLLS | 1 | 0 | 0 | 1522936 | ARU07593.1 | H9EJ42 |
| VENKAEAEWIEL | 1 | 0 | 0 | 1522937 | AST12962.1 | A0A0P0LKV0 |

| | | | | | | |
|---|---|---|---|---|---|---|
| VENKTEADWIEL | 1 | 0 | 0 | 1522938 | QEO75983.1 | A0A0P0LKV0 |
| VENNESLTWPLV | 1 | 0 | 0 | 1522941 | AUM60013.1 | K9N7C7 |
| VENSAPVPVKEY | 1 | 0 | 0 | 1522952 | AVV62524.1 | K9N7C7 |
| VENTPPHYFTLG | 1 | 0 | 0 | 1522955 | AXN92247.1 | K9N7C7 |
| VEPDLKKGPHEF | 1 | 0 | 0 | 1522958 | AVV62524.1 | K9N7C7 |
| VEPEPEPTLEEP | 1 | 0 | 0 | 1522959 | ARO76381.1 | P0C6U8 |
| VEPKNGGQFFVS | 1 | 0 | 0 | 1522963 | ARU07593.1 | H9EJ42 |
| VEPLVQDRVVEP | 1 | 0 | 0 | 1522965 | ARE29989.1 | Q4VID0 |
| VEPVRDEVSTET | 1 | 0 | 0 | 1522971 | ABB90487.1 | A0A0N7J131 |
| VEQADGVECDFS | 1 | 0 | 0 | 1522972 | ATG84679.1 | K9N5Q8 |
| VEQAEGFECDFS | 1 | 0 | 0 | 1522973 | AQZ41282.1 | K9N5Q8 |
| VEQDRNTRDVFA | 1 | 0 | 0 | 1522976 | APO40579.1 | P59594 |
| VEQEAQIDRLIN | 1 | 0 | 0 | 1522977 | AVV62526.1 | K9N5Q8 |
| VEQKVAENPKEE | 1 | 0 | 0 | 1522980 | AVP78041.1 | P0C6W2 |
| VEQLEPCVQSRL | 1 | 0 | 0 | 1522983 | AST12962.1 | A0A0P0LKV0 |
| VEQRQQAEEKKI | 1 | 0 | 0 | 1522988 | AVP78041.1 | P0C6W2 |
| VEQSFQLDTAPQ | 1 | 0 | 0 | 1522989 | ATO98120.1 | Q3LZX1 |
| VEQSQRSLNFVG | 1 | 0 | 0 | 1522990 | AVV62524.1 | K9N7C7 |
| VEQVSSDWRVYA | 1 | 0 | 0 | 1522991 | APO40579.1 | P59594 |
| VEQYNQIILNLT | 1 | 0 | 0 | 1522992 | ABB90514.1 | A0A0P0K6L9 |
| VERGLFHSSQLP | 1 | 0 | 0 | 1522995 | AJC98124.1 | P0C6X6 |
| VERGLLHSSQLP | 1 | 0 | 0 | 1522996 | AEN19363.1 | P0C6X6 |
| VERLIACENPFM | 1 | 0 | 0 | 1523001 | AXN92247.1 | K9N7C7 |
| VERLIACENSFM | 1 | 0 | 0 | 1523002 | AVN89312.1 | K9N638 |
| VERLTGGTYSTP | 1 | 0 | 0 | 1523004 | AVV62535.1 | K9N7C7 |
| VERNIVRAALLA | 1 | 0 | 0 | 1523005 | AVV62524.1 | K9N7C7 |
| VERSVSLAIDAY | 1 | 0 | 0 | 1523006 | ASU90624.1 | K9N7C7 |
| VESAAQADLQHE | 1 | 0 | 0 | 1523007 | ATQ39389.1 | K9N7C7 |
| VESASDFIAWRA | 1 | 0 | 0 | 1523009 | Q14EA9.1 | Q0ZJ84 |
| VESASEFIVWRA | 1 | 0 | 0 | 1523010 | ADN03340.1 | Q0ZJ84 |
| VESCGNYRVTKG | 1 | 0 | 0 | 1523014 | ARO76381.1 | P0C6U8 |
| VESCMVQVTCGS | 1 | 0 | 0 | 1523016 | AVV62524.1 | K9N7C7 |
| VESDIDTGDIVD | 1 | 0 | 0 | 1523019 | AVV62535.1 | K9N7C7 |
| VESDKYITTNGP | 1 | 0 | 0 | 1523020 | AVP78030.1 | P0C6W2 |
| VESGGYIPPNFA | 1 | 0 | 0 | 1523021 | AIW52740.1 | A0A0P0K6L9 |
| VESGGYIPSNFA | 1 | 0 | 0 | 1523023 | AIW52757.1 | A0A0P0K6L9 |
| VESGTEVSEFAC | 1 | 0 | 0 | 1523024 | ARO76381.1 | P0C6U8 |
| VESGVYSVSSLE | 1 | 0 | 0 | 1523027 | AIA22864.1 | K9N5Q8 |
| VESLMSTILSSC | 1 | 0 | 0 | 1523030 | AVV62535.1 | K9N7C7 |
| VESNDSVVFTYI | 1 | 0 | 0 | 1523031 | ATQ39397.1 | K9N7D2 |
| VESNVVDVNDCF | 1 | 0 | 0 | 1523033 | AGT51379.1 | H9EJ42 |
| VESPHLMGWDYP | 1 | 0 | 0 | 1523034 | AVP78041.1 | P0C6W2 |
| VESPTVDVVALQ | 1 | 0 | 0 | 1523035 | ATG84853.1 | K9N7C7 |
| VESQSKRYGFGG | 1 | 0 | 0 | 1523036 | ABB90528.1 | A0A0P0K6L9 |
| VESRLSDGVEFL | 1 | 0 | 0 | 1523037 | ARO76381.1 | P0C6U8 |
| VESTPVELPTVD | 1 | 0 | 0 | 1523040 | AVN89312.1 | K9N638 |
| VESTPVEPPTVD | 1 | 0 | 0 | 1523041 | AHI48796.1 | K9N7C7 |
| VESTPVEPSTVD | 1 | 0 | 0 | 1523042 | AIY60577.1 | K9N638 |
| VESTPVVTETSE | 1 | 0 | 0 | 1523043 | AVP78030.1 | P0C6W2 |
| VESTSVEPPTVD | 1 | 0 | 0 | 1523044 | AJG44112.1 | K9N638 |
| VESYNIFTQAGF | 1 | 0 | 0 | 1523045 | NP_073549.1 | A0A0P0LKV0 |
| VETDLKKGPHEF | 1 | 0 | 0 | 1523050 | AXN92247.1 | K9N7C7 |
| VETFYPKLQSSH | 1 | 0 | 0 | 1523053 | AVP78041.1 | P0C6W2 |
| VETGTQLEEFAC | 1 | 0 | 0 | 1523055 | APO40578.1 | P0C6X7 |
| VETKVQCSTNLF | 1 | 0 | 0 | 1523058 | AEN19363.1 | P0C6X6 |
| VETLNFQMVKEY | 1 | 0 | 0 | 1523061 | AET99208.1 | Q0ZJ80 |
| VETLVGQLEQTN | 1 | 0 | 0 | 1523063 | ASU90174.1 | K9N638 |
| VETNEIVDSVQY | 1 | 0 | 0 | 1523064 | AXN92247.1 | K9N7C7 |
| VETSNSFDALEL | 1 | 0 | 0 | 1523075 | AVP78030.1 | P0C6W2 |
| VETSNSFDALKL | 1 | 0 | 0 | 1523076 | AVP78041.1 | P0C6W2 |
| VETSNSFEVLAV | 1 | 0 | 0 | 1523078 | ACV88184.1 | P0C6X7 |
| VETSVGHYVHAR | 1 | 0 | 0 | 1523080 | AVV62524.1 | K9N7C7 |
| VETVGGLFEIQI | 1 | 0 | 0 | 1523081 | BBA20986.1 | A0A140H1H1 |
| VETVSLAGFYRD | 1 | 0 | 0 | 1523087 | ARO76381.1 | P0C6U8 |
| VEVEVNPSYFDK | 1 | 0 | 0 | 1523094 | AVV62526.1 | K9N5Q8 |
| VEVFEAQAFETP | 1 | 0 | 0 | 1523095 | AVV62535.1 | K9N7C7 |
| VEVFEVKTFETP | 1 | 0 | 0 | 1523096 | AVV62524.1 | K9N7C7 |
| VEVFTDGRLNCS | 1 | 0 | 0 | 1523098 | AVN89333.1 | K9N7C7 |
| VEVHVEDIVQVV | 1 | 0 | 0 | 1523100 | ANF29161.1 | K9N7C7 |
| VEVKDTVVQLNY | 1 | 0 | 0 | 1523102 | AVV62535.1 | K9N7C7 |
| VEVLVPSNKWTG | 1 | 0 | 0 | 1523104 | AVV62535.1 | K9N7C7 |
| VEVNALRTTGIV | 1 | 0 | 0 | 1523105 | AVV62524.1 | K9N7C7 |
| VEVNAPYYNSWQ | 1 | 0 | 0 | 1523106 | CAA83660.1 | P36334 |
| VEVNATYCNGWQ | 1 | 0 | 0 | 1523107 | ABU39950.1 | P36334 |
| VEVNPSYFDKSW | 1 | 0 | 0 | 1523109 | AVV62526.1 | K9N5Q8 |
| VEVPPQLVELPS | 1 | 0 | 0 | 1523113 | AVN89312.1 | K9N638 |
| VEVPPQVMKLPS | 1 | 0 | 0 | 1523114 | ALA50044.1 | K9N7C7 |
| VEVPPQVVELLS | 1 | 0 | 0 | 1523115 | AVN89374.1 | K9N7C7 |
| VEVPPQVVELPS | 1 | 0 | 0 | 1523116 | AVN89451.1 | K9N7C7 |
| VEVPPQVVELPY | 1 | 0 | 0 | 1523117 | ANI69888.1 | K9N638 |
| VEVPPVLQVEST | 1 | 0 | 0 | 1523121 | AVP78030.1 | P0C6W2 |
| VEVQSSGIIKPN | 1 | 0 | 0 | 1523127 | AHY22544.1 | K9N7C7 |
| VEVQSSGMIKPN | 1 | 0 | 0 | 1523129 | AXN92247.1 | K9N7C7 |
| VEVQSSGMINPN | 1 | 0 | 0 | 1523130 | ASU90746.1 | K9N7C7 |
| VEVSALRTAGVV | 1 | 0 | 0 | 1523131 | AUM60013.1 | K9N7C7 |
| VEVSPQVVKLPS | 1 | 0 | 0 | 1523132 | ASU45805.1 | K9N7C7 |
| VEVTPQVVKLPS | 1 | 0 | 0 | 1523134 | ALA50033.1 | K9N7C7 |
| VEVVCEDVTQLA | 1 | 0 | 0 | 1523135 | ATQ39389.1 | K9N7C7 |
| VEVVMGQLEETN | 1 | 0 | 0 | 1523137 | AUM60023.1 | K9N7C7 |
| VEVVMGQLEQTN | 1 | 0 | 0 | 1523138 | AUM60013.1 | K9N7C7 |
| VEWDIEYPYISH | 1 | 0 | 0 | 1523139 | AUM60013.1 | K9N7C7 |

| | | | | | | |
|---|---|---|---|---|---|---|
| VEWLTGGTYSAP | 1 | 0 | 0 | 1523142 | AVV62524.1 | K9N7C7 |
| VEYAYDQVHDEP | 1 | 0 | 0 | 1523143 | APU51933.1 | P0C6X6 |
| VEYGQFCAKINQ | 1 | 0 | 0 | 1523148 | AUM60014.1 | K9N5Q8 |
| VEYGTEDDYNGT | 1 | 0 | 0 | 1523150 | APO40578.1 | P0C6X7 |
| VEYKITQWPTAT | 1 | 0 | 0 | 1523151 | AVQ05261.1 | P0C6X6 |
| VEYLQITSKPIV | 1 | 0 | 0 | 1523152 | APT69883.1 | A0A0P0K6L9 |
| VEYLQITSTPIV | 1 | 0 | 0 | 1523154 | AIW52843.1 | A0A0P0G321 |
| VEYNIHAVLDAL | 1 | 0 | 0 | 1523156 | AVV62535.1 | K9N7C7 |
| VEYNIHAVLDTL | 1 | 0 | 0 | 1523158 | AXN92247.1 | K9N7C7 |
| VEYNLYGVSGRG | 1 | 0 | 0 | 1523160 | AVV62537.1 | K9N5Q8 |
| VEYPIIGDELKI | 1 | 0 | 0 | 1523161 | P0C6W6.1 | P0C6W2 |
| VEYSIHAVLDTL | 1 | 0 | 0 | 1523164 | ALB08320.1 | K9N7C7 |
| VEYSKNRRSRRA | 1 | 0 | 0 | 1523165 | AIX10750.1 | P36334 |
| VEYSLPIKQPFS | 1 | 0 | 0 | 1523166 | AWK59942.1 | H9EJ42 |
| VEYSLYGISGRG | 1 | 0 | 0 | 1523167 | AXN92260.1 | K9N5Q8 |
| VEYSLYGVLGRG | 1 | 0 | 0 | 1523168 | ASU90857.1 | K9N5Q8 |
| VEYSLYGVSGRG | 1 | 0 | 0 | 1523169 | AXN92228.1 | K9N5Q8 |
| VEYTYTDEVIYG | 1 | 0 | 0 | 1523170 | AEN19363.1 | P0C6X6 |
| VEYTYTDEVIYS | 1 | 0 | 0 | 1523171 | AIV41834.1 | P0C6X6 |
| VFANTVEDAVNK | 1 | 0 | 0 | 1523177 | AGT51403.1 | Q04853 |
| VFAVESGGYIPP | 1 | 0 | 0 | 1523183 | AIW52740.1 | A0A0P0K6L9 |
| VFCCLSTGCCGC | 1 | 0 | 0 | 1523189 | AIW52843.1 | A0A0P0G321 |
| VFDDEWAAAVDE | 1 | 0 | 0 | 1523201 | AQZ41284.1 | K9N7C7 |
| VFDITDAAVKAA | 1 | 0 | 0 | 1523208 | APT69868.1 | A0A0P0ILY9 |
| VFDNFNQWHEAK | 1 | 0 | 0 | 1523214 | AWK59942.1 | H9EJ42 |
| VFDSSKEFVEYK | 1 | 0 | 0 | 1523215 | AIV41984.1 | P0C6X6 |
| VFDVDKVKTPHI | 1 | 0 | 0 | 1523218 | AUM60013.1 | K9N7C7 |
| VFDVESGGYIPS | 1 | 0 | 0 | 1523219 | ARA15429.1 | A0A0P0K6L9 |
| VFDVKGVEELHD | 1 | 0 | 0 | 1523220 | AOL02451.1 | Q80872 |
| VFEAQAFETPAV | 1 | 0 | 0 | 1523222 | AVV62535.1 | K9N7C7 |
| VFEVKTFETPAG | 1 | 0 | 0 | 1523230 | AVV62524.1 | K9N7C7 |
| VFFADNNDITNT | 1 | 0 | 0 | 1523232 | ASY99807.1 | K9N4V7 |
| VFFNAFVKCNCG | 1 | 0 | 0 | 1523241 | AWK59942.1 | H9EJ42 |
| VFFNGADISDTI | 1 | 0 | 0 | 1523242 | AXN92247.1 | K9N7C7 |
| VFFNGVDISDTI | 1 | 0 | 0 | 1523243 | AQZ41321.1 | K9N7C7 |
| VFFQFENLSSED | 1 | 0 | 0 | 1523245 | AIV41978.1 | P0C6X6 |
| VFGKRGGGNVTY | 1 | 0 | 0 | 1523261 | NP_073549.1 | A0A0P0LKV0 |
| VFGYVQQGVFNK | 1 | 0 | 0 | 1523273 | ATN39862.1 | P0C6X6 |
| VFHETSDDEWTA | 1 | 0 | 0 | 1523274 | AVN89352.1 | K9N7C7 |
| VFIDAARGHAGV | 1 | 0 | 0 | 1523281 | AUM60023.1 | K9N7C7 |
| VFIDAARGPAGV | 1 | 0 | 0 | 1523282 | AUM60013.1 | K9N7C7 |
| VFIKRSDAQGTG | 1 | 0 | 0 | 1523293 | ARO76381.1 | P0C6U8 |
| VFISPYNSQNTV | 1 | 0 | 0 | 1523306 | AVP78041.1 | P0C6W2 |
| VFKADGIIYPQG | 1 | 0 | 0 | 1523312 | AHI48583.1 | K9N5Q8 |
| VFKDGCDVGFCP | 1 | 0 | 0 | 1523313 | ABE97136.1 | H9EJ42 |
| VFKDGVMHSEFP | 1 | 0 | 0 | 1523314 | AKA09651.1 | P36334 |
| VFKEGVMYSEFP | 1 | 0 | 0 | 1523315 | AMK59677.1 | P36334 |
| VFKHGVMYSEFP | 1 | 0 | 0 | 1523316 | AIL49520.1 | P36334 |
| VFKPQPAGVFTD | 1 | 0 | 0 | 1523333 | CAA83661.1 | P36334 |
| VFKPQPAGVLTD | 1 | 0 | 0 | 1523335 | QEG03776.1 | P36334 |
| VFLCAVLMYCFL | 1 | 0 | 0 | 1523346 | AVV62524.1 | K9N7C7 |
| VFNASISESKPL | 1 | 0 | 0 | 1523396 | AVL25595.1 | H9EJ69 |
| VFNASNFPSVYA | 1 | 0 | 0 | 1523397 | APO40579.1 | P59594 |
| VFNASRFPNVYA | 1 | 0 | 0 | 1523398 | ATO98181.1 | Q3LZX1 |
| VFNKATVAQIKA | 1 | 0 | 0 | 1523414 | ATN39862.1 | P0C6X6 |
| VFNLLHFTNDKT | 1 | 0 | 0 | 1523416 | AMN88692.1 | A0A140H1G9 |
| VFNNTLSRQLQP | 1 | 0 | 0 | 1523419 | CAA83660.1 | P36334 |
| VFNVTTLCYVQG | 1 | 0 | 0 | 1523427 | ARB07393.1 | A0A0P0KWE8 |
| VFNVVKQGHFIG | 1 | 0 | 0 | 1523430 | AXN92247.1 | K9N7C7 |
| VFPLDEAEDVTE | 1 | 0 | 0 | 1523435 | ASU90174.1 | K9N638 |
| VFPYDMDFGVSS | 1 | 0 | 0 | 1523448 | AHY22554.1 | K9N638 |
| VFPYDMSSDVSS | 1 | 0 | 0 | 1523451 | AUM60013.1 | K9N7C7 |
| VFQNCSAVGVRQ | 1 | 0 | 0 | 1523458 | ASU90791.1 | K9N5Q8 |
| VFQNCTAVGLRQ | 1 | 0 | 0 | 1523459 | AUM60014.1 | K9N5Q8 |
| VFQSASKIITLK | 1 | 0 | 0 | 1523464 | AVP78043.1 | Q3LZX0 |
| VFQSASKVIALH | 1 | 0 | 0 | 1523465 | ATO98194.1 | Q3LZX0 |
| VFRMPMGVYNYK | 1 | 0 | 0 | 1523485 | ARU07575.1 | A0A140H1G9 |
| VFSAVYPIDLAS | 1 | 0 | 0 | 1523497 | AUM60030.1 | K9N7A1 |
| VFSLLHFTNDKT | 1 | 0 | 0 | 1523502 | ARU07575.1 | A0A140H1G9 |
| VFSNMVETVVGQ | 1 | 0 | 0 | 1523510 | AVN89363.1 | K9N7C7 |
| VFSTGSYYCVNS | 1 | 0 | 0 | 1523513 | AUM60023.1 | K9N7C7 |
| VFSYCKKLGTTV | 1 | 0 | 0 | 1523518 | AXX83378.1 | P0C6X6 |
| VFTDGKLNCTFQ | 1 | 0 | 0 | 1523530 | AVV62524.1 | K9N7C7 |
| VFTGCVGTWDKV | 1 | 0 | 0 | 1523537 | AVV62535.1 | K9N7C7 |
| VFTKSGVNDICI | 1 | 0 | 0 | 1523543 | ANF29215.1 | K9N7C7 |
| VFVAPPIADFTP | 1 | 0 | 0 | 1523569 | AVV62528.1 | K9N4V0 |
| VFVEPKDGGHFF | 1 | 0 | 0 | 1523575 | ARE29978.1 | H9EJ42 |
| VFVEPKDGGQFF | 1 | 0 | 0 | 1523576 | AWK59942.1 | H9EJ42 |
| VFVEPKNGGQFF | 1 | 0 | 0 | 1523577 | ARU07593.1 | H9EJ42 |
| VFVNDVVYQKLL | 1 | 0 | 0 | 1523585 | AVV62535.1 | K9N7C7 |
| VFVPASRDSVPL | 1 | 0 | 0 | 1523591 | AVV62541.1 | K9N7D2 |
| VFVPATRDTVPL | 1 | 0 | 0 | 1523593 | AUM60018.1 | K9N7D2 |
| VFVPQPTGVFTN | 1 | 0 | 0 | 1523595 | AEN19366.1 | P36334 |
| VFVRYGDVDNLV | 1 | 0 | 0 | 1523601 | AWK59942.1 | H9EJ42 |
| VFVVESGGYIPS | 1 | 0 | 0 | 1523617 | AGT21353.1 | A0A0P0K6L9 |
| VFVYTDTEVCKV | 1 | 0 | 0 | 1523621 | NP_073549.1 | A0A0P0LKV0 |
| VFYCVLEPRNGT | 1 | 0 | 0 | 1523629 | AVV62526.1 | K9N5Q8 |
| VFYKETSYTSDI | 1 | 0 | 0 | 1523640 | APO40578.1 | P0C6X7 |
| VFYVKDGKYFTS | 1 | 0 | 0 | 1523647 | ALA49857.1 | K9N7C7 |
| VGAWDKVSKIAN | 1 | 0 | 0 | 1523672 | AVV62524.1 | K9N7C7 |
| VGCAMGIVNSSM | 1 | 0 | 0 | 1523674 | AUM60014.1 | K9N5Q8 |

| | | | | | | |
|---|---|---|---|---|---|---|
| VGCNNADCVACS | 1 | 0 | 0 | 1523681 | AWK59942.1 | H9EJ42 |
| VGDFQLNEEVAI | 1 | 0 | 0 | 1523698 | P0C6W6.1 | P0C6W2 |
| VGDIILKPASDG | 1 | 0 | 0 | 1523702 | AVP78030.1 | P0C6W2 |
| VGDLNCPLDPKL | 1 | 0 | 0 | 1523704 | QBP84704.1 | P36334 |
| VGDPAQLPTPRT | 1 | 0 | 0 | 1523708 | ATG84864.1 | K9N7C7 |
| VGDPQQLPAPRV | 1 | 0 | 0 | 1523709 | AWK59942.1 | H9EJ42 |
| VGEFVVNDVVIA | 1 | 0 | 0 | 1523734 | AVN89333.1 | K9N7C7 |
| VGEKPVSIINNA | 1 | 0 | 0 | 1523742 | APO40578.1 | P0C6X7 |
| VGEPTISQPSKL | 1 | 0 | 0 | 1523752 | ABE97136.1 | H9EJ42 |
| VGFDAEGAHATL | 1 | 0 | 0 | 1523773 | AEN19363.1 | P0C6X6 |
| VGFFFPYDPALE | 1 | 0 | 0 | 1523776 | AVV62535.1 | K9N7C7 |
| VGFKDVTNGTIY | 1 | 0 | 0 | 1523781 | CAA71146.1 | A0A0P0K6L9 |
| VGFKNQYCNESI | 1 | 0 | 0 | 1523783 | AXX83294.1 | P0C6X6 |
| VGGAPSWVTALY | 1 | 0 | 0 | 1523803 | AUM60013.1 | K9N7C7 |
| VGGCVASNVSPI | 1 | 0 | 0 | 1523809 | QEG03746.1 | H9EJ42 |
| VGGCVTSNVPPI | 1 | 0 | 0 | 1523810 | AWK59942.1 | H9EJ42 |
| VGGCVTSNVSPI | 1 | 0 | 0 | 1523812 | ARU07593.1 | H9EJ42 |
| VGGDFTPIDQYM | 1 | 0 | 0 | 1523815 | AVV62524.1 | K9N7C7 |
| VGGNFDNDINCP | 1 | 0 | 0 | 1523831 | AVV62524.1 | K9N7C7 |
| VGGNFTPIDQYM | 1 | 0 | 0 | 1523832 | AUM60013.1 | K9N7C7 |
| VGGTFDNDIACP | 1 | 0 | 0 | 1523839 | AUM60013.1 | K9N7C7 |
| VGGTIQILASVP | 1 | 0 | 0 | 1523842 | NP_073549.1 | A0A0P0LKV0 |
| VGHDICAQLNAK | 1 | 0 | 0 | 1523847 | ABD75519.1 | A0A140H1G9 |
| VGHDICAQLNDK | 1 | 0 | 0 | 1523848 | ARU07575.1 | A0A140H1G9 |
| VGHPAPTHYLRV | 1 | 0 | 0 | 1523851 | AVV62540.1 | K9N643 |
| VGHSDSNYDYLG | 1 | 0 | 0 | 1523852 | AVV62537.1 | K9N5Q8 |
| VGHTEFYDSQPS | 1 | 0 | 0 | 1523857 | AUM60025.1 | K9N796 |
| VGHTVCDSLNAK | 1 | 0 | 0 | 1523858 | AGT51757.1 | P0C6X6 |
| VGIADDTYVMGL | 1 | 0 | 0 | 1523870 | NP_073549.1 | A0A0P0LKV0 |
| VGIGEHCPGLGI | 1 | 0 | 0 | 1523873 | BBA20986.1 | A0A140H1H1 |
| VGIGEHCSGLAI | 1 | 0 | 0 | 1523874 | CAA83660.1 | P36334 |
| VGILTLDNQDLN | 1 | 0 | 0 | 1523880 | QVO90575.1 | P0DTD1 |
| VGIQTIEELHVR | 1 | 0 | 0 | 1523882 | AVV62524.1 | K9N7C7 |
| VGKDGVMYSEFP | 1 | 0 | 0 | 1523890 | AWW13511.1 | P36334 |
| VGKIAQWIKNTF | 1 | 0 | 0 | 1523896 | AEN19363.1 | P0C6X6 |
| VGKNGLQFAAIT | 1 | 0 | 0 | 1523898 | AUM60013.1 | K9N7C7 |
| VGKYDINHGDNG | 1 | 0 | 0 | 1523907 | AGT17757.1 | A0A140H1H0 |
| VGLLDQAWRFPC | 1 | 0 | 0 | 1523916 | AEN19363.1 | P0C6X6 |
| VGLVFVTLRSIN | 1 | 0 | 0 | 1523924 | AIV41948.1 | P0C6X6 |
| VGNFSDGKMGLF | 1 | 0 | 0 | 1523931 | ANF29217.1 | K9N5Q8 |
| VGNFSNGVSGRY | 1 | 0 | 0 | 1523933 | AVV62537.1 | K9N5Q8 |
| VGNFSNNKTGRY | 1 | 0 | 0 | 1523934 | AUM60014.1 | K9N5Q8 |
| VGNGPGIDAGYK | 1 | 0 | 0 | 1523936 | CAA83660.1 | P36334 |
| VGNLKYSTFMLN | 1 | 0 | 0 | 1523943 | ATO98193.1 | Q3LZX1 |
| VGNVILKPAADG | 1 | 0 | 0 | 1523952 | APO40578.1 | P0C6X7 |
| VGNVKFDRWSAS | 1 | 0 | 0 | 1523956 | AIW52729.1 | A0A0P0K6L9 |
| VGNVKLDRWSAS | 1 | 0 | 0 | 1523957 | AIW52740.1 | A0A0P0K6L9 |
| VGPDCAYFNGAT | 1 | 0 | 0 | 1523970 | AUM60023.1 | K9N7C7 |
| VGPDSIKSACIE | 1 | 0 | 0 | 1523975 | QBF80543.1 | K9N5Q8 |
| VGPDYAYFNGAT | 1 | 0 | 0 | 1523982 | AUM60013.1 | K9N7C7 |
| VGPMQDVNQDDD | 1 | 0 | 0 | 1523986 | AVV62533.1 | K9N4V7 |
| VGPMVDVNSEQS | 1 | 0 | 0 | 1523987 | AUM60021.1 | K9N4V7 |
| VGPNKNAGQDIS | 1 | 0 | 0 | 1523988 | APO40578.1 | P0C6X7 |
| VGPRAGKHEHSL | 1 | 0 | 0 | 1523994 | ARU07593.1 | H9EJ42 |
| VGPYVEGAINVG | 1 | 0 | 0 | 1523998 | AVV62535.1 | K9N7C7 |
| VGRPIVYETDAN | 1 | 0 | 0 | 1524018 | AQN78717.1 | P0C6X6 |
| VGRSVNSQNCTG | 1 | 0 | 0 | 1524021 | CAA33521.1 | A0A0P0KXN3 |
| VGRWSTSINTGN | 1 | 0 | 0 | 1524025 | AAQ90004.1 | A0A0P0K6L9 |
| VGSGYCVDYSTK | 1 | 0 | 0 | 1524047 | CAA83660.1 | P36334 |
| VGSIAGAGWTAG | 1 | 0 | 0 | 1524049 | AVV62526.1 | K9N5Q8 |
| VGSLIVRCSYDY | 1 | 0 | 0 | 1524053 | Q3I5I8.1 | Q3LZX5 |
| VGTCPAGTNSLT | 1 | 0 | 0 | 1524064 | QEG03757.1 | P36334 |
| VGVGNTVPMTEG | 1 | 0 | 0 | 1524100 | AVV62526.1 | K9N5Q8 |
| VGVIRISNETFL | 1 | 0 | 0 | 1524105 | BAL45640.1 | A0A0P0K6L9 |
| VGVQKYSTLQGP | 1 | 0 | 0 | 1524111 | APO40578.1 | P0C6X7 |
| VGVQSIEELHQP | 1 | 0 | 0 | 1524112 | AUM60013.1 | K9N7C7 |
| VGVQTIEDLRAR | 1 | 0 | 0 | 1524113 | AJG44112.1 | K9N638 |
| VGVQTLEDLHAR | 1 | 0 | 0 | 1524114 | ATQ39389.1 | K9N7C7 |
| VGVQTVEDLCAR | 1 | 0 | 0 | 1524115 | AKN24802.1 | K9N7C7 |
| VGVQTVEDLRVR | 1 | 0 | 0 | 1524116 | QDP16215.1 | K9N7C7 |
| VGVVDTEWGNMM | 1 | 0 | 0 | 1524124 | AVV62524.1 | K9N7C7 |
| VGWSADSCLQGD | 1 | 0 | 0 | 1524129 | BBA84088.1 | P36334 |
| VGYFFPYDPALE | 1 | 0 | 0 | 1524138 | AVV62524.1 | K9N7C7 |
| VGYGCNCDSLRQ | 1 | 0 | 0 | 1524139 | AUM60013.1 | K9N7C7 |
| VGYHSDNGNYYC | 1 | 0 | 0 | 1524141 | AUM60014.1 | K9N5Q8 |
| VGYIPTSHINVT | 1 | 0 | 0 | 1524144 | AVV62526.1 | K9N5Q8 |
| VGYNQSFVDYLM | 1 | 0 | 0 | 1524150 | ARB07605.1 | A0A140H1G9 |
| VGYQPTAHVNAT | 1 | 0 | 0 | 1524151 | AUM60014.1 | K9N5Q8 |
| VGYTKDGSVINF | 1 | 0 | 0 | 1524156 | ATG84865.1 | K9N638 |
| VGYTKEGNVINF | 1 | 0 | 0 | 1524157 | AVV62535.1 | K9N7C7 |
| VGYVPTNHVNVT | 1 | 0 | 0 | 1524162 | AVV62537.1 | K9N5Q8 |
| VHAIATIPLQAS | 1 | 0 | 0 | 1524176 | ACQ82726.1 | P59632 |
| VHARLKGGLILR | 1 | 0 | 0 | 1524181 | AGV08389.1 | K9N638 |
| VHDLNNVDPKAY | 1 | 0 | 0 | 1524194 | AVP78050.1 | Q3LZX3 |
| VHDVEPTHKVKL | 1 | 0 | 0 | 1524199 | AWK59942.1 | H9EJ42 |
| VHEPTTPISSAL | 1 | 0 | 0 | 1524205 | ALA49956.1 | K9N7C7 |
| VHEWKMVVCDKV | 1 | 0 | 0 | 1524213 | AXN92258.1 | K9N7C7 |
| VHGNGSMIEDMD | 1 | 0 | 0 | 1524232 | AXG21663.1 | K9N7C7 |
| VHGNGSMIEDVD | 1 | 0 | 0 | 1524233 | AMQ49035.1 | K9N7C7 |
| VHGNVAFYQGDV | 1 | 0 | 0 | 1524234 | ARB07424.1 | A0A0P0ILY9 |
| VHLYSSRKGDLY | 1 | 0 | 0 | 1524268 | AVV62526.1 | K9N5Q8 |

| | | | | | | |
|---|---|---|---|---|---|---|
| VHNLSGNPDEPQ | 1 | 0 | 0 | 1524278 | AHA80155.1 | P33469 |
| VHNWLNGGQSVS | 1 | 0 | 0 | 1524286 | AVN89460.1 | K9N643 |
| VHPALHLVDPQI | 1 | 0 | 0 | 1524287 | AVP78050.1 | Q3LZX3 |
| VHPVECDEYEEE | 1 | 0 | 0 | 1524291 | AVV62535.1 | K9N7C7 |
| VHQVQLIDKYCS | 1 | 0 | 0 | 1524296 | ATG84700.1 | K9N638 |
| VHSCFIERVEWD | 1 | 0 | 0 | 1524305 | AUM60013.1 | K9N7C7 |
| VHSEFNLPGCNG | 1 | 0 | 0 | 1524311 | AUM60013.1 | K9N7C7 |
| VHSLRVCVETVH | 1 | 0 | 0 | 1524314 | AVP78041.1 | P0C6W2 |
| VHSVFLLMFLLT | 1 | 0 | 0 | 1524321 | AKN24803.1 | K9N5Q8 |
| VHTIDGSSGVVN | 1 | 0 | 0 | 1524333 | APO40580.1 | P59632 |
| VHTLSVHDGYTN | 1 | 0 | 0 | 1524336 | ARO76381.1 | P0C6U8 |
| VHVDSFSCNNLD | 1 | 0 | 0 | 1524341 | BBA20986.1 | A0A140H1H1 |
| VHVGGVYDNSIT | 1 | 0 | 0 | 1524343 | ASU90174.1 | K9N638 |
| VHVMSKHTDFSSEIIG | 1 | 0 | 0 | 1524351 | P0DTC1.1 | P0DTD1 |
| VHVYVDDSIILR | 1 | 0 | 0 | 1524357 | ARA15418.1 | P0C6X6 |
| VIADTLQEIPVV | 1 | 0 | 0 | 1524380 | AQZ41284.1 | K9N7C7 |
| VIALTGTNQLCA | 1 | 0 | 0 | 1524403 | AWK59942.1 | H9EJ42 |
| VIAWNTAKQDQG | 1 | 0 | 0 | 1524412 | ATO98193.1 | Q3LZX1 |
| VIAWNTAKQDVG | 1 | 0 | 0 | 1524414 | AVP78031.1 | Q3LZX1 |
| VIDDGDGVKPFR | 1 | 0 | 0 | 1524425 | QED88039.1 | H9EJ42 |
| VIDDMHWGEGFQ | 1 | 0 | 0 | 1524428 | NP_150075.1 | Q80872 |
| VIDGVTYQLAWD | 1 | 0 | 0 | 1524431 | AWK59942.1 | H9EJ42 |
| VIDLRLDDFVMI | 1 | 0 | 0 | 1524437 | AHL18091.1 | K9N7C7 |
| VIDTPSSANPVK | 1 | 0 | 0 | 1524442 | AVV62535.1 | K9N7C7 |
| VIDVKQVGCSMR | 1 | 0 | 0 | 1524443 | AIV41894.1 | P0C6X6 |
| VIEPRNGSYCPG | 1 | 0 | 0 | 1524453 | AVV62537.1 | K9N5Q8 |
| VIEVVDKYFDCY | 1 | 0 | 0 | 1524456 | AVP78041.1 | P0C6W2 |
| VIFNAVDCKSDF | 1 | 0 | 0 | 1524463 | CAA83660.1 | P36334 |
| VIFPGQKYSSDC | 1 | 0 | 0 | 1524464 | AXN92226.1 | K9N7C7 |
| VIFSDFYLPKIG | 1 | 0 | 0 | 1524466 | AEN19363.1 | P0C6X6 |
| VIFSDPNMCLLS | 1 | 0 | 0 | 1524467 | AVN89416.1 | K9N643 |
| VIFSQSDSLGVS | 1 | 0 | 0 | 1524471 | QBP84719.1 | P0C6X6 |
| VIGDFNCTNSFI | 1 | 0 | 0 | 1524483 | BBA20986.1 | A0A140H1H1 |
| VIGDLKCPLDSR | 1 | 0 | 0 | 1524484 | AVQ05264.1 | P36334 |
| VIGDLKCPLDTR | 1 | 0 | 0 | 1524485 | AKA09647.1 | P36334 |
| VIGDLKCPLDTS | 1 | 0 | 0 | 1524486 | AKA09663.1 | P36334 |
| VIGDLKCPLESR | 1 | 0 | 0 | 1524487 | QQY99180.1 | P36334 |
| VIGDLKCTSDTS | 1 | 0 | 0 | 1524488 | CAA79896.1 | P36334 |
| VIGDLNCPLDPK | 1 | 0 | 0 | 1524491 | AWW13511.1 | P36334 |
| VIGDLNCPLDPR | 1 | 0 | 0 | 1524492 | BAM16463.1 | P36334 |
| VIGDLNCPLDSR | 1 | 0 | 0 | 1524493 | AIL49507.1 | P36334 |
| VIGDLNCPLDTR | 1 | 0 | 0 | 1524494 | AEN19366.1 | P36334 |
| VIGDLNCTLDPR | 1 | 0 | 0 | 1524495 | AWW13566.1 | P36334 |
| VIGDLNCTLHPR | 1 | 0 | 0 | 1524496 | BBA84086.1 | P36334 |
| VIGHSMQNCVLK | 1 | 0 | 0 | 1524505 | QYC64967.1 | P0DTD1 |
| VIGIVNNTVYDPLQPEL | 1 | 0 | 0 | 1524508 | P0DTC2.1 | P0DTC2 |
| VIGTTKFCGGWN | 1 | 0 | 0 | 1524518 | ADK37693.1 | H9EJ42 |
| VIGTTKFYGGWG | 1 | 0 | 0 | 1524519 | ASU45728.1 | K9N7C7 |
| VIHYNINFEENL | 1 | 0 | 0 | 1524531 | BAL45639.1 | A0A0P0K6L9 |
| VIIANRYNEVPA | 1 | 0 | 0 | 1524537 | ATN39877.2 | P0C6X6 |
| VIIANRYNEVSA | 1 | 0 | 0 | 1524538 | AEN19363.1 | P0C6X6 |
| VIKDGVMYSEFP | 1 | 0 | 0 | 1524576 | CAA79896.1 | P36334 |
| VIKFETDAYKLF | 1 | 0 | 0 | 1524583 | ARU07575.1 | A0A140H1G9 |
| VIKHGVMYSEFP | 1 | 0 | 0 | 1524586 | CAA83661.1 | P36334 |
| VILKEPCPSGTY | 1 | 0 | 0 | 1524610 | 1XAK_A | P59635 |
| VILKPAADGLKI | 1 | 0 | 0 | 1524611 | APO40578.1 | P0C6X7 |
| VILKSSDEGIKV | 1 | 0 | 0 | 1524614 | ARO76381.1 | P0C6U8 |
| VILQFGYTSRSM | 1 | 0 | 0 | 1524629 | BBM61012.1 | Q01455 |
| VILSKTDVSTPV | 1 | 0 | 0 | 1524633 | AUM60013.1 | K9N7C7 |
| VILSTCDGQKFT | 1 | 0 | 0 | 1524634 | AEN19363.1 | P0C6X6 |
| VIMTLFSPTTSS | 1 | 0 | 0 | 1524650 | APO40579.1 | P59594 |
| VINITLANFNET | 1 | 0 | 0 | 1524663 | AIW52757.1 | A0A0P0K6L9 |
| VINKALEMCIDY | 1 | 0 | 0 | 1524664 | ARO76381.1 | P0C6U8 |
| VINKGHFDGQNG | 1 | 0 | 0 | 1524666 | AVP78030.1 | P0C6W2 |
| VINNVDNEIIKS | 1 | 0 | 0 | 1524671 | AVV62535.1 | K9N7C7 |
| VINNVDNEIVRS | 1 | 0 | 0 | 1524672 | AUM60013.1 | K9N7C7 |
| VINQNAVALNTL | 1 | 0 | 0 | 1524674 | APO40579.1 | P59594 |
| VINSSAMFCATC | 1 | 0 | 0 | 1524678 | AVV62535.1 | K9N7C7 |
| VIPFNSLWQRCL | 1 | 0 | 0 | 1524692 | APO40580.1 | P59632 |
| VIPLAFMPVQSW | 1 | 0 | 0 | 1524696 | AIV41888.1 | P0C6X6 |
| VIPRNSVPVNKK | 1 | 0 | 0 | 1524700 | AMM42513.1 | A0A0N7J131 |
| VIQDVKDGTVFS | 1 | 0 | 0 | 1524714 | CAA46113.1 | A0A0P0KWE8 |
| VIQDYENVCFDF | 1 | 0 | 0 | 1524716 | QBQ01836.1 | P0C6X6 |
| VIQPSNQRDAWA | 1 | 0 | 0 | 1524727 | ATQ39390.1 | K9N5Q8 |
| VIQQLHQGFKGK | 1 | 0 | 0 | 1524728 | AUM60013.1 | K9N7C7 |
| VIRYNINFEENQ | 1 | 0 | 0 | 1524753 | ACO44679.1 | A0A0P0K6L9 |
| VISDMYDPITKN | 1 | 0 | 0 | 1524759 | AVV62535.1 | K9N7C7 |
| VISDMYDPITKT | 1 | 0 | 0 | 1524760 | AUM60013.1 | K9N7C7 |
| VISDMYDPVTKN | 1 | 0 | 0 | 1524762 | AVV62524.1 | K9N7C7 |
| VISEDLWQFVDH | 1 | 0 | 0 | 1524763 | AGT21366.1 | A0A0P0LKV0 |
| VISPSTNTVIKK | 1 | 0 | 0 | 1524770 | AUM60014.1 | K9N5Q8 |
| VISQLDENAQKF | 1 | 0 | 0 | 1524771 | AST12962.1 | A0A0P0LKV0 |
| VISQYNYKQSFA | 1 | 0 | 0 | 1524772 | AVV62537.1 | K9N5Q8 |
| VISRLGFKLDAT | 1 | 0 | 0 | 1524773 | ALA49362.1 | K9N7C7 |
| VISRMGFKLDAI | 1 | 0 | 0 | 1524774 | QBM11746.1 | K9N7C7 |
| VISTWFKLMLKA | 1 | 0 | 0 | 1524783 | ARO76381.1 | P0C6U8 |
| VITAGDGISRPK | 1 | 0 | 0 | 1524788 | ARO76383.1 | P59632 |
| VITFDNLKTILS | 1 | 0 | 0 | 1524794 | AVP78041.1 | P0C6W2 |
| VITGVPKPVEGV | 1 | 0 | 0 | 1524798 | AIW52757.1 | A0A0P0K6L9 |
| VITKKEAAAAKN | 1 | 0 | 0 | 1524803 | AXG21672.1 | K9N4V7 |
| VITKRDAAAAKN | 1 | 0 | 0 | 1524805 | ANF29235.1 | K9N4V7 |

| | | | | | | |
|---|---|---|---|---|---|---|
| VITPGTNASSEF | 1 | 0 | 0 | 1524810 | BAE93401.1 | P59594 |
| VIVDTLQETPVV | 1 | 0 | 0 | 1524831 | AVK87459.1 | K9N7C7 |
| VIVGDYVIISEK | 1 | 0 | 0 | 1524835 | AXN92247.1 | K9N7C7 |
| VIVKETNLICKA | 1 | 0 | 0 | 1524838 | AXX83348.1 | P0C6X6 |
| VIVKRDATFATH | 1 | 0 | 0 | 1524840 | AGT51337.1 | H9EJ42 |
| VIVNDDTSDTKY | 1 | 0 | 0 | 1524852 | AIV41780.1 | P0C6X6 |
| VIVNNDTSETKY | 1 | 0 | 0 | 1524853 | AOL02450.1 | P0C6X6 |
| VIVQPTHYVRVT | 1 | 0 | 0 | 1524858 | AXN92231.1 | K9N643 |
| VIVRRDATFATH | 1 | 0 | 0 | 1524861 | AWK59942.1 | H9EJ42 |
| VIVWCINFLMLF | 1 | 0 | 0 | 1524866 | ARB07436.1 | A0A140H1G9 |
| VIWDHKREAPAH | 1 | 0 | 0 | 1524870 | AVP78041.1 | P0C6W2 |
| VIWDYKRDAPAH | 1 | 0 | 0 | 1524872 | QHD43415.1 | P0DTD1 |
| VIWLSHEKASLS | 1 | 0 | 0 | 1524881 | AIV41990.1 | P0C6X6 |
| VIWLSHEQASLN | 1 | 0 | 0 | 1524882 | AVI15051.1 | P0C6X6 |
| VIYDDNTNQYAT | 1 | 0 | 0 | 1524884 | ATQ39390.1 | K9N5Q8 |
| VIYDKKTKTHAT | 1 | 0 | 0 | 1524886 | AHL18090.1 | K9N5Q8 |
| VIYDKPSDKHAT | 1 | 0 | 0 | 1524887 | AUM60014.1 | K9N5Q8 |
| VIYGFNSTETIT | 1 | 0 | 0 | 1524889 | AIL49343.1 | P30215 |
| VIYGLNFTETIT | 1 | 0 | 0 | 1524890 | ATP16766.1 | P30215 |
| VIYQLDENAQKF | 1 | 0 | 0 | 1524906 | AMM42491.1 | A0A0P0ILY9 |
| VKAALLADKFPV | 1 | 0 | 0 | 1524913 | QYC64967.1 | P0DTD1 |
| VKAEVNPDLSAY | 1 | 0 | 0 | 1524917 | AVV62535.1 | K9N7C7 |
| VKAVRNKTSACF | 1 | 0 | 0 | 1524933 | AEN19363.1 | P0C6X6 |
| VKAVTAYSVDKC | 1 | 0 | 0 | 1524934 | QEG03783.1 | A0A0P0LKV0 |
| VKAWSGICVDGI | 1 | 0 | 0 | 1524936 | AIW52843.1 | A0A0P0G321 |
| VKCEQAYQLYDA | 1 | 0 | 0 | 1524937 | AIV42008.1 | P0C6X6 |
| VKCEQFYQLYDA | 1 | 0 | 0 | 1524938 | AWW13571.1 | P0C6X6 |
| VKCHNDILTAND | 1 | 0 | 0 | 1524941 | AUM60013.1 | K9N7C7 |
| VKDFVSGLLNVQ | 1 | 0 | 0 | 1524963 | APT69882.1 | A0A0P0ILY9 |
| VKDGFSFVDDVN | 1 | 0 | 0 | 1524964 | AVV62535.1 | K9N7C7 |
| VKDGFSFVNDDT | 1 | 0 | 0 | 1524965 | AVV62524.1 | K9N7C7 |
| VKDGIVWVHEEG | 1 | 0 | 0 | 1524967 | AVV62533.1 | K9N4V7 |
| VKDGKYFTSPPP | 1 | 0 | 0 | 1524974 | AUM60013.1 | K9N7C7 |
| VKDGKYFTSTPS | 1 | 0 | 0 | 1524975 | AVV62524.1 | K9N7C7 |
| VKDIDITFLKKD | 1 | 0 | 0 | 1524977 | AVP78041.1 | P0C6W2 |
| VKDNVSFVVDDS | 1 | 0 | 0 | 1524987 | AXN92247.1 | K9N7C7 |
| VKDSLIYKYDSG | 1 | 0 | 0 | 1524991 | AVV62535.1 | K9N7C7 |
| VKDTVVQLNYCR | 1 | 0 | 0 | 1524995 | AUM60013.1 | K9N7C7 |
| VKDVDHVQIHTI | 1 | 0 | 0 | 1524998 | Q3I5J4.1 | Q3LZX0 |
| VKDVFVESSKSL | 1 | 0 | 0 | 1525000 | QEG03747.1 | H9EJ42 |
| VKDYIQTVNVIA | 1 | 0 | 0 | 1525002 | ABD75583.1 | A0A140H1G9 |
| VKEAQNANPMVI | 1 | 0 | 0 | 1525008 | ARO76381.1 | P0C6U8 |
| VKEAQNANPTVI | 1 | 0 | 0 | 1525009 | ATO98229.1 | P0C6W2 |
| VKEEPTTQKPVD | 1 | 0 | 0 | 1525012 | ARO76381.1 | P0C6U8 |
| VKEFPLLSEYPF | 1 | 0 | 0 | 1525013 | ABD75542.1 | Q0ZJ80 |
| VKEIAPVYEADT | 1 | 0 | 0 | 1525017 | ALA49362.1 | K9N7C7 |
| VKEIISIPNIIK | 1 | 0 | 0 | 1525019 | AIX09804.1 | P0C6X6 |
| VKEITRENVETL | 1 | 0 | 0 | 1525025 | AGT51393.1 | H9EJ42 |
| VKEKLNAFLVHG | 1 | 0 | 0 | 1525029 | AST12962.1 | A0A0P0LKV0 |
| VKEQLDTTQLQV | 1 | 0 | 0 | 1525032 | AHF54662.1 | P36334 |
| VKEQVVDLLVKL | 1 | 0 | 0 | 1525034 | ATQ39389.1 | K9N7C7 |
| VKETVAQFNYRR | 1 | 0 | 0 | 1525040 | QAT98908.1 | K9N7C7 |
| VKFENLTNQLPP | 1 | 0 | 0 | 1525059 | AVV62537.1 | K9N5Q8 |
| VKFQNLTNNLPP | 1 | 0 | 0 | 1525073 | AUM60024.1 | K9N5Q8 |
| VKFSATQAMEKV | 1 | 0 | 0 | 1525075 | AIL49512.1 | P36334 |
| VKFVVESNVVDV | 1 | 0 | 0 | 1525082 | AGT51379.1 | H9EJ42 |
| VKGCPAVAVHDF | 1 | 0 | 0 | 1525091 | AVP78030.1 | P0C6W2 |
| VKGDDVRQIAPA | 1 | 0 | 0 | 1525092 | APO40579.1 | P59594 |
| VKGDSCSSNCKH | 1 | 0 | 0 | 1525096 | AKN24784.1 | K9N7C7 |
| VKGLDYKTFKAI | 1 | 0 | 0 | 1525099 | ARO76381.1 | P0C6U8 |
| VKGNNEAKALFF | 1 | 0 | 0 | 1525106 | AVV62535.1 | K9N7C7 |
| VKGWVGFDVEGA | 1 | 0 | 0 | 1525115 | ABD75519.1 | A0A140H1G9 |
| VKHITNITGVSL | 1 | 0 | 0 | 1525121 | AWK59942.1 | H9EJ42 |
| VKHSSTVDAQIQ | 1 | 0 | 0 | 1525127 | AVV62535.1 | K9N7C7 |
| VKKASYDTNLNK | 1 | 0 | 0 | 1525155 | AXN92226.1 | K9N7C7 |
| VKKLSPSREDKV | 1 | 0 | 0 | 1525157 | APO40587.1 | P59636 |
| VKKPFKVEDSVI | 1 | 0 | 0 | 1525158 | AEN19363.1 | P0C6X6 |
| VKKVAFCGDQVH | 1 | 0 | 0 | 1525161 | ALT66879.1 | K9N7C7 |
| VKKVIDVTGKLS | 1 | 0 | 0 | 1525164 | AGT51687.1 | P0C6X6 |
| VKLGAALAMEKV | 1 | 0 | 0 | 1525175 | ADN03339.1 | A0A140H1H1 |
| VKLGAENKDGFY | 1 | 0 | 0 | 1525176 | AIV41955.1 | Q80872 |
| VKLNPSEEFIKH | 1 | 0 | 0 | 1525193 | ALW82730.1 | K9N7C7 |
| VKLPFEPQTIQP | 1 | 0 | 0 | 1525194 | ALW82701.1 | K9N7C7 |
| VKLPRGVGSANI | 1 | 0 | 0 | 1525195 | ARU07606.1 | P0C6X6 |
| VKLPRWSASIDT | 1 | 0 | 0 | 1525196 | QNT54756.1 | A0A0P0K6L9 |
| VKLQAGSNTELA | 1 | 0 | 0 | 1525198 | AVV62535.1 | K9N7C7 |
| VKLQNNEIKPTG | 1 | 0 | 0 | 1525201 | AUM60013.1 | K9N7C7 |
| VKMLCTHTGTGQAITV | 1 | 0 | 0 | 1525221 | P0DTC1.1 | P0DTD1 |
| VKMVNPTSKVEP | 1 | 0 | 0 | 1525231 | AEN19363.1 | P0C6X6 |
| VKNGIVLPFEDL | 1 | 0 | 0 | 1525239 | AUM60013.1 | K9N7C7 |
| VKNIPCYASAVA | 1 | 0 | 0 | 1525242 | QBP84719.1 | P0C6X6 |
| VKNLNSSEGVSD | 1 | 0 | 0 | 1525250 | AAS48454.1 | P59637 |
| VKNLNSSQGIPD | 1 | 0 | 0 | 1525251 | APO40581.1 | P59637 |
| VKNLSTLRRGAV | 1 | 0 | 0 | 1525254 | AAW33912.1 | H9EJ42 |
| VKNNKGFKKGVM | 1 | 0 | 0 | 1525255 | AIL49539.1 | P36334 |
| VKNSQSSPIIPG | 1 | 0 | 0 | 1525260 | QBF80598.1 | K9N5Q8 |
| VKNTKLYVNNIL | 1 | 0 | 0 | 1525266 | AZS52618.1 | A0A140H1H1 |
| VKNTKVFKDGVM | 1 | 0 | 0 | 1525269 | AEN19366.1 | P36334 |
| VKNTNVFKDGVM | 1 | 0 | 0 | 1525278 | BBA84088.1 | P36334 |
| VKNVTDVTGNLS | 1 | 0 | 0 | 1525283 | AVQ05261.1 | P0C6X6 |
| VKPDMADEIANL | 1 | 0 | 0 | 1525296 | ARU07581.1 | Q19U29 |

| | | | | | | |
|---|---|---|---|---|---|---|
| VKPDMADEIASL | 1 | 0 | 0 | 1525297 | AMM42394.1 | Q19U29 |
| VKPESNADQETY | 1 | 0 | 0 | 1525303 | AVV62524.1 | K9N7C7 |
| VKPFLGQAAVTT | 1 | 0 | 0 | 1525308 | P0C6W6.1 | P0C6W2 |
| VKPFLNKVVSTT | 1 | 0 | 0 | 1525310 | AVP78030.1 | P0C6W2 |
| VKPIVEEIDITV | 1 | 0 | 0 | 1525321 | APO40578.1 | P0C6X7 |
| VKPVFFNAFVKC | 1 | 0 | 0 | 1525346 | AWK59942.1 | H9EJ42 |
| VKQEQRIGLDAV | 1 | 0 | 0 | 1525354 | AVQ05261.1 | P0C6X6 |
| VKQLPEQFDSLNLSAGTQ | 1 | 0 | 0 | 1525367 | ABD75581.1 | Q19U29 |
| VKQMYKTPAIKD | 1 | 0 | 0 | 1525373 | ATO98181.1 | Q3LZX1 |
| VKQSPELFDSLN | 1 | 0 | 0 | 1525382 | ARU07581.1 | Q19U29 |
| VKQSPELLDPLN | 1 | 0 | 0 | 1525383 | ADN03343.1 | Q19U29 |
| VKQSPESFDSLN | 1 | 0 | 0 | 1525384 | AMM42397.1 | Q19U29 |
| VKRHTLSNYQHE | 1 | 0 | 0 | 1525400 | APO40578.1 | P0C6X7 |
| VKRRFSLLRPEE | 1 | 0 | 0 | 1525410 | ASU90849.1 | K9N643 |
| VKSACIKVDIQQ | 1 | 0 | 0 | 1525422 | QCQ29075.1 | K9N5Q8 |
| VKSDYCRGNSCI | 1 | 0 | 0 | 1525433 | AIX10751.1 | P36334 |
| VKSGTAILIKSP | 1 | 0 | 0 | 1525444 | ARU07575.1 | A0A140H1G9 |
| VKSIPIHHDTFS | 1 | 0 | 0 | 1525445 | AUM60027.1 | K9N643 |
| VKSKGLFKEGST | 1 | 0 | 0 | 1525446 | AET99168.1 | A0A140H1G9 |
| VKSVSPGDSSWH | 1 | 0 | 0 | 1525479 | AVL25553.1 | A0A0P0G321 |
| VKTCDLTVGSGY | 1 | 0 | 0 | 1525484 | AIL49550.1 | P36334 |
| VKTDGTLMVERS | 1 | 0 | 0 | 1525487 | ASU90624.1 | K9N7C7 |
| VKTFETPAGQFC | 1 | 0 | 0 | 1525489 | AVV62524.1 | K9N7C7 |
| VKTLQPISDLLT | 1 | 0 | 0 | 1525500 | APO40578.1 | P0C6X7 |
| VKTLQPVSDLLA | 1 | 0 | 0 | 1525501 | ARO76381.1 | P0C6U8 |
| VKVPIDLIMIEF | 1 | 0 | 0 | 1525521 | ASU90525.1 | K9N7C7 |
| VKVPIDLTFVEF | 1 | 0 | 0 | 1525522 | AVV62524.1 | K9N7C7 |
| VKVPIDLTMVEF | 1 | 0 | 0 | 1525524 | AVV62535.1 | K9N7C7 |
| VKVPKGLTFSYD | 1 | 0 | 0 | 1525525 | AVV62535.1 | K9N7C7 |
| VKVPQGLTFSYD | 1 | 0 | 0 | 1525526 | AVV62524.1 | K9N7C7 |
| VKVVRNKTSACF | 1 | 0 | 0 | 1525533 | AIV41888.1 | P0C6X6 |
| VKWADAPEPQRG | 1 | 0 | 0 | 1525534 | APT69853.1 | A0A0N7J131 |
| VKWADESEPQRG | 1 | 0 | 0 | 1525536 | ABB90491.1 | A0A0N7J131 |
| VKWEYEGGCNTI | 1 | 0 | 0 | 1525537 | AWK59942.1 | H9EJ42 |
| VKYFECPNSIYI | 1 | 0 | 0 | 1525548 | ARU07575.1 | A0A140H1G9 |
| VKYYSRVKSVRG | 1 | 0 | 0 | 1525558 | ARK08640.1 | A0A0P0LKV0 |
| VLAAYNGKPQGA | 1 | 0 | 0 | 1525572 | AEN19363.1 | P0C6X6 |
| VLAAYNGRPQGA | 1 | 0 | 0 | 1525574 | ARU07575.1 | A0A140H1G9 |
| VLACYNGAPSGV | 1 | 0 | 0 | 1525576 | APO40578.1 | P0C6X7 |
| VLAEFDTYDPIY | 1 | 0 | 0 | 1525578 | AHI48794.1 | K9N638 |
| VLAGLGESFKKV | 1 | 0 | 0 | 1525581 | AVV62535.1 | K9N7C7 |
| VLARKHETSCSQ | 1 | 0 | 0 | 1525605 | AJE26223.1 | P0C6X6 |
| VLAVVNSQDVYK | 1 | 0 | 0 | 1525618 | ANF29227.1 | K9N638 |
| VLAVVVKRLYSP | 1 | 0 | 0 | 1525619 | AVV62524.1 | K9N7C7 |
| VLAWNTNSVDSK | 1 | 0 | 0 | 1525625 | APO40579.1 | P59594 |
| VLDILLASSSLR | 1 | 0 | 0 | 1525655 | AVN89312.1 | K9N638 |
| VLDNGIIRDINA | 1 | 0 | 0 | 1525666 | AUM60013.1 | K9N7C7 |
| VLDNGIIRDINP | 1 | 0 | 0 | 1525667 | ATQ39389.1 | K9N7C7 |
| VLDQSENNCWIS | 1 | 0 | 0 | 1525672 | ABE73416.1 | H9EJ42 |
| VLDTTIPFGHSS | 1 | 0 | 0 | 1525680 | AWO68990.1 | A0A0P0G321 |
| VLDVKGLEELHY | 1 | 0 | 0 | 1525684 | AIV41836.1 | Q80872 |
| VLECTRASTSAV | 1 | 0 | 0 | 1525693 | AXN92247.1 | K9N7C7 |
| VLENFIVYNNSG | 1 | 0 | 0 | 1525705 | AWK59942.1 | H9EJ42 |
| VLEQSGGRGGSI | 1 | 0 | 0 | 1525709 | 4NJL_A | K9N5Q8 |
| VLETTDNNCWVN | 1 | 0 | 0 | 1525713 | AGT51365.1 | H9EJ42 |
| VLEWDIKQCPAK | 1 | 0 | 0 | 1525721 | AUM60013.1 | K9N7C7 |
| VLFDGRVDGQVD | 1 | 0 | 0 | 1525731 | APO40578.1 | P0C6X7 |
| VLFDGRVEGQAN | 1 | 0 | 0 | 1525732 | AVP78041.1 | P0C6W2 |
| VLFNNGTHIVIN | 1 | 0 | 0 | 1525744 | APO40579.1 | P59594 |
| VLFSGQKYSSDC | 1 | 0 | 0 | 1525753 | AQZ41321.1 | K9N7C7 |
| VLFTNSTDTVNP | 1 | 0 | 0 | 1525761 | ARB07436.1 | A0A140H1G9 |
| VLFVNDKITEFQ | 1 | 0 | 0 | 1525765 | NP_073549.1 | A0A0P0LKV0 |
| VLGCLGSECNKV | 1 | 0 | 0 | 1525775 | CAA83661.1 | P36334 |
| VLGDAIQVAKCY | 1 | 0 | 0 | 1525780 | AXN92247.1 | K9N7C7 |
| VLGLYVPIATRN | 1 | 0 | 0 | 1525799 | AXX83378.1 | P0C6X6 |
| VLGLYVPKATRN | 1 | 0 | 0 | 1525800 | AEN19363.1 | P0C6X6 |
| VLGRGVFQNCTA | 1 | 0 | 0 | 1525805 | ASU90857.1 | K9N5Q8 |
| VLHTKSAAGYGG | 1 | 0 | 0 | 1525821 | AEN19363.1 | P0C6X6 |
| VLHTKSASGYGG | 1 | 0 | 0 | 1525822 | AIV41888.1 | P0C6X6 |
| VLIVLLCLGLVT | 1 | 0 | 0 | 1525855 | ATO98193.1 | Q3LZX1 |
| VLKHCCIAAVEL | 1 | 0 | 0 | 1525865 | AUM60013.1 | K9N7C7 |
| VLKITEDGINVK | 1 | 0 | 0 | 1525869 | AWK59942.1 | H9EJ42 |
| VLKLKVDTANPK | 1 | 0 | 0 | 1525874 | AVP78041.1 | P0C6W2 |
| VLKLLTLFFYVP | 1 | 0 | 0 | 1525876 | AFD98793.1 | H9EJ24 |
| VLKQWLPSGSIL | 1 | 0 | 0 | 1525887 | ARU07574.1 | A0A140H1G9 |
| VLKTGVLQPLEQ | 1 | 0 | 0 | 1525895 | AVP78041.1 | P0C6W2 |
| VLLADKFPILHD | 1 | 0 | 0 | 1525901 | APO40578.1 | P0C6X7 |
| VLLAEFVHYDPM | 1 | 0 | 0 | 1525906 | AUM60013.1 | K9N7C7 |
| VLLAEYSTYDPI | 1 | 0 | 0 | 1525907 | AVV62535.1 | K9N7C7 |
| VLLALQQIDIKF | 1 | 0 | 0 | 1525909 | APO40578.1 | P0C6X7 |
| VLLERVYKHFNN | 1 | 0 | 0 | 1525924 | AEN19363.1 | P0C6X6 |
| VLLKQPMATSVN | 1 | 0 | 0 | 1525949 | APU51922.1 | Q4VID0 |
| VLLLYFMVDNGT | 1 | 0 | 0 | 1525955 | P30215.1 | P30215 |
| VLLRRRDASSAG | 1 | 0 | 0 | 1525971 | AVV62526.1 | K9N5Q8 |
| VLLSEPCSPNTY | 1 | 0 | 0 | 1525972 | APO40584.1 | P59635 |
| VLLSMQGAVDLN | 1 | 0 | 0 | 1525976 | AVP78041.1 | P0C6W2 |
| VLMTARTVYDDA | 1 | 0 | 0 | 1525998 | P0C6W6.1 | P0C6W2 |
| VLNASEIYCANI | 1 | 0 | 0 | 1526011 | AXN92231.1 | K9N643 |
| VLNASEIYCANL | 1 | 0 | 0 | 1526012 | QDP16209.1 | K9N643 |
| VLNASTPESKPL | 1 | 0 | 0 | 1526016 | AIW52805.1 | H9EJ69 |
| VLNAVVKQLNES | 1 | 0 | 0 | 1526017 | AVV62526.1 | K9N5Q8 |

| | | | | | | |
|---|---|---|---|---|---|---|
| VLNDPVFTGELF | 1 | 0 | 0 | 1526024 | AGT51337.1 | H9EJ42 |
| VLNDTVEVVMGQ | 1 | 0 | 0 | 1526026 | AUM60013.1 | K9N7C7 |
| VLNEVVKQLNES | 1 | 0 | 0 | 1526033 | AVV62537.1 | K9N5Q8 |
| VLNNDYYKSLPG | 1 | 0 | 0 | 1526050 | QXN66612.1 | P0DTD1 |
| VLNNKSTYRSQL | 1 | 0 | 0 | 1526059 | AZK15899.1 | K9N638 |
| VLNNPAYLAPQA | 1 | 0 | 0 | 1526062 | AIL49375.1 | P30215 |
| VLNVVCKNCGQK | 1 | 0 | 0 | 1526074 | APO40578.1 | P0C6X7 |
| VLNVVGLDARTQ | 1 | 0 | 0 | 1526077 | AIV41936.1 | P0C6X6 |
| VLPFEDLTKCRT | 1 | 0 | 0 | 1526087 | AUM60013.1 | K9N7C7 |
| VLPGAFAYSQMR | 1 | 0 | 0 | 1526094 | AXN92247.1 | K9N7C7 |
| VLPGAHAYETMK | 1 | 0 | 0 | 1526095 | AVV62535.1 | K9N7C7 |
| VLPINVIVFDGK | 1 | 0 | 0 | 1526103 | APO40578.1 | P0C6X7 |
| VLPPLLSDNQIS | 1 | 0 | 0 | 1526118 | CAA83661.1 | P36334 |
| VLPPLLSENQIN | 1 | 0 | 0 | 1526119 | ANZ78836.1 | P36334 |
| VLPPLYDPNMEA | 1 | 0 | 0 | 1526124 | AVV62526.1 | K9N5Q8 |
| VLPPLYDVNMEA | 1 | 0 | 0 | 1526126 | AUM60014.1 | K9N5Q8 |
| VLPQGYYIEGSG | 1 | 0 | 0 | 1526127 | AEN19370.1 | P33469 |
| VLPVSMTKTSVD | 1 | 0 | 0 | 1526143 | APO40579.1 | P59594 |
| VLQEDQKILAAS | 1 | 0 | 0 | 1526149 | BAL45639.1 | A0A0P0K6L9 |
| VLQLKESQVNEL | 1 | 0 | 0 | 1526156 | AUM60013.1 | K9N7C7 |
| VLQNNELMPAKL | 1 | 0 | 0 | 1526162 | AEN19363.1 | P0C6X6 |
| VLQPLEQPTNEA | 1 | 0 | 0 | 1526164 | AVP78041.1 | P0C6W2 |
| VLQQLRIESSSK | 1 | 0 | 0 | 1526170 | APO40578.1 | P0C6X7 |
| VLQYGHYMYSRF | 1 | 0 | 0 | 1526185 | AWK59934.1 | U3M6Q8 |
| VLQYGYPSRSMT | 1 | 0 | 0 | 1526186 | ASY99815.1 | K9N7A1 |
| VLRCGDCLRKPM | 1 | 0 | 0 | 1526189 | AWK59942.1 | H9EJ42 |
| VLRELYGTPDSN | 1 | 0 | 0 | 1526195 | AVV62524.1 | K9N7C7 |
| VLRIGAAANSTG | 1 | 0 | 0 | 1526196 | AHI48723.1 | K9N5Q8 |
| VLRNSNGVCIWN | 1 | 0 | 0 | 1526203 | ATG84677.1 | K9N7C7 |
| VLRPILKLPFGI | 1 | 0 | 0 | 1526205 | ATO98181.1 | Q3LZX1 |
| VLRQWLPTDAIV | 1 | 0 | 0 | 1526210 | ASY99818.1 | K9N7C7 |
| VLRVLKQKNKSM | 1 | 0 | 0 | 1526215 | ABB90496.1 | A0A0N7J131 |
| VLSDIVNGVAEG | 1 | 0 | 0 | 1526225 | APO40578.1 | P0C6X7 |
| VLSDNQKLIANK | 1 | 0 | 0 | 1526226 | ALA49671.1 | K9N5Q8 |
| VLSELYGTPDPM | 1 | 0 | 0 | 1526231 | AVV62535.1 | K9N7C7 |
| VLSKSILLVDSS | 1 | 0 | 0 | 1526267 | ATQ39389.1 | K9N7C7 |
| VLSSAVMFEDAQ | 1 | 0 | 0 | 1526289 | AXG21642.1 | K9N638 |
| VLSSDVMFEDDQ | 1 | 0 | 0 | 1526290 | AGV08522.1 | K9N7C7 |
| VLSSDVMLEDAT | 1 | 0 | 0 | 1526292 | AUM60013.1 | K9N7C7 |
| VLSSLNLPGCDG | 1 | 0 | 0 | 1526294 | APO40578.1 | P0C6X7 |
| VLSVNPYVCNAL | 1 | 0 | 0 | 1526302 | AVP78041.1 | P0C6W2 |
| VLTDVDVPFTAH | 1 | 0 | 0 | 1526316 | AGT51637.1 | P0C6X6 |
| VLTENLNAYQQQ | 1 | 0 | 0 | 1526323 | AIL49396.1 | P33469 |
| VLTFTTKVTVPK | 1 | 0 | 0 | 1526328 | AIW52712.1 | A0A0N7J131 |
| VLTIANNYSKLS | 1 | 0 | 0 | 1526333 | AST12962.1 | A0A0P0LKV0 |
| VLTSNDERLSPV | 1 | 0 | 0 | 1526354 | AVV62535.1 | K9N7C7 |
| VLTVTLQNSRTP | 1 | 0 | 0 | 1526363 | AEN19363.1 | P0C6X6 |
| VLTYSNFGVCAD | 1 | 0 | 0 | 1526366 | AIW52748.1 | A0A0P0K6L9 |
| VLVKLLEVYNGF | 1 | 0 | 0 | 1526373 | AVA26872.1 | H9EJ42 |
| VLVNGLHYPTGS | 1 | 0 | 0 | 1526375 | ALD51908.1 | K9N7D2 |
| VLVNGVHHPTGS | 1 | 0 | 0 | 1526377 | QBM11740.1 | K9N7D2 |
| VLVNGVHYPTGI | 1 | 0 | 0 | 1526379 | ALW82735.1 | K9N7D2 |
| VLVNGVHYPTGS | 1 | 0 | 0 | 1526380 | AXG21669.1 | K9N7D2 |
| VLVNGVRYPTGS | 1 | 0 | 0 | 1526381 | AXN73419.1 | K9N7D2 |
| VLVNVSQTSIAN | 1 | 0 | 0 | 1526382 | CAA71056.1 | A0A0P0K6L9 |
| VLVPHVGEVPVA | 1 | 0 | 0 | 1526388 | AVP78041.1 | P0C6W2 |
| VLVYMILPTFNM | 1 | 0 | 0 | 1526401 | AVV62535.1 | K9N7C7 |
| VLWANGVKAKGY | 1 | 0 | 0 | 1526403 | AMM42513.1 | A0A0N7J131 |
| VLWDYEAERPFT | 1 | 0 | 0 | 1526405 | AWK59942.1 | H9EJ42 |
| VLYPQQKYTSQC | 1 | 0 | 0 | 1526441 | AUM60013.1 | K9N7C7 |
| VLYQGVNCTDVS | 1 | 0 | 0 | 1526446 | ACQ82725.1 | P59594 |
| VLYVDFKPQSGG | 1 | 0 | 0 | 1526453 | CAA71056.1 | A0A0P0K6L9 |
| VLYVDGHRLASG | 1 | 0 | 0 | 1526454 | AST12967.1 | A0A0P0KXN3 |
| VMAAPAAPRAVS | 1 | 0 | 0 | 1526461 | AIZ74430.1 | K9N4V7 |
| VMCGGSLYVKPS | 1 | 0 | 0 | 1526475 | AVP78041.1 | P0C6W2 |
| VMCPADQLNDPN | 1 | 0 | 0 | 1526476 | AVV62524.1 | K9N7C7 |
| VMCPADQLSDPN | 1 | 0 | 0 | 1526477 | AXN92247.1 | K9N7C7 |
| VMDYLDSLKPRV | 1 | 0 | 0 | 1526486 | ATO98229.1 | P0C6W2 |
| VMDYLDSLKPTV | 1 | 0 | 0 | 1526487 | ARO76381.1 | P0C6U8 |
| VMEPIDYNVVTQ | 1 | 0 | 0 | 1526491 | AGT51393.1 | H9EJ42 |
| VMFDLDSTFDDI | 1 | 0 | 0 | 1526492 | ARU07575.1 | A0A140H1G9 |
| VMFEDAQGLKKF | 1 | 0 | 0 | 1526494 | AXN92247.1 | K9N7C7 |
| VMFEDAQSLKKF | 1 | 0 | 0 | 1526495 | ALA50066.1 | K9N7C7 |
| VMFEGAQGLKKF | 1 | 0 | 0 | 1526497 | ASU90140.1 | K9N7C7 |
| VMFENAQGLKKF | 1 | 0 | 0 | 1526498 | AXN92226.1 | K9N7C7 |
| VMFEPRIPTIAD | 1 | 0 | 0 | 1526499 | ALK28787.1 | A0A0P0K6L9 |
| VMFQPRLPVLSD | 1 | 0 | 0 | 1526505 | AIW52843.1 | A0A0P0G321 |
| VMGQLEETNMHS | 1 | 0 | 0 | 1526517 | AUM60023.1 | K9N7C7 |
| VMGQLEQTNMHS | 1 | 0 | 0 | 1526518 | AUM60013.1 | K9N7C7 |
| VMILKSQDLGAV | 1 | 0 | 0 | 1526524 | ANI69887.1 | K9N7C7 |
| VMISEWPLFDQQ | 1 | 0 | 0 | 1526525 | APT69896.1 | A0A0P0LKV0 |
| VMKLPSEPQTIQ | 1 | 0 | 0 | 1526536 | ALA50044.1 | K9N7C7 |
| VMLKAAMLCNRY | 1 | 0 | 0 | 1526542 | ARU07574.1 | A0A140H1G9 |
| VMLLVKHKHTFL | 1 | 0 | 0 | 1526544 | AXN92247.1 | K9N7C7 |
| VMLNTSTSNLPD | 1 | 0 | 0 | 1526553 | AMK59677.1 | P36334 |
| VMLPGEFSKKQL | 1 | 0 | 0 | 1526554 | AKK52590.1 | K9N7C7 |
| VMNDKNICLLDH | 1 | 0 | 0 | 1526566 | P0C6U5.1 | A0A140H1G9 |
| VMNFDVDTHRYR | 1 | 0 | 0 | 1526568 | ACV87155.1 | A0A140H1G9 |
| VMNMDVSLHRHR | 1 | 0 | 0 | 1526572 | AXN92247.1 | K9N7C7 |
| VMPLTAPTLVPQ | 1 | 0 | 0 | 1526580 | ATO98130.1 | P0C6W2 |
| VMPTYTVHKSDF | 1 | 0 | 0 | 1526594 | ATN39877.2 | P0C6X6 |

| | | | | | | |
|---|---|---|---|---|---|---|
| VMQFGYIDCEQT | 1 | 0 | 0 | 1526601 | AIV42002.1 | P0C6X6 |
| VMQMDIQGFVNI | 1 | 0 | 0 | 1526606 | NP_073549.1 | A0A0P0LKV0 |
| VMQNNELMPHKL | 1 | 0 | 0 | 1526607 | ARU07575.1 | A0A140H1G9 |
| VMRPNHTIKGSF | 1 | 0 | 0 | 1526621 | AUM60013.1 | K9N7C7 |
| VMSEDLWQFVDH | 1 | 0 | 0 | 1526628 | NP_073549.1 | A0A0P0LKV0 |
| VMSLKDGQINDM | 1 | 0 | 0 | 1526630 | AVP78041.1 | P0C6W2 |
| VMTSQALFDSLQ | 1 | 0 | 0 | 1526646 | AVV62524.1 | K9N7C7 |
| VMVDCRRLLKQD | 1 | 0 | 0 | 1526650 | AZP73837.1 | P0C6X6 |
| VMVDCRRLLKQE | 1 | 0 | 0 | 1526651 | AEN19363.1 | P0C6X6 |
| VMVELVAELNGI | 1 | 0 | 0 | 1526657 | APO40578.1 | P0C6X7 |
| VMWFYTYSIGEA | 1 | 0 | 0 | 1526666 | AZK15899.1 | K9N638 |
| VMYMGTLSYEQL | 1 | 0 | 0 | 1526689 | AVP78041.1 | P0C6W2 |
| VMYSEFPAIVIG | 1 | 0 | 0 | 1526692 | AIL49490.1 | P36334 |
| VNANAEAHNNLL | 1 | 0 | 0 | 1526717 | ARB66064.1 | P36334 |
| VNANGGTCFCSK | 1 | 0 | 0 | 1526720 | ARK08626.1 | H9EJ42 |
| VNANHYSPCVSI | 1 | 0 | 0 | 1526721 | AID55087.1 | K9N5Q8 |
| VNANQYSPCVPI | 1 | 0 | 0 | 1526722 | QAT98898.1 | K9N5Q8 |
| VNANQYSPCVST | 1 | 0 | 0 | 1526724 | ANC28634.1 | K9N5Q8 |
| VNAPYYNSWQNL | 1 | 0 | 0 | 1526727 | CAA83660.1 | P36334 |
| VNASYECDIPIG | 1 | 0 | 0 | 1526732 | AVP78031.1 | Q3LZX1 |
| VNATYCNGWQNL | 1 | 0 | 0 | 1526736 | ABU39950.1 | P36334 |
| VNAYNSTAISVK | 1 | 0 | 0 | 1526743 | AIL49550.1 | P36334 |
| VNCCRFQRFDEN | 1 | 0 | 0 | 1526748 | AIX10734.1 | P0C6X6 |
| VNCNGYQHHSFA | 1 | 0 | 0 | 1526762 | ABE97137.1 | A0A0P0G321 |
| VNCNGYQHHSIA | 1 | 0 | 0 | 1526763 | AVL25583.1 | A0A0P0G321 |
| VNCNGYQHHSVA | 1 | 0 | 0 | 1526764 | AIW52843.1 | A0A0P0G321 |
| VNCNGYQHNSFA | 1 | 0 | 0 | 1526765 | ALJ53431.1 | A0A0P0G321 |
| VNCNGYQHNSVA | 1 | 0 | 0 | 1526766 | AIW52841.1 | A0A0P0G321 |
| VNCNGYQHNSVV | 1 | 0 | 0 | 1526767 | YP_003767.1 | A0A0P0G321 |
| VNCTDVPTAIRA | 1 | 0 | 0 | 1526770 | ATO98181.1 | Q3LZX1 |
| VNCTDVPTTIHA | 1 | 0 | 0 | 1526773 | AVP78031.1 | Q3LZX1 |
| VNCTDVPTVIRA | 1 | 0 | 0 | 1526774 | ATO98145.1 | Q3LZX1 |
| VNDCFKKDHVVL | 1 | 0 | 0 | 1526785 | QED88038.1 | H9EJ42 |
| VNDDTGETKYVK | 1 | 0 | 0 | 1526788 | AXX83366.1 | P0C6X6 |
| VNDFTCRQISPA | 1 | 0 | 0 | 1526791 | ATG84888.1 | K9N5Q8 |
| VNDFTCSQISPP | 1 | 0 | 0 | 1526792 | AHI48528.1 | K9N5Q8 |
| VNDFTCSQISPT | 1 | 0 | 0 | 1526793 | AWF93654.1 | K9N5Q8 |
| VNDICVFENKTN | 1 | 0 | 0 | 1526798 | AUM60013.1 | K9N7C7 |
| VNDKIFTKSGFN | 1 | 0 | 0 | 1526800 | AVN89289.1 | K9N7C7 |
| VNDQKLYDQVVM | 1 | 0 | 0 | 1526816 | ARO76381.1 | P0C6U8 |
| VNDSLKPVGGLY | 1 | 0 | 0 | 1526820 | AIL49500.1 | P36334 |
| VNECVESQSKRY | 1 | 0 | 0 | 1526826 | ABB90528.1 | A0A0P0K6L9 |
| VNEFSCNGISPD | 1 | 0 | 0 | 1526837 | AVV62526.1 | K9N5Q8 |
| VNESRYGLHNLL | 1 | 0 | 0 | 1526844 | ARU07594.1 | A0A0P0G321 |
| VNETSQTGWALY | 1 | 0 | 0 | 1526847 | AFO70499.1 | U3M6Q8 |
| VNETTSQYLFST | 1 | 0 | 0 | 1526848 | ALW82700.1 | K9N7C7 |
| VNFELRRGPGKC | 1 | 0 | 0 | 1526855 | AIW52747.1 | A0A0P0K6L9 |
| VNFNFNGLKGTG | 1 | 0 | 0 | 1526866 | AVP78031.1 | Q3LZX1 |
| VNFNIDDINFSP | 1 | 0 | 0 | 1526869 | ARB66060.1 | P36334 |
| VNFNVDDINFSS | 1 | 0 | 0 | 1526874 | AVQ05264.1 | P36334 |
| VNFNVDDVNFSP | 1 | 0 | 0 | 1526875 | CAA83661.1 | P36334 |
| VNFNVTNAATIV | 1 | 0 | 0 | 1526876 | ARU07601.1 | A0A0P0K6L9 |
| VNFNVTTASATD | 1 | 0 | 0 | 1526877 | AIW52843.1 | A0A0P0G321 |
| VNFTKALAKQDA | 1 | 0 | 0 | 1526884 | AVN89336.1 | K9N4V0 |
| VNFTVDPQKAYI | 1 | 0 | 0 | 1526890 | AUM60023.1 | K9N7C7 |
| VNFTVDPQKAYM | 1 | 0 | 0 | 1526892 | AUM60013.1 | K9N7C7 |
| VNGDSEVVLKKL | 1 | 0 | 0 | 1526898 | AVP78041.1 | P0C6W2 |
| VNGELLVDNVKY | 1 | 0 | 0 | 1526900 | AUM60013.1 | K9N7C7 |
| VNGFYRLYSGDN | 1 | 0 | 0 | 1526901 | QED88017.1 | H9EJ42 |
| VNGLHYPTGSCL | 1 | 0 | 0 | 1526908 | ALD51908.1 | K9N7D2 |
| VNGMKVVDYLTR | 1 | 0 | 0 | 1526918 | AVV62535.1 | K9N7C7 |
| VNGVHYPTGGCL | 1 | 0 | 0 | 1526928 | ALA49906.1 | K9N7D2 |
| VNHNVLIKESIP | 1 | 0 | 0 | 1526948 | AWK59942.1 | H9EJ42 |
| VNIDNSANTKTL | 1 | 0 | 0 | 1526951 | AVV62524.1 | K9N7C7 |
| VNIFSSTFNVPM | 1 | 0 | 0 | 1526952 | AVP78041.1 | P0C6W2 |
| VNINIVGDFQLN | 1 | 0 | 0 | 1526958 | P0C6W6.1 | P0C6W2 |
| VNINIVGDFQLT | 1 | 0 | 0 | 1526959 | ATO98229.1 | P0C6W2 |
| VNIPADIVDRDP | 1 | 0 | 0 | 1526961 | AIX10718.1 | P33469 |
| VNITLANFNETN | 1 | 0 | 0 | 1526969 | CAA71147.1 | A0A0P0K6L9 |
| VNKHAFHKPAYD | 1 | 0 | 0 | 1526985 | AHI48661.1 | K9N7C7 |
| VNLALKNLGFDN | 1 | 0 | 0 | 1527006 | ABK63972.1 | H9EJ69 |
| VNLFQDGFVVRI | 1 | 0 | 0 | 1527011 | AUM60014.1 | K9N5Q8 |
| VNLNIPANVPYS | 1 | 0 | 0 | 1527017 | AUM60013.1 | K9N7C7 |
| VNLNLPANIPYS | 1 | 0 | 0 | 1527018 | AXN92258.1 | K9N7C7 |
| VNLNLPANVPYS | 1 | 0 | 0 | 1527019 | AXN92247.1 | K9N7C7 |
| VNLNRGLPTGLS | 1 | 0 | 0 | 1527020 | AVP78031.1 | Q3LZX1 |
| VNLRLVDCAPVV | 1 | 0 | 0 | 1527025 | AWK59942.1 | H9EJ42 |
| VNLVTVPTGYVD | 1 | 0 | 0 | 1527032 | AVP78041.1 | P0C6W2 |
| VNMEAAYTSSLV | 1 | 0 | 0 | 1527037 | AID55085.1 | K9N5Q8 |
| VNMQLMGVVMQS | 1 | 0 | 0 | 1527043 | AQZ41284.1 | K9N7C7 |
| VNNDFIEFTDSF | 1 | 0 | 0 | 1527052 | AUM60013.1 | K9N7C7 |
| VNNDYCCFKHAL | 1 | 0 | 0 | 1527053 | ARB07390.1 | A0A0P0LKV0 |
| VNNGLTCSTDLQ | 1 | 0 | 0 | 1527059 | AEN19366.1 | P36334 |
| VNNLDKSAGYPF | 1 | 0 | 0 | 1527064 | QWK72597.1 | P0DTD1 |
| VNNNAQALAKLA | 1 | 0 | 0 | 1527066 | AVK87450.1 | K9N5Q8 |
| VNNQTTYRAQLG | 1 | 0 | 0 | 1527069 | AVV62524.1 | K9N7C7 |
| VNNQTTYRTQLG | 1 | 0 | 0 | 1527070 | AVV62535.1 | K9N7C7 |
| VNNSLFDSELNK | 1 | 0 | 0 | 1527073 | AUM60013.1 | K9N7C7 |
| VNNTWMFTGSRY | 1 | 0 | 0 | 1527080 | CAA83661.1 | P36334 |
| VNNTWMHTGSGY | 1 | 0 | 0 | 1527081 | ARB66068.1 | P36334 |
| VNNYEKSAGYPF | 1 | 0 | 0 | 1527085 | AIL49570.1 | P0C6X6 |

| | | | | | | |
|---|---|---|---|---|---|---|
| VNPANMRNANGA | 1 | 0 | 0 | 1527089 | AVV62524.1 | K9N7C7 |
| VNPASMRNANGA | 1 | 0 | 0 | 1527090 | AUM60013.1 | K9N7C7 |
| VNPDLSAYYVKD | 1 | 0 | 0 | 1527092 | AVV62535.1 | K9N7C7 |
| VNPDSFNLYGYS | 1 | 0 | 0 | 1527093 | ARB07436.1 | A0A140H1G9 |
| VNPKHQIIVLYG | 1 | 0 | 0 | 1527097 | AVV62524.1 | K9N7C7 |
| VNPTNCIAPVNG | 1 | 0 | 0 | 1527103 | ATG84723.1 | K9N5Q8 |
| VNPVMDPIYDEP | 1 | 0 | 0 | 1527106 | ARO76383.1 | P59632 |
| VNQIANMFLEQT | 1 | 0 | 0 | 1527115 | QBF80541.1 | K9N7C7 |
| VNQIVLRNSNGV | 1 | 0 | 0 | 1527118 | ATG84677.1 | K9N7C7 |
| VNQQGNLLNHLN | 1 | 0 | 0 | 1527123 | BAL45637.1 | A0A0P0K6L9 |
| VNQQGNSLSHLT | 1 | 0 | 0 | 1527124 | ARA15429.1 | A0A0P0K6L9 |
| VNQSLIKSKVSA | 1 | 0 | 0 | 1527126 | ARB07393.1 | A0A0P0KWE8 |
| VNQSTNKTTFRP | 1 | 0 | 0 | 1527129 | APO40578.1 | P0C6X7 |
| VNQTVLRNSNGA | 1 | 0 | 0 | 1527131 | AVN89451.1 | K9N7C7 |
| VNRGEDIQLLKN | 1 | 0 | 0 | 1527136 | AVP78041.1 | P0C6W2 |
| VNSALEQYVFCT | 1 | 0 | 0 | 1527141 | APO40578.1 | P0C6X7 |
| VNSDFCPCADRS | 1 | 0 | 0 | 1527145 | ABD75545.1 | A0A140H1H1 |
| VNSEDIYNTLTE | 1 | 0 | 0 | 1527146 | AUM60013.1 | K9N7C7 |
| VNSGGTPLTNCV | 1 | 0 | 0 | 1527152 | AUM60013.1 | K9N7C7 |
| VNSGLLTACAFS | 1 | 0 | 0 | 1527153 | AWK59942.1 | H9EJ42 |
| VNSINDSLEPVG | 1 | 0 | 0 | 1527160 | AIL49542.1 | P36334 |
| VNSKANAIQQQF | 1 | 0 | 0 | 1527162 | AUM60013.1 | K9N7C7 |
| VNSKNDNLEPVS | 1 | 0 | 0 | 1527163 | AIL49538.1 | P36334 |
| VNSNQYSPCVSI | 1 | 0 | 0 | 1527167 | AHY22555.1 | K9N5Q8 |
| VNSQCDLTGRTP | 1 | 0 | 0 | 1527169 | AVP78042.1 | Q3LZX1 |
| VNSQCVNLTGRT | 1 | 0 | 0 | 1527170 | AVP78031.1 | Q3LZX1 |
| VNSQDVYKSLTT | 1 | 0 | 0 | 1527172 | ATQ39389.1 | K9N7C7 |
| VNSSFTEQLGVP | 1 | 0 | 0 | 1527176 | AWO68966.1 | A0A0P0G321 |
| VNSSLFVEDCNL | 1 | 0 | 0 | 1527180 | ALA49451.1 | K9N5Q8 |
| VNSSMVVDNCQL | 1 | 0 | 0 | 1527184 | AVV62537.1 | K9N5Q8 |
| VNSVFNICQAVS | 1 | 0 | 0 | 1527191 | ADO24692.1 | P0C6X6 |
| VNSVNDSLKPVG | 1 | 0 | 0 | 1527200 | AIL49500.1 | P36334 |
| VNSVSDSLEPVG | 1 | 0 | 0 | 1527204 | ARB66068.1 | P36334 |
| VNTAIPTIEDWL | 1 | 0 | 0 | 1527207 | AHI48842.1 | K9N643 |
| VNTHSIRLTITE | 1 | 0 | 0 | 1527214 | AIW52813.1 | H9EJ42 |
| VNTLLSNQLRLA | 1 | 0 | 0 | 1527217 | AWK59942.1 | H9EJ42 |
| VNTPADIDYRNP | 1 | 0 | 0 | 1527221 | AJQ24074.1 | P33469 |
| VNTPADILDRDP | 1 | 0 | 0 | 1527222 | QBP84708.1 | P33469 |
| VNTPADIVDRDP | 1 | 0 | 0 | 1527223 | AEN19370.1 | P33469 |
| VNTPADIVDRDQ | 1 | 0 | 0 | 1527224 | ANZ78879.1 | P33469 |
| VNTPADIVHRDP | 1 | 0 | 0 | 1527225 | AJQ24067.1 | P33469 |
| VNVAAWLYAAIL | 1 | 0 | 0 | 1527241 | AHI48798.1 | K9N7C7 |
| VNVALADLYHYQ | 1 | 0 | 0 | 1527246 | APO40584.1 | P59635 |
| VNVDNSANTKIL | 1 | 0 | 0 | 1527253 | AVV62535.1 | K9N7C7 |
| VNVFQTQAGCLI | 1 | 0 | 0 | 1527254 | ATO98145.1 | Q3LZX1 |
| VNVIGTRPEKFP | 1 | 0 | 0 | 1527255 | AST12962.1 | A0A0P0LKV0 |
| VNVKPKRLRKKR | 1 | 0 | 0 | 1527259 | ATG84700.1 | K9N638 |
| VNVMFDLDSTFD | 1 | 0 | 0 | 1527265 | ABD75639.1 | A0A140H1G9 |
| VNVNNTWMFTGS | 1 | 0 | 0 | 1527267 | CAA83660.1 | P36334 |
| VNVNNTWMHTGS | 1 | 0 | 0 | 1527268 | ARB66068.1 | P36334 |
| VNVPTIRKFIKF | 1 | 0 | 0 | 1527270 | AEN19363.1 | P0C6X6 |
| VNVPYLYNGSAQ | 1 | 0 | 0 | 1527271 | AVQ05263.1 | P30215 |
| VNVSHVYNGSAQ | 1 | 0 | 0 | 1527278 | QQY99179.1 | P30215 |
| VNVSYVYNGSAQ | 1 | 0 | 0 | 1527286 | AEN19365.1 | P30215 |
| VNVTAAYGLCNT | 1 | 0 | 0 | 1527287 | AVV62537.1 | K9N5Q8 |
| VNVTFFPDLNGD | 1 | 0 | 0 | 1527288 | APO40578.1 | P0C6X7 |
| VNVTKHKCDISY | 1 | 0 | 0 | 1527289 | AWW13571.1 | P0C6X6 |
| VNVTTLNGRIVN | 1 | 0 | 0 | 1527292 | AIW52843.1 | A0A0P0G321 |
| VNWNLDYPIISN | 1 | 0 | 0 | 1527307 | ABD75551.1 | A0A140H1G9 |
| VNYTKAPDVMLN | 1 | 0 | 0 | 1527319 | CAA83660.1 | P36334 |
| VPAAYKLTKLSV | 1 | 0 | 0 | 1527323 | AIW52843.1 | A0A0P0G321 |
| VPADQLADPNYD | 1 | 0 | 0 | 1527324 | AUM60013.1 | K9N7C7 |
| VPAFPETEVASY | 1 | 0 | 0 | 1527328 | AVV62530.1 | K9N7D2 |
| VPAHMPMHPMVN | 1 | 0 | 0 | 1527331 | AET99209.1 | Q0ZJ80 |
| VPAHMPMPPMVN | 1 | 0 | 0 | 1527332 | ARU07582.1 | Q0ZJ80 |
| VPAHSFTLGGKV | 1 | 0 | 0 | 1527333 | AUM60013.1 | K9N7C7 |
| VPANSLHATATI | 1 | 0 | 0 | 1527338 | APO40580.1 | P59632 |
| VPATEAKGYWYK | 1 | 0 | 0 | 1527345 | AJQ24067.1 | P33469 |
| VPCCGTGDLAEH | 1 | 0 | 0 | 1527358 | AVV62535.1 | K9N7C7 |
| VPCLYKRNFTYD | 1 | 0 | 0 | 1527361 | ALJ77695.1 | P36334 |
| VPDDAARLLHIL | 1 | 0 | 0 | 1527366 | AVV62535.1 | K9N7C7 |
| VPDDASRLIHTL | 1 | 0 | 0 | 1527367 | AVV62524.1 | K9N7C7 |
| VPDIVDVPPQVV | 1 | 0 | 0 | 1527375 | AVN89352.1 | K9N7C7 |
| VPDIVEVPPQVV | 1 | 0 | 0 | 1527376 | AVN89451.1 | K9N7C7 |
| VPDLVVEQYNQI | 1 | 0 | 0 | 1527377 | ABB90514.1 | A0A0P0K6L9 |
| VPDTVGVPPQVV | 1 | 0 | 0 | 1527382 | AHL18091.1 | K9N7C7 |
| VPDYGTYKNTCE | 1 | 0 | 0 | 1527384 | APO40578.1 | P0C6X7 |
| VPEEAPAPELQV | 1 | 0 | 0 | 1527386 | APO40578.1 | P0C6X7 |
| VPEHCQNYSGFM | 1 | 0 | 0 | 1527389 | AVN89325.1 | K9N796 |
| VPEHCQNYSGSM | 1 | 0 | 0 | 1527390 | ASU45697.1 | K9N796 |
| VPFLICSNTPAS | 1 | 0 | 0 | 1527407 | AEN19363.1 | P0C6X6 |
| VPFSPNGKPCTP | 1 | 0 | 0 | 1527413 | ABD72972.1 | P59594 |
| VPFWMTIVYVIC | 1 | 0 | 0 | 1527426 | AVP78041.1 | P0C6W2 |
| VPGDVIITSTGA | 1 | 0 | 0 | 1527428 | AWK59942.1 | H9EJ42 |
| VPGLPGTILRTI | 1 | 0 | 0 | 1527439 | AVP78041.1 | P0C6W2 |
| VPGLPGTVIRTL | 1 | 0 | 0 | 1527441 | APO40578.1 | P0C6X7 |
| VPGNVVPGDVII | 1 | 0 | 0 | 1527445 | AWK59942.1 | H9EJ42 |
| VPGYPKLFITCE | 1 | 0 | 0 | 1527458 | AQZ41330.1 | K9N7C7 |
| VPHIKEWLKVQG | 1 | 0 | 0 | 1527461 | AUM60027.1 | K9N643 |
| VPHNLTTITKPF | 1 | 0 | 0 | 1527463 | AVN89387.1 | K9N5Q8 |
| VPHNLTTITKPL | 1 | 0 | 0 | 1527464 | AXN92228.1 | K9N5Q8 |

| | | | | | | |
|---|---|---|---|---|---|---|
| VPHTRYSLADSN | 1 | 0 | 0 | 1527467 | ABD75543.1 | A0A140H1G9 |
| VPHTRYSLANSN | 1 | 0 | 0 | 1527468 | ARU07575.1 | A0A140H1G9 |
| VPIAFGIPPSEA | 1 | 0 | 0 | 1527476 | ABA60794.1 | Q19U29 |
| VPIAFGVPLSEA | 1 | 0 | 0 | 1527477 | ABD75541.1 | Q19U29 |
| VPIDEYITTYPG | 1 | 0 | 0 | 1527486 | AER30328.1 | P0C6X7 |
| VPIEYQATRVVV | 1 | 0 | 0 | 1527490 | ATO98193.1 | Q3LZX1 |
| VPIGKGSKDEQI | 1 | 0 | 0 | 1527496 | AIW52795.1 | H9EJ69 |
| VPIGTNDPIRSC | 1 | 0 | 0 | 1527497 | CAA49377.1 | A0A0P0LKV0 |
| VPIHYEMFGNYT | 1 | 0 | 0 | 1527499 | ATO98227.1 | Q3LZX5 |
| VPINMALTHVRE | 1 | 0 | 0 | 1527502 | AVV62529.1 | K9N643 |
| VPINTNSGKDDQ | 1 | 0 | 0 | 1527503 | Q3I5I7.1 | Q3LZX4 |
| VPINTNSSKDDQ | 1 | 0 | 0 | 1527505 | AVP78038.1 | Q3LZX4 |
| VPIRLAGHIRHL | 1 | 0 | 0 | 1527507 | ASU90888.1 | K9N7C7 |
| VPKGLTFSYDGL | 1 | 0 | 0 | 1527516 | AVV62535.1 | K9N7C7 |
| VPKGSITQRSRP | 1 | 0 | 0 | 1527517 | AUM60021.1 | K9N4V7 |
| VPKIFKARAIEC | 1 | 0 | 0 | 1527518 | AMM42360.1 | A0A140H1G9 |
| VPKIFKARAIEF | 1 | 0 | 0 | 1527519 | ARU07575.1 | A0A140H1G9 |
| VPKNFCPCKLNG | 1 | 0 | 0 | 1527520 | AGT51590.1 | P36334 |
| VPLAFGVPLSEA | 1 | 0 | 0 | 1527527 | ABA60796.1 | Q19U29 |
| VPLCGSGNLVEK | 1 | 0 | 0 | 1527529 | AXN92247.1 | K9N7C7 |
| VPLGIKDPILYV | 1 | 0 | 0 | 1527538 | ARU07575.1 | A0A140H1G9 |
| VPLMHKGAAWPI | 1 | 0 | 0 | 1527555 | AXN92247.1 | K9N7C7 |
| VPLNANSTPSQN | 1 | 0 | 0 | 1527558 | AQZ41279.1 | K9N4V7 |
| VPLNLSGFTKHL | 1 | 0 | 0 | 1527561 | AUM60013.1 | K9N7C7 |
| VPLNSNSTPAQN | 1 | 0 | 0 | 1527562 | AHI48825.1 | K9N4V7 |
| VPLQTIINGMHK | 1 | 0 | 0 | 1527566 | AWK59942.1 | H9EJ42 |
| VPLSAVPLCASN | 1 | 0 | 0 | 1527571 | AVV62524.1 | K9N7C7 |
| VPLSEAKGYWYR | 1 | 0 | 0 | 1527572 | ABA60796.1 | Q19U29 |
| VPLSEVYVQLRN | 1 | 0 | 0 | 1527574 | QED88043.1 | U3M6Q8 |
| VPLSFPPGQGVP | 1 | 0 | 0 | 1527575 | AUM60021.1 | K9N4V7 |
| VPLSVVYDKTTN | 1 | 0 | 0 | 1527577 | AVV62526.1 | K9N5Q8 |
| VPLTFSPGQGVP | 1 | 0 | 0 | 1527580 | AHI48535.1 | K9N4V7 |
| VPPIISEKISVM | 1 | 0 | 0 | 1527606 | AMM42546.1 | H9EJ42 |
| VPPQVMKLPSEP | 1 | 0 | 0 | 1527609 | ALA50044.1 | K9N7C7 |
| VPPQVVELPYEP | 1 | 0 | 0 | 1527612 | ANI69888.1 | K9N638 |
| VPPRCIAPVDGY | 1 | 0 | 0 | 1527617 | AUM60014.1 | K9N5Q8 |
| VPPTSPTARTTS | 1 | 0 | 0 | 1527619 | AUM60024.1 | K9N5Q8 |
| VPQAEVDWKFYD | 1 | 0 | 0 | 1527624 | AVP78041.1 | P0C6W2 |
| VPQLVNSNQYSP | 1 | 0 | 0 | 1527637 | AHY22555.1 | K9N5Q8 |
| VPQSEVDWKFYD | 1 | 0 | 0 | 1527641 | AVP78030.1 | P0C6W2 |
| VPQSVETKVQCS | 1 | 0 | 0 | 1527642 | AIV41864.1 | P0C6X6 |
| VPRASADIGANH | 1 | 0 | 0 | 1527648 | ARO76381.1 | P0C6U8 |
| VPRGSGMKETAA | 1 | 0 | 0 | 1527655 | 3R24_A | P0C6X7 |
| VPRGSLEWSHPQ | 1 | 0 | 0 | 1527656 | 5X58_A | P59594 |
| VPRNKWNGKSSL | 1 | 0 | 0 | 1527660 | AUM60013.1 | K9N7C7 |
| VPSAELTGTQSV | 1 | 0 | 0 | 1527665 | AVN89314.1 | K9N796 |
| VPSAYKLTKLSV | 1 | 0 | 0 | 1527667 | AWO68991.1 | A0A0P0G321 |
| VPSDSCNNVNVR | 1 | 0 | 0 | 1527669 | AIW52827.1 | H9EJ42 |
| VPSIVSEKISVM | 1 | 0 | 0 | 1527679 | AVA26872.1 | H9EJ42 |
| VPSLPTSGSRVA | 1 | 0 | 0 | 1527681 | ABB90513.1 | A0A0P0K6L9 |
| VPSNDNISLPFN | 1 | 0 | 0 | 1527682 | AVV62524.1 | K9N7C7 |
| VPSNKWTGKKNL | 1 | 0 | 0 | 1527683 | AVV62535.1 | K9N7C7 |
| VPSNKWTGKKTL | 1 | 0 | 0 | 1527684 | AVV62524.1 | K9N7C7 |
| VPSPLNWERKIF | 1 | 0 | 0 | 1527686 | AET99191.1 | A0A140H1H1 |
| VPSQQQNFTTAP | 1 | 0 | 0 | 1527692 | APO40579.1 | P59594 |
| VPSTEAKGYWYP | 1 | 0 | 0 | 1527698 | AJQ24074.1 | P33469 |
| VPSTLWEDGDYY | 1 | 0 | 0 | 1527699 | ALJ54501.1 | K9N5Q8 |
| VPSTSGTQVYSR | 1 | 0 | 0 | 1527703 | AXN92231.1 | K9N643 |
| VPSYVKPDSIAT | 1 | 0 | 0 | 1527714 | AVV62535.1 | K9N7C7 |
| VPTAIRADQLTP | 1 | 0 | 0 | 1527716 | ATO98181.1 | Q3LZX1 |
| VPTDNHITTYPG | 1 | 0 | 0 | 1527720 | AVP78041.1 | P0C6W2 |
| VPTGYVDTENST | 1 | 0 | 0 | 1527727 | AVP78041.1 | P0C6W2 |
| VPTGYVDTSANT | 1 | 0 | 0 | 1527729 | APO40578.1 | P0C6X7 |
| VPTKYATAKVSP | 1 | 0 | 0 | 1527739 | AQN78776.1 | P36334 |
| VPTMADFVQIEN | 1 | 0 | 0 | 1527745 | ABB90513.1 | A0A0P0K6L9 |
| VPTNFSFGITQE | 1 | 0 | 0 | 1527747 | AVV62537.1 | K9N5Q8 |
| VPTNHVNVTAAY | 1 | 0 | 0 | 1527748 | AVV62537.1 | K9N5Q8 |
| VPTNVYLVGKTL | 1 | 0 | 0 | 1527749 | AWK59942.1 | H9EJ42 |
| VPTQPLPNATYD | 1 | 0 | 0 | 1527751 | APO40578.1 | P0C6X7 |
| VPTREENVMQCF | 1 | 0 | 0 | 1527753 | AMM42619.1 | H9EJ69 |
| VPTTLAWVNNQI | 1 | 0 | 0 | 1527758 | AXN92247.1 | K9N7C7 |
| VPTTTGALVRFV | 1 | 0 | 0 | 1527760 | AGT51358.1 | H9EJ42 |
| VPTTTGILVRFV | 1 | 0 | 0 | 1527762 | AWK59942.1 | H9EJ42 |
| VPTVIRADQLTP | 1 | 0 | 0 | 1527764 | ATO98145.1 | Q3LZX1 |
| VPVAYRKVLLRK | 1 | 0 | 0 | 1527769 | AVP78041.1 | P0C6W2 |
| VPVEDIVQVVIV | 1 | 0 | 0 | 1527773 | AVK87459.1 | K9N7C7 |
| VPVEFYEDRILD | 1 | 0 | 0 | 1527775 | AXN92247.1 | K9N7C7 |
| VPVGDSFGKSLG | 1 | 0 | 0 | 1527778 | ATN39877.2 | P0C6X6 |
| VPVGESFGKSLG | 1 | 0 | 0 | 1527779 | AEN19363.1 | P0C6X6 |
| VPVGIRDPVSYV | 1 | 0 | 0 | 1527782 | ATN39862.1 | P0C6X6 |
| VPVHMPVHPMVM | 1 | 0 | 0 | 1527783 | AET99208.1 | Q0ZJ80 |
| VPVSPAFHCIES | 1 | 0 | 0 | 1527790 | ANI69893.1 | K9N7D2 |
| VPVSPVDHGVES | 1 | 0 | 0 | 1527794 | ATQ39397.1 | K9N7D2 |
| VPVSPVFHRIES | 1 | 0 | 0 | 1527795 | AVN89455.1 | K9N7D2 |
| VPVSVIYDDNTN | 1 | 0 | 0 | 1527796 | ATQ39390.1 | K9N5Q8 |
| VPVTRDSVPLHI | 1 | 0 | 0 | 1527802 | AHI48554.1 | K9N7D2 |
| VPYCYDPTVLPG | 1 | 0 | 0 | 1527812 | AVV62524.1 | K9N7C7 |
| VPYIVGDVIKEG | 1 | 0 | 0 | 1527815 | AVP78041.1 | P0C6W2 |
| VPYKVCCTLRDT | 1 | 0 | 0 | 1527816 | AUM60013.1 | K9N7C7 |
| VPYLYNGSAQAT | 1 | 0 | 0 | 1527818 | AVQ05263.1 | P30215 |
| VPYQRNNTSFPE | 1 | 0 | 0 | 1527822 | ATQ39389.1 | K9N7C7 |

| | | | | | | |
|---|---|---|---|---|---|---|
| VPYTALYYYATV | 1 | 0 | 0 | 1527829 | QBF80541.1 | K9N7C7 |
| VPYVYNGSAQST | 1 | 0 | 0 | 1527835 | P30215.1 | P30215 |
| VPYVYNGSVQAR | 1 | 0 | 0 | 1527836 | AGT51729.1 | P30215 |
| VQADTAGLYTNF | 1 | 0 | 0 | 1527841 | AHC74089.1 | K9N796 |
| VQAFGLRGSGDL | 1 | 0 | 0 | 1527844 | AUM60021.1 | K9N4V7 |
| VQALESKVLLER | 1 | 0 | 0 | 1527849 | QEG03730.1 | H9EJ42 |
| VQCMTGFNTLLV | 1 | 0 | 0 | 1527863 | ASU90554.1 | K9N5R3 |
| VQDHVDILGPLS | 1 | 0 | 0 | 1527875 | APO40578.1 | P0C6X7 |
| VQDRVVEPILAT | 1 | 0 | 0 | 1527886 | AHF54652.1 | Q4VID0 |
| VQEAVNANAQAL | 1 | 0 | 0 | 1527890 | AUM60014.1 | K9N5Q8 |
| VQEFVSYDFSIH | 1 | 0 | 0 | 1527900 | AQN78693.1 | P0C6X6 |
| VQFCDAMRNAGI | 1 | 0 | 0 | 1527918 | ATO98179.1 | P0C6W2 |
| VQFGRSGITLGV | 1 | 0 | 0 | 1527921 | ARO76381.1 | P0C6U8 |
| VQFNYCRDGQPF | 1 | 0 | 0 | 1527925 | AVK87459.1 | K9N7C7 |
| VQFQIRMIALLK | 1 | 0 | 0 | 1527930 | APO40583.1 | P59634 |
| VQGFACDCGACF | 1 | 0 | 0 | 1527936 | AVV62524.1 | K9N7C7 |
| VQGFACDCGAVF | 1 | 0 | 0 | 1527937 | AVV62535.1 | K9N7C7 |
| VQGFACECGAVY | 1 | 0 | 0 | 1527938 | AUM60013.1 | K9N7C7 |
| VQGGITQRTRTR | 1 | 0 | 0 | 1527942 | ANF29213.1 | K9N4V7 |
| VQGNVAKVANIS | 1 | 0 | 0 | 1527952 | P0C6U5.1 | A0A140H1G9 |
| VQGSITQHTRTR | 1 | 0 | 0 | 1527958 | AKN24774.1 | K9N4V7 |
| VQGSITQRIRTR | 1 | 0 | 0 | 1527960 | AKI29262.1 | K9N4V7 |
| VQIESDEYLTQR | 1 | 0 | 0 | 1527980 | AUM60013.1 | K9N7C7 |
| VQIPCMCGKQAT | 1 | 0 | 0 | 1527986 | AVP78041.1 | P0C6W2 |
| VQIQNNTVDVVV | 1 | 0 | 0 | 1527992 | AVV62524.1 | K9N7C7 |
| VQKAKQKPIYLK | 1 | 0 | 0 | 1527998 | AEN19363.1 | P0C6X6 |
| VQKHVGAPANLR | 1 | 0 | 0 | 1528011 | AVV62524.1 | K9N7C7 |
| VQKLQILIDNIN | 1 | 0 | 0 | 1528014 | ARB07425.1 | A0A0P0K6L9 |
| VQKLQSLIDNIN | 1 | 0 | 0 | 1528015 | BAL45640.1 | A0A0P0K6L9 |
| VQKRFRTRKGKR | 1 | 0 | 0 | 1528018 | AIW52723.1 | A0A0N7J131 |
| VQLEHYSSCVEC | 1 | 0 | 0 | 1528028 | NP_073549.1 | A0A0P0LKV0 |
| VQLESDAYLDAK | 1 | 0 | 0 | 1528029 | AVV62524.1 | K9N7C7 |
| VQLLIQDYIQSV | 1 | 0 | 0 | 1528034 | AAR01012.1 | P0C6X6 |
| VQLPVKDYIQTV | 1 | 0 | 0 | 1528041 | ABD75583.1 | A0A140H1G9 |
| VQLQDVQVSAQQ | 1 | 0 | 0 | 1528042 | AUM60013.1 | K9N7C7 |
| VQLQLTKLLDDS | 1 | 0 | 0 | 1528043 | ARU07593.1 | H9EJ42 |
| VQLRVVGHSMQN | 1 | 0 | 0 | 1528050 | AVP78041.1 | P0C6W2 |
| VQLTDKYCSVNV | 1 | 0 | 0 | 1528058 | AXN92247.1 | K9N7C7 |
| VQMGSNNVCTYE | 1 | 0 | 0 | 1528068 | AGT51323.1 | H9EJ42 |
| VQMIADYLAGSS | 1 | 0 | 0 | 1528070 | ATU80780.1 | A0A0P0LKV0 |
| VQMLSDTLRNLT | 1 | 0 | 0 | 1528074 | APO40578.1 | P0C6X7 |
| VQMNIAKYMQLC | 1 | 0 | 0 | 1528075 | QBF80609.1 | K9N7C7 |
| VQMTKLASTEEL | 1 | 0 | 0 | 1528076 | AAT76156.1 | Q3LZX3 |
| VQNAVPANSLHA | 1 | 0 | 0 | 1528081 | APO40580.1 | P59632 |
| VQNGVDAKGFPL | 1 | 0 | 0 | 1528082 | AMM42619.1 | H9EJ69 |
| VQNLFGNPDEPQ | 1 | 0 | 0 | 1528084 | AHA80176.1 | P33469 |
| VQNLSGNPDEPH | 1 | 0 | 0 | 1528085 | AIX10716.1 | P33469 |
| VQNTVFNASISE | 1 | 0 | 0 | 1528089 | AVL25595.1 | H9EJ69 |
| VQNTVLNASIPE | 1 | 0 | 0 | 1528090 | AIW52795.1 | H9EJ69 |
| VQNTVLNASTPE | 1 | 0 | 0 | 1528091 | AIW52805.1 | H9EJ69 |
| VQNVNLEPCQLA | 1 | 0 | 0 | 1528092 | AVV62524.1 | K9N7C7 |
| VQPIADVYRRRP | 1 | 0 | 0 | 1528109 | AHA80225.1 | P36334 |
| VQPIANVYRRKP | 1 | 0 | 0 | 1528110 | ABD43177.1 | P36334 |
| VQPLPQRQKKQP | 1 | 0 | 0 | 1528112 | APO40586.1 | P59595 |
| VQPRNVLYDSVS | 1 | 0 | 0 | 1528115 | BAL45640.1 | A0A0P0K6L9 |
| VQPRTIDSTQDG | 1 | 0 | 0 | 1528116 | QBQ01839.1 | P36334 |
| VQPRTINATQDG | 1 | 0 | 0 | 1528119 | AXX83321.1 | P36334 |
| VQPRTINLDNKL | 1 | 0 | 0 | 1528120 | ATP16757.1 | P36334 |
| VQPRTINLTQDG | 1 | 0 | 0 | 1528121 | AEN19366.1 | P36334 |
| VQPRTINSQDGV | 1 | 0 | 0 | 1528123 | QDH43726.1 | P36334 |
| VQPRTINSTQNG | 1 | 0 | 0 | 1528125 | BBA84097.1 | P36334 |
| VQPRTTQDGDNK | 1 | 0 | 0 | 1528127 | AKA09658.1 | P36334 |
| VQPTQSVVRFPN | 1 | 0 | 0 | 1528135 | AVP78031.1 | Q3LZX1 |
| VQQAQQEATHIA | 1 | 0 | 0 | 1528141 | AST12962.1 | A0A0P0LKV0 |
| VQQAQQEVTHIA | 1 | 0 | 0 | 1528143 | APT69889.1 | A0A0P0LKV0 |
| VQQDGRIPNGFS | 1 | 0 | 0 | 1528153 | AIW52843.1 | A0A0P0G321 |
| VQQDGRTPNGFS | 1 | 0 | 0 | 1528154 | AKA09627.1 | A0A0P0G321 |
| VQQDLDVILPGE | 1 | 0 | 0 | 1528155 | AXN92247.1 | K9N7C7 |
| VQQDLDVMLPGE | 1 | 0 | 0 | 1528156 | AKK52590.1 | K9N7C7 |
| VQQEFSVVLPGE | 1 | 0 | 0 | 1528157 | AVV62524.1 | K9N7C7 |
| VQQESTFVMMSA | 1 | 0 | 0 | 1528162 | APO40578.1 | P0C6X7 |
| VQQGVFNKATVA | 1 | 0 | 0 | 1528164 | ATN39862.1 | P0C6X6 |
| VQQLFYTEDIAS | 1 | 0 | 0 | 1528167 | QBF80609.1 | K9N7C7 |
| VQQLFYTEDLAS | 1 | 0 | 0 | 1528168 | AUM60013.1 | K9N7C7 |
| VQQLNNGFQGKQ | 1 | 0 | 0 | 1528170 | AVV62524.1 | K9N7C7 |
| VQQNKIRELTAE | 1 | 0 | 0 | 1528172 | AGT51784.1 | P33469 |
| VQQQQDDLFTPV | 1 | 0 | 0 | 1528176 | APO40584.1 | P59635 |
| VQQVSQWLSNLV | 1 | 0 | 0 | 1528181 | ARO76381.1 | P0C6U8 |
| VQQWGYIGNLAT | 1 | 0 | 0 | 1528183 | AVV62524.1 | K9N7C7 |
| VQQWGYVSNLAT | 1 | 0 | 0 | 1528185 | ATG84677.1 | K9N7C7 |
| VQRCTRKLNRLM | 1 | 0 | 0 | 1528189 | NP_073549.1 | A0A0P0LKV0 |
| VQRPPTLLLLFS | 1 | 0 | 0 | 1528194 | QBF80500.1 | K9N796 |
| VQSNLIRDIFVH | 1 | 0 | 0 | 1528209 | ARU07575.1 | A0A140H1G9 |
| VQSNVEVLPEGW | 1 | 0 | 0 | 1528212 | QEG03790.1 | P0C6X6 |
| VQSRADQNGYIT | 1 | 0 | 0 | 1528214 | AUM60013.1 | K9N7C7 |
| VQSSGIIKPNAL | 1 | 0 | 0 | 1528217 | AHY22544.1 | K9N7C7 |
| VQSSGMIKPNAI | 1 | 0 | 0 | 1528219 | QBF80531.1 | K9N7C7 |
| VQSSGMIKPNAL | 1 | 0 | 0 | 1528220 | AXN92247.1 | K9N7C7 |
| VQTFYPRFQASA | 1 | 0 | 0 | 1528233 | ALA49945.1 | K9N7C7 |
| VQTFYPRLQASN | 1 | 0 | 0 | 1528236 | AUM60013.1 | K9N7C7 |
| VQTIEDLRARMT | 1 | 0 | 0 | 1528239 | AJG44112.1 | K9N638 |

| | | | | | | |
|---|---|---|---|---|---|---|
| VQTLEDLHARMT | 1 | 0 | 0 | 1528241 | ATQ39389.1 | K9N7C7 |
| VQTVEDLCARMT | 1 | 0 | 0 | 1528247 | AKN24802.1 | K9N7C7 |
| VQTVEDLRVRMT | 1 | 0 | 0 | 1528248 | QDP16215.1 | K9N7C7 |
| VQTYLKSDNIVA | 1 | 0 | 0 | 1528252 | AIV42002.1 | P0C6X6 |
| VQVGQLPKYVIV | 1 | 0 | 0 | 1528260 | AWK59934.1 | U3M6Q8 |
| VQVHNLPEYMAV | 1 | 0 | 0 | 1528264 | APT69865.1 | A0A0P0KXN3 |
| VQVPIGCLDPIR | 1 | 0 | 0 | 1528270 | AWK59942.1 | H9EJ42 |
| VQVSAQQEVQLK | 1 | 0 | 0 | 1528276 | AUM60013.1 | K9N7C7 |
| VQVVDSRWNNLR | 1 | 0 | 0 | 1528283 | AGT17757.1 | A0A140H1H0 |
| VQVVIADTLQET | 1 | 0 | 0 | 1528285 | AHI48784.1 | K9N7C7 |
| VQWFHEHVGGNF | 1 | 0 | 0 | 1528287 | AVV62524.1 | K9N7C7 |
| VQWFHENVGGNF | 1 | 0 | 0 | 1528288 | AVV62535.1 | K9N7C7 |
| VQWQCKLDFLDC | 1 | 0 | 0 | 1528291 | AVV62539.1 | K9N4V0 |
| VQYGTDTISVCP | 1 | 0 | 0 | 1528304 | AGV08438.1 | K9N5Q8 |
| VQYGTDTNSACP | 1 | 0 | 0 | 1528305 | AXN92260.1 | K9N5Q8 |
| VQYGTDTNSVSP | 1 | 0 | 0 | 1528307 | AKN24803.1 | K9N5Q8 |
| VRAILKKKEKIL | 1 | 0 | 0 | 1528322 | AVV62535.1 | K9N7C7 |
| VRAYSEAAAQGF | 1 | 0 | 0 | 1528338 | AVA26872.1 | H9EJ42 |
| VRCAVTDAKWYC | 1 | 0 | 0 | 1528340 | AWK59942.1 | H9EJ42 |
| VRCEEDGQTVFY | 1 | 0 | 0 | 1528342 | AVV62535.1 | K9N7C7 |
| VRCSYYEDFLEY | 1 | 0 | 0 | 1528347 | AVP78037.1 | Q3LZX5 |
| VRDKGALEKFMS | 1 | 0 | 0 | 1528356 | AVV62535.1 | K9N7C7 |
| VRDPQTLDILDI | 1 | 0 | 0 | 1528367 | ATO98145.1 | Q3LZX1 |
| VRDTNKEVFVGD | 1 | 0 | 0 | 1528374 | AVI15051.1 | P0C6X6 |
| VREYIRYGIKPI | 1 | 0 | 0 | 1528399 | ARU07575.1 | A0A140H1G9 |
| VRFGPFNFCSET | 1 | 0 | 0 | 1528409 | NP_073549.1 | A0A0P0LKV0 |
| VRFPNITNVCPF | 1 | 0 | 0 | 1528416 | AVP78031.1 | Q3LZX1 |
| VRFSATKTVGGA | 1 | 0 | 0 | 1528417 | AVV62524.1 | K9N7C7 |
| VRFYRALYSNMA | 1 | 0 | 0 | 1528425 | ARU07576.1 | A0A140H1H0 |
| VRGFGDTVEEAI | 1 | 0 | 0 | 1528431 | APO40578.1 | P0C6X7 |
| VRGQIQVTSDNI | 1 | 0 | 0 | 1528436 | ARO76381.1 | P0C6U8 |
| VRGTTVILKEPC | 1 | 0 | 0 | 1528442 | 1XAK_A | P59635 |
| VRIGAAANATGT | 1 | 0 | 0 | 1528465 | AVK87439.1 | K9N5Q8 |
| VRIGASANSTGT | 1 | 0 | 0 | 1528469 | ATG84723.1 | K9N5Q8 |
| VRIGASSNKTGN | 1 | 0 | 0 | 1528470 | AUM60014.1 | K9N5Q8 |
| VRIGAVANSTGT | 1 | 0 | 0 | 1528471 | AGV08520.1 | K9N5Q8 |
| VRILQDKTGDFM | 1 | 0 | 0 | 1528476 | YP_009047214.1 | K9N638 |
| VRKNGHFQDHYD | 1 | 0 | 0 | 1528496 | AGT51365.1 | H9EJ42 |
| VRKRRHSPGKNL | 1 | 0 | 0 | 1528499 | ATQ39396.1 | K9N643 |
| VRLLQDKAGDFM | 1 | 0 | 0 | 1528511 | AQZ41321.1 | K9N7C7 |
| VRLLQDKTGDFT | 1 | 0 | 0 | 1528514 | AJO62148.1 | K9N7C7 |
| VRLLQGKTGDFM | 1 | 0 | 0 | 1528515 | AHI48793.1 | K9N7C7 |
| VRLQPSMALQLT | 1 | 0 | 0 | 1528520 | AVV62540.1 | K9N643 |
| VRMYNLLACLWL | 1 | 0 | 0 | 1528532 | AXN92247.1 | K9N7C7 |
| VRNLFASVKNSQ | 1 | 0 | 0 | 1528541 | QBF80598.1 | K9N5Q8 |
| VRNLFASVKSSQ | 1 | 0 | 0 | 1528542 | AXN92228.1 | K9N5Q8 |
| VRPRIVQMFADH | 1 | 0 | 0 | 1528560 | AEN19363.1 | P0C6X6 |
| VRQVAPGETGVI | 1 | 0 | 0 | 1528579 | ATO98181.1 | Q3LZX1 |
| VRRGDNFHNVLT | 1 | 0 | 0 | 1528584 | AVN89451.1 | K9N7C7 |
| VRSESAAISAQL | 1 | 0 | 0 | 1528596 | AVN89365.1 | K9N5Q8 |
| VRSIKVFTTVDN | 1 | 0 | 0 | 1528607 | AVP78041.1 | P0C6W2 |
| VRTINLGEVLIA | 1 | 0 | 0 | 1528638 | APO40578.1 | P0C6X7 |
| VRTLSTYDFNPN | 1 | 0 | 0 | 1528639 | AVP78031.1 | Q3LZX1 |
| VRTTATIPIQAT | 1 | 0 | 0 | 1528652 | AVP78032.1 | Q3LZX0 |
| VRTYSEAAAQGF | 1 | 0 | 0 | 1528655 | AWK59942.1 | H9EJ42 |
| VRVCYGSTVLNG | 1 | 0 | 0 | 1528657 | AWK59942.1 | H9EJ42 |
| VRYDMFDTNMFI | 1 | 0 | 0 | 1528681 | AUM60013.1 | K9N7C7 |
| VRYDMFESNMFI | 1 | 0 | 0 | 1528682 | AVV62524.1 | K9N7C7 |
| VSAAFHQNASSL | 1 | 0 | 0 | 1528694 | BBA20983.1 | A0A140H1H1 |
| VSAASVLGAAAK | 1 | 0 | 0 | 1528701 | AVV62535.1 | K9N7C7 |
| VSAAYHANSSEP | 1 | 0 | 0 | 1528704 | AEN19366.1 | P36334 |
| VSAGIFGVQPKI | 1 | 0 | 0 | 1528722 | AUM60013.1 | K9N7C7 |
| VSAGQEQTNCNI | 1 | 0 | 0 | 1528725 | ASU45728.1 | K9N7C7 |
| VSALIGANGNKI | 1 | 0 | 0 | 1528730 | AVN89322.1 | K9N7C7 |
| VSALKTAGVVTC | 1 | 0 | 0 | 1528731 | AVN89451.1 | K9N7C7 |
| VSAMQTMLFGMI | 1 | 0 | 0 | 1528740 | AXN92247.1 | K9N7C7 |
| VSAQQEVQLKVI | 1 | 0 | 0 | 1528744 | AUM60013.1 | K9N7C7 |
| VSARADAEGYIN | 1 | 0 | 0 | 1528745 | AVV62524.1 | K9N7C7 |
| VSARDPTIQEAI | 1 | 0 | 0 | 1528747 | AMM42397.1 | Q19U29 |
| VSATVTANLSIP | 1 | 0 | 0 | 1528755 | AAK32189.1 | A0A0P0K6L9 |
| VSAVNQSLIKSK | 1 | 0 | 0 | 1528758 | ARB07393.1 | A0A0P0KWE8 |
| VSAVNQSLSKSK | 1 | 0 | 0 | 1528760 | ABM64805.1 | A0A0P0KWE8 |
| VSAYGLCDSANP | 1 | 0 | 0 | 1528767 | AVN89334.1 | K9N5Q8 |
| VSAYGLCDVANP | 1 | 0 | 0 | 1528768 | AKM76239.1 | K9N5Q8 |
| VSCFSTCNSNAN | 1 | 0 | 0 | 1528772 | QEG59359.1 | A0A0P0G321 |
| VSCFSTCNSNAS | 1 | 0 | 0 | 1528773 | AWO68966.1 | A0A0P0G321 |
| VSCFSTRNSNAS | 1 | 0 | 0 | 1528774 | AKA09629.1 | A0A0P0G321 |
| VSCIIGSLGFYN | 1 | 0 | 0 | 1528782 | AEN19365.1 | P30215 |
| VSCLYKRNFTYD | 1 | 0 | 0 | 1528788 | CAA79896.1 | P36334 |
| VSCVCGRDATQY | 1 | 0 | 0 | 1528791 | APO40578.1 | P0C6X7 |
| VSDDEWVAAVDE | 1 | 0 | 0 | 1528802 | ASU45728.1 | K9N7C7 |
| VSDIDYVPLKSS | 1 | 0 | 0 | 1528808 | APO40578.1 | P0C6X7 |
| VSDNVRALMSCN | 1 | 0 | 0 | 1528813 | ADO24673.1 | P0C6X6 |
| VSDSLEPVGGLY | 1 | 0 | 0 | 1528817 | ARB66068.1 | P36334 |
| VSDTIPEQKQNG | 1 | 0 | 0 | 1528818 | AVV62524.1 | K9N7C7 |
| VSDTIPQQSQNG | 1 | 0 | 0 | 1528819 | AVV62535.1 | K9N7C7 |
| VSDTVEVPPHEA | 1 | 0 | 0 | 1528821 | QBF80585.1 | K9N7C7 |
| VSDTVEVPPHVV | 1 | 0 | 0 | 1528822 | AXN73447.1 | K9N638 |
| VSDTVEVPPQVV | 1 | 0 | 0 | 1528824 | AXN92247.1 | K9N7C7 |
| VSDTVVKTLQPI | 1 | 0 | 0 | 1528825 | APO40578.1 | P0C6X7 |
| VSDVWHKQLLIN | 1 | 0 | 0 | 1528831 | AEN19363.1 | P0C6X6 |

| | | | | | | |
|---|---|---|---|---|---|---|
| VSDYATKVVPFN | 1 | 0 | 0 | 1528833 | AVV62537.1 | K9N5Q8 |
| VSERVNVVPGVP | 1 | 0 | 0 | 1528855 | AFD98763.1 | H9EJ42 |
| VSFADNNDNNNQ | 1 | 0 | 0 | 1528861 | AUM60021.1 | K9N4V7 |
| VSFADNNDTTNT | 1 | 0 | 0 | 1528862 | ALW82676.1 | K9N4V7 |
| VSFDDLYVAIRG | 1 | 0 | 0 | 1528868 | AVA26874.1 | H9EJ24 |
| VSFDYLIREIET | 1 | 0 | 0 | 1528870 | ATQ39389.1 | K9N7C7 |
| VSFHEDRILDYK | 1 | 0 | 0 | 1528874 | AVV62524.1 | K9N7C7 |
| VSFNEWAMANQF | 1 | 0 | 0 | 1528885 | AVV62524.1 | K9N7C7 |
| VSFSPAFHRIES | 1 | 0 | 0 | 1528886 | AHL18095.1 | K9N7D2 |
| VSFVVDDSGTPV | 1 | 0 | 0 | 1528893 | AXN92247.1 | K9N7C7 |
| VSGCETSNVPPI | 1 | 0 | 0 | 1528898 | ACA49954.1 | H9EJ42 |
| VSGDYLWYVVDD | 1 | 0 | 0 | 1528904 | ARU07593.1 | H9EJ42 |
| VSGLVNVQKVAQ | 1 | 0 | 0 | 1528917 | AST12962.1 | A0A0P0LKV0 |
| VSGMDTALLRNN | 1 | 0 | 0 | 1528919 | AQN78779.1 | Q01455 |
| VSGTTLHPVQWQ | 1 | 0 | 0 | 1528941 | AVV62528.1 | K9N4V0 |
| VSHEVSDDEWAA | 1 | 0 | 0 | 1528955 | AXN92247.1 | K9N7C7 |
| VSHGLDLMQASQ | 1 | 0 | 0 | 1528960 | AUM60013.1 | K9N7C7 |
| VSIAPSTVWEDG | 1 | 0 | 0 | 1528982 | QEJ82215.1 | K9N5Q8 |
| VSICQYNMCEHP | 1 | 0 | 0 | 1528983 | BBO58751.1 | P36334 |
| VSIEQLLYAVQQ | 1 | 0 | 0 | 1528987 | AVV62524.1 | K9N7C7 |
| VSIPQGLTFSYD | 1 | 0 | 0 | 1529002 | ATQ39389.1 | K9N7C7 |
| VSIRTSNQGNKQ | 1 | 0 | 0 | 1529006 | AHI48842.1 | K9N643 |
| VSIVPSTVWKDG | 1 | 0 | 0 | 1529018 | ATW75480.1 | K9N5Q8 |
| VSKDGFFTYICH | 1 | 0 | 0 | 1529025 | AEN19363.1 | P0C6X6 |
| VSKIGDSIFTGC | 1 | 0 | 0 | 1529038 | ATG84700.1 | K9N638 |
| VSKLHDLDDATR | 1 | 0 | 0 | 1529044 | AVV62540.1 | K9N643 |
| VSKVVQVTIDYI | 1 | 0 | 0 | 1529065 | APO40578.1 | P0C6X7 |
| VSLEKFMSTPNA | 1 | 0 | 0 | 1529077 | ANF29193.1 | K9N7C7 |
| VSLKQGEIKDAT | 1 | 0 | 0 | 1529084 | AVP78043.1 | Q3LZX0 |
| VSLLRNTSQKSI | 1 | 0 | 0 | 1529092 | AAT76147.1 | Q3LZX1 |
| VSLNSKEELLQT | 1 | 0 | 0 | 1529100 | APO40578.1 | P0C6X7 |
| VSLTQSEMTIVK | 1 | 0 | 0 | 1529113 | ATQ39389.1 | K9N7C7 |
| VSLYNVTRDKLE | 1 | 0 | 0 | 1529135 | AXN92247.1 | K9N7C7 |
| VSLYNVTREKLD | 1 | 0 | 0 | 1529138 | AUM60013.1 | K9N7C7 |
| VSNGSNNCTTAV | 1 | 0 | 0 | 1529161 | AIW52843.1 | A0A0P0G321 |
| VSNGSNSCTTAV | 1 | 0 | 0 | 1529162 | AWO69003.1 | A0A0P0G321 |
| VSNLFADIKTQA | 1 | 0 | 0 | 1529169 | AVV62526.1 | K9N5Q8 |
| VSNLSLKQKLLY | 1 | 0 | 0 | 1529170 | QAT98908.1 | K9N7C7 |
| VSNLTKQLLELL | 1 | 0 | 0 | 1529171 | AUM60013.1 | K9N7C7 |
| VSNLTKQLLEVL | 1 | 0 | 0 | 1529172 | AUM60023.1 | K9N7C7 |
| VSNSGPLQTAVG | 1 | 0 | 0 | 1529178 | AVV62537.1 | K9N5Q8 |
| VSNVKLDRWSAS | 1 | 0 | 0 | 1529181 | AIW52741.1 | A0A0P0K6L9 |
| VSNYSHEVNLFQ | 1 | 0 | 0 | 1529182 | AUM60014.1 | K9N5Q8 |
| VSPCIKVAAMQS | 1 | 0 | 0 | 1529190 | ALA49417.1 | K9N7C7 |
| VSPDVQFEQLDN | 1 | 0 | 0 | 1529193 | AVV62526.1 | K9N5Q8 |
| VSPTSKIEPCIV | 1 | 0 | 0 | 1529217 | ARU07575.1 | A0A140H1G9 |
| VSPVFHRIESID | 1 | 0 | 0 | 1529220 | AVN89455.1 | K9N7D2 |
| VSQDSITHYPAL | 1 | 0 | 0 | 1529224 | AHL18091.1 | K9N7C7 |
| VSQDVKFENLTN | 1 | 0 | 0 | 1529225 | AVV62537.1 | K9N5Q8 |
| VSQLTKQLLDIL | 1 | 0 | 0 | 1529231 | AVV62535.1 | K9N7C7 |
| VSQQVDLVISDM | 1 | 0 | 0 | 1529235 | AQZ41341.1 | K9N7C7 |
| VSRIGDAVFTGC | 1 | 0 | 0 | 1529250 | AVV62535.1 | K9N7C7 |
| VSRIGDSIFTGC | 1 | 0 | 0 | 1529251 | AXN92247.1 | K9N7C7 |
| VSRIGDSVFTGC | 1 | 0 | 0 | 1529252 | AQZ41321.1 | K9N7C7 |
| VSRINWVTGGIA | 1 | 0 | 0 | 1529253 | AFR58690.1 | P59596 |
| VSRNSSRSSSQG | 1 | 0 | 0 | 1529259 | ASY99807.1 | K9N4V7 |
| VSRVGDSIFTGC | 1 | 0 | 0 | 1529270 | AVV62524.1 | K9N7C7 |
| VSSCDLRMGSGF | 1 | 0 | 0 | 1529283 | BBA20986.1 | A0A140H1H1 |
| VSSFEAIPSGSV | 1 | 0 | 0 | 1529293 | ALA49649.1 | K9N5Q8 |
| VSSFEVHSRGQF | 1 | 0 | 0 | 1529297 | AVV62537.1 | K9N5Q8 |
| VSSFEVRSKGEF | 1 | 0 | 0 | 1529298 | AVV62526.1 | K9N5Q8 |
| VSSFLEMKSGKQ | 1 | 0 | 0 | 1529302 | AVP78041.1 | P0C6W2 |
| VSSIKGDKGVEK | 1 | 0 | 0 | 1529315 | AWK59942.1 | H9EJ42 |
| VSSLAYYEPVQG | 1 | 0 | 0 | 1529317 | AVV62524.1 | K9N7C7 |
| VSSMLTFDGNAF | 1 | 0 | 0 | 1529324 | AVL25589.1 | A0A0P0G321 |
| VSSNGQPGGDAI | 1 | 0 | 0 | 1529325 | QAT98908.1 | K9N7C7 |
| VSSNINMLLSVP | 1 | 0 | 0 | 1529326 | AGH06535.1 | H9EJ42 |
| VSSNINRILGVN | 1 | 0 | 0 | 1529327 | AUT76985.1 | A0A0P0LKV0 |
| VSSNINRVLSVP | 1 | 0 | 0 | 1529328 | ABQ16583.1 | H9EJ42 |
| VSSNMVETLVGQ | 1 | 0 | 0 | 1529329 | ASU90174.1 | K9N638 |
| VSSQGPNFAEIS | 1 | 0 | 0 | 1529338 | AUM60014.1 | K9N5Q8 |
| VSSQGPNFQEIS | 1 | 0 | 0 | 1529339 | AVV62526.1 | K9N5Q8 |
| VSSSRRGVYYND | 1 | 0 | 0 | 1529343 | ATO98181.1 | Q3LZX1 |
| VSTAVSPDFKAF | 1 | 0 | 0 | 1529357 | AVV62535.1 | K9N7C7 |
| VSTDDCFANKHA | 1 | 0 | 0 | 1529359 | P0C6W6.1 | P0C6W2 |
| VSTDEEGDNVQI | 1 | 0 | 0 | 1529360 | AMM42619.1 | H9EJ69 |
| VSTDFLYFHFYQ | 1 | 0 | 0 | 1529361 | AGW27863.1 | A0A140H1H1 |
| VSTDVLVNTYKR | 1 | 0 | 0 | 1529366 | AVV62524.1 | K9N7C7 |
| VSTFNVPMEKLK | 1 | 0 | 0 | 1529371 | APO40578.1 | P0C6X7 |
| VSTLVYDNKLKA | 1 | 0 | 0 | 1529388 | APO40578.1 | P0C6X7 |
| VSTNFGDVNKPV | 1 | 0 | 0 | 1529390 | QED88038.1 | H9EJ42 |
| VSTPVDDFIKHV | 1 | 0 | 0 | 1529395 | AUM60013.1 | K9N7C7 |
| VSTQANAVQQEF | 1 | 0 | 0 | 1529396 | AVV62524.1 | K9N7C7 |
| VSTQPQENTIPH | 1 | 0 | 0 | 1529400 | ABD75589.1 | Q19U29 |
| VSTQPQGNSIPH | 1 | 0 | 0 | 1529403 | AMM42386.1 | Q19U29 |
| VSTSISNFGSLT | 1 | 0 | 0 | 1529412 | QBF80598.1 | K9N5Q8 |
| VSTVFVPAMRDS | 1 | 0 | 0 | 1529417 | AKN24834.1 | K9N7D2 |
| VSTVPSTVWEDG | 1 | 0 | 0 | 1529424 | ANC28634.1 | K9N5Q8 |
| VSTVSSHGFPYG | 1 | 0 | 0 | 1529425 | AKL80608.1 | K9N7D2 |
| VSTVSSHGMVPV | 1 | 0 | 0 | 1529427 | AXG21669.1 | K9N7D2 |
| VSVCQFNMCEYP | 1 | 0 | 0 | 1529432 | ANZ78837.1 | P36334 |

| | | | | | | |
|---|---|---|---|---|---|---|
| VSVCQYNMCDYP | 1 | 0 | 0 | 1529433 | ANZ78839.1 | P36334 |
| VSVDDKYKTIDE | 1 | 0 | 0 | 1529436 | ALA50066.1 | K9N7C7 |
| VSVDRESTSTHD | 1 | 0 | 0 | 1529442 | AXN92229.1 | K9N796 |
| VSVGESFGKSLG | 1 | 0 | 0 | 1529447 | QEG03773.1 | P0C6X6 |
| VSVITPGTNTSK | 1 | 0 | 0 | 1529452 | APO40579.1 | P59594 |
| VSVITPGTNTSL | 1 | 0 | 0 | 1529453 | AVP78031.1 | Q3LZX1 |
| VSVITPGTNTSS | 1 | 0 | 0 | 1529455 | AVP78042.1 | Q3LZX1 |
| VSVIYDDNTNQY | 1 | 0 | 0 | 1529456 | ATQ39390.1 | K9N5Q8 |
| VSVPLSVVYDKT | 1 | 0 | 0 | 1529466 | AVV62526.1 | K9N5Q8 |
| VSVPVSVIYDKK | 1 | 0 | 0 | 1529471 | AHL18090.1 | K9N5Q8 |
| VSVQELVQNTMD | 1 | 0 | 0 | 1529473 | ATQ39389.1 | K9N7C7 |
| VSVSRFDPSTWN | 1 | 0 | 0 | 1529474 | AIV41831.1 | P36334 |
| VSVSRFNPSTWN | 1 | 0 | 0 | 1529476 | CAA79896.1 | P36334 |
| VSVTDFSLNGTV | 1 | 0 | 0 | 1529484 | ANI69893.1 | K9N7D2 |
| VSVTEFTLNNHH | 1 | 0 | 0 | 1529485 | AVV62530.1 | K9N7D2 |
| VSVTVFSNMVET | 1 | 0 | 0 | 1529486 | AVN89363.1 | K9N7C7 |
| VSYDSVSATVTA | 1 | 0 | 0 | 1529514 | AAK32189.1 | A0A0P0K6L9 |
| VSYFYNGSAQAT | 1 | 0 | 0 | 1529521 | ARK08634.1 | P30215 |
| VSYLNSSNATIL | 1 | 0 | 0 | 1529531 | APO40578.1 | P0C6X7 |
| VSYLSEKCMIAP | 1 | 0 | 0 | 1529532 | AVV62524.1 | K9N7C7 |
| VSYVYNGSAQAT | 1 | 0 | 0 | 1529539 | AEN19365.1 | P30215 |
| VTAAYGLCNTDT | 1 | 0 | 0 | 1529547 | AVV62537.1 | K9N5Q8 |
| VTAFSKPIYVPE | 1 | 0 | 0 | 1529552 | QAT98905.1 | K9N796 |
| VTAFSKPLYVPE | 1 | 0 | 0 | 1529553 | AXN92229.1 | K9N796 |
| VTAGDGISTPKL | 1 | 0 | 0 | 1529554 | ATO98194.1 | Q3LZX0 |
| VTAGPFMDDAIN | 1 | 0 | 0 | 1529555 | AVN89322.1 | K9N7C7 |
| VTAGPFMGNAIN | 1 | 0 | 0 | 1529557 | ATQ39389.1 | K9N7C7 |
| VTAGPYVEGAIN | 1 | 0 | 0 | 1529558 | AVV62524.1 | K9N7C7 |
| VTAKGYPQFAEL | 1 | 0 | 0 | 1529563 | ABB90495.1 | A0A0N7J131 |
| VTANVCSPMACN | 1 | 0 | 0 | 1529574 | ACR82761.1 | A0A140H1G9 |
| VTAQDGCGHIST | 1 | 0 | 0 | 1529577 | ATO98193.1 | Q3LZX1 |
| VTAVVTNGYLKM | 1 | 0 | 0 | 1529587 | AVN89380.1 | K9N7A1 |
| VTDFSLNGTVST | 1 | 0 | 0 | 1529608 | ANI69893.1 | K9N7D2 |
| VTDILYPGQKYT | 1 | 0 | 0 | 1529609 | AVV62524.1 | K9N7C7 |
| VTDPSLIISSSP | 1 | 0 | 0 | 1529610 | ARE29984.1 | H9EJ42 |
| VTDVKQVGCFMR | 1 | 0 | 0 | 1529621 | AGT51518.1 | P0C6X6 |
| VTDVKQVGCSMR | 1 | 0 | 0 | 1529622 | AEN19363.1 | P0C6X6 |
| VTESVQEEAQLV | 1 | 0 | 0 | 1529634 | ATG84788.1 | K9N7C7 |
| VTESVQEKAQPV | 1 | 0 | 0 | 1529637 | AMQ49025.1 | K9N7C7 |
| VTEYHITIALLI | 1 | 0 | 0 | 1529639 | AVV62540.1 | K9N643 |
| VTFCDHMVKAGL | 1 | 0 | 0 | 1529641 | AGV08514.1 | K9N7C7 |
| VTFGDNNDNNSN | 1 | 0 | 0 | 1529648 | AVV62533.1 | K9N4V7 |
| VTFGDNNDNNTN | 1 | 0 | 0 | 1529649 | AVV62544.1 | K9N4V7 |
| VTFGEEKVHQVA | 1 | 0 | 0 | 1529654 | AVV62524.1 | K9N7C7 |
| VTFKEQPTVKEI | 1 | 0 | 0 | 1529656 | AEN19363.1 | P0C6X6 |
| VTFLNLQVEMNR | 1 | 0 | 0 | 1529664 | AGT51680.1 | P36334 |
| VTFSDPTTFGSS | 1 | 0 | 0 | 1529676 | AVN89383.1 | K9N643 |
| VTFSDSNMWYLR | 1 | 0 | 0 | 1529677 | ALA49894.1 | K9N643 |
| VTFTVDPKKAYM | 1 | 0 | 0 | 1529679 | AVV62524.1 | K9N7C7 |
| VTFVNISRVKLH | 1 | 0 | 0 | 1529682 | QEG03731.1 | A0A0P0G321 |
| VTGDNDDEDNKD | 1 | 0 | 0 | 1529685 | AGW27834.1 | A0A140H1G9 |
| VTGDNDDEDNND | 1 | 0 | 0 | 1529686 | ARB07605.1 | A0A140H1G9 |
| VTGELYHYQECI | 1 | 0 | 0 | 1529694 | AVP78047.1 | Q3LZX7 |
| VTGGAQTSTAIY | 1 | 0 | 0 | 1529700 | ARO76382.1 | P59594 |
| VTGLFKDCSRDP | 1 | 0 | 0 | 1529710 | AVV62524.1 | K9N7C7 |
| VTGNVVQQCCTA | 1 | 0 | 0 | 1529718 | ALA49384.1 | K9N7C7 |
| VTGRLIQFDNPN | 1 | 0 | 0 | 1529721 | APO40579.1 | P59594 |
| VTIADPGYMQGY | 1 | 0 | 0 | 1529737 | AXN92228.1 | K9N5Q8 |
| VTIAKTESAMVH | 1 | 0 | 0 | 1529745 | AVP78050.1 | Q3LZX3 |
| VTICDLYSIHDV | 1 | 0 | 0 | 1529748 | AAX85666.1 | P0C6X6 |
| VTICDLYSIQDV | 1 | 0 | 0 | 1529749 | AEN19363.1 | P0C6X6 |
| VTICDLYSMQDV | 1 | 0 | 0 | 1529750 | AAR01012.1 | P0C6X6 |
| VTIDCAACVCGD | 1 | 0 | 0 | 1529751 | ABD43177.1 | P36334 |
| VTIDCAAFDCGD | 1 | 0 | 0 | 1529752 | ARB66062.1 | P36334 |
| VTIDCAAFVCGD | 1 | 0 | 0 | 1529753 | CAA79896.1 | P36334 |
| VTIDCAAFVRGD | 1 | 0 | 0 | 1529754 | ANZ78843.1 | P36334 |
| VTIDCVAFVCGD | 1 | 0 | 0 | 1529756 | AIL49523.1 | P36334 |
| VTIDKLCDYLAD | 1 | 0 | 0 | 1529757 | ASU90174.1 | K9N638 |
| VTIDKLRDYLAD | 1 | 0 | 0 | 1529758 | AXN92247.1 | K9N7C7 |
| VTIELQPPCKFL | 1 | 0 | 0 | 1529763 | AVV62524.1 | K9N7C7 |
| VTINNFNPSSWN | 1 | 0 | 0 | 1529774 | BBA20986.1 | A0A140H1H1 |
| VTINNYNPSFWN | 1 | 0 | 0 | 1529775 | ABD75609.1 | A0A140H1H1 |
| VTIRGNPNNHKK | 1 | 0 | 0 | 1529780 | AVV62529.1 | K9N643 |
| VTKDIKDAPALP | 1 | 0 | 0 | 1529794 | AVV62535.1 | K9N7C7 |
| VTKGPFVEGAIN | 1 | 0 | 0 | 1529804 | AUM60023.1 | K9N7C7 |
| VTKGSFVEGAIN | 1 | 0 | 0 | 1529805 | AUM60013.1 | K9N7C7 |
| VTKHTAKEVRQK | 1 | 0 | 0 | 1529809 | AEN19370.1 | P33469 |
| VTKTSEQNNFTG | 1 | 0 | 0 | 1529838 | AVP78041.1 | P0C6W2 |
| VTLEKLRDYLAD | 1 | 0 | 0 | 1529851 | ATQ39389.1 | K9N7C7 |
| VTLEQKVKPIVE | 1 | 0 | 0 | 1529852 | APO40578.1 | P0C6X7 |
| VTLEQYSTCDMC | 1 | 0 | 0 | 1529853 | QED88039.1 | H9EJ42 |
| VTLKHFFFAQDG | 1 | 0 | 0 | 1529859 | AXN92247.1 | K9N7C7 |
| VTLKQGEIKGAT | 1 | 0 | 0 | 1529870 | AVP78032.1 | Q3LZX0 |
| VTLLDEHEFKFW | 1 | 0 | 0 | 1529871 | APT69863.1 | A0A0P0LKV0 |
| VTLLDEQESKFW | 1 | 0 | 0 | 1529873 | AIW52776.1 | A0A0P0LKV0 |
| VTLPGSDASGDC | 1 | 0 | 0 | 1529884 | ATQ39390.1 | K9N5Q8 |
| VTLRICKFGNTS | 1 | 0 | 0 | 1529889 | AWO68966.1 | A0A0P0G321 |
| VTLRSINHDLFS | 1 | 0 | 0 | 1529891 | AEN19363.1 | P0C6X6 |
| VTMTDLQSESSC | 1 | 0 | 0 | 1529911 | AST12962.1 | A0A0P0LKV0 |
| VTNAETTDFFTV | 1 | 0 | 0 | 1529914 | AAQ90006.1 | A0A0P0K6L9 |
| VTNAETTVCTVA | 1 | 0 | 0 | 1529915 | CAA71146.1 | A0A0P0K6L9 |

| | | | | | | |
|---|---|---|---|---|---|---|
| VTNGIFDGALSR | 1 | 0 | 0 | 1529919 | AVV62535.1 | K9N7C7 |
| VTNGNFDCALSR | 1 | 0 | 0 | 1529922 | AVV62524.1 | K9N7C7 |
| VTNGNLKMAGMH | 1 | 0 | 0 | 1529923 | AUM60030.1 | K9N7A1 |
| VTNMDIEKLRQY | 1 | 0 | 0 | 1529929 | AVV62524.1 | K9N7C7 |
| VTPDMADEIASL | 1 | 0 | 0 | 1529934 | AMM42457.1 | P33469 |
| VTPDMADHIASL | 1 | 0 | 0 | 1529935 | AIL49401.1 | P33469 |
| VTPDMADQIASL | 1 | 0 | 0 | 1529936 | AEN19370.1 | P33469 |
| VTPGGTSSGDAT | 1 | 0 | 0 | 1529940 | ACR82746.1 | A0A140H1G9 |
| VTPLPAADMDDF | 1 | 0 | 0 | 1529945 | AAS48456.1 | P59595 |
| VTPLTPRQYLLA | 1 | 0 | 0 | 1529950 | AWW13566.1 | P36334 |
| VTPMTDTLQMGF | 1 | 0 | 0 | 1529960 | AVV62537.1 | K9N5Q8 |
| VTPVDQYMCGVD | 1 | 0 | 0 | 1529967 | AXN92247.1 | K9N7C7 |
| VTPVMESQISTK | 1 | 0 | 0 | 1529968 | ARB07605.1 | A0A140H1G9 |
| VTQAFGRRGPGA | 1 | 0 | 0 | 1529972 | AVV62544.1 | K9N4V7 |
| VTQFAPGTQLPK | 1 | 0 | 0 | 1529976 | AVV62533.1 | K9N4V7 |
| VTQLLEKLLRGV | 1 | 0 | 0 | 1529982 | AUM60013.1 | K9N7C7 |
| VTRDIAPTDTCF | 1 | 0 | 0 | 1530002 | AVP78041.1 | P0C6W2 |
| VTRDILSTDDCF | 1 | 0 | 0 | 1530005 | ARO76381.1 | P0C6U8 |
| VTRDIVSTDDCF | 1 | 0 | 0 | 1530007 | P0C6W6.1 | P0C6W2 |
| VTRDRLDKLIAT | 1 | 0 | 0 | 1530009 | AVV62524.1 | K9N7C7 |
| VTRDRLEKLIST | 1 | 0 | 0 | 1530010 | AVV62535.1 | K9N7C7 |
| VTREKLEKLIST | 1 | 0 | 0 | 1530012 | ALD51903.1 | K9N7C7 |
| VTRGHRWDVVRP | 1 | 0 | 0 | 1530017 | QBP84719.1 | P0C6X6 |
| VTSAGEPCVASE | 1 | 0 | 0 | 1530028 | AVI15051.1 | P0C6X6 |
| VTSAGQPYVASE | 1 | 0 | 0 | 1530031 | AGT51707.1 | P0C6X6 |
| VTSDKFDFIIGH | 1 | 0 | 0 | 1530036 | AXX83348.1 | P0C6X6 |
| VTSEYSKRSAIE | 1 | 0 | 0 | 1530039 | AVV62526.1 | K9N5Q8 |
| VTSGNLFDVGYS | 1 | 0 | 0 | 1530043 | AGV08439.1 | K9N796 |
| VTSNVPPIVSEK | 1 | 0 | 0 | 1530054 | AWK59942.1 | H9EJ42 |
| VTSPPADVNTPA | 1 | 0 | 0 | 1530059 | AJQ24074.1 | P33469 |
| VTSSMTDNLQMA | 1 | 0 | 0 | 1530065 | AUM60024.1 | K9N5Q8 |
| VTSSSKSSEEHF | 1 | 0 | 0 | 1530066 | APO40578.1 | P0C6X7 |
| VTSVNFFDVGYS | 1 | 0 | 0 | 1530075 | AXN73427.1 | K9N796 |
| VTTEVKVSALKT | 1 | 0 | 0 | 1530093 | AHI48796.1 | K9N7C7 |
| VTTGDVKIMAKA | 1 | 0 | 0 | 1530099 | NP_073549.1 | A0A0P0LKV0 |
| VTTLNGRLVNYT | 1 | 0 | 0 | 1530115 | AMM42642.1 | A0A0P0G321 |
| VTTSMAPDFVAF | 1 | 0 | 0 | 1530130 | QBF80609.1 | K9N7C7 |
| VTTSQYSHRSAI | 1 | 0 | 0 | 1530131 | AUM60024.1 | K9N5Q8 |
| VTTSTEPDFVAF | 1 | 0 | 0 | 1530134 | ATG84711.1 | K9N7C7 |
| VTTSTNPEFVAF | 1 | 0 | 0 | 1530135 | AUM60013.1 | K9N7C7 |
| VTVADEPDSRAP | 1 | 0 | 0 | 1530140 | AST12968.1 | A0A0N7J131 |
| VTVAEEPDSRAP | 1 | 0 | 0 | 1530141 | AIW52721.1 | A0A0N7J131 |
| VTVAEEPDSRVP | 1 | 0 | 0 | 1530142 | APT69887.1 | A0A0N7J131 |
| VTVAVIAGIAGA | 1 | 0 | 0 | 1530146 | AVV62535.1 | K9N7C7 |
| VTVDCKQYICNG | 1 | 0 | 0 | 1530147 | AKM76239.1 | K9N5Q8 |
| VTVDGINFRTCI | 1 | 0 | 0 | 1530149 | AUM60013.1 | K9N7C7 |
| VTVDGINFRTSV | 1 | 0 | 0 | 1530150 | AVV62535.1 | K9N7C7 |
| VTVDGVNFRTCV | 1 | 0 | 0 | 1530151 | AVV62524.1 | K9N7C7 |
| VTVDTTVESAAQ | 1 | 0 | 0 | 1530154 | ATQ39389.1 | K9N7C7 |
| VTVELQPPCKFL | 1 | 0 | 0 | 1530155 | AUM60013.1 | K9N7C7 |
| VTVFTDGRLNCT | 1 | 0 | 0 | 1530161 | AUM60013.1 | K9N7C7 |
| VTVGPYVEGAIN | 1 | 0 | 0 | 1530165 | AVV62535.1 | K9N7C7 |
| VTVPHNLTTITK | 1 | 0 | 0 | 1530176 | AHI48528.1 | K9N5Q8 |
| VTVQGPPGTGKS | 1 | 0 | 0 | 1530180 | AXN92247.1 | K9N7C7 |
| VTYIPSQEKNFT | 1 | 0 | 0 | 1530198 | AVP78031.1 | Q3LZX1 |
| VTYKCCDDNYFN | 1 | 0 | 0 | 1530199 | AVP78030.1 | P0C6W2 |
| VTYKLDGVTYTE | 1 | 0 | 0 | 1530201 | APO40578.1 | P0C6X7 |
| VTYNCCDGDYFN | 1 | 0 | 0 | 1530208 | AVP78041.1 | P0C6W2 |
| VTYPSLNYAGAI | 1 | 0 | 0 | 1530209 | AUM60013.1 | K9N7C7 |
| VTYQNISINLPP | 1 | 0 | 0 | 1530212 | ATG84888.1 | K9N5Q8 |
| VTYTELNPKLDG | 1 | 0 | 0 | 1530217 | APO40578.1 | P0C6X7 |
| VTYVPSQEKNFT | 1 | 0 | 0 | 1530223 | ATO98181.1 | Q3LZX1 |
| VTYVPSQQQNFT | 1 | 0 | 0 | 1530225 | APO40579.1 | P59594 |
| VVAGGVQYYCYS | 1 | 0 | 0 | 1530243 | AVV62535.1 | K9N7C7 |
| VVANGVTAKGYP | 1 | 0 | 0 | 1530258 | ABB90495.1 | A0A0N7J131 |
| VVAQCSGVTFQL | 1 | 0 | 0 | 1530259 | 4HI3_A | P0C6U8 |
| VVAQFNYKQSFA | 1 | 0 | 0 | 1530260 | AVV62526.1 | K9N5Q8 |
| VVARTGQFYING | 1 | 0 | 0 | 1530263 | AIW52843.1 | A0A0P0G321 |
| VVAVSGSLTKKV | 1 | 0 | 0 | 1530270 | AVV62535.1 | K9N7C7 |
| VVCASVFKDGCD | 1 | 0 | 0 | 1530275 | ABE97136.1 | H9EJ42 |
| VVCATLINNISY | 1 | 0 | 0 | 1530277 | AWK59942.1 | H9EJ42 |
| VVCNNSDYASKG | 1 | 0 | 0 | 1530295 | AIX10745.1 | P0C6X6 |
| VVCYNSEYAAKG | 1 | 0 | 0 | 1530302 | AVV62535.1 | K9N7C7 |
| VVDCATYVCNGN | 1 | 0 | 0 | 1530306 | AIW52843.1 | A0A0P0G321 |
| VVDDHVGLLDLA | 1 | 0 | 0 | 1530307 | AIV41888.1 | P0C6X6 |
| VVDGFVRSFQPL | 1 | 0 | 0 | 1530322 | AFR79257.1 | A0A0P0K6L9 |
| VVDGKQIDDYRI | 1 | 0 | 0 | 1530324 | AIV41978.1 | P0C6X6 |
| VVDIKKEIAHLN | 1 | 0 | 0 | 1530325 | APO40579.1 | P59594 |
| VVDNCQLPLGQS | 1 | 0 | 0 | 1530332 | AVV62537.1 | K9N5Q8 |
| VVDQWYYTGSSF | 1 | 0 | 0 | 1530335 | AVV62537.1 | K9N5Q8 |
| VVDSRWNNARHS | 1 | 0 | 0 | 1530337 | AQN78751.1 | P30215 |
| VVDSRWNNARLS | 1 | 0 | 0 | 1530338 | AAX85668.1 | P30215 |
| VVDSRWNNLRAS | 1 | 0 | 0 | 1530342 | AGT17757.1 | A0A140H1H0 |
| VVDSVAVKNGAL | 1 | 0 | 0 | 1530343 | P0C6W6.1 | P0C6W2 |
| VVDYLTRDDMHQ | 1 | 0 | 0 | 1530361 | AVV62535.1 | K9N7C7 |
| VVEAPTTCGYLP | 1 | 0 | 0 | 1530364 | APO40578.1 | P0C6X7 |
| VVECLKLSHHSD | 1 | 0 | 0 | 1530367 | APO40578.1 | P0C6X7 |
| VVEQAEGFECDF | 1 | 0 | 0 | 1530393 | AQZ41282.1 | K9N5Q8 |
| VVESGGYIPSNF | 1 | 0 | 0 | 1530401 | AGT21353.1 | A0A0P0K6L9 |
| VVFDEVSMATNY | 1 | 0 | 0 | 1530414 | APO40578.1 | P0C6X7 |
| VVFDKNLYDKLV | 1 | 0 | 0 | 1530415 | AVP78041.1 | P0C6W2 |

| | | | | | | |
|---|---|---|---|---|---|---|
| VVFDTVEVPPQV | 1 | 0 | 0 | 1530416 | AMQ48992.1 | K9N7C7 |
| VVFGKRGGGSVT | 1 | 0 | 0 | 1530425 | ARB07390.1 | A0A0P0LKV0 |
| VVGDIILKPAID | 1 | 0 | 0 | 1530457 | AVP78041.1 | P0C6W2 |
| VVGGCVASNVPS | 1 | 0 | 0 | 1530466 | AVA26872.1 | H9EJ42 |
| VVGGCVTSNVPP | 1 | 0 | 0 | 1530467 | AWK59942.1 | H9EJ42 |
| VVGGCVTSNVSP | 1 | 0 | 0 | 1530469 | ARU07593.1 | H9EJ42 |
| VVGGTIQIFASV | 1 | 0 | 0 | 1530472 | ARB07424.1 | A0A0P0ILY9 |
| VVGKDGVMYSEF | 1 | 0 | 0 | 1530476 | AWW13511.1 | P36334 |
| VVGNVILKPAAD | 1 | 0 | 0 | 1530478 | APO40578.1 | P0C6X7 |
| VVGPRTGKHEHS | 1 | 0 | 0 | 1530492 | AWK59942.1 | H9EJ42 |
| VVGTKALAVRLT | 1 | 0 | 0 | 1530500 | ARU07575.1 | A0A140H1G9 |
| VVGVLTLDNQDL | 1 | 0 | 0 | 1530506 | QXX36617.1 | P0DTD1 |
| VVHNQDVNLHNS | 1 | 0 | 0 | 1530516 | AVP78030.1 | P0C6W2 |
| VVIADTLQEIPV | 1 | 0 | 0 | 1530521 | AQZ41284.1 | K9N7C7 |
| VVIADTLQETSV | 1 | 0 | 0 | 1530522 | ASU45706.1 | K9N7C7 |
| VVIEGLKACSYV | 1 | 0 | 0 | 1530528 | AVV62524.1 | K9N7C7 |
| VVIGDDVDDIDP | 1 | 0 | 0 | 1530529 | ABD75591.1 | A0A140H1G9 |
| VVIMTLDLLKDI | 1 | 0 | 0 | 1530544 | AQZ41284.1 | K9N7C7 |
| VVIPDFNVWNQV | 1 | 0 | 0 | 1530545 | AVV62524.1 | K9N7C7 |
| VVKAMLFKKEPL | 1 | 0 | 0 | 1530563 | ALA49373.1 | K9N7C7 |
| VVKAMLLKKEPL | 1 | 0 | 0 | 1530564 | AXN92247.1 | K9N7C7 |
| VVKDTVVQLNYC | 1 | 0 | 0 | 1530568 | AUM60013.1 | K9N7C7 |
| VVKEQVVDLLVK | 1 | 0 | 0 | 1530571 | ATQ39389.1 | K9N7C7 |
| VVKFPIDLTMIE | 1 | 0 | 0 | 1530572 | ATG84897.1 | K9N7C7 |
| VVKHIINITGVS | 1 | 0 | 0 | 1530576 | AGT51337.1 | H9EJ42 |
| VVKIYCPACHNS | 1 | 0 | 0 | 1530578 | QHD43415.1 | P0DTD1 |
| VVKLQFEPQTIQ | 1 | 0 | 0 | 1530583 | QBF80520.1 | K9N638 |
| VVKLQNNEIMPG | 1 | 0 | 0 | 1530584 | AWK59942.1 | H9EJ42 |
| VVKPTVIVNAAN | 1 | 0 | 0 | 1530598 | APO40578.1 | P0C6X7 |
| VVKRHTLSNYQH | 1 | 0 | 0 | 1530614 | APO40578.1 | P0C6X7 |
| VVKTLQPISDLL | 1 | 0 | 0 | 1530625 | APO40578.1 | P0C6X7 |
| VVKVPIDLTFVE | 1 | 0 | 0 | 1530628 | AVV62524.1 | K9N7C7 |
| VVKVPIDLTMIE | 1 | 0 | 0 | 1530630 | AXN92247.1 | K9N7C7 |
| VVKVTIDLTMIE | 1 | 0 | 0 | 1530632 | AUM60023.1 | K9N7C7 |
| VVLAEFDTYDPI | 1 | 0 | 0 | 1530635 | AHI48794.1 | K9N638 |
| VVLDEELALLRA | 1 | 0 | 0 | 1530638 | AUM60030.1 | K9N7A1 |
| VVLELDPPCKFS | 1 | 0 | 0 | 1530640 | ARU07575.1 | A0A140H1G9 |
| VVLNNKNIYRSQ | 1 | 0 | 0 | 1530666 | ASU90570.1 | K9N638 |
| VVLNNKNTYISQ | 1 | 0 | 0 | 1530668 | AVN89333.1 | K9N7C7 |
| VVLNNKNTYRSQ | 1 | 0 | 0 | 1530669 | AXN92247.1 | K9N7C7 |
| VVLPGEMAKKQI | 1 | 0 | 0 | 1530672 | AVV62535.1 | K9N7C7 |
| VVLPGEVAKKQI | 1 | 0 | 0 | 1530673 | AVV62524.1 | K9N7C7 |
| VVMEIPSLMTVK | 1 | 0 | 0 | 1530698 | ARU07575.1 | A0A140H1G9 |
| VVMGQLEETNMH | 1 | 0 | 0 | 1530700 | AUM60023.1 | K9N7C7 |
| VVMGQLEQTNMH | 1 | 0 | 0 | 1530701 | AUM60013.1 | K9N7C7 |
| VVMRPNHTIKGS | 1 | 0 | 0 | 1530706 | AUM60013.1 | K9N7C7 |
| VVMSMFSRTTSN | 1 | 0 | 0 | 1530708 | ATO98193.1 | Q3LZX1 |
| VVNAYNSTAIFV | 1 | 0 | 0 | 1530722 | AAB27260.2 | P36334 |
| VVNLNVDFKDFQ | 1 | 0 | 0 | 1530754 | AOL02450.1 | P0C6X6 |
| VVNSGSDINCNI | 1 | 0 | 0 | 1530771 | P0C6U5.1 | A0A140H1G9 |
| VVNTNPRNYSYM | 1 | 0 | 0 | 1530774 | AAA73152.1 | P30215 |
| VVNVVKAVRNKT | 1 | 0 | 0 | 1530779 | AQN78701.1 | P0C6X6 |
| VVPDIVEVPPQV | 1 | 0 | 0 | 1530785 | AVN89451.1 | K9N7C7 |
| VVPDTVGVPPQV | 1 | 0 | 0 | 1530787 | AHL18091.1 | K9N7C7 |
| VVPGLPGTILRT | 1 | 0 | 0 | 1530793 | AVP78041.1 | P0C6W2 |
| VVPGSCNFPLEA | 1 | 0 | 0 | 1530794 | AFD98750.1 | A0A0P0G321 |
| VVPGVADAERMA | 1 | 0 | 0 | 1530795 | BAL45639.1 | A0A0P0K6L9 |
| VVPLENDADKAN | 1 | 0 | 0 | 1530797 | AIV41996.1 | P0C6X6 |
| VVQFNYCRDGQP | 1 | 0 | 0 | 1530816 | AVK87459.1 | K9N7C7 |
| VVQKDGVEDTPP | 1 | 0 | 0 | 1530821 | ATQ39389.1 | K9N7C7 |
| VVQLNYTRDGQV | 1 | 0 | 0 | 1530827 | AVV62535.1 | K9N7C7 |
| VVQLPVKDYIQT | 1 | 0 | 0 | 1530829 | ABD75583.1 | A0A140H1G9 |
| VVQNTVLNASTP | 1 | 0 | 0 | 1530836 | AIW52805.1 | H9EJ69 |
| VVQPRTIDSTQD | 1 | 0 | 0 | 1530840 | QBQ01839.1 | P36334 |
| VVQPRTIISTQD | 1 | 0 | 0 | 1530841 | BBA20973.1 | P36334 |
| VVQPRTTQDGDN | 1 | 0 | 0 | 1530845 | AKA09658.1 | P36334 |
| VVQTCDLTVGSG | 1 | 0 | 0 | 1530851 | CAA83660.1 | P36334 |
| VVRKDLEEGDLP | 1 | 0 | 0 | 1530884 | ARK08633.1 | Q80872 |
| VVRLLEDKVESL | 1 | 0 | 0 | 1530887 | AVV62535.1 | K9N7C7 |
| VVRNKTSACFNF | 1 | 0 | 0 | 1530889 | AIV41888.1 | P0C6X6 |
| VVSAGQDQTTCN | 1 | 0 | 0 | 1530906 | AUM60013.1 | K9N7C7 |
| VVSAYGLCDASN | 1 | 0 | 0 | 1530909 | AHY22565.1 | K9N5Q8 |
| VVSAYGLCDLAN | 1 | 0 | 0 | 1530911 | AVN89387.1 | K9N5Q8 |
| VVSAYGLCDSAN | 1 | 0 | 0 | 1530912 | AVN89334.1 | K9N5Q8 |
| VVSDTVDVPPQV | 1 | 0 | 0 | 1530919 | ASU91261.1 | K9N638 |
| VVSDTVEVPPHE | 1 | 0 | 0 | 1530920 | QBF80585.1 | K9N7C7 |
| VVSDTVEVPPQV | 1 | 0 | 0 | 1530921 | AXN92247.1 | K9N7C7 |
| VVSDTVVKTLQP | 1 | 0 | 0 | 1530923 | APO40578.1 | P0C6X7 |
| VVSGCETSNVPP | 1 | 0 | 0 | 1530932 | ACA49954.1 | H9EJ42 |
| VVSIRSSNRGNK | 1 | 0 | 0 | 1530941 | ATG84704.1 | K9N643 |
| VVSIRTSNQGNK | 1 | 0 | 0 | 1530942 | AHI48842.1 | K9N643 |
| VVSKIGDSIFTG | 1 | 0 | 0 | 1530944 | ATG84700.1 | K9N638 |
| VVSMLLLCCCST | 1 | 0 | 0 | 1530956 | AIW52757.1 | A0A0P0K6L9 |
| VVSRLYKRNFTY | 1 | 0 | 0 | 1530964 | AHA80190.1 | P36334 |
| VVSTIGDSIFTG | 1 | 0 | 0 | 1530969 | QAT98908.1 | K9N7C7 |
| VVSTSSDPGFVA | 1 | 0 | 0 | 1530971 | AVV62524.1 | K9N7C7 |
| VVTAGDGISTPK | 1 | 0 | 0 | 1530978 | ATO98194.1 | Q3LZX0 |
| VVTDIKQVGCMM | 1 | 0 | 0 | 1530984 | ARU07575.1 | A0A140H1G9 |
| VVTGDNDDEDNK | 1 | 0 | 0 | 1530988 | AGW27834.1 | A0A140H1G9 |
| VVTGDNDQIVVT | 1 | 0 | 0 | 1530989 | ABD75623.1 | A0A140H1G9 |
| VVTGNVVKQCCI | 1 | 0 | 0 | 1530993 | ATQ39389.1 | K9N7C7 |

| | | | | | | |
|---|---|---|---|---|---|---|
| VVTGNVVQQCCT | 1 | 0 | 0 | 1530995 | ALA49384.1 | K9N7C7 |
| VVTNGYLKIAGM | 1 | 0 | 0 | 1531001 | AVV62543.1 | K9N7A1 |
| VVTRYVDNNFCG | 1 | 0 | 0 | 1531010 | ARO76381.1 | P0C6U8 |
| VVTTLKPTNQAF | 1 | 0 | 0 | 1531025 | ATO98145.1 | Q3LZX1 |
| VVVGGCVASNVP | 1 | 0 | 0 | 1531037 | AVA26872.1 | H9EJ42 |
| VVVGGCVTSNVP | 1 | 0 | 0 | 1531038 | AWK59942.1 | H9EJ42 |
| VVVGGCVTSNVS | 1 | 0 | 0 | 1531039 | ARU07593.1 | H9EJ42 |
| VVVGQLEETNMH | 1 | 0 | 0 | 1531042 | AVV62524.1 | K9N7C7 |
| VVVKRLYSPSLS | 1 | 0 | 0 | 1531046 | AVV62535.1 | K9N7C7 |
| VVVQPRTINYGV | 1 | 0 | 0 | 1531073 | AMK59677.1 | P36334 |
| VVVSGCETSNVP | 1 | 0 | 0 | 1531075 | ACA49954.1 | H9EJ42 |
| VVVTAGPFMDNA | 1 | 0 | 0 | 1531079 | AVN89363.1 | K9N7C7 |
| VVVYSLDGNMKA | 1 | 0 | 0 | 1531089 | AUM60013.1 | K9N7C7 |
| VVYQQAVAGAML | 1 | 0 | 0 | 1531123 | ADK37701.1 | A0A0P0K6L9 |
| VVYVNATGSDVN | 1 | 0 | 0 | 1531135 | ABU88826.1 | A0A0P0G321 |
| VWEEGDYYRKQL | 1 | 0 | 0 | 1531147 | ALJ76277.1 | K9N5Q8 |
| VWLESQLQKGVS | 1 | 0 | 0 | 1531172 | APO40578.1 | P0C6X7 |
| VWLLISFSFIIF | 1 | 0 | 0 | 1531183 | BBA20983.1 | A0A140H1H1 |
| VWPFYPYGRCPT | 1 | 0 | 0 | 1531195 | AGT51679.1 | P30215 |
| VWQKYTNSASTS | 1 | 0 | 0 | 1531205 | AUM60016.1 | K9N4V0 |
| VWRVIALQSVDC | 1 | 0 | 0 | 1531210 | NP_073549.1 | A0A0P0LKV0 |
| VWVHEEGATDAP | 1 | 0 | 0 | 1531222 | AVV62533.1 | K9N4V7 |
| VWVHEHGATDAP | 1 | 0 | 0 | 1531223 | AXN92267.1 | K9N4V7 |
| VWVHEYGATDAP | 1 | 0 | 0 | 1531224 | QFQ59594.1 | K9N4V7 |
| VYAGCEQSYFTK | 1 | 0 | 0 | 1531240 | AIL49582.1 | P0C6X6 |
| VYAIRNFDEQNC | 1 | 0 | 0 | 1531243 | AGT51330.1 | H9EJ42 |
| VYANDSIILRQY | 1 | 0 | 0 | 1531252 | QDH43722.1 | P0C6X6 |
| VYANLGERVRQA | 1 | 0 | 0 | 1531257 | P0C6W6.1 | P0C6W2 |
| VYAQHCFKAPKN | 1 | 0 | 0 | 1531263 | CAA79896.1 | P36334 |
| VYAQHCFKAPTN | 1 | 0 | 0 | 1531265 | CAA83660.1 | P36334 |
| VYAQHCFKVPKN | 1 | 0 | 0 | 1531267 | AGT51590.1 | P36334 |
| VYAYLGCAEGTM | 1 | 0 | 0 | 1531281 | AUM60013.1 | K9N7C7 |
| VYCGSDYMDIVK | 1 | 0 | 0 | 1531294 | AVV62535.1 | K9N7C7 |
| VYCLDGIVKAEV | 1 | 0 | 0 | 1531297 | AVV62535.1 | K9N7C7 |
| VYDAYHNLVGYY | 1 | 0 | 0 | 1531309 | ALA49649.1 | K9N5Q8 |
| VYDCFCNNINWN | 1 | 0 | 0 | 1531312 | AEN19363.1 | P0C6X6 |
| VYDDIFAESKPR | 1 | 0 | 0 | 1531316 | AGT21359.1 | A0A0P0LKV0 |
| VYDGARLFSSDL | 1 | 0 | 0 | 1531323 | P0C6U6.1 | H9EJ42 |
| VYDGFDNLVGYY | 1 | 0 | 0 | 1531326 | AVV62526.1 | K9N5Q8 |
| VYDMWLTGCKYV | 1 | 0 | 0 | 1531338 | AEN19363.1 | P0C6X6 |
| VYDNFPQGFSVL | 1 | 0 | 0 | 1531339 | APO40579.1 | P59594 |
| VYDSFDNLVGYH | 1 | 0 | 0 | 1531349 | AUM60014.1 | K9N5Q8 |
| VYDTNHGDAGFT | 1 | 0 | 0 | 1531354 | AQN78743.1 | P30215 |
| VYDVEPTHKVKL | 1 | 0 | 0 | 1531355 | AFD98798.1 | H9EJ42 |
| VYEADTEQTQSV | 1 | 0 | 0 | 1531362 | AXN92247.1 | K9N7C7 |
| VYELNGYTVQPV | 1 | 0 | 0 | 1531368 | AVR40344.1 | P36334 |
| VYENAFMPFAMG | 1 | 0 | 0 | 1531372 | AVP78041.1 | P0C6W2 |
| VYEVGDLIAQSS | 1 | 0 | 0 | 1531378 | AEN19363.1 | P0C6X6 |
| VYEVGDLMAQSS | 1 | 0 | 0 | 1531379 | YP_009555247.1 | P0C6X6 |
| VYFDNPTLPFGD | 1 | 0 | 0 | 1531385 | ATO98193.1 | Q3LZX1 |
| VYFIKMIILWLM | 1 | 0 | 0 | 1531389 | ABD75532.1 | Q0ZJ82 |
| VYFNGTGRGDCQ | 1 | 0 | 0 | 1531397 | BAL45639.1 | A0A0P0K6L9 |
| VYFSDSKPSTTE | 1 | 0 | 0 | 1531401 | ATQ39389.1 | K9N7C7 |
| VYFVARLLDEKV | 1 | 0 | 0 | 1531409 | AUM60013.1 | K9N7C7 |
| VYGDFSHQQLGG | 1 | 0 | 0 | 1531414 | APO40578.1 | P0C6X7 |
| VYGDFSHSQLGG | 1 | 0 | 0 | 1531415 | QYC64967.1 | P0DTD1 |
| VYGFSNVVYPDL | 1 | 0 | 0 | 1531416 | ARU07576.1 | A0A140H1H0 |
| VYHKHMFIVLYV | 1 | 0 | 0 | 1531426 | 6ATK_E | A0A0P0K6L9 |
| VYHRHTFIVLHV | 1 | 0 | 0 | 1531435 | AMM42532.1 | A0A0P0K6L9 |
| VYHRYRIGNYKL | 1 | 0 | 0 | 1531436 | APO40582.1 | P59596 |
| VYHTSGNVIEFH | 1 | 0 | 0 | 1531440 | APO40578.1 | P0C6X7 |
| VYIAVNDQKLYD | 1 | 0 | 0 | 1531443 | ARO76381.1 | P0C6U8 |
| VYKVYYGNVLDQ | 1 | 0 | 0 | 1531479 | APO40578.1 | P0C6X7 |
| VYLCERLTGGNY | 1 | 0 | 0 | 1531481 | AUM60013.1 | K9N7C7 |
| VYLEYIKKLYND | 1 | 0 | 0 | 1531482 | ARU07574.1 | A0A140H1G9 |
| VYLFCEHPAVGI | 1 | 0 | 0 | 1531483 | NP_073553.1 | A0A0P0KWE8 |
| VYLKHGGGVAGA | 1 | 0 | 0 | 1531497 | AVP78030.1 | P0C6W2 |
| VYMAKVGDKYYP | 1 | 0 | 0 | 1531514 | AEN19363.1 | P0C6X6 |
| VYNGSAQAISFC | 1 | 0 | 0 | 1531532 | QBP84748.1 | P30215 |
| VYNGSAQSTALC | 1 | 0 | 0 | 1531535 | P30215.1 | P30215 |
| VYNLVNVGHYDG | 1 | 0 | 0 | 1531542 | ABD75615.1 | A0A140H1G9 |
| VYNLWSTFTKVQ | 1 | 0 | 0 | 1531544 | AUM60013.1 | K9N7C7 |
| VYPDYVDVNKTL | 1 | 0 | 0 | 1531557 | AWK59979.1 | A0A0P0G321 |
| VYQNAYNCTYDR | 1 | 0 | 0 | 1531577 | ATO98193.1 | Q3LZX1 |
| VYREGYLNSTNV | 1 | 0 | 0 | 1531588 | AVP78041.1 | P0C6W2 |
| VYRRKPDPPNCN | 1 | 0 | 0 | 1531601 | AHA80201.1 | P36334 |
| VYRRRPDLPNCN | 1 | 0 | 0 | 1531603 | AHA80225.1 | P36334 |
| VYRVNTAIPTIE | 1 | 0 | 0 | 1531614 | AHI48842.1 | K9N643 |
| VYSAGHATGITP | 1 | 0 | 0 | 1531617 | ATG84701.1 | K9N5Q8 |
| VYSAGHATGSTP | 1 | 0 | 0 | 1531618 | AZU90731.1 | K9N5Q8 |
| VYSAGYATGTTP | 1 | 0 | 0 | 1531623 | ANC28634.1 | K9N5Q8 |
| VYSHKARNRSHS | 1 | 0 | 0 | 1531634 | AVN89460.1 | K9N643 |
| VYSLDGISKAEV | 1 | 0 | 0 | 1531638 | AVV62524.1 | K9N7C7 |
| VYSLDGNMKAEV | 1 | 0 | 0 | 1531639 | AUM60013.1 | K9N7C7 |
| VYSPSNHKSNTN | 1 | 0 | 0 | 1531644 | ATQ39390.1 | K9N5Q8 |
| VYSRKARNRSHS | 1 | 0 | 0 | 1531652 | ATG84704.1 | K9N643 |
| VYSSGHATGTTP | 1 | 0 | 0 | 1531657 | AVN89334.1 | K9N5Q8 |
| VYTANKWDLIIS | 1 | 0 | 0 | 1531672 | APO40578.1 | P0C6X7 |
| VYTFTTVKPGAA | 1 | 0 | 0 | 1531674 | QBF80585.1 | K9N7C7 |
| VYTNSCLVSSNG | 1 | 0 | 0 | 1531683 | QAT98908.1 | K9N7C7 |
| VYVGDPAQLPTP | 1 | 0 | 0 | 1531705 | ATG84864.1 | K9N7C7 |

| | | | | | | |
|---|---|---|---|---|---|---|
| VYVIKMIILWLM | 1 | 0 | 0 | 1531711 | AWW13547.1 | Q01455 |
| VYWVTSPPADVN | 1 | 0 | 0 | 1531732 | AJQ24074.1 | P33469 |
| VYYGNVLDQAIA | 1 | 0 | 0 | 1531738 | APO40578.1 | P0C6X7 |
| VYYPMCTNGQAV | 1 | 0 | 0 | 1531748 | AUM60013.1 | K9N7C7 |
| WAAAVDEAFPLN | 1 | 0 | 0 | 1531759 | AHI48527.1 | K9N638 |
| WAAFYVYKLQPL | 1 | 0 | 0 | 1531766 | AXN92228.1 | K9N5Q8 |
| WADESEPQRGRQ | 1 | 0 | 0 | 1531774 | ABB90491.1 | A0A0N7J131 |
| WADQSDQFRNFQ | 1 | 0 | 0 | 1531780 | AWW13570.1 | P33469 |
| WADQSDQFRSVQ | 1 | 0 | 0 | 1531782 | AIL49389.1 | P33469 |
| WADQSDQVRNVQ | 1 | 0 | 0 | 1531783 | P33469.1 | P33469 |
| WAGSKINAVIYN | 1 | 0 | 0 | 1531798 | ATQ39389.1 | K9N7C7 |
| WAMANQFTEFVG | 1 | 0 | 0 | 1531824 | AUM60013.1 | K9N7C7 |
| WAMTNGFSQVKA | 1 | 0 | 0 | 1531825 | ARB07436.1 | A0A140H1G9 |
| WARVEGKDGFVS | 1 | 0 | 0 | 1531835 | AXN92247.1 | K9N7C7 |
| WATLVDEEFPLD | 1 | 0 | 0 | 1531840 | AVV62535.1 | K9N7C7 |
| WAWNPEVNAITV | 1 | 0 | 0 | 1531844 | AST12967.1 | A0A0P0KXN3 |
| WCKDGKVQTFYP | 1 | 0 | 0 | 1531854 | ATQ39389.1 | K9N7C7 |
| WCLRCLYSTKPI | 1 | 0 | 0 | 1531862 | APO40578.1 | P0C6X7 |
| WCNVCGTRDVSI | 1 | 0 | 0 | 1531869 | AUM60013.1 | K9N7C7 |
| WCPRHVICPADQ | 1 | 0 | 0 | 1531870 | QGV13494.1 | K9N7C7 |
| WCPRHVMCPADQ | 1 | 0 | 0 | 1531871 | AXN92247.1 | K9N7C7 |
| WCPRHVMVPADQ | 1 | 0 | 0 | 1531872 | AUM60013.1 | K9N7C7 |
| WCRCSCLPDPIS | 1 | 0 | 0 | 1531873 | BBA20986.1 | A0A140H1H1 |
| WCTYLFSLIAVL | 1 | 0 | 0 | 1531880 | NP_073549.1 | A0A0P0LKV0 |
| WCVVLIVILLVC | 1 | 0 | 0 | 1531888 | NP_073554.1 | A0A0P0KH29 |
| WDDMLRPLIKDV | 1 | 0 | 0 | 1531892 | AHF54667.1 | P0C6X6 |
| WDIAVINNVDNE | 1 | 0 | 0 | 1531902 | AVV62535.1 | K9N7C7 |
| WDIKQCPAKSQT | 1 | 0 | 0 | 1531907 | AUM60013.1 | K9N7C7 |
| WDLPKCDRAMPN | 1 | 0 | 0 | 1531916 | AGH06532.1 | P0C6X6 |
| WDQAYHANCVNC | 1 | 0 | 0 | 1531919 | AVV62524.1 | K9N7C7 |
| WDQQYHANCVNC | 1 | 0 | 0 | 1531921 | AUM60013.1 | K9N7C7 |
| WDQTYHPNCSNC | 1 | 0 | 0 | 1531924 | APO40578.1 | P0C6X7 |
| WDTPTLDCINGQ | 1 | 0 | 0 | 1531936 | AUM60014.1 | K9N5Q8 |
| WDYERCNIYGSA | 1 | 0 | 0 | 1531943 | AVV62524.1 | K9N7C7 |
| WEAPHNDCTPIN | 1 | 0 | 0 | 1531962 | AVV62526.1 | K9N5Q8 |
| WEEGDYYRKQLS | 1 | 0 | 0 | 1531967 | ALJ76277.1 | K9N5Q8 |
| WEFRDYFDDNTD | 1 | 0 | 0 | 1531971 | AWK59930.1 | H9EJ42 |
| WEFRDYFDNNTD | 1 | 0 | 0 | 1531972 | AWK59942.1 | H9EJ42 |
| WEFRDYFNDNTD | 1 | 0 | 0 | 1531973 | AVA26872.1 | H9EJ42 |
| WEGLHEVLTSAM | 1 | 0 | 0 | 1531975 | AWK59942.1 | H9EJ42 |
| WEINACTYTMCE | 1 | 0 | 0 | 1531979 | ABA60802.1 | A0A140H1H1 |
| WEPDRMLKTSMA | 1 | 0 | 0 | 1531985 | AVR40343.1 | Q4VID0 |
| WERKKISNCVVD | 1 | 0 | 0 | 1531991 | ABD72995.1 | P59594 |
| WERKTFSNCNFN | 1 | 0 | 0 | 1531993 | CAA79896.1 | P36334 |
| WFDFIENPSVIG | 1 | 0 | 0 | 1532015 | AXN92226.1 | K9N7C7 |
| WFGITQDTQGVH | 1 | 0 | 0 | 1532028 | AVV62526.1 | K9N5Q8 |
| WFHEHVGGNFDN | 1 | 0 | 0 | 1532036 | AVV62524.1 | K9N7C7 |
| WFIDTFNQLTAD | 1 | 0 | 0 | 1532041 | ARB07436.1 | A0A140H1G9 |
| WFKLSYHYENNS | 1 | 0 | 0 | 1532048 | AWW13564.1 | Q80872 |
| WFKNHTSMAPNL | 1 | 0 | 0 | 1532049 | ARB07438.1 | A0A140H1H1 |
| WFKNQTSVALDL | 1 | 0 | 0 | 1532052 | ARA15421.1 | P36334 |
| WFKNQTSVAPDF | 1 | 0 | 0 | 1532053 | BBA20976.1 | P36334 |
| WFKNQTSVAPLL | 1 | 0 | 0 | 1532056 | AQN78680.1 | P36334 |
| WFKNQTSVAPVL | 1 | 0 | 0 | 1532057 | AIV41873.1 | P36334 |
| WFKNQTSVARDL | 1 | 0 | 0 | 1532058 | AIX10750.1 | P36334 |
| WFLRRFYNHVIN | 1 | 0 | 0 | 1532076 | AUM60013.1 | K9N7C7 |
| WFVKPSRTSIAS | 1 | 0 | 0 | 1532115 | AUM60013.1 | K9N7C7 |
| WFYTYSIGEASS | 1 | 0 | 0 | 1532127 | AZK15899.1 | K9N638 |
| WGNMLTGIAAHP | 1 | 0 | 0 | 1532138 | AVN89289.1 | K9N7C7 |
| WGNMMTGISARP | 1 | 0 | 0 | 1532142 | AVV62535.1 | K9N7C7 |
| WHCRSGHLIDEV | 1 | 0 | 0 | 1532150 | AUM60027.1 | K9N643 |
| WHILQFLVTGAY | 1 | 0 | 0 | 1532161 | APO40578.1 | P0C6X7 |
| WHYFDAQPLTRK | 1 | 0 | 0 | 1532203 | AXN92247.1 | K9N7C7 |
| WICANQSTSSYS | 1 | 0 | 0 | 1532210 | AKA09627.1 | A0A0P0G321 |
| WICANQSTSTYS | 1 | 0 | 0 | 1532213 | AWO68998.1 | A0A0P0G321 |
| WIFDSSKEFVEY | 1 | 0 | 0 | 1532220 | AIV41816.1 | P0C6X6 |
| WIFGSTLDNTTQ | 1 | 0 | 0 | 1532223 | APO40579.1 | P59594 |
| WIFGTTLDNTSQ | 1 | 0 | 0 | 1532225 | AVP78031.1 | Q3LZX1 |
| WIFGTTLDNTTQ | 1 | 0 | 0 | 1532226 | ATO98145.1 | Q3LZX1 |
| WIGFDVEGAHAF | 1 | 0 | 0 | 1532230 | AXN92247.1 | K9N7C7 |
| WILQEVKDNDGK | 1 | 0 | 0 | 1532245 | QRK03781.1 | A0A0P0ILY9 |
| WINPVILTPVFC | 1 | 0 | 0 | 1532253 | QQY99387.1 | H9EJ42 |
| WISPPVADYTPV | 1 | 0 | 0 | 1532260 | AVV62539.1 | K9N4V0 |
| WKCKVTDVLYPQ | 1 | 0 | 0 | 1532287 | AUM60013.1 | K9N7C7 |
| WKMVVCDKVRSF | 1 | 0 | 0 | 1532313 | AVN89451.1 | K9N7C7 |
| WKMVVCDKVRSL | 1 | 0 | 0 | 1532314 | AXN92247.1 | K9N7C7 |
| WKRQPNDDATSN | 1 | 0 | 0 | 1532320 | APT69866.1 | A0A0N7J131 |
| WKRVPTREENVI | 1 | 0 | 0 | 1532323 | AIW52805.1 | H9EJ69 |
| WLCANQSTSVYS | 1 | 0 | 0 | 1532348 | ABE73442.1 | A0A0P0G321 |
| WLDDTVYCPRAV | 1 | 0 | 0 | 1532363 | 2Q6G_A | P0C6X7 |
| WLELLEANIDAY | 1 | 0 | 0 | 1532378 | AVV62544.1 | K9N4V7 |
| WLFATLVLSYVA | 1 | 0 | 0 | 1532383 | AVV62524.1 | K9N7C7 |
| WLKGDKLFVEHY | 1 | 0 | 0 | 1532412 | ARX11394.1 | A0A0P0LKV0 |
| WLKNQTSVAPDL | 1 | 0 | 0 | 1532415 | ARB07414.1 | P36334 |
| WLKPFGGQPVSE | 1 | 0 | 0 | 1532416 | AKN24752.1 | K9N643 |
| WLLHKDGGIPDG | 1 | 0 | 0 | 1532426 | AXN92262.1 | K9N4V0 |
| WLLHKDGGVPDG | 1 | 0 | 0 | 1532427 | QBF80567.1 | K9N4V0 |
| WLMSLYSFTLKS | 1 | 0 | 0 | 1532462 | AVV62524.1 | K9N7C7 |
| WLNDKAVPSPLN | 1 | 0 | 0 | 1532469 | AIV41909.1 | P36334 |
| WLNDKLVPSPLN | 1 | 0 | 0 | 1532470 | ATN39872.1 | P36334 |
| WLNKVSVPSPLN | 1 | 0 | 0 | 1532476 | AMM42407.1 | A0A140H1H1 |

| | | | | | | |
|---|---|---|---|---|---|---|
| WLNNFNVPSPLNWERKI | 1 | 0 | 0 | 1532477 | AXT92537.1 | A0A140H1H1 |
| WLPELFMFSTLH | 1 | 0 | 0 | 1532485 | QEG03779.1 | P0C6X6 |
| WLPTDAIVIDND | 1 | 0 | 0 | 1532494 | ASY99818.1 | K9N7C7 |
| WLQPYGAKPVSE | 1 | 0 | 0 | 1532501 | AVV62529.1 | K9N643 |
| WLSAFGDVKPVT | 1 | 0 | 0 | 1532507 | AUM60027.1 | K9N643 |
| WLSNLVGSAVEK | 1 | 0 | 0 | 1532515 | ARO76381.1 | P0C6U8 |
| WLTGNAIQGFAC | 1 | 0 | 0 | 1532525 | AXN92247.1 | K9N7C7 |
| WLTNIFGTVYEK | 1 | 0 | 0 | 1532528 | AVP78041.1 | P0C6W2 |
| WLYAAILNGCNW | 1 | 0 | 0 | 1532552 | AUM60013.1 | K9N7C7 |
| WMELLDSNIDAY | 1 | 0 | 0 | 1532568 | AUM60021.1 | K9N4V7 |
| WNCNVDMYPEFS | 1 | 0 | 0 | 1532608 | AWK59942.1 | H9EJ42 |
| WNCNVDRYPANS | 1 | 0 | 0 | 1532610 | QYC64967.1 | P0DTD1 |
| WNCSAYMQLSDA | 1 | 0 | 0 | 1532613 | AUM60013.1 | K9N7C7 |
| WNCVDCDTAGIG | 1 | 0 | 0 | 1532615 | AVV62524.1 | K9N7C7 |
| WNFSWNLILTLF | 1 | 0 | 0 | 1532625 | ARE29965.1 | U3M6Q8 |
| WNGVSDLSLKQK | 1 | 0 | 0 | 1532632 | ASU90779.1 | K9N638 |
| WNIGEQSSILSP | 1 | 0 | 0 | 1532636 | AVP78030.1 | P0C6W2 |
| WNKKFGFIEDSV | 1 | 0 | 0 | 1532650 | ARB66070.1 | P36334 |
| WNKRFGFIEDSA | 1 | 0 | 0 | 1532652 | AGT51481.1 | P36334 |
| WNKYLYELIQKF | 1 | 0 | 0 | 1532659 | AGT51393.1 | H9EJ42 |
| WNKYLYELIQRF | 1 | 0 | 0 | 1532660 | AWK59942.1 | H9EJ42 |
| WNLDYPIISNEV | 1 | 0 | 0 | 1532666 | ABD75551.1 | A0A140H1G9 |
| WNNARLSDNMTA | 1 | 0 | 0 | 1532681 | AAX85668.1 | P30215 |
| WNNLRASDTSLS | 1 | 0 | 0 | 1532686 | AGT17757.1 | A0A140H1H0 |
| WNNMLRTLIDGV | 1 | 0 | 0 | 1532687 | AIW52827.1 | H9EJ42 |
| WNTAKQDQGQYY | 1 | 0 | 0 | 1532701 | ATO98193.1 | Q3LZX1 |
| WNTAKQDTGHYF | 1 | 0 | 0 | 1532702 | AVP78042.1 | Q3LZX1 |
| WNTFTKLQSLEN | 1 | 0 | 0 | 1532706 | AVP78041.1 | P0C6W2 |
| WNTKHGSGAFRE | 1 | 0 | 0 | 1532711 | ARO76381.1 | P0C6U8 |
| WNTKHSSGVTRELM | 1 | 0 | 0 | 1532716 | P0DTC1.1 | P0DTD1 |
| WNYGKPVTLPTG | 1 | 0 | 0 | 1532727 | AEN19363.1 | P0C6X6 |
| WNYLFETIPTQL | 1 | 0 | 0 | 1532729 | AXN92247.1 | K9N7C7 |
| WPFYPYGRCPTA | 1 | 0 | 0 | 1532733 | AGT51679.1 | P30215 |
| WPISNDSNCCVV | 1 | 0 | 0 | 1532737 | AGT51386.1 | H9EJ42 |
| WPLFESDKEEDL | 1 | 0 | 0 | 1532743 | APT69868.1 | A0A0P0ILY9 |
| WPLFESDKEEDV | 1 | 0 | 0 | 1532744 | AST12962.1 | A0A0P0LKV0 |
| WPLMEYDFTDYK | 1 | 0 | 0 | 1532751 | AUM60013.1 | K9N7C7 |
| WPLTIILTIFNC | 1 | 0 | 0 | 1532763 | Q01455.1 | Q01455 |
| WPLVLECTRAAA | 1 | 0 | 0 | 1532768 | AVV62524.1 | K9N7C7 |
| WPLVLECTRAAS | 1 | 0 | 0 | 1532769 | AUM60013.1 | K9N7C7 |
| WPMPINASKADG | 1 | 0 | 0 | 1532775 | AVV62537.1 | K9N5Q8 |
| WPNRTQLSESLG | 1 | 0 | 0 | 1532776 | AIZ74435.1 | K9N643 |
| WPQIAELAPSAS | 1 | 0 | 0 | 1532777 | AVV62544.1 | K9N4V7 |
| WPQIAELAPSSS | 1 | 0 | 0 | 1532778 | AVV62533.1 | K9N4V7 |
| WPQIAELVPTAS | 1 | 0 | 0 | 1532780 | AXG21672.1 | K9N4V7 |
| WPRPIDPAAADG | 1 | 0 | 0 | 1532783 | AVV62526.1 | K9N5Q8 |
| WPRPIDVFKADG | 1 | 0 | 0 | 1532784 | AHI48583.1 | K9N5Q8 |
| WPSSIALSVFSA | 1 | 0 | 0 | 1532786 | AUM60020.1 | K9N7A1 |
| WPTATGDVVLAK | 1 | 0 | 0 | 1532790 | ATN39862.1 | P0C6X6 |
| WQCKLDFLDCTG | 1 | 0 | 0 | 1532813 | AVV62539.1 | K9N4V0 |
| WQHYTNPFVNIG | 1 | 0 | 0 | 1532822 | AVV62539.1 | K9N4V0 |
| WQLHRQGKLSGF | 1 | 0 | 0 | 1532833 | AVV62539.1 | K9N4V0 |
| WQNLLYDFNGNI | 1 | 0 | 0 | 1532835 | ADN03339.1 | A0A140H1H1 |
| WQNLLYDPNGNL | 1 | 0 | 0 | 1532836 | ARB66062.1 | P36334 |
| WRIYATGTNVFQ | 1 | 0 | 0 | 1532867 | AVP78031.1 | Q3LZX1 |
| WRPEKKYLRYIN | 1 | 0 | 0 | 1532883 | YP_009555242.1 | Q04853 |
| WRVTAVHSDGIF | 1 | 0 | 0 | 1532902 | AGT51372.1 | H9EJ42 |
| WSADSCLQGNKC | 1 | 0 | 0 | 1532916 | AXX83303.1 | P36334 |
| WSASIDTGNCPF | 1 | 0 | 0 | 1532919 | ALK28781.1 | A0A0P0K6L9 |
| WSDQSDQFRNVQ | 1 | 0 | 0 | 1532924 | ARK08639.1 | P33469 |
| WSDQTVCYSGQS | 1 | 0 | 0 | 1532929 | AUM60013.1 | K9N7C7 |
| WSGISDLSLKQK | 1 | 0 | 0 | 1532946 | ALA49758.1 | K9N7C7 |
| WSGLCVDGRNGY | 1 | 0 | 0 | 1532948 | AIW52756.1 | A0A0P0K6L9 |
| WSGVSDLSLKQK | 1 | 0 | 0 | 1532952 | AXN92247.1 | K9N7C7 |
| WSIREFAVYSFY | 1 | 0 | 0 | 1532957 | AVP78042.1 | Q3LZX1 |
| WSKICDYVKPFL | 1 | 0 | 0 | 1532961 | APO40578.1 | P0C6X7 |
| WSNFTKVQGLEN | 1 | 0 | 0 | 1532973 | AVN89289.1 | K9N7C7 |
| WSRFPKSDGTGT | 1 | 0 | 0 | 1532985 | AVP78041.1 | P0C6W2 |
| WSTSINTGNCPF | 1 | 0 | 0 | 1532992 | AAQ90004.1 | A0A0P0K6L9 |
| WSVAVDEAFPLE | 1 | 0 | 0 | 1532995 | AUM60013.1 | K9N7C7 |
| WSVDAFNQFSSD | 1 | 0 | 0 | 1532996 | AEN19363.1 | P0C6X6 |
| WSYSGQNTELGV | 1 | 0 | 0 | 1533009 | APO40578.1 | P0C6X7 |
| WTAAEGVPFYLN | 1 | 0 | 0 | 1533020 | AIL49550.1 | P36334 |
| WTAAVDEAFPLD | 1 | 0 | 0 | 1533021 | AVN89451.1 | K9N7C7 |
| WTPYPELMEDLE | 1 | 0 | 0 | 1533055 | AUM60013.1 | K9N7C7 |
| WTTVLSMAVAKV | 1 | 0 | 0 | 1533062 | AEN19363.1 | P0C6X6 |
| WVAAVDEAFPLD | 1 | 0 | 0 | 1533065 | ASU45728.1 | K9N7C7 |
| WVASNQADVNIP | 1 | 0 | 0 | 1533072 | AIX10718.1 | P33469 |
| WVDQFERSHQTY | 1 | 0 | 0 | 1533076 | AZS52622.1 | Q19U29 |
| WVHEHGATDAPS | 1 | 0 | 0 | 1533100 | AXN92267.1 | K9N4V7 |
| WVHEYGATDAPS | 1 | 0 | 0 | 1533101 | QFQ59594.1 | K9N4V7 |
| WVLNNDYYKSLP | 1 | 0 | 0 | 1533120 | QXN66612.1 | P0DTD1 |
| WVLNNDYYRSIP | 1 | 0 | 0 | 1533121 | APO40578.1 | P0C6X7 |
| WVNATCIILQYL | 1 | 0 | 0 | 1533134 | AWK59942.1 | H9EJ42 |
| WVPRAAANIGSN | 1 | 0 | 0 | 1533144 | APO40578.1 | P0C6X7 |
| WVPRASADIGAN | 1 | 0 | 0 | 1533145 | ARO76381.1 | P0C6U8 |
| WVTNGNFDCALS | 1 | 0 | 0 | 1533161 | AVV62524.1 | K9N7C7 |
| WVTSPPADVNTP | 1 | 0 | 0 | 1533170 | AJQ24074.1 | P33469 |
| WWLLAKFTKLLL | 1 | 0 | 0 | 1533186 | NP_073549.1 | A0A0P0LKV0 |
| WYEFRAGRPHRL | 1 | 0 | 0 | 1533209 | ARU07575.1 | A0A140H1G9 |
| WYESTFGLSYYS | 1 | 0 | 0 | 1533211 | AEN19363.1 | P0C6X6 |

| | | | | | | |
|---|---|---|---|---|---|---|
| WYHEHIGGVYDN | 1 | 0 | 0 | 1533222 | AVN89333.1 | K9N7C7 |
| WYHEHVGGVYDS | 1 | 0 | 0 | 1533224 | AHC74096.1 | K9N7C7 |
| WYKPPFLSDFNN | 1 | 0 | 0 | 1533243 | BBA20986.1 | A0A140H1H1 |
| WYPSNRGSFSSA | 1 | 0 | 0 | 1533253 | AJQ24074.1 | P33469 |
| WYQKPFLSDFNN | 1 | 0 | 0 | 1533254 | BBA20983.1 | A0A140H1H1 |
| WYSGAIKLDPKN | 1 | 0 | 0 | 1533257 | AQZ41280.1 | K9N4V7 |
| WYVNDGRVDDDY | 1 | 0 | 0 | 1533268 | QED88011.1 | H9EJ42 |
| WYVWLLISLAGV | 1 | 0 | 0 | 1533278 | AQN78680.1 | P36334 |
| WYYSLTTNNAAT | 1 | 0 | 0 | 1533279 | AVP78031.1 | Q3LZX1 |
| YAALAAKEGLNS | 1 | 0 | 0 | 1533299 | AVV62524.1 | K9N7C7 |
| YADINGELHEDS | 1 | 0 | 0 | 1533315 | APO40578.1 | P0C6X7 |
| YADKTLQEMKTL | 1 | 0 | 0 | 1533320 | AGT51617.1 | P0C6X6 |
| YAFNSYGNMFQT | 1 | 0 | 0 | 1533351 | APO40579.1 | P59594 |
| YAHKHDIQLTNE | 1 | 0 | 0 | 1533365 | AVN89451.1 | K9N7C7 |
| YAHPAEPLVQDR | 1 | 0 | 0 | 1533370 | AHF54648.1 | Q4VID0 |
| YAIQQGFDATNS | 1 | 0 | 0 | 1533384 | AIX10750.1 | P36334 |
| YAIRNFDEQNCD | 1 | 0 | 0 | 1533388 | AGT51330.1 | H9EJ42 |
| YAISNEQHYKSW | 1 | 0 | 0 | 1533392 | ATO98193.1 | Q3LZX1 |
| YAISVKNRARTV | 1 | 0 | 0 | 1533393 | BAI44145.1 | P0C6X6 |
| YAKDQYGTDIDG | 1 | 0 | 0 | 1533400 | AIL49482.1 | P33469 |
| YANIVHEPTTPI | 1 | 0 | 0 | 1533461 | ALA49956.1 | K9N7C7 |
| YANIVYEPSTPI | 1 | 0 | 0 | 1533462 | AUM60013.1 | K9N7C7 |
| YANIVYEPSTPV | 1 | 0 | 0 | 1533463 | AVV62524.1 | K9N7C7 |
| YANVGRWSTSIN | 1 | 0 | 0 | 1533478 | AAQ90004.1 | A0A0P0K6L9 |
| YAPDEDEEEPCE | 1 | 0 | 0 | 1533482 | APO40578.1 | P0C6X7 |
| YAPEPITSFNTK | 1 | 0 | 0 | 1533483 | AVN89387.1 | K9N5Q8 |
| YAPEPITYLNTK | 1 | 0 | 0 | 1533487 | AGV08438.1 | K9N5Q8 |
| YAPNGTYNCTDA | 1 | 0 | 0 | 1533489 | ADK37704.1 | A0A0P0K6L9 |
| YAPNSRSTSRTS | 1 | 0 | 0 | 1533490 | AJQ24067.1 | P33469 |
| YAQHCFKAPTNF | 1 | 0 | 0 | 1533496 | CAA83660.1 | P36334 |
| YAQHCYKAPKNF | 1 | 0 | 0 | 1533499 | AIX10755.1 | P36334 |
| YAQLPNEITVAK | 1 | 0 | 0 | 1533500 | AUM60030.1 | K9N7A1 |
| YATGTTPQKLFV | 1 | 0 | 0 | 1533530 | ANC28634.1 | K9N5Q8 |
| YATKVVPFNNGF | 1 | 0 | 0 | 1533534 | AVV62537.1 | K9N5Q8 |
| YATLYFNNEPAF | 1 | 0 | 0 | 1533537 | AXG21664.1 | K9N638 |
| YAVQQLNNGFQG | 1 | 0 | 0 | 1533542 | AVV62524.1 | K9N7C7 |
| YAYDQVHDEPKG | 1 | 0 | 0 | 1533556 | APU51933.1 | P0C6X6 |
| YAYFTDAGVVTK | 1 | 0 | 0 | 1533564 | AIX10749.1 | P36334 |
| YAYMMPVMGMTN | 1 | 0 | 0 | 1533571 | AIW52776.1 | A0A0P0LKV0 |
| YCAVHQGAHVAS | 1 | 0 | 0 | 1533582 | AVV62524.1 | K9N7C7 |
| YCDGNLPCGICL | 1 | 0 | 0 | 1533595 | APO40578.1 | P0C6X7 |
| YCFINSTINTTP | 1 | 0 | 0 | 1533605 | AMM42635.1 | A0A0P0G321 |
| YCFNAEGDASWS | 1 | 0 | 0 | 1533612 | AXN92247.1 | K9N7C7 |
| YCFVNTTIGTET | 1 | 0 | 0 | 1533621 | AAK32191.1 | A0A0P0K6L9 |
| YCGNNSCTCQPQ | 1 | 0 | 0 | 1533631 | AEN19366.1 | P36334 |
| YCHDPTVLPGAF | 1 | 0 | 0 | 1533639 | AXN92247.1 | K9N7C7 |
| YCIHKECSIQEC | 1 | 0 | 0 | 1533641 | Q3I5I8.1 | Q3LZX5 |
| YCIIEPRNGSRC | 1 | 0 | 0 | 1533642 | AUM60014.1 | K9N5Q8 |
| YCILEPRPGNHC | 1 | 0 | 0 | 1533643 | QBF80608.1 | K9N5Q8 |
| YCILEPRSGNYC | 1 | 0 | 0 | 1533645 | AVN89387.1 | K9N5Q8 |
| YCIPYNSITDTI | 1 | 0 | 0 | 1533646 | ATO98146.1 | Q3LZX0 |
| YCMHKECSIQEC | 1 | 0 | 0 | 1533664 | ATO98227.1 | Q3LZX5 |
| YCNGWQNLLYDS | 1 | 0 | 0 | 1533668 | ABU39950.1 | P36334 |
| YCPAGNSYTSFA | 1 | 0 | 0 | 1533675 | AVN89387.1 | K9N5Q8 |
| YCPGNSNYNTFA | 1 | 0 | 0 | 1533676 | AVV62537.1 | K9N5Q8 |
| YCQVHGNAHVAS | 1 | 0 | 0 | 1533687 | ATO98118.1 | P0C6W2 |
| YCRKIGVNVCND | 1 | 0 | 0 | 1533699 | ARB07436.1 | A0A140H1G9 |
| YCSFYAPDEDEE | 1 | 0 | 0 | 1533711 | APO40578.1 | P0C6X7 |
| YCSNSSSGALDT | 1 | 0 | 0 | 1533718 | AIW52835.1 | A0A0P0G321 |
| YCSNSSSSVLDT | 1 | 0 | 0 | 1533720 | AVL25581.1 | A0A0P0G321 |
| YCTGSIPCSVCLSG | 1 | 0 | 0 | 1533728 | P0DTC1.1 | P0DTD1 |
| YCTNTTLVSGDA | 1 | 0 | 0 | 1533732 | BAL45640.1 | A0A0P0K6L9 |
| YCVDYSKNGGSG | 1 | 0 | 0 | 1533743 | 6NZK_A | P36334 |
| YCVDYSKNRLSR | 1 | 0 | 0 | 1533744 | QBP84713.1 | P36334 |
| YCVLEPRNGTRC | 1 | 0 | 0 | 1533753 | AVV62526.1 | K9N5Q8 |
| YCVRAFDGYNKD | 1 | 0 | 0 | 1533755 | AST12962.1 | A0A0P0LKV0 |
| YCVSAEANAVQQ | 1 | 0 | 0 | 1533760 | AVV62535.1 | K9N7C7 |
| YCYCGNFATCYN | 1 | 0 | 0 | 1533763 | AWK59942.1 | H9EJ42 |
| YCYDPTVLPGAH | 1 | 0 | 0 | 1533766 | AVV62524.1 | K9N7C7 |
| YCYSAKDTLPEL | 1 | 0 | 0 | 1533775 | AVV62524.1 | K9N7C7 |
| YCYTARDALDTI | 1 | 0 | 0 | 1533777 | ATQ39389.1 | K9N7C7 |
| YCYTDGLMQNAS | 1 | 0 | 0 | 1533779 | AEN19363.1 | P0C6X6 |
| YCYTEGLMQNAS | 1 | 0 | 0 | 1533780 | AGT51787.1 | P0C6X6 |
| YDANYFVCWHTN | 1 | 0 | 0 | 1533794 | Q3I5J4.1 | Q3LZX0 |
| YDAQPLRKNVQQ | 1 | 0 | 0 | 1533797 | AVV62524.1 | K9N7C7 |
| YDASNVKNVTDV | 1 | 0 | 0 | 1533800 | AVQ05261.1 | P0C6X6 |
| YDCFCNNINWNV | 1 | 0 | 0 | 1533808 | AEN19363.1 | P0C6X6 |
| YDDCMHQGPASA | 1 | 0 | 0 | 1533813 | ATG84866.1 | K9N5Q8 |
| YDDCMKQGPPSA | 1 | 0 | 0 | 1533815 | AVV62537.1 | K9N5Q8 |
| YDDCMQQGPSSA | 1 | 0 | 0 | 1533819 | AHL18090.1 | K9N5Q8 |
| YDDFLEESDVEE | 1 | 0 | 0 | 1533820 | AGT51707.1 | P0C6X6 |
| YDDIFAESKPRF | 1 | 0 | 0 | 1533822 | AGT21359.1 | A0A0P0LKV0 |
| YDDRILEFKVLD | 1 | 0 | 0 | 1533830 | ATQ39389.1 | K9N7C7 |
| YDECMQQGPQSA | 1 | 0 | 0 | 1533839 | AUM60014.1 | K9N5Q8 |
| YDFGDFVVRLPN | 1 | 0 | 0 | 1533857 | QED88024.1 | H9EJ42 |
| YDFLNIDMGVFI | 1 | 0 | 0 | 1533864 | AGT51365.1 | H9EJ42 |
| YDFLPIILQGIL | 1 | 0 | 0 | 1533866 | ARU07576.1 | A0A140H1H0 |
| YDFNGNIIGFKD | 1 | 0 | 0 | 1533867 | ADN03339.1 | A0A140H1H1 |
| YDFNKPTSKTCT | 1 | 0 | 0 | 1533868 | AUM60013.1 | K9N7C7 |
| YDFNPNVPLDYQ | 1 | 0 | 0 | 1533869 | ATO98145.1 | Q3LZX1 |
| YDFNPNVPLEYQ | 1 | 0 | 0 | 1533870 | AVP78031.1 | Q3LZX1 |

| | | | | | | |
|---|---|---|---|---|---|---|
| YDFTEERLCLFN | 1 | 0 | 0 | 1533875 | APO40578.1 | P0C6X7 |
| YDFVEHHDVRVV | 1 | 0 | 0 | 1533878 | Q3I5I8.1 | Q3LZX5 |
| YDFYPNVPIEYQ | 1 | 0 | 0 | 1533882 | ATO98120.1 | Q3LZX1 |
| YDGARLSASDLS | 1 | 0 | 0 | 1533885 | AAS58186.1 | H9EJ42 |
| YDGFDNLVGYYS | 1 | 0 | 0 | 1533889 | AVV62526.1 | K9N5Q8 |
| YDGLKGGIRKAK | 1 | 0 | 0 | 1533894 | AVV62524.1 | K9N7C7 |
| YDGSGVGVIRIS | 1 | 0 | 0 | 1533900 | BAL45639.1 | A0A0P0K6L9 |
| YDHAIGATAALP | 1 | 0 | 0 | 1533902 | ATO98145.1 | Q3LZX1 |
| YDHSQYYFNKDT | 1 | 0 | 0 | 1533906 | AEN19365.1 | P30215 |
| YDHVFGTDHKFI | 1 | 0 | 0 | 1533908 | AWK59942.1 | H9EJ42 |
| YDIEFVTGKQNI | 1 | 0 | 0 | 1533915 | ALA50044.1 | K9N7C7 |
| YDINLNKFNRAS | 1 | 0 | 0 | 1533929 | AMQ48992.1 | K9N7C7 |
| YDKLVSSFLEMKSEK | 1 | 0 | 0 | 1533945 | P0DTC1.1 | P0DTD1 |
| YDKNASLNSFKE | 1 | 0 | 0 | 1533946 | AUM60014.1 | K9N5Q8 |
| YDLVNYIRDKGC | 1 | 0 | 0 | 1533964 | AIV41836.1 | Q80872 |
| YDLYDGNMFIKF | 1 | 0 | 0 | 1533965 | AXN92247.1 | K9N7C7 |
| YDMDFGVSSFSE | 1 | 0 | 0 | 1533967 | AHY22554.1 | K9N638 |
| YDMFESNMFIKF | 1 | 0 | 0 | 1533972 | AVV62524.1 | K9N7C7 |
| YDNCYRKSNVDE | 1 | 0 | 0 | 1533976 | AIG96641.1 | A0A0P0LKV0 |
| YDNCYRNSNVGE | 1 | 0 | 0 | 1533977 | ADO24705.1 | A0A0P0LKV0 |
| YDNFTSIWPRFS | 1 | 0 | 0 | 1533983 | AGT17757.1 | A0A140H1H0 |
| YDNISSVWPLHP | 1 | 0 | 0 | 1533986 | AQN78751.1 | P30215 |
| YDNKLKAHKEKS | 1 | 0 | 0 | 1533990 | P0C6W6.1 | P0C6W2 |
| YDNVSSVWPHYP | 1 | 0 | 0 | 1533996 | AGT51769.1 | P30215 |
| YDNVSSVWPLYL | 1 | 0 | 0 | 1533997 | AGT51411.1 | P30215 |
| YDPITKTEKDTN | 1 | 0 | 0 | 1534003 | AUM60013.1 | K9N7C7 |
| YDPIYKNGAMLN | 1 | 0 | 0 | 1534004 | AVV62535.1 | K9N7C7 |
| YDPLQPELESFK | 1 | 0 | 0 | 1534012 | APO40579.1 | P59594 |
| YDPNGNLYGFRD | 1 | 0 | 0 | 1534015 | ARB66062.1 | P36334 |
| YDPSKPTSKKYT | 1 | 0 | 0 | 1534019 | AVV62535.1 | K9N7C7 |
| YDPTVLPGAHAY | 1 | 0 | 0 | 1534021 | AVV62524.1 | K9N7C7 |
| YDPTVLPGAHPY | 1 | 0 | 0 | 1534022 | AUM60013.1 | K9N7C7 |
| YDPVTKNINGSN | 1 | 0 | 0 | 1534024 | AVV62524.1 | K9N7C7 |
| YDPYLSGYYHNN | 1 | 0 | 0 | 1534025 | AVP78031.1 | Q3LZX1 |
| YDQVVMDYLDSL | 1 | 0 | 0 | 1534029 | ARO76381.1 | P0C6U8 |
| YDRLDIIQADQQ | 1 | 0 | 0 | 1534035 | AIW52757.1 | A0A0P0K6L9 |
| YDRLDPPQADQQ | 1 | 0 | 0 | 1534036 | 6U7H_A | A0A0P0K6L9 |
| YDRLSNEVTVAK | 1 | 0 | 0 | 1534043 | AMQ49021.1 | K9N7A1 |
| YDRVEQSFQLDT | 1 | 0 | 0 | 1534045 | ATO98120.1 | Q3LZX1 |
| YDSADRGQESLA | 1 | 0 | 0 | 1534046 | ATQ39389.1 | K9N7C7 |
| YDSGSLTKASDW | 1 | 0 | 0 | 1534049 | AUM60013.1 | K9N7C7 |
| YDSKESLVFPTG | 1 | 0 | 0 | 1534051 | AVV62524.1 | K9N7C7 |
| YDSNGNLYGLRD | 1 | 0 | 0 | 1534059 | ABD43179.1 | P36334 |
| YDSPCFSQQGVF | 1 | 0 | 0 | 1534060 | AAA73152.1 | P30215 |
| YDSQPSTSFYDI | 1 | 0 | 0 | 1534061 | AUM60025.1 | K9N796 |
| YDSSDHGQESLA | 1 | 0 | 0 | 1534062 | AHI48549.1 | K9N7C7 |
| YDSSGNLYGFRD | 1 | 0 | 0 | 1534066 | ARK08661.1 | P36334 |
| YDSVSAIGTANL | 1 | 0 | 0 | 1534069 | AAK32190.1 | A0A0P0K6L9 |
| YDTIKYYSIIPY | 1 | 0 | 0 | 1534075 | QBF80611.1 | K9N5Q8 |
| YDTNHGDAGFTS | 1 | 0 | 0 | 1534077 | AQN78743.1 | P30215 |
| YDTNLNKFNGAS | 1 | 0 | 0 | 1534079 | AVN89385.1 | K9N7C7 |
| YDVANNAVYDGA | 1 | 0 | 0 | 1534083 | AFD98763.1 | H9EJ42 |
| YDVAVTAGPFMG | 1 | 0 | 0 | 1534085 | ATQ39389.1 | K9N7C7 |
| YDVDFMEYHDVR | 1 | 0 | 0 | 1534087 | ATO98227.1 | Q3LZX5 |
| YDVVLSPLLSAG | 1 | 0 | 0 | 1534107 | APO40578.1 | P0C6X7 |
| YDVVVTAGPFMD | 1 | 0 | 0 | 1534109 | AVN89363.1 | K9N7C7 |
| YDYLGKLFVSDY | 1 | 0 | 0 | 1534115 | AVV62537.1 | K9N5Q8 |
| YDYSQYYFNKDT | 1 | 0 | 0 | 1534119 | AGT51679.1 | P30215 |
| YDYTQYYFNKDT | 1 | 0 | 0 | 1534121 | QEG03802.1 | P30215 |
| YEADAEIVQPAT | 1 | 0 | 0 | 1534126 | AUM60013.1 | K9N7C7 |
| YEADTEQTQSVI | 1 | 0 | 0 | 1534130 | AVN89451.1 | K9N7C7 |
| YEADTEQTQSVN | 1 | 0 | 0 | 1534131 | ATG84700.1 | K9N638 |
| YEADTEQTQSVT | 1 | 0 | 0 | 1534132 | AXN92247.1 | K9N7C7 |
| YEADVEAAEQVV | 1 | 0 | 0 | 1534133 | AVV62524.1 | K9N7C7 |
| YEASAFRDLKPL | 1 | 0 | 0 | 1534142 | AUM60013.1 | K9N7C7 |
| YECCEASYFDNK | 1 | 0 | 0 | 1534143 | AUM60013.1 | K9N7C7 |
| YEEYDLEPHKVH | 1 | 0 | 0 | 1534167 | AXN92228.1 | K9N5Q8 |
| YEFLNKHFNMMI | 1 | 0 | 0 | 1534178 | AIL49554.1 | P0C6X6 |
| YEFRAGRPHRLV | 1 | 0 | 0 | 1534179 | ARU07575.1 | A0A140H1G9 |
| YEFYNDIKPPVL | 1 | 0 | 0 | 1534180 | AOL02455.1 | Q04854 |
| YEGFITLKNNLY | 1 | 0 | 0 | 1534181 | ANI69867.1 | K9N638 |
| YEGGGTLVGVGN | 1 | 0 | 0 | 1534183 | AVV62526.1 | K9N5Q8 |
| YEGNSPYHPLAD | 1 | 0 | 0 | 1534187 | APO40584.1 | P59635 |
| YEGVNFSPHHRF | 1 | 0 | 0 | 1534191 | ABD75512.1 | A0A140H1H0 |
| YEGVNFTPYHAF | 1 | 0 | 0 | 1534192 | AEN19365.1 | P30215 |
| YEIQIPSKFTIG | 1 | 0 | 0 | 1534200 | AIL49542.1 | P36334 |
| YELEKHCYELLK | 1 | 0 | 0 | 1534208 | AVV62535.1 | K9N7C7 |
| YELNGYTVQPVA | 1 | 0 | 0 | 1534226 | AVR40344.1 | P36334 |
| YELRYNGAIRFD | 1 | 0 | 0 | 1534228 | AEN19370.1 | P33469 |
| YELSQLISGSLG | 1 | 0 | 0 | 1534232 | ABE97136.1 | H9EJ42 |
| YEMFGNYTISCE | 1 | 0 | 0 | 1534234 | ATO98227.1 | Q3LZX5 |
| YENAFLPFAMGI | 1 | 0 | 0 | 1534241 | QYC64967.1 | P0DTD1 |
| YENAFMPFAMGI | 1 | 0 | 0 | 1534244 | AVP78041.1 | P0C6W2 |
| YENDWNTKHGSG | 1 | 0 | 0 | 1534246 | APO40578.1 | P0C6X7 |
| YENRFAAIYGGD | 1 | 0 | 0 | 1534256 | AVA26874.1 | H9EJ24 |
| YENTTYECDIPI | 1 | 0 | 0 | 1534257 | APO40579.1 | P59594 |
| YENYTILQAAGL | 1 | 0 | 0 | 1534260 | QED88024.1 | H9EJ42 |
| YEQAVVNGDSEV | 1 | 0 | 0 | 1534277 | AVP78041.1 | P0C6W2 |
| YEQGSGYIPEAP | 1 | 0 | 0 | 1534279 | 6CRW_A | P59594 |
| YEQVVMDYLDSL | 1 | 0 | 0 | 1534287 | P0C6W6.1 | P0C6W2 |
| YERHPLSQFVNL | 1 | 0 | 0 | 1534302 | ARO76381.1 | P0C6U8 |

| | | | | | | |
|---|---|---|---|---|---|---|
| YERQSPFEIKSA | 1 | 0 | 0 | 1534307 | ARO76381.1 | P0C6U8 |
| YERVKDCKFVAE | 1 | 0 | 0 | 1534311 | AEN19363.1 | P0C6X6 |
| YESFDVDSGVYS | 1 | 0 | 0 | 1534312 | AUM60014.1 | K9N5Q8 |
| YESFDVDTGVYS | 1 | 0 | 0 | 1534313 | AVV62526.1 | K9N5Q8 |
| YESFDVESGVYS | 1 | 0 | 0 | 1534314 | AXN92228.1 | K9N5Q8 |
| YESFDVGSGVYS | 1 | 0 | 0 | 1534315 | AVV62537.1 | K9N5Q8 |
| YESQTPFEIKLA | 1 | 0 | 0 | 1534321 | AVP78041.1 | P0C6W2 |
| YESSAFRDLKPL | 1 | 0 | 0 | 1534322 | AUM60023.1 | K9N7C7 |
| YETMKPHVRYDM | 1 | 0 | 0 | 1534333 | AVV62535.1 | K9N7C7 |
| YEVGDLIAQSSG | 1 | 0 | 0 | 1534342 | AEN19363.1 | P0C6X6 |
| YEVGDLMAQSSG | 1 | 0 | 0 | 1534343 | YP_009555247.1 | P0C6X6 |
| YFAVRKEGEDVI | 1 | 0 | 0 | 1534367 | AVQ05261.1 | P0C6X6 |
| YFCQDPMLAVAN | 1 | 0 | 0 | 1534378 | APO40579.1 | P59594 |
| YFCYGFNEPLNV | 1 | 0 | 0 | 1534382 | ARU07576.1 | A0A140H1H0 |
| YFDKSWPRPIDP | 1 | 0 | 0 | 1534393 | AVV62526.1 | K9N5Q8 |
| YFFPYDPALETG | 1 | 0 | 0 | 1534415 | AVV62524.1 | K9N7C7 |
| YFIKGCNTLARG | 1 | 0 | 0 | 1534443 | ARU07575.1 | A0A140H1G9 |
| YFKHWSFDYHPN | 1 | 0 | 0 | 1534470 | AIW52827.1 | H9EJ42 |
| YFKNHTSPDIDL | 1 | 0 | 0 | 1534476 | AVP78031.1 | Q3LZX1 |
| YFKNTDTTCTKV | 1 | 0 | 0 | 1534480 | AVV62524.1 | K9N7C7 |
| YFLVRLLQGKTG | 1 | 0 | 0 | 1534513 | AHI48793.1 | K9N7C7 |
| YFNGTGRGACKG | 1 | 0 | 0 | 1534534 | AIW52757.1 | A0A0P0K6L9 |
| YFNGTGRGDCQG | 1 | 0 | 0 | 1534536 | BAL45639.1 | A0A0P0K6L9 |
| YFNNEPAFIMHV | 1 | 0 | 0 | 1534555 | AXG21664.1 | K9N638 |
| YFNPEPITMANT | 1 | 0 | 0 | 1534559 | AUM60014.1 | K9N5Q8 |
| YFNRPLLVDDNK | 1 | 0 | 0 | 1534560 | AEN19363.1 | P0C6X6 |
| YFPKSGANFTDL | 1 | 0 | 0 | 1534571 | AZS52618.1 | A0A140H1H1 |
| YFQGGGGSGYIP | 1 | 0 | 0 | 1534582 | 6NB6_A | P59594 |
| YFQKAGQSTYER | 1 | 0 | 0 | 1534583 | APO40578.1 | P0C6X7 |
| YFRKSCQNLDCN | 1 | 0 | 0 | 1534587 | NP_150075.1 | Q80872 |
| YFSDSKIGTTEV | 1 | 0 | 0 | 1534595 | ALD51903.1 | K9N7C7 |
| YFSLGDVEAVNF | 1 | 0 | 0 | 1534609 | AST12964.1 | A0A0P0K6L9 |
| YFSYPFSMKSDL | 1 | 0 | 0 | 1534620 | AVN89313.1 | K9N5Q8 |
| YFSYPLSMKSDL | 1 | 0 | 0 | 1534621 | AXN92228.1 | K9N5Q8 |
| YFTLGCKILTLT | 1 | 0 | 0 | 1534636 | AXN92247.1 | K9N7C7 |
| YFTNEPAFIMHL | 1 | 0 | 0 | 1534643 | AVV62524.1 | K9N7C7 |
| YFTNEPAFIMHV | 1 | 0 | 0 | 1534644 | AXN92247.1 | K9N7C7 |
| YFTQSRNLESFK | 1 | 0 | 0 | 1534648 | APO40578.1 | P0C6X7 |
| YFTSAPPVTYSP | 1 | 0 | 0 | 1534652 | AVK87459.1 | K9N7C7 |
| YFTSEPPVTYSS | 1 | 0 | 0 | 1534657 | AHI48796.1 | K9N7C7 |
| YFTSPPPITFSP | 1 | 0 | 0 | 1534659 | AUM60013.1 | K9N7C7 |
| YFYSAKDDIKSV | 1 | 0 | 0 | 1534712 | AUM60013.1 | K9N7C7 |
| YGADLKSPDLGD | 1 | 0 | 0 | 1534729 | APO40578.1 | P0C6X7 |
| YGAKPVSEYHVT | 1 | 0 | 0 | 1534736 | AVV62529.1 | K9N643 |
| YGAKPVTEYHIT | 1 | 0 | 0 | 1534737 | AVV62540.1 | K9N643 |
| YGAYQPYDHAIG | 1 | 0 | 0 | 1534749 | ATO98145.1 | Q3LZX1 |
| YGCCDASYFDNK | 1 | 0 | 0 | 1534750 | AVV62524.1 | K9N7C7 |
| YGDCTFGVPDDA | 1 | 0 | 0 | 1534770 | AVV62524.1 | K9N7C7 |
| YGDFSHSQLGGL | 1 | 0 | 0 | 1534774 | QYC64967.1 | P0DTD1 |
| YGDSAILPKGIM | 1 | 0 | 0 | 1534775 | AVP78030.1 | P0C6W2 |
| YGEPVLVNAANT | 1 | 0 | 0 | 1534786 | AVN89451.1 | K9N7C7 |
| YGFCGNGTHIFS | 1 | 0 | 0 | 1534790 | AIW52843.1 | A0A0P0G321 |
| YGFNSTETITTG | 1 | 0 | 0 | 1534796 | AIL49343.1 | P30215 |
| YGFSNVVEMPKF | 1 | 0 | 0 | 1534803 | AIW52757.1 | A0A0P0K6L9 |
| YGFTVFVNVDNS | 1 | 0 | 0 | 1534808 | AVV62535.1 | K9N7C7 |
| YGGFGCKDSFTL | 1 | 0 | 0 | 1534818 | AEN19363.1 | P0C6X6 |
| YGGVSVITPGTN | 1 | 0 | 0 | 1534825 | APO40579.1 | P59594 |
| YGIPLSEAKGYW | 1 | 0 | 0 | 1534842 | AMM42386.1 | Q19U29 |
| YGIPPSEAKGYW | 1 | 0 | 0 | 1534843 | AMM42397.1 | Q19U29 |
| YGKDAFENHSYI | 1 | 0 | 0 | 1534853 | AVV62524.1 | K9N7C7 |
| YGKMGRFFNHTL | 1 | 0 | 0 | 1534859 | AHI48802.1 | K9N5Q8 |
| YGKNAFQNHSYI | 1 | 0 | 0 | 1534860 | AVV62535.1 | K9N7C7 |
| YGKSKHYDKYFG | 1 | 0 | 0 | 1534862 | AQN78665.1 | Q04853 |
| YGKSTFNKDIYC | 1 | 0 | 0 | 1534863 | AAY32939.1 | H9EJ42 |
| YGLCDSANPTNC | 1 | 0 | 0 | 1534868 | AVN89334.1 | K9N5Q8 |
| YGLCNTDTPPRC | 1 | 0 | 0 | 1534872 | AVV62537.1 | K9N5Q8 |
| YGLEYYAFEHIV | 1 | 0 | 0 | 1534877 | QBQ01836.1 | P0C6X6 |
| YGLNSTETISTG | 1 | 0 | 0 | 1534889 | AGT51729.1 | P30215 |
| YGLNSTETITTS | 1 | 0 | 0 | 1534891 | APU51943.1 | P30215 |
| YGNCTFGSPDDA | 1 | 0 | 0 | 1534907 | AHI48794.1 | K9N638 |
| YGNMFQTQAGCL | 1 | 0 | 0 | 1534910 | APO40579.1 | P59594 |
| YGNTMYVNMAVP | 1 | 0 | 0 | 1534915 | AUM60027.1 | K9N643 |
| YGPFVDIQTAQA | 1 | 0 | 0 | 1534920 | 3ATW_A | P0C6X7 |
| YGPYKDAQVVQL | 1 | 0 | 0 | 1534931 | AEN19363.1 | P0C6X6 |
| YGQCTFGAPDDA | 1 | 0 | 0 | 1534935 | AUM60013.1 | K9N7C7 |
| YGQQFGPTYMDG | 1 | 0 | 0 | 1534940 | APO40578.1 | P0C6X7 |
| YGRCPTAADINN | 1 | 0 | 0 | 1534944 | P30215.1 | P30215 |
| YGRCPTAADINT | 1 | 0 | 0 | 1534945 | AAA73152.1 | P30215 |
| YGRGGYHYTPFH | 1 | 0 | 0 | 1534949 | AHI48797.1 | K9N638 |
| YGSGYIPEAPRD | 1 | 0 | 0 | 1534967 | 5W9H_A | K9N5Q8 |
| YGTCLNCNAPRM | 1 | 0 | 0 | 1534977 | NP_073549.1 | A0A0P0LKV0 |
| YGTDIDGIYWVA | 1 | 0 | 0 | 1534979 | AIL49434.1 | P33469 |
| YGTDTISVCPKL | 1 | 0 | 0 | 1534983 | AGV08438.1 | K9N5Q8 |
| YGTEDDYNGTPL | 1 | 0 | 0 | 1534992 | APO40578.1 | P0C6X7 |
| YGVALKDCGFVM | 1 | 0 | 0 | 1535008 | AEN19363.1 | P0C6X6 |
| YGYFLPYPDPSR | 1 | 0 | 0 | 1535026 | ALW82667.1 | K9N7C7 |
| YGYLMKRFSMMI | 1 | 0 | 0 | 1535032 | ADO24650.1 | H9EJ42 |
| YGYPNDIVGGFR | 1 | 0 | 0 | 1535035 | AWK59978.1 | H9EJ42 |
| YGYSNFVSSDLD | 1 | 0 | 0 | 1535037 | AGT17757.1 | A0A140H1H0 |
| YGYTSRSMFVYV | 1 | 0 | 0 | 1535039 | AWW13547.1 | Q01455 |
| YHAFKCNHSGSN | 1 | 0 | 0 | 1535044 | AQN78687.1 | P30215 |

| | | | | | | |
|---|---|---|---|---|---|---|
| YHCVSTQANAVQ | 1 | 0 | 0 | 1535056 | AVV62524.1 | K9N7C7 |
| YHEHIGGVYDNS | 1 | 0 | 0 | 1535061 | AVN89333.1 | K9N7C7 |
| YHFKQLGLVWNK | 1 | 0 | 0 | 1535066 | AIW52827.1 | H9EJ42 |
| YHNSQDDIMKAV | 1 | 0 | 0 | 1535109 | AAA45463.1 | A0A0N7J131 |
| YHPAHAPSFLSV | 1 | 0 | 0 | 1535115 | ARB07436.1 | A0A140H1G9 |
| YHQKHFKSIVNT | 1 | 0 | 0 | 1535125 | ADK37693.1 | H9EJ42 |
| YHQLTDNVYIKC | 1 | 0 | 0 | 1535127 | APO40578.1 | P0C6X7 |
| YHSAFADCTSCS | 1 | 0 | 0 | 1535133 | AUM60013.1 | K9N7C7 |
| YHSAFVECTGCG | 1 | 0 | 0 | 1535138 | AVV62524.1 | K9N7C7 |
| YHSGLPLHMSIS | 1 | 0 | 0 | 1535141 | AXN92231.1 | K9N643 |
| YHTASTLRSVGQ | 1 | 0 | 0 | 1535146 | ATO98181.1 | Q3LZX1 |
| YHTPATNCSDGN | 1 | 0 | 0 | 1535158 | ALA49671.1 | K9N5Q8 |
| YHTPTTDCSDGN | 1 | 0 | 0 | 1535160 | ALA49473.1 | K9N5Q8 |
| YHTSGNVIEFHL | 1 | 0 | 0 | 1535161 | APO40578.1 | P0C6X7 |
| YHYNELGIEMNM | 1 | 0 | 0 | 1535180 | AIL49563.1 | P0C6X6 |
| YHYQECARGTTV | 1 | 0 | 0 | 1535181 | APO40584.1 | P59635 |
| YHYTPFHYERNN | 1 | 0 | 0 | 1535185 | AVN89451.1 | K9N7C7 |
| YHYTPVRYERDN | 1 | 0 | 0 | 1535188 | ALA50044.1 | K9N7C7 |
| YIAPQANSGIIT | 1 | 0 | 0 | 1535195 | AEN19365.1 | P30215 |
| YIAPRANSGIIT | 1 | 0 | 0 | 1535196 | AIL49334.1 | P30215 |
| YIAVNDQKLYDQ | 1 | 0 | 0 | 1535203 | ARO76381.1 | P0C6U8 |
| YIDAGNHRSAFA | 1 | 0 | 0 | 1535217 | AKA09621.1 | A0A0P0G321 |
| YIDCEQTPCGFK | 1 | 0 | 0 | 1535221 | AIV42002.1 | P0C6X6 |
| YIDCEQYSCDFK | 1 | 0 | 0 | 1535222 | QBP84701.1 | P0C6X6 |
| YIDFVNSGGTPL | 1 | 0 | 0 | 1535225 | AUM60023.1 | K9N7C7 |
| YIECTYIGRFSK | 1 | 0 | 0 | 1535239 | ATG84705.1 | K9N7D2 |
| YIEGSGKSAPNS | 1 | 0 | 0 | 1535245 | AJQ24087.1 | P33469 |
| YIEGSGKYAPNS | 1 | 0 | 0 | 1535246 | AJQ24067.1 | P33469 |
| YIEGSGRSASNS | 1 | 0 | 0 | 1535250 | AXX83347.1 | P33469 |
| YIEGSGRSVPNS | 1 | 0 | 0 | 1535251 | AIL49415.1 | P33469 |
| YIFSTANALPEC | 1 | 0 | 0 | 1535271 | ABE97136.1 | H9EJ42 |
| YIFWRNTNPIQLSS | 1 | 0 | 0 | 1535278 | P0DTD1.1 | P0DTD1 |
| YIIMPVHTLSVH | 1 | 0 | 0 | 1535308 | ARO76381.1 | P0C6U8 |
| YIISCYWGLFSL | 1 | 0 | 0 | 1535313 | AEN19363.1 | P0C6X6 |
| YIKIKRMKQIED | 1 | 0 | 0 | 1535324 | 5X58_A | P59594 |
| YIKSGGYIPEAP | 1 | 0 | 0 | 1535337 | 6U7H_A | A0A0P0K6L9 |
| YIKWPWSHPQFE | 1 | 0 | 0 | 1535338 | 6ACC_A | P59594 |
| YIMFEPIDPIWM | 1 | 0 | 0 | 1535363 | AWU59320.1 | K9N638 |
| YINDKDTGPPPI | 1 | 0 | 0 | 1535369 | CAA79896.1 | P36334 |
| YINGFICEKLAI | 1 | 0 | 0 | 1535373 | NP_073549.1 | A0A0P0LKV0 |
| YINGYRYFSLGN | 1 | 0 | 0 | 1535380 | AIW52757.1 | A0A0P0K6L9 |
| YINIPDLPICVY | 1 | 0 | 0 | 1535383 | AEN19365.1 | P30215 |
| YINIPDLPVCVY | 1 | 0 | 0 | 1535384 | ARB07432.1 | P30215 |
| YINNKDTGPPFI | 1 | 0 | 0 | 1535396 | AKA09646.1 | P36334 |
| YINTHMIGVTLC | 1 | 0 | 0 | 1535398 | ARU07575.1 | A0A140H1G9 |
| YINVTFLNLQVE | 1 | 0 | 0 | 1535407 | AGT51680.1 | P36334 |
| YIPEAPRDGQAY | 1 | 0 | 0 | 1535412 | 6CRW_A | P59594 |
| YIPLFDMRYSFF | 1 | 0 | 0 | 1535421 | AMQ49019.1 | K9N7D2 |
| YIPLVGTPLHPV | 1 | 0 | 0 | 1535423 | ANI69891.1 | K9N4V0 |
| YIPSNFALNNWF | 1 | 0 | 0 | 1535427 | ADK37704.1 | A0A0P0K6L9 |
| YIPSQEKNFTTA | 1 | 0 | 0 | 1535428 | AVP78031.1 | Q3LZX1 |
| YIPTSHINVTAA | 1 | 0 | 0 | 1535429 | AVV62526.1 | K9N5Q8 |
| YIRMEESLMDVP | 1 | 0 | 0 | 1535442 | AIZ74435.1 | K9N643 |
| YIRRAIDCGHDD | 1 | 0 | 0 | 1535444 | AVV62526.1 | K9N5Q8 |
| YIRRSIDCGFND | 1 | 0 | 0 | 1535446 | AHZ90568.1 | K9N5Q8 |
| YISHETKLNSCC | 1 | 0 | 0 | 1535457 | ASU45783.1 | K9N7C7 |
| YISKCSRLTGDN | 1 | 0 | 0 | 1535459 | AVV62526.1 | K9N5Q8 |
| YISTCYFGVFSM | 1 | 0 | 0 | 1535470 | AUM60013.1 | K9N7C7 |
| YITANGGISFCR | 1 | 0 | 0 | 1535479 | AXN92247.1 | K9N7C7 |
| YIVGDVIKEGAL | 1 | 0 | 0 | 1535504 | AVP78041.1 | P0C6W2 |
| YIVGDVVHNGDI | 1 | 0 | 0 | 1535506 | APO40578.1 | P0C6X7 |
| YIVNISAFQQVL | 1 | 0 | 0 | 1535515 | AIL49553.1 | P0C6X6 |
| YIYKKVNNEFIN | 1 | 0 | 0 | 1535546 | AVV62524.1 | K9N7C7 |
| YIYSRVKNLNSS | 1 | 0 | 0 | 1535557 | APO40581.1 | P59637 |
| YKACSCIKDALS | 1 | 0 | 0 | 1535564 | AVV62535.1 | K9N7C7 |
| YKAGNYRRPAIV | 1 | 0 | 0 | 1535568 | AVV62543.1 | K9N7A1 |
| YKAGNYRRPPVV | 1 | 0 | 0 | 1535569 | AUM60030.1 | K9N7A1 |
| YKAIYNGVTRDI | 1 | 0 | 0 | 1535574 | APO40578.1 | P0C6X7 |
| YKCCDDNYFNKK | 1 | 0 | 0 | 1535586 | AVP78030.1 | P0C6W2 |
| YKCPQSKSIVGI | 1 | 0 | 0 | 1535589 | ARB66065.1 | P36334 |
| YKDLTGYMLDSY | 1 | 0 | 0 | 1535599 | QAT98908.1 | K9N7C7 |
| YKDSVLVNAANT | 1 | 0 | 0 | 1535601 | AVV62535.1 | K9N7C7 |
| YKDWSYSGQSTE | 1 | 0 | 0 | 1535605 | AVP78041.1 | P0C6W2 |
| YKEGFRGFTYVW | 1 | 0 | 0 | 1535610 | ARU07575.1 | A0A140H1G9 |
| YKELGLVMNMDV | 1 | 0 | 0 | 1535615 | AXN92247.1 | K9N7C7 |
| YKETNWFLHFIP | 1 | 0 | 0 | 1535618 | NP_073549.1 | A0A0P0LKV0 |
| YKETSYTSDIKP | 1 | 0 | 0 | 1535619 | APO40578.1 | P0C6X7 |
| YKGLPLEFGAST | 1 | 0 | 0 | 1535649 | P0C6W6.1 | P0C6W2 |
| YKGSITDVFYKE | 1 | 0 | 0 | 1535656 | AVP78041.1 | P0C6W2 |
| YKGTFNNKDTGP | 1 | 0 | 0 | 1535658 | AKA09658.1 | P36334 |
| YKGVITHDVSSAIN | 1 | 0 | 0 | 1535660 | P0DTD1.1 | P0DTD1 |
| YKITQWPTATGD | 1 | 0 | 0 | 1535686 | AVQ05261.1 | P0C6X6 |
| YKKDNAYYTQQP | 1 | 0 | 0 | 1535693 | APO40578.1 | P0C6X7 |
| YKKLGPIFNRAL | 1 | 0 | 0 | 1535695 | AEN19363.1 | P0C6X6 |
| YKKQQEGHIIME | 1 | 0 | 0 | 1535699 | AXN92247.1 | K9N7C7 |
| YKLDGVTYTELN | 1 | 0 | 0 | 1535712 | APO40578.1 | P0C6X7 |
| YKLEDYAFEHVV | 1 | 0 | 0 | 1535721 | AUM60013.1 | K9N7C7 |
| YKLNTDHSGSND | 1 | 0 | 0 | 1535739 | Q3I5J2.1 | Q3LZX9 |
| YKNMAVYRSLTF | 1 | 0 | 0 | 1535763 | AEN19365.1 | P30215 |
| YKNSGIGTCPAG | 1 | 0 | 0 | 1535769 | CAA83660.1 | P36334 |
| YKNTCVGSDNVA | 1 | 0 | 0 | 1535773 | APO40578.1 | P0C6X7 |

| | | | | | | |
|---|---|---|---|---|---|---|
| YKPNIWCIRCLW | 1 | 0 | 0 | 1535786 | AVP78041.1 | P0C6W2 |
| YKQQYSQLFVDT | 1 | 0 | 0 | 1535798 | AEN19363.1 | P0C6X6 |
| YKRNFTYDVNAD | 1 | 0 | 0 | 1535814 | CAA79896.1 | P36334 |
| YKRNFTYDVNAN | 1 | 0 | 0 | 1535816 | AKA09658.1 | P36334 |
| YKRNFTYDVNAT | 1 | 0 | 0 | 1535817 | AEN19366.1 | P36334 |
| YKRQQEGHIIME | 1 | 0 | 0 | 1535822 | 5YVD_A | K9N7C7 |
| YKTIDELCVNLN | 1 | 0 | 0 | 1535847 | ALA50066.1 | K9N7C7 |
| YKVCYSVKDTLA | 1 | 0 | 0 | 1535867 | AVN89451.1 | K9N7C7 |
| YKVLPPLYDPNM | 1 | 0 | 0 | 1535874 | AVV62526.1 | K9N5Q8 |
| YKVTKGKHRAGA | 1 | 0 | 0 | 1535879 | APO40578.1 | P0C6X7 |
| YKYDSGSLTKSS | 1 | 0 | 0 | 1535889 | AVV62535.1 | K9N7C7 |
| YLADNLNADETK | 1 | 0 | 0 | 1535914 | ATQ39389.1 | K9N7C7 |
| YLADNLSASEKE | 1 | 0 | 0 | 1535915 | AUM60013.1 | K9N7C7 |
| YLAHMLTDVVFE | 1 | 0 | 0 | 1535923 | AVV62540.1 | K9N643 |
| YLAVYDIEVTKG | 1 | 0 | 0 | 1535936 | AUM60023.1 | K9N7C7 |
| YLCERLTGGNYS | 1 | 0 | 0 | 1535937 | AUM60013.1 | K9N7C7 |
| YLCNFINNNLAL | 1 | 0 | 0 | 1535946 | AVV62524.1 | K9N7C7 |
| YLCNLINNNLAL | 1 | 0 | 0 | 1535947 | AXN92247.1 | K9N7C7 |
| YLDSLKPRVEAP | 1 | 0 | 0 | 1535963 | ATO98229.1 | P0C6W2 |
| YLDSLKPTVETP | 1 | 0 | 0 | 1535964 | ARO76381.1 | P0C6U8 |
| YLEAVNFNVTNA | 1 | 0 | 0 | 1535971 | AMM42536.1 | A0A0P0K6L9 |
| YLFDDAGEENLS | 1 | 0 | 0 | 1535979 | ARO76381.1 | P0C6U8 |
| YLFDEAGDEAMA | 1 | 0 | 0 | 1535980 | ARU07575.1 | A0A140H1G9 |
| YLFETIPTQLTP | 1 | 0 | 0 | 1535987 | AUM60013.1 | K9N7C7 |
| YLFQTIPTQLTP | 1 | 0 | 0 | 1535991 | AVV62524.1 | K9N7C7 |
| YLFSTINALPEI | 1 | 0 | 0 | 1535992 | ALW82729.1 | K9N7C7 |
| YLGCVVNAYNST | 1 | 0 | 0 | 1535999 | CAA79896.1 | P36334 |
| YLGDISHLINCV | 1 | 0 | 0 | 1536003 | QEG03730.1 | H9EJ42 |
| YLGDISHLVNCV | 1 | 0 | 0 | 1536004 | AWK59942.1 | H9EJ42 |
| YLGKPKEIIDGY | 1 | 0 | 0 | 1536020 | APO40578.1 | P0C6X7 |
| YLGTGPHAKDDY | 1 | 0 | 0 | 1536035 | AJQ24074.1 | P33469 |
| YLGTGPHAKDQY | 1 | 0 | 0 | 1536036 | AEN19370.1 | P33469 |
| YLGTGPHSKDQY | 1 | 0 | 0 | 1536041 | AIV41994.1 | P33469 |
| YLGTGPHSKEQY | 1 | 0 | 0 | 1536042 | ANZ78874.1 | P33469 |
| YLGTGPYAKDQY | 1 | 0 | 0 | 1536043 | AIL49482.1 | P33469 |
| YLIAYISFAPWW | 1 | 0 | 0 | 1536066 | NP_073549.1 | A0A0P0LKV0 |
| YLKNLTQTFKSM | 1 | 0 | 0 | 1536097 | ARU07575.1 | A0A140H1G9 |
| YLKRARGSSAAR | 1 | 0 | 0 | 1536105 | QDH43737.1 | H9EJ42 |
| YLLAFNQGGIIF | 1 | 0 | 0 | 1536117 | ABD43183.1 | P36334 |
| YLLDFSVDGYIR | 1 | 0 | 0 | 1536119 | AVV62526.1 | K9N5Q8 |
| YLLKFDDRGVIT | 1 | 0 | 0 | 1536129 | ADN03339.1 | A0A140H1H1 |
| YLMPILSMTDCL | 1 | 0 | 0 | 1536144 | AUM60013.1 | K9N7C7 |
| YLNNSIPNLSDF | 1 | 0 | 0 | 1536152 | BBA20986.1 | A0A140H1H1 |
| YLPATVIAGSVY | 1 | 0 | 0 | 1536174 | AVV62535.1 | K9N7C7 |
| YLQITSIPIVVD | 1 | 0 | 0 | 1536191 | APF29071.1 | A0A0P0G321 |
| YLQLATINYTQP | 1 | 0 | 0 | 1536196 | AVV62526.1 | K9N5Q8 |
| YLRHSSLSAQSE | 1 | 0 | 0 | 1536215 | AKN24751.1 | K9N4V0 |
| YLSERCNVIADP | 1 | 0 | 0 | 1536232 | AQZ41321.1 | K9N7C7 |
| YLSFNMDSATKV | 1 | 0 | 0 | 1536235 | ATO98193.1 | Q3LZX1 |
| YLSLLKRLEDLE | 1 | 0 | 0 | 1536239 | AVV62533.1 | K9N4V7 |
| YLSNVLLALQQI | 1 | 0 | 0 | 1536240 | APO40578.1 | P0C6X7 |
| YLSVDTKFKTEG | 1 | 0 | 0 | 1536244 | APO40578.1 | P0C6X7 |
| YLTCHNAAQCDC | 1 | 0 | 0 | 1536249 | CAA83660.1 | P36334 |
| YLTCHNAVQCNC | 1 | 0 | 0 | 1536251 | CAA83661.1 | P36334 |
| YLTNNISLNAPV | 1 | 0 | 0 | 1536268 | QEG59356.1 | A0A0P0G321 |
| YLTNQIRLNASV | 1 | 0 | 0 | 1536269 | AVL25553.1 | A0A0P0G321 |
| YLTSRAAQQLTI | 1 | 0 | 0 | 1536278 | AVV62535.1 | K9N7C7 |
| YLVFVTTLTSDY | 1 | 0 | 0 | 1536295 | AXN92247.1 | K9N7C7 |
| YLVLPFGNYLAI | 1 | 0 | 0 | 1536301 | AGT51619.1 | P30215 |
| YLVQQESTFVMM | 1 | 0 | 0 | 1536307 | APO40578.1 | P0C6X7 |
| YLVVYKEGFRGL | 1 | 0 | 0 | 1536311 | ARB07436.1 | A0A140H1G9 |
| YLYALVYFLQSI | 1 | 0 | 0 | 1536316 | QXJ09670.1 | P0DTC3 |
| YLYNITDDEHDE | 1 | 0 | 0 | 1536333 | AVV62526.1 | K9N5Q8 |
| YLYSAAHADPNH | 1 | 0 | 0 | 1536337 | AUM60024.1 | K9N5Q8 |
| YLYSAAHADPNR | 1 | 0 | 0 | 1536338 | AUM60014.1 | K9N5Q8 |
| YMAKSGDKYYPV | 1 | 0 | 0 | 1536341 | AXX83372.1 | P0C6X6 |
| YMCGFDGKPMLP | 1 | 0 | 0 | 1536344 | AFD98826.1 | H9EJ42 |
| YMCGSDGKPIAD | 1 | 0 | 0 | 1536347 | AUM60013.1 | K9N7C7 |
| YMCVVSSINDLS | 1 | 0 | 0 | 1536352 | AVA26872.1 | H9EJ42 |
| YMDFVNAGGAPL | 1 | 0 | 0 | 1536353 | AVV62535.1 | K9N7C7 |
| YMDFVNSGGTPL | 1 | 0 | 0 | 1536355 | AUM60013.1 | K9N7C7 |
| YMDGADVTKIKP | 1 | 0 | 0 | 1536356 | APO40578.1 | P0C6X7 |
| YMDLNPALCDSG | 1 | 0 | 0 | 1536362 | AEN19365.1 | P30215 |
| YMDLNSALCDSG | 1 | 0 | 0 | 1536364 | AIL49334.1 | P30215 |
| YMEAYNFVSASG | 1 | 0 | 0 | 1536367 | AHY22563.1 | K9N7C7 |
| YMHQMELANGTH | 1 | 0 | 0 | 1536378 | AXN92247.1 | K9N7C7 |
| YMIDPAGVCLGA | 1 | 0 | 0 | 1536380 | AEN19363.1 | P0C6X6 |
| YMIRDKLALGGS | 1 | 0 | 0 | 1536384 | AQN78757.1 | P0C6X6 |
| YMMPIMGLTICL | 1 | 0 | 0 | 1536392 | QED88024.1 | H9EJ42 |
| YMQGYDDCMHQG | 1 | 0 | 0 | 1536412 | ATG84866.1 | K9N5Q8 |
| YMQGYDDCMKQG | 1 | 0 | 0 | 1536413 | AVV62537.1 | K9N5Q8 |
| YMQGYDDCMNQG | 1 | 0 | 0 | 1536414 | AVV62526.1 | K9N5Q8 |
| YMQGYDECMQQG | 1 | 0 | 0 | 1536416 | AUM60014.1 | K9N5Q8 |
| YMQLSDAMKRQI | 1 | 0 | 0 | 1536418 | AUM60013.1 | K9N7C7 |
| YMRSVKVPAVVS | 1 | 0 | 0 | 1536422 | APO40578.1 | P0C6X7 |
| YMTHGQMEGLCL | 1 | 0 | 0 | 1536433 | AST12962.1 | A0A0P0LKV0 |
| YMVGDVINSGDI | 1 | 0 | 0 | 1536438 | ARO76381.1 | P0C6U8 |
| YMYFQKAGQSTY | 1 | 0 | 0 | 1536440 | APO40578.1 | P0C6X7 |
| YNAPCFSQQGVF | 1 | 0 | 0 | 1536448 | AGT51729.1 | P30215 |
| YNATRTFYVPAA | 1 | 0 | 0 | 1536450 | AWO68966.1 | A0A0P0G321 |
| YNDIKPPVLDVD | 1 | 0 | 0 | 1536466 | AOL02455.1 | Q04854 |

| | | | | | | |
|---|---|---|---|---|---|---|
| YNELGIEMNMDV | 1 | 0 | 0 | 1536471 | AIL49563.1 | P0C6X6 |
| YNEVPATVLQNN | 1 | 0 | 0 | 1536474 | ATN39877.2 | P0C6X6 |
| YNFLKEQHCQKA | 1 | 0 | 0 | 1536481 | ARO76392.1 | Q7TLC7 |
| YNGAIRSDSTLS | 1 | 0 | 0 | 1536499 | AHA80176.1 | P33469 |
| YNGAPSGVYQCA | 1 | 0 | 0 | 1536500 | APO40578.1 | P0C6X7 |
| YNGIMVLPGAAD | 1 | 0 | 0 | 1536509 | APT69876.1 | A0A0P0K6L9 |
| YNGKPQGAFHVT | 1 | 0 | 0 | 1536511 | AEN19363.1 | P0C6X6 |
| YNGRPQGAFHVT | 1 | 0 | 0 | 1536515 | ARU07575.1 | A0A140H1G9 |
| YNGSAQSTALCK | 1 | 0 | 0 | 1536524 | P30215.1 | P30215 |
| YNGVTRDILSTD | 1 | 0 | 0 | 1536531 | APO40578.1 | P0C6X7 |
| YNIDTGSFYGFS | 1 | 0 | 0 | 1536540 | ABD75600.1 | A0A140H1H0 |
| YNINFEENQRRG | 1 | 0 | 0 | 1536550 | ACO44679.1 | A0A0P0K6L9 |
| YNKCAYWVPRAS | 1 | 0 | 0 | 1536565 | AVP78041.1 | P0C6W2 |
| YNKDHGCGGFGH | 1 | 0 | 0 | 1536566 | AEN19363.1 | P0C6X6 |
| YNKLTEVKVPKG | 1 | 0 | 0 | 1536579 | AVV62535.1 | K9N7C7 |
| YNKLTEVKVPQG | 1 | 0 | 0 | 1536580 | AVV62524.1 | K9N7C7 |
| YNKQKANVSVLT | 1 | 0 | 0 | 1536582 | AXX83324.1 | P0C6X6 |
| YNLIKDCPAVAV | 1 | 0 | 0 | 1536598 | APO40578.1 | P0C6X7 |
| YNLNFEENISRG | 1 | 0 | 0 | 1536607 | CAA71146.1 | A0A0P0K6L9 |
| YNLPTMCDIKQM | 1 | 0 | 0 | 1536612 | AXN92247.1 | K9N7C7 |
| YNLPTMCDITQL | 1 | 0 | 0 | 1536614 | AVP78041.1 | P0C6W2 |
| YNLSSVVPSLPT | 1 | 0 | 0 | 1536619 | ABB90513.1 | A0A0P0K6L9 |
| YNLTKYTMMDLV | 1 | 0 | 0 | 1536624 | AVL25548.1 | H9EJ42 |
| YNLTMPSQVQKA | 1 | 0 | 0 | 1536625 | AAX85666.1 | P0C6X6 |
| YNLWNTFTKLQS | 1 | 0 | 0 | 1536630 | AVP78041.1 | P0C6W2 |
| YNLWQIFIETNL | 1 | 0 | 0 | 1536633 | AWK59942.1 | H9EJ42 |
| YNMCDYPHTICH | 1 | 0 | 0 | 1536640 | ANZ78839.1 | P36334 |
| YNMCEHPHTICH | 1 | 0 | 0 | 1536641 | BAM16463.1 | P36334 |
| YNMCEYPHTICH | 1 | 0 | 0 | 1536643 | AEN19366.1 | P36334 |
| YNMCEYPHTSCH | 1 | 0 | 0 | 1536644 | AKA09651.1 | P36334 |
| YNMCEYPQTICH | 1 | 0 | 0 | 1536646 | CAA79896.1 | P36334 |
| YNMDTGFAYGYS | 1 | 0 | 0 | 1536648 | AGT17757.1 | A0A140H1H0 |
| YNMLVNCFKWQV | 1 | 0 | 0 | 1536651 | AEN19363.1 | P0C6X6 |
| YNMLVNCSKWQV | 1 | 0 | 0 | 1536652 | AVI15051.1 | P0C6X6 |
| YNNCTGGAGIRD | 1 | 0 | 0 | 1536657 | CAA83661.1 | P36334 |
| YNNEGGRAFMYA | 1 | 0 | 0 | 1536661 | NP_073549.1 | A0A0P0LKV0 |
| YNNKLLAKKDLS | 1 | 0 | 0 | 1536662 | QBF80609.1 | K9N7C7 |
| YNNKLLSKKDAS | 1 | 0 | 0 | 1536667 | AUM60013.1 | K9N7C7 |
| YNNVSSVWPLYP | 1 | 0 | 0 | 1536671 | AAA73152.1 | P30215 |
| YNPPTDKLAMIM | 1 | 0 | 0 | 1536678 | NP_150075.1 | Q80872 |
| YNPSKPISKKYT | 1 | 0 | 0 | 1536680 | AVV62524.1 | K9N7C7 |
| YNQIILNLTSEI | 1 | 0 | 0 | 1536686 | ABB90514.1 | A0A0P0K6L9 |
| YNSPCFSQQGVF | 1 | 0 | 0 | 1536720 | P30215.1 | P30215 |
| YNSQNAVARSML | 1 | 0 | 0 | 1536723 | AXN92247.1 | K9N7C7 |
| YNSSCFSHQGVF | 1 | 0 | 0 | 1536733 | QBQ01838.1 | P30215 |
| YNSSCFSQQGVF | 1 | 0 | 0 | 1536734 | AEN19365.1 | P30215 |
| YNSSSSFSTFKC | 1 | 0 | 0 | 1536735 | APO40579.1 | P59594 |
| YNSTAISVQKCD | 1 | 0 | 0 | 1536738 | AEN19358.1 | P36334 |
| YNSVTDTIVVTA | 1 | 0 | 0 | 1536746 | ATO98194.1 | Q3LZX0 |
| YNSWNQAYSSPI | 1 | 0 | 0 | 1536751 | ATO98145.1 | Q3LZX1 |
| YNSWQNLLYDPN | 1 | 0 | 0 | 1536752 | ARB66062.1 | P36334 |
| YNTLTEVKIPQG | 1 | 0 | 0 | 1536766 | AUM60013.1 | K9N7C7 |
| YNTNATLNSFKE | 1 | 0 | 0 | 1536767 | AVV62537.1 | K9N5Q8 |
| YNTPCFSQQGVF | 1 | 0 | 0 | 1536768 | AGT51579.1 | P30215 |
| YNVSVMQMDIQG | 1 | 0 | 0 | 1536785 | NP_073549.1 | A0A0P0LKV0 |
| YNVVNKGHFDGD | 1 | 0 | 0 | 1536794 | APO40578.1 | P0C6X7 |
| YNWPEPIDMNKA | 1 | 0 | 0 | 1536800 | ATQ39390.1 | K9N5Q8 |
| YNYEPLTQDHLD | 1 | 0 | 0 | 1536804 | AVP78041.1 | P0C6W2 |
| YNYKQDFSNPTC | 1 | 0 | 0 | 1536814 | AUM60014.1 | K9N5Q8 |
| YNYVVSSAFWLF | 1 | 0 | 0 | 1536829 | AVN89451.1 | K9N7C7 |
| YPACINGHAVPT | 1 | 0 | 0 | 1536832 | QBF80553.1 | K9N7C7 |
| YPACTNGHALPT | 1 | 0 | 0 | 1536834 | ALL26408.1 | K9N638 |
| YPACTNGHAVPT | 1 | 0 | 0 | 1536835 | AXN92247.1 | K9N7C7 |
| YPAFMLGSSVGN | 1 | 0 | 0 | 1536838 | AXN92228.1 | K9N5Q8 |
| YPDYVDVNKTLQ | 1 | 0 | 0 | 1536856 | AWK59979.1 | A0A0P0G321 |
| YPELMEDLEADP | 1 | 0 | 0 | 1536857 | AUM60013.1 | K9N7C7 |
| YPEPINENNVVV | 1 | 0 | 0 | 1536859 | AGT51800.1 | P36334 |
| YPEPITGNNVVV | 1 | 0 | 0 | 1536862 | CAA83660.1 | P36334 |
| YPEWMDDFEADP | 1 | 0 | 0 | 1536863 | AHI48797.1 | K9N638 |
| YPGQGCAGYTLD | 1 | 0 | 0 | 1536874 | ARO76381.1 | P0C6U8 |
| YPHTICHSKLGD | 1 | 0 | 0 | 1536884 | AKA09663.1 | P36334 |
| YPHTSCHPKLGN | 1 | 0 | 0 | 1536886 | AKA09651.1 | P36334 |
| YPHTTCHPNLGN | 1 | 0 | 0 | 1536888 | BBA84088.1 | P36334 |
| YPIIGDELKINS | 1 | 0 | 0 | 1536896 | P0C6W6.1 | P0C6W2 |
| YPIILRLGSNLS | 1 | 0 | 0 | 1536899 | 2CME_A | P59636 |
| YPILSCVYAICY | 1 | 0 | 0 | 1536904 | AAR01012.1 | P0C6X6 |
| YPKCDRAMPNII | 1 | 0 | 0 | 1536908 | AKL87224.1 | P0C6X6 |
| YPKCDRVLPNMI | 1 | 0 | 0 | 1536913 | QED88038.1 | H9EJ42 |
| YPLSMKSDLSVS | 1 | 0 | 0 | 1536934 | AXN92228.1 | K9N5Q8 |
| YPLSMKSYMDPG | 1 | 0 | 0 | 1536936 | AVV62526.1 | K9N5Q8 |
| YPLTKHEDLEYQ | 1 | 0 | 0 | 1536939 | AUM60013.1 | K9N7C7 |
| YPMCTNGKVVPT | 1 | 0 | 0 | 1536946 | AVV62524.1 | K9N7C7 |
| YPMCTNGQAVPT | 1 | 0 | 0 | 1536947 | AUM60013.1 | K9N7C7 |
| YPNASFDNFRFV | 1 | 0 | 0 | 1536950 | AVP78041.1 | P0C6W2 |
| YPNDIVGGFRVL | 1 | 0 | 0 | 1536952 | AWK59978.1 | H9EJ42 |
| YPNNAIVCRFDT | 1 | 0 | 0 | 1536956 | AXN92247.1 | K9N7C7 |
| YPNTICHPNLGN | 1 | 0 | 0 | 1536958 | CAA83661.1 | P36334 |
| YPNVPIEYQATR | 1 | 0 | 0 | 1536959 | ATO98120.1 | Q3LZX1 |
| YPPEDEGEDDCE | 1 | 0 | 0 | 1536966 | AVP78041.1 | P0C6W2 |
| YPQTICHPNLGN | 1 | 0 | 0 | 1536976 | CAA79896.1 | P36334 |
| YPQVNGLTSIRW | 1 | 0 | 0 | 1536980 | AVP78041.1 | P0C6W2 |

| | | | | | | |
|---|---|---|---|---|---|---|
| YPRLQAANDWKP | 1 | 0 | 0 | 1536983 | AVV62524.1 | K9N7C7 |
| YPSNAIVCRFDT | 1 | 0 | 0 | 1536990 | AUM60013.1 | K9N7C7 |
| YPSNHIDVVSAY | 1 | 0 | 0 | 1536992 | ALX27232.1 | K9N5Q8 |
| YPSNHIEVASAY | 1 | 0 | 0 | 1536993 | ANF29261.1 | K9N5Q8 |
| YPSNQIEVVSAY | 1 | 0 | 0 | 1536995 | QBF80554.1 | K9N5Q8 |
| YPSNSIVCRFDT | 1 | 0 | 0 | 1536997 | QVU66317.1 | P0DTD1 |
| YPTGKSFSNISL | 1 | 0 | 0 | 1537005 | AVV62526.1 | K9N5Q8 |
| YPTGKSYSNISL | 1 | 0 | 0 | 1537006 | AVV62537.1 | K9N5Q8 |
| YPTGRTYSNITL | 1 | 0 | 0 | 1537007 | AUM60014.1 | K9N5Q8 |
| YPTGSCLMAFLV | 1 | 0 | 0 | 1537010 | AXG21669.1 | K9N7D2 |
| YPTSSSTYRNMA | 1 | 0 | 0 | 1537017 | AIL49538.1 | P36334 |
| YPTTLGSLELLN | 1 | 0 | 0 | 1537018 | AGT51735.1 | Q4VID0 |
| YPVASPRECNDM | 1 | 0 | 0 | 1537023 | ARO76381.1 | P0C6U8 |
| YPVCNGRVGHSG | 1 | 0 | 0 | 1537025 | ADK37701.1 | A0A0P0K6L9 |
| YPVSGIREFSNL | 1 | 0 | 0 | 1537028 | AIW52843.1 | A0A0P0G321 |
| YPYISHENKLNS | 1 | 0 | 0 | 1537034 | ALA49351.1 | K9N7C7 |
| YQDVNCTDVPTM | 1 | 0 | 0 | 1537053 | APO40579.1 | P59594 |
| YQDVNCTDVPTT | 1 | 0 | 0 | 1537055 | AVP78031.1 | Q3LZX1 |
| YQEFNDLAHTSL | 1 | 0 | 0 | 1537068 | AVA26872.1 | H9EJ42 |
| YQEFNDLVHTSL | 1 | 0 | 0 | 1537070 | AMM42571.1 | H9EJ42 |
| YQEFVIKTSHDD | 1 | 0 | 0 | 1537071 | AIV41909.1 | P36334 |
| YQGDHGDMYVYS | 1 | 0 | 0 | 1537086 | AXN92228.1 | K9N5Q8 |
| YQGLPLEFGASA | 1 | 0 | 0 | 1537093 | AEA10817.1 | P0C6X7 |
| YQHEENIYNLIK | 1 | 0 | 0 | 1537096 | APO40578.1 | P0C6X7 |
| YQHEENIYNLVK | 1 | 0 | 0 | 1537097 | AVP78041.1 | P0C6W2 |
| YQHVIKGCKDTA | 1 | 0 | 0 | 1537108 | AHI48777.1 | K9N7C7 |
| YQKWMELLDSNI | 1 | 0 | 0 | 1537123 | AUM60021.1 | K9N4V7 |
| YQKWPWYIWLGF | 1 | 0 | 0 | 1537124 | AVV62526.1 | K9N5Q8 |
| YQLDENAQKFFD | 1 | 0 | 0 | 1537126 | AMM42491.1 | A0A0P0ILY9 |
| YQLYMIDPAGVC | 1 | 0 | 0 | 1537142 | AEN19363.1 | P0C6X6 |
| YQNSSSPALLYR | 1 | 0 | 0 | 1537157 | BBA20986.1 | A0A140H1H1 |
| YQPIQQAPTGIT | 1 | 0 | 0 | 1537163 | AST12967.1 | A0A0P0KXN3 |
| YQPPQTSITSAI | 1 | 0 | 0 | 1537166 | AVP78041.1 | P0C6W2 |
| YQPTAHVNATAA | 1 | 0 | 0 | 1537168 | AUM60014.1 | K9N5Q8 |
| YQPYDHAIGATA | 1 | 0 | 0 | 1537170 | ATO98145.1 | Q3LZX1 |
| YQQQDGMVNMSP | 1 | 0 | 0 | 1537180 | AEN19370.1 | P33469 |
| YQRNNTSFPEWI | 1 | 0 | 0 | 1537182 | ATQ39389.1 | K9N7C7 |
| YQSFVQWFHEHV | 1 | 0 | 0 | 1537185 | AVV62524.1 | K9N7C7 |
| YQTNRDMQNKMA | 1 | 0 | 0 | 1537196 | AUM60024.1 | K9N5Q8 |
| YQTSNFRVQPTQ | 1 | 0 | 0 | 1537201 | AVP78031.1 | Q3LZX1 |
| YQTSNFRVQPTV | 1 | 0 | 0 | 1537202 | ATO98145.1 | Q3LZX1 |
| YRAALKSEKHQD | 1 | 0 | 0 | 1537222 | AGV08406.1 | K9N7C7 |
| YRAALNSEKRPD | 1 | 0 | 0 | 1537225 | AUM60013.1 | K9N7C7 |
| YRDLYSSPTTLQ | 1 | 0 | 0 | 1537252 | AXN92247.1 | K9N7C7 |
| YRELYDAPTTLQ | 1 | 0 | 0 | 1537259 | AVV62535.1 | K9N7C7 |
| YREYMEAYNFVS | 1 | 0 | 0 | 1537263 | AHY22563.1 | K9N7C7 |
| YRLPSTQKVSGM | 1 | 0 | 0 | 1537330 | AQN78779.1 | Q01455 |
| YRMNGIGITQQV | 1 | 0 | 0 | 1537337 | AVV62537.1 | K9N5Q8 |
| YRNDKSCPVVAA | 1 | 0 | 0 | 1537339 | ARO76381.1 | P0C6U8 |
| YRNSNVGESFVD | 1 | 0 | 0 | 1537351 | ADO24705.1 | A0A0P0LKV0 |
| YRRIPNLPDCNI | 1 | 0 | 0 | 1537371 | CAA83660.1 | P36334 |
| YRRKPDPPNCNI | 1 | 0 | 0 | 1537374 | AHA80201.1 | P36334 |
| YRRQQTSNLVAQ | 1 | 0 | 0 | 1537379 | APU51933.1 | P0C6X6 |
| YRRRPDLPNCNI | 1 | 0 | 0 | 1537382 | AHA80225.1 | P36334 |
| YRSGNGILKWAD | 1 | 0 | 0 | 1537391 | AIV41850.1 | P33469 |
| YRSHRSSKLKPF | 1 | 0 | 0 | 1537392 | ATO98145.1 | Q3LZX1 |
| YRSTSVEESFID | 1 | 0 | 0 | 1537411 | AIG96682.1 | H9EJ42 |
| YRVFPYDMDSGG | 1 | 0 | 0 | 1537419 | AHX71940.1 | K9N7C7 |
| YRVFPYDMSSDV | 1 | 0 | 0 | 1537421 | AUM60013.1 | K9N7C7 |
| YRVYPYEISDDV | 1 | 0 | 0 | 1537437 | AVV62524.1 | K9N7C7 |
| YSAAHADPNHFT | 1 | 0 | 0 | 1537448 | AUM60024.1 | K9N5Q8 |
| YSAAHADPNRFI | 1 | 0 | 0 | 1537449 | AUM60014.1 | K9N5Q8 |
| YSAGHATGTTPK | 1 | 0 | 0 | 1537456 | ALA49649.1 | K9N5Q8 |
| YSAGHATGTTSQ | 1 | 0 | 0 | 1537459 | ANF29239.1 | K9N5Q8 |
| YSAGYATGTTPQ | 1 | 0 | 0 | 1537461 | ANC28634.1 | K9N5Q8 |
| YSAKDTLPELVI | 1 | 0 | 0 | 1537463 | AVV62524.1 | K9N7C7 |
| YSALKMVQTHLS | 1 | 0 | 0 | 1537469 | ASU91185.1 | K9N7C7 |
| YSANFQHDNVLP | 1 | 0 | 0 | 1537472 | AWO68987.1 | A0A0P0G321 |
| YSCDFKGWIPGN | 1 | 0 | 0 | 1537479 | QBP84701.1 | P0C6X6 |
| YSEAAAQGFQAC | 1 | 0 | 0 | 1537502 | AWK59942.1 | H9EJ42 |
| YSEAASNGFRAC | 1 | 0 | 0 | 1537503 | NP_073549.1 | A0A0P0LKV0 |
| YSFDKAIDGFGV | 1 | 0 | 0 | 1537524 | AWK59942.1 | H9EJ42 |
| YSFEAHKDFFKD | 1 | 0 | 0 | 1537526 | ARK08657.1 | P0C6X6 |
| YSFLPKEDGDVV | 1 | 0 | 0 | 1537536 | AHI48794.1 | K9N638 |
| YSFLPKEEGDVL | 1 | 0 | 0 | 1537537 | ATQ39389.1 | K9N7C7 |
| YSFSAYSGDVPH | 1 | 0 | 0 | 1537542 | AUM60014.1 | K9N5Q8 |
| YSGDIPHYVQPG | 1 | 0 | 0 | 1537553 | AUM60024.1 | K9N5Q8 |
| YSGDVPHYVLPG | 1 | 0 | 0 | 1537556 | AUM60014.1 | K9N5Q8 |
| YSGQNTELGVEF | 1 | 0 | 0 | 1537566 | APO40578.1 | P0C6X7 |
| YSGSASEADYRC | 1 | 0 | 0 | 1537578 | AWK59942.1 | H9EJ42 |
| YSGSIRFDSTLP | 1 | 0 | 0 | 1537579 | ARU07581.1 | Q19U29 |
| YSGVVGCKSIVK | 1 | 0 | 0 | 1537582 | ARU07575.1 | A0A140H1G9 |
| YSICKNFAPNGF | 1 | 0 | 0 | 1537596 | AVV62537.1 | K9N5Q8 |
| YSIEMQLFKVGK | 1 | 0 | 0 | 1537600 | AWK59978.1 | H9EJ42 |
| YSIQDVGFKNQH | 1 | 0 | 0 | 1537615 | AWW13571.1 | P0C6X6 |
| YSKKLLNKLVGG | 1 | 0 | 0 | 1537627 | AVV62524.1 | K9N7C7 |
| YSKLPDESGDVL | 1 | 0 | 0 | 1537630 | AVV62524.1 | K9N7C7 |
| YSKLVKDVDHVQ | 1 | 0 | 0 | 1537632 | Q3I5J4.1 | Q3LZX0 |
| YSKNGGSGGAIT | 1 | 0 | 0 | 1537633 | 6NZK_A | P36334 |
| YSKRSAIEDLLF | 1 | 0 | 0 | 1537642 | AVV62526.1 | K9N5Q8 |
| YSKTRRSRRAIT | 1 | 0 | 0 | 1537644 | AVQ05264.1 | P36334 |

| | | | | | | |
|---|---|---|---|---|---|---|
| YSKYVTVQGPPG | 1 | 0 | 0 | 1537646 | AXN92247.1 | K9N7C7 |
| YSLADLPAYMSV | 1 | 0 | 0 | 1537647 | AWW13547.1 | Q01455 |
| YSLADSNGFIRF | 1 | 0 | 0 | 1537648 | ABD75543.1 | A0A140H1G9 |
| YSLDGNFRTGVD | 1 | 0 | 0 | 1537653 | ASU90030.1 | K9N7C7 |
| YSLDGNMKAEVD | 1 | 0 | 0 | 1537654 | AUM60013.1 | K9N7C7 |
| YSLKAAVHGWKM | 1 | 0 | 0 | 1537662 | 5W8T_A | K9N7C7 |
| YSLKAAVHKWKM | 1 | 0 | 0 | 1537663 | AXN92247.1 | K9N7C7 |
| YSMQDVGFKNQY | 1 | 0 | 0 | 1537691 | AAR01012.1 | P0C6X6 |
| YSNASSDVIHYN | 1 | 0 | 0 | 1537694 | BAL45639.1 | A0A0P0K6L9 |
| YSNESWFEDDYK | 1 | 0 | 0 | 1537696 | AST12962.1 | A0A0P0LKV0 |
| YSNSMDCPIVVA | 1 | 0 | 0 | 1537717 | AXX83324.1 | P0C6X6 |
| YSPATVIAGSVY | 1 | 0 | 0 | 1537728 | AVV62524.1 | K9N7C7 |
| YSPCVSIVPSTA | 1 | 0 | 0 | 1537731 | AGV08584.1 | K9N5Q8 |
| YSPEPITSLNTK | 1 | 0 | 0 | 1537734 | AHI48672.1 | K9N5Q8 |
| YSPIDIPRGLPV | 1 | 0 | 0 | 1537736 | ATO98193.1 | Q3LZX1 |
| YSPLLVDSDQPW | 1 | 0 | 0 | 1537738 | AST12968.1 | A0A0N7J131 |
| YSPQIITTDNTF | 1 | 0 | 0 | 1537745 | ATO98181.1 | Q3LZX1 |
| YSPQTITTDNTF | 1 | 0 | 0 | 1537750 | ARO76382.1 | P59594 |
| YSPQVITTDNTF | 1 | 0 | 0 | 1537751 | APO40579.1 | P59594 |
| YSPSNHKSNTND | 1 | 0 | 0 | 1537753 | ATQ39390.1 | K9N5Q8 |
| YSPVFLIVAAIV | 1 | 0 | 0 | 1537756 | QTH36422.1 | P0DTC7 |
| YSQVLSKSILLV | 1 | 0 | 0 | 1537768 | ATQ39389.1 | K9N7C7 |
| YSQYYLNKDTGV | 1 | 0 | 0 | 1537770 | AGT51729.1 | P30215 |
| YSRGDQACYERF | 1 | 0 | 0 | 1537772 | AUM60023.1 | K9N7C7 |
| YSRYCFSVNNTF | 1 | 0 | 0 | 1537798 | BBA20983.1 | A0A140H1H1 |
| YSSCVECEAKFK | 1 | 0 | 0 | 1537804 | ARB07424.1 | A0A0P0ILY9 |
| YSSNEEQAVRKF | 1 | 0 | 0 | 1537819 | QED88038.1 | H9EJ42 |
| YSSSANGILVGT | 1 | 0 | 0 | 1537830 | AVN89312.1 | K9N638 |
| YSSSANGRLVGT | 1 | 0 | 0 | 1537831 | AJG44057.1 | K9N638 |
| YSSSANGSLDGT | 1 | 0 | 0 | 1537832 | AXG21642.1 | K9N638 |
| YSSSANGSLVCT | 1 | 0 | 0 | 1537833 | AVN89333.1 | K9N7C7 |
| YSTGIDFQDELD | 1 | 0 | 0 | 1537845 | ASU91284.1 | K9N5Q8 |
| YSTKPITTSNPF | 1 | 0 | 0 | 1537850 | APO40578.1 | P0C6X7 |
| YSTSPIQPVELP | 1 | 0 | 0 | 1537860 | AIW52757.1 | A0A0P0K6L9 |
| YSTTPCELHGNG | 1 | 0 | 0 | 1537865 | ALW82645.1 | K9N7C7 |
| YSTTPCGVHGNG | 1 | 0 | 0 | 1537867 | AGV08556.1 | K9N7C7 |
| YSTYDPIYKNGA | 1 | 0 | 0 | 1537869 | AVV62535.1 | K9N7C7 |
| YSVGEYSSPIAY | 1 | 0 | 0 | 1537877 | AVV62535.1 | K9N7C7 |
| YSVSSFEAIPSG | 1 | 0 | 0 | 1537894 | ALA49649.1 | K9N5Q8 |
| YSVSSFEVRSKG | 1 | 0 | 0 | 1537899 | AVV62526.1 | K9N5Q8 |
| YSWADLPAYMTV | 1 | 0 | 0 | 1537910 | Q01455.1 | Q01455 |
| YSWFSGITQFQK | 1 | 0 | 0 | 1537911 | AEN19370.1 | P33469 |
| YSYFNKCSRLLS | 1 | 0 | 0 | 1537919 | AMW90841.1 | K9N5Q8 |
| YSYRSAIEDLLF | 1 | 0 | 0 | 1537941 | AUM60014.1 | K9N5Q8 |
| YTAATTCVTGPT | 1 | 0 | 0 | 1537949 | ATO98145.1 | Q3LZX1 |
| YTAEQCIRGAIG | 1 | 0 | 0 | 1537951 | ARB07424.1 | A0A0P0ILY9 |
| YTANDVDLQSSG | 1 | 0 | 0 | 1537961 | AUM60013.1 | K9N7C7 |
| YTARDALDTILQ | 1 | 0 | 0 | 1537964 | ATQ39389.1 | K9N7C7 |
| YTCIGYICTCYF | 1 | 0 | 0 | 1537968 | AVV62524.1 | K9N7C7 |
| YTDGLMQNASLY | 1 | 0 | 0 | 1537979 | AEN19363.1 | P0C6X6 |
| YTDGVMKNASLY | 1 | 0 | 0 | 1537980 | P0C6U5.1 | A0A140H1G9 |
| YTDIDVNPALNI | 1 | 0 | 0 | 1537983 | ANI69887.1 | K9N7C7 |
| YTDIDVNSALNM | 1 | 0 | 0 | 1537985 | AHN10810.1 | K9N7C7 |
| YTDLKFIDKLNV | 1 | 0 | 0 | 1537990 | AEN19363.1 | P0C6X6 |
| YTDNIFSVQKDG | 1 | 0 | 0 | 1537993 | AMM42643.1 | A0A0P0G321 |
| YTEDLASRFADG | 1 | 0 | 0 | 1538001 | AUM60013.1 | K9N7C7 |
| YTELNPKLDGYY | 1 | 0 | 0 | 1538011 | APO40578.1 | P0C6X7 |
| YTESAVNFTKAL | 1 | 0 | 0 | 1538015 | AVN89336.1 | K9N4V0 |
| YTEVRGSRRLAQ | 1 | 0 | 0 | 1538018 | YP_003767.1 | A0A0P0G321 |
| YTFLLRKNGRGG | 1 | 0 | 0 | 1538024 | AVV62524.1 | K9N7C7 |
| YTFTTVKPGAAF | 1 | 0 | 0 | 1538030 | AXN92247.1 | K9N7C7 |
| YTFTTVKPGASF | 1 | 0 | 0 | 1538031 | AVV62524.1 | K9N7C7 |
| YTFYGKDAFENH | 1 | 0 | 0 | 1538035 | AVV62524.1 | K9N7C7 |
| YTFYGKESFENP | 1 | 0 | 0 | 1538036 | AVN89322.1 | K9N7C7 |
| YTFYGKESLENP | 1 | 0 | 0 | 1538038 | AXN92247.1 | K9N7C7 |
| YTFYGKNAFQNH | 1 | 0 | 0 | 1538041 | AVV62535.1 | K9N7C7 |
| YTGFQGKQILGS | 1 | 0 | 0 | 1538046 | AXN92247.1 | K9N7C7 |
| YTGHQELVIKTS | 1 | 0 | 0 | 1538047 | CAA83660.1 | P36334 |
| YTGSSFFNLEPI | 1 | 0 | 0 | 1538058 | AVV62526.1 | K9N5Q8 |
| YTGSSFYSPEPI | 1 | 0 | 0 | 1538062 | AHI48672.1 | K9N5Q8 |
| YTGYQEFVIKTS | 1 | 0 | 0 | 1538070 | AIV41909.1 | P36334 |
| YTGYQESVIKTS | 1 | 0 | 0 | 1538072 | AIL49487.1 | P36334 |
| YTILPCYSGRVS | 1 | 0 | 0 | 1538087 | BBA20986.1 | A0A140H1H1 |
| YTISCEPLEINC | 1 | 0 | 0 | 1538090 | ATO98227.1 | Q3LZX5 |
| YTKAPDVMLNIS | 1 | 0 | 0 | 1538097 | CAA83660.1 | P36334 |
| YTKEGNVINFCY | 1 | 0 | 0 | 1538102 | AVV62535.1 | K9N7C7 |
| YTKIFKDDGNFV | 1 | 0 | 0 | 1538108 | AMK59674.1 | P0C6X6 |
| YTLANTTTAPLK | 1 | 0 | 0 | 1538129 | AGV08437.1 | K9N7C7 |
| YTLCYDIGNPRG | 1 | 0 | 0 | 1538133 | ARE30011.1 | P0C6X6 |
| YTLDEARTALKK | 1 | 0 | 0 | 1538134 | ARO76381.1 | P0C6U8 |
| YTLVSGDAHTPS | 1 | 0 | 0 | 1538146 | AMM42538.1 | A0A0P0K6L9 |
| YTMCEYPNTICH | 1 | 0 | 0 | 1538156 | CAA83661.1 | P36334 |
| YTMMDFVYALRH | 1 | 0 | 0 | 1538160 | ATG84732.1 | K9N7C7 |
| YTMSIGEDQSIA | 1 | 0 | 0 | 1538161 | APO40579.1 | P59594 |
| YTMSLGAENSIA | 1 | 0 | 0 | 1538163 | AVP78031.1 | Q3LZX1 |
| YTNDKACPLVAA | 1 | 0 | 0 | 1538166 | AVP78030.1 | P0C6W2 |
| YTNFRIDVPFAE | 1 | 0 | 0 | 1538173 | QAT98905.1 | K9N796 |
| YTNSCLVASDGK | 1 | 0 | 0 | 1538184 | AUM60013.1 | K9N7C7 |
| YTNSCLVSSNGQ | 1 | 0 | 0 | 1538189 | QAT98908.1 | K9N7C7 |
| YTPATDCSDGNY | 1 | 0 | 0 | 1538194 | ALA49352.1 | K9N5Q8 |
| YTPFAGYTESAV | 1 | 0 | 0 | 1538196 | ARQ84748.1 | K9N4V0 |

| | | | | | | |
|---|---|---|---|---|---|---|
| YTPIDEIPYKSF | 1 | 0 | 0 | 1538206 | AXN92247.1 | K9N7C7 |
| YTPKYTFGVVKP | 1 | 0 | 0 | 1538208 | ARB07436.1 | A0A140H1G9 |
| YTPLVGTSLHSV | 1 | 0 | 0 | 1538211 | AVN89336.1 | K9N4V0 |
| YTPVHYERDNTS | 1 | 0 | 0 | 1538221 | AXN92247.1 | K9N7C7 |
| YTQFTIGLMAAN | 1 | 0 | 0 | 1538229 | AVN89373.1 | K9N643 |
| YTQGRSCSDFEP | 1 | 0 | 0 | 1538231 | AVV62524.1 | K9N7C7 |
| YTQQPIDLVPTQ | 1 | 0 | 0 | 1538239 | APO40578.1 | P0C6X7 |
| YTSDIKPVTYKL | 1 | 0 | 0 | 1538257 | APO40578.1 | P0C6X7 |
| YTSDLLTNSVIV | 1 | 0 | 0 | 1538259 | ARO76381.1 | P0C6U8 |
| YTSSSLGSIAGA | 1 | 0 | 0 | 1538280 | AOR17480.1 | K9N5Q8 |
| YTSTSGNLVGFK | 1 | 0 | 0 | 1538282 | CAA71146.1 | A0A0P0K6L9 |
| YTTEPEFVKVLG | 1 | 0 | 0 | 1538288 | AEN19363.1 | P0C6X6 |
| YTTLPTFNTRTV | 1 | 0 | 0 | 1538291 | AUM60023.1 | K9N7C7 |
| YTTRILAKPTAL | 1 | 0 | 0 | 1538292 | AVV62540.1 | K9N643 |
| YTTTIKPVSLEH | 1 | 0 | 0 | 1538298 | 4M0W_A | P0C6U8 |
| YTVDSVTVKNGT | 1 | 0 | 0 | 1538309 | APO40578.1 | P0C6X7 |
| YTVELGTDVNEF | 1 | 0 | 0 | 1538311 | AVP78041.1 | P0C6W2 |
| YTVETGTQLEEF | 1 | 0 | 0 | 1538315 | APO40578.1 | P0C6X7 |
| YTVQKLQSLIDN | 1 | 0 | 0 | 1538325 | BAL45640.1 | A0A0P0K6L9 |
| YTWTPYPELMED | 1 | 0 | 0 | 1538341 | AUM60013.1 | K9N7C7 |
| YTYSFLPKEEGD | 1 | 0 | 0 | 1538351 | ATQ39389.1 | K9N7C7 |
| YVAALAVNVCII | 1 | 0 | 0 | 1538363 | AHI48676.1 | K9N7D2 |
| YVALRNSINADA | 1 | 0 | 0 | 1538379 | AVV62524.1 | K9N7C7 |
| YVANPTEDLCNG | 1 | 0 | 0 | 1538381 | AGT51737.1 | P0C6X6 |
| YVARLVKGDKGD | 1 | 0 | 0 | 1538389 | AST12962.1 | A0A0P0LKV0 |
| YVCLLANKEQSV | 1 | 0 | 0 | 1538405 | AUM60013.1 | K9N7C7 |
| YVCNGNPRCKNL | 1 | 0 | 0 | 1538415 | AIW52843.1 | A0A0P0G321 |
| YVCQCGGERHQQ | 1 | 0 | 0 | 1538417 | AVN89322.1 | K9N7C7 |
| YVDFKPQSGGGR | 1 | 0 | 0 | 1538425 | ABB90508.1 | A0A0P0K6L9 |
| YVDNTLYSEFST | 1 | 0 | 0 | 1538435 | ACV87129.1 | A0A140H1H1 |
| YVDVGPDSAKSV | 1 | 0 | 0 | 1538459 | QAT98898.1 | K9N5Q8 |
| YVDVGPDSFKSA | 1 | 0 | 0 | 1538460 | AQZ41296.1 | K9N5Q8 |
| YVDVGPDSVKPA | 1 | 0 | 0 | 1538463 | ASU91284.1 | K9N5Q8 |
| YVDYTFVNEYYE | 1 | 0 | 0 | 1538469 | ARU07574.1 | A0A140H1G9 |
| YVEAYNLVSASG | 1 | 0 | 0 | 1538470 | AVV62524.1 | K9N7C7 |
| YVERKDVPYTKQ | 1 | 0 | 0 | 1538481 | AUM60013.1 | K9N7C7 |
| YVFNNSLTRQLR | 1 | 0 | 0 | 1538497 | AIL49543.1 | P36334 |
| YVFNQVVSSDAM | 1 | 0 | 0 | 1538503 | AVV62524.1 | K9N7C7 |
| YVGDPAQLPTPR | 1 | 0 | 0 | 1538514 | ATG84864.1 | K9N7C7 |
| YVGIQTIEELHV | 1 | 0 | 0 | 1538515 | AVV62524.1 | K9N7C7 |
| YVGIQTLDELHA | 1 | 0 | 0 | 1538516 | AVV62535.1 | K9N7C7 |
| YVGMQTVEDLRA | 1 | 0 | 0 | 1538518 | AVN89451.1 | K9N7C7 |
| YVGSLSQNHHTF | 1 | 0 | 0 | 1538523 | AWK59942.1 | H9EJ42 |
| YVGVQSIEELHQ | 1 | 0 | 0 | 1538526 | AUM60013.1 | K9N7C7 |
| YVGVQTIEDLRA | 1 | 0 | 0 | 1538527 | AJG44112.1 | K9N638 |
| YVGVQTVEDLCA | 1 | 0 | 0 | 1538529 | AKN24802.1 | K9N7C7 |
| YVGVYDTNHGDA | 1 | 0 | 0 | 1538532 | AQN78743.1 | P30215 |
| YVHARLKCGLIL | 1 | 0 | 0 | 1538535 | AVN89290.1 | K9N7C7 |
| YVHKWFQFCRLY | 1 | 0 | 0 | 1538543 | ARU07575.1 | A0A140H1G9 |
| YVKDGKYFISEP | 1 | 0 | 0 | 1538579 | ALA49340.1 | K9N7C7 |
| YVKFQESKPPLP | 1 | 0 | 0 | 1538587 | AVV62531.1 | K9N5R3 |
| YVKPGGTGSGDA | 1 | 0 | 0 | 1538596 | ADO24677.1 | P0C6X6 |
| YVKPGGTTSGDA | 1 | 0 | 0 | 1538598 | AVP78019.1 | P0C6W2 |
| YVKPGRTSSGDA | 1 | 0 | 0 | 1538599 | AFR45548.1 | P0C6X6 |
| YVLNVDWLWFAL | 1 | 0 | 0 | 1538628 | AVV62524.1 | K9N7C7 |
| YVLPGQYTPCLY | 1 | 0 | 0 | 1538629 | AUM60014.1 | K9N5Q8 |
| YVLPNDDTLRAE | 1 | 0 | 0 | 1538631 | AVP78030.1 | P0C6W2 |
| YVLPSETPNAKE | 1 | 0 | 0 | 1538634 | P0C6W6.1 | P0C6W2 |
| YVMLNTSTSNLP | 1 | 0 | 0 | 1538650 | AMK59677.1 | P36334 |
| YVMPLTCIARRN | 1 | 0 | 0 | 1538651 | AMM42485.1 | P36334 |
| YVMPLTCIGRGN | 1 | 0 | 0 | 1538652 | AHA80218.1 | P36334 |
| YVMPLTCIRRSK | 1 | 0 | 0 | 1538657 | QEG03757.1 | P36334 |
| YVMPLTCNAISP | 1 | 0 | 0 | 1538660 | BBA20986.1 | A0A140H1H1 |
| YVMSSIYLFFTH | 1 | 0 | 0 | 1538668 | AUM60013.1 | K9N7C7 |
| YVNANGGTCFCN | 1 | 0 | 0 | 1538671 | AWK59942.1 | H9EJ42 |
| YVNATGSDVNCN | 1 | 0 | 0 | 1538673 | ABU88826.1 | A0A0P0G321 |
| YVNFEHQRGPGK | 1 | 0 | 0 | 1538675 | APT69876.1 | A0A0P0K6L9 |
| YVNMAVPHIKEW | 1 | 0 | 0 | 1538686 | AUM60027.1 | K9N643 |
| YVNNWCRCSCLP | 1 | 0 | 0 | 1538689 | ARB07438.1 | A0A140H1H1 |
| YVPEHCQNYSGC | 1 | 0 | 0 | 1538702 | AXN92229.1 | K9N796 |
| YVPEHCQNYSGS | 1 | 0 | 0 | 1538704 | ASU45697.1 | K9N796 |
| YVPLKSAVCVTA | 1 | 0 | 0 | 1538716 | AVV62524.1 | K9N7C7 |
| YVPSQEKNFTTA | 1 | 0 | 0 | 1538724 | ATO98181.1 | Q3LZX1 |
| YVPSQQQNFTTA | 1 | 0 | 0 | 1538726 | APO40579.1 | P59594 |
| YVPSYIKPESVA | 1 | 0 | 0 | 1538728 | AUM60013.1 | K9N7C7 |
| YVPTKYVTARVS | 1 | 0 | 0 | 1538732 | P36334.1 | P36334 |
| YVPTNHVNVTAA | 1 | 0 | 0 | 1538733 | AVV62537.1 | K9N5Q8 |
| YVSHGLDLMQAG | 1 | 0 | 0 | 1538769 | AVN89451.1 | K9N7C7 |
| YVSHGLDLMQAS | 1 | 0 | 0 | 1538770 | AUM60013.1 | K9N7C7 |
| YVSNGGNNCTTA | 1 | 0 | 0 | 1538781 | AWO68966.1 | A0A0P0G321 |
| YVSNGSNSCTTA | 1 | 0 | 0 | 1538783 | AWO69003.1 | A0A0P0G321 |
| YVSPDVQFEQLD | 1 | 0 | 0 | 1538786 | AVV62526.1 | K9N5Q8 |
| YVTGGLVQQVSQ | 1 | 0 | 0 | 1538805 | ARO76381.1 | P0C6U8 |
| YVTSSSKSSEEH | 1 | 0 | 0 | 1538816 | APO40578.1 | P0C6X7 |
| YVTVQGPPGTGK | 1 | 0 | 0 | 1538819 | AXN92247.1 | K9N7C7 |
| YVVDSVAVKNGA | 1 | 0 | 0 | 1538825 | P0C6W6.1 | P0C6W2 |
| YVVDVSHGLGTY | 1 | 0 | 0 | 1538827 | ARU07577.1 | A0A140H1H1 |
| YVVKNSTTFPFE | 1 | 0 | 0 | 1538834 | AXN92258.1 | K9N7C7 |
| YVYNGSAQAISF | 1 | 0 | 0 | 1538872 | QBP84748.1 | P30215 |
| YVYNGSAQATSF | 1 | 0 | 0 | 1538873 | AWW13558.1 | P30215 |
| YVYNGSAQVTSF | 1 | 0 | 0 | 1538874 | QBP84703.1 | P30215 |

| | | | | | | |
|---|---|---|---|---|---|---|
| YVYNTGRSAYVK | 1 | 0 | 0 | 1538877 | AVV62531.1 | K9N5R3 |
| YWDQTYHPNCSN | 1 | 0 | 0 | 1538896 | APO40578.1 | P0C6X7 |
| YWTVFCTNANAS | 1 | 0 | 0 | 1538935 | AVV62524.1 | K9N7C7 |
| YWVASSQADVNT | 1 | 0 | 0 | 1538938 | ANZ78879.1 | P33469 |
| YWVPRAAANIGS | 1 | 0 | 0 | 1538940 | APO40578.1 | P0C6X7 |
| YWVTSPPADVNT | 1 | 0 | 0 | 1538953 | AJQ24074.1 | P33469 |
| YYADVGMPTTFL | 1 | 0 | 0 | 1538959 | BBA20986.1 | A0A140H1H1 |
| YYCVTTKAKSIQ | 1 | 0 | 0 | 1538988 | ALA49934.1 | K9N7C7 |
| YYCYSAKDTLTE | 1 | 0 | 0 | 1538991 | AVV62535.1 | K9N7C7 |
| YYCYTSKDTLDD | 1 | 0 | 0 | 1538993 | AXN92247.1 | K9N7C7 |
| YYDDSQYYLNKD | 1 | 0 | 0 | 1538996 | AGT51480.1 | P30215 |
| YYDHSQYYFNKD | 1 | 0 | 0 | 1538999 | AEN19365.1 | P30215 |
| YYDHSQYYLNKD | 1 | 0 | 0 | 1539000 | AIX09798.1 | P30215 |
| YYDYSQYYLNKD | 1 | 0 | 0 | 1539008 | AGT51729.1 | P30215 |
| YYEFLNKHFNMM | 1 | 0 | 0 | 1539014 | AIL49554.1 | P0C6X6 |
| YYGNVLDQAIAM | 1 | 0 | 0 | 1539035 | APO40578.1 | P0C6X7 |
| YYGVFLYDNFTS | 1 | 0 | 0 | 1539037 | ARU07576.1 | A0A140H1H0 |
| YYKVEADFYLSG | 1 | 0 | 0 | 1539085 | P30215.1 | P30215 |
| YYKWPWYIWLGF | 1 | 0 | 0 | 1539087 | AHL18090.1 | K9N5Q8 |
| YYLGTGPHKDLK | 1 | 0 | 0 | 1539105 | AIW52805.1 | H9EJ69 |
| YYLPVMAAPTGI | 1 | 0 | 0 | 1539116 | ABD34826.1 | U3M6Q8 |
| YYLPVMAAPTGV | 1 | 0 | 0 | 1539117 | AWK59934.1 | U3M6Q8 |
| YYNGIMVLPGAA | 1 | 0 | 0 | 1539122 | APT69876.1 | A0A0P0K6L9 |
| YYNGIMVVPGVA | 1 | 0 | 0 | 1539124 | BAL45639.1 | A0A0P0K6L9 |
| YYNIDTGSFYGF | 1 | 0 | 0 | 1539126 | ABD75600.1 | A0A140H1H0 |
| YYNNWQNLLYDS | 1 | 0 | 0 | 1539143 | BBA20986.1 | A0A140H1H1 |
| YYPEPITGNNVV | 1 | 0 | 0 | 1539158 | CAA83660.1 | P36334 |
| YYPGTGPHKDLK | 1 | 0 | 0 | 1539159 | ABK63972.1 | H9EJ69 |
| YYPMCTNGQAVP | 1 | 0 | 0 | 1539161 | AUM60013.1 | K9N7C7 |
| YYQPIQQAPTGI | 1 | 0 | 0 | 1539183 | AST12967.1 | A0A0P0KXN3 |
| YYRITSRVMFEP | 1 | 0 | 0 | 1539187 | BAL45640.1 | A0A0P0K6L9 |
| YYSFGRCPTSSI | 1 | 0 | 0 | 1539212 | ARU07576.1 | A0A140H1H0 |
| YYSLTTNNAATK | 1 | 0 | 0 | 1539227 | AVP78031.1 | Q3LZX1 |
| YYSTTPCELHGN | 1 | 0 | 0 | 1539237 | ALW82645.1 | K9N7C7 |
| YYSYMMPVMGMT | 1 | 0 | 0 | 1539245 | NP_073549.1 | A0A0P0LKV0 |
| YYTGSSFFNPEP | 1 | 0 | 0 | 1539249 | AVV62537.1 | K9N5Q8 |
| YYVEGSGRSASN | 1 | 0 | 0 | 1539269 | ARU07581.1 | Q19U29 |
| YYVKDGKYFTSA | 1 | 0 | 0 | 1539272 | AVV62535.1 | K9N7C7 |
| YYVKPGGTSSRD | 1 | 0 | 0 | 1539275 | AWU59319.1 | K9N7C7 |
| YYVKWPWYVWLL | 1 | 0 | 0 | 1539277 | CAA79896.1 | P36334 |
| YYVMPLTCIGRR | 1 | 0 | 0 | 1539285 | AHA80213.1 | P36334 |
| YYVSNGSNSCTT | 1 | 0 | 0 | 1539299 | AWO69003.1 | A0A0P0G321 |
| YYYKVEADFYLS | 1 | 0 | 0 | 1539305 | P30215.1 | P30215 |
| YYYNMATGFAYG | 1 | 0 | 0 | 1539308 | AZS52617.1 | A0A140H1H0 |
| YYYPEPIIENNV | 1 | 0 | 0 | 1539310 | AIX09799.1 | P36334 |
| AACRKVQHMVVKAAL | 1 | 0 | 0 | 1539351 | P0DTD1.1 | P0DTD1 |
| AAETLKATEETFKLS | 1 | 0 | 0 | 1539353 | P0DTD1.1 | P0DTD1 |
| AAIFYLITPVHVMSK | 1 | 0 | 0 | 1539356 | P0DTC1.1 | P0DTD1 |
| AAIMQLFFSYFAVHF | 1 | 0 | 0 | 1539357 | P0DTC1.1 | P0DTD1 |
| AALCTFLLNKEMYLK | 1 | 0 | 0 | 1539361 | P0DTC1.1 | P0DTD1 |
| AARQGFVDSDVETKD | 1 | 0 | 0 | 1539368 | P0DTC1.1 | P0DTD1 |
| AARVVRSIFSRTLET | 1 | 0 | 0 | 1539369 | P0DTC1.1 | P0DTD1 |
| AAVNSVPWDTIANYA | 1 | 0 | 0 | 1539374 | P0DTC1.1 | P0DTD1 |
| ACEDLKPVSEEVVEN | 1 | 0 | 0 | 1539377 | P0DTC1.1 | P0DTD1 |
| ACEIVGGQIVTCAKE | 1 | 0 | 0 | 1539378 | P0DTC1.1 | P0DTD1 |
| ACHNSEVGPEHSLAE | 1 | 0 | 0 | 1539380 | P0DTC1.1 | P0DTD1 |
| ACPDGVKHVYQLRAR | 1 | 0 | 0 | 1539383 | P0DTC7.1 | P0DTC7 |
| ACPLIAAVITREVGF | 1 | 0 | 0 | 1539384 | P0DTC1.1 | P0DTD1 |
| ACVLAAECTIFKDAS | 1 | 0 | 0 | 1539388 | P0DTC1.1 | P0DTD1 |
| ACWHHSIGFDYVYNP | 1 | 0 | 0 | 1539391 | P0DTD1.1 | P0DTD1 |
| ADKYVRNLQHRLYEC | 1 | 0 | 0 | 1539397 | P0DTD1.1 | P0DTD1 |
| ADLKSFDLGDELGTD | 1 | 0 | 0 | 1539399 | P0DTC1.1 | P0DTD1 |
| ADNNCYLATALLTLQ | 1 | 0 | 0 | 1539402 | P0DTC1.1 | P0DTD1 |
| ADPAMHAASGNLLLD | 1 | 0 | 0 | 1539403 | P0DTD1.1 | P0DTD1 |
| ADPIHSLRVCVDTVR | 1 | 0 | 0 | 1539404 | P0DTC1.1 | P0DTD1 |
| ADSIIIGGAKLKALN | 1 | 0 | 0 | 1539405 | P0DTC1.1 | P0DTD1 |
| ADSKIVQLSEISMDN | 1 | 0 | 0 | 1539406 | P0DTC1.1 | P0DTD1 |
| AELEGIQYGRSGETL | 1 | 0 | 0 | 1539417 | P0DTC1.1 | P0DTD1 |
| AENVTGLFKDCSKVI | 1 | 0 | 0 | 1539418 | P0DTD1.1 | P0DTD1 |
| AFELWAKRNIKPVPE | 1 | 0 | 0 | 1539424 | P0DTD1.1 | P0DTD1 |
| AFGLVAEWFLAYILF | 1 | 0 | 0 | 1539426 | P0DTC1.1 | P0DTD1 |
| AFKQIVESCGNFKVT | 1 | 0 | 0 | 1539428 | P0DTC1.1 | P0DTD1 |
| AFLPFAMGIIAMSAF | 1 | 0 | 0 | 1539432 | P0DTC1.1 | P0DTD1 |
| AFQTVKPGNFNKDFY | 1 | 0 | 0 | 1539436 | P0DTD1.1 | P0DTD1 |
| AFVNLKQLPFFYYSD | 1 | 0 | 0 | 1539438 | P0DTD1.1 | P0DTD1 |
| AFVTNVNASSSEAFL | 1 | 0 | 0 | 1539439 | P0DTD1.1 | P0DTD1 |
| AGFPFNKWGKARLYY | 1 | 0 | 0 | 1539446 | P0DTD1.1 | P0DTD1 |
| AGKASCTLSEQLDFI | 1 | 0 | 0 | 1539448 | P0DTC1.1 | P0DTD1 |
| AGLEAPFLYLYALVY | 1 | 0 | 0 | 1539449 | P0DTC3.1 | P0DTC3 |
| AGNATEVPANSTVLS | 1 | 0 | 0 | 1539451 | P0DTC1.1 | P0DTD1 |
| AGVCVSTSGRWVLNN | 1 | 0 | 0 | 1539459 | P0DTC1.1 | P0DTD1 |
| AHAEETRKLMPVCVE | 1 | 0 | 0 | 1539461 | P0DTC1.1 | P0DTD1 |
| AHIQWMVMFTPLVPF | 1 | 0 | 0 | 1539463 | P0DTC1.1 | P0DTD1 |
| AHISTIGVCSMTDIA | 1 | 0 | 0 | 1539464 | P0DTD1.1 | P0DTD1 |
| AIATCDWTNAGDYIL | 1 | 0 | 0 | 1539466 | P0DTD1.1 | P0DTD1 |
| AIRHVRAWIGFDVEG | 1 | 0 | 0 | 1539472 | P0DTD1.1 | P0DTD1 |
| AISMWALIISVTSNY | 1 | 0 | 0 | 1539473 | P0DTC1.1 | P0DTD1 |
| AIVVTCLAYYFMRFR | 1 | 0 | 0 | 1539476 | P0DTC1.1 | P0DTD1 |
| AKSASVYYSQLMCQP | 1 | 0 | 0 | 1539487 | P0DTC1.1 | P0DTD1 |
| AKSHNIALIWNVKDF | 1 | 0 | 0 | 1539488 | P0DTC1.1 | P0DTD1 |
| AKVGILCIMSDRDLY | 1 | 0 | 0 | 1539490 | P0DTD1.1 | P0DTD1 |

| | | | | | | |
|---|---|---|---|---|---|---|
| ALDQAISMWALIISV | 1 | 0 | 0 | 1539494 | P0DTC1.1 | P0DTD1 |
| ALGGSVAIKITEHSW | 1 | 0 | 0 | 1539496 | P0DTD1.1 | P0DTD1 |
| ALITLATCELYHYQE | 1 | 0 | 0 | 1539499 | P0DTC7.1 | P0DTC7 |
| ALIWNVKDFMSLSEQ | 1 | 0 | 0 | 1539500 | P0DTC1.1 | P0DTD1 |
| ALPETTADIVVFDEI | 1 | 0 | 0 | 1539509 | P0DTD1.1 | P0DTD1 |
| ALTAESHVDTDLTKP | 1 | 0 | 0 | 1539517 | P0DTD1.1 | P0DTD1 |
| ANNSLKITEEVGHTD | 1 | 0 | 0 | 1539541 | P0DTC1.1 | P0DTD1 |
| ANYFLCWHTNCYDYC | 1 | 0 | 0 | 1539546 | P0DTC3.1 | P0DTC3 |
| APAHISTIGVCSMTD | 1 | 0 | 0 | 1539549 | P0DTD1.1 | P0DTD1 |
| APGCDVTDVTQLYLG | 1 | 0 | 0 | 1539551 | P0DTD1.1 | P0DTD1 |
| APLLSAGIFGADPIH | 1 | 0 | 0 | 1539554 | P0DTC1.1 | P0DTD1 |
| APTHLSVDTKFKTEG | 1 | 0 | 0 | 1539559 | P0DTD1.1 | P0DTD1 |
| AQAAGTDTTITVNVL | 1 | 0 | 0 | 1539560 | P0DTC1.1 | P0DTD1 |
| AQEAYEQAVANGDSE | 1 | 0 | 0 | 1539561 | P0DTC1.1 | P0DTD1 |
| AQTGIAVLDMCASLK | 1 | 0 | 0 | 1539564 | P0DTC1.1 | P0DTD1 |
| AQYELKHGTFTCASE | 1 | 0 | 0 | 1539569 | P0DTC1.1 | P0DTD1 |
| ARAGKASCTLSEQLD | 1 | 0 | 0 | 1539571 | P0DTC1.1 | P0DTD1 |
| ARFPKSDGTGTIYTE | 1 | 0 | 0 | 1539574 | P0DTC1.1 | P0DTD1 |
| ARLYYDSMSYEDQDA | 1 | 0 | 0 | 1539578 | P0DTD1.1 | P0DTD1 |
| ARQHLKDGTCGLVEV | 1 | 0 | 0 | 1539580 | P0DTC1.1 | P0DTD1 |
| ARTVAGVSICSTMTN | 1 | 0 | 0 | 1539585 | P0DTD1.1 | P0DTD1 |
| ASCCLYCRCHIDHPN | 1 | 0 | 0 | 1539590 | P0DTC1.1 | P0DTD1 |
| ASFSASTSAFVETVK | 1 | 0 | 0 | 1539592 | P0DTC1.1 | P0DTD1 |
| ASFYYVWKSYVHVVD | 1 | 0 | 0 | 1539593 | P0DTC1.1 | P0DTD1 |
| ASLPFGWLIVGVALL | 1 | 0 | 0 | 1539598 | P0DTC3.1 | P0DTC3 |
| ATCDWTNAGDYILAN | 1 | 0 | 0 | 1539606 | P0DTD1.1 | P0DTD1 |
| ATEETFKLSYGIATV | 1 | 0 | 0 | 1539608 | P0DTD1.1 | P0DTD1 |
| ATHSDKFTDGVCLFW | 1 | 0 | 0 | 1539610 | P0DTD1.1 | P0DTD1 |
| ATIPIQASLPFGWLI | 1 | 0 | 0 | 1539611 | P0DTC3.1 | P0DTC3 |
| ATNGPLKVGGSCVLS | 1 | 0 | 0 | 1539615 | P0DTC1.1 | P0DTD1 |
| ATRVECTTIVNGVRR | 1 | 0 | 0 | 1539620 | P0DTC1.1 | P0DTD1 |
| AVFDKNLYDKLVSSF | 1 | 0 | 0 | 1539628 | P0DTC1.1 | P0DTD1 |
| AVHECFVKRVDWTIE | 1 | 0 | 0 | 1539629 | P0DTD1.1 | P0DTD1 |
| AVLQSGFRKMAFPSG | 1 | 0 | 0 | 1539632 | P0DTC1.1 | P0DTD1 |
| AVVLLILMTARTVYD | 1 | 0 | 0 | 1539641 | P0DTC1.1 | P0DTD1 |
| AWIGFDVEGCHATRE | 1 | 0 | 0 | 1539642 | P0DTD1.1 | P0DTD1 |
| AYCNKTVGELGDVRE | 1 | 0 | 0 | 1539648 | P0DTC1.1 | P0DTD1 |
| AYENFNQHEVLLAPL | 1 | 0 | 0 | 1539649 | P0DTC1.1 | P0DTD1 |
| AYFNMVYMPASWVMR | 1 | 0 | 0 | 1539651 | P0DTC1.1 | P0DTD1 |
| AYPLTKHPNQEYADV | 1 | 0 | 0 | 1539654 | P0DTD1.1 | P0DTD1 |
| CALAPNMMVTNNTFT | 1 | 0 | 0 | 1539662 | P0DTC1.1 | P0DTD1 |
| CCLYCRCHIDHPNPK | 1 | 0 | 0 | 1539672 | P0DTC1.1 | P0DTD1 |
| CCNIVNVSLVKPSFY | 1 | 0 | 0 | 1539674 | YP_009724392.1 | P0DTC4 |
| CCREHEHEIAWYTER | 1 | 0 | 0 | 1539675 | P0DTC1.1 | P0DTD1 |
| CDDDYFNKKDWYDFV | 1 | 0 | 0 | 1539677 | P0DTD1.1 | P0DTD1 |
| CDGTTFTYASALWEI | 1 | 0 | 0 | 1539678 | P0DTC1.1 | P0DTD1 |
| CDRAMPNMLRIMASL | 1 | 0 | 0 | 1539681 | P0DTD1.1 | P0DTD1 |
| CDWTNAGDYILANTC | 1 | 0 | 0 | 1539682 | P0DTD1.1 | P0DTD1 |
| CEEEEFEPSTQYEYG | 1 | 0 | 0 | 1539683 | P0DTC1.1 | P0DTD1 |
| CFVKRVDWTIEYPII | 1 | 0 | 0 | 1539690 | P0DTD1.1 | P0DTD1 |
| CGFIQQKLALGGSVA | 1 | 0 | 0 | 1539693 | P0DTD1.1 | P0DTD1 |
| CGNFKVTKGKAKKGA | 1 | 0 | 0 | 1539695 | P0DTC1.1 | P0DTD1 |
| CGSVGFNIDYDCVSF | 1 | 0 | 0 | 1539697 | P0DTC1.1 | P0DTD1 |
| CGTGTSTDVVYRAFD | 1 | 0 | 0 | 1539699 | P0DTD1.1 | P0DTD1 |
| CINGLMLLEIKDTEK | 1 | 0 | 0 | 1539709 | P0DTC1.1 | P0DTD1 |
| CITRCNLGGAVCRHH | 1 | 0 | 0 | 1539711 | P0DTD1.1 | P0DTD1 |
| CLKLSHQSDIEVTGD | 1 | 0 | 0 | 1539720 | P0DTC1.1 | P0DTD1 |
| CLLNRYFRLTLGVYD | 1 | 0 | 0 | 1539721 | P0DTC1.1 | P0DTD1 |
| CLNRVCTNYMPYFFT | 1 | 0 | 0 | 1539722 | P0DTC1.1 | P0DTD1 |
| CLSTLMKCDHCGETS | 1 | 0 | 0 | 1539723 | P0DTC1.1 | P0DTD1 |
| CLYRNRDVDTDFVNE | 1 | 0 | 0 | 1539725 | P0DTD1.1 | P0DTD1 |
| CNLGGAVCRHHANEY | 1 | 0 | 0 | 1539732 | P0DTD1.1 | P0DTD1 |
| CNQMCLSTLMKCDHC | 1 | 0 | 0 | 1539734 | P0DTC1.1 | P0DTD1 |
| CNSQTSLRCGACIRR | 1 | 0 | 0 | 1539735 | P0DTD1.1 | P0DTD1 |
| CNVDRYPANSIVCRF | 1 | 0 | 0 | 1539737 | P0DTD1.1 | P0DTD1 |
| CPACHNSEVGPEHSL | 1 | 0 | 0 | 1539740 | P0DTC1.1 | P0DTD1 |
| CQEPKLGSLVVRCSF | 1 | 0 | 0 | 1539744 | P0DTC8.1 | P0DTC8 |
| CRKVQHMVVKAALLA | 1 | 0 | 0 | 1539750 | P0DTD1.1 | P0DTD1 |
| CRSKNPLLYDANYFL | 1 | 0 | 0 | 1539752 | P0DTC3.1 | P0DTC3 |
| CSFYEDFLEYHDVRV | 1 | 0 | 0 | 1539754 | P0DTC8.1 | P0DTC8 |
| CSLQSCTQHQPYVVD | 1 | 0 | 0 | 1539757 | P0DTC8.1 | P0DTC8 |
| CSLSHRFYRLANECA | 1 | 0 | 0 | 1539758 | P0DTD1.1 | P0DTD1 |
| CSVIDLLLDDFVEII | 1 | 0 | 0 | 1539761 | P0DTD1.1 | P0DTD1 |
| CTCYFGLFCLLNRYF | 1 | 0 | 0 | 1539762 | P0DTC1.1 | P0DTD1 |
| CTGSIPCSVCLSGLD | 1 | 0 | 0 | 1539764 | P0DTC1.1 | P0DTD1 |
| CTVCGMWKGYGCSCD | 1 | 0 | 0 | 1539765 | P0DTC1.1 | P0DTD1 |
| CVCSVIDLLLDDFVE | 1 | 0 | 0 | 1539767 | P0DTD1.1 | P0DTD1 |
| CVDIPGIPKDMTYRR | 1 | 0 | 0 | 1539768 | P0DTD1.1 | P0DTD1 |
| CYDGGCINANQVIVN | 1 | 0 | 0 | 1539778 | P0DTD1.1 | P0DTD1 |
| CYFGLFCLLNRYFRL | 1 | 0 | 0 | 1539780 | P0DTC1.1 | P0DTD1 |
| CYLATALLTLQQIEL | 1 | 0 | 0 | 1539783 | P0DTC1.1 | P0DTD1 |
| DAMMFTSDLATNNLV | 1 | 0 | 0 | 1539788 | P0DTC1.1 | P0DTD1 |
| DAMRNAGIVGVLTLD | 1 | 0 | 0 | 1539789 | P0DTD1.1 | P0DTD1 |
| DAPYIVGDVVQEGVL | 1 | 0 | 0 | 1539791 | P0DTC1.1 | P0DTD1 |
| DCATVHTANKWDLII | 1 | 0 | 0 | 1539800 | P0DTD1.1 | P0DTD1 |
| DCVVLHSYFTSDYYQ | 1 | 0 | 0 | 1539804 | P0DTC3.1 | P0DTC3 |
| DDFVEIIKSQDLSVV | 1 | 0 | 0 | 1539806 | P0DTD1.1 | P0DTD1 |
| DDSQQTVGQQDGSED | 1 | 0 | 0 | 1539812 | P0DTC1.1 | P0DTD1 |
| DDTLRVEAFEYYHTT | 1 | 0 | 0 | 1539813 | P0DTC1.1 | P0DTD1 |
| DDVVYCPRHVICTSE | 1 | 0 | 0 | 1539814 | P0DTC1.1 | P0DTD1 |
| DDYVYLPYPDPSRIL | 1 | 0 | 0 | 1539817 | P0DTD1.1 | P0DTD1 |

| | | | | | | |
|---|---|---|---|---|---|---|
| DEWSMATYYLFDESG | 1 | 0 | 0 | 1539829 | P0DTC1.1 | P0DTD1 |
| DFKLNEEIAIILASF | 1 | 0 | 0 | 1539833 | P0DTC1.1 | P0DTD1 |
| DFLELAMDEFIERYK | 1 | 0 | 0 | 1539834 | P0DTD1.1 | P0DTD1 |
| DFVSDADSTLIGDCA | 1 | 0 | 0 | 1539845 | P0DTD1.1 | P0DTD1 |
| DGADVTKIKPHNSHE | 1 | 0 | 0 | 1539847 | P0DTC1.1 | P0DTD1 |
| DGCNSSTCMMCYKRN | 1 | 0 | 0 | 1539848 | P0DTC1.1 | P0DTD1 |
| DGSEDNQTTTIQTIV | 1 | 0 | 0 | 1539858 | P0DTC1.1 | P0DTD1 |
| DGTCGLVEVEKGVLP | 1 | 0 | 0 | 1539860 | P0DTC1.1 | P0DTD1 |
| DGTGTIYTELEPPCR | 1 | 0 | 0 | 1539861 | P0DTC1.1 | P0DTD1 |
| DGVPFVVSTGYHFRE | 1 | 0 | 0 | 1539865 | P0DTD1.1 | P0DTD1 |
| DGWEIVKFISTCACE | 1 | 0 | 0 | 1539866 | P0DTC1.1 | P0DTD1 |
| DGYPLECIKDLLARA | 1 | 0 | 0 | 1539867 | P0DTC1.1 | P0DTD1 |
| DGYVMHANYIFWRNT | 1 | 0 | 0 | 1539868 | P0DTD1.1 | P0DTD1 |
| DISASIVAGGIVAIV | 1 | 0 | 0 | 1539887 | P0DTC1.1 | P0DTD1 |
| DKCSRIIPARARVEC | 1 | 0 | 0 | 1539894 | P0DTD1.1 | P0DTD1 |
| DKRAKVTSAMQTMLF | 1 | 0 | 0 | 1539899 | P0DTC1.1 | P0DTD1 |
| DKRTTCFSVAALTNN | 1 | 0 | 0 | 1539900 | P0DTD1.1 | P0DTD1 |
| DKSVYYTSNPTTFHL | 1 | 0 | 0 | 1539902 | P0DTC1.1 | P0DTD1 |
| DLDEWSMATYYLFDE | 1 | 0 | 0 | 1539906 | P0DTC1.1 | P0DTD1 |
| DLFRNARNGVLITEG | 1 | 0 | 0 | 1539907 | P0DTD1.1 | P0DTD1 |
| DLLLDDFVEIIKSQD | 1 | 0 | 0 | 1539911 | P0DTD1.1 | P0DTD1 |
| DLVYALRHFDEGNCD | 1 | 0 | 0 | 1539924 | P0DTD1.1 | P0DTD1 |
| DNALAYYNTTKGGRF | 1 | 0 | 0 | 1539930 | P0DTC1.1 | P0DTD1 |
| DNLKTLLSLREVRTI | 1 | 0 | 0 | 1539933 | P0DTC1.1 | P0DTD1 |
| DNTSRYWEPEFYEAM | 1 | 0 | 0 | 1539937 | P0DTD1.1 | P0DTD1 |
| DQAISMWALIISVTS | 1 | 0 | 0 | 1539940 | P0DTC1.1 | P0DTD1 |
| DRDLYDKLQFTSLEI | 1 | 0 | 0 | 1539951 | P0DTD1.1 | P0DTD1 |
| DRQTAQAAGTDTTIT | 1 | 0 | 0 | 1539953 | P0DTC1.1 | P0DTD1 |
| DSAEVAVKMFDAYVN | 1 | 0 | 0 | 1539956 | P0DTC1.1 | P0DTD1 |
| DSEVVLKKLKKSLNV | 1 | 0 | 0 | 1539961 | P0DTC1.1 | P0DTD1 |
| DSLDTYPSLETIQIT | 1 | 0 | 0 | 1539964 | P0DTC1.1 | P0DTD1 |
| DSPCESHGKQVVSDI | 1 | 0 | 0 | 1539966 | P0DTD1.1 | P0DTD1 |
| DSVTVKNGSIHLYFD | 1 | 0 | 0 | 1539971 | P0DTC1.1 | P0DTD1 |
| DTCFANKHADFDTWF | 1 | 0 | 0 | 1539972 | P0DTC1.1 | P0DTD1 |
| DTFCAGSTFISDEVA | 1 | 0 | 0 | 1539975 | P0DTC1.1 | P0DTD1 |
| DTLKEILVTYNCCDD | 1 | 0 | 0 | 1539978 | P0DTD1.1 | P0DTD1 |
| DVLECNVKTTEVVGD | 1 | 0 | 0 | 1539990 | P0DTC1.1 | P0DTD1 |
| DVVECLKLSHQSDIE | 1 | 0 | 0 | 1539997 | P0DTC1.1 | P0DTD1 |
| DYCIPYNSVTSSIVI | 1 | 0 | 0 | 1540003 | P0DTC3.1 | P0DTC3 |
| DYDYYRYNLPTMCDI | 1 | 0 | 0 | 1540005 | P0DTD1.1 | P0DTD1 |
| DYIINLIIKNLSKSL | 1 | 0 | 0 | 1540008 | P0DTC6.1 | P0DTC6 |
| DYLVSTQEFRYMNSQ | 1 | 0 | 0 | 1540010 | P0DTC1.1 | P0DTD1 |
| EASFNYLKSPNFSKL | 1 | 0 | 0 | 1540022 | P0DTC1.1 | P0DTD1 |
| EAVEAPLVGTPVCIN | 1 | 0 | 0 | 1540023 | P0DTC1.1 | P0DTD1 |
| EAVGTNLPLQLGFST | 1 | 0 | 0 | 1540024 | P0DTD1.1 | P0DTD1 |
| EAYEQAVANGDSEVV | 1 | 0 | 0 | 1540027 | P0DTC1.1 | P0DTD1 |
| ECAQVLSEMVMCGGS | 1 | 0 | 0 | 1540028 | P0DTD1.1 | P0DTD1 |
| ECTTIVNGVRRSFYV | 1 | 0 | 0 | 1540030 | P0DTC1.1 | P0DTD1 |
| ECVRGTTVLLKEPCS | 1 | 0 | 0 | 1540031 | P0DTC7.1 | P0DTC7 |
| EDDYQGKPLEFGATS | 1 | 0 | 0 | 1540032 | P0DTC1.1 | P0DTD1 |
| EDFIPMDSTVKNYFI | 1 | 0 | 0 | 1540033 | P0DTD1.1 | P0DTD1 |
| EDLLIRKSNHNFLVQ | 1 | 0 | 0 | 1540037 | P0DTC1.1 | P0DTD1 |
| EEAALCTFLLNKEMY | 1 | 0 | 0 | 1540040 | P0DTC1.1 | P0DTD1 |
| EEAIRHVRAWIGFDV | 1 | 0 | 0 | 1540042 | P0DTD1.1 | P0DTD1 |
| EEAKTVLKKCKSAFY | 1 | 0 | 0 | 1540043 | P0DTC1.1 | P0DTD1 |
| EEEFEPSTQYEYGTE | 1 | 0 | 0 | 1540045 | P0DTC1.1 | P0DTD1 |
| EEGDCEEEEFEPSTQ | 1 | 0 | 0 | 1540047 | P0DTC1.1 | P0DTD1 |
| EEHFIETISLAGSYK | 1 | 0 | 0 | 1540048 | P0DTC1.1 | P0DTD1 |
| EESSAKSASVYYSQL | 1 | 0 | 0 | 1540052 | P0DTC1.1 | P0DTD1 |
| EETGTLIVNSVLLFL | 1 | 0 | 0 | 1540053 | P0DTC4.1 | P0DTC4 |
| EETKFLTENLLLYID | 1 | 0 | 0 | 1540054 | P0DTC1.1 | P0DTD1 |
| EEVGHTDLMAAYVDN | 1 | 0 | 0 | 1540055 | P0DTC1.1 | P0DTD1 |
| EFCGTENLTKEGATT | 1 | 0 | 0 | 1540059 | P0DTC1.1 | P0DTD1 |
| EFDRDAAMQRKLEKM | 1 | 0 | 0 | 1540060 | P0DTC1.1 | P0DTD1 |
| EFIERYKLEGYAFEH | 1 | 0 | 0 | 1540062 | P0DTD1.1 | P0DTD1 |
| EFKPRSQMEIDFLEL | 1 | 0 | 0 | 1540063 | P0DTD1.1 | P0DTD1 |
| EFLRDGWEIVKFIST | 1 | 0 | 0 | 1540065 | P0DTC1.1 | P0DTD1 |
| EGKTFYVLPNDDTLR | 1 | 0 | 0 | 1540068 | P0DTC1.1 | P0DTD1 |
| EGNCDTLKEILVTYN | 1 | 0 | 0 | 1540070 | P0DTD1.1 | P0DTD1 |
| EGQINDMILSLLSKG | 1 | 0 | 0 | 1540071 | P0DTD1.1 | P0DTD1 |
| EGSVRVVTTFDSEYC | 1 | 0 | 0 | 1540074 | P0DTC1.1 | P0DTD1 |
| EGYAFEHIVYGDFSH | 1 | 0 | 0 | 1540075 | P0DTD1.1 | P0DTD1 |
| EIAIILASFSASTSA | 1 | 0 | 0 | 1540079 | P0DTC1.1 | P0DTD1 |
| EIDFLELAMDEFIER | 1 | 0 | 0 | 1540080 | P0DTD1.1 | P0DTD1 |
| EIIFLEGETLPTEVL | 1 | 0 | 0 | 1540081 | P0DTC1.1 | P0DTD1 |
| EIKESVQTFFKLVNK | 1 | 0 | 0 | 1540084 | P0DTC1.1 | P0DTD1 |
| EILQKEKVNINIVGD | 1 | 0 | 0 | 1540086 | P0DTC1.1 | P0DTD1 |
| EILVTYNCCDDDYFN | 1 | 0 | 0 | 1540087 | P0DTD1.1 | P0DTD1 |
| EKMADQAMTQMYKQA | 1 | 0 | 0 | 1540092 | P0DTC1.1 | P0DTD1 |
| EKTHVQLSLPVLQVR | 1 | 0 | 0 | 1540095 | P0DTC1.1 | P0DTD1 |
| EKWESGVKDCVVLHS | 1 | 0 | 0 | 1540096 | P0DTC3.1 | P0DTC3 |
| ELGVVHNQDVNLHSS | 1 | 0 | 0 | 1540102 | P0DTD1.1 | P0DTD1 |
| ELKHFFFAQDGNAAI | 1 | 0 | 0 | 1540105 | P0DTD1.1 | P0DTD1 |
| ELLTPLGIDLDEWSM | 1 | 0 | 0 | 1540108 | P0DTC1.1 | P0DTD1 |
| ELLVYAADPAMHAAS | 1 | 0 | 0 | 1540109 | P0DTD1.1 | P0DTD1 |
| ELMRELNGGAYTRYV | 1 | 0 | 0 | 1540110 | P0DTC1.1 | P0DTD1 |
| ELNGGAYTRYVDNNF | 1 | 0 | 0 | 1540111 | P0DTC1.1 | P0DTD1 |
| ELSRVLGLKTLATHG | 1 | 0 | 0 | 1540113 | P0DTC1.1 | P0DTD1 |
| ELTGHMLDMYSVMLT | 1 | 0 | 0 | 1540114 | P0DTD1.1 | P0DTD1 |
| ELTSMKYFVKIGPER | 1 | 0 | 0 | 1540115 | P0DTD1.1 | P0DTD1 |
| EMYLKLRSDVLLPLT | 1 | 0 | 0 | 1540119 | P0DTC1.1 | P0DTD1 |

| | | | | | | |
|---|---|---|---|---|---|---|
| ENAFLPFAMGIIAMS | 1 | 0 | 0 | 1540120 | P0DTC1.1 | P0DTD1 |
| ENDSKEGFFTYICGF | 1 | 0 | 0 | 1540121 | P0DTD1.1 | P0DTD1 |
| ENFNQHEVLLAPLLS | 1 | 0 | 0 | 1540122 | P0DTC1.1 | P0DTD1 |
| EPKLGSLVVRCSFYE | 1 | 0 | 0 | 1540129 | P0DTC8.1 | P0DTC8 |
| ERHSLSHFVNLDNLR | 1 | 0 | 0 | 1540138 | P0DTC1.1 | P0DTD1 |
| ERLKLFAAETLKATE | 1 | 0 | 0 | 1540139 | P0DTD1.1 | P0DTD1 |
| ERSEAGVCVSTSGRW | 1 | 0 | 0 | 1540141 | P0DTC1.1 | P0DTD1 |
| ERVRQALLKTVQFCD | 1 | 0 | 0 | 1540142 | P0DTD1.1 | P0DTD1 |
| ESCGNFKVTKGKAKK | 1 | 0 | 0 | 1540143 | P0DTC1.1 | P0DTD1 |
| ESELVIGAVILRGHL | 1 | 0 | 0 | 1540144 | P0DTC5.1 | P0DTC5 |
| ESGEFKLASHMYCSF | 1 | 0 | 0 | 1540145 | P0DTC1.1 | P0DTD1 |
| ESGLKTILRKGGRTI | 1 | 0 | 0 | 1540146 | P0DTC1.1 | P0DTD1 |
| ESPFELEDFIPMDST | 1 | 0 | 0 | 1540152 | P0DTD1.1 | P0DTD1 |
| ETDLTKGPHEFCSQH | 1 | 0 | 0 | 1540154 | P0DTD1.1 | P0DTD1 |
| ETGLLMPLKAPKEII | 1 | 0 | 0 | 1540155 | P0DTC1.1 | P0DTD1 |
| ETLGVLVPHVGEIPV | 1 | 0 | 0 | 1540158 | P0DTC1.1 | P0DTD1 |
| ETMSYLFQHANLDSC | 1 | 0 | 0 | 1540161 | P0DTC1.1 | P0DTD1 |
| ETNILLNVPLHGTIL | 1 | 0 | 0 | 1540162 | P0DTC5.1 | P0DTC5 |
| ETSWQTGDFVKATCE | 1 | 0 | 0 | 1540166 | P0DTC1.1 | P0DTD1 |
| EVARDLSLQFKRPIN | 1 | 0 | 0 | 1540167 | P0DTC1.1 | P0DTD1 |
| EVGKPRPPLNRNYVF | 1 | 0 | 0 | 1540168 | P0DTD1.1 | P0DTD1 |
| EVKILNNLGVDIAAN | 1 | 0 | 0 | 1540169 | P0DTD1.1 | P0DTD1 |
| EVLLAPLLSAGIFGA | 1 | 0 | 0 | 1540171 | P0DTC1.1 | P0DTD1 |
| EVNSFSGYLKLTDNV | 1 | 0 | 0 | 1540173 | P0DTC1.1 | P0DTD1 |
| EVQELYSPIFLIVAA | 1 | 0 | 0 | 1540174 | P0DTC7.1 | P0DTC7 |
| EYCPIFFITGNTLQC | 1 | 0 | 0 | 1540186 | P0DTC1.1 | P0DTD1 |
| EYCRHGTCERSEAGV | 1 | 0 | 0 | 1540187 | P0DTC1.1 | P0DTD1 |
| EYTDFATSACVLAAE | 1 | 0 | 0 | 1540195 | P0DTC1.1 | P0DTD1 |
| FAMGIIAMSAFAMMF | 1 | 0 | 0 | 1540198 | P0DTC1.1 | P0DTD1 |
| FARTRSMWSFNPETN | 1 | 0 | 0 | 1540201 | YP_009724393.1 | P0DTC5 |
| FATSACVLAAECTIF | 1 | 0 | 0 | 1540203 | P0DTC1.1 | P0DTD1 |
| FAVHFISNSWLMWLI | 1 | 0 | 0 | 1540205 | P0DTC1.1 | P0DTD1 |
| FAVSKGFFKEGSSVE | 1 | 0 | 0 | 1540206 | P0DTD1.1 | P0DTD1 |
| FAYTKRNVIPTITQM | 1 | 0 | 0 | 1540207 | P0DTD1.1 | P0DTD1 |
| FCALILAYCNKTVGE | 1 | 0 | 0 | 1540209 | P0DTC1.1 | P0DTD1 |
| FCGPDGYPLECIKDL | 1 | 0 | 0 | 1540211 | P0DTC1.1 | P0DTD1 |
| FCKLHNWNCVNCDTF | 1 | 0 | 0 | 1540213 | P0DTC1.1 | P0DTD1 |
| FDESGEFKLASHMYC | 1 | 0 | 0 | 1540221 | P0DTC1.1 | P0DTD1 |
| FDKNLYDKLVSSFLE | 1 | 0 | 0 | 1540222 | P0DTC1.1 | P0DTD1 |
| FDTWFSQRGGSYTND | 1 | 0 | 0 | 1540223 | P0DTC1.1 | P0DTD1 |
| FEHIVYGDFSHSQLG | 1 | 0 | 0 | 1540226 | P0DTD1.1 | P0DTD1 |
| FEYYHTTDPSFLGRY | 1 | 0 | 0 | 1540230 | P0DTC1.1 | P0DTD1 |
| FFDGRVDGQVDLFRN | 1 | 0 | 0 | 1540231 | P0DTD1.1 | P0DTD1 |
| FFFAQDGNAAISDYD | 1 | 0 | 0 | 1540232 | P0DTD1.1 | P0DTD1 |
| FFKEGSSVELKHFFF | 1 | 0 | 0 | 1540234 | P0DTD1.1 | P0DTD1 |
| FFSYFAVHFISNSWL | 1 | 0 | 0 | 1540238 | P0DTC1.1 | P0DTD1 |
| FGATSAALQPEEEQE | 1 | 0 | 0 | 1540242 | P0DTC1.1 | P0DTD1 |
| FGPLVRKIFVDGVPF | 1 | 0 | 0 | 1540246 | P0DTD1.1 | P0DTD1 |
| FHLVDFQVTIAEILL | 1 | 0 | 0 | 1540251 | P0DTC6.1 | P0DTC6 |
| FIKRSDARTAPHGHV | 1 | 0 | 0 | 1540254 | P0DTC1.1 | P0DTD1 |
| FIPMDSTVKNYFITD | 1 | 0 | 0 | 1540255 | P0DTD1.1 | P0DTD1 |
| FIQQKLALGGSVAIK | 1 | 0 | 0 | 1540256 | P0DTD1.1 | P0DTD1 |
| FISDEVARDLSLQFK | 1 | 0 | 0 | 1540258 | P0DTC1.1 | P0DTD1 |
| FITREEAIRHVRAWI | 1 | 0 | 0 | 1540262 | P0DTD1.1 | P0DTD1 |
| FKESPFELEDFIPMD | 1 | 0 | 0 | 1540265 | P0DTD1.1 | P0DTD1 |
| FKFRIDGDMVPHISR | 1 | 0 | 0 | 1540266 | P0DTD1.1 | P0DTD1 |
| FKLTCATTRQVVNVV | 1 | 0 | 0 | 1540270 | P0DTC1.1 | P0DTD1 |
| FKSVLYYQNNVFMSE | 1 | 0 | 0 | 1540271 | P0DTD1.1 | P0DTD1 |
| FKWDLTAFGLVAEWF | 1 | 0 | 0 | 1540274 | P0DTC1.1 | P0DTD1 |
| FLAHIQWMVMFTPLV | 1 | 0 | 0 | 1540275 | P0DTC1.1 | P0DTD1 |
| FLALITLATCELYHY | 1 | 0 | 0 | 1540276 | P0DTC7.1 | P0DTC7 |
| FLKKDAPYIVGDVVQ | 1 | 0 | 0 | 1540285 | P0DTC1.1 | P0DTD1 |
| FLPFFSNVTWFHAIH | 1 | 0 | 0 | 1540289 | YP_009724390.1 | P0DTC2 |
| FLPGVYSVIYLYLTF | 1 | 0 | 0 | 1540290 | P0DTC1.1 | P0DTD1 |
| FLTENLLLYIDINGN | 1 | 0 | 0 | 1540292 | P0DTC1.1 | P0DTD1 |
| FNEKTHVQLSLPVLQ | 1 | 0 | 0 | 1540298 | P0DTC1.1 | P0DTD1 |
| FNGECPNFVFPLNSI | 1 | 0 | 0 | 1540299 | P0DTC1.1 | P0DTD1 |
| FNQHEVLLAPLLSAG | 1 | 0 | 0 | 1540304 | P0DTC1.1 | P0DTD1 |
| FNSVCRLMKTIGPDM | 1 | 0 | 0 | 1540306 | P0DTD1.1 | P0DTD1 |
| FPKSDGTGTIYTELE | 1 | 0 | 0 | 1540313 | P0DTC1.1 | P0DTD1 |
| FPLKLRGTAVMSLKE | 1 | 0 | 0 | 1540314 | P0DTD1.1 | P0DTD1 |
| FRELGVVHNQDVNLH | 1 | 0 | 0 | 1540320 | P0DTD1.1 | P0DTD1 |
| FRIDGDMVPHISRQR | 1 | 0 | 0 | 1540321 | P0DTD1.1 | P0DTD1 |
| FRKMAFPSGKVEGCM | 1 | 0 | 0 | 1540322 | P0DTC1.1 | P0DTD1 |
| FRNARNGVLITEGSV | 1 | 0 | 0 | 1540325 | P0DTD1.1 | P0DTD1 |
| FSRTLETAQNSVRVL | 1 | 0 | 0 | 1540334 | P0DTC1.1 | P0DTD1 |
| FSSLPSYAAFATAQE | 1 | 0 | 0 | 1540337 | P0DTC1.1 | P0DTD1 |
| FSTFEEAALCTFLLN | 1 | 0 | 0 | 1540341 | P0DTC1.1 | P0DTD1 |
| FSTGVNLVAVPTGYV | 1 | 0 | 0 | 1540342 | P0DTD1.1 | P0DTD1 |
| FTDGVCLFWNCNVDR | 1 | 0 | 0 | 1540344 | P0DTD1.1 | P0DTD1 |
| FTSDLATNNLVVMAY | 1 | 0 | 0 | 1540355 | P0DTC1.1 | P0DTD1 |
| FTTTLNDFNLVAMKY | 1 | 0 | 0 | 1540357 | P0DTC1.1 | P0DTD1 |
| FVKATCEFCGTENLT | 1 | 0 | 0 | 1540364 | P0DTC1.1 | P0DTD1 |
| FVMMSAPPAQYELKH | 1 | 0 | 0 | 1540368 | P0DTC1.1 | P0DTD1 |
| FVVKRHTFSNYQHEE | 1 | 0 | 0 | 1540373 | P0DTD1.1 | P0DTD1 |
| FWITIAYIICISTKH | 1 | 0 | 0 | 1540376 | P0DTC1.1 | P0DTD1 |
| FYEDFLEYHDVRVVL | 1 | 0 | 0 | 1540379 | P0DTC8.1 | P0DTC8 |
| FYFYTSKTTVASLIN | 1 | 0 | 0 | 1540380 | P0DTC1.1 | P0DTD1 |
| FYLTNDVSFLAHIQW | 1 | 0 | 0 | 1540383 | P0DTC1.1 | P0DTD1 |
| FYRLANECAQVLSEM | 1 | 0 | 0 | 1540385 | P0DTD1.1 | P0DTD1 |
| GAKLLHKPIVWHVNN | 1 | 0 | 0 | 1540394 | P0DTC1.1 | P0DTD1 |

| | | | | | | |
|---|---|---|---|---|---|---|
| GAVCRHHANEYRLYL | 1 | 0 | 0 | 1540401 | P0DTD1.1 | P0DTD1 |
| GAYTRYVDNNFCGPD | 1 | 0 | 0 | 1540402 | P0DTC1.1 | P0DTD1 |
| GCDGGSLYVNKHAFH | 1 | 0 | 0 | 1540403 | P0DTD1.1 | P0DTD1 |
| GCNYLGKPREQIDGY | 1 | 0 | 0 | 1540409 | P0DTD1.1 | P0DTD1 |
| GDCEEEEFEPSTQYE | 1 | 0 | 0 | 1540412 | P0DTC1.1 | P0DTD1 |
| GDFVKATCEFCGTEN | 1 | 0 | 0 | 1540417 | P0DTC1.1 | P0DTD1 |
| GDRWFLNRFTTTLND | 1 | 0 | 0 | 1540422 | P0DTC1.1 | P0DTD1 |
| GEFKLASHMYCSFYP | 1 | 0 | 0 | 1540429 | P0DTC1.1 | P0DTD1 |
| GEGSEGLNDNLLEIL | 1 | 0 | 0 | 1540430 | P0DTC1.1 | P0DTD1 |
| GEYSHVVAFNTLLFL | 1 | 0 | 0 | 1540440 | P0DTC1.1 | P0DTD1 |
| GFAKFLKTNCCRFQE | 1 | 0 | 0 | 1540441 | P0DTD1.1 | P0DTD1 |
| GFSLWVYKQFDTYNL | 1 | 0 | 0 | 1540451 | P0DTD1.1 | P0DTD1 |
| GFVDSDVETKDVVEC | 1 | 0 | 0 | 1540453 | P0DTC1.1 | P0DTD1 |
| GGASCCLYCRCHIDH | 1 | 0 | 0 | 1540455 | P0DTC1.1 | P0DTD1 |
| GGIVAIVVTCLAYYF | 1 | 0 | 0 | 1540459 | P0DTC1.1 | P0DTD1 |
| GGLHLLIGLAKRFKE | 1 | 0 | 0 | 1540463 | P0DTD1.1 | P0DTD1 |
| GGSVAIKITEHSWNA | 1 | 0 | 0 | 1540470 | P0DTD1.1 | P0DTD1 |
| GGWHNMLKTVYSDVE | 1 | 0 | 0 | 1540474 | P0DTD1.1 | P0DTD1 |
| GHLRIAGHHLGRCDI | 1 | 0 | 0 | 1540477 | P0DTC5.1 | P0DTC5 |
| GHVMVELVAELEGIQ | 1 | 0 | 0 | 1540480 | P0DTC1.1 | P0DTD1 |
| GIDLDEWSMATYYLF | 1 | 0 | 0 | 1540485 | P0DTC1.1 | P0DTD1 |
| GKARLYYDSMSYEDQ | 1 | 0 | 0 | 1540494 | P0DTD1.1 | P0DTD1 |
| GLHPTQAPTHLSVDT | 1 | 0 | 0 | 1540505 | P0DTD1.1 | P0DTD1 |
| GLKTILRKGGRTIAF | 1 | 0 | 0 | 1540507 | P0DTC1.1 | P0DTD1 |
| GLLMPLKAPKEIIFL | 1 | 0 | 0 | 1540509 | P0DTC1.1 | P0DTD1 |
| GMWKGYGCSCDQLRE | 1 | 0 | 0 | 1540529 | P0DTC1.1 | P0DTD1 |
| GNPKAIKCVPQADVE | 1 | 0 | 0 | 1540537 | P0DTD1.1 | P0DTD1 |
| GNVQLRVIGHSMQNC | 1 | 0 | 0 | 1540539 | P0DTC1.1 | P0DTD1 |
| GPDGYPLECIKDLLA | 1 | 0 | 0 | 1540542 | P0DTC1.1 | P0DTD1 |
| GPKQASLNGVTLIGE | 1 | 0 | 0 | 1540547 | P0DTD1.1 | P0DTD1 |
| GPLSAQTGIAVLDMC | 1 | 0 | 0 | 1540549 | P0DTC1.1 | P0DTD1 |
| GRIRSVYPVASPNEC | 1 | 0 | 0 | 1540559 | P0DTC1.1 | P0DTD1 |
| GSEYDYVIFTQTTET | 1 | 0 | 0 | 1540566 | P0DTD1.1 | P0DTD1 |
| GSSKCVCSVIDLLLD | 1 | 0 | 0 | 1540590 | P0DTD1.1 | P0DTD1 |
| GTTTLNGLWLDDVVY | 1 | 0 | 0 | 1540614 | P0DTC1.1 | P0DTD1 |
| GVAMPNLYKMQRMLL | 1 | 0 | 0 | 1540621 | P0DTD1.1 | P0DTD1 |
| GVPVVDSYYSLLMPI | 1 | 0 | 0 | 1540633 | P0DTD1.1 | P0DTD1 |
| HANLDSCKRVLNVVC | 1 | 0 | 0 | 1540660 | P0DTC1.1 | P0DTD1 |
| HECFVKRVDWTIEYP | 1 | 0 | 0 | 1540668 | P0DTD1.1 | P0DTD1 |
| HEFCSQHTMLVKQGD | 1 | 0 | 0 | 1540670 | P0DTD1.1 | P0DTD1 |
| HEHEIAWYTERSEKS | 1 | 0 | 0 | 1540671 | P0DTC1.1 | P0DTD1 |
| HEIAWYTERSEKSYE | 1 | 0 | 0 | 1540672 | P0DTC1.1 | P0DTD1 |
| HFSMMILSDDAVVCF | 1 | 0 | 0 | 1540677 | P0DTD1.1 | P0DTD1 |
| HINAQVAKSHNIALI | 1 | 0 | 0 | 1540687 | P0DTC1.1 | P0DTD1 |
| HLLLVAAGLEAPFLY | 1 | 0 | 0 | 1540692 | P0DTC3.1 | P0DTC3 |
| HLSWEVGKPRPPLNR | 1 | 0 | 0 | 1540693 | P0DTD1.1 | P0DTD1 |
| HLYFDKAGQKTYERH | 1 | 0 | 0 | 1540694 | P0DTC1.1 | P0DTD1 |
| HNESGLKTILRKGGR | 1 | 0 | 0 | 1540698 | P0DTC1.1 | P0DTD1 |
| HPDSATLVSDIDITF | 1 | 0 | 0 | 1540703 | P0DTC1.1 | P0DTD1 |
| HQPYVVDDPCPIHFY | 1 | 0 | 0 | 1540707 | P0DTC8.1 | P0DTC8 |
| HRFYRLANECAQVLS | 1 | 0 | 0 | 1540708 | P0DTD1.1 | P0DTD1 |
| HSIGFDYVYNPFMID | 1 | 0 | 0 | 1540711 | P0DTD1.1 | P0DTD1 |
| HSKGLYRKCVKSREE | 1 | 0 | 0 | 1540712 | P0DTC1.1 | P0DTD1 |
| HSLRVCVDTVRTNVY | 1 | 0 | 0 | 1540714 | P0DTC1.1 | P0DTD1 |
| HSYGADLKSFDLGDE | 1 | 0 | 0 | 1540718 | P0DTC1.1 | P0DTD1 |
| HTTCCSLSHRFYRLA | 1 | 0 | 0 | 1540726 | P0DTD1.1 | P0DTD1 |
| HVGEIPVAYRKVLLR | 1 | 0 | 0 | 1540728 | P0DTC1.1 | P0DTD1 |
| HVICTSEDMLNPNYE | 1 | 0 | 0 | 1540729 | P0DTC1.1 | P0DTD1 |
| HVTFFIYNKIVDEPE | 1 | 0 | 0 | 1540732 | P0DTC3.1 | P0DTC3 |
| HVVAFNTLLFLMSFT | 1 | 0 | 0 | 1540733 | P0DTC1.1 | P0DTD1 |
| HYTPSFKKGAKLLHK | 1 | 0 | 0 | 1540735 | P0DTC1.1 | P0DTD1 |
| IAAVITREVGFVVPG | 1 | 0 | 0 | 1540738 | P0DTC1.1 | P0DTD1 |
| IAFGGCVFSYVGCHN | 1 | 0 | 0 | 1540739 | P0DTC1.1 | P0DTD1 |
| IAKNTVKSVGKFCLE | 1 | 0 | 0 | 1540740 | P0DTC1.1 | P0DTD1 |
| IALKGGKIVNNWLKQ | 1 | 0 | 0 | 1540741 | P0DTC1.1 | P0DTD1 |
| IASEFSSLPSYAAFA | 1 | 0 | 0 | 1540744 | P0DTC1.1 | P0DTD1 |
| IASTDTCFANKHADF | 1 | 0 | 0 | 1540745 | P0DTC1.1 | P0DTD1 |
| IATYCTGSIPCSVCL | 1 | 0 | 0 | 1540746 | P0DTC1.1 | P0DTD1 |
| IAVLDMCASLKELLQ | 1 | 0 | 0 | 1540747 | P0DTC1.1 | P0DTD1 |
| IAWYTERSEKSYELQ | 1 | 0 | 0 | 1540748 | P0DTC1.1 | P0DTD1 |
| IAYIICISTKHFYWF | 1 | 0 | 0 | 1540749 | P0DTC1.1 | P0DTD1 |
| ICISTKHFYWFFSNY | 1 | 0 | 0 | 1540750 | P0DTC1.1 | P0DTD1 |
| ICQAVTANVNALLST | 1 | 0 | 0 | 1540751 | P0DTD1.1 | P0DTD1 |
| IDGDMVPHISRQRLT | 1 | 0 | 0 | 1540756 | P0DTD1.1 | P0DTD1 |
| IDGGVTRDIASTDTC | 1 | 0 | 0 | 1540757 | P0DTC1.1 | P0DTD1 |
| IDIGNYTVSCLPFTI | 1 | 0 | 0 | 1540760 | P0DTC8.1 | P0DTC8 |
| IDSYFVVKRHTFSNY | 1 | 0 | 0 | 1540763 | P0DTD1.1 | P0DTD1 |
| IERFVSLAIDAYPLT | 1 | 0 | 0 | 1540771 | P0DTD1.1 | P0DTD1 |
| IERYKLEGYAFEHIV | 1 | 0 | 0 | 1540772 | P0DTD1.1 | P0DTD1 |
| IETISLAGSYKDWSY | 1 | 0 | 0 | 1540773 | P0DTC1.1 | P0DTD1 |
| IFFITGNTLQCIMLV | 1 | 0 | 0 | 1540774 | P0DTC1.1 | P0DTD1 |
| IFGTVYEKLKPVLDW | 1 | 0 | 0 | 1540775 | P0DTC1.1 | P0DTD1 |
| IFLEGETLPTEVLTE | 1 | 0 | 0 | 1540776 | P0DTC1.1 | P0DTD1 |
| IGALDISASIVAGGI | 1 | 0 | 0 | 1540779 | P0DTC1.1 | P0DTD1 |
| IGDELKINAACRKVQ | 1 | 0 | 0 | 1540781 | P0DTD1.1 | P0DTD1 |
| IGFDVEGCHATREAV | 1 | 0 | 0 | 1540783 | P0DTD1.1 | P0DTD1 |
| IGPDMFLGTCRRCPA | 1 | 0 | 0 | 1540786 | P0DTD1.1 | P0DTD1 |
| IGPERTCCLCDRRAT | 1 | 0 | 0 | 1540787 | P0DTD1.1 | P0DTD1 |
| IHFGAGSDKGVAPGT | 1 | 0 | 0 | 1540788 | P0DTD1.1 | P0DTD1 |
| IIAMSAFAMMFVKHK | 1 | 0 | 0 | 1540789 | P0DTC1.1 | P0DTD1 |
| IIKTIQPRVEKKKLD | 1 | 0 | 0 | 1540793 | P0DTC1.1 | P0DTD1 |

| | | | | | | |
|---|---|---|---|---|---|---|
| IINLIIKNLSKSLTE | 1 | 0 | 0 | 1540795 | P0DTC6.1 | P0DTC6 |
| IINLVQMAPISAMVR | 1 | 0 | 0 | 1540796 | P0DTC1.1 | P0DTD1 |
| IKASMPTTIAKNTVK | 1 | 0 | 0 | 1540802 | P0DTC1.1 | P0DTD1 |
| IKGTHHWLLLTILTS | 1 | 0 | 0 | 1540803 | P0DTC1.1 | P0DTD1 |
| IKNADIVEEAKKVKP | 1 | 0 | 0 | 1540806 | P0DTC1.1 | P0DTD1 |
| IKVTLVFLFVAAIFY | 1 | 0 | 0 | 1540809 | P0DTC1.1 | P0DTD1 |
| IKWDLLKYDFTEERL | 1 | 0 | 0 | 1540810 | P0DTD1.1 | P0DTD1 |
| ILFLALITLATCELY | 1 | 0 | 0 | 1540813 | P0DTC7.1 | P0DTC7 |
| ILGPLSAQTGIAVLD | 1 | 0 | 0 | 1540815 | P0DTC1.1 | P0DTD1 |
| ILKPANNSLKITEEV | 1 | 0 | 0 | 1540817 | P0DTC1.1 | P0DTD1 |
| ILLAKDTTEAFEKMV | 1 | 0 | 0 | 1540818 | P0DTC1.1 | P0DTD1 |
| ILRKGGRTIAFGGCV | 1 | 0 | 0 | 1540821 | P0DTC1.1 | P0DTD1 |
| ILTLTRALTAESHVD | 1 | 0 | 0 | 1540822 | P0DTD1.1 | P0DTD1 |
| IMASLVLARKHTTCC | 1 | 0 | 0 | 1540823 | P0DTD1.1 | P0DTD1 |
| IMMNVAKYTQLCQYL | 1 | 0 | 0 | 1540824 | P0DTD1.1 | P0DTD1 |
| IMRTFKVSIWNLDYI | 1 | 0 | 0 | 1540825 | P0DTC6.1 | P0DTC6 |
| IMTRCLAVHECFVKR | 1 | 0 | 0 | 1540826 | P0DTD1.1 | P0DTD1 |
| INAACRKVQHMVVKA | 1 | 0 | 0 | 1540827 | P0DTD1.1 | P0DTD1 |
| INRPQIGVVREFLTR | 1 | 0 | 0 | 1540831 | P0DTD1.1 | P0DTD1 |
| IPCSVCLSGLDSLDT | 1 | 0 | 0 | 1540835 | P0DTC1.1 | P0DTD1 |
| IPKDMTYRRLISMMG | 1 | 0 | 0 | 1540838 | P0DTD1.1 | P0DTD1 |
| IQFPNTYLEGSVRVV | 1 | 0 | 0 | 1540846 | P0DTC1.1 | P0DTD1 |
| IQLLKSAYENFNQHE | 1 | 0 | 0 | 1540847 | P0DTC1.1 | P0DTD1 |
| IQRKYKGIKIQEGVV | 1 | 0 | 0 | 1540849 | P0DTC1.1 | P0DTD1 |
| IQTIVEVQPQLEMEL | 1 | 0 | 0 | 1540850 | P0DTC1.1 | P0DTD1 |
| IRKLHDELTGHMLDM | 1 | 0 | 0 | 1540854 | P0DTD1.1 | P0DTD1 |
| IRKSNHNFLVQAGNV | 1 | 0 | 0 | 1540855 | P0DTC1.1 | P0DTD1 |
| IRQEEVQELYSPIFL | 1 | 0 | 0 | 1540856 | P0DTC7.1 | P0DTC7 |
| IRRPFLCCKCCYDHV | 1 | 0 | 0 | 1540859 | P0DTD1.1 | P0DTD1 |
| IRSVYPVASPNECNQ | 1 | 0 | 0 | 1540860 | P0DTC1.1 | P0DTD1 |
| ISAARQGFVDSDVET | 1 | 0 | 0 | 1540861 | P0DTC1.1 | P0DTD1 |
| ISAKNRARTVAGVSI | 1 | 0 | 0 | 1540862 | P0DTD1.1 | P0DTD1 |
| ISMDNSPNLAWPLIV | 1 | 0 | 0 | 1540867 | P0DTC1.1 | P0DTD1 |
| ISNEKQEILGTVSWN | 1 | 0 | 0 | 1540869 | P0DTC1.1 | P0DTD1 |
| ISRQRLTKYTMADLV | 1 | 0 | 0 | 1540870 | P0DTD1.1 | P0DTD1 |
| ISTCACEIVGGQIVT | 1 | 0 | 0 | 1540872 | P0DTC1.1 | P0DTD1 |
| ISTIGVCSMTDIAKK | 1 | 0 | 0 | 1540873 | P0DTD1.1 | P0DTD1 |
| ISTKHFYWFFSNYLK | 1 | 0 | 0 | 1540874 | P0DTC1.1 | P0DTD1 |
| ISVTSNYSGVVTTVM | 1 | 0 | 0 | 1540875 | P0DTC1.1 | P0DTD1 |
| ITEEVGHTDLMAAYV | 1 | 0 | 0 | 1540877 | P0DTC1.1 | P0DTD1 |
| ITGLHPTQAPTHLSV | 1 | 0 | 0 | 1540878 | P0DTD1.1 | P0DTD1 |
| ITHDVSSAINRPQIG | 1 | 0 | 0 | 1540879 | P0DTD1.1 | P0DTD1 |
| ITIAYIICISTKHFY | 1 | 0 | 0 | 1540880 | P0DTC1.1 | P0DTD1 |
| ITLATCELYHYQECV | 1 | 0 | 0 | 1540881 | P0DTC7.1 | P0DTC7 |
| ITPVHVMSKHTDFSS | 1 | 0 | 0 | 1540885 | P0DTC1.1 | P0DTD1 |
| ITTVAAFHQECSLQS | 1 | 0 | 0 | 1540889 | P0DTC8.1 | P0DTC8 |
| ITTYPGQGLNGYTVE | 1 | 0 | 0 | 1540890 | P0DTC1.1 | P0DTD1 |
| IVCRFDTRVLSNLNL | 1 | 0 | 0 | 1540894 | P0DTD1.1 | P0DTD1 |
| IVDSVTVKNGSIHLY | 1 | 0 | 0 | 1540895 | P0DTC1.1 | P0DTD1 |
| IVESCGNFKVTKGKA | 1 | 0 | 0 | 1540897 | P0DTC1.1 | P0DTD1 |
| IVFMCVEYCPIFFIT | 1 | 0 | 0 | 1540898 | P0DTC1.1 | P0DTD1 |
| IVGDVVQEGVLTAVV | 1 | 0 | 0 | 1540899 | P0DTC1.1 | P0DTD1 |
| IVKFISTCACEIVGG | 1 | 0 | 0 | 1540902 | P0DTC1.1 | P0DTD1 |
| IVMVTIMLCCMTSCC | 1 | 0 | 0 | 1540903 | P0DTC2.1 | P0DTC2 |
| IVSTIQRKYKGIKIQ | 1 | 0 | 0 | 1540908 | P0DTC1.1 | P0DTD1 |
| IVTRCLNRVCTNYMP | 1 | 0 | 0 | 1540910 | P0DTC1.1 | P0DTD1 |
| IWNVKDFMSLSEQLR | 1 | 0 | 0 | 1540913 | P0DTC1.1 | P0DTD1 |
| IYCPACHNSEVGPEH | 1 | 0 | 0 | 1540914 | P0DTC1.1 | P0DTD1 |
| IYLYLTFYLTNDVSF | 1 | 0 | 0 | 1540916 | P0DTC1.1 | P0DTD1 |
| IYSKHTPINLVRDLP | 1 | 0 | 0 | 1540918 | P0DTC2.1 | P0DTC2 |
| IYSTAALGVLMSNLG | 1 | 0 | 0 | 1540919 | P0DTC1.1 | P0DTD1 |
| KAAITILDGISQYSL | 1 | 0 | 0 | 1540920 | P0DTC1.1 | P0DTD1 |
| KAIVSTIQRKYKGIK | 1 | 0 | 0 | 1540923 | P0DTC1.1 | P0DTD1 |
| KALNDFSNSGSDVLY | 1 | 0 | 0 | 1540925 | P0DTC1.1 | P0DTD1 |
| KASCTLSEQLDFIDT | 1 | 0 | 0 | 1540927 | P0DTC1.1 | P0DTD1 |
| KATCEFCGTENLTKE | 1 | 0 | 0 | 1540928 | P0DTC1.1 | P0DTD1 |
| KATYKPNTWCIRCLW | 1 | 0 | 0 | 1540929 | P0DTC1.1 | P0DTD1 |
| KCDHCGETSWQTGDF | 1 | 0 | 0 | 1540934 | P0DTC1.1 | P0DTD1 |
| KCKSAFYILPSIISN | 1 | 0 | 0 | 1540937 | P0DTC1.1 | P0DTD1 |
| KCSAYTVELGTEVNE | 1 | 0 | 0 | 1540938 | P0DTC1.1 | P0DTD1 |
| KDFMSLSEQLRKQIR | 1 | 0 | 0 | 1540945 | P0DTC1.1 | P0DTD1 |
| KDTTEAFEKMVSLLS | 1 | 0 | 0 | 1540954 | P0DTC1.1 | P0DTD1 |
| KFLALCADSIIIGGA | 1 | 0 | 0 | 1540972 | P0DTC1.1 | P0DTD1 |
| KFLVFLGIITTVAAF | 1 | 0 | 0 | 1540973 | P0DTC8.1 | P0DTC8 |
| KGGRFVLALLSDLQD | 1 | 0 | 0 | 1540981 | P0DTC1.1 | P0DTD1 |
| KGLPWNVVRIKIVQM | 1 | 0 | 0 | 1540987 | P0DTD1.1 | P0DTD1 |
| KHCLHVVGPNVNKGE | 1 | 0 | 0 | 1540998 | P0DTC1.1 | P0DTD1 |
| KHVYQLRARSVSPKL | 1 | 0 | 0 | 1541006 | P0DTC7.1 | P0DTC7 |
| KIILFLALITLATCE | 1 | 0 | 0 | 1541011 | P0DTC7.1 | P0DTC7 |
| KIKPHNSHEGKTFYV | 1 | 0 | 0 | 1541012 | P0DTC1.1 | P0DTD1 |
| KILGLPTQTVDSSQG | 1 | 0 | 0 | 1541013 | P0DTD1.1 | P0DTD1 |
| KIQEGVVDYGARFYF | 1 | 0 | 0 | 1541015 | P0DTC1.1 | P0DTD1 |
| KIVQMLSDTLKNLSD | 1 | 0 | 0 | 1541020 | P0DTD1.1 | P0DTD1 |
| KKKLDGFMGRIRSVY | 1 | 0 | 0 | 1541029 | P0DTC1.1 | P0DTD1 |
| KKWKYPQVNGLTSIK | 1 | 0 | 0 | 1541039 | P0DTC1.1 | P0DTD1 |
| KLALGGSVAIKITEH | 1 | 0 | 0 | 1541041 | P0DTD1.1 | P0DTD1 |
| KLDGFMGRIRSVYPV | 1 | 0 | 0 | 1541044 | P0DTC1.1 | P0DTD1 |
| KLEGYAFEHIVYGDF | 1 | 0 | 0 | 1541045 | P0DTD1.1 | P0DTD1 |
| KLGSLVVRCSFYEDF | 1 | 0 | 0 | 1541047 | P0DTC8.1 | P0DTC8 |
| KLNDLCFTNVYADSFVIRGD | 1 | 0 | 0 | 1541057 | QHD43416.1 | P0DTC2 |
| KLQSSQAWQPGVAMP | 1 | 0 | 0 | 1541063 | P0DTD1.1 | P0DTD1 |

| | | | | | | |
|---|---|---|---|---|---|---|
| KLSHQSDIEVTGDSC | 1 | 0 | 0 | 1541065 | P0DTC1.1 | P0DTD1 |
| KLWAQCVQLHNDILL | 1 | 0 | 0 | 1541070 | P0DTC1.1 | P0DTD1 |
| KMQRMLLEKCDLQNY | 1 | 0 | 0 | 1541076 | P0DTD1.1 | P0DTD1 |
| KNLSDRVVFVLWAHG | 1 | 0 | 0 | 1541081 | P0DTD1.1 | P0DTD1 |
| KNVSLDNVLSTFISA | 1 | 0 | 0 | 1541089 | P0DTC1.1 | P0DTD1 |
| KPANNSLKITEEVGH | 1 | 0 | 0 | 1541091 | P0DTC1.1 | P0DTD1 |
| KPLEFGATSAALQPE | 1 | 0 | 0 | 1541096 | P0DTC1.1 | P0DTD1 |
| KPRSQMEIDFLELAM | 1 | 0 | 0 | 1541100 | P0DTD1.1 | P0DTD1 |
| KPTETICAPLTVFFD | 1 | 0 | 0 | 1541101 | P0DTD1.1 | P0DTD1 |
| KQARSEDKRAKVTSA | 1 | 0 | 0 | 1541105 | P0DTC1.1 | P0DTD1 |
| KQEILGTVSWNLREM | 1 | 0 | 0 | 1541109 | P0DTC1.1 | P0DTD1 |
| KRNRATRVECTTIVN | 1 | 0 | 0 | 1541118 | P0DTC1.1 | P0DTD1 |
| KRSDARTAPHGHVMV | 1 | 0 | 0 | 1541123 | P0DTC1.1 | P0DTD1 |
| KSDGTGTIYTELEPP | 1 | 0 | 0 | 1541132 | P0DTC1.1 | P0DTD1 |
| KSEKQVEQKIAEIPK | 1 | 0 | 0 | 1541133 | P0DTC1.1 | P0DTD1 |
| KTFYVLPNDDTLRVE | 1 | 0 | 0 | 1541143 | P0DTC1.1 | P0DTD1 |
| KVKYLYFIKGLNNLN | 1 | 0 | 0 | 1541161 | P0DTC1.1 | P0DTD1 |
| KVNINIVGDFKLNEE | 1 | 0 | 0 | 1541163 | P0DTC1.1 | P0DTD1 |
| KVVKVTIDYTEISFM | 1 | 0 | 0 | 1541169 | P0DTD1.1 | P0DTD1 |
| KWGKARLYYDSMSYE | 1 | 0 | 0 | 1541175 | P0DTD1.1 | P0DTD1 |
| KYVRNLQHRLYECLY | 1 | 0 | 0 | 1541183 | P0DTD1.1 | P0DTD1 |
| LAIDAYPLTKHPNQE | 1 | 0 | 0 | 1541189 | P0DTD1.1 | P0DTD1 |
| LAKNVSLDNVLSTFI | 1 | 0 | 0 | 1541192 | P0DTC1.1 | P0DTD1 |
| LALCADSIIIGGAKL | 1 | 0 | 0 | 1541194 | P0DTC1.1 | P0DTD1 |
| LALSKGVHFVCNLLL | 1 | 0 | 0 | 1541195 | P0DTC3.1 | P0DTC3 |
| LALYNKYKYFSGAMD | 1 | 0 | 0 | 1541196 | P0DTC1.1 | P0DTD1 |
| LANTCTERLKLFAAE | 1 | 0 | 0 | 1541198 | P0DTD1.1 | P0DTD1 |
| LARKHTTCCSLSHRF | 1 | 0 | 0 | 1541201 | P0DTD1.1 | P0DTD1 |
| LATCELYHYQECVRG | 1 | 0 | 0 | 1541205 | P0DTC7.1 | P0DTC7 |
| LATHGLAAVNSVPWD | 1 | 0 | 0 | 1541206 | P0DTC1.1 | P0DTD1 |
| LATNNLVVMAYITGG | 1 | 0 | 0 | 1541207 | P0DTC1.1 | P0DTD1 |
| LAYYFMRFRRAFGEY | 1 | 0 | 0 | 1541210 | P0DTC1.1 | P0DTD1 |
| LCADSIIIGGAKLKA | 1 | 0 | 0 | 1541211 | P0DTC1.1 | P0DTD1 |
| LCEEMLDNRATLQAI | 1 | 0 | 0 | 1541214 | P0DTC1.1 | P0DTD1 |
| LCTFLLNKEMYLKLR | 1 | 0 | 0 | 1541219 | P0DTC1.1 | P0DTD1 |
| LCTHTGTGQAITVTP | 1 | 0 | 0 | 1541220 | P0DTC1.1 | P0DTD1 |
| LCWHTNCYDYCIPYN | 1 | 0 | 0 | 1541222 | P0DTC3.1 | P0DTC3 |
| LDDRCILHCANFNVL | 1 | 0 | 0 | 1541226 | P0DTD1.1 | P0DTD1 |
| LDGVVCTEIDPKLDN | 1 | 0 | 0 | 1541230 | P0DTC1.1 | P0DTD1 |
| LDNVLSTFISAARQG | 1 | 0 | 0 | 1541234 | P0DTC1.1 | P0DTD1 |
| LDTYPSLETIQITIS | 1 | 0 | 0 | 1541238 | P0DTC1.1 | P0DTD1 |
| LEFGATSAALQPEEE | 1 | 0 | 0 | 1541244 | P0DTC1.1 | P0DTD1 |
| LEGETLPTEVLTEEV | 1 | 0 | 0 | 1541245 | P0DTC1.1 | P0DTD1 |
| LEGIQYGRSGETLGV | 1 | 0 | 0 | 1541246 | P0DTC1.1 | P0DTD1 |
| LEKCDLQNYGDSATL | 1 | 0 | 0 | 1541250 | P0DTD1.1 | P0DTD1 |
| LEMELTPVVQTIEVN | 1 | 0 | 0 | 1541252 | P0DTC1.1 | P0DTD1 |
| LEQPYVFIKRSDART | 1 | 0 | 0 | 1541256 | P0DTC1.1 | P0DTD1 |
| LETIQITISSFKWDL | 1 | 0 | 0 | 1541259 | P0DTC1.1 | P0DTD1 |
| LFAAETLKATEETFK | 1 | 0 | 0 | 1541260 | P0DTD1.1 | P0DTD1 |
| LFIRQEEVQELYSPI | 1 | 0 | 0 | 1541265 | P0DTC7.1 | P0DTC7 |
| LGAGCFVDDIVKTDG | 1 | 0 | 0 | 1541277 | P0DTD1.1 | P0DTD1 |
| LGGAVCRHHANEYRL | 1 | 0 | 0 | 1541280 | P0DTD1.1 | P0DTD1 |
| LGKYEQYIKWPWYIW | 1 | 0 | 0 | 1541283 | P0DTC2.1 | P0DTC2 |
| LGTEVNEFACVVADA | 1 | 0 | 0 | 1541288 | P0DTC1.1 | P0DTD1 |
| LGTVSWNLREMLAHA | 1 | 0 | 0 | 1541289 | P0DTC1.1 | P0DTD1 |
| LGVLVPHVGEIPVAY | 1 | 0 | 0 | 1541291 | P0DTC1.1 | P0DTD1 |
| LHFLPRVFSAVGNIC | 1 | 0 | 0 | 1541295 | P0DTC1.1 | P0DTD1 |
| LHKPIVWHVNNATNK | 1 | 0 | 0 | 1541296 | P0DTC1.1 | P0DTD1 |
| LHNDILLAKDTTEAF | 1 | 0 | 0 | 1541298 | P0DTC1.1 | P0DTD1 |
| LIDAMMFTSDLATNN | 1 | 0 | 0 | 1541301 | P0DTC1.1 | P0DTD1 |
| LIEYTDFATSACVLA | 1 | 0 | 0 | 1541302 | P0DTC1.1 | P0DTD1 |
| LIGCNYLGKPREQID | 1 | 0 | 0 | 1541303 | P0DTD1.1 | P0DTD1 |
| LILMTARTVYDDGAR | 1 | 0 | 0 | 1541306 | P0DTC1.1 | P0DTD1 |
| LIQPIGALDISASIV | 1 | 0 | 0 | 1541310 | P0DTC1.1 | P0DTD1 |
| LKDGTCGLVEVEKGV | 1 | 0 | 0 | 1541316 | P0DTC1.1 | P0DTD1 |
| LKEGQINDMILSLLS | 1 | 0 | 0 | 1541317 | P0DTD1.1 | P0DTD1 |
| LKGVEAVMYMGTLSY | 1 | 0 | 0 | 1541322 | P0DTC1.1 | P0DTD1 |
| LKINAACRKVQHMVV | 1 | 0 | 0 | 1541325 | P0DTD1.1 | P0DTD1 |
| LKLFDRYFKYWDQTY | 1 | 0 | 0 | 1541330 | P0DTD1.1 | P0DTD1 |
| LKLRGTAVMSLKEGQ | 1 | 0 | 0 | 1541332 | P0DTD1.1 | P0DTD1 |
| LKQLPFFYYSDSPCE | 1 | 0 | 0 | 1541338 | P0DTD1.1 | P0DTD1 |
| LKSFDLGDELGTDPY | 1 | 0 | 0 | 1541341 | P0DTC1.1 | P0DTD1 |
| LLARAGKASCTLSEQ | 1 | 0 | 0 | 1541351 | P0DTC1.1 | P0DTD1 |
| LLDKRTTCFSVAALT | 1 | 0 | 0 | 1541353 | P0DTD1.1 | P0DTD1 |
| LLKSAYENFNQHEVL | 1 | 0 | 0 | 1541363 | P0DTC1.1 | P0DTD1 |
| LLLLFVTVYSHLLLV | 1 | 0 | 0 | 1541369 | P0DTC3.1 | P0DTC3 |
| LLMPILTLTRALTAE | 1 | 0 | 0 | 1541371 | P0DTD1.1 | P0DTD1 |
| LLNKEMYLKLRSDVL | 1 | 0 | 0 | 1541372 | P0DTC1.1 | P0DTD1 |
| LLNVPLHGTILTRPL | 1 | 0 | 0 | 1541374 | P0DTC5.1 | P0DTC5 |
| LLPLTQYNRYLALYN | 1 | 0 | 0 | 1541376 | P0DTC1.1 | P0DTD1 |
| LLPSLATVAYFNMVY | 1 | 0 | 0 | 1541378 | P0DTC1.1 | P0DTD1 |
| LLQLCTFTRSTNSRI | 1 | 0 | 0 | 1541379 | P0DTC1.1 | P0DTD1 |
| LLSVLQQLRVESSSK | 1 | 0 | 0 | 1541386 | P0DTC1.1 | P0DTD1 |
| LLVAAGLEAPFLYLY | 1 | 0 | 0 | 1541392 | P0DTC3.1 | P0DTC3 |
| LMCQPILLLDQALVS | 1 | 0 | 0 | 1541399 | P0DTC1.1 | P0DTD1 |
| LMDGSIIQFPNTYLE | 1 | 0 | 0 | 1541400 | P0DTC1.1 | P0DTD1 |
| LMPLKAPKEIIFLEG | 1 | 0 | 0 | 1541405 | P0DTC1.1 | P0DTD1 |
| LMPVCVETKAIVSTI | 1 | 0 | 0 | 1541406 | P0DTC1.1 | P0DTD1 |
| LMWLIINLVQMAPIS | 1 | 0 | 0 | 1541409 | P0DTC1.1 | P0DTD1 |
| LNGLWLDDVVYCPRH | 1 | 0 | 0 | 1541416 | P0DTC1.1 | P0DTD1 |
| LNGYTVEEAKTVLKK | 1 | 0 | 0 | 1541420 | P0DTC1.1 | P0DTD1 |

| | | | | | | |
|---|---|---|---|---|---|---|
| LNHTKKWKYPQVNGL | 1 | 0 | 0 | 1541421 | P0DTC1.1 | P0DTD1 |
| LNNLGVDIAANTVIW | 1 | 0 | 0 | 1541425 | P0DTD1.1 | P0DTD1 |
| LNRYFRLTLGVYDYL | 1 | 0 | 0 | 1541428 | P0DTC1.1 | P0DTD1 |
| LPFAMGIIAMSAFAM | 1 | 0 | 0 | 1541432 | P0DTC1.1 | P0DTD1 |
| LPTMCDIRQLLFVVE | 1 | 0 | 0 | 1541445 | P0DTD1.1 | P0DTD1 |
| LPWNVVRIKIVQMLS | 1 | 0 | 0 | 1541449 | P0DTD1.1 | P0DTD1 |
| LQCIMLVYCFLGYFC | 1 | 0 | 0 | 1541452 | P0DTC1.1 | P0DTD1 |
| LQEFKPRSQMEIDFL | 1 | 0 | 0 | 1541454 | P0DTD1.1 | P0DTD1 |
| LQKAAITILDGISQY | 1 | 0 | 0 | 1541457 | P0DTC1.1 | P0DTD1 |
| LQKEKVNINIVGDFK | 1 | 0 | 0 | 1541459 | P0DTC1.1 | P0DTD1 |
| LQSCTQHQPYVVDDP | 1 | 0 | 0 | 1541461 | P0DTC8.1 | P0DTC8 |
| LQSINFVRIIMRLWL | 1 | 0 | 0 | 1541463 | P0DTC3.1 | P0DTC3 |
| LRDGWEIVKFISTCA | 1 | 0 | 0 | 1541466 | P0DTC1.1 | P0DTD1 |
| LREPMLQSADAQSFL | 1 | 0 | 0 | 1541467 | P0DTC1.1 | P0DTD1 |
| LREVRTIKVFTTVDN | 1 | 0 | 0 | 1541468 | P0DTC1.1 | P0DTD1 |
| LRGTAVMSLKEGQIN | 1 | 0 | 0 | 1541470 | P0DTD1.1 | P0DTD1 |
| LRIMASLVLARKHTT | 1 | 0 | 0 | 1541472 | P0DTD1.1 | P0DTD1 |
| LRVCVDTVRTNVYLA | 1 | 0 | 0 | 1541479 | P0DTC1.1 | P0DTD1 |
| LSAQTGIAVLDMCAS | 1 | 0 | 0 | 1541482 | P0DTC1.1 | P0DTD1 |
| LSDDAVVCFNSTYAS | 1 | 0 | 0 | 1541483 | P0DTD1.1 | P0DTD1 |
| LSDRVVFVLWAHGFE | 1 | 0 | 0 | 1541485 | P0DTD1.1 | P0DTD1 |
| LSHFVNLDNLRANNT | 1 | 0 | 0 | 1541489 | P0DTC1.1 | P0DTD1 |
| LSHRFYRLANECAQV | 1 | 0 | 0 | 1541490 | P0DTD1.1 | P0DTD1 |
| LSKGVHFVCNLLLLF | 1 | 0 | 0 | 1541491 | P0DTC3.1 | P0DTC3 |
| LSLLSKGRLIIRENN | 1 | 0 | 0 | 1541493 | P0DTD1.1 | P0DTD1 |
| LSLPVLQVRDVLVRG | 1 | 0 | 0 | 1541494 | P0DTC1.1 | P0DTD1 |
| LSLREVRTIKVFTTV | 1 | 0 | 0 | 1541495 | P0DTC1.1 | P0DTD1 |
| LSTDTGVEHVTFFIY | 1 | 0 | 0 | 1541499 | P0DTC3.1 | P0DTC3 |
| LSTFISAARQGFVDS | 1 | 0 | 0 | 1541500 | P0DTC1.1 | P0DTD1 |
| LTAFGLVAEWFLAYI | 1 | 0 | 0 | 1541505 | P0DTC1.1 | P0DTD1 |
| LTFYLTNDVSFLAHI | 1 | 0 | 0 | 1541513 | P0DTC1.1 | P0DTD1 |
| LTLGVYDYLVSTQEF | 1 | 0 | 0 | 1541518 | P0DTC1.1 | P0DTD1 |
| LTNDVSFLAHIQWMV | 1 | 0 | 0 | 1541519 | P0DTC1.1 | P0DTD1 |
| LTPLGIDLDEWSMAT | 1 | 0 | 0 | 1541520 | P0DTC1.1 | P0DTD1 |
| LTSIKWADNNCYLAT | 1 | 0 | 0 | 1541524 | P0DTC1.1 | P0DTD1 |
| LTSQWLTNIFGTVYE | 1 | 0 | 0 | 1541526 | P0DTC1.1 | P0DTD1 |
| LVAELEGIQYGRSGE | 1 | 0 | 0 | 1541529 | P0DTC1.1 | P0DTD1 |
| LVDFQVTIAEILLII | 1 | 0 | 0 | 1541532 | P0DTC6.1 | P0DTC6 |
| LVFLGIITTVAAFHQ | 1 | 0 | 0 | 1541534 | P0DTC8.1 | P0DTC8 |
| LVGTPVCINGLMLLE | 1 | 0 | 0 | 1541536 | P0DTC1.1 | P0DTD1 |
| LVLARKHTTCCSLSH | 1 | 0 | 0 | 1541537 | P0DTD1.1 | P0DTD1 |
| LVQSTQWSLFFFLYE | 1 | 0 | 0 | 1541542 | P0DTC1.1 | P0DTD1 |
| LVSDVGDSAEVAVKM | 1 | 0 | 0 | 1541544 | P0DTC1.1 | P0DTD1 |
| LVSSFLEMKSEKQVE | 1 | 0 | 0 | 1541545 | P0DTC1.1 | P0DTD1 |
| LVSTQEFRYMNSQGL | 1 | 0 | 0 | 1541546 | P0DTC1.1 | P0DTD1 |
| LVVRCSFYEDFLEYH | 1 | 0 | 0 | 1541548 | P0DTC8.1 | P0DTC8 |
| LWAHGFELTSMKYFV | 1 | 0 | 0 | 1541555 | P0DTD1.1 | P0DTD1 |
| LWNTFTRLQSLENVA | 1 | 0 | 0 | 1541557 | P0DTD1.1 | P0DTD1 |
| LYDKLVSSFLEMKSE | 1 | 0 | 0 | 1541561 | P0DTC1.1 | P0DTD1 |
| LYKLMGHFAWWTAFV | 1 | 0 | 0 | 1541564 | P0DTD1.1 | P0DTD1 |
| LYLTFYLTNDVSFLA | 1 | 0 | 0 | 1541567 | P0DTC1.1 | P0DTD1 |
| LYRKCVKSREETGLL | 1 | 0 | 0 | 1541570 | P0DTC1.1 | P0DTD1 |
| LYYDSMSYEDQDALF | 1 | 0 | 0 | 1541572 | P0DTD1.1 | P0DTD1 |
| MAPISAMVRMYIFFA | 1 | 0 | 0 | 1541578 | P0DTC1.1 | P0DTD1 |
| MATYYLFDESGEFKL | 1 | 0 | 0 | 1541580 | P0DTC1.1 | P0DTD1 |
| MAYITGGVVQLTSQW | 1 | 0 | 0 | 1541581 | P0DTC1.1 | P0DTD1 |
| MDEFIERYKLEGYAF | 1 | 0 | 0 | 1541584 | P0DTD1.1 | P0DTD1 |
| MDNLACEDLKPVSEE | 1 | 0 | 0 | 1541585 | P0DTC1.1 | P0DTD1 |
| MEKLKTLVATAEAEL | 1 | 0 | 0 | 1541588 | P0DTC1.1 | P0DTD1 |
| MFLARGIVFMCVEYC | 1 | 0 | 0 | 1541593 | P0DTC1.1 | P0DTD1 |
| MFLGTCRRCPAEIVD | 1 | 0 | 0 | 1541594 | P0DTD1.1 | P0DTD1 |
| MGHFAWWTAFVTNVN | 1 | 0 | 0 | 1541597 | P0DTD1.1 | P0DTD1 |
| MHANYIFWRNTNPIQ | 1 | 0 | 0 | 1541599 | P0DTD1.1 | P0DTD1 |
| MILSLLSKGRLIIRE | 1 | 0 | 0 | 1541602 | P0DTD1.1 | P0DTD1 |
| MISAGFSLWVYKQFD | 1 | 0 | 0 | 1541603 | P0DTD1.1 | P0DTD1 |
| MKYNYEPLTQDHVDI | 1 | 0 | 0 | 1541604 | P0DTC1.1 | P0DTD1 |
| MLAHAEETRKLMPVC | 1 | 0 | 0 | 1541605 | P0DTC1.1 | P0DTD1 |
| MLDNRATLQAIASEF | 1 | 0 | 0 | 1541607 | P0DTC1.1 | P0DTD1 |
| MLTNDNTSRYWEPEF | 1 | 0 | 0 | 1541614 | P0DTD1.1 | P0DTD1 |
| MLVKQGDDYVYLPYP | 1 | 0 | 0 | 1541616 | P0DTD1.1 | P0DTD1 |
| MLVYCFLGYFCTCYF | 1 | 0 | 0 | 1541617 | P0DTC1.1 | P0DTD1 |
| MMVTNNTFTLKGGAP | 1 | 0 | 0 | 1541622 | P0DTC1.1 | P0DTD1 |
| MPILTLTRALTAESH | 1 | 0 | 0 | 1541629 | P0DTD1.1 | P0DTD1 |
| MPTTIAKNTVKSVGK | 1 | 0 | 0 | 1541631 | P0DTC1.1 | P0DTD1 |
| MRIFTIGTVTLKQGE | 1 | 0 | 0 | 1541634 | P0DTC3.1 | P0DTC3 |
| MRNAGIVGVLTLDNQ | 1 | 0 | 0 | 1541637 | P0DTD1.1 | P0DTD1 |
| MSCAAGTTQTACTDD | 1 | 0 | 0 | 1541640 | P0DTC1.1 | P0DTD1 |
| MSKHTDFSSEIIGYK | 1 | 0 | 0 | 1541645 | P0DTC1.1 | P0DTD1 |
| MSLKEGQINDMILSL | 1 | 0 | 0 | 1541646 | P0DTD1.1 | P0DTD1 |
| MSMTYGQQFGPTYLD | 1 | 0 | 0 | 1541648 | P0DTC1.1 | P0DTD1 |
| MSNLGMPSYCTGYRE | 1 | 0 | 0 | 1541649 | P0DTC1.1 | P0DTD1 |
| MSYYCKSHKPPISFP | 1 | 0 | 0 | 1541651 | P0DTD1.1 | P0DTD1 |
| MVPHISRQRLTKYTM | 1 | 0 | 0 | 1541659 | P0DTD1.1 | P0DTD1 |
| MVRMYIFFASFYYVW | 1 | 0 | 0 | 1541661 | P0DTC1.1 | P0DTD1 |
| MYASAVVLLILMTAR | 1 | 0 | 0 | 1541665 | P0DTC1.1 | P0DTD1 |
| MYKGLPWNVVRIKIV | 1 | 0 | 0 | 1541668 | P0DTD1.1 | P0DTD1 |
| MYSVMLTNDNTSRYW | 1 | 0 | 0 | 1541670 | P0DTD1.1 | P0DTD1 |
| NADIVEEAKKVKPTV | 1 | 0 | 0 | 1541673 | P0DTC1.1 | P0DTD1 |
| NAGIVGVLTLDNQDL | 1 | 0 | 0 | 1541674 | P0DTD1.1 | P0DTD1 |
| NAQVAKSHNIALIWN | 1 | 0 | 0 | 1541675 | P0DTC1.1 | P0DTD1 |
| NCDTFCAGSTFISDE | 1 | 0 | 0 | 1541687 | P0DTC1.1 | P0DTD1 |

| | | | | | | |
|---|---|---|---|---|---|---|
| NCLDDRCILHCANFN | 1 | 0 | 0 | 1541689 | P0DTD1.1 | P0DTD1 |
| NDILLAKDTTEAFEK | 1 | 0 | 0 | 1541692 | P0DTC1.1 | P0DTD1 |
| NDVSFLAHIQWMVMF | 1 | 0 | 0 | 1541694 | P0DTC1.1 | P0DTD1 |
| NDYYRSLPGVFCGVD | 1 | 0 | 0 | 1541695 | P0DTC1.1 | P0DTD1 |
| NEKQEILGTVSWNLR | 1 | 0 | 0 | 1541699 | P0DTC1.1 | P0DTD1 |
| NFVFPLNSIIKTIQP | 1 | 0 | 0 | 1541713 | P0DTC1.1 | P0DTD1 |
| NGDFLHFLPRVFSAV | 1 | 0 | 0 | 1541715 | P0DTC1.1 | P0DTD1 |
| NGDSEVVLKKLKKSL | 1 | 0 | 0 | 1541716 | P0DTC1.1 | P0DTD1 |
| NGLMLLEIKDTEKYC | 1 | 0 | 0 | 1541719 | P0DTC1.1 | P0DTD1 |
| NGLTSIKWADNNCYL | 1 | 0 | 0 | 1541720 | P0DTC1.1 | P0DTD1 |
| NGNLHPDSATLVSDI | 1 | 0 | 0 | 1541722 | P0DTC1.1 | P0DTD1 |
| NHTGVVGEGSEGLND | 1 | 0 | 0 | 1541731 | P0DTC1.1 | P0DTD1 |
| NIALIWNVKDFMSLS | 1 | 0 | 0 | 1541732 | P0DTC1.1 | P0DTD1 |
| NIDYDCVSFCYMHHM | 1 | 0 | 0 | 1541734 | P0DTC1.1 | P0DTD1 |
| NIGCNHTGVVGEGSE | 1 | 0 | 0 | 1541735 | P0DTC1.1 | P0DTD1 |
| NIGEQKSILSPLYAF | 1 | 0 | 0 | 1541736 | P0DTC1.1 | P0DTD1 |
| NIIPLTTAAKLMVVI | 1 | 0 | 0 | 1541738 | P0DTC1.1 | P0DTD1 |
| NINIVGDFKLNEEIA | 1 | 0 | 0 | 1541742 | P0DTC1.1 | P0DTD1 |
| NIVGDFKLNEEIAII | 1 | 0 | 0 | 1541745 | P0DTC1.1 | P0DTD1 |
| NKDFYDFAVSKGFFK | 1 | 0 | 0 | 1541748 | P0DTD1.1 | P0DTD1 |
| NLACEDLKPVSEEVV | 1 | 0 | 0 | 1541757 | P0DTC1.1 | P0DTD1 |
| NLEEAARYMRSLKVP | 1 | 0 | 0 | 1541760 | P0DTC1.1 | P0DTD1 |
| NLGETFVTHSKGLYR | 1 | 0 | 0 | 1541761 | P0DTC1.1 | P0DTD1 |
| NLGMPSYCTGYREGY | 1 | 0 | 0 | 1541762 | P0DTC1.1 | P0DTD1 |
| NLGVDIAANTVIWDY | 1 | 0 | 0 | 1541763 | P0DTD1.1 | P0DTD1 |
| NLHPDSATLVSDIDI | 1 | 0 | 0 | 1541764 | P0DTC1.1 | P0DTD1 |
| NLHSSRLSFKELLVY | 1 | 0 | 0 | 1541765 | P0DTD1.1 | P0DTD1 |
| NLLLYIDINGNLHPD | 1 | 0 | 0 | 1541771 | P0DTC1.1 | P0DTD1 |
| NLQHRLYECLYRNRD | 1 | 0 | 0 | 1541773 | P0DTD1.1 | P0DTD1 |
| NLTKEGATTCGYLPQ | 1 | 0 | 0 | 1541777 | P0DTC1.1 | P0DTD1 |
| NMLRIMASLVLARKH | 1 | 0 | 0 | 1541781 | P0DTD1.1 | P0DTD1 |
| NNAMQVESDDYIATN | 1 | 0 | 0 | 1541785 | P0DTC1.1 | P0DTD1 |
| NNTKGSLPINVIVFD | 1 | 0 | 0 | 1541795 | P0DTC1.1 | P0DTD1 |
| NPKGFCDLKGKYVQI | 1 | 0 | 0 | 1541799 | P0DTC1.1 | P0DTD1 |
| NPTTFHLDGEVITFD | 1 | 0 | 0 | 1541805 | P0DTC1.1 | P0DTD1 |
| NRFLYIIKLIFLWLL | 1 | 0 | 0 | 1541814 | P0DTC5.1 | P0DTC5 |
| NRVCTNYMPYFFTLL | 1 | 0 | 0 | 1541818 | P0DTC1.1 | P0DTD1 |
| NSIIKTIQPRVEKKK | 1 | 0 | 0 | 1541822 | P0DTC1.1 | P0DTD1 |
| NSLKITEEVGHTDLM | 1 | 0 | 0 | 1541825 | P0DTC1.1 | P0DTD1 |
| NSPNLAWPLIVTALR | 1 | 0 | 0 | 1541826 | P0DTC1.1 | P0DTD1 |
| NSVTSSIVITSGDGT | 1 | 0 | 0 | 1541838 | P0DTC3.1 | P0DTC3 |
| NSYFTEQPIDLVPNQ | 1 | 0 | 0 | 1541839 | P0DTC1.1 | P0DTD1 |
| NTFTRLQSLENVAFN | 1 | 0 | 0 | 1541845 | P0DTD1.1 | P0DTD1 |
| NTKHSSGVTRELMRE | 1 | 0 | 0 | 1541847 | P0DTC1.1 | P0DTD1 |
| NTLQCIMLVYCFLGY | 1 | 0 | 0 | 1541849 | P0DTC1.1 | P0DTD1 |
| NTWCIRCLWSTKPVE | 1 | 0 | 0 | 1541854 | P0DTC1.1 | P0DTD1 |
| NTYLEGSVRVVTTFD | 1 | 0 | 0 | 1541856 | P0DTC1.1 | P0DTD1 |
| NVAFELWAKRNIKPV | 1 | 0 | 0 | 1541857 | P0DTD1.1 | P0DTD1 |
| NVLAWLYAAVINGDR | 1 | 0 | 0 | 1541869 | P0DTC1.1 | P0DTD1 |
| NVVTTKIALKGGKIV | 1 | 0 | 0 | 1541876 | P0DTC1.1 | P0DTD1 |
| NVYLAVFDKNLYDKL | 1 | 0 | 0 | 1541878 | P0DTC1.1 | P0DTD1 |
| NYLKSPNFSKLINII | 1 | 0 | 0 | 1541886 | P0DTC1.1 | P0DTD1 |
| NYMLTYNKVENMTPR | 1 | 0 | 0 | 1541887 | P0DTC1.1 | P0DTD1 |
| NYQCGHYKHITSKET | 1 | 0 | 0 | 1541888 | P0DTC1.1 | P0DTD1 |
| NYQVNGYPNMFITRE | 1 | 0 | 0 | 1541890 | P0DTD1.1 | P0DTD1 |
| NYTVSCLPFTINCQE | 1 | 0 | 0 | 1541891 | P0DTC8.1 | P0DTC8 |
| PASRELKVTFFPDLN | 1 | 0 | 0 | 1541897 | P0DTC1.1 | P0DTD1 |
| PDDQIGYYRRATRRI | 1 | 0 | 0 | 1541910 | P0DTC9.1 | P0DTC9 |
| PDPSRILGAGCFVDD | 1 | 0 | 0 | 1541915 | P0DTD1.1 | P0DTD1 |
| PERTCCLCDRRATCF | 1 | 0 | 0 | 1541921 | P0DTD1.1 | P0DTD1 |
| PFELEDFIPMDSTVK | 1 | 0 | 0 | 1541926 | P0DTD1.1 | P0DTD1 |
| PGQGLNGYTVEEAKT | 1 | 0 | 0 | 1541935 | P0DTC1.1 | P0DTD1 |
| PGTILRTTNGDFLHF | 1 | 0 | 0 | 1541939 | P0DTC1.1 | P0DTD1 |
| PIHSLRVCVDTVRTN | 1 | 0 | 0 | 1541945 | P0DTC1.1 | P0DTD1 |
| PILLLDQALVSDVGD | 1 | 0 | 0 | 1541947 | P0DTC1.1 | P0DTD1 |
| PINPTDQSSYIVDSV | 1 | 0 | 0 | 1541948 | P0DTC1.1 | P0DTD1 |
| PKCDRAMPNMLRIMA | 1 | 0 | 0 | 1541956 | P0DTD1.1 | P0DTD1 |
| PKEIIFLEGETLPTE | 1 | 0 | 0 | 1541958 | P0DTC1.1 | P0DTD1 |
| PKLFIRQEEVQELYS | 1 | 0 | 0 | 1541965 | P0DTC7.1 | P0DTC7 |
| PLGIDLDEWSMATYY | 1 | 0 | 0 | 1541970 | P0DTC1.1 | P0DTD1 |
| PLHGTILTRPLLESE | 1 | 0 | 0 | 1541971 | P0DTC5.1 | P0DTC5 |
| PLIAAVITREVGFVV | 1 | 0 | 0 | 1541972 | P0DTC1.1 | P0DTD1 |
| PLKAPKEIIFLEGET | 1 | 0 | 0 | 1541973 | P0DTC1.1 | P0DTD1 |
| PLLTDEMIAQYTSAL | 1 | 0 | 0 | 1541977 | P0DTC2.1 | P0DTC2 |
| PLTKHPNQEYADVFH | 1 | 0 | 0 | 1541979 | P0DTD1.1 | P0DTD1 |
| PLTQDHVDILGPLSA | 1 | 0 | 0 | 1541980 | P0DTC1.1 | P0DTD1 |
| PLTVFFDGRVDGQVD | 1 | 0 | 0 | 1541982 | P0DTD1.1 | P0DTD1 |
| PLVRKIFVDGVPFVV | 1 | 0 | 0 | 1541984 | P0DTD1.1 | P0DTD1 |
| PNDDTLRVEAFEYYH | 1 | 0 | 0 | 1541988 | P0DTC1.1 | P0DTD1 |
| PNLAWPLIVTALRAN | 1 | 0 | 0 | 1541991 | P0DTC1.1 | P0DTD1 |
| PPTSFGPLVRKIFVD | 1 | 0 | 0 | 1542004 | P0DTD1.1 | P0DTD1 |
| PQLEMELTPVVQTIE | 1 | 0 | 0 | 1542009 | P0DTC1.1 | P0DTD1 |
| PSFLGRYMSALNHTK | 1 | 0 | 0 | 1542020 | P0DTC1.1 | P0DTD1 |
| PSLATVAYFNMVYMP | 1 | 0 | 0 | 1542022 | P0DTC1.1 | P0DTD1 |
| PSRILGAGCFVDDIV | 1 | 0 | 0 | 1542024 | P0DTD1.1 | P0DTD1 |
| PTKVTFGDDTVIEVQ | 1 | 0 | 0 | 1542032 | P0DTC1.1 | P0DTD1 |
| PTNFTISVTTEILPV | 1 | 0 | 0 | 1542035 | P0DTC2.1 | P0DTC2 |
| PVALRQMSCAAGTTQ | 1 | 0 | 0 | 1542039 | P0DTC1.1 | P0DTD1 |
| PVCVETKAIVSTIQR | 1 | 0 | 0 | 1542041 | P0DTC1.1 | P0DTD1 |
| PVHVMSKHTDFSSEI | 1 | 0 | 0 | 1542044 | P0DTC1.1 | P0DTD1 |
| PVLDWLEEKFKEGVE | 1 | 0 | 0 | 1542045 | P0DTC1.1 | P0DTD1 |

| | | | | | | |
|---|---|---|---|---|---|---|
| PVVDSYYSLLMPILT | 1 | 0 | 0 | 1542051 | P0DTD1.1 | P0DTD1 |
| PYFFTLLLQLCTFTR | 1 | 0 | 0 | 1542054 | P0DTC1.1 | P0DTD1 |
| PYIKWDLLKYDFTEE | 1 | 0 | 0 | 1542056 | P0DTD1.1 | P0DTD1 |
| PYPDPSRILGAGCFV | 1 | 0 | 0 | 1542061 | P0DTD1.1 | P0DTD1 |
| QALLKTVQFCDAMRN | 1 | 0 | 0 | 1542069 | P0DTD1.1 | P0DTD1 |
| QALVSDVGDSAEVAV | 1 | 0 | 0 | 1542071 | P0DTC1.1 | P0DTD1 |
| QCFKMFYKGVITHDV | 1 | 0 | 0 | 1542077 | P0DTD1.1 | P0DTD1 |
| QCGHYKHITSKETLY | 1 | 0 | 0 | 1542078 | P0DTC1.1 | P0DTD1 |
| QELYSPIFLIVAAIV | 1 | 0 | 0 | 1542088 | P0DTC7.1 | P0DTC7 |
| QESFGGASCCLYCRC | 1 | 0 | 0 | 1542090 | P0DTC1.1 | P0DTD1 |
| QESPFVMMSAPPAQY | 1 | 0 | 0 | 1542091 | P0DTC1.1 | P0DTD1 |
| QFKHLIPLMYKGLPW | 1 | 0 | 0 | 1542095 | P0DTD1.1 | P0DTD1 |
| QGMDNLACEDLKPVS | 1 | 0 | 0 | 1542105 | P0DTC1.1 | P0DTD1 |
| QHEVLLAPLLSAGIF | 1 | 0 | 0 | 1542111 | P0DTC1.1 | P0DTD1 |
| QHLKDGTCGLVEVEK | 1 | 0 | 0 | 1542113 | P0DTC1.1 | P0DTD1 |
| QHRLYECLYRNRDVD | 1 | 0 | 0 | 1542115 | P0DTD1.1 | P0DTD1 |
| QKDVLECNVKTTEVV | 1 | 0 | 0 | 1542125 | P0DTC1.1 | P0DTD1 |
| QLFFSYFAVHFISNS | 1 | 0 | 0 | 1542132 | P0DTC1.1 | P0DTD1 |
| QLIKVTLVFLFVAAI | 1 | 0 | 0 | 1542134 | P0DTC1.1 | P0DTD1 |
| QLSEISMDNSPNLAW | 1 | 0 | 0 | 1542144 | P0DTC1.1 | P0DTD1 |
| QMCLSTLMKCDHCGE | 1 | 0 | 0 | 1542149 | P0DTC1.1 | P0DTD1 |
| QMEIDFLELAMDEFI | 1 | 0 | 0 | 1542150 | P0DTD1.1 | P0DTD1 |
| QPVSELLTPLGIDLD | 1 | 0 | 0 | 1542164 | P0DTC1.1 | P0DTD1 |
| QQDGSEDNQTTTIQT | 1 | 0 | 0 | 1542167 | P0DTC1.1 | P0DTD1 |
| QQGEVPVSIINNTVY | 1 | 0 | 0 | 1542170 | P0DTD1.1 | P0DTD1 |
| QQKLALGGSVAIKIT | 1 | 0 | 0 | 1542172 | P0DTD1.1 | P0DTD1 |
| QQWGFTGNLQSNHDL | 1 | 0 | 0 | 1542175 | P0DTD1.1 | P0DTD1 |
| QRKQDDKKIKACVEE | 1 | 0 | 0 | 1542178 | P0DTC1.1 | P0DTD1 |
| QRMLLEKCDLQNYGD | 1 | 0 | 0 | 1542179 | P0DTD1.1 | P0DTD1 |
| QSDIEVTGDSCNNYM | 1 | 0 | 0 | 1542183 | P0DTC1.1 | P0DTD1 |
| QSLENVAFNVVNKGH | 1 | 0 | 0 | 1542184 | P0DTD1.1 | P0DTD1 |
| QSRNLQEFKPRSQME | 1 | 0 | 0 | 1542185 | P0DTD1.1 | P0DTD1 |
| QSSQAWQPGVAMPNL | 1 | 0 | 0 | 1542186 | P0DTD1.1 | P0DTD1 |
| QTAQAAGTDTTITVN | 1 | 0 | 0 | 1542190 | P0DTC1.1 | P0DTD1 |
| QTTTIQTIVEVQPQL | 1 | 0 | 0 | 1542200 | P0DTC1.1 | P0DTD1 |
| QVNGYPNMFITREEA | 1 | 0 | 0 | 1542213 | P0DTD1.1 | P0DTD1 |
| QVRDVLVRGFGDSVE | 1 | 0 | 0 | 1542214 | P0DTC1.1 | P0DTD1 |
| QVTCGTTTLNGLWLD | 1 | 0 | 0 | 1542215 | P0DTC1.1 | P0DTD1 |
| QVTIAEILLIIMRTF | 1 | 0 | 0 | 1542216 | P0DTC6.1 | P0DTC6 |
| QYGRSGETLGVLVPH | 1 | 0 | 0 | 1542221 | P0DTC1.1 | P0DTD1 |
| RANNTKGSLPINVIV | 1 | 0 | 0 | 1542230 | P0DTC1.1 | P0DTD1 |
| RARSVSPKLFIRQEE | 1 | 0 | 0 | 1542233 | P0DTC7.1 | P0DTC7 |
| RDLGACIDCSARHIN | 1 | 0 | 0 | 1542240 | P0DTC1.1 | P0DTD1 |
| RDVLVRGFGDSVEEV | 1 | 0 | 0 | 1542241 | P0DTC1.1 | P0DTD1 |
| REHEHEIAWYTERSE | 1 | 0 | 0 | 1542243 | P0DTC1.1 | P0DTD1 |
| REQIDGYVMHANYIF | 1 | 0 | 0 | 1542246 | P0DTD1.1 | P0DTD1 |
| RFRRAFGEYSHVVAF | 1 | 0 | 0 | 1542251 | P0DTC1.1 | P0DTD1 |
| RGIVFMCVEYCPIFF | 1 | 0 | 0 | 1542256 | P0DTC1.1 | P0DTD1 |
| RGVYCCREHEHEIAW | 1 | 0 | 0 | 1542260 | P0DTC1.1 | P0DTD1 |
| RHFDEGNCDTLKEIL | 1 | 0 | 0 | 1542262 | P0DTD1.1 | P0DTD1 |
| RHTFSNYQHEETIYN | 1 | 0 | 0 | 1542264 | P0DTD1.1 | P0DTD1 |
| RHVRAWIGFDVEGCH | 1 | 0 | 0 | 1542265 | P0DTD1.1 | P0DTD1 |
| RKHTTCCSLSHRFYR | 1 | 0 | 0 | 1542277 | P0DTD1.1 | P0DTD1 |
| RLANECAQVLSEMVM | 1 | 0 | 0 | 1542287 | P0DTD1.1 | P0DTD1 |
| RLISMMGFKMNYQVN | 1 | 0 | 0 | 1542289 | P0DTD1.1 | P0DTD1 |
| RLQSLENVAFNVVNK | 1 | 0 | 0 | 1542292 | P0DTD1.1 | P0DTD1 |
| RLYECLYRNRDVDTD | 1 | 0 | 0 | 1542297 | P0DTD1.1 | P0DTD1 |
| RNLQEFKPRSQMEID | 1 | 0 | 0 | 1542303 | P0DTD1.1 | P0DTD1 |
| RNTNPIQLSSYSLFD | 1 | 0 | 0 | 1542305 | P0DTD1.1 | P0DTD1 |
| RNYVFTGYRVTKNSK | 1 | 0 | 0 | 1542308 | P0DTD1.1 | P0DTD1 |
| RQGFVDSDVETKDVV | 1 | 0 | 0 | 1542312 | P0DTC1.1 | P0DTD1 |
| RQVVNVVTTKIALKG | 1 | 0 | 0 | 1542315 | P0DTC1.1 | P0DTD1 |
| RQWLPTGTLLVDSDL | 1 | 0 | 0 | 1542318 | P0DTD1.1 | P0DTD1 |
| RSVSPKLFIRQEEVQ | 1 | 0 | 0 | 1542331 | P0DTC7.1 | P0DTC7 |
| RTNVYLAVFDKNLYD | 1 | 0 | 0 | 1542340 | P0DTC1.1 | P0DTD1 |
| RTTCFSVAALTNNVA | 1 | 0 | 0 | 1542341 | P0DTD1.1 | P0DTD1 |
| RVEAFEYYHTTDPSF | 1 | 0 | 0 | 1542348 | P0DTC1.1 | P0DTD1 |
| RVECTTIVNGVRRSF | 1 | 0 | 0 | 1542349 | P0DTC1.1 | P0DTD1 |
| RVIHFGAGSDKGVAP | 1 | 0 | 0 | 1542352 | P0DTD1.1 | P0DTD1 |
| RVSAKPPPGDQFKHL | 1 | 0 | 0 | 1542358 | P0DTD1.1 | P0DTD1 |
| RVVRSIFSRTLETAQ | 1 | 0 | 0 | 1542360 | P0DTC1.1 | P0DTD1 |
| RWFLNRFTTTLNDFN | 1 | 0 | 0 | 1542362 | P0DTC1.1 | P0DTD1 |
| RYFRLTLGVYDYLVS | 1 | 0 | 0 | 1542363 | P0DTC1.1 | P0DTD1 |
| RYKLEGYAFEHIVYG | 1 | 0 | 0 | 1542364 | P0DTD1.1 | P0DTD1 |
| RYWEPEFYEAMYTPH | 1 | 0 | 0 | 1542368 | P0DTD1.1 | P0DTD1 |
| SAGFSLWVYKQFDTY | 1 | 0 | 0 | 1542373 | P0DTD1.1 | P0DTD1 |
| SAINRPQIGVVREFL | 1 | 0 | 0 | 1542375 | P0DTD1.1 | P0DTD1 |
| SAKPPPGDQFKHLIP | 1 | 0 | 0 | 1542376 | P0DTD1.1 | P0DTD1 |
| SAMVRMYIFFASFYY | 1 | 0 | 0 | 1542380 | P0DTC1.1 | P0DTD1 |
| SASVYYSQLMCQPIL | 1 | 0 | 0 | 1542387 | P0DTC1.1 | P0DTD1 |
| SAVGNICYTPSKLIE | 1 | 0 | 0 | 1542388 | P0DTC1.1 | P0DTD1 |
| SAYTVELGTEVNEFA | 1 | 0 | 0 | 1542390 | P0DTC1.1 | P0DTD1 |
| SCDAIMTRCLAVHEC | 1 | 0 | 0 | 1542392 | P0DTD1.1 | P0DTD1 |
| SCLPFTINCQEPKLG | 1 | 0 | 0 | 1542394 | P0DTC8.1 | P0DTC8 |
| SCTQHQPYVVDDPCP | 1 | 0 | 0 | 1542397 | P0DTC8.1 | P0DTC8 |
| SDKAYKIEELFYSYA | 1 | 0 | 0 | 1542403 | P0DTD1.1 | P0DTD1 |
| SDLATNNLVVMAYIT | 1 | 0 | 0 | 1542405 | P0DTC1.1 | P0DTD1 |
| SDTLKNLSDRVVFVL | 1 | 0 | 0 | 1542410 | P0DTD1.1 | P0DTD1 |
| SDTYACWHHSIGFDY | 1 | 0 | 0 | 1542411 | P0DTD1.1 | P0DTD1 |
| SDVETKDVVECLKLS | 1 | 0 | 0 | 1542412 | P0DTC1.1 | P0DTD1 |
| SDVGDSAEVAVKMFD | 1 | 0 | 0 | 1542413 | P0DTC1.1 | P0DTD1 |

| | | | | | | |
|---|---|---|---|---|---|---|
| SEAARVVRSIFSRTL | 1 | 0 | 0 | 1542415 | P0DTC1.1 | P0DTD1 |
| SEARQHLKDGTCGLV | 1 | 0 | 0 | 1542419 | P0DTC1.1 | P0DTD1 |
| SEDKRAKVTSAMQTM | 1 | 0 | 0 | 1542421 | P0DTC1.1 | P0DTD1 |
| SEDNQTTTIQTIVEV | 1 | 0 | 0 | 1542422 | P0DTC1.1 | P0DTD1 |
| SEISMDNSPNLAWPL | 1 | 0 | 0 | 1542426 | P0DTC1.1 | P0DTD1 |
| SEVGPEHSLAEYHNE | 1 | 0 | 0 | 1542433 | P0DTC1.1 | P0DTD1 |
| SFGGASCCLYCRCHI | 1 | 0 | 0 | 1542439 | P0DTC1.1 | P0DTD1 |
| SFKKGAKLLHKPIVW | 1 | 0 | 0 | 1542441 | P0DTC1.1 | P0DTD1 |
| SFMLWCKDGHVETFY | 1 | 0 | 0 | 1542444 | P0DTD1.1 | P0DTD1 |
| SFNYLKSPNFSKLIN | 1 | 0 | 0 | 1542446 | P0DTC1.1 | P0DTD1 |
| SFRLFARTRSMWSFN | 1 | 0 | 0 | 1542448 | P0DTC5.1 | P0DTC5 |
| SGFRKMAFPSGKVEG | 1 | 0 | 0 | 1542457 | P0DTC1.1 | P0DTD1 |
| SGKPVPYCYDTNVLE | 1 | 0 | 0 | 1542459 | P0DTC1.1 | P0DTD1 |
| SHQSDIEVTGDSCNN | 1 | 0 | 0 | 1542483 | P0DTC1.1 | P0DTD1 |
| SIHLYFDKAGQKTYE | 1 | 0 | 0 | 1542487 | P0DTC1.1 | P0DTD1 |
| SIKNFKSVLYYQNNV | 1 | 0 | 0 | 1542492 | P0DTD1.1 | P0DTD1 |
| SINFVRIIMRLWLCW | 1 | 0 | 0 | 1542496 | P0DTC3.1 | P0DTC3 |
| SKCVCSVIDLLLDDF | 1 | 0 | 0 | 1542499 | P0DTD1.1 | P0DTD1 |
| SKFPLKLRGTAVMSL | 1 | 0 | 0 | 1542500 | P0DTD1.1 | P0DTD1 |
| SKIVQLSEISMDNSP | 1 | 0 | 0 | 1542503 | P0DTC1.1 | P0DTD1 |
| SKLIEYTDFATSACV | 1 | 0 | 0 | 1542504 | P0DTC1.1 | P0DTD1 |
| SKNPLLYDANYFLCW | 1 | 0 | 0 | 1542507 | P0DTC3.1 | P0DTC3 |
| SKTPEEHFIETISLA | 1 | 0 | 0 | 1542510 | P0DTC1.1 | P0DTD1 |
| SLIYSTAALGVLMSN | 1 | 0 | 0 | 1542513 | P0DTC1.1 | P0DTD1 |
| SLLVLVQSTQWSLFF | 1 | 0 | 0 | 1542516 | P0DTC1.1 | P0DTD1 |
| SLSEQLRKQIRSAAK | 1 | 0 | 0 | 1542521 | P0DTC1.1 | P0DTD1 |
| SLWVYKQFDTYNLWN | 1 | 0 | 0 | 1542523 | P0DTD1.1 | P0DTD1 |
| SNSWLMWLIINLVQM | 1 | 0 | 0 | 1542538 | P0DTC1.1 | P0DTD1 |
| SQLMCQPILLLDQAL | 1 | 0 | 0 | 1542558 | P0DTC1.1 | P0DTD1 |
| SQTSLRCGACIRRPF | 1 | 0 | 0 | 1542560 | P0DTD1.1 | P0DTD1 |
| SRIKASMPTTIAKNT | 1 | 0 | 0 | 1542567 | P0DTC1.1 | P0DTD1 |
| SSFKWDLTAFGLVAE | 1 | 0 | 0 | 1542578 | P0DTC1.1 | P0DTD1 |
| SSIVITSGDGTTSPI | 1 | 0 | 0 | 1542583 | P0DTC3.1 | P0DTC3 |
| SSSKTPEEHFIETIS | 1 | 0 | 0 | 1542589 | P0DTC1.1 | P0DTD1 |
| SSTCMMCYKRNRATR | 1 | 0 | 0 | 1542593 | P0DTC1.1 | P0DTD1 |
| STFISDEVARDLSLQ | 1 | 0 | 0 | 1542597 | P0DTC1.1 | P0DTD1 |
| STLEQYVFCTVNALP | 1 | 0 | 0 | 1542601 | P0DTD1.1 | P0DTD1 |
| STLMKCDHCGETSWQ | 1 | 0 | 0 | 1542602 | P0DTC1.1 | P0DTD1 |
| SVAIKITEHSWNADL | 1 | 0 | 0 | 1542612 | P0DTD1.1 | P0DTD1 |
| SVCRLMKTIGPDMFL | 1 | 0 | 0 | 1542614 | P0DTD1.1 | P0DTD1 |
| SVELKHFFFAQDGNA | 1 | 0 | 0 | 1542617 | P0DTD1.1 | P0DTD1 |
| SVFNICQAVTANVNA | 1 | 0 | 0 | 1542618 | P0DTD1.1 | P0DTD1 |
| SVGFNIDYDCVSFCY | 1 | 0 | 0 | 1542619 | P0DTC1.1 | P0DTD1 |
| SVIYLYLTFYLTNDV | 1 | 0 | 0 | 1542621 | P0DTC1.1 | P0DTD1 |
| SVLACYNGSPSGVYQ | 1 | 0 | 0 | 1542622 | P0DTC1.1 | P0DTD1 |
| SVNITFELDERIDKV | 1 | 0 | 0 | 1542630 | P0DTC1.1 | P0DTD1 |
| SVVSKVVKVTIDYTE | 1 | 0 | 0 | 1542638 | P0DTD1.1 | P0DTD1 |
| SVYYTSNPTTFHLDG | 1 | 0 | 0 | 1542641 | P0DTC1.1 | P0DTD1 |
| SWLMWLIINLVQMAP | 1 | 0 | 0 | 1542642 | P0DTC1.1 | P0DTD1 |
| SWVMRIMTWLDMVDT | 1 | 0 | 0 | 1542644 | P0DTC1.1 | P0DTD1 |
| SYFAVHFISNSWLMW | 1 | 0 | 0 | 1542649 | P0DTC1.1 | P0DTD1 |
| SYFVVKRHTFSNYQH | 1 | 0 | 0 | 1542651 | P0DTD1.1 | P0DTD1 |
| SYGIATVREVLSDRE | 1 | 0 | 0 | 1542652 | P0DTD1.1 | P0DTD1 |
| SYIVDSVTVKNGSIH | 1 | 0 | 0 | 1542653 | P0DTC1.1 | P0DTD1 |
| SYKDWSYSGQSTQLG | 1 | 0 | 0 | 1542654 | P0DTC1.1 | P0DTD1 |
| SYVGCHNKCAYWVPR | 1 | 0 | 0 | 1542659 | P0DTC1.1 | P0DTD1 |
| SYVHVVDGCNSSTCM | 1 | 0 | 0 | 1542660 | P0DTC1.1 | P0DTD1 |
| TAAKLMVVIPDYNTY | 1 | 0 | 0 | 1542662 | P0DTC1.1 | P0DTD1 |
| TAESHVDTDLTKPYI | 1 | 0 | 0 | 1542663 | P0DTD1.1 | P0DTD1 |
| TAHSCNVNRFNVAIT | 1 | 0 | 0 | 1542665 | P0DTD1.1 | P0DTD1 |
| TALRLCAYCCNIVNV | 1 | 0 | 0 | 1542667 | P0DTC4.1 | P0DTC4 |
| TANKWDLIISDMYDP | 1 | 0 | 0 | 1542668 | P0DTD1.1 | P0DTD1 |
| TAPHGHVMVELVAEL | 1 | 0 | 0 | 1542670 | P0DTC1.1 | P0DTD1 |
| TCCSLSHRFYRLANE | 1 | 0 | 0 | 1542679 | P0DTD1.1 | P0DTD1 |
| TCFSTASDTYACWHH | 1 | 0 | 0 | 1542683 | P0DTD1.1 | P0DTD1 |
| TCFSVAALTNNVAFQ | 1 | 0 | 0 | 1542684 | P0DTD1.1 | P0DTD1 |
| TCGTTTLNGLWLDDV | 1 | 0 | 0 | 1542688 | P0DTC1.1 | P0DTD1 |
| TCMMCYKRNRATRVE | 1 | 0 | 0 | 1542691 | P0DTC1.1 | P0DTD1 |
| TDFATSACVLAAECT | 1 | 0 | 0 | 1542694 | P0DTC1.1 | P0DTD1 |
| TDFNAIATCDWTNAG | 1 | 0 | 0 | 1542695 | P0DTD1.1 | P0DTD1 |
| TDGTLMIERFVSLAI | 1 | 0 | 0 | 1542699 | P0DTD1.1 | P0DTD1 |
| TDNYITTYPGQGLNG | 1 | 0 | 0 | 1542701 | P0DTC1.1 | P0DTD1 |
| TDPSFLGRYMSALNH | 1 | 0 | 0 | 1542702 | P0DTC1.1 | P0DTD1 |
| TDQSSYIVDSVTVKN | 1 | 0 | 0 | 1542703 | P0DTC1.1 | P0DTD1 |
| TDVTQLYLGGMSYYC | 1 | 0 | 0 | 1542705 | P0DTD1.1 | P0DTD1 |
| TEERLKLFDRYFKYW | 1 | 0 | 0 | 1542708 | P0DTD1.1 | P0DTD1 |
| TEKYCALAPNMMVTN | 1 | 0 | 0 | 1542715 | P0DTC1.1 | P0DTD1 |
| TEMLAKALRKVPTDN | 1 | 0 | 0 | 1542717 | P0DTC1.1 | P0DTD1 |
| TENLLLYIDINGNLH | 1 | 0 | 0 | 1542718 | P0DTC1.1 | P0DTD1 |
| TENLTKEGATTCGYL | 1 | 0 | 0 | 1542719 | P0DTC1.1 | P0DTD1 |
| TEQPIDLVPNQPYPN | 1 | 0 | 0 | 1542720 | P0DTC1.1 | P0DTD1 |
| TETICAPLTVFFDGR | 1 | 0 | 0 | 1542725 | P0DTD1.1 | P0DTD1 |
| TEVNEFACVVADAVI | 1 | 0 | 0 | 1542726 | P0DTC1.1 | P0DTD1 |
| TFEEAALCTFLLNKE | 1 | 0 | 0 | 1542730 | P0DTC1.1 | P0DTD1 |
| TFFIYNKIVDEPEEH | 1 | 0 | 0 | 1542732 | P0DTC3.1 | P0DTC3 |
| TFISAARQGFVDSDV | 1 | 0 | 0 | 1542736 | P0DTC1.1 | P0DTD1 |
| TFVTHSKGLYRKCVK | 1 | 0 | 0 | 1542745 | P0DTC1.1 | P0DTD1 |
| TGGVVQLTSQWLTNI | 1 | 0 | 0 | 1542750 | P0DTC1.1 | P0DTD1 |
| TGNTLQCIMLVYCFL | 1 | 0 | 0 | 1542755 | P0DTC1.1 | P0DTD1 |
| TGTSTDVVYRAFDIY | 1 | 0 | 0 | 1542761 | P0DTD1.1 | P0DTD1 |
| TGVEHVTFFIYNKIV | 1 | 0 | 0 | 1542762 | P0DTC3.1 | P0DTC3 |

| | | | | | | |
|---|---|---|---|---|---|---|
| TGVVGEGSEGLNDNL | 1 | 0 | 0 | 1542765 | P0DTC1.1 | P0DTD1 |
| TGYHFRELGVVHNQD | 1 | 0 | 0 | 1542766 | P0DTD1.1 | P0DTD1 |
| THLSVDTKFKTEGLC | 1 | 0 | 0 | 1542769 | P0DTD1.1 | P0DTD1 |
| THVQLSLPVLQVRDV | 1 | 0 | 0 | 1542771 | P0DTC1.1 | P0DTD1 |
| TICAPLTVFFDGRVD | 1 | 0 | 0 | 1542773 | P0DTD1.1 | P0DTD1 |
| TILDGISQYSLRLID | 1 | 0 | 0 | 1542782 | P0DTC1.1 | P0DTD1 |
| TILTRPLLESELVIG | 1 | 0 | 0 | 1542785 | YP_009724393.1 | P0DTC5 |
| TIQKDVLECNVKTTE | 1 | 0 | 0 | 1542788 | P0DTC1.1 | P0DTD1 |
| TIVEVQPQLEMELTP | 1 | 0 | 0 | 1542796 | P0DTC1.1 | P0DTD1 |
| TKFLTENLLLYIDIN | 1 | 0 | 0 | 1542804 | P0DTC1.1 | P0DTD1 |
| TKGSLPINVIVFDGK | 1 | 0 | 0 | 1542807 | P0DTC1.1 | P0DTD1 |
| TKHFYWFFSNYLKRR | 1 | 0 | 0 | 1542808 | P0DTC1.1 | P0DTD1 |
| TKHPNQEYADVFHLY | 1 | 0 | 0 | 1542809 | P0DTD1.1 | P0DTD1 |
| TKIALKGGKIVNNWL | 1 | 0 | 0 | 1542810 | P0DTC1.1 | P0DTD1 |
| TKPYIKWDLLKYDFT | 1 | 0 | 0 | 1542815 | P0DTD1.1 | P0DTD1 |
| TKYTMADLVYALRHF | 1 | 0 | 0 | 1542819 | P0DTD1.1 | P0DTD1 |
| TLTLAVPYNMRVIHF | 1 | 0 | 0 | 1542843 | P0DTD1.1 | P0DTD1 |
| TLTRALTAESHVDTD | 1 | 0 | 0 | 1542844 | P0DTD1.1 | P0DTD1 |
| TLVTMPLGYVTHGLN | 1 | 0 | 0 | 1542846 | P0DTC1.1 | P0DTD1 |
| TLYCIDGALLTKSSE | 1 | 0 | 0 | 1542847 | P0DTC1.1 | P0DTD1 |
| TMCDIRQLLFVVEVV | 1 | 0 | 0 | 1542848 | P0DTD1.1 | P0DTD1 |
| TMYICGDSTECSNLL | 1 | 0 | 0 | 1542851 | P0DTC2.1 | P0DTC2 |
| TNDNTSRYWEPEFYE | 1 | 0 | 0 | 1542854 | P0DTD1.1 | P0DTD1 |
| TNIVTRCLNRVCTNY | 1 | 0 | 0 | 1542856 | P0DTC1.1 | P0DTD1 |
| TNVLEGSVAYESLRP | 1 | 0 | 0 | 1542863 | P0DTC1.1 | P0DTD1 |
| TPCGTGTSTDVVYRA | 1 | 0 | 0 | 1542867 | P0DTD1.1 | P0DTD1 |
| TPEEHFIETISLAGS | 1 | 0 | 0 | 1542868 | P0DTC1.1 | P0DTD1 |
| TPGSGVPVVDSYYSL | 1 | 0 | 0 | 1542870 | P0DTD1.1 | P0DTD1 |
| TPHTVLQAVGACVLC | 1 | 0 | 0 | 1542873 | P0DTD1.1 | P0DTD1 |
| TPLIQPIGALDISAS | 1 | 0 | 0 | 1542874 | P0DTC1.1 | P0DTD1 |
| TPRDLGACIDCSARH | 1 | 0 | 0 | 1542876 | P0DTC1.1 | P0DTD1 |
| TQLSTDTGVEHVTFF | 1 | 0 | 0 | 1542882 | P0DTC3.1 | P0DTC3 |
| TRAKVGILCIMSDRD | 1 | 0 | 0 | 1542889 | P0DTD1.1 | P0DTD1 |
| TRALTAESHVDTDLT | 1 | 0 | 0 | 1542890 | P0DTD1.1 | P0DTD1 |
| TRCLAVHECFVKRVD | 1 | 0 | 0 | 1542891 | P0DTD1.1 | P0DTD1 |
| TRCLNRVCTNYMPYF | 1 | 0 | 0 | 1542892 | P0DTC1.1 | P0DTD1 |
| TRCNLGGAVCRHHAN | 1 | 0 | 0 | 1542893 | P0DTD1.1 | P0DTD1 |
| TSAVLQSGFRKMAFP | 1 | 0 | 0 | 1542903 | P0DTC1.1 | P0DTD1 |
| TSEAVEAPLVGTPVC | 1 | 0 | 0 | 1542904 | P0DTC1.1 | P0DTD1 |
| TSEDMLNPNYEDLLI | 1 | 0 | 0 | 1542905 | P0DTC1.1 | P0DTD1 |
| TSFGPLVRKIFVDGV | 1 | 0 | 0 | 1542906 | P0DTD1.1 | P0DTD1 |
| TSGRWVLNNDYYRSL | 1 | 0 | 0 | 1542907 | P0DTC1.1 | P0DTD1 |
| TSLRCGACIRRPFLC | 1 | 0 | 0 | 1542913 | P0DTD1.1 | P0DTD1 |
| TSNPTTFHLDGEVIT | 1 | 0 | 0 | 1542917 | P0DTC1.1 | P0DTD1 |
| TSRYWEPEFYEAMYT | 1 | 0 | 0 | 1542918 | P0DTD1.1 | P0DTD1 |
| TSSGDATTAYANSVF | 1 | 0 | 0 | 1542919 | P0DTD1.1 | P0DTD1 |
| TSVVLLSVLQQLRVE | 1 | 0 | 0 | 1542921 | P0DTC1.1 | P0DTD1 |
| TTEAFEKMVSLLSVL | 1 | 0 | 0 | 1542925 | P0DTC1.1 | P0DTD1 |
| TTEILPVSMTKTSVD | 1 | 0 | 0 | 1542927 | P0DTC2.1 | P0DTC2 |
| TTIQTIVEVQPQLEM | 1 | 0 | 0 | 1542931 | P0DTC1.1 | P0DTD1 |
| TTKGGRFVLALLSDL | 1 | 0 | 0 | 1542933 | P0DTC1.1 | P0DTD1 |
| TTLKGVEAVMYMGTL | 1 | 0 | 0 | 1542935 | P0DTC1.1 | P0DTD1 |
| TTNGDFLHFLPRVFS | 1 | 0 | 0 | 1542938 | P0DTC1.1 | P0DTD1 |
| TTQTACTDDNALAYY | 1 | 0 | 0 | 1542939 | P0DTC1.1 | P0DTD1 |
| TTRQVVNVVTTKIAL | 1 | 0 | 0 | 1542940 | P0DTC1.1 | P0DTD1 |
| TTTIKPVTYKLDGVV | 1 | 0 | 0 | 1542943 | P0DTC1.1 | P0DTD1 |
| TVAYFNMVYMPASWV | 1 | 0 | 0 | 1542952 | P0DTC1.1 | P0DTD1 |
| TVFFDGRVDGQVDLF | 1 | 0 | 0 | 1542957 | P0DTD1.1 | P0DTD1 |
| TVMFLARGIVFMCVE | 1 | 0 | 0 | 1542964 | P0DTC1.1 | P0DTD1 |
| TVRTNVYLAVFDKNL | 1 | 0 | 0 | 1542966 | P0DTC1.1 | P0DTD1 |
| TVYSDVENPHLMGWD | 1 | 0 | 0 | 1542971 | P0DTD1.1 | P0DTD1 |
| TVYTKVDGVDVELFE | 1 | 0 | 0 | 1542973 | P0DTD1.1 | P0DTD1 |
| TYACWHHSIGFDYVY | 1 | 0 | 0 | 1542975 | P0DTD1.1 | P0DTD1 |
| TYASALWEIQQVVDA | 1 | 0 | 0 | 1542976 | P0DTC1.1 | P0DTD1 |
| TYERHSLSHFVNLDN | 1 | 0 | 0 | 1542978 | P0DTC1.1 | P0DTD1 |
| TYNCCDDDYFNKKDW | 1 | 0 | 0 | 1542981 | P0DTD1.1 | P0DTD1 |
| TYPSLETIQITISSF | 1 | 0 | 0 | 1542984 | P0DTC1.1 | P0DTD1 |
| TYYLFDESGEFKLAS | 1 | 0 | 0 | 1542986 | P0DTC1.1 | P0DTD1 |
| VAEWFLAYILFTRFF | 1 | 0 | 0 | 1542989 | P0DTC1.1 | P0DTD1 |
| VAFNTLLFLMSFTVL | 1 | 0 | 0 | 1542991 | P0DTC1.1 | P0DTD1 |
| VAGFAKFLKTNCCRF | 1 | 0 | 0 | 1542993 | P0DTD1.1 | P0DTD1 |
| VAKSEFDRDAAMQRK | 1 | 0 | 0 | 1542998 | P0DTC1.1 | P0DTD1 |
| VANGDSEVVLKKLKK | 1 | 0 | 0 | 1543001 | P0DTC1.1 | P0DTD1 |
| VASPNECNQMCLSTL | 1 | 0 | 0 | 1543006 | P0DTC1.1 | P0DTD1 |
| VAVKMFDAYVNTFSS | 1 | 0 | 0 | 1543008 | P0DTC1.1 | P0DTD1 |
| VCVDTVRTNVYLAVF | 1 | 0 | 0 | 1543019 | P0DTC1.1 | P0DTD1 |
| VCVSTSGRWVLNNDY | 1 | 0 | 0 | 1543020 | P0DTC1.1 | P0DTD1 |
| VDADSKIVQLSEISM | 1 | 0 | 0 | 1543022 | P0DTC1.1 | P0DTD1 |
| VDALCEKALKYLPID | 1 | 0 | 0 | 1543023 | P0DTD1.1 | P0DTD1 |
| VDRYPANSIVCRFDT | 1 | 0 | 0 | 1543034 | P0DTD1.1 | P0DTD1 |
| VDSDVETKDVVECLK | 1 | 0 | 0 | 1543036 | P0DTC1.1 | P0DTD1 |
| VDTANPKTPKYKFVR | 1 | 0 | 0 | 1543037 | P0DTC1.1 | P0DTD1 |
| VECLKLSHQSDIEVT | 1 | 0 | 0 | 1543048 | P0DTC1.1 | P0DTD1 |
| VEHVTFFIYNKIVDE | 1 | 0 | 0 | 1543054 | P0DTC3.1 | P0DTC3 |
| VEKKKLDGFMGRIRS | 1 | 0 | 0 | 1543057 | P0DTC1.1 | P0DTD1 |
| VENMTPRDLGACIDC | 1 | 0 | 0 | 1543061 | P0DTC1.1 | P0DTD1 |
| VEQRKQDDKKIKACV | 1 | 0 | 0 | 1543064 | P0DTC1.1 | P0DTD1 |
| VETVKGLDYKAFKQI | 1 | 0 | 0 | 1543067 | P0DTC1.1 | P0DTD1 |
| VEVQPQLEMELTPVV | 1 | 0 | 0 | 1543069 | P0DTC1.1 | P0DTD1 |
| VFISPYNSQNAVASK | 1 | 0 | 0 | 1543073 | P0DTD1.1 | P0DTD1 |
| VFPLNSIIKTIQPRV | 1 | 0 | 0 | 1543078 | P0DTC1.1 | P0DTD1 |

| | | | | | | |
|---|---|---|---|---|---|---|
| VGDFKLNEEIAIILA | 1 | 0 | 0 | 1543087 | P0DTC1.1 | P0DTD1 |
| VGDSAEVAVKMFDAY | 1 | 0 | 0 | 1543089 | P0DTC1.1 | P0DTD1 |
| VGELGDVRETMSYLF | 1 | 0 | 0 | 1543090 | P0DTC1.1 | P0DTD1 |
| VGHTDLMAAYVDNSS | 1 | 0 | 0 | 1543093 | P0DTC1.1 | P0DTD1 |
| VGPEHSLAEYHNESG | 1 | 0 | 0 | 1543098 | P0DTC1.1 | P0DTD1 |
| VGQQDGSEDNQTTTI | 1 | 0 | 0 | 1543101 | P0DTC1.1 | P0DTD1 |
| VGVLTLDNQDLNGNW | 1 | 0 | 0 | 1543105 | P0DTD1.1 | P0DTD1 |
| VHFISNSWLMWLIIN | 1 | 0 | 0 | 1543109 | P0DTC1.1 | P0DTD1 |
| VHNQDVNLHSSRLSF | 1 | 0 | 0 | 1543112 | P0DTD1.1 | P0DTD1 |
| VHTANKWDLIISDMY | 1 | 0 | 0 | 1543113 | P0DTD1.1 | P0DTD1 |
| VHVVDGCNSSTCMMC | 1 | 0 | 0 | 1543114 | P0DTC1.1 | P0DTD1 |
| VIEVQGYKSVNITFE | 1 | 0 | 0 | 1543116 | P0DTC1.1 | P0DTD1 |
| VIFTQTTETAHSCNV | 1 | 0 | 0 | 1543117 | P0DTD1.1 | P0DTD1 |
| VIGFLFLTWICLLQF | 1 | 0 | 0 | 1543119 | P0DTC5.1 | P0DTC5 |
| VISTSHKLVLSVNPY | 1 | 0 | 0 | 1543126 | P0DTD1.1 | P0DTD1 |
| VKMFDAYVNTFSSTF | 1 | 0 | 0 | 1543134 | P0DTC1.1 | P0DTD1 |
| VKNGSIHLYFDKAGQ | 1 | 0 | 0 | 1543135 | P0DTC1.1 | P0DTD1 |
| VKQLSSNFGAISSVL | 1 | 0 | 0 | 1543139 | P0DTC2.1 | P0DTC2 |
| VKRHTFSNYQHEETI | 1 | 0 | 0 | 1543140 | P0DTD1.1 | P0DTD1 |
| VKTTEVVGDIILKPA | 1 | 0 | 0 | 1543145 | P0DTC1.1 | P0DTD1 |
| VKVTIDYTEISFMLW | 1 | 0 | 0 | 1543146 | P0DTD1.1 | P0DTD1 |
| VLGLAAIMQLFFSYF | 1 | 0 | 0 | 1543150 | P0DTC1.1 | P0DTD1 |
| VLNEKCSAYTVELGT | 1 | 0 | 0 | 1543156 | P0DTC1.1 | P0DTD1 |
| VLPNDDTLRVEAFEY | 1 | 0 | 0 | 1543157 | P0DTC1.1 | P0DTD1 |
| VLQVRDVLVRGFGDS | 1 | 0 | 0 | 1543160 | P0DTC1.1 | P0DTD1 |
| VLSEMVMCGGSLYVK | 1 | 0 | 0 | 1543163 | P0DTD1.1 | P0DTD1 |
| VLSNLNLPGCDGGSL | 1 | 0 | 0 | 1543165 | P0DTD1.1 | P0DTD1 |
| VLTLVYKVYYGNALD | 1 | 0 | 0 | 1543167 | P0DTC1.1 | P0DTD1 |
| VLVPHVGEIPVAYRK | 1 | 0 | 0 | 1543168 | P0DTC1.1 | P0DTD1 |
| VMRIMTWLDMVDTSL | 1 | 0 | 0 | 1543176 | P0DTC1.1 | P0DTD1 |
| VNALPETTADIVVFD | 1 | 0 | 0 | 1543180 | P0DTD1.1 | P0DTD1 |
| VNKFLALCADSIIIG | 1 | 0 | 0 | 1543186 | P0DTC1.1 | P0DTD1 |
| VPHVGEIPVAYRKVL | 1 | 0 | 0 | 1543204 | P0DTC1.1 | P0DTD1 |
| VPMEKLKTLVATAEA | 1 | 0 | 0 | 1543206 | P0DTC1.1 | P0DTD1 |
| VQLHNDILLAKDTTE | 1 | 0 | 0 | 1543215 | P0DTC1.1 | P0DTD1 |
| VQLSLPVLQVRDVLV | 1 | 0 | 0 | 1543217 | P0DTC1.1 | P0DTD1 |
| VQMAPISAMVRMYIF | 1 | 0 | 0 | 1543219 | P0DTC1.1 | P0DTD1 |
| VQPQLEMELTPVVQT | 1 | 0 | 0 | 1543221 | P0DTC1.1 | P0DTD1 |
| VQQLPETYFTQSRNL | 1 | 0 | 0 | 1543223 | P0DTD1.1 | P0DTD1 |
| VQTIEVNSFSGYLKL | 1 | 0 | 0 | 1543225 | P0DTC1.1 | P0DTD1 |
| VRCSFYEDFLEYHDV | 1 | 0 | 0 | 1543228 | P0DTC8.1 | P0DTC8 |
| VRETMSYLFQHANLD | 1 | 0 | 0 | 1543230 | P0DTC1.1 | P0DTD1 |
| VREVLSDRELHLSWE | 1 | 0 | 0 | 1543231 | P0DTD1.1 | P0DTD1 |
| VRLQAGNATEVPANS | 1 | 0 | 0 | 1543235 | P0DTC1.1 | P0DTD1 |
| VRNLQHRLYECLYRN | 1 | 0 | 0 | 1543236 | P0DTD1.1 | P0DTD1 |
| VRQCSGVTFQSAVKR | 1 | 0 | 0 | 1543238 | P0DTC1.1 | P0DTD1 |
| VSDADSTLIGDCATV | 1 | 0 | 0 | 1543245 | P0DTD1.1 | P0DTD1 |
| VSELLTPLGIDLDEW | 1 | 0 | 0 | 1543246 | P0DTC1.1 | P0DTD1 |
| VSFLAHIQWMVMFTP | 1 | 0 | 0 | 1543248 | P0DTC1.1 | P0DTD1 |
| VSFSTFEEAALCTFL | 1 | 0 | 0 | 1543249 | P0DTC1.1 | P0DTD1 |
| VSKVVKVTIDYTEIS | 1 | 0 | 0 | 1543254 | P0DTD1.1 | P0DTD1 |
| VSLDNVLSTFISAAR | 1 | 0 | 0 | 1543256 | P0DTC1.1 | P0DTD1 |
| VSPKLFIRQEEVQEL | 1 | 0 | 0 | 1543258 | P0DTC7.1 | P0DTC7 |
| VSSAINRPQIGVVRE | 1 | 0 | 0 | 1543259 | P0DTD1.1 | P0DTD1 |
| VSTGYHFRELGVVHN | 1 | 0 | 0 | 1543260 | P0DTD1.1 | P0DTD1 |
| VSTSGRWVLNNDYYR | 1 | 0 | 0 | 1543261 | P0DTC1.1 | P0DTD1 |
| VTHSKGLYRKCVKSR | 1 | 0 | 0 | 1543267 | P0DTC1.1 | P0DTD1 |
| VTMPLGYVTHGLNLE | 1 | 0 | 0 | 1543273 | P0DTC1.1 | P0DTD1 |
| VTSNYSGVVTTVMFL | 1 | 0 | 0 | 1543280 | P0DTC1.1 | P0DTD1 |
| VTTKIALKGGKIVNN | 1 | 0 | 0 | 1543283 | P0DTC1.1 | P0DTD1 |
| VTVKNGSIHLYFDKA | 1 | 0 | 0 | 1543285 | P0DTC1.1 | P0DTD1 |
| VTYKLDGVVCTEIDP | 1 | 0 | 0 | 1543286 | P0DTC1.1 | P0DTD1 |
| VVADAVIKTLQPVSE | 1 | 0 | 0 | 1543287 | P0DTC1.1 | P0DTD1 |
| VVDDPCPIHFYSKWY | 1 | 0 | 0 | 1543289 | P0DTC8.1 | P0DTC8 |
| VVGEGSEGLNDNLLE | 1 | 0 | 0 | 1543295 | P0DTC1.1 | P0DTD1 |
| VVLHSYFTSDYYQLY | 1 | 0 | 0 | 1543298 | P0DTC3.1 | P0DTC3 |
| VVNVVTTKIALKGGK | 1 | 0 | 0 | 1543303 | P0DTC1.1 | P0DTD1 |
| VYANGGKGFCKLHNW | 1 | 0 | 0 | 1543313 | P0DTC1.1 | P0DTD1 |
| VYCCREHEHEIAWYT | 1 | 0 | 0 | 1543316 | P0DTC1.1 | P0DTD1 |
| VYDDGARRVWTLMNV | 1 | 0 | 0 | 1543318 | P0DTC1.1 | P0DTD1 |
| VYIKNADIVEEAKKV | 1 | 0 | 0 | 1543323 | P0DTC1.1 | P0DTD1 |
| VYKVYYGNALDQAIS | 1 | 0 | 0 | 1543324 | P0DTC1.1 | P0DTD1 |
| VYQLRARSVSPKLFI | 1 | 0 | 0 | 1543326 | P0DTC7.1 | P0DTC7 |
| VYRGTTTYKLNVGDY | 1 | 0 | 0 | 1543327 | P0DTD1.1 | P0DTD1 |
| WDYKRDAPAHISTIG | 1 | 0 | 0 | 1543340 | P0DTD1.1 | P0DTD1 |
| WEPEFYEAMYTPHTV | 1 | 0 | 0 | 1543342 | P0DTD1.1 | P0DTD1 |
| WESGVKDCVVLHSYF | 1 | 0 | 0 | 1543343 | P0DTC3.1 | P0DTC3 |
| WHHSIGFDYVYNPFM | 1 | 0 | 0 | 1543347 | P0DTD1.1 | P0DTD1 |
| WKCRSKNPLLYDANY | 1 | 0 | 0 | 1543352 | P0DTC3.1 | P0DTC3 |
| WKYPQVNGLTSIKWA | 1 | 0 | 0 | 1543355 | P0DTC1.1 | P0DTD1 |
| WLPTGTLLVDSDLND | 1 | 0 | 0 | 1543361 | P0DTD1.1 | P0DTD1 |
| WNVVRIKIVQMLSDT | 1 | 0 | 0 | 1543370 | P0DTD1.1 | P0DTD1 |
| WRKAVFISPYNSQNA | 1 | 0 | 0 | 1543373 | P0DTD1.1 | P0DTD1 |
| WSLFFFLYENAFLPF | 1 | 0 | 0 | 1543375 | P0DTC1.1 | P0DTD1 |
| WSMATYYLFDESGEF | 1 | 0 | 0 | 1543376 | P0DTC1.1 | P0DTD1 |
| WSYSGQSTQLGIEFL | 1 | 0 | 0 | 1543378 | P0DTC1.1 | P0DTD1 |
| WTNAGDYILANTCTE | 1 | 0 | 0 | 1543381 | P0DTD1.1 | P0DTD1 |
| WVLNNDYYRSLPGVF | 1 | 0 | 0 | 1543383 | P0DTC1.1 | P0DTD1 |
| WVYKQFDTYNLWNTF | 1 | 0 | 0 | 1543384 | P0DTD1.1 | P0DTD1 |
| WYDFVENPDILRVYA | 1 | 0 | 0 | 1543386 | P0DTD1.1 | P0DTD1 |
| YAAVINGDRWFLNRF | 1 | 0 | 0 | 1543392 | P0DTC1.1 | P0DTD1 |

| | | | | | | |
|---|---|---|---|---|---|---|
| YAFEHIVYGDFSHSQ | 1 | 0 | 0 | 1543396 | P0DTD1.1 | P0DTD1 |
| YCYDTNVLEGSVAYE | 1 | 0 | 0 | 1543403 | P0DTC1.1 | P0DTD1 |
| YDAQPCSDKAYKIEE | 1 | 0 | 0 | 1543405 | P0DTD1.1 | P0DTD1 |
| YDFAVSKGFFKEGSS | 1 | 0 | 0 | 1543406 | P0DTD1.1 | P0DTD1 |
| YDSMSYEDQDALFAY | 1 | 0 | 0 | 1543409 | P0DTD1.1 | P0DTD1 |
| YDTNVLEGSVAYESL | 1 | 0 | 0 | 1543410 | P0DTC1.1 | P0DTD1 |
| YECLYRNRDVDTDFV | 1 | 0 | 0 | 1543411 | P0DTD1.1 | P0DTD1 |
| YELKHGTFTCASEYT | 1 | 0 | 0 | 1543414 | P0DTC1.1 | P0DTD1 |
| YEPLTQDHVDILGPL | 1 | 0 | 0 | 1543415 | P0DTC1.1 | P0DTD1 |
| YEQAVANGDSEVVLK | 1 | 0 | 0 | 1543416 | P0DTC1.1 | P0DTD1 |
| YFCTCYFGLFCLLNR | 1 | 0 | 0 | 1543418 | P0DTC1.1 | P0DTD1 |
| YFLQSINFVRIIMRL | 1 | 0 | 0 | 1543425 | P0DTC3.1 | P0DTC3 |
| YGQQFGPTYLDGADV | 1 | 0 | 0 | 1543433 | P0DTC1.1 | P0DTD1 |
| YHFRELGVVHNQDVN | 1 | 0 | 0 | 1543435 | P0DTD1.1 | P0DTD1 |
| YILPSIISNEKQEIL | 1 | 0 | 0 | 1543449 | P0DTC1.1 | P0DTD1 |
| YIWLGFIAGLIAIVM | 1 | 0 | 0 | 1543451 | P0DTC2.1 | P0DTC2 |
| YKAFKQIVESCGNFK | 1 | 0 | 0 | 1543452 | P0DTC1.1 | P0DTD1 |
| YKHITSKETLYCIDG | 1 | 0 | 0 | 1543457 | P0DTC1.1 | P0DTD1 |
| YKRDAPAHISTIGVC | 1 | 0 | 0 | 1543466 | P0DTD1.1 | P0DTD1 |
| YKSVNITFELDERID | 1 | 0 | 0 | 1543467 | P0DTC1.1 | P0DTD1 |
| YKYFSGAMDTTSYRE | 1 | 0 | 0 | 1543468 | P0DTC1.1 | P0DTD1 |
| YLAVFDKNLYDKLVS | 1 | 0 | 0 | 1543469 | P0DTC1.1 | P0DTD1 |
| YLEGSVRVVTTFDSE | 1 | 0 | 0 | 1543471 | P0DTC1.1 | P0DTD1 |
| YLFDESGEFKLASHM | 1 | 0 | 0 | 1543472 | P0DTC1.1 | P0DTD1 |
| YLITPVHVMSKHTDF | 1 | 0 | 0 | 1543476 | P0DTC1.1 | P0DTD1 |
| YLKRRVVFNGVSFST | 1 | 0 | 0 | 1543479 | P0DTC1.1 | P0DTD1 |
| YLNSTNVTIATYCTG | 1 | 0 | 0 | 1543480 | P0DTC1.1 | P0DTD1 |
| YLPYPDPSRILGAGC | 1 | 0 | 0 | 1543482 | P0DTD1.1 | P0DTD1 |
| YMPYFFTLLLQLCTF | 1 | 0 | 0 | 1543492 | P0DTC1.1 | P0DTD1 |
| YNLWNTFTRLQSLEN | 1 | 0 | 0 | 1543498 | P0DTD1.1 | P0DTD1 |
| YPLECIKDLLARAGK | 1 | 0 | 0 | 1543506 | P0DTC1.1 | P0DTD1 |
| YPNMFITREEAIRHV | 1 | 0 | 0 | 1543508 | P0DTD1.1 | P0DTD1 |
| YSFLPGVYSVIYLYL | 1 | 0 | 0 | 1543526 | P0DTC1.1 | P0DTD1 |
| YSHLLLVAAGLEAPF | 1 | 0 | 0 | 1543529 | P0DTC3.1 | P0DTC3 |
| YTKRNVIPTITQMNL | 1 | 0 | 0 | 1543543 | P0DTD1.1 | P0DTD1 |
| YVFIKRSDARTAPHG | 1 | 0 | 0 | 1543552 | P0DTC1.1 | P0DTD1 |
| YVWKSYVHVVDGCNS | 1 | 0 | 0 | 1543561 | P0DTC1.1 | P0DTD1 |
| YVYNPFMIDVQQWGF | 1 | 0 | 0 | 1543564 | P0DTD1.1 | P0DTD1 |
| YYFMRFRRAFGEYSH | 1 | 0 | 0 | 1543568 | P0DTC1.1 | P0DTD1 |
| YYSQLMCQPILLLDQ | 1 | 0 | 0 | 1543578 | P0DTC1.1 | P0DTD1 |
| YYTSNPTTFHLDGEV | 1 | 0 | 0 | 1543579 | P0DTC1.1 | P0DTD1 |
| ASYQTQTNSPRRARSVASQ | 1 | 0 | 0 | 1562513 | QHD43416.1 | P0DTC2 |
| ELLHAPATVCGPKKSTNLVKN | 1 | 0 | 0 | 1567893 | QHD43416.1 | P0DTC2 |
| FGAGAALQIPFAMQMAYRFNGI | 1 | 0 | 0 | 1569850 | QHD43416.1 | P0DTC2 |
| FSNVTWFHAIHVSGTNGTKRFDN | 1 | 0 | 0 | 1571025 | QHD43416.1 | P0DTC2 |
| GDEVRQIAPGQTGKIADYNYKLP | 1 | 0 | 0 | 1571473 | QHD43416.1 | P0DTC2 |
| SQCVNLTTRTQLPPAYTNSFTRGVY | 1 | 0 | 0 | 1588135 | QHD43416.1 | P0DTC2 |
| YNSASFSTFKCYGVSPTKLNDLCFT | 1 | 0 | 0 | 1594118 | QHD43416.1 | P0DTC2 |
| DFEAELSLWFKNHTS | 1 | 0 | 0 | 1594868 | Q0ZME7.1 | A0A140H1H1 |
| DFKDELEEFFKNVTS | 1 | 0 | 0 | 1594870 | QPX50158.1 | A3EX94 |
| FNATRFASVYAWNRKRISNCVADYS | 1 | 0 | 0 | 1595678 | QHD43416.1 | P0DTC2 |
| KPFERDISTEIYQAGSTPCNGVEGF | 1 | 0 | 0 | 1595939 | QHD43416.1 | P0DTC2 |
| TDEMIAQYTSALLAGTITSGW | 1 | 0 | 0 | 1596332 | QHD43416.1 | P0DTC2 |
| TMSLGAENSVAYSNNSIAIPTNFTI | 1 | 0 | 0 | 1596356 | QHD43416.1 | P0DTC2 |
| VADYSVLYNSASFSTFKCYGVSPTK | 1 | 0 | 0 | 1596399 | QHD43416.1 | P0DTC2 |
| AALQIPFAMQMAYRFNGIGVTQNVL | 1 | 0 | 0 | 1597650 | YP_009724390.1 | P0DTC2 |
| CTEVPVAIHADQLTPTWRVYSTGSN | 1 | 0 | 0 | 1597659 | YP_009724390.1 | P0DTC2 |
| DISTEIYQAGSTPCNGVEGFNCYFP | 1 | 0 | 0 | 1597664 | YP_009724390.1 | P0DTC2 |
| DSKVGGNYNYLYRLFRKSNLKPFER | 1 | 0 | 0 | 1597667 | YP_009724390.1 | P0DTC2 |
| FCGKGYHLMSFPQSAPHGVVFLHVT | 1 | 0 | 0 | 1597679 | YP_009724390.1 | P0DTC2 |
| FGMSRIGMEVTPSGTWLTYTGAIKL | 1 | 0 | 0 | 1597682 | P0DTC9-1 | P0DTC9 |
| FTNVYADSFVIRGDEVRQIAPGQTG | 1 | 0 | 0 | 1597691 | YP_009724390.1 | P0DTC2 |
| GCVIAWNSNNLDSKVGGNYNYLYR | 1 | 0 | 0 | 1597696 | YP_009724390.1 | P0DTC2 |
| GDIAARDLICAQKFNGLTVLPPLLT | 1 | 0 | 0 | 1597697 | YP_009724390.1 | P0DTC2 |
| ICASYQTQTNSPRRARSVAS | 1 | 0 | 0 | 1597705 | QHD43416.1 | P0DTC2 |
| IFGTTLDSKTQSLLIVNNAT | 1 | 0 | 0 | 1597707 | QHD43416.1 | P0DTC2 |
| KIADYNYKLPDDFTGCVIAWNSNNL | 1 | 0 | 0 | 1597719 | YP_009724390.1 | P0DTC2 |
| LALHRSYLTPGDSSSGWTAGAAAYY | 1 | 0 | 0 | 1597725 | YP_009724390.1 | P0DTC2 |
| LQSYGFQPTNGVGYQPYRVVVLSFE | 1 | 0 | 0 | 1597743 | YP_009724390.1 | P0DTC2 |
| LVKPSFYVYSRVKNLNSSRVPDL | 1 | 0 | 0 | 1597756 | P0DTC4.1 | P0DTC4 |
| NLTTRTQLPPAYTNSFTRGVYYPDK | 1 | 0 | 0 | 1597771 | YP_009724390.1 | P0DTC2 |
| PFTIYSLLLCRMNSRNYI | 1 | 0 | 0 | 1597779 | QHI42199.1 | A0A663DJA2 |
| PQGFSALEPLVDLPIGINITRFQTL | 1 | 0 | 0 | 1597782 | YP_009724390.1 | P0DTC2 |
| RDIADTTDAVRDPQTLEILDITPCS | 1 | 0 | 0 | 1597792 | YP_009724390.1 | P0DTC2 |
| RSMWSFNPETNILLNVPLHG | 1 | 0 | 0 | 1597799 | P0DTC5.1 | P0DTC5 |
| SNKKFLPFQQFGRDIADTTD | 1 | 0 | 0 | 1597808 | QHD43416.1 | P0DTC2 |
| SSNFGAISSVLNDILSRLDKVEAEV | 1 | 0 | 0 | 1597812 | YP_009724390.1 | P0DTC2 |
| VFRSSVLHSTQDLFLPFFSNVTWFH | 1 | 0 | 0 | 1597835 | YP_009724390.1 | P0DTC2 |
| VLYNSASFSTFKCYGVSPTKLNDLC | 1 | 0 | 0 | 1597840 | YP_009724390.1 | P0DTC2 |
| WLAMAVMLLLLCCCLTDMGYINVF | 1 | 0 | 0 | 1597849 | A0A663DJA2.1 | A0A663DJA2 |
| WMESEFRVYSSANNCTFEYV | 1 | 0 | 0 | 1597854 | QHD43416.1 | P0DTC2 |
| GQSKRVDFCGKGYHLMSFPQSAPHG | 1 | 0 | 0 | 1625377 | P0DTC2.1 | P0DTC2 |
| KPSFYVYSRVKNLNSSRVPDLLV | 1 | 0 | 0 | 1625395 | P0DTC4.1 | P0DTC4 |
| LFRKSNLKPFERDISTEIYQAGST | 1 | 0 | 0 | 1625406 | P0DTC2.1 | P0DTC2 |
| LGVYYHKNNKSWMESEFRVYSSA | 1 | 0 | 0 | 1625407 | P0DTC2.1 | P0DTC2 |
| RYRIGNYKLNTDHSSSSDNIA | 1 | 0 | 0 | 1625429 | P0DTC5.1 | P0DTC5 |
| VEGFNCYFPLQSYGFQPTNGVGYQP | 1 | 0 | 0 | 1625453 | P0DTC2.1 | P0DTC2 |
| AAAGIMKNPTVDGIT | 1 | 0 | 0 | 1636012 | P14340.2 | P17763 |
| AAEMEEALKGLPIRY | 1 | 0 | 0 | 1636018 | QTX93489.1 | P17763 |
| AAILAYTIGTTYFQR | 1 | 0 | 0 | 1636022 | QQW40781.1 | P17763 |
| AAILMGLDKGWPISK | 1 | 0 | 0 | 1636023 | P17763.2 | P17763 |

| | | | | | | |
|---|---|---|---|---|---|---|
| AAINQVRSLIGNEEY | 1 | 0 | 0 | 1636024 | QTX93142.1 | P17763 |
| AALLLLVAHYAIIGP | 1 | 0 | 0 | 1636028 | QTX92981.1 | P17763 |
| AAMGVPPLPLFIFSL | 1 | 0 | 0 | 1636032 | QTX93461.1 | P17763 |
| AAVLLLITHYAIIGP | 1 | 0 | 0 | 1636039 | QBQ58390.1 | P17763 |
| AAVLMGLGKGWPLHR | 1 | 0 | 0 | 1636040 | Q2YHF0.1 | P17763 |
| AAVSVSPLLLTSSQQ | 1 | 0 | 0 | 1636046 | QRG47124.1 | P17763 |
| ACLRKNGKKVIQLSR | 1 | 0 | 0 | 1636050 | P14340.2 | P17763 |
| ADDIGMGVTYLALLA | 1 | 0 | 0 | 1636051 | QTX93044.1 | P17763 |
| ADLELERAADVRWED | 1 | 0 | 0 | 1636058 | QQW40684.1 | P17763 |
| ADLELERATDVKWED | 1 | 0 | 0 | 1636059 | QTX93081.1 | P17763 |
| ADLSLEKAAEVSWEE | 1 | 0 | 0 | 1636060 | P17763.2 | P17763 |
| ADLSLEKAAEVSWEQ | 1 | 0 | 0 | 1636061 | QNH29581.1 | P17763 |
| ADLTVEKAADVTWEE | 1 | 0 | 0 | 1636062 | QTX93483.1 | P17763 |
| ADVRWEDQAEISGSS | 1 | 0 | 0 | 1636065 | QQW40684.1 | P17763 |
| ADVTWEEEAEQTGVS | 1 | 0 | 0 | 1636066 | QTX93491.1 | P17763 |
| ADWIPLALTIKGLNP | 1 | 0 | 0 | 1636067 | P14340.2 | P17763 |
| AEAIFKLTYQNKVVR | 1 | 0 | 0 | 1636071 | QEV86434.1 | P17763 |
| AEGAWRQVEKVETWA | 1 | 0 | 0 | 1636072 | QTX93479.1 | P17763 |
| AEGVITQDDMQNPKG | 1 | 0 | 0 | 1636076 | Q5UCB8.1 | P17763 |
| AEHSGASHNILVEVQ | 1 | 0 | 0 | 1636079 | P17763.2 | P17763 |
| AEHTGREIVDLMCHA | 1 | 0 | 0 | 1636080 | P14340.2 | P17763 |
| AEISGSSPILSITIS | 1 | 0 | 0 | 1636083 | P14340.2 | P17763 |
| AEPPFGESYIVIGAG | 1 | 0 | 0 | 1636088 | QFS19562.1 | P17763 |
| AEQTGVSHNLMITVD | 1 | 0 | 0 | 1636090 | QTX93491.1 | P17763 |
| AERIGEPDYEVDEDI | 1 | 0 | 0 | 1636091 | Q2YHF0.1 | P17763 |
| AETQHGTVLVQIKYE | 1 | 0 | 0 | 1636094 | QTX92739.1 | P17763 |
| AEVEPEDIDCWCNLT | 1 | 0 | 0 | 1636095 | QTX93534.1 | P17763 |
| AEVSWEEEAEHSGAS | 1 | 0 | 0 | 1636098 | P17763.2 | P17763 |
| AEVSWEQEAEHSGAS | 1 | 0 | 0 | 1636099 | QMW69345.1 | P17763 |
| AEWLWRTLGRNKKPR | 1 | 0 | 0 | 1636100 | QTX93504.1 | P17763 |
| AFFLAHAIGTSITQK | 1 | 0 | 0 | 1636103 | QTX92537.1 | P17763 |
| AFPQSNAPIQDEERD | 1 | 0 | 0 | 1636109 | QTX93489.1 | P17763 |
| AFPQSNAVIQDEERD | 1 | 0 | 0 | 1636110 | P17763.2 | P17763 |
| AGAPCKVPIEIRDVN | 1 | 0 | 0 | 1636116 | Q58HT7.1 | P17763 |
| AGATEVDSGDGNHMF | 1 | 0 | 0 | 1636118 | Q2YHF0.1 | P17763 |
| AGISYKDREWCFTGE | 1 | 0 | 0 | 1636120 | Q2YHF0.1 | P17763 |
| AGKTKRILPSIVREA | 1 | 0 | 0 | 1636121 | Q2YHF0.1 | P17763 |
| AGKTKRYLPAIVREA | 1 | 0 | 0 | 1636122 | P14340.2 | P17763 |
| AGLLFSIMKNTANTR | 1 | 0 | 0 | 1636125 | QQW40796.1 | P17763 |
| AGPLIAGGMLIACYV | 1 | 0 | 0 | 1636127 | P17763.2 | P17763 |
| AGPLVAGGLLIACYV | 1 | 0 | 0 | 1636128 | QTX93489.1 | P17763 |
| AGPMPVTAASAAQRR | 1 | 0 | 0 | 1636129 | QIC53486.1 | P17763 |
| AGPMPVTVASAAQRR | 1 | 0 | 0 | 1636130 | P17763.2 | P17763 |
| AGPWHLGKLEIDFGE | 1 | 0 | 0 | 1636131 | QTX92320.1 | P17763 |
| AGPWHLGKLELDFDL | 1 | 0 | 0 | 1636132 | P17763.2 | P17763 |
| AGPWHLGKLELDFNY | 1 | 0 | 0 | 1636133 | QTX93479.1 | P17763 |
| AGPWHLGRLEMDFDF | 1 | 0 | 0 | 1636134 | QQW40788.1 | P17763 |
| AGVLWDVPSPPPMGK | 1 | 0 | 0 | 1636140 | QTX93162.1 | P17763 |
| AGVLWDVPSPPPVEK | 1 | 0 | 0 | 1636141 | QOW96388.1 | P17763 |
| AGVSLLPVCQSSSMR | 1 | 0 | 0 | 1636144 | QIS48873.1 | P17763 |
| AGWSLRETACLGKAY | 1 | 0 | 0 | 1636147 | Q6YMS3.1 | P17763 |
| AHAIGTSITQKGIIF | 1 | 0 | 0 | 1636149 | P17763.2 | P17763 |
| AHAKKQEVVVLGSQE | 1 | 0 | 0 | 1636150 | P17763.2 | P17763 |
| AHKDGVCGIRSTTRL | 1 | 0 | 0 | 1636152 | Q2YHF0.1 | P17763 |
| AHKEGVCGIRSTTRL | 1 | 0 | 0 | 1636153 | QTX92336.1 | P17763 |
| AHYAIIGPGLQAKAT | 1 | 0 | 0 | 1636156 | P17763.2 | P17763 |
| AHYIGTSLTQKVVIF | 1 | 0 | 0 | 1636157 | QTX93479.1 | P17763 |
| AIAGAWENGVCGIRS | 1 | 0 | 0 | 1636158 | QTX93489.1 | P17763 |
| AICSAVPVDWVPTSR | 1 | 0 | 0 | 1636163 | P17763.2 | P17763 |
| AIDLDPVVYDAKFEK | 1 | 0 | 0 | 1636166 | P17763.2 | P17763 |
| AIGKAWEEGVCGIRS | 1 | 0 | 0 | 1636175 | P17763.2 | P17763 |
| AIKVLKGFKKEISNM | 1 | 0 | 0 | 1636176 | ADI44327.1 | P17763 |
| AIKVLRGFKKEISNM | 1 | 0 | 0 | 1636177 | P17763.2 | P17763 |
| AILGDTAWDFGSIGG | 1 | 0 | 0 | 1636182 | P17763.2 | P17763 |
| AILGDTAWDFGSLGG | 1 | 0 | 0 | 1636183 | P14340.2 | P17763 |
| AILGETAWDFGSVGG | 1 | 0 | 0 | 1636184 | Q2YHF0.1 | P17763 |
| AILNAHKDGVCGIRS | 1 | 0 | 0 | 1636186 | Q2YHF0.1 | P17763 |
| AILNAHKEGVCGIRS | 1 | 0 | 0 | 1636187 | QTX92334.1 | P17763 |
| AINQVRRLIGNENYL | 1 | 0 | 0 | 1636191 | P17763.2 | P17763 |
| AIQKAQEEGICGIRS | 1 | 0 | 0 | 1636194 | QTX93142.1 | P17763 |
| AKAIFKLTYQNKVVK | 1 | 0 | 0 | 1636204 | Q2YHF0.1 | P17763 |
| AKEGIKRGETDHHAV | 1 | 0 | 0 | 1636206 | P14340.2 | P17763 |
| AKEGLKRGETTHHAV | 1 | 0 | 0 | 1636207 | QTX93534.1 | P17763 |
| AKEGLKRGETTKHAV | 1 | 0 | 0 | 1636208 | QFS19562.1 | P17763 |
| AKFLCSGKITGNLVQ | 1 | 0 | 0 | 1636210 | QBA29684.1 | P17763 |
| AKFQCLELIEGKVVQ | 1 | 0 | 0 | 1636211 | QTX93368.1 | P17763 |
| AKFQCLEPIEGKVVQ | 1 | 0 | 0 | 1636212 | QTX93491.1 | P17763 |
| AKFSCSGKITGNLVQ | 1 | 0 | 0 | 1636214 | Q2YHF0.1 | P17763 |
| AKIFTPEARNSTFLI | 1 | 0 | 0 | 1636216 | Q5UCB8.1 | P17763 |
| AKIFTPETRNSTFLV | 1 | 0 | 0 | 1636217 | Q2YHF0.1 | P17763 |
| AKIIGADIQNTTFII | 1 | 0 | 0 | 1636218 | P27909.2 | P17763 |
| AKMLSTELHNHTFLI | 1 | 0 | 0 | 1636222 | QQW40788.1 | P17763 |
| AKMLSTEPHNQTFLI | 1 | 0 | 0 | 1636223 | QGW05359.1 | P17763 |
| AKMLSTESYNQTFLI | 1 | 0 | 0 | 1636224 | QTX93044.1 | P17763 |
| AKRGTAQIMEVTARW | 1 | 0 | 0 | 1636226 | P17763.2 | P17763 |
| AKRGTTQIMEVTAKW | 1 | 0 | 0 | 1636227 | QNH29581.1 | P17763 |
| AKRMAILGDTAWDFG | 1 | 0 | 0 | 1636228 | P14340.2 | P17763 |
| AKRMAILGETAWDFG | 1 | 0 | 0 | 1636229 | Q2YHF0.1 | P17763 |
| AKRQDVTVLGSQEGA | 1 | 0 | 0 | 1636230 | Q2YHF0.1 | P17763 |
| AKSALKDGSKIKHAV | 1 | 0 | 0 | 1636231 | Q58HT7.1 | P17763 |
| AKSALKDGSKIKYAV | 1 | 0 | 0 | 1636232 | Q2YHF0.1 | P17763 |

| | | | | | | |
|---|---|---|---|---|---|---|
| AKSALRDGSKIKHAV | 1 | 0 | 0 | 1636233 | AFY10039.1 | P17763 |
| ALCEALTLATGPIST | 1 | 0 | 0 | 1636240 | P14340.2 | P17763 |
| ALCEALTLATGPITT | 1 | 0 | 0 | 1636241 | QTX93491.1 | P17763 |
| ALFLAHYIGTSLTQK | 1 | 0 | 0 | 1636244 | QTX93479.1 | P17763 |
| ALKLITQFETYQLWT | 1 | 0 | 0 | 1636250 | Q6YMS3.1 | P17763 |
| ALLAKSIFKLTYQNK | 1 | 0 | 0 | 1636251 | QTX92776.1 | P17763 |
| ALLALNDMGKVRKDI | 1 | 0 | 0 | 1636252 | QTX93489.1 | P17763 |
| ALLATSIFKLTYQNK | 1 | 0 | 0 | 1636253 | P17763.2 | P17763 |
| ALLSQSTIPETILEL | 1 | 0 | 0 | 1636257 | AUN35142.1 | P17763 |
| ALLWMASVEPHWIAA | 1 | 0 | 0 | 1636261 | P17763.2 | P17763 |
| ALMVIGMAMTTTLSI | 1 | 0 | 0 | 1636265 | AMP43484.1 | P17763 |
| ALMVIGMAMTTVFSI | 1 | 0 | 0 | 1636266 | Q2YHF2.1 | P17763 |
| ALMVIGMAMTTVLSI | 1 | 0 | 0 | 1636267 | Q2YHF0.1 | P17763 |
| ALSEGVYRIMQRGLF | 1 | 0 | 0 | 1636272 | ARX96505.1 | P17763 |
| ALTALNDMGKIRKDI | 1 | 0 | 0 | 1636276 | QTX93081.1 | P17763 |
| ALTALNDMGKVRKDI | 1 | 0 | 0 | 1636277 | P17763.2 | P17763 |
| ALTGATEIQTSGGTS | 1 | 0 | 0 | 1636280 | QHR82515.1 | P17763 |
| ALTLATGPISTLWEG | 1 | 0 | 0 | 1636281 | P14340.2 | P17763 |
| ALTLATGPITTLWEG | 1 | 0 | 0 | 1636282 | QTX93489.1 | P17763 |
| AMATTLQLPEDIEQM | 1 | 0 | 0 | 1636290 | QTX93479.1 | P17763 |
| AMFTCKKNMEGKIVQ | 1 | 0 | 0 | 1636292 | BAG84557.1 | P17763 |
| AMFTCKKNMEGKVVQ | 1 | 0 | 0 | 1636293 | QQW40666.1 | P17763 |
| AMHTALTGATEIQNS | 1 | 0 | 0 | 1636294 | QTX93489.1 | P17763 |
| AMHTALTGATEIQTS | 1 | 0 | 0 | 1636295 | P17763.2 | P17763 |
| AMILSIVSLFPLCLS | 1 | 0 | 0 | 1636296 | P17763.2 | P17763 |
| AMKRYSAPFESEGVL | 1 | 0 | 0 | 1636297 | AHG06382.1 | P17763 |
| AMRRYSAHFESEGVL | 1 | 0 | 0 | 1636301 | QTX92320.1 | P17763 |
| AMYADDTAGWDTRIT | 1 | 0 | 0 | 1636308 | P14340.2 | P17763 |
| ANQATVLMGLGKGWP | 1 | 0 | 0 | 1636310 | P14340.2 | P17763 |
| ANQAVVLMGLDKGWP | 1 | 0 | 0 | 1636311 | QTX93489.1 | P17763 |
| ANVSLAAIANQAVVL | 1 | 0 | 0 | 1636313 | QTX93489.1 | P17763 |
| APHVGLGLETRTETW | 1 | 0 | 0 | 1636317 | P17763.2 | P17763 |
| APHVGMGLDTRTQTW | 1 | 0 | 0 | 1636318 | QTX93491.1 | P17763 |
| APPVNDLKYSWKTWG | 1 | 0 | 0 | 1636320 | Q2YHF0.1 | P17763 |
| AQALPVYLMTLMKGA | 1 | 0 | 0 | 1636324 | Q2YHF0.1 | P17763 |
| AQEEGICGIRSVTRL | 1 | 0 | 0 | 1636326 | QTX93142.1 | P17763 |
| AQKATLTEGVYRIMQ | 1 | 0 | 0 | 1636331 | Q2YHF0.1 | P17763 |
| AQKRAAAGIMKNPTV | 1 | 0 | 0 | 1636335 | P14340.2 | P17763 |
| AQKRTAAGIMKNPTV | 1 | 0 | 0 | 1636336 | P17763.2 | P17763 |
| AQLIRQMEGEGVFKN | 1 | 0 | 0 | 1636340 | QTX93142.1 | P17763 |
| AQLIRQMEGEGVFKS | 1 | 0 | 0 | 1636341 | P14340.2 | P17763 |
| AQNALDNLVMLHNSE | 1 | 0 | 0 | 1636344 | P17763.2 | P17763 |
| AQRIETWILRHPGFT | 1 | 0 | 0 | 1636348 | P14340.2 | P17763 |
| AQRVESWILRNPGFA | 1 | 0 | 0 | 1636349 | Q2YHF0.1 | P17763 |
| ARAAVEDEEFWKLVD | 1 | 0 | 0 | 1636354 | QTX93502.1 | P17763 |
| ARASFIEVKTCIWPK | 1 | 0 | 0 | 1636355 | P17763.2 | P17763 |
| ARDALDNLAVLHTAE | 1 | 0 | 0 | 1636356 | AHG25307.1 | P17763 |
| ARKTFVDLMRRGDLP | 1 | 0 | 0 | 1636357 | P14340.2 | P17763 |
| ARNALDNLAVLHTAE | 1 | 0 | 0 | 1636359 | QQW40770.1 | P17763 |
| ARPSFNMLKRARNRV | 1 | 0 | 0 | 1636360 | QFS19562.1 | P17763 |
| ARRMAILGDTAWDFG | 1 | 0 | 0 | 1636361 | P17763.2 | P17763 |
| ARSALKDGSKTKYAV | 1 | 0 | 0 | 1636362 | QTX92247.1 | P17763 |
| ARVSQGAGWSLRETA | 1 | 0 | 0 | 1636365 | P17763.2 | P17763 |
| ARWLWGFLSRNKKPR | 1 | 0 | 0 | 1636366 | P17763.2 | P17763 |
| ASAAQRRGRIGRNHN | 1 | 0 | 0 | 1636367 | QFS19562.1 | P17763 |
| ASAAQRRGRIGRNQN | 1 | 0 | 0 | 1636368 | P17763.2 | P17763 |
| ASAAQRRGRVGRNPQ | 1 | 0 | 0 | 1636369 | QTX93489.1 | P17763 |
| ASEMAEALKGMPIRY | 1 | 0 | 0 | 1636372 | P17763.2 | P17763 |
| ASSMVNGVVKLLTKP | 1 | 0 | 0 | 1636382 | QFS19562.1 | P17763 |
| ASSMVNGVVRLLTKP | 1 | 0 | 0 | 1636383 | P17763.2 | P17763 |
| ATDPFPQSNSPIEDI | 1 | 0 | 0 | 1636388 | Q58HT7.1 | P17763 |
| ATEEIAVQNWLARVG | 1 | 0 | 0 | 1636389 | QRG47120.1 | P17763 |
| ATEEVAVQDWLARVG | 1 | 0 | 0 | 1636390 | AHM25909.1 | P17763 |
| ATEIQTSGGTSIFAG | 1 | 0 | 0 | 1636392 | QHR82521.1 | P17763 |
| ATGPISTLWEGNPGR | 1 | 0 | 0 | 1636395 | P14340.2 | P17763 |
| ATGPITTLWEGSPGK | 1 | 0 | 0 | 1636396 | QTX93489.1 | P17763 |
| ATLLAVSGVYPLSIP | 1 | 0 | 0 | 1636398 | QFS19562.1 | P17763 |
| ATLLAVSGVYPMSIP | 1 | 0 | 0 | 1636399 | P27909.2 | P17763 |
| ATLLVWHTWQKQTQR | 1 | 0 | 0 | 1636400 | QTX93479.1 | P17763 |
| ATPPGSVEAFPQSNA | 1 | 0 | 0 | 1636403 | P17763.2 | P17763 |
| ATPPGTADAFPQSNA | 1 | 0 | 0 | 1636404 | QHR82515.1 | P17763 |
| ATQTAGPWHLGKLEI | 1 | 0 | 0 | 1636406 | QTX92333.1 | P17763 |
| ATQTVGPWHLGKLEI | 1 | 0 | 0 | 1636407 | Q2YHF0.1 | P17763 |
| ATREAQKRAAAGIMK | 1 | 0 | 0 | 1636408 | P14340.2 | P17763 |
| ATREAQKRTAAGIMK | 1 | 0 | 0 | 1636409 | P17763.2 | P17763 |
| ATTIITPMMRHTIEN | 1 | 0 | 0 | 1636411 | P17763.2 | P17763 |
| ATTILTPMLRHTIEN | 1 | 0 | 0 | 1636412 | Q2YHF0.1 | P17763 |
| ATTVITPMMRHTIEN | 1 | 0 | 0 | 1636413 | P27909.2 | P17763 |
| ATWAKNIQAAINQVR | 1 | 0 | 0 | 1636416 | QTX93142.1 | P17763 |
| ATWAQNILTAIQQVR | 1 | 0 | 0 | 1636417 | QTX93504.1 | P17763 |
| ATYGWNLVKLHSGKD | 1 | 0 | 0 | 1636420 | QFS19562.1 | P17763 |
| ATYGWNLVKLHSGVD | 1 | 0 | 0 | 1636421 | Q2YHF0.1 | P17763 |
| AVASGLLWVAEIQPQ | 1 | 0 | 0 | 1636426 | Q2YHF0.1 | P17763 |
| AVEDGRFWELVDRER | 1 | 0 | 0 | 1636427 | QRG47120.1 | P17763 |
| AVEDSGFWELVDKER | 1 | 0 | 0 | 1636428 | QQW40742.1 | P17763 |
| AVEDSRFWELVDRER | 1 | 0 | 0 | 1636430 | AGX15381.1 | P17763 |
| AVEPEVANLDIIGQR | 1 | 0 | 0 | 1636431 | P17763.2 | P17763 |
| AVFFVVTLIPLCRTS | 1 | 0 | 0 | 1636432 | ANW72070.1 | P17763 |
| AVGGITLFLGFTVHA | 1 | 0 | 0 | 1636439 | Q2YHF0.1 | P17763 |
| AVGGITLFLGFTVQA | 1 | 0 | 0 | 1636440 | Q5UCB8.1 | P17763 |
| AVGLVTLYLGVMVQA | 1 | 0 | 0 | 1636444 | QFS19562.1 | P17763 |

| | | | | | | |
|---|---|---|---|---|---|---|
| AVGMVTLYLGVMVQA | 1 | 0 | 0 | 1636446 | P17763.2 | P17763 |
| AVHADMGYWIESEKN | 1 | 0 | 0 | 1636447 | P17763.2 | P17763 |
| AVHADMGYWIESSKN | 1 | 0 | 0 | 1636448 | Q2YHF0.1 | P17763 |
| AVLHTAEAGGRAYNH | 1 | 0 | 0 | 1636454 | P14340.2 | P17763 |
| AVNDSRFWELVDKER | 1 | 0 | 0 | 1636457 | Q2YHF0.1 | P17763 |
| AVPVDWVPTSRTTWS | 1 | 0 | 0 | 1636463 | P17763.2 | P17763 |
| AVSFMTLITGNMSFR | 1 | 0 | 0 | 1636467 | QQW40674.1 | P17763 |
| AVSFVTLITGNMSFK | 1 | 0 | 0 | 1636468 | QTX93142.1 | P17763 |
| AVSGVYPLSIPATLF | 1 | 0 | 0 | 1636469 | QFS19562.1 | P17763 |
| AVTLDFKPGTSGSPI | 1 | 0 | 0 | 1636471 | Q2YHF0.1 | P17763 |
| AVVSVSPLLLTSSQQ | 1 | 0 | 0 | 1636473 | QTX93067.1 | P17763 |
| AWDFGSVGGVLNSLG | 1 | 0 | 0 | 1636475 | QTX93489.1 | P17763 |
| AWENGVCGIRSTTRM | 1 | 0 | 0 | 1636479 | QTX93489.1 | P17763 |
| AWGKAKMLSTELHNH | 1 | 0 | 0 | 1636480 | QQW40788.1 | P17763 |
| AWKHAQRIETWILRH | 1 | 0 | 0 | 1636482 | P14340.2 | P17763 |
| AWKHAQRVESWILRN | 1 | 0 | 0 | 1636483 | Q2YHF0.1 | P17763 |
| AWKHVQRIETWILRH | 1 | 0 | 0 | 1636484 | P29991.1 | P17763 |
| AWKQIQRVETWALRH | 1 | 0 | 0 | 1636485 | QFS19562.1 | P17763 |
| AWNFFEVEDYGFGMF | 1 | 0 | 0 | 1636490 | QTX92256.1 | P17763 |
| AWNFLEVEDYGFGMF | 1 | 0 | 0 | 1636491 | Q2YHF0.1 | P17763 |
| AWNIWEVEDYGFGIF | 1 | 0 | 0 | 1636492 | P17763.2 | P17763 |
| AWNSFEVEDYGFGMF | 1 | 0 | 0 | 1636493 | AKQ00032.1 | P17763 |
| AWNSLEVEDYGFGMF | 1 | 0 | 0 | 1636495 | ACN54390.1 | P17763 |
| AWNVWEVEDYGFGVF | 1 | 0 | 0 | 1636496 | QTX93489.1 | P17763 |
| AWRQVEKVETWALRH | 1 | 0 | 0 | 1636498 | QTX93479.1 | P17763 |
| AWTLYAVATTFITPM | 1 | 0 | 0 | 1636499 | Q9WDA6.1 | P17763 |
| AWTLYAVATTFVTPM | 1 | 0 | 0 | 1636500 | P14340.2 | P17763 |
| AWTLYAVATTVITPM | 1 | 0 | 0 | 1636501 | P27909.2 | P17763 |
| AWVMYGTCTQSGERR | 1 | 0 | 0 | 1636502 | Q2YHF0.1 | P17763 |
| AYMIGQTGIQRTVFF | 1 | 0 | 0 | 1636508 | Q2YHF0.1 | P17763 |
| AYTALFSGVSWIMKI | 1 | 0 | 0 | 1636510 | QTX93530.1 | P17763 |
| AYTIGTTHFQRVLIF | 1 | 0 | 0 | 1636512 | QTX93032.1 | P17763 |
| AYTIGTTYFQRVLIF | 1 | 0 | 0 | 1636513 | QQW40777.1 | P17763 |
| CAHWKEAKMLLDNIN | 1 | 0 | 0 | 1636518 | P14340.2 | P17763 |
| CAIILGGLTWMDLLR | 1 | 0 | 0 | 1636519 | Q2YHF0.1 | P17763 |
| CAVQLLLMRTSWALC | 1 | 0 | 0 | 1636521 | QTX93491.1 | P17763 |
| CDDTVTYKCPHITEV | 1 | 0 | 0 | 1636522 | QCZ25049.1 | P17763 |
| CDDTVTYKCPLIAEV | 1 | 0 | 0 | 1636523 | QTX93534.1 | P17763 |
| CDDTVTYKCPLITEV | 1 | 0 | 0 | 1636524 | QPI70484.1 | P17763 |
| CDIGESSPNPTIEAG | 1 | 0 | 0 | 1636527 | QRG47112.1 | P17763 |
| CDIGESSPNPTIEEG | 1 | 0 | 0 | 1636528 | P17763.2 | P17763 |
| CDIGESSPNPTVEAG | 1 | 0 | 0 | 1636529 | P14340.2 | P17763 |
| CDIGESSPSPTVEES | 1 | 0 | 0 | 1636530 | QTX93489.1 | P17763 |
| CDIGESSSNPTIEEG | 1 | 0 | 0 | 1636533 | ACL99188.1 | P17763 |
| CEDTITYKCPLLRQN | 1 | 0 | 0 | 1636536 | QTX93024.1 | P17763 |
| CEDTITYNCPLLRQN | 1 | 0 | 0 | 1636537 | QQW40770.1 | P17763 |
| CEDTVTYKCPLLVNT | 1 | 0 | 0 | 1636539 | Q2YHF0.1 | P17763 |
| CEPRSGIDFNEMILM | 1 | 0 | 0 | 1636540 | Q2YHF0.1 | P17763 |
| CGLFGKGGVVTCAKF | 1 | 0 | 0 | 1636543 | Q2YHF0.1 | P17763 |
| CGLFGKGSLITCAKF | 1 | 0 | 0 | 1636544 | P17763.2 | P17763 |
| CGLFGKGSLLTCAKF | 1 | 0 | 0 | 1636545 | P27909.2 | P17763 |
| CHATFTMRLLSPIRV | 1 | 0 | 0 | 1636547 | Q9WDA6.1 | P17763 |
| CHATFTTRLLSSTRV | 1 | 0 | 0 | 1636548 | Q2YHF0.1 | P17763 |
| CIAIGIITLYLGAVV | 1 | 0 | 0 | 1636549 | QCZ25049.1 | P17763 |
| CIMVGANASDKMGMG | 1 | 0 | 0 | 1636550 | P17763.2 | P17763 |
| CIMVGANASDRMGMG | 1 | 0 | 0 | 1636551 | QFS19562.1 | P17763 |
| CIMVGANVSDRMGMG | 1 | 0 | 0 | 1636552 | QIC53498.1 | P17763 |
| CIVIGIITLYLGAVV | 1 | 0 | 0 | 1636553 | QTX93512.1 | P17763 |
| CKIPFSTQDEKGVTQ | 1 | 0 | 0 | 1636554 | QFS19562.1 | P17763 |
| CKKNMEGKIVQPENL | 1 | 0 | 0 | 1636555 | BAG84557.1 | P17763 |
| CKKNMEGKVVQPENL | 1 | 0 | 0 | 1636556 | QQW40666.1 | P17763 |
| CKPLTMFLIAENKIW | 1 | 0 | 0 | 1636557 | QFS19562.1 | P17763 |
| CKPLTMFLITENKIW | 1 | 0 | 0 | 1636558 | P17763.2 | P17763 |
| CKVPIEIRDVNKEKV | 1 | 0 | 0 | 1636559 | Q2YHF0.1 | P17763 |
| CKVRMEKLRIKGMSY | 1 | 0 | 0 | 1636560 | Q2YHF0.1 | P17763 |
| CLELIEGKVVQHENL | 1 | 0 | 0 | 1636561 | QHD57348.1 | P17763 |
| CLEPIEGKVVQHENL | 1 | 0 | 0 | 1636562 | Q5UB51.1 | P17763 |
| CLESIEGKVVQHENL | 1 | 0 | 0 | 1636563 | P27915.1 | P17763 |
| CLGKSYAQMWSLMYF | 1 | 0 | 0 | 1636564 | P14340.2 | P17763 |
| CLGKSYAQMWTLMYF | 1 | 0 | 0 | 1636565 | QRG47112.1 | P17763 |
| CLKPVILKDGPERVI | 1 | 0 | 0 | 1636566 | P17763.2 | P17763 |
| CLSTTSQKTTWLPVL | 1 | 0 | 0 | 1636568 | P17763.2 | P17763 |
| CPLIAEVEPEDIDCW | 1 | 0 | 0 | 1636571 | QTX93534.1 | P17763 |
| CPLITEVEPEDIDCW | 1 | 0 | 0 | 1636572 | QHR82521.1 | P17763 |
| CPLLRQNEPEDIDCW | 1 | 0 | 0 | 1636573 | QTX92981.1 | P17763 |
| CPLLVNTEPEDIDCW | 1 | 0 | 0 | 1636574 | Q2YHF0.1 | P17763 |
| CPRITEAEPDDVDCW | 1 | 0 | 0 | 1636577 | QFS19562.1 | P17763 |
| CQSSSMRKTDWLPMA | 1 | 0 | 0 | 1636579 | AMH87199.1 | P17763 |
| CQSSSMRKTDWLPMT | 1 | 0 | 0 | 1636580 | QTX93491.1 | P17763 |
| CRLKMDKLTLKGMSY | 1 | 0 | 0 | 1636582 | P17763.2 | P17763 |
| CRLKMDKLTLKGVSY | 1 | 0 | 0 | 1636583 | QPD98750.1 | P17763 |
| CRSCTLPPLRFRGED | 1 | 0 | 0 | 1636586 | QTX92718.1 | P17763 |
| CRSCTMPPLRFLGED | 1 | 0 | 0 | 1636587 | Q2YHF0.1 | P17763 |
| CSAVPTEWFPTSRTT | 1 | 0 | 0 | 1636589 | Q2YHF0.1 | P17763 |
| CSHHFHELIMKDGRK | 1 | 0 | 0 | 1636594 | QTX93461.1 | P17763 |
| CSHHFHELIMKDGRV | 1 | 0 | 0 | 1636595 | P14340.2 | P17763 |
| CSHHFHELVMKDGRV | 1 | 0 | 0 | 1636598 | QRG47115.1 | P17763 |
| CTTLAVVSVSPLLLT | 1 | 0 | 0 | 1636602 | Q9WDA6.1 | P17763 |
| CVTKLEGKIVQYENL | 1 | 0 | 0 | 1636603 | P17763.2 | P17763 |
| CVTQVLMMRTTWALC | 1 | 0 | 0 | 1636604 | P14340.2 | P17763 |
| CVVKPIDDRFANALL | 1 | 0 | 0 | 1636605 | QTX93489.1 | P17763 |

| | | | | | | |
|---|---|---|---|---|---|---|
| CVVKPIDDRFASALT | 1 | 0 | 0 | 1636606 | QRG47115.1 | P17763 |
| CVVKPLDDRFANALT | 1 | 0 | 0 | 1636607 | QTX93060.1 | P17763 |
| CWYGMEIRPINEKEE | 1 | 0 | 0 | 1636610 | QTX93500.1 | P17763 |
| CWYGMEIRPISEKEE | 1 | 0 | 0 | 1636611 | QHJ86727.1 | P17763 |
| CWYGMEIRPLNEKEE | 1 | 0 | 0 | 1636612 | Q2YHF0.1 | P17763 |
| CWYGMEIRPLSEKEE | 1 | 0 | 0 | 1636613 | Q58HT7.1 | P17763 |
| CWYGMEIRPVKEKEE | 1 | 0 | 0 | 1636615 | P17763.2 | P17763 |
| CYVISGSSADLSLEK | 1 | 0 | 0 | 1636623 | P17763.2 | P17763 |
| CYVLTGRSADLELER | 1 | 0 | 0 | 1636625 | P14340.2 | P17763 |
| DAIDGEYRLRGEARK | 1 | 0 | 0 | 1636630 | P14340.2 | P17763 |
| DALALGMMILKIVRN | 1 | 0 | 0 | 1636631 | QQW40674.1 | P17763 |
| DALALGMMVLKIVRN | 1 | 0 | 0 | 1636632 | QTX93014.1 | P17763 |
| DARIYSDPLALKEFK | 1 | 0 | 0 | 1636637 | QTX92981.1 | P17763 |
| DARVYADPMALKDFK | 1 | 0 | 0 | 1636638 | Q2YHF0.1 | P17763 |
| DCVVKPLDERFGTSL | 1 | 0 | 0 | 1636647 | ARJ31952.1 | P17763 |
| DCVVKPLDERFSTSL | 1 | 0 | 0 | 1636648 | Q2YHF0.1 | P17763 |
| DCWCNATDTWVTYGT | 1 | 0 | 0 | 1636649 | QFS19562.1 | P17763 |
| DCWCNLTSAWVMYGT | 1 | 0 | 0 | 1636650 | Q2YHF0.1 | P17763 |
| DCWCNLTSTWVMYGT | 1 | 0 | 0 | 1636651 | Q5UCB8.1 | P17763 |
| DCWCNLTSTWVTYGT | 1 | 0 | 0 | 1636652 | QTX93489.1 | P17763 |
| DCWCNSTSTWVTYGT | 1 | 0 | 0 | 1636653 | P14340.2 | P17763 |
| DDLLNEELITEQMAP | 1 | 0 | 0 | 1636657 | Q2YHF0.1 | P17763 |
| DDLQNEELITEQMAP | 1 | 0 | 0 | 1636658 | ARJ31953.1 | P17763 |
| DDQYVFSGDPLKNDE | 1 | 0 | 0 | 1636659 | Q58HT7.1 | P17763 |
| DDQYVFSGDPLRNDE | 1 | 0 | 0 | 1636660 | Q2YHF0.1 | P17763 |
| DDTAGWDTRITEDDL | 1 | 0 | 0 | 1636661 | P17763.2 | P17763 |
| DDTAGWDTRITLEDL | 1 | 0 | 0 | 1636662 | P14340.2 | P17763 |
| DEAHFTDPASIAARG | 1 | 0 | 0 | 1636664 | P17763.2 | P17763 |
| DEAHFTDPSSIAARG | 1 | 0 | 0 | 1636665 | QFS19562.1 | P17763 |
| DETENILTVLLKTAL | 1 | 0 | 0 | 1636674 | QTX93461.1 | P17763 |
| DEVFKKRNLTIMDLH | 1 | 0 | 0 | 1636675 | P27909.2 | P17763 |
| DEVFRKRNLTIMDLH | 1 | 0 | 0 | 1636676 | P17763.2 | P17763 |
| DFGSIGGVFTSVGKL | 1 | 0 | 0 | 1636680 | P17763.2 | P17763 |
| DFGSLGGVFTSIGKA | 1 | 0 | 0 | 1636681 | P14340.2 | P17763 |
| DFGSVGGLLTSLGKA | 1 | 0 | 0 | 1636683 | Q2YHF0.1 | P17763 |
| DFKGKTVWFVPSIKA | 1 | 0 | 0 | 1636684 | P14340.2 | P17763 |
| DFKPGTSGSPIINKK | 1 | 0 | 0 | 1636685 | Q2YHF2.1 | P17763 |
| DFKPGTSGSPIINRE | 1 | 0 | 0 | 1636686 | QTX93489.1 | P17763 |
| DFKPGTSGSPIINRK | 1 | 0 | 0 | 1636687 | Q2YHF0.1 | P17763 |
| DFKPGTSGSPIVNRE | 1 | 0 | 0 | 1636688 | P17763.2 | P17763 |
| DFSPGTSGSPIIDKK | 1 | 0 | 0 | 1636691 | P14340.2 | P17763 |
| DFSPGTSGSPIVDKK | 1 | 0 | 0 | 1636692 | Q9WDA6.1 | P17763 |
| DFSPGTSGSPIVDRK | 1 | 0 | 0 | 1636693 | QRG47115.1 | P17763 |
| DGEYRLKGESRKTFV | 1 | 0 | 0 | 1636694 | QTX93489.1 | P17763 |
| DGEYRLRGEARKTFV | 1 | 0 | 0 | 1636695 | P17763.2 | P17763 |
| DGIKNNQILEENMEV | 1 | 0 | 0 | 1636699 | QHE24291.1 | P17763 |
| DGIKNNQILEENVEV | 1 | 0 | 0 | 1636700 | P14340.2 | P17763 |
| DGITVIDLEPISYDP | 1 | 0 | 0 | 1636701 | Q2YHF0.1 | P17763 |
| DGLAMGIMILKLLTD | 1 | 0 | 0 | 1636702 | QFS19562.1 | P17763 |
| DGLAMGIMMLKLLTD | 1 | 0 | 0 | 1636703 | P17763.2 | P17763 |
| DGLAMGIMMLKLLTE | 1 | 0 | 0 | 1636704 | P27909.2 | P17763 |
| DGPERVILAGPMPVT | 1 | 0 | 0 | 1636712 | P17763.2 | P17763 |
| DGPTPELEEEMFKKR | 1 | 0 | 0 | 1636713 | QTX93479.1 | P17763 |
| DGRSLVVPCRNQDEL | 1 | 0 | 0 | 1636714 | Q2YHF0.1 | P17763 |
| DGSMSIKNEEEEQTL | 1 | 0 | 0 | 1636715 | P14340.2 | P17763 |
| DGTMKIKDEERDDTL | 1 | 0 | 0 | 1636716 | P17763.2 | P17763 |
| DGTMRIKDDETENIL | 1 | 0 | 0 | 1636717 | QTX93479.1 | P17763 |
| DGTTVIVTEDCGNRG | 1 | 0 | 0 | 1636718 | QTX93067.1 | P17763 |
| DGVKNNQILEENVEV | 1 | 0 | 0 | 1636719 | P29990.1 | P17763 |
| DGVNMCTLMAIDLGE | 1 | 0 | 0 | 1636720 | QRG47115.1 | P17763 |
| DGVNMCTLMAMDLGE | 1 | 0 | 0 | 1636721 | P14340.2 | P17763 |
| DHAHWTEAKMLLDNI | 1 | 0 | 0 | 1636722 | P17763.2 | P17763 |
| DIDKKDGDLIYADDT | 1 | 0 | 0 | 1636724 | Q2YHF0.1 | P17763 |
| DIFRKKRLTIMDLHP | 1 | 0 | 0 | 1636726 | QQW40770.1 | P17763 |
| DIFRKRRLTIMDLHP | 1 | 0 | 0 | 1636727 | QTX93081.1 | P17763 |
| DIGVPLLALGCYSQV | 1 | 0 | 0 | 1636729 | P17763.2 | P17763 |
| DIISRRDQRGSGQVV | 1 | 0 | 0 | 1636731 | Q9WDA6.1 | P17763 |
| DILTEIATLPTYLSS | 1 | 0 | 0 | 1636737 | AHG06383.1 | P17763 |
| DIPMTGPLVAGGLLT | 1 | 0 | 0 | 1636738 | P14340.2 | P17763 |
| DIPQWEPSKGWNDWQ | 1 | 0 | 0 | 1636739 | P17763.2 | P17763 |
| DISEMGANFRAGRVI | 1 | 0 | 0 | 1636741 | Q2YHF0.1 | P17763 |
| DKEMAETQHGTTVVK | 1 | 0 | 0 | 1636745 | Q2YHF0.1 | P17763 |
| DKPTLDIELLKTEVT | 1 | 0 | 0 | 1636757 | P17763.2 | P17763 |
| DKTHISSWGDVPYLG | 1 | 0 | 0 | 1636759 | QTX92678.1 | P17763 |
| DKTHVSSWEEVPYLG | 1 | 0 | 0 | 1636760 | P27909.2 | P17763 |
| DLGAGTRSVSTEIEK | 1 | 0 | 0 | 1636766 | QTX92271.1 | P17763 |
| DLGAGTRSVSTETEK | 1 | 0 | 0 | 1636767 | CCM44290.1 | P17763 |
| DLGSGTRNIGIESEI | 1 | 0 | 0 | 1636768 | P14340.2 | P17763 |
| DLGSGTRNIGIESET | 1 | 0 | 0 | 1636769 | Q9WDA6.1 | P17763 |
| DLGVPLLAMGCYSQV | 1 | 0 | 0 | 1636770 | Q2YHF0.1 | P17763 |
| DLHNEEKITQQMDPE | 1 | 0 | 0 | 1636771 | QTX93491.1 | P17763 |
| DLHPGSGKTRRYLPA | 1 | 0 | 0 | 1636773 | P17763.2 | P17763 |
| DLILEIGKLPQHLTQ | 1 | 0 | 0 | 1636774 | P17763.2 | P17763 |
| DLPVWLAYKVAAEGI | 1 | 0 | 0 | 1636779 | QTX93067.1 | P17763 |
| DLPVWLSYKVASAGI | 1 | 0 | 0 | 1636780 | Q2YHF0.1 | P17763 |
| DLQNEAKITDIMEPE | 1 | 0 | 0 | 1636781 | P17763.2 | P17763 |
| DLTVVAGDVKGVLSK | 1 | 0 | 0 | 1636788 | Q2YHF0.1 | P17763 |
| DLTVVAGDVKGVLTK | 1 | 0 | 0 | 1636789 | Q5UCB8.1 | P17763 |
| DLVTEIGRVPSHLAH | 1 | 0 | 0 | 1636794 | QTX93474.1 | P17763 |
| DLWCGSLIGLSSRAT | 1 | 0 | 0 | 1636795 | Q2YHF0.1 | P17763 |
| DMGCVVSWNGKELKC | 1 | 0 | 0 | 1636798 | AXG50965.1 | P17763 |

| | | | | | | |
|---|---|---|---|---|---|---|
| DMGCVVSWSGKELKC | 1 | 0 | 0 | 1636799 | QTX92332.1 | P17763 |
| DMGCVVSWSGRELKC | 1 | 0 | 0 | 1636800 | AMP43484.1 | P17763 |
| DMGYWIESEKNETWK | 1 | 0 | 0 | 1636801 | P17763.2 | P17763 |
| DMGYWIESQKNGSWK | 1 | 0 | 0 | 1636802 | QTX93489.1 | P17763 |
| DMGYWIESSKNQTWQ | 1 | 0 | 0 | 1636803 | Q2YHF0.1 | P17763 |
| DMHNPKGLKERVEKW | 1 | 0 | 0 | 1636804 | Q2YHF0.1 | P17763 |
| DMLKVWNRVWIEDNP | 1 | 0 | 0 | 1636807 | Q2YHF0.1 | P17763 |
| DMQNPKGLKERVEKW | 1 | 0 | 0 | 1636809 | Q5UCB8.1 | P17763 |
| DMTIIGRRLQRLQEE | 1 | 0 | 0 | 1636811 | Q2YHF0.1 | P17763 |
| DMTIIGRRLQRLREE | 1 | 0 | 0 | 1636812 | AHG06381.1 | P17763 |
| DNINTPEGIIPALFE | 1 | 0 | 0 | 1636815 | P17763.2 | P17763 |
| DNPNMIDKTPVHSWE | 1 | 0 | 0 | 1636816 | Q2YHF0.1 | P17763 |
| DNPNMTDKTPVHSWE | 1 | 0 | 0 | 1636817 | Q2YHF2.1 | P17763 |
| DPEHRQLANAIFKLT | 1 | 0 | 0 | 1636821 | QTX93479.1 | P17763 |
| DPKFEKQLGQVMLLI | 1 | 0 | 0 | 1636823 | QTX93060.1 | P17763 |
| DPLKNDEDHAHWTEA | 1 | 0 | 0 | 1636828 | Q58HT7.1 | P17763 |
| DPLRNDEDHAHWTEA | 1 | 0 | 0 | 1636829 | Q2YHF0.1 | P17763 |
| DPRRCLKPVILKDGP | 1 | 0 | 0 | 1636834 | P17763.2 | P17763 |
| DPRRCLKPVILTDGP | 1 | 0 | 0 | 1636835 | QTX93489.1 | P17763 |
| DPVVYDAKFEKQLGQ | 1 | 0 | 0 | 1636837 | P17763.2 | P17763 |
| DQRGSGQVGTYGLNT | 1 | 0 | 0 | 1636843 | P17763.2 | P17763 |
| DQRGSGQVVTYGLNT | 1 | 0 | 0 | 1636844 | Q9WDA6.1 | P17763 |
| DRKEFEEYKRSGILE | 1 | 0 | 0 | 1636847 | Q2YHF0.1 | P17763 |
| DRKGKVVGLYGNGVV | 1 | 0 | 0 | 1636848 | QRG47112.1 | P17763 |
| DRSEAKEGLKRGETT | 1 | 0 | 0 | 1636852 | QFS19562.1 | P17763 |
| DRTEAKEGLKRGETT | 1 | 0 | 0 | 1636853 | AHC08451.1 | P17763 |
| DRTEAKSALKDGSKI | 1 | 0 | 0 | 1636854 | Q2YHF0.1 | P17763 |
| DRTEAKSALRDGSKI | 1 | 0 | 0 | 1636855 | AHG06381.1 | P17763 |
| DRTEARSALKDGSKT | 1 | 0 | 0 | 1636856 | QTX92247.1 | P17763 |
| DRTLAKEGIKRGETD | 1 | 0 | 0 | 1636857 | P14340.2 | P17763 |
| DSKFEKQLGQVMLLV | 1 | 0 | 0 | 1636863 | QTX93489.1 | P17763 |
| DSPKRLATAIAGAWE | 1 | 0 | 0 | 1636864 | QTX93479.1 | P17763 |
| DSPKRLSAAIGKAWE | 1 | 0 | 0 | 1636865 | P17763.2 | P17763 |
| DSYTQMCDHRLMSAA | 1 | 0 | 0 | 1636870 | P27909.2 | P17763 |
| DSYTQVCDHRLMSAA | 1 | 0 | 0 | 1636871 | P17763.2 | P17763 |
| DTAGWDTRITEDDLH | 1 | 0 | 0 | 1636872 | QPZ88403.1 | P17763 |
| DTAGWDTRITEDDLQ | 1 | 0 | 0 | 1636873 | P17763.2 | P17763 |
| DTLEVHWNHKERMVT | 1 | 0 | 0 | 1636874 | Q2YHF0.1 | P17763 |
| DTLLCDIGESSPNPT | 1 | 0 | 0 | 1636875 | P17763.2 | P17763 |
| DTLLCDIGESSSNPT | 1 | 0 | 0 | 1636878 | ACL99188.1 | P17763 |
| DTRTQTWMSAEGAWR | 1 | 0 | 0 | 1636882 | QTX93491.1 | P17763 |
| DTSEVHWNHKERMVT | 1 | 0 | 0 | 1636884 | Q2YHF2.1 | P17763 |
| DTSEVHWNYKERMVT | 1 | 0 | 0 | 1636885 | Q58HT7.1 | P17763 |
| DTTPFGQQRVFKEKV | 1 | 0 | 0 | 1636889 | P17763.2 | P17763 |
| DVDLHPASAWTLYAV | 1 | 0 | 0 | 1636893 | P17763.2 | P17763 |
| DVGKKEGGAMYADDT | 1 | 0 | 0 | 1636895 | AWK29890.1 | P17763 |
| DVIPMVTQLAMTDTT | 1 | 0 | 0 | 1636896 | Q58HT7.1 | P17763 |
| DVIPMVTQMAMTDTT | 1 | 0 | 0 | 1636897 | Q9WDA6.1 | P17763 |
| DVLPTVTQMAMTDTT | 1 | 0 | 0 | 1636899 | QTX93142.1 | P17763 |
| DVNKEKVVGRIISST | 1 | 0 | 0 | 1636900 | Q2YHF0.1 | P17763 |
| DVNKEKVVGRVISAT | 1 | 0 | 0 | 1636901 | AKQ00028.1 | P17763 |
| DVNKEKVVGRVISST | 1 | 0 | 0 | 1636902 | ARN79600.1 | P17763 |
| DVPLAGPLIAGGMLI | 1 | 0 | 0 | 1636903 | P17763.2 | P17763 |
| DVSKKEGGAMYADDT | 1 | 0 | 0 | 1636905 | P14340.2 | P17763 |
| DVTVLGSQEGAMHSA | 1 | 0 | 0 | 1636906 | Q2YHF0.1 | P17763 |
| DVVPMVTQLAMTDTT | 1 | 0 | 0 | 1636907 | Q2YHF0.1 | P17763 |
| DVVPMVTQMAMTDTT | 1 | 0 | 0 | 1636908 | P14340.2 | P17763 |
| DWDFVVTTDISEMGA | 1 | 0 | 0 | 1636909 | P14340.2 | P17763 |
| DWITDYQGKTVWFVP | 1 | 0 | 0 | 1636910 | Q2YHF0.1 | P17763 |
| DWLARVGRERLSRMA | 1 | 0 | 0 | 1636911 | QQW40703.1 | P17763 |
| DWQQVPFCSHHFHQL | 1 | 0 | 0 | 1636912 | P17763.2 | P17763 |
| DWVPTSRTTWSIHAH | 1 | 0 | 0 | 1636913 | P17763.2 | P17763 |
| DYGALTLDCSPRTGL | 1 | 0 | 0 | 1636914 | P17763.2 | P17763 |
| DYGELTLDCEPRSGI | 1 | 0 | 0 | 1636915 | Q2YHF0.1 | P17763 |
| DYGFGIFTTNIWLKL | 1 | 0 | 0 | 1636916 | P17763.2 | P17763 |
| DYGFGMFTTNIWMKF | 1 | 0 | 0 | 1636917 | Q2YHF0.1 | P17763 |
| DYMPSMKRFRKEEES | 1 | 0 | 0 | 1636919 | ABZ90961.1 | P17763 |
| DYMPSMKRFRREEEE | 1 | 0 | 0 | 1636920 | AAD20543.1 | P17763 |
| DYQGKTVWFVPSIKA | 1 | 0 | 0 | 1636921 | Q2YHF0.1 | P17763 |
| EAAAIFMTATPPGSV | 1 | 0 | 0 | 1636922 | P17763.2 | P17763 |
| EAGRTLRVLNLVENW | 1 | 0 | 0 | 1636927 | P14340.2 | P17763 |
| EAHFTDPCSVAARGY | 1 | 0 | 0 | 1636928 | AQV12653.1 | P17763 |
| EAHFTDPSSVAARGY | 1 | 0 | 0 | 1636929 | ACY70714.1 | P17763 |
| EAKHPATLRKYCIEA | 1 | 0 | 0 | 1636930 | QQW40784.1 | P17763 |
| EAKITDIMEPEHALL | 1 | 0 | 0 | 1636931 | P17763.2 | P17763 |
| EALRGLPIRYQTPAV | 1 | 0 | 0 | 1636933 | Q2YHF0.1 | P17763 |
| EAQKRTAAGIMKNPT | 1 | 0 | 0 | 1636939 | P17763.2 | P17763 |
| EARKTFVELMRRGDL | 1 | 0 | 0 | 1636940 | P17763.2 | P17763 |
| EATQLATLRKLCIEG | 1 | 0 | 0 | 1636941 | QTX93474.1 | P17763 |
| EDCDHRGPSLRTTTA | 1 | 0 | 0 | 1636949 | Q2YHF0.1 | P17763 |
| EDHWFSRENSYSGVE | 1 | 0 | 0 | 1636952 | AAX55647.1 | P17763 |
| EDIEREIPERSWNTG | 1 | 0 | 0 | 1636953 | Q2YHF0.1 | P17763 |
| EDIFRKKRLTIMDLH | 1 | 0 | 0 | 1636954 | ACO06179.1 | P17763 |
| EDKTPVESWEEIPYL | 1 | 0 | 0 | 1636955 | ADA59677.1 | P17763 |
| EDKTPVESWEEVPYL | 1 | 0 | 0 | 1636956 | ADA59900.1 | P17763 |
| EDLKNEEMVTNHMEG | 1 | 0 | 0 | 1636959 | P14340.2 | P17763 |
| EDLKNEEMVTNHMKG | 1 | 0 | 0 | 1636960 | QGW05386.1 | P17763 |
| EDNPEIEDDIFRKKR | 1 | 0 | 0 | 1636962 | QQW40770.1 | P17763 |
| EDNPEIEDDIFRKRR | 1 | 0 | 0 | 1636963 | P29990.1 | P17763 |
| EDVQVLAIEPGKNPK | 1 | 0 | 0 | 1636969 | Q2YHF0.1 | P17763 |
| EEEEQTLTILIRTGL | 1 | 0 | 0 | 1636972 | P14340.2 | P17763 |

| | | | | | | |
|---|---|---|---|---|---|---|
| EEFTKKVRSNAALGA | 1 | 0 | 0 | 1636973 | AII99332.1 | P17763 |
| EEGRTLRVLKMVEPW | 1 | 0 | 0 | 1636976 | P17763.2 | P17763 |
| EEHSSTWHYDDENPY | 1 | 0 | 0 | 1636978 | AAC64119.1 | P17763 |
| EEKITQQMDPEHRQL | 1 | 0 | 0 | 1636980 | QTX93479.1 | P17763 |
| EELGDGLAMGIMILK | 1 | 0 | 0 | 1636983 | QFS19562.1 | P17763 |
| EELGDGLAMGIMMLK | 1 | 0 | 0 | 1636984 | P17763.2 | P17763 |
| EERDDTLTILLKATL | 1 | 0 | 0 | 1636989 | P17763.2 | P17763 |
| EESRTIRVLKMVEPW | 1 | 0 | 0 | 1636994 | QTX93489.1 | P17763 |
| EEVPYLGKREDQWCG | 1 | 0 | 0 | 1636996 | P27909.2 | P17763 |
| EEVQVIAVEPGKNPK | 1 | 0 | 0 | 1636997 | P17763.2 | P17763 |
| EEVQVLALEPGKNPR | 1 | 0 | 0 | 1636998 | P14340.2 | P17763 |
| EEYTDYMPSMKRFRR | 1 | 0 | 0 | 1637000 | AAD20543.1 | P17763 |
| EFCIKVLNPYMPTVI | 1 | 0 | 0 | 1637001 | Q2YHF0.1 | P17763 |
| EFEALGFLNEDHWFG | 1 | 0 | 0 | 1637002 | Q2YHF0.1 | P17763 |
| EFEALGFLNEDHWFS | 1 | 0 | 0 | 1637003 | ABQ18243.1 | P17763 |
| EFIDGISLGLILLKM | 1 | 0 | 0 | 1637007 | Q2YHF0.1 | P17763 |
| EFTRKVRSNAAIGAV | 1 | 0 | 0 | 1637009 | P17763.2 | P17763 |
| EFWKLVDRERELHKL | 1 | 0 | 0 | 1637010 | QTX93500.1 | P17763 |
| EGDQYIYMGQPLNND | 1 | 0 | 0 | 1637011 | P17763.2 | P17763 |
| EGDQYVYMGQPLNND | 1 | 0 | 0 | 1637012 | P27909.2 | P17763 |
| EGEKKKLRPKWLDAR | 1 | 0 | 0 | 1637013 | Q2YHF0.1 | P17763 |
| EGEKKKLRPRWLDAR | 1 | 0 | 0 | 1637014 | QHR82521.1 | P17763 |
| EGERKKLKPRWLDAR | 1 | 0 | 0 | 1637015 | QTX92981.1 | P17763 |
| EGGHDLTVVAGDVKG | 1 | 0 | 0 | 1637020 | Q2YHF0.1 | P17763 |
| EGIFLPSELETPNLA | 1 | 0 | 0 | 1637021 | QMW69345.1 | P17763 |
| EGIFSPSELETPNLA | 1 | 0 | 0 | 1637022 | P17763.2 | P17763 |
| EGINKCTLIAMDLGE | 1 | 0 | 0 | 1637026 | Q2YHF0.1 | P17763 |
| EGINYADRKWCFDGI | 1 | 0 | 0 | 1637027 | QRG47120.1 | P17763 |
| EGINYADRRWCFDGI | 1 | 0 | 0 | 1637028 | P14340.2 | P17763 |
| EGINYADRRWCFDGV | 1 | 0 | 0 | 1637029 | P29990.1 | P17763 |
| EGLHKLGYILRDISK | 1 | 0 | 0 | 1637032 | P17763.2 | P17763 |
| EGLSGATWVDVVLEH | 1 | 0 | 0 | 1637033 | P17763.2 | P17763 |
| EGPLPEIEDEVFRKR | 1 | 0 | 0 | 1637038 | P17763.2 | P17763 |
| EGSSEVCDHRLMSAA | 1 | 0 | 0 | 1637039 | Q2YHF0.1 | P17763 |
| EGTTVVITENCGTRG | 1 | 0 | 0 | 1637042 | QTX93479.1 | P17763 |
| EGVLSKADLENPHLL | 1 | 0 | 0 | 1637045 | QTX93538.1 | P17763 |
| EGVSGGAWVDLVLEH | 1 | 0 | 0 | 1637047 | Q2YHF0.1 | P17763 |
| EHCGNRGPSLRTTTV | 1 | 0 | 0 | 1637048 | P17763.2 | P17763 |
| EHGTIATITPQAPTS | 1 | 0 | 0 | 1637052 | P27909.2 | P17763 |
| EHGTTATITPQAPTT | 1 | 0 | 0 | 1637053 | QFS19562.1 | P17763 |
| EHRRDKRSVALAPHV | 1 | 0 | 0 | 1637055 | P17763.2 | P17763 |
| EHRREKRSVALVPHV | 1 | 0 | 0 | 1637056 | P14340.2 | P17763 |
| EIASLPTYLSSRAKL | 1 | 0 | 0 | 1637057 | Q2YHF0.1 | P17763 |
| EIATLPTYLSSKAKL | 1 | 0 | 0 | 1637058 | AHG06382.1 | P17763 |
| EIDKKDGDLMYADDT | 1 | 0 | 0 | 1637059 | ARJ31953.1 | P17763 |
| EIEDDIFRKKRLTIM | 1 | 0 | 0 | 1637060 | QQW40770.1 | P17763 |
| EIEDDIFRKRRLTIM | 1 | 0 | 0 | 1637061 | P29990.1 | P17763 |
| EIGKLPQHLTLRAQN | 1 | 0 | 0 | 1637064 | P27909.2 | P17763 |
| EIGKLPQHLTQRAQN | 1 | 0 | 0 | 1637065 | P17763.2 | P17763 |
| EIGRVPSHLAHRTRN | 1 | 0 | 0 | 1637067 | QTX93474.1 | P17763 |
| EIKVTPQSSITEAEL | 1 | 0 | 0 | 1637070 | QTX93055.1 | P17763 |
| EIQTSGTTTIFAGHL | 1 | 0 | 0 | 1637074 | P17763.2 | P17763 |
| EISWHYDQDHPYKTW | 1 | 0 | 0 | 1637076 | AET43255.1 | P17763 |
| EITPQASTVEAILPE | 1 | 0 | 0 | 1637077 | QTX93471.1 | P17763 |
| EITPQASTVEAVLPE | 1 | 0 | 0 | 1637078 | QTX93534.1 | P17763 |
| EIWTKEGEKKKLRPR | 1 | 0 | 0 | 1637079 | QHR82521.1 | P17763 |
| EIWTKEGERKKLRPR | 1 | 0 | 0 | 1637080 | P17763.2 | P17763 |
| EKASFIEVKNCHWPK | 1 | 0 | 0 | 1637081 | QTX93151.1 | P17763 |
| EKASFIEVKTCTWPK | 1 | 0 | 0 | 1637082 | QTX93504.1 | P17763 |
| EKASLIEVKTCLWPK | 1 | 0 | 0 | 1637083 | Q2YHF0.1 | P17763 |
| EKASLIEVKTCTWPK | 1 | 0 | 0 | 1637084 | QHR82521.1 | P17763 |
| EKNETWKLARASFIE | 1 | 0 | 0 | 1637092 | P17763.2 | P17763 |
| EKQDAFCDSKLMSAA | 1 | 0 | 0 | 1637094 | QTX93081.1 | P17763 |
| EKQLGQVMLLILCVT | 1 | 0 | 0 | 1637095 | QTX93060.1 | P17763 |
| EKQLGQVMLLVLCAV | 1 | 0 | 0 | 1637096 | QTX93489.1 | P17763 |
| EKRSVALTPHSGMGL | 1 | 0 | 0 | 1637098 | Q2YHF0.1 | P17763 |
| EKRSVALVPHVGMGL | 1 | 0 | 0 | 1637099 | P14340.2 | P17763 |
| EKSIEDNPEIEDDIF | 1 | 0 | 0 | 1637100 | P14340.2 | P17763 |
| EKVDTRTPQPKPGTR | 1 | 0 | 0 | 1637102 | Q2YHF0.1 | P17763 |
| EKVDTRTQEPKEGTK | 1 | 0 | 0 | 1637103 | P14340.2 | P17763 |
| EKVETWAFRHPGFTI | 1 | 0 | 0 | 1637104 | QHR82521.1 | P17763 |
| EKVETWALRHPGFTI | 1 | 0 | 0 | 1637105 | QTX93479.1 | P17763 |
| EKVVGRIISSTPFAE | 1 | 0 | 0 | 1637106 | Q2YHF0.1 | P17763 |
| EKVVGRVISATPLAE | 1 | 0 | 0 | 1637107 | AKQ00028.1 | P17763 |
| EKVVGRVISSTPLAE | 1 | 0 | 0 | 1637108 | ACS32016.1 | P17763 |
| EKWLKECGVDRLKRM | 1 | 0 | 0 | 1637112 | Q2YHF0.1 | P17763 |
| EKWLRECGVDRLKRM | 1 | 0 | 0 | 1637113 | AMP43484.1 | P17763 |
| ELEDGAYRIKQKGIL | 1 | 0 | 0 | 1637115 | P14340.2 | P17763 |
| ELEEGVYRIKQQGIF | 1 | 0 | 0 | 1637117 | QTX93491.1 | P17763 |
| ELEKLQRKHGGNLVR | 1 | 0 | 0 | 1637118 | ARX96505.1 | P17763 |
| ELEKLQRKHGGSLVR | 1 | 0 | 0 | 1637119 | Q2YHF0.1 | P17763 |
| ELEKLQRRHGGSLVR | 1 | 0 | 0 | 1637120 | QTX92334.1 | P17763 |
| ELEKLQRRYGGSLVR | 1 | 0 | 0 | 1637121 | QTX92160.1 | P17763 |
| ELVDKERALHQEGKC | 1 | 0 | 0 | 1637129 | Q2YHF0.1 | P17763 |
| ELVDKERNLHLEGKC | 1 | 0 | 0 | 1637130 | P14340.2 | P17763 |
| ELVDRERNLHLEGKC | 1 | 0 | 0 | 1637131 | QRG47120.1 | P17763 |
| EMGANFRADRVIDPR | 1 | 0 | 0 | 1637140 | P17763.2 | P17763 |
| EMGEAAAIFMTATPP | 1 | 0 | 0 | 1637141 | Q2YHF0.1 | P17763 |
| EMGRLPTFMTQKARD | 1 | 0 | 0 | 1637142 | AHG25307.1 | P17763 |
| EMGRLPTFMTQKARN | 1 | 0 | 0 | 1637143 | QQW40770.1 | P17763 |
| EMGRLPTFMTQKTRD | 1 | 0 | 0 | 1637144 | QTX93142.1 | P17763 |

| | | | | | | |
|---|---|---|---|---|---|---|
| EMLRTRVGTKHAVLL | 1 | 0 | 0 | 1637145 | QGW05357.1 | P17763 |
| EMYTQLCDHRLMSAA | 1 | 0 | 0 | 1637147 | QHR82521.1 | P17763 |
| EMYWVSGASGNIVSS | 1 | 0 | 0 | 1637148 | Q5UCB8.1 | P17763 |
| EMYWVSGVSGNIVSS | 1 | 0 | 0 | 1637149 | Q2YHF0.1 | P17763 |
| EMYWVSNASGNIVSS | 1 | 0 | 0 | 1637150 | QTX93055.1 | P17763 |
| EMYWVSNATGNIVSS | 1 | 0 | 0 | 1637151 | QRG47112.1 | P17763 |
| ENCDHRGPSLRTTTA | 1 | 0 | 0 | 1637152 | AKQ00035.1 | P17763 |
| ENCGTRGPSLRTTTV | 1 | 0 | 0 | 1637153 | QTX93479.1 | P17763 |
| ENDMKFTVVVGDANG | 1 | 0 | 0 | 1637155 | QMW69324.1 | P17763 |
| ENDMKFTVVVGDVAG | 1 | 0 | 0 | 1637156 | QFS19562.1 | P17763 |
| ENDMKFTVVVGDVNG | 1 | 0 | 0 | 1637157 | QMT58599.1 | P17763 |
| ENDQYIFTGQPLNND | 1 | 0 | 0 | 1637158 | QTX93489.1 | P17763 |
| ENHHHAAMLDVDLHP | 1 | 0 | 0 | 1637167 | P17763.2 | P17763 |
| ENHQHATMLDVDLRP | 1 | 0 | 0 | 1637168 | QPZ88407.1 | P17763 |
| ENIKNEHKSTWHYDE | 1 | 0 | 0 | 1637170 | P27909.2 | P17763 |
| ENNIKLTVVVGDIIG | 1 | 0 | 0 | 1637172 | QTX93479.1 | P17763 |
| ENNIKLTVVVGDVIG | 1 | 0 | 0 | 1637173 | AAW66473.1 | P17763 |
| ENPYRTWAYHGSYEA | 1 | 0 | 0 | 1637174 | Q2YHF0.1 | P17763 |
| ENSLSGVEGEGLHKL | 1 | 0 | 0 | 1637175 | P17763.2 | P17763 |
| ENSSVNVSLTAIANQ | 1 | 0 | 0 | 1637176 | P14340.2 | P17763 |
| ENSTANVSLAAIANQ | 1 | 0 | 0 | 1637177 | QTX93489.1 | P17763 |
| ENSWSGVEGEGLHRL | 1 | 0 | 0 | 1637178 | Q2YHF0.1 | P17763 |
| ENWLNNNTQFCIKVL | 1 | 0 | 0 | 1637182 | P14340.2 | P17763 |
| EPDVDLGSGTRNIGI | 1 | 0 | 0 | 1637186 | P14340.2 | P17763 |
| EPEHALLAKSIFKLT | 1 | 0 | 0 | 1637187 | QTX92776.1 | P17763 |
| EPEHALLATSIFKLT | 1 | 0 | 0 | 1637188 | P17763.2 | P17763 |
| EPGKNPKHVQTKPGL | 1 | 0 | 0 | 1637189 | Q2YHF0.1 | P17763 |
| EPGKNPKNFQTMPGI | 1 | 0 | 0 | 1637190 | QTX93504.1 | P17763 |
| EPGKNPKNVQTAPGT | 1 | 0 | 0 | 1637191 | P17763.2 | P17763 |
| EPGKNPKNVQTTPGT | 1 | 0 | 0 | 1637192 | P27909.2 | P17763 |
| EPGKNPRAVQTKPGI | 1 | 0 | 0 | 1637193 | QRG47115.1 | P17763 |
| EPGKNPRAVQTKPGL | 1 | 0 | 0 | 1637194 | P14340.2 | P17763 |
| EPHMIVGIQEKGKSL | 1 | 0 | 0 | 1637195 | QTX93131.1 | P17763 |
| EPHMIVGRQEKGKSL | 1 | 0 | 0 | 1637196 | QRG47112.1 | P17763 |
| EPHMIVSKQERGKSL | 1 | 0 | 0 | 1637197 | P17763.2 | P17763 |
| EPISYDPKFEKQLGQ | 1 | 0 | 0 | 1637199 | Q2YHF0.1 | P17763 |
| EPKEGTKKLMKITAE | 1 | 0 | 0 | 1637200 | P14340.2 | P17763 |
| EPLENDEDCAHWKEA | 1 | 0 | 0 | 1637202 | P14340.2 | P17763 |
| EPLMIVAKHERGRPL | 1 | 0 | 0 | 1637203 | Q2YHF0.1 | P17763 |
| EPWLKNNQFCIKVLN | 1 | 0 | 0 | 1637206 | QTX93489.1 | P17763 |
| EPWLSSKPEFCIKVL | 1 | 0 | 0 | 1637208 | Q2YHF0.1 | P17763 |
| EQDANFVCRRTFVDR | 1 | 0 | 0 | 1637210 | QFS19562.1 | P17763 |
| EQDQNYVCKHTYVDR | 1 | 0 | 0 | 1637211 | QTX93489.1 | P17763 |
| EQDQQYICRRDVVDR | 1 | 0 | 0 | 1637213 | Q58HT7.1 | P17763 |
| EQMANGIALGLMALK | 1 | 0 | 0 | 1637214 | AHG23230.1 | P17763 |
| ERASLIEVKTCLWPK | 1 | 0 | 0 | 1637219 | QTX92247.1 | P17763 |
| ERDADLGAGTRSVST | 1 | 0 | 0 | 1637220 | AFP27206.1 | P17763 |
| ERDIPERSWNSGNEW | 1 | 0 | 0 | 1637221 | QTX93491.1 | P17763 |
| ERDIPERSWNSGYDW | 1 | 0 | 0 | 1637222 | P17763.2 | P17763 |
| ERDIPERSWNSGYEW | 1 | 0 | 0 | 1637223 | QFS19562.1 | P17763 |
| EREKSAAIDGEYRLK | 1 | 0 | 0 | 1637224 | QTX93479.1 | P17763 |
| EREKSAAIDGEYRLR | 1 | 0 | 0 | 1637225 | P17763.2 | P17763 |
| EREKSAAVDGEYRLR | 1 | 0 | 0 | 1637226 | ACW82902.1 | P17763 |
| ERELHKQGKCATCVY | 1 | 0 | 0 | 1637227 | P17763.2 | P17763 |
| ERFGTSLLFLNDMGK | 1 | 0 | 0 | 1637228 | ARJ31952.1 | P17763 |
| ERFSTSLLFLNDMGK | 1 | 0 | 0 | 1637229 | Q2YHF0.1 | P17763 |
| ERIGEPDYEVDEDIF | 1 | 0 | 0 | 1637230 | Q2YHF0.1 | P17763 |
| ERLKRMAISGDDCVV | 1 | 0 | 0 | 1637232 | P17763.2 | P17763 |
| ERLSRMAISGDDCVV | 1 | 0 | 0 | 1637233 | P14340.2 | P17763 |
| ERRREKRSVALTPHS | 1 | 0 | 0 | 1637235 | Q2YHF0.1 | P17763 |
| ERSWNSGHEWVTDFK | 1 | 0 | 0 | 1637236 | QTX92981.1 | P17763 |
| ERSWNTGFDWITDYQ | 1 | 0 | 0 | 1637237 | Q2YHF0.1 | P17763 |
| ESGRLEPSWADVRND | 1 | 0 | 0 | 1637245 | Q2YHF0.1 | P17763 |
| ESITLATGPLTTLWE | 1 | 0 | 0 | 1637246 | P17763.2 | P17763 |
| ESNIIIGIGDKALKI | 1 | 0 | 0 | 1637248 | QDG00750.1 | P17763 |
| ESNIVIGIGDKALKI | 1 | 0 | 0 | 1637249 | QTX93512.1 | P17763 |
| ESNIVIGIGDNALKI | 1 | 0 | 0 | 1637250 | QIS48873.1 | P17763 |
| ESPARLASAILNAHK | 1 | 0 | 0 | 1637251 | Q2YHF0.1 | P17763 |
| ESPSKLASAIQKAQE | 1 | 0 | 0 | 1637252 | QTX93142.1 | P17763 |
| ESRKTFVELMRRGDL | 1 | 0 | 0 | 1637254 | QTX93489.1 | P17763 |
| ESSPNPTIEAGRTLR | 1 | 0 | 0 | 1637255 | QRG47112.1 | P17763 |
| ESSPNPTIEEGRTLR | 1 | 0 | 0 | 1637256 | P17763.2 | P17763 |
| ESSPNPTVEAGRTLR | 1 | 0 | 0 | 1637257 | P14340.2 | P17763 |
| ESSSNPTIEEGRTLR | 1 | 0 | 0 | 1637262 | ACL99188.1 | P17763 |
| ETACLGKSYAQMWQL | 1 | 0 | 0 | 1637267 | P17763.2 | P17763 |
| ETAWDFGSVGGLFTS | 1 | 0 | 0 | 1637268 | Q5UCB8.1 | P17763 |
| ETDHHAVSRGSAKLR | 1 | 0 | 0 | 1637270 | P14340.2 | P17763 |
| ETGRLEPSWADVRND | 1 | 0 | 0 | 1637271 | Q58HT7.1 | P17763 |
| ETLLLLGLMILLTGG | 1 | 0 | 0 | 1637276 | QTX93489.1 | P17763 |
| ETLLLTLLATVTGG | 1 | 0 | 0 | 1637277 | P14340.2 | P17763 |
| ETLMLLALIAVLTGG | 1 | 0 | 0 | 1637278 | P17763.2 | P17763 |
| ETPNMDVIGERIKRI | 1 | 0 | 0 | 1637281 | QTX93474.1 | P17763 |
| ETPTWNKKELLVTFK | 1 | 0 | 0 | 1637282 | QHR82521.1 | P17763 |
| ETPTWNRKELLVTFK | 1 | 0 | 0 | 1637283 | QTX93491.1 | P17763 |
| ETRAETWMSSEGAWK | 1 | 0 | 0 | 1637285 | AFZ40117.1 | P17763 |
| ETTHHAVSRGSAKLQ | 1 | 0 | 0 | 1637289 | QTX93534.1 | P17763 |
| ETTKHAVSRGTAKLR | 1 | 0 | 0 | 1637290 | QFS19562.1 | P17763 |
| ETWALRHPGFTILAL | 1 | 0 | 0 | 1637291 | QTX93479.1 | P17763 |
| ETWILRHPGFTIMAA | 1 | 0 | 0 | 1637293 | P14340.2 | P17763 |
| ETWMSSEGAWKHAQR | 1 | 0 | 0 | 1637294 | P14340.2 | P17763 |
| ETWMSSEGAWKHVQR | 1 | 0 | 0 | 1637295 | P29991.1 | P17763 |

| | | | | | | |
|---|---|---|---|---|---|---|
| ETWMSSEGAWKQIQR | 1 | 0 | 0 | 1637296 | QFS19562.1 | P17763 |
| ETYQLWTALVSLTCS | 1 | 0 | 0 | 1637298 | QTX93491.1 | P17763 |
| EVANLDIIGQRIENI | 1 | 0 | 0 | 1637299 | P17763.2 | P17763 |
| EVCDHRLMSAAIKDQ | 1 | 0 | 0 | 1637300 | Q2YHF0.1 | P17763 |
| EVDEDIFRKKRLTIM | 1 | 0 | 0 | 1637301 | Q2YHF0.1 | P17763 |
| EVDSGDGNHMFAGHL | 1 | 0 | 0 | 1637302 | AHN50411.1 | P17763 |
| EVKATGSASSMINGV | 1 | 0 | 0 | 1637305 | QTX93489.1 | P17763 |
| EVKPSGSASSMVNGV | 1 | 0 | 0 | 1637306 | P17763.2 | P17763 |
| EVLTLATGPILTLWE | 1 | 0 | 0 | 1637309 | Q2YHF0.1 | P17763 |
| EVLTLATGPIMTLWE | 1 | 0 | 0 | 1637310 | ANW72071.1 | P17763 |
| EVMRGKFGKKHMIAG | 1 | 0 | 0 | 1637311 | QTX93479.1 | P17763 |
| EVMRSRWSRKMLMTG | 1 | 0 | 0 | 1637312 | P17763.2 | P17763 |
| EVQDDGTMKIKDEER | 1 | 0 | 0 | 1637313 | P17763.2 | P17763 |
| EVSETQHGTILIKVE | 1 | 0 | 0 | 1637314 | QTX93491.1 | P17763 |
| EVSKKEGGAMYADDT | 1 | 0 | 0 | 1637315 | Q9WDA6.1 | P17763 |
| EVVVLGSQEGAMHTA | 1 | 0 | 0 | 1637317 | P17763.2 | P17763 |
| EVYTQSCDHRLMSAA | 1 | 0 | 0 | 1637318 | QTX93534.1 | P17763 |
| EWCFTGERNNQILEE | 1 | 0 | 0 | 1637319 | Q2YHF0.1 | P17763 |
| EWVTDFKGKTVWFVP | 1 | 0 | 0 | 1637324 | P14340.2 | P17763 |
| EYIKTRTNDWDFVVT | 1 | 0 | 0 | 1637326 | QQW40786.1 | P17763 |
| EYPKTKLTDWDFVVT | 1 | 0 | 0 | 1637327 | Q2YHF0.1 | P17763 |
| EYVKTRANDWDFVVT | 1 | 0 | 0 | 1637329 | QRG47112.1 | P17763 |
| EYVKTRTNDWDFVVT | 1 | 0 | 0 | 1637330 | P14340.2 | P17763 |
| FAGHLKCRLKMDKLE | 1 | 0 | 0 | 1637333 | AAC63321.1 | P17763 |
| FCIKILNPYMPSVVE | 1 | 0 | 0 | 1637342 | P17763.2 | P17763 |
| FCIKVLNPYMPTVIE | 1 | 0 | 0 | 1637343 | QTX93489.1 | P17763 |
| FCSHHFHKIFMKDGR | 1 | 0 | 0 | 1637344 | Q2YHF0.1 | P17763 |
| FDFCDGTTVIVTEDC | 1 | 0 | 0 | 1637346 | QTX93067.1 | P17763 |
| FDGERNNQILEENMD | 1 | 0 | 0 | 1637347 | QTX93403.1 | P17763 |
| FDGERNNQVLEENMD | 1 | 0 | 0 | 1637348 | P17763.2 | P17763 |
| FDGQRNNQILEENMD | 1 | 0 | 0 | 1637349 | QHJ86727.1 | P17763 |
| FDLYKKSGITEVDRT | 1 | 0 | 0 | 1637350 | QTX93489.1 | P17763 |
| FEALGFLNEDHWFSR | 1 | 0 | 0 | 1637353 | ABQ18243.1 | P17763 |
| FEALGFMNEDHWFSR | 1 | 0 | 0 | 1637354 | P17763.2 | P17763 |
| FEKQLGQIMLLILCT | 1 | 0 | 0 | 1637356 | P17763.2 | P17763 |
| FEKQLGQVMLLVLCA | 1 | 0 | 0 | 1637357 | QTX93489.1 | P17763 |
| FEPEREKVDAIDGEY | 1 | 0 | 0 | 1637359 | P14340.2 | P17763 |
| FEVEDYGFGMFTTNI | 1 | 0 | 0 | 1637360 | QTX92231.1 | P17763 |
| FFIPPEKCDTLLCDI | 1 | 0 | 0 | 1637371 | QTX92722.1 | P17763 |
| FFLIVLLIPEPEKQR | 1 | 0 | 0 | 1637372 | P14340.2 | P17763 |
| FFLMVLLIPEPEKQR | 1 | 0 | 0 | 1637373 | AAD18036.1 | P17763 |
| FFMMVLLIPEPEKQR | 1 | 0 | 0 | 1637374 | QTX93489.1 | P17763 |
| FFMPPEKCDTLLCDI | 1 | 0 | 0 | 1637375 | P27909.2 | P17763 |
| FFTFVLLLSGQITWR | 1 | 0 | 0 | 1637376 | QDG00730.1 | P17763 |
| FFTPPEKCDTLLCDI | 1 | 0 | 0 | 1637377 | P17763.2 | P17763 |
| FGDSYIVIGVGNSAL | 1 | 0 | 0 | 1637381 | ARJ31953.1 | P17763 |
| FGECPGTTVTIQEDC | 1 | 0 | 0 | 1637382 | Q2YHF0.1 | P17763 |
| FGECPGTTVTIQENC | 1 | 0 | 0 | 1637383 | AKQ00035.1 | P17763 |
| FGECPGTTVTVQEDC | 1 | 0 | 0 | 1637384 | QTX92349.1 | P17763 |
| FGLGSIATQQPESNI | 1 | 0 | 0 | 1637385 | QQW40724.1 | P17763 |
| FGLGSITTQQPESNI | 1 | 0 | 0 | 1637386 | QQW40770.1 | P17763 |
| FGPEREKTQAIDGEF | 1 | 0 | 0 | 1637389 | Q2YHF0.1 | P17763 |
| FGQQRVFKEKVDTRT | 1 | 0 | 0 | 1637391 | P17763.2 | P17763 |
| FGRAKGSRAIWYMWL | 1 | 0 | 0 | 1637392 | ACN42699.1 | P17763 |
| FHELIMKDGRKLVVP | 1 | 0 | 0 | 1637393 | QTX93461.1 | P17763 |
| FHELVMKDGRVLVVP | 1 | 0 | 0 | 1637398 | QRG47120.1 | P17763 |
| FHLSTRDGEPLMIVA | 1 | 0 | 0 | 1637401 | Q2YHF0.1 | P17763 |
| FHLTSRDGEPRMIVG | 1 | 0 | 0 | 1637402 | QTX93489.1 | P17763 |
| FHLTTRNGEPHMIVG | 1 | 0 | 0 | 1637404 | QTX93024.1 | P17763 |
| FIEVKTCIWPKSHTL | 1 | 0 | 0 | 1637408 | P17763.2 | P17763 |
| FIEVKTCTWPKSHTL | 1 | 0 | 0 | 1637409 | QPZ88404.1 | P17763 |
| FIIDGPDTPECPDDQ | 1 | 0 | 0 | 1637411 | QIC53498.1 | P17763 |
| FIIDGPNTPECPDDQ | 1 | 0 | 0 | 1637412 | QFS19562.1 | P17763 |
| FIIDGPNTPECPNAS | 1 | 0 | 0 | 1637413 | QHR82521.1 | P17763 |
| FIKSTMPLVMAWRTI | 1 | 0 | 0 | 1637414 | Q2YHF0.1 | P17763 |
| FIRSTMSLVMAWRTI | 1 | 0 | 0 | 1637417 | AKQ00034.1 | P17763 |
| FIRSTTPLVMAWRTI | 1 | 0 | 0 | 1637418 | AHG06383.1 | P17763 |
| FKIQPFLALGFFLRK | 1 | 0 | 0 | 1637421 | QTX93489.1 | P17763 |
| FKLTYQNKVVKVLRP | 1 | 0 | 0 | 1637422 | Q2YHF0.1 | P17763 |
| FKLTYQNKVVRVQRP | 1 | 0 | 0 | 1637423 | P17763.2 | P17763 |
| FKMRPMFAVGLLFRR | 1 | 0 | 0 | 1637424 | QFS19562.1 | P17763 |
| FKNIQHLTVTEEVAV | 1 | 0 | 0 | 1637425 | QTX93132.1 | P17763 |
| FKSIQHLTATEEIAV | 1 | 0 | 0 | 1637426 | QRG47112.1 | P17763 |
| FKSIQHLTATEEVAV | 1 | 0 | 0 | 1637427 | ACQ44478.1 | P17763 |
| FKSIQQLTVTEEIAV | 1 | 0 | 0 | 1637428 | QDG00713.1 | P17763 |
| FKTEDGVNMCTLMAI | 1 | 0 | 0 | 1637430 | QRG47115.1 | P17763 |
| FKTEDGVNMCTLMAM | 1 | 0 | 0 | 1637431 | P14340.2 | P17763 |
| FKTENGVNMCTLMAM | 1 | 0 | 0 | 1637432 | QQW40786.1 | P17763 |
| FKVRPTFAAGLLLRK | 1 | 0 | 0 | 1637435 | P14340.2 | P17763 |
| FLDLPLPWTAGADTL | 1 | 0 | 0 | 1637437 | Q2YHF0.1 | P17763 |
| FLDLPLPWTAGADTS | 1 | 0 | 0 | 1637438 | Q2YHF2.1 | P17763 |
| FLDLPLPWTTGADTS | 1 | 0 | 0 | 1637440 | AKQ00032.1 | P17763 |
| FLEKTKKDFGLGSIA | 1 | 0 | 0 | 1637442 | QQW40724.1 | P17763 |
| FLEKTKKDLGLGSIT | 1 | 0 | 0 | 1637445 | P14340.2 | P17763 |
| FLFFMQGKGMGKLSM | 1 | 0 | 0 | 1637446 | QTX92101.1 | P17763 |
| FLFLMSGKGIGKMTL | 1 | 0 | 0 | 1637447 | QQW40770.1 | P17763 |
| FLFLMSGRGIGKMTL | 1 | 0 | 0 | 1637448 | P14340.2 | P17763 |
| FLLIMGQLTWNDLIR | 1 | 0 | 0 | 1637456 | QFS19562.1 | P17763 |
| FLNDMGKVRKDIPQW | 1 | 0 | 0 | 1637458 | Q2YHF0.1 | P17763 |
| FLVDGPDTSECPNER | 1 | 0 | 0 | 1637462 | Q2YHF0.1 | P17763 |
| FLVLWIGTNSRNTSM | 1 | 0 | 0 | 1637463 | Q58HT7.1 | P17763 |

| | | | | | | |
|---|---|---|---|---|---|---|
| FMTATPPGATDPFPQ | 1 | 0 | 0 | 1637468 | Q58HT7.1 | P17763 |
| FMTATPPGSIDPFPQ | 1 | 0 | 0 | 1637469 | AHG06382.1 | P17763 |
| FMTATPPGSRDPFPQ | 1 | 0 | 0 | 1637470 | P14340.2 | P17763 |
| FMTATPPGTTDPFPQ | 1 | 0 | 0 | 1637471 | Q2YHF0.1 | P17763 |
| FNEMILMKMKTKTWL | 1 | 0 | 0 | 1637474 | Q2YHF0.1 | P17763 |
| FNEMVLLQMEDKAWL | 1 | 0 | 0 | 1637475 | Q9WDA6.1 | P17763 |
| FNEMVLLTMKEKSWL | 1 | 0 | 0 | 1637476 | QFS19562.1 | P17763 |
| FNMLKRERNRVSTVQ | 1 | 0 | 0 | 1637480 | AAT76491.1 | P17763 |
| FPQSNAPIIDEEREI | 1 | 0 | 0 | 1637484 | QTX93142.1 | P17763 |
| FPQSNAPIMDEEREI | 1 | 0 | 0 | 1637485 | P14340.2 | P17763 |
| FPQSNSPIEDIEREI | 1 | 0 | 0 | 1637486 | Q2YHF0.1 | P17763 |
| FPTSRTTWSIHAHHQ | 1 | 0 | 0 | 1637487 | Q2YHF0.1 | P17763 |
| FQDGVFHTMWHVTRG | 1 | 0 | 0 | 1637488 | P27909.2 | P17763 |
| FQEEQGWTSASEAVN | 1 | 0 | 0 | 1637490 | Q2YHF0.1 | P17763 |
| FQEGVFHTMWHVTRG | 1 | 0 | 0 | 1637491 | P17763.2 | P17763 |
| FQENVFHTMWHVTRG | 1 | 0 | 0 | 1637492 | QTX92726.1 | P17763 |
| FQGSWNTGEEVQVIA | 1 | 0 | 0 | 1637493 | P27909.2 | P17763 |
| FQTMPGIFQTTTGEI | 1 | 0 | 0 | 1637500 | QTX93504.1 | P17763 |
| FRAGRVIDPRRCLKP | 1 | 0 | 0 | 1637503 | Q2YHF0.1 | P17763 |
| FRGSYLAGAGLAFSI | 1 | 0 | 0 | 1637506 | QTX93489.1 | P17763 |
| FRGSYLAGAGLLFSI | 1 | 0 | 0 | 1637507 | P14340.2 | P17763 |
| FRMRPMFAVGLLFRR | 1 | 0 | 0 | 1637508 | P17763.2 | P17763 |
| FRRLTSREVLLLTIG | 1 | 0 | 0 | 1637510 | QFS19562.1 | P17763 |
| FRRLTSREVLLLTVG | 1 | 0 | 0 | 1637511 | P17763.2 | P17763 |
| FSGKGPLRMVLAFIT | 1 | 0 | 0 | 1637515 | Q2YHF0.1 | P17763 |
| FSIPHDLMELIDGIS | 1 | 0 | 0 | 1637517 | Q2YHF2.1 | P17763 |
| FSKGLLNGQGPMKLV | 1 | 0 | 0 | 1637518 | AHG23230.1 | P17763 |
| FSKGLLSGQGPMKLV | 1 | 0 | 0 | 1637519 | P17763.2 | P17763 |
| FSLGMLQGRGPLKLF | 1 | 0 | 0 | 1637520 | AAT76490.1 | P17763 |
| FSPSELETPNLAERV | 1 | 0 | 0 | 1637526 | P17763.2 | P17763 |
| FSRENSLSGVEGEGL | 1 | 0 | 0 | 1637527 | P17763.2 | P17763 |
| FSRENSYSGVEGEGL | 1 | 0 | 0 | 1637528 | AAX55647.1 | P17763 |
| FSRGLLNGQGPMKLV | 1 | 0 | 0 | 1637529 | QTX93504.1 | P17763 |
| FTDENKWKSAREAVE | 1 | 0 | 0 | 1637534 | P14340.2 | P17763 |
| FTDPASIAARGYIST | 1 | 0 | 0 | 1637535 | P17763.2 | P17763 |
| FTIMAAILAYTIGTT | 1 | 0 | 0 | 1637538 | P14340.2 | P17763 |
| FTMRLLSPIRVPNYN | 1 | 0 | 0 | 1637539 | Q9WDA6.1 | P17763 |
| FTQTAGPWHLGKLEL | 1 | 0 | 0 | 1637540 | P17763.2 | P17763 |
| FTSLGKAVHQVFGSV | 1 | 0 | 0 | 1637543 | Q5UCB8.1 | P17763 |
| FTSVGKLVHQIFGTA | 1 | 0 | 0 | 1637544 | QFS19562.1 | P17763 |
| FTSVGKLVHQVFGTA | 1 | 0 | 0 | 1637545 | P27909.2 | P17763 |
| FTTRLLSSTRVPNYN | 1 | 0 | 0 | 1637546 | Q2YHF0.1 | P17763 |
| FTVIAFFLAHAIGTS | 1 | 0 | 0 | 1637547 | QTX92537.1 | P17763 |
| FTVIALFLAHAIGTS | 1 | 0 | 0 | 1637548 | P17763.2 | P17763 |
| FVDLMRRGDLPVWLA | 1 | 0 | 0 | 1637550 | P14340.2 | P17763 |
| FVELMRRGDLPVWLS | 1 | 0 | 0 | 1637552 | P17763.2 | P17763 |
| FVERNLVKPEGKVID | 1 | 0 | 0 | 1637553 | P17763.2 | P17763 |
| FVERNLVTPEGKVVD | 1 | 0 | 0 | 1637554 | Q9WDA6.1 | P17763 |
| FVERNMVIPEGRVID | 1 | 0 | 0 | 1637555 | QTX93489.1 | P17763 |
| FVERNMVTPEGKVVD | 1 | 0 | 0 | 1637556 | P14340.2 | P17763 |
| FVPSIKAGNDIANCL | 1 | 0 | 0 | 1637568 | QTX92786.1 | P17763 |
| FYKPTEQVDTLLCDI | 1 | 0 | 0 | 1637573 | Q2YHF0.1 | P17763 |
| FYLPPEKCDTLLCDI | 1 | 0 | 0 | 1637575 | QTX93489.1 | P17763 |
| GAAMTDDIGMGVTYL | 1 | 0 | 0 | 1637585 | QQW40727.1 | P17763 |
| GADIQNTTFIIDGPD | 1 | 0 | 0 | 1637587 | P27909.2 | P17763 |
| GADVQNTTFIIDGPN | 1 | 0 | 0 | 1637588 | P17763.2 | P17763 |
| GAGEKALKLSWFKKG | 1 | 0 | 0 | 1637593 | P17763.2 | P17763 |
| GAGLAFSLIKNAQTP | 1 | 0 | 0 | 1637595 | Q2YHF0.1 | P17763 |
| GAGLAFSLMKSLGGG | 1 | 0 | 0 | 1637596 | P17763.2 | P17763 |
| GAGVYKEGTFHTMWH | 1 | 0 | 0 | 1637597 | P14340.2 | P17763 |
| GAIYGAAFSGVSWTM | 1 | 0 | 0 | 1637599 | QTX92981.1 | P17763 |
| GANASDKMGMGTTYL | 1 | 0 | 0 | 1637604 | P17763.2 | P17763 |
| GANASDRMGMGTTYL | 1 | 0 | 0 | 1637605 | QFS19562.1 | P17763 |
| GANVSDRMGMGTTYL | 1 | 0 | 0 | 1637606 | QIC53498.1 | P17763 |
| GASHNILVEVQDDGT | 1 | 0 | 0 | 1637608 | P17763.2 | P17763 |
| GATMADDIGMGVTYL | 1 | 0 | 0 | 1637609 | QTX93044.1 | P17763 |
| GATTETPTWNKKELL | 1 | 0 | 0 | 1637611 | QHD57348.1 | P17763 |
| GATWVDVVLEHGGCV | 1 | 0 | 0 | 1637612 | QTX93489.1 | P17763 |
| GATWVDVVLEHGSCV | 1 | 0 | 0 | 1637613 | P17763.2 | P17763 |
| GAVFTEENQWDSARA | 1 | 0 | 0 | 1637614 | QTX93504.1 | P17763 |
| GAVFVDENQWNSAKE | 1 | 0 | 0 | 1637615 | P17763.2 | P17763 |
| GAYRIKQKGILGYSQ | 1 | 0 | 0 | 1637617 | P14340.2 | P17763 |
| GCGRGGWSYYCAGLK | 1 | 0 | 0 | 1637620 | P17763.2 | P17763 |
| GCGRGGWSYYCGGLK | 1 | 0 | 0 | 1637621 | P14340.2 | P17763 |
| GCGRGGWSYYMATLK | 1 | 0 | 0 | 1637622 | Q2YHF0.1 | P17763 |
| GCVTTMAKSKPTLDI | 1 | 0 | 0 | 1637625 | QTX93250.1 | P17763 |
| GCYSQVNPITLTAAL | 1 | 0 | 0 | 1637628 | P14340.2 | P17763 |
| GCYSQVNPLTLTAAV | 1 | 0 | 0 | 1637629 | P17763.2 | P17763 |
| GDDCVVKPIDDRFAT | 1 | 0 | 0 | 1637632 | P17763.2 | P17763 |
| GDGNHMFAGHLKCKV | 1 | 0 | 0 | 1637634 | Q2YHF0.1 | P17763 |
| GDKALKINWYKKGSS | 1 | 0 | 0 | 1637636 | QTX93504.1 | P17763 |
| GDLPVWLAHKVASEG | 1 | 0 | 0 | 1637637 | QTX93491.1 | P17763 |
| GDLPVWLSYKVASEG | 1 | 0 | 0 | 1637638 | P17763.2 | P17763 |
| GDTMSGRIGGQIHLA | 1 | 0 | 0 | 1637641 | QBA29684.1 | P17763 |
| GDTMSGRIGGQTHLA | 1 | 0 | 0 | 1637642 | AKQ00032.1 | P17763 |
| GDTMSGRMGGQIHLA | 1 | 0 | 0 | 1637643 | Q2YHF0.1 | P17763 |
| GDVPYLGKREDQWCG | 1 | 0 | 0 | 1637644 | QTX92684.1 | P17763 |
| GEDGCWYGMEIRPIN | 1 | 0 | 0 | 1637648 | QTX93500.1 | P17763 |
| GEDGCWYGMEIRPIS | 1 | 0 | 0 | 1637649 | QHJ86727.1 | P17763 |
| GEDGCWYGMEIRPLN | 1 | 0 | 0 | 1637650 | Q2YHF0.1 | P17763 |
| GEDGCWYGMEIRPLS | 1 | 0 | 0 | 1637651 | Q58HT7.1 | P17763 |

| | | | | | | |
|---|---|---|---|---|---|---|
| GEDGCWYGMEIRPVK | 1 | 0 | 0 | 1637652 | P17763.2 | P17763 |
| GEERVILAGPMPVTH | 1 | 0 | 0 | 1637656 | P14340.2 | P17763 |
| GEFRLRGEQRKTFVE | 1 | 0 | 0 | 1637657 | Q2YHF0.1 | P17763 |
| GEGIFKSIQHLTATE | 1 | 0 | 0 | 1637658 | QRG47120.1 | P17763 |
| GEGIFKSIQHLTVTE | 1 | 0 | 0 | 1637659 | AGN94888.1 | P17763 |
| GEGLHKLGYILRDVS | 1 | 0 | 0 | 1637660 | ABQ18243.1 | P17763 |
| GEGLHRLGYILEEID | 1 | 0 | 0 | 1637662 | ARJ31953.1 | P17763 |
| GEGLHRLGYILRDVG | 1 | 0 | 0 | 1637663 | AWK29891.1 | P17763 |
| GEGLHRLGYILRDVS | 1 | 0 | 0 | 1637664 | AGN94889.1 | P17763 |
| GEGVFKNIQHLTVTE | 1 | 0 | 0 | 1637667 | QTX93142.1 | P17763 |
| GEGVFKSIQHLTATE | 1 | 0 | 0 | 1637668 | QQW40772.1 | P17763 |
| GEGVFKSIQQLTVTE | 1 | 0 | 0 | 1637669 | QDG00713.1 | P17763 |
| GERIKRIKEEHSSTW | 1 | 0 | 0 | 1637679 | QTX93512.1 | P17763 |
| GEVGAIALDFKPGTS | 1 | 0 | 0 | 1637684 | P17763.2 | P17763 |
| GEYRLRGEARKTFVD | 1 | 0 | 0 | 1637690 | P14340.2 | P17763 |
| GFLNEDHWFSRENSY | 1 | 0 | 0 | 1637693 | ABQ18244.1 | P17763 |
| GFMNEDHWFSRENSL | 1 | 0 | 0 | 1637694 | P17763.2 | P17763 |
| GFTILALFLAHYIGT | 1 | 0 | 0 | 1637695 | QTX93489.1 | P17763 |
| GFTIMAAILAYTIGT | 1 | 0 | 0 | 1637696 | P14340.2 | P17763 |
| GFTVIALFLAHAIGT | 1 | 0 | 0 | 1637697 | P17763.2 | P17763 |
| GGAWVDLVLEHGGCV | 1 | 0 | 0 | 1637698 | Q2YHF0.1 | P17763 |
| GGKAYQHALNELPES | 1 | 0 | 0 | 1637704 | Q2YHF0.1 | P17763 |
| GGKAYRHAMEELPDT | 1 | 0 | 0 | 1637705 | P17763.2 | P17763 |
| GGQIHLAIMAVFKMS | 1 | 0 | 0 | 1637708 | Q2YHF0.1 | P17763 |
| GGQTHLAIMAVFKMS | 1 | 0 | 0 | 1637709 | AMP43484.1 | P17763 |
| GGRAYNHALSELPET | 1 | 0 | 0 | 1637710 | P14340.2 | P17763 |
| GGRAYQHALNELPES | 1 | 0 | 0 | 1637711 | Q5UCB8.1 | P17763 |
| GGRAYRHAMEELPDT | 1 | 0 | 0 | 1637712 | QFS19562.1 | P17763 |
| GGRAYRHAVEELPET | 1 | 0 | 0 | 1637713 | QTX93489.1 | P17763 |
| GGWKLEGEWKEGEEV | 1 | 0 | 0 | 1637720 | P14340.2 | P17763 |
| GGWRFQGSWNTGEEV | 1 | 0 | 0 | 1637721 | P27909.2 | P17763 |
| GGWRLGDKWDKEEDV | 1 | 0 | 0 | 1637722 | Q2YHF0.1 | P17763 |
| GGWRLQGSWNTGEEV | 1 | 0 | 0 | 1637723 | QTX92726.1 | P17763 |
| GGWRLRGSWNTGEEV | 1 | 0 | 0 | 1637724 | ACQ44411.1 | P17763 |
| GGWRLSAQWQKGEEV | 1 | 0 | 0 | 1637725 | QTX93491.1 | P17763 |
| GGWSYYCAGLKKVTE | 1 | 0 | 0 | 1637726 | P17763.2 | P17763 |
| GGWSYYCGGLKNVKE | 1 | 0 | 0 | 1637727 | QQW40786.1 | P17763 |
| GGWSYYCGGLKNVRE | 1 | 0 | 0 | 1637728 | P14340.2 | P17763 |
| GGWSYYMATLKNVTE | 1 | 0 | 0 | 1637729 | Q2YHF0.1 | P17763 |
| GHEEPIPMATYGWNL | 1 | 0 | 0 | 1637730 | P17763.2 | P17763 |
| GHEEPIPMSTYGWNL | 1 | 0 | 0 | 1637731 | P14340.2 | P17763 |
| GHEEPVPMSTYGWNI | 1 | 0 | 0 | 1637733 | AAC63341.1 | P17763 |
| GHGQIDNFSLGVLGM | 1 | 0 | 0 | 1637736 | P14340.2 | P17763 |
| GHLKCKVRMEKLRIK | 1 | 0 | 0 | 1637738 | Q2YHF0.1 | P17763 |
| GHLKCRLKMDKLTLK | 1 | 0 | 0 | 1637739 | P17763.2 | P17763 |
| GIGKLSMGLIAIAVA | 1 | 0 | 0 | 1637745 | Q2YHF0.1 | P17763 |
| GIGKLSMGLITIAVA | 1 | 0 | 0 | 1637746 | ARJ31953.1 | P17763 |
| GIGKMTLGMCCIITA | 1 | 0 | 0 | 1637747 | P14340.2 | P17763 |
| GIGKTSIGLICVIAS | 1 | 0 | 0 | 1637748 | QIS48873.1 | P17763 |
| GIGNRDFVEGLSGAT | 1 | 0 | 0 | 1637749 | P17763.2 | P17763 |
| GIGSRDFVEGLSGAT | 1 | 0 | 0 | 1637750 | QFS19562.1 | P17763 |
| GILGYSQIGAGVYKE | 1 | 0 | 0 | 1637752 | P14340.2 | P17763 |
| GIMKNPTVDGITVID | 1 | 0 | 0 | 1637753 | P14340.2 | P17763 |
| GIMKNPTVDGIVAID | 1 | 0 | 0 | 1637754 | P17763.2 | P17763 |
| GIMTIDLDPVIYDSK | 1 | 0 | 0 | 1637755 | QTX93489.1 | P17763 |
| GIQRTVFFVLMMLVA | 1 | 0 | 0 | 1637756 | Q58HT7.1 | P17763 |
| GISLGLILLKIVTHF | 1 | 0 | 0 | 1637758 | QBQ58394.1 | P17763 |
| GISLGLILLKMVTHF | 1 | 0 | 0 | 1637759 | Q2YHF0.1 | P17763 |
| GITVIDLDPIPYDPK | 1 | 0 | 0 | 1637760 | P14340.2 | P17763 |
| GITVIDLEPIPYDPK | 1 | 0 | 0 | 1637761 | QRG47106.1 | P17763 |
| GKALHQVFGAIYGAA | 1 | 0 | 0 | 1637763 | P14340.2 | P17763 |
| GKAVHQVFGSVYTTM | 1 | 0 | 0 | 1637764 | Q2YHF0.1 | P17763 |
| GKCESCVYNMMGKRE | 1 | 0 | 0 | 1637766 | Q2YHF0.1 | P17763 |
| GKCETCVYNMMGKRE | 1 | 0 | 0 | 1637767 | P14340.2 | P17763 |
| GKDVFFIPPEKCDTL | 1 | 0 | 0 | 1637769 | QTX92722.1 | P17763 |
| GKDVFFMPPEKCDTL | 1 | 0 | 0 | 1637770 | P27909.2 | P17763 |
| GKDVFYLPPEKCDTL | 1 | 0 | 0 | 1637771 | QTX93489.1 | P17763 |
| GKFGKKHMIAGVFFT | 1 | 0 | 0 | 1637773 | QIS48879.1 | P17763 |
| GKFGKKHMIAGVLFT | 1 | 0 | 0 | 1637774 | QTX93504.1 | P17763 |
| GKGGVVTCAKFSCSG | 1 | 0 | 0 | 1637778 | Q2YHF0.1 | P17763 |
| GKGSLITCAKFKCVT | 1 | 0 | 0 | 1637781 | P17763.2 | P17763 |
| GKGSLLTCAKFKCVT | 1 | 0 | 0 | 1637782 | P27909.2 | P17763 |
| GKGSLVTCAKFQCLE | 1 | 0 | 0 | 1637783 | QTX93489.1 | P17763 |
| GKIRKDIQQWEPSKG | 1 | 0 | 0 | 1637787 | QTX93077.1 | P17763 |
| GKLVHQVFGTAYGVL | 1 | 0 | 0 | 1637790 | P27909.2 | P17763 |
| GKMFEATARGARRMA | 1 | 0 | 0 | 1637792 | P17763.2 | P17763 |
| GKPSINMLKRVRNRV | 1 | 0 | 0 | 1637794 | QTX93479.1 | P17763 |
| GKPTLDFELIKTTAK | 1 | 0 | 0 | 1637795 | Q2YHF0.1 | P17763 |
| GKPTLDFELTKTTAK | 1 | 0 | 0 | 1637796 | Q5UCB8.1 | P17763 |
| GKREDLWCGSLIGLS | 1 | 0 | 0 | 1637797 | Q2YHF0.1 | P17763 |
| GKSEFQIYKKSGIQE | 1 | 0 | 0 | 1637800 | P14340.2 | P17763 |
| GKSEFQIYKRSGIQE | 1 | 0 | 0 | 1637801 | QFS19359.1 | P17763 |
| GKTVWFVPSIKSGND | 1 | 0 | 0 | 1637806 | P17763.2 | P17763 |
| GKVRKDIQQWEPSRG | 1 | 0 | 0 | 1637809 | P14340.2 | P17763 |
| GLDKGWPISKMDLGV | 1 | 0 | 0 | 1637816 | AAN03432.3 | P17763 |
| GLFGKTQVGVGIHME | 1 | 0 | 0 | 1637817 | Q2YHF0.1 | P17763 |
| GLGKGWPLSKMDIGV | 1 | 0 | 0 | 1637820 | P14340.2 | P17763 |
| GLGKTSIGLLCVTAS | 1 | 0 | 0 | 1637821 | P27909.2 | P17763 |
| GLGLETRTETWMSSE | 1 | 0 | 0 | 1637822 | P17763.2 | P17763 |
| GLILLKIVTHFDNTQ | 1 | 0 | 0 | 1637823 | QBQ58394.1 | P17763 |
| GLILLKIVTQFDNTQ | 1 | 0 | 0 | 1637824 | ARJ31952.1 | P17763 |

| | | | | | | |
|---|---|---|---|---|---|---|
| GLILLKMVTHFDNTQ | 1 | 0 | 0 | 1637825 | Q2YHF0.1 | P17763 |
| GLKKVTEVKGYTKGG | 1 | 0 | 0 | 1637826 | P17763.2 | P17763 |
| GLKKVTEVRGYTKGG | 1 | 0 | 0 | 1637827 | QTX93489.1 | P17763 |
| GLKNVKEVKGLTKGG | 1 | 0 | 0 | 1637828 | QQW40786.1 | P17763 |
| GLKNVREVKGLTKGG | 1 | 0 | 0 | 1637829 | P14340.2 | P17763 |
| GLLGRSQVGVGVFQD | 1 | 0 | 0 | 1637831 | P27909.2 | P17763 |
| GLLGRSQVGVGVFQE | 1 | 0 | 0 | 1637832 | P17763.2 | P17763 |
| GLLWVAEIQPQWIAA | 1 | 0 | 0 | 1637837 | Q2YHF0.1 | P17763 |
| GLNSKNTSMSFSCIA | 1 | 0 | 0 | 1637838 | QIS48879.1 | P17763 |
| GLNSKNTSMSFSCIV | 1 | 0 | 0 | 1637839 | QTX93530.1 | P17763 |
| GLQAKATREAQKRTA | 1 | 0 | 0 | 1637840 | P17763.2 | P17763 |
| GLSSRATWAKNIHTA | 1 | 0 | 0 | 1637841 | QTX92334.1 | P17763 |
| GLYGNGVVTKSGDYV | 1 | 0 | 0 | 1637844 | Q2YHF0.1 | P17763 |
| GMFTTNIWMKFREGS | 1 | 0 | 0 | 1637846 | Q2YHF0.1 | P17763 |
| GMGKLSMGLIAIAVA | 1 | 0 | 0 | 1637847 | QTX92101.1 | P17763 |
| GMGLETRAETWMSSE | 1 | 0 | 0 | 1637849 | AHG06340.1 | P17763 |
| GMGLETRTETWMSSE | 1 | 0 | 0 | 1637850 | P14340.2 | P17763 |
| GMGTTYLALMATFKM | 1 | 0 | 0 | 1637851 | QFS19562.1 | P17763 |
| GMGTTYLALMATFRM | 1 | 0 | 0 | 1637852 | P17763.2 | P17763 |
| GMGVTYLALIATFKI | 1 | 0 | 0 | 1637853 | QTX93489.1 | P17763 |
| GMGVTYLALLAAFKV | 1 | 0 | 0 | 1637854 | P14340.2 | P17763 |
| GMIKPKGKVVDLGCG | 1 | 0 | 0 | 1637855 | ARM59249.1 | P17763 |
| GMLWMADIPLQWIAS | 1 | 0 | 0 | 1637857 | QTX93520.1 | P17763 |
| GMLWMAEIPLQWIAS | 1 | 0 | 0 | 1637858 | QPI11927.1 | P17763 |
| GMVKPKGKVVDLGCG | 1 | 0 | 0 | 1637861 | Q2YHF0.1 | P17763 |
| GNDIANCLRKNGKKV | 1 | 0 | 0 | 1637863 | QTX93489.1 | P17763 |
| GNDIANCLRKNGKRV | 1 | 0 | 0 | 1637864 | P17763.2 | P17763 |
| GNSLSGVEGEGLHRL | 1 | 0 | 0 | 1637865 | QRG47112.1 | P17763 |
| GPDTPECPDDQRAWN | 1 | 0 | 0 | 1637866 | QIC53498.1 | P17763 |
| GPDTSECPNERRAWN | 1 | 0 | 0 | 1637867 | Q2YHF0.1 | P17763 |
| GPFSQHNYRQGYATQ | 1 | 0 | 0 | 1637869 | Q58HT7.1 | P17763 |
| GPIPVTPASAAQRRG | 1 | 0 | 0 | 1637870 | Q2YHF0.1 | P17763 |
| GPISQHNYRPGYFTQ | 1 | 0 | 0 | 1637872 | P17763.2 | P17763 |
| GPISQHNYRPGYHTQ | 1 | 0 | 0 | 1637873 | QIS48873.1 | P17763 |
| GPISQHNYRQGYATQ | 1 | 0 | 0 | 1637874 | Q2YHF0.1 | P17763 |
| GPLRMVLAFITFLRV | 1 | 0 | 0 | 1637875 | Q2YHF0.1 | P17763 |
| GPMPVTHSSAAQRRG | 1 | 0 | 0 | 1637876 | P14340.2 | P17763 |
| GPNTPECPNASRAWN | 1 | 0 | 0 | 1637878 | QHR82521.1 | P17763 |
| GQGTSETFSMGLLCL | 1 | 0 | 0 | 1637887 | Q2YHF0.1 | P17763 |
| GQLKKNKAIKILIGF | 1 | 0 | 0 | 1637893 | ARJ31953.1 | P17763 |
| GQLKKNKAIKILTGF | 1 | 0 | 0 | 1637894 | Q2YHF0.1 | P17763 |
| GQLKLSWFKKGSSIG | 1 | 0 | 0 | 1637895 | QQW40781.1 | P17763 |
| GQPLNNDEDHAHWTE | 1 | 0 | 0 | 1637896 | P17763.2 | P17763 |
| GQQRVFKEKVDTRTP | 1 | 0 | 0 | 1637897 | P17763.2 | P17763 |
| GQRIENIKHEHKSTW | 1 | 0 | 0 | 1637898 | QTX92739.1 | P17763 |
| GQRIENIKNEHKSTW | 1 | 0 | 0 | 1637899 | P27909.2 | P17763 |
| GQTGIQRTVFFILMM | 1 | 0 | 0 | 1637900 | Q2YHF0.1 | P17763 |
| GQTGIQRTVFFVLMM | 1 | 0 | 0 | 1637901 | Q58HT7.1 | P17763 |
| GQVGTYGLNTFTNME | 1 | 0 | 0 | 1637903 | P17763.2 | P17763 |
| GQVMLLILCVTQVLM | 1 | 0 | 0 | 1637904 | QTX93060.1 | P17763 |
| GQVMLLVLCAVQLLL | 1 | 0 | 0 | 1637905 | QTX93489.1 | P17763 |
| GRARISQGAGWSLRE | 1 | 0 | 0 | 1637910 | P14340.2 | P17763 |
| GRFWELVDRERNLHL | 1 | 0 | 0 | 1637912 | QRG47120.1 | P17763 |
| GRKLVVPCRPQDELI | 1 | 0 | 0 | 1637916 | QTX93489.1 | P17763 |
| GRPSFNMLKRARNRV | 1 | 0 | 0 | 1637921 | P17763.2 | P17763 |
| GRVISATPLAENTNS | 1 | 0 | 0 | 1637926 | AKQ00032.1 | P17763 |
| GRVISSTPLAENTNS | 1 | 0 | 0 | 1637927 | ACS32015.1 | P17763 |
| GRVLVVPCRNQDELI | 1 | 0 | 0 | 1637928 | P14340.2 | P17763 |
| GSFGCKPLTMFLITE | 1 | 0 | 0 | 1637938 | P27909.2 | P17763 |
| GSGEVDSFSLGILCV | 1 | 0 | 0 | 1637939 | P27909.2 | P17763 |
| GSGEVDSFSLGLLCI | 1 | 0 | 0 | 1637940 | P17763.2 | P17763 |
| GSGEVDSFSLGLLCV | 1 | 0 | 0 | 1637941 | QIS48856.1 | P17763 |
| GSGKTRRYLPAIVRE | 1 | 0 | 0 | 1637942 | P17763.2 | P17763 |
| GSLGCKPLTMFLIAE | 1 | 0 | 0 | 1637946 | QFS19562.1 | P17763 |
| GSLGCKPLTMFLITE | 1 | 0 | 0 | 1637947 | P17763.2 | P17763 |
| GSLIGLSSRATWAKN | 1 | 0 | 0 | 1637948 | Q2YHF0.1 | P17763 |
| GSNASDRMGMGVTYL | 1 | 0 | 0 | 1637950 | QTX93489.1 | P17763 |
| GSPIINRKGKVIGLY | 1 | 0 | 0 | 1637952 | Q2YHF0.1 | P17763 |
| GSSPILSITISEDGS | 1 | 0 | 0 | 1637958 | P14340.2 | P17763 |
| GSVEAFPQSNAVIQD | 1 | 0 | 0 | 1637961 | P17763.2 | P17763 |
| GSVYTTMFGGVSWMI | 1 | 0 | 0 | 1637963 | ARX96505.1 | P17763 |
| GSVYTTMFGGVSWMV | 1 | 0 | 0 | 1637964 | Q2YHF0.1 | P17763 |
| GTADAFPQSNAPIQD | 1 | 0 | 0 | 1637969 | QHR82515.1 | P17763 |
| GTGAQGETLGEKWKR | 1 | 0 | 0 | 1637975 | P17763.2 | P17763 |
| GTGNIGETLGEKWKS | 1 | 0 | 0 | 1637976 | P14340.2 | P17763 |
| GTGNTGETLGEKWKN | 1 | 0 | 0 | 1637977 | QQW40775.1 | P17763 |
| GTGSQGETLGEKWKK | 1 | 0 | 0 | 1637978 | QTX93489.1 | P17763 |
| GTGTTGETLGEKWKR | 1 | 0 | 0 | 1637979 | Q2YHF0.1 | P17763 |
| GTKKLMKITAEWLWK | 1 | 0 | 0 | 1637981 | P14340.2 | P17763 |
| GTLALSLTFIKSTMP | 1 | 0 | 0 | 1637982 | Q2YHF0.1 | P17763 |
| GTLALSLTFIRSTMS | 1 | 0 | 0 | 1637983 | AKQ00036.1 | P17763 |
| GTLALSLTFIRSTTP | 1 | 0 | 0 | 1637984 | AHG06383.1 | P17763 |
| GTLGLECSPRTGLDF | 1 | 0 | 0 | 1637986 | QTX93489.1 | P17763 |
| GTRMIMTTTANWLWA | 1 | 0 | 0 | 1637990 | QCZ25014.1 | P17763 |
| GTRNIGIESEIPNLD | 1 | 0 | 0 | 1637991 | P14340.2 | P17763 |
| GTRSVSTEIEKPDMT | 1 | 0 | 0 | 1637993 | QTX92271.1 | P17763 |
| GTRSVSTETEKPDMT | 1 | 0 | 0 | 1637994 | CCM44290.1 | P17763 |
| GTRVVMTTTANWLWA | 1 | 0 | 0 | 1637995 | Q2YHF0.1 | P17763 |
| GTRVVMTTTANWLWT | 1 | 0 | 0 | 1637996 | QTX92320.1 | P17763 |
| GTSGSPIIDKKGKVV | 1 | 0 | 0 | 1637997 | P14340.2 | P17763 |
| GTSGSPIINKKGKVI | 1 | 0 | 0 | 1637998 | Q2YHF2.1 | P17763 |

| | | | | | | |
|---|---|---|---|---|---|---|
| GTSGSPIINREGKVV | 1 | 0 | 0 | 1637999 | QTX93489.1 | P17763 |
| GTSGSPIVDKKGKVV | 1 | 0 | 0 | 1638001 | Q9WDA6.1 | P17763 |
| GTSGSPIVDRKGKVV | 1 | 0 | 0 | 1638002 | QRG47115.1 | P17763 |
| GTSIFAGHLKCRLKM | 1 | 0 | 0 | 1638003 | QTX93489.1 | P17763 |
| GTSITQKGIIFILLM | 1 | 0 | 0 | 1638004 | P17763.2 | P17763 |
| GTSLTQKVVIFILLM | 1 | 0 | 0 | 1638005 | QTX93461.1 | P17763 |
| GTTHFQRVLIFILLT | 1 | 0 | 0 | 1638009 | QTX93032.1 | P17763 |
| GTTYFQRVLIFILLT | 1 | 0 | 0 | 1638012 | QQW40777.1 | P17763 |
| GTYGLNTFTNMEVQL | 1 | 0 | 0 | 1638017 | P27909.2 | P17763 |
| GTYGLNTFTNMGAQL | 1 | 0 | 0 | 1638018 | QNQ18279.1 | P17763 |
| GVCGIRSTTRLENVM | 1 | 0 | 0 | 1638021 | Q2YHF0.1 | P17763 |
| GVCGIRSTTRMENLL | 1 | 0 | 0 | 1638022 | QTX93479.1 | P17763 |
| GVDVFFIPPEKCDTL | 1 | 0 | 0 | 1638024 | QTX93142.1 | P17763 |
| GVDVFFTPPEKCDTL | 1 | 0 | 0 | 1638025 | P14340.2 | P17763 |
| GVDVFYKPTEQVDTL | 1 | 0 | 0 | 1638026 | Q2YHF0.1 | P17763 |
| GVEGEGLHKLGYILR | 1 | 0 | 0 | 1638027 | P17763.2 | P17763 |
| GVEPGQLKLSWFKKG | 1 | 0 | 0 | 1638028 | QQW40781.1 | P17763 |
| GVFTTNIWLKLKERQ | 1 | 0 | 0 | 1638030 | QQW40771.1 | P17763 |
| GVFTTNIWLKLREMY | 1 | 0 | 0 | 1638031 | QHR82515.1 | P17763 |
| GVGDSALTLHWFRKG | 1 | 0 | 0 | 1638036 | Q2YHF0.1 | P17763 |
| GVGIHMEGVFHTMWH | 1 | 0 | 0 | 1638037 | Q2YHF0.1 | P17763 |
| GVGNRDFVEGVSGGA | 1 | 0 | 0 | 1638039 | Q2YHF0.1 | P17763 |
| GVGNSALTLHWFRKG | 1 | 0 | 0 | 1638040 | ARJ31953.1 | P17763 |
| GVGVFQDGVFHTMWH | 1 | 0 | 0 | 1638041 | P27909.2 | P17763 |
| GVGVFQEGVFHTMWH | 1 | 0 | 0 | 1638042 | P17763.2 | P17763 |
| GVGVFQENVFHTMWH | 1 | 0 | 0 | 1638043 | QTX92726.1 | P17763 |
| GVGVQKEGVFHTMWH | 1 | 0 | 0 | 1638045 | QTX93479.1 | P17763 |
| GVLFSGVSWTMKIGI | 1 | 0 | 0 | 1638053 | P17763.2 | P17763 |
| GVLNSLGKMVHQIFG | 1 | 0 | 0 | 1638056 | QTX93489.1 | P17763 |
| GVTAEITPQASTTEA | 1 | 0 | 0 | 1638064 | QFN25407.1 | P17763 |
| GVTAEITPQASTVEA | 1 | 0 | 0 | 1638065 | QTX93318.1 | P17763 |
| GVTVEITPQASTVEA | 1 | 0 | 0 | 1638066 | QTX93541.1 | P17763 |
| GVVKLLTKPWDVIPM | 1 | 0 | 0 | 1638071 | QFS19562.1 | P17763 |
| GVVTKNGGYVSGIAQ | 1 | 0 | 0 | 1638072 | QTX93489.1 | P17763 |
| GVVTKSGDYVSAITQ | 1 | 0 | 0 | 1638073 | Q2YHF0.1 | P17763 |
| GVVTRSGAYVSAIAQ | 1 | 0 | 0 | 1638074 | P14340.2 | P17763 |
| GVVTRSGTYVSAIAQ | 1 | 0 | 0 | 1638075 | QQW40772.1 | P17763 |
| GVYRIKQQGIFGKTQ | 1 | 0 | 0 | 1638078 | QTX93491.1 | P17763 |
| GVYRIMQRGLFGKTQ | 1 | 0 | 0 | 1638079 | Q2YHF0.1 | P17763 |
| GWDTRITEDDLLNEE | 1 | 0 | 0 | 1638081 | Q2YHF0.1 | P17763 |
| GWDTRITEDDLQNEE | 1 | 0 | 0 | 1638082 | ARX96505.1 | P17763 |
| GWDTRITLEDLKNEE | 1 | 0 | 0 | 1638083 | P14340.2 | P17763 |
| GWPISKMDLGVPLLA | 1 | 0 | 0 | 1638084 | AAN03432.3 | P17763 |
| GWPLSKMDIGVPLLA | 1 | 0 | 0 | 1638085 | P14340.2 | P17763 |
| GWSLKETACLGKSYA | 1 | 0 | 0 | 1638087 | QRG47124.1 | P17763 |
| GWSLRETACLGKSYA | 1 | 0 | 0 | 1638088 | P17763.2 | P17763 |
| GYVLGIFLRKLTSRE | 1 | 0 | 0 | 1638101 | Q2YHF0.1 | P17763 |
| GYVLGVFLRKLTSRE | 1 | 0 | 0 | 1638102 | Q5UCB8.1 | P17763 |
| HAAMLDVDLHPASAW | 1 | 0 | 0 | 1638103 | P17763.2 | P17763 |
| HAHHQWMTTEDMLSV | 1 | 0 | 0 | 1638105 | P17763.2 | P17763 |
| HAHWTEAKMLLDNIY | 1 | 0 | 0 | 1638106 | Q2YHF0.1 | P17763 |
| HAVGNDTSNHGETAT | 1 | 0 | 0 | 1638109 | AQV12768.1 | P17763 |
| HAVGNDTSNHGVTAT | 1 | 0 | 0 | 1638110 | Q2YHF0.1 | P17763 |
| HAVSRGSAKLQWFVE | 1 | 0 | 0 | 1638111 | QTX93489.1 | P17763 |
| HAVSRGSAKLRWFVE | 1 | 0 | 0 | 1638112 | P14340.2 | P17763 |
| HAVSRGSSKIRWIVE | 1 | 0 | 0 | 1638113 | Q58HT7.1 | P17763 |
| HAVSRGTAKLRWFVE | 1 | 0 | 0 | 1638114 | P17763.2 | P17763 |
| HDLMELIDGISLGLI | 1 | 0 | 0 | 1638115 | Q5UCB8.1 | P17763 |
| HDWQQVPFCSHHFHE | 1 | 0 | 0 | 1638116 | QTX93489.1 | P17763 |
| HEHKSTWHYDEDNPY | 1 | 0 | 0 | 1638118 | QTX92739.1 | P17763 |
| HERGRPLLFKTTEGI | 1 | 0 | 0 | 1638123 | Q2YHF0.1 | P17763 |
| HEWCCRSCTLPPLRF | 1 | 0 | 0 | 1638129 | P17763.2 | P17763 |
| HEWCCRSCTLPPLRY | 1 | 0 | 0 | 1638130 | AYP74696.1 | P17763 |
| HEWMTTEDMLAVWNR | 1 | 0 | 0 | 1638132 | QRG47112.1 | P17763 |
| HFHKIFMKDGRSLVV | 1 | 0 | 0 | 1638136 | Q2YHF0.1 | P17763 |
| HGAERLKRMAISGDD | 1 | 0 | 0 | 1638137 | P27909.2 | P17763 |
| HGGNLVRCPLSRNST | 1 | 0 | 0 | 1638141 | AVY51410.1 | P17763 |
| HGGSLVRCPLSRNST | 1 | 0 | 0 | 1638142 | Q2YHF0.1 | P17763 |
| HGSYEVKATGSASSM | 1 | 0 | 0 | 1638145 | QTX93489.1 | P17763 |
| HGSYEVKPSGSASSM | 1 | 0 | 0 | 1638146 | P17763.2 | P17763 |
| HGTTVVKVKYEGAGA | 1 | 0 | 0 | 1638148 | Q58HT7.1 | P17763 |
| HGTTVVKVKYEGTGA | 1 | 0 | 0 | 1638149 | Q2YHF0.1 | P17763 |
| HGTVLVQVKYEGTDA | 1 | 0 | 0 | 1638151 | P17763.2 | P17763 |
| HHQWMTTEDMLKVWN | 1 | 0 | 0 | 1638157 | Q2YHF0.1 | P17763 |
| HILLENDMKFTVVVG | 1 | 0 | 0 | 1638160 | P17763.2 | P17763 |
| HKERMVTFKVPHAKR | 1 | 0 | 0 | 1638162 | Q2YHF0.1 | P17763 |
| HKILAKAIFKLTYQN | 1 | 0 | 0 | 1638163 | Q2YHF0.1 | P17763 |
| HKLGKCGSCVYNMMG | 1 | 0 | 0 | 1638164 | QTX93489.1 | P17763 |
| HKLGYILRDVSKKEG | 1 | 0 | 0 | 1638165 | ABQ18243.1 | P17763 |
| HKLGYILREVSKKEG | 1 | 0 | 0 | 1638166 | Q9WDA6.1 | P17763 |
| HKQGKCATCVYNMMG | 1 | 0 | 0 | 1638167 | P17763.2 | P17763 |
| HKQWFLDLPLPWTAG | 1 | 0 | 0 | 1638168 | AHN50411.1 | P17763 |
| HKQWFLDLPLPWTSG | 1 | 0 | 0 | 1638169 | P17763.2 | P17763 |
| HKQWFLDLPLPWTTG | 1 | 0 | 0 | 1638170 | AFY10028.1 | P17763 |
| HKTLAKAIFKLTYQN | 1 | 0 | 0 | 1638171 | AXG50965.1 | P17763 |
| HKVASEGIKYTDRKW | 1 | 0 | 0 | 1638172 | QTX93489.1 | P17763 |
| HLAIMAVFKMSPGYV | 1 | 0 | 0 | 1638174 | Q2YHF0.1 | P17763 |
| HLGKLEIDFGECPGT | 1 | 0 | 0 | 1638179 | Q2YHF0.1 | P17763 |
| HLGKLELDFDLCEGT | 1 | 0 | 0 | 1638180 | P17763.2 | P17763 |
| HLGKLEMDFDFCDGT | 1 | 0 | 0 | 1638181 | QTX93151.1 | P17763 |
| HLGRLEMDFDFCEGT | 1 | 0 | 0 | 1638182 | QQW40788.1 | P17763 |

| | | | | | | |
|---|---|---|---|---|---|---|
| HLLEKKITQWLETKG | 1 | 0 | 0 | 1638184 | QTX93530.1 | P17763 |
| HMEGVFHTMWHVTRG | 1 | 0 | 0 | 1638188 | Q2YHF0.1 | P17763 |
| HMFAGHLKCKVRMEK | 1 | 0 | 0 | 1638189 | Q2YHF0.1 | P17763 |
| HNGRLITANPVVTKK | 1 | 0 | 0 | 1638194 | QTX93489.1 | P17763 |
| HNHTFLIDGPETAEC | 1 | 0 | 0 | 1638195 | QQW40786.1 | P17763 |
| HPASAWTLYAVATTV | 1 | 0 | 0 | 1638196 | AHI43742.1 | P17763 |
| HQLIMKDGREIVVPC | 1 | 0 | 0 | 1638206 | P17763.2 | P17763 |
| HQVGNDTQGVTVEIT | 1 | 0 | 0 | 1638212 | QTX93541.1 | P17763 |
| HQVGNESTEHGTTAT | 1 | 0 | 0 | 1638213 | QFS19562.1 | P17763 |
| HQVGNETTEHGTIAT | 1 | 0 | 0 | 1638215 | P27909.2 | P17763 |
| HQWMTTEDMLTVWNR | 1 | 0 | 0 | 1638216 | QTX93489.1 | P17763 |
| HRGPSLRTTTASGKL | 1 | 0 | 0 | 1638217 | Q2YHF0.1 | P17763 |
| HRLGYILEDIDKKDG | 1 | 0 | 0 | 1638222 | Q2YHF0.1 | P17763 |
| HRLGYILEEIDKKDG | 1 | 0 | 0 | 1638223 | ARJ31953.1 | P17763 |
| HRLGYILRDVGKKEG | 1 | 0 | 0 | 1638224 | AWK29891.1 | P17763 |
| HRLGYILRDVSKKEG | 1 | 0 | 0 | 1638225 | AGN94889.1 | P17763 |
| HRLMSAAIKDERAVH | 1 | 0 | 0 | 1638226 | QTX93541.1 | P17763 |
| HRLMSAAIKDQKAVH | 1 | 0 | 0 | 1638227 | Q2YHF0.1 | P17763 |
| HRLMSAAIKDSKAVH | 1 | 0 | 0 | 1638228 | P17763.2 | P17763 |
| HRRDLRLASMAICSA | 1 | 0 | 0 | 1638229 | Q2YHF0.1 | P17763 |
| HSALAGATEVDSGDG | 1 | 0 | 0 | 1638230 | Q2YHF0.1 | P17763 |
| HTLWSNGVLESQMLI | 1 | 0 | 0 | 1638240 | Q2YHF0.1 | P17763 |
| HTQTAGPWHLGRLEM | 1 | 0 | 0 | 1638245 | QQW40786.1 | P17763 |
| HVAAENHQHATMLDV | 1 | 0 | 0 | 1638246 | ARQ80406.1 | P17763 |
| HVAVENHHHAAMLDV | 1 | 0 | 0 | 1638248 | QTX92726.1 | P17763 |
| HYAIIGPGLQAKATR | 1 | 0 | 0 | 1638256 | P17763.2 | P17763 |
| HYDHENPYRTWAYHG | 1 | 0 | 0 | 1638260 | Q2YHF0.1 | P17763 |
| HYDQENPYRTWAYHG | 1 | 0 | 0 | 1638262 | Q58HT7.1 | P17763 |
| IAASIILEFFLIVLL | 1 | 0 | 0 | 1638268 | P14340.2 | P17763 |
| IAGGMLIACYVISGS | 1 | 0 | 0 | 1638270 | P17763.2 | P17763 |
| IAGVLFTFVLLLSGQ | 1 | 0 | 0 | 1638271 | QTX93487.1 | P17763 |
| IALFLAHAIGTSITQ | 1 | 0 | 0 | 1638276 | P17763.2 | P17763 |
| IAMDLGELCEDTMTY | 1 | 0 | 0 | 1638280 | P17763.2 | P17763 |
| IAMDLGEMCDDTVTY | 1 | 0 | 0 | 1638281 | QTX93489.1 | P17763 |
| IAMDLGEMCEDTVTY | 1 | 0 | 0 | 1638282 | Q2YHF0.1 | P17763 |
| IANQAAILMGLDKGW | 1 | 0 | 0 | 1638286 | P17763.2 | P17763 |
| IANQAAVLMGLGKGW | 1 | 0 | 0 | 1638287 | Q2YHF0.1 | P17763 |
| IAQAKASQEGPLPEI | 1 | 0 | 0 | 1638289 | P17763.2 | P17763 |
| IAQTEKSIEDNPEIE | 1 | 0 | 0 | 1638290 | P14340.2 | P17763 |
| IAQTNAEPDGPTPEL | 1 | 0 | 0 | 1638291 | QTX93489.1 | P17763 |
| IASSGMLWMADIPLQ | 1 | 0 | 0 | 1638294 | AQV12569.1 | P17763 |
| IASSGMLWMAEIPLQ | 1 | 0 | 0 | 1638295 | QPI11927.1 | P17763 |
| IATITPQAPTSEIQL | 1 | 0 | 0 | 1638296 | P27909.2 | P17763 |
| IAVKNWLVRVGRERL | 1 | 0 | 0 | 1638297 | QQW40772.1 | P17763 |
| IAVLTGGVTLFFLSG | 1 | 0 | 0 | 1638298 | P17763.2 | P17763 |
| IAVSMANIFRGSYLA | 1 | 0 | 0 | 1638300 | P17763.2 | P17763 |
| IDEEREIPERSWNSG | 1 | 0 | 0 | 1638308 | QTX93142.1 | P17763 |
| IDLDPIPYDPKFEKQ | 1 | 0 | 0 | 1638309 | P14340.2 | P17763 |
| IDLDPVIYDSKFEKQ | 1 | 0 | 0 | 1638310 | QTX93489.1 | P17763 |
| IDLEPIPYDPKFEKQ | 1 | 0 | 0 | 1638311 | QRG47106.1 | P17763 |
| IEAEPPFGESNIIIG | 1 | 0 | 0 | 1638313 | QQD79912.1 | P17763 |
| IEAKLTNTTTASRCP | 1 | 0 | 0 | 1638315 | QQW40788.1 | P17763 |
| IEAKLTNTTTESRCP | 1 | 0 | 0 | 1638316 | QTX93024.1 | P17763 |
| IEALISNITTATRCP | 1 | 0 | 0 | 1638317 | AKQ00032.1 | P17763 |
| IEASISNITTATRCP | 1 | 0 | 0 | 1638318 | Q2YHF0.1 | P17763 |
| IEENPWMEDKTHISS | 1 | 0 | 0 | 1638322 | QTX92698.1 | P17763 |
| IEENPWMEDKTHVSS | 1 | 0 | 0 | 1638323 | P17763.2 | P17763 |
| IEGKITNVTTDSRCP | 1 | 0 | 0 | 1638324 | QHR82515.1 | P17763 |
| IEIRDVNKEKVVGRV | 1 | 0 | 0 | 1638325 | ARN79603.1 | P17763 |
| IEKIKQEHEISWHYD | 1 | 0 | 0 | 1638326 | AFK65749.1 | P17763 |
| IEKIKQEHETSWHYD | 1 | 0 | 0 | 1638327 | P14340.2 | P17763 |
| IEKPDMTIIGRRLQR | 1 | 0 | 0 | 1638328 | QTX92271.1 | P17763 |
| IENTTANISLTAIAN | 1 | 0 | 0 | 1638329 | P17763.2 | P17763 |
| IEPSWADVRKDLISY | 1 | 0 | 0 | 1638330 | QGW51895.1 | P17763 |
| IFGSAYTALFSGVSW | 1 | 0 | 0 | 1638337 | QTX93489.1 | P17763 |
| IFMKDGRSLVVPCRN | 1 | 0 | 0 | 1638338 | Q2YHF0.1 | P17763 |
| IFMTATPPGSVEAFP | 1 | 0 | 0 | 1638339 | P17763.2 | P17763 |
| IFPYSIPATLLVWHT | 1 | 0 | 0 | 1638342 | QTX93483.1 | P17763 |
| IGAVFQEEQGWTSAS | 1 | 0 | 0 | 1638345 | Q2YHF0.1 | P17763 |
| IGIESETPNLDIIGK | 1 | 0 | 0 | 1638353 | Q9WDA6.1 | P17763 |
| IGIGILLTWLGLNSR | 1 | 0 | 0 | 1638354 | P17763.2 | P17763 |
| IGIGVLLTWLGLNSR | 1 | 0 | 0 | 1638355 | QFS19562.1 | P17763 |
| IGKRIEKIKQEHEIS | 1 | 0 | 0 | 1638356 | AFK65749.1 | P17763 |
| IGKRIEKIKQEHETS | 1 | 0 | 0 | 1638357 | P14340.2 | P17763 |
| IGMAMTTTLSIPHDL | 1 | 0 | 0 | 1638360 | AMP43484.1 | P17763 |
| IGMAMTTVFSIPHDL | 1 | 0 | 0 | 1638361 | Q2YHF2.1 | P17763 |
| IGMAMTTVLSIPHDL | 1 | 0 | 0 | 1638362 | Q2YHF0.1 | P17763 |
| IGNENYLDYMTSMKR | 1 | 0 | 0 | 1638363 | AAG14999.1 | P17763 |
| IGPGLQAKATREAQK | 1 | 0 | 0 | 1638364 | P17763.2 | P17763 |
| IGQTGIQRTVFFVLM | 1 | 0 | 0 | 1638366 | Q58HT7.1 | P17763 |
| IGRNPAQEDDQYVFS | 1 | 0 | 0 | 1638367 | Q2YHF0.1 | P17763 |
| IGRRLQRLQEEHKET | 1 | 0 | 0 | 1638369 | Q2YHF0.1 | P17763 |
| IGRRLQRLREEHKET | 1 | 0 | 0 | 1638370 | AHG06383.1 | P17763 |
| IGTSITQKGIIFILL | 1 | 0 | 0 | 1638373 | P17763.2 | P17763 |
| IGTSLTQKVVIFILL | 1 | 0 | 0 | 1638374 | QTX93461.1 | P17763 |
| IGTTYFQRVLIFILL | 1 | 0 | 0 | 1638375 | QQW40777.1 | P17763 |
| IGVLLTWIGLNSKNT | 1 | 0 | 0 | 1638377 | QTX93491.1 | P17763 |
| IGVPLLAIGCYSQVN | 1 | 0 | 0 | 1638378 | P14340.2 | P17763 |
| IHAHHQWMTTEDMLT | 1 | 0 | 0 | 1638379 | QTX93489.1 | P17763 |
| IHAKHEWMTTEDMLA | 1 | 0 | 0 | 1638380 | QRG47115.1 | P17763 |
| IHAKHEWMTTEDMLT | 1 | 0 | 0 | 1638381 | P14340.2 | P17763 |

| | | | | | | |
|---|---|---|---|---|---|---|
| IHARHEWMTTEDMLT | 1 | 0 | 0 | 1638382 | QNQ18279.1 | P17763 |
| IHATHEWMTTEDMLT | 1 | 0 | 0 | 1638383 | QQW40786.1 | P17763 |
| IIGIGDKALKINWYK | 1 | 0 | 0 | 1638392 | QDG00750.1 | P17763 |
| IILEFFLIVLLIPEP | 1 | 0 | 0 | 1638398 | P14340.2 | P17763 |
| IILEFFLMVLLIPEP | 1 | 0 | 0 | 1638399 | P17763.2 | P17763 |
| IIPALFEPEREKSAA | 1 | 0 | 0 | 1638402 | P17763.2 | P17763 |
| IKILIGFRKEIGRML | 1 | 0 | 0 | 1638407 | ARJ31953.1 | P17763 |
| IKILTGFRKEIGRML | 1 | 0 | 0 | 1638408 | Q2YHF0.1 | P17763 |
| IKQKGILGYSQIGAG | 1 | 0 | 0 | 1638411 | P14340.2 | P17763 |
| IKQQGIFGKTQVGVG | 1 | 0 | 0 | 1638413 | QTX93491.1 | P17763 |
| IKQRGILGYSQIGAG | 1 | 0 | 0 | 1638414 | QRG47115.1 | P17763 |
| IKRGETDHHAVSRGS | 1 | 0 | 0 | 1638415 | P14340.2 | P17763 |
| IKRKLRTLVLAPTRV | 1 | 0 | 0 | 1638416 | QMT58602.1 | P17763 |
| ILARWGSFKKNGAIK | 1 | 0 | 0 | 1638427 | P17763.2 | P17763 |
| ILARWGTFKKSGAIK | 1 | 0 | 0 | 1638428 | QTX93247.1 | P17763 |
| ILARWSSFKKNGAIK | 1 | 0 | 0 | 1638429 | AXX75195.1 | P17763 |
| ILAVVITLCAIILGG | 1 | 0 | 0 | 1638430 | AKQ00036.1 | P17763 |
| ILEENMDVEIWTKEG | 1 | 0 | 0 | 1638431 | QTX93345.1 | P17763 |
| ILEVDRTEARSALKD | 1 | 0 | 0 | 1638434 | QTX92258.1 | P17763 |
| ILFEEVMRGKFGKKH | 1 | 0 | 0 | 1638435 | QTX93489.1 | P17763 |
| ILIGFLVLWIGTNSR | 1 | 0 | 0 | 1638438 | Q58HT7.1 | P17763 |
| ILIGLLVLWIGTNSR | 1 | 0 | 0 | 1638439 | Q2YHF0.1 | P17763 |
| ILIGVVITWIGMNSR | 1 | 0 | 0 | 1638441 | QQW40770.1 | P17763 |
| ILIRTGLLVISGLFP | 1 | 0 | 0 | 1638442 | P14340.2 | P17763 |
| ILKIVRNMEKYQLAV | 1 | 0 | 0 | 1638443 | QQW40674.1 | P17763 |
| ILLKATLLAVSGVYP | 1 | 0 | 0 | 1638445 | QFS19562.1 | P17763 |
| ILLWYAQIQPHWIAA | 1 | 0 | 0 | 1638447 | P14340.2 | P17763 |
| ILMKMKKKTWLVHKQ | 1 | 0 | 0 | 1638448 | Q58HT7.1 | P17763 |
| ILMKMKTKTWLVHKQ | 1 | 0 | 0 | 1638449 | Q2YHF0.1 | P17763 |
| ILNPYMPSVVETLEQ | 1 | 0 | 0 | 1638452 | P17763.2 | P17763 |
| ILPEEQDQNYVCKHT | 1 | 0 | 0 | 1638453 | QTX93510.1 | P17763 |
| ILQRGLLGRSQVGVG | 1 | 0 | 0 | 1638454 | P17763.2 | P17763 |
| ILRDISKIPGGAMYA | 1 | 0 | 0 | 1638455 | AAX55647.1 | P17763 |
| ILRDISKIPGGNMYA | 1 | 0 | 0 | 1638456 | P17763.2 | P17763 |
| ILTDGEERVILAGPM | 1 | 0 | 0 | 1638457 | P14340.2 | P17763 |
| ILTDGPERVILAGPI | 1 | 0 | 0 | 1638458 | ACL99055.1 | P17763 |
| ILVVVITFCAIILGG | 1 | 0 | 0 | 1638463 | ANW72070.1 | P17763 |
| ILVVVTTLCAIILGG | 1 | 0 | 0 | 1638464 | Q2YHF0.1 | P17763 |
| IMAISCVPNAVILQN | 1 | 0 | 0 | 1638466 | QRG47124.1 | P17763 |
| IMEVDRSEAKEGLKR | 1 | 0 | 0 | 1638469 | QFS19562.1 | P17763 |
| IMKDGRVLVVPCRNQ | 1 | 0 | 0 | 1638470 | P14340.2 | P17763 |
| IMKNPTVDGIMTIDL | 1 | 0 | 0 | 1638471 | QTX93489.1 | P17763 |
| IMKNPTVDGITVIDL | 1 | 0 | 0 | 1638472 | P14340.2 | P17763 |
| IMQRGLFGKTQVGVG | 1 | 0 | 0 | 1638473 | Q2YHF0.1 | P17763 |
| IMQRGLLGRSQVGVG | 1 | 0 | 0 | 1638474 | QTX92726.1 | P17763 |
| IMTTTANWLWALLGK | 1 | 0 | 0 | 1638476 | QCZ25011.1 | P17763 |
| INEKEENMVKSLVSA | 1 | 0 | 0 | 1638477 | QTX93512.1 | P17763 |
| IPFSTEDGQGKAHNG | 1 | 0 | 0 | 1638483 | QTX93500.1 | P17763 |
| IPLCRTSCLQKQSHW | 1 | 0 | 0 | 1638489 | Q2YHF0.1 | P17763 |
| IPSMFEPEREKVDAI | 1 | 0 | 0 | 1638491 | P14340.2 | P17763 |
| IPTLFGPEREKTQAI | 1 | 0 | 0 | 1638492 | Q2YHF0.1 | P17763 |
| IQDEERDIPERSWNS | 1 | 0 | 0 | 1638497 | P17763.2 | P17763 |
| IQEVDRTLAKEGIKR | 1 | 0 | 0 | 1638498 | P14340.2 | P17763 |
| IQRVETWALRHPGFT | 1 | 0 | 0 | 1638505 | QFS19562.1 | P17763 |
| IRSTTRLENVMWKQI | 1 | 0 | 0 | 1638512 | Q2YHF0.1 | P17763 |
| IRVPNYNLIIMDEAH | 1 | 0 | 0 | 1638514 | Q9WDA6.1 | P17763 |
| IRYQTTAVKSEHTGK | 1 | 0 | 0 | 1638515 | P17763.2 | P17763 |
| IRYQTTAVKSEHTGR | 1 | 0 | 0 | 1638516 | P27909.2 | P17763 |
| ISGDDCVVKPLDERF | 1 | 0 | 0 | 1638519 | Q2YHF0.1 | P17763 |
| ISKIPGGAMYADDTA | 1 | 0 | 0 | 1638524 | QTX93489.1 | P17763 |
| ISKIPGGNMYADDTA | 1 | 0 | 0 | 1638525 | P17763.2 | P17763 |
| ISKMDIGVPLLALGC | 1 | 0 | 0 | 1638526 | P17763.2 | P17763 |
| ISNITTATRCPTQGE | 1 | 0 | 0 | 1638531 | Q2YHF0.1 | P17763 |
| ISQGAGWSLRETACL | 1 | 0 | 0 | 1638534 | P14340.2 | P17763 |
| ISRKDQRGSGQVGTY | 1 | 0 | 0 | 1638535 | QRG47124.1 | P17763 |
| ISRRDQRGSGQVGTY | 1 | 0 | 0 | 1638537 | P17763.2 | P17763 |
| ISTLWEGNPGRFWNT | 1 | 0 | 0 | 1638539 | P14340.2 | P17763 |
| ISTRVEMGEAAAIFM | 1 | 0 | 0 | 1638540 | Q2YHF0.1 | P17763 |
| ISTRVGMGEAAAIFM | 1 | 0 | 0 | 1638541 | P17763.2 | P17763 |
| ISYGGGWKLEGEWKE | 1 | 0 | 0 | 1638542 | P14340.2 | P17763 |
| ISYGGGWRFQGSWNT | 1 | 0 | 0 | 1638543 | P27909.2 | P17763 |
| ISYGGGWRLGDKWDK | 1 | 0 | 0 | 1638544 | Q2YHF0.1 | P17763 |
| ISYGGGWRLQGSWNT | 1 | 0 | 0 | 1638545 | QTX92726.1 | P17763 |
| ISYGGGWRLRGSWNT | 1 | 0 | 0 | 1638546 | ACQ44411.1 | P17763 |
| ITASILLWYAQIQPH | 1 | 0 | 0 | 1638549 | P14340.2 | P17763 |
| ITEDDLHNEEKITQQ | 1 | 0 | 0 | 1638550 | QTX93491.1 | P17763 |
| ITEDDLQNEAKITDI | 1 | 0 | 0 | 1638551 | P17763.2 | P17763 |
| ITEQMAPHHKILAKA | 1 | 0 | 0 | 1638552 | Q2YHF0.1 | P17763 |
| ITEQMAPHHKTLAKA | 1 | 0 | 0 | 1638553 | QBA29684.1 | P17763 |
| ITEVDRTEAKEGLKR | 1 | 0 | 0 | 1638554 | QTX93489.1 | P17763 |
| ITGNLVQIENLEYTV | 1 | 0 | 0 | 1638557 | Q2YHF0.1 | P17763 |
| ITLTAALLLLVAHYA | 1 | 0 | 0 | 1638561 | QTX92981.1 | P17763 |
| ITPHSGEENAVGNDT | 1 | 0 | 0 | 1638564 | QQW40798.1 | P17763 |
| ITPMMRHTIENTTAN | 1 | 0 | 0 | 1638565 | P17763.2 | P17763 |
| ITQAERIGEPDYEVD | 1 | 0 | 0 | 1638566 | Q2YHF0.1 | P17763 |
| ITQDDMHNPKGLKER | 1 | 0 | 0 | 1638567 | Q58HT7.1 | P17763 |
| ITQDDMQNPKGLKER | 1 | 0 | 0 | 1638568 | Q5UCB8.1 | P17763 |
| ITQFETYQLWTALVS | 1 | 0 | 0 | 1638569 | QTX93491.1 | P17763 |
| ITRDDMHNPKGLKER | 1 | 0 | 0 | 1638572 | Q2YHF0.1 | P17763 |
| ITTLWEGSPGKFWNT | 1 | 0 | 0 | 1638575 | QTX93489.1 | P17763 |
| ITVHTGDQHQVGNDT | 1 | 0 | 0 | 1638576 | QTX93534.1 | P17763 |

| | | | | | | |
|---|---|---|---|---|---|---|
| ITYKCPLLRQNEPED | 1 | 0 | 0 | 1638577 | QTX93024.1 | P17763 |
| ITYNCPLLRQNEPED | 1 | 0 | 0 | 1638578 | QQW40770.1 | P17763 |
| IVAKHERGRPLLFKT | 1 | 0 | 0 | 1638580 | Q2YHF0.1 | P17763 |
| IVERGMIKPKGKVVD | 1 | 0 | 0 | 1638582 | ARM59249.1 | P17763 |
| IVERGMVKPKGKVVD | 1 | 0 | 0 | 1638583 | Q2YHF0.1 | P17763 |
| IVGIQEKGKSLLFKT | 1 | 0 | 0 | 1638584 | QTX93131.1 | P17763 |
| IVGKNERGKSLLFKT | 1 | 0 | 0 | 1638585 | QTX93489.1 | P17763 |
| IVGRQEKGKSLLFKT | 1 | 0 | 0 | 1638586 | QRG47112.1 | P17763 |
| IVKLMSGKDVFYLPP | 1 | 0 | 0 | 1638590 | AAC63341.1 | P17763 |
| IVLEFFMMVLLIPEP | 1 | 0 | 0 | 1638591 | QTX93489.1 | P17763 |
| IVMDEAHFTDPCSVA | 1 | 0 | 0 | 1638593 | AQV12653.1 | P17763 |
| IVMDEAHFTDPSSVA | 1 | 0 | 0 | 1638594 | Q2YHF0.1 | P17763 |
| IVQPENLEYTIVITP | 1 | 0 | 0 | 1638599 | BAG84557.1 | P17763 |
| IVSGIFPYSIPATLL | 1 | 0 | 0 | 1638600 | QTX93489.1 | P17763 |
| IVSKQERGKSLLFKT | 1 | 0 | 0 | 1638601 | P17763.2 | P17763 |
| IVTCAMFTCKKNMEG | 1 | 0 | 0 | 1638604 | BAG84557.1 | P17763 |
| IVTDKEKPVNIEAEP | 1 | 0 | 0 | 1638605 | P17763.2 | P17763 |
| IVTDKEKPVNIETEP | 1 | 0 | 0 | 1638606 | P27909.2 | P17763 |
| IWTKEGERKKLKPRW | 1 | 0 | 0 | 1638609 | P14340.2 | P17763 |
| IWTREGEKKKLRPKW | 1 | 0 | 0 | 1638610 | Q2YHF0.1 | P17763 |
| IWTREGEKKKLRPRW | 1 | 0 | 0 | 1638611 | AFY10053.1 | P17763 |
| IWYMWLGARFLEFEA | 1 | 0 | 0 | 1638612 | P17763.2 | P17763 |
| IYMGEPLENDEDCAH | 1 | 0 | 0 | 1638617 | P14340.2 | P17763 |
| KALKLSWFKKGSSIG | 1 | 0 | 0 | 1638630 | P17763.2 | P17763 |
| KASQEGPLPEIEDEV | 1 | 0 | 0 | 1638634 | P17763.2 | P17763 |
| KATREAQKRTAAGIM | 1 | 0 | 0 | 1638636 | P17763.2 | P17763 |
| KATYEPDVDLGSGTR | 1 | 0 | 0 | 1638637 | P14340.2 | P17763 |
| KAYAGPISQHNYRQG | 1 | 0 | 0 | 1638639 | Q2YHF0.1 | P17763 |
| KCATCVYNMMGKREK | 1 | 0 | 0 | 1638640 | P17763.2 | P17763 |
| KCTLIAMDLGEMCED | 1 | 0 | 0 | 1638647 | Q2YHF0.1 | P17763 |
| KDERAVHADMGYWIE | 1 | 0 | 0 | 1638648 | QTX93489.1 | P17763 |
| KDGDLIYADDTAGWD | 1 | 0 | 0 | 1638649 | Q2YHF0.1 | P17763 |
| KDGDLMYADDTAGWD | 1 | 0 | 0 | 1638650 | Q5UCB8.1 | P17763 |
| KDIPQWQPSKGWHDW | 1 | 0 | 0 | 1638651 | QTX93489.1 | P17763 |
| KDIQQWEPSKGWNDW | 1 | 0 | 0 | 1638652 | QTX93077.1 | P17763 |
| KDQKAVHADMGYWIE | 1 | 0 | 0 | 1638660 | Q2YHF0.1 | P17763 |
| KDSKAVHADMGYWIE | 1 | 0 | 0 | 1638662 | P17763.2 | P17763 |
| KDVDLGAGTRHVNAE | 1 | 0 | 0 | 1638663 | QTX93489.1 | P17763 |
| KEAKMLLDNINTPEG | 1 | 0 | 0 | 1638666 | P14340.2 | P17763 |
| KEEYVDYMPAMKRYS | 1 | 0 | 0 | 1638669 | QTX92249.1 | P17763 |
| KEEYVDYMPAMRRYS | 1 | 0 | 0 | 1638670 | QTX92339.1 | P17763 |
| KEGGAMYADDTAGWD | 1 | 0 | 0 | 1638672 | ASK38693.1 | P17763 |
| KEIGRMLNILNGRKR | 1 | 0 | 0 | 1638673 | Q2YHF0.1 | P17763 |
| KEISNMLSIINRRKK | 1 | 0 | 0 | 1638674 | QDG00742.1 | P17763 |
| KEKPVNIETEPPFGE | 1 | 0 | 0 | 1638676 | P27909.2 | P17763 |
| KEPGVVSPTSYLDVD | 1 | 0 | 0 | 1638678 | QTX93479.1 | P17763 |
| KERALHQEGKCESCV | 1 | 0 | 0 | 1638679 | Q2YHF0.1 | P17763 |
| KERNLHLEGKCETCV | 1 | 0 | 0 | 1638680 | P14340.2 | P17763 |
| KERVEKWLKECGVDR | 1 | 0 | 0 | 1638681 | Q2YHF0.1 | P17763 |
| KERVEKWLRECGVDR | 1 | 0 | 0 | 1638682 | AMP43484.1 | P17763 |
| KETACLGKAYAQMWS | 1 | 0 | 0 | 1638683 | QTX93541.1 | P17763 |
| KETACLGKSYAQMWT | 1 | 0 | 0 | 1638684 | QRG47112.1 | P17763 |
| KETWHYDHENPYRTW | 1 | 0 | 0 | 1638686 | Q2YHF0.1 | P17763 |
| KETWHYDQENPYRTW | 1 | 0 | 0 | 1638687 | Q58HT7.1 | P17763 |
| KFSIDKEMAETQHGT | 1 | 0 | 0 | 1638693 | Q2YHF0.1 | P17763 |
| KFTVVVGDANGILAQ | 1 | 0 | 0 | 1638695 | QMW69339.1 | P17763 |
| KFTVVVGDVAGILAQ | 1 | 0 | 0 | 1638696 | QFS19562.1 | P17763 |
| KFTVVVGDVNGILAQ | 1 | 0 | 0 | 1638697 | QHE24256.1 | P17763 |
| KGEDAPCKIPFSTED | 1 | 0 | 0 | 1638700 | QTX93489.1 | P17763 |
| KGEDVPCKIPFSTED | 1 | 0 | 0 | 1638701 | AMH87199.1 | P17763 |
| KGGPGHEEPIPMATY | 1 | 0 | 0 | 1638704 | P17763.2 | P17763 |
| KGGPGHEEPIPMSTY | 1 | 0 | 0 | 1638705 | P14340.2 | P17763 |
| KGGPGHEEPVPMSTY | 1 | 0 | 0 | 1638707 | AAC63341.1 | P17763 |
| KGKELKCGSGIFVTN | 1 | 0 | 0 | 1638708 | QTX93489.1 | P17763 |
| KGKRIEPSWADVKKD | 1 | 0 | 0 | 1638709 | P14340.2 | P17763 |
| KGKRIEPSWADVRKD | 1 | 0 | 0 | 1638710 | QGW51895.1 | P17763 |
| KGLTKGGPGHEEPIP | 1 | 0 | 0 | 1638711 | P14340.2 | P17763 |
| KGMSYAMCLNAFVLK | 1 | 0 | 0 | 1638712 | AHG06373.1 | P17763 |
| KGMSYAMCSNAFVLK | 1 | 0 | 0 | 1638713 | QHR82521.1 | P17763 |
| KGMSYAMCTNTFVLK | 1 | 0 | 0 | 1638714 | QHJ86727.1 | P17763 |
| KGSRAIWYMWLGARF | 1 | 0 | 0 | 1638719 | P17763.2 | P17763 |
| KGWNDWQQVPFCSHH | 1 | 0 | 0 | 1638722 | P17763.2 | P17763 |
| KGWPISKMDIGVPLL | 1 | 0 | 0 | 1638723 | P17763.2 | P17763 |
| KGWPLHRMDLGVPLL | 1 | 0 | 0 | 1638724 | Q2YHF0.1 | P17763 |
| KGYTKGGPGHEEPIP | 1 | 0 | 0 | 1638725 | P17763.2 | P17763 |
| KHGKEIKVTPQSSIT | 1 | 0 | 0 | 1638727 | QTX93055.1 | P17763 |
| KIGEFGKAKGSRAIW | 1 | 0 | 0 | 1638733 | QEI46448.1 | P17763 |
| KIKDEERDDTLTILL | 1 | 0 | 0 | 1638736 | P17763.2 | P17763 |
| KIRWIVERGMIKPKG | 1 | 0 | 0 | 1638745 | ARM59249.1 | P17763 |
| KIRWIVERGMVKPKG | 1 | 0 | 0 | 1638746 | Q2YHF0.1 | P17763 |
| KIVGLYGNGVVTTSG | 1 | 0 | 0 | 1638747 | P17763.2 | P17763 |
| KIYGGPISQHNYRPG | 1 | 0 | 0 | 1638749 | P17763.2 | P17763 |
| KKDLISYGGGWKLEG | 1 | 0 | 0 | 1638752 | P14340.2 | P17763 |
| KKDLISYGGGWRFQG | 1 | 0 | 0 | 1638753 | P17763.2 | P17763 |
| KKDLISYGGGWRLQG | 1 | 0 | 0 | 1638754 | QTX92726.1 | P17763 |
| KKDLISYGGGWRLRG | 1 | 0 | 0 | 1638755 | ACQ44459.1 | P17763 |
| KKEISNMLNIMNRRK | 1 | 0 | 0 | 1638756 | P17763.2 | P17763 |
| KKEISNMLSIINKRK | 1 | 0 | 0 | 1638757 | ADI44327.1 | P17763 |
| KKEISNMLSIINRRK | 1 | 0 | 0 | 1638758 | QDG00742.1 | P17763 |
| KKGSSIGQMFETTMR | 1 | 0 | 0 | 1638760 | QTX92981.1 | P17763 |
| KKHMIAGVLFTFVLL | 1 | 0 | 0 | 1638761 | QTX93504.1 | P17763 |

| | | | | | | |
|---|---|---|---|---|---|---|
| KKITQWLETKGVERL | 1 | 0 | 0 | 1638762 | QTX93479.1 | P17763 |
| KKLKPRWLDARIYSD | 1 | 0 | 0 | 1638764 | QTX92981.1 | P17763 |
| KKLRPKWLDARVYAD | 1 | 0 | 0 | 1638765 | Q2YHF0.1 | P17763 |
| KKLRPRWLDARVYAD | 1 | 0 | 0 | 1638766 | AQV12649.1 | P17763 |
| KKMIRPQPMEHKYSW | 1 | 0 | 0 | 1638768 | P17763.2 | P17763 |
| KKRLTIMDLHPGAGK | 1 | 0 | 0 | 1638769 | QQW40770.1 | P17763 |
| KKSGAIKVLKGFKKE | 1 | 0 | 0 | 1638771 | QBM00871.1 | P17763 |
| KKSGAIKVLRGFKKE | 1 | 0 | 0 | 1638772 | UAV34166.1 | P17763 |
| KKSGIQEVDRTLAKE | 1 | 0 | 0 | 1638773 | P14340.2 | P17763 |
| KKSGITEVDRTEAKE | 1 | 0 | 0 | 1638774 | QTX93489.1 | P17763 |
| KKVRSNAALGAIFTD | 1 | 0 | 0 | 1638777 | AII99332.1 | P17763 |
| KKVVRPPFNMLKRER | 1 | 0 | 0 | 1638779 | Q58HT7.1 | P17763 |
| KLASAIQKAQEEGIC | 1 | 0 | 0 | 1638780 | QTX93142.1 | P17763 |
| KLELKGMSYAMCLNA | 1 | 0 | 0 | 1638785 | AHG06373.1 | P17763 |
| KLELKGMSYAMCLNT | 1 | 0 | 0 | 1638786 | AAC63321.1 | P17763 |
| KLELKGMSYAMCSNA | 1 | 0 | 0 | 1638787 | QHR82521.1 | P17763 |
| KLELKGMSYAMCTNT | 1 | 0 | 0 | 1638788 | QHJ86727.1 | P17763 |
| KLGEFGKAKGSRAIW | 1 | 0 | 0 | 1638791 | P17763.2 | P17763 |
| KLGEFGRAKGSRAIW | 1 | 0 | 0 | 1638792 | ACN42699.1 | P17763 |
| KLHSGKDVFFIPPEK | 1 | 0 | 0 | 1638796 | QTX92698.1 | P17763 |
| KLHSGKDVFFMPPEK | 1 | 0 | 0 | 1638797 | P27909.2 | P17763 |
| KLHSGVDVFYKPTEQ | 1 | 0 | 0 | 1638798 | Q2YHF0.1 | P17763 |
| KLLTKPWDVIPMVTQ | 1 | 0 | 0 | 1638803 | QFS19562.1 | P17763 |
| KLLTKPWDVVPMVTQ | 1 | 0 | 0 | 1638804 | QFS19207.1 | P17763 |
| KLMSGKDVFYLPPEK | 1 | 0 | 0 | 1638805 | AAC63341.1 | P17763 |
| KLQWFVERNMVIPEG | 1 | 0 | 0 | 1638806 | QTX93489.1 | P17763 |
| KLRWFVERNLVKPEG | 1 | 0 | 0 | 1638808 | P17763.2 | P17763 |
| KLRWFVERNLVTPEG | 1 | 0 | 0 | 1638809 | Q9WDA6.1 | P17763 |
| KLRWFVERNMVTPEG | 1 | 0 | 0 | 1638810 | P14340.2 | P17763 |
| KLTYQNKVVKVQRPT | 1 | 0 | 0 | 1638814 | QTX93489.1 | P17763 |
| KLTYQNKVVRVQRPA | 1 | 0 | 0 | 1638815 | P17763.2 | P17763 |
| KLVMAFIAFLRFLAI | 1 | 0 | 0 | 1638816 | P17763.2 | P17763 |
| KMEALQRKYGGALVR | 1 | 0 | 0 | 1638819 | P14340.2 | P17763 |
| KMEALQRQYGGALVR | 1 | 0 | 0 | 1638820 | ACW82894.1 | P17763 |
| KMLLDNINTPEGIIP | 1 | 0 | 0 | 1638825 | P17763.2 | P17763 |
| KMSPGYVLGIFLRKL | 1 | 0 | 0 | 1638826 | Q2YHF0.1 | P17763 |
| KMSPGYVLGVFLRKL | 1 | 0 | 0 | 1638827 | Q5UCB8.1 | P17763 |
| KNENDQYIYMGEPLE | 1 | 0 | 0 | 1638831 | P14340.2 | P17763 |
| KNGGYVSGIAQTNAE | 1 | 0 | 0 | 1638834 | QTX93489.1 | P17763 |
| KNGKRVIQLSRKTFD | 1 | 0 | 0 | 1638835 | QFS19562.1 | P17763 |
| KNIQAAINQVRSLIG | 1 | 0 | 0 | 1638840 | QTX93142.1 | P17763 |
| KNLAGPVSQHNYRPG | 1 | 0 | 0 | 1638842 | QTX93151.1 | P17763 |
| KNNQFCIKVLNPYMP | 1 | 0 | 0 | 1638845 | QTX93489.1 | P17763 |
| KNPRLCTREEFISKV | 1 | 0 | 0 | 1638846 | Q2YHF0.1 | P17763 |
| KNTPFNMLKRERNRV | 1 | 0 | 0 | 1638848 | P29990.1 | P17763 |
| KNTSMSFSCIAIGII | 1 | 0 | 0 | 1638849 | QIS48873.1 | P17763 |
| KNTSMSFSCIVIGII | 1 | 0 | 0 | 1638850 | QTX93530.1 | P17763 |
| KPLDERFSTSLLFLN | 1 | 0 | 0 | 1638853 | Q2YHF0.1 | P17763 |
| KPRLCTREEFTKKVR | 1 | 0 | 0 | 1638855 | QTX93504.1 | P17763 |
| KPTYEKDADLGAGTR | 1 | 0 | 0 | 1638858 | Q2YHF0.1 | P17763 |
| KPTYEKDVDLGAGTR | 1 | 0 | 0 | 1638859 | CCM44290.1 | P17763 |
| KPTYERDADLGAGTR | 1 | 0 | 0 | 1638860 | AFP27206.1 | P17763 |
| KPWDVIPMVTQIAMT | 1 | 0 | 0 | 1638861 | P17763.2 | P17763 |
| KPWDVVPMVTQMAMT | 1 | 0 | 0 | 1638862 | P14340.2 | P17763 |
| KQEHEISWHYDQDHP | 1 | 0 | 0 | 1638864 | AET43255.1 | P17763 |
| KQEHETSWHYDQDHP | 1 | 0 | 0 | 1638865 | P14340.2 | P17763 |
| KQEVVVLGSQEGAMH | 1 | 0 | 0 | 1638866 | P17763.2 | P17763 |
| KQISNELNHILLEND | 1 | 0 | 0 | 1638868 | P17763.2 | P17763 |
| KQITNELNYVLWEGG | 1 | 0 | 0 | 1638869 | Q2YHF0.1 | P17763 |
| KQITPELNHILAENE | 1 | 0 | 0 | 1638870 | QTX93142.1 | P17763 |
| KRALAPPVNDLKYSW | 1 | 0 | 0 | 1638882 | Q2YHF0.1 | P17763 |
| KRALTPPASDLKYSW | 1 | 0 | 0 | 1638884 | ALJ53458.1 | P17763 |
| KRALTPPVNDLKYSW | 1 | 0 | 0 | 1638885 | Q2YHF2.1 | P17763 |
| KRALTPPVSDLKYSW | 1 | 0 | 0 | 1638886 | AEW50182.1 | P17763 |
| KRARNRVSTGSQLAK | 1 | 0 | 0 | 1638887 | AAT76483.1 | P17763 |
| KRARNRVSTVSQLAK | 1 | 0 | 0 | 1638888 | P17763.2 | P17763 |
| KREKKIGEFGKAKGS | 1 | 0 | 0 | 1638893 | QEI46448.1 | P17763 |
| KREKKLGEFGKAKGS | 1 | 0 | 0 | 1638894 | P17763.2 | P17763 |
| KREKKLGEFGRAKGS | 1 | 0 | 0 | 1638895 | ACN42699.1 | P17763 |
| KRGLRTLILAPTRVV | 1 | 0 | 0 | 1638899 | P14340.2 | P17763 |
| KRIKEEHSSTWHYDD | 1 | 0 | 0 | 1638900 | QTX93500.1 | P17763 |
| KRILPSIVREALKRR | 1 | 0 | 0 | 1638901 | Q2YHF0.1 | P17763 |
| KRMAISGDDCVVKPL | 1 | 0 | 0 | 1638902 | AAW28113.1 | P17763 |
| KRRLRTLILAPTRVV | 1 | 0 | 0 | 1638905 | ACA48751.1 | P17763 |
| KRRLTIMDLHPGAGK | 1 | 0 | 0 | 1638906 | QTX93081.1 | P17763 |
| KRSGILEVDRTEAKS | 1 | 0 | 0 | 1638907 | Q2YHF0.1 | P17763 |
| KRSGILEVDRTEARS | 1 | 0 | 0 | 1638908 | QTX92258.1 | P17763 |
| KRSGIMEVDRSEAKE | 1 | 0 | 0 | 1638909 | QFS19562.1 | P17763 |
| KRSGIQEVDRTLAKE | 1 | 0 | 0 | 1638910 | QFS19360.1 | P17763 |
| KRTLTPQPMELKYSW | 1 | 0 | 0 | 1638913 | QTX93479.1 | P17763 |
| KRVIQLSRKTFDTEY | 1 | 0 | 0 | 1638914 | QFS19562.1 | P17763 |
| KRVRNRVSTGSQLAK | 1 | 0 | 0 | 1638916 | QTX93345.1 | P17763 |
| KSEHTGKEIVDLMCH | 1 | 0 | 0 | 1638920 | P17763.2 | P17763 |
| KSGKKVIQLSRKTFD | 1 | 0 | 0 | 1638921 | Q2YHF0.1 | P17763 |
| KSGKRVIQLSRKTFD | 1 | 0 | 0 | 1638922 | AQV12350.1 | P17763 |
| KSIFKLTYQNKVVRV | 1 | 0 | 0 | 1638923 | QTX92776.1 | P17763 |
| KSLAGPISQHNHRPG | 1 | 0 | 0 | 1638924 | QTX93500.1 | P17763 |
| KSLAGPISQHNYRPG | 1 | 0 | 0 | 1638925 | QHR82521.1 | P17763 |
| KSLLFKTEDGVNMCT | 1 | 0 | 0 | 1638927 | P14340.2 | P17763 |
| KSLLFKTENGVNMCT | 1 | 0 | 0 | 1638928 | QQW40784.1 | P17763 |
| KSPRLCTREEFISKV | 1 | 0 | 0 | 1638931 | QFS19537.1 | P17763 |

| | | | | | | |
|---|---|---|---|---|---|---|
| KSYAGPFSQHNYRQG | 1 | 0 | 0 | 1638935 | ARJ31953.1 | P17763 |
| KTAHAKKQEVVVLGS | 1 | 0 | 0 | 1638936 | P17763.2 | P17763 |
| KTCIWPKSHTLWSNG | 1 | 0 | 0 | 1638940 | P17763.2 | P17763 |
| KTCLWPKTHTLWSNG | 1 | 0 | 0 | 1638941 | Q2YHF0.1 | P17763 |
| KTCTWPKSHTLWSNG | 1 | 0 | 0 | 1638942 | QPZ88404.1 | P17763 |
| KTFDTEYQKTKNNDW | 1 | 0 | 0 | 1638948 | P17763.2 | P17763 |
| KTKLNDWDFVVTTDI | 1 | 0 | 0 | 1638956 | AHG23230.1 | P17763 |
| KTKNNDWDYVVTTDI | 1 | 0 | 0 | 1638957 | P17763.2 | P17763 |
| KTLTGEIGAVTLDFK | 1 | 0 | 0 | 1638965 | Q2YHF0.1 | P17763 |
| KTNTGTIGAVSLDFS | 1 | 0 | 0 | 1638966 | QTX92981.1 | P17763 |
| KTPRMCTREEFTKKV | 1 | 0 | 0 | 1638967 | AII99332.1 | P17763 |
| KTQVGVGIHMEGVFH | 1 | 0 | 0 | 1638970 | Q2YHF0.1 | P17763 |
| KTQVGVGVQKEGVFH | 1 | 0 | 0 | 1638971 | QTX93479.1 | P17763 |
| KTVWFVPSIKAGNDI | 1 | 0 | 0 | 1638974 | QTX92786.1 | P17763 |
| KVAAEGINYADRKWC | 1 | 0 | 0 | 1638975 | QRG47120.1 | P17763 |
| KVAAEGINYADRRWC | 1 | 0 | 0 | 1638976 | QTX93142.1 | P17763 |
| KVASAGISYKDREWC | 1 | 0 | 0 | 1638977 | Q2YHF0.1 | P17763 |
| KVDTRTPKAKRGTAQ | 1 | 0 | 0 | 1638978 | P17763.2 | P17763 |
| KVDTRTPRAKRGTTQ | 1 | 0 | 0 | 1638979 | QNH29581.1 | P17763 |
| KVDTRTPRPMPGTRK | 1 | 0 | 0 | 1638980 | QTX93500.1 | P17763 |
| KVEYKGEDVPCKIPF | 1 | 0 | 0 | 1638981 | AMH87199.1 | P17763 |
| KVIQLSRKTFDSEYI | 1 | 0 | 0 | 1638987 | QQW40786.1 | P17763 |
| KVIQLSRKTFDTEYP | 1 | 0 | 0 | 1638988 | Q2YHF0.1 | P17763 |
| KVLNPYMPSVIEKME | 1 | 0 | 0 | 1638993 | P14340.2 | P17763 |
| KVPHAKRQDVTVLGS | 1 | 0 | 0 | 1638995 | Q2YHF0.1 | P17763 |
| KVRKDIPQWEPSKGW | 1 | 0 | 0 | 1638998 | P17763.2 | P17763 |
| KVRTNAAMGAVFTEE | 1 | 0 | 0 | 1638999 | QTX93489.1 | P17763 |
| KVVGLYGNGVVTKNG | 1 | 0 | 0 | 1639000 | QTX93489.1 | P17763 |
| KYEGTDAPCKIPFST | 1 | 0 | 0 | 1639005 | QFS19562.1 | P17763 |
| KYEGTGAPCKVPIEI | 1 | 0 | 0 | 1639006 | Q2YHF0.1 | P17763 |
| KYTDRKWCFDGERNN | 1 | 0 | 0 | 1639009 | QTX93504.1 | P17763 |
| LAAIANQAVVLMGLD | 1 | 0 | 0 | 1639013 | QTX93489.1 | P17763 |
| LAANAICSAVPVDWV | 1 | 0 | 0 | 1639015 | P17763.2 | P17763 |
| LAENTNSVTNIELEP | 1 | 0 | 0 | 1639018 | ARJ31952.1 | P17763 |
| LAGFMAYMIGQTGIQ | 1 | 0 | 0 | 1639021 | Q2YHF0.1 | P17763 |
| LAHAIGTSITQKGII | 1 | 0 | 0 | 1639024 | P17763.2 | P17763 |
| LAHRTRNALDNLVML | 1 | 0 | 0 | 1639025 | QTX93474.1 | P17763 |
| LAHYIGTSLTQKVVI | 1 | 0 | 0 | 1639026 | QTX93479.1 | P17763 |
| LAIPPTAGILARWGS | 1 | 0 | 0 | 1639028 | P17763.2 | P17763 |
| LAIPPTAGILARWGT | 1 | 0 | 0 | 1639029 | QTX93247.1 | P17763 |
| LAIPPTAGILARWSS | 1 | 0 | 0 | 1639030 | AXX75195.1 | P17763 |
| LAIPPTAGVLARWGT | 1 | 0 | 0 | 1639031 | AYA22932.1 | P17763 |
| LAKRFSKGLLNGQGP | 1 | 0 | 0 | 1639035 | AHG23230.1 | P17763 |
| LAKRFSRGLLNGQGP | 1 | 0 | 0 | 1639037 | QTX93504.1 | P17763 |
| LALFLAHYIGTSLTQ | 1 | 0 | 0 | 1639040 | QTX93479.1 | P17763 |
| LAMTDTTPFGQQRVF | 1 | 0 | 0 | 1639047 | Q2YHF0.1 | P17763 |
| LASSLLKNDIPMTGP | 1 | 0 | 0 | 1639053 | P14340.2 | P17763 |
| LASSLLRNDVPMAGP | 1 | 0 | 0 | 1639054 | QTX93489.1 | P17763 |
| LATGPILTLWEGNPG | 1 | 0 | 0 | 1639055 | Q2YHF0.1 | P17763 |
| LATGPIMTLWEGNPG | 1 | 0 | 0 | 1639056 | ANW72071.1 | P17763 |
| LATGPLTTLWEGSPG | 1 | 0 | 0 | 1639057 | P17763.2 | P17763 |
| LATLRKLCIEGKITN | 1 | 0 | 0 | 1639058 | QTX93489.1 | P17763 |
| LATVTGGIFLFLMSG | 1 | 0 | 0 | 1639060 | P14340.2 | P17763 |
| LAVFFLLIMGQLTWN | 1 | 0 | 0 | 1639061 | QTX92695.1 | P17763 |
| LAVFLLLTMGQLTWN | 1 | 0 | 0 | 1639062 | P17763.2 | P17763 |
| LAVTIMAISCVPNAV | 1 | 0 | 0 | 1639065 | QRG47112.1 | P17763 |
| LAYTIGTTYFQRVLI | 1 | 0 | 0 | 1639069 | QQW40777.1 | P17763 |
| LCISIMIEEVMRSRW | 1 | 0 | 0 | 1639072 | P17763.2 | P17763 |
| LCLAILFEEVMRGKF | 1 | 0 | 0 | 1639073 | QTX93474.1 | P17763 |
| LCVGQLLLMRTTWAL | 1 | 0 | 0 | 1639075 | AQV12653.1 | P17763 |
| LDARTYSDPLALKEF | 1 | 0 | 0 | 1639078 | QTX93461.1 | P17763 |
| LDARTYSDPLALREF | 1 | 0 | 0 | 1639079 | P17763.2 | P17763 |
| LDFELIKTEAKHPAT | 1 | 0 | 0 | 1639080 | QQW40763.1 | P17763 |
| LDFELIKTTAKEVAL | 1 | 0 | 0 | 1639081 | Q2YHF0.1 | P17763 |
| LDFELTKTTAKEVAL | 1 | 0 | 0 | 1639082 | Q5UCB8.1 | P17763 |
| LDFNEMILLTMKNKA | 1 | 0 | 0 | 1639083 | QTX93489.1 | P17763 |
| LDIIGQRIENIKHEH | 1 | 0 | 0 | 1639084 | QTX92739.1 | P17763 |
| LDIIGQRIENIKNEH | 1 | 0 | 0 | 1639085 | P27909.2 | P17763 |
| LDNIVMLHTTEKGGR | 1 | 0 | 0 | 1639093 | Q5UCB8.1 | P17763 |
| LDNIVMLHTTERGGK | 1 | 0 | 0 | 1639094 | Q2YHF0.1 | P17763 |
| LDNIVMLHTTERGGR | 1 | 0 | 0 | 1639095 | AUN35144.1 | P17763 |
| LDNLVMLHNSEQGGK | 1 | 0 | 0 | 1639096 | P17763.2 | P17763 |
| LDNLVMLHNSEQGGR | 1 | 0 | 0 | 1639097 | QFS19562.1 | P17763 |
| LDNLVMLHTSEHGGR | 1 | 0 | 0 | 1639098 | QTX93489.1 | P17763 |
| LDVDLHPASAWTLYA | 1 | 0 | 0 | 1639105 | P17763.2 | P17763 |
| LDVDLRPASAWTLYA | 1 | 0 | 0 | 1639106 | QPZ88407.1 | P17763 |
| LDYAWKTMAMVLSIV | 1 | 0 | 0 | 1639107 | QHF19278.1 | P17763 |
| LDYAWKTTAMALSIV | 1 | 0 | 0 | 1639108 | AHI43747.1 | P17763 |
| LEENMEVEIWTKEGE | 1 | 0 | 0 | 1639113 | QNH29582.1 | P17763 |
| LEENVEVEIWTKEGE | 1 | 0 | 0 | 1639115 | P14340.2 | P17763 |
| LEGEWKEGEEVQVLA | 1 | 0 | 0 | 1639118 | P14340.2 | P17763 |
| LEGKIVQYENLKYSV | 1 | 0 | 0 | 1639119 | P17763.2 | P17763 |
| LEHGGCVTTMAKNKP | 1 | 0 | 0 | 1639123 | QTX93489.1 | P17763 |
| LEHGGCVTTMAQGKP | 1 | 0 | 0 | 1639125 | Q2YHF0.1 | P17763 |
| LEHGSCVTTMAKDKP | 1 | 0 | 0 | 1639126 | P17763.2 | P17763 |
| LEIDFGECPGTTVTI | 1 | 0 | 0 | 1639127 | Q2YHF0.1 | P17763 |
| LEIDFGECPGTTVTV | 1 | 0 | 0 | 1639128 | QTX92349.1 | P17763 |
| LEKAAEVSWEEEAEH | 1 | 0 | 0 | 1639130 | P17763.2 | P17763 |
| LEKAAEVSWEQEAEH | 1 | 0 | 0 | 1639131 | QNH29581.1 | P17763 |
| LELDFNYCEGTTVVI | 1 | 0 | 0 | 1639134 | QTX93489.1 | P17763 |
| LELTDALALGMMILK | 1 | 0 | 0 | 1639135 | QQW40684.1 | P17763 |

| | | | | | | |
|---|---|---|---|---|---|---|
| LELTDALALGMMVLK | 1 | 0 | 0 | 1639136 | P14340.2 | P17763 |
| LEMDFDFCDGTTVIV | 1 | 0 | 0 | 1639137 | QTX93067.1 | P17763 |
| LENPHLLEKKITQWL | 1 | 0 | 0 | 1639138 | QTX93539.1 | P17763 |
| LEPNWASVKKDLISY | 1 | 0 | 0 | 1639140 | QPZ88403.1 | P17763 |
| LEPPFGDSYIVIGVG | 1 | 0 | 0 | 1639141 | Q2YHF0.1 | P17763 |
| LEPSWADVRNDMISY | 1 | 0 | 0 | 1639142 | Q2YHF0.1 | P17763 |
| LEPSWASVKKDLISY | 1 | 0 | 0 | 1639143 | P17763.2 | P17763 |
| LEQMQRKHGGMLVRN | 1 | 0 | 0 | 1639145 | P17763.2 | P17763 |
| LERAADVRWEDQAEI | 1 | 0 | 0 | 1639147 | QQW40684.1 | P17763 |
| LERATDVKWEDQAEI | 1 | 0 | 0 | 1639148 | QTX93081.1 | P17763 |
| LERLQRKHGGMLVRN | 1 | 0 | 0 | 1639149 | QQD79919.1 | P17763 |
| LESEMIIPKIYGGPI | 1 | 0 | 0 | 1639150 | P17763.2 | P17763 |
| LESEMIIPKNLAGPV | 1 | 0 | 0 | 1639151 | QTX93151.1 | P17763 |
| LESQMLIPKAYAGPI | 1 | 0 | 0 | 1639152 | Q2YHF0.1 | P17763 |
| LESQMLIPKSYAGPF | 1 | 0 | 0 | 1639153 | ARJ31953.1 | P17763 |
| LESQMLIPRSYAGPF | 1 | 0 | 0 | 1639154 | QBA29684.1 | P17763 |
| LETPNLAERVLDWLE | 1 | 0 | 0 | 1639157 | QFS19562.1 | P17763 |
| LEVEDYGFGMFTTNI | 1 | 0 | 0 | 1639159 | ACN54390.1 | P17763 |
| LFEPEREKSAAIDGE | 1 | 0 | 0 | 1639167 | P17763.2 | P17763 |
| LFEPEREKSAAVDGE | 1 | 0 | 0 | 1639168 | ACW82902.1 | P17763 |
| LFPISIPITAAAWYL | 1 | 0 | 0 | 1639178 | QTX93036.1 | P17763 |
| LFPLCLSTTSQKTTW | 1 | 0 | 0 | 1639179 | P17763.2 | P17763 |
| LFSGVSWIMKIGIGV | 1 | 0 | 0 | 1639182 | QPC96841.1 | P17763 |
| LFVEECLRRRVTRKH | 1 | 0 | 0 | 1639183 | Q2YHF0.1 | P17763 |
| LGAGTRHVAVEPEVA | 1 | 0 | 0 | 1639184 | P17763.2 | P17763 |
| LGAGTRHVNAEPETP | 1 | 0 | 0 | 1639185 | QTX93489.1 | P17763 |
| LGAIFTDENKWKSAR | 1 | 0 | 0 | 1639186 | P14340.2 | P17763 |
| LGAMTAGIFLFFMQG | 1 | 0 | 0 | 1639187 | Q2YHF0.1 | P17763 |
| LGDKWDKEEDVQVLA | 1 | 0 | 0 | 1639188 | Q2YHF0.1 | P17763 |
| LGDTAWDFGSVGGVL | 1 | 0 | 0 | 1639189 | QTX93489.1 | P17763 |
| LGEKWKKKLNQLSRK | 1 | 0 | 0 | 1639191 | QTX93479.1 | P17763 |
| LGEKWKNRLNALGKS | 1 | 0 | 0 | 1639192 | Q9WDA6.1 | P17763 |
| LGEKWKNRLNSLGKS | 1 | 0 | 0 | 1639193 | QCZ24978.1 | P17763 |
| LGEKWKRQLNQLSKS | 1 | 0 | 0 | 1639194 | P17763.2 | P17763 |
| LGEKWKRQLNSLDRK | 1 | 0 | 0 | 1639195 | Q2YHF0.1 | P17763 |
| LGEKWKSRLNALGKS | 1 | 0 | 0 | 1639196 | P14340.2 | P17763 |
| LGELCEDTITYKCPL | 1 | 0 | 0 | 1639197 | QTX93024.1 | P17763 |
| LGELCEDTITYNCPL | 1 | 0 | 0 | 1639198 | QQW40770.1 | P17763 |
| LGELCEDTMTYKCPR | 1 | 0 | 0 | 1639199 | P17763.2 | P17763 |
| LGEMCDDTVTYKCPH | 1 | 0 | 0 | 1639200 | QDG00730.1 | P17763 |
| LGEMCDDTVTYKCPL | 1 | 0 | 0 | 1639201 | QTX93530.1 | P17763 |
| LGEMCEDTVTYKCPL | 1 | 0 | 0 | 1639202 | Q2YHF0.1 | P17763 |
| LGFLNEDHWFGRENS | 1 | 0 | 0 | 1639203 | Q2YHF0.1 | P17763 |
| LGFLNEDHWFSRENS | 1 | 0 | 0 | 1639204 | ABQ18243.1 | P17763 |
| LGFLNEDHWFSRGNS | 1 | 0 | 0 | 1639205 | QRG47115.1 | P17763 |
| LGGVFTSIGKALHQV | 1 | 0 | 0 | 1639206 | P14340.2 | P17763 |
| LGIGHVAAENHQHAT | 1 | 0 | 0 | 1639207 | AVY51407.1 | P17763 |
| LGIGHVAVENHHHAA | 1 | 0 | 0 | 1639208 | QTX92739.1 | P17763 |
| LGKKKNPRLCTREEF | 1 | 0 | 0 | 1639210 | Q58HT7.1 | P17763 |
| LGKKKSPRLCTREEF | 1 | 0 | 0 | 1639211 | ARM59249.1 | P17763 |
| LGKKKTPRMCTREEF | 1 | 0 | 0 | 1639212 | P14340.2 | P17763 |
| LGKSYAQMWQLMYFH | 1 | 0 | 0 | 1639213 | P17763.2 | P17763 |
| LGLETRTETWMSSEG | 1 | 0 | 0 | 1639218 | P17763.2 | P17763 |
| LGLGSIATQQPESNI | 1 | 0 | 0 | 1639219 | P29990.1 | P17763 |
| LGLGSITTQESESNI | 1 | 0 | 0 | 1639220 | AWI47685.1 | P17763 |
| LGMSKEPGVVSPTSY | 1 | 0 | 0 | 1639223 | QTX93479.1 | P17763 |
| LGQVMLLVLCAGQLL | 1 | 0 | 0 | 1639229 | Q2YHF0.1 | P17763 |
| LGRKKNPRLCTREEF | 1 | 0 | 0 | 1639232 | Q2YHF0.1 | P17763 |
| LGRVMVMVGAAMTDD | 1 | 0 | 0 | 1639233 | QQW40724.1 | P17763 |
| LGRVMVMVGATMTDD | 1 | 0 | 0 | 1639234 | P14340.2 | P17763 |
| LGRVVVMVGATMADD | 1 | 0 | 0 | 1639235 | QTX93044.1 | P17763 |
| LGSQEGAMHSALAGA | 1 | 0 | 0 | 1639236 | Q2YHF0.1 | P17763 |
| LGVPLLALGCYSQVN | 1 | 0 | 0 | 1639239 | AAN03432.3 | P17763 |
| LHLEGKCETCVYNMM | 1 | 0 | 0 | 1639241 | P14340.2 | P17763 |
| LHPASAWTLYAVATT | 1 | 0 | 0 | 1639242 | P17763.2 | P17763 |
| LHPGAGKTKRILPSI | 1 | 0 | 0 | 1639244 | Q2YHF0.1 | P17763 |
| LHPGAGKTKRYLPAI | 1 | 0 | 0 | 1639245 | P14340.2 | P17763 |
| LHQEGKCESCVYNMM | 1 | 0 | 0 | 1639246 | Q2YHF0.1 | P17763 |
| LHRMDLGVPLLAMGC | 1 | 0 | 0 | 1639247 | Q2YHF0.1 | P17763 |
| LHYAWKTMAMILSIV | 1 | 0 | 0 | 1639248 | P17763.2 | P17763 |
| LHYAWKTMAMVLSIV | 1 | 0 | 0 | 1639249 | QTX92698.1 | P17763 |
| LIAIAVASGLLWVAE | 1 | 0 | 0 | 1639251 | Q2YHF0.1 | P17763 |
| LICVIASSGMLWMAD | 1 | 0 | 0 | 1639252 | P27915.1 | P17763 |
| LICVIASSGMLWMAE | 1 | 0 | 0 | 1639253 | QHJ86727.1 | P17763 |
| LIEKTKTDFGFYQVK | 1 | 0 | 0 | 1639254 | Q5UCB8.1 | P17763 |
| LIEVKTCLWPKTHTL | 1 | 0 | 0 | 1639255 | Q2YHF0.1 | P17763 |
| LIEVKTCTWPKSHTL | 1 | 0 | 0 | 1639256 | QIS48873.1 | P17763 |
| LIGLTARATWATNIQ | 1 | 0 | 0 | 1639261 | P17763.2 | P17763 |
| LIGNEEYTDYMPSMK | 1 | 0 | 0 | 1639262 | AAG15003.1 | P17763 |
| LIGSEEYTDYMPSMK | 1 | 0 | 0 | 1639264 | QGW51895.1 | P17763 |
| LIKTEAKHPATLRKY | 1 | 0 | 0 | 1639267 | QQW40763.1 | P17763 |
| LIKTEAKQPATLRKY | 1 | 0 | 0 | 1639268 | QTX92984.1 | P17763 |
| LIKTTAKEVALLRTY | 1 | 0 | 0 | 1639269 | Q2YHF0.1 | P17763 |
| LILGAQALPVYLMTL | 1 | 0 | 0 | 1639272 | Q2YHF0.1 | P17763 |
| LIMLGDTMSGRIGGQ | 1 | 0 | 0 | 1639273 | Q5UCB8.1 | P17763 |
| LIMLGDTMSGRMGGQ | 1 | 0 | 0 | 1639274 | Q2YHF0.1 | P17763 |
| LIRLCIMVGANASDK | 1 | 0 | 0 | 1639275 | P17763.2 | P17763 |
| LIRLCIMVGANASDR | 1 | 0 | 0 | 1639276 | QFS19562.1 | P17763 |
| LIRLCIMVGANVSDR | 1 | 0 | 0 | 1639277 | QIS48854.1 | P17763 |
| LITANPVVTKKEEPV | 1 | 0 | 0 | 1639282 | QTX93489.1 | P17763 |
| LITCAKFKCVTKLEG | 1 | 0 | 0 | 1639283 | P17763.2 | P17763 |

| | | | | | | |
|---|---|---|---|---|---|---|
| LITIAVASGLLWVAE | 1 | 0 | 0 | 1639284 | ARJ31953.1 | P17763 |
| LIYADDTAGWDTRIT | 1 | 0 | 0 | 1639290 | Q2YHF0.1 | P17763 |
| LKCGSGIFVIDNVHT | 1 | 0 | 0 | 1639294 | Q2YHF0.1 | P17763 |
| LKCGSGIFVTNEVHT | 1 | 0 | 0 | 1639295 | P17763.2 | P17763 |
| LKCGSGIFVVDNVHT | 1 | 0 | 0 | 1639296 | AXB26598.1 | P17763 |
| LKDGSKIKHAVSRGS | 1 | 0 | 0 | 1639298 | Q58HT7.1 | P17763 |
| LKDGSKIKYAVSRGT | 1 | 0 | 0 | 1639299 | Q2YHF0.1 | P17763 |
| LKDGSKTKYAVSRGT | 1 | 0 | 0 | 1639300 | QTX92247.1 | P17763 |
| LKIVTHFDNTQVGTL | 1 | 0 | 0 | 1639304 | QBQ58394.1 | P17763 |
| LKIVTQFDNTQVGTL | 1 | 0 | 0 | 1639305 | ARJ31952.1 | P17763 |
| LKLKEKQDAFCDSKL | 1 | 0 | 0 | 1639310 | QTX93117.1 | P17763 |
| LKLKERQDVFCDSKL | 1 | 0 | 0 | 1639311 | QQW40722.1 | P17763 |
| LKLRDSYTQMCDHRL | 1 | 0 | 0 | 1639312 | P27909.2 | P17763 |
| LKLRDSYTQVCDHRL | 1 | 0 | 0 | 1639313 | P17763.2 | P17763 |
| LKLREMYTQLCDHRL | 1 | 0 | 0 | 1639314 | QHR82521.1 | P17763 |
| LKMDKLELKGMSYAM | 1 | 0 | 0 | 1639316 | AAC63321.1 | P17763 |
| LKMVEPWLKNNQFCI | 1 | 0 | 0 | 1639317 | QTX93489.1 | P17763 |
| LKMVEPWLRGNQFCI | 1 | 0 | 0 | 1639318 | P17763.2 | P17763 |
| LKMVEPWLSSKPEFC | 1 | 0 | 0 | 1639319 | Q2YHF0.1 | P17763 |
| LKPVILTDGPERVIL | 1 | 0 | 0 | 1639321 | QTX93489.1 | P17763 |
| LKRGETTHHAVSRGS | 1 | 0 | 0 | 1639322 | QTX93534.1 | P17763 |
| LKRGETTKHAVSRGT | 1 | 0 | 0 | 1639323 | QFS19562.1 | P17763 |
| LLAIGCYSQVNPITL | 1 | 0 | 0 | 1639329 | P14340.2 | P17763 |
| LLALGCYSQVNPLTL | 1 | 0 | 0 | 1639330 | P17763.2 | P17763 |
| LLALIAVLTGGVTLF | 1 | 0 | 0 | 1639331 | P17763.2 | P17763 |
| LLCVTASSALLWMAS | 1 | 0 | 0 | 1639337 | P27909.2 | P17763 |
| LLETTKKDLGIGHVA | 1 | 0 | 0 | 1639339 | QFS19562.1 | P17763 |
| LLETTKRDLGMSKEP | 1 | 0 | 0 | 1639340 | QTX93489.1 | P17763 |
| LLGLMILLTGGAMLF | 1 | 0 | 0 | 1639344 | QTX93489.1 | P17763 |
| LLIACYVITGTSADL | 1 | 0 | 0 | 1639345 | QTX93489.1 | P17763 |
| LLITHYAIIGPGLQA | 1 | 0 | 0 | 1639347 | QBQ58390.1 | P17763 |
| LLKNDIPMTGPLVAG | 1 | 0 | 0 | 1639348 | P14340.2 | P17763 |
| LLKNDVPLAGPLIAG | 1 | 0 | 0 | 1639349 | P17763.2 | P17763 |
| LLLTMGQLTWNDLIR | 1 | 0 | 0 | 1639350 | P17763.2 | P17763 |
| LLNGQGPMKLVMAFI | 1 | 0 | 0 | 1639352 | QBM00871.1 | P17763 |
| LLPVCQSSSMRKTDW | 1 | 0 | 0 | 1639354 | AHG23230.1 | P17763 |
| LLRALIMLGDTMSGR | 1 | 0 | 0 | 1639358 | Q2YHF0.1 | P17763 |
| LLSGQGPMKLVMAFI | 1 | 0 | 0 | 1639360 | P17763.2 | P17763 |
| LLSLTFIKTTLSLDY | 1 | 0 | 0 | 1639361 | P27909.2 | P17763 |
| LLSLTFVKTTFSLHY | 1 | 0 | 0 | 1639362 | P17763.2 | P17763 |
| LLSSLLKNDVPLAGP | 1 | 0 | 0 | 1639363 | P17763.2 | P17763 |
| LLSSTRVPNYNLIVM | 1 | 0 | 0 | 1639365 | Q2YHF0.1 | P17763 |
| LLTCAKFKCVTKLEG | 1 | 0 | 0 | 1639366 | P27909.2 | P17763 |
| LLTLLATVTGGIFLF | 1 | 0 | 0 | 1639367 | P14340.2 | P17763 |
| LLTSSQQKADWIPLA | 1 | 0 | 0 | 1639368 | Q9WDA6.1 | P17763 |
| LLTSSQQKTDWIPLA | 1 | 0 | 0 | 1639369 | QTX93135.1 | P17763 |
| LLVAHYAIIGPGLQA | 1 | 0 | 0 | 1639371 | P14340.2 | P17763 |
| LLVLWIGTNSRNTSM | 1 | 0 | 0 | 1639372 | Q2YHF0.1 | P17763 |
| LMATFKMRPMFAVGL | 1 | 0 | 0 | 1639374 | QFS19562.1 | P17763 |
| LMATFRMRPMFAVGL | 1 | 0 | 0 | 1639375 | P17763.2 | P17763 |
| LMKITAEWLWKELGK | 1 | 0 | 0 | 1639379 | P14340.2 | P17763 |
| LMMATIGIALLSQST | 1 | 0 | 0 | 1639380 | QRG47115.1 | P17763 |
| LMMTTIGIVLLSQST | 1 | 0 | 0 | 1639381 | P14340.2 | P17763 |
| LMRRGDLPVWLAHKV | 1 | 0 | 0 | 1639382 | QTX93491.1 | P17763 |
| LMRRGDLPVWLSYKV | 1 | 0 | 0 | 1639383 | P17763.2 | P17763 |
| LMYADDTAGWDTRIT | 1 | 0 | 0 | 1639385 | Q5UCB8.1 | P17763 |
| LMYFHRRDLRLASMA | 1 | 0 | 0 | 1639387 | Q2YHF0.1 | P17763 |
| LNAFVLKKEVSETQH | 1 | 0 | 0 | 1639388 | AHG06373.1 | P17763 |
| LNALGKSEFQIYKKS | 1 | 0 | 0 | 1639389 | P14340.2 | P17763 |
| LNDMGKIRKDIQQWE | 1 | 0 | 0 | 1639391 | QTX93117.1 | P17763 |
| LNDMGKVRKDIPQWQ | 1 | 0 | 0 | 1639392 | QTX93489.1 | P17763 |
| LNDMGKVRKDIQQWE | 1 | 0 | 0 | 1639393 | P14340.2 | P17763 |
| LNEKEENMVKSQVSA | 1 | 0 | 0 | 1639395 | Q2YHF0.1 | P17763 |
| LNELPESLETLMLVA | 1 | 0 | 0 | 1639396 | Q2YHF0.1 | P17763 |
| LNLVENWLNNNTQFC | 1 | 0 | 0 | 1639399 | P14340.2 | P17763 |
| LNPTAIFLTTLSRTN | 1 | 0 | 0 | 1639400 | P14340.2 | P17763 |
| LNQLSKSEFNTYKRS | 1 | 0 | 0 | 1639401 | P17763.2 | P17763 |
| LNQLSRKEFDLYKKS | 1 | 0 | 0 | 1639402 | QTX93474.1 | P17763 |
| LNSLDRKEFEEYKRS | 1 | 0 | 0 | 1639403 | Q2YHF0.1 | P17763 |
| LNTFTNMGAQLIRQM | 1 | 0 | 0 | 1639405 | QQW40674.1 | P17763 |
| LNTFVLKKEVSETQH | 1 | 0 | 0 | 1639406 | QTX93530.1 | P17763 |
| LNWFKKGSSIGQMFE | 1 | 0 | 0 | 1639408 | QTX93024.1 | P17763 |
| LPEYGTLGLECSPRT | 1 | 0 | 0 | 1639418 | QTX93489.1 | P17763 |
| LPLFIFSLKDTLKRR | 1 | 0 | 0 | 1639421 | AHG23230.1 | P17763 |
| LPQHLTLRAQNALDN | 1 | 0 | 0 | 1639430 | P27909.2 | P17763 |
| LPQHLTQRAQNALDN | 1 | 0 | 0 | 1639431 | P17763.2 | P17763 |
| LPSELETPNLAERVL | 1 | 0 | 0 | 1639433 | QMT58612.1 | P17763 |
| LPTFMTQKTRDALDN | 1 | 0 | 0 | 1639436 | QTX93142.1 | P17763 |
| LPTYLSSKAKLALDN | 1 | 0 | 0 | 1639438 | ARM59249.1 | P17763 |
| LPVYLMTLMKGASKR | 1 | 0 | 0 | 1639440 | Q2YHF0.1 | P17763 |
| LQAKATREAQKRAAA | 1 | 0 | 0 | 1639443 | P14340.2 | P17763 |
| LQAKATREAQKRTAA | 1 | 0 | 0 | 1639444 | P17763.2 | P17763 |
| LQGSWNTGEEVQVIA | 1 | 0 | 0 | 1639447 | QFS19562.1 | P17763 |
| LQKTEATQLATLRKL | 1 | 0 | 0 | 1639450 | QTX93474.1 | P17763 |
| LQNAWKVSCTILAAV | 1 | 0 | 0 | 1639451 | QRG47133.1 | P17763 |
| LQNAWKVSCTILAVV | 1 | 0 | 0 | 1639452 | P14340.2 | P17763 |
| LQNAWKVSCTTLAVV | 1 | 0 | 0 | 1639453 | Q9WDA6.1 | P17763 |
| LQRKHGGNLVRCPLS | 1 | 0 | 0 | 1639455 | ARX96505.1 | P17763 |
| LQRKHGGSLVRCPLS | 1 | 0 | 0 | 1639456 | Q2YHF0.1 | P17763 |
| LQRLQEEHKETWHYD | 1 | 0 | 0 | 1639458 | Q2YHF0.1 | P17763 |
| LQRLREEHKETWHYD | 1 | 0 | 0 | 1639459 | AHG06383.1 | P17763 |

| | | | | | | |
|---|---|---|---|---|---|---|
| LQRQYGGALVRNPLS | 1 | 0 | 0 | 1639460 | ACW82894.1 | P17763 |
| LQRRHGGSLVRCPLS | 1 | 0 | 0 | 1639461 | QTX92334.1 | P17763 |
| LQRRYGGSLVRCPLS | 1 | 0 | 0 | 1639462 | QTX92160.1 | P17763 |
| LRDGSKIKHAVSRGS | 1 | 0 | 0 | 1639469 | AFY10039.1 | P17763 |
| LRETACLGKAYAQMW | 1 | 0 | 0 | 1639471 | Q6YMS3.1 | P17763 |
| LRFLGEDGCWYGMEI | 1 | 0 | 0 | 1639472 | AFZ40118.1 | P17763 |
| LRFRGEDGCWYGMEI | 1 | 0 | 0 | 1639473 | QTX92718.1 | P17763 |
| LRGEARKTFVDLMRR | 1 | 0 | 0 | 1639474 | P14340.2 | P17763 |
| LRGEQRKTFVELMRR | 1 | 0 | 0 | 1639475 | Q2YHF0.1 | P17763 |
| LRGSWNTGEEVQVIA | 1 | 0 | 0 | 1639476 | ACQ44411.1 | P17763 |
| LRHPGFTILALFLAH | 1 | 0 | 0 | 1639477 | QTX93479.1 | P17763 |
| LRHTIENTSANLSLA | 1 | 0 | 0 | 1639478 | Q2YHF0.1 | P17763 |
| LRKLTSKELMMTTIG | 1 | 0 | 0 | 1639481 | P14340.2 | P17763 |
| LRLASMAICSAVPTE | 1 | 0 | 0 | 1639487 | Q2YHF0.1 | P17763 |
| LRNPGFALLAGFMAY | 1 | 0 | 0 | 1639489 | Q2YHF0.1 | P17763 |
| LRPASAWTLYAVATT | 1 | 0 | 0 | 1639490 | QTX93045.1 | P17763 |
| LRPTPKGAVMDIISR | 1 | 0 | 0 | 1639491 | Q2YHF0.1 | P17763 |
| LRPTPRGAVMDIISR | 1 | 0 | 0 | 1639492 | Q5UCB8.1 | P17763 |
| LRTLVLAPTRVVASE | 1 | 0 | 0 | 1639494 | QMT58604.1 | P17763 |
| LRVLSIPPTAGILKR | 1 | 0 | 0 | 1639495 | Q2YHF0.1 | P17763 |
| LSAQWQKGEEVQVIA | 1 | 0 | 0 | 1639497 | QTX93491.1 | P17763 |
| LSEKEENMVKSQVTA | 1 | 0 | 0 | 1639499 | Q5UCB8.1 | P17763 |
| LSELPETLETLLLLT | 1 | 0 | 0 | 1639500 | P14340.2 | P17763 |
| LSGKGLGKTSIGLLC | 1 | 0 | 0 | 1639503 | P27909.2 | P17763 |
| LSGRGLGKTSIGLLC | 1 | 0 | 0 | 1639504 | P17763.2 | P17763 |
| LSIPHDLMEFIDGIS | 1 | 0 | 0 | 1639506 | Q2YHF0.1 | P17763 |
| LSIPHDLMELIDGIS | 1 | 0 | 0 | 1639507 | Q5UCB8.1 | P17763 |
| LSKGKRALAPPVNDL | 1 | 0 | 0 | 1639508 | Q2YHF0.1 | P17763 |
| LSLTFIKSTMPLVMA | 1 | 0 | 0 | 1639509 | Q2YHF0.1 | P17763 |
| LSMGLIAIAVASGLL | 1 | 0 | 0 | 1639511 | Q2YHF0.1 | P17763 |
| LSMGLITIAVASGLL | 1 | 0 | 0 | 1639512 | ARJ31953.1 | P17763 |
| LSRKTFDSEYIKTRT | 1 | 0 | 0 | 1639515 | QQW40786.1 | P17763 |
| LSRKTFDSEYVKTRA | 1 | 0 | 0 | 1639516 | QRG47112.1 | P17763 |
| LSRKTFDSEYVKTRT | 1 | 0 | 0 | 1639517 | P14340.2 | P17763 |
| LSRKTFDTEYPKTKL | 1 | 0 | 0 | 1639518 | Q2YHF0.1 | P17763 |
| LSRNSTHEMYWISNG | 1 | 0 | 0 | 1639519 | QTX93489.1 | P17763 |
| LSRNSTHEMYWVSCG | 1 | 0 | 0 | 1639520 | P17763.2 | P17763 |
| LSSKAKLALDNIVML | 1 | 0 | 0 | 1639522 | AQV12653.1 | P17763 |
| LSSRAKLALDNIVML | 1 | 0 | 0 | 1639523 | Q2YHF0.1 | P17763 |
| LSVWNRVWIEENPWM | 1 | 0 | 0 | 1639524 | P17763.2 | P17763 |
| LSWFKKGSSIGKMFE | 1 | 0 | 0 | 1639525 | P17763.2 | P17763 |
| LSWFKKGSSIGQMFE | 1 | 0 | 0 | 1639526 | QQW40781.1 | P17763 |
| LTAIANQATVLMGLG | 1 | 0 | 0 | 1639530 | P14340.2 | P17763 |
| LTALNDMGKVRKDIP | 1 | 0 | 0 | 1639531 | P17763.2 | P17763 |
| LTDFQSHQLWATLLS | 1 | 0 | 0 | 1639532 | P17763.2 | P17763 |
| LTDFQSHQLWTTLLS | 1 | 0 | 0 | 1639533 | QIC53498.1 | P17763 |
| LTEFQPHQLWTTLLS | 1 | 0 | 0 | 1639535 | P27909.2 | P17763 |
| LTIMDLHPGSGKTRR | 1 | 0 | 0 | 1639540 | P17763.2 | P17763 |
| LTIPPTAGILKRWGT | 1 | 0 | 0 | 1639541 | P14340.2 | P17763 |
| LTKGKRALTPPASDL | 1 | 0 | 0 | 1639542 | ALJ53458.1 | P17763 |
| LTKGKRALTPPVNDL | 1 | 0 | 0 | 1639543 | Q5UCB8.1 | P17763 |
| LTKGKRALTPPVSDL | 1 | 0 | 0 | 1639544 | AEW50182.1 | P17763 |
| LTKRFSLGMLQGRGP | 1 | 0 | 0 | 1639546 | AAT76490.1 | P17763 |
| LTKTTAKEVALLRTY | 1 | 0 | 0 | 1639547 | Q5UCB8.1 | P17763 |
| LTLDCEPRSGIDFNE | 1 | 0 | 0 | 1639549 | Q2YHF0.1 | P17763 |
| LTLDCSPRTGLDFNE | 1 | 0 | 0 | 1639550 | P17763.2 | P17763 |
| LTLRAQNALDNLVML | 1 | 0 | 0 | 1639555 | P27909.2 | P17763 |
| LTLTAAVLLLITHYA | 1 | 0 | 0 | 1639556 | QBQ58390.1 | P17763 |
| LTMFLIAENKIWGRK | 1 | 0 | 0 | 1639559 | QFS19562.1 | P17763 |
| LTMFLITENKIWGRK | 1 | 0 | 0 | 1639560 | P17763.2 | P17763 |
| LTNTTTASRCPTQGE | 1 | 0 | 0 | 1639561 | QQW40784.1 | P17763 |
| LTNTTTESRCPTQGE | 1 | 0 | 0 | 1639562 | QTX93024.1 | P17763 |
| LTPMLRHTIENTSAN | 1 | 0 | 0 | 1639565 | Q2YHF0.1 | P17763 |
| LTQRAQNALDNLVML | 1 | 0 | 0 | 1639569 | P17763.2 | P17763 |
| LTSLGKAVHQVFGSV | 1 | 0 | 0 | 1639571 | Q2YHF0.1 | P17763 |
| LTSRATWAKNIQAAI | 1 | 0 | 0 | 1639572 | QTX93142.1 | P17763 |
| LTSRATWAKNIQTAI | 1 | 0 | 0 | 1639573 | P14340.2 | P17763 |
| LTSRATWAQNILTAI | 1 | 0 | 0 | 1639574 | QTX93504.1 | P17763 |
| LVDRERELHKLGKCG | 1 | 0 | 0 | 1639581 | QTX93479.1 | P17763 |
| LVGRARVSQGAGWSL | 1 | 0 | 0 | 1639586 | P17763.2 | P17763 |
| LVGVVTLYLGVMVQA | 1 | 0 | 0 | 1639592 | AAB33855.1 | P17763 |
| LVHRERELHKQGKCA | 1 | 0 | 0 | 1639593 | P17763.2 | P17763 |
| LVKLHSGKDVFFIPP | 1 | 0 | 0 | 1639598 | QTX92722.1 | P17763 |
| LVKRFSTGLFSGKGP | 1 | 0 | 0 | 1639599 | Q2YHF0.1 | P17763 |
| LVQIENLEYTVVVTV | 1 | 0 | 0 | 1639606 | Q2YHF0.1 | P17763 |
| LVQIKYEGTDAPCKI | 1 | 0 | 0 | 1639607 | QFS19562.1 | P17763 |
| LVQVKYEGTDAPCKI | 1 | 0 | 0 | 1639609 | P17763.2 | P17763 |
| LVRCPLSRNSTHEMY | 1 | 0 | 0 | 1639610 | Q2YHF0.1 | P17763 |
| LVRNPLSRNSTHEMY | 1 | 0 | 0 | 1639615 | P17763.2 | P17763 |
| LVSLTCSNTIFTLTV | 1 | 0 | 0 | 1639616 | QTX93504.1 | P17763 |
| LVTCAKFQCLELIEG | 1 | 0 | 0 | 1639620 | QTX93368.1 | P17763 |
| LVTCAKFQCLESIEG | 1 | 0 | 0 | 1639622 | P27915.1 | P17763 |
| LVVPCRNQDELIGRA | 1 | 0 | 0 | 1639625 | P14340.2 | P17763 |
| LWALLGKKKNPRLCT | 1 | 0 | 0 | 1639629 | Q58HT7.1 | P17763 |
| LWALLGKKKSPRLCT | 1 | 0 | 0 | 1639630 | AQV12653.1 | P17763 |
| LWALLGRKKNPRLCT | 1 | 0 | 0 | 1639631 | Q2YHF0.1 | P17763 |
| LWATLLSLTFIKTTF | 1 | 0 | 0 | 1639632 | QFS19562.1 | P17763 |
| LWATLLSLTFVKTTF | 1 | 0 | 0 | 1639633 | P17763.2 | P17763 |
| LWEGNPGRFWNTTIA | 1 | 0 | 0 | 1639637 | P14340.2 | P17763 |
| LWEGSPGKFWNTTIA | 1 | 0 | 0 | 1639638 | P17763.2 | P17763 |
| LWKELGKKKTPRMCT | 1 | 0 | 0 | 1639639 | P14340.2 | P17763 |

| | | | | | | |
|---|---|---|---|---|---|---|
| LWTLLGRKKNPRLCT | 1 | 0 | 0 | 1639640 | QTX92332.1 | P17763 |
| LWTTLLSLTFIKTTL | 1 | 0 | 0 | 1639641 | P27909.2 | P17763 |
| LYAVATTIITPMMRH | 1 | 0 | 0 | 1639642 | P17763.2 | P17763 |
| LYAVATTVITPMMRH | 1 | 0 | 0 | 1639643 | P27909.2 | P17763 |
| LYGNGVVTKNGGYVS | 1 | 0 | 0 | 1639645 | QTX93489.1 | P17763 |
| LYGNGVVTKSGDYVS | 1 | 0 | 0 | 1639646 | Q2YHF0.1 | P17763 |
| LYGNGVVTRSGAYVS | 1 | 0 | 0 | 1639647 | P14340.2 | P17763 |
| LYGNGVVTRSGTYVS | 1 | 0 | 0 | 1639648 | QQW40772.1 | P17763 |
| LYGNGVVTTSGTYVS | 1 | 0 | 0 | 1639649 | P17763.2 | P17763 |
| MAAILAYTIGTTHFQ | 1 | 0 | 0 | 1639655 | P14340.2 | P17763 |
| MAAILAYTIGTTYFQ | 1 | 0 | 0 | 1639656 | QQW40781.1 | P17763 |
| MADIPLQWIASAIVL | 1 | 0 | 0 | 1639657 | QTX93520.1 | P17763 |
| MAIDLGELCEDTITY | 1 | 0 | 0 | 1639658 | QRG47115.1 | P17763 |
| MAIGIVSILLSSLLK | 1 | 0 | 0 | 1639659 | P27909.2 | P17763 |
| MAISGDDCVVKPIDD | 1 | 0 | 0 | 1639661 | P17763.2 | P17763 |
| MAMDLGELCEDTITY | 1 | 0 | 0 | 1639666 | P14340.2 | P17763 |
| MANIFRGSYLAGAGL | 1 | 0 | 0 | 1639669 | P17763.2 | P17763 |
| MAPHHKILAKAIFKL | 1 | 0 | 0 | 1639670 | Q2YHF0.1 | P17763 |
| MAPHHKTLAKAIFKL | 1 | 0 | 0 | 1639671 | AXG50965.1 | P17763 |
| MASVEPHWIAASIIL | 1 | 0 | 0 | 1639673 | P17763.2 | P17763 |
| MAVGLVSILASSLLR | 1 | 0 | 0 | 1639676 | QTX93489.1 | P17763 |
| MCCIITASILLWYAQ | 1 | 0 | 0 | 1639677 | P14340.2 | P17763 |
| MCHATFTMRLLSPVR | 1 | 0 | 0 | 1639678 | P17763.2 | P17763 |
| MCTLIAMDLGELCED | 1 | 0 | 0 | 1639680 | P17763.2 | P17763 |
| MCTLMAIDLGELCED | 1 | 0 | 0 | 1639682 | QRG47115.1 | P17763 |
| MCTLMAMDLGELCED | 1 | 0 | 0 | 1639683 | P14340.2 | P17763 |
| MCTREEFTKKVRSNA | 1 | 0 | 0 | 1639684 | AII99332.1 | P17763 |
| MDEEREIPERSWNSG | 1 | 0 | 0 | 1639691 | Q9WDA6.1 | P17763 |
| MDIISRKDQRGSGQV | 1 | 0 | 0 | 1639693 | QRG47124.1 | P17763 |
| MDKLTLKGMSYVMCT | 1 | 0 | 0 | 1639694 | P17763.2 | P17763 |
| MDKLTLKGVSYVMCT | 1 | 0 | 0 | 1639695 | QPD98750.1 | P17763 |
| MDVIGERIKRIKEEH | 1 | 0 | 0 | 1639696 | QTX93474.1 | P17763 |
| MDVISRRDQRGSGQV | 1 | 0 | 0 | 1639697 | P17763.2 | P17763 |
| MEDKAWLVHRQWFLD | 1 | 0 | 0 | 1639699 | Q9WDA6.1 | P17763 |
| MEELPDTIETLMLLA | 1 | 0 | 0 | 1639700 | P17763.2 | P17763 |
| MEGEHKKLAEAIFKL | 1 | 0 | 0 | 1639702 | P14340.2 | P17763 |
| MEGKIVQPENLEYTI | 1 | 0 | 0 | 1639703 | BAG84557.1 | P17763 |
| MEGKVVQPENLEYTI | 1 | 0 | 0 | 1639704 | QQW40666.1 | P17763 |
| MEHKYSWKSWGKAKI | 1 | 0 | 0 | 1639705 | P17763.2 | P17763 |
| MEIRPINEKEENMVK | 1 | 0 | 0 | 1639707 | QTX93500.1 | P17763 |
| MEIRPISEKEENMVK | 1 | 0 | 0 | 1639708 | QHJ86727.1 | P17763 |
| MEIRPLNEKEENMVK | 1 | 0 | 0 | 1639710 | Q2YHF0.1 | P17763 |
| MEIRPVKEKEENLVK | 1 | 0 | 0 | 1639716 | P17763.2 | P17763 |
| MEIRPVKEKEENLVR | 1 | 0 | 0 | 1639717 | P27909.2 | P17763 |
| MEITAEWLWRTLGRN | 1 | 0 | 0 | 1639718 | QTX93500.1 | P17763 |
| MEKLRIKGMSYTMCS | 1 | 0 | 0 | 1639719 | Q2YHF0.1 | P17763 |
| MELKYSWKTWGKAKI | 1 | 0 | 0 | 1639720 | QTX93489.1 | P17763 |
| MEVEIWTKEGERKKL | 1 | 0 | 0 | 1639725 | QNH29582.1 | P17763 |
| MEVTAKWLWGFLSRN | 1 | 0 | 0 | 1639727 | QPD98749.1 | P17763 |
| MFESTYRGAKRMAIL | 1 | 0 | 0 | 1639733 | Q2YHF0.1 | P17763 |
| MFETTMRGAKRMAIL | 1 | 0 | 0 | 1639734 | QTX92984.1 | P17763 |
| MGANFRAGRVIDPRR | 1 | 0 | 0 | 1639739 | Q2YHF0.1 | P17763 |
| MGIMILKLLTDFQSH | 1 | 0 | 0 | 1639748 | QFS19562.1 | P17763 |
| MGIMMLKLLTDFQSH | 1 | 0 | 0 | 1639749 | P17763.2 | P17763 |
| MGIMMLKLLTEFQPH | 1 | 0 | 0 | 1639750 | P27909.2 | P17763 |
| MGKAELEDGAYRIKQ | 1 | 0 | 0 | 1639751 | QTX93162.1 | P17763 |
| MGLDKGWPISKMDIG | 1 | 0 | 0 | 1639752 | P17763.2 | P17763 |
| MGLDTRAQTWMSAEG | 1 | 0 | 0 | 1639753 | QHR82515.1 | P17763 |
| MGLETRAETWMSSEG | 1 | 0 | 0 | 1639755 | AHG06340.1 | P17763 |
| MGLETRTETWMSSEG | 1 | 0 | 0 | 1639756 | P14340.2 | P17763 |
| MGLGKGWPLHRMDLG | 1 | 0 | 0 | 1639757 | Q2YHF0.1 | P17763 |
| MGYWIESQKNGSWKL | 1 | 0 | 0 | 1639761 | QTX93489.1 | P17763 |
| MIDKTPVHSWEDIPY | 1 | 0 | 0 | 1639764 | Q2YHF0.1 | P17763 |
| MIIMDEAHFTDPSSI | 1 | 0 | 0 | 1639767 | QFS19562.1 | P17763 |
| MIIPKNLAGPVSQHN | 1 | 0 | 0 | 1639768 | QTX93151.1 | P17763 |
| MILLTGGAMLFLISG | 1 | 0 | 0 | 1639769 | QTX93489.1 | P17763 |
| MKDGREIVVPCRNQD | 1 | 0 | 0 | 1639773 | P17763.2 | P17763 |
| MKEKSWLVHKQWFLD | 1 | 0 | 0 | 1639774 | QFS19562.1 | P17763 |
| MKFREGSSEVCDHRL | 1 | 0 | 0 | 1639775 | Q2YHF0.1 | P17763 |
| MKGEHKKLAEAIFKL | 1 | 0 | 0 | 1639776 | QRG47100.1 | P17763 |
| MKHKKATYEPDVDLG | 1 | 0 | 0 | 1639777 | QRG47107.1 | P17763 |
| MKKKTWLVHKQWFLD | 1 | 0 | 0 | 1639779 | AHN50411.1 | P17763 |
| MKPVILTDGEERVIL | 1 | 0 | 0 | 1639780 | P14340.2 | P17763 |
| MKRFKNESDPEGALW | 1 | 0 | 0 | 1639781 | P17763.2 | P17763 |
| MKRFRKEEESEGAIW | 1 | 0 | 0 | 1639782 | ABZ90961.1 | P17763 |
| MKTKTWLVHKQWFLD | 1 | 0 | 0 | 1639785 | Q2YHF0.1 | P17763 |
| MLAVWNRVWIQENPW | 1 | 0 | 0 | 1639787 | QRG47107.1 | P17763 |
| MLFLISGKGIGKTSI | 1 | 0 | 0 | 1639793 | QTX93491.1 | P17763 |
| MLIPKAYAGPISQHN | 1 | 0 | 0 | 1639796 | Q2YHF0.1 | P17763 |
| MLIPKSYAGPFSQHN | 1 | 0 | 0 | 1639797 | ARJ31953.1 | P17763 |
| MLIPRSYAGPFSQHN | 1 | 0 | 0 | 1639798 | QBA29684.1 | P17763 |
| MLKLLTDFQSHQLWA | 1 | 0 | 0 | 1639799 | P17763.2 | P17763 |
| MLKLLTDFQSHQLWT | 1 | 0 | 0 | 1639800 | QIC53498.1 | P17763 |
| MLKLLTEFQPHQLWT | 1 | 0 | 0 | 1639801 | P27909.2 | P17763 |
| MLLDNINTPEGIIPS | 1 | 0 | 0 | 1639802 | P14340.2 | P17763 |
| MLLDNIYTPEGIIPT | 1 | 0 | 0 | 1639803 | Q2YHF0.1 | P17763 |
| MLLVLCAGQLLLMRT | 1 | 0 | 0 | 1639806 | Q2YHF0.1 | P17763 |
| MLLVLCVGQLLLMRT | 1 | 0 | 0 | 1639807 | AXG50965.1 | P17763 |
| MLQGRGPLKLFMALV | 1 | 0 | 0 | 1639808 | AAT76490.1 | P17763 |
| MLTVWNRVWIEDNPW | 1 | 0 | 0 | 1639809 | QTX93489.1 | P17763 |
| MLTVWNRVWIQENPW | 1 | 0 | 0 | 1639810 | ADA59809.1 | P17763 |

| | | | | | | |
|---|---|---|---|---|---|---|
| MLTVWNRVWIRENPW | 1 | 0 | 0 | 1639811 | ADA59630.1 | P17763 |
| MMGKREKKLGEFGKA | 1 | 0 | 0 | 1639821 | P17763.2 | P17763 |
| MNNQRKKAKNTPFNM | 1 | 0 | 0 | 1639832 | QTX93078.1 | P17763 |
| MNNQRKKARSTPFNM | 1 | 0 | 0 | 1639834 | QRG47115.1 | P17763 |
| MNNQRKKTARPSFNM | 1 | 0 | 0 | 1639835 | QFS19562.1 | P17763 |
| MNNQRKKTGKPSINM | 1 | 0 | 0 | 1639836 | QTX93483.1 | P17763 |
| MNNQRKKTGRPSFNM | 1 | 0 | 0 | 1639837 | P17763.2 | P17763 |
| MNQRKKVVRPPFNML | 1 | 0 | 0 | 1639838 | Q58HT7.1 | P17763 |
| MQGKGIGKLSMGLIA | 1 | 0 | 0 | 1639853 | Q2YHF0.1 | P17763 |
| MQGKGIGKLSMGLIT | 1 | 0 | 0 | 1639854 | ARJ31953.1 | P17763 |
| MQGKGMGKLSMGLIA | 1 | 0 | 0 | 1639855 | QTX92101.1 | P17763 |
| MRCVGIGNRDFVEGL | 1 | 0 | 0 | 1639862 | P17763.2 | P17763 |
| MRCVGIGSRDFVEGL | 1 | 0 | 0 | 1639863 | QFS19562.1 | P17763 |
| MRCVGVGNRDFVEGV | 1 | 0 | 0 | 1639864 | Q2YHF0.1 | P17763 |
| MRHKKATYEPDVDLG | 1 | 0 | 0 | 1639865 | P14340.2 | P17763 |
| MRHTIENTTANISLT | 1 | 0 | 0 | 1639866 | P17763.2 | P17763 |
| MRRGDLPVWLAYKVA | 1 | 0 | 0 | 1639868 | QTX93131.1 | P17763 |
| MRRGDLPVWLAYRVA | 1 | 0 | 0 | 1639869 | P14340.2 | P17763 |
| MRRGDLPVWLSYKVA | 1 | 0 | 0 | 1639870 | P17763.2 | P17763 |
| MRTTWAFCEVLTLAT | 1 | 0 | 0 | 1639874 | Q2YHF0.1 | P17763 |
| MRTTWALCESITLAT | 1 | 0 | 0 | 1639875 | P17763.2 | P17763 |
| MRTTWALCEVLTLAT | 1 | 0 | 0 | 1639876 | AQV12525.1 | P17763 |
| MSCIAVGGITLFLGF | 1 | 0 | 0 | 1639877 | Q2YHF0.1 | P17763 |
| MSFSCIAIGIITLYL | 1 | 0 | 0 | 1639878 | QHJ86727.1 | P17763 |
| MSFSCIVIGIITLYL | 1 | 0 | 0 | 1639879 | QTX93530.1 | P17763 |
| MSGKGIGKMTLGMCC | 1 | 0 | 0 | 1639880 | QQW40770.1 | P17763 |
| MSGRGIGKMTLGMCC | 1 | 0 | 0 | 1639881 | P14340.2 | P17763 |
| MSYTMCSGKFSIDKE | 1 | 0 | 0 | 1639888 | Q2YHF0.1 | P17763 |
| MSYVMCTGSFKLEKE | 1 | 0 | 0 | 1639889 | P17763.2 | P17763 |
| MTCIAVGGITLFLGF | 1 | 0 | 0 | 1639890 | Q58HT7.1 | P17763 |
| MTCIAVGLVTLYLGV | 1 | 0 | 0 | 1639891 | QFS19562.1 | P17763 |
| MTCIAVGMVTLYLGV | 1 | 0 | 0 | 1639892 | P17763.2 | P17763 |
| MTDKTPVHSWEDIPY | 1 | 0 | 0 | 1639893 | Q2YHF2.1 | P17763 |
| MTGTLAVFFLLIMGQ | 1 | 0 | 0 | 1639896 | QTX92698.1 | P17763 |
| MTGTLAVFLLLIMGQ | 1 | 0 | 0 | 1639897 | P27909.2 | P17763 |
| MTGTLAVFLLLTMGQ | 1 | 0 | 0 | 1639898 | P17763.2 | P17763 |
| MTLITGNMSFRDLGR | 1 | 0 | 0 | 1639899 | QQW40674.1 | P17763 |
| MTQKARDALDNLAVL | 1 | 0 | 0 | 1639900 | AHG25307.1 | P17763 |
| MTQKARNALDNLAVL | 1 | 0 | 0 | 1639901 | QQW40770.1 | P17763 |
| MTTEDMLKVWNRVWI | 1 | 0 | 0 | 1639903 | Q2YHF0.1 | P17763 |
| MTTTLSIPHDLMELI | 1 | 0 | 0 | 1639904 | AMP43484.1 | P17763 |
| MTTVFSIPHDLMELI | 1 | 0 | 0 | 1639905 | Q2YHF2.1 | P17763 |
| MTTVLSIPHDLMEFI | 1 | 0 | 0 | 1639906 | Q2YHF0.1 | P17763 |
| MTYKCPRITEAEPDD | 1 | 0 | 0 | 1639908 | QFS19562.1 | P17763 |
| MVHQIFGSAYTALFS | 1 | 0 | 0 | 1639911 | QTX93489.1 | P17763 |
| MVLAFITFLRVLSIP | 1 | 0 | 0 | 1639919 | Q2YHF0.1 | P17763 |
| MVMVGAAMTDDIGMG | 1 | 0 | 0 | 1639920 | QQW40684.1 | P17763 |
| MVMVGATMTDDIGMG | 1 | 0 | 0 | 1639921 | P14340.2 | P17763 |
| MVTFKVPHAKRQDVT | 1 | 0 | 0 | 1639925 | Q2YHF0.1 | P17763 |
| MVTQLAMTDTTPFGQ | 1 | 0 | 0 | 1639927 | Q2YHF0.1 | P17763 |
| MVTQMAMTDTTPFGQ | 1 | 0 | 0 | 1639928 | P14340.2 | P17763 |
| MWHVTRGAVLMHKGK | 1 | 0 | 0 | 1639933 | P14340.2 | P17763 |
| MWHVTRGAVLMYQGK | 1 | 0 | 0 | 1639934 | P17763.2 | P17763 |
| MWHVTRGAVLTHNGK | 1 | 0 | 0 | 1639935 | QTX93491.1 | P17763 |
| MWHVTRGAVLTYNGK | 1 | 0 | 0 | 1639936 | QDG00739.1 | P17763 |
| MWHVTRGSVICHESG | 1 | 0 | 0 | 1639937 | Q2YHF0.1 | P17763 |
| MWHVTRGSVICHETG | 1 | 0 | 0 | 1639938 | Q58HT7.1 | P17763 |
| MWLGARFLEFEALGF | 1 | 0 | 0 | 1639939 | P17763.2 | P17763 |
| MWSLMYFHRRDLRLA | 1 | 0 | 0 | 1639941 | P14340.2 | P17763 |
| MWTLMYFHRRDLRLA | 1 | 0 | 0 | 1639942 | QRG47112.1 | P17763 |
| MYADDTAGWDTRITE | 1 | 0 | 0 | 1639943 | P17763.2 | P17763 |
| MYFHRRDLRLAANAI | 1 | 0 | 0 | 1639947 | P17763.2 | P17763 |
| MYWVSCGTGNIVSAV | 1 | 0 | 0 | 1639950 | P17763.2 | P17763 |
| NAAIGAVFVDENQWN | 1 | 0 | 0 | 1639951 | P17763.2 | P17763 |
| NAAMGAVFTEENQWD | 1 | 0 | 0 | 1639952 | QTX93489.1 | P17763 |
| NAEPDGPTPELEEEM | 1 | 0 | 0 | 1639955 | QTX93489.1 | P17763 |
| NAEPETPNMDVIGER | 1 | 0 | 0 | 1639956 | QTX93461.1 | P17763 |
| NAICSAVPSHWVPTS | 1 | 0 | 0 | 1639958 | P14340.2 | P17763 |
| NAICSAVPVHWVPTS | 1 | 0 | 0 | 1639959 | QTX93489.1 | P17763 |
| NAPIIDEEREIPERS | 1 | 0 | 0 | 1639963 | QTX93142.1 | P17763 |
| NAPIMDEEREIPERS | 1 | 0 | 0 | 1639964 | P14340.2 | P17763 |
| NASRAWNVWEVEDYG | 1 | 0 | 0 | 1639965 | QHR82521.1 | P17763 |
| NATDTWVTYGTCSQT | 1 | 0 | 0 | 1639966 | QFS19562.1 | P17763 |
| NAVGNDTGKHGKEIK | 1 | 0 | 0 | 1639969 | QQW40777.1 | P17763 |
| NAVILQNAWKVSCTT | 1 | 0 | 0 | 1639971 | Q9WDA6.1 | P17763 |
| NCLRKSGKKVIQLSR | 1 | 0 | 0 | 1639973 | Q2YHF0.1 | P17763 |
| NCLRKSGKRVIQLSR | 1 | 0 | 0 | 1639974 | AQV12350.1 | P17763 |
| NDEDCAHWKEAKMLL | 1 | 0 | 0 | 1639976 | P14340.2 | P17763 |
| NDEDHAHWTEAKMLL | 1 | 0 | 0 | 1639977 | P17763.2 | P17763 |
| NDIAACLRKNGKKVI | 1 | 0 | 0 | 1639978 | P14340.2 | P17763 |
| NDIANCLRKSGKKVI | 1 | 0 | 0 | 1639979 | Q2YHF0.1 | P17763 |
| NDIANCLRKSGKRVI | 1 | 0 | 0 | 1639980 | AQV12350.1 | P17763 |
| NDLKYSWKTWGKAKI | 1 | 0 | 0 | 1639981 | Q2YHF0.1 | P17763 |
| NDMGKVRKDIPQWEP | 1 | 0 | 0 | 1639982 | P17763.2 | P17763 |
| NDQYIYMGEPLENDE | 1 | 0 | 0 | 1639985 | P14340.2 | P17763 |
| NDTGKHGKEIKVTPQ | 1 | 0 | 0 | 1639986 | QTX93107.1 | P17763 |
| NDTQGVTVEITPQAS | 1 | 0 | 0 | 1639987 | QTX93541.1 | P17763 |
| NDTSNHGETATITPR | 1 | 0 | 0 | 1639988 | AQV12768.1 | P17763 |
| NDTSNHGVTATITPR | 1 | 0 | 0 | 1639989 | Q2YHF0.1 | P17763 |
| NDWDYVVTTDISEMG | 1 | 0 | 0 | 1639990 | P17763.2 | P17763 |
| NDWTQVPFCSHHFHE | 1 | 0 | 0 | 1639992 | P14340.2 | P17763 |

| | | | | | | |
|---|---|---|---|---|---|---|
| NEAIMAVGMVSILAS | 1 | 0 | 0 | 1639993 | P14340.2 | P17763 |
| NEAVMAVGMVSILAS | 1 | 0 | 0 | 1639994 | AUN35140.1 | P17763 |
| NEDHWFGRENSWSGV | 1 | 0 | 0 | 1639996 | Q2YHF0.1 | P17763 |
| NEDHWFSRENSLSGV | 1 | 0 | 0 | 1639997 | P17763.2 | P17763 |
| NEDHWFSRENSWSGV | 1 | 0 | 0 | 1639998 | AXG50965.1 | P17763 |
| NEELITEQMAPHHKI | 1 | 0 | 0 | 1639999 | Q2YHF0.1 | P17763 |
| NEELITEQMAPHHKT | 1 | 0 | 0 | 1640000 | QBA29684.1 | P17763 |
| NEEMVTNHMEGEHKK | 1 | 0 | 0 | 1640001 | P14340.2 | P17763 |
| NEEMVTNHMKGEHKK | 1 | 0 | 0 | 1640002 | QGW05389.1 | P17763 |
| NEGVMAVGLVSILAS | 1 | 0 | 0 | 1640003 | QTX93489.1 | P17763 |
| NEHKSTWHYDEDNPY | 1 | 0 | 0 | 1640004 | P27909.2 | P17763 |
| NELNYVLWEGGHDLT | 1 | 0 | 0 | 1640010 | Q2YHF0.1 | P17763 |
| NEMGFLEKTKKDFGL | 1 | 0 | 0 | 1640012 | QQW40742.1 | P17763 |
| NEMGFLEKTKKDLGL | 1 | 0 | 0 | 1640013 | P14340.2 | P17763 |
| NEMGLIEKTKTDFGF | 1 | 0 | 0 | 1640014 | Q2YHF0.1 | P17763 |
| NEMGLLETTKKDLGI | 1 | 0 | 0 | 1640015 | P17763.2 | P17763 |
| NEMGLLETTKRDLGM | 1 | 0 | 0 | 1640016 | QTX93489.1 | P17763 |
| NERRAWNFFEVEDYG | 1 | 0 | 0 | 1640018 | QTX92256.1 | P17763 |
| NERRAWNFLEVEDYG | 1 | 0 | 0 | 1640019 | Q2YHF0.1 | P17763 |
| NERRAWNSFEVEDYG | 1 | 0 | 0 | 1640020 | AKQ00032.1 | P17763 |
| NERRAWNSLEVEDYG | 1 | 0 | 0 | 1640021 | AHK05342.1 | P17763 |
| NESTEHGTTATITPQ | 1 | 0 | 0 | 1640024 | QFS19562.1 | P17763 |
| NETTEHGTIATITPQ | 1 | 0 | 0 | 1640026 | P27909.2 | P17763 |
| NFRADRVIDPRRCLK | 1 | 0 | 0 | 1640033 | P17763.2 | P17763 |
| NGDTHAVGNDTSNHG | 1 | 0 | 0 | 1640035 | Q2YHF0.1 | P17763 |
| NGIALGLMALKLITQ | 1 | 0 | 0 | 1640036 | Q6YMS3.1 | P17763 |
| NGKELKCGSGIFVVD | 1 | 0 | 0 | 1640037 | ARM59249.1 | P17763 |
| NGVNMCTLMAMDLGE | 1 | 0 | 0 | 1640044 | QQW40786.1 | P17763 |
| NGVVKLLTKPWDVIP | 1 | 0 | 0 | 1640045 | QFS19562.1 | P17763 |
| NGVVKLLTKPWDVVP | 1 | 0 | 0 | 1640046 | QFS19207.1 | P17763 |
| NGVVRLLTKPWDVIP | 1 | 0 | 0 | 1640047 | P17763.2 | P17763 |
| NHGETATITPRSPSV | 1 | 0 | 0 | 1640048 | AQV12768.1 | P17763 |
| NHGVTATITPRSPSV | 1 | 0 | 0 | 1640049 | Q2YHF0.1 | P17763 |
| NHNKEGDQYIYMGQP | 1 | 0 | 0 | 1640050 | QFS19562.1 | P17763 |
| NILVEVQDDGTMKIK | 1 | 0 | 0 | 1640053 | P17763.2 | P17763 |
| NINTPEGIIPSMFEP | 1 | 0 | 0 | 1640056 | P14340.2 | P17763 |
| NIYTPEGIIPTLFGP | 1 | 0 | 0 | 1640059 | Q2YHF0.1 | P17763 |
| NKKGKVIGLYGNGVV | 1 | 0 | 0 | 1640060 | Q2YHF2.1 | P17763 |
| NKPTLDFELIKTEAK | 1 | 0 | 0 | 1640063 | QTX92981.1 | P17763 |
| NKPTLDIELLKTEVT | 1 | 0 | 0 | 1640064 | P27909.2 | P17763 |
| NKWKSAREAVEDGRF | 1 | 0 | 0 | 1640066 | QTE05819.1 | P17763 |
| NKWKSAREAVEDSGF | 1 | 0 | 0 | 1640067 | QQW40742.1 | P17763 |
| NLAERVLDWLEKHGA | 1 | 0 | 0 | 1640069 | P27909.2 | P17763 |
| NLAERVLDWLEKYGV | 1 | 0 | 0 | 1640070 | QFS19562.1 | P17763 |
| NLDIIGKRIEKIKQE | 1 | 0 | 0 | 1640073 | P14340.2 | P17763 |
| NLIGKEEYVDYMPAM | 1 | 0 | 0 | 1640075 | QTX92339.1 | P17763 |
| NLITEMGRLPTFMTQ | 1 | 0 | 0 | 1640077 | P14340.2 | P17763 |
| NLLLGVGLAMATTLQ | 1 | 0 | 0 | 1640081 | QTX93489.1 | P17763 |
| NLTSTWVMYGTCTQS | 1 | 0 | 0 | 1640084 | Q5UCB8.1 | P17763 |
| NLVKPEGKVIDLGCG | 1 | 0 | 0 | 1640086 | P17763.2 | P17763 |
| NLVTPEGKVVDLGCG | 1 | 0 | 0 | 1640087 | Q9WDA6.1 | P17763 |
| NMDVEIWTKEGEKKK | 1 | 0 | 0 | 1640089 | QHR82521.1 | P17763 |
| NMDVEIWTKEGERKK | 1 | 0 | 0 | 1640090 | P17763.2 | P17763 |
| NMEAQLIRQMEGEGV | 1 | 0 | 0 | 1640091 | P14340.2 | P17763 |
| NMEVQLIRQMESEGI | 1 | 0 | 0 | 1640092 | P27909.2 | P17763 |
| NMLKRERNRVSTPQG | 1 | 0 | 0 | 1640095 | Q2YHF0.1 | P17763 |
| NMMGKREKKIGEFGK | 1 | 0 | 0 | 1640096 | QEI46448.1 | P17763 |
| NMMGKREKKLGEFGK | 1 | 0 | 0 | 1640097 | P17763.2 | P17763 |
| NMMGKREKKLGEFGR | 1 | 0 | 0 | 1640098 | ACN42699.1 | P17763 |
| NMVIPEGRVIDLGCG | 1 | 0 | 0 | 1640102 | QTX93489.1 | P17763 |
| NMVTPEGKVVDLGCG | 1 | 0 | 0 | 1640103 | P14340.2 | P17763 |
| NNDEDHAHWTEAKML | 1 | 0 | 0 | 1640104 | P17763.2 | P17763 |
| NNNTQFCIKVLNPYM | 1 | 0 | 0 | 1640105 | P14340.2 | P17763 |
| NNQILEENMEVEIWT | 1 | 0 | 0 | 1640106 | QNH29582.1 | P17763 |
| NNQILEENVEVEIWT | 1 | 0 | 0 | 1640107 | P14340.2 | P17763 |
| NPGRFWNTTIAVSTA | 1 | 0 | 0 | 1640109 | Q2YHF0.1 | P17763 |
| NPKHVQTKPGLFKTL | 1 | 0 | 0 | 1640116 | Q2YHF0.1 | P17763 |
| NPKNVQTAPGTFKTP | 1 | 0 | 0 | 1640117 | P17763.2 | P17763 |
| NPKNVQTTPGTFKTP | 1 | 0 | 0 | 1640118 | P27909.2 | P17763 |
| NPRAVQTKPGIFKTN | 1 | 0 | 0 | 1640123 | QRG47115.1 | P17763 |
| NPRAVQTKPGLFKTN | 1 | 0 | 0 | 1640124 | P14340.2 | P17763 |
| NPTIEAGRTLRVLNL | 1 | 0 | 0 | 1640127 | QRG47112.1 | P17763 |
| NPTIEEGRTLRVLKM | 1 | 0 | 0 | 1640128 | P17763.2 | P17763 |
| NPTVEAGRTLRVLNL | 1 | 0 | 0 | 1640130 | P14340.2 | P17763 |
| NPWMEDKTPVESWEE | 1 | 0 | 0 | 1640131 | ADA59546.1 | P17763 |
| NPWMEDKTPVTTWED | 1 | 0 | 0 | 1640132 | QTX93491.1 | P17763 |
| NPYKTWAYHGSYEVK | 1 | 0 | 0 | 1640133 | P17763.2 | P17763 |
| NQAGEHRRDKRSVAL | 1 | 0 | 0 | 1640134 | QTX93489.1 | P17763 |
| NQDELVGRARVSQGA | 1 | 0 | 0 | 1640135 | P17763.2 | P17763 |
| NQNKEGDQYIYMGQP | 1 | 0 | 0 | 1640142 | P17763.2 | P17763 |
| NQNKEGDQYVYMGQP | 1 | 0 | 0 | 1640143 | P27909.2 | P17763 |
| NRFTMKHKKATYEPD | 1 | 0 | 0 | 1640147 | QRG47107.1 | P17763 |
| NRFTMRHKKATYEPD | 1 | 0 | 0 | 1640148 | P14340.2 | P17763 |
| NRFTTRHRKPTYEKD | 1 | 0 | 0 | 1640149 | AKA62184.1 | P17763 |
| NRFTTRHRKPTYERD | 1 | 0 | 0 | 1640150 | AFP27206.1 | P17763 |
| NRGPSLRTTTVTGKI | 1 | 0 | 0 | 1640151 | QFS19562.1 | P17763 |
| NRKGKVIGLYGNGVV | 1 | 0 | 0 | 1640153 | Q2YHF0.1 | P17763 |
| NRVSTGSQLAKRFSK | 1 | 0 | 0 | 1640156 | AAT76483.1 | P17763 |
| NRVSTGSQLAKRFSR | 1 | 0 | 0 | 1640157 | AFZ40116.1 | P17763 |
| NRVSTVQQLTKRFSL | 1 | 0 | 0 | 1640158 | AAT76491.1 | P17763 |
| NRVSTVSQLAKRFSK | 1 | 0 | 0 | 1640159 | P17763.2 | P17763 |

| | | | | | | |
|---|---|---|---|---|---|---|
| NRVWIEENPWMEDKT | 1 | 0 | 0 | 1640160 | P17763.2 | P17763 |
| NSEQGGKAYRHAMEE | 1 | 0 | 0 | 1640161 | P17763.2 | P17763 |
| NSEQGGRAYRHAMEE | 1 | 0 | 0 | 1640162 | QFS19562.1 | P17763 |
| NSGHEWVTDFKGKTV | 1 | 0 | 0 | 1640164 | QTX92981.1 | P17763 |
| NSLSGVEGEGLHKLG | 1 | 0 | 0 | 1640165 | P17763.2 | P17763 |
| NSPIEDIEREIPERS | 1 | 0 | 0 | 1640166 | Q2YHF0.1 | P17763 |
| NSRSTSLSMTCIAVG | 1 | 0 | 0 | 1640168 | P17763.2 | P17763 |
| NSTHEMYWVSGASGN | 1 | 0 | 0 | 1640170 | Q5UCB8.1 | P17763 |
| NSTHEMYWVSGVSGN | 1 | 0 | 0 | 1640171 | Q2YHF0.1 | P17763 |
| NSTHEMYWVSNASGN | 1 | 0 | 0 | 1640172 | QTX93055.1 | P17763 |
| NSTHEMYWVSNATGN | 1 | 0 | 0 | 1640173 | QRG47112.1 | P17763 |
| NSTSTWVTYGTCTAT | 1 | 0 | 0 | 1640174 | QQW40770.1 | P17763 |
| NSTSTWVTYGTCTTT | 1 | 0 | 0 | 1640175 | P14340.2 | P17763 |
| NSYSGVEGEGLHKLG | 1 | 0 | 0 | 1640176 | AAX55647.1 | P17763 |
| NTFTNMEVQLIRQME | 1 | 0 | 0 | 1640178 | P27909.2 | P17763 |
| NTGFDWITDYQGKTV | 1 | 0 | 0 | 1640179 | Q2YHF0.1 | P17763 |
| NTISKMLLNRFTTRH | 1 | 0 | 0 | 1640184 | ARM59249.1 | P17763 |
| NTQVGTLALSLTFIK | 1 | 0 | 0 | 1640185 | Q2YHF0.1 | P17763 |
| NTQVGTLALSLTFIR | 1 | 0 | 0 | 1640186 | P09866.2 | P17763 |
| NTTSKMLLNRFTTRH | 1 | 0 | 0 | 1640188 | CCM44290.1 | P17763 |
| NWLARVGRERLSRMA | 1 | 0 | 0 | 1640202 | P14340.2 | P17763 |
| NWLVRVGRERLSRMA | 1 | 0 | 0 | 1640203 | QQW40771.1 | P17763 |
| NYNLIVMDEAHFTDP | 1 | 0 | 0 | 1640205 | AFY10047.1 | P17763 |
| NYVCKHTYVDRGWGN | 1 | 0 | 0 | 1640206 | QTX93489.1 | P17763 |
| PAIKTEHTGREIVDL | 1 | 0 | 0 | 1640212 | QNG98968.1 | P17763 |
| PAIRAEHTGREIVDL | 1 | 0 | 0 | 1640213 | P14340.2 | P17763 |
| PAIVREAIKRGLRTL | 1 | 0 | 0 | 1640214 | P14340.2 | P17763 |
| PAMKRYSAPFESEGV | 1 | 0 | 0 | 1640216 | AHG06382.1 | P17763 |
| PAMRRYSAHFESEGV | 1 | 0 | 0 | 1640217 | QTX92320.1 | P17763 |
| PAQEDDQYVFSGDPL | 1 | 0 | 0 | 1640218 | Q2YHF0.1 | P17763 |
| PAVKSEHTGREIVDL | 1 | 0 | 0 | 1640220 | Q2YHF0.1 | P17763 |
| PDTIETLMLLALIAV | 1 | 0 | 0 | 1640230 | P17763.2 | P17763 |
| PECPNASRAWNVWEV | 1 | 0 | 0 | 1640233 | QHR82521.1 | P17763 |
| PECPSASRAWNVWEV | 1 | 0 | 0 | 1640234 | QTX93479.1 | P17763 |
| PEDIDCWCNLTSAWV | 1 | 0 | 0 | 1640235 | ACL99241.1 | P17763 |
| PEDIDCWCNLTSTWV | 1 | 0 | 0 | 1640236 | QTX93489.1 | P17763 |
| PEDIDCWCNSTSTWV | 1 | 0 | 0 | 1640237 | P14340.2 | P17763 |
| PEGIIPSMFEPEREK | 1 | 0 | 0 | 1640238 | P14340.2 | P17763 |
| PEGIIPTLFGPEREK | 1 | 0 | 0 | 1640239 | Q2YHF0.1 | P17763 |
| PEGKVIDLGCGRGGW | 1 | 0 | 0 | 1640240 | P17763.2 | P17763 |
| PEGKVVDLGCGRGGW | 1 | 0 | 0 | 1640241 | P14340.2 | P17763 |
| PEGRVIDLGCGRGGW | 1 | 0 | 0 | 1640242 | QTX92874.1 | P17763 |
| PEIEDEVFKKRNLTI | 1 | 0 | 0 | 1640243 | P27909.2 | P17763 |
| PEIEDEVFRKRNLTI | 1 | 0 | 0 | 1640244 | P17763.2 | P17763 |
| PEKCDTLLCDIGESS | 1 | 0 | 0 | 1640245 | P17763.2 | P17763 |
| PELNHILAENEVKLT | 1 | 0 | 0 | 1640246 | QTX93142.1 | P17763 |
| PERSWNSGNEWITDF | 1 | 0 | 0 | 1640248 | QTX93491.1 | P17763 |
| PERSWNSGYDWITDF | 1 | 0 | 0 | 1640249 | P17763.2 | P17763 |
| PERSWNSGYEWITDF | 1 | 0 | 0 | 1640250 | QFS19562.1 | P17763 |
| PESLETLMLVALLGA | 1 | 0 | 0 | 1640251 | Q2YHF0.1 | P17763 |
| PETLETLLLLTLLAT | 1 | 0 | 0 | 1640254 | P14340.2 | P17763 |
| PETVLELTDALALGM | 1 | 0 | 0 | 1640255 | QTX93044.1 | P17763 |
| PFGQQRVFKEKVDTR | 1 | 0 | 0 | 1640263 | P17763.2 | P17763 |
| PGGAMYADDTAGWDT | 1 | 0 | 0 | 1640270 | QTX93489.1 | P17763 |
| PGGNMYADDTAGWDT | 1 | 0 | 0 | 1640271 | P17763.2 | P17763 |
| PGIFKTNTGTIGAVS | 1 | 0 | 0 | 1640273 | QRG47115.1 | P17763 |
| PGKFWNTTIAVSMAN | 1 | 0 | 0 | 1640274 | P17763.2 | P17763 |
| PGLFKTNTGTIGAVS | 1 | 0 | 0 | 1640275 | QTX93055.1 | P17763 |
| PGRFWNTTIAVSMAN | 1 | 0 | 0 | 1640277 | QHE24278.1 | P17763 |
| PGTFKTPEGEVGAIA | 1 | 0 | 0 | 1640278 | P17763.2 | P17763 |
| PGTTVTIQEDCDHRG | 1 | 0 | 0 | 1640285 | Q2YHF0.1 | P17763 |
| PGTTVTIQENCDHRG | 1 | 0 | 0 | 1640286 | AKQ00035.1 | P17763 |
| PGTTVTVQEDCDHRG | 1 | 0 | 0 | 1640287 | QTX92349.1 | P17763 |
| PHSGMGLETRAETWM | 1 | 0 | 0 | 1640290 | Q2YHF0.1 | P17763 |
| PHVGLGLETRTETWM | 1 | 0 | 0 | 1640291 | P17763.2 | P17763 |
| PHVGMGLDTRAQTWM | 1 | 0 | 0 | 1640292 | QHR82515.1 | P17763 |
| PHVGMGLDTRTQTWM | 1 | 0 | 0 | 1640293 | QTX93491.1 | P17763 |
| PHVGMGLETRTETWM | 1 | 0 | 0 | 1640294 | P14340.2 | P17763 |
| PIDDRFANALLALND | 1 | 0 | 0 | 1640295 | QTX93489.1 | P17763 |
| PIDDRFASALTALND | 1 | 0 | 0 | 1640296 | QRG47115.1 | P17763 |
| PILTLWEGNPGRFWN | 1 | 0 | 0 | 1640298 | Q2YHF0.1 | P17763 |
| PIMTLWEGNPGRFWN | 1 | 0 | 0 | 1640299 | ANW72071.1 | P17763 |
| PINEKEENMVKSLVS | 1 | 0 | 0 | 1640300 | QTX93512.1 | P17763 |
| PIPMATYGWNLVKLH | 1 | 0 | 0 | 1640301 | QTX92516.1 | P17763 |
| PIPMSTYGWNLVRLQ | 1 | 0 | 0 | 1640302 | P14340.2 | P17763 |
| PIPYDPKFEKQLGQV | 1 | 0 | 0 | 1640303 | P14340.2 | P17763 |
| PISEKEENMVKSLVS | 1 | 0 | 0 | 1640305 | QHJ86727.1 | P17763 |
| PKGAVMDIISRKDQR | 1 | 0 | 0 | 1640312 | Q2YHF0.1 | P17763 |
| PKGKVVDLGCGRGGW | 1 | 0 | 0 | 1640313 | Q2YHF0.1 | P17763 |
| PKGLKERVEKWLKEC | 1 | 0 | 0 | 1640314 | Q2YHF0.1 | P17763 |
| PKGLKERVEKWLREC | 1 | 0 | 0 | 1640315 | AMP43484.1 | P17763 |
| PKNENDQYIYMGEPL | 1 | 0 | 0 | 1640317 | P14340.2 | P17763 |
| PLALKEFKEFAAGRK | 1 | 0 | 0 | 1640320 | P14340.2 | P17763 |
| PLALREFKEFAAGRR | 1 | 0 | 0 | 1640321 | P17763.2 | P17763 |
| PLALTIKGLNPTAIF | 1 | 0 | 0 | 1640322 | P14340.2 | P17763 |
| PLDDRFANALTALND | 1 | 0 | 0 | 1640323 | QTX93060.1 | P17763 |
| PLLALGCYSQVNPLT | 1 | 0 | 0 | 1640326 | P17763.2 | P17763 |
| PLLAMGCYSQVNPTT | 1 | 0 | 0 | 1640327 | Q2YHF0.1 | P17763 |
| PLNEKEENMVKSQVS | 1 | 0 | 0 | 1640328 | Q2YHF0.1 | P17763 |
| PLPWTAGADTLEVHW | 1 | 0 | 0 | 1640329 | Q2YHF0.1 | P17763 |
| PLPWTAGADTSEVHW | 1 | 0 | 0 | 1640330 | Q2YHF2.1 | P17763 |

| | | | | | | |
|---|---|---|---|---|---|---|
| PLPWTSGASTSQETW | 1 | 0 | 0 | 1640331 | P17763.2 | P17763 |
| PLPWTTGADTSEVHW | 1 | 0 | 0 | 1640332 | AKQ00032.1 | P17763 |
| PLSEKEENMVKSQVT | 1 | 0 | 0 | 1640333 | Q5UCB8.1 | P17763 |
| PLSRNSTHEMYWVSG | 1 | 0 | 0 | 1640334 | Q2YHF0.1 | P17763 |
| PLSRNSTHEMYWVSN | 1 | 0 | 0 | 1640335 | QTX92981.1 | P17763 |
| PLTLTAAVLMLVAHY | 1 | 0 | 0 | 1640336 | QFS19562.1 | P17763 |
| PLTTLWEGSPGKFWN | 1 | 0 | 0 | 1640337 | P17763.2 | P17763 |
| PMALKDFKEFASGRK | 1 | 0 | 0 | 1640339 | Q2YHF0.1 | P17763 |
| PMAVAAMGVPPLPLF | 1 | 0 | 0 | 1640340 | AMH87199.1 | P17763 |
| PMFAVGLLFRRLTSR | 1 | 0 | 0 | 1640341 | P17763.2 | P17763 |
| PMPGTRKVMEITAEW | 1 | 0 | 0 | 1640349 | QTX93491.1 | P17763 |
| PNSLEELGDGLAMGI | 1 | 0 | 0 | 1640356 | P17763.2 | P17763 |
| PPFGESNIIIGIGDK | 1 | 0 | 0 | 1640358 | QHD57351.1 | P17763 |
| PPFGESNIVIGIGDK | 1 | 0 | 0 | 1640359 | QTX93512.1 | P17763 |
| PPFGESNIVIGIGDN | 1 | 0 | 0 | 1640360 | QIS48873.1 | P17763 |
| PPTAGVLARWGTFKK | 1 | 0 | 0 | 1640362 | ADI44327.1 | P17763 |
| PQGLVKRFSTGLFSG | 1 | 0 | 0 | 1640363 | Q2YHF0.1 | P17763 |
| PQWEPSKGWKNWQEV | 1 | 0 | 0 | 1640367 | Q2YHF0.1 | P17763 |
| PRGAVMDIISRKDQR | 1 | 0 | 0 | 1640370 | Q5UCB8.1 | P17763 |
| PRGTVMDIISRRDQR | 1 | 0 | 0 | 1640371 | P14340.2 | P17763 |
| PRRCLKPVILTDGPE | 1 | 0 | 0 | 1640373 | QTX93489.1 | P17763 |
| PRRCMKPVILTDGEE | 1 | 0 | 0 | 1640374 | P14340.2 | P17763 |
| PRTGLDFNEMILLTM | 1 | 0 | 0 | 1640376 | QTX93489.1 | P17763 |
| PRWLDARIYSDPLAL | 1 | 0 | 0 | 1640378 | QTX92981.1 | P17763 |
| PSIVREALKRRLRTL | 1 | 0 | 0 | 1640385 | Q2YHF0.1 | P17763 |
| PSKGWKNWQEVPFCS | 1 | 0 | 0 | 1640386 | Q2YHF0.1 | P17763 |
| PSVEVKLPDYGELTL | 1 | 0 | 0 | 1640390 | Q2YHF0.1 | P17763 |
| PTAGILARWGSFKKN | 1 | 0 | 0 | 1640392 | P17763.2 | P17763 |
| PTAGILARWGTFKKS | 1 | 0 | 0 | 1640393 | QTX93247.1 | P17763 |
| PTAGILARWSSFKKN | 1 | 0 | 0 | 1640394 | AXX75195.1 | P17763 |
| PTAGILKRWGTIKKS | 1 | 0 | 0 | 1640395 | P14340.2 | P17763 |
| PTAGVLARWGTFKKS | 1 | 0 | 0 | 1640396 | AYA22932.1 | P17763 |
| PTEWFPTSRTTWSIH | 1 | 0 | 0 | 1640397 | Q2YHF0.1 | P17763 |
| PTFAAGLLLRKLTSK | 1 | 0 | 0 | 1640398 | P14340.2 | P17763 |
| PTIEKDVDLGAGTRH | 1 | 0 | 0 | 1640401 | QTX93489.1 | P17763 |
| PTSEIQLTDYGALTL | 1 | 0 | 0 | 1640409 | P17763.2 | P17763 |
| PTSRTTWSIHAHHQW | 1 | 0 | 0 | 1640410 | P17763.2 | P17763 |
| PTSRTTWSIHAKHEW | 1 | 0 | 0 | 1640411 | P14340.2 | P17763 |
| PTSRTTWSIHARHEW | 1 | 0 | 0 | 1640412 | QNQ18279.1 | P17763 |
| PTSRTTWSIHATHEW | 1 | 0 | 0 | 1640413 | QQW40786.1 | P17763 |
| PTTEIQLTDYGALTL | 1 | 0 | 0 | 1640414 | QFS19562.1 | P17763 |
| PTVDGIMTIDLDPVI | 1 | 0 | 0 | 1640415 | QTX93489.1 | P17763 |
| PTYERDVDLGAGTRH | 1 | 0 | 0 | 1640416 | P17763.2 | P17763 |
| PVESWEEIPYLGKRE | 1 | 0 | 0 | 1640418 | ADA59677.1 | P17763 |
| PVESWEEVPYLGKRE | 1 | 0 | 0 | 1640419 | ADA59901.1 | P17763 |
| PVIYDSKFEKQLGQV | 1 | 0 | 0 | 1640422 | QTX93489.1 | P17763 |
| PVKEKEENLVKSMVS | 1 | 0 | 0 | 1640424 | P17763.2 | P17763 |
| PVKEKEENLVRSMVS | 1 | 0 | 0 | 1640425 | P27909.2 | P17763 |
| PVLLGSFGCKPLTMF | 1 | 0 | 0 | 1640426 | P27909.2 | P17763 |
| PVLLGSLGCKPLTMF | 1 | 0 | 0 | 1640427 | P17763.2 | P17763 |
| PVMKRYSAHFESEGV | 1 | 0 | 0 | 1640428 | Q2YHF0.1 | P17763 |
| PVMKRYSAPFESEGV | 1 | 0 | 0 | 1640429 | Q58HT7.1 | P17763 |
| PVPMSTYGWNIVKLM | 1 | 0 | 0 | 1640432 | AAC63341.1 | P17763 |
| PVRVPNYNLIIMDEA | 1 | 0 | 0 | 1640433 | P14340.2 | P17763 |
| PVTAASAAQRRGRVG | 1 | 0 | 0 | 1640435 | QTX93491.1 | P17763 |
| PVTTWEDVPYLGKRE | 1 | 0 | 0 | 1640439 | QTX93491.1 | P17763 |
| PVTVASAAQRRGRIG | 1 | 0 | 0 | 1640440 | P17763.2 | P17763 |
| PWMEDKTHISSWGDV | 1 | 0 | 0 | 1640443 | QTX92678.1 | P17763 |
| PWMEDKTHVSSWEEV | 1 | 0 | 0 | 1640444 | P27909.2 | P17763 |
| PWTSGATTETPTWNK | 1 | 0 | 0 | 1640445 | QHD57350.1 | P17763 |
| PWTSGATTETPTWNR | 1 | 0 | 0 | 1640446 | QTX93500.1 | P17763 |
| PYLGKREDQWCGSLI | 1 | 0 | 0 | 1640448 | ABZ90961.1 | P17763 |
| PYMPTVIEELEKLQR | 1 | 0 | 0 | 1640451 | Q2YHF0.1 | P17763 |
| QAIDGEFRLRGEQRK | 1 | 0 | 0 | 1640455 | Q2YHF0.1 | P17763 |
| QASTTEAILPEYGTL | 1 | 0 | 0 | 1640458 | QIS48873.1 | P17763 |
| QASTVEAILPEYGTL | 1 | 0 | 0 | 1640459 | QTX93471.1 | P17763 |
| QASTVEAVLPEYGTL | 1 | 0 | 0 | 1640460 | QTX93534.1 | P17763 |
| QEEHKETWHYDHENP | 1 | 0 | 0 | 1640472 | Q2YHF0.1 | P17763 |
| QEEHKETWHYDQENP | 1 | 0 | 0 | 1640473 | Q58HT7.1 | P17763 |
| QEKGKSLLFKTEDGV | 1 | 0 | 0 | 1640476 | P14340.2 | P17763 |
| QEKGKSLLFKTENGV | 1 | 0 | 0 | 1640477 | QQW40784.1 | P17763 |
| QERGKSLLFKTSAGV | 1 | 0 | 0 | 1640482 | P17763.2 | P17763 |
| QESESNILDIDLRPA | 1 | 0 | 0 | 1640483 | QRG47112.1 | P17763 |
| QEVPFCSHHFHKIFM | 1 | 0 | 0 | 1640484 | Q2YHF0.1 | P17763 |
| QFCIKVLNPYMPSVI | 1 | 0 | 0 | 1640487 | P14340.2 | P17763 |
| QGAGWSLRETACLGK | 1 | 0 | 0 | 1640488 | P17763.2 | P17763 |
| QGEAILPEEQDQNYV | 1 | 0 | 0 | 1640489 | QTX93510.1 | P17763 |
| QGEATLPEEQDQNYV | 1 | 0 | 0 | 1640490 | QTX93534.1 | P17763 |
| QGEATLVEEQDANFV | 1 | 0 | 0 | 1640491 | QFS19562.1 | P17763 |
| QGEAVLPEEQDQNYV | 1 | 0 | 0 | 1640492 | Q6YMS3.1 | P17763 |
| QGEPYLKEEQDQQYI | 1 | 0 | 0 | 1640495 | Q2YHF0.1 | P17763 |
| QGETLGEKWKKKLNQ | 1 | 0 | 0 | 1640496 | QTX93479.1 | P17763 |
| QGETLGEKWKRQLNQ | 1 | 0 | 0 | 1640497 | P17763.2 | P17763 |
| QGKRLEPSWASVKKD | 1 | 0 | 0 | 1640499 | P17763.2 | P17763 |
| QGPMKLVMAFIAFLR | 1 | 0 | 0 | 1640505 | P17763.2 | P17763 |
| QGWTSASEAVNDSRF | 1 | 0 | 0 | 1640509 | Q2YHF0.1 | P17763 |
| QHLTATEEIAVQNWL | 1 | 0 | 0 | 1640510 | QRG47120.1 | P17763 |
| QHLTATEEVAVQDWL | 1 | 0 | 0 | 1640511 | AHM25909.1 | P17763 |
| QHLTVTEEVAVQNWL | 1 | 0 | 0 | 1640512 | QTX93135.1 | P17763 |
| QHNHRPGYHTQTAGP | 1 | 0 | 0 | 1640513 | QTX93500.1 | P17763 |
| QHNYRPGYFTQTAGP | 1 | 0 | 0 | 1640514 | P17763.2 | P17763 |

| | | | | | | |
|---|---|---|---|---|---|---|
| QHNYRQGYATQTAGP | 1 | 0 | 0 | 1640516 | QTX92333.1 | P17763 |
| QHNYRQGYATQTVGP | 1 | 0 | 0 | 1640517 | Q2YHF0.1 | P17763 |
| QILLMRTTWALCESI | 1 | 0 | 0 | 1640524 | P17763.2 | P17763 |
| QKEGVFHTMWHVTRG | 1 | 0 | 0 | 1640529 | QTX93479.1 | P17763 |
| QKNGSWKLEKASFIE | 1 | 0 | 0 | 1640531 | QTX93504.1 | P17763 |
| QKNGSWKLEKASLIE | 1 | 0 | 0 | 1640532 | QHR82521.1 | P17763 |
| QKVETWALRHPGFTV | 1 | 0 | 0 | 1640536 | P17763.2 | P17763 |
| QLCDHRLMSAAVKDE | 1 | 0 | 0 | 1640537 | QDG00742.1 | P17763 |
| QLIRQMEGEGVLSKA | 1 | 0 | 0 | 1640540 | AXB26602.1 | P17763 |
| QLIRQMESEGIFLPS | 1 | 0 | 0 | 1640541 | QPB40126.1 | P17763 |
| QLIRQMESEGIFSPS | 1 | 0 | 0 | 1640542 | P17763.2 | P17763 |
| QLLLMRTTWALCEVL | 1 | 0 | 0 | 1640546 | QBB90024.1 | P17763 |
| QLSRKTFDTEYQKTK | 1 | 0 | 0 | 1640550 | P17763.2 | P17763 |
| QMCDHRLMSAAIKDS | 1 | 0 | 0 | 1640552 | P27909.2 | P17763 |
| QMEGEGVLSKADLEN | 1 | 0 | 0 | 1640555 | QTX93500.1 | P17763 |
| QMESEGIFLPSELET | 1 | 0 | 0 | 1640556 | QNH29581.1 | P17763 |
| QMESEGIFSPSELET | 1 | 0 | 0 | 1640557 | P17763.2 | P17763 |
| QNKVVKVQRPTPKGT | 1 | 0 | 0 | 1640563 | QTX93504.1 | P17763 |
| QNSGGTSIFAGHLKC | 1 | 0 | 0 | 1640564 | QTX93489.1 | P17763 |
| QNTTFIIDGPDTPEC | 1 | 0 | 0 | 1640568 | P27909.2 | P17763 |
| QPHQLWTTLLSLTFI | 1 | 0 | 0 | 1640573 | AKQ00039.1 | P17763 |
| QPKPGTRMIMTTTAN | 1 | 0 | 0 | 1640574 | QCZ25014.1 | P17763 |
| QPKPGTRMVMTTTAN | 1 | 0 | 0 | 1640575 | AUY62013.1 | P17763 |
| QPKPGTRVVMTTTAN | 1 | 0 | 0 | 1640576 | Q2YHF0.1 | P17763 |
| QQLTVTEEIAVKNWL | 1 | 0 | 0 | 1640582 | QBQ58389.1 | P17763 |
| QQPESNILDIDLRPA | 1 | 0 | 0 | 1640584 | P14340.2 | P17763 |
| QRGSGQVGTYGLNTF | 1 | 0 | 0 | 1640585 | P17763.2 | P17763 |
| QRIETWILRHPGFTI | 1 | 0 | 0 | 1640586 | P14340.2 | P17763 |
| QRKHGGMLVRNPLSR | 1 | 0 | 0 | 1640587 | P17763.2 | P17763 |
| QRKTFVELMRRGDLP | 1 | 0 | 0 | 1640588 | Q2YHF0.1 | P17763 |
| QRPTPRGTVMDIISR | 1 | 0 | 0 | 1640589 | P14340.2 | P17763 |
| QRRGRIGRNHNKEGD | 1 | 0 | 0 | 1640590 | QFS19562.1 | P17763 |
| QRRGRIGRNQNKEGD | 1 | 0 | 0 | 1640591 | P17763.2 | P17763 |
| QRRGRVGRNPQKEND | 1 | 0 | 0 | 1640592 | QTX93489.1 | P17763 |
| QRVESWILRNPGFAL | 1 | 0 | 0 | 1640595 | Q2YHF0.1 | P17763 |
| QSHQLWTTLLSLTFI | 1 | 0 | 0 | 1640601 | QIC53498.1 | P17763 |
| QSTIPETVLELTDAL | 1 | 0 | 0 | 1640607 | QTX93044.1 | P17763 |
| QTGSASSMVNGVVRL | 1 | 0 | 0 | 1640614 | QTX92981.1 | P17763 |
| QTSGGTSIFAGHLKC | 1 | 0 | 0 | 1640615 | QHR82521.1 | P17763 |
| QTWMSAEGAWRQVEK | 1 | 0 | 0 | 1640617 | QTX93479.1 | P17763 |
| QVCDHRLMSAAIKDS | 1 | 0 | 0 | 1640619 | P17763.2 | P17763 |
| QVKTETTILDVDLRP | 1 | 0 | 0 | 1640624 | Q58HT7.1 | P17763 |
| QVNPITLTAALLLLV | 1 | 0 | 0 | 1640628 | QTX92981.1 | P17763 |
| QVNPLTLTAAVLLLI | 1 | 0 | 0 | 1640629 | QBQ58390.1 | P17763 |
| QVRSLIGNEEYTDYM | 1 | 0 | 0 | 1640631 | AAG15003.1 | P17763 |
| QVRSLIGSEEYTDYM | 1 | 0 | 0 | 1640632 | QGW51895.1 | P17763 |
| QWCGSLIGLTSRATW | 1 | 0 | 0 | 1640639 | ABZ90963.1 | P17763 |
| QWDSARAAVEDEEFW | 1 | 0 | 0 | 1640640 | QTX93504.1 | P17763 |
| QWEPSKGWNDWTQVP | 1 | 0 | 0 | 1640641 | QTX93077.1 | P17763 |
| QWFFDLPLPWTSGAT | 1 | 0 | 0 | 1640643 | QTX93489.1 | P17763 |
| QWLETKGVERLKRMA | 1 | 0 | 0 | 1640644 | QTX93491.1 | P17763 |
| QWMTTEDMLSVWNRV | 1 | 0 | 0 | 1640645 | P17763.2 | P17763 |
| QWNSAKEAVEDERFW | 1 | 0 | 0 | 1640646 | P17763.2 | P17763 |
| QWQPSKGWHDWQQVP | 1 | 0 | 0 | 1640647 | QTX93489.1 | P17763 |
| QYICRRDMVDRGWGN | 1 | 0 | 0 | 1640649 | Q2YHF0.1 | P17763 |
| QYICRRDVVDRGWGN | 1 | 0 | 0 | 1640650 | Q58HT7.1 | P17763 |
| QYSDRRWCFDGERNN | 1 | 0 | 0 | 1640656 | P17763.2 | P17763 |
| RARISQGAGWSLKET | 1 | 0 | 0 | 1640668 | QRG47124.1 | P17763 |
| RARISQGAGWSLRET | 1 | 0 | 0 | 1640669 | P14340.2 | P17763 |
| RATWAKNIHTAITQV | 1 | 0 | 0 | 1640671 | QTX92334.1 | P17763 |
| RCPTQGEATLPEEQD | 1 | 0 | 0 | 1640675 | QTX93534.1 | P17763 |
| RDFVEGLSGATWVDV | 1 | 0 | 0 | 1640679 | P17763.2 | P17763 |
| RDFVEGVSGGAWVDL | 1 | 0 | 0 | 1640680 | Q2YHF0.1 | P17763 |
| RDVDLGAGTRHVAVE | 1 | 0 | 0 | 1640689 | P17763.2 | P17763 |
| REALKRRLRTLILAP | 1 | 0 | 0 | 1640692 | Q2YHF0.1 | P17763 |
| RECGVDRLKRMAISG | 1 | 0 | 0 | 1640693 | AMP43484.1 | P17763 |
| REEHKETWHYDQENP | 1 | 0 | 0 | 1640695 | AHG06383.1 | P17763 |
| REIPERSWNSGHEWV | 1 | 0 | 0 | 1640697 | QTX92981.1 | P17763 |
| REIPERSWNTGFDWI | 1 | 0 | 0 | 1640698 | Q2YHF0.1 | P17763 |
| REIVVPCRNQDELVG | 1 | 0 | 0 | 1640699 | P17763.2 | P17763 |
| REKKLGEFGKAKGSR | 1 | 0 | 0 | 1640700 | P17763.2 | P17763 |
| REKKLGEFGRAKGSR | 1 | 0 | 0 | 1640701 | ACN42699.1 | P17763 |
| REKTQAIDGEFRLRG | 1 | 0 | 0 | 1640702 | Q2YHF0.1 | P17763 |
| REKVDAIDGEYRLRG | 1 | 0 | 0 | 1640703 | P14340.2 | P17763 |
| RERNLHLEGKCETCV | 1 | 0 | 0 | 1640708 | QRG47115.1 | P17763 |
| RERNRVSTPQGLVKR | 1 | 0 | 0 | 1640709 | Q2YHF0.1 | P17763 |
| RETACLGKSYAQMWS | 1 | 0 | 0 | 1640710 | P14340.2 | P17763 |
| RFANALLALNDMGKV | 1 | 0 | 0 | 1640713 | QTX93489.1 | P17763 |
| RFANALTALNDMGKI | 1 | 0 | 0 | 1640714 | QTX93094.1 | P17763 |
| RFASALTALNDMGKV | 1 | 0 | 0 | 1640715 | P14340.2 | P17763 |
| RFTMAHRKPTYERDV | 1 | 0 | 0 | 1640720 | P17763.2 | P17763 |
| RFTMTHRRPTIEKDV | 1 | 0 | 0 | 1640721 | QTX93489.1 | P17763 |
| RFWDLVHRERELHKQ | 1 | 0 | 0 | 1640722 | P17763.2 | P17763 |
| RGARRMAILGDTAWD | 1 | 0 | 0 | 1640723 | P17763.2 | P17763 |
| RGKRIEPSWADVKKD | 1 | 0 | 0 | 1640728 | QRG47112.1 | P17763 |
| RGLPIRYQTPAVKSE | 1 | 0 | 0 | 1640729 | Q2YHF0.1 | P17763 |
| RGSAKLQWFVERNMV | 1 | 0 | 0 | 1640734 | QTX93489.1 | P17763 |
| RGSAKLRWFVERNLV | 1 | 0 | 0 | 1640735 | Q9WDA6.1 | P17763 |
| RGSAKLRWFVERNMV | 1 | 0 | 0 | 1640736 | P14340.2 | P17763 |
| RGSSKIRWIVERGMI | 1 | 0 | 0 | 1640738 | ANC57614.1 | P17763 |
| RGSSKIRWIVERGMV | 1 | 0 | 0 | 1640739 | Q58HT7.1 | P17763 |

| | | | | | | |
|---|---|---|---|---|---|---|
| RGTSKIRWIVERGMV | 1 | 0 | 0 | 1640741 | Q2YHF0.1 | P17763 |
| RGYISTRVEMGEAAA | 1 | 0 | 0 | 1640744 | Q2YHF0.1 | P17763 |
| RGYISTRVEMGEAAG | 1 | 0 | 0 | 1640745 | P14340.2 | P17763 |
| RGYTKGGPGHEEPVP | 1 | 0 | 0 | 1640747 | QTX93489.1 | P17763 |
| RHPGFTIMAAILAYT | 1 | 0 | 0 | 1640751 | P14340.2 | P17763 |
| RHPGFTVIAFFLAHA | 1 | 0 | 0 | 1640752 | QTX92537.1 | P17763 |
| RHPGFTVIALFLAHA | 1 | 0 | 0 | 1640753 | P17763.2 | P17763 |
| RHSIENSSVNVSLTA | 1 | 0 | 0 | 1640754 | P14340.2 | P17763 |
| RIGRNHNKEGDQYIY | 1 | 0 | 0 | 1640758 | QFS19562.1 | P17763 |
| RIGRNQNKEGDQYIY | 1 | 0 | 0 | 1640759 | P17763.2 | P17763 |
| RIGRNQNKEGDQYVY | 1 | 0 | 0 | 1640760 | P27909.2 | P17763 |
| RIKDDETENILTVLL | 1 | 0 | 0 | 1640763 | QTX93474.1 | P17763 |
| RIKGMSYTMCSGKFS | 1 | 0 | 0 | 1640764 | Q2YHF0.1 | P17763 |
| RITEDDLLNEELITE | 1 | 0 | 0 | 1640772 | Q2YHF0.1 | P17763 |
| RITEDDLQNEELITE | 1 | 0 | 0 | 1640773 | ARJ31953.1 | P17763 |
| RITLEDLKNEEMVTN | 1 | 0 | 0 | 1640774 | P14340.2 | P17763 |
| RKDIPQWEPSKGWKN | 1 | 0 | 0 | 1640779 | Q2YHF0.1 | P17763 |
| RKDLISYGGGWKLEG | 1 | 0 | 0 | 1640780 | QGW51895.1 | P17763 |
| RKEIGRMLNILNRRR | 1 | 0 | 0 | 1640781 | P14340.2 | P17763 |
| RKGSSIGKMFESTYR | 1 | 0 | 0 | 1640783 | Q2YHF0.1 | P17763 |
| RKHMILAVVITLCAI | 1 | 0 | 0 | 1640784 | AKQ00036.1 | P17763 |
| RKHMILVVVITFCAI | 1 | 0 | 0 | 1640785 | ANW72070.1 | P17763 |
| RKHMILVVVTTLCAI | 1 | 0 | 0 | 1640786 | Q2YHF0.1 | P17763 |
| RKKAKNTPFNMLKRE | 1 | 0 | 0 | 1640787 | P29990.1 | P17763 |
| RKKARSTPFNMLKRE | 1 | 0 | 0 | 1640789 | QRG47115.1 | P17763 |
| RKKLRPRWLDARTYS | 1 | 0 | 0 | 1640790 | P17763.2 | P17763 |
| RKKTARPSFNMLKRA | 1 | 0 | 0 | 1640791 | QFS19562.1 | P17763 |
| RKKTGKPSINMLKRV | 1 | 0 | 0 | 1640792 | QTX93483.1 | P17763 |
| RKKTGRPSFNMLKRA | 1 | 0 | 0 | 1640793 | P17763.2 | P17763 |
| RKLCIEGKITNITTD | 1 | 0 | 0 | 1640795 | QTX93491.1 | P17763 |
| RKLCIEGKITNVTTD | 1 | 0 | 0 | 1640796 | QHR82515.1 | P17763 |
| RKLTSRETALMVIGM | 1 | 0 | 0 | 1640797 | Q2YHF0.1 | P17763 |
| RKMLMTGTLAVFFLL | 1 | 0 | 0 | 1640798 | QTX92695.1 | P17763 |
| RKRNLTIMDLHPGSG | 1 | 0 | 0 | 1640801 | P17763.2 | P17763 |
| RKWCFDGERNNQILE | 1 | 0 | 0 | 1640806 | QTX93403.1 | P17763 |
| RKWCFDGQRNNQILE | 1 | 0 | 0 | 1640807 | QIS48873.1 | P17763 |
| RKYCIEAKLTNTTTA | 1 | 0 | 0 | 1640808 | QQW40788.1 | P17763 |
| RKYCIEAKLTNTTTE | 1 | 0 | 0 | 1640809 | QTX93024.1 | P17763 |
| RLAANAICSAVPSHW | 1 | 0 | 0 | 1640812 | P14340.2 | P17763 |
| RLASAILNAHKDGVC | 1 | 0 | 0 | 1640813 | Q2YHF0.1 | P17763 |
| RLASAILNAHKEGVC | 1 | 0 | 0 | 1640814 | QTX92334.1 | P17763 |
| RLKGESRKTFVELMR | 1 | 0 | 0 | 1640819 | QTX93489.1 | P17763 |
| RLKRMAISGDDCVVK | 1 | 0 | 0 | 1640820 | P17763.2 | P17763 |
| RLLSPVRVPNYNLII | 1 | 0 | 0 | 1640824 | P14340.2 | P17763 |
| RLLTKPWDVIPMVTQ | 1 | 0 | 0 | 1640825 | P17763.2 | P17763 |
| RLQSGVDVFFIPPEK | 1 | 0 | 0 | 1640829 | QTX93142.1 | P17763 |
| RLQSGVDVFFTPPEK | 1 | 0 | 0 | 1640830 | P14340.2 | P17763 |
| RLRGEARKTFVELMR | 1 | 0 | 0 | 1640831 | P17763.2 | P17763 |
| RMAILGDTAWDFGSV | 1 | 0 | 0 | 1640836 | ACY70637.1 | P17763 |
| RMAISGDDCVVKPID | 1 | 0 | 0 | 1640837 | P17763.2 | P17763 |
| RMAISGDDCVVKPLD | 1 | 0 | 0 | 1640838 | P14340.2 | P17763 |
| RMLINRFTMKHKKAT | 1 | 0 | 0 | 1640842 | QRG47107.1 | P17763 |
| RNDMISYGGGWRLGD | 1 | 0 | 0 | 1640847 | Q2YHF0.1 | P17763 |
| RNNQILEENMDVEIW | 1 | 0 | 0 | 1640851 | QTX93345.1 | P17763 |
| RNNQVLEENMDVEIW | 1 | 0 | 0 | 1640852 | P17763.2 | P17763 |
| RNSTFLVDGPDTSEC | 1 | 0 | 0 | 1640855 | Q2YHF0.1 | P17763 |
| RPASAWTLYAVATTF | 1 | 0 | 0 | 1640858 | P14340.2 | P17763 |
| RPGYFTQTAGPWHLG | 1 | 0 | 0 | 1640861 | P17763.2 | P17763 |
| RPLLFKTTEGINKCT | 1 | 0 | 0 | 1640863 | Q2YHF0.1 | P17763 |
| RPQDELIGRARISQG | 1 | 0 | 0 | 1640864 | QTX93489.1 | P17763 |
| RPQPTELKYSWKAWG | 1 | 0 | 0 | 1640866 | QQW40796.1 | P17763 |
| RPTPKGTVMDIISRK | 1 | 0 | 0 | 1640867 | QTX93504.1 | P17763 |
| RQGYATQTAGPWHLG | 1 | 0 | 0 | 1640868 | QTX92333.1 | P17763 |
| RQGYATQTVGPWHLG | 1 | 0 | 0 | 1640869 | Q2YHF0.1 | P17763 |
| RQLANAIFKLTYQNK | 1 | 0 | 0 | 1640871 | QTX93479.1 | P17763 |
| RQMEAEGVITQDDMH | 1 | 0 | 0 | 1640873 | Q58HT7.1 | P17763 |
| RQMEAEGVITRDDMH | 1 | 0 | 0 | 1640875 | Q2YHF0.1 | P17763 |
| RQMEGEGIFKSIQHL | 1 | 0 | 0 | 1640876 | QRG47106.1 | P17763 |
| RQMEGEGVFKNIQHL | 1 | 0 | 0 | 1640877 | QTX93142.1 | P17763 |
| RQMEGEGVFKSIQHL | 1 | 0 | 0 | 1640878 | P14340.2 | P17763 |
| RQMEGEGVFKSIQQL | 1 | 0 | 0 | 1640879 | QQW40688.1 | P17763 |
| RQNEPEDIDCWCNST | 1 | 0 | 0 | 1640880 | QTX92981.1 | P17763 |
| RRDMVDRGWGNGCGL | 1 | 0 | 0 | 1640882 | Q2YHF0.1 | P17763 |
| RRGDLPVWLSYKVAS | 1 | 0 | 0 | 1640885 | P17763.2 | P17763 |
| RRGRIGRNPAQEDDQ | 1 | 0 | 0 | 1640886 | Q2YHF0.1 | P17763 |
| RRGRVGRNPKNENDQ | 1 | 0 | 0 | 1640888 | QQW40788.1 | P17763 |
| RRTFVDRGWGNGCGL | 1 | 0 | 0 | 1640898 | P17763.2 | P17763 |
| RRVTRKHMILAVVIT | 1 | 0 | 0 | 1640899 | AMP43484.1 | P17763 |
| RRVTRKHMILVVVIT | 1 | 0 | 0 | 1640900 | Q5UCB8.1 | P17763 |
| RRVTRKHMILVVVTT | 1 | 0 | 0 | 1640901 | Q2YHF0.1 | P17763 |
| RRWCFDGERNNQVLE | 1 | 0 | 0 | 1640902 | P17763.2 | P17763 |
| RSGAYVSAIAQTEKS | 1 | 0 | 0 | 1640904 | P14340.2 | P17763 |
| RSGTYVSAIAQTEKS | 1 | 0 | 0 | 1640906 | QQW40770.1 | P17763 |
| RSQVGVGVFQDGVFH | 1 | 0 | 0 | 1640915 | P27909.2 | P17763 |
| RSQVGVGVFQEGVFH | 1 | 0 | 0 | 1640916 | P17763.2 | P17763 |
| RSTPFNMLKRERNRV | 1 | 0 | 0 | 1640920 | QRG47112.1 | P17763 |
| RSYAGPFSQHNYRQG | 1 | 0 | 0 | 1640922 | QBA29684.1 | P17763 |
| RTAAGIMKNPTVDGI | 1 | 0 | 0 | 1640923 | P17763.2 | P17763 |
| RTPKAKRGTAQIMEV | 1 | 0 | 0 | 1640939 | P17763.2 | P17763 |
| RTPRAKRGTTQIMEV | 1 | 0 | 0 | 1640940 | QNH29581.1 | P17763 |
| RTPRPMPGTRKVMEI | 1 | 0 | 0 | 1640941 | QTX93496.1 | P17763 |

| | | | | | | |
|---|---|---|---|---|---|---|
| RTSCLQKQSHWVEIT | 1 | 0 | 0 | 1640943 | Q2YHF0.1 | P17763 |
| RTSWALCEALTLATG | 1 | 0 | 0 | 1640945 | QTX93491.1 | P17763 |
| RTTWALCEALTLATG | 1 | 0 | 0 | 1640946 | P14340.2 | P17763 |
| RTTWSIHAHHQWMTT | 1 | 0 | 0 | 1640947 | P17763.2 | P17763 |
| RTWAYHGSYEAPSTG | 1 | 0 | 0 | 1640950 | Q2YHF0.1 | P17763 |
| RTYCIEALISNITTA | 1 | 0 | 0 | 1640952 | AKQ00034.1 | P17763 |
| RTYCIEASISNITTA | 1 | 0 | 0 | 1640953 | Q2YHF0.1 | P17763 |
| RVAAEGINYADRRWC | 1 | 0 | 0 | 1640954 | P14340.2 | P17763 |
| RVFKEKVDTRTPQPK | 1 | 0 | 0 | 1640962 | Q2YHF0.1 | P17763 |
| RVFKEKVDTRTQEPK | 1 | 0 | 0 | 1640963 | P14340.2 | P17763 |
| RVGRERLSRMAISGD | 1 | 0 | 0 | 1640964 | P14340.2 | P17763 |
| RVGRNPQKENDQYIF | 1 | 0 | 0 | 1640965 | QTX93489.1 | P17763 |
| RVIDPRRCLKPVILT | 1 | 0 | 0 | 1640967 | QTX93489.1 | P17763 |
| RVILAGPMPVTAASA | 1 | 0 | 0 | 1640971 | QIC53486.1 | P17763 |
| RVILAGPMPVTVASA | 1 | 0 | 0 | 1640972 | P17763.2 | P17763 |
| RVIQLSRKTFDTEYP | 1 | 0 | 0 | 1640973 | QBA29684.1 | P17763 |
| RVLDWLEKHGAERLK | 1 | 0 | 0 | 1640978 | P27909.2 | P17763 |
| RVLDWLEKYGVERLK | 1 | 0 | 0 | 1640979 | QFS19562.1 | P17763 |
| RVSTPQGLVKRFSTG | 1 | 0 | 0 | 1640989 | Q2YHF0.1 | P17763 |
| RVVAAEMEEALRGLP | 1 | 0 | 0 | 1640991 | P14340.2 | P17763 |
| RWCFDGIKNNQILEE | 1 | 0 | 0 | 1640998 | P14340.2 | P17763 |
| RWCFDGVKNNQILEE | 1 | 0 | 0 | 1640999 | P29990.1 | P17763 |
| RWGSFKKNGAIKVLR | 1 | 0 | 0 | 1641000 | P17763.2 | P17763 |
| RYQTPAIKTEHTGRE | 1 | 0 | 0 | 1641004 | QNG98968.1 | P17763 |
| RYQTPAVKSEHTGRE | 1 | 0 | 0 | 1641006 | Q2YHF0.1 | P17763 |
| SAAIDGEYRLKGESR | 1 | 0 | 0 | 1641007 | QTX93479.1 | P17763 |
| SAAIKDERAVHADMG | 1 | 0 | 0 | 1641009 | QTX93541.1 | P17763 |
| SAAIKDQKAVHADMG | 1 | 0 | 0 | 1641010 | Q2YHF0.1 | P17763 |
| SAAQRRGRIGRNPAQ | 1 | 0 | 0 | 1641012 | Q2YHF0.1 | P17763 |
| SAAQRRGRVGRNPKN | 1 | 0 | 0 | 1641014 | QQW40786.1 | P17763 |
| SAAVDGEYRLRGEAR | 1 | 0 | 0 | 1641018 | ACW82995.1 | P17763 |
| SAAVKDERAVHADMG | 1 | 0 | 0 | 1641019 | Q6YMS3.1 | P17763 |
| SAEGAWRQVEKVETW | 1 | 0 | 0 | 1641020 | QTX93479.1 | P17763 |
| SALTLHWFRKGSSIG | 1 | 0 | 0 | 1641030 | Q2YHF0.1 | P17763 |
| SAREAVEDGRFWELV | 1 | 0 | 0 | 1641033 | QTE05819.1 | P17763 |
| SAREAVEDSGFWELV | 1 | 0 | 0 | 1641034 | QQW40742.1 | P17763 |
| SASEAVNDSRFWELV | 1 | 0 | 0 | 1641035 | Q2YHF0.1 | P17763 |
| SASRAWNVWEVEDYG | 1 | 0 | 0 | 1641036 | QTX93479.1 | P17763 |
| SATPLAENTNSVTNI | 1 | 0 | 0 | 1641038 | AKQ00034.1 | P17763 |
| SAVNMTSRMLLNRFT | 1 | 0 | 0 | 1641039 | P17763.2 | P17763 |
| SAVPSHWVPTSRTTW | 1 | 0 | 0 | 1641040 | P14340.2 | P17763 |
| SAVPVHWVPTSRTTW | 1 | 0 | 0 | 1641043 | QTX93489.1 | P17763 |
| SAWTLYAVATTIITP | 1 | 0 | 0 | 1641044 | P17763.2 | P17763 |
| SAWTLYAVATTILTP | 1 | 0 | 0 | 1641045 | Q2YHF0.1 | P17763 |
| SAWTLYAVATTVITP | 1 | 0 | 0 | 1641046 | P27909.2 | P17763 |
| SCGTGNIVSAVNMTS | 1 | 0 | 0 | 1641048 | P17763.2 | P17763 |
| SCVTTMAKDKPTLDI | 1 | 0 | 0 | 1641049 | P17763.2 | P17763 |
| SCVTTMAKNKPTLDI | 1 | 0 | 0 | 1641050 | P27909.2 | P17763 |
| SDKMGMGTTYLALMA | 1 | 0 | 0 | 1641054 | P17763.2 | P17763 |
| SDLKYSWKTWGKAKI | 1 | 0 | 0 | 1641055 | ANK35834.1 | P17763 |
| SDRMGMGTTYLALMA | 1 | 0 | 0 | 1641056 | QFS19562.1 | P17763 |
| SDRMGMGVTYLALIA | 1 | 0 | 0 | 1641057 | QTX93489.1 | P17763 |
| SECPNERRAWNFFEV | 1 | 0 | 0 | 1641060 | QTX92256.1 | P17763 |
| SECPNERRAWNFLEV | 1 | 0 | 0 | 1641061 | Q2YHF0.1 | P17763 |
| SECPNERRAWNSFEV | 1 | 0 | 0 | 1641062 | AKQ00032.1 | P17763 |
| SECPNERRAWNSLEV | 1 | 0 | 0 | 1641063 | AHK05342.1 | P17763 |
| SEGAWKHAQRIETWI | 1 | 0 | 0 | 1641065 | P14340.2 | P17763 |
| SEGAWKHAQRVESWI | 1 | 0 | 0 | 1641066 | Q2YHF0.1 | P17763 |
| SEGAWKHVQRIETWI | 1 | 0 | 0 | 1641067 | P29991.1 | P17763 |
| SEGAWKQIQKVETWA | 1 | 0 | 0 | 1641068 | P17763.2 | P17763 |
| SEGAWKQIQRVETWA | 1 | 0 | 0 | 1641069 | QFS19562.1 | P17763 |
| SEGFQYSDRRWCFDG | 1 | 0 | 0 | 1641071 | P17763.2 | P17763 |
| SEGIKYTDRKWCFDG | 1 | 0 | 0 | 1641072 | QTX93489.1 | P17763 |
| SEHTGREIVDLMCHA | 1 | 0 | 0 | 1641074 | P27909.2 | P17763 |
| SEIPNLDIIGKRIEK | 1 | 0 | 0 | 1641076 | P14340.2 | P17763 |
| SETFSMGLLCLTLFV | 1 | 0 | 0 | 1641080 | Q2YHF0.1 | P17763 |
| SETPNLDIIGKRIEK | 1 | 0 | 0 | 1641081 | Q9WDA6.1 | P17763 |
| SFKDLGRVVVMVGAT | 1 | 0 | 0 | 1641085 | QTX93151.1 | P17763 |
| SFKLEKEVAETQHGT | 1 | 0 | 0 | 1641086 | P17763.2 | P17763 |
| SFRDLGRVMVMVGAA | 1 | 0 | 0 | 1641089 | QQD36453.1 | P17763 |
| SFRDLGRVMVMVGAT | 1 | 0 | 0 | 1641090 | P14340.2 | P17763 |
| SGALWDVPSPAAAQK | 1 | 0 | 0 | 1641095 | Q2YHF0.1 | P17763 |
| SGALWDVPSPAATQK | 1 | 0 | 0 | 1641096 | Q5UCB8.1 | P17763 |
| SGDDCVVKPIDDRFA | 1 | 0 | 0 | 1641098 | P17763.2 | P17763 |
| SGDDCVVKPLDDRFA | 1 | 0 | 0 | 1641099 | P14340.2 | P17763 |
| SGEENAVGNDTGKHG | 1 | 0 | 0 | 1641105 | QQW40784.1 | P17763 |
| SGFWELVDKERNLHL | 1 | 0 | 0 | 1641106 | QQW40742.1 | P17763 |
| SGIDFNEMILMKMKT | 1 | 0 | 0 | 1641110 | Q2YHF0.1 | P17763 |
| SGIFVIDNVHTWTEQ | 1 | 0 | 0 | 1641111 | Q2YHF0.1 | P17763 |
| SGIFVTNEVHTWTEQ | 1 | 0 | 0 | 1641112 | P17763.2 | P17763 |
| SGIFVVDNVHTWTEQ | 1 | 0 | 0 | 1641113 | AXB26598.1 | P17763 |
| SGINMCTLIAMDLGE | 1 | 0 | 0 | 1641114 | QTX93489.1 | P17763 |
| SGKELKCGSGIFVID | 1 | 0 | 0 | 1641115 | Q2YHF0.1 | P17763 |
| SGNIVSSVNTISKML | 1 | 0 | 0 | 1641120 | AXG50965.1 | P17763 |
| SGQITWRDMAHTLIM | 1 | 0 | 0 | 1641121 | QTX93489.1 | P17763 |
| SGQVGTYGLNTFTNM | 1 | 0 | 0 | 1641122 | P17763.2 | P17763 |
| SGQVVTYGLNTFTNM | 1 | 0 | 0 | 1641123 | Q9WDA6.1 | P17763 |
| SGRELKCGSGIFVVD | 1 | 0 | 0 | 1641124 | AKQ00034.1 | P17763 |
| SGRIGGQIHLAIMAV | 1 | 0 | 0 | 1641125 | QPJ20200.1 | P17763 |
| SGRIGGQTHLAIMAV | 1 | 0 | 0 | 1641126 | AKQ00032.1 | P17763 |
| SGRMGGQIHLAIMAV | 1 | 0 | 0 | 1641127 | QTX92385.1 | P17763 |

| | | | | | | |
|---|---|---|---|---|---|---|
| SGSASSMVNGVVKLL | 1 | 0 | 0 | 1641128 | QFS19562.1 | P17763 |
| SGSASSMVNGVVRLL | 1 | 0 | 0 | 1641129 | P17763.2 | P17763 |
| SGSSADLSLEKAAEV | 1 | 0 | 0 | 1641131 | P17763.2 | P17763 |
| SGTTTIFAGHLKCRL | 1 | 0 | 0 | 1641135 | P17763.2 | P17763 |
| SGVEGEGLHKLGYIL | 1 | 0 | 0 | 1641136 | P17763.2 | P17763 |
| SGVEGEGLHRLGYIL | 1 | 0 | 0 | 1641137 | QTX92694.1 | P17763 |
| SGVLWDTPSPPEVER | 1 | 0 | 0 | 1641138 | P17763.2 | P17763 |
| SGVLWDTPSPPKVER | 1 | 0 | 0 | 1641139 | QTX92784.1 | P17763 |
| SGVLWDVPSPPETQK | 1 | 0 | 0 | 1641140 | QTX93593.1 | P17763 |
| SHWVPTSRTTWSIHA | 1 | 0 | 0 | 1641146 | QTX93208.1 | P17763 |
| SIALDLVTEIGRVPS | 1 | 0 | 0 | 1641147 | QTX93593.1 | P17763 |
| SIATQQPESNILDID | 1 | 0 | 0 | 1641148 | QTX93077.1 | P17763 |
| SIDPFPQSNSPIEDI | 1 | 0 | 0 | 1641149 | ANW72071.1 | P17763 |
| SIGKMFEATARGARR | 1 | 0 | 0 | 1641150 | P17763.2 | P17763 |
| SIGKMFESTYRGAKR | 1 | 0 | 0 | 1641151 | QTX92385.1 | P17763 |
| SIGQMFETTMRGAKR | 1 | 0 | 0 | 1641152 | QTX93024.1 | P17763 |
| SIHAHHQWMTTEDML | 1 | 0 | 0 | 1641153 | P17763.2 | P17763 |
| SIKNEEEEQTLTILI | 1 | 0 | 0 | 1641159 | QTX93024.1 | P17763 |
| SIKSGNDIANCLRKN | 1 | 0 | 0 | 1641160 | P17763.2 | P17763 |
| SIPATLFVWYFWQKK | 1 | 0 | 0 | 1641164 | P17763.2 | P17763 |
| SIPATLLVWHTWQKQ | 1 | 0 | 0 | 1641165 | QTX93491.1 | P17763 |
| SIPITAAAWYLWEVK | 1 | 0 | 0 | 1641167 | QTX93024.1 | P17763 |
| SIPPTAGILKRWGQL | 1 | 0 | 0 | 1641168 | QBH74475.1 | P17763 |
| SITLDILTEIASLPT | 1 | 0 | 0 | 1641171 | QTX92385.1 | P17763 |
| SITLDILTEIATLPT | 1 | 0 | 0 | 1641172 | ANW72071.1 | P17763 |
| SITTQESESNILDID | 1 | 0 | 0 | 1641174 | QTX93073.1 | P17763 |
| SITTQQPESNILDID | 1 | 0 | 0 | 1641175 | QTX93112.1 | P17763 |
| SIVSLFPLCLSTTSQ | 1 | 0 | 0 | 1641176 | P17763.2 | P17763 |
| SKADLENPHLLEKKI | 1 | 0 | 0 | 1641178 | QTX93551.1 | P17763 |
| SKGWHDWQQVPFCSH | 1 | 0 | 0 | 1641181 | QTX93593.1 | P17763 |
| SKGWNDWTQVPFCSH | 1 | 0 | 0 | 1641183 | QTX93077.1 | P17763 |
| SKIKHAVSRGSSKIR | 1 | 0 | 0 | 1641185 | QPJ20200.1 | P17763 |
| SKIKYAVSRGTSKIR | 1 | 0 | 0 | 1641187 | QTX92385.1 | P17763 |
| SKMDIGVPLLAIGCY | 1 | 0 | 0 | 1641189 | QTX93024.1 | P17763 |
| SKNQTWQIEKASLIE | 1 | 0 | 0 | 1641192 | QTX92385.1 | P17763 |
| SKPTLDIELQKTEAT | 1 | 0 | 0 | 1641194 | QTX93521.1 | P17763 |
| SKSEFNTYKRSGIME | 1 | 0 | 0 | 1641195 | QFS19562.1 | P17763 |
| SKTKYAVSRGTSKIR | 1 | 0 | 0 | 1641196 | QTX92372.1 | P17763 |
| SKVRSNAAIGAVFQE | 1 | 0 | 0 | 1641197 | QTX92385.1 | P17763 |
| SLAAIANQAAVLMGL | 1 | 0 | 0 | 1641198 | QTX92385.1 | P17763 |
| SLGLLCVSIMIEEVM | 1 | 0 | 0 | 1641203 | QPB40130.1 | P17763 |
| SLIGLTSRATWAKNI | 1 | 0 | 0 | 1641204 | QTX93024.1 | P17763 |
| SLIGLTSRATWAQNI | 1 | 0 | 0 | 1641205 | ABZ90965.1 | P17763 |
| SLRTTTASGKLVTQW | 1 | 0 | 0 | 1641215 | QTX92385.1 | P17763 |
| SLRTTTVTGKIIHEW | 1 | 0 | 0 | 1641216 | QFS19562.1 | P17763 |
| SLTAIANQAAILMGL | 1 | 0 | 0 | 1641217 | P17763.2 | P17763 |
| SLTLNLITEMGRLPT | 1 | 0 | 0 | 1641219 | QTX93024.1 | P17763 |
| SLVASVELPNSLEEL | 1 | 0 | 0 | 1641220 | P17763.2 | P17763 |
| SMAICSAVPTEWFPT | 1 | 0 | 0 | 1641224 | QTX92385.1 | P17763 |
| SMANIFRGSYLAGAG | 1 | 0 | 0 | 1641227 | P17763.2 | P17763 |
| SMGLLCLTLFVEECL | 1 | 0 | 0 | 1641231 | QTX92385.1 | P17763 |
| SMKRFRKEEESEGAI | 1 | 0 | 0 | 1641232 | ABZ90965.1 | P17763 |
| SMRKTDWLPMAVAAM | 1 | 0 | 0 | 1641239 | AXX75610.1 | P17763 |
| SMRKTDWLPMTVAAM | 1 | 0 | 0 | 1641240 | QTX93593.1 | P17763 |
| SNAAIGAVFQEEQGW | 1 | 0 | 0 | 1641241 | QTX92385.1 | P17763 |
| SNAALGAIFTDENKW | 1 | 0 | 0 | 1641242 | QTX93024.1 | P17763 |
| SNAFVLKKEVSETQH | 1 | 0 | 0 | 1641243 | QHR82521.1 | P17763 |
| SNAPIQDEERDIPER | 1 | 0 | 0 | 1641244 | QTX93593.1 | P17763 |
| SNAVIQDEERDIPER | 1 | 0 | 0 | 1641245 | P17763.2 | P17763 |
| SNGTGNIVSSVNMVS | 1 | 0 | 0 | 1641249 | QTX93593.1 | P17763 |
| SNGVLESEMIIPKIY | 1 | 0 | 0 | 1641250 | P17763.2 | P17763 |
| SNGVLESEMIIPKNL | 1 | 0 | 0 | 1641251 | QTX93201.1 | P17763 |
| SNGVLESQMLIPKAY | 1 | 0 | 0 | 1641252 | QTX92385.1 | P17763 |
| SNGVLESQMLIPKSY | 1 | 0 | 0 | 1641253 | QDZ58857.1 | P17763 |
| SNGVLESQMLIPRSY | 1 | 0 | 0 | 1641254 | QPJ20200.1 | P17763 |
| SPAAAQKATLTEGVY | 1 | 0 | 0 | 1641259 | QTX92385.1 | P17763 |
| SPAATQKAALSEGVY | 1 | 0 | 0 | 1641260 | QDZ58857.1 | P17763 |
| SPAATQKATLSEGVY | 1 | 0 | 0 | 1641261 | QPJ20200.1 | P17763 |
| SPGKFWNTTIAVSMA | 1 | 0 | 0 | 1641264 | P17763.2 | P17763 |
| SPIIDKKGKVVGLYG | 1 | 0 | 0 | 1641268 | QTX93201.1 | P17763 |
| SPIINKKGKVIGLYG | 1 | 0 | 0 | 1641269 | QPJ20200.1 | P17763 |
| SPIINRKGKVIGLYG | 1 | 0 | 0 | 1641270 | QTX92385.1 | P17763 |
| SPIVDKKGKVVGLYG | 1 | 0 | 0 | 1641272 | QQW40772.1 | P17763 |
| SPIVDRKGKVVGLYG | 1 | 0 | 0 | 1641273 | QTX93073.1 | P17763 |
| SPIVNREGKIVGLYG | 1 | 0 | 0 | 1641274 | P17763.2 | P17763 |
| SPPETQKAELEEGVY | 1 | 0 | 0 | 1641277 | QTX93593.1 | P17763 |
| SPPEVERAVLDDGIY | 1 | 0 | 0 | 1641278 | P17763.2 | P17763 |
| SPPKVERAVLDDGIY | 1 | 0 | 0 | 1641279 | QTX92784.1 | P17763 |
| SPPPMGKAELEDGAY | 1 | 0 | 0 | 1641280 | QTX93201.1 | P17763 |
| SPPPVEKAELEDGAY | 1 | 0 | 0 | 1641281 | QUD37232.1 | P17763 |
| SPSELETPNLAERVL | 1 | 0 | 0 | 1641287 | P17763.2 | P17763 |
| SPTVEESRTIRVLKM | 1 | 0 | 0 | 1641289 | QTX93593.1 | P17763 |
| SQGAGWSLKETACLG | 1 | 0 | 0 | 1641297 | QTX92829.1 | P17763 |
| SQGAGWSLRETACLG | 1 | 0 | 0 | 1641298 | P17763.2 | P17763 |
| SQQKADWIPLALTIK | 1 | 0 | 0 | 1641309 | QTX93213.1 | P17763 |
| SQQKTDWIPLALTIK | 1 | 0 | 0 | 1641310 | QTX93201.1 | P17763 |
| SQTGEHRRDKRSVAL | 1 | 0 | 0 | 1641314 | P17763.2 | P17763 |
| SQVNPLTLTAAVLML | 1 | 0 | 0 | 1641320 | QFS19562.1 | P17763 |
| SQVNPTTLTASLVML | 1 | 0 | 0 | 1641321 | QTX92385.1 | P17763 |
| SRDGEPRMIVGKNER | 1 | 0 | 0 | 1641326 | QTX93593.1 | P17763 |
| SRDPFPQSNAPIIDE | 1 | 0 | 0 | 1641327 | QTX93201.1 | P17763 |

| | | | | | | |
|---|---|---|---|---|---|---|
| SRDPFPQSNAPIMDE | 1 | 0 | 0 | 1641328 | QTX93213.1 | P17763 |
| SRFWELVDKERALHQ | 1 | 0 | 0 | 1641329 | QTX92385.1 | P17763 |
| SRFWELVDKERNLHL | 1 | 0 | 0 | 1641330 | QTX93201.1 | P17763 |
| SRFWELVDRERNLHL | 1 | 0 | 0 | 1641331 | QGW05386.1 | P17763 |
| SRKDQRGSGQVGTYG | 1 | 0 | 0 | 1641340 | QTX92977.1 | P17763 |
| SRKEFDLYKKSGITE | 1 | 0 | 0 | 1641341 | QTX93474.1 | P17763 |
| SRNKKPRICTREEFT | 1 | 0 | 0 | 1641345 | P17763.2 | P17763 |
| SRRDQRGSGQVGTYG | 1 | 0 | 0 | 1641346 | P17763.2 | P17763 |
| SRWSRKMLMTGTLAV | 1 | 0 | 0 | 1641348 | P17763.2 | P17763 |
| SSEGAWKHAQRIETW | 1 | 0 | 0 | 1641352 | QTX93201.1 | P17763 |
| SSEGAWKHAQRVESW | 1 | 0 | 0 | 1641353 | QTX92385.1 | P17763 |
| SSEGAWKHVQRIETW | 1 | 0 | 0 | 1641354 | QTX93073.1 | P17763 |
| SSEGAWKQIQRVETW | 1 | 0 | 0 | 1641355 | QFS19562.1 | P17763 |
| SSIAARGYISTRVGM | 1 | 0 | 0 | 1641358 | QFS19562.1 | P17763 |
| SSKPEFCIKVLNPYM | 1 | 0 | 0 | 1641359 | QTX92385.1 | P17763 |
| SSMINGVVKLLTKPW | 1 | 0 | 0 | 1641365 | QTX93593.1 | P17763 |
| SSMVNGVVKLLTKPW | 1 | 0 | 0 | 1641366 | QFS19562.1 | P17763 |
| SSMVNGVVRLLTKPW | 1 | 0 | 0 | 1641367 | P17763.2 | P17763 |
| SSTPFAENTNSVTNI | 1 | 0 | 0 | 1641372 | QTX92385.1 | P17763 |
| STANIFRGSYLAGAG | 1 | 0 | 0 | 1641378 | QTX92385.1 | P17763 |
| STELHNHTFLIDGPE | 1 | 0 | 0 | 1641381 | QQW40798.1 | P17763 |
| STEPHNQTFLIDGPE | 1 | 0 | 0 | 1641382 | QGW51895.1 | P17763 |
| STESYNQTFLIDGPE | 1 | 0 | 0 | 1641383 | QTX93077.1 | P17763 |
| STGLFSGKGPLRMVL | 1 | 0 | 0 | 1641384 | QBA83313.1 | P17763 |
| STGSASSMVNGVVKL | 1 | 0 | 0 | 1641385 | QTX92385.1 | P17763 |
| STHEMYWISNGTGNI | 1 | 0 | 0 | 1641386 | QTX93593.1 | P17763 |
| STHEMYWVSCGTGNI | 1 | 0 | 0 | 1641387 | P17763.2 | P17763 |
| STSQETWNRQDLLVT | 1 | 0 | 0 | 1641396 | P17763.2 | P17763 |
| STWHYDDENPYKTWA | 1 | 0 | 0 | 1641399 | AAC64119.1 | P17763 |
| STWHYDEDNPYKTWA | 1 | 0 | 0 | 1641400 | P17763.2 | P17763 |
| STYGWNIVKLMSGKD | 1 | 0 | 0 | 1641401 | AAC63341.1 | P17763 |
| STYGWNLVRLQSGVD | 1 | 0 | 0 | 1641402 | QTX93024.1 | P17763 |
| SVAARGYISTRVEMG | 1 | 0 | 0 | 1641403 | QTX92385.1 | P17763 |
| SVALAPHVGMGLDTR | 1 | 0 | 0 | 1641405 | QTX93593.1 | P17763 |
| SVALTPHSGMGLETR | 1 | 0 | 0 | 1641406 | QTX92385.1 | P17763 |
| SVELPNSLEELGDGL | 1 | 0 | 0 | 1641409 | P17763.2 | P17763 |
| SVIEKMEALQRKYGG | 1 | 0 | 0 | 1641420 | QTX93201.1 | P17763 |
| SVIEKMEALQRQYGG | 1 | 0 | 0 | 1641421 | ACW82894.1 | P17763 |
| SVIEKMETLQRKYGG | 1 | 0 | 0 | 1641422 | QTX93189.1 | P17763 |
| SVSGDLILEIGKLPQ | 1 | 0 | 0 | 1641434 | P17763.2 | P17763 |
| SWEDIPYLGKREDLW | 1 | 0 | 0 | 1641438 | QTX92385.1 | P17763 |
| SWGKAKIIGADVQNT | 1 | 0 | 0 | 1641441 | P17763.2 | P17763 |
| SWILRNPGFALLAGF | 1 | 0 | 0 | 1641442 | QTX92385.1 | P17763 |
| SWKLEKASFIEVKTC | 1 | 0 | 0 | 1641443 | QTX93593.1 | P17763 |
| SWPLNEAVMAVGMVS | 1 | 0 | 0 | 1641447 | QTX93073.1 | P17763 |
| SWPLNEGIMAIGIVS | 1 | 0 | 0 | 1641448 | QUD37222.1 | P17763 |
| SWPLNEGIMAVGIVS | 1 | 0 | 0 | 1641449 | P17763.2 | P17763 |
| SWPLNEGVMAVGLVS | 1 | 0 | 0 | 1641450 | QTX93593.1 | P17763 |
| SYLAGAGLAFSLIKN | 1 | 0 | 0 | 1641466 | QTX92385.1 | P17763 |
| SYLAGAGLAFSLMKS | 1 | 0 | 0 | 1641467 | P17763.2 | P17763 |
| TAAGIMKNPTVDGIM | 1 | 0 | 0 | 1641484 | QTX93593.1 | P17763 |
| TAAVLMLVAHYAIIG | 1 | 0 | 0 | 1641485 | QFS19562.1 | P17763 |
| TAEAGGRAYNHALSE | 1 | 0 | 0 | 1641487 | QTX93024.1 | P17763 |
| TAETQNSSFIIDGPN | 1 | 0 | 0 | 1641488 | QTX93593.1 | P17763 |
| TAEWLWKELGKKKTP | 1 | 0 | 0 | 1641489 | QTX93024.1 | P17763 |
| TAGADTLEVHWNHKE | 1 | 0 | 0 | 1641490 | QTX92372.1 | P17763 |
| TAGADTSEVHWNHKE | 1 | 0 | 0 | 1641491 | QTX92321.1 | P17763 |
| TAGADTSEVHWNYKE | 1 | 0 | 0 | 1641492 | QPJ20200.1 | P17763 |
| TAGIFLFFMQGKGIG | 1 | 0 | 0 | 1641493 | QTX92372.1 | P17763 |
| TAGIFLFFMQGKGMG | 1 | 0 | 0 | 1641494 | QTX92385.1 | P17763 |
| TAGILKRWGQLKKNK | 1 | 0 | 0 | 1641495 | QBH74475.1 | P17763 |
| TAIFLTTLSRTNKKR | 1 | 0 | 0 | 1641496 | QTX93201.1 | P17763 |
| TAINQVRSLIGNEEY | 1 | 0 | 0 | 1641497 | AAG15002.1 | P17763 |
| TAINQVRSLIGSEEY | 1 | 0 | 0 | 1641498 | QGW51895.1 | P17763 |
| TAIQQVRSLIGNEEF | 1 | 0 | 0 | 1641499 | ABZ90965.1 | P17763 |
| TAKEVALLRTYCIEA | 1 | 0 | 0 | 1641500 | QTX92385.1 | P17763 |
| TALLIVSGIFPYSIP | 1 | 0 | 0 | 1641504 | QTX93593.1 | P17763 |
| TANISLTAIANQAAI | 1 | 0 | 0 | 1641506 | P17763.2 | P17763 |
| TANPIVTDKEKPVNI | 1 | 0 | 0 | 1641507 | P17763.2 | P17763 |
| TANWLWALLGKKKNP | 1 | 0 | 0 | 1641508 | QOV09272.1 | P17763 |
| TANWLWALLGKKKSP | 1 | 0 | 0 | 1641509 | QPJ20200.1 | P17763 |
| TANWLWALLGRKKNP | 1 | 0 | 0 | 1641510 | QTX92391.1 | P17763 |
| TANWLWTLLGRKKNP | 1 | 0 | 0 | 1641511 | QTX92385.1 | P17763 |
| TAQIMEVTARWLWGF | 1 | 0 | 0 | 1641512 | P17763.2 | P17763 |
| TARATWATNIQVAIN | 1 | 0 | 0 | 1641513 | P17763.2 | P17763 |
| TARGARRMAILGDTA | 1 | 0 | 0 | 1641514 | P17763.2 | P17763 |
| TASLVMLLVHYAIIG | 1 | 0 | 0 | 1641516 | QTX92385.1 | P17763 |
| TASSALLWMASVEPH | 1 | 0 | 0 | 1641517 | QUD37222.1 | P17763 |
| TATGEHRREKRSVAL | 1 | 0 | 0 | 1641519 | QQW40772.1 | P17763 |
| TATITPQAPTTEIQL | 1 | 0 | 0 | 1641520 | QFS19562.1 | P17763 |
| TATITPRSPSVEVKL | 1 | 0 | 0 | 1641521 | QTX92385.1 | P17763 |
| TCSNTIFTLTVAWRT | 1 | 0 | 0 | 1641528 | QTX93593.1 | P17763 |
| TCVYNMMGKREKKIG | 1 | 0 | 0 | 1641529 | QGW51895.1 | P17763 |
| TDAPCKIPFSTQDEK | 1 | 0 | 0 | 1641530 | QFS19562.1 | P17763 |
| TDFPGKTVWFVPSIK | 1 | 0 | 0 | 1641533 | P17763.2 | P17763 |
| TDIMEPEHALLAKSI | 1 | 0 | 0 | 1641537 | QTX92825.1 | P17763 |
| TDIMEPEHALLATSI | 1 | 0 | 0 | 1641538 | P17763.2 | P17763 |
| TDISEMGANFRADRV | 1 | 0 | 0 | 1641539 | P17763.2 | P17763 |
| TDPCSVAARGYISTR | 1 | 0 | 0 | 1641543 | QPJ20200.1 | P17763 |
| TDPSSVAARGYISTR | 1 | 0 | 0 | 1641544 | ACY70714.1 | P17763 |
| TDVKWEDQAEISGSS | 1 | 0 | 0 | 1641546 | QTX93201.1 | P17763 |

| | | | | | | |
|---|---|---|---|---|---|---|
| TDWIPLALTIKGLNP | 1 | 0 | 0 | 1641549 | QTX93201.1 | P17763 |
| TDWLPMAVAAMGVPP | 1 | 0 | 0 | 1641550 | AXX75610.1 | P17763 |
| TDWLPMTVAAMGVPP | 1 | 0 | 0 | 1641551 | QTX93491.1 | P17763 |
| TEAILPEYGTLGLEC | 1 | 0 | 0 | 1641553 | AHG23230.1 | P17763 |
| TEDGQGKAHNGRLIT | 1 | 0 | 0 | 1641555 | QTX93593.1 | P17763 |
| TEDMLSVWNRVWIEE | 1 | 0 | 0 | 1641556 | P17763.2 | P17763 |
| TEENQWDSARAAVED | 1 | 0 | 0 | 1641557 | QTX93593.1 | P17763 |
| TEHTGREIVDLMCHA | 1 | 0 | 0 | 1641561 | QGW05344.1 | P17763 |
| TEKPDMTIIGRRLQR | 1 | 0 | 0 | 1641562 | CCM44290.1 | P17763 |
| TELKYSWKAWGKAKM | 1 | 0 | 0 | 1641563 | QQW40798.1 | P17763 |
| TEPPFGESYIVVGAG | 1 | 0 | 0 | 1641564 | QUD37222.1 | P17763 |
| TEQVDTLLCDIGESS | 1 | 0 | 0 | 1641566 | QTX92385.1 | P17763 |
| TEYQKTKNNDWDYVV | 1 | 0 | 0 | 1641572 | P17763.2 | P17763 |
| TFDSEYIKTRTNDWD | 1 | 0 | 0 | 1641573 | QSV27725.1 | P17763 |
| TFDSEYVKTRANDWD | 1 | 0 | 0 | 1641574 | QTX93073.1 | P17763 |
| TFDSEYVKTRTNDWD | 1 | 0 | 0 | 1641575 | QTX93201.1 | P17763 |
| TFDTEYPKTKLTDWD | 1 | 0 | 0 | 1641576 | QTX92385.1 | P17763 |
| TFIKTTFSLHYAWKT | 1 | 0 | 0 | 1641579 | QFS19562.1 | P17763 |
| TFIKTTLSLDYAWKT | 1 | 0 | 0 | 1641580 | QUD37222.1 | P17763 |
| TFVELMRRGDLPVWL | 1 | 0 | 0 | 1641592 | P17763.2 | P17763 |
| TFVKTTFSLHYAWKT | 1 | 0 | 0 | 1641593 | P17763.2 | P17763 |
| TGAPCKVPIEIRDVN | 1 | 0 | 0 | 1641594 | QTX92385.1 | P17763 |
| TGATEIQTSGTTTIF | 1 | 0 | 0 | 1641596 | P17763.2 | P17763 |
| TGDQHQVGNDTQGVT | 1 | 0 | 0 | 1641598 | QTX93593.1 | P17763 |
| TGDQHQVGNESTEHG | 1 | 0 | 0 | 1641599 | QFS19562.1 | P17763 |
| TGDQHQVGNETQGVT | 1 | 0 | 0 | 1641600 | QHR82521.1 | P17763 |
| TGDQHQVGNETTEHG | 1 | 0 | 0 | 1641601 | P17763.2 | P17763 |
| TGERNNQILEENMEV | 1 | 0 | 0 | 1641604 | QTX92385.1 | P17763 |
| TGETLGEKWKNRLNA | 1 | 0 | 0 | 1641605 | QQW40798.1 | P17763 |
| TGETLGEKWKNRLNS | 1 | 0 | 0 | 1641606 | QGW51895.1 | P17763 |
| TGETLGEKWKRQLNS | 1 | 0 | 0 | 1641607 | QTX92385.1 | P17763 |
| TGFRKEIGRMLNILN | 1 | 0 | 0 | 1641608 | QBH74438.1 | P17763 |
| TGGAMLFLISGKGIG | 1 | 0 | 0 | 1641609 | QTX93593.1 | P17763 |
| TGGIFLFLMSGKGIG | 1 | 0 | 0 | 1641610 | QTX93213.1 | P17763 |
| TGGIFLFLMSGRGIG | 1 | 0 | 0 | 1641611 | QTX93201.1 | P17763 |
| TGGVTLFFLSGKGLG | 1 | 0 | 0 | 1641612 | QFS19562.1 | P17763 |
| TGGVTLFFLSGRGLG | 1 | 0 | 0 | 1641613 | P17763.2 | P17763 |
| TGKEIVDLMCHATFT | 1 | 0 | 0 | 1641615 | P17763.2 | P17763 |
| TGLLVISGLFPISIP | 1 | 0 | 0 | 1641620 | QTX93077.1 | P17763 |
| TGLLVISGLFPVSIP | 1 | 0 | 0 | 1641621 | QTX93201.1 | P17763 |
| TGLLVISGVFPVSIP | 1 | 0 | 0 | 1641622 | QTX93073.1 | P17763 |
| TGNMSFKDLGRVVVM | 1 | 0 | 0 | 1641624 | QTX93201.1 | P17763 |
| TGNMSFRDLGRVMVM | 1 | 0 | 0 | 1641625 | QTX93213.1 | P17763 |
| TGPLVAGGLLTVCYV | 1 | 0 | 0 | 1641626 | QTX93024.1 | P17763 |
| TGRSADLELERATDV | 1 | 0 | 0 | 1641628 | QTX93201.1 | P17763 |
| TGSASSMINGVVKLL | 1 | 0 | 0 | 1641629 | QTX93593.1 | P17763 |
| TGSQLAKRFSKGLLN | 1 | 0 | 0 | 1641630 | AHG23230.1 | P17763 |
| TGSQLAKRFSKGLLS | 1 | 0 | 0 | 1641631 | AAT76483.1 | P17763 |
| TGSQLAKRFSRGLLN | 1 | 0 | 0 | 1641632 | QTX93593.1 | P17763 |
| TGTSADLTVEKAADV | 1 | 0 | 0 | 1641633 | QTX93593.1 | P17763 |
| THFDNTQVGTLALSL | 1 | 0 | 0 | 1641637 | QTX92372.1 | P17763 |
| THRRPTIEKDVDLGA | 1 | 0 | 0 | 1641639 | QTX93491.1 | P17763 |
| TIAVSMANIFRGSYL | 1 | 0 | 0 | 1641643 | P17763.2 | P17763 |
| TIAVSTANIFRGSYL | 1 | 0 | 0 | 1641644 | QTX92385.1 | P17763 |
| TIFAGHLKCRLKMDK | 1 | 0 | 0 | 1641648 | P17763.2 | P17763 |
| TIFTLTVAWRTATLI | 1 | 0 | 0 | 1641649 | QTX93593.1 | P17763 |
| TIGIALLSQSTIPET | 1 | 0 | 0 | 1641651 | QTX93073.1 | P17763 |
| TIGLSLVASVELPNS | 1 | 0 | 0 | 1641652 | QFS19562.1 | P17763 |
| TIKGLNPTAIFLTTL | 1 | 0 | 0 | 1641655 | QTX93024.1 | P17763 |
| TILIKVEYKGEDVPC | 1 | 0 | 0 | 1641660 | QUD37256.1 | P17763 |
| TIMDLHPGAGKTKRI | 1 | 0 | 0 | 1641661 | QTX92385.1 | P17763 |
| TIMDLHPGAGKTKRY | 1 | 0 | 0 | 1641662 | QTX93024.1 | P17763 |
| TISEDGSMSIKNEEE | 1 | 0 | 0 | 1641665 | QTX93024.1 | P17763 |
| TKKDFGLGSIATQQP | 1 | 0 | 0 | 1641675 | QQW40724.1 | P17763 |
| TKKDFGLGSITTQQP | 1 | 0 | 0 | 1641676 | QQW40798.1 | P17763 |
| TKKDLGIGHVAAENH | 1 | 0 | 0 | 1641677 | QFS19562.1 | P17763 |
| TKKDLGIGHVAVENH | 1 | 0 | 0 | 1641678 | QTX92825.1 | P17763 |
| TKKDLGLGSIATQQP | 1 | 0 | 0 | 1641679 | QTX93223.1 | P17763 |
| TKKDLGLGSITTQES | 1 | 0 | 0 | 1641680 | QTX93073.1 | P17763 |
| TKLTDWDFVVTTDIS | 1 | 0 | 0 | 1641682 | QTX92385.1 | P17763 |
| TKPWDVIPMVTQLAM | 1 | 0 | 0 | 1641686 | QTX92631.1 | P17763 |
| TKPWDVIPMVTQMAM | 1 | 0 | 0 | 1641687 | QQW40772.1 | P17763 |
| TKPWDVLPTVTQMAM | 1 | 0 | 0 | 1641688 | QTX93201.1 | P17763 |
| TKPWDVVPMVTQLAM | 1 | 0 | 0 | 1641689 | QTX92385.1 | P17763 |
| TKPWDVVPMVTQMAM | 1 | 0 | 0 | 1641690 | QTX93073.1 | P17763 |
| TKRDLGMSKEPGVVS | 1 | 0 | 0 | 1641691 | QTX93593.1 | P17763 |
| TLFFLSGRGLGKTSI | 1 | 0 | 0 | 1641697 | P17763.2 | P17763 |
| TLILAGVSLLPVCQS | 1 | 0 | 0 | 1641702 | QTX93374.1 | P17763 |
| TLKGMSYVMCTGSFK | 1 | 0 | 0 | 1641704 | P17763.2 | P17763 |
| TLKGVSYVMCTGSFK | 1 | 0 | 0 | 1641705 | QPD98754.1 | P17763 |
| TLKNVTEVKGYTKGG | 1 | 0 | 0 | 1641706 | QTX92372.1 | P17763 |
| TLPEEQDQNYVCKHT | 1 | 0 | 0 | 1641708 | QTX93593.1 | P17763 |
| TLPPLRFKGEDGCWY | 1 | 0 | 0 | 1641709 | P17763.2 | P17763 |
| TLPPLRFRGEDGCWY | 1 | 0 | 0 | 1641710 | QUD37222.1 | P17763 |
| TLPPLRYMGEDGCWY | 1 | 0 | 0 | 1641711 | QTX93593.1 | P17763 |
| TLQLPEDIEQMANGI | 1 | 0 | 0 | 1641712 | QTX93593.1 | P17763 |
| TLRVLKMVEPWLRGN | 1 | 0 | 0 | 1641715 | P17763.2 | P17763 |
| TLRVLKMVEPWLSSK | 1 | 0 | 0 | 1641716 | QTX92385.1 | P17763 |
| TLSEGVYRIMQRGLF | 1 | 0 | 0 | 1641717 | ANW72071.1 | P17763 |
| TLTEGVYRIMQRGLF | 1 | 0 | 0 | 1641719 | QTX92385.1 | P17763 |
| TLVEEQDANFVCRRT | 1 | 0 | 0 | 1641722 | QFS19562.1 | P17763 |

| | | | | | | |
|---|---|---|---|---|---|---|
| TMAKDKPTLDIELLK | 1 | 0 | 0 | 1641726 | P17763.2 | P17763 |
| TMAKNKPTLDFELIK | 1 | 0 | 0 | 1641727 | QTX93024.1 | P17763 |
| TMAKNKPTLDIELLK | 1 | 0 | 0 | 1641728 | QUD37222.1 | P17763 |
| TMAKNKPTLDIELQK | 1 | 0 | 0 | 1641729 | QTX93593.1 | P17763 |
| TMAKSKPTLDIELQK | 1 | 0 | 0 | 1641730 | QTX93521.1 | P17763 |
| TMAQGKPTLDFELIK | 1 | 0 | 0 | 1641732 | QTX92385.1 | P17763 |
| TMAQGKPTLDFELTK | 1 | 0 | 0 | 1641733 | QPJ20200.1 | P17763 |
| TMPPLRFLGEDGCWY | 1 | 0 | 0 | 1641736 | QTX92385.1 | P17763 |
| TMRGAKRMAILGDTA | 1 | 0 | 0 | 1641738 | QTX93024.1 | P17763 |
| TNIELEPPFGDSYIV | 1 | 0 | 0 | 1641743 | QTX92385.1 | P17763 |
| TNIWLKLKEKQDAFC | 1 | 0 | 0 | 1641744 | QTX93201.1 | P17763 |
| TNIWLKLKERQDVFC | 1 | 0 | 0 | 1641745 | QQW40741.1 | P17763 |
| TNIWLKLRDSYTQMC | 1 | 0 | 0 | 1641746 | QUD37222.1 | P17763 |
| TNIWLKLRDSYTQVC | 1 | 0 | 0 | 1641747 | P17763.2 | P17763 |
| TNIWLKLREMYTQLC | 1 | 0 | 0 | 1641748 | QHR82521.1 | P17763 |
| TNIWLKLREVYTQLC | 1 | 0 | 0 | 1641749 | QBK46952.1 | P17763 |
| TNIWLKLREVYTQSC | 1 | 0 | 0 | 1641750 | QTX93593.1 | P17763 |
| TNIWMKFREGSSEVC | 1 | 0 | 0 | 1641751 | QTX92385.1 | P17763 |
| TNMEAQLIRQMEGEG | 1 | 0 | 0 | 1641754 | QTX93189.1 | P17763 |
| TNMEVQLIRQMEAEG | 1 | 0 | 0 | 1641755 | QTX92385.1 | P17763 |
| TNMGAQLIRQMEGEG | 1 | 0 | 0 | 1641756 | QSV27725.1 | P17763 |
| TPEARNSTFLIDGPD | 1 | 0 | 0 | 1641757 | QPJ20200.1 | P17763 |
| TPEGIIPALFEPERE | 1 | 0 | 0 | 1641758 | P17763.2 | P17763 |
| TPETRNSTFLVDGPD | 1 | 0 | 0 | 1641759 | QTX92385.1 | P17763 |
| TPHSGMGLETRAETW | 1 | 0 | 0 | 1641761 | QTX92385.1 | P17763 |
| TPMLRHSIENSSVNV | 1 | 0 | 0 | 1641764 | QTX93024.1 | P17763 |
| TPMLRHTIENSTANV | 1 | 0 | 0 | 1641765 | QTX93593.1 | P17763 |
| TPPASDLKYSWKTWG | 1 | 0 | 0 | 1641768 | QDZ58857.1 | P17763 |
| TPPGATDPFPQSNSP | 1 | 0 | 0 | 1641769 | QDZ58857.1 | P17763 |
| TPPGSIDPFPQSNSP | 1 | 0 | 0 | 1641770 | ANW72071.1 | P17763 |
| TPPGSRDPFPQSNAP | 1 | 0 | 0 | 1641771 | QTX93024.1 | P17763 |
| TPPGTTDPFPQSNSP | 1 | 0 | 0 | 1641772 | QTX92385.1 | P17763 |
| TPPVNDLKYSWKTWG | 1 | 0 | 0 | 1641773 | QPJ20200.1 | P17763 |
| TPPVSDLKYSWKTWG | 1 | 0 | 0 | 1641774 | AFX65870.1 | P17763 |
| TPQAPTSEIQLTDYG | 1 | 0 | 0 | 1641775 | P17763.2 | P17763 |
| TPQAPTTEIQLTDYG | 1 | 0 | 0 | 1641776 | QFS19562.1 | P17763 |
| TPQPMELKYSWKTWG | 1 | 0 | 0 | 1641777 | QTX93593.1 | P17763 |
| TPRSPSVEVKLPDYG | 1 | 0 | 0 | 1641778 | QTX92385.1 | P17763 |
| TPVHSWEDIPYLGKR | 1 | 0 | 0 | 1641779 | QTX92385.1 | P17763 |
| TQFDNTQVGTLALSL | 1 | 0 | 0 | 1641783 | QDZ58857.1 | P17763 |
| TQHGTILIKVEYKGE | 1 | 0 | 0 | 1641784 | QTX93593.1 | P17763 |
| TQKAALSEGVYRIMQ | 1 | 0 | 0 | 1641785 | QDZ58857.1 | P17763 |
| TQKAELEEGVYRIKQ | 1 | 0 | 0 | 1641786 | QTX93593.1 | P17763 |
| TQKATLSEGVYRIMQ | 1 | 0 | 0 | 1641787 | QPJ20200.1 | P17763 |
| TQQMDPEHRQLANAI | 1 | 0 | 0 | 1641795 | QTX93593.1 | P17763 |
| TQSGERRREKRSVAL | 1 | 0 | 0 | 1641796 | QTX92385.1 | P17763 |
| TQWCCRSCTMPPLRF | 1 | 0 | 0 | 1641799 | QTX92385.1 | P17763 |
| TRAETWMSSEGAWKH | 1 | 0 | 0 | 1641801 | AHG06340.1 | P17763 |
| TRANDWDFVVTTDIS | 1 | 0 | 0 | 1641802 | QTX93073.1 | P17763 |
| TRAQTWMSAEGAWRQ | 1 | 0 | 0 | 1641803 | QHR82515.1 | P17763 |
| TRDALDNLAVLHTAE | 1 | 0 | 0 | 1641806 | QTX93201.1 | P17763 |
| TRDGEPLMIVAKHER | 1 | 0 | 0 | 1641807 | QTX92385.1 | P17763 |
| TRGAVLMHKGKRIEP | 1 | 0 | 0 | 1641808 | QTX93201.1 | P17763 |
| TRGAVLMHRGKRIEP | 1 | 0 | 0 | 1641809 | QTX93073.1 | P17763 |
| TRGAVLTHNGKRLEP | 1 | 0 | 0 | 1641811 | QTX93491.1 | P17763 |
| TRGGEPHMIVSKQER | 1 | 0 | 0 | 1641812 | P17763.2 | P17763 |
| TRGSVICHESGRLEP | 1 | 0 | 0 | 1641814 | QTX92385.1 | P17763 |
| TRGSVICHETGRLEP | 1 | 0 | 0 | 1641815 | QPJ20200.1 | P17763 |
| TRHRKPTYEKDADLG | 1 | 0 | 0 | 1641816 | QTX92372.1 | P17763 |
| TRHRKPTYEKDVDLG | 1 | 0 | 0 | 1641817 | CCM44290.1 | P17763 |
| TRHRKPTYERDADLG | 1 | 0 | 0 | 1641818 | AFP27206.1 | P17763 |
| TRHVAVEPEVANLDI | 1 | 0 | 0 | 1641819 | P17763.2 | P17763 |
| TRHVNAEPETPNMDV | 1 | 0 | 0 | 1641820 | QTX93593.1 | P17763 |
| TRKVMEITAEWLWRT | 1 | 0 | 0 | 1641824 | QTX93593.1 | P17763 |
| TRLENVMWKQITNEL | 1 | 0 | 0 | 1641826 | QTX92385.1 | P17763 |
| TRMENLLWKQIANEL | 1 | 0 | 0 | 1641827 | QTX93593.1 | P17763 |
| TRNALDNLVMLHTSE | 1 | 0 | 0 | 1641828 | QTX93593.1 | P17763 |
| TRNGEPHMIVGIQEK | 1 | 0 | 0 | 1641829 | QTX93131.1 | P17763 |
| TRNGEPHMIVGRQEK | 1 | 0 | 0 | 1641830 | QTX93073.1 | P17763 |
| TRRYLPAIVREAIKR | 1 | 0 | 0 | 1641835 | QFS19562.1 | P17763 |
| TRTETWMSSEGAWKH | 1 | 0 | 0 | 1641836 | QTX93024.1 | P17763 |
| TRTNDWDFVVTTDIS | 1 | 0 | 0 | 1641838 | QTX93201.1 | P17763 |
| TRTPQPKPGTRMIMT | 1 | 0 | 0 | 1641839 | QPJ20200.1 | P17763 |
| TRTPQPKPGTRMVMT | 1 | 0 | 0 | 1641840 | QDZ58857.1 | P17763 |
| TRTPQPKPGTRVVMT | 1 | 0 | 0 | 1641841 | QTX92372.1 | P17763 |
| TRTQEPKEGTKKLMK | 1 | 0 | 0 | 1641842 | QTX93024.1 | P17763 |
| TRTQTWMSAEGAWRQ | 1 | 0 | 0 | 1641843 | QTX93593.1 | P17763 |
| TRVGTKHAVLLVAVS | 1 | 0 | 0 | 1641847 | QGW51895.1 | P17763 |
| TRVPNYNLIVMDEAH | 1 | 0 | 0 | 1641848 | QTX92385.1 | P17763 |
| TRVVAAEMEEALKGL | 1 | 0 | 0 | 1641849 | QTX93593.1 | P17763 |
| TRVVAAEMEEALRGL | 1 | 0 | 0 | 1641850 | QTX93024.1 | P17763 |
| TRVVASEMAEALKGM | 1 | 0 | 0 | 1641851 | P17763.2 | P17763 |
| TSEHGGRAYRHAVEE | 1 | 0 | 0 | 1641856 | QTX93593.1 | P17763 |
| TSGASTSQETWNRQD | 1 | 0 | 0 | 1641857 | P17763.2 | P17763 |
| TSGTYVSAIAQAKAS | 1 | 0 | 0 | 1641861 | P17763.2 | P17763 |
| TSIGLICVIASSGML | 1 | 0 | 0 | 1641862 | AHG23230.1 | P17763 |
| TSIGLLCVMASSVLL | 1 | 0 | 0 | 1641863 | QFS19562.1 | P17763 |
| TSIGLLCVTASSALL | 1 | 0 | 0 | 1641864 | QUD37222.1 | P17763 |
| TSKELMMATIGIALL | 1 | 0 | 0 | 1641866 | QTX93073.1 | P17763 |
| TSKELMMTTIGIVLL | 1 | 0 | 0 | 1641867 | QTX93201.1 | P17763 |
| TSLLFLNDMGKVRKD | 1 | 0 | 0 | 1641870 | QTX92385.1 | P17763 |

| | | | | | | |
|---|---|---|---|---|---|---|
| TSLSMTCIAVGLVTL | 1 | 0 | 0 | 1641871 | QFS19562.1 | P17763 |
| TSLSMTCIAVGMVTL | 1 | 0 | 0 | 1641872 | P17763.2 | P17763 |
| TSMAMSCIAVGGITL | 1 | 0 | 0 | 1641873 | QTX92385.1 | P17763 |
| TSMAMTCIAVGGITL | 1 | 0 | 0 | 1641874 | QPJ20200.1 | P17763 |
| TSQKTTWLPVLLGSF | 1 | 0 | 0 | 1641875 | QUD37222.1 | P17763 |
| TSQKTTWLPVLLGSL | 1 | 0 | 0 | 1641876 | P17763.2 | P17763 |
| TSRENLLLGVGLAMA | 1 | 0 | 0 | 1641877 | QTX93593.1 | P17763 |
| TSREVLLLTVGLSLV | 1 | 0 | 0 | 1641878 | P17763.2 | P17763 |
| TSRTTWSIHAHHQWM | 1 | 0 | 0 | 1641879 | P17763.2 | P17763 |
| TSYLDVDLHPASAWT | 1 | 0 | 0 | 1641883 | QTX93593.1 | P17763 |
| TTASRCPTQGEPSLN | 1 | 0 | 0 | 1641886 | QQW40772.1 | P17763 |
| TTATRCPTQGEPYLK | 1 | 0 | 0 | 1641887 | QTX92385.1 | P17763 |
| TTAVKSEHTGKEIVD | 1 | 0 | 0 | 1641888 | P17763.2 | P17763 |
| TTAVKSEHTGREIVD | 1 | 0 | 0 | 1641889 | QUD37222.1 | P17763 |
| TTDPFPQSNSPIEDI | 1 | 0 | 0 | 1641891 | QTX92385.1 | P17763 |
| TTDSRCPTQGEAILP | 1 | 0 | 0 | 1641894 | QTX93551.1 | P17763 |
| TTDSRCPTQGEATLP | 1 | 0 | 0 | 1641895 | QTX93593.1 | P17763 |
| TTDSRCPTQGEATLV | 1 | 0 | 0 | 1641896 | P17763.2 | P17763 |
| TTDSRCPTQGEAVLP | 1 | 0 | 0 | 1641897 | AHG23230.1 | P17763 |
| TTEDMLAVWNRVWIQ | 1 | 0 | 0 | 1641898 | QRG47112.1 | P17763 |
| TTEDMLTVWNRVWIE | 1 | 0 | 0 | 1641902 | QTX93593.1 | P17763 |
| TTEDMLTVWNRVWIQ | 1 | 0 | 0 | 1641903 | ADA59898.1 | P17763 |
| TTEDMLTVWNRVWIR | 1 | 0 | 0 | 1641904 | ADA59742.1 | P17763 |
| TTEKGGRAYQHALNE | 1 | 0 | 0 | 1641905 | QPJ20200.1 | P17763 |
| TTERGGKAYQHALNE | 1 | 0 | 0 | 1641906 | QTX92385.1 | P17763 |
| TTERGGRAYQHALNE | 1 | 0 | 0 | 1641907 | QDZ58857.1 | P17763 |
| TTESRCPTQGEPSLN | 1 | 0 | 0 | 1641908 | QTX93073.1 | P17763 |
| TTFITPMLRHSIENS | 1 | 0 | 0 | 1641909 | QSV27725.1 | P17763 |
| TTFSLHYAWKTMAMI | 1 | 0 | 0 | 1641910 | P17763.2 | P17763 |
| TTFVTPMLRHSIENS | 1 | 0 | 0 | 1641911 | QTX93189.1 | P17763 |
| TTGADTSEVHWNYKE | 1 | 0 | 0 | 1641912 | AKQ00032.1 | P17763 |
| TTLSLDYAWKTMAMV | 1 | 0 | 0 | 1641918 | QPB40130.1 | P17763 |
| TTMFGGVSWMIRILI | 1 | 0 | 0 | 1641920 | ARX96505.1 | P17763 |
| TTPLVMAWRTIMAVF | 1 | 0 | 0 | 1641921 | ANW72071.1 | P17763 |
| TTQIMEVTAKWLWGF | 1 | 0 | 0 | 1641922 | QUD37222.1 | P17763 |
| TTTGEHRREKRSVAL | 1 | 0 | 0 | 1641925 | QTX93189.1 | P17763 |
| TTVITPMLRHTIENS | 1 | 0 | 0 | 1641927 | QTX93593.1 | P17763 |
| TTVSGKLIHEWCCRS | 1 | 0 | 0 | 1641928 | QTX93593.1 | P17763 |
| TTWLPVLLGSFGCKP | 1 | 0 | 0 | 1641929 | QUD37222.1 | P17763 |
| TTWLPVLLGSLGCKP | 1 | 0 | 0 | 1641930 | P17763.2 | P17763 |
| TTWSIHAHHQWMTTE | 1 | 0 | 0 | 1641931 | P17763.2 | P17763 |
| TTWSIHAKHEWMTTE | 1 | 0 | 0 | 1641932 | QTX93024.1 | P17763 |
| TTWSIHARHEWMTTE | 1 | 0 | 0 | 1641933 | QNQ18279.1 | P17763 |
| TTWSIHATHEWMTTE | 1 | 0 | 0 | 1641934 | QQW40798.1 | P17763 |
| TVDDDGTMRIKDDET | 1 | 0 | 0 | 1641940 | QTX93491.1 | P17763 |
| TVFFILMMLVAPSYG | 1 | 0 | 0 | 1641941 | QTX92385.1 | P17763 |
| TVFFVLMMLVAPSYG | 1 | 0 | 0 | 1641942 | QPJ20200.1 | P17763 |
| TVGLSLVASVELPNS | 1 | 0 | 0 | 1641944 | P17763.2 | P17763 |
| TVIEELEKLQRKHGG | 1 | 0 | 0 | 1641948 | QTX92321.1 | P17763 |
| TVIEELEKLQRRHGG | 1 | 0 | 0 | 1641949 | QTX92385.1 | P17763 |
| TVIEELEKLQRRYGG | 1 | 0 | 0 | 1641950 | QTX92177.1 | P17763 |
| TVLMGLGKGWPLSKM | 1 | 0 | 0 | 1641960 | QTX93024.1 | P17763 |
| TVQQLTKRFSLGMLQ | 1 | 0 | 0 | 1641964 | QCF42071.2 | P17763 |
| TVSQLAKRFSKGLLS | 1 | 0 | 0 | 1641967 | P17763.2 | P17763 |
| TVTQMAMTDTTPFGQ | 1 | 0 | 0 | 1641968 | QTX93201.1 | P17763 |
| TWAFRHPGFTILALF | 1 | 0 | 0 | 1641972 | QHR82521.1 | P17763 |
| TWALRHPGFTILALF | 1 | 0 | 0 | 1641973 | QTX93593.1 | P17763 |
| TWALRHPGFTVIAFF | 1 | 0 | 0 | 1641974 | QUD37222.1 | P17763 |
| TWALRHPGFTVIALF | 1 | 0 | 0 | 1641975 | P17763.2 | P17763 |
| TWATNIQVAINQVRR | 1 | 0 | 0 | 1641976 | P17763.2 | P17763 |
| TWAYHGSYEVKATGS | 1 | 0 | 0 | 1641977 | QTX93593.1 | P17763 |
| TWAYHGSYEVKPSGS | 1 | 0 | 0 | 1641978 | P17763.2 | P17763 |
| TWGKAKIFTPEARNS | 1 | 0 | 0 | 1641979 | QPJ20200.1 | P17763 |
| TWGKAKIFTPETRNS | 1 | 0 | 0 | 1641980 | QTX92385.1 | P17763 |
| TWGKAKMLSTEPHNQ | 1 | 0 | 0 | 1641981 | QGW51895.1 | P17763 |
| TWGKAKMLSTESYNQ | 1 | 0 | 0 | 1641982 | QTX93077.1 | P17763 |
| TWILRHPGFTIMAAI | 1 | 0 | 0 | 1641985 | QTX93024.1 | P17763 |
| TWKIEKASFIEVKNC | 1 | 0 | 0 | 1641986 | QTX93201.1 | P17763 |
| TWKLARASFIEVKTC | 1 | 0 | 0 | 1641987 | P17763.2 | P17763 |
| TWKMEKASFIEVKSC | 1 | 0 | 0 | 1641988 | QTX93073.1 | P17763 |
| TWLVHKQWFLDLPLP | 1 | 0 | 0 | 1641990 | AHN50411.1 | P17763 |
| TWMDLLRALIMLGDT | 1 | 0 | 0 | 1641991 | QTX92385.1 | P17763 |
| TWMSAEGAWRQVEKV | 1 | 0 | 0 | 1641992 | QTX93593.1 | P17763 |
| TWMSSEGAWKHAQRI | 1 | 0 | 0 | 1641993 | QTX93201.1 | P17763 |
| TWMSSEGAWKHAQRV | 1 | 0 | 0 | 1641994 | QTX92385.1 | P17763 |
| TWMSSEGAWKHVQRI | 1 | 0 | 0 | 1641995 | QTX93073.1 | P17763 |
| TWMSSEGAWKQIQKV | 1 | 0 | 0 | 1641996 | P17763.2 | P17763 |
| TWMSSEGAWKQIQRV | 1 | 0 | 0 | 1641997 | QFS19562.1 | P17763 |
| TWQIEKASLIEVKTC | 1 | 0 | 0 | 1641998 | QTX92385.1 | P17763 |
| TWSIHAHHQWMTTED | 1 | 0 | 0 | 1642002 | P17763.2 | P17763 |
| TWVMYGTCTQSGERR | 1 | 0 | 0 | 1642003 | QTX92346.1 | P17763 |
| TWVTYGTCNQAGEHR | 1 | 0 | 0 | 1642004 | QTX93593.1 | P17763 |
| TWVTYGTCSQTGEHR | 1 | 0 | 0 | 1642005 | P17763.2 | P17763 |
| TWVTYGTCTATGEHR | 1 | 0 | 0 | 1642006 | QQW40772.1 | P17763 |
| TWVTYGTCTTTGEHR | 1 | 0 | 0 | 1642007 | QTX93189.1 | P17763 |
| TYLALMATFKMRPMF | 1 | 0 | 0 | 1642009 | QFS19562.1 | P17763 |
| TYLALMATFRMRPMF | 1 | 0 | 0 | 1642010 | P17763.2 | P17763 |
| TYRGAKRMAILGETA | 1 | 0 | 0 | 1642014 | QTX92385.1 | P17763 |
| TYSDPLALKEFKDFA | 1 | 0 | 0 | 1642015 | QTX93593.1 | P17763 |
| TYSDPLALREFKEFA | 1 | 0 | 0 | 1642016 | P17763.2 | P17763 |
| VAEIQPQWIAASIIL | 1 | 0 | 0 | 1642019 | QTX92385.1 | P17763 |

| | | | | | | |
|---|---|---|---|---|---|---|
| VAGDVKGVLSKGKRA | 1 | 0 | 0 | 1642021 | QTX92385.1 | P17763 |
| VAGDVKGVLTKGKRA | 1 | 0 | 0 | 1642022 | QPJ20200.1 | P17763 |
| VAGGLLIACYVITGT | 1 | 0 | 0 | 1642023 | QTX93593.1 | P17763 |
| VAGGLLTVCYVLTGR | 1 | 0 | 0 | 1642024 | QTX93024.1 | P17763 |
| VAGILAQGKKMIRPQ | 1 | 0 | 0 | 1642025 | QFS19562.1 | P17763 |
| VALAPHVGMGLDTRA | 1 | 0 | 0 | 1642029 | QHR82515.1 | P17763 |
| VALAPHVGMGLDTRT | 1 | 0 | 0 | 1642030 | QTX93593.1 | P17763 |
| VALLRTYCIEALISN | 1 | 0 | 0 | 1642033 | AKQ00034.1 | P17763 |
| VALLRTYCIEASISN | 1 | 0 | 0 | 1642034 | QTX92385.1 | P17763 |
| VALTPHSGMGLETRA | 1 | 0 | 0 | 1642035 | QTX92385.1 | P17763 |
| VALVPHVGMGLETRT | 1 | 0 | 0 | 1642036 | QTX93024.1 | P17763 |
| VAVQDWLARVGRERL | 1 | 0 | 0 | 1642043 | QGW51895.1 | P17763 |
| VDENQWNSAKEAVED | 1 | 0 | 0 | 1642051 | P17763.2 | P17763 |
| VDLMCHATFTMRLLS | 1 | 0 | 0 | 1642054 | P17763.2 | P17763 |
| VDLMCHATFTTRLLS | 1 | 0 | 0 | 1642055 | ADU76222.1 | P17763 |
| VDLVLEHGGCVTTMA | 1 | 0 | 0 | 1642056 | QTX92385.1 | P17763 |
| VDNFSLGVLGMALFL | 1 | 0 | 0 | 1642057 | QTX93201.1 | P17763 |
| VDNFTMGVLCLAILF | 1 | 0 | 0 | 1642058 | QTX93474.1 | P17763 |
| VDRLKRMAISGDDCV | 1 | 0 | 0 | 1642061 | ACY70845.1 | P17763 |
| VDSFSLGILCVSIMI | 1 | 0 | 0 | 1642062 | QOW96385.1 | P17763 |
| VDSFSLGLLCISIMI | 1 | 0 | 0 | 1642063 | P17763.2 | P17763 |
| VDSFSLGLLCVSIMI | 1 | 0 | 0 | 1642064 | QPB40130.1 | P17763 |
| VDVVLEHGSCVTTMA | 1 | 0 | 0 | 1642066 | P17763.2 | P17763 |
| VDYMPAMKRYSAPFE | 1 | 0 | 0 | 1642067 | ANW72071.1 | P17763 |
| VDYMPAMRRYSAHFE | 1 | 0 | 0 | 1642068 | QTX92385.1 | P17763 |
| VDYMPVMKRYSAHFE | 1 | 0 | 0 | 1642069 | QTX92391.1 | P17763 |
| VEAILPEYGTLGLEC | 1 | 0 | 0 | 1642070 | QTX93474.1 | P17763 |
| VEAVLPEYGTLGLEC | 1 | 0 | 0 | 1642071 | QTX93593.1 | P17763 |
| VEDEEFWKLVDRERE | 1 | 0 | 0 | 1642073 | QTX93593.1 | P17763 |
| VEDERFWDLVHRERE | 1 | 0 | 0 | 1642074 | P17763.2 | P17763 |
| VEELPETMETLLLLG | 1 | 0 | 0 | 1642075 | QTX93593.1 | P17763 |
| VEKAADVTWEEEAEQ | 1 | 0 | 0 | 1642077 | QTX93593.1 | P17763 |
| VEKAELEDGAYRIKQ | 1 | 0 | 0 | 1642078 | QUD37232.1 | P17763 |
| VEKVETWALRHPGFT | 1 | 0 | 0 | 1642080 | QTX93593.1 | P17763 |
| VERAVLDDGIYRILQ | 1 | 0 | 0 | 1642084 | P17763.2 | P17763 |
| VERAVLDDGIYRIMQ | 1 | 0 | 0 | 1642085 | QFS19562.1 | P17763 |
| VEVEIWTKEGERKKL | 1 | 0 | 0 | 1642086 | QTX93208.1 | P17763 |
| VFHTMWHVTRGAVLM | 1 | 0 | 0 | 1642091 | P17763.2 | P17763 |
| VFHTMWHVTRGAVLT | 1 | 0 | 0 | 1642092 | QTX93350.1 | P17763 |
| VFHTMWHVTRGSVIC | 1 | 0 | 0 | 1642093 | QTX92385.1 | P17763 |
| VFKEKVDTRTPKAKR | 1 | 0 | 0 | 1642094 | P17763.2 | P17763 |
| VFKEKVDTRTPRAKR | 1 | 0 | 0 | 1642095 | QUD37222.1 | P17763 |
| VFKEKVDTRTPRPMP | 1 | 0 | 0 | 1642096 | QTX93593.1 | P17763 |
| VFPVSIPITAAAWYL | 1 | 0 | 0 | 1642099 | QTX93073.1 | P17763 |
| VFSGDPLKNDEDHAH | 1 | 0 | 0 | 1642102 | Q58HT7.1 | P17763 |
| VFSGDPLRNDEDHAH | 1 | 0 | 0 | 1642103 | QTX92385.1 | P17763 |
| VGGLLTSLGKAVHQV | 1 | 0 | 0 | 1642107 | QTX92385.1 | P17763 |
| VGPWHLGKLEIDFGE | 1 | 0 | 0 | 1642113 | QTX92387.1 | P17763 |
| VGRNPKNENDQYIYM | 1 | 0 | 0 | 1642114 | QQW40798.1 | P17763 |
| VHTWTEQYKFQADSP | 1 | 0 | 0 | 1642125 | P17763.2 | P17763 |
| VHWNYKERMVTFKVP | 1 | 0 | 0 | 1642127 | QPJ20200.1 | P17763 |
| VHYAIIGPGLQAKAT | 1 | 0 | 0 | 1642129 | QTX92607.1 | P17763 |
| VIAVEPGKNPKNFQT | 1 | 0 | 0 | 1642130 | QTX93593.1 | P17763 |
| VIAVEPGKNPKNVQT | 1 | 0 | 0 | 1642131 | P17763.2 | P17763 |
| VICHESGRLEPSWAD | 1 | 0 | 0 | 1642132 | QTX92385.1 | P17763 |
| VICHETGRLEPSWAD | 1 | 0 | 0 | 1642133 | QPJ20200.1 | P17763 |
| VIDLEPISYDPKFEK | 1 | 0 | 0 | 1642134 | QTX92385.1 | P17763 |
| VIDLGCGRGGWSYYC | 1 | 0 | 0 | 1642136 | P17763.2 | P17763 |
| VIDNVHTWTEQYKFQ | 1 | 0 | 0 | 1642137 | QTX92385.1 | P17763 |
| VIGIGDNALKINWYK | 1 | 0 | 0 | 1642140 | AHG23230.1 | P17763 |
| VIGIITLYLGAVVQA | 1 | 0 | 0 | 1642141 | QTX93593.1 | P17763 |
| VIGVLEQGKRTLTPQ | 1 | 0 | 0 | 1642142 | AFY10047.1 | P17763 |
| VILAGPIPVTPASAA | 1 | 0 | 0 | 1642149 | QTX92385.1 | P17763 |
| VILAGPMPVTHSSAA | 1 | 0 | 0 | 1642150 | QTX93024.1 | P17763 |
| VINWKGRELKCGSGI | 1 | 0 | 0 | 1642154 | P17763.2 | P17763 |
| VIQLSRKTFDTEYQK | 1 | 0 | 0 | 1642155 | QFS19562.1 | P17763 |
| VISGLFPISIPITAA | 1 | 0 | 0 | 1642157 | QTX93077.1 | P17763 |
| VISGVFPVSIPITAA | 1 | 0 | 0 | 1642159 | QTX93073.1 | P17763 |
| VITFCAIILGGLTWM | 1 | 0 | 0 | 1642160 | ALJ53458.1 | P17763 |
| VITLCAIILGGLTWM | 1 | 0 | 0 | 1642161 | QDZ58857.1 | P17763 |
| VIVTEDCGNRGPSLR | 1 | 0 | 0 | 1642162 | QTX93077.1 | P17763 |
| VKEKEENLVKSMVSA | 1 | 0 | 0 | 1642166 | P17763.2 | P17763 |
| VKEVKGLTKGGPGHE | 1 | 0 | 0 | 1642168 | QQW40798.1 | P17763 |
| VKGVLSKGKRALAPP | 1 | 0 | 0 | 1642171 | QTX92385.1 | P17763 |
| VKGVLTKGKRALTPP | 1 | 0 | 0 | 1642172 | QPJ20200.1 | P17763 |
| VKLLTKPWDVIPMVT | 1 | 0 | 0 | 1642174 | QFS19562.1 | P17763 |
| VKLLTKPWDVVPMVT | 1 | 0 | 0 | 1642175 | QFS19207.1 | P17763 |
| VKLPDYGELTLDCEP | 1 | 0 | 0 | 1642176 | QTX92385.1 | P17763 |
| VKRFSTGLFSGKGPL | 1 | 0 | 0 | 1642179 | QBA83313.1 | P17763 |
| VKVQRPTPKGTVMDI | 1 | 0 | 0 | 1642181 | QTX93593.1 | P17763 |
| VLAIEPGKNPKHVQT | 1 | 0 | 0 | 1642183 | QTX92385.1 | P17763 |
| VLALEPGKNPRAVQT | 1 | 0 | 0 | 1642184 | QTX93024.1 | P17763 |
| VLARWGTFKKSGAIK | 1 | 0 | 0 | 1642186 | QBM00871.1 | P17763 |
| VLDDGIYRIMQRGLL | 1 | 0 | 0 | 1642187 | QFS19562.1 | P17763 |
| VLEENMDVEIWTKEG | 1 | 0 | 0 | 1642188 | P17763.2 | P17763 |
| VLKIVRNMEKYQLAV | 1 | 0 | 0 | 1642193 | QTX93213.1 | P17763 |
| VLLKTALLIVSGIFP | 1 | 0 | 0 | 1642197 | QTX93593.1 | P17763 |
| VLLLSGQITWRDMAH | 1 | 0 | 0 | 1642198 | QTX93593.1 | P17763 |
| VLLLTIGLSLVASVE | 1 | 0 | 0 | 1642199 | QFS19562.1 | P17763 |
| VLLLTVGLSLVASVE | 1 | 0 | 0 | 1642200 | P17763.2 | P17763 |
| VLLQMEDKAWLVHRQ | 1 | 0 | 0 | 1642201 | QRG47133.1 | P17763 |

| | | | | | | |
|---|---|---|---|---|---|---|
| VLLSQSTIPETILEL | 1 | 0 | 0 | 1642202 | QTX93201.1 | P17763 |
| VLLSQSTIPETVLEL | 1 | 0 | 0 | 1642203 | QTX93198.1 | P17763 |
| VLLTWLGLNSRSTSL | 1 | 0 | 0 | 1642205 | QFS19562.1 | P17763 |
| VLLVAVSFMTLITGN | 1 | 0 | 0 | 1642206 | QGW51895.1 | P17763 |
| VLLWMASVEPHWIAA | 1 | 0 | 0 | 1642207 | QFS19562.1 | P17763 |
| VLMHRGKRIEPSWAD | 1 | 0 | 0 | 1642208 | QTX93073.1 | P17763 |
| VLMMRTTWALCEALT | 1 | 0 | 0 | 1642209 | QTX93024.1 | P17763 |
| VLMYQGKRLEPSWAS | 1 | 0 | 0 | 1642210 | P17763.2 | P17763 |
| VLNPYMPTVIEHLER | 1 | 0 | 0 | 1642211 | QTX93593.1 | P17763 |
| VLPEEQDQNYVCKHT | 1 | 0 | 0 | 1642214 | AHG23230.1 | P17763 |
| VLTHNGKRLEPNWAS | 1 | 0 | 0 | 1642218 | QTX93491.1 | P17763 |
| VMAWRTIMAVFFVVT | 1 | 0 | 0 | 1642227 | ANW72071.1 | P17763 |
| VMDIISRRDQRGSGQ | 1 | 0 | 0 | 1642229 | QTX93024.1 | P17763 |
| VMKRYSAHFESEGVL | 1 | 0 | 0 | 1642235 | QTX92391.1 | P17763 |
| VMKRYSAPFESEGVL | 1 | 0 | 0 | 1642236 | QOV09272.1 | P17763 |
| VMLHNSEQGGKAYRH | 1 | 0 | 0 | 1642239 | P17763.2 | P17763 |
| VMLHNSEQGGRAYRH | 1 | 0 | 0 | 1642240 | QFS19562.1 | P17763 |
| VMLHTSEHGGRAYRH | 1 | 0 | 0 | 1642241 | QTX93593.1 | P17763 |
| VMLHTTEKGGRAYQH | 1 | 0 | 0 | 1642242 | QPJ20200.1 | P17763 |
| VMLHTTERGGKAYQH | 1 | 0 | 0 | 1642243 | QTX92385.1 | P17763 |
| VMLHTTERGGRAYQH | 1 | 0 | 0 | 1642244 | QDZ58857.1 | P17763 |
| VMLLVHYAIIGPGLQ | 1 | 0 | 0 | 1642245 | QTX92385.1 | P17763 |
| VMTTTANWLWALLGK | 1 | 0 | 0 | 1642252 | QDZ58857.1 | P17763 |
| VMTTTANWLWALLGR | 1 | 0 | 0 | 1642253 | QTX92391.1 | P17763 |
| VMTTTANWLWTLLGR | 1 | 0 | 0 | 1642254 | QTX92386.1 | P17763 |
| VNGILAQGKKMIRPQ | 1 | 0 | 0 | 1642257 | QMT58614.1 | P17763 |
| VNGVVKLLTKPWDVI | 1 | 0 | 0 | 1642258 | QFS19562.1 | P17763 |
| VNGVVKLLTKPWDVV | 1 | 0 | 0 | 1642259 | QFS19207.1 | P17763 |
| VNGVVRLLTKPWDVL | 1 | 0 | 0 | 1642260 | QTX93201.1 | P17763 |
| VNIEAEPPFGESYIV | 1 | 0 | 0 | 1642261 | P17763.2 | P17763 |
| VNIETEPPFGESYIV | 1 | 0 | 0 | 1642262 | QUD37222.1 | P17763 |
| VNTEPEDIDCWCNLT | 1 | 0 | 0 | 1642269 | QTX92385.1 | P17763 |
| VNVSLTAIANQATVL | 1 | 0 | 0 | 1642272 | QTX93024.1 | P17763 |
| VPCKIPFSTEDGQGK | 1 | 0 | 0 | 1642276 | QUD37256.1 | P17763 |
| VPCRNQDELVGRARV | 1 | 0 | 0 | 1642277 | P17763.2 | P17763 |
| VPFCSHHFHQLIMKD | 1 | 0 | 0 | 1642278 | P17763.2 | P17763 |
| VPHVGMGLETRTETW | 1 | 0 | 0 | 1642280 | QTX93024.1 | P17763 |
| VPPLPLFIFSLKDTL | 1 | 0 | 0 | 1642283 | QTX93374.1 | P17763 |
| VQLIRQMEAEGVITQ | 1 | 0 | 0 | 1642297 | QTX92367.1 | P17763 |
| VQLIRQMEAEGVITR | 1 | 0 | 0 | 1642298 | QTX92385.1 | P17763 |
| VQRIETWILRHPGFT | 1 | 0 | 0 | 1642302 | QTX93073.1 | P17763 |
| VQTAPGTFKTPEGEV | 1 | 0 | 0 | 1642304 | P17763.2 | P17763 |
| VQTKPGIFKTNTGTI | 1 | 0 | 0 | 1642305 | QRG47133.1 | P17763 |
| VQTKPGLFKTLTGEI | 1 | 0 | 0 | 1642306 | QTX92385.1 | P17763 |
| VQTKPGLFKTNTGTI | 1 | 0 | 0 | 1642307 | QTX93201.1 | P17763 |
| VQTTPGTFKTPEGEV | 1 | 0 | 0 | 1642308 | QTX92659.1 | P17763 |
| VREVKGLTKGGPGHE | 1 | 0 | 0 | 1642311 | QTX93024.1 | P17763 |
| VRLLTKPWDVIPMVT | 1 | 0 | 0 | 1642318 | P17763.2 | P17763 |
| VRLLTKPWDVLPTVT | 1 | 0 | 0 | 1642319 | QTX93201.1 | P17763 |
| VRLLTKPWDVVPMVT | 1 | 0 | 0 | 1642320 | QTX93073.1 | P17763 |
| VRNPLSRNSTHEMYW | 1 | 0 | 0 | 1642325 | P17763.2 | P17763 |
| VRRLIGNENYLDYMT | 1 | 0 | 0 | 1642327 | AAG14999.1 | P17763 |
| VRVQRPAKNGTVMDV | 1 | 0 | 0 | 1642329 | P17763.2 | P17763 |
| VSGASGNIVSSVNTI | 1 | 0 | 0 | 1642336 | QPJ20200.1 | P17763 |
| VSGASGNIVSSVNTT | 1 | 0 | 0 | 1642337 | QDZ58857.1 | P17763 |
| VSGVSGNIVSSVNTT | 1 | 0 | 0 | 1642341 | QTX92385.1 | P17763 |
| VSLVLVGVVTLYLGV | 1 | 0 | 0 | 1642347 | AAB33855.1 | P17763 |
| VSNASGNIVSSVNMI | 1 | 0 | 0 | 1642350 | QTX93201.1 | P17763 |
| VSNATGNIVSSVNMI | 1 | 0 | 0 | 1642351 | QTX93073.1 | P17763 |
| VSPLLLTSSQQKADW | 1 | 0 | 0 | 1642352 | QTX93213.1 | P17763 |
| VSSVNTISKMLLNRF | 1 | 0 | 0 | 1642355 | QPJ20200.1 | P17763 |
| VSSVNTTSKMLLNRF | 1 | 0 | 0 | 1642356 | CCM44290.1 | P17763 |
| VSSWEEVPYLGKRED | 1 | 0 | 0 | 1642357 | QUD37222.1 | P17763 |
| VSTEIEKPDMTIIGR | 1 | 0 | 0 | 1642358 | QTX92372.1 | P17763 |
| VSTETEKPDMTIIGR | 1 | 0 | 0 | 1642359 | CCM44290.1 | P17763 |
| VSWIMKIGIGVLLTW | 1 | 0 | 0 | 1642363 | QPC96841.1 | P17763 |
| VSYVMCTGSFKLEKE | 1 | 0 | 0 | 1642367 | QPD98754.1 | P17763 |
| VTEEIAVQNWLARVG | 1 | 0 | 0 | 1642373 | QTX93201.1 | P17763 |
| VTEVKGYTKGGPGHE | 1 | 0 | 0 | 1642375 | P17763.2 | P17763 |
| VTEVRGYTKGGPGHE | 1 | 0 | 0 | 1642376 | QTX93593.1 | P17763 |
| VTHSSAAQRRGRIGR | 1 | 0 | 0 | 1642378 | QTX93024.1 | P17763 |
| VTHSSAAQRRGRVGR | 1 | 0 | 0 | 1642379 | QQW40798.1 | P17763 |
| VTIQEDCDHRGPSLR | 1 | 0 | 0 | 1642381 | QTX92389.1 | P17763 |
| VTIQENCDHRGPSLR | 1 | 0 | 0 | 1642382 | QPJ20200.1 | P17763 |
| VTLITGNMSFKDLGR | 1 | 0 | 0 | 1642384 | QTX93201.1 | P17763 |
| VTLITGNMSFRDLGR | 1 | 0 | 0 | 1642385 | QTX93213.1 | P17763 |
| VTNHMEGEHKKLAEA | 1 | 0 | 0 | 1642389 | QEV86434.1 | P17763 |
| VTNHMKGEHKKLAEA | 1 | 0 | 0 | 1642390 | QTX93073.1 | P17763 |
| VTPASAAQRRGRIGR | 1 | 0 | 0 | 1642392 | QTX92385.1 | P17763 |
| VTQIAMTDTTPFGQQ | 1 | 0 | 0 | 1642395 | P17763.2 | P17763 |
| VTQMAMTDTTPFGQQ | 1 | 0 | 0 | 1642396 | QTX93024.1 | P17763 |
| VTQNGRLITANPIVT | 1 | 0 | 0 | 1642397 | P17763.2 | P17763 |
| VTTLCAIILGGLTWM | 1 | 0 | 0 | 1642400 | QTX92385.1 | P17763 |
| VTVHNGDTHAVGNDT | 1 | 0 | 0 | 1642401 | QTX92385.1 | P17763 |
| VTVHTGDQHQVGNES | 1 | 0 | 0 | 1642402 | QFS19562.1 | P17763 |
| VTVHTGDQHQVGNET | 1 | 0 | 0 | 1642403 | P17763.2 | P17763 |
| VTVQEDCDHRGPSLR | 1 | 0 | 0 | 1642404 | QTX92385.1 | P17763 |
| VTYGLNTFTNMEAQL | 1 | 0 | 0 | 1642405 | QTX93213.1 | P17763 |
| VTYKCPHITEVEPED | 1 | 0 | 0 | 1642406 | QFS19534.1 | P17763 |
| VTYKCPLIAEVEPED | 1 | 0 | 0 | 1642407 | QTX93593.1 | P17763 |
| VTYKCPLITEVEPED | 1 | 0 | 0 | 1642408 | QTX93318.1 | P17763 |

| | | | | | | |
|---|---|---|---|---|---|---|
| VTYKCPLLVNTEPED | 1 | 0 | 0 | 1642409 | QTX92385.1 | P17763 |
| VVDLGCGRGGWSYYC | 1 | 0 | 0 | 1642416 | QTX93024.1 | P17763 |
| VVDLGCGRGGWSYYM | 1 | 0 | 0 | 1642417 | QTX92385.1 | P17763 |
| VVDNVHTWTEQYKFQ | 1 | 0 | 0 | 1642418 | QCE20767.1 | P17763 |
| VVETLEQMQRKHGGM | 1 | 0 | 0 | 1642419 | P17763.2 | P17763 |
| VVGDANGILAQGKKM | 1 | 0 | 0 | 1642425 | QUD37222.1 | P17763 |
| VVGDIIGVLEQGKRT | 1 | 0 | 0 | 1642426 | QTX93551.1 | P17763 |
| VVGDVNGILAQGKKM | 1 | 0 | 0 | 1642429 | QMT58614.1 | P17763 |
| VVITWIGMNSRSTSL | 1 | 0 | 0 | 1642438 | QQW40772.1 | P17763 |
| VVKPIDDRFATALTA | 1 | 0 | 0 | 1642441 | P17763.2 | P17763 |
| VVKVKYEGAGAPCKV | 1 | 0 | 0 | 1642443 | QOV09272.1 | P17763 |
| VVKVKYEGTGAPCKV | 1 | 0 | 0 | 1642444 | QTX92385.1 | P17763 |
| VVKVLRPTPKGAVMD | 1 | 0 | 0 | 1642445 | QTX92385.1 | P17763 |
| VVKVLRPTPRGAVMD | 1 | 0 | 0 | 1642446 | QDZ58857.1 | P17763 |
| VVLGSQEGAMHTALT | 1 | 0 | 0 | 1642450 | P17763.2 | P17763 |
| VVLMGLDKGWPISKM | 1 | 0 | 0 | 1642452 | QTX93593.1 | P17763 |
| VVMVGATMADDIGMG | 1 | 0 | 0 | 1642455 | QTX93045.1 | P17763 |
| VVPCRNQDELIGRAR | 1 | 0 | 0 | 1642462 | QTX93024.1 | P17763 |
| VVPCRPQDELIGRAR | 1 | 0 | 0 | 1642463 | QTX93593.1 | P17763 |
| VVPMVTQMAMTDTTP | 1 | 0 | 0 | 1642465 | QTX93073.1 | P17763 |
| VVQHENLKYTVIITV | 1 | 0 | 0 | 1642469 | QTX93474.1 | P17763 |
| VVRVQRPTPRGTVMD | 1 | 0 | 0 | 1642471 | QTX93024.1 | P17763 |
| VVSPTSYLDVDLHPA | 1 | 0 | 0 | 1642472 | QTX93593.1 | P17763 |
| VVSWNGKELKCGSGI | 1 | 0 | 0 | 1642473 | QPJ20200.1 | P17763 |
| VVSWSGKELKCGSGI | 1 | 0 | 0 | 1642474 | QTX92385.1 | P17763 |
| VVSWSGRELKCGSGI | 1 | 0 | 0 | 1642475 | QDZ58857.1 | P17763 |
| VVTCAKFLCSGKITG | 1 | 0 | 0 | 1642476 | QPJ20200.1 | P17763 |
| VVTCAKFSCSGKITG | 1 | 0 | 0 | 1642477 | QTX92372.1 | P17763 |
| VVTTDISEMGANFKA | 1 | 0 | 0 | 1642479 | QTX93024.1 | P17763 |
| VVTTDISEMGANFRA | 1 | 0 | 0 | 1642480 | P17763.2 | P17763 |
| VVVDEHCGNRGPSLR | 1 | 0 | 0 | 1642482 | P17763.2 | P17763 |
| VWIEDNPNMIDKTPV | 1 | 0 | 0 | 1642487 | QTX92385.1 | P17763 |
| VWIEDNPNMTDKTPV | 1 | 0 | 0 | 1642488 | QPJ20200.1 | P17763 |
| VWLAHKVASEGIKYT | 1 | 0 | 0 | 1642490 | QTX93593.1 | P17763 |
| VWLSYKVASEGFQYS | 1 | 0 | 0 | 1642492 | P17763.2 | P17763 |
| VWNRVWIEDNPNMID | 1 | 0 | 0 | 1642493 | QTX92385.1 | P17763 |
| VWNRVWIEDNPNMTD | 1 | 0 | 0 | 1642494 | QPJ20200.1 | P17763 |
| VYPLSIPATLFVWYF | 1 | 0 | 0 | 1642500 | QTX92825.1 | P17763 |
| VYPMSIPATLFVWYF | 1 | 0 | 0 | 1642501 | P17763.2 | P17763 |
| WADVKKDLISYGGGW | 1 | 0 | 0 | 1642507 | QTX93189.1 | P17763 |
| WADVRKDLISYGGGW | 1 | 0 | 0 | 1642508 | QSV27725.1 | P17763 |
| WADVRNDMISYGGGW | 1 | 0 | 0 | 1642509 | QTX92385.1 | P17763 |
| WALCESITLATGPLT | 1 | 0 | 0 | 1642511 | P17763.2 | P17763 |
| WALCEVLTLATGPIM | 1 | 0 | 0 | 1642512 | ANW72071.1 | P17763 |
| WDKEEDVQVLAIEPG | 1 | 0 | 0 | 1642521 | QTX92385.1 | P17763 |
| WDTPSPPEVERAVLD | 1 | 0 | 0 | 1642524 | P17763.2 | P17763 |
| WDTPSPPKVERAVLD | 1 | 0 | 0 | 1642525 | QTX92784.1 | P17763 |
| WDTRITEDDLHNEEK | 1 | 0 | 0 | 1642526 | QTX93593.1 | P17763 |
| WDTRITEDDLQNEAK | 1 | 0 | 0 | 1642527 | P17763.2 | P17763 |
| WDVPSPAAAQKATLT | 1 | 0 | 0 | 1642529 | QTX92385.1 | P17763 |
| WDVPSPAATQKAALS | 1 | 0 | 0 | 1642530 | QDZ58857.1 | P17763 |
| WDVPSPAATQKATLS | 1 | 0 | 0 | 1642531 | QPJ20200.1 | P17763 |
| WDVPSPPETQKAELE | 1 | 0 | 0 | 1642532 | QTX93593.1 | P17763 |
| WDVPSPPPMGKAELE | 1 | 0 | 0 | 1642533 | QTX93201.1 | P17763 |
| WDVPSPPPVEKAELE | 1 | 0 | 0 | 1642534 | QUD37232.1 | P17763 |
| WEDVPYLGKREDQWC | 1 | 0 | 0 | 1642537 | QFS19562.1 | P17763 |
| WEEEAEHSGASHNIL | 1 | 0 | 0 | 1642538 | P17763.2 | P17763 |
| WEEEAEQTGVSHNLM | 1 | 0 | 0 | 1642541 | QTX93593.1 | P17763 |
| WEEIPYLGKREDQWC | 1 | 0 | 0 | 1642542 | ADA59677.1 | P17763 |
| WEEVPYLGKREDQWC | 1 | 0 | 0 | 1642544 | ADA59905.1 | P17763 |
| WEGNPGRFWNTTIAV | 1 | 0 | 0 | 1642545 | QTX93024.1 | P17763 |
| WEGSPGKFWNTTIAV | 1 | 0 | 0 | 1642546 | P17763.2 | P17763 |
| WEQEAEHSGASHNIL | 1 | 0 | 0 | 1642549 | QPZ88407.1 | P17763 |
| WEVEDYGFGVFTTNI | 1 | 0 | 0 | 1642551 | QIC53463.1 | P17763 |
| WFGRENSWSGVEGEG | 1 | 0 | 0 | 1642552 | QTX92385.1 | P17763 |
| WFSRENSLSGVEGEG | 1 | 0 | 0 | 1642556 | P17763.2 | P17763 |
| WFSRENSWSGVEGEG | 1 | 0 | 0 | 1642557 | QCE20767.1 | P17763 |
| WFSRGNSLSGVEGEG | 1 | 0 | 0 | 1642558 | QTX93073.1 | P17763 |
| WFVPSIKAGNDIANC | 1 | 0 | 0 | 1642559 | QTX92871.1 | P17763 |
| WFVPSIKSGNDIANC | 1 | 0 | 0 | 1642560 | P17763.2 | P17763 |
| WGFLSRNKKPRICTR | 1 | 0 | 0 | 1642563 | P17763.2 | P17763 |
| WGNGCGLFGKGGVVT | 1 | 0 | 0 | 1642564 | ACN94866.1 | P17763 |
| WGNGCGLFGKGSLIT | 1 | 0 | 0 | 1642565 | P17763.2 | P17763 |
| WGNGCGLFGKGSLLT | 1 | 0 | 0 | 1642566 | QUD37222.1 | P17763 |
| WIEDNPWMEDKTPVT | 1 | 0 | 0 | 1642569 | QTX93593.1 | P17763 |
| WIESALNDTWKMEKA | 1 | 0 | 0 | 1642570 | QTX93073.1 | P17763 |
| WIESQKNGSWKLEKA | 1 | 0 | 0 | 1642571 | QTX93593.1 | P17763 |
| WIESSKNQTWQIEKA | 1 | 0 | 0 | 1642572 | QTX92385.1 | P17763 |
| WIESSKNQTWQIERA | 1 | 0 | 0 | 1642573 | QTX92257.1 | P17763 |
| WIGTNSRNTSMAMSC | 1 | 0 | 0 | 1642575 | QTX92385.1 | P17763 |
| WIGTNSRNTSMAMTC | 1 | 0 | 0 | 1642576 | QCE20767.1 | P17763 |
| WIQENPWMEDKTPVE | 1 | 0 | 0 | 1642578 | ADA59846.1 | P17763 |
| WIRENPWMEDKTPVE | 1 | 0 | 0 | 1642579 | ADA59643.1 | P17763 |
| WKEGEEVQVLALEPG | 1 | 0 | 0 | 1642584 | QTX93024.1 | P17763 |
| WKHAQRIETWILRHP | 1 | 0 | 0 | 1642585 | QTX93201.1 | P17763 |
| WKHAQRVESWILRNP | 1 | 0 | 0 | 1642586 | QTX92385.1 | P17763 |
| WKHVQRIETWILRHP | 1 | 0 | 0 | 1642587 | QTX93073.1 | P17763 |
| WKKKLNQLSRKEFDL | 1 | 0 | 0 | 1642588 | QTX93474.1 | P17763 |
| WKNRLNALGKSEFQI | 1 | 0 | 0 | 1642592 | QQW40772.1 | P17763 |
| WKNRLNSLGKSEFQI | 1 | 0 | 0 | 1642593 | QGW51895.1 | P17763 |
| WKNWQEVPFCSHHFH | 1 | 0 | 0 | 1642594 | QTX92385.1 | P17763 |

| | | | | | | |
|---|---|---|---|---|---|---|
| WKQIQKVETWALRHP | 1 | 0 | 0 | 1642595 | P17763.2 | P17763 |
| WKQIQRVETWALRHP | 1 | 0 | 0 | 1642596 | QFS19562.1 | P17763 |
| WKRQLNQLSKSEFNT | 1 | 0 | 0 | 1642597 | P17763.2 | P17763 |
| WKRQLNSLDRKEFEE | 1 | 0 | 0 | 1642598 | QTX92385.1 | P17763 |
| WKSRLNALGKSEFQI | 1 | 0 | 0 | 1642599 | QTX93189.1 | P17763 |
| WKTMAMILSIVSLFP | 1 | 0 | 0 | 1642600 | P17763.2 | P17763 |
| WKTTAMALSIVSLFP | 1 | 0 | 0 | 1642601 | ANC57575.1 | P17763 |
| WKVSCTTLAVVSVSP | 1 | 0 | 0 | 1642602 | QQW40695.1 | P17763 |
| WLAYKVAAEGINYAD | 1 | 0 | 0 | 1642604 | QTX93189.1 | P17763 |
| WLAYRVAAEGINYAD | 1 | 0 | 0 | 1642605 | QTX93213.1 | P17763 |
| WLEKHGAERLKRMAI | 1 | 0 | 0 | 1642606 | QUD37222.1 | P17763 |
| WLEKYGVERLKRMAI | 1 | 0 | 0 | 1642607 | QFS19562.1 | P17763 |
| WLGARFLEFEALGFM | 1 | 0 | 0 | 1642614 | P17763.2 | P17763 |
| WLGARYLEFEALGFL | 1 | 0 | 0 | 1642615 | ACJ09459.1 | P17763 |
| WLGLNSRSTSLSMTC | 1 | 0 | 0 | 1642616 | P17763.2 | P17763 |
| WLSYKVASAGISYKD | 1 | 0 | 0 | 1642621 | QTX92385.1 | P17763 |
| WMIRILIGFLVLWIG | 1 | 0 | 0 | 1642626 | ARX96505.1 | P17763 |
| WMVRILIGFLVLWIG | 1 | 0 | 0 | 1642629 | QPJ20200.1 | P17763 |
| WMVRILIGLLVLWIG | 1 | 0 | 0 | 1642630 | QTX92385.1 | P17763 |
| WNKKELLVTFKNAHA | 1 | 0 | 0 | 1642637 | QHR82521.1 | P17763 |
| WNLVKLHSGKDVFFI | 1 | 0 | 0 | 1642638 | QTX92726.1 | P17763 |
| WNLVKLHSGKDVFFM | 1 | 0 | 0 | 1642639 | QUD37222.1 | P17763 |
| WNLVKLHSGVDVFYK | 1 | 0 | 0 | 1642640 | QTX92385.1 | P17763 |
| WNRVWIEDNPWMEDK | 1 | 0 | 0 | 1642641 | QTX93593.1 | P17763 |
| WNRVWIQENPWMEDK | 1 | 0 | 0 | 1642642 | ADA59898.1 | P17763 |
| WNRVWIRENPWMEDK | 1 | 0 | 0 | 1642643 | ADA59811.1 | P17763 |
| WNSGYDWITDFPGKT | 1 | 0 | 0 | 1642647 | P17763.2 | P17763 |
| WNSGYEWITDFPGKT | 1 | 0 | 0 | 1642648 | QFS19562.1 | P17763 |
| WNTTIAVSMANIFRG | 1 | 0 | 0 | 1642649 | P17763.2 | P17763 |
| WPKTHTLWSNGVLES | 1 | 0 | 0 | 1642653 | QHD26414.1 | P17763 |
| WRQVEKVETWAFRHP | 1 | 0 | 0 | 1642663 | QTX93318.1 | P17763 |
| WRQVEKVETWALRHP | 1 | 0 | 0 | 1642664 | QTX93593.1 | P17763 |
| WRTLGRNKKPRLCTR | 1 | 0 | 0 | 1642665 | QTX93593.1 | P17763 |
| WSLRETACLGKSYAQ | 1 | 0 | 0 | 1642667 | P17763.2 | P17763 |
| WTEAKMLLDNINTPE | 1 | 0 | 0 | 1642669 | P17763.2 | P17763 |
| WTMKIGIGILLTWLG | 1 | 0 | 0 | 1642671 | P17763.2 | P17763 |
| WTMKIGIGVLLTWLG | 1 | 0 | 0 | 1642672 | QFS19562.1 | P17763 |
| WTMKILIGVIITWIG | 1 | 0 | 0 | 1642673 | QTX93189.1 | P17763 |
| WTMKILIGVVITWIG | 1 | 0 | 0 | 1642674 | QQW40772.1 | P17763 |
| WYKKGSSIGKMFEAT | 1 | 0 | 0 | 1642683 | QTX92565.1 | P17763 |
| YADPMALKDFKEFAS | 1 | 0 | 0 | 1642685 | QTX92385.1 | P17763 |
| YADRKWCFDGIKNNQ | 1 | 0 | 0 | 1642686 | QYU75493.1 | P17763 |
| YADRRWCFDGVKNNQ | 1 | 0 | 0 | 1642687 | QQW40798.1 | P17763 |
| YAMCLNAFVLKKEVS | 1 | 0 | 0 | 1642688 | AHG06373.1 | P17763 |
| YAMCLNTFVLKKEVS | 1 | 0 | 0 | 1642689 | QTX93593.1 | P17763 |
| YAMCSNAFVLKKEVS | 1 | 0 | 0 | 1642690 | QHR82521.1 | P17763 |
| YAMCTNTFVLKKEVS | 1 | 0 | 0 | 1642691 | AHG23230.1 | P17763 |
| YAQMWQLMYFHRRDL | 1 | 0 | 0 | 1642694 | P17763.2 | P17763 |
| YAVATTFITPMLRHS | 1 | 0 | 0 | 1642696 | QSV27725.1 | P17763 |
| YAVATTFVTPMLRHS | 1 | 0 | 0 | 1642697 | QTX93189.1 | P17763 |
| YAVATTVITPMLRHT | 1 | 0 | 0 | 1642698 | QTX93350.1 | P17763 |
| YAVSRGTSKIRWIVE | 1 | 0 | 0 | 1642699 | QTX92385.1 | P17763 |
| YDAKFEKQLGQIMLL | 1 | 0 | 0 | 1642700 | P17763.2 | P17763 |
| YDDENPYKTWAYHGS | 1 | 0 | 0 | 1642701 | AAC64119.1 | P17763 |
| YDEDNPYKTWAYHGS | 1 | 0 | 0 | 1642702 | P17763.2 | P17763 |
| YDPKFEKQLGQVMLL | 1 | 0 | 0 | 1642703 | QTX93024.1 | P17763 |
| YDWITDFPGKTVWFV | 1 | 0 | 0 | 1642704 | P17763.2 | P17763 |
| YEAPSTGSASSMVNG | 1 | 0 | 0 | 1642705 | QTX92385.1 | P17763 |
| YETKQTGSASSMVNG | 1 | 0 | 0 | 1642708 | QTX93024.1 | P17763 |
| YEWITDFPGKTVWFV | 1 | 0 | 0 | 1642709 | QFS19562.1 | P17763 |
| YFQRVLIFILLTAVA | 1 | 0 | 0 | 1642714 | QSV27725.1 | P17763 |
| YGGALVRNPLSRNST | 1 | 0 | 0 | 1642715 | QTX93024.1 | P17763 |
| YGGSLVRCPLSRNST | 1 | 0 | 0 | 1642717 | QTX92177.1 | P17763 |
| YGTCNQAGEHRRDKR | 1 | 0 | 0 | 1642724 | QTX93593.1 | P17763 |
| YGTCSQTGEHRRDKR | 1 | 0 | 0 | 1642725 | P17763.2 | P17763 |
| YGTCTATGEHRREKR | 1 | 0 | 0 | 1642726 | QQW40772.1 | P17763 |
| YGTCTQSGERRREKR | 1 | 0 | 0 | 1642727 | QTX92385.1 | P17763 |
| YGTCTTTGEHRREKR | 1 | 0 | 0 | 1642728 | QTX93189.1 | P17763 |
| YGVERLKRMAISGDD | 1 | 0 | 0 | 1642729 | QFS19562.1 | P17763 |
| YHGSYETKQTGSASS | 1 | 0 | 0 | 1642733 | QTX93024.1 | P17763 |
| YIFTGQPLNNDEDHA | 1 | 0 | 0 | 1642734 | QTX93593.1 | P17763 |
| YIIIGVEPGQLKLSW | 1 | 0 | 0 | 1642735 | QSV27725.1 | P17763 |
| YILEDIDKKDGDLIY | 1 | 0 | 0 | 1642736 | QTX92385.1 | P17763 |
| YILEEIDKKDGDLMY | 1 | 0 | 0 | 1642737 | QDZ58857.1 | P17763 |
| YILRDVGKKEGGAMY | 1 | 0 | 0 | 1642738 | QBQ87236.1 | P17763 |
| YILRDVSKKEGGAMY | 1 | 0 | 0 | 1642739 | ABQ18243.1 | P17763 |
| YILREVSKKEGGAMY | 1 | 0 | 0 | 1642740 | QSX24591.1 | P17763 |
| YILWENNIKLTVVVG | 1 | 0 | 0 | 1642741 | QTX93593.1 | P17763 |
| YIVIGVGDSALTLHW | 1 | 0 | 0 | 1642742 | QTX92385.1 | P17763 |
| YIVIGVGNSALTLHW | 1 | 0 | 0 | 1642743 | QDZ58857.1 | P17763 |
| YIVVGAGEKALKLSW | 1 | 0 | 0 | 1642744 | P17763.2 | P17763 |
| YIYMGQPLNNDEDHA | 1 | 0 | 0 | 1642745 | P17763.2 | P17763 |
| YKDREWCFTGERNNQ | 1 | 0 | 0 | 1642747 | QTX92385.1 | P17763 |
| YKEGTFHTMWHVTRG | 1 | 0 | 0 | 1642749 | QTX93024.1 | P17763 |
| YKERMVTFKVPHAKR | 1 | 0 | 0 | 1642750 | QPJ20200.1 | P17763 |
| YKVASEGFQYSDRRW | 1 | 0 | 0 | 1642752 | P17763.2 | P17763 |
| YLAGAGLAFSIMKSV | 1 | 0 | 0 | 1642753 | QTX93593.1 | P17763 |
| YLAGAGLLFSIMKNT | 1 | 0 | 0 | 1642754 | QTX93024.1 | P17763 |
| YLGKREDQWCGSLIG | 1 | 0 | 0 | 1642757 | ABZ90963.1 | P17763 |
| YLKEEQDQQYICRRD | 1 | 0 | 0 | 1642758 | QTX92385.1 | P17763 |
| YMPTVIEHLERLQRK | 1 | 0 | 0 | 1642767 | QTX93593.1 | P17763 |

| | | | | | | |
|---|---|---|---|---|---|---|
| YMTSMKRFKNESDPE | 1 | 0 | 0 | 1642771 | ABZ81545.1 | P17763 |
| YMWLGARYLEFEALG | 1 | 0 | 0 | 1642774 | ACJ09459.1 | P17763 |
| YNHALSELPETLETL | 1 | 0 | 0 | 1642775 | QTX93024.1 | P17763 |
| YNQTFLIDGPETAEC | 1 | 0 | 0 | 1642776 | QTX93077.1 | P17763 |
| YQHALNELPESLETL | 1 | 0 | 0 | 1642784 | QTX92385.1 | P17763 |
| YQNKVVKVLRPTPRG | 1 | 0 | 0 | 1642788 | QDZ58857.1 | P17763 |
| YRHAMEELPDTIETL | 1 | 0 | 0 | 1642794 | P17763.2 | P17763 |
| YRHAVEELPETMETL | 1 | 0 | 0 | 1642795 | QTX93593.1 | P17763 |
| YRQGYATQTVGPWHL | 1 | 0 | 0 | 1642797 | QTX92387.1 | P17763 |
| YSDPLALKEFKEFAA | 1 | 0 | 0 | 1642803 | QTX93024.1 | P17763 |
| YSQIGAGVYKEGTFH | 1 | 0 | 0 | 1642806 | QTX93024.1 | P17763 |
| YSVIVTVHTGDQHQV | 1 | 0 | 0 | 1642809 | P17763.2 | P17763 |
| YSWKAWGKAKMLSTE | 1 | 0 | 0 | 1642811 | QQW40798.1 | P17763 |
| YSWKSWGKAKIIGAD | 1 | 0 | 0 | 1642812 | P17763.2 | P17763 |
| YSWKTWGKAKIFTPE | 1 | 0 | 0 | 1642813 | QTX92385.1 | P17763 |
| YSWKTWGKAKIVTAE | 1 | 0 | 0 | 1642814 | QTX93593.1 | P17763 |
| YTIVITPHSGEENAV | 1 | 0 | 0 | 1642820 | QQW40798.1 | P17763 |
| YTVIITVHTGDQHQV | 1 | 0 | 0 | 1642822 | QTX93593.1 | P17763 |
| YTVVVTVHNGDTHAV | 1 | 0 | 0 | 1642823 | QTX92385.1 | P17763 |
| YVLWEGGHDLTVVAG | 1 | 0 | 0 | 1642826 | QTX92385.1 | P17763 |
| YVSAIAQAKASQEGP | 1 | 0 | 0 | 1642827 | P17763.2 | P17763 |
| YVSAIAQTEKSIEDN | 1 | 0 | 0 | 1642828 | QTX93024.1 | P17763 |
| YVSAITQAERIGEPD | 1 | 0 | 0 | 1642829 | QTX92385.1 | P17763 |
| YVSGIAQTNAEPDGP | 1 | 0 | 0 | 1642834 | QTX93593.1 | P17763 |
| YVVTTDISEMGANFR | 1 | 0 | 0 | 1642835 | P17763.2 | P17763 |
| YVYMGQPLNNDEDHA | 1 | 0 | 0 | 1642836 | QUD37222.1 | P17763 |
| YYCAGLKKVTEVKGY | 1 | 0 | 0 | 1642841 | P17763.2 | P17763 |
| YYCAGLKKVTEVRGY | 1 | 0 | 0 | 1642842 | AAC63341.1 | P17763 |
| YYCGGLKNVREVKGL | 1 | 0 | 0 | 1642844 | QTX93024.1 | P17763 |
| AAALKSLGFDKPQEKD | 1 | 0 | 0 | 1642919 | NP_073556.1 | A0A0N7J131 |
| AAAVDEAFPLDEAEDV | 1 | 0 | 0 | 1642924 | YP_009047202.1 | K9N7C7 |
| AACCHLAKALNDFSNS | 1 | 0 | 0 | 1642933 | YP_009724389.1 | P0DTD1 |
| AACHLAKALQTYSETG | 1 | 0 | 0 | 1642935 | YP_009047202.1 | K9N7C7 |
| AADLGVLIQNGAKHVQ | 1 | 0 | 0 | 1642942 | YP_173236.1 | A0A140H1G9 |
| AADLGVLIQNSAKHVQ | 1 | 0 | 0 | 1642943 | YP_009555238.1 | P0C6X6 |
| AADMDDFSRQLQNSMS | 1 | 0 | 0 | 1642945 | NP_828858.1 | P59595 |
| AADPALHVASASALYD | 1 | 0 | 0 | 1642946 | YP_009555238.1 | P0C6X6 |
| AADPAMHIASSNAFLD | 1 | 0 | 0 | 1642948 | YP_009047202.1 | K9N7C7 |
| AAECTIFKDAAGKPVP | 1 | 0 | 0 | 1642951 | QIQ54046.1 | P0C6U8 |
| AAECTIFKDASGKPVP | 1 | 0 | 0 | 1642953 | YP_009724389.1 | P0DTD1 |
| AAEFKYMVANGLHAPH | 1 | 0 | 0 | 1642955 | YP_003766.2 | H9EJ42 |
| AAESHLDADLTKPYVK | 1 | 0 | 0 | 1642958 | QIQ54047.1 | P0C6X7 |
| AAESKAFVDTIVPPCP | 1 | 0 | 0 | 1642960 | NP_073549.1 | A0A0P0LKV0 |
| AAETHRDCDFNKPLIE | 1 | 0 | 0 | 1642961 | YP_009047202.1 | K9N7C7 |
| AAETQKATEEAFKQSY | 1 | 0 | 0 | 1642965 | YP_009555238.1 | P0C6X6 |
| AAETVKAKEESVKSSY | 1 | 0 | 0 | 1642966 | NP_073549.1 | A0A0P0LKV0 |
| AAFATAQEAYEQAVAN | 1 | 0 | 0 | 1642968 | YP_009724389.1 | P0DTD1 |
| AAFDFSLDVFKIGDVK | 1 | 0 | 0 | 1642969 | YP_003766.2 | H9EJ42 |
| AAFHANSSEPALLFRN | 1 | 0 | 0 | 1642974 | YP_009555241.1 | P36334 |
| AAFHQECSLQSCTQHQ | 1 | 0 | 0 | 1642975 | YP_009724396.1 | P0DTC8 |
| AAGGKVSFSDDVEVKD | 1 | 0 | 0 | 1642983 | NP_073549.1 | A0A0P0LKV0 |
| AAGLEFTDENYNNLVP | 1 | 0 | 0 | 1642990 | YP_173236.1 | A0A140H1G9 |
| AAIDFVQYEVDRRVLF | 1 | 0 | 0 | 1643004 | YP_173236.1 | A0A140H1G9 |
| AAIELSKELKRPVNPT | 1 | 0 | 0 | 1643005 | YP_173236.1 | A0A140H1G9 |
| AAIHYLNSKIVDLAQH | 1 | 0 | 0 | 1643007 | YP_009555238.1 | P0C6X6 |
| AAIITREIGFIVPGLP | 1 | 0 | 0 | 1643008 | NP_828849.2 | P0C6X7 |
| AAIKLVPTTTGVLVRF | 1 | 0 | 0 | 1643011 | YP_003766.2 | H9EJ42 |
| AAIKLYNPKAIHDIGN | 1 | 0 | 0 | 1643012 | NP_073549.1 | A0A0P0LKV0 |
| AAILNGCAWFVKPNRT | 1 | 0 | 0 | 1643015 | YP_009047202.1 | K9N7C7 |
| AAISDYDYYRYNLPTM | 1 | 0 | 0 | 1643025 | YP_009724389.1 | P0DTD1 |
| AAITAPYVVLTGLGES | 1 | 0 | 0 | 1643026 | YP_009047202.1 | K9N7C7 |
| AAITDYNYYKYNLPTM | 1 | 0 | 0 | 1643027 | YP_009555238.1 | P0C6X6 |
| AAITDYNYYSYNLPTM | 1 | 0 | 0 | 1643028 | YP_009047202.1 | K9N7C7 |
| AAITILDGISQYSLRL | 1 | 0 | 0 | 1643029 | YP_009724389.1 | P0DTD1 |
| AAITSGIKFQTVKPGN | 1 | 0 | 0 | 1643030 | YP_173236.1 | A0A140H1G9 |
| AAITSGVKFQTVKPGN | 1 | 0 | 0 | 1643031 | YP_009555238.1 | P0C6X6 |
| AAITTSNCAKRLAQRV | 1 | 0 | 0 | 1643032 | NP_828849.2 | P0C6X7 |
| AAIYGGDHYVVLGGET | 1 | 0 | 0 | 1643035 | YP_003768.1 | H9EJ24 |
| AAKAYKDYLASGGQPI | 1 | 0 | 0 | 1643036 | YP_009724389.1 | P0DTD1 |
| AAKGYIAGIQNFKETL | 1 | 0 | 0 | 1643037 | YP_009047202.1 | K9N7C7 |
| AAKKNNLPFKLTCATT | 1 | 0 | 0 | 1643042 | YP_009724389.1 | P0DTD1 |
| AAKLMVVVPDYGTYKN | 1 | 0 | 0 | 1643044 | NP_828849.2 | P0C6X7 |
| AAKLMVVVPDYNTYKN | 1 | 0 | 0 | 1643045 | QIQ54046.1 | P0C6U8 |
| AAKRVLGLQTQTVDSA | 1 | 0 | 0 | 1643047 | YP_009555238.1 | P0C6X6 |
| AALAMEKVNECVKSQS | 1 | 0 | 0 | 1643051 | YP_173238.1 | A0A140H1H1 |
| AALAVNVCLIPLLLLL | 1 | 0 | 0 | 1643052 | YP_009047208.1 | K9N7D2 |
| AALDLSKELKRPIQPT | 1 | 0 | 0 | 1643054 | YP_009555238.1 | P0C6X6 |
| AALDYLATVKTVPAGN | 1 | 0 | 0 | 1643055 | YP_003766.2 | H9EJ42 |
| AALGVLMSNLGMPSYC | 1 | 0 | 0 | 1643060 | YP_009724389.1 | P0DTD1 |
| AALKSLGFDKPQEKDK | 1 | 0 | 0 | 1643064 | NP_073556.1 | A0A0N7J131 |
| AALLAGSFDKVYDIGN | 1 | 0 | 0 | 1643066 | YP_009047202.1 | K9N7C7 |
| AALNAFVSQVLNKYTE | 1 | 0 | 0 | 1643068 | YP_003767.1 | A0A0P0G321 |
| AALPQSKDSNFLNESG | 1 | 0 | 0 | 1643072 | YP_009047203.1 | K9N7C7 |
| AALPQSKDSNFLNRVR | 1 | 0 | 0 | 1643073 | YP_009047202.1 | K9N7C7 |
| AALQPEEEQEEDWLDD | 1 | 0 | 0 | 1643075 | YP_009724389.1 | P0DTD1 |
| AALSAQLAKDKVNECV | 1 | 0 | 0 | 1643077 | YP_009047204.1 | K9N5Q8 |
| AALSTGLTSQTVKPGH | 1 | 0 | 0 | 1643080 | NP_073549.1 | A0A0P0LKV0 |
| AAMLFDSHIVSKESGN | 1 | 0 | 0 | 1643090 | NP_073556.1 | A0A0N7J131 |
| AANSSEWAYCVDLHNK | 1 | 0 | 0 | 1643109 | YP_003766.2 | H9EJ42 |
| AANTHLKHGGGIAGAI | 1 | 0 | 0 | 1643112 | YP_009047202.1 | K9N7C7 |
| AANTLNIIVPDKSVYD | 1 | 0 | 0 | 1643113 | YP_009555238.1 | P0C6X6 |

| | | | | | | |
|---|---|---|---|---|---|---|
| AAPRAVSFADNNDITN | 1 | 0 | 0 | 1643125 | YP_009047211.1 | K9N4V7 |
| AAQKAYQEAMDSGDTS | 1 | 0 | 0 | 1643130 | YP_009047202.1 | K9N7C7 |
| AARLLGLQTQTVDSAQ | 1 | 0 | 0 | 1643139 | NP_073549.1 | A0A0P0LKV0 |
| AARQGVVDTDVDTKDV | 1 | 0 | 0 | 1643146 | NP_828849.2 | P0C6X7 |
| AASFNKAINNIVASFS | 1 | 0 | 0 | 1643153 | YP_003767.1 | A0A0P0G321 |
| AASFNKAMTNIVDAFT | 1 | 0 | 0 | 1643154 | NP_073551.1 | A0A0P0K6L9 |
| AASYNKYKYYSGSASE | 1 | 0 | 0 | 1643164 | YP_003766.2 | H9EJ42 |
| AASYNRYKYYSGNANE | 1 | 0 | 0 | 1643165 | NP_073549.1 | A0A0P0LKV0 |
| AATLPKGIMMNVAKYT | 1 | 0 | 0 | 1643170 | QIQ54047.1 | P0C6X7 |
| AATRGATVVIGTSKFY | 1 | 0 | 0 | 1643173 | YP_009724389.1 | P0DTD1 |
| AATVAAMFPPWSAAAG | 1 | 0 | 0 | 1643176 | YP_173238.1 | A0A140H1H1 |
| AATVRLQAGNATEVPA | 1 | 0 | 0 | 1643178 | YP_009724389.1 | P0DTD1 |
| AATVRLQAGSNTEFAS | 1 | 0 | 0 | 1643179 | YP_009047202.1 | K9N7C7 |
| AAVDEAFPLDEAEDVT | 1 | 0 | 0 | 1643183 | YP_009047202.1 | K9N7C7 |
| AAVDLSMDKLRRVDWK | 1 | 0 | 0 | 1643184 | YP_009047202.1 | K9N7C7 |
| AAVHGWKMVVCDKVRS | 1 | 0 | 0 | 1643190 | YP_009047202.1 | K9N7C7 |
| AAVITRDVGFVVPGLP | 1 | 0 | 0 | 1643193 | QIQ54046.1 | P0C6U8 |
| AAVKAAESKAFVDTIV | 1 | 0 | 0 | 1643195 | NP_073549.1 | A0A0P0LKV0 |
| AAVKDFDFYRYNKPTI | 1 | 0 | 0 | 1643196 | YP_003766.2 | H9EJ42 |
| AAVTILDGISEQSLRL | 1 | 0 | 0 | 1643203 | NP_828849.2 | P0C6X7 |
| AAVTLALKNLGFDNQS | 1 | 0 | 0 | 1643204 | YP_003771.1 | H9EJ69 |
| AAYATAQEAYEQAVAN | 1 | 0 | 0 | 1643209 | NP_828849.2 | P0C6X7 |
| AAYLDAVKQGAKPVGN | 1 | 0 | 0 | 1643213 | NP_073549.1 | A0A0P0LKV0 |
| AAYNRYRIGNYKLNTD | 1 | 0 | 0 | 1643218 | NP_828855.1 | P59596 |
| AAYVDNSSLTIKKPNE | 1 | 0 | 0 | 1643225 | YP_009724389.1 | P0DTD1 |
| AAYVDNTSLTIKKPNE | 1 | 0 | 0 | 1643226 | QIQ54046.1 | P0C6U8 |
| ACAQYYNGIMVLPGVA | 1 | 0 | 0 | 1643235 | NP_073551.1 | A0A0P0K6L9 |
| ACCHLAKALNDFSNSG | 1 | 0 | 0 | 1643237 | YP_009724389.1 | P0DTD1 |
| ACDGLASAYRANSFDV | 1 | 0 | 0 | 1643242 | YP_009047202.1 | K9N7C7 |
| ACEHISSTMSQYSRST | 1 | 0 | 0 | 1643247 | YP_009047204.1 | K9N5Q8 |
| ACEVVVTNLNKSAGWP | 1 | 0 | 0 | 1643251 | YP_003766.2 | H9EJ42 |
| ACFSKCHNCCDEYGGH | 1 | 0 | 0 | 1643254 | YP_173238.1 | A0A140H1H1 |
| ACGCGASYTANDVEVQ | 1 | 0 | 0 | 1643259 | YP_009047202.1 | K9N7C7 |
| ACGSPGYNLKNGEVEF | 1 | 0 | 0 | 1643260 | NP_073549.1 | A0A0P0LKV0 |
| ACGSPGYNVRNDGTVE | 1 | 0 | 0 | 1643261 | YP_003766.2 | H9EJ42 |
| ACGTNVPLQLGFSTGV | 1 | 0 | 0 | 1643262 | YP_009047202.1 | K9N7C7 |
| ACHNPEVGPEHSLAEY | 1 | 0 | 0 | 1643267 | QHR63299.1 | P0C6W2 |
| ACIKTAQADTAGLYTN | 1 | 0 | 0 | 1643274 | YP_009047205.1 | K9N796 |
| ACKCGVKQEQRTGLDA | 1 | 0 | 0 | 1643278 | YP_009555238.1 | P0C6X6 |
| ACKTIEDALRNSARLE | 1 | 0 | 0 | 1643282 | NP_073551.1 | A0A0P0K6L9 |
| ACNEDNALAYYNTSKG | 1 | 0 | 0 | 1643290 | QIQ54046.1 | P0C6U8 |
| ACNIAKSAYERDRAVA | 1 | 0 | 0 | 1643294 | YP_009555238.1 | P0C6X6 |
| ACNIAKSVYERDKAVA | 1 | 0 | 0 | 1643295 | YP_173236.1 | A0A140H1G9 |
| ACNLGGAVCRKHATEY | 1 | 0 | 0 | 1643296 | YP_009047202.1 | K9N7C7 |
| ACQDPEIGPEHSVADY | 1 | 0 | 0 | 1643304 | NP_828849.2 | P0C6X7 |
| ACQLPYCYFRNSSANY | 1 | 0 | 0 | 1643306 | YP_173237.1 | A0A140H1H0 |
| ACQPPYCYFRNSTTNY | 1 | 0 | 0 | 1643308 | YP_009555240.1 | P30215 |
| ACSCGSCCKFDEDDSE | 1 | 0 | 0 | 1643318 | NP_828851.1 | P59594 |
| ACTDDNALAYYNNSKG | 1 | 0 | 0 | 1643329 | NP_828849.2 | P0C6X7 |
| ACTDDNALAYYNTTKG | 1 | 0 | 0 | 1643330 | YP_009724389.1 | P0DTD1 |
| ACVEEVTTTLEETKFL | 1 | 0 | 0 | 1643337 | YP_009724389.1 | P0DTD1 |
| ACVKDFDFKFYDAQPI | 1 | 0 | 0 | 1643339 | YP_009555238.1 | P0C6X6 |
| ACVLAAECTIFKDAAG | 1 | 0 | 0 | 1643342 | QIQ54046.1 | P0C6U8 |
| ACVSVPVSVIYDKETK | 1 | 0 | 0 | 1643346 | YP_009047204.1 | K9N5Q8 |
| ACYAYLAKAMLDFSRD | 1 | 0 | 0 | 1643360 | NP_073549.1 | A0A0P0LKV0 |
| ACYDGCAQGVFGVNMR | 1 | 0 | 0 | 1643361 | NP_073549.1 | A0A0P0LKV0 |
| ACYEGIASGVFGVNLR | 1 | 0 | 0 | 1643362 | YP_003766.2 | H9EJ42 |
| ADAGFMKQYGECLGDI | 1 | 0 | 0 | 1643368 | NP_828851.1 | P59594 |
| ADDLNQLTGYKKPASR | 1 | 0 | 0 | 1643386 | YP_009724389.1 | P0DTD1 |
| ADDLNQMSGYKKPASR | 1 | 0 | 0 | 1643387 | QIQ54046.1 | P0C6U8 |
| ADDLNQMTGFTKPASR | 1 | 0 | 0 | 1643388 | NP_828849.2 | P0C6X7 |
| ADEIANLVLAKLGKDS | 1 | 0 | 0 | 1643395 | YP_173242.1 | Q19U29 |
| ADEIIGYKAIEDGVTR | 1 | 0 | 0 | 1643396 | QIQ54046.1 | P0C6U8 |
| ADGELPVAISGDKVFV | 1 | 0 | 0 | 1643407 | NP_073549.1 | A0A0P0LKV0 |
| ADGLEAYADKTLQEMK | 1 | 0 | 0 | 1643413 | YP_009555238.1 | P0C6X6 |
| ADGSLIPVRPRNSSDN | 1 | 0 | 0 | 1643417 | YP_003767.1 | A0A0P0G321 |
| ADICYKFVLWDYERSN | 1 | 0 | 0 | 1643423 | YP_009047202.1 | K9N7C7 |
| ADINGKLYHDSQNMLR | 1 | 0 | 0 | 1643426 | NP_828849.2 | P0C6X7 |
| ADINGNLYPDSTSLVE | 1 | 0 | 0 | 1643427 | QIQ54046.1 | P0C6U8 |
| ADINIPDLPICVYDPL | 1 | 0 | 0 | 1643428 | YP_009555240.1 | P30215 |
| ADISAVQSSFGFDQQQ | 1 | 0 | 0 | 1643431 | YP_173236.1 | A0A140H1G9 |
| ADIVDRDPSSDEAIPT | 1 | 0 | 0 | 1643434 | YP_009555245.1 | P33469 |
| ADIVEEAKKVKPTIVV | 1 | 0 | 0 | 1643435 | QHR63299.1 | P0C6W2 |
| ADIVEEAKKVKPTVVV | 1 | 0 | 0 | 1643436 | YP_009724389.1 | P0DTD1 |
| ADIVVFDEISMATNYD | 1 | 0 | 0 | 1643437 | YP_009724389.1 | P0DTD1 |
| ADIVVVDEVSMCTNYD | 1 | 0 | 0 | 1643438 | NP_073549.1 | A0A0P0LKV0 |
| ADKDKDLSGAVPSDLN | 1 | 0 | 0 | 1643440 | NP_073549.1 | A0A0P0LKV0 |
| ADKILYQYENLSLADI | 1 | 0 | 0 | 1643443 | YP_173236.1 | A0A140H1G9 |
| ADKPNHFINFPLTHFQ | 1 | 0 | 0 | 1643446 | YP_009555239.1 | Q80872 |
| ADLALTNMYKEARIND | 1 | 0 | 0 | 1643452 | YP_009555238.1 | P0C6X6 |
| ADLGVLIQNGAKHVQG | 1 | 0 | 0 | 1643453 | YP_173236.1 | A0A140H1G9 |
| ADLGVLIQNSAKHVQG | 1 | 0 | 0 | 1643454 | YP_009555238.1 | P0C6X6 |
| ADLNGVIVDKVGELKV | 1 | 0 | 0 | 1643459 | YP_173236.1 | A0A140H1G9 |
| ADLTKPYVKWDLLKYD | 1 | 0 | 0 | 1643464 | QIQ54047.1 | P0C6X7 |
| ADMDDFSRQLQNSMSG | 1 | 0 | 0 | 1643470 | NP_828858.1 | P59595 |
| ADNNDITNTNLSRGRG | 1 | 0 | 0 | 1643481 | YP_009047211.1 | K9N4V7 |
| ADPAMHIASSNAFLDL | 1 | 0 | 0 | 1643484 | YP_009047202.1 | K9N7C7 |
| ADQAMTSMYKQARAED | 1 | 0 | 0 | 1643493 | YP_009047202.1 | K9N7C7 |
| ADSFTCNNIDAAKIYG | 1 | 0 | 0 | 1643506 | YP_009555241.1 | P36334 |
| ADSSIAYSNNTIAIPT | 1 | 0 | 0 | 1643520 | NP_828851.1 | P59594 |
| ADSYYSYMMPMLTMCH | 1 | 0 | 0 | 1643523 | YP_173236.1 | A0A140H1G9 |

| | | | | | | |
|---|---|---|---|---|---|---|
| ADTAGLYTNFRIDVPS | 1 | 0 | 0 | 1643524 | YP_009047205.1 | K9N796 |
| ADTAHANNINRFNVAI | 1 | 0 | 0 | 1643525 | YP_009047202.1 | K9N7C7 |
| ADTDWKFYDAQPCSDK | 1 | 0 | 0 | 1643526 | QIQ54047.1 | P0C6X7 |
| ADTEQTQNVTVKPKRL | 1 | 0 | 0 | 1643527 | YP_009047202.1 | K9N7C7 |
| ADTLMQGVTLSSNLNT | 1 | 0 | 0 | 1643528 | YP_173238.1 | A0A140H1H1 |
| ADTLQETPVVPDTVEV | 1 | 0 | 0 | 1643529 | YP_009047202.1 | K9N7C7 |
| ADTSLSGYRLKDCVMY | 1 | 0 | 0 | 1643531 | NP_828849.2 | P0C6X7 |
| ADTSTPSDVSSRDPTT | 1 | 0 | 0 | 1643532 | YP_173242.1 | Q19U29 |
| ADVDDPKLMGWDYPKC | 1 | 0 | 0 | 1643536 | NP_073549.1 | A0A0P0LKV0 |
| ADVIVNPANGHMLHGG | 1 | 0 | 0 | 1643540 | YP_173236.1 | A0A140H1G9 |
| ADVSEMLTFDKKAFTL | 1 | 0 | 0 | 1643545 | NP_073551.1 | A0A0P0K6L9 |
| ADVVKEQVSDLLVKLL | 1 | 0 | 0 | 1643550 | YP_009047202.1 | K9N7C7 |
| ADVYRRKPNLPNCNIE | 1 | 0 | 0 | 1643551 | YP_009555241.1 | P36334 |
| ADYDVAVTAGPFMDNA | 1 | 0 | 0 | 1643553 | YP_009047202.1 | K9N7C7 |
| ADYKKCTKGLSIADLA | 1 | 0 | 0 | 1643556 | NP_073551.1 | A0A0P0K6L9 |
| AEAELAKNVSLDNVLS | 1 | 0 | 0 | 1643567 | YP_009724389.1 | P0DTD1 |
| AEAFEYYHTTDPSFLG | 1 | 0 | 0 | 1643568 | QHR63299.1 | P0C6W2 |
| AEAQIDRLINGRLTAL | 1 | 0 | 0 | 1643572 | YP_009555241.1 | P36334 |
| AEAVIKTLQPVSELLT | 1 | 0 | 0 | 1643575 | QIQ54046.1 | P0C6U8 |
| AEAVVKTLQPVSDLLT | 1 | 0 | 0 | 1643576 | NP_828849.2 | P0C6X7 |
| AECKRALGLSISDHEF | 1 | 0 | 0 | 1643579 | NP_073549.1 | A0A0P0LKV0 |
| AECTIFKDAAGKPVPY | 1 | 0 | 0 | 1643580 | QIQ54046.1 | P0C6U8 |
| AECTIFKDAMGKPVPY | 1 | 0 | 0 | 1643581 | NP_828849.2 | P0C6X7 |
| AECTIFKDASGKPVPY | 1 | 0 | 0 | 1643582 | YP_009724389.1 | P0DTD1 |
| AEDIFGPCWSALASAL | 1 | 0 | 0 | 1643584 | NP_073549.1 | A0A0P0LKV0 |
| AEDMLNPNYEDLLIRK | 1 | 0 | 0 | 1643587 | NP_828849.2 | P0C6X7 |
| AEDTLTKLSKYLANEA | 1 | 0 | 0 | 1643588 | NP_073549.1 | A0A0P0LKV0 |
| AEDVVDQVEEVNSIFD | 1 | 0 | 0 | 1643591 | NP_073549.1 | A0A0P0LKV0 |
| AEEALSKLSEYLISDS | 1 | 0 | 0 | 1643592 | YP_003766.2 | H9EJ42 |
| AEETRKLMPICMDTKA | 1 | 0 | 0 | 1643598 | QIQ54046.1 | P0C6U8 |
| AEETRKLMPICMDVRA | 1 | 0 | 0 | 1643599 | NP_828849.2 | P0C6X7 |
| AEETRKLMPVCMETKA | 1 | 0 | 0 | 1643600 | QHR63299.1 | P0C6W2 |
| AEETRKLMPVCVETKA | 1 | 0 | 0 | 1643601 | YP_009724389.1 | P0DTD1 |
| AEEYCNYLESYNIVTT | 1 | 0 | 0 | 1643602 | YP_173236.1 | A0A140H1G9 |
| AEEYREYLESYNTATT | 1 | 0 | 0 | 1643603 | YP_009555238.1 | P0C6X6 |
| AEFDRESSVQKKINRM | 1 | 0 | 0 | 1643604 | NP_073549.1 | A0A0P0LKV0 |
| AEFDTYDPIYKNGAMY | 1 | 0 | 0 | 1643605 | YP_009047202.1 | K9N7C7 |
| AEFKYMVANGLHAPHG | 1 | 0 | 0 | 1643606 | YP_003766.2 | H9EJ42 |
| AEFKYMVANGLNAPNG | 1 | 0 | 0 | 1643607 | NP_073549.1 | A0A0P0LKV0 |
| AEFVEPGCGGILAVID | 1 | 0 | 0 | 1643608 | NP_073549.1 | A0A0P0LKV0 |
| AEGAHATRDSIGTNFP | 1 | 0 | 0 | 1643609 | YP_009555238.1 | P0C6X6 |
| AEHDFFTFDIDGSRVP | 1 | 0 | 0 | 1643616 | YP_173236.1 | A0A140H1G9 |
| AEHDFFTFDVEGSRVP | 1 | 0 | 0 | 1643617 | YP_009555238.1 | P0C6X6 |
| AEHDFFTWKDGRVIYG | 1 | 0 | 0 | 1643618 | YP_003766.2 | H9EJ42 |
| AEHVNNSYECDIPVGA | 1 | 0 | 0 | 1643620 | QIQ54048.1 | P59594 |
| AEIAFMLWCKDGHVET | 1 | 0 | 0 | 1643621 | QIQ54047.1 | P0C6X7 |
| AEIPKEEVKPFITESK | 1 | 0 | 0 | 1643625 | YP_009724389.1 | P0DTD1 |
| AEISFMLWCKDGHVET | 1 | 0 | 0 | 1643629 | NP_828849.2 | P0C6X7 |
| AEIVDTVSALVYDNKL | 1 | 0 | 0 | 1643630 | YP_009724389.1 | P0DTD1 |
| AEIVNTVSELVYENKF | 1 | 0 | 0 | 1643631 | NP_073549.1 | A0A0P0LKV0 |
| AELAKNVSLDNVLSTF | 1 | 0 | 0 | 1643635 | YP_009724389.1 | P0DTD1 |
| AELAPTPGAFFFGSKL | 1 | 0 | 0 | 1643638 | YP_173242.1 | Q19U29 |
| AELDGVQYGRSGETLG | 1 | 0 | 0 | 1643640 | QIQ54046.1 | P0C6U8 |
| AELIPNQAALFFDSEV | 1 | 0 | 0 | 1643646 | YP_003771.1 | H9EJ69 |
| AELKCSVKSFEIDKGI | 1 | 0 | 0 | 1643647 | NP_828851.1 | P59594 |
| AELVPSTAAMLFDSHI | 1 | 0 | 0 | 1643652 | NP_073556.1 | A0A0N7J131 |
| AEMDGIQYGRSGITLG | 1 | 0 | 0 | 1643654 | NP_828849.2 | P0C6X7 |
| AEMLKLGTNDPQFPIL | 1 | 0 | 0 | 1643655 | YP_173242.1 | Q19U29 |
| AEMTSNVRRQVASGVI | 1 | 0 | 0 | 1643656 | NP_073549.1 | A0A0P0LKV0 |
| AEPKGKYSQKAYALLR | 1 | 0 | 0 | 1643660 | YP_173236.1 | A0A140H1G9 |
| AEPVRDEVSIETDIID | 1 | 0 | 0 | 1643661 | NP_073556.1 | A0A0N7J131 |
| AEQAAAAMYKEARAVN | 1 | 0 | 0 | 1643662 | NP_073549.1 | A0A0P0LKV0 |
| AEQAATQMYKEARSVN | 1 | 0 | 0 | 1643663 | YP_003766.2 | H9EJ42 |
| AERDCYTFKKTVAKAP | 1 | 0 | 0 | 1643667 | YP_173236.1 | A0A140H1G9 |
| AERMAMYTGSLIGGIA | 1 | 0 | 0 | 1643668 | NP_073551.1 | A0A0P0K6L9 |
| AESHVDTDLTKPYIKW | 1 | 0 | 0 | 1643674 | YP_009724389.1 | P0DTD1 |
| AESKAFVDTIVPPCPS | 1 | 0 | 0 | 1643675 | NP_073549.1 | A0A0P0LKV0 |
| AESKPWFVRKAEDIFG | 1 | 0 | 0 | 1643676 | NP_073549.1 | A0A0P0LKV0 |
| AESTGTQSVSVDLEST | 1 | 0 | 0 | 1643677 | YP_009047205.1 | K9N796 |
| AESVIVEDDVIENVKS | 1 | 0 | 0 | 1643678 | YP_173236.1 | A0A140H1G9 |
| AETELAKNVSLDNVLS | 1 | 0 | 0 | 1643680 | QIQ54046.1 | P0C6U8 |
| AETHRDCDFNKPLIEW | 1 | 0 | 0 | 1643681 | YP_009047202.1 | K9N7C7 |
| AETIKAKEESVKSSYA | 1 | 0 | 0 | 1643682 | YP_003766.2 | H9EJ42 |
| AETLKATEETFKLSYG | 1 | 0 | 0 | 1643683 | YP_009724389.1 | P0DTD1 |
| AETLRATEEASKQSYA | 1 | 0 | 0 | 1643684 | YP_009047202.1 | K9N7C7 |
| AETVKAKEESVKSSYA | 1 | 0 | 0 | 1643686 | NP_073549.1 | A0A0P0LKV0 |
| AETVRVEEEEEEDWLD | 1 | 0 | 0 | 1643687 | NP_828849.2 | P0C6X7 |
| AEVEWKFYDAQPCSDK | 1 | 0 | 0 | 1643688 | NP_828849.2 | P0C6X7 |
| AEVSRQVIQDVKDGTV | 1 | 0 | 0 | 1643692 | NP_073552.1 | A0A0P0KWE8 |
| AEVVEQEVTPTEQEEE | 1 | 0 | 0 | 1643693 | QIQ54046.1 | P0C6U8 |
| AEVVKQMYGVNLQSGK | 1 | 0 | 0 | 1643694 | YP_003766.2 | H9EJ42 |
| AEVVVNPANGHMAHGG | 1 | 0 | 0 | 1643695 | YP_009555238.1 | P0C6X6 |
| AEWIELFPHNDRIKSF | 1 | 0 | 0 | 1643698 | NP_073549.1 | A0A0P0LKV0 |
| AEYHNESGIKTTLRKG | 1 | 0 | 0 | 1643703 | QIQ54046.1 | P0C6U8 |
| AEYHNESGLKPILRKG | 1 | 0 | 0 | 1643704 | QHR63299.1 | P0C6W2 |
| AEYHNESGLKTILRKG | 1 | 0 | 0 | 1643705 | YP_009724389.1 | P0DTD1 |
| AFACPDGTKHTFQLRA | 1 | 0 | 0 | 1643709 | QIQ54053.1 | P59635 |
| AFANSVFNICQAVSAN | 1 | 0 | 0 | 1643715 | YP_009555238.1 | P0C6X6 |
| AFATAQEAYEQAVANG | 1 | 0 | 0 | 1643720 | YP_009724389.1 | P0DTD1 |
| AFATLKEVVGPKELLL | 1 | 0 | 0 | 1643722 | YP_003766.2 | H9EJ42 |
| AFAVIGDLKCTSDNIN | 1 | 0 | 0 | 1643726 | YP_009555241.1 | P36334 |

| | | | | | | |
|---|---|---|---|---|---|---|
| AFDFSLDVFKIGDVKF | 1 | 0 | 0 | 1643734 | YP_003766.2 | H9EJ42 |
| AFDKSAFTNLKQLPFF | 1 | 0 | 0 | 1643746 | NP_828849.2 | P0C6X7 |
| AFDKSAFVNLKQLPFF | 1 | 0 | 0 | 1643747 | YP_009724389.1 | P0DTD1 |
| AFDVYNKDASFIGKNL | 1 | 0 | 0 | 1643750 | NP_073549.1 | A0A0P0LKV0 |
| AFELFAKRKMGLTPPL | 1 | 0 | 0 | 1643761 | YP_003766.2 | H9EJ42 |
| AFELFAKRKVGLTPPL | 1 | 0 | 0 | 1643762 | NP_073549.1 | A0A0P0LKV0 |
| AFEYYHTLDESFLGRY | 1 | 0 | 0 | 1643767 | NP_828849.2 | P0C6X7 |
| AFFFGSKLDLVKRDSE | 1 | 0 | 0 | 1643770 | YP_173242.1 | Q19U29 |
| AFGGDQVHEVAAVRSV | 1 | 0 | 0 | 1643783 | YP_009047202.1 | K9N7C7 |
| AFGGVSVITPGTNASS | 1 | 0 | 0 | 1643784 | NP_828851.1 | P59594 |
| AFGLQDCVTGINDDDY | 1 | 0 | 0 | 1643786 | YP_003766.2 | H9EJ42 |
| AFGLRGPGDLQGNFGD | 1 | 0 | 0 | 1643787 | YP_009047211.1 | K9N4V7 |
| AFGRRGPEQTQGNFGD | 1 | 0 | 0 | 1643793 | YP_009724397.2 | P0DTC9 |
| AFHANSSEPALLFRNI | 1 | 0 | 0 | 1643797 | YP_009555241.1 | P36334 |
| AFHQECSLQSCAQHQP | 1 | 0 | 0 | 1643798 | QHR63307.1 | Q3LZX5 |
| AFHQECSLQSCTQHQP | 1 | 0 | 0 | 1643799 | YP_009724396.1 | P0DTC8 |
| AFHTPAFDKSAFTNLK | 1 | 0 | 0 | 1643804 | NP_828849.2 | P0C6X7 |
| AFHTPAFDKSAFVNLK | 1 | 0 | 0 | 1643805 | YP_009724389.1 | P0DTD1 |
| AFISVLDITDAAVKAA | 1 | 0 | 0 | 1643813 | NP_073549.1 | A0A0P0LKV0 |
| AFLDTVVSTVKIGGLT | 1 | 0 | 0 | 1643836 | NP_073549.1 | A0A0P0LKV0 |
| AFLGASVTEDVKFAAS | 1 | 0 | 0 | 1643838 | YP_003766.2 | H9EJ42 |
| AFLVHDNVAFYQGDVD | 1 | 0 | 0 | 1643859 | NP_073549.1 | A0A0P0LKV0 |
| AFMDKQVHQVQLTDKY | 1 | 0 | 0 | 1643863 | YP_009047202.1 | K9N7C7 |
| AFMKHKGGDSTDFIAL | 1 | 0 | 0 | 1643865 | YP_009047202.1 | K9N7C7 |
| AFMLGSSVGNFSDGKM | 1 | 0 | 0 | 1643866 | YP_009047204.1 | K9N5Q8 |
| AFNHPIQVDQLNSSYF | 1 | 0 | 0 | 1643871 | YP_009047204.1 | K9N5Q8 |
| AFNNWFLLTNTSSVVD | 1 | 0 | 0 | 1643874 | NP_073551.1 | A0A0P0K6L9 |
| AFNQDGIIFNAVDCMS | 1 | 0 | 0 | 1643876 | YP_009555241.1 | P36334 |
| AFNVINKGHFDGQQGE | 1 | 0 | 0 | 1643882 | QIQ54047.1 | P0C6X7 |
| AFNVNVMGTRPEKFPT | 1 | 0 | 0 | 1643883 | NP_073549.1 | A0A0P0LKV0 |
| AFNVSDAYANLVPYYQ | 1 | 0 | 0 | 1643884 | YP_003766.2 | H9EJ42 |
| AFNVVNKGHFDGQQGE | 1 | 0 | 0 | 1643887 | YP_009724389.1 | P0DTD1 |
| AFPLDEAEDVTESVQE | 1 | 0 | 0 | 1643892 | YP_009047202.1 | K9N7C7 |
| AFQKVQDAVNNNAQAL | 1 | 0 | 0 | 1643899 | YP_009047204.1 | K9N5Q8 |
| AFRKARNGVFISTEKL | 1 | 0 | 0 | 1643908 | YP_173236.1 | A0A140H1G9 |
| AFSASLFKPVQLVPVS | 1 | 0 | 0 | 1643910 | YP_009047208.1 | K9N7D2 |
| AFSLDVSEKSGNFKHL | 1 | 0 | 0 | 1643916 | NP_828851.1 | P59594 |
| AFSVDPKKTYLDFIQQ | 1 | 0 | 0 | 1643919 | YP_009555238.1 | P0C6X6 |
| AFSVDPKKTYLDYIQQ | 1 | 0 | 0 | 1643920 | YP_173236.1 | A0A140H1G9 |
| AFTKAAGGKVSFSDDV | 1 | 0 | 0 | 1643924 | NP_073549.1 | A0A0P0LKV0 |
| AFTKLAGGKISFSDDV | 1 | 0 | 0 | 1643925 | YP_003766.2 | H9EJ42 |
| AFTREMQQHPLLNPSA | 1 | 0 | 0 | 1643931 | NP_073556.1 | A0A0N7J131 |
| AFTTELKDSVKKFFKL | 1 | 0 | 0 | 1643932 | QIQ54046.1 | P0C6U8 |
| AFVDTIVPPCPSILKV | 1 | 0 | 0 | 1643937 | NP_073549.1 | A0A0P0LKV0 |
| AFVETVKGLDYKAFKQ | 1 | 0 | 0 | 1643939 | YP_009724389.1 | P0DTD1 |
| AFVETVKGLDYKSFKQ | 1 | 0 | 0 | 1643940 | QIQ54046.1 | P0C6U8 |
| AFVETVKGLDYKTFKQ | 1 | 0 | 0 | 1643941 | QHR63299.1 | P0C6W2 |
| AFVSQVLNKYTEVRGS | 1 | 0 | 0 | 1643950 | YP_003767.1 | A0A0P0G321 |
| AFYQGDVDTVVNGVDF | 1 | 0 | 0 | 1643966 | NP_073549.1 | A0A0P0LKV0 |
| AFYVKDGKYFTSEPPV | 1 | 0 | 0 | 1643968 | YP_009047202.1 | K9N7C7 |
| AGAINAASKGAVQKES | 1 | 0 | 0 | 1643977 | YP_009047202.1 | K9N7C7 |
| AGALNKATNGAMQKES | 1 | 0 | 0 | 1643980 | NP_828849.2 | P0C6X7 |
| AGALNKATNNAMQIES | 1 | 0 | 0 | 1643981 | QIQ54046.1 | P0C6U8 |
| AGALNKATNNAMQVES | 1 | 0 | 0 | 1643982 | YP_009724389.1 | P0DTD1 |
| AGEAPVSIINNAVYTK | 1 | 0 | 0 | 1644005 | NP_828849.2 | P0C6X7 |
| AGEDIQLLKAAYENFN | 1 | 0 | 0 | 1644006 | NP_828849.2 | P0C6X7 |
| AGEMPCAIINDKVVAK | 1 | 0 | 0 | 1644009 | YP_009555238.1 | P0C6X6 |
| AGFMKQYGECLGDINA | 1 | 0 | 0 | 1644017 | NP_828851.1 | P59594 |
| AGGHSFGIDLKSYDLG | 1 | 0 | 0 | 1644028 | QIQ54046.1 | P0C6U8 |
| AGGHSYGADLKSFDLG | 1 | 0 | 0 | 1644029 | YP_009724389.1 | P0DTD1 |
| AGGHSYGIDLKSYDLG | 1 | 0 | 0 | 1644030 | NP_828849.2 | P0C6X7 |
| AGGITYVSNSGNLLGF | 1 | 0 | 0 | 1644032 | YP_003767.1 | A0A0P0G321 |
| AGHATGTTPQKLFVAN | 1 | 0 | 0 | 1644041 | YP_009047204.1 | K9N5Q8 |
| AGHSLGRCDIKDLPKE | 1 | 0 | 0 | 1644043 | NP_828855.1 | P59596 |
| AGICASYQTQTNSRSV | 1 | 0 | 0 | 1644049 | QHR63300.2 | Q3LZX1 |
| AGINYVKSPKFSKLFT | 1 | 0 | 0 | 1644064 | NP_828849.2 | P0C6X7 |
| AGIQNFKETLYYQNNV | 1 | 0 | 0 | 1644067 | YP_009047202.1 | K9N7C7 |
| AGKVKVGTGVMVECDS | 1 | 0 | 0 | 1644078 | NP_073549.1 | A0A0P0LKV0 |
| AGLEFTDENYNNLVPT | 1 | 0 | 0 | 1644082 | YP_173236.1 | A0A140H1G9 |
| AGLPYGANKEGIIWVA | 1 | 0 | 0 | 1644090 | QIQ54056.1 | P59595 |
| AGLVGVLTLDNQDLNG | 1 | 0 | 0 | 1644094 | YP_009047202.1 | K9N7C7 |
| AGMFGLYDDILTNNKP | 1 | 0 | 0 | 1644100 | YP_003766.2 | H9EJ42 |
| AGMHFGACDYDRLPNE | 1 | 0 | 0 | 1644101 | YP_009047210.1 | K9N7A1 |
| AGNIFGLYKNSATGSL | 1 | 0 | 0 | 1644105 | YP_003766.2 | H9EJ42 |
| AGNSYTSFATYHTPAT | 1 | 0 | 0 | 1644109 | YP_009047204.1 | K9N5Q8 |
| AGNVIGNDALTRFTIF | 1 | 0 | 0 | 1644113 | YP_173236.1 | A0A140H1G9 |
| AGNVWQIQTIQDSDGT | 1 | 0 | 0 | 1644115 | YP_009555238.1 | P0C6X6 |
| AGPFMDNAINVGGTGL | 1 | 0 | 0 | 1644117 | YP_009047202.1 | K9N7C7 |
| AGRKVNFNEKPVVMEI | 1 | 0 | 0 | 1644128 | YP_173236.1 | A0A140H1G9 |
| AGSDKGIAPGTSVLRQ | 1 | 0 | 0 | 1644135 | YP_009047202.1 | K9N7C7 |
| AGSDKGVAPGTTVLKR | 1 | 0 | 0 | 1644137 | YP_003766.2 | H9EJ42 |
| AGSFDKVYDIGNPKGI | 1 | 0 | 0 | 1644140 | YP_009047202.1 | K9N7C7 |
| AGSKSPIQYIDIGNYT | 1 | 0 | 0 | 1644143 | YP_009724396.1 | P0DTC8 |
| AGSRSRANSGNRTPTS | 1 | 0 | 0 | 1644147 | YP_009555245.1 | P33469 |
| AGSVWHIQTVQDADGI | 1 | 0 | 0 | 1644155 | YP_173236.1 | A0A140H1G9 |
| AGSYKDWSYSGQSTQL | 1 | 0 | 0 | 1644158 | YP_009724389.1 | P0DTD1 |
| AGTAVINLRADQINDM | 1 | 0 | 0 | 1644161 | YP_009555238.1 | P0C6X6 |
| AGTDLEGKFYGPFVDR | 1 | 0 | 0 | 1644163 | NP_828849.2 | P0C6X7 |
| AGTDLEGNFYGPFVDR | 1 | 0 | 0 | 1644164 | YP_009724389.1 | P0DTD1 |
| AGTDLEGTFYGPFVDR | 1 | 0 | 0 | 1644165 | QHR63299.1 | P0C6W2 |
| AGTILVDNDLYPFVSD | 1 | 0 | 0 | 1644174 | YP_009555238.1 | P0C6X6 |

| | | | | | | |
|---|---|---|---|---|---|---|
| AGTNYLTCDNLCTPDP | 1 | 0 | 0 | 1644177 | YP_009555241.1 | P36334 |
| AGVLLKQLMVNKSSCY | 1 | 0 | 0 | 1644197 | YP_173243.1 | Q0ZJ80 |
| AGVNITLANFNETKGP | 1 | 0 | 0 | 1644200 | NP_073551.1 | A0A0P0K6L9 |
| AGVVANGVKAKGYPQF | 1 | 0 | 0 | 1644207 | NP_073556.1 | A0A0N7J131 |
| AGVVTGNVVKQCCTAA | 1 | 0 | 0 | 1644208 | YP_009047202.1 | K9N7C7 |
| AGVVWTLNDVKDNDGR | 1 | 0 | 0 | 1644209 | YP_003766.2 | H9EJ42 |
| AGVVWTLQEVKDNDGK | 1 | 0 | 0 | 1644210 | NP_073549.1 | A0A0P0LKV0 |
| AGWPLNKFGKAGLYYE | 1 | 0 | 0 | 1644211 | NP_073549.1 | A0A0P0LKV0 |
| AGYKVLPPLMDVNMEA | 1 | 0 | 0 | 1644216 | YP_009047204.1 | K9N5Q8 |
| AGYTESAVNSTKALAK | 1 | 0 | 0 | 1644218 | YP_009047206.1 | K9N4V0 |
| AGYWRRQDRKINTGNG | 1 | 0 | 0 | 1644220 | YP_009047211.1 | K9N4V7 |
| AHAEETRKLMPICMDV | 1 | 0 | 0 | 1644224 | NP_828849.2 | P0C6X7 |
| AHAEETRKLMPVCMET | 1 | 0 | 0 | 1644225 | QHR63299.1 | P0C6W2 |
| AHAPSFLAVDDKYKAT | 1 | 0 | 0 | 1644228 | YP_009555238.1 | P0C6X6 |
| AHAPSFLAVDDKYKVN | 1 | 0 | 0 | 1644229 | YP_173236.1 | A0A140H1G9 |
| AHATRDSIGTNFPLQL | 1 | 0 | 0 | 1644232 | YP_009555238.1 | P0C6X6 |
| AHATRENIGTNFPLQI | 1 | 0 | 0 | 1644233 | YP_173236.1 | A0A140H1G9 |
| AHGGGLAKALDVYTKG | 1 | 0 | 0 | 1644238 | NP_073549.1 | A0A0P0LKV0 |
| AHKHDIQITNESYNNY | 1 | 0 | 0 | 1644248 | YP_009047202.1 | K9N7C7 |
| AHSELAKGVALDGVLS | 1 | 0 | 0 | 1644259 | NP_828849.2 | P0C6X7 |
| AHVCGDNIGTNVPLQV | 1 | 0 | 0 | 1644270 | YP_003766.2 | H9EJ42 |
| AIATVKSEDGNIKNIN | 1 | 0 | 0 | 1644291 | NP_073549.1 | A0A0P0LKV0 |
| AIAVAAGKKFSKETAA | 1 | 0 | 0 | 1644292 | YP_173236.1 | A0A140H1G9 |
| AICHEGKAHFPREGVF | 1 | 0 | 0 | 1644297 | QIQ54048.1 | P59594 |
| AIDAYPLSKHPKPEYR | 1 | 0 | 0 | 1644304 | NP_073549.1 | A0A0P0LKV0 |
| AIDGFGVFDIKNSSVN | 1 | 0 | 0 | 1644313 | YP_003766.2 | H9EJ42 |
| AIDILTEGQLQSLSKD | 1 | 0 | 0 | 1644315 | YP_003766.2 | H9EJ42 |
| AIDYDHEYSIMRLHNF | 1 | 0 | 0 | 1644320 | NP_073549.1 | A0A0P0LKV0 |
| AIDYKHYTPSFKKGAK | 1 | 0 | 0 | 1644321 | YP_009724389.1 | P0DTD1 |
| AIDYRHYSASFKKGAK | 1 | 0 | 0 | 1644322 | NP_828849.2 | P0C6X7 |
| AIEDGVTRDISSNDNC | 1 | 0 | 0 | 1644323 | QIQ54046.1 | P0C6U8 |
| AIEDILFSKLVTSGLG | 1 | 0 | 0 | 1644324 | NP_073551.1 | A0A0P0K6L9 |
| AIEDLLFDKVKLSDVG | 1 | 0 | 0 | 1644325 | YP_009555241.1 | P36334 |
| AIEDLLFDKVTIADPG | 1 | 0 | 0 | 1644326 | YP_009047204.1 | K9N5Q8 |
| AIEDVEDVISSEAVED | 1 | 0 | 0 | 1644327 | YP_173236.1 | A0A140H1G9 |
| AIEEIEYVHGDALHTL | 1 | 0 | 0 | 1644328 | NP_073549.1 | A0A0P0LKV0 |
| AIELSKELKRPVNPTD | 1 | 0 | 0 | 1644331 | YP_173236.1 | A0A140H1G9 |
| AIFNDHHLNRPGVYCG | 1 | 0 | 0 | 1644340 | YP_009047202.1 | K9N7C7 |
| AIGKKIKHEGDWDSFC | 1 | 0 | 0 | 1644347 | NP_073549.1 | A0A0P0LKV0 |
| AIGTNLPLQLGFSTGV | 1 | 0 | 0 | 1644353 | QHR63299.1 | P0C6W2 |
| AIHAEQLTPAWRVYSA | 1 | 0 | 0 | 1644356 | QIQ54048.1 | P59594 |
| AIIAVELLLLDFKTAV | 1 | 0 | 0 | 1644363 | YP_003766.2 | H9EJ42 |
| AIIIDNDINDYVSDAD | 1 | 0 | 0 | 1644365 | YP_003766.2 | H9EJ42 |
| AIIIDNDLNEFVSDAD | 1 | 0 | 0 | 1644366 | YP_009047202.1 | K9N7C7 |
| AIINDKVVAKIDKEDV | 1 | 0 | 0 | 1644373 | YP_009555238.1 | P0C6X6 |
| AIITANLSIPSNWTTS | 1 | 0 | 0 | 1644376 | YP_003767.1 | A0A0P0G321 |
| AIKCVDIVKEAQSANP | 1 | 0 | 0 | 1644380 | NP_828849.2 | P0C6X7 |
| AIKCVPQADVEWKFYD | 1 | 0 | 0 | 1644382 | YP_009724389.1 | P0DTD1 |
| AIKCVPQAEVEWKFYD | 1 | 0 | 0 | 1644383 | NP_828849.2 | P0C6X7 |
| AIKDCVDINKAIPSGN | 1 | 0 | 0 | 1644384 | NP_073549.1 | A0A0P0LKV0 |
| AIKITEFSWNADLYKL | 1 | 0 | 0 | 1644388 | YP_173236.1 | A0A140H1G9 |
| AIKITEHSWNADLYKL | 1 | 0 | 0 | 1644390 | YP_009724389.1 | P0DTD1 |
| AIKLDDKDPQFKDNVI | 1 | 0 | 0 | 1644393 | NP_828858.1 | P59595 |
| AIKLDDKDPSFKDNVI | 1 | 0 | 0 | 1644394 | QIQ54056.1 | P59595 |
| AIKLDPKNPNYNKWLE | 1 | 0 | 0 | 1644395 | YP_009047211.1 | K9N4V7 |
| AIKLNFGMLNGNAIAT | 1 | 0 | 0 | 1644396 | NP_073549.1 | A0A0P0LKV0 |
| AIKVTEYSWNKKLYEL | 1 | 0 | 0 | 1644404 | NP_073549.1 | A0A0P0LKV0 |
| AILEDEFTPFDVVRQC | 1 | 0 | 0 | 1644407 | QIQ54046.1 | P0C6U8 |
| AILMSEDVKLNPSEDF | 1 | 0 | 0 | 1644410 | YP_009047202.1 | K9N7C7 |
| AILNTVKFCDHMVKAG | 1 | 0 | 0 | 1644415 | YP_009047202.1 | K9N7C7 |
| AIMNDKVVVKINNVDT | 1 | 0 | 0 | 1644428 | YP_173236.1 | A0A140H1G9 |
| AINAASKGAVQKESDE | 1 | 0 | 0 | 1644439 | YP_009047202.1 | K9N7C7 |
| AINVTINNYNPSSWNR | 1 | 0 | 0 | 1644448 | YP_173238.1 | A0A140H1H1 |
| AIPNGRKVDLQLGNLG | 1 | 0 | 0 | 1644452 | YP_009555241.1 | P36334 |
| AIPSGNLIKFDIGSVV | 1 | 0 | 0 | 1644454 | NP_073549.1 | A0A0P0LKV0 |
| AIPSLAANTLNIIVPD | 1 | 0 | 0 | 1644455 | YP_009555238.1 | P0C6X6 |
| AIPTNFTISVTTEILP | 1 | 0 | 0 | 1644459 | YP_009724390.1 | P0DTC2 |
| AIQEGFDATNSALVKI | 1 | 0 | 0 | 1644464 | YP_009555241.1 | P36334 |
| AIQSFDNSYLNESGAL | 1 | 0 | 0 | 1644469 | NP_073550.1 | A0A0P0ILY9 |
| AIRNVRGWLGMDVESA | 1 | 0 | 0 | 1644478 | YP_003766.2 | H9EJ42 |
| AISDYDYYRYNLPTMC | 1 | 0 | 0 | 1644482 | YP_009724389.1 | P0DTD1 |
| AISGDKVFVRDGNTDN | 1 | 0 | 0 | 1644483 | NP_073549.1 | A0A0P0LKV0 |
| AISLSFNNLLGFDSSK | 1 | 0 | 0 | 1644485 | YP_009047202.1 | K9N7C7 |
| AISNSIQAIYDRLDSI | 1 | 0 | 0 | 1644491 | YP_003767.1 | A0A0P0G321 |
| AISSNTDNETLQYWVT | 1 | 0 | 0 | 1644495 | YP_173238.1 | A0A140H1H1 |
| AISVQTCDLTVGSGYC | 1 | 0 | 0 | 1644500 | YP_009555241.1 | P36334 |
| AITDYNYYKYNLPTMV | 1 | 0 | 0 | 1644503 | YP_009555238.1 | P0C6X6 |
| AITDYNYYSYNLPTMC | 1 | 0 | 0 | 1644504 | YP_009047202.1 | K9N7C7 |
| AITPYVCNTSGCNVND | 1 | 0 | 0 | 1644510 | NP_073549.1 | A0A0P0LKV0 |
| AITQTSQALQTVATAL | 1 | 0 | 0 | 1644512 | NP_073551.1 | A0A0P0K6L9 |
| AITSGIKFQTVKPGNF | 1 | 0 | 0 | 1644521 | YP_173236.1 | A0A140H1G9 |
| AITSGVKFQTVKPGNF | 1 | 0 | 0 | 1644522 | YP_009555238.1 | P0C6X6 |
| AITTGYRFTNFEPFTV | 1 | 0 | 0 | 1644523 | YP_009555241.1 | P36334 |
| AITVTPEANMDQESFG | 1 | 0 | 0 | 1644527 | YP_009724389.1 | P0DTD1 |
| AIVCASVKRDGVQVGY | 1 | 0 | 0 | 1644529 | NP_073549.1 | A0A0P0LKV0 |
| AIVQASESKVLLERFL | 1 | 0 | 0 | 1644546 | YP_003766.2 | H9EJ42 |
| AIVTANLSIPSNWTTS | 1 | 0 | 0 | 1644548 | NP_073551.1 | A0A0P0K6L9 |
| AIWDAFVAAIKLVPTT | 1 | 0 | 0 | 1644554 | YP_003766.2 | H9EJ42 |
| AIYDRLDSIQADQQVD | 1 | 0 | 0 | 1644559 | YP_003767.1 | A0A0P0G321 |
| AIYDRLDTIQADQQVD | 1 | 0 | 0 | 1644560 | NP_073551.1 | A0A0P0K6L9 |
| AIYNLLKDCPAVAVHD | 1 | 0 | 0 | 1644563 | QIQ54047.1 | P0C6X7 |

| | | | | | | |
|---|---|---|---|---|---|---|
| AKAGDKYYPVVVDDHV | 1 | 0 | 0 | 1644569 | YP_009555238.1 | P0C6X6 |
| AKAMDTFTNNNGSDVL | 1 | 0 | 0 | 1644578 | YP_009555238.1 | P0C6X6 |
| AKAMETFNHNNGNDVL | 1 | 0 | 0 | 1644579 | YP_173236.1 | A0A140H1G9 |
| AKAMLDFSRDHNDILY | 1 | 0 | 0 | 1644580 | NP_073549.1 | A0A0P0LKV0 |
| AKAMLDYAKDHNDMLY | 1 | 0 | 0 | 1644581 | YP_003766.2 | H9EJ42 |
| AKAYKDYLASGGQPIT | 1 | 0 | 0 | 1644589 | YP_009724389.1 | P0DTD1 |
| AKCWTEEDLSIGPHEF | 1 | 0 | 0 | 1644591 | NP_073549.1 | A0A0P0LKV0 |
| AKCWTETDLTKGPHEF | 1 | 0 | 0 | 1644592 | YP_009724389.1 | P0DTD1 |
| AKDTTEAFEKXVSLLS | 1 | 0 | 0 | 1644603 | QIQ54046.1 | P0C6U8 |
| AKEESVKSSYAYATLK | 1 | 0 | 0 | 1644607 | NP_073549.1 | A0A0P0LKV0 |
| AKEGAKTVNTSLGNRK | 1 | 0 | 0 | 1644608 | YP_003771.1 | H9EJ69 |
| AKFKNSVASINSAIVC | 1 | 0 | 0 | 1644620 | NP_073549.1 | A0A0P0LKV0 |
| AKFLKTNCCRFQEKDE | 1 | 0 | 0 | 1644621 | YP_009724389.1 | P0DTD1 |
| AKFPLKLAGTAVINLR | 1 | 0 | 0 | 1644623 | YP_009555238.1 | P0C6X6 |
| AKGDSCSSNCKHSVAQ | 1 | 0 | 0 | 1644628 | YP_009047202.1 | K9N7C7 |
| AKGFPQLAELIPNQAA | 1 | 0 | 0 | 1644630 | YP_003771.1 | H9EJ69 |
| AKGGFKFGDPADSRDF | 1 | 0 | 0 | 1644631 | YP_009555238.1 | P0C6X6 |
| AKGHFKFDEPSDATDF | 1 | 0 | 0 | 1644632 | YP_173236.1 | A0A140H1G9 |
| AKGYIAGIQNFKETLY | 1 | 0 | 0 | 1644639 | YP_009047202.1 | K9N7C7 |
| AKGYPQFAELVPSTAA | 1 | 0 | 0 | 1644640 | NP_073556.1 | A0A0N7J131 |
| AKHDFFKFRIDGDMVP | 1 | 0 | 0 | 1644644 | YP_009724389.1 | P0DTD1 |
| AKHIVYVGDPAQLPAP | 1 | 0 | 0 | 1644647 | YP_009047202.1 | K9N7C7 |
| AKICIVDNVYMAKAGD | 1 | 0 | 0 | 1644652 | YP_009555238.1 | P0C6X6 |
| AKIGILCIMSDRDLYD | 1 | 0 | 0 | 1644654 | NP_828849.2 | P0C6X7 |
| AKIQSVVNSNAQALNS | 1 | 0 | 0 | 1644658 | YP_173238.1 | A0A140H1H1 |
| AKIYGMCFSSITIDKF | 1 | 0 | 0 | 1644661 | YP_009555241.1 | P36334 |
| AKKCLHVVGPNLNAGE | 1 | 0 | 0 | 1644662 | NP_828849.2 | P0C6X7 |
| AKKFDNFTGECPNFVF | 1 | 0 | 0 | 1644664 | QIQ54046.1 | P0C6U8 |
| AKKFDTFNGECPNFVF | 1 | 0 | 0 | 1644666 | YP_009724389.1 | P0DTD1 |
| AKKGAWNIGEQKSILS | 1 | 0 | 0 | 1644667 | YP_009724389.1 | P0DTD1 |
| AKKLERMADLALTNMY | 1 | 0 | 0 | 1644674 | YP_009555238.1 | P0C6X6 |
| AKKNNLPFKLTCATTR | 1 | 0 | 0 | 1644678 | YP_009724389.1 | P0DTD1 |
| AKKQVVLVSNNQEDFD | 1 | 0 | 0 | 1644682 | YP_009555238.1 | P0C6X6 |
| AKKSMYDGDRFVKHDL | 1 | 0 | 0 | 1644683 | NP_073549.1 | A0A0P0LKV0 |
| AKLGFDSSKEFVEYKI | 1 | 0 | 0 | 1644688 | YP_009555238.1 | P0C6X6 |
| AKLKIQVVNSGPDQTC | 1 | 0 | 0 | 1644694 | YP_009555238.1 | P0C6X6 |
| AKLMVVIPDYNTYKNT | 1 | 0 | 0 | 1644699 | YP_009724389.1 | P0DTD1 |
| AKLMVVVPDYGTYKNT | 1 | 0 | 0 | 1644700 | NP_828849.2 | P0C6X7 |
| AKLMVVVPDYNTYKNT | 1 | 0 | 0 | 1644701 | QIQ54046.1 | P0C6U8 |
| AKLSAGVEFLKDAWEI | 1 | 0 | 0 | 1644703 | NP_828849.2 | P0C6X7 |
| AKNAYEKDKAVARKLE | 1 | 0 | 0 | 1644706 | YP_009047202.1 | K9N7C7 |
| AKNCLHVVGPNVNRGE | 1 | 0 | 0 | 1644708 | QHR63299.1 | P0C6W2 |
| AKPEEKLSAYDLACCM | 1 | 0 | 0 | 1644723 | NP_073549.1 | A0A0P0LKV0 |
| AKPFLNKVTSVAASGV | 1 | 0 | 0 | 1644724 | QIQ54046.1 | P0C6U8 |
| AKPFLNKVVSTTTNIV | 1 | 0 | 0 | 1644725 | YP_009724389.1 | P0DTD1 |
| AKQDVSLLSKCYKAMN | 1 | 0 | 0 | 1644737 | YP_009047202.1 | K9N7C7 |
| AKQGAGDAGHSLTWLW | 1 | 0 | 0 | 1644738 | NP_073549.1 | A0A0P0LKV0 |
| AKQKPIYLKGSGSDFS | 1 | 0 | 0 | 1644740 | YP_009555238.1 | P0C6X6 |
| AKQVDYVPLKSATCIT | 1 | 0 | 0 | 1644741 | YP_009555238.1 | P0C6X6 |
| AKRAVRSHPDFKLLHN | 1 | 0 | 0 | 1644742 | YP_009047202.1 | K9N7C7 |
| AKRFKESPFELEDFIP | 1 | 0 | 0 | 1644743 | YP_009724389.1 | P0DTD1 |
| AKRKMGLTPPLSILKN | 1 | 0 | 0 | 1644746 | YP_003766.2 | H9EJ42 |
| AKRKVGLTPPLSILKN | 1 | 0 | 0 | 1644747 | NP_073549.1 | A0A0P0LKV0 |
| AKRNIKPVPEIKILNN | 1 | 0 | 0 | 1644749 | NP_828849.2 | P0C6X7 |
| AKRNIKPVPEVKILNN | 1 | 0 | 0 | 1644750 | YP_009724389.1 | P0DTD1 |
| AKRSKDSPLELEDFIP | 1 | 0 | 0 | 1644751 | QIQ54047.1 | P0C6X7 |
| AKRSKESPLELEDFIP | 1 | 0 | 0 | 1644752 | QHR63299.1 | P0C6W2 |
| AKRVLGLQTQTVDSAQ | 1 | 0 | 0 | 1644755 | YP_009555238.1 | P0C6X6 |
| AKRVLGVQTQTVDSAQ | 1 | 0 | 0 | 1644756 | YP_173236.1 | A0A140H1G9 |
| AKTDLFNGYKIGCNCA | 1 | 0 | 0 | 1644775 | YP_173236.1 | A0A140H1G9 |
| AKTGTPKPSRNQSPAS | 1 | 0 | 0 | 1644777 | NP_073556.1 | A0A0N7J131 |
| AKVEISVTPIKNTVDT | 1 | 0 | 0 | 1644787 | NP_073549.1 | A0A0P0LKV0 |
| AKVKNTKVIKDRVMYS | 1 | 0 | 0 | 1644792 | YP_009555241.1 | P36334 |
| AKVSAEVSRQVIQDVK | 1 | 0 | 0 | 1644797 | NP_073552.1 | A0A0P0KWE8 |
| AKWYCYDKNPINSNVK | 1 | 0 | 0 | 1644804 | NP_073549.1 | A0A0P0LKV0 |
| ALALLLLDRLNALESK | 1 | 0 | 0 | 1644822 | QIQ54056.1 | P59595 |
| ALANQFTEFVGTQSVD | 1 | 0 | 0 | 1644825 | YP_009047202.1 | K9N7C7 |
| ALASALKQLKVTTGEL | 1 | 0 | 0 | 1644827 | NP_073549.1 | A0A0P0LKV0 |
| ALASDIFDTPSVLQAT | 1 | 0 | 0 | 1644828 | YP_009047202.1 | K9N7C7 |
| ALASMTGVSIETLLAA | 1 | 0 | 0 | 1644829 | YP_173236.1 | A0A140H1G9 |
| ALASMTGVSLETLLAA | 1 | 0 | 0 | 1644830 | YP_009555238.1 | P0C6X6 |
| ALCFDKSKQFVPVQVV | 1 | 0 | 0 | 1644843 | YP_173237.1 | A0A140H1H0 |
| ALDLSKELKRPIQPTD | 1 | 0 | 0 | 1644857 | YP_009555238.1 | P0C6X6 |
| ALEAQVQIDRLINGRL | 1 | 0 | 0 | 1644870 | YP_173238.1 | A0A140H1H1 |
| ALEDLLFSKVVTSGLG | 1 | 0 | 0 | 1644871 | YP_003767.1 | A0A0P0G321 |
| ALEFNPSQTSPATAEP | 1 | 0 | 0 | 1644872 | NP_073556.1 | A0A0N7J131 |
| ALFASLDVKPYGYPND | 1 | 0 | 0 | 1644881 | YP_003766.2 | H9EJ42 |
| ALFFDSEVSTDEVGDN | 1 | 0 | 0 | 1644884 | YP_003771.1 | H9EJ69 |
| ALFVSYGSSVETETYF | 1 | 0 | 0 | 1644896 | YP_173236.1 | A0A140H1G9 |
| ALGAMQTGFTTTNEAF | 1 | 0 | 0 | 1644898 | YP_009047204.1 | K9N5Q8 |
| ALGKVVANAMLKCVAF | 1 | 0 | 0 | 1644910 | NP_073549.1 | A0A0P0LKV0 |
| ALGSMDIDVFNKLSTS | 1 | 0 | 0 | 1644914 | NP_073549.1 | A0A0P0LKV0 |
| ALGVLMSNLGMPSYCT | 1 | 0 | 0 | 1644915 | YP_009724389.1 | P0DTD1 |
| ALIATAHSSIKQGTQI | 1 | 0 | 0 | 1644925 | YP_009555238.1 | P0C6X6 |
| ALIAVANWLAPTNAYM | 1 | 0 | 0 | 1644926 | YP_009047202.1 | K9N7C7 |
| ALILAYCNKTVGELGD | 1 | 0 | 0 | 1644928 | YP_009724389.1 | P0DTD1 |
| ALKKCKSAFYVLPSEA | 1 | 0 | 0 | 1644947 | NP_828849.2 | P0C6X7 |
| ALKSLGFDKPQEKDKK | 1 | 0 | 0 | 1644959 | NP_073556.1 | A0A0N7J131 |
| ALKTAGVVTGNVVKQC | 1 | 0 | 0 | 1644960 | YP_009047202.1 | K9N7C7 |
| ALKTQFNYYKKVNGVV | 1 | 0 | 0 | 1644961 | QHR63299.1 | P0C6W2 |
| ALLADKFPVLHDIGNP | 1 | 0 | 0 | 1644963 | YP_009724389.1 | P0DTD1 |

| | | | | | | |
|---|---|---|---|---|---|---|
| ALLAGSFDKVYDIGNP | 1 | 0 | 0 | 1644966 | YP_009047202.1 | K9N7C7 |
| ALLDWVKHLNKTLNEG | 1 | 0 | 0 | 1644972 | NP_073549.1 | A0A0P0LKV0 |
| ALLEDEFTPFDVVRQC | 1 | 0 | 0 | 1644973 | YP_009724389.1 | P0DTD1 |
| ALLHIAGCQTTNGLNT | 1 | 0 | 0 | 1644977 | NP_073551.1 | A0A0P0K6L9 |
| ALLLLDRLNALESKMS | 1 | 0 | 0 | 1644983 | QIQ54056.1 | P59595 |
| ALLNLTDEDLARDFSP | 1 | 0 | 0 | 1644986 | YP_009047207.1 | K9N643 |
| ALLSKTSEYKGPVTDV | 1 | 0 | 0 | 1644995 | QIQ54046.1 | P0C6U8 |
| ALLSTDGNKIADKHVR | 1 | 0 | 0 | 1644996 | QHR63299.1 | P0C6W2 |
| ALLSTDGNKIADKYVR | 1 | 0 | 0 | 1644997 | YP_009724389.1 | P0DTD1 |
| ALLSTDGNKIGDKYIR | 1 | 0 | 0 | 1644998 | QIQ54047.1 | P0C6X7 |
| ALLTKSSEYKGPITDI | 1 | 0 | 0 | 1644999 | QHR63299.1 | P0C6W2 |
| ALLTKSSEYKGPITDV | 1 | 0 | 0 | 1645000 | YP_009724389.1 | P0DTD1 |
| ALMGANGNKIVDKEVK | 1 | 0 | 0 | 1645005 | YP_009047202.1 | K9N7C7 |
| ALNAFVSQVLNKYTEV | 1 | 0 | 0 | 1645008 | YP_003767.1 | A0A0P0G321 |
| ALNAYVSQQLSDISLV | 1 | 0 | 0 | 1645009 | YP_173238.1 | A0A140H1H1 |
| ALNAYVSQQLSDSTLV | 1 | 0 | 0 | 1645010 | YP_009555241.1 | P36334 |
| ALNDFSNSGADVLYQP | 1 | 0 | 0 | 1645012 | NP_828849.2 | P0C6X7 |
| ALNDFSNSGSDVLYQP | 1 | 0 | 0 | 1645013 | YP_009724389.1 | P0DTD1 |
| ALNESYIDLKELGNYT | 1 | 0 | 0 | 1645014 | YP_009047204.1 | K9N5Q8 |
| ALNKATNGAMQKESDD | 1 | 0 | 0 | 1645021 | NP_828849.2 | P0C6X7 |
| ALNKATNNAMQIESDD | 1 | 0 | 0 | 1645022 | QIQ54046.1 | P0C6U8 |
| ALNKATNNAMQVESDD | 1 | 0 | 0 | 1645023 | YP_009724389.1 | P0DTD1 |
| ALNKATNNAMQVESDH | 1 | 0 | 0 | 1645024 | QHR63299.1 | P0C6W2 |
| ALNKIQDVVNQQGNSL | 1 | 0 | 0 | 1645025 | NP_073551.1 | A0A0P0K6L9 |
| ALNKIQDVVNQQGSAL | 1 | 0 | 0 | 1645026 | YP_003767.1 | A0A0P0G321 |
| ALNNIINNARDGCVPL | 1 | 0 | 0 | 1645031 | YP_009724389.1 | P0DTD1 |
| ALNSEKHQDHVSLTVP | 1 | 0 | 0 | 1645035 | YP_009047202.1 | K9N7C7 |
| ALNSILDNAVKGCVPL | 1 | 0 | 0 | 1645036 | YP_009555238.1 | P0C6X6 |
| ALNTICIKDADYNAKV | 1 | 0 | 0 | 1645038 | NP_073549.1 | A0A0P0LKV0 |
| ALNTPKDHIGTRNPAN | 1 | 0 | 0 | 1645040 | YP_009724397.2 | P0DTC9 |
| ALPDTPSTLTPRSVRS | 1 | 0 | 0 | 1645043 | YP_009047204.1 | K9N5Q8 |
| ALPIYYQHTDINFTAT | 1 | 0 | 0 | 1645052 | YP_003767.1 | A0A0P0G321 |
| ALPKGFYAEGSRGGSQ | 1 | 0 | 0 | 1645053 | QIQ54056.1 | P59595 |
| ALPQSKDSNFLNESGV | 1 | 0 | 0 | 1645057 | YP_009047203.1 | K9N7C7 |
| ALPQSKDSNFLNRVRG | 1 | 0 | 0 | 1645058 | YP_009047202.1 | K9N7C7 |
| ALQDAYYRARAGDAAN | 1 | 0 | 0 | 1645060 | QIQ54046.1 | P0C6U8 |
| ALQDAYYRARAGEAAN | 1 | 0 | 0 | 1645061 | YP_009724389.1 | P0DTD1 |
| ALQEAYYRARAGDAAN | 1 | 0 | 0 | 1645062 | NP_828849.2 | P0C6X7 |
| ALQHAFMKHKGGDSTD | 1 | 0 | 0 | 1645065 | YP_009047202.1 | K9N7C7 |
| ALQPEEEQEEDWLDDD | 1 | 0 | 0 | 1645070 | YP_009724389.1 | P0DTD1 |
| ALQTDVLQENQKILAA | 1 | 0 | 0 | 1645076 | NP_073551.1 | A0A0P0K6L9 |
| ALQTVATALNKIQDVV | 1 | 0 | 0 | 1645078 | NP_073551.1 | A0A0P0K6L9 |
| ALRANSAVKLQNNELS | 1 | 0 | 0 | 1645081 | YP_009724389.1 | P0DTD1 |
| ALRMGSGFCVDYNSPS | 1 | 0 | 0 | 1645095 | YP_173238.1 | A0A140H1H1 |
| ALRNFDEKDCEVFKEI | 1 | 0 | 0 | 1645096 | NP_073549.1 | A0A0P0LKV0 |
| ALRNSLTNDAYSRFLG | 1 | 0 | 0 | 1645098 | YP_009047202.1 | K9N7C7 |
| ALSEAREHLKNGTCGL | 1 | 0 | 0 | 1645103 | NP_828849.2 | P0C6X7 |
| ALSFEEQDEIYAYTKR | 1 | 0 | 0 | 1645106 | YP_009555238.1 | P0C6X6 |
| ALSFEEQNEIYAYTKR | 1 | 0 | 0 | 1645107 | YP_173236.1 | A0A140H1G9 |
| ALSGVFCGVDAMNLIA | 1 | 0 | 0 | 1645109 | NP_828849.2 | P0C6X7 |
| ALSIFDTWANWDSNWA | 1 | 0 | 0 | 1645113 | NP_073555.1 | A0A0P0KXN3 |
| ALSTGLTSQTVKPGHF | 1 | 0 | 0 | 1645127 | NP_073549.1 | A0A0P0LKV0 |
| ALVATAHSELAKGVAL | 1 | 0 | 0 | 1645146 | NP_828849.2 | P0C6X7 |
| ALVKIQAVVNANAEAL | 1 | 0 | 0 | 1645154 | YP_009555241.1 | P36334 |
| ALVLAKGGFKFGDPAD | 1 | 0 | 0 | 1645157 | YP_009555238.1 | P0C6X6 |
| ALVYDNKLRAHKDKSQ | 1 | 0 | 0 | 1645171 | QIQ54047.1 | P0C6X7 |
| ALYAIQEGFDATNSAL | 1 | 0 | 0 | 1645179 | YP_009555241.1 | P36334 |
| ALYEQVVMDYLDNLKP | 1 | 0 | 0 | 1645181 | NP_828849.2 | P0C6X7 |
| ALYLDYVQCKCGNLEQ | 1 | 0 | 0 | 1645184 | YP_173236.1 | A0A140H1G9 |
| ALYLDYVQHKCGNLHQ | 1 | 0 | 0 | 1645185 | YP_009555238.1 | P0C6X6 |
| ALYNKYKYFSGAMDTT | 1 | 0 | 0 | 1645189 | YP_009724389.1 | P0DTD1 |
| ALYVTWSEGNSITGVP | 1 | 0 | 0 | 1645195 | YP_003767.1 | A0A0P0G321 |
| AMANGYTSVSSVECYS | 1 | 0 | 0 | 1645204 | YP_003766.2 | H9EJ42 |
| AMDEFIQRYKLEGYAF | 1 | 0 | 0 | 1645209 | NP_828849.2 | P0C6X7 |
| AMDTFTNNNGSDVLYQ | 1 | 0 | 0 | 1645212 | YP_009555238.1 | P0C6X6 |
| AMEPIYDEPTTTTSVP | 1 | 0 | 0 | 1645216 | QHR63301.1 | Q3LZX0 |
| AMETFNHNNGNDVLYQ | 1 | 0 | 0 | 1645219 | YP_173236.1 | A0A140H1G9 |
| AMFTKVVVGPTTEVKF | 1 | 0 | 0 | 1645223 | YP_003766.2 | H9EJ42 |
| AMGQGQNSADPKVYPI | 1 | 0 | 0 | 1645227 | NP_828859.1 | P59636 |
| AMHIASSNAFLDLRTS | 1 | 0 | 0 | 1645230 | YP_009047202.1 | K9N7C7 |
| AMLDFSRDHNDILYTP | 1 | 0 | 0 | 1645239 | NP_073549.1 | A0A0P0LKV0 |
| AMLFDSHIVSKESGNT | 1 | 0 | 0 | 1645241 | NP_073556.1 | A0A0N7J131 |
| AMLGAEMTSNVRRQVA | 1 | 0 | 0 | 1645243 | NP_073549.1 | A0A0P0LKV0 |
| AMLSENFTSYGFSNVV | 1 | 0 | 0 | 1645249 | NP_073551.1 | A0A0P0K6L9 |
| AMNGEDAFSILAAKTG | 1 | 0 | 0 | 1645257 | NP_073549.1 | A0A0P0LKV0 |
| AMNVAKAEFDRESSVQ | 1 | 0 | 0 | 1645260 | NP_073549.1 | A0A0P0LKV0 |
| AMQIESDDYIARNGPL | 1 | 0 | 0 | 1645274 | QIQ54046.1 | P0C6U8 |
| AMQKESDDYIKLNGPL | 1 | 0 | 0 | 1645275 | NP_828849.2 | P0C6X7 |
| AMQTGFTTTNEAFQKV | 1 | 0 | 0 | 1645279 | YP_009047204.1 | K9N5Q8 |
| AMQVESDDYIATNGPL | 1 | 0 | 0 | 1645282 | YP_009724389.1 | P0DTD1 |
| AMRHVRGWLGMDVEGA | 1 | 0 | 0 | 1645285 | NP_073549.1 | A0A0P0LKV0 |
| AMRPNHTIKGSFLNGS | 1 | 0 | 0 | 1645291 | NP_828849.2 | P0C6X7 |
| AMTAVNESRYGLQNLL | 1 | 0 | 0 | 1645295 | YP_003767.1 | A0A0P0G321 |
| AMTQMYKQARSEDKRA | 1 | 0 | 0 | 1645302 | YP_009724389.1 | P0DTD1 |
| AMTSMYKQARAEDKKA | 1 | 0 | 0 | 1645303 | YP_009047202.1 | K9N7C7 |
| ANAKGFIRFPEVLREG | 1 | 0 | 0 | 1645324 | YP_009555238.1 | P0C6X6 |
| ANALSRAVNVPTIRKF | 1 | 0 | 0 | 1645326 | YP_009555238.1 | P0C6X6 |
| ANASSSEGFLLGINYL | 1 | 0 | 0 | 1645329 | YP_009047202.1 | K9N7C7 |
| ANAYYKYDNGNFIKLP | 1 | 0 | 0 | 1645332 | NP_073549.1 | A0A0P0LKV0 |
| ANDPVGFTLKNTVCTV | 1 | 0 | 0 | 1645340 | YP_009724389.1 | P0DTD1 |
| ANDPVGFTLRNTVCTV | 1 | 0 | 0 | 1645341 | NP_828849.2 | P0C6X7 |

| | | | | | | |
|---|---|---|---|---|---|---|
| ANECAQVLSEYVLCGG | 1 | 0 | 0 | 1645349 | YP_009047202.1 | K9N7C7 |
| ANENAINKGGRTVQSH | 1 | 0 | 0 | 1645354 | NP_073549.1 | A0A0P0LKV0 |
| ANENLAHGGGLAKALD | 1 | 0 | 0 | 1645355 | NP_073549.1 | A0A0P0LKV0 |
| ANEYRLYLDAYNMMIS | 1 | 0 | 0 | 1645357 | YP_009724389.1 | P0DTD1 |
| ANEYRQYLDAYNMMIS | 1 | 0 | 0 | 1645358 | NP_828849.2 | P0C6X7 |
| ANFLDDNVLPETYVAL | 1 | 0 | 0 | 1645364 | YP_003767.1 | A0A0P0G321 |
| ANFNETKGPLCVDTSH | 1 | 0 | 0 | 1645367 | NP_073551.1 | A0A0P0K6L9 |
| ANGLNAPNGPFDALFL | 1 | 0 | 0 | 1645391 | NP_073549.1 | A0A0P0LKV0 |
| ANGNKIVDKEVKDMQF | 1 | 0 | 0 | 1645395 | YP_009047202.1 | K9N7C7 |
| ANGTHTGSAFDGTMYG | 1 | 0 | 0 | 1645401 | YP_009047202.1 | K9N7C7 |
| ANGVKAKGYPQFAELV | 1 | 0 | 0 | 1645402 | NP_073556.1 | A0A0N7J131 |
| ANHQADTSTPSDVSSR | 1 | 0 | 0 | 1645405 | YP_173242.1 | Q19U29 |
| ANIFIGDKVGHYVHVK | 1 | 0 | 0 | 1645409 | YP_009555238.1 | P0C6X6 |
| ANIIYCNSVINRLRCD | 1 | 0 | 0 | 1645414 | NP_073551.1 | A0A0P0K6L9 |
| ANIVYEPTTPISSALI | 1 | 0 | 0 | 1645419 | YP_009047202.1 | K9N7C7 |
| ANKEGIIWVATEGALN | 1 | 0 | 0 | 1645421 | QIQ54056.1 | P59595 |
| ANKFEQFDQWYESTFG | 1 | 0 | 0 | 1645423 | YP_009555238.1 | P0C6X6 |
| ANKFFQFDQWYESTFG | 1 | 0 | 0 | 1645424 | YP_173236.1 | A0A140H1G9 |
| ANKFNQALGAMQTGFT | 1 | 0 | 0 | 1645425 | YP_009047204.1 | K9N5Q8 |
| ANKHADFDTWFSQRGG | 1 | 0 | 0 | 1645426 | YP_009724389.1 | P0DTD1 |
| ANKHVGFDSWFSQRGG | 1 | 0 | 0 | 1645429 | QIQ54046.1 | P0C6U8 |
| ANKIVGGAPTWFNALR | 1 | 0 | 0 | 1645430 | YP_009047202.1 | K9N7C7 |
| ANKLILINDVAFVSTF | 1 | 0 | 0 | 1645431 | YP_009555238.1 | P0C6X6 |
| ANKSVGIISMPLGYVS | 1 | 0 | 0 | 1645432 | YP_009047202.1 | K9N7C7 |
| ANKWDLIISDMYDPKT | 1 | 0 | 0 | 1645433 | YP_009724389.1 | P0DTD1 |
| ANKWDLIISDMYDPRT | 1 | 0 | 0 | 1645434 | NP_828849.2 | P0C6X7 |
| ANLKLPPHDVTVLRDN | 1 | 0 | 0 | 1645440 | YP_003768.1 | H9EJ24 |
| ANLRQDDSVRNLFASV | 1 | 0 | 0 | 1645444 | YP_009047204.1 | K9N5Q8 |
| ANLRVVGHAMQGTLLK | 1 | 0 | 0 | 1645445 | YP_009047202.1 | K9N7C7 |
| ANLSAPTLVPQENYSS | 1 | 0 | 0 | 1645446 | YP_009555238.1 | P0C6X6 |
| ANLVLAKLGKDSKPQQ | 1 | 0 | 0 | 1645449 | YP_173242.1 | Q19U29 |
| ANMSLAECKRALGLSI | 1 | 0 | 0 | 1645460 | NP_073549.1 | A0A0P0LKV0 |
| ANNLTAPRNSWEAMAL | 1 | 0 | 0 | 1645468 | YP_009047202.1 | K9N7C7 |
| ANNYSKLFDEGYTVVI | 1 | 0 | 0 | 1645473 | NP_073549.1 | A0A0P0LKV0 |
| ANPSTPAYTFTTVKPG | 1 | 0 | 0 | 1645481 | YP_009047202.1 | K9N7C7 |
| ANPTNCIAPVNGYFIK | 1 | 0 | 0 | 1645483 | YP_009047204.1 | K9N5Q8 |
| ANQFNKAISQIQESLT | 1 | 0 | 0 | 1645487 | NP_828851.1 | P59594 |
| ANQFNSAIGKIQDSLS | 1 | 0 | 0 | 1645488 | YP_009724390.1 | P0DTC2 |
| ANQFTEFVGTQSVDML | 1 | 0 | 0 | 1645489 | YP_009047202.1 | K9N7C7 |
| ANQQQLKQLEKACNIA | 1 | 0 | 0 | 1645491 | YP_009555238.1 | P0C6X6 |
| ANQVIVNNLDKSAGFP | 1 | 0 | 0 | 1645493 | YP_009724389.1 | P0DTD1 |
| ANRYNEVANAVMQNNE | 1 | 0 | 0 | 1645499 | YP_173236.1 | A0A140H1G9 |
| ANSAVKLQNNELSPVA | 1 | 0 | 0 | 1645501 | YP_009724389.1 | P0DTD1 |
| ANSGNRTPTSGVTPDM | 1 | 0 | 0 | 1645505 | YP_009555245.1 | P33469 |
| ANSNGFIRFPDVISEG | 1 | 0 | 0 | 1645511 | YP_173236.1 | A0A140H1G9 |
| ANSSEPALLFRNIKCN | 1 | 0 | 0 | 1645513 | YP_009555241.1 | P36334 |
| ANSSEWAYCVDLHNKI | 1 | 0 | 0 | 1645514 | YP_003766.2 | H9EJ42 |
| ANSTGTVIISPSTSAT | 1 | 0 | 0 | 1645516 | YP_009047204.1 | K9N5Q8 |
| ANSTPAQNAGYWRRQD | 1 | 0 | 0 | 1645517 | YP_009047211.1 | K9N4V7 |
| ANSVFNICQAVSANVC | 1 | 0 | 0 | 1645521 | YP_009555238.1 | P0C6X6 |
| ANSVFNICQAVTANVN | 1 | 0 | 0 | 1645523 | YP_009724389.1 | P0DTD1 |
| ANTGSVCYTPIDEIPY | 1 | 0 | 0 | 1645528 | YP_009047202.1 | K9N7C7 |
| ANTHLKHGGGIAGAIN | 1 | 0 | 0 | 1645530 | YP_009047202.1 | K9N7C7 |
| ANTLNIIVPDKSVYDQ | 1 | 0 | 0 | 1645533 | YP_009555238.1 | P0C6X6 |
| ANTNQDSYGGASICLY | 1 | 0 | 0 | 1645535 | YP_003766.2 | H9EJ42 |
| ANTVEDAVNKLVFLAV | 1 | 0 | 0 | 1645537 | YP_009555242.1 | Q04853 |
| ANVGRVIKFETDAYKL | 1 | 0 | 0 | 1645544 | YP_173236.1 | A0A140H1G9 |
| ANVGRWSASINTGNCP | 1 | 0 | 0 | 1645545 | NP_073551.1 | A0A0P0K6L9 |
| ANVNALLSTDGNKIAD | 1 | 0 | 0 | 1645549 | YP_009724389.1 | P0DTD1 |
| ANVNALLSTDGNKIGD | 1 | 0 | 0 | 1645550 | QIQ54047.1 | P0C6X7 |
| ANVPAIDVLKKLLSSL | 1 | 0 | 0 | 1645551 | YP_003766.2 | H9EJ42 |
| ANVSALMGANGNKIVD | 1 | 0 | 0 | 1645553 | YP_009047202.1 | K9N7C7 |
| ANWLAPTNAYMRTTHT | 1 | 0 | 0 | 1645563 | YP_009047202.1 | K9N7C7 |
| ANYLGKPKEQIDGYTM | 1 | 0 | 0 | 1645574 | NP_828849.2 | P0C6X7 |
| ANYQKVGMQKYSTLQG | 1 | 0 | 0 | 1645576 | YP_009724389.1 | P0DTD1 |
| ANYQKVGMQXYSTLQG | 1 | 0 | 0 | 1645577 | QIQ54047.1 | P0C6X7 |
| ANYVGKYDINHGDSGF | 1 | 0 | 0 | 1645580 | YP_173237.1 | A0A140H1H0 |
| APALQEAYYRARAGDA | 1 | 0 | 0 | 1645585 | NP_828849.2 | P0C6X7 |
| APGCDVNDVTKLYLGG | 1 | 0 | 0 | 1645606 | YP_009555238.1 | P0C6X6 |
| APIDIDAFPVKPICSS | 1 | 0 | 0 | 1645630 | YP_003766.2 | H9EJ42 |
| APIELENKFTPLSVES | 1 | 0 | 0 | 1645631 | YP_009047202.1 | K9N7C7 |
| APIKGVTFGEDTVWEV | 1 | 0 | 0 | 1645632 | NP_828849.2 | P0C6X7 |
| APKEESTDQMSEPPKE | 1 | 0 | 0 | 1645634 | YP_009047211.1 | K9N4V7 |
| APKEVTFLEGDSHDTV | 1 | 0 | 0 | 1645637 | NP_828849.2 | P0C6X7 |
| APKSGYFVNVNNTWMY | 1 | 0 | 0 | 1645644 | YP_009555241.1 | P36334 |
| APLIELCVDEAGSKSP | 1 | 0 | 0 | 1645648 | YP_009724396.1 | P0DTC8 |
| APLRAPTMANQEKYST | 1 | 0 | 0 | 1645657 | NP_073549.1 | A0A0P0LKV0 |
| APLTVFFDGRVNGQVD | 1 | 0 | 0 | 1645660 | QHR63299.1 | P0C6W2 |
| APLVYLNHSVPKLSDF | 1 | 0 | 0 | 1645662 | YP_173238.1 | A0A140H1H1 |
| APNSRSTSRTSSRASS | 1 | 0 | 0 | 1645671 | YP_009555245.1 | P33469 |
| APQTIQPEVKEVAPVY | 1 | 0 | 0 | 1645683 | YP_009047202.1 | K9N7C7 |
| APQVTYQNISTNLPPP | 1 | 0 | 0 | 1645684 | YP_009047204.1 | K9N5Q8 |
| APRAVSFADNNDITNT | 1 | 0 | 0 | 1645685 | YP_009047211.1 | K9N4V7 |
| APRITFGGPSDSTDNN | 1 | 0 | 0 | 1645686 | QIQ54056.1 | P59595 |
| APRTLLTKGTLEPEYF | 1 | 0 | 0 | 1645690 | YP_009724389.1 | P0DTD1 |
| APRVLISKGVMEPIDY | 1 | 0 | 0 | 1645692 | NP_073549.1 | A0A0P0LKV0 |
| APRVMITKGVMEPVDY | 1 | 0 | 0 | 1645695 | YP_003766.2 | H9EJ42 |
| APRWYFYYTGTGPEAA | 1 | 0 | 0 | 1645696 | YP_009047211.1 | K9N4V7 |
| APSDVYQQLAGVKLQS | 1 | 0 | 0 | 1645698 | YP_173236.1 | A0A140H1G9 |
| APSFLAVDDKYKATGD | 1 | 0 | 0 | 1645699 | YP_009555238.1 | P0C6X6 |
| APSFLAVDDKYKVNEN | 1 | 0 | 0 | 1645700 | YP_173236.1 | A0A140H1G9 |

| | | | | | | |
|---|---|---|---|---|---|---|
| APSRSQSRSQSRGRGE | 1 | 0 | 0 | 1645701 | NP_073556.1 | A0A0N7J131 |
| APSTFGTRNPNNDSAI | 1 | 0 | 0 | 1645702 | YP_009047211.1 | K9N4V7 |
| APSYCNGVRELYLNSS | 1 | 0 | 0 | 1645704 | NP_828849.2 | P0C6X7 |
| APTLVPQEHYVRITGL | 1 | 0 | 0 | 1645714 | YP_009724389.1 | P0DTD1 |
| APTLVPQENYASIRFS | 1 | 0 | 0 | 1645715 | YP_173236.1 | A0A140H1G9 |
| APTLVPQENYSSIRFA | 1 | 0 | 0 | 1645716 | YP_009555238.1 | P0C6X6 |
| APTNAYMRTTHTDIGV | 1 | 0 | 0 | 1645718 | YP_009047202.1 | K9N7C7 |
| APTPGAFFFGSKLDLV | 1 | 0 | 0 | 1645719 | YP_173242.1 | Q19U29 |
| APTYVSVDDKYKTSDE | 1 | 0 | 0 | 1645722 | YP_009047202.1 | K9N7C7 |
| APVEFYEDRILDFKVL | 1 | 0 | 0 | 1645723 | YP_009047202.1 | K9N7C7 |
| APVNTVKPKPVINQLD | 1 | 0 | 0 | 1645726 | NP_073549.1 | A0A0P0LKV0 |
| APVSIINNAVYTKVDG | 1 | 0 | 0 | 1645728 | NP_828849.2 | P0C6X7 |
| APVYEADTEQTQNVTV | 1 | 0 | 0 | 1645730 | YP_009047202.1 | K9N7C7 |
| APVYIWTADEAIKFLK | 1 | 0 | 0 | 1645731 | YP_009555244.1 | Q01455 |
| APYIVGDIITSGNLTT | 1 | 0 | 0 | 1645735 | QIQ54046.1 | P0C6U8 |
| APYVMLNTSIPNLPDF | 1 | 0 | 0 | 1645739 | YP_009555241.1 | P36334 |
| AQADTAGLYTNFRIDV | 1 | 0 | 0 | 1645744 | YP_009047205.1 | K9N796 |
| AQALSKLASELSNTFG | 1 | 0 | 0 | 1645747 | YP_009047204.1 | K9N5Q8 |
| AQFKTFEKVDGVYTNF | 1 | 0 | 0 | 1645770 | YP_009555238.1 | P0C6X6 |
| AQGARGTYRAALNSEK | 1 | 0 | 0 | 1645773 | YP_009047202.1 | K9N7C7 |
| AQIDRLINGRLTALNA | 1 | 0 | 0 | 1645789 | YP_009555241.1 | P36334 |
| AQIDRLINGRLTTLNA | 1 | 0 | 0 | 1645790 | YP_009047204.1 | K9N5Q8 |
| AQIIAIIKDWNFAWSL | 1 | 0 | 0 | 1645791 | YP_009047210.1 | K9N7A1 |
| AQIKALFLDKVDILLT | 1 | 0 | 0 | 1645792 | YP_009555238.1 | P0C6X6 |
| AQLPAPRTLLTKGTLE | 1 | 0 | 0 | 1645803 | YP_009724389.1 | P0DTD1 |
| AQLPAPRVLLSKGSLE | 1 | 0 | 0 | 1645805 | YP_173236.1 | A0A140H1G9 |
| AQLPAPRVLLSKGTLE | 1 | 0 | 0 | 1645806 | YP_009555238.1 | P0C6X6 |
| AQNAGYWRRQDRKINT | 1 | 0 | 0 | 1645811 | YP_009047211.1 | K9N4V7 |
| AQNTVLNASIPESKPL | 1 | 0 | 0 | 1645817 | YP_003771.1 | H9EJ69 |
| AQPCSDKAYKIEELFY | 1 | 0 | 0 | 1645818 | YP_009724389.1 | P0DTD1 |
| AQPKQTATSQQPSGGN | 1 | 0 | 0 | 1645822 | YP_009555245.1 | P33469 |
| AQPLPQRQKKQPTVTL | 1 | 0 | 0 | 1645823 | NP_828858.1 | P59595 |
| AQPVKIDNASPASTVH | 1 | 0 | 0 | 1645830 | NP_828852.2 | P59632 |
| AQTSDTAHACNANRFN | 1 | 0 | 0 | 1645851 | NP_073549.1 | A0A0P0LKV0 |
| AQVLTEVVYSNGGFYF | 1 | 0 | 0 | 1645862 | NP_073549.1 | A0A0P0LKV0 |
| AQVQIDRLINGRLTAL | 1 | 0 | 0 | 1645863 | YP_173238.1 | A0A140H1H1 |
| AQVVIADTLQETPVVP | 1 | 0 | 0 | 1645865 | YP_009047202.1 | K9N7C7 |
| AQVVQLPVKDYVQTVN | 1 | 0 | 0 | 1645867 | YP_173236.1 | A0A140H1G9 |
| AQYIFSTVNALPECNA | 1 | 0 | 0 | 1645871 | YP_003766.2 | H9EJ42 |
| AQYVFSTVNALPEVNA | 1 | 0 | 0 | 1645874 | NP_073549.1 | A0A0P0LKV0 |
| AQYYNGIMVLPGVADA | 1 | 0 | 0 | 1645875 | NP_073551.1 | A0A0P0K6L9 |
| ARAEDKKAKIVSAMQT | 1 | 0 | 0 | 1645879 | YP_009047202.1 | K9N7C7 |
| ARAIDILTEGQLQSLS | 1 | 0 | 0 | 1645886 | YP_003766.2 | H9EJ42 |
| ARAMLKCVALCDAMVA | 1 | 0 | 0 | 1645890 | YP_003766.2 | H9EJ42 |
| ARARVECFDKFKVNST | 1 | 0 | 0 | 1645892 | YP_009724389.1 | P0DTD1 |
| ARDGVRRGDNFHSVLT | 1 | 0 | 0 | 1645902 | YP_009047202.1 | K9N7C7 |
| ARDGVVPLSVIPATSA | 1 | 0 | 0 | 1645903 | YP_003766.2 | H9EJ42 |
| ARDLSLQFKRPINPTD | 1 | 0 | 0 | 1645905 | YP_009724389.1 | P0DTD1 |
| ARGTYRAALNSEKHQD | 1 | 0 | 0 | 1645921 | YP_009047202.1 | K9N7C7 |
| ARIAGNGGDAALALLL | 1 | 0 | 0 | 1645927 | QIQ54056.1 | P59595 |
| ARIRAKHYVYIGDPAQ | 1 | 0 | 0 | 1645934 | YP_009555238.1 | P0C6X6 |
| ARIVFTACSHAAVDSL | 1 | 0 | 0 | 1645935 | NP_073549.1 | A0A0P0LKV0 |
| ARIVFVACAHAAVDSL | 1 | 0 | 0 | 1645936 | YP_003766.2 | H9EJ42 |
| ARIVYTACSHAAVDAL | 1 | 0 | 0 | 1645937 | YP_009724389.1 | P0DTD1 |
| ARKLERMADLALTNMY | 1 | 0 | 0 | 1645944 | YP_173236.1 | A0A140H1G9 |
| ARKSAPLIELCVDEAG | 1 | 0 | 0 | 1645947 | YP_009724396.1 | P0DTC8 |
| ARLEIVKLGAEKKDGF | 1 | 0 | 0 | 1645950 | YP_009555239.1 | Q80872 |
| ARLEPCNGTDIDKCVR | 1 | 0 | 0 | 1645951 | YP_003766.2 | H9EJ42 |
| ARLKGGLILKFDSGTV | 1 | 0 | 0 | 1645956 | YP_009047202.1 | K9N7C7 |
| ARLKRFPVNTIVNGVQ | 1 | 0 | 0 | 1645957 | NP_073549.1 | A0A0P0LKV0 |
| ARLKRVPLQTIINGMH | 1 | 0 | 0 | 1645958 | YP_003766.2 | H9EJ42 |
| ARLLGLQTQTVDSAQG | 1 | 0 | 0 | 1645959 | NP_073549.1 | A0A0P0LKV0 |
| ARLRAKHYVYIGDPAQ | 1 | 0 | 0 | 1645962 | YP_009724389.1 | P0DTD1 |
| ARLYYEALSFEEQDEI | 1 | 0 | 0 | 1645969 | YP_009555238.1 | P0C6X6 |
| ARNGVVPLSVIPATSA | 1 | 0 | 0 | 1645981 | NP_073549.1 | A0A0P0LKV0 |
| ARNPIDLPPSHATTYL | 1 | 0 | 0 | 1645983 | NP_073549.1 | A0A0P0LKV0 |
| ARPKQRRPQGLPNNTA | 1 | 0 | 0 | 1645984 | QHR63308.1 | Q3LZX4 |
| ARQAYEDAIANGSSSQ | 1 | 0 | 0 | 1645986 | YP_003766.2 | H9EJ42 |
| ARQSDNMTAVACQPPY | 1 | 0 | 0 | 1645994 | YP_009555240.1 | P30215 |
| ARVECFDKFKVNSTLE | 1 | 0 | 0 | 1646024 | YP_009724389.1 | P0DTD1 |
| ARVECYDRFKVNETNS | 1 | 0 | 0 | 1646025 | YP_009047202.1 | K9N7C7 |
| ARVECYSGFKPNNNSA | 1 | 0 | 0 | 1646026 | NP_073549.1 | A0A0P0LKV0 |
| ARVECYSGFKPNNTSA | 1 | 0 | 0 | 1646027 | YP_003766.2 | H9EJ42 |
| ARVEHPDVDGICKLRG | 1 | 0 | 0 | 1646029 | YP_173236.1 | A0A140H1G9 |
| ARVSPGLCIAGDRGIA | 1 | 0 | 0 | 1646031 | YP_009555241.1 | P36334 |
| ARVVYTAASHAAVDAL | 1 | 0 | 0 | 1646034 | YP_009555238.1 | P0C6X6 |
| ARVVYTACSHAAVDAL | 1 | 0 | 0 | 1646035 | YP_009047202.1 | K9N7C7 |
| ARYMRSLKVPATVSVS | 1 | 0 | 0 | 1646039 | YP_009724389.1 | P0DTD1 |
| ASAETVRVEEEEEEDW | 1 | 0 | 0 | 1646044 | NP_828849.2 | P0C6X7 |
| ASALKQLKVTTGELVR | 1 | 0 | 0 | 1646049 | NP_073549.1 | A0A0P0LKV0 |
| ASALWEIQQVVDADSK | 1 | 0 | 0 | 1646052 | YP_009724389.1 | P0DTD1 |
| ASANIGCNHTGVVGEG | 1 | 0 | 0 | 1646054 | YP_009724389.1 | P0DTD1 |
| ASATIQEIVSDREVIL | 1 | 0 | 0 | 1646061 | YP_173236.1 | A0A140H1G9 |
| ASATIQEIVSERELIL | 1 | 0 | 0 | 1646062 | YP_009555238.1 | P0C6X6 |
| ASAYRANSFDVPTFCA | 1 | 0 | 0 | 1646065 | YP_009047202.1 | K9N7C7 |
| ASCPIGTNYRSCESTT | 1 | 0 | 0 | 1646072 | YP_173238.1 | A0A140H1H1 |
| ASCTLSEQLDFIDTKR | 1 | 0 | 0 | 1646073 | YP_009724389.1 | P0DTD1 |
| ASDDLYVKRYFKGCET | 1 | 0 | 0 | 1646074 | YP_173236.1 | A0A140H1G9 |
| ASDKFYRLSNELAQVL | 1 | 0 | 0 | 1646078 | NP_073549.1 | A0A0P0LKV0 |
| ASECFMKSDIFGQDFK | 1 | 0 | 0 | 1646091 | NP_073549.1 | A0A0P0LKV0 |
| ASECFVKSDIFGSDFK | 1 | 0 | 0 | 1646092 | YP_003766.2 | H9EJ42 |

| | | | | | | |
|---|---|---|---|---|---|---|
| ASESKVLLERFLPKCP | 1 | 0 | 0 | 1646100 | YP_003766.2 | H9EJ42 |
| ASEVVVNNLDKSAGHP | 1 | 0 | 0 | 1646101 | YP_009047202.1 | K9N7C7 |
| ASFDNFKFVCDNIKFA | 1 | 0 | 0 | 1646103 | YP_009724389.1 | P0DTD1 |
| ASFDNFKFVCDNIKLA | 1 | 0 | 0 | 1646104 | QHR63299.1 | P0C6W2 |
| ASFDNFKLTCSNTKFA | 1 | 0 | 0 | 1646105 | NP_828849.2 | P0C6X7 |
| ASFKVIDNGVVRDISV | 1 | 0 | 0 | 1646112 | YP_173236.1 | A0A140H1G9 |
| ASFLEMKSETKVQDHF | 1 | 0 | 0 | 1646113 | QIQ54046.1 | P0C6U8 |
| ASFVEYEVAKKNLADA | 1 | 0 | 0 | 1646125 | YP_173236.1 | A0A140H1G9 |
| ASFVEYEVAKKNLDEA | 1 | 0 | 0 | 1646126 | YP_009555238.1 | P0C6X6 |
| ASIITKLNSLNEPLVT | 1 | 0 | 0 | 1646162 | NP_828849.2 | P0C6X7 |
| ASIPESKPLADDDSAI | 1 | 0 | 0 | 1646167 | YP_003771.1 | H9EJ69 |
| ASIQHLHEGFGGKNIL | 1 | 0 | 0 | 1646168 | YP_003766.2 | H9EJ42 |
| ASKAYRDYLASGGQPI | 1 | 0 | 0 | 1646172 | QIQ54046.1 | P0C6U8 |
| ASKFASVYAWNRKRIS | 1 | 0 | 0 | 1646173 | QIQ54048.1 | P59594 |
| ASKGAVQKESDEYILA | 1 | 0 | 0 | 1646175 | YP_009047202.1 | K9N7C7 |
| ASKLTIVSPDLESYSK | 1 | 0 | 0 | 1646184 | YP_003766.2 | H9EJ42 |
| ASKSASVYYSQLMCQP | 1 | 0 | 0 | 1646188 | NP_828849.2 | P0C6X7 |
| ASLDVKPYGYPNDFVG | 1 | 0 | 0 | 1646189 | YP_003766.2 | H9EJ42 |
| ASLFKPVQLVPVSPAF | 1 | 0 | 0 | 1646191 | YP_009047208.1 | K9N7D2 |
| ASLINTLNDLNETLVT | 1 | 0 | 0 | 1646198 | YP_009724389.1 | P0DTD1 |
| ASLKELLQNGMNGRTI | 1 | 0 | 0 | 1646199 | YP_009724389.1 | P0DTD1 |
| ASLKELLQNGVNGRTI | 1 | 0 | 0 | 1646200 | QIQ54046.1 | P0C6U8 |
| ASLNGVTLIGEAVKTQ | 1 | 0 | 0 | 1646202 | YP_009724389.1 | P0DTD1 |
| ASLQEILSRLDALEAE | 1 | 0 | 0 | 1646211 | YP_009555241.1 | P36334 |
| ASLRQIFDVAPIELEN | 1 | 0 | 0 | 1646213 | YP_009047202.1 | K9N7C7 |
| ASLSAPTLVPQENYAS | 1 | 0 | 0 | 1646214 | YP_173236.1 | A0A140H1G9 |
| ASLVLAKLGKDATKPQ | 1 | 0 | 0 | 1646215 | YP_009555245.1 | P33469 |
| ASNKPHVLEDPCKVQH | 1 | 0 | 0 | 1646232 | NP_849176.1 | Q7TFA0 |
| ASNQADVNTPADIVDR | 1 | 0 | 0 | 1646233 | YP_009555245.1 | P33469 |
| ASNQVAVLYQDVNCTE | 1 | 0 | 0 | 1646234 | QHR63300.2 | Q3LZX1 |
| ASNSRPGSRSQSRGPN | 1 | 0 | 0 | 1646238 | YP_173242.1 | Q19U29 |
| ASNVKKVTDVTGKLSD | 1 | 0 | 0 | 1646243 | YP_009555238.1 | P0C6X6 |
| ASPMKSDEGGLCPSTG | 1 | 0 | 0 | 1646246 | YP_173236.1 | A0A140H1G9 |
| ASPYDLEVLLQDALQS | 1 | 0 | 0 | 1646251 | YP_009555238.1 | P0C6X6 |
| ASQRVGTDSGFAAYNR | 1 | 0 | 0 | 1646257 | NP_828855.1 | P59596 |
| ASQVIVNNYDKSAGYP | 1 | 0 | 0 | 1646258 | YP_009555238.1 | P0C6X6 |
| ASREFKVTFFPDLNGD | 1 | 0 | 0 | 1646259 | QHR63299.1 | P0C6W2 |
| ASRFLGLQIQTVDSSQ | 1 | 0 | 0 | 1646265 | YP_003766.2 | H9EJ42 |
| ASRMYCTFAIEDVEDV | 1 | 0 | 0 | 1646267 | YP_173236.1 | A0A140H1G9 |
| ASSEKPTKEEPKKVKP | 1 | 0 | 0 | 1646276 | QIQ54046.1 | P0C6U8 |
| ASSFVGMPSFVAYETA | 1 | 0 | 0 | 1646278 | NP_073549.1 | A0A0P0LKV0 |
| ASSGNRSGNGILKWAD | 1 | 0 | 0 | 1646280 | YP_009555245.1 | P33469 |
| ASSIRGCCESTKLPYY | 1 | 0 | 0 | 1646283 | NP_073551.1 | A0A0P0K6L9 |
| ASSKPLYVPEHCQNYS | 1 | 0 | 0 | 1646284 | YP_009047205.1 | K9N796 |
| ASSLSRNSSRSSSQGS | 1 | 0 | 0 | 1646286 | YP_009047211.1 | K9N4V7 |
| ASSPALVDKRTVCFSV | 1 | 0 | 0 | 1646288 | NP_073549.1 | A0A0P0LKV0 |
| ASSQTSAKSLARSQSS | 1 | 0 | 0 | 1646291 | NP_073556.1 | A0A0N7J131 |
| ASSRSSTRNNSRDSSR | 1 | 0 | 0 | 1646295 | YP_003771.1 | H9EJ69 |
| ASSSYDALLHRNSLDP | 1 | 0 | 0 | 1646303 | YP_003766.2 | H9EJ42 |
| ASSVQQQEEQEEDWLE | 1 | 0 | 0 | 1646305 | QIQ54046.1 | P0C6U8 |
| ASTSAFVETVKGLDYK | 1 | 0 | 0 | 1646314 | YP_009724389.1 | P0DTD1 |
| ASTSAVKLQNNEIKPS | 1 | 0 | 0 | 1646315 | YP_009047202.1 | K9N7C7 |
| ASVKRDGVQVGYCVHG | 1 | 0 | 0 | 1646325 | NP_073549.1 | A0A0P0LKV0 |
| ASVKSSQSSPIIPGFG | 1 | 0 | 0 | 1646326 | YP_009047204.1 | K9N5Q8 |
| ASVPEKFLNAFDVFVT | 1 | 0 | 0 | 1646331 | NP_073549.1 | A0A0P0LKV0 |
| ASVTEDVKFAASTGVI | 1 | 0 | 0 | 1646334 | YP_003766.2 | H9EJ42 |
| ASVYSVLETFQNNVVN | 1 | 0 | 0 | 1646338 | YP_009555238.1 | P0C6X6 |
| ASVYYSQLMCQPILLL | 1 | 0 | 0 | 1646339 | YP_009724389.1 | P0DTD1 |
| ASYNKYKYYSGSASEA | 1 | 0 | 0 | 1646353 | YP_003766.2 | H9EJ42 |
| ASYNRYKYYSGNANEA | 1 | 0 | 0 | 1646354 | NP_073549.1 | A0A0P0LKV0 |
| ASYQTQTNSRSVASQS | 1 | 0 | 0 | 1646356 | QHR63300.2 | Q3LZX1 |
| ASYRFVTFEPFNVSFV | 1 | 0 | 0 | 1646357 | YP_173238.1 | A0A140H1H1 |
| ASYTANDVEVQSSGMI | 1 | 0 | 0 | 1646358 | YP_009047202.1 | K9N7C7 |
| ATAEAELAKNVSLDNV | 1 | 0 | 0 | 1646360 | YP_009724389.1 | P0DTD1 |
| ATAEPVRDEVSIETDI | 1 | 0 | 0 | 1646361 | NP_073556.1 | A0A0N7J131 |
| ATAETELAKNVSLDNV | 1 | 0 | 0 | 1646362 | QIQ54046.1 | P0C6U8 |
| ATAFNNALLSIQNGFS | 1 | 0 | 0 | 1646363 | YP_173238.1 | A0A140H1H1 |
| ATAHSELAKGVALDGV | 1 | 0 | 0 | 1646366 | NP_828849.2 | P0C6X7 |
| ATALNKIQDVVNQQGN | 1 | 0 | 0 | 1646370 | NP_073551.1 | A0A0P0K6L9 |
| ATAQEAYEQAVANGDS | 1 | 0 | 0 | 1646372 | YP_009724389.1 | P0DTD1 |
| ATAVKTGGKSLPAIKL | 1 | 0 | 0 | 1646376 | NP_073549.1 | A0A0P0LKV0 |
| ATCDWTESGDYTLANT | 1 | 0 | 0 | 1646377 | YP_009047202.1 | K9N7C7 |
| ATCPFAKGDSCSSNCK | 1 | 0 | 0 | 1646391 | YP_009047202.1 | K9N7C7 |
| ATDCSDGNYNRNASLN | 1 | 0 | 0 | 1646398 | YP_009047204.1 | K9N5Q8 |
| ATEFSRVSAKPPPGDQ | 1 | 0 | 0 | 1646412 | QIQ54047.1 | P0C6X7 |
| ATELGIEFLKRGDKVV | 1 | 0 | 0 | 1646415 | QIQ54046.1 | P0C6U8 |
| ATFAELEAAQKAYQEA | 1 | 0 | 0 | 1646420 | YP_009047202.1 | K9N7C7 |
| ATFLDLYYEMNLIQES | 1 | 0 | 0 | 1646427 | YP_173238.1 | A0A140H1H1 |
| ATFVDVLVNVSATNIQ | 1 | 0 | 0 | 1646431 | YP_003767.1 | A0A0P0G321 |
| ATFYPQLQSAEWKCGY | 1 | 0 | 0 | 1646435 | NP_073549.1 | A0A0P0LKV0 |
| ATGDLAVCLGIGDSAV | 1 | 0 | 0 | 1646438 | YP_009555238.1 | P0C6X6 |
| ATGDVVLASDDLYVKR | 1 | 0 | 0 | 1646439 | YP_173236.1 | A0A140H1G9 |
| ATGDVVLATDDLYVKR | 1 | 0 | 0 | 1646440 | YP_009555238.1 | P0C6X6 |
| ATGSDVNCNGYQHNSV | 1 | 0 | 0 | 1646443 | YP_003767.1 | A0A0P0G321 |
| ATGSLDVEVFNRLATS | 1 | 0 | 0 | 1646444 | YP_003766.2 | H9EJ42 |
| ATGTTPQKLFVANYSQ | 1 | 0 | 0 | 1646445 | YP_009047204.1 | K9N5Q8 |
| ATHGIAAINSVPWSKI | 1 | 0 | 0 | 1646447 | NP_828849.2 | P0C6X7 |
| ATHHDKFTDGVCLFWN | 1 | 0 | 0 | 1646450 | NP_828849.2 | P0C6X7 |
| ATIGVCKYTDIDVNSA | 1 | 0 | 0 | 1646456 | YP_009047202.1 | K9N7C7 |
| ATIPYKVCNSVKDTLA | 1 | 0 | 0 | 1646464 | YP_009047202.1 | K9N7C7 |
| ATIQRKYKGIKIQEGI | 1 | 0 | 0 | 1646466 | NP_828849.2 | P0C6X7 |

| | | | | | | |
|---|---|---|---|---|---|---|
| ATIRLQAGKQTELAVN | 1 | 0 | 0 | 1646468 | YP_003766.2 | H9EJ42 |
| ATKGEGDGGITSEGNA | 1 | 0 | 0 | 1646472 | NP_073549.1 | A0A0P0LKV0 |
| ATKMFDSFVNSFVSLY | 1 | 0 | 0 | 1646475 | YP_009047202.1 | K9N7C7 |
| ATKQYNVTQAFGRRGP | 1 | 0 | 0 | 1646478 | QHR63308.1 | Q3LZX4 |
| ATKYLVQQESPFVMMS | 1 | 0 | 0 | 1646480 | YP_009724389.1 | P0DTD1 |
| ATLIATLNSLGETLVT | 1 | 0 | 0 | 1646484 | QIQ54046.1 | P0C6U8 |
| ATLMTFSGNVDLDALA | 1 | 0 | 0 | 1646490 | YP_009047202.1 | K9N7C7 |
| ATLNSLGETLVTMPLG | 1 | 0 | 0 | 1646491 | QIQ54046.1 | P0C6U8 |
| ATLTAPTIVNQERYVK | 1 | 0 | 0 | 1646498 | YP_009047202.1 | K9N7C7 |
| ATLVNDIDITFLKKDA | 1 | 0 | 0 | 1646500 | QHR63299.1 | P0C6W2 |
| ATLVSDIDITFLKKDA | 1 | 0 | 0 | 1646501 | YP_009724389.1 | P0DTD1 |
| ATLYYQNGVFMSTAKC | 1 | 0 | 0 | 1646504 | NP_073549.1 | A0A0P0LKV0 |
| ATNGAMQKESDDYIKL | 1 | 0 | 0 | 1646510 | NP_828849.2 | P0C6X7 |
| ATNGPLKVGGGCVLSG | 1 | 0 | 0 | 1646511 | QHR63299.1 | P0C6W2 |
| ATNKATYKPNTWCIRC | 1 | 0 | 0 | 1646516 | YP_009724389.1 | P0DTD1 |
| ATNNAMQIESDDYIAR | 1 | 0 | 0 | 1646517 | QIQ54046.1 | P0C6U8 |
| ATNNAMQVESDDYIAT | 1 | 0 | 0 | 1646518 | YP_009724389.1 | P0DTD1 |
| ATNNAMQVESDHYIAT | 1 | 0 | 0 | 1646519 | QHR63299.1 | P0C6W2 |
| ATNSALVKIQAVVNAN | 1 | 0 | 0 | 1646523 | YP_009555241.1 | P36334 |
| ATPSDFVRATATIPIQ | 1 | 0 | 0 | 1646528 | YP_009724391.1 | P0DTC3 |
| ATQYLVQQESSFVMMS | 1 | 0 | 0 | 1646534 | NP_828849.2 | P0C6X7 |
| ATRDAVGTNLPLQLGF | 1 | 0 | 0 | 1646535 | NP_828849.2 | P0C6X7 |
| ATRDSIGTNFPLQLGF | 1 | 0 | 0 | 1646536 | YP_009555238.1 | P0C6X6 |
| ATREAIGTNLPLQLGF | 1 | 0 | 0 | 1646538 | QHR63299.1 | P0C6W2 |
| ATRENIGTNFPLQIGF | 1 | 0 | 0 | 1646540 | YP_173236.1 | A0A140H1G9 |
| ATRRVRGGDGKMKDLS | 1 | 0 | 0 | 1646551 | QIQ54056.1 | P59595 |
| ATRRVRGGDGKMKELS | 1 | 0 | 0 | 1646552 | NP_828858.1 | P59595 |
| ATSAALQPEEEQEEDW | 1 | 0 | 0 | 1646559 | YP_009724389.1 | P0DTD1 |
| ATSASKLTIVSPDLES | 1 | 0 | 0 | 1646562 | YP_003766.2 | H9EJ42 |
| ATSASLFPPWTAAAGV | 1 | 0 | 0 | 1646563 | YP_009555241.1 | P36334 |
| ATSCQLYYNLPAANVS | 1 | 0 | 0 | 1646564 | YP_009555241.1 | P36334 |
| ATSDWTDVRDYKLAND | 1 | 0 | 0 | 1646566 | YP_003766.2 | H9EJ42 |
| ATSIVAKQGAGDAGHS | 1 | 0 | 0 | 1646567 | NP_073549.1 | A0A0P0LKV0 |
| ATSQQPSGGNVVPYYS | 1 | 0 | 0 | 1646569 | YP_009555245.1 | P33469 |
| ATSVTPQPEEELEEDW | 1 | 0 | 0 | 1646572 | QHR63299.1 | P0C6W2 |
| ATTANVSALMGANGNK | 1 | 0 | 0 | 1646575 | YP_009047202.1 | K9N7C7 |
| ATTFASVYAWNRKRIS | 1 | 0 | 0 | 1646582 | QHR63300.2 | Q3LZX1 |
| ATTYLEAVKHGAKPVS | 1 | 0 | 0 | 1646592 | YP_003766.2 | H9EJ42 |
| ATVCGPKLSTDLIKNQ | 1 | 0 | 0 | 1646597 | NP_828851.1 | P59594 |
| ATVHTANKWDLIISDM | 1 | 0 | 0 | 1646602 | YP_009724389.1 | P0DTD1 |
| ATVIRGHLYIQGVKLG | 1 | 0 | 0 | 1646603 | YP_173241.1 | Q0ZJ82 |
| ATVKSEDGNIKNINWF | 1 | 0 | 0 | 1646604 | NP_073549.1 | A0A0P0LKV0 |
| ATVNQVCVLLAKKIDV | 1 | 0 | 0 | 1646610 | YP_173236.1 | A0A140H1G9 |
| ATVQSKMSDVKCTSVV | 1 | 0 | 0 | 1646613 | YP_009724389.1 | P0DTD1 |
| ATVREVLSDRELHLSW | 1 | 0 | 0 | 1646614 | YP_009724389.1 | P0DTD1 |
| ATVREVLSDRELYLSW | 1 | 0 | 0 | 1646615 | QIQ54047.1 | P0C6X7 |
| ATVRLQAGKQTEFVSN | 1 | 0 | 0 | 1646616 | NP_073549.1 | A0A0P0LKV0 |
| ATVRLQAGNATEVPAN | 1 | 0 | 0 | 1646617 | YP_009724389.1 | P0DTD1 |
| ATVSVSSPDAVTAYNG | 1 | 0 | 0 | 1646619 | YP_009724389.1 | P0DTD1 |
| ATVVVTLKDSDVNDMV | 1 | 0 | 0 | 1646627 | YP_003766.2 | H9EJ42 |
| ATYHTPATDCSDGNYN | 1 | 0 | 0 | 1646633 | YP_009047204.1 | K9N5Q8 |
| ATYYNSWQNLLYDSNG | 1 | 0 | 0 | 1646641 | YP_009555241.1 | P36334 |
| AVAAALKSLGFDKPQE | 1 | 0 | 0 | 1646642 | NP_073556.1 | A0A0N7J131 |
| AVAAGLEFTDENYNNL | 1 | 0 | 0 | 1646644 | YP_173236.1 | A0A140H1G9 |
| AVADSYYSYMMPMLTM | 1 | 0 | 0 | 1646648 | YP_173236.1 | A0A140H1G9 |
| AVAKHDFFKFRIDGDM | 1 | 0 | 0 | 1646650 | YP_009724389.1 | P0DTD1 |
| AVAKHDFFTWHEGRTI | 1 | 0 | 0 | 1646651 | NP_073549.1 | A0A0P0LKV0 |
| AVANAHRYDVLLSDLS | 1 | 0 | 0 | 1646654 | YP_003766.2 | H9EJ42 |
| AVANWLAPTNAYMRTT | 1 | 0 | 0 | 1646657 | YP_009047202.1 | K9N7C7 |
| AVARKLERMADLALTN | 1 | 0 | 0 | 1646660 | YP_173236.1 | A0A140H1G9 |
| AVASKILGLPTQTVDS | 1 | 0 | 0 | 1646664 | YP_009724389.1 | P0DTD1 |
| AVATFYPQLQSAEWKC | 1 | 0 | 0 | 1646665 | NP_073549.1 | A0A0P0LKV0 |
| AVAVHDFFKFRVDGDM | 1 | 0 | 0 | 1646666 | NP_828849.2 | P0C6X7 |
| AVAYADKPNHFINFPL | 1 | 0 | 0 | 1646668 | YP_009555239.1 | Q80872 |
| AVCLKHAEEYREYLES | 1 | 0 | 0 | 1646674 | YP_009555238.1 | P0C6X6 |
| AVCSKHANLYQKYVEA | 1 | 0 | 0 | 1646679 | YP_003766.2 | H9EJ42 |
| AVDAAKAYKDYLASGG | 1 | 0 | 0 | 1646682 | YP_009724389.1 | P0DTD1 |
| AVDASKAYRDYLASGG | 1 | 0 | 0 | 1646686 | QIQ54046.1 | P0C6U8 |
| AVDDKYKVNENLAVNL | 1 | 0 | 0 | 1646694 | YP_173236.1 | A0A140H1G9 |
| AVDEAFPLDEAEDVTE | 1 | 0 | 0 | 1646695 | YP_009047202.1 | K9N7C7 |
| AVDINKLCEEMLDNRA | 1 | 0 | 0 | 1646698 | YP_009724389.1 | P0DTD1 |
| AVDINRLCEEMLDNRA | 1 | 0 | 0 | 1646699 | NP_828849.2 | P0C6X7 |
| AVDPAAAYLDAVKQGA | 1 | 0 | 0 | 1646701 | NP_073549.1 | A0A0P0LKV0 |
| AVDPAKAYKDYLASGG | 1 | 0 | 0 | 1646702 | NP_828849.2 | P0C6X7 |
| AVDVVSIRSSNQGNKQ | 1 | 0 | 0 | 1646708 | YP_009047207.1 | K9N643 |
| AVEDTIDGVVEDTIND | 1 | 0 | 0 | 1646710 | YP_173236.1 | A0A140H1G9 |
| AVEDTQGMDNLACESQ | 1 | 0 | 0 | 1646711 | NP_828849.2 | P0C6X7 |
| AVESGGYIPSDFAFNN | 1 | 0 | 0 | 1646718 | NP_073551.1 | A0A0P0K6L9 |
| AVFISPYNSQNAVAAK | 1 | 0 | 0 | 1646724 | QIQ54047.1 | P0C6X7 |
| AVFISPYNSQNFAAKR | 1 | 0 | 0 | 1646727 | YP_009555238.1 | P0C6X6 |
| AVFISPYNSQNYVAKR | 1 | 0 | 0 | 1646729 | YP_173236.1 | A0A140H1G9 |
| AVGIVNTDFKLEIH | 1 | 0 | 0 | 1646741 | NP_073553.1 | A0A0P0KWE8 |
| AVGNICYTPSKLIEYS | 1 | 0 | 0 | 1646742 | NP_828849.2 | P0C6X7 |
| AVGTNLPLQLGFSTGV | 1 | 0 | 0 | 1646746 | YP_009724389.1 | P0DTD1 |
| AVGVRQQRFVYDAYQN | 1 | 0 | 0 | 1646747 | YP_009047204.1 | K9N5Q8 |
| AVHDFFKFRVDGDMVP | 1 | 0 | 0 | 1646749 | NP_828849.2 | P0C6X7 |
| AVIDQDFGSTVFNVPT | 1 | 0 | 0 | 1646758 | YP_009555238.1 | P0C6X6 |
| AVIGDFNCTNFAINDL | 1 | 0 | 0 | 1646760 | YP_173238.1 | A0A140H1H1 |
| AVIGDLKCTSDNINDK | 1 | 0 | 0 | 1646761 | YP_009555241.1 | P36334 |
| AVIKTLQPVSELLIPL | 1 | 0 | 0 | 1646763 | QHR63299.1 | P0C6W2 |
| AVIKTLQPVSELLTPL | 1 | 0 | 0 | 1646764 | YP_009724389.1 | P0DTD1 |

| | | | | | | |
|---|---|---|---|---|---|---|
| AVIMTLDLLKDIKFVI | 1 | 0 | 0 | 1646768 | YP_009047202.1 | K9N7C7 |
| AVINLRADQINDMVYS | 1 | 0 | 0 | 1646771 | YP_009555238.1 | P0C6X6 |
| AVITRDVGFVVPGLPG | 1 | 0 | 0 | 1646773 | QIQ54046.1 | P0C6U8 |
| AVITTQQAGAGIKYFC | 1 | 0 | 0 | 1646775 | NP_073549.1 | A0A0P0LKV0 |
| AVIYSGRESLIFPSGT | 1 | 0 | 0 | 1646778 | YP_009047202.1 | K9N7C7 |
| AVKAAESKAFVDTIVP | 1 | 0 | 0 | 1646779 | NP_073549.1 | A0A0P0LKV0 |
| AVKDFDFYRYNKPTIL | 1 | 0 | 0 | 1646780 | YP_003766.2 | H9EJ42 |
| AVKGCVPLNAIPSLAA | 1 | 0 | 0 | 1646782 | YP_009555238.1 | P0C6X6 |
| AVKGCVPLSAIPALAA | 1 | 0 | 0 | 1646783 | YP_173236.1 | A0A140H1G9 |
| AVKITEHSWNADLYKL | 1 | 0 | 0 | 1646786 | NP_828849.2 | P0C6X7 |
| AVKQGAKPVGNCVKML | 1 | 0 | 0 | 1646792 | NP_073549.1 | A0A0P0LKV0 |
| AVKRVRAWVGFDAEGA | 1 | 0 | 0 | 1646794 | YP_009555238.1 | P0C6X6 |
| AVKSDYCGGNSCTCRP | 1 | 0 | 0 | 1646795 | YP_009555241.1 | P36334 |
| AVKTQFNYYKKVDGVV | 1 | 0 | 0 | 1646799 | YP_009724389.1 | P0DTD1 |
| AVLDMCAALKELLQNG | 1 | 0 | 0 | 1646800 | NP_828849.2 | P0C6X7 |
| AVLFSATAVKTGGKSL | 1 | 0 | 0 | 1646804 | NP_073549.1 | A0A0P0LKV0 |
| AVLFSTVVIKNLTPIK | 1 | 0 | 0 | 1646805 | YP_003766.2 | H9EJ42 |
| AVLSDVDGLKYTKIMK | 1 | 0 | 0 | 1646819 | YP_173236.1 | A0A140H1G9 |
| AVLTIANNYSKLFDEG | 1 | 0 | 0 | 1646820 | NP_073549.1 | A0A0P0LKV0 |
| AVMSLKENQINDMIYS | 1 | 0 | 0 | 1646835 | NP_828849.2 | P0C6X7 |
| AVMYMGTLSYDNLKTG | 1 | 0 | 0 | 1646838 | NP_828849.2 | P0C6X7 |
| AVMYMGTLSYEQFKKG | 1 | 0 | 0 | 1646839 | YP_009724389.1 | P0DTD1 |
| AVNESRYGLQNLLQLP | 1 | 0 | 0 | 1646843 | YP_003767.1 | A0A0P0G321 |
| AVNFNVTTAETTDFCT | 1 | 0 | 0 | 1646844 | NP_073551.1 | A0A0P0K6L9 |
| AVNIAKNAYEKDKAVA | 1 | 0 | 0 | 1646847 | YP_009047202.1 | K9N7C7 |
| AVNNNAQALSKLASEL | 1 | 0 | 0 | 1646853 | YP_009047204.1 | K9N5Q8 |
| AVNQSLSNAKVSAEVS | 1 | 0 | 0 | 1646854 | NP_073552.1 | A0A0P0KWE8 |
| AVNSVPWDTIANYAKP | 1 | 0 | 0 | 1646858 | YP_009724389.1 | P0DTD1 |
| AVPTGYVDTENNTEFT | 1 | 0 | 0 | 1646870 | NP_828849.2 | P0C6X7 |
| AVPTGYVDTPNNTDFS | 1 | 0 | 0 | 1646872 | YP_009724389.1 | P0DTD1 |
| AVQIGNNNVCTYEHVI | 1 | 0 | 0 | 1646877 | NP_073549.1 | A0A0P0LKV0 |
| AVQIGSNNVCTYEHVI | 1 | 0 | 0 | 1646878 | YP_003766.2 | H9EJ42 |
| AVQKESDEYILAKGPL | 1 | 0 | 0 | 1646879 | YP_009047202.1 | K9N7C7 |
| AVQSKDLNFLNRVRGT | 1 | 0 | 0 | 1646882 | YP_173236.1 | A0A140H1G9 |
| AVRKEGQDVIFSQFDS | 1 | 0 | 0 | 1646885 | YP_009555238.1 | P0C6X6 |
| AVRQVRSWIGFDVEGA | 1 | 0 | 0 | 1646888 | YP_009047202.1 | K9N7C7 |
| AVRSHPDFKLLHNLQA | 1 | 0 | 0 | 1646890 | YP_009047202.1 | K9N7C7 |
| AVRSSFNFDQKELLAY | 1 | 0 | 0 | 1646891 | YP_009555238.1 | P0C6X6 |
| AVSCGHYQTNIYSQNL | 1 | 0 | 0 | 1646897 | NP_073549.1 | A0A0P0LKV0 |
| AVSFADNNDITNTNLS | 1 | 0 | 0 | 1646898 | YP_009047211.1 | K9N4V7 |
| AVSFDYLIREAKTRVL | 1 | 0 | 0 | 1646899 | YP_009047202.1 | K9N7C7 |
| AVSRSMLGLTTQTVDS | 1 | 0 | 0 | 1646907 | YP_009047202.1 | K9N7C7 |
| AVSSNINCVLSVNSSN | 1 | 0 | 0 | 1646909 | NP_073549.1 | A0A0P0LKV0 |
| AVSSNINRLLSVPSDS | 1 | 0 | 0 | 1646910 | YP_003766.2 | H9EJ42 |
| AVSSPMSNMTENERLL | 1 | 0 | 0 | 1646911 | NP_073555.1 | A0A0P0KXN3 |
| AVTANVCSLMACNGHK | 1 | 0 | 0 | 1646916 | YP_173236.1 | A0A140H1G9 |
| AVTANVNALLSTDGNK | 1 | 0 | 0 | 1646917 | YP_009724389.1 | P0DTD1 |
| AVTAYNGYLTSSSKTP | 1 | 0 | 0 | 1646918 | YP_009724389.1 | P0DTD1 |
| AVTDAKWYCYDKNPIN | 1 | 0 | 0 | 1646920 | NP_073549.1 | A0A0P0LKV0 |
| AVTDAKWYCYDKQPVN | 1 | 0 | 0 | 1646921 | YP_003766.2 | H9EJ42 |
| AVTRYVDNNFCGPDGY | 1 | 0 | 0 | 1646927 | NP_828849.2 | P0C6X7 |
| AVTTYNGYLTSSSKTS | 1 | 0 | 0 | 1646929 | NP_828849.2 | P0C6X7 |
| AVVCDAIENALNSCKE | 1 | 0 | 0 | 1646932 | YP_173236.1 | A0A140H1G9 |
| AVVCFNSTYASQGLVA | 1 | 0 | 0 | 1646933 | YP_009724389.1 | P0DTD1 |
| AVVCYNSNYAAQGLVA | 1 | 0 | 0 | 1646935 | NP_828849.2 | P0C6X7 |
| AVVGAMLSENFTSYGF | 1 | 0 | 0 | 1646938 | NP_073551.1 | A0A0P0K6L9 |
| AVVIPTKKAGGTTEML | 1 | 0 | 0 | 1646941 | YP_009724389.1 | P0DTD1 |
| AVVKTLQPVSDLLTNM | 1 | 0 | 0 | 1646946 | NP_828849.2 | P0C6X7 |
| AVVLLERYVSLAIDAY | 1 | 0 | 0 | 1646947 | YP_003766.2 | H9EJ42 |
| AVVNANAEALNNLLQQ | 1 | 0 | 0 | 1646949 | YP_009555241.1 | P36334 |
| AVVNDKIFTKSGVNDI | 1 | 0 | 0 | 1646950 | YP_009047202.1 | K9N7C7 |
| AVVNLRPDQLNDLVYS | 1 | 0 | 0 | 1646953 | YP_173236.1 | A0A140H1G9 |
| AVVSVSSPDAVTTYNG | 1 | 0 | 0 | 1646955 | NP_828849.2 | P0C6X7 |
| AVYANVGRWSASINTG | 1 | 0 | 0 | 1646959 | NP_073551.1 | A0A0P0K6L9 |
| AVYQQSIIGAMTAVNE | 1 | 0 | 0 | 1646966 | YP_003767.1 | A0A0P0G321 |
| AVYTKVDGIDVEIFEN | 1 | 0 | 0 | 1646971 | NP_828849.2 | P0C6X7 |
| AWDVIRKDLSYEQQNV | 1 | 0 | 0 | 1646979 | YP_003766.2 | H9EJ42 |
| AWFTERSDKSYEHQTP | 1 | 0 | 0 | 1646985 | NP_828849.2 | P0C6X7 |
| AWHVVRDPRYVMRLQS | 1 | 0 | 0 | 1646990 | YP_009555238.1 | P0C6X6 |
| AWHVVRDSRFVMRLQT | 1 | 0 | 0 | 1646991 | YP_173236.1 | A0A140H1G9 |
| AWIGFDVEGCHATRDA | 1 | 0 | 0 | 1646992 | NP_828849.2 | P0C6X7 |
| AWLNDKSVPSPLNWER | 1 | 0 | 0 | 1647000 | YP_009555241.1 | P36334 |
| AWNSKHIDAKEGGNFN | 1 | 0 | 0 | 1647013 | QHR63300.2 | Q3LZX1 |
| AWNSVKQDALTGGNYG | 1 | 0 | 0 | 1647015 | QIQ54048.1 | P59594 |
| AWNTRNIDATSTGNYN | 1 | 0 | 0 | 1647016 | NP_828851.1 | P59594 |
| AWRIYSTGNNVFQTQA | 1 | 0 | 0 | 1647021 | NP_828851.1 | P59594 |
| AWSGLCVDGTNGYVLR | 1 | 0 | 0 | 1647026 | NP_073551.1 | A0A0P0K6L9 |
| AWVGFDAEGAHATRDS | 1 | 0 | 0 | 1647029 | YP_009555238.1 | P0C6X6 |
| AWWTAFVTNVNASSSE | 1 | 0 | 0 | 1647033 | YP_009724389.1 | P0DTD1 |
| AYADKPNHFINFPLTH | 1 | 0 | 0 | 1647039 | YP_009555239.1 | Q80872 |
| AYANSVFNICQAVTAN | 1 | 0 | 0 | 1647051 | YP_009724389.1 | P0DTD1 |
| AYANSVFNILQATTAN | 1 | 0 | 0 | 1647053 | YP_009047202.1 | K9N7C7 |
| AYAPTYVSVDDKYKTS | 1 | 0 | 0 | 1647055 | YP_009047202.1 | K9N7C7 |
| AYATAQEAYEQAVANG | 1 | 0 | 0 | 1647057 | NP_828849.2 | P0C6X7 |
| AYCVDLHNKINLCDDP | 1 | 0 | 0 | 1647064 | YP_003766.2 | H9EJ42 |
| AYDKRAMAKLKPAPFF | 1 | 0 | 0 | 1647068 | NP_073549.1 | A0A0P0LKV0 |
| AYDVSAFRDLKPLPFF | 1 | 0 | 0 | 1647072 | YP_009047202.1 | K9N7C7 |
| AYEDAIANGSSSQLIK | 1 | 0 | 0 | 1647073 | YP_003766.2 | H9EJ42 |
| AYEKDKAVARKLERMA | 1 | 0 | 0 | 1647074 | YP_009047202.1 | K9N7C7 |
| AYENARQAYEDAIANG | 1 | 0 | 0 | 1647075 | YP_003766.2 | H9EJ42 |
| AYENFNHHEKLLAPLL | 1 | 0 | 0 | 1647076 | QIQ54046.1 | P0C6U8 |

| | | | | | | |
|---|---|---|---|---|---|---|
| AYENFNSQDILLAPLL | 1 | 0 | 0 | 1647079 | NP_828849.2 | P0C6X7 |
| AYESLRPDTRYVLMDG | 1 | 0 | 0 | 1647083 | YP_009724389.1 | P0DTD1 |
| AYETARQEYENAVANG | 1 | 0 | 0 | 1647084 | NP_073549.1 | A0A0P0LKV0 |
| AYFNGAIIRDSDVVKQ | 1 | 0 | 0 | 1647088 | YP_009047202.1 | K9N7C7 |
| AYFSKKHVEVFTDGKL | 1 | 0 | 0 | 1647092 | YP_009047202.1 | K9N7C7 |
| AYFTEQPIDLVPTQPY | 1 | 0 | 0 | 1647094 | QIQ54046.1 | P0C6U8 |
| AYGNCTFGAPDDASRL | 1 | 0 | 0 | 1647100 | YP_009047202.1 | K9N7C7 |
| AYIDVLRNSFGKDLNA | 1 | 0 | 0 | 1647103 | NP_073549.1 | A0A0P0LKV0 |
| AYIKKLYNDLGNQILD | 1 | 0 | 0 | 1647105 | YP_009555238.1 | P0C6X6 |
| AYISNKADLKFVKWEY | 1 | 0 | 0 | 1647108 | YP_003766.2 | H9EJ42 |
| AYKDYLASGGQPITNC | 1 | 0 | 0 | 1647113 | YP_009724389.1 | P0DTD1 |
| AYKFLNINDCTRIVPA | 1 | 0 | 0 | 1647115 | YP_009555238.1 | P0C6X6 |
| AYLAKAMLDFSRDHND | 1 | 0 | 0 | 1647124 | NP_073549.1 | A0A0P0LKV0 |
| AYLDAVKQGAKPVGNC | 1 | 0 | 0 | 1647125 | NP_073549.1 | A0A0P0LKV0 |
| AYLDFVNAGGAPLTNC | 1 | 0 | 0 | 1647126 | YP_009047202.1 | K9N7C7 |
| AYNGYLTSSSKTPEEH | 1 | 0 | 0 | 1647144 | YP_009724389.1 | P0DTD1 |
| AYNPDLSQYYCDNGKY | 1 | 0 | 0 | 1647151 | YP_173236.1 | A0A140H1G9 |
| AYNRYRIGNYKLNTDH | 1 | 0 | 0 | 1647153 | NP_828855.1 | P59596 |
| AYNSILFENGIPLMPL | 1 | 0 | 0 | 1647154 | YP_003766.2 | H9EJ42 |
| AYNTINNEQGTPLTPI | 1 | 0 | 0 | 1647157 | NP_073549.1 | A0A0P0LKV0 |
| AYNVVNKGHFDGHAGE | 1 | 0 | 0 | 1647159 | NP_828849.2 | P0C6X7 |
| AYPLTKHEDIEYQNVF | 1 | 0 | 0 | 1647163 | YP_009047202.1 | K9N7C7 |
| AYPLVHHENEEYQKVF | 1 | 0 | 0 | 1647165 | YP_173236.1 | A0A140H1G9 |
| AYPLVYHENEEYQKVF | 1 | 0 | 0 | 1647167 | YP_009555238.1 | P0C6X6 |
| AYQDYMQIAPVPAEVL | 1 | 0 | 0 | 1647168 | YP_003769.1 | H9EJA2 |
| AYQNLVGYYSDDGNYY | 1 | 0 | 0 | 1647171 | YP_009047204.1 | K9N5Q8 |
| AYQVAARYFDCYEGGC | 1 | 0 | 0 | 1647173 | NP_073549.1 | A0A0P0LKV0 |
| AYRDYLASGGQPITNC | 1 | 0 | 0 | 1647175 | QIQ54046.1 | P0C6U8 |
| AYRKILLRKNGNKGAG | 1 | 0 | 0 | 1647181 | QIQ54046.1 | P0C6U8 |
| AYRKVLLRKNGNKGAG | 1 | 0 | 0 | 1647182 | YP_009724389.1 | P0DTD1 |
| AYRNVLLRKNGNKGAG | 1 | 0 | 0 | 1647184 | NP_828849.2 | P0C6X7 |
| AYSLFDMAKFPLKLAG | 1 | 0 | 0 | 1647187 | YP_009555238.1 | P0C6X6 |
| AYSNKTVGELGDVRET | 1 | 0 | 0 | 1647189 | NP_828849.2 | P0C6X7 |
| AYSPQLTLVFPEVKMI | 1 | 0 | 0 | 1647193 | YP_009047202.1 | K9N7C7 |
| AYSQMRPHVRYDLYDG | 1 | 0 | 0 | 1647194 | YP_009047202.1 | K9N7C7 |
| AYSSPLQSYACWTHSC | 1 | 0 | 0 | 1647196 | YP_009047202.1 | K9N7C7 |
| AYSSSANGSLVGTTLQ | 1 | 0 | 0 | 1647197 | YP_009047202.1 | K9N7C7 |
| AYSTMRLHNFSVSHNG | 1 | 0 | 0 | 1647198 | YP_003766.2 | H9EJ42 |
| AYSVDKCTRIIPARAR | 1 | 0 | 0 | 1647199 | NP_073549.1 | A0A0P0LKV0 |
| AYTAFSGPVDKGHYTV | 1 | 0 | 0 | 1647201 | NP_073549.1 | A0A0P0LKV0 |
| AYTMSLGADSSIAYSN | 1 | 0 | 0 | 1647207 | NP_828851.1 | P59594 |
| AYTNSSTRGVYYPDKV | 1 | 0 | 0 | 1647209 | QHR63300.2 | Q3LZX1 |
| AYTRYVDNNFCGPDGY | 1 | 0 | 0 | 1647210 | YP_009724389.1 | P0DTD1 |
| AYTTFSGSFDNGHYVV | 1 | 0 | 0 | 1647212 | YP_003766.2 | H9EJ42 |
| AYTVELGTEVNEFACV | 1 | 0 | 0 | 1647213 | YP_009724389.1 | P0DTD1 |
| AYVANPTEDLCDGSVV | 1 | 0 | 0 | 1647215 | YP_009555238.1 | P0C6X6 |
| AYVDNSSLTIKKPNEL | 1 | 0 | 0 | 1647216 | YP_009724389.1 | P0DTD1 |
| AYVDNTSLTIKKPNEL | 1 | 0 | 0 | 1647217 | QIQ54046.1 | P0C6U8 |
| AYVDTFSATFSVPMEK | 1 | 0 | 0 | 1647218 | NP_828849.2 | P0C6X7 |
| AYVDVLCNSFFKELTA | 1 | 0 | 0 | 1647219 | YP_003766.2 | H9EJ42 |
| AYVENTSITIKKPNEL | 1 | 0 | 0 | 1647220 | NP_828849.2 | P0C6X7 |
| AYVIIDKVDFVNGFYR | 1 | 0 | 0 | 1647223 | YP_003766.2 | H9EJ42 |
| AYVMIDKVEFENGFYR | 1 | 0 | 0 | 1647225 | NP_073549.1 | A0A0P0LKV0 |
| AYVNSNQNTDSDSLSS | 1 | 0 | 0 | 1647226 | YP_173242.1 | Q19U29 |
| AYVNTFSSTFNVPMDK | 1 | 0 | 0 | 1647227 | QIQ54046.1 | P0C6U8 |
| AYVNTFSSTFNVPMEK | 1 | 0 | 0 | 1647228 | YP_009724389.1 | P0DTD1 |
| AYVSQQLSDISLVKFG | 1 | 0 | 0 | 1647229 | YP_173238.1 | A0A140H1H1 |
| AYVSQQLSDSTLVKFS | 1 | 0 | 0 | 1647230 | YP_009555241.1 | P36334 |
| AYVTGGVVQMTTQWLT | 1 | 0 | 0 | 1647232 | QIQ54046.1 | P0C6U8 |
| AYYEPVQGRKMLMALL | 1 | 0 | 0 | 1647245 | YP_009047202.1 | K9N7C7 |
| AYYKYDAKNYVRFPEI | 1 | 0 | 0 | 1647248 | YP_003766.2 | H9EJ42 |
| AYYKYDNGNFIKLPEV | 1 | 0 | 0 | 1647249 | NP_073549.1 | A0A0P0LKV0 |
| AYYTEQPIDLVPTQPL | 1 | 0 | 0 | 1647258 | NP_828849.2 | P0C6X7 |
| CADSIVIGGAKLKALN | 1 | 0 | 0 | 1647279 | QIQ54046.1 | P0C6U8 |
| CAFGGVSVITPGTNAS | 1 | 0 | 0 | 1647285 | NP_828851.1 | P59594 |
| CAFSVDPKKTYLDYIQ | 1 | 0 | 0 | 1647290 | YP_173236.1 | A0A140H1G9 |
| CAGNVIGNDALTRFTI | 1 | 0 | 0 | 1647294 | YP_173236.1 | A0A140H1G9 |
| CAGRKVNFNEKPVVME | 1 | 0 | 0 | 1647296 | YP_173236.1 | A0A140H1G9 |
| CAGRRVTFKEQPTVKE | 1 | 0 | 0 | 1647297 | YP_009555238.1 | P0C6X6 |
| CAIINDKVVAKIDKED | 1 | 0 | 0 | 1647302 | YP_009555238.1 | P0C6X6 |
| CAIVQASESKVLLERF | 1 | 0 | 0 | 1647307 | YP_003766.2 | H9EJ42 |
| CAKEIKESVQTFFKLV | 1 | 0 | 0 | 1647310 | YP_009724389.1 | P0DTD1 |
| CALILAYCNKTVGELG | 1 | 0 | 0 | 1647322 | YP_009724389.1 | P0DTD1 |
| CALPDTPSTLTPRSVR | 1 | 0 | 0 | 1647325 | YP_009047204.1 | K9N5Q8 |
| CAMRPNHTIKGSFLNG | 1 | 0 | 0 | 1647331 | NP_828849.2 | P0C6X7 |
| CANDPVGFTLKNTVCT | 1 | 0 | 0 | 1647332 | YP_009724389.1 | P0DTD1 |
| CANDPVGFTLRNTVCT | 1 | 0 | 0 | 1647333 | NP_828849.2 | P0C6X7 |
| CAPVVCPKGKIVVIAG | 1 | 0 | 0 | 1647349 | YP_003766.2 | H9EJ42 |
| CAQVLSEYVLCGGGYY | 1 | 0 | 0 | 1647357 | YP_009047202.1 | K9N7C7 |
| CARNPIDLPPSHATTY | 1 | 0 | 0 | 1647361 | NP_073549.1 | A0A0P0LKV0 |
| CASVKRDGVQVGYCVH | 1 | 0 | 0 | 1647368 | NP_073549.1 | A0A0P0LKV0 |
| CASYQTQTNSRSVASQ | 1 | 0 | 0 | 1647373 | QHR63300.2 | Q3LZX1 |
| CATCPFAKGDSCSSNC | 1 | 0 | 0 | 1647374 | YP_009047202.1 | K9N7C7 |
| CATTRQVVNVVTTKIA | 1 | 0 | 0 | 1647379 | YP_009724389.1 | P0DTD1 |
| CAWDYHVTKVLAEKFD | 1 | 0 | 0 | 1647390 | NP_073549.1 | A0A0P0LKV0 |
| CAYCCNIVNVSLVKPS | 1 | 0 | 0 | 1647395 | YP_009724392.1 | P0DTC4 |
| CCDCGSTKLPYYEFEK | 1 | 0 | 0 | 1647406 | YP_003767.1 | A0A0P0G321 |
| CCDDYTGYQELVIKTS | 1 | 0 | 0 | 1647408 | YP_009555241.1 | P36334 |
| CCDEYGGHHDFVIKTS | 1 | 0 | 0 | 1647409 | YP_173238.1 | A0A140H1H1 |
| CCESTKLPYYDVEKIH | 1 | 0 | 0 | 1647413 | NP_073551.1 | A0A0P0K6L9 |
| CCFSVAAITSGVKFQT | 1 | 0 | 0 | 1647417 | YP_009555238.1 | P0C6X6 |

| | | | | | | |
|---|---|---|---|---|---|---|
| CCHLAKALNDFSNSGA | 1 | 0 | 0 | 1647421 | NP_828849.2 | P0C6X7 |
| CCHLAKALNDFSNSGS | 1 | 0 | 0 | 1647422 | YP_009724389.1 | P0DTD1 |
| CCLKCGFSFDLNGLDA | 1 | 0 | 0 | 1647436 | YP_173236.1 | A0A140H1G9 |
| CCRFQEKDEDDNLIDS | 1 | 0 | 0 | 1647453 | YP_009724389.1 | P0DTD1 |
| CCRFQEKDEDGNLIDS | 1 | 0 | 0 | 1647454 | QIQ54047.1 | P0C6X7 |
| CCRFQEKDEEGNLLDS | 1 | 0 | 0 | 1647455 | NP_828849.2 | P0C6X7 |
| CCRFQRIDDDGNKLDK | 1 | 0 | 0 | 1647456 | YP_173236.1 | A0A140H1G9 |
| CCRFQRVDENGDKLDQ | 1 | 0 | 0 | 1647457 | YP_009555238.1 | P0C6X6 |
| CCTASDKFYRLSNELA | 1 | 0 | 0 | 1647469 | NP_073549.1 | A0A0P0LKV0 |
| CCTVYDSSEVKAQSSG | 1 | 0 | 0 | 1647479 | YP_173236.1 | A0A140H1G9 |
| CCYDHVIATPHKMVLS | 1 | 0 | 0 | 1647481 | YP_009047202.1 | K9N7C7 |
| CCYDHVMATDHKYVLS | 1 | 0 | 0 | 1647483 | YP_009555238.1 | P0C6X6 |
| CCYDHVMATNHKYVLS | 1 | 0 | 0 | 1647484 | YP_173236.1 | A0A140H1G9 |
| CCYVGVQTVEDLRARM | 1 | 0 | 0 | 1647486 | YP_009047202.1 | K9N7C7 |
| CDAIENALNSCKEHPV | 1 | 0 | 0 | 1647488 | YP_173236.1 | A0A140H1G9 |
| CDAKFKNSVASINSAI | 1 | 0 | 0 | 1647490 | NP_073549.1 | A0A0P0LKV0 |
| CDDCNGYTDNIFSVQQ | 1 | 0 | 0 | 1647499 | YP_003767.1 | A0A0P0G321 |
| CDDYTGYQELVIKTSH | 1 | 0 | 0 | 1647504 | YP_009555241.1 | P36334 |
| CDEECSEVEASDLEEG | 1 | 0 | 0 | 1647505 | YP_009047202.1 | K9N7C7 |
| CDEFTLAEVVKQMYGV | 1 | 0 | 0 | 1647506 | YP_003766.2 | H9EJ42 |
| CDEVSWQTCDFLKATC | 1 | 0 | 0 | 1647511 | NP_828849.2 | P0C6X7 |
| CDEYGGHHDFVIKTSH | 1 | 0 | 0 | 1647512 | YP_173238.1 | A0A140H1H1 |
| CDFEIACKCGVKQEQR | 1 | 0 | 0 | 1647515 | YP_009555238.1 | P0C6X6 |
| CDFNKPLIEWPLTEYD | 1 | 0 | 0 | 1647518 | YP_009047202.1 | K9N7C7 |
| CDFYGWVSGNMMDGFS | 1 | 0 | 0 | 1647520 | YP_173236.1 | A0A140H1G9 |
| CDGLASAYRANSFDVP | 1 | 0 | 0 | 1647526 | YP_009047202.1 | K9N7C7 |
| CDGLKFTEESFYKNMY | 1 | 0 | 0 | 1647527 | YP_173236.1 | A0A140H1G9 |
| CDGSLGCKMCLFGYQE | 1 | 0 | 0 | 1647531 | NP_073549.1 | A0A0P0LKV0 |
| CDGVNVTKHKCDINYK | 1 | 0 | 0 | 1647535 | YP_009555238.1 | P0C6X6 |
| CDICKSTVVEVKSAIV | 1 | 0 | 0 | 1647540 | YP_003766.2 | H9EJ42 |
| CDIDKWLNNFNVPSPL | 1 | 0 | 0 | 1647541 | YP_173238.1 | A0A140H1H1 |
| CDINYKGKVFFQFDNL | 1 | 0 | 0 | 1647545 | YP_009555238.1 | P0C6X6 |
| CDKRATCFSTASDTYA | 1 | 0 | 0 | 1647553 | QHR63299.1 | P0C6W2 |
| CDKVRSLKLSDNNCYL | 1 | 0 | 0 | 1647556 | YP_009047202.1 | K9N7C7 |
| CDLQNYGENAVIPKGI | 1 | 0 | 0 | 1647561 | NP_828849.2 | P0C6X7 |
| CDNIKLADDLNQLTGY | 1 | 0 | 0 | 1647569 | QHR63299.1 | P0C6W2 |
| CDNINSILDEVNGLLD | 1 | 0 | 0 | 1647571 | YP_173238.1 | A0A140H1H1 |
| CDNPFFAVSKPMGTQT | 1 | 0 | 0 | 1647573 | NP_828851.1 | P59594 |
| CDNSYFDSKGWYDPVE | 1 | 0 | 0 | 1647574 | YP_003766.2 | H9EJ42 |
| CDQLSFDVPDGFYSTS | 1 | 0 | 0 | 1647578 | NP_073551.1 | A0A0P0K6L9 |
| CDRRATCFSTASDTYA | 1 | 0 | 0 | 1647585 | YP_009724389.1 | P0DTD1 |
| CDRTAIQSFDNSYLNE | 1 | 0 | 0 | 1647586 | NP_073550.1 | A0A0P0ILY9 |
| CDRTAIQSFDNSYLNR | 1 | 0 | 0 | 1647587 | NP_073549.1 | A0A0P0LKV0 |
| CDSFGPGNTFINGDIA | 1 | 0 | 0 | 1647591 | YP_003766.2 | H9EJ42 |
| CDSLNAKLGFDSSKEF | 1 | 0 | 0 | 1647593 | YP_009555238.1 | P0C6X6 |
| CDSPFEKLQCEHLQFG | 1 | 0 | 0 | 1647596 | YP_003767.1 | A0A0P0G321 |
| CDVKTTEVVGDVILKP | 1 | 0 | 0 | 1647606 | QIQ54046.1 | P0C6U8 |
| CDVNDVTKLYLGGMSY | 1 | 0 | 0 | 1647610 | YP_009555238.1 | P0C6X6 |
| CDVSDVTKLYLGGMSY | 1 | 0 | 0 | 1647611 | YP_173236.1 | A0A140H1G9 |
| CDVVQEQVNYVPLRAS | 1 | 0 | 0 | 1647616 | YP_003766.2 | H9EJ42 |
| CDWTESGDYTLANTTT | 1 | 0 | 0 | 1647618 | YP_009047202.1 | K9N7C7 |
| CEEEDFEPPTQYEYGT | 1 | 0 | 0 | 1647634 | QHR63299.1 | P0C6W2 |
| CEEGTMYFVPRAKSVV | 1 | 0 | 0 | 1647637 | YP_009047202.1 | K9N7C7 |
| CEEVANDLTTALRRPI | 1 | 0 | 0 | 1647640 | YP_009047202.1 | K9N7C7 |
| CEFCGTENQVKEGPTT | 1 | 0 | 0 | 1647642 | QIQ54046.1 | P0C6U8 |
| CEHPAVGIVNTDFKLE | 1 | 0 | 0 | 1647652 | NP_073553.1 | A0A0P0KWE8 |
| CEILCEYADCKESYFS | 1 | 0 | 0 | 1647654 | YP_173236.1 | A0A140H1G9 |
| CEIVGGQIVTCAKEIK | 1 | 0 | 0 | 1647655 | YP_009724389.1 | P0DTD1 |
| CEKAIGKKIKHEGDWD | 1 | 0 | 0 | 1647657 | NP_073549.1 | A0A0P0LKV0 |
| CEKNLQLMRKVDLYNG | 1 | 0 | 0 | 1647662 | YP_003768.1 | H9EJ24 |
| CELCHSGFDMLDTYAA | 1 | 0 | 0 | 1647665 | YP_173236.1 | A0A140H1G9 |
| CELFVNDSYRQFDLVQ | 1 | 0 | 0 | 1647667 | YP_173236.1 | A0A140H1G9 |
| CELYVNNAYRLFDLVQ | 1 | 0 | 0 | 1647670 | YP_009555238.1 | P0C6X6 |
| CENLESAVLFVNDKIT | 1 | 0 | 0 | 1647672 | NP_073549.1 | A0A0P0LKV0 |
| CEQSYQLYDASNVKKV | 1 | 0 | 0 | 1647680 | YP_009555238.1 | P0C6X6 |
| CESHGKQVVSDIDYVP | 1 | 0 | 0 | 1647685 | YP_009724389.1 | P0DTD1 |
| CESTKLPYYDVEKIHI | 1 | 0 | 0 | 1647687 | NP_073551.1 | A0A0P0K6L9 |
| CETFWRYNFDITESKY | 1 | 0 | 0 | 1647690 | NP_073549.1 | A0A0P0LKV0 |
| CEVVHDQVNYVPLRAT | 1 | 0 | 0 | 1647693 | NP_073549.1 | A0A0P0LKV0 |
| CEVVVTNLNKSAGWPL | 1 | 0 | 0 | 1647694 | YP_003766.2 | H9EJ42 |
| CFALRNFDEKDCEVFK | 1 | 0 | 0 | 1647701 | NP_073549.1 | A0A0P0LKV0 |
| CFANKFEQFDQWYEST | 1 | 0 | 0 | 1647702 | YP_009555238.1 | P0C6X6 |
| CFDKFKVNSTLEQYVF | 1 | 0 | 0 | 1647714 | YP_009724389.1 | P0DTD1 |
| CFDKSKQFVPVQVVDS | 1 | 0 | 0 | 1647715 | YP_173237.1 | A0A140H1H0 |
| CFGKRGPNQNFGGGEM | 1 | 0 | 0 | 1647726 | YP_009555245.1 | P33469 |
| CFGKRGPSQNFGNAEM | 1 | 0 | 0 | 1647727 | YP_173242.1 | Q19U29 |
| CFKNDNVVLKITEDGI | 1 | 0 | 0 | 1647749 | YP_003766.2 | H9EJ42 |
| CFKVFFKGNVQVDNGS | 1 | 0 | 0 | 1647751 | YP_003766.2 | H9EJ42 |
| CFKVYFKGQTTHESSS | 1 | 0 | 0 | 1647752 | YP_173236.1 | A0A140H1G9 |
| CFMKSDIFGQDFKTFD | 1 | 0 | 0 | 1647762 | NP_073549.1 | A0A0P0LKV0 |
| CFSKCHNCCDEYGGHH | 1 | 0 | 0 | 1647778 | YP_173238.1 | A0A140H1H1 |
| CFSVAAITSGVKFQTV | 1 | 0 | 0 | 1647788 | YP_009555238.1 | P0C6X6 |
| CFSVAALSTGLTSQTV | 1 | 0 | 0 | 1647790 | NP_073549.1 | A0A0P0LKV0 |
| CFTGVDGELPVAVVND | 1 | 0 | 0 | 1647796 | YP_003766.2 | H9EJ42 |
| CFVKNVDWSITYPMIA | 1 | 0 | 0 | 1647808 | NP_073549.1 | A0A0P0LKV0 |
| CFVKSDIFGSDFKTFD | 1 | 0 | 0 | 1647812 | YP_003766.2 | H9EJ42 |
| CFVNTTIGNETTSAFV | 1 | 0 | 0 | 1647818 | NP_073551.1 | A0A0P0K6L9 |
| CGADGKPVMSEDLWQF | 1 | 0 | 0 | 1647829 | NP_073549.1 | A0A0P0LKV0 |
| CGASYTANDVEVQSSG | 1 | 0 | 0 | 1647830 | YP_009047202.1 | K9N7C7 |
| CGCGASYTANDVEVQS | 1 | 0 | 0 | 1647832 | YP_009047202.1 | K9N7C7 |
| CGDIVSGYEKSTFNKD | 1 | 0 | 0 | 1647839 | YP_003766.2 | H9EJ42 |

| | | | | | | |
|---|---|---|---|---|---|---|
| CGDKFFPIVMNDKNIC | 1 | 0 | 0 | 1647840 | YP_173236.1 | A0A140H1G9 |
| CGDNIGTNVPLQVGFS | 1 | 0 | 0 | 1647841 | YP_003766.2 | H9EJ42 |
| CGDSIECSNLLLQYGS | 1 | 0 | 0 | 1647843 | QIQ54048.1 | P59594 |
| CGDYAACKSQLVEYGS | 1 | 0 | 0 | 1647846 | YP_009555241.1 | P36334 |
| CGEPWFLTDYNAILQS | 1 | 0 | 0 | 1647847 | YP_003766.2 | H9EJ42 |
| CGGCCDDYTGYQELVI | 1 | 0 | 0 | 1647862 | YP_009555241.1 | P36334 |
| CGGCYYVKPGGTSSGD | 1 | 0 | 0 | 1647863 | YP_009555238.1 | P0C6X6 |
| CGGGYYVKPGGTSSGD | 1 | 0 | 0 | 1647865 | YP_009047202.1 | K9N7C7 |
| CGGSLYVKPGGTSSGD | 1 | 0 | 0 | 1647869 | YP_009724389.1 | P0DTD1 |
| CGHYQTNIYSQNLCVD | 1 | 0 | 0 | 1647874 | NP_073549.1 | A0A0P0LKV0 |
| CGKGYHLMSFPQAAPH | 1 | 0 | 0 | 1647882 | NP_828851.1 | P59594 |
| CGKQATKYLVQQESPF | 1 | 0 | 0 | 1647886 | YP_009724389.1 | P0DTD1 |
| CGKQATQYLVQQESPF | 1 | 0 | 0 | 1647887 | QHR63299.1 | P0C6W2 |
| CGLFKDCARNPIDLPP | 1 | 0 | 0 | 1647888 | NP_073549.1 | A0A0P0LKV0 |
| CGLVELEKGVLPQLEQ | 1 | 0 | 0 | 1647891 | NP_828849.2 | P0C6X7 |
| CGNDSWLTGNAIQGFA | 1 | 0 | 0 | 1647896 | YP_009047202.1 | K9N7C7 |
| CGNLHQRELLGVSDVW | 1 | 0 | 0 | 1647905 | YP_009555238.1 | P0C6X6 |
| CGQKTTTLTGVEAVMY | 1 | 0 | 0 | 1647915 | NP_828849.2 | P0C6X7 |
| CGQQQTTLKGVEAVMY | 1 | 0 | 0 | 1647916 | YP_009724389.1 | P0DTD1 |
| CGRQATQYLVQQESSF | 1 | 0 | 0 | 1647921 | QIQ54046.1 | P0C6U8 |
| CGSCGNDSWLTGNAIQ | 1 | 0 | 0 | 1647923 | YP_009047202.1 | K9N7C7 |
| CGSCGSVGYTKEGSVI | 1 | 0 | 0 | 1647924 | YP_009047202.1 | K9N7C7 |
| CGSGNLVEKLSPWFMD | 1 | 0 | 0 | 1647930 | YP_009047202.1 | K9N7C7 |
| CGSPGYNLKNGEVEFV | 1 | 0 | 0 | 1647932 | NP_073549.1 | A0A0P0LKV0 |
| CGSPGYNVRNDGTVEF | 1 | 0 | 0 | 1647933 | YP_003766.2 | H9EJ42 |
| CGSSSRSFFEDLLFDK | 1 | 0 | 0 | 1647938 | YP_173238.1 | A0A140H1H1 |
| CGSVGYVIMGDCVKFV | 1 | 0 | 0 | 1647942 | YP_009555238.1 | P0C6X6 |
| CGSVGYVLTGDSVKFV | 1 | 0 | 0 | 1647943 | YP_173236.1 | A0A140H1G9 |
| CGTENQVKEGPTTCGY | 1 | 0 | 0 | 1647947 | QIQ54046.1 | P0C6U8 |
| CGVCLSGLDSLDTYPS | 1 | 0 | 0 | 1647960 | QHR63299.1 | P0C6W2 |
| CGVDAVNLLTNMFTPL | 1 | 0 | 0 | 1647962 | YP_009724389.1 | P0DTD1 |
| CGVDSVNLLTNMFTPL | 1 | 0 | 0 | 1647964 | QIQ54046.1 | P0C6U8 |
| CGVSDVTKLYLGGMSY | 1 | 0 | 0 | 1647973 | YP_009047202.1 | K9N7C7 |
| CGYTDFAEDVCTCYDN | 1 | 0 | 0 | 1647984 | NP_073549.1 | A0A0P0LKV0 |
| CHALDCELYVNNAYRL | 1 | 0 | 0 | 1647985 | YP_009555238.1 | P0C6X6 |
| CHATREAIGTNLPLQL | 1 | 0 | 0 | 1647987 | QHR63299.1 | P0C6W2 |
| CHIDHPNPKGFCDLKG | 1 | 0 | 0 | 1647997 | YP_009724389.1 | P0DTD1 |
| CHIDHPNPKGYCELKG | 1 | 0 | 0 | 1647998 | QIQ54046.1 | P0C6U8 |
| CHLAKALNDFSNSGAD | 1 | 0 | 0 | 1648000 | NP_828849.2 | P0C6X7 |
| CHLAKALNDFSNSGSD | 1 | 0 | 0 | 1648001 | YP_009724389.1 | P0DTD1 |
| CHMFCNRTVYGPIKNV | 1 | 0 | 0 | 1648004 | NP_073554.1 | A0A0P0KH29 |
| CHNCCDEYGGHHDFVI | 1 | 0 | 0 | 1648006 | YP_173238.1 | A0A140H1H1 |
| CHSVAVIGDEQIDGKF | 1 | 0 | 0 | 1648017 | YP_173236.1 | A0A140H1G9 |
| CHVLDCELFVNDSYRQ | 1 | 0 | 0 | 1648020 | YP_173236.1 | A0A140H1G9 |
| CIDGALLTKSSEYKGP | 1 | 0 | 0 | 1648035 | YP_009724389.1 | P0DTD1 |
| CIDMKGKMFVRPVIED | 1 | 0 | 0 | 1648036 | YP_173241.1 | Q0ZJ82 |
| CIDMKGTMYVRPIIED | 1 | 0 | 0 | 1648037 | YP_009555244.1 | Q01455 |
| CIEEVTTTLEETKFLT | 1 | 0 | 0 | 1648040 | QIQ54046.1 | P0C6U8 |
| CIFNGKFLSNTKYYDD | 1 | 0 | 0 | 1648044 | YP_009555240.1 | P30215 |
| CIKACEVVVTNLNKSA | 1 | 0 | 0 | 1648054 | YP_003766.2 | H9EJ42 |
| CIKDADYNAKVEISVT | 1 | 0 | 0 | 1648056 | NP_073549.1 | A0A0P0LKV0 |
| CIKISTVQSKLTDLKC | 1 | 0 | 0 | 1648060 | YP_003766.2 | H9EJ42 |
| CIKTAQADTAGLYTNF | 1 | 0 | 0 | 1648064 | YP_009047205.1 | K9N796 |
| CIKVATVQSKMSDVKC | 1 | 0 | 0 | 1648066 | YP_009724389.1 | P0DTD1 |
| CIMSDRDLYDKLQFTS | 1 | 0 | 0 | 1648077 | YP_009724389.1 | P0DTD1 |
| CINANQVIVNNLDKSA | 1 | 0 | 0 | 1648081 | YP_009724389.1 | P0DTD1 |
| CIPASQVIVNNYDKSA | 1 | 0 | 0 | 1648085 | YP_009555238.1 | P0C6X6 |
| CIRCLWNTKPVETSNS | 1 | 0 | 0 | 1648091 | QHR63299.1 | P0C6W2 |
| CITTNGSWAIFNDHHL | 1 | 0 | 0 | 1648115 | YP_009047202.1 | K9N7C7 |
| CIVDNVYMAKAGDKYY | 1 | 0 | 0 | 1648117 | YP_009555238.1 | P0C6X6 |
| CIVGNKFYNVSTGLIP | 1 | 0 | 0 | 1648120 | YP_173236.1 | A0A140H1G9 |
| CKCGVKQEQRTGLDAV | 1 | 0 | 0 | 1648146 | YP_009555238.1 | P0C6X6 |
| CKDGHVETFYPKLQAS | 1 | 0 | 0 | 1648148 | NP_828849.2 | P0C6X7 |
| CKDGHVETFYPKLQSS | 1 | 0 | 0 | 1648149 | YP_009724389.1 | P0DTD1 |
| CKDGQVQTFYPRLQAS | 1 | 0 | 0 | 1648150 | YP_009047202.1 | K9N7C7 |
| CKDNAVATFYPQLQSA | 1 | 0 | 0 | 1648151 | NP_073549.1 | A0A0P0LKV0 |
| CKFKGKCVQVPIGCLD | 1 | 0 | 0 | 1648161 | YP_003766.2 | H9EJ42 |
| CKHCGQKTTTLTGVEA | 1 | 0 | 0 | 1648169 | NP_828849.2 | P0C6X7 |
| CKLPLGQSLCALPDTP | 1 | 0 | 0 | 1648185 | YP_009047204.1 | K9N5Q8 |
| CKPHQVNISLNGNTSV | 1 | 0 | 0 | 1648193 | YP_003767.1 | A0A0P0G321 |
| CKPKVEIDGNVMHANY | 1 | 0 | 0 | 1648194 | YP_009555238.1 | P0C6X6 |
| CKRALGLSISDHEFTS | 1 | 0 | 0 | 1648196 | NP_073549.1 | A0A0P0LKV0 |
| CKSHKPPSASCPIGTN | 1 | 0 | 0 | 1648203 | YP_173238.1 | A0A140H1H1 |
| CKSKGSSRNESWHFDK | 1 | 0 | 0 | 1648206 | YP_173238.1 | A0A140H1H1 |
| CKSQLVEYGSFCDNIN | 1 | 0 | 0 | 1648208 | YP_009555241.1 | P36334 |
| CKSVNWNLEYPIISNE | 1 | 0 | 0 | 1648210 | YP_173236.1 | A0A140H1G9 |
| CKTCGQQQTTLKGVEA | 1 | 0 | 0 | 1648212 | YP_009724389.1 | P0DTD1 |
| CKTIEDALRNSARLES | 1 | 0 | 0 | 1648217 | NP_073551.1 | A0A0P0K6L9 |
| CKTILNIVGPDARQDG | 1 | 0 | 0 | 1648218 | YP_173236.1 | A0A140H1G9 |
| CKTKSLLPNTGVYDLS | 1 | 0 | 0 | 1648220 | YP_173238.1 | A0A140H1H1 |
| CKTVLNVVGPDARTQG | 1 | 0 | 0 | 1648223 | YP_009555238.1 | P0C6X6 |
| CKVTDVLFPGQKYSSD | 1 | 0 | 0 | 1648227 | YP_009047202.1 | K9N7C7 |
| CLASECFMKSDIFGQD | 1 | 0 | 0 | 1648246 | NP_073549.1 | A0A0P0LKV0 |
| CLASECFVKSDIFGSD | 1 | 0 | 0 | 1648247 | YP_003766.2 | H9EJ42 |
| CLCDKRATCFSTASDT | 1 | 0 | 0 | 1648257 | QHR63299.1 | P0C6W2 |
| CLCDKRATCFSTSSDT | 1 | 0 | 0 | 1648258 | NP_828849.2 | P0C6X7 |
| CLGPHCGSSSRSFFED | 1 | 0 | 0 | 1648295 | YP_173238.1 | A0A140H1H1 |
| CLHVVGPNLNAGEDIQ | 1 | 0 | 0 | 1648301 | NP_828849.2 | P0C6X7 |
| CLHVVGPNLNKGEDIQ | 1 | 0 | 0 | 1648302 | QIQ54046.1 | P0C6U8 |
| CLHVVGPNVNKGEDIQ | 1 | 0 | 0 | 1648303 | YP_009724389.1 | P0DTD1 |
| CLHVVGPNVNRGEDIQ | 1 | 0 | 0 | 1648304 | QHR63299.1 | P0C6W2 |

| | | | | | | |
|---|---|---|---|---|---|---|
| CLKCGFSFDLNGLDAL | 1 | 0 | 0 | 1648315 | YP_173236.1 | A0A140H1G9 |
| CLKHAEEYREYLESYN | 1 | 0 | 0 | 1648318 | YP_009555238.1 | P0C6X6 |
| CLNASEVVVNNLDKSA | 1 | 0 | 0 | 1648345 | YP_009047202.1 | K9N7C7 |
| CLPDPITAYDPRSCSQ | 1 | 0 | 0 | 1648351 | YP_173238.1 | A0A140H1H1 |
| CLPTFSMAPVEFYEDR | 1 | 0 | 0 | 1648353 | YP_009047202.1 | K9N7C7 |
| CLQHLHVASNSKLWHY | 1 | 0 | 0 | 1648356 | YP_009555238.1 | P0C6X6 |
| CLQHLHVASNSKLWQY | 1 | 0 | 0 | 1648357 | YP_173236.1 | A0A140H1G9 |
| CLRCLWSTKPVETSNI | 1 | 0 | 0 | 1648360 | QIQ54046.1 | P0C6U8 |
| CLSGLDSLDSYPALET | 1 | 0 | 0 | 1648366 | NP_828849.2 | P0C6X7 |
| CLSGLDSLDTYPSLET | 1 | 0 | 0 | 1648367 | YP_009724389.1 | P0DTD1 |
| CLSSMNIAANSSEWAY | 1 | 0 | 0 | 1648371 | YP_003766.2 | H9EJ42 |
| CLTDCLYLKNLTQTFT | 1 | 0 | 0 | 1648376 | YP_173236.1 | A0A140H1G9 |
| CLVGAEHVNNSYECDI | 1 | 0 | 0 | 1648383 | QIQ54048.1 | P59594 |
| CLVSSDGQPGGDAISL | 1 | 0 | 0 | 1648386 | YP_009047202.1 | K9N7C7 |
| CLWNTKPVETSNSFDV | 1 | 0 | 0 | 1648388 | QHR63299.1 | P0C6W2 |
| CLWSTKPVDTSNSFEV | 1 | 0 | 0 | 1648391 | NP_828849.2 | P0C6X7 |
| CLWSTKPVETSNIFDV | 1 | 0 | 0 | 1648392 | QIQ54046.1 | P0C6U8 |
| CLWSTKPVETSNSFDV | 1 | 0 | 0 | 1648393 | YP_009724389.1 | P0DTD1 |
| CLYLKNLTQTFTSMLT | 1 | 0 | 0 | 1648403 | YP_173236.1 | A0A140H1G9 |
| CMFENKTTLPTNIAFE | 1 | 0 | 0 | 1648412 | YP_009047202.1 | K9N7C7 |
| CMLRACIKTAQADTAG | 1 | 0 | 0 | 1648417 | YP_009047205.1 | K9N796 |
| CNAASVNQIVLRNSNG | 1 | 0 | 0 | 1648438 | YP_009047202.1 | K9N7C7 |
| CNADIVVVDEVSMCTN | 1 | 0 | 0 | 1648439 | YP_003766.2 | H9EJ42 |
| CNAISSNTDNETLQYW | 1 | 0 | 0 | 1648440 | YP_173238.1 | A0A140H1H1 |
| CNAPGCDVNDVTKLYL | 1 | 0 | 0 | 1648442 | YP_009555238.1 | P0C6X6 |
| CNASGCGVSDVKKLYL | 1 | 0 | 0 | 1648449 | YP_003766.2 | H9EJ42 |
| CNAVAKHDFFTWHEGR | 1 | 0 | 0 | 1648451 | NP_073549.1 | A0A0P0LKV0 |
| CNDEKVMTFYPRLQAA | 1 | 0 | 0 | 1648457 | YP_009555238.1 | P0C6X6 |
| CNEDNALAYYNTSKGG | 1 | 0 | 0 | 1648460 | QIQ54046.1 | P0C6U8 |
| CNGHKIEDLSIRNLQK | 1 | 0 | 0 | 1648471 | YP_173236.1 | A0A140H1G9 |
| CNGNPRCKNLLKQYTS | 1 | 0 | 0 | 1648474 | YP_003767.1 | A0A0P0G321 |
| CNGNVRCVELLKQYTS | 1 | 0 | 0 | 1648475 | NP_073551.1 | A0A0P0K6L9 |
| CNGVRELYLNSSNVTT | 1 | 0 | 0 | 1648484 | NP_828849.2 | P0C6X7 |
| CNGYTDNIFSVQQDGR | 1 | 0 | 0 | 1648486 | YP_003767.1 | A0A0P0G321 |
| CNHINNLKIKNFDYLD | 1 | 0 | 0 | 1648488 | YP_173237.1 | A0A140H1H0 |
| CNIAKSAYERDRAVAK | 1 | 0 | 0 | 1648492 | YP_009555238.1 | P0C6X6 |
| CNIEAWLNDKSVPSPL | 1 | 0 | 0 | 1648494 | YP_009555241.1 | P36334 |
| CNIGGAVCSKHANLYQ | 1 | 0 | 0 | 1648497 | YP_003766.2 | H9EJ42 |
| CNIGGAVCSKHANLYR | 1 | 0 | 0 | 1648498 | NP_073549.1 | A0A0P0LKV0 |
| CNLGGAVCRKHATEYR | 1 | 0 | 0 | 1648511 | YP_009047202.1 | K9N7C7 |
| CNLYNYGAGIKLPSGI | 1 | 0 | 0 | 1648518 | YP_003766.2 | H9EJ42 |
| CNNIDAAKIYGMCFSS | 1 | 0 | 0 | 1648526 | YP_009555241.1 | P36334 |
| CNNINWNVEYPIISNE | 1 | 0 | 0 | 1648527 | YP_009555238.1 | P0C6X6 |
| CNNLVPTYLKSDNIVA | 1 | 0 | 0 | 1648528 | YP_009555238.1 | P0C6X6 |
| CNQPDQVAVYQQSIIG | 1 | 0 | 0 | 1648536 | YP_003767.1 | A0A0P0G321 |
| CNRCCDRYEEYDLEPH | 1 | 0 | 0 | 1648537 | YP_009047204.1 | K9N5Q8 |
| CNRSGSNDIWMQNKGL | 1 | 0 | 0 | 1648539 | YP_009555240.1 | P30215 |
| CNSFFKELTANMSMAE | 1 | 0 | 0 | 1648545 | YP_003766.2 | H9EJ42 |
| CNSNANLSMLQLGVPD | 1 | 0 | 0 | 1648547 | YP_003767.1 | A0A0P0G321 |
| CNSVINRLRCDQLSFD | 1 | 0 | 0 | 1648550 | NP_073551.1 | A0A0P0K6L9 |
| CNTPTQCYYNNSNNGK | 1 | 0 | 0 | 1648559 | YP_009555238.1 | P0C6X6 |
| CNTSGCNVNDVTKLYL | 1 | 0 | 0 | 1648560 | NP_073549.1 | A0A0P0LKV0 |
| CNVHRNEHVASGDAIM | 1 | 0 | 0 | 1648573 | NP_073549.1 | A0A0P0LKV0 |
| CNVLDDFIVFNNNGTN | 1 | 0 | 0 | 1648577 | NP_073549.1 | A0A0P0LKV0 |
| CNVSFVGELKAAEAKV | 1 | 0 | 0 | 1648585 | NP_073549.1 | A0A0P0LKV0 |
| CNYLESYNIVTTAGFT | 1 | 0 | 0 | 1648593 | YP_173236.1 | A0A140H1G9 |
| CPACHNPEVGPEHSLA | 1 | 0 | 0 | 1648598 | QHR63299.1 | P0C6W2 |
| CPACQDPEIGPEHSVA | 1 | 0 | 0 | 1648600 | NP_828849.2 | P0C6X7 |
| CPAEIVDTVSALVYDN | 1 | 0 | 0 | 1648602 | YP_009724389.1 | P0DTD1 |
| CPAVAKHDFFKFRIDG | 1 | 0 | 0 | 1648606 | YP_009724389.1 | P0DTD1 |
| CPCKLNGSCVGSGPGK | 1 | 0 | 0 | 1648609 | YP_009555241.1 | P36334 |
| CPEWMDDFEADPKGKY | 1 | 0 | 0 | 1648614 | YP_009047202.1 | K9N7C7 |
| CPFAKGDSCSSNCKHS | 1 | 0 | 0 | 1648615 | YP_009047202.1 | K9N7C7 |
| CPFGEVFNASKFASVY | 1 | 0 | 0 | 1648616 | QIQ54048.1 | P59594 |
| CPIGTNYRSCESTTVL | 1 | 0 | 0 | 1648624 | YP_173238.1 | A0A140H1H1 |
| CPKEIVDTVSALVYEN | 1 | 0 | 0 | 1648632 | YP_009555238.1 | P0C6X6 |
| CPKEIVETVSALVYDN | 1 | 0 | 0 | 1648633 | YP_173236.1 | A0A140H1G9 |
| CPKEIVSTVSALVYNN | 1 | 0 | 0 | 1648634 | YP_009047202.1 | K9N7C7 |
| CPKFVFPLNSKVKVIQ | 1 | 0 | 0 | 1648635 | NP_828849.2 | P0C6X7 |
| CPKLEFANDTKIASQL | 1 | 0 | 0 | 1648637 | YP_009047204.1 | K9N5Q8 |
| CPLCCTVYDSSEVKAQ | 1 | 0 | 0 | 1648638 | YP_173236.1 | A0A140H1G9 |
| CPLIPGSSTTSTGPC | 1 | 0 | 0 | 1648640 | P03138.3 | Q76R62 |
| CPSGTYEGNSPFHPLA | 1 | 0 | 0 | 1648654 | NP_828857.1 | P59635 |
| CPTAADINIPDLPICV | 1 | 0 | 0 | 1648657 | YP_009555240.1 | P30215 |
| CQAVSANVCALMSCNG | 1 | 0 | 0 | 1648661 | YP_009555238.1 | P0C6X6 |
| CQAVTANVNALLSTDG | 1 | 0 | 0 | 1648663 | YP_009724389.1 | P0DTD1 |
| CQLPYCYFRNSSANYV | 1 | 0 | 0 | 1648675 | YP_173237.1 | A0A140H1H0 |
| CQLSFAGYTESAVNST | 1 | 0 | 0 | 1648676 | YP_009047206.1 | K9N4V0 |
| CQTADTAHANNINRFN | 1 | 0 | 0 | 1648686 | YP_009047202.1 | K9N7C7 |
| CQTTNGLNTSYSVCNG | 1 | 0 | 0 | 1648688 | NP_073551.1 | A0A0P0K6L9 |
| CQYKGKWVQVPIGTND | 1 | 0 | 0 | 1648694 | NP_073549.1 | A0A0P0LKV0 |
| CRCHIDHPNPKGFCDL | 1 | 0 | 0 | 1648708 | YP_009724389.1 | P0DTD1 |
| CRCHIDHPNPKGYCEL | 1 | 0 | 0 | 1648709 | QIQ54046.1 | P0C6U8 |
| CRFDTRVLNNLNLPGC | 1 | 0 | 0 | 1648720 | YP_009555238.1 | P0C6X6 |
| CRFQEKDEDDNLIDSY | 1 | 0 | 0 | 1648725 | YP_009724389.1 | P0DTD1 |
| CRFQEKDEDGNLIDSY | 1 | 0 | 0 | 1648726 | QIQ54047.1 | P0C6X7 |
| CRFQEKDEEGNLLDSY | 1 | 0 | 0 | 1648727 | NP_828849.2 | P0C6X7 |
| CRFQRIDDDGNKLDKF | 1 | 0 | 0 | 1648728 | YP_173236.1 | A0A140H1G9 |
| CRFQRVDENGDKLDQF | 1 | 0 | 0 | 1648729 | YP_009555238.1 | P0C6X6 |
| CRHHANEYRQYLDAYN | 1 | 0 | 0 | 1648739 | NP_828849.2 | P0C6X7 |
| CRKHATEYREYMEAYN | 1 | 0 | 0 | 1648746 | YP_009047202.1 | K9N7C7 |

| | | | | | | |
|---|---|---|---|---|---|---|
| CRKIGVNVCSDSTFEE | 1 | 0 | 0 | 1648747 | YP_173236.1 | A0A140H1G9 |
| CRLAMLGAEMTSNVRR | 1 | 0 | 0 | 1648753 | NP_073549.1 | A0A0P0LKV0 |
| CRLKNSVSDVAYNRYL | 1 | 0 | 0 | 1648754 | YP_173236.1 | A0A140H1G9 |
| CRLYIEESDLIISNFK | 1 | 0 | 0 | 1648757 | YP_173236.1 | A0A140H1G9 |
| CRMNSRNYIAQVDVVN | 1 | 0 | 0 | 1648758 | YP_009725255.1 | A0A663DJA2 |
| CRPIKLVDSELLSTLS | 1 | 0 | 0 | 1648760 | NP_073549.1 | A0A0P0LKV0 |
| CRQYSNESWFEDDYRA | 1 | 0 | 0 | 1648763 | NP_073549.1 | A0A0P0LKV0 |
| CRRCPAEIVDTVSALV | 1 | 0 | 0 | 1648764 | YP_009724389.1 | P0DTD1 |
| CRTDMEKDFIALDQDV | 1 | 0 | 0 | 1648769 | YP_009555238.1 | P0C6X6 |
| CRVGECVESNAGVCFG | 1 | 0 | 0 | 1648773 | NP_073549.1 | A0A0P0LKV0 |
| CSASDMTNPDYTNLLC | 1 | 0 | 0 | 1648784 | YP_009555238.1 | P0C6X6 |
| CSCVGSSVAVQSKDLN | 1 | 0 | 0 | 1648796 | YP_173236.1 | A0A140H1G9 |
| CSCVSTDTTVQSKDTN | 1 | 0 | 0 | 1648797 | YP_009555238.1 | P0C6X6 |
| CSDFLPLSDMEKDFLS | 1 | 0 | 0 | 1648799 | YP_009047202.1 | K9N7C7 |
| CSDGNYNRNASLNSFK | 1 | 0 | 0 | 1648801 | YP_009047204.1 | K9N5Q8 |
| CSETVNGYAKSNFVKD | 1 | 0 | 0 | 1648806 | NP_073549.1 | A0A0P0LKV0 |
| CSFGGVSVITPGTNAS | 1 | 0 | 0 | 1648810 | QHR63300.2 | Q3LZX1 |
| CSFYPPDEDEEEGDCE | 1 | 0 | 0 | 1648816 | YP_009724389.1 | P0DTD1 |
| CSFYPPDEDYEEEYSE | 1 | 0 | 0 | 1648817 | QIQ54046.1 | P0C6U8 |
| CSFYPPDEEEEDDAEC | 1 | 0 | 0 | 1648818 | NP_828849.2 | P0C6X7 |
| CSGLAVKSDYCGGNSC | 1 | 0 | 0 | 1648819 | YP_009555241.1 | P36334 |
| CSGVMWLYTYSIGEAS | 1 | 0 | 0 | 1648820 | YP_009047202.1 | K9N7C7 |
| CSGVTFQSAVKRTIKG | 1 | 0 | 0 | 1648823 | YP_009724389.1 | P0DTD1 |
| CSICLSGLDSLDSYPA | 1 | 0 | 0 | 1648828 | NP_828849.2 | P0C6X7 |
| CSKHAEEYCNYLESYN | 1 | 0 | 0 | 1648837 | YP_173236.1 | A0A140H1G9 |
| CSKHANLYQKYVEAYN | 1 | 0 | 0 | 1648838 | YP_003766.2 | H9EJ42 |
| CSKSCLGYHPAHAPSF | 1 | 0 | 0 | 1648845 | YP_173236.1 | A0A140H1G9 |
| CSKSYSGYHPAHAPSF | 1 | 0 | 0 | 1648847 | YP_009555238.1 | P0C6X6 |
| CSLSEQLDFIDTKRGV | 1 | 0 | 0 | 1648858 | QIQ54046.1 | P0C6U8 |
| CSNDLLQPNTEVFTDV | 1 | 0 | 0 | 1648866 | YP_173238.1 | A0A140H1H1 |
| CSNSSSGVLDTTIPFG | 1 | 0 | 0 | 1648870 | YP_003767.1 | A0A0P0G321 |
| CSNTKFADDLNQMTGF | 1 | 0 | 0 | 1648871 | NP_828849.2 | P0C6X7 |
| CSQHTLYIKDGDDGYF | 1 | 0 | 0 | 1648881 | YP_009047202.1 | K9N7C7 |
| CSQHTMLVKMDGDDVY | 1 | 0 | 0 | 1648884 | YP_009555238.1 | P0C6X6 |
| CSQHTMLVKQGDDYVY | 1 | 0 | 0 | 1648885 | YP_009724389.1 | P0DTD1 |
| CSQHTMQIVDENGKYY | 1 | 0 | 0 | 1648886 | NP_073549.1 | A0A0P0LKV0 |
| CSQHTMQIVDKDGTYY | 1 | 0 | 0 | 1648887 | YP_003766.2 | H9EJ42 |
| CSRETSGLSPAYAPTY | 1 | 0 | 0 | 1648895 | YP_009047202.1 | K9N7C7 |
| CSRLLSDDRTEVPQLV | 1 | 0 | 0 | 1648898 | YP_009047204.1 | K9N5Q8 |
| CSSGTYEGNSPFHPLA | 1 | 0 | 0 | 1648900 | YP_009724395.1 | P0DTC7 |
| CSSHYWSPQGNLGGNC | 1 | 0 | 0 | 1648901 | YP_173236.1 | A0A140H1G9 |
| CSTNLFKDCSKSYSGY | 1 | 0 | 0 | 1648920 | YP_009555238.1 | P0C6X6 |
| CSTRVNVDGFNEWAMA | 1 | 0 | 0 | 1648921 | YP_003766.2 | H9EJ42 |
| CSVCLSGLDSLDTYPS | 1 | 0 | 0 | 1648926 | YP_009724389.1 | P0DTD1 |
| CSVCLSGMDSLDAYPA | 1 | 0 | 0 | 1648927 | QIQ54046.1 | P0C6U8 |
| CSVKSFEIDKGIYQTS | 1 | 0 | 0 | 1648934 | NP_828851.1 | P59594 |
| CSVYTVESGTEVTEFA | 1 | 0 | 0 | 1648939 | NP_828849.2 | P0C6X7 |
| CTARVVYTAASHAAVD | 1 | 0 | 0 | 1648944 | YP_009555238.1 | P0C6X6 |
| CTASDKFYRLSNELAQ | 1 | 0 | 0 | 1648945 | NP_073549.1 | A0A0P0LKV0 |
| CTCDRTAIQSFDNSYL | 1 | 0 | 0 | 1648948 | NP_073549.1 | A0A0P0LKV0 |
| CTCGKQATKYLVQQES | 1 | 0 | 0 | 1648949 | YP_009724389.1 | P0DTD1 |
| CTCGKQATQYLVQQES | 1 | 0 | 0 | 1648950 | QHR63299.1 | P0C6W2 |
| CTDDNALAYYNNSKGG | 1 | 0 | 0 | 1648958 | NP_828849.2 | P0C6X7 |
| CTDDNALAYYNTTKGG | 1 | 0 | 0 | 1648959 | YP_009724389.1 | P0DTD1 |
| CTDDYFNKKDWYDFVE | 1 | 0 | 0 | 1648961 | QIQ54047.1 | P0C6X7 |
| CTDNSANTKVLRNKGV | 1 | 0 | 0 | 1648962 | YP_009047202.1 | K9N7C7 |
| CTEIDPKLDGYYKKDN | 1 | 0 | 0 | 1648965 | QIQ54046.1 | P0C6U8 |
| CTEIDPKLDNYYKKDN | 1 | 0 | 0 | 1648966 | YP_009724389.1 | P0DTD1 |
| CTFEDELAPSDVYQQL | 1 | 0 | 0 | 1648972 | YP_173236.1 | A0A140H1G9 |
| CTFTKSTNSRIRASLP | 1 | 0 | 0 | 1648980 | NP_828849.2 | P0C6X7 |
| CTGSIPCGVCLSGLDS | 1 | 0 | 0 | 1648990 | QHR63299.1 | P0C6W2 |
| CTGSPYIDDFNKIASC | 1 | 0 | 0 | 1648994 | YP_173236.1 | A0A140H1G9 |
| CTGYREGYLNSTNVTI | 1 | 0 | 0 | 1648998 | YP_009724389.1 | P0DTD1 |
| CTGYREGYLNSTNVTT | 1 | 0 | 0 | 1648999 | QHR63299.1 | P0C6W2 |
| CTIDSNTTQDTYGGAS | 1 | 0 | 0 | 1649003 | NP_073549.1 | A0A0P0LKV0 |
| CTIFKDASGKPVPYCY | 1 | 0 | 0 | 1649006 | YP_009724389.1 | P0DTD1 |
| CTKGLSIADLACAQYY | 1 | 0 | 0 | 1649011 | NP_073551.1 | A0A0P0K6L9 |
| CTKYNIYDVSGVGVIR | 1 | 0 | 0 | 1649016 | NP_073551.1 | A0A0P0K6L9 |
| CTKYNIYDYVGTGIIR | 1 | 0 | 0 | 1649017 | YP_003767.1 | A0A0P0G321 |
| CTLSEQLDFIDTKRGV | 1 | 0 | 0 | 1649025 | YP_009724389.1 | P0DTD1 |
| CTLSEQLDYIESKRGV | 1 | 0 | 0 | 1649026 | NP_828849.2 | P0C6X7 |
| CTMTDIAKKPTESACS | 1 | 0 | 0 | 1649031 | NP_828849.2 | P0C6X7 |
| CTNFAINDLNTTVPRI | 1 | 0 | 0 | 1649037 | YP_173238.1 | A0A140H1H1 |
| CTNNTLVSGDAHIPFG | 1 | 0 | 0 | 1649040 | NP_073551.1 | A0A0P0K6L9 |
| CTPNVTINCQDPAGGA | 1 | 0 | 0 | 1649050 | NP_849177.1 | Q80H93 |
| CTPTKKAFPYGTCLNC | 1 | 0 | 0 | 1649052 | NP_073549.1 | A0A0P0LKV0 |
| CTQLNRALSGIAAEQD | 1 | 0 | 0 | 1649054 | NP_828851.1 | P59594 |
| CTRASTSAVKLQNNEI | 1 | 0 | 0 | 1649056 | YP_009047202.1 | K9N7C7 |
| CTRLEGLGGNNVYCYN | 1 | 0 | 0 | 1649064 | NP_073549.1 | A0A0P0LKV0 |
| CTSDNINDKDTGPPPI | 1 | 0 | 0 | 1649066 | YP_009555241.1 | P36334 |
| CTSYRDGYLNSTNVTT | 1 | 0 | 0 | 1649077 | QIQ54046.1 | P0C6U8 |
| CTTFDDVQAPNYTQHT | 1 | 0 | 0 | 1649082 | NP_828851.1 | P59594 |
| CTVQGPPGTGKSHLAI | 1 | 0 | 0 | 1649095 | YP_009555238.1 | P0C6X6 |
| CTVQQCFGKRGPNQNF | 1 | 0 | 0 | 1649096 | YP_009555245.1 | P33469 |
| CTVYDSSEVKAQSSGV | 1 | 0 | 0 | 1649101 | YP_173236.1 | A0A140H1G9 |
| CTYKRAFLDKLDVNSG | 1 | 0 | 0 | 1649107 | YP_173241.1 | Q0ZJ82 |
| CTYMDLLLDDFVSVLK | 1 | 0 | 0 | 1649110 | NP_073549.1 | A0A0P0LKV0 |
| CVDGFGVNKIQPWTND | 1 | 0 | 0 | 1649132 | NP_073549.1 | A0A0P0LKV0 |
| CVDINKAIPSGNLIKF | 1 | 0 | 0 | 1649137 | NP_073549.1 | A0A0P0LKV0 |
| CVDIVKEAQSANPMVI | 1 | 0 | 0 | 1649139 | NP_828849.2 | P0C6X7 |
| CVDLHNKINLCDDPEK | 1 | 0 | 0 | 1649140 | YP_003766.2 | H9EJ42 |

| | | | | | | |
|---|---|---|---|---|---|---|
| CVECDAKFKNSVASIN | 1 | 0 | 0 | 1649147 | NP_073549.1 | A0A0P0LKV0 |
| CVEEVTTTLEETKFLT | 1 | 0 | 0 | 1649148 | YP_009724389.1 | P0DTD1 |
| CVEMHNKINLCDDPET | 1 | 0 | 0 | 1649150 | NP_073549.1 | A0A0P0LKV0 |
| CVFYFAVRKEGQDVIF | 1 | 0 | 0 | 1649167 | YP_009555238.1 | P0C6X6 |
| CVGSSVAVQSKDLNFL | 1 | 0 | 0 | 1649171 | YP_173236.1 | A0A140H1G9 |
| CVGYTIEEAKTALKKS | 1 | 0 | 0 | 1649173 | QIQ54046.1 | P0C6U8 |
| CVIANVPAIDVLKKLL | 1 | 0 | 0 | 1649175 | YP_003766.2 | H9EJ42 |
| CVIAWNSKHIDAKEGG | 1 | 0 | 0 | 1649176 | QHR63300.2 | Q3LZX1 |
| CVIAWNSVKQDALTGG | 1 | 0 | 0 | 1649178 | QIQ54048.1 | P59594 |
| CVKDFDFKFYDAQPIV | 1 | 0 | 0 | 1649187 | YP_009555238.1 | P0C6X6 |
| CVKDYEFKFYDAFPVA | 1 | 0 | 0 | 1649188 | YP_173236.1 | A0A140H1G9 |
| CVKPKEETGLLMPLKA | 1 | 0 | 0 | 1649195 | QHR63299.1 | P0C6W2 |
| CVKSQSPRINFCGNGN | 1 | 0 | 0 | 1649198 | YP_173238.1 | A0A140H1H1 |
| CVLAAECTIFKDAAGK | 1 | 0 | 0 | 1649203 | QIQ54046.1 | P0C6U8 |
| CVLAAECTIFKDASGK | 1 | 0 | 0 | 1649205 | YP_009724389.1 | P0DTD1 |
| CVLSVNSSNCNNFNVK | 1 | 0 | 0 | 1649222 | NP_073549.1 | A0A0P0LKV0 |
| CVMSNMQLFESLNFIT | 1 | 0 | 0 | 1649226 | YP_173236.1 | A0A140H1G9 |
| CVNCDSFGPGNTFING | 1 | 0 | 0 | 1649230 | YP_003766.2 | H9EJ42 |
| CVPHNMRVLHYGAGSD | 1 | 0 | 0 | 1649242 | YP_003766.2 | H9EJ42 |
| CVPLSAIPALAANTLT | 1 | 0 | 0 | 1649245 | YP_173236.1 | A0A140H1G9 |
| CVPLYVRDYVDKWDDF | 1 | 0 | 0 | 1649246 | NP_073549.1 | A0A0P0LKV0 |
| CVPQADTDWKFYDAQP | 1 | 0 | 0 | 1649247 | QIQ54047.1 | P0C6X7 |
| CVPQADVEWKFYDAQP | 1 | 0 | 0 | 1649248 | YP_009724389.1 | P0DTD1 |
| CVPQAEVEWKFYDAQP | 1 | 0 | 0 | 1649249 | NP_828849.2 | P0C6X7 |
| CVSTDTTVQSKDTNFL | 1 | 0 | 0 | 1649280 | YP_009555238.1 | P0C6X6 |
| CVSVPVSVIYDKETKT | 1 | 0 | 0 | 1649283 | YP_009047204.1 | K9N5Q8 |
| CVVIPSKKAGGTTEML | 1 | 0 | 0 | 1649305 | NP_828849.2 | P0C6X7 |
| CVVPGNVKPGDAVITT | 1 | 0 | 0 | 1649311 | NP_073549.1 | A0A0P0LKV0 |
| CVVPSINDLSFDKNLG | 1 | 0 | 0 | 1649314 | YP_003766.2 | H9EJ42 |
| CVYVDGLESKQVDYVP | 1 | 0 | 0 | 1649328 | YP_173236.1 | A0A140H1G9 |
| CWKCKSKNPLLYDANY | 1 | 0 | 0 | 1649341 | NP_828852.2 | P59632 |
| CWTETDLTKGPHEFCS | 1 | 0 | 0 | 1649356 | YP_009724389.1 | P0DTD1 |
| CWVEEDLTKGPHEFCS | 1 | 0 | 0 | 1649358 | YP_003766.2 | H9EJ42 |
| CYCGNSATCYNSVSNE | 1 | 0 | 0 | 1649373 | YP_003766.2 | H9EJ42 |
| CYDFDGVTTSDKCIFN | 1 | 0 | 0 | 1649375 | YP_003766.2 | H9EJ42 |
| CYDGCAQGVFGVNMRT | 1 | 0 | 0 | 1649376 | NP_073549.1 | A0A0P0LKV0 |
| CYDKFKINDTTCKYVF | 1 | 0 | 0 | 1649385 | YP_173236.1 | A0A140H1G9 |
| CYDKFKINDTTRKYVF | 1 | 0 | 0 | 1649386 | YP_009555238.1 | P0C6X6 |
| CYDKNPINSNVKTLEY | 1 | 0 | 0 | 1649387 | NP_073549.1 | A0A0P0LKV0 |
| CYDKQPVNSNVKLLDY | 1 | 0 | 0 | 1649388 | YP_003766.2 | H9EJ42 |
| CYDRFKVNETNSQYLF | 1 | 0 | 0 | 1649391 | YP_009047202.1 | K9N7C7 |
| CYEGIASGVFGVNLRT | 1 | 0 | 0 | 1649398 | YP_003766.2 | H9EJ42 |
| CYFRNSSANYVGKYDI | 1 | 0 | 0 | 1649406 | YP_173237.1 | A0A140H1H0 |
| CYNSDYASKGYIANIS | 1 | 0 | 0 | 1649439 | YP_009555238.1 | P0C6X6 |
| CYNTALMEGSLPYSSI | 1 | 0 | 0 | 1649444 | NP_073549.1 | A0A0P0LKV0 |
| CYNTDLIEGSKPYSTL | 1 | 0 | 0 | 1649445 | YP_003766.2 | H9EJ42 |
| CYPAGVNITLANFNET | 1 | 0 | 0 | 1649447 | NP_073551.1 | A0A0P0K6L9 |
| CYQLYSTQLSTDTGVE | 1 | 0 | 0 | 1649448 | QIQ54049.1 | P59632 |
| CYRCPKEIVDTVSALV | 1 | 0 | 0 | 1649450 | YP_009555238.1 | P0C6X6 |
| CYRCPKEIVETVSALV | 1 | 0 | 0 | 1649451 | YP_173236.1 | A0A140H1G9 |
| CYRCPKEIVSTVSALV | 1 | 0 | 0 | 1649452 | YP_009047202.1 | K9N7C7 |
| CYSDGVMKNASLYTSL | 1 | 0 | 0 | 1649456 | YP_173236.1 | A0A140H1G9 |
| CYSGRVSAAFHANSSE | 1 | 0 | 0 | 1649459 | YP_009555241.1 | P36334 |
| CYSILAAKTGVSVEQL | 1 | 0 | 0 | 1649460 | YP_003766.2 | H9EJ42 |
| CYTPSKLIEYSDFATS | 1 | 0 | 0 | 1649468 | NP_828849.2 | P0C6X7 |
| CYTPSKLIEYTDFATS | 1 | 0 | 0 | 1649469 | YP_009724389.1 | P0DTD1 |
| CYVCKPHQVNISLNGN | 1 | 0 | 0 | 1649471 | YP_003767.1 | A0A0P0G321 |
| CYVGVQTVEDLRARMT | 1 | 0 | 0 | 1649472 | YP_009047202.1 | K9N7C7 |
| CYYVKPGGTSSGDATT | 1 | 0 | 0 | 1649487 | YP_009555238.1 | P0C6X6 |
| DAAGKPVPYCYDTNVL | 1 | 0 | 0 | 1649490 | QIQ54046.1 | P0C6U8 |
| DAAIKDFDYYRYNRPT | 1 | 0 | 0 | 1649491 | NP_073549.1 | A0A0P0LKV0 |
| DAAKAYKDYLASGGQP | 1 | 0 | 0 | 1649492 | YP_009724389.1 | P0DTD1 |
| DAALALLLLDRLNALE | 1 | 0 | 0 | 1649494 | QIQ54056.1 | P59595 |
| DAAMQRKLEKMADQAM | 1 | 0 | 0 | 1649496 | YP_009724389.1 | P0DTD1 |
| DAANHSIPDLQRAAVT | 1 | 0 | 0 | 1649499 | NP_828849.2 | P0C6X7 |
| DAASNIFVINKDTYAA | 1 | 0 | 0 | 1649506 | YP_009047202.1 | K9N7C7 |
| DAAVKAAESKAFVDTI | 1 | 0 | 0 | 1649507 | NP_073549.1 | A0A0P0LKV0 |
| DAAVKDFDFYRYNKPT | 1 | 0 | 0 | 1649508 | YP_003766.2 | H9EJ42 |
| DADGINKQLTDISVDS | 1 | 0 | 0 | 1649514 | YP_173236.1 | A0A140H1G9 |
| DADKANFLNAAVFYAQ | 1 | 0 | 0 | 1649516 | YP_009555238.1 | P0C6X6 |
| DADLAKPLIKWDLLKY | 1 | 0 | 0 | 1649517 | NP_828849.2 | P0C6X7 |
| DADYNAKVEISVTPIK | 1 | 0 | 0 | 1649525 | NP_073549.1 | A0A0P0LKV0 |
| DAEDTLTKLSKYLANE | 1 | 0 | 0 | 1649527 | NP_073549.1 | A0A0P0LKV0 |
| DAEEALSKLSEYLISD | 1 | 0 | 0 | 1649528 | YP_003766.2 | H9EJ42 |
| DAEGAHATRDSIGTNF | 1 | 0 | 0 | 1649531 | YP_009555238.1 | P0C6X6 |
| DAERMAMYTGSLIGGI | 1 | 0 | 0 | 1649533 | NP_073551.1 | A0A0P0K6L9 |
| DAERMAMYTGSLIGGM | 1 | 0 | 0 | 1649534 | YP_003767.1 | A0A0P0G321 |
| DAFSLDVSEKSGNFKH | 1 | 0 | 0 | 1649546 | NP_828851.1 | P59594 |
| DAFVAAIKLVPTTTGV | 1 | 0 | 0 | 1649548 | YP_003766.2 | H9EJ42 |
| DAGFMKQYGECLGDIN | 1 | 0 | 0 | 1649551 | NP_828851.1 | P59594 |
| DAIENALNSCKEHPVV | 1 | 0 | 0 | 1649559 | YP_173236.1 | A0A140H1G9 |
| DAIFSLTKRNILPTMT | 1 | 0 | 0 | 1649560 | NP_073549.1 | A0A0P0LKV0 |
| DAIIIDNDINDYVSDA | 1 | 0 | 0 | 1649562 | YP_003766.2 | H9EJ42 |
| DAIIIDNDLNEFVSDA | 1 | 0 | 0 | 1649563 | YP_009047202.1 | K9N7C7 |
| DAITQTSQALQTVATA | 1 | 0 | 0 | 1649573 | NP_073551.1 | A0A0P0K6L9 |
| DAIVVDNDVVDYVSDA | 1 | 0 | 0 | 1649575 | NP_073549.1 | A0A0P0LKV0 |
| DAKFKNSVASINSAIV | 1 | 0 | 0 | 1649579 | NP_073549.1 | A0A0P0LKV0 |
| DAKGFPQLAELIPNQA | 1 | 0 | 0 | 1649581 | YP_003771.1 | H9EJ69 |
| DAKQVDYVPLKSATCI | 1 | 0 | 0 | 1649585 | YP_009555238.1 | P0C6X6 |
| DAKWYCYDKNPINSNV | 1 | 0 | 0 | 1649586 | NP_073549.1 | A0A0P0LKV0 |
| DAKWYCYDKQPVNSNV | 1 | 0 | 0 | 1649587 | YP_003766.2 | H9EJ42 |

| | | | | | | |
|---|---|---|---|---|---|---|
| DALASMTGVSIETLLA | 1 | 0 | 0 | 1649589 | YP_173236.1 | A0A140H1G9 |
| DALEAEAQIDRLINGR | 1 | 0 | 0 | 1649595 | YP_009555241.1 | P36334 |
| DALEAQVQIDRLINGR | 1 | 0 | 0 | 1649596 | YP_173238.1 | A0A140H1H1 |
| DALFVSYGSSVETETY | 1 | 0 | 0 | 1649602 | YP_173236.1 | A0A140H1G9 |
| DALLHRNSLDPFCFDV | 1 | 0 | 0 | 1649606 | YP_003766.2 | H9EJ42 |
| DALNNIINNARDGCVP | 1 | 0 | 0 | 1649608 | YP_009724389.1 | P0DTD1 |
| DALNTICIKDADYNAK | 1 | 0 | 0 | 1649609 | NP_073549.1 | A0A0P0LKV0 |
| DALRLSAHLETNDVSS | 1 | 0 | 0 | 1649611 | YP_003767.1 | A0A0P0G321 |
| DALSTNHGHKVVELVA | 1 | 0 | 0 | 1649614 | NP_828849.2 | P0C6X7 |
| DAMGQGQNSADPKVYP | 1 | 0 | 0 | 1649620 | NP_828859.1 | P59636 |
| DANKLILINDVAFVST | 1 | 0 | 0 | 1649629 | YP_009555238.1 | P0C6X6 |
| DAPYIVGDIITSGNLT | 1 | 0 | 0 | 1649637 | QIQ54046.1 | P0C6U8 |
| DAQIDRLINGRLTTLN | 1 | 0 | 0 | 1649642 | YP_009047204.1 | K9N5Q8 |
| DAQVVQLPVKDYVQTV | 1 | 0 | 0 | 1649649 | YP_173236.1 | A0A140H1G9 |
| DARTAPHGHVMVELVA | 1 | 0 | 0 | 1649654 | YP_009724389.1 | P0DTD1 |
| DASFIGKNLKSNCVRF | 1 | 0 | 0 | 1649660 | NP_073549.1 | A0A0P0LKV0 |
| DASKAYRDYLASGGQP | 1 | 0 | 0 | 1649664 | QIQ54046.1 | P0C6U8 |
| DATKPQQVTKHTAKEV | 1 | 0 | 0 | 1649674 | YP_009555245.1 | P33469 |
| DATPSDFVRATATIPI | 1 | 0 | 0 | 1649676 | YP_009724391.1 | P0DTC3 |
| DAVIKTLQPVSELLIP | 1 | 0 | 0 | 1649692 | QHR63299.1 | P0C6W2 |
| DAVIKTLQPVSELLTP | 1 | 0 | 0 | 1649693 | YP_009724389.1 | P0DTD1 |
| DAVILLERYVSLAIDA | 1 | 0 | 0 | 1649694 | NP_073549.1 | A0A0P0LKV0 |
| DAVKQGAKPVGNCVKM | 1 | 0 | 0 | 1649696 | NP_073549.1 | A0A0P0LKV0 |
| DAVMHFGTLAKTDLFN | 1 | 0 | 0 | 1649698 | YP_173236.1 | A0A140H1G9 |
| DAVNNNAQALSKLASE | 1 | 0 | 0 | 1649703 | YP_009047204.1 | K9N5Q8 |
| DAVTAYNGYLTSSSKT | 1 | 0 | 0 | 1649705 | YP_009724389.1 | P0DTD1 |
| DAVTTYNGYLTSSSKT | 1 | 0 | 0 | 1649707 | NP_828849.2 | P0C6X7 |
| DAVVCYNSNYAAQGLV | 1 | 0 | 0 | 1649709 | NP_828849.2 | P0C6X7 |
| DAVVLLERYVSLAIDA | 1 | 0 | 0 | 1649710 | YP_003766.2 | H9EJ42 |
| DAYAEVHAEPKGKYSQ | 1 | 0 | 0 | 1649715 | YP_173236.1 | A0A140H1G9 |
| DAYDQVHDEPKGKFSK | 1 | 0 | 0 | 1649717 | YP_009555238.1 | P0C6X6 |
| DAYPLSKHPKPEYRKV | 1 | 0 | 0 | 1649725 | NP_073549.1 | A0A0P0LKV0 |
| DAYPLSKHPNSEYRKV | 1 | 0 | 0 | 1649726 | YP_003766.2 | H9EJ42 |
| DAYPLVHHENEEYQKV | 1 | 0 | 0 | 1649729 | YP_173236.1 | A0A140H1G9 |
| DAYPLVYHENEEYQKV | 1 | 0 | 0 | 1649730 | YP_009555238.1 | P0C6X6 |
| DAYQNLVGYYSDDGNY | 1 | 0 | 0 | 1649731 | YP_009047204.1 | K9N5Q8 |
| DCARNPIDLPPSHATT | 1 | 0 | 0 | 1649740 | NP_073549.1 | A0A0P0LKV0 |
| DCDFNKPLIEWPLTEY | 1 | 0 | 0 | 1649747 | YP_009047202.1 | K9N7C7 |
| DCDIDKWLNNFNVPSP | 1 | 0 | 0 | 1649748 | YP_173238.1 | A0A140H1H1 |
| DCDVVQEQVNYVPLRA | 1 | 0 | 0 | 1649753 | YP_003766.2 | H9EJ42 |
| DCFKNDNVVLKITEDG | 1 | 0 | 0 | 1649763 | YP_003766.2 | H9EJ42 |
| DCKLPLGQSLCALPDT | 1 | 0 | 0 | 1649790 | YP_009047204.1 | K9N5Q8 |
| DCLAAETHRDCDFNKP | 1 | 0 | 0 | 1649792 | YP_009047202.1 | K9N7C7 |
| DCLYLKNLTQTFTSML | 1 | 0 | 0 | 1649798 | YP_173236.1 | A0A140H1G9 |
| DCNGYTDNIFSVQQDG | 1 | 0 | 0 | 1649806 | YP_003767.1 | A0A0P0G321 |
| DCNHINNLKIKNFDYL | 1 | 0 | 0 | 1649807 | YP_173237.1 | A0A140H1H0 |
| DCNMYICGDSTECANL | 1 | 0 | 0 | 1649809 | NP_828851.1 | P59594 |
| DCPAVAKHDFFKFRID | 1 | 0 | 0 | 1649812 | YP_009724389.1 | P0DTD1 |
| DCSDGNYNRNASLNSF | 1 | 0 | 0 | 1649821 | YP_009047204.1 | K9N5Q8 |
| DCSILCEILCEYADCK | 1 | 0 | 0 | 1649822 | YP_173236.1 | A0A140H1G9 |
| DCSKSCLGYHPAHAPS | 1 | 0 | 0 | 1649824 | YP_173236.1 | A0A140H1G9 |
| DCSKSYSGYHPAHAPS | 1 | 0 | 0 | 1649825 | YP_009555238.1 | P0C6X6 |
| DCSRETSGLSPAYAPT | 1 | 0 | 0 | 1649830 | YP_009047202.1 | K9N7C7 |
| DCVDINKAIPSGNLIK | 1 | 0 | 0 | 1649839 | NP_073549.1 | A0A0P0LKV0 |
| DCVFYFAVRKEGQDVI | 1 | 0 | 0 | 1649841 | YP_009555238.1 | P0C6X6 |
| DCYDKFKINDTTCKYV | 1 | 0 | 0 | 1649855 | YP_173236.1 | A0A140H1G9 |
| DDAARRVWTLMNVITL | 1 | 0 | 0 | 1649862 | NP_828849.2 | P0C6X7 |
| DDARTFVQSNVDVVPE | 1 | 0 | 0 | 1649866 | YP_009555238.1 | P0C6X6 |
| DDASRLLHTVLAKAEL | 1 | 0 | 0 | 1649867 | YP_009047202.1 | K9N7C7 |
| DDAVVCYNSNYAAQGL | 1 | 0 | 0 | 1649869 | NP_828849.2 | P0C6X7 |
| DDCNGYTDNIFSVQQD | 1 | 0 | 0 | 1649874 | YP_003767.1 | A0A0P0G321 |
| DDDFVSAVANAHRYDV | 1 | 0 | 0 | 1649879 | YP_003766.2 | H9EJ42 |
| DDDSAIIEIVNEVLH | 1 | 0 | 0 | 1649881 | YP_003771.1 | H9EJ69 |
| DDDSQQTVVQEDDSEV | 1 | 0 | 0 | 1649883 | QHR63299.1 | P0C6W2 |
| DDDYFNKKDWYDFVEN | 1 | 0 | 0 | 1649884 | YP_009724389.1 | P0DTD1 |
| DDEDVVTGDNDDEDVV | 1 | 0 | 0 | 1649886 | YP_173236.1 | A0A140H1G9 |
| DDEDVVTGDNNDEEIV | 1 | 0 | 0 | 1649887 | YP_173236.1 | A0A140H1G9 |
| DDEWAAAVDEAFPLDE | 1 | 0 | 0 | 1649889 | YP_009047202.1 | K9N7C7 |
| DDFCRQYSNESWFEDD | 1 | 0 | 0 | 1649890 | NP_073549.1 | A0A0P0LKV0 |
| DDFIVFNNNGTNVTQV | 1 | 0 | 0 | 1649893 | NP_073549.1 | A0A0P0LKV0 |
| DDFQVNWNTKHGSGVT | 1 | 0 | 0 | 1649898 | QIQ54046.1 | P0C6U8 |
| DDFSRQLQNSMSGASA | 1 | 0 | 0 | 1649900 | NP_828858.1 | P59595 |
| DDGNKLDKFFVVKRTN | 1 | 0 | 0 | 1649920 | YP_173236.1 | A0A140H1G9 |
| DDGVVCYNSDYAAKGY | 1 | 0 | 0 | 1649925 | YP_009047202.1 | K9N7C7 |
| DDGVVCYNSDYASKGY | 1 | 0 | 0 | 1649926 | YP_009555238.1 | P0C6X6 |
| DDIFAESKPWFVRKAE | 1 | 0 | 0 | 1649934 | NP_073549.1 | A0A0P0LKV0 |
| DDILQQANKSVGIISM | 1 | 0 | 0 | 1649936 | YP_009047202.1 | K9N7C7 |
| DDITKTDAVILLERYV | 1 | 0 | 0 | 1649940 | NP_073549.1 | A0A0P0LKV0 |
| DDIYYPASCNGVLPVA | 1 | 0 | 0 | 1649943 | YP_003766.2 | H9EJ42 |
| DDKKIKACVEEVTTTL | 1 | 0 | 0 | 1649946 | YP_009724389.1 | P0DTD1 |
| DDLLKTDSVLLIERFV | 1 | 0 | 0 | 1649953 | YP_009555238.1 | P0C6X6 |
| DDLNQLTGYKKPASRE | 1 | 0 | 0 | 1649954 | YP_009724389.1 | P0DTD1 |
| DDLNQMSGYKKPASRE | 1 | 0 | 0 | 1649955 | QIQ54046.1 | P0C6U8 |
| DDLNQMTGFTKPASRE | 1 | 0 | 0 | 1649956 | NP_828849.2 | P0C6X7 |
| DDLVNYIRDKGCVADQ | 1 | 0 | 0 | 1649957 | YP_009555239.1 | Q80872 |
| DDMLRHLIKDVDNPVL | 1 | 0 | 0 | 1649966 | YP_173236.1 | A0A140H1G9 |
| DDMLRRLIKDVDNPVL | 1 | 0 | 0 | 1649967 | YP_009555238.1 | P0C6X6 |
| DDNVLPETYVALPIYY | 1 | 0 | 0 | 1649976 | YP_003767.1 | A0A0P0G321 |
| DDPKLMGWDYPKCDRA | 1 | 0 | 0 | 1649980 | NP_073549.1 | A0A0P0LKV0 |
| DDQIVVTGDDVDDIES | 1 | 0 | 0 | 1649985 | YP_173236.1 | A0A140H1G9 |
| DDQPSLQVESANLMLS | 1 | 0 | 0 | 1649986 | YP_003766.2 | H9EJ42 |

| | | | | | | |
|---|---|---|---|---|---|---|
| DDSQQTVVQEDDSEVN | 1 | 0 | 0 | 1650001 | QHR63299.1 | P0C6W2 |
| DDSQYYFNKDTGVIYG | 1 | 0 | 0 | 1650002 | YP_009555240.1 | P30215 |
| DDTSETKYVKSLSIVD | 1 | 0 | 0 | 1650011 | YP_009555238.1 | P0C6X6 |
| DDVDVVKPFRVEGNFS | 1 | 0 | 0 | 1650021 | YP_003766.2 | H9EJ42 |
| DDVNASLFVDYSNLLH | 1 | 0 | 0 | 1650029 | YP_009555238.1 | P0C6X6 |
| DDVQAPNYTQHTSSMR | 1 | 0 | 0 | 1650030 | NP_828851.1 | P59594 |
| DDVVKTDAVVLLERYV | 1 | 0 | 0 | 1650035 | YP_003766.2 | H9EJ42 |
| DDYCDGSLGCKMCLFG | 1 | 0 | 0 | 1650041 | NP_073549.1 | A0A0P0LKV0 |
| DDYIARNGPLNVGGSC | 1 | 0 | 0 | 1650044 | QIQ54046.1 | P0C6U8 |
| DDYIATNGPLKVGGSC | 1 | 0 | 0 | 1650045 | YP_009724389.1 | P0DTD1 |
| DDYQGKPLEFGATSAA | 1 | 0 | 0 | 1650048 | YP_009724389.1 | P0DTD1 |
| DDYQGKSLEFGATSVT | 1 | 0 | 0 | 1650049 | QHR63299.1 | P0C6W2 |
| DDYTGYQELVIKTSHD | 1 | 0 | 0 | 1650052 | YP_009555241.1 | P36334 |
| DEAFPLDEAEDVTESV | 1 | 0 | 0 | 1650059 | YP_009047202.1 | K9N7C7 |
| DEAIPTRFPPGTVLPQ | 1 | 0 | 0 | 1650064 | YP_009555245.1 | P33469 |
| DEAQPLPQRQKKQPTV | 1 | 0 | 0 | 1650066 | NP_828858.1 | P59595 |
| DECRLAMLGAEMTSNV | 1 | 0 | 0 | 1650069 | NP_073549.1 | A0A0P0LKV0 |
| DEDDNLIDSYFVVKRH | 1 | 0 | 0 | 1650071 | YP_009724389.1 | P0DTD1 |
| DEDEEEGDCEEEDFEP | 1 | 0 | 0 | 1650072 | QHR63299.1 | P0C6W2 |
| DEDEEEGDCEEEEFEP | 1 | 0 | 0 | 1650073 | YP_009724389.1 | P0DTD1 |
| DEDVVTGDNDDEDVVT | 1 | 0 | 0 | 1650077 | YP_173236.1 | A0A140H1G9 |
| DEDVVTGDNNDEEIVT | 1 | 0 | 0 | 1650078 | YP_173236.1 | A0A140H1G9 |
| DEEIVTGDNDDQIVVT | 1 | 0 | 0 | 1650087 | YP_173236.1 | A0A140H1G9 |
| DEFFKNVSTSIPNFGS | 1 | 0 | 0 | 1650089 | YP_009047204.1 | K9N5Q8 |
| DEFSINEVVKQMFGVN | 1 | 0 | 0 | 1650092 | NP_073549.1 | A0A0P0LKV0 |
| DEFSSNVANYQKVGMQ | 1 | 0 | 0 | 1650093 | YP_009724389.1 | P0DTD1 |
| DEIANLVLAKLGKDSK | 1 | 0 | 0 | 1650110 | YP_173242.1 | Q19U29 |
| DEISMATNYDLSVVNA | 1 | 0 | 0 | 1650116 | YP_009724389.1 | P0DTD1 |
| DEKDCEVFKEILVLTG | 1 | 0 | 0 | 1650119 | NP_073549.1 | A0A0P0LKV0 |
| DEKQNGHSLYLADNLT | 1 | 0 | 0 | 1650121 | YP_009047202.1 | K9N7C7 |
| DELAPSDVYQQLAGVK | 1 | 0 | 0 | 1650125 | YP_173236.1 | A0A140H1G9 |
| DELDEFFKNVSTSIPN | 1 | 0 | 0 | 1650127 | YP_009047204.1 | K9N5Q8 |
| DELFAMTKRNVIPTMT | 1 | 0 | 0 | 1650128 | YP_009047202.1 | K9N7C7 |
| DENGTITDAVDCSQNP | 1 | 0 | 0 | 1650142 | NP_828851.1 | P59594 |
| DENYNNLVPTYLKSDN | 1 | 0 | 0 | 1650144 | YP_173236.1 | A0A140H1G9 |
| DEPEDHVQIHTIDGSS | 1 | 0 | 0 | 1650145 | QIQ54049.1 | P59632 |
| DEPEEHVQIHTIDGSS | 1 | 0 | 0 | 1650146 | YP_009724391.1 | P0DTC3 |
| DEPQKDVYELRYNGAI | 1 | 0 | 0 | 1650148 | YP_009555245.1 | P33469 |
| DEQIDGKFVTKFSGDK | 1 | 0 | 0 | 1650150 | YP_173236.1 | A0A140H1G9 |
| DEQNCDVLKEVLVLTG | 1 | 0 | 0 | 1650152 | YP_003766.2 | H9EJ42 |
| DERIDKVLNEKCSAYT | 1 | 0 | 0 | 1650153 | YP_009724389.1 | P0DTD1 |
| DERIDKVLNEKCSTYT | 1 | 0 | 0 | 1650154 | QHR63299.1 | P0C6W2 |
| DERVDKVLNEKCSNYT | 1 | 0 | 0 | 1650155 | QIQ54046.1 | P0C6U8 |
| DERVDKVLNEKCSVYT | 1 | 0 | 0 | 1650156 | NP_828849.2 | P0C6X7 |
| DESASKSASVYYSQLM | 1 | 0 | 0 | 1650157 | NP_828849.2 | P0C6X7 |
| DESCNNLVPTYLKSDN | 1 | 0 | 0 | 1650158 | YP_009555238.1 | P0C6X6 |
| DEVGSKSPIQYIDIGN | 1 | 0 | 0 | 1650172 | QHR63307.1 | Q3LZX5 |
| DEVNGLLDTTQLHVAD | 1 | 0 | 0 | 1650175 | YP_173238.1 | A0A140H1H1 |
| DEVTTTLEETKFLTNK | 1 | 0 | 0 | 1650183 | NP_828849.2 | P0C6X7 |
| DEWAAAVDEAFPLDEA | 1 | 0 | 0 | 1650184 | YP_009047202.1 | K9N7C7 |
| DEWSVATFYLFDDAGE | 1 | 0 | 0 | 1650188 | NP_828849.2 | P0C6X7 |
| DEWSYTGSSFYAPEPI | 1 | 0 | 0 | 1650189 | YP_009047204.1 | K9N5Q8 |
| DEYGGHHDFVIKTSHD | 1 | 0 | 0 | 1650190 | YP_173238.1 | A0A140H1H1 |
| DEYILAKGPLQVGDSV | 1 | 0 | 0 | 1650191 | YP_009047202.1 | K9N7C7 |
| DFADEIIGYKAIEDGV | 1 | 0 | 0 | 1650195 | QIQ54046.1 | P0C6U8 |
| DFAFNNWFLLTNTSSV | 1 | 0 | 0 | 1650197 | NP_073551.1 | A0A0P0K6L9 |
| DFAQGPFIDGNIIHAN | 1 | 0 | 0 | 1650200 | NP_073549.1 | A0A0P0LKV0 |
| DFCRQYSNESWFEDDY | 1 | 0 | 0 | 1650208 | NP_073549.1 | A0A0P0LKV0 |
| DFDGVTTSDKCIFNSA | 1 | 0 | 0 | 1650214 | YP_003766.2 | H9EJ42 |
| DFDIIQKCQITSVVGT | 1 | 0 | 0 | 1650215 | YP_173236.1 | A0A140H1G9 |
| DFDITESKYSCKEVLK | 1 | 0 | 0 | 1650216 | YP_003766.2 | H9EJ42 |
| DFDIVVAWHVVRDSRF | 1 | 0 | 0 | 1650217 | YP_173236.1 | A0A140H1G9 |
| DFDLISKCQITAVEGT | 1 | 0 | 0 | 1650219 | YP_009555238.1 | P0C6X6 |
| DFEADPKGKYAQNLLK | 1 | 0 | 0 | 1650224 | YP_009047202.1 | K9N7C7 |
| DFEIACKCGVKQEQRT | 1 | 0 | 0 | 1650225 | YP_009555238.1 | P0C6X6 |
| DFFKFRIDGDMVPHIS | 1 | 0 | 0 | 1650229 | YP_009724389.1 | P0DTD1 |
| DFFKFRVDGDMVPHIS | 1 | 0 | 0 | 1650230 | NP_828849.2 | P0C6X7 |
| DFFTWHEGRTIYGNVS | 1 | 0 | 0 | 1650234 | NP_073549.1 | A0A0P0LKV0 |
| DFGGFNFLQILPDPSK | 1 | 0 | 0 | 1650245 | QIQ54048.1 | P59594 |
| DFGLGDLVDSYFENDS | 1 | 0 | 0 | 1650247 | NP_073549.1 | A0A0P0LKV0 |
| DFHSVEIEAGEVKPFA | 1 | 0 | 0 | 1650253 | YP_003766.2 | H9EJ42 |
| DFIDTKRGVYCCREHE | 1 | 0 | 0 | 1650259 | YP_009724389.1 | P0DTD1 |
| DFINGIFAKVKNTKVI | 1 | 0 | 0 | 1650262 | YP_009555241.1 | P36334 |
| DFIVNAANENLAHGGG | 1 | 0 | 0 | 1650275 | NP_073549.1 | A0A0P0LKV0 |
| DFKPQSGGGKCFNCYP | 1 | 0 | 0 | 1650285 | NP_073551.1 | A0A0P0K6L9 |
| DFKPRSQMETDFLELA | 1 | 0 | 0 | 1650286 | NP_828849.2 | P0C6X7 |
| DFLARAGKSMCTLSEQ | 1 | 0 | 0 | 1650297 | NP_828849.2 | P0C6X7 |
| DFMLKTLYKDVDNPHL | 1 | 0 | 0 | 1650317 | YP_009047202.1 | K9N7C7 |
| DFNCTNFAINDLNTTV | 1 | 0 | 0 | 1650322 | YP_173238.1 | A0A140H1H1 |
| DFNKPLIEWPLTEYDF | 1 | 0 | 0 | 1650330 | YP_009047202.1 | K9N7C7 |
| DFNTILNTACGVFEVD | 1 | 0 | 0 | 1650337 | YP_009555238.1 | P0C6X6 |
| DFQENWNTKHSSGVTR | 1 | 0 | 0 | 1650342 | YP_009724389.1 | P0DTD1 |
| DFQVNWNTKHGSGVTR | 1 | 0 | 0 | 1650347 | QIQ54046.1 | P0C6U8 |
| DFSAVSSPMSNMTENE | 1 | 0 | 0 | 1650350 | NP_073555.1 | A0A0P0KXN3 |
| DFSLADSVVEVVTTSL | 1 | 0 | 0 | 1650360 | YP_009555238.1 | P0C6X6 |
| DFSLDVFKIGDVKFKR | 1 | 0 | 0 | 1650361 | YP_003766.2 | H9EJ42 |
| DFSNSGADVLYQPPQT | 1 | 0 | 0 | 1650363 | NP_828849.2 | P0C6X7 |
| DFSNSGSDVLYQPPQT | 1 | 0 | 0 | 1650364 | YP_009724389.1 | P0DTD1 |
| DFSRDHNDILYTPPTV | 1 | 0 | 0 | 1650367 | NP_073549.1 | A0A0P0LKV0 |
| DFSSEIIGYKAIDGGV | 1 | 0 | 0 | 1650369 | YP_009724389.1 | P0DTD1 |
| DFTDYKLELFNKYFKH | 1 | 0 | 0 | 1650375 | YP_009555238.1 | P0C6X6 |

| | | | | | | |
|---|---|---|---|---|---|---|
| DFTDYKLELFNKYFKY | 1 | 0 | 0 | 1650376 | YP_173236.1 | A0A140H1G9 |
| DFTGSVYGNFDDQPSL | 1 | 0 | 0 | 1650387 | YP_003766.2 | H9EJ42 |
| DFTPRSDMEYDFLNMD | 1 | 0 | 0 | 1650391 | YP_003766.2 | H9EJ42 |
| DFTPVQVVDSRWNNAR | 1 | 0 | 0 | 1650392 | YP_009555240.1 | P30215 |
| DFVDLESVIQDYENVC | 1 | 0 | 0 | 1650399 | YP_009555238.1 | P0C6X6 |
| DFVENPDIINIYKKLG | 1 | 0 | 0 | 1650403 | YP_173236.1 | A0A140H1G9 |
| DFVENPDIINVYKKLG | 1 | 0 | 0 | 1650404 | YP_009555238.1 | P0C6X6 |
| DFVTASDTTLRCCTVT | 1 | 0 | 0 | 1650427 | YP_003766.2 | H9EJ42 |
| DFVVNAANENLLHGGG | 1 | 0 | 0 | 1650432 | YP_003766.2 | H9EJ42 |
| DFYASMYENSTILQAA | 1 | 0 | 0 | 1650438 | YP_003766.2 | H9EJ42 |
| DFYEFVKSKGLFKEGS | 1 | 0 | 0 | 1650442 | YP_173236.1 | A0A140H1G9 |
| DFYGWVSGNMMDGFSC | 1 | 0 | 0 | 1650444 | YP_173236.1 | A0A140H1G9 |
| DFYSKLGVGFTSHFCN | 1 | 0 | 0 | 1650454 | YP_173236.1 | A0A140H1G9 |
| DGALLSKTSEYKGPVT | 1 | 0 | 0 | 1650460 | QIQ54046.1 | P0C6U8 |
| DGALLTKSSEYKGPIT | 1 | 0 | 0 | 1650461 | YP_009724389.1 | P0DTD1 |
| DGCAQGVFGVNMRTNW | 1 | 0 | 0 | 1650465 | NP_073549.1 | A0A0P0LKV0 |
| DGCVPLNIIPLTTAAK | 1 | 0 | 0 | 1650472 | YP_009724389.1 | P0DTD1 |
| DGDRFVKHDLSLLSVT | 1 | 0 | 0 | 1650480 | NP_073549.1 | A0A0P0LKV0 |
| DGDVLLAEFDTYDPIY | 1 | 0 | 0 | 1650481 | YP_009047202.1 | K9N7C7 |
| DGDYYRKQLSPLEGGG | 1 | 0 | 0 | 1650485 | YP_009047204.1 | K9N5Q8 |
| DGELPVAISGDKVFVR | 1 | 0 | 0 | 1650486 | NP_073549.1 | A0A0P0LKV0 |
| DGELPVAVVNDKVFVR | 1 | 0 | 0 | 1650487 | YP_003766.2 | H9EJ42 |
| DGEVITFDNLKTLLSL | 1 | 0 | 0 | 1650494 | YP_009724389.1 | P0DTD1 |
| DGFGVFDIKNSSVNTV | 1 | 0 | 0 | 1650502 | YP_003766.2 | H9EJ42 |
| DGFGVNKIQPWTNDAL | 1 | 0 | 0 | 1650503 | NP_073549.1 | A0A0P0LKV0 |
| DGFYSANFLDDNVLPE | 1 | 0 | 0 | 1650515 | YP_003767.1 | A0A0P0G321 |
| DGGCINANQVIVNNLD | 1 | 0 | 0 | 1650520 | YP_009724389.1 | P0DTD1 |
| DGGCLNASEVVVNNLD | 1 | 0 | 0 | 1650522 | YP_009047202.1 | K9N7C7 |
| DGGDSSESGAKETKEI | 1 | 0 | 0 | 1650523 | YP_009555238.1 | P0C6X6 |
| DGGITSEGNALYNNEG | 1 | 0 | 0 | 1650525 | NP_073549.1 | A0A0P0LKV0 |
| DGGKIWNGVIKNVNSV | 1 | 0 | 0 | 1650526 | NP_073549.1 | A0A0P0LKV0 |
| DGHRLASGVQVHNLPE | 1 | 0 | 0 | 1650535 | NP_073555.1 | A0A0P0KXN3 |
| DGHVETFYPKLQASQA | 1 | 0 | 0 | 1650536 | NP_828849.2 | P0C6X7 |
| DGHVETFYPKLQSSQA | 1 | 0 | 0 | 1650537 | YP_009724389.1 | P0DTD1 |
| DGIDVEIFENKTTLPV | 1 | 0 | 0 | 1650540 | NP_828849.2 | P0C6X7 |
| DGIIWVATEGALNTPK | 1 | 0 | 0 | 1650544 | YP_009724397.2 | P0DTC9 |
| DGISQYSLRLIDAMLF | 1 | 0 | 0 | 1650550 | QIQ54046.1 | P0C6U8 |
| DGISQYSLRLIDAMMF | 1 | 0 | 0 | 1650551 | YP_009724389.1 | P0DTD1 |
| DGISTPKLKEDYQIGG | 1 | 0 | 0 | 1650553 | NP_828852.2 | P59632 |
| DGKFVTKFSGDKFDFI | 1 | 0 | 0 | 1650561 | YP_173236.1 | A0A140H1G9 |
| DGKLNCSFQDAASNIF | 1 | 0 | 0 | 1650562 | YP_009047202.1 | K9N7C7 |
| DGKMKDLSPRWYFYYL | 1 | 0 | 0 | 1650564 | YP_009724397.2 | P0DTC9 |
| DGLASAYRANSFDVPT | 1 | 0 | 0 | 1650576 | YP_009047202.1 | K9N7C7 |
| DGLCMFWNCNVDKYPS | 1 | 0 | 0 | 1650581 | YP_173236.1 | A0A140H1G9 |
| DGLEAYADKTLQEMKA | 1 | 0 | 0 | 1650583 | YP_009555238.1 | P0C6X6 |
| DGLESKQVDYVPLRSA | 1 | 0 | 0 | 1650584 | YP_173236.1 | A0A140H1G9 |
| DGLKFTEESFYKNMYL | 1 | 0 | 0 | 1650588 | YP_173236.1 | A0A140H1G9 |
| DGLKITEEVGHTDLMA | 1 | 0 | 0 | 1650589 | QHR63299.1 | P0C6W2 |
| DGLKYTKILKDDGNFV | 1 | 0 | 0 | 1650590 | YP_009555238.1 | P0C6X6 |
| DGLKYTKIMKDDGNCV | 1 | 0 | 0 | 1650591 | YP_173236.1 | A0A140H1G9 |
| DGLRGAIRKAKDYGFT | 1 | 0 | 0 | 1650593 | YP_009047202.1 | K9N7C7 |
| DGMDAKQVDYVPLKSA | 1 | 0 | 0 | 1650595 | YP_009555238.1 | P0C6X6 |
| DGMMNMSPKPQRQRGH | 1 | 0 | 0 | 1650597 | YP_009555245.1 | P33469 |
| DGNAAISDYDYYRYNL | 1 | 0 | 0 | 1650598 | YP_009724389.1 | P0DTD1 |
| DGNAAITDYNYYKYNL | 1 | 0 | 0 | 1650599 | YP_009555238.1 | P0C6X6 |
| DGNKLDKFFVVKRTNL | 1 | 0 | 0 | 1650610 | YP_173236.1 | A0A140H1G9 |
| DGNYNRNASLNSFKEY | 1 | 0 | 0 | 1650617 | YP_009047204.1 | K9N5Q8 |
| DGQKFTDESFYKNMYL | 1 | 0 | 0 | 1650622 | YP_009555238.1 | P0C6X6 |
| DGQRVYDDVDASLFVD | 1 | 0 | 0 | 1650629 | YP_173236.1 | A0A140H1G9 |
| DGRDNGALEAFKRSNN | 1 | 0 | 0 | 1650632 | YP_009555238.1 | P0C6X6 |
| DGRDTGALEAFRKARN | 1 | 0 | 0 | 1650633 | YP_173236.1 | A0A140H1G9 |
| DGRVEGQVDLFRNARN | 1 | 0 | 0 | 1650643 | NP_828849.2 | P0C6X7 |
| DGSCEVVHDQVNYVPL | 1 | 0 | 0 | 1650646 | NP_073549.1 | A0A0P0LKV0 |
| DGSCSCVSTDTTVQSK | 1 | 0 | 0 | 1650648 | YP_009555238.1 | P0C6X6 |
| DGSLGCKMCLFGYQEL | 1 | 0 | 0 | 1650653 | NP_073549.1 | A0A0P0LKV0 |
| DGSLIPVRPRNSSDNG | 1 | 0 | 0 | 1650654 | YP_003767.1 | A0A0P0G321 |
| DGSSGVANPAMDPIYD | 1 | 0 | 0 | 1650657 | NP_828852.2 | P59632 |
| DGSSGVVNPAMDPIYD | 1 | 0 | 0 | 1650658 | QIQ54049.1 | P59632 |
| DGSSGVVNPAMEPIYD | 1 | 0 | 0 | 1650659 | QHR63301.1 | Q3LZX0 |
| DGSSGVVNPVMEPIYD | 1 | 0 | 0 | 1650660 | YP_009724391.1 | P0DTC3 |
| DGTCGIIDVEKGVLPQ | 1 | 0 | 0 | 1650664 | QIQ54046.1 | P0C6U8 |
| DGTNKQLNEISDDCNW | 1 | 0 | 0 | 1650673 | YP_009555238.1 | P0C6X6 |
| DGTTSPISEHDYQIGG | 1 | 0 | 0 | 1650678 | YP_009724391.1 | P0DTC3 |
| DGVDSFEGVLPINTDT | 1 | 0 | 0 | 1650683 | YP_003766.2 | H9EJ42 |
| DGVDVELFENKTILPV | 1 | 0 | 0 | 1650684 | QIQ54047.1 | P0C6X7 |
| DGVDVELFENKTTLPV | 1 | 0 | 0 | 1650685 | YP_009724389.1 | P0DTD1 |
| DGVENPMLMGWDYPKC | 1 | 0 | 0 | 1650686 | YP_003766.2 | H9EJ42 |
| DGVKHVYQLRARSVSP | 1 | 0 | 0 | 1650689 | YP_009724395.1 | P0DTC7 |
| DGVMYGGFEDQPNLQV | 1 | 0 | 0 | 1650694 | NP_073549.1 | A0A0P0LKV0 |
| DGVNFNVDDINFSPVL | 1 | 0 | 0 | 1650696 | YP_009555241.1 | P36334 |
| DGVNFTNRFVPVGESF | 1 | 0 | 0 | 1650698 | YP_009555238.1 | P0C6X6 |
| DGVNVTKHKCDINYKG | 1 | 0 | 0 | 1650700 | YP_009555238.1 | P0C6X6 |
| DGVTTSDKCIFNSACT | 1 | 0 | 0 | 1650714 | YP_003766.2 | H9EJ42 |
| DGVVCYNKDYAELGYI | 1 | 0 | 0 | 1650718 | YP_003766.2 | H9EJ42 |
| DGVVCYNSDYAAKGYI | 1 | 0 | 0 | 1650719 | YP_009047202.1 | K9N7C7 |
| DGVVCYNSDYASKGYI | 1 | 0 | 0 | 1650720 | YP_009555238.1 | P0C6X6 |
| DGVVPLSVIPATSASK | 1 | 0 | 0 | 1650722 | YP_003766.2 | H9EJ42 |
| DGVVWVAKEGAKTVNT | 1 | 0 | 0 | 1650725 | YP_003771.1 | H9EJ69 |
| DGVYYPSNGTNILPVA | 1 | 0 | 0 | 1650730 | NP_073549.1 | A0A0P0LKV0 |
| DGYCKLFITKEEAVKR | 1 | 0 | 0 | 1650734 | YP_009555238.1 | P0C6X6 |
| DGYLNSTNVTTTAYCT | 1 | 0 | 0 | 1650742 | QIQ54046.1 | P0C6U8 |

| | | | | | | |
|---|---|---|---|---|---|---|
| DGYSKLFITKDEAIKR | 1 | 0 | 0 | 1650747 | YP_173236.1 | A0A140H1G9 |
| DGYTNEIIGYKAIQDG | 1 | 0 | 0 | 1650749 | NP_828849.2 | P0C6X7 |
| DGYYKKDNAYFTEQPI | 1 | 0 | 0 | 1650751 | QIQ54046.1 | P0C6U8 |
| DGYYKKDNAYYTEQPI | 1 | 0 | 0 | 1650752 | NP_828849.2 | P0C6X7 |
| DHDYQIGGYTEKWESG | 1 | 0 | 0 | 1650760 | QIQ54049.1 | P59632 |
| DHEHEIAWFTERSDKS | 1 | 0 | 0 | 1650763 | NP_828849.2 | P0C6X7 |
| DHFDVVETKVEITPEE | 1 | 0 | 0 | 1650767 | QIQ54046.1 | P0C6U8 |
| DHGDMYVYSAGHATGT | 1 | 0 | 0 | 1650770 | YP_009047204.1 | K9N5Q8 |
| DHIGTRNPNNNAAIVL | 1 | 0 | 0 | 1650775 | QIQ54056.1 | P59595 |
| DHIGTRNPNNNAATVL | 1 | 0 | 0 | 1650776 | NP_828858.1 | P59595 |
| DHNDILYTPPTVSYGS | 1 | 0 | 0 | 1650785 | NP_073549.1 | A0A0P0LKV0 |
| DHQDLKWARFPKSDGT | 1 | 0 | 0 | 1650791 | NP_828849.2 | P0C6X7 |
| DHSLLATLDDPYVEDS | 1 | 0 | 0 | 1650795 | YP_173242.1 | Q19U29 |
| DHVDILGPLSAQTGIA | 1 | 0 | 0 | 1650799 | YP_009724389.1 | P0DTD1 |
| DHVQIHTIDGSSGVVN | 1 | 0 | 0 | 1650806 | QIQ54049.1 | P59632 |
| DHYIATNGPLKVGGGC | 1 | 0 | 0 | 1650811 | QHR63299.1 | P0C6W2 |
| DIAEDVVDQVEEVNSI | 1 | 0 | 0 | 1650818 | NP_073549.1 | A0A0P0LKV0 |
| DIAQVVIADTLQETPV | 1 | 0 | 0 | 1650824 | YP_009047202.1 | K9N7C7 |
| DICMFENKTTLPTNIA | 1 | 0 | 0 | 1650829 | YP_009047202.1 | K9N7C7 |
| DIDITFLKKDAPYIVG | 1 | 0 | 0 | 1650839 | YP_009724389.1 | P0DTD1 |
| DIDKWLNNFNVPSPLN | 1 | 0 | 0 | 1650841 | YP_173238.1 | A0A140H1H1 |
| DIDVFNKLSTSDWSDI | 1 | 0 | 0 | 1650842 | NP_073549.1 | A0A0P0LKV0 |
| DIDYCVRAFDVYNKDA | 1 | 0 | 0 | 1650844 | NP_073549.1 | A0A0P0LKV0 |
| DIEEVELLDAEFVEPG | 1 | 0 | 0 | 1650847 | NP_073549.1 | A0A0P0LKV0 |
| DIEYPYISHEKKLNSC | 1 | 0 | 0 | 1650853 | YP_009047202.1 | K9N7C7 |
| DIFAESKPWFVRKAED | 1 | 0 | 0 | 1650855 | NP_073549.1 | A0A0P0LKV0 |
| DIFGPCWSALASALKQ | 1 | 0 | 0 | 1650857 | NP_073549.1 | A0A0P0LKV0 |
| DIFGQDFKTFDLLKYD | 1 | 0 | 0 | 1650858 | NP_073549.1 | A0A0P0LKV0 |
| DIFGSDFKTFDLLKYD | 1 | 0 | 0 | 1650859 | YP_003766.2 | H9EJ42 |
| DIFVHEDLHVVEVLTK | 1 | 0 | 0 | 1650865 | YP_173236.1 | A0A140H1G9 |
| DIFYKENSYITTIKPV | 1 | 0 | 0 | 1650867 | QHR63299.1 | P0C6W2 |
| DIHLPAYASFKVIDNG | 1 | 0 | 0 | 1650888 | YP_173236.1 | A0A140H1G9 |
| DIIQKCQITSVVGTKA | 1 | 0 | 0 | 1650902 | YP_173236.1 | A0A140H1G9 |
| DIITSGNLTTVVIPTK | 1 | 0 | 0 | 1650905 | QIQ54046.1 | P0C6U8 |
| DIKNSSVNTVCFVDVD | 1 | 0 | 0 | 1650909 | YP_003766.2 | H9EJ42 |
| DILQQANKSVGIISMP | 1 | 0 | 0 | 1650924 | YP_009047202.1 | K9N7C7 |
| DILSLNNPIFINYSKE | 1 | 0 | 0 | 1650928 | YP_173237.1 | A0A140H1H0 |
| DINAYANWCCLKCGFS | 1 | 0 | 0 | 1650938 | YP_173236.1 | A0A140H1G9 |
| DINDYVSDADFSITGD | 1 | 0 | 0 | 1650939 | YP_003766.2 | H9EJ42 |
| DINGKLYHDSQNMLRG | 1 | 0 | 0 | 1650943 | NP_828849.2 | P0C6X7 |
| DINGNLYPDSTSLVEX | 1 | 0 | 0 | 1650946 | QIQ54046.1 | P0C6U8 |
| DINKAIPSGNLIKFDI | 1 | 0 | 0 | 1650950 | NP_073549.1 | A0A0P0LKV0 |
| DINKLCEEMLDNRATL | 1 | 0 | 0 | 1650951 | YP_009724389.1 | P0DTD1 |
| DINRLCEEMLDNRATL | 1 | 0 | 0 | 1650955 | NP_828849.2 | P0C6X7 |
| DINYKGKVFFQFDNLS | 1 | 0 | 0 | 1650957 | YP_009555238.1 | P0C6X6 |
| DIPGGCAMPIVANWAY | 1 | 0 | 0 | 1650958 | NP_073551.1 | A0A0P0K6L9 |
| DIPGIPKDMTYRRLIS | 1 | 0 | 0 | 1650959 | YP_009724389.1 | P0DTD1 |
| DIPYNVSLSKFNSCKS | 1 | 0 | 0 | 1650965 | YP_173237.1 | A0A140H1H0 |
| DIQITNESYNNYVPSY | 1 | 0 | 0 | 1650967 | YP_009047202.1 | K9N7C7 |
| DIQLLKAAYENFNSQD | 1 | 0 | 0 | 1650968 | NP_828849.2 | P0C6X7 |
| DIQQTFFDKTWPRPID | 1 | 0 | 0 | 1650974 | YP_009047204.1 | K9N5Q8 |
| DIRDYKLANDAKESLR | 1 | 0 | 0 | 1650980 | NP_073549.1 | A0A0P0LKV0 |
| DISAVQSSFGFDQQQL | 1 | 0 | 0 | 1650987 | YP_173236.1 | A0A140H1G9 |
| DISDTTDAVRDPQTLE | 1 | 0 | 0 | 1650989 | QIQ54048.1 | P59594 |
| DISSNDNCFANKHVGF | 1 | 0 | 0 | 1650999 | QIQ54046.1 | P0C6U8 |
| DITDAAVKAAESKAFV | 1 | 0 | 0 | 1651007 | NP_073549.1 | A0A0P0LKV0 |
| DITESKYSCKEVFKNC | 1 | 0 | 0 | 1651008 | NP_073549.1 | A0A0P0LKV0 |
| DITESKYSCKEVLKNC | 1 | 0 | 0 | 1651009 | YP_003766.2 | H9EJ42 |
| DITFLKKDAPYIVGDV | 1 | 0 | 0 | 1651010 | YP_009724389.1 | P0DTD1 |
| DITKTDAVILLERYVS | 1 | 0 | 0 | 1651011 | NP_073549.1 | A0A0P0LKV0 |
| DITNTNLSRGRGRNPK | 1 | 0 | 0 | 1651015 | YP_009047211.1 | K9N4V7 |
| DIVCKVENKTEAEWIE | 1 | 0 | 0 | 1651018 | NP_073549.1 | A0A0P0LKV0 |
| DIVKEAQSANPMVIVN | 1 | 0 | 0 | 1651024 | NP_828849.2 | P0C6X7 |
| DIVKGQIQVASDNIKD | 1 | 0 | 0 | 1651025 | NP_828849.2 | P0C6X7 |
| DIVKTDGTLMVERFVS | 1 | 0 | 0 | 1651027 | YP_009047202.1 | K9N7C7 |
| DIVSGYEKSTFNKDIY | 1 | 0 | 0 | 1651031 | YP_003766.2 | H9EJ42 |
| DIVSSFEGYDFKYIEN | 1 | 0 | 0 | 1651033 | NP_073549.1 | A0A0P0LKV0 |
| DIVVAWHVVRDSRFVM | 1 | 0 | 0 | 1651035 | YP_173236.1 | A0A140H1G9 |
| DKCIFNSACTRLEGLG | 1 | 0 | 0 | 1651064 | YP_003766.2 | H9EJ42 |
| DKCTKYNIYDVSGVGV | 1 | 0 | 0 | 1651069 | NP_073551.1 | A0A0P0K6L9 |
| DKDKDLSGAVPSDLNT | 1 | 0 | 0 | 1651075 | NP_073549.1 | A0A0P0LKV0 |
| DKEDVVIFINNTTYPT | 1 | 0 | 0 | 1651082 | YP_009555238.1 | P0C6X6 |
| DKFAIPNGRKVDLQLG | 1 | 0 | 0 | 1651087 | YP_009555241.1 | P36334 |
| DKFDFIIGHGMSFSMT | 1 | 0 | 0 | 1651089 | YP_009555238.1 | P0C6X6 |
| DKFDLLISDMYDGRIK | 1 | 0 | 0 | 1651091 | YP_003766.2 | H9EJ42 |
| DKFDLLISDMYDGRTK | 1 | 0 | 0 | 1651092 | NP_073549.1 | A0A0P0LKV0 |
| DKFKINDTTCKYVFTT | 1 | 0 | 0 | 1651097 | YP_173236.1 | A0A140H1G9 |
| DKFKINDTTRKYVFTT | 1 | 0 | 0 | 1651098 | YP_009555238.1 | P0C6X6 |
| DKFKVNSTLEQYVFCT | 1 | 0 | 0 | 1651099 | YP_009724389.1 | P0DTD1 |
| DKFYRLSNELAQVLTE | 1 | 0 | 0 | 1651104 | NP_073549.1 | A0A0P0LKV0 |
| DKGDAEDTLTKLSKYL | 1 | 0 | 0 | 1651106 | NP_073549.1 | A0A0P0LKV0 |
| DKGIAPGTSVLRQWLP | 1 | 0 | 0 | 1651108 | YP_009047202.1 | K9N7C7 |
| DKGIYQTSNFRVVPSG | 1 | 0 | 0 | 1651109 | NP_828851.1 | P59594 |
| DKGVAPGTTVLKRWLP | 1 | 0 | 0 | 1651111 | YP_003766.2 | H9EJ42 |
| DKITEFQLDYSIDVID | 1 | 0 | 0 | 1651119 | NP_073549.1 | A0A0P0LKV0 |
| DKKAFTLANVSSFGDY | 1 | 0 | 0 | 1651121 | NP_073551.1 | A0A0P0K6L9 |
| DKKIKACVEEVTTTLE | 1 | 0 | 0 | 1651125 | YP_009724389.1 | P0DTD1 |
| DKLDTGAQKFFQFGDF | 1 | 0 | 0 | 1651138 | YP_003766.2 | H9EJ42 |
| DKLEKLISTARDGVRR | 1 | 0 | 0 | 1651140 | YP_009047202.1 | K9N7C7 |
| DKLGFNVDLPFVEYKV | 1 | 0 | 0 | 1651142 | YP_173236.1 | A0A140H1G9 |
| DKLRDYLADYDVAVTA | 1 | 0 | 0 | 1651149 | YP_009047202.1 | K9N7C7 |

| | | | | | | |
|---|---|---|---|---|---|---|
| DKLSVYDIACCMRAGS | 1 | 0 | 0 | 1651153 | YP_003766.2 | H9EJ42 |
| DKNLITTANTGTSVTE | 1 | 0 | 0 | 1651160 | YP_009555238.1 | P0C6X6 |
| DKNPINSNVKTLEYDY | 1 | 0 | 0 | 1651163 | NP_073549.1 | A0A0P0LKV0 |
| DKPNHFINFPLTHFQG | 1 | 0 | 0 | 1651166 | YP_009555239.1 | Q80872 |
| DKPQEKDKKSAKTGTP | 1 | 0 | 0 | 1651167 | NP_073556.1 | A0A0N7J131 |
| DKQGQLLDTKLNGILQ | 1 | 0 | 0 | 1651170 | YP_003766.2 | H9EJ42 |
| DKQPVNSNVKLLDYDY | 1 | 0 | 0 | 1651171 | YP_003766.2 | H9EJ42 |
| DKRATCFSTASDTYAC | 1 | 0 | 0 | 1651179 | QHR63299.1 | P0C6W2 |
| DKRATCFSTSSDTYAC | 1 | 0 | 0 | 1651180 | NP_828849.2 | P0C6X7 |
| DKSAFTNLKQLPFFYY | 1 | 0 | 0 | 1651184 | NP_828849.2 | P0C6X7 |
| DKSAFVNLKQLPFFYY | 1 | 0 | 0 | 1651185 | YP_009724389.1 | P0DTD1 |
| DKSAGHPFNKFGKARV | 1 | 0 | 0 | 1651187 | YP_009047202.1 | K9N7C7 |
| DKSEPLCLFKKNFTYN | 1 | 0 | 0 | 1651192 | YP_173238.1 | A0A140H1H1 |
| DKSFEQQVGVIADKDK | 1 | 0 | 0 | 1651193 | NP_073549.1 | A0A0P0LKV0 |
| DKSYEHQTPFEIKSAK | 1 | 0 | 0 | 1651201 | NP_828849.2 | P0C6X7 |
| DKVDFVNGFYRLYSGD | 1 | 0 | 0 | 1651210 | YP_003766.2 | H9EJ42 |
| DKVDILLTVDGVNFTN | 1 | 0 | 0 | 1651211 | YP_009555238.1 | P0C6X6 |
| DKVDSTFVTEYYEFLN | 1 | 0 | 0 | 1651212 | YP_009555238.1 | P0C6X6 |
| DKVFGYVQQGIFNKAT | 1 | 0 | 0 | 1651216 | YP_009555238.1 | P0C6X6 |
| DKVFVRDGNTDNLVFV | 1 | 0 | 0 | 1651220 | NP_073549.1 | A0A0P0LKV0 |
| DKVGELKVEFWFAMRK | 1 | 0 | 0 | 1651223 | YP_173236.1 | A0A140H1G9 |
| DKVGHYVHVKCEQSYQ | 1 | 0 | 0 | 1651224 | YP_009555238.1 | P0C6X6 |
| DKVKLSDVGFVEAYNN | 1 | 0 | 0 | 1651225 | YP_009555241.1 | P36334 |
| DKVLNEKCSVYTVESG | 1 | 0 | 0 | 1651230 | NP_828849.2 | P0C6X7 |
| DKVRSLKLSDNNCYLN | 1 | 0 | 0 | 1651233 | YP_009047202.1 | K9N7C7 |
| DKVVYHTTSKPITFHM | 1 | 0 | 0 | 1651238 | QIQ54046.1 | P0C6U8 |
| DKVYCPRHVICSASDM | 1 | 0 | 0 | 1651239 | YP_009555238.1 | P0C6X6 |
| DKVYCPRHVICSSSNM | 1 | 0 | 0 | 1651240 | YP_173236.1 | A0A140H1G9 |
| DKWFVNDGRVANGYVC | 1 | 0 | 0 | 1651243 | NP_073549.1 | A0A0P0LKV0 |
| DKWLNNFNVPSPLNWE | 1 | 0 | 0 | 1651244 | YP_173238.1 | A0A140H1H1 |
| DKYFDCYDGGCINANQ | 1 | 0 | 0 | 1651247 | YP_009724389.1 | P0DTD1 |
| DLALTNMYKEARINDK | 1 | 0 | 0 | 1651266 | YP_009555238.1 | P0C6X6 |
| DLAVCLGIGDSAVTYS | 1 | 0 | 0 | 1651271 | YP_009555238.1 | P0C6X6 |
| DLAVQIGNNNVCTYEH | 1 | 0 | 0 | 1651272 | NP_073549.1 | A0A0P0LKV0 |
| DLAVQIGSNNVCTYEH | 1 | 0 | 0 | 1651273 | YP_003766.2 | H9EJ42 |
| DLDHNFGSAGVVANGV | 1 | 0 | 0 | 1651292 | NP_073556.1 | A0A0N7J131 |
| DLDVILPGEFSKKQLG | 1 | 0 | 0 | 1651296 | YP_009047202.1 | K9N7C7 |
| DLEDFKPRSQMETDFL | 1 | 0 | 0 | 1651297 | NP_828849.2 | P0C6X7 |
| DLEGNFYGPFVDRQTA | 1 | 0 | 0 | 1651301 | YP_009724389.1 | P0DTD1 |
| DLEIGYTVDCSCGKKL | 1 | 0 | 0 | 1651303 | YP_009555238.1 | P0C6X6 |
| DLERDFIDMDDNLFIA | 1 | 0 | 0 | 1651304 | YP_173236.1 | A0A140H1G9 |
| DLESVIQDYENVCFEF | 1 | 0 | 0 | 1651306 | YP_009555238.1 | P0C6X6 |
| DLEVLLQDALQSREAV | 1 | 0 | 0 | 1651308 | YP_009555238.1 | P0C6X6 |
| DLGVISKVHEVIIDNK | 1 | 0 | 0 | 1651337 | YP_003766.2 | H9EJ42 |
| DLGVLIQNGAKHVQGN | 1 | 0 | 0 | 1651338 | YP_173236.1 | A0A140H1G9 |
| DLGVLIQNSAKHVQGN | 1 | 0 | 0 | 1651339 | YP_009555238.1 | P0C6X6 |
| DLHNKINLCDDPEKAQ | 1 | 0 | 0 | 1651341 | YP_003766.2 | H9EJ42 |
| DLHVVEVLTKTAVKSG | 1 | 0 | 0 | 1651346 | YP_173236.1 | A0A140H1G9 |
| DLIEGSKPYSTLQPNA | 1 | 0 | 0 | 1651350 | YP_003766.2 | H9EJ42 |
| DLIISDMYDPKTKNVT | 1 | 0 | 0 | 1651353 | YP_009724389.1 | P0DTD1 |
| DLISKCQITAVEGTKK | 1 | 0 | 0 | 1651359 | YP_009555238.1 | P0C6X6 |
| DLKCTSDNINDKDTGP | 1 | 0 | 0 | 1651365 | YP_009555241.1 | P36334 |
| DLKFIDKLNVLFDGRD | 1 | 0 | 0 | 1651369 | YP_009555238.1 | P0C6X6 |
| DLLCLRDDNQTLLVPK | 1 | 0 | 0 | 1651387 | YP_173236.1 | A0A140H1G9 |
| DLLFDKVTIADPGYMQ | 1 | 0 | 0 | 1651392 | YP_009047204.1 | K9N5Q8 |
| DLLIKAYNTINNEQGT | 1 | 0 | 0 | 1651396 | NP_073549.1 | A0A0P0LKV0 |
| DLLISDMYDGRTKAID | 1 | 0 | 0 | 1651400 | NP_073549.1 | A0A0P0LKV0 |
| DLLKDIKFVIPALQHA | 1 | 0 | 0 | 1651401 | YP_009047202.1 | K9N7C7 |
| DLLKTDSVLLIERFVS | 1 | 0 | 0 | 1651402 | YP_009555238.1 | P0C6X6 |
| DLLKYDFTEERLKLFD | 1 | 0 | 0 | 1651404 | YP_009724389.1 | P0DTD1 |
| DLLKYDFTEHKENLFN | 1 | 0 | 0 | 1651406 | YP_003766.2 | H9EJ42 |
| DLNANMSLAECKRALG | 1 | 0 | 0 | 1651432 | NP_073549.1 | A0A0P0LKV0 |
| DLNDFVSDADSTLIGD | 1 | 0 | 0 | 1651434 | YP_009724389.1 | P0DTD1 |
| DLNEFVSDADITLFGD | 1 | 0 | 0 | 1651435 | YP_009047202.1 | K9N7C7 |
| DLNETLVTMPLGYVTH | 1 | 0 | 0 | 1651436 | YP_009724389.1 | P0DTD1 |
| DLNGDVVAIDYKHYTP | 1 | 0 | 0 | 1651438 | YP_009724389.1 | P0DTD1 |
| DLNGDVVAIDYRHYSA | 1 | 0 | 0 | 1651439 | NP_828849.2 | P0C6X7 |
| DLNGVIVDKVGELKVE | 1 | 0 | 0 | 1651447 | YP_173236.1 | A0A140H1G9 |
| DLNQLTGYKKPASREF | 1 | 0 | 0 | 1651450 | QHR63299.1 | P0C6W2 |
| DLPDDVVAANMFMGVG | 1 | 0 | 0 | 1651459 | YP_173236.1 | A0A140H1G9 |
| DLPKEITVATSRTLSY | 1 | 0 | 0 | 1651464 | YP_009724393.1 | P0DTC5 |
| DLQDLKWARFPKSDGT | 1 | 0 | 0 | 1651473 | YP_009724389.1 | P0DTD1 |
| DLQGNFGDLQLNKLGT | 1 | 0 | 0 | 1651475 | YP_009047211.1 | K9N4V7 |
| DLQLGSSGFLQSSNYK | 1 | 0 | 0 | 1651479 | YP_173238.1 | A0A140H1H1 |
| DLQLNKLGTEDPRWPQ | 1 | 0 | 0 | 1651480 | YP_009047211.1 | K9N4V7 |
| DLQNYGDSATLPKGIM | 1 | 0 | 0 | 1651481 | YP_009724389.1 | P0DTD1 |
| DLQNYGENAVIPKGIM | 1 | 0 | 0 | 1651483 | NP_828849.2 | P0C6X7 |
| DLRFPRGQGVPINTNS | 1 | 0 | 0 | 1651490 | QIQ54056.1 | P59595 |
| DLSAFYVKDGKYFTSE | 1 | 0 | 0 | 1651495 | YP_009047202.1 | K9N7C7 |
| DLSKELKRPIQPTDVA | 1 | 0 | 0 | 1651508 | YP_009555238.1 | P0C6X6 |
| DLSKFQLKLKGTPVLQ | 1 | 0 | 0 | 1651511 | YP_009047202.1 | K9N7C7 |
| DLSQYYCDNGKYYTKP | 1 | 0 | 0 | 1651524 | YP_173236.1 | A0A140H1G9 |
| DLSVVNARLRAKHYVY | 1 | 0 | 0 | 1651535 | YP_009724389.1 | P0DTD1 |
| DLTLDGYSKLFITKDE | 1 | 0 | 0 | 1651552 | YP_173236.1 | A0A140H1G9 |
| DLTVVSKVHEVIIDNK | 1 | 0 | 0 | 1651560 | NP_073549.1 | A0A0P0LKV0 |
| DLVDSYFENDSILQSV | 1 | 0 | 0 | 1651563 | NP_073549.1 | A0A0P0LKV0 |
| DLVPNQPYPNASFDNF | 1 | 0 | 0 | 1651569 | YP_009724389.1 | P0DTD1 |
| DLVPTQPLPNASFDNF | 1 | 0 | 0 | 1651571 | NP_828849.2 | P0C6X7 |
| DLVPTQPYPNSNFDNF | 1 | 0 | 0 | 1651572 | QIQ54046.1 | P0C6U8 |
| DLVQNGVDAKGFPQLA | 1 | 0 | 0 | 1651573 | YP_003771.1 | H9EJ69 |
| DLWQFVDHFGENEEII | 1 | 0 | 0 | 1651583 | NP_073549.1 | A0A0P0LKV0 |

| | | | | | | |
|---|---|---|---|---|---|---|
| DLYDKLQFTSLEIPRR | 1 | 0 | 0 | 1651588 | YP_009724389.1 | P0DTD1 |
| DLYDKLQFTSLEVPRR | 1 | 0 | 0 | 1651589 | QIQ54047.1 | P0C6X7 |
| DLYVALRGRIDKDLSL | 1 | 0 | 0 | 1651603 | NP_073553.1 | A0A0P0KWE8 |
| DLYVNVYRSTSPDPKF | 1 | 0 | 0 | 1651606 | YP_009047202.1 | K9N7C7 |
| DMAKFPLKLAGTAVIN | 1 | 0 | 0 | 1651610 | YP_009555238.1 | P0C6X6 |
| DMCAALKELLQNGMNG | 1 | 0 | 0 | 1651612 | NP_828849.2 | P0C6X7 |
| DMCASLKELLQNGMNG | 1 | 0 | 0 | 1651613 | YP_009724389.1 | P0DTD1 |
| DMCASLKELLQNGVNG | 1 | 0 | 0 | 1651614 | QIQ54046.1 | P0C6U8 |
| DMEKDFLSFDSDVFIK | 1 | 0 | 0 | 1651622 | YP_009047202.1 | K9N7C7 |
| DMEYDFLNMDMGVFIN | 1 | 0 | 0 | 1651623 | YP_003766.2 | H9EJ42 |
| DMGVFINKYGLEDFNF | 1 | 0 | 0 | 1651625 | YP_003766.2 | H9EJ42 |
| DMKGKMFVRPVIEDYH | 1 | 0 | 0 | 1651631 | YP_173241.1 | Q0ZJ82 |
| DMKGTMYVRPIIEDYH | 1 | 0 | 0 | 1651632 | YP_009555244.1 | Q01455 |
| DMLNPNYEDLLIRKSN | 1 | 0 | 0 | 1651636 | YP_009724389.1 | P0DTD1 |
| DMLRHLIKDVDNPVLM | 1 | 0 | 0 | 1651637 | YP_173236.1 | A0A140H1G9 |
| DMLRRLIKDVDNPVLM | 1 | 0 | 0 | 1651638 | YP_009555238.1 | P0C6X6 |
| DMLYSPPTISYNSTLQ | 1 | 0 | 0 | 1651639 | YP_003766.2 | H9EJ42 |
| DMSSVDTILNMARNGV | 1 | 0 | 0 | 1651650 | NP_073549.1 | A0A0P0LKV0 |
| DMTKFSLKLAGTAVVN | 1 | 0 | 0 | 1651653 | YP_173236.1 | A0A140H1G9 |
| DMTNPDYTNLLCRVTS | 1 | 0 | 0 | 1651654 | YP_009555238.1 | P0C6X6 |
| DMVDTSLSGFKLKDCV | 1 | 0 | 0 | 1651656 | YP_009724389.1 | P0DTD1 |
| DMVDTSLSGYKLKDCI | 1 | 0 | 0 | 1651657 | QIQ54046.1 | P0C6U8 |
| DMYDGRTKAIDGENVS | 1 | 0 | 0 | 1651664 | NP_073549.1 | A0A0P0LKV0 |
| DMYDPITKNIGEYNVS | 1 | 0 | 0 | 1651665 | YP_009555238.1 | P0C6X6 |
| DMYDPKTKNVTKENDS | 1 | 0 | 0 | 1651667 | YP_009724389.1 | P0DTD1 |
| DMYDPLTKNIGDYNVS | 1 | 0 | 0 | 1651668 | YP_173236.1 | A0A140H1G9 |
| DMYDPRTKHVTKENDS | 1 | 0 | 0 | 1651669 | NP_828849.2 | P0C6X7 |
| DMYDPTTKNVTGSNES | 1 | 0 | 0 | 1651670 | YP_009047202.1 | K9N7C7 |
| DNALSGVLSAAIKDCV | 1 | 0 | 0 | 1651681 | NP_073549.1 | A0A0P0LKV0 |
| DNASPASTVHATATIP | 1 | 0 | 0 | 1651683 | NP_828852.2 | P59632 |
| DNAVATFYPQLQSAEW | 1 | 0 | 0 | 1651684 | NP_073549.1 | A0A0P0LKV0 |
| DNAVKGCVPLNAIPSL | 1 | 0 | 0 | 1651685 | YP_009555238.1 | P0C6X6 |
| DNAVKGCVPLSAIPAL | 1 | 0 | 0 | 1651686 | YP_173236.1 | A0A140H1G9 |
| DNAYLKWARVEGKDGF | 1 | 0 | 0 | 1651688 | YP_009047202.1 | K9N7C7 |
| DNCDFYGWVSGNMMDG | 1 | 0 | 0 | 1651690 | YP_173236.1 | A0A140H1G9 |
| DNCSLEKFMSTPNAIF | 1 | 0 | 0 | 1651693 | YP_009047202.1 | K9N7C7 |
| DNDALNNIINNARDGC | 1 | 0 | 0 | 1651697 | YP_009724389.1 | P0DTD1 |
| DNDDEDVVTGDNDDED | 1 | 0 | 0 | 1651698 | YP_173236.1 | A0A140H1G9 |
| DNDDEDVVTGDNNDEE | 1 | 0 | 0 | 1651699 | YP_173236.1 | A0A140H1G9 |
| DNDDQIVVTGDDVDDI | 1 | 0 | 0 | 1651700 | YP_173236.1 | A0A140H1G9 |
| DNDFNTILNTACGVFE | 1 | 0 | 0 | 1651701 | YP_009555238.1 | P0C6X6 |
| DNDGRPVHVKEITKEN | 1 | 0 | 0 | 1651703 | YP_003766.2 | H9EJ42 |
| DNDINDYVSDADFSIT | 1 | 0 | 0 | 1651704 | YP_003766.2 | H9EJ42 |
| DNFTGECPNFVFPLNS | 1 | 0 | 0 | 1651723 | QIQ54046.1 | P0C6U8 |
| DNGALEAFKRSNNGVY | 1 | 0 | 0 | 1651726 | YP_009555238.1 | P0C6X6 |
| DNGKYYTKPIIKAQFK | 1 | 0 | 0 | 1651731 | YP_173236.1 | A0A140H1G9 |
| DNGSSINRKQLEIVKL | 1 | 0 | 0 | 1651736 | YP_003766.2 | H9EJ42 |
| DNGSSINRRQLDVVKR | 1 | 0 | 0 | 1651737 | NP_073549.1 | A0A0P0LKV0 |
| DNGVIRDVSVEDVCFA | 1 | 0 | 0 | 1651741 | YP_009555238.1 | P0C6X6 |
| DNHLSTFYPQLQSAEW | 1 | 0 | 0 | 1651743 | YP_003766.2 | H9EJ42 |
| DNIGTNVPLQVGFSNG | 1 | 0 | 0 | 1651747 | YP_003766.2 | H9EJ42 |
| DNIKFADDLNQLTGYK | 1 | 0 | 0 | 1651749 | YP_009724389.1 | P0DTD1 |
| DNIKLADDLNQLTGYK | 1 | 0 | 0 | 1651750 | QHR63299.1 | P0C6W2 |
| DNINAILTEVNELLDT | 1 | 0 | 0 | 1651752 | YP_009555241.1 | P36334 |
| DNINLHTQVVDMSMTY | 1 | 0 | 0 | 1651755 | YP_009724389.1 | P0DTD1 |
| DNINSILDEVNGLLDT | 1 | 0 | 0 | 1651756 | YP_173238.1 | A0A140H1H1 |
| DNINSTLVDLKWLNRV | 1 | 0 | 0 | 1651757 | NP_073551.1 | A0A0P0K6L9 |
| DNISVAVPKSRVQQNK | 1 | 0 | 0 | 1651758 | YP_009555245.1 | P33469 |
| DNIVAADLGVLIQNSA | 1 | 0 | 0 | 1651760 | YP_009555238.1 | P0C6X6 |
| DNKFDVLKVDDVDDGG | 1 | 0 | 0 | 1651764 | YP_009555238.1 | P0C6X6 |
| DNKLKAHKDKSAQCFK | 1 | 0 | 0 | 1651766 | YP_009724389.1 | P0DTD1 |
| DNKPYRWMLWCKDNHL | 1 | 0 | 0 | 1651772 | YP_003766.2 | H9EJ42 |
| DNLKPRVEAPKQEEPP | 1 | 0 | 0 | 1651780 | NP_828849.2 | P0C6X7 |
| DNLLEILQKEKVNINI | 1 | 0 | 0 | 1651785 | YP_009724389.1 | P0DTD1 |
| DNLRANNTKGSIPINV | 1 | 0 | 0 | 1651787 | QIQ54046.1 | P0C6U8 |
| DNLRANNTKGSLPINV | 1 | 0 | 0 | 1651788 | YP_009724389.1 | P0DTD1 |
| DNLSSEDLKAVRSSFN | 1 | 0 | 0 | 1651790 | YP_009555238.1 | P0C6X6 |
| DNLVFTNKTTLPTNVA | 1 | 0 | 0 | 1651793 | YP_003766.2 | H9EJ42 |
| DNNDEEIVTGDNDDQI | 1 | 0 | 0 | 1651805 | YP_173236.1 | A0A140H1G9 |
| DNNDITNTNLSRGRGR | 1 | 0 | 0 | 1651806 | YP_009047211.1 | K9N4V7 |
| DNNQNGDRSGARPKQR | 1 | 0 | 0 | 1651809 | QIQ54056.1 | P59595 |
| DNPHLMGWDYPKCDRA | 1 | 0 | 0 | 1651813 | YP_009047202.1 | K9N7C7 |
| DNPSSSEVDMICSTTA | 1 | 0 | 0 | 1651815 | YP_009555238.1 | P0C6X6 |
| DNPVLMGWDYPKCDRA | 1 | 0 | 0 | 1651816 | YP_009555238.1 | P0C6X6 |
| DNQALNSILDNAVKGC | 1 | 0 | 0 | 1651820 | YP_009555238.1 | P0C6X6 |
| DNQDLNGKWYDFGDFV | 1 | 0 | 0 | 1651821 | YP_009047202.1 | K9N7C7 |
| DNQDLNGKWYDFGDYV | 1 | 0 | 0 | 1651822 | YP_009555238.1 | P0C6X6 |
| DNQSKSPSSSGTSTPK | 1 | 0 | 0 | 1651828 | YP_003771.1 | H9EJ69 |
| DNQTTTIQTIVEVQPQ | 1 | 0 | 0 | 1651830 | YP_009724389.1 | P0DTD1 |
| DNSAKFWEEAFYRDLY | 1 | 0 | 0 | 1651833 | YP_009047202.1 | K9N7C7 |
| DNSANTKVLRNKGVDY | 1 | 0 | 0 | 1651834 | YP_009047202.1 | K9N7C7 |
| DNSSLTIKKPNELSRV | 1 | 0 | 0 | 1651841 | YP_009724389.1 | P0DTD1 |
| DNSSTLQSVASSFVSM | 1 | 0 | 0 | 1651843 | YP_003766.2 | H9EJ42 |
| DNSTTVSVTVSSNMVE | 1 | 0 | 0 | 1651844 | YP_009047202.1 | K9N7C7 |
| DNSYFDSKGWYDPVEN | 1 | 0 | 0 | 1651845 | YP_003766.2 | H9EJ42 |
| DNTKFADDLNQMSGYK | 1 | 0 | 0 | 1651852 | QIQ54046.1 | P0C6U8 |
| DNTNLHTQLVDMSMTY | 1 | 0 | 0 | 1651853 | NP_828849.2 | P0C6X7 |
| DNTSCPEWMDDFEADP | 1 | 0 | 0 | 1651854 | YP_009047202.1 | K9N7C7 |
| DNVAFYQGDVDTVVNG | 1 | 0 | 0 | 1651859 | NP_073549.1 | A0A0P0LKV0 |
| DNVLPETYVALPIYYQ | 1 | 0 | 0 | 1651864 | YP_003767.1 | A0A0P0G321 |
| DNVSIKNVDIVSEAKN | 1 | 0 | 0 | 1651869 | QIQ54046.1 | P0C6U8 |

| | | | | | | |
|---|---|---|---|---|---|---|
| DNVSSVWPLYPYGRCP | 1 | 0 | 0 | 1651870 | YP_009555240.1 | P30215 |
| DNVTDFNAIATCDWTN | 1 | 0 | 0 | 1651871 | YP_009724389.1 | P0DTD1 |
| DNVYCYNTDLIEGSKP | 1 | 0 | 0 | 1651874 | YP_003766.2 | H9EJ42 |
| DNVYIKNADIVEEAKK | 1 | 0 | 0 | 1651875 | YP_009724389.1 | P0DTD1 |
| DNVYMAKAGDKYYPVV | 1 | 0 | 0 | 1651876 | YP_009555238.1 | P0C6X6 |
| DNYYKKDNSYFTEQPI | 1 | 0 | 0 | 1651882 | YP_009724389.1 | P0DTD1 |
| DPAAAYLDAVKQGAKP | 1 | 0 | 0 | 1651883 | NP_073549.1 | A0A0P0LKV0 |
| DPAKAYKDYLASGGQP | 1 | 0 | 0 | 1651887 | NP_828849.2 | P0C6X7 |
| DPAMHIASSNAFLDLR | 1 | 0 | 0 | 1651890 | YP_009047202.1 | K9N7C7 |
| DPAPIDIDAFPVKPIC | 1 | 0 | 0 | 1651892 | YP_003766.2 | H9EJ42 |
| DPAQLPAPRTLLTKGT | 1 | 0 | 0 | 1651893 | YP_009724389.1 | P0DTD1 |
| DPAQLPAPRVLLSKGS | 1 | 0 | 0 | 1651895 | YP_173236.1 | A0A140H1G9 |
| DPAQLPAPRVLLSKGT | 1 | 0 | 0 | 1651896 | YP_009555238.1 | P0C6X6 |
| DPDLSAFYVKDGKYFT | 1 | 0 | 0 | 1651899 | YP_009047202.1 | K9N7C7 |
| DPGYMQGYDDCMQQGP | 1 | 0 | 0 | 1651906 | YP_009047204.1 | K9N5Q8 |
| DPIDDFQVNWNTKHGS | 1 | 0 | 0 | 1651907 | QIQ54046.1 | P0C6U8 |
| DPIEDYEQNWNTKHGS | 1 | 0 | 0 | 1651908 | NP_828849.2 | P0C6X7 |
| DPIRFCLENNVCNVCG | 1 | 0 | 0 | 1651913 | YP_003766.2 | H9EJ42 |
| DPIRFCLENTVCKVCG | 1 | 0 | 0 | 1651914 | NP_073549.1 | A0A0P0LKV0 |
| DPITAYDPRSCSQKKS | 1 | 0 | 0 | 1651915 | YP_173238.1 | A0A140H1H1 |
| DPKFVDKYYAFLNKHF | 1 | 0 | 0 | 1651919 | YP_009047202.1 | K9N7C7 |
| DPKKTYLDFIQQGGTP | 1 | 0 | 0 | 1651921 | YP_009555238.1 | P0C6X6 |
| DPKKTYLDYIQQGGVP | 1 | 0 | 0 | 1651922 | YP_173236.1 | A0A140H1G9 |
| DPKLDGYYKKDNAYFT | 1 | 0 | 0 | 1651923 | QIQ54046.1 | P0C6U8 |
| DPKLDNYYKKDNSYFT | 1 | 0 | 0 | 1651924 | YP_009724389.1 | P0DTD1 |
| DPKLMGWDYPKCDRAM | 1 | 0 | 0 | 1651925 | NP_073549.1 | A0A0P0LKV0 |
| DPKNPNYNKWLELLEQ | 1 | 0 | 0 | 1651926 | YP_009047211.1 | K9N4V7 |
| DPKTKNVTKENDSKEG | 1 | 0 | 0 | 1651929 | YP_009724389.1 | P0DTD1 |
| DPLKPTKRSFIEDLLF | 1 | 0 | 0 | 1651934 | NP_828851.1 | P59594 |
| DPLTKNIGDYNVSKDG | 1 | 0 | 0 | 1651940 | YP_173236.1 | A0A140H1G9 |
| DPNQTNVVPPALHLVD | 1 | 0 | 0 | 1651943 | NP_828859.1 | P59636 |
| DPPCKFTVQDAKGLKI | 1 | 0 | 0 | 1651946 | YP_009555238.1 | P0C6X6 |
| DPPNVQIHTIDGSSGV | 1 | 0 | 0 | 1651947 | NP_828852.2 | P59632 |
| DPQKAYLDFVNAGGAP | 1 | 0 | 0 | 1651952 | YP_009047202.1 | K9N7C7 |
| DPRSCSQKKSLVGVGE | 1 | 0 | 0 | 1651955 | YP_173238.1 | A0A140H1H1 |
| DPRTKHVTKENDSKEG | 1 | 0 | 0 | 1651956 | NP_828849.2 | P0C6X7 |
| DPRWPQIAELAPTASA | 1 | 0 | 0 | 1651958 | YP_009047211.1 | K9N4V7 |
| DPSFKDNVILLNKHID | 1 | 0 | 0 | 1651961 | QIQ54056.1 | P59595 |
| DPSSDEAIPTRFPPGT | 1 | 0 | 0 | 1651969 | YP_009555245.1 | P33469 |
| DPTTKNVTGSNESKAL | 1 | 0 | 0 | 1651977 | YP_009047202.1 | K9N7C7 |
| DPVGFCLSNTPCNVCQ | 1 | 0 | 0 | 1651983 | YP_009047202.1 | K9N7C7 |
| DPVGFTLRNTVCTVCG | 1 | 0 | 0 | 1651985 | NP_828849.2 | P0C6X7 |
| DPVVDWHYFDAQPLTR | 1 | 0 | 0 | 1651988 | YP_009047202.1 | K9N7C7 |
| DPYEDFQENWNTKHSS | 1 | 0 | 0 | 1651989 | YP_009724389.1 | P0DTD1 |
| DQAMTQMYKQARSEDK | 1 | 0 | 0 | 1651996 | YP_009724389.1 | P0DTD1 |
| DQAMTSMYKQARAEDK | 1 | 0 | 0 | 1651997 | YP_009047202.1 | K9N7C7 |
| DQDALFAYTKRNVIPT | 1 | 0 | 0 | 1652001 | YP_009724389.1 | P0DTD1 |
| DQDFGSTVFNVPTKVL | 1 | 0 | 0 | 1652002 | YP_009555238.1 | P0C6X6 |
| DQDLIRQGTDYKHWPQ | 1 | 0 | 0 | 1652003 | NP_828858.1 | P59595 |
| DQFKTTGDLAVQIGSN | 1 | 0 | 0 | 1652012 | YP_003766.2 | H9EJ42 |
| DQIVVTGDDVDDIESI | 1 | 0 | 0 | 1652024 | YP_173236.1 | A0A140H1G9 |
| DQLNDLVYSLIERGKL | 1 | 0 | 0 | 1652026 | YP_173236.1 | A0A140H1G9 |
| DQLNSSYFKLSIPTNF | 1 | 0 | 0 | 1652027 | YP_009047204.1 | K9N5Q8 |
| DQLSFDVPDGFYSTSP | 1 | 0 | 0 | 1652031 | NP_073551.1 | A0A0P0K6L9 |
| DQMSEPPKEQRVQGSI | 1 | 0 | 0 | 1652034 | YP_009047211.1 | K9N4V7 |
| DQPSLQVESANLMLSD | 1 | 0 | 0 | 1652037 | YP_003766.2 | H9EJ42 |
| DQSSYIVDSVAVKNGA | 1 | 0 | 0 | 1652046 | NP_828849.2 | P0C6X7 |
| DQSSYVVDSVAVKNGS | 1 | 0 | 0 | 1652048 | QIQ54046.1 | P0C6U8 |
| DQTPIVWHAKDFNSLS | 1 | 0 | 0 | 1652051 | NP_073549.1 | A0A0P0LKV0 |
| DQTYHPNCINCLDDRC | 1 | 0 | 0 | 1652053 | NP_828849.2 | P0C6X7 |
| DQTYHPNCVNCLDDRC | 1 | 0 | 0 | 1652054 | YP_009724389.1 | P0DTD1 |
| DQVAVYQQSIIGAMTA | 1 | 0 | 0 | 1652055 | YP_003767.1 | A0A0P0G321 |
| DQVEEVNSIFDIETVD | 1 | 0 | 0 | 1652056 | NP_073549.1 | A0A0P0LKV0 |
| DQYGCDYSGKLADCLQ | 1 | 0 | 0 | 1652069 | YP_173236.1 | A0A140H1G9 |
| DQYLCGADGKPVMSED | 1 | 0 | 0 | 1652072 | NP_073549.1 | A0A0P0LKV0 |
| DQYMCGVDGKPISAYA | 1 | 0 | 0 | 1652073 | YP_009047202.1 | K9N7C7 |
| DRCIKISTVQSKLTDL | 1 | 0 | 0 | 1652084 | YP_003766.2 | H9EJ42 |
| DRDGQRTYDDVNASLF | 1 | 0 | 0 | 1652089 | YP_009555238.1 | P0C6X6 |
| DRDGQRVYDDVDASLF | 1 | 0 | 0 | 1652090 | YP_173236.1 | A0A140H1G9 |
| DREVILCWETGKVKPP | 1 | 0 | 0 | 1652098 | YP_173236.1 | A0A140H1G9 |
| DRFKTSGDLAVQIGNN | 1 | 0 | 0 | 1652099 | NP_073549.1 | A0A0P0LKV0 |
| DRFKVNETNSQYLFST | 1 | 0 | 0 | 1652100 | YP_009047202.1 | K9N7C7 |
| DRGIAPKSGYFVNVNN | 1 | 0 | 0 | 1652104 | YP_009555241.1 | P36334 |
| DRKIKKYPCMVGPDYA | 1 | 0 | 0 | 1652114 | YP_009047202.1 | K9N7C7 |
| DRLNALESKMSGKGSQ | 1 | 0 | 0 | 1652124 | QIQ54056.1 | P59595 |
| DRLNEVAKNLNESLID | 1 | 0 | 0 | 1652125 | YP_009724390.1 | P0DTC2 |
| DRNHNSQDDIMKAVAA | 1 | 0 | 0 | 1652132 | NP_073556.1 | A0A0N7J131 |
| DRNTREVFAQVKQMYK | 1 | 0 | 0 | 1652133 | NP_828851.1 | P59594 |
| DRSGARPKQRRPQGLP | 1 | 0 | 0 | 1652142 | QIQ54056.1 | P59595 |
| DRSNNSSRASSRSSTR | 1 | 0 | 0 | 1652144 | YP_003771.1 | H9EJ69 |
| DRTAIQSFDNSYLNES | 1 | 0 | 0 | 1652145 | NP_073550.1 | A0A0P0ILY9 |
| DRYFKYWDQTYHPNCI | 1 | 0 | 0 | 1652163 | NP_828849.2 | P0C6X7 |
| DRYFKYWDQTYHPNCV | 1 | 0 | 0 | 1652164 | YP_009724389.1 | P0DTD1 |
| DSATLPKGIMMNVAKY | 1 | 0 | 0 | 1652172 | YP_009724389.1 | P0DTD1 |
| DSATLVNDIDITFLKK | 1 | 0 | 0 | 1652173 | QHR63299.1 | P0C6W2 |
| DSATLVSDIDITFLKK | 1 | 0 | 0 | 1652174 | YP_009724389.1 | P0DTD1 |
| DSCNNVNVRDLQRRLY | 1 | 0 | 0 | 1652180 | YP_003766.2 | H9EJ42 |
| DSCNNYMLTYNKVENM | 1 | 0 | 0 | 1652181 | YP_009724389.1 | P0DTD1 |
| DSDCDVVQEQVNYVPL | 1 | 0 | 0 | 1652183 | YP_003766.2 | H9EJ42 |
| DSDGTNKQLNEISDDC | 1 | 0 | 0 | 1652184 | YP_009555238.1 | P0C6X6 |
| DSDINEMVLSLVRSGK | 1 | 0 | 0 | 1652185 | NP_073549.1 | A0A0P0LKV0 |

| | | | | | | |
|---|---|---|---|---|---|---|
| DSDLNDFVSDADSTLI | 1 | 0 | 0 | 1652186 | YP_009724389.1 | P0DTD1 |
| DSDSLSSKPQRKRGVK | 1 | 0 | 0 | 1652188 | YP_173242.1 | Q19U29 |
| DSDVETRFITKSMISA | 1 | 0 | 0 | 1652192 | YP_173236.1 | A0A140H1G9 |
| DSDVFIKKYGLENYAF | 1 | 0 | 0 | 1652193 | YP_009047202.1 | K9N7C7 |
| DSEISANGCSTIAQAV | 1 | 0 | 0 | 1652197 | NP_073549.1 | A0A0P0LKV0 |
| DSEISGFGFAIPSVAV | 1 | 0 | 0 | 1652198 | YP_003766.2 | H9EJ42 |
| DSELLSTLSVDFNGVL | 1 | 0 | 0 | 1652199 | NP_073549.1 | A0A0P0LKV0 |
| DSFEGVLPINTDTVLS | 1 | 0 | 0 | 1652207 | YP_003766.2 | H9EJ42 |
| DSFGPGNTFINGDIAR | 1 | 0 | 0 | 1652208 | YP_003766.2 | H9EJ42 |
| DSFNILACYEGIASGV | 1 | 0 | 0 | 1652212 | YP_003766.2 | H9EJ42 |
| DSFSCNNFDESKIYGS | 1 | 0 | 0 | 1652214 | YP_173238.1 | A0A140H1H1 |
| DSFTCNNIDAAKIYGM | 1 | 0 | 0 | 1652215 | YP_009555241.1 | P36334 |
| DSFTNGAIVGTPVCVN | 1 | 0 | 0 | 1652217 | NP_828849.2 | P0C6X7 |
| DSFVITGDEVRQIAPG | 1 | 0 | 0 | 1652218 | QHR63300.2 | Q3LZX1 |
| DSFVKMMVDGFVHYAG | 1 | 0 | 0 | 1652219 | NP_073549.1 | A0A0P0LKV0 |
| DSFVNSFVSLYNVTRD | 1 | 0 | 0 | 1652220 | YP_009047202.1 | K9N7C7 |
| DSFVVKGDDVRQIAPG | 1 | 0 | 0 | 1652221 | NP_828851.1 | P59594 |
| DSFVVKGDEVRQIAPG | 1 | 0 | 0 | 1652222 | QIQ54048.1 | P59594 |
| DSGFAAYNRYRIGNYK | 1 | 0 | 0 | 1652225 | NP_828855.1 | P59596 |
| DSGTVSKTSDWKCKVT | 1 | 0 | 0 | 1652232 | YP_009047202.1 | K9N7C7 |
| DSHDTVLTSEEVVLKN | 1 | 0 | 0 | 1652235 | NP_828849.2 | P0C6X7 |
| DSHIVSKESGNTVVLT | 1 | 0 | 0 | 1652236 | NP_073556.1 | A0A0N7J131 |
| DSIEMQLFKVGKVDSI | 1 | 0 | 0 | 1652240 | YP_003766.2 | H9EJ42 |
| DSIESVGGLYEIKIPT | 1 | 0 | 0 | 1652241 | YP_173238.1 | A0A140H1H1 |
| DSIGTNFPLQLGFSTG | 1 | 0 | 0 | 1652244 | YP_009555238.1 | P0C6X6 |
| DSIRATATIPIQASLP | 1 | 0 | 0 | 1652249 | QIQ54049.1 | P59632 |
| DSKFQVGEFVFEKVDY | 1 | 0 | 0 | 1652264 | NP_073549.1 | A0A0P0LKV0 |
| DSKGWYDPVENEDIHR | 1 | 0 | 0 | 1652265 | YP_003766.2 | H9EJ42 |
| DSKTQSLLIVNNATNV | 1 | 0 | 0 | 1652274 | YP_009724390.1 | P0DTC2 |
| DSLCSSHYWSPQGNLG | 1 | 0 | 0 | 1652278 | YP_173236.1 | A0A140H1G9 |
| DSLNAKLGFDSSKEFV | 1 | 0 | 0 | 1652289 | YP_009555238.1 | P0C6X6 |
| DSLNLSAGTQHISNDF | 1 | 0 | 0 | 1652290 | YP_173242.1 | Q19U29 |
| DSLRQAALPQSKDSNF | 1 | 0 | 0 | 1652292 | YP_009047202.1 | K9N7C7 |
| DSMSYEDQDALFAYTK | 1 | 0 | 0 | 1652298 | YP_009724389.1 | P0DTD1 |
| DSNGCVVEASTDFHQL | 1 | 0 | 0 | 1652302 | YP_003766.2 | H9EJ42 |
| DSNGNIIGFKDFVTNK | 1 | 0 | 0 | 1652303 | YP_173238.1 | A0A140H1H1 |
| DSNGNLYGFRDYITNR | 1 | 0 | 0 | 1652304 | YP_009555241.1 | P36334 |
| DSNTTQDTYGGASVCI | 1 | 0 | 0 | 1652306 | NP_073549.1 | A0A0P0LKV0 |
| DSPFEKLQCEHLQFGL | 1 | 0 | 0 | 1652311 | YP_003767.1 | A0A0P0G321 |
| DSQQTVVQEDDSEVNQ | 1 | 0 | 0 | 1652317 | QHR63299.1 | P0C6W2 |
| DSRSDCNHINNLKIKN | 1 | 0 | 0 | 1652323 | YP_173237.1 | A0A140H1H0 |
| DSRWNNARQSDNMTAV | 1 | 0 | 0 | 1652326 | YP_009555240.1 | P30215 |
| DSSESGAKETKEINII | 1 | 0 | 0 | 1652330 | YP_009555238.1 | P0C6X6 |
| DSSEVKAQSSGVIPEN | 1 | 0 | 0 | 1652331 | YP_173236.1 | A0A140H1G9 |
| DSSRSTSRQQSRTRSD | 1 | 0 | 0 | 1652341 | YP_003771.1 | H9EJ69 |
| DSTDNNQNGDRSGARP | 1 | 0 | 0 | 1652345 | QIQ54056.1 | P59595 |
| DSTFVTEYYEFLNKHF | 1 | 0 | 0 | 1652353 | YP_009555238.1 | P0C6X6 |
| DSTGSNQNGERSGARP | 1 | 0 | 0 | 1652354 | QHR63308.1 | Q3LZX4 |
| DSTLIGDCATVHTANK | 1 | 0 | 0 | 1652356 | YP_009724389.1 | P0DTD1 |
| DSTLVKFSAAQAMEKV | 1 | 0 | 0 | 1652359 | YP_009555241.1 | P36334 |
| DSTYGPLQTPVGCVLG | 1 | 0 | 0 | 1652365 | YP_009047204.1 | K9N5Q8 |
| DSVFYVVKNSTAFPFE | 1 | 0 | 0 | 1652374 | YP_009047202.1 | K9N7C7 |
| DSVIVNDDTSETKYVK | 1 | 0 | 0 | 1652375 | YP_009555238.1 | P0C6X6 |
| DSVLLIERFVSLAIDA | 1 | 0 | 0 | 1652379 | YP_009555238.1 | P0C6X6 |
| DSVNLLTNMFTPLIQP | 1 | 0 | 0 | 1652382 | QIQ54046.1 | P0C6U8 |
| DSVQYAHKHDIQITNE | 1 | 0 | 0 | 1652385 | YP_009047202.1 | K9N7C7 |
| DSVRATATIPIQASLP | 1 | 0 | 0 | 1652386 | QHR63301.1 | Q3LZX0 |
| DSVRDPKTSEILDISP | 1 | 0 | 0 | 1652387 | NP_828851.1 | P59594 |
| DSVREEVDIIEQPFEE | 1 | 0 | 0 | 1652388 | YP_003766.2 | H9EJ42 |
| DSVVEVVTTSLTPCGY | 1 | 0 | 0 | 1652396 | YP_009555238.1 | P0C6X6 |
| DSYCKLKNSLSDVAFN | 1 | 0 | 0 | 1652400 | YP_009555238.1 | P0C6X6 |
| DSYCRLKNSVSDVAYN | 1 | 0 | 0 | 1652401 | YP_173236.1 | A0A140H1G9 |
| DSYFDNSSTLQSVASS | 1 | 0 | 0 | 1652402 | YP_003766.2 | H9EJ42 |
| DTAAYREAACSQLAKA | 1 | 0 | 0 | 1652423 | YP_009555238.1 | P0C6X6 |
| DTAGVLLKQLMVNKSS | 1 | 0 | 0 | 1652427 | YP_173243.1 | Q0ZJ80 |
| DTAHANNINRFNVAIT | 1 | 0 | 0 | 1652430 | YP_009047202.1 | K9N7C7 |
| DTAKKSMYDGDRFVKH | 1 | 0 | 0 | 1652431 | NP_073549.1 | A0A0P0LKV0 |
| DTENNTEFTRVNAKPP | 1 | 0 | 0 | 1652447 | NP_828849.2 | P0C6X7 |
| DTEQTQNVTVKPKRLR | 1 | 0 | 0 | 1652448 | YP_009047202.1 | K9N7C7 |
| DTFKGECPKFVFPLNS | 1 | 0 | 0 | 1652454 | NP_828849.2 | P0C6X7 |
| DTFLSMFDVDKKSLNA | 1 | 0 | 0 | 1652457 | YP_009555238.1 | P0C6X6 |
| DTFNGECPNFVFPLNS | 1 | 0 | 0 | 1652459 | YP_009724389.1 | P0DTD1 |
| DTFSATFSVPMEKLKA | 1 | 0 | 0 | 1652460 | NP_828849.2 | P0C6X7 |
| DTFTNNNGSDVLYQPP | 1 | 0 | 0 | 1652461 | YP_009555238.1 | P0C6X6 |
| DTGPPPISTDTVDVTN | 1 | 0 | 0 | 1652467 | YP_009555241.1 | P36334 |
| DTIANYAKPFLNKVVS | 1 | 0 | 0 | 1652476 | YP_009724389.1 | P0DTD1 |
| DTINDDEDVVTGDNDD | 1 | 0 | 0 | 1652481 | YP_173236.1 | A0A140H1G9 |
| DTIVTYAKPFLNKVTS | 1 | 0 | 0 | 1652485 | QIQ54046.1 | P0C6U8 |
| DTIVVTEGDGISTPKL | 1 | 0 | 0 | 1652486 | NP_828852.2 | P59632 |
| DTKLNGILQQAALDYL | 1 | 0 | 0 | 1652494 | YP_003766.2 | H9EJ42 |
| DTKRGVYCCREHEHEI | 1 | 0 | 0 | 1652496 | YP_009724389.1 | P0DTD1 |
| DTLDDILQQANKSVGI | 1 | 0 | 0 | 1652499 | YP_009047202.1 | K9N7C7 |
| DTLMQGVTLSSNLNTN | 1 | 0 | 0 | 1652506 | YP_173238.1 | A0A140H1H1 |
| DTLQETPVVPDTVEVP | 1 | 0 | 0 | 1652507 | YP_009047202.1 | K9N7C7 |
| DTLTKLSKYLANEAQV | 1 | 0 | 0 | 1652515 | NP_073549.1 | A0A0P0LKV0 |
| DTNSVCPKLEFANDTK | 1 | 0 | 0 | 1652523 | YP_009047204.1 | K9N5Q8 |
| DTPNATEFSRVSAKPP | 1 | 0 | 0 | 1652530 | QIQ54047.1 | P0C6X7 |
| DTPNNTDFSRVSAKPP | 1 | 0 | 0 | 1652531 | YP_009724389.1 | P0DTD1 |
| DTRTRSTLNLEGVNGG | 1 | 0 | 0 | 1652539 | NP_073549.1 | A0A0P0LKV0 |
| DTRTRSVFNLEGVNGG | 1 | 0 | 0 | 1652540 | YP_003766.2 | H9EJ42 |
| DTRVLNKLNLPGCNGG | 1 | 0 | 0 | 1652542 | YP_173236.1 | A0A140H1G9 |

| | | | | | | |
|---|---|---|---|---|---|---|
| DTRVLNNLNLPGCNGG | 1 | 0 | 0 | 1652543 | YP_009555238.1 | P0C6X6 |
| DTRVLSNLNLPGCDGG | 1 | 0 | 0 | 1652544 | YP_009724389.1 | P0DTD1 |
| DTSLSGFKLKDCVMYA | 1 | 0 | 0 | 1652549 | YP_009724389.1 | P0DTD1 |
| DTSLSGYKLKDCIMYA | 1 | 0 | 0 | 1652550 | QIQ54046.1 | P0C6U8 |
| DTSNSFEVLAVEDTQG | 1 | 0 | 0 | 1652553 | NP_828849.2 | P0C6X7 |
| DTSPQVLKALQKAVNI | 1 | 0 | 0 | 1652554 | YP_009047202.1 | K9N7C7 |
| DTSTPSDVSSRDPTTQ | 1 | 0 | 0 | 1652555 | YP_173242.1 | Q19U29 |
| DTTATSCQLYYNLPAA | 1 | 0 | 0 | 1652556 | YP_009555241.1 | P36334 |
| DTTPKEEFVVKEKLNA | 1 | 0 | 0 | 1652565 | NP_073549.1 | A0A0P0LKV0 |
| DTTQLHVADTLMQGVT | 1 | 0 | 0 | 1652566 | YP_173238.1 | A0A140H1H1 |
| DTVLTSEEVVLKNGEL | 1 | 0 | 0 | 1652580 | NP_828849.2 | P0C6X7 |
| DTVSALVYENKLKAKN | 1 | 0 | 0 | 1652585 | YP_009555238.1 | P0C6X6 |
| DTVVSTVKIGGLTFKT | 1 | 0 | 0 | 1652591 | NP_073549.1 | A0A0P0LKV0 |
| DVAVTAGPFMDNAINV | 1 | 0 | 0 | 1652614 | YP_009047202.1 | K9N7C7 |
| DVDDIESIYDFDTYKA | 1 | 0 | 0 | 1652630 | YP_173236.1 | A0A140H1G9 |
| DVDDPKLMGWDYPKCD | 1 | 0 | 0 | 1652631 | NP_073549.1 | A0A0P0LKV0 |
| DVDFHSVEIEAGEVKP | 1 | 0 | 0 | 1652633 | YP_003766.2 | H9EJ42 |
| DVDGLKYTKILKDDGN | 1 | 0 | 0 | 1652636 | YP_009555238.1 | P0C6X6 |
| DVDGLKYTKIMKDDGN | 1 | 0 | 0 | 1652637 | YP_173236.1 | A0A140H1G9 |
| DVDNINFKSLVGCLGP | 1 | 0 | 0 | 1652642 | YP_173238.1 | A0A140H1H1 |
| DVDNLVFTNKTTLPTN | 1 | 0 | 0 | 1652643 | YP_003766.2 | H9EJ42 |
| DVDNPHLMGWDYPKCD | 1 | 0 | 0 | 1652644 | YP_009047202.1 | K9N7C7 |
| DVDNPVLMGWDYPKCD | 1 | 0 | 0 | 1652645 | YP_009555238.1 | P0C6X6 |
| DVDTVVNGVDFDFIVN | 1 | 0 | 0 | 1652652 | NP_073549.1 | A0A0P0LKV0 |
| DVEGAHATRENIGTNF | 1 | 0 | 0 | 1652663 | YP_173236.1 | A0A140H1G9 |
| DVEGCHATREAIGTNL | 1 | 0 | 0 | 1652666 | QHR63299.1 | P0C6W2 |
| DVEIFENKTTLPVNVA | 1 | 0 | 0 | 1652671 | NP_828849.2 | P0C6X7 |
| DVEKGVLPQLEQPYVF | 1 | 0 | 0 | 1652673 | QIQ54046.1 | P0C6U8 |
| DVELFENKTILPVNVA | 1 | 0 | 0 | 1652674 | QIQ54047.1 | P0C6X7 |
| DVELFENKTTLPVNVA | 1 | 0 | 0 | 1652675 | YP_009724389.1 | P0DTD1 |
| DVENPHLMGWDYPKCD | 1 | 0 | 0 | 1652678 | YP_009724389.1 | P0DTD1 |
| DVESAHVCGDNIGTNV | 1 | 0 | 0 | 1652680 | YP_003766.2 | H9EJ42 |
| DVESGVYSVSSFEAKP | 1 | 0 | 0 | 1652681 | YP_009047204.1 | K9N5Q8 |
| DVETPHLMGWDYPKCD | 1 | 0 | 0 | 1652684 | NP_828849.2 | P0C6X7 |
| DVEWKFYDAQPCSDKA | 1 | 0 | 0 | 1652689 | YP_009724389.1 | P0DTD1 |
| DVFKIGDVKFKRLGDY | 1 | 0 | 0 | 1652695 | YP_003766.2 | H9EJ42 |
| DVFYKENSYTTTIKPI | 1 | 0 | 0 | 1652701 | QIQ54046.1 | P0C6U8 |
| DVFYKENSYTTTIKPV | 1 | 0 | 0 | 1652702 | YP_009724389.1 | P0DTD1 |
| DVFYKETSYTTTIKPV | 1 | 0 | 0 | 1652703 | NP_828849.2 | P0C6X7 |
| DVGFKNQYCNGSIACQ | 1 | 0 | 0 | 1652708 | YP_009555238.1 | P0C6X6 |
| DVGFVEAYNNCTGGAE | 1 | 0 | 0 | 1652709 | YP_009555241.1 | P36334 |
| DVGFVEAYNNCTGGSE | 1 | 0 | 0 | 1652710 | YP_173238.1 | A0A140H1H1 |
| DVHKDNFKDGLCMFWN | 1 | 0 | 0 | 1652717 | YP_173236.1 | A0A140H1G9 |
| DVIFSQFDSLGVSSNQ | 1 | 0 | 0 | 1652722 | YP_009555238.1 | P0C6X6 |
| DVIRKDLSYEQQNVLA | 1 | 0 | 0 | 1652730 | YP_003766.2 | H9EJ42 |
| DVIRYNLNFEENLRRG | 1 | 0 | 0 | 1652731 | NP_073551.1 | A0A0P0K6L9 |
| DVISSEAVEDTIDGVV | 1 | 0 | 0 | 1652733 | YP_173236.1 | A0A140H1G9 |
| DVIVAWHVVRDPRYVM | 1 | 0 | 0 | 1652735 | YP_009555238.1 | P0C6X6 |
| DVIVNPANGHMLHGGG | 1 | 0 | 0 | 1652738 | YP_173236.1 | A0A140H1G9 |
| DVKPKIKACIDEVTTT | 1 | 0 | 0 | 1652755 | NP_828849.2 | P0C6X7 |
| DVKPYGYPNDFVGGFR | 1 | 0 | 0 | 1652757 | YP_003766.2 | H9EJ42 |
| DVKTTEVVGNVILKPS | 1 | 0 | 0 | 1652761 | NP_828849.2 | P0C6X7 |
| DVLCNSFFKELTANMS | 1 | 0 | 0 | 1652763 | YP_003766.2 | H9EJ42 |
| DVLFYCSNSSSGVLDT | 1 | 0 | 0 | 1652767 | YP_003767.1 | A0A0P0G321 |
| DVLKSEDAQGMDNLAC | 1 | 0 | 0 | 1652771 | YP_009724389.1 | P0DTD1 |
| DVLKSEDTQGMDNLAC | 1 | 0 | 0 | 1652772 | QHR63299.1 | P0C6W2 |
| DVLQENQKILAASFNK | 1 | 0 | 0 | 1652783 | NP_073551.1 | A0A0P0K6L9 |
| DVLRNSFGKDLNANMS | 1 | 0 | 0 | 1652785 | NP_073549.1 | A0A0P0LKV0 |
| DVLSDVIRYNLNFEEN | 1 | 0 | 0 | 1652786 | NP_073551.1 | A0A0P0K6L9 |
| DVLSQNQKLIANAFNN | 1 | 0 | 0 | 1652788 | YP_009555241.1 | P36334 |
| DVLVRGFGDSVEEVLS | 1 | 0 | 0 | 1652793 | YP_009724389.1 | P0DTD1 |
| DVMRYNLNFSANSLDN | 1 | 0 | 0 | 1652797 | YP_003767.1 | A0A0P0G321 |
| DVNDVTKLYLGGMSYY | 1 | 0 | 0 | 1652806 | YP_009555238.1 | P0C6X6 |
| DVNKTLQEFAQNLPKY | 1 | 0 | 0 | 1652808 | YP_003767.1 | A0A0P0G321 |
| DVNKTLQELSYKLPNY | 1 | 0 | 0 | 1652809 | NP_073551.1 | A0A0P0K6L9 |
| DVNMQIMGVVMQSGVR | 1 | 0 | 0 | 1652812 | YP_009047202.1 | K9N7C7 |
| DVPDGFYSTSPIQSVE | 1 | 0 | 0 | 1652823 | NP_073551.1 | A0A0P0K6L9 |
| DVPYPKQSIFTINSVV | 1 | 0 | 0 | 1652831 | YP_009047202.1 | K9N7C7 |
| DVQLRAFDICNTNRAG | 1 | 0 | 0 | 1652833 | YP_173236.1 | A0A140H1G9 |
| DVRDYKLANDVKDTLR | 1 | 0 | 0 | 1652838 | YP_003766.2 | H9EJ42 |
| DVRDYLLSKMTSLPKD | 1 | 0 | 0 | 1652839 | YP_173236.1 | A0A140H1G9 |
| DVRETMNYLFQHANLD | 1 | 0 | 0 | 1652840 | QHR63299.1 | P0C6W2 |
| DVSEMLTFDKKAFTLA | 1 | 0 | 0 | 1652850 | NP_073551.1 | A0A0P0K6L9 |
| DVSKADGIIYPQGRTY | 1 | 0 | 0 | 1652857 | YP_009047204.1 | K9N5Q8 |
| DVSLTYLLGTAKKQVV | 1 | 0 | 0 | 1652863 | YP_009555238.1 | P0C6X6 |
| DVSPFEMPFEELNGLK | 1 | 0 | 0 | 1652866 | NP_073549.1 | A0A0P0LKV0 |
| DVSVEDVCFANKFEQF | 1 | 0 | 0 | 1652871 | YP_009555238.1 | P0C6X6 |
| DVTVLRDNLKPVTTLS | 1 | 0 | 0 | 1652894 | YP_003768.1 | H9EJ24 |
| DVTVTTDKSFEQQVGV | 1 | 0 | 0 | 1652895 | NP_073549.1 | A0A0P0LKV0 |
| DVVAANMFMGVGVGHY | 1 | 0 | 0 | 1652897 | YP_173236.1 | A0A140H1G9 |
| DVVAIDYKHYTPSFKK | 1 | 0 | 0 | 1652898 | YP_009724389.1 | P0DTD1 |
| DVVDQVEEVNSIFDIE | 1 | 0 | 0 | 1652902 | NP_073549.1 | A0A0P0LKV0 |
| DVVETKVEITPEESAS | 1 | 0 | 0 | 1652907 | QIQ54046.1 | P0C6U8 |
| DVVKEQVSDLLVKLLR | 1 | 0 | 0 | 1652915 | YP_009047202.1 | K9N7C7 |
| DVVKTDAVVLLERYVS | 1 | 0 | 0 | 1652919 | YP_003766.2 | H9EJ42 |
| DVVLASDDLYVKRYFK | 1 | 0 | 0 | 1652921 | YP_173236.1 | A0A140H1G9 |
| DVVLATDDLYVKRYER | 1 | 0 | 0 | 1652922 | YP_009555238.1 | P0C6X6 |
| DVVNQQGNSLNHLTSQ | 1 | 0 | 0 | 1652926 | NP_073551.1 | A0A0P0K6L9 |
| DVVNQQGSALNHLTSQ | 1 | 0 | 0 | 1652927 | YP_003767.1 | A0A0P0G321 |
| DVVQYEADRRAFVDYT | 1 | 0 | 0 | 1652931 | YP_009555238.1 | P0C6X6 |
| DVVSIRSSNQGNKQIV | 1 | 0 | 0 | 1652939 | YP_009047207.1 | K9N643 |

| | | | | | | |
|---|---|---|---|---|---|---|
| DVVTGDNDDEDVVTGD | 1 | 0 | 0 | 1652940 | YP_173236.1 | A0A140H1G9 |
| DVVTGDNNDEEIVTGD | 1 | 0 | 0 | 1652941 | YP_173236.1 | A0A140H1G9 |
| DVVWKHITDPLFSNMQ | 1 | 0 | 0 | 1652944 | NP_073549.1 | A0A0P0LKV0 |
| DVYDDIFAESKPWFVR | 1 | 0 | 0 | 1652954 | NP_073549.1 | A0A0P0LKV0 |
| DVYNDALFVSYGSSVE | 1 | 0 | 0 | 1652960 | YP_173236.1 | A0A140H1G9 |
| DVYNKDASFIGKNLKS | 1 | 0 | 0 | 1652962 | NP_073549.1 | A0A0P0LKV0 |
| DVYRRKPNLPNCNIEA | 1 | 0 | 0 | 1652966 | YP_009555241.1 | P36334 |
| DWDIEYPYISHEKKLN | 1 | 0 | 0 | 1652971 | YP_009047202.1 | K9N7C7 |
| DWKFYDAQPCSDKAYK | 1 | 0 | 0 | 1652979 | QIQ54047.1 | P0C6X7 |
| DWLDDDSQQTVGQQDG | 1 | 0 | 0 | 1652983 | YP_009724389.1 | P0DTD1 |
| DWLDDDSQQTVVQEDD | 1 | 0 | 0 | 1652984 | QHR63299.1 | P0C6W2 |
| DWLEEKFKEGIEFLKD | 1 | 0 | 0 | 1652986 | QIQ54046.1 | P0C6U8 |
| DWLEEKFKEGVEFLRD | 1 | 0 | 0 | 1652987 | YP_009724389.1 | P0DTD1 |
| DWRLINKFDVINGVKT | 1 | 0 | 0 | 1652996 | YP_173236.1 | A0A140H1G9 |
| DWSDIRDYKLANDAKE | 1 | 0 | 0 | 1652998 | NP_073549.1 | A0A0P0LKV0 |
| DWSITYPMIANENAIN | 1 | 0 | 0 | 1652999 | NP_073549.1 | A0A0P0LKV0 |
| DWSVEYPIIGDELRVN | 1 | 0 | 0 | 1653000 | NP_828849.2 | P0C6X7 |
| DWTDVRDYKLANDVKD | 1 | 0 | 0 | 1653004 | YP_003766.2 | H9EJ42 |
| DWTESGDYTLANTTTE | 1 | 0 | 0 | 1653005 | YP_009047202.1 | K9N7C7 |
| DWTIEYPIIGDELKIN | 1 | 0 | 0 | 1653007 | YP_009724389.1 | P0DTD1 |
| DWTVTYPFIANEKFIN | 1 | 0 | 0 | 1653009 | YP_003766.2 | H9EJ42 |
| DWVKHLNKNLNEGVLE | 1 | 0 | 0 | 1653013 | YP_003766.2 | H9EJ42 |
| DWVKHLNKTLNEGVLE | 1 | 0 | 0 | 1653014 | NP_073549.1 | A0A0P0LKV0 |
| DWYDFVENPDIINIYK | 1 | 0 | 0 | 1653015 | YP_173236.1 | A0A140H1G9 |
| DWYDFVENPDIINVYK | 1 | 0 | 0 | 1653016 | YP_009555238.1 | P0C6X6 |
| DYAAKGYIAGIQNFKE | 1 | 0 | 0 | 1653020 | YP_009047202.1 | K9N7C7 |
| DYAELGYIADISAFKA | 1 | 0 | 0 | 1653024 | YP_003766.2 | H9EJ42 |
| DYAFEHIVYGNFNQKI | 1 | 0 | 0 | 1653026 | YP_009555238.1 | P0C6X6 |
| DYAYFNGAIIRDSDVV | 1 | 0 | 0 | 1653035 | YP_009047202.1 | K9N7C7 |
| DYCDGSLGCKMCLFGY | 1 | 0 | 0 | 1653037 | NP_073549.1 | A0A0P0LKV0 |
| DYCVRAFDVYNKDASF | 1 | 0 | 0 | 1653043 | NP_073549.1 | A0A0P0LKV0 |
| DYDVAVTAGPFMDNAI | 1 | 0 | 0 | 1653050 | YP_009047202.1 | K9N7C7 |
| DYEFKFYDAFPVAKSV | 1 | 0 | 0 | 1653056 | YP_173236.1 | A0A140H1G9 |
| DYEQNWNTKHGSGALR | 1 | 0 | 0 | 1653059 | NP_828849.2 | P0C6X7 |
| DYFNKKDWYDFVENPD | 1 | 0 | 0 | 1653062 | YP_009724389.1 | P0DTD1 |
| DYGSDTVTYKSTATTK | 1 | 0 | 0 | 1653071 | NP_073549.1 | A0A0P0LKV0 |
| DYHVTKVLAEKFDYNV | 1 | 0 | 0 | 1653082 | NP_073549.1 | A0A0P0LKV0 |
| DYIARNGPLNVGGSCL | 1 | 0 | 0 | 1653083 | QIQ54046.1 | P0C6U8 |
| DYIATNGPLKVGGSCV | 1 | 0 | 0 | 1653084 | YP_009724389.1 | P0DTD1 |
| DYIESKRGVYCCRDHE | 1 | 0 | 0 | 1653085 | NP_828849.2 | P0C6X7 |
| DYINVTFLDLQVEMNR | 1 | 0 | 0 | 1653092 | YP_009555241.1 | P36334 |
| DYISSNGPLKVGAGVM | 1 | 0 | 0 | 1653095 | YP_003766.2 | H9EJ42 |
| DYKHWPQIAQFAPSAS | 1 | 0 | 0 | 1653100 | YP_009724397.2 | P0DTC9 |
| DYKKCTKGLSIADLAC | 1 | 0 | 0 | 1653102 | NP_073551.1 | A0A0P0K6L9 |
| DYKLANDAKESLRLFA | 1 | 0 | 0 | 1653103 | NP_073549.1 | A0A0P0LKV0 |
| DYKLANDVKDTLRLFA | 1 | 0 | 0 | 1653104 | YP_003766.2 | H9EJ42 |
| DYKLELFNKYFKHWSM | 1 | 0 | 0 | 1653105 | YP_009555238.1 | P0C6X6 |
| DYKRQNNLAIEEIEYV | 1 | 0 | 0 | 1653110 | NP_073549.1 | A0A0P0LKV0 |
| DYKSCTKGLSIADLAC | 1 | 0 | 0 | 1653111 | YP_003767.1 | A0A0P0G321 |
| DYKSFKQIVESCGNFK | 1 | 0 | 0 | 1653112 | QIQ54046.1 | P0C6U8 |
| DYLDIHPSLCNNGKIS | 1 | 0 | 0 | 1653120 | YP_173237.1 | A0A140H1H0 |
| DYLDNLKPRVEAPKQE | 1 | 0 | 0 | 1653121 | NP_828849.2 | P0C6X7 |
| DYNAILQSNNPQCAIV | 1 | 0 | 0 | 1653134 | YP_003766.2 | H9EJ42 |
| DYNAKVEISVTPIKNT | 1 | 0 | 0 | 1653135 | NP_073549.1 | A0A0P0LKV0 |
| DYNLSSVIPSLPTSGS | 1 | 0 | 0 | 1653136 | NP_073551.1 | A0A0P0K6L9 |
| DYNLSSVLPQRNIRSS | 1 | 0 | 0 | 1653137 | YP_003767.1 | A0A0P0G321 |
| DYNSPSSSSSRRKRRS | 1 | 0 | 0 | 1653139 | YP_173238.1 | A0A140H1H1 |
| DYNYYKYNLPTMVDIK | 1 | 0 | 0 | 1653147 | YP_009555238.1 | P0C6X6 |
| DYQGKSLEFGATSVTP | 1 | 0 | 0 | 1653156 | QHR63299.1 | P0C6W2 |
| DYQIGGYSEDRHSGVK | 1 | 0 | 0 | 1653159 | NP_828852.2 | P59632 |
| DYSDAVSYKSSTTYKL | 1 | 0 | 0 | 1653166 | YP_009047202.1 | K9N7C7 |
| DYSIDVIDNEIIVKPN | 1 | 0 | 0 | 1653168 | NP_073549.1 | A0A0P0LKV0 |
| DYSNLLHSKVKSVPNM | 1 | 0 | 0 | 1653170 | YP_009555238.1 | P0C6X6 |
| DYTGSLADGLEAYADK | 1 | 0 | 0 | 1653177 | YP_009555238.1 | P0C6X6 |
| DYTGYQELVIKTSHDD | 1 | 0 | 0 | 1653179 | YP_009555241.1 | P36334 |
| DYTLANTTTEPLKLFA | 1 | 0 | 0 | 1653182 | YP_009047202.1 | K9N7C7 |
| DYVDKWDDFCRQYSNE | 1 | 0 | 0 | 1653188 | NP_073549.1 | A0A0P0LKV0 |
| DYVDVNKTLQEFAQNL | 1 | 0 | 0 | 1653189 | YP_003767.1 | A0A0P0G321 |
| DYVGTGIIRSSNQSLA | 1 | 0 | 0 | 1653191 | YP_003767.1 | A0A0P0G321 |
| DYYQLYSTQLSTDTGV | 1 | 0 | 0 | 1653225 | YP_009724391.1 | P0DTC3 |
| EAACCHLAKALNDFSN | 1 | 0 | 0 | 1653236 | YP_009724389.1 | P0DTD1 |
| EAAIELSKELKRPVNP | 1 | 0 | 0 | 1653239 | YP_173236.1 | A0A140H1G9 |
| EAALDLSKELKRPIQP | 1 | 0 | 0 | 1653241 | YP_009555238.1 | P0C6X6 |
| EAARCMRSLKAPAVVS | 1 | 0 | 0 | 1653245 | NP_828849.2 | P0C6X7 |
| EAARYMRSLKVPATVS | 1 | 0 | 0 | 1653247 | YP_009724389.1 | P0DTD1 |
| EADEGICFNFNGSWVL | 1 | 0 | 0 | 1653253 | YP_009555238.1 | P0C6X6 |
| EADTEQTQNVTVKPKR | 1 | 0 | 0 | 1653258 | YP_009047202.1 | K9N7C7 |
| EAEAQIDRLINGRLTA | 1 | 0 | 0 | 1653263 | YP_009555241.1 | P36334 |
| EAELAKNVSLDNVLST | 1 | 0 | 0 | 1653266 | YP_009724389.1 | P0DTD1 |
| EAFEYYHTLDESFLGR | 1 | 0 | 0 | 1653275 | NP_828849.2 | P0C6X7 |
| EAFKRSNNGVYISTTK | 1 | 0 | 0 | 1653279 | YP_009555238.1 | P0C6X6 |
| EAFPLDEAEDVTESVQ | 1 | 0 | 0 | 1653283 | YP_009047202.1 | K9N7C7 |
| EAFQKVQDAVNNNAQA | 1 | 0 | 0 | 1653284 | YP_009047204.1 | K9N5Q8 |
| EAFRKARNGVFISTEK | 1 | 0 | 0 | 1653285 | YP_173236.1 | A0A140H1G9 |
| EAGLPYGANKEGIIWV | 1 | 0 | 0 | 1653294 | QIQ54056.1 | P59595 |
| EAIGTNLPLQLGFSTG | 1 | 0 | 0 | 1653297 | QHR63299.1 | P0C6W2 |
| EAILMSEDVKLNPSED | 1 | 0 | 0 | 1653304 | YP_009047202.1 | K9N7C7 |
| EAIPTRFPPGTILPQG | 1 | 0 | 0 | 1653305 | YP_173242.1 | Q19U29 |
| EAIPTRFPPGTVLPQG | 1 | 0 | 0 | 1653306 | YP_009555245.1 | P33469 |
| EAKCWTETDLTKGPHE | 1 | 0 | 0 | 1653309 | YP_009724389.1 | P0DTD1 |
| EAKVITIKVTEDGVNV | 1 | 0 | 0 | 1653324 | NP_073549.1 | A0A0P0LKV0 |

| | | | | | | |
|---|---|---|---|---|---|---|
| EALKTQFNYYKKVNGV | 1 | 0 | 0 | 1653326 | QHR63299.1 | P0C6W2 |
| EALNNLLQQLSNRFGA | 1 | 0 | 0 | 1653328 | YP_009555241.1 | P36334 |
| EALSFEEQDEIYAYTK | 1 | 0 | 0 | 1653333 | YP_009555238.1 | P0C6X6 |
| EALSFEEQNEIYAYTK | 1 | 0 | 0 | 1653334 | YP_173236.1 | A0A140H1G9 |
| EAPLKCVRNVFENFED | 1 | 0 | 0 | 1653347 | NP_073549.1 | A0A0P0LKV0 |
| EAPVSIINNAVYTKVD | 1 | 0 | 0 | 1653350 | NP_828849.2 | P0C6X7 |
| EAQIDRLINGRLTALN | 1 | 0 | 0 | 1653353 | YP_009555241.1 | P36334 |
| EAQPLPQRQKKQPTVT | 1 | 0 | 0 | 1653355 | NP_828858.1 | P59595 |
| EAQVQIDRLINGRLTA | 1 | 0 | 0 | 1653358 | YP_173238.1 | A0A140H1H1 |
| EARAFQSTPIVVQMTK | 1 | 0 | 0 | 1653360 | NP_828859.1 | P59636 |
| EARFSGSANQQQLKQL | 1 | 0 | 0 | 1653363 | YP_009555238.1 | P0C6X6 |
| EARSVNRKSKVISAMH | 1 | 0 | 0 | 1653367 | YP_003766.2 | H9EJ42 |
| EARTVLKKCKSAFYIL | 1 | 0 | 0 | 1653368 | QHR63299.1 | P0C6W2 |
| EATGLFADRDGYSFKK | 1 | 0 | 0 | 1653385 | YP_009555238.1 | P0C6X6 |
| EATGLFAERDCYTFKK | 1 | 0 | 0 | 1653386 | YP_173236.1 | A0A140H1G9 |
| EAVEDTIDGVVEDTIN | 1 | 0 | 0 | 1653391 | YP_173236.1 | A0A140H1G9 |
| EAVIKTLQPVSELLTP | 1 | 0 | 0 | 1653393 | QIQ54046.1 | P0C6U8 |
| EAVKTQFNYYKKVDGV | 1 | 0 | 0 | 1653396 | YP_009724389.1 | P0DTD1 |
| EAVMYMGTLSYDNLKT | 1 | 0 | 0 | 1653398 | NP_828849.2 | P0C6X7 |
| EAVMYMGTLSYEQFKK | 1 | 0 | 0 | 1653399 | YP_009724389.1 | P0DTD1 |
| EAWLNDKSVPSPLNWE | 1 | 0 | 0 | 1653409 | YP_009555241.1 | P36334 |
| EAYADKTLQEMKALFP | 1 | 0 | 0 | 1653412 | YP_009555238.1 | P0C6X6 |
| EAYNSILFENGIPLMP | 1 | 0 | 0 | 1653418 | YP_003766.2 | H9EJ42 |
| EAYYRARAGDAANFCA | 1 | 0 | 0 | 1653420 | NP_828849.2 | P0C6X7 |
| ECAQVLSEYVLCGGGY | 1 | 0 | 0 | 1653424 | YP_009047202.1 | K9N7C7 |
| ECDEECSEVEASDLEE | 1 | 0 | 0 | 1653428 | YP_009047202.1 | K9N7C7 |
| ECDNFTGYLKLTDNVS | 1 | 0 | 0 | 1653431 | QIQ54046.1 | P0C6U8 |
| ECDVKTTEVVGDVILK | 1 | 0 | 0 | 1653433 | QIQ54046.1 | P0C6U8 |
| ECEEEEIDETCEHEYG | 1 | 0 | 0 | 1653435 | NP_828849.2 | P0C6X7 |
| ECFDKFKVNSTLEQYV | 1 | 0 | 0 | 1653437 | YP_009724389.1 | P0DTD1 |
| ECFMKSDIFGQDFKTF | 1 | 0 | 0 | 1653438 | NP_073549.1 | A0A0P0LKV0 |
| ECFVKSDIFGSDFKTF | 1 | 0 | 0 | 1653441 | YP_003766.2 | H9EJ42 |
| ECGSCGNDSWLTGNAI | 1 | 0 | 0 | 1653442 | YP_009047202.1 | K9N7C7 |
| ECKRALGLSISDHEFT | 1 | 0 | 0 | 1653452 | NP_073549.1 | A0A0P0LKV0 |
| ECNADIVVVDEVSMCT | 1 | 0 | 0 | 1653459 | YP_003766.2 | H9EJ42 |
| ECPKFVFPLNSKVKVI | 1 | 0 | 0 | 1653464 | NP_828849.2 | P0C6X7 |
| ECRLAMLGAEMTSNVR | 1 | 0 | 0 | 1653469 | NP_073549.1 | A0A0P0LKV0 |
| ECTIFKDAAGKPVPYC | 1 | 0 | 0 | 1653478 | QIQ54046.1 | P0C6U8 |
| ECTIFKDAMGKPVPYC | 1 | 0 | 0 | 1653479 | NP_828849.2 | P0C6X7 |
| ECTIFKDASGKPVPYC | 1 | 0 | 0 | 1653480 | YP_009724389.1 | P0DTD1 |
| ECTRASTSAVKLQNNE | 1 | 0 | 0 | 1653482 | YP_009047202.1 | K9N7C7 |
| ECYDRFKVNETNSQYL | 1 | 0 | 0 | 1653500 | YP_009047202.1 | K9N7C7 |
| ECYQLYSTQLSTDTGV | 1 | 0 | 0 | 1653501 | QIQ54049.1 | P59632 |
| ECYSGFKPNNNSAQYV | 1 | 0 | 0 | 1653502 | NP_073549.1 | A0A0P0LKV0 |
| ECYSGFKPNNTSAQYI | 1 | 0 | 0 | 1653503 | YP_003766.2 | H9EJ42 |
| ECYSILAAKTGVSVEQ | 1 | 0 | 0 | 1653504 | YP_003766.2 | H9EJ42 |
| EDAEEKLDNPSSSEVD | 1 | 0 | 0 | 1653506 | YP_009555238.1 | P0C6X6 |
| EDAMGQGQNSADPKVY | 1 | 0 | 0 | 1653512 | NP_828859.1 | P59636 |
| EDAYAEVHAEPKGKYS | 1 | 0 | 0 | 1653517 | YP_173236.1 | A0A140H1G9 |
| EDCKLPLGQSLCALPD | 1 | 0 | 0 | 1653519 | YP_009047204.1 | K9N5Q8 |
| EDDAECEEEEIDETCE | 1 | 0 | 0 | 1653520 | NP_828849.2 | P0C6X7 |
| EDDSEVNQTTITQSIA | 1 | 0 | 0 | 1653523 | QHR63299.1 | P0C6W2 |
| EDDYQGKSLEFGATSV | 1 | 0 | 0 | 1653528 | QHR63299.1 | P0C6W2 |
| EDEEEGDCEEEDFEPP | 1 | 0 | 0 | 1653531 | QHR63299.1 | P0C6W2 |
| EDEEEGDCEEEEFEPS | 1 | 0 | 0 | 1653532 | YP_009724389.1 | P0DTD1 |
| EDEFTPEDVNMQIMGV | 1 | 0 | 0 | 1653533 | YP_009047202.1 | K9N7C7 |
| EDEFTPFDVVRQCSGV | 1 | 0 | 0 | 1653534 | YP_009724389.1 | P0DTD1 |
| EDEILEWFGITQTAQG | 1 | 0 | 0 | 1653535 | YP_009047204.1 | K9N5Q8 |
| EDEKLVDVCEKAIGKK | 1 | 0 | 0 | 1653536 | NP_073549.1 | A0A0P0LKV0 |
| EDELAPSDVYQQLAGV | 1 | 0 | 0 | 1653537 | YP_173236.1 | A0A140H1G9 |
| EDELTPSDVYQQLAGI | 1 | 0 | 0 | 1653538 | YP_009555238.1 | P0C6X6 |
| EDFDLISKCQITAVEG | 1 | 0 | 0 | 1653540 | YP_009555238.1 | P0C6X6 |
| EDFKPRSQMETDFLEL | 1 | 0 | 0 | 1653544 | NP_828849.2 | P0C6X7 |
| EDFQENWNTKHSSGVT | 1 | 0 | 0 | 1653551 | YP_009724389.1 | P0DTD1 |
| EDFYASMYENSTILQA | 1 | 0 | 0 | 1653553 | YP_003766.2 | H9EJ42 |
| EDGDVLLAEFDTYDPI | 1 | 0 | 0 | 1653555 | YP_009047202.1 | K9N7C7 |
| EDGVTRDISSNDNCFA | 1 | 0 | 0 | 1653563 | QIQ54046.1 | P0C6U8 |
| EDHSLLATLDDPYVED | 1 | 0 | 0 | 1653565 | YP_173242.1 | Q19U29 |
| EDHVQIHTIDGSSGVV | 1 | 0 | 0 | 1653566 | QIQ54049.1 | P59632 |
| EDIAQVVIADTLQETP | 1 | 0 | 0 | 1653567 | YP_009047202.1 | K9N7C7 |
| EDIFGPCWSALASALK | 1 | 0 | 0 | 1653570 | NP_073549.1 | A0A0P0LKV0 |
| EDIHRVYAALGKVVAN | 1 | 0 | 0 | 1653572 | NP_073549.1 | A0A0P0LKV0 |
| EDILFSKLVTSGLGTV | 1 | 0 | 0 | 1653574 | NP_073551.1 | A0A0P0K6L9 |
| EDIQLLKAAYENFNSQ | 1 | 0 | 0 | 1653575 | NP_828849.2 | P0C6X7 |
| EDIQLLKSAYENFNQH | 1 | 0 | 0 | 1653576 | YP_009724389.1 | P0DTD1 |
| EDKFDLLISDMYDGRI | 1 | 0 | 0 | 1653579 | YP_003766.2 | H9EJ42 |
| EDKFDLLISDMYDGRT | 1 | 0 | 0 | 1653580 | NP_073549.1 | A0A0P0LKV0 |
| EDKLSVYDIACCMRAG | 1 | 0 | 0 | 1653583 | YP_003766.2 | H9EJ42 |
| EDLEIGYTVDCSCGKK | 1 | 0 | 0 | 1653588 | YP_009555238.1 | P0C6X6 |
| EDLKAVRSSFNFDQKE | 1 | 0 | 0 | 1653591 | YP_009555238.1 | P0C6X6 |
| EDLKPVSEEVVENPTI | 1 | 0 | 0 | 1653593 | YP_009724389.1 | P0DTD1 |
| EDLLEILSRERVNINI | 1 | 0 | 0 | 1653595 | NP_828849.2 | P0C6X7 |
| EDLLFDKVKLSDVGFV | 1 | 0 | 0 | 1653596 | YP_009555241.1 | P36334 |
| EDLLFDKVTIADPGYM | 1 | 0 | 0 | 1653597 | YP_009047204.1 | K9N5Q8 |
| EDLLFNKVTLADAGFM | 1 | 0 | 0 | 1653599 | NP_828851.1 | P59594 |
| EDLLFSKVVTSGLGTV | 1 | 0 | 0 | 1653600 | YP_003767.1 | A0A0P0G321 |
| EDLMAAYVDNTSLTIK | 1 | 0 | 0 | 1653603 | QIQ54046.1 | P0C6U8 |
| EDLRARMTYVCQCGGE | 1 | 0 | 0 | 1653605 | YP_009047202.1 | K9N7C7 |
| EDLRFPRGQGVPINTN | 1 | 0 | 0 | 1653606 | QIQ54056.1 | P59595 |
| EDLSIRNLQKRLYSNV | 1 | 0 | 0 | 1653609 | YP_173236.1 | A0A140H1G9 |
| EDLWQFVDHFGENEEI | 1 | 0 | 0 | 1653612 | NP_073549.1 | A0A0P0LKV0 |

| | | | | | | |
|---|---|---|---|---|---|---|
| EDMLNPNYEDLLIRKS | 1 | 0 | 0 | 1653614 | YP_009724389.1 | P0DTD1 |
| EDNALAYYNTSKGGRF | 1 | 0 | 0 | 1653616 | QIQ54046.1 | P0C6U8 |
| EDQDALFAYTKRNVIP | 1 | 0 | 0 | 1653620 | YP_009724389.1 | P0DTD1 |
| EDRSNNSSRASSRSST | 1 | 0 | 0 | 1653624 | YP_003771.1 | H9EJ69 |
| EDTINDDEDVVTGDND | 1 | 0 | 0 | 1653630 | YP_173236.1 | A0A140H1G9 |
| EDTKLVTEEVVETPTI | 1 | 0 | 0 | 1653631 | QIQ54046.1 | P0C6U8 |
| EDTLTKLSKYLANEAQ | 1 | 0 | 0 | 1653633 | NP_073549.1 | A0A0P0LKV0 |
| EDTVWEVQGYKNVRIT | 1 | 0 | 0 | 1653637 | NP_828849.2 | P0C6X7 |
| EDVEDVISSEAVEDTI | 1 | 0 | 0 | 1653641 | YP_173236.1 | A0A140H1G9 |
| EDVKFAASTGVIDISA | 1 | 0 | 0 | 1653644 | YP_003766.2 | H9EJ42 |
| EDVTESVQEEAQPVEV | 1 | 0 | 0 | 1653648 | YP_009047202.1 | K9N7C7 |
| EDVVDQVEEVNSIFDI | 1 | 0 | 0 | 1653650 | NP_073549.1 | A0A0P0LKV0 |
| EDVVIFINNTTYPTNV | 1 | 0 | 0 | 1653651 | YP_009555238.1 | P0C6X6 |
| EDVVTGDNDDEDVVTG | 1 | 0 | 0 | 1653652 | YP_173236.1 | A0A140H1G9 |
| EDVVTGDNNDEEIVTG | 1 | 0 | 0 | 1653653 | YP_173236.1 | A0A140H1G9 |
| EDWLDDDSQQTVGQQD | 1 | 0 | 0 | 1653655 | YP_009724389.1 | P0DTD1 |
| EDWLDDDSQQTVVQED | 1 | 0 | 0 | 1653656 | QHR63299.1 | P0C6W2 |
| EDWLDDTTEQSEIEPE | 1 | 0 | 0 | 1653657 | NP_828849.2 | P0C6X7 |
| EDYAFDHIVYGSFNHK | 1 | 0 | 0 | 1653659 | YP_173236.1 | A0A140H1G9 |
| EDYAFEHIVYGNFNQK | 1 | 0 | 0 | 1653660 | YP_009555238.1 | P0C6X6 |
| EDYEEEYSEEEQPEQP | 1 | 0 | 0 | 1653661 | QIQ54046.1 | P0C6U8 |
| EDYEQNWNTKHGSGAL | 1 | 0 | 0 | 1653662 | NP_828849.2 | P0C6X7 |
| EEAARYMRSLKVPATV | 1 | 0 | 0 | 1653669 | YP_009724389.1 | P0DTD1 |
| EEADEGICFNFNGSWV | 1 | 0 | 0 | 1653670 | YP_009555238.1 | P0C6X6 |
| EEAKKVKPTVVVNAAN | 1 | 0 | 0 | 1653677 | YP_009724389.1 | P0DTD1 |
| EEALSEAREHLKNGTC | 1 | 0 | 0 | 1653681 | NP_828849.2 | P0C6X7 |
| EEALSEARQHLKDGTC | 1 | 0 | 0 | 1653682 | QHR63299.1 | P0C6W2 |
| EEALSKLSEYLISDSI | 1 | 0 | 0 | 1653684 | YP_003766.2 | H9EJ42 |
| EEDDAECEEEEIDETC | 1 | 0 | 0 | 1653692 | NP_828849.2 | P0C6X7 |
| EEDGFAFYPACTNGHA | 1 | 0 | 0 | 1653696 | YP_009047202.1 | K9N7C7 |
| EEDWLDDDSQQTVGQQ | 1 | 0 | 0 | 1653701 | YP_009724389.1 | P0DTD1 |
| EEDWLDDDSQQTVVQE | 1 | 0 | 0 | 1653702 | QHR63299.1 | P0C6W2 |
| EEDWLDDTTEQSEIEP | 1 | 0 | 0 | 1653703 | NP_828849.2 | P0C6X7 |
| EEDWLETEAEVVEQEV | 1 | 0 | 0 | 1653704 | QIQ54046.1 | P0C6U8 |
| EEEDWLDDTTEQSEIE | 1 | 0 | 0 | 1653707 | NP_828849.2 | P0C6X7 |
| EEEEDWLDDTTEQSEI | 1 | 0 | 0 | 1653709 | NP_828849.2 | P0C6X7 |
| EEEEEDWLDDTTEQSE | 1 | 0 | 0 | 1653710 | NP_828849.2 | P0C6X7 |
| EEEEEEDWLDDTTEQS | 1 | 0 | 0 | 1653711 | NP_828849.2 | P0C6X7 |
| EEELEEDWLDDDSQQT | 1 | 0 | 0 | 1653720 | QHR63299.1 | P0C6W2 |
| EEEQEEDWLDDDSQQT | 1 | 0 | 0 | 1653722 | YP_009724389.1 | P0DTD1 |
| EEEQPEQPTQYEYGTE | 1 | 0 | 0 | 1653723 | QIQ54046.1 | P0C6U8 |
| EEFASHVANFQKSGYS | 1 | 0 | 0 | 1653726 | YP_009047202.1 | K9N7C7 |
| EEFIQTNSPKVTIDCS | 1 | 0 | 0 | 1653729 | YP_173238.1 | A0A140H1H1 |
| EEFVVKEKLNAFLVHD | 1 | 0 | 0 | 1653731 | NP_073549.1 | A0A0P0LKV0 |
| EEFYNFLRLRGFFDEG | 1 | 0 | 0 | 1653733 | YP_003766.2 | H9EJ42 |
| EEHVQIHTIDGSSGVV | 1 | 0 | 0 | 1653746 | YP_009724391.1 | P0DTC3 |
| EEIDETCEHEYGTEDD | 1 | 0 | 0 | 1653748 | NP_828849.2 | P0C6X7 |
| EEIEYVHGDALHTLRN | 1 | 0 | 0 | 1653749 | NP_073549.1 | A0A0P0LKV0 |
| EEILGTVSWNLREMLA | 1 | 0 | 0 | 1653751 | NP_828849.2 | P0C6X7 |
| EEILGTVSWNLREMLT | 1 | 0 | 0 | 1653752 | QIQ54046.1 | P0C6U8 |
| EEIVTGDNDDQIVVTG | 1 | 0 | 0 | 1653754 | YP_173236.1 | A0A140H1G9 |
| EEKFKEGIEFLKDGWE | 1 | 0 | 0 | 1653756 | QIQ54046.1 | P0C6U8 |
| EEKFKEGVEFLRDGWE | 1 | 0 | 0 | 1653757 | YP_009724389.1 | P0DTD1 |
| EEKSVVQKPVDVKPKI | 1 | 0 | 0 | 1653761 | NP_828849.2 | P0C6X7 |
| EELDQWFKNQTSVAPD | 1 | 0 | 0 | 1653764 | YP_009555241.1 | P36334 |
| EELEEDWLDDDSQQTV | 1 | 0 | 0 | 1653765 | QHR63299.1 | P0C6W2 |
| EELFYSYATHHDKFTD | 1 | 0 | 0 | 1653766 | NP_828849.2 | P0C6X7 |
| EELFYSYATHSDKFTD | 1 | 0 | 0 | 1653768 | YP_009724389.1 | P0DTD1 |
| EELHDDLVNYIRDKGC | 1 | 0 | 0 | 1653769 | YP_009555239.1 | Q80872 |
| EELRFPRGQGVPINTN | 1 | 0 | 0 | 1653776 | NP_828858.1 | P59595 |
| EELVASFLEMKSETKV | 1 | 0 | 0 | 1653777 | QIQ54046.1 | P0C6U8 |
| EENLNAYVNSNQNTDS | 1 | 0 | 0 | 1653782 | YP_173242.1 | Q19U29 |
| EEPKKVKPCIEEVTTT | 1 | 0 | 0 | 1653786 | QIQ54046.1 | P0C6U8 |
| EEQDAIFSLTKRNILP | 1 | 0 | 0 | 1653792 | NP_073549.1 | A0A0P0LKV0 |
| EEQDEIYAYTKRNVLP | 1 | 0 | 0 | 1653794 | YP_009555238.1 | P0C6X6 |
| EEQEEDWLDDDSQQTV | 1 | 0 | 0 | 1653795 | YP_009724389.1 | P0DTD1 |
| EEQEEDWLETEAEVVE | 1 | 0 | 0 | 1653796 | QIQ54046.1 | P0C6U8 |
| EEQNEIYAYTKRNVLP | 1 | 0 | 0 | 1653797 | YP_173236.1 | A0A140H1G9 |
| EEQPEQPTQYEYGTES | 1 | 0 | 0 | 1653798 | QIQ54046.1 | P0C6U8 |
| EESASSEKPTKEEPKK | 1 | 0 | 0 | 1653802 | QIQ54046.1 | P0C6U8 |
| EESDLIISNFKFDDYD | 1 | 0 | 0 | 1653803 | YP_173236.1 | A0A140H1G9 |
| EESFYKNMYLKSAVMQ | 1 | 0 | 0 | 1653806 | YP_173236.1 | A0A140H1G9 |
| EESGAVLFCTPTKKAF | 1 | 0 | 0 | 1653807 | NP_073549.1 | A0A0P0LKV0 |
| EESQGNVVTSVMESQI | 1 | 0 | 0 | 1653809 | YP_173236.1 | A0A140H1G9 |
| EESVKSSYAFATLKEV | 1 | 0 | 0 | 1653813 | YP_003766.2 | H9EJ42 |
| EESVKSSYAYATLKEI | 1 | 0 | 0 | 1653814 | NP_073549.1 | A0A0P0LKV0 |
| EETDLTVTSAGQPCVA | 1 | 0 | 0 | 1653815 | YP_009555238.1 | P0C6X6 |
| EETIYNLVKDCPAVAV | 1 | 0 | 0 | 1653820 | NP_828849.2 | P0C6X7 |
| EETKFLTENLLLYADI | 1 | 0 | 0 | 1653821 | QIQ54046.1 | P0C6U8 |
| EETRKLMPICMDTKAI | 1 | 0 | 0 | 1653824 | QIQ54046.1 | P0C6U8 |
| EETRKLMPICMDVRAI | 1 | 0 | 0 | 1653825 | NP_828849.2 | P0C6X7 |
| EETRKLMPVCMETKAI | 1 | 0 | 0 | 1653826 | QHR63299.1 | P0C6W2 |
| EETRKLMPVCVETKAI | 1 | 0 | 0 | 1653827 | YP_009724389.1 | P0DTD1 |
| EEVDIIEQPFEEVEHV | 1 | 0 | 0 | 1653832 | YP_003766.2 | H9EJ42 |
| EEVILKTGELQPLEEP | 1 | 0 | 0 | 1653837 | QIQ54046.1 | P0C6U8 |
| EEVLSEARQHLKDGTC | 1 | 0 | 0 | 1653842 | YP_009724389.1 | P0DTD1 |
| EEVNSIFDIETVDVKH | 1 | 0 | 0 | 1653843 | NP_073549.1 | A0A0P0LKV0 |
| EEVQQELYSPLFLIVA | 1 | 0 | 0 | 1653847 | NP_828857.1 | P59635 |
| EEVTTTLEETKFLTEN | 1 | 0 | 0 | 1653848 | YP_009724389.1 | P0DTD1 |
| EEVVENPTIQKDILEC | 1 | 0 | 0 | 1653849 | QHR63299.1 | P0C6W2 |
| EEVVENPTIQKEVIEC | 1 | 0 | 0 | 1653851 | NP_828849.2 | P0C6X7 |

| | | | | | | |
|---|---|---|---|---|---|---|
| EEVVLKTGDLQPLEQP | 1 | 0 | 0 | 1653854 | YP_009724389.1 | P0DTD1 |
| EEYCNYLESYNIVTTA | 1 | 0 | 0 | 1653855 | YP_173236.1 | A0A140H1G9 |
| EEYREYLESYNTATTA | 1 | 0 | 0 | 1653859 | YP_009555238.1 | P0C6X6 |
| EFANDTKIASQLGNCV | 1 | 0 | 0 | 1653866 | YP_009047204.1 | K9N5Q8 |
| EFATRTFTATTAVGSC | 1 | 0 | 0 | 1653870 | YP_009047202.1 | K9N7C7 |
| EFCGTENQVKEGPTTC | 1 | 0 | 0 | 1653873 | QIQ54046.1 | P0C6U8 |
| EFCSQHTMLVKMDGDD | 1 | 0 | 0 | 1653876 | YP_009555238.1 | P0C6X6 |
| EFCSQHTMLVKQGDDY | 1 | 0 | 0 | 1653877 | YP_009724389.1 | P0DTD1 |
| EFCSQHTMQIVDENGK | 1 | 0 | 0 | 1653878 | NP_073549.1 | A0A0P0LKV0 |
| EFCSQHTMQIVDKDGT | 1 | 0 | 0 | 1653879 | YP_003766.2 | H9EJ42 |
| EFDTYDPIYKNGAMYK | 1 | 0 | 0 | 1653884 | YP_009047202.1 | K9N7C7 |
| EFEDRSNNSSRASSRS | 1 | 0 | 0 | 1653888 | YP_003771.1 | H9EJ69 |
| EFEFVEGQGVPIAPGV | 1 | 0 | 0 | 1653889 | YP_009555245.1 | P33469 |
| EFEPSTQYEYGTEDDY | 1 | 0 | 0 | 1653891 | YP_009724389.1 | P0DTD1 |
| EFHLDGEVLSLDKLKS | 1 | 0 | 0 | 1653900 | NP_828849.2 | P0C6X7 |
| EFIQTNSPKVTIDCSL | 1 | 0 | 0 | 1653905 | YP_173238.1 | A0A140H1H1 |
| EFIQTSSPKVTIDCAA | 1 | 0 | 0 | 1653906 | YP_009555241.1 | P36334 |
| EFKVTFFPDLNGDVVA | 1 | 0 | 0 | 1653911 | QHR63299.1 | P0C6W2 |
| EFKYMVANGLHAPHGP | 1 | 0 | 0 | 1653912 | YP_003766.2 | H9EJ42 |
| EFKYMVANGLNAPNGP | 1 | 0 | 0 | 1653913 | NP_073549.1 | A0A0P0LKV0 |
| EFLKDAWEILKFLITG | 1 | 0 | 0 | 1653917 | NP_828849.2 | P0C6X7 |
| EFLKRGDKIVYHTLES | 1 | 0 | 0 | 1653919 | NP_828849.2 | P0C6X7 |
| EFLKRGDKSVYYTSNP | 1 | 0 | 0 | 1653920 | YP_009724389.1 | P0DTD1 |
| EFLKRGDKVVYHTTSK | 1 | 0 | 0 | 1653921 | QIQ54046.1 | P0C6U8 |
| EFMLWCKDGQVQTFYP | 1 | 0 | 0 | 1653927 | YP_009047202.1 | K9N7C7 |
| EFNPSQTSPATAEPVR | 1 | 0 | 0 | 1653930 | NP_073556.1 | A0A0N7J131 |
| EFNRLATCDWTESGDY | 1 | 0 | 0 | 1653931 | YP_009047202.1 | K9N7C7 |
| EFPLLSEYPLLKQKDI | 1 | 0 | 0 | 1653933 | YP_173243.1 | Q0ZJ80 |
| EFQLDYSIDVIDNEII | 1 | 0 | 0 | 1653936 | NP_073549.1 | A0A0P0LKV0 |
| EFRFMTANNLTAPRNS | 1 | 0 | 0 | 1653939 | YP_009047202.1 | K9N7C7 |
| EFRVYSSANNCTFEYI | 1 | 0 | 0 | 1653941 | QIQ54048.1 | P59594 |
| EFRYMNSQGLLPPKNS | 1 | 0 | 0 | 1653944 | YP_009724389.1 | P0DTD1 |
| EFRYMNSQGLLPPKSS | 1 | 0 | 0 | 1653945 | NP_828849.2 | P0C6X7 |
| EFSHLATFAELEAAQK | 1 | 0 | 0 | 1653946 | YP_009047202.1 | K9N7C7 |
| EFSKKQLGLLQPTDNS | 1 | 0 | 0 | 1653950 | YP_009047202.1 | K9N7C7 |
| EFSRVSAKPPPGDQFK | 1 | 0 | 0 | 1653952 | QIQ54047.1 | P0C6X7 |
| EFSSNVANYQKVGMQK | 1 | 0 | 0 | 1653955 | YP_009724389.1 | P0DTD1 |
| EFSWNADLYKLMSCFA | 1 | 0 | 0 | 1653958 | YP_173236.1 | A0A140H1G9 |
| EFTDENYNNLVPTYLK | 1 | 0 | 0 | 1653960 | YP_173236.1 | A0A140H1G9 |
| EFTRVNAKPPPGDQFK | 1 | 0 | 0 | 1653966 | NP_828849.2 | P0C6X7 |
| EFVEGQGVPIAPGVPA | 1 | 0 | 0 | 1653970 | YP_009555245.1 | P33469 |
| EFVVKEKLNAFLVHDN | 1 | 0 | 0 | 1653984 | NP_073549.1 | A0A0P0LKV0 |
| EFWFAMRKDGDDVIFS | 1 | 0 | 0 | 1653987 | YP_173236.1 | A0A140H1G9 |
| EFYEDRILDFKVLDNG | 1 | 0 | 0 | 1653992 | YP_009047202.1 | K9N7C7 |
| EFYNDVKPPVLDVDDV | 1 | 0 | 0 | 1653995 | YP_009555243.1 | Q04854 |
| EGAHASRNACGTNVPL | 1 | 0 | 0 | 1653998 | YP_009047202.1 | K9N7C7 |
| EGAHATRDSIGTNFPL | 1 | 0 | 0 | 1653999 | YP_009555238.1 | P0C6X6 |
| EGAHATRENIGTNFPL | 1 | 0 | 0 | 1654000 | YP_173236.1 | A0A140H1G9 |
| EGATTCGYLPQNAVVK | 1 | 0 | 0 | 1654004 | YP_009724389.1 | P0DTD1 |
| EGCHATREAIGTNLPL | 1 | 0 | 0 | 1654006 | QHR63299.1 | P0C6W2 |
| EGCHATREAVGTNLPL | 1 | 0 | 0 | 1654007 | YP_009724389.1 | P0DTD1 |
| EGDGGITSEGNALYNN | 1 | 0 | 0 | 1654014 | NP_073549.1 | A0A0P0LKV0 |
| EGDQIIFYEGVNFNPY | 1 | 0 | 0 | 1654020 | YP_173237.1 | A0A140H1H0 |
| EGEETDLTVTSAGQPC | 1 | 0 | 0 | 1654023 | YP_009555238.1 | P0C6X6 |
| EGELPVAVVNDKIFTK | 1 | 0 | 0 | 1654025 | YP_009047202.1 | K9N7C7 |
| EGFLLGINYLGTIKEN | 1 | 0 | 0 | 1654036 | YP_009047202.1 | K9N7C7 |
| EGGCNTIELDSPCRFM | 1 | 0 | 0 | 1654043 | YP_003766.2 | H9EJ42 |
| EGGKTFMYAYISNKAD | 1 | 0 | 0 | 1654045 | YP_003766.2 | H9EJ42 |
| EGGLCPSTGQAMESVG | 1 | 0 | 0 | 1654046 | YP_173236.1 | A0A140H1G9 |
| EGGRAFMYAYVTTKPG | 1 | 0 | 0 | 1654049 | NP_073549.1 | A0A0P0LKV0 |
| EGHIIMEEMLKGSSTI | 1 | 0 | 0 | 1654052 | YP_009047202.1 | K9N7C7 |
| EGKFYGPFVDRQTAQA | 1 | 0 | 0 | 1654065 | NP_828849.2 | P0C6X7 |
| EGKQGNFKNLREFVFK | 1 | 0 | 0 | 1654067 | YP_009724390.1 | P0DTC2 |
| EGLNDNLLEILQKEKV | 1 | 0 | 0 | 1654077 | YP_009724389.1 | P0DTD1 |
| EGNALYNNEGGRAFMY | 1 | 0 | 0 | 1654080 | NP_073549.1 | A0A0P0LKV0 |
| EGNSPFHPLADNKFAL | 1 | 0 | 0 | 1654087 | YP_009724395.1 | P0DTC7 |
| EGQDVIFSQFDSLGVS | 1 | 0 | 0 | 1654092 | YP_009555238.1 | P0C6X6 |
| EGQLQSLSKDYISSNG | 1 | 0 | 0 | 1654095 | YP_003766.2 | H9EJ42 |
| EGRTIYGNVSRQDLTK | 1 | 0 | 0 | 1654100 | NP_073549.1 | A0A0P0LKV0 |
| EGSFPCSICLSGLDSL | 1 | 0 | 0 | 1654105 | NP_828849.2 | P0C6X7 |
| EGSGRSASNSRPGSRS | 1 | 0 | 0 | 1654107 | YP_173242.1 | Q19U29 |
| EGSISYSELRPDTRYV | 1 | 0 | 0 | 1654108 | NP_828849.2 | P0C6X7 |
| EGSKPYSTLQPNAYYK | 1 | 0 | 0 | 1654109 | YP_003766.2 | H9EJ42 |
| EGSLPYSSIQANAYYK | 1 | 0 | 0 | 1654110 | NP_073549.1 | A0A0P0LKV0 |
| EGSRVPHIVRKDLTKY | 1 | 0 | 0 | 1654112 | YP_009555238.1 | P0C6X6 |
| EGSVAYESLRPDTRYV | 1 | 0 | 0 | 1654118 | YP_009724389.1 | P0DTD1 |
| EGSVKGLQPSVGPKQA | 1 | 0 | 0 | 1654120 | YP_009724389.1 | P0DTD1 |
| EGSVKGLTPSKGPAQA | 1 | 0 | 0 | 1654121 | NP_828849.2 | P0C6X7 |
| EGSVPYESLRPDTRYV | 1 | 0 | 0 | 1654122 | QIQ54046.1 | P0C6U8 |
| EGSVRVVTTFDAEYCR | 1 | 0 | 0 | 1654123 | NP_828849.2 | P0C6X7 |
| EGTGGNSQSSSRASSL | 1 | 0 | 0 | 1654126 | YP_009047211.1 | K9N4V7 |
| EGTKKLAARLSFNVGR | 1 | 0 | 0 | 1654127 | YP_009555238.1 | P0C6X6 |
| EGVCITTNGSWAIFND | 1 | 0 | 0 | 1654130 | YP_009047202.1 | K9N7C7 |
| EGVFWVANHQADTSTP | 1 | 0 | 0 | 1654137 | YP_173242.1 | Q19U29 |
| EGVGAKVSAFLQKLED | 1 | 0 | 0 | 1654138 | YP_009555238.1 | P0C6X6 |
| EGVKVTQELGHEDLMA | 1 | 0 | 0 | 1654139 | NP_828849.2 | P0C6X7 |
| EGVLPINTDTVLSVAP | 1 | 0 | 0 | 1654142 | YP_003766.2 | H9EJ42 |
| EGVNGGSLYVNKHAFH | 1 | 0 | 0 | 1654147 | YP_003766.2 | H9EJ42 |
| EGVNGGSLYVNNHAFH | 1 | 0 | 0 | 1654148 | NP_073549.1 | A0A0P0LKV0 |
| EGYDFKYIENGQLKNF | 1 | 0 | 0 | 1654159 | NP_073549.1 | A0A0P0LKV0 |
| EGYLNSTNVTIATYCT | 1 | 0 | 0 | 1654160 | YP_009724389.1 | P0DTD1 |

| | | | | | | |
|---|---|---|---|---|---|---|
| EGYLNSTNVTTAIYCT | 1 | 0 | 0 | 1654161 | QHR63299.1 | P0C6W2 |
| EHESKFWDESFYASMY | 1 | 0 | 0 | 1654181 | NP_073549.1 | A0A0P0LKV0 |
| EHPAVGIVNTDFKLEI | 1 | 0 | 0 | 1654199 | NP_073553.1 | A0A0P0KWE8 |
| EHSLAEYHNESGIKTT | 1 | 0 | 0 | 1654205 | QIQ54046.1 | P0C6U8 |
| EHSLAEYHNESGLKPI | 1 | 0 | 0 | 1654206 | QHR63299.1 | P0C6W2 |
| EHSLAEYHNESGLKTI | 1 | 0 | 0 | 1654207 | YP_009724389.1 | P0DTD1 |
| EHSLLVEAYNSILFEN | 1 | 0 | 0 | 1654208 | YP_003766.2 | H9EJ42 |
| EHSWNADLYKLMGHFA | 1 | 0 | 0 | 1654210 | YP_009724389.1 | P0DTD1 |
| EHSWNADLYKLMGHFS | 1 | 0 | 0 | 1654211 | NP_828849.2 | P0C6X7 |
| EHVNNSYECDIPVGAG | 1 | 0 | 0 | 1654223 | QIQ54048.1 | P59594 |
| EHVQIHTIDGSSGVVN | 1 | 0 | 0 | 1654225 | YP_009724391.1 | P0DTC3 |
| EHVTSVNLFDVGYSVN | 1 | 0 | 0 | 1654227 | YP_009047205.1 | K9N796 |
| EHYNEWAQANGFTAMN | 1 | 0 | 0 | 1654230 | NP_073549.1 | A0A0P0LKV0 |
| EIACKCGVKQEQRTGL | 1 | 0 | 0 | 1654235 | YP_009555238.1 | P0C6X6 |
| EIANLVLAKLGKDSKP | 1 | 0 | 0 | 1654238 | YP_173242.1 | Q19U29 |
| EIATKMFDSFVNSFVS | 1 | 0 | 0 | 1654242 | YP_009047202.1 | K9N7C7 |
| EIDGNVMHANYLFWRN | 1 | 0 | 0 | 1654247 | YP_009555238.1 | P0C6X6 |
| EIDPKLDGYYKKDNAY | 1 | 0 | 0 | 1654249 | QIQ54046.1 | P0C6U8 |
| EIDPKLDNYYKKDNSY | 1 | 0 | 0 | 1654250 | YP_009724389.1 | P0DTD1 |
| EIEAGEVKPFAVYKNV | 1 | 0 | 0 | 1654251 | YP_003766.2 | H9EJ42 |
| EIEPEPEPTPEEPVNQ | 1 | 0 | 0 | 1654252 | NP_828849.2 | P0C6X7 |
| EIEPKLDGYYKKDNAY | 1 | 0 | 0 | 1654253 | NP_828849.2 | P0C6X7 |
| EIEYVHGDALHTLRNG | 1 | 0 | 0 | 1654254 | NP_073549.1 | A0A0P0LKV0 |
| EIFENKTTLPVNVAFE | 1 | 0 | 0 | 1654255 | NP_828849.2 | P0C6X7 |
| EIIGGQLVAFTTELKD | 1 | 0 | 0 | 1654264 | QIQ54046.1 | P0C6U8 |
| EIIGYKAIDGGVTRDI | 1 | 0 | 0 | 1654265 | YP_009724389.1 | P0DTD1 |
| EIIGYKAIEDGVTRDI | 1 | 0 | 0 | 1654266 | QIQ54046.1 | P0C6U8 |
| EIIGYKAIQDGVTRDI | 1 | 0 | 0 | 1654267 | NP_828849.2 | P0C6X7 |
| EIIVKPNISLCVPLYV | 1 | 0 | 0 | 1654272 | NP_073549.1 | A0A0P0LKV0 |
| EIKDATPSDFVRATAT | 1 | 0 | 0 | 1654274 | YP_009724391.1 | P0DTC3 |
| EIKDATPSDSVRATAT | 1 | 0 | 0 | 1654275 | QHR63301.1 | Q3LZX0 |
| EIKDTEKYCALAPNMM | 1 | 0 | 0 | 1654277 | YP_009724389.1 | P0DTD1 |
| EIKLAKKFDNFTGECP | 1 | 0 | 0 | 1654282 | QIQ54046.1 | P0C6U8 |
| EIKLAKKFDTFNGECP | 1 | 0 | 0 | 1654283 | YP_009724389.1 | P0DTD1 |
| EILCEYADCKESYFSK | 1 | 0 | 0 | 1654289 | YP_173236.1 | A0A140H1G9 |
| EIPHFNQKLPNGVTVV | 1 | 0 | 0 | 1654317 | NP_073556.1 | A0A0N7J131 |
| EIPKEEVKPFITESKP | 1 | 0 | 0 | 1654318 | YP_009724389.1 | P0DTD1 |
| EIPRRNVATLQAENVT | 1 | 0 | 0 | 1654320 | YP_009724389.1 | P0DTD1 |
| EIPYKSFSDSGCILPS | 1 | 0 | 0 | 1654323 | YP_009047202.1 | K9N7C7 |
| EISCNVCTKRATVYNS | 1 | 0 | 0 | 1654333 | YP_009555238.1 | P0C6X6 |
| EISMATNYDLSVVNAR | 1 | 0 | 0 | 1654337 | YP_009724389.1 | P0DTD1 |
| EISVQKKINRMAEQAA | 1 | 0 | 0 | 1654342 | YP_003766.2 | H9EJ42 |
| EISVTPIKNTVDTTPK | 1 | 0 | 0 | 1654343 | NP_073549.1 | A0A0P0LKV0 |
| EITKPLIEAEKPNLNS | 1 | 0 | 0 | 1654346 | YP_173243.1 | Q0ZJ80 |
| EIVDTVSALVYDNKLK | 1 | 0 | 0 | 1654353 | YP_009724389.1 | P0DTD1 |
| EIVDTVSALVYDNKLR | 1 | 0 | 0 | 1654354 | QIQ54047.1 | P0C6X7 |
| EIVDTVSALVYENKLK | 1 | 0 | 0 | 1654355 | YP_009555238.1 | P0C6X6 |
| EIVETVSALVYDNKLK | 1 | 0 | 0 | 1654356 | YP_173236.1 | A0A140H1G9 |
| EIVGGQIVTCAKEIKE | 1 | 0 | 0 | 1654358 | YP_009724389.1 | P0DTD1 |
| EIVGPKELLLLWESGK | 1 | 0 | 0 | 1654359 | NP_073549.1 | A0A0P0LKV0 |
| EIVKLGAEKKDGFYET | 1 | 0 | 0 | 1654363 | YP_009555239.1 | Q80872 |
| EIVKSSDVVDSNENLT | 1 | 0 | 0 | 1654364 | YP_009047202.1 | K9N7C7 |
| EIVSDREVILCWETGK | 1 | 0 | 0 | 1654371 | YP_173236.1 | A0A140H1G9 |
| EIVSERELILSWEIGK | 1 | 0 | 0 | 1654372 | YP_009555238.1 | P0C6X6 |
| EIVTGDNDDQIVVTGD | 1 | 0 | 0 | 1654374 | YP_173236.1 | A0A140H1G9 |
| EIVWYTERSDKSYELQ | 1 | 0 | 0 | 1654376 | QIQ54046.1 | P0C6U8 |
| EKACNIAKSAYERDRA | 1 | 0 | 0 | 1654385 | YP_009555238.1 | P0C6X6 |
| EKACNIAKSVYERDKA | 1 | 0 | 0 | 1654386 | YP_173236.1 | A0A140H1G9 |
| EKAIGKKIKHEGDWDS | 1 | 0 | 0 | 1654388 | NP_073549.1 | A0A0P0LKV0 |
| EKASLNSLTYFNRPSL | 1 | 0 | 0 | 1654392 | YP_009555238.1 | P0C6X6 |
| EKCDLQNYGDSATLPK | 1 | 0 | 0 | 1654395 | YP_009724389.1 | P0DTD1 |
| EKCSNYTVELGTNIDE | 1 | 0 | 0 | 1654400 | QIQ54046.1 | P0C6U8 |
| EKCSVYTVESGTEVTE | 1 | 0 | 0 | 1654402 | NP_828849.2 | P0C6X7 |
| EKDEDDNLIDSYFVVK | 1 | 0 | 0 | 1654406 | YP_009724389.1 | P0DTD1 |
| EKDEDGNLIDSYFIVK | 1 | 0 | 0 | 1654407 | QIQ54047.1 | P0C6X7 |
| EKDFIALDQDVFIQKY | 1 | 0 | 0 | 1654409 | YP_009555238.1 | P0C6X6 |
| EKDFLSFDSDVFIKKY | 1 | 0 | 0 | 1654410 | YP_009047202.1 | K9N7C7 |
| EKFINGCGRNVQGHVV | 1 | 0 | 0 | 1654415 | YP_003766.2 | H9EJ42 |
| EKFKEGIEFLKDGWEI | 1 | 0 | 0 | 1654416 | QIQ54046.1 | P0C6U8 |
| EKFKEGVEFLRDGWEI | 1 | 0 | 0 | 1654417 | YP_009724389.1 | P0DTD1 |
| EKGAYNKDHGRGGFGH | 1 | 0 | 0 | 1654425 | YP_009555238.1 | P0C6X6 |
| EKGDYGDAVVYRGTTT | 1 | 0 | 0 | 1654426 | YP_009724389.1 | P0DTD1 |
| EKGIYQTSNFRVQPTD | 1 | 0 | 0 | 1654427 | QHR63300.2 | Q3LZX1 |
| EKGIYQTSNFRVQPTI | 1 | 0 | 0 | 1654429 | QIQ54048.1 | P59594 |
| EKGKLLIRDTNKEVFV | 1 | 0 | 0 | 1654430 | YP_009555238.1 | P0C6X6 |
| EKGRLIIRENNRVVVS | 1 | 0 | 0 | 1654431 | NP_828849.2 | P0C6X7 |
| EKGVLPQLEQPYVFIK | 1 | 0 | 0 | 1654433 | YP_009724389.1 | P0DTD1 |
| EKGVLPQLEQPYVFVK | 1 | 0 | 0 | 1654434 | QIQ54046.1 | P0C6U8 |
| EKHQDHVSLTVPLCGS | 1 | 0 | 0 | 1654437 | YP_009047202.1 | K9N7C7 |
| EKICIIDNMYMGKCGD | 1 | 0 | 0 | 1654438 | YP_173236.1 | A0A140H1G9 |
| EKKKTEGFMGRIRSVY | 1 | 0 | 0 | 1654444 | NP_828849.2 | P0C6X7 |
| EKLFVEHYNEWAQANG | 1 | 0 | 0 | 1654453 | NP_073549.1 | A0A0P0LKV0 |
| EKLISTARDGVRRGDN | 1 | 0 | 0 | 1654456 | YP_009047202.1 | K9N7C7 |
| EKLKALVATAHSELAK | 1 | 0 | 0 | 1654457 | NP_828849.2 | P0C6X7 |
| EKLKPILDWLEEKFKE | 1 | 0 | 0 | 1654459 | QIQ54046.1 | P0C6U8 |
| EKLKPVLDWLEEKFKE | 1 | 0 | 0 | 1654460 | YP_009724389.1 | P0DTD1 |
| EKLKTLVATAEAELAK | 1 | 0 | 0 | 1654462 | YP_009724389.1 | P0DTD1 |
| EKLSAYDLACCMRAGA | 1 | 0 | 0 | 1654467 | NP_073549.1 | A0A0P0LKV0 |
| EKLSPWFMDGENAYEV | 1 | 0 | 0 | 1654469 | YP_009047202.1 | K9N7C7 |
| EKLSRLSMIKGPQRAD | 1 | 0 | 0 | 1654470 | YP_173236.1 | A0A140H1G9 |
| EKPTKEEPKKVKPCIE | 1 | 0 | 0 | 1654481 | QIQ54046.1 | P0C6U8 |

| | | | | | | |
|---|---|---|---|---|---|---|
| EKQNGHSLYLADNLTA | 1 | 0 | 0 | 1654485 | YP_009047202.1 | K9N7C7 |
| EKQVEQKIAEIPKEEV | 1 | 0 | 0 | 1654486 | YP_009724389.1 | P0DTD1 |
| EKSGNFKHLREFVFKN | 1 | 0 | 0 | 1654490 | NP_828851.1 | P59594 |
| EKSTFNKDIYCGNSMV | 1 | 0 | 0 | 1654493 | YP_003766.2 | H9EJ42 |
| EKSVVQKPVDVKPKIK | 1 | 0 | 0 | 1654495 | NP_828849.2 | P0C6X7 |
| EKSYELQTPFEIKLAK | 1 | 0 | 0 | 1654496 | YP_009724389.1 | P0DTD1 |
| EKVDYGSDTVTYKSTA | 1 | 0 | 0 | 1654501 | NP_073549.1 | A0A0P0LKV0 |
| EKVMTFYPRLQAASDW | 1 | 0 | 0 | 1654504 | YP_009555238.1 | P0C6X6 |
| EKYCALAPNMMVTNNT | 1 | 0 | 0 | 1654512 | YP_009724389.1 | P0DTD1 |
| EKYFKYWDQTYHANCV | 1 | 0 | 0 | 1654513 | YP_009047202.1 | K9N7C7 |
| EKYSTIYKLHPSFNVS | 1 | 0 | 0 | 1654515 | NP_073549.1 | A0A0P0LKV0 |
| ELAKGVALDGVLSTFV | 1 | 0 | 0 | 1654519 | NP_828849.2 | P0C6X7 |
| ELAMDEFIERYKLEGY | 1 | 0 | 0 | 1654522 | YP_009724389.1 | P0DTD1 |
| ELAMDEFIQRYKLEGY | 1 | 0 | 0 | 1654523 | NP_828849.2 | P0C6X7 |
| ELAPSDVYQQLAGVKL | 1 | 0 | 0 | 1654526 | YP_173236.1 | A0A140H1G9 |
| ELAPTPGAFFFGSKLD | 1 | 0 | 0 | 1654529 | YP_173242.1 | Q19U29 |
| ELAQVLTEVVYSNGGF | 1 | 0 | 0 | 1654530 | NP_073549.1 | A0A0P0LKV0 |
| ELCDNPFFAVSKPMGT | 1 | 0 | 0 | 1654535 | NP_828851.1 | P59594 |
| ELDEFFKNVSTSIPNF | 1 | 0 | 0 | 1654543 | YP_009047204.1 | K9N5Q8 |
| ELDERIDKVLNEKCSA | 1 | 0 | 0 | 1654544 | YP_009724389.1 | P0DTD1 |
| ELDERIDKVLNEKCST | 1 | 0 | 0 | 1654545 | QHR63299.1 | P0C6W2 |
| ELDERVDKVLNEKCSN | 1 | 0 | 0 | 1654546 | QIQ54046.1 | P0C6U8 |
| ELDERVDKVLNEKCSV | 1 | 0 | 0 | 1654547 | NP_828849.2 | P0C6X7 |
| ELDGVQYGRSGETLGV | 1 | 0 | 0 | 1654548 | QIQ54046.1 | P0C6U8 |
| ELDNDFNTILNTACGV | 1 | 0 | 0 | 1654550 | YP_009555238.1 | P0C6X6 |
| ELDPPCKFSIQDVKGL | 1 | 0 | 0 | 1654552 | YP_173236.1 | A0A140H1G9 |
| ELDQWFKNQTSVAPDL | 1 | 0 | 0 | 1654554 | YP_009555241.1 | P36334 |
| ELEEDWLDDDSQQTVV | 1 | 0 | 0 | 1654561 | QHR63299.1 | P0C6W2 |
| ELEKGVLPQLEQPYVF | 1 | 0 | 0 | 1654565 | NP_828849.2 | P0C6X7 |
| ELEPPCRFVTDTPKGP | 1 | 0 | 0 | 1654570 | YP_009724389.1 | P0DTD1 |
| ELFAKRKMGLTPPLSI | 1 | 0 | 0 | 1654571 | YP_003766.2 | H9EJ42 |
| ELFAKRKVGLTPPLSI | 1 | 0 | 0 | 1654572 | NP_073549.1 | A0A0P0LKV0 |
| ELFENKTILPVNVAFE | 1 | 0 | 0 | 1654575 | QIQ54047.1 | P0C6X7 |
| ELFENKTTLPVNVAFE | 1 | 0 | 0 | 1654576 | YP_009724389.1 | P0DTD1 |
| ELFYSYATHHDKFTDG | 1 | 0 | 0 | 1654585 | NP_828849.2 | P0C6X7 |
| ELFYSYATHSDKFKDG | 1 | 0 | 0 | 1654586 | QIQ54047.1 | P0C6X7 |
| ELFYSYATHSDKFTDG | 1 | 0 | 0 | 1654587 | YP_009724389.1 | P0DTD1 |
| ELGIEFLKRGDKVVYH | 1 | 0 | 0 | 1654593 | QIQ54046.1 | P0C6U8 |
| ELGSGSHVGSSFDGVM | 1 | 0 | 0 | 1654602 | NP_073549.1 | A0A0P0LKV0 |
| ELGVEFLKRGDKIVYH | 1 | 0 | 0 | 1654608 | NP_828849.2 | P0C6X7 |
| ELHDDLVNYIRDKGCV | 1 | 0 | 0 | 1654614 | YP_009555239.1 | Q80872 |
| ELHTVIPDYVDVNKTL | 1 | 0 | 0 | 1654617 | YP_003767.1 | A0A0P0G321 |
| ELHWAPEFPWMFEDAE | 1 | 0 | 0 | 1654618 | YP_009555238.1 | P0C6X6 |
| ELILSWEIGKVKPPLN | 1 | 0 | 0 | 1654621 | YP_009555238.1 | P0C6X6 |
| ELKAAEAKVITIKVTE | 1 | 0 | 0 | 1654628 | NP_073549.1 | A0A0P0LKV0 |
| ELKFNPPALQDAYYRA | 1 | 0 | 0 | 1654632 | YP_009724389.1 | P0DTD1 |
| ELKITFFPDLNGDVVA | 1 | 0 | 0 | 1654639 | QIQ54046.1 | P0C6U8 |
| ELKQLEAKTASLFQTT | 1 | 0 | 0 | 1654642 | YP_003767.1 | A0A0P0G321 |
| ELKVTFFPDLNGDVVA | 1 | 0 | 0 | 1654647 | YP_009724389.1 | P0DTD1 |
| ELLDAEFVEPGCGGIL | 1 | 0 | 0 | 1654649 | NP_073549.1 | A0A0P0LKV0 |
| ELLDTTQLQVANSLMN | 1 | 0 | 0 | 1654650 | YP_009555241.1 | P36334 |
| ELLEQNIDAYKTFPKK | 1 | 0 | 0 | 1654652 | YP_009047211.1 | K9N4V7 |
| ELLLLWESGKAKPPLN | 1 | 0 | 0 | 1654663 | NP_073549.1 | A0A0P0LKV0 |
| ELLLSWESGKVKPPLN | 1 | 0 | 0 | 1654664 | YP_003766.2 | H9EJ42 |
| ELLNAPATVCGPKLST | 1 | 0 | 0 | 1654667 | NP_828851.1 | P59594 |
| ELLTPLGIDLDEWGVA | 1 | 0 | 0 | 1654677 | QIQ54046.1 | P0C6U8 |
| ELNGGAVTRYVDNNFC | 1 | 0 | 0 | 1654690 | NP_828849.2 | P0C6X7 |
| ELNGLKILKQLDNNCW | 1 | 0 | 0 | 1654693 | NP_073549.1 | A0A0P0LKV0 |
| ELNGVVVDKVGDTDCV | 1 | 0 | 0 | 1654694 | YP_009555238.1 | P0C6X6 |
| ELPKFFYASNGTYNCT | 1 | 0 | 0 | 1654698 | NP_073551.1 | A0A0P0K6L9 |
| ELPTGVHAGTDLEGKF | 1 | 0 | 0 | 1654701 | NP_828849.2 | P0C6X7 |
| ELPTGVHAGTDLEGNF | 1 | 0 | 0 | 1654702 | YP_009724389.1 | P0DTD1 |
| ELPTGVHAGTDLEGTF | 1 | 0 | 0 | 1654703 | QHR63299.1 | P0C6W2 |
| ELPVAISGDKVFVRDG | 1 | 0 | 0 | 1654704 | NP_073549.1 | A0A0P0LKV0 |
| ELPVAVVNDKIFTKSG | 1 | 0 | 0 | 1654705 | YP_009047202.1 | K9N7C7 |
| ELPVAVVNDKVFVRYG | 1 | 0 | 0 | 1654706 | YP_003766.2 | H9EJ42 |
| ELQTIVPEYIDVNKTL | 1 | 0 | 0 | 1654712 | NP_073551.1 | A0A0P0K6L9 |
| ELQTPFEIKLAKKFDT | 1 | 0 | 0 | 1654713 | YP_009724389.1 | P0DTD1 |
| ELRFPRGQGVPINTNS | 1 | 0 | 0 | 1654714 | NP_828858.1 | P59595 |
| ELRPDTRYVLMDGSII | 1 | 0 | 0 | 1654715 | NP_828849.2 | P0C6X7 |
| ELRYMNANGLRPPKNS | 1 | 0 | 0 | 1654717 | YP_009555238.1 | P0C6X6 |
| ELRYMNANGLRPPRNS | 1 | 0 | 0 | 1654718 | YP_173236.1 | A0A140H1G9 |
| ELSKELKRPVNPTDAS | 1 | 0 | 0 | 1654725 | YP_173236.1 | A0A140H1G9 |
| ELSNTFGAISASIGDI | 1 | 0 | 0 | 1654728 | YP_009047204.1 | K9N5Q8 |
| ELSTGCHTGTDFNGDF | 1 | 0 | 0 | 1654737 | YP_009555238.1 | P0C6X6 |
| ELSTGCHTGTDFTGNF | 1 | 0 | 0 | 1654738 | YP_173236.1 | A0A140H1G9 |
| ELSVTFFPDLNGDVVA | 1 | 0 | 0 | 1654742 | NP_828849.2 | P0C6X7 |
| ELSYKLPNYTVPDLVV | 1 | 0 | 0 | 1654744 | NP_073551.1 | A0A0P0K6L9 |
| ELTAEDISLLKKMDEP | 1 | 0 | 0 | 1654745 | YP_009555245.1 | P33469 |
| ELTLKHFFFTQKGDAA | 1 | 0 | 0 | 1654754 | NP_073549.1 | A0A0P0LKV0 |
| ELTPSDVYQQLAGIKL | 1 | 0 | 0 | 1654756 | YP_009555238.1 | P0C6X6 |
| ELTPVVQTIEVNSFSG | 1 | 0 | 0 | 1654757 | YP_009724389.1 | P0DTD1 |
| ELTRELNGGAVTRYVD | 1 | 0 | 0 | 1654758 | NP_828849.2 | P0C6X7 |
| ELTSASYFCKIGKEQK | 1 | 0 | 0 | 1654759 | YP_009047202.1 | K9N7C7 |
| ELVAELNGIQYGRSGE | 1 | 0 | 0 | 1654766 | QHR63299.1 | P0C6W2 |
| ELVASFLEMKSETKVQ | 1 | 0 | 0 | 1654768 | QIQ54046.1 | P0C6U8 |
| ELVTDIVVVDEVSMLT | 1 | 0 | 0 | 1654776 | YP_173236.1 | A0A140H1G9 |
| ELVYENKFVPVKEASK | 1 | 0 | 0 | 1654778 | NP_073549.1 | A0A0P0LKV0 |
| ELVYENKFVPVKPASK | 1 | 0 | 0 | 1654779 | YP_003766.2 | H9EJ42 |
| ELWAKRNIKPVPEIKI | 1 | 0 | 0 | 1654780 | NP_828849.2 | P0C6X7 |
| ELWAKRNIKPVPEVKI | 1 | 0 | 0 | 1654781 | YP_009724389.1 | P0DTD1 |

| | | | | | | |
|---|---|---|---|---|---|---|
| ELYAKRAVRSHPDFKL | 1 | 0 | 0 | 1654783 | YP_009047202.1 | K9N7C7 |
| ELYLNSSNVTTMDFCE | 1 | 0 | 0 | 1654788 | NP_828849.2 | P0C6X7 |
| ELYVNNAYRLFDLVQY | 1 | 0 | 0 | 1654795 | YP_009555238.1 | P0C6X6 |
| EMAKEVKTSSKVVLSD | 1 | 0 | 0 | 1654796 | NP_073549.1 | A0A0P0LKV0 |
| EMDGIQYGRSGITLGV | 1 | 0 | 0 | 1654800 | NP_828849.2 | P0C6X7 |
| EMHNKINLCDDPETAQ | 1 | 0 | 0 | 1654804 | NP_073549.1 | A0A0P0LKV0 |
| EMKSEKQVEQKIAEIP | 1 | 0 | 0 | 1654808 | YP_009724389.1 | P0DTD1 |
| EMKSEKQVEQKTAEIP | 1 | 0 | 0 | 1654809 | QHR63299.1 | P0C6W2 |
| EMKSETKVQDHFDVVE | 1 | 0 | 0 | 1654810 | QIQ54046.1 | P0C6U8 |
| EMLKLGTNDPQFPILA | 1 | 0 | 0 | 1654817 | YP_173242.1 | Q19U29 |
| EMLSLQQVVKALNESY | 1 | 0 | 0 | 1654819 | YP_009047204.1 | K9N5Q8 |
| EMLSRALKKVPVDEYI | 1 | 0 | 0 | 1654820 | NP_828849.2 | P0C6X7 |
| EMLTFDKKAFTLANVS | 1 | 0 | 0 | 1654821 | NP_073551.1 | A0A0P0K6L9 |
| EMLTKALRKVPTDNYI | 1 | 0 | 0 | 1654823 | QHR63299.1 | P0C6W2 |
| EMPCAIINDKVVAKID | 1 | 0 | 0 | 1654826 | YP_009555238.1 | P0C6X6 |
| EMQKPRWKRQPNDDVT | 1 | 0 | 0 | 1654828 | NP_073556.1 | A0A0N7J131 |
| EMQQHPLLNPSALEFN | 1 | 0 | 0 | 1654830 | NP_073556.1 | A0A0N7J131 |
| EMSCDCGKKFDDQVGC | 1 | 0 | 0 | 1654833 | YP_003766.2 | H9EJ42 |
| EMTSNVRRQVASGVID | 1 | 0 | 0 | 1654836 | NP_073549.1 | A0A0P0LKV0 |
| ENAVANGSSPQIIKQL | 1 | 0 | 0 | 1654853 | NP_073549.1 | A0A0P0LKV0 |
| ENDITNGPHEFCSQHT | 1 | 0 | 0 | 1654859 | YP_173236.1 | A0A140H1G9 |
| ENEEIIINGHTYVCAW | 1 | 0 | 0 | 1654866 | NP_073549.1 | A0A0P0LKV0 |
| ENEFRVYSSANNCTFE | 1 | 0 | 0 | 1654869 | QIQ54048.1 | P59594 |
| ENFNHHEKLLAPLLSA | 1 | 0 | 0 | 1654872 | QIQ54046.1 | P0C6U8 |
| ENIDYFNMRRAKFSLE | 1 | 0 | 0 | 1654894 | YP_009555238.1 | P0C6X6 |
| ENIGTNFPLQIGFSTG | 1 | 0 | 0 | 1654895 | YP_173236.1 | A0A140H1G9 |
| ENKSAELNYTVQKLQT | 1 | 0 | 0 | 1654902 | NP_073551.1 | A0A0P0K6L9 |
| ENKTILPVNVAFELWA | 1 | 0 | 0 | 1654904 | QIQ54047.1 | P0C6X7 |
| ENKTTLPTNIAFELYA | 1 | 0 | 0 | 1654905 | YP_009047202.1 | K9N7C7 |
| ENKTTLPVNVAFELWA | 1 | 0 | 0 | 1654906 | YP_009724389.1 | P0DTD1 |
| ENKYSQLDEEQPMEID | 1 | 0 | 0 | 1654907 | YP_009724394.1 | P0DTC6 |
| ENLNAYVNSNQNTDSD | 1 | 0 | 0 | 1654917 | YP_173242.1 | Q19U29 |
| ENMTPRDLGACIDCNA | 1 | 0 | 0 | 1654924 | NP_828849.2 | P0C6X7 |
| ENNTEFTRVNAKPPPG | 1 | 0 | 0 | 1654930 | NP_828849.2 | P0C6X7 |
| ENPDIINIYKKLGPIF | 1 | 0 | 0 | 1654934 | YP_173236.1 | A0A140H1G9 |
| ENPDIINVYKKLGPIF | 1 | 0 | 0 | 1654935 | YP_009555238.1 | P0C6X6 |
| ENPFMVNQLAYSSSAN | 1 | 0 | 0 | 1654937 | YP_009047202.1 | K9N7C7 |
| ENPHLMGWDYPKCDRA | 1 | 0 | 0 | 1654939 | YP_009724389.1 | P0DTD1 |
| ENPMLMGWDYPKCDRA | 1 | 0 | 0 | 1654940 | YP_003766.2 | H9EJ42 |
| ENPTIQKDILECNVKT | 1 | 0 | 0 | 1654942 | QHR63299.1 | P0C6W2 |
| ENPTIQKDVLECNVKT | 1 | 0 | 0 | 1654943 | YP_009724389.1 | P0DTD1 |
| ENQINDMIYSLLEKGR | 1 | 0 | 0 | 1654949 | NP_828849.2 | P0C6X7 |
| ENQISGYTLAATSASL | 1 | 0 | 0 | 1654950 | YP_009555241.1 | P36334 |
| ENQKILAASFNKAINN | 1 | 0 | 0 | 1654951 | YP_003767.1 | A0A0P0G321 |
| ENQKILAASFNKAMTN | 1 | 0 | 0 | 1654952 | NP_073551.1 | A0A0P0K6L9 |
| ENQKLIANKFNQALGA | 1 | 0 | 0 | 1654953 | YP_009047204.1 | K9N5Q8 |
| ENQKQIANQFNKAISQ | 1 | 0 | 0 | 1654954 | NP_828851.1 | P59594 |
| ENRYSVLSVDSVSEES | 1 | 0 | 0 | 1654957 | YP_173236.1 | A0A140H1G9 |
| ENSYITTIKPVIYKLD | 1 | 0 | 0 | 1654962 | QHR63299.1 | P0C6W2 |
| ENSYTTTIKPIVYKLD | 1 | 0 | 0 | 1654963 | QIQ54046.1 | P0C6U8 |
| ENSYTTTIKPVTYKLD | 1 | 0 | 0 | 1654964 | YP_009724389.1 | P0DTD1 |
| ENTSITIKKPNELSLA | 1 | 0 | 0 | 1654966 | NP_828849.2 | P0C6X7 |
| ENVAFNVVNKGHFDGQ | 1 | 0 | 0 | 1654969 | YP_009724389.1 | P0DTD1 |
| ENVFAVESGGYIPSDF | 1 | 0 | 0 | 1654974 | NP_073551.1 | A0A0P0K6L9 |
| ENYNNLVPTYLKSDNI | 1 | 0 | 0 | 1654990 | YP_173236.1 | A0A140H1G9 |
| EPCNLYNYGAGIKLPS | 1 | 0 | 0 | 1654999 | YP_003766.2 | H9EJ42 |
| EPEDHVQIHTIDGSSG | 1 | 0 | 0 | 1655007 | QIQ54049.1 | P59632 |
| EPEEHVQIHTIDGSSG | 1 | 0 | 0 | 1655008 | YP_009724391.1 | P0DTC3 |
| EPEIPHFNQKLPNGVT | 1 | 0 | 0 | 1655011 | NP_073556.1 | A0A0N7J131 |
| EPEPEPTPEEPVNQFT | 1 | 0 | 0 | 1655013 | NP_828849.2 | P0C6X7 |
| EPEPTPEEPVNQFTGY | 1 | 0 | 0 | 1655014 | NP_828849.2 | P0C6X7 |
| EPFNVSFVNDSIESVG | 1 | 0 | 0 | 1655016 | YP_173238.1 | A0A140H1H1 |
| EPFTVNSVNDSLEPVG | 1 | 0 | 0 | 1655017 | YP_009555241.1 | P36334 |
| EPITSLNTKYVAPQVT | 1 | 0 | 0 | 1655024 | YP_009047204.1 | K9N5Q8 |
| EPKGKYSQKAYALLRQ | 1 | 0 | 0 | 1655028 | YP_173236.1 | A0A140H1G9 |
| EPKKVKPCIEEVTTTL | 1 | 0 | 0 | 1655031 | QIQ54046.1 | P0C6U8 |
| EPKLDGYYKKDNAYYT | 1 | 0 | 0 | 1655032 | NP_828849.2 | P0C6X7 |
| EPMLQSADAQSFLNRV | 1 | 0 | 0 | 1655047 | YP_009724389.1 | P0DTD1 |
| EPPCRFVTDTPKGPKV | 1 | 0 | 0 | 1655049 | YP_009724389.1 | P0DTD1 |
| EPPKEQRVQGSITQRT | 1 | 0 | 0 | 1655050 | YP_009047211.1 | K9N4V7 |
| EPQIITTDNTFVSGSC | 1 | 0 | 0 | 1655056 | QHR63300.2 | Q3LZX1 |
| EPQKDVYELRYNGAIR | 1 | 0 | 0 | 1655057 | YP_009555245.1 | P33469 |
| EPTGYGVRRKNSEPEI | 1 | 0 | 0 | 1655066 | NP_073556.1 | A0A0N7J131 |
| EPTPVVQTEVNSFSGY | 1 | 0 | 0 | 1655069 | QHR63299.1 | P0C6W2 |
| EPVASIITKLNSLNEP | 1 | 0 | 0 | 1655071 | NP_828849.2 | P0C6X7 |
| EPVRDEVSIETDIIDE | 1 | 0 | 0 | 1655080 | NP_073556.1 | A0A0N7J131 |
| EQAATQMYKEARSVNR | 1 | 0 | 0 | 1655086 | YP_003766.2 | H9EJ42 |
| EQDAIFSLTKRNILPT | 1 | 0 | 0 | 1655091 | NP_073549.1 | A0A0P0LKV0 |
| EQDAQIDRLINGRLTT | 1 | 0 | 0 | 1655093 | YP_009047204.1 | K9N5Q8 |
| EQDEIYAYTKRNVLPT | 1 | 0 | 0 | 1655094 | YP_009555238.1 | P0C6X6 |
| EQDELFAMTKRNVIPT | 1 | 0 | 0 | 1655095 | YP_009047202.1 | K9N7C7 |
| EQEAILMSEDVKLNPS | 1 | 0 | 0 | 1655099 | YP_009047202.1 | K9N7C7 |
| EQEEDWLDDDSQQTVG | 1 | 0 | 0 | 1655100 | YP_009724389.1 | P0DTD1 |
| EQEEDWLETEAEVVEQ | 1 | 0 | 0 | 1655101 | QIQ54046.1 | P0C6U8 |
| EQFDSLNLSAGTQHIS | 1 | 0 | 0 | 1655107 | YP_173242.1 | Q19U29 |
| EQFVGPLTVNEKRRL | 1 | 0 | 0 | 1655113 | P03155.1 | Q69028 |
| EQIDGKFVTKFSGDKF | 1 | 0 | 0 | 1655115 | YP_173236.1 | A0A140H1G9 |
| EQIDGYTMHANYIFWR | 1 | 0 | 0 | 1655116 | NP_828849.2 | P0C6X7 |
| EQIDGYVMHANYIFWR | 1 | 0 | 0 | 1655117 | YP_009724389.1 | P0DTD1 |
| EQISAFTKPSSIKEMQ | 1 | 0 | 0 | 1655119 | YP_003771.1 | H9EJ69 |
| EQKIAEIPKEEVKPFI | 1 | 0 | 0 | 1655122 | YP_009724389.1 | P0DTD1 |

| | | | | | | |
|---|---|---|---|---|---|---|
| EQLDFIDTKRGVYCCR | 1 | 0 | 0 | 1655125 | YP_009724389.1 | P0DTD1 |
| EQLDYIESKRGVYCCR | 1 | 0 | 0 | 1655126 | NP_828849.2 | P0C6X7 |
| EQLEPCAQSRLLSGVA | 1 | 0 | 0 | 1655127 | NP_073549.1 | A0A0P0LKV0 |
| EQLKPVLDWLEEKFKE | 1 | 0 | 0 | 1655130 | QHR63299.1 | P0C6W2 |
| EQNEIYAYTKRNVLPT | 1 | 0 | 0 | 1655142 | YP_173236.1 | A0A140H1G9 |
| EQNIDAYKTFPKKEKK | 1 | 0 | 0 | 1655144 | YP_009047211.1 | K9N4V7 |
| EQPEQPTQYEYGTESD | 1 | 0 | 0 | 1655146 | QIQ54046.1 | P0C6U8 |
| EQPIDLVPNQPYPNAS | 1 | 0 | 0 | 1655148 | YP_009724389.1 | P0DTD1 |
| EQPIDLVPTQPLPNAS | 1 | 0 | 0 | 1655149 | NP_828849.2 | P0C6X7 |
| EQPKIEPKPVSVIKVA | 1 | 0 | 0 | 1655151 | NP_073549.1 | A0A0P0LKV0 |
| EQPTSEAVEAPLVGTP | 1 | 0 | 0 | 1655153 | YP_009724389.1 | P0DTD1 |
| EQPWKVIPRNLVPINK | 1 | 0 | 0 | 1655155 | NP_073556.1 | A0A0N7J131 |
| EQQVGVIADKDKDLSG | 1 | 0 | 0 | 1655161 | NP_073549.1 | A0A0P0LKV0 |
| EQRVQGSITQRTRTRP | 1 | 0 | 0 | 1655165 | YP_009047211.1 | K9N4V7 |
| EQSEIEPEPEPTPEEP | 1 | 0 | 0 | 1655166 | NP_828849.2 | P0C6X7 |
| EQSMYNLLKGCNAVAK | 1 | 0 | 0 | 1655170 | NP_073549.1 | A0A0P0LKV0 |
| EQSYFTKKDWYDFVEN | 1 | 0 | 0 | 1655172 | YP_009555238.1 | P0C6X6 |
| EQTNCNTSSLAYYEPV | 1 | 0 | 0 | 1655174 | YP_009047202.1 | K9N7C7 |
| EQTQNVTVKPKRLRKK | 1 | 0 | 0 | 1655179 | YP_009047202.1 | K9N7C7 |
| EQVSDLLVKLLRGMPI | 1 | 0 | 0 | 1655183 | YP_009047202.1 | K9N7C7 |
| EQVVMDYLDNLKPRVE | 1 | 0 | 0 | 1655184 | NP_828849.2 | P0C6X7 |
| EQYNQTILNLTSEIST | 1 | 0 | 0 | 1655190 | NP_073551.1 | A0A0P0K6L9 |
| ERCELANYKQSIPMPR | 1 | 0 | 0 | 1655198 | YP_009047202.1 | K9N7C7 |
| ERDFIDMDDNLFIAKY | 1 | 0 | 0 | 1655204 | YP_173236.1 | A0A140H1G9 |
| ERDLLIKAYNTINNEQ | 1 | 0 | 0 | 1655209 | NP_073549.1 | A0A0P0LKV0 |
| ERELILSWEIGKVKPP | 1 | 0 | 0 | 1655212 | YP_009555238.1 | P0C6X6 |
| ERGKLLVRDTRKEIFV | 1 | 0 | 0 | 1655222 | YP_173236.1 | A0A140H1G9 |
| ERGSVQVDNGSSINRR | 1 | 0 | 0 | 1655225 | NP_073549.1 | A0A0P0LKV0 |
| ERIDKVLNEKCSAYTV | 1 | 0 | 0 | 1655230 | YP_009724389.1 | P0DTD1 |
| ERIDKVLNEKCSTYTV | 1 | 0 | 0 | 1655231 | QHR63299.1 | P0C6W2 |
| ERIDYSDAVSYKSSTT | 1 | 0 | 0 | 1655232 | YP_009047202.1 | K9N7C7 |
| ERLCLFDRYFKYWDQT | 1 | 0 | 0 | 1655239 | NP_828849.2 | P0C6X7 |
| ERLIACENPFMVNQLA | 1 | 0 | 0 | 1655240 | YP_009047202.1 | K9N7C7 |
| ERLKLFDRYFKYWDQT | 1 | 0 | 0 | 1655244 | YP_009724389.1 | P0DTD1 |
| ERLNLFNRYFKYWDQT | 1 | 0 | 0 | 1655246 | QIQ54047.1 | P0C6X7 |
| ERMADLALTNMYKEAR | 1 | 0 | 0 | 1655247 | YP_009555238.1 | P0C6X6 |
| ERQLLLVWEAGKSKPP | 1 | 0 | 0 | 1655256 | YP_009047202.1 | K9N7C7 |
| ERSEKSYELQTPFEIK | 1 | 0 | 0 | 1655261 | YP_009724389.1 | P0DTD1 |
| ERSGARPKQRRPQGLP | 1 | 0 | 0 | 1655263 | QHR63308.1 | Q3LZX4 |
| ERTCCLCDKRATCFST | 1 | 0 | 0 | 1655267 | QHR63299.1 | P0C6W2 |
| ERTCCLCDRRATCFST | 1 | 0 | 0 | 1655268 | YP_009724389.1 | P0DTD1 |
| ERVDKVLNEKCSNYTV | 1 | 0 | 0 | 1655269 | QIQ54046.1 | P0C6U8 |
| ERVDWDIEYPYISHEK | 1 | 0 | 0 | 1655271 | YP_009047202.1 | K9N7C7 |
| ESAALSAQLAKDKVNE | 1 | 0 | 0 | 1655291 | YP_009047204.1 | K9N5Q8 |
| ESADVSEMLTFDKKAF | 1 | 0 | 0 | 1655293 | NP_073551.1 | A0A0P0K6L9 |
| ESAHVCGDNIGTNVPL | 1 | 0 | 0 | 1655294 | YP_003766.2 | H9EJ42 |
| ESASKSASVYYSQLMC | 1 | 0 | 0 | 1655299 | NP_828849.2 | P0C6X7 |
| ESASSEKPTKEEPKKV | 1 | 0 | 0 | 1655300 | QIQ54046.1 | P0C6U8 |
| ESAVNSTKALAKQDAA | 1 | 0 | 0 | 1655302 | YP_009047206.1 | K9N4V0 |
| ESAYMPIADPTHFDIE | 1 | 0 | 0 | 1655304 | NP_073549.1 | A0A0P0LKV0 |
| ESCNNLVPTYLKSDNI | 1 | 0 | 0 | 1655308 | YP_009555238.1 | P0C6X6 |
| ESDADRANFLNAVVFY | 1 | 0 | 0 | 1655310 | YP_173236.1 | A0A140H1G9 |
| ESDDYIARNGPLNVGG | 1 | 0 | 0 | 1655311 | QIQ54046.1 | P0C6U8 |
| ESDDYIATNGPLKVGG | 1 | 0 | 0 | 1655312 | YP_009724389.1 | P0DTD1 |
| ESDDYIKLNGPLTVGG | 1 | 0 | 0 | 1655313 | NP_828849.2 | P0C6X7 |
| ESDEYILAKGPLQVGD | 1 | 0 | 0 | 1655314 | YP_009047202.1 | K9N7C7 |
| ESDHYIATNGPLKVGG | 1 | 0 | 0 | 1655315 | QHR63299.1 | P0C6W2 |
| ESDIYFVKNIPRYASA | 1 | 0 | 0 | 1655316 | YP_009555238.1 | P0C6X6 |
| ESDLIISNFKFDDYDF | 1 | 0 | 0 | 1655317 | YP_173236.1 | A0A140H1G9 |
| ESDYKGLPLEFGASSV | 1 | 0 | 0 | 1655320 | QIQ54046.1 | P0C6U8 |
| ESFIDDYYGYLRKHFS | 1 | 0 | 0 | 1655329 | YP_003766.2 | H9EJ42 |
| ESFKKVATIPYKVCNS | 1 | 0 | 0 | 1655330 | YP_009047202.1 | K9N7C7 |
| ESFSVTLLDNQEDKFW | 1 | 0 | 0 | 1655333 | YP_003766.2 | H9EJ42 |
| ESFVDDFYGYLQKHFS | 1 | 0 | 0 | 1655334 | NP_073549.1 | A0A0P0LKV0 |
| ESFYKNMYLKSAVMQS | 1 | 0 | 0 | 1655336 | YP_173236.1 | A0A140H1G9 |
| ESGAKETKEINIIKLS | 1 | 0 | 0 | 1655338 | YP_009555238.1 | P0C6X6 |
| ESGDYTLANTTTEPLK | 1 | 0 | 0 | 1655341 | YP_009047202.1 | K9N7C7 |
| ESGGYIPSDFAFNNWF | 1 | 0 | 0 | 1655344 | NP_073551.1 | A0A0P0K6L9 |
| ESGKAKPPLNRNSVFT | 1 | 0 | 0 | 1655346 | NP_073549.1 | A0A0P0LKV0 |
| ESGKVKPPLNRNSVFT | 1 | 0 | 0 | 1655347 | YP_003766.2 | H9EJ42 |
| ESGLKPILRKGGRTIA | 1 | 0 | 0 | 1655349 | QHR63299.1 | P0C6W2 |
| ESGVKDCVVLHSYFTS | 1 | 0 | 0 | 1655354 | YP_009724391.1 | P0DTC3 |
| ESGVYSVSSFEAKPSG | 1 | 0 | 0 | 1655355 | YP_009047204.1 | K9N5Q8 |
| ESHGKQVVSDIDYVPL | 1 | 0 | 0 | 1655357 | YP_009724389.1 | P0DTD1 |
| ESHLDADLTKPYVKWD | 1 | 0 | 0 | 1655358 | QIQ54047.1 | P0C6X7 |
| ESHVDTDLTKPYIKWD | 1 | 0 | 0 | 1655360 | YP_009724389.1 | P0DTD1 |
| ESIHYLGTTGHTLKSG | 1 | 0 | 0 | 1655362 | YP_003766.2 | H9EJ42 |
| ESIKSLNNSYINLKDI | 1 | 0 | 0 | 1655363 | YP_173238.1 | A0A140H1H1 |
| ESIPIVWGVKDFNTLS | 1 | 0 | 0 | 1655364 | YP_003766.2 | H9EJ42 |
| ESKAFVDTIVPPCPSI | 1 | 0 | 0 | 1655369 | NP_073549.1 | A0A0P0LKV0 |
| ESKFWDESFYASMYEK | 1 | 0 | 0 | 1655373 | NP_073549.1 | A0A0P0LKV0 |
| ESKIYGSCFKSIVLDK | 1 | 0 | 0 | 1655374 | YP_173238.1 | A0A140H1H1 |
| ESKLSVEQRQQVDKKI | 1 | 0 | 0 | 1655375 | QHR63299.1 | P0C6W2 |
| ESKPQSRNPSSDRNHN | 1 | 0 | 0 | 1655379 | NP_073556.1 | A0A0N7J131 |
| ESKPSVEQRKQDDKKI | 1 | 0 | 0 | 1655380 | YP_009724389.1 | P0DTD1 |
| ESKPWFVRKAEDIFGP | 1 | 0 | 0 | 1655381 | NP_073549.1 | A0A0P0LKV0 |
| ESKQVDYVPLRSATCI | 1 | 0 | 0 | 1655382 | YP_173236.1 | A0A140H1G9 |
| ESKRGVYCCRDHEHEI | 1 | 0 | 0 | 1655383 | NP_828849.2 | P0C6X7 |
| ESKVLLERFLPKCPEI | 1 | 0 | 0 | 1655384 | YP_003766.2 | H9EJ42 |
| ESKVSGKGQQQQGQTV | 1 | 0 | 0 | 1655385 | NP_828858.1 | P59595 |
| ESKYSCKEVFKNCNVL | 1 | 0 | 0 | 1655386 | NP_073549.1 | A0A0P0LKV0 |

| | | | | | | |
|---|---|---|---|---|---|---|
| ESKYSCKEVLKNCNVL | 1 | 0 | 0 | 1655387 | YP_003766.2 | H9EJ42 |
| ESLNDNLLEILTKEKV | 1 | 0 | 0 | 1655395 | QIQ54046.1 | P0C6U8 |
| ESLRLFAAETVKAKEE | 1 | 0 | 0 | 1655397 | NP_073549.1 | A0A0P0LKV0 |
| ESLRPDTRYVLMDGSI | 1 | 0 | 0 | 1655398 | YP_009724389.1 | P0DTD1 |
| ESLVPGFNEKTHVQLS | 1 | 0 | 0 | 1655401 | YP_009724389.1 | P0DTD1 |
| ESMSYQEQDELFAMTK | 1 | 0 | 0 | 1655402 | YP_009047202.1 | K9N7C7 |
| ESPFVMMSAPPAQYEL | 1 | 0 | 0 | 1655407 | YP_009724389.1 | P0DTD1 |
| ESQGNVVTSVMESQIS | 1 | 0 | 0 | 1655410 | YP_173236.1 | A0A140H1G9 |
| ESRYGLQNLLQLPNFY | 1 | 0 | 0 | 1655418 | YP_003767.1 | A0A0P0G321 |
| ESSCGLFKDCARNPID | 1 | 0 | 0 | 1655421 | NP_073549.1 | A0A0P0LKV0 |
| ESSKSLGKQLGVVSDG | 1 | 0 | 0 | 1655424 | YP_003766.2 | H9EJ42 |
| ESSRKLLGLTNLSEIT | 1 | 0 | 0 | 1655425 | YP_173243.1 | Q0ZJ80 |
| ESSSKLWAQCVQLHND | 1 | 0 | 0 | 1655428 | YP_009724389.1 | P0DTD1 |
| ESSVQKKINRMAEQAA | 1 | 0 | 0 | 1655430 | NP_073549.1 | A0A0P0LKV0 |
| ESTADQETYGGASVCL | 1 | 0 | 0 | 1655431 | YP_009047202.1 | K9N7C7 |
| ESTDQMSEPPKEQRVQ | 1 | 0 | 0 | 1655432 | YP_009047211.1 | K9N4V7 |
| ESTDSIVFTYIPASGY | 1 | 0 | 0 | 1655433 | YP_009047208.1 | K9N7D2 |
| ESTFGLSYYSNSMACP | 1 | 0 | 0 | 1655434 | YP_009555238.1 | P0C6X6 |
| ESTFGSVYYHNSMDCP | 1 | 0 | 0 | 1655435 | YP_173236.1 | A0A140H1G9 |
| ESTGTQSVSVDLESTS | 1 | 0 | 0 | 1655436 | YP_009047205.1 | K9N796 |
| ESTINKSVANLKLPPH | 1 | 0 | 0 | 1655437 | YP_003768.1 | H9EJ24 |
| ESTKLPYYDVEKIHIQ | 1 | 0 | 0 | 1655438 | NP_073551.1 | A0A0P0K6L9 |
| ESTQITTDTGIENATF | 1 | 0 | 0 | 1655441 | NP_828852.2 | P59632 |
| ESTSTHDGPTEHVTSV | 1 | 0 | 0 | 1655442 | YP_009047205.1 | K9N796 |
| ESTTVLDHTDWCRCSC | 1 | 0 | 0 | 1655443 | YP_173238.1 | A0A140H1H1 |
| ESVKTQFNYFKKVDGI | 1 | 0 | 0 | 1655451 | NP_828849.2 | P0C6X7 |
| ESYIDLKELGNYTYYN | 1 | 0 | 0 | 1655459 | YP_009047204.1 | K9N5Q8 |
| ESYNNYVPSYVKPDSV | 1 | 0 | 0 | 1655462 | YP_009047202.1 | K9N7C7 |
| ETAQNSVRALQKAAIT | 1 | 0 | 0 | 1655475 | QHR63299.1 | P0C6W2 |
| ETARQEYENAVANGSS | 1 | 0 | 0 | 1655477 | NP_073549.1 | A0A0P0LKV0 |
| ETCTIAGKAFDKVFDY | 1 | 0 | 0 | 1655481 | NP_073549.1 | A0A0P0LKV0 |
| ETDFLELAMDEFIQRY | 1 | 0 | 0 | 1655484 | NP_828849.2 | P0C6X7 |
| ETDGICPENHVMVDCR | 1 | 0 | 0 | 1655485 | YP_009555238.1 | P0C6X6 |
| ETDLKKGPHEFCSQHT | 1 | 0 | 0 | 1655486 | YP_009047202.1 | K9N7C7 |
| ETELAKNVSLDNVLST | 1 | 0 | 0 | 1655490 | QIQ54046.1 | P0C6U8 |
| ETETYFKVNGLWSPTI | 1 | 0 | 0 | 1655491 | YP_173236.1 | A0A140H1G9 |
| ETFNHNNGNDVLYQPP | 1 | 0 | 0 | 1655494 | YP_173236.1 | A0A140H1G9 |
| ETFYPKLQASQAWQPG | 1 | 0 | 0 | 1655502 | NP_828849.2 | P0C6X7 |
| ETFYPKLQSSQAWQPG | 1 | 0 | 0 | 1655503 | YP_009724389.1 | P0DTD1 |
| ETGSDLLYQPPNCSIT | 1 | 0 | 0 | 1655507 | YP_009047202.1 | K9N7C7 |
| ETHRDCDFNKPLIEWP | 1 | 0 | 0 | 1655509 | YP_009047202.1 | K9N7C7 |
| ETIKAKEESVKSSYAF | 1 | 0 | 0 | 1655511 | YP_003766.2 | H9EJ42 |
| ETIMKVLNENLNAYQQ | 1 | 0 | 0 | 1655513 | YP_009555245.1 | P33469 |
| ETKALKELYGPVDPTF | 1 | 0 | 0 | 1655526 | YP_009047202.1 | K9N7C7 |
| ETKCTLKSLTVEKGIY | 1 | 0 | 0 | 1655528 | QIQ54048.1 | P59594 |
| ETKDVVECLKLSHQSD | 1 | 0 | 0 | 1655529 | YP_009724389.1 | P0DTD1 |
| ETKEINIIKLSGVKKP | 1 | 0 | 0 | 1655530 | YP_009555238.1 | P0C6X6 |
| ETKEQKHEMQKPRWKR | 1 | 0 | 0 | 1655531 | NP_073556.1 | A0A0N7J131 |
| ETKFLTENLLLYADIN | 1 | 0 | 0 | 1655532 | QIQ54046.1 | P0C6U8 |
| ETKFLTNKLLLFADIN | 1 | 0 | 0 | 1655534 | NP_828849.2 | P0C6X7 |
| ETKGPLCVDTSHFTTK | 1 | 0 | 0 | 1655535 | NP_073551.1 | A0A0P0K6L9 |
| ETKVEITPEESASSEK | 1 | 0 | 0 | 1655537 | QIQ54046.1 | P0C6U8 |
| ETKVQCSTNLFKDCSK | 1 | 0 | 0 | 1655538 | YP_009555238.1 | P0C6X6 |
| ETKVQDHFDVVETKVE | 1 | 0 | 0 | 1655539 | QIQ54046.1 | P0C6U8 |
| ETLPTEVLSEEVILKT | 1 | 0 | 0 | 1655551 | QIQ54046.1 | P0C6U8 |
| ETLPTEVLTEEVVLKT | 1 | 0 | 0 | 1655552 | YP_009724389.1 | P0DTD1 |
| ETMNYLFQHANLDSCK | 1 | 0 | 0 | 1655564 | QHR63299.1 | P0C6W2 |
| ETNNLMCIDMKGTMYV | 1 | 0 | 0 | 1655577 | YP_009555244.1 | Q01455 |
| ETNSQYLFSTINALPE | 1 | 0 | 0 | 1655578 | YP_009047202.1 | K9N7C7 |
| ETPHLMGWDYPKCDRA | 1 | 0 | 0 | 1655581 | NP_828849.2 | P0C6X7 |
| ETPVSIVNNTVYTKVD | 1 | 0 | 0 | 1655587 | QIQ54047.1 | P0C6X7 |
| ETPVVPDTVEVPPQVV | 1 | 0 | 0 | 1655588 | YP_009047202.1 | K9N7C7 |
| ETRKLMPVCMETKAIV | 1 | 0 | 0 | 1655594 | QHR63299.1 | P0C6W2 |
| ETSNIFDVLQSEDTQG | 1 | 0 | 0 | 1655603 | QIQ54046.1 | P0C6U8 |
| ETSNSFDVLKSEDTQG | 1 | 0 | 0 | 1655605 | QHR63299.1 | P0C6W2 |
| ETSYTTTIKPVSYKLD | 1 | 0 | 0 | 1655610 | NP_828849.2 | P0C6X7 |
| ETTADIVVFDEISMAT | 1 | 0 | 0 | 1655611 | YP_009724389.1 | P0DTD1 |
| ETTDFCTVALASYADV | 1 | 0 | 0 | 1655612 | NP_073551.1 | A0A0P0K6L9 |
| ETTDMVKSKGVCATGD | 1 | 0 | 0 | 1655613 | YP_009555238.1 | P0C6X6 |
| ETTSAFVGALPKTVRE | 1 | 0 | 0 | 1655614 | NP_073551.1 | A0A0P0K6L9 |
| ETVKAKEESVKSSYAY | 1 | 0 | 0 | 1655618 | NP_073549.1 | A0A0P0LKV0 |
| ETVKGLDYKAFKQIVE | 1 | 0 | 0 | 1655619 | YP_009724389.1 | P0DTD1 |
| ETVKGLDYKSFKQIVE | 1 | 0 | 0 | 1655620 | QIQ54046.1 | P0C6U8 |
| ETVKGLDYKTFKQIVE | 1 | 0 | 0 | 1655621 | QHR63299.1 | P0C6W2 |
| ETVNGYAKSNFVKDDY | 1 | 0 | 0 | 1655623 | NP_073549.1 | A0A0P0LKV0 |
| ETVRVEEEEEEDWLDD | 1 | 0 | 0 | 1655626 | NP_828849.2 | P0C6X7 |
| ETYFTQSRDLEDFKPR | 1 | 0 | 0 | 1655633 | NP_828849.2 | P0C6X7 |
| ETYIKWPWWVWLCISV | 1 | 0 | 0 | 1655637 | NP_073551.1 | A0A0P0K6L9 |
| ETYVALPIYYQHTDIN | 1 | 0 | 0 | 1655638 | YP_003767.1 | A0A0P0G321 |
| EVDIIEQPFEEVEHVL | 1 | 0 | 0 | 1655656 | YP_003766.2 | H9EJ42 |
| EVDIQQTFFDKTWPRP | 1 | 0 | 0 | 1655657 | YP_009047204.1 | K9N5Q8 |
| EVDPDLSAFYVKDGKY | 1 | 0 | 0 | 1655659 | YP_009047202.1 | K9N7C7 |
| EVEKGVLPQLEQPYVF | 1 | 0 | 0 | 1655666 | YP_009724389.1 | P0DTD1 |
| EVEWKFYDAQPCSDKA | 1 | 0 | 0 | 1655670 | NP_828849.2 | P0C6X7 |
| EVFKNCNVLDDFIVFN | 1 | 0 | 0 | 1655676 | NP_073549.1 | A0A0P0LKV0 |
| EVFNATKFPSVYAWER | 1 | 0 | 0 | 1655678 | NP_828851.1 | P59594 |
| EVFNRLATSDWTDVRD | 1 | 0 | 0 | 1655681 | YP_003766.2 | H9EJ42 |
| EVFTDGKLNCSFQDAA | 1 | 0 | 0 | 1655682 | YP_009047202.1 | K9N7C7 |
| EVHGNGSMIEDIDYVP | 1 | 0 | 0 | 1655698 | YP_009047202.1 | K9N7C7 |
| EVIECDVKTTEVVGNV | 1 | 0 | 0 | 1655701 | NP_828849.2 | P0C6X7 |
| EVILCWETGKVKPPLN | 1 | 0 | 0 | 1655705 | YP_173236.1 | A0A140H1G9 |

| | | | | | | |
|---|---|---|---|---|---|---|
| EVILKTGELQPLEEPT | 1 | 0 | 0 | 1655706 | QIQ54046.1 | P0C6U8 |
| EVITFDNLKTLLSLRE | 1 | 0 | 0 | 1655709 | YP_009724389.1 | P0DTD1 |
| EVKAQSSGVIPENPVL | 1 | 0 | 0 | 1655711 | YP_173236.1 | A0A140H1G9 |
| EVKDNDGKNVHLKDVT | 1 | 0 | 0 | 1655714 | NP_073549.1 | A0A0P0LKV0 |
| EVKFNAPALQEAYYRA | 1 | 0 | 0 | 1655716 | NP_828849.2 | P0C6X7 |
| EVKPFITESKPSVEQR | 1 | 0 | 0 | 1655722 | YP_009724389.1 | P0DTD1 |
| EVKTIKVFTTVDNTNL | 1 | 0 | 0 | 1655727 | NP_828849.2 | P0C6X7 |
| EVLLDVCNTKEVKVFV | 1 | 0 | 0 | 1655738 | NP_073549.1 | A0A0P0LKV0 |
| EVLSDRELHLSWEVGK | 1 | 0 | 0 | 1655742 | YP_009724389.1 | P0DTD1 |
| EVLSDRELYLSWEVGK | 1 | 0 | 0 | 1655743 | QIQ54047.1 | P0C6X7 |
| EVLSEEVILKTGELQP | 1 | 0 | 0 | 1655745 | QIQ54046.1 | P0C6U8 |
| EVLTEEVVLKTGDLQP | 1 | 0 | 0 | 1655747 | YP_009724389.1 | P0DTD1 |
| EVNELLDTTQLQVANS | 1 | 0 | 0 | 1655757 | YP_009555241.1 | P36334 |
| EVNGLLDTTQLHVADT | 1 | 0 | 0 | 1655758 | YP_173238.1 | A0A140H1H1 |
| EVNSIFDIETVDVKHD | 1 | 0 | 0 | 1655762 | NP_073549.1 | A0A0P0LKV0 |
| EVPPQVVKLPSAPQTI | 1 | 0 | 0 | 1655767 | YP_009047202.1 | K9N7C7 |
| EVPRRNVATLQAENVT | 1 | 0 | 0 | 1655769 | QIQ54047.1 | P0C6X7 |
| EVPVSIINNTVYTKVD | 1 | 0 | 0 | 1655771 | YP_009724389.1 | P0DTD1 |
| EVQGYKNVNITFELDE | 1 | 0 | 0 | 1655775 | QIQ54046.1 | P0C6U8 |
| EVRASRQLAQQKVNEC | 1 | 0 | 0 | 1655783 | NP_073551.1 | A0A0P0K6L9 |
| EVRGSRRLAQQKINEC | 1 | 0 | 0 | 1655785 | YP_003767.1 | A0A0P0G321 |
| EVRQKILNKPRQKRSP | 1 | 0 | 0 | 1655787 | YP_009555245.1 | P33469 |
| EVSMCTNYDLSIINAR | 1 | 0 | 0 | 1655795 | YP_009047202.1 | K9N7C7 |
| EVSQFQSKLTDVKCAN | 1 | 0 | 0 | 1655798 | YP_009555238.1 | P0C6X6 |
| EVSQIQSKLTDVKCAN | 1 | 0 | 0 | 1655799 | YP_173236.1 | A0A140H1G9 |
| EVSRELGNITKTNVQP | 1 | 0 | 0 | 1655800 | NP_073549.1 | A0A0P0LKV0 |
| EVSRQVIQDVKDGTVT | 1 | 0 | 0 | 1655801 | NP_073552.1 | A0A0P0KWE8 |
| EVTFLEGDSHDTVLTS | 1 | 0 | 0 | 1655807 | NP_828849.2 | P0C6X7 |
| EVTGDSCNNFMLTYNK | 1 | 0 | 0 | 1655808 | NP_828849.2 | P0C6X7 |
| EVTGDSCNNYMLTYNK | 1 | 0 | 0 | 1655809 | YP_009724389.1 | P0DTD1 |
| EVTTTLEETKFLTENL | 1 | 0 | 0 | 1655813 | YP_009724389.1 | P0DTD1 |
| EVTTTLEETKFLTNKL | 1 | 0 | 0 | 1655814 | NP_828849.2 | P0C6X7 |
| EVVENPTIQKDILECN | 1 | 0 | 0 | 1655817 | QHR63299.1 | P0C6W2 |
| EVVENPTIQKEVIECD | 1 | 0 | 0 | 1655819 | NP_828849.2 | P0C6X7 |
| EVVGDVILKPVQDGVK | 1 | 0 | 0 | 1655825 | QIQ54046.1 | P0C6U8 |
| EVVGPKELLLSWESGK | 1 | 0 | 0 | 1655827 | YP_003766.2 | H9EJ42 |
| EVVHDQVNYVPLRATN | 1 | 0 | 0 | 1655828 | NP_073549.1 | A0A0P0LKV0 |
| EVVLKTGDLQPLEQPT | 1 | 0 | 0 | 1655834 | YP_009724389.1 | P0DTD1 |
| EVVVNNLDKSAGHPFN | 1 | 0 | 0 | 1655839 | YP_009047202.1 | K9N7C7 |
| EVVVNPANGHMAHGGG | 1 | 0 | 0 | 1655840 | YP_009555238.1 | P0C6X6 |
| EVVVTNLNKSAGWPLN | 1 | 0 | 0 | 1655841 | NP_073549.1 | A0A0P0LKV0 |
| EVVYSNGGFYFKPGGT | 1 | 0 | 0 | 1655843 | NP_073549.1 | A0A0P0LKV0 |
| EVYVHLRNWNFSWNLI | 1 | 0 | 0 | 1655847 | YP_003770.1 | U3M6Q8 |
| EWAAAVDEAFPLDEAE | 1 | 0 | 0 | 1655849 | YP_009047202.1 | K9N7C7 |
| EWAMANGYTSVSSVEC | 1 | 0 | 0 | 1655851 | YP_003766.2 | H9EJ42 |
| EWAYCVDLHNKINLCD | 1 | 0 | 0 | 1655853 | YP_003766.2 | H9EJ42 |
| EWIELFPHNDRIKSFS | 1 | 0 | 0 | 1655864 | NP_073549.1 | A0A0P0LKV0 |
| EWKFYDAQPCSDKAYK | 1 | 0 | 0 | 1655867 | YP_009724389.1 | P0DTD1 |
| EWMDDFEADPKGKYAQ | 1 | 0 | 0 | 1655868 | YP_009047202.1 | K9N7C7 |
| EWPLSVQQAQQEATLP | 1 | 0 | 0 | 1655872 | NP_073549.1 | A0A0P0LKV0 |
| EWPLTEYDFTDYKVQL | 1 | 0 | 0 | 1655873 | YP_009047202.1 | K9N7C7 |
| EWSYTGSSFYAPEPIT | 1 | 0 | 0 | 1655877 | YP_009047204.1 | K9N5Q8 |
| EYDFLNMDMGVFINKY | 1 | 0 | 0 | 1655895 | YP_003766.2 | H9EJ42 |
| EYDFTDYKVQLFEKYF | 1 | 0 | 0 | 1655896 | YP_009047202.1 | K9N7C7 |
| EYDYVIFTQTTETAHS | 1 | 0 | 0 | 1655899 | YP_009724389.1 | P0DTD1 |
| EYDYVIYSQTAETAHS | 1 | 0 | 0 | 1655901 | YP_009555238.1 | P0C6X6 |
| EYENAVANGSSPQIIK | 1 | 0 | 0 | 1655903 | NP_073549.1 | A0A0P0LKV0 |
| EYEVAKKNLADAKNSG | 1 | 0 | 0 | 1655904 | YP_173236.1 | A0A140H1G9 |
| EYGGHHDFVIKTSHDD | 1 | 0 | 0 | 1655908 | YP_173238.1 | A0A140H1H1 |
| EYGSDTVTYKSTVTTK | 1 | 0 | 0 | 1655910 | YP_003766.2 | H9EJ42 |
| EYGTESDYKGLPLEFG | 1 | 0 | 0 | 1655915 | QIQ54046.1 | P0C6U8 |
| EYGTFCDNINSILDEV | 1 | 0 | 0 | 1655916 | YP_173238.1 | A0A140H1H1 |
| EYHNESGIKTTLRKGG | 1 | 0 | 0 | 1655918 | QIQ54046.1 | P0C6U8 |
| EYHNESGLKPILRKGG | 1 | 0 | 0 | 1655919 | QHR63299.1 | P0C6W2 |
| EYHNESGLKTILRKGG | 1 | 0 | 0 | 1655920 | YP_009724389.1 | P0DTD1 |
| EYIDVNKTLQELSYKL | 1 | 0 | 0 | 1655921 | NP_073551.1 | A0A0P0K6L9 |
| EYIFERIDYSDAVSYK | 1 | 0 | 0 | 1655922 | YP_009047202.1 | K9N7C7 |
| EYIKKLYNDLGTQILD | 1 | 0 | 0 | 1655923 | YP_173236.1 | A0A140H1G9 |
| EYIQTTIQKVTVDCKQ | 1 | 0 | 0 | 1655925 | YP_009047204.1 | K9N5Q8 |
| EYKGPITDIFYKENSY | 1 | 0 | 0 | 1655931 | QHR63299.1 | P0C6W2 |
| EYKGPITDVFYKENSY | 1 | 0 | 0 | 1655932 | YP_009724389.1 | P0DTD1 |
| EYKGPVTDVFYKENSY | 1 | 0 | 0 | 1655933 | QIQ54046.1 | P0C6U8 |
| EYKGPVTDVFYKETSY | 1 | 0 | 0 | 1655934 | NP_828849.2 | P0C6X7 |
| EYKHWPQIAQFAPSAS | 1 | 0 | 0 | 1655935 | QIQ54056.1 | P59595 |
| EYLESYNTATTAGFTF | 1 | 0 | 0 | 1655941 | YP_009555238.1 | P0C6X6 |
| EYNFIIYDSSDRGQES | 1 | 0 | 0 | 1655948 | YP_009047202.1 | K9N7C7 |
| EYPIIGDELKINAACR | 1 | 0 | 0 | 1655953 | YP_009724389.1 | P0DTD1 |
| EYPIIGDELRVNSACR | 1 | 0 | 0 | 1655954 | NP_828849.2 | P0C6X7 |
| EYPIISNELSINTSCR | 1 | 0 | 0 | 1655955 | YP_009555238.1 | P0C6X6 |
| EYPIISNEVSINTSCR | 1 | 0 | 0 | 1655956 | YP_173236.1 | A0A140H1G9 |
| EYPLLKQKDIGIDTAG | 1 | 0 | 0 | 1655957 | YP_173243.1 | Q0ZJ80 |
| EYREYLESYNTATTAG | 1 | 0 | 0 | 1655964 | YP_009555238.1 | P0C6X6 |
| EYRLYLDAYNMMISAG | 1 | 0 | 0 | 1655968 | YP_009724389.1 | P0DTD1 |
| EYRQYLDAYNMMISAG | 1 | 0 | 0 | 1655969 | NP_828849.2 | P0C6X7 |
| EYSEEEQPEQPTQYEY | 1 | 0 | 0 | 1655971 | QIQ54046.1 | P0C6U8 |
| EYSWNKKLYELVQRFS | 1 | 0 | 0 | 1655975 | NP_073549.1 | A0A0P0LKV0 |
| EYTFEKGDYGDAVVYR | 1 | 0 | 0 | 1655978 | YP_009724389.1 | P0DTD1 |
| EYTGNYQCGHYKHITS | 1 | 0 | 0 | 1655979 | YP_009724389.1 | P0DTD1 |
| EYTGNYQCGHYTHITA | 1 | 0 | 0 | 1655980 | NP_828849.2 | P0C6X7 |
| EYYHTLDESFLGRYMS | 1 | 0 | 0 | 1655994 | NP_828849.2 | P0C6X7 |
| FAAETIKAKEESVKSS | 1 | 0 | 0 | 1655999 | YP_003766.2 | H9EJ42 |

| | | | | | | |
|---|---|---|---|---|---|---|
| FAAETVKAKEESVKSS | 1 | 0 | 0 | 1656003 | NP_073549.1 | A0A0P0LKV0 |
| FAAKRVLGLQTQTVDS | 1 | 0 | 0 | 1656008 | YP_009555238.1 | P0C6X6 |
| FAAYNRYRIGNYKLNT | 1 | 0 | 0 | 1656012 | NP_828855.1 | P59596 |
| FACGCGASYTANDVEV | 1 | 0 | 0 | 1656014 | YP_009047202.1 | K9N7C7 |
| FADDLNQMSGYKKPAS | 1 | 0 | 0 | 1656023 | QIQ54046.1 | P0C6U8 |
| FADEIIGYKAIEDGVT | 1 | 0 | 0 | 1656025 | QIQ54046.1 | P0C6U8 |
| FADINGKLYHDSQNML | 1 | 0 | 0 | 1656028 | NP_828849.2 | P0C6X7 |
| FADNNDITNTNLSRGR | 1 | 0 | 0 | 1656030 | YP_009047211.1 | K9N4V7 |
| FAEDVCTCYDNSIQGS | 1 | 0 | 0 | 1656035 | NP_073549.1 | A0A0P0LKV0 |
| FAELEAAQKAYQEAMD | 1 | 0 | 0 | 1656037 | YP_009047202.1 | K9N7C7 |
| FAGYTESAVNSTKALA | 1 | 0 | 0 | 1656054 | YP_009047206.1 | K9N4V0 |
| FAINDLNTTVPRISEY | 1 | 0 | 0 | 1656060 | YP_173238.1 | A0A140H1H1 |
| FAIPNGRKVDLQLGNL | 1 | 0 | 0 | 1656061 | YP_009555241.1 | P36334 |
| FAIPSVAVRTYSEAAA | 1 | 0 | 0 | 1656063 | YP_003766.2 | H9EJ42 |
| FAKGDSCSSNCKHSVA | 1 | 0 | 0 | 1656067 | YP_009047202.1 | K9N7C7 |
| FAKRKMGLTPPLSILK | 1 | 0 | 0 | 1656070 | YP_003766.2 | H9EJ42 |
| FAKRKVGLTPPLSILK | 1 | 0 | 0 | 1656071 | NP_073549.1 | A0A0P0LKV0 |
| FALRNFDEKDCEVFKE | 1 | 0 | 0 | 1656085 | NP_073549.1 | A0A0P0LKV0 |
| FAMRHVRGWLGMDVEG | 1 | 0 | 0 | 1656095 | NP_073549.1 | A0A0P0LKV0 |
| FANDTKIASQLGNCVE | 1 | 0 | 0 | 1656099 | YP_009047204.1 | K9N5Q8 |
| FANFILHDVNSGLTCS | 1 | 0 | 0 | 1656100 | YP_009555241.1 | P36334 |
| FANGFVVRIGAAANST | 1 | 0 | 0 | 1656101 | YP_009047204.1 | K9N5Q8 |
| FANKFEQFDQWYESTF | 1 | 0 | 0 | 1656102 | YP_009555238.1 | P0C6X6 |
| FANKFFQFDQWYESTF | 1 | 0 | 0 | 1656103 | YP_173236.1 | A0A140H1G9 |
| FANSVFNICQAVSANV | 1 | 0 | 0 | 1656111 | YP_009555238.1 | P0C6X6 |
| FANSVFNICQAVTANV | 1 | 0 | 0 | 1656112 | YP_173236.1 | A0A140H1G9 |
| FANTGSVCYTPIDEIP | 1 | 0 | 0 | 1656113 | YP_009047202.1 | K9N7C7 |
| FAQDGNAAISDYDYYR | 1 | 0 | 0 | 1656118 | YP_009724389.1 | P0DTD1 |
| FAQTSDTAHACNANRF | 1 | 0 | 0 | 1656124 | NP_073549.1 | A0A0P0LKV0 |
| FASLDVKPYGYPNDFV | 1 | 0 | 0 | 1656139 | YP_003766.2 | H9EJ42 |
| FASVKSSQSSPIIPGF | 1 | 0 | 0 | 1656145 | YP_009047204.1 | K9N5Q8 |
| FATAQEAYEQAVANGD | 1 | 0 | 0 | 1656147 | YP_009724389.1 | P0DTD1 |
| FATLMTFSGNVDLDAL | 1 | 0 | 0 | 1656152 | YP_009047202.1 | K9N7C7 |
| FATYHTPATDCSDGNY | 1 | 0 | 0 | 1656161 | YP_009047204.1 | K9N5Q8 |
| FAVDPAAAYLDAVKQG | 1 | 0 | 0 | 1656165 | NP_073549.1 | A0A0P0LKV0 |
| FAVDPAKAYKDYLASG | 1 | 0 | 0 | 1656166 | NP_828849.2 | P0C6X7 |
| FAVESGGYIPSDFAFN | 1 | 0 | 0 | 1656167 | NP_073551.1 | A0A0P0K6L9 |
| FAVIGDLKCTSDNIND | 1 | 0 | 0 | 1656170 | YP_009555241.1 | P36334 |
| FAYESAVVNGIRVLKT | 1 | 0 | 0 | 1656180 | NP_073549.1 | A0A0P0LKV0 |
| FCDGVNVTKHKCDINY | 1 | 0 | 0 | 1656208 | YP_009555238.1 | P0C6X6 |
| FCDNINSILDEVNGLL | 1 | 0 | 0 | 1656212 | YP_173238.1 | A0A140H1H1 |
| FCEHPAVGIVNTDFKL | 1 | 0 | 0 | 1656215 | NP_073553.1 | A0A0P0KWE8 |
| FCGTENQVKEGPTTCG | 1 | 0 | 0 | 1656231 | QIQ54046.1 | P0C6U8 |
| FCGVDAVNLLTNMFTP | 1 | 0 | 0 | 1656233 | YP_009724389.1 | P0DTD1 |
| FCGVDSVNLLTNMFTP | 1 | 0 | 0 | 1656234 | QIQ54046.1 | P0C6U8 |
| FCQTADTAHANNINRF | 1 | 0 | 0 | 1656273 | YP_009047202.1 | K9N7C7 |
| FCQYKGKWVQVPIGTN | 1 | 0 | 0 | 1656274 | NP_073549.1 | A0A0P0LKV0 |
| FCRQYSNESWFEDDYR | 1 | 0 | 0 | 1656276 | NP_073549.1 | A0A0P0LKV0 |
| FCSETVNGYAKSNFVK | 1 | 0 | 0 | 1656278 | NP_073549.1 | A0A0P0LKV0 |
| FCSQHTLYIKDGDDGY | 1 | 0 | 0 | 1656282 | YP_009047202.1 | K9N7C7 |
| FCSQHTMLVKQGDDYV | 1 | 0 | 0 | 1656285 | YP_009724389.1 | P0DTD1 |
| FCSQHTMQIVDENGKY | 1 | 0 | 0 | 1656286 | NP_073549.1 | A0A0P0LKV0 |
| FCSQHTMQIVDKDGTY | 1 | 0 | 0 | 1656287 | YP_003766.2 | H9EJ42 |
| FCTPTKKAFPYGTCLN | 1 | 0 | 0 | 1656298 | NP_073549.1 | A0A0P0LKV0 |
| FCTQLNRALSGIAAEQ | 1 | 0 | 0 | 1656299 | NP_828851.1 | P59594 |
| FDAEGAHATRDSIGTN | 1 | 0 | 0 | 1656324 | YP_009555238.1 | P0C6X6 |
| FDATNSALVKIQAVVN | 1 | 0 | 0 | 1656331 | YP_009555241.1 | P36334 |
| FDAYVDTFSATFSVPM | 1 | 0 | 0 | 1656333 | NP_828849.2 | P0C6X7 |
| FDCYDGGCINANQVIV | 1 | 0 | 0 | 1656342 | YP_009724389.1 | P0DTD1 |
| FDDQPSLQVESANLML | 1 | 0 | 0 | 1656349 | YP_003766.2 | H9EJ42 |
| FDDSDCDVVQEQVNYV | 1 | 0 | 0 | 1656351 | YP_003766.2 | H9EJ42 |
| FDDVQAPNYTQHTSSM | 1 | 0 | 0 | 1656352 | NP_828851.1 | P59594 |
| FDEAGEEVLAPKLYCA | 1 | 0 | 0 | 1656355 | YP_009555238.1 | P0C6X6 |
| FDEGNCDTLKEILVTY | 1 | 0 | 0 | 1656356 | YP_009724389.1 | P0DTD1 |
| FDEISMATNYDLSVVN | 1 | 0 | 0 | 1656361 | YP_009724389.1 | P0DTD1 |
| FDEQNCDVLKEVLVLT | 1 | 0 | 0 | 1656364 | YP_003766.2 | H9EJ42 |
| FDFSLDVFKIGDVKFK | 1 | 0 | 0 | 1656374 | YP_003766.2 | H9EJ42 |
| FDGKSKCDESASKSAS | 1 | 0 | 0 | 1656382 | NP_828849.2 | P0C6X7 |
| FDGKSKCDESSARAAS | 1 | 0 | 0 | 1656383 | QIQ54046.1 | P0C6U8 |
| FDGKSKCEESSAKSAS | 1 | 0 | 0 | 1656384 | YP_009724389.1 | P0DTD1 |
| FDGRDNGALEAFKRSN | 1 | 0 | 0 | 1656387 | YP_009555238.1 | P0C6X6 |
| FDGVMYGGFEDQPNLQ | 1 | 0 | 0 | 1656394 | NP_073549.1 | A0A0P0LKV0 |
| FDGVTTSDKCIFNSAC | 1 | 0 | 0 | 1656395 | YP_003766.2 | H9EJ42 |
| FDIETVDVKHDVSPFE | 1 | 0 | 0 | 1656402 | NP_073549.1 | A0A0P0LKV0 |
| FDIIQKCQITSVVGTK | 1 | 0 | 0 | 1656404 | YP_173236.1 | A0A140H1G9 |
| FDIKNSSVNTVCFVDV | 1 | 0 | 0 | 1656405 | YP_003766.2 | H9EJ42 |
| FDIVKGQIQVASDNIK | 1 | 0 | 0 | 1656412 | NP_828849.2 | P0C6X7 |
| FDIVVAWHVVRDSRFV | 1 | 0 | 0 | 1656413 | YP_173236.1 | A0A140H1G9 |
| FDKFKVNSTLEQYVFC | 1 | 0 | 0 | 1656423 | YP_009724389.1 | P0DTD1 |
| FDKPQEKDKKSAKTGT | 1 | 0 | 0 | 1656428 | NP_073556.1 | A0A0N7J131 |
| FDKVKLSDVGFVEAYN | 1 | 0 | 0 | 1656437 | YP_009555241.1 | P36334 |
| FDKVVDNVYVTYAGSV | 1 | 0 | 0 | 1656439 | YP_173236.1 | A0A140H1G9 |
| FDKVYDIGNPKGIPIV | 1 | 0 | 0 | 1656440 | YP_009047202.1 | K9N7C7 |
| FDLISKCQITAVEGTK | 1 | 0 | 0 | 1656448 | YP_009555238.1 | P0C6X6 |
| FDLSKFQLKLKGTPVL | 1 | 0 | 0 | 1656456 | YP_009047202.1 | K9N7C7 |
| FDLYVNVYRSTSPDPK | 1 | 0 | 0 | 1656459 | YP_009047202.1 | K9N7C7 |
| FDMAKFPLKLAGTAVI | 1 | 0 | 0 | 1656460 | YP_009555238.1 | P0C6X6 |
| FDNQSKSPSSSGTSTP | 1 | 0 | 0 | 1656479 | YP_003771.1 | H9EJ69 |
| FDNSYLNESGALVPLD | 1 | 0 | 0 | 1656483 | NP_073550.1 | A0A0P0ILY9 |
| FDPIENEDIHRVYAAL | 1 | 0 | 0 | 1656485 | NP_073549.1 | A0A0P0LKV0 |
| FDSEYCRHGTCEKSEA | 1 | 0 | 0 | 1656500 | QIQ54046.1 | P0C6U8 |

| | | | | | | |
|---|---|---|---|---|---|---|
| FDSEYCRHGTCERSEA | 1 | 0 | 0 | 1656501 | YP_009724389.1 | P0DTD1 |
| FDSGTVSKTSDWKCKV | 1 | 0 | 0 | 1656503 | YP_009047202.1 | K9N7C7 |
| FDSHIVSKESGNTVVL | 1 | 0 | 0 | 1656504 | NP_073556.1 | A0A0N7J131 |
| FDSKGWYDPVENEDIH | 1 | 0 | 0 | 1656505 | YP_003766.2 | H9EJ42 |
| FDSLNLSAGTQHISND | 1 | 0 | 0 | 1656508 | YP_173242.1 | Q19U29 |
| FDTFNGECPNFVFPLN | 1 | 0 | 0 | 1656518 | YP_009724389.1 | P0DTD1 |
| FDTRVLNKLNLPGCNG | 1 | 0 | 0 | 1656523 | YP_173236.1 | A0A140H1G9 |
| FDTRVLNNLNLPGCNG | 1 | 0 | 0 | 1656524 | YP_009555238.1 | P0C6X6 |
| FDVAPIELENKFTPLS | 1 | 0 | 0 | 1656531 | YP_009047202.1 | K9N7C7 |
| FDVDKKSLNALIATAH | 1 | 0 | 0 | 1656532 | YP_009555238.1 | P0C6X6 |
| FDVEGAHATRENIGTN | 1 | 0 | 0 | 1656537 | YP_173236.1 | A0A140H1G9 |
| FDVEGCHATREAIGTN | 1 | 0 | 0 | 1656539 | QHR63299.1 | P0C6W2 |
| FDVESGVYSVSSFEAK | 1 | 0 | 0 | 1656542 | YP_009047204.1 | K9N5Q8 |
| FDVINGVKTVKYFECP | 1 | 0 | 0 | 1656545 | YP_173236.1 | A0A140H1G9 |
| FDVIVAWHVVRDPRYV | 1 | 0 | 0 | 1656546 | YP_009555238.1 | P0C6X6 |
| FDVLKSEDAQGMDNLA | 1 | 0 | 0 | 1656548 | YP_009724389.1 | P0DTD1 |
| FDVLKSEDTQGMDNLA | 1 | 0 | 0 | 1656549 | QHR63299.1 | P0C6W2 |
| FDVPDGFYSTSPIQSV | 1 | 0 | 0 | 1656553 | NP_073551.1 | A0A0P0K6L9 |
| FDVYNKDASFIGKNLK | 1 | 0 | 0 | 1656564 | NP_073549.1 | A0A0P0LKV0 |
| FDVYVDTFLSMFDVDK | 1 | 0 | 0 | 1656566 | YP_009555238.1 | P0C6X6 |
| FDVYVDTFMSHFDVDR | 1 | 0 | 0 | 1656567 | YP_173236.1 | A0A140H1G9 |
| FDYHPNCSDCYDDMCV | 1 | 0 | 0 | 1656568 | YP_003766.2 | H9EJ42 |
| FDYLDIHPSLCNNGKI | 1 | 0 | 0 | 1656569 | YP_173237.1 | A0A140H1H0 |
| FEDDVVTSLCKKSFGK | 1 | 0 | 0 | 1656589 | YP_003766.2 | H9EJ42 |
| FEDELAPSDVYQQLAG | 1 | 0 | 0 | 1656592 | YP_173236.1 | A0A140H1G9 |
| FEDELTPSDVYQQLAG | 1 | 0 | 0 | 1656593 | YP_009555238.1 | P0C6X6 |
| FEDLLFDKVKLSDVGF | 1 | 0 | 0 | 1656594 | YP_173238.1 | A0A140H1H1 |
| FEDRSNNSSRASSRSS | 1 | 0 | 0 | 1656596 | YP_003771.1 | H9EJ69 |
| FEEQDEIYAYTKRNVL | 1 | 0 | 0 | 1656602 | YP_009555238.1 | P0C6X6 |
| FEFVEGQGVPIAPGVP | 1 | 0 | 0 | 1656608 | YP_009555245.1 | P33469 |
| FEGVLPINTDTVLSVA | 1 | 0 | 0 | 1656613 | YP_003766.2 | H9EJ42 |
| FEGYDFKYIENGQLKN | 1 | 0 | 0 | 1656614 | NP_073549.1 | A0A0P0LKV0 |
| FEIACKCGVKQEQRTG | 1 | 0 | 0 | 1656623 | YP_009555238.1 | P0C6X6 |
| FEIAQLYGSCITPNVC | 1 | 0 | 0 | 1656624 | YP_009555238.1 | P0C6X6 |
| FEIDGNVMHANYLFWR | 1 | 0 | 0 | 1656625 | YP_173236.1 | A0A140H1G9 |
| FEIDKGIYQTSNFRVV | 1 | 0 | 0 | 1656626 | NP_828851.1 | P59594 |
| FEIKHADLHSSQVCGL | 1 | 0 | 0 | 1656627 | YP_003766.2 | H9EJ42 |
| FEIKLAKKFDNFTGEC | 1 | 0 | 0 | 1656628 | QIQ54046.1 | P0C6U8 |
| FEIKLAKKFDTFNGEC | 1 | 0 | 0 | 1656629 | YP_009724389.1 | P0DTD1 |
| FEIYDGGCIPASQVIV | 1 | 0 | 0 | 1656632 | YP_009555238.1 | P0C6X6 |
| FEIYDGGCLNASEVVV | 1 | 0 | 0 | 1656633 | YP_009047202.1 | K9N7C7 |
| FEKGDYGDAVVYRGTT | 1 | 0 | 0 | 1656636 | YP_009724389.1 | P0DTD1 |
| FEKVDYGSDTVTYKST | 1 | 0 | 0 | 1656641 | NP_073549.1 | A0A0P0LKV0 |
| FEKVEYGSDTVTYKST | 1 | 0 | 0 | 1656642 | YP_003766.2 | H9EJ42 |
| FEKYFKYWDQTYHANC | 1 | 0 | 0 | 1656644 | YP_009047202.1 | K9N7C7 |
| FELDERIDKVLNEKCS | 1 | 0 | 0 | 1656646 | YP_009724389.1 | P0DTD1 |
| FELDERVDKVLNEKCS | 1 | 0 | 0 | 1656647 | NP_828849.2 | P0C6X7 |
| FELFAKRKMGLTPPLS | 1 | 0 | 0 | 1656649 | YP_003766.2 | H9EJ42 |
| FELLNAPATVCGPKLS | 1 | 0 | 0 | 1656654 | NP_828851.1 | P59594 |
| FELWAKRNIKPVPEIK | 1 | 0 | 0 | 1656660 | NP_828849.2 | P0C6X7 |
| FELWAKRNIKPVPEVK | 1 | 0 | 0 | 1656661 | YP_009724389.1 | P0DTD1 |
| FELYAKRAVRSHPDFK | 1 | 0 | 0 | 1656662 | YP_009047202.1 | K9N7C7 |
| FEMPFEELNGLKILKQ | 1 | 0 | 0 | 1656664 | NP_073549.1 | A0A0P0LKV0 |
| FENKTILPVNVAFELW | 1 | 0 | 0 | 1656671 | QIQ54047.1 | P0C6X7 |
| FENKTTLPTNIAFELY | 1 | 0 | 0 | 1656672 | YP_009047202.1 | K9N7C7 |
| FENKTTLPVNVAFELW | 1 | 0 | 0 | 1656673 | YP_009724389.1 | P0DTD1 |
| FENYIKWPWWVWLIIS | 1 | 0 | 0 | 1656676 | YP_003767.1 | A0A0P0G321 |
| FEPFNVSFVNDSIESV | 1 | 0 | 0 | 1656677 | YP_173238.1 | A0A140H1H1 |
| FEPFTVNSVNDSLEPV | 1 | 0 | 0 | 1656678 | YP_009555241.1 | P36334 |
| FEPSTQYEYGTEDDYQ | 1 | 0 | 0 | 1656682 | YP_009724389.1 | P0DTD1 |
| FEQQVGVIADKDKDLS | 1 | 0 | 0 | 1656684 | NP_073549.1 | A0A0P0LKV0 |
| FERGSVQVDNGSSINR | 1 | 0 | 0 | 1656689 | NP_073549.1 | A0A0P0LKV0 |
| FERIDYSDAVSYKSST | 1 | 0 | 0 | 1656690 | YP_009047202.1 | K9N7C7 |
| FETIMKVLEENLNAYV | 1 | 0 | 0 | 1656700 | YP_173242.1 | Q19U29 |
| FETIMKVLNENLNAYQ | 1 | 0 | 0 | 1656701 | YP_009555245.1 | P33469 |
| FEVLAVEDTQGMDNLA | 1 | 0 | 0 | 1656706 | NP_828849.2 | P0C6X7 |
| FEYTSPTDKLAMIMGY | 1 | 0 | 0 | 1656710 | YP_009555239.1 | Q80872 |
| FEYYHTLDESFLGRYM | 1 | 0 | 0 | 1656712 | NP_828849.2 | P0C6X7 |
| FFEDLLFDKVKLSDVG | 1 | 0 | 0 | 1656734 | YP_173238.1 | A0A140H1H1 |
| FFEIKHADLHSSQVCG | 1 | 0 | 0 | 1656736 | YP_003766.2 | H9EJ42 |
| FFEITMTDLQSESSCG | 1 | 0 | 0 | 1656737 | NP_073549.1 | A0A0P0LKV0 |
| FFFTQKGDAAIKDFDY | 1 | 0 | 0 | 1656745 | NP_073549.1 | A0A0P0LKV0 |
| FFIFNKLVKDPPNVQI | 1 | 0 | 0 | 1656761 | NP_828852.2 | P59632 |
| FFIYNKIVDEPEEHVQ | 1 | 0 | 0 | 1656767 | YP_009724391.1 | P0DTC3 |
| FFIYSRIVDEPEDHVQ | 1 | 0 | 0 | 1656768 | QIQ54049.1 | P59632 |
| FFKCTMGVYDFKVSAA | 1 | 0 | 0 | 1656769 | YP_003766.2 | H9EJ42 |
| FFKELTANMSMAECKA | 1 | 0 | 0 | 1656772 | YP_003766.2 | H9EJ42 |
| FFKFRVDGDMVPHISR | 1 | 0 | 0 | 1656774 | NP_828849.2 | P0C6X7 |
| FFKGNVQVDNGSSINR | 1 | 0 | 0 | 1656775 | YP_003766.2 | H9EJ42 |
| FFLSKHSDFGLGDLVD | 1 | 0 | 0 | 1656782 | NP_073549.1 | A0A0P0LKV0 |
| FFPDLNGDVVAIDYKH | 1 | 0 | 0 | 1656792 | YP_009724389.1 | P0DTD1 |
| FFPDLNGDVVAIDYRH | 1 | 0 | 0 | 1656793 | NP_828849.2 | P0C6X7 |
| FFSPESVFSVFGDVKL | 1 | 0 | 0 | 1656810 | YP_003766.2 | H9EJ42 |
| FFSPQIITTDNTFVSG | 1 | 0 | 0 | 1656811 | NP_828851.1 | P59594 |
| FFTCNSNANLSMLQLG | 1 | 0 | 0 | 1656816 | YP_003767.1 | A0A0P0G321 |
| FFTQDGNAAITDYNYY | 1 | 0 | 0 | 1656823 | YP_009555238.1 | P0C6X6 |
| FFTQKGDAAIKDFDYY | 1 | 0 | 0 | 1656824 | NP_073549.1 | A0A0P0LKV0 |
| FGACDYDRLPNEVTVA | 1 | 0 | 0 | 1656866 | YP_009047210.1 | K9N7A1 |
| FGAGSDKGIAPGTSVL | 1 | 0 | 0 | 1656870 | YP_009047202.1 | K9N7C7 |
| FGAISASIGDIIQRLD | 1 | 0 | 0 | 1656873 | YP_009047204.1 | K9N5Q8 |
| FGAPSYCNGVRELYLN | 1 | 0 | 0 | 1656880 | NP_828849.2 | P0C6X7 |

| | | | | | | |
|---|---|---|---|---|---|---|
| FGASSVQQQEEQEEDW | 1 | 0 | 0 | 1656883 | QIQ54046.1 | P0C6U8 |
| FGDDTVIEVQGYKSVN | 1 | 0 | 0 | 1656893 | YP_009724389.1 | P0DTD1 |
| FGDLQLNKLGTEDPRW | 1 | 0 | 0 | 1656903 | YP_009047211.1 | K9N4V7 |
| FGDQELIRQGTEYKHW | 1 | 0 | 0 | 1656910 | QIQ54056.1 | P59595 |
| FGDSRSDCNHINNLKI | 1 | 0 | 0 | 1656911 | YP_173237.1 | A0A140H1H0 |
| FGDSVEEVLSEARQHL | 1 | 0 | 0 | 1656915 | YP_009724389.1 | P0DTD1 |
| FGDYNLSSVLPQRNIR | 1 | 0 | 0 | 1656919 | YP_003767.1 | A0A0P0G321 |
| FGEDTVWEVQGYKNVR | 1 | 0 | 0 | 1656924 | NP_828849.2 | P0C6X7 |
| FGEVFNATKFPSVYAW | 1 | 0 | 0 | 1656927 | NP_828851.1 | P59594 |
| FGGDQVHEVAAVRSVT | 1 | 0 | 0 | 1656947 | YP_009047202.1 | K9N7C7 |
| FGGFNFLQILPDPSKP | 1 | 0 | 0 | 1656948 | QIQ54048.1 | P59594 |
| FGGFNFSQILPDPLKP | 1 | 0 | 0 | 1656949 | NP_828851.1 | P59594 |
| FGGKQILGYSSLNDEF | 1 | 0 | 0 | 1656953 | NP_073549.1 | A0A0P0LKV0 |
| FGGPSDSTDNNQNGDR | 1 | 0 | 0 | 1656955 | QIQ54056.1 | P59595 |
| FGGPSDSTGSNQNGER | 1 | 0 | 0 | 1656956 | YP_009724397.2 | P0DTC9 |
| FGGTTVVRPLVEDSTS | 1 | 0 | 0 | 1656959 | YP_009047210.1 | K9N7A1 |
| FGGVSVITPGTNASNQ | 1 | 0 | 0 | 1656961 | QHR63300.2 | Q3LZX1 |
| FGGVSVITPGTNASSE | 1 | 0 | 0 | 1656962 | NP_828851.1 | P59594 |
| FGHVYDFKVEDAYDQV | 1 | 0 | 0 | 1656965 | YP_009555238.1 | P0C6X6 |
| FGKDLNANMSLAECKR | 1 | 0 | 0 | 1656977 | NP_073549.1 | A0A0P0LKV0 |
| FGKRGPNQNFGGGEML | 1 | 0 | 0 | 1656983 | YP_009555245.1 | P33469 |
| FGKRGPSQNFGNAEML | 1 | 0 | 0 | 1656984 | YP_173242.1 | Q19U29 |
| FGLGDLVDSYFENDSI | 1 | 0 | 0 | 1656990 | NP_073549.1 | A0A0P0LKV0 |
| FGLQDCVTGINDDDYV | 1 | 0 | 0 | 1656993 | YP_003766.2 | H9EJ42 |
| FGLRGPGDLQGNFGDL | 1 | 0 | 0 | 1656995 | YP_009047211.1 | K9N4V7 |
| FGLVTICDFYSKLGVG | 1 | 0 | 0 | 1656999 | YP_173236.1 | A0A140H1G9 |
| FGLYKNSATGSLDVEV | 1 | 0 | 0 | 1657002 | YP_003766.2 | H9EJ42 |
| FGLYKQSCTGSPYIDD | 1 | 0 | 0 | 1657004 | YP_009555238.1 | P0C6X6 |
| FGMIKKLDNDVLNGII | 1 | 0 | 0 | 1657007 | YP_009047202.1 | K9N7C7 |
| FGMLNGNAIATVKSED | 1 | 0 | 0 | 1657009 | NP_073549.1 | A0A0P0LKV0 |
| FGPFNFCSETVNGYAK | 1 | 0 | 0 | 1657022 | NP_073549.1 | A0A0P0LKV0 |
| FGPGNTFINGDIAREL | 1 | 0 | 0 | 1657023 | YP_003766.2 | H9EJ42 |
| FGPRDLDHNFGSAGVV | 1 | 0 | 0 | 1657031 | NP_073556.1 | A0A0N7J131 |
| FGPTYLDGADVTKIKP | 1 | 0 | 0 | 1657033 | YP_009724389.1 | P0DTD1 |
| FGRDISDTTDAVRDPQ | 1 | 0 | 0 | 1657037 | QIQ54048.1 | P59594 |
| FGRRGPEQTQGNFGDQ | 1 | 0 | 0 | 1657039 | YP_009724397.2 | P0DTC9 |
| FGSAGVVANGVKAKGY | 1 | 0 | 0 | 1657040 | NP_073556.1 | A0A0N7J131 |
| FGSRLELAKVQNLSGN | 1 | 0 | 0 | 1657051 | YP_009555245.1 | P33469 |
| FGSVACEHISSTMSQY | 1 | 0 | 0 | 1657054 | YP_009047204.1 | K9N5Q8 |
| FGTVYEKLKPVLDWLE | 1 | 0 | 0 | 1657066 | YP_009724389.1 | P0DTD1 |
| FGTVYEQLKPVLDWLE | 1 | 0 | 0 | 1657067 | QHR63299.1 | P0C6W2 |
| FGVFDIKNSSVNTVCF | 1 | 0 | 0 | 1657073 | YP_003766.2 | H9EJ42 |
| FGVNKIQPWTNDALNT | 1 | 0 | 0 | 1657078 | NP_073549.1 | A0A0P0LKV0 |
| FGVTNLEDLCLKYNYC | 1 | 0 | 0 | 1657085 | YP_173239.1 | Q0ZJ84 |
| FGVTQEYIQTTIQKVT | 1 | 0 | 0 | 1657086 | YP_009047204.1 | K9N5Q8 |
| FGWIKISADNKVIYTT | 1 | 0 | 0 | 1657091 | YP_009555238.1 | P0C6X6 |
| FGYIDCEQDSCDFKGW | 1 | 0 | 0 | 1657098 | YP_009555238.1 | P0C6X6 |
| FGYVQQGIFNKATVAQ | 1 | 0 | 0 | 1657103 | YP_009555238.1 | P0C6X6 |
| FHANSSEPALLFRNIK | 1 | 0 | 0 | 1657106 | YP_009555241.1 | P36334 |
| FHIEGTGGNSQSSSRA | 1 | 0 | 0 | 1657113 | YP_009047211.1 | K9N4V7 |
| FHLDGEVITFDNLKTL | 1 | 0 | 0 | 1657115 | YP_009724389.1 | P0DTD1 |
| FHLDGEVLSLDKLKSL | 1 | 0 | 0 | 1657116 | NP_828849.2 | P0C6X7 |
| FHMDGEVINIDSLKTL | 1 | 0 | 0 | 1657121 | QIQ54046.1 | P0C6U8 |
| FHQECSLQSCAQHQPY | 1 | 0 | 0 | 1657126 | QHR63307.1 | Q3LZX5 |
| FHQKLLKSIAATRGAT | 1 | 0 | 0 | 1657131 | YP_009724389.1 | P0DTD1 |
| FHSVEIEAGEVKPFAV | 1 | 0 | 0 | 1657137 | YP_003766.2 | H9EJ42 |
| FHSVLTTFIDAARGPA | 1 | 0 | 0 | 1657138 | YP_009047202.1 | K9N7C7 |
| FHTPAFDKSAFTNLKQ | 1 | 0 | 0 | 1657143 | NP_828849.2 | P0C6X7 |
| FHTPAFDKSAFVNLKQ | 1 | 0 | 0 | 1657144 | YP_009724389.1 | P0DTD1 |
| FHYERDNTSCPEWMDD | 1 | 0 | 0 | 1657151 | YP_009047202.1 | K9N7C7 |
| FIANEKFINGCGRNVQ | 1 | 0 | 0 | 1657161 | YP_003766.2 | H9EJ42 |
| FIAQSKGLYRQCIRGK | 1 | 0 | 0 | 1657162 | NP_828849.2 | P0C6X7 |
| FICEEVANDLTTALRR | 1 | 0 | 0 | 1657169 | YP_009047202.1 | K9N7C7 |
| FIDAARGPAGVESDVE | 1 | 0 | 0 | 1657175 | YP_009047202.1 | K9N7C7 |
| FIDTKRGVYCCREHEH | 1 | 0 | 0 | 1657188 | YP_009724389.1 | P0DTD1 |
| FIDVVNKALEMCIDQV | 1 | 0 | 0 | 1657189 | NP_828849.2 | P0C6X7 |
| FIFEKVEYGSDTVTYK | 1 | 0 | 0 | 1657201 | YP_003766.2 | H9EJ42 |
| FIFNKLVKDPPNVQIH | 1 | 0 | 0 | 1657203 | NP_828852.2 | P59632 |
| FIGATIRLQAGKQTEL | 1 | 0 | 0 | 1657205 | YP_003766.2 | H9EJ42 |
| FIGVEGELPVAVVNDK | 1 | 0 | 0 | 1657210 | YP_009047202.1 | K9N7C7 |
| FIIYDSSDRGQESLAR | 1 | 0 | 0 | 1657222 | YP_009047202.1 | K9N7C7 |
| FIKGLNNLNRGMVLGS | 1 | 0 | 0 | 1657228 | YP_009724389.1 | P0DTD1 |
| FIKTNNTRIVDEWSYT | 1 | 0 | 0 | 1657239 | YP_009047204.1 | K9N5Q8 |
| FINGACGSPGYNLKNG | 1 | 0 | 0 | 1657253 | NP_073549.1 | A0A0P0LKV0 |
| FINGACGSPGYNVRND | 1 | 0 | 0 | 1657254 | YP_003766.2 | H9EJ42 |
| FINGIFAKVKNTKVIK | 1 | 0 | 0 | 1657257 | YP_009555241.1 | P36334 |
| FINKYGLEDFNFEHVV | 1 | 0 | 0 | 1657258 | YP_003766.2 | H9EJ42 |
| FINNTTYPTNVAVELF | 1 | 0 | 0 | 1657260 | YP_009555238.1 | P0C6X6 |
| FIQTAPGFGVAVADSY | 1 | 0 | 0 | 1657273 | YP_173236.1 | A0A140H1G9 |
| FIQTTPGSGVPIVDSY | 1 | 0 | 0 | 1657276 | QHR63299.1 | P0C6W2 |
| FIRQEEVQELYSPLFL | 1 | 0 | 0 | 1657283 | QIQ54053.1 | P59635 |
| FIRQEEVQQELYSPLF | 1 | 0 | 0 | 1657284 | NP_828857.1 | P59635 |
| FISPYNSQNAVAAKIL | 1 | 0 | 0 | 1657297 | QIQ54047.1 | P0C6X7 |
| FISPYNSQNAVASKIL | 1 | 0 | 0 | 1657298 | YP_009724389.1 | P0DTD1 |
| FISPYNSQNFAAKRVL | 1 | 0 | 0 | 1657300 | YP_009555238.1 | P0C6X6 |
| FISPYNSQNYVAKRVL | 1 | 0 | 0 | 1657302 | YP_173236.1 | A0A140H1G9 |
| FISVLDITDAAVKAAE | 1 | 0 | 0 | 1657308 | NP_073549.1 | A0A0P0LKV0 |
| FITESKPSVEQRKQDD | 1 | 0 | 0 | 1657315 | YP_009724389.1 | P0DTD1 |
| FITKDEAIKRVRGWVG | 1 | 0 | 0 | 1657321 | YP_173236.1 | A0A140H1G9 |
| FITMSSKGQKGDAEEA | 1 | 0 | 0 | 1657329 | YP_003766.2 | H9EJ42 |
| FITPEVSRELGNITKT | 1 | 0 | 0 | 1657331 | NP_073549.1 | A0A0P0LKV0 |

| | | | | | | |
|---|---|---|---|---|---|---|
| FITQRNFYEPQIITTD | 1 | 0 | 0 | 1657334 | QIQ54048.1 | P59594 |
| FITREEAIKHVRAWVG | 1 | 0 | 0 | 1657335 | QIQ54047.1 | P0C6X7 |
| FITTCSCEIIGGQLVA | 1 | 0 | 0 | 1657339 | QIQ54046.1 | P0C6U8 |
| FITVEAAIELSKELKR | 1 | 0 | 0 | 1657340 | YP_173236.1 | A0A140H1G9 |
| FITVEAALDLSKELKR | 1 | 0 | 0 | 1657341 | YP_009555238.1 | P0C6X6 |
| FIVNAANENLAHGGGL | 1 | 0 | 0 | 1657354 | NP_073549.1 | A0A0P0LKV0 |
| FIYNKIVDEPEEHVQI | 1 | 0 | 0 | 1657364 | YP_009724391.1 | P0DTC3 |
| FIYSRIVDEPEDHVQI | 1 | 0 | 0 | 1657365 | QIQ54049.1 | P59632 |
| FKCTMGVYDFKVSAAE | 1 | 0 | 0 | 1657377 | YP_003766.2 | H9EJ42 |
| FKCYGVSATKLNDLCF | 1 | 0 | 0 | 1657378 | NP_828851.1 | P59594 |
| FKDAAGKPVPYCYDTN | 1 | 0 | 0 | 1657380 | QIQ54046.1 | P0C6U8 |
| FKDAMGKPVPYCYDTN | 1 | 0 | 0 | 1657381 | NP_828849.2 | P0C6X7 |
| FKDASGKPVPYCYDTN | 1 | 0 | 0 | 1657382 | YP_009724389.1 | P0DTD1 |
| FKDCARNPIDLPPSHA | 1 | 0 | 0 | 1657384 | NP_073549.1 | A0A0P0LKV0 |
| FKDCSKSCLGYHPAHA | 1 | 0 | 0 | 1657386 | YP_173236.1 | A0A140H1G9 |
| FKDCSKSYSGYHPAHA | 1 | 0 | 0 | 1657387 | YP_009555238.1 | P0C6X6 |
| FKDCSRETSGLSPAYA | 1 | 0 | 0 | 1657389 | YP_009047202.1 | K9N7C7 |
| FKDFQFMLWCNDNKIM | 1 | 0 | 0 | 1657392 | YP_173236.1 | A0A140H1G9 |
| FKDGIYFAATEKSNVV | 1 | 0 | 0 | 1657394 | NP_828851.1 | P59594 |
| FKEGIEFLKDGWEIVK | 1 | 0 | 0 | 1657403 | QIQ54046.1 | P0C6U8 |
| FKEGVEFLRDGWEIVK | 1 | 0 | 0 | 1657407 | YP_009724389.1 | P0DTD1 |
| FKELLVYAADPAMHAA | 1 | 0 | 0 | 1657409 | YP_009724389.1 | P0DTD1 |
| FKELTANMSMAECKAT | 1 | 0 | 0 | 1657410 | YP_003766.2 | H9EJ42 |
| FKFSDGQGVPIAFGVP | 1 | 0 | 0 | 1657425 | YP_173242.1 | Q19U29 |
| FKFVCDNIKLADDLNQ | 1 | 0 | 0 | 1657427 | QHR63299.1 | P0C6W2 |
| FKFVCDNTKFADDLNQ | 1 | 0 | 0 | 1657428 | QIQ54046.1 | P0C6U8 |
| FKGNVQVDNGSSINRK | 1 | 0 | 0 | 1657435 | YP_003766.2 | H9EJ42 |
| FKGWIPGNMIDGFACT | 1 | 0 | 0 | 1657438 | YP_009555238.1 | P0C6X6 |
| FKHWSMPYHPNTVDCQ | 1 | 0 | 0 | 1657448 | YP_009555238.1 | P0C6X6 |
| FKIGDVKFKRLGDYVL | 1 | 0 | 0 | 1657450 | YP_003766.2 | H9EJ42 |
| FKKCGGCCDDYTGYQE | 1 | 0 | 0 | 1657462 | YP_009555241.1 | P36334 |
| FKKVDGIIQQLPETYF | 1 | 0 | 0 | 1657471 | NP_828849.2 | P0C6X7 |
| FKKVETLNFQMVKEFP | 1 | 0 | 0 | 1657472 | YP_173243.1 | Q0ZJ80 |
| FKLKISTQLFEGDKFI | 1 | 0 | 0 | 1657482 | YP_003766.2 | H9EJ42 |
| FKLLGIGGVPIIEVSQ | 1 | 0 | 0 | 1657484 | YP_009555238.1 | P0C6X6 |
| FKLTCSNTKFADDLNQ | 1 | 0 | 0 | 1657499 | NP_828849.2 | P0C6X7 |
| FKMNYQVNGYPNMFIT | 1 | 0 | 0 | 1657510 | YP_009724389.1 | P0DTD1 |
| FKNDNVVLKITEDGIN | 1 | 0 | 0 | 1657515 | YP_003766.2 | H9EJ42 |
| FKNVDKDDAFYIVKRC | 1 | 0 | 0 | 1657526 | NP_073549.1 | A0A0P0LKV0 |
| FKPGGTTSGDASTAYA | 1 | 0 | 0 | 1657530 | YP_003766.2 | H9EJ42 |
| FKPQSGGGKCFNCYPA | 1 | 0 | 0 | 1657540 | NP_073551.1 | A0A0P0K6L9 |
| FKPRSQMETDFLELAM | 1 | 0 | 0 | 1657543 | NP_828849.2 | P0C6X7 |
| FKPVQLVPVSPAFHRI | 1 | 0 | 0 | 1657544 | YP_009047208.1 | K9N7D2 |
| FKQANNNCFVNVSCLM | 1 | 0 | 0 | 1657546 | YP_009555238.1 | P0C6X6 |
| FKRGDGTPHPYCYSDG | 1 | 0 | 0 | 1657551 | YP_173236.1 | A0A140H1G9 |
| FKSENRYSVLSVDSVS | 1 | 0 | 0 | 1657560 | YP_173236.1 | A0A140H1G9 |
| FKSISLTVGEVFGKIL | 1 | 0 | 0 | 1657563 | YP_173236.1 | A0A140H1G9 |
| FKSNFSQYYYNIDTGS | 1 | 0 | 0 | 1657567 | YP_173237.1 | A0A140H1H0 |
| FKTADGNQRQLLPRWY | 1 | 0 | 0 | 1657571 | YP_009555245.1 | P33469 |
| FKTADGQQKQLLPRWY | 1 | 0 | 0 | 1657572 | YP_173242.1 | Q19U29 |
| FKTFDLLKYDFTEHKE | 1 | 0 | 0 | 1657575 | NP_073549.1 | A0A0P0LKV0 |
| FKTFEKVDGVYTNFKL | 1 | 0 | 0 | 1657576 | YP_009555238.1 | P0C6X6 |
| FKTLKPGDSFNILACY | 1 | 0 | 0 | 1657580 | YP_003766.2 | H9EJ42 |
| FKTSGDLAVQIGNNNV | 1 | 0 | 0 | 1657582 | NP_073549.1 | A0A0P0LKV0 |
| FKTTGDLAVQIGSNNV | 1 | 0 | 0 | 1657584 | YP_003766.2 | H9EJ42 |
| FKVEDAYDQVHDEPKG | 1 | 0 | 0 | 1657587 | YP_009555238.1 | P0C6X6 |
| FKVFFKGNVQVDNGSS | 1 | 0 | 0 | 1657589 | YP_003766.2 | H9EJ42 |
| FKVGKVDSIVQKCYEL | 1 | 0 | 0 | 1657590 | YP_003766.2 | H9EJ42 |
| FKVLDNGIIRDVNPDD | 1 | 0 | 0 | 1657592 | YP_009047202.1 | K9N7C7 |
| FKVNETNSQYLFSTIN | 1 | 0 | 0 | 1657593 | YP_009047202.1 | K9N7C7 |
| FKVQNVNLERCELANY | 1 | 0 | 0 | 1657596 | YP_009047202.1 | K9N7C7 |
| FKVYFKGQTTHESSSA | 1 | 0 | 0 | 1657603 | YP_173236.1 | A0A140H1G9 |
| FKWAPEFRWLLPDAAE | 1 | 0 | 0 | 1657605 | YP_173236.1 | A0A140H1G9 |
| FKYMVANGLNAPNGPF | 1 | 0 | 0 | 1657616 | NP_073549.1 | A0A0P0LKV0 |
| FKYWDQTYHANCVNCT | 1 | 0 | 0 | 1657618 | YP_009047202.1 | K9N7C7 |
| FKYWDQTYHPNCINCL | 1 | 0 | 0 | 1657619 | NP_828849.2 | P0C6X7 |
| FKYWDQTYHPNCVNCL | 1 | 0 | 0 | 1657620 | YP_009724389.1 | P0DTD1 |
| FKYWGMKYHPNTVDCD | 1 | 0 | 0 | 1657621 | YP_173236.1 | A0A140H1G9 |
| FKYWGQDYHPDCVDCH | 1 | 0 | 0 | 1657622 | NP_073549.1 | A0A0P0LKV0 |
| FLAVDDKYKATGDLAV | 1 | 0 | 0 | 1657643 | YP_009555238.1 | P0C6X6 |
| FLAVDDKYKVNENLAV | 1 | 0 | 0 | 1657644 | YP_173236.1 | A0A140H1G9 |
| FLDDNVLPETYVALPI | 1 | 0 | 0 | 1657665 | YP_003767.1 | A0A0P0G321 |
| FLDDVEMVAYNPDLSQ | 1 | 0 | 0 | 1657666 | YP_173236.1 | A0A140H1G9 |
| FLDKVDILLTVDGVNF | 1 | 0 | 0 | 1657670 | YP_009555238.1 | P0C6X6 |
| FLDLYYEMNLIQESIK | 1 | 0 | 0 | 1657673 | YP_173238.1 | A0A140H1H1 |
| FLDTVVSTVKIGGLTF | 1 | 0 | 0 | 1657676 | NP_073549.1 | A0A0P0LKV0 |
| FLEELNAFTREMQQHP | 1 | 0 | 0 | 1657677 | NP_073556.1 | A0A0N7J131 |
| FLELAMDEFIERYKLE | 1 | 0 | 0 | 1657682 | YP_009724389.1 | P0DTD1 |
| FLELAMDEFIQRYKLE | 1 | 0 | 0 | 1657683 | NP_828849.2 | P0C6X7 |
| FLEMKSEKQVEQKIAE | 1 | 0 | 0 | 1657685 | YP_009724389.1 | P0DTD1 |
| FLEMKSEKQVEQKTAE | 1 | 0 | 0 | 1657686 | QHR63299.1 | P0C6W2 |
| FLEMKSETKVQDHFDV | 1 | 0 | 0 | 1657687 | QIQ54046.1 | P0C6U8 |
| FLFDEAGEEVLAPKLY | 1 | 0 | 0 | 1657693 | YP_009555238.1 | P0C6X6 |
| FLFGDSRSDCNHINNL | 1 | 0 | 0 | 1657695 | YP_173237.1 | A0A140H1H0 |
| FLFGDSRSDCNHIVNI | 1 | 0 | 0 | 1657696 | YP_009555240.1 | P30215 |
| FLGASVTEDVKFAAST | 1 | 0 | 0 | 1657709 | YP_003766.2 | H9EJ42 |
| FLGLQIQTVDSSQGSE | 1 | 0 | 0 | 1657715 | YP_003766.2 | H9EJ42 |
| FLGVYYHKNNKSWMES | 1 | 0 | 0 | 1657724 | YP_009724390.1 | P0DTC2 |
| FLHTVLLPTQYKDVEA | 1 | 0 | 0 | 1657738 | NP_073551.1 | A0A0P0K6L9 |
| FLKRGDKIVYHTLESP | 1 | 0 | 0 | 1657771 | NP_828849.2 | P0C6X7 |
| FLKRGDKSVYYTSNPI | 1 | 0 | 0 | 1657772 | QHR63299.1 | P0C6W2 |

| | | | | | | |
|---|---|---|---|---|---|---|
| FLKRGDKSVYYTSNPT | 1 | 0 | 0 | 1657773 | YP_009724389.1 | P0DTD1 |
| FLKRGDKVVYHTTSKP | 1 | 0 | 0 | 1657774 | QIQ54046.1 | P0C6U8 |
| FLKTNCCRFQEKDEDG | 1 | 0 | 0 | 1657776 | QIQ54047.1 | P0C6X7 |
| FLKTNCCRFQEKDEEG | 1 | 0 | 0 | 1657777 | NP_828849.2 | P0C6X7 |
| FLLGINYLGTIKENID | 1 | 0 | 0 | 1657788 | YP_009047202.1 | K9N7C7 |
| FLLSYNQNGTITDAVD | 1 | 0 | 0 | 1657803 | QIQ54048.1 | P59594 |
| FLLTNTSSVVDGVVRS | 1 | 0 | 0 | 1657807 | NP_073551.1 | A0A0P0K6L9 |
| FLNGITYTSTSGNLLG | 1 | 0 | 0 | 1657820 | NP_073551.1 | A0A0P0K6L9 |
| FLNGSCGSVGFNIDYD | 1 | 0 | 0 | 1657821 | YP_009724389.1 | P0DTD1 |
| FLNRFTTTLNDFNLVA | 1 | 0 | 0 | 1657831 | YP_009724389.1 | P0DTD1 |
| FLNRVRGTSVDARLVP | 1 | 0 | 0 | 1657834 | YP_009555238.1 | P0C6X6 |
| FLNRYTTTLNDFNLVA | 1 | 0 | 0 | 1657836 | QIQ54046.1 | P0C6U8 |
| FLPFQQFGRDISDTTD | 1 | 0 | 0 | 1657841 | QIQ54048.1 | P59594 |
| FLPKEDGDVLLAEFDT | 1 | 0 | 0 | 1657848 | YP_009047202.1 | K9N7C7 |
| FLPQFTSDQAVTFLKE | 1 | 0 | 0 | 1657850 | YP_173241.1 | Q0ZJ82 |
| FLQSSNYKIDTTSSSC | 1 | 0 | 0 | 1657862 | YP_173238.1 | A0A140H1H1 |
| FLRLIDDNGIVLNSIL | 1 | 0 | 0 | 1657868 | YP_003769.1 | H9EJA2 |
| FLSDFINGIFAKVKNT | 1 | 0 | 0 | 1657876 | YP_009555241.1 | P36334 |
| FLSFDSDVFIKKYGLE | 1 | 0 | 0 | 1657878 | YP_009047202.1 | K9N7C7 |
| FLSGDDCFVSNSSVIQ | 1 | 0 | 0 | 1657880 | YP_173236.1 | A0A140H1G9 |
| FLSKHSDFGLGDLVDS | 1 | 0 | 0 | 1657882 | NP_073549.1 | A0A0P0LKV0 |
| FLSLSDQFKTTGDLAV | 1 | 0 | 0 | 1657885 | YP_003766.2 | H9EJ42 |
| FLSMFDVDKKSLNALI | 1 | 0 | 0 | 1657888 | YP_009555238.1 | P0C6X6 |
| FLSNTKYYDDSQYYFN | 1 | 0 | 0 | 1657890 | YP_009555240.1 | P30215 |
| FLVHDNVAFYQGDVDT | 1 | 0 | 0 | 1657913 | NP_073549.1 | A0A0P0LKV0 |
| FMDNAINVGGTGLQYA | 1 | 0 | 0 | 1657951 | YP_009047202.1 | K9N7C7 |
| FMGRIRSVYPVASPQE | 1 | 0 | 0 | 1657957 | NP_828849.2 | P0C6X7 |
| FMKHKGGDSTDFIALI | 1 | 0 | 0 | 1657968 | YP_009047202.1 | K9N7C7 |
| FMKSDIFGQDFKTFDL | 1 | 0 | 0 | 1657970 | NP_073549.1 | A0A0P0LKV0 |
| FMLGSSVGNFSDGKMG | 1 | 0 | 0 | 1657972 | YP_009047204.1 | K9N5Q8 |
| FMLKTLYKDVDNPHLM | 1 | 0 | 0 | 1657974 | YP_009047202.1 | K9N7C7 |
| FMLKYDENGTITDAVD | 1 | 0 | 0 | 1657975 | NP_828851.1 | P59594 |
| FMLWCKDGQVQTFYPR | 1 | 0 | 0 | 1657981 | YP_009047202.1 | K9N7C7 |
| FMNVTLDKCTKYNIYD | 1 | 0 | 0 | 1657988 | NP_073551.1 | A0A0P0K6L9 |
| FMSEAKCWTETDLTKG | 1 | 0 | 0 | 1658000 | YP_009724389.1 | P0DTD1 |
| FMSHFDVDRKSFNNFV | 1 | 0 | 0 | 1658005 | YP_173236.1 | A0A140H1G9 |
| FMSTSKCWVEEDLTKG | 1 | 0 | 0 | 1658010 | YP_003766.2 | H9EJ42 |
| FMVNQLAYSSSANGSL | 1 | 0 | 0 | 1658016 | YP_009047202.1 | K9N7C7 |
| FNAPALQEAYYRARAG | 1 | 0 | 0 | 1658028 | NP_828849.2 | P0C6X7 |
| FNATGSDVNCNGYQHN | 1 | 0 | 0 | 1658030 | YP_003767.1 | A0A0P0G321 |
| FNAVDCMSDFMSEIKC | 1 | 0 | 0 | 1658034 | YP_009555241.1 | P36334 |
| FNCFYALNNAFLAFSI | 1 | 0 | 0 | 1658035 | YP_173241.1 | Q0ZJ82 |
| FNCTNFAINDLNTTVP | 1 | 0 | 0 | 1658039 | YP_173238.1 | A0A140H1H1 |
| FNEDVCVCFDNSIQGS | 1 | 0 | 0 | 1658051 | YP_003766.2 | H9EJ42 |
| FNETKGPLCVDTSHFT | 1 | 0 | 0 | 1658056 | NP_073551.1 | A0A0P0K6L9 |
| FNFDQKELLAYYNMLV | 1 | 0 | 0 | 1658060 | YP_009555238.1 | P0C6X6 |
| FNFNGLTGTGVLTPSS | 1 | 0 | 0 | 1658067 | NP_828851.1 | P59594 |
| FNGAIIRDSDVVKQPV | 1 | 0 | 0 | 1658076 | YP_009047202.1 | K9N7C7 |
| FNGIGVTQNVLYENQK | 1 | 0 | 0 | 1658080 | YP_009724390.1 | P0DTC2 |
| FNGKFLSNTKYYDDSQ | 1 | 0 | 0 | 1658082 | YP_009555240.1 | P30215 |
| FNGLTGTGVLTESNKK | 1 | 0 | 0 | 1658083 | YP_009724390.1 | P0DTC2 |
| FNGTGRGDCKGFSSDV | 1 | 0 | 0 | 1658089 | NP_073551.1 | A0A0P0K6L9 |
| FNHHEKLLAPLLSAGI | 1 | 0 | 0 | 1658097 | QIQ54046.1 | P0C6U8 |
| FNHNNGNDVLYQPPTA | 1 | 0 | 0 | 1658100 | YP_173236.1 | A0A140H1G9 |
| FNHPIQVDQLNSSYFK | 1 | 0 | 0 | 1658101 | YP_009047204.1 | K9N5Q8 |
| FNICQAVSANVCALMS | 1 | 0 | 0 | 1658104 | YP_009555238.1 | P0C6X6 |
| FNKAIDGFGVFDIKNS | 1 | 0 | 0 | 1658117 | YP_003766.2 | H9EJ42 |
| FNKDTGVIYGLNSTET | 1 | 0 | 0 | 1658124 | YP_009555240.1 | P30215 |
| FNKEFYDFLRSQGFFD | 1 | 0 | 0 | 1658125 | NP_073549.1 | A0A0P0LKV0 |
| FNKKDWYDFVENPDIL | 1 | 0 | 0 | 1658130 | YP_009724389.1 | P0DTD1 |
| FNKLVKDPPNVQIHTI | 1 | 0 | 0 | 1658132 | NP_828852.2 | P59632 |
| FNKPLIEWPLTEYDFT | 1 | 0 | 0 | 1658133 | YP_009047202.1 | K9N7C7 |
| FNKVTLADAGFIKQYG | 1 | 0 | 0 | 1658136 | YP_009724390.1 | P0DTC2 |
| FNKYFKHWSMPYHPNT | 1 | 0 | 0 | 1658141 | YP_009555238.1 | P0C6X6 |
| FNKYKYFSGAMETAAY | 1 | 0 | 0 | 1658144 | YP_009047202.1 | K9N7C7 |
| FNNALYAIQEGFDATN | 1 | 0 | 0 | 1658164 | YP_009555241.1 | P36334 |
| FNNFFISYAKPEDKLS | 1 | 0 | 0 | 1658165 | YP_003766.2 | H9EJ42 |
| FNNFVSSYAKPEEKLS | 1 | 0 | 0 | 1658168 | NP_073549.1 | A0A0P0LKV0 |
| FNNLLGFDSSKPVTKK | 1 | 0 | 0 | 1658170 | YP_009047202.1 | K9N7C7 |
| FNNNGTNVTQVKNASV | 1 | 0 | 0 | 1658171 | NP_073549.1 | A0A0P0LKV0 |
| FNNSLTRQLQPINYFD | 1 | 0 | 0 | 1658173 | YP_009555241.1 | P36334 |
| FNNWFLLTNTSSVVDG | 1 | 0 | 0 | 1658175 | NP_073551.1 | A0A0P0K6L9 |
| FNPETNCLLNVPFGGT | 1 | 0 | 0 | 1658179 | YP_009047210.1 | K9N7A1 |
| FNPETNILLNVPLHGT | 1 | 0 | 0 | 1658180 | YP_009724393.1 | P0DTC5 |
| FNPETNILLNVPLRGT | 1 | 0 | 0 | 1658181 | NP_828855.1 | P59596 |
| FNPSTWNKRFGFIEDS | 1 | 0 | 0 | 1658186 | YP_009555241.1 | P36334 |
| FNQALGAMQTGFTTTN | 1 | 0 | 0 | 1658188 | YP_009047204.1 | K9N5Q8 |
| FNQDGIIFNAVDCMSD | 1 | 0 | 0 | 1658192 | YP_009555241.1 | P36334 |
| FNRALVSATEFADKLV | 1 | 0 | 0 | 1658202 | YP_009555238.1 | P0C6X6 |
| FNRGRQFYEFYNDVKP | 1 | 0 | 0 | 1658206 | YP_009555243.1 | Q04854 |
| FNRLATCDWTESGDYT | 1 | 0 | 0 | 1658208 | YP_009047202.1 | K9N7C7 |
| FNRPSLVDDNKFDVLK | 1 | 0 | 0 | 1658210 | YP_009555238.1 | P0C6X6 |
| FNSCKSDILSLNNPIF | 1 | 0 | 0 | 1658214 | YP_173237.1 | A0A140H1H0 |
| FNTILNTACGVFEVDD | 1 | 0 | 0 | 1658226 | YP_009555238.1 | P0C6X6 |
| FNTMLDASAPATGWLL | 1 | 0 | 0 | 1658233 | YP_173236.1 | A0A140H1G9 |
| FNVDDINFSPVLGCLG | 1 | 0 | 0 | 1658241 | YP_009555241.1 | P36334 |
| FNVINKGHFDGQQGET | 1 | 0 | 0 | 1658247 | QIQ54047.1 | P0C6X7 |
| FNVPMDKLKTLIATAE | 1 | 0 | 0 | 1658253 | QIQ54046.1 | P0C6U8 |
| FNVSDAYANLVPYYQL | 1 | 0 | 0 | 1658259 | NP_073549.1 | A0A0P0LKV0 |
| FNVSFVNDSIESVGGL | 1 | 0 | 0 | 1658261 | YP_173238.1 | A0A140H1H1 |
| FNVVNKGHFDGQQGEV | 1 | 0 | 0 | 1658270 | YP_009724389.1 | P0DTD1 |

| | | | | | | |
|---|---|---|---|---|---|---|
| FNYEPLTQDHVDILGP | 1 | 0 | 0 | 1658276 | QIQ54046.1 | P0C6U8 |
| FNYFKKVDGIIQQLPE | 1 | 0 | 0 | 1658277 | NP_828849.2 | P0C6X7 |
| FNYNVSVLQMDVQGLV | 1 | 0 | 0 | 1658284 | YP_003766.2 | H9EJ42 |
| FNYYKKVDGVVQQLPE | 1 | 0 | 0 | 1658289 | YP_009724389.1 | P0DTD1 |
| FNYYKKVNGVVQQLPE | 1 | 0 | 0 | 1658290 | QHR63299.1 | P0C6W2 |
| FPCAGRKVNFNEKPVV | 1 | 0 | 0 | 1658293 | YP_173236.1 | A0A140H1G9 |
| FPDLNGDVVAIDYKHY | 1 | 0 | 0 | 1658298 | YP_009724389.1 | P0DTD1 |
| FPDLNGDVVAIDYRHY | 1 | 0 | 0 | 1658299 | NP_828849.2 | P0C6X7 |
| FPHNDRIKSFSTFESA | 1 | 0 | 0 | 1658310 | NP_073549.1 | A0A0P0LKV0 |
| FPLNSIIKTIQPRVEK | 1 | 0 | 0 | 1658332 | YP_009724389.1 | P0DTD1 |
| FPLNSKVKVIQPRVEK | 1 | 0 | 0 | 1658333 | NP_828849.2 | P0C6X7 |
| FPLNSTIKTIQPRVEK | 1 | 0 | 0 | 1658334 | QHR63299.1 | P0C6W2 |
| FPPGQGVPLNANSTPA | 1 | 0 | 0 | 1658342 | YP_009047211.1 | K9N4V7 |
| FPQVNGLTSIKWADNN | 1 | 0 | 0 | 1658355 | QIQ54046.1 | P0C6U8 |
| FPRGQGVPINTNSSPD | 1 | 0 | 0 | 1658360 | YP_009724397.2 | P0DTC9 |
| FPRGQGVPINTNSTKD | 1 | 0 | 0 | 1658361 | QIQ54056.1 | P59595 |
| FQDAASNIFVINKDTY | 1 | 0 | 0 | 1658386 | YP_009047202.1 | K9N7C7 |
| FQEKDEDDNLIDSYFV | 1 | 0 | 0 | 1658390 | YP_009724389.1 | P0DTD1 |
| FQEKDEDGNLIDSYFI | 1 | 0 | 0 | 1658391 | QIQ54047.1 | P0C6X7 |
| FQENWNTKHSSGVTRD | 1 | 0 | 0 | 1658393 | QHR63299.1 | P0C6W2 |
| FQENWNTKHSSGVTRE | 1 | 0 | 0 | 1658394 | YP_009724389.1 | P0DTD1 |
| FQKCEQLLREYGQFCS | 1 | 0 | 0 | 1658419 | YP_009047204.1 | K9N5Q8 |
| FQKGKEFEFVEGQGVP | 1 | 0 | 0 | 1658422 | YP_009555245.1 | P33469 |
| FQKGRDFKFSDGQGVP | 1 | 0 | 0 | 1658423 | YP_173242.1 | Q19U29 |
| FQKVQDAVNNNAQALS | 1 | 0 | 0 | 1658425 | YP_009047204.1 | K9N5Q8 |
| FQLKLKGTPVLQLKES | 1 | 0 | 0 | 1658428 | YP_009047202.1 | K9N7C7 |
| FQQFGRDISDTTDAVR | 1 | 0 | 0 | 1658442 | QIQ54048.1 | P59594 |
| FQRIDDDGNKLDKFFV | 1 | 0 | 0 | 1658445 | YP_173236.1 | A0A140H1G9 |
| FQRVDENGDKLDQFFV | 1 | 0 | 0 | 1658446 | YP_009555238.1 | P0C6X6 |
| FQTRAGCLIGAEHVNN | 1 | 0 | 0 | 1658460 | YP_009724390.1 | P0DTC2 |
| FQTRAGCLVGAEHVNN | 1 | 0 | 0 | 1658461 | QIQ54048.1 | P59594 |
| FQTTVELQGLIDQINS | 1 | 0 | 0 | 1658462 | YP_003767.1 | A0A0P0G321 |
| FQTVKPGNFNQDFYEF | 1 | 0 | 0 | 1658465 | YP_173236.1 | A0A140H1G9 |
| FQVNWNTKHGSGVTRE | 1 | 0 | 0 | 1658468 | QIQ54046.1 | P0C6U8 |
| FRDAEGRMTPYCHDPT | 1 | 0 | 0 | 1658480 | YP_009047202.1 | K9N7C7 |
| FRDELGVRVLDQSDNN | 1 | 0 | 0 | 1658481 | YP_003766.2 | H9EJ42 |
| FRDYFNNNTDSIVIGG | 1 | 0 | 0 | 1658485 | YP_003766.2 | H9EJ42 |
| FRIDVPSAESTGTQSV | 1 | 0 | 0 | 1658508 | YP_009047205.1 | K9N796 |
| FRKANLKPFERDISTE | 1 | 0 | 0 | 1658511 | QHR63300.2 | Q3LZX1 |
| FRKARNGVFISTEKLS | 1 | 0 | 0 | 1658512 | YP_173236.1 | A0A140H1G9 |
| FRKSKLKPFERDISTE | 1 | 0 | 0 | 1658514 | QIQ54048.1 | P59594 |
| FRNSSANYVGKYDINH | 1 | 0 | 0 | 1658537 | YP_173237.1 | A0A140H1H0 |
| FRNSTTNYVGVYDINH | 1 | 0 | 0 | 1658538 | YP_009555240.1 | P30215 |
| FRSVNQRSIIAYTMSL | 1 | 0 | 0 | 1658557 | QIQ54048.1 | P59594 |
| FRYMNSQGLLPPKNSI | 1 | 0 | 0 | 1658583 | YP_009724389.1 | P0DTD1 |
| FRYMNSQGLLPPKSSI | 1 | 0 | 0 | 1658584 | NP_828849.2 | P0C6X7 |
| FSASLFKPVQLVPVSP | 1 | 0 | 0 | 1658588 | YP_009047208.1 | K9N7D2 |
| FSASTSAFVETVKGLD | 1 | 0 | 0 | 1658590 | YP_009724389.1 | P0DTD1 |
| FSAVSSPMSNMTENER | 1 | 0 | 0 | 1658595 | NP_073555.1 | A0A0P0KXN3 |
| FSCNNFDESKIYGSCF | 1 | 0 | 0 | 1658601 | YP_173238.1 | A0A140H1H1 |
| FSCPLCCTVYDSSEVK | 1 | 0 | 0 | 1658602 | YP_173236.1 | A0A140H1G9 |
| FSDGQGVPIAFGVPPS | 1 | 0 | 0 | 1658609 | YP_173242.1 | Q19U29 |
| FSFEQSHNNCYVNVAC | 1 | 0 | 0 | 1658620 | YP_173236.1 | A0A140H1G9 |
| FSFGVTQEYIQTTIQK | 1 | 0 | 0 | 1658624 | YP_009047204.1 | K9N5Q8 |
| FSFRDELGVRVLDQSD | 1 | 0 | 0 | 1658627 | YP_003766.2 | H9EJ42 |
| FSGFKLDGFNHQFVTA | 1 | 0 | 0 | 1658637 | YP_003766.2 | H9EJ42 |
| FSGNVDLDALASDIFD | 1 | 0 | 0 | 1658640 | YP_009047202.1 | K9N7C7 |
| FSGVASQEGVLSEREK | 1 | 0 | 0 | 1658644 | YP_003770.1 | U3M6Q8 |
| FSIVNSAPDGLLFLHT | 1 | 0 | 0 | 1658668 | YP_003767.1 | A0A0P0G321 |
| FSKCHNCCDEYGGHHD | 1 | 0 | 0 | 1658669 | YP_173238.1 | A0A140H1H1 |
| FSKKDWYDFVENPDII | 1 | 0 | 0 | 1658674 | YP_173236.1 | A0A140H1G9 |
| FSKKHVEVFTDGKLNC | 1 | 0 | 0 | 1658675 | YP_009047202.1 | K9N7C7 |
| FSKKQLGLLQPTDNST | 1 | 0 | 0 | 1658676 | YP_009047202.1 | K9N7C7 |
| FSLADSVVEVVTTSLT | 1 | 0 | 0 | 1658684 | YP_009555238.1 | P0C6X6 |
| FSLELQDHNETCHA | 1 | 0 | 0 | 1658690 | YP_009725318.1 | P0DTD8 |
| FSLKDIPGGCAMPIVA | 1 | 0 | 0 | 1658698 | NP_073551.1 | A0A0P0K6L9 |
| FSLKLAGTAVVNLRPD | 1 | 0 | 0 | 1658701 | YP_173236.1 | A0A140H1G9 |
| FSMAPVEFYEDRILDF | 1 | 0 | 0 | 1658720 | YP_009047202.1 | K9N7C7 |
| FSMVRKLDNQALNSIL | 1 | 0 | 0 | 1658729 | YP_009555238.1 | P0C6X6 |
| FSNCNFNMSSLMSFIQ | 1 | 0 | 0 | 1658731 | YP_009555241.1 | P36334 |
| FSNGVDFVAQPEGCVL | 1 | 0 | 0 | 1658733 | NP_073549.1 | A0A0P0LKV0 |
| FSNLVLNNCTKYNIYD | 1 | 0 | 0 | 1658735 | YP_003767.1 | A0A0P0G321 |
| FSNSGADVLYQPPQTS | 1 | 0 | 0 | 1658738 | NP_828849.2 | P0C6X7 |
| FSNSGSDVLYQPPQTS | 1 | 0 | 0 | 1658739 | YP_009724389.1 | P0DTD1 |
| FSNSNYVLQDFDVVFG | 1 | 0 | 0 | 1658741 | YP_003766.2 | H9EJ42 |
| FSNVVELPKFFYASNG | 1 | 0 | 0 | 1658744 | NP_073551.1 | A0A0P0K6L9 |
| FSNVYADSFVVKGDDV | 1 | 0 | 0 | 1658746 | NP_828851.1 | P59594 |
| FSNYQHEEAIYNLLKD | 1 | 0 | 0 | 1658749 | QIQ54047.1 | P0C6X7 |
| FSNYQHEETIYNLLKD | 1 | 0 | 0 | 1658750 | YP_009724389.1 | P0DTD1 |
| FSPQIITTDNTFVSGN | 1 | 0 | 0 | 1658758 | NP_828851.1 | P59594 |
| FSQFDSLGVSSNQSPQ | 1 | 0 | 0 | 1658761 | YP_009555238.1 | P0C6X6 |
| FSRDHNDILYTPPTVS | 1 | 0 | 0 | 1658774 | NP_073549.1 | A0A0P0LKV0 |
| FSRNTTFDFLSNASSS | 1 | 0 | 0 | 1658775 | YP_003767.1 | A0A0P0G321 |
| FSRQLQNSMSGASADS | 1 | 0 | 0 | 1658776 | NP_828858.1 | P59595 |
| FSSEIIGYKAIDGGVT | 1 | 0 | 0 | 1658787 | YP_009724389.1 | P0DTD1 |
| FSSRMYCSFYPPDEEE | 1 | 0 | 0 | 1658794 | NP_828849.2 | P0C6X7 |
| FSSTFNVPMDKLKTLI | 1 | 0 | 0 | 1658798 | QIQ54046.1 | P0C6U8 |
| FSSVNDAITQTAEAIH | 1 | 0 | 0 | 1658800 | YP_003767.1 | A0A0P0G321 |
| FSTFESAYMPIADPTH | 1 | 0 | 0 | 1658804 | NP_073549.1 | A0A0P0LKV0 |
| FSTNDLYVALRGRIDK | 1 | 0 | 0 | 1658814 | NP_073553.1 | A0A0P0KWE8 |
| FSTVNALPEVNADIVV | 1 | 0 | 0 | 1658820 | NP_073549.1 | A0A0P0LKV0 |

| | | | | | | |
|---|---|---|---|---|---|---|
| FSTVVIKNLTPIKLNF | 1 | 0 | 0 | 1658821 | YP_003766.2 | H9EJ42 |
| FSVAAITSGVKFQTVK | 1 | 0 | 0 | 1658823 | YP_009555238.1 | P0C6X6 |
| FSVAALSTGLTSQTVK | 1 | 0 | 0 | 1658825 | NP_073549.1 | A0A0P0LKV0 |
| FSVDPATTYLEAVKHG | 1 | 0 | 0 | 1658830 | YP_003766.2 | H9EJ42 |
| FSVDPKKTYLDFIQQG | 1 | 0 | 0 | 1658831 | YP_009555238.1 | P0C6X6 |
| FSVDPKKTYLDYIQQG | 1 | 0 | 0 | 1658832 | YP_173236.1 | A0A140H1G9 |
| FSVNDFTCSQISPAAI | 1 | 0 | 0 | 1658839 | YP_009047204.1 | K9N5Q8 |
| FSVNNTFCPCAKPSFA | 1 | 0 | 0 | 1658840 | YP_173238.1 | A0A140H1H1 |
| FSVQKHIGAPANLRVV | 1 | 0 | 0 | 1658844 | YP_009047202.1 | K9N7C7 |
| FSWNADLYKLMSCFAF | 1 | 0 | 0 | 1658852 | YP_173236.1 | A0A140H1G9 |
| FSYCRKIGVNVCSDST | 1 | 0 | 0 | 1658856 | YP_173236.1 | A0A140H1G9 |
| FSYDGLRGAIRKAKDY | 1 | 0 | 0 | 1658858 | YP_009047202.1 | K9N7C7 |
| FTACSHAAVDSLCAKA | 1 | 0 | 0 | 1658868 | NP_073549.1 | A0A0P0LKV0 |
| FTAFANTVEDAVNKLV | 1 | 0 | 0 | 1658869 | YP_009555242.1 | Q04853 |
| FTANKIVGGAPTWFNA | 1 | 0 | 0 | 1658875 | YP_009047202.1 | K9N7C7 |
| FTCNNIDAAKIYGMCF | 1 | 0 | 0 | 1658883 | YP_009555241.1 | P36334 |
| FTCNSNANLSMLQLGV | 1 | 0 | 0 | 1658884 | YP_003767.1 | A0A0P0G321 |
| FTCRTDMEKDFIALDQ | 1 | 0 | 0 | 1658885 | YP_009555238.1 | P0C6X6 |
| FTDGKLNCSFQDAASN | 1 | 0 | 0 | 1658890 | YP_009047202.1 | K9N7C7 |
| FTDYKLELFNKYFKHW | 1 | 0 | 0 | 1658898 | YP_009555238.1 | P0C6X6 |
| FTEESFYKNMYLKSAV | 1 | 0 | 0 | 1658905 | YP_173236.1 | A0A140H1G9 |
| FTFDIDGSRVPHIVRR | 1 | 0 | 0 | 1658916 | YP_173236.1 | A0A140H1G9 |
| FTFDVEGSRVPHIVRK | 1 | 0 | 0 | 1658917 | YP_009555238.1 | P0C6X6 |
| FTFKQANNNCFVNVSC | 1 | 0 | 0 | 1658918 | YP_009555238.1 | P0C6X6 |
| FTGSSYYYPEPISDKN | 1 | 0 | 0 | 1658932 | YP_173238.1 | A0A140H1H1 |
| FTGSVYGNFDDQPSLQ | 1 | 0 | 0 | 1658933 | YP_003766.2 | H9EJ42 |
| FTGVDGELPVAVVNDK | 1 | 0 | 0 | 1658935 | YP_003766.2 | H9EJ42 |
| FTGYHFTKNGKTVLGE | 1 | 0 | 0 | 1658938 | YP_009555238.1 | P0C6X6 |
| FTGYHFTSTGKTVLGE | 1 | 0 | 0 | 1658939 | YP_173236.1 | A0A140H1G9 |
| FTGYHITKNSKVQLGE | 1 | 0 | 0 | 1658940 | YP_009047202.1 | K9N7C7 |
| FTGYRVTKNSKTQIGE | 1 | 0 | 0 | 1658943 | QIQ54047.1 | P0C6X7 |
| FTGYRVTKNSKVQIGE | 1 | 0 | 0 | 1658944 | YP_009724389.1 | P0DTD1 |
| FTIGTVTLKQGEIKDA | 1 | 0 | 0 | 1658951 | YP_009724391.1 | P0DTC3 |
| FTIVGQEEFIQTNSPK | 1 | 0 | 0 | 1658961 | YP_173238.1 | A0A140H1H1 |
| FTKAAGGKVSFSDDVE | 1 | 0 | 0 | 1658963 | NP_073549.1 | A0A0P0LKV0 |
| FTKKDWYDFVENPDII | 1 | 0 | 0 | 1658966 | YP_009555238.1 | P0C6X6 |
| FTKLAGGKISFSDDVI | 1 | 0 | 0 | 1658967 | YP_003766.2 | H9EJ42 |
| FTKNGKTVLGEYVFDK | 1 | 0 | 0 | 1658970 | YP_009555238.1 | P0C6X6 |
| FTKSGVNDICMFENKT | 1 | 0 | 0 | 1658975 | YP_009047202.1 | K9N7C7 |
| FTKSTNSRIRASLPTT | 1 | 0 | 0 | 1658976 | NP_828849.2 | P0C6X7 |
| FTKSVCGYTDFAEDVC | 1 | 0 | 0 | 1658977 | NP_073549.1 | A0A0P0LKV0 |
| FTKVVVGPTTEVKFSV | 1 | 0 | 0 | 1658979 | YP_003766.2 | H9EJ42 |
| FTLAEVVKQMYGVNLQ | 1 | 0 | 0 | 1658980 | YP_003766.2 | H9EJ42 |
| FTLANVSSFGDYNLSS | 1 | 0 | 0 | 1658981 | NP_073551.1 | A0A0P0K6L9 |
| FTLGTVTLKQGEIKDA | 1 | 0 | 0 | 1658986 | QHR63301.1 | Q3LZX0 |
| FTLQLVSAVNQSLSNA | 1 | 0 | 0 | 1658998 | NP_073552.1 | A0A0P0KWE8 |
| FTMLRKLDNDALNNII | 1 | 0 | 0 | 1659009 | YP_009724389.1 | P0DTD1 |
| FTNDKTIFYTTEIASK | 1 | 0 | 0 | 1659011 | YP_173236.1 | A0A140H1G9 |
| FTNDVSFLAHLQWFAM | 1 | 0 | 0 | 1659012 | NP_828849.2 | P0C6X7 |
| FTNEPAFIMHVSWYIM | 1 | 0 | 0 | 1659013 | YP_009047202.1 | K9N7C7 |
| FTNFEPFTVNSVNDSL | 1 | 0 | 0 | 1659014 | YP_009555241.1 | P36334 |
| FTNKTTLPTNVAFELF | 1 | 0 | 0 | 1659016 | YP_003766.2 | H9EJ42 |
| FTNNNGSDVLYQPPTA | 1 | 0 | 0 | 1659019 | YP_009555238.1 | P0C6X6 |
| FTNSILMLDKQGQLLD | 1 | 0 | 0 | 1659021 | YP_003766.2 | H9EJ42 |
| FTNSTDTVNHDSFNLY | 1 | 0 | 0 | 1659022 | YP_173236.1 | A0A140H1G9 |
| FTNVYADSFVITGDEV | 1 | 0 | 0 | 1659024 | QHR63300.2 | Q3LZX1 |
| FTNVYADSFVVKGDEV | 1 | 0 | 0 | 1659025 | QIQ54048.1 | P59594 |
| FTPEDVNMQIMGVVMQ | 1 | 0 | 0 | 1659028 | YP_009047202.1 | K9N7C7 |
| FTPGKQSSSRASSGNR | 1 | 0 | 0 | 1659030 | YP_009555245.1 | P33469 |
| FTPRSDMEYDFLNMDM | 1 | 0 | 0 | 1659037 | YP_003766.2 | H9EJ42 |
| FTPYHAFKCNRSGSND | 1 | 0 | 0 | 1659039 | YP_009555240.1 | P30215 |
| FTQDGNAAITDYNYYK | 1 | 0 | 0 | 1659043 | YP_009555238.1 | P0C6X6 |
| FTQKGDAAIKDFDYYR | 1 | 0 | 0 | 1659044 | NP_073549.1 | A0A0P0LKV0 |
| FTQSRDLEDFKPRSQM | 1 | 0 | 0 | 1659045 | NP_828849.2 | P0C6X7 |
| FTREMQQHPLLNPSAL | 1 | 0 | 0 | 1659052 | NP_073556.1 | A0A0N7J131 |
| FTSDQAVTFLKEWNFS | 1 | 0 | 0 | 1659068 | YP_173241.1 | Q0ZJ82 |
| FTSDYYQLYSTQLSTD | 1 | 0 | 0 | 1659069 | YP_009724391.1 | P0DTC3 |
| FTSECYQLYSTQLSTD | 1 | 0 | 0 | 1659070 | QIQ54049.1 | P59632 |
| FTSEPPVTYSPATILA | 1 | 0 | 0 | 1659072 | YP_009047202.1 | K9N7C7 |
| FTSLEIPRRNVATLQA | 1 | 0 | 0 | 1659076 | YP_009724389.1 | P0DTD1 |
| FTSLEVPRRNVATLQA | 1 | 0 | 0 | 1659077 | QIQ54047.1 | P0C6X7 |
| FTSTGKTVLGEYVFDK | 1 | 0 | 0 | 1659080 | YP_173236.1 | A0A140H1G9 |
| FTTELKDSVKKFFKLV | 1 | 0 | 0 | 1659086 | QIQ54046.1 | P0C6U8 |
| FTTTNEAFQKVQDAVN | 1 | 0 | 0 | 1659097 | YP_009047204.1 | K9N5Q8 |
| FTTVDNINLHTQVVDM | 1 | 0 | 0 | 1659098 | YP_009724389.1 | P0DTD1 |
| FTTVDNTNLHTQLVDM | 1 | 0 | 0 | 1659099 | NP_828849.2 | P0C6X7 |
| FTVDPQKAYLDFVNAG | 1 | 0 | 0 | 1659104 | YP_009047202.1 | K9N7C7 |
| FTVNSVNDSLEPVGGL | 1 | 0 | 0 | 1659111 | YP_009555241.1 | P36334 |
| FTVSTQPQGNTIPHYS | 1 | 0 | 0 | 1659113 | YP_173242.1 | Q19U29 |
| FVANYSQDVKQFANGF | 1 | 0 | 0 | 1659154 | YP_009047204.1 | K9N5Q8 |
| FVAYETARQEYENAVA | 1 | 0 | 0 | 1659164 | NP_073549.1 | A0A0P0LKV0 |
| FVCDNIKFADDLNQLT | 1 | 0 | 0 | 1659165 | YP_009724389.1 | P0DTD1 |
| FVCDNIKLADDLNQLT | 1 | 0 | 0 | 1659166 | QHR63299.1 | P0C6W2 |
| FVCDNTKFADDLNQMS | 1 | 0 | 0 | 1659167 | QIQ54046.1 | P0C6U8 |
| FVCTDNSANTKVLRNK | 1 | 0 | 0 | 1659172 | YP_009047202.1 | K9N7C7 |
| FVDHFGENEEIIINGH | 1 | 0 | 0 | 1659186 | NP_073549.1 | A0A0P0LKV0 |
| FVDQYGCDYSGKLADC | 1 | 0 | 0 | 1659192 | YP_173236.1 | A0A140H1G9 |
| FVDRQTAQAAGTDTTI | 1 | 0 | 0 | 1659194 | YP_009724389.1 | P0DTD1 |
| FVDYSNLLHSKVKSVP | 1 | 0 | 0 | 1659206 | YP_009555238.1 | P0C6X6 |
| FVEAYNNCTGGAEIRD | 1 | 0 | 0 | 1659208 | YP_009555241.1 | P36334 |
| FVEAYNNCTGGSEIRD | 1 | 0 | 0 | 1659209 | YP_173238.1 | A0A140H1H1 |

| | | | | | | |
|---|---|---|---|---|---|---|
| FVEDCKLPLGQSLCAL | 1 | 0 | 0 | 1659210 | YP_009047204.1 | K9N5Q8 |
| FVEHYNEWAQANGFTA | 1 | 0 | 0 | 1659213 | NP_073549.1 | A0A0P0LKV0 |
| FVENPDIINIYKKLGP | 1 | 0 | 0 | 1659219 | YP_173236.1 | A0A140H1G9 |
| FVEYEVAKKNLADAKN | 1 | 0 | 0 | 1659231 | YP_173236.1 | A0A140H1G9 |
| FVFEKVDYGSDTVTYK | 1 | 0 | 0 | 1659235 | NP_073549.1 | A0A0P0LKV0 |
| FVFPLNSKVKVIQPRV | 1 | 0 | 0 | 1659243 | NP_828849.2 | P0C6X7 |
| FVGADGELPVAISGDK | 1 | 0 | 0 | 1659248 | NP_073549.1 | A0A0P0LKV0 |
| FVGLSPTVWLSVIWM | 1 | 0 | 0 | 1659256 | P03138.3 | Q76R62 |
| FVGMPSFVAYETARQE | 1 | 0 | 0 | 1659257 | NP_073549.1 | A0A0P0LKV0 |
| FVGTQSVDMLAVKTGV | 1 | 0 | 0 | 1659261 | YP_009047202.1 | K9N7C7 |
| FVHEDLHVVEVLTKTA | 1 | 0 | 0 | 1659264 | YP_173236.1 | A0A140H1G9 |
| FVHYAGVVWTLQEVKD | 1 | 0 | 0 | 1659265 | NP_073549.1 | A0A0P0LKV0 |
| FVKDDYCDGSLGCKMC | 1 | 0 | 0 | 1659280 | NP_073549.1 | A0A0P0LKV0 |
| FVKRVDWSVEYPIIGD | 1 | 0 | 0 | 1659305 | NP_828849.2 | P0C6X7 |
| FVKSDIFGSDFKTFDL | 1 | 0 | 0 | 1659307 | YP_003766.2 | H9EJ42 |
| FVKSKVGNYRLPSSKP | 1 | 0 | 0 | 1659311 | YP_173241.1 | Q0ZJ82 |
| FVMQFGYIDCEQDSCD | 1 | 0 | 0 | 1659356 | YP_009555238.1 | P0C6X6 |
| FVMQYGYVDCFNDNCD | 1 | 0 | 0 | 1659357 | YP_173236.1 | A0A140H1G9 |
| FVNAGGAPLTNCVKML | 1 | 0 | 0 | 1659359 | YP_009047202.1 | K9N7C7 |
| FVNDGRVANGYVCGTG | 1 | 0 | 0 | 1659360 | NP_073549.1 | A0A0P0LKV0 |
| FVNDKITEFQLDYSID | 1 | 0 | 0 | 1659361 | NP_073549.1 | A0A0P0LKV0 |
| FVNDSIESVGGLYEIK | 1 | 0 | 0 | 1659362 | YP_173238.1 | A0A140H1H1 |
| FVNDSYRQFDLVQYDF | 1 | 0 | 0 | 1659363 | YP_173236.1 | A0A140H1G9 |
| FVNEFYAYLRKHFSMM | 1 | 0 | 0 | 1659364 | YP_009724389.1 | P0DTD1 |
| FVNEYYEFLCKHFSMM | 1 | 0 | 0 | 1659365 | YP_173236.1 | A0A140H1G9 |
| FVNGFYRLYSGDTFWR | 1 | 0 | 0 | 1659367 | YP_003766.2 | H9EJ42 |
| FVNIAHASLREGVQLE | 1 | 0 | 0 | 1659369 | YP_173236.1 | A0A140H1G9 |
| FVNISRSELQTIVPEY | 1 | 0 | 0 | 1659371 | NP_073551.1 | A0A0P0K6L9 |
| FVNISRVELHTVIPDY | 1 | 0 | 0 | 1659372 | YP_003767.1 | A0A0P0G321 |
| FVNKTSLPTNIAFELF | 1 | 0 | 0 | 1659373 | NP_073549.1 | A0A0P0LKV0 |
| FVNLDNLRANNTKGSL | 1 | 0 | 0 | 1659375 | YP_009724389.1 | P0DTD1 |
| FVNLDNLRASNTKGSL | 1 | 0 | 0 | 1659376 | QHR63299.1 | P0C6W2 |
| FVNLKQLPFFYYSDSP | 1 | 0 | 0 | 1659377 | YP_009724389.1 | P0DTD1 |
| FVNMASFVEYEVAKKN | 1 | 0 | 0 | 1659378 | YP_009555238.1 | P0C6X6 |
| FVNSFVSLYNVTRDKL | 1 | 0 | 0 | 1659380 | YP_009047202.1 | K9N7C7 |
| FVNTTIGNETTSAFVG | 1 | 0 | 0 | 1659384 | NP_073551.1 | A0A0P0K6L9 |
| FVNVNNTWMYTGSGYY | 1 | 0 | 0 | 1659386 | YP_009555241.1 | P36334 |
| FVNVPYVYNGSAQATA | 1 | 0 | 0 | 1659387 | YP_009555240.1 | P30215 |
| FVQDPAPIDIDAFPVK | 1 | 0 | 0 | 1659398 | YP_003766.2 | H9EJ42 |
| FVQIPAQCVRDPVGFC | 1 | 0 | 0 | 1659401 | YP_009047202.1 | K9N7C7 |
| FVQNDVCSTEDFNVWA | 1 | 0 | 0 | 1659403 | YP_173236.1 | A0A140H1G9 |
| FVQQKPEPVNPVSFVV | 1 | 0 | 0 | 1659404 | NP_073549.1 | A0A0P0LKV0 |
| FVQSDKCSVEDFNVWA | 1 | 0 | 0 | 1659405 | YP_009555238.1 | P0C6X6 |
| FVQSNVDVVPEGWRVV | 1 | 0 | 0 | 1659407 | YP_009555238.1 | P0C6X6 |
| FVQVAPGCGVPIVDSY | 1 | 0 | 0 | 1659408 | NP_828849.2 | P0C6X7 |
| FVQWFVGLSPTVWLS | 1 | 0 | 0 | 1659411 | P03138.3 | Q76R62 |
| FVRHIIVGCNNADCVA | 1 | 0 | 0 | 1659415 | YP_003766.2 | H9EJ42 |
| FVRPVIEDYHTLTATV | 1 | 0 | 0 | 1659421 | YP_173241.1 | Q0ZJ82 |
| FVRYGDVDNLVFTNKT | 1 | 0 | 0 | 1659423 | YP_003766.2 | H9EJ42 |
| FVSQVLNKYTEVRGSR | 1 | 0 | 0 | 1659460 | YP_003767.1 | A0A0P0G321 |
| FVSSYAKPEEKLSAYD | 1 | 0 | 0 | 1659463 | NP_073549.1 | A0A0P0LKV0 |
| FVSYGSSVETETYFKV | 1 | 0 | 0 | 1659468 | YP_173236.1 | A0A140H1G9 |
| FVTEYYEFLNKHFSMM | 1 | 0 | 0 | 1659477 | YP_009555238.1 | P0C6X6 |
| FVTFEPFNVSFVNDSI | 1 | 0 | 0 | 1659478 | YP_173238.1 | A0A140H1H1 |
| FVTQRNFYEPQIITTD | 1 | 0 | 0 | 1659491 | YP_009724390.1 | P0DTC2 |
| FVVDKSLSIEEFADVV | 1 | 0 | 0 | 1659499 | YP_009047202.1 | K9N7C7 |
| FVVKEKLNAFLVHDNV | 1 | 0 | 0 | 1659508 | NP_073549.1 | A0A0P0LKV0 |
| FVVNAANENLLHGGGV | 1 | 0 | 0 | 1659520 | YP_003766.2 | H9EJ42 |
| FVVQPVGVVDTEWGNM | 1 | 0 | 0 | 1659525 | YP_009047202.1 | K9N7C7 |
| FVYDAYQNLVGYYSDD | 1 | 0 | 0 | 1659538 | YP_009047204.1 | K9N5Q8 |
| FVYFNATGSDVNCNGY | 1 | 0 | 0 | 1659543 | YP_003767.1 | A0A0P0G321 |
| FVYSSNEEQAVLKFLD | 1 | 0 | 0 | 1659548 | YP_003766.2 | H9EJ42 |
| FWFAMRKDGDDVIFSR | 1 | 0 | 0 | 1659566 | YP_173236.1 | A0A140H1G9 |
| FWSFNPETNNLMCIDM | 1 | 0 | 0 | 1659593 | YP_009555244.1 | Q01455 |
| FWTLFCTSVNTSSSEA | 1 | 0 | 0 | 1659594 | YP_003766.2 | H9EJ42 |
| FWTMFCTSVNTSSSEA | 1 | 0 | 0 | 1659595 | NP_073549.1 | A0A0P0LKV0 |
| FWTVFCTNANASSSEG | 1 | 0 | 0 | 1659597 | YP_009555238.1 | P0C6X6 |
| FWTVFCTNVNASSSEG | 1 | 0 | 0 | 1659598 | YP_173236.1 | A0A140H1G9 |
| FWVANHQADTSTPSDV | 1 | 0 | 0 | 1659599 | YP_173242.1 | Q19U29 |
| FYAPEPITSLNTKYVA | 1 | 0 | 0 | 1659609 | YP_009047204.1 | K9N5Q8 |
| FYASNGTYNCTDAVLT | 1 | 0 | 0 | 1659616 | NP_073551.1 | A0A0P0K6L9 |
| FYCFVNTTIGNETTSA | 1 | 0 | 0 | 1659622 | NP_073551.1 | A0A0P0K6L9 |
| FYCSNSSSGVLDTTIP | 1 | 0 | 0 | 1659624 | YP_003767.1 | A0A0P0G321 |
| FYCTNNTLVSGDAHIP | 1 | 0 | 0 | 1659626 | NP_073551.1 | A0A0P0K6L9 |
| FYDRDGQRTYDDVNAS | 1 | 0 | 0 | 1659636 | YP_009555238.1 | P0C6X6 |
| FYDRDGQRVYDDVDAS | 1 | 0 | 0 | 1659637 | YP_173236.1 | A0A140H1G9 |
| FYEFYNDVKPPVLDVD | 1 | 0 | 0 | 1659644 | YP_009555243.1 | Q04854 |
| FYELDNDFNTILNTAC | 1 | 0 | 0 | 1659647 | YP_009555238.1 | P0C6X6 |
| FYFAVRKEGQDVIFSQ | 1 | 0 | 0 | 1659652 | YP_009555238.1 | P0C6X6 |
| FYFDDSDCDVVQEQVN | 1 | 0 | 0 | 1659654 | YP_003766.2 | H9EJ42 |
| FYGGWDNMLKNLMADV | 1 | 0 | 0 | 1659675 | NP_073549.1 | A0A0P0LKV0 |
| FYGWVSGNMMDGFSCP | 1 | 0 | 0 | 1659685 | YP_173236.1 | A0A140H1G9 |
| FYIYDEEGGYDVSKPV | 1 | 0 | 0 | 1659696 | YP_003766.2 | H9EJ42 |
| FYKENSYITTIKPVIY | 1 | 0 | 0 | 1659698 | QHR63299.1 | P0C6W2 |
| FYKENSYTTTIKPIVY | 1 | 0 | 0 | 1659699 | QIQ54046.1 | P0C6U8 |
| FYKENSYTTTIKPVTY | 1 | 0 | 0 | 1659700 | YP_009724389.1 | P0DTD1 |
| FYKETSYTTTIKPVSY | 1 | 0 | 0 | 1659701 | NP_828849.2 | P0C6X7 |
| FYKKDGVYYPSNGTNI | 1 | 0 | 0 | 1659704 | NP_073549.1 | A0A0P0LKV0 |
| FYKNMYLKSAVMQSVG | 1 | 0 | 0 | 1659705 | YP_173236.1 | A0A140H1G9 |
| FYLFDDAGEENFSSRM | 1 | 0 | 0 | 1659709 | NP_828849.2 | P0C6X7 |
| FYMEADTVVNAFYGVA | 1 | 0 | 0 | 1659724 | YP_009555238.1 | P0C6X6 |

| | | | | | | |
|---|---|---|---|---|---|---|
| FYPKLQASQAWQPGVA | 1 | 0 | 0 | 1659737 | NP_828849.2 | P0C6X7 |
| FYPKLQSSQAWQPGVA | 1 | 0 | 0 | 1659738 | YP_009724389.1 | P0DTD1 |
| FYPPDEDEEEGDCEEE | 1 | 0 | 0 | 1659742 | YP_009724389.1 | P0DTD1 |
| FYPPDEDYEEEYSEEE | 1 | 0 | 0 | 1659743 | QIQ54046.1 | P0C6U8 |
| FYPPDEEEEDDAECEE | 1 | 0 | 0 | 1659744 | NP_828849.2 | P0C6X7 |
| FYPRLQAASDWKPGYS | 1 | 0 | 0 | 1659747 | YP_009555238.1 | P0C6X6 |
| FYQGDVDTVVNGVDFD | 1 | 0 | 0 | 1659753 | NP_073549.1 | A0A0P0LKV0 |
| FYRLANECAQVLSEYV | 1 | 0 | 0 | 1659761 | YP_009047202.1 | K9N7C7 |
| FYRLSNELAQVLTEVV | 1 | 0 | 0 | 1659764 | NP_073549.1 | A0A0P0LKV0 |
| FYSANFLDDNVLPETY | 1 | 0 | 0 | 1659768 | YP_003767.1 | A0A0P0G321 |
| FYSYATHHDKFTDGVC | 1 | 0 | 0 | 1659776 | NP_828849.2 | P0C6X7 |
| FYSYATHSDKFKDGVC | 1 | 0 | 0 | 1659777 | QIQ54047.1 | P0C6X7 |
| FYSYATHSDKFTDGVC | 1 | 0 | 0 | 1659778 | YP_009724389.1 | P0DTD1 |
| FYTSKTTVASLINTLN | 1 | 0 | 0 | 1659787 | YP_009724389.1 | P0DTD1 |
| FYVKDGKYFTSEPPVT | 1 | 0 | 0 | 1659791 | YP_009047202.1 | K9N7C7 |
| FYVRVKHGDFSAVSSP | 1 | 0 | 0 | 1659804 | NP_073555.1 | A0A0P0KXN3 |
| FYVVKNSTAFPFETLS | 1 | 0 | 0 | 1659806 | YP_009047202.1 | K9N7C7 |
| FYVYDVHKDNFKDGLC | 1 | 0 | 0 | 1659809 | YP_173236.1 | A0A140H1G9 |
| GAAIHYLNSKIVDLAQ | 1 | 0 | 0 | 1659836 | YP_009555238.1 | P0C6X6 |
| GAALAMEKVNECVKSQ | 1 | 0 | 0 | 1659837 | YP_173238.1 | A0A140H1H1 |
| GACDYDRLPNEVTVAK | 1 | 0 | 0 | 1659839 | YP_009047210.1 | K9N7A1 |
| GACGSPGYNLKNGEVE | 1 | 0 | 0 | 1659841 | NP_073549.1 | A0A0P0LKV0 |
| GACGSPGYNVRNDGTV | 1 | 0 | 0 | 1659842 | YP_003766.2 | H9EJ42 |
| GACIDCNARHINAQVA | 1 | 0 | 0 | 1659843 | NP_828849.2 | P0C6X7 |
| GACIDCSARHINAQVA | 1 | 0 | 0 | 1659844 | YP_009724389.1 | P0DTD1 |
| GACSCGSCCKFDEDDS | 1 | 0 | 0 | 1659848 | NP_828851.1 | P59594 |
| GADGELPVAISGDKVF | 1 | 0 | 0 | 1659851 | NP_073549.1 | A0A0P0LKV0 |
| GADISDTIPDEKQNGH | 1 | 0 | 0 | 1659853 | YP_009047202.1 | K9N7C7 |
| GADSSIAYSNNTIAIP | 1 | 0 | 0 | 1659857 | NP_828851.1 | P59594 |
| GADVLYQPPQTSITSA | 1 | 0 | 0 | 1659858 | NP_828849.2 | P0C6X7 |
| GADVTKIKPHASHDSK | 1 | 0 | 0 | 1659859 | QIQ54046.1 | P0C6U8 |
| GADVTKIKPHNSHEGK | 1 | 0 | 0 | 1659860 | YP_009724389.1 | P0DTD1 |
| GADVTKIKPHVNHEGK | 1 | 0 | 0 | 1659861 | NP_828849.2 | P0C6X7 |
| GAEHVNNSYECDIPVG | 1 | 0 | 0 | 1659864 | QIQ54048.1 | P59594 |
| GAEMTSNVRRQVASGV | 1 | 0 | 0 | 1659867 | NP_073549.1 | A0A0P0LKV0 |
| GAFFFGSKLDLVKRDS | 1 | 0 | 0 | 1659871 | YP_173242.1 | Q19U29 |
| GAFGPGFTPPHGGLL | 1 | 0 | 0 | 1659873 | P12934.3 | Q76R62 |
| GAFHVTMRSSYTIKGS | 1 | 0 | 0 | 1659874 | YP_009555238.1 | P0C6X6 |
| GAGGHSFGIDLKSYDL | 1 | 0 | 0 | 1659881 | QIQ54046.1 | P0C6U8 |
| GAGGHSYGIDLKSYDL | 1 | 0 | 0 | 1659883 | NP_828849.2 | P0C6X7 |
| GAGICASYQTQTNSRS | 1 | 0 | 0 | 1659887 | QHR63300.2 | Q3LZX1 |
| GAGNGQAITTSVDANT | 1 | 0 | 0 | 1659891 | YP_003766.2 | H9EJ42 |
| GAGSDKGIAPGTSVLR | 1 | 0 | 0 | 1659894 | YP_009047202.1 | K9N7C7 |
| GAGSDKGVAPGTTVLK | 1 | 0 | 0 | 1659896 | YP_003766.2 | H9EJ42 |
| GAGSEKGVAPGSAVLR | 1 | 0 | 0 | 1659898 | YP_009555238.1 | P0C6X6 |
| GAGVMLECEKFNVFNV | 1 | 0 | 0 | 1659900 | YP_003766.2 | H9EJ42 |
| GAHATRDSIGTNFPLQ | 1 | 0 | 0 | 1659902 | YP_009555238.1 | P0C6X6 |
| GAHATRENIGTNFPLQ | 1 | 0 | 0 | 1659903 | YP_173236.1 | A0A140H1G9 |
| GAHLTKMSEYKGPVTD | 1 | 0 | 0 | 1659904 | NP_828849.2 | P0C6X7 |
| GAIIRDSDVVKQPVKF | 1 | 0 | 0 | 1659913 | YP_009047202.1 | K9N7C7 |
| GAIKLDDKDPQFKDNV | 1 | 0 | 0 | 1659915 | NP_828858.1 | P59595 |
| GAIKLDDKDPSFKDNV | 1 | 0 | 0 | 1659916 | QIQ54056.1 | P59595 |
| GAINAASKGAVQKESD | 1 | 0 | 0 | 1659921 | YP_009047202.1 | K9N7C7 |
| GAISASIGDIIQRLDV | 1 | 0 | 0 | 1659924 | YP_009047204.1 | K9N5Q8 |
| GAITTGYRFTNFEPFT | 1 | 0 | 0 | 1659928 | YP_009555241.1 | P36334 |
| GAKETKEINIIKLSGV | 1 | 0 | 0 | 1659931 | YP_009555238.1 | P0C6X6 |
| GALDISASIVAGGIVA | 1 | 0 | 0 | 1659947 | YP_009724389.1 | P0DTD1 |
| GALDISASIVAGGLVA | 1 | 0 | 0 | 1659948 | QIQ54046.1 | P0C6U8 |
| GALDTTSYREAACCHL | 1 | 0 | 0 | 1659949 | NP_828849.2 | P0C6X7 |
| GALLSKTSEYKGPVTD | 1 | 0 | 0 | 1659955 | QIQ54046.1 | P0C6U8 |
| GALLTKSSEYKGPITD | 1 | 0 | 0 | 1659956 | YP_009724389.1 | P0DTD1 |
| GALNKATNGAMQKESD | 1 | 0 | 0 | 1659957 | NP_828849.2 | P0C6X7 |
| GALNKATNNAMQIESD | 1 | 0 | 0 | 1659958 | QIQ54046.1 | P0C6U8 |
| GALNKATNNAMQVESD | 1 | 0 | 0 | 1659959 | YP_009724389.1 | P0DTD1 |
| GALRELTRELNGGAVT | 1 | 0 | 0 | 1659963 | NP_828849.2 | P0C6X7 |
| GAMDTTSYREAACCHL | 1 | 0 | 0 | 1659966 | YP_009724389.1 | P0DTD1 |
| GAMLSENFTSYGFSNV | 1 | 0 | 0 | 1659970 | NP_073551.1 | A0A0P0K6L9 |
| GAMQKESDDYIKLNGP | 1 | 0 | 0 | 1659971 | NP_828849.2 | P0C6X7 |
| GAMQTGFTTTNEAFQK | 1 | 0 | 0 | 1659972 | YP_009047204.1 | K9N5Q8 |
| GAMTAVNESRYGLQNL | 1 | 0 | 0 | 1659973 | YP_003767.1 | A0A0P0G321 |
| GANFRDLSLKGTTYLS | 1 | 0 | 0 | 1659975 | YP_173238.1 | A0A140H1H1 |
| GANGNKIVDKEVKDMQ | 1 | 0 | 0 | 1659976 | YP_009047202.1 | K9N7C7 |
| GANLRQDDSVRNLFAS | 1 | 0 | 0 | 1659979 | YP_009047204.1 | K9N5Q8 |
| GANYLGKPKEQIDGYT | 1 | 0 | 0 | 1659981 | NP_828849.2 | P0C6X7 |
| GAPIKGVTFGEDTVWE | 1 | 0 | 0 | 1659984 | NP_828849.2 | P0C6X7 |
| GAPSYCNGVRELYLNS | 1 | 0 | 0 | 1659987 | NP_828849.2 | P0C6X7 |
| GAPVKKVAFGGDQVHE | 1 | 0 | 0 | 1659990 | YP_009047202.1 | K9N7C7 |
| GARGTYRAALNSEKHQ | 1 | 0 | 0 | 1659994 | YP_009047202.1 | K9N7C7 |
| GARIPDVPTTLAWVNN | 1 | 0 | 0 | 1659995 | YP_009047202.1 | K9N7C7 |
| GARKSAPLIELCVDEA | 1 | 0 | 0 | 1659998 | YP_009724396.1 | P0DTC8 |
| GARPKQRRPQGLPNNT | 1 | 0 | 0 | 1660001 | QHR63308.1 | Q3LZX4 |
| GASQRVGTDSGFAAYN | 1 | 0 | 0 | 1660006 | NP_828855.1 | P59596 |
| GASSVQQQEEQEEDWL | 1 | 0 | 0 | 1660007 | QIQ54046.1 | P0C6U8 |
| GASVTEDVKFAASTGV | 1 | 0 | 0 | 1660011 | YP_003766.2 | H9EJ42 |
| GATIRLQAGKQTELAV | 1 | 0 | 0 | 1660015 | YP_003766.2 | H9EJ42 |
| GATVRLQAGKQTEFVS | 1 | 0 | 0 | 1660020 | NP_073549.1 | A0A0P0LKV0 |
| GAVCLKHAEEYREYLE | 1 | 0 | 0 | 1660023 | YP_009555238.1 | P0C6X6 |
| GAVCSKHANLYQKYVE | 1 | 0 | 0 | 1660028 | YP_003766.2 | H9EJ42 |
| GAVDINKLCEEMLDNR | 1 | 0 | 0 | 1660030 | YP_009724389.1 | P0DTD1 |
| GAVDINRLCEEMLDNR | 1 | 0 | 0 | 1660031 | NP_828849.2 | P0C6X7 |
| GAVLFCTPTKKAFPYG | 1 | 0 | 0 | 1660038 | NP_073549.1 | A0A0P0LKV0 |

| | | | | | | |
|---|---|---|---|---|---|---|
| GAVQKESDEYILAKGP | 1 | 0 | 0 | 1660042 | YP_009047202.1 | K9N7C7 |
| GAVSCGHYQTNIYSQN | 1 | 0 | 0 | 1660043 | NP_073549.1 | A0A0P0LKV0 |
| GAVTRYVDNNFCGPDG | 1 | 0 | 0 | 1660044 | NP_828849.2 | P0C6X7 |
| GAWDGYLSSCCGTPAK | 1 | 0 | 0 | 1660045 | YP_003766.2 | H9EJ42 |
| GAYSLFDMAKFPLKLA | 1 | 0 | 0 | 1660050 | YP_009555238.1 | P0C6X6 |
| GCACDRTTIQSVDISY | 1 | 0 | 0 | 1660053 | YP_003766.2 | H9EJ42 |
| GCAQGVFGVNMRTNWT | 1 | 0 | 0 | 1660056 | NP_073549.1 | A0A0P0LKV0 |
| GCCDCGSTKLPYYEFE | 1 | 0 | 0 | 1660058 | YP_003767.1 | A0A0P0G321 |
| GCCDDYTGYQELVIKT | 1 | 0 | 0 | 1660059 | YP_009555241.1 | P36334 |
| GCCDNSYFDSKGWYDP | 1 | 0 | 0 | 1660060 | YP_003766.2 | H9EJ42 |
| GCCESTKLPYYDVEKI | 1 | 0 | 0 | 1660062 | NP_073551.1 | A0A0P0K6L9 |
| GCDVNDVTKLYLGGMS | 1 | 0 | 0 | 1660071 | YP_009555238.1 | P0C6X6 |
| GCDYSGKLADCLQAYG | 1 | 0 | 0 | 1660073 | YP_173236.1 | A0A140H1G9 |
| GCEEGTMYFVPRAKSV | 1 | 0 | 0 | 1660078 | YP_009047202.1 | K9N7C7 |
| GCGASYTANDVEVQSS | 1 | 0 | 0 | 1660087 | YP_009047202.1 | K9N7C7 |
| GCGTNCMGKLKCNRCC | 1 | 0 | 0 | 1660092 | YP_009047204.1 | K9N5Q8 |
| GCGVSDVTKLYLGGMS | 1 | 0 | 0 | 1660098 | YP_009047202.1 | K9N7C7 |
| GCHATREAIGTNLPLQ | 1 | 0 | 0 | 1660100 | QHR63299.1 | P0C6W2 |
| GCHATREAVGTNLPLQ | 1 | 0 | 0 | 1660101 | YP_009724389.1 | P0DTD1 |
| GCIKACEVVVTNLNKS | 1 | 0 | 0 | 1660106 | YP_003766.2 | H9EJ42 |
| GCINANQVIVNNLDKS | 1 | 0 | 0 | 1660109 | YP_009724389.1 | P0DTD1 |
| GCIPASQVIVNNYDKS | 1 | 0 | 0 | 1660110 | YP_009555238.1 | P0C6X6 |
| GCITSREVVVTNLNKS | 1 | 0 | 0 | 1660113 | NP_073549.1 | A0A0P0LKV0 |
| GCLNASEVVVNNLDKS | 1 | 0 | 0 | 1660130 | YP_009047202.1 | K9N7C7 |
| GCLSSMNIAANSSEWA | 1 | 0 | 0 | 1660131 | YP_003766.2 | H9EJ42 |
| GCLVGAEHVNNSYECD | 1 | 0 | 0 | 1660133 | QIQ54048.1 | P59594 |
| GCMLRACIKTAQADTA | 1 | 0 | 0 | 1660134 | YP_009047205.1 | K9N796 |
| GCNVNDVTKLYLGGLN | 1 | 0 | 0 | 1660152 | NP_073549.1 | A0A0P0LKV0 |
| GCQTTNGLNTSYSVCN | 1 | 0 | 0 | 1660155 | NP_073551.1 | A0A0P0K6L9 |
| GCSTIAQAVRRYSEAA | 1 | 0 | 0 | 1660163 | NP_073549.1 | A0A0P0LKV0 |
| GCTCDRTAIQSFDNSY | 1 | 0 | 0 | 1660165 | NP_073549.1 | A0A0P0LKV0 |
| GCVGYTIEEAKTALKK | 1 | 0 | 0 | 1660177 | QIQ54046.1 | P0C6U8 |
| GCVIAWNSVKQDALTG | 1 | 0 | 0 | 1660180 | QIQ54048.1 | P59594 |
| GCVPLSAIPALAANTL | 1 | 0 | 0 | 1660189 | YP_173236.1 | A0A140H1G9 |
| GCYVVKPGGTSSGDAT | 1 | 0 | 0 | 1660202 | YP_009555238.1 | P0C6X6 |
| GDAAIKDFDYYRYNRP | 1 | 0 | 0 | 1660203 | NP_073549.1 | A0A0P0LKV0 |
| GDAALALLLLDRLNAL | 1 | 0 | 0 | 1660204 | QIQ54056.1 | P59595 |
| GDAAVKDFDFYRYNKP | 1 | 0 | 0 | 1660207 | YP_003766.2 | H9EJ42 |
| GDAEDTLTKLSKYLAN | 1 | 0 | 0 | 1660208 | NP_073549.1 | A0A0P0LKV0 |
| GDAEEALSKLSEYLIS | 1 | 0 | 0 | 1660209 | YP_003766.2 | H9EJ42 |
| GDCEEEDFEPPTQYEY | 1 | 0 | 0 | 1660228 | QHR63299.1 | P0C6W2 |
| GDDCFVSNSSVIQEVL | 1 | 0 | 0 | 1660239 | YP_173236.1 | A0A140H1G9 |
| GDDVRQIAPGQTGVIA | 1 | 0 | 0 | 1660245 | NP_828851.1 | P59594 |
| GDEVRQIAPGQTGVIA | 1 | 0 | 0 | 1660255 | QIQ54048.1 | P59594 |
| GDFADEIIGYKAIEDG | 1 | 0 | 0 | 1660256 | QIQ54046.1 | P0C6U8 |
| GDFNCTNFAINDLNTT | 1 | 0 | 0 | 1660270 | YP_173238.1 | A0A140H1H1 |
| GDFSAVSSPMSNMTEN | 1 | 0 | 0 | 1660272 | NP_073555.1 | A0A0P0KXN3 |
| GDGFVPFLLDGLIPRS | 1 | 0 | 0 | 1660288 | YP_173236.1 | A0A140H1G9 |
| GDGGITSEGNALYNNE | 1 | 0 | 0 | 1660289 | NP_073549.1 | A0A0P0LKV0 |
| GDGKMKELSPRWYFYY | 1 | 0 | 0 | 1660293 | NP_828858.1 | P59595 |
| GDGNALYNTEGGKTFM | 1 | 0 | 0 | 1660295 | YP_003766.2 | H9EJ42 |
| GDGTTSPISEHDYQIG | 1 | 0 | 0 | 1660298 | YP_009724391.1 | P0DTC3 |
| GDIILKPANNSLKITE | 1 | 0 | 0 | 1660305 | YP_009724389.1 | P0DTD1 |
| GDIIQRLDVLEQDAQI | 1 | 0 | 0 | 1660307 | YP_009047204.1 | K9N5Q8 |
| GDIITSGNLTTVVIPT | 1 | 0 | 0 | 1660308 | QIQ54046.1 | P0C6U8 |
| GDINARDLICAQKFNG | 1 | 0 | 0 | 1660309 | NP_828851.1 | P59594 |
| GDIVSGYEKSTFNKDI | 1 | 0 | 0 | 1660314 | YP_003766.2 | H9EJ42 |
| GDKGDAEDTLTKLSKY | 1 | 0 | 0 | 1660324 | NP_073549.1 | A0A0P0LKV0 |
| GDKVFVRDGNTDNLVF | 1 | 0 | 0 | 1660330 | NP_073549.1 | A0A0P0LKV0 |
| GDKVVYHTTSKPITFH | 1 | 0 | 0 | 1660332 | QIQ54046.1 | P0C6U8 |
| GDLAVCLGIGDSAVTY | 1 | 0 | 0 | 1660335 | YP_009555238.1 | P0C6X6 |
| GDLAVQIGNNNVCTYE | 1 | 0 | 0 | 1660336 | NP_073549.1 | A0A0P0LKV0 |
| GDLAVQIGSNNVCTYE | 1 | 0 | 0 | 1660337 | YP_003766.2 | H9EJ42 |
| GDLFYTIKFSGFKLDG | 1 | 0 | 0 | 1660338 | YP_003766.2 | H9EJ42 |
| GDLKCTSDNINDKDTG | 1 | 0 | 0 | 1660339 | YP_009555241.1 | P36334 |
| GDLQGNFGDLQLNKLG | 1 | 0 | 0 | 1660343 | YP_009047211.1 | K9N4V7 |
| GDLQLNKLGTEDPRWP | 1 | 0 | 0 | 1660344 | YP_009047211.1 | K9N4V7 |
| GDLVDSYFENDSILQS | 1 | 0 | 0 | 1660348 | NP_073549.1 | A0A0P0LKV0 |
| GDNDDEDVVTGDNDDE | 1 | 0 | 0 | 1660351 | YP_173236.1 | A0A140H1G9 |
| GDNDDEDVVTGDNNDE | 1 | 0 | 0 | 1660352 | YP_173236.1 | A0A140H1G9 |
| GDNDDQIVVTGDDVDD | 1 | 0 | 0 | 1660353 | YP_173236.1 | A0A140H1G9 |
| GDNIGTNVPLQVGFSN | 1 | 0 | 0 | 1660355 | YP_003766.2 | H9EJ42 |
| GDNNDEEIVTGDNDDQ | 1 | 0 | 0 | 1660357 | YP_173236.1 | A0A140H1G9 |
| GDNSAKFWEEAFYRDL | 1 | 0 | 0 | 1660358 | YP_009047202.1 | K9N7C7 |
| GDNVGTNVPLQVGFSN | 1 | 0 | 0 | 1660360 | NP_073549.1 | A0A0P0LKV0 |
| GDNVYCYNTDLIEGSK | 1 | 0 | 0 | 1660362 | YP_003766.2 | H9EJ42 |
| GDPAQLPAPRTLLTKG | 1 | 0 | 0 | 1660366 | YP_009724389.1 | P0DTD1 |
| GDPAQLPAPRVLLSKG | 1 | 0 | 0 | 1660368 | YP_009555238.1 | P0C6X6 |
| GDQDLIRQGTDYKHWP | 1 | 0 | 0 | 1660371 | NP_828858.1 | P59595 |
| GDQELIRQGTDYKHWP | 1 | 0 | 0 | 1660372 | YP_009724397.2 | P0DTC9 |
| GDQELIRQGTEYKHWP | 1 | 0 | 0 | 1660373 | QIQ54056.1 | P59595 |
| GDRCIKISTVQSKLTD | 1 | 0 | 0 | 1660378 | YP_003766.2 | H9EJ42 |
| GDRGIAPKSGYFVNVN | 1 | 0 | 0 | 1660381 | YP_009555241.1 | P36334 |
| GDRSGARPKQRRPQGL | 1 | 0 | 0 | 1660383 | QIQ54056.1 | P59595 |
| GDSCNNYMLTYNKVEN | 1 | 0 | 0 | 1660391 | YP_009724389.1 | P0DTD1 |
| GDSGKLLSELLKEKYT | 1 | 0 | 0 | 1660398 | YP_003766.2 | H9EJ42 |
| GDSHDTVLTSEEVVLK | 1 | 0 | 0 | 1660400 | NP_828849.2 | P0C6X7 |
| GDSSESGAKETKEINI | 1 | 0 | 0 | 1660407 | YP_009555238.1 | P0C6X6 |
| GDSSSGWTAGAAAYYV | 1 | 0 | 0 | 1660408 | YP_009724390.1 | P0DTC2 |
| GDSSWHIYLKSGTCPF | 1 | 0 | 0 | 1660409 | YP_003767.1 | A0A0P0G321 |
| GDSVEEALSEAREHLK | 1 | 0 | 0 | 1660415 | NP_828849.2 | P0C6X7 |

| | | | | | | |
|---|---|---|---|---|---|---|
| GDSVEEALSEARQHLK | 1 | 0 | 0 | 1660416 | QHR63299.1 | P0C6W2 |
| GDSVEEALSEARQHLL | 1 | 0 | 0 | 1660417 | QIQ54046.1 | P0C6U8 |
| GDSVEEVLSEARQHLK | 1 | 0 | 0 | 1660418 | YP_009724389.1 | P0DTD1 |
| GDVDNLVFTNKTTLPT | 1 | 0 | 0 | 1660426 | YP_003766.2 | H9EJ42 |
| GDVEACMVQVTCGSMT | 1 | 0 | 0 | 1660428 | YP_009047202.1 | K9N7C7 |
| GDVGIAPKQGYFIKHN | 1 | 0 | 0 | 1660432 | YP_173238.1 | A0A140H1H1 |
| GDVIKPVCAKSPPGEQ | 1 | 0 | 0 | 1660436 | YP_003766.2 | H9EJ42 |
| GDVILKPVQDGVKITE | 1 | 0 | 0 | 1660437 | QIQ54046.1 | P0C6U8 |
| GDVITSGDITCVVIPS | 1 | 0 | 0 | 1660438 | NP_828849.2 | P0C6X7 |
| GDVKFKRLGDYVLTEN | 1 | 0 | 0 | 1660439 | YP_003766.2 | H9EJ42 |
| GDVLLAEFDTYDPIYK | 1 | 0 | 0 | 1660442 | YP_009047202.1 | K9N7C7 |
| GDVVAIDYKHYTPSFK | 1 | 0 | 0 | 1660449 | YP_009724389.1 | P0DTD1 |
| GDVVAIDYRHYSASFK | 1 | 0 | 0 | 1660450 | NP_828849.2 | P0C6X7 |
| GDVVLASDDLYVKRYF | 1 | 0 | 0 | 1660452 | YP_173236.1 | A0A140H1G9 |
| GDVVLATDDLYVKRYE | 1 | 0 | 0 | 1660453 | YP_009555238.1 | P0C6X6 |
| GDVVQEGVLTAVVIPT | 1 | 0 | 0 | 1660454 | YP_009724389.1 | P0DTD1 |
| GDYNLSSVLPQRNIRS | 1 | 0 | 0 | 1660468 | YP_003767.1 | A0A0P0G321 |
| GDYYRKQLSPLEGGGW | 1 | 0 | 0 | 1660478 | YP_009047204.1 | K9N5Q8 |
| GEALKTQFNYYKKVNG | 1 | 0 | 0 | 1660482 | QHR63299.1 | P0C6W2 |
| GEAPVSIINNAVYTKV | 1 | 0 | 0 | 1660483 | NP_828849.2 | P0C6X7 |
| GEAVKTQFNYYKKVDG | 1 | 0 | 0 | 1660485 | YP_009724389.1 | P0DTD1 |
| GECPKFVFPLNSKVKV | 1 | 0 | 0 | 1660488 | NP_828849.2 | P0C6X7 |
| GEDFLGWIVPFGFMPS | 1 | 0 | 0 | 1660495 | YP_173236.1 | A0A140H1G9 |
| GEDIQLLKAAYENFNS | 1 | 0 | 0 | 1660496 | NP_828849.2 | P0C6X7 |
| GEDIQLLKSAYENFNQ | 1 | 0 | 0 | 1660497 | YP_009724389.1 | P0DTD1 |
| GEFSKKQLGLLQPTDN | 1 | 0 | 0 | 1660506 | YP_009047202.1 | K9N7C7 |
| GEGDGGITSEGNALYN | 1 | 0 | 0 | 1660509 | NP_073549.1 | A0A0P0LKV0 |
| GEIKDATPSDSVRATA | 1 | 0 | 0 | 1660518 | QHR63301.1 | Q3LZX0 |
| GEKGAYNKDHGRGGFG | 1 | 0 | 0 | 1660521 | YP_009555238.1 | P0C6X6 |
| GEKLFVEHYNEWAQAN | 1 | 0 | 0 | 1660522 | NP_073549.1 | A0A0P0LKV0 |
| GELGDVRETMNYLFQH | 1 | 0 | 0 | 1660524 | QHR63299.1 | P0C6W2 |
| GELGDVRETMSHLFQH | 1 | 0 | 0 | 1660525 | QIQ54046.1 | P0C6U8 |
| GELGDVRETMSYLFQH | 1 | 0 | 0 | 1660526 | YP_009724389.1 | P0DTD1 |
| GELGDVRETMTHLLQH | 1 | 0 | 0 | 1660527 | NP_828849.2 | P0C6X7 |
| GELKVEFWFAMRKDGD | 1 | 0 | 0 | 1660529 | YP_173236.1 | A0A140H1G9 |
| GELPVAISGDKVFVRD | 1 | 0 | 0 | 1660532 | NP_073549.1 | A0A0P0LKV0 |
| GELPVAVVNDKIFTKS | 1 | 0 | 0 | 1660533 | YP_009047202.1 | K9N7C7 |
| GELPVAVVNDKVFVRY | 1 | 0 | 0 | 1660534 | YP_003766.2 | H9EJ42 |
| GELQPLEEPTAQAVEV | 1 | 0 | 0 | 1660535 | QIQ54046.1 | P0C6U8 |
| GEMPCAIINDKVVAKI | 1 | 0 | 0 | 1660538 | YP_009555238.1 | P0C6X6 |
| GENAVIPKGIMMNVAK | 1 | 0 | 0 | 1660540 | NP_828849.2 | P0C6X7 |
| GENDNISVAVPKSRVQ | 1 | 0 | 0 | 1660542 | YP_009555245.1 | P33469 |
| GERQLLLVWEAGKSKP | 1 | 0 | 0 | 1660556 | YP_009047202.1 | K9N7C7 |
| GERSGARPKQRRPQGL | 1 | 0 | 0 | 1660557 | QHR63308.1 | Q3LZX4 |
| GERVRQAILNTVKFCD | 1 | 0 | 0 | 1660558 | YP_009047202.1 | K9N7C7 |
| GESVKTQFNYFKKVDG | 1 | 0 | 0 | 1660567 | NP_828849.2 | P0C6X7 |
| GETLPTEVLSEEVILK | 1 | 0 | 0 | 1660578 | QIQ54046.1 | P0C6U8 |
| GETLPTEVLTEEVVLK | 1 | 0 | 0 | 1660579 | YP_009724389.1 | P0DTD1 |
| GETPVSIVNNTVYTKV | 1 | 0 | 0 | 1660583 | QIQ54047.1 | P0C6X7 |
| GEVFNATKFPSVYAWE | 1 | 0 | 0 | 1660591 | NP_828851.1 | P59594 |
| GEVPVSIINNTVYTKV | 1 | 0 | 0 | 1660599 | YP_009724389.1 | P0DTD1 |
| GEYIFERIDYSDAVSY | 1 | 0 | 0 | 1660601 | YP_009047202.1 | K9N7C7 |
| GFAAYNRYRIGNYKLN | 1 | 0 | 0 | 1660609 | NP_828855.1 | P59596 |
| GFACGCGASYTANDVE | 1 | 0 | 0 | 1660610 | YP_009047202.1 | K9N7C7 |
| GFAIPSVAVRTYSEAA | 1 | 0 | 0 | 1660613 | YP_003766.2 | H9EJ42 |
| GFCDLKGKYVQIPTTC | 1 | 0 | 0 | 1660617 | YP_009724389.1 | P0DTD1 |
| GFDAWFSQRGGSYKND | 1 | 0 | 0 | 1660632 | NP_828849.2 | P0C6X7 |
| GFDKPQEKDKKSAKTG | 1 | 0 | 0 | 1660634 | NP_073556.1 | A0A0N7J131 |
| GFDMLDNYKAIDVVQY | 1 | 0 | 0 | 1660638 | YP_009555238.1 | P0C6X6 |
| GFDNQSKSPSSSGTST | 1 | 0 | 0 | 1660640 | YP_003771.1 | H9EJ69 |
| GFDSSKEFVEYKITEW | 1 | 0 | 0 | 1660642 | YP_009555238.1 | P0C6X6 |
| GFDSSKPVTKKYTYSF | 1 | 0 | 0 | 1660643 | YP_009047202.1 | K9N7C7 |
| GFDYVYNPFMIDVQQW | 1 | 0 | 0 | 1660650 | YP_009724389.1 | P0DTD1 |
| GFEDQPNLQVESANQM | 1 | 0 | 0 | 1660651 | NP_073549.1 | A0A0P0LKV0 |
| GFETIMKVLNENLNAY | 1 | 0 | 0 | 1660656 | YP_009555245.1 | P33469 |
| GFGDSVEEALSEAREH | 1 | 0 | 0 | 1660672 | NP_828849.2 | P0C6X7 |
| GFGDSVEEALSEARQH | 1 | 0 | 0 | 1660673 | QHR63299.1 | P0C6W2 |
| GFGDSVEEVLSEARQH | 1 | 0 | 0 | 1660674 | YP_009724389.1 | P0DTD1 |
| GFGGKQILGYSSLNDE | 1 | 0 | 0 | 1660679 | NP_073549.1 | A0A0P0LKV0 |
| GFGITVQYGTDTNSVC | 1 | 0 | 0 | 1660681 | YP_009047204.1 | K9N5Q8 |
| GFGVAVADSYYSYMMP | 1 | 0 | 0 | 1660683 | YP_173236.1 | A0A140H1G9 |
| GFGVDEEKCGVLDGSY | 1 | 0 | 0 | 1660684 | YP_173238.1 | A0A140H1H1 |
| GFGVFDIKNSSVNTVC | 1 | 0 | 0 | 1660685 | YP_003766.2 | H9EJ42 |
| GFGVNKIQPWTNDALN | 1 | 0 | 0 | 1660686 | NP_073549.1 | A0A0P0LKV0 |
| GFIEDSVFKPRPAGVL | 1 | 0 | 0 | 1660696 | YP_009555241.1 | P36334 |
| GFKDAVTFATVDHSAF | 1 | 0 | 0 | 1660707 | NP_073549.1 | A0A0P0LKV0 |
| GFKDFVTNKTYNIFPC | 1 | 0 | 0 | 1660708 | YP_173238.1 | A0A140H1H1 |
| GFKFGDPADSRDFLRV | 1 | 0 | 0 | 1660710 | YP_009555238.1 | P0C6X6 |
| GFKMNYQVNGYPNMFI | 1 | 0 | 0 | 1660717 | YP_009724389.1 | P0DTD1 |
| GFLQSSNYKIDTTSSS | 1 | 0 | 0 | 1660736 | YP_173238.1 | A0A140H1H1 |
| GFNDLSQLHCSYESFD | 1 | 0 | 0 | 1660748 | YP_009047204.1 | K9N5Q8 |
| GFNEWAMANGYTSVSS | 1 | 0 | 0 | 1660751 | YP_003766.2 | H9EJ42 |
| GFNFLQILPDPSKPSK | 1 | 0 | 0 | 1660752 | QIQ54048.1 | P59594 |
| GFNFSQILPDPLKPTK | 1 | 0 | 0 | 1660753 | NP_828851.1 | P59594 |
| GFNLEEAARCMRSLKA | 1 | 0 | 0 | 1660759 | NP_828849.2 | P0C6X7 |
| GFNVSELEDACFKFNY | 1 | 0 | 0 | 1660764 | YP_009555242.1 | Q04853 |
| GFPQLAELIPNQAALF | 1 | 0 | 0 | 1660768 | YP_003771.1 | H9EJ69 |
| GFQGRQILGSCTFEDE | 1 | 0 | 0 | 1660773 | YP_173236.1 | A0A140H1G9 |
| GFQGRQIMGSCSFEDE | 1 | 0 | 0 | 1660774 | YP_009555238.1 | P0C6X6 |
| GFQKCEQLLREYGQFC | 1 | 0 | 0 | 1660776 | YP_009047204.1 | K9N5Q8 |
| GFRDYITNRTFMIRSC | 1 | 0 | 0 | 1660779 | YP_009555241.1 | P36334 |

| | | | | | | |
|---|---|---|---|---|---|---|
| GFSCPLCCTVYDSSEV | 1 | 0 | 0 | 1660790 | YP_173236.1 | A0A140H1G9 |
| GFSNGVDFVAQPEGCV | 1 | 0 | 0 | 1660796 | NP_073549.1 | A0A0P0LKV0 |
| GFSNVVELPKFFYASN | 1 | 0 | 0 | 1660798 | NP_073551.1 | A0A0P0K6L9 |
| GFTAFANTVEDAVNKL | 1 | 0 | 0 | 1660807 | YP_009555242.1 | Q04853 |
| GFTAMNGEDAFSILAA | 1 | 0 | 0 | 1660808 | NP_073549.1 | A0A0P0LKV0 |
| GFTTTNEAFQKVQDAV | 1 | 0 | 0 | 1660821 | YP_009047204.1 | K9N5Q8 |
| GFTVKPVATVHRRIPD | 1 | 0 | 0 | 1660823 | YP_173238.1 | A0A140H1H1 |
| GFVVRIGAAANSTGTV | 1 | 0 | 0 | 1660843 | YP_009047204.1 | K9N5Q8 |
| GFVYFNATGSDVNCNG | 1 | 0 | 0 | 1660846 | YP_003767.1 | A0A0P0G321 |
| GFWRDGSCSCVGSSVA | 1 | 0 | 0 | 1660848 | YP_173236.1 | A0A140H1G9 |
| GFYFKPGGTTSGDAST | 1 | 0 | 0 | 1660852 | YP_003766.2 | H9EJ42 |
| GFYSANFLDDNVLPET | 1 | 0 | 0 | 1660859 | YP_003767.1 | A0A0P0G321 |
| GGAKLKALNLGETFVA | 1 | 0 | 0 | 1660865 | QIQ54046.1 | P0C6U8 |
| GGAKLKALNLGETFVT | 1 | 0 | 0 | 1660866 | YP_009724389.1 | P0DTD1 |
| GGAPIKGVTFGEDTVW | 1 | 0 | 0 | 1660869 | NP_828849.2 | P0C6X7 |
| GGAPVKKVAFGGDQVH | 1 | 0 | 0 | 1660873 | YP_009047202.1 | K9N7C7 |
| GGAVCLKHAEEYREYL | 1 | 0 | 0 | 1660879 | YP_009555238.1 | P0C6X6 |
| GGAVCSKHANLYQKYV | 1 | 0 | 0 | 1660884 | YP_003766.2 | H9EJ42 |
| GGAVTRYVDNNFCGPD | 1 | 0 | 0 | 1660887 | NP_828849.2 | P0C6X7 |
| GGAYSLFDMAKFPLKL | 1 | 0 | 0 | 1660888 | YP_009555238.1 | P0C6X6 |
| GGAYSLFDMTKFSLKL | 1 | 0 | 0 | 1660889 | YP_173236.1 | A0A140H1G9 |
| GGCCDDYTGYQELVIK | 1 | 0 | 0 | 1660892 | YP_009555241.1 | P36334 |
| GGCIKACEVVVTNLNK | 1 | 0 | 0 | 1660893 | YP_003766.2 | H9EJ42 |
| GGCINANQVIVNNLDK | 1 | 0 | 0 | 1660894 | YP_009724389.1 | P0DTD1 |
| GGCIPASQVIVNNYDK | 1 | 0 | 0 | 1660895 | YP_009555238.1 | P0C6X6 |
| GGCITSREVVVTNLNK | 1 | 0 | 0 | 1660896 | NP_073549.1 | A0A0P0LKV0 |
| GGCLNASEVVVNNLDK | 1 | 0 | 0 | 1660897 | YP_009047202.1 | K9N7C7 |
| GGCYYVKPGGTSSGDA | 1 | 0 | 0 | 1660905 | YP_009555238.1 | P0C6X6 |
| GGDAALALLLLDRLNA | 1 | 0 | 0 | 1660906 | QIQ54056.1 | P59595 |
| GGDAISLSFNNLLGFD | 1 | 0 | 0 | 1660907 | YP_009047202.1 | K9N7C7 |
| GGDFNLTLLEPVSIST | 1 | 0 | 0 | 1660908 | YP_009047204.1 | K9N5Q8 |
| GGDLLYLDLLNRLQAL | 1 | 0 | 0 | 1660910 | YP_009047211.1 | K9N4V7 |
| GGDNVYCYNTDLIEGS | 1 | 0 | 0 | 1660911 | YP_003766.2 | H9EJ42 |
| GGDRCIKISTVQSKLT | 1 | 0 | 0 | 1660914 | YP_003766.2 | H9EJ42 |
| GGDSSESGAKETKEIN | 1 | 0 | 0 | 1660916 | YP_009555238.1 | P0C6X6 |
| GGEMLKLGTSDPQFPI | 1 | 0 | 0 | 1660919 | YP_009555245.1 | P33469 |
| GGERHRQLVEHTTPWL | 1 | 0 | 0 | 1660920 | YP_009047202.1 | K9N7C7 |
| GGFNFSQILPDPLKPT | 1 | 0 | 0 | 1660928 | NP_828851.1 | P59594 |
| GGFYRFMVDPTTVLND | 1 | 0 | 0 | 1660934 | YP_003766.2 | H9EJ42 |
| GGGIAGAINAASKGAV | 1 | 0 | 0 | 1660938 | YP_009047202.1 | K9N7C7 |
| GGGKCFNCYPAGVNIT | 1 | 0 | 0 | 1660941 | NP_073551.1 | A0A0P0K6L9 |
| GGGLAKALDVYTKGKL | 1 | 0 | 0 | 1660942 | NP_073549.1 | A0A0P0LKV0 |
| GGGVAGALNKATNGAM | 1 | 0 | 0 | 1660945 | NP_828849.2 | P0C6X7 |
| GGGVAGALNKATNNAM | 1 | 0 | 0 | 1660946 | YP_009724389.1 | P0DTD1 |
| GGGYYVKPGGTSSGDA | 1 | 0 | 0 | 1660951 | YP_009047202.1 | K9N7C7 |
| GGHSFGIDLKSYDLGD | 1 | 0 | 0 | 1660953 | QIQ54046.1 | P0C6U8 |
| GGHSYGADLKSFDLGD | 1 | 0 | 0 | 1660954 | YP_009724389.1 | P0DTD1 |
| GGHSYGIDLKSYDLGD | 1 | 0 | 0 | 1660955 | NP_828849.2 | P0C6X7 |
| GGIAGAINAASKGAVQ | 1 | 0 | 0 | 1660956 | YP_009047202.1 | K9N7C7 |
| GGITSEGNALYNNEGG | 1 | 0 | 0 | 1660966 | NP_073549.1 | A0A0P0LKV0 |
| GGITYVSNSGNLLGFK | 1 | 0 | 0 | 1660967 | YP_003767.1 | A0A0P0G321 |
| GGKCFNCYPAGVNITL | 1 | 0 | 0 | 1660969 | NP_073551.1 | A0A0P0K6L9 |
| GGKPCIKVATVQSKMS | 1 | 0 | 0 | 1660979 | YP_009724389.1 | P0DTD1 |
| GGKQILGYSSLNDEFS | 1 | 0 | 0 | 1660980 | NP_073549.1 | A0A0P0LKV0 |
| GGLAKALDVYTKGKLQ | 1 | 0 | 0 | 1660986 | NP_073549.1 | A0A0P0LKV0 |
| GGLCPSTGQAMESVGF | 1 | 0 | 0 | 1660987 | YP_173236.1 | A0A140H1G9 |
| GGMIRYYDVMANGGTG | 1 | 0 | 0 | 1661010 | YP_009555238.1 | P0C6X6 |
| GGMSYYCEDHKPQYSF | 1 | 0 | 0 | 1661012 | YP_009555238.1 | P0C6X6 |
| GGNCAGNVIGNDALTR | 1 | 0 | 0 | 1661017 | YP_173236.1 | A0A140H1G9 |
| GGNSQSSSRASSLSRN | 1 | 0 | 0 | 1661026 | YP_009047211.1 | K9N4V7 |
| GGPRTIKVSTVQSKLT | 1 | 0 | 0 | 1661031 | NP_073549.1 | A0A0P0LKV0 |
| GGPSDSTDNNQNGDRS | 1 | 0 | 0 | 1661032 | QIQ54056.1 | P59595 |
| GGQIVTCAKEIKESVQ | 1 | 0 | 0 | 1661036 | YP_009724389.1 | P0DTD1 |
| GGRNGARPKQRRPQGL | 1 | 0 | 0 | 1661044 | NP_828858.1 | P59595 |
| GGSIAIKVTEYSWNKK | 1 | 0 | 0 | 1661054 | NP_073549.1 | A0A0P0LKV0 |
| GGSLYVKPGGTSSGDA | 1 | 0 | 0 | 1661058 | YP_009724389.1 | P0DTD1 |
| GGSQASSRSSSRSRGN | 1 | 0 | 0 | 1661064 | NP_828858.1 | P59595 |
| GGTPVCLGQNSQSPT | 1 | 0 | 0 | 1661082 | C7DN83.1 | Q76R62 |
| GGTSSGDATTAFANSV | 1 | 0 | 0 | 1661083 | YP_009555238.1 | P0C6X6 |
| GGTSSGDATTAYANSV | 1 | 0 | 0 | 1661084 | YP_009724389.1 | P0DTD1 |
| GGTTSGDASTAYANSI | 1 | 0 | 0 | 1661089 | YP_003766.2 | H9EJ42 |
| GGTTSGDATTAYANSV | 1 | 0 | 0 | 1661090 | NP_073549.1 | A0A0P0LKV0 |
| GGTTVVRPLVEDSTSV | 1 | 0 | 0 | 1661091 | YP_009047210.1 | K9N7A1 |
| GGVAGALNKATNGAMQ | 1 | 0 | 0 | 1661092 | NP_828849.2 | P0C6X7 |
| GGVAGALNKATNNAMQ | 1 | 0 | 0 | 1661093 | YP_009724389.1 | P0DTD1 |
| GGVLGDGNALYNTEGG | 1 | 0 | 0 | 1661099 | YP_003766.2 | H9EJ42 |
| GGVPVIEVSQIQSKLT | 1 | 0 | 0 | 1661102 | YP_173236.1 | A0A140H1G9 |
| GGVSVITPGTNASNQV | 1 | 0 | 0 | 1661105 | QHR63300.2 | Q3LZX1 |
| GGVSVITPGTNASSEV | 1 | 0 | 0 | 1661106 | NP_828851.1 | P59594 |
| GGVVQMTTQWLTNIFG | 1 | 0 | 0 | 1661112 | QIQ54046.1 | P0C6U8 |
| GGWDFMLKTLYKDVDN | 1 | 0 | 0 | 1661116 | YP_009047202.1 | K9N7C7 |
| GGWDNMLKNLMADVDD | 1 | 0 | 0 | 1661117 | NP_073549.1 | A0A0P0LKV0 |
| GGYHYTPFHYERDNTS | 1 | 0 | 0 | 1661124 | YP_009047202.1 | K9N7C7 |
| GGYYVKPGGTSSGDAT | 1 | 0 | 0 | 1661134 | YP_009047202.1 | K9N7C7 |
| GHAMPSLFKVQNVNLE | 1 | 0 | 0 | 1661136 | YP_009047202.1 | K9N7C7 |
| GHEDLMAAYVDNTSLT | 1 | 0 | 0 | 1661142 | QIQ54046.1 | P0C6U8 |
| GHEDLMAAYVENTSIT | 1 | 0 | 0 | 1661143 | NP_828849.2 | P0C6X7 |
| GHFNKEFYDFLRSQGF | 1 | 0 | 0 | 1661151 | NP_073549.1 | A0A0P0LKV0 |
| GHKNGQGENDNISVAV | 1 | 0 | 0 | 1661164 | YP_009555245.1 | P33469 |
| GHLYIQGIKLGTGYSL | 1 | 0 | 0 | 1661170 | YP_009555244.1 | Q01455 |
| GHLYIQGVKLGTGYTL | 1 | 0 | 0 | 1661171 | YP_173241.1 | Q0ZJ82 |

| | | | | | | |
|---|---|---|---|---|---|---|
| GHRLASGVQVHNLPEY | 1 | 0 | 0 | 1661187 | NP_073555.1 | A0A0P0KXN3 |
| GHSMQNCLLRLKVDTS | 1 | 0 | 0 | 1661194 | NP_828849.2 | P0C6X7 |
| GHSMQNCVLKLKVDTA | 1 | 0 | 0 | 1661196 | YP_009724389.1 | P0DTD1 |
| GHTDLMAAYVDNSSLT | 1 | 0 | 0 | 1661199 | YP_009724389.1 | P0DTD1 |
| GHTVCDSLNAKLGFDS | 1 | 0 | 0 | 1661203 | YP_009555238.1 | P0C6X6 |
| GHVETFYPKLQASQAW | 1 | 0 | 0 | 1661205 | NP_828849.2 | P0C6X7 |
| GHVETFYPKLQSSQAW | 1 | 0 | 0 | 1661206 | YP_009724389.1 | P0DTD1 |
| GHYAGSRSSSGNRSGI | 1 | 0 | 0 | 1661213 | YP_173242.1 | Q19U29 |
| GHYQTNIYSQNLCVDG | 1 | 0 | 0 | 1661219 | NP_073549.1 | A0A0P0LKV0 |
| GHYTGQAGEMPCAIIN | 1 | 0 | 0 | 1661222 | YP_009555238.1 | P0C6X6 |
| GHYTVYDTAKKSMYDG | 1 | 0 | 0 | 1661225 | NP_073549.1 | A0A0P0LKV0 |
| GIAAEQDRNTREVFAQ | 1 | 0 | 0 | 1661229 | NP_828851.1 | P59594 |
| GIAAINSVPWSKILAY | 1 | 0 | 0 | 1661230 | NP_828849.2 | P0C6X7 |
| GIADDTYVMGLHGNQT | 1 | 0 | 0 | 1661232 | NP_073549.1 | A0A0P0LKV0 |
| GIASGVFGVNLRTNFT | 1 | 0 | 0 | 1661242 | YP_003766.2 | H9EJ42 |
| GIATVREVLSDRELHL | 1 | 0 | 0 | 1661243 | YP_009724389.1 | P0DTD1 |
| GIATVREVLSDRELYL | 1 | 0 | 0 | 1661244 | QIQ54047.1 | P0C6X7 |
| GICADGSLIPVRPRNS | 1 | 0 | 0 | 1661248 | YP_003767.1 | A0A0P0G321 |
| GICASYQTQTNSRSVA | 1 | 0 | 0 | 1661252 | QHR63300.2 | Q3LZX1 |
| GICPENHVMVDCRRLL | 1 | 0 | 0 | 1661257 | YP_009555238.1 | P0C6X6 |
| GIDICSQDKVFGYVQQ | 1 | 0 | 0 | 1661262 | YP_009555238.1 | P0C6X6 |
| GIDVEIFENKTTLPVN | 1 | 0 | 0 | 1661269 | NP_828849.2 | P0C6X7 |
| GIDVTKLKCSDFYADK | 1 | 0 | 0 | 1661270 | YP_173236.1 | A0A140H1G9 |
| GIEFLKDGWEIVKFIT | 1 | 0 | 0 | 1661271 | QIQ54046.1 | P0C6U8 |
| GIEFLKRGDKSVYYTS | 1 | 0 | 0 | 1661272 | YP_009724389.1 | P0DTD1 |
| GIEFLKRGDKVVYHTT | 1 | 0 | 0 | 1661273 | QIQ54046.1 | P0C6U8 |
| GIENATFFIFNKLVKD | 1 | 0 | 0 | 1661274 | NP_828852.2 | P59632 |
| GIETAVGHYVHARLKG | 1 | 0 | 0 | 1661275 | YP_009047202.1 | K9N7C7 |
| GIGAVGGDLLYLDLLN | 1 | 0 | 0 | 1661294 | YP_009047211.1 | K9N4V7 |
| GIGDSAVTYSRLISLM | 1 | 0 | 0 | 1661295 | YP_009555238.1 | P0C6X6 |
| GIGGDRCIKISTVQSK | 1 | 0 | 0 | 1661297 | YP_003766.2 | H9EJ42 |
| GIGGKPCIKVATVQSK | 1 | 0 | 0 | 1661299 | NP_828849.2 | P0C6X7 |
| GIGGVPIIEVSQFQSK | 1 | 0 | 0 | 1661302 | YP_009555238.1 | P0C6X6 |
| GIGGVPVIEVSQIQSK | 1 | 0 | 0 | 1661303 | YP_173236.1 | A0A140H1G9 |
| GIGTCPAGTNYLTCDN | 1 | 0 | 0 | 1661308 | YP_009555241.1 | P36334 |
| GIGVTQNVLYENQKLI | 1 | 0 | 0 | 1661309 | YP_009724390.1 | P0DTC2 |
| GIIDVEKGVLPQLEQP | 1 | 0 | 0 | 1661317 | QIQ54046.1 | P0C6U8 |
| GIIRDVNPDDKCFANK | 1 | 0 | 0 | 1661322 | YP_009047202.1 | K9N7C7 |
| GIIRSSNQSLAGGITY | 1 | 0 | 0 | 1661323 | YP_003767.1 | A0A0P0G321 |
| GIISNARNGCIPLSVI | 1 | 0 | 0 | 1661325 | YP_009047202.1 | K9N7C7 |
| GIIWVATEGALNTPKD | 1 | 0 | 0 | 1661327 | YP_009724397.2 | P0DTC9 |
| GIKDVVLQGLKACCYV | 1 | 0 | 0 | 1661331 | YP_009047202.1 | K9N7C7 |
| GIKFQTVKPGNFNQDF | 1 | 0 | 0 | 1661332 | YP_173236.1 | A0A140H1G9 |
| GIKLGTGYSLADLPAY | 1 | 0 | 0 | 1661337 | YP_009555244.1 | Q01455 |
| GIKRFDNPVLPFNDGV | 1 | 0 | 0 | 1661344 | QHR63300.2 | Q3LZX1 |
| GIKVLPPLLSENQISG | 1 | 0 | 0 | 1661347 | YP_009555241.1 | P36334 |
| GIKYVAQPTEDVVDGD | 1 | 0 | 0 | 1661349 | YP_173236.1 | A0A140H1G9 |
| GILCIMSDRDLYDKLQ | 1 | 0 | 0 | 1661352 | YP_009724389.1 | P0DTD1 |
| GILDGDYAMQFFKMGR | 1 | 0 | 0 | 1661355 | NP_073549.1 | A0A0P0LKV0 |
| GILKWADQSDQFRNVQ | 1 | 0 | 0 | 1661359 | YP_009555245.1 | P33469 |
| GILLEAYNAFLDTVVS | 1 | 0 | 0 | 1661361 | NP_073549.1 | A0A0P0LKV0 |
| GILQQAALDYLATVKT | 1 | 0 | 0 | 1661365 | YP_003766.2 | H9EJ42 |
| GILSNMNIASNSKEWA | 1 | 0 | 0 | 1661366 | NP_073549.1 | A0A0P0LKV0 |
| GINASVVNIQKEIDRL | 1 | 0 | 0 | 1661376 | YP_009724390.1 | P0DTC2 |
| GINDDDYVIALTGTNQ | 1 | 0 | 0 | 1661377 | YP_003766.2 | H9EJ42 |
| GINSGTTCSNDLLQPN | 1 | 0 | 0 | 1661381 | YP_173238.1 | A0A140H1H1 |
| GINYLGDFAQGPFIDG | 1 | 0 | 0 | 1661384 | NP_073549.1 | A0A0P0LKV0 |
| GINYLGDFIQGPFIAG | 1 | 0 | 0 | 1661385 | YP_003766.2 | H9EJ42 |
| GINYLGTIKENIDGGA | 1 | 0 | 0 | 1661387 | YP_009047202.1 | K9N7C7 |
| GIQYGRSGETLGVLVP | 1 | 0 | 0 | 1661399 | YP_009724389.1 | P0DTD1 |
| GIREFSNLVLNNCTKY | 1 | 0 | 0 | 1661402 | YP_003767.1 | A0A0P0G321 |
| GIRFEYTSPTDKLAMI | 1 | 0 | 0 | 1661403 | YP_009555239.1 | Q80872 |
| GISEQSLRLVDAMVYT | 1 | 0 | 0 | 1661406 | NP_828849.2 | P0C6X7 |
| GISFCRRHNWNCVDCD | 1 | 0 | 0 | 1661407 | YP_009047202.1 | K9N7C7 |
| GISQYSLRLIDAMLFT | 1 | 0 | 0 | 1661408 | QIQ54046.1 | P0C6U8 |
| GISQYSLRLIDAMMFT | 1 | 0 | 0 | 1661409 | YP_009724389.1 | P0DTD1 |
| GISTPKLKEDYQIGGY | 1 | 0 | 0 | 1661412 | NP_828852.2 | P59632 |
| GITGDNVETLNEDLLE | 1 | 0 | 0 | 1661414 | NP_828849.2 | P0C6X7 |
| GITQFQKGKEFEFVEG | 1 | 0 | 0 | 1661421 | YP_009555245.1 | P33469 |
| GITQFQKGRDFKFSDG | 1 | 0 | 0 | 1661422 | YP_173242.1 | Q19U29 |
| GITSEGNALYNNEGGR | 1 | 0 | 0 | 1661425 | NP_073549.1 | A0A0P0LKV0 |
| GITVQYGTDTNSVCPK | 1 | 0 | 0 | 1661426 | YP_009047204.1 | K9N5Q8 |
| GITYTSTSGNLLGFKD | 1 | 0 | 0 | 1661428 | NP_073551.1 | A0A0P0K6L9 |
| GITYVSNSGNLLGFKN | 1 | 0 | 0 | 1661429 | YP_003767.1 | A0A0P0G321 |
| GIVDYGVRFFFYTSKE | 1 | 0 | 0 | 1661432 | NP_828849.2 | P0C6X7 |
| GIVYSSQLYEDVPSIN | 1 | 0 | 0 | 1661444 | YP_003768.1 | H9EJ24 |
| GIYQTSNFRVQPTDSI | 1 | 0 | 0 | 1661448 | QHR63300.2 | Q3LZX1 |
| GIYQTSNFRVQPTISI | 1 | 0 | 0 | 1661450 | QIQ54048.1 | P59594 |
| GKAGLYYESISYEEQD | 1 | 0 | 0 | 1661453 | NP_073549.1 | A0A0P0LKV0 |
| GKAKKGAWNIGEQKSI | 1 | 0 | 0 | 1661455 | YP_009724389.1 | P0DTD1 |
| GKCGDKFFPIVMNDKN | 1 | 0 | 0 | 1661465 | YP_173236.1 | A0A140H1G9 |
| GKCLKMNCVRFKNADL | 1 | 0 | 0 | 1661466 | YP_003766.2 | H9EJ42 |
| GKDLNANMSLAECKRA | 1 | 0 | 0 | 1661474 | NP_073549.1 | A0A0P0LKV0 |
| GKEDLKFPRGQGVPIN | 1 | 0 | 0 | 1661476 | YP_009724397.2 | P0DTC9 |
| GKEDLRFPRGQGVPIN | 1 | 0 | 0 | 1661477 | QIQ54056.1 | P59595 |
| GKEELRFPRGQGVPIN | 1 | 0 | 0 | 1661478 | NP_828858.1 | P59595 |
| GKEFEFVEGQGVPIAP | 1 | 0 | 0 | 1661479 | YP_009555245.1 | P33469 |
| GKFKKNAWNIGEPKSI | 1 | 0 | 0 | 1661489 | QIQ54046.1 | P0C6U8 |
| GKFLEELNAFTREMQQ | 1 | 0 | 0 | 1661490 | NP_073556.1 | A0A0N7J131 |
| GKFLSNTKYYDDSQYY | 1 | 0 | 0 | 1661491 | YP_009555240.1 | P30215 |
| GKFYGPFVDRQTAQAA | 1 | 0 | 0 | 1661499 | NP_828849.2 | P0C6X7 |

| | | | | | | |
|---|---|---|---|---|---|---|
| GKGFCKLHNWNCVNCD | 1 | 0 | 0 | 1661500 | YP_009724389.1 | P0DTD1 |
| GKGNKDEQIGYWNVQE | 1 | 0 | 0 | 1661501 | YP_003771.1 | H9EJ69 |
| GKGYHLMSFPQAAPHG | 1 | 0 | 0 | 1661505 | NP_828851.1 | P59594 |
| GKHEHSLLVEAYNSIL | 1 | 0 | 0 | 1661507 | YP_003766.2 | H9EJ42 |
| GKHERDLLIKAYNTIN | 1 | 0 | 0 | 1661508 | NP_073549.1 | A0A0P0LKV0 |
| GKIQDSLSSTASALGK | 1 | 0 | 0 | 1661513 | YP_009724390.1 | P0DTC2 |
| GKIWNGVIKNVNSVRD | 1 | 0 | 0 | 1661522 | NP_073549.1 | A0A0P0LKV0 |
| GKKIKHEGDWDSFCKT | 1 | 0 | 0 | 1661525 | NP_073549.1 | A0A0P0LKV0 |
| GKLKCNRCCDRYEEYD | 1 | 0 | 0 | 1661532 | YP_009047204.1 | K9N5Q8 |
| GKLLIRDTNKEVFVGD | 1 | 0 | 0 | 1661536 | YP_009555238.1 | P0C6X6 |
| GKLNCSFQDAASNIFV | 1 | 0 | 0 | 1661540 | YP_009047202.1 | K9N7C7 |
| GKLQDVVNQNAQALNT | 1 | 0 | 0 | 1661541 | YP_009724390.1 | P0DTC2 |
| GKLRPFERDISNVPFS | 1 | 0 | 0 | 1661543 | NP_828851.1 | P59594 |
| GKMDTAAYREAACSQL | 1 | 0 | 0 | 1661546 | YP_009555238.1 | P0C6X6 |
| GKMFVRPVIEDYHTLT | 1 | 0 | 0 | 1661547 | YP_173241.1 | Q0ZJ82 |
| GKPCIKVATVQSKMSD | 1 | 0 | 0 | 1661556 | YP_009724389.1 | P0DTD1 |
| GKPGNVGGNDALSIST | 1 | 0 | 0 | 1661558 | YP_009555238.1 | P0C6X6 |
| GKPILWVNKASYDTNL | 1 | 0 | 0 | 1661560 | YP_009047202.1 | K9N7C7 |
| GKPKEQIDGYTMHANY | 1 | 0 | 0 | 1661563 | NP_828849.2 | P0C6X7 |
| GKPREQIDGYVMHANY | 1 | 0 | 0 | 1661566 | YP_009724389.1 | P0DTD1 |
| GKQATKYLVQQESPFV | 1 | 0 | 0 | 1661577 | YP_009724389.1 | P0DTD1 |
| GKQATQYLVQQESPFV | 1 | 0 | 0 | 1661578 | QHR63299.1 | P0C6W2 |
| GKQGNFKNLREFVFKN | 1 | 0 | 0 | 1661579 | YP_009724390.1 | P0DTC2 |
| GKQIDDHRITSITSDK | 1 | 0 | 0 | 1661580 | YP_009555238.1 | P0C6X6 |
| GKQSSSRASSGNRSGN | 1 | 0 | 0 | 1661587 | YP_009555245.1 | P33469 |
| GKQVVSDIDYVPLKSA | 1 | 0 | 0 | 1661591 | YP_009724389.1 | P0DTD1 |
| GKRGPNQNFGGGEMLK | 1 | 0 | 0 | 1661593 | YP_009555245.1 | P33469 |
| GKSIIYTGDWEGLHEV | 1 | 0 | 0 | 1661603 | YP_003766.2 | H9EJ42 |
| GKSKCDESASKSASVY | 1 | 0 | 0 | 1661604 | NP_828849.2 | P0C6X7 |
| GKSLEFGATSVTPQPE | 1 | 0 | 0 | 1661609 | QHR63299.1 | P0C6W2 |
| GKSLGNVFCDGVNVTK | 1 | 0 | 0 | 1661610 | YP_009555238.1 | P0C6X6 |
| GKTVLGEYVFDKSELT | 1 | 0 | 0 | 1661623 | YP_009555238.1 | P0C6X6 |
| GKVDSIVQKCYELSHL | 1 | 0 | 0 | 1661625 | YP_003766.2 | H9EJ42 |
| GKVFFQFDNLSSEDLK | 1 | 0 | 0 | 1661627 | YP_009555238.1 | P0C6X6 |
| GKVKVGTGVMVECDSL | 1 | 0 | 0 | 1661632 | NP_073549.1 | A0A0P0LKV0 |
| GKWVQVPIGTNDPIRF | 1 | 0 | 0 | 1661639 | NP_073549.1 | A0A0P0LKV0 |
| GKYAQNLLKKLIGGDV | 1 | 0 | 0 | 1661642 | YP_009047202.1 | K9N7C7 |
| GKYFTFKQANNNCFVN | 1 | 0 | 0 | 1661646 | YP_009555238.1 | P0C6X6 |
| GKYSQKAYALLRQYRG | 1 | 0 | 0 | 1661648 | YP_173236.1 | A0A140H1G9 |
| GKYYTKPIIKAQFKPF | 1 | 0 | 0 | 1661653 | YP_173236.1 | A0A140H1G9 |
| GLAAVNSVPWDTIANY | 1 | 0 | 0 | 1661658 | YP_009724389.1 | P0DTD1 |
| GLAGKVKVGTGVMVEC | 1 | 0 | 0 | 1661661 | NP_073549.1 | A0A0P0LKV0 |
| GLAKRFKESPFELEDF | 1 | 0 | 0 | 1661664 | YP_009724389.1 | P0DTD1 |
| GLAKRSKDSPLELEDF | 1 | 0 | 0 | 1661665 | QIQ54047.1 | P0C6X7 |
| GLAKRSKESPLELEDF | 1 | 0 | 0 | 1661666 | QHR63299.1 | P0C6W2 |
| GLALDNALSGVLSAAI | 1 | 0 | 0 | 1661668 | NP_073549.1 | A0A0P0LKV0 |
| GLASAYRANSFDVPTF | 1 | 0 | 0 | 1661671 | YP_009047202.1 | K9N7C7 |
| GLCDAANPTNCIAPVN | 1 | 0 | 0 | 1661677 | YP_009047204.1 | K9N5Q8 |
| GLDAAWNKFVLGDVEI | 1 | 0 | 0 | 1661692 | NP_073549.1 | A0A0P0LKV0 |
| GLDGLIDSYFDNSSTL | 1 | 0 | 0 | 1661696 | YP_003766.2 | H9EJ42 |
| GLDSLDSYPALETIQV | 1 | 0 | 0 | 1661699 | NP_828849.2 | P0C6X7 |
| GLDSLDTYPSLETIQI | 1 | 0 | 0 | 1661700 | YP_009724389.1 | P0DTD1 |
| GLDYKAFKQIVESCGN | 1 | 0 | 0 | 1661701 | YP_009724389.1 | P0DTD1 |
| GLDYKSFKQIVESCGN | 1 | 0 | 0 | 1661702 | QIQ54046.1 | P0C6U8 |
| GLDYKTFKQIVESCGN | 1 | 0 | 0 | 1661703 | QHR63299.1 | P0C6W2 |
| GLEAYADKTLQEMKAL | 1 | 0 | 0 | 1661707 | YP_009555238.1 | P0C6X6 |
| GLEDFNFEHVVYGDVS | 1 | 0 | 0 | 1661708 | NP_073549.1 | A0A0P0LKV0 |
| GLEDYAFDHIVYGSFN | 1 | 0 | 0 | 1661709 | YP_173236.1 | A0A140H1G9 |
| GLEDYAFEHIVYGNFN | 1 | 0 | 0 | 1661710 | YP_009555238.1 | P0C6X6 |
| GLEFTDENYNNLVPTY | 1 | 0 | 0 | 1661711 | YP_173236.1 | A0A140H1G9 |
| GLELHWAPEFPWMFED | 1 | 0 | 0 | 1661712 | YP_009555238.1 | P0C6X6 |
| GLELTDESCNNLVPTY | 1 | 0 | 0 | 1661713 | YP_009555238.1 | P0C6X6 |
| GLENIAFNVVKQGHFI | 1 | 0 | 0 | 1661715 | YP_009047202.1 | K9N7C7 |
| GLENIAFNVVNKGSFV | 1 | 0 | 0 | 1661716 | NP_073549.1 | A0A0P0LKV0 |
| GLESKQVDYVPLRSAT | 1 | 0 | 0 | 1661718 | YP_173236.1 | A0A140H1G9 |
| GLFDAIWDAFVAAIKL | 1 | 0 | 0 | 1661722 | YP_003766.2 | H9EJ42 |
| GLFKDCARNPIDLPPS | 1 | 0 | 0 | 1661724 | NP_073549.1 | A0A0P0LKV0 |
| GLFKDCSRETSGLSPA | 1 | 0 | 0 | 1661727 | YP_009047202.1 | K9N7C7 |
| GLFTLQLVSAVNQSLS | 1 | 0 | 0 | 1661738 | NP_073552.1 | A0A0P0KWE8 |
| GLGDLVDSYFENDSIL | 1 | 0 | 0 | 1661741 | NP_073549.1 | A0A0P0LKV0 |
| GLGTVDVDYKSCTKGL | 1 | 0 | 0 | 1661748 | YP_003767.1 | A0A0P0G321 |
| GLGVTMDVLNKNQKLI | 1 | 0 | 0 | 1661751 | YP_173238.1 | A0A140H1H1 |
| GLGVTMDVLSQNQKLI | 1 | 0 | 0 | 1661752 | YP_009555241.1 | P36334 |
| GLGYIADISAFKATLY | 1 | 0 | 0 | 1661753 | NP_073549.1 | A0A0P0LKV0 |
| GLHLLIGLYRRQQTSN | 1 | 0 | 0 | 1661763 | YP_009555238.1 | P0C6X6 |
| GLIDLLVNVLSIFSSS | 1 | 0 | 0 | 1661773 | YP_003766.2 | H9EJ42 |
| GLIDQINSTYVDLKLL | 1 | 0 | 0 | 1661774 | YP_003767.1 | A0A0P0G321 |
| GLIDSYFDNSSTLQSV | 1 | 0 | 0 | 1661775 | YP_003766.2 | H9EJ42 |
| GLKFTEESFYKNMYLK | 1 | 0 | 0 | 1661780 | YP_173236.1 | A0A140H1G9 |
| GLKITEEVGHTDLMAA | 1 | 0 | 0 | 1661784 | QHR63299.1 | P0C6W2 |
| GLKSSTGFVYFNATGS | 1 | 0 | 0 | 1661793 | YP_003767.1 | A0A0P0G321 |
| GLKYTKILKDDGNFVV | 1 | 0 | 0 | 1661802 | YP_009555238.1 | P0C6X6 |
| GLKYTKIMKDDGNCVV | 1 | 0 | 0 | 1661803 | YP_173236.1 | A0A140H1G9 |
| GLLDTTQLHVADTLMQ | 1 | 0 | 0 | 1661806 | YP_173238.1 | A0A140H1H1 |
| GLLEVSVCQYNMCEYP | 1 | 0 | 0 | 1661807 | YP_009555241.1 | P36334 |
| GLLHSSQLPLDVYDLT | 1 | 0 | 0 | 1661813 | YP_009555238.1 | P0C6X6 |
| GLLPPKNSIDAFKLNV | 1 | 0 | 0 | 1661820 | QIQ54046.1 | P0C6U8 |
| GLLPPKSSIDAFKLNI | 1 | 0 | 0 | 1661821 | NP_828849.2 | P0C6X7 |
| GLLTACAFSVDPATTY | 1 | 0 | 0 | 1661826 | YP_003766.2 | H9EJ42 |
| GLMLLEIKDKEQYCAL | 1 | 0 | 0 | 1661831 | NP_828849.2 | P0C6X7 |
| GLMLLEIKDTEKYCAL | 1 | 0 | 0 | 1661832 | YP_009724389.1 | P0DTD1 |

| | | | | | | |
|---|---|---|---|---|---|---|
| GLNDNLLEILQKEKVN | 1 | 0 | 0 | 1661840 | YP_009724389.1 | P0DTD1 |
| GLNGYTVEEARTVLKK | 1 | 0 | 0 | 1661842 | QHR63299.1 | P0C6W2 |
| GLNKPFVKDNVSFVAD | 1 | 0 | 0 | 1661843 | YP_009047202.1 | K9N7C7 |
| GLNLEEAARYMRSLKV | 1 | 0 | 0 | 1661845 | YP_009724389.1 | P0DTD1 |
| GLNNLNRGMVLGSLAA | 1 | 0 | 0 | 1661846 | YP_009724389.1 | P0DTD1 |
| GLPTQTVDSSQGSEYD | 1 | 0 | 0 | 1661859 | YP_009724389.1 | P0DTD1 |
| GLPYGANKEGIIWVAT | 1 | 0 | 0 | 1661862 | QIQ54056.1 | P59595 |
| GLQDCVTGINDDDYVI | 1 | 0 | 0 | 1661863 | YP_003766.2 | H9EJ42 |
| GLQFQILDEGVDCKIQ | 1 | 0 | 0 | 1661865 | YP_009555239.1 | Q80872 |
| GLQIQTVDSSQGSEYD | 1 | 0 | 0 | 1661866 | YP_003766.2 | H9EJ42 |
| GLQPSVGPKQASLNGV | 1 | 0 | 0 | 1661869 | YP_009724389.1 | P0DTD1 |
| GLQTQTVDSAQGSEYD | 1 | 0 | 0 | 1661870 | NP_073549.1 | A0A0P0LKV0 |
| GLRGAIRKAKDYGFTV | 1 | 0 | 0 | 1661873 | YP_009047202.1 | K9N7C7 |
| GLRGPGDLQGNFGDLQ | 1 | 0 | 0 | 1661874 | YP_009047211.1 | K9N4V7 |
| GLSPAYAPTYVSVDDK | 1 | 0 | 0 | 1661885 | YP_009047202.1 | K9N7C7 |
| GLSRYVARLSSNSRI | 1 | 0 | 0 | 1661886 | P03155.1 | Q69028 |
| GLTNCLASECFVKSDI | 1 | 0 | 0 | 1661900 | YP_003766.2 | H9EJ42 |
| GLTNLSEITKPLIEAE | 1 | 0 | 0 | 1661901 | YP_173243.1 | Q0ZJ80 |
| GLTNQVVKPGHFNEEF | 1 | 0 | 0 | 1661902 | YP_003766.2 | H9EJ42 |
| GLTPSKGPAQASVNGV | 1 | 0 | 0 | 1661904 | NP_828849.2 | P0C6X7 |
| GLTSQTVKPGHFNKEF | 1 | 0 | 0 | 1661910 | NP_073549.1 | A0A0P0LKV0 |
| GLTTQTVDSSQGSEYQ | 1 | 0 | 0 | 1661911 | YP_009047202.1 | K9N7C7 |
| GLVELEKGVLPQLEQP | 1 | 0 | 0 | 1661922 | NP_828849.2 | P0C6X7 |
| GLVEVEKGVLPQLEQP | 1 | 0 | 0 | 1661923 | YP_009724389.1 | P0DTD1 |
| GLVGVLTLDNQDLNGK | 1 | 0 | 0 | 1661927 | YP_009047202.1 | K9N7C7 |
| GLVNVQKVEQPKIEPK | 1 | 0 | 0 | 1661933 | NP_073549.1 | A0A0P0LKV0 |
| GLVTICDFYSKLGVGF | 1 | 0 | 0 | 1661937 | YP_173236.1 | A0A140H1G9 |
| GLYDDILTNNKPWFVR | 1 | 0 | 0 | 1661950 | YP_003766.2 | H9EJ42 |
| GLYEIQIPSEFTIGNM | 1 | 0 | 0 | 1661952 | YP_009555241.1 | P36334 |
| GLYKNSATGSLDVEVF | 1 | 0 | 0 | 1661958 | YP_003766.2 | H9EJ42 |
| GLYKQSCTGSPYIDDF | 1 | 0 | 0 | 1661960 | YP_009555238.1 | P0C6X6 |
| GLYPTLNISDEFSSNV | 1 | 0 | 0 | 1661964 | YP_009724389.1 | P0DTD1 |
| GLYRKCVKPKEETGLL | 1 | 0 | 0 | 1661965 | QHR63299.1 | P0C6W2 |
| GLYRRQQTSNLVVQEF | 1 | 0 | 0 | 1661968 | YP_009555238.1 | P0C6X6 |
| GMDAKQVDYVPLKSAT | 1 | 0 | 0 | 1661980 | YP_009555238.1 | P0C6X6 |
| GMETLACEDTKLVTEE | 1 | 0 | 0 | 1661988 | QIQ54046.1 | P0C6U8 |
| GMFGLYDDILTNNKPW | 1 | 0 | 0 | 1661992 | YP_003766.2 | H9EJ42 |
| GMHFGACDYDRLPNEV | 1 | 0 | 0 | 1661995 | YP_009047210.1 | K9N7A1 |
| GMIKKLDNDVLNGIIS | 1 | 0 | 0 | 1661998 | YP_009047202.1 | K9N7C7 |
| GMIRYYDVMANGGTGF | 1 | 0 | 0 | 1662000 | YP_009555238.1 | P0C6X6 |
| GMLNGNAIATVKSEDG | 1 | 0 | 0 | 1662008 | NP_073549.1 | A0A0P0LKV0 |
| GMPIPDFDLDDFIDAP | 1 | 0 | 0 | 1662014 | YP_009047202.1 | K9N7C7 |
| GMPSFVAYETARQEYE | 1 | 0 | 0 | 1662015 | NP_073549.1 | A0A0P0LKV0 |
| GMPSYCTGYREGYLNS | 1 | 0 | 0 | 1662016 | YP_009724389.1 | P0DTD1 |
| GMPSYCTSYRDGYLNS | 1 | 0 | 0 | 1662017 | QIQ54046.1 | P0C6U8 |
| GMQXYSTLQGPPGTGK | 1 | 0 | 0 | 1662023 | QIQ54047.1 | P0C6X7 |
| GMSYYCEDHKPQYSFK | 1 | 0 | 0 | 1662031 | YP_009555238.1 | P0C6X6 |
| GNAAITDYNYYKYNLP | 1 | 0 | 0 | 1662046 | YP_009555238.1 | P0C6X6 |
| GNAIATVKSEDGNIKN | 1 | 0 | 0 | 1662051 | NP_073549.1 | A0A0P0LKV0 |
| GNCAGNVIGNDALTRF | 1 | 0 | 0 | 1662059 | YP_173236.1 | A0A140H1G9 |
| GNCYRCPKEIVETVSA | 1 | 0 | 0 | 1662070 | YP_173236.1 | A0A140H1G9 |
| GNDVLYQPPTASVSTS | 1 | 0 | 0 | 1662075 | YP_173236.1 | A0A140H1G9 |
| GNELAQVLTEVVYSNG | 1 | 0 | 0 | 1662076 | YP_003766.2 | H9EJ42 |
| GNFDDQPSLQVESANL | 1 | 0 | 0 | 1662078 | YP_003766.2 | H9EJ42 |
| GNFGDLQLNKLGTEDP | 1 | 0 | 0 | 1662079 | YP_009047211.1 | K9N4V7 |
| GNFGDQELIRQGTEYK | 1 | 0 | 0 | 1662082 | QIQ54056.1 | P59595 |
| GNFNQDFYEFVKSKGL | 1 | 0 | 0 | 1662092 | YP_173236.1 | A0A140H1G9 |
| GNFTGLYSSTVPVFN | 1 | 0 | 0 | 1662098 | P03155.1 | Q69028 |
| GNFYCFVNTTIGNETT | 1 | 0 | 0 | 1662100 | NP_073551.1 | A0A0P0K6L9 |
| GNFYGPFVDRQTAQAA | 1 | 0 | 0 | 1662103 | YP_009724389.1 | P0DTD1 |
| GNGILKWADQSDQFRN | 1 | 0 | 0 | 1662106 | YP_009555245.1 | P33469 |
| GNGSMIEDIDYVPLKS | 1 | 0 | 0 | 1662112 | YP_009047202.1 | K9N7C7 |
| GNICYTPSKLIEYSDF | 1 | 0 | 0 | 1662120 | NP_828849.2 | P0C6X7 |
| GNICYTPSKLIEYTDF | 1 | 0 | 0 | 1662121 | YP_009724389.1 | P0DTD1 |
| GNIFGLYKNSATGSLD | 1 | 0 | 0 | 1662122 | YP_003766.2 | H9EJ42 |
| GNIFIVTPCNQPDQVA | 1 | 0 | 0 | 1662123 | YP_003767.1 | A0A0P0G321 |
| GNIIGFKDFVTNKTYN | 1 | 0 | 0 | 1662124 | YP_173238.1 | A0A140H1H1 |
| GNKDEQIGYWNVQERW | 1 | 0 | 0 | 1662128 | YP_003771.1 | H9EJ69 |
| GNKFYNVSTGLIPGFV | 1 | 0 | 0 | 1662129 | YP_173236.1 | A0A140H1G9 |
| GNKIVDKEVKDMQFDL | 1 | 0 | 0 | 1662137 | YP_009047202.1 | K9N7C7 |
| GNKQIVHSYPILHHPG | 1 | 0 | 0 | 1662139 | YP_009047207.1 | K9N643 |
| GNLGSNGKPGNVGGND | 1 | 0 | 0 | 1662143 | YP_009555238.1 | P0C6X6 |
| GNLHQRELLGVSDVWH | 1 | 0 | 0 | 1662147 | YP_009555238.1 | P0C6X6 |
| GNLVEKLSPWFMDGEN | 1 | 0 | 0 | 1662159 | YP_009047202.1 | K9N7C7 |
| GNMFQFATLPVYDTIK | 1 | 0 | 0 | 1662164 | YP_009047204.1 | K9N5Q8 |
| GNMVEFIQTSSPKVTI | 1 | 0 | 0 | 1662169 | YP_009555241.1 | P36334 |
| GNPDEPQKDVYELRYN | 1 | 0 | 0 | 1662175 | YP_009555245.1 | P33469 |
| GNPKGIPIVDDPVVDW | 1 | 0 | 0 | 1662180 | YP_009047202.1 | K9N7C7 |
| GNRKRNQKPLEPKFSI | 1 | 0 | 0 | 1662190 | YP_003771.1 | H9EJ69 |
| GNRSGNGILKWADQSD | 1 | 0 | 0 | 1662192 | YP_009555245.1 | P33469 |
| GNSPFHPLADNKFALA | 1 | 0 | 0 | 1662201 | QIQ54053.1 | P59635 |
| GNSPFHPLADNKFALT | 1 | 0 | 0 | 1662202 | YP_009724395.1 | P0DTC7 |
| GNSQSSSRASSLSRNS | 1 | 0 | 0 | 1662203 | YP_009047211.1 | K9N4V7 |
| GNSRNSTPGSSRGNSP | 1 | 0 | 0 | 1662204 | NP_828858.1 | P59595 |
| GNSTRGTSPGPSGIGA | 1 | 0 | 0 | 1662206 | YP_009047211.1 | K9N4V7 |
| GNSYTSFATYHTPATD | 1 | 0 | 0 | 1662207 | YP_009047204.1 | K9N5Q8 |
| GNVDLDALASDIFDTP | 1 | 0 | 0 | 1662225 | YP_009047202.1 | K9N7C7 |
| GNVEAVNFNVTTAETT | 1 | 0 | 0 | 1662226 | NP_073551.1 | A0A0P0K6L9 |
| GNVFCDGVNVTKHKCD | 1 | 0 | 0 | 1662228 | YP_009555238.1 | P0C6X6 |
| GNVILKPSDEGVKVTQ | 1 | 0 | 0 | 1662233 | NP_828849.2 | P0C6X7 |
| GNVKGLQPSVGPKQAS | 1 | 0 | 0 | 1662234 | QIQ54047.1 | P0C6X7 |

| | | | | | | |
|---|---|---|---|---|---|---|
| GNVKPGDAVITTQQAG | 1 | 0 | 0 | 1662235 | NP_073549.1 | A0A0P0LKV0 |
| GNVQVDNGSSINRKQL | 1 | 0 | 0 | 1662241 | YP_003766.2 | H9EJ42 |
| GNVSRHNLTKYTMMDL | 1 | 0 | 0 | 1662243 | YP_003766.2 | H9EJ42 |
| GNVSRQDLTKYTMMDL | 1 | 0 | 0 | 1662244 | NP_073549.1 | A0A0P0LKV0 |
| GNVSTVFVPATRDSVP | 1 | 0 | 0 | 1662245 | YP_009047208.1 | K9N7D2 |
| GNVVKTAVQPTAPAYV | 1 | 0 | 0 | 1662249 | YP_003766.2 | H9EJ42 |
| GNVVTSVMESQISTKE | 1 | 0 | 0 | 1662252 | YP_173236.1 | A0A140H1G9 |
| GNVWQIQTIQDSDGTN | 1 | 0 | 0 | 1662253 | YP_009555238.1 | P0C6X6 |
| GNYKLNTDHAGSNDNI | 1 | 0 | 0 | 1662257 | NP_828855.1 | P59596 |
| GNYKLNTDHSNSSDNI | 1 | 0 | 0 | 1662258 | QIQ54051.1 | P59596 |
| GNYNRNASLNSFKEYF | 1 | 0 | 0 | 1662262 | YP_009047204.1 | K9N5Q8 |
| GNYRLPSSKPSGMDTA | 1 | 0 | 0 | 1662266 | YP_173241.1 | Q0ZJ82 |
| GNYRLPSTQKGSGMDT | 1 | 0 | 0 | 1662267 | YP_009555244.1 | Q01455 |
| GPAGVESDVETNEIVD | 1 | 0 | 0 | 1662274 | YP_009047202.1 | K9N7C7 |
| GPASARDLICAQYVAG | 1 | 0 | 0 | 1662276 | YP_009047204.1 | K9N5Q8 |
| GPAYVMIDKVEFENGF | 1 | 0 | 0 | 1662278 | NP_073549.1 | A0A0P0LKV0 |
| GPDVFLHKCYRCPAEI | 1 | 0 | 0 | 1662293 | NP_073549.1 | A0A0P0LKV0 |
| GPEAALPFRAVKDGIV | 1 | 0 | 0 | 1662295 | YP_009047211.1 | K9N4V7 |
| GPEAGLPYGANKDGII | 1 | 0 | 0 | 1662296 | YP_009724397.2 | P0DTC9 |
| GPEAGLPYGANKEGII | 1 | 0 | 0 | 1662297 | QIQ54056.1 | P59595 |
| GPEASLPYGANKEGIV | 1 | 0 | 0 | 1662298 | NP_828858.1 | P59595 |
| GPEHSLAEYHNESGIK | 1 | 0 | 0 | 1662299 | QIQ54046.1 | P0C6U8 |
| GPEHSLAEYHNESGLK | 1 | 0 | 0 | 1662300 | YP_009724389.1 | P0DTD1 |
| GPFMDNAINVGGTGLQ | 1 | 0 | 0 | 1662312 | YP_009047202.1 | K9N7C7 |
| GPFNFCSETVNGYAKS | 1 | 0 | 0 | 1662313 | NP_073549.1 | A0A0P0LKV0 |
| GPGDLQGNFGDLQLNK | 1 | 0 | 0 | 1662316 | YP_009047211.1 | K9N4V7 |
| GPGNTFINGDIARELG | 1 | 0 | 0 | 1662318 | YP_003766.2 | H9EJ42 |
| GPHEFCSQHTMQIVDK | 1 | 0 | 0 | 1662326 | YP_003766.2 | H9EJ42 |
| GPISQFNYKQSFSNPT | 1 | 0 | 0 | 1662333 | YP_009047204.1 | K9N5Q8 |
| GPITDIFYKENSYITT | 1 | 0 | 0 | 1662334 | QHR63299.1 | P0C6W2 |
| GPITDVFYKENSYTTT | 1 | 0 | 0 | 1662335 | YP_009724389.1 | P0DTD1 |
| GPKELLLLWESGKAKP | 1 | 0 | 0 | 1662338 | NP_073549.1 | A0A0P0LKV0 |
| GPKELLLSWESGKVKP | 1 | 0 | 0 | 1662339 | YP_003766.2 | H9EJ42 |
| GPLKVGAGVMLECEKF | 1 | 0 | 0 | 1662349 | YP_003766.2 | H9EJ42 |
| GPNLNAGEDIQLLKAA | 1 | 0 | 0 | 1662360 | NP_828849.2 | P0C6X7 |
| GPNLNKGEDIQLLKVA | 1 | 0 | 0 | 1662361 | QIQ54046.1 | P0C6U8 |
| GPNNRSLSRSNSNFRH | 1 | 0 | 0 | 1662362 | YP_173242.1 | Q19U29 |
| GPNVNKGEDIQLLKSA | 1 | 0 | 0 | 1662364 | YP_009724389.1 | P0DTD1 |
| GPNVNRGEDIQLLKSA | 1 | 0 | 0 | 1662365 | QHR63299.1 | P0C6W2 |
| GPPPISTDTVDVTNGL | 1 | 0 | 0 | 1662371 | YP_009555241.1 | P36334 |
| GPPRAELNGVVVDKVG | 1 | 0 | 0 | 1662372 | YP_009555238.1 | P0C6X6 |
| GPQRADLNGVIVDKVG | 1 | 0 | 0 | 1662375 | YP_173236.1 | A0A140H1G9 |
| GPQSNQRSAPRITFGG | 1 | 0 | 0 | 1662376 | NP_828858.1 | P59595 |
| GPRDLDHNFGSAGVVA | 1 | 0 | 0 | 1662380 | NP_073556.1 | A0A0N7J131 |
| GPRTIKVSTVQSKLTD | 1 | 0 | 0 | 1662383 | NP_073549.1 | A0A0P0LKV0 |
| GPSDSTDNNQNGDRSG | 1 | 0 | 0 | 1662385 | QIQ54056.1 | P59595 |
| GPSQNFGNAEMLKLGT | 1 | 0 | 0 | 1662388 | YP_173242.1 | Q19U29 |
| GPSSQPYYCFINSTIN | 1 | 0 | 0 | 1662389 | YP_003767.1 | A0A0P0G321 |
| GPTEHVTSVNLFDVGY | 1 | 0 | 0 | 1662392 | YP_009047205.1 | K9N796 |
| GPTYLDGADVTKIKPH | 1 | 0 | 0 | 1662396 | YP_009724389.1 | P0DTD1 |
| GPVDKGHYTVYDTAKK | 1 | 0 | 0 | 1662397 | NP_073549.1 | A0A0P0LKV0 |
| GPVDPTFLHRFYSLKA | 1 | 0 | 0 | 1662398 | YP_009047202.1 | K9N7C7 |
| GPVTDVFYKENSYTTT | 1 | 0 | 0 | 1662399 | QIQ54046.1 | P0C6U8 |
| GPVTDVFYKETSYTTT | 1 | 0 | 0 | 1662400 | NP_828849.2 | P0C6X7 |
| GPYKDAQVVQLLIQDY | 1 | 0 | 0 | 1662402 | YP_009555238.1 | P0C6X6 |
| GPYRDAQVVQLPVKDY | 1 | 0 | 0 | 1662403 | YP_173236.1 | A0A140H1G9 |
| GQAGEMPCAIINDKVV | 1 | 0 | 0 | 1662407 | YP_009555238.1 | P0C6X6 |
| GQAITVTPEANMDQES | 1 | 0 | 0 | 1662410 | YP_009724389.1 | P0DTD1 |
| GQATELGIEFLKRGDK | 1 | 0 | 0 | 1662413 | QIQ54046.1 | P0C6U8 |
| GQDFKTFDLLKYDFTE | 1 | 0 | 0 | 1662415 | NP_073549.1 | A0A0P0LKV0 |
| GQDVIFSQFDSLGVSS | 1 | 0 | 0 | 1662416 | YP_009555238.1 | P0C6X6 |
| GQEEFIQTNSPKVTID | 1 | 0 | 0 | 1662418 | YP_173238.1 | A0A140H1H1 |
| GQGCAGYTLEEAKTAL | 1 | 0 | 0 | 1662427 | NP_828849.2 | P0C6X7 |
| GQGCVGYTIEEAKTAL | 1 | 0 | 0 | 1662428 | QIQ54046.1 | P0C6U8 |
| GQGENDNISVAVPKSR | 1 | 0 | 0 | 1662429 | YP_009555245.1 | P33469 |
| GQGLNGYTVEEAKTVL | 1 | 0 | 0 | 1662432 | YP_009724389.1 | P0DTD1 |
| GQGQNSADPKVYPIIL | 1 | 0 | 0 | 1662433 | NP_828859.1 | P59636 |
| GQGVPINTNSSPDDQI | 1 | 0 | 0 | 1662438 | YP_009724397.2 | P0DTC9 |
| GQGVPLNANSTPAQNA | 1 | 0 | 0 | 1662440 | YP_009047211.1 | K9N4V7 |
| GQIVTCAKEIKESVQT | 1 | 0 | 0 | 1662447 | YP_009724389.1 | P0DTD1 |
| GQKGDAEEALSKLSEY | 1 | 0 | 0 | 1662449 | YP_003766.2 | H9EJ42 |
| GQKTTTLTGVEAVMYM | 1 | 0 | 0 | 1662450 | NP_828849.2 | P0C6X7 |
| GQLLDTKLNGILQQAA | 1 | 0 | 0 | 1662461 | YP_003766.2 | H9EJ42 |
| GQLQSLSKDYISSNGP | 1 | 0 | 0 | 1662463 | YP_003766.2 | H9EJ42 |
| GQNSADPKVYPIILRL | 1 | 0 | 0 | 1662466 | NP_828859.1 | P59636 |
| GQQFGPTYLDGADVTK | 1 | 0 | 0 | 1662475 | YP_009724389.1 | P0DTD1 |
| GQQQQSQTVTKKSAAE | 1 | 0 | 0 | 1662482 | QHR63308.1 | Q3LZX4 |
| GQQQTTLKGVEAVMYM | 1 | 0 | 0 | 1662483 | YP_009724389.1 | P0DTD1 |
| GQQVDLVISDMYDPTT | 1 | 0 | 0 | 1662485 | YP_009047202.1 | K9N7C7 |
| GQRTELGVEFLKRGDK | 1 | 0 | 0 | 1662486 | NP_828849.2 | P0C6X7 |
| GQRVYDDVDASLFVDI | 1 | 0 | 0 | 1662489 | YP_173236.1 | A0A140H1G9 |
| GQSLCALPDTPSTLTP | 1 | 0 | 0 | 1662491 | YP_009047204.1 | K9N5Q8 |
| GQSTQLGIEFLKRGDK | 1 | 0 | 0 | 1662492 | YP_009724389.1 | P0DTD1 |
| GQTGVIADYNYKLPDD | 1 | 0 | 0 | 1662496 | NP_828851.1 | P59594 |
| GQVDLFRNARNGVLIT | 1 | 0 | 0 | 1662500 | YP_009724389.1 | P0DTD1 |
| GRAIIVSVEQLEPCAQ | 1 | 0 | 0 | 1662509 | NP_073549.1 | A0A0P0LKV0 |
| GRCIVLDVKGVEELHD | 1 | 0 | 0 | 1662510 | YP_009555239.1 | Q80872 |
| GRCPTAADINIPDLPI | 1 | 0 | 0 | 1662511 | YP_009555240.1 | P30215 |
| GRDATQYLVQQESSFV | 1 | 0 | 0 | 1662515 | NP_828849.2 | P0C6X7 |
| GRDFKFSDGQGVPIAF | 1 | 0 | 0 | 1662516 | YP_173242.1 | Q19U29 |
| GRDISDTTDAVRDPQT | 1 | 0 | 0 | 1662518 | QIQ54048.1 | P59594 |

| | | | | | | |
|---|---|---|---|---|---|---|
| GRDNGALEAFKRSNNG | 1 | 0 | 0 | 1662520 | YP_009555238.1 | P0C6X6 |
| GRDTGALEAFRKARNG | 1 | 0 | 0 | 1662521 | YP_173236.1 | A0A140H1G9 |
| GREISCNVCTKRATVY | 1 | 0 | 0 | 1662524 | YP_009555238.1 | P0C6X6 |
| GRGDCKGFSSDVLSDV | 1 | 0 | 0 | 1662533 | NP_073551.1 | A0A0P0K6L9 |
| GRGESKPQSRNPSSDR | 1 | 0 | 0 | 1662534 | NP_073556.1 | A0A0N7J131 |
| GRGGYHYTPFHYERDN | 1 | 0 | 0 | 1662537 | YP_009047202.1 | K9N7C7 |
| GRKMLMALLSDNAYLK | 1 | 0 | 0 | 1662552 | YP_009047202.1 | K9N7C7 |
| GRKVDLQLGNLGYLQS | 1 | 0 | 0 | 1662554 | YP_009555241.1 | P36334 |
| GRKVNFNEKPVVMEIP | 1 | 0 | 0 | 1662555 | YP_173236.1 | A0A140H1G9 |
| GRLIIRENNRVVISSD | 1 | 0 | 0 | 1662561 | YP_009724389.1 | P0DTD1 |
| GRLIIRENNRVVVSSD | 1 | 0 | 0 | 1662562 | NP_828849.2 | P0C6X7 |
| GRNGARPKQRRPQGLP | 1 | 0 | 0 | 1662571 | NP_828858.1 | P59595 |
| GRNLSDFTPRSDMEYD | 1 | 0 | 0 | 1662573 | YP_003766.2 | H9EJ42 |
| GRQATQYLVQQESSFV | 1 | 0 | 0 | 1662581 | QIQ54046.1 | P0C6U8 |
| GRQFYEFYNDVKPPVL | 1 | 0 | 0 | 1662582 | YP_009555243.1 | Q04854 |
| GRQIMGSCSFEDELTP | 1 | 0 | 0 | 1662585 | YP_009555238.1 | P0C6X6 |
| GRRGPEQTQGNFGDQE | 1 | 0 | 0 | 1662589 | YP_009724397.2 | P0DTC9 |
| GRSVYVKFQDSKPPLP | 1 | 0 | 0 | 1662602 | YP_009047209.1 | K9N5R3 |
| GRTGELPCAIMNDKVV | 1 | 0 | 0 | 1662603 | YP_173236.1 | A0A140H1G9 |
| GRTILGSALLEDEFTP | 1 | 0 | 0 | 1662606 | YP_009724389.1 | P0DTD1 |
| GRTILGSTILEDEFTP | 1 | 0 | 0 | 1662607 | NP_828849.2 | P0C6X7 |
| GRVSAAFHANSSEPAL | 1 | 0 | 0 | 1662623 | YP_009555241.1 | P36334 |
| GSAGVVANGVKAKGYP | 1 | 0 | 0 | 1662635 | NP_073556.1 | A0A0N7J131 |
| GSAILEDEFTPFDVVR | 1 | 0 | 0 | 1662636 | QIQ54046.1 | P0C6U8 |
| GSALLEDEFTPFDVVR | 1 | 0 | 0 | 1662637 | YP_009724389.1 | P0DTD1 |
| GSCGSVGFNIDYDCVS | 1 | 0 | 0 | 1662651 | YP_009724389.1 | P0DTD1 |
| GSCGSVGYTKEGSVIN | 1 | 0 | 0 | 1662652 | YP_009047202.1 | K9N7C7 |
| GSCSFEDELTPSDVYQ | 1 | 0 | 0 | 1662665 | YP_009555238.1 | P0C6X6 |
| GSCTFEDELAPSDVYQ | 1 | 0 | 0 | 1662666 | YP_173236.1 | A0A140H1G9 |
| GSCVGSGPGKNNGIGT | 1 | 0 | 0 | 1662667 | YP_009555241.1 | P36334 |
| GSCYVCKPHQVNISLN | 1 | 0 | 0 | 1662670 | YP_003767.1 | A0A0P0G321 |
| GSDAALALLLLDRLNQ | 1 | 0 | 0 | 1662671 | QHR63308.1 | Q3LZX4 |
| GSDFSLADSVVEVVTT | 1 | 0 | 0 | 1662674 | YP_009555238.1 | P0C6X6 |
| GSDFTGSVYGNFDDQP | 1 | 0 | 0 | 1662675 | YP_003766.2 | H9EJ42 |
| GSDKGIAPGTSVLRQW | 1 | 0 | 0 | 1662677 | YP_009047202.1 | K9N7C7 |
| GSDKGVAPGTTVLKRW | 1 | 0 | 0 | 1662679 | YP_003766.2 | H9EJ42 |
| GSDVLYQPPQTSITSA | 1 | 0 | 0 | 1662686 | YP_009724389.1 | P0DTD1 |
| GSDVLYQPPTASVSTS | 1 | 0 | 0 | 1662687 | YP_009555238.1 | P0C6X6 |
| GSECSKASSRSAIEDL | 1 | 0 | 0 | 1662690 | YP_009555241.1 | P36334 |
| GSEKGVAPGSAVLRQW | 1 | 0 | 0 | 1662694 | YP_009555238.1 | P0C6X6 |
| GSFCTQLNRALSGIAA | 1 | 0 | 0 | 1662705 | NP_828851.1 | P59594 |
| GSFDNGHYVVYDAANN | 1 | 0 | 0 | 1662708 | YP_003766.2 | H9EJ42 |
| GSFINGACGSPGYNLK | 1 | 0 | 0 | 1662710 | NP_073549.1 | A0A0P0LKV0 |
| GSFLNGSCGSVGFNID | 1 | 0 | 0 | 1662716 | YP_009724389.1 | P0DTD1 |
| GSGALRELTRELNGGA | 1 | 0 | 0 | 1662723 | NP_828849.2 | P0C6X7 |
| GSGFCVDYNSPSSSSS | 1 | 0 | 0 | 1662724 | YP_173238.1 | A0A140H1H1 |
| GSGNLVEKLSPWFMDG | 1 | 0 | 0 | 1662730 | YP_009047202.1 | K9N7C7 |
| GSGRSASNSRPGSRSQ | 1 | 0 | 0 | 1662734 | YP_173242.1 | Q19U29 |
| GSGVAIVDSYYSYLMP | 1 | 0 | 0 | 1662738 | YP_009047202.1 | K9N7C7 |
| GSGVTRELMRELNGGA | 1 | 0 | 0 | 1662741 | QIQ54046.1 | P0C6U8 |
| GSGYYYPEPITENNVV | 1 | 0 | 0 | 1662743 | YP_009555241.1 | P36334 |
| GSIAIKVTEYSWNKKL | 1 | 0 | 0 | 1662749 | NP_073549.1 | A0A0P0LKV0 |
| GSIPINVIVFDGKSKC | 1 | 0 | 0 | 1662759 | QIQ54046.1 | P0C6U8 |
| GSIRFDSTLPGFETIM | 1 | 0 | 0 | 1662760 | YP_173242.1 | Q19U29 |
| GSISYSELRPDTRYVL | 1 | 0 | 0 | 1662761 | NP_828849.2 | P0C6X7 |
| GSITQRTRTRPSVQPG | 1 | 0 | 0 | 1662762 | YP_009047211.1 | K9N4V7 |
| GSLAATVRLQAGNATE | 1 | 0 | 0 | 1662774 | YP_009724389.1 | P0DTD1 |
| GSLADGLEAYADKTLQ | 1 | 0 | 0 | 1662775 | YP_009555238.1 | P0C6X6 |
| GSLDVEVFNRLATSDW | 1 | 0 | 0 | 1662776 | YP_003766.2 | H9EJ42 |
| GSLIGGMVLGGLTSAA | 1 | 0 | 0 | 1662786 | YP_003767.1 | A0A0P0G321 |
| GSLIPVRPRNSSDNGI | 1 | 0 | 0 | 1662787 | YP_003767.1 | A0A0P0G321 |
| GSLPINVIVFDGKSKC | 1 | 0 | 0 | 1662791 | YP_009724389.1 | P0DTD1 |
| GSLYVKPGGTSSGDAT | 1 | 0 | 0 | 1662803 | YP_009724389.1 | P0DTD1 |
| GSMDIDVFNKLSTSDW | 1 | 0 | 0 | 1662810 | NP_073549.1 | A0A0P0LKV0 |
| GSMIEDIDYVPLKSAV | 1 | 0 | 0 | 1662811 | YP_009047202.1 | K9N7C7 |
| GSNDIWMQNKGLFYTQ | 1 | 0 | 0 | 1662815 | YP_009555240.1 | P30215 |
| GSNGKPGNVGGNDALS | 1 | 0 | 0 | 1662818 | YP_009555238.1 | P0C6X6 |
| GSNQNGERSGARPKQR | 1 | 0 | 0 | 1662824 | QHR63308.1 | Q3LZX4 |
| GSPGSLNFKKVETLNF | 1 | 0 | 0 | 1662829 | YP_173243.1 | Q0ZJ80 |
| GSPGYNLKNGEVEFVY | 1 | 0 | 0 | 1662830 | NP_073549.1 | A0A0P0LKV0 |
| GSPGYNVRNDGTVEFC | 1 | 0 | 0 | 1662831 | YP_003766.2 | H9EJ42 |
| GSPYIDDFNKIASCKW | 1 | 0 | 0 | 1662837 | YP_173236.1 | A0A140H1G9 |
| GSQASSRSSSRSRGNS | 1 | 0 | 0 | 1662840 | NP_828858.1 | P59595 |
| GSQQQSQTVTKKSAAE | 1 | 0 | 0 | 1662843 | QIQ54056.1 | P59595 |
| GSRSGNSTRGTSPGPS | 1 | 0 | 0 | 1662850 | YP_009047211.1 | K9N4V7 |
| GSRSRANSGNRTPTSG | 1 | 0 | 0 | 1662852 | YP_009555245.1 | P33469 |
| GSRVPHIVRKDLTKYT | 1 | 0 | 0 | 1662855 | YP_009555238.1 | P0C6X6 |
| GSSFDGVMYGGFEDQP | 1 | 0 | 0 | 1662859 | NP_073549.1 | A0A0P0LKV0 |
| GSSFYAPEPITSLNTK | 1 | 0 | 0 | 1662860 | YP_009047204.1 | K9N5Q8 |
| GSSGFLQSSNYKIDTT | 1 | 0 | 0 | 1662861 | YP_173238.1 | A0A140H1H1 |
| GSSGVANPAMDPIYDE | 1 | 0 | 0 | 1662864 | NP_828852.2 | P59632 |
| GSSGVVNPAMDPIYDE | 1 | 0 | 0 | 1662865 | QIQ54049.1 | P59632 |
| GSSGVVNPAMEPIYDE | 1 | 0 | 0 | 1662866 | QHR63301.1 | Q3LZX0 |
| GSSGVVNPVMEPIYDE | 1 | 0 | 0 | 1662867 | YP_009724391.1 | P0DTC3 |
| GSSRGNSPARMASGGG | 1 | 0 | 0 | 1662873 | NP_828858.1 | P59595 |
| GSSRGTSPARIAGNGG | 1 | 0 | 0 | 1662874 | QIQ54056.1 | P59595 |
| GSSRGTSPARMAGNGG | 1 | 0 | 0 | 1662875 | YP_009724397.2 | P0DTC9 |
| GSSRGTSPARMAGNGS | 1 | 0 | 0 | 1662876 | QHR63308.1 | Q3LZX4 |
| GSSRNESWHFDKSEPL | 1 | 0 | 0 | 1662877 | YP_173238.1 | A0A140H1H1 |
| GSSSRSFFEDLLFDKV | 1 | 0 | 0 | 1662880 | YP_173238.1 | A0A140H1H1 |
| GSSVETETYFKVNGLW | 1 | 0 | 0 | 1662885 | YP_173236.1 | A0A140H1G9 |

| | | | | | | |
|---|---|---|---|---|---|---|
| GSSVGNFSDGKMGRFF | 1 | 0 | 0 | 1662886 | YP_009047204.1 | K9N5Q8 |
| GSSYYYPEPISDKNVV | 1 | 0 | 0 | 1662888 | YP_173238.1 | A0A140H1H1 |
| GSTILEDEFTPFDVVR | 1 | 0 | 0 | 1662894 | NP_828849.2 | P0C6X7 |
| GSTMNNKSQSVIIINN | 1 | 0 | 0 | 1662900 | NP_828851.1 | P59594 |
| GSVACEHISSTMSQYS | 1 | 0 | 0 | 1662908 | YP_009047204.1 | K9N5Q8 |
| GSVAYESLRPDTRYVL | 1 | 0 | 0 | 1662914 | YP_009724389.1 | P0DTD1 |
| GSVGYVLTGDSVKFVY | 1 | 0 | 0 | 1662921 | YP_173236.1 | A0A140H1G9 |
| GSVKGLQPSVGPKQAS | 1 | 0 | 0 | 1662924 | YP_009724389.1 | P0DTD1 |
| GSVKGLTPSKGPAQAS | 1 | 0 | 0 | 1662925 | NP_828849.2 | P0C6X7 |
| GSVLEMAKEVKTSSKV | 1 | 0 | 0 | 1662926 | NP_073549.1 | A0A0P0LKV0 |
| GSVPYESLRPDTRYVL | 1 | 0 | 0 | 1662930 | QIQ54046.1 | P0C6U8 |
| GSVQVDNGSSINRRQL | 1 | 0 | 0 | 1662931 | NP_073549.1 | A0A0P0LKV0 |
| GSVRVVTTFDAEYCRH | 1 | 0 | 0 | 1662932 | NP_828849.2 | P0C6X7 |
| GSVRVVTTFDSEYCRH | 1 | 0 | 0 | 1662933 | YP_009724389.1 | P0DTD1 |
| GSVVEQAEGVECDFSP | 1 | 0 | 0 | 1662935 | YP_009047204.1 | K9N5Q8 |
| GSVVFVDKYMCGFDGK | 1 | 0 | 0 | 1662936 | YP_003766.2 | H9EJ42 |
| GSVVIKEPVHVYADDS | 1 | 0 | 0 | 1662937 | YP_009555238.1 | P0C6X6 |
| GSVVKPVRARAPPGEQ | 1 | 0 | 0 | 1662938 | NP_073549.1 | A0A0P0LKV0 |
| GSVWHIQTVQDADGIN | 1 | 0 | 0 | 1662941 | YP_173236.1 | A0A140H1G9 |
| GSVYGNFDDQPSLQVE | 1 | 0 | 0 | 1662943 | YP_003766.2 | H9EJ42 |
| GSVYTNSCLVSSDGQP | 1 | 0 | 0 | 1662944 | YP_009047202.1 | K9N7C7 |
| GSVYYHNSMDCPIVVA | 1 | 0 | 0 | 1662945 | YP_173236.1 | A0A140H1G9 |
| GSWAIFNDHHLNRPGV | 1 | 0 | 0 | 1662946 | YP_009047202.1 | K9N7C7 |
| GSYKDWSYSGQATELG | 1 | 0 | 0 | 1662953 | QIQ54046.1 | P0C6U8 |
| GSYRDWSYSGQRTELG | 1 | 0 | 0 | 1662958 | NP_828849.2 | P0C6X7 |
| GTAKKQVVLVSNNQED | 1 | 0 | 0 | 1662965 | YP_009555238.1 | P0C6X6 |
| GTALPKGFYAEGSRGG | 1 | 0 | 0 | 1662966 | QIQ54056.1 | P59595 |
| GTAVINLRADQINDMV | 1 | 0 | 0 | 1662970 | YP_009555238.1 | P0C6X6 |
| GTAVMSLKENQINDMI | 1 | 0 | 0 | 1662974 | NP_828849.2 | P0C6X7 |
| GTAVVNLRPDQLNDLV | 1 | 0 | 0 | 1662975 | YP_173236.1 | A0A140H1G9 |
| GTCGIIDVEKGVLPQL | 1 | 0 | 0 | 1662982 | QIQ54046.1 | P0C6U8 |
| GTCPFSFSKLNNFQKF | 1 | 0 | 0 | 1662988 | YP_003767.1 | A0A0P0G321 |
| GTCRRCPAEIVDTVSA | 1 | 0 | 0 | 1662989 | YP_009724389.1 | P0DTD1 |
| GTCYRCPKEIVDTVSA | 1 | 0 | 0 | 1662990 | YP_009555238.1 | P0C6X6 |
| GTDFNGDFYGPYKDAQ | 1 | 0 | 0 | 1662991 | YP_009555238.1 | P0C6X6 |
| GTDFTGNFYGPYRDAQ | 1 | 0 | 0 | 1662992 | YP_173236.1 | A0A140H1G9 |
| GTDLEGKFYGPFVDRQ | 1 | 0 | 0 | 1662997 | NP_828849.2 | P0C6X7 |
| GTDLEGNFYGPFVDRQ | 1 | 0 | 0 | 1662998 | YP_009724389.1 | P0DTD1 |
| GTDLEGTFYGPFVDRQ | 1 | 0 | 0 | 1662999 | QHR63299.1 | P0C6W2 |
| GTDPIDDFQVNWNTKH | 1 | 0 | 0 | 1663000 | QIQ54046.1 | P0C6U8 |
| GTDPIEDYEQNWNTKH | 1 | 0 | 0 | 1663001 | NP_828849.2 | P0C6X7 |
| GTDSGFAAYNRYRIGN | 1 | 0 | 0 | 1663003 | NP_828855.1 | P59596 |
| GTDTNSVCPKLEFAND | 1 | 0 | 0 | 1663004 | YP_009047204.1 | K9N5Q8 |
| GTEDDYQGKPLEFGAT | 1 | 0 | 0 | 1663009 | YP_009724389.1 | P0DTD1 |
| GTEDDYQGKSLEFGAT | 1 | 0 | 0 | 1663010 | QHR63299.1 | P0C6W2 |
| GTEDDYQGLPLEFGAS | 1 | 0 | 0 | 1663011 | NP_828849.2 | P0C6X7 |
| GTEDPRWPQIAELAPT | 1 | 0 | 0 | 1663012 | YP_009047211.1 | K9N4V7 |
| GTENQVKEGPTTCGYL | 1 | 0 | 0 | 1663015 | QIQ54046.1 | P0C6U8 |
| GTESDYKGLPLEFGAS | 1 | 0 | 0 | 1663016 | QIQ54046.1 | P0C6U8 |
| GTFCDNINSILDEVNG | 1 | 0 | 0 | 1663020 | YP_173238.1 | A0A140H1H1 |
| GTFTVVMRPNYTIKGS | 1 | 0 | 0 | 1663030 | YP_009047202.1 | K9N7C7 |
| GTFYGPFVDRQTAQAA | 1 | 0 | 0 | 1663035 | QHR63299.1 | P0C6W2 |
| GTGGNSQSSSRASSLS | 1 | 0 | 0 | 1663038 | YP_009047211.1 | K9N4V7 |
| GTGIIRSSNQSLAGGI | 1 | 0 | 0 | 1663040 | YP_003767.1 | A0A0P0G321 |
| GTGMAITVKPDATTSQ | 1 | 0 | 0 | 1663048 | YP_009555238.1 | P0C6X6 |
| GTGPEAALPFRAVKDG | 1 | 0 | 0 | 1663049 | YP_009047211.1 | K9N4V7 |
| GTGPEAGLPYGANKEG | 1 | 0 | 0 | 1663051 | QIQ54056.1 | P59595 |
| GTGPEASLPYGANKEG | 1 | 0 | 0 | 1663052 | NP_828858.1 | P59595 |
| GTGPYANASYGESLEG | 1 | 0 | 0 | 1663056 | YP_173242.1 | Q19U29 |
| GTGQAITVTPEANMDQ | 1 | 0 | 0 | 1663057 | YP_009724389.1 | P0DTD1 |
| GTGRGDCKGFSSDVLS | 1 | 0 | 0 | 1663058 | NP_073551.1 | A0A0P0K6L9 |
| GTGTIYTELEPPCRFV | 1 | 0 | 0 | 1663059 | YP_009724389.1 | P0DTD1 |
| GTGVLTTSKKQFLPFQ | 1 | 0 | 0 | 1663064 | QIQ54048.1 | P59594 |
| GTGYSLADLPAYMTVA | 1 | 0 | 0 | 1663066 | YP_009555244.1 | Q01455 |
| GTIKENIDGGAMHANY | 1 | 0 | 0 | 1663078 | YP_009047202.1 | K9N7C7 |
| GTIYTELEPPCRFVTD | 1 | 0 | 0 | 1663094 | YP_009724389.1 | P0DTD1 |
| GTLLVDSDLNDFVSDA | 1 | 0 | 0 | 1663111 | YP_009724389.1 | P0DTD1 |
| GTMYVRPIIEDYHTLT | 1 | 0 | 0 | 1663123 | YP_009555244.1 | Q01455 |
| GTNASNQVAVLYQDVN | 1 | 0 | 0 | 1663124 | QHR63300.2 | Q3LZX1 |
| GTNKQLNEISDDCNWP | 1 | 0 | 0 | 1663136 | YP_009555238.1 | P0C6X6 |
| GTNTSNQVAVLYQDVN | 1 | 0 | 0 | 1663141 | YP_009724390.1 | P0DTC2 |
| GTNYLTCDNLCTPDPI | 1 | 0 | 0 | 1663146 | YP_009555241.1 | P36334 |
| GTPAKKLCVVPGNVVP | 1 | 0 | 0 | 1663148 | YP_003766.2 | H9EJ42 |
| GTPKPSRNQSPASSQT | 1 | 0 | 0 | 1663152 | NP_073556.1 | A0A0N7J131 |
| GTPVCINGLMLLEIKD | 1 | 0 | 0 | 1663157 | YP_009724389.1 | P0DTD1 |
| GTPVCVNGLMLLEIKD | 1 | 0 | 0 | 1663158 | NP_828849.2 | P0C6X7 |
| GTPVVEYLSKEDLHTL | 1 | 0 | 0 | 1663160 | YP_009047202.1 | K9N7C7 |
| GTQHISNDFTPEDHSL | 1 | 0 | 0 | 1663161 | YP_173242.1 | Q19U29 |
| GTRNPANNAAIVLQLP | 1 | 0 | 0 | 1663170 | YP_009724397.2 | P0DTC9 |
| GTRNPNNNAAIVLQLP | 1 | 0 | 0 | 1663172 | QIQ54056.1 | P59595 |
| GTRNPNNNAATVLQLP | 1 | 0 | 0 | 1663173 | NP_828858.1 | P59595 |
| GTSPARIAGNGGDAAL | 1 | 0 | 0 | 1663183 | QIQ54056.1 | P59595 |
| GTSPGPSGIGAVGGDL | 1 | 0 | 0 | 1663186 | YP_009047211.1 | K9N4V7 |
| GTSTPKKPNKPLSQPR | 1 | 0 | 0 | 1663191 | YP_003771.1 | H9EJ69 |
| GTTCSNDLLQPNTEVF | 1 | 0 | 0 | 1663199 | YP_173238.1 | A0A140H1H1 |
| GTTKFYGGWDDMLRRL | 1 | 0 | 0 | 1663208 | YP_009555238.1 | P0C6X6 |
| GTTSPISEHDYQIGGY | 1 | 0 | 0 | 1663222 | YP_009724391.1 | P0DTC3 |
| GTTVLLKEPCPSGTYE | 1 | 0 | 0 | 1663230 | NP_828857.1 | P59635 |
| GTTVVRPLVEDSTSVT | 1 | 0 | 0 | 1663232 | YP_009047210.1 | K9N7A1 |
| GTVSKTSDWKCKVTDV | 1 | 0 | 0 | 1663244 | YP_009047202.1 | K9N7C7 |
| GTVSWNLREMLAHAEE | 1 | 0 | 0 | 1663245 | YP_009724389.1 | P0DTD1 |

| | | | | | | |
|---|---|---|---|---|---|---|
| GTVSWNLREMLTHAEE | 1 | 0 | 0 | 1663246 | QIQ54046.1 | P0C6U8 |
| GTVTLKQGEIKDATPS | 1 | 0 | 0 | 1663248 | YP_009724391.1 | P0DTC3 |
| GTVYEKLKPILDWLEE | 1 | 0 | 0 | 1663249 | QIQ54046.1 | P0C6U8 |
| GTVYEKLKPVLDWLEE | 1 | 0 | 0 | 1663250 | YP_009724389.1 | P0DTD1 |
| GTVYEQLKPVLDWLEE | 1 | 0 | 0 | 1663251 | QHR63299.1 | P0C6W2 |
| GTWLTYHGAIKLDDKD | 1 | 0 | 0 | 1663252 | NP_828858.1 | P59595 |
| GTWLTYTGAIKLDDKD | 1 | 0 | 0 | 1663253 | YP_009724397.2 | P0DTC9 |
| GTYEGNSPFHPLADNK | 1 | 0 | 0 | 1663254 | YP_009724395.1 | P0DTC7 |
| GTYYCVTTKAKSVQQD | 1 | 0 | 0 | 1663265 | YP_009047202.1 | K9N7C7 |
| GVAFLAVFQSATKIIA | 1 | 0 | 0 | 1663272 | NP_828852.2 | P59632 |
| GVAGALNKATNNAMQI | 1 | 0 | 0 | 1663275 | QIQ54046.1 | P0C6U8 |
| GVAGALNKATNNAMQV | 1 | 0 | 0 | 1663276 | YP_009724389.1 | P0DTD1 |
| GVALKDCGFVMQFGYI | 1 | 0 | 0 | 1663286 | YP_009555238.1 | P0C6X6 |
| GVARAIDILTEGQLQS | 1 | 0 | 0 | 1663295 | YP_003766.2 | H9EJ42 |
| GVAVADSYYSYMMPML | 1 | 0 | 0 | 1663300 | YP_173236.1 | A0A140H1G9 |
| GVAYTAFSGPVDKGHY | 1 | 0 | 0 | 1663302 | NP_073549.1 | A0A0P0LKV0 |
| GVCITTNGSWAIFNDH | 1 | 0 | 0 | 1663308 | YP_009047202.1 | K9N7C7 |
| GVCKYTDIDVNSALNI | 1 | 0 | 0 | 1663310 | YP_009047202.1 | K9N7C7 |
| GVCSMTDIAKKPTENI | 1 | 0 | 0 | 1663317 | QHR63299.1 | P0C6W2 |
| GVCSMTDIAKKPTETI | 1 | 0 | 0 | 1663318 | YP_009724389.1 | P0DTD1 |
| GVCTMTDIAKKPTESA | 1 | 0 | 0 | 1663319 | NP_828849.2 | P0C6X7 |
| GVCYDFDGVTTSDKCI | 1 | 0 | 0 | 1663325 | YP_003766.2 | H9EJ42 |
| GVDAKGFPQLAELIPN | 1 | 0 | 0 | 1663327 | YP_003771.1 | H9EJ69 |
| GVDAMNLIANIFTPLV | 1 | 0 | 0 | 1663328 | NP_828849.2 | P0C6X7 |
| GVDAVNLLTNMFTPLI | 1 | 0 | 0 | 1663330 | YP_009724389.1 | P0DTD1 |
| GVDFDFIVNAANENLA | 1 | 0 | 0 | 1663333 | NP_073549.1 | A0A0P0LKV0 |
| GVDFVAQPEGCVLTNT | 1 | 0 | 0 | 1663334 | NP_073549.1 | A0A0P0LKV0 |
| GVDGELPVAVVNDKVF | 1 | 0 | 0 | 1663336 | YP_003766.2 | H9EJ42 |
| GVDIAANTVIWDYKRE | 1 | 0 | 0 | 1663339 | NP_828849.2 | P0C6X7 |
| GVDSFEGVLPINTDTV | 1 | 0 | 0 | 1663340 | YP_003766.2 | H9EJ42 |
| GVDSVNLLTNMFTPLI | 1 | 0 | 0 | 1663341 | QIQ54046.1 | P0C6U8 |
| GVDVELFENKTILPVN | 1 | 0 | 0 | 1663342 | QIQ54047.1 | P0C6X7 |
| GVDVELFENKTTLPVN | 1 | 0 | 0 | 1663343 | YP_009724389.1 | P0DTD1 |
| GVDYTKKFLTVDGVQY | 1 | 0 | 0 | 1663344 | YP_009047202.1 | K9N7C7 |
| GVEAVMYMGTLSYDNL | 1 | 0 | 0 | 1663345 | NP_828849.2 | P0C6X7 |
| GVECDFSPLLSGTPPQ | 1 | 0 | 0 | 1663348 | YP_009047204.1 | K9N5Q8 |
| GVEELHDDLVNYIRDK | 1 | 0 | 0 | 1663349 | YP_009555239.1 | Q80872 |
| GVEFLKRGDKIVYHTL | 1 | 0 | 0 | 1663351 | NP_828849.2 | P0C6X7 |
| GVEFLRDGWEIVKFIS | 1 | 0 | 0 | 1663352 | YP_009724389.1 | P0DTD1 |
| GVEGELPVAVVNDKIF | 1 | 0 | 0 | 1663353 | YP_009047202.1 | K9N7C7 |
| GVENPMLMGWDYPKCD | 1 | 0 | 0 | 1663358 | YP_003766.2 | H9EJ42 |
| GVESDVETNEIVDSVQ | 1 | 0 | 0 | 1663360 | YP_009047202.1 | K9N7C7 |
| GVFCGVDAVNLLTNMF | 1 | 0 | 0 | 1663362 | YP_009724389.1 | P0DTD1 |
| GVFCGVDSVNLLTNMF | 1 | 0 | 0 | 1663363 | QIQ54046.1 | P0C6U8 |
| GVFDIKNSSVNTVCFV | 1 | 0 | 0 | 1663364 | YP_003766.2 | H9EJ42 |
| GVFDIVKGQIQVASDN | 1 | 0 | 0 | 1663365 | NP_828849.2 | P0C6X7 |
| GVFMSTAKCWTEEDLS | 1 | 0 | 0 | 1663375 | NP_073549.1 | A0A0P0LKV0 |
| GVFVQDPAPIDIDAFP | 1 | 0 | 0 | 1663384 | YP_003766.2 | H9EJ42 |
| GVFWVANHQADTSTPS | 1 | 0 | 0 | 1663387 | YP_173242.1 | Q19U29 |
| GVGGKPCIKVATVQSK | 1 | 0 | 0 | 1663391 | YP_009724389.1 | P0DTD1 |
| GVGNTFICEEVANDLT | 1 | 0 | 0 | 1663395 | YP_009047202.1 | K9N7C7 |
| GVGSANIFIGDKVGHY | 1 | 0 | 0 | 1663396 | YP_009555238.1 | P0C6X6 |
| GVHAGTDLEGKFYGPF | 1 | 0 | 0 | 1663401 | NP_828849.2 | P0C6X7 |
| GVHAGTDLEGNFYGPF | 1 | 0 | 0 | 1663402 | YP_009724389.1 | P0DTD1 |
| GVHAGTDLEGTFYGPF | 1 | 0 | 0 | 1663403 | QHR63299.1 | P0C6W2 |
| GVIADKDKDLSGAVPS | 1 | 0 | 0 | 1663409 | NP_073549.1 | A0A0P0LKV0 |
| GVIADYNYKLPDDFMG | 1 | 0 | 0 | 1663410 | NP_828851.1 | P59594 |
| GVIADYNYKLPDDFTG | 1 | 0 | 0 | 1663411 | QIQ54048.1 | P59594 |
| GVIDISAGMFGLYDDI | 1 | 0 | 0 | 1663412 | YP_003766.2 | H9EJ42 |
| GVIIMCADVPDAFQSV | 1 | 0 | 0 | 1663414 | YP_003766.2 | H9EJ42 |
| GVIKNVNSVRDWLKSL | 1 | 0 | 0 | 1663415 | NP_073549.1 | A0A0P0LKV0 |
| GVIRDVSVEDVCFANK | 1 | 0 | 0 | 1663420 | YP_009555238.1 | P0C6X6 |
| GVIRVSNDTFLNGITY | 1 | 0 | 0 | 1663423 | NP_073551.1 | A0A0P0K6L9 |
| GVIRYYDITANGGTGF | 1 | 0 | 0 | 1663424 | YP_173236.1 | A0A140H1G9 |
| GVISKVHEVIIDNKPY | 1 | 0 | 0 | 1663425 | YP_003766.2 | H9EJ42 |
| GVKAKGYPQFAELVPS | 1 | 0 | 0 | 1663431 | NP_073556.1 | A0A0N7J131 |
| GVKDCVVLHSYFTSDY | 1 | 0 | 0 | 1663433 | YP_009724391.1 | P0DTC3 |
| GVKDFNTLSQEGKKYL | 1 | 0 | 0 | 1663434 | YP_003766.2 | H9EJ42 |
| GVKDYVVVHGYFTEVY | 1 | 0 | 0 | 1663435 | NP_828852.2 | P59632 |
| GVKFQTVKPGNFNQDF | 1 | 0 | 0 | 1663436 | YP_009555238.1 | P0C6X6 |
| GVKITEEVGHEDLMAA | 1 | 0 | 0 | 1663439 | QIQ54046.1 | P0C6U8 |
| GVKLGTGYTLSDLPVY | 1 | 0 | 0 | 1663441 | YP_173241.1 | Q0ZJ82 |
| GVKQEQRTGLDAVMHF | 1 | 0 | 0 | 1663445 | YP_009555238.1 | P0C6X6 |
| GVKQLPEQFDSLNLSA | 1 | 0 | 0 | 1663446 | YP_173242.1 | Q19U29 |
| GVKVTQELGHEDLMAA | 1 | 0 | 0 | 1663449 | NP_828849.2 | P0C6X7 |
| GVLDGSYNVSCLCSTD | 1 | 0 | 0 | 1663451 | YP_173238.1 | A0A140H1H1 |
| GVLDTTIPFGPSSQPY | 1 | 0 | 0 | 1663452 | YP_003767.1 | A0A0P0G321 |
| GVLESFSVTLLDNQED | 1 | 0 | 0 | 1663455 | YP_003766.2 | H9EJ42 |
| GVLGCKAIVKETNLIC | 1 | 0 | 0 | 1663456 | YP_009555238.1 | P0C6X6 |
| GVLGDGNALYNTEGGK | 1 | 0 | 0 | 1663457 | YP_003766.2 | H9EJ42 |
| GVLIQNGAKHVQGNVA | 1 | 0 | 0 | 1663460 | YP_173236.1 | A0A140H1G9 |
| GVLITEGNVKGLQPSV | 1 | 0 | 0 | 1663462 | QIQ54047.1 | P0C6X7 |
| GVLITEGSVKGLQPSV | 1 | 0 | 0 | 1663463 | YP_009724389.1 | P0DTD1 |
| GVLLKQLMVNKSSCYR | 1 | 0 | 0 | 1663467 | YP_173243.1 | Q0ZJ80 |
| GVLPINTDTVLSVAPE | 1 | 0 | 0 | 1663471 | YP_003766.2 | H9EJ42 |
| GVLQSGLVKMSHPSGD | 1 | 0 | 0 | 1663477 | YP_009047202.1 | K9N7C7 |
| GVLSAAIKDCVDINKA | 1 | 0 | 0 | 1663478 | NP_073549.1 | A0A0P0LKV0 |
| GVLTLDNQDLNGKWYD | 1 | 0 | 0 | 1663482 | YP_009047202.1 | K9N7C7 |
| GVLTTSKKQFLPFQQF | 1 | 0 | 0 | 1663488 | QIQ54048.1 | P59594 |
| GVMKNASLYTSLVPHT | 1 | 0 | 0 | 1663497 | YP_173236.1 | A0A140H1G9 |
| GVMWLYTYSIGEASSP | 1 | 0 | 0 | 1663500 | YP_009047202.1 | K9N7C7 |

| | | | | | | |
|---|---|---|---|---|---|---|
| GVMYGGFEDQPNLQVE | 1 | 0 | 0 | 1663501 | NP_073549.1 | A0A0P0LKV0 |
| GVNDAITQTSQALQTV | 1 | 0 | 0 | 1663503 | NP_073551.1 | A0A0P0K6L9 |
| GVNDICMFENKTTLPT | 1 | 0 | 0 | 1663504 | YP_009047202.1 | K9N7C7 |
| GVNFKSISLTVGEVFG | 1 | 0 | 0 | 1663506 | YP_173236.1 | A0A140H1G9 |
| GVNFNPYHRFKCFPNG | 1 | 0 | 0 | 1663507 | YP_173237.1 | A0A140H1H0 |
| GVNFNVDDINFSPVLG | 1 | 0 | 0 | 1663508 | YP_009555241.1 | P36334 |
| GVNFRTVVLNNKNTYR | 1 | 0 | 0 | 1663509 | YP_009047202.1 | K9N7C7 |
| GVNFTNRFVPVGESFG | 1 | 0 | 0 | 1663510 | YP_009555238.1 | P0C6X6 |
| GVNFTPYHAFKCNRSG | 1 | 0 | 0 | 1663511 | YP_009555240.1 | P30215 |
| GVNFVVQPVGVVDTEW | 1 | 0 | 0 | 1663512 | YP_009047202.1 | K9N7C7 |
| GVNITLANFNETKGPL | 1 | 0 | 0 | 1663517 | NP_073551.1 | A0A0P0K6L9 |
| GVNKIQPWTNDALNTI | 1 | 0 | 0 | 1663518 | NP_073549.1 | A0A0P0LKV0 |
| GVNLRTNFTIKGSFIN | 1 | 0 | 0 | 1663521 | YP_003766.2 | H9EJ42 |
| GVNVTKHKCDINYKGK | 1 | 0 | 0 | 1663528 | YP_009555238.1 | P0C6X6 |
| GVPDNSSTIVTGLLPT | 1 | 0 | 0 | 1663532 | YP_003767.1 | A0A0P0G321 |
| GVPIAFGVPPSEAKGY | 1 | 0 | 0 | 1663537 | YP_173242.1 | Q19U29 |
| GVPIINCVKMLCDHAG | 1 | 0 | 0 | 1663540 | YP_173236.1 | A0A140H1G9 |
| GVPINTNSGPDDQIGY | 1 | 0 | 0 | 1663541 | NP_828858.1 | P59595 |
| GVPINTNSTKDDQIGY | 1 | 0 | 0 | 1663543 | QIQ54056.1 | P59595 |
| GVPLNANSTPAQNAGY | 1 | 0 | 0 | 1663545 | YP_009047211.1 | K9N4V7 |
| GVPVIEVSQIQSKLTD | 1 | 0 | 0 | 1663550 | YP_173236.1 | A0A140H1G9 |
| GVQIPCTCGKQATKYL | 1 | 0 | 0 | 1663556 | YP_009724389.1 | P0DTD1 |
| GVQIPCTCGKQATQYL | 1 | 0 | 0 | 1663557 | QHR63299.1 | P0C6W2 |
| GVQTVEDLRARMTYVC | 1 | 0 | 0 | 1663561 | YP_009047202.1 | K9N7C7 |
| GVQYGRSGETLGVLVP | 1 | 0 | 0 | 1663564 | QIQ54046.1 | P0C6U8 |
| GVRDARIKKAMFTKVV | 1 | 0 | 0 | 1663567 | YP_003766.2 | H9EJ42 |
| GVRELYLNSSNVTTMD | 1 | 0 | 0 | 1663568 | NP_828849.2 | P0C6X7 |
| GVRQQRFVYDAYQNLV | 1 | 0 | 0 | 1663575 | YP_009047204.1 | K9N5Q8 |
| GVRRKNSEPEIPHFNQ | 1 | 0 | 0 | 1663577 | NP_073556.1 | A0A0N7J131 |
| GVSATKLNDLCFSNVY | 1 | 0 | 0 | 1663582 | NP_828851.1 | P59594 |
| GVSDLSLKQKLLYTFY | 1 | 0 | 0 | 1663584 | YP_009047202.1 | K9N7C7 |
| GVSDVKKLYLGGLNYY | 1 | 0 | 0 | 1663585 | YP_003766.2 | H9EJ42 |
| GVSDVWHKQLLLNRGV | 1 | 0 | 0 | 1663587 | YP_009555238.1 | P0C6X6 |
| GVSEIVNTVAGIPSNV | 1 | 0 | 0 | 1663588 | NP_073549.1 | A0A0P0LKV0 |
| GVSERINVVPGVPTNV | 1 | 0 | 0 | 1663589 | YP_003766.2 | H9EJ42 |
| GVSGCLTDCLYLKNLT | 1 | 0 | 0 | 1663592 | YP_173236.1 | A0A140H1G9 |
| GVSGRGVFQNCTAVGV | 1 | 0 | 0 | 1663593 | YP_009047204.1 | K9N5Q8 |
| GVSKPNAIDVEHLELK | 1 | 0 | 0 | 1663599 | YP_003766.2 | H9EJ42 |
| GVSSNQSPQGNLGSNG | 1 | 0 | 0 | 1663608 | YP_009555238.1 | P0C6X6 |
| GVSVITPGTNASNQVA | 1 | 0 | 0 | 1663610 | QHR63300.2 | Q3LZX1 |
| GVSVITPGTNASSEVA | 1 | 0 | 0 | 1663611 | NP_828851.1 | P59594 |
| GVTFGEDTVWEVQGYK | 1 | 0 | 0 | 1663615 | NP_828849.2 | P0C6X7 |
| GVTLDKLRDYLADYDV | 1 | 0 | 0 | 1663622 | YP_009047202.1 | K9N7C7 |
| GVTLSSNLNTNLHFDV | 1 | 0 | 0 | 1663627 | YP_173238.1 | A0A140H1H1 |
| GVTLSTKLKDGVNFNV | 1 | 0 | 0 | 1663628 | YP_009555241.1 | P36334 |
| GVTMDVLNKNQKLIAT | 1 | 0 | 0 | 1663630 | YP_173238.1 | A0A140H1H1 |
| GVTNLEDLCLKYNYCQ | 1 | 0 | 0 | 1663633 | YP_173239.1 | Q0ZJ84 |
| GVTQNVLYENQKLIAN | 1 | 0 | 0 | 1663636 | YP_009724390.1 | P0DTC2 |
| GVTQNVLYENQKQIAN | 1 | 0 | 0 | 1663637 | NP_828851.1 | P59594 |
| GVTRDIASTDTCFANK | 1 | 0 | 0 | 1663638 | YP_009724389.1 | P0DTD1 |
| GVTRDIISTDDCFANK | 1 | 0 | 0 | 1663639 | NP_828849.2 | P0C6X7 |
| GVTRDISSNDNCFANK | 1 | 0 | 0 | 1663640 | QIQ54046.1 | P0C6U8 |
| GVTRDLMRELNGGAYT | 1 | 0 | 0 | 1663641 | QHR63299.1 | P0C6W2 |
| GVTRELMRELNGGAYT | 1 | 0 | 0 | 1663642 | YP_009724389.1 | P0DTD1 |
| GVTTSDKCIFNSACTR | 1 | 0 | 0 | 1663645 | YP_003766.2 | H9EJ42 |
| GVTYQLAWDVIRKDLS | 1 | 0 | 0 | 1663649 | YP_003766.2 | H9EJ42 |
| GVTYTEIEPKLDGYYK | 1 | 0 | 0 | 1663650 | NP_828849.2 | P0C6X7 |
| GVVANGVKAKGYPQFA | 1 | 0 | 0 | 1663652 | NP_073556.1 | A0A0N7J131 |
| GVVCTEIDPKLDNYYK | 1 | 0 | 0 | 1663655 | YP_009724389.1 | P0DTD1 |
| GVVCYNKDYAELGYIA | 1 | 0 | 0 | 1663656 | YP_003766.2 | H9EJ42 |
| GVVCYNSDYAAKGYIA | 1 | 0 | 0 | 1663657 | YP_009047202.1 | K9N7C7 |
| GVVCYNSDYASKGYIA | 1 | 0 | 0 | 1663658 | YP_009555238.1 | P0C6X6 |
| GVVDKQVILVSNNKED | 1 | 0 | 0 | 1663659 | YP_173236.1 | A0A140H1G9 |
| GVVDTEWGNMLTGIAA | 1 | 0 | 0 | 1663661 | YP_009047202.1 | K9N7C7 |
| GVVDYGARFYFYTSKT | 1 | 0 | 0 | 1663662 | YP_009724389.1 | P0DTD1 |
| GVVDYGVRFYFYTSKT | 1 | 0 | 0 | 1663663 | QIQ54046.1 | P0C6U8 |
| GVVEDTINDDEDVVTG | 1 | 0 | 0 | 1663664 | YP_173236.1 | A0A140H1G9 |
| GVVGEGSESLNDNLLE | 1 | 0 | 0 | 1663670 | QIQ54046.1 | P0C6U8 |
| GVVLKRSLKYNLKASA | 1 | 0 | 0 | 1663676 | NP_073549.1 | A0A0P0LKV0 |
| GVVPLSVIPATSAARL | 1 | 0 | 0 | 1663682 | NP_073549.1 | A0A0P0LKV0 |
| GVVPLSVIPATSASKL | 1 | 0 | 0 | 1663683 | YP_003766.2 | H9EJ42 |
| GVVRDISVNDLCFANK | 1 | 0 | 0 | 1663686 | YP_173236.1 | A0A140H1G9 |
| GVVREFLTRNPAWRKA | 1 | 0 | 0 | 1663687 | YP_009724389.1 | P0DTD1 |
| GVVREFLTRNPTWRKA | 1 | 0 | 0 | 1663688 | QHR63299.1 | P0C6W2 |
| GVVTGNVVKQCCTAAV | 1 | 0 | 0 | 1663693 | YP_009047202.1 | K9N7C7 |
| GVVWTLNDVKDNDGRP | 1 | 0 | 0 | 1663702 | YP_003766.2 | H9EJ42 |
| GVVWVAKEGAKTVNTS | 1 | 0 | 0 | 1663704 | YP_003771.1 | H9EJ69 |
| GVYDFCVSPAEFKYMV | 1 | 0 | 0 | 1663711 | NP_073549.1 | A0A0P0LKV0 |
| GVYDFKVSAAEFKYMV | 1 | 0 | 0 | 1663712 | YP_003766.2 | H9EJ42 |
| GVYDFKVSTQEFRFMT | 1 | 0 | 0 | 1663713 | YP_009047202.1 | K9N7C7 |
| GVYDYLVSTQEFRYMN | 1 | 0 | 0 | 1663717 | YP_009724389.1 | P0DTD1 |
| GVYKPLLENIDYFNMR | 1 | 0 | 0 | 1663723 | YP_009555238.1 | P0C6X6 |
| GVYQCAMRPNFTIKGS | 1 | 0 | 0 | 1663725 | YP_009724389.1 | P0DTD1 |
| GVYQCAMRPNHTIKGS | 1 | 0 | 0 | 1663726 | NP_828849.2 | P0C6X7 |
| GVYSVSSFEAKPSGSV | 1 | 0 | 0 | 1663730 | YP_009047204.1 | K9N5Q8 |
| GVYWVASNQADVNTPA | 1 | 0 | 0 | 1663733 | YP_009555245.1 | P33469 |
| GVYYHNNNKTWVENEF | 1 | 0 | 0 | 1663735 | QIQ54048.1 | P59594 |
| GWDDMLRHLIKDVDNP | 1 | 0 | 0 | 1663744 | YP_173236.1 | A0A140H1G9 |
| GWDDMLRRLIKDVDNP | 1 | 0 | 0 | 1663745 | YP_009555238.1 | P0C6X6 |
| GWDFMLKTLYKDVDNP | 1 | 0 | 0 | 1663746 | YP_009047202.1 | K9N7C7 |
| GWDNMLKNLMADVDDP | 1 | 0 | 0 | 1663747 | NP_073549.1 | A0A0P0LKV0 |

| | | | | | | |
|---|---|---|---|---|---|---|
| GWDYPKCDRALPNMIR | 1 | 0 | 0 | 1663748 | YP_003766.2 | H9EJ42 |
| GWDYPKCDRAMPNLLR | 1 | 0 | 0 | 1663750 | YP_009555238.1 | P0C6X6 |
| GWDYPKCDRAMPNMCR | 1 | 0 | 0 | 1663751 | YP_009047202.1 | K9N7C7 |
| GWDYPKCDRAMPNMLR | 1 | 0 | 0 | 1663752 | YP_009724389.1 | P0DTD1 |
| GWDYPKCDRAMPSMIR | 1 | 0 | 0 | 1663753 | NP_073549.1 | A0A0P0LKV0 |
| GWFDVYDDIFAESKPW | 1 | 0 | 0 | 1663756 | NP_073549.1 | A0A0P0LKV0 |
| GWIKISADNKVIYTTE | 1 | 0 | 0 | 1663762 | YP_009555238.1 | P0C6X6 |
| GWLGMDVEGAHVTGDN | 1 | 0 | 0 | 1663766 | NP_073549.1 | A0A0P0LKV0 |
| GWLGMDVESAHVCGDN | 1 | 0 | 0 | 1663767 | YP_003766.2 | H9EJ42 |
| GWTTGAAAYYVGYLQQ | 1 | 0 | 0 | 1663788 | QIQ54048.1 | P59594 |
| GWVFYVRVKHGDFSAV | 1 | 0 | 0 | 1663790 | NP_073555.1 | A0A0P0KXN3 |
| GWVGFDVEGAHATREN | 1 | 0 | 0 | 1663791 | YP_173236.1 | A0A140H1G9 |
| GWVSGNMMDGFSCPLC | 1 | 0 | 0 | 1663793 | YP_173236.1 | A0A140H1G9 |
| GWYDPVENEDIHRVYA | 1 | 0 | 0 | 1663796 | YP_003766.2 | H9EJ42 |
| GYCVDYSKNRRSRGAI | 1 | 0 | 0 | 1663810 | YP_009555241.1 | P36334 |
| GYDDCMQQGPASARDL | 1 | 0 | 0 | 1663812 | YP_009047204.1 | K9N5Q8 |
| GYDFKYIENGQLKNFE | 1 | 0 | 0 | 1663813 | NP_073549.1 | A0A0P0LKV0 |
| GYEKSTFNKDIYCGNS | 1 | 0 | 0 | 1663817 | YP_003766.2 | H9EJ42 |
| GYFIKTNNTRIVDEWS | 1 | 0 | 0 | 1663823 | YP_009047204.1 | K9N5Q8 |
| GYFVNVNNTWMYTGSG | 1 | 0 | 0 | 1663832 | YP_009555241.1 | P36334 |
| GYGSTFITPEVSRELG | 1 | 0 | 0 | 1663838 | NP_073549.1 | A0A0P0LKV0 |
| GYGVRRKNSEPEIPHF | 1 | 0 | 0 | 1663839 | NP_073556.1 | A0A0N7J131 |
| GYHPAHAPSFLAVDDK | 1 | 0 | 0 | 1663847 | YP_009555238.1 | P0C6X6 |
| GYHYTPFHYERDNTSC | 1 | 0 | 0 | 1663852 | YP_009047202.1 | K9N7C7 |
| GYIDCEQDSCDFKGWI | 1 | 0 | 0 | 1663858 | YP_009555238.1 | P0C6X6 |
| GYIGATVRLQAGKQTE | 1 | 0 | 0 | 1663859 | NP_073549.1 | A0A0P0LKV0 |
| GYKVLPPLMDVNMEAA | 1 | 0 | 0 | 1663878 | YP_009047204.1 | K9N5Q8 |
| GYLKLTDNVYIKNADI | 1 | 0 | 0 | 1663881 | YP_009724389.1 | P0DTD1 |
| GYLNSTNVTTAYCTG | 1 | 0 | 0 | 1663885 | QIQ54046.1 | P0C6U8 |
| GYLQQRTFLLSYNQNG | 1 | 0 | 0 | 1663891 | QIQ54048.1 | P59594 |
| GYLTSSSKTPEEHFIE | 1 | 0 | 0 | 1663895 | YP_009724389.1 | P0DTD1 |
| GYLTSSSKTSEEHFVE | 1 | 0 | 0 | 1663896 | NP_828849.2 | P0C6X7 |
| GYNLKNGEVEFVYMHQ | 1 | 0 | 0 | 1663900 | NP_073549.1 | A0A0P0LKV0 |
| GYNVRNDGTVEFCYLH | 1 | 0 | 0 | 1663903 | YP_003766.2 | H9EJ42 |
| GYPQFAELVPSTAAML | 1 | 0 | 0 | 1663912 | NP_073556.1 | A0A0N7J131 |
| GYQHNSVVDVMRYNLN | 1 | 0 | 0 | 1663916 | YP_003767.1 | A0A0P0G321 |
| GYQPIDVVRDLPSGFN | 1 | 0 | 0 | 1663917 | NP_828851.1 | P59594 |
| GYQVRAFLNKLNENVV | 1 | 0 | 0 | 1663920 | YP_173236.1 | A0A140H1G9 |
| GYREGYLNSTNVTIAT | 1 | 0 | 0 | 1663921 | YP_009724389.1 | P0DTD1 |
| GYREGYLNSTNVTTAI | 1 | 0 | 0 | 1663922 | QHR63299.1 | P0C6W2 |
| GYRFTNFEPFTVNSVN | 1 | 0 | 0 | 1663923 | YP_009555241.1 | P36334 |
| GYRVTKNSKTQIGEYT | 1 | 0 | 0 | 1663926 | QIQ54047.1 | P0C6X7 |
| GYRVTKNSKVQIGEYT | 1 | 0 | 0 | 1663927 | YP_009724389.1 | P0DTD1 |
| GYSKLFITKDEAIKRV | 1 | 0 | 0 | 1663933 | YP_173236.1 | A0A140H1G9 |
| GYSSLCDEFTLAEVVK | 1 | 0 | 0 | 1663940 | YP_003766.2 | H9EJ42 |
| GYTDFAEDVCTCYDNS | 1 | 0 | 0 | 1663943 | NP_073549.1 | A0A0P0LKV0 |
| GYTESAVNSTKALAKQ | 1 | 0 | 0 | 1663947 | YP_009047206.1 | K9N4V0 |
| GYTLAATSASLFPPWT | 1 | 0 | 0 | 1663951 | YP_009555241.1 | P36334 |
| GYTNEIIGYKAIQDGV | 1 | 0 | 0 | 1663955 | NP_828849.2 | P0C6X7 |
| GYTNSSTRGVYYPDKV | 1 | 0 | 0 | 1663956 | QIQ54048.1 | P59594 |
| GYTVEEAKTVLKKCKS | 1 | 0 | 0 | 1663962 | YP_009724389.1 | P0DTD1 |
| GYTVEEARTVLKKCKS | 1 | 0 | 0 | 1663963 | QHR63299.1 | P0C6W2 |
| GYVDCFNDNCDFYGWV | 1 | 0 | 0 | 1663971 | YP_173236.1 | A0A140H1G9 |
| GYVDTENNTEFTRVNA | 1 | 0 | 0 | 1663972 | NP_828849.2 | P0C6X7 |
| GYVDTPNNTDFSRVSA | 1 | 0 | 0 | 1663974 | YP_009724389.1 | P0DTD1 |
| GYVLTGDSVKFVYMHQ | 1 | 0 | 0 | 1663982 | YP_173236.1 | A0A140H1G9 |
| GYVQQGIFNKATVAQI | 1 | 0 | 0 | 1663985 | YP_009555238.1 | P0C6X6 |
| GYVTHGFNLEEAARCM | 1 | 0 | 0 | 1663987 | NP_828849.2 | P0C6X7 |
| GYVTHGLNLEEAARYM | 1 | 0 | 0 | 1663988 | YP_009724389.1 | P0DTD1 |
| GYWRRQDRKINTGNGI | 1 | 0 | 0 | 1663992 | YP_009047211.1 | K9N4V7 |
| GYYKKDNAYFTEQPID | 1 | 0 | 0 | 1664000 | QIQ54046.1 | P0C6U8 |
| GYYVKPGGTSSGDATT | 1 | 0 | 0 | 1664009 | YP_009047202.1 | K9N7C7 |
| HAEETRKLMPVCMETK | 1 | 0 | 0 | 1664024 | QHR63299.1 | P0C6W2 |
| HAEEYREYLESYNTAT | 1 | 0 | 0 | 1664027 | YP_009555238.1 | P0C6X6 |
| HAEPKGKYSQKAYALL | 1 | 0 | 0 | 1664028 | YP_173236.1 | A0A140H1G9 |
| HAFHTPAFDKSAFTNL | 1 | 0 | 0 | 1664032 | NP_828849.2 | P0C6X7 |
| HAFMKHKGGDSTDFIA | 1 | 0 | 0 | 1664039 | YP_009047202.1 | K9N7C7 |
| HAGTDLEGKFYGPFVD | 1 | 0 | 0 | 1664044 | NP_828849.2 | P0C6X7 |
| HAGTDLEGNFYGPFVD | 1 | 0 | 0 | 1664045 | YP_009724389.1 | P0DTD1 |
| HAGTDLEGTFYGPFVD | 1 | 0 | 0 | 1664046 | QHR63299.1 | P0C6W2 |
| HALDCELYVNNAYRLF | 1 | 0 | 0 | 1664056 | YP_009555238.1 | P0C6X6 |
| HANEYRLYLDAYNMMI | 1 | 0 | 0 | 1664066 | YP_009724389.1 | P0DTD1 |
| HANEYRQYLDAYNMMI | 1 | 0 | 0 | 1664067 | NP_828849.2 | P0C6X7 |
| HANSSEPALLFRNIKC | 1 | 0 | 0 | 1664073 | YP_009555241.1 | P36334 |
| HAPSFLAVDDKYKATG | 1 | 0 | 0 | 1664082 | YP_009555238.1 | P0C6X6 |
| HAPSFLAVDDKYKVNE | 1 | 0 | 0 | 1664083 | YP_173236.1 | A0A140H1G9 |
| HARLKGGLILKFDSGT | 1 | 0 | 0 | 1664084 | YP_009047202.1 | K9N7C7 |
| HATGTTPQKLFVANYS | 1 | 0 | 0 | 1664090 | YP_009047204.1 | K9N5Q8 |
| HATRDSIGTNFPLQLG | 1 | 0 | 0 | 1664093 | YP_009555238.1 | P0C6X6 |
| HATREAIGTNLPLQLG | 1 | 0 | 0 | 1664094 | QHR63299.1 | P0C6W2 |
| HATRENIGTNFPLQIG | 1 | 0 | 0 | 1664096 | YP_173236.1 | A0A140H1G9 |
| HCGETSWQTGDFVKAT | 1 | 0 | 0 | 1664110 | YP_009724389.1 | P0DTD1 |
| HCGSSSRSFFEDLLFD | 1 | 0 | 0 | 1664112 | YP_173238.1 | A0A140H1H1 |
| HCLHVVGPNVNKGEDI | 1 | 0 | 0 | 1664115 | YP_009724389.1 | P0DTD1 |
| HDAIVVDNDVVDYVSD | 1 | 0 | 0 | 1664136 | NP_073549.1 | A0A0P0LKV0 |
| HDASSAINRPQLTFVK | 1 | 0 | 0 | 1664138 | YP_009047202.1 | K9N7C7 |
| HDDLVNYIRDKGCVAD | 1 | 0 | 0 | 1664140 | YP_009555239.1 | Q80872 |
| HDFFKFRIDGDMVPHI | 1 | 0 | 0 | 1664146 | YP_009724389.1 | P0DTD1 |
| HDFFKFRVDGDMVPHI | 1 | 0 | 0 | 1664147 | NP_828849.2 | P0C6X7 |
| HDGYTNEIIGYKAIQD | 1 | 0 | 0 | 1664154 | NP_828849.2 | P0C6X7 |
| HDICAQLNDKLGFNVD | 1 | 0 | 0 | 1664156 | YP_173236.1 | A0A140H1G9 |

| | | | | | | |
|---|---|---|---|---|---|---|
| HDIQITNESYNNYVPS | 1 | 0 | 0 | 1664165 | YP_009047202.1 | K9N7C7 |
| HDNVAFYQGDVDTVVN | 1 | 0 | 0 | 1664174 | NP_073549.1 | A0A0P0LKV0 |
| HDTVLTSEEVVLKNGE | 1 | 0 | 0 | 1664186 | NP_828849.2 | P0C6X7 |
| HDVSPFEMPFEELNGL | 1 | 0 | 0 | 1664192 | NP_073549.1 | A0A0P0LKV0 |
| HDVTVLRDNLKPVTTL | 1 | 0 | 0 | 1664194 | YP_003768.1 | H9EJ24 |
| HDVTVTTDKSFEQQVG | 1 | 0 | 0 | 1664195 | NP_073549.1 | A0A0P0LKV0 |
| HEAKFGVVTTNSDKCP | 1 | 0 | 0 | 1664199 | YP_003766.2 | H9EJ42 |
| HEDLHVVEVLTKTAVK | 1 | 0 | 0 | 1664204 | YP_173236.1 | A0A140H1G9 |
| HEDLMAAYVDNTSLTI | 1 | 0 | 0 | 1664205 | QIQ54046.1 | P0C6U8 |
| HEETIYNLVKDCPAVA | 1 | 0 | 0 | 1664210 | NP_828849.2 | P0C6X7 |
| HEFCSQHTMQIVDKDG | 1 | 0 | 0 | 1664218 | YP_003766.2 | H9EJ42 |
| HEGRTIYGNVSRQDLT | 1 | 0 | 0 | 1664228 | NP_073549.1 | A0A0P0LKV0 |
| HEHSLLVEAYNSILFE | 1 | 0 | 0 | 1664232 | YP_003766.2 | H9EJ42 |
| HEMQKPRWKRQPNDDV | 1 | 0 | 0 | 1664241 | NP_073556.1 | A0A0N7J131 |
| HEQSMYNLLKGCNAVA | 1 | 0 | 0 | 1664244 | NP_073549.1 | A0A0P0LKV0 |
| HERDLLIKAYNTINNE | 1 | 0 | 0 | 1664246 | NP_073549.1 | A0A0P0LKV0 |
| HFDVDRKSFNNFVNIA | 1 | 0 | 0 | 1664278 | YP_173236.1 | A0A140H1G9 |
| HFDVVETKVEITPEES | 1 | 0 | 0 | 1664279 | QIQ54046.1 | P0C6U8 |
| HFFFTQKGDAAIKDFD | 1 | 0 | 0 | 1664284 | NP_073549.1 | A0A0P0LKV0 |
| HFGACDYDRLPNEVTV | 1 | 0 | 0 | 1664287 | YP_009047210.1 | K9N7A1 |
| HFGAGSDKGIAPGTSV | 1 | 0 | 0 | 1664288 | YP_009047202.1 | K9N7C7 |
| HFIGVEGELPVAVVND | 1 | 0 | 0 | 1664295 | YP_009047202.1 | K9N7C7 |
| HFNKEFYDFLRSQGFF | 1 | 0 | 0 | 1664308 | NP_073549.1 | A0A0P0LKV0 |
| HFNQKLPNGVTVVEEP | 1 | 0 | 0 | 1664309 | NP_073556.1 | A0A0N7J131 |
| HFVNLDNLRASNTKGS | 1 | 0 | 0 | 1664337 | QHR63299.1 | P0C6W2 |
| HGAIKLDDKDPQFKDN | 1 | 0 | 0 | 1664349 | NP_828858.1 | P59595 |
| HGCACDRTTIQSVDIS | 1 | 0 | 0 | 1664353 | YP_003766.2 | H9EJ42 |
| HGCTCDRTAIQSFDNS | 1 | 0 | 0 | 1664354 | NP_073549.1 | A0A0P0LKV0 |
| HGDFADEIIGYKAIED | 1 | 0 | 0 | 1664357 | QIQ54046.1 | P0C6U8 |
| HGFNLEEAARCMRSLK | 1 | 0 | 0 | 1664366 | NP_828849.2 | P0C6X7 |
| HGGGLAKALDVYTKGK | 1 | 0 | 0 | 1664368 | NP_073549.1 | A0A0P0LKV0 |
| HGGGVAGALNKATNNA | 1 | 0 | 0 | 1664370 | YP_009724389.1 | P0DTD1 |
| HGIAAINSVPWSKILA | 1 | 0 | 0 | 1664378 | NP_828849.2 | P0C6X7 |
| HGKEDLKFPRGQGVPI | 1 | 0 | 0 | 1664380 | YP_009724397.2 | P0DTC9 |
| HGKLRPFERDISNVPF | 1 | 0 | 0 | 1664382 | NP_828851.1 | P59594 |
| HGKQVVSDIDYVPLKS | 1 | 0 | 0 | 1664383 | YP_009724389.1 | P0DTD1 |
| HGNGSMIEDIDYVPLK | 1 | 0 | 0 | 1664392 | YP_009047202.1 | K9N7C7 |
| HGSGALRELTRELNGG | 1 | 0 | 0 | 1664401 | NP_828849.2 | P0C6X7 |
| HHANEYRLYLDAYNMM | 1 | 0 | 0 | 1664422 | YP_009724389.1 | P0DTD1 |
| HHANEYRQYLDAYNMM | 1 | 0 | 0 | 1664423 | NP_828849.2 | P0C6X7 |
| HHMELPTGVHAGTDLE | 1 | 0 | 0 | 1664431 | YP_009724389.1 | P0DTD1 |
| HHPELKLFRNLNIDVC | 1 | 0 | 0 | 1664433 | YP_009555238.1 | P0C6X6 |
| HIAGCQTTNGLNTSYS | 1 | 0 | 0 | 1664441 | NP_073551.1 | A0A0P0K6L9 |
| HIASSNAFLDLRTSCF | 1 | 0 | 0 | 1664442 | YP_009047202.1 | K9N7C7 |
| HIDHPNPKGFCDLKGK | 1 | 0 | 0 | 1664446 | YP_009724389.1 | P0DTD1 |
| HIGAPANLRVVGHAMQ | 1 | 0 | 0 | 1664453 | YP_009047202.1 | K9N7C7 |
| HIGLAGKVKVGTGVMV | 1 | 0 | 0 | 1664454 | NP_073549.1 | A0A0P0LKV0 |
| HIGTRNPNNNAAIVLQ | 1 | 0 | 0 | 1664457 | QIQ54056.1 | P59595 |
| HIGTRNPNNNAATVLQ | 1 | 0 | 0 | 1664458 | NP_828858.1 | P59595 |
| HIIPGSPGSLNFKKVE | 1 | 0 | 0 | 1664460 | YP_173243.1 | Q0ZJ80 |
| HILGRCIVLDVKGVEE | 1 | 0 | 0 | 1664464 | YP_009555239.1 | Q80872 |
| HINQATTKTTFKPNTW | 1 | 0 | 0 | 1664474 | NP_828849.2 | P0C6X7 |
| HIQTVQDADGINKQLT | 1 | 0 | 0 | 1664478 | YP_173236.1 | A0A140H1G9 |
| HIVSKESGNTVVLTFT | 1 | 0 | 0 | 1664502 | NP_073556.1 | A0A0N7J131 |
| HKAVFISPYNSQNFAA | 1 | 0 | 0 | 1664515 | YP_009555238.1 | P0C6X6 |
| HKAYIDVLRNSFGKDL | 1 | 0 | 0 | 1664516 | NP_073549.1 | A0A0P0LKV0 |
| HKHDIQITNESYNNYV | 1 | 0 | 0 | 1664540 | YP_009047202.1 | K9N7C7 |
| HKPIVWHVNNTINKAT | 1 | 0 | 0 | 1664567 | QIQ54046.1 | P0C6U8 |
| HKPPSASCPIGTNYRS | 1 | 0 | 0 | 1664569 | YP_173238.1 | A0A140H1H1 |
| HLAKALNDFSNSGADV | 1 | 0 | 0 | 1664587 | NP_828849.2 | P0C6X7 |
| HLAKALNDFSNSGSDV | 1 | 0 | 0 | 1664588 | YP_009724389.1 | P0DTD1 |
| HLATFAELEAAQKAYQ | 1 | 0 | 0 | 1664591 | YP_009047202.1 | K9N7C7 |
| HLDGEVITFDNLKTLL | 1 | 0 | 0 | 1664595 | YP_009724389.1 | P0DTD1 |
| HLDGEVLSLDKLKSLL | 1 | 0 | 0 | 1664596 | NP_828849.2 | P0C6X7 |
| HLHVASNSKLWQYCSV | 1 | 0 | 0 | 1664614 | YP_173236.1 | A0A140H1G9 |
| HLIKDVDNPVLMGWDY | 1 | 0 | 0 | 1664616 | YP_173236.1 | A0A140H1G9 |
| HLKHGGGIAGAINAAS | 1 | 0 | 0 | 1664627 | YP_009047202.1 | K9N7C7 |
| HLKVNCCRFQRVDENG | 1 | 0 | 0 | 1664634 | YP_009555238.1 | P0C6X6 |
| HLLDGTCGIIDVEKGV | 1 | 0 | 0 | 1664635 | QIQ54046.1 | P0C6U8 |
| HLLIGLAKRSKESPLE | 1 | 0 | 0 | 1664638 | QHR63299.1 | P0C6W2 |
| HLMGWDYPKCDRAMPN | 1 | 0 | 0 | 1664650 | YP_009724389.1 | P0DTD1 |
| HLNKTLNEGVLESFSV | 1 | 0 | 0 | 1664657 | NP_073549.1 | A0A0P0LKV0 |
| HLPAYASFKVIDNGVV | 1 | 0 | 0 | 1664662 | YP_173236.1 | A0A140H1G9 |
| HLRIAGHHLGRCDIKD | 1 | 0 | 0 | 1664669 | YP_009724393.1 | P0DTC5 |
| HLSMTTSTTNFGEDFL | 1 | 0 | 0 | 1664675 | YP_173236.1 | A0A140H1G9 |
| HLSTFYPQLQSAEWKC | 1 | 0 | 0 | 1664676 | YP_003766.2 | H9EJ42 |
| HLYIQGIKLGTGYSLA | 1 | 0 | 0 | 1664697 | YP_009555244.1 | Q01455 |
| HLYIQGVKLGTGYTLS | 1 | 0 | 0 | 1664698 | YP_173241.1 | Q0ZJ82 |
| HMELPTGVHAGTDLEG | 1 | 0 | 0 | 1664707 | YP_009724389.1 | P0DTD1 |
| HMYCSFYPPDEDEEEG | 1 | 0 | 0 | 1664722 | YP_009724389.1 | P0DTD1 |
| HNCCDEYGGHHDFVIK | 1 | 0 | 0 | 1664723 | YP_173238.1 | A0A140H1H1 |
| HNDMLYSPPTISYNST | 1 | 0 | 0 | 1664728 | YP_003766.2 | H9EJ42 |
| HNMRVLHYGAGSDKGV | 1 | 0 | 0 | 1664765 | YP_003766.2 | H9EJ42 |
| HNNGNDVLYQPPTASV | 1 | 0 | 0 | 1664768 | YP_173236.1 | A0A140H1G9 |
| HNPEVGPEHSLAEYHN | 1 | 0 | 0 | 1664771 | QHR63299.1 | P0C6W2 |
| HNQDVNLHSSRLSFKE | 1 | 0 | 0 | 1664773 | YP_009724389.1 | P0DTD1 |
| HNSEVGPEHSLAEYHN | 1 | 0 | 0 | 1664775 | YP_009724389.1 | P0DTD1 |
| HPAFNVSDAYANLVPY | 1 | 0 | 0 | 1664794 | YP_003766.2 | H9EJ42 |
| HPAHAPSFLAVDDKYK | 1 | 0 | 0 | 1664795 | YP_009555238.1 | P0C6X6 |
| HPDFKLLHNLQADICY | 1 | 0 | 0 | 1664798 | YP_009047202.1 | K9N7C7 |
| HPIQVDQLNSSYFKLS | 1 | 0 | 0 | 1664809 | YP_009047204.1 | K9N5Q8 |

| | | | | | | |
|---|---|---|---|---|---|---|
| HPLLNPSALEFNPSQT | 1 | 0 | 0 | 1664814 | NP_073556.1 | A0A0N7J131 |
| HPNCINCLDDRCILHC | 1 | 0 | 0 | 1664817 | NP_828849.2 | P0C6X7 |
| HPNCVNCLDDRCILHC | 1 | 0 | 0 | 1664819 | YP_009724389.1 | P0DTD1 |
| HPSFNVSDAYANLVPY | 1 | 0 | 0 | 1664827 | NP_073549.1 | A0A0P0LKV0 |
| HPSLCNNGKISSSAGD | 1 | 0 | 0 | 1664829 | YP_173237.1 | A0A140H1H0 |
| HPYCYSDGVMKNASLY | 1 | 0 | 0 | 1664839 | YP_173236.1 | A0A140H1G9 |
| HQADTSTPSDVSSRDP | 1 | 0 | 0 | 1664840 | YP_173242.1 | Q19U29 |
| HQDLKWARFPKSDGTG | 1 | 0 | 0 | 1664842 | NP_828849.2 | P0C6X7 |
| HQKLLKSIAATRGATV | 1 | 0 | 0 | 1664853 | YP_009724389.1 | P0DTD1 |
| HQKMLKSMAATRGATC | 1 | 0 | 0 | 1664854 | YP_009047202.1 | K9N7C7 |
| HQTPFEIKSAKKFDTF | 1 | 0 | 0 | 1664868 | NP_828849.2 | P0C6X7 |
| HQVNISLNGNTSVCVR | 1 | 0 | 0 | 1664869 | YP_003767.1 | A0A0P0G321 |
| HRDCDFNKPLIEWPLT | 1 | 0 | 0 | 1664874 | YP_009047202.1 | K9N7C7 |
| HRLASGVQVHNLPEYM | 1 | 0 | 0 | 1664887 | NP_073555.1 | A0A0P0KXN3 |
| HRLYECLYRNRDVDHE | 1 | 0 | 0 | 1664890 | NP_828849.2 | P0C6X7 |
| HRSYLTPGKLESGWTT | 1 | 0 | 0 | 1664903 | QIQ54048.1 | P59594 |
| HRYKAGNYRSPPITAD | 1 | 0 | 0 | 1664910 | YP_009047210.1 | K9N7A1 |
| HSDKFKDGVCLFWNCN | 1 | 0 | 0 | 1664920 | QIQ54047.1 | P0C6X7 |
| HSELAKGVALDGVLST | 1 | 0 | 0 | 1664925 | NP_828849.2 | P0C6X7 |
| HSFSVQKHIGAPANLR | 1 | 0 | 0 | 1664931 | YP_009047202.1 | K9N7C7 |
| HSLAEYHNESGIKTTL | 1 | 0 | 0 | 1664945 | QIQ54046.1 | P0C6U8 |
| HSLAEYHNESGLKPIL | 1 | 0 | 0 | 1664946 | QHR63299.1 | P0C6W2 |
| HSLAEYHNESGLKTIL | 1 | 0 | 0 | 1664947 | YP_009724389.1 | P0DTD1 |
| HSLLATLDDPYVEDSV | 1 | 0 | 0 | 1664953 | YP_173242.1 | Q19U29 |
| HSMAVVDGKQIDDHRI | 1 | 0 | 0 | 1664963 | YP_009555238.1 | P0C6X6 |
| HSSGVTRDLMRELNGG | 1 | 0 | 0 | 1664978 | QHR63299.1 | P0C6W2 |
| HSSQLPLDVYDLTMPS | 1 | 0 | 0 | 1664982 | YP_009555238.1 | P0C6X6 |
| HSVASLSAPTLVPQEN | 1 | 0 | 0 | 1664993 | YP_173236.1 | A0A140H1G9 |
| HSVAVIGDEQIDGKFV | 1 | 0 | 0 | 1664995 | YP_173236.1 | A0A140H1G9 |
| HSVEIEAGEVKPFAVY | 1 | 0 | 0 | 1664996 | YP_003766.2 | H9EJ42 |
| HSVLTTFIDAARGPAG | 1 | 0 | 0 | 1665000 | YP_009047202.1 | K9N7C7 |
| HSWNADLYKLMGHFAW | 1 | 0 | 0 | 1665008 | YP_009724389.1 | P0DTD1 |
| HSWNADLYKLMGHFSW | 1 | 0 | 0 | 1665009 | NP_828849.2 | P0C6X7 |
| HTDFSSEIIGYKAIDG | 1 | 0 | 0 | 1665020 | YP_009724389.1 | P0DTD1 |
| HTDLMAAYVDNSSLTI | 1 | 0 | 0 | 1665023 | YP_009724389.1 | P0DTD1 |
| HTFGNPVIPFKDGIYF | 1 | 0 | 0 | 1665027 | NP_828851.1 | P59594 |
| HTIDGSSGVANPAMDP | 1 | 0 | 0 | 1665047 | NP_828852.2 | P59632 |
| HTIDGSSGVVNPAMDP | 1 | 0 | 0 | 1665048 | QIQ54049.1 | P59632 |
| HTIDGSSGVVNPAMEP | 1 | 0 | 0 | 1665049 | QHR63301.1 | Q3LZX0 |
| HTIDGSSGVVNPVMEP | 1 | 0 | 0 | 1665050 | YP_009724391.1 | P0DTC3 |
| HTLDESFLGRYMSALN | 1 | 0 | 0 | 1665061 | NP_828849.2 | P0C6X7 |
| HTLRNGSVLEMAKEVK | 1 | 0 | 0 | 1665064 | NP_073549.1 | A0A0P0LKV0 |
| HTMQIVDENGKYYLPY | 1 | 0 | 0 | 1665077 | NP_073549.1 | A0A0P0LKV0 |
| HTMQIVDKDGTYYLPY | 1 | 0 | 0 | 1665078 | YP_003766.2 | H9EJ42 |
| HTPAFDKSAFTNLKQL | 1 | 0 | 0 | 1665085 | NP_828849.2 | P0C6X7 |
| HTPAFDKSAFVNLKQL | 1 | 0 | 0 | 1665086 | YP_009724389.1 | P0DTD1 |
| HTPATDCSDGNYNRNA | 1 | 0 | 0 | 1665087 | YP_009047204.1 | K9N5Q8 |
| HTPIDLVRDLPRGFAA | 1 | 0 | 0 | 1665091 | QIQ54048.1 | P59594 |
| HTPKHVFKTLKPGDSF | 1 | 0 | 0 | 1665094 | YP_003766.2 | H9EJ42 |
| HTVCDSLNAKLGFDSS | 1 | 0 | 0 | 1665109 | YP_009555238.1 | P0C6X6 |
| HVADTLMQGVTLSSNL | 1 | 0 | 0 | 1665124 | YP_173238.1 | A0A140H1H1 |
| HVAQNTVLNASIPESK | 1 | 0 | 0 | 1665127 | YP_003771.1 | H9EJ69 |
| HVCGDNIGTNVPLQVG | 1 | 0 | 0 | 1665140 | YP_003766.2 | H9EJ42 |
| HVDILGPLSAQTGIAV | 1 | 0 | 0 | 1665142 | YP_009724389.1 | P0DTD1 |
| HVETFYPKLQASQAWQ | 1 | 0 | 0 | 1665146 | NP_828849.2 | P0C6X7 |
| HVETFYPKLQSSQAWQ | 1 | 0 | 0 | 1665147 | YP_009724389.1 | P0DTD1 |
| HVEVFTDGKLNCSFQD | 1 | 0 | 0 | 1665148 | YP_009047202.1 | K9N7C7 |
| HVFKTLKPGDSFNILA | 1 | 0 | 0 | 1665150 | YP_003766.2 | H9EJ42 |
| HVIASNTTSAIDYDHE | 1 | 0 | 0 | 1665164 | NP_073549.1 | A0A0P0LKV0 |
| HVLDCELFVNDSYRQF | 1 | 0 | 0 | 1665178 | YP_173236.1 | A0A140H1G9 |
| HVLTKHGDFADEIIGY | 1 | 0 | 0 | 1665184 | QIQ54046.1 | P0C6U8 |
| HVNNATNKATYKPNTW | 1 | 0 | 0 | 1665198 | YP_009724389.1 | P0DTD1 |
| HVNNSYECDIPVGAGI | 1 | 0 | 0 | 1665200 | QIQ54048.1 | P59594 |
| HVNNTINKATFKPNTW | 1 | 0 | 0 | 1665201 | QIQ54046.1 | P0C6U8 |
| HVPHPSMDGYCKFKGK | 1 | 0 | 0 | 1665203 | YP_003766.2 | H9EJ42 |
| HVQIHTIDGSSGVVNP | 1 | 0 | 0 | 1665206 | YP_009724391.1 | P0DTC3 |
| HVTKENDSKEGFFTYL | 1 | 0 | 0 | 1665226 | NP_828849.2 | P0C6X7 |
| HVTKVLAEKFDYNVSV | 1 | 0 | 0 | 1665227 | NP_073549.1 | A0A0P0LKV0 |
| HVVGPDARAKQDVSLL | 1 | 0 | 0 | 1665238 | YP_009047202.1 | K9N7C7 |
| HVVGPNLNAGEDIQLL | 1 | 0 | 0 | 1665239 | NP_828849.2 | P0C6X7 |
| HVVGPNVNKGEDIQLL | 1 | 0 | 0 | 1665241 | YP_009724389.1 | P0DTD1 |
| HVVGPNVNRGEDIQLL | 1 | 0 | 0 | 1665242 | QHR63299.1 | P0C6W2 |
| HVVRDPRYVMRLQSAA | 1 | 0 | 0 | 1665244 | YP_009555238.1 | P0C6X6 |
| HVVRDSRFVMRLQTIA | 1 | 0 | 0 | 1665245 | YP_173236.1 | A0A140H1G9 |
| HVYDFKVEDAYDQVHD | 1 | 0 | 0 | 1665250 | YP_009555238.1 | P0C6X6 |
| HWAPEFPWMFEDAEEK | 1 | 0 | 0 | 1665255 | YP_009555238.1 | P0C6X6 |
| HWMGIRFEYTSPTDKL | 1 | 0 | 0 | 1665269 | YP_009555239.1 | Q80872 |
| HWSFDYHPNCSDCYDD | 1 | 0 | 0 | 1665273 | YP_003766.2 | H9EJ42 |
| HWSMPYHPNTVDCQDD | 1 | 0 | 0 | 1665275 | YP_009555238.1 | P0C6X6 |
| HYAGSRSSSGNRSGIL | 1 | 0 | 0 | 1665276 | YP_173242.1 | Q19U29 |
| HYAGVVWTLNDVKDND | 1 | 0 | 0 | 1665277 | YP_003766.2 | H9EJ42 |
| HYDGRTGELPCAIMND | 1 | 0 | 0 | 1665284 | YP_173236.1 | A0A140H1G9 |
| HYGAGSDKGVAPGTTV | 1 | 0 | 0 | 1665292 | YP_003766.2 | H9EJ42 |
| HYKELGVVMNLDVDTH | 1 | 0 | 0 | 1665297 | YP_173236.1 | A0A140H1G9 |
| HYNEWAQANGFTAMNG | 1 | 0 | 0 | 1665304 | NP_073549.1 | A0A0P0LKV0 |
| HYQTNIYSQNLCVDGF | 1 | 0 | 0 | 1665309 | NP_073549.1 | A0A0P0LKV0 |
| HYQTNLYSFNKAIDGF | 1 | 0 | 0 | 1665310 | YP_003766.2 | H9EJ42 |
| HYRALSGVFCGVDAMN | 1 | 0 | 0 | 1665311 | NP_828849.2 | P0C6X7 |
| HYTGQAGEMPCAIIND | 1 | 0 | 0 | 1665318 | YP_009555238.1 | P0C6X6 |
| HYTPFHYERDNTSCPE | 1 | 0 | 0 | 1665321 | YP_009047202.1 | K9N7C7 |
| HYTVYDTAKKSMYDGD | 1 | 0 | 0 | 1665324 | NP_073549.1 | A0A0P0LKV0 |

| | | | | | | |
|---|---|---|---|---|---|---|
| HYVHARLKGGLILKFD | 1 | 0 | 0 | 1665325 | YP_009047202.1 | K9N7C7 |
| HYWSPQGNLGGNCAGN | 1 | 0 | 0 | 1665336 | YP_173236.1 | A0A140H1G9 |
| IAANTVIWDYKRDAPA | 1 | 0 | 0 | 1665348 | YP_009724389.1 | P0DTD1 |
| IAANTVIWDYKREAPA | 1 | 0 | 0 | 1665349 | NP_828849.2 | P0C6X7 |
| IAARPPPGEQFKHLVP | 1 | 0 | 0 | 1665351 | YP_009047202.1 | K9N7C7 |
| IAATRGATVVIGTSKF | 1 | 0 | 0 | 1665352 | YP_009724389.1 | P0DTD1 |
| IAAYTAALVSGTATAG | 1 | 0 | 0 | 1665356 | NP_828851.1 | P59594 |
| IACKCGVKQEQRTGLD | 1 | 0 | 0 | 1665359 | YP_009555238.1 | P0C6X6 |
| IACVKDFDFKFYDAQP | 1 | 0 | 0 | 1665364 | YP_009555238.1 | P0C6X6 |
| IADKDKDLSGAVPSDL | 1 | 0 | 0 | 1665369 | NP_073549.1 | A0A0P0LKV0 |
| IADTLQETPVVPDTVE | 1 | 0 | 0 | 1665377 | YP_009047202.1 | K9N7C7 |
| IAEDVVDQVEEVNSIF | 1 | 0 | 0 | 1665381 | NP_073549.1 | A0A0P0LKV0 |
| IAEIPKEEVKPFITES | 1 | 0 | 0 | 1665385 | YP_009724389.1 | P0DTD1 |
| IAFELFAKRKVGLTPP | 1 | 0 | 0 | 1665389 | NP_073549.1 | A0A0P0LKV0 |
| IAFNHPIQVDQLNSSY | 1 | 0 | 0 | 1665396 | YP_009047204.1 | K9N5Q8 |
| IAGCQTTNGLNTSYSV | 1 | 0 | 0 | 1665403 | NP_073551.1 | A0A0P0K6L9 |
| IAGHHLGRCDIKDLPK | 1 | 0 | 0 | 1665405 | YP_009724393.1 | P0DTC5 |
| IAGIQNFKETLYYQNN | 1 | 0 | 0 | 1665406 | YP_009047202.1 | K9N7C7 |
| IAGNGGDAALALLLLD | 1 | 0 | 0 | 1665412 | QIQ54056.1 | P59595 |
| IAGVAGARIPDVPTTL | 1 | 0 | 0 | 1665417 | YP_009047202.1 | K9N7C7 |
| IAHASLREGVQLEKVL | 1 | 0 | 0 | 1665420 | YP_173236.1 | A0A140H1G9 |
| IAIKVTEYSWNKKLYE | 1 | 0 | 0 | 1665427 | NP_073549.1 | A0A0P0LKV0 |
| IALNKIQDVVNQQGSA | 1 | 0 | 0 | 1665460 | YP_003767.1 | A0A0P0G321 |
| IANENAINKGGRTVQS | 1 | 0 | 0 | 1665475 | NP_073549.1 | A0A0P0LKV0 |
| IANKFNQALGAMQTGF | 1 | 0 | 0 | 1665482 | YP_009047204.1 | K9N5Q8 |
| IANLVLAKLGKDSKPQ | 1 | 0 | 0 | 1665483 | YP_173242.1 | Q19U29 |
| IANMFLEQTQHSLNFV | 1 | 0 | 0 | 1665484 | YP_009047202.1 | K9N7C7 |
| IANNYSKLFDEGYTVV | 1 | 0 | 0 | 1665485 | NP_073549.1 | A0A0P0LKV0 |
| IANQFNSAIGKIQDSL | 1 | 0 | 0 | 1665488 | YP_009724390.1 | P0DTC2 |
| IANRYNEVANAVMQNN | 1 | 0 | 0 | 1665489 | YP_173236.1 | A0A140H1G9 |
| IANRYNEVSATVLQNN | 1 | 0 | 0 | 1665490 | YP_009555238.1 | P0C6X6 |
| IANVPAIDVLKKLLSS | 1 | 0 | 0 | 1665491 | YP_003766.2 | H9EJ42 |
| IAPKSGYFVNVNNTWM | 1 | 0 | 0 | 1665496 | YP_009555241.1 | P36334 |
| IAPVNGYFIKTNNTRI | 1 | 0 | 0 | 1665501 | YP_009047204.1 | K9N5Q8 |
| IAQVVIADTLQETPVV | 1 | 0 | 0 | 1665508 | YP_009047202.1 | K9N7C7 |
| IASLVLAKLGKDATKP | 1 | 0 | 0 | 1665523 | YP_009555245.1 | P33469 |
| IASNSKEWAYCVEMHN | 1 | 0 | 0 | 1665525 | NP_073549.1 | A0A0P0LKV0 |
| IASNTTSAIDYDHEYS | 1 | 0 | 0 | 1665526 | NP_073549.1 | A0A0P0LKV0 |
| IATAETELAKNVSLDN | 1 | 0 | 0 | 1665532 | QIQ54046.1 | P0C6U8 |
| IATAFNNALLSIQNGF | 1 | 0 | 0 | 1665533 | YP_173238.1 | A0A140H1H1 |
| IATKMFDSFVNSFVSL | 1 | 0 | 0 | 1665541 | YP_009047202.1 | K9N7C7 |
| IATLNSLGETLVTMPL | 1 | 0 | 0 | 1665543 | QIQ54046.1 | P0C6U8 |
| IATNGPLKVGGGCVLS | 1 | 0 | 0 | 1665544 | QHR63299.1 | P0C6W2 |
| IATVKSEDGNIKNINW | 1 | 0 | 0 | 1665548 | NP_073549.1 | A0A0P0LKV0 |
| IAVAAGKKFSKETAAM | 1 | 0 | 0 | 1665552 | YP_173236.1 | A0A140H1G9 |
| IAVANWLAPTNAYMRT | 1 | 0 | 0 | 1665554 | YP_009047202.1 | K9N7C7 |
| IAVKITEHSWNADLYK | 1 | 0 | 0 | 1665560 | NP_828849.2 | P0C6X7 |
| IAWFTERSDKSYEHQT | 1 | 0 | 0 | 1665568 | NP_828849.2 | P0C6X7 |
| IAWNSKHIDAKEGGNF | 1 | 0 | 0 | 1665571 | QHR63300.2 | Q3LZX1 |
| IAWNSVKQDALTGGNY | 1 | 0 | 0 | 1665573 | QIQ54048.1 | P59594 |
| IAYENARQAYEDAIAN | 1 | 0 | 0 | 1665577 | YP_003766.2 | H9EJ42 |
| IAYRNVLLRKNGNKGA | 1 | 0 | 0 | 1665582 | NP_828849.2 | P0C6X7 |
| ICAAFDFSLDVFKIGD | 1 | 0 | 0 | 1665585 | YP_003766.2 | H9EJ42 |
| ICASYQTQTNSRSVAS | 1 | 0 | 0 | 1665596 | QHR63300.2 | Q3LZX1 |
| ICIKDADYNAKVEISV | 1 | 0 | 0 | 1665631 | NP_073549.1 | A0A0P0LKV0 |
| ICIVDNVYMAKAGDKY | 1 | 0 | 0 | 1665634 | YP_009555238.1 | P0C6X6 |
| ICMFENKTTLPTNIAF | 1 | 0 | 0 | 1665655 | YP_009047202.1 | K9N7C7 |
| ICNTNRAGIGLYYKVN | 1 | 0 | 0 | 1665659 | YP_173236.1 | A0A140H1G9 |
| ICQAVSANVCALMSCN | 1 | 0 | 0 | 1665665 | YP_009555238.1 | P0C6X6 |
| ICSASDMTNPDYTNLL | 1 | 0 | 0 | 1665669 | YP_009555238.1 | P0C6X6 |
| ICSNTPLSKDLPDDVV | 1 | 0 | 0 | 1665671 | YP_173236.1 | A0A140H1G9 |
| ICSSSNMNEPDYSALL | 1 | 0 | 0 | 1665674 | YP_173236.1 | A0A140H1G9 |
| ICTSEDMLNPNYEDLL | 1 | 0 | 0 | 1665679 | YP_009724389.1 | P0DTD1 |
| IDAYPLSKHPKPEYRK | 1 | 0 | 0 | 1665711 | NP_073549.1 | A0A0P0LKV0 |
| IDAYPLSKHPNSEYRK | 1 | 0 | 0 | 1665712 | YP_003766.2 | H9EJ42 |
| IDAYPLTKHEDIEYQN | 1 | 0 | 0 | 1665713 | YP_009047202.1 | K9N7C7 |
| IDDFQVNWNTKHGSGV | 1 | 0 | 0 | 1665729 | QIQ54046.1 | P0C6U8 |
| IDEHVFYKKDGVYYPS | 1 | 0 | 0 | 1665736 | NP_073549.1 | A0A0P0LKV0 |
| IDGALLSKTSEYKGPV | 1 | 0 | 0 | 1665749 | QIQ54046.1 | P0C6U8 |
| IDGFGVFDIKNSSVNT | 1 | 0 | 0 | 1665755 | YP_003766.2 | H9EJ42 |
| IDGGKIWNGVIKNVNS | 1 | 0 | 0 | 1665757 | NP_073549.1 | A0A0P0LKV0 |
| IDGKFVTKFSGDKFDF | 1 | 0 | 0 | 1665759 | YP_173236.1 | A0A140H1G9 |
| IDGSSGVANPAMDPIY | 1 | 0 | 0 | 1665764 | NP_828852.2 | P59632 |
| IDGSSGVVNPAMDPIY | 1 | 0 | 0 | 1665765 | QIQ54049.1 | P59632 |
| IDGSSGVVNPAMEPIY | 1 | 0 | 0 | 1665766 | QHR63301.1 | Q3LZX0 |
| IDGSSGVVNPVMEPIY | 1 | 0 | 0 | 1665767 | YP_009724391.1 | P0DTC3 |
| IDGVENPMLMGWDYPK | 1 | 0 | 0 | 1665768 | YP_003766.2 | H9EJ42 |
| IDHPNPKGFCDLKGKY | 1 | 0 | 0 | 1665775 | YP_009724389.1 | P0DTD1 |
| IDHPNPKGYCELKGKY | 1 | 0 | 0 | 1665776 | QIQ54046.1 | P0C6U8 |
| IDIDAFPVKPICSSVY | 1 | 0 | 0 | 1665779 | YP_003766.2 | H9EJ42 |
| IDILTEGQLQSLSKDY | 1 | 0 | 0 | 1665783 | YP_003766.2 | H9EJ42 |
| IDINGNLHPDSATLVN | 1 | 0 | 0 | 1665784 | QHR63299.1 | P0C6W2 |
| IDITFLKKDAPYIVGD | 1 | 0 | 0 | 1665789 | YP_009724389.1 | P0DTD1 |
| IDKEDVVIFINNTTYP | 1 | 0 | 0 | 1665795 | YP_009555238.1 | P0C6X6 |
| IDKFAIPNGRKVDLQL | 1 | 0 | 0 | 1665796 | YP_009555241.1 | P36334 |
| IDKLNVLFDGRDNGAL | 1 | 0 | 0 | 1665798 | YP_009555238.1 | P0C6X6 |
| IDKWLNNFNVPSPLNW | 1 | 0 | 0 | 1665803 | YP_173238.1 | A0A140H1H1 |
| IDLLCLRDDNQTLLVP | 1 | 0 | 0 | 1665813 | YP_173236.1 | A0A140H1G9 |
| IDLLNLREIKPAVNVV | 1 | 0 | 0 | 1665816 | YP_009555238.1 | P0C6X6 |
| IDLVPNQPYPNASFDN | 1 | 0 | 0 | 1665822 | YP_009724389.1 | P0DTD1 |
| IDLVPTQPLPNASFDN | 1 | 0 | 0 | 1665823 | NP_828849.2 | P0C6X7 |

| | | | | | | |
|---|---|---|---|---|---|---|
| IDLVPTQPYPNSNFDN | 1 | 0 | 0 | 1665824 | QIQ54046.1 | P0C6U8 |
| IDMKGKMFVRPVIEDY | 1 | 0 | 0 | 1665827 | YP_173241.1 | Q0ZJ82 |
| IDMKGTMYVRPIIEDY | 1 | 0 | 0 | 1665828 | YP_009555244.1 | Q01455 |
| IDNDINDYVSDADFSI | 1 | 0 | 0 | 1665832 | YP_003766.2 | H9EJ42 |
| IDNINSTLVDLKWLNR | 1 | 0 | 0 | 1665836 | NP_073551.1 | A0A0P0K6L9 |
| IDPKLDGYYKKDNAYF | 1 | 0 | 0 | 1665841 | QIQ54046.1 | P0C6U8 |
| IDPKLDNYYKKDNSYF | 1 | 0 | 0 | 1665842 | YP_009724389.1 | P0DTD1 |
| IDQDFGSTVFNVPTKV | 1 | 0 | 0 | 1665844 | YP_009555238.1 | P0C6X6 |
| IDQINSTYVDLKLLNR | 1 | 0 | 0 | 1665845 | YP_003767.1 | A0A0P0G321 |
| IDQVTIAGAKLRSLNL | 1 | 0 | 0 | 1665846 | NP_828849.2 | P0C6X7 |
| IDSDVDTKCLADSVMS | 1 | 0 | 0 | 1665851 | YP_009555238.1 | P0C6X6 |
| IDSDVETRFITKSMIS | 1 | 0 | 0 | 1665852 | YP_173236.1 | A0A140H1G9 |
| IDSLKTLLALREVKTI | 1 | 0 | 0 | 1665853 | QIQ54046.1 | P0C6U8 |
| IDSNTTQDTYGGASVC | 1 | 0 | 0 | 1665854 | NP_073549.1 | A0A0P0LKV0 |
| IDTKRGVYCCREHEHE | 1 | 0 | 0 | 1665861 | YP_009724389.1 | P0DTD1 |
| IDVEIFENKTTLPVNV | 1 | 0 | 0 | 1665869 | NP_828849.2 | P0C6X7 |
| IDVEKGVLPQLEQPYV | 1 | 0 | 0 | 1665870 | QIQ54046.1 | P0C6U8 |
| IDVFNKLSTSDWSDIR | 1 | 0 | 0 | 1665871 | NP_073549.1 | A0A0P0LKV0 |
| IDVIDNEIIVKPNISL | 1 | 0 | 0 | 1665873 | NP_073549.1 | A0A0P0LKV0 |
| IDVLLTVDGVNFKSIS | 1 | 0 | 0 | 1665875 | YP_173236.1 | A0A140H1G9 |
| IDVLRNSFGKDLNANM | 1 | 0 | 0 | 1665876 | NP_073549.1 | A0A0P0LKV0 |
| IDVNKTLQELSYKLPN | 1 | 0 | 0 | 1665877 | NP_073551.1 | A0A0P0K6L9 |
| IDVNSALNICFDIRDN | 1 | 0 | 0 | 1665878 | YP_009047202.1 | K9N7C7 |
| IDVPSAESTGTQSVSV | 1 | 0 | 0 | 1665879 | YP_009047205.1 | K9N796 |
| IDVSKADGIIYPQGRT | 1 | 0 | 0 | 1665881 | YP_009047204.1 | K9N5Q8 |
| IDVVNKALEMCIDQVT | 1 | 0 | 0 | 1665883 | NP_828849.2 | P0C6X7 |
| IDVVQYEADRRAFVDY | 1 | 0 | 0 | 1665884 | YP_009555238.1 | P0C6X6 |
| IDVVRDLPSGFNTLKP | 1 | 0 | 0 | 1665885 | NP_828851.1 | P59594 |
| IDYCVRAFDVYNKDAS | 1 | 0 | 0 | 1665889 | NP_073549.1 | A0A0P0LKV0 |
| IEAGEVKPFAVYKNVK | 1 | 0 | 0 | 1665903 | YP_003766.2 | H9EJ42 |
| IECGSCGNDSWLTGNA | 1 | 0 | 0 | 1665908 | YP_009047202.1 | K9N7C7 |
| IEDAIIVNDENSSIKV | 1 | 0 | 0 | 1665912 | YP_173236.1 | A0A140H1G9 |
| IEDALRLSAHLETNDV | 1 | 0 | 0 | 1665913 | YP_003767.1 | A0A0P0G321 |
| IEDIDYVPLKSAVCIT | 1 | 0 | 0 | 1665917 | YP_009047202.1 | K9N7C7 |
| IEDILFSKLVTSGLGT | 1 | 0 | 0 | 1665918 | NP_073551.1 | A0A0P0K6L9 |
| IEDLLFDKVKLSDVGF | 1 | 0 | 0 | 1665919 | YP_009555241.1 | P36334 |
| IEDLLFDKVTIADPGY | 1 | 0 | 0 | 1665920 | YP_009047204.1 | K9N5Q8 |
| IEDLSIRNLQKRLYSN | 1 | 0 | 0 | 1665923 | YP_173236.1 | A0A140H1G9 |
| IEDSVFKPRPAGVLTN | 1 | 0 | 0 | 1665924 | YP_009555241.1 | P36334 |
| IEDYEQNWNTKHGSGA | 1 | 0 | 0 | 1665926 | NP_828849.2 | P0C6X7 |
| IEEFADVVKEQVSDLL | 1 | 0 | 0 | 1665930 | YP_009047202.1 | K9N7C7 |
| IEELFYSYATHHDKFT | 1 | 0 | 0 | 1665933 | NP_828849.2 | P0C6X7 |
| IEELFYSYATHSDKFK | 1 | 0 | 0 | 1665934 | QIQ54047.1 | P0C6X7 |
| IEESDLIISNFKFDDY | 1 | 0 | 0 | 1665936 | YP_173236.1 | A0A140H1G9 |
| IEEVELLDAEFVEPGC | 1 | 0 | 0 | 1665938 | NP_073549.1 | A0A0P0LKV0 |
| IEFLKRGDKSVYYTSN | 1 | 0 | 0 | 1665941 | YP_009724389.1 | P0DTD1 |
| IEFLKRGDKVVYHTTS | 1 | 0 | 0 | 1665942 | QIQ54046.1 | P0C6U8 |
| IEFMLWCKDGQVQTFY | 1 | 0 | 0 | 1665943 | YP_009047202.1 | K9N7C7 |
| IEGSGRSAPNSRSTSR | 1 | 0 | 0 | 1665947 | YP_009555245.1 | P33469 |
| IEGSKPYSTLQPNAYY | 1 | 0 | 0 | 1665948 | YP_003766.2 | H9EJ42 |
| IEGTGGNSQSSSRASS | 1 | 0 | 0 | 1665949 | YP_009047211.1 | K9N4V7 |
| IEKLYKDLTGHMLDSY | 1 | 0 | 0 | 1665953 | YP_009047202.1 | K9N7C7 |
| IELGSGSHVGSSFDGV | 1 | 0 | 0 | 1665962 | NP_073549.1 | A0A0P0LKV0 |
| IELKFNPPALQDAYYR | 1 | 0 | 0 | 1665964 | YP_009724389.1 | P0DTD1 |
| IELSKELKRPVNPTDA | 1 | 0 | 0 | 1665965 | YP_173236.1 | A0A140H1G9 |
| IEMQLFKVGKVDSIVQ | 1 | 0 | 0 | 1665968 | YP_003766.2 | H9EJ42 |
| IEPEPEPTPEEPVNQF | 1 | 0 | 0 | 1665982 | NP_828849.2 | P0C6X7 |
| IEPKLDGYYKKDNAYY | 1 | 0 | 0 | 1665983 | NP_828849.2 | P0C6X7 |
| IEPVYRVKLCFEFEDE | 1 | 0 | 0 | 1665985 | NP_073549.1 | A0A0P0LKV0 |
| IERGKLLVRDTRKEIF | 1 | 0 | 0 | 1665993 | YP_173236.1 | A0A140H1G9 |
| IERVDWDIEYPYISHE | 1 | 0 | 0 | 1665995 | YP_009047202.1 | K9N7C7 |
| IESDDYIARNGPLNVG | 1 | 0 | 0 | 1665997 | QIQ54046.1 | P0C6U8 |
| IESIHYLGTTGHTLKS | 1 | 0 | 0 | 1665998 | YP_003766.2 | H9EJ42 |
| IESKRGVYCCRDHEHE | 1 | 0 | 0 | 1666000 | NP_828849.2 | P0C6X7 |
| IEVDIQQTFFDKTWPR | 1 | 0 | 0 | 1666010 | YP_009047204.1 | K9N5Q8 |
| IEVQGYKNVNITFELD | 1 | 0 | 0 | 1666013 | QIQ54046.1 | P0C6U8 |
| IEVQGYKSVNITFELD | 1 | 0 | 0 | 1666014 | YP_009724389.1 | P0DTD1 |
| IEVSQFQSKLTDVKCA | 1 | 0 | 0 | 1666015 | YP_009555238.1 | P0C6X6 |
| IEVSQIQSKLTDVKCA | 1 | 0 | 0 | 1666016 | YP_173236.1 | A0A140H1G9 |
| IEVTGDSCNNYMLTYN | 1 | 0 | 0 | 1666017 | YP_009724389.1 | P0DTD1 |
| IEWPLTEYDFTDYKVQ | 1 | 0 | 0 | 1666019 | YP_009047202.1 | K9N7C7 |
| IEYPIIGDELKINAAC | 1 | 0 | 0 | 1666021 | YP_009724389.1 | P0DTD1 |
| IFADTLVEVGLVGVLT | 1 | 0 | 0 | 1666027 | YP_173236.1 | A0A140H1G9 |
| IFAQTSDTAHACNANR | 1 | 0 | 0 | 1666032 | NP_073549.1 | A0A0P0LKV0 |
| IFCQTADTAHANNINR | 1 | 0 | 0 | 1666040 | YP_009047202.1 | K9N7C7 |
| IFDIETVDVKHDVSPF | 1 | 0 | 0 | 1666045 | NP_073549.1 | A0A0P0LKV0 |
| IFDVAPIELENKFTPL | 1 | 0 | 0 | 1666049 | YP_009047202.1 | K9N7C7 |
| IFENKTTLPVNVAFEL | 1 | 0 | 0 | 1666054 | NP_828849.2 | P0C6X7 |
| IFERGSVQVDNGSSIN | 1 | 0 | 0 | 1666055 | NP_073549.1 | A0A0P0LKV0 |
| IFERIDYSDAVSYKSS | 1 | 0 | 0 | 1666056 | YP_009047202.1 | K9N7C7 |
| IFGLYKNSATGSLDVE | 1 | 0 | 0 | 1666075 | YP_003766.2 | H9EJ42 |
| IFGQDFKTFDLLKYDF | 1 | 0 | 0 | 1666078 | NP_073549.1 | A0A0P0LKV0 |
| IFIETNLQSLENIAFN | 1 | 0 | 0 | 1666090 | YP_003766.2 | H9EJ42 |
| IFINNTTYPTNVAVEL | 1 | 0 | 0 | 1666094 | YP_009555238.1 | P0C6X6 |
| IFKDAAGKPVPYCYDT | 1 | 0 | 0 | 1666102 | QIQ54046.1 | P0C6U8 |
| IFKDAMGKPVPYCYDT | 1 | 0 | 0 | 1666103 | NP_828849.2 | P0C6X7 |
| IFKDASGKPVPYCYDT | 1 | 0 | 0 | 1666104 | YP_009724389.1 | P0DTD1 |
| IFKNNTSFPTNIAVEL | 1 | 0 | 0 | 1666107 | YP_173236.1 | A0A140H1G9 |
| IFNAVDCMSDFMSEIK | 1 | 0 | 0 | 1666125 | YP_009555241.1 | P36334 |
| IFNCFYALNNAFLAFS | 1 | 0 | 0 | 1666126 | YP_173241.1 | Q0ZJ82 |
| IFNGKFLSNTKYYDDS | 1 | 0 | 0 | 1666129 | YP_009555240.1 | P30215 |

| | | | | | | |
|---|---|---|---|---|---|---|
| IFNKATVAQIKALFLD | 1 | 0 | 0 | 1666131 | YP_009555238.1 | P0C6X6 |
| IFNKLVKDPPNVQIHT | 1 | 0 | 0 | 1666132 | NP_828852.2 | P59632 |
| IFNSACTRLEGLGGDN | 1 | 0 | 0 | 1666136 | YP_003766.2 | H9EJ42 |
| IFSQFDSLGVSSNQSP | 1 | 0 | 0 | 1666158 | YP_009555238.1 | P0C6X6 |
| IFSTVNALPECNADIV | 1 | 0 | 0 | 1666166 | YP_003766.2 | H9EJ42 |
| IFSVQQDGRIPNGFPF | 1 | 0 | 0 | 1666169 | YP_003767.1 | A0A0P0G321 |
| IFTIGTVTLKQGEIKD | 1 | 0 | 0 | 1666171 | YP_009724391.1 | P0DTC3 |
| IFTINSVVQKDGVENT | 1 | 0 | 0 | 1666172 | YP_009047202.1 | K9N7C7 |
| IFTLGTVTLKQGEIKD | 1 | 0 | 0 | 1666174 | QHR63301.1 | Q3LZX0 |
| IFVHEDLHVVEVLTKT | 1 | 0 | 0 | 1666190 | YP_173236.1 | A0A140H1G9 |
| IFVQQKPEPVNPVSFV | 1 | 0 | 0 | 1666194 | NP_073549.1 | A0A0P0LKV0 |
| IFYKENSYITTIKPVI | 1 | 0 | 0 | 1666207 | QHR63299.1 | P0C6W2 |
| IGAGICASYQTQTNSP | 1 | 0 | 0 | 1666215 | YP_009724390.1 | P0DTC2 |
| IGAGICASYQTQTNSR | 1 | 0 | 0 | 1666216 | QHR63300.2 | Q3LZX1 |
| IGAMTAVNESRYGLQN | 1 | 0 | 0 | 1666221 | YP_003767.1 | A0A0P0G321 |
| IGANYLGKPKEQIDGY | 1 | 0 | 0 | 1666222 | NP_828849.2 | P0C6X7 |
| IGAPANLRVVGHAMQG | 1 | 0 | 0 | 1666223 | YP_009047202.1 | K9N7C7 |
| IGATIRLQAGKQTELA | 1 | 0 | 0 | 1666224 | YP_003766.2 | H9EJ42 |
| IGATVRLQAGKQTEFV | 1 | 0 | 0 | 1666225 | NP_073549.1 | A0A0P0LKV0 |
| IGAVGGDLLYLDLLNR | 1 | 0 | 0 | 1666226 | YP_009047211.1 | K9N4V7 |
| IGDCATVHTANKWDLI | 1 | 0 | 0 | 1666234 | YP_009724389.1 | P0DTD1 |
| IGDFNCTNFAINDLNT | 1 | 0 | 0 | 1666238 | YP_173238.1 | A0A140H1H1 |
| IGDIIQRLDVLEQDAQ | 1 | 0 | 0 | 1666239 | YP_009047204.1 | K9N5Q8 |
| IGDLKCTSDNINDKDT | 1 | 0 | 0 | 1666242 | YP_009555241.1 | P36334 |
| IGDPAQLPAPRTLLTK | 1 | 0 | 0 | 1666243 | YP_009724389.1 | P0DTD1 |
| IGDVKFKRLGDYVLTE | 1 | 0 | 0 | 1666250 | YP_003766.2 | H9EJ42 |
| IGEALKTQFNYYKKVN | 1 | 0 | 0 | 1666252 | QHR63299.1 | P0C6W2 |
| IGEAVKTQFNYYKKVD | 1 | 0 | 0 | 1666254 | YP_009724389.1 | P0DTD1 |
| IGEHCSGLAVKSDYCG | 1 | 0 | 0 | 1666256 | YP_009555241.1 | P36334 |
| IGEPKSILSPLYAFPS | 1 | 0 | 0 | 1666257 | QIQ54046.1 | P0C6U8 |
| IGESVKTQFNYFKKVD | 1 | 0 | 0 | 1666259 | NP_828849.2 | P0C6X7 |
| IGEYTFEKGDYGDAVV | 1 | 0 | 0 | 1666261 | YP_009724389.1 | P0DTD1 |
| IGFKDFVTNKTYNIFP | 1 | 0 | 0 | 1666268 | YP_173238.1 | A0A140H1H1 |
| IGGAVCSKHANLYQKY | 1 | 0 | 0 | 1666274 | YP_003766.2 | H9EJ42 |
| IGGDRCIKISTVQSKL | 1 | 0 | 0 | 1666276 | YP_003766.2 | H9EJ42 |
| IGGKPCIKVATVQSKM | 1 | 0 | 0 | 1666280 | NP_828849.2 | P0C6X7 |
| IGGVPIIEVSQFQSKL | 1 | 0 | 0 | 1666290 | YP_009555238.1 | P0C6X6 |
| IGGVPVIEVSQIQSKL | 1 | 0 | 0 | 1666291 | YP_173236.1 | A0A140H1G9 |
| IGHTVCDSLNAKLGFD | 1 | 0 | 0 | 1666300 | YP_009555238.1 | P0C6X6 |
| IGIDTAGVLLKQLMVN | 1 | 0 | 0 | 1666301 | YP_173243.1 | Q0ZJ80 |
| IGINYLCKPKVEIDGN | 1 | 0 | 0 | 1666307 | YP_009555238.1 | P0C6X6 |
| IGINYLGKSSFEIDGN | 1 | 0 | 0 | 1666309 | YP_173236.1 | A0A140H1G9 |
| IGKGNKDEQIGYWNVQ | 1 | 0 | 0 | 1666313 | YP_003771.1 | H9EJ69 |
| IGLAGKVKVGTGVMVE | 1 | 0 | 0 | 1666321 | NP_073549.1 | A0A0P0LKV0 |
| IGLAKRFKESPFELED | 1 | 0 | 0 | 1666323 | YP_009724389.1 | P0DTD1 |
| IGLAKRSKDSPLELED | 1 | 0 | 0 | 1666324 | QIQ54047.1 | P0C6X7 |
| IGLAKRSKESPLELED | 1 | 0 | 0 | 1666325 | QHR63299.1 | P0C6W2 |
| IGLYRRQQTSNLVVQE | 1 | 0 | 0 | 1666339 | YP_009555238.1 | P0C6X6 |
| IGNMVEFIQTSSPKVT | 1 | 0 | 0 | 1666347 | YP_009555241.1 | P36334 |
| IGNNNVCTYEHVISYM | 1 | 0 | 0 | 1666348 | NP_073549.1 | A0A0P0LKV0 |
| IGNPKGIPIVDDPVVD | 1 | 0 | 0 | 1666353 | YP_009047202.1 | K9N7C7 |
| IGNPKGIRCAVTDAKW | 1 | 0 | 0 | 1666354 | NP_073549.1 | A0A0P0LKV0 |
| IGNYKLNTDHAGSNDN | 1 | 0 | 0 | 1666357 | NP_828855.1 | P59596 |
| IGNYKLNTDHSNSSDN | 1 | 0 | 0 | 1666358 | QIQ54051.1 | P59596 |
| IGPDVFLHKCYRCPAE | 1 | 0 | 0 | 1666363 | NP_073549.1 | A0A0P0LKV0 |
| IGSNNVCTYEHVISFM | 1 | 0 | 0 | 1666378 | YP_003766.2 | H9EJ42 |
| IGTNDPIRFCLENTVC | 1 | 0 | 0 | 1666385 | NP_073549.1 | A0A0P0LKV0 |
| IGTNFPLQIGFSTGVD | 1 | 0 | 0 | 1666386 | YP_173236.1 | A0A140H1G9 |
| IGTNFPLQLGFSTGID | 1 | 0 | 0 | 1666387 | YP_009555238.1 | P0C6X6 |
| IGTNLPLQLGFSTGVN | 1 | 0 | 0 | 1666388 | QHR63299.1 | P0C6W2 |
| IGTNVPLQVGFSNGVN | 1 | 0 | 0 | 1666389 | YP_003766.2 | H9EJ42 |
| IGTNYRSCESTTVLDH | 1 | 0 | 0 | 1666390 | YP_173238.1 | A0A140H1H1 |
| IGTRNPANNAAIVLQL | 1 | 0 | 0 | 1666392 | YP_009724397.2 | P0DTC9 |
| IGTVTLKQGEIKDATP | 1 | 0 | 0 | 1666401 | YP_009724391.1 | P0DTC3 |
| IGVCKYTDIDVNSALN | 1 | 0 | 0 | 1666405 | YP_009047202.1 | K9N7C7 |
| IGVCTMTDIAKKPTES | 1 | 0 | 0 | 1666408 | NP_828849.2 | P0C6X7 |
| IGVEGELPVAVVNDKI | 1 | 0 | 0 | 1666410 | YP_009047202.1 | K9N7C7 |
| IGVNVCSDSTFEETSL | 1 | 0 | 0 | 1666412 | YP_173236.1 | A0A140H1G9 |
| IGVTQNVLYENQKLIA | 1 | 0 | 0 | 1666415 | YP_009724390.1 | P0DTC2 |
| IGYHYKELGVVMNLDV | 1 | 0 | 0 | 1666424 | YP_173236.1 | A0A140H1G9 |
| IGYVTHGFNLEEAARC | 1 | 0 | 0 | 1666433 | NP_828849.2 | P0C6X7 |
| IHDGYTNEIIGYKAIQ | 1 | 0 | 0 | 1666445 | NP_828849.2 | P0C6X7 |
| IHFGAGSDKGIAPGTS | 1 | 0 | 0 | 1666448 | YP_009047202.1 | K9N7C7 |
| IHLPAYASFKVIDNGV | 1 | 0 | 0 | 1666457 | YP_173236.1 | A0A140H1G9 |
| IHTIDGSSGVANPAMD | 1 | 0 | 0 | 1666472 | NP_828852.2 | P59632 |
| IHTIDGSSGVVNPAMD | 1 | 0 | 0 | 1666473 | QIQ54049.1 | P59632 |
| IHTIDGSSGVVNPAME | 1 | 0 | 0 | 1666474 | QHR63301.1 | Q3LZX0 |
| IHTIDGSSGVVNPVME | 1 | 0 | 0 | 1666475 | YP_009724391.1 | P0DTC3 |
| IIANRYNEVANAVMQN | 1 | 0 | 0 | 1666491 | YP_173236.1 | A0A140H1G9 |
| IIAVIYSGRESLIFPS | 1 | 0 | 0 | 1666497 | YP_009047202.1 | K9N7C7 |
| IIDNDINDYVSDADFS | 1 | 0 | 0 | 1666502 | YP_003766.2 | H9EJ42 |
| IIDVEKGVLPQLEQPY | 1 | 0 | 0 | 1666507 | QIQ54046.1 | P0C6U8 |
| IIEVSQFQSKLTDVKC | 1 | 0 | 0 | 1666510 | YP_009555238.1 | P0C6X6 |
| IIFNAVDCMSDFMSEI | 1 | 0 | 0 | 1666517 | YP_009555241.1 | P36334 |
| IIFVQQKPEPVNPVSF | 1 | 0 | 0 | 1666520 | NP_073549.1 | A0A0P0LKV0 |
| IIGAMTAVNESRYGLQ | 1 | 0 | 0 | 1666525 | YP_003767.1 | A0A0P0G321 |
| IIGSLGFYNPPTNVVS | 1 | 0 | 0 | 1666533 | YP_009555240.1 | P30215 |
| IIIDNDINDYVSDADF | 1 | 0 | 0 | 1666540 | YP_003766.2 | H9EJ42 |
| IIIDNDLNEFVSDADI | 1 | 0 | 0 | 1666541 | YP_009047202.1 | K9N7C7 |
| IIIGGAKLKALNLGET | 1 | 0 | 0 | 1666542 | YP_009724389.1 | P0DTD1 |
| IIINNSTNVVIRACNF | 1 | 0 | 0 | 1666546 | NP_828851.1 | P59594 |

| | | | | | | |
|---|---|---|---|---|---|---|
| IIKAQFKTFEKVDGVY | 1 | 0 | 0 | 1666549 | YP_009555238.1 | P0C6X6 |
| IIKNLSKPLTENKYSQ | 1 | 0 | 0 | 1666556 | QIQ54052.1 | P59634 |
| IIKNLSKSLTENKYSQ | 1 | 0 | 0 | 1666557 | YP_009724394.1 | P0DTC6 |
| IIKSSVKSYDDLVYSG | 1 | 0 | 0 | 1666561 | YP_173236.1 | A0A140H1G9 |
| IINARIKAKHIVYVGD | 1 | 0 | 0 | 1666597 | YP_009047202.1 | K9N7C7 |
| IINDKVVAKIDKEDVV | 1 | 0 | 0 | 1666599 | YP_009555238.1 | P0C6X6 |
| IINNARDGCVPLNIIP | 1 | 0 | 0 | 1666606 | YP_009724389.1 | P0DTD1 |
| IINNAVYTKVDGIDVE | 1 | 0 | 0 | 1666607 | NP_828849.2 | P0C6X7 |
| IINNTVYTKVDGVDVE | 1 | 0 | 0 | 1666610 | YP_009724389.1 | P0DTD1 |
| IIPARARVECFDKFKV | 1 | 0 | 0 | 1666615 | YP_009724389.1 | P0DTD1 |
| IIPGSPGSLNFKKVET | 1 | 0 | 0 | 1666618 | YP_173243.1 | Q0ZJ80 |
| IIRDVNPDDKCFANKH | 1 | 0 | 0 | 1666629 | YP_009047202.1 | K9N7C7 |
| IIRENNRVVISSDVLV | 1 | 0 | 0 | 1666631 | YP_009724389.1 | P0DTD1 |
| IIRENNRVVVSSDILV | 1 | 0 | 0 | 1666632 | NP_828849.2 | P0C6X7 |
| IIRGHLYIQGIKLGTG | 1 | 0 | 0 | 1666634 | YP_009555244.1 | Q01455 |
| IIRSSNQSLAGGITYV | 1 | 0 | 0 | 1666637 | YP_003767.1 | A0A0P0G321 |
| IISDMYDPRTKHVTKE | 1 | 0 | 0 | 1666646 | NP_828849.2 | P0C6X7 |
| IISNFKFDDYDFSVED | 1 | 0 | 0 | 1666660 | YP_173236.1 | A0A140H1G9 |
| IITSGNLTTVVIPTKK | 1 | 0 | 0 | 1666679 | QIQ54046.1 | P0C6U8 |
| IITSTSAGCGVKYYAG | 1 | 0 | 0 | 1666680 | YP_003766.2 | H9EJ42 |
| IITTDNTFVSGNCDVV | 1 | 0 | 0 | 1666681 | YP_009724390.1 | P0DTC2 |
| IITTDNTFVSGSCDVV | 1 | 0 | 0 | 1666682 | QHR63300.2 | Q3LZX1 |
| IIVKPNISLCVPLYVR | 1 | 0 | 0 | 1666687 | NP_073549.1 | A0A0P0LKV0 |
| IIVPDKSVYDQVVDNV | 1 | 0 | 0 | 1666690 | YP_009555238.1 | P0C6X6 |
| IIWVATEGALNTPKDH | 1 | 0 | 0 | 1666695 | YP_009724397.2 | P0DTC9 |
| IIYDSSDRGQESLARS | 1 | 0 | 0 | 1666697 | YP_009047202.1 | K9N7C7 |
| IIYTGDWEGLHEVLTS | 1 | 0 | 0 | 1666700 | YP_003766.2 | H9EJ42 |
| IKACEVVVTNLNKSAG | 1 | 0 | 0 | 1666701 | YP_003766.2 | H9EJ42 |
| IKACVEEVTTTLEETK | 1 | 0 | 0 | 1666703 | YP_009724389.1 | P0DTD1 |
| IKAQFKTFEKVDGVYT | 1 | 0 | 0 | 1666709 | YP_009555238.1 | P0C6X6 |
| IKAYNTINNEQGTPLT | 1 | 0 | 0 | 1666712 | NP_073549.1 | A0A0P0LKV0 |
| IKCVDIVKEAQSANPM | 1 | 0 | 0 | 1666716 | NP_828849.2 | P0C6X7 |
| IKCVPQADVEWKFYDA | 1 | 0 | 0 | 1666718 | YP_009724389.1 | P0DTD1 |
| IKCVPQAEVEWKFYDA | 1 | 0 | 0 | 1666719 | NP_828849.2 | P0C6X7 |
| IKDADYNAKVEISVTP | 1 | 0 | 0 | 1666720 | NP_073549.1 | A0A0P0LKV0 |
| IKDATPSDFVRATATI | 1 | 0 | 0 | 1666721 | YP_009724391.1 | P0DTC3 |
| IKDATPSDSVRATATI | 1 | 0 | 0 | 1666722 | QHR63301.1 | Q3LZX0 |
| IKDCVDINKAIPSGNL | 1 | 0 | 0 | 1666723 | NP_073549.1 | A0A0P0LKV0 |
| IKDFDYYRYNRPTMLD | 1 | 0 | 0 | 1666725 | NP_073549.1 | A0A0P0LKV0 |
| IKDFGGFNFLQILPDP | 1 | 0 | 0 | 1666726 | QIQ54048.1 | P59594 |
| IKDGDDGYFLPYPDPS | 1 | 0 | 0 | 1666729 | YP_009047202.1 | K9N7C7 |
| IKDRVMYSEFPAITIG | 1 | 0 | 0 | 1666735 | YP_009555241.1 | P36334 |
| IKDTEKYCALAPNMMV | 1 | 0 | 0 | 1666736 | YP_009724389.1 | P0DTD1 |
| IKDVDNPVLMGWDYPK | 1 | 0 | 0 | 1666737 | YP_009555238.1 | P0C6X6 |
| IKENVDFINSMQFNYK | 1 | 0 | 0 | 1666744 | YP_009555239.1 | Q80872 |
| IKEPCSSGTYEGNSPF | 1 | 0 | 0 | 1666745 | QIQ54053.1 | P59635 |
| IKESIPIVWGVKDFNT | 1 | 0 | 0 | 1666747 | YP_003766.2 | H9EJ42 |
| IKFADDLNQLTGYKKP | 1 | 0 | 0 | 1666750 | YP_009724389.1 | P0DTD1 |
| IKFCDGENVSKDGFFT | 1 | 0 | 0 | 1666751 | YP_003766.2 | H9EJ42 |
| IKFKTEGLCVDIPGIP | 1 | 0 | 0 | 1666756 | NP_828849.2 | P0C6X7 |
| IKFQTVKPGNFNQDFY | 1 | 0 | 0 | 1666759 | YP_173236.1 | A0A140H1G9 |
| IKFSGFKLDGFNHQFV | 1 | 0 | 0 | 1666760 | YP_003766.2 | H9EJ42 |
| IKGLNNLNRGMVLGSL | 1 | 0 | 0 | 1666765 | YP_009724389.1 | P0DTD1 |
| IKGPQRADLNGVIVDK | 1 | 0 | 0 | 1666766 | YP_173236.1 | A0A140H1G9 |
| IKGVTFGEDTVWEVQG | 1 | 0 | 0 | 1666772 | NP_828849.2 | P0C6X7 |
| IKILNNLGVDIAANTV | 1 | 0 | 0 | 1666779 | NP_828849.2 | P0C6X7 |
| IKIPTNFTIVGQEEFI | 1 | 0 | 0 | 1666780 | YP_173238.1 | A0A140H1H1 |
| IKISADNKVIYTTEIA | 1 | 0 | 0 | 1666784 | YP_009555238.1 | P0C6X6 |
| IKISTVQSKLTDLKCT | 1 | 0 | 0 | 1666785 | YP_003766.2 | H9EJ42 |
| IKITEFSWNADLYKLM | 1 | 0 | 0 | 1666786 | YP_173236.1 | A0A140H1G9 |
| IKITEFSWNAELYKLM | 1 | 0 | 0 | 1666787 | YP_009555238.1 | P0C6X6 |
| IKITEHSWNADLYKLM | 1 | 0 | 0 | 1666788 | YP_009724389.1 | P0DTD1 |
| IKKLDNDVLNGIISNA | 1 | 0 | 0 | 1666795 | YP_009047202.1 | K9N7C7 |
| IKKLYNDLGNQILDSY | 1 | 0 | 0 | 1666796 | YP_009555238.1 | P0C6X6 |
| IKLADDLNQLTGYKKP | 1 | 0 | 0 | 1666803 | QHR63299.1 | P0C6W2 |
| IKLAKKFDNFTGECPN | 1 | 0 | 0 | 1666804 | QIQ54046.1 | P0C6U8 |
| IKLDDKDPQFKDNVIL | 1 | 0 | 0 | 1666808 | NP_828858.1 | P59595 |
| IKLDDKDPSFKDNVIL | 1 | 0 | 0 | 1666809 | QIQ54056.1 | P59595 |
| IKLDPKNPNYNKWLEL | 1 | 0 | 0 | 1666810 | YP_009047211.1 | K9N4V7 |
| IKLGTGYSLADLPAYM | 1 | 0 | 0 | 1666813 | YP_009555244.1 | Q01455 |
| IKLLGIGGKPCIKVAT | 1 | 0 | 0 | 1666820 | NP_828849.2 | P0C6X7 |
| IKLLGVGGKPCIKVAT | 1 | 0 | 0 | 1666821 | YP_009724389.1 | P0DTD1 |
| IKLNFGMLNGMPVSSI | 1 | 0 | 0 | 1666822 | YP_003766.2 | H9EJ42 |
| IKLNFGMLNGNAIATV | 1 | 0 | 0 | 1666823 | NP_073549.1 | A0A0P0LKV0 |
| IKLNGPLTVGGSCLLS | 1 | 0 | 0 | 1666824 | NP_828849.2 | P0C6X7 |
| IKLPQFYIYDEEGGYD | 1 | 0 | 0 | 1666826 | YP_003766.2 | H9EJ42 |
| IKLPSGIMLNVVKYTQ | 1 | 0 | 0 | 1666827 | YP_003766.2 | H9EJ42 |
| IKNLSKPLTENKYSQL | 1 | 0 | 0 | 1666844 | QIQ54052.1 | P59634 |
| IKNLTPIKLNFGMLNG | 1 | 0 | 0 | 1666846 | YP_003766.2 | H9EJ42 |
| IKNVDIVSEAKNVKPT | 1 | 0 | 0 | 1666851 | QIQ54046.1 | P0C6U8 |
| IKNVNSVRDWLKSLKL | 1 | 0 | 0 | 1666852 | NP_073549.1 | A0A0P0LKV0 |
| IKPVPEIKILNNLGVD | 1 | 0 | 0 | 1666866 | NP_828849.2 | P0C6X7 |
| IKPVPEVKILNNLGVD | 1 | 0 | 0 | 1666867 | YP_009724389.1 | P0DTD1 |
| IKPVSYKLDGVTYTEI | 1 | 0 | 0 | 1666868 | NP_828849.2 | P0C6X7 |
| IKQLKKAMNVAKAEFD | 1 | 0 | 0 | 1666876 | NP_073549.1 | A0A0P0LKV0 |
| IKQLKRAMNIAKSEFD | 1 | 0 | 0 | 1666877 | YP_003766.2 | H9EJ42 |
| IKQPFSFSFRDELGVR | 1 | 0 | 0 | 1666880 | YP_003766.2 | H9EJ42 |
| IKQYGDCLGDIAARDL | 1 | 0 | 0 | 1666882 | YP_009724390.1 | P0DTC2 |
| IKRFDNPVLPFNDGVY | 1 | 0 | 0 | 1666884 | QHR63300.2 | Q3LZX1 |
| IKRSDALSTNHGHKVV | 1 | 0 | 0 | 1666887 | NP_828849.2 | P0C6X7 |
| IKRVRGWVGFDVEGAH | 1 | 0 | 0 | 1666890 | YP_173236.1 | A0A140H1G9 |

| | | | | | | |
|---|---|---|---|---|---|---|
| IKSFSTFESAYMPIAD | 1 | 0 | 0 | 1666892 | NP_073549.1 | A0A0P0LKV0 |
| IKSLNNSYINLKDIGT | 1 | 0 | 0 | 1666896 | YP_173238.1 | A0A140H1H1 |
| IKSSVKSYDDLVYSGV | 1 | 0 | 0 | 1666900 | YP_173236.1 | A0A140H1G9 |
| IKSVMDHEQSMYNLLK | 1 | 0 | 0 | 1666901 | NP_073549.1 | A0A0P0LKV0 |
| IKTAQADTAGLYTNFR | 1 | 0 | 0 | 1666903 | YP_009047205.1 | K9N796 |
| IKTIQPRVEKKKLEGF | 1 | 0 | 0 | 1666905 | QIQ54046.1 | P0C6U8 |
| IKTLQPVSELLIPLGI | 1 | 0 | 0 | 1666906 | QHR63299.1 | P0C6W2 |
| IKTLQPVSELLTPLGI | 1 | 0 | 0 | 1666907 | YP_009724389.1 | P0DTD1 |
| IKTNNTRIVDEWSYTG | 1 | 0 | 0 | 1666908 | YP_009047204.1 | K9N5Q8 |
| IKVATVQSKMSDVKCT | 1 | 0 | 0 | 1666913 | YP_009724389.1 | P0DTD1 |
| IKVFTTVDNINLHTQV | 1 | 0 | 0 | 1666916 | YP_009724389.1 | P0DTD1 |
| IKVFTTVDNTNLHTQL | 1 | 0 | 0 | 1666917 | NP_828849.2 | P0C6X7 |
| IKVFYELDNDFNTILN | 1 | 0 | 0 | 1666918 | YP_009555238.1 | P0C6X6 |
| IKVLPPLLSENQISGY | 1 | 0 | 0 | 1666923 | YP_009555241.1 | P36334 |
| IKVTEDGVNVHDVTVT | 1 | 0 | 0 | 1666928 | NP_073549.1 | A0A0P0LKV0 |
| IKVTEYSWNKKLYELV | 1 | 0 | 0 | 1666929 | NP_073549.1 | A0A0P0LKV0 |
| ILAASFNKAINNIVAS | 1 | 0 | 0 | 1666953 | YP_003767.1 | A0A0P0G321 |
| ILAASFNKAMTNIVDA | 1 | 0 | 0 | 1666954 | NP_073551.1 | A0A0P0K6L9 |
| ILASVPEKFLNAFDVF | 1 | 0 | 0 | 1666971 | NP_073549.1 | A0A0P0LKV0 |
| ILAYCNKTVGELGDVR | 1 | 0 | 0 | 1666976 | YP_009724389.1 | P0DTD1 |
| ILAYSNKTVGELGDVR | 1 | 0 | 0 | 1666977 | NP_828849.2 | P0C6X7 |
| ILCEYADCKESYFSKK | 1 | 0 | 0 | 1666981 | YP_173236.1 | A0A140H1G9 |
| ILCIMSDRDLYDKLQF | 1 | 0 | 0 | 1666982 | YP_009724389.1 | P0DTD1 |
| ILCWETGKVKPPLNKN | 1 | 0 | 0 | 1666987 | YP_173236.1 | A0A140H1G9 |
| ILDEVNGLLDTTQLHV | 1 | 0 | 0 | 1666989 | YP_173238.1 | A0A140H1H1 |
| ILDFKVLDNGIIRDVN | 1 | 0 | 0 | 1666990 | YP_009047202.1 | K9N7C7 |
| ILDGISQYSLRLIDAM | 1 | 0 | 0 | 1666993 | YP_009724389.1 | P0DTD1 |
| ILDNAVKGCVPLNAIP | 1 | 0 | 0 | 1666997 | YP_009555238.1 | P0C6X6 |
| ILDNAVKGCVPLSAIP | 1 | 0 | 0 | 1666998 | YP_173236.1 | A0A140H1G9 |
| ILDWLEEKFKEGIEFL | 1 | 0 | 0 | 1667002 | QIQ54046.1 | P0C6U8 |
| ILECNVKTTEVVGDII | 1 | 0 | 0 | 1667005 | QHR63299.1 | P0C6W2 |
| ILEDEFTPFDVVRQCS | 1 | 0 | 0 | 1667006 | NP_828849.2 | P0C6X7 |
| ILGFRKIPMGVGLSP | 1 | 0 | 0 | 1667026 | P03155.1 | Q69028 |
| ILGLPTQTVDSSQGSE | 1 | 0 | 0 | 1667029 | YP_009724389.1 | P0DTD1 |
| ILGRCIVLDVKGVEEL | 1 | 0 | 0 | 1667034 | YP_009555239.1 | Q80872 |
| ILGSAILEDEFTPFDV | 1 | 0 | 0 | 1667035 | QIQ54046.1 | P0C6U8 |
| ILGSALLEDEFTPFDV | 1 | 0 | 0 | 1667036 | YP_009724389.1 | P0DTD1 |
| ILGSCTFEDELAPSDV | 1 | 0 | 0 | 1667037 | YP_173236.1 | A0A140H1G9 |
| ILGSTILEDEFTPFDV | 1 | 0 | 0 | 1667039 | NP_828849.2 | P0C6X7 |
| ILGYSSLNDEFSINEV | 1 | 0 | 0 | 1667046 | NP_073549.1 | A0A0P0LKV0 |
| ILKFDSGTVSKTSDWK | 1 | 0 | 0 | 1667063 | YP_009047202.1 | K9N7C7 |
| ILKTGELQPLEEPTAQ | 1 | 0 | 0 | 1667075 | QIQ54046.1 | P0C6U8 |
| ILKWADQSDQFRNVQT | 1 | 0 | 0 | 1667077 | YP_009555245.1 | P33469 |
| ILLERYVSLAIDAYPL | 1 | 0 | 0 | 1667085 | NP_073549.1 | A0A0P0LKV0 |
| ILLLDQALVSDVGDSA | 1 | 0 | 0 | 1667096 | YP_009724389.1 | P0DTD1 |
| ILLRKNGNKGAGGHSF | 1 | 0 | 0 | 1667106 | QIQ54046.1 | P0C6U8 |
| ILLVDKKLITTACNGI | 1 | 0 | 0 | 1667114 | YP_173236.1 | A0A140H1G9 |
| ILMLDKQGQLLDTKLN | 1 | 0 | 0 | 1667118 | YP_003766.2 | H9EJ42 |
| ILMSEDVKLNPSEDFI | 1 | 0 | 0 | 1667119 | YP_009047202.1 | K9N7C7 |
| ILMVDKNLITTANTGT | 1 | 0 | 0 | 1667123 | YP_009555238.1 | P0C6X6 |
| ILNIVGPDARQDGRQS | 1 | 0 | 0 | 1667128 | YP_173236.1 | A0A140H1G9 |
| ILNLTSEISTLENKSA | 1 | 0 | 0 | 1667130 | NP_073551.1 | A0A0P0K6L9 |
| ILNTVKFCDHMVKAGL | 1 | 0 | 0 | 1667136 | YP_009047202.1 | K9N7C7 |
| ILPDPLKPTKRSFIED | 1 | 0 | 0 | 1667137 | NP_828851.1 | P59594 |
| ILPGEFSKKQLGLLQP | 1 | 0 | 0 | 1667139 | YP_009047202.1 | K9N7C7 |
| ILPTMTQLNLKYAISG | 1 | 0 | 0 | 1667146 | NP_073549.1 | A0A0P0LKV0 |
| ILPVAFTKAAGGKVSF | 1 | 0 | 0 | 1667148 | NP_073549.1 | A0A0P0LKV0 |
| ILQQANKSVGIISMPL | 1 | 0 | 0 | 1667161 | YP_009047202.1 | K9N7C7 |
| ILQSNNPQCAIVQASE | 1 | 0 | 0 | 1667162 | YP_003766.2 | H9EJ42 |
| ILSCGIFGIKLETSLE | 1 | 0 | 0 | 1667178 | NP_073549.1 | A0A0P0LKV0 |
| ILSDDAVVCYNSNYAA | 1 | 0 | 0 | 1667182 | NP_828849.2 | P0C6X7 |
| ILSDDGVVCYNKDYAE | 1 | 0 | 0 | 1667183 | YP_003766.2 | H9EJ42 |
| ILSDDGVVCYNSDYAA | 1 | 0 | 0 | 1667184 | YP_009047202.1 | K9N7C7 |
| ILSDDGVVCYNSDYAS | 1 | 0 | 0 | 1667185 | YP_009555238.1 | P0C6X6 |
| ILSDDSVVCYNKTYAG | 1 | 0 | 0 | 1667186 | NP_073549.1 | A0A0P0LKV0 |
| ILSLNNPIFINYSKEV | 1 | 0 | 0 | 1667197 | YP_173237.1 | A0A140H1H0 |
| ILSNMNIASNSKEWAY | 1 | 0 | 0 | 1667200 | NP_073549.1 | A0A0P0LKV0 |
| ILSTSDLSVAFDKLAQ | 1 | 0 | 0 | 1667211 | YP_173236.1 | A0A140H1G9 |
| ILSWEIGKVKPPLNKN | 1 | 0 | 0 | 1667213 | YP_009555238.1 | P0C6X6 |
| ILTEVNELLDTTQLQV | 1 | 0 | 0 | 1667218 | YP_009555241.1 | P36334 |
| ILTNNKPWFVRKASGL | 1 | 0 | 0 | 1667229 | YP_003766.2 | H9EJ42 |
| ILVDNDLYPFVSDSVA | 1 | 0 | 0 | 1667240 | YP_009555238.1 | P0C6X6 |
| ILVSNNKEDFDIIQKC | 1 | 0 | 0 | 1667250 | YP_173236.1 | A0A140H1G9 |
| ILWVNKASYDTNLNKF | 1 | 0 | 0 | 1667260 | YP_009047202.1 | K9N7C7 |
| IMCADVPDAFQSVYRT | 1 | 0 | 0 | 1667278 | YP_003766.2 | H9EJ42 |
| IMGSCSFEDELTPSDV | 1 | 0 | 0 | 1667289 | YP_009555238.1 | P0C6X6 |
| IMKDDGNCVVLELDPP | 1 | 0 | 0 | 1667296 | YP_173236.1 | A0A140H1G9 |
| IMKVLNENLNAYQQQD | 1 | 0 | 0 | 1667300 | YP_009555245.1 | P33469 |
| IMNDKVVVKINNVDTV | 1 | 0 | 0 | 1667311 | YP_173236.1 | A0A140H1G9 |
| IMSDRDLYDKLQFTSL | 1 | 0 | 0 | 1667328 | YP_009724389.1 | P0DTD1 |
| INAASKGAVQKESDEY | 1 | 0 | 0 | 1667349 | YP_009047202.1 | K9N7C7 |
| INANQVIVNNLDKSAG | 1 | 0 | 0 | 1667355 | YP_009724389.1 | P0DTD1 |
| INARIRAKHYVYIGDP | 1 | 0 | 0 | 1667362 | YP_009555238.1 | P0C6X6 |
| INASVVNIQKEIDRLN | 1 | 0 | 0 | 1667363 | YP_009724390.1 | P0DTC2 |
| INCVLSVNSSNCNNFN | 1 | 0 | 0 | 1667371 | NP_073549.1 | A0A0P0LKV0 |
| INDDEDVVTGDNDDED | 1 | 0 | 0 | 1667375 | YP_173236.1 | A0A140H1G9 |
| INDKVVAKIDKEDVVI | 1 | 0 | 0 | 1667378 | YP_009555238.1 | P0C6X6 |
| INEVVKQMFGVNLQSG | 1 | 0 | 0 | 1667392 | NP_073549.1 | A0A0P0LKV0 |
| INGACGSPGYNLKNGE | 1 | 0 | 0 | 1667402 | NP_073549.1 | A0A0P0LKV0 |
| INGACGSPGYNVRNDG | 1 | 0 | 0 | 1667403 | YP_003766.2 | H9EJ42 |
| INGFNVSELEDACFKF | 1 | 0 | 0 | 1667412 | YP_009555242.1 | Q04853 |

| | | | | | | |
|---|---|---|---|---|---|---|
| INGLGVTMDVLNKNQK | 1 | 0 | 0 | 1667417 | YP_173238.1 | A0A140H1H1 |
| INGMHKSFYVNANGGT | 1 | 0 | 0 | 1667421 | YP_003766.2 | H9EJ42 |
| INHTFGNPVIPFKDGI | 1 | 0 | 0 | 1667434 | NP_828851.1 | P59594 |
| INKAIPSGNLIKFDIG | 1 | 0 | 0 | 1667445 | NP_073549.1 | A0A0P0LKV0 |
| INKATFKPNTWCLRCL | 1 | 0 | 0 | 1667446 | QIQ54046.1 | P0C6U8 |
| INKCSRLLSDDRTEVP | 1 | 0 | 0 | 1667447 | YP_009047204.1 | K9N5Q8 |
| INKFLKANPLWHKAVF | 1 | 0 | 0 | 1667450 | YP_009555238.1 | P0C6X6 |
| INKGHFDGQQGETPVS | 1 | 0 | 0 | 1667452 | QIQ54047.1 | P0C6X7 |
| INKQLTDISVDSNWPL | 1 | 0 | 0 | 1667456 | YP_173236.1 | A0A140H1G9 |
| INKSVANLKLPPHDVT | 1 | 0 | 0 | 1667457 | YP_003768.1 | H9EJ24 |
| INKYGLEDFNFEHVVY | 1 | 0 | 0 | 1667459 | YP_003766.2 | H9EJ42 |
| INKYGLELHWAPEFPW | 1 | 0 | 0 | 1667460 | YP_009555238.1 | P0C6X6 |
| INLHTQVVDMSMTYGQ | 1 | 0 | 0 | 1667463 | YP_009724389.1 | P0DTD1 |
| INLRADQINDMVYSLL | 1 | 0 | 0 | 1667470 | YP_009555238.1 | P0C6X6 |
| INNAVYTKVDGIDVEI | 1 | 0 | 0 | 1667476 | NP_828849.2 | P0C6X7 |
| INNLKIKNFDYLDIHP | 1 | 0 | 0 | 1667481 | YP_173237.1 | A0A140H1H0 |
| INNTVYDPLQPELDSF | 1 | 0 | 0 | 1667487 | NP_828851.1 | P59594 |
| INNTVYTKVDGVDVEL | 1 | 0 | 0 | 1667488 | YP_009724389.1 | P0DTD1 |
| INQATTKTTFKPNTWC | 1 | 0 | 0 | 1667499 | NP_828849.2 | P0C6X7 |
| INQLDEKAQKFFDFGD | 1 | 0 | 0 | 1667501 | NP_073549.1 | A0A0P0LKV0 |
| INSAIVCASVKRDGVQ | 1 | 0 | 0 | 1667515 | NP_073549.1 | A0A0P0LKV0 |
| INSILDEVNGLLDTTQ | 1 | 0 | 0 | 1667518 | YP_173238.1 | A0A140H1H1 |
| INSNVKTLEYDYMTHG | 1 | 0 | 0 | 1667520 | NP_073549.1 | A0A0P0LKV0 |
| INSTLVDLKWLNRVET | 1 | 0 | 0 | 1667522 | NP_073551.1 | A0A0P0K6L9 |
| INSTQDGDNKLQGLLE | 1 | 0 | 0 | 1667523 | YP_009555241.1 | P36334 |
| INTISPEKLKNYAASY | 1 | 0 | 0 | 1667531 | YP_003766.2 | H9EJ42 |
| INTLNDLNETLVTMPL | 1 | 0 | 0 | 1667532 | YP_009724389.1 | P0DTD1 |
| INTNSGPDDQIGYYRR | 1 | 0 | 0 | 1667533 | NP_828858.1 | P59595 |
| INTNSTKDDQIGYYRR | 1 | 0 | 0 | 1667535 | QIQ54056.1 | P59595 |
| INVIVFDGKSKCDESA | 1 | 0 | 0 | 1667545 | NP_828849.2 | P0C6X7 |
| INVIVFDGKSKCDESS | 1 | 0 | 0 | 1667546 | QIQ54046.1 | P0C6U8 |
| INVIVFDGKSKCEESS | 1 | 0 | 0 | 1667547 | YP_009724389.1 | P0DTD1 |
| INWNVEYPIISNELSI | 1 | 0 | 0 | 1667560 | YP_009555238.1 | P0C6X6 |
| INYKGKVFFQFDNLSS | 1 | 0 | 0 | 1667563 | YP_009555238.1 | P0C6X6 |
| INYLGTIKENIDGGAM | 1 | 0 | 0 | 1667568 | YP_009047202.1 | K9N7C7 |
| IPAKVRVDCYDKFKIN | 1 | 0 | 0 | 1667573 | YP_173236.1 | A0A140H1G9 |
| IPARARVECFDKFKVN | 1 | 0 | 0 | 1667578 | YP_009724389.1 | P0DTD1 |
| IPASQVIVNNYDKSAG | 1 | 0 | 0 | 1667581 | YP_009555238.1 | P0C6X6 |
| IPATSASKLTIVSPDL | 1 | 0 | 0 | 1667583 | YP_003766.2 | H9EJ42 |
| IPATSIVAKQGAGDAG | 1 | 0 | 0 | 1667584 | NP_073549.1 | A0A0P0LKV0 |
| IPCSVCLSGMDSLDAY | 1 | 0 | 0 | 1667587 | QIQ54046.1 | P0C6U8 |
| IPCTCGKQATKYLVQQ | 1 | 0 | 0 | 1667588 | YP_009724389.1 | P0DTD1 |
| IPCTCGKQATQYLVQQ | 1 | 0 | 0 | 1667589 | QHR63299.1 | P0C6W2 |
| IPDFDLDDFIDAPCYC | 1 | 0 | 0 | 1667592 | YP_009047202.1 | K9N7C7 |
| IPDKQVFDKVVDNVYV | 1 | 0 | 0 | 1667595 | YP_173236.1 | A0A140H1G9 |
| IPDYNTYKNTCDGTTF | 1 | 0 | 0 | 1667601 | YP_009724389.1 | P0DTD1 |
| IPDYVDVNKTLQEFAQ | 1 | 0 | 0 | 1667602 | YP_003767.1 | A0A0P0G321 |
| IPESKPLADDDSAIIE | 1 | 0 | 0 | 1667604 | YP_003771.1 | H9EJ69 |
| IPFEIKLAKKFDNFTG | 1 | 0 | 0 | 1667609 | QIQ54046.1 | P0C6U8 |
| IPFKDGIYFAATEKSN | 1 | 0 | 0 | 1667612 | NP_828851.1 | P59594 |
| IPFRLTCATTRQVVNV | 1 | 0 | 0 | 1667613 | NP_828849.2 | P0C6X7 |
| IPGSPGSLNFKKVETL | 1 | 0 | 0 | 1667624 | YP_173243.1 | Q0ZJ80 |
| IPHFNQKLPNGVTVVE | 1 | 0 | 0 | 1667626 | NP_073556.1 | A0A0N7J131 |
| IPIDLLNLREIKPAVN | 1 | 0 | 0 | 1667630 | YP_009555238.1 | P0C6X6 |
| IPINVIVFDGKSKCDE | 1 | 0 | 0 | 1667635 | QIQ54046.1 | P0C6U8 |
| IPIVWGVKDFNTLSQE | 1 | 0 | 0 | 1667638 | YP_003766.2 | H9EJ42 |
| IPKEEVKPFITESKPS | 1 | 0 | 0 | 1667641 | YP_009724389.1 | P0DTD1 |
| IPKEEVKSSITESKLS | 1 | 0 | 0 | 1667642 | QHR63299.1 | P0C6W2 |
| IPLTTAAKLMVVIPDY | 1 | 0 | 0 | 1667659 | YP_009724389.1 | P0DTD1 |
| IPLTTAAKLMVVVPDY | 1 | 0 | 0 | 1667660 | NP_828849.2 | P0C6X7 |
| IPMDSTVKNYFITDAQ | 1 | 0 | 0 | 1667662 | YP_009724389.1 | P0DTD1 |
| IPNFGSLTQINTTLLD | 1 | 0 | 0 | 1667664 | YP_009047204.1 | K9N5Q8 |
| IPNGRKVDLQLGNLGY | 1 | 0 | 0 | 1667666 | YP_009555241.1 | P36334 |
| IPRRNVATLQAENVTG | 1 | 0 | 0 | 1667678 | YP_009724389.1 | P0DTD1 |
| IPSGNLIKFDIGSVVV | 1 | 0 | 0 | 1667683 | NP_073549.1 | A0A0P0LKV0 |
| IPSKKAGGTTEMLSRA | 1 | 0 | 0 | 1667684 | NP_828849.2 | P0C6X7 |
| IPSLAANTLNIIVPDK | 1 | 0 | 0 | 1667685 | YP_009555238.1 | P0C6X6 |
| IPTITQMNLKYAISAK | 1 | 0 | 0 | 1667692 | YP_009724389.1 | P0DTD1 |
| IPTMTQMNLKYAISAK | 1 | 0 | 0 | 1667696 | YP_009047202.1 | K9N7C7 |
| IPTNFTIVGQEEFIQT | 1 | 0 | 0 | 1667700 | YP_173238.1 | A0A140H1H1 |
| IPTQCYYNNGSSGRIV | 1 | 0 | 0 | 1667701 | YP_173236.1 | A0A140H1G9 |
| IPTTCANDPVGFTLKN | 1 | 0 | 0 | 1667705 | YP_009724389.1 | P0DTD1 |
| IPTTCANDPVGFTLRN | 1 | 0 | 0 | 1667706 | NP_828849.2 | P0C6X7 |
| IPVAYRKVLLRKNGNK | 1 | 0 | 0 | 1667707 | YP_009724389.1 | P0DTD1 |
| IPVRPRNSSDNGISAI | 1 | 0 | 0 | 1667710 | YP_003767.1 | A0A0P0G321 |
| IPYKSFSDSGCILPSE | 1 | 0 | 0 | 1667713 | YP_009047202.1 | K9N7C7 |
| IPYNVSLSKFNSCKSD | 1 | 0 | 0 | 1667718 | YP_173237.1 | A0A140H1H0 |
| IQADSFTCNNIDAAKI | 1 | 0 | 0 | 1667721 | YP_009555241.1 | P36334 |
| IQAIYDRLDTIQADQQ | 1 | 0 | 0 | 1667723 | NP_073551.1 | A0A0P0K6L9 |
| IQANAYYKYDNGNFIK | 1 | 0 | 0 | 1667724 | NP_073549.1 | A0A0P0LKV0 |
| IQCKTKSLLPNTGVYD | 1 | 0 | 0 | 1667730 | YP_173238.1 | A0A140H1H1 |
| IQDSDGTNKQLNEISD | 1 | 0 | 0 | 1667732 | YP_009555238.1 | P0C6X6 |
| IQDVGFKNQYCNGSIA | 1 | 0 | 0 | 1667733 | YP_009555238.1 | P0C6X6 |
| IQDVKDGTVTFNLLAY | 1 | 0 | 0 | 1667734 | NP_073552.1 | A0A0P0KWE8 |
| IQDVKGLKIKYLYFIK | 1 | 0 | 0 | 1667735 | YP_173236.1 | A0A140H1G9 |
| IQDVVNQQGNSLNHLT | 1 | 0 | 0 | 1667736 | NP_073551.1 | A0A0P0K6L9 |
| IQDVVNQQGSALNHLT | 1 | 0 | 0 | 1667737 | YP_003767.1 | A0A0P0G321 |
| IQEGFDATNSALVKIQ | 1 | 0 | 0 | 1667741 | YP_009555241.1 | P36334 |
| IQGFACGCGASYTAND | 1 | 0 | 0 | 1667754 | YP_009047202.1 | K9N7C7 |
| IQGIKLGTGYSLADLP | 1 | 0 | 0 | 1667756 | YP_009555244.1 | Q01455 |
| IQGVKLGTGYTLSDLP | 1 | 0 | 0 | 1667762 | YP_173241.1 | Q0ZJ82 |

| | | | | | | |
|---|---|---|---|---|---|---|
| IQILASVPEKFLNAFD | 1 | 0 | 0 | 1667764 | NP_073549.1 | A0A0P0LKV0 |
| IQISYNDFYASGCVLS | 1 | 0 | 0 | 1667767 | YP_173236.1 | A0A140H1G9 |
| IQITNESYNNYVPSYV | 1 | 0 | 0 | 1667769 | YP_009047202.1 | K9N7C7 |
| IQKDIVECDVKTTEVV | 1 | 0 | 0 | 1667772 | QIQ54046.1 | P0C6U8 |
| IQKEVIECDVKTTEVV | 1 | 0 | 0 | 1667775 | NP_828849.2 | P0C6X7 |
| IQKVTVDCKQYVCNGF | 1 | 0 | 0 | 1667776 | YP_009047204.1 | K9N5Q8 |
| IQKYGLEDFNFEHVVY | 1 | 0 | 0 | 1667777 | NP_073549.1 | A0A0P0LKV0 |
| IQLLKAAYENFNSQDI | 1 | 0 | 0 | 1667781 | NP_828849.2 | P0C6X7 |
| IQLLKSAYENFNQHDV | 1 | 0 | 0 | 1667782 | QHR63299.1 | P0C6W2 |
| IQLLKVAYENFNHHEK | 1 | 0 | 0 | 1667784 | QIQ54046.1 | P0C6U8 |
| IQLNNDVRDYLLSKMT | 1 | 0 | 0 | 1667786 | YP_173236.1 | A0A140H1G9 |
| IQLSSYSLFDMSKFPL | 1 | 0 | 0 | 1667788 | YP_009724389.1 | P0DTD1 |
| IQNGAKHVQGNVAKAA | 1 | 0 | 0 | 1667793 | YP_173236.1 | A0A140H1G9 |
| IQNLLYCDSPFEKLQC | 1 | 0 | 0 | 1667795 | YP_003767.1 | A0A0P0G321 |
| IQNSAKHVQGNVAKIA | 1 | 0 | 0 | 1667797 | YP_009555238.1 | P0C6X6 |
| IQPRVEKKKLDGFMGR | 1 | 0 | 0 | 1667802 | YP_009724389.1 | P0DTD1 |
| IQPRVEKKKLEGFMGR | 1 | 0 | 0 | 1667803 | QIQ54046.1 | P0C6U8 |
| IQPRVEKKKTEGFMGR | 1 | 0 | 0 | 1667804 | NP_828849.2 | P0C6X7 |
| IQPWTNDALNTICIKD | 1 | 0 | 0 | 1667806 | NP_073549.1 | A0A0P0LKV0 |
| IQSFDNSYLNESGALV | 1 | 0 | 0 | 1667828 | NP_073550.1 | A0A0P0ILY9 |
| IQSLKVCIETVRTQVF | 1 | 0 | 0 | 1667832 | QIQ54046.1 | P0C6U8 |
| IQSVVNSNAQALNSLL | 1 | 0 | 0 | 1667836 | YP_173238.1 | A0A140H1H1 |
| IQTIQDSDGTNKQLNE | 1 | 0 | 0 | 1667839 | YP_009555238.1 | P0C6X6 |
| IQTNSPKVTIDCSLFV | 1 | 0 | 0 | 1667841 | YP_173238.1 | A0A140H1H1 |
| IQTTIQKVTVDCKQYV | 1 | 0 | 0 | 1667843 | YP_009047204.1 | K9N5Q8 |
| IQTVQDADGINKQLTD | 1 | 0 | 0 | 1667848 | YP_173236.1 | A0A140H1G9 |
| IQVAKCYGESVLVNAA | 1 | 0 | 0 | 1667849 | YP_009047202.1 | K9N7C7 |
| IQVDQLNSSYFKLSIP | 1 | 0 | 0 | 1667851 | YP_009047204.1 | K9N5Q8 |
| IQVVNSGPDQTCNTPT | 1 | 0 | 0 | 1667855 | YP_009555238.1 | P0C6X6 |
| IQVVNSGSDMNCNIPT | 1 | 0 | 0 | 1667856 | YP_173236.1 | A0A140H1G9 |
| IRCLWNTKPVETSNSF | 1 | 0 | 0 | 1667870 | QHR63299.1 | P0C6W2 |
| IRCLWSTKPVETSNSF | 1 | 0 | 0 | 1667871 | YP_009724389.1 | P0DTD1 |
| IRDIVCKVENKTEAEW | 1 | 0 | 0 | 1667873 | NP_073549.1 | A0A0P0LKV0 |
| IRDKGCVADQSRKWIG | 1 | 0 | 0 | 1667874 | YP_009555239.1 | Q80872 |
| IRDVSVEDVCFANKFE | 1 | 0 | 0 | 1667885 | YP_009555238.1 | P0C6X6 |
| IRDYKLANDAKESLRL | 1 | 0 | 0 | 1667886 | NP_073549.1 | A0A0P0LKV0 |
| IRENNRVVISSDVLVN | 1 | 0 | 0 | 1667893 | YP_009724389.1 | P0DTD1 |
| IRENNRVVVSSDILVN | 1 | 0 | 0 | 1667894 | NP_828849.2 | P0C6X7 |
| IRFEYTSPTDKLAMIM | 1 | 0 | 0 | 1667902 | YP_009555239.1 | Q80872 |
| IRGCCESTKLPYYDVE | 1 | 0 | 0 | 1667908 | NP_073551.1 | A0A0P0K6L9 |
| IRGDEVRQIAPGQTGK | 1 | 0 | 0 | 1667909 | YP_009724390.1 | P0DTC2 |
| IRGHLYIQGIKLGTGY | 1 | 0 | 0 | 1667911 | YP_009555244.1 | Q01455 |
| IRGHLYIQGVKLGTGY | 1 | 0 | 0 | 1667912 | YP_173241.1 | Q0ZJ82 |
| IRGPPRAELNGVVVDK | 1 | 0 | 0 | 1667914 | YP_009555238.1 | P0C6X6 |
| IRKDLSYEQQNVLAIE | 1 | 0 | 0 | 1667927 | YP_003766.2 | H9EJ42 |
| IRLQAGVATEYAANSS | 1 | 0 | 0 | 1667940 | YP_173236.1 | A0A140H1G9 |
| IRQEEVQQELYSPLFL | 1 | 0 | 0 | 1667951 | NP_828857.1 | P59635 |
| IRQLLFVVEVVDKYFD | 1 | 0 | 0 | 1667954 | YP_009724389.1 | P0DTD1 |
| IRRAIDCGFNDLSQLH | 1 | 0 | 0 | 1667957 | YP_009047204.1 | K9N5Q8 |
| IRSAAKKNNLPFKLTC | 1 | 0 | 0 | 1667961 | YP_009724389.1 | P0DTD1 |
| IRSSNQGNKQIVHSYP | 1 | 0 | 0 | 1667966 | YP_009047207.1 | K9N643 |
| IRSSNQSLAGGITYVS | 1 | 0 | 0 | 1667967 | YP_003767.1 | A0A0P0G321 |
| IRVGARKSAPLIELCV | 1 | 0 | 0 | 1667977 | YP_009724396.1 | P0DTC8 |
| ISAGIFSVPADVSLTY | 1 | 0 | 0 | 1667998 | YP_173236.1 | A0A140H1G9 |
| ISAGIFSVPSDVSLTY | 1 | 0 | 0 | 1667999 | YP_009555238.1 | P0C6X6 |
| ISDEFSSNVANYQKVG | 1 | 0 | 0 | 1668022 | YP_009724389.1 | P0DTD1 |
| ISDMYDGRTKAIDGEN | 1 | 0 | 0 | 1668027 | NP_073549.1 | A0A0P0LKV0 |
| ISDMYDPKTKNVTKEN | 1 | 0 | 0 | 1668030 | YP_009724389.1 | P0DTD1 |
| ISDMYDPLTKNIGDYN | 1 | 0 | 0 | 1668031 | YP_173236.1 | A0A140H1G9 |
| ISDMYDPRTKHVTKEN | 1 | 0 | 0 | 1668032 | NP_828849.2 | P0C6X7 |
| ISDMYDPTTKNVTGSN | 1 | 0 | 0 | 1668033 | YP_009047202.1 | K9N7C7 |
| ISDTTDAVRDPQTLEI | 1 | 0 | 0 | 1668038 | QIQ54048.1 | P59594 |
| ISEHDYQIGGYTEKWE | 1 | 0 | 0 | 1668042 | YP_009724391.1 | P0DTC3 |
| ISEWPLSVQQAQQEAT | 1 | 0 | 0 | 1668046 | NP_073549.1 | A0A0P0LKV0 |
| ISGDKVFVRDGNTDNL | 1 | 0 | 0 | 1668057 | NP_073549.1 | A0A0P0LKV0 |
| ISGYTLAATSASLFPP | 1 | 0 | 0 | 1668066 | YP_009555241.1 | P36334 |
| ISKDSKFQIGEFIFEK | 1 | 0 | 0 | 1668081 | YP_003766.2 | H9EJ42 |
| ISLSFNNLLGFDSSKP | 1 | 0 | 0 | 1668106 | YP_009047202.1 | K9N7C7 |
| ISLTMLDIQPEDYKSV | 1 | 0 | 0 | 1668108 | YP_009555239.1 | Q80872 |
| ISLTVGEVFGKILGNV | 1 | 0 | 0 | 1668110 | YP_173236.1 | A0A140H1G9 |
| ISLVSVTDFSLNGNVS | 1 | 0 | 0 | 1668113 | YP_009047208.1 | K9N7D2 |
| ISMATNYDLSVVNARL | 1 | 0 | 0 | 1668114 | YP_009724389.1 | P0DTD1 |
| ISMMGFKMNYQVNGYP | 1 | 0 | 0 | 1668116 | YP_009724389.1 | P0DTD1 |
| ISNCVADYSVLYNSAS | 1 | 0 | 0 | 1668126 | YP_009724390.1 | P0DTC2 |
| ISNDSNGCVVEASTDF | 1 | 0 | 0 | 1668130 | YP_003766.2 | H9EJ42 |
| ISNFKFDDYDFSVEDA | 1 | 0 | 0 | 1668136 | YP_173236.1 | A0A140H1G9 |
| ISNKADLKFVKWEYEG | 1 | 0 | 0 | 1668137 | YP_003766.2 | H9EJ42 |
| ISNKLCVVPGNVKPGD | 1 | 0 | 0 | 1668138 | NP_073549.1 | A0A0P0LKV0 |
| ISPYNSQNAVAAKILG | 1 | 0 | 0 | 1668150 | QIQ54047.1 | P0C6X7 |
| ISPYNSQNAVASKILG | 1 | 0 | 0 | 1668151 | YP_009724389.1 | P0DTD1 |
| ISPYNSQNAVSRSMLG | 1 | 0 | 0 | 1668152 | YP_009047202.1 | K9N7C7 |
| ISPYNSQNFAAKRVLG | 1 | 0 | 0 | 1668153 | YP_009555238.1 | P0C6X6 |
| ISQDSITHYPALKMVQ | 1 | 0 | 0 | 1668157 | YP_009047202.1 | K9N7C7 |
| ISQPFLMDLEGKQGNF | 1 | 0 | 0 | 1668162 | QIQ54048.1 | P59594 |
| ISQWPISNDSNGCVVE | 1 | 0 | 0 | 1668165 | YP_003766.2 | H9EJ42 |
| ISQYSLRLIDAMMFTS | 1 | 0 | 0 | 1668167 | YP_009724389.1 | P0DTD1 |
| ISRMGFKLDATVPGYP | 1 | 0 | 0 | 1668169 | YP_009047202.1 | K9N7C7 |
| ISRSELQTIVPEYIDV | 1 | 0 | 0 | 1668171 | NP_073551.1 | A0A0P0K6L9 |
| ISSNDNCFANKHVGFD | 1 | 0 | 0 | 1668183 | QIQ54046.1 | P0C6U8 |
| ISSNGPLKVGAGVMLE | 1 | 0 | 0 | 1668184 | YP_003766.2 | H9EJ42 |
| ISSNTDNETLQYWVTP | 1 | 0 | 0 | 1668185 | YP_173238.1 | A0A140H1H1 |

| | | | | | | |
|---|---|---|---|---|---|---|
| ISTARDGVRRGDNFHS | 1 | 0 | 0 | 1668193 | YP_009047202.1 | K9N7C7 |
| ISTDDCFANKHAGFDA | 1 | 0 | 0 | 1668195 | NP_828849.2 | P0C6X7 |
| ISTEKLSRLSMIKGPQ | 1 | 0 | 0 | 1668200 | YP_173236.1 | A0A140H1G9 |
| ISTGSRSARSAIEDLL | 1 | 0 | 0 | 1668203 | YP_009047204.1 | K9N5Q8 |
| ISTLENKSAELNYTVQ | 1 | 0 | 0 | 1668211 | NP_073551.1 | A0A0P0K6L9 |
| ISTPKLKEDYQIGGYS | 1 | 0 | 0 | 1668213 | NP_828852.2 | P59632 |
| ISTQLFEGDKFIGTFE | 1 | 0 | 0 | 1668214 | YP_003766.2 | H9EJ42 |
| ISTVQSKLTDLKCTNV | 1 | 0 | 0 | 1668219 | YP_003766.2 | H9EJ42 |
| ISVLDITDAAVKAAES | 1 | 0 | 0 | 1668229 | NP_073549.1 | A0A0P0LKV0 |
| ISVMDKLDTGAQKFFQ | 1 | 0 | 0 | 1668230 | YP_003766.2 | H9EJ42 |
| ISVNDLCFANKFFQFD | 1 | 0 | 0 | 1668231 | YP_173236.1 | A0A140H1G9 |
| ISVTPIKNTVDTTPKE | 1 | 0 | 0 | 1668235 | NP_073549.1 | A0A0P0LKV0 |
| ISYSELRPDTRYVLMD | 1 | 0 | 0 | 1668250 | NP_828849.2 | P0C6X7 |
| ITAKETLYRIDGAHLT | 1 | 0 | 0 | 1668259 | NP_828849.2 | P0C6X7 |
| ITAPYVVLTGLGESFK | 1 | 0 | 0 | 1668264 | YP_009047202.1 | K9N7C7 |
| ITAQPVKIDNASPAST | 1 | 0 | 0 | 1668265 | NP_828852.2 | P59632 |
| ITDAAVKAAESKAFVD | 1 | 0 | 0 | 1668272 | NP_073549.1 | A0A0P0LKV0 |
| ITDHDYQIGGYTEKWE | 1 | 0 | 0 | 1668278 | QIQ54049.1 | P59632 |
| ITDIFYKENSYITTIK | 1 | 0 | 0 | 1668279 | QHR63299.1 | P0C6W2 |
| ITDVFYKENSYTTTIK | 1 | 0 | 0 | 1668282 | YP_009724389.1 | P0DTD1 |
| ITDYNYYKYNLPTMVD | 1 | 0 | 0 | 1668283 | YP_009555238.1 | P0C6X6 |
| ITEFQLDYSIDVIDNE | 1 | 0 | 0 | 1668290 | NP_073549.1 | A0A0P0LKV0 |
| ITEFSWNADLYKLMSC | 1 | 0 | 0 | 1668291 | YP_173236.1 | A0A140H1G9 |
| ITEGSVKGLTPSKGPA | 1 | 0 | 0 | 1668295 | NP_828849.2 | P0C6X7 |
| ITEHSWNADLYKLMGH | 1 | 0 | 0 | 1668296 | YP_009724389.1 | P0DTD1 |
| ITESKPSVEQRKQDDK | 1 | 0 | 0 | 1668303 | YP_009724389.1 | P0DTD1 |
| ITFELDERIDKVLNEK | 1 | 0 | 0 | 1668311 | YP_009724389.1 | P0DTD1 |
| ITFELDERVDKVLNEK | 1 | 0 | 0 | 1668312 | NP_828849.2 | P0C6X7 |
| ITFGGPSDSTDNNQNG | 1 | 0 | 0 | 1668315 | QIQ54056.1 | P59595 |
| ITFHLDGEVITFDNLK | 1 | 0 | 0 | 1668318 | QHR63299.1 | P0C6W2 |
| ITFHMDGEVINIDSLK | 1 | 0 | 0 | 1668319 | QIQ54046.1 | P0C6U8 |
| ITGDEVRQIAPGQTGK | 1 | 0 | 0 | 1668325 | QHR63300.2 | Q3LZX1 |
| ITGLYPTLNISDEFSS | 1 | 0 | 0 | 1668333 | YP_009724389.1 | P0DTD1 |
| ITIKVTEDGVNVHDVT | 1 | 0 | 0 | 1668356 | NP_073549.1 | A0A0P0LKV0 |
| ITITYQGLFPYQGDHG | 1 | 0 | 0 | 1668362 | YP_009047204.1 | K9N5Q8 |
| ITKDSKFQVGEFVFEK | 1 | 0 | 0 | 1668367 | NP_073549.1 | A0A0P0LKV0 |
| ITKENVETLTWPLILN | 1 | 0 | 0 | 1668370 | YP_003766.2 | H9EJ42 |
| ITKPLIEAEKPNLNSL | 1 | 0 | 0 | 1668379 | YP_173243.1 | Q0ZJ80 |
| ITKTDAVILLERYVSL | 1 | 0 | 0 | 1668383 | NP_073549.1 | A0A0P0LKV0 |
| ITLANFNETKGPLCVD | 1 | 0 | 0 | 1668386 | NP_073551.1 | A0A0P0K6L9 |
| ITLKCCTVTYLNDPSS | 1 | 0 | 0 | 1668392 | NP_073549.1 | A0A0P0LKV0 |
| ITMTDLQSESSCGLFK | 1 | 0 | 0 | 1668407 | NP_073549.1 | A0A0P0LKV0 |
| ITNCVKMLCTHTGTGQ | 1 | 0 | 0 | 1668410 | YP_009724389.1 | P0DTD1 |
| ITNESYNNYVPSYVKP | 1 | 0 | 0 | 1668411 | YP_009047202.1 | K9N7C7 |
| ITNLCPFGEVFNASKF | 1 | 0 | 0 | 1668416 | QIQ54048.1 | P59594 |
| ITNLCPFGEVFNATKF | 1 | 0 | 0 | 1668417 | NP_828851.1 | P59594 |
| ITNTNLSRGRGRNPKP | 1 | 0 | 0 | 1668421 | YP_009047211.1 | K9N4V7 |
| ITPEVSRELGNITKTN | 1 | 0 | 0 | 1668426 | NP_073549.1 | A0A0P0LKV0 |
| ITPGTNASNQVAVLYQ | 1 | 0 | 0 | 1668427 | QHR63300.2 | Q3LZX1 |
| ITPNVCFVKGDIINVA | 1 | 0 | 0 | 1668431 | YP_173236.1 | A0A140H1G9 |
| ITQFQKGKEFEFVEGQ | 1 | 0 | 0 | 1668437 | YP_009555245.1 | P33469 |
| ITQMNLKYAISAKNRA | 1 | 0 | 0 | 1668440 | YP_009724389.1 | P0DTD1 |
| ITQQVLSENQKLIANK | 1 | 0 | 0 | 1668442 | YP_009047204.1 | K9N5Q8 |
| ITQRNFYEPQIITTDN | 1 | 0 | 0 | 1668444 | QIQ54048.1 | P59594 |
| ITQRTRTRPSVQPGPM | 1 | 0 | 0 | 1668445 | YP_009047211.1 | K9N4V7 |
| ITQTSQALQTVATALN | 1 | 0 | 0 | 1668449 | NP_073551.1 | A0A0P0K6L9 |
| ITRDVGFVVPGLPGTI | 1 | 0 | 0 | 1668461 | QIQ54046.1 | P0C6U8 |
| ITREEAVRQVRSWIGF | 1 | 0 | 0 | 1668462 | YP_009047202.1 | K9N7C7 |
| ITREIGFIVPGLPGTV | 1 | 0 | 0 | 1668463 | NP_828849.2 | P0C6X7 |
| ITREVGFVVPGLPGTI | 1 | 0 | 0 | 1668464 | YP_009724389.1 | P0DTD1 |
| ITSEGNALYNNEGGRA | 1 | 0 | 0 | 1668472 | NP_073549.1 | A0A0P0LKV0 |
| ITSGDGTTSPISEHDY | 1 | 0 | 0 | 1668474 | YP_009724391.1 | P0DTC3 |
| ITSGIKFQTVKPGNFN | 1 | 0 | 0 | 1668476 | YP_173236.1 | A0A140H1G9 |
| ITSGNLTTVVIPTKKA | 1 | 0 | 0 | 1668477 | QIQ54046.1 | P0C6U8 |
| ITSGVKFQTVKPGNFN | 1 | 0 | 0 | 1668478 | YP_009555238.1 | P0C6X6 |
| ITSLNTKYVAPQVTYQ | 1 | 0 | 0 | 1668484 | YP_009047204.1 | K9N5Q8 |
| ITSREVVVTNLNKSAG | 1 | 0 | 0 | 1668485 | NP_073549.1 | A0A0P0LKV0 |
| ITTDNTFVSGNCDVVI | 1 | 0 | 0 | 1668495 | YP_009724390.1 | P0DTC2 |
| ITTDNTFVSGSCDVVI | 1 | 0 | 0 | 1668496 | QHR63300.2 | Q3LZX1 |
| ITTGYRFTNFEPFTVN | 1 | 0 | 0 | 1668500 | YP_009555241.1 | P36334 |
| ITTNGSWAIFNDHHLN | 1 | 0 | 0 | 1668504 | YP_009047202.1 | K9N7C7 |
| ITTSVDANTNQDSYGG | 1 | 0 | 0 | 1668507 | YP_003766.2 | H9EJ42 |
| ITVTPEANMDQESFGG | 1 | 0 | 0 | 1668521 | YP_009724389.1 | P0DTD1 |
| ITYPMIANENAINKGG | 1 | 0 | 0 | 1668525 | NP_073549.1 | A0A0P0LKV0 |
| ITYTSTSGNLLGFKDV | 1 | 0 | 0 | 1668527 | NP_073551.1 | A0A0P0K6L9 |
| ITYTYKMLVAKDNKNL | 1 | 0 | 0 | 1668528 | YP_003771.1 | H9EJ69 |
| ITYVSNSGNLLGFKNV | 1 | 0 | 0 | 1668529 | YP_003767.1 | A0A0P0G321 |
| IVAKQGAGDAGHSLTW | 1 | 0 | 0 | 1668542 | NP_073549.1 | A0A0P0LKV0 |
| IVAWHVVRDPRYVMRL | 1 | 0 | 0 | 1668551 | YP_009555238.1 | P0C6X6 |
| IVCASVKRDGVQVGYC | 1 | 0 | 0 | 1668554 | NP_073549.1 | A0A0P0LKV0 |
| IVDENGKYYLPYPDPS | 1 | 0 | 0 | 1668569 | NP_073549.1 | A0A0P0LKV0 |
| IVDEPEDHVQIHTIDG | 1 | 0 | 0 | 1668570 | QIQ54049.1 | P59632 |
| IVDEWSYTGSSFYAPE | 1 | 0 | 0 | 1668571 | YP_009047204.1 | K9N5Q8 |
| IVDKDGTYYLPYPDPS | 1 | 0 | 0 | 1668575 | YP_003766.2 | H9EJ42 |
| IVDKEVKDMQFDLYVN | 1 | 0 | 0 | 1668576 | YP_009047202.1 | K9N7C7 |
| IVDNVYMAKAGDKYYP | 1 | 0 | 0 | 1668579 | YP_009555238.1 | P0C6X6 |
| IVDQVHVLNHVDPIIV | 1 | 0 | 0 | 1668580 | YP_173243.1 | Q0ZJ80 |
| IVDTVSALVYDNKLKA | 1 | 0 | 0 | 1668587 | YP_009724389.1 | P0DTD1 |
| IVDTVSALVYDNKLRA | 1 | 0 | 0 | 1668588 | QIQ54047.1 | P0C6X7 |
| IVDTVSALVYENKLKA | 1 | 0 | 0 | 1668589 | YP_009555238.1 | P0C6X6 |
| IVEDDVIENVKSSLSS | 1 | 0 | 0 | 1668593 | YP_173236.1 | A0A140H1G9 |

| | | | | | | |
|---|---|---|---|---|---|---|
| IVETVSALVYDNKLKA | 1 | 0 | 0 | 1668603 | YP_173236.1 | A0A140H1G9 |
| IVFDGKSKCDESASKS | 1 | 0 | 0 | 1668605 | NP_828849.2 | P0C6X7 |
| IVFDGKSKCDESSARA | 1 | 0 | 0 | 1668606 | QIQ54046.1 | P0C6U8 |
| IVFDGKSKCEESSAKS | 1 | 0 | 0 | 1668607 | YP_009724389.1 | P0DTD1 |
| IVFNNNGTNVTQVKNA | 1 | 0 | 0 | 1668615 | NP_073549.1 | A0A0P0LKV0 |
| IVFTACSHAAVDSLCA | 1 | 0 | 0 | 1668617 | NP_073549.1 | A0A0P0LKV0 |
| IVGERQLLLVWEAGKS | 1 | 0 | 0 | 1668633 | YP_009047202.1 | K9N7C7 |
| IVGGQIVTCAKEIKES | 1 | 0 | 0 | 1668636 | YP_009724389.1 | P0DTD1 |
| IVGNKFYNVSTGLIPG | 1 | 0 | 0 | 1668641 | YP_173236.1 | A0A140H1G9 |
| IVGPDARQDGRQSYVL | 1 | 0 | 0 | 1668642 | YP_173236.1 | A0A140H1G9 |
| IVGPKELLLLWESGKA | 1 | 0 | 0 | 1668643 | NP_073549.1 | A0A0P0LKV0 |
| IVGQEEFIQTNSPKVT | 1 | 0 | 0 | 1668644 | YP_173238.1 | A0A140H1H1 |
| IVIGGAKLKALNLGET | 1 | 0 | 0 | 1668656 | QIQ54046.1 | P0C6U8 |
| IVIPDKQVFDKVVDNV | 1 | 0 | 0 | 1668663 | YP_173236.1 | A0A140H1G9 |
| IVITSGDGTTSPISEH | 1 | 0 | 0 | 1668666 | YP_009724391.1 | P0DTC3 |
| IVKCKGLNKPFVKDNV | 1 | 0 | 0 | 1668668 | YP_009047202.1 | K9N7C7 |
| IVKEAQSANPMVIVNA | 1 | 0 | 0 | 1668669 | NP_828849.2 | P0C6X7 |
| IVKGQIQVASDNIKDC | 1 | 0 | 0 | 1668675 | NP_828849.2 | P0C6X7 |
| IVKLGAEKKDGFYETI | 1 | 0 | 0 | 1668678 | YP_009555239.1 | Q80872 |
| IVKSSDVVDSNENLTW | 1 | 0 | 0 | 1668687 | YP_009047202.1 | K9N7C7 |
| IVLDKFAIPNSRRSDL | 1 | 0 | 0 | 1668692 | YP_173238.1 | A0A140H1H1 |
| IVLDVKGVEELHDDLV | 1 | 0 | 0 | 1668693 | YP_009555239.1 | Q80872 |
| IVNAANENLAHGGGLA | 1 | 0 | 0 | 1668721 | NP_073549.1 | A0A0P0LKV0 |
| IVNAANIHLKHGGGVA | 1 | 0 | 0 | 1668722 | NP_828849.2 | P0C6X7 |
| IVNARIEPCSSGLSTD | 1 | 0 | 0 | 1668724 | YP_009047202.1 | K9N7C7 |
| IVNDDTSETKYVKSLS | 1 | 0 | 0 | 1668725 | YP_009555238.1 | P0C6X6 |
| IVNNLDKSAGFPFNKW | 1 | 0 | 0 | 1668734 | YP_009724389.1 | P0DTD1 |
| IVNNTVYTKVDGVDVE | 1 | 0 | 0 | 1668736 | QIQ54047.1 | P0C6X7 |
| IVNNYDKSAGYPFNKF | 1 | 0 | 0 | 1668738 | YP_009555238.1 | P0C6X6 |
| IVNPANGHMLHGGGVA | 1 | 0 | 0 | 1668739 | YP_173236.1 | A0A140H1G9 |
| IVNTVSELVYENKFVP | 1 | 0 | 0 | 1668745 | NP_073549.1 | A0A0P0LKV0 |
| IVPDKSVYDQVVDNVY | 1 | 0 | 0 | 1668750 | YP_009555238.1 | P0C6X6 |
| IVPEYIDVNKTLQELS | 1 | 0 | 0 | 1668751 | NP_073551.1 | A0A0P0K6L9 |
| IVPLCIFNGKFLSNTK | 1 | 0 | 0 | 1668756 | YP_009555240.1 | P30215 |
| IVPSTVWEDGDYYRKQ | 1 | 0 | 0 | 1668760 | YP_009047204.1 | K9N5Q8 |
| IVQASESKVLLERFLP | 1 | 0 | 0 | 1668763 | YP_003766.2 | H9EJ42 |
| IVQESSRKLLGLTNLS | 1 | 0 | 0 | 1668764 | YP_173243.1 | Q0ZJ80 |
| IVQLSEINMDNSPNLA | 1 | 0 | 0 | 1668767 | NP_828849.2 | P0C6X7 |
| IVQLSEISMDNSPNLA | 1 | 0 | 0 | 1668768 | YP_009724389.1 | P0DTD1 |
| IVQLSEITMDNSPNIA | 1 | 0 | 0 | 1668769 | QIQ54046.1 | P0C6U8 |
| IVQMISDYLSNLSDIL | 1 | 0 | 0 | 1668773 | YP_003766.2 | H9EJ42 |
| IVQMLSDTLKNLSDRV | 1 | 0 | 0 | 1668776 | YP_009724389.1 | P0DTD1 |
| IVQMLSDYLLDLSDSV | 1 | 0 | 0 | 1668777 | YP_173236.1 | A0A140H1G9 |
| IVSDREVILCWETGKV | 1 | 0 | 0 | 1668796 | YP_173236.1 | A0A140H1G9 |
| IVSEAKNVKPTIVVNA | 1 | 0 | 0 | 1668797 | QIQ54046.1 | P0C6U8 |
| IVSEKISVMDKLDTGA | 1 | 0 | 0 | 1668798 | YP_003766.2 | H9EJ42 |
| IVSERELILSWEIGKV | 1 | 0 | 0 | 1668799 | YP_009555238.1 | P0C6X6 |
| IVSGYEKSTFNKDIYC | 1 | 0 | 0 | 1668801 | YP_003766.2 | H9EJ42 |
| IVSKESGNTVVLTFTT | 1 | 0 | 0 | 1668804 | NP_073556.1 | A0A0N7J131 |
| IVSSFEGYDFKYIENG | 1 | 0 | 0 | 1668808 | NP_073549.1 | A0A0P0LKV0 |
| IVTALRANSAVKLQNN | 1 | 0 | 0 | 1668818 | YP_009724389.1 | P0DTD1 |
| IVTANLSIPSNWTTSV | 1 | 0 | 0 | 1668819 | NP_073551.1 | A0A0P0K6L9 |
| IVTCAKEIKESVQTFF | 1 | 0 | 0 | 1668820 | YP_009724389.1 | P0DTD1 |
| IVTGDNDDQIVVTGDD | 1 | 0 | 0 | 1668821 | YP_173236.1 | A0A140H1G9 |
| IVTGLFKDCSRETSGL | 1 | 0 | 0 | 1668822 | YP_009047202.1 | K9N7C7 |
| IVTLEQYSTCDICKST | 1 | 0 | 0 | 1668825 | YP_003766.2 | H9EJ42 |
| IVVAVMDEDIGSTMFN | 1 | 0 | 0 | 1668837 | YP_173236.1 | A0A140H1G9 |
| IVVAWHVVRDSRFVMR | 1 | 0 | 0 | 1668838 | YP_173236.1 | A0A140H1G9 |
| IVVFDEISMATNYDLS | 1 | 0 | 0 | 1668843 | YP_009724389.1 | P0DTD1 |
| IVVTGDDVDDIESIYD | 1 | 0 | 0 | 1668859 | YP_173236.1 | A0A140H1G9 |
| IVWGVKDFNTLSQEGK | 1 | 0 | 0 | 1668867 | YP_003766.2 | H9EJ42 |
| IVWHAKDFNSLSAEGR | 1 | 0 | 0 | 1668868 | NP_073549.1 | A0A0P0LKV0 |
| IVWHINQATTKTTFKP | 1 | 0 | 0 | 1668869 | NP_828849.2 | P0C6X7 |
| IVWHVNNATNKATYKP | 1 | 0 | 0 | 1668870 | YP_009724389.1 | P0DTD1 |
| IVWHVNNTINKATFKP | 1 | 0 | 0 | 1668871 | QIQ54046.1 | P0C6U8 |
| IVYAVLSDVDGLKYTK | 1 | 0 | 0 | 1668876 | YP_173236.1 | A0A140H1G9 |
| IVYHTLESPVEFHLDG | 1 | 0 | 0 | 1668887 | NP_828849.2 | P0C6X7 |
| IVYKAACVVDVNDSHS | 1 | 0 | 0 | 1668888 | YP_009555238.1 | P0C6X6 |
| IVYNNSGSNITQIKNA | 1 | 0 | 0 | 1668890 | YP_003766.2 | H9EJ42 |
| IWNGVIKNVNSVRDWL | 1 | 0 | 0 | 1668921 | NP_073549.1 | A0A0P0LKV0 |
| IWNVKDYMSLSEQLRK | 1 | 0 | 0 | 1668926 | NP_828849.2 | P0C6X7 |
| IWTADEAIKFLKEWNF | 1 | 0 | 0 | 1668931 | YP_009555244.1 | Q01455 |
| IWVATEGALNTPKDHI | 1 | 0 | 0 | 1668932 | YP_009724397.2 | P0DTC9 |
| IYAQTSDTAHACNVNR | 1 | 0 | 0 | 1668940 | YP_003766.2 | H9EJ42 |
| IYCNSVINRLRCDQLS | 1 | 0 | 0 | 1668945 | NP_073551.1 | A0A0P0K6L9 |
| IYDEEGGYDVSKPVMI | 1 | 0 | 0 | 1668954 | YP_003766.2 | H9EJ42 |
| IYDRLDSIQADQQVDR | 1 | 0 | 0 | 1668960 | YP_003767.1 | A0A0P0G321 |
| IYDRLDTIQADQQVDR | 1 | 0 | 0 | 1668961 | NP_073551.1 | A0A0P0K6L9 |
| IYDSSDRGQESLARSA | 1 | 0 | 0 | 1668962 | YP_009047202.1 | K9N7C7 |
| IYFAATEKSNVVRGWV | 1 | 0 | 0 | 1668967 | NP_828851.1 | P59594 |
| IYFITMSSKGQKGDAE | 1 | 0 | 0 | 1668969 | YP_003766.2 | H9EJ42 |
| IYKLDGVVCTEIDPKL | 1 | 0 | 0 | 1668987 | QHR63299.1 | P0C6W2 |
| IYKTPPIKDFGGFNFL | 1 | 0 | 0 | 1668993 | QIQ54048.1 | P59594 |
| IYLKGSGSDFSLADSV | 1 | 0 | 0 | 1668998 | YP_009555238.1 | P0C6X6 |
| IYMCVVPSINDLSFDK | 1 | 0 | 0 | 1669002 | YP_003766.2 | H9EJ42 |
| IYNKIVDEPEEHVQIH | 1 | 0 | 0 | 1669007 | YP_009724391.1 | P0DTC3 |
| IYNRVKSGSPGDSSWH | 1 | 0 | 0 | 1669015 | YP_003767.1 | A0A0P0G321 |
| IYQTSNFRVQPTDSIV | 1 | 0 | 0 | 1669024 | QHR63300.2 | Q3LZX1 |
| IYQTSNFRVQPTISIV | 1 | 0 | 0 | 1669026 | QIQ54048.1 | P59594 |
| IYSQNLCVDGFGVNKI | 1 | 0 | 0 | 1669036 | NP_073549.1 | A0A0P0LKV0 |
| IYSQTAETAHSVNVNR | 1 | 0 | 0 | 1669037 | YP_009555238.1 | P0C6X6 |

| | | | | | | |
|---|---|---|---|---|---|---|
| IYSRIVDEPEDHVQIH | 1 | 0 | 0 | 1669038 | QIQ54049.1 | P59632 |
| IYSTGNNVFQTQAGCL | 1 | 0 | 0 | 1669042 | NP_828851.1 | P59594 |
| IYTELEPPCRFVTDTP | 1 | 0 | 0 | 1669043 | YP_009724389.1 | P0DTD1 |
| IYTGDWEGLHEVLTSA | 1 | 0 | 0 | 1669044 | YP_003766.2 | H9EJ42 |
| IYVRKNGQFQDHYDGF | 1 | 0 | 0 | 1669052 | YP_003766.2 | H9EJ42 |
| IYVTNEIGLNASVTLK | 1 | 0 | 0 | 1669053 | YP_003767.1 | A0A0P0G321 |
| KAACVVDVNDSHSMAV | 1 | 0 | 0 | 1669059 | YP_009555238.1 | P0C6X6 |
| KAAESKAFVDTIVPPC | 1 | 0 | 0 | 1669061 | NP_073549.1 | A0A0P0LKV0 |
| KAAGGKVSFSDDVEVK | 1 | 0 | 0 | 1669062 | NP_073549.1 | A0A0P0LKV0 |
| KAALLADKFPVLHDIG | 1 | 0 | 0 | 1669065 | YP_009724389.1 | P0DTD1 |
| KAAYENFNSQDILLAP | 1 | 0 | 0 | 1669070 | NP_828849.2 | P0C6X7 |
| KACEVVVTNLNKSAGW | 1 | 0 | 0 | 1669073 | YP_003766.2 | H9EJ42 |
| KACNIAKSAYERDRAV | 1 | 0 | 0 | 1669075 | YP_009555238.1 | P0C6X6 |
| KACNIAKSVYERDKAV | 1 | 0 | 0 | 1669076 | YP_173236.1 | A0A140H1G9 |
| KACVEEVTTTLEETKF | 1 | 0 | 0 | 1669079 | YP_009724389.1 | P0DTD1 |
| KADDFVTASDTTLRCC | 1 | 0 | 0 | 1669081 | YP_003766.2 | H9EJ42 |
| KADGIIYPQGRTYSNI | 1 | 0 | 0 | 1669083 | YP_009047204.1 | K9N5Q8 |
| KAEVVVNPANGHMAHG | 1 | 0 | 0 | 1669092 | YP_009555238.1 | P0C6X6 |
| KAGDKYYPVVVDDHVG | 1 | 0 | 0 | 1669101 | YP_009555238.1 | P0C6X6 |
| KAIACVKDFDFKFYDA | 1 | 0 | 0 | 1669114 | YP_009555238.1 | P0C6X6 |
| KAIAVAAGKKFSKETA | 1 | 0 | 0 | 1669115 | YP_173236.1 | A0A140H1G9 |
| KAIDVDWVEFYGFKDA | 1 | 0 | 0 | 1669121 | NP_073549.1 | A0A0P0LKV0 |
| KAIGKKIKHEGDWDSF | 1 | 0 | 0 | 1669124 | NP_073549.1 | A0A0P0LKV0 |
| KAIPSGNLIKFDIGSV | 1 | 0 | 0 | 1669133 | NP_073549.1 | A0A0P0LKV0 |
| KAKEESVKSSYAFATL | 1 | 0 | 0 | 1669140 | YP_003766.2 | H9EJ42 |
| KAKGYPQFAELVPSTA | 1 | 0 | 0 | 1669144 | NP_073556.1 | A0A0N7J131 |
| KAKKGAWNIGEQKSIL | 1 | 0 | 0 | 1669148 | YP_009724389.1 | P0DTD1 |
| KAKQKPIYLKGSGSDF | 1 | 0 | 0 | 1669152 | YP_009555238.1 | P0C6X6 |
| KALCFDKSKQFVPVQV | 1 | 0 | 0 | 1669157 | YP_173237.1 | A0A140H1H0 |
| KALEMCIDQVTIAGAK | 1 | 0 | 0 | 1669159 | NP_828849.2 | P0C6X7 |
| KALNESYIDLKELGNY | 1 | 0 | 0 | 1669170 | YP_009047204.1 | K9N5Q8 |
| KALVATAHSELAKGVA | 1 | 0 | 0 | 1669177 | NP_828849.2 | P0C6X7 |
| KALYEQVVMDYLDNLK | 1 | 0 | 0 | 1669178 | NP_828849.2 | P0C6X7 |
| KAMDTFTNNNGSDVLY | 1 | 0 | 0 | 1669180 | YP_009555238.1 | P0C6X6 |
| KAMETFNHNNGNDVLY | 1 | 0 | 0 | 1669181 | YP_173236.1 | A0A140H1G9 |
| KAMFTKVVVGPTTEVK | 1 | 0 | 0 | 1669182 | YP_003766.2 | H9EJ42 |
| KAMLDFSRDHNDILYT | 1 | 0 | 0 | 1669183 | NP_073549.1 | A0A0P0LKV0 |
| KAMLDYAKDHNDMLYS | 1 | 0 | 0 | 1669184 | YP_003766.2 | H9EJ42 |
| KAMNVAKAEFDRESSV | 1 | 0 | 0 | 1669187 | NP_073549.1 | A0A0P0LKV0 |
| KANPLWHKAVFISPYN | 1 | 0 | 0 | 1669192 | YP_009555238.1 | P0C6X6 |
| KAPKEVTFLEGDSHDT | 1 | 0 | 0 | 1669199 | NP_828849.2 | P0C6X7 |
| KAPKNFCPCKLNGSCV | 1 | 0 | 0 | 1669200 | YP_009555241.1 | P36334 |
| KAPLVYLNHSVPKLSD | 1 | 0 | 0 | 1669202 | YP_173238.1 | A0A140H1H1 |
| KAPYVMLNTSIPNLPD | 1 | 0 | 0 | 1669207 | YP_009555241.1 | P36334 |
| KAQFKPFAKVDGVYTN | 1 | 0 | 0 | 1669209 | YP_173236.1 | A0A140H1G9 |
| KAQFKTFEKVDGVYTN | 1 | 0 | 0 | 1669210 | YP_009555238.1 | P0C6X6 |
| KAQSSGVIPENPVLFT | 1 | 0 | 0 | 1669214 | YP_173236.1 | A0A140H1G9 |
| KARLYYEALSFEEQDE | 1 | 0 | 0 | 1669219 | YP_009555238.1 | P0C6X6 |
| KARNGVFISTEKLSRL | 1 | 0 | 0 | 1669221 | YP_173236.1 | A0A140H1G9 |
| KARVECYDRFKVNETN | 1 | 0 | 0 | 1669223 | YP_009047202.1 | K9N7C7 |
| KARVYYESMSYQEQDE | 1 | 0 | 0 | 1669224 | YP_009047202.1 | K9N7C7 |
| KASYDTNLNKFNRASL | 1 | 0 | 0 | 1669232 | YP_009047202.1 | K9N7C7 |
| KATGDLAVCLGIGDSA | 1 | 0 | 0 | 1669240 | YP_009555238.1 | P0C6X6 |
| KATNNAMQIESDDYIA | 1 | 0 | 0 | 1669248 | QIQ54046.1 | P0C6U8 |
| KATNNAMQVESDDYIA | 1 | 0 | 0 | 1669249 | YP_009724389.1 | P0DTD1 |
| KATNNAMQVESDHYIA | 1 | 0 | 0 | 1669250 | QHR63299.1 | P0C6W2 |
| KAVAAALKSLGFDKPQ | 1 | 0 | 0 | 1669258 | NP_073556.1 | A0A0N7J131 |
| KAVARKLERMADQAMT | 1 | 0 | 0 | 1669261 | YP_009047202.1 | K9N7C7 |
| KAVFISPYNSQNAVAA | 1 | 0 | 0 | 1669263 | QIQ54047.1 | P0C6X7 |
| KAVFISPYNSQNAVAS | 1 | 0 | 0 | 1669264 | YP_009724389.1 | P0DTD1 |
| KAVFISPYNSQNFAAK | 1 | 0 | 0 | 1669266 | YP_009555238.1 | P0C6X6 |
| KAVNIAKNAYEKDKAV | 1 | 0 | 0 | 1669270 | YP_009047202.1 | K9N7C7 |
| KAVRSSFNFDQKELLA | 1 | 0 | 0 | 1669272 | YP_009555238.1 | P0C6X6 |
| KAYIDVLRNSFGKDLN | 1 | 0 | 0 | 1669279 | NP_073549.1 | A0A0P0LKV0 |
| KAYKDYLASGGQPITN | 1 | 0 | 0 | 1669280 | YP_009724389.1 | P0DTD1 |
| KAYLDFVNAGGAPLTN | 1 | 0 | 0 | 1669285 | YP_009047202.1 | K9N7C7 |
| KAYNTINNEQGTPLTP | 1 | 0 | 0 | 1669286 | NP_073549.1 | A0A0P0LKV0 |
| KAYNVTQAFGRRGPEQ | 1 | 0 | 0 | 1669287 | YP_009724397.2 | P0DTC9 |
| KAYRDYLASGGQPITN | 1 | 0 | 0 | 1669289 | QIQ54046.1 | P0C6U8 |
| KCCSIDSDVETRFITK | 1 | 0 | 0 | 1669297 | YP_173236.1 | A0A140H1G9 |
| KCDINYKGKVFFQFDN | 1 | 0 | 0 | 1669306 | YP_009555238.1 | P0C6X6 |
| KCDLQNYGDSATLPKG | 1 | 0 | 0 | 1669308 | YP_009724389.1 | P0DTD1 |
| KCDLQNYGENAVIPKG | 1 | 0 | 0 | 1669310 | NP_828849.2 | P0C6X7 |
| KCFNCYPAGVNITLAN | 1 | 0 | 0 | 1669323 | NP_073551.1 | A0A0P0K6L9 |
| KCGDKFFPIVMNDKNI | 1 | 0 | 0 | 1669325 | YP_173236.1 | A0A140H1G9 |
| KCGFSFDLNGLDALFF | 1 | 0 | 0 | 1669327 | YP_173236.1 | A0A140H1G9 |
| KCGGCCDDYTGYQELV | 1 | 0 | 0 | 1669328 | YP_009555241.1 | P36334 |
| KCGNLEQNHILGVNNS | 1 | 0 | 0 | 1669330 | YP_173236.1 | A0A140H1G9 |
| KCGNLHQRELLGVSDV | 1 | 0 | 0 | 1669331 | YP_009555238.1 | P0C6X6 |
| KCGVKQEQRTGLDAVM | 1 | 0 | 0 | 1669333 | YP_009555238.1 | P0C6X6 |
| KCHNCCDEYGGHHDFV | 1 | 0 | 0 | 1669337 | YP_173238.1 | A0A140H1H1 |
| KCKGLNKPFVKDNVSF | 1 | 0 | 0 | 1669342 | YP_009047202.1 | K9N7C7 |
| KCKTQSIAPPTGVYEL | 1 | 0 | 0 | 1669346 | YP_009555241.1 | P36334 |
| KCKVTDVLFPGQKYSS | 1 | 0 | 0 | 1669347 | YP_009047202.1 | K9N7C7 |
| KCLHVVGPNLNAGEDI | 1 | 0 | 0 | 1669350 | NP_828849.2 | P0C6X7 |
| KCNCGSESWSVGAWDG | 1 | 0 | 0 | 1669356 | YP_003766.2 | H9EJ42 |
| KCNIGGAVCSKHANLY | 1 | 0 | 0 | 1669359 | NP_073549.1 | A0A0P0LKV0 |
| KCNRCCDRYEEYDLEP | 1 | 0 | 0 | 1669361 | YP_009047204.1 | K9N5Q8 |
| KCNRSGSNDIWMQNKG | 1 | 0 | 0 | 1669362 | YP_009555240.1 | P30215 |
| KCSRLLSDDRTEVPQL | 1 | 0 | 0 | 1669376 | YP_009047204.1 | K9N5Q8 |
| KCSVYTVESGTEVTEF | 1 | 0 | 0 | 1669382 | NP_828849.2 | P0C6X7 |

| | | | | | | |
|---|---|---|---|---|---|---|
| KCTKGLSIADLACAQY | 1 | 0 | 0 | 1669384 | NP_073551.1 | A0A0P0K6L9 |
| KCTKYNIYDVSGVGVI | 1 | 0 | 0 | 1669385 | NP_073551.1 | A0A0P0K6L9 |
| KCTSDNINDKDTGPPP | 1 | 0 | 0 | 1669393 | YP_009555241.1 | P36334 |
| KCVDIVKEAQSANPMV | 1 | 0 | 0 | 1669399 | NP_828849.2 | P0C6X7 |
| KCVKPKEETGLLMPLK | 1 | 0 | 0 | 1669400 | QHR63299.1 | P0C6W2 |
| KCVPQADTDWKFYDAQ | 1 | 0 | 0 | 1669403 | QIQ54047.1 | P0C6X7 |
| KCVPQADVEWKFYDAQ | 1 | 0 | 0 | 1669404 | YP_009724389.1 | P0DTD1 |
| KCVPQAEVEWKFYDAQ | 1 | 0 | 0 | 1669405 | NP_828849.2 | P0C6X7 |
| KCWTEEDLSIGPHEFC | 1 | 0 | 0 | 1669411 | NP_073549.1 | A0A0P0LKV0 |
| KCWTETDLTKGPHEFC | 1 | 0 | 0 | 1669412 | YP_009724389.1 | P0DTD1 |
| KCWVEEDLTKGPHEFC | 1 | 0 | 0 | 1669413 | YP_003766.2 | H9EJ42 |
| KCYGVSATKLNDLCFS | 1 | 0 | 0 | 1669422 | NP_828851.1 | P59594 |
| KDAAGKPVPYCYDTNV | 1 | 0 | 0 | 1669427 | QIQ54046.1 | P0C6U8 |
| KDADYNAKVEISVTPI | 1 | 0 | 0 | 1669428 | NP_073549.1 | A0A0P0LKV0 |
| KDAPYIVGDIITSGNL | 1 | 0 | 0 | 1669431 | QIQ54046.1 | P0C6U8 |
| KDASFIGKNLKSNCVR | 1 | 0 | 0 | 1669435 | NP_073549.1 | A0A0P0LKV0 |
| KDASGKPVPYCYDTNV | 1 | 0 | 0 | 1669436 | YP_009724389.1 | P0DTD1 |
| KDATKPQQVTKHTAKE | 1 | 0 | 0 | 1669437 | YP_009555245.1 | P33469 |
| KDATPSDSVRATATIP | 1 | 0 | 0 | 1669438 | QHR63301.1 | Q3LZX0 |
| KDCARNPIDLPPSHAT | 1 | 0 | 0 | 1669441 | NP_073549.1 | A0A0P0LKV0 |
| KDCSKSYSGYHPAHAP | 1 | 0 | 0 | 1669450 | YP_009555238.1 | P0C6X6 |
| KDCVDINKAIPSGNLI | 1 | 0 | 0 | 1669453 | NP_073549.1 | A0A0P0LKV0 |
| KDDGNCVVLELDPPCK | 1 | 0 | 0 | 1669461 | YP_173236.1 | A0A140H1G9 |
| KDDGNFVVLELDPPCK | 1 | 0 | 0 | 1669462 | YP_009555238.1 | P0C6X6 |
| KDDYCDGSLGCKMCLF | 1 | 0 | 0 | 1669464 | NP_073549.1 | A0A0P0LKV0 |
| KDFDKFYDAQPIVKS | 1 | 0 | 0 | 1669469 | YP_009555238.1 | P0C6X6 |
| KDFDFYRYNKPTILDI | 1 | 0 | 0 | 1669470 | YP_003766.2 | H9EJ42 |
| KDFGGFNFLQILPDPS | 1 | 0 | 0 | 1669472 | QIQ54048.1 | P59594 |
| KDGDDGYFLPYPDPSR | 1 | 0 | 0 | 1669487 | YP_009047202.1 | K9N7C7 |
| KDGFLYVYKGYQPIDV | 1 | 0 | 0 | 1669492 | NP_828851.1 | P59594 |
| KDGHVETFYPKLQASQ | 1 | 0 | 0 | 1669497 | NP_828849.2 | P0C6X7 |
| KDGHVETFYPKLQSSQ | 1 | 0 | 0 | 1669498 | YP_009724389.1 | P0DTD1 |
| KDGIYFAATEKSNVVR | 1 | 0 | 0 | 1669501 | NP_828851.1 | P59594 |
| KDGKYFTSEPPVTYSP | 1 | 0 | 0 | 1669503 | YP_009047202.1 | K9N7C7 |
| KDGQVQTFYPRLQASA | 1 | 0 | 0 | 1669505 | YP_009047202.1 | K9N7C7 |
| KDGVNFNVDDINFSPV | 1 | 0 | 0 | 1669512 | YP_009555241.1 | P36334 |
| KDGVYYPSNGTNILPV | 1 | 0 | 0 | 1669513 | NP_073549.1 | A0A0P0LKV0 |
| KDHIGTRNPNNNAAIV | 1 | 0 | 0 | 1669518 | QIQ54056.1 | P59595 |
| KDHIGTRNPNNNAATV | 1 | 0 | 0 | 1669519 | NP_828858.1 | P59595 |
| KDIVECDVKTTEVVGD | 1 | 0 | 0 | 1669529 | QIQ54046.1 | P0C6U8 |
| KDKCVNFNFNGLTGTG | 1 | 0 | 0 | 1669532 | QIQ54048.1 | P59594 |
| KDKDLSGAVPSDLNTS | 1 | 0 | 0 | 1669533 | NP_073549.1 | A0A0P0LKV0 |
| KDKKKKTDESQPLPQR | 1 | 0 | 0 | 1669537 | QIQ54056.1 | P59595 |
| KDLNANMSLAECKRAL | 1 | 0 | 0 | 1669553 | NP_073549.1 | A0A0P0LKV0 |
| KDLNFLNRVRGTSVNA | 1 | 0 | 0 | 1669554 | YP_173236.1 | A0A140H1G9 |
| KDNAVATFYPQLQSAE | 1 | 0 | 0 | 1669566 | NP_073549.1 | A0A0P0LKV0 |
| KDNAYFTEQPIDLVPT | 1 | 0 | 0 | 1669567 | QIQ54046.1 | P0C6U8 |
| KDNAYYTEQPIDLVPT | 1 | 0 | 0 | 1669568 | NP_828849.2 | P0C6X7 |
| KDNDGKNVHLKDVTKE | 1 | 0 | 0 | 1669569 | NP_073549.1 | A0A0P0LKV0 |
| KDNFKDGLCMFWNCNV | 1 | 0 | 0 | 1669571 | YP_173236.1 | A0A140H1G9 |
| KDNHLSTFYPQLQSAE | 1 | 0 | 0 | 1669572 | YP_003766.2 | H9EJ42 |
| KDNKNLPKFIEQISAF | 1 | 0 | 0 | 1669573 | YP_003771.1 | H9EJ69 |
| KDNSYFTEQPIDLVPN | 1 | 0 | 0 | 1669574 | YP_009724389.1 | P0DTD1 |
| KDNTVLQALQSEFVNM | 1 | 0 | 0 | 1669575 | YP_009555238.1 | P0C6X6 |
| KDNVILLNKHIDAYKT | 1 | 0 | 0 | 1669576 | NP_828858.1 | P59595 |
| KDPPNVQIHTIDGSSG | 1 | 0 | 0 | 1669581 | NP_828852.2 | P59632 |
| KDPSFKDNVILLNKHI | 1 | 0 | 0 | 1669583 | QIQ54056.1 | P59595 |
| KDQVILLNKHIDAYKT | 1 | 0 | 0 | 1669586 | YP_009724397.2 | P0DTC9 |
| KDSFKDGLCMFWNCNV | 1 | 0 | 0 | 1669591 | YP_009555238.1 | P0C6X6 |
| KDSNFLNRVRGSIVNA | 1 | 0 | 0 | 1669598 | YP_009047202.1 | K9N7C7 |
| KDSYCRLKNSVSDVAY | 1 | 0 | 0 | 1669603 | YP_173236.1 | A0A140H1G9 |
| KDTLDDILQQANKSVG | 1 | 0 | 0 | 1669610 | YP_009047202.1 | K9N7C7 |
| KDTNFLNRVRGTSVDA | 1 | 0 | 0 | 1669612 | YP_009555238.1 | P0C6X6 |
| KDTWPDANKVGAGAF | 1 | 0 | 0 | 1669615 | P03138.3 | Q76R62 |
| KDVDNPHLMGWDYPKC | 1 | 0 | 0 | 1669617 | YP_009047202.1 | K9N7C7 |
| KDVDNPVLMGWDYPKC | 1 | 0 | 0 | 1669618 | YP_009555238.1 | P0C6X6 |
| KDWRLINKFDVINGVK | 1 | 0 | 0 | 1669635 | YP_173236.1 | A0A140H1G9 |
| KDWYDFVENPDIINIY | 1 | 0 | 0 | 1669638 | YP_173236.1 | A0A140H1G9 |
| KDWYDFVENPDIINVY | 1 | 0 | 0 | 1669639 | YP_009555238.1 | P0C6X6 |
| KDWYDFVENPDILRVY | 1 | 0 | 0 | 1669640 | YP_009724389.1 | P0DTD1 |
| KDYEFKFYDAFPVAKS | 1 | 0 | 0 | 1669642 | YP_173236.1 | A0A140H1G9 |
| KDYISSNGPLKVGAGV | 1 | 0 | 0 | 1669644 | YP_003766.2 | H9EJ42 |
| KEAQSANPMVIVNAAN | 1 | 0 | 0 | 1669649 | NP_828849.2 | P0C6X7 |
| KEDGDVLLAEFDTYDP | 1 | 0 | 0 | 1669655 | YP_009047202.1 | K9N7C7 |
| KEDLRFPRGQGVPINT | 1 | 0 | 0 | 1669658 | QIQ54056.1 | P59595 |
| KEDVVIFINNTTYPTN | 1 | 0 | 0 | 1669659 | YP_009555238.1 | P0C6X6 |
| KEDYQIGGYSEDRHSG | 1 | 0 | 0 | 1669660 | NP_828852.2 | P59632 |
| KEEFVVKEKLNAFLVH | 1 | 0 | 0 | 1669663 | NP_073549.1 | A0A0P0LKV0 |
| KEELRFPRGQGVPINT | 1 | 0 | 0 | 1669666 | NP_828858.1 | P59595 |
| KEEPKKVKPCIEEVTT | 1 | 0 | 0 | 1669667 | QIQ54046.1 | P0C6U8 |
| KEESVKSSYAFATLKE | 1 | 0 | 0 | 1669669 | YP_003766.2 | H9EJ42 |
| KEESVKSSYAYATLKE | 1 | 0 | 0 | 1669670 | NP_073549.1 | A0A0P0LKV0 |
| KEETGLLMPLKAPKEI | 1 | 0 | 0 | 1669671 | QHR63299.1 | P0C6W2 |
| KEFEFVEGQGVPIAPG | 1 | 0 | 0 | 1669675 | YP_009555245.1 | P33469 |
| KEFPLLSEYPLLKQKD | 1 | 0 | 0 | 1669677 | YP_173243.1 | Q0ZJ80 |
| KEGAKTVNTSLGNRKR | 1 | 0 | 0 | 1669680 | YP_003771.1 | H9EJ69 |
| KEGIEFLKDGWEIVKF | 1 | 0 | 0 | 1669687 | QIQ54046.1 | P0C6U8 |
| KEGIVWVATEGALNTP | 1 | 0 | 0 | 1669689 | NP_828858.1 | P59595 |
| KEGVEFLRDGWEIVKF | 1 | 0 | 0 | 1669699 | YP_009724389.1 | P0DTD1 |
| KEILSHIIPGSPGSLN | 1 | 0 | 0 | 1669707 | YP_173243.1 | Q0ZJ80 |
| KEIVDTVSALVYENKL | 1 | 0 | 0 | 1669715 | YP_009555238.1 | P0C6X6 |

| | | | | | | |
|---|---|---|---|---|---|---|
| KEIVETVSALVYDNKL | 1 | 0 | 0 | 1669716 | YP_173236.1 | A0A140H1G9 |
| KEKKQKAPKEESTDQM | 1 | 0 | 0 | 1669720 | YP_009047211.1 | K9N4V7 |
| KEKTYYELTKSCGVVA | 1 | 0 | 0 | 1669722 | YP_173236.1 | A0A140H1G9 |
| KELGIVMNMDVDTHRY | 1 | 0 | 0 | 1669728 | YP_009555238.1 | P0C6X6 |
| KELGVVMNLDVDTHRY | 1 | 0 | 0 | 1669731 | YP_173236.1 | A0A140H1G9 |
| KELLLLWESGKAKPPL | 1 | 0 | 0 | 1669736 | NP_073549.1 | A0A0P0LKV0 |
| KELLLSWESGKVKPPL | 1 | 0 | 0 | 1669737 | YP_003766.2 | H9EJ42 |
| KELLQNGMNGRTILGS | 1 | 0 | 0 | 1669738 | YP_009724389.1 | P0DTD1 |
| KELLQNGVNGRTILGS | 1 | 0 | 0 | 1669739 | QIQ54046.1 | P0C6U8 |
| KENQINDMIYSLLEKG | 1 | 0 | 0 | 1669755 | NP_828849.2 | P0C6X7 |
| KENSYITTIKPVIYKL | 1 | 0 | 0 | 1669756 | QHR63299.1 | P0C6W2 |
| KENSYTTTIKPIVYKL | 1 | 0 | 0 | 1669757 | QIQ54046.1 | P0C6U8 |
| KENSYTTTIKPVTYKL | 1 | 0 | 0 | 1669758 | YP_009724389.1 | P0DTD1 |
| KENVDFINSMQFNYKI | 1 | 0 | 0 | 1669759 | YP_009555239.1 | Q80872 |
| KEQRVQGSITQRTRTR | 1 | 0 | 0 | 1669772 | YP_009047211.1 | K9N4V7 |
| KESDDYIKLNGPLTVG | 1 | 0 | 0 | 1669779 | NP_828849.2 | P0C6X7 |
| KESDEYILAKGPLQVG | 1 | 0 | 0 | 1669780 | YP_009047202.1 | K9N7C7 |
| KESYFSKKDWYDFVEN | 1 | 0 | 0 | 1669789 | YP_173236.1 | A0A140H1G9 |
| KETSYTTTIKPVSYKL | 1 | 0 | 0 | 1669801 | NP_828849.2 | P0C6X7 |
| KEVCKTTTGIPEYNFI | 1 | 0 | 0 | 1669807 | YP_009047202.1 | K9N7C7 |
| KEVIECDVKTTEVVGN | 1 | 0 | 0 | 1669810 | NP_828849.2 | P0C6X7 |
| KEVRTIKVFTTVDNIN | 1 | 0 | 0 | 1669821 | QHR63299.1 | P0C6W2 |
| KEVTFLEGDSHDTVLT | 1 | 0 | 0 | 1669823 | NP_828849.2 | P0C6X7 |
| KFADDLNQMSGYKKPA | 1 | 0 | 0 | 1669832 | QIQ54046.1 | P0C6U8 |
| KFADDLNQMTGFTKPA | 1 | 0 | 0 | 1669833 | NP_828849.2 | P0C6X7 |
| KFAIPNGRKVDLQLGN | 1 | 0 | 0 | 1669834 | YP_009555241.1 | P36334 |
| KFASVYAWNRKRISNC | 1 | 0 | 0 | 1669840 | QIQ54048.1 | P59594 |
| KFDNFTGECPNFVFPL | 1 | 0 | 0 | 1669855 | QIQ54046.1 | P0C6U8 |
| KFDNPVLPXNDGVYFA | 1 | 0 | 0 | 1669856 | QIQ54048.1 | P59594 |
| KFDNRGVITNAVDCSS | 1 | 0 | 0 | 1669857 | YP_173238.1 | A0A140H1H1 |
| KFDSGTVSKTSDWKCK | 1 | 0 | 0 | 1669858 | YP_009047202.1 | K9N7C7 |
| KFFYASNGTYNCTDAV | 1 | 0 | 0 | 1669876 | NP_073551.1 | A0A0P0K6L9 |
| KFIDKLNVLFDGRDNG | 1 | 0 | 0 | 1669890 | YP_009555238.1 | P0C6X6 |
| KFKEGIEFLKDGWEIV | 1 | 0 | 0 | 1669903 | QIQ54046.1 | P0C6U8 |
| KFKEVCKTTTGIPEYN | 1 | 0 | 0 | 1669905 | YP_009047202.1 | K9N7C7 |
| KFKGKCVQVPIGCLDP | 1 | 0 | 0 | 1669906 | YP_003766.2 | H9EJ42 |
| KFKKNAWNIGEPKSIL | 1 | 0 | 0 | 1669911 | QIQ54046.1 | P0C6U8 |
| KFKNSVASINSAIVCA | 1 | 0 | 0 | 1669912 | NP_073549.1 | A0A0P0LKV0 |
| KFLALCADSIVIGGAK | 1 | 0 | 0 | 1669920 | QIQ54046.1 | P0C6U8 |
| KFLEELNAFTREMQQH | 1 | 0 | 0 | 1669922 | NP_073556.1 | A0A0N7J131 |
| KFLKTNCCRFQEKDEE | 1 | 0 | 0 | 1669931 | NP_828849.2 | P0C6X7 |
| KFLSNTKYYDDSQYYF | 1 | 0 | 0 | 1669936 | YP_009555240.1 | P30215 |
| KFNQALGAMQTGFTTT | 1 | 0 | 0 | 1669951 | YP_009047204.1 | K9N5Q8 |
| KFNSCKSDILSLNNPI | 1 | 0 | 0 | 1669953 | YP_173237.1 | A0A140H1H0 |
| KFNYQFPKVGYCRVPS | 1 | 0 | 0 | 1669956 | YP_009555242.1 | Q04853 |
| KFPLKLAGTAVINLRA | 1 | 0 | 0 | 1669958 | YP_009555238.1 | P0C6X6 |
| KFPQVNGLTSIKWADN | 1 | 0 | 0 | 1669962 | QIQ54046.1 | P0C6U8 |
| KFQTVKPGNFNQDFYD | 1 | 0 | 0 | 1669970 | YP_009555238.1 | P0C6X6 |
| KFQTVKPGNFNQDFYE | 1 | 0 | 0 | 1669971 | YP_173236.1 | A0A140H1G9 |
| KFRRVFGEYNHVVAAN | 1 | 0 | 0 | 1669977 | NP_828849.2 | P0C6X7 |
| KFSGFKLDGFNHQFVT | 1 | 0 | 0 | 1669983 | YP_003766.2 | H9EJ42 |
| KFTEESFYKNMYLKSA | 1 | 0 | 0 | 1669996 | YP_173236.1 | A0A140H1G9 |
| KFVCDNIKFADDLNQL | 1 | 0 | 0 | 1670006 | YP_009724389.1 | P0DTD1 |
| KFVCDNIKLADDLNQL | 1 | 0 | 0 | 1670007 | QHR63299.1 | P0C6W2 |
| KFVCDNTKFADDLNQM | 1 | 0 | 0 | 1670008 | QIQ54046.1 | P0C6U8 |
| KFVFPLNSKVKVIQPR | 1 | 0 | 0 | 1670010 | NP_828849.2 | P0C6X7 |
| KFVVESNVMDVNDCFK | 1 | 0 | 0 | 1670029 | YP_003766.2 | H9EJ42 |
| KFYDAFPVAKSVKQLF | 1 | 0 | 0 | 1670034 | YP_173236.1 | A0A140H1G9 |
| KFYDAQPCSDKAYKIE | 1 | 0 | 0 | 1670035 | YP_009724389.1 | P0DTD1 |
| KFYGGWDDMLRHLIKD | 1 | 0 | 0 | 1670037 | YP_173236.1 | A0A140H1G9 |
| KFYGGWDNMLKNLMAD | 1 | 0 | 0 | 1670040 | NP_073549.1 | A0A0P0LKV0 |
| KFYGGWHNMLKTVYSD | 1 | 0 | 0 | 1670041 | YP_009724389.1 | P0DTD1 |
| KFYGGWNNMLKTVYSD | 1 | 0 | 0 | 1670042 | QIQ54047.1 | P0C6X7 |
| KFYLYKKVNNEFIDPT | 1 | 0 | 0 | 1670047 | YP_009047202.1 | K9N7C7 |
| KFYQHVINGCKDTACL | 1 | 0 | 0 | 1670049 | YP_009047202.1 | K9N7C7 |
| KGAGGHSFGIDLKSYD | 1 | 0 | 0 | 1670055 | QIQ54046.1 | P0C6U8 |
| KGAGGHSYGIDLKSYD | 1 | 0 | 0 | 1670057 | NP_828849.2 | P0C6X7 |
| KGAHVASADAIMTRCL | 1 | 0 | 0 | 1670058 | YP_173236.1 | A0A140H1G9 |
| KGAHVASSDAIMTRCL | 1 | 0 | 0 | 1670059 | YP_009555238.1 | P0C6X6 |
| KGAVQKESDEYILAKG | 1 | 0 | 0 | 1670062 | YP_009047202.1 | K9N7C7 |
| KGAWNIGEQKSILSPL | 1 | 0 | 0 | 1670063 | YP_009724389.1 | P0DTD1 |
| KGAYNKDHGRGGFGHV | 1 | 0 | 0 | 1670065 | YP_009555238.1 | P0C6X6 |
| KGCCSCGSCCKFDEDD | 1 | 0 | 0 | 1670066 | YP_009724390.1 | P0DTC2 |
| KGCFTGVDGELPVAVV | 1 | 0 | 0 | 1670068 | YP_003766.2 | H9EJ42 |
| KGCNAVAKHDFFTWHE | 1 | 0 | 0 | 1670069 | NP_073549.1 | A0A0P0LKV0 |
| KGCVADQSRKWIGHCT | 1 | 0 | 0 | 1670072 | YP_009555239.1 | Q80872 |
| KGCVPLNAIPSLAANT | 1 | 0 | 0 | 1670073 | YP_009555238.1 | P0C6X6 |
| KGCVPLSAIPALAANT | 1 | 0 | 0 | 1670074 | YP_173236.1 | A0A140H1G9 |
| KGDAAIKDFDYYRYNR | 1 | 0 | 0 | 1670075 | NP_073549.1 | A0A0P0LKV0 |
| KGDAEDTLTKLSKYLA | 1 | 0 | 0 | 1670076 | NP_073549.1 | A0A0P0LKV0 |
| KGDAEEALSKLSEYLI | 1 | 0 | 0 | 1670077 | YP_003766.2 | H9EJ42 |
| KGDEVRQIAPGQTGVI | 1 | 0 | 0 | 1670079 | QIQ54048.1 | P59594 |
| KGDSCSSNCKHSVAQL | 1 | 0 | 0 | 1670084 | YP_009047202.1 | K9N7C7 |
| KGDYGDAVVYRGTTTY | 1 | 0 | 0 | 1670086 | YP_009724389.1 | P0DTD1 |
| KGECPKFVFPLNSKVK | 1 | 0 | 0 | 1670088 | NP_828849.2 | P0C6X7 |
| KGEDIQLLKSAYENFN | 1 | 0 | 0 | 1670089 | YP_009724389.1 | P0DTD1 |
| KGEDIQLLKVAYENFN | 1 | 0 | 0 | 1670090 | QIQ54046.1 | P0C6U8 |
| KGEGDGGITSEGNALY | 1 | 0 | 0 | 1670091 | NP_073549.1 | A0A0P0LKV0 |
| KGFCDLKGKYVQIPTT | 1 | 0 | 0 | 1670093 | YP_009724389.1 | P0DTD1 |
| KGFCKLHNWNCVNCDT | 1 | 0 | 0 | 1670094 | YP_009724389.1 | P0DTD1 |
| KGFFKEGSSVELKHFF | 1 | 0 | 0 | 1670095 | YP_009724389.1 | P0DTD1 |

| | | | | | | |
|---|---|---|---|---|---|---|
| KGFIRFPEVLREGLVR | 1 | 0 | 0 | 1670097 | YP_009555238.1 | P0C6X6 |
| KGFPQLAELIPNQAAL | 1 | 0 | 0 | 1670099 | YP_003771.1 | H9EJ69 |
| KGGAPIKGVTFGEDTV | 1 | 0 | 0 | 1670103 | NP_828849.2 | P0C6X7 |
| KGGAPVKKVAFGGDQV | 1 | 0 | 0 | 1670105 | YP_009047202.1 | K9N7C7 |
| KGGFKFGDPADSRDFL | 1 | 0 | 0 | 1670108 | YP_009555238.1 | P0C6X6 |
| KGGLILKFDSGTVSKT | 1 | 0 | 0 | 1670112 | YP_009047202.1 | K9N7C7 |
| KGHFDGQQGETPVSIV | 1 | 0 | 0 | 1670121 | QIQ54047.1 | P0C6X7 |
| KGHFKFDEPSDATDFI | 1 | 0 | 0 | 1670123 | YP_173236.1 | A0A140H1G9 |
| KGHYTVYDTAKKSMYD | 1 | 0 | 0 | 1670124 | NP_073549.1 | A0A0P0LKV0 |
| KGIAPGTSVLRQWLPT | 1 | 0 | 0 | 1670125 | YP_009047202.1 | K9N7C7 |
| KGIFCVMCDKTLFDSL | 1 | 0 | 0 | 1670127 | YP_003766.2 | H9EJ42 |
| KGIKIQEGIVDYGVRF | 1 | 0 | 0 | 1670130 | NP_828849.2 | P0C6X7 |
| KGIKIQEGVVDYGARF | 1 | 0 | 0 | 1670131 | YP_009724389.1 | P0DTD1 |
| KGIKIQEGVVDYGVRF | 1 | 0 | 0 | 1670132 | QIQ54046.1 | P0C6U8 |
| KGIKVLPPLLSENQIS | 1 | 0 | 0 | 1670133 | YP_009555241.1 | P36334 |
| KGIMMNVAKYTQLCQY | 1 | 0 | 0 | 1670137 | YP_009724389.1 | P0DTD1 |
| KGIYQTSNFRVQPTDS | 1 | 0 | 0 | 1670140 | QHR63300.2 | Q3LZX1 |
| KGKAKKGAWNIGEQKS | 1 | 0 | 0 | 1670143 | YP_009724389.1 | P0DTD1 |
| KGKEFEFVEGQGVPIA | 1 | 0 | 0 | 1670145 | YP_009555245.1 | P33469 |
| KGKFKKNAWNIGEPKS | 1 | 0 | 0 | 1670146 | QIQ54046.1 | P0C6U8 |
| KGKFVQIPAQCVRDPV | 1 | 0 | 0 | 1670148 | YP_009047202.1 | K9N7C7 |
| KGKHERDLLIKAYNTI | 1 | 0 | 0 | 1670149 | NP_073549.1 | A0A0P0LKV0 |
| KGKLLIRDTNKEVFVG | 1 | 0 | 0 | 1670151 | YP_009555238.1 | P0C6X6 |
| KGKLQRLSKEHIGLAG | 1 | 0 | 0 | 1670152 | NP_073549.1 | A0A0P0LKV0 |
| KGKMFVRPVIEDYHTL | 1 | 0 | 0 | 1670153 | YP_173241.1 | Q0ZJ82 |
| KGKPILWVNKASYDTN | 1 | 0 | 0 | 1670154 | YP_009047202.1 | K9N7C7 |
| KGKVFFQFDNLSSEDL | 1 | 0 | 0 | 1670157 | YP_009555238.1 | P0C6X6 |
| KGKWVQVPIGTNDPIR | 1 | 0 | 0 | 1670158 | NP_073549.1 | A0A0P0LKV0 |
| KGKYAQNLLKKLIGGD | 1 | 0 | 0 | 1670159 | YP_009047202.1 | K9N7C7 |
| KGKYVQIPTTCANDPV | 1 | 0 | 0 | 1670161 | YP_009724389.1 | P0DTD1 |
| KGLACVKDYEFKFYDA | 1 | 0 | 0 | 1670162 | YP_173236.1 | A0A140H1G9 |
| KGLDYKSFKQIVESCG | 1 | 0 | 0 | 1670166 | QIQ54046.1 | P0C6U8 |
| KGLFKEGSTVDLKHFF | 1 | 0 | 0 | 1670168 | YP_173236.1 | A0A140H1G9 |
| KGLLKEGSSVDLKHFF | 1 | 0 | 0 | 1670172 | YP_009555238.1 | P0C6X6 |
| KGLNNLNRGMVLGSLA | 1 | 0 | 0 | 1670174 | YP_009724389.1 | P0DTD1 |
| KGLNVLWNKFVTGDVG | 1 | 0 | 0 | 1670175 | YP_003766.2 | H9EJ42 |
| KGLSIADLACAQYYNG | 1 | 0 | 0 | 1670182 | NP_073551.1 | A0A0P0K6L9 |
| KGLYRKCVKPKEETGL | 1 | 0 | 0 | 1670186 | QHR63299.1 | P0C6W2 |
| KGLYRKCVKSREETGL | 1 | 0 | 0 | 1670187 | YP_009724389.1 | P0DTD1 |
| KGLYRQCIRGKEQLQL | 1 | 0 | 0 | 1670188 | NP_828849.2 | P0C6X7 |
| KGNKDEQIGYWNVQER | 1 | 0 | 0 | 1670190 | YP_003771.1 | H9EJ69 |
| KGNVTHDASSAINRPQ | 1 | 0 | 0 | 1670192 | YP_009047202.1 | K9N7C7 |
| KGPITDIFYKENSYIT | 1 | 0 | 0 | 1670198 | QHR63299.1 | P0C6W2 |
| KGPITDVFYKENSYTT | 1 | 0 | 0 | 1670199 | YP_009724389.1 | P0DTD1 |
| KGPLCVDTSHFTTKYV | 1 | 0 | 0 | 1670201 | NP_073551.1 | A0A0P0K6L9 |
| KGPQRADLNGVIVDKV | 1 | 0 | 0 | 1670203 | YP_173236.1 | A0A140H1G9 |
| KGPVTDVFYKENSYTT | 1 | 0 | 0 | 1670204 | QIQ54046.1 | P0C6U8 |
| KGPVTDVFYKETSYTT | 1 | 0 | 0 | 1670205 | NP_828849.2 | P0C6X7 |
| KGQIQVASDNIKDCVK | 1 | 0 | 0 | 1670206 | NP_828849.2 | P0C6X7 |
| KGQKGDAEEALSKLSE | 1 | 0 | 0 | 1670207 | YP_003766.2 | H9EJ42 |
| KGQTTHESSSAVNIQQ | 1 | 0 | 0 | 1670213 | YP_173236.1 | A0A140H1G9 |
| KGRDFKFSDGQGVPIA | 1 | 0 | 0 | 1670214 | YP_173242.1 | Q19U29 |
| KGRLIIRENNKVVISS | 1 | 0 | 0 | 1670215 | QIQ54047.1 | P0C6X7 |
| KGRLIIRENNRVVISS | 1 | 0 | 0 | 1670216 | YP_009724389.1 | P0DTD1 |
| KGRLIIRENNRVVVSS | 1 | 0 | 0 | 1670217 | NP_828849.2 | P0C6X7 |
| KGSFINGACGSPGYNV | 1 | 0 | 0 | 1670218 | YP_003766.2 | H9EJ42 |
| KGSFLCGSCGSVGYTK | 1 | 0 | 0 | 1670219 | YP_009047202.1 | K9N7C7 |
| KGSFLNGSCGSVGFNI | 1 | 0 | 0 | 1670222 | YP_009724389.1 | P0DTD1 |
| KGSFVGADGELPVAIS | 1 | 0 | 0 | 1670223 | NP_073549.1 | A0A0P0LKV0 |
| KGSGSDFSLADSVVEV | 1 | 0 | 0 | 1670226 | YP_009555238.1 | P0C6X6 |
| KGSIPINVIVFDGKSK | 1 | 0 | 0 | 1670227 | QIQ54046.1 | P0C6U8 |
| KGSSRNESWHFDKSEP | 1 | 0 | 0 | 1670233 | YP_173238.1 | A0A140H1H1 |
| KGTGVFVQDPAPIDID | 1 | 0 | 0 | 1670236 | YP_003766.2 | H9EJ42 |
| KGTIYSITPCNPPDQL | 1 | 0 | 0 | 1670239 | NP_073551.1 | A0A0P0K6L9 |
| KGTLEPEYFNSVCRLM | 1 | 0 | 0 | 1670240 | YP_009724389.1 | P0DTD1 |
| KGTMYVRPIIEDYHTL | 1 | 0 | 0 | 1670242 | YP_009555244.1 | Q01455 |
| KGTPVLQLKESQINEL | 1 | 0 | 0 | 1670243 | YP_009047202.1 | K9N7C7 |
| KGVAPGTTVLKRWLPP | 1 | 0 | 0 | 1670249 | YP_003766.2 | H9EJ42 |
| KGVCATGDCYVSTGGK | 1 | 0 | 0 | 1670250 | YP_009555238.1 | P0C6X6 |
| KGVCFGFDKWYVNDGR | 1 | 0 | 0 | 1670251 | YP_003766.2 | H9EJ42 |
| KGVCQVGDCYVSTGGK | 1 | 0 | 0 | 1670252 | YP_173236.1 | A0A140H1G9 |
| KGVDYTKKFLTVDGVQ | 1 | 0 | 0 | 1670253 | YP_009047202.1 | K9N7C7 |
| KGVEELHDDLVNYIRD | 1 | 0 | 0 | 1670255 | YP_009555239.1 | Q80872 |
| KGVGSANIFIGDKVGH | 1 | 0 | 0 | 1670257 | YP_009555238.1 | P0C6X6 |
| KGVITHDVSSAINRPQ | 1 | 0 | 0 | 1670260 | YP_009724389.1 | P0DTD1 |
| KGVLPQLEQPYVFIKR | 1 | 0 | 0 | 1670261 | YP_009724389.1 | P0DTD1 |
| KGVLPQLEQPYVFVKR | 1 | 0 | 0 | 1670262 | QIQ54046.1 | P0C6U8 |
| KGVMEPVDYNVVTQRM | 1 | 0 | 0 | 1670264 | YP_003766.2 | H9EJ42 |
| KGVQIPCTCGKQATKY | 1 | 0 | 0 | 1670265 | YP_009724389.1 | P0DTD1 |
| KGVQIPCTCGKQATQY | 1 | 0 | 0 | 1670266 | QHR63299.1 | P0C6W2 |
| KGVTLDKLRDYLADYD | 1 | 0 | 0 | 1670271 | YP_009047202.1 | K9N7C7 |
| KGVTTHESSSAVNMQQ | 1 | 0 | 0 | 1670272 | YP_009555238.1 | P0C6X6 |
| KGWIPGNMIDGFACTT | 1 | 0 | 0 | 1670275 | YP_009555238.1 | P0C6X6 |
| KGWYDPVENEDIHRVY | 1 | 0 | 0 | 1670277 | YP_003766.2 | H9EJ42 |
| KGYCELKGKYVQIPTT | 1 | 0 | 0 | 1670278 | QIQ54046.1 | P0C6U8 |
| KGYGCSCDQLREPLMQ | 1 | 0 | 0 | 1670279 | NP_828849.2 | P0C6X7 |
| KGYGCSCDQLREPMLQ | 1 | 0 | 0 | 1670280 | YP_009724389.1 | P0DTD1 |
| KGYHLMSFPQAAPHGV | 1 | 0 | 0 | 1670281 | NP_828851.1 | P59594 |
| KGYIAGIQNFKETLYY | 1 | 0 | 0 | 1670283 | YP_009047202.1 | K9N7C7 |
| KGYPQFAELVPSTAAM | 1 | 0 | 0 | 1670286 | NP_073556.1 | A0A0N7J131 |
| KGYQPIDVVRDLPSGF | 1 | 0 | 0 | 1670287 | NP_828851.1 | P59594 |

| | | | | | | |
|---|---|---|---|---|---|---|
| KHAFHTPAFDKSAFTN | 1 | 0 | 0 | 1670298 | NP_828849.2 | P0C6X7 |
| KHAFHTPAFDKSAFVN | 1 | 0 | 0 | 1670299 | YP_009724389.1 | P0DTD1 |
| KHDFFKFRIDGDMVPH | 1 | 0 | 0 | 1670316 | YP_009724389.1 | P0DTD1 |
| KHDIQITNESYNNYVP | 1 | 0 | 0 | 1670319 | YP_009047202.1 | K9N7C7 |
| KHDVSPFEMPFEELNG | 1 | 0 | 0 | 1670321 | NP_073549.1 | A0A0P0LKV0 |
| KHEHSLLVEAYNSILF | 1 | 0 | 0 | 1670325 | YP_003766.2 | H9EJ42 |
| KHERDLLIKAYNTINN | 1 | 0 | 0 | 1670327 | NP_073549.1 | A0A0P0LKV0 |
| KHFFFTQKGDAAIKDF | 1 | 0 | 0 | 1670333 | NP_073549.1 | A0A0P0LKV0 |
| KHGDFADEIIGYKAIE | 1 | 0 | 0 | 1670341 | QIQ54046.1 | P0C6U8 |
| KHGDFSAVSSPMSNMT | 1 | 0 | 0 | 1670342 | NP_073555.1 | A0A0P0KXN3 |
| KHGGGVAGALNKATNN | 1 | 0 | 0 | 1670346 | YP_009724389.1 | P0DTD1 |
| KHGSGVTRELMRELNG | 1 | 0 | 0 | 1670348 | QIQ54046.1 | P0C6U8 |
| KHGTFTCASEYTGNYQ | 1 | 0 | 0 | 1670352 | YP_009724389.1 | P0DTD1 |
| KHIGAPANLRVVGHAM | 1 | 0 | 0 | 1670355 | YP_009047202.1 | K9N7C7 |
| KHKATVVVQLKDSDIN | 1 | 0 | 0 | 1670363 | NP_073549.1 | A0A0P0LKV0 |
| KHLNKTLNEGVLESFS | 1 | 0 | 0 | 1670381 | NP_073549.1 | A0A0P0LKV0 |
| KHTDFSSEIIGYKAID | 1 | 0 | 0 | 1670405 | YP_009724389.1 | P0DTD1 |
| KHVTKENDSKEGFFTY | 1 | 0 | 0 | 1670427 | NP_828849.2 | P0C6X7 |
| KHWSMPYHPNTVDCQD | 1 | 0 | 0 | 1670432 | YP_009555238.1 | P0C6X6 |
| KIAEIPKEEVKPFITE | 1 | 0 | 0 | 1670439 | YP_009724389.1 | P0DTD1 |
| KICIVDNVYMAKAGDK | 1 | 0 | 0 | 1670448 | YP_009555238.1 | P0C6X6 |
| KIDGDYVYLPYPDPSR | 1 | 0 | 0 | 1670451 | YP_173236.1 | A0A140H1G9 |
| KIDNASPASTVHATAT | 1 | 0 | 0 | 1670453 | NP_828852.2 | P59632 |
| KIDTTSSSCQLYYSLP | 1 | 0 | 0 | 1670454 | YP_173238.1 | A0A140H1H1 |
| KIDVLLTVDGVNFKSI | 1 | 0 | 0 | 1670455 | YP_173236.1 | A0A140H1G9 |
| KIEDAIIVNDENSSIK | 1 | 0 | 0 | 1670456 | YP_173236.1 | A0A140H1G9 |
| KIEELFYSYATHHDKF | 1 | 0 | 0 | 1670460 | NP_828849.2 | P0C6X7 |
| KIFERGSVQVDNGSSI | 1 | 0 | 0 | 1670466 | NP_073549.1 | A0A0P0LKV0 |
| KIGDVKFKRLGDYVLT | 1 | 0 | 0 | 1670477 | YP_003766.2 | H9EJ42 |
| KIGGLTFKTYAFDKPY | 1 | 0 | 0 | 1670479 | NP_073549.1 | A0A0P0LKV0 |
| KIKACVEEVTTTLEET | 1 | 0 | 0 | 1670500 | YP_009724389.1 | P0DTD1 |
| KIKKYPCMVGPDYAYF | 1 | 0 | 0 | 1670502 | YP_009047202.1 | K9N7C7 |
| KILAASFNKAINNIVA | 1 | 0 | 0 | 1670510 | YP_003767.1 | A0A0P0G321 |
| KILAASFNKAMTNIVD | 1 | 0 | 0 | 1670511 | NP_073551.1 | A0A0P0K6L9 |
| KILKDDGNFVVLELDP | 1 | 0 | 0 | 1670516 | YP_009555238.1 | P0C6X6 |
| KILLRKNGNKGAGGHS | 1 | 0 | 0 | 1670518 | QIQ54046.1 | P0C6U8 |
| KINKYGLELHWAPEFP | 1 | 0 | 0 | 1670538 | YP_009555238.1 | P0C6X6 |
| KIPANIVLPQGGYVAD | 1 | 0 | 0 | 1670546 | YP_009555238.1 | P0C6X6 |
| KIPTNFTIVGQEEFIQ | 1 | 0 | 0 | 1670547 | YP_173238.1 | A0A140H1H1 |
| KIQAVVNANAEALNNL | 1 | 0 | 0 | 1670548 | YP_009555241.1 | P36334 |
| KIQDVVNQQGNSLNHL | 1 | 0 | 0 | 1670550 | NP_073551.1 | A0A0P0K6L9 |
| KIQDVVNQQGSALNHL | 1 | 0 | 0 | 1670551 | YP_003767.1 | A0A0P0G321 |
| KIQPWTNDALNTICIK | 1 | 0 | 0 | 1670556 | NP_073549.1 | A0A0P0LKV0 |
| KIQVVNSGPDQTCNTP | 1 | 0 | 0 | 1670559 | YP_009555238.1 | P0C6X6 |
| KISADNKVIYTTEIAS | 1 | 0 | 0 | 1670562 | YP_009555238.1 | P0C6X6 |
| KISTQLFEGDKFIGTF | 1 | 0 | 0 | 1670571 | YP_003766.2 | H9EJ42 |
| KISTVQSKLTDLKCTN | 1 | 0 | 0 | 1670572 | YP_003766.2 | H9EJ42 |
| KITEFQLDYSIDVIDN | 1 | 0 | 0 | 1670579 | NP_073549.1 | A0A0P0LKV0 |
| KITEFSWNADLYKLMS | 1 | 0 | 0 | 1670580 | YP_173236.1 | A0A140H1G9 |
| KITEHSWNADLYKLMG | 1 | 0 | 0 | 1670582 | YP_009724389.1 | P0DTD1 |
| KITFFPDLNGDVVAID | 1 | 0 | 0 | 1670586 | QIQ54046.1 | P0C6U8 |
| KIVDKEVKDMQFDLYV | 1 | 0 | 0 | 1670593 | YP_009047202.1 | K9N7C7 |
| KIVGGAPTWFNALRDF | 1 | 0 | 0 | 1670595 | YP_009047202.1 | K9N7C7 |
| KIVYAILSDVDGLKYT | 1 | 0 | 0 | 1670612 | YP_009555238.1 | P0C6X6 |
| KIVYHTLESPVEFHLD | 1 | 0 | 0 | 1670613 | NP_828849.2 | P0C6X7 |
| KIWNGVIKNVNSVRDW | 1 | 0 | 0 | 1670614 | NP_073549.1 | A0A0P0LKV0 |
| KIYCPACHNPEVGPEH | 1 | 0 | 0 | 1670616 | QHR63299.1 | P0C6W2 |
| KKAKIVSAMQTMLFGM | 1 | 0 | 0 | 1670632 | YP_009047202.1 | K9N7C7 |
| KKAMFTKVVVGPTTEV | 1 | 0 | 0 | 1670633 | YP_003766.2 | H9EJ42 |
| KKAMNVAKAEFDRESS | 1 | 0 | 0 | 1670634 | NP_073549.1 | A0A0P0LKV0 |
| KKCGGCCDDYTGYQEL | 1 | 0 | 0 | 1670637 | YP_009555241.1 | P36334 |
| KKCKSAFYVLPSEAPN | 1 | 0 | 0 | 1670639 | NP_828849.2 | P0C6X7 |
| KKCLHVVGPNLNAGED | 1 | 0 | 0 | 1670640 | NP_828849.2 | P0C6X7 |
| KKCTKGLSIADLACAQ | 1 | 0 | 0 | 1670641 | NP_073551.1 | A0A0P0K6L9 |
| KKCVKSRGDSGLLMPL | 1 | 0 | 0 | 1670642 | QIQ54046.1 | P0C6U8 |
| KKDAPYIVGDIITSGN | 1 | 0 | 0 | 1670644 | QIQ54046.1 | P0C6U8 |
| KKDGFYETIVSHWMGI | 1 | 0 | 0 | 1670646 | YP_009555239.1 | Q80872 |
| KKDGVYYPSNGTNILP | 1 | 0 | 0 | 1670647 | NP_073549.1 | A0A0P0LKV0 |
| KKDKKKKTDESQPLPQ | 1 | 0 | 0 | 1670651 | QIQ54056.1 | P59595 |
| KKDNAYFTEQPIDLVP | 1 | 0 | 0 | 1670653 | QIQ54046.1 | P0C6U8 |
| KKDNAYYTEQPIDLVP | 1 | 0 | 0 | 1670654 | NP_828849.2 | P0C6X7 |
| KKDNSYFTEQPIDLVP | 1 | 0 | 0 | 1670655 | YP_009724389.1 | P0DTD1 |
| KKDWYDFVENPDIINI | 1 | 0 | 0 | 1670656 | YP_173236.1 | A0A140H1G9 |
| KKDWYDFVENPDIINV | 1 | 0 | 0 | 1670657 | YP_009555238.1 | P0C6X6 |
| KKDWYDFVENPDILRV | 1 | 0 | 0 | 1670658 | YP_009724389.1 | P0DTD1 |
| KKEKKQKAPKEESTDQ | 1 | 0 | 0 | 1670660 | YP_009047211.1 | K9N4V7 |
| KKFDNFTGECPNFVFP | 1 | 0 | 0 | 1670664 | QIQ54046.1 | P0C6U8 |
| KKFDNPVLPXNDGVYF | 1 | 0 | 0 | 1670665 | QIQ54048.1 | P59594 |
| KKFDTFNGECPNFVFP | 1 | 0 | 0 | 1670667 | YP_009724389.1 | P0DTD1 |
| KKGCFTGVDGELPVAV | 1 | 0 | 0 | 1670676 | YP_003766.2 | H9EJ42 |
| KKGVQIPCTCGKQATK | 1 | 0 | 0 | 1670682 | YP_009724389.1 | P0DTD1 |
| KKIDVLLTVDGVNFKS | 1 | 0 | 0 | 1670686 | YP_173236.1 | A0A140H1G9 |
| KKIKACVEEVTTTLEE | 1 | 0 | 0 | 1670689 | YP_009724389.1 | P0DTD1 |
| KKIKHEGDWDSFCKTI | 1 | 0 | 0 | 1670690 | NP_073549.1 | A0A0P0LKV0 |
| KKKKTDESQPLPQRQK | 1 | 0 | 0 | 1670698 | QIQ54056.1 | P59595 |
| KKKTDESQPLPQRQKK | 1 | 0 | 0 | 1670702 | QIQ54056.1 | P59595 |
| KKKTEGFMGRIRSVYP | 1 | 0 | 0 | 1670703 | NP_828849.2 | P0C6X7 |
| KKLCVVPGNVVPGDVI | 1 | 0 | 0 | 1670705 | YP_003766.2 | H9EJ42 |
| KKLDNDVLNGIISNAR | 1 | 0 | 0 | 1670707 | YP_009047202.1 | K9N7C7 |
| KKLERMADLALTNMYK | 1 | 0 | 0 | 1670709 | YP_009555238.1 | P0C6X6 |
| KKLGPIFNRALVSATE | 1 | 0 | 0 | 1670711 | YP_009555238.1 | P0C6X6 |

| | | | | | | |
|---|---|---|---|---|---|---|
| KKLIGGDVTPVDQYMC | 1 | 0 | 0 | 1670712 | YP_009047202.1 | K9N7C7 |
| KKLKKSLNVAKSEFDR | 1 | 0 | 0 | 1670715 | YP_009724389.1 | P0DTD1 |
| KKLQRQLYDNCYRNSN | 1 | 0 | 0 | 1670720 | NP_073549.1 | A0A0P0LKV0 |
| KKLYNDLGNQILDSYS | 1 | 0 | 0 | 1670725 | YP_009555238.1 | P0C6X6 |
| KKLYNDLGTQILDSYS | 1 | 0 | 0 | 1670726 | YP_173236.1 | A0A140H1G9 |
| KKMAQPSGCVERCVVR | 1 | 0 | 0 | 1670727 | YP_003766.2 | H9EJ42 |
| KKNAWNIGEPKSILSP | 1 | 0 | 0 | 1670729 | QIQ54046.1 | P0C6U8 |
| KKNLADAKNSGSVNQQ | 1 | 0 | 0 | 1670731 | YP_173236.1 | A0A140H1G9 |
| KKNLDEARFSGSANQQ | 1 | 0 | 0 | 1670732 | YP_009555238.1 | P0C6X6 |
| KKPNKPLSQPRADKPS | 1 | 0 | 0 | 1670744 | YP_003771.1 | H9EJ69 |
| KKPRQKRTATKAYNVT | 1 | 0 | 0 | 1670745 | YP_009724397.2 | P0DTC9 |
| KKQFLPFQQFGRDISD | 1 | 0 | 0 | 1670751 | QIQ54048.1 | P59594 |
| KKQVVLVSNNQEDFDL | 1 | 0 | 0 | 1670756 | YP_009555238.1 | P0C6X6 |
| KKRNVDPLSNFEHKVI | 1 | 0 | 0 | 1670757 | YP_009047202.1 | K9N7C7 |
| KKSAKTGTPKPSRNQS | 1 | 0 | 0 | 1670762 | NP_073556.1 | A0A0N7J131 |
| KKSKSAYYVLPSIIPN | 1 | 0 | 0 | 1670764 | QIQ54046.1 | P0C6U8 |
| KKSKVVSALQTMLFSM | 1 | 0 | 0 | 1670765 | YP_009555238.1 | P0C6X6 |
| KKSLNVAKSEFDRDAA | 1 | 0 | 0 | 1670767 | YP_009724389.1 | P0DTD1 |
| KKSMYDGDRFVKHDLS | 1 | 0 | 0 | 1670769 | NP_073549.1 | A0A0P0LKV0 |
| KKSNLLIQEFLQYDSS | 1 | 0 | 0 | 1670770 | YP_173236.1 | A0A140H1G9 |
| KKTDESQPLPQRQKKQ | 1 | 0 | 0 | 1670774 | QIQ54056.1 | P59595 |
| KKTEGFMGRIRSVYPV | 1 | 0 | 0 | 1670775 | NP_828849.2 | P0C6X7 |
| KKTSWADQSERNYQTF | 1 | 0 | 0 | 1670776 | YP_173242.1 | Q19U29 |
| KKTYLDYIQQGGVPII | 1 | 0 | 0 | 1670780 | YP_173236.1 | A0A140H1G9 |
| KKVDGIIQQLPETYFT | 1 | 0 | 0 | 1670783 | NP_828849.2 | P0C6X7 |
| KKVDGVVQQLPETYFT | 1 | 0 | 0 | 1670784 | YP_009724389.1 | P0DTD1 |
| KKVKPTIVVNAANVYL | 1 | 0 | 0 | 1670787 | QHR63299.1 | P0C6W2 |
| KKVNGVVQQLPETYFT | 1 | 0 | 0 | 1670789 | QHR63299.1 | P0C6W2 |
| KKVNNEFIDPTECIYT | 1 | 0 | 0 | 1670790 | YP_009047202.1 | K9N7C7 |
| KKVPVDEYITTYPGQG | 1 | 0 | 0 | 1670791 | NP_828849.2 | P0C6X7 |
| KKYPCMVGPDYAYFNG | 1 | 0 | 0 | 1670800 | YP_009047202.1 | K9N7C7 |
| KKYTYSFLPKEDGDVL | 1 | 0 | 0 | 1670802 | YP_009047202.1 | K9N7C7 |
| KLADCLQAYGHYSLQD | 1 | 0 | 0 | 1670804 | YP_173236.1 | A0A140H1G9 |
| KLAGGKISFSDDVIVH | 1 | 0 | 0 | 1670807 | YP_003766.2 | H9EJ42 |
| KLAGTAVINLRADQIN | 1 | 0 | 0 | 1670808 | YP_009555238.1 | P0C6X6 |
| KLAGTAVVNLRPDQLN | 1 | 0 | 0 | 1670809 | YP_173236.1 | A0A140H1G9 |
| KLAKKFDNFTGECPNF | 1 | 0 | 0 | 1670812 | QIQ54046.1 | P0C6U8 |
| KLANDAKESLRLFAAE | 1 | 0 | 0 | 1670818 | NP_073549.1 | A0A0P0LKV0 |
| KLANDVKDTLRLFAAE | 1 | 0 | 0 | 1670819 | YP_003766.2 | H9EJ42 |
| KLASELSNTFGAISAS | 1 | 0 | 0 | 1670822 | YP_009047204.1 | K9N5Q8 |
| KLCVVPGNVKPGDAVI | 1 | 0 | 0 | 1670835 | NP_073549.1 | A0A0P0LKV0 |
| KLDGFNHQFVTASSAT | 1 | 0 | 0 | 1670840 | YP_003766.2 | H9EJ42 |
| KLDGVTYTEIEPKLDG | 1 | 0 | 0 | 1670841 | NP_828849.2 | P0C6X7 |
| KLDGYYKKDNAYFTEQ | 1 | 0 | 0 | 1670844 | QIQ54046.1 | P0C6U8 |
| KLDGYYKKDNAYYTEQ | 1 | 0 | 0 | 1670845 | NP_828849.2 | P0C6X7 |
| KLDNDALNNIINNARD | 1 | 0 | 0 | 1670850 | YP_009724389.1 | P0DTD1 |
| KLDNDVLNGIISNARN | 1 | 0 | 0 | 1670851 | YP_009047202.1 | K9N7C7 |
| KLDNQALNSILDNAVK | 1 | 0 | 0 | 1670853 | YP_009555238.1 | P0C6X6 |
| KLDNYYKKDNSYFTEQ | 1 | 0 | 0 | 1670854 | YP_009724389.1 | P0DTD1 |
| KLDTGAQKFFQFGDFV | 1 | 0 | 0 | 1670857 | YP_003766.2 | H9EJ42 |
| KLEFANDTKIASQLGN | 1 | 0 | 0 | 1670862 | YP_009047204.1 | K9N5Q8 |
| KLEKLISTARDGVRRG | 1 | 0 | 0 | 1670865 | YP_009047202.1 | K9N7C7 |
| KLERMADLALTNMYKE | 1 | 0 | 0 | 1670869 | YP_009555238.1 | P0C6X6 |
| KLERMADQAMTSMYKQ | 1 | 0 | 0 | 1670870 | YP_009047202.1 | K9N7C7 |
| KLFITKDEAIKRVRGW | 1 | 0 | 0 | 1670882 | YP_173236.1 | A0A140H1G9 |
| KLFLSGDDCFVSNSSV | 1 | 0 | 0 | 1670886 | YP_173236.1 | A0A140H1G9 |
| KLFVESCTIYMCVVPS | 1 | 0 | 0 | 1670894 | YP_003766.2 | H9EJ42 |
| KLGAEKKDGFYETIVS | 1 | 0 | 0 | 1670896 | YP_009555239.1 | Q80872 |
| KLGASQRVAGDSGFAA | 1 | 0 | 0 | 1670897 | YP_009724393.1 | P0DTC5 |
| KLGASQRVGTDSGFAA | 1 | 0 | 0 | 1670898 | NP_828855.1 | P59596 |
| KLGFDSSKEFVEYKIT | 1 | 0 | 0 | 1670900 | YP_009555238.1 | P0C6X6 |
| KLGPIFNRALVSATEF | 1 | 0 | 0 | 1670905 | YP_009555238.1 | P0C6X6 |
| KLGTGYSLADLPAYMT | 1 | 0 | 0 | 1670910 | YP_009555244.1 | Q01455 |
| KLHDELTGHMLDMYSV | 1 | 0 | 0 | 1670918 | YP_009724389.1 | P0DTD1 |
| KLHPAFNVSDAYANLV | 1 | 0 | 0 | 1670921 | YP_003766.2 | H9EJ42 |
| KLHPSFNVSDAYANLV | 1 | 0 | 0 | 1670922 | NP_073549.1 | A0A0P0LKV0 |
| KLIANQFNSAIGKIQD | 1 | 0 | 0 | 1670925 | YP_009724390.1 | P0DTC2 |
| KLISTARDGVRRGDNF | 1 | 0 | 0 | 1670942 | YP_009047202.1 | K9N7C7 |
| KLKAHKDKSAQCFKMF | 1 | 0 | 0 | 1670946 | YP_009724389.1 | P0DTD1 |
| KLKAKNDNSSLCFKVY | 1 | 0 | 0 | 1670947 | YP_173236.1 | A0A140H1G9 |
| KLKAKNESSSLCFKVY | 1 | 0 | 0 | 1670948 | YP_009555238.1 | P0C6X6 |
| KLKCNRCCDRYEEYDL | 1 | 0 | 0 | 1670952 | YP_009047204.1 | K9N5Q8 |
| KLKDGVNFNVDDINFS | 1 | 0 | 0 | 1670956 | YP_009555241.1 | P36334 |
| KLKEDYQIGGYSEDRH | 1 | 0 | 0 | 1670957 | NP_828852.2 | P59632 |
| KLKIQVVNSGSDMNCN | 1 | 0 | 0 | 1670965 | YP_173236.1 | A0A140H1G9 |
| KLKISTQLFEGDKFIG | 1 | 0 | 0 | 1670966 | YP_003766.2 | H9EJ42 |
| KLKPFERDISTEIYQA | 1 | 0 | 0 | 1670974 | QIQ54048.1 | P59594 |
| KLKPILDWLEEKFKEG | 1 | 0 | 0 | 1670975 | QIQ54046.1 | P0C6U8 |
| KLKPMPFFYFDDSDCD | 1 | 0 | 0 | 1670976 | YP_003766.2 | H9EJ42 |
| KLKPVLDWLEEKFKEG | 1 | 0 | 0 | 1670977 | YP_009724389.1 | P0DTD1 |
| KLKVSTNLFEGDKFVG | 1 | 0 | 0 | 1670988 | NP_073549.1 | A0A0P0LKV0 |
| KLLHKPIVWHVNNTIN | 1 | 0 | 0 | 1671006 | QIQ54046.1 | P0C6U8 |
| KLLHWSDQTICYKDSV | 1 | 0 | 0 | 1671008 | YP_009047202.1 | K9N7C7 |
| KLLIRDTNKEVFVGDS | 1 | 0 | 0 | 1671010 | YP_009555238.1 | P0C6X6 |
| KLLKSIAATRGATVVI | 1 | 0 | 0 | 1671012 | YP_009724389.1 | P0DTD1 |
| KLMGWDYPKCDRAMPS | 1 | 0 | 0 | 1671032 | NP_073549.1 | A0A0P0LKV0 |
| KLMVVVPDYGTYKNTC | 1 | 0 | 0 | 1671041 | NP_828849.2 | P0C6X7 |
| KLMVVVPDYNTYKNTC | 1 | 0 | 0 | 1671042 | QIQ54046.1 | P0C6U8 |
| KLNFGMLNGNAIATVK | 1 | 0 | 0 | 1671050 | NP_073549.1 | A0A0P0LKV0 |
| KLNGILQQAALDYLAT | 1 | 0 | 0 | 1671051 | YP_003766.2 | H9EJ42 |
| KLNTDHSNSSDNIALL | 1 | 0 | 0 | 1671065 | QIQ54051.1 | P59596 |

| | | | | | | |
|---|---|---|---|---|---|---|
| KLPCQFKDKNLQDLWV | 1 | 0 | 0 | 1671070 | YP_009555238.1 | P0C6X6 |
| KLPFKFKDLAIENMWL | 1 | 0 | 0 | 1671076 | YP_173236.1 | A0A140H1G9 |
| KLPKGVGSANIFIGDK | 1 | 0 | 0 | 1671077 | YP_009555238.1 | P0C6X6 |
| KLPLGQSLCALPDTPS | 1 | 0 | 0 | 1671080 | YP_009047204.1 | K9N5Q8 |
| KLPNYTVPDLVVEQYN | 1 | 0 | 0 | 1671082 | NP_073551.1 | A0A0P0K6L9 |
| KLPPHDVTVLRDNLKP | 1 | 0 | 0 | 1671083 | YP_003768.1 | H9EJ24 |
| KLPQFYIYDEEGGYDV | 1 | 0 | 0 | 1671084 | YP_003766.2 | H9EJ42 |
| KLPSAPQTIQPEVKEV | 1 | 0 | 0 | 1671085 | YP_009047202.1 | K9N7C7 |
| KLQASQAWQPGVAMPN | 1 | 0 | 0 | 1671091 | NP_828849.2 | P0C6X7 |
| KLQFTSLEIPRRNVAT | 1 | 0 | 0 | 1671094 | YP_009724389.1 | P0DTD1 |
| KLQFTSLEVPRRNVAT | 1 | 0 | 0 | 1671095 | QIQ54047.1 | P0C6X7 |
| KLQTLIDNINSTLVDL | 1 | 0 | 0 | 1671111 | NP_073551.1 | A0A0P0K6L9 |
| KLRAHKDKSQQCFKMF | 1 | 0 | 0 | 1671112 | QIQ54047.1 | P0C6X7 |
| KLRDYLADYDVAVTAG | 1 | 0 | 0 | 1671115 | YP_009047202.1 | K9N7C7 |
| KLRPFERDISNVPFSP | 1 | 0 | 0 | 1671120 | NP_828851.1 | P59594 |
| KLRSDVLLPLTQYNRY | 1 | 0 | 0 | 1671123 | YP_009724389.1 | P0DTD1 |
| KLSAGVEFLKDAWEIL | 1 | 0 | 0 | 1671129 | NP_828849.2 | P0C6X7 |
| KLSDNNCYLNAVIMTL | 1 | 0 | 0 | 1671134 | YP_009047202.1 | K9N7C7 |
| KLSDVGFVEAYNNCTG | 1 | 0 | 0 | 1671135 | YP_009555241.1 | P36334 |
| KLSPWFMDGENAYEVV | 1 | 0 | 0 | 1671147 | YP_009047202.1 | K9N7C7 |
| KLSRLSMIKGPQRADL | 1 | 0 | 0 | 1671148 | YP_173236.1 | A0A140H1G9 |
| KLSTDLIKNQCVNFNF | 1 | 0 | 0 | 1671149 | NP_828851.1 | P59594 |
| KLTCATTRQVVNVVTT | 1 | 0 | 0 | 1671158 | YP_009724389.1 | P0DTD1 |
| KLTCSNTKFADDLNQM | 1 | 0 | 0 | 1671160 | NP_828849.2 | P0C6X7 |
| KLTDLKCTNVVLMGIL | 1 | 0 | 0 | 1671162 | NP_073549.1 | A0A0P0LKV0 |
| KLTDNVYIKNADIVEE | 1 | 0 | 0 | 1671166 | YP_009724389.1 | P0DTD1 |
| KLTDVKCANVVLLNCL | 1 | 0 | 0 | 1671167 | YP_009555238.1 | P0C6X6 |
| KLTFNKQEASVPILTT | 1 | 0 | 0 | 1671169 | YP_173236.1 | A0A140H1G9 |
| KLVKDPPNVQIHTIDG | 1 | 0 | 0 | 1671190 | NP_828852.2 | P59632 |
| KLVSMKGTGVFVQDPA | 1 | 0 | 0 | 1671202 | YP_003766.2 | H9EJ42 |
| KLVTEEVVETPTIQKD | 1 | 0 | 0 | 1671204 | QIQ54046.1 | P0C6U8 |
| KLYCAFTAPEDDDFLE | 1 | 0 | 0 | 1671213 | YP_009555238.1 | P0C6X6 |
| KLYLGGMSYYCEDHKP | 1 | 0 | 0 | 1671222 | YP_009555238.1 | P0C6X6 |
| KMAFPSGKVEGCMVQV | 1 | 0 | 0 | 1671230 | YP_009724389.1 | P0DTD1 |
| KMDGDDVYLPYPNPSR | 1 | 0 | 0 | 1671239 | YP_009555238.1 | P0C6X6 |
| KMDTAAYREAACSQLA | 1 | 0 | 0 | 1671240 | YP_009555238.1 | P0C6X6 |
| KMFDSFVNSFVSLYNV | 1 | 0 | 0 | 1671243 | YP_009047202.1 | K9N7C7 |
| KMFVRPVIEDYHTLTA | 1 | 0 | 0 | 1671245 | YP_173241.1 | Q0ZJ82 |
| KMGLTPPLSILKNLGV | 1 | 0 | 0 | 1671248 | YP_003766.2 | H9EJ42 |
| KMGVLKADDFVTASDT | 1 | 0 | 0 | 1671251 | YP_003766.2 | H9EJ42 |
| KMLKSMAATRGATCVI | 1 | 0 | 0 | 1671266 | YP_009047202.1 | K9N7C7 |
| KMLVAKDNKNLPKFIE | 1 | 0 | 0 | 1671271 | YP_003771.1 | H9EJ69 |
| KMNCVRFKNADLKDGY | 1 | 0 | 0 | 1671275 | YP_003766.2 | H9EJ42 |
| KMNYQVNGYPNMFITR | 1 | 0 | 0 | 1671276 | YP_009724389.1 | P0DTD1 |
| KMQRMLLDKCDLQNYG | 1 | 0 | 0 | 1671279 | QIQ54047.1 | P0C6X7 |
| KMSEYKGPVTDVFYKE | 1 | 0 | 0 | 1671285 | NP_828849.2 | P0C6X7 |
| KMTSLPKDWRLINKFD | 1 | 0 | 0 | 1671290 | YP_173236.1 | A0A140H1G9 |
| KMVKRQSYGTNSGVAI | 1 | 0 | 0 | 1671291 | YP_009047210.1 | K9N7A1 |
| KMVNWFNTMLDASAPA | 1 | 0 | 0 | 1671294 | YP_173236.1 | A0A140H1G9 |
| KNASLYTSLVPHTRYS | 1 | 0 | 0 | 1671305 | YP_173236.1 | A0A140H1G9 |
| KNAYEKDKAVARKLER | 1 | 0 | 0 | 1671308 | YP_009047202.1 | K9N7C7 |
| KNCLHVVGPNVNRGED | 1 | 0 | 0 | 1671310 | QHR63299.1 | P0C6W2 |
| KNFHIEGTGGNSQSSS | 1 | 0 | 0 | 1671322 | YP_009047211.1 | K9N4V7 |
| KNFTTAPAICHDGKAH | 1 | 0 | 0 | 1671327 | YP_009724390.1 | P0DTC2 |
| KNFTTAPAICHEGKAH | 1 | 0 | 0 | 1671328 | QIQ54048.1 | P59594 |
| KNGFQGRQIMGSCSFE | 1 | 0 | 0 | 1671334 | YP_009555238.1 | P0C6X6 |
| KNGKTVLGEYVFDKSE | 1 | 0 | 0 | 1671335 | YP_009555238.1 | P0C6X6 |
| KNGNKGAGGHSFGIDL | 1 | 0 | 0 | 1671336 | QIQ54046.1 | P0C6U8 |
| KNGNKGAGGHSYGADL | 1 | 0 | 0 | 1671337 | YP_009724389.1 | P0DTD1 |
| KNGNKGAGGHSYGIDL | 1 | 0 | 0 | 1671338 | NP_828849.2 | P0C6X7 |
| KNGQFQDHYDGFYTQG | 1 | 0 | 0 | 1671339 | YP_003766.2 | H9EJ42 |
| KNGTCGLVELEKGVLP | 1 | 0 | 0 | 1671343 | NP_828849.2 | P0C6X7 |
| KNIGDYNVSKDGFFTY | 1 | 0 | 0 | 1671348 | YP_173236.1 | A0A140H1G9 |
| KNIGEYNVSKDGFFTY | 1 | 0 | 0 | 1671349 | YP_009555238.1 | P0C6X6 |
| KNLDEARFSGSANQQQ | 1 | 0 | 0 | 1671359 | YP_009555238.1 | P0C6X6 |
| KNLGFDNQSKSPSSSG | 1 | 0 | 0 | 1671360 | YP_003771.1 | H9EJ69 |
| KNLKSNCVRFKNVDKD | 1 | 0 | 0 | 1671365 | NP_073549.1 | A0A0P0LKV0 |
| KNLSKPLTENKYSQLD | 1 | 0 | 0 | 1671383 | QIQ54052.1 | P59634 |
| KNLSKSLTENKYSQLD | 1 | 0 | 0 | 1671384 | YP_009724394.1 | P0DTC6 |
| KNLTPIKLNFGMLNGM | 1 | 0 | 0 | 1671385 | YP_003766.2 | H9EJ42 |
| KNMCTGSPSIVEFNRL | 1 | 0 | 0 | 1671389 | YP_009047202.1 | K9N7C7 |
| KNMWEFRDYFNNNTDS | 1 | 0 | 0 | 1671390 | YP_003766.2 | H9EJ42 |
| KNMYLKSAVMQSVGAC | 1 | 0 | 0 | 1671391 | YP_173236.1 | A0A140H1G9 |
| KNNTSFPTNIAVELFT | 1 | 0 | 0 | 1671398 | YP_173236.1 | A0A140H1G9 |
| KNPINSNVKTLEYDYM | 1 | 0 | 0 | 1671399 | NP_073549.1 | A0A0P0LKV0 |
| KNQCVNFNFNGLTGTG | 1 | 0 | 0 | 1671404 | NP_828851.1 | P59594 |
| KNQKLIATAFNNALLS | 1 | 0 | 0 | 1671405 | YP_173238.1 | A0A140H1H1 |
| KNSATGSLDVEVFNRL | 1 | 0 | 0 | 1671412 | YP_003766.2 | H9EJ42 |
| KNSIDAFKLNIKLLGV | 1 | 0 | 0 | 1671416 | YP_009724389.1 | P0DTD1 |
| KNSIDAFKLNVKLLGI | 1 | 0 | 0 | 1671417 | QIQ54046.1 | P0C6U8 |
| KNSSVNTVCFVDVDFH | 1 | 0 | 0 | 1671422 | YP_003766.2 | H9EJ42 |
| KNSVSDVAYNRYLSLY | 1 | 0 | 0 | 1671427 | YP_173236.1 | A0A140H1G9 |
| KNTCVGSDNVTDFNAI | 1 | 0 | 0 | 1671431 | YP_009724389.1 | P0DTD1 |
| KNTKLYVNKTLYSEFS | 1 | 0 | 0 | 1671433 | YP_173238.1 | A0A140H1H1 |
| KNVDIVSEAKNVKPTI | 1 | 0 | 0 | 1671441 | QIQ54046.1 | P0C6U8 |
| KNVDWSITYPMIANEN | 1 | 0 | 0 | 1671443 | NP_073549.1 | A0A0P0LKV0 |
| KNVDWTVTYPFIANEK | 1 | 0 | 0 | 1671444 | YP_003766.2 | H9EJ42 |
| KNVHLKDVTKENQEIL | 1 | 0 | 0 | 1671445 | NP_073549.1 | A0A0P0LKV0 |
| KNVTGSNESKALFFTY | 1 | 0 | 0 | 1671456 | YP_009047202.1 | K9N7C7 |
| KNVTKENDSKEGFFTY | 1 | 0 | 0 | 1671457 | YP_009724389.1 | P0DTD1 |
| KNWFDPIENEDIHRVY | 1 | 0 | 0 | 1671460 | NP_073549.1 | A0A0P0LKV0 |

| | | | | | | |
|---|---|---|---|---|---|---|
| KNYFITDAQTGSSKCV | 1 | 0 | 0 | 1671463 | YP_009724389.1 | P0DTD1 |
| KPCIKVATVQSKMSDV | 1 | 0 | 0 | 1671479 | YP_009724389.1 | P0DTD1 |
| KPCTPPALNCYWPLND | 1 | 0 | 0 | 1671481 | NP_828851.1 | P59594 |
| KPDLSQYYCDGGKYYT | 1 | 0 | 0 | 1671483 | YP_009555238.1 | P0C6X6 |
| KPEEKLSAYDLACCMR | 1 | 0 | 0 | 1671488 | NP_073549.1 | A0A0P0LKV0 |
| KPFARAAFEHLKPMPF | 1 | 0 | 0 | 1671493 | YP_009555238.1 | P0C6X6 |
| KPFITESKPSVEQRKQ | 1 | 0 | 0 | 1671496 | YP_009724389.1 | P0DTD1 |
| KPFKVEDSVIVNDDTS | 1 | 0 | 0 | 1671497 | YP_009555238.1 | P0C6X6 |
| KPFLGQAAITTSNCAK | 1 | 0 | 0 | 1671498 | NP_828849.2 | P0C6X7 |
| KPFLNKVTSVAASGVA | 1 | 0 | 0 | 1671499 | QIQ54046.1 | P0C6U8 |
| KPFLNKVVSTTTNIVT | 1 | 0 | 0 | 1671500 | YP_009724389.1 | P0DTD1 |
| KPGDAVITTQQAGAGI | 1 | 0 | 0 | 1671506 | NP_073549.1 | A0A0P0LKV0 |
| KPGDSFNILACYEGIA | 1 | 0 | 0 | 1671507 | YP_003766.2 | H9EJ42 |
| KPGGTSSGDATTAFAN | 1 | 0 | 0 | 1671510 | YP_009555238.1 | P0C6X6 |
| KPGGTSSGDATTAYAN | 1 | 0 | 0 | 1671511 | YP_009724389.1 | P0DTD1 |
| KPGHFNEEFYNFLRLR | 1 | 0 | 0 | 1671515 | YP_003766.2 | H9EJ42 |
| KPGHFNKEFYDFLRSQ | 1 | 0 | 0 | 1671516 | NP_073549.1 | A0A0P0LKV0 |
| KPGNFNKDFYDFAVSK | 1 | 0 | 0 | 1671519 | YP_009724389.1 | P0DTD1 |
| KPGNFNQDFYDFVLSK | 1 | 0 | 0 | 1671520 | YP_009555238.1 | P0C6X6 |
| KPGNFNQDFYEFVKSK | 1 | 0 | 0 | 1671521 | YP_173236.1 | A0A140H1G9 |
| KPHQVNISLNGNTSVC | 1 | 0 | 0 | 1671532 | YP_003767.1 | A0A0P0G321 |
| KPIIKAQFKPFAKVDG | 1 | 0 | 0 | 1671539 | YP_173236.1 | A0A140H1G9 |
| KPILWVNKASYDTNLN | 1 | 0 | 0 | 1671542 | YP_009047202.1 | K9N7C7 |
| KPITFHMDGEVINIDS | 1 | 0 | 0 | 1671546 | QIQ54046.1 | P0C6U8 |
| KPIVWHVNNATNKATY | 1 | 0 | 0 | 1671550 | YP_009724389.1 | P0DTD1 |
| KPIVWHVNNTINKATF | 1 | 0 | 0 | 1671551 | QIQ54046.1 | P0C6U8 |
| KPIVYKLDGVVCTEID | 1 | 0 | 0 | 1671552 | QIQ54046.1 | P0C6U8 |
| KPIYLKGSGSDFSLAD | 1 | 0 | 0 | 1671553 | YP_009555238.1 | P0C6X6 |
| KPKEETGLLMPLKAPK | 1 | 0 | 0 | 1671554 | QHR63299.1 | P0C6W2 |
| KPKEQIDGYTMHANYI | 1 | 0 | 0 | 1671555 | NP_828849.2 | P0C6X7 |
| KPKPVINQLDEKAQKF | 1 | 0 | 0 | 1671557 | NP_073549.1 | A0A0P0LKV0 |
| KPKRLRKKRNVDPLSN | 1 | 0 | 0 | 1671558 | YP_009047202.1 | K9N7C7 |
| KPKVEIDGNVMHANYL | 1 | 0 | 0 | 1671559 | YP_009555238.1 | P0C6X6 |
| KPLEPKFSIALPPELS | 1 | 0 | 0 | 1671563 | YP_003771.1 | H9EJ69 |
| KPLFAFNVNVMGTRPE | 1 | 0 | 0 | 1671564 | NP_073549.1 | A0A0P0LKV0 |
| KPLIEAEKPNLNSLCL | 1 | 0 | 0 | 1671566 | YP_173243.1 | Q0ZJ80 |
| KPLIEWPLTEYDFTDY | 1 | 0 | 0 | 1671567 | YP_009047202.1 | K9N7C7 |
| KPLLENIDYFNMRRAK | 1 | 0 | 0 | 1671571 | YP_009555238.1 | P0C6X6 |
| KPLQVFVYSSNEEQAV | 1 | 0 | 0 | 1671575 | YP_003766.2 | H9EJ42 |
| KPLTENKYSQLDEEQP | 1 | 0 | 0 | 1671578 | QIQ54052.1 | P59634 |
| KPNRTSVVSFNEWALA | 1 | 0 | 0 | 1671601 | YP_009047202.1 | K9N7C7 |
| KPPSASCPIGTNYRSC | 1 | 0 | 0 | 1671614 | YP_173238.1 | A0A140H1H1 |
| KPQEKDKKSAKTGTPK | 1 | 0 | 0 | 1671615 | NP_073556.1 | A0A0N7J131 |
| KPQRKRGVKQLPEQFD | 1 | 0 | 0 | 1671620 | YP_173242.1 | Q19U29 |
| KPQRQRGHKNGQGEND | 1 | 0 | 0 | 1671621 | YP_009555245.1 | P33469 |
| KPQSGGGKCFNCYPAG | 1 | 0 | 0 | 1671622 | NP_073551.1 | A0A0P0K6L9 |
| KPREQIDGYVMHANYI | 1 | 0 | 0 | 1671626 | YP_009724389.1 | P0DTD1 |
| KPRQKRTATKAYNVTQ | 1 | 0 | 0 | 1671630 | YP_009724397.2 | P0DTC9 |
| KPRSQMETDFLELAMD | 1 | 0 | 0 | 1671634 | NP_828849.2 | P0C6X7 |
| KPRVEAPKQEEPPNTE | 1 | 0 | 0 | 1671635 | NP_828849.2 | P0C6X7 |
| KPRWKRQPNDDVTSNV | 1 | 0 | 0 | 1671636 | NP_073556.1 | A0A0N7J131 |
| KPSSIKEMQSQSSHVA | 1 | 0 | 0 | 1671647 | YP_003771.1 | H9EJ69 |
| KPTKEEPKKVKPCIEE | 1 | 0 | 0 | 1671658 | QIQ54046.1 | P0C6U8 |
| KPVATVHRRIPDLPDC | 1 | 0 | 0 | 1671663 | YP_173238.1 | A0A140H1H1 |
| KPVDTSNSFEVLAVED | 1 | 0 | 0 | 1671667 | NP_828849.2 | P0C6X7 |
| KPVDVKPKIKACIDEV | 1 | 0 | 0 | 1671668 | NP_828849.2 | P0C6X7 |
| KPVETSNIFDVLQSED | 1 | 0 | 0 | 1671669 | QIQ54046.1 | P0C6U8 |
| KPVETSNSFDVLKSED | 1 | 0 | 0 | 1671670 | YP_009724389.1 | P0DTD1 |
| KPVMISQWPISNDSNG | 1 | 0 | 0 | 1671682 | YP_003766.2 | H9EJ42 |
| KPVPEIKILNNLGVDI | 1 | 0 | 0 | 1671684 | NP_828849.2 | P0C6X7 |
| KPVPEVKILNNLGVDI | 1 | 0 | 0 | 1671685 | YP_009724389.1 | P0DTD1 |
| KPVRARAPPGEQFTHI | 1 | 0 | 0 | 1671690 | NP_073549.1 | A0A0P0LKV0 |
| KPVSEEVVENPTIQKD | 1 | 0 | 0 | 1671691 | YP_009724389.1 | P0DTD1 |
| KPYSTLQPNAYYKYDA | 1 | 0 | 0 | 1671710 | YP_003766.2 | H9EJ42 |
| KQANNNCFVNVSCLML | 1 | 0 | 0 | 1671713 | YP_009555238.1 | P0C6X6 |
| KQARAEDKKAKIVSAM | 1 | 0 | 0 | 1671714 | YP_009047202.1 | K9N7C7 |
| KQATKYLVQQESPFVM | 1 | 0 | 0 | 1671718 | YP_009724389.1 | P0DTD1 |
| KQATQYLVQQESPFVM | 1 | 0 | 0 | 1671719 | QHR63299.1 | P0C6W2 |
| KQCTVQQCFGKRGPNQ | 1 | 0 | 0 | 1671723 | YP_009555245.1 | P33469 |
| KQDALTGGNYGYLYRL | 1 | 0 | 0 | 1671725 | QIQ54048.1 | P59594 |
| KQDDKKIKACVEEVTT | 1 | 0 | 0 | 1671726 | YP_009724389.1 | P0DTD1 |
| KQDVSLLSKCYKAMNA | 1 | 0 | 0 | 1671727 | YP_009047202.1 | K9N7C7 |
| KQEASVPILTTPFSLK | 1 | 0 | 0 | 1671728 | YP_173236.1 | A0A140H1G9 |
| KQFLPFQQFGRDISDT | 1 | 0 | 0 | 1671736 | QIQ54048.1 | P59594 |
| KQGAGDAGHSLTWLWL | 1 | 0 | 0 | 1671739 | NP_073549.1 | A0A0P0LKV0 |
| KQGAKPVGNCVKMLTN | 1 | 0 | 0 | 1671741 | NP_073549.1 | A0A0P0LKV0 |
| KQGDDYVYLPYPDPSR | 1 | 0 | 0 | 1671742 | YP_009724389.1 | P0DTD1 |
| KQGEIKDATPSDSVRA | 1 | 0 | 0 | 1671743 | QHR63301.1 | Q3LZX0 |
| KQGNFKNLREFVFKNI | 1 | 0 | 0 | 1671745 | YP_009724390.1 | P0DTC2 |
| KQGNFKNLREFVFKNV | 1 | 0 | 0 | 1671746 | QIQ54048.1 | P59594 |
| KQGQLLDTKLNGILQQ | 1 | 0 | 0 | 1671747 | YP_003766.2 | H9EJ42 |
| KQIANQFNKAISQIQE | 1 | 0 | 0 | 1671751 | NP_828851.1 | P59594 |
| KQIDDHRITSITSDKF | 1 | 0 | 0 | 1671752 | YP_009555238.1 | P0C6X6 |
| KQIRSAAKKNNLPFKL | 1 | 0 | 0 | 1671756 | YP_009724389.1 | P0DTD1 |
| KQKPIYLKGSGSDFSL | 1 | 0 | 0 | 1671764 | YP_009555238.1 | P0C6X6 |
| KQLFYVYDVHKDNFKD | 1 | 0 | 0 | 1671773 | YP_173236.1 | A0A140H1G9 |
| KQLKKAMNVAKAEFDR | 1 | 0 | 0 | 1671779 | NP_073549.1 | A0A0P0LKV0 |
| KQLKRAMNIAKSEFDH | 1 | 0 | 0 | 1671780 | YP_003766.2 | H9EJ42 |
| KQLNEISDDCNWPLVI | 1 | 0 | 0 | 1671787 | YP_009555238.1 | P0C6X6 |
| KQLSSNFGAISSVLND | 1 | 0 | 0 | 1671791 | YP_009724390.1 | P0DTC2 |
| KQNGHSLYLADNLTAD | 1 | 0 | 0 | 1671799 | YP_009047202.1 | K9N7C7 |

| | | | | | | |
|---|---|---|---|---|---|---|
| KQPFSFSFRDELGVRV | 1 | 0 | 0 | 1671801 | YP_003766.2 | H9EJ42 |
| KQPVNSNVKLLDYDYA | 1 | 0 | 0 | 1671804 | YP_003766.2 | H9EJ42 |
| KQSCTGSPYIDDFNKI | 1 | 0 | 0 | 1671809 | YP_173236.1 | A0A140H1G9 |
| KQSCTGSPYIDDFNRI | 1 | 0 | 0 | 1671810 | YP_009555238.1 | P0C6X6 |
| KQSIFTINSVVQKDGV | 1 | 0 | 0 | 1671812 | YP_009047202.1 | K9N7C7 |
| KQSSSRASSGNRSGNG | 1 | 0 | 0 | 1671815 | YP_009555245.1 | P33469 |
| KQTATSQQPSGGNVVP | 1 | 0 | 0 | 1671821 | YP_009555245.1 | P33469 |
| KQVEQKIAEIPKEEVK | 1 | 0 | 0 | 1671827 | YP_009724389.1 | P0DTD1 |
| KQVEQKTAEIPKEEVK | 1 | 0 | 0 | 1671828 | QHR63299.1 | P0C6W2 |
| KQVGCMMRLFYDRDGQ | 1 | 0 | 0 | 1671830 | YP_173236.1 | A0A140H1G9 |
| KQVGCSMRLFYDRDGQ | 1 | 0 | 0 | 1671831 | YP_009555238.1 | P0C6X6 |
| KQVLDLLNKGMQLLHT | 1 | 0 | 0 | 1671834 | YP_009047202.1 | K9N7C7 |
| KQVVLVSNNQEDFDLI | 1 | 0 | 0 | 1671835 | YP_009555238.1 | P0C6X6 |
| KQVVSDIDYVPLKSAT | 1 | 0 | 0 | 1671836 | YP_009724389.1 | P0DTD1 |
| KQYGDCLGDIAARDLI | 1 | 0 | 0 | 1671838 | YP_009724390.1 | P0DTC2 |
| KQYGECLGDINARDLI | 1 | 0 | 0 | 1671839 | NP_828851.1 | P59594 |
| KQYNVTQAFGRRGPEQ | 1 | 0 | 0 | 1671840 | QHR63308.1 | Q3LZX4 |
| KRAFLDKLDVNSGFAV | 1 | 0 | 0 | 1671847 | YP_173241.1 | Q0ZJ82 |
| KRALGLSISDHEFTSA | 1 | 0 | 0 | 1671850 | NP_073549.1 | A0A0P0LKV0 |
| KRAMAKLKPAPFFYYD | 1 | 0 | 0 | 1671851 | NP_073549.1 | A0A0P0LKV0 |
| KRAMNIAKSEFDHEIS | 1 | 0 | 0 | 1671852 | YP_003766.2 | H9EJ42 |
| KRAVRSHPDFKLLHNL | 1 | 0 | 0 | 1671857 | YP_009047202.1 | K9N7C7 |
| KRCIKSVMDHEQSMYN | 1 | 0 | 0 | 1671859 | NP_073549.1 | A0A0P0LKV0 |
| KRCTKSVMEHEQSMYN | 1 | 0 | 0 | 1671860 | YP_003766.2 | H9EJ42 |
| KRDAPAHISTIGVCSM | 1 | 0 | 0 | 1671861 | YP_009724389.1 | P0DTD1 |
| KRDGVQVGYCVHGIKY | 1 | 0 | 0 | 1671862 | NP_073549.1 | A0A0P0LKV0 |
| KRDSEADSPVKDVFEL | 1 | 0 | 0 | 1671863 | YP_173242.1 | Q19U29 |
| KRFGFIEDSVFKPRPA | 1 | 0 | 0 | 1671866 | YP_009555241.1 | P36334 |
| KRFKESPFELEDFIPM | 1 | 0 | 0 | 1671869 | YP_009724389.1 | P0DTD1 |
| KRGDKSVYYTSNPTTF | 1 | 0 | 0 | 1671879 | YP_009724389.1 | P0DTD1 |
| KRGDKVVYHTTSKPIT | 1 | 0 | 0 | 1671880 | QIQ54046.1 | P0C6U8 |
| KRGEKGAYNKDHGRGG | 1 | 0 | 0 | 1671881 | YP_009555238.1 | P0C6X6 |
| KRGGGNVTYTDQYLCG | 1 | 0 | 0 | 1671883 | NP_073549.1 | A0A0P0LKV0 |
| KRGMQLYKSYSEQVIP | 1 | 0 | 0 | 1671884 | YP_173240.1 | Q0ZJ83 |
| KRGPNQNFGGGEMLKL | 1 | 0 | 0 | 1671885 | YP_009555245.1 | P33469 |
| KRGVKQLPEQFDSLNL | 1 | 0 | 0 | 1671887 | YP_173242.1 | Q19U29 |
| KRGVTVPCVCGRQATQ | 1 | 0 | 0 | 1671888 | QIQ54046.1 | P0C6U8 |
| KRGVYCCRDHEHEIAW | 1 | 0 | 0 | 1671889 | NP_828849.2 | P0C6X7 |
| KRIVQMIADFLAGSSD | 1 | 0 | 0 | 1671898 | NP_073549.1 | A0A0P0LKV0 |
| KRIVYKAACVVDVNDS | 1 | 0 | 0 | 1671899 | YP_009555238.1 | P0C6X6 |
| KRKMGLTPPLSILKNL | 1 | 0 | 0 | 1671901 | YP_003766.2 | H9EJ42 |
| KRKPLDYKRQNNLAIE | 1 | 0 | 0 | 1671902 | NP_073549.1 | A0A0P0LKV0 |
| KRKVGLTPPLSILKNL | 1 | 0 | 0 | 1671903 | NP_073549.1 | A0A0P0LKV0 |
| KRLGDYVLTENALVRL | 1 | 0 | 0 | 1671905 | YP_003766.2 | H9EJ42 |
| KRLKNGFQGRQIMGSC | 1 | 0 | 0 | 1671906 | YP_009555238.1 | P0C6X6 |
| KRLRKKRNVDPLSNFE | 1 | 0 | 0 | 1671907 | YP_009047202.1 | K9N7C7 |
| KRLVFTNCNYNLTKLL | 1 | 0 | 0 | 1671908 | YP_009047204.1 | K9N5Q8 |
| KRLYSHVYRSDKVDST | 1 | 0 | 0 | 1671911 | YP_009555238.1 | P0C6X6 |
| KRNFTYDVNADYLYFH | 1 | 0 | 0 | 1671914 | YP_009555241.1 | P36334 |
| KRNIKPVPEIKILNNL | 1 | 0 | 0 | 1671915 | NP_828849.2 | P0C6X7 |
| KRNIKPVPEVKILNNL | 1 | 0 | 0 | 1671916 | YP_009724389.1 | P0DTD1 |
| KRNILPTMTQLNLKYA | 1 | 0 | 0 | 1671917 | NP_073549.1 | A0A0P0LKV0 |
| KRNQKPLEPKFSIALP | 1 | 0 | 0 | 1671918 | YP_003771.1 | H9EJ69 |
| KRNRSLRVKCSTIVGG | 1 | 0 | 0 | 1671921 | YP_009555238.1 | P0C6X6 |
| KRNVDPLSNFEHKVIT | 1 | 0 | 0 | 1671922 | YP_009047202.1 | K9N7C7 |
| KRNVIPTITQMNLKYA | 1 | 0 | 0 | 1671923 | YP_009724389.1 | P0DTD1 |
| KRNVIPTMTQMNLKYA | 1 | 0 | 0 | 1671924 | YP_009047202.1 | K9N7C7 |
| KRNVLPTLTQMNLKYA | 1 | 0 | 0 | 1671925 | YP_009555238.1 | P0C6X6 |
| KRNVLPTMTQLNLKYA | 1 | 0 | 0 | 1671926 | YP_003766.2 | H9EJ42 |
| KRPINPTDQSSYIVDS | 1 | 0 | 0 | 1671927 | YP_009724389.1 | P0DTD1 |
| KRPINPTDQSSYVVDS | 1 | 0 | 0 | 1671928 | QIQ54046.1 | P0C6U8 |
| KRPIQPTDVAYHTVTD | 1 | 0 | 0 | 1671930 | YP_009555238.1 | P0C6X6 |
| KRPVNPTDASHYVVTD | 1 | 0 | 0 | 1671931 | YP_173236.1 | A0A140H1G9 |
| KRQIRIACRKCNLAFR | 1 | 0 | 0 | 1671932 | YP_009047202.1 | K9N7C7 |
| KRQNNLAIEEIEYVHG | 1 | 0 | 0 | 1671933 | NP_073549.1 | A0A0P0LKV0 |
| KRQPNDDVTSNVTQCF | 1 | 0 | 0 | 1671934 | NP_073556.1 | A0A0N7J131 |
| KRQSYGTNSGVAIYHR | 1 | 0 | 0 | 1671935 | YP_009047210.1 | K9N7A1 |
| KRQYLLKFDNRGVITN | 1 | 0 | 0 | 1671936 | YP_173238.1 | A0A140H1H1 |
| KRRDSTYGPLQTPVGC | 1 | 0 | 0 | 1671938 | YP_009047204.1 | K9N5Q8 |
| KRRFSLLRHEDLSVIV | 1 | 0 | 0 | 1671939 | YP_009047207.1 | K9N643 |
| KRRSISASYRFVTFEP | 1 | 0 | 0 | 1671941 | YP_173238.1 | A0A140H1H1 |
| KRRVVFNGVSFSTFEE | 1 | 0 | 0 | 1671942 | YP_009724389.1 | P0DTD1 |
| KRSDALSTNHGHKVVE | 1 | 0 | 0 | 1671943 | NP_828849.2 | P0C6X7 |
| KRSIRHHPELKILRNL | 1 | 0 | 0 | 1671948 | YP_173236.1 | A0A140H1G9 |
| KRSKDSPLELEDFIPM | 1 | 0 | 0 | 1671949 | QIQ54047.1 | P0C6X7 |
| KRSKESPLELEDFIPM | 1 | 0 | 0 | 1671950 | QHR63299.1 | P0C6W2 |
| KRSVRHHPELKLFRNL | 1 | 0 | 0 | 1671955 | YP_009555238.1 | P0C6X6 |
| KRTDLTIYNREMKCYE | 1 | 0 | 0 | 1671958 | YP_009555238.1 | P0C6X6 |
| KRTNLEVYNKEKTYYE | 1 | 0 | 0 | 1671962 | YP_173236.1 | A0A140H1G9 |
| KRTVCFSVAALSTGLT | 1 | 0 | 0 | 1671968 | NP_073549.1 | A0A0P0LKV0 |
| KRTVKGTHHWLLLTIL | 1 | 0 | 0 | 1671969 | QIQ54046.1 | P0C6U8 |
| KRVDWSVEYPIIGDEL | 1 | 0 | 0 | 1671973 | NP_828849.2 | P0C6X7 |
| KRVDWTIEYPIIGDEL | 1 | 0 | 0 | 1671974 | YP_009724389.1 | P0DTD1 |
| KRVLGLQTQTVDSAQG | 1 | 0 | 0 | 1671976 | YP_009555238.1 | P0C6X6 |
| KRVLGVQTQTVDSAQG | 1 | 0 | 0 | 1671977 | YP_173236.1 | A0A140H1G9 |
| KRVLNVVCKTCGQQQT | 1 | 0 | 0 | 1671979 | YP_009724389.1 | P0DTD1 |
| KRVPTREENVIQCFGP | 1 | 0 | 0 | 1671982 | YP_003771.1 | H9EJ69 |
| KRVRAWVGFDAEGAHA | 1 | 0 | 0 | 1671983 | YP_009555238.1 | P0C6X6 |
| KRVRGWVGFDVEGAHA | 1 | 0 | 0 | 1671984 | YP_173236.1 | A0A140H1G9 |
| KRWLGILLEAYNAFLD | 1 | 0 | 0 | 1671985 | NP_073549.1 | A0A0P0LKV0 |
| KRWLPHDAIVVDNDVV | 1 | 0 | 0 | 1671986 | NP_073549.1 | A0A0P0LKV0 |

| | | | | | | |
|---|---|---|---|---|---|---|
| KRWLPPDAIIIDNDIN | 1 | 0 | 0 | 1671987 | YP_003766.2 | H9EJ42 |
| KRYCTVQGPPGTGKSH | 1 | 0 | 0 | 1671992 | YP_009555238.1 | P0C6X6 |
| KRYVVHHLSMTTSTTN | 1 | 0 | 0 | 1671996 | YP_173236.1 | A0A140H1G9 |
| KSAELNYTVQKLQTLI | 1 | 0 | 0 | 1671999 | NP_073551.1 | A0A0P0K6L9 |
| KSAFTNLKQLPFFYYS | 1 | 0 | 0 | 1672000 | NP_828849.2 | P0C6X7 |
| KSAFVNLKQLPFFYYS | 1 | 0 | 0 | 1672001 | YP_009724389.1 | P0DTD1 |
| KSAFYILPSIISNEKQ | 1 | 0 | 0 | 1672002 | YP_009724389.1 | P0DTD1 |
| KSAGWPLNKFGKAGLY | 1 | 0 | 0 | 1672006 | NP_073549.1 | A0A0P0LKV0 |
| KSAKKFDTFKGECPKF | 1 | 0 | 0 | 1672010 | NP_828849.2 | P0C6X7 |
| KSAKTGTPKPSRNQSP | 1 | 0 | 0 | 1672011 | NP_073556.1 | A0A0N7J131 |
| KSALLADKFPVLHDIG | 1 | 0 | 0 | 1672012 | NP_828849.2 | P0C6X7 |
| KSARLKRFPVNTIVNG | 1 | 0 | 0 | 1672015 | NP_073549.1 | A0A0P0LKV0 |
| KSAYENFNQHDVLLAP | 1 | 0 | 0 | 1672021 | QHR63299.1 | P0C6W2 |
| KSAYENFNQHEVLLAP | 1 | 0 | 0 | 1672022 | YP_009724389.1 | P0DTD1 |
| KSAYERDRAVAKKLER | 1 | 0 | 0 | 1672023 | YP_009555238.1 | P0C6X6 |
| KSCLGYHPAHAPSFLA | 1 | 0 | 0 | 1672026 | YP_173236.1 | A0A140H1G9 |
| KSCTKGLSIADLACAQ | 1 | 0 | 0 | 1672029 | YP_003767.1 | A0A0P0G321 |
| KSDEGGLCPSTGQAME | 1 | 0 | 0 | 1672030 | YP_173236.1 | A0A140H1G9 |
| KSDIFGQDFKTFDLLK | 1 | 0 | 0 | 1672033 | NP_073549.1 | A0A0P0LKV0 |
| KSDIFGSDFKTFDLLK | 1 | 0 | 0 | 1672034 | YP_003766.2 | H9EJ42 |
| KSDIHLPAYASFKVID | 1 | 0 | 0 | 1672035 | YP_173236.1 | A0A140H1G9 |
| KSDILSLNNPIFINYS | 1 | 0 | 0 | 1672036 | YP_173237.1 | A0A140H1H0 |
| KSEDAQGMDNLACEDL | 1 | 0 | 0 | 1672042 | YP_009724389.1 | P0DTD1 |
| KSEDTQGMDNLACEDL | 1 | 0 | 0 | 1672044 | QHR63299.1 | P0C6W2 |
| KSENRYSVLSVDSVSE | 1 | 0 | 0 | 1672050 | YP_173236.1 | A0A140H1G9 |
| KSETKVQDHFDVVETK | 1 | 0 | 0 | 1672052 | QIQ54046.1 | P0C6U8 |
| KSFEIDKGIYQTSNFR | 1 | 0 | 0 | 1672054 | NP_828851.1 | P59594 |
| KSFEQQVGVIADKDKD | 1 | 0 | 0 | 1672055 | NP_073549.1 | A0A0P0LKV0 |
| KSFKQIVESCGNFKVT | 1 | 0 | 0 | 1672059 | QIQ54046.1 | P0C6U8 |
| KSFKTIVESCGNYKVT | 1 | 0 | 0 | 1672060 | NP_828849.2 | P0C6X7 |
| KSFLPQFTSDQAVTFL | 1 | 0 | 0 | 1672061 | YP_173241.1 | Q0ZJ82 |
| KSFSTFESAYMPIADP | 1 | 0 | 0 | 1672065 | NP_073549.1 | A0A0P0LKV0 |
| KSGCKLTNAKPPKYSS | 1 | 0 | 0 | 1672070 | YP_003766.2 | H9EJ42 |
| KSGSLVLNNPAYIAPQ | 1 | 0 | 0 | 1672076 | YP_009555240.1 | P30215 |
| KSGSPGDSSWHIYLKS | 1 | 0 | 0 | 1672077 | YP_003767.1 | A0A0P0G321 |
| KSGTAILIKSPLHSLG | 1 | 0 | 0 | 1672078 | YP_173236.1 | A0A140H1G9 |
| KSGVNDICMFENKTTL | 1 | 0 | 0 | 1672081 | YP_009047202.1 | K9N7C7 |
| KSGYFVNVNNTWMYTG | 1 | 0 | 0 | 1672082 | YP_009555241.1 | P36334 |
| KSHKPPSASCPIGTNY | 1 | 0 | 0 | 1672089 | YP_173238.1 | A0A140H1H1 |
| KSIAATRGATVVIGTS | 1 | 0 | 0 | 1672095 | YP_009724389.1 | P0DTD1 |
| KSIIYTGDWEGLHEVL | 1 | 0 | 0 | 1672099 | YP_003766.2 | H9EJ42 |
| KSISLTVGEVFGKILG | 1 | 0 | 0 | 1672104 | YP_173236.1 | A0A140H1G9 |
| KSKELLDVSLDSLGAA | 1 | 0 | 0 | 1672114 | YP_009555238.1 | P0C6X6 |
| KSKGLFKEGSTVDLKH | 1 | 0 | 0 | 1672115 | YP_173236.1 | A0A140H1G9 |
| KSKGLNVLWNKFVTGD | 1 | 0 | 0 | 1672116 | YP_003766.2 | H9EJ42 |
| KSKGSSRNESWHFDKS | 1 | 0 | 0 | 1672117 | YP_173238.1 | A0A140H1H1 |
| KSKGVCQVGDCYVSTG | 1 | 0 | 0 | 1672119 | YP_173236.1 | A0A140H1G9 |
| KSKSAYYVLPSIIPNK | 1 | 0 | 0 | 1672125 | QIQ54046.1 | P0C6U8 |
| KSKVGNYRLPSSKPSG | 1 | 0 | 0 | 1672126 | YP_173241.1 | Q0ZJ82 |
| KSKVGNYRLPSTQKGS | 1 | 0 | 0 | 1672127 | YP_009555244.1 | Q01455 |
| KSLEFGATSVTPQPEE | 1 | 0 | 0 | 1672136 | QHR63299.1 | P0C6W2 |
| KSLGFDKPQEKDKKSA | 1 | 0 | 0 | 1672137 | NP_073556.1 | A0A0N7J131 |
| KSLKLNLTQQGLLGTC | 1 | 0 | 0 | 1672140 | NP_073549.1 | A0A0P0LKV0 |
| KSLLPNTGVYDLSGFT | 1 | 0 | 0 | 1672141 | YP_173238.1 | A0A140H1H1 |
| KSLNVAKSEFDRDAAM | 1 | 0 | 0 | 1672147 | YP_009724389.1 | P0DTD1 |
| KSLTENKYSQLDEEQP | 1 | 0 | 0 | 1672153 | YP_009724394.1 | P0DTC6 |
| KSLTVEKGIYQTSNFR | 1 | 0 | 0 | 1672155 | QIQ54048.1 | P59594 |
| KSMAATRGATCVIGTT | 1 | 0 | 0 | 1672159 | YP_009047202.1 | K9N7C7 |
| KSMYDGDRFVKHDLSL | 1 | 0 | 0 | 1672162 | NP_073549.1 | A0A0P0LKV0 |
| KSNCVRFKNVDKDDAF | 1 | 0 | 0 | 1672163 | NP_073549.1 | A0A0P0LKV0 |
| KSNFSQYYYNIDTGSV | 1 | 0 | 0 | 1672164 | YP_173237.1 | A0A140H1H0 |
| KSNLLIQEFLQYDSSI | 1 | 0 | 0 | 1672172 | YP_173236.1 | A0A140H1G9 |
| KSPSSSGTSTPKKPNK | 1 | 0 | 0 | 1672180 | YP_003771.1 | H9EJ69 |
| KSQLVEYGSFCDNINA | 1 | 0 | 0 | 1672185 | YP_009555241.1 | P36334 |
| KSQSPRINFCGNGNHI | 1 | 0 | 0 | 1672189 | YP_173238.1 | A0A140H1H1 |
| KSQSSRINFCGNGNHI | 1 | 0 | 0 | 1672190 | YP_009555241.1 | P36334 |
| KSQSVIIINNSTNVVI | 1 | 0 | 0 | 1672191 | NP_828851.1 | P59594 |
| KSREETGLLMPLKAPK | 1 | 0 | 0 | 1672192 | YP_009724389.1 | P0DTD1 |
| KSRGDSGLLMPLKAPK | 1 | 0 | 0 | 1672194 | QIQ54046.1 | P0C6U8 |
| KSRVQQNKSRELTAED | 1 | 0 | 0 | 1672195 | YP_009555245.1 | P33469 |
| KSSALGSMDIDVFNKL | 1 | 0 | 0 | 1672196 | NP_073549.1 | A0A0P0LKV0 |
| KSSDVVDSNENLTWPL | 1 | 0 | 0 | 1672200 | YP_009047202.1 | K9N7C7 |
| KSSEYKGPITDIFYKE | 1 | 0 | 0 | 1672201 | QHR63299.1 | P0C6W2 |
| KSSEYKGPITDVFYKE | 1 | 0 | 0 | 1672202 | YP_009724389.1 | P0DTD1 |
| KSSFEIDGNVMHANYL | 1 | 0 | 0 | 1672203 | YP_173236.1 | A0A140H1G9 |
| KSSIDAFKLNIKLLGI | 1 | 0 | 0 | 1672204 | NP_828849.2 | P0C6X7 |
| KSSITESKLSVEQRQQ | 1 | 0 | 0 | 1672205 | QHR63299.1 | P0C6W2 |
| KSSTGFVYFNATGSDV | 1 | 0 | 0 | 1672209 | YP_003767.1 | A0A0P0G321 |
| KSTFNKDIYCGNSMVC | 1 | 0 | 0 | 1672217 | YP_003766.2 | H9EJ42 |
| KSTNSRIRASLPTTIA | 1 | 0 | 0 | 1672220 | NP_828849.2 | P0C6X7 |
| KSVANLKLPPHDVTVL | 1 | 0 | 0 | 1672225 | YP_003768.1 | H9EJ24 |
| KSVMDHEQSMYNLLKG | 1 | 0 | 0 | 1672239 | NP_073549.1 | A0A0P0LKV0 |
| KSVMEHEQSMYNLLNF | 1 | 0 | 0 | 1672240 | YP_003766.2 | H9EJ42 |
| KSVNWNLEYPIISNEV | 1 | 0 | 0 | 1672242 | YP_173236.1 | A0A140H1G9 |
| KSVVQKPVDVKPKIKA | 1 | 0 | 0 | 1672246 | NP_828849.2 | P0C6X7 |
| KSVYERDKAVARKLER | 1 | 0 | 0 | 1672250 | YP_173236.1 | A0A140H1G9 |
| KSWMESEFRVYSSANN | 1 | 0 | 0 | 1672253 | YP_009724390.1 | P0DTC2 |
| KSYAASYNRYKYYSGN | 1 | 0 | 0 | 1672255 | NP_073549.1 | A0A0P0LKV0 |
| KSYELQTPFEIKLAKK | 1 | 0 | 0 | 1672262 | YP_009724389.1 | P0DTD1 |
| KSYSGYHPAHAPSFLA | 1 | 0 | 0 | 1672265 | YP_009555238.1 | P0C6X6 |
| KTADGNQRQLLPRWYF | 1 | 0 | 0 | 1672269 | YP_009555245.1 | P33469 |

| | | | | | | |
|---|---|---|---|---|---|---|
| KTAGVVTGNVVKQCCT | 1 | 0 | 0 | 1672272 | YP_009047202.1 | K9N7C7 |
| KTALKKCKSAFYVLPS | 1 | 0 | 0 | 1672273 | NP_828849.2 | P0C6X7 |
| KTALKKSKSAYYVLPS | 1 | 0 | 0 | 1672274 | QIQ54046.1 | P0C6U8 |
| KTCGQQQTTLKGVEAV | 1 | 0 | 0 | 1672282 | YP_009724389.1 | P0DTD1 |
| KTCVIANVPAIDVLKK | 1 | 0 | 0 | 1672284 | YP_003766.2 | H9EJ42 |
| KTDAVILLERYVSLAI | 1 | 0 | 0 | 1672285 | NP_073549.1 | A0A0P0LKV0 |
| KTDAVVLLERYVSLAI | 1 | 0 | 0 | 1672286 | YP_003766.2 | H9EJ42 |
| KTDESQPLPQRQKKQQ | 1 | 0 | 0 | 1672288 | QIQ54056.1 | P59595 |
| KTDLFNGYKIGCNCAG | 1 | 0 | 0 | 1672291 | YP_173236.1 | A0A140H1G9 |
| KTDSVLLIERFVSLAI | 1 | 0 | 0 | 1672292 | YP_009555238.1 | P0C6X6 |
| KTEAEWIELFPHNDRI | 1 | 0 | 0 | 1672293 | NP_073549.1 | A0A0P0LKV0 |
| KTEGFMGRIRSVYPVA | 1 | 0 | 0 | 1672295 | NP_828849.2 | P0C6X7 |
| KTFDLLKYDFTEHKEN | 1 | 0 | 0 | 1672300 | YP_003766.2 | H9EJ42 |
| KTFDLLKYDFTEHKEV | 1 | 0 | 0 | 1672301 | NP_073549.1 | A0A0P0LKV0 |
| KTGDLQPLEQPTSEAV | 1 | 0 | 0 | 1672316 | YP_009724389.1 | P0DTD1 |
| KTGELQPLEEPTAQAV | 1 | 0 | 0 | 1672317 | QIQ54046.1 | P0C6U8 |
| KTGHYTGQAGEMPCAI | 1 | 0 | 0 | 1672319 | YP_009555238.1 | P0C6X6 |
| KTGTPKPSRNQSPASS | 1 | 0 | 0 | 1672323 | NP_073556.1 | A0A0N7J131 |
| KTILNIVGPDARQDGR | 1 | 0 | 0 | 1672339 | YP_173236.1 | A0A140H1G9 |
| KTKNVTKENDSKEGFF | 1 | 0 | 0 | 1672348 | YP_009724389.1 | P0DTD1 |
| KTKSLLPNTGVYDLSG | 1 | 0 | 0 | 1672351 | YP_173238.1 | A0A140H1H1 |
| KTLKPGDSFNILACYE | 1 | 0 | 0 | 1672359 | YP_003766.2 | H9EJ42 |
| KTLLYSFEAHKDSFKD | 1 | 0 | 0 | 1672362 | YP_009555238.1 | P0C6X6 |
| KTLQEFAQNLPKYVKP | 1 | 0 | 0 | 1672364 | YP_003767.1 | A0A0P0G321 |
| KTLQELSYKLPNYTVP | 1 | 0 | 0 | 1672365 | NP_073551.1 | A0A0P0K6L9 |
| KTLQEMKALFPTWSQE | 1 | 0 | 0 | 1672366 | YP_009555238.1 | P0C6X6 |
| KTLQPVSELLIPLGID | 1 | 0 | 0 | 1672368 | QHR63299.1 | P0C6W2 |
| KTLQPVSELLTPLGID | 1 | 0 | 0 | 1672369 | YP_009724389.1 | P0DTD1 |
| KTLYKDVDNPHLMGWD | 1 | 0 | 0 | 1672375 | YP_009047202.1 | K9N7C7 |
| KTNCCRFQEKDEDGNL | 1 | 0 | 0 | 1672381 | QIQ54047.1 | P0C6X7 |
| KTNCCRFQEKDEEGNL | 1 | 0 | 0 | 1672382 | NP_828849.2 | P0C6X7 |
| KTNNTRIVDEWSYTGS | 1 | 0 | 0 | 1672383 | YP_009047204.1 | K9N5Q8 |
| KTPPIKDFGGFNFLQI | 1 | 0 | 0 | 1672390 | QIQ54048.1 | P59594 |
| KTPTLKYFGGFNFSQI | 1 | 0 | 0 | 1672391 | NP_828851.1 | P59594 |
| KTPVATLIATLNSLGE | 1 | 0 | 0 | 1672392 | QIQ54046.1 | P0C6U8 |
| KTQFNYFKKVDGIIQQ | 1 | 0 | 0 | 1672393 | NP_828849.2 | P0C6X7 |
| KTQFNYYKKVDGVVQQ | 1 | 0 | 0 | 1672394 | YP_009724389.1 | P0DTD1 |
| KTQFNYYKKVNGVVQQ | 1 | 0 | 0 | 1672395 | QHR63299.1 | P0C6W2 |
| KTQIGEYTFEKGDYGD | 1 | 0 | 0 | 1672396 | QIQ54047.1 | P0C6X7 |
| KTSEYKGPVTDVFYKE | 1 | 0 | 0 | 1672407 | QIQ54046.1 | P0C6U8 |
| KTSGDLAVQIGNNNVC | 1 | 0 | 0 | 1672408 | NP_073549.1 | A0A0P0LKV0 |
| KTSWADQSERNYQTFN | 1 | 0 | 0 | 1672417 | YP_173242.1 | Q19U29 |
| KTTGDLAVQIGSNNVC | 1 | 0 | 0 | 1672424 | YP_003766.2 | H9EJ42 |
| KTTPAPVYIWTADEAI | 1 | 0 | 0 | 1672433 | YP_009555244.1 | Q01455 |
| KTTTLTGVEAVMYMGT | 1 | 0 | 0 | 1672436 | NP_828849.2 | P0C6X7 |
| KTTVASLINTLNDLNE | 1 | 0 | 0 | 1672437 | YP_009724389.1 | P0DTD1 |
| KTVGELGDVRETMNYL | 1 | 0 | 0 | 1672440 | QHR63299.1 | P0C6W2 |
| KTVKIEDAIIVNDENS | 1 | 0 | 0 | 1672441 | YP_173236.1 | A0A140H1G9 |
| KTWPRPIDVSKADGII | 1 | 0 | 0 | 1672455 | YP_009047204.1 | K9N5Q8 |
| KTWVENEFRVYSSANN | 1 | 0 | 0 | 1672456 | QIQ54048.1 | P59594 |
| KTYLDFIQQGGTPIAN | 1 | 0 | 0 | 1672461 | YP_009555238.1 | P0C6X6 |
| KTYLDYIQQGGVPIIN | 1 | 0 | 0 | 1672462 | YP_173236.1 | A0A140H1G9 |
| KVAFGGDQVHEVAAVR | 1 | 0 | 0 | 1672468 | YP_009047202.1 | K9N7C7 |
| KVATVQSKMSDVKCTS | 1 | 0 | 0 | 1672473 | YP_009724389.1 | P0DTD1 |
| KVAYENFNHHEKLLAP | 1 | 0 | 0 | 1672474 | QIQ54046.1 | P0C6U8 |
| KVDGIDVEIFENKTTL | 1 | 0 | 0 | 1672485 | NP_828849.2 | P0C6X7 |
| KVDGIIQQLPETYFTQ | 1 | 0 | 0 | 1672486 | NP_828849.2 | P0C6X7 |
| KVDGVDVELFENKTIL | 1 | 0 | 0 | 1672487 | QIQ54047.1 | P0C6X7 |
| KVDGVDVELFENKTTL | 1 | 0 | 0 | 1672488 | YP_009724389.1 | P0DTD1 |
| KVDGVVQQLPETYFTQ | 1 | 0 | 0 | 1672489 | YP_009724389.1 | P0DTD1 |
| KVDLQLGNLGYLQSFN | 1 | 0 | 0 | 1672493 | YP_009555241.1 | P36334 |
| KVDSTFVTEYYEFLNK | 1 | 0 | 0 | 1672496 | YP_009555238.1 | P0C6X6 |
| KVDTSNPKTPKYKFVR | 1 | 0 | 0 | 1672499 | NP_828849.2 | P0C6X7 |
| KVDYGSDTVTYKSTAT | 1 | 0 | 0 | 1672500 | NP_073549.1 | A0A0P0LKV0 |
| KVEFWFAMRKDGDDVI | 1 | 0 | 0 | 1672506 | YP_173236.1 | A0A140H1G9 |
| KVEISVTPIKNTVDTT | 1 | 0 | 0 | 1672509 | NP_073549.1 | A0A0P0LKV0 |
| KVEQPKIEPKPVSVIK | 1 | 0 | 0 | 1672515 | NP_073549.1 | A0A0P0LKV0 |
| KVETLNFQMVKEFPLL | 1 | 0 | 0 | 1672516 | YP_173243.1 | Q0ZJ80 |
| KVFFKGNVQVDNGSSI | 1 | 0 | 0 | 1672520 | YP_003766.2 | H9EJ42 |
| KVFFQFDNLSSEDLKA | 1 | 0 | 0 | 1672521 | YP_009555238.1 | P0C6X6 |
| KVFGYVQQGIFNKATV | 1 | 0 | 0 | 1672523 | YP_009555238.1 | P0C6X6 |
| KVFTTVDNINLHTQVV | 1 | 0 | 0 | 1672531 | YP_009724389.1 | P0DTD1 |
| KVFTTVDNTNLHTQLV | 1 | 0 | 0 | 1672532 | NP_828849.2 | P0C6X7 |
| KVGAGVMLECEKFNVF | 1 | 0 | 0 | 1672540 | YP_003766.2 | H9EJ42 |
| KVGELKVEFWFAMRKD | 1 | 0 | 0 | 1672542 | YP_173236.1 | A0A140H1G9 |
| KVGILCIMSDRDLYDK | 1 | 0 | 0 | 1672546 | YP_009724389.1 | P0DTD1 |
| KVGKVDSIVQKCYELS | 1 | 0 | 0 | 1672547 | YP_003766.2 | H9EJ42 |
| KVGMQKYSTLQGPPGT | 1 | 0 | 0 | 1672549 | YP_009724389.1 | P0DTD1 |
| KVGMQXYSTLQGPPGT | 1 | 0 | 0 | 1672550 | QIQ54047.1 | P0C6X7 |
| KVGNYRLPSSKPSGMD | 1 | 0 | 0 | 1672551 | YP_173241.1 | Q0ZJ82 |
| KVGNYRLPSTQKGSGM | 1 | 0 | 0 | 1672552 | YP_009555244.1 | Q01455 |
| KVGTGVMVECDSLRIF | 1 | 0 | 0 | 1672554 | NP_073549.1 | A0A0P0LKV0 |
| KVIQPRVEKKKTEGFM | 1 | 0 | 0 | 1672569 | NP_828849.2 | P0C6X7 |
| KVKATKGEGDGGITSE | 1 | 0 | 0 | 1672578 | NP_073549.1 | A0A0P0LKV0 |
| KVKLIFEFEDDVVTSL | 1 | 0 | 0 | 1672580 | YP_003766.2 | H9EJ42 |
| KVKLSDVGFVEAYNNC | 1 | 0 | 0 | 1672582 | YP_009555241.1 | P36334 |
| KVKVGTGVMVECDSLR | 1 | 0 | 0 | 1672594 | NP_073549.1 | A0A0P0LKV0 |
| KVKVIQPRVEKKKTEG | 1 | 0 | 0 | 1672595 | NP_828849.2 | P0C6X7 |
| KVLAEKFDYNVSVMQM | 1 | 0 | 0 | 1672599 | NP_073549.1 | A0A0P0LKV0 |
| KVLDNGIIRDVNPDDK | 1 | 0 | 0 | 1672600 | YP_009047202.1 | K9N7C7 |
| KVLEENLNAYVNSNQN | 1 | 0 | 0 | 1672604 | YP_173242.1 | Q19U29 |

| | | | | | | |
|---|---|---|---|---|---|---|
| KVLLERFLPKCPEILL | 1 | 0 | 0 | 1672607 | YP_003766.2 | H9EJ42 |
| KVLLRKNGNKGAGGHS | 1 | 0 | 0 | 1672608 | YP_009724389.1 | P0DTD1 |
| KVLNEKCSVYTVESGT | 1 | 0 | 0 | 1672612 | NP_828849.2 | P0C6X7 |
| KVLNENLNAYQQQDGM | 1 | 0 | 0 | 1672613 | YP_009555245.1 | P33469 |
| KVLPPLLSENQISGYT | 1 | 0 | 0 | 1672617 | YP_009555241.1 | P36334 |
| KVNCCRFQRIDDDGNK | 1 | 0 | 0 | 1672626 | YP_173236.1 | A0A140H1G9 |
| KVNCCRFQRVDENGDK | 1 | 0 | 0 | 1672627 | YP_009555238.1 | P0C6X6 |
| KVNECVKSQSPRINFC | 1 | 0 | 0 | 1672630 | YP_173238.1 | A0A140H1H1 |
| KVNETNSQYLFSTINA | 1 | 0 | 0 | 1672633 | YP_009047202.1 | K9N7C7 |
| KVNFNEKPVVMEIPSL | 1 | 0 | 0 | 1672634 | YP_173236.1 | A0A140H1G9 |
| KVNGVVQQLPETYFTQ | 1 | 0 | 0 | 1672636 | QHR63299.1 | P0C6W2 |
| KVNNEFIDPTECIYTQ | 1 | 0 | 0 | 1672639 | YP_009047202.1 | K9N7C7 |
| KVPVDEYITTYPGQGC | 1 | 0 | 0 | 1672648 | NP_828849.2 | P0C6X7 |
| KVQCSTNLFKDCSKSY | 1 | 0 | 0 | 1672649 | YP_009555238.1 | P0C6X6 |
| KVQDAVNNNAQALSKL | 1 | 0 | 0 | 1672650 | YP_009047204.1 | K9N5Q8 |
| KVQIGEYTFEKGDYGD | 1 | 0 | 0 | 1672655 | YP_009724389.1 | P0DTD1 |
| KVQNLSGNPDEPQKDV | 1 | 0 | 0 | 1672658 | YP_009555245.1 | P33469 |
| KVQNVNLERCELANYK | 1 | 0 | 0 | 1672659 | YP_009047202.1 | K9N7C7 |
| KVRSLKLSDNNCYLNA | 1 | 0 | 0 | 1672663 | YP_009047202.1 | K9N7C7 |
| KVSAEVSRQVIQDVKD | 1 | 0 | 0 | 1672667 | NP_073552.1 | A0A0P0KWE8 |
| KVSALKTAGVVTGNVV | 1 | 0 | 0 | 1672669 | YP_009047202.1 | K9N7C7 |
| KVSTNLFEGDKFVGTF | 1 | 0 | 0 | 1672678 | NP_073549.1 | A0A0P0LKV0 |
| KVSTQEFRFMTANNLT | 1 | 0 | 0 | 1672679 | YP_009047202.1 | K9N7C7 |
| KVSTVQSKLTDLKCTN | 1 | 0 | 0 | 1672681 | NP_073549.1 | A0A0P0LKV0 |
| KVTDVLFPGQKYSSDC | 1 | 0 | 0 | 1672684 | YP_009047202.1 | K9N7C7 |
| KVTEDGVNVHDVTVTT | 1 | 0 | 0 | 1672686 | NP_073549.1 | A0A0P0LKV0 |
| KVTEYSWNKKLYELVQ | 1 | 0 | 0 | 1672687 | NP_073549.1 | A0A0P0LKV0 |
| KVTFFPDLNGDVVAID | 1 | 0 | 0 | 1672688 | YP_009724389.1 | P0DTD1 |
| KVTFGDDTVIEVQGYK | 1 | 0 | 0 | 1672689 | YP_009724389.1 | P0DTD1 |
| KVTLADAGFIKQYGDC | 1 | 0 | 0 | 1672700 | YP_009724390.1 | P0DTC2 |
| KVTLADAGFMKQYGEC | 1 | 0 | 0 | 1672701 | NP_828851.1 | P59594 |
| KVVAKIDKEDVVIFIN | 1 | 0 | 0 | 1672711 | YP_009555238.1 | P0C6X6 |
| KVVNANVLTKDQTPIV | 1 | 0 | 0 | 1672726 | NP_073549.1 | A0A0P0LKV0 |
| KVVNINVDFKDFQFML | 1 | 0 | 0 | 1672728 | YP_173236.1 | A0A140H1G9 |
| KVVNVNVDFKDFQFML | 1 | 0 | 0 | 1672729 | YP_009555238.1 | P0C6X6 |
| KVVVGPTTEVKFSVIE | 1 | 0 | 0 | 1672735 | YP_003766.2 | H9EJ42 |
| KVVYHTTSKPITFHMD | 1 | 0 | 0 | 1672737 | QIQ54046.1 | P0C6U8 |
| KVYFKGQTTHESSSAV | 1 | 0 | 0 | 1672741 | YP_173236.1 | A0A140H1G9 |
| KWADASEPQRGRQGRI | 1 | 0 | 0 | 1672745 | NP_073556.1 | A0A0N7J131 |
| KWADQSDQFRNVQTRG | 1 | 0 | 0 | 1672748 | YP_009555245.1 | P33469 |
| KWDDFCRQYSNESWFE | 1 | 0 | 0 | 1672753 | NP_073549.1 | A0A0P0LKV0 |
| KWDLIISDMYDPKTKH | 1 | 0 | 0 | 1672755 | QIQ54047.1 | P0C6X7 |
| KWDLIISDMYDPKTKN | 1 | 0 | 0 | 1672756 | YP_009724389.1 | P0DTD1 |
| KWDLIISDMYDPRTKH | 1 | 0 | 0 | 1672757 | NP_828849.2 | P0C6X7 |
| KWDLLKYDFTEERLCL | 1 | 0 | 0 | 1672758 | NP_828849.2 | P0C6X7 |
| KWDLLKYDFTEERLKL | 1 | 0 | 0 | 1672759 | YP_009724389.1 | P0DTD1 |
| KWDLLKYDFTEERLNL | 1 | 0 | 0 | 1672760 | QIQ54047.1 | P0C6X7 |
| KWLELLEQNIDAYKTF | 1 | 0 | 0 | 1672774 | YP_009047211.1 | K9N4V7 |
| KWLNNFNVPSPLNWER | 1 | 0 | 0 | 1672777 | YP_173238.1 | A0A140H1H1 |
| KWVQVPIGTNDPIRFC | 1 | 0 | 0 | 1672796 | NP_073549.1 | A0A0P0LKV0 |
| KWYCYDKNPINSNVKT | 1 | 0 | 0 | 1672798 | NP_073549.1 | A0A0P0LKV0 |
| KXTCVGSDNVTDFNAI | 1 | 0 | 0 | 1672804 | QIQ54047.1 | P0C6X7 |
| KYATVVVGSTEEVKSS | 1 | 0 | 0 | 1672809 | NP_073549.1 | A0A0P0LKV0 |
| KYDAKNYVRFPEILAR | 1 | 0 | 0 | 1672814 | YP_003766.2 | H9EJ42 |
| KYDENGTITDAVDCSQ | 1 | 0 | 0 | 1672815 | NP_828851.1 | P59594 |
| KYDFTEERLKLFDRYF | 1 | 0 | 0 | 1672817 | YP_009724389.1 | P0DTD1 |
| KYDFTEHKENLFNKYF | 1 | 0 | 0 | 1672819 | YP_003766.2 | H9EJ42 |
| KYDNGNFIKLPEVIAQ | 1 | 0 | 0 | 1672822 | NP_073549.1 | A0A0P0LKV0 |
| KYFDCYDGGCINANQV | 1 | 0 | 0 | 1672826 | YP_009724389.1 | P0DTD1 |
| KYFEIYDGGCIPASQV | 1 | 0 | 0 | 1672830 | YP_009555238.1 | P0C6X6 |
| KYFEIYDGGCLNASEV | 1 | 0 | 0 | 1672831 | YP_009047202.1 | K9N7C7 |
| KYFKHWSMPYHPNTVD | 1 | 0 | 0 | 1672835 | YP_009555238.1 | P0C6X6 |
| KYFSGALDTTSYREAA | 1 | 0 | 0 | 1672841 | NP_828849.2 | P0C6X7 |
| KYFSGAMDTTSYREAA | 1 | 0 | 0 | 1672842 | YP_009724389.1 | P0DTD1 |
| KYFSGAMETAAYREAA | 1 | 0 | 0 | 1672843 | YP_009047202.1 | K9N7C7 |
| KYIENGQLKNFEAPLK | 1 | 0 | 0 | 1672857 | NP_073549.1 | A0A0P0LKV0 |
| KYKATGDLAVCLGIGD | 1 | 0 | 0 | 1672861 | YP_009555238.1 | P0C6X6 |
| KYKGIKIQEGVVDYGA | 1 | 0 | 0 | 1672864 | YP_009724389.1 | P0DTD1 |
| KYKTSDELCVNLNLPA | 1 | 0 | 0 | 1672867 | YP_009047202.1 | K9N7C7 |
| KYKYFSGALDTTSYRE | 1 | 0 | 0 | 1672869 | NP_828849.2 | P0C6X7 |
| KYLANEAQVQLEHYSS | 1 | 0 | 0 | 1672873 | NP_073549.1 | A0A0P0LKV0 |
| KYLVQQESPFVMMSAP | 1 | 0 | 0 | 1672882 | YP_009724389.1 | P0DTD1 |
| KYMVANGLHAPHGPFD | 1 | 0 | 0 | 1672891 | YP_003766.2 | H9EJ42 |
| KYMVANGLNAPNGPFD | 1 | 0 | 0 | 1672892 | NP_073549.1 | A0A0P0LKV0 |
| KYNIYDVSGVGVIRVS | 1 | 0 | 0 | 1672895 | NP_073551.1 | A0A0P0K6L9 |
| KYNIYDYVGTGIIRSS | 1 | 0 | 0 | 1672896 | YP_003767.1 | A0A0P0G321 |
| KYPCMVGPDYAYFNGA | 1 | 0 | 0 | 1672901 | YP_009047202.1 | K9N7C7 |
| KYPQVNGLTSIKWADN | 1 | 0 | 0 | 1672904 | YP_009724389.1 | P0DTD1 |
| KYSYINKCSRLLSDDR | 1 | 0 | 0 | 1672919 | YP_009047204.1 | K9N5Q8 |
| KYTEVRASRQLAQQKV | 1 | 0 | 0 | 1672924 | NP_073551.1 | A0A0P0K6L9 |
| KYTFGNVKPGETFTVL | 1 | 0 | 0 | 1672926 | YP_173236.1 | A0A140H1G9 |
| KYTKILKDDGNFVVLE | 1 | 0 | 0 | 1672929 | YP_009555238.1 | P0C6X6 |
| KYTKIMKDDGNCVVLE | 1 | 0 | 0 | 1672930 | YP_173236.1 | A0A140H1G9 |
| KYWDQTYHANCVNCTD | 1 | 0 | 0 | 1672960 | YP_009047202.1 | K9N7C7 |
| KYWDQTYHPNCINCLD | 1 | 0 | 0 | 1672961 | NP_828849.2 | P0C6X7 |
| KYWDQTYHPNCVNCLD | 1 | 0 | 0 | 1672962 | YP_009724389.1 | P0DTD1 |
| KYWGQDYHPDCVDCHD | 1 | 0 | 0 | 1672964 | NP_073549.1 | A0A0P0LKV0 |
| KYYDDSQYYFNKDTGV | 1 | 0 | 0 | 1672968 | YP_009555240.1 | P30215 |
| KYYSGNANEADYRCAC | 1 | 0 | 0 | 1672972 | NP_073549.1 | A0A0P0LKV0 |
| LAAECTIFKDAAGKPV | 1 | 0 | 0 | 1672979 | QIQ54046.1 | P0C6U8 |
| LAAECTIFKDAMGKPV | 1 | 0 | 0 | 1672980 | NP_828849.2 | P0C6X7 |

| | | | | | | |
|---|---|---|---|---|---|---|
| LAAECTIFKDASGKPV | 1 | 0 | 0 | 1672981 | YP_009724389.1 | P0DTD1 |
| LAANTLNIIVPDKSVY | 1 | 0 | 0 | 1672995 | YP_009555238.1 | P0C6X6 |
| LAASFNKAINNIVASF | 1 | 0 | 0 | 1672998 | YP_003767.1 | A0A0P0G321 |
| LAASFNKAMTNIVDAF | 1 | 0 | 0 | 1672999 | NP_073551.1 | A0A0P0K6L9 |
| LAATSASLFPPWTAAA | 1 | 0 | 0 | 1673002 | YP_009555241.1 | P36334 |
| LAATVRLQAGNATEVP | 1 | 0 | 0 | 1673003 | YP_009724389.1 | P0DTD1 |
| LADAGFMKQYGECLGD | 1 | 0 | 0 | 1673027 | NP_828851.1 | P59594 |
| LADDDSAIIEIVNEVL | 1 | 0 | 0 | 1673030 | YP_003771.1 | H9EJ69 |
| LADGLEAYADKTLQEM | 1 | 0 | 0 | 1673032 | YP_009555238.1 | P0C6X6 |
| LADSVVEVVTTSLTPC | 1 | 0 | 0 | 1673042 | YP_009555238.1 | P0C6X6 |
| LADYDVAVTAGPFMDN | 1 | 0 | 0 | 1673046 | YP_009047202.1 | K9N7C7 |
| LAECKRALGLSISDHE | 1 | 0 | 0 | 1673047 | NP_073549.1 | A0A0P0LKV0 |
| LAEFDTYDPIYKNGAM | 1 | 0 | 0 | 1673048 | YP_009047202.1 | K9N7C7 |
| LAELKCSVKSFEIDKG | 1 | 0 | 0 | 1673054 | NP_828851.1 | P59594 |
| LAEYHNESGIKTTLRK | 1 | 0 | 0 | 1673056 | QIQ54046.1 | P0C6U8 |
| LAEYHNESGLKPILRK | 1 | 0 | 0 | 1673057 | QHR63299.1 | P0C6W2 |
| LAEYHNESGLKTILRK | 1 | 0 | 0 | 1673058 | YP_009724389.1 | P0DTD1 |
| LAFLGASVTEDVKFAA | 1 | 0 | 0 | 1673063 | YP_003766.2 | H9EJ42 |
| LAFNQDGIIFNAVDCM | 1 | 0 | 0 | 1673067 | YP_009555241.1 | P36334 |
| LAFSYMDDVVLGAKS | 1 | 0 | 0 | 1673070 | P03155.1 | Q69028 |
| LAGFDMLDNYKAIDVV | 1 | 0 | 0 | 1673072 | YP_009555238.1 | P0C6X6 |
| LAGGITYVSNSGNLLG | 1 | 0 | 0 | 1673073 | YP_003767.1 | A0A0P0G321 |
| LAGKVKVGTGVMVECD | 1 | 0 | 0 | 1673077 | NP_073549.1 | A0A0P0LKV0 |
| LAGSYKDWSYSGQSTQ | 1 | 0 | 0 | 1673083 | YP_009724389.1 | P0DTD1 |
| LAGTITSGWTFGAGAA | 1 | 0 | 0 | 1673087 | YP_009724390.1 | P0DTC2 |
| LAHGGGLAKALDVYTK | 1 | 0 | 0 | 1673094 | NP_073549.1 | A0A0P0LKV0 |
| LAIDAYPLSKHPKPEY | 1 | 0 | 0 | 1673099 | NP_073549.1 | A0A0P0LKV0 |
| LAIDAYPLTKHEDIEY | 1 | 0 | 0 | 1673101 | YP_009047202.1 | K9N7C7 |
| LAIEEIEYVHGDALHT | 1 | 0 | 0 | 1673105 | NP_073549.1 | A0A0P0LKV0 |
| LAKAMDTFTNNNGSDV | 1 | 0 | 0 | 1673130 | YP_009555238.1 | P0C6X6 |
| LAKAMETFNHNNGNDV | 1 | 0 | 0 | 1673131 | YP_173236.1 | A0A140H1G9 |
| LAKAMLDFSRDHNDIL | 1 | 0 | 0 | 1673132 | NP_073549.1 | A0A0P0LKV0 |
| LAKGGFKFGDPADSRD | 1 | 0 | 0 | 1673138 | YP_009555238.1 | P0C6X6 |
| LAKGHFKFDEPSDATD | 1 | 0 | 0 | 1673139 | YP_173236.1 | A0A140H1G9 |
| LAKKFDNFTGECPNFV | 1 | 0 | 0 | 1673146 | QIQ54046.1 | P0C6U8 |
| LAKKFDTFNGECPNFV | 1 | 0 | 0 | 1673147 | YP_009724389.1 | P0DTD1 |
| LAKKIDVLLTVDGVNF | 1 | 0 | 0 | 1673148 | YP_173236.1 | A0A140H1G9 |
| LAKLGKDATKPQQVTK | 1 | 0 | 0 | 1673149 | YP_009555245.1 | P33469 |
| LAKRFKESPFELEDFI | 1 | 0 | 0 | 1673157 | YP_009724389.1 | P0DTD1 |
| LAKRSKDSPLELEDFI | 1 | 0 | 0 | 1673158 | QIQ54047.1 | P0C6X7 |
| LAKRSKESPLELEDFI | 1 | 0 | 0 | 1673159 | QHR63299.1 | P0C6W2 |
| LALHRSYLTPGKLESG | 1 | 0 | 0 | 1673174 | QIQ54048.1 | P59594 |
| LALLLDRLNALESKM | 1 | 0 | 0 | 1673181 | QIQ54056.1 | P59595 |
| LALQQLEVKFNAPALQ | 1 | 0 | 0 | 1673188 | NP_828849.2 | P0C6X7 |
| LALTNMYKEARINDKK | 1 | 0 | 0 | 1673195 | YP_009555238.1 | P0C6X6 |
| LAMDEFIERYKLEGYA | 1 | 0 | 0 | 1673202 | YP_009724389.1 | P0DTD1 |
| LAMDEFIQRYKLEGYA | 1 | 0 | 0 | 1673203 | NP_828849.2 | P0C6X7 |
| LAMLGAEMTSNVRRQV | 1 | 0 | 0 | 1673206 | NP_073549.1 | A0A0P0LKV0 |
| LANECAQVLSEYVLCG | 1 | 0 | 0 | 1673217 | YP_009047202.1 | K9N7C7 |
| LANFNETKGPLCVDTS | 1 | 0 | 0 | 1673220 | NP_073551.1 | A0A0P0K6L9 |
| LANKEQEAILMSEDVK | 1 | 0 | 0 | 1673222 | YP_009047202.1 | K9N7C7 |
| LANQFTEFVGTQSVDM | 1 | 0 | 0 | 1673224 | YP_009047202.1 | K9N7C7 |
| LANSNGFIRFPDVISE | 1 | 0 | 0 | 1673226 | YP_173236.1 | A0A140H1G9 |
| LANTTTEPLKLFAAET | 1 | 0 | 0 | 1673228 | YP_009047202.1 | K9N7C7 |
| LAPSDVYQQLAGVKLQ | 1 | 0 | 0 | 1673239 | YP_173236.1 | A0A140H1G9 |
| LAPTNAYMRTTHTDIG | 1 | 0 | 0 | 1673242 | YP_009047202.1 | K9N7C7 |
| LAPTPGAFFFGSKLDL | 1 | 0 | 0 | 1673243 | YP_173242.1 | Q19U29 |
| LAQPVDFLALTASSIA | 1 | 0 | 0 | 1673246 | YP_009555238.1 | P0C6X6 |
| LARDGVVPLSVIPATS | 1 | 0 | 0 | 1673257 | YP_003766.2 | H9EJ42 |
| LASALKQLKVTTGELV | 1 | 0 | 0 | 1673270 | NP_073549.1 | A0A0P0LKV0 |
| LASAYRANSFDVPTFC | 1 | 0 | 0 | 1673271 | YP_009047202.1 | K9N7C7 |
| LASDDLYVKRYFKGCE | 1 | 0 | 0 | 1673273 | YP_173236.1 | A0A140H1G9 |
| LASECFMKSDIFGQDF | 1 | 0 | 0 | 1673275 | NP_073549.1 | A0A0P0LKV0 |
| LASECFVKSDIFGSDF | 1 | 0 | 0 | 1673276 | YP_003766.2 | H9EJ42 |
| LASHMYCSFYPPDEDE | 1 | 0 | 0 | 1673282 | YP_009724389.1 | P0DTD1 |
| LASMTGVSIETLLAAI | 1 | 0 | 0 | 1673287 | YP_173236.1 | A0A140H1G9 |
| LASVPEKFLNAFDVFV | 1 | 0 | 0 | 1673292 | NP_073549.1 | A0A0P0LKV0 |
| LATFAELEAAQKAYQE | 1 | 0 | 0 | 1673298 | YP_009047202.1 | K9N7C7 |
| LAVDDKYKVNENLAVN | 1 | 0 | 0 | 1673322 | YP_173236.1 | A0A140H1G9 |
| LAVDVVSIRSSNQGNK | 1 | 0 | 0 | 1673324 | YP_009047207.1 | K9N643 |
| LAVQIGNNNVCTYEHV | 1 | 0 | 0 | 1673345 | NP_073549.1 | A0A0P0LKV0 |
| LAVQIGSNNVCTYEHV | 1 | 0 | 0 | 1673346 | YP_003766.2 | H9EJ42 |
| LAVSGQAFCDYADKLC | 1 | 0 | 0 | 1673348 | YP_009555238.1 | P0C6X6 |
| LAWDVIRKDLSYEQQN | 1 | 0 | 0 | 1673354 | YP_003766.2 | H9EJ42 |
| LAWNTRNIDATSTGNY | 1 | 0 | 0 | 1673357 | NP_828851.1 | P59594 |
| LAYFSKKHVEVFTDGK | 1 | 0 | 0 | 1673362 | YP_009047202.1 | K9N7C7 |
| LAYQDYMQIAPVPAEV | 1 | 0 | 0 | 1673367 | YP_003769.1 | H9EJA2 |
| LCALPDTPSTLTPRSV | 1 | 0 | 0 | 1673392 | YP_009047204.1 | K9N5Q8 |
| LCATCPFAKGDSCSSN | 1 | 0 | 0 | 1673399 | YP_009047202.1 | K9N7C7 |
| LCCTVYDSSEVKAQSS | 1 | 0 | 0 | 1673412 | YP_173236.1 | A0A140H1G9 |
| LCDEFTLAEVVKQMYG | 1 | 0 | 0 | 1673418 | YP_003766.2 | H9EJ42 |
| LCDNPFFAVSKPMGTQ | 1 | 0 | 0 | 1673426 | NP_828851.1 | P59594 |
| LCDRRATCFSTASDTY | 1 | 0 | 0 | 1673427 | YP_009724389.1 | P0DTD1 |
| LCDSGKISSKAGNSIF | 1 | 0 | 0 | 1673428 | YP_009555240.1 | P30215 |
| LCEILCEYADCKESYF | 1 | 0 | 0 | 1673432 | YP_173236.1 | A0A140H1G9 |
| LCFDKSKQFVPVQVVD | 1 | 0 | 0 | 1673440 | YP_173237.1 | A0A140H1H0 |
| LCFKVYFKGQTTHESS | 1 | 0 | 0 | 1673443 | YP_173236.1 | A0A140H1G9 |
| LCGADGKPVMSEDLWQ | 1 | 0 | 0 | 1673456 | NP_073549.1 | A0A0P0LKV0 |
| LCGDIVSGYEKSTFNK | 1 | 0 | 0 | 1673457 | YP_003766.2 | H9EJ42 |
| LCIFNGKFLSNTKYYD | 1 | 0 | 0 | 1673473 | YP_009555240.1 | P30215 |
| LCIMSDRDLYDKLQFT | 1 | 0 | 0 | 1673474 | YP_009724389.1 | P0DTD1 |

| | | | | | | |
|---|---|---|---|---|---|---|
| LCKTVLNVVGPDARTQ | 1 | 0 | 0 | 1673489 | YP_009555238.1 | P0C6X6 |
| LCLRDDNQTLLVPKIF | 1 | 0 | 0 | 1673501 | YP_173236.1 | A0A140H1G9 |
| LCMFWNCNVDKYPSNS | 1 | 0 | 0 | 1673507 | YP_173236.1 | A0A140H1G9 |
| LCNLINNNLALGGSVA | 1 | 0 | 0 | 1673511 | YP_009047202.1 | K9N7C7 |
| LCNNGKISSSAGDSIF | 1 | 0 | 0 | 1673512 | YP_173237.1 | A0A140H1H0 |
| LCPFGEVFNASKFASV | 1 | 0 | 0 | 1673517 | QIQ54048.1 | P59594 |
| LCRMNSRNYIAQVDVV | 1 | 0 | 0 | 1673530 | YP_009725255.1 | A0A663DJA2 |
| LCSSHYWSPQGNLGGN | 1 | 0 | 0 | 1673538 | YP_173236.1 | A0A140H1G9 |
| LCSTRVNVDGFNEWAM | 1 | 0 | 0 | 1673540 | YP_003766.2 | H9EJ42 |
| LCTFTKSTNSRIRASL | 1 | 0 | 0 | 1673542 | NP_828849.2 | P0C6X7 |
| LCVDGFGVNKIQPWTN | 1 | 0 | 0 | 1673555 | NP_073549.1 | A0A0P0LKV0 |
| LCVDGTNGYVLRQPNL | 1 | 0 | 0 | 1673556 | NP_073551.1 | A0A0P0K6L9 |
| LCVVPGNVKPGDAVIT | 1 | 0 | 0 | 1673573 | NP_073549.1 | A0A0P0LKV0 |
| LCWETGKVKPPLNKNY | 1 | 0 | 0 | 1673575 | YP_173236.1 | A0A140H1G9 |
| LDAANHSIPDLQRAAV | 1 | 0 | 0 | 1673585 | NP_828849.2 | P0C6X7 |
| LDADLTKPYVKWDLLK | 1 | 0 | 0 | 1673587 | QIQ54047.1 | P0C6X7 |
| LDALASMTGVSIETLL | 1 | 0 | 0 | 1673591 | YP_173236.1 | A0A140H1G9 |
| LDALEAEAQIDRLING | 1 | 0 | 0 | 1673592 | YP_009555241.1 | P36334 |
| LDALEAQVQIDRLING | 1 | 0 | 0 | 1673593 | YP_173238.1 | A0A140H1H1 |
| LDAVKQGAKPVGNCVK | 1 | 0 | 0 | 1673599 | NP_073549.1 | A0A0P0LKV0 |
| LDDFIVFNNNGTNVTQ | 1 | 0 | 0 | 1673613 | NP_073549.1 | A0A0P0LKV0 |
| LDDILQQANKSVGIIS | 1 | 0 | 0 | 1673622 | YP_009047202.1 | K9N7C7 |
| LDDKDPQFKDNVILLN | 1 | 0 | 0 | 1673623 | NP_828858.1 | P59595 |
| LDDKDPSFKDNVILLN | 1 | 0 | 0 | 1673624 | QIQ54056.1 | P59595 |
| LDDNVLPETYVALPIY | 1 | 0 | 0 | 1673628 | YP_003767.1 | A0A0P0G321 |
| LDDVKKIEYKPDLSQY | 1 | 0 | 0 | 1673636 | YP_009555238.1 | P0C6X6 |
| LDERIDKVLNEKCSAY | 1 | 0 | 0 | 1673646 | YP_009724389.1 | P0DTD1 |
| LDERVDKVLNEKCSNY | 1 | 0 | 0 | 1673648 | QIQ54046.1 | P0C6U8 |
| LDERVDKVLNEKCSVY | 1 | 0 | 0 | 1673649 | NP_828849.2 | P0C6X7 |
| LDFIDTKRGVYCCREH | 1 | 0 | 0 | 1673656 | YP_009724389.1 | P0DTD1 |
| LDFSRDHNDILYTPPT | 1 | 0 | 0 | 1673660 | NP_073549.1 | A0A0P0LKV0 |
| LDFVNAGGAPLTNCVK | 1 | 0 | 0 | 1673662 | YP_009047202.1 | K9N7C7 |
| LDGDYAMQFFKMGRVA | 1 | 0 | 0 | 1673667 | NP_073549.1 | A0A0P0LKV0 |
| LDGEVLSLDKLKSLLS | 1 | 0 | 0 | 1673670 | NP_828849.2 | P0C6X7 |
| LDGISQYSLRLIDAML | 1 | 0 | 0 | 1673674 | QIQ54046.1 | P0C6U8 |
| LDGISQYSLRLIDAMM | 1 | 0 | 0 | 1673675 | YP_009724389.1 | P0DTD1 |
| LDGTCGIIDVEKGVLP | 1 | 0 | 0 | 1673682 | QIQ54046.1 | P0C6U8 |
| LDGYSKLFITKDEAIK | 1 | 0 | 0 | 1673689 | YP_173236.1 | A0A140H1G9 |
| LDGYYKKDNAYFTEQP | 1 | 0 | 0 | 1673690 | QIQ54046.1 | P0C6U8 |
| LDGYYKKDNAYYTEQP | 1 | 0 | 0 | 1673691 | NP_828849.2 | P0C6X7 |
| LDIHPSLCNNGKISSS | 1 | 0 | 0 | 1673697 | YP_173237.1 | A0A140H1H0 |
| LDITDAAVKAAESKAF | 1 | 0 | 0 | 1673703 | NP_073549.1 | A0A0P0LKV0 |
| LDKCTKYNIYDVSGVG | 1 | 0 | 0 | 1673706 | NP_073551.1 | A0A0P0K6L9 |
| LDKQGQLLDTKLNGIL | 1 | 0 | 0 | 1673716 | YP_003766.2 | H9EJ42 |
| LDKVDILLTVDGVNFT | 1 | 0 | 0 | 1673720 | YP_009555238.1 | P0C6X6 |
| LDKVEAEVQIDRLITG | 1 | 0 | 0 | 1673721 | YP_009724390.1 | P0DTC2 |
| LDLLNRLQALESGKVK | 1 | 0 | 0 | 1673731 | YP_009047211.1 | K9N4V7 |
| LDLSKELKRPIQPTDV | 1 | 0 | 0 | 1673738 | YP_009555238.1 | P0C6X6 |
| LDLTLDGYSKLFITKD | 1 | 0 | 0 | 1673742 | YP_173236.1 | A0A140H1G9 |
| LDMCAALKELLQNGMN | 1 | 0 | 0 | 1673748 | NP_828849.2 | P0C6X7 |
| LDMVDTSLSGFKLKDC | 1 | 0 | 0 | 1673754 | YP_009724389.1 | P0DTD1 |
| LDMVDTSLSGYKLKDC | 1 | 0 | 0 | 1673755 | QIQ54046.1 | P0C6U8 |
| LDNAVKGCVPLNAIPS | 1 | 0 | 0 | 1673760 | YP_009555238.1 | P0C6X6 |
| LDNDALNNIINNARDG | 1 | 0 | 0 | 1673762 | YP_009724389.1 | P0DTD1 |
| LDNDFNTILNTACGVF | 1 | 0 | 0 | 1673763 | YP_009555238.1 | P0C6X6 |
| LDNDVLNGIISNARNG | 1 | 0 | 0 | 1673764 | YP_009047202.1 | K9N7C7 |
| LDNGVIRDVSVEDVCF | 1 | 0 | 0 | 1673767 | YP_009555238.1 | P0C6X6 |
| LDNLKPRVEAPKQEEP | 1 | 0 | 0 | 1673768 | NP_828849.2 | P0C6X7 |
| LDNLRANNTKGSIPIN | 1 | 0 | 0 | 1673770 | QIQ54046.1 | P0C6U8 |
| LDNLRANNTKGSLPIN | 1 | 0 | 0 | 1673771 | YP_009724389.1 | P0DTD1 |
| LDNLRASNTKGSLPIN | 1 | 0 | 0 | 1673772 | QHR63299.1 | P0C6W2 |
| LDNQDLNGKWYDFGDF | 1 | 0 | 0 | 1673777 | YP_009047202.1 | K9N7C7 |
| LDNQDLNGKWYDFGDY | 1 | 0 | 0 | 1673778 | YP_009555238.1 | P0C6X6 |
| LDNYYKKDNSYFTEQP | 1 | 0 | 0 | 1673787 | YP_009724389.1 | P0DTD1 |
| LDPFCFDVNTLLSNQL | 1 | 0 | 0 | 1673788 | YP_003766.2 | H9EJ42 |
| LDPKNPNYNKWLELLE | 1 | 0 | 0 | 1673790 | YP_009047211.1 | K9N4V7 |
| LDPPCKFSIQDVKGLK | 1 | 0 | 0 | 1673794 | YP_173236.1 | A0A140H1G9 |
| LDQALVSDVGDSAEVA | 1 | 0 | 0 | 1673798 | YP_009724389.1 | P0DTD1 |
| LDQALVSDVGDSTEVA | 1 | 0 | 0 | 1673799 | QHR63299.1 | P0C6W2 |
| LDQALVSDVGDSTEVS | 1 | 0 | 0 | 1673800 | NP_828849.2 | P0C6X7 |
| LDQDVFIQKYGLEDYA | 1 | 0 | 0 | 1673803 | YP_009555238.1 | P0C6X6 |
| LDRLNALESKMSGKGS | 1 | 0 | 0 | 1673809 | QIQ54056.1 | P59595 |
| LDRLNQLESKVSGKGQ | 1 | 0 | 0 | 1673811 | NP_828858.1 | P59595 |
| LDSIQADQQVDRLITG | 1 | 0 | 0 | 1673816 | YP_003767.1 | A0A0P0G321 |
| LDTKLNGILQQAALDY | 1 | 0 | 0 | 1673840 | YP_003766.2 | H9EJ42 |
| LDTTQLHVADTLMQGV | 1 | 0 | 0 | 1673843 | YP_173238.1 | A0A140H1H1 |
| LDTVVSTVKIGGLTFK | 1 | 0 | 0 | 1673846 | NP_073549.1 | A0A0P0LKV0 |
| LDVFKIGDVKFKRLGD | 1 | 0 | 0 | 1673852 | YP_003766.2 | H9EJ42 |
| LDVKGVEELHDDLVNY | 1 | 0 | 0 | 1673855 | YP_009555239.1 | Q80872 |
| LDVKPYGYPNDFVGGF | 1 | 0 | 0 | 1673856 | YP_003766.2 | H9EJ42 |
| LDWLEEKFKEGIEFLK | 1 | 0 | 0 | 1673869 | QIQ54046.1 | P0C6U8 |
| LDWLEEKFKEGVEFLR | 1 | 0 | 0 | 1673870 | YP_009724389.1 | P0DTD1 |
| LDWVKHLNKNLNEGVL | 1 | 0 | 0 | 1673871 | YP_003766.2 | H9EJ42 |
| LDWVKHLNKTLNEGVL | 1 | 0 | 0 | 1673872 | NP_073549.1 | A0A0P0LKV0 |
| LDYAKDHNDMLYSPPT | 1 | 0 | 0 | 1673873 | YP_003766.2 | H9EJ42 |
| LDYIESKRGVYCCRDH | 1 | 0 | 0 | 1673876 | NP_828849.2 | P0C6X7 |
| LDYKAFKQIVESCGNF | 1 | 0 | 0 | 1673881 | YP_009724389.1 | P0DTD1 |
| LDYKRQNNLAIEEIEY | 1 | 0 | 0 | 1673882 | NP_073549.1 | A0A0P0LKV0 |
| LDYKSFKQIVESCGNF | 1 | 0 | 0 | 1673883 | QIQ54046.1 | P0C6U8 |
| LDYKTFKQIVESCGNF | 1 | 0 | 0 | 1673885 | QHR63299.1 | P0C6W2 |
| LDYSIDVIDNEIIVKP | 1 | 0 | 0 | 1673887 | NP_073549.1 | A0A0P0LKV0 |

| | | | | | | |
|---|---|---|---|---|---|---|
| LEALETPVDSFTNGAI | 1 | 0 | 0 | 1673900 | NP_828849.2 | P0C6X7 |
| LEAQVQIDRLINGRLT | 1 | 0 | 0 | 1673904 | YP_173238.1 | A0A140H1H1 |
| LEAYADKTLQEMKALF | 1 | 0 | 0 | 1673910 | YP_009555238.1 | P0C6X6 |
| LEDEFTPEDVNMQIMG | 1 | 0 | 0 | 1673921 | YP_009047202.1 | K9N7C7 |
| LEDEFTPFDVVRQCSG | 1 | 0 | 0 | 1673922 | YP_009724389.1 | P0DTD1 |
| LEDFKPRSQMETDFLE | 1 | 0 | 0 | 1673924 | NP_828849.2 | P0C6X7 |
| LEDFNFEHVVYGDVSK | 1 | 0 | 0 | 1673925 | NP_073549.1 | A0A0P0LKV0 |
| LEDKFDLLISDMYDGR | 1 | 0 | 0 | 1673926 | NP_073549.1 | A0A0P0LKV0 |
| LEDTLDDGPSVETSDS | 1 | 0 | 0 | 1673929 | YP_009555238.1 | P0C6X6 |
| LEEAARYMRSLKVPAT | 1 | 0 | 0 | 1673933 | YP_009724389.1 | P0DTD1 |
| LEEDWLDDDSQQTVVQ | 1 | 0 | 0 | 1673936 | QHR63299.1 | P0C6W2 |
| LEEFDVVFGKRGGGNV | 1 | 0 | 0 | 1673938 | NP_073549.1 | A0A0P0LKV0 |
| LEEKFKEGIEFLKDGW | 1 | 0 | 0 | 1673941 | QIQ54046.1 | P0C6U8 |
| LEEKFKEGVEFLRDGW | 1 | 0 | 0 | 1673942 | YP_009724389.1 | P0DTD1 |
| LEENLNAYVNSNQNTD | 1 | 0 | 0 | 1673944 | YP_173242.1 | Q19U29 |
| LEEPTAQAVEVPLVGT | 1 | 0 | 0 | 1673945 | QIQ54046.1 | P0C6U8 |
| LEFANDTKIASQLGNC | 1 | 0 | 0 | 1673951 | YP_009047204.1 | K9N5Q8 |
| LEFNPSQTSPATAEPV | 1 | 0 | 0 | 1673956 | NP_073556.1 | A0A0N7J131 |
| LEFTDENYNNLVPTYL | 1 | 0 | 0 | 1673958 | YP_173236.1 | A0A140H1G9 |
| LEGKQGNFKNLREFVF | 1 | 0 | 0 | 1673966 | YP_009724390.1 | P0DTC2 |
| LEGLGGNNVVYCYNTAL | 1 | 0 | 0 | 1673968 | NP_073549.1 | A0A0P0LKV0 |
| LEGNFYGPFVDRQTAQ | 1 | 0 | 0 | 1673969 | YP_009724389.1 | P0DTD1 |
| LEGSISYSELRPDTRY | 1 | 0 | 0 | 1673970 | NP_828849.2 | P0C6X7 |
| LEGSVAYESLRPDTRY | 1 | 0 | 0 | 1673971 | YP_009724389.1 | P0DTD1 |
| LEGSVPYESLRPDTRY | 1 | 0 | 0 | 1673972 | QIQ54046.1 | P0C6U8 |
| LEGSVRVVTTFDAEYC | 1 | 0 | 0 | 1673973 | NP_828849.2 | P0C6X7 |
| LEIGYTVDCSCGKKLI | 1 | 0 | 0 | 1673982 | YP_009555238.1 | P0C6X6 |
| LEIKDKEQYCALSPGL | 1 | 0 | 0 | 1673983 | NP_828849.2 | P0C6X7 |
| LEIKDTEKYCALAPNM | 1 | 0 | 0 | 1673984 | YP_009724389.1 | P0DTD1 |
| LEILDITPCSFGGVSV | 1 | 0 | 0 | 1673985 | YP_009724390.1 | P0DTC2 |
| LEILSRERVNINIVGD | 1 | 0 | 0 | 1673987 | NP_828849.2 | P0C6X7 |
| LEIPRRNVATLQAENV | 1 | 0 | 0 | 1673989 | YP_009724389.1 | P0DTD1 |
| LEITACQYTMCEYPHT | 1 | 0 | 0 | 1673991 | YP_173238.1 | A0A140H1H1 |
| LEIVKLGAEKKDGFYE | 1 | 0 | 0 | 1673993 | YP_009555239.1 | Q80872 |
| LEIVQESSRKLLGLTN | 1 | 0 | 0 | 1673994 | YP_173243.1 | Q0ZJ80 |
| LEKACNIAKSAYERDR | 1 | 0 | 0 | 1673995 | YP_009555238.1 | P0C6X6 |
| LEKGKLLIRDTNKEVF | 1 | 0 | 0 | 1674001 | YP_009555238.1 | P0C6X6 |
| LEKGRLIIRENNRVVV | 1 | 0 | 0 | 1674002 | NP_828849.2 | P0C6X7 |
| LEKLISTARDGVRRGD | 1 | 0 | 0 | 1674005 | YP_009047202.1 | K9N7C7 |
| LELADTSLSGYRLKDC | 1 | 0 | 0 | 1674009 | NP_828849.2 | P0C6X7 |
| LELAMDEFIERYKLEG | 1 | 0 | 0 | 1674011 | YP_009724389.1 | P0DTD1 |
| LELAMDEFIQRYKLEG | 1 | 0 | 0 | 1674012 | NP_828849.2 | P0C6X7 |
| LELHWAPEFPWMFEDA | 1 | 0 | 0 | 1674019 | YP_009555238.1 | P0C6X6 |
| LELLEQNIDAYKTFPK | 1 | 0 | 0 | 1674022 | YP_009047211.1 | K9N4V7 |
| LELLPNVFKLKISTQL | 1 | 0 | 0 | 1674023 | YP_003766.2 | H9EJ42 |
| LELSTGCHTGTDFTGN | 1 | 0 | 0 | 1674026 | YP_173236.1 | A0A140H1G9 |
| LEMAKEVKTSSKVVLS | 1 | 0 | 0 | 1674030 | NP_073549.1 | A0A0P0LKV0 |
| LEMKSEKQVEQKIAEI | 1 | 0 | 0 | 1674034 | YP_009724389.1 | P0DTD1 |
| LEMKSETKVQDHFDVV | 1 | 0 | 0 | 1674036 | QIQ54046.1 | P0C6U8 |
| LENIDYFNMRRAKFSL | 1 | 0 | 0 | 1674041 | YP_009555238.1 | P0C6X6 |
| LENKSAELNYTVQKLQ | 1 | 0 | 0 | 1674044 | NP_073551.1 | A0A0P0K6L9 |
| LENVAFNVVNKGHFDG | 1 | 0 | 0 | 1674049 | YP_009724389.1 | P0DTD1 |
| LENVAYNVVNKGHFDG | 1 | 0 | 0 | 1674050 | NP_828849.2 | P0C6X7 |
| LEPCNLYNYGAGIKLP | 1 | 0 | 0 | 1674057 | YP_003766.2 | H9EJ42 |
| LEPKFSIALPPELSVV | 1 | 0 | 0 | 1674061 | YP_003771.1 | H9EJ69 |
| LEPPCRFVIDTPTGPQ | 1 | 0 | 0 | 1674064 | NP_073549.1 | A0A0P0LKV0 |
| LEQDAQIDRLINGRLT | 1 | 0 | 0 | 1674071 | YP_009047204.1 | K9N5Q8 |
| LEQNIDAYKTFPKKEK | 1 | 0 | 0 | 1674073 | YP_009047211.1 | K9N4V7 |
| LEQPTSEAVEAPLVGT | 1 | 0 | 0 | 1674074 | YP_009724389.1 | P0DTD1 |
| LEQYVFCTVNALPEAT | 1 | 0 | 0 | 1674084 | QIQ54047.1 | P0C6X7 |
| LEQYVFCTVNALPETT | 1 | 0 | 0 | 1674085 | YP_009724389.1 | P0DTD1 |
| LERCELANYKQSIPMP | 1 | 0 | 0 | 1674086 | YP_009047202.1 | K9N7C7 |
| LERDFIDMDDNLFIAK | 1 | 0 | 0 | 1674087 | YP_173236.1 | A0A140H1G9 |
| LERVSLWNYGKPINLP | 1 | 0 | 0 | 1674091 | YP_173236.1 | A0A140H1G9 |
| LERVYKHLNNYDCVVT | 1 | 0 | 0 | 1674092 | YP_009555238.1 | P0C6X6 |
| LERYGFHPTTGVNYQP | 1 | 0 | 0 | 1674093 | QIQ54048.1 | P59594 |
| LESADVSEMLTFDKKA | 1 | 0 | 0 | 1674095 | NP_073551.1 | A0A0P0K6L9 |
| LESFSVTLLDNQEDKF | 1 | 0 | 0 | 1674100 | YP_003766.2 | H9EJ42 |
| LESKQVDYVPLRSATC | 1 | 0 | 0 | 1674105 | YP_173236.1 | A0A140H1G9 |
| LESKVSGKGQQQQGQT | 1 | 0 | 0 | 1674106 | NP_828858.1 | P59595 |
| LESTSTHDGPTEHVTS | 1 | 0 | 0 | 1674110 | YP_009047205.1 | K9N796 |
| LETDGICPENHVMVDC | 1 | 0 | 0 | 1674118 | YP_009555238.1 | P0C6X6 |
| LETFQNNVVNYQHIGM | 1 | 0 | 0 | 1674120 | YP_009555238.1 | P0C6X6 |
| LEVKFNAPALQEAYYR | 1 | 0 | 0 | 1674132 | NP_828849.2 | P0C6X7 |
| LEVLLDVCNTKEVKVF | 1 | 0 | 0 | 1674133 | NP_073549.1 | A0A0P0LKV0 |
| LEVLLQDALQSREAVL | 1 | 0 | 0 | 1674134 | YP_009555238.1 | P0C6X6 |
| LEVPRRNVATLQAENV | 1 | 0 | 0 | 1674135 | QIQ54047.1 | P0C6X7 |
| LEVTGDSCNNFMLTYN | 1 | 0 | 0 | 1674137 | NP_828849.2 | P0C6X7 |
| LEVYNKEKTYYELTKS | 1 | 0 | 0 | 1674139 | YP_173236.1 | A0A140H1G9 |
| LEVYVHLRNWNFSWNL | 1 | 0 | 0 | 1674140 | YP_003770.1 | U3M6Q8 |
| LEYIKKLYNDLGTQIL | 1 | 0 | 0 | 1674144 | YP_173236.1 | A0A140H1G9 |
| LEYPIISNEVSINTSC | 1 | 0 | 0 | 1674145 | YP_173236.1 | A0A140H1G9 |
| LFAAETIKAKEESVKS | 1 | 0 | 0 | 1674147 | YP_003766.2 | H9EJ42 |
| LFAAETVKAKEESVKS | 1 | 0 | 0 | 1674151 | NP_073549.1 | A0A0P0LKV0 |
| LFADINGKLYHDSQNM | 1 | 0 | 0 | 1674153 | NP_828849.2 | P0C6X7 |
| LFAKRKMGLTPPLSIL | 1 | 0 | 0 | 1674159 | YP_003766.2 | H9EJ42 |
| LFAKRKVGLTPPLSIL | 1 | 0 | 0 | 1674160 | NP_073549.1 | A0A0P0LKV0 |
| LFASVKSSQSSPIIPG | 1 | 0 | 0 | 1674171 | YP_009047204.1 | K9N5Q8 |
| LFATLMTFSGNVDLDA | 1 | 0 | 0 | 1674172 | YP_009047202.1 | K9N7C7 |
| LFCEHPAVGIVNTDFK | 1 | 0 | 0 | 1674178 | NP_073553.1 | A0A0P0KWE8 |
| LFDEAGEEVLAPKLYC | 1 | 0 | 0 | 1674198 | YP_009555238.1 | P0C6X6 |

| | | | | | | |
|---|---|---|---|---|---|---|
| LFDGRDNGALEAFKRS | 1 | 0 | 0 | 1674202 | YP_009555238.1 | P0C6X6 |
| LFDLSKFQLKLKGTPV | 1 | 0 | 0 | 1674208 | YP_009047202.1 | K9N7C7 |
| LFDRYFKYWDQTYHPN | 1 | 0 | 0 | 1674215 | YP_009724389.1 | P0DTD1 |
| LFDSHIVSKESGNTVV | 1 | 0 | 0 | 1674216 | NP_073556.1 | A0A0N7J131 |
| LFDVIVAWHVVRDPRY | 1 | 0 | 0 | 1674218 | YP_009555238.1 | P0C6X6 |
| LFEGDKFIGTFESAAA | 1 | 0 | 0 | 1674221 | YP_003766.2 | H9EJ42 |
| LFEGDKFVGTFESAAA | 1 | 0 | 0 | 1674222 | NP_073549.1 | A0A0P0LKV0 |
| LFENGIPLMPLLSCGI | 1 | 0 | 0 | 1674224 | YP_003766.2 | H9EJ42 |
| LFENKTILPVNVAFEL | 1 | 0 | 0 | 1674225 | QIQ54047.1 | P0C6X7 |
| LFENKTTLPVNVAFEL | 1 | 0 | 0 | 1674226 | YP_009724389.1 | P0DTD1 |
| LFGDSRSDCNHINNLK | 1 | 0 | 0 | 1674247 | YP_173237.1 | A0A140H1H0 |
| LFGMIKKLDNDVLNGI | 1 | 0 | 0 | 1674249 | YP_009047202.1 | K9N7C7 |
| LFITKDEAIKRVRGWV | 1 | 0 | 0 | 1674272 | YP_173236.1 | A0A140H1G9 |
| LFKDCARNPIDLPPSH | 1 | 0 | 0 | 1674280 | NP_073549.1 | A0A0P0LKV0 |
| LFKDCSKSCLGYHPAH | 1 | 0 | 0 | 1674282 | YP_173236.1 | A0A140H1G9 |
| LFKDCSKSYSGYHPAH | 1 | 0 | 0 | 1674283 | YP_009555238.1 | P0C6X6 |
| LFKPLTKKNYSELDDE | 1 | 0 | 0 | 1674293 | NP_828856.1 | P59634 |
| LFKPVQLVPVSPAFHR | 1 | 0 | 0 | 1674294 | YP_009047208.1 | K9N7D2 |
| LFKVGKVDSIVQKCYE | 1 | 0 | 0 | 1674297 | YP_003766.2 | H9EJ42 |
| LFLFDEAGEEVLAPKL | 1 | 0 | 0 | 1674303 | YP_009555238.1 | P0C6X6 |
| LFLSGDDCFVSNSSVI | 1 | 0 | 0 | 1674320 | YP_173236.1 | A0A140H1G9 |
| LFLTLTSGSDLDRCTT | 1 | 0 | 0 | 1674323 | NP_828851.1 | P59594 |
| LFNKYKYFSGAMETAA | 1 | 0 | 0 | 1674349 | YP_009047202.1 | K9N7C7 |
| LFPHNDRIKSFSTFES | 1 | 0 | 0 | 1674354 | NP_073549.1 | A0A0P0LKV0 |
| LFSMVRKLDNQALNSI | 1 | 0 | 0 | 1674394 | YP_009555238.1 | P0C6X6 |
| LFSTVVIKNLTPIKLN | 1 | 0 | 0 | 1674400 | YP_003766.2 | H9EJ42 |
| LFSVNDFTCSQISPAA | 1 | 0 | 0 | 1674402 | YP_009047204.1 | K9N5Q8 |
| LFTLQLVSAVNQSLSN | 1 | 0 | 0 | 1674413 | NP_073552.1 | A0A0P0KWE8 |
| LFTMLRKLDNDALNNI | 1 | 0 | 0 | 1674414 | YP_009724389.1 | P0DTD1 |
| LFTNSILMLDKQGQLL | 1 | 0 | 0 | 1674415 | YP_003766.2 | H9EJ42 |
| LFTNSTDTVNHDSFNL | 1 | 0 | 0 | 1674416 | YP_173236.1 | A0A140H1G9 |
| LFVANYSQDVKQFANG | 1 | 0 | 0 | 1674422 | YP_009047204.1 | K9N5Q8 |
| LFVDTLVNKIPANIVL | 1 | 0 | 0 | 1674426 | YP_009555238.1 | P0C6X6 |
| LFVEDCKLPLGQSLCA | 1 | 0 | 0 | 1674428 | YP_009047204.1 | K9N5Q8 |
| LFVEHYNEWAQANGFT | 1 | 0 | 0 | 1674429 | NP_073549.1 | A0A0P0LKV0 |
| LFVSYGSSVETETYFK | 1 | 0 | 0 | 1674443 | YP_173236.1 | A0A140H1G9 |
| LFYCSNSSSGVLDTTI | 1 | 0 | 0 | 1674465 | YP_003767.1 | A0A0P0G321 |
| LFYDRDGQRTYDDVNA | 1 | 0 | 0 | 1674466 | YP_009555238.1 | P0C6X6 |
| LFYSYATHHDKFTDGV | 1 | 0 | 0 | 1674471 | NP_828849.2 | P0C6X7 |
| LFYSYATHSDKFKDGV | 1 | 0 | 0 | 1674472 | QIQ54047.1 | P0C6X7 |
| LFYSYATHSDKFTDGV | 1 | 0 | 0 | 1674473 | YP_009724389.1 | P0DTD1 |
| LFYVYDVHKDNFKDGL | 1 | 0 | 0 | 1674478 | YP_173236.1 | A0A140H1G9 |
| LGAAIHYLNSKIVDLA | 1 | 0 | 0 | 1674480 | YP_009555238.1 | P0C6X6 |
| LGAEMTSNVRRQVASG | 1 | 0 | 0 | 1674485 | NP_073549.1 | A0A0P0LKV0 |
| LGAMQTGFTTTNEAFQ | 1 | 0 | 0 | 1674494 | YP_009047204.1 | K9N5Q8 |
| LGASQRVGTDSGFAAY | 1 | 0 | 0 | 1674496 | NP_828855.1 | P59596 |
| LGASVTEDVKFAASTG | 1 | 0 | 0 | 1674497 | YP_003766.2 | H9EJ42 |
| LGCLSSMNIAANSSEW | 1 | 0 | 0 | 1674504 | YP_003766.2 | H9EJ42 |
| LGDELGTDPIDDFQVN | 1 | 0 | 0 | 1674510 | QIQ54046.1 | P0C6U8 |
| LGDELGTDPIEDYEQN | 1 | 0 | 0 | 1674511 | NP_828849.2 | P0C6X7 |
| LGDFIQGPFIAGNTVH | 1 | 0 | 0 | 1674514 | YP_003766.2 | H9EJ42 |
| LGEYIFERIDYSDAVS | 1 | 0 | 0 | 1674539 | YP_009047202.1 | K9N7C7 |
| LGEYVFDKSELTNGVY | 1 | 0 | 0 | 1674540 | YP_009555238.1 | P0C6X6 |
| LGFDKPQEKDKKSAKT | 1 | 0 | 0 | 1674541 | NP_073556.1 | A0A0N7J131 |
| LGFDNQSKSPSSSGTS | 1 | 0 | 0 | 1674542 | YP_003771.1 | H9EJ69 |
| LGFDSSKEFVEYKITE | 1 | 0 | 0 | 1674543 | YP_009555238.1 | P0C6X6 |
| LGFDSSKPVTKKYTYS | 1 | 0 | 0 | 1674544 | YP_009047202.1 | K9N7C7 |
| LGHGCACDRTTIQSVD | 1 | 0 | 0 | 1674594 | YP_003766.2 | H9EJ42 |
| LGIEFLKRGDKSVYYT | 1 | 0 | 0 | 1674599 | YP_009724389.1 | P0DTD1 |
| LGIEFLKRGDKVVYHT | 1 | 0 | 0 | 1674600 | QIQ54046.1 | P0C6U8 |
| LGIGGDRCIKISTVQS | 1 | 0 | 0 | 1674602 | YP_003766.2 | H9EJ42 |
| LGIGGKPCIKVATVQS | 1 | 0 | 0 | 1674604 | NP_828849.2 | P0C6X7 |
| LGINYLGTIKENIDGG | 1 | 0 | 0 | 1674618 | YP_009047202.1 | K9N7C7 |
| LGKFLEELNAFTREMQ | 1 | 0 | 0 | 1674625 | NP_073556.1 | A0A0N7J131 |
| LGKPKEQIDGYTMHAN | 1 | 0 | 0 | 1674628 | NP_828849.2 | P0C6X7 |
| LGKPREQIDGYVMHAN | 1 | 0 | 0 | 1674629 | YP_009724389.1 | P0DTD1 |
| LGLPTQTVDSSQGSEY | 1 | 0 | 0 | 1674647 | YP_009724389.1 | P0DTD1 |
| LGLQIQTVDSSQGSEY | 1 | 0 | 0 | 1674648 | YP_003766.2 | H9EJ42 |
| LGLTNLSEITKPLIEA | 1 | 0 | 0 | 1674653 | YP_173243.1 | Q0ZJ80 |
| LGLTTQTVDSSQGSEY | 1 | 0 | 0 | 1674654 | YP_009047202.1 | K9N7C7 |
| LGMSLEACYVRGCNPK | 1 | 0 | 0 | 1674667 | YP_009555238.1 | P0C6X6 |
| LGNCYRCPKEIVETVS | 1 | 0 | 0 | 1674670 | YP_173236.1 | A0A140H1G9 |
| LGNELAQVLTEVVYSN | 1 | 0 | 0 | 1674671 | YP_003766.2 | H9EJ42 |
| LGNRKRNQKPLEPKFS | 1 | 0 | 0 | 1674677 | YP_003771.1 | H9EJ69 |
| LGNVEAVNFNVTTAET | 1 | 0 | 0 | 1674679 | NP_073551.1 | A0A0P0K6L9 |
| LGNVFCDGVNVTKHKC | 1 | 0 | 0 | 1674681 | YP_009555238.1 | P0C6X6 |
| LGNVVKTAVQPTAPAY | 1 | 0 | 0 | 1674682 | YP_003766.2 | H9EJ42 |
| LGPDIFLGNCYRCPKE | 1 | 0 | 0 | 1674684 | YP_173236.1 | A0A140H1G9 |
| LGPDIFLGTCYRCPKE | 1 | 0 | 0 | 1674685 | YP_009555238.1 | P0C6X6 |
| LGPDIFLSMCYRCPKE | 1 | 0 | 0 | 1674686 | YP_009047202.1 | K9N7C7 |
| LGPHCGSSSRSFFEDL | 1 | 0 | 0 | 1674687 | YP_173238.1 | A0A140H1H1 |
| LGQSLCALPDTPSTLT | 1 | 0 | 0 | 1674695 | YP_009047204.1 | K9N5Q8 |
| LGRCIVLDVKGVEELH | 1 | 0 | 0 | 1674697 | YP_009555239.1 | Q80872 |
| LGRELNCNVCSNRATC | 1 | 0 | 0 | 1674699 | YP_173236.1 | A0A140H1G9 |
| LGSAILEDEFTPFDVV | 1 | 0 | 0 | 1674704 | QIQ54046.1 | P0C6U8 |
| LGSALLEDEFTPFDVV | 1 | 0 | 0 | 1674705 | YP_009724389.1 | P0DTD1 |
| LGSCTFEDELAPSDVY | 1 | 0 | 0 | 1674706 | YP_173236.1 | A0A140H1G9 |
| LGSECSKASSRSAIED | 1 | 0 | 0 | 1674707 | YP_009555241.1 | P36334 |
| LGSLIDCNAASVNQIV | 1 | 0 | 0 | 1674715 | YP_009047202.1 | K9N7C7 |
| LGSMDIDVFNKLSTSD | 1 | 0 | 0 | 1674720 | NP_073549.1 | A0A0P0LKV0 |
| LGSNGKPGNVGGNDAL | 1 | 0 | 0 | 1674721 | YP_009555238.1 | P0C6X6 |

| | | | | | | |
|---|---|---|---|---|---|---|
| LGSSGFLQSSNYKIDT | 1 | 0 | 0 | 1674723 | YP_173238.1 | A0A140H1H1 |
| LGSSVGNFSDGKMGRF | 1 | 0 | 0 | 1674724 | YP_009047204.1 | K9N5Q8 |
| LGSTILEDEFTPFDVV | 1 | 0 | 0 | 1674725 | NP_828849.2 | P0C6X7 |
| LGTAKKQVVLVSNNQE | 1 | 0 | 0 | 1674728 | YP_009555238.1 | P0C6X6 |
| LGTCRRCPAEIVDTVS | 1 | 0 | 0 | 1674730 | YP_009724389.1 | P0DTD1 |
| LGTCYRCPKEIVDTVS | 1 | 0 | 0 | 1674731 | YP_009555238.1 | P0C6X6 |
| LGTDPIDDFQVNWNTK | 1 | 0 | 0 | 1674732 | QIQ54046.1 | P0C6U8 |
| LGTDPIEDYEQNWNTK | 1 | 0 | 0 | 1674733 | NP_828849.2 | P0C6X7 |
| LGTGPEAGLPYGANKE | 1 | 0 | 0 | 1674738 | QIQ54056.1 | P59595 |
| LGTGPEASLPYGANKE | 1 | 0 | 0 | 1674739 | NP_828858.1 | P59595 |
| LGTGPHAKDQYGTDID | 1 | 0 | 0 | 1674740 | YP_009555245.1 | P33469 |
| LGTGYSLADLPAYMTV | 1 | 0 | 0 | 1674744 | YP_009555244.1 | Q01455 |
| LGTIKENIDGGAMHAN | 1 | 0 | 0 | 1674746 | YP_009047202.1 | K9N7C7 |
| LGTNDPQFPILAELAP | 1 | 0 | 0 | 1674748 | YP_173242.1 | Q19U29 |
| LGTVTLKQGEIKDATP | 1 | 0 | 0 | 1674762 | QHR63301.1 | Q3LZX0 |
| LGVEFLKRGDKIVYHT | 1 | 0 | 0 | 1674769 | NP_828849.2 | P0C6X7 |
| LGVGGKPCIKVATVQS | 1 | 0 | 0 | 1674771 | YP_009724389.1 | P0DTD1 |
| LGVSSNQSPQGNLGSN | 1 | 0 | 0 | 1674789 | YP_009555238.1 | P0C6X6 |
| LGVTMDVLNKNQKLIA | 1 | 0 | 0 | 1674791 | YP_173238.1 | A0A140H1H1 |
| LGVTMDVLSQNQKLIA | 1 | 0 | 0 | 1674792 | YP_009555241.1 | P36334 |
| LGVYYHNNNKTWVENE | 1 | 0 | 0 | 1674806 | QIQ54048.1 | P59594 |
| LGYHPAHAPSFLAVDD | 1 | 0 | 0 | 1674812 | YP_173236.1 | A0A140H1G9 |
| LGYVTHGLNLEEAARY | 1 | 0 | 0 | 1674820 | YP_009724389.1 | P0DTD1 |
| LHDDLVNYIRDKGCVA | 1 | 0 | 0 | 1674831 | YP_009555239.1 | Q80872 |
| LHIAGCQTTNGLNTSY | 1 | 0 | 0 | 1674856 | NP_073551.1 | A0A0P0K6L9 |
| LHKAYIDVLRNSFGKD | 1 | 0 | 0 | 1674857 | NP_073549.1 | A0A0P0LKV0 |
| LHLKVNCCRFQRVDEN | 1 | 0 | 0 | 1674869 | YP_009555238.1 | P0C6X6 |
| LHPAFNVSDAYANLVP | 1 | 0 | 0 | 1674895 | YP_003766.2 | H9EJ42 |
| LHPSFNVSDAYANLVP | 1 | 0 | 0 | 1674898 | NP_073549.1 | A0A0P0LKV0 |
| LHQRELLGVSDVWHKQ | 1 | 0 | 0 | 1674905 | YP_009555238.1 | P0C6X6 |
| LHRSYLTPGKLESGWT | 1 | 0 | 0 | 1674911 | QIQ54048.1 | P59594 |
| LHSSQLPLDVYDLTMP | 1 | 0 | 0 | 1674915 | YP_009555238.1 | P0C6X6 |
| LHTQLVDMSMTYGQQF | 1 | 0 | 0 | 1674930 | NP_828849.2 | P0C6X7 |
| LHTQVVDMSMTYGQQF | 1 | 0 | 0 | 1674931 | YP_009724389.1 | P0DTD1 |
| LHVADTLMQGVTLSSN | 1 | 0 | 0 | 1674939 | YP_173238.1 | A0A140H1H1 |
| LHVVGPDARAKQDVSL | 1 | 0 | 0 | 1674946 | YP_009047202.1 | K9N7C7 |
| LHVVGPNLNAGEDIQL | 1 | 0 | 0 | 1674947 | NP_828849.2 | P0C6X7 |
| LHVVGPNVNKGEDIQL | 1 | 0 | 0 | 1674949 | YP_009724389.1 | P0DTD1 |
| LHVVGPNVNRGEDIQL | 1 | 0 | 0 | 1674950 | QHR63299.1 | P0C6W2 |
| LHYGAGSDKGVAPGTT | 1 | 0 | 0 | 1674955 | YP_003766.2 | H9EJ42 |
| LIANAFNNALYAIQEG | 1 | 0 | 0 | 1674966 | YP_009555241.1 | P36334 |
| LIANKFNQALGAMQTG | 1 | 0 | 0 | 1674968 | YP_009047204.1 | K9N5Q8 |
| LIANQFNSAIGKIQDS | 1 | 0 | 0 | 1674969 | YP_009724390.1 | P0DTC2 |
| LIARCWYLHEGHQTAA | 1 | 0 | 0 | 1674970 | NP_849177.1 | Q80H93 |
| LIATAETELAKNVSLD | 1 | 0 | 0 | 1674971 | QIQ54046.1 | P0C6U8 |
| LIATLNSLGETLVTMP | 1 | 0 | 0 | 1674974 | QIQ54046.1 | P0C6U8 |
| LIAVANWLAPTNAYMR | 1 | 0 | 0 | 1674975 | YP_009047202.1 | K9N7C7 |
| LIDGVENPMLMGWDYP | 1 | 0 | 0 | 1674997 | YP_003766.2 | H9EJ42 |
| LIDNINSTLVDLKWLN | 1 | 0 | 0 | 1675001 | NP_073551.1 | A0A0P0K6L9 |
| LIDQINSTYVDLKLLN | 1 | 0 | 0 | 1675002 | YP_003767.1 | A0A0P0G321 |
| LIDSYFDNSSTLQSVA | 1 | 0 | 0 | 1675003 | YP_003766.2 | H9EJ42 |
| LIEGSKPYSTLQPNAY | 1 | 0 | 0 | 1675009 | YP_003766.2 | H9EJ42 |
| LIERFVSLAIDAYPLV | 1 | 0 | 0 | 1675013 | YP_009555238.1 | P0C6X6 |
| LIERGKLLVRDTRKEI | 1 | 0 | 0 | 1675014 | YP_173236.1 | A0A140H1G9 |
| LIEWPLTEYDFTDYKV | 1 | 0 | 0 | 1675015 | YP_009047202.1 | K9N7C7 |
| LIGANYLGKPKEQIDG | 1 | 0 | 0 | 1675031 | NP_828849.2 | P0C6X7 |
| LIGDCATVHTANKWDL | 1 | 0 | 0 | 1675033 | YP_009724389.1 | P0DTD1 |
| LIGESVKTQFNYFKKV | 1 | 0 | 0 | 1675036 | NP_828849.2 | P0C6X7 |
| LIGINYLCKPKVEIDG | 1 | 0 | 0 | 1675042 | YP_009555238.1 | P0C6X6 |
| LIGLAKRFKESPFELE | 1 | 0 | 0 | 1675047 | YP_009724389.1 | P0DTD1 |
| LIGLAKRSKDSPLELE | 1 | 0 | 0 | 1675048 | QIQ54047.1 | P0C6X7 |
| LIGLAKRSKESPLELE | 1 | 0 | 0 | 1675049 | QHR63299.1 | P0C6W2 |
| LIGLYRRQQTSNLVVQ | 1 | 0 | 0 | 1675053 | YP_009555238.1 | P0C6X6 |
| LIIKNLSKPLTENKYS | 1 | 0 | 0 | 1675066 | QIQ54052.1 | P59634 |
| LIIRENNRVVISSDVL | 1 | 0 | 0 | 1675072 | YP_009724389.1 | P0DTD1 |
| LIIRENNRVVVSSDIL | 1 | 0 | 0 | 1675073 | NP_828849.2 | P0C6X7 |
| LIISDMYDPITKNIGE | 1 | 0 | 0 | 1675074 | YP_009555238.1 | P0C6X6 |
| LIKAYNTINNEQGTPL | 1 | 0 | 0 | 1675082 | NP_073549.1 | A0A0P0LKV0 |
| LIKDVDNPVLMGWDYP | 1 | 0 | 0 | 1675083 | YP_009555238.1 | P0C6X6 |
| LIKEPCSSGTYEGNSP | 1 | 0 | 0 | 1675084 | QIQ54053.1 | P59635 |
| LIKESIPIVWGVKDFN | 1 | 0 | 0 | 1675085 | YP_003766.2 | H9EJ42 |
| LIKQLKRAMNIAKSEF | 1 | 0 | 0 | 1675092 | YP_003766.2 | H9EJ42 |
| LILAYCNKTVGELGDV | 1 | 0 | 0 | 1675099 | YP_009724389.1 | P0DTD1 |
| LILSWEIGKVKPPLNK | 1 | 0 | 0 | 1675111 | YP_009555238.1 | P0C6X6 |
| LIMAYGNCTFGAPDDA | 1 | 0 | 0 | 1675115 | YP_009047202.1 | K9N7C7 |
| LINKFDVINGVKTVKY | 1 | 0 | 0 | 1675123 | YP_173236.1 | A0A140H1G9 |
| LINKFLKANPLWHKAV | 1 | 0 | 0 | 1675124 | YP_009555238.1 | P0C6X6 |
| LINTLNDLNETLVTMP | 1 | 0 | 0 | 1675128 | YP_009724389.1 | P0DTD1 |
| LIPGFVLPCNAQEQQI | 1 | 0 | 0 | 1675130 | YP_173236.1 | A0A140H1G9 |
| LIPVRPRNSSDNGISA | 1 | 0 | 0 | 1675141 | YP_003767.1 | A0A0P0G321 |
| LIQESIKSLNNSYINL | 1 | 0 | 0 | 1675145 | YP_173238.1 | A0A140H1H1 |
| LIQNGAKHVQGNVAKA | 1 | 0 | 0 | 1675148 | YP_173236.1 | A0A140H1G9 |
| LISAGIFSVPADVSLT | 1 | 0 | 0 | 1675168 | YP_173236.1 | A0A140H1G9 |
| LISAGIFSVPSDVSLT | 1 | 0 | 0 | 1675169 | YP_009555238.1 | P0C6X6 |
| LISDMYDGRTKAIDGE | 1 | 0 | 0 | 1675171 | NP_073549.1 | A0A0P0LKV0 |
| LISMMGFKMNYQVNGY | 1 | 0 | 0 | 1675184 | YP_009724389.1 | P0DTD1 |
| LISQDSITHYPALKMV | 1 | 0 | 0 | 1675187 | YP_009047202.1 | K9N7C7 |
| LISTARDGVRRGDNFH | 1 | 0 | 0 | 1675190 | YP_009047202.1 | K9N7C7 |
| LITEGNVKGLQPSVGP | 1 | 0 | 0 | 1675193 | QIQ54047.1 | P0C6X7 |
| LITEGSVKGLTPSKGP | 1 | 0 | 0 | 1675195 | NP_828849.2 | P0C6X7 |
| LITGVFDIVKGQIQVA | 1 | 0 | 0 | 1675199 | NP_828849.2 | P0C6X7 |

| | | | | | | |
|---|---|---|---|---|---|---|
| LIVDQVHVLNHVDPII | 1 | 0 | 0 | 1675212 | YP_173243.1 | Q0ZJ80 |
| LKAAVHGWKMVVCDKV | 1 | 0 | 0 | 1675239 | YP_009047202.1 | K9N7C7 |
| LKAAYENFNSQDILLA | 1 | 0 | 0 | 1675240 | NP_828849.2 | P0C6X7 |
| LKADDFVTASDTTLRC | 1 | 0 | 0 | 1675242 | YP_003766.2 | H9EJ42 |
| LKALVATAHSELAKGV | 1 | 0 | 0 | 1675253 | NP_828849.2 | P0C6X7 |
| LKAPKEVTFLEGDSHD | 1 | 0 | 0 | 1675259 | NP_828849.2 | P0C6X7 |
| LKAVRSSFNFDQKELL | 1 | 0 | 0 | 1675267 | YP_009555238.1 | P0C6X6 |
| LKCGFSFDLNGLDALF | 1 | 0 | 0 | 1675269 | YP_173236.1 | A0A140H1G9 |
| LKCGSPYQHYDACSVK | 1 | 0 | 0 | 1675270 | YP_173236.1 | A0A140H1G9 |
| LKCNRCCDRYEEYDLE | 1 | 0 | 0 | 1675271 | YP_009047204.1 | K9N5Q8 |
| LKCTSDNINDKDTGPP | 1 | 0 | 0 | 1675277 | YP_009555241.1 | P36334 |
| LKDDGNFVVLELDPPC | 1 | 0 | 0 | 1675289 | YP_009555238.1 | P0C6X6 |
| LKDGVNFNVDDINFSP | 1 | 0 | 0 | 1675292 | YP_009555241.1 | P36334 |
| LKDIKFVIPALQHAFM | 1 | 0 | 0 | 1675297 | YP_009047202.1 | K9N7C7 |
| LKDLLLYAADPALHVA | 1 | 0 | 0 | 1675301 | YP_009555238.1 | P0C6X6 |
| LKDLLLYAADPAMHVA | 1 | 0 | 0 | 1675302 | YP_173236.1 | A0A140H1G9 |
| LKDSDINEMVLSLVRS | 1 | 0 | 0 | 1675305 | NP_073549.1 | A0A0P0LKV0 |
| LKDSDVNDMVLSLIKS | 1 | 0 | 0 | 1675306 | YP_003766.2 | H9EJ42 |
| LKEDYQIGGYSEDRHS | 1 | 0 | 0 | 1675309 | NP_828852.2 | P59632 |
| LKEIVGPKELLLLWES | 1 | 0 | 0 | 1675315 | NP_073549.1 | A0A0P0LKV0 |
| LKELLQNGMNGRTILG | 1 | 0 | 0 | 1675318 | YP_009724389.1 | P0DTD1 |
| LKELLQNGVNGRTILG | 1 | 0 | 0 | 1675319 | QIQ54046.1 | P0C6U8 |
| LKELYGPVDPTFLHRF | 1 | 0 | 0 | 1675321 | YP_009047202.1 | K9N7C7 |
| LKENQINDMIYSLLEK | 1 | 0 | 0 | 1675322 | NP_828849.2 | P0C6X7 |
| LKEVVGPKELLLSWES | 1 | 0 | 0 | 1675329 | YP_003766.2 | H9EJ42 |
| LKEWNFSLGIILLFIT | 1 | 0 | 0 | 1675330 | YP_009555244.1 | Q01455 |
| LKFDNRGVITNAVDCS | 1 | 0 | 0 | 1675332 | YP_173238.1 | A0A140H1H1 |
| LKFDSGTVSKTSDWKC | 1 | 0 | 0 | 1675333 | YP_009047202.1 | K9N7C7 |
| LKFIDKLNVLFDGRDN | 1 | 0 | 0 | 1675336 | YP_009555238.1 | P0C6X6 |
| LKFTEESFYKNMYLKS | 1 | 0 | 0 | 1675345 | YP_173236.1 | A0A140H1G9 |
| LKGGAPIKGVTFGEDT | 1 | 0 | 0 | 1675352 | NP_828849.2 | P0C6X7 |
| LKGGAPVKKVAFGGDQ | 1 | 0 | 0 | 1675354 | YP_009047202.1 | K9N7C7 |
| LKGGLILKFDSGTVSK | 1 | 0 | 0 | 1675359 | YP_009047202.1 | K9N7C7 |
| LKGKYVQIPTTCANDP | 1 | 0 | 0 | 1675361 | YP_009724389.1 | P0DTD1 |
| LKGSGSDFSLADSVVE | 1 | 0 | 0 | 1675364 | YP_009555238.1 | P0C6X6 |
| LKGVTLDKLRDYLADY | 1 | 0 | 0 | 1675372 | YP_009047202.1 | K9N7C7 |
| LKHAEEYREYLESYNT | 1 | 0 | 0 | 1675375 | YP_009555238.1 | P0C6X6 |
| LKHFFFTQKGDAAIKD | 1 | 0 | 0 | 1675381 | NP_073549.1 | A0A0P0LKV0 |
| LKHGTFTCASEYTGNY | 1 | 0 | 0 | 1675386 | YP_009724389.1 | P0DTD1 |
| LKISTQLFEGDKFIGT | 1 | 0 | 0 | 1675402 | YP_003766.2 | H9EJ42 |
| LKKAMNVAKAEFDRES | 1 | 0 | 0 | 1675409 | NP_073549.1 | A0A0P0LKV0 |
| LKKCKSAFYVLPSEAP | 1 | 0 | 0 | 1675411 | NP_828849.2 | P0C6X7 |
| LKKLIGGDVTPVDQYM | 1 | 0 | 0 | 1675418 | YP_009047202.1 | K9N7C7 |
| LKKLKKSLNVAKSEFD | 1 | 0 | 0 | 1675419 | YP_009724389.1 | P0DTD1 |
| LKKSLNVAKSEFDRDA | 1 | 0 | 0 | 1675430 | YP_009724389.1 | P0DTD1 |
| LKLAGTAVINLRADQI | 1 | 0 | 0 | 1675433 | YP_009555238.1 | P0C6X6 |
| LKLAGTAVVNLRPDQL | 1 | 0 | 0 | 1675434 | YP_173236.1 | A0A140H1G9 |
| LKLGTNDPQFPILAEL | 1 | 0 | 0 | 1675442 | YP_173242.1 | Q19U29 |
| LKLKVSTNLFEGDKFV | 1 | 0 | 0 | 1675451 | NP_073549.1 | A0A0P0LKV0 |
| LKLPPHDVTVLRDNLK | 1 | 0 | 0 | 1675456 | YP_003768.1 | H9EJ24 |
| LKLRSDVLLPLTQYNR | 1 | 0 | 0 | 1675462 | YP_009724389.1 | P0DTD1 |
| LKLSDNNCYLNAVIMT | 1 | 0 | 0 | 1675464 | YP_009047202.1 | K9N7C7 |
| LKLTDNVYIKNADIVE | 1 | 0 | 0 | 1675469 | YP_009724389.1 | P0DTD1 |
| LKLTFNKQEASVPILT | 1 | 0 | 0 | 1675470 | YP_173236.1 | A0A140H1G9 |
| LKMVKRQSYGTNSGVA | 1 | 0 | 0 | 1675482 | YP_009047210.1 | K9N7A1 |
| LKNLGFDNQSKSPSSS | 1 | 0 | 0 | 1675492 | YP_003771.1 | H9EJ69 |
| LKNLTQTFTSMLTNYF | 1 | 0 | 0 | 1675497 | YP_173236.1 | A0A140H1G9 |
| LKNNLYRLVWHVERKD | 1 | 0 | 0 | 1675498 | YP_009047202.1 | K9N7C7 |
| LKNSVSDVAYNRYLSL | 1 | 0 | 0 | 1675500 | YP_173236.1 | A0A140H1G9 |
| LKNYAASYNKYKYYSG | 1 | 0 | 0 | 1675503 | YP_003766.2 | H9EJ42 |
| LKPGDSFNILACYEGI | 1 | 0 | 0 | 1675508 | YP_003766.2 | H9EJ42 |
| LKPILDWLEEKFKEGI | 1 | 0 | 0 | 1675511 | QIQ54046.1 | P0C6U8 |
| LKPMPFFYFDDSDCDV | 1 | 0 | 0 | 1675514 | YP_003766.2 | H9EJ42 |
| LKPRVEAPKQEEPPNT | 1 | 0 | 0 | 1675516 | NP_828849.2 | P0C6X7 |
| LKPVLDWLEEKFKEGV | 1 | 0 | 0 | 1675520 | YP_009724389.1 | P0DTD1 |
| LKPVSEEVVENPTIQK | 1 | 0 | 0 | 1675522 | YP_009724389.1 | P0DTD1 |
| LKQGEIKDATPSDFVR | 1 | 0 | 0 | 1675527 | YP_009724391.1 | P0DTC3 |
| LKQGEIKDATPSDSVR | 1 | 0 | 0 | 1675528 | QHR63301.1 | Q3LZX0 |
| LKQKDIGIDTAGVLLK | 1 | 0 | 0 | 1675529 | YP_173243.1 | Q0ZJ80 |
| LKQLMVNKSSCYRDGI | 1 | 0 | 0 | 1675538 | YP_173243.1 | Q0ZJ80 |
| LKRAMNIAKSEFDHEI | 1 | 0 | 0 | 1675544 | YP_003766.2 | H9EJ42 |
| LKRGDKIVYHTLESPV | 1 | 0 | 0 | 1675546 | NP_828849.2 | P0C6X7 |
| LKRGDKSVYYTSNPIT | 1 | 0 | 0 | 1675547 | QHR63299.1 | P0C6W2 |
| LKRGDKSVYYTSNPTT | 1 | 0 | 0 | 1675548 | YP_009724389.1 | P0DTD1 |
| LKRGDKVVYHTTSKPI | 1 | 0 | 0 | 1675549 | QIQ54046.1 | P0C6U8 |
| LKRRDSTYGPLQTPVG | 1 | 0 | 0 | 1675554 | YP_009047204.1 | K9N5Q8 |
| LKSAYENFNQHDVLLA | 1 | 0 | 0 | 1675563 | QHR63299.1 | P0C6W2 |
| LKSAYENFNQHEVLLA | 1 | 0 | 0 | 1675564 | YP_009724389.1 | P0DTD1 |
| LKSEDAQGMDNLACED | 1 | 0 | 0 | 1675566 | YP_009724389.1 | P0DTD1 |
| LKSEDTQGMDNLACED | 1 | 0 | 0 | 1675567 | QHR63299.1 | P0C6W2 |
| LKSIAATRGATVVIGT | 1 | 0 | 0 | 1675573 | YP_009724389.1 | P0DTD1 |
| LKSIVNTRNATVVIGT | 1 | 0 | 0 | 1675576 | YP_003766.2 | H9EJ42 |
| LKSLGFDKPQEKDKKS | 1 | 0 | 0 | 1675580 | NP_073556.1 | A0A0N7J131 |
| LKSMAATRGATCVIGT | 1 | 0 | 0 | 1675583 | YP_009047202.1 | K9N7C7 |
| LKSSTGFVYFNATGSD | 1 | 0 | 0 | 1675589 | YP_003767.1 | A0A0P0G321 |
| LKTAGVVTGNVVKQCC | 1 | 0 | 0 | 1675594 | YP_009047202.1 | K9N7C7 |
| LKTCVIANVPAIDVLK | 1 | 0 | 0 | 1675595 | YP_003766.2 | H9EJ42 |
| LKTDSVLLIERFVSLA | 1 | 0 | 0 | 1675596 | YP_009555238.1 | P0C6X6 |
| LKTGDLQPLEQPTSEA | 1 | 0 | 0 | 1675597 | YP_009724389.1 | P0DTD1 |
| LKTGELQPLEEPTAQA | 1 | 0 | 0 | 1675598 | QIQ54046.1 | P0C6U8 |
| LKTNCCRFQEKDEDDN | 1 | 0 | 0 | 1675616 | YP_009724389.1 | P0DTD1 |

| | | | | | | |
|---|---|---|---|---|---|---|
| LKTNCCRFQEKDEDGN | 1 | 0 | 0 | 1675617 | QIQ54047.1 | P0C6X7 |
| LKTNCCRFQEKDEEGN | 1 | 0 | 0 | 1675618 | NP_828849.2 | P0C6X7 |
| LKTQFNYYKKVNGVVQ | 1 | 0 | 0 | 1675619 | QHR63299.1 | P0C6W2 |
| LKVAYENFNHHEKLLA | 1 | 0 | 0 | 1675626 | QIQ54046.1 | P0C6U8 |
| LKVDTANPKTPKYKFV | 1 | 0 | 0 | 1675630 | YP_009724389.1 | P0DTD1 |
| LKVEFWFAMRKDGDDV | 1 | 0 | 0 | 1675633 | YP_173236.1 | A0A140H1G9 |
| LKVGAGVMLECEKFNV | 1 | 0 | 0 | 1675635 | YP_003766.2 | H9EJ42 |
| LKVNCCRFQRVDENGD | 1 | 0 | 0 | 1675639 | YP_009555238.1 | P0C6X6 |
| LKVSTNLFEGDKFVGT | 1 | 0 | 0 | 1675641 | NP_073549.1 | A0A0P0LKV0 |
| LKWADQSDQFRNVQTR | 1 | 0 | 0 | 1675645 | YP_009555245.1 | P33469 |
| LKYDFTEERLKLFDRY | 1 | 0 | 0 | 1675654 | YP_009724389.1 | P0DTD1 |
| LKYDFTEERLNLFNRY | 1 | 0 | 0 | 1675655 | QIQ54047.1 | P0C6X7 |
| LKYSYINKCSRLLSDD | 1 | 0 | 0 | 1675663 | YP_009047204.1 | K9N5Q8 |
| LKYTKILKDDGNFVVL | 1 | 0 | 0 | 1675664 | YP_009555238.1 | P0C6X6 |
| LKYTKIMKDDGNCVVL | 1 | 0 | 0 | 1675665 | YP_173236.1 | A0A140H1G9 |
| LLAFNQDGIIFNAVDC | 1 | 0 | 0 | 1675674 | YP_009555241.1 | P36334 |
| LLAGSFDKVYDIGNPK | 1 | 0 | 0 | 1675675 | YP_009047202.1 | K9N7C7 |
| LLAGTITSGWTFGAGA | 1 | 0 | 0 | 1675676 | YP_009724390.1 | P0DTC2 |
| LLAPLLSAGIFGADPI | 1 | 0 | 0 | 1675690 | YP_009724389.1 | P0DTD1 |
| LLCATCPFAKGDSCSS | 1 | 0 | 0 | 1675707 | YP_009047202.1 | K9N7C7 |
| LLCLRDDNQTLLVPKI | 1 | 0 | 0 | 1675717 | YP_173236.1 | A0A140H1G9 |
| LLCRMNSRNYIAQVDV | 1 | 0 | 0 | 1675718 | YP_009725255.1 | A0A663DJA2 |
| LLDGTCGIIDVEKGVL | 1 | 0 | 0 | 1675740 | QIQ54046.1 | P0C6U8 |
| LLDNQEDKFWCEDFYA | 1 | 0 | 0 | 1675749 | YP_003766.2 | H9EJ42 |
| LLDQALVSDVGDSAEV | 1 | 0 | 0 | 1675750 | YP_009724389.1 | P0DTD1 |
| LLDQALVSDVGDSTEV | 1 | 0 | 0 | 1675751 | QHR63299.1 | P0C6W2 |
| LLDQAWRFPCAGRKVN | 1 | 0 | 0 | 1675752 | YP_173236.1 | A0A140H1G9 |
| LLDQAWRVPCAGRRVT | 1 | 0 | 0 | 1675753 | YP_009555238.1 | P0C6X6 |
| LLDRLNALESKMSGKG | 1 | 0 | 0 | 1675754 | QIQ54056.1 | P59595 |
| LLDTKLNGILQQAALD | 1 | 0 | 0 | 1675758 | YP_003766.2 | H9EJ42 |
| LLDTTQLHVADTLMQG | 1 | 0 | 0 | 1675759 | YP_173238.1 | A0A140H1H1 |
| LLDWVKHLNKNLNEGV | 1 | 0 | 0 | 1675764 | YP_003766.2 | H9EJ42 |
| LLDWVKHLNKTLNEGV | 1 | 0 | 0 | 1675765 | NP_073549.1 | A0A0P0LKV0 |
| LLEDEFTPFDVVRQCS | 1 | 0 | 0 | 1675769 | YP_009724389.1 | P0DTD1 |
| LLEGSISYSELRPDTR | 1 | 0 | 0 | 1675771 | NP_828849.2 | P0C6X7 |
| LLEIKDTEKYCALAPN | 1 | 0 | 0 | 1675773 | YP_009724389.1 | P0DTD1 |
| LLEKGKLLIRDTNKEV | 1 | 0 | 0 | 1675779 | YP_009555238.1 | P0C6X6 |
| LLEKGRLIIRENNRVV | 1 | 0 | 0 | 1675780 | NP_828849.2 | P0C6X7 |
| LLENIDYFNMRRAKFS | 1 | 0 | 0 | 1675781 | YP_009555238.1 | P0C6X6 |
| LLEQNIDAYKTFPKKE | 1 | 0 | 0 | 1675783 | YP_009047211.1 | K9N4V7 |
| LLFADINGKLYHDSQN | 1 | 0 | 0 | 1675792 | NP_828849.2 | P0C6X7 |
| LLGCLSSMNIAANSSE | 1 | 0 | 0 | 1675831 | YP_003766.2 | H9EJ42 |
| LLGFDSSKPVTKKYTY | 1 | 0 | 0 | 1675833 | YP_009047202.1 | K9N7C7 |
| LLGIGGKPCIKVATVQ | 1 | 0 | 0 | 1675839 | NP_828849.2 | P0C6X7 |
| LLGINYLGTIKENIDG | 1 | 0 | 0 | 1675844 | YP_009047202.1 | K9N7C7 |
| LLGLTNLSEITKPLIE | 1 | 0 | 0 | 1675847 | YP_173243.1 | Q0ZJ80 |
| LLGTAKKQVVLVSNNQ | 1 | 0 | 0 | 1675853 | YP_009555238.1 | P0C6X6 |
| LLGVGGKPCIKVATVQ | 1 | 0 | 0 | 1675858 | YP_009724389.1 | P0DTD1 |
| LLHIAGCQTTNGLNTS | 1 | 0 | 0 | 1675865 | NP_073551.1 | A0A0P0K6L9 |
| LLHSSQLPLDVYDLTM | 1 | 0 | 0 | 1675874 | YP_009555238.1 | P0C6X6 |
| LLIERFVSLAIDAYPL | 1 | 0 | 0 | 1675880 | YP_009555238.1 | P0C6X6 |
| LLIGLAKRFKESPFEL | 1 | 0 | 0 | 1675883 | YP_009724389.1 | P0DTD1 |
| LLIGLAKRSKDSPLEL | 1 | 0 | 0 | 1675884 | QIQ54047.1 | P0C6X7 |
| LLIGLAKRSKESPLEL | 1 | 0 | 0 | 1675885 | QHR63299.1 | P0C6W2 |
| LLIKAYNTINNEQGTP | 1 | 0 | 0 | 1675891 | NP_073549.1 | A0A0P0LKV0 |
| LLIQEFLQYDSSIHSY | 1 | 0 | 0 | 1675899 | YP_173236.1 | A0A140H1G9 |
| LLISDMYDGRTKAIDG | 1 | 0 | 0 | 1675906 | NP_073549.1 | A0A0P0LKV0 |
| LLKAAYENFNSQDILL | 1 | 0 | 0 | 1675920 | NP_828849.2 | P0C6X7 |
| LLKDCPAVAKHDFFKF | 1 | 0 | 0 | 1675922 | YP_009724389.1 | P0DTD1 |
| LLKDIKFVIPALQHAF | 1 | 0 | 0 | 1675924 | YP_009047202.1 | K9N7C7 |
| LLKEPCSSGTYEGNSP | 1 | 0 | 0 | 1675929 | YP_009724395.1 | P0DTC7 |
| LLKFDNRGVITNAVDC | 1 | 0 | 0 | 1675930 | YP_173238.1 | A0A140H1H1 |
| LLKLKVSTNLFEGDKF | 1 | 0 | 0 | 1675937 | NP_073549.1 | A0A0P0LKV0 |
| LLKSIAATRGATVVIG | 1 | 0 | 0 | 1675947 | YP_009724389.1 | P0DTD1 |
| LLKTDSVLLIERFVSL | 1 | 0 | 0 | 1675948 | YP_009555238.1 | P0C6X6 |
| LLKVAYENFNHHEKLL | 1 | 0 | 0 | 1675952 | QIQ54046.1 | P0C6U8 |
| LLKYDFTEERLCLFDR | 1 | 0 | 0 | 1675954 | NP_828849.2 | P0C6X7 |
| LLKYDFTEERLKLFDR | 1 | 0 | 0 | 1675955 | YP_009724389.1 | P0DTD1 |
| LLKYDFTEERLNLFNR | 1 | 0 | 0 | 1675956 | QIQ54047.1 | P0C6X7 |
| LLKYDFTEHKENLFNK | 1 | 0 | 0 | 1675957 | YP_003766.2 | H9EJ42 |
| LLKYDFTEHKEVLFNK | 1 | 0 | 0 | 1675958 | NP_073549.1 | A0A0P0LKV0 |
| LLLCRMNSRNYIAQVD | 1 | 0 | 0 | 1675965 | YP_009725255.1 | A0A663DJA2 |
| LLLDDFVEIIKSQDLS | 1 | 0 | 0 | 1675966 | YP_009724389.1 | P0DTD1 |
| LLLDDFVSVLKSLDLT | 1 | 0 | 0 | 1675968 | NP_073549.1 | A0A0P0LKV0 |
| LLLDQALVSDVGDSAE | 1 | 0 | 0 | 1675972 | YP_009724389.1 | P0DTD1 |
| LLLDQALVSDVGDSTE | 1 | 0 | 0 | 1675973 | QHR63299.1 | P0C6W2 |
| LLLDRLNALESKMSGK | 1 | 0 | 0 | 1675974 | QIQ54056.1 | P59595 |
| LLLLDRLNALESKMSG | 1 | 0 | 0 | 1675987 | QIQ54056.1 | P59595 |
| LLLLWESGKAKPPLNR | 1 | 0 | 0 | 1675995 | NP_073549.1 | A0A0P0LKV0 |
| LLLNRGVYKPLLENID | 1 | 0 | 0 | 1676000 | YP_009555238.1 | P0C6X6 |
| LLLRHDIQLNNDVRDY | 1 | 0 | 0 | 1676005 | YP_173236.1 | A0A140H1G9 |
| LLLSWESGKVKPPLNR | 1 | 0 | 0 | 1676012 | YP_003766.2 | H9EJ42 |
| LLLWESGKAKPPLNRN | 1 | 0 | 0 | 1676021 | NP_073549.1 | A0A0P0LKV0 |
| LLLYADINGNLYPDST | 1 | 0 | 0 | 1676024 | QIQ54046.1 | P0C6U8 |
| LLMPLKAPKEVTFLEG | 1 | 0 | 0 | 1676036 | NP_828849.2 | P0C6X7 |
| LLNCLQHLHVASNSKL | 1 | 0 | 0 | 1676040 | YP_009555238.1 | P0C6X6 |
| LLNGIAYTTFSGSFDN | 1 | 0 | 0 | 1676045 | YP_003766.2 | H9EJ42 |
| LLNLTDEDLARDFSPI | 1 | 0 | 0 | 1676058 | YP_009047207.1 | K9N643 |
| LLNPSALEFNPSQTSP | 1 | 0 | 0 | 1676060 | NP_073556.1 | A0A0N7J131 |
| LLNRGDYNMLLKNIDL | 1 | 0 | 0 | 1676063 | YP_173236.1 | A0A140H1G9 |
| LLNRGVYKPLLENIDY | 1 | 0 | 0 | 1676064 | YP_009555238.1 | P0C6X6 |

| | | | | | | |
|---|---|---|---|---|---|---|
| LLPPKNSIDAFKLNVK | 1 | 0 | 0 | 1676081 | QIQ54046.1 | P0C6U8 |
| LLPPKSSIDAFKLNIK | 1 | 0 | 0 | 1676082 | NP_828849.2 | P0C6X7 |
| LLQLPNFYYVSNGGNN | 1 | 0 | 0 | 1676106 | YP_003767.1 | A0A0P0G321 |
| LLRKNGNKGAGGHSFG | 1 | 0 | 0 | 1676129 | QIQ54046.1 | P0C6U8 |
| LLRKNGNKGAGGHSYG | 1 | 0 | 0 | 1676130 | YP_009724389.1 | P0DTD1 |
| LLSDDRTEVPQLVNAN | 1 | 0 | 0 | 1676147 | YP_009047204.1 | K9N5Q8 |
| LLSEYPLLKQKDIGID | 1 | 0 | 0 | 1676156 | YP_173243.1 | Q0ZJ80 |
| LLSKGRLIIRENNRVV | 1 | 0 | 0 | 1676169 | YP_009724389.1 | P0DTD1 |
| LLSKGSLEPRHFNSIT | 1 | 0 | 0 | 1676170 | YP_173236.1 | A0A140H1G9 |
| LLSKTSEYKGPVTDVF | 1 | 0 | 0 | 1676173 | QIQ54046.1 | P0C6U8 |
| LLSMQGAVDINKLCEE | 1 | 0 | 0 | 1676181 | YP_009724389.1 | P0DTD1 |
| LLSMQGAVDINRLCEE | 1 | 0 | 0 | 1676182 | NP_828849.2 | P0C6X7 |
| LLSNFGAPSYCNGVRE | 1 | 0 | 0 | 1676183 | NP_828849.2 | P0C6X7 |
| LLSNQLRLAFLGASVT | 1 | 0 | 0 | 1676184 | YP_003766.2 | H9EJ42 |
| LLSQGKLLIRDNDTLS | 1 | 0 | 0 | 1676185 | YP_009047202.1 | K9N7C7 |
| LLSTDGNKIADKHVRN | 1 | 0 | 0 | 1676190 | QHR63299.1 | P0C6W2 |
| LLSTDGNKIADKYVRN | 1 | 0 | 0 | 1676191 | YP_009724389.1 | P0DTD1 |
| LLSTDGNKIGDKYIRN | 1 | 0 | 0 | 1676192 | QIQ54047.1 | P0C6X7 |
| LLSVLLSMQGAVDINK | 1 | 0 | 0 | 1676197 | YP_009724389.1 | P0DTD1 |
| LLSVLLSMQGAVDINR | 1 | 0 | 0 | 1676198 | NP_828849.2 | P0C6X7 |
| LLSWESGKVKPPLNRN | 1 | 0 | 0 | 1676203 | YP_003766.2 | H9EJ42 |
| LLSYNQNGTITDAVDC | 1 | 0 | 0 | 1676205 | QIQ54048.1 | P59594 |
| LLTFNDNQAITQVPAT | 1 | 0 | 0 | 1676210 | YP_003766.2 | H9EJ42 |
| LLTKGTLEPEYFNSVC | 1 | 0 | 0 | 1676215 | YP_009724389.1 | P0DTD1 |
| LLTKSSEYKGPITDVF | 1 | 0 | 0 | 1676217 | YP_009724389.1 | P0DTD1 |
| LLTNMGIDLDEWSVAT | 1 | 0 | 0 | 1676222 | NP_828849.2 | P0C6X7 |
| LLTNTSSVVDGVVRSF | 1 | 0 | 0 | 1676224 | NP_073551.1 | A0A0P0K6L9 |
| LLTRGTLEPENFNSVT | 1 | 0 | 0 | 1676228 | YP_009047202.1 | K9N7C7 |
| LLVDKKLITTACNGIS | 1 | 0 | 0 | 1676243 | YP_173236.1 | A0A140H1G9 |
| LLVDSDLNDFVSDADS | 1 | 0 | 0 | 1676244 | YP_009724389.1 | P0DTD1 |
| LLVTASSKPLYVPEHC | 1 | 0 | 0 | 1676276 | YP_009047205.1 | K9N796 |
| LLWESGKAKPPLNRNS | 1 | 0 | 0 | 1676284 | NP_073549.1 | A0A0P0LKV0 |
| LLYADINGNLYPDSTS | 1 | 0 | 0 | 1676291 | QIQ54046.1 | P0C6U8 |
| LLYCDSPFEKLQCEHL | 1 | 0 | 0 | 1676295 | YP_003767.1 | A0A0P0G321 |
| LLYDSNGNIIGFKDFV | 1 | 0 | 0 | 1676298 | YP_173238.1 | A0A140H1H1 |
| LLYDSNGNLYGFRDYI | 1 | 0 | 0 | 1676299 | YP_009555241.1 | P36334 |
| LLYIDINGNLHPDSAT | 1 | 0 | 0 | 1676305 | YP_009724389.1 | P0DTD1 |
| LLYLDLLNRLQALESG | 1 | 0 | 0 | 1676307 | YP_009047211.1 | K9N4V7 |
| LLYQPPNCSITSGVLQ | 1 | 0 | 0 | 1676312 | YP_009047202.1 | K9N7C7 |
| LLYSFEAHKDSFKDGL | 1 | 0 | 0 | 1676314 | YP_009555238.1 | P0C6X6 |
| LMAAYVDNSSLTIKKP | 1 | 0 | 0 | 1676321 | YP_009724389.1 | P0DTD1 |
| LMADVDDPKLMGWDYP | 1 | 0 | 0 | 1676325 | NP_073549.1 | A0A0P0LKV0 |
| LMEGSLPYSSIQANAY | 1 | 0 | 0 | 1676340 | NP_073549.1 | A0A0P0LKV0 |
| LMGANGNKIVDKEVKD | 1 | 0 | 0 | 1676344 | YP_009047202.1 | K9N7C7 |
| LMGFKLDLTLDGYSKL | 1 | 0 | 0 | 1676345 | YP_173236.1 | A0A140H1G9 |
| LMGILSNMNIASNSKE | 1 | 0 | 0 | 1676350 | NP_073549.1 | A0A0P0LKV0 |
| LMGWDYPKCDRALPNM | 1 | 0 | 0 | 1676352 | YP_003766.2 | H9EJ42 |
| LMGWDYPKCDRAMPNI | 1 | 0 | 0 | 1676353 | YP_173236.1 | A0A140H1G9 |
| LMGWDYPKCDRAMPNL | 1 | 0 | 0 | 1676354 | YP_009555238.1 | P0C6X6 |
| LMGWDYPKCDRAMPNM | 1 | 0 | 0 | 1676355 | YP_009724389.1 | P0DTD1 |
| LMGWDYPKCDRAMPSM | 1 | 0 | 0 | 1676356 | NP_073549.1 | A0A0P0LKV0 |
| LMLDKQGQLLDTKLNG | 1 | 0 | 0 | 1676370 | YP_003766.2 | H9EJ42 |
| LMLLEIKDTEKYCALA | 1 | 0 | 0 | 1676374 | YP_009724389.1 | P0DTD1 |
| LMPHKLKIQVVNSGSD | 1 | 0 | 0 | 1676388 | YP_173236.1 | A0A140H1G9 |
| LMPLKAPKEVTFLEGD | 1 | 0 | 0 | 1676395 | NP_828849.2 | P0C6X7 |
| LMQGVTLSSNLNTNLH | 1 | 0 | 0 | 1676402 | YP_173238.1 | A0A140H1H1 |
| LMSEDVKLNPSEDFIK | 1 | 0 | 0 | 1676410 | YP_009047202.1 | K9N7C7 |
| LMTFSGNVDLDALASD | 1 | 0 | 0 | 1676424 | YP_009047202.1 | K9N7C7 |
| LMVDKNLITTANTGTS | 1 | 0 | 0 | 1676427 | YP_009555238.1 | P0C6X6 |
| LMVVVPDYGTYKNTCD | 1 | 0 | 0 | 1676431 | NP_828849.2 | P0C6X7 |
| LMVVVPDYNTYKNTCE | 1 | 0 | 0 | 1676432 | QIQ54046.1 | P0C6U8 |
| LNAFVAQQLVRSESAA | 1 | 0 | 0 | 1676446 | YP_009047204.1 | K9N5Q8 |
| LNAGEDIQLLKAAYEN | 1 | 0 | 0 | 1676448 | NP_828849.2 | P0C6X7 |
| LNAKLGFDSSKEFVEY | 1 | 0 | 0 | 1676450 | YP_009555238.1 | P0C6X6 |
| LNALIATAHSSIKQGT | 1 | 0 | 0 | 1676452 | YP_009555238.1 | P0C6X6 |
| LNANMSLAECKRALGL | 1 | 0 | 0 | 1676453 | NP_073549.1 | A0A0P0LKV0 |
| LNASEVVVNNLDKSAG | 1 | 0 | 0 | 1676459 | YP_009047202.1 | K9N7C7 |
| LNASVTLKICKFSRNT | 1 | 0 | 0 | 1676461 | YP_003767.1 | A0A0P0G321 |
| LNAYVNSNQNTDSDSL | 1 | 0 | 0 | 1676465 | YP_173242.1 | Q19U29 |
| LNAYVSQQLSDSTLVK | 1 | 0 | 0 | 1676467 | YP_009555241.1 | P36334 |
| LNCERVVKLQNNEIMP | 1 | 0 | 0 | 1676470 | YP_003766.2 | H9EJ42 |
| LNCLQHLHVASNSKLW | 1 | 0 | 0 | 1676471 | YP_009555238.1 | P0C6X6 |
| LNCSFQDAASNIFVIN | 1 | 0 | 0 | 1676475 | YP_009047202.1 | K9N7C7 |
| LNDFSNSGADVLYQPP | 1 | 0 | 0 | 1676484 | NP_828849.2 | P0C6X7 |
| LNDFSNSGSDVLYQPP | 1 | 0 | 0 | 1676485 | YP_009724389.1 | P0DTD1 |
| LNDLNETLVTMPLGYV | 1 | 0 | 0 | 1676493 | YP_009724389.1 | P0DTD1 |
| LNDNLLEILQKEKVNI | 1 | 0 | 0 | 1676495 | YP_009724389.1 | P0DTD1 |
| LNDPSSKTVCTYMDLL | 1 | 0 | 0 | 1676497 | NP_073549.1 | A0A0P0LKV0 |
| LNDVKDNDGRPVHVKE | 1 | 0 | 0 | 1676499 | YP_003766.2 | H9EJ42 |
| LNEKCSVYTVESGTEV | 1 | 0 | 0 | 1676511 | NP_828849.2 | P0C6X7 |
| LNENLNAYQQQDGMMN | 1 | 0 | 0 | 1676513 | YP_009555245.1 | P33469 |
| LNESYIDLKELGNYTY | 1 | 0 | 0 | 1676517 | YP_009047204.1 | K9N5Q8 |
| LNETLVTMPLGYVTHG | 1 | 0 | 0 | 1676518 | YP_009724389.1 | P0DTD1 |
| LNFGMLNGNAIATVKS | 1 | 0 | 0 | 1676522 | NP_073549.1 | A0A0P0LKV0 |
| LNFKKVETLNFQMVKE | 1 | 0 | 0 | 1676524 | YP_173243.1 | Q0ZJ80 |
| LNGDVVAIDYKHYTPS | 1 | 0 | 0 | 1676536 | YP_009724389.1 | P0DTD1 |
| LNGGAVTRYVDNNFCG | 1 | 0 | 0 | 1676539 | NP_828849.2 | P0C6X7 |
| LNGIISNARNGCIPLS | 1 | 0 | 0 | 1676543 | YP_009047202.1 | K9N7C7 |
| LNGITYTSTSGNLLGF | 1 | 0 | 0 | 1676547 | NP_073551.1 | A0A0P0K6L9 |
| LNGNAIATVKSEDGNI | 1 | 0 | 0 | 1676559 | NP_073549.1 | A0A0P0LKV0 |
| LNGNVSTVFVPATRDS | 1 | 0 | 0 | 1676563 | YP_009047208.1 | K9N7D2 |

| | | | | | | |
|---|---|---|---|---|---|---|
| LNGSCGSVGFNIDYDC | 1 | 0 | 0 | 1676568 | YP_009724389.1 | P0DTD1 |
| LNGVIVDKVGELKVEF | 1 | 0 | 0 | 1676572 | YP_173236.1 | A0A140H1G9 |
| LNGVTLIGEAVKTQFN | 1 | 0 | 0 | 1676574 | YP_009724389.1 | P0DTD1 |
| LNGYTVEEARTVLKKC | 1 | 0 | 0 | 1676578 | QHR63299.1 | P0C6W2 |
| LNHGCTCDRTAIQSFD | 1 | 0 | 0 | 1676582 | NP_073549.1 | A0A0P0LKV0 |
| LNHSVPKLSDFESELS | 1 | 0 | 0 | 1676586 | YP_173238.1 | A0A140H1H1 |
| LNICFDIRDNCSLEKF | 1 | 0 | 0 | 1676592 | YP_009047202.1 | K9N7C7 |
| LNIIVPDKSVYDQVVD | 1 | 0 | 0 | 1676598 | YP_009555238.1 | P0C6X6 |
| LNILFDGRDTGALEAF | 1 | 0 | 0 | 1676601 | YP_173236.1 | A0A140H1G9 |
| LNISDEFSSNVANYQK | 1 | 0 | 0 | 1676604 | YP_009724389.1 | P0DTD1 |
| LNIVGPDARQDGRQSY | 1 | 0 | 0 | 1676605 | YP_173236.1 | A0A140H1G9 |
| LNKATNGAMQKESDDY | 1 | 0 | 0 | 1676606 | NP_828849.2 | P0C6X7 |
| LNKATNNAMQIESDDY | 1 | 0 | 0 | 1676607 | QIQ54046.1 | P0C6U8 |
| LNKATNNAMQVESDDY | 1 | 0 | 0 | 1676608 | YP_009724389.1 | P0DTD1 |
| LNKATNNAMQVESDHY | 1 | 0 | 0 | 1676609 | QHR63299.1 | P0C6W2 |
| LNKFGKASLYYESISY | 1 | 0 | 0 | 1676613 | YP_003766.2 | H9EJ42 |
| LNKFNRASLRQIFDVA | 1 | 0 | 0 | 1676614 | YP_009047202.1 | K9N7C7 |
| LNKHFSMMILSDDGVV | 1 | 0 | 0 | 1676617 | YP_009047202.1 | K9N7C7 |
| LNKIQDVVNQQGNSLN | 1 | 0 | 0 | 1676619 | NP_073551.1 | A0A0P0K6L9 |
| LNKIQDVVNQQGSALN | 1 | 0 | 0 | 1676620 | YP_003767.1 | A0A0P0G321 |
| LNKLGTEDPRWPQIAE | 1 | 0 | 0 | 1676621 | YP_009047211.1 | K9N4V7 |
| LNKLLHWSDQTICYKD | 1 | 0 | 0 | 1676622 | YP_009047202.1 | K9N7C7 |
| LNKLNLPGCNGGSLYV | 1 | 0 | 0 | 1676624 | YP_173236.1 | A0A140H1G9 |
| LNKNLNEGVLESFSVT | 1 | 0 | 0 | 1676625 | YP_003766.2 | H9EJ42 |
| LNKNQKLIATAFNNAL | 1 | 0 | 0 | 1676626 | YP_173238.1 | A0A140H1H1 |
| LNKSAGWPLNKFGKAG | 1 | 0 | 0 | 1676634 | NP_073549.1 | A0A0P0LKV0 |
| LNKSAGWPLNKFGKAS | 1 | 0 | 0 | 1676635 | YP_003766.2 | H9EJ42 |
| LNKTLNEGVLESFSVT | 1 | 0 | 0 | 1676636 | NP_073549.1 | A0A0P0LKV0 |
| LNLSAGTQHISNDFTP | 1 | 0 | 0 | 1676660 | YP_173242.1 | Q19U29 |
| LNLSSELKQLEAKTAS | 1 | 0 | 0 | 1676662 | YP_003767.1 | A0A0P0G321 |
| LNLTDEDLARDFSPIA | 1 | 0 | 0 | 1676663 | YP_009047207.1 | K9N643 |
| LNMDMGVFINKYGLED | 1 | 0 | 0 | 1676668 | YP_003766.2 | H9EJ42 |
| LNNCTKYNIYDYVGTG | 1 | 0 | 0 | 1676672 | YP_003767.1 | A0A0P0G321 |
| LNNIINNARDGCVPLN | 1 | 0 | 0 | 1676681 | YP_009724389.1 | P0DTD1 |
| LNNLNRGMVLGSLAAT | 1 | 0 | 0 | 1676688 | YP_009724389.1 | P0DTD1 |
| LNNSYINLKDIGTYEM | 1 | 0 | 0 | 1676692 | YP_173238.1 | A0A140H1H1 |
| LNPNKTKRWGYSLNF | 1 | 0 | 0 | 1676698 | P03155.1 | Q69028 |
| LNPSALEFNPSQTSPA | 1 | 0 | 0 | 1676701 | NP_073556.1 | A0A0N7J131 |
| LNPSEDFIKHVRTNGG | 1 | 0 | 0 | 1676702 | YP_009047202.1 | K9N7C7 |
| LNQSYINLKDIGTYEY | 1 | 0 | 0 | 1676710 | YP_009555241.1 | P36334 |
| LNRARGSSAARLEPCN | 1 | 0 | 0 | 1676713 | YP_003766.2 | H9EJ42 |
| LNRGDYNMLLKNIDLF | 1 | 0 | 0 | 1676716 | YP_173236.1 | A0A140H1G9 |
| LNRGVYKPLLENIDYF | 1 | 0 | 0 | 1676718 | YP_009555238.1 | P0C6X6 |
| LNRYTTTLNDFNLVAM | 1 | 0 | 0 | 1676736 | QIQ54046.1 | P0C6U8 |
| LNSCCRIVERNVVRAA | 1 | 0 | 0 | 1676737 | YP_009047202.1 | K9N7C7 |
| LNSEKHQDHVSLTVPL | 1 | 0 | 0 | 1676739 | YP_009047202.1 | K9N7C7 |
| LNSILDNAVKGCVPLN | 1 | 0 | 0 | 1676742 | YP_009555238.1 | P0C6X6 |
| LNSILDNAVKGCVPLS | 1 | 0 | 0 | 1676743 | YP_173236.1 | A0A140H1G9 |
| LNSKVKVIQPRVEKKK | 1 | 0 | 0 | 1676746 | NP_828849.2 | P0C6X7 |
| LNSLGETLVTMPLGYV | 1 | 0 | 0 | 1676749 | QIQ54046.1 | P0C6U8 |
| LNSLNEPLVTMPIGYV | 1 | 0 | 0 | 1676751 | NP_828849.2 | P0C6X7 |
| LNSLTYFNKPSFKSEN | 1 | 0 | 0 | 1676752 | YP_173236.1 | A0A140H1G9 |
| LNSLTYFNRPSLVDDN | 1 | 0 | 0 | 1676753 | YP_009555238.1 | P0C6X6 |
| LNSSNVTTMDFCEGSF | 1 | 0 | 0 | 1676756 | NP_828849.2 | P0C6X7 |
| LNSTIKTIQPRVEKKK | 1 | 0 | 0 | 1676759 | QHR63299.1 | P0C6W2 |
| LNSTNVTTTAYCTGSI | 1 | 0 | 0 | 1676762 | QIQ54046.1 | P0C6U8 |
| LNTDHSNSSDNIALLV | 1 | 0 | 0 | 1676769 | QIQ54051.1 | P59596 |
| LNTICIKDADYNAKVE | 1 | 0 | 0 | 1676770 | NP_073549.1 | A0A0P0LKV0 |
| LNTNLHFDVDNINFKS | 1 | 0 | 0 | 1676774 | YP_173238.1 | A0A140H1H1 |
| LNTPKDHIGTRNPANN | 1 | 0 | 0 | 1676775 | YP_009724397.2 | P0DTC9 |
| LNTSIPNLPDFKEELD | 1 | 0 | 0 | 1676778 | YP_009555241.1 | P36334 |
| LNTTLFLNGYYPTSGS | 1 | 0 | 0 | 1676781 | YP_009555241.1 | P36334 |
| LNVAKSEFDRDAAMQR | 1 | 0 | 0 | 1676786 | YP_009724389.1 | P0DTD1 |
| LNVQYRINGLGVTMDV | 1 | 0 | 0 | 1676799 | YP_009555241.1 | P36334 |
| LNVVCKTCGQQQTTLK | 1 | 0 | 0 | 1676801 | YP_009724389.1 | P0DTD1 |
| LNVVGPDARTQGKQSY | 1 | 0 | 0 | 1676802 | YP_009555238.1 | P0C6X6 |
| LNYTVQKLQTLIDNIN | 1 | 0 | 0 | 1676811 | NP_073551.1 | A0A0P0K6L9 |
| LPAIKLNFGMLNGNAI | 1 | 0 | 0 | 1676820 | NP_073549.1 | A0A0P0LKV0 |
| LPAINVTINNYNPSSW | 1 | 0 | 0 | 1676821 | YP_173238.1 | A0A140H1H1 |
| LPAPRTLLTKGTLEPE | 1 | 0 | 0 | 1676823 | YP_009724389.1 | P0DTD1 |
| LPAPRVLLSKGSLEPR | 1 | 0 | 0 | 1676826 | YP_173236.1 | A0A140H1G9 |
| LPAPRVLLSKGTLEPK | 1 | 0 | 0 | 1676827 | YP_009555238.1 | P0C6X6 |
| LPAYASFKVIDNGVVR | 1 | 0 | 0 | 1676829 | YP_173236.1 | A0A140H1G9 |
| LPCQFKDKNLQDLWVL | 1 | 0 | 0 | 1676833 | YP_009555238.1 | P0C6X6 |
| LPDCDIDKWLNNFNVP | 1 | 0 | 0 | 1676836 | YP_173238.1 | A0A140H1H1 |
| LPDPLKPTKRSFIEDL | 1 | 0 | 0 | 1676847 | NP_828851.1 | P59594 |
| LPEQFDSLNLSAGTQH | 1 | 0 | 0 | 1676856 | YP_173242.1 | Q19U29 |
| LPETYFTQSRDLEDFK | 1 | 0 | 0 | 1676859 | NP_828849.2 | P0C6X7 |
| LPETYFTQSRNLQEFK | 1 | 0 | 0 | 1676861 | YP_009724389.1 | P0DTD1 |
| LPETYVALPIYYQHTD | 1 | 0 | 0 | 1676862 | YP_003767.1 | A0A0P0G321 |
| LPFKFKDLAIENMWLS | 1 | 0 | 0 | 1676878 | YP_173236.1 | A0A140H1G9 |
| LPFKLTCATTRQVVNV | 1 | 0 | 0 | 1676879 | YP_009724389.1 | P0DTD1 |
| LPFNDGVYFASTEKSN | 1 | 0 | 0 | 1676880 | YP_009724390.1 | P0DTC2 |
| LPFQQFGRDISDTTDA | 1 | 0 | 0 | 1676881 | QIQ54048.1 | P59594 |
| LPGEFSKKQLGLLQPT | 1 | 0 | 0 | 1676892 | YP_009047202.1 | K9N7C7 |
| LPGVFCGVDAVNLLTN | 1 | 0 | 0 | 1676901 | YP_009724389.1 | P0DTD1 |
| LPGVFCGVDSVNLLTN | 1 | 0 | 0 | 1676902 | QIQ54046.1 | P0C6U8 |
| LPHDAIVVDNDVVDYV | 1 | 0 | 0 | 1676906 | NP_073549.1 | A0A0P0LKV0 |
| LPINTDTVLSVAPEVD | 1 | 0 | 0 | 1676911 | YP_003766.2 | H9EJ42 |
| LPINVIVFDGKSKCDE | 1 | 0 | 0 | 1676912 | NP_828849.2 | P0C6X7 |
| LPINVIVFDGKSKCEE | 1 | 0 | 0 | 1676913 | YP_009724389.1 | P0DTD1 |

| | | | | | | |
|---|---|---|---|---|---|---|
| LPIYYQHTDINFTATA | 1 | 0 | 0 | 1676915 | YP_003767.1 | A0A0P0G321 |
| LPKDWRLINKFDVING | 1 | 0 | 0 | 1676917 | YP_173236.1 | A0A140H1G9 |
| LPKEDGDVLLAEFDTY | 1 | 0 | 0 | 1676918 | YP_009047202.1 | K9N7C7 |
| LPKFFYASNGTYNCTD | 1 | 0 | 0 | 1676919 | NP_073551.1 | A0A0P0K6L9 |
| LPKFIEQISAFTKPSS | 1 | 0 | 0 | 1676920 | YP_003771.1 | H9EJ69 |
| LPKGVGSANIFIGDKV | 1 | 0 | 0 | 1676923 | YP_009555238.1 | P0C6X6 |
| LPLAYFSDSKLGTTEV | 1 | 0 | 0 | 1676931 | YP_009047202.1 | K9N7C7 |
| LPLTCNAISSNTDNET | 1 | 0 | 0 | 1676942 | YP_173238.1 | A0A140H1H1 |
| LPNCNIEAWLNDKSVP | 1 | 0 | 0 | 1676948 | YP_009555241.1 | P36334 |
| LPNDDTLRAEAFEYYH | 1 | 0 | 0 | 1676949 | QHR63299.1 | P0C6W2 |
| LPNDDTLRSEAFEYYH | 1 | 0 | 0 | 1676950 | QIQ54046.1 | P0C6U8 |
| LPNFYYVSNGGNNCTT | 1 | 0 | 0 | 1676953 | YP_003767.1 | A0A0P0G321 |
| LPNYTVPDLVVEQYNQ | 1 | 0 | 0 | 1676962 | NP_073551.1 | A0A0P0K6L9 |
| LPPDAIIIDNDINDYV | 1 | 0 | 0 | 1676965 | YP_003766.2 | H9EJ42 |
| LPPELSVVEFEDRSNN | 1 | 0 | 0 | 1676966 | YP_003771.1 | H9EJ69 |
| LPPHDVTVLRDNLKPV | 1 | 0 | 0 | 1676968 | YP_003768.1 | H9EJ24 |
| LPPKNSIDAFKLNIKL | 1 | 0 | 0 | 1676970 | YP_009724389.1 | P0DTD1 |
| LPPKNSIDAFKLNVKL | 1 | 0 | 0 | 1676971 | QIQ54046.1 | P0C6U8 |
| LPPKSSIDAFKLNIKL | 1 | 0 | 0 | 1676972 | NP_828849.2 | P0C6X7 |
| LPPLLSENQISGYTLA | 1 | 0 | 0 | 1676974 | YP_009555241.1 | P36334 |
| LPQFTSDQAVTFLKEW | 1 | 0 | 0 | 1676983 | YP_173241.1 | Q0ZJ82 |
| LPQFYIYDEEGGYDVS | 1 | 0 | 0 | 1676984 | YP_003766.2 | H9EJ42 |
| LPQRNIRSSRIAGRSA | 1 | 0 | 0 | 1676995 | YP_003767.1 | A0A0P0G321 |
| LPQSKDSNFLNESGVL | 1 | 0 | 0 | 1676997 | YP_009047203.1 | K9N7C7 |
| LPQSKDSNFLNRVRGS | 1 | 0 | 0 | 1676998 | YP_009047202.1 | K9N7C7 |
| LPSAPQTIQPEVKEVA | 1 | 0 | 0 | 1677006 | YP_009047202.1 | K9N7C7 |
| LPSTQKGSGMDTALLR | 1 | 0 | 0 | 1677022 | YP_009555244.1 | Q01455 |
| LPSYAAFATAQEAYEQ | 1 | 0 | 0 | 1677023 | YP_009724389.1 | P0DTD1 |
| LPSYAAYATAQEAYEQ | 1 | 0 | 0 | 1677024 | NP_828849.2 | P0C6X7 |
| LPTAFAVIGDLKCTSD | 1 | 0 | 0 | 1677025 | YP_009555241.1 | P36334 |
| LPTDAIIIDNDLNEFV | 1 | 0 | 0 | 1677026 | YP_009047202.1 | K9N7C7 |
| LPTEVLSEEVILKTGE | 1 | 0 | 0 | 1677028 | QIQ54046.1 | P0C6U8 |
| LPTEVLTEEVVLKTGD | 1 | 0 | 0 | 1677029 | YP_009724389.1 | P0DTD1 |
| LPTFSMAPVEFYEDRI | 1 | 0 | 0 | 1677030 | YP_009047202.1 | K9N7C7 |
| LPTGTLLVDSDLNDFV | 1 | 0 | 0 | 1677033 | YP_009724389.1 | P0DTD1 |
| LPTGVHAGTDLEGKFY | 1 | 0 | 0 | 1677034 | NP_828849.2 | P0C6X7 |
| LPTGVHAGTDLEGNFY | 1 | 0 | 0 | 1677035 | YP_009724389.1 | P0DTD1 |
| LPTGVHAGTDLEGTFY | 1 | 0 | 0 | 1677036 | QHR63299.1 | P0C6W2 |
| LPTKALCFDKSKQFVP | 1 | 0 | 0 | 1677038 | YP_173237.1 | A0A140H1H0 |
| LPTLTQMNLKYAISAK | 1 | 0 | 0 | 1677039 | YP_009555238.1 | P0C6X6 |
| LPTQTVDSSQGSEYDY | 1 | 0 | 0 | 1677048 | YP_009724389.1 | P0DTD1 |
| LPTSGSRVAGRSAIED | 1 | 0 | 0 | 1677050 | NP_073551.1 | A0A0P0K6L9 |
| LPVAFTKAAGGKVSFS | 1 | 0 | 0 | 1677056 | NP_073549.1 | A0A0P0LKV0 |
| LPXNDGVYFASTEKSN | 1 | 0 | 0 | 1677079 | QIQ54048.1 | P59594 |
| LPYCYFRNSSANYVGK | 1 | 0 | 0 | 1677080 | YP_173237.1 | A0A140H1H0 |
| LQAASDWKPGYSMPVL | 1 | 0 | 0 | 1677094 | YP_009555238.1 | P0C6X6 |
| LQAGLRKMAQPSGFVE | 1 | 0 | 0 | 1677101 | NP_073549.1 | A0A0P0LKV0 |
| LQAGNATEVPANSTVL | 1 | 0 | 0 | 1677102 | YP_009724389.1 | P0DTD1 |
| LQAGSNTEFASNSSVL | 1 | 0 | 0 | 1677103 | YP_009047202.1 | K9N7C7 |
| LQAGTATEYASNSSIL | 1 | 0 | 0 | 1677104 | YP_009555238.1 | P0C6X6 |
| LQAIASEFSSLPSYAA | 1 | 0 | 0 | 1677106 | YP_009724389.1 | P0DTD1 |
| LQALQSEFVNMASFVE | 1 | 0 | 0 | 1677108 | YP_009555238.1 | P0C6X6 |
| LQASADWKPGHAMPSL | 1 | 0 | 0 | 1677110 | YP_009047202.1 | K9N7C7 |
| LQASQAWQPGVAMPNL | 1 | 0 | 0 | 1677112 | NP_828849.2 | P0C6X7 |
| LQDAYYRARAGDAANF | 1 | 0 | 0 | 1677124 | QIQ54046.1 | P0C6U8 |
| LQDAYYRARAGEAANF | 1 | 0 | 0 | 1677125 | YP_009724389.1 | P0DTD1 |
| LQDCIVGIADDTYVMG | 1 | 0 | 0 | 1677126 | NP_073549.1 | A0A0P0LKV0 |
| LQDCVTGINDDDYVIA | 1 | 0 | 0 | 1677127 | YP_003766.2 | H9EJ42 |
| LQDFLPRSTMEEDFLN | 1 | 0 | 0 | 1677129 | NP_073549.1 | A0A0P0LKV0 |
| LQDLKWARFPKSDGTG | 1 | 0 | 0 | 1677132 | YP_009724389.1 | P0DTD1 |
| LQEAYYRARAGDAANF | 1 | 0 | 0 | 1677138 | NP_828849.2 | P0C6X7 |
| LQELSYKLPNYTVPDL | 1 | 0 | 0 | 1677145 | NP_073551.1 | A0A0P0K6L9 |
| LQENQKILAASFNKAI | 1 | 0 | 0 | 1677147 | YP_003767.1 | A0A0P0G321 |
| LQENQKILAASFNKAM | 1 | 0 | 0 | 1677148 | NP_073551.1 | A0A0P0K6L9 |
| LQETPVVPDTVEVPPQ | 1 | 0 | 0 | 1677149 | YP_009047202.1 | K9N7C7 |
| LQEVKDNDGKNVHLKD | 1 | 0 | 0 | 1677150 | NP_073549.1 | A0A0P0LKV0 |
| LQFGLQDGFYSANFLD | 1 | 0 | 0 | 1677155 | YP_003767.1 | A0A0P0G321 |
| LQFKRPINPTDQSSYI | 1 | 0 | 0 | 1677158 | YP_009724389.1 | P0DTD1 |
| LQFTSLEIPRRNVATL | 1 | 0 | 0 | 1677161 | YP_009724389.1 | P0DTD1 |
| LQFTSLEVPRRNVATL | 1 | 0 | 0 | 1677162 | QIQ54047.1 | P0C6X7 |
| LQGLENIAFNVVNKGS | 1 | 0 | 0 | 1677170 | NP_073549.1 | A0A0P0LKV0 |
| LQGLIDQINSTYVDLK | 1 | 0 | 0 | 1677171 | YP_003767.1 | A0A0P0G321 |
| LQGNFGDLQLNKLGTE | 1 | 0 | 0 | 1677174 | YP_009047211.1 | K9N4V7 |
| LQHAFMKHKGGDSTDF | 1 | 0 | 0 | 1677177 | YP_009047202.1 | K9N7C7 |
| LQHLHVASNSKLWQYC | 1 | 0 | 0 | 1677182 | YP_173236.1 | A0A140H1G9 |
| LQIQTVDSSQGSEYDY | 1 | 0 | 0 | 1677189 | YP_003766.2 | H9EJ42 |
| LQKRLYSHVYRSDKVD | 1 | 0 | 0 | 1677198 | YP_009555238.1 | P0C6X6 |
| LQLGSSGFLQSSNYKI | 1 | 0 | 0 | 1677206 | YP_173238.1 | A0A140H1H1 |
| LQLNKLGTEDPRWPQI | 1 | 0 | 0 | 1677212 | YP_009047211.1 | K9N4V7 |
| LQLPNFYYVSNGGNNC | 1 | 0 | 0 | 1677213 | YP_003767.1 | A0A0P0G321 |
| LQLVSAVNQSLSNAKV | 1 | 0 | 0 | 1677216 | NP_073552.1 | A0A0P0KWE8 |
| LQMGFGITVQYGTDTN | 1 | 0 | 0 | 1677218 | YP_009047204.1 | K9N5Q8 |
| LQNNEIKPSGLKTMVV | 1 | 0 | 0 | 1677223 | YP_009047202.1 | K9N7C7 |
| LQNYGDSATLPKGIMM | 1 | 0 | 0 | 1677230 | YP_009724389.1 | P0DTD1 |
| LQPEEEQEEDWLDDDS | 1 | 0 | 0 | 1677233 | YP_009724389.1 | P0DTD1 |
| LQQEMTIVKCKGLNKP | 1 | 0 | 0 | 1677250 | YP_009047202.1 | K9N7C7 |
| LQQGTFLCANEYTGNY | 1 | 0 | 0 | 1677251 | NP_828849.2 | P0C6X7 |
| LQQRTFLLSYNQNGTI | 1 | 0 | 0 | 1677258 | QIQ54048.1 | P59594 |
| LQQVVKALNESYIDLK | 1 | 0 | 0 | 1677259 | YP_009047204.1 | K9N5Q8 |
| LQSFNYRIDTTATSCQ | 1 | 0 | 0 | 1677278 | YP_009555241.1 | P36334 |
| LQSLSKDYISSNGPLK | 1 | 0 | 0 | 1677298 | YP_003766.2 | H9EJ42 |

| | | | | | | |
|---|---|---|---|---|---|---|
| LQSNNPQCAIVQASES | 1 | 0 | 0 | 1677303 | YP_003766.2 | H9EJ42 |
| LQSSNYKIDTTSSSCQ | 1 | 0 | 0 | 1677306 | YP_173238.1 | A0A140H1H1 |
| LQTIVPEYIDVNKTLQ | 1 | 0 | 0 | 1677318 | NP_073551.1 | A0A0P0K6L9 |
| LQTLIDNINSTLVDLK | 1 | 0 | 0 | 1677319 | NP_073551.1 | A0A0P0K6L9 |
| LQTPFEIKLAKKFDTF | 1 | 0 | 0 | 1677322 | YP_009724389.1 | P0DTD1 |
| LQTVATALNKIQDVVN | 1 | 0 | 0 | 1677325 | NP_073551.1 | A0A0P0K6L9 |
| LQVESANLMLSDNVVA | 1 | 0 | 0 | 1677331 | YP_003766.2 | H9EJ42 |
| LQVESANQMLTVNVVA | 1 | 0 | 0 | 1677332 | NP_073549.1 | A0A0P0LKV0 |
| LQVFVYSSNEEQAVLK | 1 | 0 | 0 | 1677335 | YP_003766.2 | H9EJ42 |
| LQVGFSNGVDFVAQPE | 1 | 0 | 0 | 1677337 | NP_073549.1 | A0A0P0LKV0 |
| LQYDSSIHSYFITDQE | 1 | 0 | 0 | 1677345 | YP_173236.1 | A0A140H1G9 |
| LQYDVLFYCSNSSSGV | 1 | 0 | 0 | 1677346 | YP_003767.1 | A0A0P0G321 |
| LQYSKPHFISQGLDAA | 1 | 0 | 0 | 1677354 | NP_073549.1 | A0A0P0LKV0 |
| LRACIKTAQADTAGLY | 1 | 0 | 0 | 1677356 | YP_009047205.1 | K9N796 |
| LRAFDIYNASVAGIGL | 1 | 0 | 0 | 1677361 | YP_009555238.1 | P0C6X6 |
| LRAKHYVYIGDPAQLP | 1 | 0 | 0 | 1677365 | YP_009724389.1 | P0DTD1 |
| LRANSAVKLQNNELSP | 1 | 0 | 0 | 1677369 | YP_009724389.1 | P0DTD1 |
| LRAPTMANQEKYSTIY | 1 | 0 | 0 | 1677372 | NP_073549.1 | A0A0P0LKV0 |
| LRASNTKGSLPINVIV | 1 | 0 | 0 | 1677375 | QHR63299.1 | P0C6W2 |
| LRATEEASKQSYAIAT | 1 | 0 | 0 | 1677377 | YP_009047202.1 | K9N7C7 |
| LRCLWSTKPVDTSNSF | 1 | 0 | 0 | 1677387 | NP_828849.2 | P0C6X7 |
| LRDNLKPVTTLSTITA | 1 | 0 | 0 | 1677393 | YP_003768.1 | H9EJ24 |
| LRDYLADYDVAVTAGP | 1 | 0 | 0 | 1677394 | YP_009047202.1 | K9N7C7 |
| LREMLAHAEETRKLMP | 1 | 0 | 0 | 1677403 | YP_009724389.1 | P0DTD1 |
| LRFPRGQGVPINTNSG | 1 | 0 | 0 | 1677412 | NP_828858.1 | P59595 |
| LRFPRGQGVPINTNST | 1 | 0 | 0 | 1677413 | QIQ54056.1 | P59595 |
| LRGEDMSFLEKDAPYM | 1 | 0 | 0 | 1677418 | NP_828849.2 | P0C6X7 |
| LRGMPIPDFDLDDFID | 1 | 0 | 0 | 1677424 | YP_009047202.1 | K9N7C7 |
| LRGPGDLQGNFGDLQL | 1 | 0 | 0 | 1677425 | YP_009047211.1 | K9N4V7 |
| LRGRIDKDLSLSRKVE | 1 | 0 | 0 | 1677426 | NP_073553.1 | A0A0P0KWE8 |
| LRGTAVMSLKENQIND | 1 | 0 | 0 | 1677428 | NP_828849.2 | P0C6X7 |
| LRGTIVTRPLMESELV | 1 | 0 | 0 | 1677429 | NP_828855.1 | P59596 |
| LRHGKLRPFERDISNV | 1 | 0 | 0 | 1677440 | NP_828851.1 | P59594 |
| LRHLIKDVDNPVLMGW | 1 | 0 | 0 | 1677441 | YP_173236.1 | A0A140H1G9 |
| LRIAGHHLGRCDIKDL | 1 | 0 | 0 | 1677444 | YP_009724393.1 | P0DTC5 |
| LRKGGRTIAFGGCVFS | 1 | 0 | 0 | 1677455 | YP_009724389.1 | P0DTD1 |
| LRKGGRTKAFGGCVFS | 1 | 0 | 0 | 1677456 | QIQ54046.1 | P0C6U8 |
| LRKGGRTRCFGGCVFA | 1 | 0 | 0 | 1677457 | NP_828849.2 | P0C6X7 |
| LRKKRNVDPLSNFEHK | 1 | 0 | 0 | 1677462 | YP_009047202.1 | K9N7C7 |
| LRKLDNDALNNIINNA | 1 | 0 | 0 | 1677463 | YP_009724389.1 | P0DTD1 |
| LRKNGNKGAGGHSFGI | 1 | 0 | 0 | 1677466 | QIQ54046.1 | P0C6U8 |
| LRKNGNKGAGGHSYGA | 1 | 0 | 0 | 1677467 | YP_009724389.1 | P0DTD1 |
| LRKNGNKGAGGHSYGI | 1 | 0 | 0 | 1677468 | NP_828849.2 | P0C6X7 |
| LRLAFLGASVTEDVKF | 1 | 0 | 0 | 1677479 | YP_003766.2 | H9EJ42 |
| LRLHTVESGDINVVAA | 1 | 0 | 0 | 1677484 | YP_009047202.1 | K9N7C7 |
| LRLIDAMLFTSELTTD | 1 | 0 | 0 | 1677485 | QIQ54046.1 | P0C6U8 |
| LRLIDAMMFTSDLATN | 1 | 0 | 0 | 1677486 | YP_009724389.1 | P0DTD1 |
| LRLIDAMMFTSDLVTN | 1 | 0 | 0 | 1677487 | QHR63299.1 | P0C6W2 |
| LRLIDDNGIVLNSILW | 1 | 0 | 0 | 1677488 | YP_003769.1 | H9EJA2 |
| LRLVDAMVYTSDLLTN | 1 | 0 | 0 | 1677495 | NP_828849.2 | P0C6X7 |
| LRLVDCAPVVCPKGKI | 1 | 0 | 0 | 1677496 | YP_003766.2 | H9EJ42 |
| LRMGSGFCVDYNSPSS | 1 | 0 | 0 | 1677500 | YP_173238.1 | A0A140H1H1 |
| LRNCTFMYTYNITEDE | 1 | 0 | 0 | 1677502 | YP_009047204.1 | K9N5Q8 |
| LRNFDEKDCEVFKEIL | 1 | 0 | 0 | 1677503 | NP_073549.1 | A0A0P0LKV0 |
| LRNGSVLEMAKEVKTS | 1 | 0 | 0 | 1677504 | NP_073549.1 | A0A0P0LKV0 |
| LRNSFGKDLNANMSLA | 1 | 0 | 0 | 1677509 | NP_073549.1 | A0A0P0LKV0 |
| LRPDTRYVLMDGSIIQ | 1 | 0 | 0 | 1677516 | YP_009724389.1 | P0DTD1 |
| LRPFERDISNVPFSPD | 1 | 0 | 0 | 1677517 | NP_828851.1 | P59594 |
| LRPPKNSFEALMLNFK | 1 | 0 | 0 | 1677519 | YP_009555238.1 | P0C6X6 |
| LRQAALPQSKDSNFLN | 1 | 0 | 0 | 1677521 | YP_009047202.1 | K9N7C7 |
| LRQIFDVAPIELENKF | 1 | 0 | 0 | 1677524 | YP_009047202.1 | K9N7C7 |
| LRRLDMSSVDTILNMA | 1 | 0 | 0 | 1677539 | NP_073549.1 | A0A0P0LKV0 |
| LRRLDMSSVETVLNLA | 1 | 0 | 0 | 1677540 | YP_003766.2 | H9EJ42 |
| LRRLIKDVDNPVLMGW | 1 | 0 | 0 | 1677541 | YP_009555238.1 | P0C6X6 |
| LRSDVLLPLTQYNRYL | 1 | 0 | 0 | 1677548 | YP_009724389.1 | P0DTD1 |
| LRSITAQPVKIDNASP | 1 | 0 | 0 | 1677555 | NP_828852.2 | P59632 |
| LRSQGFFDEGSELTLK | 1 | 0 | 0 | 1677557 | NP_073549.1 | A0A0P0LKV0 |
| LRTNFTIKGSFINGAC | 1 | 0 | 0 | 1677567 | YP_003766.2 | H9EJ42 |
| LSAGIFGADPIHSLRV | 1 | 0 | 0 | 1677601 | YP_009724389.1 | P0DTD1 |
| LSAGVEFLKDAWEILK | 1 | 0 | 0 | 1677606 | NP_828849.2 | P0C6X7 |
| LSAHLETNDVSSMLTF | 1 | 0 | 0 | 1677608 | YP_003767.1 | A0A0P0G321 |
| LSAPTLVPQEHYVRIT | 1 | 0 | 0 | 1677612 | YP_009724389.1 | P0DTD1 |
| LSAPTLVPQENYASIR | 1 | 0 | 0 | 1677613 | YP_173236.1 | A0A140H1G9 |
| LSAPTLVPQENYSSIR | 1 | 0 | 0 | 1677614 | YP_009555238.1 | P0C6X6 |
| LSCGIFGIKLETSLEV | 1 | 0 | 0 | 1677621 | NP_073549.1 | A0A0P0LKV0 |
| LSDDGVVCYNKDYAEL | 1 | 0 | 0 | 1677632 | YP_003766.2 | H9EJ42 |
| LSDDGVVCYNSDYAAK | 1 | 0 | 0 | 1677633 | YP_009047202.1 | K9N7C7 |
| LSDDGVVCYNSDYASK | 1 | 0 | 0 | 1677634 | YP_009555238.1 | P0C6X6 |
| LSDFINGIFAKVKNTK | 1 | 0 | 0 | 1677638 | YP_009555241.1 | P36334 |
| LSDFTPRSDMEYDFLN | 1 | 0 | 0 | 1677640 | YP_003766.2 | H9EJ42 |
| LSDLQDLKWARFPKSD | 1 | 0 | 0 | 1677646 | YP_009724389.1 | P0DTD1 |
| LSDMEKDFLSFDSDVF | 1 | 0 | 0 | 1677649 | YP_009047202.1 | K9N7C7 |
| LSDNAYLKWARVEGKD | 1 | 0 | 0 | 1677650 | YP_009047202.1 | K9N7C7 |
| LSDQFKTTGDLAVQIG | 1 | 0 | 0 | 1677654 | YP_003766.2 | H9EJ42 |
| LSDRELHLSWEVGKPR | 1 | 0 | 0 | 1677655 | YP_009724389.1 | P0DTD1 |
| LSDRFKTSGDLAVQIG | 1 | 0 | 0 | 1677657 | NP_073549.1 | A0A0P0LKV0 |
| LSDVDGLKYTKILKDD | 1 | 0 | 0 | 1677666 | YP_009555238.1 | P0C6X6 |
| LSDVDGLKYTKIMKDD | 1 | 0 | 0 | 1677667 | YP_173236.1 | A0A140H1G9 |
| LSDVGFVEAYNNCTGG | 1 | 0 | 0 | 1677668 | YP_009555241.1 | P36334 |
| LSDVIRYNLNFEENLR | 1 | 0 | 0 | 1677669 | NP_073551.1 | A0A0P0K6L9 |
| LSDYLLDLSDSVVFIT | 1 | 0 | 0 | 1677671 | YP_173236.1 | A0A140H1G9 |

| | | | | | | |
|---|---|---|---|---|---|---|
| LSEAREHLKNGTCGLV | 1 | 0 | 0 | 1677672 | NP_828849.2 | P0C6X7 |
| LSEEVILKTGELQPLE | 1 | 0 | 0 | 1677675 | QIQ54046.1 | P0C6U8 |
| LSEFSHLATFAELEAA | 1 | 0 | 0 | 1677676 | YP_009047202.1 | K9N7C7 |
| LSEITKPLIEAEKPNL | 1 | 0 | 0 | 1677679 | YP_173243.1 | Q0ZJ80 |
| LSENQKLIANKFNQAL | 1 | 0 | 0 | 1677686 | YP_009047204.1 | K9N5Q8 |
| LSEQLDFIDTKRGVYC | 1 | 0 | 0 | 1677687 | YP_009724389.1 | P0DTD1 |
| LSEQLDYIESKRGVYC | 1 | 0 | 0 | 1677688 | NP_828849.2 | P0C6X7 |
| LSERCNVIADSKSFTL | 1 | 0 | 0 | 1677690 | YP_009047202.1 | K9N7C7 |
| LSESQISGYTTAATVA | 1 | 0 | 0 | 1677691 | YP_173238.1 | A0A140H1H1 |
| LSEYGTFCDNINSILD | 1 | 0 | 0 | 1677694 | YP_173238.1 | A0A140H1H1 |
| LSEYPLLKQKDIGIDT | 1 | 0 | 0 | 1677696 | YP_173243.1 | Q0ZJ80 |
| LSFAGYTESAVNSTKA | 1 | 0 | 0 | 1677698 | YP_009047206.1 | K9N4V0 |
| LSFDSDVFIKKYGLEN | 1 | 0 | 0 | 1677703 | YP_009047202.1 | K9N7C7 |
| LSFDVPDGFYSTSPIQ | 1 | 0 | 0 | 1677704 | NP_073551.1 | A0A0P0K6L9 |
| LSFEEQDEIYAYTKRN | 1 | 0 | 0 | 1677706 | YP_009555238.1 | P0C6X6 |
| LSFEEQNEIYAYTKRN | 1 | 0 | 0 | 1677707 | YP_173236.1 | A0A140H1G9 |
| LSFKELLVYAADPAMH | 1 | 0 | 0 | 1677711 | YP_009724389.1 | P0DTD1 |
| LSFNNFVSSYAKPEEK | 1 | 0 | 0 | 1677716 | NP_073549.1 | A0A0P0LKV0 |
| LSFNNLLGFDSSKPVT | 1 | 0 | 0 | 1677717 | YP_009047202.1 | K9N7C7 |
| LSGAVPSDLNTSELLT | 1 | 0 | 0 | 1677723 | NP_073549.1 | A0A0P0LKV0 |
| LSGDDCFVSNSSVIQE | 1 | 0 | 0 | 1677725 | YP_173236.1 | A0A140H1G9 |
| LSGFETIMKVLNENLN | 1 | 0 | 0 | 1677728 | YP_009555245.1 | P33469 |
| LSGNPDEPQKDVYELR | 1 | 0 | 0 | 1677740 | YP_009555245.1 | P33469 |
| LSGVFCGVDAMNLIAN | 1 | 0 | 0 | 1677747 | NP_828849.2 | P0C6X7 |
| LSIDDGHLWNLFVEKF | 1 | 0 | 0 | 1677767 | YP_003766.2 | H9EJ42 |
| LSKDLPDDVVAANMFM | 1 | 0 | 0 | 1677791 | YP_173236.1 | A0A140H1G9 |
| LSKDYISSNGPLKVGA | 1 | 0 | 0 | 1677792 | YP_003766.2 | H9EJ42 |
| LSKELKRPIQPTDVAY | 1 | 0 | 0 | 1677795 | YP_009555238.1 | P0C6X6 |
| LSKELKRPVNPTDASH | 1 | 0 | 0 | 1677796 | YP_173236.1 | A0A140H1G9 |
| LSKFQLKLKGTPVLQL | 1 | 0 | 0 | 1677800 | YP_009047202.1 | K9N7C7 |
| LSKGRLIIRENNRVVI | 1 | 0 | 0 | 1677804 | YP_009724389.1 | P0DTD1 |
| LSKGSLEPRHFNSITK | 1 | 0 | 0 | 1677805 | YP_173236.1 | A0A140H1G9 |
| LSKHSDFGLGDLVDSY | 1 | 0 | 0 | 1677812 | NP_073549.1 | A0A0P0LKV0 |
| LSKLASELSNTFGAIS | 1 | 0 | 0 | 1677813 | YP_009047204.1 | K9N5Q8 |
| LSKMGVLKADDFVTAS | 1 | 0 | 0 | 1677816 | YP_003766.2 | H9EJ42 |
| LSKPLTENKYSQLDEE | 1 | 0 | 0 | 1677818 | QIQ54052.1 | P59634 |
| LSKSLTENKYSQLDEE | 1 | 0 | 0 | 1677820 | YP_009724394.1 | P0DTC6 |
| LSKTSEYKGPVTDVFY | 1 | 0 | 0 | 1677821 | QIQ54046.1 | P0C6U8 |
| LSLCAFSVDPKKTYLD | 1 | 0 | 0 | 1677827 | YP_009555238.1 | P0C6X6 |
| LSLKELMMYAADPAMH | 1 | 0 | 0 | 1677838 | YP_009047202.1 | K9N7C7 |
| LSLQQVVKALNESYID | 1 | 0 | 0 | 1677855 | YP_009047204.1 | K9N5Q8 |
| LSLSDRFKTSGDLAVQ | 1 | 0 | 0 | 1677860 | NP_073549.1 | A0A0P0LKV0 |
| LSLSMARRNLDSLEAR | 1 | 0 | 0 | 1677861 | NP_828859.1 | P59636 |
| LSLVNFTVDPQKAYLD | 1 | 0 | 0 | 1677867 | YP_009047202.1 | K9N7C7 |
| LSLYNKYRYYSGKMDT | 1 | 0 | 0 | 1677870 | YP_009555238.1 | P0C6X6 |
| LSMARRNLDSLEARAF | 1 | 0 | 0 | 1677871 | NP_828859.1 | P59636 |
| LSMFDVDKKSLNALIA | 1 | 0 | 0 | 1677875 | YP_009555238.1 | P0C6X6 |
| LSMIKGPQRADLNGVI | 1 | 0 | 0 | 1677879 | YP_173236.1 | A0A140H1G9 |
| LSMIRGPPRAELNGVV | 1 | 0 | 0 | 1677880 | YP_009555238.1 | P0C6X6 |
| LSMLQLGVPDNSSTIV | 1 | 0 | 0 | 1677882 | YP_003767.1 | A0A0P0G321 |
| LSMQGAVDINKLCEEM | 1 | 0 | 0 | 1677883 | YP_009724389.1 | P0DTD1 |
| LSMQGAVDINRLCEEM | 1 | 0 | 0 | 1677884 | NP_828849.2 | P0C6X7 |
| LSMTTSTTNFGEDFLG | 1 | 0 | 0 | 1677886 | YP_173236.1 | A0A140H1G9 |
| LSNAKVSAEVSRQVIQ | 1 | 0 | 0 | 1677887 | NP_073552.1 | A0A0P0KWE8 |
| LSNELAQVLTEVVYSN | 1 | 0 | 0 | 1677889 | NP_073549.1 | A0A0P0LKV0 |
| LSNMNIASNSKEWAYC | 1 | 0 | 0 | 1677899 | NP_073549.1 | A0A0P0LKV0 |
| LSNQLRLAFLGASVTE | 1 | 0 | 0 | 1677900 | YP_003766.2 | H9EJ42 |
| LSNTKYYDDSQYYFNK | 1 | 0 | 0 | 1677903 | YP_009555240.1 | P30215 |
| LSPAYAPTYVSVDDKY | 1 | 0 | 0 | 1677906 | YP_009047202.1 | K9N7C7 |
| LSPWFMDGENAYEVVK | 1 | 0 | 0 | 1677916 | YP_009047202.1 | K9N7C7 |
| LSQPRADKPSQLKKPR | 1 | 0 | 0 | 1677923 | YP_003771.1 | H9EJ69 |
| LSRALKKVPVDEYITT | 1 | 0 | 0 | 1677926 | NP_828849.2 | P0C6X7 |
| LSRLDALEAEAQIDRL | 1 | 0 | 0 | 1677932 | YP_009555241.1 | P36334 |
| LSRNSSRSSSQGSRSG | 1 | 0 | 0 | 1677939 | YP_009047211.1 | K9N4V7 |
| LSRSNSNFRHSDSIVK | 1 | 0 | 0 | 1677940 | YP_173242.1 | Q19U29 |
| LSSACTMFTMADGSPQ | 1 | 0 | 0 | 1677943 | YP_009555238.1 | P0C6X6 |
| LSSNLNTNLHFDVDNI | 1 | 0 | 0 | 1677960 | YP_173238.1 | A0A140H1H1 |
| LSSVIPSLPTSGSRVA | 1 | 0 | 0 | 1677964 | NP_073551.1 | A0A0P0K6L9 |
| LSSVLPQRNIRSSRIA | 1 | 0 | 0 | 1677966 | YP_003767.1 | A0A0P0G321 |
| LSSYEYCQPPKSVEKI | 1 | 0 | 0 | 1677968 | YP_173236.1 | A0A140H1G9 |
| LSTCDGLKFTEESFYK | 1 | 0 | 0 | 1677969 | YP_173236.1 | A0A140H1G9 |
| LSTDGNKIADKHVRNL | 1 | 0 | 0 | 1677971 | QHR63299.1 | P0C6W2 |
| LSTDGNKIADKYVRNL | 1 | 0 | 0 | 1677972 | YP_009724389.1 | P0DTD1 |
| LSTDGNKIGDKYIRNL | 1 | 0 | 0 | 1677973 | QIQ54047.1 | P0C6X7 |
| LSTGLTSQTVKPGHFN | 1 | 0 | 0 | 1677987 | NP_073549.1 | A0A0P0LKV0 |
| LSTKLKDGVNFNVDDI | 1 | 0 | 0 | 1677989 | YP_009555241.1 | P36334 |
| LSTTLVKDKCVNFNFN | 1 | 0 | 0 | 1678009 | QIQ54048.1 | P59594 |
| LSVDIKFKTEGLCVDI | 1 | 0 | 0 | 1678020 | NP_828849.2 | P0C6X7 |
| LSVDSVSEESQGNVVT | 1 | 0 | 0 | 1678021 | YP_173236.1 | A0A140H1G9 |
| LSVDTKFKTEGLCVDI | 1 | 0 | 0 | 1678022 | YP_009724389.1 | P0DTD1 |
| LSVLLSMQGAVDINKL | 1 | 0 | 0 | 1678037 | YP_009724389.1 | P0DTD1 |
| LSVLLSMQGAVDINRL | 1 | 0 | 0 | 1678038 | NP_828849.2 | P0C6X7 |
| LSVNSSNCNNFNVKKL | 1 | 0 | 0 | 1678045 | NP_073549.1 | A0A0P0LKV0 |
| LSVQQAQQEATLPDIA | 1 | 0 | 0 | 1678047 | NP_073549.1 | A0A0P0LKV0 |
| LSVSPYVCNAPGCGVS | 1 | 0 | 0 | 1678050 | YP_009047202.1 | K9N7C7 |
| LSVVEFEDRSNNSSRA | 1 | 0 | 0 | 1678056 | YP_003771.1 | H9EJ69 |
| LSWEIGKVKPPLNKNY | 1 | 0 | 0 | 1678062 | YP_009555238.1 | P0C6X6 |
| LSWESGKVKPPLNRNS | 1 | 0 | 0 | 1678063 | YP_003766.2 | H9EJ42 |
| LSXELLNGPATVCGPK | 1 | 0 | 0 | 1678065 | QIQ54048.1 | P59594 |
| LSYEQFKKGVQIPCTC | 1 | 0 | 0 | 1678067 | YP_009724389.1 | P0DTD1 |
| LSYEQLKKGVQIPCTC | 1 | 0 | 0 | 1678068 | QHR63299.1 | P0C6W2 |

| | | | | | | |
|---|---|---|---|---|---|---|
| LSYHYENNSWFFRHVY | 1 | 0 | 0 | 1678076 | YP_009555239.1 | Q80872 |
| LSYKLPNYTVPDLVVE | 1 | 0 | 0 | 1678078 | NP_073551.1 | A0A0P0K6L9 |
| LSYNQNGTITDAVDCS | 1 | 0 | 0 | 1678081 | QIQ54048.1 | P59594 |
| LSYYVPSVEYTYTDEV | 1 | 0 | 0 | 1678086 | YP_009555238.1 | P0C6X6 |
| LTADETKALKELYGPV | 1 | 0 | 0 | 1678088 | YP_009047202.1 | K9N7C7 |
| LTAEDISLLKKMDEPY | 1 | 0 | 0 | 1678090 | YP_009555245.1 | P33469 |
| LTAFSPAQDIWGTSAA | 1 | 0 | 0 | 1678094 | NP_828851.1 | P59594 |
| LTALNAYVSQQLSDIS | 1 | 0 | 0 | 1678095 | YP_173238.1 | A0A140H1H1 |
| LTALNAYVSQQLSDST | 1 | 0 | 0 | 1678096 | YP_009555241.1 | P36334 |
| LTANMSMAECKATLGL | 1 | 0 | 0 | 1678098 | YP_003766.2 | H9EJ42 |
| LTCNAISSNTDNETLQ | 1 | 0 | 0 | 1678118 | YP_173238.1 | A0A140H1H1 |
| LTCSNTKFADDLNQMT | 1 | 0 | 0 | 1678121 | NP_828849.2 | P0C6X7 |
| LTDAALSIKENVDFIN | 1 | 0 | 0 | 1678124 | YP_009555239.1 | Q80872 |
| LTDCLYLKNLTQTFTS | 1 | 0 | 0 | 1678126 | YP_173236.1 | A0A140H1G9 |
| LTDESCNNLVPTYLKS | 1 | 0 | 0 | 1678130 | YP_009555238.1 | P0C6X6 |
| LTDLKCTNVVLLGCLS | 1 | 0 | 0 | 1678133 | YP_003766.2 | H9EJ42 |
| LTDLKCTNVVLMGILS | 1 | 0 | 0 | 1678134 | NP_073549.1 | A0A0P0LKV0 |
| LTDNVYIKNADIVEEA | 1 | 0 | 0 | 1678138 | YP_009724389.1 | P0DTD1 |
| LTDVKCANVVLLNCLQ | 1 | 0 | 0 | 1678139 | YP_009555238.1 | P0C6X6 |
| LTDYNAILQSNNPQCA | 1 | 0 | 0 | 1678140 | YP_003766.2 | H9EJ42 |
| LTEEVVLKTGDLQPLE | 1 | 0 | 0 | 1678144 | YP_009724389.1 | P0DTD1 |
| LTENKYSQLDEEQPME | 1 | 0 | 0 | 1678147 | YP_009724394.1 | P0DTC6 |
| LTEVNELLDTTQLQVA | 1 | 0 | 0 | 1678151 | YP_009555241.1 | P36334 |
| LTFNDNQAITQVPATS | 1 | 0 | 0 | 1678164 | YP_003766.2 | H9EJ42 |
| LTFNKQEASVPILTTP | 1 | 0 | 0 | 1678165 | YP_173236.1 | A0A140H1G9 |
| LTFPPGQGVPLNANST | 1 | 0 | 0 | 1678166 | YP_009047211.1 | K9N4V7 |
| LTFSYDGLRGAIRKAK | 1 | 0 | 0 | 1678168 | YP_009047202.1 | K9N7C7 |
| LTFVNVPYVYNGSAQA | 1 | 0 | 0 | 1678172 | YP_009555240.1 | P30215 |
| LTGCCDNSYFDSKGWY | 1 | 0 | 0 | 1678178 | YP_003766.2 | H9EJ42 |
| LTGDSVKFVYMHQLEL | 1 | 0 | 0 | 1678183 | YP_173236.1 | A0A140H1G9 |
| LTGILDGDYAMQFFKM | 1 | 0 | 0 | 1678189 | NP_073549.1 | A0A0P0LKV0 |
| LTGTNQLCAKILPFSD | 1 | 0 | 0 | 1678196 | YP_003766.2 | H9EJ42 |
| LTGVEAVMYMGTLSYD | 1 | 0 | 0 | 1678197 | NP_828849.2 | P0C6X7 |
| LTIANNYSKLFDEGYT | 1 | 0 | 0 | 1678206 | NP_073549.1 | A0A0P0LKV0 |
| LTIVIPDKQVFDKVVD | 1 | 0 | 0 | 1678220 | YP_173236.1 | A0A140H1G9 |
| LTKALRKVPTDNYITT | 1 | 0 | 0 | 1678224 | QHR63299.1 | P0C6W2 |
| LTKGTLEPEYFNSVCR | 1 | 0 | 0 | 1678230 | YP_009724389.1 | P0DTD1 |
| LTKHGDFADEIIGYKA | 1 | 0 | 0 | 1678232 | QIQ54046.1 | P0C6U8 |
| LTKLSKYLANEAQVQL | 1 | 0 | 0 | 1678237 | NP_073549.1 | A0A0P0LKV0 |
| LTKMSEYKGPVTDVFY | 1 | 0 | 0 | 1678239 | NP_828849.2 | P0C6X7 |
| LTKSSEYKGPITDIFY | 1 | 0 | 0 | 1678248 | QHR63299.1 | P0C6W2 |
| LTKSSEYKGPITDVFY | 1 | 0 | 0 | 1678249 | YP_009724389.1 | P0DTD1 |
| LTLDDRGLALDNALSG | 1 | 0 | 0 | 1678258 | NP_073549.1 | A0A0P0LKV0 |
| LTLDGYSKLFITKDEA | 1 | 0 | 0 | 1678259 | YP_173236.1 | A0A140H1G9 |
| LTLDNQDLNGKWYDFG | 1 | 0 | 0 | 1678261 | YP_009047202.1 | K9N7C7 |
| LTLTSGSDLDRCTTFD | 1 | 0 | 0 | 1678286 | NP_828851.1 | P59594 |
| LTMIEFMLWCKDGQVQ | 1 | 0 | 0 | 1678295 | YP_009047202.1 | K9N7C7 |
| LTMLDIQPEDYKSVDV | 1 | 0 | 0 | 1678297 | YP_009555239.1 | Q80872 |
| LTNHDVVYAQHCFKAP | 1 | 0 | 0 | 1678312 | YP_009555241.1 | P36334 |
| LTNLSEITKPLIEAEK | 1 | 0 | 0 | 1678317 | YP_173243.1 | Q0ZJ80 |
| LTNQVVKPGHFNEEFY | 1 | 0 | 0 | 1678323 | YP_003766.2 | H9EJ42 |
| LTNTGSVVKPVRARAP | 1 | 0 | 0 | 1678325 | NP_073549.1 | A0A0P0LKV0 |
| LTNTSSVVDGVVRSFQ | 1 | 0 | 0 | 1678326 | NP_073551.1 | A0A0P0K6L9 |
| LTPCGYSEPPKVAAKI | 1 | 0 | 0 | 1678332 | YP_009555238.1 | P0C6X6 |
| LTPGKLESGWTTGAAA | 1 | 0 | 0 | 1678336 | QIQ54048.1 | P59594 |
| LTPIKLNFGMLNGMPV | 1 | 0 | 0 | 1678338 | YP_003766.2 | H9EJ42 |
| LTPILSCGIFGIKLET | 1 | 0 | 0 | 1678339 | NP_073549.1 | A0A0P0LKV0 |
| LTPSDVYQQLAGIKLQ | 1 | 0 | 0 | 1678347 | YP_009555238.1 | P0C6X6 |
| LTPTESYVDVGPDSVK | 1 | 0 | 0 | 1678350 | YP_009047204.1 | K9N5Q8 |
| LTPVVQTIEVNSFSGY | 1 | 0 | 0 | 1678352 | YP_009724389.1 | P0DTD1 |
| LTQDHVDILGPLSAQT | 1 | 0 | 0 | 1678355 | YP_009724389.1 | P0DTD1 |
| LTQMNLKYAISAKNRA | 1 | 0 | 0 | 1678363 | YP_009555238.1 | P0C6X6 |
| LTRELNGGAVTRYVDN | 1 | 0 | 0 | 1678370 | NP_828849.2 | P0C6X7 |
| LTRGTLEPENFNSVTR | 1 | 0 | 0 | 1678372 | YP_009047202.1 | K9N7C7 |
| LTSEEVVLKNGELEAL | 1 | 0 | 0 | 1678386 | NP_828849.2 | P0C6X7 |
| LTSEISTLENKSAELN | 1 | 0 | 0 | 1678387 | NP_073551.1 | A0A0P0K6L9 |
| LTSHSVANLSAPTLVP | 1 | 0 | 0 | 1678392 | YP_009555238.1 | P0C6X6 |
| LTSHSVASLSAPTLVP | 1 | 0 | 0 | 1678393 | YP_173236.1 | A0A140H1G9 |
| LTSHTVMPLSAPTLVP | 1 | 0 | 0 | 1678395 | YP_009724389.1 | P0DTD1 |
| LTSQTVKPGHFNKEFY | 1 | 0 | 0 | 1678407 | NP_073549.1 | A0A0P0LKV0 |
| LTSSSKTPEEHFIETI | 1 | 0 | 0 | 1678412 | YP_009724389.1 | P0DTD1 |
| LTSSSKTSEEHFVETV | 1 | 0 | 0 | 1678413 | NP_828849.2 | P0C6X7 |
| LTTAAKLMVVIPDYNT | 1 | 0 | 0 | 1678415 | YP_009724389.1 | P0DTD1 |
| LTTAAKLMVVVPDYGT | 1 | 0 | 0 | 1678416 | NP_828849.2 | P0C6X7 |
| LTTAAKLMVVVPDYNT | 1 | 0 | 0 | 1678417 | QIQ54046.1 | P0C6U8 |
| LTTFIDAARGPAGVES | 1 | 0 | 0 | 1678422 | YP_009047202.1 | K9N7C7 |
| LTTQTVDSSQGSEYQY | 1 | 0 | 0 | 1678435 | YP_009047202.1 | K9N7C7 |
| LTTRTGIPPGYTNSST | 1 | 0 | 0 | 1678436 | QIQ54048.1 | P59594 |
| LTTRTQLPPAYTNSST | 1 | 0 | 0 | 1678438 | QHR63300.2 | Q3LZX1 |
| LTTSKKQFLPFQQFGR | 1 | 0 | 0 | 1678439 | QIQ54048.1 | P59594 |
| LTTYYLDDVKKIEYKP | 1 | 0 | 0 | 1678448 | YP_009555238.1 | P0C6X6 |
| LTVDGVNFTNRFVPVG | 1 | 0 | 0 | 1678452 | YP_009555238.1 | P0C6X6 |
| LTVDVANPSTPAYTFT | 1 | 0 | 0 | 1678454 | YP_009047202.1 | K9N7C7 |
| LTVEKGIYQTSNFRVQ | 1 | 0 | 0 | 1678455 | QIQ54048.1 | P59594 |
| LTVVSKVHEVIIDNKP | 1 | 0 | 0 | 1678473 | NP_073549.1 | A0A0P0LKV0 |
| LTYFNKPSFKSENRYS | 1 | 0 | 0 | 1678484 | YP_173236.1 | A0A140H1G9 |
| LTYFNRPSLVDDNKFD | 1 | 0 | 0 | 1678485 | YP_009555238.1 | P0C6X6 |
| LTYNKQMANVSVLTTP | 1 | 0 | 0 | 1678490 | YP_009555238.1 | P0C6X6 |
| LTYNKVENMTPRDLGA | 1 | 0 | 0 | 1678491 | YP_009724389.1 | P0DTD1 |
| LTYTGAIKLDDKDPSF | 1 | 0 | 0 | 1678495 | QIQ54056.1 | P59595 |
| LVAFTTELKDSVKKFF | 1 | 0 | 0 | 1678505 | QIQ54046.1 | P0C6U8 |

| | | | | | | |
|---|---|---|---|---|---|---|
| LVALANMLPAHVFMRF | 1 | 0 | 0 | 1678510 | YP_009555238.1 | P0C6X6 |
| LVAMKFNYEPLTQDHV | 1 | 0 | 0 | 1678513 | QIQ54046.1 | P0C6U8 |
| LVAMKYNYEPLTQDHV | 1 | 0 | 0 | 1678514 | YP_009724389.1 | P0DTD1 |
| LVASFLEMKSETKVQD | 1 | 0 | 0 | 1678515 | QIQ54046.1 | P0C6U8 |
| LVATAHSELAKGVALD | 1 | 0 | 0 | 1678520 | NP_828849.2 | P0C6X7 |
| LVAVPTGYVDTENNTE | 1 | 0 | 0 | 1678521 | NP_828849.2 | P0C6X7 |
| LVAVPTGYVDTPNNTD | 1 | 0 | 0 | 1678523 | YP_009724389.1 | P0DTD1 |
| LVDGVSRLYQPLRLTC | 1 | 0 | 0 | 1678541 | YP_003767.1 | A0A0P0G321 |
| LVDLKWLNRVETYIKW | 1 | 0 | 0 | 1678545 | NP_073551.1 | A0A0P0K6L9 |
| LVDMSMTYGQQFGPTY | 1 | 0 | 0 | 1678547 | NP_828849.2 | P0C6X7 |
| LVDNDLYPFVSDSVAT | 1 | 0 | 0 | 1678549 | YP_009555238.1 | P0C6X6 |
| LVDQRTICFSVAALST | 1 | 0 | 0 | 1678551 | YP_003766.2 | H9EJ42 |
| LVDSDLNDFVSDADST | 1 | 0 | 0 | 1678552 | YP_009724389.1 | P0DTD1 |
| LVDSSLVTSVGDSSEI | 1 | 0 | 0 | 1678555 | YP_009047202.1 | K9N7C7 |
| LVDSYFENDSILQSVA | 1 | 0 | 0 | 1678556 | NP_073549.1 | A0A0P0LKV0 |
| LVEKLSPWFMDGENAY | 1 | 0 | 0 | 1678563 | YP_009047202.1 | K9N7C7 |
| LVEVEKGVLPQLEQPY | 1 | 0 | 0 | 1678568 | YP_009724389.1 | P0DTD1 |
| LVEXVDVTFLKKDAPY | 1 | 0 | 0 | 1678570 | QIQ54046.1 | P0C6U8 |
| LVFQNNVANYQHIGMK | 1 | 0 | 0 | 1678586 | YP_173236.1 | A0A140H1G9 |
| LVFSNSNYLLEEFDVV | 1 | 0 | 0 | 1678588 | NP_073549.1 | A0A0P0LKV0 |
| LVFTNKTTLPTNVAFE | 1 | 0 | 0 | 1678591 | YP_003766.2 | H9EJ42 |
| LVFVNKTSLPTNIAFE | 1 | 0 | 0 | 1678594 | NP_073549.1 | A0A0P0LKV0 |
| LVGVLTLDNQDLNGKW | 1 | 0 | 0 | 1678610 | YP_009047202.1 | K9N7C7 |
| LVHDNVAFYQGDVDTV | 1 | 0 | 0 | 1678613 | NP_073549.1 | A0A0P0LKV0 |
| LVHHENEEYQKVFRVY | 1 | 0 | 0 | 1678614 | YP_173236.1 | A0A140H1G9 |
| LVIIANRYNEVANAVM | 1 | 0 | 0 | 1678625 | YP_173236.1 | A0A140H1G9 |
| LVKDKCVNFNFNGLTG | 1 | 0 | 0 | 1678637 | QIQ54048.1 | P59594 |
| LVKDPPNVQIHTIDGS | 1 | 0 | 0 | 1678638 | NP_828852.2 | P59632 |
| LVKIQAVVNANAEALN | 1 | 0 | 0 | 1678646 | YP_009555241.1 | P36334 |
| LVKMDGDDVYLPYPNP | 1 | 0 | 0 | 1678652 | YP_009555238.1 | P0C6X6 |
| LVKQGDDYVYLPYPDP | 1 | 0 | 0 | 1678659 | YP_009724389.1 | P0DTD1 |
| LVKTGHYTGQAGEMPC | 1 | 0 | 0 | 1678665 | YP_009555238.1 | P0C6X6 |
| LVLAKGGFKFGDPADS | 1 | 0 | 0 | 1678668 | YP_009555238.1 | P0C6X6 |
| LVLAKGHFKFDEPSDA | 1 | 0 | 0 | 1678669 | YP_173236.1 | A0A140H1G9 |
| LVLAKLGKDATKPQQV | 1 | 0 | 0 | 1678670 | YP_009555245.1 | P33469 |
| LVLAKLGKDSKPQQVT | 1 | 0 | 0 | 1678671 | YP_173242.1 | Q19U29 |
| LVLGVNEKTHVQLSLP | 1 | 0 | 0 | 1678683 | NP_828849.2 | P0C6X7 |
| LVLNNCTKYNIYDYVG | 1 | 0 | 0 | 1678695 | YP_003767.1 | A0A0P0G321 |
| LVMQKLPFKFKDLAIE | 1 | 0 | 0 | 1678722 | YP_173236.1 | A0A140H1G9 |
| LVNAANTHLKHGGGIA | 1 | 0 | 0 | 1678725 | YP_009047202.1 | K9N7C7 |
| LVNANQYSPCVSIVPS | 1 | 0 | 0 | 1678726 | YP_009047204.1 | K9N5Q8 |
| LVNDIDITFLKKDAPY | 1 | 0 | 0 | 1678729 | QHR63299.1 | P0C6W2 |
| LVNFTVDPQKAYLDFV | 1 | 0 | 0 | 1678730 | YP_009047202.1 | K9N7C7 |
| LVNHYFQTRHYLHTL | 1 | 0 | 0 | 1678732 | P03155.1 | Q69028 |
| LVNKMVNWFNTMLDAS | 1 | 0 | 0 | 1678737 | YP_173236.1 | A0A140H1G9 |
| LVNSSLFVEDCKLPLG | 1 | 0 | 0 | 1678739 | YP_009047204.1 | K9N5Q8 |
| LVNVGHYDGRTGELPC | 1 | 0 | 0 | 1678740 | YP_173236.1 | A0A140H1G9 |
| LVNVQKVEQPKIEPKP | 1 | 0 | 0 | 1678743 | NP_073549.1 | A0A0P0LKV0 |
| LVPCASGLSTDVQLRA | 1 | 0 | 0 | 1678749 | YP_009555238.1 | P0C6X6 |
| LVPGFNEKTHVQLSLP | 1 | 0 | 0 | 1678752 | YP_009724389.1 | P0DTD1 |
| LVPIGKGNKDEQIGYW | 1 | 0 | 0 | 1678760 | YP_003771.1 | H9EJ69 |
| LVPNQPYPNASFDNFK | 1 | 0 | 0 | 1678765 | YP_009724389.1 | P0DTD1 |
| LVPQEHYVRITGLYPT | 1 | 0 | 0 | 1678766 | YP_009724389.1 | P0DTD1 |
| LVPTQPLPNASFDNFK | 1 | 0 | 0 | 1678771 | NP_828849.2 | P0C6X7 |
| LVPTQPYPNSNFDNFK | 1 | 0 | 0 | 1678772 | QIQ54046.1 | P0C6U8 |
| LVPTYLKSDNIVAADL | 1 | 0 | 0 | 1678774 | YP_009555238.1 | P0C6X6 |
| LVQNGVDAKGFPQLAE | 1 | 0 | 0 | 1678788 | YP_003771.1 | H9EJ69 |
| LVQQESSFVMMSAPPA | 1 | 0 | 0 | 1678792 | NP_828849.2 | P0C6X7 |
| LVQQTSQWLSNLLGTT | 1 | 0 | 0 | 1678793 | NP_828849.2 | P0C6X7 |
| LVQYDFTDYKLELFNK | 1 | 0 | 0 | 1678797 | YP_009555238.1 | P0C6X6 |
| LVRGFGDSVEEVLSEA | 1 | 0 | 0 | 1678806 | YP_009724389.1 | P0DTD1 |
| LVSAVNQSLSNAKVSA | 1 | 0 | 0 | 1678820 | NP_073552.1 | A0A0P0KWE8 |
| LVSDIDITFLKKDAPY | 1 | 0 | 0 | 1678822 | YP_009724389.1 | P0DTD1 |
| LVSGDAHIPFGTVLGN | 1 | 0 | 0 | 1678826 | NP_073551.1 | A0A0P0K6L9 |
| LVSGTATAGWTFGAGA | 1 | 0 | 0 | 1678827 | NP_828851.1 | P59594 |
| LVSNNKEDFDIIQKCQ | 1 | 0 | 0 | 1678831 | YP_173236.1 | A0A140H1G9 |
| LVSNNQEDFDLISKCQ | 1 | 0 | 0 | 1678832 | YP_009555238.1 | P0C6X6 |
| LVSSDGQPGGDAISLS | 1 | 0 | 0 | 1678834 | YP_009047202.1 | K9N7C7 |
| LVSVTDFSLNGNVSTV | 1 | 0 | 0 | 1678841 | YP_009047208.1 | K9N7D2 |
| LVTASSKPLYVPEHCQ | 1 | 0 | 0 | 1678843 | YP_009047205.1 | K9N796 |
| LVTDIVVVDEVSMLTN | 1 | 0 | 0 | 1678845 | YP_173236.1 | A0A140H1G9 |
| LVTEEVVETPTIQKDI | 1 | 0 | 0 | 1678846 | QIQ54046.1 | P0C6U8 |
| LVTICDFYSKLGVGFT | 1 | 0 | 0 | 1678850 | YP_173236.1 | A0A140H1G9 |
| LVTYNCCDDDYFNKKD | 1 | 0 | 0 | 1678872 | YP_009724389.1 | P0DTD1 |
| LVTYNCCTDDYFNKKD | 1 | 0 | 0 | 1678873 | QIQ54047.1 | P0C6X7 |
| LVVEQYNQTILNLTSE | 1 | 0 | 0 | 1678876 | NP_073551.1 | A0A0P0K6L9 |
| LVVVVPDHDSFVKMMV | 1 | 0 | 0 | 1678886 | NP_073549.1 | A0A0P0LKV0 |
| LVVYQQAVVGAMLSEN | 1 | 0 | 0 | 1678890 | NP_073551.1 | A0A0P0K6L9 |
| LVWEAGKSKPPLNRNY | 1 | 0 | 0 | 1678891 | YP_009047202.1 | K9N7C7 |
| LVWHVERKDVPYPKQS | 1 | 0 | 0 | 1678892 | YP_009047202.1 | K9N7C7 |
| LVYAADPAMHAASGNL | 1 | 0 | 0 | 1678897 | YP_009724389.1 | P0DTD1 |
| LVYALRHFDQNSEVLK | 1 | 0 | 0 | 1678899 | YP_009047202.1 | K9N7C7 |
| LVYDNKLKAHKDKSAQ | 1 | 0 | 0 | 1678903 | YP_009724389.1 | P0DTD1 |
| LVYDNKLKAKNDNSSL | 1 | 0 | 0 | 1678904 | YP_173236.1 | A0A140H1G9 |
| LVYDNKLRAHKDKSQQ | 1 | 0 | 0 | 1678905 | QIQ54047.1 | P0C6X7 |
| LVYENKFVPVKPASKQ | 1 | 0 | 0 | 1678907 | YP_003766.2 | H9EJ42 |
| LVYHENEEYQKVFRVY | 1 | 0 | 0 | 1678911 | YP_009555238.1 | P0C6X6 |
| LVYLNHSVPKLSDFES | 1 | 0 | 0 | 1678913 | YP_173238.1 | A0A140H1H1 |
| LWAKRNIKPVPEIKIL | 1 | 0 | 0 | 1678922 | NP_828849.2 | P0C6X7 |
| LWAKRNIKPVPEVKIL | 1 | 0 | 0 | 1678923 | YP_009724389.1 | P0DTD1 |
| LWCKDGHVETFYPKLQ | 1 | 0 | 0 | 1678926 | YP_009724389.1 | P0DTD1 |

| | | | | | | |
|---|---|---|---|---|---|---|
| LWCKDGQVQTFYPRLQ | 1 | 0 | 0 | 1678927 | YP_009047202.1 | K9N7C7 |
| LWDITVINNVDNEIVK | 1 | 0 | 0 | 1678936 | YP_009047202.1 | K9N7C7 |
| LWEIQQVVDADSKIVQ | 1 | 0 | 0 | 1678941 | YP_009724389.1 | P0DTD1 |
| LWESGKAKPPLNRNSV | 1 | 0 | 0 | 1678942 | NP_073549.1 | A0A0P0LKV0 |
| LWIPIIKSSVKSYDDL | 1 | 0 | 0 | 1678952 | YP_173236.1 | A0A140H1G9 |
| LWNCQLSFAGYTESAV | 1 | 0 | 0 | 1678971 | YP_009047206.1 | K9N4V0 |
| LWNTKPVETSNSFDVL | 1 | 0 | 0 | 1678979 | QHR63299.1 | P0C6W2 |
| LWQFVDHFGENEEIII | 1 | 0 | 0 | 1678988 | NP_073549.1 | A0A0P0LKV0 |
| LWSTKPVDTSNSFEVL | 1 | 0 | 0 | 1678996 | NP_828849.2 | P0C6X7 |
| LWSTKPVETSNIFDVL | 1 | 0 | 0 | 1678997 | QIQ54046.1 | P0C6U8 |
| LWSTKPVETSNSFDVL | 1 | 0 | 0 | 1678998 | YP_009724389.1 | P0DTD1 |
| LWVNKASYDTNLNKFN | 1 | 0 | 0 | 1679002 | YP_009047202.1 | K9N7C7 |
| LYADINGNLYPDSTSL | 1 | 0 | 0 | 1679017 | QIQ54046.1 | P0C6U8 |
| LYAIQEGFDATNSALV | 1 | 0 | 0 | 1679021 | YP_009555241.1 | P36334 |
| LYAKRAVRSHPDFKLL | 1 | 0 | 0 | 1679024 | YP_009047202.1 | K9N7C7 |
| LYCAFTAPEDDDFLEE | 1 | 0 | 0 | 1679028 | YP_009555238.1 | P0C6X6 |
| LYCDSPFEKLQCEHLQ | 1 | 0 | 0 | 1679029 | YP_003767.1 | A0A0P0G321 |
| LYCRAHIEHPDVSGVC | 1 | 0 | 0 | 1679032 | YP_009047202.1 | K9N7C7 |
| LYDASNVKKVTDVTGK | 1 | 0 | 0 | 1679039 | YP_009555238.1 | P0C6X6 |
| LYDGNMFIKFPEVVFE | 1 | 0 | 0 | 1679041 | YP_009047202.1 | K9N7C7 |
| LYDNCYRLTSVEESFI | 1 | 0 | 0 | 1679047 | YP_003766.2 | H9EJ42 |
| LYDSNGNIIGFKDFVT | 1 | 0 | 0 | 1679050 | YP_173238.1 | A0A140H1H1 |
| LYDSNGNLYGFRDYIT | 1 | 0 | 0 | 1679051 | YP_009555241.1 | P36334 |
| LYEQVVMDYLDNLKPR | 1 | 0 | 0 | 1679064 | NP_828849.2 | P0C6X7 |
| LYIDINGNLHPDSATL | 1 | 0 | 0 | 1679104 | YP_009724389.1 | P0DTD1 |
| LYIKDGDDGYFLPYPD | 1 | 0 | 0 | 1679110 | YP_009047202.1 | K9N7C7 |
| LYIQGIKLGTGYSLAD | 1 | 0 | 0 | 1679114 | YP_009555244.1 | Q01455 |
| LYIQGVKLGTGYTLSD | 1 | 0 | 0 | 1679115 | YP_173241.1 | Q0ZJ82 |
| LYKDVDNPHLMGWDYP | 1 | 0 | 0 | 1679117 | YP_009047202.1 | K9N7C7 |
| LYKKVNNEFIDPTECI | 1 | 0 | 0 | 1679123 | YP_009047202.1 | K9N7C7 |
| LYKMQRMLLDKCDLQN | 1 | 0 | 0 | 1679128 | QIQ54047.1 | P0C6X7 |
| LYKNMCTGSPSIVEFN | 1 | 0 | 0 | 1679131 | YP_009047202.1 | K9N7C7 |
| LYKNSATGSLDVEVFN | 1 | 0 | 0 | 1679132 | YP_003766.2 | H9EJ42 |
| LYKNTCVGSDNVTDFN | 1 | 0 | 0 | 1679133 | YP_009724389.1 | P0DTD1 |
| LYKQSCTGSPYIDDFN | 1 | 0 | 0 | 1679136 | YP_009555238.1 | P0C6X6 |
| LYKSSALGSMDIDVFN | 1 | 0 | 0 | 1679138 | NP_073549.1 | A0A0P0LKV0 |
| LYKSYSEQVIPPTSDY | 1 | 0 | 0 | 1679139 | YP_173240.1 | Q0ZJ83 |
| LYKXTCVGSDNVTDFN | 1 | 0 | 0 | 1679142 | QIQ54047.1 | P0C6X7 |
| LYKYLNSPMERVSLWN | 1 | 0 | 0 | 1679143 | YP_009555238.1 | P0C6X6 |
| LYLDAYNMMISAGFSL | 1 | 0 | 0 | 1679146 | YP_009724389.1 | P0DTD1 |
| LYLDLLNRLQALESGK | 1 | 0 | 0 | 1679147 | YP_009047211.1 | K9N4V7 |
| LYLDYVQCKCGNLEQN | 1 | 0 | 0 | 1679148 | YP_173236.1 | A0A140H1G9 |
| LYLKNLTQTFTSMLTN | 1 | 0 | 0 | 1679159 | YP_173236.1 | A0A140H1G9 |
| LYLNSSNVTTMDFCEG | 1 | 0 | 0 | 1679160 | NP_828849.2 | P0C6X7 |
| LYNKYKYFSGALDTTS | 1 | 0 | 0 | 1679182 | NP_828849.2 | P0C6X7 |
| LYNKYKYFSGAMDTTS | 1 | 0 | 0 | 1679183 | YP_009724389.1 | P0DTD1 |
| LYNKYRYYSGKMDTAA | 1 | 0 | 0 | 1679184 | YP_009555238.1 | P0C6X6 |
| LYNTGRSVYVKFQDSK | 1 | 0 | 0 | 1679195 | YP_009047209.1 | K9N5R3 |
| LYNYGAGIKLPSGIML | 1 | 0 | 0 | 1679199 | YP_003766.2 | H9EJ42 |
| LYPDSTSLVEXVDVTF | 1 | 0 | 0 | 1679201 | QIQ54046.1 | P0C6U8 |
| LYPTITVPEEFASHVA | 1 | 0 | 0 | 1679203 | YP_009047202.1 | K9N7C7 |
| LYPTLNISDEFSSNVA | 1 | 0 | 0 | 1679204 | YP_009724389.1 | P0DTD1 |
| LYPYGRCPTAADINIP | 1 | 0 | 0 | 1679205 | YP_009555240.1 | P30215 |
| LYQPPTASVSTSFLQS | 1 | 0 | 0 | 1679213 | YP_009555238.1 | P0C6X6 |
| LYRKCVKPKEETGLLM | 1 | 0 | 0 | 1679218 | QHR63299.1 | P0C6W2 |
| LYRLFRKSNLKPFERD | 1 | 0 | 0 | 1679222 | YP_009724390.1 | P0DTC2 |
| LYRNRDVDTDFVNEFY | 1 | 0 | 0 | 1679226 | YP_009724389.1 | P0DTD1 |
| LYRRQQTSNLVVQEFV | 1 | 0 | 0 | 1679229 | YP_009555238.1 | P0C6X6 |
| LYSFEAHKDSFKDGLC | 1 | 0 | 0 | 1679235 | YP_009555238.1 | P0C6X6 |
| LYSNVYRTDYVDYTFV | 1 | 0 | 0 | 1679241 | YP_173236.1 | A0A140H1G9 |
| LYSPPTISYNSTLQSG | 1 | 0 | 0 | 1679247 | YP_003766.2 | H9EJ42 |
| LYSTQLSTDTGVEHTT | 1 | 0 | 0 | 1679250 | QIQ54049.1 | P59632 |
| LYSTQLSTDTGVEHVT | 1 | 0 | 0 | 1679251 | YP_009724391.1 | P0DTC3 |
| LYVALRGRIDKDLSLS | 1 | 0 | 0 | 1679266 | NP_073553.1 | A0A0P0KWE8 |
| LYVDFKPQSGGGKCFN | 1 | 0 | 0 | 1679267 | NP_073551.1 | A0A0P0K6L9 |
| LYVIDGALLSKTSEYK | 1 | 0 | 0 | 1679272 | QIQ54046.1 | P0C6U8 |
| LYVNVYRSTSPDPKFV | 1 | 0 | 0 | 1679285 | YP_009047202.1 | K9N7C7 |
| LYVSWSDGDGITGVPQ | 1 | 0 | 0 | 1679290 | NP_073551.1 | A0A0P0K6L9 |
| LYVVVVESDADRANFL | 1 | 0 | 0 | 1679292 | YP_173236.1 | A0A140H1G9 |
| LYYKVNCCRFQRIDDD | 1 | 0 | 0 | 1679302 | YP_173236.1 | A0A140H1G9 |
| MAATRGATCVIGTTKF | 1 | 0 | 0 | 1679314 | YP_009047202.1 | K9N7C7 |
| MAAYVDNSSLTIKKPN | 1 | 0 | 0 | 1679315 | YP_009724389.1 | P0DTD1 |
| MAAYVDNTSLTIKKPN | 1 | 0 | 0 | 1679316 | QIQ54046.1 | P0C6U8 |
| MADLALTNMYKEARIN | 1 | 0 | 0 | 1679327 | YP_009555238.1 | P0C6X6 |
| MADVDDPKLMGWDYPK | 1 | 0 | 0 | 1679333 | NP_073549.1 | A0A0P0LKV0 |
| MAEQAAAAMYKEARAV | 1 | 0 | 0 | 1679335 | NP_073549.1 | A0A0P0LKV0 |
| MAFSASLFKPVQLVPV | 1 | 0 | 0 | 1679340 | YP_009047208.1 | K9N7D2 |
| MAKAGDKYYPVVVDDH | 1 | 0 | 0 | 1679352 | YP_009555238.1 | P0C6X6 |
| MAKEVKTSSKVVLSDA | 1 | 0 | 0 | 1679355 | NP_073549.1 | A0A0P0LKV0 |
| MAKFPLKLAGTAVINL | 1 | 0 | 0 | 1679356 | YP_009555238.1 | P0C6X6 |
| MANQEKYSTIYKLHPS | 1 | 0 | 0 | 1679372 | NP_073549.1 | A0A0P0LKV0 |
| MAPVEFYEDRILDFKV | 1 | 0 | 0 | 1679376 | YP_009047202.1 | K9N7C7 |
| MARNGVVPLSVIPATS | 1 | 0 | 0 | 1679381 | NP_073549.1 | A0A0P0LKV0 |
| MARRNLDSLEARAFQS | 1 | 0 | 0 | 1679382 | NP_828859.1 | P59636 |
| MASFVEYEVAKKNLAD | 1 | 0 | 0 | 1679384 | YP_173236.1 | A0A140H1G9 |
| MASVNWADDRAARKKF | 1 | 0 | 0 | 1679394 | YP_003771.1 | H9EJ69 |
| MATVKWADASEPQRGR | 1 | 0 | 0 | 1679399 | NP_073556.1 | A0A0N7J131 |
| MAVAYADKPNHFINFP | 1 | 0 | 0 | 1679402 | YP_009555239.1 | Q80872 |
| MAYVTGGVVQMTTQWL | 1 | 0 | 0 | 1679409 | QIQ54046.1 | P0C6U8 |
| MCDIKQMLFCMEVVNK | 1 | 0 | 0 | 1679417 | YP_009047202.1 | K9N7C7 |
| MCDIRQLLFVVEVVDK | 1 | 0 | 0 | 1679418 | YP_009724389.1 | P0DTD1 |

| | | | | | | |
|---|---|---|---|---|---|---|
| MCIDMKGTMYVRPIIE | 1 | 0 | 0 | 1679432 | YP_009555244.1 | Q01455 |
| MCLEPCNLYNYGAGIK | 1 | 0 | 0 | 1679437 | YP_003766.2 | H9EJ42 |
| MCLEPCNLYNYGAGLK | 1 | 0 | 0 | 1679438 | NP_073549.1 | A0A0P0LKV0 |
| MCTGSPSIVEFNRLAT | 1 | 0 | 0 | 1679453 | YP_009047202.1 | K9N7C7 |
| MCTLSEQLDYIESKRG | 1 | 0 | 0 | 1679455 | NP_828849.2 | P0C6X7 |
| MCVVPSINDLSFDKNL | 1 | 0 | 0 | 1679464 | YP_003766.2 | H9EJ42 |
| MCYRCPKEIVSTVSAL | 1 | 0 | 0 | 1679466 | YP_009047202.1 | K9N7C7 |
| MDADLAKPLIKWDLLK | 1 | 0 | 0 | 1679467 | NP_828849.2 | P0C6X7 |
| MDAKQVDYVPLKSATC | 1 | 0 | 0 | 1679468 | YP_009555238.1 | P0C6X6 |
| MDGFSCPLCCTVYDSS | 1 | 0 | 0 | 1679483 | YP_173236.1 | A0A140H1G9 |
| MDGMDAKQVDYVPLKS | 1 | 0 | 0 | 1679486 | YP_009555238.1 | P0C6X6 |
| MDIDVFNKLSTSDWSD | 1 | 0 | 0 | 1679490 | NP_073549.1 | A0A0P0LKV0 |
| MDKLDTGAQKFFQFGD | 1 | 0 | 0 | 1679494 | YP_003766.2 | H9EJ42 |
| MDLNPVLCDSGKISSK | 1 | 0 | 0 | 1679506 | YP_009555240.1 | P30215 |
| MDLVYALRHFDQNSEV | 1 | 0 | 0 | 1679507 | YP_009047202.1 | K9N7C7 |
| MDMGVFINKYGLEDFN | 1 | 0 | 0 | 1679509 | YP_003766.2 | H9EJ42 |
| MDNAINVGGTGLQYAA | 1 | 0 | 0 | 1679510 | YP_009047202.1 | K9N7C7 |
| MDPNQTNVVPPALHLV | 1 | 0 | 0 | 1679516 | NP_828859.1 | P59636 |
| MDTFTNNNGSDVLYQP | 1 | 0 | 0 | 1679525 | YP_009555238.1 | P0C6X6 |
| MDVLSQNQKLIANAFN | 1 | 0 | 0 | 1679532 | YP_009555241.1 | P36334 |
| MDYLDNLKPRVEAPKQ | 1 | 0 | 0 | 1679541 | NP_828849.2 | P0C6X7 |
| MEDAMGQGQNSADPKV | 1 | 0 | 0 | 1679547 | NP_828859.1 | P59636 |
| MEKDFIALDQDVFIQK | 1 | 0 | 0 | 1679556 | YP_009555238.1 | P0C6X6 |
| MEKDFLSFDSDVFIKK | 1 | 0 | 0 | 1679557 | YP_009047202.1 | K9N7C7 |
| MEKVNECVKSQSSRIN | 1 | 0 | 0 | 1679561 | YP_009555241.1 | P36334 |
| MELPTGVHAGTDLEGK | 1 | 0 | 0 | 1679564 | NP_828849.2 | P0C6X7 |
| MELPTGVHAGTDLEGN | 1 | 0 | 0 | 1679565 | YP_009724389.1 | P0DTD1 |
| MELPTGVHAGTDLEGT | 1 | 0 | 0 | 1679566 | QHR63299.1 | P0C6W2 |
| MELTPVVQTIEVNSFS | 1 | 0 | 0 | 1679568 | YP_009724389.1 | P0DTD1 |
| MEPTPVVQTEVNSFSG | 1 | 0 | 0 | 1679571 | QHR63299.1 | P0C6W2 |
| MESQISTKEVKLKGVR | 1 | 0 | 0 | 1679578 | YP_173236.1 | A0A140H1G9 |
| METDFLELAMDEFIQR | 1 | 0 | 0 | 1679581 | NP_828849.2 | P0C6X7 |
| METFNHNNGNDVLYQP | 1 | 0 | 0 | 1679582 | YP_173236.1 | A0A140H1G9 |
| MEYDFLNMDMGVFINK | 1 | 0 | 0 | 1679588 | YP_003766.2 | H9EJ42 |
| MFDAYVDTFSATFSVP | 1 | 0 | 0 | 1679592 | NP_828849.2 | P0C6X7 |
| MFDAYVNTFSSTFNVP | 1 | 0 | 0 | 1679593 | YP_009724389.1 | P0DTD1 |
| MFDLDSTFDDILGKVC | 1 | 0 | 0 | 1679594 | YP_173236.1 | A0A140H1G9 |
| MFDSFVNSFVSLYNVT | 1 | 0 | 0 | 1679595 | YP_009047202.1 | K9N7C7 |
| MFDVDKKSLNALIATA | 1 | 0 | 0 | 1679596 | YP_009555238.1 | P0C6X6 |
| MFDVYVDTFLSMFDVD | 1 | 0 | 0 | 1679597 | YP_009555238.1 | P0C6X6 |
| MFENKTTLPTNIAFEL | 1 | 0 | 0 | 1679601 | YP_009047202.1 | K9N7C7 |
| MFEPRIPTMADFVQIE | 1 | 0 | 0 | 1679602 | NP_073551.1 | A0A0P0K6L9 |
| MFGLYDDILTNNKPWF | 1 | 0 | 0 | 1679606 | YP_003766.2 | H9EJ42 |
| MFITREEAIKHVRAWV | 1 | 0 | 0 | 1679614 | QIQ54047.1 | P0C6X7 |
| MFKRGDGTPHPYCYSD | 1 | 0 | 0 | 1679617 | YP_173236.1 | A0A140H1G9 |
| MFLRLIDDNGIVLNSI | 1 | 0 | 0 | 1679630 | YP_003769.1 | H9EJA2 |
| MFTGSSYYYPEPISDK | 1 | 0 | 0 | 1679647 | YP_173238.1 | A0A140H1H1 |
| MFTKVVVGPTTEVKFS | 1 | 0 | 0 | 1679648 | YP_003766.2 | H9EJ42 |
| MFVRPVIEDYHTLTAT | 1 | 0 | 0 | 1679662 | YP_173241.1 | Q0ZJ82 |
| MGANGNKIVDKEVKDM | 1 | 0 | 0 | 1679672 | YP_009047202.1 | K9N7C7 |
| MGFKMNYQVNGYPNMF | 1 | 0 | 0 | 1679681 | YP_009724389.1 | P0DTD1 |
| MGILSNMNIASNSKEW | 1 | 0 | 0 | 1679692 | NP_073549.1 | A0A0P0LKV0 |
| MGIRFEYTSPTDKLAM | 1 | 0 | 0 | 1679694 | YP_009555239.1 | Q80872 |
| MGKCGDKFFPIVMNDK | 1 | 0 | 0 | 1679695 | YP_173236.1 | A0A140H1G9 |
| MGLHGNQTLFCNIMKF | 1 | 0 | 0 | 1679701 | NP_073549.1 | A0A0P0LKV0 |
| MGQGQNSADPKVYPII | 1 | 0 | 0 | 1679707 | NP_828859.1 | P59636 |
| MGSCSFEDELTPSDVY | 1 | 0 | 0 | 1679712 | YP_009555238.1 | P0C6X6 |
| MGSGFCVDYNSPSSSS | 1 | 0 | 0 | 1679713 | YP_173238.1 | A0A140H1H1 |
| MGTLSYDNLKTGVSIP | 1 | 0 | 0 | 1679714 | NP_828849.2 | P0C6X7 |
| MGTLSYEQLKKGVQIP | 1 | 0 | 0 | 1679716 | QHR63299.1 | P0C6W2 |
| MGVFINKYGLEDFNFE | 1 | 0 | 0 | 1679720 | YP_003766.2 | H9EJ42 |
| MGVYDFKVSAAEFKYM | 1 | 0 | 0 | 1679725 | YP_003766.2 | H9EJ42 |
| MGWDYPKCDRALPNMI | 1 | 0 | 0 | 1679728 | YP_003766.2 | H9EJ42 |
| MGWDYPKCDRAMPNIL | 1 | 0 | 0 | 1679729 | YP_173236.1 | A0A140H1G9 |
| MGWDYPKCDRAMPNLL | 1 | 0 | 0 | 1679730 | YP_009555238.1 | P0C6X6 |
| MGWDYPKCDRAMPNMC | 1 | 0 | 0 | 1679731 | YP_009047202.1 | K9N7C7 |
| MGWDYPKCDRAMPNML | 1 | 0 | 0 | 1679732 | YP_009724389.1 | P0DTD1 |
| MGWDYPKCDRAMPSMI | 1 | 0 | 0 | 1679733 | NP_073549.1 | A0A0P0LKV0 |
| MHFGACDYDRLPNEVT | 1 | 0 | 0 | 1679741 | YP_009047210.1 | K9N7A1 |
| MHFGTLAKTDLFNGYK | 1 | 0 | 0 | 1679742 | YP_173236.1 | A0A140H1G9 |
| MHIASSNAFLDLRTSC | 1 | 0 | 0 | 1679748 | YP_009047202.1 | K9N7C7 |
| MHIDPFPKRVIDF | 1 | 0 | 0 | 1679749 | NP_073554.1 | A0A0P0KH29 |
| MHLSTLMKCNHCDEVS | 1 | 0 | 0 | 1679755 | NP_828849.2 | P0C6X7 |
| MHPMVNPSKGSSGLLI | 1 | 0 | 0 | 1679757 | YP_173243.1 | Q0ZJ80 |
| MIANENAINKGGRTVQ | 1 | 0 | 0 | 1679773 | NP_073549.1 | A0A0P0LKV0 |
| MIEFMLWCKDGQVQTF | 1 | 0 | 0 | 1679782 | YP_009047202.1 | K9N7C7 |
| MIFWFALTLQDDDECC | 1 | 0 | 0 | 1679790 | QIQ54054.1 | Q7TFA1 |
| MIKGPQRADLNGVIVD | 1 | 0 | 0 | 1679796 | YP_173236.1 | A0A140H1G9 |
| MIKKLDNDVLNGIISN | 1 | 0 | 0 | 1679797 | YP_009047202.1 | K9N7C7 |
| MILSDDGVVCYNKDYA | 1 | 0 | 0 | 1679807 | YP_003766.2 | H9EJ42 |
| MILSDDGVVCYNSDYA | 1 | 0 | 0 | 1679808 | YP_009047202.1 | K9N7C7 |
| MIRGPPRAELNGVVVD | 1 | 0 | 0 | 1679814 | YP_009555238.1 | P0C6X6 |
| MISEWPLSVQQAQQEA | 1 | 0 | 0 | 1679824 | NP_073549.1 | A0A0P0LKV0 |
| MISQWPISNDSNGCVV | 1 | 0 | 0 | 1679825 | YP_003766.2 | H9EJ42 |
| MKAVAAALKSLGFDKP | 1 | 0 | 0 | 1679834 | NP_073556.1 | A0A0N7J131 |
| MKDDGNCVVLELDPPC | 1 | 0 | 0 | 1679839 | YP_173236.1 | A0A140H1G9 |
| MKGKMFVRPVIEDYHT | 1 | 0 | 0 | 1679848 | YP_173241.1 | Q0ZJ82 |
| MKGTGVFVQDPAPIDI | 1 | 0 | 0 | 1679849 | YP_003766.2 | H9EJ42 |
| MKGTMYVRPIIEDYHT | 1 | 0 | 0 | 1679850 | YP_009555244.1 | Q01455 |
| MKQYGECLGDINARDL | 1 | 0 | 0 | 1679864 | NP_828851.1 | P59594 |
| MKSDEGGLCPSTGQAM | 1 | 0 | 0 | 1679867 | YP_173236.1 | A0A140H1G9 |

| | | | | | | |
|---|---|---|---|---|---|---|
| MKSDIFGQDFKTFDLL | 1 | 0 | 0 | 1679868 | NP_073549.1 | A0A0P0LKV0 |
| MKVKATKGEGDGGITS | 1 | 0 | 0 | 1679874 | NP_073549.1 | A0A0P0LKV0 |
| MKVLNENLNAYQQQDG | 1 | 0 | 0 | 1679876 | YP_009555245.1 | P33469 |
| MLDFSRDHNDILYTPP | 1 | 0 | 0 | 1679900 | NP_073549.1 | A0A0P0LKV0 |
| MLDKQGQLLDTKLNGI | 1 | 0 | 0 | 1679903 | YP_003766.2 | H9EJ42 |
| MLDYAKDHNDMLYSPP | 1 | 0 | 0 | 1679911 | YP_003766.2 | H9EJ42 |
| MLFDSHIVSKESGNTV | 1 | 0 | 0 | 1679917 | NP_073556.1 | A0A0N7J131 |
| MLFGMIKKLDNDVLNG | 1 | 0 | 0 | 1679918 | YP_009047202.1 | K9N7C7 |
| MLFSMVRKLDNQALNS | 1 | 0 | 0 | 1679922 | YP_009555238.1 | P0C6X6 |
| MLFTMLRKLDNDALNN | 1 | 0 | 0 | 1679924 | YP_009724389.1 | P0DTD1 |
| MLGAEMTSNVRRQVAS | 1 | 0 | 0 | 1679928 | NP_073549.1 | A0A0P0LKV0 |
| MLGLTTQTVDSSQGSE | 1 | 0 | 0 | 1679931 | YP_009047202.1 | K9N7C7 |
| MLGSSVGNFSDGKMGR | 1 | 0 | 0 | 1679932 | YP_009047204.1 | K9N5Q8 |
| MLKRRDSTYGPLQTPV | 1 | 0 | 0 | 1679948 | YP_009047204.1 | K9N5Q8 |
| MLKSMAATRGATCVIG | 1 | 0 | 0 | 1679949 | YP_009047202.1 | K9N7C7 |
| MLMGWDYPKCDRALPN | 1 | 0 | 0 | 1679978 | YP_003766.2 | H9EJ42 |
| MLNGNAIATVKSEDGN | 1 | 0 | 0 | 1679982 | NP_073549.1 | A0A0P0LKV0 |
| MLNTSIPNLPDFKEEL | 1 | 0 | 0 | 1679985 | YP_009555241.1 | P36334 |
| MLPLAYFSDSKLGTTE | 1 | 0 | 0 | 1679990 | YP_009047202.1 | K9N7C7 |
| MLRACIKTAQADTAGL | 1 | 0 | 0 | 1679998 | YP_009047205.1 | K9N796 |
| MLRGEDMSFLEKDAPY | 1 | 0 | 0 | 1679999 | NP_828849.2 | P0C6X7 |
| MLRHLIKDVDNPVLMG | 1 | 0 | 0 | 1680000 | YP_173236.1 | A0A140H1G9 |
| MLRKLDNDALNNIINN | 1 | 0 | 0 | 1680004 | YP_009724389.1 | P0DTD1 |
| MLRLHTVESGDINVVA | 1 | 0 | 0 | 1680005 | YP_009047202.1 | K9N7C7 |
| MLRRLIKDVDNPVLMG | 1 | 0 | 0 | 1680008 | YP_009555238.1 | P0C6X6 |
| MLSDTLKNLSDRVVFV | 1 | 0 | 0 | 1680014 | YP_009724389.1 | P0DTD1 |
| MLSENFTSYGFSNVVE | 1 | 0 | 0 | 1680016 | NP_073551.1 | A0A0P0K6L9 |
| MLSLQQVVKALNESYI | 1 | 0 | 0 | 1680017 | YP_009047204.1 | K9N5Q8 |
| MLSRALKKVPVDEYIT | 1 | 0 | 0 | 1680019 | NP_828849.2 | P0C6X7 |
| MLTFDKKAFTLANVSS | 1 | 0 | 0 | 1680021 | NP_073551.1 | A0A0P0K6L9 |
| MLTLDDRGLALDNALS | 1 | 0 | 0 | 1680027 | NP_073549.1 | A0A0P0LKV0 |
| MLTYNKVENMTPRDLG | 1 | 0 | 0 | 1680038 | YP_009724389.1 | P0DTD1 |
| MLVKMDGDDVYLPYPN | 1 | 0 | 0 | 1680041 | YP_009555238.1 | P0C6X6 |
| MLWCKDGQVQTFYPRL | 1 | 0 | 0 | 1680050 | YP_009047202.1 | K9N7C7 |
| MLYSPPTISYNSTLQS | 1 | 0 | 0 | 1680055 | YP_003766.2 | H9EJ42 |
| MMILSDDGVVCYNKDY | 1 | 0 | 0 | 1680068 | YP_003766.2 | H9EJ42 |
| MMRLFYDRDGQRVYDD | 1 | 0 | 0 | 1680081 | YP_173236.1 | A0A140H1G9 |
| MNASPYDLEVLLQDAL | 1 | 0 | 0 | 1680092 | YP_009555238.1 | P0C6X6 |
| MNGRTILGSALLEDEF | 1 | 0 | 0 | 1680103 | YP_009724389.1 | P0DTD1 |
| MNKSFLPQFTSDQAVT | 1 | 0 | 0 | 1680110 | YP_173241.1 | Q0ZJ82 |
| MNLSTYSLFDLSKFQL | 1 | 0 | 0 | 1680115 | YP_009047202.1 | K9N7C7 |
| MNNKSQSVIIINNSTN | 1 | 0 | 0 | 1680120 | NP_828851.1 | P59594 |
| MNSQGLLPPKNSIDAF | 1 | 0 | 0 | 1680123 | YP_009724389.1 | P0DTD1 |
| MNSQGLLPPKSSIDAF | 1 | 0 | 0 | 1680124 | NP_828849.2 | P0C6X7 |
| MNSRNYIAQVDVVNFN | 1 | 0 | 0 | 1680125 | YP_009725255.1 | A0A663DJA2 |
| MNVTLDKCTKYNIYDV | 1 | 0 | 0 | 1680133 | NP_073551.1 | A0A0P0K6L9 |
| MPCAIINDKVVAKIDK | 1 | 0 | 0 | 1680139 | YP_009555238.1 | P0C6X6 |
| MPCPACQDPEIGPEHS | 1 | 0 | 0 | 1680141 | NP_828849.2 | P0C6X7 |
| MPFEELNGLKILKQLD | 1 | 0 | 0 | 1680142 | NP_073549.1 | A0A0P0LKV0 |
| MPFFYFDDSDCDVVQE | 1 | 0 | 0 | 1680143 | YP_003766.2 | H9EJ42 |
| MPHKLKIQVVNSGSDM | 1 | 0 | 0 | 1680154 | YP_173236.1 | A0A140H1G9 |
| MPIADPTHFDIEEVEL | 1 | 0 | 0 | 1680155 | NP_073549.1 | A0A0P0LKV0 |
| MPIGYVTHGFNLEEAA | 1 | 0 | 0 | 1680158 | NP_828849.2 | P0C6X7 |
| MPIPDFDLDDFIDAPC | 1 | 0 | 0 | 1680164 | YP_009047202.1 | K9N7C7 |
| MPKIIKVFYELDNDFN | 1 | 0 | 0 | 1680166 | YP_009555238.1 | P0C6X6 |
| MPLGYVTHGLNLEEAA | 1 | 0 | 0 | 1680170 | YP_009724389.1 | P0DTD1 |
| MPLKAPKEVTFLEGDS | 1 | 0 | 0 | 1680173 | NP_828849.2 | P0C6X7 |
| MPLSAPTLVPQEHYVR | 1 | 0 | 0 | 1680176 | YP_009724389.1 | P0DTD1 |
| MPNLYKMQRMLLDKCD | 1 | 0 | 0 | 1680185 | QIQ54047.1 | P0C6X7 |
| MPNLYKMQRMLLEKCD | 1 | 0 | 0 | 1680186 | YP_009724389.1 | P0DTD1 |
| MPSFVAYETARQEYEN | 1 | 0 | 0 | 1680191 | NP_073549.1 | A0A0P0LKV0 |
| MPSLFKVQNVNLERCE | 1 | 0 | 0 | 1680192 | YP_009047202.1 | K9N7C7 |
| MPYHPNTVDCQDDRCI | 1 | 0 | 0 | 1680215 | YP_009555238.1 | P0C6X6 |
| MPYTKLFQKGECCICH | 1 | 0 | 0 | 1680216 | YP_003766.2 | H9EJ42 |
| MQGAVDINKLCEEMLD | 1 | 0 | 0 | 1680225 | YP_009724389.1 | P0DTD1 |
| MQGAVDINRLCEEMLD | 1 | 0 | 0 | 1680226 | NP_828849.2 | P0C6X7 |
| MQGVTLSSNLNTNLHF | 1 | 0 | 0 | 1680230 | YP_173238.1 | A0A140H1H1 |
| MQIESDDYIARNGPLN | 1 | 0 | 0 | 1680233 | QIQ54046.1 | P0C6U8 |
| MQIVDENGKYYLPYPD | 1 | 0 | 0 | 1680235 | NP_073549.1 | A0A0P0LKV0 |
| MQIVDKDGTYYLPYPD | 1 | 0 | 0 | 1680236 | YP_003766.2 | H9EJ42 |
| MQKESDDYIKLNGPLT | 1 | 0 | 0 | 1680237 | NP_828849.2 | P0C6X7 |
| MQKLPCQFKDKNLQDL | 1 | 0 | 0 | 1680238 | YP_009555238.1 | P0C6X6 |
| MQKLPFKFKDLAIENM | 1 | 0 | 0 | 1680239 | YP_173236.1 | A0A140H1G9 |
| MQKPRWKRQPNDDVTS | 1 | 0 | 0 | 1680240 | NP_073556.1 | A0A0N7J131 |
| MQKYSTLQGPPGTGKS | 1 | 0 | 0 | 1680241 | YP_009724389.1 | P0DTD1 |
| MQNCLLRLKVDTSNPK | 1 | 0 | 0 | 1680255 | NP_828849.2 | P0C6X7 |
| MQNCVLKLKVDAANPK | 1 | 0 | 0 | 1680256 | QIQ54046.1 | P0C6U8 |
| MQQHPLLNPSALEFNP | 1 | 0 | 0 | 1680263 | NP_073556.1 | A0A0N7J131 |
| MQRMLLDKCDLQNYGE | 1 | 0 | 0 | 1680266 | QIQ54047.1 | P0C6X7 |
| MQTGFTTTNEAFQKVQ | 1 | 0 | 0 | 1680275 | YP_009047204.1 | K9N5Q8 |
| MQVESDHYIATNGPLK | 1 | 0 | 0 | 1680280 | QHR63299.1 | P0C6W2 |
| MQXYSTLQGPPGTGKS | 1 | 0 | 0 | 1680282 | QIQ54047.1 | P0C6X7 |
| MRAGSKVVNHNVLIKE | 1 | 0 | 0 | 1680286 | YP_003766.2 | H9EJ42 |
| MRDAGIVGVLTLDNQD | 1 | 0 | 0 | 1680288 | QHR63299.1 | P0C6W2 |
| MRELNGGAYTRYVDNN | 1 | 0 | 0 | 1680289 | YP_009724389.1 | P0DTD1 |
| MRGCCDCGSTKLPYYE | 1 | 0 | 0 | 1680293 | YP_003767.1 | A0A0P0G321 |
| MRHVRGWLGMDVEGAH | 1 | 0 | 0 | 1680297 | NP_073549.1 | A0A0P0LKV0 |
| MRKVDLYNGAVIYIFA | 1 | 0 | 0 | 1680304 | YP_003768.1 | H9EJ24 |
| MRLFYDRDGQRTYDDV | 1 | 0 | 0 | 1680306 | YP_009555238.1 | P0C6X6 |
| MRLFYDRDGQRVYDDV | 1 | 0 | 0 | 1680307 | YP_173236.1 | A0A140H1G9 |
| MRNFDEQNCDVLKEVL | 1 | 0 | 0 | 1680314 | YP_003766.2 | H9EJ42 |

| | | | | | | |
|---|---|---|---|---|---|---|
| MRPNHTIKGSFLNGSC | 1 | 0 | 0 | 1680317 | NP_828849.2 | P0C6X7 |
| MRRGQRVDLPPKVHFY | 1 | 0 | 0 | 1680321 | YP_003771.1 | H9EJ69 |
| MRVLHLGAGSDKEVAP | 1 | 0 | 0 | 1680336 | YP_173236.1 | A0A140H1G9 |
| MRVLHYGAGSDKGVAP | 1 | 0 | 0 | 1680339 | YP_003766.2 | H9EJ42 |
| MRYNLNFSANSLDNLK | 1 | 0 | 0 | 1680343 | YP_003767.1 | A0A0P0G321 |
| MSAPPAEYKLKHGTFL | 1 | 0 | 0 | 1680349 | QIQ54046.1 | P0C6U8 |
| MSCDCGKKFDDQVGCL | 1 | 0 | 0 | 1680356 | YP_003766.2 | H9EJ42 |
| MSDFMSEIKCKTQSIA | 1 | 0 | 0 | 1680359 | YP_009555241.1 | P36334 |
| MSDRDLYDKLQFTSLE | 1 | 0 | 0 | 1680363 | YP_009724389.1 | P0DTD1 |
| MSEAKCWTETDLTKGP | 1 | 0 | 0 | 1680366 | YP_009724389.1 | P0DTD1 |
| MSEDLWQFVDHFGENE | 1 | 0 | 0 | 1680369 | NP_073549.1 | A0A0P0LKV0 |
| MSEDVKLNPSEDFIKH | 1 | 0 | 0 | 1680370 | YP_009047202.1 | K9N7C7 |
| MSEPPKEQRVQGSITQ | 1 | 0 | 0 | 1680374 | YP_009047211.1 | K9N4V7 |
| MSESKCWVEHDINNGP | 1 | 0 | 0 | 1680375 | YP_009555238.1 | P0C6X6 |
| MSESKCWVENDITNGP | 1 | 0 | 0 | 1680376 | YP_173236.1 | A0A140H1G9 |
| MSEYKGPVTDVFYKET | 1 | 0 | 0 | 1680377 | NP_828849.2 | P0C6X7 |
| MSFTPGKQSSSRASSG | 1 | 0 | 0 | 1680385 | YP_009555245.1 | P33469 |
| MSHFDVDRKSFNNFVN | 1 | 0 | 0 | 1680394 | YP_173236.1 | A0A140H1G9 |
| MSKINKYGLELHWAPE | 1 | 0 | 0 | 1680403 | YP_009555238.1 | P0C6X6 |
| MSLAECKRALGLSISD | 1 | 0 | 0 | 1680404 | NP_073549.1 | A0A0P0LKV0 |
| MSLGADSSIAYSNNTI | 1 | 0 | 0 | 1680407 | NP_828851.1 | P59594 |
| MSLKENQINDMIYSLL | 1 | 0 | 0 | 1680410 | NP_828849.2 | P0C6X7 |
| MSLYMAISPKFTTSLS | 1 | 0 | 0 | 1680417 | NP_828853.1 | P59633 |
| MSNMQLFESLNFITLP | 1 | 0 | 0 | 1680424 | YP_173236.1 | A0A140H1G9 |
| MSNSSVPLLEVYVHLR | 1 | 0 | 0 | 1680427 | YP_003770.1 | U3M6Q8 |
| MSNYQHEETIYNLVKD | 1 | 0 | 0 | 1680428 | NP_828849.2 | P0C6X7 |
| MSQFKLTHQNNDDHGN | 1 | 0 | 0 | 1680430 | YP_009047211.1 | K9N4V7 |
| MSSVDTILNMARNGVV | 1 | 0 | 0 | 1680440 | NP_073549.1 | A0A0P0LKV0 |
| MSSVETVLNLARDGVV | 1 | 0 | 0 | 1680441 | YP_003766.2 | H9EJ42 |
| MSTSKCWVEEDLTKGP | 1 | 0 | 0 | 1680446 | YP_003766.2 | H9EJ42 |
| MSYTPGHYAGSRSSSG | 1 | 0 | 0 | 1680455 | YP_173242.1 | Q19U29 |
| MTAVNESRYGLQNLLQ | 1 | 0 | 0 | 1680464 | YP_003767.1 | A0A0P0G321 |
| MTDIAKKPTENICAPL | 1 | 0 | 0 | 1680468 | QHR63299.1 | P0C6W2 |
| MTDIAKKPTESACSSL | 1 | 0 | 0 | 1680469 | NP_828849.2 | P0C6X7 |
| MTDIAKKPTETICAPL | 1 | 0 | 0 | 1680470 | YP_009724389.1 | P0DTD1 |
| MTFSGNVDLDALASDI | 1 | 0 | 0 | 1680476 | YP_009047202.1 | K9N7C7 |
| MTFYPRLQAASDWKPG | 1 | 0 | 0 | 1680478 | YP_009555238.1 | P0C6X6 |
| MTIVKCKGLNKPFVKD | 1 | 0 | 0 | 1680487 | YP_009047202.1 | K9N7C7 |
| MTNHSFSVQKHIGAPA | 1 | 0 | 0 | 1680501 | YP_009047202.1 | K9N7C7 |
| MTQLNLKYAISGKERA | 1 | 0 | 0 | 1680509 | NP_073549.1 | A0A0P0LKV0 |
| MTQMNLKYAISAKNRA | 1 | 0 | 0 | 1680511 | YP_009047202.1 | K9N7C7 |
| MTQMYKQARSEDKRAK | 1 | 0 | 0 | 1680512 | YP_009724389.1 | P0DTD1 |
| MTSLPKDWRLINKFDV | 1 | 0 | 0 | 1680522 | YP_173236.1 | A0A140H1G9 |
| MTSMYKQARAEDKKAK | 1 | 0 | 0 | 1680523 | YP_009047202.1 | K9N7C7 |
| MTWLDMVDTSLSGFKL | 1 | 0 | 0 | 1680537 | YP_009724389.1 | P0DTD1 |
| MTWLDMVDTSLSGYKL | 1 | 0 | 0 | 1680538 | QIQ54046.1 | P0C6U8 |
| MVANGLHAPHGPFDAL | 1 | 0 | 0 | 1680546 | YP_003766.2 | H9EJ42 |
| MVANGLNAPNGPFDAL | 1 | 0 | 0 | 1680547 | NP_073549.1 | A0A0P0LKV0 |
| MVDTSLSGFKLKDCVM | 1 | 0 | 0 | 1680558 | YP_009724389.1 | P0DTD1 |
| MVDTSLSGYKLKDCIM | 1 | 0 | 0 | 1680559 | QIQ54046.1 | P0C6U8 |
| MVKEFPLLSEYPLLKQ | 1 | 0 | 0 | 1680576 | YP_173243.1 | Q0ZJ80 |
| MVKSKGVCATGDCYVS | 1 | 0 | 0 | 1680578 | YP_009555238.1 | P0C6X6 |
| MVKSKGVCQVGDCYVS | 1 | 0 | 0 | 1680579 | YP_173236.1 | A0A140H1G9 |
| MVNWFNTMLDASAPAT | 1 | 0 | 0 | 1680601 | YP_173236.1 | A0A140H1G9 |
| MVQTCTPNVTINCQDP | 1 | 0 | 0 | 1680606 | NP_849177.1 | Q80H93 |
| MVRKLDNQALNSILDN | 1 | 0 | 0 | 1680610 | YP_009555238.1 | P0C6X6 |
| MVTDIVVVDEVSMLTN | 1 | 0 | 0 | 1680615 | YP_009555238.1 | P0C6X6 |
| MVTNNTFTLKGGAPTK | 1 | 0 | 0 | 1680618 | YP_009724389.1 | P0DTD1 |
| MVVCDKVRSLKLSDNN | 1 | 0 | 0 | 1680619 | YP_009047202.1 | K9N7C7 |
| MVVIPDYNTYKNTCDG | 1 | 0 | 0 | 1680620 | YP_009724389.1 | P0DTD1 |
| MVVVPDYGTYKNTCDG | 1 | 0 | 0 | 1680624 | NP_828849.2 | P0C6X7 |
| MVVVPDYNTYKNTCEG | 1 | 0 | 0 | 1680625 | QIQ54046.1 | P0C6U8 |
| MWEFRDYFNNNTDSIV | 1 | 0 | 0 | 1680632 | YP_003766.2 | H9EJ42 |
| MWKGYGCSCDQLREPL | 1 | 0 | 0 | 1680637 | NP_828849.2 | P0C6X7 |
| MWKGYGCSCDQLREPM | 1 | 0 | 0 | 1680638 | YP_009724389.1 | P0DTD1 |
| MWLYTYSIGEASSPIA | 1 | 0 | 0 | 1680646 | YP_009047202.1 | K9N7C7 |
| MYAADPAMHIASSNAF | 1 | 0 | 0 | 1680653 | YP_009047202.1 | K9N7C7 |
| MYCSFYPPDEDEEEGD | 1 | 0 | 0 | 1680659 | YP_009724389.1 | P0DTD1 |
| MYCSFYPPDEDYEEEY | 1 | 0 | 0 | 1680660 | QIQ54046.1 | P0C6U8 |
| MYCSFYPPDEEEEDDA | 1 | 0 | 0 | 1680661 | NP_828849.2 | P0C6X7 |
| MYCTFAIEDVEDVISS | 1 | 0 | 0 | 1680662 | YP_173236.1 | A0A140H1G9 |
| MYDGDRFVKHDLSLLS | 1 | 0 | 0 | 1680663 | NP_073549.1 | A0A0P0LKV0 |
| MYDGRTKAIDGENVSK | 1 | 0 | 0 | 1680665 | NP_073549.1 | A0A0P0LKV0 |
| MYDPITKNIGEYNVSK | 1 | 0 | 0 | 1680667 | YP_009555238.1 | P0C6X6 |
| MYDPKTKNVTKENDSK | 1 | 0 | 0 | 1680669 | YP_009724389.1 | P0DTD1 |
| MYDPLTKNIGDYNVSK | 1 | 0 | 0 | 1680670 | YP_173236.1 | A0A140H1G9 |
| MYDPRTKHVTKENDSK | 1 | 0 | 0 | 1680671 | NP_828849.2 | P0C6X7 |
| MYDPTTKNVTGSNESK | 1 | 0 | 0 | 1680672 | YP_009047202.1 | K9N7C7 |
| MYGGFEDQPNLQVESA | 1 | 0 | 0 | 1680680 | NP_073549.1 | A0A0P0LKV0 |
| MYKEARSVNRKSKVIS | 1 | 0 | 0 | 1680695 | YP_003766.2 | H9EJ42 |
| MYKQARAEDKKAKIVS | 1 | 0 | 0 | 1680699 | YP_009047202.1 | K9N7C7 |
| MYLRKRGEKGAYNKDH | 1 | 0 | 0 | 1680709 | YP_009555238.1 | P0C6X6 |
| MYMGTLSYDNLKTGVS | 1 | 0 | 0 | 1680713 | NP_828849.2 | P0C6X7 |
| MYMGTLSYEQFKKGVQ | 1 | 0 | 0 | 1680714 | YP_009724389.1 | P0DTD1 |
| MYNLLKGCNAVAKHDF | 1 | 0 | 0 | 1680719 | NP_073549.1 | A0A0P0LKV0 |
| MYVRPIIEDYHTLTVT | 1 | 0 | 0 | 1680734 | YP_009555244.1 | Q01455 |
| MYVYSAGHATGTTPQK | 1 | 0 | 0 | 1680736 | YP_009047204.1 | K9N5Q8 |
| NAAITDYNYYKYNLPT | 1 | 0 | 0 | 1680741 | YP_009555238.1 | P0C6X6 |
| NAANTHLKHGGGIAGA | 1 | 0 | 0 | 1680747 | YP_009047202.1 | K9N7C7 |
| NAASVNQIVLRNSNGA | 1 | 0 | 0 | 1680752 | YP_009047202.1 | K9N7C7 |
| NADIVVVDEVSMCTNY | 1 | 0 | 0 | 1680763 | NP_073549.1 | A0A0P0LKV0 |

| | | | | | | |
|---|---|---|---|---|---|---|
| NAFLVHDNVAFYQGDV | 1 | 0 | 0 | 1680782 | NP_073549.1 | A0A0P0LKV0 |
| NAFTREMQQHPLLNPS | 1 | 0 | 0 | 1680787 | NP_073556.1 | A0A0N7J131 |
| NAFVSQVLNKYTEVRG | 1 | 0 | 0 | 1680790 | YP_003767.1 | A0A0P0G321 |
| NAGEDIQLLKAAYENF | 1 | 0 | 0 | 1680793 | NP_828849.2 | P0C6X7 |
| NAGYWRRQDRKINTGN | 1 | 0 | 0 | 1680798 | YP_009047211.1 | K9N4V7 |
| NAIATVKSEDGNIKNI | 1 | 0 | 0 | 1680803 | NP_073549.1 | A0A0P0LKV0 |
| NAILTEVNELLDTTQL | 1 | 0 | 0 | 1680808 | YP_009555241.1 | P36334 |
| NAIPSLAANTLNIIVP | 1 | 0 | 0 | 1680811 | YP_009555238.1 | P0C6X6 |
| NAISSNTDNETLQYWV | 1 | 0 | 0 | 1680813 | YP_173238.1 | A0A140H1H1 |
| NAKLGFDSSKEFVEYK | 1 | 0 | 0 | 1680820 | YP_009555238.1 | P0C6X6 |
| NAKVEISVTPIKNTVD | 1 | 0 | 0 | 1680823 | NP_073549.1 | A0A0P0LKV0 |
| NAKVSAEVSRQVIQDV | 1 | 0 | 0 | 1680824 | NP_073552.1 | A0A0P0KWE8 |
| NALIATAHSSIKQGTQ | 1 | 0 | 0 | 1680832 | YP_009555238.1 | P0C6X6 |
| NALLSIQNGFSATNSA | 1 | 0 | 0 | 1680835 | YP_173238.1 | A0A140H1H1 |
| NALLSTDGNKIADKHV | 1 | 0 | 0 | 1680836 | QHR63299.1 | P0C6W2 |
| NALLSTDGNKIADKYV | 1 | 0 | 0 | 1680837 | YP_009724389.1 | P0DTD1 |
| NALLSTDGNKIGDKYI | 1 | 0 | 0 | 1680838 | QIQ54047.1 | P0C6X7 |
| NALSRAVNVPTIRKFI | 1 | 0 | 0 | 1680849 | YP_009555238.1 | P0C6X6 |
| NALYAIQEGFDATNSA | 1 | 0 | 0 | 1680852 | YP_009555241.1 | P36334 |
| NAMQVESDHYIATNGP | 1 | 0 | 0 | 1680858 | QHR63299.1 | P0C6W2 |
| NANMSLAECKRALGLS | 1 | 0 | 0 | 1680868 | NP_073549.1 | A0A0P0LKV0 |
| NANQVIVNNLDKSAGF | 1 | 0 | 0 | 1680869 | YP_009724389.1 | P0DTD1 |
| NAPGCDVNDVTKLYLG | 1 | 0 | 0 | 1680877 | YP_009555238.1 | P0C6X6 |
| NAQALSKLASELSNTF | 1 | 0 | 0 | 1680887 | YP_009047204.1 | K9N5Q8 |
| NARIRAKHYVYIGDPA | 1 | 0 | 0 | 1680898 | YP_009555238.1 | P0C6X6 |
| NARLRAKHYVYIGDPA | 1 | 0 | 0 | 1680899 | YP_009724389.1 | P0DTD1 |
| NARQAYEDAIANGSSS | 1 | 0 | 0 | 1680904 | YP_003766.2 | H9EJ42 |
| NASEVVVNNLDKSAGH | 1 | 0 | 0 | 1680907 | YP_009047202.1 | K9N7C7 |
| NASGCGVSDVKKLYLG | 1 | 0 | 0 | 1680911 | YP_003766.2 | H9EJ42 |
| NASKFASVYAWNRKRI | 1 | 0 | 0 | 1680913 | QIQ54048.1 | P59594 |
| NASLNSFKEYFNLRNC | 1 | 0 | 0 | 1680915 | YP_009047204.1 | K9N5Q8 |
| NASLYTSLVPHTRYSL | 1 | 0 | 0 | 1680917 | YP_173236.1 | A0A140H1G9 |
| NASNQVAVLYQDVNCT | 1 | 0 | 0 | 1680918 | QHR63300.2 | Q3LZX1 |
| NASPYDLEVLLQDALQ | 1 | 0 | 0 | 1680920 | YP_009555238.1 | P0C6X6 |
| NASSSEAFLIGANYLG | 1 | 0 | 0 | 1680923 | NP_828849.2 | P0C6X7 |
| NATEFSRVSAKPPPGD | 1 | 0 | 0 | 1680935 | QIQ54047.1 | P0C6X7 |
| NATGCDVTDVTQLYLG | 1 | 0 | 0 | 1680939 | QIQ54047.1 | P0C6X7 |
| NATGSDVNCNGYQHNS | 1 | 0 | 0 | 1680940 | YP_003767.1 | A0A0P0G321 |
| NATKFPSVYAWERKKI | 1 | 0 | 0 | 1680941 | NP_828851.1 | P59594 |
| NATNKATYKPNTWCIR | 1 | 0 | 0 | 1680942 | YP_009724389.1 | P0DTD1 |
| NAVAKHDFFTWHEGRT | 1 | 0 | 0 | 1680952 | NP_073549.1 | A0A0P0LKV0 |
| NAVATFYPQLQSAEWK | 1 | 0 | 0 | 1680955 | NP_073549.1 | A0A0P0LKV0 |
| NAVDCMSDFMSEIKCK | 1 | 0 | 0 | 1680957 | YP_009555241.1 | P36334 |
| NAVDCSSSFFSEIQCK | 1 | 0 | 0 | 1680958 | YP_173238.1 | A0A140H1H1 |
| NAVKGCVPLNAIPSLA | 1 | 0 | 0 | 1680961 | YP_009555238.1 | P0C6X6 |
| NAVSRSMLGLTTQTVD | 1 | 0 | 0 | 1680966 | YP_009047202.1 | K9N7C7 |
| NAVYTKVDGIDVEIFE | 1 | 0 | 0 | 1680975 | NP_828849.2 | P0C6X7 |
| NAYANWCCLKCGFSFD | 1 | 0 | 0 | 1680977 | YP_173236.1 | A0A140H1G9 |
| NAYEKDKAVARKLERM | 1 | 0 | 0 | 1680978 | YP_009047202.1 | K9N7C7 |
| NAYVNSNQNTDSDSLS | 1 | 0 | 0 | 1680987 | YP_173242.1 | Q19U29 |
| NAYVSQQLSDSTLVKF | 1 | 0 | 0 | 1680989 | YP_009555241.1 | P36334 |
| NAYYKYDAKNYVRFPE | 1 | 0 | 0 | 1680991 | YP_003766.2 | H9EJ42 |
| NAYYKYDNGNFIKLPE | 1 | 0 | 0 | 1680992 | NP_073549.1 | A0A0P0LKV0 |
| NCAGNVIGNDALTRFT | 1 | 0 | 0 | 1680994 | YP_173236.1 | A0A140H1G9 |
| NCCDDDYFNKKDWYDF | 1 | 0 | 0 | 1681001 | YP_009724389.1 | P0DTD1 |
| NCCDEYGGHHDFVIKT | 1 | 0 | 0 | 1681002 | YP_173238.1 | A0A140H1H1 |
| NCCRFQEKDEDDNLID | 1 | 0 | 0 | 1681003 | YP_009724389.1 | P0DTD1 |
| NCCRFQEKDEDGNLID | 1 | 0 | 0 | 1681004 | QIQ54047.1 | P0C6X7 |
| NCCRFQRIDDDGNKLD | 1 | 0 | 0 | 1681006 | YP_173236.1 | A0A140H1G9 |
| NCCRFQRVDENGDKLD | 1 | 0 | 0 | 1681007 | YP_009555238.1 | P0C6X6 |
| NCCTDDYFNKKDWYDF | 1 | 0 | 0 | 1681009 | QIQ54047.1 | P0C6X7 |
| NCDFYGWVSGNMMDGF | 1 | 0 | 0 | 1681011 | YP_173236.1 | A0A140H1G9 |
| NCDSFGPGNTFINGDI | 1 | 0 | 0 | 1681012 | YP_003766.2 | H9EJ42 |
| NCDVSDVTKLYLGGMS | 1 | 0 | 0 | 1681020 | YP_173236.1 | A0A140H1G9 |
| NCIDCDSYKPGNTFIT | 1 | 0 | 0 | 1681033 | YP_009555238.1 | P0C6X6 |
| NCLHVVGPNVNRGEDI | 1 | 0 | 0 | 1681044 | QHR63299.1 | P0C6W2 |
| NCLQHLHVASNSKLWH | 1 | 0 | 0 | 1681048 | YP_009555238.1 | P0C6X6 |
| NCLQHLHVASNSKLWQ | 1 | 0 | 0 | 1681049 | YP_173236.1 | A0A140H1G9 |
| NCNIPTQCYYNNGSSG | 1 | 0 | 0 | 1681058 | YP_173236.1 | A0A140H1G9 |
| NCQLSFAGYTESAVNS | 1 | 0 | 0 | 1681077 | YP_009047206.1 | K9N4V0 |
| NCSFQDAASNIFVINK | 1 | 0 | 0 | 1681079 | YP_009047202.1 | K9N7C7 |
| NCTKYNIYDYVGTGII | 1 | 0 | 0 | 1681096 | YP_003767.1 | A0A0P0G321 |
| NCTNFAINDLNTTVPR | 1 | 0 | 0 | 1681097 | YP_173238.1 | A0A140H1H1 |
| NCVKMLTNGSGSGQAI | 1 | 0 | 0 | 1681107 | NP_073549.1 | A0A0P0LKV0 |
| NCVLSVNSSNCNNFNV | 1 | 0 | 0 | 1681113 | NP_073549.1 | A0A0P0LKV0 |
| NCYRCPKEIVETVSAL | 1 | 0 | 0 | 1681136 | YP_173236.1 | A0A140H1G9 |
| NDAITQTSQALQTVAT | 1 | 0 | 0 | 1681148 | NP_073551.1 | A0A0P0K6L9 |
| NDDDAWFKLSYHYENN | 1 | 0 | 0 | 1681161 | YP_009555239.1 | Q80872 |
| NDDEDVVTGDNDDEDV | 1 | 0 | 0 | 1681163 | YP_173236.1 | A0A140H1G9 |
| NDDEDVVTGDNNDEEI | 1 | 0 | 0 | 1681164 | YP_173236.1 | A0A140H1G9 |
| NDDQIVVTGDDVDDIE | 1 | 0 | 0 | 1681166 | YP_173236.1 | A0A140H1G9 |
| NDDTSETKYVKSLSIV | 1 | 0 | 0 | 1681170 | YP_009555238.1 | P0C6X6 |
| NDDVTSNVTQCFGPRD | 1 | 0 | 0 | 1681171 | NP_073556.1 | A0A0N7J131 |
| NDEEIVTGDNDDQIVV | 1 | 0 | 0 | 1681172 | YP_173236.1 | A0A140H1G9 |
| NDEKVMTFYPRLQAAS | 1 | 0 | 0 | 1681174 | YP_009555238.1 | P0C6X6 |
| NDFNTILNTACGVFEV | 1 | 0 | 0 | 1681178 | YP_009555238.1 | P0C6X6 |
| NDFSNSGADVLYQPPQ | 1 | 0 | 0 | 1681179 | NP_828849.2 | P0C6X7 |
| NDICMFENKTTLPTNI | 1 | 0 | 0 | 1681194 | YP_009047202.1 | K9N7C7 |
| NDIDITFLKKDAPYIV | 1 | 0 | 0 | 1681195 | QHR63299.1 | P0C6W2 |
| NDINDYVSDADFSITG | 1 | 0 | 0 | 1681201 | YP_003766.2 | H9EJ42 |
| NDITNTNLSRGRGRNP | 1 | 0 | 0 | 1681204 | YP_009047211.1 | K9N4V7 |

| | | | | | | |
|---|---|---|---|---|---|---|
| NDLNETLVTMPLGYVT | 1 | 0 | 0 | 1681233 | YP_009724389.1 | P0DTD1 |
| NDLYVALRGRIDKDLS | 1 | 0 | 0 | 1681242 | NP_073553.1 | A0A0P0KWE8 |
| NDMLYSPPTISYNSTL | 1 | 0 | 0 | 1681245 | YP_003766.2 | H9EJ42 |
| NDNCDFYGWVSGNMMD | 1 | 0 | 0 | 1681248 | YP_173236.1 | A0A140H1G9 |
| NDNLLEILQKEKVNIN | 1 | 0 | 0 | 1681254 | YP_009724389.1 | P0DTD1 |
| NDSNGCVVEASTDFHQ | 1 | 0 | 0 | 1681276 | YP_003766.2 | H9EJ42 |
| NDTFLNGITYTSTSGN | 1 | 0 | 0 | 1681281 | NP_073551.1 | A0A0P0K6L9 |
| NEAFQKVQDAVNNNAQ | 1 | 0 | 0 | 1681314 | YP_009047204.1 | K9N5Q8 |
| NECAQVLSEYVLCGGG | 1 | 0 | 0 | 1681318 | YP_009047202.1 | K9N7C7 |
| NEDLLEILSRERVNIN | 1 | 0 | 0 | 1681330 | NP_828849.2 | P0C6X7 |
| NEDNALAYYNTSKGGR | 1 | 0 | 0 | 1681331 | QIQ54046.1 | P0C6U8 |
| NEFRVYSSANNCTFEY | 1 | 0 | 0 | 1681343 | QIQ54048.1 | P59594 |
| NEIIGYKAIQDGVTRD | 1 | 0 | 0 | 1681353 | NP_828849.2 | P0C6X7 |
| NEIIVKPNISLCVPLY | 1 | 0 | 0 | 1681354 | NP_073549.1 | A0A0P0LKV0 |
| NEIKPSGLKTMVVSAG | 1 | 0 | 0 | 1681355 | YP_009047202.1 | K9N7C7 |
| NEKCSVYTVESGTEVT | 1 | 0 | 0 | 1681367 | NP_828849.2 | P0C6X7 |
| NELAQVLTEVVYSNGG | 1 | 0 | 0 | 1681374 | NP_073549.1 | A0A0P0LKV0 |
| NELLDTTQLQVANSLM | 1 | 0 | 0 | 1681375 | YP_009555241.1 | P36334 |
| NENLNAYQQQDGMMNM | 1 | 0 | 0 | 1681395 | YP_009555245.1 | P33469 |
| NESRYGLQNLLQLPNF | 1 | 0 | 0 | 1681412 | YP_003767.1 | A0A0P0G321 |
| NESWHFDKSEPLCLFK | 1 | 0 | 0 | 1681415 | YP_173238.1 | A0A140H1H1 |
| NESYNNYVPSYVKPDS | 1 | 0 | 0 | 1681417 | YP_009047202.1 | K9N7C7 |
| NETLQYWVTPLSKRQY | 1 | 0 | 0 | 1681419 | YP_173238.1 | A0A140H1H1 |
| NETLVTMPLGYVTHGL | 1 | 0 | 0 | 1681420 | YP_009724389.1 | P0DTD1 |
| NETNSQYLFSTINALP | 1 | 0 | 0 | 1681421 | YP_009047202.1 | K9N7C7 |
| NEVVKQMFGVNLQSGK | 1 | 0 | 0 | 1681429 | NP_073549.1 | A0A0P0LKV0 |
| NEYRLYLDAYNMMISA | 1 | 0 | 0 | 1681435 | YP_009724389.1 | P0DTD1 |
| NEYRQYLDAYNMMISA | 1 | 0 | 0 | 1681436 | NP_828849.2 | P0C6X7 |
| NFAAKRVLGLQTQTVD | 1 | 0 | 0 | 1681439 | YP_009555238.1 | P0C6X6 |
| NFAINDLNTTVPRISE | 1 | 0 | 0 | 1681440 | YP_173238.1 | A0A140H1H1 |
| NFDDQPSLQVESANLM | 1 | 0 | 0 | 1681453 | YP_003766.2 | H9EJ42 |
| NFDEQNCDVLKEVLVL | 1 | 0 | 0 | 1681455 | YP_003766.2 | H9EJ42 |
| NFDQKELLAYYNMLVN | 1 | 0 | 0 | 1681461 | YP_009555238.1 | P0C6X6 |
| NFDYLDIHPSLCNNGK | 1 | 0 | 0 | 1681463 | YP_173237.1 | A0A140H1H0 |
| NFEPFTVNSVNDSLEP | 1 | 0 | 0 | 1681470 | YP_009555241.1 | P36334 |
| NFFISYAKPEDKLSVY | 1 | 0 | 0 | 1681474 | YP_003766.2 | H9EJ42 |
| NFFSPQIITTDNTFVS | 1 | 0 | 0 | 1681475 | NP_828851.1 | P59594 |
| NFGDLQLNKLGTEDPR | 1 | 0 | 0 | 1681479 | YP_009047211.1 | K9N4V7 |
| NFGDQELIRQGTEYKH | 1 | 0 | 0 | 1681482 | QIQ54056.1 | P59595 |
| NFGMLNGNAIATVKSE | 1 | 0 | 0 | 1681488 | NP_073549.1 | A0A0P0LKV0 |
| NFIIYDSSDRGQESLA | 1 | 0 | 0 | 1681496 | YP_009047202.1 | K9N7C7 |
| NFKFVCDNIKFADDLN | 1 | 0 | 0 | 1681510 | YP_009724389.1 | P0DTD1 |
| NFKFVCDNIKLADDLN | 1 | 0 | 0 | 1681511 | QHR63299.1 | P0C6W2 |
| NFKFVCDNTKFADDLN | 1 | 0 | 0 | 1681512 | QIQ54046.1 | P0C6U8 |
| NFKKVETLNFQMVKEF | 1 | 0 | 0 | 1681514 | YP_173243.1 | Q0ZJ80 |
| NFKLTCSNTKFADDLN | 1 | 0 | 0 | 1681518 | NP_828849.2 | P0C6X7 |
| NFKSISLTVGEVFGKI | 1 | 0 | 0 | 1681523 | YP_173236.1 | A0A140H1G9 |
| NFLDDNVLPETYVALP | 1 | 0 | 0 | 1681528 | YP_003767.1 | A0A0P0G321 |
| NFLQILPDPSKPSKRS | 1 | 0 | 0 | 1681538 | QIQ54048.1 | P59594 |
| NFNEKPVVMEIPSLMT | 1 | 0 | 0 | 1681547 | YP_173236.1 | A0A140H1G9 |
| NFNETKGPLCVDTSHF | 1 | 0 | 0 | 1681548 | NP_073551.1 | A0A0P0K6L9 |
| NFNGLTGTGVLTESNK | 1 | 0 | 0 | 1681553 | YP_009724390.1 | P0DTC2 |
| NFNHHEKLLAPLLSAG | 1 | 0 | 0 | 1681557 | QIQ54046.1 | P0C6U8 |
| NFNTLFATTIPNTAFG | 1 | 0 | 0 | 1681575 | NP_073549.1 | A0A0P0LKV0 |
| NFNVDDINFSPVLGCL | 1 | 0 | 0 | 1681576 | YP_009555241.1 | P36334 |
| NFRIDVPSAESTGTQS | 1 | 0 | 0 | 1681597 | YP_009047205.1 | K9N796 |
| NFSDGKMGRFFNHTLV | 1 | 0 | 0 | 1681605 | YP_009047204.1 | K9N5Q8 |
| NFSQILPDPLKPTKRS | 1 | 0 | 0 | 1681617 | NP_828851.1 | P59594 |
| NFSQYYYNIDTGSVYG | 1 | 0 | 0 | 1681618 | YP_173237.1 | A0A140H1H0 |
| NFTIVGQEEFIQTNSP | 1 | 0 | 0 | 1681628 | YP_173238.1 | A0A140H1H1 |
| NFTVDPQKAYLDFVNA | 1 | 0 | 0 | 1681637 | YP_009047202.1 | K9N7C7 |
| NFVKDDYCDGSLGCKM | 1 | 0 | 0 | 1681647 | NP_073549.1 | A0A0P0LKV0 |
| NFVSSYAKPEEKLSAY | 1 | 0 | 0 | 1681651 | NP_073549.1 | A0A0P0LKV0 |
| NFVVQPVGVVDTEWGN | 1 | 0 | 0 | 1681654 | YP_009047202.1 | K9N7C7 |
| NFYCFVNTTIGNETTS | 1 | 0 | 0 | 1681657 | NP_073551.1 | A0A0P0K6L9 |
| NGACGSPGYNLKNGEV | 1 | 0 | 0 | 1681664 | NP_073549.1 | A0A0P0LKV0 |
| NGACGSPGYNVRNDGT | 1 | 0 | 0 | 1681665 | YP_003766.2 | H9EJ42 |
| NGAIIRDSDVVKQPVK | 1 | 0 | 0 | 1681669 | YP_009047202.1 | K9N7C7 |
| NGALEAFKRSNNGVYI | 1 | 0 | 0 | 1681673 | YP_009555238.1 | P0C6X6 |
| NGARPKQRRPQGLPNN | 1 | 0 | 0 | 1681677 | NP_828858.1 | P59595 |
| NGDAAVKDFDFYRYNK | 1 | 0 | 0 | 1681688 | YP_003766.2 | H9EJ42 |
| NGDRSGARPKQRRPQG | 1 | 0 | 0 | 1681693 | QIQ54056.1 | P59595 |
| NGDVVAIDYKHYTPSF | 1 | 0 | 0 | 1681697 | YP_009724389.1 | P0DTD1 |
| NGECPNFVFPLNSTIK | 1 | 0 | 0 | 1681701 | QHR63299.1 | P0C6W2 |
| NGERSGARPKQRRPQG | 1 | 0 | 0 | 1681705 | QHR63308.1 | Q3LZX4 |
| NGFNVSELEDACFKFN | 1 | 0 | 0 | 1681713 | YP_009555242.1 | Q04853 |
| NGGAVTRYVDNNFCGP | 1 | 0 | 0 | 1681726 | NP_828849.2 | P0C6X7 |
| NGGAYSLFDMAKFPLK | 1 | 0 | 0 | 1681727 | YP_009555238.1 | P0C6X6 |
| NGGAYSLFDMTKFSLK | 1 | 0 | 0 | 1681728 | YP_173236.1 | A0A140H1G9 |
| NGGRNGARPKQRRPQG | 1 | 0 | 0 | 1681735 | NP_828858.1 | P59595 |
| NGHKIEDLSIRNLQKR | 1 | 0 | 0 | 1681746 | YP_173236.1 | A0A140H1G9 |
| NGIAYTTFSGSFDNGH | 1 | 0 | 0 | 1681754 | YP_003766.2 | H9EJ42 |
| NGIGVTQNVLYENQKL | 1 | 0 | 0 | 1681758 | YP_009724390.1 | P0DTC2 |
| NGIGVTQNVLYENQKQ | 1 | 0 | 0 | 1681759 | NP_828851.1 | P59594 |
| NGIIRDVNPDDKCFAN | 1 | 0 | 0 | 1681760 | YP_009047202.1 | K9N7C7 |
| NGIISNARNGCIPLSV | 1 | 0 | 0 | 1681761 | YP_009047202.1 | K9N7C7 |
| NGITYTSTSGNLLGFK | 1 | 0 | 0 | 1681774 | NP_073551.1 | A0A0P0K6L9 |
| NGKFLSNTKYYDDSQY | 1 | 0 | 0 | 1681777 | YP_009555240.1 | P30215 |
| NGKLYHDSQNMLRGED | 1 | 0 | 0 | 1681780 | NP_828849.2 | P0C6X7 |
| NGKYYTKPIIKAQFKP | 1 | 0 | 0 | 1681788 | YP_173236.1 | A0A140H1G9 |
| NGLGVTMDVLNKNQKL | 1 | 0 | 0 | 1681792 | YP_173238.1 | A0A140H1H1 |

| | | | | | | |
|---|---|---|---|---|---|---|
| NGLLDTTQLHVADTLM | 1 | 0 | 0 | 1681796 | YP_173238.1 | A0A140H1H1 |
| NGNAIATVKSEDGNIK | 1 | 0 | 0 | 1681827 | NP_073549.1 | A0A0P0LKV0 |
| NGNKIVDKEVKDMQFD | 1 | 0 | 0 | 1681839 | YP_009047202.1 | K9N7C7 |
| NGNVRCVELLKQYTSA | 1 | 0 | 0 | 1681846 | NP_073551.1 | A0A0P0K6L9 |
| NGNVSTVFVPATRDSV | 1 | 0 | 0 | 1681847 | YP_009047208.1 | K9N7D2 |
| NGNWYDFGDFIQTTPG | 1 | 0 | 0 | 1681848 | YP_009724389.1 | P0DTD1 |
| NGPLKVGAGVMLECEK | 1 | 0 | 0 | 1681855 | YP_003766.2 | H9EJ42 |
| NGQAITTSVDANTNQD | 1 | 0 | 0 | 1681864 | YP_003766.2 | H9EJ42 |
| NGRKVDLQLGNLGYLQ | 1 | 0 | 0 | 1681875 | YP_009555241.1 | P36334 |
| NGSCGSVGFNIDYDCV | 1 | 0 | 0 | 1681886 | YP_009724389.1 | P0DTD1 |
| NGSDAALALLLLDRLN | 1 | 0 | 0 | 1681888 | QHR63308.1 | Q3LZX4 |
| NGSDVLYQPPTASVST | 1 | 0 | 0 | 1681889 | YP_009555238.1 | P0C6X6 |
| NGSGSGQAITCTIDSN | 1 | 0 | 0 | 1681891 | NP_073549.1 | A0A0P0LKV0 |
| NGSMIEDIDYVPLKSA | 1 | 0 | 0 | 1681896 | YP_009047202.1 | K9N7C7 |
| NGSVLEMAKEVKTSSK | 1 | 0 | 0 | 1681907 | NP_073549.1 | A0A0P0LKV0 |
| NGSWAIFNDHHLNRPG | 1 | 0 | 0 | 1681908 | YP_009047202.1 | K9N7C7 |
| NGTDIDKCVRAFDIYN | 1 | 0 | 0 | 1681911 | YP_003766.2 | H9EJ42 |
| NGTGRGDCKGFSSDVL | 1 | 0 | 0 | 1681913 | NP_073551.1 | A0A0P0K6L9 |
| NGVDAKGFPQLAELIP | 1 | 0 | 0 | 1681929 | YP_003771.1 | H9EJ69 |
| NGVDFDFIVNAANENL | 1 | 0 | 0 | 1681930 | NP_073549.1 | A0A0P0LKV0 |
| NGVFMSTAKCWTEEDL | 1 | 0 | 0 | 1681935 | NP_073549.1 | A0A0P0LKV0 |
| NGVIKNVNSVRDWLKS | 1 | 0 | 0 | 1681942 | NP_073549.1 | A0A0P0LKV0 |
| NGVIRDVSVEDVCFAN | 1 | 0 | 0 | 1681943 | YP_009555238.1 | P0C6X6 |
| NGVIVDKVGELKVEFW | 1 | 0 | 0 | 1681945 | YP_173236.1 | A0A140H1G9 |
| NGVKAKGYPQFAELVP | 1 | 0 | 0 | 1681946 | NP_073556.1 | A0A0N7J131 |
| NGVLITEGNVKGLQPS | 1 | 0 | 0 | 1681951 | QIQ54047.1 | P0C6X7 |
| NGVRELYLNSSNVTTM | 1 | 0 | 0 | 1681959 | NP_828849.2 | P0C6X7 |
| NGVTLIGEAVKTQFNY | 1 | 0 | 0 | 1681965 | YP_009724389.1 | P0DTD1 |
| NGVTLSTKLKDGVNFN | 1 | 0 | 0 | 1681967 | YP_009555241.1 | P36334 |
| NGVVPLSVIPATSAAR | 1 | 0 | 0 | 1681969 | NP_073549.1 | A0A0P0LKV0 |
| NGYFIKTNNTRIVDEW | 1 | 0 | 0 | 1681978 | YP_009047204.1 | K9N5Q8 |
| NGYLTSSSKTPEEHFI | 1 | 0 | 0 | 1681980 | YP_009724389.1 | P0DTD1 |
| NGYLTSSSKTSEEHFV | 1 | 0 | 0 | 1681981 | NP_828849.2 | P0C6X7 |
| NGYTDNIFSVQQDGRI | 1 | 0 | 0 | 1681987 | YP_003767.1 | A0A0P0G321 |
| NHHEKLLAPLLSAGIF | 1 | 0 | 0 | 1682018 | QIQ54046.1 | P0C6U8 |
| NHLSTFYPQLQSAEWK | 1 | 0 | 0 | 1682031 | YP_003766.2 | H9EJ42 |
| NHNNGNDVLYQPPTAS | 1 | 0 | 0 | 1682037 | YP_173236.1 | A0A140H1G9 |
| NHPIQVDQLNSSYFKL | 1 | 0 | 0 | 1682040 | YP_009047204.1 | K9N5Q8 |
| NHQADTSTPSDVSSRD | 1 | 0 | 0 | 1682041 | YP_173242.1 | Q19U29 |
| NHSFSVQKHIGAPANL | 1 | 0 | 0 | 1682046 | YP_009047202.1 | K9N7C7 |
| NHSVPKLSDFESELSH | 1 | 0 | 0 | 1682051 | YP_173238.1 | A0A140H1H1 |
| NIAFELFAKRKVGLTP | 1 | 0 | 0 | 1682065 | NP_073549.1 | A0A0P0LKV0 |
| NICQAVSANVCALMSC | 1 | 0 | 0 | 1682085 | YP_009555238.1 | P0C6X6 |
| NICYTPSKLIEYTDFA | 1 | 0 | 0 | 1682089 | YP_009724389.1 | P0DTD1 |
| NIEAWLNDKSVPSPLN | 1 | 0 | 0 | 1682103 | YP_009555241.1 | P36334 |
| NIFDVLQSEDTQGMET | 1 | 0 | 0 | 1682108 | QIQ54046.1 | P0C6U8 |
| NIFGLYKNSATGSLDV | 1 | 0 | 0 | 1682109 | YP_003766.2 | H9EJ42 |
| NIFVINKDTYAALRNS | 1 | 0 | 0 | 1682123 | YP_009047202.1 | K9N7C7 |
| NIGGAVCSKHANLYQK | 1 | 0 | 0 | 1682130 | YP_003766.2 | H9EJ42 |
| NIGGAVCSKHANLYRA | 1 | 0 | 0 | 1682131 | NP_073549.1 | A0A0P0LKV0 |
| NIGTNFPLQIGFSTGV | 1 | 0 | 0 | 1682133 | YP_173236.1 | A0A140H1G9 |
| NIGTNVPLQVGFSNGV | 1 | 0 | 0 | 1682134 | YP_003766.2 | H9EJ42 |
| NIIVPDKSVYDQVVDN | 1 | 0 | 0 | 1682149 | YP_009555238.1 | P0C6X6 |
| NIKFADDLNQLTGYKK | 1 | 0 | 0 | 1682154 | YP_009724389.1 | P0DTD1 |
| NIKPVPEIKILNNLGV | 1 | 0 | 0 | 1682159 | NP_828849.2 | P0C6X7 |
| NIKPVPEVKILNNLGV | 1 | 0 | 0 | 1682160 | YP_009724389.1 | P0DTD1 |
| NILHVVGPDARAKQDV | 1 | 0 | 0 | 1682167 | YP_009047202.1 | K9N7C7 |
| NINAILTEVNELLDTT | 1 | 0 | 0 | 1682179 | YP_009555241.1 | P36334 |
| NINLHTQVVDMSMTYG | 1 | 0 | 0 | 1682188 | YP_009724389.1 | P0DTD1 |
| NINPHNYSYMDLNPVL | 1 | 0 | 0 | 1682189 | YP_009555240.1 | P30215 |
| NINSTLVDLKWLNRVE | 1 | 0 | 0 | 1682193 | NP_073551.1 | A0A0P0K6L9 |
| NINWNVEYPIISNELS | 1 | 0 | 0 | 1682196 | YP_009555238.1 | P0C6X6 |
| NIPTQCYYNNGSSGRI | 1 | 0 | 0 | 1682200 | YP_173236.1 | A0A140H1G9 |
| NIQNLLYCDSPFEKLQ | 1 | 0 | 0 | 1682201 | YP_003767.1 | A0A0P0G321 |
| NISDEFSSNVANYQKV | 1 | 0 | 0 | 1682207 | YP_009724389.1 | P0DTD1 |
| NISLNGNTSVCVRTSH | 1 | 0 | 0 | 1682210 | YP_003767.1 | A0A0P0G321 |
| NITFELDERIDKVLNE | 1 | 0 | 0 | 1682220 | YP_009724389.1 | P0DTD1 |
| NITFELDERVDKVLNE | 1 | 0 | 0 | 1682221 | QIQ54046.1 | P0C6U8 |
| NITNLCPFGEVFNASK | 1 | 0 | 0 | 1682227 | QIQ54048.1 | P59594 |
| NITNLCPFGEVFNATK | 1 | 0 | 0 | 1682228 | NP_828851.1 | P59594 |
| NITSGFLGPLLVLQA | 1 | 0 | 0 | 1682233 | P03138.3 | Q76R62 |
| NIVAADLGVLIQNSAK | 1 | 0 | 0 | 1682235 | YP_009555238.1 | P0C6X6 |
| NIVGPDARQDGRQSYV | 1 | 0 | 0 | 1682241 | YP_173236.1 | A0A140H1G9 |
| NIYDYVGTGIIRSSNQ | 1 | 0 | 0 | 1682254 | YP_003767.1 | A0A0P0G321 |
| NIYSQNLCVDGFGVNK | 1 | 0 | 0 | 1682257 | NP_073549.1 | A0A0P0LKV0 |
| NKAIDGFGVFDIKNSS | 1 | 0 | 0 | 1682260 | YP_003766.2 | H9EJ42 |
| NKAIPSGNLIKFDIGS | 1 | 0 | 0 | 1682262 | NP_073549.1 | A0A0P0LKV0 |
| NKASYDTNLNKFNRAS | 1 | 0 | 0 | 1682267 | YP_009047202.1 | K9N7C7 |
| NKATNGAMQKESDDYI | 1 | 0 | 0 | 1682269 | NP_828849.2 | P0C6X7 |
| NKATNNAMQIESDDYI | 1 | 0 | 0 | 1682270 | QIQ54046.1 | P0C6U8 |
| NKATNNAMQVESDDYI | 1 | 0 | 0 | 1682271 | YP_009724389.1 | P0DTD1 |
| NKATNNAMQVESDHYI | 1 | 0 | 0 | 1682272 | QHR63299.1 | P0C6W2 |
| NKCSRLLSDDRTEVPQ | 1 | 0 | 0 | 1682278 | YP_009047204.1 | K9N5Q8 |
| NKDASFIGKNLKSNCV | 1 | 0 | 0 | 1682279 | NP_073549.1 | A0A0P0LKV0 |
| NKDGFLYVYKGYQPID | 1 | 0 | 0 | 1682282 | NP_828851.1 | P59594 |
| NKEDFDIIQKCQITSV | 1 | 0 | 0 | 1682291 | YP_173236.1 | A0A140H1G9 |
| NKEGIVWVATEGALNT | 1 | 0 | 0 | 1682294 | NP_828858.1 | P59595 |
| NKEQEAILMSEDVKLN | 1 | 0 | 0 | 1682298 | YP_009047202.1 | K9N7C7 |
| NKFNQALGAMQTGFTT | 1 | 0 | 0 | 1682315 | YP_009047204.1 | K9N5Q8 |
| NKFVPVKPASKQCFKV | 1 | 0 | 0 | 1682320 | YP_003766.2 | H9EJ42 |
| NKGAGGHSFGIDLKSY | 1 | 0 | 0 | 1682325 | QIQ54046.1 | P0C6U8 |

| | | | | | | |
|---|---|---|---|---|---|---|
| NKGAGGHSYGIDLKSY | 1 | 0 | 0 | 1682327 | NP_828849.2 | P0C6X7 |
| NKGEDIQLLKVAYENF | 1 | 0 | 0 | 1682328 | QIQ54046.1 | P0C6U8 |
| NKGHFDGQQGETPVSI | 1 | 0 | 0 | 1682331 | QIQ54047.1 | P0C6X7 |
| NKGHFDGQQGEVPVSI | 1 | 0 | 0 | 1682332 | YP_009724389.1 | P0DTD1 |
| NKHAFHTPAFDKSAFT | 1 | 0 | 0 | 1682340 | NP_828849.2 | P0C6X7 |
| NKHCNVQQCFGKRGPS | 1 | 0 | 0 | 1682345 | YP_173242.1 | Q19U29 |
| NKINLCDDPEKAQSML | 1 | 0 | 0 | 1682358 | YP_003766.2 | H9EJ42 |
| NKINLCDDPETAQELL | 1 | 0 | 0 | 1682359 | NP_073549.1 | A0A0P0LKV0 |
| NKIQDVVNQQGNSLNH | 1 | 0 | 0 | 1682361 | NP_073551.1 | A0A0P0K6L9 |
| NKIQDVVNQQGSALNH | 1 | 0 | 0 | 1682362 | YP_003767.1 | A0A0P0G321 |
| NKIQPWTNDALNTICI | 1 | 0 | 0 | 1682363 | NP_073549.1 | A0A0P0LKV0 |
| NKIVDKEVKDMQFDLY | 1 | 0 | 0 | 1682366 | YP_009047202.1 | K9N7C7 |
| NKIVGGAPTWFNALRD | 1 | 0 | 0 | 1682367 | YP_009047202.1 | K9N7C7 |
| NKKDWYDFVENPDILR | 1 | 0 | 0 | 1682369 | YP_009724389.1 | P0DTD1 |
| NKLCVVPGNVKPGDAV | 1 | 0 | 0 | 1682373 | NP_073549.1 | A0A0P0LKV0 |
| NKLKAHKDKSAQCFKM | 1 | 0 | 0 | 1682378 | YP_009724389.1 | P0DTD1 |
| NKLLHWSDQTICYKDS | 1 | 0 | 0 | 1682382 | YP_009047202.1 | K9N7C7 |
| NKLNLPGCNGGSLYVN | 1 | 0 | 0 | 1682385 | YP_173236.1 | A0A140H1G9 |
| NKLSTSDWSDIRDYKL | 1 | 0 | 0 | 1682389 | NP_073549.1 | A0A0P0LKV0 |
| NKLVKDPPNVQIHTID | 1 | 0 | 0 | 1682391 | NP_828852.2 | P59632 |
| NKMVNWFNTMLDASAP | 1 | 0 | 0 | 1682394 | YP_173236.1 | A0A140H1G9 |
| NKPLIEWPLTEYDFTD | 1 | 0 | 0 | 1682404 | YP_009047202.1 | K9N7C7 |
| NKPLSQPRADKPSQLK | 1 | 0 | 0 | 1682406 | YP_003771.1 | H9EJ69 |
| NKPRQKRSPNKQCTVQ | 1 | 0 | 0 | 1682407 | YP_009555245.1 | P33469 |
| NKPWRWMLWCKDNAVA | 1 | 0 | 0 | 1682411 | NP_073549.1 | A0A0P0LKV0 |
| NKQEASVPILTTPFSL | 1 | 0 | 0 | 1682414 | YP_173236.1 | A0A140H1G9 |
| NKQLNEISDDCNWPLV | 1 | 0 | 0 | 1682416 | YP_009555238.1 | P0C6X6 |
| NKQMANVSVLTTPFSL | 1 | 0 | 0 | 1682418 | YP_009555238.1 | P0C6X6 |
| NKRFGFIEDSVFKPRP | 1 | 0 | 0 | 1682421 | YP_009555241.1 | P36334 |
| NKSAGWPLNKFGKAGL | 1 | 0 | 0 | 1682425 | NP_073549.1 | A0A0P0LKV0 |
| NKSFLPQFTSDQAVTF | 1 | 0 | 0 | 1682427 | YP_173241.1 | Q0ZJ82 |
| NKSVANLKLPPHDVTV | 1 | 0 | 0 | 1682431 | YP_003768.1 | H9EJ24 |
| NKSWMESEFRVYSSAN | 1 | 0 | 0 | 1682433 | YP_009724390.1 | P0DTC2 |
| NKTLQEFAQNLPKYVK | 1 | 0 | 0 | 1682437 | YP_003767.1 | A0A0P0G321 |
| NKTVGELGDVRETMNY | 1 | 0 | 0 | 1682444 | QHR63299.1 | P0C6W2 |
| NKTVGELGDVRETMSY | 1 | 0 | 0 | 1682446 | YP_009724389.1 | P0DTD1 |
| NKTWVENEFRVYSSAN | 1 | 0 | 0 | 1682448 | QIQ54048.1 | P59594 |
| NKVIYTTEIASKLTCK | 1 | 0 | 0 | 1682452 | YP_009555238.1 | P0C6X6 |
| NKVTLADAGFIKQYGD | 1 | 0 | 0 | 1682456 | YP_009724390.1 | P0DTC2 |
| NKVTLADAGFMKQYGE | 1 | 0 | 0 | 1682457 | NP_828851.1 | P59594 |
| NKYFKHWSMPYHPNTV | 1 | 0 | 0 | 1682471 | YP_009555238.1 | P0C6X6 |
| NKYKYYSGSASEADYR | 1 | 0 | 0 | 1682480 | YP_003766.2 | H9EJ42 |
| NLAIEEIEYVHGDALH | 1 | 0 | 0 | 1682491 | NP_073549.1 | A0A0P0LKV0 |
| NLARDGVVPLSVIPAT | 1 | 0 | 0 | 1682500 | YP_003766.2 | H9EJ42 |
| NLCPFGEVFNASKFAS | 1 | 0 | 0 | 1682506 | QIQ54048.1 | P59594 |
| NLCPFGEVFNATKFPS | 1 | 0 | 0 | 1682507 | NP_828851.1 | P59594 |
| NLCVDGFGVNKIQPWT | 1 | 0 | 0 | 1682512 | NP_073549.1 | A0A0P0LKV0 |
| NLDEARFSGSANQQQL | 1 | 0 | 0 | 1682515 | YP_009555238.1 | P0C6X6 |
| NLDNLRANNTKGSIPI | 1 | 0 | 0 | 1682519 | QIQ54046.1 | P0C6U8 |
| NLDNLRANNTKGSLPI | 1 | 0 | 0 | 1682520 | YP_009724389.1 | P0DTD1 |
| NLDNLRASNTKGSLPI | 1 | 0 | 0 | 1682521 | QHR63299.1 | P0C6W2 |
| NLDSLEARAFQSTPIV | 1 | 0 | 0 | 1682524 | NP_828859.1 | P59636 |
| NLEDLCLKYNYCQPIV | 1 | 0 | 0 | 1682529 | YP_173239.1 | Q0ZJ84 |
| NLEYPIISNEVSINTS | 1 | 0 | 0 | 1682542 | YP_173236.1 | A0A140H1G9 |
| NLFEGDKFVGTFESAA | 1 | 0 | 0 | 1682545 | NP_073549.1 | A0A0P0LKV0 |
| NLFKDCSKSCLGYHPA | 1 | 0 | 0 | 1682548 | YP_173236.1 | A0A140H1G9 |
| NLFKDCSKSYSGYHPA | 1 | 0 | 0 | 1682549 | YP_009555238.1 | P0C6X6 |
| NLGFDNQSKSPSSSGT | 1 | 0 | 0 | 1682558 | YP_003771.1 | H9EJ69 |
| NLGSNGKPGNVGGNDA | 1 | 0 | 0 | 1682570 | YP_009555238.1 | P0C6X6 |
| NLHFDVDNINFKSLVG | 1 | 0 | 0 | 1682575 | YP_173238.1 | A0A140H1H1 |
| NLHTQLVDMSMTYGQQ | 1 | 0 | 0 | 1682582 | NP_828849.2 | P0C6X7 |
| NLHTQVVDMSMTYGQQ | 1 | 0 | 0 | 1682583 | YP_009724389.1 | P0DTD1 |
| NLIIKNLSKPLTENKY | 1 | 0 | 0 | 1682588 | QIQ54052.1 | P59634 |
| NLKLPPHDVTVLRDNL | 1 | 0 | 0 | 1682605 | YP_003768.1 | H9EJ24 |
| NLKPRVEAPKQEEPPN | 1 | 0 | 0 | 1682609 | NP_828849.2 | P0C6X7 |
| NLKSNCVRFKNVDKDD | 1 | 0 | 0 | 1682614 | NP_073549.1 | A0A0P0LKV0 |
| NLLEGSISYSELRPDT | 1 | 0 | 0 | 1682623 | NP_828849.2 | P0C6X7 |
| NLLEILQKEKVNINIV | 1 | 0 | 0 | 1682624 | YP_009724389.1 | P0DTD1 |
| NLLGFDSSKPVTKKYT | 1 | 0 | 0 | 1682627 | YP_009047202.1 | K9N7C7 |
| NLLIQEFLQYDSSIHS | 1 | 0 | 0 | 1682634 | YP_173236.1 | A0A140H1G9 |
| NLLYCDSPFEKLQCEH | 1 | 0 | 0 | 1682651 | YP_003767.1 | A0A0P0G321 |
| NLLYDSNGNIIGFKDF | 1 | 0 | 0 | 1682652 | YP_173238.1 | A0A140H1H1 |
| NLNAGEDIQLLKAAYE | 1 | 0 | 0 | 1682659 | NP_828849.2 | P0C6X7 |
| NLNAYVNSNQNTDSDS | 1 | 0 | 0 | 1682661 | YP_173242.1 | Q19U29 |
| NLNFSANSLDNLKSGV | 1 | 0 | 0 | 1682666 | YP_003767.1 | A0A0P0G321 |
| NLNKGEDIQLLKVAYE | 1 | 0 | 0 | 1682672 | QIQ54046.1 | P0C6U8 |
| NLNTNLHFDVDNINFK | 1 | 0 | 0 | 1682683 | YP_173238.1 | A0A140H1H1 |
| NLPKFIEQISAFTKPS | 1 | 0 | 0 | 1682691 | YP_003771.1 | H9EJ69 |
| NLPLQLGFSTGVNLVA | 1 | 0 | 0 | 1682693 | YP_009724389.1 | P0DTD1 |
| NLPTYYIYDEEGGNDL | 1 | 0 | 0 | 1682701 | NP_073549.1 | A0A0P0LKV0 |
| NLQSNHDLYCQVHGNA | 1 | 0 | 0 | 1682713 | YP_009724389.1 | P0DTD1 |
| NLQSNHDQHCQVHGNA | 1 | 0 | 0 | 1682714 | NP_828849.2 | P0C6X7 |
| NLQSNHDQYCQVHGNA | 1 | 0 | 0 | 1682715 | QIQ54047.1 | P0C6X7 |
| NLQSNHDRYCQVHGNA | 1 | 0 | 0 | 1682716 | QHR63299.1 | P0C6W2 |
| NLQVESANQMLTVNVV | 1 | 0 | 0 | 1682717 | NP_073549.1 | A0A0P0LKV0 |
| NLRANNTKGSIPINVI | 1 | 0 | 0 | 1682719 | QIQ54046.1 | P0C6U8 |
| NLRQDDSVRNLFASVK | 1 | 0 | 0 | 1682731 | YP_009047204.1 | K9N5Q8 |
| NLRVVGHAMQGTLLKL | 1 | 0 | 0 | 1682735 | YP_009047202.1 | K9N7C7 |
| NLSAGTQHISNDFTPE | 1 | 0 | 0 | 1682736 | YP_173242.1 | Q19U29 |
| NLSAPTLVPQENYSSI | 1 | 0 | 0 | 1682737 | YP_009555238.1 | P0C6X6 |
| NLSEITKPLIEAEKPN | 1 | 0 | 0 | 1682743 | YP_173243.1 | Q0ZJ80 |

| | | | | | | |
|---|---|---|---|---|---|---|
| NLSGNPDEPQKDVYEL | 1 | 0 | 0 | 1682744 | YP_009555245.1 | P33469 |
| NLSKPLTENKYSQLDE | 1 | 0 | 0 | 1682747 | QIQ54052.1 | P59634 |
| NLSKSLTENKYSQLDE | 1 | 0 | 0 | 1682748 | YP_009724394.1 | P0DTC6 |
| NLSMLQLGVPDNSSTI | 1 | 0 | 0 | 1682751 | YP_003767.1 | A0A0P0G321 |
| NLSSVLPQRNIRSSRI | 1 | 0 | 0 | 1682757 | YP_003767.1 | A0A0P0G321 |
| NLTDEDLARDFSPIAL | 1 | 0 | 0 | 1682762 | YP_009047207.1 | K9N643 |
| NLTPIKLNFGMLNGMP | 1 | 0 | 0 | 1682769 | YP_003766.2 | H9EJ42 |
| NLTTRTGIPPGYTNSS | 1 | 0 | 0 | 1682774 | QIQ54048.1 | P59594 |
| NLTTRTQLPPAYTNSS | 1 | 0 | 0 | 1682776 | QHR63300.2 | Q3LZX1 |
| NLVAMKFNYEPLTQDH | 1 | 0 | 0 | 1682780 | QIQ54046.1 | P0C6U8 |
| NLVAMKYNYEPLTQDH | 1 | 0 | 0 | 1682781 | YP_009724389.1 | P0DTD1 |
| NLVAVPTGYVDTPNNT | 1 | 0 | 0 | 1682784 | YP_009724389.1 | P0DTD1 |
| NLVEKLSPWFMDGENA | 1 | 0 | 0 | 1682786 | YP_009047202.1 | K9N7C7 |
| NLVFTNKTTLPTNVAF | 1 | 0 | 0 | 1682788 | YP_003766.2 | H9EJ42 |
| NLVFVNKTSLPTNIAF | 1 | 0 | 0 | 1682789 | NP_073549.1 | A0A0P0LKV0 |
| NLVKNKCVNFNFNGLT | 1 | 0 | 0 | 1682796 | YP_009724390.1 | P0DTC2 |
| NLVKTGHYTGQAGEMP | 1 | 0 | 0 | 1682797 | YP_009555238.1 | P0C6X6 |
| NLVLAKLGKDSKPQQV | 1 | 0 | 0 | 1682798 | YP_173242.1 | Q19U29 |
| NLVLNNCTKYNIYDYV | 1 | 0 | 0 | 1682799 | YP_003767.1 | A0A0P0G321 |
| NLVNVGHYDGRTGELP | 1 | 0 | 0 | 1682801 | YP_173236.1 | A0A140H1G9 |
| NLVPIGKGNKDEQIGY | 1 | 0 | 0 | 1682802 | YP_003771.1 | H9EJ69 |
| NLVPTYLKSDNIVAAD | 1 | 0 | 0 | 1682804 | YP_009555238.1 | P0C6X6 |
| NLYKMQRMLLDKCDLQ | 1 | 0 | 0 | 1682823 | QIQ54047.1 | P0C6X7 |
| NLYNYGAGIKLPSGIM | 1 | 0 | 0 | 1682825 | YP_003766.2 | H9EJ42 |
| NLYPDSTSLVEXVDVT | 1 | 0 | 0 | 1682827 | QIQ54046.1 | P0C6U8 |
| NLYRLVWHVERKDVPY | 1 | 0 | 0 | 1682830 | YP_009047202.1 | K9N7C7 |
| NLYVVVVESDADRANF | 1 | 0 | 0 | 1682832 | YP_173236.1 | A0A140H1G9 |
| NMARNGVVPLSVIPAT | 1 | 0 | 0 | 1682835 | NP_073549.1 | A0A0P0LKV0 |
| NMASFVEYEVAKKNLA | 1 | 0 | 0 | 1682836 | YP_173236.1 | A0A140H1G9 |
| NMDMGVFINKYGLEDF | 1 | 0 | 0 | 1682843 | YP_003766.2 | H9EJ42 |
| NMFITREEAIKHVRAW | 1 | 0 | 0 | 1682850 | QIQ54047.1 | P0C6X7 |
| NMFQFATLPVYDTIKY | 1 | 0 | 0 | 1682855 | YP_009047204.1 | K9N5Q8 |
| NMNIASNSKEWAYCVE | 1 | 0 | 0 | 1682884 | NP_073549.1 | A0A0P0LKV0 |
| NMRVIHFGAGSDKGIA | 1 | 0 | 0 | 1682892 | YP_009047202.1 | K9N7C7 |
| NMRVIHFGAGSDKGVA | 1 | 0 | 0 | 1682893 | YP_009724389.1 | P0DTD1 |
| NMRVLHLGAGSDKEVA | 1 | 0 | 0 | 1682894 | YP_173236.1 | A0A140H1G9 |
| NMRVLHLGAGSDYGVA | 1 | 0 | 0 | 1682895 | NP_073549.1 | A0A0P0LKV0 |
| NMRVLHLGAGSEKGVA | 1 | 0 | 0 | 1682896 | YP_009555238.1 | P0C6X6 |
| NMRVLHYGAGSDKGVA | 1 | 0 | 0 | 1682897 | YP_003766.2 | H9EJ42 |
| NMSLAECKRALGLSIS | 1 | 0 | 0 | 1682898 | NP_073549.1 | A0A0P0LKV0 |
| NMSPKPQRQRGHKNGQ | 1 | 0 | 0 | 1682900 | YP_009555245.1 | P33469 |
| NMVEFIQTSSPKVTID | 1 | 0 | 0 | 1682908 | YP_009555241.1 | P36334 |
| NMVETVVGQLEQTNMH | 1 | 0 | 0 | 1682909 | YP_009047202.1 | K9N7C7 |
| NMWEFRDYFNNNTDSI | 1 | 0 | 0 | 1682912 | YP_003766.2 | H9EJ42 |
| NNAAIVLQLPQGTALP | 1 | 0 | 0 | 1682921 | QIQ54056.1 | P59595 |
| NNALYAIQEGFDATNS | 1 | 0 | 0 | 1682927 | YP_009555241.1 | P36334 |
| NNAMQVESDHYIATNG | 1 | 0 | 0 | 1682930 | QHR63299.1 | P0C6W2 |
| NNARQSDNMTAVACQP | 1 | 0 | 0 | 1682933 | YP_009555240.1 | P30215 |
| NNATNKATYKPNTWCI | 1 | 0 | 0 | 1682934 | YP_009724389.1 | P0DTD1 |
| NNAVYTKVDGIDVEIF | 1 | 0 | 0 | 1682937 | NP_828849.2 | P0C6X7 |
| NNCTKYNIYDYVGTGI | 1 | 0 | 0 | 1682946 | YP_003767.1 | A0A0P0G321 |
| NNDEEIVTGDNDDQIV | 1 | 0 | 0 | 1682958 | YP_173236.1 | A0A140H1G9 |
| NNDITNTNLSRGRGRN | 1 | 0 | 0 | 1682959 | YP_009047211.1 | K9N4V7 |
| NNEFIDPTECIYTQSR | 1 | 0 | 0 | 1682965 | YP_009047202.1 | K9N7C7 |
| NNEIKPSGLKTMVVSA | 1 | 0 | 0 | 1682968 | YP_009047202.1 | K9N7C7 |
| NNELMPAKLKIQVVNS | 1 | 0 | 0 | 1682971 | YP_009555238.1 | P0C6X6 |
| NNELSPVALRQMSCAA | 1 | 0 | 0 | 1682973 | YP_009724389.1 | P0DTD1 |
| NNFVSSYAKPEEKLSA | 1 | 0 | 0 | 1682988 | NP_073549.1 | A0A0P0LKV0 |
| NNGNDVLYQPPTASVS | 1 | 0 | 0 | 1682994 | YP_173236.1 | A0A140H1G9 |
| NNHAFHTPAYDKRAMA | 1 | 0 | 0 | 1683002 | NP_073549.1 | A0A0P0LKV0 |
| NNIINNARDGCVPLNI | 1 | 0 | 0 | 1683004 | YP_009724389.1 | P0DTD1 |
| NNISLTTQPYFDSYLG | 1 | 0 | 0 | 1683008 | YP_173238.1 | A0A140H1H1 |
| NNKEDFDIIQKCQITS | 1 | 0 | 0 | 1683012 | YP_173236.1 | A0A140H1G9 |
| NNKPWFVRKASGLFDA | 1 | 0 | 0 | 1683015 | YP_003766.2 | H9EJ42 |
| NNLAIEEIEYVHGDAL | 1 | 0 | 0 | 1683020 | NP_073549.1 | A0A0P0LKV0 |
| NNLKIKNFDYLDIHPS | 1 | 0 | 0 | 1683027 | YP_173237.1 | A0A140H1H0 |
| NNLLGFDSSKPVTKKY | 1 | 0 | 0 | 1683028 | YP_009047202.1 | K9N7C7 |
| NNLMCIDMKGTMYVRP | 1 | 0 | 0 | 1683032 | YP_009555244.1 | Q01455 |
| NNLNRGMVLGSLAATV | 1 | 0 | 0 | 1683034 | YP_009724389.1 | P0DTD1 |
| NNLTAPRNSWEAMALN | 1 | 0 | 0 | 1683037 | YP_009047202.1 | K9N7C7 |
| NNLVPTYLKSDNIVAA | 1 | 0 | 0 | 1683038 | YP_009555238.1 | P0C6X6 |
| NNNGSDVLYQPPTASV | 1 | 0 | 0 | 1683048 | YP_009555238.1 | P0C6X6 |
| NNNGTNVTQVKNASVY | 1 | 0 | 0 | 1683049 | NP_073549.1 | A0A0P0LKV0 |
| NNQNGDRSGARPKQRR | 1 | 0 | 0 | 1683060 | QIQ54056.1 | P59595 |
| NNRSLSRSNSNFRHSD | 1 | 0 | 0 | 1683063 | YP_173242.1 | Q19U29 |
| NNSAQYVFSTVNALPE | 1 | 0 | 0 | 1683066 | NP_073549.1 | A0A0P0LKV0 |
| NNSGSNITQIKNACVY | 1 | 0 | 0 | 1683067 | YP_003766.2 | H9EJ42 |
| NNSLTRQLQPINYFDS | 1 | 0 | 0 | 1683071 | YP_009555241.1 | P36334 |
| NNSRDSSRSTSRQQSR | 1 | 0 | 0 | 1683073 | YP_003771.1 | H9EJ69 |
| NNSSRASSRSSTRNNS | 1 | 0 | 0 | 1683074 | YP_003771.1 | H9EJ69 |
| NNSTNVVIRACNFELC | 1 | 0 | 0 | 1683075 | NP_828851.1 | P59594 |
| NNTDFSRVSAKPPPGD | 1 | 0 | 0 | 1683081 | YP_009724389.1 | P0DTD1 |
| NNTEFTRVNAKPPPGD | 1 | 0 | 0 | 1683083 | NP_828849.2 | P0C6X7 |
| NNTINKATFKPNTWCL | 1 | 0 | 0 | 1683087 | QIQ54046.1 | P0C6U8 |
| NNTLVSGDAHIPFGTV | 1 | 0 | 0 | 1683090 | NP_073551.1 | A0A0P0K6L9 |
| NNTRIVDEWSYTGSSF | 1 | 0 | 0 | 1683091 | YP_009047204.1 | K9N5Q8 |
| NNTSAQYIFSTVNALP | 1 | 0 | 0 | 1683092 | YP_003766.2 | H9EJ42 |
| NNVAFQTVKPGNFNKD | 1 | 0 | 0 | 1683099 | YP_009724389.1 | P0DTD1 |
| NNVDTVIFKNNTSFPT | 1 | 0 | 0 | 1683105 | YP_173236.1 | A0A140H1G9 |
| NNVFMSEAKCWTETDL | 1 | 0 | 0 | 1683106 | YP_009724389.1 | P0DTD1 |
| NNVFMSEAKCWVETDL | 1 | 0 | 0 | 1683107 | YP_009047202.1 | K9N7C7 |

| | | | | | | |
|---|---|---|---|---|---|---|
| NNVFMSESKCWVEHDI | 1 | 0 | 0 | 1683108 | YP_009555238.1 | P0C6X6 |
| NNVFMSESKCWVENDI | 1 | 0 | 0 | 1683109 | YP_173236.1 | A0A140H1G9 |
| NNVYCYNTALMEGSLP | 1 | 0 | 0 | 1683118 | NP_073549.1 | A0A0P0LKV0 |
| NNWFLLTNTSSVVDGV | 1 | 0 | 0 | 1683121 | NP_073551.1 | A0A0P0K6L9 |
| NNYDKSAGYPFNKFGK | 1 | 0 | 0 | 1683125 | YP_009555238.1 | P0C6X6 |
| NNYVPSYVKPDSVSTS | 1 | 0 | 0 | 1683132 | YP_009047202.1 | K9N7C7 |
| NPANNAAIVLQLPQGT | 1 | 0 | 0 | 1683139 | YP_009724397.2 | P0DTC9 |
| NPDDKCFANKHRSFTQ | 1 | 0 | 0 | 1683144 | YP_009047202.1 | K9N7C7 |
| NPDEPQKDVYELRYNG | 1 | 0 | 0 | 1683145 | YP_009555245.1 | P33469 |
| NPDIINIYKKLGPIFN | 1 | 0 | 0 | 1683146 | YP_173236.1 | A0A140H1G9 |
| NPDIINVYKKLGPIFN | 1 | 0 | 0 | 1683147 | YP_009555238.1 | P0C6X6 |
| NPETNCLLNVPFGGTT | 1 | 0 | 0 | 1683153 | YP_009047210.1 | K9N7A1 |
| NPETNILLNVPLHGTI | 1 | 0 | 0 | 1683154 | YP_009724393.1 | P0DTC5 |
| NPETNILLNVPLRGTI | 1 | 0 | 0 | 1683155 | NP_828855.1 | P59596 |
| NPEVGPEHSLAEYHNE | 1 | 0 | 0 | 1683158 | QHR63299.1 | P0C6W2 |
| NPEVNAITVTTVLGQT | 1 | 0 | 0 | 1683159 | NP_073555.1 | A0A0P0KXN3 |
| NPFMVNQLAYSSSANG | 1 | 0 | 0 | 1683163 | YP_009047202.1 | K9N7C7 |
| NPFTRTVFENLKPMPF | 1 | 0 | 0 | 1683164 | YP_173236.1 | A0A140H1G9 |
| NPHLMGWDYPKCDRAM | 1 | 0 | 0 | 1683168 | YP_009724389.1 | P0DTD1 |
| NPIDLPPSHATTYLSL | 1 | 0 | 0 | 1683170 | NP_073549.1 | A0A0P0LKV0 |
| NPINSNVKTLEYDYMT | 1 | 0 | 0 | 1683172 | NP_073549.1 | A0A0P0LKV0 |
| NPIQLSSYSLFDMSKF | 1 | 0 | 0 | 1683173 | YP_009724389.1 | P0DTD1 |
| NPITFHLDGEVITFDN | 1 | 0 | 0 | 1683174 | QHR63299.1 | P0C6W2 |
| NPKGIPIVDDPVVDWH | 1 | 0 | 0 | 1683181 | YP_009047202.1 | K9N7C7 |
| NPMLMGWDYPKCDRAL | 1 | 0 | 0 | 1683197 | YP_003766.2 | H9EJ42 |
| NPNNNAAIVLQLPQGT | 1 | 0 | 0 | 1683200 | QIQ54056.1 | P59595 |
| NPNNNAATVLQLPQGT | 1 | 0 | 0 | 1683201 | NP_828858.1 | P59595 |
| NPNYEDLLIRKSNHNF | 1 | 0 | 0 | 1683202 | YP_009724389.1 | P0DTD1 |
| NPNYEDLLIRKSNHSF | 1 | 0 | 0 | 1683203 | NP_828849.2 | P0C6X7 |
| NPSALEFNPSQTSPAT | 1 | 0 | 0 | 1683211 | NP_073556.1 | A0A0N7J131 |
| NPSEDFIKHVRTNGGY | 1 | 0 | 0 | 1683212 | YP_009047202.1 | K9N7C7 |
| NPSSSEVDMICSTTAQ | 1 | 0 | 0 | 1683219 | YP_009555238.1 | P0C6X6 |
| NPSTPAYTFTTVKPGA | 1 | 0 | 0 | 1683221 | YP_009047202.1 | K9N7C7 |
| NPSTWNKRFGFIEDSV | 1 | 0 | 0 | 1683222 | YP_009555241.1 | P36334 |
| NPTDQSSYIVDSVAVK | 1 | 0 | 0 | 1683227 | NP_828849.2 | P0C6X7 |
| NPTDQSSYVVDSVAVK | 1 | 0 | 0 | 1683229 | QIQ54046.1 | P0C6U8 |
| NPTEDLCDGSVVIKEP | 1 | 0 | 0 | 1683230 | YP_009555238.1 | P0C6X6 |
| NPVLFTNSTDTVNHDS | 1 | 0 | 0 | 1683242 | YP_173236.1 | A0A140H1G9 |
| NPVLMGWDYPKCDRAM | 1 | 0 | 0 | 1683244 | YP_009555238.1 | P0C6X6 |
| NPVLPFNDGVYFASTE | 1 | 0 | 0 | 1683245 | YP_009724390.1 | P0DTC2 |
| NPVLPXNDGVYFASTE | 1 | 0 | 0 | 1683246 | QIQ54048.1 | P59594 |
| NPVSFVVKPVCSSIFR | 1 | 0 | 0 | 1683247 | NP_073549.1 | A0A0P0LKV0 |
| NPYHRFKCFPNGSNDV | 1 | 0 | 0 | 1683251 | YP_173237.1 | A0A140H1H0 |
| NPYTPKYTFGNVKPGE | 1 | 0 | 0 | 1683252 | YP_173236.1 | A0A140H1G9 |
| NQALGAMQTGFTTTNE | 1 | 0 | 0 | 1683259 | YP_009047204.1 | K9N5Q8 |
| NQALNSILDNAVKGCV | 1 | 0 | 0 | 1683261 | YP_009555238.1 | P0C6X6 |
| NQATTKTTFKPNTWCL | 1 | 0 | 0 | 1683262 | NP_828849.2 | P0C6X7 |
| NQDVNIHSSRLSFKEL | 1 | 0 | 0 | 1683278 | QIQ54047.1 | P0C6X7 |
| NQDVNLHSSRLSFKEL | 1 | 0 | 0 | 1683279 | YP_009724389.1 | P0DTD1 |
| NQEKYSTIYKLHPSFN | 1 | 0 | 0 | 1683284 | NP_073549.1 | A0A0P0LKV0 |
| NQFTEFVGTQSVDMLA | 1 | 0 | 0 | 1683291 | YP_009047202.1 | K9N7C7 |
| NQKILAASFNKAINNI | 1 | 0 | 0 | 1683304 | YP_003767.1 | A0A0P0G321 |
| NQKILAASFNKAMTNI | 1 | 0 | 0 | 1683305 | NP_073551.1 | A0A0P0K6L9 |
| NQKLIANKFNQALGAM | 1 | 0 | 0 | 1683307 | YP_009047204.1 | K9N5Q8 |
| NQKPLEPKFSIALPPE | 1 | 0 | 0 | 1683311 | YP_003771.1 | H9EJ69 |
| NQKQIANQFNKAISQI | 1 | 0 | 0 | 1683312 | NP_828851.1 | P59594 |
| NQLAYSSSANGSLVGT | 1 | 0 | 0 | 1683313 | YP_009047202.1 | K9N7C7 |
| NQLESKVSGKGQQQQG | 1 | 0 | 0 | 1683317 | NP_828858.1 | P59595 |
| NQLTADLQHKLKKACV | 1 | 0 | 0 | 1683319 | YP_173236.1 | A0A140H1G9 |
| NQNGDRSGARPKQRRP | 1 | 0 | 0 | 1683328 | QIQ54056.1 | P59595 |
| NQNGERSGARPKQRRP | 1 | 0 | 0 | 1683329 | QHR63308.1 | Q3LZX4 |
| NQNGTITDAVDCSLDP | 1 | 0 | 0 | 1683332 | QIQ54048.1 | P59594 |
| NQPDQVAVYQQSIIGA | 1 | 0 | 0 | 1683334 | YP_003767.1 | A0A0P0G321 |
| NQPYPNASFDNFKFVC | 1 | 0 | 0 | 1683335 | YP_009724389.1 | P0DTD1 |
| NQQGNSLNHLTSQLRQ | 1 | 0 | 0 | 1683336 | NP_073551.1 | A0A0P0K6L9 |
| NQQGSALNHLTSQLRH | 1 | 0 | 0 | 1683337 | YP_003767.1 | A0A0P0G321 |
| NQSKSPSSSGTSTPKK | 1 | 0 | 0 | 1683346 | YP_003771.1 | H9EJ69 |
| NQSLAGGITYVSNSGN | 1 | 0 | 0 | 1683347 | YP_003767.1 | A0A0P0G321 |
| NQSLSNAKVSAEVSRQ | 1 | 0 | 0 | 1683348 | NP_073552.1 | A0A0P0KWE8 |
| NQSPASSQTSAKSLAR | 1 | 0 | 0 | 1683349 | NP_073556.1 | A0A0N7J131 |
| NQSPQGNLGSNGKPGN | 1 | 0 | 0 | 1683350 | YP_009555238.1 | P0C6X6 |
| NQTNVVPPALHLVDPQ | 1 | 0 | 0 | 1683358 | NP_828859.1 | P59636 |
| NQVIVNNLDKSAGFPF | 1 | 0 | 0 | 1683365 | YP_009724389.1 | P0DTD1 |
| NQVVKPGHFNEEFYNF | 1 | 0 | 0 | 1683369 | YP_003766.2 | H9EJ42 |
| NQWHEAKFGVVTTNSD | 1 | 0 | 0 | 1683371 | YP_003766.2 | H9EJ42 |
| NQYYIYVTNEIGLNAS | 1 | 0 | 0 | 1683374 | YP_003767.1 | A0A0P0G321 |
| NRALSGIAAEQDRNTR | 1 | 0 | 0 | 1683377 | NP_828851.1 | P59594 |
| NRALTGIAVEQDKNTQ | 1 | 0 | 0 | 1683378 | YP_009724390.1 | P0DTC2 |
| NRALVSATEFADKLVE | 1 | 0 | 0 | 1683379 | YP_009555238.1 | P0C6X6 |
| NRATRVECTTIVNGVR | 1 | 0 | 0 | 1683387 | YP_009724389.1 | P0DTD1 |
| NRGEDIQLLKSAYENF | 1 | 0 | 0 | 1683406 | QHR63299.1 | P0C6W2 |
| NRGRQFYEFYNDVKPP | 1 | 0 | 0 | 1683409 | YP_009555243.1 | Q04854 |
| NRGVYKPLLENIDYFN | 1 | 0 | 0 | 1683411 | YP_009555238.1 | P0C6X6 |
| NRLLSVPSDSCNNVNV | 1 | 0 | 0 | 1683423 | YP_003766.2 | H9EJ42 |
| NRNASLNSFKEYFNLR | 1 | 0 | 0 | 1683433 | YP_009047204.1 | K9N5Q8 |
| NRRAAAYSSPLQSYAC | 1 | 0 | 0 | 1683447 | YP_009047202.1 | K9N7C7 |
| NRRQLDVVKRFIHKNS | 1 | 0 | 0 | 1683448 | NP_073549.1 | A0A0P0LKV0 |
| NRRSFKTADGNQRQLL | 1 | 0 | 0 | 1683449 | YP_009555245.1 | P33469 |
| NRRSRGAITTGYRFTN | 1 | 0 | 0 | 1683450 | YP_009555241.1 | P36334 |
| NRSGSNDIWMQNKGLF | 1 | 0 | 0 | 1683455 | YP_009555240.1 | P30215 |
| NRSLSRSNSNFRHSDS | 1 | 0 | 0 | 1683457 | YP_173242.1 | Q19U29 |

| | | | | | | |
|---|---|---|---|---|---|---|
| NRVKSGSPGDSSWHIY | 1 | 0 | 0 | 1683465 | YP_003767.1 | A0A0P0G321 |
| NRYKYYSGNANEADYR | 1 | 0 | 0 | 1683476 | NP_073549.1 | A0A0P0LKV0 |
| NRYNEVANAVMQNNEL | 1 | 0 | 0 | 1683479 | YP_173236.1 | A0A140H1G9 |
| NRYRIGNYKLNTDHAG | 1 | 0 | 0 | 1683482 | NP_828855.1 | P59596 |
| NRYSVLSVDSVSEESQ | 1 | 0 | 0 | 1683483 | YP_173236.1 | A0A140H1G9 |
| NSAIVCASVKRDGVQV | 1 | 0 | 0 | 1683490 | NP_073549.1 | A0A0P0LKV0 |
| NSALAKIQSVVNSNAQ | 1 | 0 | 0 | 1683493 | YP_173238.1 | A0A140H1H1 |
| NSALVKIQAVVNANAE | 1 | 0 | 0 | 1683495 | YP_009555241.1 | P36334 |
| NSAQYVFSTVNALPEV | 1 | 0 | 0 | 1683498 | NP_073549.1 | A0A0P0LKV0 |
| NSATGSLDVEVFNRLA | 1 | 0 | 0 | 1683501 | YP_003766.2 | H9EJ42 |
| NSAVFISPYNSQNYVA | 1 | 0 | 0 | 1683503 | YP_173236.1 | A0A140H1G9 |
| NSAVKLQNNELSPVAL | 1 | 0 | 0 | 1683504 | YP_009724389.1 | P0DTD1 |
| NSCLVSSDGQPGGDAI | 1 | 0 | 0 | 1683508 | YP_009047202.1 | K9N7C7 |
| NSEKHQDHVSLTVPLC | 1 | 0 | 0 | 1683515 | YP_009047202.1 | K9N7C7 |
| NSFDVLKSEDTQGMDN | 1 | 0 | 0 | 1683522 | QHR63299.1 | P0C6W2 |
| NSFDVPTFCANRSAMC | 1 | 0 | 0 | 1683523 | YP_009047202.1 | K9N7C7 |
| NSFFKELTANMSMAEC | 1 | 0 | 0 | 1683527 | YP_003766.2 | H9EJ42 |
| NSFGKDLNANMSLAEC | 1 | 0 | 0 | 1683528 | NP_073549.1 | A0A0P0LKV0 |
| NSFKEYFNLRNCTFMY | 1 | 0 | 0 | 1683529 | YP_009047204.1 | K9N5Q8 |
| NSGNLLGFKNVSTGNI | 1 | 0 | 0 | 1683540 | YP_003767.1 | A0A0P0G321 |
| NSGNRTPTSGVTPDMA | 1 | 0 | 0 | 1683541 | YP_009555245.1 | P33469 |
| NSGSDMNCNIPTQCYY | 1 | 0 | 0 | 1683544 | YP_173236.1 | A0A140H1G9 |
| NSGSDVLYQPPQTSIT | 1 | 0 | 0 | 1683545 | YP_009724389.1 | P0DTD1 |
| NSGSNITQIKNACVYF | 1 | 0 | 0 | 1683546 | YP_003766.2 | H9EJ42 |
| NSIFDIETVDVKHDVS | 1 | 0 | 0 | 1683555 | NP_073549.1 | A0A0P0LKV0 |
| NSILDEVNGLLDTTQL | 1 | 0 | 0 | 1683559 | YP_173238.1 | A0A140H1H1 |
| NSILDNAVKGCVPLNA | 1 | 0 | 0 | 1683560 | YP_009555238.1 | P0C6X6 |
| NSILDNAVKGCVPLSA | 1 | 0 | 0 | 1683561 | YP_173236.1 | A0A140H1G9 |
| NSILMLDKQGQLLDTK | 1 | 0 | 0 | 1683563 | YP_003766.2 | H9EJ42 |
| NSISPEKLKSYAASYN | 1 | 0 | 0 | 1683570 | NP_073549.1 | A0A0P0LKV0 |
| NSKHIDAKEGGNFNYL | 1 | 0 | 0 | 1683580 | QHR63300.2 | Q3LZX1 |
| NSKIVDLAQHFSDFGT | 1 | 0 | 0 | 1683581 | YP_009555238.1 | P0C6X6 |
| NSKVKVIQPRVEKKKT | 1 | 0 | 0 | 1683587 | NP_828849.2 | P0C6X7 |
| NSKVQIGEYTFEKGDY | 1 | 0 | 0 | 1683588 | YP_009724389.1 | P0DTD1 |
| NSLGETLVTMPLGYVT | 1 | 0 | 0 | 1683595 | QIQ54046.1 | P0C6U8 |
| NSLTYFNRPSLVDDNK | 1 | 0 | 0 | 1683609 | YP_009555238.1 | P0C6X6 |
| NSNANLSMLQLGVPDN | 1 | 0 | 0 | 1683615 | YP_003767.1 | A0A0P0G321 |
| NSNNPDWDFNPNKDT | 1 | 0 | 0 | 1683623 | A0A096ZU61.1 | Q76R62 |
| NSNQNTDSDSLSSKPQ | 1 | 0 | 0 | 1683624 | YP_173242.1 | Q19U29 |
| NSPFHPLADNKFALAC | 1 | 0 | 0 | 1683633 | QIQ54053.1 | P59635 |
| NSPFHPLADNKFALTC | 1 | 0 | 0 | 1683634 | YP_009724395.1 | P0DTC7 |
| NSQDDIMKAVAAALKS | 1 | 0 | 0 | 1683643 | NP_073556.1 | A0A0N7J131 |
| NSQGLLPPKNSIDAFK | 1 | 0 | 0 | 1683646 | YP_009724389.1 | P0DTD1 |
| NSQGLLPPKSSIDAFK | 1 | 0 | 0 | 1683647 | NP_828849.2 | P0C6X7 |
| NSQNAVAAKILGLPTQ | 1 | 0 | 0 | 1683648 | QIQ54047.1 | P0C6X7 |
| NSQNAVSRSMLGLTTQ | 1 | 0 | 0 | 1683650 | YP_009047202.1 | K9N7C7 |
| NSQNFAAKRVLGLQTQ | 1 | 0 | 0 | 1683651 | YP_009555238.1 | P0C6X6 |
| NSQNYVAKRVLGVQTQ | 1 | 0 | 0 | 1683654 | YP_173236.1 | A0A140H1G9 |
| NSQSSSRASSLSRNSS | 1 | 0 | 0 | 1683656 | YP_009047211.1 | K9N4V7 |
| NSQYLFSTINALPETS | 1 | 0 | 0 | 1683658 | YP_009047202.1 | K9N7C7 |
| NSRIRASLPTTIAKNS | 1 | 0 | 0 | 1683662 | NP_828849.2 | P0C6X7 |
| NSRNSTPGSSRGNSPA | 1 | 0 | 0 | 1683663 | NP_828858.1 | P59595 |
| NSRSTSRTSSRASSAG | 1 | 0 | 0 | 1683668 | YP_009555245.1 | P33469 |
| NSSANYVGKYDINHGD | 1 | 0 | 0 | 1683673 | YP_173237.1 | A0A140H1H0 |
| NSSDNGISAIITANLS | 1 | 0 | 0 | 1683674 | YP_003767.1 | A0A0P0G321 |
| NSSEWAYCVDLHNKIN | 1 | 0 | 0 | 1683676 | YP_003766.2 | H9EJ42 |
| NSSRASSRSSTRNNSR | 1 | 0 | 0 | 1683685 | YP_003771.1 | H9EJ69 |
| NSSRNSTPGSSRGTSP | 1 | 0 | 0 | 1683686 | YP_009724397.2 | P0DTC9 |
| NSSRSSSQGSRSGNST | 1 | 0 | 0 | 1683687 | YP_009047211.1 | K9N4V7 |
| NSSSGVLDTTIPFGPS | 1 | 0 | 0 | 1683688 | YP_003767.1 | A0A0P0G321 |
| NSSVPLLEVYVHLRNW | 1 | 0 | 0 | 1683696 | YP_003770.1 | U3M6Q8 |
| NSTDTVNHDSFNLYGY | 1 | 0 | 0 | 1683701 | YP_173236.1 | A0A140H1G9 |
| NSTPGSSRGTSPARIA | 1 | 0 | 0 | 1683722 | QIQ54056.1 | P59595 |
| NSTPGSSRGTSPARMA | 1 | 0 | 0 | 1683723 | YP_009724397.2 | P0DTC9 |
| NSTQDGDNKLQGLLEV | 1 | 0 | 0 | 1683725 | YP_009555241.1 | P36334 |
| NSTRGTSPGPSGIGAV | 1 | 0 | 0 | 1683726 | YP_009047211.1 | K9N4V7 |
| NSTTMCVPHNMRVLHY | 1 | 0 | 0 | 1683729 | YP_003766.2 | H9EJ42 |
| NSTTNYVGVYDINHGD | 1 | 0 | 0 | 1683730 | YP_009555240.1 | P30215 |
| NSTTVSVTVSSNMVET | 1 | 0 | 0 | 1683731 | YP_009047202.1 | K9N7C7 |
| NSVCPKLEFANDTKIA | 1 | 0 | 0 | 1683743 | YP_009047204.1 | K9N5Q8 |
| NSVFNICQAVSANVCA | 1 | 0 | 0 | 1683745 | YP_009555238.1 | P0C6X6 |
| NSVINRLRCDQLSFDV | 1 | 0 | 0 | 1683754 | NP_073551.1 | A0A0P0K6L9 |
| NSVKQDALTGGNYGYL | 1 | 0 | 0 | 1683756 | QIQ54048.1 | P59594 |
| NSVNDSLEPVGGLYEI | 1 | 0 | 0 | 1683763 | YP_009555241.1 | P36334 |
| NSVVLSRKYTSFPWL | 1 | 0 | 0 | 1683778 | P12900.1 | Q69028 |
| NSWQNLLYDSNGNIIG | 1 | 0 | 0 | 1683786 | YP_173238.1 | A0A140H1H1 |
| NSWQNLLYDSNGNLYG | 1 | 0 | 0 | 1683787 | YP_009555241.1 | P36334 |
| NSYFDSKGWYDPVENE | 1 | 0 | 0 | 1683790 | YP_003766.2 | H9EJ42 |
| NSYITTIKPVIYKLDG | 1 | 0 | 0 | 1683794 | QHR63299.1 | P0C6W2 |
| NSYTSFATYHTPATDC | 1 | 0 | 0 | 1683798 | YP_009047204.1 | K9N5Q8 |
| NSYTTTIKPIVYKLDG | 1 | 0 | 0 | 1683799 | QIQ54046.1 | P0C6U8 |
| NSYTTTIKPVTYKLDG | 1 | 0 | 0 | 1683800 | YP_009724389.1 | P0DTD1 |
| NTDFSRVSAKPPPGDQ | 1 | 0 | 0 | 1683819 | YP_009724389.1 | P0DTD1 |
| NTDHSNSSDNIALLVQ | 1 | 0 | 0 | 1683820 | QIQ54051.1 | P59596 |
| NTDLIEGSKPYSTLQP | 1 | 0 | 0 | 1683822 | YP_003766.2 | H9EJ42 |
| NTEFTRVNAKPPPGDQ | 1 | 0 | 0 | 1683830 | NP_828849.2 | P0C6X7 |
| NTFSSTFNVPMDKLKT | 1 | 0 | 0 | 1683840 | QIQ54046.1 | P0C6U8 |
| NTGRSVYVKFQDSKPP | 1 | 0 | 0 | 1683855 | YP_009047209.1 | K9N5R3 |
| NTGSVCYTPIDEIPYK | 1 | 0 | 0 | 1683856 | YP_009047202.1 | K9N7C7 |
| NTGSVVKPVRARAPPG | 1 | 0 | 0 | 1683857 | NP_073549.1 | A0A0P0LKV0 |
| NTGVCYDFDGVTTSDK | 1 | 0 | 0 | 1683859 | YP_003766.2 | H9EJ42 |

| | | | | | | |
|---|---|---|---|---|---|---|
| NTGVYDLSGFTVKPVA | 1 | 0 | 0 | 1683860 | YP_173238.1 | A0A140H1H1 |
| NTHLKHGGGIAGAINA | 1 | 0 | 0 | 1683861 | YP_009047202.1 | K9N7C7 |
| NTICIKDADYNAKVEI | 1 | 0 | 0 | 1683866 | NP_073549.1 | A0A0P0LKV0 |
| NTILNTACGVFEVDDT | 1 | 0 | 0 | 1683869 | YP_009555238.1 | P0C6X6 |
| NTINKATFKPNTWCLR | 1 | 0 | 0 | 1683870 | QIQ54046.1 | P0C6U8 |
| NTISPEKLKNYAASYN | 1 | 0 | 0 | 1683873 | YP_003766.2 | H9EJ42 |
| NTKFADDLNQMTGFTK | 1 | 0 | 0 | 1683878 | NP_828849.2 | P0C6X7 |
| NTKHGSGVTRELMREL | 1 | 0 | 0 | 1683882 | QIQ54046.1 | P0C6U8 |
| NTKPVETSNSFDVLKS | 1 | 0 | 0 | 1683886 | QHR63299.1 | P0C6W2 |
| NTKVIKDRVMYSEFPA | 1 | 0 | 0 | 1683888 | YP_009555241.1 | P36334 |
| NTKYVAPQVTYQNIST | 1 | 0 | 0 | 1683890 | YP_009047204.1 | K9N5Q8 |
| NTKYYDDSQYYFNKDT | 1 | 0 | 0 | 1683891 | YP_009555240.1 | P30215 |
| NTLNDLNETLVTMPLG | 1 | 0 | 0 | 1683900 | YP_009724389.1 | P0DTD1 |
| NTLNIIVPDKSVYDQV | 1 | 0 | 0 | 1683901 | YP_009555238.1 | P0C6X6 |
| NTLVSGDAHIPFGTVL | 1 | 0 | 0 | 1683910 | NP_073551.1 | A0A0P0K6L9 |
| NTNLHFDVDNINFKSL | 1 | 0 | 0 | 1683912 | YP_173238.1 | A0A140H1H1 |
| NTNLHTQLVDMSMTYG | 1 | 0 | 0 | 1683913 | NP_828849.2 | P0C6X7 |
| NTNLSRGRGRNPKPRA | 1 | 0 | 0 | 1683914 | YP_009047211.1 | K9N4V7 |
| NTNSTKDDQIGYYRRA | 1 | 0 | 0 | 1683920 | QIQ54056.1 | P59595 |
| NTPKDHIGTRNPANNA | 1 | 0 | 0 | 1683925 | YP_009724397.2 | P0DTC9 |
| NTPTQCYYNNSNNGKI | 1 | 0 | 0 | 1683929 | YP_009555238.1 | P0C6X6 |
| NTREVFAQVKQMYKTP | 1 | 0 | 0 | 1683932 | NP_828851.1 | P59594 |
| NTRNIDATSTGNYNYK | 1 | 0 | 0 | 1683936 | NP_828851.1 | P59594 |
| NTSAQYIFSTVNALPE | 1 | 0 | 0 | 1683937 | YP_003766.2 | H9EJ42 |
| NTSCPEWMDDFEADPK | 1 | 0 | 0 | 1683938 | YP_009047202.1 | K9N7C7 |
| NTSGCNVNDVTKLYLG | 1 | 0 | 0 | 1683943 | NP_073549.1 | A0A0P0LKV0 |
| NTSIPNLPDFKEELDQ | 1 | 0 | 0 | 1683944 | YP_009555241.1 | P36334 |
| NTSLGNRKRNQKPLEP | 1 | 0 | 0 | 1683947 | YP_003771.1 | H9EJ69 |
| NTVDTTPKEEFVVKEK | 1 | 0 | 0 | 1683977 | NP_073549.1 | A0A0P0LKV0 |
| NTVKPKPVINQLDEKA | 1 | 0 | 0 | 1683984 | NP_073549.1 | A0A0P0LKV0 |
| NTVLNASIPESKPLAD | 1 | 0 | 0 | 1683986 | YP_003771.1 | H9EJ69 |
| NTVVLTFTTRVTVPKD | 1 | 0 | 0 | 1683992 | NP_073556.1 | A0A0N7J131 |
| NTVYDPLQPELDSFKE | 1 | 0 | 0 | 1683994 | YP_009724390.1 | P0DTC2 |
| NVAFELFAKRKMGLTP | 1 | 0 | 0 | 1684009 | YP_003766.2 | H9EJ42 |
| NVAFQTVKPGNFNKDF | 1 | 0 | 0 | 1684013 | YP_009724389.1 | P0DTD1 |
| NVANYQKVGMQKYSTL | 1 | 0 | 0 | 1684030 | YP_009724389.1 | P0DTD1 |
| NVANYQKVGMQXYSTL | 1 | 0 | 0 | 1684031 | QIQ54047.1 | P0C6X7 |
| NVCALMSCNGNKIEDL | 1 | 0 | 0 | 1684037 | YP_009555238.1 | P0C6X6 |
| NVCHSVAVIGDEQIDG | 1 | 0 | 0 | 1684043 | YP_173236.1 | A0A140H1G9 |
| NVDDINFSPVLGCLGS | 1 | 0 | 0 | 1684053 | YP_009555241.1 | P36334 |
| NVDESFVDDFYGYLQK | 1 | 0 | 0 | 1684054 | NP_073549.1 | A0A0P0LKV0 |
| NVDGFNEWAMANGYTS | 1 | 0 | 0 | 1684058 | YP_003766.2 | H9EJ42 |
| NVDIVSEAKNVKPTIV | 1 | 0 | 0 | 1684060 | QIQ54046.1 | P0C6U8 |
| NVDLDALASDIFDTPS | 1 | 0 | 0 | 1684066 | YP_009047202.1 | K9N7C7 |
| NVDPLSNFEHKVITEC | 1 | 0 | 0 | 1684070 | YP_009047202.1 | K9N7C7 |
| NVDWSITYPMIANENA | 1 | 0 | 0 | 1684075 | NP_073549.1 | A0A0P0LKV0 |
| NVEAVNFNVTTAETTD | 1 | 0 | 0 | 1684078 | NP_073551.1 | A0A0P0K6L9 |
| NVEYPIISNELSINTS | 1 | 0 | 0 | 1684081 | YP_009555238.1 | P0C6X6 |
| NVFAVESGGYIPSDFA | 1 | 0 | 0 | 1684083 | NP_073551.1 | A0A0P0K6L9 |
| NVFCDGVNVTKHKCDI | 1 | 0 | 0 | 1684085 | YP_009555238.1 | P0C6X6 |
| NVFKLKISTQLFEGDK | 1 | 0 | 0 | 1684089 | YP_003766.2 | H9EJ42 |
| NVFMSEAKCWTETDLT | 1 | 0 | 0 | 1684090 | YP_009724389.1 | P0DTD1 |
| NVFMSEAKCWVETDLK | 1 | 0 | 0 | 1684091 | YP_009047202.1 | K9N7C7 |
| NVFMSESKCWVEHDIN | 1 | 0 | 0 | 1684092 | YP_009555238.1 | P0C6X6 |
| NVFMSESKCWVENDIT | 1 | 0 | 0 | 1684093 | YP_173236.1 | A0A140H1G9 |
| NVGHYDGRTGELPCAI | 1 | 0 | 0 | 1684107 | YP_173236.1 | A0A140H1G9 |
| NVGRWSASINTGNCPF | 1 | 0 | 0 | 1684110 | NP_073551.1 | A0A0P0K6L9 |
| NVHDVTVTTDKSFEQQ | 1 | 0 | 0 | 1684112 | NP_073549.1 | A0A0P0LKV0 |
| NVHRNEHVASGDAIMT | 1 | 0 | 0 | 1684117 | NP_073549.1 | A0A0P0LKV0 |
| NVHTPKHVFKTLKPGD | 1 | 0 | 0 | 1684118 | YP_003766.2 | H9EJ42 |
| NVIVFDGKSKCDESAS | 1 | 0 | 0 | 1684135 | NP_828849.2 | P0C6X7 |
| NVIVFDGKSKCDESSA | 1 | 0 | 0 | 1684136 | QIQ54046.1 | P0C6U8 |
| NVIVFDGKSKCEESSA | 1 | 0 | 0 | 1684137 | YP_009724389.1 | P0DTD1 |
| NVKDYMSLSEQLRKQI | 1 | 0 | 0 | 1684143 | NP_828849.2 | P0C6X7 |
| NVKLLDYDYATHGQLD | 1 | 0 | 0 | 1684148 | YP_003766.2 | H9EJ42 |
| NVKPGDAVITTQQAGA | 1 | 0 | 0 | 1684150 | NP_073549.1 | A0A0P0LKV0 |
| NVKSSLSSYEYCQPPK | 1 | 0 | 0 | 1684153 | YP_173236.1 | A0A140H1G9 |
| NVLDDFIVFNNNGTNV | 1 | 0 | 0 | 1684158 | NP_073549.1 | A0A0P0LKV0 |
| NVLEGSVAYESLRPDT | 1 | 0 | 0 | 1684159 | YP_009724389.1 | P0DTD1 |
| NVLEGSVPYESLRPDT | 1 | 0 | 0 | 1684160 | QIQ54046.1 | P0C6U8 |
| NVLFDGRDNGALEAFK | 1 | 0 | 0 | 1684163 | YP_009555238.1 | P0C6X6 |
| NVLIKESIPIVWGVKD | 1 | 0 | 0 | 1684166 | YP_003766.2 | H9EJ42 |
| NVLLRKNGNKGAGGHS | 1 | 0 | 0 | 1684167 | NP_828849.2 | P0C6X7 |
| NVLPETYVALPIYYQH | 1 | 0 | 0 | 1684169 | YP_003767.1 | A0A0P0G321 |
| NVLQDVLSLRHDIALD | 1 | 0 | 0 | 1684172 | YP_009555238.1 | P0C6X6 |
| NVLYENQKQIANQFNK | 1 | 0 | 0 | 1684181 | NP_828851.1 | P59594 |
| NVMDVNDCFKNDNVVL | 1 | 0 | 0 | 1684183 | YP_003766.2 | H9EJ42 |
| NVNKGEDIQLLKSAYE | 1 | 0 | 0 | 1684194 | YP_009724389.1 | P0DTD1 |
| NVPAIDVLKKLLSSLT | 1 | 0 | 0 | 1684205 | YP_003766.2 | H9EJ42 |
| NVPLQVGFSNGVDFVA | 1 | 0 | 0 | 1684213 | NP_073549.1 | A0A0P0LKV0 |
| NVQIHTIDGSSGVANP | 1 | 0 | 0 | 1684229 | NP_828852.2 | P59632 |
| NVQKVEQPKIEPKPVS | 1 | 0 | 0 | 1684232 | NP_073549.1 | A0A0P0LKV0 |
| NVQVDNGSSINRKQLE | 1 | 0 | 0 | 1684240 | YP_003766.2 | H9EJ42 |
| NVQYRINGLGVTMDVL | 1 | 0 | 0 | 1684241 | YP_009555241.1 | P36334 |
| NVSALMGANGNKIVDK | 1 | 0 | 0 | 1684250 | YP_009047202.1 | K9N7C7 |
| NVSDAYANLVPYYQLI | 1 | 0 | 0 | 1684255 | NP_073549.1 | A0A0P0LKV0 |
| NVSFVGELKAAEAKVI | 1 | 0 | 0 | 1684259 | NP_073549.1 | A0A0P0LKV0 |
| NVSFVNDSIESVGGLY | 1 | 0 | 0 | 1684260 | YP_173238.1 | A0A140H1H1 |
| NVSIKNVDIVSEAKNV | 1 | 0 | 0 | 1684261 | QIQ54046.1 | P0C6U8 |
| NVSSFGDYNLSSVIPS | 1 | 0 | 0 | 1684274 | NP_073551.1 | A0A0P0K6L9 |
| NVTDFNAIATCDWTNA | 1 | 0 | 0 | 1684287 | YP_009724389.1 | P0DTD1 |

| | | | | | | |
|---|---|---|---|---|---|---|
| NVTGSNESKALFFTYL | 1 | 0 | 0 | 1684294 | YP_009047202.1 | K9N7C7 |
| NVTINCQDPAGGALIA | 1 | 0 | 0 | 1684297 | NP_849177.1 | Q80H93 |
| NVTKENDSKEGFFTYI | 1 | 0 | 0 | 1684299 | YP_009724389.1 | P0DTD1 |
| NVTKHKCDINYKGKVF | 1 | 0 | 0 | 1684300 | YP_009555238.1 | P0C6X6 |
| NVTLDKCTKYNIYDVS | 1 | 0 | 0 | 1684302 | NP_073551.1 | A0A0P0K6L9 |
| NVVCKHCGQKTTTLTG | 1 | 0 | 0 | 1684324 | NP_828849.2 | P0C6X7 |
| NVVCKTCGQQQTTLKG | 1 | 0 | 0 | 1684325 | YP_009724389.1 | P0DTD1 |
| NVVELPKFFYASNGTY | 1 | 0 | 0 | 1684326 | NP_073551.1 | A0A0P0K6L9 |
| NVVGPDARTQGKQSYV | 1 | 0 | 0 | 1684328 | YP_009555238.1 | P0C6X6 |
| NVVNKGHFDGQQGEVP | 1 | 0 | 0 | 1684346 | YP_009724389.1 | P0DTD1 |
| NVVPPALHLVDPQIQL | 1 | 0 | 0 | 1684351 | NP_828859.1 | P59636 |
| NVVRYSLDGNFRTEVD | 1 | 0 | 0 | 1684356 | YP_009047202.1 | K9N7C7 |
| NVVSHVNGDWFLFGDS | 1 | 0 | 0 | 1684358 | YP_009555240.1 | P30215 |
| NVWQIQTIQDSDGTNK | 1 | 0 | 0 | 1684372 | YP_009555238.1 | P0C6X6 |
| NVYCYNTDLIEGSKPY | 1 | 0 | 0 | 1684378 | YP_003766.2 | H9EJ42 |
| NVYMAKAGDKYYPVVV | 1 | 0 | 0 | 1684388 | YP_009555238.1 | P0C6X6 |
| NVYRSTSPDPKFVDKY | 1 | 0 | 0 | 1684389 | YP_009047202.1 | K9N7C7 |
| NVYRTDYVDYTFVNEY | 1 | 0 | 0 | 1684390 | YP_173236.1 | A0A140H1G9 |
| NWFDPIENEDIHRVYA | 1 | 0 | 0 | 1684401 | NP_073549.1 | A0A0P0LKV0 |
| NWFLLTNTSSVVDGVV | 1 | 0 | 0 | 1684405 | NP_073551.1 | A0A0P0K6L9 |
| NWFNTMLDASAPATGW | 1 | 0 | 0 | 1684406 | YP_173236.1 | A0A140H1G9 |
| NWLAPTNAYMRTTHTD | 1 | 0 | 0 | 1684411 | YP_009047202.1 | K9N7C7 |
| NWLKPYGGQPVSEYHI | 1 | 0 | 0 | 1684412 | YP_009047207.1 | K9N643 |
| NWNLEYPIISNEVSIN | 1 | 0 | 0 | 1684422 | YP_173236.1 | A0A140H1G9 |
| NWNVEYPIISNELSIN | 1 | 0 | 0 | 1684427 | YP_009555238.1 | P0C6X6 |
| NYAKPFLNKVVSTTTN | 1 | 0 | 0 | 1684441 | YP_009724389.1 | P0DTD1 |
| NYDLSVVNARLRAKHY | 1 | 0 | 0 | 1684452 | YP_009724389.1 | P0DTD1 |
| NYFKKVDGIIQQLPET | 1 | 0 | 0 | 1684465 | NP_828849.2 | P0C6X7 |
| NYGAGIKLPSGIMLNV | 1 | 0 | 0 | 1684469 | YP_003766.2 | H9EJ42 |
| NYGDSATLPKGIMMNV | 1 | 0 | 0 | 1684471 | YP_009724389.1 | P0DTD1 |
| NYKIDTTSSSCQLYYS | 1 | 0 | 0 | 1684489 | YP_173238.1 | A0A140H1H1 |
| NYKLNTDHAGSNDNIA | 1 | 0 | 0 | 1684492 | NP_828855.1 | P59596 |
| NYKLNTDHSNSSDNIA | 1 | 0 | 0 | 1684493 | QIQ54051.1 | P59596 |
| NYLGKPKEQIDGYTMH | 1 | 0 | 0 | 1684511 | NP_828849.2 | P0C6X7 |
| NYLGTIKENIDGGAMH | 1 | 0 | 0 | 1684514 | YP_009047202.1 | K9N7C7 |
| NYNNLVPTYLKSDNIV | 1 | 0 | 0 | 1684533 | YP_173236.1 | A0A140H1G9 |
| NYNRNASLNSFKEYFN | 1 | 0 | 0 | 1684535 | YP_009047204.1 | K9N5Q8 |
| NYNVSVLQMDVQGLVN | 1 | 0 | 0 | 1684536 | YP_003766.2 | H9EJ42 |
| NYQKVGMQKYSTLQGP | 1 | 0 | 0 | 1684546 | YP_009724389.1 | P0DTD1 |
| NYQKVGMQXYSTLQGP | 1 | 0 | 0 | 1684547 | QIQ54047.1 | P0C6X7 |
| NYRLPSTQKGSGMDTA | 1 | 0 | 0 | 1684553 | YP_009555244.1 | Q01455 |
| NYTVCDDCNGYTDNIF | 1 | 0 | 0 | 1684572 | YP_003767.1 | A0A0P0G321 |
| NYTVPDLVVEQYNQTI | 1 | 0 | 0 | 1684574 | NP_073551.1 | A0A0P0K6L9 |
| NYTVQKLQTLIDNINS | 1 | 0 | 0 | 1684575 | NP_073551.1 | A0A0P0K6L9 |
| NYVAKRVLGVQTQTVD | 1 | 0 | 0 | 1684580 | YP_173236.1 | A0A140H1G9 |
| NYVGKYDINHGDSGFI | 1 | 0 | 0 | 1684590 | YP_173237.1 | A0A140H1H0 |
| NYVGVYDINHGDAGFT | 1 | 0 | 0 | 1684591 | YP_009555240.1 | P30215 |
| NYVPSYVKPDSVSTSD | 1 | 0 | 0 | 1684596 | YP_009047202.1 | K9N7C7 |
| NYYKKDNSYFTEQPID | 1 | 0 | 0 | 1684601 | YP_009724389.1 | P0DTD1 |
| NYYKKVDGVVQQLPET | 1 | 0 | 0 | 1684602 | YP_009724389.1 | P0DTD1 |
| NYYKKVNGVVQQLPET | 1 | 0 | 0 | 1684603 | QHR63299.1 | P0C6W2 |
| PAAAYLDAVKQGAKPV | 1 | 0 | 0 | 1684608 | NP_073549.1 | A0A0P0LKV0 |
| PAAPRAVSFADNNDIT | 1 | 0 | 0 | 1684613 | YP_009047211.1 | K9N4V7 |
| PAAVDTKCLASIDEVS | 1 | 0 | 0 | 1684615 | YP_173236.1 | A0A140H1G9 |
| PACHNPEVGPEHSLAE | 1 | 0 | 0 | 1684618 | QHR63299.1 | P0C6W2 |
| PACQDPEIGPEHSVAD | 1 | 0 | 0 | 1684620 | NP_828849.2 | P0C6X7 |
| PADSRDFLRVVFSQVD | 1 | 0 | 0 | 1684624 | YP_009555238.1 | P0C6X6 |
| PAEFKYMVANGLNAPN | 1 | 0 | 0 | 1684627 | NP_073549.1 | A0A0P0LKV0 |
| PAEIVDTVSALVYDNK | 1 | 0 | 0 | 1684628 | YP_009724389.1 | P0DTD1 |
| PAFDKSAFTNLKQLPF | 1 | 0 | 0 | 1684632 | NP_828849.2 | P0C6X7 |
| PAFDKSAFVNLKQLPF | 1 | 0 | 0 | 1684633 | YP_009724389.1 | P0DTD1 |
| PAFMLGSSVGNFSDGK | 1 | 0 | 0 | 1684636 | YP_009047204.1 | K9N5Q8 |
| PAFNVSDAYANLVPYY | 1 | 0 | 0 | 1684637 | YP_003766.2 | H9EJ42 |
| PAGNSYTSFATYHTPA | 1 | 0 | 0 | 1684641 | YP_009047204.1 | K9N5Q8 |
| PAGTILVDNDLYPFVS | 1 | 0 | 0 | 1684642 | YP_009555238.1 | P0C6X6 |
| PAGVNITLANFNETKG | 1 | 0 | 0 | 1684647 | NP_073551.1 | A0A0P0K6L9 |
| PAHAPSFLAVDDKYKA | 1 | 0 | 0 | 1684648 | YP_009555238.1 | P0C6X6 |
| PAHAPSFLAVDDKYKV | 1 | 0 | 0 | 1684649 | YP_173236.1 | A0A140H1G9 |
| PAIKLNFGMLNGNAIA | 1 | 0 | 0 | 1684658 | NP_073549.1 | A0A0P0LKV0 |
| PAINVTINNYNPSSWN | 1 | 0 | 0 | 1684659 | YP_173238.1 | A0A140H1H1 |
| PAKAYKDYLASGGQPI | 1 | 0 | 0 | 1684662 | NP_828849.2 | P0C6X7 |
| PAKLKIQVVNSGPDQT | 1 | 0 | 0 | 1684664 | YP_009555238.1 | P0C6X6 |
| PAKVRVDCYDKFKIND | 1 | 0 | 0 | 1684665 | YP_173236.1 | A0A140H1G9 |
| PAKVRVECYDKFKIND | 1 | 0 | 0 | 1684666 | YP_009555238.1 | P0C6X6 |
| PALAANTLTIVIPDKQ | 1 | 0 | 0 | 1684667 | YP_173236.1 | A0A140H1G9 |
| PALQDAYYRARAGDAA | 1 | 0 | 0 | 1684676 | QIQ54046.1 | P0C6U8 |
| PALQDAYYRARAGEAA | 1 | 0 | 0 | 1684677 | YP_009724389.1 | P0DTD1 |
| PALQEAYYRARAGDAA | 1 | 0 | 0 | 1684678 | NP_828849.2 | P0C6X7 |
| PALQHAFMKHKGGDST | 1 | 0 | 0 | 1684679 | YP_009047202.1 | K9N7C7 |
| PAMHIASSNAFLDLRT | 1 | 0 | 0 | 1684686 | YP_009047202.1 | K9N7C7 |
| PAPIDIDAFPVKPICS | 1 | 0 | 0 | 1684702 | YP_003766.2 | H9EJ42 |
| PAPRTLLTKGTLEPEY | 1 | 0 | 0 | 1684703 | YP_009724389.1 | P0DTD1 |
| PAPRTLLTRGTLEPEN | 1 | 0 | 0 | 1684704 | YP_009047202.1 | K9N7C7 |
| PAPRVLISKGVMEPID | 1 | 0 | 0 | 1684705 | NP_073549.1 | A0A0P0LKV0 |
| PAPRVLLSKGSLEPRH | 1 | 0 | 0 | 1684706 | YP_173236.1 | A0A140H1G9 |
| PAPRVLLSKGTLEPKY | 1 | 0 | 0 | 1684707 | YP_009555238.1 | P0C6X6 |
| PAPVYIWTADEAIKFL | 1 | 0 | 0 | 1684709 | YP_009555244.1 | Q01455 |
| PAQNAGYWRRQDRKIN | 1 | 0 | 0 | 1684720 | YP_009047211.1 | K9N4V7 |
| PARARVECFDKFKVNS | 1 | 0 | 0 | 1684723 | YP_009724389.1 | P0DTD1 |
| PARIAGNGGDAALALL | 1 | 0 | 0 | 1684726 | QIQ54056.1 | P59595 |
| PARLEIVKLGAEKKDG | 1 | 0 | 0 | 1684727 | YP_009555239.1 | Q80872 |

| | | | | | | |
|---|---|---|---|---|---|---|
| PARMASGGGETALALL | 1 | 0 | 0 | 1684730 | NP_828858.1 | P59595 |
| PASSQTSAKSLARSQS | 1 | 0 | 0 | 1684740 | NP_073556.1 | A0A0N7J131 |
| PASTVHATATIPLQAS | 1 | 0 | 0 | 1684741 | NP_828852.2 | P59632 |
| PATDCSDGNYNRNASL | 1 | 0 | 0 | 1684746 | YP_009047204.1 | K9N5Q8 |
| PATSASKLTIVSPDLE | 1 | 0 | 0 | 1684752 | YP_003766.2 | H9EJ42 |
| PATSIVAKQGAGDAGH | 1 | 0 | 0 | 1684753 | NP_073549.1 | A0A0P0LKV0 |
| PATVCGPKLSTDLIKN | 1 | 0 | 0 | 1684757 | NP_828851.1 | P59594 |
| PATVSVSSPDAVTAYN | 1 | 0 | 0 | 1684759 | YP_009724389.1 | P0DTD1 |
| PAVAKHDFFKFRIDGD | 1 | 0 | 0 | 1684760 | YP_009724389.1 | P0DTD1 |
| PAVAVHDFFKFRVDGD | 1 | 0 | 0 | 1684761 | NP_828849.2 | P0C6X7 |
| PAVVSVSSPDAVTTYN | 1 | 0 | 0 | 1684768 | NP_828849.2 | P0C6X7 |
| PAYAPTYVSVDDKYKT | 1 | 0 | 0 | 1684773 | YP_009047202.1 | K9N7C7 |
| PAYTNSSTRGVYYPDK | 1 | 0 | 0 | 1684783 | QHR63300.2 | Q3LZX1 |
| PCAFGGVSVITPGTNA | 1 | 0 | 0 | 1684786 | NP_828851.1 | P59594 |
| PCAIINDKVVAKIDKE | 1 | 0 | 0 | 1684789 | YP_009555238.1 | P0C6X6 |
| PCESHGKQVVSDIDYV | 1 | 0 | 0 | 1684795 | YP_009724389.1 | P0DTD1 |
| PCEVHGNGSMIEDIDY | 1 | 0 | 0 | 1684796 | YP_009047202.1 | K9N7C7 |
| PCGYSEPPKVAAKICI | 1 | 0 | 0 | 1684801 | YP_009555238.1 | P0C6X6 |
| PCIKVATVQSKMSDVK | 1 | 0 | 0 | 1684804 | YP_009724389.1 | P0DTD1 |
| PCKFTVQDAKGLKIKY | 1 | 0 | 0 | 1684809 | YP_009555238.1 | P0C6X6 |
| PCNAQEQQIYFFEGVA | 1 | 0 | 0 | 1684812 | YP_173236.1 | A0A140H1G9 |
| PCNLYNYGAGIKLPSG | 1 | 0 | 0 | 1684818 | YP_003766.2 | H9EJ42 |
| PCPACQDPEIGPEHSV | 1 | 0 | 0 | 1684823 | NP_828849.2 | P0C6X7 |
| PCPSGTYEGNSPFHPL | 1 | 0 | 0 | 1684825 | NP_828857.1 | P59635 |
| PCSSGTYEGNSPFHPL | 1 | 0 | 0 | 1684835 | YP_009724395.1 | P0DTC7 |
| PCTCGKQATKYLVQQE | 1 | 0 | 0 | 1684838 | YP_009724389.1 | P0DTD1 |
| PCTCGKQATQYLVQQE | 1 | 0 | 0 | 1684839 | QHR63299.1 | P0C6W2 |
| PCVSIVPSTVWEDGDY | 1 | 0 | 0 | 1684846 | YP_009047204.1 | K9N5Q8 |
| PCVYVDGLESKQVDYV | 1 | 0 | 0 | 1684849 | YP_173236.1 | A0A140H1G9 |
| PDAIIIDNDINDYVSD | 1 | 0 | 0 | 1684855 | YP_003766.2 | H9EJ42 |
| PDARAKQDVSLLSKCY | 1 | 0 | 0 | 1684857 | YP_009047202.1 | K9N7C7 |
| PDAVTAYNGYLTSSSK | 1 | 0 | 0 | 1684861 | YP_009724389.1 | P0DTD1 |
| PDAVTTYNGYLTSSSK | 1 | 0 | 0 | 1684862 | NP_828849.2 | P0C6X7 |
| PDDASRLLHTVLAKAE | 1 | 0 | 0 | 1684866 | YP_009047202.1 | K9N7C7 |
| PDEDEEEGDCEEEDFE | 1 | 0 | 0 | 1684875 | QHR63299.1 | P0C6W2 |
| PDEPQKDVYELRYNGA | 1 | 0 | 0 | 1684881 | YP_009555245.1 | P33469 |
| PDIINIYKKLGPIFNR | 1 | 0 | 0 | 1684901 | YP_173236.1 | A0A140H1G9 |
| PDIINVYKKLGPIFNR | 1 | 0 | 0 | 1684902 | YP_009555238.1 | P0C6X6 |
| PDKQVFDKVVDNVYVT | 1 | 0 | 0 | 1684904 | YP_173236.1 | A0A140H1G9 |
| PDKSVYDQVVDNVYVT | 1 | 0 | 0 | 1684905 | YP_009555238.1 | P0C6X6 |
| PDLNGDVVAIDYKHYT | 1 | 0 | 0 | 1684912 | YP_009724389.1 | P0DTD1 |
| PDLNGDVVAIDYRHYS | 1 | 0 | 0 | 1684913 | NP_828849.2 | P0C6X7 |
| PDLPDCDIDKWLNNFN | 1 | 0 | 0 | 1684914 | YP_173238.1 | A0A140H1H1 |
| PDLSQYYCDNGKYYTK | 1 | 0 | 0 | 1684920 | YP_173236.1 | A0A140H1G9 |
| PDNSSTIVTGLLPTHW | 1 | 0 | 0 | 1684925 | YP_003767.1 | A0A0P0G321 |
| PDPKFVDKYYAFLNKH | 1 | 0 | 0 | 1684928 | YP_009047202.1 | K9N7C7 |
| PDPLKPTKRSFIEDLL | 1 | 0 | 0 | 1684929 | NP_828851.1 | P59594 |
| PDQTCNTPTQCYYNNS | 1 | 0 | 0 | 1684937 | YP_009555238.1 | P0C6X6 |
| PDQVAVYQQSIIGAMT | 1 | 0 | 0 | 1684938 | YP_003767.1 | A0A0P0G321 |
| PDSTSLVEXVDVTFLK | 1 | 0 | 0 | 1684942 | QIQ54046.1 | P0C6U8 |
| PDWDFNPNKDTWPDA | 1 | 0 | 0 | 1684955 | P03138.3 | Q76R62 |
| PDYAYFNGAIIRDSDV | 1 | 0 | 0 | 1684957 | YP_009047202.1 | K9N7C7 |
| PDYNTYKNTCDGTTFT | 1 | 0 | 0 | 1684959 | YP_009724389.1 | P0DTD1 |
| PDYNTYKNTCEGSTFT | 1 | 0 | 0 | 1684960 | QIQ54046.1 | P0C6U8 |
| PDYTNLLCRVTSSDFT | 1 | 0 | 0 | 1684962 | YP_009555238.1 | P0C6X6 |
| PDYVDVNKTLQEFAQN | 1 | 0 | 0 | 1684963 | YP_003767.1 | A0A0P0G321 |
| PEAALPFRAVKDGIVW | 1 | 0 | 0 | 1684964 | YP_009047211.1 | K9N4V7 |
| PEAGLPYGANKEGIIW | 1 | 0 | 0 | 1684965 | QIQ54056.1 | P59595 |
| PEASLPYGANKEGIVW | 1 | 0 | 0 | 1684967 | NP_828858.1 | P59595 |
| PEDHSLLATLDDPYVE | 1 | 0 | 0 | 1684972 | YP_173242.1 | Q19U29 |
| PEDHVQIHTIDGSSGV | 1 | 0 | 0 | 1684973 | QIQ54049.1 | P59632 |
| PEEELEEDWLDDDSQQ | 1 | 0 | 0 | 1684977 | QHR63299.1 | P0C6W2 |
| PEEEQEEDWLDDDSQQ | 1 | 0 | 0 | 1684978 | YP_009724389.1 | P0DTD1 |
| PEEFASHVANFQKSGY | 1 | 0 | 0 | 1684979 | YP_009047202.1 | K9N7C7 |
| PEEHVQIHTIDGSSGV | 1 | 0 | 0 | 1684981 | YP_009724391.1 | P0DTC3 |
| PEHSLAEYHNESGIKT | 1 | 0 | 0 | 1684995 | QIQ54046.1 | P0C6U8 |
| PEHSLAEYHNESGLKP | 1 | 0 | 0 | 1684996 | QHR63299.1 | P0C6W2 |
| PEHSLAEYHNESGLKT | 1 | 0 | 0 | 1684997 | YP_009724389.1 | P0DTD1 |
| PEIKILNNLGVDIAAN | 1 | 0 | 0 | 1685000 | NP_828849.2 | P0C6X7 |
| PEIPHFNQKLPNGVTV | 1 | 0 | 0 | 1685003 | NP_073556.1 | A0A0N7J131 |
| PEPEPTPEEPVNQFTG | 1 | 0 | 0 | 1685023 | NP_828849.2 | P0C6X7 |
| PEPISDKNVVFMNTCS | 1 | 0 | 0 | 1685024 | YP_173238.1 | A0A140H1H1 |
| PEQFDSLNLSAGTQHI | 1 | 0 | 0 | 1685029 | YP_173242.1 | Q19U29 |
| PEQPTQYEYGTESDYK | 1 | 0 | 0 | 1685030 | QIQ54046.1 | P0C6U8 |
| PESKPLADDDSAIIEI | 1 | 0 | 0 | 1685036 | YP_003771.1 | H9EJ69 |
| PETNNLMCIDMKGKMF | 1 | 0 | 0 | 1685044 | YP_173241.1 | Q0ZJ82 |
| PETNNLMCIDMKGTMY | 1 | 0 | 0 | 1685045 | YP_009555244.1 | Q01455 |
| PETTADIVVFDEISMA | 1 | 0 | 0 | 1685047 | YP_009724389.1 | P0DTD1 |
| PETYFTQSRDLEDFKP | 1 | 0 | 0 | 1685049 | NP_828849.2 | P0C6X7 |
| PEVSRELGNITKTNVQ | 1 | 0 | 0 | 1685062 | NP_073549.1 | A0A0P0LKV0 |
| PEWMDDFEADPKGKYA | 1 | 0 | 0 | 1685064 | YP_009047202.1 | K9N7C7 |
| PEYIDVNKTLQELSYK | 1 | 0 | 0 | 1685066 | NP_073551.1 | A0A0P0K6L9 |
| PFEIKLAKKFDNFTGE | 1 | 0 | 0 | 1685090 | QIQ54046.1 | P0C6U8 |
| PFFSFEQSHNNCYVNV | 1 | 0 | 0 | 1685101 | YP_173236.1 | A0A140H1G9 |
| PFGEVFNASKFASVYA | 1 | 0 | 0 | 1685111 | QIQ54048.1 | P59594 |
| PFGEVFNATKFPSVYA | 1 | 0 | 0 | 1685112 | NP_828851.1 | P59594 |
| PFGGTTVVRPLVEDST | 1 | 0 | 0 | 1685117 | YP_009047210.1 | K9N7A1 |
| PFHYERDNTSCPEWMD | 1 | 0 | 0 | 1685128 | YP_009047202.1 | K9N7C7 |
| PFIANEKFINGCGRNV | 1 | 0 | 0 | 1685130 | YP_003766.2 | H9EJ42 |
| PFITESKPSVEQRKQD | 1 | 0 | 0 | 1685132 | YP_009724389.1 | P0DTD1 |
| PFKDGIYFAATEKSNV | 1 | 0 | 0 | 1685135 | NP_828851.1 | P59594 |

| | | | | | | |
|---|---|---|---|---|---|---|
| PFMDNAINVGGTGLQY | 1 | 0 | 0 | 1685155 | YP_009047202.1 | K9N7C7 |
| PFMVNQLAYSSSANGS | 1 | 0 | 0 | 1685159 | YP_009047202.1 | K9N7C7 |
| PFNVSFVNDSIESVGG | 1 | 0 | 0 | 1685168 | YP_173238.1 | A0A140H1H1 |
| PFQQFGRDISDTTDAV | 1 | 0 | 0 | 1685169 | QIQ54048.1 | P59594 |
| PFTINCQEPKLGSLVV | 1 | 0 | 0 | 1685186 | YP_009724396.1 | P0DTC8 |
| PFTVNSVNDSLEPVGG | 1 | 0 | 0 | 1685191 | YP_009555241.1 | P36334 |
| PGAFFFGSKLDLVKRD | 1 | 0 | 0 | 1685215 | YP_173242.1 | Q19U29 |
| PGCDVNDVTKLYLGGM | 1 | 0 | 0 | 1685219 | YP_009555238.1 | P0C6X6 |
| PGDLQGNFGDLQLNKL | 1 | 0 | 0 | 1685228 | YP_009047211.1 | K9N4V7 |
| PGEFSKKQLGLLQPTD | 1 | 0 | 0 | 1685234 | YP_009047202.1 | K9N7C7 |
| PGFGVAVADSYYSYMM | 1 | 0 | 0 | 1685245 | YP_173236.1 | A0A140H1G9 |
| PGGDAISLSFNNLLGF | 1 | 0 | 0 | 1685252 | YP_009047202.1 | K9N7C7 |
| PGGTSSGDATTAFANS | 1 | 0 | 0 | 1685253 | YP_009555238.1 | P0C6X6 |
| PGGTSSGDATTAYANS | 1 | 0 | 0 | 1685254 | YP_009724389.1 | P0DTD1 |
| PGHFNKEFYDFLRSQG | 1 | 0 | 0 | 1685260 | NP_073549.1 | A0A0P0LKV0 |
| PGHYAGSRSSSGNRSG | 1 | 0 | 0 | 1685261 | YP_173242.1 | Q19U29 |
| PGKQSSSRASSGNRSG | 1 | 0 | 0 | 1685269 | YP_009555245.1 | P33469 |
| PGNFNKDFYDFAVSKG | 1 | 0 | 0 | 1685283 | YP_009724389.1 | P0DTD1 |
| PGNFNQDFYDFVLSKG | 1 | 0 | 0 | 1685284 | YP_009555238.1 | P0C6X6 |
| PGNFNQDFYDFVVSKG | 1 | 0 | 0 | 1685285 | YP_009047202.1 | K9N7C7 |
| PGNFNQDFYEFVKSKG | 1 | 0 | 0 | 1685286 | YP_173236.1 | A0A140H1G9 |
| PGNTFINGDIARELGN | 1 | 0 | 0 | 1685288 | YP_003766.2 | H9EJ42 |
| PGNVKPGDAVITTQQA | 1 | 0 | 0 | 1685292 | NP_073549.1 | A0A0P0LKV0 |
| PGQGCVGYTIEEAKTA | 1 | 0 | 0 | 1685297 | QIQ54046.1 | P0C6U8 |
| PGQGLNGYTVEEARTV | 1 | 0 | 0 | 1685299 | QHR63299.1 | P0C6W2 |
| PGQGVPLNANSTPAQN | 1 | 0 | 0 | 1685300 | YP_009047211.1 | K9N4V7 |
| PGQTGVIADYNYKLPD | 1 | 0 | 0 | 1685304 | NP_828851.1 | P59594 |
| PGSLNFKKVETLNFQM | 1 | 0 | 0 | 1685315 | YP_173243.1 | Q0ZJ80 |
| PGSSRGTSPARIAGNG | 1 | 0 | 0 | 1685319 | QIQ54056.1 | P59595 |
| PGSSRGTSPARMAGNG | 1 | 0 | 0 | 1685320 | YP_009724397.2 | P0DTC9 |
| PGTNASNQVAVLYQDV | 1 | 0 | 0 | 1685336 | QHR63300.2 | Q3LZX1 |
| PGTNTSNQVAVLYQDV | 1 | 0 | 0 | 1685338 | YP_009724390.1 | P0DTC2 |
| PGVFCGVDAVNLLTNM | 1 | 0 | 0 | 1685345 | YP_009724389.1 | P0DTD1 |
| PGVFCGVDSVNLLTNM | 1 | 0 | 0 | 1685346 | QIQ54046.1 | P0C6U8 |
| PGYMQGYDDCMQQGPA | 1 | 0 | 0 | 1685353 | YP_009047204.1 | K9N5Q8 |
| PGYNLKNGEVEFVYMH | 1 | 0 | 0 | 1685354 | NP_073549.1 | A0A0P0LKV0 |
| PGYTNSSTRGVYYPDK | 1 | 0 | 0 | 1685359 | QIQ54048.1 | P59594 |
| PHCGSSSRSFFEDLLF | 1 | 0 | 0 | 1685363 | YP_173238.1 | A0A140H1H1 |
| PHDAIVVDNDVVDYVS | 1 | 0 | 0 | 1685364 | NP_073549.1 | A0A0P0LKV0 |
| PHDVTVLRDNLKPVTT | 1 | 0 | 0 | 1685366 | YP_003768.1 | H9EJ24 |
| PHEFCSQHTLYIKDGD | 1 | 0 | 0 | 1685367 | YP_009047202.1 | K9N7C7 |
| PHEFCSQHTMQIVDKD | 1 | 0 | 0 | 1685372 | YP_003766.2 | H9EJ42 |
| PHKLKIQVVNSGSDMN | 1 | 0 | 0 | 1685390 | YP_173236.1 | A0A140H1G9 |
| PHLMGWDYPKCDRAMP | 1 | 0 | 0 | 1685393 | YP_009724389.1 | P0DTD1 |
| PHPYCYSDGVMKNASL | 1 | 0 | 0 | 1685403 | YP_173236.1 | A0A140H1G9 |
| PHQVNISLNGNTSVCV | 1 | 0 | 0 | 1685404 | YP_003767.1 | A0A0P0G321 |
| PIADPTHFDIEEVELL | 1 | 0 | 0 | 1685422 | NP_073549.1 | A0A0P0LKV0 |
| PIAFGVPPSEAKGYWY | 1 | 0 | 0 | 1685424 | YP_173242.1 | Q19U29 |
| PIAYRNVLLRKNGNKG | 1 | 0 | 0 | 1685429 | NP_828849.2 | P0C6X7 |
| PIDDFQVNWNTKHGSG | 1 | 0 | 0 | 1685436 | QIQ54046.1 | P0C6U8 |
| PIDIDAFPVKPICSSV | 1 | 0 | 0 | 1685439 | YP_003766.2 | H9EJ42 |
| PIDLVPNQPYPNASFD | 1 | 0 | 0 | 1685446 | YP_009724389.1 | P0DTD1 |
| PIDLVPTQPLPNASFD | 1 | 0 | 0 | 1685447 | NP_828849.2 | P0C6X7 |
| PIDLVPTQPYPNSNFD | 1 | 0 | 0 | 1685448 | QIQ54046.1 | P0C6U8 |
| PIDYNVVTQRMCAIGP | 1 | 0 | 0 | 1685452 | NP_073549.1 | A0A0P0LKV0 |
| PIEDYEQNWNTKHGSG | 1 | 0 | 0 | 1685453 | NP_828849.2 | P0C6X7 |
| PIGKGNKDEQIGYWNV | 1 | 0 | 0 | 1685472 | YP_003771.1 | H9EJ69 |
| PIGMFFPYDIELVTGK | 1 | 0 | 0 | 1685473 | YP_009047202.1 | K9N7C7 |
| PIGTNDPIRFCLENTV | 1 | 0 | 0 | 1685474 | NP_073549.1 | A0A0P0LKV0 |
| PIGTNYRSCESTTVLD | 1 | 0 | 0 | 1685475 | YP_173238.1 | A0A140H1H1 |
| PIGYVTHGFNLEEAAR | 1 | 0 | 0 | 1685476 | NP_828849.2 | P0C6X7 |
| PIIEVSQFQSKLTDVK | 1 | 0 | 0 | 1685480 | YP_009555238.1 | P0C6X6 |
| PIIKAQFKPFAKVDGV | 1 | 0 | 0 | 1685483 | YP_173236.1 | A0A140H1G9 |
| PIIKSSVKSYDDLVYS | 1 | 0 | 0 | 1685484 | YP_173236.1 | A0A140H1G9 |
| PIKDFGGFNFLQILPD | 1 | 0 | 0 | 1685492 | QIQ54048.1 | P59594 |
| PIKGVTFGEDTVWEVQ | 1 | 0 | 0 | 1685493 | NP_828849.2 | P0C6X7 |
| PILKSSVKSYDSLVYT | 1 | 0 | 0 | 1685506 | YP_009555238.1 | P0C6X6 |
| PILMVDKNLITTANTG | 1 | 0 | 0 | 1685509 | YP_009555238.1 | P0C6X6 |
| PILSCGIFGIKLETSL | 1 | 0 | 0 | 1685512 | NP_073549.1 | A0A0P0LKV0 |
| PILWVNKASYDTNLNK | 1 | 0 | 0 | 1685519 | YP_009047202.1 | K9N7C7 |
| PINATDRSHYYVDSVT | 1 | 0 | 0 | 1685522 | YP_009047202.1 | K9N7C7 |
| PINSNVKTLEYDYMTH | 1 | 0 | 0 | 1685530 | NP_073549.1 | A0A0P0LKV0 |
| PINTDTVLSVAPEVDW | 1 | 0 | 0 | 1685531 | YP_003766.2 | H9EJ42 |
| PINTNSGPDDQIGYYR | 1 | 0 | 0 | 1685532 | NP_828858.1 | P59595 |
| PINTNSTKDDQIGYYR | 1 | 0 | 0 | 1685534 | QIQ54056.1 | P59595 |
| PINVIVFDGKSKCDES | 1 | 0 | 0 | 1685535 | NP_828849.2 | P0C6X7 |
| PINVIVFDGKSKCEES | 1 | 0 | 0 | 1685536 | YP_009724389.1 | P0DTD1 |
| PIPDFDLDDFIDAPCY | 1 | 0 | 0 | 1685538 | YP_009047202.1 | K9N7C7 |
| PIQLSSYSLFDMSKFP | 1 | 0 | 0 | 1685541 | YP_009724389.1 | P0DTD1 |
| PIQPTDVAYHTVTDVK | 1 | 0 | 0 | 1685542 | YP_009555238.1 | P0C6X6 |
| PIQSLKVCIETVRTQV | 1 | 0 | 0 | 1685544 | QIQ54046.1 | P0C6U8 |
| PIQYIDIGNYTVSCSP | 1 | 0 | 0 | 1685548 | QHR63307.1 | Q3LZX5 |
| PISEHDYQIGGYTEKW | 1 | 0 | 0 | 1685555 | YP_009724391.1 | P0DTC3 |
| PISNDSNGCVVEASTD | 1 | 0 | 0 | 1685558 | YP_003766.2 | H9EJ42 |
| PITDHDYQIGGYTEKW | 1 | 0 | 0 | 1685564 | QIQ54049.1 | P59632 |
| PITENNVVVMSTCAVN | 1 | 0 | 0 | 1685568 | YP_009555241.1 | P36334 |
| PITFHMDGEVINIDSL | 1 | 0 | 0 | 1685570 | QIQ54046.1 | P0C6U8 |
| PITKNIGEYNVSKDGF | 1 | 0 | 0 | 1685573 | YP_009555238.1 | P0C6X6 |
| PITSLNTKYVAPQVTY | 1 | 0 | 0 | 1685575 | YP_009047204.1 | K9N5Q8 |
| PIVMNDKNICLLDQAW | 1 | 0 | 0 | 1685584 | YP_173236.1 | A0A140H1G9 |
| PIVWGVKDFNTLSQEG | 1 | 0 | 0 | 1685596 | YP_003766.2 | H9EJ42 |

| | | | | | | |
|---|---|---|---|---|---|---|
| PIVWHAKDFNSLSAEG | 1 | 0 | 0 | 1685597 | NP_073549.1 | A0A0P0LKV0 |
| PIVWHVNNATNKATYK | 1 | 0 | 0 | 1685599 | YP_009724389.1 | P0DTD1 |
| PIVWHVNNTINKATFK | 1 | 0 | 0 | 1685600 | QIQ54046.1 | P0C6U8 |
| PIVYKLDGVVCTEIDP | 1 | 0 | 0 | 1685601 | QIQ54046.1 | P0C6U8 |
| PIYKNGAMYKGKPILW | 1 | 0 | 0 | 1685603 | YP_009047202.1 | K9N7C7 |
| PIYLKGSGSDFSLADS | 1 | 0 | 0 | 1685604 | YP_009555238.1 | P0C6X6 |
| PKDHIGTRNPNNNAAI | 1 | 0 | 0 | 1685622 | QIQ54056.1 | P59595 |
| PKDHIGTRNPNNNAAT | 1 | 0 | 0 | 1685623 | NP_828858.1 | P59595 |
| PKDWRLINKFDVINGV | 1 | 0 | 0 | 1685627 | YP_173236.1 | A0A140H1G9 |
| PKEDGDVLLAEFDTYD | 1 | 0 | 0 | 1685628 | YP_009047202.1 | K9N7C7 |
| PKEEFVVKEKLNAFLV | 1 | 0 | 0 | 1685629 | NP_073549.1 | A0A0P0LKV0 |
| PKEETGLLMPLKAPKE | 1 | 0 | 0 | 1685631 | QHR63299.1 | P0C6W2 |
| PKEEVKPFITESKPSV | 1 | 0 | 0 | 1685632 | YP_009724389.1 | P0DTD1 |
| PKEEVKSSITESKLSV | 1 | 0 | 0 | 1685633 | QHR63299.1 | P0C6W2 |
| PKEIVDTVSALVYENK | 1 | 0 | 0 | 1685635 | YP_009555238.1 | P0C6X6 |
| PKEIVETVSALVYDNK | 1 | 0 | 0 | 1685636 | YP_173236.1 | A0A140H1G9 |
| PKELLLLWESGKAKPP | 1 | 0 | 0 | 1685638 | NP_073549.1 | A0A0P0LKV0 |
| PKELLLSWESGKVKPP | 1 | 0 | 0 | 1685639 | YP_003766.2 | H9EJ42 |
| PKEQIDGYTMHANYIF | 1 | 0 | 0 | 1685640 | NP_828849.2 | P0C6X7 |
| PKEQRVQGSITQRTRT | 1 | 0 | 0 | 1685641 | YP_009047211.1 | K9N4V7 |
| PKEVTFLEGDSHDTVL | 1 | 0 | 0 | 1685643 | NP_828849.2 | P0C6X7 |
| PKFFYASNGTYNCTDA | 1 | 0 | 0 | 1685644 | NP_073551.1 | A0A0P0K6L9 |
| PKFIEQISAFTKPSSI | 1 | 0 | 0 | 1685645 | YP_003771.1 | H9EJ69 |
| PKFVFPLNSKVKVIQP | 1 | 0 | 0 | 1685651 | NP_828849.2 | P0C6X7 |
| PKGKYSQKAYALLRQY | 1 | 0 | 0 | 1685660 | YP_173236.1 | A0A140H1G9 |
| PKGVGSANIFIGDKVG | 1 | 0 | 0 | 1685664 | YP_009555238.1 | P0C6X6 |
| PKHVFKTLKPGDSFNI | 1 | 0 | 0 | 1685669 | YP_003766.2 | H9EJ42 |
| PKIIKVFYELDNDFNT | 1 | 0 | 0 | 1685673 | YP_009555238.1 | P0C6X6 |
| PKKDKKKKTDESQPLP | 1 | 0 | 0 | 1685677 | QIQ54056.1 | P59595 |
| PKKEKKQKAPKEESTD | 1 | 0 | 0 | 1685678 | YP_009047211.1 | K9N4V7 |
| PKKPNKPLSQPRADKP | 1 | 0 | 0 | 1685680 | YP_003771.1 | H9EJ69 |
| PKKTYLDYIQQGGVPI | 1 | 0 | 0 | 1685683 | YP_173236.1 | A0A140H1G9 |
| PKKVKPCIEEVTTTLE | 1 | 0 | 0 | 1685684 | QIQ54046.1 | P0C6U8 |
| PKLDGYYKKDNAYFTE | 1 | 0 | 0 | 1685685 | QIQ54046.1 | P0C6U8 |
| PKLDGYYKKDNAYYTE | 1 | 0 | 0 | 1685686 | NP_828849.2 | P0C6X7 |
| PKLDNYYKKDNSYFTE | 1 | 0 | 0 | 1685687 | YP_009724389.1 | P0DTD1 |
| PKLEFANDTKIASQLG | 1 | 0 | 0 | 1685688 | YP_009047204.1 | K9N5Q8 |
| PKLKEDYQIGGYSEDR | 1 | 0 | 0 | 1685694 | NP_828852.2 | P59632 |
| PKLMGWDYPKCDRAMP | 1 | 0 | 0 | 1685695 | NP_073549.1 | A0A0P0LKV0 |
| PKLQASQAWQPGVAMP | 1 | 0 | 0 | 1685696 | NP_828849.2 | P0C6X7 |
| PKLSTDLIKNQCVNFN | 1 | 0 | 0 | 1685699 | NP_828851.1 | P59594 |
| PKNFCPCKLNGSCVGS | 1 | 0 | 0 | 1685703 | YP_009555241.1 | P36334 |
| PKNSFEALMLNFKLLG | 1 | 0 | 0 | 1685707 | YP_009555238.1 | P0C6X6 |
| PKNSIDAFKLNIKLLG | 1 | 0 | 0 | 1685708 | YP_009724389.1 | P0DTD1 |
| PKNSIDAFKLNVKLLG | 1 | 0 | 0 | 1685709 | QIQ54046.1 | P0C6U8 |
| PKQSIFTINSVVQKDG | 1 | 0 | 0 | 1685725 | YP_009047202.1 | K9N7C7 |
| PKRLRKKRNVDPLSNF | 1 | 0 | 0 | 1685727 | YP_009047202.1 | K9N7C7 |
| PKSGYFVNVNNTWMYT | 1 | 0 | 0 | 1685731 | YP_009555241.1 | P36334 |
| PKSILSPLYAFPSEAA | 1 | 0 | 0 | 1685732 | QIQ54046.1 | P0C6U8 |
| PKSRVQQNKSRELTAE | 1 | 0 | 0 | 1685733 | YP_009555245.1 | P33469 |
| PKSSIDAFKLNIKLLG | 1 | 0 | 0 | 1685734 | NP_828849.2 | P0C6X7 |
| PKTKNVTKENDSKEGF | 1 | 0 | 0 | 1685739 | YP_009724389.1 | P0DTD1 |
| PKVEIDGNVMHANYLF | 1 | 0 | 0 | 1685744 | YP_009555238.1 | P0C6X6 |
| PKYTFGNVKPGETFTV | 1 | 0 | 0 | 1685757 | YP_173236.1 | A0A140H1G9 |
| PLAELKCSVKSFEIDK | 1 | 0 | 0 | 1685765 | NP_828851.1 | P59594 |
| PLAVDVVSIRSSNQGN | 1 | 0 | 0 | 1685768 | YP_009047207.1 | K9N643 |
| PLCASNKLRVVIPDFT | 1 | 0 | 0 | 1685774 | YP_009047202.1 | K9N7C7 |
| PLCGSGNLVEKLSPWF | 1 | 0 | 0 | 1685777 | YP_009047202.1 | K9N7C7 |
| PLCIFNGKFLSNTKYY | 1 | 0 | 0 | 1685778 | YP_009555240.1 | P30215 |
| PLCSAGNIFGLYKNSA | 1 | 0 | 0 | 1685784 | YP_003766.2 | H9EJ42 |
| PLDYKRQNNLAIEEIE | 1 | 0 | 0 | 1685791 | NP_073549.1 | A0A0P0LKV0 |
| PLEEPTAQAVEVPLVG | 1 | 0 | 0 | 1685795 | QIQ54046.1 | P0C6U8 |
| PLEGGGWLVASGSTVA | 1 | 0 | 0 | 1685799 | YP_009047204.1 | K9N5Q8 |
| PLEPKFSIALPPELSV | 1 | 0 | 0 | 1685802 | YP_003771.1 | H9EJ69 |
| PLEQPTSEAVEAPLVG | 1 | 0 | 0 | 1685803 | YP_009724389.1 | P0DTD1 |
| PLERYGFHPTTGVNYQ | 1 | 0 | 0 | 1685805 | QIQ54048.1 | P59594 |
| PLFLFDEAGEEVLAPK | 1 | 0 | 0 | 1685811 | YP_009555238.1 | P0C6X6 |
| PLGMSLEACYVRGCNP | 1 | 0 | 0 | 1685820 | YP_009555238.1 | P0C6X6 |
| PLGYVTHGLNLEEAAR | 1 | 0 | 0 | 1685824 | YP_009724389.1 | P0DTD1 |
| PLHSLGGFPKGYVMGL | 1 | 0 | 0 | 1685828 | YP_173236.1 | A0A140H1G9 |
| PLIELCVDEAGSKSPI | 1 | 0 | 0 | 1685831 | YP_009724396.1 | P0DTC8 |
| PLIEWPLTEYDFTDYK | 1 | 0 | 0 | 1685834 | YP_009047202.1 | K9N7C7 |
| PLKAPKEVTFLEGDSH | 1 | 0 | 0 | 1685844 | NP_828849.2 | P0C6X7 |
| PLKLAGTAVINLRADQ | 1 | 0 | 0 | 1685846 | YP_009555238.1 | P0C6X6 |
| PLKVGAGVMLECEKFN | 1 | 0 | 0 | 1685854 | YP_003766.2 | H9EJ42 |
| PLLEVYVHLRNWNFSW | 1 | 0 | 0 | 1685862 | YP_003770.1 | U3M6Q8 |
| PLLKQKDIGIDTAGVL | 1 | 0 | 0 | 1685866 | YP_173243.1 | Q0ZJ80 |
| PLLNPSALEFNPSQTS | 1 | 0 | 0 | 1685869 | NP_073556.1 | A0A0N7J131 |
| PLLSAGIFGADPIHSL | 1 | 0 | 0 | 1685872 | YP_009724389.1 | P0DTD1 |
| PLLSENQISGYTLAAT | 1 | 0 | 0 | 1685877 | YP_009555241.1 | P36334 |
| PLLSEYPLLKQKDIGI | 1 | 0 | 0 | 1685878 | YP_173243.1 | Q0ZJ80 |
| PLNAIPSLAANTLNII | 1 | 0 | 0 | 1685901 | YP_009555238.1 | P0C6X6 |
| PLNANSTPAQNAGYWR | 1 | 0 | 0 | 1685902 | YP_009047211.1 | K9N4V7 |
| PLNSIIKTIQPRVEKK | 1 | 0 | 0 | 1685916 | YP_009724389.1 | P0DTD1 |
| PLNSKVKVIQPRVEKK | 1 | 0 | 0 | 1685917 | NP_828849.2 | P0C6X7 |
| PLNSTIKTIQPRVEKK | 1 | 0 | 0 | 1685918 | QHR63299.1 | P0C6W2 |
| PLPNASFDNFKLTCSN | 1 | 0 | 0 | 1685925 | NP_828849.2 | P0C6X7 |
| PLQASLPFGWLVIGVA | 1 | 0 | 0 | 1685929 | NP_828852.2 | P59632 |
| PLRAPTIANQEKYSSI | 1 | 0 | 0 | 1685943 | YP_003766.2 | H9EJ42 |
| PLRAPTMANQEKYSTI | 1 | 0 | 0 | 1685944 | NP_073549.1 | A0A0P0LKV0 |
| PLRASSCVTRCNIGGA | 1 | 0 | 0 | 1685945 | YP_003766.2 | H9EJ42 |

| | | | | | | |
|---|---|---|---|---|---|---|
| PLRATNCITKCNIGGA | 1 | 0 | 0 | 1685946 | NP_073549.1 | A0A0P0LKV0 |
| PLRGTIVTRPLMESEL | 1 | 0 | 0 | 1685947 | NP_828855.1 | P59596 |
| PLSAPTLVPQEHYVRI | 1 | 0 | 0 | 1685952 | YP_009724389.1 | P0DTD1 |
| PLSAQTGIAVLDMCAA | 1 | 0 | 0 | 1685953 | NP_828849.2 | P0C6X7 |
| PLSDMEKDFLSFDSDV | 1 | 0 | 0 | 1685955 | YP_009047202.1 | K9N7C7 |
| PLSKDLPDDVVAANMF | 1 | 0 | 0 | 1685960 | YP_173236.1 | A0A140H1G9 |
| PLSQPRADKPSQLKKP | 1 | 0 | 0 | 1685966 | YP_003771.1 | H9EJ69 |
| PLSVQQAQQEATLPDI | 1 | 0 | 0 | 1685971 | NP_073549.1 | A0A0P0LKV0 |
| PLTCNAISSNTDNETL | 1 | 0 | 0 | 1685973 | YP_173238.1 | A0A140H1H1 |
| PLTENKYSQLDEEQPM | 1 | 0 | 0 | 1685975 | QIQ54052.1 | P59634 |
| PLTKKNYSELDDEEPM | 1 | 0 | 0 | 1685985 | NP_828856.1 | P59634 |
| PLTKNIGDYNVSKDGF | 1 | 0 | 0 | 1685986 | YP_173236.1 | A0A140H1G9 |
| PLTPILSCGIFGIKLE | 1 | 0 | 0 | 1685988 | NP_073549.1 | A0A0P0LKV0 |
| PLVYHENEEYQKVFRV | 1 | 0 | 0 | 1686024 | YP_009555238.1 | P0C6X6 |
| PLVYLNHSVPKLSDFE | 1 | 0 | 0 | 1686025 | YP_173238.1 | A0A140H1H1 |
| PLYPYGRCPTAADINI | 1 | 0 | 0 | 1686030 | YP_009555240.1 | P30215 |
| PMHPMVNPSKGSSGLL | 1 | 0 | 0 | 1686046 | YP_173243.1 | Q0ZJ80 |
| PMIANENAINKGGRTV | 1 | 0 | 0 | 1686047 | NP_073549.1 | A0A0P0LKV0 |
| PMLMGWDYPKCDRALP | 1 | 0 | 0 | 1686051 | YP_003766.2 | H9EJ42 |
| PMVNPSKGSSGLLITK | 1 | 0 | 0 | 1686063 | YP_173243.1 | Q0ZJ80 |
| PNALLCATCPFAKGDS | 1 | 0 | 0 | 1686067 | YP_009047202.1 | K9N7C7 |
| PNCINCLDDRCILHCA | 1 | 0 | 0 | 1686074 | NP_828849.2 | P0C6X7 |
| PNDDVTSNVTQCFGPR | 1 | 0 | 0 | 1686082 | NP_073556.1 | A0A0N7J131 |
| PNHTIKGSFLNGSCGS | 1 | 0 | 0 | 1686110 | NP_828849.2 | P0C6X7 |
| PNITNLCPFGEVFNAS | 1 | 0 | 0 | 1686115 | QIQ54048.1 | P59594 |
| PNITNLCPFGEVFNAT | 1 | 0 | 0 | 1686116 | YP_009724390.1 | P0DTC2 |
| PNKHCNVQQCFGKRGP | 1 | 0 | 0 | 1686117 | YP_173242.1 | Q19U29 |
| PNKPLSQPRADKPSQL | 1 | 0 | 0 | 1686119 | YP_003771.1 | H9EJ69 |
| PNLNAGEDIQLLKAAY | 1 | 0 | 0 | 1686125 | NP_828849.2 | P0C6X7 |
| PNLNSLCLLNHKEILS | 1 | 0 | 0 | 1686127 | YP_173243.1 | Q0ZJ80 |
| PNLYKMQRMLLDKCDL | 1 | 0 | 0 | 1686133 | QIQ54047.1 | P0C6X7 |
| PNLYVVVVESDADRAN | 1 | 0 | 0 | 1686135 | YP_173236.1 | A0A140H1G9 |
| PNMFITREEAIKHVRA | 1 | 0 | 0 | 1686138 | QIQ54047.1 | P0C6X7 |
| PNNDSAIVTQFAPGTK | 1 | 0 | 0 | 1686148 | YP_009047211.1 | K9N4V7 |
| PNNNAATVLQLPQGTT | 1 | 0 | 0 | 1686150 | NP_828858.1 | P59595 |
| PNNNSAQYVFSTVNAL | 1 | 0 | 0 | 1686151 | NP_073549.1 | A0A0P0LKV0 |
| PNNRSLSRSNSNFRHS | 1 | 0 | 0 | 1686152 | YP_173242.1 | Q19U29 |
| PNNTDFSRVSAKPPPG | 1 | 0 | 0 | 1686154 | YP_009724389.1 | P0DTD1 |
| PNQPYPNASFDNFKFV | 1 | 0 | 0 | 1686163 | YP_009724389.1 | P0DTD1 |
| PNSNFDNFKFVCDNTK | 1 | 0 | 0 | 1686168 | QIQ54046.1 | P0C6U8 |
| PNSRSTSRTSSRASSA | 1 | 0 | 0 | 1686170 | YP_009555245.1 | P33469 |
| PNTVDCDNDRCIIHCA | 1 | 0 | 0 | 1686177 | YP_173236.1 | A0A140H1G9 |
| PNTVDCQDDRCIIHCA | 1 | 0 | 0 | 1686178 | YP_009555238.1 | P0C6X6 |
| PNVNKGEDIQLLKSAY | 1 | 0 | 0 | 1686188 | YP_009724389.1 | P0DTD1 |
| PNVNRGEDIQLLKSAY | 1 | 0 | 0 | 1686189 | QHR63299.1 | P0C6W2 |
| PNVQIHTIDGSSGVAN | 1 | 0 | 0 | 1686190 | NP_828852.2 | P59632 |
| PNVTINCQDPAGGALI | 1 | 0 | 0 | 1686191 | NP_849177.1 | Q80H93 |
| PNYTVPDLVVEQYNQT | 1 | 0 | 0 | 1686199 | NP_073551.1 | A0A0P0K6L9 |
| PPALHLVDPQIQLTIT | 1 | 0 | 0 | 1686202 | NP_828859.1 | P59636 |
| PPALQDAYYRARAGDA | 1 | 0 | 0 | 1686203 | QIQ54046.1 | P0C6U8 |
| PPALQDAYYRARAGEA | 1 | 0 | 0 | 1686204 | YP_009724389.1 | P0DTD1 |
| PPAQYELKHGTFTCAS | 1 | 0 | 0 | 1686206 | YP_009724389.1 | P0DTD1 |
| PPAYTNSSTRGVYYPD | 1 | 0 | 0 | 1686208 | QHR63300.2 | Q3LZX1 |
| PPCKFTVQDAKGLKIK | 1 | 0 | 0 | 1686211 | YP_009555238.1 | P0C6X6 |
| PPDAIIIDNDINDYVS | 1 | 0 | 0 | 1686215 | YP_003766.2 | H9EJ42 |
| PPDEDEEEGDCEEEEF | 1 | 0 | 0 | 1686217 | YP_009724389.1 | P0DTD1 |
| PPGQGVPLNANSTPAQ | 1 | 0 | 0 | 1686231 | YP_009047211.1 | K9N4V7 |
| PPGYTNSSTRGVYYPD | 1 | 0 | 0 | 1686240 | QIQ54048.1 | P59594 |
| PPHDVTVLRDNLKPVT | 1 | 0 | 0 | 1686241 | YP_003768.1 | H9EJ24 |
| PPIKDFGGFNFLQILP | 1 | 0 | 0 | 1686244 | QIQ54048.1 | P59594 |
| PPILSESQISGYTTAA | 1 | 0 | 0 | 1686245 | YP_173238.1 | A0A140H1H1 |
| PPISFPLCANGQVFGL | 1 | 0 | 0 | 1686246 | YP_009724389.1 | P0DTD1 |
| PPKEQRVQGSITQRTR | 1 | 0 | 0 | 1686249 | YP_009047211.1 | K9N4V7 |
| PPKNSIDAFKLNIKLL | 1 | 0 | 0 | 1686251 | YP_009724389.1 | P0DTD1 |
| PPKNSIDAFKLNVKLL | 1 | 0 | 0 | 1686252 | QIQ54046.1 | P0C6U8 |
| PPKSSIDAFKLNIKLL | 1 | 0 | 0 | 1686253 | NP_828849.2 | P0C6X7 |
| PPLLSENQISGYTLAA | 1 | 0 | 0 | 1686259 | YP_009555241.1 | P36334 |
| PPLLTDDMIAAYTAAL | 1 | 0 | 0 | 1686260 | NP_828851.1 | P59594 |
| PPNVQIHTIDGSSGVA | 1 | 0 | 0 | 1686274 | NP_828852.2 | P59632 |
| PPPGEQFKHLVPLMHK | 1 | 0 | 0 | 1686276 | YP_009047202.1 | K9N7C7 |
| PPQVVKLPSAPQTIQP | 1 | 0 | 0 | 1686281 | YP_009047202.1 | K9N7C7 |
| PPRAELNGVVVDKVGD | 1 | 0 | 0 | 1686283 | YP_009555238.1 | P0C6X6 |
| PPSASCPIGTNYRSCE | 1 | 0 | 0 | 1686285 | YP_173238.1 | A0A140H1H1 |
| PPTEPKKDKKKKTDEA | 1 | 0 | 0 | 1686291 | NP_828858.1 | P59595 |
| PPTEPKKDKKKKTDES | 1 | 0 | 0 | 1686292 | QIQ54056.1 | P59595 |
| PPWTAAAGVPFYLNVQ | 1 | 0 | 0 | 1686306 | YP_009555241.1 | P36334 |
| PQADTDWKFYDAQPCS | 1 | 0 | 0 | 1686309 | QIQ54047.1 | P0C6X7 |
| PQADVEWKFYDAQPCS | 1 | 0 | 0 | 1686310 | YP_009724389.1 | P0DTD1 |
| PQAEVEWKFYDAQPCS | 1 | 0 | 0 | 1686311 | NP_828849.2 | P0C6X7 |
| PQAMQWNSTTFHQTL | 1 | 0 | 0 | 1686313 | P03138.3 | Q76R62 |
| PQEKDKKSAKTGTPKP | 1 | 0 | 0 | 1686319 | NP_073556.1 | A0A0N7J131 |
| PQFTSDQAVTFLKEWN | 1 | 0 | 0 | 1686326 | YP_173241.1 | Q0ZJ82 |
| PQFYIYDEEGGYDVSK | 1 | 0 | 0 | 1686327 | YP_003766.2 | H9EJ42 |
| PQGNLGGNCAGNVIGN | 1 | 0 | 0 | 1686333 | YP_173236.1 | A0A140H1G9 |
| PQGNLGSNGKPGNVGG | 1 | 0 | 0 | 1686334 | YP_009555238.1 | P0C6X6 |
| PQGTALPKGFYAEGSR | 1 | 0 | 0 | 1686337 | QIQ54056.1 | P59595 |
| PQGYYIEGSGRSAPNS | 1 | 0 | 0 | 1686338 | YP_009555245.1 | P33469 |
| PQIITTDNTFVSGSCD | 1 | 0 | 0 | 1686346 | QHR63300.2 | Q3LZX1 |
| PQKAYLDFVNAGGAPL | 1 | 0 | 0 | 1686351 | YP_009047202.1 | K9N7C7 |
| PQNQRNAPRITFGGPS | 1 | 0 | 0 | 1686370 | YP_009724397.2 | P0DTC9 |
| PQPEEELEEDWLDDDS | 1 | 0 | 0 | 1686371 | QHR63299.1 | P0C6W2 |

| | | | | | | |
|---|---|---|---|---|---|---|
| PQRADLNGVIVDKVGE | 1 | 0 | 0 | 1686379 | YP_173236.1 | A0A140H1G9 |
| PQRKRGVKQLPEQFDS | 1 | 0 | 0 | 1686381 | YP_173242.1 | Q19U29 |
| PQRQRGHKNGQGENDN | 1 | 0 | 0 | 1686385 | YP_009555245.1 | P33469 |
| PQSGGGKCFNCYPAGV | 1 | 0 | 0 | 1686388 | NP_073551.1 | A0A0P0K6L9 |
| PQSKDSNFLNESGVLL | 1 | 0 | 0 | 1686389 | YP_009047203.1 | K9N7C7 |
| PQSKDSNFLNRVRGSI | 1 | 0 | 0 | 1686390 | YP_009047202.1 | K9N7C7 |
| PQSNQRSAPRITFGGP | 1 | 0 | 0 | 1686392 | NP_828858.1 | P59595 |
| PQVNGLTSIKWADNNC | 1 | 0 | 0 | 1686402 | YP_009724389.1 | P0DTD1 |
| PQVTYQNISTNLPPPL | 1 | 0 | 0 | 1686403 | YP_009047204.1 | K9N5Q8 |
| PQVVKLPSAPQTIQPE | 1 | 0 | 0 | 1686404 | YP_009047202.1 | K9N7C7 |
| PQXRAPRITFGGPSDS | 1 | 0 | 0 | 1686406 | QIQ54056.1 | P59595 |
| PRAELNGVVVDKVGDT | 1 | 0 | 0 | 1686411 | YP_009555238.1 | P0C6X6 |
| PRASANIGCNHTGVVG | 1 | 0 | 0 | 1686414 | YP_009724389.1 | P0DTD1 |
| PRAVSFADNNDITNTN | 1 | 0 | 0 | 1686415 | YP_009047211.1 | K9N4V7 |
| PRGQGVPINTNSSPDD | 1 | 0 | 0 | 1686429 | YP_009724397.2 | P0DTC9 |
| PRGQGVPINTNSTKDD | 1 | 0 | 0 | 1686430 | QIQ54056.1 | P59595 |
| PRHVICSSSNMNEPDY | 1 | 0 | 0 | 1686437 | YP_173236.1 | A0A140H1G9 |
| PRHVICTAEDMLNPNY | 1 | 0 | 0 | 1686438 | NP_828849.2 | P0C6X7 |
| PRITFGGPSDSTDNNQ | 1 | 0 | 0 | 1686444 | QIQ54056.1 | P59595 |
| PRLQAASDWKPGYSMP | 1 | 0 | 0 | 1686451 | YP_009555238.1 | P0C6X6 |
| PRLQASADWKPGHAMP | 1 | 0 | 0 | 1686452 | YP_009047202.1 | K9N7C7 |
| PRNSSDNGISAIITAN | 1 | 0 | 0 | 1686458 | YP_003767.1 | A0A0P0G321 |
| PRNSWEAMALNFKLIG | 1 | 0 | 0 | 1686459 | YP_009047202.1 | K9N7C7 |
| PRPIDVSKADGIIYPQ | 1 | 0 | 0 | 1686463 | YP_009047204.1 | K9N5Q8 |
| PRQKRTATKAYNVTQA | 1 | 0 | 0 | 1686466 | YP_009724397.2 | P0DTC9 |
| PRQKRTATKQYNVTQA | 1 | 0 | 0 | 1686467 | QHR63308.1 | Q3LZX4 |
| PRRNVATLQAENVTGL | 1 | 0 | 0 | 1686470 | YP_009724389.1 | P0DTD1 |
| PRSDMEYDFLNMDMGV | 1 | 0 | 0 | 1686473 | YP_003766.2 | H9EJ42 |
| PRSQMETDFLELAMDE | 1 | 0 | 0 | 1686476 | NP_828849.2 | P0C6X7 |
| PRTGKHEHSLLVEAYN | 1 | 0 | 0 | 1686481 | YP_003766.2 | H9EJ42 |
| PRTIKVSTVQSKLTDL | 1 | 0 | 0 | 1686482 | NP_073549.1 | A0A0P0LKV0 |
| PRTINSTQDGDNKLQG | 1 | 0 | 0 | 1686483 | YP_009555241.1 | P36334 |
| PRTKHVTKENDSKEGF | 1 | 0 | 0 | 1686484 | NP_828849.2 | P0C6X7 |
| PRTLLTKGTLEPEYFN | 1 | 0 | 0 | 1686485 | YP_009724389.1 | P0DTD1 |
| PRVEAPKQEEPPNTED | 1 | 0 | 0 | 1686487 | NP_828849.2 | P0C6X7 |
| PRVEKKKLDGFMGRIR | 1 | 0 | 0 | 1686488 | YP_009724389.1 | P0DTD1 |
| PRVEKKKLEGFMGRIR | 1 | 0 | 0 | 1686489 | QIQ54046.1 | P0C6U8 |
| PRVEKKKTEGFMGRIR | 1 | 0 | 0 | 1686490 | NP_828849.2 | P0C6X7 |
| PRWKRQPNDDVTSNVT | 1 | 0 | 0 | 1686497 | NP_073556.1 | A0A0N7J131 |
| PSALEFNPSQTSPATA | 1 | 0 | 0 | 1686508 | NP_073556.1 | A0A0N7J131 |
| PSARIVYTACSHAAVD | 1 | 0 | 0 | 1686510 | YP_009724389.1 | P0DTD1 |
| PSASAFFGMSRIGMEV | 1 | 0 | 0 | 1686511 | YP_009724397.2 | P0DTC9 |
| PSASCPIGTNYRSCES | 1 | 0 | 0 | 1686512 | YP_173238.1 | A0A140H1H1 |
| PSDFAFNNWFLLTNTS | 1 | 0 | 0 | 1686518 | NP_073551.1 | A0A0P0K6L9 |
| PSDSTDNNQNGDRSGA | 1 | 0 | 0 | 1686524 | QIQ54056.1 | P59595 |
| PSDVSLTYLLGTAKKQ | 1 | 0 | 0 | 1686527 | YP_009555238.1 | P0C6X6 |
| PSEDFIKHVRTNGGYN | 1 | 0 | 0 | 1686536 | YP_009047202.1 | K9N7C7 |
| PSEKDKHLDNNVQRCT | 1 | 0 | 0 | 1686540 | NP_073549.1 | A0A0P0LKV0 |
| PSFASSCKSHKPPSAS | 1 | 0 | 0 | 1686541 | YP_173238.1 | A0A140H1H1 |
| PSFKDNVILLNKHIDA | 1 | 0 | 0 | 1686542 | QIQ54056.1 | P59595 |
| PSFLAVDDKYKATGDL | 1 | 0 | 0 | 1686545 | YP_009555238.1 | P0C6X6 |
| PSFLAVDDKYKVNENL | 1 | 0 | 0 | 1686546 | YP_173236.1 | A0A140H1G9 |
| PSFNVSDAYANLVPYY | 1 | 0 | 0 | 1686548 | NP_073549.1 | A0A0P0LKV0 |
| PSFVAYETARQEYENA | 1 | 0 | 0 | 1686549 | NP_073549.1 | A0A0P0LKV0 |
| PSGIGAVGGDLLYLDL | 1 | 0 | 0 | 1686558 | YP_009047211.1 | K9N4V7 |
| PSGTYEGNSPFHPLAD | 1 | 0 | 0 | 1686569 | NP_828857.1 | P59635 |
| PSGVYQCAMRPNHTIK | 1 | 0 | 0 | 1686572 | NP_828849.2 | P0C6X7 |
| PSIISNEKQEILGTVS | 1 | 0 | 0 | 1686576 | YP_009724389.1 | P0DTD1 |
| PSINDLSFDKNLGRCV | 1 | 0 | 0 | 1686579 | YP_003766.2 | H9EJ42 |
| PSKLLNGIAYTTFSGS | 1 | 0 | 0 | 1686587 | YP_003766.2 | H9EJ42 |
| PSLAANTLNIIVPDKS | 1 | 0 | 0 | 1686588 | YP_009555238.1 | P0C6X6 |
| PSLCNNGKISSSAGDS | 1 | 0 | 0 | 1686590 | YP_173237.1 | A0A140H1H0 |
| PSMDGYCKFKGKCVQV | 1 | 0 | 0 | 1686604 | YP_003766.2 | H9EJ42 |
| PSRSQSRSQSRGRGES | 1 | 0 | 0 | 1686627 | NP_073556.1 | A0A0N7J131 |
| PSSDEAIPTRFPPGTV | 1 | 0 | 0 | 1686628 | YP_009555245.1 | P33469 |
| PSSIKEMQSQSSHVAQ | 1 | 0 | 0 | 1686630 | YP_003771.1 | H9EJ69 |
| PSSSGTSTPKKPNKPL | 1 | 0 | 0 | 1686638 | YP_003771.1 | H9EJ69 |
| PSTFGTRNPNNDSAIV | 1 | 0 | 0 | 1686642 | YP_009047211.1 | K9N4V7 |
| PSTLTPRSVRSVPGEM | 1 | 0 | 0 | 1686644 | YP_009047204.1 | K9N5Q8 |
| PSTPAYTFTTVKPGAA | 1 | 0 | 0 | 1686645 | YP_009047202.1 | K9N7C7 |
| PSTQKGSGMDTALLRN | 1 | 0 | 0 | 1686646 | YP_009555244.1 | Q01455 |
| PSTTVLIDYDHAYSTM | 1 | 0 | 0 | 1686652 | YP_003766.2 | H9EJ42 |
| PSTVWEDGDYYRKQLS | 1 | 0 | 0 | 1686653 | YP_009047204.1 | K9N5Q8 |
| PSTWNKRFGFIEDSVF | 1 | 0 | 0 | 1686654 | YP_009555241.1 | P36334 |
| PSYAAFATAQEAYEQA | 1 | 0 | 0 | 1686666 | YP_009724389.1 | P0DTD1 |
| PSYAAYATAQEAYEQA | 1 | 0 | 0 | 1686667 | NP_828849.2 | P0C6X7 |
| PSYCNGVRELYLNSSN | 1 | 0 | 0 | 1686668 | NP_828849.2 | P0C6X7 |
| PSYCTGYREGYLNSTN | 1 | 0 | 0 | 1686669 | YP_009724389.1 | P0DTD1 |
| PSYCTSYRDGYLNSTN | 1 | 0 | 0 | 1686670 | QIQ54046.1 | P0C6U8 |
| PSYIAYENARQAYEDA | 1 | 0 | 0 | 1686671 | YP_003766.2 | H9EJ42 |
| PTAADINIPDLPICVY | 1 | 0 | 0 | 1686675 | YP_009555240.1 | P30215 |
| PTARVVYTACSHAAVD | 1 | 0 | 0 | 1686681 | YP_009047202.1 | K9N7C7 |
| PTDNSTTVSVTVSSNM | 1 | 0 | 0 | 1686691 | YP_009047202.1 | K9N7C7 |
| PTDQSSYVVDSVAVKN | 1 | 0 | 0 | 1686695 | QIQ54046.1 | P0C6U8 |
| PTEHVTSVNLFDVGYS | 1 | 0 | 0 | 1686703 | YP_009047205.1 | K9N796 |
| PTEVLSEEVILKTGEL | 1 | 0 | 0 | 1686713 | QIQ54046.1 | P0C6U8 |
| PTEVLTEEVVLKTGDL | 1 | 0 | 0 | 1686714 | YP_009724389.1 | P0DTD1 |
| PTFSMAPVEFYEDRIL | 1 | 0 | 0 | 1686718 | YP_009047202.1 | K9N7C7 |
| PTGTFTVVMRPNYTIK | 1 | 0 | 0 | 1686725 | YP_009047202.1 | K9N7C7 |
| PTGTLLVDSDLNDFVS | 1 | 0 | 0 | 1686726 | YP_009724389.1 | P0DTD1 |
| PTGVHAGTDLEGKFYG | 1 | 0 | 0 | 1686727 | NP_828849.2 | P0C6X7 |

| | | | | | | |
|---|---|---|---|---|---|---|
| PTGVHAGTDLEGNFYG | 1 | 0 | 0 | 1686728 | YP_009724389.1 | P0DTD1 |
| PTGVHAGTDLEGTFYG | 1 | 0 | 0 | 1686729 | QHR63299.1 | P0C6W2 |
| PTGYGVRRKNSEPEIP | 1 | 0 | 0 | 1686732 | NP_073556.1 | A0A0N7J131 |
| PTIQKEVIECDVKTTE | 1 | 0 | 0 | 1686749 | NP_828849.2 | P0C6X7 |
| PTITQMNLKYAISAKN | 1 | 0 | 0 | 1686754 | YP_009724389.1 | P0DTD1 |
| PTIVSEKISVMDKLDT | 1 | 0 | 0 | 1686757 | YP_003766.2 | H9EJ42 |
| PTKALCFDKSKQFVPV | 1 | 0 | 0 | 1686761 | YP_173237.1 | A0A140H1H0 |
| PTKEEPKKVKPCIEEV | 1 | 0 | 0 | 1686762 | QIQ54046.1 | P0C6U8 |
| PTLNISDEFSSNVANY | 1 | 0 | 0 | 1686777 | YP_009724389.1 | P0DTD1 |
| PTLTQMNLKYAISAKN | 1 | 0 | 0 | 1686778 | YP_009555238.1 | P0C6X6 |
| PTLVPQENYASIRFSS | 1 | 0 | 0 | 1686780 | YP_173236.1 | A0A140H1G9 |
| PTLVPQENYSSIRFAS | 1 | 0 | 0 | 1686781 | YP_009555238.1 | P0C6X6 |
| PTMLDIGQARVAYQVA | 1 | 0 | 0 | 1686787 | NP_073549.1 | A0A0P0LKV0 |
| PTMTQLNLKYAISGKE | 1 | 0 | 0 | 1686788 | NP_073549.1 | A0A0P0LKV0 |
| PTMTQMNLKYAISAKN | 1 | 0 | 0 | 1686789 | YP_009047202.1 | K9N7C7 |
| PTNAYMRTTHTDIGVY | 1 | 0 | 0 | 1686792 | YP_009047202.1 | K9N7C7 |
| PTNFTIVGQEEFIQTN | 1 | 0 | 0 | 1686797 | YP_173238.1 | A0A140H1H1 |
| PTNIAFELFAKRKVGL | 1 | 0 | 0 | 1686799 | NP_073549.1 | A0A0P0LKV0 |
| PTNVAFELFAKRKMGL | 1 | 0 | 0 | 1686802 | YP_003766.2 | H9EJ42 |
| PTPEEPVNQFTGYLKL | 1 | 0 | 0 | 1686806 | NP_828849.2 | P0C6X7 |
| PTPGAFFFGSKLDLVK | 1 | 0 | 0 | 1686807 | YP_173242.1 | Q19U29 |
| PTPVVQTEVNSFSGYL | 1 | 0 | 0 | 1686809 | QHR63299.1 | P0C6W2 |
| PTQCYYNNSNNGKIVY | 1 | 0 | 0 | 1686814 | YP_009555238.1 | P0C6X6 |
| PTQPLPNASFDNFKLT | 1 | 0 | 0 | 1686816 | NP_828849.2 | P0C6X7 |
| PTQTVDSSQGSEYDYV | 1 | 0 | 0 | 1686818 | YP_009724389.1 | P0DTD1 |
| PTQYEYGTEDDYQGKS | 1 | 0 | 0 | 1686819 | QHR63299.1 | P0C6W2 |
| PTQYKDVEAWSGLCVD | 1 | 0 | 0 | 1686821 | NP_073551.1 | A0A0P0K6L9 |
| PTSEEVVENPTIQKEV | 1 | 0 | 0 | 1686826 | NP_828849.2 | P0C6X7 |
| PTSGSRVAGRSAIEDI | 1 | 0 | 0 | 1686828 | NP_073551.1 | A0A0P0K6L9 |
| PTTCANDPVGFTLKNT | 1 | 0 | 0 | 1686834 | YP_009724389.1 | P0DTD1 |
| PTTCANDPVGFTLRNT | 1 | 0 | 0 | 1686835 | NP_828849.2 | P0C6X7 |
| PTTKNVTGSNESKALF | 1 | 0 | 0 | 1686846 | YP_009047202.1 | K9N7C7 |
| PTVKEIISMPKIIKVF | 1 | 0 | 0 | 1686858 | YP_009555238.1 | P0C6X6 |
| PTYLDGADVTKIKPHA | 1 | 0 | 0 | 1686870 | QIQ54046.1 | P0C6U8 |
| PTYLDGADVTKIKPHN | 1 | 0 | 0 | 1686871 | YP_009724389.1 | P0DTD1 |
| PTYLDGADVTKIKPHV | 1 | 0 | 0 | 1686872 | NP_828849.2 | P0C6X7 |
| PTYLKSDNIVAADLGV | 1 | 0 | 0 | 1686873 | YP_009555238.1 | P0C6X6 |
| PTYTVHKSDFQLPVYA | 1 | 0 | 0 | 1686876 | YP_009555238.1 | P0C6X6 |
| PTYVSVDDKYKTSDEL | 1 | 0 | 0 | 1686877 | YP_009047202.1 | K9N7C7 |
| PVAFTKAAGGKVSFSD | 1 | 0 | 0 | 1686879 | NP_073549.1 | A0A0P0LKV0 |
| PVAFTKLAGGKISFSD | 1 | 0 | 0 | 1686880 | YP_003766.2 | H9EJ42 |
| PVASIITKLNSLNEPL | 1 | 0 | 0 | 1686885 | NP_828849.2 | P0C6X7 |
| PVATGDVVLASDDLYV | 1 | 0 | 0 | 1686889 | YP_173236.1 | A0A140H1G9 |
| PVATLIATLNSLGETL | 1 | 0 | 0 | 1686890 | QIQ54046.1 | P0C6U8 |
| PVAVVNDKIFTKSGVN | 1 | 0 | 0 | 1686892 | YP_009047202.1 | K9N7C7 |
| PVAYRKILLRKNGNKG | 1 | 0 | 0 | 1686894 | QIQ54046.1 | P0C6U8 |
| PVAYRKVLLRKNGNKG | 1 | 0 | 0 | 1686895 | YP_009724389.1 | P0DTD1 |
| PVDEYITTYPGQGCAG | 1 | 0 | 0 | 1686903 | NP_828849.2 | P0C6X7 |
| PVDQYMCGVDGKPISA | 1 | 0 | 0 | 1686908 | YP_009047202.1 | K9N7C7 |
| PVDTSNSFEVLAVEDT | 1 | 0 | 0 | 1686910 | NP_828849.2 | P0C6X7 |
| PVDVKPKIKACIDEVT | 1 | 0 | 0 | 1686911 | NP_828849.2 | P0C6X7 |
| PVEFHLDGEVLSLDKL | 1 | 0 | 0 | 1686915 | NP_828849.2 | P0C6X7 |
| PVEFYEDRILDFKVLD | 1 | 0 | 0 | 1686916 | YP_009047202.1 | K9N7C7 |
| PVETSNIFDVLQSEDT | 1 | 0 | 0 | 1686920 | QIQ54046.1 | P0C6U8 |
| PVETSNSFDVLKSEDA | 1 | 0 | 0 | 1686921 | YP_009724389.1 | P0DTD1 |
| PVETSNSFDVLKSEDT | 1 | 0 | 0 | 1686922 | QHR63299.1 | P0C6W2 |
| PVGNCVKMLTNGSGSG | 1 | 0 | 0 | 1686937 | NP_073549.1 | A0A0P0LKV0 |
| PVGVVDTEWGNMLTGI | 1 | 0 | 0 | 1686939 | YP_009047202.1 | K9N7C7 |
| PVHVKEITKENVETLT | 1 | 0 | 0 | 1686943 | YP_003766.2 | H9EJ42 |
| PVHVLTKHGDFADEII | 1 | 0 | 0 | 1686944 | QIQ54046.1 | P0C6U8 |
| PVIEVSQIQSKLTDVK | 1 | 0 | 0 | 1686949 | YP_173236.1 | A0A140H1G9 |
| PVKDYVQTVNVIAWLY | 1 | 0 | 0 | 1686959 | YP_173236.1 | A0A140H1G9 |
| PVKEASKQCFKIFERG | 1 | 0 | 0 | 1686960 | NP_073549.1 | A0A0P0LKV0 |
| PVKIDNASPASTVHAT | 1 | 0 | 0 | 1686963 | NP_828852.2 | P59632 |
| PVKKVAFGGDQVHEVA | 1 | 0 | 0 | 1686964 | YP_009047202.1 | K9N7C7 |
| PVKPICSSVYLGVKGS | 1 | 0 | 0 | 1686966 | YP_003766.2 | H9EJ42 |
| PVLKTINGLVCIVGNK | 1 | 0 | 0 | 1686976 | YP_173236.1 | A0A140H1G9 |
| PVLMGWDYPKCDRAMP | 1 | 0 | 0 | 1686977 | YP_009555238.1 | P0C6X6 |
| PVLPFNDGVYFASTEK | 1 | 0 | 0 | 1686978 | YP_009724390.1 | P0DTC2 |
| PVLPKNMWEFRDYFNN | 1 | 0 | 0 | 1686979 | YP_003766.2 | H9EJ42 |
| PVLPXNDGVYFASTEK | 1 | 0 | 0 | 1686980 | QIQ54048.1 | P59594 |
| PVMISEWPLSVQQAQQ | 1 | 0 | 0 | 1686992 | NP_073549.1 | A0A0P0LKV0 |
| PVMISQWPISNDSNGC | 1 | 0 | 0 | 1686993 | YP_003766.2 | H9EJ42 |
| PVMTNGSNILEAFTKP | 1 | 0 | 0 | 1686995 | NP_073549.1 | A0A0P0LKV0 |
| PVNSNVKLLDYDYATH | 1 | 0 | 0 | 1687003 | YP_003766.2 | H9EJ42 |
| PVNTVKPKPVINQLDE | 1 | 0 | 0 | 1687005 | NP_073549.1 | A0A0P0LKV0 |
| PVPYCYDTNVLEGSVA | 1 | 0 | 0 | 1687012 | YP_009724389.1 | P0DTD1 |
| PVRDEVSIETDIIDEV | 1 | 0 | 0 | 1687021 | NP_073556.1 | A0A0N7J131 |
| PVRPRNSSDNGISAII | 1 | 0 | 0 | 1687022 | YP_003767.1 | A0A0P0G321 |
| PVSEEVVENPTIQKDI | 1 | 0 | 0 | 1687025 | QHR63299.1 | P0C6W2 |
| PVSEEVVENPTIQKDV | 1 | 0 | 0 | 1687026 | YP_009724389.1 | P0DTD1 |
| PVSIINNAVYTKVDGI | 1 | 0 | 0 | 1687032 | NP_828849.2 | P0C6X7 |
| PVSVIYDKETKTHATL | 1 | 0 | 0 | 1687043 | YP_009047204.1 | K9N5Q8 |
| PVTKKYTYSFLPKEDG | 1 | 0 | 0 | 1687048 | YP_009047202.1 | K9N7C7 |
| PVVEYLSKEDLHTLYV | 1 | 0 | 0 | 1687059 | YP_009047202.1 | K9N7C7 |
| PVVGIVYSSQLYEDVP | 1 | 0 | 0 | 1687060 | YP_003768.1 | H9EJ24 |
| PVVPDTVEVPPQVVKL | 1 | 0 | 0 | 1687064 | YP_009047202.1 | K9N7C7 |
| PVVVDDHVGLLDQAWR | 1 | 0 | 0 | 1687067 | YP_009555238.1 | P0C6X6 |
| PVVYRKVLLRKNGNKG | 1 | 0 | 0 | 1687068 | QHR63299.1 | P0C6W2 |
| PVYASYKVLDNGVIRD | 1 | 0 | 0 | 1687070 | YP_009555238.1 | P0C6X6 |
| PVYEADTEQTQNVTVK | 1 | 0 | 0 | 1687072 | YP_009047202.1 | K9N7C7 |

| | | | | | | |
|---|---|---|---|---|---|---|
| PVYIWTADEAIKFLKE | 1 | 0 | 0 | 1687075 | YP_009555244.1 | Q01455 |
| PWDTIANYAKPFLNKV | 1 | 0 | 0 | 1687082 | YP_009724389.1 | P0DTD1 |
| PWFMDGENAYEVVKAM | 1 | 0 | 0 | 1687085 | YP_009047202.1 | K9N7C7 |
| PWMFEDAEEKLDNPSS | 1 | 0 | 0 | 1687091 | YP_009555238.1 | P0C6X6 |
| PWTNDALNTICIKDAD | 1 | 0 | 0 | 1687099 | NP_073549.1 | A0A0P0LKV0 |
| PXNDGVYFASTEKSNI | 1 | 0 | 0 | 1687111 | QIQ54048.1 | P59594 |
| PYCYFRNSSANYVGKY | 1 | 0 | 0 | 1687119 | YP_173237.1 | A0A140H1H0 |
| PYCYSDGVMKNASLYT | 1 | 0 | 0 | 1687121 | YP_173236.1 | A0A140H1G9 |
| PYDLEVLLQDALQSRE | 1 | 0 | 0 | 1687124 | YP_009555238.1 | P0C6X6 |
| PYEDFQENWNTKHSSG | 1 | 0 | 0 | 1687126 | YP_009724389.1 | P0DTD1 |
| PYESLRPDTRYVLMDG | 1 | 0 | 0 | 1687127 | QIQ54046.1 | P0C6U8 |
| PYGRCPTAADINIPDL | 1 | 0 | 0 | 1687136 | YP_009555240.1 | P30215 |
| PYHPNTVDCQDDRCII | 1 | 0 | 0 | 1687140 | YP_009555238.1 | P0C6X6 |
| PYHRFKCFPNGSNDVW | 1 | 0 | 0 | 1687141 | YP_173237.1 | A0A140H1H0 |
| PYIDDFNRIASCKWTD | 1 | 0 | 0 | 1687143 | YP_009555238.1 | P0C6X6 |
| PYIVGDIITSGNLTTV | 1 | 0 | 0 | 1687147 | QIQ54046.1 | P0C6U8 |
| PYKVCNSVKDTLAYYA | 1 | 0 | 0 | 1687152 | YP_009047202.1 | K9N7C7 |
| PYNSQNAVAAKILGLP | 1 | 0 | 0 | 1687156 | QIQ54047.1 | P0C6X7 |
| PYNSQNAVASKILGLP | 1 | 0 | 0 | 1687157 | YP_009724389.1 | P0DTD1 |
| PYNSQNAVSRSMLGLT | 1 | 0 | 0 | 1687158 | YP_009047202.1 | K9N7C7 |
| PYNSQNFAAKRVLGLQ | 1 | 0 | 0 | 1687159 | YP_009555238.1 | P0C6X6 |
| PYNSQNYVAARLLGLQ | 1 | 0 | 0 | 1687160 | NP_073549.1 | A0A0P0LKV0 |
| PYNVSLSKFNSCKSDI | 1 | 0 | 0 | 1687165 | YP_173237.1 | A0A140H1H0 |
| PYPNASFDNFKFVCDN | 1 | 0 | 0 | 1687171 | YP_009724389.1 | P0DTD1 |
| PYPNSNFDNFKFVCDN | 1 | 0 | 0 | 1687173 | QIQ54046.1 | P0C6U8 |
| PYQHYDACSVKKYTGV | 1 | 0 | 0 | 1687176 | YP_173236.1 | A0A140H1G9 |
| PYTPKYTFGNVKPGET | 1 | 0 | 0 | 1687189 | YP_173236.1 | A0A140H1G9 |
| PYVCNAPGCDVNDVTK | 1 | 0 | 0 | 1687192 | YP_009555238.1 | P0C6X6 |
| PYVCNASGCGVSDVKK | 1 | 0 | 0 | 1687196 | YP_003766.2 | H9EJ42 |
| PYVMLNTSIPNLPDFK | 1 | 0 | 0 | 1687207 | YP_009555241.1 | P36334 |
| QAALDYLATVKTVPAG | 1 | 0 | 0 | 1687226 | YP_003766.2 | H9EJ42 |
| QAALPQSKDSNFLNES | 1 | 0 | 0 | 1687228 | YP_009047203.1 | K9N7C7 |
| QAALPQSKDSNFLNRV | 1 | 0 | 0 | 1687229 | YP_009047202.1 | K9N7C7 |
| QAASDWKPGYSMPVLY | 1 | 0 | 0 | 1687231 | YP_009555238.1 | P0C6X6 |
| QADQQVDRLITGRLAA | 1 | 0 | 0 | 1687236 | NP_073551.1 | A0A0P0K6L9 |
| QADSFTCNNIDAAKIY | 1 | 0 | 0 | 1687237 | YP_009555241.1 | P36334 |
| QADTAGLYTNFRIDVP | 1 | 0 | 0 | 1687238 | YP_009047205.1 | K9N796 |
| QADTSTPSDVSSRDPT | 1 | 0 | 0 | 1687240 | YP_173242.1 | Q19U29 |
| QADVEWKFYDAQPCSD | 1 | 0 | 0 | 1687241 | YP_009724389.1 | P0DTD1 |
| QAEVEWKFYDAQPCSD | 1 | 0 | 0 | 1687246 | NP_828849.2 | P0C6X7 |
| QAFGLRGPGDLQGNFG | 1 | 0 | 0 | 1687249 | YP_009047211.1 | K9N4V7 |
| QAFTFSPTYKAFLCK | 1 | 0 | 0 | 1687253 | P03155.1 | Q69028 |
| QAGEMPCAIINDKVVA | 1 | 0 | 0 | 1687256 | YP_009555238.1 | P0C6X6 |
| QAIASEFSSLPSYAAF | 1 | 0 | 0 | 1687272 | YP_009724389.1 | P0DTD1 |
| QAIASEFSSLPSYAAY | 1 | 0 | 0 | 1687273 | NP_828849.2 | P0C6X7 |
| QAITVTPEANMDQESF | 1 | 0 | 0 | 1687283 | YP_009724389.1 | P0DTD1 |
| QAIYDRLDSIQADQQV | 1 | 0 | 0 | 1687284 | YP_003767.1 | A0A0P0G321 |
| QAIYDRLDTIQADQQV | 1 | 0 | 0 | 1687285 | NP_073551.1 | A0A0P0K6L9 |
| QALGAMQTGFTTTNEA | 1 | 0 | 0 | 1687288 | YP_009047204.1 | K9N5Q8 |
| QALHGANLRQDDSVRN | 1 | 0 | 0 | 1687289 | YP_009047204.1 | K9N5Q8 |
| QALNSILDNAVKGCVP | 1 | 0 | 0 | 1687292 | YP_009555238.1 | P0C6X6 |
| QALQTVATALNKIQDV | 1 | 0 | 0 | 1687296 | NP_073551.1 | A0A0P0K6L9 |
| QALSKLASELSNTFGA | 1 | 0 | 0 | 1687297 | YP_009047204.1 | K9N5Q8 |
| QAMTQMYKQARSEDKR | 1 | 0 | 0 | 1687303 | YP_009724389.1 | P0DTD1 |
| QAMTSMYKQARAEDKK | 1 | 0 | 0 | 1687304 | YP_009047202.1 | K9N7C7 |
| QANAYYKYDNGNFIKL | 1 | 0 | 0 | 1687305 | NP_073549.1 | A0A0P0LKV0 |
| QANNNCFVNVSCLMLQ | 1 | 0 | 0 | 1687309 | YP_009555238.1 | P0C6X6 |
| QASADWKPGHAMPSLF | 1 | 0 | 0 | 1687322 | YP_009047202.1 | K9N7C7 |
| QASESKVLLERFLPKC | 1 | 0 | 0 | 1687323 | YP_003766.2 | H9EJ42 |
| QASLNGVTLIGEAVKT | 1 | 0 | 0 | 1687325 | YP_009724389.1 | P0DTD1 |
| QASSRSSSRSRGNSRN | 1 | 0 | 0 | 1687331 | NP_828858.1 | P59595 |
| QATELGIEFLKRGDKV | 1 | 0 | 0 | 1687335 | QIQ54046.1 | P0C6U8 |
| QATKYLVQQESPFVMM | 1 | 0 | 0 | 1687337 | YP_009724389.1 | P0DTD1 |
| QATNDWKPGYSMPVLY | 1 | 0 | 0 | 1687339 | YP_173236.1 | A0A140H1G9 |
| QATQYLVQQESSFVMM | 1 | 0 | 0 | 1687341 | QIQ54046.1 | P0C6U8 |
| QATTANVSALMGANGN | 1 | 0 | 0 | 1687342 | YP_009047202.1 | K9N7C7 |
| QATTKTTFKPNTWCLR | 1 | 0 | 0 | 1687343 | NP_828849.2 | P0C6X7 |
| QAVSSNINRLLSVPSD | 1 | 0 | 0 | 1687353 | YP_003766.2 | H9EJ42 |
| QAVVGAMLSENFTSYG | 1 | 0 | 0 | 1687358 | NP_073551.1 | A0A0P0K6L9 |
| QAWRFPCAGRKVNFNE | 1 | 0 | 0 | 1687361 | YP_173236.1 | A0A140H1G9 |
| QAYEDAIANGSSSQLI | 1 | 0 | 0 | 1687363 | YP_003766.2 | H9EJ42 |
| QCAMRPNHTIKGSFLN | 1 | 0 | 0 | 1687368 | NP_828849.2 | P0C6X7 |
| QCFGKRGPNQNFGGGE | 1 | 0 | 0 | 1687372 | YP_009555245.1 | P33469 |
| QCFGKRGPSQNFGNAE | 1 | 0 | 0 | 1687373 | YP_173242.1 | Q19U29 |
| QCKTKSLLPNTGVYDL | 1 | 0 | 0 | 1687391 | YP_173238.1 | A0A140H1H1 |
| QCSGVTFQSAIKRTVK | 1 | 0 | 0 | 1687394 | QIQ54046.1 | P0C6U8 |
| QCSGVTFQSAVKRTIK | 1 | 0 | 0 | 1687395 | YP_009724389.1 | P0DTD1 |
| QCSTNLFKDCSKSYSG | 1 | 0 | 0 | 1687396 | YP_009555238.1 | P0C6X6 |
| QCTVQQCFGKRGPNQN | 1 | 0 | 0 | 1687398 | YP_009555245.1 | P33469 |
| QCYYNNSNNGKIVYAI | 1 | 0 | 0 | 1687408 | YP_009555238.1 | P0C6X6 |
| QDADGINKQLTDISVD | 1 | 0 | 0 | 1687411 | YP_173236.1 | A0A140H1G9 |
| QDAQIDRLINGRLTTL | 1 | 0 | 0 | 1687418 | YP_009047204.1 | K9N5Q8 |
| QDAVNNNAQALSKLAS | 1 | 0 | 0 | 1687419 | YP_009047204.1 | K9N5Q8 |
| QDAYYRARAGEAANFC | 1 | 0 | 0 | 1687421 | YP_009724389.1 | P0DTD1 |
| QDDIMKAVAAALKSLG | 1 | 0 | 0 | 1687424 | NP_073556.1 | A0A0N7J131 |
| QDDKKIKACVEEVTTT | 1 | 0 | 0 | 1687425 | YP_009724389.1 | P0DTD1 |
| QDELFAMTKRNVIPTM | 1 | 0 | 0 | 1687430 | YP_009047202.1 | K9N7C7 |
| QDFGSTVFNVPTKVLR | 1 | 0 | 0 | 1687432 | YP_009555238.1 | P0C6X6 |
| QDGFYSANFLDDNVLP | 1 | 0 | 0 | 1687439 | YP_003767.1 | A0A0P0G321 |
| QDGNAAITDYNYYKYN | 1 | 0 | 0 | 1687443 | YP_009555238.1 | P0C6X6 |
| QDGVKITEEVGHEDLM | 1 | 0 | 0 | 1687448 | QIQ54046.1 | P0C6U8 |

| | | | | | | |
|---|---|---|---|---|---|---|
| QDHVDILGPLSAQTGI | 1 | 0 | 0 | 1687451 | YP_009724389.1 | P0DTD1 |
| QDKVFGYVQQGIFNKA | 1 | 0 | 0 | 1687458 | YP_009555238.1 | P0C6X6 |
| QDLIRQGTDYKHWPQI | 1 | 0 | 0 | 1687464 | NP_828858.1 | P59595 |
| QDLNGKWYDFGDFVIT | 1 | 0 | 0 | 1687466 | YP_009047202.1 | K9N7C7 |
| QDLNGKWYDFGDYVIA | 1 | 0 | 0 | 1687467 | YP_009555238.1 | P0C6X6 |
| QDLNGNWYDFGDFIQT | 1 | 0 | 0 | 1687470 | YP_009724389.1 | P0DTD1 |
| QDLNGNWYDFGDFVQV | 1 | 0 | 0 | 1687471 | NP_828849.2 | P0C6X7 |
| QDPAPIDIDAFPVKPI | 1 | 0 | 0 | 1687480 | YP_003766.2 | H9EJ42 |
| QDPEIGPEHSVADYHN | 1 | 0 | 0 | 1687481 | NP_828849.2 | P0C6X7 |
| QDRNTREVFAQVKQMY | 1 | 0 | 0 | 1687483 | NP_828851.1 | P59594 |
| QDSPLKLEDFIPMDST | 1 | 0 | 0 | 1687489 | NP_828849.2 | P0C6X7 |
| QDVFIQKYGLEDYAFE | 1 | 0 | 0 | 1687495 | YP_009555238.1 | P0C6X6 |
| QDVGFKNQYCNGSIAC | 1 | 0 | 0 | 1687496 | YP_009555238.1 | P0C6X6 |
| QDVIFSQFDSLGVSSN | 1 | 0 | 0 | 1687497 | YP_009555238.1 | P0C6X6 |
| QDVNLHSSRLSFKELL | 1 | 0 | 0 | 1687504 | YP_009724389.1 | P0DTD1 |
| QDVVNQQGNSLNHLTS | 1 | 0 | 0 | 1687507 | NP_073551.1 | A0A0P0K6L9 |
| QDVVNQQGSALNHLTS | 1 | 0 | 0 | 1687508 | YP_003767.1 | A0A0P0G321 |
| QDVYKSLTIVDIPQSL | 1 | 0 | 0 | 1687509 | YP_009047202.1 | K9N7C7 |
| QDYMQIAPVPAEVLNV | 1 | 0 | 0 | 1687513 | YP_003769.1 | H9EJA2 |
| QEAILMSEDVKLNPSE | 1 | 0 | 0 | 1687515 | YP_009047202.1 | K9N7C7 |
| QEAYYRARAGDAANFC | 1 | 0 | 0 | 1687523 | NP_828849.2 | P0C6X7 |
| QEDDSEVNQTTITQSI | 1 | 0 | 0 | 1687533 | QHR63299.1 | P0C6W2 |
| QEDFDLISKCQITAVE | 1 | 0 | 0 | 1687534 | YP_009555238.1 | P0C6X6 |
| QEEDWLDDDSQQTVGQ | 1 | 0 | 0 | 1687537 | YP_009724389.1 | P0DTD1 |
| QEEDWLETEAEVVEQE | 1 | 0 | 0 | 1687538 | QIQ54046.1 | P0C6U8 |
| QEEELPITEIVPAVEQ | 1 | 0 | 0 | 1687539 | QIQ54046.1 | P0C6U8 |
| QEEFIQTNSPKVTIDC | 1 | 0 | 0 | 1687540 | YP_173238.1 | A0A140H1H1 |
| QEEQEEDWLETEAEVV | 1 | 0 | 0 | 1687542 | QIQ54046.1 | P0C6U8 |
| QEFAQNLPKYVKPNFD | 1 | 0 | 0 | 1687548 | YP_003767.1 | A0A0P0G321 |
| QEFRYMNSQGLLPPKN | 1 | 0 | 0 | 1687553 | YP_009724389.1 | P0DTD1 |
| QEFRYMNSQGLLPPKS | 1 | 0 | 0 | 1687554 | NP_828849.2 | P0C6X7 |
| QEGFDATNSALVKIQA | 1 | 0 | 0 | 1687556 | YP_009555241.1 | P36334 |
| QEHYVRITGLYPTLNI | 1 | 0 | 0 | 1687567 | YP_009724389.1 | P0DTD1 |
| QEKDEDGNLIDSYFIV | 1 | 0 | 0 | 1687575 | QIQ54047.1 | P0C6X7 |
| QEKDKKSAKTGTPKPS | 1 | 0 | 0 | 1687577 | NP_073556.1 | A0A0N7J131 |
| QELSYKLPNYTVPDLV | 1 | 0 | 0 | 1687593 | NP_073551.1 | A0A0P0K6L9 |
| QENQKILAASFNKAIN | 1 | 0 | 0 | 1687600 | YP_003767.1 | A0A0P0G321 |
| QENQKILAASFNKAMT | 1 | 0 | 0 | 1687601 | NP_073551.1 | A0A0P0K6L9 |
| QENWNTKHSSGVTRDL | 1 | 0 | 0 | 1687602 | QHR63299.1 | P0C6W2 |
| QERWRMRRGQRVDLPP | 1 | 0 | 0 | 1687615 | YP_003771.1 | H9EJ69 |
| QESIKSLNNSYINLKD | 1 | 0 | 0 | 1687618 | YP_173238.1 | A0A140H1H1 |
| QESLARSACVYYSQVL | 1 | 0 | 0 | 1687619 | YP_009047202.1 | K9N7C7 |
| QETPVVPDTVEVPPQV | 1 | 0 | 0 | 1687625 | YP_009047202.1 | K9N7C7 |
| QEVIDDMHWGDGFQIK | 1 | 0 | 0 | 1687627 | YP_009555239.1 | Q80872 |
| QEVKDNDGKNVHLKDV | 1 | 0 | 0 | 1687628 | NP_073549.1 | A0A0P0LKV0 |
| QEVLLLRHDIQLNNDV | 1 | 0 | 0 | 1687629 | YP_173236.1 | A0A140H1G9 |
| QEYIQTTIQKVTVDCK | 1 | 0 | 0 | 1687635 | YP_009047204.1 | K9N5Q8 |
| QFCDAMRNAGIVGVLT | 1 | 0 | 0 | 1687648 | YP_009724389.1 | P0DTD1 |
| QFDINAYANWCCLKCG | 1 | 0 | 0 | 1687656 | YP_173236.1 | A0A140H1G9 |
| QFDNLSSEDLKAVRSS | 1 | 0 | 0 | 1687659 | YP_009555238.1 | P0C6X6 |
| QFDSLGVSSNQSPQGN | 1 | 0 | 0 | 1687662 | YP_009555238.1 | P0C6X6 |
| QFDSLNLSAGTQHISN | 1 | 0 | 0 | 1687663 | YP_173242.1 | Q19U29 |
| QFGPTYLDGADVTKIK | 1 | 0 | 0 | 1687672 | YP_009724389.1 | P0DTD1 |
| QFGRDISDTTDAVRDP | 1 | 0 | 0 | 1687674 | QIQ54048.1 | P59594 |
| QFKTFEKVDGVYTNFK | 1 | 0 | 0 | 1687694 | YP_009555238.1 | P0C6X6 |
| QFKTTGDLAVQIGSNN | 1 | 0 | 0 | 1687695 | YP_003766.2 | H9EJ42 |
| QFLPFQQFGRDISDTT | 1 | 0 | 0 | 1687696 | QIQ54048.1 | P59594 |
| QFNYFKKVDGIIQQLP | 1 | 0 | 0 | 1687706 | NP_828849.2 | P0C6X7 |
| QFNYYKKVDGVVQQLP | 1 | 0 | 0 | 1687710 | YP_009724389.1 | P0DTD1 |
| QFNYYKKVNGVVQQLP | 1 | 0 | 0 | 1687711 | QHR63299.1 | P0C6W2 |
| QFQILDEGVDCKIQTA | 1 | 0 | 0 | 1687717 | YP_009555239.1 | Q80872 |
| QFQKGKEFEFVEGQGV | 1 | 0 | 0 | 1687718 | YP_009555245.1 | P33469 |
| QFTEFVGTQSVDMLAV | 1 | 0 | 0 | 1687728 | YP_009047202.1 | K9N7C7 |
| QFTSDQAVTFLKEWNF | 1 | 0 | 0 | 1687731 | YP_173241.1 | Q0ZJ82 |
| QFTSLEIPRRNVATLQ | 1 | 0 | 0 | 1687732 | YP_009724389.1 | P0DTD1 |
| QFTSLEVPRRNVATLQ | 1 | 0 | 0 | 1687733 | QIQ54047.1 | P0C6X7 |
| QFVDHFGENEEIIING | 1 | 0 | 0 | 1687735 | NP_073549.1 | A0A0P0LKV0 |
| QFYEFYNDVKPPVLDV | 1 | 0 | 0 | 1687742 | YP_009555243.1 | Q04854 |
| QFYIYDEEGGYDVSKP | 1 | 0 | 0 | 1687744 | YP_003766.2 | H9EJ42 |
| QGAKPVGNCVKMLTNG | 1 | 0 | 0 | 1687750 | NP_073549.1 | A0A0P0LKV0 |
| QGARGTYRAALNSEKH | 1 | 0 | 0 | 1687751 | YP_009047202.1 | K9N7C7 |
| QGAVDINKLCEEMLDN | 1 | 0 | 0 | 1687752 | YP_009724389.1 | P0DTD1 |
| QGAVDINRLCEEMLDN | 1 | 0 | 0 | 1687753 | NP_828849.2 | P0C6X7 |
| QGCAGYTLEEAKTALK | 1 | 0 | 0 | 1687754 | NP_828849.2 | P0C6X7 |
| QGCVGYTIEEAKTALK | 1 | 0 | 0 | 1687756 | QIQ54046.1 | P0C6U8 |
| QGDKCNIFANFILHDV | 1 | 0 | 0 | 1687759 | YP_009555241.1 | P36334 |
| QGEIKDATPSDFVRAT | 1 | 0 | 0 | 1687761 | YP_009724391.1 | P0DTC3 |
| QGEIKDATPSDSVRAT | 1 | 0 | 0 | 1687762 | QHR63301.1 | Q3LZX0 |
| QGETPVSIVNNTVYTK | 1 | 0 | 0 | 1687764 | QIQ54047.1 | P0C6X7 |
| QGEVPVSIINNTVYTK | 1 | 0 | 0 | 1687765 | YP_009724389.1 | P0DTD1 |
| QGFACGCGASYTANDV | 1 | 0 | 0 | 1687766 | YP_009047202.1 | K9N7C7 |
| QGGFKKFDNPVLPXND | 1 | 0 | 0 | 1687777 | QIQ54048.1 | P59594 |
| QGIKLGTGYSLADLPA | 1 | 0 | 0 | 1687790 | YP_009555244.1 | Q01455 |
| QGKSLEFGATSVTPQP | 1 | 0 | 0 | 1687801 | QHR63299.1 | P0C6W2 |
| QGLIDQINSTYVDLKL | 1 | 0 | 0 | 1687806 | YP_003767.1 | A0A0P0G321 |
| QGLLPPKSSIDAFKLN | 1 | 0 | 0 | 1687812 | NP_828849.2 | P0C6X7 |
| QGLNGYTVEEAKTVLK | 1 | 0 | 0 | 1687813 | YP_009724389.1 | P0DTD1 |
| QGLNGYTVEEARTVLK | 1 | 0 | 0 | 1687814 | QHR63299.1 | P0C6W2 |
| QGMETLACEDTKLVTE | 1 | 0 | 0 | 1687822 | QIQ54046.1 | P0C6U8 |
| QGNFGDLQLNKLGTED | 1 | 0 | 0 | 1687823 | YP_009047211.1 | K9N4V7 |
| QGNFGDQELIRQGTEY | 1 | 0 | 0 | 1687826 | QIQ54056.1 | P59595 |

| | | | | | | |
|---|---|---|---|---|---|---|
| QGNFKNLREFVFKNVD | 1 | 0 | 0 | 1687828 | QIQ54048.1 | P59594 |
| QGNKQIVHSYPILHHP | 1 | 0 | 0 | 1687829 | YP_009047207.1 | K9N643 |
| QGNLGGNCAGNVIGND | 1 | 0 | 0 | 1687830 | YP_173236.1 | A0A140H1G9 |
| QGNLGSNGKPGNVGGN | 1 | 0 | 0 | 1687831 | YP_009555238.1 | P0C6X6 |
| QGNVVTSVMESQISTK | 1 | 0 | 0 | 1687836 | YP_173236.1 | A0A140H1G9 |
| QGPASARDLICAQYVA | 1 | 0 | 0 | 1687837 | YP_009047204.1 | K9N5Q8 |
| QGQLLDTKLNGILQQA | 1 | 0 | 0 | 1687844 | YP_003766.2 | H9EJ42 |
| QGQNSADPKVYPIILR | 1 | 0 | 0 | 1687845 | NP_828859.1 | P59636 |
| QGSITQRTRTRPSVQP | 1 | 0 | 0 | 1687860 | YP_009047211.1 | K9N4V7 |
| QGSRSGNSTRGTSPGP | 1 | 0 | 0 | 1687861 | YP_009047211.1 | K9N4V7 |
| QGVKLGTGYTLSDLPV | 1 | 0 | 0 | 1687875 | YP_173241.1 | Q0ZJ82 |
| QGVPIAFGVPPSEAKG | 1 | 0 | 0 | 1687876 | YP_173242.1 | Q19U29 |
| QGVPINTNSGPDDQIG | 1 | 0 | 0 | 1687878 | NP_828858.1 | P59595 |
| QGVPINTNSSPDDQIG | 1 | 0 | 0 | 1687879 | YP_009724397.2 | P0DTC9 |
| QGVPINTNSTKDDQIG | 1 | 0 | 0 | 1687880 | QIQ54056.1 | P59595 |
| QGVPLNANSTPAQNAG | 1 | 0 | 0 | 1687881 | YP_009047211.1 | K9N4V7 |
| QGVTLSSNLNTNLHFD | 1 | 0 | 0 | 1687882 | YP_173238.1 | A0A140H1H1 |
| QHAFMKHKGGDSTDFI | 1 | 0 | 0 | 1687891 | YP_009047202.1 | K9N7C7 |
| QHEETIYNLVKDCPAV | 1 | 0 | 0 | 1687899 | NP_828849.2 | P0C6X7 |
| QHGKEDLKFPRGQGVP | 1 | 0 | 0 | 1687902 | YP_009724397.2 | P0DTC9 |
| QHGKEDLRFPRGQGVP | 1 | 0 | 0 | 1687903 | QIQ54056.1 | P59595 |
| QHGKEELRFPRGQGVP | 1 | 0 | 0 | 1687904 | NP_828858.1 | P59595 |
| QHGKVPLTFPPGQGVP | 1 | 0 | 0 | 1687905 | YP_009047211.1 | K9N4V7 |
| QHIGMKRYCTVQGPPG | 1 | 0 | 0 | 1687906 | YP_009555238.1 | P0C6X6 |
| QHLHVASNSKLWQYCS | 1 | 0 | 0 | 1687915 | YP_173236.1 | A0A140H1G9 |
| QHPLLNPSALEFNPSQ | 1 | 0 | 0 | 1687921 | NP_073556.1 | A0A0N7J131 |
| QHTLYIKDGDDGYFLP | 1 | 0 | 0 | 1687930 | YP_009047202.1 | K9N7C7 |
| QHTMQIVDENGKYYLP | 1 | 0 | 0 | 1687934 | NP_073549.1 | A0A0P0LKV0 |
| QHTMQIVDKDGTYYLP | 1 | 0 | 0 | 1687935 | YP_003766.2 | H9EJ42 |
| QIAPGQTGVIADYNYK | 1 | 0 | 0 | 1687943 | NP_828851.1 | P59594 |
| QIDGKFVTKFSGDKFD | 1 | 0 | 0 | 1687949 | YP_173236.1 | A0A140H1G9 |
| QIESDDYIARNGPLNV | 1 | 0 | 0 | 1687959 | QIQ54046.1 | P0C6U8 |
| QIFDVAPIELENKFTP | 1 | 0 | 0 | 1687960 | YP_009047202.1 | K9N7C7 |
| QIGEYTFEKGDYGDAV | 1 | 0 | 0 | 1687964 | YP_009724389.1 | P0DTD1 |
| QIGSNNVCTYEHVISF | 1 | 0 | 0 | 1687969 | YP_003766.2 | H9EJ42 |
| QIHTIDGSSGVANPAM | 1 | 0 | 0 | 1687975 | NP_828852.2 | P59632 |
| QIHTIDGSSGVVNPAM | 1 | 0 | 0 | 1687976 | QHR63301.1 | Q3LZX0 |
| QIHTIDGSSGVVNPVM | 1 | 0 | 0 | 1687977 | YP_009724391.1 | P0DTC3 |
| QIITTDNTFVSGSCDV | 1 | 0 | 0 | 1687986 | QHR63300.2 | Q3LZX1 |
| QILASVPEKFLNAFDV | 1 | 0 | 0 | 1687993 | NP_073549.1 | A0A0P0LKV0 |
| QILGSCTFEDELAPSD | 1 | 0 | 0 | 1687997 | YP_173236.1 | A0A140H1G9 |
| QILGYSSLNDEFSINE | 1 | 0 | 0 | 1687999 | NP_073549.1 | A0A0P0LKV0 |
| QILPDPLKPTKRSFIE | 1 | 0 | 0 | 1688001 | NP_828851.1 | P59594 |
| QIMGSCSFEDELTPSD | 1 | 0 | 0 | 1688004 | YP_009555238.1 | P0C6X6 |
| QINDMVYSLLEKGKLL | 1 | 0 | 0 | 1688008 | YP_009555238.1 | P0C6X6 |
| QINSTYVDLKLLNRFE | 1 | 0 | 0 | 1688011 | YP_003767.1 | A0A0P0G321 |
| QIPATSIVAKQGAGDA | 1 | 0 | 0 | 1688014 | NP_073549.1 | A0A0P0LKV0 |
| QIPCTCGKQATKYLVQ | 1 | 0 | 0 | 1688015 | YP_009724389.1 | P0DTD1 |
| QIPCTCGKQATQYLVQ | 1 | 0 | 0 | 1688016 | QHR63299.1 | P0C6W2 |
| QIPFEIKLAKKFDNFT | 1 | 0 | 0 | 1688018 | QIQ54046.1 | P0C6U8 |
| QIPTTCANDPVGFTLK | 1 | 0 | 0 | 1688020 | YP_009724389.1 | P0DTD1 |
| QIPTTCANDPVGFTLR | 1 | 0 | 0 | 1688021 | NP_828849.2 | P0C6X7 |
| QIQSKLTDVKCANVVL | 1 | 0 | 0 | 1688026 | YP_173236.1 | A0A140H1G9 |
| QIQTIQDSDGTNKQLN | 1 | 0 | 0 | 1688027 | YP_009555238.1 | P0C6X6 |
| QIQTVDSSQGSEYDYV | 1 | 0 | 0 | 1688028 | YP_003766.2 | H9EJ42 |
| QIRSAAKKNNLPFKLT | 1 | 0 | 0 | 1688032 | YP_009724389.1 | P0DTD1 |
| QITKDSKFQVGEFVFE | 1 | 0 | 0 | 1688043 | NP_073549.1 | A0A0P0LKV0 |
| QITNESYNNYVPSYVK | 1 | 0 | 0 | 1688044 | YP_009047202.1 | K9N7C7 |
| QITYTYKMLVAKDNKN | 1 | 0 | 0 | 1688049 | YP_003771.1 | H9EJ69 |
| QIVDENGKYYLPYPDP | 1 | 0 | 0 | 1688050 | NP_073549.1 | A0A0P0LKV0 |
| QIVDKDGTYYLPYPDP | 1 | 0 | 0 | 1688051 | YP_003766.2 | H9EJ42 |
| QIVTCAKEIKESVQTF | 1 | 0 | 0 | 1688056 | YP_009724389.1 | P0DTD1 |
| QIVVTGDDVDDIESIY | 1 | 0 | 0 | 1688058 | YP_173236.1 | A0A140H1G9 |
| QKAKQKPIYLKGSGSD | 1 | 0 | 0 | 1688067 | YP_009555238.1 | P0C6X6 |
| QKAYLDFVNAGGAPLT | 1 | 0 | 0 | 1688073 | YP_009047202.1 | K9N7C7 |
| QKDIGIDTAGVLLKQL | 1 | 0 | 0 | 1688082 | YP_173243.1 | Q0ZJ80 |
| QKDIVECDVKTTEVVG | 1 | 0 | 0 | 1688084 | QIQ54046.1 | P0C6U8 |
| QKELLAYYNNMLVNCFK | 1 | 0 | 0 | 1688089 | YP_009555238.1 | P0C6X6 |
| QKESDDYIKLNGPLTV | 1 | 0 | 0 | 1688090 | NP_828849.2 | P0C6X7 |
| QKESDEYILAKGPLQV | 1 | 0 | 0 | 1688091 | YP_009047202.1 | K9N7C7 |
| QKEVIECDVKTTEVVG | 1 | 0 | 0 | 1688092 | NP_828849.2 | P0C6X7 |
| QKGDAAIKDFDYYRYN | 1 | 0 | 0 | 1688099 | NP_073549.1 | A0A0P0LKV0 |
| QKGDAEEALSKLSEYL | 1 | 0 | 0 | 1688100 | YP_003766.2 | H9EJ42 |
| QKGKEFEFVEGQGVPI | 1 | 0 | 0 | 1688103 | YP_009555245.1 | P33469 |
| QKHIGAPANLRVVGHA | 1 | 0 | 0 | 1688108 | YP_009047202.1 | K9N7C7 |
| QKIAEIPKEEVKPFIT | 1 | 0 | 0 | 1688110 | YP_009724389.1 | P0DTD1 |
| QKILAASFNKAINNIV | 1 | 0 | 0 | 1688112 | YP_003767.1 | A0A0P0G321 |
| QKILAASFNKAMTNIV | 1 | 0 | 0 | 1688113 | NP_073551.1 | A0A0P0K6L9 |
| QKLETDGICPENHVMV | 1 | 0 | 0 | 1688124 | YP_009555238.1 | P0C6X6 |
| QKLIANKFNQALGAMQ | 1 | 0 | 0 | 1688127 | YP_009047204.1 | K9N5Q8 |
| QKLLKSIAATRGATVV | 1 | 0 | 0 | 1688130 | YP_009724389.1 | P0DTD1 |
| QKLPCQFKDKNLQDLW | 1 | 0 | 0 | 1688132 | YP_009555238.1 | P0C6X6 |
| QKLPFKFKDLAIENMW | 1 | 0 | 0 | 1688133 | YP_173236.1 | A0A140H1G9 |
| QKLPNGVTVVEEPDSR | 1 | 0 | 0 | 1688134 | NP_073556.1 | A0A0N7J131 |
| QKLQTLIDNINSTLVD | 1 | 0 | 0 | 1688135 | NP_073551.1 | A0A0P0K6L9 |
| QKMLKSMAATRGATCV | 1 | 0 | 0 | 1688136 | YP_009047202.1 | K9N7C7 |
| QKPIYLKGSGSDFSLA | 1 | 0 | 0 | 1688139 | YP_009555238.1 | P0C6X6 |
| QKPLEPKFSIALPPEL | 1 | 0 | 0 | 1688140 | YP_003771.1 | H9EJ69 |
| QKPRWKRQPNDDVTSN | 1 | 0 | 0 | 1688142 | NP_073556.1 | A0A0N7J131 |
| QKPVDVKPKIKACIDE | 1 | 0 | 0 | 1688143 | NP_828849.2 | P0C6X7 |
| QKQIANQFNKAISQIQ | 1 | 0 | 0 | 1688144 | NP_828851.1 | P59594 |

| | | | | | | |
|---|---|---|---|---|---|---|
| QKRLYSHVYRSDKVDS | 1 | 0 | 0 | 1688149 | YP_009555238.1 | P0C6X6 |
| QKRTATKAYNVTQAFG | 1 | 0 | 0 | 1688153 | YP_009724397.2 | P0DTC9 |
| QKRTATKQYNVTQAFG | 1 | 0 | 0 | 1688154 | QHR63308.1 | Q3LZX4 |
| QKRTPNKHCNVQQCFG | 1 | 0 | 0 | 1688155 | YP_173242.1 | Q19U29 |
| QKSILSPLYAFASEAA | 1 | 0 | 0 | 1688157 | YP_009724389.1 | P0DTD1 |
| QKTTTLTGVEAVMYMG | 1 | 0 | 0 | 1688160 | NP_828849.2 | P0C6X7 |
| QKVEQPKIEPKPVSVI | 1 | 0 | 0 | 1688163 | NP_073549.1 | A0A0P0LKV0 |
| QKVGMQKYSTLQGPPG | 1 | 0 | 0 | 1688166 | YP_009724389.1 | P0DTD1 |
| QKVGMQXYSTLQGPPG | 1 | 0 | 0 | 1688167 | QIQ54047.1 | P0C6X7 |
| QKVQDAVNNNAQALSK | 1 | 0 | 0 | 1688169 | YP_009047204.1 | K9N5Q8 |
| QKVTVDCKQYVCNGFQ | 1 | 0 | 0 | 1688172 | YP_009047204.1 | K9N5Q8 |
| QKYVEAYNTFTQAGFN | 1 | 0 | 0 | 1688180 | YP_003766.2 | H9EJ42 |
| QLCTFTKSTNSRIRAS | 1 | 0 | 0 | 1688203 | NP_828849.2 | P0C6X7 |
| QLDEKAQKFFDFGDFL | 1 | 0 | 0 | 1688205 | NP_073549.1 | A0A0P0LKV0 |
| QLDFIDTKRGVYCCRE | 1 | 0 | 0 | 1688206 | YP_009724389.1 | P0DTD1 |
| QLDYIESKRGVYCCRD | 1 | 0 | 0 | 1688211 | NP_828849.2 | P0C6X7 |
| QLDYSIDVIDNEIIVK | 1 | 0 | 0 | 1688212 | NP_073549.1 | A0A0P0LKV0 |
| QLEKACNIAKSAYERD | 1 | 0 | 0 | 1688217 | YP_009555238.1 | P0C6X6 |
| QLESKMSGKGQQQQSQ | 1 | 0 | 0 | 1688230 | QHR63308.1 | Q3LZX4 |
| QLESKVSGKGQQQQGQ | 1 | 0 | 0 | 1688231 | NP_828858.1 | P59595 |
| QLEVKFNAPALQEAYY | 1 | 0 | 0 | 1688233 | NP_828849.2 | P0C6X7 |
| QLFEGDKFIGTFESAA | 1 | 0 | 0 | 1688235 | YP_003766.2 | H9EJ42 |
| QLFKPLTKKNYSELDD | 1 | 0 | 0 | 1688241 | NP_828856.1 | P59634 |
| QLFYTEDMASRFADGL | 1 | 0 | 0 | 1688247 | YP_009047202.1 | K9N7C7 |
| QLFYVYDVHKDNFKDG | 1 | 0 | 0 | 1688248 | YP_173236.1 | A0A140H1G9 |
| QLGEYIFERIDYSDAV | 1 | 0 | 0 | 1688251 | YP_009047202.1 | K9N7C7 |
| QLGIEFLKRGDKSVYY | 1 | 0 | 0 | 1688258 | YP_009724389.1 | P0DTD1 |
| QLGLVWNKDVNTHSTR | 1 | 0 | 0 | 1688260 | NP_073549.1 | A0A0P0LKV0 |
| QLGSSGFLQSSNYKID | 1 | 0 | 0 | 1688264 | YP_173238.1 | A0A140H1H1 |
| QLHVADTLMQGVTLSS | 1 | 0 | 0 | 1688270 | YP_173238.1 | A0A140H1H1 |
| QLIGKQKITTIQGPPG | 1 | 0 | 0 | 1688271 | YP_003766.2 | H9EJ42 |
| QLIGKQRITTIQGPPG | 1 | 0 | 0 | 1688272 | NP_073549.1 | A0A0P0LKV0 |
| QLKKAMNVAKAEFDRE | 1 | 0 | 0 | 1688278 | NP_073549.1 | A0A0P0LKV0 |
| QLKRAMNIAKSEFDHE | 1 | 0 | 0 | 1688285 | YP_003766.2 | H9EJ42 |
| QLLDTKLNGILQQAAL | 1 | 0 | 0 | 1688293 | YP_003766.2 | H9EJ42 |
| QLLKAAYENFNSQDIL | 1 | 0 | 0 | 1688303 | NP_828849.2 | P0C6X7 |
| QLLKSAYENFNQHDVL | 1 | 0 | 0 | 1688305 | QHR63299.1 | P0C6W2 |
| QLLKVAYENFNHHEKL | 1 | 0 | 0 | 1688307 | QIQ54046.1 | P0C6U8 |
| QLLLVWEAGKSKPPLN | 1 | 0 | 0 | 1688311 | YP_009047202.1 | K9N7C7 |
| QLLMPLKAPKEVTFLE | 1 | 0 | 0 | 1688312 | NP_828849.2 | P0C6X7 |
| QLMVNKSSCYRDGIST | 1 | 0 | 0 | 1688320 | YP_173243.1 | Q0ZJ80 |
| QLNKLGTEDPRWPQIA | 1 | 0 | 0 | 1688324 | YP_009047211.1 | K9N4V7 |
| QLPAPRTLLTKGTLEP | 1 | 0 | 0 | 1688330 | YP_009724389.1 | P0DTD1 |
| QLPAPRVLLSKGSLEP | 1 | 0 | 0 | 1688333 | YP_173236.1 | A0A140H1G9 |
| QLPAPRVLLSKGTLEP | 1 | 0 | 0 | 1688334 | YP_009555238.1 | P0C6X6 |
| QLPETYFTQSRNLKEF | 1 | 0 | 0 | 1688338 | QHR63299.1 | P0C6W2 |
| QLPETYFTQSRNLQEF | 1 | 0 | 0 | 1688339 | YP_009724389.1 | P0DTD1 |
| QLPLDVYDLTMPSQVQ | 1 | 0 | 0 | 1688342 | YP_009555238.1 | P0C6X6 |
| QLPNFYYVSNGGNNCT | 1 | 0 | 0 | 1688343 | YP_003767.1 | A0A0P0G321 |
| QLPVKDYVQTVNVIAW | 1 | 0 | 0 | 1688347 | YP_173236.1 | A0A140H1G9 |
| QLPYCYFRNSSANYVG | 1 | 0 | 0 | 1688349 | YP_173237.1 | A0A140H1H0 |
| QLQSLSKDYISSNGPL | 1 | 0 | 0 | 1688359 | YP_003766.2 | H9EJ42 |
| QLRLAFLGASVTEDVK | 1 | 0 | 0 | 1688369 | YP_003766.2 | H9EJ42 |
| QLSFAGYTESAVNSTK | 1 | 0 | 0 | 1688380 | YP_009047206.1 | K9N4V0 |
| QLSFDVPDGFYSTSPI | 1 | 0 | 0 | 1688381 | NP_073551.1 | A0A0P0K6L9 |
| QLSLSMARRNLDSLEA | 1 | 0 | 0 | 1688385 | NP_828859.1 | P59636 |
| QLSSYSLFDMSKFPLK | 1 | 0 | 0 | 1688389 | YP_009724389.1 | P0DTD1 |
| QLTGILDGDYAMQFFK | 1 | 0 | 0 | 1688398 | NP_073549.1 | A0A0P0LKV0 |
| QLTLESTINKSVANLK | 1 | 0 | 0 | 1688404 | YP_003768.1 | H9EJ24 |
| QLVAFTTELKDSVKKF | 1 | 0 | 0 | 1688410 | QIQ54046.1 | P0C6U8 |
| QLVPVSPAFHRIESTD | 1 | 0 | 0 | 1688416 | YP_009047208.1 | K9N7D2 |
| QLVSAVNQSLSNAKVS | 1 | 0 | 0 | 1688418 | NP_073552.1 | A0A0P0KWE8 |
| QLYSTQLSTDTGVEHT | 1 | 0 | 0 | 1688429 | QIQ54049.1 | P59632 |
| QLYSTQLSTDTGVEHV | 1 | 0 | 0 | 1688430 | YP_009724391.1 | P0DTC3 |
| QMETDFLELAMDEFIQ | 1 | 0 | 0 | 1688444 | NP_828849.2 | P0C6X7 |
| QMLSDTLKNLSDRVVF | 1 | 0 | 0 | 1688455 | YP_009724389.1 | P0DTD1 |
| QMNLKYAISAKNRART | 1 | 0 | 0 | 1688458 | YP_009724389.1 | P0DTD1 |
| QMRPHVRYDLYDGNMF | 1 | 0 | 0 | 1688461 | YP_009047202.1 | K9N7C7 |
| QMYKEARSVNRKSKVI | 1 | 0 | 0 | 1688471 | YP_003766.2 | H9EJ42 |
| QNAGYWRRQDRKINTG | 1 | 0 | 0 | 1688474 | YP_009047211.1 | K9N4V7 |
| QNAKEIRHKILTKPRQ | 1 | 0 | 0 | 1688475 | YP_173242.1 | Q19U29 |
| QNAVAAKILGLPTQTV | 1 | 0 | 0 | 1688481 | QIQ54047.1 | P0C6X7 |
| QNAVASKILGLPTQTV | 1 | 0 | 0 | 1688482 | YP_009724389.1 | P0DTD1 |
| QNAVSRSMLGLTTQTV | 1 | 0 | 0 | 1688483 | YP_009047202.1 | K9N7C7 |
| QNDVCSTEDFNVWAMA | 1 | 0 | 0 | 1688494 | YP_173236.1 | A0A140H1G9 |
| QNFAAKRVLGLQTQTV | 1 | 0 | 0 | 1688496 | YP_009555238.1 | P0C6X6 |
| QNGAKHVQGNVAKAAN | 1 | 0 | 0 | 1688501 | YP_173236.1 | A0A140H1G9 |
| QNGDAAVKDFDFYRYN | 1 | 0 | 0 | 1688502 | YP_003766.2 | H9EJ42 |
| QNGDRSGARPKQRRPQ | 1 | 0 | 0 | 1688503 | QIQ54056.1 | P59595 |
| QNGVDAKGFPQLAELI | 1 | 0 | 0 | 1688510 | YP_003771.1 | H9EJ69 |
| QNGVFMSTAKCWTEED | 1 | 0 | 0 | 1688511 | NP_073549.1 | A0A0P0LKV0 |
| QNIDAYKTFPKKEKKQ | 1 | 0 | 0 | 1688515 | YP_009047211.1 | K9N4V7 |
| QNISTNLPPPLLGNST | 1 | 0 | 0 | 1688517 | YP_009047204.1 | K9N5Q8 |
| QNKGLFYTQVYKNMAV | 1 | 0 | 0 | 1688518 | YP_009555240.1 | P30215 |
| QNKSRELTAEDISLLK | 1 | 0 | 0 | 1688519 | YP_009555245.1 | P33469 |
| QNLCVDGFGVNKIQPW | 1 | 0 | 0 | 1688520 | NP_073549.1 | A0A0P0LKV0 |
| QNLLYCDSPFEKLQCE | 1 | 0 | 0 | 1688525 | YP_003767.1 | A0A0P0G321 |
| QNLLYDSNGNIIGFKD | 1 | 0 | 0 | 1688526 | YP_173238.1 | A0A140H1H1 |
| QNLLYDSNGNLYGFRD | 1 | 0 | 0 | 1688527 | YP_009555241.1 | P36334 |
| QNLSGNPDEPQKDVYE | 1 | 0 | 0 | 1688529 | YP_009555245.1 | P33469 |
| QNNEIKPSGLKTMVVS | 1 | 0 | 0 | 1688534 | YP_009047202.1 | K9N7C7 |

| | | | | | | |
|---|---|---|---|---|---|---|
| QNNELSPVALRQMSCA | 1 | 0 | 0 | 1688539 | YP_009724389.1 | P0DTD1 |
| QNNLAIEEIEYVHGDA | 1 | 0 | 0 | 1688540 | NP_073549.1 | A0A0P0LKV0 |
| QNNVFMSEAKCWTETD | 1 | 0 | 0 | 1688542 | YP_009724389.1 | P0DTD1 |
| QNNVFMSEAKCWVETD | 1 | 0 | 0 | 1688543 | YP_009047202.1 | K9N7C7 |
| QNNVFMSESKCWVEHD | 1 | 0 | 0 | 1688544 | YP_009555238.1 | P0C6X6 |
| QNNVFMSESKCWVEND | 1 | 0 | 0 | 1688545 | YP_173236.1 | A0A140H1G9 |
| QNPYTPKYTFGNVKPG | 1 | 0 | 0 | 1688549 | YP_173236.1 | A0A140H1G9 |
| QNSADPKVYPIILRLG | 1 | 0 | 0 | 1688552 | NP_828859.1 | P59636 |
| QNTVLNASIPESKPLA | 1 | 0 | 0 | 1688562 | YP_003771.1 | H9EJ69 |
| QNVLYENQKQIANQFN | 1 | 0 | 0 | 1688565 | NP_828851.1 | P59594 |
| QNVTVKPKRLRKKRNV | 1 | 0 | 0 | 1688567 | YP_009047202.1 | K9N7C7 |
| QNYGDSATLPKGIMMN | 1 | 0 | 0 | 1688569 | YP_009724389.1 | P0DTD1 |
| QNYGENAVIPKGIMMN | 1 | 0 | 0 | 1688571 | NP_828849.2 | P0C6X7 |
| QNYVAKRVLGVQTQTV | 1 | 0 | 0 | 1688574 | YP_173236.1 | A0A140H1G9 |
| QPCSDKAYKIEELFYS | 1 | 0 | 0 | 1688577 | YP_009724389.1 | P0DTD1 |
| QPDQVAVYQQSIIGAM | 1 | 0 | 0 | 1688579 | YP_003767.1 | A0A0P0G321 |
| QPEEELEEDWLDDDSQ | 1 | 0 | 0 | 1688581 | QHR63299.1 | P0C6W2 |
| QPEEEQEEDWLDDDSQ | 1 | 0 | 0 | 1688582 | YP_009724389.1 | P0DTD1 |
| QPEQPTQYEYGTESDY | 1 | 0 | 0 | 1688584 | QIQ54046.1 | P0C6U8 |
| QPGQTFSVLACYNGSP | 1 | 0 | 0 | 1688595 | YP_009724389.1 | P0DTD1 |
| QPIDLVPNQPYPNASF | 1 | 0 | 0 | 1688600 | YP_009724389.1 | P0DTD1 |
| QPIDLVPTQPLPNASF | 1 | 0 | 0 | 1688601 | NP_828849.2 | P0C6X7 |
| QPIDLVPTQPYPNSNF | 1 | 0 | 0 | 1688602 | QIQ54046.1 | P0C6U8 |
| QPIQSLKVCIETVRTQ | 1 | 0 | 0 | 1688608 | QIQ54046.1 | P0C6U8 |
| QPITNCVKMLCTHTGT | 1 | 0 | 0 | 1688609 | YP_009724389.1 | P0DTD1 |
| QPKIEPKPVSVIKVAP | 1 | 0 | 0 | 1688614 | NP_073549.1 | A0A0P0LKV0 |
| QPLEEPTAQAVEVPLV | 1 | 0 | 0 | 1688617 | QIQ54046.1 | P0C6U8 |
| QPLPNASFDNFKLTCS | 1 | 0 | 0 | 1688620 | NP_828849.2 | P0C6X7 |
| QPNDDVTSNVTQCFGP | 1 | 0 | 0 | 1688627 | NP_073556.1 | A0A0N7J131 |
| QPPKSVEKICIIDNMY | 1 | 0 | 0 | 1688633 | YP_173236.1 | A0A140H1G9 |
| QPPTASVSTSFLQSGI | 1 | 0 | 0 | 1688636 | YP_009555238.1 | P0C6X6 |
| QPRADKPSQLKKPRWK | 1 | 0 | 0 | 1688641 | YP_003771.1 | H9EJ69 |
| QPRTINSTQDGDNKLQ | 1 | 0 | 0 | 1688645 | YP_009555241.1 | P36334 |
| QPRVEKKKLDGFMGRI | 1 | 0 | 0 | 1688646 | YP_009724389.1 | P0DTD1 |
| QPRVEKKKLEGFMGRI | 1 | 0 | 0 | 1688647 | QIQ54046.1 | P0C6U8 |
| QPRVEKKKTEGFMGRI | 1 | 0 | 0 | 1688648 | NP_828849.2 | P0C6X7 |
| QPSLQVESANLMLSDN | 1 | 0 | 0 | 1688653 | YP_003766.2 | H9EJ42 |
| QPTDNSTTVSVTVSSN | 1 | 0 | 0 | 1688656 | YP_009047202.1 | K9N7C7 |
| QPTSEAVEAPLVGTPV | 1 | 0 | 0 | 1688666 | YP_009724389.1 | P0DTD1 |
| QPTSEEVVENPTIQKE | 1 | 0 | 0 | 1688667 | NP_828849.2 | P0C6X7 |
| QPVGVVDTEWGNMLTG | 1 | 0 | 0 | 1688674 | YP_009047202.1 | K9N7C7 |
| QPVKIDNASPASTVHA | 1 | 0 | 0 | 1688676 | NP_828852.2 | P59632 |
| QPVNSNVKLLDYDYAT | 1 | 0 | 0 | 1688677 | YP_003766.2 | H9EJ42 |
| QPVSDLLTNMGIDLDE | 1 | 0 | 0 | 1688678 | NP_828849.2 | P0C6X7 |
| QPYPNASFDNFKFVCD | 1 | 0 | 0 | 1688689 | YP_009724389.1 | P0DTD1 |
| QQAALDYLATVKTVPA | 1 | 0 | 0 | 1688698 | YP_003766.2 | H9EJ42 |
| QQAVVGAMLSENFTSY | 1 | 0 | 0 | 1688703 | NP_073551.1 | A0A0P0K6L9 |
| QQCFGKRGPNQNFGGG | 1 | 0 | 0 | 1688704 | YP_009555245.1 | P33469 |
| QQCFGKRGPSQNFGNA | 1 | 0 | 0 | 1688705 | YP_173242.1 | Q19U29 |
| QQDGRIPNGFPFNNWF | 1 | 0 | 0 | 1688708 | YP_003767.1 | A0A0P0G321 |
| QQEEQEEDWLETEAEV | 1 | 0 | 0 | 1688712 | QIQ54046.1 | P0C6U8 |
| QQFGPTYLDGADVTKI | 1 | 0 | 0 | 1688719 | YP_009724389.1 | P0DTD1 |
| QQGIFNKATVAQIKAL | 1 | 0 | 0 | 1688727 | YP_009555238.1 | P0C6X6 |
| QQGNSLNHLTSQLRQN | 1 | 0 | 0 | 1688729 | NP_073551.1 | A0A0P0K6L9 |
| QQGTFLCANEYTGNYQ | 1 | 0 | 0 | 1688733 | NP_828849.2 | P0C6X7 |
| QQHPLLNPSALEFNPS | 1 | 0 | 0 | 1688735 | NP_073556.1 | A0A0N7J131 |
| QQLFYTEDMASRFADG | 1 | 0 | 0 | 1688751 | YP_009047202.1 | K9N7C7 |
| QQLPETYFTQSRNLKE | 1 | 0 | 0 | 1688759 | QHR63299.1 | P0C6W2 |
| QQLPETYFTQSRNLQE | 1 | 0 | 0 | 1688760 | YP_009724389.1 | P0DTD1 |
| QQPTSEEVVENPTIQK | 1 | 0 | 0 | 1688772 | NP_828849.2 | P0C6X7 |
| QQQEEQEEDWLETEAE | 1 | 0 | 0 | 1688774 | QIQ54046.1 | P0C6U8 |
| QQQQSQTVTKKSAAEA | 1 | 0 | 0 | 1688779 | QHR63308.1 | Q3LZX4 |
| QQQSQTVTKKSAAEAS | 1 | 0 | 0 | 1688780 | QHR63308.1 | Q3LZX4 |
| QQQTTLKGVEAVMYMG | 1 | 0 | 0 | 1688781 | YP_009724389.1 | P0DTD1 |
| QQRFVYDAYQNLVGYY | 1 | 0 | 0 | 1688782 | YP_009047204.1 | K9N5Q8 |
| QQRTFLLSYNQNGTIT | 1 | 0 | 0 | 1688784 | QIQ54048.1 | P59594 |
| QQSIIGAMTAVNESRY | 1 | 0 | 0 | 1688785 | YP_003767.1 | A0A0P0G321 |
| QQSRTRSDSNQSSSDL | 1 | 0 | 0 | 1688787 | YP_003771.1 | H9EJ69 |
| QQTSNLVVQEFVSYDS | 1 | 0 | 0 | 1688789 | YP_009555238.1 | P0C6X6 |
| QQTTLKGVEAVMYMGT | 1 | 0 | 0 | 1688791 | YP_009724389.1 | P0DTD1 |
| QQTVVQEDDSEVNQTT | 1 | 0 | 0 | 1688793 | QHR63299.1 | P0C6W2 |
| QQVDKKIKACVEEVTT | 1 | 0 | 0 | 1688794 | QHR63299.1 | P0C6W2 |
| QQVDLVISDMYDPTTK | 1 | 0 | 0 | 1688795 | YP_009047202.1 | K9N7C7 |
| QQVGVIADKDKDLSGA | 1 | 0 | 0 | 1688798 | NP_073549.1 | A0A0P0LKV0 |
| QQVLSENQKLIANKFN | 1 | 0 | 0 | 1688799 | YP_009047204.1 | K9N5Q8 |
| QQVLYYQNNVFMSESK | 1 | 0 | 0 | 1688800 | YP_009555238.1 | P0C6X6 |
| QQVVDADSKIVQLSEI | 1 | 0 | 0 | 1688803 | YP_009724389.1 | P0DTD1 |
| QQVVKALNESYIDLKE | 1 | 0 | 0 | 1688804 | YP_009047204.1 | K9N5Q8 |
| QRADLNGVIVDKVGEL | 1 | 0 | 0 | 1688817 | YP_173236.1 | A0A140H1G9 |
| QRFVYDAYQNLVGYYS | 1 | 0 | 0 | 1688825 | YP_009047204.1 | K9N5Q8 |
| QRGGSYTNDKACPLIA | 1 | 0 | 0 | 1688827 | YP_009724389.1 | P0DTD1 |
| QRGGSYTNDKTCPIVA | 1 | 0 | 0 | 1688828 | QIQ54046.1 | P0C6U8 |
| QRGHKNGQGENDNISV | 1 | 0 | 0 | 1688829 | YP_009555245.1 | P33469 |
| QRIDDDGNKLDKFFVV | 1 | 0 | 0 | 1688832 | YP_173236.1 | A0A140H1G9 |
| QRKLEKMADQAMTQMY | 1 | 0 | 0 | 1688836 | YP_009724389.1 | P0DTD1 |
| QRKRGVKQLPEQFDSL | 1 | 0 | 0 | 1688838 | YP_173242.1 | Q19U29 |
| QRKYKGIKIQEGVVDY | 1 | 0 | 0 | 1688840 | YP_009724389.1 | P0DTD1 |
| QRLDVLEQDAQIDRLI | 1 | 0 | 0 | 1688841 | YP_009047204.1 | K9N5Q8 |
| QRMLLDKCDLQNYGEA | 1 | 0 | 0 | 1688848 | QIQ54047.1 | P0C6X7 |
| QRQRGHKNGQGENDNI | 1 | 0 | 0 | 1688859 | YP_009555245.1 | P33469 |
| QRSVLTPLCGFPSQAA | 1 | 0 | 0 | 1688865 | NP_828849.2 | P0C6X7 |

| | | | | | | |
|---|---|---|---|---|---|---|
| QRTELGVEFLKRGDKI | 1 | 0 | 0 | 1688866 | NP_828849.2 | P0C6X7 |
| QRTFLLSYNQNGTITD | 1 | 0 | 0 | 1688867 | QIQ54048.1 | P59594 |
| QRVDENGDKLDQFFVV | 1 | 0 | 0 | 1688873 | YP_009555238.1 | P0C6X6 |
| QRVGTDSGFAAYNRYR | 1 | 0 | 0 | 1688876 | NP_828855.1 | P59596 |
| QRVQGSITQRTRTRPS | 1 | 0 | 0 | 1688879 | YP_009047211.1 | K9N4V7 |
| QRVYDDVDASLFVDIN | 1 | 0 | 0 | 1688880 | YP_173236.1 | A0A140H1G9 |
| QSATKIIALNKRWQLA | 1 | 0 | 0 | 1688895 | NP_828852.2 | P59632 |
| QSCTGSPYIDDFNKIA | 1 | 0 | 0 | 1688900 | YP_173236.1 | A0A140H1G9 |
| QSCTGSPYIDDFNRIA | 1 | 0 | 0 | 1688901 | YP_009555238.1 | P0C6X6 |
| QSEIEPEPEPTPEEPV | 1 | 0 | 0 | 1688912 | NP_828849.2 | P0C6X7 |
| QSESSCGLFKDCARNP | 1 | 0 | 0 | 1688914 | NP_073549.1 | A0A0P0LKV0 |
| QSFDNSYLNESGALVP | 1 | 0 | 0 | 1688915 | NP_073550.1 | A0A0P0ILY9 |
| QSGGGKCFNCYPAGVN | 1 | 0 | 0 | 1688924 | NP_073551.1 | A0A0P0K6L9 |
| QSGLVKMSHPSGDVEA | 1 | 0 | 0 | 1688931 | YP_009047202.1 | K9N7C7 |
| QSIFTINSVVQKDGVE | 1 | 0 | 0 | 1688937 | YP_009047202.1 | K9N7C7 |
| QSIIGAMTAVNESRYG | 1 | 0 | 0 | 1688939 | YP_003767.1 | A0A0P0G321 |
| QSKDSNFLNRVRGSIV | 1 | 0 | 0 | 1688945 | YP_009047202.1 | K9N7C7 |
| QSKSPSSSGTSTPKKP | 1 | 0 | 0 | 1688957 | YP_003771.1 | H9EJ69 |
| QSLCALPDTPSTLTPR | 1 | 0 | 0 | 1688961 | YP_009047204.1 | K9N5Q8 |
| QSLENVAYNVVNKGHF | 1 | 0 | 0 | 1688965 | NP_828849.2 | P0C6X7 |
| QSLENVIYNLVNVGHY | 1 | 0 | 0 | 1688966 | YP_173236.1 | A0A140H1G9 |
| QSLSKDYISSNGPLKV | 1 | 0 | 0 | 1688976 | YP_003766.2 | H9EJ42 |
| QSLSNAKVSAEVSRQV | 1 | 0 | 0 | 1688977 | NP_073552.1 | A0A0P0KWE8 |
| QSLTFSYDGLRGAIRK | 1 | 0 | 0 | 1688978 | YP_009047202.1 | K9N7C7 |
| QSNVDVVPEGWRVVNK | 1 | 0 | 0 | 1688991 | YP_009555238.1 | P0C6X6 |
| QSPASSQTSAKSLARS | 1 | 0 | 0 | 1688993 | NP_073556.1 | A0A0N7J131 |
| QSPQGNLGSNGKPGNV | 1 | 0 | 0 | 1688994 | YP_009555238.1 | P0C6X6 |
| QSPRRRRRQSRESQC | 1 | 0 | 0 | 1688996 | SRC279982 | Q69028 |
| QSQSSHVAQNTVLNAS | 1 | 0 | 0 | 1688999 | YP_003771.1 | H9EJ69 |
| QSRDLEDFKPRSQMET | 1 | 0 | 0 | 1689001 | NP_828849.2 | P0C6X7 |
| QSRGPNNRSLSRSNSN | 1 | 0 | 0 | 1689003 | YP_173242.1 | Q19U29 |
| QSRGRGESKPQSRNPS | 1 | 0 | 0 | 1689004 | NP_073556.1 | A0A0N7J131 |
| QSRTRSDSNQSSSDLV | 1 | 0 | 0 | 1689013 | YP_003771.1 | H9EJ69 |
| QSSHVAQNTVLNASIP | 1 | 0 | 0 | 1689021 | YP_003771.1 | H9EJ69 |
| QSSNYKIDTTSSSCQL | 1 | 0 | 0 | 1689023 | YP_173238.1 | A0A140H1H1 |
| QSSPIIPGFGGDFNLT | 1 | 0 | 0 | 1689024 | YP_009047204.1 | K9N5Q8 |
| QSSSRASSGNRSGNGI | 1 | 0 | 0 | 1689028 | YP_009555245.1 | P33469 |
| QSSSRASSLSRNSSRS | 1 | 0 | 0 | 1689029 | YP_009047211.1 | K9N4V7 |
| QSSYIVDSVAVKNGAL | 1 | 0 | 0 | 1689030 | NP_828849.2 | P0C6X7 |
| QSSYVVDSVAVKNGSL | 1 | 0 | 0 | 1689032 | QIQ54046.1 | P0C6U8 |
| QSTQLGIEFLKRGDKS | 1 | 0 | 0 | 1689035 | YP_009724389.1 | P0DTD1 |
| QSVIIINNSTNVVIRA | 1 | 0 | 0 | 1689047 | NP_828851.1 | P59594 |
| QSVVNSNAQALNSLLQ | 1 | 0 | 0 | 1689051 | YP_173238.1 | A0A140H1H1 |
| QSYQLYDASNVKKVTD | 1 | 0 | 0 | 1689064 | YP_009555238.1 | P0C6X6 |
| QTAAFRDVLVVLNKRT | 1 | 0 | 0 | 1689067 | NP_849177.1 | Q80H93 |
| QTADTAHANNINRFNV | 1 | 0 | 0 | 1689071 | YP_009047202.1 | K9N7C7 |
| QTATSQQPSGGNVVPY | 1 | 0 | 0 | 1689080 | YP_009555245.1 | P33469 |
| QTCNTPTQCYYNNSNN | 1 | 0 | 0 | 1689084 | YP_009555238.1 | P0C6X6 |
| QTCTPNVTINCQDPAG | 1 | 0 | 0 | 1689085 | NP_849177.1 | Q80H93 |
| QTFKSVLTTYYLDDVK | 1 | 0 | 0 | 1689094 | YP_009555238.1 | P0C6X6 |
| QTFYPRLQASADWKPG | 1 | 0 | 0 | 1689100 | YP_009047202.1 | K9N7C7 |
| QTGFTTTNEAFQKVQD | 1 | 0 | 0 | 1689103 | YP_009047204.1 | K9N5Q8 |
| QTGVIADYNYKLPDDF | 1 | 0 | 0 | 1689109 | NP_828851.1 | P59594 |
| QTIQDSDGTNKQLNEI | 1 | 0 | 0 | 1689119 | YP_009555238.1 | P0C6X6 |
| QTIVPEYIDVNKTLQE | 1 | 0 | 0 | 1689122 | NP_073551.1 | A0A0P0K6L9 |
| QTLIDNINSTLVDLKW | 1 | 0 | 0 | 1689126 | NP_073551.1 | A0A0P0K6L9 |
| QTLQDPRVRGLYFPA | 1 | 0 | 0 | 1689131 | P03138.3 | Q76R62 |
| QTNIYSQNLCVDGFGV | 1 | 0 | 0 | 1689138 | NP_073549.1 | A0A0P0LKV0 |
| QTNLYSFNKAIDGFGV | 1 | 0 | 0 | 1689139 | YP_003766.2 | H9EJ42 |
| QTNVVPPALHLVDPQI | 1 | 0 | 0 | 1689145 | NP_828859.1 | P59636 |
| QTPIVWHAKDFNSLSA | 1 | 0 | 0 | 1689148 | NP_073549.1 | A0A0P0LKV0 |
| QTRGRRAQPKQTATSQ | 1 | 0 | 0 | 1689159 | YP_009555245.1 | P33469 |
| QTSDTAHACNANRFNV | 1 | 0 | 0 | 1689161 | NP_073549.1 | A0A0P0LKV0 |
| QTSNFRVQPTDSIVRF | 1 | 0 | 0 | 1689170 | QHR63300.2 | Q3LZX1 |
| QTSNFRVQPTISIVRF | 1 | 0 | 0 | 1689172 | QIQ54048.1 | P59594 |
| QTSNLVVQEFVSYDSS | 1 | 0 | 0 | 1689174 | YP_009555238.1 | P0C6X6 |
| QTTPGSGVPIVDSYYS | 1 | 0 | 0 | 1689187 | QHR63299.1 | P0C6W2 |
| QTTPGSGVPVVDSYYS | 1 | 0 | 0 | 1689188 | YP_009724389.1 | P0DTD1 |
| QTVATALNKIQDVVNQ | 1 | 0 | 0 | 1689191 | NP_073551.1 | A0A0P0K6L9 |
| QTVDSSQGSEYQYVIF | 1 | 0 | 0 | 1689196 | YP_009047202.1 | K9N7C7 |
| QTVKPGHFNKEFYDFL | 1 | 0 | 0 | 1689200 | NP_073549.1 | A0A0P0LKV0 |
| QTVKPGNFNKDFYDFA | 1 | 0 | 0 | 1689201 | YP_009724389.1 | P0DTD1 |
| QTVKPGNFNQDFYDFV | 1 | 0 | 0 | 1689202 | YP_009555238.1 | P0C6X6 |
| QTVKPGNFNQDFYEFV | 1 | 0 | 0 | 1689203 | YP_173236.1 | A0A140H1G9 |
| QTVQDADGINKQLTDI | 1 | 0 | 0 | 1689207 | YP_173236.1 | A0A140H1G9 |
| QTVRPGNFNQDFYDFV | 1 | 0 | 0 | 1689208 | YP_009047202.1 | K9N7C7 |
| QTVVQEDDSEVNQTTI | 1 | 0 | 0 | 1689211 | QHR63299.1 | P0C6W2 |
| QTYHPNCINCLDDRCI | 1 | 0 | 0 | 1689214 | NP_828849.2 | P0C6X7 |
| QTYHPNCVNCLDDRCI | 1 | 0 | 0 | 1689215 | YP_009724389.1 | P0DTD1 |
| QVAVLYQDVNCTEVPM | 1 | 0 | 0 | 1689229 | QIQ54048.1 | P59594 |
| QVAVYQQSIIGAMTAV | 1 | 0 | 0 | 1689231 | YP_003767.1 | A0A0P0G321 |
| QVDKKIKACVEEVTTT | 1 | 0 | 0 | 1689236 | QHR63299.1 | P0C6W2 |
| QVDLVISDMYDPTTKN | 1 | 0 | 0 | 1689239 | YP_009047202.1 | K9N7C7 |
| QVDNGSSINRKQLEIV | 1 | 0 | 0 | 1689240 | YP_003766.2 | H9EJ42 |
| QVDNGSSINRRQLDVV | 1 | 0 | 0 | 1689241 | NP_073549.1 | A0A0P0LKV0 |
| QVDQLNSSYFKLSIPT | 1 | 0 | 0 | 1689242 | YP_009047204.1 | K9N5Q8 |
| QVEEVNSIFDIETVDV | 1 | 0 | 0 | 1689250 | NP_073549.1 | A0A0P0LKV0 |
| QVEQKIAEIPKEEVKP | 1 | 0 | 0 | 1689252 | YP_009724389.1 | P0DTD1 |
| QVESANLMLSDNVVAF | 1 | 0 | 0 | 1689254 | YP_003766.2 | H9EJ42 |
| QVESANQMLTVNVVAF | 1 | 0 | 0 | 1689255 | NP_073549.1 | A0A0P0LKV0 |
| QVESDHYIATNGPLKV | 1 | 0 | 0 | 1689257 | QHR63299.1 | P0C6W2 |

| | | | | | | |
|---|---|---|---|---|---|---|
| QVFDKVVDNVYVTYAG | 1 | 0 | 0 | 1689259 | YP_173236.1 | A0A140H1G9 |
| QVGFSNGVDFVAQPEG | 1 | 0 | 0 | 1689273 | NP_073549.1 | A0A0P0LKV0 |
| QVGVIADKDKDLSGAV | 1 | 0 | 0 | 1689278 | NP_073549.1 | A0A0P0LKV0 |
| QVIVNNLDKSAGFPFN | 1 | 0 | 0 | 1689290 | YP_009724389.1 | P0DTD1 |
| QVIVNNYDKSAGYPFN | 1 | 0 | 0 | 1689291 | YP_009555238.1 | P0C6X6 |
| QVLSENQKLIANKFNQ | 1 | 0 | 0 | 1689309 | YP_009047204.1 | K9N5Q8 |
| QVNISLNGNTSVCVRT | 1 | 0 | 0 | 1689317 | YP_003767.1 | A0A0P0G321 |
| QVNWNTKHGSGVTREL | 1 | 0 | 0 | 1689318 | QIQ54046.1 | P0C6U8 |
| QVPATSIVAKQGAGFK | 1 | 0 | 0 | 1689321 | YP_003766.2 | H9EJ42 |
| QVPIGTNDPIRFCLEN | 1 | 0 | 0 | 1689323 | NP_073549.1 | A0A0P0LKV0 |
| QVPVGIKDPVSYVLTH | 1 | 0 | 0 | 1689325 | YP_009555238.1 | P0C6X6 |
| QVRSWIGFDVEGAHAS | 1 | 0 | 0 | 1689334 | YP_009047202.1 | K9N7C7 |
| QVTIAGAKLRSLNLGE | 1 | 0 | 0 | 1689341 | NP_828849.2 | P0C6X7 |
| QVTYQNISTNLPPPLL | 1 | 0 | 0 | 1689347 | YP_009047204.1 | K9N5Q8 |
| QVVDSRWNNARQSDNM | 1 | 0 | 0 | 1689353 | YP_009555240.1 | P30215 |
| QVVIADTLQETPVVPD | 1 | 0 | 0 | 1689355 | YP_009047202.1 | K9N7C7 |
| QVVKALNESYIDLKEL | 1 | 0 | 0 | 1689356 | YP_009047204.1 | K9N5Q8 |
| QVVKLPSAPQTIQPEV | 1 | 0 | 0 | 1689357 | YP_009047202.1 | K9N7C7 |
| QVVKPGHFNEEFYNFL | 1 | 0 | 0 | 1689358 | YP_003766.2 | H9EJ42 |
| QVVLVSNNQEDFDLIS | 1 | 0 | 0 | 1689359 | YP_009555238.1 | P0C6X6 |
| QVVMDYLDNLKPRVEA | 1 | 0 | 0 | 1689360 | NP_828849.2 | P0C6X7 |
| QVVQLPVKDYVQTVNV | 1 | 0 | 0 | 1689366 | YP_173236.1 | A0A140H1G9 |
| QVVTYPSLNYAGALWD | 1 | 0 | 0 | 1689367 | YP_009047202.1 | K9N7C7 |
| QVYIAVNDKALYEQVV | 1 | 0 | 0 | 1689370 | NP_828849.2 | P0C6X7 |
| QWDLIISDMYDPITKN | 1 | 0 | 0 | 1689376 | YP_009555238.1 | P0C6X6 |
| QWLPAGTILVDNDLYP | 1 | 0 | 0 | 1689393 | YP_009555238.1 | P0C6X6 |
| QWLPTDAIIIDNDLNE | 1 | 0 | 0 | 1689395 | YP_009047202.1 | K9N7C7 |
| QWPISNDSNGCVVEAS | 1 | 0 | 0 | 1689407 | YP_003766.2 | H9EJ42 |
| QWYHEHVGGVYDNSIT | 1 | 0 | 0 | 1689415 | YP_009047202.1 | K9N7C7 |
| QXRAPRITFGGPSDST | 1 | 0 | 0 | 1689416 | QIQ54056.1 | P59595 |
| QXYSTLQGPPGTGKSH | 1 | 0 | 0 | 1689417 | QIQ54047.1 | P0C6X7 |
| QYAHKHDIQITNESYN | 1 | 0 | 0 | 1689419 | YP_009047202.1 | K9N7C7 |
| QYDFTDYKLELFNKYF | 1 | 0 | 0 | 1689425 | YP_009555238.1 | P0C6X6 |
| QYDVLFYCSNSSSGVL | 1 | 0 | 0 | 1689427 | YP_003767.1 | A0A0P0G321 |
| QYENLSLADISAVQSS | 1 | 0 | 0 | 1689431 | YP_173236.1 | A0A140H1G9 |
| QYEYGTEDDYQGKPLE | 1 | 0 | 0 | 1689433 | YP_009724389.1 | P0DTD1 |
| QYEYGTESDYKGLPLE | 1 | 0 | 0 | 1689435 | QIQ54046.1 | P0C6U8 |
| QYGCDYSGKLADCLQA | 1 | 0 | 0 | 1689437 | YP_173236.1 | A0A140H1G9 |
| QYGTDTNSVCPKLEFA | 1 | 0 | 0 | 1689448 | YP_009047204.1 | K9N5Q8 |
| QYIFSTVNALPECNAD | 1 | 0 | 0 | 1689457 | YP_003766.2 | H9EJ42 |
| QYKGKWVQVPIGTNDP | 1 | 0 | 0 | 1689462 | NP_073549.1 | A0A0P0LKV0 |
| QYLCGADGKPVMSEDL | 1 | 0 | 0 | 1689463 | NP_073549.1 | A0A0P0LKV0 |
| QYLDAYNMMISAGFSL | 1 | 0 | 0 | 1689464 | NP_828849.2 | P0C6X7 |
| QYLFSTINALPETSAD | 1 | 0 | 0 | 1689465 | YP_009047202.1 | K9N7C7 |
| QYLLAFNQDGIIFNAV | 1 | 0 | 0 | 1689467 | YP_009555241.1 | P36334 |
| QYLLKFDNRGVITNAV | 1 | 0 | 0 | 1689468 | YP_173238.1 | A0A140H1H1 |
| QYMCGVDGKPISAYAF | 1 | 0 | 0 | 1689475 | YP_009047202.1 | K9N7C7 |
| QYRINGLGVTMDVLNK | 1 | 0 | 0 | 1689482 | YP_173238.1 | A0A140H1H1 |
| QYRINGLGVTMDVLSQ | 1 | 0 | 0 | 1689483 | YP_009555241.1 | P36334 |
| QYSPCVSIVPSTVWED | 1 | 0 | 0 | 1689489 | YP_009047204.1 | K9N5Q8 |
| QYSTCDICKSTVVEVK | 1 | 0 | 0 | 1689492 | YP_003766.2 | H9EJ42 |
| QYTSACKTIEDALRNS | 1 | 0 | 0 | 1689495 | NP_073551.1 | A0A0P0K6L9 |
| QYVAGYKVLPPLMDVN | 1 | 0 | 0 | 1689497 | YP_009047204.1 | K9N5Q8 |
| QYVFSTVNALPEVNAD | 1 | 0 | 0 | 1689501 | NP_073549.1 | A0A0P0LKV0 |
| QYYFNKDTGVIYGLNS | 1 | 0 | 0 | 1689508 | YP_009555240.1 | P30215 |
| QYYIYVTNEIGLNASV | 1 | 0 | 0 | 1689509 | YP_003767.1 | A0A0P0G321 |
| RACIKTAQADTAGLYT | 1 | 0 | 0 | 1689524 | YP_009047205.1 | K9N796 |
| RADLNGVIVDKVGELK | 1 | 0 | 0 | 1689531 | YP_173236.1 | A0A140H1G9 |
| RADQINDMVYSLLEKG | 1 | 0 | 0 | 1689532 | YP_009555238.1 | P0C6X6 |
| RAELNGVVVDKVGDTD | 1 | 0 | 0 | 1689535 | YP_009555238.1 | P0C6X6 |
| RAFDVYNKDASFIGKN | 1 | 0 | 0 | 1689543 | NP_073549.1 | A0A0P0LKV0 |
| RAFISVLDITDAAVKA | 1 | 0 | 0 | 1689548 | NP_073549.1 | A0A0P0LKV0 |
| RAGCLVGAEHVNNSYE | 1 | 0 | 0 | 1689558 | QIQ54048.1 | P59594 |
| RAIDCGFNDLSQLHCS | 1 | 0 | 0 | 1689572 | YP_009047204.1 | K9N5Q8 |
| RAIIVSVEQLEPCAQS | 1 | 0 | 0 | 1689575 | NP_073549.1 | A0A0P0LKV0 |
| RAKHYVYIGDPAQLPA | 1 | 0 | 0 | 1689582 | YP_009724389.1 | P0DTD1 |
| RALAAESHLDADLTKP | 1 | 0 | 0 | 1689592 | QIQ54047.1 | P0C6X7 |
| RALAAESHMDADLAKP | 1 | 0 | 0 | 1689593 | NP_828849.2 | P0C6X7 |
| RALSGIAAEQDRNTRE | 1 | 0 | 0 | 1689600 | NP_828851.1 | P59594 |
| RALSGVFCGVDAMNLI | 1 | 0 | 0 | 1689601 | NP_828849.2 | P0C6X7 |
| RAMNIAKSEFDHEISV | 1 | 0 | 0 | 1689608 | YP_003766.2 | H9EJ42 |
| RANSAVKLQNNELSPV | 1 | 0 | 0 | 1689618 | YP_009724389.1 | P0DTD1 |
| RANSGNRTPTSGVTPD | 1 | 0 | 0 | 1689620 | YP_009555245.1 | P33469 |
| RAPRITFGGPSDSTDN | 1 | 0 | 0 | 1689623 | QIQ54056.1 | P59595 |
| RAPTMANQEKYSTIYK | 1 | 0 | 0 | 1689626 | NP_073549.1 | A0A0P0LKV0 |
| RARVECFDKFKVNSTL | 1 | 0 | 0 | 1689643 | YP_009724389.1 | P0DTD1 |
| RARVEHPDVDGLCKLR | 1 | 0 | 0 | 1689647 | YP_009555238.1 | P0C6X6 |
| RASLRQIFDVAPIELE | 1 | 0 | 0 | 1689653 | YP_009047202.1 | K9N7C7 |
| RASSAGSRSRANSGNR | 1 | 0 | 0 | 1689656 | YP_009555245.1 | P33469 |
| RASSGNRSGNGILKWA | 1 | 0 | 0 | 1689658 | YP_009555245.1 | P33469 |
| RASSLSRNSSRSSSQG | 1 | 0 | 0 | 1689659 | YP_009047211.1 | K9N4V7 |
| RASSRSSTRNNSRDSS | 1 | 0 | 0 | 1689660 | YP_003771.1 | H9EJ69 |
| RATATIPIQASLPFGW | 1 | 0 | 0 | 1689662 | YP_009724391.1 | P0DTC3 |
| RATRRVRGGDGKMKDL | 1 | 0 | 0 | 1689671 | QIQ54056.1 | P59595 |
| RATRRVRGGDGKMKEL | 1 | 0 | 0 | 1689672 | NP_828858.1 | P59595 |
| RAVRSHPDFKLLHNLQ | 1 | 0 | 0 | 1689682 | YP_009047202.1 | K9N7C7 |
| RAVSFADNNDITNTNL | 1 | 0 | 0 | 1689683 | YP_009047211.1 | K9N4V7 |
| RAWAFCVKCHNDILAA | 1 | 0 | 0 | 1689684 | YP_009047202.1 | K9N7C7 |
| RAWVGFDAEGAHATRD | 1 | 0 | 0 | 1689687 | YP_009555238.1 | P0C6X6 |
| RCHIDHPNPKGFCDLK | 1 | 0 | 0 | 1689714 | YP_009724389.1 | P0DTD1 |
| RCHIDHPNPKGYCELK | 1 | 0 | 0 | 1689715 | QIQ54046.1 | P0C6U8 |

| | | | | | | |
|---|---|---|---|---|---|---|
| RCIKISTVQSKLTDLK | 1 | 0 | 0 | 1689717 | YP_003766.2 | H9EJ42 |
| RCIVLDVKGVEELHDD | 1 | 0 | 0 | 1689721 | YP_009555239.1 | Q80872 |
| RCLWNTKPVETSNSFD | 1 | 0 | 0 | 1689732 | QHR63299.1 | P0C6W2 |
| RCLWSTKPVDTSNSFE | 1 | 0 | 0 | 1689733 | NP_828849.2 | P0C6X7 |
| RCLWSTKPVETSNIFD | 1 | 0 | 0 | 1689734 | QIQ54046.1 | P0C6U8 |
| RCLWSTKPVETSNSFD | 1 | 0 | 0 | 1689735 | YP_009724389.1 | P0DTD1 |
| RCNIGGAVCSKHANLY | 1 | 0 | 0 | 1689738 | YP_003766.2 | H9EJ42 |
| RCPAEIVDTVSALVYD | 1 | 0 | 0 | 1689743 | YP_009724389.1 | P0DTD1 |
| RCPKEIVDTVSALVYE | 1 | 0 | 0 | 1689745 | YP_009555238.1 | P0C6X6 |
| RCPTAADINIPDLPIC | 1 | 0 | 0 | 1689748 | YP_009555240.1 | P30215 |
| RCWYLHEGHQTAAFRD | 1 | 0 | 0 | 1689764 | NP_849177.1 | Q80H93 |
| RDAQVVQLPVKDYVQT | 1 | 0 | 0 | 1689770 | YP_173236.1 | A0A140H1G9 |
| RDATQYLVQQESSFVM | 1 | 0 | 0 | 1689773 | NP_828849.2 | P0C6X7 |
| RDAVGTNLPLQLGFST | 1 | 0 | 0 | 1689774 | NP_828849.2 | P0C6X7 |
| RDCDFNKPLIEWPLTE | 1 | 0 | 0 | 1689776 | YP_009047202.1 | K9N7C7 |
| RDELGVRVLDQSDNNC | 1 | 0 | 0 | 1689779 | YP_003766.2 | H9EJ42 |
| RDGCVPLNIIPLTTAA | 1 | 0 | 0 | 1689789 | YP_009724389.1 | P0DTD1 |
| RDGQRVYDDVDASLFV | 1 | 0 | 0 | 1689794 | YP_173236.1 | A0A140H1G9 |
| RDGVRRGDNFHSVLTT | 1 | 0 | 0 | 1689798 | YP_009047202.1 | K9N7C7 |
| RDGVVPLSVIPATSAS | 1 | 0 | 0 | 1689799 | YP_003766.2 | H9EJ42 |
| RDGYLNSTNVTTTAYC | 1 | 0 | 0 | 1689801 | QIQ54046.1 | P0C6U8 |
| RDHNDILYTPPTVSYG | 1 | 0 | 0 | 1689804 | NP_073549.1 | A0A0P0LKV0 |
| RDISDTTDAVRDPQTL | 1 | 0 | 0 | 1689809 | QIQ54048.1 | P59594 |
| RDKAVARKLERMADLA | 1 | 0 | 0 | 1689816 | YP_173236.1 | A0A140H1G9 |
| RDKLEKLISTARDGVR | 1 | 0 | 0 | 1689819 | YP_009047202.1 | K9N7C7 |
| RDLEDFKPRSQMETDF | 1 | 0 | 0 | 1689822 | NP_828849.2 | P0C6X7 |
| RDLLIKAYNTINNEQG | 1 | 0 | 0 | 1689832 | NP_073549.1 | A0A0P0LKV0 |
| RDLSLQFKRPINPTDQ | 1 | 0 | 0 | 1689840 | YP_009724389.1 | P0DTD1 |
| RDLYDKLQFTSLEIPR | 1 | 0 | 0 | 1689841 | YP_009724389.1 | P0DTD1 |
| RDLYDKLQFTSLEVPR | 1 | 0 | 0 | 1689842 | QIQ54047.1 | P0C6X7 |
| RDNGALEAFKRSNNGV | 1 | 0 | 0 | 1689846 | YP_009555238.1 | P0C6X6 |
| RDNLKPVTTLSTITAY | 1 | 0 | 0 | 1689847 | YP_003768.1 | H9EJ24 |
| RDPRYVMRLQSAATIR | 1 | 0 | 0 | 1689851 | YP_009555238.1 | P0C6X6 |
| RDPSSDEAIPTRFPPG | 1 | 0 | 0 | 1689852 | YP_009555245.1 | P33469 |
| RDRAVAKKLERMADLA | 1 | 0 | 0 | 1689855 | YP_009555238.1 | P0C6X6 |
| RDSHKGVCFGFDKWYV | 1 | 0 | 0 | 1689859 | YP_003766.2 | H9EJ42 |
| RDSIGTNFPLQLGFST | 1 | 0 | 0 | 1689860 | YP_009555238.1 | P0C6X6 |
| RDSRFVMRLQTIATIC | 1 | 0 | 0 | 1689861 | YP_173236.1 | A0A140H1G9 |
| RDSSRSTSRQQSRTRS | 1 | 0 | 0 | 1689862 | YP_003771.1 | H9EJ69 |
| RDSTYGPLQTPVGCVL | 1 | 0 | 0 | 1689863 | YP_009047204.1 | K9N5Q8 |
| RDSVPLHIIAPSSLIV | 1 | 0 | 0 | 1689864 | YP_009047208.1 | K9N7D2 |
| RDVDTDFVNEFYAYLR | 1 | 0 | 0 | 1689870 | YP_009724389.1 | P0DTD1 |
| RDVGFVVPGLPGTIFR | 1 | 0 | 0 | 1689872 | QIQ54046.1 | P0C6U8 |
| RDVSVEDVCFANKFEQ | 1 | 0 | 0 | 1689877 | YP_009555238.1 | P0C6X6 |
| RDYKLANDAKESLRLF | 1 | 0 | 0 | 1689882 | NP_073549.1 | A0A0P0LKV0 |
| RDYKLANDVKDTLRLF | 1 | 0 | 0 | 1689883 | YP_003766.2 | H9EJ42 |
| REAACCHLAKALNDFS | 1 | 0 | 0 | 1689889 | YP_009724389.1 | P0DTD1 |
| REAIGTNLPLQLGFST | 1 | 0 | 0 | 1689892 | QHR63299.1 | P0C6W2 |
| REEVDIIEQPFEEVEH | 1 | 0 | 0 | 1689904 | YP_003766.2 | H9EJ42 |
| REFKVTFFPDLNGDVV | 1 | 0 | 0 | 1689906 | QHR63299.1 | P0C6W2 |
| REGYLNSTNVTIATYC | 1 | 0 | 0 | 1689918 | YP_009724389.1 | P0DTD1 |
| REGYLNSTNVTTAIYC | 1 | 0 | 0 | 1689919 | QHR63299.1 | P0C6W2 |
| REISCNVCTKRATVYN | 1 | 0 | 0 | 1689926 | YP_009555238.1 | P0C6X6 |
| RELILSWEIGKVKPPL | 1 | 0 | 0 | 1689935 | YP_009555238.1 | P0C6X6 |
| RELKITFFPDLNGDVV | 1 | 0 | 0 | 1689936 | QIQ54046.1 | P0C6U8 |
| RELKVTFFPDLNGDVV | 1 | 0 | 0 | 1689937 | YP_009724389.1 | P0DTD1 |
| RELNGGAVTRYVDNNF | 1 | 0 | 0 | 1689941 | NP_828849.2 | P0C6X7 |
| RELSVTFFPDLNGDVV | 1 | 0 | 0 | 1689943 | NP_828849.2 | P0C6X7 |
| RELTAEDISLLKKMDE | 1 | 0 | 0 | 1689944 | YP_009555245.1 | P33469 |
| RELYLNSSNVTTMDFC | 1 | 0 | 0 | 1689946 | NP_828849.2 | P0C6X7 |
| REMQQHPLLNPSALEF | 1 | 0 | 0 | 1689952 | NP_073556.1 | A0A0N7J131 |
| RENIGTNFPLQIGFST | 1 | 0 | 0 | 1689954 | YP_173236.1 | A0A140H1G9 |
| RENNRVVISSDVLVNN | 1 | 0 | 0 | 1689956 | YP_009724389.1 | P0DTD1 |
| RENNRVVVSSDILVNN | 1 | 0 | 0 | 1689957 | NP_828849.2 | P0C6X7 |
| REPMLQSADAQSFLNR | 1 | 0 | 0 | 1689961 | YP_009724389.1 | P0DTD1 |
| RERVNINIVGDFHLNE | 1 | 0 | 0 | 1689967 | NP_828849.2 | P0C6X7 |
| RETMNYLFQHANLDSC | 1 | 0 | 0 | 1689971 | QHR63299.1 | P0C6W2 |
| REVFAQVKQMYKTPTL | 1 | 0 | 0 | 1689977 | NP_828851.1 | P59594 |
| REVILCWETGKVKPPL | 1 | 0 | 0 | 1689979 | YP_173236.1 | A0A140H1G9 |
| REVKTIKVFTTVDNTN | 1 | 0 | 0 | 1689981 | NP_828849.2 | P0C6X7 |
| REVRTIKVFTTVDNIN | 1 | 0 | 0 | 1689984 | YP_009724389.1 | P0DTD1 |
| REVVVTNLNKSAGWPL | 1 | 0 | 0 | 1689985 | NP_073549.1 | A0A0P0LKV0 |
| REYLESYNTATTAGFT | 1 | 0 | 0 | 1689989 | YP_009555238.1 | P0C6X6 |
| RFDTRVLNNLNLPGCN | 1 | 0 | 0 | 1690010 | YP_009555238.1 | P0C6X6 |
| RFEYTSPTDKLAMIMG | 1 | 0 | 0 | 1690016 | YP_009555239.1 | Q80872 |
| RFGFIEDSVFKPRPAG | 1 | 0 | 0 | 1690025 | YP_009555241.1 | P36334 |
| RFKTSGDLAVQIGNNN | 1 | 0 | 0 | 1690044 | NP_073549.1 | A0A0P0LKV0 |
| RFKVNETNSQYLFSTI | 1 | 0 | 0 | 1690045 | YP_009047202.1 | K9N7C7 |
| RFLGLQIQTVDSSQGS | 1 | 0 | 0 | 1690047 | YP_003766.2 | H9EJ42 |
| RFMTANNLTAPRNSWE | 1 | 0 | 0 | 1690051 | YP_009047202.1 | K9N7C7 |
| RFNGIGVTQNVLYENQ | 1 | 0 | 0 | 1690054 | YP_009724390.1 | P0DTC2 |
| RFNPSTWNKRFGFIED | 1 | 0 | 0 | 1690055 | YP_009555241.1 | P36334 |
| RFPCAGRKVNFNEKPV | 1 | 0 | 0 | 1690062 | YP_173236.1 | A0A140H1G9 |
| RFPRGQGVPINTNSTK | 1 | 0 | 0 | 1690072 | QIQ54056.1 | P59595 |
| RFQEKDEDDNLIDSYF | 1 | 0 | 0 | 1690074 | YP_009724389.1 | P0DTD1 |
| RFQEKDEDGNLIDSYF | 1 | 0 | 0 | 1690075 | QIQ54047.1 | P0C6X7 |
| RFQEKDEEGNLLDSYF | 1 | 0 | 0 | 1690076 | NP_828849.2 | P0C6X7 |
| RFQRIDDDGNKLDKFF | 1 | 0 | 0 | 1690078 | YP_173236.1 | A0A140H1G9 |
| RFQRVDENGDKLDQFF | 1 | 0 | 0 | 1690079 | YP_009555238.1 | P0C6X6 |
| RFTNFEPFTVNSVNDS | 1 | 0 | 0 | 1690095 | YP_009555241.1 | P36334 |
| RFVTFEPFNVSFVNDS | 1 | 0 | 0 | 1690114 | YP_173238.1 | A0A140H1H1 |

| | | | | | | |
|---|---|---|---|---|---|---|
| RFVYDAYQNLVGYYSD | 1 | 0 | 0 | 1690115 | YP_009047204.1 | K9N5Q8 |
| RFYRLANECAQVLSEY | 1 | 0 | 0 | 1690124 | YP_009047202.1 | K9N7C7 |
| RGAITTGYRFTNFEPF | 1 | 0 | 0 | 1690129 | YP_009555241.1 | P36334 |
| RGCCDCGSTKLPYYEF | 1 | 0 | 0 | 1690136 | YP_003767.1 | A0A0P0G321 |
| RGCCESTKLPYYDVEK | 1 | 0 | 0 | 1690137 | NP_073551.1 | A0A0P0K6L9 |
| RGDEVRQIAPGQTGKI | 1 | 0 | 0 | 1690142 | YP_009724390.1 | P0DTC2 |
| RGDKVVYHTTSKPITF | 1 | 0 | 0 | 1690147 | QIQ54046.1 | P0C6U8 |
| RGDYNMLLKNIDLFVK | 1 | 0 | 0 | 1690150 | YP_173236.1 | A0A140H1G9 |
| RGEDIQLLKSAYENFN | 1 | 0 | 0 | 1690151 | QHR63299.1 | P0C6W2 |
| RGEKGAYNKDHGRGGF | 1 | 0 | 0 | 1690153 | YP_009555238.1 | P0C6X6 |
| RGFGDSVEEVLSEARQ | 1 | 0 | 0 | 1690160 | YP_009724389.1 | P0DTD1 |
| RGGDGKMKELSPRWYF | 1 | 0 | 0 | 1690165 | NP_828858.1 | P59595 |
| RGGSQASSRSSSRSRG | 1 | 0 | 0 | 1690168 | NP_828858.1 | P59595 |
| RGGSYKNDKSCPVVAA | 1 | 0 | 0 | 1690170 | NP_828849.2 | P0C6X7 |
| RGGSYTNDKACPLIAA | 1 | 0 | 0 | 1690171 | YP_009724389.1 | P0DTD1 |
| RGGSYTNDKTCPIVAA | 1 | 0 | 0 | 1690172 | QIQ54046.1 | P0C6U8 |
| RGGYHYTPFHYERDNT | 1 | 0 | 0 | 1690173 | YP_009047202.1 | K9N7C7 |
| RGHKNGQGENDNISVA | 1 | 0 | 0 | 1690174 | YP_009555245.1 | P33469 |
| RGHLYIQGIKLGTGYS | 1 | 0 | 0 | 1690177 | YP_009555244.1 | Q01455 |
| RGHLYIQGVKLGTGYT | 1 | 0 | 0 | 1690178 | YP_173241.1 | Q0ZJ82 |
| RGIAPKSGYFVNVNNT | 1 | 0 | 0 | 1690180 | YP_009555241.1 | P36334 |
| RGKFVQVPLGIKDPIL | 1 | 0 | 0 | 1690185 | YP_173236.1 | A0A140H1G9 |
| RGKFVQVPVGIKDPVS | 1 | 0 | 0 | 1690186 | YP_009555238.1 | P0C6X6 |
| RGKLLVRDTRKEIFVG | 1 | 0 | 0 | 1690187 | YP_173236.1 | A0A140H1G9 |
| RGLALDNALSGVLSAA | 1 | 0 | 0 | 1690188 | NP_073549.1 | A0A0P0LKV0 |
| RGLLHSSQLPLDVYDL | 1 | 0 | 0 | 1690190 | YP_009555238.1 | P0C6X6 |
| RGMPIPDFDLDDFIDA | 1 | 0 | 0 | 1690194 | YP_009047202.1 | K9N7C7 |
| RGNGKCLSFSNHLVST | 1 | 0 | 0 | 1690197 | NP_073549.1 | A0A0P0LKV0 |
| RGPGDLQGNFGDLQLN | 1 | 0 | 0 | 1690203 | YP_009047211.1 | K9N4V7 |
| RGPNQNFGGGEMLKLG | 1 | 0 | 0 | 1690205 | YP_009555245.1 | P33469 |
| RGPPRAELNGVVVDKV | 1 | 0 | 0 | 1690206 | YP_009555238.1 | P0C6X6 |
| RGQGVPINTNSSPDDQ | 1 | 0 | 0 | 1690210 | YP_009724397.2 | P0DTC9 |
| RGQGVPINTNSTKDDQ | 1 | 0 | 0 | 1690211 | QIQ54056.1 | P59595 |
| RGRAIIVSVEQLEPCA | 1 | 0 | 0 | 1690215 | NP_073549.1 | A0A0P0LKV0 |
| RGRIDKDLSLSRKVEL | 1 | 0 | 0 | 1690218 | NP_073553.1 | A0A0P0KWE8 |
| RGRQFYEFYNDVKPPV | 1 | 0 | 0 | 1690221 | YP_009555243.1 | Q04854 |
| RGSIVNARIEPCSSGL | 1 | 0 | 0 | 1690227 | YP_009047202.1 | K9N7C7 |
| RGSVQVDNGSSINRRQ | 1 | 0 | 0 | 1690230 | NP_073549.1 | A0A0P0LKV0 |
| RGTAVMSLKENQINDM | 1 | 0 | 0 | 1690232 | NP_828849.2 | P0C6X7 |
| RGTLEPENFNSVTRLM | 1 | 0 | 0 | 1690236 | YP_009047202.1 | K9N7C7 |
| RGTSPARIAGNGGDAA | 1 | 0 | 0 | 1690238 | QIQ54056.1 | P59595 |
| RGTSPARMAGNGSDAA | 1 | 0 | 0 | 1690240 | QHR63308.1 | Q3LZX4 |
| RGTSPGPSGIGAVGGD | 1 | 0 | 0 | 1690241 | YP_009047211.1 | K9N4V7 |
| RGTTVLLKEPCPSGTY | 1 | 0 | 0 | 1690246 | NP_828857.1 | P59635 |
| RGTYRAALNSEKHQDH | 1 | 0 | 0 | 1690248 | YP_009047202.1 | K9N7C7 |
| RGVKQLPEQFDSLNLS | 1 | 0 | 0 | 1690253 | YP_173242.1 | Q19U29 |
| RGVYKPLLENIDYFNM | 1 | 0 | 0 | 1690260 | YP_009555238.1 | P0C6X6 |
| RGWLGMDVEGAHVTGD | 1 | 0 | 0 | 1690265 | NP_073549.1 | A0A0P0LKV0 |
| RGWLGMDVESAHVCGD | 1 | 0 | 0 | 1690266 | YP_003766.2 | H9EJ42 |
| RGWVFGSTMNNKSQSV | 1 | 0 | 0 | 1690268 | NP_828851.1 | P59594 |
| RGWVGFDVEGAHATRE | 1 | 0 | 0 | 1690269 | YP_173236.1 | A0A140H1G9 |
| RHGKLRPFERDISNVP | 1 | 0 | 0 | 1690285 | NP_828851.1 | P59594 |
| RHHANEYRLYLDAYNM | 1 | 0 | 0 | 1690289 | YP_009724389.1 | P0DTD1 |
| RHHANEYRQYLDAYNM | 1 | 0 | 0 | 1690290 | NP_828849.2 | P0C6X7 |
| RHLIKDVDNPVLMGWD | 1 | 0 | 0 | 1690302 | YP_173236.1 | A0A140H1G9 |
| RIAGHHLGRCDIKDLP | 1 | 0 | 0 | 1690340 | YP_009724393.1 | P0DTC5 |
| RIDDDGNKLDKFFVVK | 1 | 0 | 0 | 1690350 | YP_173236.1 | A0A140H1G9 |
| RIDVPSAESTGTQSVS | 1 | 0 | 0 | 1690357 | YP_009047205.1 | K9N796 |
| RIFTIGTVTLKQGEIK | 1 | 0 | 0 | 1690366 | YP_009724391.1 | P0DTC3 |
| RIFTLGTVTLKQGEIK | 1 | 0 | 0 | 1690367 | QHR63301.1 | Q3LZX0 |
| RIGNYKLNTDHAGSND | 1 | 0 | 0 | 1690373 | NP_828855.1 | P59596 |
| RIGNYKLNTDHSNSSD | 1 | 0 | 0 | 1690374 | QIQ54051.1 | P59596 |
| RINGLGVTMDVLNKNQ | 1 | 0 | 0 | 1690409 | YP_173238.1 | A0A140H1H1 |
| RIRAKHYVVIGDPAQL | 1 | 0 | 0 | 1690421 | YP_009555238.1 | P0C6X6 |
| RIRSVYPVASPQECNN | 1 | 0 | 0 | 1690428 | NP_828849.2 | P0C6X7 |
| RITFELDERVDKVLNE | 1 | 0 | 0 | 1690434 | NP_828849.2 | P0C6X7 |
| RITFGGPSDSTDNNQN | 1 | 0 | 0 | 1690435 | QIQ54056.1 | P59595 |
| RITGLYPTLNISDEFS | 1 | 0 | 0 | 1690437 | YP_009724389.1 | P0DTD1 |
| RIVDEPEDHVQIHTID | 1 | 0 | 0 | 1690442 | QIQ54049.1 | P59632 |
| RIVDEWSYTGSSFYAP | 1 | 0 | 0 | 1690443 | YP_009047204.1 | K9N5Q8 |
| RIVFTACSHAAVDSLC | 1 | 0 | 0 | 1690445 | NP_073549.1 | A0A0P0LKV0 |
| RIVQMLSDTLDKLSDY | 1 | 0 | 0 | 1690453 | YP_009047202.1 | K9N7C7 |
| RIVYAVLSDVDGLKYT | 1 | 0 | 0 | 1690461 | YP_173236.1 | A0A140H1G9 |
| RKARNGVFISTEKLSR | 1 | 0 | 0 | 1690470 | YP_173236.1 | A0A140H1G9 |
| RKCVKPKEETGLLMPL | 1 | 0 | 0 | 1690476 | QHR63299.1 | P0C6W2 |
| RKCVKSREETGLLMPL | 1 | 0 | 0 | 1690477 | YP_009724389.1 | P0DTD1 |
| RKDLSYEQQNVLAIES | 1 | 0 | 0 | 1690481 | YP_003766.2 | H9EJ42 |
| RKGKHERDLLIKAYNT | 1 | 0 | 0 | 1690495 | NP_073549.1 | A0A0P0LKV0 |
| RKIKKYPCMVGPDYAY | 1 | 0 | 0 | 1690512 | YP_009047202.1 | K9N7C7 |
| RKILLRKNGNKGAGGH | 1 | 0 | 0 | 1690513 | QIQ54046.1 | P0C6U8 |
| RKKKSNLLIQEFLQYD | 1 | 0 | 0 | 1690519 | YP_173236.1 | A0A140H1G9 |
| RKKRNVDPLSNFEHKV | 1 | 0 | 0 | 1690520 | YP_009047202.1 | K9N7C7 |
| RKLDNDALNNIINNAR | 1 | 0 | 0 | 1690521 | YP_009724389.1 | P0DTD1 |
| RKLDNQALNSILDNAV | 1 | 0 | 0 | 1690522 | YP_009555238.1 | P0C6X6 |
| RKLEKMADQAMTQMYK | 1 | 0 | 0 | 1690523 | YP_009724389.1 | P0DTD1 |
| RKLERMADLALTNMYK | 1 | 0 | 0 | 1690524 | YP_173236.1 | A0A140H1G9 |
| RKLERMADQAMTSMYK | 1 | 0 | 0 | 1690525 | YP_009047202.1 | K9N7C7 |
| RKLHDELTGHMLDMYS | 1 | 0 | 0 | 1690527 | YP_009724389.1 | P0DTD1 |
| RKNGNKGAGGHSFGID | 1 | 0 | 0 | 1690540 | QIQ54046.1 | P0C6U8 |
| RKNGNKGAGGHSYGAD | 1 | 0 | 0 | 1690541 | YP_009724389.1 | P0DTD1 |
| RKNGNKGAGGHSYGID | 1 | 0 | 0 | 1690542 | NP_828849.2 | P0C6X7 |

| | | | | | | |
|---|---|---|---|---|---|---|
| RKNGQFQDHYDGFYTQ | 1 | 0 | 0 | 1690543 | YP_003766.2 | H9EJ42 |
| RKPLDYKRQNNLAIEE | 1 | 0 | 0 | 1690545 | NP_073549.1 | A0A0P0LKV0 |
| RKPNLPNCNIEAWLND | 1 | 0 | 0 | 1690548 | YP_009555241.1 | P36334 |
| RKQDDKKIKACVEEVT | 1 | 0 | 0 | 1690549 | YP_009724389.1 | P0DTD1 |
| RKQIRSAAKKNNIPFR | 1 | 0 | 0 | 1690550 | NP_828849.2 | P0C6X7 |
| RKQIRSAAKKNNLPFK | 1 | 0 | 0 | 1690551 | YP_009724389.1 | P0DTD1 |
| RKRGEKGAYNKDHGRG | 1 | 0 | 0 | 1690554 | YP_009555238.1 | P0C6X6 |
| RKRGVKQLPEQFDSLN | 1 | 0 | 0 | 1690555 | YP_173242.1 | Q19U29 |
| RKRNQKPLEPKFSIAL | 1 | 0 | 0 | 1690559 | YP_003771.1 | H9EJ69 |
| RKRRSISASYRFVTFE | 1 | 0 | 0 | 1690560 | YP_173238.1 | A0A140H1H1 |
| RKSAPLIELCVDEAGS | 1 | 0 | 0 | 1690563 | YP_009724396.1 | P0DTC8 |
| RKSKLKPFERDISTEI | 1 | 0 | 0 | 1690568 | QIQ54048.1 | P59594 |
| RKVLLRKNGNKGAGGH | 1 | 0 | 0 | 1690588 | YP_009724389.1 | P0DTD1 |
| RKVPTDNYITTYPGQG | 1 | 0 | 0 | 1690591 | YP_009724389.1 | P0DTD1 |
| RKYKGIKIQEGIVDYG | 1 | 0 | 0 | 1690599 | NP_828849.2 | P0C6X7 |
| RKYKGIKIQEGVVDYG | 1 | 0 | 0 | 1690600 | YP_009724389.1 | P0DTD1 |
| RLAFLGASVTEDVKFA | 1 | 0 | 0 | 1690604 | YP_003766.2 | H9EJ42 |
| RLAMLGAEMTSNVRRQ | 1 | 0 | 0 | 1690606 | NP_073549.1 | A0A0P0LKV0 |
| RLANECAQVLSEYVLC | 1 | 0 | 0 | 1690609 | YP_009047202.1 | K9N7C7 |
| RLASGVQVHNLPEYMT | 1 | 0 | 0 | 1690612 | NP_073555.1 | A0A0P0KXN3 |
| RLDALEAQVQIDRLIN | 1 | 0 | 0 | 1690623 | YP_173238.1 | A0A140H1H1 |
| RLDMSSVDTILNMARN | 1 | 0 | 0 | 1690625 | NP_073549.1 | A0A0P0LKV0 |
| RLEGLGGDNVYCYNTD | 1 | 0 | 0 | 1690631 | YP_003766.2 | H9EJ42 |
| RLEGLGGNNVYCYNTA | 1 | 0 | 0 | 1690632 | NP_073549.1 | A0A0P0LKV0 |
| RLEIVKLGAEKKDGFY | 1 | 0 | 0 | 1690633 | YP_009555239.1 | Q80872 |
| RLEPCNGTDIDKCVRA | 1 | 0 | 0 | 1690635 | YP_003766.2 | H9EJ42 |
| RLESADVSEMLTFDKK | 1 | 0 | 0 | 1690637 | NP_073551.1 | A0A0P0K6L9 |
| RLFARTRSMWSFNPET | 1 | 0 | 0 | 1690640 | YP_009724393.1 | P0DTC5 |
| RLFRKANLKPFERDIS | 1 | 0 | 0 | 1690647 | QHR63300.2 | Q3LZX1 |
| RLFYDRDGQRTYDDVN | 1 | 0 | 0 | 1690652 | YP_009555238.1 | P0C6X6 |
| RLFYDRDGQRVYDDVD | 1 | 0 | 0 | 1690653 | YP_173236.1 | A0A140H1G9 |
| RLIIRENNRVVISSDV | 1 | 0 | 0 | 1690669 | YP_009724389.1 | P0DTD1 |
| RLIIRENNRVVVSSDI | 1 | 0 | 0 | 1690670 | NP_828849.2 | P0C6X7 |
| RLIKDVDNPVLMGWDY | 1 | 0 | 0 | 1690671 | YP_009555238.1 | P0C6X6 |
| RLINKFDVINGVKTVK | 1 | 0 | 0 | 1690674 | YP_173236.1 | A0A140H1G9 |
| RLKGGAPIKGVTFGED | 1 | 0 | 0 | 1690686 | NP_828849.2 | P0C6X7 |
| RLKGGLILKFDSGTVS | 1 | 0 | 0 | 1690688 | YP_009047202.1 | K9N7C7 |
| RLKNSVSDVAYNRYLS | 1 | 0 | 0 | 1690694 | YP_173236.1 | A0A140H1G9 |
| RLLGLQTQTVDSAQGS | 1 | 0 | 0 | 1690700 | NP_073549.1 | A0A0P0LKV0 |
| RLLSDDRTEVPQLVNA | 1 | 0 | 0 | 1690708 | YP_009047204.1 | K9N5Q8 |
| RLNALESKMSGKGSQQ | 1 | 0 | 0 | 1690719 | QIQ54056.1 | P59595 |
| RLPSTQKGSGMDTALL | 1 | 0 | 0 | 1690729 | YP_009555244.1 | Q01455 |
| RLQAASDWKPGYSMPV | 1 | 0 | 0 | 1690731 | YP_009555238.1 | P0C6X6 |
| RLQAGNATEVPANSTV | 1 | 0 | 0 | 1690734 | YP_009724389.1 | P0DTD1 |
| RLQAGSNTEFASNSSV | 1 | 0 | 0 | 1690735 | YP_009047202.1 | K9N7C7 |
| RLQAGTATEYASNSSI | 1 | 0 | 0 | 1690736 | YP_009555238.1 | P0C6X6 |
| RLQAGVATEYAANSSI | 1 | 0 | 0 | 1690737 | YP_173236.1 | A0A140H1G9 |
| RLQASADWKPGHAMPS | 1 | 0 | 0 | 1690739 | YP_009047202.1 | K9N7C7 |
| RLRAKHYVYIGDPAQL | 1 | 0 | 0 | 1690747 | YP_009724389.1 | P0DTD1 |
| RLRKKRNVDPLSNFEH | 1 | 0 | 0 | 1690751 | YP_009047202.1 | K9N7C7 |
| RLSAHLETNDVSSMLT | 1 | 0 | 0 | 1690752 | YP_003767.1 | A0A0P0G321 |
| RLSKMGILKAEEFVAA | 1 | 0 | 0 | 1690756 | NP_073549.1 | A0A0P0LKV0 |
| RLSLKELMMYAADPAM | 1 | 0 | 0 | 1690760 | YP_009047202.1 | K9N7C7 |
| RLTANVGRVIKFETDA | 1 | 0 | 0 | 1690768 | YP_173236.1 | A0A140H1G9 |
| RLVDAMVYTSDLLTNS | 1 | 0 | 0 | 1690783 | NP_828849.2 | P0C6X7 |
| RLYDNCYRLTSVEESF | 1 | 0 | 0 | 1690799 | YP_003766.2 | H9EJ42 |
| RLYLDAYNMMISAGFS | 1 | 0 | 0 | 1690804 | YP_009724389.1 | P0DTD1 |
| RLYSHVYRSDKVDSTF | 1 | 0 | 0 | 1690810 | YP_009555238.1 | P0C6X6 |
| RMADLALTNMYKEARI | 1 | 0 | 0 | 1690815 | YP_009555238.1 | P0C6X6 |
| RMEDAMGQGQNSADPK | 1 | 0 | 0 | 1690832 | NP_828859.1 | P59636 |
| RMGFKLDATVPGYPKL | 1 | 0 | 0 | 1690835 | YP_009047202.1 | K9N7C7 |
| RMGSGFCVDYNSPSSS | 1 | 0 | 0 | 1690836 | YP_173238.1 | A0A140H1H1 |
| RMLLDKCDLQNYGEAA | 1 | 0 | 0 | 1690838 | QIQ54047.1 | P0C6X7 |
| RMNSRNYIAQVDVVNF | 1 | 0 | 0 | 1690843 | YP_009725255.1 | A0A663DJA2 |
| RMRRGQRVDLPPKVHF | 1 | 0 | 0 | 1690846 | YP_003771.1 | H9EJ69 |
| RMYCSFYPPDEDYEEE | 1 | 0 | 0 | 1690851 | QIQ54046.1 | P0C6U8 |
| RMYCSFYPPDEEEEDD | 1 | 0 | 0 | 1690852 | NP_828849.2 | P0C6X7 |
| RMYCTFAIEDVEDVIS | 1 | 0 | 0 | 1690853 | YP_173236.1 | A0A140H1G9 |
| RNASLNSFKEYFNLRN | 1 | 0 | 0 | 1690861 | YP_009047204.1 | K9N5Q8 |
| RNDCMLLCDILSIYAG | 1 | 0 | 0 | 1690866 | YP_009555238.1 | P0C6X6 |
| RNDCSILCEILCEYAD | 1 | 0 | 0 | 1690867 | YP_173236.1 | A0A140H1G9 |
| RNESWHFDKSEPLCLF | 1 | 0 | 0 | 1690871 | YP_173238.1 | A0A140H1H1 |
| RNFDEQNCDVLKEVLV | 1 | 0 | 0 | 1690873 | YP_003766.2 | H9EJ42 |
| RNGARPKQRRPQGLPN | 1 | 0 | 0 | 1690879 | NP_828858.1 | P59595 |
| RNGSVLEMAKEVKTSS | 1 | 0 | 0 | 1690882 | NP_073549.1 | A0A0P0LKV0 |
| RNGVVPLSVIPATSAA | 1 | 0 | 0 | 1690887 | NP_073549.1 | A0A0P0LKV0 |
| RNHNSQDDIMKAVAAA | 1 | 0 | 0 | 1690888 | NP_073556.1 | A0A0N7J131 |
| RNIDATSTGNYNYKYR | 1 | 0 | 0 | 1690889 | NP_828851.1 | P59594 |
| RNIKPVPEIKILNNLG | 1 | 0 | 0 | 1690891 | NP_828849.2 | P0C6X7 |
| RNIKPVPEVKILNNLG | 1 | 0 | 0 | 1690892 | YP_009724389.1 | P0DTD1 |
| RNLDSLEARAFQSTPI | 1 | 0 | 0 | 1690899 | NP_828859.1 | P59636 |
| RNLKEFKPRSQMEIDF | 1 | 0 | 0 | 1690902 | QHR63299.1 | P0C6W2 |
| RNPANNAAIVLQLPQG | 1 | 0 | 0 | 1690917 | YP_009724397.2 | P0DTC9 |
| RNPAWRKAVFISPYNS | 1 | 0 | 0 | 1690918 | YP_009724389.1 | P0DTD1 |
| RNPNNNAATVLQLPQG | 1 | 0 | 0 | 1690923 | NP_828858.1 | P59595 |
| RNPSSDRNHNSQDDIM | 1 | 0 | 0 | 1690924 | NP_073556.1 | A0A0N7J131 |
| RNPTWRKAVFISPYNS | 1 | 0 | 0 | 1690925 | QHR63299.1 | P0C6W2 |
| RNRDVDTDFVNEFYAY | 1 | 0 | 0 | 1690932 | YP_009724389.1 | P0DTD1 |
| RNSFGKDLNANMSLAE | 1 | 0 | 0 | 1690939 | NP_073549.1 | A0A0P0LKV0 |
| RNSSANYVGKYDINHG | 1 | 0 | 0 | 1690944 | YP_173237.1 | A0A140H1H0 |
| RNSSDNGISAIITANL | 1 | 0 | 0 | 1690945 | YP_003767.1 | A0A0P0G321 |

| | | | | | | |
|---|---|---|---|---|---|---|
| RNSSRNSTPGSSRGTS | 1 | 0 | 0 | 1690946 | YP_009724397.2 | P0DTC9 |
| RNSSRSSSQGSRSGNS | 1 | 0 | 0 | 1690947 | YP_009047211.1 | K9N4V7 |
| RNSTPGSSRGTSPARM | 1 | 0 | 0 | 1690951 | YP_009724397.2 | P0DTC9 |
| RNSTTNYVGVYDINHG | 1 | 0 | 0 | 1690953 | YP_009555240.1 | P30215 |
| RNTREVFAQVKQMYKT | 1 | 0 | 0 | 1690962 | NP_828851.1 | P59594 |
| RNVFENFEDWHYAKFG | 1 | 0 | 0 | 1690968 | NP_073549.1 | A0A0P0LKV0 |
| RNVLLRKNGNKGAGGH | 1 | 0 | 0 | 1690971 | NP_828849.2 | P0C6X7 |
| RPDTRYVLMDGSIIQF | 1 | 0 | 0 | 1690989 | YP_009724389.1 | P0DTD1 |
| RPGNFNQDFYDFVVSK | 1 | 0 | 0 | 1690998 | YP_009047202.1 | K9N7C7 |
| RPIDVSKADGIIYPQG | 1 | 0 | 0 | 1691004 | YP_009047204.1 | K9N5Q8 |
| RPIIEDYHTLTVTIIR | 1 | 0 | 0 | 1691006 | YP_009555244.1 | Q01455 |
| RPNHTIKGSFLNGSCG | 1 | 0 | 0 | 1691024 | NP_828849.2 | P0C6X7 |
| RPRNSSDNGISAIITA | 1 | 0 | 0 | 1691038 | YP_003767.1 | A0A0P0G321 |
| RPVIEDYHTLTATVIR | 1 | 0 | 0 | 1691046 | YP_173241.1 | Q0ZJ82 |
| RQAALPQSKDSNFLNE | 1 | 0 | 0 | 1691049 | YP_009047203.1 | K9N7C7 |
| RQAALPQSKDSNFLNR | 1 | 0 | 0 | 1691050 | YP_009047202.1 | K9N7C7 |
| RQATQYLVQQESSFVM | 1 | 0 | 0 | 1691054 | QIQ54046.1 | P0C6U8 |
| RQAYEDAIANGSSSQL | 1 | 0 | 0 | 1691055 | YP_003766.2 | H9EJ42 |
| RQFYEFYNDVKPPVLD | 1 | 0 | 0 | 1691073 | YP_009555243.1 | Q04854 |
| RQIAPGQTGVIADYNY | 1 | 0 | 0 | 1691084 | NP_828851.1 | P59594 |
| RQIFDVAPIELENKFT | 1 | 0 | 0 | 1691085 | YP_009047202.1 | K9N7C7 |
| RQILGSCTFEDELAPS | 1 | 0 | 0 | 1691087 | YP_173236.1 | A0A140H1G9 |
| RQIMGSCSFEDELTPS | 1 | 0 | 0 | 1691088 | YP_009555238.1 | P0C6X6 |
| RQKQSVWLANCDFDIV | 1 | 0 | 0 | 1691092 | YP_173236.1 | A0A140H1G9 |
| RQKRTATKAYNVTQAF | 1 | 0 | 0 | 1691095 | YP_009724397.2 | P0DTC9 |
| RQKRTATKQYNVTQAF | 1 | 0 | 0 | 1691096 | QHR63308.1 | Q3LZX4 |
| RQLFKPLTKKNYSELD | 1 | 0 | 0 | 1691100 | NP_828856.1 | P59634 |
| RQLQNSMSGASADSTQ | 1 | 0 | 0 | 1691108 | NP_828858.1 | P59595 |
| RQLYDNCYRNSNVDES | 1 | 0 | 0 | 1691111 | NP_073549.1 | A0A0P0LKV0 |
| RQNNLAIEEIEYVHGD | 1 | 0 | 0 | 1691115 | NP_073549.1 | A0A0P0LKV0 |
| RQPNDDVTSNVTQCFG | 1 | 0 | 0 | 1691116 | NP_073556.1 | A0A0N7J131 |
| RQQRFVYDAYQNLVGY | 1 | 0 | 0 | 1691119 | YP_009047204.1 | K9N5Q8 |
| RQQSRTRSDSNQSSSD | 1 | 0 | 0 | 1691120 | YP_003771.1 | H9EJ69 |
| RQQTSNLVVQEFVSYD | 1 | 0 | 0 | 1691121 | YP_009555238.1 | P0C6X6 |
| RQQVDKKIKACVEEVT | 1 | 0 | 0 | 1691122 | QHR63299.1 | P0C6W2 |
| RQRGHKNGQGENDNIS | 1 | 0 | 0 | 1691123 | YP_009555245.1 | P33469 |
| RQSDNMTAVACQPPYC | 1 | 0 | 0 | 1691127 | YP_009555240.1 | P30215 |
| RQVIQDVKDGTVTFNL | 1 | 0 | 0 | 1691135 | NP_073552.1 | A0A0P0KWE8 |
| RQVRSWIGFDVEGAHA | 1 | 0 | 0 | 1691136 | YP_009047202.1 | K9N7C7 |
| RQWLPTDAIIIDNDLN | 1 | 0 | 0 | 1691141 | YP_009047202.1 | K9N7C7 |
| RQYLDAYNMMISAGFS | 1 | 0 | 0 | 1691145 | NP_828849.2 | P0C6X7 |
| RQYLLAFNQDGIIFNA | 1 | 0 | 0 | 1691146 | YP_009555241.1 | P36334 |
| RQYSNESWFEDDYRAF | 1 | 0 | 0 | 1691150 | NP_073549.1 | A0A0P0LKV0 |
| RRAIDCGFNDLSQLHC | 1 | 0 | 0 | 1691155 | YP_009047204.1 | K9N5Q8 |
| RRCPAEIVDTVSALVY | 1 | 0 | 0 | 1691164 | YP_009724389.1 | P0DTD1 |
| RRDSTYGPLQTPVGCV | 1 | 0 | 0 | 1691167 | YP_009047204.1 | K9N5Q8 |
| RRGQRVDLPPKVHFYY | 1 | 0 | 0 | 1691176 | YP_003771.1 | H9EJ69 |
| RRIVQMLSDTLDKLSD | 1 | 0 | 0 | 1691184 | YP_009047202.1 | K9N7C7 |
| RRKKKSNLLIQEFLQY | 1 | 0 | 0 | 1691185 | YP_173236.1 | A0A140H1G9 |
| RRKNSEPEIPHFNQKL | 1 | 0 | 0 | 1691186 | NP_073556.1 | A0A0N7J131 |
| RRKPNLPNCNIEAWLN | 1 | 0 | 0 | 1691187 | YP_009555241.1 | P36334 |
| RRLIKDVDNPVLMGWD | 1 | 0 | 0 | 1691193 | YP_009555238.1 | P0C6X6 |
| RRLYDNCYRLTSVEES | 1 | 0 | 0 | 1691199 | YP_003766.2 | H9EJ42 |
| RRNLDSLEARAFQSTP | 1 | 0 | 0 | 1691201 | NP_828859.1 | P59636 |
| RRQQTSNLVVQEFVSY | 1 | 0 | 0 | 1691210 | YP_009555238.1 | P0C6X6 |
| RRSFKTADGQQKQLLP | 1 | 0 | 0 | 1691220 | YP_173242.1 | Q19U29 |
| RRSISASYRFVTFEPF | 1 | 0 | 0 | 1691223 | YP_173238.1 | A0A140H1H1 |
| RRSRGAITTGYRFTNF | 1 | 0 | 0 | 1691224 | YP_009555241.1 | P36334 |
| RRVRGGDGKMKDLSPR | 1 | 0 | 0 | 1691233 | QIQ54056.1 | P59595 |
| RSAAKKNNLPFKLTCA | 1 | 0 | 0 | 1691243 | YP_009724389.1 | P0DTD1 |
| RSAPNSRSTSRTSSRA | 1 | 0 | 0 | 1691252 | YP_009555245.1 | P33469 |
| RSDCNHINNLKIKNFD | 1 | 0 | 0 | 1691264 | YP_173237.1 | A0A140H1H0 |
| RSDGVVWVAKEGAKTV | 1 | 0 | 0 | 1691267 | YP_003771.1 | H9EJ69 |
| RSDKSYEHQTPFEIKS | 1 | 0 | 0 | 1691268 | NP_828849.2 | P0C6X7 |
| RSDLQLGSSGFLQSSN | 1 | 0 | 0 | 1691272 | YP_173238.1 | A0A140H1H1 |
| RSDMEYDFLNMDMGVF | 1 | 0 | 0 | 1691273 | YP_003766.2 | H9EJ42 |
| RSDSNQSSSDLVAAVT | 1 | 0 | 0 | 1691274 | YP_003771.1 | H9EJ69 |
| RSDVLLPLTQYNRYLA | 1 | 0 | 0 | 1691276 | YP_009724389.1 | P0DTD1 |
| RSEAFEYYHTLDESFL | 1 | 0 | 0 | 1691277 | NP_828849.2 | P0C6X7 |
| RSEKSYELQTPFEIKL | 1 | 0 | 0 | 1691281 | YP_009724389.1 | P0DTD1 |
| RSELQTIVPEYIDVNK | 1 | 0 | 0 | 1691282 | NP_073551.1 | A0A0P0K6L9 |
| RSETLLPLTQYNRYLA | 1 | 0 | 0 | 1691284 | NP_828849.2 | P0C6X7 |
| RSFFEDLLFDKVKLSD | 1 | 0 | 0 | 1691286 | YP_173238.1 | A0A140H1H1 |
| RSFHFTDFYNYTGEGQ | 1 | 0 | 0 | 1691287 | YP_009555240.1 | P30215 |
| RSFKTADGNQRQLLPR | 1 | 0 | 0 | 1691288 | YP_009555245.1 | P33469 |
| RSGARPKQRRPQGLPN | 1 | 0 | 0 | 1691296 | QHR63308.1 | Q3LZX4 |
| RSGNGILKWADQSDQF | 1 | 0 | 0 | 1691305 | YP_009555245.1 | P33469 |
| RSGNSTRGTSPGPSGI | 1 | 0 | 0 | 1691307 | YP_009047211.1 | K9N4V7 |
| RSHPDFKLLHNLQADI | 1 | 0 | 0 | 1691312 | YP_009047202.1 | K9N7C7 |
| RSIRHHPELKILRNLN | 1 | 0 | 0 | 1691323 | YP_173236.1 | A0A140H1G9 |
| RSISASYRFVTFEPFN | 1 | 0 | 0 | 1691324 | YP_173238.1 | A0A140H1H1 |
| RSKDSPLELEDFIPMD | 1 | 0 | 0 | 1691327 | QIQ54047.1 | P0C6X7 |
| RSLKAPAVVSVSSPDA | 1 | 0 | 0 | 1691332 | NP_828849.2 | P0C6X7 |
| RSLPGVFCGVDAVNLL | 1 | 0 | 0 | 1691338 | YP_009724389.1 | P0DTD1 |
| RSLPGVFCGVDSVNLL | 1 | 0 | 0 | 1691339 | QIQ54046.1 | P0C6U8 |
| RSLSRSNSNFRHSDSI | 1 | 0 | 0 | 1691342 | YP_173242.1 | Q19U29 |
| RSMLKRRDSTYGPLQT | 1 | 0 | 0 | 1691347 | YP_009047204.1 | K9N5Q8 |
| RSNSNFRHSDSIVKPD | 1 | 0 | 0 | 1691355 | YP_173242.1 | Q19U29 |
| RSQMEIDFLELAMDEF | 1 | 0 | 0 | 1691361 | YP_009724389.1 | P0DTD1 |
| RSQMETDFLELAMDEF | 1 | 0 | 0 | 1691362 | NP_828849.2 | P0C6X7 |
| RSRGAITTGYRFTNFE | 1 | 0 | 0 | 1691368 | YP_009555241.1 | P36334 |

| | | | | | | |
|---|---|---|---|---|---|---|
| RSRNSSRNSTPGSSRG | 1 | 0 | 0 | 1691371 | YP_009724397.2 | P0DTC9 |
| RSSFNFDQKELLAYYN | 1 | 0 | 0 | 1691373 | YP_009555238.1 | P0C6X6 |
| RSSSQGSRSGNSTRGT | 1 | 0 | 0 | 1691379 | YP_009047211.1 | K9N4V7 |
| RSSSRSRGNSRNSTPG | 1 | 0 | 0 | 1691380 | NP_828858.1 | P59595 |
| RSSTRNNSRDSSRSTS | 1 | 0 | 0 | 1691382 | YP_003771.1 | H9EJ69 |
| RSTRSMLKRRDSTYGP | 1 | 0 | 0 | 1691390 | YP_009047204.1 | K9N5Q8 |
| RSTSPDPKFVDKYYAF | 1 | 0 | 0 | 1691391 | YP_009047202.1 | K9N7C7 |
| RSTSQKSIVAYTMSLG | 1 | 0 | 0 | 1691392 | NP_828851.1 | P59594 |
| RSTSRQQSRTRSDSNQ | 1 | 0 | 0 | 1691393 | YP_003771.1 | H9EJ69 |
| RSTSRTSSRASSAGSR | 1 | 0 | 0 | 1691394 | YP_009555245.1 | P33469 |
| RSVFNLEGVNGGSLYV | 1 | 0 | 0 | 1691397 | YP_003766.2 | H9EJ42 |
| RSVNQRSIIAYTMSLG | 1 | 0 | 0 | 1691402 | QIQ54048.1 | P59594 |
| RSVRGRAIIVSVEQLE | 1 | 0 | 0 | 1691406 | NP_073549.1 | A0A0P0LKV0 |
| RSVRHHPELKLFRNLN | 1 | 0 | 0 | 1691407 | YP_009555238.1 | P0C6X6 |
| RSVRSVPGEMRLASIA | 1 | 0 | 0 | 1691408 | YP_009047204.1 | K9N5Q8 |
| RSVYPVASPNECNPMH | 1 | 0 | 0 | 1691412 | QIQ54046.1 | P0C6U8 |
| RSVYPVASPNECNQMC | 1 | 0 | 0 | 1691413 | YP_009724389.1 | P0DTD1 |
| RSVYPVASPQECNNMH | 1 | 0 | 0 | 1691414 | NP_828849.2 | P0C6X7 |
| RSVYVKFQDSKPPLPP | 1 | 0 | 0 | 1691415 | YP_009047209.1 | K9N5R3 |
| RSWIGFDVEGAHASRN | 1 | 0 | 0 | 1691416 | YP_009047202.1 | K9N7C7 |
| RSYLTPGKLESGWTTG | 1 | 0 | 0 | 1691421 | QIQ54048.1 | P59594 |
| RTAIQSFDNSYLNESG | 1 | 0 | 0 | 1691424 | NP_073550.1 | A0A0P0ILY9 |
| RTDMEKDFIALDQDVF | 1 | 0 | 0 | 1691437 | YP_009555238.1 | P0C6X6 |
| RTDSLCSSHYWSPQGN | 1 | 0 | 0 | 1691438 | YP_173236.1 | A0A140H1G9 |
| RTELGVEFLKRGDKIV | 1 | 0 | 0 | 1691441 | NP_828849.2 | P0C6X7 |
| RTEVPQLVNANQYSPC | 1 | 0 | 0 | 1691443 | YP_009047204.1 | K9N5Q8 |
| RTFLLSYNQNGTITDA | 1 | 0 | 0 | 1691447 | QIQ54048.1 | P59594 |
| RTFVVDKSLSIEEFAD | 1 | 0 | 0 | 1691453 | YP_009047202.1 | K9N7C7 |
| RTGIPPGYTNSSTRGV | 1 | 0 | 0 | 1691459 | QIQ54048.1 | P59594 |
| RTGKHEHSLLVEAYNS | 1 | 0 | 0 | 1691460 | YP_003766.2 | H9EJ42 |
| RTILGSAILEDEFTPF | 1 | 0 | 0 | 1691477 | QIQ54046.1 | P0C6U8 |
| RTILGSALLEDEFTPF | 1 | 0 | 0 | 1691478 | YP_009724389.1 | P0DTD1 |
| RTILGSTILEDEFTPF | 1 | 0 | 0 | 1691479 | NP_828849.2 | P0C6X7 |
| RTINSTQDGDNKLQGL | 1 | 0 | 0 | 1691480 | YP_009555241.1 | P36334 |
| RTKHVTKENDSKEGFF | 1 | 0 | 0 | 1691485 | NP_828849.2 | P0C6X7 |
| RTLLTKGTLEPEYFNS | 1 | 0 | 0 | 1691494 | YP_009724389.1 | P0DTD1 |
| RTNGGYNSWHLVEGEL | 1 | 0 | 0 | 1691502 | YP_009047202.1 | K9N7C7 |
| RTNLEVYNKEKTYYEL | 1 | 0 | 0 | 1691503 | YP_173236.1 | A0A140H1G9 |
| RTPNKHCNVQQCFGKR | 1 | 0 | 0 | 1691508 | YP_173242.1 | Q19U29 |
| RTPTSGVTPDMADQIA | 1 | 0 | 0 | 1691509 | YP_009555245.1 | P33469 |
| RTRSDSNQSSSDLVAA | 1 | 0 | 0 | 1691520 | YP_003771.1 | H9EJ69 |
| RTSSRASSAGSRSRAN | 1 | 0 | 0 | 1691531 | YP_009555245.1 | P33469 |
| RTVCFSVAALSTGLTS | 1 | 0 | 0 | 1691542 | NP_073549.1 | A0A0P0LKV0 |
| RTVFENLKPMPFFYYS | 1 | 0 | 0 | 1691543 | YP_173236.1 | A0A140H1G9 |
| RTVYDDAARRVWTLMN | 1 | 0 | 0 | 1691553 | NP_828849.2 | P0C6X7 |
| RTVYDDGARRVWTLMN | 1 | 0 | 0 | 1691554 | YP_009724389.1 | P0DTD1 |
| RTYDDVNASLFVDYSN | 1 | 0 | 0 | 1691556 | YP_009555238.1 | P0C6X6 |
| RVDENGDKLDQFFVVK | 1 | 0 | 0 | 1691576 | YP_009555238.1 | P0C6X6 |
| RVDWDIEYPYISHEKK | 1 | 0 | 0 | 1691585 | YP_009047202.1 | K9N7C7 |
| RVDWSVEYPIIGDELR | 1 | 0 | 0 | 1691587 | NP_828849.2 | P0C6X7 |
| RVDWTIEYPIIGDELK | 1 | 0 | 0 | 1691588 | YP_009724389.1 | P0DTD1 |
| RVEAPKQEEPPNTEDS | 1 | 0 | 0 | 1691590 | NP_828849.2 | P0C6X7 |
| RVECFDKFKVNSTLEQ | 1 | 0 | 0 | 1691592 | YP_009724389.1 | P0DTD1 |
| RVECYDRFKVNETNSQ | 1 | 0 | 0 | 1691596 | YP_009047202.1 | K9N7C7 |
| RVECYSGFKPNNNSAQ | 1 | 0 | 0 | 1691597 | NP_073549.1 | A0A0P0LKV0 |
| RVECYSGFKPNNTSAQ | 1 | 0 | 0 | 1691598 | YP_003766.2 | H9EJ42 |
| RVEEEEEEDWLDDTTE | 1 | 0 | 0 | 1691599 | NP_828849.2 | P0C6X7 |
| RVEKKKTEGFMGRIRS | 1 | 0 | 0 | 1691608 | NP_828849.2 | P0C6X7 |
| RVFPYDMDSGVSSFSE | 1 | 0 | 0 | 1691617 | YP_009047202.1 | K9N7C7 |
| RVGLCEEADEGICFNF | 1 | 0 | 0 | 1691623 | YP_009555238.1 | P0C6X6 |
| RVGRSVNETSQTGWAF | 1 | 0 | 0 | 1691625 | YP_003770.1 | U3M6Q8 |
| RVGRSVNSQNSTGWVF | 1 | 0 | 0 | 1691626 | NP_073555.1 | A0A0P0KXN3 |
| RVHSEFNLPGCDGGSL | 1 | 0 | 0 | 1691628 | YP_009047202.1 | K9N7C7 |
| RVIRRERCDMTFPCVS | 1 | 0 | 0 | 1691637 | YP_003766.2 | H9EJ42 |
| RVISRMGFKLDATVPG | 1 | 0 | 0 | 1691638 | YP_009047202.1 | K9N7C7 |
| RVKHGDFSAVSSPMSN | 1 | 0 | 0 | 1691645 | NP_073555.1 | A0A0P0KXN3 |
| RVKNTKLYVNKTLYSE | 1 | 0 | 0 | 1691649 | YP_173238.1 | A0A140H1H1 |
| RVKSGSPGDSSWHIYL | 1 | 0 | 0 | 1691650 | YP_003767.1 | A0A0P0G321 |
| RVLDQSDNNCWISTTL | 1 | 0 | 0 | 1691652 | YP_003766.2 | H9EJ42 |
| RVLGLQTQTVDSAQGS | 1 | 0 | 0 | 1691656 | YP_009555238.1 | P0C6X6 |
| RVLGVQTQTVDSAQGS | 1 | 0 | 0 | 1691658 | YP_173236.1 | A0A140H1G9 |
| RVLHLGAGSDKEVAPG | 1 | 0 | 0 | 1691659 | YP_173236.1 | A0A140H1G9 |
| RVLNNLNLPGCNGGSL | 1 | 0 | 0 | 1691668 | YP_009555238.1 | P0C6X6 |
| RVLNVVCKTCGQQQTT | 1 | 0 | 0 | 1691670 | YP_009724389.1 | P0DTD1 |
| RVMFNGVTFSTFEEAA | 1 | 0 | 0 | 1691677 | NP_828849.2 | P0C6X7 |
| RVNAKPPPGDQFKHLI | 1 | 0 | 0 | 1691682 | NP_828849.2 | P0C6X7 |
| RVNVDGFNEWAMANGY | 1 | 0 | 0 | 1691687 | YP_003766.2 | H9EJ42 |
| RVNVPYVCLLANKEQE | 1 | 0 | 0 | 1691688 | YP_009047202.1 | K9N7C7 |
| RVPHIVRKDLTKYTML | 1 | 0 | 0 | 1691690 | YP_009555238.1 | P0C6X6 |
| RVQGSITQRTRTRPSV | 1 | 0 | 0 | 1691695 | YP_009047211.1 | K9N4V7 |
| RVQPTDSIVRFPNITN | 1 | 0 | 0 | 1691696 | QHR63300.2 | Q3LZX1 |
| RVQPTESIVRFPNITN | 1 | 0 | 0 | 1691697 | YP_009724390.1 | P0DTC2 |
| RVQQNKSRELTAEDIS | 1 | 0 | 0 | 1691699 | YP_009555245.1 | P33469 |
| RVRAWVGFDAEGAHAT | 1 | 0 | 0 | 1691702 | YP_009555238.1 | P0C6X6 |
| RVRGGDGKMKDLSPRW | 1 | 0 | 0 | 1691703 | QIQ54056.1 | P59595 |
| RVRGSIVNARIEPCSS | 1 | 0 | 0 | 1691705 | YP_009047202.1 | K9N7C7 |
| RVRGSSAARLEPCNGT | 1 | 0 | 0 | 1691706 | NP_073549.1 | A0A0P0LKV0 |
| RVRGWVGFDVEGAHAT | 1 | 0 | 0 | 1691709 | YP_173236.1 | A0A140H1G9 |
| RVRSVRGRAIIVSVEQ | 1 | 0 | 0 | 1691714 | NP_073549.1 | A0A0P0LKV0 |
| RVSAAFHANSSEPALL | 1 | 0 | 0 | 1691715 | YP_009555241.1 | P36334 |
| RVTKNSKTQIGEYTFE | 1 | 0 | 0 | 1691728 | QIQ54047.1 | P0C6X7 |

| | | | | | | |
|---|---|---|---|---|---|---|
| RVTKNSKVQIGEYTFE | 1 | 0 | 0 | 1691729 | YP_009724389.1 | P0DTD1 |
| RVTVPKDHPHLGKFLE | 1 | 0 | 0 | 1691734 | NP_073556.1 | A0A0N7J131 |
| RVVFNGTSFSTFEEAA | 1 | 0 | 0 | 1691736 | QIQ54046.1 | P0C6U8 |
| RVVFNGVSFSTFEEAA | 1 | 0 | 0 | 1691737 | YP_009724389.1 | P0DTD1 |
| RVVPSGDVVRFPNITN | 1 | 0 | 0 | 1691749 | NP_828851.1 | P59594 |
| RVVTTFDAEYCRHGTC | 1 | 0 | 0 | 1691752 | NP_828849.2 | P0C6X7 |
| RVVTTFDSEYCRHGTC | 1 | 0 | 0 | 1691753 | YP_009724389.1 | P0DTD1 |
| RVVYTAASHAAVDALC | 1 | 0 | 0 | 1691757 | YP_009555238.1 | P0C6X6 |
| RVYDDVDASLFVDINN | 1 | 0 | 0 | 1691765 | YP_173236.1 | A0A140H1G9 |
| RVYSSANNCTFEYISQ | 1 | 0 | 0 | 1691773 | QIQ54048.1 | P59594 |
| RVYYESMSYQEQDELF | 1 | 0 | 0 | 1691776 | YP_009047202.1 | K9N7C7 |
| RWHTMVQTCTPNVTIN | 1 | 0 | 0 | 1691780 | NP_849177.1 | Q80H93 |
| RWKRQPNDDVTSNVTQ | 1 | 0 | 0 | 1691781 | NP_073556.1 | A0A0N7J131 |
| RWLLPDAAEELASPMK | 1 | 0 | 0 | 1691784 | YP_173236.1 | A0A140H1G9 |
| RWLPHDAIVVDNDVVD | 1 | 0 | 0 | 1691785 | NP_073549.1 | A0A0P0LKV0 |
| RWLPPDAIIIDNDIND | 1 | 0 | 0 | 1691786 | YP_003766.2 | H9EJ42 |
| RWNNERASDISLSVAC | 1 | 0 | 0 | 1691790 | YP_173237.1 | A0A140H1H0 |
| RWPQIAELAPTASAFM | 1 | 0 | 0 | 1691791 | YP_009047211.1 | K9N4V7 |
| RWRMRRGQRVDLPPKV | 1 | 0 | 0 | 1691796 | YP_003771.1 | H9EJ69 |
| RYFKYWDQTYHPNCIN | 1 | 0 | 0 | 1691826 | NP_828849.2 | P0C6X7 |
| RYFKYWDQTYHPNCVN | 1 | 0 | 0 | 1691827 | YP_009724389.1 | P0DTD1 |
| RYGDVDNLVFTNKTTL | 1 | 0 | 0 | 1691832 | YP_003766.2 | H9EJ42 |
| RYIYNRVKSGSPGDSS | 1 | 0 | 0 | 1691842 | YP_003767.1 | A0A0P0G321 |
| RYKAGNYRSPPITADI | 1 | 0 | 0 | 1691843 | YP_009047210.1 | K9N7A1 |
| RYKYYSGNANEADYRC | 1 | 0 | 0 | 1691845 | NP_073549.1 | A0A0P0LKV0 |
| RYLRHGKLRPFERDIS | 1 | 0 | 0 | 1691848 | NP_828851.1 | P59594 |
| RYMNANGLRPPKNSFE | 1 | 0 | 0 | 1691852 | YP_009555238.1 | P0C6X6 |
| RYMNANGLRPPRNSFE | 1 | 0 | 0 | 1691853 | YP_173236.1 | A0A140H1G9 |
| RYMNSQGLLPPKNSID | 1 | 0 | 0 | 1691854 | YP_009724389.1 | P0DTD1 |
| RYMNSQGLLPPKSSID | 1 | 0 | 0 | 1691855 | NP_828849.2 | P0C6X7 |
| RYNEVANAVMQNNELM | 1 | 0 | 0 | 1691859 | YP_173236.1 | A0A140H1G9 |
| RYNFDITESKYSCKEV | 1 | 0 | 0 | 1691861 | NP_073549.1 | A0A0P0LKV0 |
| RYNLNFSANSLDNLKS | 1 | 0 | 0 | 1691867 | YP_003767.1 | A0A0P0G321 |
| RYNRPTMLDIGQARVA | 1 | 0 | 0 | 1691869 | NP_073549.1 | A0A0P0LKV0 |
| RYRIGNYKLNTDHAGS | 1 | 0 | 0 | 1691873 | NP_828855.1 | P59596 |
| RYRIGNYKLNTDHSNS | 1 | 0 | 0 | 1691874 | QIQ54051.1 | P59596 |
| RYSVLSVDSVSEESQG | 1 | 0 | 0 | 1691881 | YP_173236.1 | A0A140H1G9 |
| RYTTTLNDFNLVAMKF | 1 | 0 | 0 | 1691883 | QIQ54046.1 | P0C6U8 |
| RYVDNNFCGPDGYPLE | 1 | 0 | 0 | 1691886 | YP_009724389.1 | P0DTD1 |
| RYYSGKMDTAAYREAA | 1 | 0 | 0 | 1691896 | YP_009555238.1 | P0C6X6 |
| SAAEFKYMVANGLHAP | 1 | 0 | 0 | 1691902 | YP_003766.2 | H9EJ42 |
| SAAFHANSSEPALLFR | 1 | 0 | 0 | 1691903 | YP_009555241.1 | P36334 |
| SAAIKDCVDINKAIPS | 1 | 0 | 0 | 1691906 | NP_073549.1 | A0A0P0LKV0 |
| SAAKKNNLPFKLTCAT | 1 | 0 | 0 | 1691908 | YP_009724389.1 | P0DTD1 |
| SAALQPEEEQEEDWLD | 1 | 0 | 0 | 1691910 | YP_009724389.1 | P0DTD1 |
| SAALSAQLAKDKVNEC | 1 | 0 | 0 | 1691911 | YP_009047204.1 | K9N5Q8 |
| SAAQAMEKVNECVKSQ | 1 | 0 | 0 | 1691912 | YP_009555241.1 | P36334 |
| SAARLEPCNGTDIDKC | 1 | 0 | 0 | 1691913 | YP_003766.2 | H9EJ42 |
| SAARLTPCGTGTSTDV | 1 | 0 | 0 | 1691915 | YP_009724389.1 | P0DTD1 |
| SAARLVVVVPDHDSFV | 1 | 0 | 0 | 1691916 | NP_073549.1 | A0A0P0LKV0 |
| SAAVLKSKWWLLAKFT | 1 | 0 | 0 | 1691921 | NP_073549.1 | A0A0P0LKV0 |
| SACDGLASAYRANSFD | 1 | 0 | 0 | 1691922 | YP_009047202.1 | K9N7C7 |
| SACKTIEDALRNSARL | 1 | 0 | 0 | 1691926 | NP_073551.1 | A0A0P0K6L9 |
| SACVLAAECTIFKDAA | 1 | 0 | 0 | 1691933 | QIQ54046.1 | P0C6U8 |
| SADAIVLKLPSLMKVM | 1 | 0 | 0 | 1691938 | YP_173236.1 | A0A140H1G9 |
| SADASTFLNRVCGVSA | 1 | 0 | 0 | 1691942 | NP_828849.2 | P0C6X7 |
| SADIGSGHTGITGDNV | 1 | 0 | 0 | 1691944 | NP_828849.2 | P0C6X7 |
| SADSCLQGDKCNIFAN | 1 | 0 | 0 | 1691948 | YP_009555241.1 | P36334 |
| SADVSEMLTFDKKAFT | 1 | 0 | 0 | 1691949 | NP_073551.1 | A0A0P0K6L9 |
| SADWKPGHAMPSLFKV | 1 | 0 | 0 | 1691950 | YP_009047202.1 | K9N7C7 |
| SAETVRVEEEEEEDWL | 1 | 0 | 0 | 1691954 | NP_828849.2 | P0C6X7 |
| SAEVAVKMFDAYVNTF | 1 | 0 | 0 | 1691955 | YP_009724389.1 | P0DTD1 |
| SAEVSRQVIQDVKDGT | 1 | 0 | 0 | 1691956 | NP_073552.1 | A0A0P0KWE8 |
| SAFVETVKGLDYKSFK | 1 | 0 | 0 | 1691976 | QIQ54046.1 | P0C6U8 |
| SAFYVKDGKYFTSEPP | 1 | 0 | 0 | 1691983 | YP_009047202.1 | K9N7C7 |
| SAGHATGTTPQKLFVA | 1 | 0 | 0 | 1691992 | YP_009047204.1 | K9N5Q8 |
| SAGIFGADPVHSLRVC | 1 | 0 | 0 | 1691995 | QHR63299.1 | P0C6W2 |
| SAGLELTDESCNNLVP | 1 | 0 | 0 | 1692001 | YP_009555238.1 | P0C6X6 |
| SAGMFGLYDDILTNNK | 1 | 0 | 0 | 1692002 | YP_003766.2 | H9EJ42 |
| SAGNIFGLYKNSATGS | 1 | 0 | 0 | 1692003 | YP_003766.2 | H9EJ42 |
| SAGSRSRANSGNRTPT | 1 | 0 | 0 | 1692008 | YP_009555245.1 | P33469 |
| SAGTQHISNDFTPEDH | 1 | 0 | 0 | 1692009 | YP_173242.1 | Q19U29 |
| SAGVEFLKDAWEILKF | 1 | 0 | 0 | 1692010 | NP_828849.2 | P0C6X7 |
| SAGVVANGVKAKGYPQ | 1 | 0 | 0 | 1692013 | NP_073556.1 | A0A0N7J131 |
| SAGWPLNKFGKAGLYY | 1 | 0 | 0 | 1692014 | NP_073549.1 | A0A0P0LKV0 |
| SAHLETNDVSSMLTFD | 1 | 0 | 0 | 1692017 | YP_003767.1 | A0A0P0G321 |
| SAHVCGDNIGTNVPLQ | 1 | 0 | 0 | 1692018 | YP_003766.2 | H9EJ42 |
| SAIDYDHEYSIMRLHN | 1 | 0 | 0 | 1692019 | NP_073549.1 | A0A0P0LKV0 |
| SAIEDLLFDKVTIADP | 1 | 0 | 0 | 1692022 | YP_009047204.1 | K9N5Q8 |
| SAIITANLSIPSNWTT | 1 | 0 | 0 | 1692025 | YP_003767.1 | A0A0P0G321 |
| SAILEDEFTPFDVVRQ | 1 | 0 | 0 | 1692027 | QIQ54046.1 | P0C6U8 |
| SAIVCASVKRDGVQVG | 1 | 0 | 0 | 1692033 | NP_073549.1 | A0A0P0LKV0 |
| SAIVCASVLKDGCDVG | 1 | 0 | 0 | 1692034 | YP_003766.2 | H9EJ42 |
| SAIVTANLSIPSNWTT | 1 | 0 | 0 | 1692035 | NP_073551.1 | A0A0P0K6L9 |
| SAIVTQFAPGTKLPKN | 1 | 0 | 0 | 1692036 | YP_009047211.1 | K9N4V7 |
| SAKSLARSQSSETKEQ | 1 | 0 | 0 | 1692045 | NP_073556.1 | A0A0N7J131 |
| SAKTGTPKPSRNQSPA | 1 | 0 | 0 | 1692046 | NP_073556.1 | A0A0N7J131 |
| SALASALKQLKVTTGE | 1 | 0 | 0 | 1692048 | NP_073549.1 | A0A0P0LKV0 |
| SALEFNPSQTSPATAE | 1 | 0 | 0 | 1692050 | NP_073556.1 | A0A0N7J131 |
| SALGSMDIDVFNKLST | 1 | 0 | 0 | 1692054 | NP_073549.1 | A0A0P0LKV0 |
| SALIAVANWLAPTNAY | 1 | 0 | 0 | 1692055 | YP_009047202.1 | K9N7C7 |

| | | | | | | |
|---|---|---|---|---|---|---|
| SALKQLKVTTGELVRF | 1 | 0 | 0 | 1692056 | NP_073549.1 | A0A0P0LKV0 |
| SALKTAGVVTGNVVKQ | 1 | 0 | 0 | 1692057 | YP_009047202.1 | K9N7C7 |
| SALLADKFPVLHDIGN | 1 | 0 | 0 | 1692058 | NP_828849.2 | P0C6X7 |
| SALLEDEFTPFDVVRQ | 1 | 0 | 0 | 1692062 | YP_009724389.1 | P0DTD1 |
| SALMGANGNKIVDKEV | 1 | 0 | 0 | 1692063 | YP_009047202.1 | K9N7C7 |
| SALVKIQAVVNANAEA | 1 | 0 | 0 | 1692071 | YP_009555241.1 | P36334 |
| SALVYDNKLRAHKDKS | 1 | 0 | 0 | 1692076 | QIQ54047.1 | P0C6X7 |
| SALWEIQQVVDADSKI | 1 | 0 | 0 | 1692079 | YP_009724389.1 | P0DTD1 |
| SANFLDDNVLPETYVA | 1 | 0 | 0 | 1692091 | YP_003767.1 | A0A0P0G321 |
| SANNCTFEYISQPFLM | 1 | 0 | 0 | 1692099 | QIQ54048.1 | P59594 |
| SANNCTFEYVSQPFLM | 1 | 0 | 0 | 1692100 | YP_009724390.1 | P0DTC2 |
| SANYVGKYDINHGDSG | 1 | 0 | 0 | 1692108 | YP_173237.1 | A0A140H1H0 |
| SAPLIELCVDEAGSKS | 1 | 0 | 0 | 1692112 | YP_009724396.1 | P0DTC8 |
| SAPNSRSTSRTSSRAS | 1 | 0 | 0 | 1692115 | YP_009555245.1 | P33469 |
| SAPQTIQPEVKEVAPV | 1 | 0 | 0 | 1692120 | YP_009047202.1 | K9N7C7 |
| SAPTLVPQEHYVRITG | 1 | 0 | 0 | 1692122 | YP_009724389.1 | P0DTD1 |
| SAPTLVPQENYASIRF | 1 | 0 | 0 | 1692123 | YP_173236.1 | A0A140H1G9 |
| SAPTLVPQENYSSIRF | 1 | 0 | 0 | 1692124 | YP_009555238.1 | P0C6X6 |
| SAQYIFSTVNALPECN | 1 | 0 | 0 | 1692132 | YP_003766.2 | H9EJ42 |
| SAQYVFSTVNALPEVN | 1 | 0 | 0 | 1692133 | NP_073549.1 | A0A0P0LKV0 |
| SARAASVYYSQLMCQP | 1 | 0 | 0 | 1692134 | QIQ54046.1 | P0C6U8 |
| SARIVYTACSHAAVDA | 1 | 0 | 0 | 1692138 | YP_009724389.1 | P0DTD1 |
| SASAFFGMSRIGMEVT | 1 | 0 | 0 | 1692144 | YP_009724397.2 | P0DTC9 |
| SASCPIGTNYRSCEST | 1 | 0 | 0 | 1692147 | YP_173238.1 | A0A140H1H1 |
| SASDMTNPDYTNLLCR | 1 | 0 | 0 | 1692148 | YP_009555238.1 | P0C6X6 |
| SASEADYRCACYAHLA | 1 | 0 | 0 | 1692149 | YP_003766.2 | H9EJ42 |
| SASINTGNCPFSFGKV | 1 | 0 | 0 | 1692155 | NP_073551.1 | A0A0P0K6L9 |
| SASKLTIVSPDLESYS | 1 | 0 | 0 | 1692160 | YP_003766.2 | H9EJ42 |
| SASLFKPVQLVPVSPA | 1 | 0 | 0 | 1692162 | YP_009047208.1 | K9N7D2 |
| SASLQEILSRLDALEA | 1 | 0 | 0 | 1692164 | YP_009555241.1 | P36334 |
| SASNSRPGSRSQSRGP | 1 | 0 | 0 | 1692165 | YP_173242.1 | Q19U29 |
| SASSEKPTKEEPKKVK | 1 | 0 | 0 | 1692166 | QIQ54046.1 | P0C6U8 |
| SASTSAFVETVKGLDY | 1 | 0 | 0 | 1692168 | YP_009724389.1 | P0DTD1 |
| SASYRFVTFEPFNVSF | 1 | 0 | 0 | 1692172 | YP_173238.1 | A0A140H1H1 |
| SATGSLDVEVFNRLAT | 1 | 0 | 0 | 1692182 | YP_003766.2 | H9EJ42 |
| SATKIIALNKRWQLAL | 1 | 0 | 0 | 1692186 | NP_828852.2 | P59632 |
| SATKLNDLCFSNVYAD | 1 | 0 | 0 | 1692187 | NP_828851.1 | P59594 |
| SATLVNDIDITFLKKD | 1 | 0 | 0 | 1692189 | QHR63299.1 | P0C6W2 |
| SATLVSDIDITFLKKD | 1 | 0 | 0 | 1692190 | YP_009724389.1 | P0DTD1 |
| SAVAVVGGTIQILASV | 1 | 0 | 0 | 1692197 | NP_073549.1 | A0A0P0LKV0 |
| SAVFISPYNSQNYVAK | 1 | 0 | 0 | 1692199 | YP_173236.1 | A0A140H1G9 |
| SAVKLQNNELSPVALR | 1 | 0 | 0 | 1692202 | YP_009724389.1 | P0DTD1 |
| SAVNQSLSNAKVSAEV | 1 | 0 | 0 | 1692211 | NP_073552.1 | A0A0P0KWE8 |
| SAVNSTKALAKQDAAQ | 1 | 0 | 0 | 1692212 | YP_009047206.1 | K9N4V0 |
| SAVSSPMSNMTENERL | 1 | 0 | 0 | 1692217 | NP_073555.1 | A0A0P0KXN3 |
| SAVVNGIRVLKTSDNN | 1 | 0 | 0 | 1692220 | NP_073549.1 | A0A0P0LKV0 |
| SAYGLCDAANPTNCIA | 1 | 0 | 0 | 1692228 | YP_009047204.1 | K9N5Q8 |
| SAYRANSFDVPTFCAN | 1 | 0 | 0 | 1692230 | YP_009047202.1 | K9N7C7 |
| SCDCGKKFDDQVGCLF | 1 | 0 | 0 | 1692246 | YP_003766.2 | H9EJ42 |
| SCEKNLQLMRKVDLYN | 1 | 0 | 0 | 1692253 | YP_003768.1 | H9EJ24 |
| SCEVVHDQVNYVPLRA | 1 | 0 | 0 | 1692257 | NP_073549.1 | A0A0P0LKV0 |
| SCGHYQTNIYSQNLCV | 1 | 0 | 0 | 1692267 | NP_073549.1 | A0A0P0LKV0 |
| SCGLFKDCARNPIDLP | 1 | 0 | 0 | 1692271 | NP_073549.1 | A0A0P0LKV0 |
| SCGNDSWLTGNAIQGF | 1 | 0 | 0 | 1692272 | YP_009047202.1 | K9N7C7 |
| SCIIGSLGFYNPPTNV | 1 | 0 | 0 | 1692284 | YP_009555240.1 | P30215 |
| SCLVSSDGQPGGDAIS | 1 | 0 | 0 | 1692313 | YP_009047202.1 | K9N7C7 |
| SCNNLVPTYLKSDNIV | 1 | 0 | 0 | 1692321 | YP_009555238.1 | P0C6X6 |
| SCNNVNVRDLQRRLYD | 1 | 0 | 0 | 1692322 | YP_003766.2 | H9EJ42 |
| SCPEWMDDFEADPKGK | 1 | 0 | 0 | 1692327 | YP_009047202.1 | K9N7C7 |
| SCPIGTNYRSCESTTV | 1 | 0 | 0 | 1692328 | YP_173238.1 | A0A140H1H1 |
| SCPLCCTVYDSSEVKA | 1 | 0 | 0 | 1692330 | YP_173236.1 | A0A140H1G9 |
| SCSCVSTDTTVQSKDT | 1 | 0 | 0 | 1692339 | YP_009555238.1 | P0C6X6 |
| SCSDFLPLSDMEKDFL | 1 | 0 | 0 | 1692340 | YP_009047202.1 | K9N7C7 |
| SCSFEDELTPSDVYQQ | 1 | 0 | 0 | 1692341 | YP_009555238.1 | P0C6X6 |
| SCSLSEQLDFIDTKRG | 1 | 0 | 0 | 1692343 | QIQ54046.1 | P0C6U8 |
| SCSSNCKHSVAQLVSY | 1 | 0 | 0 | 1692347 | YP_009047202.1 | K9N7C7 |
| SCTFEDELAPSDVYQQ | 1 | 0 | 0 | 1692349 | YP_173236.1 | A0A140H1G9 |
| SCTGSPYIDDFNKIAS | 1 | 0 | 0 | 1692350 | YP_173236.1 | A0A140H1G9 |
| SCTGSPYIDDFNRIAS | 1 | 0 | 0 | 1692351 | YP_009555238.1 | P0C6X6 |
| SCTKGLSIADLACAQY | 1 | 0 | 0 | 1692353 | YP_003767.1 | A0A0P0G321 |
| SCTLSEQLDFIDTKRG | 1 | 0 | 0 | 1692354 | YP_009724389.1 | P0DTD1 |
| SCVGSSVAVQSKDLNF | 1 | 0 | 0 | 1692360 | YP_173236.1 | A0A140H1G9 |
| SCVSTDTTVQSKDTNF | 1 | 0 | 0 | 1692363 | YP_009555238.1 | P0C6X6 |
| SCYSGRVSAAFHANSS | 1 | 0 | 0 | 1692369 | YP_009555241.1 | P36334 |
| SDAALALLLLDRLNQL | 1 | 0 | 0 | 1692374 | QHR63308.1 | Q3LZX4 |
| SDAFSLDVSEKSGNFK | 1 | 0 | 0 | 1692380 | NP_828851.1 | P59594 |
| SDCDVVQEQVNYVPLR | 1 | 0 | 0 | 1692389 | YP_003766.2 | H9EJ42 |
| SDCNHINNLKIKNFDY | 1 | 0 | 0 | 1692391 | YP_173237.1 | A0A140H1H0 |
| SDDAVVCYNSNYAAQG | 1 | 0 | 0 | 1692397 | NP_828849.2 | P0C6X7 |
| SDDDFVSAVANAHRYD | 1 | 0 | 0 | 1692399 | YP_003766.2 | H9EJ42 |
| SDDEWAAAVDEAFPLD | 1 | 0 | 0 | 1692400 | YP_009047202.1 | K9N7C7 |
| SDDGVVCYNKDYAELG | 1 | 0 | 0 | 1692402 | YP_003766.2 | H9EJ42 |
| SDDGVVCYNSDYASKG | 1 | 0 | 0 | 1692404 | YP_009555238.1 | P0C6X6 |
| SDDYIARNGPLNVGGS | 1 | 0 | 0 | 1692411 | QIQ54046.1 | P0C6U8 |
| SDDYIATNGPLKVGGS | 1 | 0 | 0 | 1692412 | YP_009724389.1 | P0DTD1 |
| SDDYIKLNGPLTVGGS | 1 | 0 | 0 | 1692413 | NP_828849.2 | P0C6X7 |
| SDEAIPTRFPPGTVLP | 1 | 0 | 0 | 1692415 | YP_009555245.1 | P33469 |
| SDEYILAKGPLQVGDS | 1 | 0 | 0 | 1692421 | YP_009047202.1 | K9N7C7 |
| SDFAFNNWFLLTNTSS | 1 | 0 | 0 | 1692422 | NP_073551.1 | A0A0P0K6L9 |
| SDFINGIFAKVKNTKV | 1 | 0 | 0 | 1692429 | YP_009555241.1 | P36334 |
| SDFLPLSDMEKDFLSF | 1 | 0 | 0 | 1692431 | YP_009047202.1 | K9N7C7 |

| | | | | | | |
|---|---|---|---|---|---|---|
| SDFNNGIFSRVKNTKL | 1 | 0 | 0 | 1692433 | YP_173238.1 | A0A140H1H1 |
| SDFSLADSVVEVVTTS | 1 | 0 | 0 | 1692436 | YP_009555238.1 | P0C6X6 |
| SDFTGSVYGNFDDQPS | 1 | 0 | 0 | 1692438 | YP_003766.2 | H9EJ42 |
| SDFTPRSDMEYDFLNM | 1 | 0 | 0 | 1692439 | YP_003766.2 | H9EJ42 |
| SDFVDLESVIQDYENV | 1 | 0 | 0 | 1692441 | YP_009555238.1 | P0C6X6 |
| SDGNYNRNASLNSFKE | 1 | 0 | 0 | 1692451 | YP_009047204.1 | K9N5Q8 |
| SDGTNKQLNEISDDCN | 1 | 0 | 0 | 1692456 | YP_009555238.1 | P0C6X6 |
| SDGVDSFEGVLPINTD | 1 | 0 | 0 | 1692457 | YP_003766.2 | H9EJ42 |
| SDGVMKNASLYTSLVP | 1 | 0 | 0 | 1692458 | YP_173236.1 | A0A140H1G9 |
| SDGVVWVAKEGAKTVN | 1 | 0 | 0 | 1692459 | YP_003771.1 | H9EJ69 |
| SDHYIATNGPLKVGGG | 1 | 0 | 0 | 1692463 | QHR63299.1 | P0C6W2 |
| SDIDITFLKKDAPYIV | 1 | 0 | 0 | 1692464 | YP_009724389.1 | P0DTD1 |
| SDIDYVPLKSATCITR | 1 | 0 | 0 | 1692465 | YP_009724389.1 | P0DTD1 |
| SDIFGQDFKTFDLLKY | 1 | 0 | 0 | 1692468 | NP_073549.1 | A0A0P0LKV0 |
| SDIFGSDFKTFDLLKY | 1 | 0 | 0 | 1692469 | YP_003766.2 | H9EJ42 |
| SDIHLPAYASFKVIDN | 1 | 0 | 0 | 1692470 | YP_173236.1 | A0A140H1G9 |
| SDILSLNNPIFINYSK | 1 | 0 | 0 | 1692472 | YP_173237.1 | A0A140H1H0 |
| SDIRDYKLANDAKESL | 1 | 0 | 0 | 1692475 | NP_073549.1 | A0A0P0LKV0 |
| SDISLSVACQLPYCYF | 1 | 0 | 0 | 1692476 | YP_173237.1 | A0A140H1H0 |
| SDISLVKFGAALAMEK | 1 | 0 | 0 | 1692477 | YP_173238.1 | A0A140H1H1 |
| SDITLKCCTVTYLNDP | 1 | 0 | 0 | 1692478 | NP_073549.1 | A0A0P0LKV0 |
| SDIYFVKNIPRYASAV | 1 | 0 | 0 | 1692479 | YP_009555238.1 | P0C6X6 |
| SDKFDFIIGHGMSFSM | 1 | 0 | 0 | 1692486 | YP_009555238.1 | P0C6X6 |
| SDKFKDGVCLFWNCNV | 1 | 0 | 0 | 1692487 | QIQ54047.1 | P0C6X7 |
| SDKFYRLSNELAQVLT | 1 | 0 | 0 | 1692489 | NP_073549.1 | A0A0P0LKV0 |
| SDLIISNFKFDDYDFS | 1 | 0 | 0 | 1692503 | YP_173236.1 | A0A140H1G9 |
| SDLNDFVSDADSTLIG | 1 | 0 | 0 | 1692508 | YP_009724389.1 | P0DTD1 |
| SDLNTSELLTKAIDVD | 1 | 0 | 0 | 1692509 | NP_073549.1 | A0A0P0LKV0 |
| SDLQDLKWARFPKSDG | 1 | 0 | 0 | 1692511 | YP_009724389.1 | P0DTD1 |
| SDLQLGSSGFLQSSNY | 1 | 0 | 0 | 1692512 | YP_173238.1 | A0A140H1H1 |
| SDLVQNGVDAKGFPQL | 1 | 0 | 0 | 1692522 | YP_003771.1 | H9EJ69 |
| SDMEKDFLSFDSDVFI | 1 | 0 | 0 | 1692525 | YP_009047202.1 | K9N7C7 |
| SDMEYDFLNMDMGVFI | 1 | 0 | 0 | 1692526 | YP_003766.2 | H9EJ42 |
| SDMYDGRTKAIDGENV | 1 | 0 | 0 | 1692530 | NP_073549.1 | A0A0P0LKV0 |
| SDMYDPKTKNVTKEND | 1 | 0 | 0 | 1692533 | YP_009724389.1 | P0DTD1 |
| SDMYDPLTKNIGDYNV | 1 | 0 | 0 | 1692534 | YP_173236.1 | A0A140H1G9 |
| SDMYDPRTKHVTKEND | 1 | 0 | 0 | 1692535 | NP_828849.2 | P0C6X7 |
| SDMYDPTTKNVTGSNE | 1 | 0 | 0 | 1692536 | YP_009047202.1 | K9N7C7 |
| SDNINDKDTGPPPIST | 1 | 0 | 0 | 1692544 | YP_009555241.1 | P36334 |
| SDNVTDFNAIATCDWT | 1 | 0 | 0 | 1692551 | YP_009724389.1 | P0DTD1 |
| SDPQFPILAELAPTAG | 1 | 0 | 0 | 1692555 | YP_009555245.1 | P33469 |
| SDQFKTTGDLAVQIGS | 1 | 0 | 0 | 1692557 | YP_003766.2 | H9EJ42 |
| SDRELHLSWEVGKPRP | 1 | 0 | 0 | 1692562 | YP_009724389.1 | P0DTD1 |
| SDREVILCWETGKVKP | 1 | 0 | 0 | 1692564 | YP_173236.1 | A0A140H1G9 |
| SDRFKTSGDLAVQIGN | 1 | 0 | 0 | 1692565 | NP_073549.1 | A0A0P0LKV0 |
| SDSCNNVNVRDLQRRL | 1 | 0 | 0 | 1692576 | YP_003766.2 | H9EJ42 |
| SDSGCILPSECTMFRD | 1 | 0 | 0 | 1692579 | YP_009047202.1 | K9N7C7 |
| SDSIRATATIPIQASL | 1 | 0 | 0 | 1692580 | QIQ54049.1 | P59632 |
| SDSNQSSSDLVAAVTL | 1 | 0 | 0 | 1692586 | YP_003771.1 | H9EJ69 |
| SDSTDNNQNGDRSGAR | 1 | 0 | 0 | 1692589 | QIQ54056.1 | P59595 |
| SDSTLVKFSAAQAMEK | 1 | 0 | 0 | 1692592 | YP_009555241.1 | P36334 |
| SDSVATYFGDCITLPF | 1 | 0 | 0 | 1692593 | YP_009555238.1 | P0C6X6 |
| SDSVQCYTPHIQISYN | 1 | 0 | 0 | 1692594 | YP_173236.1 | A0A140H1G9 |
| SDSVRATATIPIQASL | 1 | 0 | 0 | 1692595 | QHR63301.1 | Q3LZX0 |
| SDTAHACNANRFNVAI | 1 | 0 | 0 | 1692597 | NP_073549.1 | A0A0P0LKV0 |
| SDTTDAVRDPQTLEIL | 1 | 0 | 0 | 1692607 | QIQ54048.1 | P59594 |
| SDTVTYKSTATTKLVP | 1 | 0 | 0 | 1692609 | NP_073549.1 | A0A0P0LKV0 |
| SDVDGLKYTKILKDDG | 1 | 0 | 0 | 1692616 | YP_009555238.1 | P0C6X6 |
| SDVDGLKYTKIMKDDG | 1 | 0 | 0 | 1692617 | YP_173236.1 | A0A140H1G9 |
| SDVENPHLMGWDYPKC | 1 | 0 | 0 | 1692621 | YP_009724389.1 | P0DTD1 |
| SDVETPHLMGWDYPKC | 1 | 0 | 0 | 1692624 | NP_828849.2 | P0C6X7 |
| SDVFIKKYGLENYAFE | 1 | 0 | 0 | 1692626 | YP_009047202.1 | K9N7C7 |
| SDVGFVEAYNNCTGGA | 1 | 0 | 0 | 1692630 | YP_009555241.1 | P36334 |
| SDVGFVEAYNNCTGGS | 1 | 0 | 0 | 1692631 | YP_173238.1 | A0A140H1H1 |
| SDVIRYNLNFEENLRR | 1 | 0 | 0 | 1692632 | NP_073551.1 | A0A0P0K6L9 |
| SDVLSDVIRYNLNFEE | 1 | 0 | 0 | 1692636 | NP_073551.1 | A0A0P0K6L9 |
| SDVSLTYLLGTAKKQV | 1 | 0 | 0 | 1692643 | YP_009555238.1 | P0C6X6 |
| SDVSSRDPTTQEAIPT | 1 | 0 | 0 | 1692644 | YP_173242.1 | Q19U29 |
| SDWSDIRDYKLANDAK | 1 | 0 | 0 | 1692654 | NP_073549.1 | A0A0P0LKV0 |
| SDWTDVRDYKLANDVK | 1 | 0 | 0 | 1692655 | YP_003766.2 | H9EJ42 |
| SDYKGLPLEFGASSVQ | 1 | 0 | 0 | 1692664 | QIQ54046.1 | P0C6U8 |
| SDYYQLYSTQLSTDTG | 1 | 0 | 0 | 1692668 | YP_009724391.1 | P0DTC3 |
| SEADYRCACYAHLAKA | 1 | 0 | 0 | 1692673 | YP_003766.2 | H9EJ42 |
| SEAFEYYHTLDESFLG | 1 | 0 | 0 | 1692675 | NP_828849.2 | P0C6X7 |
| SEAKCWTETDLTKGPH | 1 | 0 | 0 | 1692683 | YP_009724389.1 | P0DTD1 |
| SEAVEDTIDGVVEDTI | 1 | 0 | 0 | 1692692 | YP_173236.1 | A0A140H1G9 |
| SECFMKSDIFGQDFKT | 1 | 0 | 0 | 1692694 | NP_073549.1 | A0A0P0LKV0 |
| SECFVKSDIFGSDFKT | 1 | 0 | 0 | 1692695 | YP_003766.2 | H9EJ42 |
| SECSKASSRSAIEDLL | 1 | 0 | 0 | 1692697 | YP_009555241.1 | P36334 |
| SECYQLYSTQLSTDTG | 1 | 0 | 0 | 1692700 | QIQ54049.1 | P59632 |
| SEDFIKHVRTNGGYNS | 1 | 0 | 0 | 1692702 | YP_009047202.1 | K9N7C7 |
| SEDLKAVRSSFNFDQK | 1 | 0 | 0 | 1692705 | YP_009555238.1 | P0C6X6 |
| SEDLWQFVDHFGENEE | 1 | 0 | 0 | 1692706 | NP_073549.1 | A0A0P0LKV0 |
| SEDVKLNPSEDFIKHV | 1 | 0 | 0 | 1692712 | YP_009047202.1 | K9N7C7 |
| SEEEQPEQPTQYEYGT | 1 | 0 | 0 | 1692713 | QIQ54046.1 | P0C6U8 |
| SEESQGNVVTSVMESQ | 1 | 0 | 0 | 1692715 | YP_173236.1 | A0A140H1G9 |
| SEEVILKTGELQPLEE | 1 | 0 | 0 | 1692717 | QIQ54046.1 | P0C6U8 |
| SEEVVENPTIQKDILE | 1 | 0 | 0 | 1692718 | QHR63299.1 | P0C6W2 |
| SEEVVENPTIQKEVIE | 1 | 0 | 0 | 1692720 | NP_828849.2 | P0C6X7 |
| SEFNLPGCDGGSLYVN | 1 | 0 | 0 | 1692726 | YP_009047202.1 | K9N7C7 |
| SEFSSLPSYAAFATAQ | 1 | 0 | 0 | 1692730 | YP_009724389.1 | P0DTD1 |

| | | | | | | |
|---|---|---|---|---|---|---|
| SEFSSLPSYAAYATAQ | 1 | 0 | 0 | 1692731 | NP_828849.2 | P0C6X7 |
| SEGFLLGINYLGTIKE | 1 | 0 | 0 | 1692738 | YP_009047202.1 | K9N7C7 |
| SEGNALYNNEGGRAFM | 1 | 0 | 0 | 1692741 | NP_073549.1 | A0A0P0LKV0 |
| SEHDYQIGGYTEKWES | 1 | 0 | 0 | 1692743 | YP_009724391.1 | P0DTC3 |
| SEIATKMFDSFVNSFV | 1 | 0 | 0 | 1692744 | YP_009047202.1 | K9N7C7 |
| SEIEPEPEPTPEEPVN | 1 | 0 | 0 | 1692745 | NP_828849.2 | P0C6X7 |
| SEIIGYKAIDGGVTRD | 1 | 0 | 0 | 1692746 | YP_009724389.1 | P0DTD1 |
| SEIKCKTQSIAPPTGV | 1 | 0 | 0 | 1692747 | YP_009555241.1 | P36334 |
| SEILDISPCAFGGVSV | 1 | 0 | 0 | 1692748 | NP_828851.1 | P59594 |
| SEIRDLLCVQSFNGIK | 1 | 0 | 0 | 1692751 | YP_173238.1 | A0A140H1H1 |
| SEISANGCSTIAQAVR | 1 | 0 | 0 | 1692752 | NP_073549.1 | A0A0P0LKV0 |
| SEISGFGFAIPSVAVR | 1 | 0 | 0 | 1692753 | YP_003766.2 | H9EJ42 |
| SEISVIMHLQWIVMYG | 1 | 0 | 0 | 1692756 | YP_173236.1 | A0A140H1G9 |
| SEISVIMHLQWLVMYG | 1 | 0 | 0 | 1692757 | YP_009555238.1 | P0C6X6 |
| SEITKPLIEAEKPNLN | 1 | 0 | 0 | 1692758 | YP_173243.1 | Q0ZJ80 |
| SEIVMCGGCYYVKPGG | 1 | 0 | 0 | 1692761 | YP_009555238.1 | P0C6X6 |
| SEIVNTVAGIPSNVYL | 1 | 0 | 0 | 1692762 | NP_073549.1 | A0A0P0LKV0 |
| SEIYCANIHRFKPVYR | 1 | 0 | 0 | 1692763 | YP_009047207.1 | K9N643 |
| SEKHQDHVSLTVPLCG | 1 | 0 | 0 | 1692766 | YP_009047202.1 | K9N7C7 |
| SEKISVMDKLDTGAQK | 1 | 0 | 0 | 1692767 | YP_003766.2 | H9EJ42 |
| SEKPTKEEPKKVKPCI | 1 | 0 | 0 | 1692769 | QIQ54046.1 | P0C6U8 |
| SEKQVEQKIAEIPKEE | 1 | 0 | 0 | 1692770 | YP_009724389.1 | P0DTD1 |
| SEKSGNFKHLREFVFK | 1 | 0 | 0 | 1692772 | NP_828851.1 | P59594 |
| SEKSYELQTPFEIKLA | 1 | 0 | 0 | 1692773 | YP_009724389.1 | P0DTD1 |
| SELAKGVALDGVLSTF | 1 | 0 | 0 | 1692774 | NP_828849.2 | P0C6X7 |
| SELKQLEAKTASLFQT | 1 | 0 | 0 | 1692777 | YP_003767.1 | A0A0P0G321 |
| SELLKEKYTCSITFEM | 1 | 0 | 0 | 1692780 | YP_003766.2 | H9EJ42 |
| SELQTIVPEYIDVNKT | 1 | 0 | 0 | 1692785 | NP_073551.1 | A0A0P0K6L9 |
| SELRPDTRYVLMDGSI | 1 | 0 | 0 | 1692786 | NP_828849.2 | P0C6X7 |
| SELVYENKFVPVKEAS | 1 | 0 | 0 | 1692796 | NP_073549.1 | A0A0P0LKV0 |
| SELVYENKFVPVKPAS | 1 | 0 | 0 | 1692797 | YP_003766.2 | H9EJ42 |
| SEMLTFDKKAFTLANV | 1 | 0 | 0 | 1692798 | NP_073551.1 | A0A0P0K6L9 |
| SENQISGYTLAATSAS | 1 | 0 | 0 | 1692801 | YP_009555241.1 | P36334 |
| SENQKLIANKFNQALG | 1 | 0 | 0 | 1692802 | YP_009047204.1 | K9N5Q8 |
| SENRYSVLSVDSVSEE | 1 | 0 | 0 | 1692803 | YP_173236.1 | A0A140H1G9 |
| SEPPKEQRVQGSITQR | 1 | 0 | 0 | 1692808 | YP_009047211.1 | K9N4V7 |
| SEQLDFIDTKRGVYCC | 1 | 0 | 0 | 1692813 | YP_009724389.1 | P0DTD1 |
| SEQLDYIESKRGVYCC | 1 | 0 | 0 | 1692814 | NP_828849.2 | P0C6X7 |
| SEQLRKQIRSAAKKNN | 1 | 0 | 0 | 1692815 | YP_009724389.1 | P0DTD1 |
| SEQSLRLVDAMVYTSD | 1 | 0 | 0 | 1692817 | NP_828849.2 | P0C6X7 |
| SERELILSWEIGKVKP | 1 | 0 | 0 | 1692820 | YP_009555238.1 | P0C6X6 |
| SESGAKETKEINIIKL | 1 | 0 | 0 | 1692824 | YP_009555238.1 | P0C6X6 |
| SESKVLLERFLPKCPE | 1 | 0 | 0 | 1692827 | YP_003766.2 | H9EJ42 |
| SESQISGYTTAATVAA | 1 | 0 | 0 | 1692829 | YP_173238.1 | A0A140H1H1 |
| SESSCGLFKDCARNPI | 1 | 0 | 0 | 1692830 | NP_073549.1 | A0A0P0LKV0 |
| SETSWQTGDFLKATCE | 1 | 0 | 0 | 1692840 | QIQ54046.1 | P0C6U8 |
| SETVNGYAKSNFVKDD | 1 | 0 | 0 | 1692841 | NP_073549.1 | A0A0P0LKV0 |
| SEVKAQSSGVIPENPV | 1 | 0 | 0 | 1692847 | YP_173236.1 | A0A140H1G9 |
| SEVVVNNLDKSAGHPF | 1 | 0 | 0 | 1692853 | YP_009047202.1 | K9N7C7 |
| SEWAYCVDLHNKINLC | 1 | 0 | 0 | 1692854 | YP_003766.2 | H9EJ42 |
| SEYGTFCDNINSILDE | 1 | 0 | 0 | 1692862 | YP_173238.1 | A0A140H1H1 |
| SEYKGPITDIFYKENS | 1 | 0 | 0 | 1692864 | QHR63299.1 | P0C6W2 |
| SEYKGPITDVFYKENS | 1 | 0 | 0 | 1692865 | YP_009724389.1 | P0DTD1 |
| SEYKGPVTDVFYKENS | 1 | 0 | 0 | 1692866 | QIQ54046.1 | P0C6U8 |
| SEYKGPVTDVFYKETS | 1 | 0 | 0 | 1692867 | NP_828849.2 | P0C6X7 |
| SEYPLLKQKDIGIDTA | 1 | 0 | 0 | 1692869 | YP_173243.1 | Q0ZJ80 |
| SFADNNDITNTNLSRG | 1 | 0 | 0 | 1692877 | YP_009047211.1 | K9N4V7 |
| SFAGYTESAVNSTKAL | 1 | 0 | 0 | 1692878 | YP_009047206.1 | K9N4V0 |
| SFASSCKSHKPPSASC | 1 | 0 | 0 | 1692881 | YP_173238.1 | A0A140H1H1 |
| SFATYHTPATDCSDGN | 1 | 0 | 0 | 1692882 | YP_009047204.1 | K9N5Q8 |
| SFAVDPAAAYLDAVKQ | 1 | 0 | 0 | 1692883 | NP_073549.1 | A0A0P0LKV0 |
| SFCDNINAILTEVNEL | 1 | 0 | 0 | 1692887 | YP_009555241.1 | P36334 |
| SFCTQLNRALSGIAAE | 1 | 0 | 0 | 1692890 | NP_828851.1 | P59594 |
| SFDDLYVAIRGSCEKN | 1 | 0 | 0 | 1692894 | YP_003768.1 | H9EJ24 |
| SFDGVMYGGFEDQPNL | 1 | 0 | 0 | 1692896 | NP_073549.1 | A0A0P0LKV0 |
| SFDNFKLTCSNTKFAD | 1 | 0 | 0 | 1692903 | NP_828849.2 | P0C6X7 |
| SFDNSYLNESGALVPL | 1 | 0 | 0 | 1692905 | NP_073550.1 | A0A0P0ILY9 |
| SFDNSYLNRVRGSSAA | 1 | 0 | 0 | 1692906 | NP_073549.1 | A0A0P0LKV0 |
| SFDSDVFIKKYGLENY | 1 | 0 | 0 | 1692907 | YP_009047202.1 | K9N7C7 |
| SFDVLKSEDTQGMDNL | 1 | 0 | 0 | 1692910 | QHR63299.1 | P0C6W2 |
| SFDVPDGFYSTSPIQS | 1 | 0 | 0 | 1692911 | NP_073551.1 | A0A0P0K6L9 |
| SFDVPTFCANRSAMCN | 1 | 0 | 0 | 1692912 | YP_009047202.1 | K9N7C7 |
| SFDYHPNCSDCYDDMC | 1 | 0 | 0 | 1692914 | YP_003766.2 | H9EJ42 |
| SFEAHKDSFKDGLCMF | 1 | 0 | 0 | 1692916 | YP_009555238.1 | P0C6X6 |
| SFEDELTPSDVYQQLA | 1 | 0 | 0 | 1692920 | YP_009555238.1 | P0C6X6 |
| SFEEQDEIYAYTKRNV | 1 | 0 | 0 | 1692921 | YP_009555238.1 | P0C6X6 |
| SFEEQNEIYAYTKRNV | 1 | 0 | 0 | 1692922 | YP_173236.1 | A0A140H1G9 |
| SFEGVLPINTDTVLSV | 1 | 0 | 0 | 1692923 | YP_003766.2 | H9EJ42 |
| SFEGYDFKYIENGQLK | 1 | 0 | 0 | 1692924 | NP_073549.1 | A0A0P0LKV0 |
| SFEIDKGIYQTSNFRV | 1 | 0 | 0 | 1692926 | NP_828851.1 | P59594 |
| SFEQQVGVIADKDKDL | 1 | 0 | 0 | 1692933 | NP_073549.1 | A0A0P0LKV0 |
| SFEVLAVEDTQGMDNL | 1 | 0 | 0 | 1692935 | NP_828849.2 | P0C6X7 |
| SFFEDLLFDKVKLSDV | 1 | 0 | 0 | 1692937 | YP_173238.1 | A0A140H1H1 |
| SFFKELTANMSMAECK | 1 | 0 | 0 | 1692938 | YP_003766.2 | H9EJ42 |
| SFGDYNLSSVIPSLPT | 1 | 0 | 0 | 1692941 | NP_073551.1 | A0A0P0K6L9 |
| SFGDYNLSSVLPQRNI | 1 | 0 | 0 | 1692942 | YP_003767.1 | A0A0P0G321 |
| SFGGVSVITPGTNASN | 1 | 0 | 0 | 1692946 | QHR63300.2 | Q3LZX1 |
| SFGKDLNANMSLAECK | 1 | 0 | 0 | 1692949 | NP_073549.1 | A0A0P0LKV0 |
| SFGPGNTFINGDIARE | 1 | 0 | 0 | 1692953 | YP_003766.2 | H9EJ42 |
| SFGVTQEYIQTTIQKV | 1 | 0 | 0 | 1692958 | YP_009047204.1 | K9N5Q8 |
| SFIDYTTTVTSFHGYD | 1 | 0 | 0 | 1692965 | YP_003766.2 | H9EJ42 |

| | | | | | | |
|---|---|---|---|---|---|---|
| SFINGACGSPGYNLKN | 1 | 0 | 0 | 1692971 | NP_073549.1 | A0A0P0LKV0 |
| SFIQADSFTCNNIDAA | 1 | 0 | 0 | 1692973 | YP_009555241.1 | P36334 |
| SFKDNVILLNKHIDAY | 1 | 0 | 0 | 1692976 | QIQ54056.1 | P59595 |
| SFKELLVYAADPAMHA | 1 | 0 | 0 | 1692977 | YP_009724389.1 | P0DTD1 |
| SFKEYFNLRNCTFMYT | 1 | 0 | 0 | 1692978 | YP_009047204.1 | K9N5Q8 |
| SFKQIVESCGNFKVTK | 1 | 0 | 0 | 1692987 | QIQ54046.1 | P0C6U8 |
| SFKSENRYSVLSVDSV | 1 | 0 | 0 | 1692988 | YP_173236.1 | A0A140H1G9 |
| SFKTADGQQKQLLPRW | 1 | 0 | 0 | 1692990 | YP_173242.1 | Q19U29 |
| SFLAVDDKYKATGDLA | 1 | 0 | 0 | 1693000 | YP_009555238.1 | P0C6X6 |
| SFLAVDDKYKVNENLA | 1 | 0 | 0 | 1693001 | YP_173236.1 | A0A140H1G9 |
| SFLEMKSEKQVEQKIA | 1 | 0 | 0 | 1693006 | YP_009724389.1 | P0DTD1 |
| SFLEMKSEKQVEQKTA | 1 | 0 | 0 | 1693007 | QHR63299.1 | P0C6W2 |
| SFLEMKSETKVQDHFD | 1 | 0 | 0 | 1693008 | QIQ54046.1 | P0C6U8 |
| SFLNGSCGSVGFNIDY | 1 | 0 | 0 | 1693013 | YP_009724389.1 | P0DTD1 |
| SFLPKEDGDVLLAEFD | 1 | 0 | 0 | 1693017 | YP_009047202.1 | K9N7C7 |
| SFLPQFTSDQAVTFLK | 1 | 0 | 0 | 1693018 | YP_173241.1 | Q0ZJ82 |
| SFNEWALANQFTEFVG | 1 | 0 | 0 | 1693026 | YP_009047202.1 | K9N7C7 |
| SFNFDQKELLAYYNML | 1 | 0 | 0 | 1693027 | YP_009555238.1 | P0C6X6 |
| SFNNFVSSYAKPEEKL | 1 | 0 | 0 | 1693038 | NP_073549.1 | A0A0P0LKV0 |
| SFNNLLGFDSSKPVTK | 1 | 0 | 0 | 1693039 | YP_009047202.1 | K9N7C7 |
| SFNPETNCLLNVPFGG | 1 | 0 | 0 | 1693040 | YP_009047210.1 | K9N7A1 |
| SFNPETNILLNVPLRG | 1 | 0 | 0 | 1693042 | NP_828855.1 | P59596 |
| SFNVSDAYANLVPYYQ | 1 | 0 | 0 | 1693045 | NP_073549.1 | A0A0P0LKV0 |
| SFQDAASNIFVINKDT | 1 | 0 | 0 | 1693059 | YP_009047202.1 | K9N7C7 |
| SFRDELGVRVLDQSDN | 1 | 0 | 0 | 1693061 | YP_003766.2 | H9EJ42 |
| SFSCNNFDESKIYGSC | 1 | 0 | 0 | 1693071 | YP_173238.1 | A0A140H1H1 |
| SFSFRDELGVRVLDQS | 1 | 0 | 0 | 1693077 | YP_003766.2 | H9EJ42 |
| SFSTFESAYMPIADPT | 1 | 0 | 0 | 1693085 | NP_073549.1 | A0A0P0LKV0 |
| SFSVQKHIGAPANLRV | 1 | 0 | 0 | 1693088 | YP_009047202.1 | K9N7C7 |
| SFTCNNIDAAKIYGMC | 1 | 0 | 0 | 1693092 | YP_009555241.1 | P36334 |
| SFTCRTDMEKDFIALD | 1 | 0 | 0 | 1693093 | YP_009555238.1 | P0C6X6 |
| SFTPGKQSSSRASSGN | 1 | 0 | 0 | 1693100 | YP_009555245.1 | P33469 |
| SFTTYGNTVSEAVSRL | 1 | 0 | 0 | 1693103 | YP_173239.1 | Q0ZJ84 |
| SFVAYETARQEYENAV | 1 | 0 | 0 | 1693108 | NP_073549.1 | A0A0P0LKV0 |
| SFVDDFYGYLQKHFSM | 1 | 0 | 0 | 1693109 | NP_073549.1 | A0A0P0LKV0 |
| SFVDYLLTTIPKAIVL | 1 | 0 | 0 | 1693110 | YP_173236.1 | A0A140H1G9 |
| SFVEYEVAKKNLADAK | 1 | 0 | 0 | 1693111 | YP_173236.1 | A0A140H1G9 |
| SFVEYEVAKKNLDEAR | 1 | 0 | 0 | 1693112 | YP_009555238.1 | P0C6X6 |
| SFVGELKAAEAKVITI | 1 | 0 | 0 | 1693114 | NP_073549.1 | A0A0P0LKV0 |
| SFVGMPSFVAYETARQ | 1 | 0 | 0 | 1693115 | NP_073549.1 | A0A0P0LKV0 |
| SFVNDSIESVGGLYEI | 1 | 0 | 0 | 1693122 | YP_173238.1 | A0A140H1H1 |
| SFVSMPSYIAYENARQ | 1 | 0 | 0 | 1693128 | YP_003766.2 | H9EJ42 |
| SFVVKGDDVRQIAPGQ | 1 | 0 | 0 | 1693129 | NP_828851.1 | P59594 |
| SFYASMYEKSTVLQAA | 1 | 0 | 0 | 1693136 | NP_073549.1 | A0A0P0LKV0 |
| SFYMPLLVSSDKAPYR | 1 | 0 | 0 | 1693144 | YP_003771.1 | H9EJ69 |
| SFYPPDEDEEEGDCEE | 1 | 0 | 0 | 1693145 | YP_009724389.1 | P0DTD1 |
| SFYPPDEDYEEEYSEE | 1 | 0 | 0 | 1693146 | QIQ54046.1 | P0C6U8 |
| SFYPPDEEEEDDAECE | 1 | 0 | 0 | 1693147 | NP_828849.2 | P0C6X7 |
| SGAIKLDPKNPNYNKW | 1 | 0 | 0 | 1693160 | YP_009047211.1 | K9N4V7 |
| SGAKETKEINIIKLSG | 1 | 0 | 0 | 1693161 | YP_009555238.1 | P0C6X6 |
| SGALRELTRELNGGAV | 1 | 0 | 0 | 1693164 | NP_828849.2 | P0C6X7 |
| SGARPKQRRPQGLPNN | 1 | 0 | 0 | 1693168 | QHR63308.1 | Q3LZX4 |
| SGAVLFCTPTKKAFPY | 1 | 0 | 0 | 1693169 | NP_073549.1 | A0A0P0LKV0 |
| SGCMLRACIKTAQADT | 1 | 0 | 0 | 1693177 | YP_009047205.1 | K9N796 |
| SGCNVNDVTKLYLGGL | 1 | 0 | 0 | 1693178 | NP_073549.1 | A0A0P0LKV0 |
| SGDAHIPFGTVLGNFY | 1 | 0 | 0 | 1693182 | NP_073551.1 | A0A0P0K6L9 |
| SGDDCFVSNSSVIQEV | 1 | 0 | 0 | 1693190 | YP_173236.1 | A0A140H1G9 |
| SGDGTTSPISEHDYQI | 1 | 0 | 0 | 1693194 | YP_009724391.1 | P0DTC3 |
| SGDKVFVRDGNTDNLV | 1 | 0 | 0 | 1693198 | NP_073549.1 | A0A0P0LKV0 |
| SGDLAVQIGNNNVCTY | 1 | 0 | 0 | 1693199 | NP_073549.1 | A0A0P0LKV0 |
| SGDYTLANTTTEPLKL | 1 | 0 | 0 | 1693206 | YP_009047202.1 | K9N7C7 |
| SGFAAYNRYRIGNYKL | 1 | 0 | 0 | 1693216 | NP_828855.1 | P59596 |
| SGFETIMKVLNENLNA | 1 | 0 | 0 | 1693222 | YP_009555245.1 | P33469 |
| SGFKLDGFNHQFVTAS | 1 | 0 | 0 | 1693225 | YP_003766.2 | H9EJ42 |
| SGFKPNNTSAQYIFST | 1 | 0 | 0 | 1693228 | YP_003766.2 | H9EJ42 |
| SGFLQSSNYKIDTTSS | 1 | 0 | 0 | 1693229 | YP_173238.1 | A0A140H1H1 |
| SGGGKCFNCYPAGVNI | 1 | 0 | 0 | 1693238 | NP_073551.1 | A0A0P0K6L9 |
| SGGYIPSDFAFNNWFL | 1 | 0 | 0 | 1693242 | NP_073551.1 | A0A0P0K6L9 |
| SGHYQTNLYSFNKAID | 1 | 0 | 0 | 1693249 | YP_003766.2 | H9EJ42 |
| SGIGAVGGDLLYLDLL | 1 | 0 | 0 | 1693253 | YP_009047211.1 | K9N4V7 |
| SGIKFQTVKPGNFNQD | 1 | 0 | 0 | 1693254 | YP_173236.1 | A0A140H1G9 |
| SGITQFQKGKEFEFVE | 1 | 0 | 0 | 1693262 | YP_009555245.1 | P33469 |
| SGLSPAYAPTYVSVDD | 1 | 0 | 0 | 1693305 | YP_009047202.1 | K9N7C7 |
| SGLVNVQKVEQPKIEP | 1 | 0 | 0 | 1693311 | NP_073549.1 | A0A0P0LKV0 |
| SGNGILKWADQSDQFR | 1 | 0 | 0 | 1693319 | YP_009555245.1 | P33469 |
| SGNLLGFKDVTKGTIY | 1 | 0 | 0 | 1693322 | NP_073551.1 | A0A0P0K6L9 |
| SGNLLGFKNVSTGNIF | 1 | 0 | 0 | 1693323 | YP_003767.1 | A0A0P0G321 |
| SGNLVEKLSPWFMDGE | 1 | 0 | 0 | 1693326 | YP_009047202.1 | K9N7C7 |
| SGNPDEPQKDVYELRY | 1 | 0 | 0 | 1693328 | YP_009555245.1 | P33469 |
| SGNRSGNGILKWADQS | 1 | 0 | 0 | 1693330 | YP_009555245.1 | P33469 |
| SGNSTRGTSPGPSGIG | 1 | 0 | 0 | 1693332 | YP_009047211.1 | K9N4V7 |
| SGNVDLDALASDIFDT | 1 | 0 | 0 | 1693334 | YP_009047202.1 | K9N7C7 |
| SGPVDKGHYTVYDTAK | 1 | 0 | 0 | 1693339 | NP_073549.1 | A0A0P0LKV0 |
| SGRSASNSRPGSRSQS | 1 | 0 | 0 | 1693357 | YP_173242.1 | Q19U29 |
| SGRVSAAFHANSSEPA | 1 | 0 | 0 | 1693358 | YP_009555241.1 | P36334 |
| SGSHVGSSFDGVMYGG | 1 | 0 | 0 | 1693370 | NP_073549.1 | A0A0P0LKV0 |
| SGSLVLNNPAYIAPQA | 1 | 0 | 0 | 1693375 | YP_009555240.1 | P30215 |
| SGSNDIWMQNKGLFYT | 1 | 0 | 0 | 1693376 | YP_009555240.1 | P30215 |
| SGTPVVEYLSKEDLHT | 1 | 0 | 0 | 1693393 | YP_009047202.1 | K9N7C7 |
| SGTSTPKKPNKPLSQP | 1 | 0 | 0 | 1693395 | YP_003771.1 | H9EJ69 |
| SGTTCSNDLLQPNTEV | 1 | 0 | 0 | 1693396 | YP_173238.1 | A0A140H1H1 |

| | | | | | | |
|---|---|---|---|---|---|---|
| SGTVSKTSDWKCKVTD | 1 | 0 | 0 | 1693398 | YP_009047202.1 | K9N7C7 |
| SGTWLTYHGAIKLDDK | 1 | 0 | 0 | 1693399 | NP_828858.1 | P59595 |
| SGTWLTYTGAIKLDDK | 1 | 0 | 0 | 1693400 | YP_009724397.2 | P0DTC9 |
| SGTYEGNSPFHPLADN | 1 | 0 | 0 | 1693401 | YP_009724395.1 | P0DTC7 |
| SGVFCGVDAMNLIANI | 1 | 0 | 0 | 1693410 | NP_828849.2 | P0C6X7 |
| SGVFGVNLRTNFTIKG | 1 | 0 | 0 | 1693411 | YP_003766.2 | H9EJ42 |
| SGVKFQTVKPGNFNQD | 1 | 0 | 0 | 1693419 | YP_009555238.1 | P0C6X6 |
| SGVLDTTIPFGPSSQP | 1 | 0 | 0 | 1693421 | YP_003767.1 | A0A0P0G321 |
| SGVLQSGLVKMSHPSG | 1 | 0 | 0 | 1693424 | YP_009047202.1 | K9N7C7 |
| SGVMWLYTYSIGEASS | 1 | 0 | 0 | 1693427 | YP_009047202.1 | K9N7C7 |
| SGVNDICMFENKTTLP | 1 | 0 | 0 | 1693428 | YP_009047202.1 | K9N7C7 |
| SGVTRDLMRELNGGAY | 1 | 0 | 0 | 1693439 | QHR63299.1 | P0C6W2 |
| SGVTRELMRELNGGAY | 1 | 0 | 0 | 1693440 | YP_009724389.1 | P0DTD1 |
| SGVVNPAMDPIYDEPT | 1 | 0 | 0 | 1693442 | QIQ54049.1 | P59632 |
| SGVYQCAMRPNHTIKG | 1 | 0 | 0 | 1693448 | NP_828849.2 | P0C6X7 |
| SGVYSVSSFEAKPSGS | 1 | 0 | 0 | 1693449 | YP_009047204.1 | K9N5Q8 |
| SGYEKSTFNKDIYCGN | 1 | 0 | 0 | 1693454 | YP_003766.2 | H9EJ42 |
| SGYHPAHAPSFLAVDD | 1 | 0 | 0 | 1693456 | YP_009555238.1 | P0C6X6 |
| SHDTVLTSEEVVLKNG | 1 | 0 | 0 | 1693476 | NP_828849.2 | P0C6X7 |
| SHGKQVVSDIDYVPLK | 1 | 0 | 0 | 1693494 | YP_009724389.1 | P0DTD1 |
| SHIVSKESGNTVVLTF | 1 | 0 | 0 | 1693503 | NP_073556.1 | A0A0N7J131 |
| SHKPPSASCPIGTNYR | 1 | 0 | 0 | 1693508 | YP_173238.1 | A0A140H1H1 |
| SHLATFAELEAAQKAY | 1 | 0 | 0 | 1693511 | YP_009047202.1 | K9N7C7 |
| SHLDADLTKPYVKWDL | 1 | 0 | 0 | 1693512 | QIQ54047.1 | P0C6X7 |
| SHMYCSFYPPDEDEEE | 1 | 0 | 0 | 1693523 | YP_009724389.1 | P0DTD1 |
| SHPDFKLLHNLQADIC | 1 | 0 | 0 | 1693531 | YP_009047202.1 | K9N7C7 |
| SHVDTDLTKPYIKWDL | 1 | 0 | 0 | 1693554 | YP_009724389.1 | P0DTD1 |
| SHVNGDWFLFGDSRSD | 1 | 0 | 0 | 1693556 | YP_009555240.1 | P30215 |
| SHWMGIRFEYTSPTDK | 1 | 0 | 0 | 1693560 | YP_009555239.1 | Q80872 |
| SHYWSPQGNLGGNCAG | 1 | 0 | 0 | 1693563 | YP_173236.1 | A0A140H1G9 |
| SIAATRGATVVIGTSK | 1 | 0 | 0 | 1693568 | YP_009724389.1 | P0DTD1 |
| SIADLACAQYYNGIMV | 1 | 0 | 0 | 1693571 | NP_073551.1 | A0A0P0K6L9 |
| SIAFNHPIQVDQLNSS | 1 | 0 | 0 | 1693573 | YP_009047204.1 | K9N5Q8 |
| SIAGVGWTAGLSSFAA | 1 | 0 | 0 | 1693575 | YP_009047204.1 | K9N5Q8 |
| SIAIKVTEYSWNKKLY | 1 | 0 | 0 | 1693576 | NP_073549.1 | A0A0P0LKV0 |
| SIAIPTNFTISVTTEI | 1 | 0 | 0 | 1693577 | YP_009724390.1 | P0DTC2 |
| SIDEVSDDYVQDSTVL | 1 | 0 | 0 | 1693593 | YP_173236.1 | A0A140H1G9 |
| SIDVIDNEIIVKPNIS | 1 | 0 | 0 | 1693597 | NP_073549.1 | A0A0P0LKV0 |
| SIESVGGLYEIKIPTN | 1 | 0 | 0 | 1693602 | YP_173238.1 | A0A140H1H1 |
| SIFDIETVDVKHDVSP | 1 | 0 | 0 | 1693607 | NP_073549.1 | A0A0P0LKV0 |
| SIFSRTLETAQHSVRV | 1 | 0 | 0 | 1693616 | QIQ54046.1 | P0C6U8 |
| SIFSRTLETAQNSVRA | 1 | 0 | 0 | 1693617 | QHR63299.1 | P0C6W2 |
| SIFSRTLETAQNSVRV | 1 | 0 | 0 | 1693618 | YP_009724389.1 | P0DTD1 |
| SIFTINSVVQKDGVEN | 1 | 0 | 0 | 1693621 | YP_009047202.1 | K9N7C7 |
| SIGTNFPLQLGFSTGI | 1 | 0 | 0 | 1693631 | YP_009555238.1 | P0C6X6 |
| SIHDGYTNEIIGYKAI | 1 | 0 | 0 | 1693634 | NP_828849.2 | P0C6X7 |
| SIHSYFITDEKSGGSK | 1 | 0 | 0 | 1693636 | YP_009555238.1 | P0C6X6 |
| SIIAVQPRNVSYDSVS | 1 | 0 | 0 | 1693639 | NP_073551.1 | A0A0P0K6L9 |
| SIIGAMTAVNESRYGL | 1 | 0 | 0 | 1693640 | YP_003767.1 | A0A0P0G321 |
| SIINNAVYTKVDGIDV | 1 | 0 | 0 | 1693646 | NP_828849.2 | P0C6X7 |
| SIINNTVYTKVDGVDV | 1 | 0 | 0 | 1693647 | YP_009724389.1 | P0DTD1 |
| SIIQFPNTYLEGSVRV | 1 | 0 | 0 | 1693650 | YP_009724389.1 | P0DTD1 |
| SIIYTGDWEGLHEVLT | 1 | 0 | 0 | 1693657 | YP_003766.2 | H9EJ42 |
| SIKENVDFINSMQFNY | 1 | 0 | 0 | 1693659 | YP_009555239.1 | Q80872 |
| SIKGDKGVEKLVNWYI | 1 | 0 | 0 | 1693660 | YP_003766.2 | H9EJ42 |
| SIKNVDIVSEAKNVKP | 1 | 0 | 0 | 1693663 | QIQ54046.1 | P0C6U8 |
| SIKSLNNSYINLKDIG | 1 | 0 | 0 | 1693667 | YP_173238.1 | A0A140H1H1 |
| SILDEVNGLLDTTQLH | 1 | 0 | 0 | 1693674 | YP_173238.1 | A0A140H1H1 |
| SILDNAVKGCVPLNAI | 1 | 0 | 0 | 1693675 | YP_009555238.1 | P0C6X6 |
| SILDNAVKGCVPLSAI | 1 | 0 | 0 | 1693676 | YP_173236.1 | A0A140H1G9 |
| SILMLDKQGQLLDTKL | 1 | 0 | 0 | 1693682 | YP_003766.2 | H9EJ42 |
| SINDLSFDKNLGRCVR | 1 | 0 | 0 | 1693696 | YP_003766.2 | H9EJ42 |
| SINSAIVCASVKRDGV | 1 | 0 | 0 | 1693703 | NP_073549.1 | A0A0P0LKV0 |
| SINTGNCPFSFGKVNN | 1 | 0 | 0 | 1693704 | NP_073551.1 | A0A0P0K6L9 |
| SIPESKPLADDDSAII | 1 | 0 | 0 | 1693712 | YP_003771.1 | H9EJ69 |
| SIPIDLLNLREIKPAV | 1 | 0 | 0 | 1693715 | YP_009555238.1 | P0C6X6 |
| SIPINVIVFDGKSKCD | 1 | 0 | 0 | 1693716 | QIQ54046.1 | P0C6U8 |
| SIPIVWGVKDFNTLSQ | 1 | 0 | 0 | 1693717 | YP_003766.2 | H9EJ42 |
| SIPNFGSLTQINTTLL | 1 | 0 | 0 | 1693720 | YP_009047204.1 | K9N5Q8 |
| SIQAIYDRLDTIQADQ | 1 | 0 | 0 | 1693726 | NP_073551.1 | A0A0P0K6L9 |
| SIQANAYYKYDNGNFI | 1 | 0 | 0 | 1693727 | NP_073549.1 | A0A0P0LKV0 |
| SIQDVGFKNQYCNGSI | 1 | 0 | 0 | 1693728 | YP_009555238.1 | P0C6X6 |
| SIRATATIPIQASLPF | 1 | 0 | 0 | 1693737 | QIQ54049.1 | P59632 |
| SIRFDSTLPGFETIMK | 1 | 0 | 0 | 1693739 | YP_173242.1 | Q19U29 |
| SIRGCCESTKLPYYDV | 1 | 0 | 0 | 1693741 | NP_073551.1 | A0A0P0K6L9 |
| SIRHHPELKILRNLNI | 1 | 0 | 0 | 1693742 | YP_173236.1 | A0A140H1G9 |
| SIRSSNQGNKQIVHSY | 1 | 0 | 0 | 1693748 | YP_009047207.1 | K9N643 |
| SISASYRFVTFEPFNV | 1 | 0 | 0 | 1693750 | YP_173238.1 | A0A140H1H1 |
| SISLTVGEVFGKILGN | 1 | 0 | 0 | 1693755 | YP_173236.1 | A0A140H1G9 |
| SISTGSRSARSAIEDL | 1 | 0 | 0 | 1693757 | YP_009047204.1 | K9N5Q8 |
| SISYSELRPDTRYVLM | 1 | 0 | 0 | 1693761 | NP_828849.2 | P0C6X7 |
| SITAQPVKIDNASPAS | 1 | 0 | 0 | 1693762 | NP_828852.2 | P59632 |
| SITESKLSVEQRQQVD | 1 | 0 | 0 | 1693764 | QHR63299.1 | P0C6W2 |
| SITFEMSCDCGKKFDD | 1 | 0 | 0 | 1693766 | YP_003766.2 | H9EJ42 |
| SITKIMCCLGPDIFLG | 1 | 0 | 0 | 1693772 | YP_173236.1 | A0A140H1G9 |
| SITQRTRTRPSVQPGP | 1 | 0 | 0 | 1693774 | YP_009047211.1 | K9N4V7 |
| SITSDKFDFIIGHGMS | 1 | 0 | 0 | 1693776 | YP_009555238.1 | P0C6X6 |
| SITYPMIANENAINKG | 1 | 0 | 0 | 1693780 | NP_073549.1 | A0A0P0LKV0 |
| SIVAKQGAGDAGHSLT | 1 | 0 | 0 | 1693783 | NP_073549.1 | A0A0P0LKV0 |
| SIVIGGAKLKALNLGE | 1 | 0 | 0 | 1693796 | QIQ54046.1 | P0C6U8 |
| SIVNNTVYTKVDGVDV | 1 | 0 | 0 | 1693807 | QIQ54047.1 | P0C6X7 |

| | | | | | | |
|---|---|---|---|---|---|---|
| SIVPSTVWEDGDYYRK | 1 | 0 | 0 | 1693810 | YP_009047204.1 | K9N5Q8 |
| SIVYETDANKLILIND | 1 | 0 | 0 | 1693818 | YP_009555238.1 | P0C6X6 |
| SKADGIIYPQGRTYSN | 1 | 0 | 0 | 1693827 | YP_009047204.1 | K9N5Q8 |
| SKAFVDTIVPPCPSIL | 1 | 0 | 0 | 1693828 | NP_073549.1 | A0A0P0LKV0 |
| SKAYRDYLASGGQPIT | 1 | 0 | 0 | 1693835 | QIQ54046.1 | P0C6U8 |
| SKCHNCCDEYGGHHDF | 1 | 0 | 0 | 1693839 | YP_173238.1 | A0A140H1H1 |
| SKCQITAVEGTKKLAA | 1 | 0 | 0 | 1693841 | YP_009555238.1 | P0C6X6 |
| SKCWVEEDLTKGPHEF | 1 | 0 | 0 | 1693843 | YP_003766.2 | H9EJ42 |
| SKDSNFLNRVRGSIVN | 1 | 0 | 0 | 1693854 | YP_009047202.1 | K9N7C7 |
| SKDSPLELEDFIPMDS | 1 | 0 | 0 | 1693855 | QIQ54047.1 | P0C6X7 |
| SKDTLDDILQQANKSV | 1 | 0 | 0 | 1693856 | YP_009047202.1 | K9N7C7 |
| SKDTNFLNRVRGTSVD | 1 | 0 | 0 | 1693857 | YP_009555238.1 | P0C6X6 |
| SKDYISSNGPLKVGAG | 1 | 0 | 0 | 1693858 | YP_003766.2 | H9EJ42 |
| SKELKRPIQPTDVAYH | 1 | 0 | 0 | 1693866 | YP_009555238.1 | P0C6X6 |
| SKELKRPVNPTDASHY | 1 | 0 | 0 | 1693867 | YP_173236.1 | A0A140H1G9 |
| SKEPVASIITKLNSLN | 1 | 0 | 0 | 1693869 | NP_828849.2 | P0C6X7 |
| SKFASVYAWNRKRISN | 1 | 0 | 0 | 1693878 | QIQ54048.1 | P59594 |
| SKFNSCKSDILSLNNP | 1 | 0 | 0 | 1693883 | YP_173237.1 | A0A140H1H0 |
| SKGAVQKESDEYILAK | 1 | 0 | 0 | 1693892 | YP_009047202.1 | K9N7C7 |
| SKGRLIIRENNRVVIS | 1 | 0 | 0 | 1693908 | YP_009724389.1 | P0DTD1 |
| SKGSLEPRHFNSITKI | 1 | 0 | 0 | 1693909 | YP_173236.1 | A0A140H1G9 |
| SKGSSRNESWHFDKSE | 1 | 0 | 0 | 1693911 | YP_173238.1 | A0A140H1H1 |
| SKGWYDPVENEDIHRV | 1 | 0 | 0 | 1693918 | YP_003766.2 | H9EJ42 |
| SKHAEEYCNYLESYNI | 1 | 0 | 0 | 1693921 | YP_173236.1 | A0A140H1G9 |
| SKHANLYQKYVEAYNT | 1 | 0 | 0 | 1693922 | YP_003766.2 | H9EJ42 |
| SKHIDAKEGGNFNYLY | 1 | 0 | 0 | 1693924 | QHR63300.2 | Q3LZX1 |
| SKHSDFGLGDLVDSYF | 1 | 0 | 0 | 1693929 | NP_073549.1 | A0A0P0LKV0 |
| SKHTDFSSEIIGYKAI | 1 | 0 | 0 | 1693930 | YP_009724389.1 | P0DTD1 |
| SKIITGLHPTQAPTHL | 1 | 0 | 0 | 1693939 | NP_828849.2 | P0C6X7 |
| SKINKYGLELHWAPEF | 1 | 0 | 0 | 1693944 | YP_009555238.1 | P0C6X6 |
| SKISLVSVTDFSLNGN | 1 | 0 | 0 | 1693946 | YP_009047208.1 | K9N7D2 |
| SKIVDLAQHFSDFGTS | 1 | 0 | 0 | 1693948 | YP_009555238.1 | P0C6X6 |
| SKIVQLSEINMDNSPN | 1 | 0 | 0 | 1693950 | NP_828849.2 | P0C6X7 |
| SKIYGSCFKSIVLDKF | 1 | 0 | 0 | 1693953 | YP_173238.1 | A0A140H1H1 |
| SKKDWYDFVENPDIIN | 1 | 0 | 0 | 1693956 | YP_173236.1 | A0A140H1G9 |
| SKKHVEVFTDGKLNCS | 1 | 0 | 0 | 1693957 | YP_009047202.1 | K9N7C7 |
| SKKQFLPFQQFGRDIS | 1 | 0 | 0 | 1693959 | QIQ54048.1 | P59594 |
| SKLASELSNTFGAISA | 1 | 0 | 0 | 1693963 | YP_009047204.1 | K9N5Q8 |
| SKLFITKDEAIKRVRG | 1 | 0 | 0 | 1693966 | YP_173236.1 | A0A140H1G9 |
| SKLLDTCFEATEATFN | 1 | 0 | 0 | 1693976 | YP_009047202.1 | K9N7C7 |
| SKLTIVSPDLESYSKI | 1 | 0 | 0 | 1693987 | YP_003766.2 | H9EJ42 |
| SKPITFHMDGEVINID | 1 | 0 | 0 | 1694005 | QIQ54046.1 | P0C6U8 |
| SKPLTENKYSQLDEEQ | 1 | 0 | 0 | 1694007 | QIQ54052.1 | P59634 |
| SKPQSRNPSSDRNHNS | 1 | 0 | 0 | 1694014 | NP_073556.1 | A0A0N7J131 |
| SKPVMISQWPISNDSN | 1 | 0 | 0 | 1694017 | YP_003766.2 | H9EJ42 |
| SKPYSTLQPNAYYKYD | 1 | 0 | 0 | 1694020 | YP_003766.2 | H9EJ42 |
| SKRGVYCCRDHEHEIA | 1 | 0 | 0 | 1694028 | NP_828849.2 | P0C6X7 |
| SKSCLGYHPAHAPSFL | 1 | 0 | 0 | 1694038 | YP_173236.1 | A0A140H1G9 |
| SKSLTENKYSQLDEEQ | 1 | 0 | 0 | 1694043 | YP_009724394.1 | P0DTC6 |
| SKSPSSSGTSTPKKPN | 1 | 0 | 0 | 1694044 | YP_003771.1 | H9EJ69 |
| SKSYSGYHPAHAPSFL | 1 | 0 | 0 | 1694047 | YP_009555238.1 | P0C6X6 |
| SKTQIGEYTFEKGDYG | 1 | 0 | 0 | 1694054 | QIQ54047.1 | P0C6X7 |
| SKTQSLLIVNNATNVV | 1 | 0 | 0 | 1694055 | YP_009724390.1 | P0DTC2 |
| SKTSDWKCKVTDVLFP | 1 | 0 | 0 | 1694056 | YP_009047202.1 | K9N7C7 |
| SKTSEYKGPVTDVFYK | 1 | 0 | 0 | 1694058 | QIQ54046.1 | P0C6U8 |
| SKTTVASLINTLNDLN | 1 | 0 | 0 | 1694062 | YP_009724389.1 | P0DTD1 |
| SKVGNYRLPSSKPSGM | 1 | 0 | 0 | 1694066 | YP_173241.1 | Q0ZJ82 |
| SKVGNYRLPSTQKGSG | 1 | 0 | 0 | 1694067 | YP_009555244.1 | Q01455 |
| SKVITGLHPTQAPTYL | 1 | 0 | 0 | 1694072 | QIQ54047.1 | P0C6X7 |
| SKVKVIQPRVEKKKTE | 1 | 0 | 0 | 1694074 | NP_828849.2 | P0C6X7 |
| SKVKVVPNLYVVVVES | 1 | 0 | 0 | 1694075 | YP_173236.1 | A0A140H1G9 |
| SKVLLERFLPKCPEIL | 1 | 0 | 0 | 1694076 | YP_003766.2 | H9EJ42 |
| SKVQIGEYTFEKGDYG | 1 | 0 | 0 | 1694077 | YP_009724389.1 | P0DTD1 |
| SKVVLSDALDKLYKVF | 1 | 0 | 0 | 1694085 | NP_073549.1 | A0A0P0LKV0 |
| SKVVNINVDFKDFQFM | 1 | 0 | 0 | 1694088 | YP_173236.1 | A0A140H1G9 |
| SKVVNVNVDFKDFQFM | 1 | 0 | 0 | 1694089 | YP_009555238.1 | P0C6X6 |
| SKYLANEAQVQLEHYS | 1 | 0 | 0 | 1694096 | NP_073549.1 | A0A0P0LKV0 |
| SLAANTLNIIVPDKSV | 1 | 0 | 0 | 1694102 | YP_009555238.1 | P0C6X6 |
| SLADGLEAYADKTLQE | 1 | 0 | 0 | 1694104 | YP_009555238.1 | P0C6X6 |
| SLADISAVQSSFGFDQ | 1 | 0 | 0 | 1694105 | YP_173236.1 | A0A140H1G9 |
| SLADSVVEVVTTSLTP | 1 | 0 | 0 | 1694107 | YP_009555238.1 | P0C6X6 |
| SLAECKRALGLSISDH | 1 | 0 | 0 | 1694108 | NP_073549.1 | A0A0P0LKV0 |
| SLAEYHNESGIKTTLR | 1 | 0 | 0 | 1694109 | QIQ54046.1 | P0C6U8 |
| SLAEYHNESGLKPILR | 1 | 0 | 0 | 1694110 | QHR63299.1 | P0C6W2 |
| SLAEYHNESGLKTILR | 1 | 0 | 0 | 1694111 | YP_009724389.1 | P0DTD1 |
| SLAGSYKDWSYSGQST | 1 | 0 | 0 | 1694114 | YP_009724389.1 | P0DTD1 |
| SLAIDAYPLTKHEDIE | 1 | 0 | 0 | 1694119 | YP_009047202.1 | K9N7C7 |
| SLANSNGFIRFPDVIS | 1 | 0 | 0 | 1694129 | YP_173236.1 | A0A140H1G9 |
| SLCAFSVDPKKTYLDF | 1 | 0 | 0 | 1694135 | YP_009555238.1 | P0C6X6 |
| SLCAFSVDPKKTYLDY | 1 | 0 | 0 | 1694136 | YP_173236.1 | A0A140H1G9 |
| SLCALPDTPSTLTPRS | 1 | 0 | 0 | 1694139 | YP_009047204.1 | K9N5Q8 |
| SLCDEFTLAEVVKQMY | 1 | 0 | 0 | 1694140 | YP_003766.2 | H9EJ42 |
| SLCFKVYFKGQTTHES | 1 | 0 | 0 | 1694141 | YP_173236.1 | A0A140H1G9 |
| SLCNNGKISSSAGDSI | 1 | 0 | 0 | 1694147 | YP_173237.1 | A0A140H1H0 |
| SLCSSHYWSPQGNLGG | 1 | 0 | 0 | 1694149 | YP_173236.1 | A0A140H1G9 |
| SLDLQDCIVGIADDTY | 1 | 0 | 0 | 1694156 | NP_073549.1 | A0A0P0LKV0 |
| SLDPFCFDVNTLLSNQ | 1 | 0 | 0 | 1694160 | YP_003766.2 | H9EJ42 |
| SLDVEVFNRLATSDWT | 1 | 0 | 0 | 1694165 | YP_003766.2 | H9EJ42 |
| SLDVFKIGDVKFKRLG | 1 | 0 | 0 | 1694166 | YP_003766.2 | H9EJ42 |
| SLDVKPYGYPNDFVGG | 1 | 0 | 0 | 1694167 | YP_003766.2 | H9EJ42 |
| SLEFGATSVTPQPEEE | 1 | 0 | 0 | 1694173 | QHR63299.1 | P0C6W2 |

| | | | | | | |
|---|---|---|---|---|---|---|
| SLEGVFWVANHQADTS | 1 | 0 | 0 | 1694175 | YP_173242.1 | Q19U29 |
| SLEIPRRNVATLQAEN | 1 | 0 | 0 | 1694176 | YP_009724389.1 | P0DTD1 |
| SLEVLLDVCNTKEVKV | 1 | 0 | 0 | 1694193 | NP_073549.1 | A0A0P0LKV0 |
| SLEVPRRNVATLQAEN | 1 | 0 | 0 | 1694194 | QIQ54047.1 | P0C6X7 |
| SLFAQSEEEEPFSN | 1 | 0 | 0 | 1694196 | YP_009047206.1 | K9N4V0 |
| SLFATLMTFSGNVDLD | 1 | 0 | 0 | 1694197 | YP_009047202.1 | K9N7C7 |
| SLFKPVQLVPVSPAFH | 1 | 0 | 0 | 1694212 | YP_009047208.1 | K9N7D2 |
| SLFVEDCKLPLGQSLC | 1 | 0 | 0 | 1694224 | YP_009047204.1 | K9N5Q8 |
| SLGFDKPQEKDKKSAK | 1 | 0 | 0 | 1694233 | NP_073556.1 | A0A0N7J131 |
| SLGNVFCDGVNVTKHK | 1 | 0 | 0 | 1694241 | YP_009555238.1 | P0C6X6 |
| SLGVSSNQSPQGNLGS | 1 | 0 | 0 | 1694244 | YP_009555238.1 | P0C6X6 |
| SLIDCNAASVNQIVLR | 1 | 0 | 0 | 1694255 | YP_009047202.1 | K9N7C7 |
| SLIERGKLLVRDTRKE | 1 | 0 | 0 | 1694258 | YP_173236.1 | A0A140H1G9 |
| SLIGGMVLGGLTSAAA | 1 | 0 | 0 | 1694263 | YP_003767.1 | A0A0P0G321 |
| SLIIASSPALVDQRTI | 1 | 0 | 0 | 1694264 | YP_003766.2 | H9EJ42 |
| SLINTLNDLNETLVTM | 1 | 0 | 0 | 1694268 | YP_009724389.1 | P0DTD1 |
| SLIPVRPRNSSDNGIS | 1 | 0 | 0 | 1694269 | YP_003767.1 | A0A0P0G321 |
| SLKDLLLYAADPALHV | 1 | 0 | 0 | 1694281 | YP_009555238.1 | P0C6X6 |
| SLKDLLLYAADPAMHV | 1 | 0 | 0 | 1694282 | YP_173236.1 | A0A140H1G9 |
| SLKELLQNGMNGRTIL | 1 | 0 | 0 | 1694284 | YP_009724389.1 | P0DTD1 |
| SLKITEEVGHTDLMAA | 1 | 0 | 0 | 1694295 | YP_009724389.1 | P0DTD1 |
| SLKLAGTAVVNLRPDQ | 1 | 0 | 0 | 1694296 | YP_173236.1 | A0A140H1G9 |
| SLKLSDNNCYLNAVIM | 1 | 0 | 0 | 1694298 | YP_009047202.1 | K9N7C7 |
| SLLEKGRLIIRENNRV | 1 | 0 | 0 | 1694309 | NP_828849.2 | P0C6X7 |
| SLLKKMDEPYTEDTSE | 1 | 0 | 0 | 1694317 | YP_009555245.1 | P33469 |
| SLLKLKVSTNLFEGDK | 1 | 0 | 0 | 1694318 | NP_073549.1 | A0A0P0LKV0 |
| SLLLLYKTFGRKLHL | 1 | 0 | 0 | 1694323 | A0A161J4U8.1 | Q69028 |
| SLLNQIWQKYLNSPYT | 1 | 0 | 0 | 1694327 | YP_009047206.1 | K9N4V0 |
| SLLSKGRLIIRENNRV | 1 | 0 | 0 | 1694336 | YP_009724389.1 | P0DTD1 |
| SLLVEAYNSILFENGI | 1 | 0 | 0 | 1694343 | YP_003766.2 | H9EJ42 |
| SLMKVMTHMDDFSIKS | 1 | 0 | 0 | 1694354 | YP_173236.1 | A0A140H1G9 |
| SLNAKLGFDSSKEFVE | 1 | 0 | 0 | 1694360 | YP_009555238.1 | P0C6X6 |
| SLNALIATAHSSIKQG | 1 | 0 | 0 | 1694361 | YP_009555238.1 | P0C6X6 |
| SLNDEFSINEVVKQMF | 1 | 0 | 0 | 1694362 | NP_073549.1 | A0A0P0LKV0 |
| SLNDNLLEILTKEKVN | 1 | 0 | 0 | 1694363 | QIQ54046.1 | P0C6U8 |
| SLNEPLVTMPIGYVTH | 1 | 0 | 0 | 1694364 | NP_828849.2 | P0C6X7 |
| SLNGNVSTVFVPATRD | 1 | 0 | 0 | 1694369 | YP_009047208.1 | K9N7D2 |
| SLNGVTLIGEAVKTQF | 1 | 0 | 0 | 1694371 | YP_009724389.1 | P0DTD1 |
| SLNLSCVSKVVNINVD | 1 | 0 | 0 | 1694376 | YP_173236.1 | A0A140H1G9 |
| SLNNPIFINYSKEVYF | 1 | 0 | 0 | 1694377 | YP_173237.1 | A0A140H1H0 |
| SLNNSYINLKDIGTYE | 1 | 0 | 0 | 1694378 | YP_173238.1 | A0A140H1H1 |
| SLNVAKSEFDRDAAMQ | 1 | 0 | 0 | 1694383 | YP_009724389.1 | P0DTD1 |
| SLNVQYRINGLGVTMD | 1 | 0 | 0 | 1694384 | YP_173238.1 | A0A140H1H1 |
| SLPAIKLNFGMLNGNA | 1 | 0 | 0 | 1694386 | NP_073549.1 | A0A0P0LKV0 |
| SLPAINVTINNYNPSS | 1 | 0 | 0 | 1694387 | YP_173238.1 | A0A140H1H1 |
| SLPGVFCGVDAVNLLT | 1 | 0 | 0 | 1694392 | YP_009724389.1 | P0DTD1 |
| SLPGVFCGVDSVNLLT | 1 | 0 | 0 | 1694393 | QIQ54046.1 | P0C6U8 |
| SLPINVIVFDGKSKCD | 1 | 0 | 0 | 1694394 | NP_828849.2 | P0C6X7 |
| SLPINVIVFDGKSKCE | 1 | 0 | 0 | 1694395 | YP_009724389.1 | P0DTD1 |
| SLPKDWRLINKFDVIN | 1 | 0 | 0 | 1694396 | YP_173236.1 | A0A140H1G9 |
| SLPSYAAFATAQEAYE | 1 | 0 | 0 | 1694398 | YP_009724389.1 | P0DTD1 |
| SLPSYAAYATAQEAYE | 1 | 0 | 0 | 1694399 | NP_828849.2 | P0C6X7 |
| SLPYGANKEGIVWVAT | 1 | 0 | 0 | 1694408 | NP_828858.1 | P59595 |
| SLQQVVKALNESYIDL | 1 | 0 | 0 | 1694417 | YP_009047204.1 | K9N5Q8 |
| SLQSCAQHQPYVVDDP | 1 | 0 | 0 | 1694418 | QHR63307.1 | Q3LZX5 |
| SLQVESANLMLSDNVV | 1 | 0 | 0 | 1694423 | YP_003766.2 | H9EJ42 |
| SLQVYGHNYYLPVMAA | 1 | 0 | 0 | 1694424 | YP_003770.1 | U3M6Q8 |
| SLRPDTRYVLMDGSII | 1 | 0 | 0 | 1694438 | YP_009724389.1 | P0DTD1 |
| SLRQIFDVAPIELENK | 1 | 0 | 0 | 1694440 | YP_009047202.1 | K9N7C7 |
| SLSAPTLVPQENYASI | 1 | 0 | 0 | 1694447 | YP_173236.1 | A0A140H1G9 |
| SLSDRFKTSGDLAVQI | 1 | 0 | 0 | 1694449 | NP_073549.1 | A0A0P0LKV0 |
| SLSEQLDFIDTKRGVY | 1 | 0 | 0 | 1694452 | QIQ54046.1 | P0C6U8 |
| SLSFNNLLGFDSSKPV | 1 | 0 | 0 | 1694454 | YP_009047202.1 | K9N7C7 |
| SLSGFKLKDCVMYASA | 1 | 0 | 0 | 1694455 | YP_009724389.1 | P0DTD1 |
| SLSIEEFADVVKEQVS | 1 | 0 | 0 | 1694462 | YP_009047202.1 | K9N7C7 |
| SLSKDYISSNGPLKVG | 1 | 0 | 0 | 1694464 | YP_003766.2 | H9EJ42 |
| SLSKFNSCKSDILSLN | 1 | 0 | 0 | 1694465 | YP_173237.1 | A0A140H1H0 |
| SLSMARRNLDSLEARA | 1 | 0 | 0 | 1694469 | NP_828859.1 | P59636 |
| SLSMIRGPPRAELNGV | 1 | 0 | 0 | 1694470 | YP_009555238.1 | P0C6X6 |
| SLSNAKVSAEVSRQVI | 1 | 0 | 0 | 1694471 | NP_073552.1 | A0A0P0KWE8 |
| SLSRNSSRSSSQGSRS | 1 | 0 | 0 | 1694475 | YP_009047211.1 | K9N4V7 |
| SLSRSNSNFRHSDSIV | 1 | 0 | 0 | 1694476 | YP_173242.1 | Q19U29 |
| SLSSKPQRKRGVKQLP | 1 | 0 | 0 | 1694477 | YP_173242.1 | Q19U29 |
| SLSSNHDLYCSVHKGA | 1 | 0 | 0 | 1694478 | YP_009555238.1 | P0C6X6 |
| SLSSYEYCQPPKSVEK | 1 | 0 | 0 | 1694480 | YP_173236.1 | A0A140H1G9 |
| SLTENKYSQLDEEQPM | 1 | 0 | 0 | 1694487 | YP_009724394.1 | P0DTC6 |
| SLTFSYDGLRGAIRKA | 1 | 0 | 0 | 1694488 | YP_009047202.1 | K9N7C7 |
| SLTFVNVPYVYNGSAQ | 1 | 0 | 0 | 1694489 | YP_009555240.1 | P30215 |
| SLTMLDIQPEDYKSVD | 1 | 0 | 0 | 1694495 | YP_009555239.1 | Q80872 |
| SLTPCGYSEPPKVAAK | 1 | 0 | 0 | 1694497 | YP_009555238.1 | P0C6X6 |
| SLTQINTTLLDLTYEM | 1 | 0 | 0 | 1694498 | YP_009047204.1 | K9N5Q8 |
| SLTVEKGIYQTSNFRV | 1 | 0 | 0 | 1694506 | QIQ54048.1 | P59594 |
| SLVGTTLQGKPIGMFF | 1 | 0 | 0 | 1694523 | YP_009047202.1 | K9N7C7 |
| SLVLAKLGKDATKPQQ | 1 | 0 | 0 | 1694531 | YP_009555245.1 | P33469 |
| SLVNFTVDPQKAYLDF | 1 | 0 | 0 | 1694541 | YP_009047202.1 | K9N7C7 |
| SLVSVTDFSLNGNVST | 1 | 0 | 0 | 1694548 | YP_009047208.1 | K9N7D2 |
| SLWRVTAVHSDGMFVA | 1 | 0 | 0 | 1694559 | YP_003766.2 | H9EJ42 |
| SLYNKYRYYSGKMDTA | 1 | 0 | 0 | 1694570 | YP_009555238.1 | P0C6X6 |
| SLYVNNHAFHTPAYDK | 1 | 0 | 0 | 1694584 | NP_073549.1 | A0A0P0LKV0 |
| SLYVSWSDGDGITGVP | 1 | 0 | 0 | 1694585 | NP_073551.1 | A0A0P0K6L9 |
| SMAATRGATCVIGTTK | 1 | 0 | 0 | 1694587 | YP_009047202.1 | K9N7C7 |

| | | | | | | |
|---|---|---|---|---|---|---|
| SMAPVEFYEDRILDFK | 1 | 0 | 0 | 1694592 | YP_009047202.1 | K9N7C7 |
| SMARRNLDSLEARAFQ | 1 | 0 | 0 | 1694593 | NP_828859.1 | P59636 |
| SMATNYDLSVVNARLR | 1 | 0 | 0 | 1694594 | YP_009724389.1 | P0DTD1 |
| SMCTLSEQLDYIESKR | 1 | 0 | 0 | 1694598 | NP_828849.2 | P0C6X7 |
| SMCYRCPKEIVSTVSA | 1 | 0 | 0 | 1694602 | YP_009047202.1 | K9N7C7 |
| SMDIDVFNKLSTSDWS | 1 | 0 | 0 | 1694605 | NP_073549.1 | A0A0P0LKV0 |
| SMFDVDKKSLNALIAT | 1 | 0 | 0 | 1694608 | YP_009555238.1 | P0C6X6 |
| SMIKGPQRADLNGVIV | 1 | 0 | 0 | 1694616 | YP_173236.1 | A0A140H1G9 |
| SMIRGPPRAELNGVVV | 1 | 0 | 0 | 1694617 | YP_009555238.1 | P0C6X6 |
| SMKGTGVFVQDPAPID | 1 | 0 | 0 | 1694620 | YP_003766.2 | H9EJ42 |
| SMKSDLSVSSAGPISQ | 1 | 0 | 0 | 1694622 | YP_009047204.1 | K9N5Q8 |
| SMLGLTTQTVDSSQGS | 1 | 0 | 0 | 1694624 | YP_009047202.1 | K9N7C7 |
| SMLKRRDSTYGPLQTP | 1 | 0 | 0 | 1694625 | YP_009047204.1 | K9N5Q8 |
| SMPGIYKTQRMCLEPC | 1 | 0 | 0 | 1694642 | NP_073549.1 | A0A0P0LKV0 |
| SMPKIIKVFYELDNDF | 1 | 0 | 0 | 1694645 | YP_009555238.1 | P0C6X6 |
| SMPSYIAYENARQAYE | 1 | 0 | 0 | 1694647 | YP_003766.2 | H9EJ42 |
| SMPYHPNTVDCQDDRC | 1 | 0 | 0 | 1694651 | YP_009555238.1 | P0C6X6 |
| SMQGAVDINKLCEEML | 1 | 0 | 0 | 1694653 | YP_009724389.1 | P0DTD1 |
| SMQGAVDINRLCEEML | 1 | 0 | 0 | 1694654 | NP_828849.2 | P0C6X7 |
| SMRLFYDRDGQRTYDD | 1 | 0 | 0 | 1694661 | YP_009555238.1 | P0C6X6 |
| SMSSFELPQLYGLCIT | 1 | 0 | 0 | 1694663 | YP_173236.1 | A0A140H1G9 |
| SMSSFRSVNQRSIIAY | 1 | 0 | 0 | 1694664 | QIQ54048.1 | P59594 |
| SMSYEDQDALFAYTKR | 1 | 0 | 0 | 1694666 | YP_009724389.1 | P0DTD1 |
| SMTDIAKKPTENICAP | 1 | 0 | 0 | 1694669 | QHR63299.1 | P0C6W2 |
| SMTDIAKKPTETICAP | 1 | 0 | 0 | 1694670 | YP_009724389.1 | P0DTD1 |
| SMVLIDVDVPFTAHFA | 1 | 0 | 0 | 1694684 | YP_009555238.1 | P0C6X6 |
| SMVRKLDNQALNSILD | 1 | 0 | 0 | 1694686 | YP_009555238.1 | P0C6X6 |
| SMYDGDRFVKHDLSLL | 1 | 0 | 0 | 1694691 | NP_073549.1 | A0A0P0LKV0 |
| SMYKQARAEDKKAKIV | 1 | 0 | 0 | 1694694 | YP_009047202.1 | K9N7C7 |
| SNAKVSAEVSRQVIQD | 1 | 0 | 0 | 1694701 | NP_073552.1 | A0A0P0KWE8 |
| SNANLSMLQLGVPDNS | 1 | 0 | 0 | 1694702 | YP_003767.1 | A0A0P0G321 |
| SNCVADYSVLYNSASF | 1 | 0 | 0 | 1694711 | YP_009724390.1 | P0DTC2 |
| SNCVRFKNVDKDDAFY | 1 | 0 | 0 | 1694714 | NP_073549.1 | A0A0P0LKV0 |
| SNDIWMQNKGLFYTQV | 1 | 0 | 0 | 1694718 | YP_009555240.1 | P30215 |
| SNDLLQPNTEVFTDVC | 1 | 0 | 0 | 1694719 | YP_173238.1 | A0A140H1H1 |
| SNDNCFANKHVGFDSW | 1 | 0 | 0 | 1694720 | QIQ54046.1 | P0C6U8 |
| SNDNCTGDIVTHLKNW | 1 | 0 | 0 | 1694721 | NP_073555.1 | A0A0P0KXN3 |
| SNDSNGCVVEASTDFH | 1 | 0 | 0 | 1694722 | YP_003766.2 | H9EJ42 |
| SNDYCCFKHALGCDYV | 1 | 0 | 0 | 1694725 | NP_073549.1 | A0A0P0LKV0 |
| SNEEQAVLKFLDGLDL | 1 | 0 | 0 | 1694726 | YP_003766.2 | H9EJ42 |
| SNELAQVLTEVVYSNG | 1 | 0 | 0 | 1694728 | NP_073549.1 | A0A0P0LKV0 |
| SNESWFEDDYRAFISV | 1 | 0 | 0 | 1694733 | NP_073549.1 | A0A0P0LKV0 |
| SNFDNFKFVCDNTKFA | 1 | 0 | 0 | 1694738 | QIQ54046.1 | P0C6U8 |
| SNFGICADGSLIPVRP | 1 | 0 | 0 | 1694742 | YP_003767.1 | A0A0P0G321 |
| SNFRVQPTESIVRFPN | 1 | 0 | 0 | 1694749 | YP_009724390.1 | P0DTC2 |
| SNFSQYYYNIDTGSVY | 1 | 0 | 0 | 1694752 | YP_173237.1 | A0A140H1H0 |
| SNFVKDDYCDGSLGCK | 1 | 0 | 0 | 1694753 | NP_073549.1 | A0A0P0LKV0 |
| SNFYASGCVLSSACTM | 1 | 0 | 0 | 1694754 | YP_009555238.1 | P0C6X6 |
| SNGCVVEASTDFHQLE | 1 | 0 | 0 | 1694757 | YP_003766.2 | H9EJ42 |
| SNGNIIGFKDFVTNKT | 1 | 0 | 0 | 1694765 | YP_173238.1 | A0A140H1H1 |
| SNGPLKVGAGVMLECE | 1 | 0 | 0 | 1694767 | YP_003766.2 | H9EJ42 |
| SNHDIICNVHKGAHVA | 1 | 0 | 0 | 1694776 | YP_173236.1 | A0A140H1G9 |
| SNHDLYCQVHGNAHVA | 1 | 0 | 0 | 1694777 | YP_009724389.1 | P0DTD1 |
| SNHDLYCSVHKGAHVA | 1 | 0 | 0 | 1694778 | YP_009555238.1 | P0C6X6 |
| SNHDQHCQVHGNAHVA | 1 | 0 | 0 | 1694779 | NP_828849.2 | P0C6X7 |
| SNHDQYCQVHGNAHVA | 1 | 0 | 0 | 1694780 | QIQ54047.1 | P0C6X7 |
| SNHDRYCQVHGNAHVA | 1 | 0 | 0 | 1694781 | QHR63299.1 | P0C6W2 |
| SNIFDVLQSEDTQGME | 1 | 0 | 0 | 1694787 | QIQ54046.1 | P0C6U8 |
| SNIFVINKDTYAALRN | 1 | 0 | 0 | 1694788 | YP_009047202.1 | K9N7C7 |
| SNINCVLSVNSSNCNN | 1 | 0 | 0 | 1694792 | NP_073549.1 | A0A0P0LKV0 |
| SNITITYQGLFPYQGD | 1 | 0 | 0 | 1694794 | YP_009047204.1 | K9N5Q8 |
| SNKLCVVPGNVKPGDA | 1 | 0 | 0 | 1694799 | NP_073549.1 | A0A0P0LKV0 |
| SNLNLPGCDGGSLYVN | 1 | 0 | 0 | 1694813 | YP_009724389.1 | P0DTD1 |
| SNLNTNLHFDVDNINF | 1 | 0 | 0 | 1694814 | YP_173238.1 | A0A140H1H1 |
| SNLSWLSLDVSAAFY | 1 | 0 | 0 | 1694816 | P03155.1 | Q69028 |
| SNLVLNNCTKYNIYDY | 1 | 0 | 0 | 1694817 | YP_003767.1 | A0A0P0G321 |
| SNMNIASNSKEWAYCV | 1 | 0 | 0 | 1694821 | NP_073549.1 | A0A0P0LKV0 |
| SNMVETVVGQLEQTNM | 1 | 0 | 0 | 1694827 | YP_009047202.1 | K9N7C7 |
| SNNGVYISTTKVKSLS | 1 | 0 | 0 | 1694830 | YP_009555238.1 | P0C6X6 |
| SNNKEDFDIIQKCQIT | 1 | 0 | 0 | 1694831 | YP_173236.1 | A0A140H1G9 |
| SNNSSRASSRSSTRNN | 1 | 0 | 0 | 1694837 | YP_003771.1 | H9EJ69 |
| SNQADVNTPADIVDRD | 1 | 0 | 0 | 1694844 | YP_009555245.1 | P33469 |
| SNQNGERSGARPKQRR | 1 | 0 | 0 | 1694847 | QHR63308.1 | Q3LZX4 |
| SNQNTDSDSLSSKPQR | 1 | 0 | 0 | 1694849 | YP_173242.1 | Q19U29 |
| SNQSPQGNLGSNGKPG | 1 | 0 | 0 | 1694852 | YP_009555238.1 | P0C6X6 |
| SNSFDVLKSEDAQGMD | 1 | 0 | 0 | 1694859 | YP_009724389.1 | P0DTD1 |
| SNSFDVLKSEDTQGMD | 1 | 0 | 0 | 1694860 | QHR63299.1 | P0C6W2 |
| SNSGADVLYQPPQTSI | 1 | 0 | 0 | 1694862 | NP_828849.2 | P0C6X7 |
| SNSGNLLGFKNVSTGN | 1 | 0 | 0 | 1694863 | YP_003767.1 | A0A0P0G321 |
| SNSGSDVLYQPPQTSI | 1 | 0 | 0 | 1694864 | YP_009724389.1 | P0DTD1 |
| SNSRPGSRSQSRGPNN | 1 | 0 | 0 | 1694875 | YP_173242.1 | Q19U29 |
| SNSSILSLCAFSVDPK | 1 | 0 | 0 | 1694877 | YP_009555238.1 | P0C6X6 |
| SNSSSGVLDTTIPFGP | 1 | 0 | 0 | 1694878 | YP_003767.1 | A0A0P0G321 |
| SNSSVIQEVLLLRHDI | 1 | 0 | 0 | 1694879 | YP_173236.1 | A0A140H1G9 |
| SNSSVLSLVNFTVDPQ | 1 | 0 | 0 | 1694880 | YP_009047202.1 | K9N7C7 |
| SNSSVPLLEVYVHLRN | 1 | 0 | 0 | 1694881 | YP_003770.1 | U3M6Q8 |
| SNTDNETLQYWVTPLS | 1 | 0 | 0 | 1694884 | YP_173238.1 | A0A140H1H1 |
| SNTKFADDLNQMTGFT | 1 | 0 | 0 | 1694887 | NP_828849.2 | P0C6X7 |
| SNTKYYDDSQYYFNKD | 1 | 0 | 0 | 1694889 | YP_009555240.1 | P30215 |
| SNVANYQKVGMQKYST | 1 | 0 | 0 | 1694896 | YP_009724389.1 | P0DTD1 |
| SNVANYQKVGMQXYST | 1 | 0 | 0 | 1694897 | QIQ54047.1 | P0C6X7 |

| | | | | | | |
|---|---|---|---|---|---|---|
| SNVDESFVDDFYGYLQ | 1 | 0 | 0 | 1694898 | NP_073549.1 | A0A0P0LKV0 |
| SNVDVVPEGWRVVNKF | 1 | 0 | 0 | 1694899 | YP_009555238.1 | P0C6X6 |
| SNVKKVTDVTGKLSDC | 1 | 0 | 0 | 1694902 | YP_009555238.1 | P0C6X6 |
| SNVKLLDYDYATHGQL | 1 | 0 | 0 | 1694903 | YP_003766.2 | H9EJ42 |
| SNVKTLEYDYMTHGQM | 1 | 0 | 0 | 1694904 | NP_073549.1 | A0A0P0LKV0 |
| SNVVELPKFFYASNGT | 1 | 0 | 0 | 1694914 | NP_073551.1 | A0A0P0K6L9 |
| SNVVYPDLDCIYISLK | 1 | 0 | 0 | 1694916 | YP_173237.1 | A0A140H1H0 |
| SNVYRTDYVDYTFVNE | 1 | 0 | 0 | 1694919 | YP_173236.1 | A0A140H1G9 |
| SNYKIDTTSSSCQLYY | 1 | 0 | 0 | 1694926 | YP_173238.1 | A0A140H1H1 |
| SNYLLEEFDVVFGKRG | 1 | 0 | 0 | 1694928 | NP_073549.1 | A0A0P0LKV0 |
| SPAAPRAVSFADNNDI | 1 | 0 | 0 | 1694938 | YP_009047211.1 | K9N4V7 |
| SPAEFKYMVANGLNAP | 1 | 0 | 0 | 1694939 | NP_073549.1 | A0A0P0LKV0 |
| SPARIAGNGGDAALAL | 1 | 0 | 0 | 1694944 | QIQ54056.1 | P59595 |
| SPARLEIVKLGAEKKD | 1 | 0 | 0 | 1694945 | YP_009555239.1 | Q80872 |
| SPARMAGNGGDAALAL | 1 | 0 | 0 | 1694946 | YP_009724397.2 | P0DTC9 |
| SPARMAGNGSDAALAL | 1 | 0 | 0 | 1694947 | QHR63308.1 | Q3LZX4 |
| SPARMASGGGETALAL | 1 | 0 | 0 | 1694948 | NP_828858.1 | P59595 |
| SPAYAPTYVSVDDKYK | 1 | 0 | 0 | 1694952 | YP_009047202.1 | K9N7C7 |
| SPCKELEGVGAKVSAF | 1 | 0 | 0 | 1694956 | YP_009555238.1 | P0C6X6 |
| SPCVSIVPSTVWEDGD | 1 | 0 | 0 | 1694958 | YP_009047204.1 | K9N5Q8 |
| SPDPKFVDKYYAFLNK | 1 | 0 | 0 | 1694964 | YP_009047202.1 | K9N7C7 |
| SPFEKLQCEHLQFGLQ | 1 | 0 | 0 | 1694969 | YP_003767.1 | A0A0P0G321 |
| SPFEMPFEELNGLKIL | 1 | 0 | 0 | 1694971 | NP_073549.1 | A0A0P0LKV0 |
| SPFHPLADNKFALACT | 1 | 0 | 0 | 1694972 | QIQ54053.1 | P59635 |
| SPFHPLADNKFALTCF | 1 | 0 | 0 | 1694973 | YP_009724395.1 | P0DTC7 |
| SPFHPLADNKFALTCT | 1 | 0 | 0 | 1694974 | NP_828857.1 | P59635 |
| SPFTINCQEPKLGSLV | 1 | 0 | 0 | 1694977 | QHR63307.1 | Q3LZX5 |
| SPGSLNFKKVETLNFQ | 1 | 0 | 0 | 1694984 | YP_173243.1 | Q0ZJ80 |
| SPGYNLKNGEVEFVYM | 1 | 0 | 0 | 1694985 | NP_073549.1 | A0A0P0LKV0 |
| SPGYNVRNDGTVEFCY | 1 | 0 | 0 | 1694986 | YP_003766.2 | H9EJ42 |
| SPIIPGFGGDFNLTLL | 1 | 0 | 0 | 1694991 | YP_009047204.1 | K9N5Q8 |
| SPIQSVELPVSIVSLP | 1 | 0 | 0 | 1694992 | NP_073551.1 | A0A0P0K6L9 |
| SPIQYIDIGNYTVSCS | 1 | 0 | 0 | 1694994 | QHR63307.1 | Q3LZX5 |
| SPISEHDYQIGGYTEK | 1 | 0 | 0 | 1694995 | YP_009724391.1 | P0DTC3 |
| SPITDHDYQIGGYTEK | 1 | 0 | 0 | 1694996 | QIQ54049.1 | P59632 |
| SPLKLEDFIPMDSTVK | 1 | 0 | 0 | 1695011 | NP_828849.2 | P0C6X7 |
| SPMKSDEGGLCPSTGQ | 1 | 0 | 0 | 1695021 | YP_173236.1 | A0A140H1G9 |
| SPMSNMTENERLLHFF | 1 | 0 | 0 | 1695022 | NP_073555.1 | A0A0P0KXN3 |
| SPNKQCTVQQCFGKRG | 1 | 0 | 0 | 1695029 | YP_009555245.1 | P33469 |
| SPQGNLGGNCAGNVIG | 1 | 0 | 0 | 1695037 | YP_173236.1 | A0A140H1G9 |
| SPQGNLGSNGKPGNVG | 1 | 0 | 0 | 1695038 | YP_009555238.1 | P0C6X6 |
| SPQIITTDNTFVSGNC | 1 | 0 | 0 | 1695040 | NP_828851.1 | P59594 |
| SPSSSGTSTPKKPNKP | 1 | 0 | 0 | 1695054 | YP_003771.1 | H9EJ69 |
| SPVEFHLDGEVLSLDK | 1 | 0 | 0 | 1695067 | NP_828849.2 | P0C6X7 |
| SPVKDVFELHYSGSIR | 1 | 0 | 0 | 1695068 | YP_173242.1 | Q19U29 |
| SPVMTNGSNILEAFTK | 1 | 0 | 0 | 1695070 | NP_073549.1 | A0A0P0LKV0 |
| SPYDLEVLLQDALQSR | 1 | 0 | 0 | 1695072 | YP_009555238.1 | P0C6X6 |
| SPYIDDFNKIASCKWT | 1 | 0 | 0 | 1695074 | YP_173236.1 | A0A140H1G9 |
| SPYIDDFNRIASCKWT | 1 | 0 | 0 | 1695075 | YP_009555238.1 | P0C6X6 |
| SPYNSQNAVAAKILGL | 1 | 0 | 0 | 1695076 | QIQ54047.1 | P0C6X7 |
| SPYNSQNFAAKRVLGL | 1 | 0 | 0 | 1695079 | YP_009555238.1 | P0C6X6 |
| SPYVCNAPGCDVNDVT | 1 | 0 | 0 | 1695085 | YP_009555238.1 | P0C6X6 |
| SQASSRSSSRSRNSSR | 1 | 0 | 0 | 1695093 | YP_009724397.2 | P0DTC9 |
| SQDDIMKAVAAALKSL | 1 | 0 | 0 | 1695098 | NP_073556.1 | A0A0N7J131 |
| SQDKVFGYVQQGIFNK | 1 | 0 | 0 | 1695100 | YP_009555238.1 | P0C6X6 |
| SQDSITHYPALKMVQT | 1 | 0 | 0 | 1695104 | YP_009047202.1 | K9N7C7 |
| SQDSPLKLEDFIPMDS | 1 | 0 | 0 | 1695105 | NP_828849.2 | P0C6X7 |
| SQDVKQFANGFVVRIG | 1 | 0 | 0 | 1695107 | YP_009047204.1 | K9N5Q8 |
| SQDVYKSLTIVDIPQS | 1 | 0 | 0 | 1695108 | YP_009047202.1 | K9N7C7 |
| SQEGKKYLVKTTKAKG | 1 | 0 | 0 | 1695109 | YP_003766.2 | H9EJ42 |
| SQERNFTTAPAICHEG | 1 | 0 | 0 | 1695112 | NP_828851.1 | P59594 |
| SQFDSLGVSSNQSPQG | 1 | 0 | 0 | 1695114 | YP_009555238.1 | P0C6X6 |
| SQGLLPPKNSIDAFKL | 1 | 0 | 0 | 1695126 | YP_009724389.1 | P0DTD1 |
| SQGLLPPKSSIDAFKL | 1 | 0 | 0 | 1695127 | NP_828849.2 | P0C6X7 |
| SQGNVVTSVMESQIST | 1 | 0 | 0 | 1695129 | YP_173236.1 | A0A140H1G9 |
| SQGSRSGNSTRGTSPG | 1 | 0 | 0 | 1695133 | YP_009047211.1 | K9N4V7 |
| SQHTLYIKDGDDGYFL | 1 | 0 | 0 | 1695134 | YP_009047202.1 | K9N7C7 |
| SQHTMQIVDENGKYYL | 1 | 0 | 0 | 1695138 | NP_073549.1 | A0A0P0LKV0 |
| SQHTMQIVDKDGTYYL | 1 | 0 | 0 | 1695139 | YP_003766.2 | H9EJ42 |
| SQILPDPLKPTKRSFI | 1 | 0 | 0 | 1695141 | NP_828851.1 | P59594 |
| SQIQSKLTDVKCANVV | 1 | 0 | 0 | 1695146 | YP_173236.1 | A0A140H1G9 |
| SQISGYTTAATVAAMF | 1 | 0 | 0 | 1695147 | YP_173238.1 | A0A140H1H1 |
| SQISPAAIASNCYSSL | 1 | 0 | 0 | 1695148 | YP_009047204.1 | K9N5Q8 |
| SQISYSNFYASGCVLS | 1 | 0 | 0 | 1695150 | YP_009555238.1 | P0C6X6 |
| SQIVTGLFKDCSRETS | 1 | 0 | 0 | 1695151 | YP_009047202.1 | K9N7C7 |
| SQLFVDTLVNKIPANI | 1 | 0 | 0 | 1695157 | YP_009555238.1 | P0C6X6 |
| SQLPLDVYDLTMPSQV | 1 | 0 | 0 | 1695167 | YP_009555238.1 | P0C6X6 |
| SQLSLSMARRNLDSLE | 1 | 0 | 0 | 1695170 | NP_828859.1 | P59636 |
| SQMETDFLELAMDEFI | 1 | 0 | 0 | 1695173 | NP_828849.2 | P0C6X7 |
| SQNAVAAKILGLPTQT | 1 | 0 | 0 | 1695175 | QIQ54047.1 | P0C6X7 |
| SQNAVSRSMLGLTTQT | 1 | 0 | 0 | 1695177 | YP_009047202.1 | K9N7C7 |
| SQNFAAKRVLGLQTQT | 1 | 0 | 0 | 1695178 | YP_009555238.1 | P0C6X6 |
| SQNFGNAEMLKLGTND | 1 | 0 | 0 | 1695179 | YP_173242.1 | Q19U29 |
| SQNYVAKRVLGVQTQT | 1 | 0 | 0 | 1695187 | YP_173236.1 | A0A140H1G9 |
| SQPLPQRQKKQQTVTL | 1 | 0 | 0 | 1695190 | QIQ54056.1 | P59595 |
| SQPRADKPSQLKKPRW | 1 | 0 | 0 | 1695191 | YP_003771.1 | H9EJ69 |
| SQQGVFRYDNVSSVWP | 1 | 0 | 0 | 1695196 | YP_009555240.1 | P30215 |
| SQQQSQTVTKKSAAEA | 1 | 0 | 0 | 1695201 | QIQ54056.1 | P59595 |
| SQQTVGQQDGSEDNQT | 1 | 0 | 0 | 1695202 | YP_009724389.1 | P0DTD1 |
| SQQTVVQEDDSEVNQT | 1 | 0 | 0 | 1695203 | QHR63299.1 | P0C6W2 |
| SQRVGTDSGFAAYNRY | 1 | 0 | 0 | 1695207 | NP_828855.1 | P59596 |

| | | | | | | |
|---|---|---|---|---|---|---|
| SQSRGRGESKPQSRNP | 1 | 0 | 0 | 1695214 | NP_073556.1 | A0A0N7J131 |
| SQSRSQSRGRGESKPQ | 1 | 0 | 0 | 1695215 | NP_073556.1 | A0A0N7J131 |
| SQSSETKEQKHEMQKP | 1 | 0 | 0 | 1695216 | NP_073556.1 | A0A0N7J131 |
| SQSSHVAQNTVLNASI | 1 | 0 | 0 | 1695217 | YP_003771.1 | H9EJ69 |
| SQSSPIIPGFGGDFNL | 1 | 0 | 0 | 1695218 | YP_009047204.1 | K9N5Q8 |
| SQSSRINFCGNGNHII | 1 | 0 | 0 | 1695219 | YP_009555241.1 | P36334 |
| SQSSSRASSLSRNSSR | 1 | 0 | 0 | 1695220 | YP_009047211.1 | K9N4V7 |
| SQSVIIINNSTNVVIR | 1 | 0 | 0 | 1695221 | NP_828851.1 | P59594 |
| SQTSAKSLARSQSSET | 1 | 0 | 0 | 1695225 | NP_073556.1 | A0A0N7J131 |
| SQTVKPGHFNKEFYDF | 1 | 0 | 0 | 1695231 | NP_073549.1 | A0A0P0LKV0 |
| SQWPISNDSNGCVVEA | 1 | 0 | 0 | 1695247 | YP_003766.2 | H9EJ42 |
| SQYLFSTINALPETSA | 1 | 0 | 0 | 1695248 | YP_009047202.1 | K9N7C7 |
| SQYYFNKDTGVIYGLN | 1 | 0 | 0 | 1695254 | YP_009555240.1 | P30215 |
| SRANSGNRTPTSGVTP | 1 | 0 | 0 | 1695257 | YP_009555245.1 | P33469 |
| SRASSAGSRSRANSGN | 1 | 0 | 0 | 1695259 | YP_009555245.1 | P33469 |
| SRASSRSSTRNNSRDS | 1 | 0 | 0 | 1695260 | YP_003771.1 | H9EJ69 |
| SRDHNDILYTPPTVSY | 1 | 0 | 0 | 1695265 | NP_073549.1 | A0A0P0LKV0 |
| SRDLEDFKPRSQMETD | 1 | 0 | 0 | 1695266 | NP_828849.2 | P0C6X7 |
| SRDPTTQEAIPTRFPP | 1 | 0 | 0 | 1695267 | YP_173242.1 | Q19U29 |
| SRDSSRSTSRQQSRTR | 1 | 0 | 0 | 1695268 | YP_003771.1 | H9EJ69 |
| SRELKVTFFPDLNGDV | 1 | 0 | 0 | 1695275 | YP_009724389.1 | P0DTD1 |
| SRELTAEDISLLKKMD | 1 | 0 | 0 | 1695277 | YP_009555245.1 | P33469 |
| SRERVNINIVGDFHLN | 1 | 0 | 0 | 1695278 | NP_828849.2 | P0C6X7 |
| SREVVVTNLNKSAGWP | 1 | 0 | 0 | 1695280 | NP_073549.1 | A0A0P0LKV0 |
| SRFLGLQIQTVDSSQG | 1 | 0 | 0 | 1695283 | YP_003766.2 | H9EJ42 |
| SRGAITTGYRFTNFEP | 1 | 0 | 0 | 1695287 | YP_009555241.1 | P36334 |
| SRGLYKKCVKSRGDSG | 1 | 0 | 0 | 1695289 | QIQ54046.1 | P0C6U8 |
| SRGNSRNSTPGSSRGN | 1 | 0 | 0 | 1695291 | NP_828858.1 | P59595 |
| SRGPNNRSLSRSNSNF | 1 | 0 | 0 | 1695292 | YP_173242.1 | Q19U29 |
| SRGTSPARIAGNGGDA | 1 | 0 | 0 | 1695295 | QIQ54056.1 | P59595 |
| SRIMFEPRIPTMADFV | 1 | 0 | 0 | 1695310 | NP_073551.1 | A0A0P0K6L9 |
| SRINFCGNGNHIISLV | 1 | 0 | 0 | 1695311 | YP_009555241.1 | P36334 |
| SRLDALEAEAQIDRLI | 1 | 0 | 0 | 1695318 | YP_009555241.1 | P36334 |
| SRLDALEAQVQIDRLI | 1 | 0 | 0 | 1695319 | YP_173238.1 | A0A140H1H1 |
| SRLELAKVQNLSGNPD | 1 | 0 | 0 | 1695321 | YP_009555245.1 | P33469 |
| SRLLSDDRTEVPQLVN | 1 | 0 | 0 | 1695327 | YP_009047204.1 | K9N5Q8 |
| SRMGFKLDATVPGYPK | 1 | 0 | 0 | 1695336 | YP_009047202.1 | K9N7C7 |
| SRMYCSFYPPDEDYEE | 1 | 0 | 0 | 1695337 | QIQ54046.1 | P0C6U8 |
| SRMYCSFYPPDEEEED | 1 | 0 | 0 | 1695338 | NP_828849.2 | P0C6X7 |
| SRNESWHFDKSEPLCL | 1 | 0 | 0 | 1695341 | YP_173238.1 | A0A140H1H1 |
| SRNPSSDRNHNSQDDI | 1 | 0 | 0 | 1695344 | NP_073556.1 | A0A0N7J131 |
| SRNSSRNSTPGSSRGT | 1 | 0 | 0 | 1695346 | YP_009724397.2 | P0DTC9 |
| SRNSSRSSSQGSRSGN | 1 | 0 | 0 | 1695347 | YP_009047211.1 | K9N4V7 |
| SRNSTPGSSRGTSPAR | 1 | 0 | 0 | 1695349 | YP_009724397.2 | P0DTC9 |
| SRNTTFDFLSNASSSF | 1 | 0 | 0 | 1695350 | YP_003767.1 | A0A0P0G321 |
| SRQLQNSMSGASADST | 1 | 0 | 0 | 1695356 | NP_828858.1 | P59595 |
| SRQVIQDVKDGTVTFN | 1 | 0 | 0 | 1695359 | NP_073552.1 | A0A0P0KWE8 |
| SRRKRRSISASYRFVT | 1 | 0 | 0 | 1695361 | YP_173238.1 | A0A140H1H1 |
| SRSGNSTRGTSPGPSG | 1 | 0 | 0 | 1695372 | YP_009047211.1 | K9N4V7 |
| SRSRANSGNRTPTSGV | 1 | 0 | 0 | 1695381 | YP_009555245.1 | P33469 |
| SRSRNSSRNSTPGSSR | 1 | 0 | 0 | 1695383 | YP_009724397.2 | P0DTC9 |
| SRSSSQGSRSGNSTRG | 1 | 0 | 0 | 1695385 | YP_009047211.1 | K9N4V7 |
| SRSSSRSRGNSRNSTP | 1 | 0 | 0 | 1695386 | NP_828858.1 | P59595 |
| SRSSTRNNSRDSSRST | 1 | 0 | 0 | 1695388 | YP_003771.1 | H9EJ69 |
| SRSTRSMLKRRDSTYG | 1 | 0 | 0 | 1695389 | YP_009047204.1 | K9N5Q8 |
| SRSTSRTSSRASSAGS | 1 | 0 | 0 | 1695391 | YP_009555245.1 | P33469 |
| SRSVASQSIIAYTMSL | 1 | 0 | 0 | 1695392 | QHR63300.2 | Q3LZX1 |
| SRTRSDSNQSSSDLVA | 1 | 0 | 0 | 1695404 | YP_003771.1 | H9EJ69 |
| SRVGRSVNSQNSTGWV | 1 | 0 | 0 | 1695411 | NP_073555.1 | A0A0P0KXN3 |
| SRVISRMGFKLDATVP | 1 | 0 | 0 | 1695415 | YP_009047202.1 | K9N7C7 |
| SRVLGLKTLATHGLAA | 1 | 0 | 0 | 1695421 | YP_009724389.1 | P0DTD1 |
| SRVLSSFEPRSDLERD | 1 | 0 | 0 | 1695423 | YP_173236.1 | A0A140H1G9 |
| SRVPHIVRKDLTKYTM | 1 | 0 | 0 | 1695425 | YP_009555238.1 | P0C6X6 |
| SRWNNARQSDNMTAVA | 1 | 0 | 0 | 1695430 | YP_009555240.1 | P30215 |
| SRWNNERASDISLSVA | 1 | 0 | 0 | 1695431 | YP_173237.1 | A0A140H1H0 |
| SRYRIGNYKLNTDHSN | 1 | 0 | 0 | 1695435 | QIQ54051.1 | P59596 |
| SSAGSRSRANSGNRTP | 1 | 0 | 0 | 1695446 | YP_009555245.1 | P33469 |
| SSALGSMDIDVFNKLS | 1 | 0 | 0 | 1695450 | NP_073549.1 | A0A0P0LKV0 |
| SSANYVGKYDINHGDS | 1 | 0 | 0 | 1695455 | YP_173237.1 | A0A140H1H0 |
| SSCGLFKDCARNPIDL | 1 | 0 | 0 | 1695463 | NP_073549.1 | A0A0P0LKV0 |
| SSCSLSEQLDFIDTKR | 1 | 0 | 0 | 1695466 | QIQ54046.1 | P0C6U8 |
| SSCVECDAKFKNSVAS | 1 | 0 | 0 | 1695467 | NP_073549.1 | A0A0P0LKV0 |
| SSDEAIPTRFPPGTVL | 1 | 0 | 0 | 1695472 | YP_009555245.1 | P33469 |
| SSDGQPGGDAISLSFN | 1 | 0 | 0 | 1695475 | YP_009047202.1 | K9N7C7 |
| SSEGFLLGINYLGTIK | 1 | 0 | 0 | 1695497 | YP_009047202.1 | K9N7C7 |
| SSEIIGYKAIDGGVTR | 1 | 0 | 0 | 1695499 | YP_009724389.1 | P0DTD1 |
| SSEKPTKEEPKKVKPC | 1 | 0 | 0 | 1695500 | QIQ54046.1 | P0C6U8 |
| SSELKQLEAKTASLFQ | 1 | 0 | 0 | 1695501 | YP_003767.1 | A0A0P0G321 |
| SSESGAKETKEINIIK | 1 | 0 | 0 | 1695503 | YP_009555238.1 | P0C6X6 |
| SSETKEQKHEMQKPRW | 1 | 0 | 0 | 1695504 | NP_073556.1 | A0A0N7J131 |
| SSEVKAQSSGVIPENP | 1 | 0 | 0 | 1695506 | YP_173236.1 | A0A140H1G9 |
| SSEWAYCVDLHNKINL | 1 | 0 | 0 | 1695508 | YP_003766.2 | H9EJ42 |
| SSEYKGPITDIFYKEN | 1 | 0 | 0 | 1695509 | QHR63299.1 | P0C6W2 |
| SSEYKGPITDVFYKEN | 1 | 0 | 0 | 1695510 | YP_009724389.1 | P0DTD1 |
| SSFDGVMYGGFEDQPN | 1 | 0 | 0 | 1695513 | NP_073549.1 | A0A0P0LKV0 |
| SSFEAKPSGSVVEQAE | 1 | 0 | 0 | 1695514 | YP_009047204.1 | K9N5Q8 |
| SSFEGYDFKYIENGQL | 1 | 0 | 0 | 1695515 | NP_073549.1 | A0A0P0LKV0 |
| SSFGDYNLSSVIPSLP | 1 | 0 | 0 | 1695520 | NP_073551.1 | A0A0P0K6L9 |
| SSFLEMKSEKQVEQKI | 1 | 0 | 0 | 1695524 | YP_009724389.1 | P0DTD1 |
| SSFLEMKSEKQVEQKT | 1 | 0 | 0 | 1695525 | QHR63299.1 | P0C6W2 |
| SSFNFDQKELLAYYNM | 1 | 0 | 0 | 1695527 | YP_009555238.1 | P0C6X6 |

| | | | | | | |
|---|---|---|---|---|---|---|
| SSFVGMPSFVAYETAR | 1 | 0 | 0 | 1695533 | NP_073549.1 | A0A0P0LKV0 |
| SSGFLQSSNYKIDTTS | 1 | 0 | 0 | 1695540 | YP_173238.1 | A0A140H1H1 |
| SSGNRSGNGILKWADQ | 1 | 0 | 0 | 1695545 | YP_009555245.1 | P33469 |
| SSGTSTPKKPNKPLSQ | 1 | 0 | 0 | 1695547 | YP_003771.1 | H9EJ69 |
| SSGTYEGNSPFHPLAD | 1 | 0 | 0 | 1695548 | YP_009724395.1 | P0DTC7 |
| SSGVLDTTIPFGPSSQ | 1 | 0 | 0 | 1695551 | YP_003767.1 | A0A0P0G321 |
| SSGVTRDLMRELNGGA | 1 | 0 | 0 | 1695553 | QHR63299.1 | P0C6W2 |
| SSGVTRELMRELNGGA | 1 | 0 | 0 | 1695554 | YP_009724389.1 | P0DTD1 |
| SSGVVNPAMDPIYDEP | 1 | 0 | 0 | 1695555 | QIQ54049.1 | P59632 |
| SSGVVNPAMEPIYDEP | 1 | 0 | 0 | 1695556 | QHR63301.1 | Q3LZX0 |
| SSHYWSPQGNLGGNCA | 1 | 0 | 0 | 1695561 | YP_173236.1 | A0A140H1G9 |
| SSIAYSNNTIAIPTNF | 1 | 0 | 0 | 1695563 | NP_828851.1 | P59594 |
| SSIKEMQSQSSHVAQN | 1 | 0 | 0 | 1695570 | YP_003771.1 | H9EJ69 |
| SSIKGDKGVEKLVNWY | 1 | 0 | 0 | 1695571 | YP_003766.2 | H9EJ42 |
| SSIQAIYDRLDTIQAD | 1 | 0 | 0 | 1695578 | NP_073551.1 | A0A0P0K6L9 |
| SSIQANAYYKYDNGNF | 1 | 0 | 0 | 1695579 | NP_073549.1 | A0A0P0LKV0 |
| SSIRGCCESTKLPYYD | 1 | 0 | 0 | 1695581 | NP_073551.1 | A0A0P0K6L9 |
| SSITIDKFAIPNGRKV | 1 | 0 | 0 | 1695583 | YP_009555241.1 | P36334 |
| SSIVITSGDGTPSPIT | 1 | 0 | 0 | 1695584 | QIQ54049.1 | P59632 |
| SSIYKLHPAFNVSDAY | 1 | 0 | 0 | 1695587 | YP_003766.2 | H9EJ42 |
| SSKLWAQCVQLHNDIL | 1 | 0 | 0 | 1695592 | YP_009724389.1 | P0DTD1 |
| SSKPLYVPEHCQNYSG | 1 | 0 | 0 | 1695593 | YP_009047205.1 | K9N796 |
| SSKSLGKQLGVVSDGV | 1 | 0 | 0 | 1695599 | YP_003766.2 | H9EJ42 |
| SSKTSEEHFVETVSLA | 1 | 0 | 0 | 1695602 | NP_828849.2 | P0C6X7 |
| SSKVVLSDALDKLYKV | 1 | 0 | 0 | 1695606 | NP_073549.1 | A0A0P0LKV0 |
| SSLCDEFTLAEVVKQM | 1 | 0 | 0 | 1695610 | YP_003766.2 | H9EJ42 |
| SSLCFKVYFKGQTTHE | 1 | 0 | 0 | 1695611 | YP_173236.1 | A0A140H1G9 |
| SSLCTMFKRGDGTPHP | 1 | 0 | 0 | 1695613 | YP_173236.1 | A0A140H1G9 |
| SSLFVEDCKLPLGQSL | 1 | 0 | 0 | 1695615 | YP_009047204.1 | K9N5Q8 |
| SSLNDEFSINEVVKQM | 1 | 0 | 0 | 1695621 | NP_073549.1 | A0A0P0LKV0 |
| SSLPSYAAYATAQEAY | 1 | 0 | 0 | 1695623 | NP_828849.2 | P0C6X7 |
| SSLQEILSRLDALEAQ | 1 | 0 | 0 | 1695624 | YP_173238.1 | A0A140H1H1 |
| SSLSRNSSRSSSQGSR | 1 | 0 | 0 | 1695626 | YP_009047211.1 | K9N4V7 |
| SSLSSYEYCQPPKSVE | 1 | 0 | 0 | 1695627 | YP_173236.1 | A0A140H1G9 |
| SSLTCNVSFVGELKAA | 1 | 0 | 0 | 1695628 | NP_073549.1 | A0A0P0LKV0 |
| SSNDNCFANKHVGFDS | 1 | 0 | 0 | 1695646 | QIQ54046.1 | P0C6U8 |
| SSNEEQAVLKFLDGLD | 1 | 0 | 0 | 1695647 | YP_003766.2 | H9EJ42 |
| SSNGPLKVGAGVMLEC | 1 | 0 | 0 | 1695649 | YP_003766.2 | H9EJ42 |
| SSNINCVLSVNSSNCN | 1 | 0 | 0 | 1695651 | NP_073549.1 | A0A0P0LKV0 |
| SSNINRLLSVPSDSCN | 1 | 0 | 0 | 1695652 | YP_003766.2 | H9EJ42 |
| SSNLNTNLHFDVDNIN | 1 | 0 | 0 | 1695653 | YP_173238.1 | A0A140H1H1 |
| SSNMVETVVGQLEQTN | 1 | 0 | 0 | 1695655 | YP_009047202.1 | K9N7C7 |
| SSNQGNKQIVHSYPIL | 1 | 0 | 0 | 1695656 | YP_009047207.1 | K9N643 |
| SSNQSPQGNLGSNGKP | 1 | 0 | 0 | 1695658 | YP_009555238.1 | P0C6X6 |
| SSNTDNETLQYWVTPL | 1 | 0 | 0 | 1695659 | YP_173238.1 | A0A140H1H1 |
| SSNVANYQKVGMQKYS | 1 | 0 | 0 | 1695660 | YP_009724389.1 | P0DTD1 |
| SSNVANYQKVGMQXYS | 1 | 0 | 0 | 1695661 | QIQ54047.1 | P0C6X7 |
| SSNYKIDTTSSSCQLY | 1 | 0 | 0 | 1695663 | YP_173238.1 | A0A140H1H1 |
| SSPARLEIVKLGAEKK | 1 | 0 | 0 | 1695666 | YP_009555239.1 | Q80872 |
| SSPMSNMTENERLLHF | 1 | 0 | 0 | 1695674 | NP_073555.1 | A0A0P0KXN3 |
| SSQGSRSGNSTRGTSP | 1 | 0 | 0 | 1695683 | YP_009047211.1 | K9N4V7 |
| SSQLPLDVYDLTMPSQ | 1 | 0 | 0 | 1695685 | YP_009555238.1 | P0C6X6 |
| SSQPYYCFINSTINTT | 1 | 0 | 0 | 1695686 | YP_003767.1 | A0A0P0G321 |
| SSQSSPIIPGFGGDFN | 1 | 0 | 0 | 1695687 | YP_009047204.1 | K9N5Q8 |
| SSQTSAKSLARSQSSE | 1 | 0 | 0 | 1695688 | NP_073556.1 | A0A0N7J131 |
| SSRASSAGSRSRANSG | 1 | 0 | 0 | 1695691 | YP_009555245.1 | P33469 |
| SSRASSGNRSGNGILK | 1 | 0 | 0 | 1695692 | YP_009555245.1 | P33469 |
| SSRASSLSRNSSRSSS | 1 | 0 | 0 | 1695693 | YP_009047211.1 | K9N4V7 |
| SSRASSRSSTRNNSRD | 1 | 0 | 0 | 1695694 | YP_003771.1 | H9EJ69 |
| SSRGNSPARMASGGGE | 1 | 0 | 0 | 1695696 | NP_828858.1 | P59595 |
| SSRGTSPARIAGNGGD | 1 | 0 | 0 | 1695697 | QIQ54056.1 | P59595 |
| SSRMYCSFYPPDEDYE | 1 | 0 | 0 | 1695704 | QIQ54046.1 | P0C6U8 |
| SSRMYCSFYPPDEEEE | 1 | 0 | 0 | 1695705 | NP_828849.2 | P0C6X7 |
| SSRNESWHFDKSEPLC | 1 | 0 | 0 | 1695706 | YP_173238.1 | A0A140H1H1 |
| SSRNSTPGSSRGTSPA | 1 | 0 | 0 | 1695708 | YP_009724397.2 | P0DTC9 |
| SSRRKRRSISASYRFV | 1 | 0 | 0 | 1695709 | YP_173238.1 | A0A140H1H1 |
| SSRSFFEDLLFDKVKL | 1 | 0 | 0 | 1695711 | YP_173238.1 | A0A140H1H1 |
| SSRSRGNSRNSTPGSS | 1 | 0 | 0 | 1695712 | NP_828858.1 | P59595 |
| SSRSRNSSRNSTPGSS | 1 | 0 | 0 | 1695714 | YP_009724397.2 | P0DTC9 |
| SSRSSSQGSRSGNSTR | 1 | 0 | 0 | 1695715 | YP_009047211.1 | K9N4V7 |
| SSRSSTRNNSRDSSRS | 1 | 0 | 0 | 1695719 | YP_003771.1 | H9EJ69 |
| SSSAVNIQQIYLISKF | 1 | 0 | 0 | 1695726 | YP_173236.1 | A0A140H1G9 |
| SSSAVNMQQIYLINKF | 1 | 0 | 0 | 1695727 | YP_009555238.1 | P0C6X6 |
| SSSEVDMICSTTAQKL | 1 | 0 | 0 | 1695737 | YP_009555238.1 | P0C6X6 |
| SSSGTSTPKKPNKPLS | 1 | 0 | 0 | 1695743 | YP_003771.1 | H9EJ69 |
| SSSGVLDTTIPFGPSS | 1 | 0 | 0 | 1695744 | YP_003767.1 | A0A0P0G321 |
| SSSKTSEEHFVETVSL | 1 | 0 | 0 | 1695749 | NP_828849.2 | P0C6X7 |
| SSSLQEILSRLDALEA | 1 | 0 | 0 | 1695751 | YP_173238.1 | A0A140H1H1 |
| SSSQGSRSGNSTRGTS | 1 | 0 | 0 | 1695755 | YP_009047211.1 | K9N4V7 |
| SSSRASSGNRSGNGIL | 1 | 0 | 0 | 1695757 | YP_009555245.1 | P33469 |
| SSSRASSLSRNSSRSS | 1 | 0 | 0 | 1695758 | YP_009047211.1 | K9N4V7 |
| SSSRRKRRSISASYRF | 1 | 0 | 0 | 1695759 | YP_173238.1 | A0A140H1H1 |
| SSSRSFFEDLLFDKVK | 1 | 0 | 0 | 1695760 | YP_173238.1 | A0A140H1H1 |
| SSSRSRGNSRNSTPGS | 1 | 0 | 0 | 1695761 | NP_828858.1 | P59595 |
| SSSRSRNSSRNSTPGS | 1 | 0 | 0 | 1695762 | YP_009724397.2 | P0DTC9 |
| SSSSRRKRRSISASYR | 1 | 0 | 0 | 1695763 | YP_173238.1 | A0A140H1H1 |
| SSSYDALLHRNSLDPF | 1 | 0 | 0 | 1695765 | YP_003766.2 | H9EJ42 |
| SSTFNVPMDKLKTLIA | 1 | 0 | 0 | 1695768 | QIQ54046.1 | P0C6U8 |
| SSTFNVPMEKLKTLVA | 1 | 0 | 0 | 1695769 | YP_009724389.1 | P0DTD1 |
| SSTFVEEEHVNRMDTF | 1 | 0 | 0 | 1695770 | NP_073549.1 | A0A0P0LKV0 |
| SSTGFVYFNATGSDVN | 1 | 0 | 0 | 1695771 | YP_003767.1 | A0A0P0G321 |

| | | | | | | |
|---|---|---|---|---|---|---|
| SSTIHNYFITETNTAA | 1 | 0 | 0 | 1695772 | YP_009047202.1 | K9N7C7 |
| SSTIRLQAGVATEYAA | 1 | 0 | 0 | 1695773 | YP_173236.1 | A0A140H1G9 |
| SSTRGVYYPDKVFRSS | 1 | 0 | 0 | 1695778 | QHR63300.2 | Q3LZX1 |
| SSTRNNSRDSSRSTSR | 1 | 0 | 0 | 1695779 | YP_003771.1 | H9EJ69 |
| SSTVRLQAGTATEYAS | 1 | 0 | 0 | 1695781 | YP_009555238.1 | P0C6X6 |
| SSVAVQSKDLNFLNRV | 1 | 0 | 0 | 1695784 | YP_173236.1 | A0A140H1G9 |
| SSVDLKHFFFTQDGNA | 1 | 0 | 0 | 1695785 | YP_009555238.1 | P0C6X6 |
| SSVDTILNMARNGVVP | 1 | 0 | 0 | 1695786 | NP_073549.1 | A0A0P0LKV0 |
| SSVETETYFKVNGLWS | 1 | 0 | 0 | 1695789 | YP_173236.1 | A0A140H1G9 |
| SSVETVLNLARDGVVP | 1 | 0 | 0 | 1695790 | YP_003766.2 | H9EJ42 |
| SSVKSYDDLVYSGVVG | 1 | 0 | 0 | 1695794 | YP_173236.1 | A0A140H1G9 |
| SSVLPQRNIRSSRIAG | 1 | 0 | 0 | 1695800 | YP_003767.1 | A0A0P0G321 |
| SSVNDAITQTAEAIHT | 1 | 0 | 0 | 1695802 | YP_003767.1 | A0A0P0G321 |
| SSVPLLEVYVHLRNWN | 1 | 0 | 0 | 1695804 | YP_003770.1 | U3M6Q8 |
| SSVQQQEEQEEDWLET | 1 | 0 | 0 | 1695806 | QIQ54046.1 | P0C6U8 |
| SSVWPLYPYGRCPTAA | 1 | 0 | 0 | 1695809 | YP_009555240.1 | P30215 |
| SSYAKPEEKLSAYDLA | 1 | 0 | 0 | 1695817 | NP_073549.1 | A0A0P0LKV0 |
| SSYDALLHRNSLDPFC | 1 | 0 | 0 | 1695819 | YP_003766.2 | H9EJ42 |
| SSYIVDSVAVKNGALH | 1 | 0 | 0 | 1695822 | NP_828849.2 | P0C6X7 |
| SSYSLFDMSKFPLKLR | 1 | 0 | 0 | 1695825 | YP_009724389.1 | P0DTD1 |
| SSYVVDSVAVKNGSLH | 1 | 0 | 0 | 1695827 | QIQ54046.1 | P0C6U8 |
| SSYYYPEPISDKNVVF | 1 | 0 | 0 | 1695828 | YP_173238.1 | A0A140H1H1 |
| STALGKLQDVVNQNAQ | 1 | 0 | 0 | 1695836 | NP_828851.1 | P59594 |
| STARDGVRRGDNFHSV | 1 | 0 | 0 | 1695839 | YP_009047202.1 | K9N7C7 |
| STAVLTIANNYSKLFD | 1 | 0 | 0 | 1695842 | NP_073549.1 | A0A0P0LKV0 |
| STCDGLKFTEESFYKN | 1 | 0 | 0 | 1695847 | YP_173236.1 | A0A140H1G9 |
| STDDCFANKHAGFDAW | 1 | 0 | 0 | 1695853 | NP_828849.2 | P0C6X7 |
| STDGNKIADKYVRNLQ | 1 | 0 | 0 | 1695858 | YP_009724389.1 | P0DTD1 |
| STDGNKIGDKYIRNLQ | 1 | 0 | 0 | 1695859 | QIQ54047.1 | P0C6X7 |
| STDNNQNGDRSGARPK | 1 | 0 | 0 | 1695862 | QIQ54056.1 | P59595 |
| STDTVNHDSFNLYGYS | 1 | 0 | 0 | 1695870 | YP_173236.1 | A0A140H1G9 |
| STEDFNVWAMANGFSQ | 1 | 0 | 0 | 1695881 | YP_173236.1 | A0A140H1G9 |
| STEKLSRLSMIKGPQR | 1 | 0 | 0 | 1695886 | YP_173236.1 | A0A140H1G9 |
| STFITPEVSRELGNIT | 1 | 0 | 0 | 1695904 | NP_073549.1 | A0A0P0LKV0 |
| STFKCYGVSATKLNDL | 1 | 0 | 0 | 1695905 | NP_828851.1 | P59594 |
| STFNKDIYCGNSMVCK | 1 | 0 | 0 | 1695910 | YP_003766.2 | H9EJ42 |
| STFVTEYYEFLNKHFS | 1 | 0 | 0 | 1695920 | YP_009555238.1 | P0C6X6 |
| STFYPQLQSAEWKCGY | 1 | 0 | 0 | 1695921 | YP_003766.2 | H9EJ42 |
| STGFVYFNATGSDVNC | 1 | 0 | 0 | 1695926 | YP_003767.1 | A0A0P0G321 |
| STGKTVLGEYVFDKSE | 1 | 0 | 0 | 1695931 | YP_173236.1 | A0A140H1G9 |
| STGLTSQTVKPGHFNK | 1 | 0 | 0 | 1695934 | NP_073549.1 | A0A0P0LKV0 |
| STGSNQNGERSGARPK | 1 | 0 | 0 | 1695938 | QHR63308.1 | Q3LZX4 |
| STGTQSVSVDLESTST | 1 | 0 | 0 | 1695940 | YP_009047205.1 | K9N796 |
| STIGVCSMTDIAKKPT | 1 | 0 | 0 | 1695953 | YP_009724389.1 | P0DTD1 |
| STIGVCTMTDIAKKPT | 1 | 0 | 0 | 1695954 | NP_828849.2 | P0C6X7 |
| STIGVCTMTDIAKKST | 1 | 0 | 0 | 1695955 | QIQ54047.1 | P0C6X7 |
| STIKTIQPRVEKKKLE | 1 | 0 | 0 | 1695959 | QIQ54046.1 | P0C6U8 |
| STILEDEFTPFDVVRQ | 1 | 0 | 0 | 1695960 | NP_828849.2 | P0C6X7 |
| STINTTHVSTFVGILP | 1 | 0 | 0 | 1695965 | YP_003767.1 | A0A0P0G321 |
| STIRLQAGVATEYAAN | 1 | 0 | 0 | 1695967 | YP_173236.1 | A0A140H1G9 |
| STIVTGLLPTHWFCAN | 1 | 0 | 0 | 1695973 | YP_003767.1 | A0A0P0G321 |
| STIYKLHPSFNVSDAY | 1 | 0 | 0 | 1695974 | NP_073549.1 | A0A0P0LKV0 |
| STKALAKQDAAQRIAW | 1 | 0 | 0 | 1695975 | YP_009047206.1 | K9N4V0 |
| STKLKDGVNFNVDDIN | 1 | 0 | 0 | 1695980 | YP_009555241.1 | P36334 |
| STKPVDTSNSFEVLAV | 1 | 0 | 0 | 1695983 | NP_828849.2 | P0C6X7 |
| STKPVETSNIFDVLQS | 1 | 0 | 0 | 1695984 | QIQ54046.1 | P0C6U8 |
| STKPVETSNSFDVLKS | 1 | 0 | 0 | 1695985 | YP_009724389.1 | P0DTD1 |
| STLENKSAELNYTVQK | 1 | 0 | 0 | 1695988 | NP_073551.1 | A0A0P0K6L9 |
| STLQPNAYYKYDAKNY | 1 | 0 | 0 | 1696003 | YP_003766.2 | H9EJ42 |
| STNDLYVALRGRIDKD | 1 | 0 | 0 | 1696031 | NP_073553.1 | A0A0P0KWE8 |
| STNLFEGDKFVGTFES | 1 | 0 | 0 | 1696035 | NP_073549.1 | A0A0P0LKV0 |
| STNLPPPLLGNSTGID | 1 | 0 | 0 | 1696037 | YP_009047204.1 | K9N5Q8 |
| STPAQNAGYWRRQDRK | 1 | 0 | 0 | 1696045 | YP_009047211.1 | K9N4V7 |
| STPGSSRGNSPARMAS | 1 | 0 | 0 | 1696049 | NP_828858.1 | P59595 |
| STPGSSRGTSPARIAG | 1 | 0 | 0 | 1696050 | QIQ54056.1 | P59595 |
| STPGSSRGTSPARMAG | 1 | 0 | 0 | 1696051 | YP_009724397.2 | P0DTC9 |
| STPKKPNKPLSQPRAD | 1 | 0 | 0 | 1696055 | YP_003771.1 | H9EJ69 |
| STPKLKEDYQIGGYSE | 1 | 0 | 0 | 1696056 | NP_828852.2 | P59632 |
| STPSDVSSRDPTTQEA | 1 | 0 | 0 | 1696059 | YP_173242.1 | Q19U29 |
| STQDGDNKLQGLLEVS | 1 | 0 | 0 | 1696061 | YP_009555241.1 | P36334 |
| STQFAFACPDGVKHVY | 1 | 0 | 0 | 1696065 | YP_009724395.1 | P0DTC7 |
| STQKGSGMDTALLRNN | 1 | 0 | 0 | 1696067 | YP_009555244.1 | Q01455 |
| STQLGIEFLKRGDKSV | 1 | 0 | 0 | 1696069 | YP_009724389.1 | P0DTD1 |
| STRGTSPGPSGIGAVG | 1 | 0 | 0 | 1696077 | YP_009047211.1 | K9N4V7 |
| STRSMLKRRDSTYGPL | 1 | 0 | 0 | 1696082 | YP_009047204.1 | K9N5Q8 |
| STRVNVDGFNEWAMAN | 1 | 0 | 0 | 1696083 | YP_003766.2 | H9EJ42 |
| STSAFVETVKGLDYKA | 1 | 0 | 0 | 1696085 | YP_009724389.1 | P0DTD1 |
| STSAFVETVKGLDYKS | 1 | 0 | 0 | 1696086 | QIQ54046.1 | P0C6U8 |
| STSAFVETVKGLDYKT | 1 | 0 | 0 | 1696087 | QHR63299.1 | P0C6W2 |
| STSAVKLQNNEIKPSG | 1 | 0 | 0 | 1696090 | YP_009047202.1 | K9N7C7 |
| STSDLSVAFDKLAQLL | 1 | 0 | 0 | 1696092 | YP_173236.1 | A0A140H1G9 |
| STSDWSDIRDYKLAND | 1 | 0 | 0 | 1696093 | NP_073549.1 | A0A0P0LKV0 |
| STSFLQSGIVKMVNPT | 1 | 0 | 0 | 1696094 | YP_009555238.1 | P0C6X6 |
| STSGNLLGFKDVTKGT | 1 | 0 | 0 | 1696097 | NP_073551.1 | A0A0P0K6L9 |
| STSIPNFGSLTQINTT | 1 | 0 | 0 | 1696102 | YP_009047204.1 | K9N5Q8 |
| STSKCWVEEDLTKGPH | 1 | 0 | 0 | 1696103 | YP_003766.2 | H9EJ42 |
| STSLVEXVDVTFLKKD | 1 | 0 | 0 | 1696104 | QIQ54046.1 | P0C6U8 |
| STSPDPKFVDKYYAFL | 1 | 0 | 0 | 1696105 | YP_009047202.1 | K9N7C7 |
| STSRQQSRTRSDSNQS | 1 | 0 | 0 | 1696108 | YP_003771.1 | H9EJ69 |
| STSRTSSRASSAGSRS | 1 | 0 | 0 | 1696109 | YP_009555245.1 | P33469 |
| STSSDTYACWNHSVGF | 1 | 0 | 0 | 1696111 | NP_828849.2 | P0C6X7 |

| | | | | | | |
|---|---|---|---|---|---|---|
| STSTHDGPTEHVTSVN | 1 | 0 | 0 | 1696112 | YP_009047205.1 | K9N796 |
| STTLVKDKCVNFNFNG | 1 | 0 | 0 | 1696122 | QIQ54048.1 | P59594 |
| STTNYVGVYDINHGDA | 1 | 0 | 0 | 1696126 | YP_009555240.1 | P30215 |
| STTPCEVHGNGSMIED | 1 | 0 | 0 | 1696127 | YP_009047202.1 | K9N7C7 |
| STTVSVTVSSNMVETV | 1 | 0 | 0 | 1696131 | YP_009047202.1 | K9N7C7 |
| STTWNGGAYSLFDMTK | 1 | 0 | 0 | 1696132 | YP_173236.1 | A0A140H1G9 |
| STVLNGVWLGDTVTCP | 1 | 0 | 0 | 1696144 | YP_003766.2 | H9EJ42 |
| STVVIKNLTPIKLNFG | 1 | 0 | 0 | 1696162 | YP_003766.2 | H9EJ42 |
| STVWEDGDYYRKQLSP | 1 | 0 | 0 | 1696163 | YP_009047204.1 | K9N5Q8 |
| STVWNGGAYSLFDMAK | 1 | 0 | 0 | 1696164 | YP_009555238.1 | P0C6X6 |
| STWNKRFGFIEDSVFK | 1 | 0 | 0 | 1696165 | YP_009555241.1 | P36334 |
| STYGVCKYTDLKFIEN | 1 | 0 | 0 | 1696170 | YP_173236.1 | A0A140H1G9 |
| STYVDLKLLNRFENYI | 1 | 0 | 0 | 1696176 | YP_003767.1 | A0A0P0G321 |
| SVAAITSGVKFQTVKP | 1 | 0 | 0 | 1696179 | YP_009555238.1 | P0C6X6 |
| SVAALSTGLTSQTVKP | 1 | 0 | 0 | 1696181 | NP_073549.1 | A0A0P0LKV0 |
| SVACEHISSTMSQYSR | 1 | 0 | 0 | 1696187 | YP_009047204.1 | K9N5Q8 |
| SVADYHNHSNIETRLR | 1 | 0 | 0 | 1696189 | NP_828849.2 | P0C6X7 |
| SVANLSAPTLVPQENY | 1 | 0 | 0 | 1696201 | YP_009555238.1 | P0C6X6 |
| SVAVIGDEQIDGKFVT | 1 | 0 | 0 | 1696210 | YP_173236.1 | A0A140H1G9 |
| SVAYESLRPDTRYVLM | 1 | 0 | 0 | 1696216 | YP_009724389.1 | P0DTD1 |
| SVCPKLEFANDTKIAS | 1 | 0 | 0 | 1696233 | YP_009047204.1 | K9N5Q8 |
| SVDAFNQFSSDFQHKL | 1 | 0 | 0 | 1696240 | YP_009555238.1 | P0C6X6 |
| SVDIKFKTEGLCVDIP | 1 | 0 | 0 | 1696251 | NP_828849.2 | P0C6X7 |
| SVDISYLNRARGSSAA | 1 | 0 | 0 | 1696252 | YP_003766.2 | H9EJ42 |
| SVDLKHFFFTQDGNAA | 1 | 0 | 0 | 1696255 | YP_009555238.1 | P0C6X6 |
| SVDPATTYLEAVKHGA | 1 | 0 | 0 | 1696257 | YP_003766.2 | H9EJ42 |
| SVDPKKTYLDFIQQGG | 1 | 0 | 0 | 1696258 | YP_009555238.1 | P0C6X6 |
| SVDPKKTYLDYIQQGG | 1 | 0 | 0 | 1696259 | YP_173236.1 | A0A140H1G9 |
| SVDSVSEESQGNVVTS | 1 | 0 | 0 | 1696261 | YP_173236.1 | A0A140H1G9 |
| SVECYSILAAKTGVSV | 1 | 0 | 0 | 1696265 | YP_003766.2 | H9EJ42 |
| SVEDFNVWALSNGFSQ | 1 | 0 | 0 | 1696267 | YP_009555238.1 | P0C6X6 |
| SVEDVCFANKFEQFDQ | 1 | 0 | 0 | 1696268 | YP_009555238.1 | P0C6X6 |
| SVEEALSEAREHLKNG | 1 | 0 | 0 | 1696269 | NP_828849.2 | P0C6X7 |
| SVEIEAGEVKPFAVYK | 1 | 0 | 0 | 1696274 | YP_003766.2 | H9EJ42 |
| SVEQRQQVDKKIKACV | 1 | 0 | 0 | 1696284 | QHR63299.1 | P0C6W2 |
| SVETETYFKVNGLWSP | 1 | 0 | 0 | 1696286 | YP_173236.1 | A0A140H1G9 |
| SVETVLNLARDGVVPL | 1 | 0 | 0 | 1696288 | YP_003766.2 | H9EJ42 |
| SVEYPIIGDELRVNSA | 1 | 0 | 0 | 1696289 | NP_828849.2 | P0C6X7 |
| SVFNICQAVSANVCAL | 1 | 0 | 0 | 1696296 | YP_009555238.1 | P0C6X6 |
| SVFNLEGVNGGSLYVN | 1 | 0 | 0 | 1696301 | YP_003766.2 | H9EJ42 |
| SVGYVLTGDSVKFVYM | 1 | 0 | 0 | 1696328 | YP_173236.1 | A0A140H1G9 |
| SVHQGAHVASNDAIMT | 1 | 0 | 0 | 1696331 | YP_009047202.1 | K9N7C7 |
| SVIDLKLDDFVMILKS | 1 | 0 | 0 | 1696333 | YP_009047202.1 | K9N7C7 |
| SVIDLLLDDFVEIIKS | 1 | 0 | 0 | 1696334 | YP_009724389.1 | P0DTD1 |
| SVIELATVNLRLVDCA | 1 | 0 | 0 | 1696335 | YP_003766.2 | H9EJ42 |
| SVIKVAPKPYRVDGKF | 1 | 0 | 0 | 1696340 | NP_073549.1 | A0A0P0LKV0 |
| SVILSTCDGLKFTEES | 1 | 0 | 0 | 1696341 | YP_173236.1 | A0A140H1G9 |
| SVINRLRCDQLSFDVP | 1 | 0 | 0 | 1696350 | NP_073551.1 | A0A0P0K6L9 |
| SVIPATSASKLTIVSP | 1 | 0 | 0 | 1696352 | YP_003766.2 | H9EJ42 |
| SVITPGTNASNQVAVL | 1 | 0 | 0 | 1696358 | QHR63300.2 | Q3LZX1 |
| SVIVEDDVIENVKSSL | 1 | 0 | 0 | 1696361 | YP_173236.1 | A0A140H1G9 |
| SVIVNDDTSETKYVKS | 1 | 0 | 0 | 1696362 | YP_009555238.1 | P0C6X6 |
| SVIYDKETKTHATLFG | 1 | 0 | 0 | 1696365 | YP_009047204.1 | K9N5Q8 |
| SVKGLQPSVGPKQASL | 1 | 0 | 0 | 1696370 | YP_009724389.1 | P0DTD1 |
| SVKGLTPSKGPAQASV | 1 | 0 | 0 | 1696371 | NP_828849.2 | P0C6X7 |
| SVKQDALTGGNYGYLY | 1 | 0 | 0 | 1696377 | QIQ54048.1 | P59594 |
| SVKRDGVQVGYCVHGI | 1 | 0 | 0 | 1696379 | NP_073549.1 | A0A0P0LKV0 |
| SVKSFEIDKGIYQTSN | 1 | 0 | 0 | 1696381 | NP_828851.1 | P59594 |
| SVKSSQSSPIIPGFGG | 1 | 0 | 0 | 1696382 | YP_009047204.1 | K9N5Q8 |
| SVKTQFNYFKKVDGII | 1 | 0 | 0 | 1696389 | NP_828849.2 | P0C6X7 |
| SVLDITDAAVKAAESK | 1 | 0 | 0 | 1696393 | NP_073549.1 | A0A0P0LKV0 |
| SVLEMAKEVKTSSKVV | 1 | 0 | 0 | 1696396 | NP_073549.1 | A0A0P0LKV0 |
| SVLETFQNNVVNYQHI | 1 | 0 | 0 | 1696397 | YP_009555238.1 | P0C6X6 |
| SVLLIERFVSLAIDAY | 1 | 0 | 0 | 1696405 | YP_009555238.1 | P0C6X6 |
| SVLLSMQGAVDINKLC | 1 | 0 | 0 | 1696407 | YP_009724389.1 | P0DTD1 |
| SVLLSMQGAVDINRLC | 1 | 0 | 0 | 1696408 | NP_828849.2 | P0C6X7 |
| SVLPQRNIRSSRIAGR | 1 | 0 | 0 | 1696412 | YP_003767.1 | A0A0P0G321 |
| SVLSVDSVSEESQGNV | 1 | 0 | 0 | 1696421 | YP_173236.1 | A0A140H1G9 |
| SVLTTFIDAARGPAGV | 1 | 0 | 0 | 1696424 | YP_009047202.1 | K9N7C7 |
| SVLTTYYLDDVKKIEY | 1 | 0 | 0 | 1696426 | YP_009555238.1 | P0C6X6 |
| SVMESQISTKEVKLKG | 1 | 0 | 0 | 1696436 | YP_173236.1 | A0A140H1G9 |
| SVMLTNDNTSRYWEPE | 1 | 0 | 0 | 1696440 | YP_009724389.1 | P0DTD1 |
| SVNDAITQTAEAIHTV | 1 | 0 | 0 | 1696446 | YP_003767.1 | A0A0P0G321 |
| SVNDFTCSQISPAAIA | 1 | 0 | 0 | 1696447 | YP_009047204.1 | K9N5Q8 |
| SVNDLCFANKFFQFDQ | 1 | 0 | 0 | 1696448 | YP_173236.1 | A0A140H1G9 |
| SVNDSLEPVGGLYEIQ | 1 | 0 | 0 | 1696449 | YP_009555241.1 | P36334 |
| SVPEKFLNAFDVFVTA | 1 | 0 | 0 | 1696475 | NP_073549.1 | A0A0P0LKV0 |
| SVPLLEVYVHLRNWNF | 1 | 0 | 0 | 1696479 | YP_003770.1 | U3M6Q8 |
| SVPLVFQNNVANYQHI | 1 | 0 | 0 | 1696480 | YP_173236.1 | A0A140H1G9 |
| SVPVSVIYDKETKTHA | 1 | 0 | 0 | 1696487 | YP_009047204.1 | K9N5Q8 |
| SVPYESLRPDTRYVLM | 1 | 0 | 0 | 1696491 | QIQ54046.1 | P0C6U8 |
| SVQKHIGAPANLRVVG | 1 | 0 | 0 | 1696495 | YP_009047202.1 | K9N7C7 |
| SVQQAQQEATLPDIAE | 1 | 0 | 0 | 1696498 | NP_073549.1 | A0A0P0LKV0 |
| SVQQQEEQEEDWLETE | 1 | 0 | 0 | 1696501 | QIQ54046.1 | P0C6U8 |
| SVQVDNGSSINRRQLD | 1 | 0 | 0 | 1696504 | NP_073549.1 | A0A0P0LKV0 |
| SVQYAHKHDIQITNES | 1 | 0 | 0 | 1696506 | YP_009047202.1 | K9N7C7 |
| SVRDPKTSEILDISPC | 1 | 0 | 0 | 1696509 | NP_828851.1 | P59594 |
| SVRGRAIIVSVEQLEP | 1 | 0 | 0 | 1696513 | NP_073549.1 | A0A0P0LKV0 |
| SVRHHPELKLFRNLNI | 1 | 0 | 0 | 1696514 | YP_009555238.1 | P0C6X6 |
| SVRVVTTFDAEYCRHG | 1 | 0 | 0 | 1696521 | NP_828849.2 | P0C6X7 |
| SVRVVTTFDSEYCRHG | 1 | 0 | 0 | 1696522 | YP_009724389.1 | P0DTD1 |

| | | | | | | |
|---|---|---|---|---|---|---|
| SVSAIVTANLSIPSNW | 1 | 0 | 0 | 1696523 | NP_073551.1 | A0A0P0K6L9 |
| SVSNDYCCFKHALGCD | 1 | 0 | 0 | 1696528 | NP_073549.1 | A0A0P0LKV0 |
| SVSRFNPSTWNKRFGF | 1 | 0 | 0 | 1696535 | YP_009555241.1 | P36334 |
| SVSSAGPISQFNYKQS | 1 | 0 | 0 | 1696536 | YP_009047204.1 | K9N5Q8 |
| SVSSCALRMGSGFCVD | 1 | 0 | 0 | 1696537 | YP_173238.1 | A0A140H1H1 |
| SVSSFEAKPSGSVVEQ | 1 | 0 | 0 | 1696538 | YP_009047204.1 | K9N5Q8 |
| SVSSPDAVTTYNGYLT | 1 | 0 | 0 | 1696540 | NP_828849.2 | P0C6X7 |
| SVSSVECYSILAAKTG | 1 | 0 | 0 | 1696541 | YP_003766.2 | H9EJ42 |
| SVSTSDLGSLIDCNAA | 1 | 0 | 0 | 1696543 | YP_009047202.1 | K9N7C7 |
| SVTAVVTNGHLKMAGM | 1 | 0 | 0 | 1696547 | YP_009047210.1 | K9N7A1 |
| SVTDTIVVTEGDGIST | 1 | 0 | 0 | 1696551 | NP_828852.2 | P59632 |
| SVTEDVKFAASTGVID | 1 | 0 | 0 | 1696552 | YP_003766.2 | H9EJ42 |
| SVTFFPDLNGDVVAID | 1 | 0 | 0 | 1696554 | NP_828849.2 | P0C6X7 |
| SVTPIKNTVDTTPKEE | 1 | 0 | 0 | 1696562 | NP_073549.1 | A0A0P0LKV0 |
| SVTPQPEEELEEDWLD | 1 | 0 | 0 | 1696563 | QHR63299.1 | P0C6W2 |
| SVTRLMCNLGPDIFLS | 1 | 0 | 0 | 1696565 | YP_009047202.1 | K9N7C7 |
| SVTSSIVITSGDGTTS | 1 | 0 | 0 | 1696568 | YP_009724391.1 | P0DTC3 |
| SVTVSSNMVETVVGQL | 1 | 0 | 0 | 1696574 | YP_009047202.1 | K9N7C7 |
| SVVDVMRYNLNFSANS | 1 | 0 | 0 | 1696581 | YP_003767.1 | A0A0P0G321 |
| SVVEFEDRSNNSSRAS | 1 | 0 | 0 | 1696582 | YP_003771.1 | H9EJ69 |
| SVVEVVTTSLTPCGYS | 1 | 0 | 0 | 1696584 | YP_009555238.1 | P0C6X6 |
| SVVKPVRARAPPGEQF | 1 | 0 | 0 | 1696591 | NP_073549.1 | A0A0P0LKV0 |
| SVVNIQKEIDRLNEVA | 1 | 0 | 0 | 1696599 | YP_009724390.1 | P0DTC2 |
| SVVNSNAQALNSLLQQ | 1 | 0 | 0 | 1696600 | YP_173238.1 | A0A140H1H1 |
| SVVQKPVDVKPKIKAC | 1 | 0 | 0 | 1696602 | NP_828849.2 | P0C6X7 |
| SVVSFNEWALANQFTE | 1 | 0 | 0 | 1696606 | YP_009047202.1 | K9N7C7 |
| SVVVQPRTINSTQDGD | 1 | 0 | 0 | 1696612 | YP_009555241.1 | P36334 |
| SVVVYMCVVPSEKDKH | 1 | 0 | 0 | 1696613 | NP_073549.1 | A0A0P0LKV0 |
| SVWHIQTVQDADGINK | 1 | 0 | 0 | 1696615 | YP_173236.1 | A0A140H1G9 |
| SVYDIACCMRAGSKVV | 1 | 0 | 0 | 1696621 | YP_003766.2 | H9EJ42 |
| SVYDQVVDNVYVTYAG | 1 | 0 | 0 | 1696622 | YP_009555238.1 | P0C6X6 |
| SVYERDKAVARKLERM | 1 | 0 | 0 | 1696623 | YP_173236.1 | A0A140H1G9 |
| SVYGNFDDQPSLQVES | 1 | 0 | 0 | 1696626 | YP_003766.2 | H9EJ42 |
| SVYKSDIHLPAYASFK | 1 | 0 | 0 | 1696628 | YP_173236.1 | A0A140H1G9 |
| SVYLGVKGSGHYQTNL | 1 | 0 | 0 | 1696629 | YP_003766.2 | H9EJ42 |
| SVYSVLETFQNNVVNY | 1 | 0 | 0 | 1696635 | YP_009555238.1 | P0C6X6 |
| SVYTNSCLVSSDGQPG | 1 | 0 | 0 | 1696637 | YP_009047202.1 | K9N7C7 |
| SWADQSERNYQTFNRG | 1 | 0 | 0 | 1696644 | YP_173242.1 | Q19U29 |
| SWAIFNDHHLNRPGVY | 1 | 0 | 0 | 1696647 | YP_009047202.1 | K9N7C7 |
| SWESGKVKPPLNRNSV | 1 | 0 | 0 | 1696651 | YP_003766.2 | H9EJ42 |
| SWFTALTQHGKEDLKF | 1 | 0 | 0 | 1696659 | YP_009724397.2 | P0DTC9 |
| SWFTALTQHGKEDLRF | 1 | 0 | 0 | 1696660 | QIQ54056.1 | P59595 |
| SWIGFDVEGAHASRNA | 1 | 0 | 0 | 1696665 | YP_009047202.1 | K9N7C7 |
| SWMESEFRVYSSANNC | 1 | 0 | 0 | 1696669 | YP_009724390.1 | P0DTC2 |
| SWNADLYKLMGHFAWW | 1 | 0 | 0 | 1696670 | YP_009724389.1 | P0DTD1 |
| SWNLREMLAHAEETRK | 1 | 0 | 0 | 1696678 | YP_009724389.1 | P0DTD1 |
| SWNLREMLTHAEETRK | 1 | 0 | 0 | 1696679 | QIQ54046.1 | P0C6U8 |
| SWQNLLYDSNGNIIGF | 1 | 0 | 0 | 1696681 | YP_173238.1 | A0A140H1H1 |
| SWQNLLYDSNGNLYGF | 1 | 0 | 0 | 1696682 | YP_009555241.1 | P36334 |
| SWWTAFVTNVNASSSE | 1 | 0 | 0 | 1696698 | NP_828849.2 | P0C6X7 |
| SYAAFATAQEAYEQAV | 1 | 0 | 0 | 1696702 | YP_009724389.1 | P0DTD1 |
| SYAASYNRYKYYSGNA | 1 | 0 | 0 | 1696703 | NP_073549.1 | A0A0P0LKV0 |
| SYAAYATAQEAYEQAV | 1 | 0 | 0 | 1696704 | NP_828849.2 | P0C6X7 |
| SYAKPEEKLSAYDLAC | 1 | 0 | 0 | 1696710 | NP_073549.1 | A0A0P0LKV0 |
| SYATHHDKFTDGVCLF | 1 | 0 | 0 | 1696714 | NP_828849.2 | P0C6X7 |
| SYATHSDKFKDGVCLF | 1 | 0 | 0 | 1696715 | QIQ54047.1 | P0C6X7 |
| SYCNGVRELYLNSSNV | 1 | 0 | 0 | 1696719 | NP_828849.2 | P0C6X7 |
| SYCRLKNSVSDVAYNR | 1 | 0 | 0 | 1696722 | YP_173236.1 | A0A140H1G9 |
| SYCTGYREGYLNSTNV | 1 | 0 | 0 | 1696724 | YP_009724389.1 | P0DTD1 |
| SYCTSYRDGYLNSTNV | 1 | 0 | 0 | 1696725 | QIQ54046.1 | P0C6U8 |
| SYDGLRGAIRKAKDYG | 1 | 0 | 0 | 1696728 | YP_009047202.1 | K9N7C7 |
| SYDNLKTGVSIPCVCG | 1 | 0 | 0 | 1696731 | NP_828849.2 | P0C6X7 |
| SYEDQDALFAYTKRNV | 1 | 0 | 0 | 1696740 | YP_009724389.1 | P0DTD1 |
| SYEEQDAIFSLTKRNI | 1 | 0 | 0 | 1696741 | NP_073549.1 | A0A0P0LKV0 |
| SYEHQTPFEIKSAKKF | 1 | 0 | 0 | 1696743 | NP_828849.2 | P0C6X7 |
| SYELQTPFEIKLAKKF | 1 | 0 | 0 | 1696746 | YP_009724389.1 | P0DTD1 |
| SYEQFKKGVQIPCTCG | 1 | 0 | 0 | 1696747 | YP_009724389.1 | P0DTD1 |
| SYFCKIGKEQKCCMCN | 1 | 0 | 0 | 1696757 | YP_009047202.1 | K9N7C7 |
| SYFDSKGWYDPVENED | 1 | 0 | 0 | 1696761 | YP_003766.2 | H9EJ42 |
| SYFTEDLLCVADDKPI | 1 | 0 | 0 | 1696770 | NP_073549.1 | A0A0P0LKV0 |
| SYFTEQPIDLVPNQPY | 1 | 0 | 0 | 1696771 | YP_009724389.1 | P0DTD1 |
| SYGSSVETETYFKVNG | 1 | 0 | 0 | 1696790 | YP_173236.1 | A0A140H1G9 |
| SYHTVSLLRSTSQKSI | 1 | 0 | 0 | 1696797 | NP_828851.1 | P59594 |
| SYIAYENARQAYEDAI | 1 | 0 | 0 | 1696799 | YP_003766.2 | H9EJ42 |
| SYIDLKELGNYTYYNK | 1 | 0 | 0 | 1696800 | YP_009047204.1 | K9N5Q8 |
| SYIHVVDGCTSSTCMM | 1 | 0 | 0 | 1696801 | QIQ54046.1 | P0C6U8 |
| SYITTIKPVIYKLDGV | 1 | 0 | 0 | 1696807 | QHR63299.1 | P0C6W2 |
| SYIVDSVAVKNGALHL | 1 | 0 | 0 | 1696808 | NP_828849.2 | P0C6X7 |
| SYKLDGVTYTEIEPKL | 1 | 0 | 0 | 1696815 | NP_828849.2 | P0C6X7 |
| SYKLPNYTVPDLVVEQ | 1 | 0 | 0 | 1696817 | NP_073551.1 | A0A0P0K6L9 |
| SYKNDKSCPVVAAIIT | 1 | 0 | 0 | 1696818 | NP_828849.2 | P0C6X7 |
| SYLSERCNVIADSKSF | 1 | 0 | 0 | 1696834 | YP_009047202.1 | K9N7C7 |
| SYLTPGKLESGWTTGA | 1 | 0 | 0 | 1696836 | QIQ54048.1 | P59594 |
| SYNDFYASGCVLSSLC | 1 | 0 | 0 | 1696843 | YP_173236.1 | A0A140H1G9 |
| SYNNYVPSYVKPDSVS | 1 | 0 | 0 | 1696848 | YP_009047202.1 | K9N7C7 |
| SYNQNGTITDAVDCSL | 1 | 0 | 0 | 1696849 | QIQ54048.1 | P59594 |
| SYNRYKYYSGNANEAD | 1 | 0 | 0 | 1696850 | NP_073549.1 | A0A0P0LKV0 |
| SYQLYDASNVKKVTDV | 1 | 0 | 0 | 1696861 | YP_009555238.1 | P0C6X6 |
| SYQTQTNSRSVASQSI | 1 | 0 | 0 | 1696864 | QHR63300.2 | Q3LZX1 |
| SYRDGYLNSTNVTTTA | 1 | 0 | 0 | 1696865 | QIQ54046.1 | P0C6U8 |
| SYREAACCHLAKALND | 1 | 0 | 0 | 1696867 | YP_009724389.1 | P0DTD1 |

| | | | | | | |
|---|---|---|---|---|---|---|
| SYRFVTFEPFNVSFVN | 1 | 0 | 0 | 1696868 | YP_173238.1 | A0A140H1H1 |
| SYSELRPDTRYVLMDG | 1 | 0 | 0 | 1696870 | NP_828849.2 | P0C6X7 |
| SYSGYHPAHAPSFLAV | 1 | 0 | 0 | 1696875 | YP_009555238.1 | P0C6X6 |
| SYSKIVCDGSVHYAGV | 1 | 0 | 0 | 1696876 | YP_003766.2 | H9EJ42 |
| SYSLFDMSKFPLKLRG | 1 | 0 | 0 | 1696877 | YP_009724389.1 | P0DTD1 |
| SYSNFYASGCVLSSAC | 1 | 0 | 0 | 1696878 | YP_009555238.1 | P0C6X6 |
| SYTGSSFYAPEPITSL | 1 | 0 | 0 | 1696886 | YP_009047204.1 | K9N5Q8 |
| SYTKSVCKYTDFNEDV | 1 | 0 | 0 | 1696891 | YP_003766.2 | H9EJ42 |
| SYTNDKACPLIAAVIT | 1 | 0 | 0 | 1696892 | YP_009724389.1 | P0DTD1 |
| SYTNDKTCPIVAAVIT | 1 | 0 | 0 | 1696893 | QIQ54046.1 | P0C6U8 |
| SYTPGHYAGSRSSSGN | 1 | 0 | 0 | 1696894 | YP_173242.1 | Q19U29 |
| SYTTTIKPIVYKLDGV | 1 | 0 | 0 | 1696896 | QIQ54046.1 | P0C6U8 |
| SYTTTIKPVSYKLDGV | 1 | 0 | 0 | 1696897 | NP_828849.2 | P0C6X7 |
| SYTTTIKPVTYKLDGV | 1 | 0 | 0 | 1696898 | YP_009724389.1 | P0DTD1 |
| SYVLNNISLTTQPYFD | 1 | 0 | 0 | 1696909 | YP_173238.1 | A0A140H1H1 |
| SYVVDSVAVKNGSLHL | 1 | 0 | 0 | 1696912 | QIQ54046.1 | P0C6U8 |
| TAAFRDVLVVLNKRTN | 1 | 0 | 0 | 1696934 | NP_849177.1 | Q80H93 |
| TAAMLFDSHIVSKESG | 1 | 0 | 0 | 1696940 | NP_073556.1 | A0A0N7J131 |
| TAATVAAMFPPWSAAA | 1 | 0 | 0 | 1696943 | YP_173238.1 | A0A140H1H1 |
| TACAFSVDPATTYLEA | 1 | 0 | 0 | 1696947 | YP_003766.2 | H9EJ42 |
| TACNEDNALAYYNTSK | 1 | 0 | 0 | 1696950 | QIQ54046.1 | P0C6U8 |
| TACTDDNALAYYNNSK | 1 | 0 | 0 | 1696958 | NP_828849.2 | P0C6X7 |
| TACTDDNALAYYNTTK | 1 | 0 | 0 | 1696959 | YP_009724389.1 | P0DTD1 |
| TADEAIKFLKEWNFSL | 1 | 0 | 0 | 1696960 | YP_009555244.1 | Q01455 |
| TADTAHANNINRFNVA | 1 | 0 | 0 | 1696967 | YP_009047202.1 | K9N7C7 |
| TAEAELAKNVSLDNVL | 1 | 0 | 0 | 1696968 | YP_009724389.1 | P0DTD1 |
| TAEDISLLKKMDEPYT | 1 | 0 | 0 | 1696970 | YP_009555245.1 | P33469 |
| TAEPVRDEVSIETDII | 1 | 0 | 0 | 1696973 | NP_073556.1 | A0A0N7J131 |
| TAETAHSVNVNRFNVA | 1 | 0 | 0 | 1696976 | YP_009555238.1 | P0C6X6 |
| TAETELAKNVSLDNVL | 1 | 0 | 0 | 1696977 | QIQ54046.1 | P0C6U8 |
| TAFANTVEDAVNKLVF | 1 | 0 | 0 | 1696981 | YP_009555242.1 | Q04853 |
| TAFHQECSLQSCAQHQ | 1 | 0 | 0 | 1696986 | QHR63307.1 | Q3LZX5 |
| TAFNNALLSIQNGFSA | 1 | 0 | 0 | 1696989 | YP_173238.1 | A0A140H1H1 |
| TAFSPAQDIWGTSAAA | 1 | 0 | 0 | 1696993 | NP_828851.1 | P59594 |
| TAGVVTGNVVKQCCTA | 1 | 0 | 0 | 1697006 | YP_009047202.1 | K9N7C7 |
| TAHSELAKGVALDGVL | 1 | 0 | 0 | 1697014 | NP_828849.2 | P0C6X7 |
| TAIQSFDNSYLNESGA | 1 | 0 | 0 | 1697021 | NP_073550.1 | A0A0P0ILY9 |
| TAKKQVVLVSNNQEDF | 1 | 0 | 0 | 1697031 | YP_009555238.1 | P0C6X6 |
| TAKKSMYDGDRFVKHD | 1 | 0 | 0 | 1697032 | NP_073549.1 | A0A0P0LKV0 |
| TALNAYVSQQLSDISL | 1 | 0 | 0 | 1697043 | YP_173238.1 | A0A140H1H1 |
| TALNAYVSQQLSDSTL | 1 | 0 | 0 | 1697044 | YP_009555241.1 | P36334 |
| TALNKIQDVVNQQGNS | 1 | 0 | 0 | 1697045 | NP_073551.1 | A0A0P0K6L9 |
| TALPKGFYAEGSRGGS | 1 | 0 | 0 | 1697046 | QIQ54056.1 | P59595 |
| TALRANSAVKLQNNEL | 1 | 0 | 0 | 1697048 | YP_009724389.1 | P0DTD1 |
| TANKIVGGAPTWFNAL | 1 | 0 | 0 | 1697063 | YP_009047202.1 | K9N7C7 |
| TANVGRVIKFETDAYK | 1 | 0 | 0 | 1697074 | YP_173236.1 | A0A140H1G9 |
| TANVNALLSTDGNKIA | 1 | 0 | 0 | 1697075 | YP_009724389.1 | P0DTD1 |
| TANVNALLSTDGNKIG | 1 | 0 | 0 | 1697076 | QIQ54047.1 | P0C6X7 |
| TANVSALMGANGNKIV | 1 | 0 | 0 | 1697077 | YP_009047202.1 | K9N7C7 |
| TAPRNSWEAMALNFKL | 1 | 0 | 0 | 1697089 | YP_009047202.1 | K9N7C7 |
| TAPTIVNQERYVKITG | 1 | 0 | 0 | 1697090 | YP_009047202.1 | K9N7C7 |
| TAPYVVLTGLGESFKK | 1 | 0 | 0 | 1697091 | YP_009047202.1 | K9N7C7 |
| TAQADTAGLYTNFRID | 1 | 0 | 0 | 1697095 | YP_009047205.1 | K9N796 |
| TAQPVKIDNASPASTV | 1 | 0 | 0 | 1697106 | NP_828852.2 | P59632 |
| TARDGVRRGDNFHSVL | 1 | 0 | 0 | 1697107 | YP_009047202.1 | K9N7C7 |
| TARVVYTAASHAAVDA | 1 | 0 | 0 | 1697113 | YP_009555238.1 | P0C6X6 |
| TARVVYTACSHAAVDA | 1 | 0 | 0 | 1697114 | YP_009047202.1 | K9N7C7 |
| TASDKFYRLSNELAQV | 1 | 0 | 0 | 1697118 | NP_073549.1 | A0A0P0LKV0 |
| TATKQYNVTQAFGRRG | 1 | 0 | 0 | 1697140 | QHR63308.1 | Q3LZX4 |
| TATSQQPSGGNVVPYY | 1 | 0 | 0 | 1697143 | YP_009555245.1 | P33469 |
| TAVINLRADQINDMVY | 1 | 0 | 0 | 1697155 | YP_009555238.1 | P0C6X6 |
| TAVLTIANNYSKLFDE | 1 | 0 | 0 | 1697160 | NP_073549.1 | A0A0P0LKV0 |
| TAVMSLKENQINDMIY | 1 | 0 | 0 | 1697162 | NP_828849.2 | P0C6X7 |
| TAVNESRYGLQNLLQL | 1 | 0 | 0 | 1697164 | YP_003767.1 | A0A0P0G321 |
| TAVVIPTKKAGGTTEM | 1 | 0 | 0 | 1697168 | YP_009724389.1 | P0DTD1 |
| TAVVNLRPDQLNDLVY | 1 | 0 | 0 | 1697169 | YP_173236.1 | A0A140H1G9 |
| TAYNGYLTSSSKTPEE | 1 | 0 | 0 | 1697183 | YP_009724389.1 | P0DTD1 |
| TCAKEIKESVQTFFKL | 1 | 0 | 0 | 1697187 | YP_009724389.1 | P0DTD1 |
| TCANDPVGFTLKNTVC | 1 | 0 | 0 | 1697189 | YP_009724389.1 | P0DTD1 |
| TCANDPVGFTLRNTVC | 1 | 0 | 0 | 1697190 | NP_828849.2 | P0C6X7 |
| TCATTRQVVNVVTTKI | 1 | 0 | 0 | 1697193 | YP_009724389.1 | P0DTD1 |
| TCDGLKFTEESFYKNM | 1 | 0 | 0 | 1697207 | YP_173236.1 | A0A140H1G9 |
| TCDICKSTVVEVKSAI | 1 | 0 | 0 | 1697211 | YP_003766.2 | H9EJ42 |
| TCDRTAIQSFDNSYLN | 1 | 0 | 0 | 1697214 | NP_073549.1 | A0A0P0LKV0 |
| TCEFCGTENQVKEGPT | 1 | 0 | 0 | 1697220 | QIQ54046.1 | P0C6U8 |
| TCFSTQFAFACPDGVK | 1 | 0 | 0 | 1697242 | YP_009724395.1 | P0DTC7 |
| TCGIIDVEKGVLPQLE | 1 | 0 | 0 | 1697246 | QIQ54046.1 | P0C6U8 |
| TCGKQATKYLVQQESP | 1 | 0 | 0 | 1697247 | YP_009724389.1 | P0DTD1 |
| TCGKQATQYLVQQESP | 1 | 0 | 0 | 1697248 | QHR63299.1 | P0C6W2 |
| TCGLVELEKGVLPQLE | 1 | 0 | 0 | 1697249 | NP_828849.2 | P0C6X7 |
| TCGQQQTTLKGVEAVM | 1 | 0 | 0 | 1697251 | YP_009724389.1 | P0DTD1 |
| TCNAISSNTDNETLQY | 1 | 0 | 0 | 1697279 | YP_173238.1 | A0A140H1H1 |
| TCNNIDAAKIYGMCFS | 1 | 0 | 0 | 1697280 | YP_009555241.1 | P36334 |
| TCNSNANLSMLQLGVP | 1 | 0 | 0 | 1697282 | YP_003767.1 | A0A0P0G321 |
| TCNTPTQCYYNNSNNG | 1 | 0 | 0 | 1697283 | YP_009555238.1 | P0C6X6 |
| TCNVSFVGELKAAEAK | 1 | 0 | 0 | 1697284 | NP_073549.1 | A0A0P0LKV0 |
| TCRRCPAEIVDTVSAL | 1 | 0 | 0 | 1697293 | YP_009724389.1 | P0DTD1 |
| TCRTDMEKDFIALDQD | 1 | 0 | 0 | 1697295 | YP_009555238.1 | P0C6X6 |
| TCSNDLLQPNTEVFTD | 1 | 0 | 0 | 1697298 | YP_173238.1 | A0A140H1H1 |
| TCSNTKFADDLNQMTG | 1 | 0 | 0 | 1697299 | NP_828849.2 | P0C6X7 |
| TCTERLKLFAAETLKA | 1 | 0 | 0 | 1697303 | YP_009724389.1 | P0DTD1 |

| | | | | | | |
|---|---|---|---|---|---|---|
| TCTIAGKAFDKVFDYV | 1 | 0 | 0 | 1697304 | NP_073549.1 | A0A0P0LKV0 |
| TCTPNVTINCQDPAGG | 1 | 0 | 0 | 1697307 | NP_849177.1 | Q80H93 |
| TCVIANVPAIDVLKKL | 1 | 0 | 0 | 1697311 | YP_003766.2 | H9EJ42 |
| TCVVIPSKKAGGTTEM | 1 | 0 | 0 | 1697315 | NP_828849.2 | P0C6X7 |
| TCYRCPKEIVDTVSAL | 1 | 0 | 0 | 1697323 | YP_009555238.1 | P0C6X6 |
| TDAAVKAAESKAFVDT | 1 | 0 | 0 | 1697325 | NP_073549.1 | A0A0P0LKV0 |
| TDAIIIDNDLNEFVSD | 1 | 0 | 0 | 1697328 | YP_009047202.1 | K9N7C7 |
| TDAKWYCYDKNPINSN | 1 | 0 | 0 | 1697329 | NP_073549.1 | A0A0P0LKV0 |
| TDAKWYCYDKQPVNSN | 1 | 0 | 0 | 1697330 | YP_003766.2 | H9EJ42 |
| TDANKLILINDVAFVS | 1 | 0 | 0 | 1697331 | YP_009555238.1 | P0C6X6 |
| TDAVILLERYVSLAID | 1 | 0 | 0 | 1697338 | NP_073549.1 | A0A0P0LKV0 |
| TDAVVLLERYVSLAID | 1 | 0 | 0 | 1697340 | YP_003766.2 | H9EJ42 |
| TDCLAAETHRDCDFNK | 1 | 0 | 0 | 1697342 | YP_009047202.1 | K9N7C7 |
| TDCLYLKNLTQTFTSM | 1 | 0 | 0 | 1697343 | YP_173236.1 | A0A140H1G9 |
| TDCSDGNYNRNASLNS | 1 | 0 | 0 | 1697344 | YP_009047204.1 | K9N5Q8 |
| TDCVFYFAVRKEGQDV | 1 | 0 | 0 | 1697345 | YP_009555238.1 | P0C6X6 |
| TDDNALAYYNNSKGGR | 1 | 0 | 0 | 1697349 | NP_828849.2 | P0C6X7 |
| TDDYFNKKDWYDFVEN | 1 | 0 | 0 | 1697352 | QIQ54047.1 | P0C6X7 |
| TDECRLAMLGAEMTSN | 1 | 0 | 0 | 1697353 | NP_073549.1 | A0A0P0LKV0 |
| TDENYNNLVPTYLKSD | 1 | 0 | 0 | 1697356 | YP_173236.1 | A0A140H1G9 |
| TDESQPLPQRQKKQQT | 1 | 0 | 0 | 1697360 | QIQ54056.1 | P59595 |
| TDFSRVSAKPPPGDQF | 1 | 0 | 0 | 1697374 | YP_009724389.1 | P0DTD1 |
| TDFSSEIIGYKAIDGG | 1 | 0 | 0 | 1697375 | YP_009724389.1 | P0DTD1 |
| TDFVNEFYAYLRKHFS | 1 | 0 | 0 | 1697377 | YP_009724389.1 | P0DTD1 |
| TDGKLNCSFQDAASNI | 1 | 0 | 0 | 1697381 | YP_009047202.1 | K9N7C7 |
| TDHDYQIGGYTEKWES | 1 | 0 | 0 | 1697389 | QIQ54049.1 | P59632 |
| TDIAKKPTENICAPLT | 1 | 0 | 0 | 1697395 | QHR63299.1 | P0C6W2 |
| TDIAKKPTESACSSLT | 1 | 0 | 0 | 1697396 | NP_828849.2 | P0C6X7 |
| TDIAKKPTETICAPLT | 1 | 0 | 0 | 1697397 | YP_009724389.1 | P0DTD1 |
| TDIDYCVRAFDVYNKD | 1 | 0 | 0 | 1697402 | NP_073549.1 | A0A0P0LKV0 |
| TDIFYKENSYITTIKP | 1 | 0 | 0 | 1697403 | QHR63299.1 | P0C6W2 |
| TDIVVVDEVSMLTNYE | 1 | 0 | 0 | 1697410 | YP_009555238.1 | P0C6X6 |
| TDKSFEQQVGVIADKD | 1 | 0 | 0 | 1697412 | NP_073549.1 | A0A0P0LKV0 |
| TDLEGKFYGPFVDRQT | 1 | 0 | 0 | 1697414 | NP_828849.2 | P0C6X7 |
| TDLEGNFYGPFVDRQT | 1 | 0 | 0 | 1697415 | YP_009724389.1 | P0DTD1 |
| TDLEGTFYGPFVDRQT | 1 | 0 | 0 | 1697416 | QHR63299.1 | P0C6W2 |
| TDLMAAYVDNSSLTIK | 1 | 0 | 0 | 1697426 | YP_009724389.1 | P0DTD1 |
| TDLQSESSCGLFKDCA | 1 | 0 | 0 | 1697428 | NP_073549.1 | A0A0P0LKV0 |
| TDMVKSKGVCATGDCY | 1 | 0 | 0 | 1697434 | YP_009555238.1 | P0C6X6 |
| TDNLVFVNKTSLPTNI | 1 | 0 | 0 | 1697437 | NP_073549.1 | A0A0P0LKV0 |
| TDNNQNGDRSGARPKQ | 1 | 0 | 0 | 1697439 | QIQ54056.1 | P59595 |
| TDNSANTKVLRNKGVD | 1 | 0 | 0 | 1697441 | YP_009047202.1 | K9N7C7 |
| TDNSTTVSVTVSSNMV | 1 | 0 | 0 | 1697442 | YP_009047202.1 | K9N7C7 |
| TDNTFVSGNCDVVIGI | 1 | 0 | 0 | 1697443 | YP_009724390.1 | P0DTC2 |
| TDNVSIKNVDIVSEAK | 1 | 0 | 0 | 1697446 | QIQ54046.1 | P0C6U8 |
| TDNVYIKNADIVEEAK | 1 | 0 | 0 | 1697447 | YP_009724389.1 | P0DTD1 |
| TDPIDDFQVNWNTKHG | 1 | 0 | 0 | 1697450 | QIQ54046.1 | P0C6U8 |
| TDPIEDYEQNWNTKHG | 1 | 0 | 0 | 1697451 | NP_828849.2 | P0C6X7 |
| TDPYEDFQENWNTKHS | 1 | 0 | 0 | 1697457 | YP_009724389.1 | P0DTD1 |
| TDQSSYIVDSVAVKNG | 1 | 0 | 0 | 1697460 | NP_828849.2 | P0C6X7 |
| TDQSSYVVDSVAVKNG | 1 | 0 | 0 | 1697462 | QIQ54046.1 | P0C6U8 |
| TDQYLCGADGKPVMSE | 1 | 0 | 0 | 1697463 | NP_073549.1 | A0A0P0LKV0 |
| TDSDSLSSKPQRKRGV | 1 | 0 | 0 | 1697466 | YP_173242.1 | Q19U29 |
| TDSFSCNNFDESKIYG | 1 | 0 | 0 | 1697467 | YP_173238.1 | A0A140H1H1 |
| TDSGFAAYNRYRIGNY | 1 | 0 | 0 | 1697468 | NP_828855.1 | P59596 |
| TDSIVFTYIPASGYVA | 1 | 0 | 0 | 1697469 | YP_009047208.1 | K9N7D2 |
| TDSIVRFPNITNLCPF | 1 | 0 | 0 | 1697471 | QHR63300.2 | Q3LZX1 |
| TDSIVVMAYVTGGVVQ | 1 | 0 | 0 | 1697472 | QIQ54046.1 | P0C6U8 |
| TDSLCSSHYWSPQGNL | 1 | 0 | 0 | 1697473 | YP_173236.1 | A0A140H1G9 |
| TDSMLTLDDRGLALDN | 1 | 0 | 0 | 1697474 | NP_073549.1 | A0A0P0LKV0 |
| TDSVLLIERFVSLAID | 1 | 0 | 0 | 1697476 | YP_009555238.1 | P0C6X6 |
| TDTIVVTEGDGISTPK | 1 | 0 | 0 | 1697484 | NP_828852.2 | P59632 |
| TDTNSVCPKLEFANDT | 1 | 0 | 0 | 1697485 | YP_009047204.1 | K9N5Q8 |
| TDTVNHDSFNLYGYSV | 1 | 0 | 0 | 1697493 | YP_173236.1 | A0A140H1G9 |
| TDVAYHTVTDVKQVGC | 1 | 0 | 0 | 1697494 | YP_009555238.1 | P0C6X6 |
| TDVFYKENSYTTTIKP | 1 | 0 | 0 | 1697498 | YP_009724389.1 | P0DTD1 |
| TDVFYKETSYTTTIKP | 1 | 0 | 0 | 1697499 | NP_828849.2 | P0C6X7 |
| TDVLFPGQKYSSDCNV | 1 | 0 | 0 | 1697502 | YP_009047202.1 | K9N7C7 |
| TDVLQENQKILAASFN | 1 | 0 | 0 | 1697503 | NP_073551.1 | A0A0P0K6L9 |
| TDVQLRAFDIYNASVA | 1 | 0 | 0 | 1697506 | YP_009555238.1 | P0C6X6 |
| TDVRDYKLANDVKDTL | 1 | 0 | 0 | 1697507 | YP_003766.2 | H9EJ42 |
| TDVVYRAFDIYNDKVA | 1 | 0 | 0 | 1697512 | YP_009724389.1 | P0DTD1 |
| TDVVYRAFDIYNEKVA | 1 | 0 | 0 | 1697513 | NP_828849.2 | P0C6X7 |
| TDYKLELFNKYFKHWS | 1 | 0 | 0 | 1697520 | YP_009555238.1 | P0C6X6 |
| TDYKLELFNKYFKYWG | 1 | 0 | 0 | 1697521 | YP_173236.1 | A0A140H1G9 |
| TDYNAILQSNNPQCAI | 1 | 0 | 0 | 1697524 | YP_003766.2 | H9EJ42 |
| TDYNYYKYNLPTMVDI | 1 | 0 | 0 | 1697525 | YP_009555238.1 | P0C6X6 |
| TEAEWIELFPHNDRIK | 1 | 0 | 0 | 1697530 | NP_073549.1 | A0A0P0LKV0 |
| TEDDYQGKSLEFGATS | 1 | 0 | 0 | 1697541 | QHR63299.1 | P0C6W2 |
| TEDPRWPQIAELAPTA | 1 | 0 | 0 | 1697550 | YP_009047211.1 | K9N4V7 |
| TEDVKFAASTGVIDIS | 1 | 0 | 0 | 1697552 | YP_003766.2 | H9EJ42 |
| TEEKSVVQKPVDVKPK | 1 | 0 | 0 | 1697558 | NP_828849.2 | P0C6X7 |
| TEESFYKNMYLKSAVM | 1 | 0 | 0 | 1697563 | YP_173236.1 | A0A140H1G9 |
| TEEVKSSRVERSTAVL | 1 | 0 | 0 | 1697567 | NP_073549.1 | A0A0P0LKV0 |
| TEEVVLKTGDLQPLEQ | 1 | 0 | 0 | 1697569 | YP_009724389.1 | P0DTD1 |
| TEFQLDYSIDVIDNEI | 1 | 0 | 0 | 1697573 | NP_073549.1 | A0A0P0LKV0 |
| TEFSRVSAKPPPGDQF | 1 | 0 | 0 | 1697574 | QIQ54047.1 | P0C6X7 |
| TEFSWNADLYKLMSCF | 1 | 0 | 0 | 1697575 | YP_173236.1 | A0A140H1G9 |
| TEFTRVNAKPPPGDQF | 1 | 0 | 0 | 1697577 | NP_828849.2 | P0C6X7 |
| TEGCVSTNFGDVIKPV | 1 | 0 | 0 | 1697581 | YP_003766.2 | H9EJ42 |
| TEGLMQNASLYSSLVP | 1 | 0 | 0 | 1697586 | YP_009555238.1 | P0C6X6 |

| | | | | | | |
|---|---|---|---|---|---|---|
| TEGQLQSLSKDYISSN | 1 | 0 | 0 | 1697588 | YP_003766.2 | H9EJ42 |
| TEGSVKGLTPSKGPAQ | 1 | 0 | 0 | 1697590 | NP_828849.2 | P0C6X7 |
| TEHSWNADLYKLMGHF | 1 | 0 | 0 | 1697593 | YP_009724389.1 | P0DTD1 |
| TEHVTSVNLFDVGYSV | 1 | 0 | 0 | 1697595 | YP_009047205.1 | K9N796 |
| TEIDPKLDGYYKKDNA | 1 | 0 | 0 | 1697598 | QIQ54046.1 | P0C6U8 |
| TEIDPKLDNYYKKDNS | 1 | 0 | 0 | 1697599 | YP_009724389.1 | P0DTD1 |
| TEIEPKLDGYYKKDNA | 1 | 0 | 0 | 1697600 | NP_828849.2 | P0C6X7 |
| TEKLSRLSMIKGPQRA | 1 | 0 | 0 | 1697608 | YP_173236.1 | A0A140H1G9 |
| TELAKNVSLDNVLSTF | 1 | 0 | 0 | 1697614 | QIQ54046.1 | P0C6U8 |
| TELEPPCRFVTDTPKG | 1 | 0 | 0 | 1697616 | YP_009724389.1 | P0DTD1 |
| TELGIEFLKRGDKVVY | 1 | 0 | 0 | 1697617 | QIQ54046.1 | P0C6U8 |
| TELGVEFLKRGDKIVY | 1 | 0 | 0 | 1697618 | NP_828849.2 | P0C6X7 |
| TELKDSVKKFFKLVNK | 1 | 0 | 0 | 1697619 | QIQ54046.1 | P0C6U8 |
| TEMLSRALKKVPVDEY | 1 | 0 | 0 | 1697625 | NP_828849.2 | P0C6X7 |
| TENKYSQLDEEQPMEI | 1 | 0 | 0 | 1697629 | YP_009724394.1 | P0DTC6 |
| TENNTEFTRVNAKPPP | 1 | 0 | 0 | 1697634 | NP_828849.2 | P0C6X7 |
| TEPKKDKKKKTDEAQP | 1 | 0 | 0 | 1697639 | NP_828858.1 | P59595 |
| TEQPIDLVPTQPLPNA | 1 | 0 | 0 | 1697647 | NP_828849.2 | P0C6X7 |
| TEQSEIEPEPEPTPEE | 1 | 0 | 0 | 1697649 | NP_828849.2 | P0C6X7 |
| TEQTQNVTVKPKRLRK | 1 | 0 | 0 | 1697650 | YP_009047202.1 | K9N7C7 |
| TERSDKSYEHQTPFEI | 1 | 0 | 0 | 1697654 | NP_828849.2 | P0C6X7 |
| TERSEKSYELQTPFEI | 1 | 0 | 0 | 1697656 | YP_009724389.1 | P0DTD1 |
| TESAVNSTKALAKQDA | 1 | 0 | 0 | 1697658 | YP_009047206.1 | K9N4V0 |
| TESDYKGLPLEFGASS | 1 | 0 | 0 | 1697659 | QIQ54046.1 | P0C6U8 |
| TESKPSVEQRKQDDKK | 1 | 0 | 0 | 1697663 | YP_009724389.1 | P0DTD1 |
| TEVDDYVLANECIERL | 1 | 0 | 0 | 1697678 | YP_173236.1 | A0A140H1G9 |
| TEVLSEEVILKTGELQ | 1 | 0 | 0 | 1697683 | QIQ54046.1 | P0C6U8 |
| TEVLTEEVVLKTGDLQ | 1 | 0 | 0 | 1697684 | YP_009724389.1 | P0DTD1 |
| TEVNELLDTTQLQVAN | 1 | 0 | 0 | 1697687 | YP_009555241.1 | P36334 |
| TEVPMAIHAEQLTPAW | 1 | 0 | 0 | 1697691 | QIQ54048.1 | P59594 |
| TEVRGSRRLAQQKINE | 1 | 0 | 0 | 1697694 | YP_003767.1 | A0A0P0G321 |
| TEVVGDIILKPANDGL | 1 | 0 | 0 | 1697697 | QHR63299.1 | P0C6W2 |
| TEVVGDIILKPANNSL | 1 | 0 | 0 | 1697698 | YP_009724389.1 | P0DTD1 |
| TEVVGDVILKPVQDGV | 1 | 0 | 0 | 1697699 | QIQ54046.1 | P0C6U8 |
| TEVVYSNGGFYFKPGG | 1 | 0 | 0 | 1697702 | NP_073549.1 | A0A0P0LKV0 |
| TEYDFTDYKVQLFEKY | 1 | 0 | 0 | 1697708 | YP_009047202.1 | K9N7C7 |
| TEYSWNKKLYELVQRF | 1 | 0 | 0 | 1697711 | NP_073549.1 | A0A0P0LKV0 |
| TEYYEFLNKHFSMMIL | 1 | 0 | 0 | 1697714 | YP_009555238.1 | P0C6X6 |
| TFAELEAAQKAYQEAM | 1 | 0 | 0 | 1697715 | YP_009047202.1 | K9N7C7 |
| TFASVYAWNRKRISNC | 1 | 0 | 0 | 1697717 | QHR63300.2 | Q3LZX1 |
| TFCDNINSILDEVNGL | 1 | 0 | 0 | 1697724 | YP_173238.1 | A0A140H1H1 |
| TFDDVQAPNYTQHTSS | 1 | 0 | 0 | 1697734 | NP_828851.1 | P59594 |
| TFDSEYCRHGTCEKSE | 1 | 0 | 0 | 1697744 | QIQ54046.1 | P0C6U8 |
| TFDSEYCRHGTCERSE | 1 | 0 | 0 | 1697745 | YP_009724389.1 | P0DTD1 |
| TFEDELAPSDVYQQLA | 1 | 0 | 0 | 1697748 | YP_173236.1 | A0A140H1G9 |
| TFEKGDYGDAVVYRGT | 1 | 0 | 0 | 1697753 | YP_009724389.1 | P0DTD1 |
| TFELDERIDKVLNEKC | 1 | 0 | 0 | 1697755 | YP_009724389.1 | P0DTD1 |
| TFELDERVDKVLNEKC | 1 | 0 | 0 | 1697756 | NP_828849.2 | P0C6X7 |
| TFEMSCDCGKKFDDQV | 1 | 0 | 0 | 1697757 | YP_003766.2 | H9EJ42 |
| TFEPFNVSFVNDSIES | 1 | 0 | 0 | 1697758 | YP_173238.1 | A0A140H1H1 |
| TFFIFNKLVKDPPNVQ | 1 | 0 | 0 | 1697765 | NP_828852.2 | P59632 |
| TFFIYSRIVDEPEDHV | 1 | 0 | 0 | 1697767 | QIQ54049.1 | P59632 |
| TFFPDLNGDVVAIDYK | 1 | 0 | 0 | 1697770 | YP_009724389.1 | P0DTD1 |
| TFFPDLNGDVVAIDYR | 1 | 0 | 0 | 1697771 | NP_828849.2 | P0C6X7 |
| TFGAISASIGDIIQRL | 1 | 0 | 0 | 1697776 | YP_009047204.1 | K9N5Q8 |
| TFGDDTVIEVQGYKNV | 1 | 0 | 0 | 1697778 | QIQ54046.1 | P0C6U8 |
| TFGDDTVIEVQGYKSV | 1 | 0 | 0 | 1697779 | YP_009724389.1 | P0DTD1 |
| TFGEDTVWEVQGYKNV | 1 | 0 | 0 | 1697780 | NP_828849.2 | P0C6X7 |
| TFGGPSDSTDNNQNGD | 1 | 0 | 0 | 1697781 | QIQ54056.1 | P59595 |
| TFGGPSDSTGSNQNGE | 1 | 0 | 0 | 1697782 | YP_009724397.2 | P0DTC9 |
| TFGNVKPGETFTVLAA | 1 | 0 | 0 | 1697789 | YP_173236.1 | A0A140H1G9 |
| TFGVVKPGETFTVLAA | 1 | 0 | 0 | 1697793 | YP_009555238.1 | P0C6X6 |
| TFHLDGEVITFDNLKI | 1 | 0 | 0 | 1697794 | QHR63299.1 | P0C6W2 |
| TFHLDGEVITFDNLKT | 1 | 0 | 0 | 1697795 | YP_009724389.1 | P0DTD1 |
| TFHMDGEVINIDSLKT | 1 | 0 | 0 | 1697796 | QIQ54046.1 | P0C6U8 |
| TFIDAARGPAGVESDV | 1 | 0 | 0 | 1697800 | YP_009047202.1 | K9N7C7 |
| TFITPEVSRELGNITK | 1 | 0 | 0 | 1697807 | NP_073549.1 | A0A0P0LKV0 |
| TFITVEAAIELSKELK | 1 | 0 | 0 | 1697808 | YP_173236.1 | A0A140H1G9 |
| TFITVEAALDLSKELK | 1 | 0 | 0 | 1697809 | YP_009555238.1 | P0C6X6 |
| TFKCYGVSATKLNDLC | 1 | 0 | 0 | 1697812 | NP_828851.1 | P59594 |
| TFKQANNNCFVNVSCL | 1 | 0 | 0 | 1697823 | YP_009555238.1 | P0C6X6 |
| TFLDLQVEMNRLQEAI | 1 | 0 | 0 | 1697833 | YP_009555241.1 | P36334 |
| TFLLSYNQNGTITDAV | 1 | 0 | 0 | 1697847 | QIQ54048.1 | P59594 |
| TFLLTFNDNQAITQVP | 1 | 0 | 0 | 1697848 | YP_003766.2 | H9EJ42 |
| TFLNGITYTSTSGNLL | 1 | 0 | 0 | 1697850 | NP_073551.1 | A0A0P0K6L9 |
| TFLSLSDQFKTTGDLA | 1 | 0 | 0 | 1697853 | YP_003766.2 | H9EJ42 |
| TFLSMFDVDKKSLNAL | 1 | 0 | 0 | 1697854 | YP_009555238.1 | P0C6X6 |
| TFMLKYDENGTITDAV | 1 | 0 | 0 | 1697860 | NP_828851.1 | P59594 |
| TFNDNQAITQVPATSI | 1 | 0 | 0 | 1697864 | YP_003766.2 | H9EJ42 |
| TFNHNNGNDVLYQPPT | 1 | 0 | 0 | 1697868 | YP_173236.1 | A0A140H1G9 |
| TFNKDIYCGNSMVCKM | 1 | 0 | 0 | 1697869 | YP_003766.2 | H9EJ42 |
| TFNKQEASVPILTTPF | 1 | 0 | 0 | 1697870 | YP_173236.1 | A0A140H1G9 |
| TFNVPMDKLKTLIATA | 1 | 0 | 0 | 1697875 | QIQ54046.1 | P0C6U8 |
| TFPPGQGVPLNANSTP | 1 | 0 | 0 | 1697879 | YP_009047211.1 | K9N4V7 |
| TFSATFSVPMEKLKAL | 1 | 0 | 0 | 1697891 | NP_828849.2 | P0C6X7 |
| TFSGNVDLDALASDIF | 1 | 0 | 0 | 1697893 | YP_009047202.1 | K9N7C7 |
| TFSGSFDNGHYVVYDA | 1 | 0 | 0 | 1697894 | YP_003766.2 | H9EJ42 |
| TFSMAPVEFYEDRILD | 1 | 0 | 0 | 1697896 | YP_009047202.1 | K9N7C7 |
| TFSSTFNVPMDKLKTL | 1 | 0 | 0 | 1697900 | QIQ54046.1 | P0C6U8 |
| TFSYDGLRGAIRKAKD | 1 | 0 | 0 | 1697905 | YP_009047202.1 | K9N7C7 |
| TFTKSTNSRIRASLPT | 1 | 0 | 0 | 1697911 | NP_828849.2 | P0C6X7 |

| | | | | | | |
|---|---|---|---|---|---|---|
| TFTNNNGSDVLYQPPT | 1 | 0 | 0 | 1697915 | YP_009555238.1 | P0C6X6 |
| TFVIDMRSYEKLANSI | 1 | 0 | 0 | 1697940 | NP_073549.1 | A0A0P0LKV0 |
| TFVNEYYEFLCKHFSM | 1 | 0 | 0 | 1697943 | YP_173236.1 | A0A140H1G9 |
| TFVSAARQGVVDTDVD | 1 | 0 | 0 | 1697949 | NP_828849.2 | P0C6X7 |
| TFVVDKSLSIEEFADV | 1 | 0 | 0 | 1697957 | YP_009047202.1 | K9N7C7 |
| TFYLFDDAGEENFSSR | 1 | 0 | 0 | 1697973 | NP_828849.2 | P0C6X7 |
| TFYPKLQASQAWQPGV | 1 | 0 | 0 | 1697976 | NP_828849.2 | P0C6X7 |
| TFYPKLQSSQAWQPGV | 1 | 0 | 0 | 1697977 | YP_009724389.1 | P0DTD1 |
| TFYPRLQAASDWKPGY | 1 | 0 | 0 | 1697981 | YP_009555238.1 | P0C6X6 |
| TGAICDFEIACKCGVK | 1 | 0 | 0 | 1697988 | YP_009555238.1 | P0C6X6 |
| TGAIKLDDKDPSFKDN | 1 | 0 | 0 | 1697990 | QIQ54056.1 | P59595 |
| TGCCDNSYFDSKGWYD | 1 | 0 | 0 | 1697993 | YP_003766.2 | H9EJ42 |
| TGCVIAWNSVKQDALT | 1 | 0 | 0 | 1698010 | QIQ54048.1 | P59594 |
| TGDLAVCLGIGDSAVT | 1 | 0 | 0 | 1698019 | YP_009555238.1 | P0C6X6 |
| TGDLAVQIGSNNVCTY | 1 | 0 | 0 | 1698020 | YP_003766.2 | H9EJ42 |
| TGDLFYTIKFSGFKLD | 1 | 0 | 0 | 1698021 | YP_003766.2 | H9EJ42 |
| TGDNDDEDVVTGDNDD | 1 | 0 | 0 | 1698023 | YP_173236.1 | A0A140H1G9 |
| TGDNDDEDVVTGDNND | 1 | 0 | 0 | 1698024 | YP_173236.1 | A0A140H1G9 |
| TGDNNDEEIVTGDNDD | 1 | 0 | 0 | 1698026 | YP_173236.1 | A0A140H1G9 |
| TGDSVKFVYMHQLELS | 1 | 0 | 0 | 1698031 | YP_173236.1 | A0A140H1G9 |
| TGDVVLASDDLYVKRY | 1 | 0 | 0 | 1698034 | YP_173236.1 | A0A140H1G9 |
| TGDVVLATDDLYVKRY | 1 | 0 | 0 | 1698035 | YP_009555238.1 | P0C6X6 |
| TGECPNFVFPLNSTIK | 1 | 0 | 0 | 1698037 | QIQ54046.1 | P0C6U8 |
| TGELQPLEEPTAQAVE | 1 | 0 | 0 | 1698041 | QIQ54046.1 | P0C6U8 |
| TGFTTTNEAFQKVQDA | 1 | 0 | 0 | 1698052 | YP_009047204.1 | K9N5Q8 |
| TGFVYFNATGSDVNCN | 1 | 0 | 0 | 1698053 | YP_003767.1 | A0A0P0G321 |
| TGGLVQQTSQWLSNLL | 1 | 0 | 0 | 1698063 | NP_828849.2 | P0C6X7 |
| TGGNSQSSSRASSLSR | 1 | 0 | 0 | 1698064 | YP_009047211.1 | K9N4V7 |
| TGGVVQMTTQWLTNIF | 1 | 0 | 0 | 1698068 | QIQ54046.1 | P0C6U8 |
| TGHYTGQAGEMPCAII | 1 | 0 | 0 | 1698074 | YP_009555238.1 | P0C6X6 |
| TGIIRSSNQSLAGGIT | 1 | 0 | 0 | 1698083 | YP_003767.1 | A0A0P0G321 |
| TGILDGDYAMQFFKMG | 1 | 0 | 0 | 1698084 | NP_073549.1 | A0A0P0LKV0 |
| TGKTVLGEYVFDKSEL | 1 | 0 | 0 | 1698098 | YP_173236.1 | A0A140H1G9 |
| TGLFKDCSRETSGLSP | 1 | 0 | 0 | 1698106 | YP_009047202.1 | K9N7C7 |
| TGLYPTLNISDEFSSN | 1 | 0 | 0 | 1698126 | YP_009724389.1 | P0DTD1 |
| TGNYQCGHYTHITAKE | 1 | 0 | 0 | 1698143 | NP_828849.2 | P0C6X7 |
| TGPEAALPFRAVKDGI | 1 | 0 | 0 | 1698146 | YP_009047211.1 | K9N4V7 |
| TGPEAGLPYGANKEGI | 1 | 0 | 0 | 1698148 | QIQ54056.1 | P59595 |
| TGPEASLPYGANKEGI | 1 | 0 | 0 | 1698149 | NP_828858.1 | P59595 |
| TGPPPISTDTVDVTNG | 1 | 0 | 0 | 1698153 | YP_009555241.1 | P36334 |
| TGQAGEMPCAIINDKV | 1 | 0 | 0 | 1698156 | YP_009555238.1 | P0C6X6 |
| TGQAITVTPEANMDQE | 1 | 0 | 0 | 1698157 | YP_009724389.1 | P0DTD1 |
| TGRGDCKGFSSDVLSD | 1 | 0 | 0 | 1698162 | NP_073551.1 | A0A0P0K6L9 |
| TGRSVYVKFQDSKPPL | 1 | 0 | 0 | 1698167 | YP_009047209.1 | K9N5R3 |
| TGSLADGLEAYADKTL | 1 | 0 | 0 | 1698177 | YP_009555238.1 | P0C6X6 |
| TGSNQNGERSGARPKQ | 1 | 0 | 0 | 1698183 | QHR63308.1 | Q3LZX4 |
| TGSPYIDDFNKIASCK | 1 | 0 | 0 | 1698186 | YP_173236.1 | A0A140H1G9 |
| TGSPYIDDFNRIASCK | 1 | 0 | 0 | 1698187 | YP_009555238.1 | P0C6X6 |
| TGSSFYAPEPITSLNT | 1 | 0 | 0 | 1698189 | YP_009047204.1 | K9N5Q8 |
| TGSVVKPVRARAPPGE | 1 | 0 | 0 | 1698194 | NP_073549.1 | A0A0P0LKV0 |
| TGSVYGNFDDQPSLQV | 1 | 0 | 0 | 1698196 | YP_003766.2 | H9EJ42 |
| TGTDFNGDFYGPYKDA | 1 | 0 | 0 | 1698200 | YP_009555238.1 | P0C6X6 |
| TGTFTVVMRPNYTIKG | 1 | 0 | 0 | 1698202 | YP_009047202.1 | K9N7C7 |
| TGTGPEAALPFRAVKD | 1 | 0 | 0 | 1698204 | YP_009047211.1 | K9N4V7 |
| TGTGQAITVTPEANMD | 1 | 0 | 0 | 1698205 | YP_009724389.1 | P0DTD1 |
| TGTGVLTTSKKQFLPF | 1 | 0 | 0 | 1698208 | QIQ54048.1 | P59594 |
| TGTLLVDSDLNDFVSD | 1 | 0 | 0 | 1698211 | YP_009724389.1 | P0DTD1 |
| TGTPKPSRNQSPASSQ | 1 | 0 | 0 | 1698213 | NP_073556.1 | A0A0N7J131 |
| TGVCYDFDGVTTSDKC | 1 | 0 | 0 | 1698222 | YP_003766.2 | H9EJ42 |
| TGVDGELPVAVVNDKV | 1 | 0 | 0 | 1698224 | YP_003766.2 | H9EJ42 |
| TGVEAVMYMGTLSYDN | 1 | 0 | 0 | 1698225 | NP_828849.2 | P0C6X7 |
| TGVFDIVKGQIQVASD | 1 | 0 | 0 | 1698228 | NP_828849.2 | P0C6X7 |
| TGVHAGTDLEGKFYGP | 1 | 0 | 0 | 1698230 | NP_828849.2 | P0C6X7 |
| TGVHAGTDLEGNFYGP | 1 | 0 | 0 | 1698231 | YP_009724389.1 | P0DTD1 |
| TGVHAGTDLEGTFYGP | 1 | 0 | 0 | 1698232 | QHR63299.1 | P0C6W2 |
| TGVIADYNYKLPDDFM | 1 | 0 | 0 | 1698233 | NP_828851.1 | P59594 |
| TGVIADYNYKLPDDFT | 1 | 0 | 0 | 1698234 | QIQ54048.1 | P59594 |
| TGVLTTSKKQFLPFQQ | 1 | 0 | 0 | 1698241 | QIQ54048.1 | P59594 |
| TGVVLKRSLKYNLKAS | 1 | 0 | 0 | 1698260 | NP_073549.1 | A0A0P0LKV0 |
| TGYGVRRKNSEPEIPH | 1 | 0 | 0 | 1698270 | NP_073556.1 | A0A0N7J131 |
| TGYHFTKNGKTVLGEY | 1 | 0 | 0 | 1698272 | YP_009555238.1 | P0C6X6 |
| TGYHFTSTGKTVLGEY | 1 | 0 | 0 | 1698273 | YP_173236.1 | A0A140H1G9 |
| TGYHITKNSKVQLGEY | 1 | 0 | 0 | 1698274 | YP_009047202.1 | K9N7C7 |
| TGYREGYLNSTNVTIA | 1 | 0 | 0 | 1698280 | YP_009724389.1 | P0DTD1 |
| TGYREGYLNSTNVTTA | 1 | 0 | 0 | 1698281 | QHR63299.1 | P0C6W2 |
| TGYRFTNFEPFTVNSV | 1 | 0 | 0 | 1698282 | YP_009555241.1 | P36334 |
| TGYRVTKNSKTQIGEY | 1 | 0 | 0 | 1698283 | QIQ54047.1 | P0C6X7 |
| TGYRVTKNSKVQIGEY | 1 | 0 | 0 | 1698284 | YP_009724389.1 | P0DTD1 |
| TGYSLADLPAYMTVAK | 1 | 0 | 0 | 1698285 | YP_009555244.1 | Q01455 |
| TGYVDTENNTEFTRVN | 1 | 0 | 0 | 1698287 | NP_828849.2 | P0C6X7 |
| TGYVDTPNNTDFSRVS | 1 | 0 | 0 | 1698289 | YP_009724389.1 | P0DTD1 |
| THGFNLEEAARCMRSL | 1 | 0 | 0 | 1698309 | NP_828849.2 | P0C6X7 |
| THGIAAINSVPWSKIL | 1 | 0 | 0 | 1698310 | NP_828849.2 | P0C6X7 |
| THHDKFTDGVCLFWNC | 1 | 0 | 0 | 1698315 | NP_828849.2 | P0C6X7 |
| THLKHGGGIAGAINAA | 1 | 0 | 0 | 1698328 | YP_009047202.1 | K9N7C7 |
| THRDCDFNKPLIEWPL | 1 | 0 | 0 | 1698340 | YP_009047202.1 | K9N7C7 |
| THSDKFKDGVCLFWNC | 1 | 0 | 0 | 1698344 | QIQ54047.1 | P0C6X7 |
| THWFCANQSTSVYSAN | 1 | 0 | 0 | 1698360 | YP_003767.1 | A0A0P0G321 |
| TIALNKIQDVVNQQGS | 1 | 0 | 0 | 1698377 | YP_003767.1 | A0A0P0G321 |
| TIANNYSKLFDEGYTV | 1 | 0 | 0 | 1698379 | NP_073549.1 | A0A0P0LKV0 |
| TICDFYSKLGVGFTSH | 1 | 0 | 0 | 1698390 | YP_173236.1 | A0A140H1G9 |

| | | | | | | |
|---|---|---|---|---|---|---|
| TICGIKYVAQPTEDVV | 1 | 0 | 0 | 1698394 | YP_173236.1 | A0A140H1G9 |
| TICIKDADYNAKVEIS | 1 | 0 | 0 | 1698396 | NP_073549.1 | A0A0P0LKV0 |
| TIDGSSGVANPAMDPI | 1 | 0 | 0 | 1698401 | NP_828852.2 | P59632 |
| TIDGSSGVVNPAMDPI | 1 | 0 | 0 | 1698402 | QIQ54049.1 | P59632 |
| TIDGSSGVVNPAMEPI | 1 | 0 | 0 | 1698403 | QHR63301.1 | Q3LZX0 |
| TIDGSSGVVNPVMEPI | 1 | 0 | 0 | 1698404 | YP_009724391.1 | P0DTC3 |
| TIDKFAIPNGRKVDLQ | 1 | 0 | 0 | 1698406 | YP_009555241.1 | P36334 |
| TIDSNTTQDTYGGASV | 1 | 0 | 0 | 1698407 | NP_073549.1 | A0A0P0LKV0 |
| TIEDALRNSARLESAD | 1 | 0 | 0 | 1698412 | NP_073551.1 | A0A0P0K6L9 |
| TIEYPIIGDELKINAA | 1 | 0 | 0 | 1698417 | YP_009724389.1 | P0DTD1 |
| TIFKDAAGKPVPYCYD | 1 | 0 | 0 | 1698419 | QIQ54046.1 | P0C6U8 |
| TIFKDAMGKPVPYCYD | 1 | 0 | 0 | 1698420 | NP_828849.2 | P0C6X7 |
| TIFKDASGKPVPYCYD | 1 | 0 | 0 | 1698421 | YP_009724389.1 | P0DTD1 |
| TIFNCFYALNNAFLAF | 1 | 0 | 0 | 1698424 | YP_173241.1 | Q0ZJ82 |
| TIGNETTSAFVGALPK | 1 | 0 | 0 | 1698430 | NP_073551.1 | A0A0P0K6L9 |
| TIGVCTMTDIAKKPTE | 1 | 0 | 0 | 1698437 | NP_828849.2 | P0C6X7 |
| TIIFVQQKPEPVNPVS | 1 | 0 | 0 | 1698441 | NP_073549.1 | A0A0P0LKV0 |
| TIKAKEESVKSSYAFA | 1 | 0 | 0 | 1698452 | YP_003766.2 | H9EJ42 |
| TIKENIDGGAMHANYI | 1 | 0 | 0 | 1698455 | YP_009047202.1 | K9N7C7 |
| TIKFSGFKLDGFNHQF | 1 | 0 | 0 | 1698456 | YP_003766.2 | H9EJ42 |
| TIKPVSYKLDGVTYTE | 1 | 0 | 0 | 1698468 | NP_828849.2 | P0C6X7 |
| TIKTIQPRVEKKKLDG | 1 | 0 | 0 | 1698471 | QHR63299.1 | P0C6W2 |
| TIKTIQPRVEKKKLEG | 1 | 0 | 0 | 1698472 | QIQ54046.1 | P0C6U8 |
| TIKVFTTVDNINLHTQ | 1 | 0 | 0 | 1698473 | YP_009724389.1 | P0DTD1 |
| TIKVFTTVDNTNLHTQ | 1 | 0 | 0 | 1698474 | NP_828849.2 | P0C6X7 |
| TIKVSTVQSKLTDLKC | 1 | 0 | 0 | 1698475 | NP_073549.1 | A0A0P0LKV0 |
| TIKVTEDGVNVHDVTV | 1 | 0 | 0 | 1698476 | NP_073549.1 | A0A0P0LKV0 |
| TILEDEFTPFDVVRQC | 1 | 0 | 0 | 1698483 | NP_828849.2 | P0C6X7 |
| TILGSAILEDEFTPFD | 1 | 0 | 0 | 1698488 | QIQ54046.1 | P0C6U8 |
| TILGSALLEDEFTPFD | 1 | 0 | 0 | 1698489 | YP_009724389.1 | P0DTD1 |
| TILGSTILEDEFTPFD | 1 | 0 | 0 | 1698490 | NP_828849.2 | P0C6X7 |
| TILNIVGPDARQDGRQ | 1 | 0 | 0 | 1698493 | YP_173236.1 | A0A140H1G9 |
| TILNTACGVFEVDDTV | 1 | 0 | 0 | 1698496 | YP_009555238.1 | P0C6X6 |
| TIMKVLNENLNAYQQQ | 1 | 0 | 0 | 1698508 | YP_009555245.1 | P33469 |
| TINDDEDVVTGDNDDE | 1 | 0 | 0 | 1698519 | YP_173236.1 | A0A140H1G9 |
| TINKATFKPNTWCLRC | 1 | 0 | 0 | 1698523 | QIQ54046.1 | P0C6U8 |
| TINSVVQKDGVENTPP | 1 | 0 | 0 | 1698530 | YP_009047202.1 | K9N7C7 |
| TIQDSDGTNKQLNEIS | 1 | 0 | 0 | 1698546 | YP_009555238.1 | P0C6X6 |
| TIQKDIVECDVKTTEV | 1 | 0 | 0 | 1698553 | QIQ54046.1 | P0C6U8 |
| TIQKEVIECDVKTTEV | 1 | 0 | 0 | 1698555 | NP_828849.2 | P0C6X7 |
| TIQPRVEKKKLDGFMG | 1 | 0 | 0 | 1698558 | YP_009724389.1 | P0DTD1 |
| TIRLQAGKQTELAVNS | 1 | 0 | 0 | 1698570 | YP_003766.2 | H9EJ42 |
| TIRLQAGVATEYAANS | 1 | 0 | 0 | 1698571 | YP_173236.1 | A0A140H1G9 |
| TITQMNLKYAISAKNR | 1 | 0 | 0 | 1698596 | YP_009724389.1 | P0DTD1 |
| TIVDIPQSLTFSYDGL | 1 | 0 | 0 | 1698608 | YP_009047202.1 | K9N7C7 |
| TIVGQEEFIQTNSPKV | 1 | 0 | 0 | 1698615 | YP_173238.1 | A0A140H1H1 |
| TIVIPDKQVFDKVVDN | 1 | 0 | 0 | 1698617 | YP_173236.1 | A0A140H1G9 |
| TIVKCKGLNKPFVKDN | 1 | 0 | 0 | 1698618 | YP_009047202.1 | K9N7C7 |
| TIVPEYIDVNKTLQEL | 1 | 0 | 0 | 1698625 | NP_073551.1 | A0A0P0K6L9 |
| TIVSEKISVMDKLDTG | 1 | 0 | 0 | 1698627 | YP_003766.2 | H9EJ42 |
| TIYGNVSRQDLTKYTM | 1 | 0 | 0 | 1698643 | NP_073549.1 | A0A0P0LKV0 |
| TIYMCVVPSINDLSFD | 1 | 0 | 0 | 1698645 | YP_003766.2 | H9EJ42 |
| TIYNLLKDCPAVAKHD | 1 | 0 | 0 | 1698646 | YP_009724389.1 | P0DTD1 |
| TKAAGGKVSFSDDVEV | 1 | 0 | 0 | 1698654 | NP_073549.1 | A0A0P0LKV0 |
| TKAIDGENVSKEGFFT | 1 | 0 | 0 | 1698657 | NP_073549.1 | A0A0P0LKV0 |
| TKALCFDKSKQFVPVQ | 1 | 0 | 0 | 1698665 | YP_173237.1 | A0A140H1H0 |
| TKALKELYGPVDPTFL | 1 | 0 | 0 | 1698666 | YP_009047202.1 | K9N7C7 |
| TKAPLVYLNHSVPKLS | 1 | 0 | 0 | 1698668 | YP_173238.1 | A0A140H1H1 |
| TKAPYVMLNTSIPNLP | 1 | 0 | 0 | 1698669 | YP_009555241.1 | P36334 |
| TKCNIGGAVCSKHANL | 1 | 0 | 0 | 1698673 | NP_073549.1 | A0A0P0LKV0 |
| TKDEAIKRVRGWVGFD | 1 | 0 | 0 | 1698677 | YP_173236.1 | A0A140H1G9 |
| TKDSKFQVGEFVFEKV | 1 | 0 | 0 | 1698679 | NP_073549.1 | A0A0P0LKV0 |
| TKDSYCRLKNSVSDVA | 1 | 0 | 0 | 1698681 | YP_173236.1 | A0A140H1G9 |
| TKDVIECLKLSHHSDL | 1 | 0 | 0 | 1698682 | NP_828849.2 | P0C6X7 |
| TKEEPKKVKPCIEEVT | 1 | 0 | 0 | 1698686 | QIQ54046.1 | P0C6U8 |
| TKENDSKEGFFTYICG | 1 | 0 | 0 | 1698691 | YP_009724389.1 | P0DTD1 |
| TKENDSKEGFFTYLCG | 1 | 0 | 0 | 1698692 | NP_828849.2 | P0C6X7 |
| TKFADDLNQMSGYKKP | 1 | 0 | 0 | 1698698 | QIQ54046.1 | P0C6U8 |
| TKFADDLNQMTGFTKP | 1 | 0 | 0 | 1698699 | NP_828849.2 | P0C6X7 |
| TKFYGGWDNMLKNLMA | 1 | 0 | 0 | 1698715 | NP_073549.1 | A0A0P0LKV0 |
| TKGEGDGGITSEGNAL | 1 | 0 | 0 | 1698717 | NP_073549.1 | A0A0P0LKV0 |
| TKGKFKKNAWNIGEPK | 1 | 0 | 0 | 1698721 | QIQ54046.1 | P0C6U8 |
| TKGLSIADLACAQYYN | 1 | 0 | 0 | 1698724 | NP_073551.1 | A0A0P0K6L9 |
| TKHGDFADEIIGYKAI | 1 | 0 | 0 | 1698736 | QIQ54046.1 | P0C6U8 |
| TKHGSGALRELTRELN | 1 | 0 | 0 | 1698737 | NP_828849.2 | P0C6X7 |
| TKHGSGVTRELMRELN | 1 | 0 | 0 | 1698738 | QIQ54046.1 | P0C6U8 |
| TKHKCDINYKGKVFFQ | 1 | 0 | 0 | 1698739 | YP_009555238.1 | P0C6X6 |
| TKHSSGVTRDLMRELN | 1 | 0 | 0 | 1698741 | QHR63299.1 | P0C6W2 |
| TKHSSGVTRELMRELN | 1 | 0 | 0 | 1698742 | YP_009724389.1 | P0DTD1 |
| TKHVTKENDSKEGFFT | 1 | 0 | 0 | 1698745 | NP_828849.2 | P0C6X7 |
| TKHVTRENDSKEGFFT | 1 | 0 | 0 | 1698746 | QIQ54047.1 | P0C6X7 |
| TKKAFPYGTCLNCNAP | 1 | 0 | 0 | 1698758 | NP_073549.1 | A0A0P0LKV0 |
| TKKDWYDFVENPDIIN | 1 | 0 | 0 | 1698762 | YP_009555238.1 | P0C6X6 |
| TKKYTYSFLPKEDGDV | 1 | 0 | 0 | 1698771 | YP_009047202.1 | K9N7C7 |
| TKLAGGKISFSDDVIV | 1 | 0 | 0 | 1698773 | YP_003766.2 | H9EJ42 |
| TKLKDGVNFNVDDINF | 1 | 0 | 0 | 1698778 | YP_009555241.1 | P36334 |
| TKLNDLCFSNVYADSF | 1 | 0 | 0 | 1698784 | NP_828851.1 | P59594 |
| TKLNGILQQAALDYLA | 1 | 0 | 0 | 1698786 | YP_003766.2 | H9EJ42 |
| TKLSKYLANEAQVQLE | 1 | 0 | 0 | 1698793 | NP_073549.1 | A0A0P0LKV0 |
| TKLVTEEVVETPTIQK | 1 | 0 | 0 | 1698798 | QIQ54046.1 | P0C6U8 |
| TKMFDSFVNSFVSLYN | 1 | 0 | 0 | 1698805 | YP_009047202.1 | K9N7C7 |

| | | | | | | |
|---|---|---|---|---|---|---|
| TKMSEYKGPVTDVFYK | 1 | 0 | 0 | 1698807 | NP_828849.2 | P0C6X7 |
| TKNGKTVLGEYVFDKS | 1 | 0 | 0 | 1698808 | YP_009555238.1 | P0C6X6 |
| TKNIGDYNVSKDGFFT | 1 | 0 | 0 | 1698809 | YP_173236.1 | A0A140H1G9 |
| TKNIGEYNVSKDGFFT | 1 | 0 | 0 | 1698810 | YP_009555238.1 | P0C6X6 |
| TKNVTGSNESKALFFT | 1 | 0 | 0 | 1698814 | YP_009047202.1 | K9N7C7 |
| TKNVTKENDSKEGFFT | 1 | 0 | 0 | 1698815 | YP_009724389.1 | P0DTD1 |
| TKPIIKAQFKPFAKVD | 1 | 0 | 0 | 1698819 | YP_173236.1 | A0A140H1G9 |
| TKPLIEAEKPNLNSLC | 1 | 0 | 0 | 1698821 | YP_173243.1 | Q0ZJ80 |
| TKPVDTSNSFEVLAVE | 1 | 0 | 0 | 1698826 | NP_828849.2 | P0C6X7 |
| TKPVETSNIFDVLQSE | 1 | 0 | 0 | 1698827 | QIQ54046.1 | P0C6U8 |
| TKPVETSNSFDVLKSE | 1 | 0 | 0 | 1698828 | YP_009724389.1 | P0DTD1 |
| TKQYNVTQAFGRRGPE | 1 | 0 | 0 | 1698833 | QHR63308.1 | Q3LZX4 |
| TKRNILPTMTQLNLKY | 1 | 0 | 0 | 1698841 | NP_073549.1 | A0A0P0LKV0 |
| TKRNVIPTMTQMNLKY | 1 | 0 | 0 | 1698843 | YP_009047202.1 | K9N7C7 |
| TKRNVLPTLTQMNLKY | 1 | 0 | 0 | 1698844 | YP_009555238.1 | P0C6X6 |
| TKRNVLPTMTQLNLKY | 1 | 0 | 0 | 1698845 | YP_003766.2 | H9EJ42 |
| TKSGVNDICMFENKTT | 1 | 0 | 0 | 1698852 | YP_009047202.1 | K9N7C7 |
| TKSLLPNTGVYDLSGF | 1 | 0 | 0 | 1698853 | YP_173238.1 | A0A140H1H1 |
| TKSSEYKGPITDVFYK | 1 | 0 | 0 | 1698857 | YP_009724389.1 | P0DTD1 |
| TKSTNSRIRASLPTTI | 1 | 0 | 0 | 1698858 | NP_828849.2 | P0C6X7 |
| TKTDAVILLERYVSLA | 1 | 0 | 0 | 1698864 | NP_073549.1 | A0A0P0LKV0 |
| TKVDGIDVEIFENKTT | 1 | 0 | 0 | 1698871 | NP_828849.2 | P0C6X7 |
| TKVDGVDVELFENKTI | 1 | 0 | 0 | 1698872 | QIQ54047.1 | P0C6X7 |
| TKVDGVDVELFENKTT | 1 | 0 | 0 | 1698873 | YP_009724389.1 | P0DTD1 |
| TKVLAEKFDYNVSVMQ | 1 | 0 | 0 | 1698877 | NP_073549.1 | A0A0P0LKV0 |
| TKVVVGPTTEVKFSVI | 1 | 0 | 0 | 1698886 | YP_003766.2 | H9EJ42 |
| TKYLVQQESPFVMMSA | 1 | 0 | 0 | 1698888 | YP_009724389.1 | P0DTD1 |
| TKYNIYDVSGVGVIRV | 1 | 0 | 0 | 1698889 | NP_073551.1 | A0A0P0K6L9 |
| TKYNIYDYVGTGIIRS | 1 | 0 | 0 | 1698890 | YP_003767.1 | A0A0P0G321 |
| TKYTEVRASRQLAQQK | 1 | 0 | 0 | 1698891 | NP_073551.1 | A0A0P0K6L9 |
| TKYVAPQVTYQNISTN | 1 | 0 | 0 | 1698896 | YP_009047204.1 | K9N5Q8 |
| TKYYDDSQYYFNKDTG | 1 | 0 | 0 | 1698900 | YP_009555240.1 | P30215 |
| TLAATSASLFPPWTAA | 1 | 0 | 0 | 1698901 | YP_009555241.1 | P36334 |
| TLADAGFMKQYGECLG | 1 | 0 | 0 | 1698906 | NP_828851.1 | P59594 |
| TLAKTDLFNGYKIGCN | 1 | 0 | 0 | 1698909 | YP_173236.1 | A0A140H1G9 |
| TLANFNETKGPLCVDT | 1 | 0 | 0 | 1698912 | NP_073551.1 | A0A0P0K6L9 |
| TLDAANHSIPDLQRAA | 1 | 0 | 0 | 1698935 | NP_828849.2 | P0C6X7 |
| TLDDILQQANKSVGII | 1 | 0 | 0 | 1698938 | YP_009047202.1 | K9N7C7 |
| TLDDRGLALDNALSGV | 1 | 0 | 0 | 1698940 | NP_073549.1 | A0A0P0LKV0 |
| TLDESFLGRYMSALNH | 1 | 0 | 0 | 1698941 | NP_828849.2 | P0C6X7 |
| TLDGYSKLFITKDEAI | 1 | 0 | 0 | 1698943 | YP_173236.1 | A0A140H1G9 |
| TLDKCTKYNIYDVSGV | 1 | 0 | 0 | 1698944 | NP_073551.1 | A0A0P0K6L9 |
| TLDNQDLNGKWYDFGD | 1 | 0 | 0 | 1698949 | YP_009047202.1 | K9N7C7 |
| TLENKSAELNYTVQKL | 1 | 0 | 0 | 1698960 | NP_073551.1 | A0A0P0K6L9 |
| TLESTINKSVANLKLP | 1 | 0 | 0 | 1698968 | YP_003768.1 | H9EJ24 |
| TLETAQNSVRALQKAA | 1 | 0 | 0 | 1698970 | QHR63299.1 | P0C6W2 |
| TLFGDCVTVRVGQQVD | 1 | 0 | 0 | 1698981 | YP_009047202.1 | K9N7C7 |
| TLGNVEAVNFNVTTAE | 1 | 0 | 0 | 1698997 | NP_073551.1 | A0A0P0K6L9 |
| TLGTVTLKQGEIKDAT | 1 | 0 | 0 | 1698998 | QHR63301.1 | Q3LZX0 |
| TLIATLNSLGETLVTM | 1 | 0 | 0 | 1699012 | QIQ54046.1 | P0C6U8 |
| TLIDNINSTLVDLKWL | 1 | 0 | 0 | 1699016 | NP_073551.1 | A0A0P0K6L9 |
| TLIGDCATVHTANKWD | 1 | 0 | 0 | 1699019 | YP_009724389.1 | P0DTD1 |
| TLKATEETFKLSYGIA | 1 | 0 | 0 | 1699032 | YP_009724389.1 | P0DTD1 |
| TLKEILVTYNCCDDDY | 1 | 0 | 0 | 1699035 | YP_009724389.1 | P0DTD1 |
| TLKEVVGPKELLLSWE | 1 | 0 | 0 | 1699038 | YP_003766.2 | H9EJ42 |
| TLKGGAPTKVTFGDDT | 1 | 0 | 0 | 1699040 | YP_009724389.1 | P0DTD1 |
| TLKPGDSFNILACYEG | 1 | 0 | 0 | 1699057 | YP_003766.2 | H9EJ42 |
| TLKQGEIKDATPSDFV | 1 | 0 | 0 | 1699059 | YP_009724391.1 | P0DTC3 |
| TLKQGEIKDATPSDSV | 1 | 0 | 0 | 1699060 | QHR63301.1 | Q3LZX0 |
| TLKSLTVEKGIYQTSN | 1 | 0 | 0 | 1699063 | QIQ54048.1 | P59594 |
| TLLALHRSYLTPGDSS | 1 | 0 | 0 | 1699067 | YP_009724390.1 | P0DTC2 |
| TLLTKGTLEPEYFNSV | 1 | 0 | 0 | 1699095 | YP_009724389.1 | P0DTD1 |
| TLLVDSDLNDFVSDAD | 1 | 0 | 0 | 1699098 | YP_009724389.1 | P0DTD1 |
| TLMQGVTLSSNLNTNL | 1 | 0 | 0 | 1699109 | YP_173238.1 | A0A140H1H1 |
| TLMTFSGNVDLDALAS | 1 | 0 | 0 | 1699111 | YP_009047202.1 | K9N7C7 |
| TLNDFNLVAMKFNYEP | 1 | 0 | 0 | 1699114 | QIQ54046.1 | P0C6U8 |
| TLNDFNLVAMKYNYEP | 1 | 0 | 0 | 1699115 | YP_009724389.1 | P0DTD1 |
| TLNDLNETLVTMPLGY | 1 | 0 | 0 | 1699116 | YP_009724389.1 | P0DTD1 |
| TLNDVKDNDGRPVHVK | 1 | 0 | 0 | 1699117 | YP_003766.2 | H9EJ42 |
| TLNEDLLEILSRERVN | 1 | 0 | 0 | 1699118 | NP_828849.2 | P0C6X7 |
| TLNIIVPDKSVYDQVV | 1 | 0 | 0 | 1699125 | YP_009555238.1 | P0C6X6 |
| TLNISDEFSSNVANYQ | 1 | 0 | 0 | 1699126 | YP_009724389.1 | P0DTD1 |
| TLNLEGVNGGSLYVNN | 1 | 0 | 0 | 1699127 | NP_073549.1 | A0A0P0LKV0 |
| TLNSLGETLVTMPLGY | 1 | 0 | 0 | 1699129 | QIQ54046.1 | P0C6U8 |
| TLPTEVLSEEVILKTG | 1 | 0 | 0 | 1699139 | QIQ54046.1 | P0C6U8 |
| TLPTEVLTEEVVLKTG | 1 | 0 | 0 | 1699140 | YP_009724389.1 | P0DTD1 |
| TLQEFAQNLPKYVKPN | 1 | 0 | 0 | 1699154 | YP_003767.1 | A0A0P0G321 |
| TLQELSYKLPNYTVPD | 1 | 0 | 0 | 1699155 | NP_073551.1 | A0A0P0K6L9 |
| TLQEMKALFPTWSQEL | 1 | 0 | 0 | 1699156 | YP_009555238.1 | P0C6X6 |
| TLQETPVVPDTVEVPP | 1 | 0 | 0 | 1699157 | YP_009047202.1 | K9N7C7 |
| TLQLVSAVNQSLSNAK | 1 | 0 | 0 | 1699161 | NP_073552.1 | A0A0P0KWE8 |
| TLQPNAYYKYDAKNYV | 1 | 0 | 0 | 1699163 | YP_003766.2 | H9EJ42 |
| TLQPVSELLTPLGIDL | 1 | 0 | 0 | 1699166 | YP_009724389.1 | P0DTD1 |
| TLQQIELKFNPPALQD | 1 | 0 | 0 | 1699167 | YP_009724389.1 | P0DTD1 |
| TLQSLENVIYNLVNVG | 1 | 0 | 0 | 1699169 | YP_173236.1 | A0A140H1G9 |
| TLRCCTVTYLNELSSK | 1 | 0 | 0 | 1699175 | YP_003766.2 | H9EJ42 |
| TLREILVTYNCCDDDY | 1 | 0 | 0 | 1699176 | QHR63299.1 | P0C6W2 |
| TLRNGSVLEMAKEVKT | 1 | 0 | 0 | 1699182 | NP_073549.1 | A0A0P0LKV0 |
| TLRSITAQPVKIDNAS | 1 | 0 | 0 | 1699189 | NP_828852.2 | P59632 |
| TLSEQLDFIDTKRGVY | 1 | 0 | 0 | 1699194 | YP_009724389.1 | P0DTD1 |
| TLSEQLDYIESKRGVY | 1 | 0 | 0 | 1699195 | NP_828849.2 | P0C6X7 |

| | | | | | | |
|---|---|---|---|---|---|---|
| TLSSNLNTNLHFDVDN | 1 | 0 | 0 | 1699201 | YP_173238.1 | A0A140H1H1 |
| TLSTKLKDGVNFNVDD | 1 | 0 | 0 | 1699205 | YP_009555241.1 | P36334 |
| TLSYEQFKKGVQIPCT | 1 | 0 | 0 | 1699212 | YP_009724389.1 | P0DTD1 |
| TLSYEQLKKGVQIPCT | 1 | 0 | 0 | 1699213 | QHR63299.1 | P0C6W2 |
| TLTAPTIVNQERYVKI | 1 | 0 | 0 | 1699216 | YP_009047202.1 | K9N7C7 |
| TLTKLSKYLANEAQVQ | 1 | 0 | 0 | 1699221 | NP_073549.1 | A0A0P0LKV0 |
| TLTPRSVRSVPGEMRL | 1 | 0 | 0 | 1699228 | YP_009047204.1 | K9N5Q8 |
| TLTQMNLKYAISAKNR | 1 | 0 | 0 | 1699229 | YP_009555238.1 | P0C6X6 |
| TLVDLKWLNRVETYIK | 1 | 0 | 0 | 1699242 | NP_073551.1 | A0A0P0K6L9 |
| TLVKDKCVNFNFNGLT | 1 | 0 | 0 | 1699247 | QIQ54048.1 | P59594 |
| TLVNDIDITFLKKDAP | 1 | 0 | 0 | 1699256 | QHR63299.1 | P0C6W2 |
| TLVNKMVNWFNTMLDA | 1 | 0 | 0 | 1699258 | YP_173236.1 | A0A140H1G9 |
| TLVSDIDITFLKKDAP | 1 | 0 | 0 | 1699263 | YP_009724389.1 | P0DTD1 |
| TLVSGDAHIPFGTVLG | 1 | 0 | 0 | 1699264 | NP_073551.1 | A0A0P0K6L9 |
| TLYIKDGDDGYFLPYP | 1 | 0 | 0 | 1699277 | YP_009047202.1 | K9N7C7 |
| TMFTMADGSPQPYCYT | 1 | 0 | 0 | 1699317 | YP_009555238.1 | P0C6X6 |
| TMGVYDFKVSAAEFKY | 1 | 0 | 0 | 1699320 | YP_003766.2 | H9EJ42 |
| TMIEFMLWCKDGQVQT | 1 | 0 | 0 | 1699327 | YP_009047202.1 | K9N7C7 |
| TMLDIGQARVAYQVAA | 1 | 0 | 0 | 1699332 | NP_073549.1 | A0A0P0LKV0 |
| TMLDLCYALRHFDRND | 1 | 0 | 0 | 1699334 | YP_009555238.1 | P0C6X6 |
| TMLFGMIKKLDNDVLN | 1 | 0 | 0 | 1699336 | YP_009047202.1 | K9N7C7 |
| TMLFSMVRKLDNQALN | 1 | 0 | 0 | 1699337 | YP_009555238.1 | P0C6X6 |
| TMLFTMLRKLDNDALN | 1 | 0 | 0 | 1699338 | YP_009724389.1 | P0DTD1 |
| TMLRKLDNDALNNIIN | 1 | 0 | 0 | 1699339 | YP_009724389.1 | P0DTD1 |
| TMLVKMDGDDVYLPYP | 1 | 0 | 0 | 1699341 | YP_009555238.1 | P0C6X6 |
| TMMDLCFALRNFDEKD | 1 | 0 | 0 | 1699343 | NP_073549.1 | A0A0P0LKV0 |
| TMPIGYVTHGFNLEEA | 1 | 0 | 0 | 1699349 | NP_828849.2 | P0C6X7 |
| TMQIVDENGKYYLPYP | 1 | 0 | 0 | 1699353 | NP_073549.1 | A0A0P0LKV0 |
| TMQIVDKDGTYYLPYP | 1 | 0 | 0 | 1699354 | YP_003766.2 | H9EJ42 |
| TMSLGADSSIAYSNNT | 1 | 0 | 0 | 1699360 | NP_828851.1 | P59594 |
| TMTDIAKKPTESACSS | 1 | 0 | 0 | 1699367 | NP_828849.2 | P0C6X7 |
| TMTQLNLKYAISGKER | 1 | 0 | 0 | 1699374 | NP_073549.1 | A0A0P0LKV0 |
| TMTQMNLKYAISAKNR | 1 | 0 | 0 | 1699375 | YP_009047202.1 | K9N7C7 |
| TMVQTCTPNVTINCQD | 1 | 0 | 0 | 1699382 | NP_849177.1 | Q80H93 |
| TMYVRPIIEDYHTLTV | 1 | 0 | 0 | 1699390 | YP_009555244.1 | Q01455 |
| TNAKPPKYSSKVVLSG | 1 | 0 | 0 | 1699393 | YP_003766.2 | H9EJ42 |
| TNANASSSEGFLIGIN | 1 | 0 | 0 | 1699394 | YP_009555238.1 | P0C6X6 |
| TNANASSSEGFLLGIN | 1 | 0 | 0 | 1699395 | YP_009047202.1 | K9N7C7 |
| TNASNQVAVLYQDVNC | 1 | 0 | 0 | 1699396 | QHR63300.2 | Q3LZX1 |
| TNCCRFQEKDEDGNLI | 1 | 0 | 0 | 1699404 | QIQ54047.1 | P0C6X7 |
| TNCCRFQEKDEEGNLL | 1 | 0 | 0 | 1699405 | NP_828849.2 | P0C6X7 |
| TNCVKMLCTHTGTGQA | 1 | 0 | 0 | 1699414 | YP_009724389.1 | P0DTD1 |
| TNDLYVALRGRIDKDL | 1 | 0 | 0 | 1699424 | NP_073553.1 | A0A0P0KWE8 |
| TNEAFQKVQDAVNNNA | 1 | 0 | 0 | 1699434 | YP_009047204.1 | K9N5Q8 |
| TNEIGLNASVTLKICK | 1 | 0 | 0 | 1699435 | YP_003767.1 | A0A0P0G321 |
| TNEIIGYKAIQDGVTR | 1 | 0 | 0 | 1699436 | NP_828849.2 | P0C6X7 |
| TNESYNNYVPSYVKPD | 1 | 0 | 0 | 1699439 | YP_009047202.1 | K9N7C7 |
| TNFAINDLNTTVPRIS | 1 | 0 | 0 | 1699440 | YP_173238.1 | A0A140H1H1 |
| TNFEPFTVNSVNDSLE | 1 | 0 | 0 | 1699441 | YP_009555241.1 | P36334 |
| TNFTIVGQEEFIQTNS | 1 | 0 | 0 | 1699455 | YP_173238.1 | A0A140H1H1 |
| TNGIKRFDNPVLPFND | 1 | 0 | 0 | 1699462 | QHR63300.2 | Q3LZX1 |
| TNGSGSGQAITCTIDS | 1 | 0 | 0 | 1699469 | NP_073549.1 | A0A0P0LKV0 |
| TNGSWAIFNDHHLNRP | 1 | 0 | 0 | 1699472 | YP_009047202.1 | K9N7C7 |
| TNGTKRFDNPVLPFND | 1 | 0 | 0 | 1699473 | YP_009724390.1 | P0DTC2 |
| TNHDRYCSVHQGAHVA | 1 | 0 | 0 | 1699478 | YP_009047202.1 | K9N7C7 |
| TNHHAICNVHRNEHVA | 1 | 0 | 0 | 1699481 | NP_073549.1 | A0A0P0LKV0 |
| TNHSFSVQKHIGAPAN | 1 | 0 | 0 | 1699484 | YP_009047202.1 | K9N7C7 |
| TNIAFELFAKRKVGLT | 1 | 0 | 0 | 1699485 | NP_073549.1 | A0A0P0LKV0 |
| TNIFGTVYEKLKPILD | 1 | 0 | 0 | 1699489 | QIQ54046.1 | P0C6U8 |
| TNIQNLLYCDSPFEKL | 1 | 0 | 0 | 1699495 | YP_003767.1 | A0A0P0G321 |
| TNIYSQNLCVDGFGVN | 1 | 0 | 0 | 1699499 | NP_073549.1 | A0A0P0LKV0 |
| TNKQLNEISDDCNWPL | 1 | 0 | 0 | 1699504 | YP_009555238.1 | P0C6X6 |
| TNLCPFGEVFNASKFA | 1 | 0 | 0 | 1699507 | QIQ54048.1 | P59594 |
| TNLCPFGEVFNATKFP | 1 | 0 | 0 | 1699508 | NP_828851.1 | P59594 |
| TNLHFDVDNINFKSLV | 1 | 0 | 0 | 1699518 | YP_173238.1 | A0A140H1H1 |
| TNLHTQLVDMSMTYGQ | 1 | 0 | 0 | 1699519 | NP_828849.2 | P0C6X7 |
| TNLSEITKPLIEAEKP | 1 | 0 | 0 | 1699529 | YP_173243.1 | Q0ZJ80 |
| TNLSRGRGRNPKPRAA | 1 | 0 | 0 | 1699530 | YP_009047211.1 | K9N4V7 |
| TNNAMQIESDDYIARN | 1 | 0 | 0 | 1699539 | QIQ54046.1 | P0C6U8 |
| TNNTLVSGDAHIPFGT | 1 | 0 | 0 | 1699548 | NP_073551.1 | A0A0P0K6L9 |
| TNNTRIVDEWSYTGSS | 1 | 0 | 0 | 1699549 | YP_009047204.1 | K9N5Q8 |
| TNPFTRTVFENLKPMP | 1 | 0 | 0 | 1699553 | YP_173236.1 | A0A140H1G9 |
| TNPIQLSSYSLFDMSK | 1 | 0 | 0 | 1699554 | YP_009724389.1 | P0DTD1 |
| TNQVVKPGHFNEEFYN | 1 | 0 | 0 | 1699557 | YP_003766.2 | H9EJ42 |
| TNRQSGRQPTPLSPP | 1 | 0 | 0 | 1699561 | P03138.3 | Q76R62 |
| TNSALAKIQSVVNSNA | 1 | 0 | 0 | 1699564 | YP_173238.1 | A0A140H1H1 |
| TNSALVKIQAVVNANA | 1 | 0 | 0 | 1699565 | YP_009555241.1 | P36334 |
| TNSCLVSSDGQPGGDA | 1 | 0 | 0 | 1699566 | YP_009047202.1 | K9N7C7 |
| TNSILMLDKQGQLLDT | 1 | 0 | 0 | 1699571 | YP_003766.2 | H9EJ42 |
| TNSQYLFSTINALPET | 1 | 0 | 0 | 1699574 | YP_009047202.1 | K9N7C7 |
| TNSTDTVNHDSFNLYG | 1 | 0 | 0 | 1699580 | YP_173236.1 | A0A140H1G9 |
| TNSVCPKLEFANDTKI | 1 | 0 | 0 | 1699582 | YP_009047204.1 | K9N5Q8 |
| TNTGSVVKPVRARAPP | 1 | 0 | 0 | 1699586 | NP_073549.1 | A0A0P0LKV0 |
| TNTNLSRGRGRNPKPR | 1 | 0 | 0 | 1699588 | YP_009047211.1 | K9N4V7 |
| TNTSNQVAVLYQDVNC | 1 | 0 | 0 | 1699589 | YP_009724390.1 | P0DTC2 |
| TNTSSVVDGVVRSFQP | 1 | 0 | 0 | 1699590 | NP_073551.1 | A0A0P0K6L9 |
| TNVAFELFAKRKMGLT | 1 | 0 | 0 | 1699591 | YP_003766.2 | H9EJ42 |
| TNVNASSSEAFLIGAN | 1 | 0 | 0 | 1699597 | NP_828849.2 | P0C6X7 |
| TNVVLLGCLSSMNIAA | 1 | 0 | 0 | 1699611 | YP_003766.2 | H9EJ42 |
| TNVVPPALHLVDPQIQ | 1 | 0 | 0 | 1699613 | NP_828859.1 | P59636 |
| TNYVGVYDINHGDAGF | 1 | 0 | 0 | 1699634 | YP_009555240.1 | P30215 |

| | | | | | | |
|---|---|---|---|---|---|---|
| TPAFDKSAFTNLKQLP | 1 | 0 | 0 | 1699636 | NP_828849.2 | P0C6X7 |
| TPAFDKSAFVNLKQLP | 1 | 0 | 0 | 1699637 | YP_009724389.1 | P0DTD1 |
| TPAKKLCVVPGNVVPG | 1 | 0 | 0 | 1699638 | YP_003766.2 | H9EJ42 |
| TPAPVYIWTADEAIKF | 1 | 0 | 0 | 1699639 | YP_009555244.1 | Q01455 |
| TPAQNAGYWRRQDRKI | 1 | 0 | 0 | 1699640 | YP_009047211.1 | K9N4V7 |
| TPATDCSDGNYNRNAS | 1 | 0 | 0 | 1699642 | YP_009047204.1 | K9N5Q8 |
| TPAWRIYSTGNNVFQT | 1 | 0 | 0 | 1699643 | NP_828851.1 | P59594 |
| TPCEVHGNGSMIEDID | 1 | 0 | 0 | 1699649 | YP_009047202.1 | K9N7C7 |
| TPCGYSEPPKVAAKIC | 1 | 0 | 0 | 1699651 | YP_009555238.1 | P0C6X6 |
| TPCNQPDQVAVYQQSI | 1 | 0 | 0 | 1699656 | YP_003767.1 | A0A0P0G321 |
| TPCVYVDGLESKQVDY | 1 | 0 | 0 | 1699660 | YP_173236.1 | A0A140H1G9 |
| TPEVSRELGNITKTNV | 1 | 0 | 0 | 1699670 | NP_073549.1 | A0A0P0LKV0 |
| TPFHYERDNTSCPEWM | 1 | 0 | 0 | 1699676 | YP_009047202.1 | K9N7C7 |
| TPGAFFFGSKLDLVKR | 1 | 0 | 0 | 1699681 | YP_173242.1 | Q19U29 |
| TPGHYAGSRSSSGNRS | 1 | 0 | 0 | 1699683 | YP_173242.1 | Q19U29 |
| TPGKQSSSRASSGNRS | 1 | 0 | 0 | 1699685 | YP_009555245.1 | P33469 |
| TPGSSRGNSPARMASG | 1 | 0 | 0 | 1699688 | NP_828858.1 | P59595 |
| TPGSSRGTSPARMAGN | 1 | 0 | 0 | 1699690 | YP_009724397.2 | P0DTC9 |
| TPGTNASNQVAVLYQD | 1 | 0 | 0 | 1699691 | QHR63300.2 | Q3LZX1 |
| TPGTNTSNQVAVLYQD | 1 | 0 | 0 | 1699693 | YP_009724390.1 | P0DTC2 |
| TPHLMGWDYPKCDRAM | 1 | 0 | 0 | 1699697 | NP_828849.2 | P0C6X7 |
| TPHPYCYSDGVMKNAS | 1 | 0 | 0 | 1699698 | YP_173236.1 | A0A140H1G9 |
| TPIAYRNVLLRKNGNK | 1 | 0 | 0 | 1699702 | NP_828849.2 | P0C6X7 |
| TPIDLVRDLPRGFAAL | 1 | 0 | 0 | 1699704 | QIQ54048.1 | P59594 |
| TPIKLNFGMLNGMPVS | 1 | 0 | 0 | 1699706 | YP_003766.2 | H9EJ42 |
| TPIKNTVDTTPKEEFV | 1 | 0 | 0 | 1699707 | NP_073549.1 | A0A0P0LKV0 |
| TPILSCGIFGIKLETS | 1 | 0 | 0 | 1699709 | NP_073549.1 | A0A0P0LKV0 |
| TPINLVRDLPPGFSAL | 1 | 0 | 0 | 1699710 | QHR63300.2 | Q3LZX1 |
| TPIVVDCSTYVCNGNV | 1 | 0 | 0 | 1699714 | NP_073551.1 | A0A0P0K6L9 |
| TPIVWHAKDFNSLSAE | 1 | 0 | 0 | 1699716 | NP_073549.1 | A0A0P0LKV0 |
| TPKDHIGTRNPNNNAA | 1 | 0 | 0 | 1699718 | NP_828858.1 | P59595 |
| TPKEEFVVKEKLNAFL | 1 | 0 | 0 | 1699719 | NP_073549.1 | A0A0P0LKV0 |
| TPKHVFKTLKPGDSFN | 1 | 0 | 0 | 1699721 | YP_003766.2 | H9EJ42 |
| TPKKPNKPLSQPRADK | 1 | 0 | 0 | 1699723 | YP_003771.1 | H9EJ69 |
| TPKLKEDYQIGGYSED | 1 | 0 | 0 | 1699724 | NP_828852.2 | P59632 |
| TPKYTFGNVKPGETFT | 1 | 0 | 0 | 1699728 | YP_173236.1 | A0A140H1G9 |
| TPLTSRQYLLAFNQDG | 1 | 0 | 0 | 1699742 | YP_009555241.1 | P36334 |
| TPNATEFSRVSAKPPP | 1 | 0 | 0 | 1699750 | QIQ54047.1 | P0C6X7 |
| TPNKHCNVQQCFGKRG | 1 | 0 | 0 | 1699753 | YP_173242.1 | Q19U29 |
| TPNNTDFSRVSAKPPP | 1 | 0 | 0 | 1699754 | YP_009724389.1 | P0DTD1 |
| TPNVTINCQDPAGGAL | 1 | 0 | 0 | 1699757 | NP_849177.1 | Q80H93 |
| TPPIKDFGGFNFLQIL | 1 | 0 | 0 | 1699761 | QIQ54048.1 | P59594 |
| TPQKLFVANYSQDVKQ | 1 | 0 | 0 | 1699765 | YP_009047204.1 | K9N5Q8 |
| TPQPEEELEEDWLDDD | 1 | 0 | 0 | 1699766 | QHR63299.1 | P0C6W2 |
| TPRSDMEYDFLNMDMG | 1 | 0 | 0 | 1699771 | YP_003766.2 | H9EJ42 |
| TPRSVRSVPGEMRLAS | 1 | 0 | 0 | 1699772 | YP_009047204.1 | K9N5Q8 |
| TPSDVSSRDPTTQEAI | 1 | 0 | 0 | 1699776 | YP_173242.1 | Q19U29 |
| TPSDVYQQLAGIKLQS | 1 | 0 | 0 | 1699777 | YP_009555238.1 | P0C6X6 |
| TPSKGPAQASVNGVTL | 1 | 0 | 0 | 1699781 | NP_828849.2 | P0C6X7 |
| TPSTLTPRSVRSVPGE | 1 | 0 | 0 | 1699786 | YP_009047204.1 | K9N5Q8 |
| TPTESYVDVGPDSVKS | 1 | 0 | 0 | 1699790 | YP_009047204.1 | K9N5Q8 |
| TPTKKAFPYGTCLNCN | 1 | 0 | 0 | 1699793 | NP_073549.1 | A0A0P0LKV0 |
| TPTQCYYNNSNNGKIV | 1 | 0 | 0 | 1699795 | YP_009555238.1 | P0C6X6 |
| TPVATLIATLNSLGET | 1 | 0 | 0 | 1699797 | QIQ54046.1 | P0C6U8 |
| TPVAYRKILLRKNGNK | 1 | 0 | 0 | 1699798 | QIQ54046.1 | P0C6U8 |
| TPVHVLTKHGDFADEI | 1 | 0 | 0 | 1699805 | QIQ54046.1 | P0C6U8 |
| TPVQVVDSRWNNARQS | 1 | 0 | 0 | 1699810 | YP_009555240.1 | P30215 |
| TPVVEYLSKEDLHTLY | 1 | 0 | 0 | 1699812 | YP_009047202.1 | K9N7C7 |
| TPVVPDTVEVPPQVVK | 1 | 0 | 0 | 1699813 | YP_009047202.1 | K9N7C7 |
| TPVVQTEVNSFSGYLK | 1 | 0 | 0 | 1699814 | QHR63299.1 | P0C6W2 |
| TPVVQTIEVNSFSGYL | 1 | 0 | 0 | 1699815 | YP_009724389.1 | P0DTD1 |
| TPVVYRKVLLRKNGNK | 1 | 0 | 0 | 1699816 | QHR63299.1 | P0C6W2 |
| TPYHAFKCNRSGSNDI | 1 | 0 | 0 | 1699821 | YP_009555240.1 | P30215 |
| TPYVCNASGCGVSDVK | 1 | 0 | 0 | 1699822 | YP_003766.2 | H9EJ42 |
| TPYVCNTSGCNVNDVT | 1 | 0 | 0 | 1699823 | NP_073549.1 | A0A0P0LKV0 |
| TQCYYNNSNNGKIVYA | 1 | 0 | 0 | 1699834 | YP_009555238.1 | P0C6X6 |
| TQDGDNKLQGLLEVSV | 1 | 0 | 0 | 1699835 | YP_009555241.1 | P36334 |
| TQDGNAAITDYNYYKY | 1 | 0 | 0 | 1699836 | YP_009555238.1 | P0C6X6 |
| TQDHVDILGPLSAQTG | 1 | 0 | 0 | 1699837 | YP_009724389.1 | P0DTD1 |
| TQEFRFMTANNLTAPR | 1 | 0 | 0 | 1699843 | YP_009047202.1 | K9N7C7 |
| TQEFRYMNSQGLLPPK | 1 | 0 | 0 | 1699844 | YP_009724389.1 | P0DTD1 |
| TQEYIQTTIQKVTVDC | 1 | 0 | 0 | 1699846 | YP_009047204.1 | K9N5Q8 |
| TQFAFACPDGVKHVYQ | 1 | 0 | 0 | 1699847 | YP_009724395.1 | P0DTC7 |
| TQFNYFKKVDGIIQQL | 1 | 0 | 0 | 1699850 | NP_828849.2 | P0C6X7 |
| TQFNYYKKVDGVVQQL | 1 | 0 | 0 | 1699851 | YP_009724389.1 | P0DTD1 |
| TQFNYYKKVNGVVQQL | 1 | 0 | 0 | 1699852 | QHR63299.1 | P0C6W2 |
| TQFQKGKEFEFVEGQG | 1 | 0 | 0 | 1699853 | YP_009555245.1 | P33469 |
| TQGNFGDQELIRQGTE | 1 | 0 | 0 | 1699861 | QIQ54056.1 | P59595 |
| TQGRNLQDFLPRSTME | 1 | 0 | 0 | 1699862 | NP_073549.1 | A0A0P0LKV0 |
| TQIPATSIVAKQGAGD | 1 | 0 | 0 | 1699877 | NP_073549.1 | A0A0P0LKV0 |
| TQKGDAAIKDFDYYRY | 1 | 0 | 0 | 1699881 | NP_073549.1 | A0A0P0LKV0 |
| TQKGSGMDTALLRNNI | 1 | 0 | 0 | 1699882 | YP_009555244.1 | Q01455 |
| TQLGIEFLKRGDKSVY | 1 | 0 | 0 | 1699889 | YP_009724389.1 | P0DTD1 |
| TQLHVADTLMQGVTLS | 1 | 0 | 0 | 1699890 | YP_173238.1 | A0A140H1H1 |
| TQLNRALSGIAAEQDR | 1 | 0 | 0 | 1699892 | NP_828851.1 | P59594 |
| TQLPPAYTNSSTRGVY | 1 | 0 | 0 | 1699894 | QHR63300.2 | Q3LZX1 |
| TQMNLKYAISAKNRAR | 1 | 0 | 0 | 1699902 | YP_009724389.1 | P0DTD1 |
| TQMYKEARSVNRKSKV | 1 | 0 | 0 | 1699903 | YP_003766.2 | H9EJ42 |
| TQMYKQARSEDKRAKV | 1 | 0 | 0 | 1699904 | YP_009724389.1 | P0DTD1 |
| TQNVLYENQKQIANQF | 1 | 0 | 0 | 1699908 | NP_828851.1 | P59594 |
| TQNVTVKPKRLRKKRN | 1 | 0 | 0 | 1699909 | YP_009047202.1 | K9N7C7 |

| | | | | | | |
|---|---|---|---|---|---|---|
| TQPKFTVSTQPQGNTI | 1 | 0 | 0 | 1699911 | YP_173242.1 | Q19U29 |
| TQPLPNASFDNFKLTC | 1 | 0 | 0 | 1699912 | NP_828849.2 | P0C6X7 |
| TQQVLSENQKLIANKF | 1 | 0 | 0 | 1699920 | YP_009047204.1 | K9N5Q8 |
| TQRMCAIGPDVFLHKC | 1 | 0 | 0 | 1699922 | NP_073549.1 | A0A0P0LKV0 |
| TQRNFYEPQIITTDNT | 1 | 0 | 0 | 1699925 | YP_009724390.1 | P0DTC2 |
| TQSRDLEDFKPRSQME | 1 | 0 | 0 | 1699930 | NP_828849.2 | P0C6X7 |
| TQTSQALQTVATALNK | 1 | 0 | 0 | 1699949 | NP_073551.1 | A0A0P0K6L9 |
| TQTVDSSQGSEYQYVI | 1 | 0 | 0 | 1699953 | YP_009047202.1 | K9N7C7 |
| TQWYHEHVGGVYDNSI | 1 | 0 | 0 | 1699965 | YP_009047202.1 | K9N7C7 |
| TQYEYGTESDYKGLPL | 1 | 0 | 0 | 1699969 | QIQ54046.1 | P0C6U8 |
| TRAGCLVGAEHVNNSY | 1 | 0 | 0 | 1699975 | QIQ54048.1 | P59594 |
| TRCNIGGAVCSKHANL | 1 | 0 | 0 | 1699995 | YP_003766.2 | H9EJ42 |
| TRDAVGTNLPLQLGFS | 1 | 0 | 0 | 1699999 | NP_828849.2 | P0C6X7 |
| TRDFAIRNVRGWLGMD | 1 | 0 | 0 | 1700000 | YP_003766.2 | H9EJ42 |
| TRDFAMRHVRGWLGMD | 1 | 0 | 0 | 1700001 | NP_073549.1 | A0A0P0LKV0 |
| TRDIASTDTCFANKHA | 1 | 0 | 0 | 1700002 | YP_009724389.1 | P0DTD1 |
| TRDIISTDDCFANKHA | 1 | 0 | 0 | 1700003 | NP_828849.2 | P0C6X7 |
| TRDISSNDNCFANKHV | 1 | 0 | 0 | 1700004 | QIQ54046.1 | P0C6U8 |
| TRDKLEKLISTARDGV | 1 | 0 | 0 | 1700005 | YP_009047202.1 | K9N7C7 |
| TRDLMRELNGGAYTRY | 1 | 0 | 0 | 1700006 | QHR63299.1 | P0C6W2 |
| TRDRFYRLANECAQVL | 1 | 0 | 0 | 1700007 | YP_009047202.1 | K9N7C7 |
| TRDSIGTNFPLQLGFS | 1 | 0 | 0 | 1700008 | YP_009555238.1 | P0C6X6 |
| TRDSVPLHIIAPSSLI | 1 | 0 | 0 | 1700009 | YP_009047208.1 | K9N7D2 |
| TRDVGFVVPGLPGTIF | 1 | 0 | 0 | 1700010 | QIQ54046.1 | P0C6U8 |
| TREAIGTNLPLQLGFS | 1 | 0 | 0 | 1700011 | QHR63299.1 | P0C6W2 |
| TREAVGTNLPLQLGFS | 1 | 0 | 0 | 1700012 | YP_009724389.1 | P0DTD1 |
| TREEAIKHVRAWVGFD | 1 | 0 | 0 | 1700013 | QIQ54047.1 | P0C6X7 |
| TREEAVRHVRAWIGFD | 1 | 0 | 0 | 1700015 | QHR63299.1 | P0C6W2 |
| TREEAVRQVRSWIGFD | 1 | 0 | 0 | 1700016 | YP_009047202.1 | K9N7C7 |
| TREENVIQCFGPRDFN | 1 | 0 | 0 | 1700017 | YP_003771.1 | H9EJ69 |
| TREIGFIVPGLPGTVL | 1 | 0 | 0 | 1700018 | NP_828849.2 | P0C6X7 |
| TRELMRELNGGAYTRY | 1 | 0 | 0 | 1700019 | YP_009724389.1 | P0DTD1 |
| TRELNGGAVTRYVDNN | 1 | 0 | 0 | 1700020 | NP_828849.2 | P0C6X7 |
| TREMQQHPLLNPSALE | 1 | 0 | 0 | 1700021 | NP_073556.1 | A0A0N7J131 |
| TRENDSKEGFFTYICG | 1 | 0 | 0 | 1700022 | QIQ54047.1 | P0C6X7 |
| TRENIGTNFPLQIGFS | 1 | 0 | 0 | 1700023 | YP_173236.1 | A0A140H1G9 |
| TREVFAQVKQMYKTPT | 1 | 0 | 0 | 1700024 | NP_828851.1 | P59594 |
| TREVGFVVPGLPGTIL | 1 | 0 | 0 | 1700025 | YP_009724389.1 | P0DTD1 |
| TRGRRAQPKQTATSQQ | 1 | 0 | 0 | 1700037 | YP_009555245.1 | P33469 |
| TRLEGLGGDNVYCYNT | 1 | 0 | 0 | 1700060 | YP_003766.2 | H9EJ42 |
| TRLEGLGGNNVYCYNT | 1 | 0 | 0 | 1700061 | NP_073549.1 | A0A0P0LKV0 |
| TRMEDAMGQGQNSADP | 1 | 0 | 0 | 1700069 | NP_828859.1 | P59636 |
| TRNIDATSTGNYNYKY | 1 | 0 | 0 | 1700071 | NP_828851.1 | P59594 |
| TRNPANNAAIVLQLPQ | 1 | 0 | 0 | 1700074 | YP_009724397.2 | P0DTC9 |
| TRNPNNDSAIVTQFAP | 1 | 0 | 0 | 1700076 | YP_009047211.1 | K9N4V7 |
| TRNPNNNAAIVLQLPQ | 1 | 0 | 0 | 1700077 | QIQ54056.1 | P59595 |
| TRNPNNNAATVLQLPQ | 1 | 0 | 0 | 1700078 | NP_828858.1 | P59595 |
| TRRVRGGDGKMKDLSP | 1 | 0 | 0 | 1700091 | QIQ54056.1 | P59595 |
| TRSDSNQSSSDLVAAV | 1 | 0 | 0 | 1700093 | YP_003771.1 | H9EJ69 |
| TRSMLKRRDSTYGPLQ | 1 | 0 | 0 | 1700096 | YP_009047204.1 | K9N5Q8 |
| TRSTNSRIKASMPTTI | 1 | 0 | 0 | 1700101 | YP_009724389.1 | P0DTD1 |
| TRTGIPPGYTNSSTRG | 1 | 0 | 0 | 1700105 | QIQ54048.1 | P59594 |
| TRTRSVFNLEGVNGGS | 1 | 0 | 0 | 1700112 | YP_003766.2 | H9EJ42 |
| TRVLNNLNLPGCNGGS | 1 | 0 | 0 | 1700120 | YP_009555238.1 | P0C6X6 |
| TRVLSNLNLPGCDGGS | 1 | 0 | 0 | 1700121 | YP_009724389.1 | P0DTD1 |
| TRVNAKPPPGDQFKHL | 1 | 0 | 0 | 1700123 | NP_828849.2 | P0C6X7 |
| TRVNVDGFNEWAMANG | 1 | 0 | 0 | 1700124 | YP_003766.2 | H9EJ42 |
| TRVTVPKDHPHLGKFL | 1 | 0 | 0 | 1700126 | NP_073556.1 | A0A0N7J131 |
| TSAALQPEEEQEEDWL | 1 | 0 | 0 | 1700133 | YP_009724389.1 | P0DTD1 |
| TSACKTIEDALRNSAR | 1 | 0 | 0 | 1700136 | NP_073551.1 | A0A0P0K6L9 |
| TSAKSLARSQSSETKE | 1 | 0 | 0 | 1700150 | NP_073556.1 | A0A0N7J131 |
| TSASKLTIVSPDLESY | 1 | 0 | 0 | 1700155 | YP_003766.2 | H9EJ42 |
| TSASLFPPWTAAAGVP | 1 | 0 | 0 | 1700156 | YP_009555241.1 | P36334 |
| TSCPEWMDDFEADPKG | 1 | 0 | 0 | 1700167 | YP_009047202.1 | K9N7C7 |
| TSDELCVNLNLPANVP | 1 | 0 | 0 | 1700174 | YP_009047202.1 | K9N7C7 |
| TSDKCIFNSACTRLEG | 1 | 0 | 0 | 1700175 | YP_003766.2 | H9EJ42 |
| TSDNINDKDTGPPPIS | 1 | 0 | 0 | 1700183 | YP_009555241.1 | P36334 |
| TSDQAVTFLKEWNFSL | 1 | 0 | 0 | 1700186 | YP_173241.1 | Q0ZJ82 |
| TSDTAHACNANRFNVA | 1 | 0 | 0 | 1700188 | NP_073549.1 | A0A0P0LKV0 |
| TSDTAHACNVNRFNVA | 1 | 0 | 0 | 1700189 | YP_003766.2 | H9EJ42 |
| TSDYYQLYSTQLSTDT | 1 | 0 | 0 | 1700194 | YP_009724391.1 | P0DTC3 |
| TSECYQLYSTQLSTDT | 1 | 0 | 0 | 1700196 | QIQ54049.1 | P59632 |
| TSEEVVENPTIQKEVI | 1 | 0 | 0 | 1700199 | NP_828849.2 | P0C6X7 |
| TSEEVVLKNGELEALE | 1 | 0 | 0 | 1700200 | NP_828849.2 | P0C6X7 |
| TSEGNALYNNEGGRAF | 1 | 0 | 0 | 1700201 | NP_073549.1 | A0A0P0LKV0 |
| TSEYKGPVTDVFYKEN | 1 | 0 | 0 | 1700208 | QIQ54046.1 | P0C6U8 |
| TSFATYHTPATDCSDG | 1 | 0 | 0 | 1700209 | YP_009047204.1 | K9N5Q8 |
| TSFTTYGNTVSEAVSR | 1 | 0 | 0 | 1700219 | YP_173239.1 | Q0ZJ84 |
| TSGCNVNDVTKLYLGG | 1 | 0 | 0 | 1700221 | NP_073549.1 | A0A0P0LKV0 |
| TSGDGTTSPISEHDYQ | 1 | 0 | 0 | 1700225 | YP_009724391.1 | P0DTC3 |
| TSGDLAVQIGNNNVCT | 1 | 0 | 0 | 1700227 | NP_073549.1 | A0A0P0LKV0 |
| TSGIKFQTVKPGNFNQ | 1 | 0 | 0 | 1700229 | YP_173236.1 | A0A140H1G9 |
| TSGLGTVDVDYKSCTK | 1 | 0 | 0 | 1700231 | YP_003767.1 | A0A0P0G321 |
| TSGLSPAYAPTYVSVD | 1 | 0 | 0 | 1700232 | YP_009047202.1 | K9N7C7 |
| TSGNLLGFKDVTKGTI | 1 | 0 | 0 | 1700233 | NP_073551.1 | A0A0P0K6L9 |
| TSGSDLDRCTTFDDVQ | 1 | 0 | 0 | 1700238 | NP_828851.1 | P59594 |
| TSGSRVAGRSAIEDIL | 1 | 0 | 0 | 1700239 | NP_073551.1 | A0A0P0K6L9 |
| TSGVKFQTVKPGNFNQ | 1 | 0 | 0 | 1700242 | YP_009555238.1 | P0C6X6 |
| TSIPNFGSLTQINTTL | 1 | 0 | 0 | 1700265 | YP_009047204.1 | K9N5Q8 |
| TSIVAKQGAGDAGHSL | 1 | 0 | 0 | 1700270 | NP_073549.1 | A0A0P0LKV0 |
| TSKCWVEEDLTKGPHE | 1 | 0 | 0 | 1700274 | YP_003766.2 | H9EJ42 |

| | | | | | | |
|---|---|---|---|---|---|---|
| TSKDTLDDILQQANKS | 1 | 0 | 0 | 1700275 | YP_009047202.1 | K9N7C7 |
| TSKKQFLPFQQFGRDI | 1 | 0 | 0 | 1700283 | QIQ54048.1 | P59594 |
| TSKPITFHMDGEVINI | 1 | 0 | 0 | 1700285 | QIQ54046.1 | P0C6U8 |
| TSKTTVASLINTLNDL | 1 | 0 | 0 | 1700287 | YP_009724389.1 | P0DTD1 |
| TSLEIPRRNVATLQAE | 1 | 0 | 0 | 1700291 | YP_009724389.1 | P0DTD1 |
| TSLEVPRRNVATLQAE | 1 | 0 | 0 | 1700293 | QIQ54047.1 | P0C6X7 |
| TSLGNRKRNQKPLEPK | 1 | 0 | 0 | 1700294 | YP_003771.1 | H9EJ69 |
| TSLPKDWRLINKFDVI | 1 | 0 | 0 | 1700301 | YP_173236.1 | A0A140H1G9 |
| TSLSGFKLKDCVMYAS | 1 | 0 | 0 | 1700306 | YP_009724389.1 | P0DTD1 |
| TSLSGYKLKDCIMYAS | 1 | 0 | 0 | 1700307 | QIQ54046.1 | P0C6U8 |
| TSLTPCGYSEPPKVAA | 1 | 0 | 0 | 1700311 | YP_009555238.1 | P0C6X6 |
| TSMLTNYFLDDVEMVA | 1 | 0 | 0 | 1700319 | YP_173236.1 | A0A140H1G9 |
| TSMYKQARAEDKKAKI | 1 | 0 | 0 | 1700320 | YP_009047202.1 | K9N7C7 |
| TSNIFDVLQSEDTQGM | 1 | 0 | 0 | 1700328 | QIQ54046.1 | P0C6U8 |
| TSNSFDVLKSEDAQGM | 1 | 0 | 0 | 1700334 | YP_009724389.1 | P0DTD1 |
| TSNSFDVLKSEDTQGM | 1 | 0 | 0 | 1700335 | QHR63299.1 | P0C6W2 |
| TSNSFEVLAVEDTQGM | 1 | 0 | 0 | 1700336 | NP_828849.2 | P0C6X7 |
| TSPARIAGNGGDAALA | 1 | 0 | 0 | 1700342 | QIQ54056.1 | P59595 |
| TSPARMAGNGSDAALA | 1 | 0 | 0 | 1700344 | QHR63308.1 | Q3LZX4 |
| TSPDPKFVDKYYAFLN | 1 | 0 | 0 | 1700346 | YP_009047202.1 | K9N7C7 |
| TSPGPSGIGAVGGDLL | 1 | 0 | 0 | 1700348 | YP_009047211.1 | K9N4V7 |
| TSPISEHDYQIGGYTE | 1 | 0 | 0 | 1700350 | YP_009724391.1 | P0DTC3 |
| TSQALQTVATALNKIQ | 1 | 0 | 0 | 1700354 | NP_073551.1 | A0A0P0K6L9 |
| TSQLRHNFQAISNSIQ | 1 | 0 | 0 | 1700358 | YP_003767.1 | A0A0P0G321 |
| TSQTVKPGHFNKEFYD | 1 | 0 | 0 | 1700363 | NP_073549.1 | A0A0P0LKV0 |
| TSREVVVTNLNKSAGW | 1 | 0 | 0 | 1700367 | NP_073549.1 | A0A0P0LKV0 |
| TSSKVVLSDALDKLYK | 1 | 0 | 0 | 1700383 | NP_073549.1 | A0A0P0LKV0 |
| TSSRASSAGSRSRANS | 1 | 0 | 0 | 1700389 | YP_009555245.1 | P33469 |
| TSTALGKLQDVVNQNA | 1 | 0 | 0 | 1700398 | NP_828851.1 | P59594 |
| TSTDVVYRAFDIYNDK | 1 | 0 | 0 | 1700400 | YP_009724389.1 | P0DTD1 |
| TSTDVVYRAFDIYNEK | 1 | 0 | 0 | 1700401 | NP_828849.2 | P0C6X7 |
| TSTGKTVLGEYVFDKS | 1 | 0 | 0 | 1700403 | YP_173236.1 | A0A140H1G9 |
| TSTHDGPTEHVTSVNL | 1 | 0 | 0 | 1700405 | YP_009047205.1 | K9N796 |
| TSTPKKPNKPLSQPRA | 1 | 0 | 0 | 1700411 | YP_003771.1 | H9EJ69 |
| TSTPSDVSSRDPTTQE | 1 | 0 | 0 | 1700412 | YP_173242.1 | Q19U29 |
| TSTSGNLLGFKDVTKG | 1 | 0 | 0 | 1700414 | NP_073551.1 | A0A0P0K6L9 |
| TSVDANTNQDSYGGAS | 1 | 0 | 0 | 1700420 | YP_003766.2 | H9EJ42 |
| TSVMESQISTKEVKLK | 1 | 0 | 0 | 1700428 | YP_173236.1 | A0A140H1G9 |
| TSVTPQPEEELEEDWL | 1 | 0 | 0 | 1700438 | QHR63299.1 | P0C6W2 |
| TSWADQSERNYQTFNR | 1 | 0 | 0 | 1700447 | YP_173242.1 | Q19U29 |
| TSYECDIPIGAGICAS | 1 | 0 | 0 | 1700451 | NP_828851.1 | P59594 |
| TSYRDGYLNSTNVTTT | 1 | 0 | 0 | 1700454 | QIQ54046.1 | P0C6U8 |
| TSYSVCNGCVGYSENV | 1 | 0 | 0 | 1700456 | NP_073551.1 | A0A0P0K6L9 |
| TSYTTTIKPVSYKLDG | 1 | 0 | 0 | 1700460 | NP_828849.2 | P0C6X7 |
| TTAATVAAMFPPWSAA | 1 | 0 | 0 | 1700466 | YP_173238.1 | A0A140H1H1 |
| TTANVSALMGANGNKI | 1 | 0 | 0 | 1700478 | YP_009047202.1 | K9N7C7 |
| TTCANDPVGFTLKNTV | 1 | 0 | 0 | 1700493 | YP_009724389.1 | P0DTD1 |
| TTCANDPVGFTLRNTV | 1 | 0 | 0 | 1700494 | NP_828849.2 | P0C6X7 |
| TTCSNDLLQPNTEVFT | 1 | 0 | 0 | 1700504 | YP_173238.1 | A0A140H1H1 |
| TTDKSFEQQVGVIADK | 1 | 0 | 0 | 1700507 | NP_073549.1 | A0A0P0LKV0 |
| TTDMVKSKGVCATGDC | 1 | 0 | 0 | 1700508 | YP_009555238.1 | P0C6X6 |
| TTDNTFVSGNCDVVIG | 1 | 0 | 0 | 1700510 | YP_009724390.1 | P0DTC2 |
| TTDNTFVSGSCDVVIG | 1 | 0 | 0 | 1700511 | QHR63300.2 | Q3LZX1 |
| TTELKDSVKKFFKLVN | 1 | 0 | 0 | 1700520 | QIQ54046.1 | P0C6U8 |
| TTETAHSCNVNRFNVA | 1 | 0 | 0 | 1700528 | YP_009724389.1 | P0DTD1 |
| TTEVVGDIILKPANDG | 1 | 0 | 0 | 1700532 | QHR63299.1 | P0C6W2 |
| TTEVVGDIILKPANNS | 1 | 0 | 0 | 1700533 | YP_009724389.1 | P0DTD1 |
| TTFASVYAWNRKRISN | 1 | 0 | 0 | 1700537 | QHR63300.2 | Q3LZX1 |
| TTFDSEYCRHGTCEKS | 1 | 0 | 0 | 1700542 | QIQ54046.1 | P0C6U8 |
| TTFDSEYCRHGTCERS | 1 | 0 | 0 | 1700543 | YP_009724389.1 | P0DTD1 |
| TTFHLDGEVITFDNLK | 1 | 0 | 0 | 1700546 | YP_009724389.1 | P0DTD1 |
| TTFIDAARGPAGVESD | 1 | 0 | 0 | 1700547 | YP_009047202.1 | K9N7C7 |
| TTFMLKYDENGTITDA | 1 | 0 | 0 | 1700552 | NP_828851.1 | P59594 |
| TTGDLAVQIGSNNVCT | 1 | 0 | 0 | 1700556 | YP_003766.2 | H9EJ42 |
| TTGYRFTNFEPFTVNS | 1 | 0 | 0 | 1700567 | YP_009555241.1 | P36334 |
| TTIGNETTSAFVGALP | 1 | 0 | 0 | 1700575 | NP_073551.1 | A0A0P0K6L9 |
| TTIKPVSYKLDGVTYT | 1 | 0 | 0 | 1700579 | NP_828849.2 | P0C6X7 |
| TTKFYGGWDNMLKNLM | 1 | 0 | 0 | 1700608 | NP_073549.1 | A0A0P0LKV0 |
| TTKNVTGSNESKALFF | 1 | 0 | 0 | 1700618 | YP_009047202.1 | K9N7C7 |
| TTLNAFVAQQLVRSES | 1 | 0 | 0 | 1700638 | YP_009047204.1 | K9N5Q8 |
| TTLNDFNLVAMKFNYE | 1 | 0 | 0 | 1700639 | QIQ54046.1 | P0C6U8 |
| TTLNDFNLVAMKYNYE | 1 | 0 | 0 | 1700640 | YP_009724389.1 | P0DTD1 |
| TTLQSLENVIYNLVNV | 1 | 0 | 0 | 1700649 | YP_173236.1 | A0A140H1G9 |
| TTLVKDKCVNFNFNGL | 1 | 0 | 0 | 1700656 | QIQ54048.1 | P59594 |
| TTNEAFQKVQDAVNNN | 1 | 0 | 0 | 1700665 | YP_009047204.1 | K9N5Q8 |
| TTNGSWAIFNDHHLNR | 1 | 0 | 0 | 1700669 | YP_009047202.1 | K9N7C7 |
| TTNYVGVYDINHGDAG | 1 | 0 | 0 | 1700676 | YP_009555240.1 | P30215 |
| TTPAPVYIWTADEAIK | 1 | 0 | 0 | 1700677 | YP_009555244.1 | Q01455 |
| TTPCEVHGNGSMIEDI | 1 | 0 | 0 | 1700678 | YP_009047202.1 | K9N7C7 |
| TTPKEEFVVKEKLNAF | 1 | 0 | 0 | 1700685 | NP_073549.1 | A0A0P0LKV0 |
| TTQLHVADTLMQGVTL | 1 | 0 | 0 | 1700691 | YP_173238.1 | A0A140H1H1 |
| TTQTVDSSQGSEYQYV | 1 | 0 | 0 | 1700698 | YP_009047202.1 | K9N7C7 |
| TTRTGIPPGYTNSSTR | 1 | 0 | 0 | 1700706 | QIQ54048.1 | P59594 |
| TTRVTVPKDHPHLGKF | 1 | 0 | 0 | 1700710 | NP_073556.1 | A0A0N7J131 |
| TTSDKCIFNSACTRLE | 1 | 0 | 0 | 1700713 | YP_003766.2 | H9EJ42 |
| TTSKKQFLPFQQFGRD | 1 | 0 | 0 | 1700716 | QIQ54048.1 | P59594 |
| TTSKPITFHMDGEVIN | 1 | 0 | 0 | 1700718 | QIQ54046.1 | P0C6U8 |
| TTSLTPCGYSEPPKVA | 1 | 0 | 0 | 1700720 | YP_009555238.1 | P0C6X6 |
| TTSPISEHDYQIGGYT | 1 | 0 | 0 | 1700724 | YP_009724391.1 | P0DTC3 |
| TTSTALGKLQDVVNQN | 1 | 0 | 0 | 1700728 | NP_828851.1 | P59594 |
| TTTIKPVSYKLDGVTY | 1 | 0 | 0 | 1700740 | NP_828849.2 | P0C6X7 |

| | | | | | | |
|---|---|---|---|---|---|---|
| TTTLEETKFLTNKLLL | 1 | 0 | 0 | 1700745 | NP_828849.2 | P0C6X7 |
| TTTLNDFNLVAMKFNY | 1 | 0 | 0 | 1700746 | QIQ54046.1 | P0C6U8 |
| TTTLNDFNLVAMKYNY | 1 | 0 | 0 | 1700747 | YP_009724389.1 | P0DTD1 |
| TTVASLINTLNDLNET | 1 | 0 | 0 | 1700760 | YP_009724389.1 | P0DTD1 |
| TTVDNTNLHTQLVDMS | 1 | 0 | 0 | 1700761 | NP_828849.2 | P0C6X7 |
| TTVLGQTYYQPIQQAP | 1 | 0 | 0 | 1700767 | NP_073555.1 | A0A0P0KXN3 |
| TTVSVTVSSNMVETVV | 1 | 0 | 0 | 1700778 | YP_009047202.1 | K9N7C7 |
| TTVVIPTKKAGGTTEM | 1 | 0 | 0 | 1700782 | QIQ54046.1 | P0C6U8 |
| TTVVRPLVEDSTSVTA | 1 | 0 | 0 | 1700783 | YP_009047210.1 | K9N7A1 |
| TTWNGGAYSLFDMTKF | 1 | 0 | 0 | 1700787 | YP_173236.1 | A0A140H1G9 |
| TTYLEAVKHGAKPVSN | 1 | 0 | 0 | 1700793 | YP_003766.2 | H9EJ42 |
| TTYNGYLTSSSKTSEE | 1 | 0 | 0 | 1700796 | NP_828849.2 | P0C6X7 |
| TTYYLDDVKKIEYKPD | 1 | 0 | 0 | 1700802 | YP_009555238.1 | P0C6X6 |
| TVASLINTLNDLNETL | 1 | 0 | 0 | 1700817 | YP_009724389.1 | P0DTD1 |
| TVATALNKIQDVVNQQ | 1 | 0 | 0 | 1700818 | NP_073551.1 | A0A0P0K6L9 |
| TVCDSLNAKLGFDSSK | 1 | 0 | 0 | 1700826 | YP_009555238.1 | P0C6X6 |
| TVCFSVAALSTGLTSQ | 1 | 0 | 0 | 1700827 | NP_073549.1 | A0A0P0LKV0 |
| TVCGPKLSTDLIKNQC | 1 | 0 | 0 | 1700831 | NP_828851.1 | P59594 |
| TVDCKQYVCNGFQKCE | 1 | 0 | 0 | 1700841 | YP_009047204.1 | K9N5Q8 |
| TVDGVNFKSISLTVGE | 1 | 0 | 0 | 1700845 | YP_173236.1 | A0A140H1G9 |
| TVDGVNFRTVVLNNKN | 1 | 0 | 0 | 1700846 | YP_009047202.1 | K9N7C7 |
| TVDGVNFTNRFVPVGE | 1 | 0 | 0 | 1700847 | YP_009555238.1 | P0C6X6 |
| TVDLKHFFFTQDGNAA | 1 | 0 | 0 | 1700850 | YP_173236.1 | A0A140H1G9 |
| TVDNINLHTQVVDMSM | 1 | 0 | 0 | 1700852 | YP_009724389.1 | P0DTD1 |
| TVDNTNLHTQLVDMSM | 1 | 0 | 0 | 1700853 | NP_828849.2 | P0C6X7 |
| TVDPQKAYLDFVNAGG | 1 | 0 | 0 | 1700854 | YP_009047202.1 | K9N7C7 |
| TVDSAQGSEYDYVIFA | 1 | 0 | 0 | 1700855 | NP_073549.1 | A0A0P0LKV0 |
| TVDSSQGSEYDYVIFT | 1 | 0 | 0 | 1700857 | YP_009724389.1 | P0DTD1 |
| TVDSSQGSEYDYVIYA | 1 | 0 | 0 | 1700858 | YP_003766.2 | H9EJ42 |
| TVDSSQGSEYQYVIFC | 1 | 0 | 0 | 1700859 | YP_009047202.1 | K9N7C7 |
| TVDTTPKEEFVVKEKL | 1 | 0 | 0 | 1700860 | NP_073549.1 | A0A0P0LKV0 |
| TVDVANPSTPAYTFTT | 1 | 0 | 0 | 1700861 | YP_009047202.1 | K9N7C7 |
| TVDVDYKSCTKGLSIA | 1 | 0 | 0 | 1700862 | YP_003767.1 | A0A0P0G321 |
| TVEAAIELSKELKRPV | 1 | 0 | 0 | 1700867 | YP_173236.1 | A0A140H1G9 |
| TVEAALDLSKELKRPI | 1 | 0 | 0 | 1700868 | YP_009555238.1 | P0C6X6 |
| TVEDLRARMTYVCQCG | 1 | 0 | 0 | 1700870 | YP_009047202.1 | K9N7C7 |
| TVGEVFGKILGNVFCD | 1 | 0 | 0 | 1700903 | YP_173236.1 | A0A140H1G9 |
| TVIDILLDDFVALVKS | 1 | 0 | 0 | 1700917 | YP_009555238.1 | P0C6X6 |
| TVIPDYVDVNKTLQEF | 1 | 0 | 0 | 1700925 | YP_003767.1 | A0A0P0G321 |
| TVIRGHLYIQGVKLGT | 1 | 0 | 0 | 1700926 | YP_173241.1 | Q0ZJ82 |
| TVKAKEESVKSSYAYA | 1 | 0 | 0 | 1700931 | NP_073549.1 | A0A0P0LKV0 |
| TVKGLDYKTFKQIVES | 1 | 0 | 0 | 1700938 | QHR63299.1 | P0C6W2 |
| TVKIEDAIIVNDENSS | 1 | 0 | 0 | 1700940 | YP_173236.1 | A0A140H1G9 |
| TVKIGGLTFKTYAFDK | 1 | 0 | 0 | 1700941 | NP_073549.1 | A0A0P0LKV0 |
| TVKPGHFNKEFYDFLR | 1 | 0 | 0 | 1700946 | NP_073549.1 | A0A0P0LKV0 |
| TVKPGNFNKDFYDFAV | 1 | 0 | 0 | 1700947 | YP_009724389.1 | P0DTD1 |
| TVKPGNFNQDFYDFVL | 1 | 0 | 0 | 1700948 | YP_009555238.1 | P0C6X6 |
| TVKPGNFNQDFYEFVK | 1 | 0 | 0 | 1700949 | YP_173236.1 | A0A140H1G9 |
| TVKPKPVINQLDEKAQ | 1 | 0 | 0 | 1700950 | NP_073549.1 | A0A0P0LKV0 |
| TVKPVATVHRRIPDLP | 1 | 0 | 0 | 1700952 | YP_173238.1 | A0A140H1H1 |
| TVKSEDGNIKNINWFV | 1 | 0 | 0 | 1700953 | NP_073549.1 | A0A0P0LKV0 |
| TVKWADASEPQRGRQG | 1 | 0 | 0 | 1700957 | NP_073556.1 | A0A0N7J131 |
| TVLGEYVFDKSELTNG | 1 | 0 | 0 | 1700967 | YP_009555238.1 | P0C6X6 |
| TVLIKEPCSSGTYEGN | 1 | 0 | 0 | 1700971 | QIQ54053.1 | P59635 |
| TVLLKEPCSSGTYEGN | 1 | 0 | 0 | 1700976 | YP_009724395.1 | P0DTC7 |
| TVLLPTQYKDVEAWSG | 1 | 0 | 0 | 1700978 | NP_073551.1 | A0A0P0K6L9 |
| TVLNDPVFTGDLFYTI | 1 | 0 | 0 | 1700980 | YP_003766.2 | H9EJ42 |
| TVLNVVGPDARTQGKQ | 1 | 0 | 0 | 1700984 | YP_009555238.1 | P0C6X6 |
| TVLRDNLKPVTTLSTI | 1 | 0 | 0 | 1700997 | YP_003768.1 | H9EJ24 |
| TVLTSEEVVLKNGELE | 1 | 0 | 0 | 1701003 | NP_828849.2 | P0C6X7 |
| TVMPLSAPTLVPQEHY | 1 | 0 | 0 | 1701007 | YP_009724389.1 | P0DTD1 |
| TVNGYAKSNFVKDDYC | 1 | 0 | 0 | 1701014 | NP_073549.1 | A0A0P0LKV0 |
| TVNSVNDSLEPVGGLY | 1 | 0 | 0 | 1701020 | YP_009555241.1 | P36334 |
| TVPDLVVEQYNQTILN | 1 | 0 | 0 | 1701028 | NP_073551.1 | A0A0P0K6L9 |
| TVPEEFASHVANFQKS | 1 | 0 | 0 | 1701029 | YP_009047202.1 | K9N7C7 |
| TVQDADGINKQLTDIS | 1 | 0 | 0 | 1701036 | YP_173236.1 | A0A140H1G9 |
| TVQDAKGLKIKYLYFV | 1 | 0 | 0 | 1701037 | YP_009555238.1 | P0C6X6 |
| TVQGPPGTGKSHLAIG | 1 | 0 | 0 | 1701041 | YP_009555238.1 | P0C6X6 |
| TVQKLQTLIDNINSTL | 1 | 0 | 0 | 1701042 | NP_073551.1 | A0A0P0K6L9 |
| TVQQCFGKRGPNQNFG | 1 | 0 | 0 | 1701044 | YP_009555245.1 | P33469 |
| TVQYGTDTNSVCPKLE | 1 | 0 | 0 | 1701050 | YP_009047204.1 | K9N5Q8 |
| TVRLQAGKQTEFVSNS | 1 | 0 | 0 | 1701057 | NP_073549.1 | A0A0P0LKV0 |
| TVRVEEEEEEDWLDDT | 1 | 0 | 0 | 1701066 | NP_828849.2 | P0C6X7 |
| TVSALVYENKLKAKNE | 1 | 0 | 0 | 1701072 | YP_009555238.1 | P0C6X6 |
| TVSALVYNNKLLAKKE | 1 | 0 | 0 | 1701073 | YP_009047202.1 | K9N7C7 |
| TVSDDDFVSAVANAHR | 1 | 0 | 0 | 1701076 | YP_003766.2 | H9EJ42 |
| TVSKTSDWKCKVTDVL | 1 | 0 | 0 | 1701080 | YP_009047202.1 | K9N7C7 |
| TVSSNMVETVVGQLEQ | 1 | 0 | 0 | 1701087 | YP_009047202.1 | K9N7C7 |
| TVSVSSPDAVTAYNGY | 1 | 0 | 0 | 1701089 | YP_009724389.1 | P0DTD1 |
| TVSVTVSSNMVETVVG | 1 | 0 | 0 | 1701090 | YP_009047202.1 | K9N7C7 |
| TVSWNLREMLTHAEET | 1 | 0 | 0 | 1701092 | QIQ54046.1 | P0C6U8 |
| TVTIALNKIQDVVNQQ | 1 | 0 | 0 | 1701100 | YP_003767.1 | A0A0P0G321 |
| TVTLKQGEIKDATPSD | 1 | 0 | 0 | 1701103 | YP_009724391.1 | P0DTC3 |
| TVTPEANMDQESFGGA | 1 | 0 | 0 | 1701107 | YP_009724389.1 | P0DTD1 |
| TVTTDKSFEQQVGVIA | 1 | 0 | 0 | 1701110 | NP_073549.1 | A0A0P0LKV0 |
| TVTYPFIANEKFINGC | 1 | 0 | 0 | 1701118 | YP_003766.2 | H9EJ42 |
| TVVIKNLTPIKLNFGM | 1 | 0 | 0 | 1701130 | YP_003766.2 | H9EJ42 |
| TVVIPTKKAGGTTEML | 1 | 0 | 0 | 1701131 | QIQ54046.1 | P0C6U8 |
| TVVQEDDSEVNQTTIT | 1 | 0 | 0 | 1701137 | QHR63299.1 | P0C6W2 |
| TVVQRCASNKPHVLED | 1 | 0 | 0 | 1701139 | NP_849176.1 | Q7TFA0 |
| TVVSKVHEVIIDNKPW | 1 | 0 | 0 | 1701141 | NP_073549.1 | A0A0P0LKV0 |

| | | | | | | |
|---|---|---|---|---|---|---|
| TVVSTVKIGGLTFKTY | 1 | 0 | 0 | 1701142 | NP_073549.1 | A0A0P0LKV0 |
| TVVVTLKDSDVNDMVL | 1 | 0 | 0 | 1701148 | YP_003766.2 | H9EJ42 |
| TVWEDGDYYRKQLSPL | 1 | 0 | 0 | 1701151 | YP_009047204.1 | K9N5Q8 |
| TVWNGGAYSLFDMAKF | 1 | 0 | 0 | 1701154 | YP_009555238.1 | P0C6X6 |
| TVYDSSEVKAQSSGVI | 1 | 0 | 0 | 1701162 | YP_173236.1 | A0A140H1G9 |
| TVYDTAKKSMYDGDRF | 1 | 0 | 0 | 1701163 | NP_073549.1 | A0A0P0LKV0 |
| TVYEKLKPILDWLEEK | 1 | 0 | 0 | 1701164 | QIQ54046.1 | P0C6U8 |
| TVYEKLKPVLDWLEEK | 1 | 0 | 0 | 1701165 | YP_009724389.1 | P0DTD1 |
| TVYEQLKPVLDWLEEK | 1 | 0 | 0 | 1701166 | QHR63299.1 | P0C6W2 |
| TWFSQRGGSYTNDKAC | 1 | 0 | 0 | 1701191 | YP_009724389.1 | P0DTD1 |
| TWHEGRTIYGNVSRQD | 1 | 0 | 0 | 1701192 | NP_073549.1 | A0A0P0LKV0 |
| TWLDMVDTSLSGFKLK | 1 | 0 | 0 | 1701196 | YP_009724389.1 | P0DTD1 |
| TWLDMVDTSLSGYKLK | 1 | 0 | 0 | 1701197 | QIQ54046.1 | P0C6U8 |
| TWLTYHGAIKLDDKDP | 1 | 0 | 0 | 1701203 | NP_828858.1 | P59595 |
| TWLTYTGAIKLDDKDP | 1 | 0 | 0 | 1701205 | YP_009724397.2 | P0DTC9 |
| TWNGGAYSLFDMTKFS | 1 | 0 | 0 | 1701208 | YP_173236.1 | A0A140H1G9 |
| TWNKRFGFIEDSVFKP | 1 | 0 | 0 | 1701209 | YP_009555241.1 | P36334 |
| TWPRPIDVSKADGIIY | 1 | 0 | 0 | 1701212 | YP_009047204.1 | K9N5Q8 |
| TWSEGNSITGVPYPVS | 1 | 0 | 0 | 1701218 | YP_003767.1 | A0A0P0G321 |
| TWVENEFRVYSSANNC | 1 | 0 | 0 | 1701223 | QIQ54048.1 | P59594 |
| TYAALRNSLTNDAYSR | 1 | 0 | 0 | 1701226 | YP_009047202.1 | K9N7C7 |
| TYAGNVWQIQTIQDSD | 1 | 0 | 0 | 1701233 | YP_009555238.1 | P0C6X6 |
| TYAGSVWHIQTVQDAD | 1 | 0 | 0 | 1701234 | YP_173236.1 | A0A140H1G9 |
| TYAKPFLNKVTSVAAS | 1 | 0 | 0 | 1701236 | QIQ54046.1 | P0C6U8 |
| TYEGNSPFHPLADNKF | 1 | 0 | 0 | 1701245 | YP_009724395.1 | P0DTC7 |
| TYFNRPSLVDDNKFDV | 1 | 0 | 0 | 1701260 | YP_009555238.1 | P0C6X6 |
| TYFTQSRDLEDFKPRS | 1 | 0 | 0 | 1701262 | NP_828849.2 | P0C6X7 |
| TYGNMTLNGLWLDDKV | 1 | 0 | 0 | 1701268 | YP_009555238.1 | P0C6X6 |
| TYHGAIKLDDKDPQFK | 1 | 0 | 0 | 1701277 | NP_828858.1 | P59595 |
| TYHPNCINCLDDRCIL | 1 | 0 | 0 | 1701278 | NP_828849.2 | P0C6X7 |
| TYHPNCVNCLDDRCIL | 1 | 0 | 0 | 1701280 | YP_009724389.1 | P0DTD1 |
| TYHTPATDCSDGNYNR | 1 | 0 | 0 | 1701281 | YP_009047204.1 | K9N5Q8 |
| TYKLDGVVCTEIDPKL | 1 | 0 | 0 | 1701296 | YP_009724389.1 | P0DTD1 |
| TYKMLVAKDNKNLPKF | 1 | 0 | 0 | 1701302 | YP_003771.1 | H9EJ69 |
| TYKSTATTKLVPGMLF | 1 | 0 | 0 | 1701310 | NP_073549.1 | A0A0P0LKV0 |
| TYKSTVTTKLVPGMIF | 1 | 0 | 0 | 1701311 | YP_003766.2 | H9EJ42 |
| TYLDFIQQGGTPIANC | 1 | 0 | 0 | 1701314 | YP_009555238.1 | P0C6X6 |
| TYLDGADVTKIKPHAS | 1 | 0 | 0 | 1701315 | QIQ54046.1 | P0C6U8 |
| TYLDGADVTKIKPHNS | 1 | 0 | 0 | 1701316 | YP_009724389.1 | P0DTD1 |
| TYLDYIQQGGVPIINC | 1 | 0 | 0 | 1701318 | YP_173236.1 | A0A140H1G9 |
| TYLKSDNIVAADLGVL | 1 | 0 | 0 | 1701323 | YP_009555238.1 | P0C6X6 |
| TYLNDPSSKTVCTYMD | 1 | 0 | 0 | 1701326 | NP_073549.1 | A0A0P0LKV0 |
| TYLNLSSELKQLEAKT | 1 | 0 | 0 | 1701329 | YP_003767.1 | A0A0P0G321 |
| TYNGYLTSSSKTSEEH | 1 | 0 | 0 | 1701339 | NP_828849.2 | P0C6X7 |
| TYNKVENMTPRDLGAC | 1 | 0 | 0 | 1701343 | YP_009724389.1 | P0DTD1 |
| TYPFIANEKFINGCGR | 1 | 0 | 0 | 1701346 | YP_003766.2 | H9EJ42 |
| TYPGQGCVGYTIEEAK | 1 | 0 | 0 | 1701348 | QIQ54046.1 | P0C6U8 |
| TYPGQGLNGYTVEEAR | 1 | 0 | 0 | 1701350 | QHR63299.1 | P0C6W2 |
| TYPMIANENAINKGGR | 1 | 0 | 0 | 1701351 | NP_073549.1 | A0A0P0LKV0 |
| TYQNISTNLPPPLLGN | 1 | 0 | 0 | 1701359 | YP_009047204.1 | K9N5Q8 |
| TYSFLPKEDGDVLLAE | 1 | 0 | 0 | 1701366 | YP_009047202.1 | K9N7C7 |
| TYTDQYLCGADGKPVM | 1 | 0 | 0 | 1701378 | NP_073549.1 | A0A0P0LKV0 |
| TYTEIEPKLDGYYKKD | 1 | 0 | 0 | 1701379 | NP_828849.2 | P0C6X7 |
| TYTGAIKLDDKDPSFK | 1 | 0 | 0 | 1701381 | QIQ54056.1 | P59595 |
| TYTSTSGNLLGFKDVT | 1 | 0 | 0 | 1701382 | NP_073551.1 | A0A0P0K6L9 |
| TYTYKMLVAKDNKNLP | 1 | 0 | 0 | 1701387 | YP_003771.1 | H9EJ69 |
| TYVCNGNVRCVELLKQ | 1 | 0 | 0 | 1701392 | NP_073551.1 | A0A0P0K6L9 |
| TYVSNSGNLLGFKNVS | 1 | 0 | 0 | 1701400 | YP_003767.1 | A0A0P0G321 |
| TYYCVTTKAKSVQQDL | 1 | 0 | 0 | 1701406 | YP_009047202.1 | K9N7C7 |
| TYYIYDEEGGNDLSLP | 1 | 0 | 0 | 1701409 | NP_073549.1 | A0A0P0LKV0 |
| TYYLDDVKKIEYKPDL | 1 | 0 | 0 | 1701410 | YP_009555238.1 | P0C6X6 |
| TYYNSWQNLLYDSNGN | 1 | 0 | 0 | 1701415 | YP_009555241.1 | P36334 |
| VAAALKSLGFDKPQEK | 1 | 0 | 0 | 1701418 | NP_073556.1 | A0A0N7J131 |
| VAADLGVLIQNSAKHV | 1 | 0 | 0 | 1701420 | YP_009555238.1 | P0C6X6 |
| VAAFHQECSLQSCTQH | 1 | 0 | 0 | 1701421 | YP_009724396.1 | P0DTC8 |
| VAAITSGVKFQTVKPG | 1 | 0 | 0 | 1701433 | YP_009555238.1 | P0C6X6 |
| VAALSTGLTSQTVKPG | 1 | 0 | 0 | 1701439 | NP_073549.1 | A0A0P0LKV0 |
| VACEHISSTMSQYSRS | 1 | 0 | 0 | 1701460 | YP_009047204.1 | K9N5Q8 |
| VACQLPYCYFRNSSAN | 1 | 0 | 0 | 1701463 | YP_173237.1 | A0A140H1H0 |
| VACQPPYCYFRNSTTN | 1 | 0 | 0 | 1701464 | YP_009555240.1 | P30215 |
| VADAVIKTLQPVSELL | 1 | 0 | 0 | 1701467 | YP_009724389.1 | P0DTD1 |
| VADQSRKWIGHCTIAQ | 1 | 0 | 0 | 1701472 | YP_009555239.1 | Q80872 |
| VADTLMQGVTLSSNLN | 1 | 0 | 0 | 1701474 | YP_173238.1 | A0A140H1H1 |
| VAEAVIKTLQPVSELL | 1 | 0 | 0 | 1701479 | QIQ54046.1 | P0C6U8 |
| VAFGGDQVHEVAAVRS | 1 | 0 | 0 | 1701494 | YP_009047202.1 | K9N7C7 |
| VAFGLQDCVTGINDDD | 1 | 0 | 0 | 1701495 | YP_003766.2 | H9EJ42 |
| VAFTKAAGGKVSFSDD | 1 | 0 | 0 | 1701510 | NP_073549.1 | A0A0P0LKV0 |
| VAFTKLAGGKISFSDD | 1 | 0 | 0 | 1701511 | YP_003766.2 | H9EJ42 |
| VAFTTELKDSVKKFFK | 1 | 0 | 0 | 1701512 | QIQ54046.1 | P0C6U8 |
| VAFYQGDVDTVVNGVD | 1 | 0 | 0 | 1701516 | NP_073549.1 | A0A0P0LKV0 |
| VAGALNKATNGAMQKE | 1 | 0 | 0 | 1701517 | NP_828849.2 | P0C6X7 |
| VAGALNKATNNAMQIE | 1 | 0 | 0 | 1701518 | QIQ54046.1 | P0C6U8 |
| VAGALNKATNNAMQVE | 1 | 0 | 0 | 1701519 | YP_009724389.1 | P0DTD1 |
| VAGYKVLPPLMDVNME | 1 | 0 | 0 | 1701533 | YP_009047204.1 | K9N5Q8 |
| VAIADSYYSYIMPMLT | 1 | 0 | 0 | 1701536 | YP_009555238.1 | P0C6X6 |
| VAIDYKHYTPSFKKGA | 1 | 0 | 0 | 1701539 | YP_009724389.1 | P0DTD1 |
| VAIKCVDIVKEAQSAN | 1 | 0 | 0 | 1701544 | NP_828849.2 | P0C6X7 |
| VAIKITEHSWNADLYK | 1 | 0 | 0 | 1701547 | YP_009724389.1 | P0DTD1 |
| VAIQEVIDDMHWGDGF | 1 | 0 | 0 | 1701550 | YP_009555239.1 | Q80872 |
| VAKAIAVAAGKKFSKE | 1 | 0 | 0 | 1701566 | YP_173236.1 | A0A140H1G9 |
| VAKEGAKTVNTSLGNR | 1 | 0 | 0 | 1701572 | YP_003771.1 | H9EJ69 |

| | | | | | | |
|---|---|---|---|---|---|---|
| VAKHDFFKFRIDGDMV | 1 | 0 | 0 | 1701574 | YP_009724389.1 | P0DTD1 |
| VAKQGAGDAGHSLTWL | 1 | 0 | 0 | 1701585 | NP_073549.1 | A0A0P0LKV0 |
| VAKRVLGVQTQTVDSA | 1 | 0 | 0 | 1701587 | YP_173236.1 | A0A140H1G9 |
| VALRGRIDKDLSLSRK | 1 | 0 | 0 | 1701613 | NP_073553.1 | A0A0P0KWE8 |
| VANGLNAPNGPFDALF | 1 | 0 | 0 | 1701636 | NP_073549.1 | A0A0P0LKV0 |
| VANGVKAKGYPQFAEL | 1 | 0 | 0 | 1701638 | NP_073556.1 | A0A0N7J131 |
| VANHQADTSTPSDVSS | 1 | 0 | 0 | 1701640 | YP_173242.1 | Q19U29 |
| VANLKLPPHDVTVLRD | 1 | 0 | 0 | 1701642 | YP_003768.1 | H9EJ24 |
| VANLSAPTLVPQENYS | 1 | 0 | 0 | 1701643 | YP_009555238.1 | P0C6X6 |
| VANPSTPAYTFTTVKP | 1 | 0 | 0 | 1701646 | YP_009047202.1 | K9N7C7 |
| VANWLAPTNAYMRTTH | 1 | 0 | 0 | 1701650 | YP_009047202.1 | K9N7C7 |
| VANYQKVGMQKYSTLQ | 1 | 0 | 0 | 1701653 | YP_009724389.1 | P0DTD1 |
| VANYQKVGMQXYSTLQ | 1 | 0 | 0 | 1701654 | QIQ54047.1 | P0C6X7 |
| VAPGTAVLKRWLPHDA | 1 | 0 | 0 | 1701661 | NP_073549.1 | A0A0P0LKV0 |
| VAPIELENKFTPLSVE | 1 | 0 | 0 | 1701665 | YP_009047202.1 | K9N7C7 |
| VAPQVTYQNISTNLPP | 1 | 0 | 0 | 1701668 | YP_009047204.1 | K9N5Q8 |
| VAPVNTVKPKPVINQL | 1 | 0 | 0 | 1701669 | NP_073549.1 | A0A0P0LKV0 |
| VAPVYEADTEQTQNVT | 1 | 0 | 0 | 1701670 | YP_009047202.1 | K9N7C7 |
| VAQNTVLNASIPESKP | 1 | 0 | 0 | 1701676 | YP_003771.1 | H9EJ69 |
| VARLMKGDKGDAEDTL | 1 | 0 | 0 | 1701687 | NP_073549.1 | A0A0P0LKV0 |
| VASDSEISANGCSTIA | 1 | 0 | 0 | 1701695 | NP_073549.1 | A0A0P0LKV0 |
| VASDSEISGFGFAIPS | 1 | 0 | 0 | 1701696 | YP_003766.2 | H9EJ42 |
| VASFLEMKSETKVQDH | 1 | 0 | 0 | 1701698 | QIQ54046.1 | P0C6U8 |
| VASIITKLNSLNEPLV | 1 | 0 | 0 | 1701704 | NP_828849.2 | P0C6X7 |
| VASINSAIVCASVKRD | 1 | 0 | 0 | 1701707 | NP_073549.1 | A0A0P0LKV0 |
| VASLINTLNDLNETLV | 1 | 0 | 0 | 1701709 | YP_009724389.1 | P0DTD1 |
| VASLSAPTLVPQENYA | 1 | 0 | 0 | 1701710 | YP_173236.1 | A0A140H1G9 |
| VASNQADVNTPADIVD | 1 | 0 | 0 | 1701712 | YP_009555245.1 | P33469 |
| VASNSKLWHYCSTLHN | 1 | 0 | 0 | 1701713 | YP_009555238.1 | P0C6X6 |
| VASNSKLWQYCSVLHN | 1 | 0 | 0 | 1701714 | YP_173236.1 | A0A140H1G9 |
| VASRFLGLQIQTVDSS | 1 | 0 | 0 | 1701719 | YP_003766.2 | H9EJ42 |
| VASSFVGMPSFVAYET | 1 | 0 | 0 | 1701721 | NP_073549.1 | A0A0P0LKV0 |
| VASSSYDALLHRNSLD | 1 | 0 | 0 | 1701724 | YP_003766.2 | H9EJ42 |
| VATAHSELAKGVALDG | 1 | 0 | 0 | 1701729 | NP_828849.2 | P0C6X7 |
| VATALNKIQDVVNQQG | 1 | 0 | 0 | 1701730 | NP_073551.1 | A0A0P0K6L9 |
| VATFYPQLQSAEWKCG | 1 | 0 | 0 | 1701736 | NP_073549.1 | A0A0P0LKV0 |
| VATIPYKVCNSVKDTL | 1 | 0 | 0 | 1701737 | YP_009047202.1 | K9N7C7 |
| VATLIATLNSLGETLV | 1 | 0 | 0 | 1701738 | QIQ54046.1 | P0C6U8 |
| VATRNATVVIGTTKFY | 1 | 0 | 0 | 1701742 | NP_073549.1 | A0A0P0LKV0 |
| VATVQSKMSDVKCTSV | 1 | 0 | 0 | 1701746 | YP_009724389.1 | P0DTD1 |
| VAVADSYYSYMMPMLT | 1 | 0 | 0 | 1701750 | YP_173236.1 | A0A140H1G9 |
| VAVDGAKTEPTGYGVR | 1 | 0 | 0 | 1701751 | NP_073556.1 | A0A0N7J131 |
| VAVHDFFKFRVDGDMV | 1 | 0 | 0 | 1701753 | NP_828849.2 | P0C6X7 |
| VAVIGDEQIDGKFVTK | 1 | 0 | 0 | 1701756 | YP_173236.1 | A0A140H1G9 |
| VAVPTGYVDTENNTEF | 1 | 0 | 0 | 1701768 | NP_828849.2 | P0C6X7 |
| VAVPTGYVDTPNNTDF | 1 | 0 | 0 | 1701770 | YP_009724389.1 | P0DTD1 |
| VAVVNDKIFTKSGVND | 1 | 0 | 0 | 1701776 | YP_009047202.1 | K9N7C7 |
| VAVVNDKVFVRYGDVD | 1 | 0 | 0 | 1701777 | YP_003766.2 | H9EJ42 |
| VAVYANVGRWSASINT | 1 | 0 | 0 | 1701778 | NP_073551.1 | A0A0P0K6L9 |
| VAVYQQSIIGAMTAVN | 1 | 0 | 0 | 1701779 | YP_003767.1 | A0A0P0G321 |
| VAWHVVRDPRYVMRLQ | 1 | 0 | 0 | 1701780 | YP_009555238.1 | P0C6X6 |
| VAWHVVRDSRFVMRLQ | 1 | 0 | 0 | 1701781 | YP_173236.1 | A0A140H1G9 |
| VAYADKPNHFINFPLT | 1 | 0 | 0 | 1701784 | YP_009555239.1 | Q80872 |
| VAYENFNHHEKLLAPL | 1 | 0 | 0 | 1701787 | QIQ54046.1 | P0C6U8 |
| VAYESLRPDTRYVLMD | 1 | 0 | 0 | 1701788 | YP_009724389.1 | P0DTD1 |
| VAYRKILLRKNGNKGA | 1 | 0 | 0 | 1701800 | QIQ54046.1 | P0C6U8 |
| VAYRKVLLRKNGNKGA | 1 | 0 | 0 | 1701801 | YP_009724389.1 | P0DTD1 |
| VAYTMSLGADSSIAYS | 1 | 0 | 0 | 1701805 | NP_828851.1 | P59594 |
| VCALMSCNGNKIEDLS | 1 | 0 | 0 | 1701813 | YP_009555238.1 | P0C6X6 |
| VCASVKRDGVQVGYCV | 1 | 0 | 0 | 1701814 | NP_073549.1 | A0A0P0LKV0 |
| VCDAIENALNSCKEHP | 1 | 0 | 0 | 1701819 | YP_173236.1 | A0A140H1G9 |
| VCDDCNGYTDNIFSVQ | 1 | 0 | 0 | 1701820 | YP_003767.1 | A0A0P0G321 |
| VCDNIKLADDLNQLTG | 1 | 0 | 0 | 1701826 | QHR63299.1 | P0C6W2 |
| VCDSLNAKLGFDSSKE | 1 | 0 | 0 | 1701829 | YP_009555238.1 | P0C6X6 |
| VCFGFDKWFVNDGRVA | 1 | 0 | 0 | 1701836 | NP_073549.1 | A0A0P0LKV0 |
| VCFSVAALSTGLTSQT | 1 | 0 | 0 | 1701843 | NP_073549.1 | A0A0P0LKV0 |
| VCGDNIGTNVPLQVGF | 1 | 0 | 0 | 1701852 | YP_003766.2 | H9EJ42 |
| VCGYTDFAEDVCTCYD | 1 | 0 | 0 | 1701868 | NP_073549.1 | A0A0P0LKV0 |
| VCHSVAVIGDEQIDGK | 1 | 0 | 0 | 1701870 | YP_173236.1 | A0A140H1G9 |
| VCITTNGSWAIFNDHH | 1 | 0 | 0 | 1701877 | YP_009047202.1 | K9N7C7 |
| VCKHCGQKTTTLTGVE | 1 | 0 | 0 | 1701882 | NP_828849.2 | P0C6X7 |
| VCKPHQVNISLNGNTS | 1 | 0 | 0 | 1701885 | YP_003767.1 | A0A0P0G321 |
| VCKTCGQQQTTLKGVE | 1 | 0 | 0 | 1701886 | YP_009724389.1 | P0DTD1 |
| VCKYTDIDVNSALNIC | 1 | 0 | 0 | 1701894 | YP_009047202.1 | K9N7C7 |
| VCLGQNSQSPTSNHS | 1 | 0 | 0 | 1701902 | P03138.3 | Q76R62 |
| VCLKHAEEYREYLESY | 1 | 0 | 0 | 1701907 | YP_009555238.1 | P0C6X6 |
| VCNAPGCDVNDVTKLY | 1 | 0 | 0 | 1701915 | YP_009555238.1 | P0C6X6 |
| VCNASGCGVSDVKKLY | 1 | 0 | 0 | 1701919 | YP_003766.2 | H9EJ42 |
| VCNGNVRCVELLKQYT | 1 | 0 | 0 | 1701924 | NP_073551.1 | A0A0P0K6L9 |
| VCNTSGCNVNDVTKLY | 1 | 0 | 0 | 1701929 | NP_073549.1 | A0A0P0LKV0 |
| VCPKLEFANDTKIASQ | 1 | 0 | 0 | 1701932 | YP_009047204.1 | K9N5Q8 |
| VCRHHANEYRQYLDAY | 1 | 0 | 0 | 1701945 | NP_828849.2 | P0C6X7 |
| VCRLMKTIGPDMFLGT | 1 | 0 | 0 | 1701947 | YP_009724389.1 | P0DTD1 |
| VCSKHAEEYCNYLESY | 1 | 0 | 0 | 1701952 | YP_173236.1 | A0A140H1G9 |
| VCSKHANLYQKYVEAY | 1 | 0 | 0 | 1701953 | YP_003766.2 | H9EJ42 |
| VCSMTDIAKKPTENIC | 1 | 0 | 0 | 1701957 | QHR63299.1 | P0C6W2 |
| VCSMTDIAKKPTETIC | 1 | 0 | 0 | 1701958 | YP_009724389.1 | P0DTD1 |
| VCTDNSANTKVLRNKG | 1 | 0 | 0 | 1701968 | YP_009047202.1 | K9N7C7 |
| VCTEIDPKLDGYYKKD | 1 | 0 | 0 | 1701969 | QIQ54046.1 | P0C6U8 |
| VCTEIDPKLDNYYKKD | 1 | 0 | 0 | 1701970 | YP_009724389.1 | P0DTD1 |
| VCTMTDIAKKPTESAC | 1 | 0 | 0 | 1701973 | NP_828849.2 | P0C6X7 |

| | | | | | | |
|---|---|---|---|---|---|---|
| VCYNSDYASKGYIANI | 1 | 0 | 0 | 1702011 | YP_009555238.1 | P0C6X6 |
| VDAAKAYKDYLASGGQ | 1 | 0 | 0 | 1702014 | YP_009724389.1 | P0DTD1 |
| VDAFNQFSSDFQHKLK | 1 | 0 | 0 | 1702020 | YP_009555238.1 | P0C6X6 |
| VDAKGFPQLAELIPNQ | 1 | 0 | 0 | 1702022 | YP_003771.1 | H9EJ69 |
| VDALCEKAYKFLNIND | 1 | 0 | 0 | 1702025 | YP_009555238.1 | P0C6X6 |
| VDANTNQDSYGGASIC | 1 | 0 | 0 | 1702028 | YP_003766.2 | H9EJ42 |
| VDASKAYRDYLASGGQ | 1 | 0 | 0 | 1702030 | QIQ54046.1 | P0C6U8 |
| VDAVNLLTNMFTPLIQ | 1 | 0 | 0 | 1702033 | YP_009724389.1 | P0DTD1 |
| VDCSLDPLSETKCTLK | 1 | 0 | 0 | 1702049 | QIQ54048.1 | P59594 |
| VDCSTYVCNGNVRCVE | 1 | 0 | 0 | 1702052 | NP_073551.1 | A0A0P0K6L9 |
| VDDIESIYDFDTYKAL | 1 | 0 | 0 | 1702062 | YP_173236.1 | A0A140H1G9 |
| VDDINFSPVLGCLGSE | 1 | 0 | 0 | 1702063 | YP_009555241.1 | P36334 |
| VDDIYYPASCNGVLPV | 1 | 0 | 0 | 1702067 | YP_003766.2 | H9EJ42 |
| VDDLLKTDSVLLIERF | 1 | 0 | 0 | 1702071 | YP_009555238.1 | P0C6X6 |
| VDDPKLMGWDYPKCDR | 1 | 0 | 0 | 1702074 | NP_073549.1 | A0A0P0LKV0 |
| VDDVVKTDAVVLLERY | 1 | 0 | 0 | 1702079 | YP_003766.2 | H9EJ42 |
| VDEAFPLDEAEDVTES | 1 | 0 | 0 | 1702082 | YP_009047202.1 | K9N7C7 |
| VDENGDKLDQFFVVKR | 1 | 0 | 0 | 1702085 | YP_009555238.1 | P0C6X6 |
| VDENGKYYLPYPDPSR | 1 | 0 | 0 | 1702086 | NP_073549.1 | A0A0P0LKV0 |
| VDEPEDHVQIHTIDGS | 1 | 0 | 0 | 1702087 | QIQ54049.1 | P59632 |
| VDEPEEHVQIHTIDGS | 1 | 0 | 0 | 1702088 | YP_009724391.1 | P0DTC3 |
| VDEVGSKSPIQYIDIG | 1 | 0 | 0 | 1702090 | QHR63307.1 | Q3LZX5 |
| VDEVSMCTNYDLSIIN | 1 | 0 | 0 | 1702092 | YP_009047202.1 | K9N7C7 |
| VDEVSMCTNYDLSVIN | 1 | 0 | 0 | 1702093 | NP_073549.1 | A0A0P0LKV0 |
| VDEWSYTGSSFYAPEP | 1 | 0 | 0 | 1702095 | YP_009047204.1 | K9N5Q8 |
| VDFHSVEIEAGEVKPF | 1 | 0 | 0 | 1702100 | YP_003766.2 | H9EJ42 |
| VDFINSMQFNYKITIN | 1 | 0 | 0 | 1702101 | YP_009555239.1 | Q80872 |
| VDFKPQSGGGKCFNCY | 1 | 0 | 0 | 1702105 | NP_073551.1 | A0A0P0K6L9 |
| VDGELPVAVVNDKVFV | 1 | 0 | 0 | 1702120 | YP_003766.2 | H9EJ42 |
| VDGFGVNKIQPWTNDA | 1 | 0 | 0 | 1702121 | NP_073549.1 | A0A0P0LKV0 |
| VDGHRLASGVQVHNLP | 1 | 0 | 0 | 1702125 | NP_073555.1 | A0A0P0KXN3 |
| VDGIDVEIFENKTTLP | 1 | 0 | 0 | 1702127 | NP_828849.2 | P0C6X7 |
| VDGLESKQVDYVPLRS | 1 | 0 | 0 | 1702134 | YP_173236.1 | A0A140H1G9 |
| VDGLKYTKILKDDGNF | 1 | 0 | 0 | 1702135 | YP_009555238.1 | P0C6X6 |
| VDGLKYTKIMKDDGNC | 1 | 0 | 0 | 1702136 | YP_173236.1 | A0A140H1G9 |
| VDGVDVELFENKTILP | 1 | 0 | 0 | 1702139 | QIQ54047.1 | P0C6X7 |
| VDGVDVELFENKTTLP | 1 | 0 | 0 | 1702140 | YP_009724389.1 | P0DTD1 |
| VDGVNFKSISLTVGEV | 1 | 0 | 0 | 1702141 | YP_173236.1 | A0A140H1G9 |
| VDGVNFRTVVLNNKNT | 1 | 0 | 0 | 1702142 | YP_009047202.1 | K9N7C7 |
| VDGVNFTNRFVPVGES | 1 | 0 | 0 | 1702143 | YP_009555238.1 | P0C6X6 |
| VDGVSRLYQPLRLTCL | 1 | 0 | 0 | 1702148 | YP_003767.1 | A0A0P0G321 |
| VDHFGENEEIIINGHT | 1 | 0 | 0 | 1702156 | NP_073549.1 | A0A0P0LKV0 |
| VDIAANTVIWDYKRDA | 1 | 0 | 0 | 1702161 | YP_009724389.1 | P0DTD1 |
| VDINKAIPSGNLIKFD | 1 | 0 | 0 | 1702169 | NP_073549.1 | A0A0P0LKV0 |
| VDINKLCEEMLDNRAT | 1 | 0 | 0 | 1702170 | YP_009724389.1 | P0DTD1 |
| VDINRLCEEMLDNRAT | 1 | 0 | 0 | 1702172 | NP_828849.2 | P0C6X7 |
| VDIQQTFFDKTWPRPI | 1 | 0 | 0 | 1702176 | YP_009047204.1 | K9N5Q8 |
| VDIVKEAQSANPMVIV | 1 | 0 | 0 | 1702180 | NP_828849.2 | P0C6X7 |
| VDIVSEAKNVKPTIVV | 1 | 0 | 0 | 1702181 | QIQ54046.1 | P0C6U8 |
| VDKEVKDMQFDLYVNV | 1 | 0 | 0 | 1702185 | YP_009047202.1 | K9N7C7 |
| VDKKIKACVEEVTTTL | 1 | 0 | 0 | 1702188 | QHR63299.1 | P0C6W2 |
| VDKVLNEKCSVYTVES | 1 | 0 | 0 | 1702199 | NP_828849.2 | P0C6X7 |
| VDKYMCGFDGKPVLPK | 1 | 0 | 0 | 1702202 | YP_003766.2 | H9EJ42 |
| VDLHNKINLCDDPEKA | 1 | 0 | 0 | 1702213 | YP_003766.2 | H9EJ42 |
| VDLYGGNMFQFATLPV | 1 | 0 | 0 | 1702226 | YP_009047204.1 | K9N5Q8 |
| VDNDLYPFVSDSVATY | 1 | 0 | 0 | 1702234 | YP_009555238.1 | P0C6X6 |
| VDNGSSINRRQLDVVK | 1 | 0 | 0 | 1702238 | NP_073549.1 | A0A0P0LKV0 |
| VDNINLHTQVVDMSMT | 1 | 0 | 0 | 1702240 | YP_009724389.1 | P0DTD1 |
| VDNLVFTNKTTLPTNV | 1 | 0 | 0 | 1702241 | YP_003766.2 | H9EJ42 |
| VDNNFCGPDGYPLDCI | 1 | 0 | 0 | 1702242 | NP_828849.2 | P0C6X7 |
| VDNNFCGPDGYPLECI | 1 | 0 | 0 | 1702243 | YP_009724389.1 | P0DTD1 |
| VDNPHLMGWDYPKCDR | 1 | 0 | 0 | 1702244 | YP_009047202.1 | K9N7C7 |
| VDNPVLMGWDYPKCDR | 1 | 0 | 0 | 1702245 | YP_009555238.1 | P0C6X6 |
| VDNSSLTIKKPNELSR | 1 | 0 | 0 | 1702246 | YP_009724389.1 | P0DTD1 |
| VDNTNLHTQLVDMSMT | 1 | 0 | 0 | 1702247 | NP_828849.2 | P0C6X7 |
| VDNTSLTIKKPNELSV | 1 | 0 | 0 | 1702248 | QIQ54046.1 | P0C6U8 |
| VDNVYMAKAGDKYYPV | 1 | 0 | 0 | 1702249 | YP_009555238.1 | P0C6X6 |
| VDPAAAYLDAVKQGAK | 1 | 0 | 0 | 1702252 | NP_073549.1 | A0A0P0LKV0 |
| VDPAKAYKDYLASGGQ | 1 | 0 | 0 | 1702253 | NP_828849.2 | P0C6X7 |
| VDPATTYLEAVKHGAK | 1 | 0 | 0 | 1702254 | YP_003766.2 | H9EJ42 |
| VDPDLSAFYVKDGKYF | 1 | 0 | 0 | 1702255 | YP_009047202.1 | K9N7C7 |
| VDPKKTYLDFIQQGGT | 1 | 0 | 0 | 1702256 | YP_009555238.1 | P0C6X6 |
| VDPKKTYLDYIQQGGV | 1 | 0 | 0 | 1702257 | YP_173236.1 | A0A140H1G9 |
| VDPKYQVIVLKDNVLS | 1 | 0 | 0 | 1702258 | YP_009047202.1 | K9N7C7 |
| VDPQKAYLDFVNAGGA | 1 | 0 | 0 | 1702261 | YP_009047202.1 | K9N7C7 |
| VDPTTVLNDPVFTGDL | 1 | 0 | 0 | 1702263 | YP_003766.2 | H9EJ42 |
| VDQLNSSYFKLSIPTN | 1 | 0 | 0 | 1702264 | YP_009047204.1 | K9N5Q8 |
| VDQVEEVNSIFDIETV | 1 | 0 | 0 | 1702266 | NP_073549.1 | A0A0P0LKV0 |
| VDQYMCGVDGKPISAY | 1 | 0 | 0 | 1702270 | YP_009047202.1 | K9N7C7 |
| VDSAQGSEYDYVIFAQ | 1 | 0 | 0 | 1702281 | NP_073549.1 | A0A0P0LKV0 |
| VDSAQGSEYDYVIYSQ | 1 | 0 | 0 | 1702282 | YP_009555238.1 | P0C6X6 |
| VDSDLNDFVSDADSTL | 1 | 0 | 0 | 1702283 | YP_009724389.1 | P0DTD1 |
| VDSEQPWKVIPRNLVP | 1 | 0 | 0 | 1702287 | NP_073556.1 | A0A0N7J131 |
| VDSFEGVLPINTDTVL | 1 | 0 | 0 | 1702288 | YP_003766.2 | H9EJ42 |
| VDSFTNGAIVGTPVCV | 1 | 0 | 0 | 1702289 | NP_828849.2 | P0C6X7 |
| VDSLCAKAVTAYSVDK | 1 | 0 | 0 | 1702292 | NP_073549.1 | A0A0P0LKV0 |
| VDSNWPLVIIANRYNE | 1 | 0 | 0 | 1702294 | YP_173236.1 | A0A140H1G9 |
| VDSRWNNARQSDNMTA | 1 | 0 | 0 | 1702295 | YP_009555240.1 | P30215 |
| VDSRWNNERASDISLS | 1 | 0 | 0 | 1702296 | YP_173237.1 | A0A140H1H0 |
| VDSSQGSEYDYVIFTQ | 1 | 0 | 0 | 1702298 | YP_009724389.1 | P0DTD1 |
| VDSSQGSEYDYVIYAQ | 1 | 0 | 0 | 1702299 | YP_003766.2 | H9EJ42 |

| | | | | | | |
|---|---|---|---|---|---|---|
| VDSVNLLTNMFTPLIQ | 1 | 0 | 0 | 1702303 | QIQ54046.1 | P0C6U8 |
| VDSVQYAHKHDIQITN | 1 | 0 | 0 | 1702304 | YP_009047202.1 | K9N7C7 |
| VDSYFENDSILQSVAS | 1 | 0 | 0 | 1702308 | NP_073549.1 | A0A0P0LKV0 |
| VDTANTKTPKYKFVRI | 1 | 0 | 0 | 1702312 | QHR63299.1 | P0C6W2 |
| VDTDFVNEFYAYLRKH | 1 | 0 | 0 | 1702313 | YP_009724389.1 | P0DTD1 |
| VDTDLTKPYIKWDLLK | 1 | 0 | 0 | 1702314 | YP_009724389.1 | P0DTD1 |
| VDTENNTEFTRVNAKP | 1 | 0 | 0 | 1702316 | NP_828849.2 | P0C6X7 |
| VDTFLSMFDVDKKSLN | 1 | 0 | 0 | 1702318 | YP_009555238.1 | P0C6X6 |
| VDTFMSHFDVDRKSFN | 1 | 0 | 0 | 1702319 | YP_173236.1 | A0A140H1G9 |
| VDTFSATFSVPMEKLK | 1 | 0 | 0 | 1702320 | NP_828849.2 | P0C6X7 |
| VDTILNMARNGVVPLS | 1 | 0 | 0 | 1702322 | NP_073549.1 | A0A0P0LKV0 |
| VDTLVNKIPANIVLPQ | 1 | 0 | 0 | 1702329 | YP_009555238.1 | P0C6X6 |
| VDTPNNTDFSRVSAKP | 1 | 0 | 0 | 1702331 | YP_009724389.1 | P0DTD1 |
| VDTSLSGFKLKDCVMY | 1 | 0 | 0 | 1702333 | YP_009724389.1 | P0DTD1 |
| VDTSLSGYKLKDCIMY | 1 | 0 | 0 | 1702334 | QIQ54046.1 | P0C6U8 |
| VDTSNSFEVLAVEDTQ | 1 | 0 | 0 | 1702336 | NP_828849.2 | P0C6X7 |
| VDTTPKEEFVVKEKLN | 1 | 0 | 0 | 1702338 | NP_073549.1 | A0A0P0LKV0 |
| VDTVIFKNNTSFPTNI | 1 | 0 | 0 | 1702339 | YP_173236.1 | A0A140H1G9 |
| VDTVRTNVYLAVFDKN | 1 | 0 | 0 | 1702340 | YP_009724389.1 | P0DTD1 |
| VDTVSALVYDNKLKAH | 1 | 0 | 0 | 1702341 | YP_009724389.1 | P0DTD1 |
| VDTVSALVYDNKLRAH | 1 | 0 | 0 | 1702342 | QIQ54047.1 | P0C6X7 |
| VDTVSALVYENKLKAK | 1 | 0 | 0 | 1702343 | YP_009555238.1 | P0C6X6 |
| VDTVVNGVDFDFIVNA | 1 | 0 | 0 | 1702344 | NP_073549.1 | A0A0P0LKV0 |
| VDVAIQEVIDDMHWGD | 1 | 0 | 0 | 1702345 | YP_009555239.1 | Q80872 |
| VDVANPSTPAYTFTTV | 1 | 0 | 0 | 1702346 | YP_009047202.1 | K9N7C7 |
| VDVDYKSCTKGLSIAD | 1 | 0 | 0 | 1702349 | YP_003767.1 | A0A0P0G321 |
| VDVELFENKTILPVNV | 1 | 0 | 0 | 1702350 | QIQ54047.1 | P0C6X7 |
| VDVELFENKTTLPVNV | 1 | 0 | 0 | 1702351 | YP_009724389.1 | P0DTD1 |
| VDVGPDSVKSACIEVD | 1 | 0 | 0 | 1702352 | YP_009047204.1 | K9N5Q8 |
| VDVLKDMPVLKTINGL | 1 | 0 | 0 | 1702356 | YP_173236.1 | A0A140H1G9 |
| VDVMRYNLNFSANSLD | 1 | 0 | 0 | 1702358 | YP_003767.1 | A0A0P0G321 |
| VDVNDSHSMAVVDGKQ | 1 | 0 | 0 | 1702359 | YP_009555238.1 | P0C6X6 |
| VDVNKTLQEFAQNLPK | 1 | 0 | 0 | 1702360 | YP_003767.1 | A0A0P0G321 |
| VDVSHETSDDEWAAAV | 1 | 0 | 0 | 1702364 | YP_009047202.1 | K9N7C7 |
| VDVTFLKKDAPYIVGD | 1 | 0 | 0 | 1702366 | QIQ54046.1 | P0C6U8 |
| VDVTNGLGTYYVLDRV | 1 | 0 | 0 | 1702367 | YP_009555241.1 | P36334 |
| VDVVALQQEMTIVKCK | 1 | 0 | 0 | 1702368 | YP_009047202.1 | K9N7C7 |
| VDVVPEGWRVVNKFYQ | 1 | 0 | 0 | 1702370 | YP_009555238.1 | P0C6X6 |
| VDVVSIRSSNQGNKQI | 1 | 0 | 0 | 1702371 | YP_009047207.1 | K9N643 |
| VDWDIEYPYISHEKKL | 1 | 0 | 0 | 1702374 | YP_009047202.1 | K9N7C7 |
| VDWSITYPMIANENAI | 1 | 0 | 0 | 1702378 | NP_073549.1 | A0A0P0LKV0 |
| VDWSVEYPIIGDELRV | 1 | 0 | 0 | 1702379 | NP_828849.2 | P0C6X7 |
| VDWTIEYPIIGDELKI | 1 | 0 | 0 | 1702380 | YP_009724389.1 | P0DTD1 |
| VDYKSCTKGLSIADLA | 1 | 0 | 0 | 1702391 | YP_003767.1 | A0A0P0G321 |
| VDYNSPSSSSSRRKRR | 1 | 0 | 0 | 1702393 | YP_173238.1 | A0A140H1H1 |
| VDYSNLLHSKVKSVPN | 1 | 0 | 0 | 1702396 | YP_009555238.1 | P0C6X6 |
| VDYTFVNEYYEFLCKH | 1 | 0 | 0 | 1702397 | YP_173236.1 | A0A140H1G9 |
| VEAAIELSKELKRPVN | 1 | 0 | 0 | 1702403 | YP_173236.1 | A0A140H1G9 |
| VEAALDLSKELKRPIQ | 1 | 0 | 0 | 1702404 | YP_009555238.1 | P0C6X6 |
| VEAPKQEEPPNTEDSK | 1 | 0 | 0 | 1702410 | NP_828849.2 | P0C6X7 |
| VEAVMYMGTLSYDNLK | 1 | 0 | 0 | 1702418 | NP_828849.2 | P0C6X7 |
| VEAVMYMGTLSYEQFK | 1 | 0 | 0 | 1702419 | YP_009724389.1 | P0DTD1 |
| VEAVNFNVTTAETTDF | 1 | 0 | 0 | 1702421 | NP_073551.1 | A0A0P0K6L9 |
| VEAYNSILFENGIPLM | 1 | 0 | 0 | 1702425 | YP_003766.2 | H9EJ42 |
| VECFDKFKVNSTLEQY | 1 | 0 | 0 | 1702432 | YP_009724389.1 | P0DTD1 |
| VECYDRFKVNETNSQY | 1 | 0 | 0 | 1702438 | YP_009047202.1 | K9N7C7 |
| VECYSGFKPNNNSAQY | 1 | 0 | 0 | 1702439 | NP_073549.1 | A0A0P0LKV0 |
| VECYSGFKPNNTSAQY | 1 | 0 | 0 | 1702440 | YP_003766.2 | H9EJ42 |
| VECYSILAAKTGVSVE | 1 | 0 | 0 | 1702441 | YP_003766.2 | H9EJ42 |
| VEDAYAEVHAEPKGKY | 1 | 0 | 0 | 1702443 | YP_173236.1 | A0A140H1G9 |
| VEDAYDQVHDEPKGKF | 1 | 0 | 0 | 1702444 | YP_009555238.1 | P0C6X6 |
| VEDCKLPLGQSLCALP | 1 | 0 | 0 | 1702445 | YP_009047204.1 | K9N5Q8 |
| VEDIAQVVIADTLQET | 1 | 0 | 0 | 1702448 | YP_009047202.1 | K9N7C7 |
| VEDTIDGVVEDTINDD | 1 | 0 | 0 | 1702452 | YP_173236.1 | A0A140H1G9 |
| VEDTINDDEDVVTGDN | 1 | 0 | 0 | 1702453 | YP_173236.1 | A0A140H1G9 |
| VEEAKKVKPTVVVNAA | 1 | 0 | 0 | 1702458 | YP_009724389.1 | P0DTD1 |
| VEEALSEAREHLKNGT | 1 | 0 | 0 | 1702460 | NP_828849.2 | P0C6X7 |
| VEEALSEARQHLKDGT | 1 | 0 | 0 | 1702461 | QHR63299.1 | P0C6W2 |
| VEEALSEARQHLLDGT | 1 | 0 | 0 | 1702462 | QIQ54046.1 | P0C6U8 |
| VEEDDVEGEETDLTVT | 1 | 0 | 0 | 1702464 | YP_009555238.1 | P0C6X6 |
| VEEDLTKGPHEFCSQH | 1 | 0 | 0 | 1702465 | YP_003766.2 | H9EJ42 |
| VEEDVEMSDFVDLESV | 1 | 0 | 0 | 1702466 | YP_009555238.1 | P0C6X6 |
| VEEEEEEDWLDDTTEQ | 1 | 0 | 0 | 1702467 | NP_828849.2 | P0C6X7 |
| VEEEHVNRMDTFCFNV | 1 | 0 | 0 | 1702468 | NP_073549.1 | A0A0P0LKV0 |
| VEELHDDLVNYIRDKG | 1 | 0 | 0 | 1702469 | YP_009555239.1 | Q80872 |
| VEELKKLLEQWNLVIG | 1 | 0 | 0 | 1702470 | YP_009724393.1 | P0DTC5 |
| VEEVLSEARQHLKDGT | 1 | 0 | 0 | 1702474 | YP_009724389.1 | P0DTD1 |
| VEEVNSIFDIETVDVK | 1 | 0 | 0 | 1702475 | NP_073549.1 | A0A0P0LKV0 |
| VEFEDRSNNSSRASSR | 1 | 0 | 0 | 1702478 | YP_003771.1 | H9EJ69 |
| VEFHLDGEVLSLDKLK | 1 | 0 | 0 | 1702480 | NP_828849.2 | P0C6X7 |
| VEFLKRGDKIVYHTLE | 1 | 0 | 0 | 1702484 | NP_828849.2 | P0C6X7 |
| VEFNRLATCDWTESGD | 1 | 0 | 0 | 1702486 | YP_009047202.1 | K9N7C7 |
| VEFWFAMRKDGDDVIF | 1 | 0 | 0 | 1702488 | YP_173236.1 | A0A140H1G9 |
| VEFYEDRILDFKVLDN | 1 | 0 | 0 | 1702489 | YP_009047202.1 | K9N7C7 |
| VEGAHATRENIGTNFP | 1 | 0 | 0 | 1702492 | YP_173236.1 | A0A140H1G9 |
| VEGAHVTGDNVGTNVP | 1 | 0 | 0 | 1702493 | NP_073549.1 | A0A0P0LKV0 |
| VEGCHATREAIGTNLP | 1 | 0 | 0 | 1702495 | QHR63299.1 | P0C6W2 |
| VEGELPVAVVNDKIFT | 1 | 0 | 0 | 1702500 | YP_009047202.1 | K9N7C7 |
| VEGSGRSASNSRPGSR | 1 | 0 | 0 | 1702506 | YP_173242.1 | Q19U29 |
| VEHLELKETVFVEPKD | 1 | 0 | 0 | 1702512 | YP_003766.2 | H9EJ42 |
| VEIDGNVMHANYLFWR | 1 | 0 | 0 | 1702520 | YP_009555238.1 | P0C6X6 |

| | | | | | | |
|---|---|---|---|---|---|---|
| VEIEAGEVKPFAVYKN | 1 | 0 | 0 | 1702521 | YP_003766.2 | H9EJ42 |
| VEIFENKTTLPVNVAF | 1 | 0 | 0 | 1702522 | NP_828849.2 | P0C6X7 |
| VEISVTPIKNTVDTTP | 1 | 0 | 0 | 1702526 | NP_073549.1 | A0A0P0LKV0 |
| VEKGVLPQLEQPYVFI | 1 | 0 | 0 | 1702530 | YP_009724389.1 | P0DTD1 |
| VEKGVLPQLEQPYVFV | 1 | 0 | 0 | 1702531 | QIQ54046.1 | P0C6U8 |
| VEKKKTEGFMGRIRSV | 1 | 0 | 0 | 1702536 | NP_828849.2 | P0C6X7 |
| VEKLSPWFMDGENAYE | 1 | 0 | 0 | 1702538 | YP_009047202.1 | K9N7C7 |
| VELFENKTILPVNVAF | 1 | 0 | 0 | 1702545 | QIQ54047.1 | P0C6X7 |
| VELFENKTTLPVNVAF | 1 | 0 | 0 | 1702546 | YP_009724389.1 | P0DTD1 |
| VELPKFFYASNGTYNC | 1 | 0 | 0 | 1702555 | NP_073551.1 | A0A0P0K6L9 |
| VELVAELDGVQYGRSG | 1 | 0 | 0 | 1702559 | QIQ54046.1 | P0C6U8 |
| VELVAELEGIQYGRSG | 1 | 0 | 0 | 1702560 | YP_009724389.1 | P0DTD1 |
| VELVAELNGIQYGRSG | 1 | 0 | 0 | 1702561 | QHR63299.1 | P0C6W2 |
| VELVAEMDGIQYGRSG | 1 | 0 | 0 | 1702562 | NP_828849.2 | P0C6X7 |
| VEMHNKINLCDDPETA | 1 | 0 | 0 | 1702566 | NP_073549.1 | A0A0P0LKV0 |
| VEMVAYNPDLSQYYCD | 1 | 0 | 0 | 1702569 | YP_173236.1 | A0A140H1G9 |
| VENEFRVYSSANNCTF | 1 | 0 | 0 | 1702573 | QIQ54048.1 | P59594 |
| VENPDIINIYKKLGPI | 1 | 0 | 0 | 1702577 | YP_173236.1 | A0A140H1G9 |
| VENPDIINVYKKLGPI | 1 | 0 | 0 | 1702578 | YP_009555238.1 | P0C6X6 |
| VENPDILRVYANLGER | 1 | 0 | 0 | 1702579 | YP_009724389.1 | P0DTD1 |
| VENPHLMGWDYPKCDR | 1 | 0 | 0 | 1702580 | YP_009724389.1 | P0DTD1 |
| VENPMLMGWDYPKCDR | 1 | 0 | 0 | 1702581 | YP_003766.2 | H9EJ42 |
| VENPTIQKDILECNVK | 1 | 0 | 0 | 1702583 | QHR63299.1 | P0C6W2 |
| VENPTIQKDVLECNVK | 1 | 0 | 0 | 1702584 | YP_009724389.1 | P0DTD1 |
| VENTSITIKKPNELSL | 1 | 0 | 0 | 1702587 | NP_828849.2 | P0C6X7 |
| VEQPKIEPKPVSVIKV | 1 | 0 | 0 | 1702602 | NP_073549.1 | A0A0P0LKV0 |
| VEQYNQTILNLTSEIS | 1 | 0 | 0 | 1702606 | NP_073551.1 | A0A0P0K6L9 |
| VESAHVCGDNIGTNVP | 1 | 0 | 0 | 1702615 | YP_003766.2 | H9EJ42 |
| VESANLMLSDNVVAFL | 1 | 0 | 0 | 1702616 | YP_003766.2 | H9EJ42 |
| VESANQMLTVNVVAFL | 1 | 0 | 0 | 1702617 | NP_073549.1 | A0A0P0LKV0 |
| VESDDYIATNGPLKVG | 1 | 0 | 0 | 1702624 | YP_009724389.1 | P0DTD1 |
| VESDHYIATNGPLKVG | 1 | 0 | 0 | 1702625 | QHR63299.1 | P0C6W2 |
| VESGGYIPSDFAFNNW | 1 | 0 | 0 | 1702629 | NP_073551.1 | A0A0P0K6L9 |
| VESGVYSVSSFEAKPS | 1 | 0 | 0 | 1702631 | YP_009047204.1 | K9N5Q8 |
| VESSKSLGKQLGVVSD | 1 | 0 | 0 | 1702634 | YP_003766.2 | H9EJ42 |
| VETCTIAGKAFDKVFD | 1 | 0 | 0 | 1702638 | NP_073549.1 | A0A0P0LKV0 |
| VETETYFKVNGLWSPT | 1 | 0 | 0 | 1702640 | YP_173236.1 | A0A140H1G9 |
| VETFYPKLQASQAWQP | 1 | 0 | 0 | 1702641 | NP_828849.2 | P0C6X7 |
| VETFYPKLQSSQAWQP | 1 | 0 | 0 | 1702642 | YP_009724389.1 | P0DTD1 |
| VETKVEITPEESASSE | 1 | 0 | 0 | 1702645 | QIQ54046.1 | P0C6U8 |
| VETLNEDLLEILSRER | 1 | 0 | 0 | 1702647 | NP_828849.2 | P0C6X7 |
| VETLNFQMVKEFPLLS | 1 | 0 | 0 | 1702648 | YP_173243.1 | Q0ZJ80 |
| VETPHLMGWDYPKCDR | 1 | 0 | 0 | 1702651 | NP_828849.2 | P0C6X7 |
| VETSNIFDVLQSEDTQ | 1 | 0 | 0 | 1702656 | QIQ54046.1 | P0C6U8 |
| VETSNSFDVLKSEDAQ | 1 | 0 | 0 | 1702657 | YP_009724389.1 | P0DTD1 |
| VETSNSFDVLKSEDTQ | 1 | 0 | 0 | 1702658 | QHR63299.1 | P0C6W2 |
| VETVLNLARDGVVPLS | 1 | 0 | 0 | 1702662 | YP_003766.2 | H9EJ42 |
| VETVSALVYDNKLKAK | 1 | 0 | 0 | 1702663 | YP_173236.1 | A0A140H1G9 |
| VETYIKWPWWVWLCIS | 1 | 0 | 0 | 1702665 | NP_073551.1 | A0A0P0K6L9 |
| VEVEKGVLPQLEQPYV | 1 | 0 | 0 | 1702666 | YP_009724389.1 | P0DTD1 |
| VEVFNRLATSDWTDVR | 1 | 0 | 0 | 1702667 | YP_003766.2 | H9EJ42 |
| VEVFTDGKLNCSFQDA | 1 | 0 | 0 | 1702668 | YP_009047202.1 | K9N7C7 |
| VEVNATYYNSWQNLLY | 1 | 0 | 0 | 1702673 | YP_009555241.1 | P36334 |
| VEVVDKYFDCYDGGCI | 1 | 0 | 0 | 1702680 | YP_009724389.1 | P0DTD1 |
| VEWKFYDAQPCSDKAY | 1 | 0 | 0 | 1702682 | YP_009724389.1 | P0DTD1 |
| VEXVDVTFLKKDAPYI | 1 | 0 | 0 | 1702683 | QIQ54046.1 | P0C6U8 |
| VEYEVAKKNLADAKNS | 1 | 0 | 0 | 1702686 | YP_173236.1 | A0A140H1G9 |
| VEYPIIGDELRVNSAC | 1 | 0 | 0 | 1702696 | NP_828849.2 | P0C6X7 |
| VEYPIISNELSINTSC | 1 | 0 | 0 | 1702697 | YP_009555238.1 | P0C6X6 |
| VFAVESGGYIPSDFAF | 1 | 0 | 0 | 1702707 | NP_073551.1 | A0A0P0K6L9 |
| VFCDGVNVTKHKCDIN | 1 | 0 | 0 | 1702711 | YP_009555238.1 | P0C6X6 |
| VFCGVDAVNLLTNMFT | 1 | 0 | 0 | 1702713 | YP_009724389.1 | P0DTD1 |
| VFCGVDSVNLLTNMFT | 1 | 0 | 0 | 1702714 | QIQ54046.1 | P0C6U8 |
| VFDEISMATNYDLSVV | 1 | 0 | 0 | 1702722 | YP_009724389.1 | P0DTD1 |
| VFDGKSKCDESASKSA | 1 | 0 | 0 | 1702723 | NP_828849.2 | P0C6X7 |
| VFDGKSKCDESSARAA | 1 | 0 | 0 | 1702724 | QIQ54046.1 | P0C6U8 |
| VFDGKSKCEESSAKSA | 1 | 0 | 0 | 1702725 | YP_009724389.1 | P0DTD1 |
| VFDIKNSSVNTVCFVD | 1 | 0 | 0 | 1702726 | YP_003766.2 | H9EJ42 |
| VFDIVKGQIQVASDNI | 1 | 0 | 0 | 1702727 | NP_828849.2 | P0C6X7 |
| VFDKDLYEELVASFLE | 1 | 0 | 0 | 1702728 | QIQ54046.1 | P0C6U8 |
| VFDKVVDNVYVTYAGS | 1 | 0 | 0 | 1702731 | YP_173236.1 | A0A140H1G9 |
| VFENFEDWHYAKFGFT | 1 | 0 | 0 | 1702738 | NP_073549.1 | A0A0P0LKV0 |
| VFEVDDTVDMEEFYAV | 1 | 0 | 0 | 1702741 | YP_009555238.1 | P0C6X6 |
| VFFKGNVQVDNGSSIN | 1 | 0 | 0 | 1702747 | YP_003766.2 | H9EJ42 |
| VFFQFDNLSSEDLKAV | 1 | 0 | 0 | 1702750 | YP_009555238.1 | P0C6X6 |
| VFFSPESVFSVFGDVK | 1 | 0 | 0 | 1702752 | YP_003766.2 | H9EJ42 |
| VFGYVQQGIFNKATVA | 1 | 0 | 0 | 1702771 | YP_009555238.1 | P0C6X6 |
| VFINKYGLEDFNFEHV | 1 | 0 | 0 | 1702779 | YP_003766.2 | H9EJ42 |
| VFIQKYGLEDFNFEHV | 1 | 0 | 0 | 1702781 | NP_073549.1 | A0A0P0LKV0 |
| VFISPYNSQNAVAAKI | 1 | 0 | 0 | 1702784 | QIQ54047.1 | P0C6X7 |
| VFISPYNSQNFAAKRV | 1 | 0 | 0 | 1702787 | YP_009555238.1 | P0C6X6 |
| VFISPYNSQNYVAKRV | 1 | 0 | 0 | 1702789 | YP_173236.1 | A0A140H1G9 |
| VFKIGDVKFKRLGDYV | 1 | 0 | 0 | 1702795 | YP_003766.2 | H9EJ42 |
| VFKTLKPGDSFNILAC | 1 | 0 | 0 | 1702804 | YP_003766.2 | H9EJ42 |
| VFLVDKNPHNTTESR | 1 | 0 | 0 | 1702828 | A0A161J4U8.1 | Q69028 |
| VFMSEAKCWTETDLTK | 1 | 0 | 0 | 1702837 | YP_009724389.1 | P0DTD1 |
| VFMSEAKCWVETDLKK | 1 | 0 | 0 | 1702838 | YP_009047202.1 | K9N7C7 |
| VFMSESKCWVEHDINN | 1 | 0 | 0 | 1702839 | YP_009555238.1 | P0C6X6 |
| VFMSESKCWVENDITN | 1 | 0 | 0 | 1702840 | YP_173236.1 | A0A140H1G9 |
| VFMSTSKCWVEEDLTK | 1 | 0 | 0 | 1702842 | YP_003766.2 | H9EJ42 |
| VFNATTFASVYAWNRK | 1 | 0 | 0 | 1702847 | QHR63300.2 | Q3LZX1 |

| | | | | | | |
|---|---|---|---|---|---|---|
| VFNDVYNDALFVSYGS | 1 | 0 | 0 | 1702848 | YP_173236.1 | A0A140H1G9 |
| VFNICQAVSANVCALM | 1 | 0 | 0 | 1702852 | YP_009555238.1 | P0C6X6 |
| VFNNNGTNVTQVKNAS | 1 | 0 | 0 | 1702860 | NP_073549.1 | A0A0P0LKV0 |
| VFPLNSKVKVIQPRVE | 1 | 0 | 0 | 1702871 | NP_828849.2 | P0C6X7 |
| VFPYDMDSGVSSFSEL | 1 | 0 | 0 | 1702874 | YP_009047202.1 | K9N7C7 |
| VFRAFDICNYKAKVAG | 1 | 0 | 0 | 1702886 | YP_009047202.1 | K9N7C7 |
| VFSTVNALPEVNADIV | 1 | 0 | 0 | 1702904 | NP_073549.1 | A0A0P0LKV0 |
| VFTACSHAAVDSLCAK | 1 | 0 | 0 | 1702911 | NP_073549.1 | A0A0P0LKV0 |
| VFTDGKLNCSFQDAAS | 1 | 0 | 0 | 1702914 | YP_009047202.1 | K9N7C7 |
| VFTGYHITKNSKVQLG | 1 | 0 | 0 | 1702919 | YP_009047202.1 | K9N7C7 |
| VFTNKTTLPTNVAFEL | 1 | 0 | 0 | 1702927 | YP_003766.2 | H9EJ42 |
| VFTTINALPEMVTDIV | 1 | 0 | 0 | 1702931 | YP_009555238.1 | P0C6X6 |
| VFTTVDNINLHTQVVD | 1 | 0 | 0 | 1702932 | YP_009724389.1 | P0DTD1 |
| VFTTVDNTNLHTQLVD | 1 | 0 | 0 | 1702933 | NP_828849.2 | P0C6X7 |
| VFVCTDNSANTKVLRN | 1 | 0 | 0 | 1702938 | YP_009047202.1 | K9N7C7 |
| VFVDDITKTDAVILLE | 1 | 0 | 0 | 1702940 | NP_073549.1 | A0A0P0LKV0 |
| VFVDDVVKTDAVVLLE | 1 | 0 | 0 | 1702941 | YP_003766.2 | H9EJ42 |
| VFVDKYMCGFDGKPVL | 1 | 0 | 0 | 1702942 | YP_003766.2 | H9EJ42 |
| VFVNKTSLPTNIAFEL | 1 | 0 | 0 | 1702953 | NP_073549.1 | A0A0P0LKV0 |
| VFWVANHQADTSTPSD | 1 | 0 | 0 | 1702971 | YP_173242.1 | Q19U29 |
| VFYCTNNTLVSGDAHI | 1 | 0 | 0 | 1702977 | NP_073551.1 | A0A0P0K6L9 |
| VFYELDNDFNTILNTA | 1 | 0 | 0 | 1702978 | YP_009555238.1 | P0C6X6 |
| VFYFAVRKEGQDVIFS | 1 | 0 | 0 | 1702979 | YP_009555238.1 | P0C6X6 |
| VFYFITPVHVLTKHGD | 1 | 0 | 0 | 1702980 | QIQ54046.1 | P0C6U8 |
| VFYKENSYTTTIKPIV | 1 | 0 | 0 | 1702982 | QIQ54046.1 | P0C6U8 |
| VFYKENSYTTTIKPVT | 1 | 0 | 0 | 1702983 | YP_009724389.1 | P0DTD1 |
| VFYKETSYTTTIKPVS | 1 | 0 | 0 | 1702984 | NP_828849.2 | P0C6X7 |
| VFYVRVKHGDFSAVSS | 1 | 0 | 0 | 1702989 | NP_073555.1 | A0A0P0KXN3 |
| VGADGELPVAISGDKV | 1 | 0 | 0 | 1702995 | NP_073549.1 | A0A0P0LKV0 |
| VGAEHVNNSYECDIPV | 1 | 0 | 0 | 1702996 | QIQ54048.1 | P59594 |
| VGAMLSENFTSYGFSN | 1 | 0 | 0 | 1703003 | NP_073551.1 | A0A0P0K6L9 |
| VGAWDGYLSSCCGTPA | 1 | 0 | 0 | 1703006 | YP_003766.2 | H9EJ42 |
| VGFSNGVDFVAQPEGC | 1 | 0 | 0 | 1703077 | NP_073549.1 | A0A0P0LKV0 |
| VGFVEAYNNCTGGAEI | 1 | 0 | 0 | 1703082 | YP_009555241.1 | P36334 |
| VGFVEAYNNCTGGSEI | 1 | 0 | 0 | 1703083 | YP_173238.1 | A0A140H1H1 |
| VGGKPCIKVATVQSKM | 1 | 0 | 0 | 1703093 | YP_009724389.1 | P0DTD1 |
| VGGMIRYYDVMANGGT | 1 | 0 | 0 | 1703097 | YP_009555238.1 | P0C6X6 |
| VGGQIVTCAKEIKESV | 1 | 0 | 0 | 1703099 | YP_009724389.1 | P0DTD1 |
| VGGVYDNSITCPLTVA | 1 | 0 | 0 | 1703108 | YP_009047202.1 | K9N7C7 |
| VGHYDGRTGELPCAIM | 1 | 0 | 0 | 1703115 | YP_173236.1 | A0A140H1G9 |
| VGINYLGDFAQGPFID | 1 | 0 | 0 | 1703127 | NP_073549.1 | A0A0P0LKV0 |
| VGITQQVLSENQKLIA | 1 | 0 | 0 | 1703128 | YP_009047204.1 | K9N5Q8 |
| VGIVYSSQLYEDVPSI | 1 | 0 | 0 | 1703130 | YP_003768.1 | H9EJ24 |
| VGLVGVLTLDNQDLNG | 1 | 0 | 0 | 1703145 | YP_009555238.1 | P0C6X6 |
| VGMPSFVAYETARQEY | 1 | 0 | 0 | 1703147 | NP_073549.1 | A0A0P0LKV0 |
| VGNCVKMLTNGSGSGQ | 1 | 0 | 0 | 1703151 | NP_073549.1 | A0A0P0LKV0 |
| VGNICYTPSKLIEYSD | 1 | 0 | 0 | 1703154 | NP_828849.2 | P0C6X7 |
| VGNICYTPSKLIEYTD | 1 | 0 | 0 | 1703155 | YP_009724389.1 | P0DTD1 |
| VGNKFYNVSTGLIPGF | 1 | 0 | 0 | 1703156 | YP_173236.1 | A0A140H1G9 |
| VGNYRLPSSKPSGMDT | 1 | 0 | 0 | 1703160 | YP_173241.1 | Q0ZJ82 |
| VGNYRLPSTQKGSGMD | 1 | 0 | 0 | 1703161 | YP_009555244.1 | Q01455 |
| VGPDARAKQDVSLLSK | 1 | 0 | 0 | 1703162 | YP_009047202.1 | K9N7C7 |
| VGPDARTQGKQSYVLL | 1 | 0 | 0 | 1703164 | YP_009555238.1 | P0C6X6 |
| VGPKELLLLWESGKAK | 1 | 0 | 0 | 1703169 | NP_073549.1 | A0A0P0LKV0 |
| VGPNLNAGEDIQLLKA | 1 | 0 | 0 | 1703172 | NP_828849.2 | P0C6X7 |
| VGPNLNKGEDIQLLKV | 1 | 0 | 0 | 1703173 | QIQ54046.1 | P0C6U8 |
| VGPNVNKGEDIQLLKS | 1 | 0 | 0 | 1703174 | YP_009724389.1 | P0DTD1 |
| VGPNVNRGEDIQLLKS | 1 | 0 | 0 | 1703175 | QHR63299.1 | P0C6W2 |
| VGQEEFIQTNSPKVTI | 1 | 0 | 0 | 1703179 | YP_173238.1 | A0A140H1H1 |
| VGRSVNETSQTGWAFY | 1 | 0 | 0 | 1703190 | YP_003770.1 | U3M6Q8 |
| VGRSVNSQNSTGWVFY | 1 | 0 | 0 | 1703191 | NP_073555.1 | A0A0P0KXN3 |
| VGRWSASINTGNCPFS | 1 | 0 | 0 | 1703193 | NP_073551.1 | A0A0P0K6L9 |
| VGSANIFIGDKVGHYV | 1 | 0 | 0 | 1703194 | YP_009555238.1 | P0C6X6 |
| VGSSFDGVMYGGFEDQ | 1 | 0 | 0 | 1703205 | NP_073549.1 | A0A0P0LKV0 |
| VGSSVAVQSKDLNFLN | 1 | 0 | 0 | 1703206 | YP_173236.1 | A0A140H1G9 |
| VGTDSGFAAYNRYRIG | 1 | 0 | 0 | 1703208 | NP_828855.1 | P59596 |
| VGTGIIRSSNQSLAGG | 1 | 0 | 0 | 1703210 | YP_003767.1 | A0A0P0G321 |
| VGTNVPLQVGFSNGVD | 1 | 0 | 0 | 1703216 | NP_073549.1 | A0A0P0LKV0 |
| VGTQSVDMLAVKTGVA | 1 | 0 | 0 | 1703219 | YP_009047202.1 | K9N7C7 |
| VGVIADKDKDLSGAVP | 1 | 0 | 0 | 1703229 | NP_073549.1 | A0A0P0LKV0 |
| VGVRQQRFVYDAYQNL | 1 | 0 | 0 | 1703236 | YP_009047204.1 | K9N5Q8 |
| VGVSERINVVPGVPTN | 1 | 0 | 0 | 1703238 | YP_003766.2 | H9EJ42 |
| VGVVDTEWGNMLTGIA | 1 | 0 | 0 | 1703240 | YP_009047202.1 | K9N7C7 |
| VGVYDINHGDAGFTSI | 1 | 0 | 0 | 1703241 | YP_009555240.1 | P30215 |
| VGYQVRAFLNKLNENV | 1 | 0 | 0 | 1703253 | YP_173236.1 | A0A140H1G9 |
| VGYTIEEAKTALKKSK | 1 | 0 | 0 | 1703255 | QIQ54046.1 | P0C6U8 |
| VHAEPKGKYSQKAYAL | 1 | 0 | 0 | 1703262 | YP_173236.1 | A0A140H1G9 |
| VHAGTDLEGKFYGPFV | 1 | 0 | 0 | 1703263 | NP_828849.2 | P0C6X7 |
| VHAGTDLEGNFYGPFV | 1 | 0 | 0 | 1703264 | YP_009724389.1 | P0DTD1 |
| VHAGTDLEGTFYGPFV | 1 | 0 | 0 | 1703265 | QHR63299.1 | P0C6W2 |
| VHARLKGGLILKFDSG | 1 | 0 | 0 | 1703267 | YP_009047202.1 | K9N7C7 |
| VHDFFKFRVDGDMVPH | 1 | 0 | 0 | 1703272 | NP_828849.2 | P0C6X7 |
| VHDNVAFYQGDVDTVV | 1 | 0 | 0 | 1703273 | NP_073549.1 | A0A0P0LKV0 |
| VHKDNFKDGLCMFWNC | 1 | 0 | 0 | 1703298 | YP_173236.1 | A0A140H1G9 |
| VHNWLKPYGGQPVSEY | 1 | 0 | 0 | 1703313 | YP_009047207.1 | K9N643 |
| VHRNEHVASGDAIMTR | 1 | 0 | 0 | 1703317 | NP_073549.1 | A0A0P0LKV0 |
| VHTPKHVFKTLKPGDS | 1 | 0 | 0 | 1703326 | YP_003766.2 | H9EJ42 |
| VHVKCEQSYQLYDASN | 1 | 0 | 0 | 1703327 | YP_009555238.1 | P0C6X6 |
| VHVLTKHGDFADEIIG | 1 | 0 | 0 | 1703330 | QIQ54046.1 | P0C6U8 |
| VIADKDKDLSGAVPSD | 1 | 0 | 0 | 1703339 | NP_073549.1 | A0A0P0LKV0 |
| VIADTLQETPVVPDTV | 1 | 0 | 0 | 1703341 | YP_009047202.1 | K9N7C7 |

| | | | | | | |
|---|---|---|---|---|---|---|
| VIADYNYKLPDDFMGC | 1 | 0 | 0 | 1703342 | NP_828851.1 | P59594 |
| VIADYNYKLPDDFTGC | 1 | 0 | 0 | 1703343 | QIQ54048.1 | P59594 |
| VIANVPAIDVLKKLLS | 1 | 0 | 0 | 1703351 | YP_003766.2 | H9EJ42 |
| VIASNTTSAIDYDHEY | 1 | 0 | 0 | 1703354 | NP_073549.1 | A0A0P0LKV0 |
| VIAWNSKHIDAKEGGN | 1 | 0 | 0 | 1703360 | QHR63300.2 | Q3LZX1 |
| VIAWNSVKQDALTGGN | 1 | 0 | 0 | 1703362 | QIQ54048.1 | P59594 |
| VICSASDMTNPDYTNL | 1 | 0 | 0 | 1703366 | YP_009555238.1 | P0C6X6 |
| VICSSSNMNEPDYSAL | 1 | 0 | 0 | 1703367 | YP_173236.1 | A0A140H1G9 |
| VIDDVDVVKPFRVEGN | 1 | 0 | 0 | 1703373 | YP_003766.2 | H9EJ42 |
| VIDGALLSKTSEYKGP | 1 | 0 | 0 | 1703375 | QIQ54046.1 | P0C6U8 |
| VIDMRSYEKLANSISP | 1 | 0 | 0 | 1703384 | NP_073549.1 | A0A0P0LKV0 |
| VIDQDFGSTVFNVPTK | 1 | 0 | 0 | 1703387 | YP_009555238.1 | P0C6X6 |
| VIEVSQIQSKLTDVKC | 1 | 0 | 0 | 1703398 | YP_173236.1 | A0A140H1G9 |
| VIFAQTSDTAHACNAN | 1 | 0 | 0 | 1703400 | NP_073549.1 | A0A0P0LKV0 |
| VIFCQTADTAHANNIN | 1 | 0 | 0 | 1703401 | YP_009047202.1 | K9N7C7 |
| VIFINNTTYPTNVAVE | 1 | 0 | 0 | 1703404 | YP_009555238.1 | P0C6X6 |
| VIFKNNTSFPTNIAVE | 1 | 0 | 0 | 1703405 | YP_173236.1 | A0A140H1G9 |
| VIFSQFDSLGVSSNQS | 1 | 0 | 0 | 1703409 | YP_009555238.1 | P0C6X6 |
| VIGDFNCTNFAINDLN | 1 | 0 | 0 | 1703415 | YP_173238.1 | A0A140H1H1 |
| VIGDLKCTSDNINDKD | 1 | 0 | 0 | 1703416 | YP_009555241.1 | P36334 |
| VIIANRYNEVANAVMQ | 1 | 0 | 0 | 1703444 | YP_173236.1 | A0A140H1G9 |
| VIIINNSTNVVIRACN | 1 | 0 | 0 | 1703449 | NP_828851.1 | P59594 |
| VIKNLTPIKLNFGMLN | 1 | 0 | 0 | 1703463 | YP_003766.2 | H9EJ42 |
| VIKNVNSVRDWLKSLK | 1 | 0 | 0 | 1703464 | NP_073549.1 | A0A0P0LKV0 |
| VIKPVCAKSPPGEQFR | 1 | 0 | 0 | 1703465 | YP_003766.2 | H9EJ42 |
| VIKTLQPVSELLIPLG | 1 | 0 | 0 | 1703468 | QHR63299.1 | P0C6W2 |
| VIKTLQPVSELLTPLG | 1 | 0 | 0 | 1703469 | YP_009724389.1 | P0DTD1 |
| VIKVAPKPYRVDGKFS | 1 | 0 | 0 | 1703470 | NP_073549.1 | A0A0P0LKV0 |
| VILCWETGKVKPPLNK | 1 | 0 | 0 | 1703474 | YP_173236.1 | A0A140H1G9 |
| VILKTGELQPLEEPTA | 1 | 0 | 0 | 1703479 | QIQ54046.1 | P0C6U8 |
| VILLERYVSLAIDAYP | 1 | 0 | 0 | 1703480 | NP_073549.1 | A0A0P0LKV0 |
| VILPGEFSKKQLGLLQ | 1 | 0 | 0 | 1703487 | YP_009047202.1 | K9N7C7 |
| VILVSNNKEDFDIIQK | 1 | 0 | 0 | 1703497 | YP_173236.1 | A0A140H1G9 |
| VINARIRAKHYVYIGD | 1 | 0 | 0 | 1703503 | YP_009555238.1 | P0C6X6 |
| VINKGHFDGQQGETPV | 1 | 0 | 0 | 1703512 | QIQ54047.1 | P0C6X7 |
| VINLRADQINDMVYSL | 1 | 0 | 0 | 1703513 | YP_009555238.1 | P0C6X6 |
| VINQLDEKAQKFFDFG | 1 | 0 | 0 | 1703516 | NP_073549.1 | A0A0P0LKV0 |
| VIPATSASKLTIVSPD | 1 | 0 | 0 | 1703523 | YP_003766.2 | H9EJ42 |
| VIPDFTVWNQVVTYPS | 1 | 0 | 0 | 1703524 | YP_009047202.1 | K9N7C7 |
| VIPDKQVFDKVVDNVY | 1 | 0 | 0 | 1703525 | YP_173236.1 | A0A140H1G9 |
| VIPDYNTYKNTCDGTT | 1 | 0 | 0 | 1703526 | YP_009724389.1 | P0DTD1 |
| VIPDYVDVNKTLQEFA | 1 | 0 | 0 | 1703527 | YP_003767.1 | A0A0P0G321 |
| VIPFKDGIYFAATEKS | 1 | 0 | 0 | 1703529 | NP_828851.1 | P59594 |
| VIPSKKAGGTTEMLSR | 1 | 0 | 0 | 1703535 | NP_828849.2 | P0C6X7 |
| VIPTKKAGGTTEMLTK | 1 | 0 | 0 | 1703539 | QHR63299.1 | P0C6W2 |
| VIQDVKDGTVTFNLLA | 1 | 0 | 0 | 1703542 | NP_073552.1 | A0A0P0KWE8 |
| VIRDVSVEDVCFANKF | 1 | 0 | 0 | 1703548 | YP_009555238.1 | P0C6X6 |
| VIRGHLYIQGVKLGTG | 1 | 0 | 0 | 1703553 | YP_173241.1 | Q0ZJ82 |
| VIRKDLSYEQQNVLAI | 1 | 0 | 0 | 1703554 | YP_003766.2 | H9EJ42 |
| VIRYNLNFEENLRRGT | 1 | 0 | 0 | 1703558 | NP_073551.1 | A0A0P0K6L9 |
| VISRMGFKLDATVPGY | 1 | 0 | 0 | 1703571 | YP_009047202.1 | K9N7C7 |
| VITNVGEPIISQPSKL | 1 | 0 | 0 | 1703590 | YP_003766.2 | H9EJ42 |
| VITPGTNASNQVAVLY | 1 | 0 | 0 | 1703591 | QHR63300.2 | Q3LZX1 |
| VITRDVGFVVPGLPGT | 1 | 0 | 0 | 1703594 | QIQ54046.1 | P0C6U8 |
| VITREVGFVVPGLPGT | 1 | 0 | 0 | 1703595 | YP_009724389.1 | P0DTD1 |
| VITSGDGTTSPISEHD | 1 | 0 | 0 | 1703597 | YP_009724391.1 | P0DTC3 |
| VIVAWHVVRDPRYVMR | 1 | 0 | 0 | 1703602 | YP_009555238.1 | P0C6X6 |
| VIVEDDVIENVKSSLS | 1 | 0 | 0 | 1703604 | YP_173236.1 | A0A140H1G9 |
| VIVFDGKSKCDESASK | 1 | 0 | 0 | 1703605 | NP_828849.2 | P0C6X7 |
| VIVFDGKSKCDESSAR | 1 | 0 | 0 | 1703606 | QIQ54046.1 | P0C6U8 |
| VIVFDGKSKCEESSAK | 1 | 0 | 0 | 1703607 | YP_009724389.1 | P0DTD1 |
| VIVNDDTSETKYVKSL | 1 | 0 | 0 | 1703614 | YP_009555238.1 | P0C6X6 |
| VIVNNLDKSAGFPFNK | 1 | 0 | 0 | 1703615 | YP_009724389.1 | P0DTD1 |
| VIVNNYDKSAGYPFNK | 1 | 0 | 0 | 1703616 | YP_009555238.1 | P0C6X6 |
| VIYDKETKTHATLFGS | 1 | 0 | 0 | 1703631 | YP_009047204.1 | K9N5Q8 |
| VIYSQTAETAHSVNVN | 1 | 0 | 0 | 1703640 | YP_009555238.1 | P0C6X6 |
| VKAAESKAFVDTIVPP | 1 | 0 | 0 | 1703642 | NP_073549.1 | A0A0P0LKV0 |
| VKAKGYPQFAELVPST | 1 | 0 | 0 | 1703649 | NP_073556.1 | A0A0N7J131 |
| VKALNESYIDLKELGN | 1 | 0 | 0 | 1703650 | YP_009047204.1 | K9N5Q8 |
| VKAQSSGVIPENPVLF | 1 | 0 | 0 | 1703653 | YP_173236.1 | A0A140H1G9 |
| VKATKGEGDGGITSEG | 1 | 0 | 0 | 1703655 | NP_073549.1 | A0A0P0LKV0 |
| VKCEQSYQLYDASNVK | 1 | 0 | 0 | 1703659 | YP_009555238.1 | P0C6X6 |
| VKCKGLNKPFVKDNVS | 1 | 0 | 0 | 1703662 | YP_009047202.1 | K9N7C7 |
| VKDDYCDGSLGCKMCL | 1 | 0 | 0 | 1703672 | NP_073549.1 | A0A0P0LKV0 |
| VKDFDFKFYDAQPIVK | 1 | 0 | 0 | 1703673 | YP_009555238.1 | P0C6X6 |
| VKDFDFYRYNKPTILD | 1 | 0 | 0 | 1703674 | YP_003766.2 | H9EJ42 |
| VKDFNTLSQEGKKYLV | 1 | 0 | 0 | 1703676 | YP_003766.2 | H9EJ42 |
| VKDFVSGLVNVQKVEQ | 1 | 0 | 0 | 1703677 | NP_073549.1 | A0A0P0LKV0 |
| VKDGKYFTSEPPVTYS | 1 | 0 | 0 | 1703679 | YP_009047202.1 | K9N7C7 |
| VKDKCVNFNFNGLTGT | 1 | 0 | 0 | 1703682 | QIQ54048.1 | P59594 |
| VKDNDGKNVHLKDVTK | 1 | 0 | 0 | 1703684 | NP_073549.1 | A0A0P0LKV0 |
| VKDNVSFVADDSGTPV | 1 | 0 | 0 | 1703686 | YP_009047202.1 | K9N7C7 |
| VKDPPNVQIHTIDGSS | 1 | 0 | 0 | 1703687 | NP_828852.2 | P59632 |
| VKDYMSLSEQLRKQIR | 1 | 0 | 0 | 1703694 | NP_828849.2 | P0C6X7 |
| VKEAQSANPMVIVNAA | 1 | 0 | 0 | 1703697 | NP_828849.2 | P0C6X7 |
| VKFGAALAMEKVNECV | 1 | 0 | 0 | 1703712 | YP_173238.1 | A0A140H1H1 |
| VKFQDSKPPLPPDEWV | 1 | 0 | 0 | 1703719 | YP_009047209.1 | K9N5R3 |
| VKFQTVKPGNFNQDFY | 1 | 0 | 0 | 1703720 | YP_009555238.1 | P0C6X6 |
| VKGQIQVASDNIKDCV | 1 | 0 | 0 | 1703742 | NP_828849.2 | P0C6X7 |
| VKHDVSPFEMPFEELN | 1 | 0 | 0 | 1703749 | NP_073549.1 | A0A0P0LKV0 |
| VKHGDFSAVSSPMSNM | 1 | 0 | 0 | 1703751 | NP_073555.1 | A0A0P0KXN3 |

| | | | | | | |
|---|---|---|---|---|---|---|
| VKHLNKNLNEGVLESF | 1 | 0 | 0 | 1703758 | YP_003766.2 | H9EJ42 |
| VKHLNKTLNEGVLESF | 1 | 0 | 0 | 1703759 | NP_073549.1 | A0A0P0LKV0 |
| VKIDGDYVYLPYPDPS | 1 | 0 | 0 | 1703763 | YP_173236.1 | A0A140H1G9 |
| VKIDNASPASTVHATA | 1 | 0 | 0 | 1703764 | NP_828852.2 | P59632 |
| VKIEDAIIVNDENSSI | 1 | 0 | 0 | 1703765 | YP_173236.1 | A0A140H1G9 |
| VKIGAVKHCQCGTVAT | 1 | 0 | 0 | 1703767 | NP_073549.1 | A0A0P0LKV0 |
| VKIGGLTFKTYAFDKP | 1 | 0 | 0 | 1703768 | NP_073549.1 | A0A0P0LKV0 |
| VKIGPERTCCLCDKRA | 1 | 0 | 0 | 1703769 | QHR63299.1 | P0C6W2 |
| VKIGPERTCCLCDRRA | 1 | 0 | 0 | 1703770 | YP_009724389.1 | P0DTD1 |
| VKIGPIKYCYCGNSAT | 1 | 0 | 0 | 1703771 | YP_003766.2 | H9EJ42 |
| VKILNNLGVDIAANTV | 1 | 0 | 0 | 1703772 | YP_009724389.1 | P0DTD1 |
| VKINNVDTVIFKNNTS | 1 | 0 | 0 | 1703774 | YP_173236.1 | A0A140H1G9 |
| VKIQAVVNANAEALNN | 1 | 0 | 0 | 1703775 | YP_009555241.1 | P36334 |
| VKITEHSWNADLYKLM | 1 | 0 | 0 | 1703777 | NP_828849.2 | P0C6X7 |
| VKIYCPACHNPEVGPE | 1 | 0 | 0 | 1703779 | QHR63299.1 | P0C6W2 |
| VKKGCFTGVDGELPVA | 1 | 0 | 0 | 1703782 | YP_003766.2 | H9EJ42 |
| VKKVAFGGDQVHEVAA | 1 | 0 | 0 | 1703787 | YP_009047202.1 | K9N7C7 |
| VKLGAEKKDGFYETIV | 1 | 0 | 0 | 1703793 | YP_009555239.1 | Q80872 |
| VKLGTGYTLSDLPVYV | 1 | 0 | 0 | 1703794 | YP_173241.1 | Q0ZJ82 |
| VKLPSAPQTIQPEVKE | 1 | 0 | 0 | 1703806 | YP_009047202.1 | K9N7C7 |
| VKLQNNELSPVALRQM | 1 | 0 | 0 | 1703810 | YP_009724389.1 | P0DTD1 |
| VKLSDVGFVEAYNNCT | 1 | 0 | 0 | 1703812 | YP_009555241.1 | P36334 |
| VKMDGDDVYLPYPNPS | 1 | 0 | 0 | 1703815 | YP_009555238.1 | P0C6X6 |
| VKMPCPACQDPEIGPE | 1 | 0 | 0 | 1703824 | NP_828849.2 | P0C6X7 |
| VKNSTAFPFETLSACR | 1 | 0 | 0 | 1703840 | YP_009047202.1 | K9N7C7 |
| VKNTKLYVNKTLYSEF | 1 | 0 | 0 | 1703841 | YP_173238.1 | A0A140H1H1 |
| VKPFITESKPSVEQRK | 1 | 0 | 0 | 1703854 | YP_009724389.1 | P0DTD1 |
| VKPGDAVITTQQAGAG | 1 | 0 | 0 | 1703858 | NP_073549.1 | A0A0P0LKV0 |
| VKPGGTSSGDATTAFA | 1 | 0 | 0 | 1703861 | YP_009555238.1 | P0C6X6 |
| VKPGHFNEEFYNFLRL | 1 | 0 | 0 | 1703863 | YP_003766.2 | H9EJ42 |
| VKPGHFNKEFYDFLRS | 1 | 0 | 0 | 1703864 | NP_073549.1 | A0A0P0LKV0 |
| VKPGNFNKDFYDFAVS | 1 | 0 | 0 | 1703865 | YP_009724389.1 | P0DTD1 |
| VKPGNFNQDFYDFVLS | 1 | 0 | 0 | 1703866 | YP_009555238.1 | P0C6X6 |
| VKPGNFNQDFYEFVKS | 1 | 0 | 0 | 1703867 | YP_173236.1 | A0A140H1G9 |
| VKPKEETGLLMPLKAP | 1 | 0 | 0 | 1703869 | QHR63299.1 | P0C6W2 |
| VKPKIKACIDEVTTTL | 1 | 0 | 0 | 1703870 | NP_828849.2 | P0C6X7 |
| VKPKPVINQLDEKAQK | 1 | 0 | 0 | 1703871 | NP_073549.1 | A0A0P0LKV0 |
| VKPKRLRKKRNVDPLS | 1 | 0 | 0 | 1703872 | YP_009047202.1 | K9N7C7 |
| VKPNRTSVVSFNEWAL | 1 | 0 | 0 | 1703876 | YP_009047202.1 | K9N7C7 |
| VKPVATVHRRIPDLPD | 1 | 0 | 0 | 1703884 | YP_173238.1 | A0A140H1H1 |
| VKPVRARAPPGEQFTH | 1 | 0 | 0 | 1703887 | NP_073549.1 | A0A0P0LKV0 |
| VKQDALTGGNYGYLYR | 1 | 0 | 0 | 1703890 | QIQ54048.1 | P59594 |
| VKQGAKPVGNCVKMLT | 1 | 0 | 0 | 1703893 | NP_073549.1 | A0A0P0LKV0 |
| VKQGDDYVYLPYPDPS | 1 | 0 | 0 | 1703894 | YP_009724389.1 | P0DTD1 |
| VKQGHFIGVEGELPVA | 1 | 0 | 0 | 1703895 | YP_009047202.1 | K9N7C7 |
| VKQLFYVYDVHKDNFK | 1 | 0 | 0 | 1703896 | YP_173236.1 | A0A140H1G9 |
| VKRDGVQVGYCVHGIK | 1 | 0 | 0 | 1703908 | NP_073549.1 | A0A0P0LKV0 |
| VKRRFSLLRHEDLSVI | 1 | 0 | 0 | 1703918 | YP_009047207.1 | K9N643 |
| VKRTDLTIYNREMKCY | 1 | 0 | 0 | 1703920 | YP_009555238.1 | P0C6X6 |
| VKRTNLEVYNKEKTYY | 1 | 0 | 0 | 1703922 | YP_173236.1 | A0A140H1G9 |
| VKRVDWSVEYPIIGDE | 1 | 0 | 0 | 1703923 | NP_828849.2 | P0C6X7 |
| VKRVDWTIEYPIIGDE | 1 | 0 | 0 | 1703924 | YP_009724389.1 | P0DTD1 |
| VKRVRAWVGFDAEGAH | 1 | 0 | 0 | 1703925 | YP_009555238.1 | P0C6X6 |
| VKSALLADKFPVLHDI | 1 | 0 | 0 | 1703930 | NP_828849.2 | P0C6X7 |
| VKSDIFGSDFKTFDLL | 1 | 0 | 0 | 1703931 | YP_003766.2 | H9EJ42 |
| VKSFEIDKGIYQTSNF | 1 | 0 | 0 | 1703935 | NP_828851.1 | P59594 |
| VKSKVGNYRLPSSKPS | 1 | 0 | 0 | 1703943 | YP_173241.1 | Q0ZJ82 |
| VKSKVGNYRLPSTQKG | 1 | 0 | 0 | 1703944 | YP_009555244.1 | Q01455 |
| VKSLSMIRGPPRAELN | 1 | 0 | 0 | 1703949 | YP_009555238.1 | P0C6X6 |
| VKSREETGLLMPLKAP | 1 | 0 | 0 | 1703956 | YP_009724389.1 | P0DTD1 |
| VKSSDVVDSNENLTWP | 1 | 0 | 0 | 1703958 | YP_009047202.1 | K9N7C7 |
| VKSSITESKLSVEQRQ | 1 | 0 | 0 | 1703959 | QHR63299.1 | P0C6W2 |
| VKSSLSSYEYCQPPKS | 1 | 0 | 0 | 1703960 | YP_173236.1 | A0A140H1G9 |
| VKSSQSSPIIPGFGGD | 1 | 0 | 0 | 1703961 | YP_009047204.1 | K9N5Q8 |
| VKTDAVVLLERYVSLA | 1 | 0 | 0 | 1703972 | YP_003766.2 | H9EJ42 |
| VKTGHYTGQAGEMPCA | 1 | 0 | 0 | 1703977 | YP_009555238.1 | P0C6X6 |
| VKTQFNYFKKVDGIIQ | 1 | 0 | 0 | 1703986 | NP_828849.2 | P0C6X7 |
| VKTQFNYYKKVDGVVQ | 1 | 0 | 0 | 1703987 | YP_009724389.1 | P0DTD1 |
| VKVGTGVMVECDSLRI | 1 | 0 | 0 | 1703998 | NP_073549.1 | A0A0P0LKV0 |
| VKVIQPRVEKKKTEGF | 1 | 0 | 0 | 1703999 | NP_828849.2 | P0C6X7 |
| VKWADASEPQRGRQGR | 1 | 0 | 0 | 1704009 | NP_073556.1 | A0A0N7J131 |
| VLAAECTIFKDAAGKP | 1 | 0 | 0 | 1704026 | QIQ54046.1 | P0C6U8 |
| VLAAECTIFKDAMGKP | 1 | 0 | 0 | 1704027 | NP_828849.2 | P0C6X7 |
| VLAAECTIFKDASGKP | 1 | 0 | 0 | 1704028 | YP_009724389.1 | P0DTD1 |
| VLAIESIHYLGTTGHT | 1 | 0 | 0 | 1704037 | YP_003766.2 | H9EJ42 |
| VLAKGGFKFGDPADSR | 1 | 0 | 0 | 1704040 | YP_009555238.1 | P0C6X6 |
| VLAKGHFKFDEPSDAT | 1 | 0 | 0 | 1704041 | YP_173236.1 | A0A140H1G9 |
| VLAKLGKDATKPQQVT | 1 | 0 | 0 | 1704042 | YP_009555245.1 | P33469 |
| VLASDDLYVKRYFKGC | 1 | 0 | 0 | 1704055 | YP_173236.1 | A0A140H1G9 |
| VLATDDLYVKRYERGC | 1 | 0 | 0 | 1704056 | YP_009555238.1 | P0C6X6 |
| VLAYFSKKHVEVFTDG | 1 | 0 | 0 | 1704062 | YP_009047202.1 | K9N7C7 |
| VLDDFIVFNNNGTNVT | 1 | 0 | 0 | 1704077 | NP_073549.1 | A0A0P0LKV0 |
| VLDITDAAVKAAESKA | 1 | 0 | 0 | 1704080 | NP_073549.1 | A0A0P0LKV0 |
| VLDQSDNNCWISTTLV | 1 | 0 | 0 | 1704091 | YP_003766.2 | H9EJ42 |
| VLDVKGVEELHDDLVN | 1 | 0 | 0 | 1704098 | YP_009555239.1 | Q80872 |
| VLEDTLDDGPSVETSD | 1 | 0 | 0 | 1704103 | YP_009555238.1 | P0C6X6 |
| VLEENLNAYVNSNQNT | 1 | 0 | 0 | 1704104 | YP_173242.1 | Q19U29 |
| VLEGSVAYESLRPDTR | 1 | 0 | 0 | 1704105 | YP_009724389.1 | P0DTD1 |
| VLEGSVPYESLRPDTR | 1 | 0 | 0 | 1704106 | QIQ54046.1 | P0C6U8 |
| VLEMAKEVKTSSKVVL | 1 | 0 | 0 | 1704110 | NP_073549.1 | A0A0P0LKV0 |
| VLEQDAQIDRLINGRL | 1 | 0 | 0 | 1704112 | YP_009047204.1 | K9N5Q8 |

| | | | | | | |
|---|---|---|---|---|---|---|
| VLESFSVTLLDNQEDK | 1 | 0 | 0 | 1704114 | YP_003766.2 | H9EJ42 |
| VLETFQNNVVNYQHIG | 1 | 0 | 0 | 1704115 | YP_009555238.1 | P0C6X6 |
| VLFDGRDNGALEAFKR | 1 | 0 | 0 | 1704122 | YP_009555238.1 | P0C6X6 |
| VLFSTVVIKNLTPIKL | 1 | 0 | 0 | 1704136 | YP_003766.2 | H9EJ42 |
| VLFTNSTDTVNHDSFN | 1 | 0 | 0 | 1704139 | YP_173236.1 | A0A140H1G9 |
| VLFYCSNSSSGVLDTT | 1 | 0 | 0 | 1704143 | YP_003767.1 | A0A0P0G321 |
| VLGEYVFDKSELTNGV | 1 | 0 | 0 | 1704149 | YP_009555238.1 | P0C6X6 |
| VLGLQTQTVDSAQGSE | 1 | 0 | 0 | 1704159 | YP_009555238.1 | P0C6X6 |
| VLGVNEKTHVQLSLPV | 1 | 0 | 0 | 1704168 | NP_828849.2 | P0C6X7 |
| VLHLGAGSDKEVAPGS | 1 | 0 | 0 | 1704178 | YP_173236.1 | A0A140H1G9 |
| VLHYGAGSDKGVAPGT | 1 | 0 | 0 | 1704186 | YP_003766.2 | H9EJ42 |
| VLIDYDHAYSTMRLHN | 1 | 0 | 0 | 1704189 | YP_003766.2 | H9EJ42 |
| VLIKEPCSSGTYEGNS | 1 | 0 | 0 | 1704191 | QIQ54053.1 | P59635 |
| VLIQNGAKHVQGNVAK | 1 | 0 | 0 | 1704196 | YP_173236.1 | A0A140H1G9 |
| VLITEGNVKGLQPSVG | 1 | 0 | 0 | 1704199 | QIQ54047.1 | P0C6X7 |
| VLITEGSVKGLTPSKG | 1 | 0 | 0 | 1704201 | NP_828849.2 | P0C6X7 |
| VLKLPSLMKVMTHMDD | 1 | 0 | 0 | 1704221 | YP_173236.1 | A0A140H1G9 |
| VLKSEDAQGMDNLACE | 1 | 0 | 0 | 1704229 | YP_009724389.1 | P0DTD1 |
| VLKSEDTQGMDNLACE | 1 | 0 | 0 | 1704230 | QHR63299.1 | P0C6W2 |
| VLKTGDLQPLEQPTSE | 1 | 0 | 0 | 1704233 | YP_009724389.1 | P0DTD1 |
| VLLALQQLEVKFNAPA | 1 | 0 | 0 | 1704240 | NP_828849.2 | P0C6X7 |
| VLLDWVKHLNKNLNEG | 1 | 0 | 0 | 1704247 | YP_003766.2 | H9EJ42 |
| VLLERYVSLAIDAYPL | 1 | 0 | 0 | 1704250 | YP_003766.2 | H9EJ42 |
| VLLIERFVSLAIDAYP | 1 | 0 | 0 | 1704260 | YP_009555238.1 | P0C6X6 |
| VLLKEPCSSGTYEGNS | 1 | 0 | 0 | 1704264 | YP_009724395.1 | P0DTC7 |
| VLLNCLQHLHVASNSK | 1 | 0 | 0 | 1704269 | YP_009555238.1 | P0C6X6 |
| VLLRKNGNKGAGGHSY | 1 | 0 | 0 | 1704279 | YP_009724389.1 | P0DTD1 |
| VLLSKGSLEPRHFNSI | 1 | 0 | 0 | 1704282 | YP_173236.1 | A0A140H1G9 |
| VLLSMQGAVDINKLCE | 1 | 0 | 0 | 1704285 | YP_009724389.1 | P0DTD1 |
| VLLSMQGAVDINRLCE | 1 | 0 | 0 | 1704286 | NP_828849.2 | P0C6X7 |
| VLMGILSNMNIASNSK | 1 | 0 | 0 | 1704298 | NP_073549.1 | A0A0P0LKV0 |
| VLMGWDYPKCDRAMPN | 1 | 0 | 0 | 1704299 | YP_009555238.1 | P0C6X6 |
| VLNEKCSVYTVESGTE | 1 | 0 | 0 | 1704310 | NP_828849.2 | P0C6X7 |
| VLNENLNAYQQQDGMM | 1 | 0 | 0 | 1704311 | YP_009555245.1 | P33469 |
| VLNNCTKYNIYDYVGT | 1 | 0 | 0 | 1704320 | YP_003767.1 | A0A0P0G321 |
| VLNNISLTTQPYFDSY | 1 | 0 | 0 | 1704326 | YP_173238.1 | A0A140H1H1 |
| VLNNLNLPGCNGGSLY | 1 | 0 | 0 | 1704328 | YP_009555238.1 | P0C6X6 |
| VLNQSYINLKDIGTYE | 1 | 0 | 0 | 1704330 | YP_009555241.1 | P36334 |
| VLNVVCKTCGQQQTTL | 1 | 0 | 0 | 1704334 | YP_009724389.1 | P0DTD1 |
| VLNVVGPDARTQGKQS | 1 | 0 | 0 | 1704335 | YP_009555238.1 | P0C6X6 |
| VLPETYVALPIYYQHT | 1 | 0 | 0 | 1704338 | YP_003767.1 | A0A0P0G321 |
| VLPFNDGVYFASTEKS | 1 | 0 | 0 | 1704340 | YP_009724390.1 | P0DTC2 |
| VLPINTDTVLSVAPEV | 1 | 0 | 0 | 1704343 | YP_003766.2 | H9EJ42 |
| VLPLTCNAISSNTDNE | 1 | 0 | 0 | 1704346 | YP_173238.1 | A0A140H1H1 |
| VLPPLLSENQISGYTL | 1 | 0 | 0 | 1704353 | YP_009555241.1 | P36334 |
| VLPPLMDVNMEAAYTS | 1 | 0 | 0 | 1704356 | YP_009047204.1 | K9N5Q8 |
| VLPQRNIRSSRIAGRS | 1 | 0 | 0 | 1704362 | YP_003767.1 | A0A0P0G321 |
| VLPSIIPNKKEEILGT | 1 | 0 | 0 | 1704366 | QIQ54046.1 | P0C6U8 |
| VLPXNDGVYFASTEKS | 1 | 0 | 0 | 1704371 | QIQ54048.1 | P59594 |
| VLQENQKILAASFNKA | 1 | 0 | 0 | 1704378 | NP_073551.1 | A0A0P0K6L9 |
| VLRDNLKPVTTLSTIT | 1 | 0 | 0 | 1704397 | YP_003768.1 | H9EJ24 |
| VLRNSFGKDLNANMSL | 1 | 0 | 0 | 1704403 | NP_073549.1 | A0A0P0LKV0 |
| VLSAAIKDCVDINKAI | 1 | 0 | 0 | 1704412 | NP_073549.1 | A0A0P0LKV0 |
| VLSDRELHLSWEVGKP | 1 | 0 | 0 | 1704415 | YP_009724389.1 | P0DTD1 |
| VLSDVDGLKYTKIMKD | 1 | 0 | 0 | 1704417 | YP_173236.1 | A0A140H1G9 |
| VLSDVIRYNLNFEENL | 1 | 0 | 0 | 1704418 | NP_073551.1 | A0A0P0K6L9 |
| VLSEEVILKTGELQPL | 1 | 0 | 0 | 1704420 | QIQ54046.1 | P0C6U8 |
| VLSENQKLIANKFNQA | 1 | 0 | 0 | 1704423 | YP_009047204.1 | K9N5Q8 |
| VLSLRHDIALDDDART | 1 | 0 | 0 | 1704440 | YP_009555238.1 | P0C6X6 |
| VLSLVNFTVDPQKAYL | 1 | 0 | 0 | 1704442 | YP_009047202.1 | K9N7C7 |
| VLSMTDCLAAETHRDC | 1 | 0 | 0 | 1704445 | YP_009047202.1 | K9N7C7 |
| VLSSLCTMFKRGDGTP | 1 | 0 | 0 | 1704453 | YP_173236.1 | A0A140H1G9 |
| VLSVDSVSEESQGNVV | 1 | 0 | 0 | 1704459 | YP_173236.1 | A0A140H1G9 |
| VLSVNPYVCNAPGCDV | 1 | 0 | 0 | 1704460 | YP_009724389.1 | P0DTD1 |
| VLSVNPYVCNATGCDV | 1 | 0 | 0 | 1704461 | QIQ54047.1 | P0C6X7 |
| VLSVNSSNCNNFNVKK | 1 | 0 | 0 | 1704462 | NP_073549.1 | A0A0P0LKV0 |
| VLSXELLNGPATVCGP | 1 | 0 | 0 | 1704466 | QIQ54048.1 | P59594 |
| VLTEEVVLKTGDLQPL | 1 | 0 | 0 | 1704469 | YP_009724389.1 | P0DTD1 |
| VLTENALVRLTTEVVR | 1 | 0 | 0 | 1704470 | YP_003766.2 | H9EJ42 |
| VLTGCCDNSYFDSKGW | 1 | 0 | 0 | 1704473 | YP_003766.2 | H9EJ42 |
| VLTGDSVKFVYMHQLE | 1 | 0 | 0 | 1704475 | YP_173236.1 | A0A140H1G9 |
| VLTIANNYSKLFDEGY | 1 | 0 | 0 | 1704479 | NP_073549.1 | A0A0P0LKV0 |
| VLTKHGDFADEIIGYK | 1 | 0 | 0 | 1704481 | QIQ54046.1 | P0C6U8 |
| VLTLDNQDLNGKWYDF | 1 | 0 | 0 | 1704483 | YP_009047202.1 | K9N7C7 |
| VLTSEEVVLKNGELEA | 1 | 0 | 0 | 1704495 | NP_828849.2 | P0C6X7 |
| VLTTFIDAARGPAGVE | 1 | 0 | 0 | 1704502 | YP_009047202.1 | K9N7C7 |
| VLTTSKKQFLPFQQFG | 1 | 0 | 0 | 1704504 | QIQ54048.1 | P59594 |
| VLVRGFGDSVEEVLSE | 1 | 0 | 0 | 1704531 | YP_009724389.1 | P0DTD1 |
| VLVSNNQEDFDLISKC | 1 | 0 | 0 | 1704532 | YP_009555238.1 | P0C6X6 |
| VLVSPGLCISGDVGIA | 1 | 0 | 0 | 1704533 | YP_173238.1 | A0A140H1H1 |
| VLWDYEAERPLTSFTK | 1 | 0 | 0 | 1704545 | NP_073549.1 | A0A0P0LKV0 |
| VMAYVTGGVVQMTTQW | 1 | 0 | 0 | 1704581 | QIQ54046.1 | P0C6U8 |
| VMDVNDCFKNDNVVLK | 1 | 0 | 0 | 1704589 | YP_003766.2 | H9EJ42 |
| VMDYLDNLKPRVEAPK | 1 | 0 | 0 | 1704591 | NP_828849.2 | P0C6X7 |
| VMESQISTKEVKLKGV | 1 | 0 | 0 | 1704596 | YP_173236.1 | A0A140H1G9 |
| VMGLHGNQTLFCNIMK | 1 | 0 | 0 | 1704607 | NP_073549.1 | A0A0P0LKV0 |
| VMISEWPLSVQQAQQE | 1 | 0 | 0 | 1704618 | NP_073549.1 | A0A0P0LKV0 |
| VMISQWPISNDSNGCV | 1 | 0 | 0 | 1704619 | YP_003766.2 | H9EJ42 |
| VMLNTSIPNLPDFKEE | 1 | 0 | 0 | 1704628 | YP_009555241.1 | P36334 |
| VMPLSAPTLVPQEHYV | 1 | 0 | 0 | 1704643 | YP_009724389.1 | P0DTD1 |
| VMQFGYIDCEQDSCDF | 1 | 0 | 0 | 1704647 | YP_009555238.1 | P0C6X6 |

| | | | | | | |
|---|---|---|---|---|---|---|
| VMQKLPFKFKDLAIEN | 1 | 0 | 0 | 1704649 | YP_173236.1 | A0A140H1G9 |
| VMRYNLNFSANSLDNL | 1 | 0 | 0 | 1704660 | YP_003767.1 | A0A0P0G321 |
| VMSLKENQINDMIYSL | 1 | 0 | 0 | 1704665 | NP_828849.2 | P0C6X7 |
| VMSLSYNSVLDLSKFN | 1 | 0 | 0 | 1704666 | NP_073549.1 | A0A0P0LKV0 |
| VMTFYPRLQAASDWKP | 1 | 0 | 0 | 1704671 | YP_009555238.1 | P0C6X6 |
| VMWLYTYSIGEASSPI | 1 | 0 | 0 | 1704687 | YP_009047202.1 | K9N7C7 |
| VMYGGFEDQPNLQVES | 1 | 0 | 0 | 1704694 | NP_073549.1 | A0A0P0LKV0 |
| VMYMGTLSYEQFKKGV | 1 | 0 | 0 | 1704698 | YP_009724389.1 | P0DTD1 |
| VNAANTHLKHGGGIAG | 1 | 0 | 0 | 1704705 | YP_009047202.1 | K9N7C7 |
| VNADIVVVDEVSMCTN | 1 | 0 | 0 | 1704710 | NP_073549.1 | A0A0P0LKV0 |
| VNAGGAPLTNCVKMLT | 1 | 0 | 0 | 1704713 | YP_009047202.1 | K9N7C7 |
| VNALLSTDGNKIADKY | 1 | 0 | 0 | 1704718 | YP_009724389.1 | P0DTD1 |
| VNALLSTDGNKIGDKY | 1 | 0 | 0 | 1704719 | QIQ54047.1 | P0C6X7 |
| VNANGGSKFCKKHRFF | 1 | 0 | 0 | 1704725 | NP_073549.1 | A0A0P0LKV0 |
| VNARLRAKHYVYIGDP | 1 | 0 | 0 | 1704731 | YP_009724389.1 | P0DTD1 |
| VNASSSEAFLIGANYL | 1 | 0 | 0 | 1704734 | NP_828849.2 | P0C6X7 |
| VNCCRFQRIDDDGNKL | 1 | 0 | 0 | 1704742 | YP_173236.1 | A0A140H1G9 |
| VNCCRFQRVDENGDKL | 1 | 0 | 0 | 1704743 | YP_009555238.1 | P0C6X6 |
| VNCDSFGPGNTFINGD | 1 | 0 | 0 | 1704745 | YP_003766.2 | H9EJ42 |
| VNDAITQTSQALQTVA | 1 | 0 | 0 | 1704755 | NP_073551.1 | A0A0P0K6L9 |
| VNDDTSETKYVKSLSI | 1 | 0 | 0 | 1704757 | YP_009555238.1 | P0C6X6 |
| VNDICMFENKTTLPTN | 1 | 0 | 0 | 1704762 | YP_009047202.1 | K9N7C7 |
| VNDIDITFLKKDAPYI | 1 | 0 | 0 | 1704763 | QHR63299.1 | P0C6W2 |
| VNDKITEFQLDYSIDV | 1 | 0 | 0 | 1704766 | NP_073549.1 | A0A0P0LKV0 |
| VNDLCFANKFFQFDQW | 1 | 0 | 0 | 1704768 | YP_173236.1 | A0A140H1G9 |
| VNDSYRQFDLVQYDFT | 1 | 0 | 0 | 1704773 | YP_173236.1 | A0A140H1G9 |
| VNEKRRLKLIMPARF | 1 | 0 | 0 | 1704783 | P03155.1 | Q69028 |
| VNELLDTTQLQVANSL | 1 | 0 | 0 | 1704785 | YP_009555241.1 | P36334 |
| VNESRYGLQNLLQLPN | 1 | 0 | 0 | 1704787 | YP_003767.1 | A0A0P0G321 |
| VNETNSQYLFSTINAL | 1 | 0 | 0 | 1704788 | YP_009047202.1 | K9N7C7 |
| VNFKSISLTVGEVFGK | 1 | 0 | 0 | 1704792 | YP_173236.1 | A0A140H1G9 |
| VNFNVDDINFSPVLGC | 1 | 0 | 0 | 1704799 | YP_009555241.1 | P36334 |
| VNFTVDPQKAYLDFVN | 1 | 0 | 0 | 1704807 | YP_009047202.1 | K9N7C7 |
| VNFVVQPVGVVDTEWG | 1 | 0 | 0 | 1704810 | YP_009047202.1 | K9N7C7 |
| VNGDWFLFGDSRSDCN | 1 | 0 | 0 | 1704812 | YP_009555240.1 | P30215 |
| VNGLLDTTQLHVADTL | 1 | 0 | 0 | 1704819 | YP_173238.1 | A0A140H1H1 |
| VNGVDFDFIVNAANEN | 1 | 0 | 0 | 1704828 | NP_073549.1 | A0A0P0LKV0 |
| VNGYAKSNFVKDDYCD | 1 | 0 | 0 | 1704834 | NP_073549.1 | A0A0P0LKV0 |
| VNGYFIKTNNTRIVDE | 1 | 0 | 0 | 1704835 | YP_009047204.1 | K9N5Q8 |
| VNIAKNAYEKDKAVAR | 1 | 0 | 0 | 1704843 | YP_009047202.1 | K9N7C7 |
| VNINPHNYSYMDLNPV | 1 | 0 | 0 | 1704850 | YP_009555240.1 | P30215 |
| VNISLNGNTSVCVRTS | 1 | 0 | 0 | 1704854 | YP_003767.1 | A0A0P0G321 |
| VNITFELDERIDKVLN | 1 | 0 | 0 | 1704857 | YP_009724389.1 | P0DTD1 |
| VNITLANFNETKGPLC | 1 | 0 | 0 | 1704859 | NP_073551.1 | A0A0P0K6L9 |
| VNKASYDTNLNKFNRA | 1 | 0 | 0 | 1704861 | YP_009047202.1 | K9N7C7 |
| VNKFLALCADSIVIGG | 1 | 0 | 0 | 1704863 | QIQ54046.1 | P0C6U8 |
| VNKGEDIQLLKSAYEN | 1 | 0 | 0 | 1704866 | YP_009724389.1 | P0DTD1 |
| VNKGHFDGHAGEAPVS | 1 | 0 | 0 | 1704867 | NP_828849.2 | P0C6X7 |
| VNKGHFDGQQGEVPVS | 1 | 0 | 0 | 1704868 | YP_009724389.1 | P0DTD1 |
| VNKGSFVGADGELPVA | 1 | 0 | 0 | 1704869 | NP_073549.1 | A0A0P0LKV0 |
| VNKHAFHTPAFDKSAF | 1 | 0 | 0 | 1704872 | YP_009724389.1 | P0DTD1 |
| VNKIQPWTNDALNTIC | 1 | 0 | 0 | 1704877 | NP_073549.1 | A0A0P0LKV0 |
| VNKMVNWFNTMLDASA | 1 | 0 | 0 | 1704879 | YP_173236.1 | A0A140H1G9 |
| VNKTLQEFAQNLPKYV | 1 | 0 | 0 | 1704881 | YP_003767.1 | A0A0P0G321 |
| VNKTLQELSYKLPNYT | 1 | 0 | 0 | 1704882 | NP_073551.1 | A0A0P0K6L9 |
| VNKTLYSEFSTIVIGS | 1 | 0 | 0 | 1704883 | YP_173238.1 | A0A140H1H1 |
| VNKYFEIYDGGCLNAS | 1 | 0 | 0 | 1704885 | YP_009047202.1 | K9N7C7 |
| VNLDNLRANNTKGSIP | 1 | 0 | 0 | 1704887 | QIQ54046.1 | P0C6U8 |
| VNLDNLRANNTKGSLP | 1 | 0 | 0 | 1704888 | YP_009724389.1 | P0DTD1 |
| VNLDNLRASNTKGSLP | 1 | 0 | 0 | 1704889 | QHR63299.1 | P0C6W2 |
| VNMQQIYLINKFLKAN | 1 | 0 | 0 | 1704912 | YP_009555238.1 | P0C6X6 |
| VNMRVLHLGAGSDKEV | 1 | 0 | 0 | 1704914 | YP_173236.1 | A0A140H1G9 |
| VNMRVLHLGAGSEKGV | 1 | 0 | 0 | 1704915 | YP_009555238.1 | P0C6X6 |
| VNNATNKATYKPNTWC | 1 | 0 | 0 | 1704916 | YP_009724389.1 | P0DTD1 |
| VNNHAFHTPAYDKRAM | 1 | 0 | 0 | 1704921 | NP_073549.1 | A0A0P0LKV0 |
| VNNSYECDIPVGAGIC | 1 | 0 | 0 | 1704927 | QIQ54048.1 | P59594 |
| VNNTINKATFKPNTWC | 1 | 0 | 0 | 1704929 | QIQ54046.1 | P0C6U8 |
| VNNTVYDPLQPELDSF | 1 | 0 | 0 | 1704930 | YP_009724390.1 | P0DTC2 |
| VNNYDKSAGYPFNKFG | 1 | 0 | 0 | 1704934 | YP_009555238.1 | P0C6X6 |
| VNPAMDPIYDEPTTTT | 1 | 0 | 0 | 1704935 | QIQ54049.1 | P59632 |
| VNPDDKCFANKHRSFT | 1 | 0 | 0 | 1704939 | YP_009047202.1 | K9N7C7 |
| VNQLAYSSSANGSLVG | 1 | 0 | 0 | 1704951 | YP_009047202.1 | K9N7C7 |
| VNQQGNSLNHLTSQLR | 1 | 0 | 0 | 1704953 | NP_073551.1 | A0A0P0K6L9 |
| VNQQGSALNHLTSQLR | 1 | 0 | 0 | 1704954 | YP_003767.1 | A0A0P0G321 |
| VNQSLSNAKVSAEVSR | 1 | 0 | 0 | 1704957 | NP_073552.1 | A0A0P0KWE8 |
| VNRGEDIQLLKSAYEN | 1 | 0 | 0 | 1704964 | QHR63299.1 | P0C6W2 |
| VNSIFDIETVDVKHDV | 1 | 0 | 0 | 1704981 | NP_073549.1 | A0A0P0LKV0 |
| VNSNAQALNSLLQQLF | 1 | 0 | 0 | 1704984 | YP_173238.1 | A0A140H1H1 |
| VNSNQNTDSDSLSSKP | 1 | 0 | 0 | 1704985 | YP_173242.1 | Q19U29 |
| VNSNVKLLDYDYATHG | 1 | 0 | 0 | 1704986 | YP_003766.2 | H9EJ42 |
| VNSVNDSLEPVGGLYE | 1 | 0 | 0 | 1704996 | YP_009555241.1 | P36334 |
| VNTFSSTFNVPMDKLK | 1 | 0 | 0 | 1705001 | QIQ54046.1 | P0C6U8 |
| VNTFSSTFNVPMEKLK | 1 | 0 | 0 | 1705002 | YP_009724389.1 | P0DTD1 |
| VNTSLGNRKRNQKPLE | 1 | 0 | 0 | 1705009 | YP_003771.1 | H9EJ69 |
| VNTTIGNETTSAFVGA | 1 | 0 | 0 | 1705013 | NP_073551.1 | A0A0P0K6L9 |
| VNTVAGIPSNVYLVGK | 1 | 0 | 0 | 1705014 | NP_073549.1 | A0A0P0LKV0 |
| VNTVCFVDVDFHSVEI | 1 | 0 | 0 | 1705015 | YP_003766.2 | H9EJ42 |
| VNTVKPKPVINQLDEK | 1 | 0 | 0 | 1705016 | NP_073549.1 | A0A0P0LKV0 |
| VNTVSELVYENKFVPV | 1 | 0 | 0 | 1705017 | NP_073549.1 | A0A0P0LKV0 |
| VNVDGFNEWAMANGYT | 1 | 0 | 0 | 1705024 | YP_003766.2 | H9EJ42 |
| VNVGHYDGRTGELPCA | 1 | 0 | 0 | 1705025 | YP_173236.1 | A0A140H1G9 |

| | | | | | | |
|---|---|---|---|---|---|---|
| VNVHDVTVTTDKSFEQ | 1 | 0 | 0 | 1705026 | NP_073549.1 | A0A0P0LKV0 |
| VNVNNTWMYTGSGYYY | 1 | 0 | 0 | 1705036 | YP_009555241.1 | P36334 |
| VNVNRFNVAITRAKKG | 1 | 0 | 0 | 1705037 | YP_009555238.1 | P0C6X6 |
| VNVNVDFKDFQFMLWC | 1 | 0 | 0 | 1705038 | YP_009555238.1 | P0C6X6 |
| VNVQKVEQPKIEPKPV | 1 | 0 | 0 | 1705043 | NP_073549.1 | A0A0P0LKV0 |
| VNVSATNIQNLLYCDS | 1 | 0 | 0 | 1705045 | YP_003767.1 | A0A0P0G321 |
| VNVTKHKCDINYKGKV | 1 | 0 | 0 | 1705050 | YP_009555238.1 | P0C6X6 |
| VNVYRSTSPDPKFVDK | 1 | 0 | 0 | 1705057 | YP_009047202.1 | K9N7C7 |
| VNWADDRAARKKFPPP | 1 | 0 | 0 | 1705058 | YP_003771.1 | H9EJ69 |
| VNWFNTMLDASAPATG | 1 | 0 | 0 | 1705059 | YP_173236.1 | A0A140H1G9 |
| VNWNLEYPIISNEVSI | 1 | 0 | 0 | 1705060 | YP_173236.1 | A0A140H1G9 |
| VNWNTKHGSGVTRELM | 1 | 0 | 0 | 1705061 | QIQ54046.1 | P0C6U8 |
| VNYTVCDDCNGYTDNI | 1 | 0 | 0 | 1705070 | YP_003767.1 | A0A0P0G321 |
| VPATRDSVPLHIIAPS | 1 | 0 | 0 | 1705083 | YP_009047208.1 | K9N7D2 |
| VPDGFYSTSPIQSVEL | 1 | 0 | 0 | 1705094 | NP_073551.1 | A0A0P0K6L9 |
| VPDHDSFVKMMVDGFV | 1 | 0 | 0 | 1705095 | NP_073549.1 | A0A0P0LKV0 |
| VPDKSVYDQVVDNVYV | 1 | 0 | 0 | 1705096 | YP_009555238.1 | P0C6X6 |
| VPDLVVEQYNQTILNL | 1 | 0 | 0 | 1705097 | NP_073551.1 | A0A0P0K6L9 |
| VPDTVEVPPQVVKLPS | 1 | 0 | 0 | 1705099 | YP_009047202.1 | K9N7C7 |
| VPDYNTYKNTCEGSTF | 1 | 0 | 0 | 1705101 | QIQ54046.1 | P0C6U8 |
| VPEEFASHVANFQKSG | 1 | 0 | 0 | 1705102 | YP_009047202.1 | K9N7C7 |
| VPEKFLNAFDVFVTAI | 1 | 0 | 0 | 1705106 | NP_073549.1 | A0A0P0LKV0 |
| VPEYIDVNKTLQELSY | 1 | 0 | 0 | 1705108 | NP_073551.1 | A0A0P0K6L9 |
| VPFGGTTVVRPLVEDS | 1 | 0 | 0 | 1705111 | YP_009047210.1 | K9N7A1 |
| VPGNVKPGDAVITTQQ | 1 | 0 | 0 | 1705134 | NP_073549.1 | A0A0P0LKV0 |
| VPHIVRKDLTKYTMLD | 1 | 0 | 0 | 1705140 | YP_009555238.1 | P0C6X6 |
| VPHNMRVLHYGAGSDK | 1 | 0 | 0 | 1705144 | YP_003766.2 | H9EJ42 |
| VPIAFGVPPSEAKGYW | 1 | 0 | 0 | 1705151 | YP_173242.1 | Q19U29 |
| VPIGKGNKDEQIGYWN | 1 | 0 | 0 | 1705155 | YP_003771.1 | H9EJ69 |
| VPIGTNDPIRFCLENT | 1 | 0 | 0 | 1705156 | NP_073549.1 | A0A0P0LKV0 |
| VPIINCVKMLCDHAGT | 1 | 0 | 0 | 1705158 | YP_173236.1 | A0A140H1G9 |
| VPINTNSGPDDQIGYY | 1 | 0 | 0 | 1705161 | NP_828858.1 | P59595 |
| VPINTNSTKDDQIGYY | 1 | 0 | 0 | 1705163 | QIQ54056.1 | P59595 |
| VPLCIFNGKFLSNTKY | 1 | 0 | 0 | 1705174 | YP_009555240.1 | P30215 |
| VPLLEVYVHLRNWNFS | 1 | 0 | 0 | 1705182 | YP_003770.1 | U3M6Q8 |
| VPLNANSTPAQNAGYW | 1 | 0 | 0 | 1705188 | YP_009047211.1 | K9N4V7 |
| VPLRASSCVTRCNIGG | 1 | 0 | 0 | 1705195 | YP_003766.2 | H9EJ42 |
| VPLRGTIVTRPLMESE | 1 | 0 | 0 | 1705197 | NP_828855.1 | P59596 |
| VPLSAIPALAANTLTI | 1 | 0 | 0 | 1705199 | YP_173236.1 | A0A140H1G9 |
| VPLSVIPATSASKLTI | 1 | 0 | 0 | 1705201 | YP_003766.2 | H9EJ42 |
| VPLYVRDYVDKWDDFC | 1 | 0 | 0 | 1705206 | NP_073549.1 | A0A0P0LKV0 |
| VPNQPYPNASFDNFKF | 1 | 0 | 0 | 1705213 | YP_009724389.1 | P0DTD1 |
| VPPALHLVDPQIQLTI | 1 | 0 | 0 | 1705214 | NP_828859.1 | P59636 |
| VPQADTDWKFYDAQPC | 1 | 0 | 0 | 1705218 | QIQ54047.1 | P0C6X7 |
| VPQADVEWKFYDAQPC | 1 | 0 | 0 | 1705219 | YP_009724389.1 | P0DTD1 |
| VPQAEVEWKFYDAQPC | 1 | 0 | 0 | 1705220 | NP_828849.2 | P0C6X7 |
| VPQEHYVRITGLYPTL | 1 | 0 | 0 | 1705222 | YP_009724389.1 | P0DTD1 |
| VPRASANIGCNHTGVV | 1 | 0 | 0 | 1705230 | YP_009724389.1 | P0DTD1 |
| VPRRNVATLQAENVTG | 1 | 0 | 0 | 1705233 | QIQ54047.1 | P0C6X7 |
| VPSINDLSFDKNLGRC | 1 | 0 | 0 | 1705241 | YP_003766.2 | H9EJ42 |
| VPSTVWEDGDYYRKQL | 1 | 0 | 0 | 1705248 | YP_009047204.1 | K9N5Q8 |
| VPSYVKPDSVSTSDLG | 1 | 0 | 0 | 1705250 | YP_009047202.1 | K9N7C7 |
| VPTDNYITTYPGQGLN | 1 | 0 | 0 | 1705252 | YP_009724389.1 | P0DTD1 |
| VPTGYVDTENNTEFTR | 1 | 0 | 0 | 1705254 | NP_828849.2 | P0C6X7 |
| VPTGYVDTPNNTDFSR | 1 | 0 | 0 | 1705256 | YP_009724389.1 | P0DTD1 |
| VPTQPLPNASFDNFKL | 1 | 0 | 0 | 1705265 | NP_828849.2 | P0C6X7 |
| VPTQPYPNSNFDNFKF | 1 | 0 | 0 | 1705266 | QIQ54046.1 | P0C6U8 |
| VPTYLKSDNIVAADLG | 1 | 0 | 0 | 1705271 | YP_009555238.1 | P0C6X6 |
| VPVDEYITTYPGQGCA | 1 | 0 | 0 | 1705272 | NP_828849.2 | P0C6X7 |
| VPVEDIAQVVIADTLQ | 1 | 0 | 0 | 1705273 | YP_009047202.1 | K9N7C7 |
| VPVIEVSQIQSKLTDV | 1 | 0 | 0 | 1705276 | YP_173236.1 | A0A140H1G9 |
| VPVQVVDSRWNNERAS | 1 | 0 | 0 | 1705282 | YP_173237.1 | A0A140H1H0 |
| VPVSPAFHRIESTDSI | 1 | 0 | 0 | 1705284 | YP_009047208.1 | K9N7D2 |
| VPVSVIYDKETKTHAT | 1 | 0 | 0 | 1705285 | YP_009047204.1 | K9N5Q8 |
| VPWDTIANYAKPFLNK | 1 | 0 | 0 | 1705289 | YP_009724389.1 | P0DTD1 |
| VPYESLRPDTRYVLMD | 1 | 0 | 0 | 1705296 | QIQ54046.1 | P0C6U8 |
| VPYPKQSIFTINSVVQ | 1 | 0 | 0 | 1705299 | YP_009047202.1 | K9N7C7 |
| VQASESKVLLERFLPK | 1 | 0 | 0 | 1705314 | YP_003766.2 | H9EJ42 |
| VQDADGINKQLTDISV | 1 | 0 | 0 | 1705321 | YP_173236.1 | A0A140H1G9 |
| VQDAKGLKIKYLYFVK | 1 | 0 | 0 | 1705322 | YP_009555238.1 | P0C6X6 |
| VQDAVNNNAQALSKLA | 1 | 0 | 0 | 1705323 | YP_009047204.1 | K9N5Q8 |
| VQDGVKITEEVGHEDL | 1 | 0 | 0 | 1705324 | QIQ54046.1 | P0C6U8 |
| VQDPAPIDIDAFPVKP | 1 | 0 | 0 | 1705327 | YP_003766.2 | H9EJ42 |
| VQEDDSEVNQTTITQS | 1 | 0 | 0 | 1705329 | QHR63299.1 | P0C6W2 |
| VQGSITQRTRTRPSVQ | 1 | 0 | 0 | 1705355 | YP_009047211.1 | K9N4V7 |
| VQIGEYTFEKGDYGDA | 1 | 0 | 0 | 1705364 | YP_009724389.1 | P0DTD1 |
| VQIGNNNVCTYEHVIS | 1 | 0 | 0 | 1705365 | NP_073549.1 | A0A0P0LKV0 |
| VQIGSNNVCTYEHVIS | 1 | 0 | 0 | 1705366 | YP_003766.2 | H9EJ42 |
| VQIHTIDGSSGVANPA | 1 | 0 | 0 | 1705367 | NP_828852.2 | P59632 |
| VQIHTIDGSSGVVNPA | 1 | 0 | 0 | 1705368 | QHR63301.1 | Q3LZX0 |
| VQIHTIDGSSGVVNPV | 1 | 0 | 0 | 1705369 | YP_009724391.1 | P0DTC3 |
| VQIPCTCGKQATKYLV | 1 | 0 | 0 | 1705371 | YP_009724389.1 | P0DTD1 |
| VQIPCTCGKQATQYLV | 1 | 0 | 0 | 1705372 | QHR63299.1 | P0C6W2 |
| VQIPTTCANDPVGFTL | 1 | 0 | 0 | 1705373 | YP_009724389.1 | P0DTD1 |
| VQITYTYKMLVAKDNK | 1 | 0 | 0 | 1705374 | YP_003771.1 | H9EJ69 |
| VQKESDEYILAKGPLQ | 1 | 0 | 0 | 1705379 | YP_009047202.1 | K9N7C7 |
| VQKHIGAPANLRVVGH | 1 | 0 | 0 | 1705380 | YP_009047202.1 | K9N7C7 |
| VQKLQTLIDNINSTLV | 1 | 0 | 0 | 1705383 | NP_073551.1 | A0A0P0K6L9 |
| VQKPVDVKPKIKACID | 1 | 0 | 0 | 1705384 | NP_828849.2 | P0C6X7 |
| VQKVEQPKIEPKPVSV | 1 | 0 | 0 | 1705386 | NP_073549.1 | A0A0P0LKV0 |
| VQLFEKYFKYWDQTYH | 1 | 0 | 0 | 1705389 | YP_009047202.1 | K9N7C7 |

| | | | | | | |
|---|---|---|---|---|---|---|
| VQLPVKDYVQTVNVIA | 1 | 0 | 0 | 1705394 | YP_173236.1 | A0A140H1G9 |
| VQLRAFDICNTNRAGI | 1 | 0 | 0 | 1705396 | YP_173236.1 | A0A140H1G9 |
| VQNDVCSTEDFNVWAM | 1 | 0 | 0 | 1705420 | YP_173236.1 | A0A140H1G9 |
| VQNGVDAKGFPQLAEL | 1 | 0 | 0 | 1705421 | YP_003771.1 | H9EJ69 |
| VQNLSGNPDEPQKDVY | 1 | 0 | 0 | 1705422 | YP_009555245.1 | P33469 |
| VQNVNLERCELANYKQ | 1 | 0 | 0 | 1705423 | YP_009047202.1 | K9N7C7 |
| VQPRTINSTQDGDNKL | 1 | 0 | 0 | 1705431 | YP_009555241.1 | P36334 |
| VQPVGVVDTEWGNMLT | 1 | 0 | 0 | 1705439 | YP_009047202.1 | K9N7C7 |
| VQQAQQEATLPDIAED | 1 | 0 | 0 | 1705440 | NP_073549.1 | A0A0P0LKV0 |
| VQQCFGKRGPNQNFGG | 1 | 0 | 0 | 1705441 | YP_009555245.1 | P33469 |
| VQQCFGKRGPSQNFGN | 1 | 0 | 0 | 1705442 | YP_173242.1 | Q19U29 |
| VQQESPFVMMSAPPAQ | 1 | 0 | 0 | 1705446 | YP_009724389.1 | P0DTD1 |
| VQQGIFNKATVAQIKA | 1 | 0 | 0 | 1705448 | YP_009555238.1 | P0C6X6 |
| VQQLFYTEDMASRFAD | 1 | 0 | 0 | 1705450 | YP_009047202.1 | K9N7C7 |
| VQQQEEQEEDWLETEA | 1 | 0 | 0 | 1705454 | QIQ54046.1 | P0C6U8 |
| VQRCASNKPHVLEDPC | 1 | 0 | 0 | 1705460 | NP_849176.1 | Q7TFA0 |
| VQTCTPNVTINCQDPA | 1 | 0 | 0 | 1705485 | NP_849177.1 | Q80H93 |
| VQTFYPRLQASADWKP | 1 | 0 | 0 | 1705489 | YP_009047202.1 | K9N7C7 |
| VQTVNVIAWLYAAILN | 1 | 0 | 0 | 1705495 | YP_173236.1 | A0A140H1G9 |
| VQVDNGSSINRKQLEI | 1 | 0 | 0 | 1705498 | YP_003766.2 | H9EJ42 |
| VQVDNGSSINRRQLDV | 1 | 0 | 0 | 1705499 | NP_073549.1 | A0A0P0LKV0 |
| VQVHNLPEYMTVAVPS | 1 | 0 | 0 | 1705503 | NP_073555.1 | A0A0P0KXN3 |
| VQYAHKHDIQITNESY | 1 | 0 | 0 | 1705515 | YP_009047202.1 | K9N7C7 |
| VQYDFTDYKLELFNKY | 1 | 0 | 0 | 1705516 | YP_009555238.1 | P0C6X6 |
| VQYGTDTNSVCPKLEF | 1 | 0 | 0 | 1705520 | YP_009047204.1 | K9N5Q8 |
| VQYRINGLGVTMDVLN | 1 | 0 | 0 | 1705521 | YP_173238.1 | A0A140H1H1 |
| VQYRINGLGVTMDVLS | 1 | 0 | 0 | 1705522 | YP_009555241.1 | P36334 |
| VRAFDVYNKDASFIGK | 1 | 0 | 0 | 1705527 | NP_073549.1 | A0A0P0LKV0 |
| VRASRQLAQQKVNECV | 1 | 0 | 0 | 1705533 | NP_073551.1 | A0A0P0K6L9 |
| VRATATIPIQASLPFG | 1 | 0 | 0 | 1705534 | YP_009724391.1 | P0DTC3 |
| VRAWVGFDAEGAHATR | 1 | 0 | 0 | 1705536 | YP_009555238.1 | P0C6X6 |
| VRDEVSIETDIIDEVN | 1 | 0 | 0 | 1705544 | NP_073556.1 | A0A0N7J131 |
| VRDGNTDNLVFVNKTS | 1 | 0 | 0 | 1705545 | NP_073549.1 | A0A0P0LKV0 |
| VRDPKTSEILDISPCA | 1 | 0 | 0 | 1705552 | NP_828851.1 | P59594 |
| VRDPRYVMRLQSAATI | 1 | 0 | 0 | 1705553 | YP_009555238.1 | P0C6X6 |
| VRDSRFVMRLQTIATI | 1 | 0 | 0 | 1705555 | YP_173236.1 | A0A140H1G9 |
| VRDYKLANDVKDTLRL | 1 | 0 | 0 | 1705560 | YP_003766.2 | H9EJ42 |
| VRDYLLSKMTSLPKDW | 1 | 0 | 0 | 1705561 | YP_173236.1 | A0A140H1G9 |
| VRDYVDKWDDFCRQYS | 1 | 0 | 0 | 1705562 | NP_073549.1 | A0A0P0LKV0 |
| VREEVDIIEQPFEEVE | 1 | 0 | 0 | 1705563 | YP_003766.2 | H9EJ42 |
| VRELYLNSSNVTTMDF | 1 | 0 | 0 | 1705568 | NP_828849.2 | P0C6X7 |
| VRETMNYLFQHANLDS | 1 | 0 | 0 | 1705570 | QHR63299.1 | P0C6W2 |
| VRGFGDSVEEVLSEAR | 1 | 0 | 0 | 1705593 | YP_009724389.1 | P0DTD1 |
| VRGRAIIVSVEQLEPC | 1 | 0 | 0 | 1705595 | NP_073549.1 | A0A0P0LKV0 |
| VRGSIVNARIEPCSSG | 1 | 0 | 0 | 1705596 | YP_009047202.1 | K9N7C7 |
| VRGSRRLAQQKINECV | 1 | 0 | 0 | 1705597 | YP_003767.1 | A0A0P0G321 |
| VRGSSAARLEPCNGTD | 1 | 0 | 0 | 1705598 | NP_073549.1 | A0A0P0LKV0 |
| VRGWLGMDVESAHVCG | 1 | 0 | 0 | 1705606 | YP_003766.2 | H9EJ42 |
| VRGWVGFDVEGAHATR | 1 | 0 | 0 | 1705608 | YP_173236.1 | A0A140H1G9 |
| VRHHPELKLFRNLNID | 1 | 0 | 0 | 1705609 | YP_009555238.1 | P0C6X6 |
| VRKDGKPVDHYDGFYT | 1 | 0 | 0 | 1705629 | NP_073549.1 | A0A0P0LKV0 |
| VRKEGQDVIFSQFDSL | 1 | 0 | 0 | 1705631 | YP_009555238.1 | P0C6X6 |
| VRKLDNQALNSILDNA | 1 | 0 | 0 | 1705635 | YP_009555238.1 | P0C6X6 |
| VRKNGQFQDHYDGFYT | 1 | 0 | 0 | 1705637 | YP_003766.2 | H9EJ42 |
| VRLQAGKQTEFVSNSH | 1 | 0 | 0 | 1705641 | NP_073549.1 | A0A0P0LKV0 |
| VRLQAGSNTEFASNSS | 1 | 0 | 0 | 1705643 | YP_009047202.1 | K9N7C7 |
| VRLQAGTATEYASNSS | 1 | 0 | 0 | 1705644 | YP_009555238.1 | P0C6X6 |
| VRLSKMGILKAEEFVA | 1 | 0 | 0 | 1705645 | NP_073549.1 | A0A0P0LKV0 |
| VRNDGTVEFCYLHQIE | 1 | 0 | 0 | 1705652 | YP_003766.2 | H9EJ42 |
| VRPIIEDYHTLTVTII | 1 | 0 | 0 | 1705660 | YP_009555244.1 | Q01455 |
| VRPRNSSDNGISAIIT | 1 | 0 | 0 | 1705663 | YP_003767.1 | A0A0P0G321 |
| VRPVIEDYHTLTATVI | 1 | 0 | 0 | 1705664 | YP_173241.1 | Q0ZJ82 |
| VRQQRFVYDAYQNLVG | 1 | 0 | 0 | 1705675 | YP_009047204.1 | K9N5Q8 |
| VRQVRSWIGFDVEGAH | 1 | 0 | 0 | 1705678 | YP_009047202.1 | K9N7C7 |
| VRRVNVPYVCLLANKE | 1 | 0 | 0 | 1705688 | YP_009047202.1 | K9N7C7 |
| VRSHPDFKLLHNLQAD | 1 | 0 | 0 | 1705694 | YP_009047202.1 | K9N7C7 |
| VRSLKLSDNNCYLNAV | 1 | 0 | 0 | 1705698 | YP_009047202.1 | K9N7C7 |
| VRSSFNFDQKELLAYY | 1 | 0 | 0 | 1705699 | YP_009555238.1 | P0C6X6 |
| VRSVRGRAIIVSVEQL | 1 | 0 | 0 | 1705701 | NP_073549.1 | A0A0P0LKV0 |
| VRSWIGFDVEGAHASR | 1 | 0 | 0 | 1705703 | YP_009047202.1 | K9N7C7 |
| VRTANALSRAVNVPTI | 1 | 0 | 0 | 1705704 | YP_009555238.1 | P0C6X6 |
| VRTNGGYNSWHLVEGE | 1 | 0 | 0 | 1705707 | YP_009047202.1 | K9N7C7 |
| VRVEEEEEEDWLDDTT | 1 | 0 | 0 | 1705720 | NP_828849.2 | P0C6X7 |
| VRVKHGDFSAVSSPMS | 1 | 0 | 0 | 1705723 | NP_073555.1 | A0A0P0KXN3 |
| VRVVTTFDAEYCRHGT | 1 | 0 | 0 | 1705727 | NP_828849.2 | P0C6X7 |
| VRVVTTFDSEYCRHGT | 1 | 0 | 0 | 1705728 | YP_009724389.1 | P0DTD1 |
| VRYGDVDNLVFTNKTT | 1 | 0 | 0 | 1705731 | YP_003766.2 | H9EJ42 |
| VSAAEFKYMVANGLHA | 1 | 0 | 0 | 1705735 | YP_003766.2 | H9EJ42 |
| VSAAFHANSSEPALLF | 1 | 0 | 0 | 1705736 | YP_009555241.1 | P36334 |
| VSAEVSRQVIQDVKDG | 1 | 0 | 0 | 1705740 | NP_073552.1 | A0A0P0KWE8 |
| VSALKTAGVVTGNVVK | 1 | 0 | 0 | 1705748 | YP_009047202.1 | K9N7C7 |
| VSALMGANGNKIVDKE | 1 | 0 | 0 | 1705749 | YP_009047202.1 | K9N7C7 |
| VSALVYENKLKAKNES | 1 | 0 | 0 | 1705754 | YP_009555238.1 | P0C6X6 |
| VSALVYNNKLLAKKEL | 1 | 0 | 0 | 1705755 | YP_009047202.1 | K9N7C7 |
| VSANVCALMSCNGNKI | 1 | 0 | 0 | 1705760 | YP_009555238.1 | P0C6X6 |
| VSATKLNDLCFSNVYA | 1 | 0 | 0 | 1705764 | NP_828851.1 | P59594 |
| VSAVNQSLSNAKVSAE | 1 | 0 | 0 | 1705768 | NP_073552.1 | A0A0P0KWE8 |
| VSCGHYQTNIYSQNLC | 1 | 0 | 0 | 1705771 | NP_073549.1 | A0A0P0LKV0 |
| VSDAYANLVPYYQLIG | 1 | 0 | 0 | 1705788 | NP_073549.1 | A0A0P0LKV0 |
| VSDDDFVSAVANAHRY | 1 | 0 | 0 | 1705789 | YP_003766.2 | H9EJ42 |
| VSDIDITFLKKDAPYI | 1 | 0 | 0 | 1705794 | YP_009724389.1 | P0DTD1 |

| | | | | | | |
|---|---|---|---|---|---|---|
| VSDIDYVPLKSATCIT | 1 | 0 | 0 | 1705795 | YP_009724389.1 | P0DTD1 |
| VSDREVILCWETGKVK | 1 | 0 | 0 | 1705800 | YP_173236.1 | A0A140H1G9 |
| VSEESQGNVVTSVMES | 1 | 0 | 0 | 1705813 | YP_173236.1 | A0A140H1G9 |
| VSEEVVENPTIQKDIL | 1 | 0 | 0 | 1705815 | QHR63299.1 | P0C6W2 |
| VSEEVVENPTIQKDVL | 1 | 0 | 0 | 1705816 | YP_009724389.1 | P0DTD1 |
| VSEKISVMDKLDTGAQ | 1 | 0 | 0 | 1705818 | YP_003766.2 | H9EJ42 |
| VSELVYENKFVPVKEA | 1 | 0 | 0 | 1705824 | NP_073549.1 | A0A0P0LKV0 |
| VSELVYENKFVPVKPA | 1 | 0 | 0 | 1705825 | YP_003766.2 | H9EJ42 |
| VSEMLTFDKKAFTLAN | 1 | 0 | 0 | 1705826 | NP_073551.1 | A0A0P0K6L9 |
| VSERINVVPGVPTNVY | 1 | 0 | 0 | 1705828 | YP_003766.2 | H9EJ42 |
| VSFADNNDITNTNLSR | 1 | 0 | 0 | 1705830 | YP_009047211.1 | K9N4V7 |
| VSFVGELKAAEAKVIT | 1 | 0 | 0 | 1705846 | NP_073549.1 | A0A0P0LKV0 |
| VSFVNDSIESVGGLYE | 1 | 0 | 0 | 1705847 | YP_173238.1 | A0A140H1H1 |
| VSGDAHIPFGTVLGNF | 1 | 0 | 0 | 1705851 | NP_073551.1 | A0A0P0K6L9 |
| VSGLVNVQKVEQPKIE | 1 | 0 | 0 | 1705857 | NP_073549.1 | A0A0P0LKV0 |
| VSGTATAGWTFGAGAA | 1 | 0 | 0 | 1705864 | NP_828851.1 | P59594 |
| VSGYEKSTFNKDIYCG | 1 | 0 | 0 | 1705869 | YP_003766.2 | H9EJ42 |
| VSHVCKCGHNMTLIAA | 1 | 0 | 0 | 1705877 | YP_173236.1 | A0A140H1G9 |
| VSIINNAVYTKVDGID | 1 | 0 | 0 | 1705883 | NP_828849.2 | P0C6X7 |
| VSIINNTVYTKVDGVD | 1 | 0 | 0 | 1705884 | YP_009724389.1 | P0DTD1 |
| VSIKNVDIVSEAKNVK | 1 | 0 | 0 | 1705885 | QIQ54046.1 | P0C6U8 |
| VSIPIDLLNLREIKPA | 1 | 0 | 0 | 1705891 | YP_009555238.1 | P0C6X6 |
| VSIRSSNQGNKQIVHS | 1 | 0 | 0 | 1705892 | YP_009047207.1 | K9N643 |
| VSISTGSRSARSAIED | 1 | 0 | 0 | 1705893 | YP_009047204.1 | K9N5Q8 |
| VSIVNNTVYTKVDGVD | 1 | 0 | 0 | 1705895 | QIQ54047.1 | P0C6X7 |
| VSIVPSTVWEDGDYYR | 1 | 0 | 0 | 1705896 | YP_009047204.1 | K9N5Q8 |
| VSKADGIIYPQGRTYS | 1 | 0 | 0 | 1705899 | YP_009047204.1 | K9N5Q8 |
| VSKESGNTVVLTFTTR | 1 | 0 | 0 | 1705904 | NP_073556.1 | A0A0N7J131 |
| VSKTSDWKCKVTDVLF | 1 | 0 | 0 | 1705914 | YP_009047202.1 | K9N7C7 |
| VSKVVNINVDFKDFQF | 1 | 0 | 0 | 1705921 | YP_173236.1 | A0A140H1G9 |
| VSLAIDAYPLSKHPNS | 1 | 0 | 0 | 1705925 | YP_003766.2 | H9EJ42 |
| VSLDSLGAAIHYLNSK | 1 | 0 | 0 | 1705932 | YP_009555238.1 | P0C6X6 |
| VSLETLLAAIKRLKNG | 1 | 0 | 0 | 1705933 | YP_009555238.1 | P0C6X6 |
| VSLIWNVKDYMSLSEQ | 1 | 0 | 0 | 1705939 | NP_828849.2 | P0C6X7 |
| VSLTYLLGTAKKQVVL | 1 | 0 | 0 | 1705952 | YP_009555238.1 | P0C6X6 |
| VSMKGTGVFVQDPAPI | 1 | 0 | 0 | 1705965 | YP_003766.2 | H9EJ42 |
| VSNCIKMLSNGAGNGQ | 1 | 0 | 0 | 1705971 | YP_003766.2 | H9EJ42 |
| VSNGGNNCTTAVMTYS | 1 | 0 | 0 | 1705975 | YP_003767.1 | A0A0P0G321 |
| VSNNKEDFDIIQKCQI | 1 | 0 | 0 | 1705980 | YP_173236.1 | A0A140H1G9 |
| VSNNQEDFDLISKCQI | 1 | 0 | 0 | 1705981 | YP_009555238.1 | P0C6X6 |
| VSNSGNLLGFKNVSTG | 1 | 0 | 0 | 1705983 | YP_003767.1 | A0A0P0G321 |
| VSPFEMPFEELNGLKI | 1 | 0 | 0 | 1705989 | NP_073549.1 | A0A0P0LKV0 |
| VSPYVCNAPGCDVNDV | 1 | 0 | 0 | 1705996 | YP_009555238.1 | P0C6X6 |
| VSQIQSKLTDVKCANV | 1 | 0 | 0 | 1706002 | YP_173236.1 | A0A140H1G9 |
| VSRQVIQDVKDGTVTF | 1 | 0 | 0 | 1706017 | NP_073552.1 | A0A0P0KWE8 |
| VSSAFWLFTHIPMAGL | 1 | 0 | 0 | 1706022 | YP_009047202.1 | K9N7C7 |
| VSSFEGYDFKYIENGQ | 1 | 0 | 0 | 1706029 | NP_073549.1 | A0A0P0LKV0 |
| VSSFLEMKSEKQVEQK | 1 | 0 | 0 | 1706031 | YP_009724389.1 | P0DTD1 |
| VSSNINCVLSVNSSNC | 1 | 0 | 0 | 1706039 | NP_073549.1 | A0A0P0LKV0 |
| VSSNINRLLSVPSDSC | 1 | 0 | 0 | 1706040 | YP_003766.2 | H9EJ42 |
| VSSNMVETVVGQLEQT | 1 | 0 | 0 | 1706041 | YP_009047202.1 | K9N7C7 |
| VSSNQSPQGNLGSNGK | 1 | 0 | 0 | 1706042 | YP_009555238.1 | P0C6X6 |
| VSSPMSNMTENERLLH | 1 | 0 | 0 | 1706045 | NP_073555.1 | A0A0P0KXN3 |
| VSSVECYSILAAKTGV | 1 | 0 | 0 | 1706049 | YP_003766.2 | H9EJ42 |
| VSSYAKPEEKLSAYDL | 1 | 0 | 0 | 1706051 | NP_073549.1 | A0A0P0LKV0 |
| VSTDTTVQSKDTNFLN | 1 | 0 | 0 | 1706055 | YP_009555238.1 | P0C6X6 |
| VSTFNVLQDVLSLRHD | 1 | 0 | 0 | 1706058 | YP_009555238.1 | P0C6X6 |
| VSTIGVCTMTDIAKKP | 1 | 0 | 0 | 1706065 | NP_828849.2 | P0C6X7 |
| VSTNLFEGDKFVGTFE | 1 | 0 | 0 | 1706072 | NP_073549.1 | A0A0P0LKV0 |
| VSTQEFRFMTANNLTA | 1 | 0 | 0 | 1706073 | YP_009047202.1 | K9N7C7 |
| VSTSIPNFGSLTQINT | 1 | 0 | 0 | 1706080 | YP_009047204.1 | K9N5Q8 |
| VSVEDVCFANKFEQFD | 1 | 0 | 0 | 1706093 | YP_009555238.1 | P0C6X6 |
| VSVIKVAPKPYRVDGK | 1 | 0 | 0 | 1706095 | NP_073549.1 | A0A0P0LKV0 |
| VSVITPGTNASNQVAV | 1 | 0 | 0 | 1706096 | QHR63300.2 | Q3LZX1 |
| VSVKMFDAYVDTFSAT | 1 | 0 | 0 | 1706100 | NP_828849.2 | P0C6X7 |
| VSVPVSVIYDKETKTH | 1 | 0 | 0 | 1706105 | YP_009047204.1 | K9N5Q8 |
| VSVSSPDAVTAYNGYL | 1 | 0 | 0 | 1706107 | YP_009724389.1 | P0DTD1 |
| VSVSSPDAVTTYNGYL | 1 | 0 | 0 | 1706108 | NP_828849.2 | P0C6X7 |
| VSVTVSSNMVETVVGQ | 1 | 0 | 0 | 1706110 | YP_009047202.1 | K9N7C7 |
| VSWNLREMLTHAEETR | 1 | 0 | 0 | 1706116 | QIQ54046.1 | P0C6U8 |
| VSYGSSVETETYFKVN | 1 | 0 | 0 | 1706127 | YP_173236.1 | A0A140H1G9 |
| VSYKLDGVTYTEIEPK | 1 | 0 | 0 | 1706130 | NP_828849.2 | P0C6X7 |
| VSYLSERCNVIADSKS | 1 | 0 | 0 | 1706132 | YP_009047202.1 | K9N7C7 |
| VTAFHQECSLQSCAQH | 1 | 0 | 0 | 1706136 | QHR63307.1 | Q3LZX5 |
| VTALRANSAVKLQNNE | 1 | 0 | 0 | 1706141 | YP_009724389.1 | P0DTD1 |
| VTANVNALLSTDGNKI | 1 | 0 | 0 | 1706145 | YP_009724389.1 | P0DTD1 |
| VTASDTTLRCCTVTYL | 1 | 0 | 0 | 1706148 | YP_003766.2 | H9EJ42 |
| VTASSKPLYVPEHCQN | 1 | 0 | 0 | 1706150 | YP_009047205.1 | K9N796 |
| VTAYNGYLTSSSKTPE | 1 | 0 | 0 | 1706153 | YP_009724389.1 | P0DTD1 |
| VTCAKEIKESVQTFFK | 1 | 0 | 0 | 1706156 | YP_009724389.1 | P0DTD1 |
| VTCCTASDKFYRLSNE | 1 | 0 | 0 | 1706157 | NP_073549.1 | A0A0P0LKV0 |
| VTDAKWYCYDKNPINS | 1 | 0 | 0 | 1706165 | NP_073549.1 | A0A0P0LKV0 |
| VTDAKWYCYDKQPVNS | 1 | 0 | 0 | 1706166 | YP_003766.2 | H9EJ42 |
| VTDECRLAMLGAEMTS | 1 | 0 | 0 | 1706167 | NP_073549.1 | A0A0P0LKV0 |
| VTDIVVVDEVSMLTNY | 1 | 0 | 0 | 1706171 | YP_009555238.1 | P0C6X6 |
| VTDTIVVTEGDGISTP | 1 | 0 | 0 | 1706174 | NP_828852.2 | P59632 |
| VTDVFYKENSYTTTIK | 1 | 0 | 0 | 1706176 | QIQ54046.1 | P0C6U8 |
| VTDVFYKETSYTTTIK | 1 | 0 | 0 | 1706177 | NP_828849.2 | P0C6X7 |
| VTDVLFPGQKYSSDCN | 1 | 0 | 0 | 1706179 | YP_009047202.1 | K9N7C7 |
| VTEDVKFAASTGVIDI | 1 | 0 | 0 | 1706184 | YP_003766.2 | H9EJ42 |
| VTEYSWNKKLYELVQR | 1 | 0 | 0 | 1706190 | NP_073549.1 | A0A0P0LKV0 |

| | | | | | | |
|---|---|---|---|---|---|---|
| VTEYYEFLNKHFSMMI | 1 | 0 | 0 | 1706191 | YP_009555238.1 | P0C6X6 |
| VTFEPFNVSFVNDSIE | 1 | 0 | 0 | 1706194 | YP_173238.1 | A0A140H1H1 |
| VTFGDDTVIEVQGYKN | 1 | 0 | 0 | 1706198 | QIQ54046.1 | P0C6U8 |
| VTFGDDTVIEVQGYKS | 1 | 0 | 0 | 1706199 | YP_009724389.1 | P0DTD1 |
| VTFGEDTVWEVQGYKN | 1 | 0 | 0 | 1706200 | NP_828849.2 | P0C6X7 |
| VTFLDLQVEMNRLQEA | 1 | 0 | 0 | 1706204 | YP_009555241.1 | P36334 |
| VTGDDVDDIESIYDFD | 1 | 0 | 0 | 1706217 | YP_173236.1 | A0A140H1G9 |
| VTGDNDDEDVVTGDND | 1 | 0 | 0 | 1706218 | YP_173236.1 | A0A140H1G9 |
| VTGDNDDEDVVTGDNN | 1 | 0 | 0 | 1706219 | YP_173236.1 | A0A140H1G9 |
| VTGDNNDEEIVTGDND | 1 | 0 | 0 | 1706221 | YP_173236.1 | A0A140H1G9 |
| VTGGVFLVDKNPHNT | 1 | 0 | 0 | 1706229 | P03155.1 | Q69028 |
| VTGGVVQMTTQWLTNI | 1 | 0 | 0 | 1706230 | QIQ54046.1 | P0C6U8 |
| VTGINDDDYVIALTGT | 1 | 0 | 0 | 1706231 | YP_003766.2 | H9EJ42 |
| VTHGFNLEEAARCMRS | 1 | 0 | 0 | 1706241 | NP_828849.2 | P0C6X7 |
| VTIAGAKLRSLNLGEV | 1 | 0 | 0 | 1706251 | NP_828849.2 | P0C6X7 |
| VTIALNKIQDVVNQQG | 1 | 0 | 0 | 1706252 | YP_003767.1 | A0A0P0G321 |
| VTICDFYSKLGVGFTS | 1 | 0 | 0 | 1706254 | YP_173236.1 | A0A140H1G9 |
| VTIDYAEIAFMLWCKD | 1 | 0 | 0 | 1706256 | QIQ54047.1 | P0C6X7 |
| VTIDYAEISFMLWCKD | 1 | 0 | 0 | 1706257 | NP_828849.2 | P0C6X7 |
| VTIDYTEISFMLWCKD | 1 | 0 | 0 | 1706258 | YP_009724389.1 | P0DTD1 |
| VTKENDSKEGFFTYIC | 1 | 0 | 0 | 1706271 | YP_009724389.1 | P0DTD1 |
| VTKENDSKEGFFTYLC | 1 | 0 | 0 | 1706272 | NP_828849.2 | P0C6X7 |
| VTKFSGDKFDFIVGYG | 1 | 0 | 0 | 1706277 | YP_173236.1 | A0A140H1G9 |
| VTKGKFKKNAWNIGEP | 1 | 0 | 0 | 1706280 | QIQ54046.1 | P0C6U8 |
| VTKKYTYSFLPKEDGD | 1 | 0 | 0 | 1706289 | YP_009047202.1 | K9N7C7 |
| VTKVLAEKFDYNVSVM | 1 | 0 | 0 | 1706300 | NP_073549.1 | A0A0P0LKV0 |
| VTLADAGFMKQYGECL | 1 | 0 | 0 | 1706305 | NP_828851.1 | P59594 |
| VTLDKCTKYNIYDVSG | 1 | 0 | 0 | 1706313 | NP_073551.1 | A0A0P0K6L9 |
| VTLIGEALKTQFNYYK | 1 | 0 | 0 | 1706316 | QHR63299.1 | P0C6W2 |
| VTLIGESVKTQFNYFK | 1 | 0 | 0 | 1706318 | NP_828849.2 | P0C6X7 |
| VTLKQGEIKDATPSDF | 1 | 0 | 0 | 1706324 | YP_009724391.1 | P0DTC3 |
| VTLKQGEIKDATPSDS | 1 | 0 | 0 | 1706325 | QHR63301.1 | Q3LZX0 |
| VTLLDEHESKFWDESF | 1 | 0 | 0 | 1706326 | NP_073549.1 | A0A0P0LKV0 |
| VTLSSNLNTNLHFDVD | 1 | 0 | 0 | 1706333 | YP_173238.1 | A0A140H1H1 |
| VTLSTKLKDGVNFNVD | 1 | 0 | 0 | 1706334 | YP_009555241.1 | P36334 |
| VTMDVLNKNQKLIATA | 1 | 0 | 0 | 1706339 | YP_173238.1 | A0A140H1H1 |
| VTMDVLSQNQKLIANA | 1 | 0 | 0 | 1706340 | YP_009555241.1 | P36334 |
| VTMPIGYVTHGFNLEE | 1 | 0 | 0 | 1706343 | NP_828849.2 | P0C6X7 |
| VTNEIGLNASVTLKIC | 1 | 0 | 0 | 1706346 | YP_003767.1 | A0A0P0G321 |
| VTNLNKSAGWPLNKFG | 1 | 0 | 0 | 1706351 | NP_073549.1 | A0A0P0LKV0 |
| VTNVNASSSEAFLIGA | 1 | 0 | 0 | 1706354 | NP_828849.2 | P0C6X7 |
| VTNVNASSSEAFLIGC | 1 | 0 | 0 | 1706355 | YP_009724389.1 | P0DTD1 |
| VTPCNQPDQVAVYQQS | 1 | 0 | 0 | 1706357 | YP_003767.1 | A0A0P0G321 |
| VTPEANMDQESFGGAS | 1 | 0 | 0 | 1706359 | YP_009724389.1 | P0DTD1 |
| VTPIKNTVDTTPKEEF | 1 | 0 | 0 | 1706361 | NP_073549.1 | A0A0P0LKV0 |
| VTPQPEEELEEDWLDD | 1 | 0 | 0 | 1706366 | QHR63299.1 | P0C6W2 |
| VTQEYIQTTIQKVTVD | 1 | 0 | 0 | 1706374 | YP_009047204.1 | K9N5Q8 |
| VTQNVLYENQKQIANQ | 1 | 0 | 0 | 1706384 | NP_828851.1 | P59594 |
| VTQRMCAIGPDVFLHK | 1 | 0 | 0 | 1706386 | NP_073549.1 | A0A0P0LKV0 |
| VTQRNFYEPQIITTDN | 1 | 0 | 0 | 1706387 | YP_009724390.1 | P0DTC2 |
| VTRDKLEKLISTARDG | 1 | 0 | 0 | 1706395 | YP_009047202.1 | K9N7C7 |
| VTSFTTYGNTVSEAVS | 1 | 0 | 0 | 1706408 | YP_173239.1 | Q0ZJ84 |
| VTSVMESQISTKEVKL | 1 | 0 | 0 | 1706422 | YP_173236.1 | A0A140H1G9 |
| VTTFDSEYCRHGTCEK | 1 | 0 | 0 | 1706433 | QIQ54046.1 | P0C6U8 |
| VTTFDSEYCRHGTCER | 1 | 0 | 0 | 1706434 | YP_009724389.1 | P0DTD1 |
| VTTKAKSVQQDLDVIL | 1 | 0 | 0 | 1706440 | YP_009047202.1 | K9N7C7 |
| VTTNSDKCPIVVGVSE | 1 | 0 | 0 | 1706451 | YP_003766.2 | H9EJ42 |
| VTTTLEETKFLTNKLL | 1 | 0 | 0 | 1706458 | NP_828849.2 | P0C6X7 |
| VTTYNGYLTSSSKTSE | 1 | 0 | 0 | 1706461 | NP_828849.2 | P0C6X7 |
| VTVDGVNFRTVVLNNK | 1 | 0 | 0 | 1706465 | YP_009047202.1 | K9N7C7 |
| VTVKETVVQFNYRRDG | 1 | 0 | 0 | 1706468 | YP_009047202.1 | K9N7C7 |
| VTVLRDNLKPVTTLST | 1 | 0 | 0 | 1706471 | YP_003768.1 | H9EJ24 |
| VTVSSNMVETVVGQLE | 1 | 0 | 0 | 1706478 | YP_009047202.1 | K9N7C7 |
| VTYAGNVWQIQTIQDS | 1 | 0 | 0 | 1706488 | YP_009555238.1 | P0C6X6 |
| VTYAGSVWHIQTVQDA | 1 | 0 | 0 | 1706489 | YP_173236.1 | A0A140H1G9 |
| VTYGNMTLNGLWLDDK | 1 | 0 | 0 | 1706494 | YP_009555238.1 | P0C6X6 |
| VTYGSMTLNGLWLDDK | 1 | 0 | 0 | 1706495 | YP_173236.1 | A0A140H1G9 |
| VTYKSTATTKLVPGML | 1 | 0 | 0 | 1706499 | NP_073549.1 | A0A0P0LKV0 |
| VTYLNELSSKVVCTYM | 1 | 0 | 0 | 1706502 | YP_003766.2 | H9EJ42 |
| VTYPFIANEKFINGCG | 1 | 0 | 0 | 1706505 | YP_003766.2 | H9EJ42 |
| VTYQNISTNLPPPLLG | 1 | 0 | 0 | 1706508 | YP_009047204.1 | K9N5Q8 |
| VTYTDQYLCGADGKPV | 1 | 0 | 0 | 1706511 | NP_073549.1 | A0A0P0LKV0 |
| VTYTEIEPKLDGYYKK | 1 | 0 | 0 | 1706512 | NP_828849.2 | P0C6X7 |
| VVAIDYKHYTPSFKKG | 1 | 0 | 0 | 1706531 | YP_009724389.1 | P0DTD1 |
| VVANGVKAKGYPQFAE | 1 | 0 | 0 | 1706538 | NP_073556.1 | A0A0N7J131 |
| VVAVIDQDFGSTVFNV | 1 | 0 | 0 | 1706543 | YP_009555238.1 | P0C6X6 |
| VVAWHVVRDSRFVMRL | 1 | 0 | 0 | 1706546 | YP_173236.1 | A0A140H1G9 |
| VVCDAIENALNSCKEH | 1 | 0 | 0 | 1706550 | YP_173236.1 | A0A140H1G9 |
| VVCDKVRSLKLSDNNC | 1 | 0 | 0 | 1706551 | YP_009047202.1 | K9N7C7 |
| VVCKHCGQKTTTLTGV | 1 | 0 | 0 | 1706556 | NP_828849.2 | P0C6X7 |
| VVCKTCGQQQTTLKGV | 1 | 0 | 0 | 1706557 | YP_009724389.1 | P0DTD1 |
| VVCTEIDPKLDGYYKK | 1 | 0 | 0 | 1706561 | QIQ54046.1 | P0C6U8 |
| VVCTEIDPKLDNYYKK | 1 | 0 | 0 | 1706562 | YP_009724389.1 | P0DTD1 |
| VVCYNSDYASKGYIAN | 1 | 0 | 0 | 1706567 | YP_009555238.1 | P0C6X6 |
| VVDDHVGLLDQAWRVP | 1 | 0 | 0 | 1706574 | YP_009555238.1 | P0C6X6 |
| VVDDIYYPASCNGVLP | 1 | 0 | 0 | 1706575 | YP_003766.2 | H9EJ42 |
| VVDEVSMCTNYDLSII | 1 | 0 | 0 | 1706577 | YP_009047202.1 | K9N7C7 |
| VVDEVSMCTNYDLSVI | 1 | 0 | 0 | 1706578 | NP_073549.1 | A0A0P0LKV0 |
| VVDEVSMLTNYELSVI | 1 | 0 | 0 | 1706579 | YP_009555238.1 | P0C6X6 |
| VVDNDVVDYVSDADFS | 1 | 0 | 0 | 1706591 | NP_073549.1 | A0A0P0LKV0 |
| VVDQVEEVNSIFDIET | 1 | 0 | 0 | 1706594 | NP_073549.1 | A0A0P0LKV0 |

| | | | | | | |
|---|---|---|---|---|---|---|
| VVDSVAVKNGSLHLYF | 1 | 0 | 0 | 1706598 | QIQ54046.1 | P0C6U8 |
| VVDVMRYNLNFSANSL | 1 | 0 | 0 | 1706602 | YP_003767.1 | A0A0P0G321 |
| VVDVNDSHSMAVVDGK | 1 | 0 | 0 | 1706603 | YP_009555238.1 | P0C6X6 |
| VVDYVSDADFSVTGDC | 1 | 0 | 0 | 1706608 | NP_073549.1 | A0A0P0LKV0 |
| VVEDTINDDEDVVTGD | 1 | 0 | 0 | 1706614 | YP_173236.1 | A0A140H1G9 |
| VVEFEDRSNNSSRASS | 1 | 0 | 0 | 1706616 | YP_003771.1 | H9EJ69 |
| VVELPKFFYASNGTYN | 1 | 0 | 0 | 1706617 | NP_073551.1 | A0A0P0K6L9 |
| VVENDADKANFLNAAV | 1 | 0 | 0 | 1706619 | YP_009555238.1 | P0C6X6 |
| VVENPTIQKDILECNV | 1 | 0 | 0 | 1706620 | QHR63299.1 | P0C6W2 |
| VVENPTIQKDVLECNV | 1 | 0 | 0 | 1706621 | YP_009724389.1 | P0DTD1 |
| VVENPTIQKEVIECDV | 1 | 0 | 0 | 1706622 | NP_828849.2 | P0C6X7 |
| VVEQYNQTILNLTSEI | 1 | 0 | 0 | 1706625 | NP_073551.1 | A0A0P0K6L9 |
| VVETKVEITPEESASS | 1 | 0 | 0 | 1706630 | QIQ54046.1 | P0C6U8 |
| VVEVVTTSLTPCGYSE | 1 | 0 | 0 | 1706635 | YP_009555238.1 | P0C6X6 |
| VVEYLSKEDLHTLYVD | 1 | 0 | 0 | 1706636 | YP_009047202.1 | K9N7C7 |
| VVFDEISMATNYDLSV | 1 | 0 | 0 | 1706637 | YP_009724389.1 | P0DTD1 |
| VVFVDKYMCGFDGKPV | 1 | 0 | 0 | 1706652 | YP_003766.2 | H9EJ42 |
| VVGAMLSENFTSYGFS | 1 | 0 | 0 | 1706659 | NP_073551.1 | A0A0P0K6L9 |
| VVGDVILKPVQDGVKI | 1 | 0 | 0 | 1706664 | QIQ54046.1 | P0C6U8 |
| VVGIVYSSQLYEDVPS | 1 | 0 | 0 | 1706672 | YP_003768.1 | H9EJ24 |
| VVGPDARAKQDVSLLS | 1 | 0 | 0 | 1706674 | YP_009047202.1 | K9N7C7 |
| VVGPDARTQGKQSYVL | 1 | 0 | 0 | 1706675 | YP_009555238.1 | P0C6X6 |
| VVGPKELLLSWESGKV | 1 | 0 | 0 | 1706676 | YP_003766.2 | H9EJ42 |
| VVGPNLNAGEDIQLLK | 1 | 0 | 0 | 1706677 | NP_828849.2 | P0C6X7 |
| VVGPNLNKGEDIQLLK | 1 | 0 | 0 | 1706678 | QIQ54046.1 | P0C6U8 |
| VVGPNVNKGEDIQLLK | 1 | 0 | 0 | 1706679 | YP_009724389.1 | P0DTD1 |
| VVGPNVNRGEDIQLLK | 1 | 0 | 0 | 1706680 | QHR63299.1 | P0C6W2 |
| VVGTLSSTIRLQAGVA | 1 | 0 | 0 | 1706688 | YP_173236.1 | A0A140H1G9 |
| VVGVSERINVVPGVPT | 1 | 0 | 0 | 1706691 | YP_003766.2 | H9EJ42 |
| VVIADTLQETPVVPDT | 1 | 0 | 0 | 1706700 | YP_009047202.1 | K9N7C7 |
| VVIFINNTTYPTNVAV | 1 | 0 | 0 | 1706703 | YP_009555238.1 | P0C6X6 |
| VVIKNLTPIKLNFGML | 1 | 0 | 0 | 1706713 | YP_003766.2 | H9EJ42 |
| VVIPDYNTYKNTCDGT | 1 | 0 | 0 | 1706719 | YP_009724389.1 | P0DTD1 |
| VVIPSKKAGGTTEMLS | 1 | 0 | 0 | 1706720 | NP_828849.2 | P0C6X7 |
| VVIPTKKAGGTTEMLA | 1 | 0 | 0 | 1706721 | YP_009724389.1 | P0DTD1 |
| VVIPTKKAGGTTEMLT | 1 | 0 | 0 | 1706722 | QHR63299.1 | P0C6W2 |
| VVITNVGEPIISQPSK | 1 | 0 | 0 | 1706726 | YP_003766.2 | H9EJ42 |
| VVKALNESYIDLKELG | 1 | 0 | 0 | 1706728 | YP_009047204.1 | K9N5Q8 |
| VVKLPSAPQTIQPEVK | 1 | 0 | 0 | 1706742 | YP_009047202.1 | K9N7C7 |
| VVKNSTAFPFETLSAC | 1 | 0 | 0 | 1706746 | YP_009047202.1 | K9N7C7 |
| VVKPGHFNEEFYNFLR | 1 | 0 | 0 | 1706749 | YP_003766.2 | H9EJ42 |
| VVKPVRARAPPGEQFT | 1 | 0 | 0 | 1706751 | NP_073549.1 | A0A0P0LKV0 |
| VVKQGHFIGVEGELPV | 1 | 0 | 0 | 1706753 | YP_009047202.1 | K9N7C7 |
| VVKTDAVVLLERYVSL | 1 | 0 | 0 | 1706767 | YP_003766.2 | H9EJ42 |
| VVLASDDLYVKRYFKG | 1 | 0 | 0 | 1706776 | YP_173236.1 | A0A140H1G9 |
| VVLATDDLYVKRYERG | 1 | 0 | 0 | 1706777 | YP_009555238.1 | P0C6X6 |
| VVLKTGDLQPLEQPTS | 1 | 0 | 0 | 1706790 | YP_009724389.1 | P0DTD1 |
| VVLLERYVSLAIDAYP | 1 | 0 | 0 | 1706792 | YP_003766.2 | H9EJ42 |
| VVLSDALDKLYKVFGS | 1 | 0 | 0 | 1706804 | NP_073549.1 | A0A0P0LKV0 |
| VVLSXELLNGPATVCG | 1 | 0 | 0 | 1706809 | QIQ54048.1 | P59594 |
| VVLVSNNQEDFDLISK | 1 | 0 | 0 | 1706813 | YP_009555238.1 | P0C6X6 |
| VVMDYLDNLKPRVEAP | 1 | 0 | 0 | 1706817 | NP_828849.2 | P0C6X7 |
| VVNAANENLLHGGGVA | 1 | 0 | 0 | 1706824 | YP_003766.2 | H9EJ42 |
| VVNAANVHLKHGGGVA | 1 | 0 | 0 | 1706825 | QIQ54046.1 | P0C6U8 |
| VVNAANVYLKHGGGVA | 1 | 0 | 0 | 1706826 | YP_009724389.1 | P0DTD1 |
| VVNANAEALNNLLQQL | 1 | 0 | 0 | 1706828 | YP_009555241.1 | P36334 |
| VVNARLRAKHYVYIGD | 1 | 0 | 0 | 1706831 | YP_009724389.1 | P0DTD1 |
| VVNDKIFTKSGVNDIC | 1 | 0 | 0 | 1706834 | YP_009047202.1 | K9N7C7 |
| VVNDKVFVRYGDVDNL | 1 | 0 | 0 | 1706835 | YP_003766.2 | H9EJ42 |
| VVNGKYFTFKQANNNC | 1 | 0 | 0 | 1706839 | YP_009555238.1 | P0C6X6 |
| VVNGVDFDFIVNAANE | 1 | 0 | 0 | 1706841 | NP_073549.1 | A0A0P0LKV0 |
| VVNINVDFKDFQFMLW | 1 | 0 | 0 | 1706843 | YP_173236.1 | A0A140H1G9 |
| VVNKGHFDGQQGEVPV | 1 | 0 | 0 | 1706848 | YP_009724389.1 | P0DTD1 |
| VVNNLDKSAGHPFNKF | 1 | 0 | 0 | 1706852 | YP_009047202.1 | K9N7C7 |
| VVNPANGHMAHGGGVA | 1 | 0 | 0 | 1706855 | YP_009555238.1 | P0C6X6 |
| VVNQQGNSLNHLTSQL | 1 | 0 | 0 | 1706857 | NP_073551.1 | A0A0P0K6L9 |
| VVNQQGSALNHLTSQL | 1 | 0 | 0 | 1706858 | YP_003767.1 | A0A0P0G321 |
| VVNSNAQALNSLLQQL | 1 | 0 | 0 | 1706861 | YP_173238.1 | A0A140H1H1 |
| VVNVNVDFKDFQFMLW | 1 | 0 | 0 | 1706865 | YP_009555238.1 | P0C6X6 |
| VVNYTVCDDCNGYTDN | 1 | 0 | 0 | 1706869 | YP_003767.1 | A0A0P0G321 |
| VVPDHDSFVKMMVDGF | 1 | 0 | 0 | 1706870 | NP_073549.1 | A0A0P0LKV0 |
| VVPDTVEVPPQVVKLP | 1 | 0 | 0 | 1706871 | YP_009047202.1 | K9N7C7 |
| VVPDYNTYKNTCEGST | 1 | 0 | 0 | 1706873 | QIQ54046.1 | P0C6U8 |
| VVPGNVKPGDAVITTQ | 1 | 0 | 0 | 1706878 | NP_073549.1 | A0A0P0LKV0 |
| VVPLSVIPATSASKLT | 1 | 0 | 0 | 1706882 | YP_003766.2 | H9EJ42 |
| VVPPALHLVDPQIQLT | 1 | 0 | 0 | 1706884 | NP_828859.1 | P59636 |
| VVPSINDLSFDKNLGR | 1 | 0 | 0 | 1706886 | YP_003766.2 | H9EJ42 |
| VVQEDDSEVNQTTITQ | 1 | 0 | 0 | 1706888 | QHR63299.1 | P0C6W2 |
| VVQKPVDVKPKIKACI | 1 | 0 | 0 | 1706894 | NP_828849.2 | P0C6X7 |
| VVQLPVKDYVQTVNVI | 1 | 0 | 0 | 1706897 | YP_173236.1 | A0A140H1G9 |
| VVQPRTINSTQDGDNK | 1 | 0 | 0 | 1706902 | YP_009555241.1 | P36334 |
| VVQPVGVVDTEWGNML | 1 | 0 | 0 | 1706903 | YP_009047202.1 | K9N7C7 |
| VVQRCASNKPHVLEDP | 1 | 0 | 0 | 1706905 | NP_849176.1 | Q7TFA0 |
| VVQYEADRRAFVDYTG | 1 | 0 | 0 | 1706909 | YP_009555238.1 | P0C6X6 |
| VVRDISVNDLCFANKF | 1 | 0 | 0 | 1706914 | YP_173236.1 | A0A140H1G9 |
| VVRDPRYVMRLQSAAT | 1 | 0 | 0 | 1706916 | YP_009555238.1 | P0C6X6 |
| VVRDSRFVMRLQTIAT | 1 | 0 | 0 | 1706917 | YP_173236.1 | A0A140H1G9 |
| VVRIGAAANSTGTVII | 1 | 0 | 0 | 1706924 | YP_009047204.1 | K9N5Q8 |
| VVRTANALSRAVNVPT | 1 | 0 | 0 | 1706937 | YP_009555238.1 | P0C6X6 |
| VVSDIDYVPLKSATCI | 1 | 0 | 0 | 1706949 | YP_009724389.1 | P0DTD1 |
| VVSFNEWALANQFTEF | 1 | 0 | 0 | 1706950 | YP_009047202.1 | K9N7C7 |

| | | | | | | |
|---|---|---|---|---|---|---|
| VVSHICKCGESMVLID | 1 | 0 | 0 | 1706951 | YP_009555238.1 | P0C6X6 |
| VVSIGYHYKELGVVMN | 1 | 0 | 0 | 1706955 | YP_173236.1 | A0A140H1G9 |
| VVSIRSSNQGNKQIVH | 1 | 0 | 0 | 1706956 | YP_009047207.1 | K9N643 |
| VVSTVKIGGLTFKTYA | 1 | 0 | 0 | 1706970 | NP_073549.1 | A0A0P0LKV0 |
| VVSVSSPDAVTTYNGY | 1 | 0 | 0 | 1706971 | NP_828849.2 | P0C6X7 |
| VVTGDDVDDIESIYDF | 1 | 0 | 0 | 1706978 | YP_173236.1 | A0A140H1G9 |
| VVTGDNDDEDVVTGDN | 1 | 0 | 0 | 1706979 | YP_173236.1 | A0A140H1G9 |
| VVTGDNNDEEIVTGDN | 1 | 0 | 0 | 1706980 | YP_173236.1 | A0A140H1G9 |
| VVTGNVVKQCCTAAVD | 1 | 0 | 0 | 1706981 | YP_009047202.1 | K9N7C7 |
| VVTQRMCAIGPDVFLH | 1 | 0 | 0 | 1706988 | NP_073549.1 | A0A0P0LKV0 |
| VVTSVMESQISTKEVK | 1 | 0 | 0 | 1706992 | YP_173236.1 | A0A140H1G9 |
| VVTTFDAEYCRHGTCE | 1 | 0 | 0 | 1706993 | NP_828849.2 | P0C6X7 |
| VVTTFDSEYCRHGTCE | 1 | 0 | 0 | 1706994 | YP_009724389.1 | P0DTD1 |
| VVTTNSDKCPIVVGVS | 1 | 0 | 0 | 1706999 | YP_003766.2 | H9EJ42 |
| VVTYPSLNYAGALWDI | 1 | 0 | 0 | 1707003 | YP_009047202.1 | K9N7C7 |
| VVVDEVSMCTNYDLSV | 1 | 0 | 0 | 1707006 | NP_073549.1 | A0A0P0LKV0 |
| VVVDEVSMLTNYELSV | 1 | 0 | 0 | 1707007 | YP_009555238.1 | P0C6X6 |
| VVVMSTCAVNYTKAPY | 1 | 0 | 0 | 1707023 | YP_009555241.1 | P36334 |
| VVVNGKYFTFKQANNN | 1 | 0 | 0 | 1707026 | YP_009555238.1 | P0C6X6 |
| VVVNGPFFSFEQSHNN | 1 | 0 | 0 | 1707027 | YP_173236.1 | A0A140H1G9 |
| VVVNNLDKSAGHPFNK | 1 | 0 | 0 | 1707028 | YP_009047202.1 | K9N7C7 |
| VVVNSQDVYKSLTIVD | 1 | 0 | 0 | 1707030 | YP_009047202.1 | K9N7C7 |
| VVVPDHDSFVKMMVDG | 1 | 0 | 0 | 1707031 | NP_073549.1 | A0A0P0LKV0 |
| VVVPDYGTYKNTCDGN | 1 | 0 | 0 | 1707032 | NP_828849.2 | P0C6X7 |
| VVVPDYNTYKNTCEGS | 1 | 0 | 0 | 1707033 | QIQ54046.1 | P0C6U8 |
| VVVVENDADKANFLNA | 1 | 0 | 0 | 1707039 | YP_009555238.1 | P0C6X6 |
| VVVVESDADRANFLNA | 1 | 0 | 0 | 1707040 | YP_173236.1 | A0A140H1G9 |
| VVVVPDHDSFVKMMVD | 1 | 0 | 0 | 1707041 | NP_073549.1 | A0A0P0LKV0 |
| VVWTLNDVKDNDGRPV | 1 | 0 | 0 | 1707046 | YP_003766.2 | H9EJ42 |
| VVWTLQEVKDNDGKNV | 1 | 0 | 0 | 1707047 | NP_073549.1 | A0A0P0LKV0 |
| VVWVAKEGAKTVNTSL | 1 | 0 | 0 | 1707048 | YP_003771.1 | H9EJ69 |
| VVWVAVDGAKTEPTGY | 1 | 0 | 0 | 1707050 | NP_073556.1 | A0A0N7J131 |
| VVYHTTSKPITFHMDG | 1 | 0 | 0 | 1707059 | QIQ54046.1 | P0C6U8 |
| VVYQQAVVGAMLSENF | 1 | 0 | 0 | 1707068 | NP_073551.1 | A0A0P0K6L9 |
| VVYRKVLLRKNGNKGA | 1 | 0 | 0 | 1707072 | QHR63299.1 | P0C6W2 |
| VVYSNGGFYFKPGGTT | 1 | 0 | 0 | 1707073 | NP_073549.1 | A0A0P0LKV0 |
| VWEAGKSKPPLNRNYV | 1 | 0 | 0 | 1707084 | YP_009047202.1 | K9N7C7 |
| VWGVKDFNTLSQEGKK | 1 | 0 | 0 | 1707090 | YP_003766.2 | H9EJ42 |
| VWHINQATTKTTFKPN | 1 | 0 | 0 | 1707092 | NP_828849.2 | P0C6X7 |
| VWHIQTVQDADGINKQ | 1 | 0 | 0 | 1707093 | YP_173236.1 | A0A140H1G9 |
| VWHVERKDVPYPKQSI | 1 | 0 | 0 | 1707095 | YP_009047202.1 | K9N7C7 |
| VWHVNNATNKATYKPN | 1 | 0 | 0 | 1707096 | YP_009724389.1 | P0DTD1 |
| VWHVNNTINKATFKPN | 1 | 0 | 0 | 1707097 | QIQ54046.1 | P0C6U8 |
| VWNGGAYSLFDMAKFP | 1 | 0 | 0 | 1707112 | YP_009555238.1 | P0C6X6 |
| VWQIQTIQDSDGTNKQ | 1 | 0 | 0 | 1707120 | YP_009555238.1 | P0C6X6 |
| VWTLNDVKDNDGRPVH | 1 | 0 | 0 | 1707126 | YP_003766.2 | H9EJ42 |
| VWTLQEVKDNDGKNVH | 1 | 0 | 0 | 1707127 | NP_073549.1 | A0A0P0LKV0 |
| VWVAKEGAKTVNTSLG | 1 | 0 | 0 | 1707128 | YP_003771.1 | H9EJ69 |
| VWVANELSRLVKSPTV | 1 | 0 | 0 | 1707129 | YP_173236.1 | A0A140H1G9 |
| VWVAVDGAKTEPTGYG | 1 | 0 | 0 | 1707131 | NP_073556.1 | A0A0N7J131 |
| VWVHEDGATDAPSTFG | 1 | 0 | 0 | 1707132 | YP_009047211.1 | K9N4V7 |
| VYAADPAMHAASGNLL | 1 | 0 | 0 | 1707135 | YP_009724389.1 | P0DTD1 |
| VYADDSIILRQYNLVD | 1 | 0 | 0 | 1707138 | YP_009555238.1 | P0C6X6 |
| VYADSFVVKGDEVRQI | 1 | 0 | 0 | 1707142 | QIQ54048.1 | P59594 |
| VYALRHFDEGNCDTLK | 1 | 0 | 0 | 1707146 | YP_009724389.1 | P0DTD1 |
| VYALRHFDEGNCDTLR | 1 | 0 | 0 | 1707147 | QHR63299.1 | P0C6W2 |
| VYALRHFDQNSEVLKA | 1 | 0 | 0 | 1707148 | YP_009047202.1 | K9N7C7 |
| VYAMRNFDEQNCDVLK | 1 | 0 | 0 | 1707149 | YP_003766.2 | H9EJ42 |
| VYANVGRWSASINTGN | 1 | 0 | 0 | 1707154 | NP_073551.1 | A0A0P0K6L9 |
| VYAVLSDVDGLKYTKI | 1 | 0 | 0 | 1707158 | YP_173236.1 | A0A140H1G9 |
| VYCPRHVIASNTTSAI | 1 | 0 | 0 | 1707168 | NP_073549.1 | A0A0P0LKV0 |
| VYCPRHVICSSSNMNE | 1 | 0 | 0 | 1707170 | YP_173236.1 | A0A140H1G9 |
| VYCPRHVICTAEDMLN | 1 | 0 | 0 | 1707171 | NP_828849.2 | P0C6X7 |
| VYCPRHVICTSEDMLN | 1 | 0 | 0 | 1707172 | YP_009724389.1 | P0DTD1 |
| VYCYNTDLIEGSKPYS | 1 | 0 | 0 | 1707174 | YP_003766.2 | H9EJ42 |
| VYDAANNAVYDGARLF | 1 | 0 | 0 | 1707175 | YP_003766.2 | H9EJ42 |
| VYDAYQNLVGYYSDDG | 1 | 0 | 0 | 1707176 | YP_009047204.1 | K9N5Q8 |
| VYDDVDASLFVDINNL | 1 | 0 | 0 | 1707182 | YP_173236.1 | A0A140H1G9 |
| VYDFCVSPAEFKYMVA | 1 | 0 | 0 | 1707183 | NP_073549.1 | A0A0P0LKV0 |
| VYDFKVEDAYDQVHDE | 1 | 0 | 0 | 1707184 | YP_009555238.1 | P0C6X6 |
| VYDFKVSAAEFKYMVA | 1 | 0 | 0 | 1707185 | YP_003766.2 | H9EJ42 |
| VYDIACCMRAGSKVVN | 1 | 0 | 0 | 1707189 | YP_003766.2 | H9EJ42 |
| VYDIGNPKGIPIVDDP | 1 | 0 | 0 | 1707190 | YP_009047202.1 | K9N7C7 |
| VYDNKLKAHKDKSAQC | 1 | 0 | 0 | 1707196 | YP_009724389.1 | P0DTD1 |
| VYDNKLRAHKDKSQQC | 1 | 0 | 0 | 1707198 | QIQ54047.1 | P0C6X7 |
| VYDQVVDNVYVTYAGN | 1 | 0 | 0 | 1707202 | YP_009555238.1 | P0C6X6 |
| VYDSSEVKAQSSGVIP | 1 | 0 | 0 | 1707203 | YP_173236.1 | A0A140H1G9 |
| VYDTAKKSMYDGDRFV | 1 | 0 | 0 | 1707204 | NP_073549.1 | A0A0P0LKV0 |
| VYDVHKDNFKDGLCMF | 1 | 0 | 0 | 1707206 | YP_173236.1 | A0A140H1G9 |
| VYEADTEQTQNVTVKP | 1 | 0 | 0 | 1707208 | YP_009047202.1 | K9N7C7 |
| VYEKLKPILDWLEEKF | 1 | 0 | 0 | 1707209 | QIQ54046.1 | P0C6U8 |
| VYEKLKPVLDWLEEKF | 1 | 0 | 0 | 1707210 | YP_009724389.1 | P0DTD1 |
| VYENKFVPVKEASKQC | 1 | 0 | 0 | 1707216 | NP_073549.1 | A0A0P0LKV0 |
| VYENKFVPVKPASKQC | 1 | 0 | 0 | 1707217 | YP_003766.2 | H9EJ42 |
| VYEQLKPVLDWLEEKF | 1 | 0 | 0 | 1707220 | QHR63299.1 | P0C6W2 |
| VYERDKAVARKLERMA | 1 | 0 | 0 | 1707221 | YP_173236.1 | A0A140H1G9 |
| VYFNATGSDVNCNGYQ | 1 | 0 | 0 | 1707232 | YP_003767.1 | A0A0P0G321 |
| VYGNFDDQPSLQVESA | 1 | 0 | 0 | 1707247 | YP_003766.2 | H9EJ42 |
| VYHTTSKPITFHMDGE | 1 | 0 | 0 | 1707258 | QIQ54046.1 | P0C6U8 |
| VYIGDPAQLPAPRTLL | 1 | 0 | 0 | 1707260 | YP_009724389.1 | P0DTD1 |
| VYIWTADEAIKFLKEW | 1 | 0 | 0 | 1707265 | YP_009555244.1 | Q01455 |

| | | | | | | |
|---|---|---|---|---|---|---|
| VYKLDGVVCTEIDPKL | 1 | 0 | 0 | 1707273 | QIQ54046.1 | P0C6U8 |
| VYKSDIHLPAYASFKV | 1 | 0 | 0 | 1707282 | YP_173236.1 | A0A140H1G9 |
| VYLEDKFDLLISDMYD | 1 | 0 | 0 | 1707289 | NP_073549.1 | A0A0P0LKV0 |
| VYLNHSVPKLSDFESE | 1 | 0 | 0 | 1707299 | YP_173238.1 | A0A140H1H1 |
| VYMAKAGDKYYPVVVD | 1 | 0 | 0 | 1707308 | YP_009555238.1 | P0C6X6 |
| VYNDALFVSYGSSVET | 1 | 0 | 0 | 1707315 | YP_173236.1 | A0A140H1G9 |
| VYNKDASFIGKNLKSN | 1 | 0 | 0 | 1707319 | NP_073549.1 | A0A0P0LKV0 |
| VYNLVKTGHYTGQAGE | 1 | 0 | 0 | 1707321 | YP_009555238.1 | P0C6X6 |
| VYNNSGSNITQIKNAC | 1 | 0 | 0 | 1707324 | YP_003766.2 | H9EJ42 |
| VYQCAMRPNHTIKGSF | 1 | 0 | 0 | 1707341 | NP_828849.2 | P0C6X7 |
| VYQQAVVGAMLSENFT | 1 | 0 | 0 | 1707343 | NP_073551.1 | A0A0P0K6L9 |
| VYQQSIIGAMTAVNES | 1 | 0 | 0 | 1707346 | YP_003767.1 | A0A0P0G321 |
| VYRKVLLRKNGNKGAG | 1 | 0 | 0 | 1707355 | QHR63299.1 | P0C6W2 |
| VYRRKPNLPNCNIEAW | 1 | 0 | 0 | 1707356 | YP_009555241.1 | P36334 |
| VYRSTSPDPKFVDKYY | 1 | 0 | 0 | 1707359 | YP_009047202.1 | K9N7C7 |
| VYSAGHATGTTPQKLF | 1 | 0 | 0 | 1707365 | YP_009047204.1 | K9N5Q8 |
| VYSDVENPHLMGWDYP | 1 | 0 | 0 | 1707367 | YP_009724389.1 | P0DTD1 |
| VYSDVETPHLMGWDYP | 1 | 0 | 0 | 1707368 | NP_828849.2 | P0C6X7 |
| VYSNGGFYFKPGGTTS | 1 | 0 | 0 | 1707377 | NP_073549.1 | A0A0P0LKV0 |
| VYSRKARKRSHSPTKK | 1 | 0 | 0 | 1707378 | YP_009047207.1 | K9N643 |
| VYSRVKNLNSSEGVPD | 1 | 0 | 0 | 1707379 | NP_828854.1 | P59637 |
| VYSSANNCTFEYISQP | 1 | 0 | 0 | 1707382 | QIQ54048.1 | P59594 |
| VYSSNEEQAVLKFLDG | 1 | 0 | 0 | 1707384 | YP_003766.2 | H9EJ42 |
| VYSVLETFQNNVVNYQ | 1 | 0 | 0 | 1707389 | YP_009555238.1 | P0C6X6 |
| VYSVSSFEAKPSGSVV | 1 | 0 | 0 | 1707391 | YP_009047204.1 | K9N5Q8 |
| VYTDTEVCKVKDFVSG | 1 | 0 | 0 | 1707395 | NP_073549.1 | A0A0P0LKV0 |
| VYTKVDGIDVEIFENK | 1 | 0 | 0 | 1707398 | NP_828849.2 | P0C6X7 |
| VYTNSCLVSSDGQPGG | 1 | 0 | 0 | 1707402 | YP_009047202.1 | K9N7C7 |
| VYTSDLLTNSVIIMAY | 1 | 0 | 0 | 1707403 | NP_828849.2 | P0C6X7 |
| VYVDGLESKQVDYVPL | 1 | 0 | 0 | 1707409 | YP_173236.1 | A0A140H1G9 |
| VYVDTFMSHFDVDRKS | 1 | 0 | 0 | 1707411 | YP_173236.1 | A0A140H1G9 |
| VYVGDPAQLPAPRTLL | 1 | 0 | 0 | 1707413 | YP_009047202.1 | K9N7C7 |
| VYVKFQDSKPPLPPDE | 1 | 0 | 0 | 1707420 | YP_009047209.1 | K9N5R3 |
| VYVKSKVGNYRLPSTQ | 1 | 0 | 0 | 1707421 | YP_009555244.1 | Q01455 |
| VYVRKDGKPVDHYDGF | 1 | 0 | 0 | 1707424 | NP_073549.1 | A0A0P0LKV0 |
| VYYGNALDQAISMWAL | 1 | 0 | 0 | 1707432 | YP_009724389.1 | P0DTD1 |
| VYYHNNNKTWVENEFR | 1 | 0 | 0 | 1707434 | QIQ54048.1 | P59594 |
| VYYKGVTTHESSSAVN | 1 | 0 | 0 | 1707436 | YP_009555238.1 | P0C6X6 |
| VYYNSWQNLLYDSNGN | 1 | 0 | 0 | 1707437 | YP_173238.1 | A0A140H1H1 |
| VYYVARLMKGDKGDAE | 1 | 0 | 0 | 1707451 | NP_073549.1 | A0A0P0LKV0 |
| WAAAVDEAFPLDEAED | 1 | 0 | 0 | 1707453 | YP_009047202.1 | K9N7C7 |
| WADQSERNYQTFNRGR | 1 | 0 | 0 | 1707461 | YP_173242.1 | Q19U29 |
| WAFYVRAKHGDFSGVA | 1 | 0 | 0 | 1707467 | YP_003770.1 | U3M6Q8 |
| WAIFNDHHLNRPGVYC | 1 | 0 | 0 | 1707474 | YP_009047202.1 | K9N7C7 |
| WAKRNIKPVPEIKILN | 1 | 0 | 0 | 1707475 | NP_828849.2 | P0C6X7 |
| WAKRNIKPVPEVKILN | 1 | 0 | 0 | 1707476 | YP_009724389.1 | P0DTD1 |
| WAPEFPWMFEDAEEKL | 1 | 0 | 0 | 1707485 | YP_009555238.1 | P0C6X6 |
| WAYCVDLHNKINLCDD | 1 | 0 | 0 | 1707492 | YP_003766.2 | H9EJ42 |
| WCKDGHVETFYPKLQA | 1 | 0 | 0 | 1707501 | NP_828849.2 | P0C6X7 |
| WCKDGHVETFYPKLQS | 1 | 0 | 0 | 1707502 | YP_009724389.1 | P0DTD1 |
| WCKDGQVQTFYPRLQA | 1 | 0 | 0 | 1707503 | YP_009047202.1 | K9N7C7 |
| WCKDNAVATFYPQLQS | 1 | 0 | 0 | 1707504 | NP_073549.1 | A0A0P0LKV0 |
| WDDFCRQYSNESWFED | 1 | 0 | 0 | 1707526 | NP_073549.1 | A0A0P0LKV0 |
| WDDMLRHLIKDVDNPV | 1 | 0 | 0 | 1707527 | YP_173236.1 | A0A140H1G9 |
| WDDMLRRLIKDVDNPV | 1 | 0 | 0 | 1707528 | YP_009555238.1 | P0C6X6 |
| WDFMLKTLYKDVDNPH | 1 | 0 | 0 | 1707531 | YP_009047202.1 | K9N7C7 |
| WDIEYPYISHEKKLNS | 1 | 0 | 0 | 1707533 | YP_009047202.1 | K9N7C7 |
| WDITVINNVDNEIVKS | 1 | 0 | 0 | 1707534 | YP_009047202.1 | K9N7C7 |
| WDLLKYDFTEERLKLF | 1 | 0 | 0 | 1707542 | YP_009724389.1 | P0DTD1 |
| WDQTYHPNCINCLDDR | 1 | 0 | 0 | 1707549 | NP_828849.2 | P0C6X7 |
| WDQTYHPNCVNCLDDR | 1 | 0 | 0 | 1707550 | YP_009724389.1 | P0DTD1 |
| WDTIANYAKPFLNKVV | 1 | 0 | 0 | 1707553 | YP_009724389.1 | P0DTD1 |
| WDVIRKDLSYEQQNVL | 1 | 0 | 0 | 1707555 | YP_003766.2 | H9EJ42 |
| WDYHVTKVLAEKFDYN | 1 | 0 | 0 | 1707561 | NP_073549.1 | A0A0P0LKV0 |
| WDYPKCDRALPNMIRM | 1 | 0 | 0 | 1707564 | YP_003766.2 | H9EJ42 |
| WDYPKCDRAMPNMCRI | 1 | 0 | 0 | 1707567 | YP_009047202.1 | K9N7C7 |
| WEAGKSKPPLNRNYVF | 1 | 0 | 0 | 1707571 | YP_009047202.1 | K9N7C7 |
| WEDGDYYRKQLSPLEG | 1 | 0 | 0 | 1707573 | YP_009047204.1 | K9N5Q8 |
| WESGKAKPPLNRNSVF | 1 | 0 | 0 | 1707588 | NP_073549.1 | A0A0P0LKV0 |
| WESGKVKPPLNRNSVF | 1 | 0 | 0 | 1707589 | YP_003766.2 | H9EJ42 |
| WFALTLQDDDECCQV | 1 | 0 | 0 | 1707598 | QIQ54054.1 | Q7TFA1 |
| WFAMRKDGDDVIFSRT | 1 | 0 | 0 | 1707600 | YP_173236.1 | A0A140H1G9 |
| WFDPIENEDIHRVYAA | 1 | 0 | 0 | 1707606 | NP_073549.1 | A0A0P0LKV0 |
| WFKNQTSVAPDLSLDY | 1 | 0 | 0 | 1707622 | YP_009555241.1 | P36334 |
| WFLLTNTSSVVDGVVR | 1 | 0 | 0 | 1707634 | NP_073551.1 | A0A0P0K6L9 |
| WFMDGENAYEVVKAML | 1 | 0 | 0 | 1707640 | YP_009047202.1 | K9N7C7 |
| WFNTMLDASAPATGWL | 1 | 0 | 0 | 1707646 | YP_173236.1 | A0A140H1G9 |
| WFSLEIQDLEEPCTKV | 1 | 0 | 0 | 1707651 | NP_849175.1 | Q7TFA1 |
| WFTERSDKSYEHQTPF | 1 | 0 | 0 | 1707659 | NP_828849.2 | P0C6X7 |
| WGMKYHPNTVDCDNDR | 1 | 0 | 0 | 1707682 | YP_173236.1 | A0A140H1G9 |
| WGVKDFNTLSQEGKKY | 1 | 0 | 0 | 1707687 | YP_003766.2 | H9EJ42 |
| WHEGRTIYGNVSRQDL | 1 | 0 | 0 | 1707696 | NP_073549.1 | A0A0P0LKV0 |
| WHINQATTKTTFKPNT | 1 | 0 | 0 | 1707701 | NP_828849.2 | P0C6X7 |
| WHIQTVQDADGINKQL | 1 | 0 | 0 | 1707702 | YP_173236.1 | A0A140H1G9 |
| WHVERKDVPYPKQSIF | 1 | 0 | 0 | 1707714 | YP_009047202.1 | K9N7C7 |
| WHVNNATNKATYKPNT | 1 | 0 | 0 | 1707715 | YP_009724389.1 | P0DTD1 |
| WHVNNTINKATFKPNT | 1 | 0 | 0 | 1707716 | QIQ54046.1 | P0C6U8 |
| WHVVRDPRYVMRLQSA | 1 | 0 | 0 | 1707718 | YP_009555238.1 | P0C6X6 |
| WIELFPHNDRIKSFST | 1 | 0 | 0 | 1707726 | NP_073549.1 | A0A0P0LKV0 |
| WIGFDVEGAHASRNAC | 1 | 0 | 0 | 1707729 | YP_009047202.1 | K9N7C7 |
| WKRQPNDDVTSNVTQC | 1 | 0 | 0 | 1707784 | NP_073556.1 | A0A0N7J131 |

| | | | | | | |
|---|---|---|---|---|---|---|
| WLAPTNAYMRTTHTDI | 1 | 0 | 0 | 1707794 | YP_009047202.1 | K9N7C7 |
| WLDDDSQQTVGQQDGS | 1 | 0 | 0 | 1707801 | YP_009724389.1 | P0DTD1 |
| WLDMVDTSLSGFKLKD | 1 | 0 | 0 | 1707809 | YP_009724389.1 | P0DTD1 |
| WLDMVDTSLSGYKLKD | 1 | 0 | 0 | 1707810 | QIQ54046.1 | P0C6U8 |
| WLEEKFKEGIEFLKDG | 1 | 0 | 0 | 1707812 | QIQ54046.1 | P0C6U8 |
| WLEEKFKEGVEFLRDG | 1 | 0 | 0 | 1707813 | YP_009724389.1 | P0DTD1 |
| WLELADTSLSGYRLKD | 1 | 0 | 0 | 1707815 | NP_828849.2 | P0C6X7 |
| WLGDTVTCPRHVIAPS | 1 | 0 | 0 | 1707824 | YP_003766.2 | H9EJ42 |
| WLKPYGGQPVSEYHIT | 1 | 0 | 0 | 1707840 | YP_009047207.1 | K9N643 |
| WLNDKSVPSPLNWERK | 1 | 0 | 0 | 1707869 | YP_009555241.1 | P36334 |
| WLPHDAIVVDNDVVDY | 1 | 0 | 0 | 1707877 | NP_073549.1 | A0A0P0LKV0 |
| WLPPDAIIIDNDINDY | 1 | 0 | 0 | 1707878 | YP_003766.2 | H9EJ42 |
| WLPTDAIIIDNDLNEF | 1 | 0 | 0 | 1707880 | YP_009047202.1 | K9N7C7 |
| WLTYTGAIKLDDKDPN | 1 | 0 | 0 | 1707901 | YP_009724397.2 | P0DTC9 |
| WLTYTGAIKLDDKDPS | 1 | 0 | 0 | 1707902 | QIQ54056.1 | P59595 |
| WLYTYSIGEASSPIAY | 1 | 0 | 0 | 1707915 | YP_009047202.1 | K9N7C7 |
| WMDDFEADPKGKYAQN | 1 | 0 | 0 | 1707916 | YP_009047202.1 | K9N7C7 |
| WMFEDAEEKLDNPSSS | 1 | 0 | 0 | 1707918 | YP_009555238.1 | P0C6X6 |
| WMGIRFEYTSPTDKLA | 1 | 0 | 0 | 1707920 | YP_009555239.1 | Q80872 |
| WNCQLSFAGYTESAVN | 1 | 0 | 0 | 1707944 | YP_009047206.1 | K9N4V0 |
| WNGGAYSLFDMAKFPL | 1 | 0 | 0 | 1707952 | YP_009555238.1 | P0C6X6 |
| WNGGAYSLFDMTKFSL | 1 | 0 | 0 | 1707953 | YP_173236.1 | A0A140H1G9 |
| WNKDVNTHSTRLTITE | 1 | 0 | 0 | 1707961 | NP_073549.1 | A0A0P0LKV0 |
| WNKRFGFIEDSVFKPR | 1 | 0 | 0 | 1707966 | YP_009555241.1 | P36334 |
| WNKVTQIANMFLEQTQ | 1 | 0 | 0 | 1707967 | YP_009047202.1 | K9N7C7 |
| WNLEYPIISNEVSINT | 1 | 0 | 0 | 1707971 | YP_173236.1 | A0A140H1G9 |
| WNLREMLAHAEETRKL | 1 | 0 | 0 | 1707974 | YP_009724389.1 | P0DTD1 |
| WNSKHIDAKEGGNFNY | 1 | 0 | 0 | 1707988 | QHR63300.2 | Q3LZX1 |
| WNSVKQDALTGGNYGY | 1 | 0 | 0 | 1707990 | QIQ54048.1 | P59594 |
| WNTKHGSGVTRELMRE | 1 | 0 | 0 | 1707996 | QIQ54046.1 | P0C6U8 |
| WNTKPVETSNSFDVLK | 1 | 0 | 0 | 1707999 | QHR63299.1 | P0C6W2 |
| WNTRNIDATSTGNYNY | 1 | 0 | 0 | 1708000 | NP_828851.1 | P59594 |
| WNVEYPIISNELSINT | 1 | 0 | 0 | 1708001 | YP_009555238.1 | P0C6X6 |
| WPISNDSNGCVVEAST | 1 | 0 | 0 | 1708011 | YP_003766.2 | H9EJ42 |
| WPLSVQQAQQEATLPD | 1 | 0 | 0 | 1708018 | NP_073549.1 | A0A0P0LKV0 |
| WPLYPYGRCPTAADIN | 1 | 0 | 0 | 1708027 | YP_009555240.1 | P30215 |
| WPRPIDVSKADGIIYP | 1 | 0 | 0 | 1708031 | YP_009047204.1 | K9N5Q8 |
| WPVATGDVVLASDDLY | 1 | 0 | 0 | 1708034 | YP_173236.1 | A0A140H1G9 |
| WQFVDHFGENEEIIIN | 1 | 0 | 0 | 1708049 | NP_073549.1 | A0A0P0LKV0 |
| WQIQTIQDSDGTNKQL | 1 | 0 | 0 | 1708051 | YP_009555238.1 | P0C6X6 |
| WQNLLYDSNGNIIGFK | 1 | 0 | 0 | 1708056 | YP_173238.1 | A0A140H1H1 |
| WRFPCAGRKVNFNEKP | 1 | 0 | 0 | 1708070 | YP_173236.1 | A0A140H1G9 |
| WRLINKFDVINGVKTV | 1 | 0 | 0 | 1708076 | YP_173236.1 | A0A140H1G9 |
| WRMRRGQRVDLPPKVH | 1 | 0 | 0 | 1708077 | YP_003771.1 | H9EJ69 |
| WRRQDRKINTGNGIKQ | 1 | 0 | 0 | 1708087 | YP_009047211.1 | K9N4V7 |
| WRYDFDITESKYSCKE | 1 | 0 | 0 | 1708097 | YP_003766.2 | H9EJ42 |
| WRYNFDITESKYSCKE | 1 | 0 | 0 | 1708098 | NP_073549.1 | A0A0P0LKV0 |
| WSADSCLQGDKCNIFA | 1 | 0 | 0 | 1708100 | YP_009555241.1 | P36334 |
| WSASINTGNCPFSFGK | 1 | 0 | 0 | 1708103 | NP_073551.1 | A0A0P0K6L9 |
| WSDIRDYKLANDAKES | 1 | 0 | 0 | 1708105 | NP_073549.1 | A0A0P0LKV0 |
| WSEGNSITGVPYPVSG | 1 | 0 | 0 | 1708107 | YP_003767.1 | A0A0P0G321 |
| WSFDYHPNCSDCYDDM | 1 | 0 | 0 | 1708108 | YP_003766.2 | H9EJ42 |
| WSFNPETNNLMCIDMK | 1 | 0 | 0 | 1708112 | YP_009555244.1 | Q01455 |
| WSITYPMIANENAINK | 1 | 0 | 0 | 1708118 | NP_073549.1 | A0A0P0LKV0 |
| WSMPYHPNTVDCQDDR | 1 | 0 | 0 | 1708127 | YP_009555238.1 | P0C6X6 |
| WSPQGNLGGNCAGNVI | 1 | 0 | 0 | 1708131 | YP_173236.1 | A0A140H1G9 |
| WSTKPVDTSNSFEVLA | 1 | 0 | 0 | 1708138 | NP_828849.2 | P0C6X7 |
| WSTKPVETSNIFDVLQ | 1 | 0 | 0 | 1708139 | QIQ54046.1 | P0C6U8 |
| WSTKPVETSNSFDVLK | 1 | 0 | 0 | 1708140 | YP_009724389.1 | P0DTD1 |
| WSVEYPIIGDELRVNS | 1 | 0 | 0 | 1708144 | NP_828849.2 | P0C6X7 |
| WSYTGSSFYAPEPITS | 1 | 0 | 0 | 1708154 | YP_009047204.1 | K9N5Q8 |
| WTAFVTNVNASSSEAF | 1 | 0 | 0 | 1708156 | YP_009724389.1 | P0DTD1 |
| WTDVRDYKLANDVKDT | 1 | 0 | 0 | 1708160 | YP_003766.2 | H9EJ42 |
| WTESGDYTLANTTTEP | 1 | 0 | 0 | 1708162 | YP_009047202.1 | K9N7C7 |
| WTETDLTKGPHEFCSQ | 1 | 0 | 0 | 1708163 | YP_009724389.1 | P0DTD1 |
| WTIEYPIIGDELKINA | 1 | 0 | 0 | 1708168 | YP_009724389.1 | P0DTD1 |
| WTLNDVKDNDGRPVHV | 1 | 0 | 0 | 1708174 | YP_003766.2 | H9EJ42 |
| WTLQEVKDNDGKNVHL | 1 | 0 | 0 | 1708175 | NP_073549.1 | A0A0P0LKV0 |
| WTMFCTSVNTSSSEAF | 1 | 0 | 0 | 1708176 | NP_073549.1 | A0A0P0LKV0 |
| WTNDALNTICIKDADY | 1 | 0 | 0 | 1708180 | NP_073549.1 | A0A0P0LKV0 |
| WTVFCTNANASSSEGF | 1 | 0 | 0 | 1708186 | YP_009047202.1 | K9N7C7 |
| WTVFCTNVNASSSEGF | 1 | 0 | 0 | 1708187 | YP_173236.1 | A0A140H1G9 |
| WTVTYPFIANEKFING | 1 | 0 | 0 | 1708188 | YP_003766.2 | H9EJ42 |
| WVAKEGAKTVNTSLGN | 1 | 0 | 0 | 1708190 | YP_003771.1 | H9EJ69 |
| WVANHQADTSTPSDVS | 1 | 0 | 0 | 1708192 | YP_173242.1 | Q19U29 |
| WVASNQADVNTPADIV | 1 | 0 | 0 | 1708193 | YP_009555245.1 | P33469 |
| WVATEGALNTPKDHIG | 1 | 0 | 0 | 1708194 | YP_009724397.2 | P0DTC9 |
| WVAVDGAKTEPTGYGV | 1 | 0 | 0 | 1708195 | NP_073556.1 | A0A0N7J131 |
| WVEEDLTKGPHEFCSQ | 1 | 0 | 0 | 1708197 | YP_003766.2 | H9EJ42 |
| WVENDITNGPHEFCSQ | 1 | 0 | 0 | 1708200 | YP_173236.1 | A0A140H1G9 |
| WVENEFRVYSSANNCT | 1 | 0 | 0 | 1708201 | QIQ54048.1 | P59594 |
| WVFYVRVKHGDFSAVS | 1 | 0 | 0 | 1708206 | NP_073555.1 | A0A0P0KXN3 |
| WVGFDAEGAHATRDSI | 1 | 0 | 0 | 1708207 | YP_009555238.1 | P0C6X6 |
| WVGFDVEGAHATRENI | 1 | 0 | 0 | 1708208 | YP_173236.1 | A0A140H1G9 |
| WVKHLNKNLNEGVLES | 1 | 0 | 0 | 1708212 | YP_003766.2 | H9EJ42 |
| WVKHLNKTLNEGVLES | 1 | 0 | 0 | 1708213 | NP_073549.1 | A0A0P0LKV0 |
| WVNKASYDTNLNKFNR | 1 | 0 | 0 | 1708229 | YP_009047202.1 | K9N7C7 |
| WVSGNMMDGFSCPLCC | 1 | 0 | 0 | 1708239 | YP_173236.1 | A0A140H1G9 |
| WWTAFVTNVNASSSEA | 1 | 0 | 0 | 1708258 | YP_009724389.1 | P0DTD1 |
| WWTVFCTNANASSSEG | 1 | 0 | 0 | 1708260 | YP_009047202.1 | K9N7C7 |
| WYCYDKNPINSNVKTL | 1 | 0 | 0 | 1708263 | NP_073549.1 | A0A0P0LKV0 |

| | | | | | | |
|---|---|---|---|---|---|---|
| WYDFVENPDIINVYKK | 1 | 0 | 0 | 1708271 | YP_009555238.1 | P0C6X6 |
| WYDPVENEDIHRVYAS | 1 | 0 | 0 | 1708273 | YP_003766.2 | H9EJ42 |
| WYKGSNLEEEILLMLA | 1 | 0 | 0 | 1708294 | YP_009555238.1 | P0C6X6 |
| WYRHSRRSFKTADGQQ | 1 | 0 | 0 | 1708300 | YP_173242.1 | Q19U29 |
| WYTERSEKSYELQTPF | 1 | 0 | 0 | 1708302 | YP_009724389.1 | P0DTD1 |
| WYVNDGRVDDGYICGD | 1 | 0 | 0 | 1708306 | YP_003766.2 | H9EJ42 |
| XELLNGPATVCGPKLS | 1 | 0 | 0 | 1708312 | QIQ54048.1 | P59594 |
| XNDGVYFASTEKSNII | 1 | 0 | 0 | 1708314 | QIQ54048.1 | P59594 |
| XRAPRITFGGPSDSTD | 1 | 0 | 0 | 1708315 | QIQ54056.1 | P59595 |
| XTCVGSDNVTDFNAIA | 1 | 0 | 0 | 1708317 | QIQ54047.1 | P0C6X7 |
| XVDVTFLKKDAPYIVG | 1 | 0 | 0 | 1708318 | QIQ54046.1 | P0C6U8 |
| XYSTLQGPPGTGKSHF | 1 | 0 | 0 | 1708320 | QIQ54047.1 | P0C6X7 |
| YAADPAMHIASSNAFL | 1 | 0 | 0 | 1708325 | YP_009047202.1 | K9N7C7 |
| YAAFATAQEAYEQAVA | 1 | 0 | 0 | 1708327 | YP_009724389.1 | P0DTD1 |
| YAAILNNCAWFVQNDV | 1 | 0 | 0 | 1708331 | YP_173236.1 | A0A140H1G9 |
| YAAILNNCNWFVQSDK | 1 | 0 | 0 | 1708332 | YP_009555238.1 | P0C6X6 |
| YAAKGYIAGIQNFKET | 1 | 0 | 0 | 1708334 | YP_009047202.1 | K9N7C7 |
| YAASYNRYKYYSGNAN | 1 | 0 | 0 | 1708341 | NP_073549.1 | A0A0P0LKV0 |
| YAAYATAQEAYEQAVA | 1 | 0 | 0 | 1708344 | NP_828849.2 | P0C6X7 |
| YADINGNLYPDSTSLV | 1 | 0 | 0 | 1708352 | QIQ54046.1 | P0C6U8 |
| YADKPNHFINFPLTHF | 1 | 0 | 0 | 1708355 | YP_009555239.1 | Q80872 |
| YADKTLQEMKALFPTW | 1 | 0 | 0 | 1708356 | YP_009555238.1 | P0C6X6 |
| YADSFVVKGDEVRQIA | 1 | 0 | 0 | 1708360 | QIQ54048.1 | P59594 |
| YAFATLKEVVGPKELL | 1 | 0 | 0 | 1708372 | YP_003766.2 | H9EJ42 |
| YAGNVWQIQTIQDSDG | 1 | 0 | 0 | 1708388 | YP_009555238.1 | P0C6X6 |
| YAGSRSSSGNRSGILK | 1 | 0 | 0 | 1708390 | YP_173242.1 | Q19U29 |
| YAGSVWHIQTVQDADG | 1 | 0 | 0 | 1708391 | YP_173236.1 | A0A140H1G9 |
| YAGVVWTLNDVKDNDG | 1 | 0 | 0 | 1708392 | YP_003766.2 | H9EJ42 |
| YAHKHDIQITNESYNN | 1 | 0 | 0 | 1708394 | YP_009047202.1 | K9N7C7 |
| YAIQEGFDATNSALVK | 1 | 0 | 0 | 1708403 | YP_009555241.1 | P36334 |
| YAISGKERARTVGGVS | 1 | 0 | 0 | 1708406 | NP_073549.1 | A0A0P0LKV0 |
| YAKPEEKLSAYDLACC | 1 | 0 | 0 | 1708413 | NP_073549.1 | A0A0P0LKV0 |
| YAKPFLNKVTSVAASG | 1 | 0 | 0 | 1708414 | QIQ54046.1 | P0C6U8 |
| YAKPFLNKVVSTTTNI | 1 | 0 | 0 | 1708415 | YP_009724389.1 | P0DTD1 |
| YAKRAVRSHPDFKLLH | 1 | 0 | 0 | 1708416 | YP_009047202.1 | K9N7C7 |
| YAKSNFVKDDYCDGSL | 1 | 0 | 0 | 1708417 | NP_073549.1 | A0A0P0LKV0 |
| YALLDWVKHLNKTLNE | 1 | 0 | 0 | 1708420 | NP_073549.1 | A0A0P0LKV0 |
| YALLHIAGCQTTNGLN | 1 | 0 | 0 | 1708421 | NP_073551.1 | A0A0P0K6L9 |
| YALRHFDEGNCDTLKE | 1 | 0 | 0 | 1708425 | YP_009724389.1 | P0DTD1 |
| YALRHFDQNSEVLKAI | 1 | 0 | 0 | 1708427 | YP_009047202.1 | K9N7C7 |
| YANVGRWSASINTGNC | 1 | 0 | 0 | 1708450 | NP_073551.1 | A0A0P0K6L9 |
| YAPEPITSLNTKYVAP | 1 | 0 | 0 | 1708453 | YP_009047204.1 | K9N5Q8 |
| YAPTYVSVDDKYKTSD | 1 | 0 | 0 | 1708454 | YP_009047202.1 | K9N7C7 |
| YATAQEAYEQAVANGD | 1 | 0 | 0 | 1708482 | NP_828849.2 | P0C6X7 |
| YATHHDKFTDGVCLFW | 1 | 0 | 0 | 1708485 | NP_828849.2 | P0C6X7 |
| YAYATLKEIVGPKELL | 1 | 0 | 0 | 1708501 | NP_073549.1 | A0A0P0LKV0 |
| YAYFNGAIIRDSDVVK | 1 | 0 | 0 | 1708503 | YP_009047202.1 | K9N7C7 |
| YCAFTAPEDDDFLEES | 1 | 0 | 0 | 1708512 | YP_009555238.1 | P0C6X6 |
| YCDGSLGCKMCLFGYQ | 1 | 0 | 0 | 1708525 | NP_073549.1 | A0A0P0LKV0 |
| YCDNGKYYTKPIIKAQ | 1 | 0 | 0 | 1708526 | YP_173236.1 | A0A140H1G9 |
| YCDSPFEKLQCEHLQF | 1 | 0 | 0 | 1708527 | YP_003767.1 | A0A0P0G321 |
| YCFSVNNTFCPCAKPS | 1 | 0 | 0 | 1708535 | YP_173238.1 | A0A140H1H1 |
| YCFVNTTIGNETTSAF | 1 | 0 | 0 | 1708536 | NP_073551.1 | A0A0P0K6L9 |
| YCGNSATCYNSVSNEY | 1 | 0 | 0 | 1708538 | YP_003766.2 | H9EJ42 |
| YCILEPRSGNHCPAGN | 1 | 0 | 0 | 1708543 | YP_009047204.1 | K9N5Q8 |
| YCNGVRELYLNSSNVT | 1 | 0 | 0 | 1708555 | NP_828849.2 | P0C6X7 |
| YCPACHNPEVGPEHSL | 1 | 0 | 0 | 1708559 | QHR63299.1 | P0C6W2 |
| YCPRHVIASNTTSAID | 1 | 0 | 0 | 1708563 | NP_073549.1 | A0A0P0LKV0 |
| YCPRHVICSSSNMNEP | 1 | 0 | 0 | 1708565 | YP_173236.1 | A0A140H1G9 |
| YCRCHIDHPNPKGFCD | 1 | 0 | 0 | 1708576 | YP_009724389.1 | P0DTD1 |
| YCRCHIDHPNPKGYCE | 1 | 0 | 0 | 1708577 | QIQ54046.1 | P0C6U8 |
| YCRKIGVNVCSDSTFE | 1 | 0 | 0 | 1708582 | YP_173236.1 | A0A140H1G9 |
| YCRVGECVESNAGVCF | 1 | 0 | 0 | 1708587 | NP_073549.1 | A0A0P0LKV0 |
| YCSFYPPDEDEEEGDC | 1 | 0 | 0 | 1708590 | YP_009724389.1 | P0DTD1 |
| YCSFYPPDEDYEEEYS | 1 | 0 | 0 | 1708591 | QIQ54046.1 | P0C6U8 |
| YCSFYPPDEEEEDDAE | 1 | 0 | 0 | 1708592 | NP_828849.2 | P0C6X7 |
| YCSNSSSGVLDTTIPF | 1 | 0 | 0 | 1708593 | YP_003767.1 | A0A0P0G321 |
| YCTFAIEDVEDVISSE | 1 | 0 | 0 | 1708600 | YP_173236.1 | A0A140H1G9 |
| YCTGYREGYLNSTNVT | 1 | 0 | 0 | 1708605 | YP_009724389.1 | P0DTD1 |
| YCTNNTLVSGDAHIPF | 1 | 0 | 0 | 1708607 | NP_073551.1 | A0A0P0K6L9 |
| YCTSYRDGYLNSTNVT | 1 | 0 | 0 | 1708608 | QIQ54046.1 | P0C6U8 |
| YCTVQGPPGTGKSHLA | 1 | 0 | 0 | 1708610 | YP_009555238.1 | P0C6X6 |
| YCVDLHNKINLCDDPE | 1 | 0 | 0 | 1708612 | YP_003766.2 | H9EJ42 |
| YCVEMHNKINLCDDPE | 1 | 0 | 0 | 1708614 | NP_073549.1 | A0A0P0LKV0 |
| YCVRAFDVYNKDASFI | 1 | 0 | 0 | 1708616 | NP_073549.1 | A0A0P0LKV0 |
| YCVTTKAKSVQQDLDV | 1 | 0 | 0 | 1708617 | YP_009047202.1 | K9N7C7 |
| YCYCGNSATCYNSVSN | 1 | 0 | 0 | 1708618 | YP_003766.2 | H9EJ42 |
| YCYDKNPINSNVKTLE | 1 | 0 | 0 | 1708619 | NP_073549.1 | A0A0P0LKV0 |
| YCYDKQPVNSNVKLLD | 1 | 0 | 0 | 1708620 | YP_003766.2 | H9EJ42 |
| YCYFRNSSANYVGKYD | 1 | 0 | 0 | 1708624 | YP_173237.1 | A0A140H1H0 |
| YCYFRNSTTNYVGVYD | 1 | 0 | 0 | 1708625 | YP_009555240.1 | P30215 |
| YCYNTDLIEGSKPYST | 1 | 0 | 0 | 1708627 | YP_003766.2 | H9EJ42 |
| YCYSDGVMKNASLYTS | 1 | 0 | 0 | 1708628 | YP_173236.1 | A0A140H1G9 |
| YDALLHRNSLDPFCFD | 1 | 0 | 0 | 1708635 | YP_003766.2 | H9EJ42 |
| YDAYQNLVGYYSDDGN | 1 | 0 | 0 | 1708643 | YP_009047204.1 | K9N5Q8 |
| YDDGSCEVVHDQVNYV | 1 | 0 | 0 | 1708653 | NP_073549.1 | A0A0P0LKV0 |
| YDDSQYYFNKDTGVIY | 1 | 0 | 0 | 1708658 | YP_009555240.1 | P30215 |
| YDDVDASLFVDINNLL | 1 | 0 | 0 | 1708659 | YP_173236.1 | A0A140H1G9 |
| YDEEGGYDVSKPVMIS | 1 | 0 | 0 | 1708662 | YP_003766.2 | H9EJ42 |
| YDENGTITDAVDCSQN | 1 | 0 | 0 | 1708663 | NP_828851.1 | P59594 |
| YDFDGVTTSDKCIFNS | 1 | 0 | 0 | 1708666 | YP_003766.2 | H9EJ42 |

| | | | | | | |
|---|---|---|---|---|---|---|
| YDFGDFIQTAPGFGVA | 1 | 0 | 0 | 1708669 | YP_173236.1 | A0A140H1G9 |
| YDFGDFIQTTPGSGVP | 1 | 0 | 0 | 1708670 | YP_009724389.1 | P0DTD1 |
| YDFGDFVITQPGSGVA | 1 | 0 | 0 | 1708671 | YP_009047202.1 | K9N7C7 |
| YDFGDFVQVAPGCGVP | 1 | 0 | 0 | 1708673 | NP_828849.2 | P0C6X7 |
| YDFGDFVVSLPNMGVP | 1 | 0 | 0 | 1708674 | YP_003766.2 | H9EJ42 |
| YDFGDYVIAAPGCGVA | 1 | 0 | 0 | 1708675 | YP_009555238.1 | P0C6X6 |
| YDFKVEDAYDQVHDEP | 1 | 0 | 0 | 1708676 | YP_009555238.1 | P0C6X6 |
| YDFLNMDMGVFINKYG | 1 | 0 | 0 | 1708681 | YP_003766.2 | H9EJ42 |
| YDFTDYKLELFNKYFK | 1 | 0 | 0 | 1708685 | YP_009555238.1 | P0C6X6 |
| YDFTEERLKLFDRYFK | 1 | 0 | 0 | 1708688 | YP_009724389.1 | P0DTD1 |
| YDFVENPDIINVYKKL | 1 | 0 | 0 | 1708693 | YP_009555238.1 | P0C6X6 |
| YDGCAQGVFGVNMRTN | 1 | 0 | 0 | 1708698 | NP_073549.1 | A0A0P0LKV0 |
| YDGDRFVKHDLSLLSV | 1 | 0 | 0 | 1708699 | NP_073549.1 | A0A0P0LKV0 |
| YDGGCLNASEVVVNNL | 1 | 0 | 0 | 1708704 | YP_009047202.1 | K9N7C7 |
| YDGLRGAIRKAKDYGF | 1 | 0 | 0 | 1708705 | YP_009047202.1 | K9N7C7 |
| YDGNMFIKFPEVVFES | 1 | 0 | 0 | 1708706 | YP_009047202.1 | K9N7C7 |
| YDGRTGELPCAIMNDK | 1 | 0 | 0 | 1708708 | YP_173236.1 | A0A140H1G9 |
| YDHVISTSHKLVLSVN | 1 | 0 | 0 | 1708714 | YP_009724389.1 | P0DTD1 |
| YDIGNPKGIPIVDDPV | 1 | 0 | 0 | 1708721 | YP_009047202.1 | K9N7C7 |
| YDKFKINDTTCKYVFT | 1 | 0 | 0 | 1708728 | YP_173236.1 | A0A140H1G9 |
| YDKFKINDTTRKYVFT | 1 | 0 | 0 | 1708729 | YP_009555238.1 | P0C6X6 |
| YDKNPINSNVKTLEYD | 1 | 0 | 0 | 1708733 | NP_073549.1 | A0A0P0LKV0 |
| YDKQPVNSNVKLLDYD | 1 | 0 | 0 | 1708734 | YP_003766.2 | H9EJ42 |
| YDLEVLLQDALQSREA | 1 | 0 | 0 | 1708740 | YP_009555238.1 | P0C6X6 |
| YDLSVVNARLRAKHYV | 1 | 0 | 0 | 1708749 | YP_009724389.1 | P0DTD1 |
| YDNGNFIKLPEVIAQG | 1 | 0 | 0 | 1708761 | NP_073549.1 | A0A0P0LKV0 |
| YDNKLKAHKDKSAQCF | 1 | 0 | 0 | 1708763 | YP_009724389.1 | P0DTD1 |
| YDNKLRAHKDKSQQCF | 1 | 0 | 0 | 1708765 | QIQ54047.1 | P0C6X7 |
| YDPITKNIGEYNVSKD | 1 | 0 | 0 | 1708770 | YP_009555238.1 | P0C6X6 |
| YDPKTKHVTRENDSKE | 1 | 0 | 0 | 1708772 | QIQ54047.1 | P0C6X7 |
| YDPKTKNVTKENDSKE | 1 | 0 | 0 | 1708773 | YP_009724389.1 | P0DTD1 |
| YDPLTKNIGDYNVSKD | 1 | 0 | 0 | 1708776 | YP_173236.1 | A0A140H1G9 |
| YDPRTKHVTKENDSKE | 1 | 0 | 0 | 1708778 | NP_828849.2 | P0C6X7 |
| YDPTTKNVTGSNESKA | 1 | 0 | 0 | 1708779 | YP_009047202.1 | K9N7C7 |
| YDPVENEDIHRVYASL | 1 | 0 | 0 | 1708780 | YP_003766.2 | H9EJ42 |
| YDQVVDNVYVTYAGNV | 1 | 0 | 0 | 1708782 | YP_009555238.1 | P0C6X6 |
| YDRDGQRTYDDVNASL | 1 | 0 | 0 | 1708783 | YP_009555238.1 | P0C6X6 |
| YDRDGQRVYDDVDASL | 1 | 0 | 0 | 1708784 | YP_173236.1 | A0A140H1G9 |
| YDRFKVNETNSQYLFS | 1 | 0 | 0 | 1708785 | YP_009047202.1 | K9N7C7 |
| YDRLDSIQADQQVDRL | 1 | 0 | 0 | 1708786 | YP_003767.1 | A0A0P0G321 |
| YDRLDTIQADQQVDRL | 1 | 0 | 0 | 1708787 | NP_073551.1 | A0A0P0K6L9 |
| YDRLPNEVTVAKPNVL | 1 | 0 | 0 | 1708788 | YP_009047210.1 | K9N7A1 |
| YDSNGNIIGFKDFVTN | 1 | 0 | 0 | 1708791 | YP_173238.1 | A0A140H1H1 |
| YDSSEVKAQSSGVIPE | 1 | 0 | 0 | 1708794 | YP_173236.1 | A0A140H1G9 |
| YDTAKKSMYDGDRFVK | 1 | 0 | 0 | 1708798 | NP_073549.1 | A0A0P0LKV0 |
| YDVAVTAGPFMDNAIN | 1 | 0 | 0 | 1708805 | YP_009047202.1 | K9N7C7 |
| YDVHKDNFKDGLCMFW | 1 | 0 | 0 | 1708806 | YP_173236.1 | A0A140H1G9 |
| YDVLFYCSNSSSGVLD | 1 | 0 | 0 | 1708807 | YP_003767.1 | A0A0P0G321 |
| YDYVGTGIIRSSNQSL | 1 | 0 | 0 | 1708820 | YP_003767.1 | A0A0P0G321 |
| YEADTEQTQNVTVKPK | 1 | 0 | 0 | 1708829 | YP_009047202.1 | K9N7C7 |
| YEDAIANGSSSQLIKQ | 1 | 0 | 0 | 1708840 | YP_003766.2 | H9EJ42 |
| YEDFQENWNTKHSSGV | 1 | 0 | 0 | 1708842 | YP_009724389.1 | P0DTD1 |
| YEELVASFLEMKSETK | 1 | 0 | 0 | 1708849 | QIQ54046.1 | P0C6U8 |
| YEEQDAIFSLTKRNIL | 1 | 0 | 0 | 1708850 | NP_073549.1 | A0A0P0LKV0 |
| YEGGCIKACEVVVTNL | 1 | 0 | 0 | 1708860 | YP_003766.2 | H9EJ42 |
| YEGGCITSREVVVTNL | 1 | 0 | 0 | 1708861 | NP_073549.1 | A0A0P0LKV0 |
| YEGNSPFHPLADNKFA | 1 | 0 | 0 | 1708864 | YP_009724395.1 | P0DTC7 |
| YEHQTPFEIKSAKKFD | 1 | 0 | 0 | 1708867 | NP_828849.2 | P0C6X7 |
| YEKDKAVARKLERMAD | 1 | 0 | 0 | 1708872 | YP_009047202.1 | K9N7C7 |
| YEKLKPILDWLEEKFK | 1 | 0 | 0 | 1708874 | QIQ54046.1 | P0C6U8 |
| YEKLKPVLDWLEEKFK | 1 | 0 | 0 | 1708875 | YP_009724389.1 | P0DTD1 |
| YEKSTFNKDIYCGNSM | 1 | 0 | 0 | 1708876 | YP_003766.2 | H9EJ42 |
| YELDNDFNTILNTACG | 1 | 0 | 0 | 1708878 | YP_009555238.1 | P0C6X6 |
| YELQTPFEIKLAKKFD | 1 | 0 | 0 | 1708886 | YP_009724389.1 | P0DTD1 |
| YEMLSLQQVVKALNES | 1 | 0 | 0 | 1708893 | YP_009047204.1 | K9N5Q8 |
| YEMYVKWPWYVWLLIS | 1 | 0 | 0 | 1708895 | YP_173238.1 | A0A140H1H1 |
| YENARQAYEDAIANGS | 1 | 0 | 0 | 1708899 | YP_003766.2 | H9EJ42 |
| YENAVANGSSPQIIKQ | 1 | 0 | 0 | 1708900 | NP_073549.1 | A0A0P0LKV0 |
| YENFNHHEKLLAPLLS | 1 | 0 | 0 | 1708901 | QIQ54046.1 | P0C6U8 |
| YENKFVPVKEASKQCF | 1 | 0 | 0 | 1708905 | NP_073549.1 | A0A0P0LKV0 |
| YENKFVPVKPASKQCF | 1 | 0 | 0 | 1708906 | YP_003766.2 | H9EJ42 |
| YENLSLADISAVQSSF | 1 | 0 | 0 | 1708908 | YP_173236.1 | A0A140H1G9 |
| YENQKQIANQFNKAIS | 1 | 0 | 0 | 1708911 | NP_828851.1 | P59594 |
| YEPQIITTDNTFVSGS | 1 | 0 | 0 | 1708917 | QHR63300.2 | Q3LZX1 |
| YEQLKPVLDWLEEKFK | 1 | 0 | 0 | 1708923 | QHR63299.1 | P0C6W2 |
| YEQNWNTKHGSGALRE | 1 | 0 | 0 | 1708925 | NP_828849.2 | P0C6X7 |
| YEQVVMDYLDNLKPRV | 1 | 0 | 0 | 1708927 | NP_828849.2 | P0C6X7 |
| YEQYIKWPWYVWLGFI | 1 | 0 | 0 | 1708929 | NP_828851.1 | P59594 |
| YERDKAVARKLERMAD | 1 | 0 | 0 | 1708930 | YP_173236.1 | A0A140H1G9 |
| YERDNTSCPEWMDDFE | 1 | 0 | 0 | 1708931 | YP_009047202.1 | K9N7C7 |
| YERDRAVAKKLERMAD | 1 | 0 | 0 | 1708932 | YP_009555238.1 | P0C6X6 |
| YESLRPDTRYVLMDGS | 1 | 0 | 0 | 1708946 | YP_009724389.1 | P0DTD1 |
| YETARQEYENAVANGS | 1 | 0 | 0 | 1708950 | NP_073549.1 | A0A0P0LKV0 |
| YEVAKKNLDEARFSGS | 1 | 0 | 0 | 1708954 | YP_009555238.1 | P0C6X6 |
| YEYGTEDDYQGKPLEF | 1 | 0 | 0 | 1708961 | YP_009724389.1 | P0DTD1 |
| YEYGTESDYKGLPLEF | 1 | 0 | 0 | 1708963 | QIQ54046.1 | P0C6U8 |
| YFAKLGRELNCNVCSN | 1 | 0 | 0 | 1708967 | YP_173236.1 | A0A140H1G9 |
| YFAKVGREISCNVCTK | 1 | 0 | 0 | 1708968 | YP_009555238.1 | P0C6X6 |
| YFAVRKEGQDVIFSQF | 1 | 0 | 0 | 1708975 | YP_009555238.1 | P0C6X6 |
| YFDKAGQKTYERHPLS | 1 | 0 | 0 | 1708987 | NP_828849.2 | P0C6X7 |
| YFDKAGQKTYERHSLS | 1 | 0 | 0 | 1708988 | YP_009724389.1 | P0DTD1 |

| | | | | | | |
|---|---|---|---|---|---|---|
| YFDSKGWYDPVENEDI | 1 | 0 | 0 | 1708991 | YP_003766.2 | H9EJ42 |
| YFEIYDGGCIPASQVI | 1 | 0 | 0 | 1708996 | YP_009555238.1 | P0C6X6 |
| YFEIYDGGCLNASEVV | 1 | 0 | 0 | 1708997 | YP_009047202.1 | K9N7C7 |
| YFGGFNFSQILPDPLK | 1 | 0 | 0 | 1709010 | NP_828851.1 | P59594 |
| YFIKGLNNLNRGMVLG | 1 | 0 | 0 | 1709020 | YP_009724389.1 | P0DTD1 |
| YFIKTNNTRIVDEWSY | 1 | 0 | 0 | 1709022 | YP_009047204.1 | K9N5Q8 |
| YFITMSSKGQKGDAEE | 1 | 0 | 0 | 1709027 | YP_003766.2 | H9EJ42 |
| YFKHWSMPYHPNTVDC | 1 | 0 | 0 | 1709034 | YP_009555238.1 | P0C6X6 |
| YFKKVDGIIQQLPETY | 1 | 0 | 0 | 1709036 | NP_828849.2 | P0C6X7 |
| YFKPGGTTSGDASTAY | 1 | 0 | 0 | 1709038 | YP_003766.2 | H9EJ42 |
| YFKYWDQTYHPNCINC | 1 | 0 | 0 | 1709042 | NP_828849.2 | P0C6X7 |
| YFKYWDQTYHPNCVNC | 1 | 0 | 0 | 1709043 | YP_009724389.1 | P0DTD1 |
| YFKYWGQDYHPDCVDC | 1 | 0 | 0 | 1709045 | NP_073549.1 | A0A0P0LKV0 |
| YFLDDVEMVAYNPDLS | 1 | 0 | 0 | 1709047 | YP_173236.1 | A0A140H1G9 |
| YFMVDNVTRLHDA | 1 | 0 | 0 | 1709059 | YP_009555240.1 | P30215 |
| YFMYDIVSSFEGYDFK | 1 | 0 | 0 | 1709060 | NP_073549.1 | A0A0P0LKV0 |
| YFNATGSDVNCNGYQH | 1 | 0 | 0 | 1709061 | YP_003767.1 | A0A0P0G321 |
| YFNGAIIRDSDVVKQP | 1 | 0 | 0 | 1709062 | YP_009047202.1 | K9N7C7 |
| YFNGTGRGDCKGFSSD | 1 | 0 | 0 | 1709063 | NP_073551.1 | A0A0P0K6L9 |
| YFNKKDWYDFVENPDI | 1 | 0 | 0 | 1709066 | YP_009724389.1 | P0DTD1 |
| YFNRPSLVDDNKFDVL | 1 | 0 | 0 | 1709072 | YP_009555238.1 | P0C6X6 |
| YFRNSSANYVGKYDIN | 1 | 0 | 0 | 1709085 | YP_173237.1 | A0A140H1H0 |
| YFRNSTTNYVGVYDIN | 1 | 0 | 0 | 1709086 | YP_009555240.1 | P30215 |
| YFSGAMETAAYREAAA | 1 | 0 | 0 | 1709090 | YP_009047202.1 | K9N7C7 |
| YFSKKDWYDFVENPDI | 1 | 0 | 0 | 1709091 | YP_173236.1 | A0A140H1G9 |
| YFSKKHVEVFTDGKLN | 1 | 0 | 0 | 1709092 | YP_009047202.1 | K9N7C7 |
| YFSQLLCRPIKLVDSE | 1 | 0 | 0 | 1709095 | NP_073549.1 | A0A0P0LKV0 |
| YFTKKDWYDFVENPDI | 1 | 0 | 0 | 1709106 | YP_009555238.1 | P0C6X6 |
| YFTQSRDLEDFKPRSQ | 1 | 0 | 0 | 1709112 | NP_828849.2 | P0C6X7 |
| YFVNVNNTWMYTGSGY | 1 | 0 | 0 | 1709138 | YP_009555241.1 | P36334 |
| YGADLKSFDLGDELGT | 1 | 0 | 0 | 1709157 | YP_009724389.1 | P0DTD1 |
| YGAFMDKQVHQVQLTD | 1 | 0 | 0 | 1709158 | YP_009047202.1 | K9N7C7 |
| YGAGSDKGVAPGTTVL | 1 | 0 | 0 | 1709161 | YP_003766.2 | H9EJ42 |
| YGCDYSGKLADCLQAY | 1 | 0 | 0 | 1709168 | YP_173236.1 | A0A140H1G9 |
| YGDSATLPKGIMMNVA | 1 | 0 | 0 | 1709186 | YP_009724389.1 | P0DTD1 |
| YGDVDNLVFTNKTTLP | 1 | 0 | 0 | 1709187 | YP_003766.2 | H9EJ42 |
| YGENAVIPKGIMMNVA | 1 | 0 | 0 | 1709192 | NP_828849.2 | P0C6X7 |
| YGGFEDQPNLQVESAN | 1 | 0 | 0 | 1709216 | NP_073549.1 | A0A0P0LKV0 |
| YGGWDDMLRRLIKDVD | 1 | 0 | 0 | 1709222 | YP_009555238.1 | P0C6X6 |
| YGGWDFMLKTLYKDVD | 1 | 0 | 0 | 1709223 | YP_009047202.1 | K9N7C7 |
| YGGWDNMLKNLMADVD | 1 | 0 | 0 | 1709224 | NP_073549.1 | A0A0P0LKV0 |
| YGIATVREVLSDRELH | 1 | 0 | 0 | 1709231 | YP_009724389.1 | P0DTD1 |
| YGKPINLPTGCMMNVA | 1 | 0 | 0 | 1709238 | YP_173236.1 | A0A140H1G9 |
| YGKPVTLPTGCMMNVA | 1 | 0 | 0 | 1709239 | YP_009555238.1 | P0C6X6 |
| YGLNSTETITTGFDLN | 1 | 0 | 0 | 1709254 | YP_009555240.1 | P30215 |
| YGNFDDQPSLQVESAN | 1 | 0 | 0 | 1709263 | YP_003766.2 | H9EJ42 |
| YGNMTLNGLWLDDKVY | 1 | 0 | 0 | 1709265 | YP_009555238.1 | P0C6X6 |
| YGNVSRHNLTKYTMMD | 1 | 0 | 0 | 1709268 | YP_003766.2 | H9EJ42 |
| YGPYRDAQVVQLPVKD | 1 | 0 | 0 | 1709275 | YP_173236.1 | A0A140H1G9 |
| YGRCPTAADINIPDLP | 1 | 0 | 0 | 1709280 | YP_009555240.1 | P30215 |
| YGSSVETETYFKVNGL | 1 | 0 | 0 | 1709294 | YP_173236.1 | A0A140H1G9 |
| YGSTVLNGVWLGDTVT | 1 | 0 | 0 | 1709297 | YP_003766.2 | H9EJ42 |
| YGTDTNSVCPKLEFAN | 1 | 0 | 0 | 1709302 | YP_009047204.1 | K9N5Q8 |
| YGTEDDYQGKPLEFGA | 1 | 0 | 0 | 1709303 | YP_009724389.1 | P0DTD1 |
| YGTEDDYQGKSLEFGA | 1 | 0 | 0 | 1709304 | QHR63299.1 | P0C6W2 |
| YGTESDYKGLPLEFGA | 1 | 0 | 0 | 1709306 | QIQ54046.1 | P0C6U8 |
| YGTFCDNINSILDEVN | 1 | 0 | 0 | 1709307 | YP_173238.1 | A0A140H1H1 |
| YGVCKYTDLKFIENLN | 1 | 0 | 0 | 1709313 | YP_173236.1 | A0A140H1G9 |
| YGVCMYTDLKFIDKLN | 1 | 0 | 0 | 1709314 | YP_009555238.1 | P0C6X6 |
| YGVRRKNSEPEIPHFN | 1 | 0 | 0 | 1709320 | NP_073556.1 | A0A0N7J131 |
| YGWVSGNMMDGFSCPL | 1 | 0 | 0 | 1709325 | YP_173236.1 | A0A140H1G9 |
| YHAFKCNRSGSNDIWM | 1 | 0 | 0 | 1709336 | YP_009555240.1 | P30215 |
| YHGAIKLDDKDPQFKD | 1 | 0 | 0 | 1709346 | NP_828858.1 | P59595 |
| YHNESGIKTTLRKGGR | 1 | 0 | 0 | 1709357 | QIQ54046.1 | P0C6U8 |
| YHNESGLKPILRKGGR | 1 | 0 | 0 | 1709358 | QHR63299.1 | P0C6W2 |
| YHPAHAPSFLAVDDKY | 1 | 0 | 0 | 1709363 | YP_009555238.1 | P0C6X6 |
| YHPNCINCLDDRCILH | 1 | 0 | 0 | 1709365 | NP_828849.2 | P0C6X7 |
| YHPNCVNCLDDRCILH | 1 | 0 | 0 | 1709367 | YP_009724389.1 | P0DTD1 |
| YHQKMLKSMAATRGAT | 1 | 0 | 0 | 1709371 | YP_009047202.1 | K9N7C7 |
| YHTPATDCSDGNYNRN | 1 | 0 | 0 | 1709381 | YP_009047204.1 | K9N5Q8 |
| YHTTSKPITFHMDGEV | 1 | 0 | 0 | 1709384 | QIQ54046.1 | P0C6U8 |
| YHVTKVLAEKFDYNVS | 1 | 0 | 0 | 1709388 | NP_073549.1 | A0A0P0LKV0 |
| YHYKELGVVMNLDVDT | 1 | 0 | 0 | 1709392 | YP_173236.1 | A0A140H1G9 |
| YHYTPFHYERDNTSCP | 1 | 0 | 0 | 1709395 | YP_009047202.1 | K9N7C7 |
| YIAYENARQAYEDAIA | 1 | 0 | 0 | 1709407 | YP_003766.2 | H9EJ42 |
| YIDINGNLHPDSATLV | 1 | 0 | 0 | 1709420 | YP_009724389.1 | P0DTD1 |
| YIDVLRNSFGKDLNAN | 1 | 0 | 0 | 1709423 | NP_073549.1 | A0A0P0LKV0 |
| YIDVNKTLQELSYKLP | 1 | 0 | 0 | 1709424 | NP_073551.1 | A0A0P0K6L9 |
| YIEKLYKDLTGHMLDS | 1 | 0 | 0 | 1709429 | YP_009047202.1 | K9N7C7 |
| YIESKRGVYCCRDHEH | 1 | 0 | 0 | 1709432 | NP_828849.2 | P0C6X7 |
| YIFERIDYSDAVSYKS | 1 | 0 | 0 | 1709434 | YP_009047202.1 | K9N7C7 |
| YIGATVRLQAGKQTEF | 1 | 0 | 0 | 1709444 | NP_073549.1 | A0A0P0LKV0 |
| YIGDPAQLPAPRVLLS | 1 | 0 | 0 | 1709446 | YP_009555238.1 | P0C6X6 |
| YIKDGDDGYFLPYPDP | 1 | 0 | 0 | 1709460 | YP_009047202.1 | K9N7C7 |
| YIKKLYNDLGNQILDS | 1 | 0 | 0 | 1709462 | YP_009555238.1 | P0C6X6 |
| YIKKLYNDLGTQILDS | 1 | 0 | 0 | 1709463 | YP_173236.1 | A0A140H1G9 |
| YILANECTERLKLFAA | 1 | 0 | 0 | 1709473 | YP_009555238.1 | P0C6X6 |
| YINGFNVSELEDACFK | 1 | 0 | 0 | 1709485 | YP_009555242.1 | Q04853 |
| YIQGIKLGTGYSLADL | 1 | 0 | 0 | 1709499 | YP_009555244.1 | Q01455 |
| YIQGVKLGTGYTLSDL | 1 | 0 | 0 | 1709500 | YP_173241.1 | Q0ZJ82 |
| YISSNGPLKVGAGVML | 1 | 0 | 0 | 1709519 | YP_003766.2 | H9EJ42 |

| | | | | | | |
|---|---|---|---|---|---|---|
| YIVDSVAVKNGALHLY | 1 | 0 | 0 | 1709530 | NP_828849.2 | P0C6X7 |
| YIWTADEAIKFLKEWN | 1 | 0 | 0 | 1709547 | YP_009555244.1 | Q01455 |
| YIYDEEGGNDLSLPVM | 1 | 0 | 0 | 1709549 | NP_073549.1 | A0A0P0LKV0 |
| YIYDEEGGYDVSKPVM | 1 | 0 | 0 | 1709550 | YP_003766.2 | H9EJ42 |
| YIYNRVKSGSPGDSSW | 1 | 0 | 0 | 1709552 | YP_003767.1 | A0A0P0G321 |
| YIYVRKNGQFQDHYDG | 1 | 0 | 0 | 1709553 | YP_003766.2 | H9EJ42 |
| YIYVTNEIGLNASVTL | 1 | 0 | 0 | 1709554 | YP_003767.1 | A0A0P0G321 |
| YKAACVVDVNDSHSMA | 1 | 0 | 0 | 1709555 | YP_009555238.1 | P0C6X6 |
| YKATGDLAVCLGIGDS | 1 | 0 | 0 | 1709565 | YP_009555238.1 | P0C6X6 |
| YKDSVFYVVKNSTAFP | 1 | 0 | 0 | 1709571 | YP_009047202.1 | K9N7C7 |
| YKDVDNPHLMGWDYPK | 1 | 0 | 0 | 1709572 | YP_009047202.1 | K9N7C7 |
| YKDWSYSGQSTQLGIE | 1 | 0 | 0 | 1709575 | YP_009724389.1 | P0DTD1 |
| YKDYLASGGQPITNCV | 1 | 0 | 0 | 1709576 | YP_009724389.1 | P0DTD1 |
| YKELGVVMNLDVDTHR | 1 | 0 | 0 | 1709584 | YP_173236.1 | A0A140H1G9 |
| YKENSYITTIKPVIYK | 1 | 0 | 0 | 1709585 | QHR63299.1 | P0C6W2 |
| YKENSYTTTIKPIVYK | 1 | 0 | 0 | 1709586 | QIQ54046.1 | P0C6U8 |
| YKENSYTTTIKPVTYK | 1 | 0 | 0 | 1709587 | YP_009724389.1 | P0DTD1 |
| YKETSYTTTIKPVSYK | 1 | 0 | 0 | 1709589 | NP_828849.2 | P0C6X7 |
| YKGIKIQEGIVDYGVR | 1 | 0 | 0 | 1709598 | NP_828849.2 | P0C6X7 |
| YKGIKIQEGVVDYGAR | 1 | 0 | 0 | 1709599 | YP_009724389.1 | P0DTD1 |
| YKGIKIQEGVVDYGVR | 1 | 0 | 0 | 1709600 | QIQ54046.1 | P0C6U8 |
| YKGKWVQVPIGTNDPI | 1 | 0 | 0 | 1709605 | NP_073549.1 | A0A0P0LKV0 |
| YKGNVTHDASSAINRP | 1 | 0 | 0 | 1709610 | YP_009047202.1 | K9N7C7 |
| YKGPITDIFYKENSYI | 1 | 0 | 0 | 1709611 | QHR63299.1 | P0C6W2 |
| YKGPITDVFYKENSYT | 1 | 0 | 0 | 1709612 | YP_009724389.1 | P0DTD1 |
| YKGPVTDVFYKENSYT | 1 | 0 | 0 | 1709613 | QIQ54046.1 | P0C6U8 |
| YKGPVTDVFYKETSYT | 1 | 0 | 0 | 1709614 | NP_828849.2 | P0C6X7 |
| YKGYQPIDVVRDLPSG | 1 | 0 | 0 | 1709618 | NP_828851.1 | P59594 |
| YKIDTTSSSCQLYYSL | 1 | 0 | 0 | 1709627 | YP_173238.1 | A0A140H1H1 |
| YKIEELFYSYATHHDK | 1 | 0 | 0 | 1709628 | NP_828849.2 | P0C6X7 |
| YKKCTKGLSIADLACA | 1 | 0 | 0 | 1709636 | NP_073551.1 | A0A0P0K6L9 |
| YKKDGVYYPSNGTNIL | 1 | 0 | 0 | 1709638 | NP_073549.1 | A0A0P0LKV0 |
| YKKDNAYFTEQPIDLV | 1 | 0 | 0 | 1709639 | QIQ54046.1 | P0C6U8 |
| YKKDNAYYTEQPIDLV | 1 | 0 | 0 | 1709640 | NP_828849.2 | P0C6X7 |
| YKKDNSYFTEQPIDLV | 1 | 0 | 0 | 1709641 | YP_009724389.1 | P0DTD1 |
| YKKVDGVVQQLPETYF | 1 | 0 | 0 | 1709648 | YP_009724389.1 | P0DTD1 |
| YKKVNGVVQQLPETYF | 1 | 0 | 0 | 1709649 | QHR63299.1 | P0C6W2 |
| YKKVNNEFIDPTECIY | 1 | 0 | 0 | 1709650 | YP_009047202.1 | K9N7C7 |
| YKLANDAKESLRLFAA | 1 | 0 | 0 | 1709651 | NP_073549.1 | A0A0P0LKV0 |
| YKLANDVKDTLRLFAA | 1 | 0 | 0 | 1709652 | YP_003766.2 | H9EJ42 |
| YKLDGVTYTEIEPKLD | 1 | 0 | 0 | 1709653 | NP_828849.2 | P0C6X7 |
| YKLFLSGDDCFVSNSS | 1 | 0 | 0 | 1709659 | YP_173236.1 | A0A140H1G9 |
| YKLHPAFNVSDAYANL | 1 | 0 | 0 | 1709661 | YP_003766.2 | H9EJ42 |
| YKLHPSFNVSDAYANL | 1 | 0 | 0 | 1709662 | NP_073549.1 | A0A0P0LKV0 |
| YKLNTDHAGSNDNIAL | 1 | 0 | 0 | 1709669 | NP_828855.1 | P59596 |
| YKLNTDHSNSSDNIAL | 1 | 0 | 0 | 1709670 | QIQ54051.1 | P59596 |
| YKLNTDHSSSSDNIAL | 1 | 0 | 0 | 1709671 | YP_009724393.1 | P0DTC5 |
| YKLPNYTVPDLVVEQY | 1 | 0 | 0 | 1709675 | NP_073551.1 | A0A0P0K6L9 |
| YKMLVAKDNKNLPKFI | 1 | 0 | 0 | 1709685 | YP_003771.1 | H9EJ69 |
| YKMQRMLLDKCDLQNY | 1 | 0 | 0 | 1709686 | QIQ54047.1 | P0C6X7 |
| YKNMCTGSPSIVEFNR | 1 | 0 | 0 | 1709693 | YP_009047202.1 | K9N7C7 |
| YKNMYLKSAVMQSVGA | 1 | 0 | 0 | 1709694 | YP_173236.1 | A0A140H1G9 |
| YKNSATGSLDVEVFNR | 1 | 0 | 0 | 1709696 | YP_003766.2 | H9EJ42 |
| YKNTCVGSDNVTDFNA | 1 | 0 | 0 | 1709700 | YP_009724389.1 | P0DTD1 |
| YKNVRITFELDERVDK | 1 | 0 | 0 | 1709704 | NP_828849.2 | P0C6X7 |
| YKPLLENIDYFNMRRA | 1 | 0 | 0 | 1709709 | YP_009555238.1 | P0C6X6 |
| YKQARAEDKKAKIVSA | 1 | 0 | 0 | 1709713 | YP_009047202.1 | K9N7C7 |
| YKQSCTGSPYIDDFNK | 1 | 0 | 0 | 1709717 | YP_173236.1 | A0A140H1G9 |
| YKQSCTGSPYIDDFNR | 1 | 0 | 0 | 1709718 | YP_009555238.1 | P0C6X6 |
| YKRAFLDKLDVNSGFA | 1 | 0 | 0 | 1709721 | YP_173241.1 | Q0ZJ82 |
| YKRQNNLAIEEIEYVH | 1 | 0 | 0 | 1709731 | NP_073549.1 | A0A0P0LKV0 |
| YKSCTKGLSIADLACA | 1 | 0 | 0 | 1709732 | YP_003767.1 | A0A0P0G321 |
| YKSSALGSMDIDVFNK | 1 | 0 | 0 | 1709738 | NP_073549.1 | A0A0P0LKV0 |
| YKTPPIKDFGGFNFLQ | 1 | 0 | 0 | 1709751 | QIQ54048.1 | P59594 |
| YKVLPPLMDVNMEAAY | 1 | 0 | 0 | 1709762 | YP_009047204.1 | K9N5Q8 |
| YKVNCCRFQRIDDDGN | 1 | 0 | 0 | 1709763 | YP_173236.1 | A0A140H1G9 |
| YKVNENLAVNLNICEP | 1 | 0 | 0 | 1709764 | YP_173236.1 | A0A140H1G9 |
| YKXTCVGSDNVTDFNA | 1 | 0 | 0 | 1709771 | QIQ54047.1 | P0C6X7 |
| YKYDAKNYVRFPEILA | 1 | 0 | 0 | 1709772 | YP_003766.2 | H9EJ42 |
| YKYDNGNFIKLPEVIA | 1 | 0 | 0 | 1709773 | NP_073549.1 | A0A0P0LKV0 |
| YKYLNSPMERVSLWNY | 1 | 0 | 0 | 1709778 | YP_009555238.1 | P0C6X6 |
| YKYYSGNANEADYRCA | 1 | 0 | 0 | 1709784 | NP_073549.1 | A0A0P0LKV0 |
| YLADYDVAVTAGPFMD | 1 | 0 | 0 | 1709788 | YP_009047202.1 | K9N7C7 |
| YLAKAMLDFSRDHNDI | 1 | 0 | 0 | 1709790 | NP_073549.1 | A0A0P0LKV0 |
| YLAYIKKLYNDLGNQI | 1 | 0 | 0 | 1709799 | YP_009555238.1 | P0C6X6 |
| YLCGADGKPVMSEDLW | 1 | 0 | 0 | 1709802 | NP_073549.1 | A0A0P0LKV0 |
| YLDAVKQGAKPVGNCV | 1 | 0 | 0 | 1709807 | NP_073549.1 | A0A0P0LKV0 |
| YLDDVKKIEYKPDLSQ | 1 | 0 | 0 | 1709809 | YP_009555238.1 | P0C6X6 |
| YLDFIQQGGTPIANCV | 1 | 0 | 0 | 1709810 | YP_009555238.1 | P0C6X6 |
| YLDFVNAGGAPLTNCV | 1 | 0 | 0 | 1709811 | YP_009047202.1 | K9N7C7 |
| YLDIHPSLCNNGKISS | 1 | 0 | 0 | 1709815 | YP_173237.1 | A0A140H1H0 |
| YLDLLNRLQALESGKV | 1 | 0 | 0 | 1709816 | YP_009047211.1 | K9N4V7 |
| YLDNLKPRVEAPKQEE | 1 | 0 | 0 | 1709817 | NP_828849.2 | P0C6X7 |
| YLEDKFDLLISDMYDG | 1 | 0 | 0 | 1709824 | NP_073549.1 | A0A0P0LKV0 |
| YLEGSVRVVTTFDAEY | 1 | 0 | 0 | 1709825 | NP_828849.2 | P0C6X7 |
| YLEYIKKLYNDLGTQI | 1 | 0 | 0 | 1709830 | YP_173236.1 | A0A140H1G9 |
| YLFDDAGEENFSSRMY | 1 | 0 | 0 | 1709832 | NP_828849.2 | P0C6X7 |
| YLFSTINALPETSADI | 1 | 0 | 0 | 1709842 | YP_009047202.1 | K9N7C7 |
| YLGDFIQGPFIAGNTV | 1 | 0 | 0 | 1709848 | YP_003766.2 | H9EJ42 |
| YLGKPKEQIDGYTMHA | 1 | 0 | 0 | 1709858 | NP_828849.2 | P0C6X7 |
| YLGTIKENIDGGAMHA | 1 | 0 | 0 | 1709868 | YP_009047202.1 | K9N7C7 |

| | | | | | | |
|---|---|---|---|---|---|---|
| YLISKFLKANPVWNSA | 1 | 0 | 0 | 1709883 | YP_173236.1 | A0A140H1G9 |
| YLKGSGSDFSLADSVV | 1 | 0 | 0 | 1709885 | YP_009555238.1 | P0C6X6 |
| YLKNLTQTFTSMLTNY | 1 | 0 | 0 | 1709893 | YP_173236.1 | A0A140H1G9 |
| YLKPTTFMLKYDENGT | 1 | 0 | 0 | 1709895 | NP_828851.1 | P59594 |
| YLLAFNQDGIIFNAVD | 1 | 0 | 0 | 1709904 | YP_009555241.1 | P36334 |
| YLLEEFDVVFGKRGGG | 1 | 0 | 0 | 1709906 | NP_073549.1 | A0A0P0LKV0 |
| YLLKFDNRGVITNAVD | 1 | 0 | 0 | 1709912 | YP_173238.1 | A0A140H1H1 |
| YLNDPSSKTVCTYMDL | 1 | 0 | 0 | 1709918 | NP_073549.1 | A0A0P0LKV0 |
| YLNHSVPKLSDFESEL | 1 | 0 | 0 | 1709922 | YP_173238.1 | A0A140H1H1 |
| YLNLSSELKQLEAKTA | 1 | 0 | 0 | 1709924 | YP_003767.1 | A0A0P0G321 |
| YLNSSNVTTMDFCEGS | 1 | 0 | 0 | 1709930 | NP_828849.2 | P0C6X7 |
| YLNSTNVTTTAYCTGS | 1 | 0 | 0 | 1709933 | QIQ54046.1 | P0C6U8 |
| YLNVQYRINGLGVTMD | 1 | 0 | 0 | 1709941 | YP_009555241.1 | P36334 |
| YLQQRTFLLSYNQNGT | 1 | 0 | 0 | 1709957 | QIQ54048.1 | P59594 |
| YLRHGKLRPFERDISN | 1 | 0 | 0 | 1709963 | NP_828851.1 | P59594 |
| YLRYINGFNVSELEDA | 1 | 0 | 0 | 1709972 | YP_009555242.1 | Q04853 |
| YLSERCNVIADSKSFT | 1 | 0 | 0 | 1709973 | YP_009047202.1 | K9N7C7 |
| YLSLSDRFKTSGDLAV | 1 | 0 | 0 | 1709976 | NP_073549.1 | A0A0P0LKV0 |
| YLTPGKLESGWTTGAA | 1 | 0 | 0 | 1709995 | QIQ54048.1 | P59594 |
| YLTSSSKTPEEHFIET | 1 | 0 | 0 | 1709998 | YP_009724389.1 | P0DTD1 |
| YLTSSSKTSEEHFVET | 1 | 0 | 0 | 1709999 | NP_828849.2 | P0C6X7 |
| YLVQQESSFVMMSAPP | 1 | 0 | 0 | 1710009 | NP_828849.2 | P0C6X7 |
| YLYKKVNNEFIDPTEC | 1 | 0 | 0 | 1710025 | YP_009047202.1 | K9N7C7 |
| YLYNTGRSVYVKFQDS | 1 | 0 | 0 | 1710030 | YP_009047209.1 | K9N5R3 |
| YMAKAGDKYYPVVVDD | 1 | 0 | 0 | 1710035 | YP_009555238.1 | P0C6X6 |
| YMCVVPSINDLSFDKN | 1 | 0 | 0 | 1710039 | YP_003766.2 | H9EJ42 |
| YMDGMDAKQVDYVPLK | 1 | 0 | 0 | 1710040 | YP_009555238.1 | P0C6X6 |
| YMGKCGDKFFPIVMND | 1 | 0 | 0 | 1710048 | YP_173236.1 | A0A140H1G9 |
| YMGTLSYDNLKTGVSI | 1 | 0 | 0 | 1710049 | NP_828849.2 | P0C6X7 |
| YMGTLSYEQFKKGVQI | 1 | 0 | 0 | 1710050 | YP_009724389.1 | P0DTD1 |
| YMNANGLRPPKNSFEA | 1 | 0 | 0 | 1710067 | YP_009555238.1 | P0C6X6 |
| YMNSQGLLPPKNSIDA | 1 | 0 | 0 | 1710069 | YP_009724389.1 | P0DTD1 |
| YMNSQGLLPPKSSIDA | 1 | 0 | 0 | 1710070 | NP_828849.2 | P0C6X7 |
| YMVANGLNAPNGPFDA | 1 | 0 | 0 | 1710090 | NP_073549.1 | A0A0P0LKV0 |
| YNAFLDTVVSTVKIGG | 1 | 0 | 0 | 1710092 | NP_073549.1 | A0A0P0LKV0 |
| YNAKVEISVTPIKNTV | 1 | 0 | 0 | 1710094 | NP_073549.1 | A0A0P0LKV0 |
| YNCCDDDYFNKKDWYD | 1 | 0 | 0 | 1710096 | YP_009724389.1 | P0DTD1 |
| YNCCTDDYFNKKDWYD | 1 | 0 | 0 | 1710097 | QIQ54047.1 | P0C6X7 |
| YNEWAQANGFTAMNGE | 1 | 0 | 0 | 1710111 | NP_073549.1 | A0A0P0LKV0 |
| YNFIIYDSSDRGQESL | 1 | 0 | 0 | 1710113 | YP_009047202.1 | K9N7C7 |
| YNGSPSGVYQCAMRPN | 1 | 0 | 0 | 1710128 | YP_009724389.1 | P0DTD1 |
| YNGYLTSSSKTPEEHF | 1 | 0 | 0 | 1710129 | YP_009724389.1 | P0DTD1 |
| YNGYLTSSSKTSEEHF | 1 | 0 | 0 | 1710130 | NP_828849.2 | P0C6X7 |
| YNKDASFIGKNLKSNC | 1 | 0 | 0 | 1710142 | NP_073549.1 | A0A0P0LKV0 |
| YNKDYAELGYIADISA | 1 | 0 | 0 | 1710144 | YP_003766.2 | H9EJ42 |
| YNKIVDEPEEHVQIHT | 1 | 0 | 0 | 1710146 | YP_009724391.1 | P0DTC3 |
| YNKVENMTPRDLGACI | 1 | 0 | 0 | 1710152 | YP_009724389.1 | P0DTD1 |
| YNKWLELLEQNIDAYK | 1 | 0 | 0 | 1710153 | YP_009047211.1 | K9N4V7 |
| YNKYKYFSGAMDTTSY | 1 | 0 | 0 | 1710156 | YP_009724389.1 | P0DTD1 |
| YNLKNGEVEFVYMHQI | 1 | 0 | 0 | 1710162 | NP_073549.1 | A0A0P0LKV0 |
| YNLLKDCPAVAKHDFF | 1 | 0 | 0 | 1710164 | YP_009724389.1 | P0DTD1 |
| YNLLKGCNAVAKHDFF | 1 | 0 | 0 | 1710166 | NP_073549.1 | A0A0P0LKV0 |
| YNLNFSANSLDNLKSG | 1 | 0 | 0 | 1710169 | YP_003767.1 | A0A0P0G321 |
| YNLSSVLPQRNIRSSR | 1 | 0 | 0 | 1710175 | YP_003767.1 | A0A0P0G321 |
| YNLVKTGHYTGQAGEM | 1 | 0 | 0 | 1710179 | YP_009555238.1 | P0C6X6 |
| YNLVNVGHYDGRTGEL | 1 | 0 | 0 | 1710180 | YP_173236.1 | A0A140H1G9 |
| YNNLVPTYLKSDNIVA | 1 | 0 | 0 | 1710199 | YP_173236.1 | A0A140H1G9 |
| YNNSGSNITQIKNACV | 1 | 0 | 0 | 1710200 | YP_003766.2 | H9EJ42 |
| YNNSNNGKIVYAILSD | 1 | 0 | 0 | 1710202 | YP_009555238.1 | P0C6X6 |
| YNNYVPSYVKPDSVST | 1 | 0 | 0 | 1710204 | YP_009047202.1 | K9N7C7 |
| YNQNGTITDAVDCSLD | 1 | 0 | 0 | 1710216 | QIQ54048.1 | P59594 |
| YNRNASLNSFKEYFNL | 1 | 0 | 0 | 1710221 | YP_009047204.1 | K9N5Q8 |
| YNRVKSGSPGDSSWHI | 1 | 0 | 0 | 1710223 | YP_003767.1 | A0A0P0G321 |
| YNRYKYYSGNANEADY | 1 | 0 | 0 | 1710224 | NP_073549.1 | A0A0P0LKV0 |
| YNRYRIGNYKLNTDHA | 1 | 0 | 0 | 1710227 | NP_828855.1 | P59596 |
| YNSDYASKGYIANISA | 1 | 0 | 0 | 1710230 | YP_009555238.1 | P0C6X6 |
| YNSQNAVAAKILGLPT | 1 | 0 | 0 | 1710237 | QIQ54047.1 | P0C6X7 |
| YNSQNAVSRSMLGLTT | 1 | 0 | 0 | 1710239 | YP_009047202.1 | K9N7C7 |
| YNSQNFAAKRVLGLQT | 1 | 0 | 0 | 1710240 | YP_009555238.1 | P0C6X6 |
| YNSQNYVAKRVLGVQT | 1 | 0 | 0 | 1710242 | YP_173236.1 | A0A140H1G9 |
| YNSVSNDYCCFKHALG | 1 | 0 | 0 | 1710252 | NP_073549.1 | A0A0P0LKV0 |
| YNSVSNEYCCFKHALG | 1 | 0 | 0 | 1710253 | YP_003766.2 | H9EJ42 |
| YNSWQNLLYDSNGNII | 1 | 0 | 0 | 1710257 | YP_173238.1 | A0A140H1H1 |
| YNSWQNLLYDSNGNLY | 1 | 0 | 0 | 1710258 | YP_009555241.1 | P36334 |
| YNTDLIEGSKPYSTLQ | 1 | 0 | 0 | 1710261 | YP_003766.2 | H9EJ42 |
| YNTGRSVYVKFQDSKP | 1 | 0 | 0 | 1710264 | YP_009047209.1 | K9N5R3 |
| YNVRNDGTVEFCYLHQ | 1 | 0 | 0 | 1710272 | YP_003766.2 | H9EJ42 |
| YNVSLSKFNSCKSDIL | 1 | 0 | 0 | 1710277 | YP_173237.1 | A0A140H1H0 |
| YNYGAGIKLPSGIMLN | 1 | 0 | 0 | 1710291 | YP_003766.2 | H9EJ42 |
| YNYYKYNLPTMVDIKQ | 1 | 0 | 0 | 1710299 | YP_009555238.1 | P0C6X6 |
| YPAGVNITLANFNETK | 1 | 0 | 0 | 1710303 | NP_073551.1 | A0A0P0K6L9 |
| YPCMVGPDYAYFNGAI | 1 | 0 | 0 | 1710310 | YP_009047202.1 | K9N7C7 |
| YPFIANEKFINGCGRN | 1 | 0 | 0 | 1710323 | YP_003766.2 | H9EJ42 |
| YPGQGCVGYTIEEAKT | 1 | 0 | 0 | 1710329 | QIQ54046.1 | P0C6U8 |
| YPGQGLNGYTVEEART | 1 | 0 | 0 | 1710331 | QHR63299.1 | P0C6W2 |
| YPIISNEVSINTSCRL | 1 | 0 | 0 | 1710339 | YP_173236.1 | A0A140H1G9 |
| YPKLQASQAWQPGVAM | 1 | 0 | 0 | 1710348 | NP_828849.2 | P0C6X7 |
| YPKQSIFTINSVVQKD | 1 | 0 | 0 | 1710351 | YP_009047202.1 | K9N7C7 |
| YPLLKQKDIGIDTAGV | 1 | 0 | 0 | 1710359 | YP_173243.1 | Q0ZJ80 |
| YPLVYHENEEYQKVFR | 1 | 0 | 0 | 1710367 | YP_009555238.1 | P0C6X6 |
| YPMIANENAINKGGRT | 1 | 0 | 0 | 1710369 | NP_073549.1 | A0A0P0LKV0 |

| | | | | | | |
|---|---|---|---|---|---|---|
| YPNSNFDNFKFVCDNT | 1 | 0 | 0 | 1710377 | QIQ54046.1 | P0C6U8 |
| YPPDEDEEEGDCEEED | 1 | 0 | 0 | 1710378 | QHR63299.1 | P0C6W2 |
| YPPDEDEEEGDCEEEE | 1 | 0 | 0 | 1710379 | YP_009724389.1 | P0DTD1 |
| YPPDEDYEEEYSEEEQ | 1 | 0 | 0 | 1710380 | QIQ54046.1 | P0C6U8 |
| YPPDEEEEDDAECEEE | 1 | 0 | 0 | 1710381 | NP_828849.2 | P0C6X7 |
| YPQVNGLTSIKWADNN | 1 | 0 | 0 | 1710389 | YP_009724389.1 | P0DTD1 |
| YPRLQAASDWKPGYSM | 1 | 0 | 0 | 1710390 | YP_009555238.1 | P0C6X6 |
| YPRLQASADWKPGHAM | 1 | 0 | 0 | 1710391 | YP_009047202.1 | K9N7C7 |
| YPTLNISDEFSSNVAN | 1 | 0 | 0 | 1710403 | YP_009724389.1 | P0DTD1 |
| YPYGRCPTAADINIPD | 1 | 0 | 0 | 1710411 | YP_009555240.1 | P30215 |
| YQCAMRPNHTIKGSFL | 1 | 0 | 0 | 1710417 | NP_828849.2 | P0C6X7 |
| YQDVNCTEVPVAIHAD | 1 | 0 | 0 | 1710422 | YP_009724390.1 | P0DTC2 |
| YQDYMQIAPVPAEVLN | 1 | 0 | 0 | 1710423 | YP_003769.1 | H9EJA2 |
| YQGDVDTVVNGVDFDF | 1 | 0 | 0 | 1710435 | NP_073549.1 | A0A0P0LKV0 |
| YQGKSLEFGATSVTPQ | 1 | 0 | 0 | 1710438 | QHR63299.1 | P0C6W2 |
| YQHIGMKRYCTVQGPP | 1 | 0 | 0 | 1710444 | YP_009555238.1 | P0C6X6 |
| YQKVGMQKYSTLQGPP | 1 | 0 | 0 | 1710455 | YP_009724389.1 | P0DTD1 |
| YQKVGMQXYSTLQGPP | 1 | 0 | 0 | 1710456 | QIQ54047.1 | P0C6X7 |
| YQLESTQITTDTGIEN | 1 | 0 | 0 | 1710459 | NP_828852.2 | P59632 |
| YQLIGKQKITTIQGPP | 1 | 0 | 0 | 1710461 | YP_003766.2 | H9EJ42 |
| YQLYSTQLSTDTGVEH | 1 | 0 | 0 | 1710467 | YP_009724391.1 | P0DTC3 |
| YQNGVFMSTAKCWTEE | 1 | 0 | 0 | 1710470 | NP_073549.1 | A0A0P0LKV0 |
| YQNISTNLPPPLLGNS | 1 | 0 | 0 | 1710471 | YP_009047204.1 | K9N5Q8 |
| YQNNVFMSEAKCWTET | 1 | 0 | 0 | 1710473 | YP_009724389.1 | P0DTD1 |
| YQNNVFMSEAKCWVET | 1 | 0 | 0 | 1710474 | YP_009047202.1 | K9N7C7 |
| YQNNVFMSESKCWVEN | 1 | 0 | 0 | 1710476 | YP_173236.1 | A0A140H1G9 |
| YQPPTASVSTSFLQSG | 1 | 0 | 0 | 1710485 | YP_009555238.1 | P0C6X6 |
| YQQAVVGAMLSENFTS | 1 | 0 | 0 | 1710489 | NP_073551.1 | A0A0P0K6L9 |
| YQQSIIGAMTAVNESR | 1 | 0 | 0 | 1710493 | YP_003767.1 | A0A0P0G321 |
| YQTNIYSQNLCVDGFG | 1 | 0 | 0 | 1710496 | NP_073549.1 | A0A0P0LKV0 |
| YQTNLYSFNKAIDGFG | 1 | 0 | 0 | 1710497 | YP_003766.2 | H9EJ42 |
| YQTQTNSRSVASQSII | 1 | 0 | 0 | 1710499 | QHR63300.2 | Q3LZX1 |
| YQTSNFRVQPTDSIVR | 1 | 0 | 0 | 1710500 | QHR63300.2 | Q3LZX1 |
| YQTSNFRVQPTISIVR | 1 | 0 | 0 | 1710502 | QIQ54048.1 | P59594 |
| YRAFDIYNDKVAGFAK | 1 | 0 | 0 | 1710511 | YP_009724389.1 | P0DTD1 |
| YRAFDIYNEKVAGFAK | 1 | 0 | 0 | 1710512 | NP_828849.2 | P0C6X7 |
| YRAFISVLDITDAAVK | 1 | 0 | 0 | 1710514 | NP_073549.1 | A0A0P0LKV0 |
| YRALSGVFCGVDAMNL | 1 | 0 | 0 | 1710515 | NP_828849.2 | P0C6X7 |
| YRCPKEIVDTVSALVY | 1 | 0 | 0 | 1710526 | YP_009555238.1 | P0C6X6 |
| YRCPKEIVETVSALVY | 1 | 0 | 0 | 1710527 | YP_173236.1 | A0A140H1G9 |
| YRDGISTISVPAHMPM | 1 | 0 | 0 | 1710530 | YP_173243.1 | Q0ZJ80 |
| YRDGYLNSTNVTTTAY | 1 | 0 | 0 | 1710531 | QIQ54046.1 | P0C6U8 |
| YRDWSYSGQRTELGVE | 1 | 0 | 0 | 1710533 | NP_828849.2 | P0C6X7 |
| YREAACCHLAKALNDF | 1 | 0 | 0 | 1710536 | YP_009724389.1 | P0DTD1 |
| YREAACSQLAKAMDTF | 1 | 0 | 0 | 1710537 | YP_009555238.1 | P0C6X6 |
| YREGYLNSTNVTTAIY | 1 | 0 | 0 | 1710540 | QHR63299.1 | P0C6W2 |
| YREYLESYNTATTAGF | 1 | 0 | 0 | 1710541 | YP_009555238.1 | P0C6X6 |
| YRFTNFEPFTVNSVND | 1 | 0 | 0 | 1710545 | YP_009555241.1 | P36334 |
| YRFVTFEPFNVSFVND | 1 | 0 | 0 | 1710546 | YP_173238.1 | A0A140H1H1 |
| YRIGNYKLNTDHAGSN | 1 | 0 | 0 | 1710557 | NP_828855.1 | P59596 |
| YRIGNYKLNTDHSNSS | 1 | 0 | 0 | 1710558 | QIQ54051.1 | P59596 |
| YRINGLGVTMDVLNKN | 1 | 0 | 0 | 1710560 | YP_173238.1 | A0A140H1H1 |
| YRINGLGVTMDVLSQN | 1 | 0 | 0 | 1710561 | YP_009555241.1 | P36334 |
| YRKCVKPKEETGLLMP | 1 | 0 | 0 | 1710567 | QHR63299.1 | P0C6W2 |
| YRKILLRKNGNKGAGG | 1 | 0 | 0 | 1710569 | QIQ54046.1 | P0C6U8 |
| YRKQLSPLEGGGWLVA | 1 | 0 | 0 | 1710570 | YP_009047204.1 | K9N5Q8 |
| YRKVLLRKNGNKGAGG | 1 | 0 | 0 | 1710574 | YP_009724389.1 | P0DTD1 |
| YRLANECAQVLSEYVL | 1 | 0 | 0 | 1710577 | YP_009047202.1 | K9N7C7 |
| YRLFRKSNLKPFERDI | 1 | 0 | 0 | 1710581 | YP_009724390.1 | P0DTC2 |
| YRLPSSKPSGMDTALL | 1 | 0 | 0 | 1710586 | YP_173241.1 | Q0ZJ82 |
| YRLPSTQKGSGMDTAL | 1 | 0 | 0 | 1710587 | YP_009555244.1 | Q01455 |
| YRLYLDAYNMMISAGF | 1 | 0 | 0 | 1710592 | YP_009724389.1 | P0DTD1 |
| YRNRDVDTDFVNEFYA | 1 | 0 | 0 | 1710598 | YP_009724389.1 | P0DTD1 |
| YRNVLLRKNGNKGAGG | 1 | 0 | 0 | 1710600 | NP_828849.2 | P0C6X7 |
| YRQYLDAYNMMISAGF | 1 | 0 | 0 | 1710605 | NP_828849.2 | P0C6X7 |
| YRRATRRVRGGDGKMK | 1 | 0 | 0 | 1710607 | NP_828858.1 | P59595 |
| YRRQQTSNLVVQEFVS | 1 | 0 | 0 | 1710611 | YP_009555238.1 | P0C6X6 |
| YRSLPGVFCGVDAVNL | 1 | 0 | 0 | 1710615 | YP_009724389.1 | P0DTD1 |
| YRSLPGVFCGVDSVNL | 1 | 0 | 0 | 1710616 | QIQ54046.1 | P0C6U8 |
| YRSTSPDPKFVDKYYA | 1 | 0 | 0 | 1710621 | YP_009047202.1 | K9N7C7 |
| YSAGHATGTTPQKLFV | 1 | 0 | 0 | 1710642 | YP_009047204.1 | K9N5Q8 |
| YSANFLDDNVLPETYV | 1 | 0 | 0 | 1710644 | YP_003767.1 | A0A0P0G321 |
| YSDVENPHLMGWDYPK | 1 | 0 | 0 | 1710659 | YP_009724389.1 | P0DTD1 |
| YSDVETPHLMGWDYPK | 1 | 0 | 0 | 1710660 | NP_828849.2 | P0C6X7 |
| YSEEEQPEQPTQYEYG | 1 | 0 | 0 | 1710664 | QIQ54046.1 | P0C6U8 |
| YSELRPDTRYVLMDGS | 1 | 0 | 0 | 1710667 | NP_828849.2 | P0C6X7 |
| YSENVFAVESGGYIPS | 1 | 0 | 0 | 1710668 | NP_073551.1 | A0A0P0K6L9 |
| YSFLPKEDGDVLLAEF | 1 | 0 | 0 | 1710679 | YP_009047202.1 | K9N7C7 |
| YSGDTFWRYDFDITES | 1 | 0 | 0 | 1710685 | YP_003766.2 | H9EJ42 |
| YSGFKPNNTSAQYIFS | 1 | 0 | 0 | 1710687 | YP_003766.2 | H9EJ42 |
| YSGNANEADYRCACYA | 1 | 0 | 0 | 1710691 | NP_073549.1 | A0A0P0LKV0 |
| YSGRVSAAFHANSSEP | 1 | 0 | 0 | 1710696 | YP_009555241.1 | P36334 |
| YSGYHPAHAPSFLAVD | 1 | 0 | 0 | 1710702 | YP_009555238.1 | P0C6X6 |
| YSIDVIDNEIIVKPNI | 1 | 0 | 0 | 1710709 | NP_073549.1 | A0A0P0LKV0 |
| YSILAAKTGVSVEQLL | 1 | 0 | 0 | 1710712 | YP_003766.2 | H9EJ42 |
| YSIQDVGFKNQYCNGS | 1 | 0 | 0 | 1710714 | YP_009555238.1 | P0C6X6 |
| YSKHTPIDLVRDLPRG | 1 | 0 | 0 | 1710717 | QIQ54048.1 | P59594 |
| YSKLFITKDEAIKRVR | 1 | 0 | 0 | 1710722 | YP_173236.1 | A0A140H1G9 |
| YSLFDMAKFPLKLAGT | 1 | 0 | 0 | 1710733 | YP_009555238.1 | P0C6X6 |
| YSLPAINVTINNYNPS | 1 | 0 | 0 | 1710743 | YP_173238.1 | A0A140H1H1 |
| YSNFYASGCVLSSACT | 1 | 0 | 0 | 1710756 | YP_009555238.1 | P0C6X6 |

| | | | | | | |
|---|---|---|---|---|---|---|
| YSNKTVGELGDVRETM | 1 | 0 | 0 | 1710759 | NP_828849.2 | P0C6X7 |
| YSNVYRTDYVDYTFVN | 1 | 0 | 0 | 1710767 | YP_173236.1 | A0A140H1G9 |
| YSPCVSIVPSTVWEDG | 1 | 0 | 0 | 1710769 | YP_009047204.1 | K9N5Q8 |
| YSQMRPHVRYDLYDGN | 1 | 0 | 0 | 1710782 | YP_009047202.1 | K9N7C7 |
| YSQNLCVDGFGVNKIQ | 1 | 0 | 0 | 1710783 | NP_073549.1 | A0A0P0LKV0 |
| YSRIVDEPEDHVQIHT | 1 | 0 | 0 | 1710787 | QIQ54049.1 | P59632 |
| YSRSTRSMLKRRDSTY | 1 | 0 | 0 | 1710793 | YP_009047204.1 | K9N5Q8 |
| YSRVISRMGFKLDATV | 1 | 0 | 0 | 1710795 | YP_009047202.1 | K9N7C7 |
| YSRVKNLNSSEGVPDL | 1 | 0 | 0 | 1710796 | NP_828854.1 | P59637 |
| YSSCVECDAKFKNSVA | 1 | 0 | 0 | 1710803 | NP_073549.1 | A0A0P0LKV0 |
| YSSLCDEFTLAEVVKQ | 1 | 0 | 0 | 1710810 | YP_003766.2 | H9EJ42 |
| YSSNEEQAVLKFLDGL | 1 | 0 | 0 | 1710814 | YP_003766.2 | H9EJ42 |
| YSTGNNVFQTQAGCLI | 1 | 0 | 0 | 1710822 | NP_828851.1 | P59594 |
| YSTLQGPPGTGKSHFA | 1 | 0 | 0 | 1710824 | YP_009724389.1 | P0DTD1 |
| YSTTPCEVHGNGSMIE | 1 | 0 | 0 | 1710830 | YP_009047202.1 | K9N7C7 |
| YSVILSTCDGLKFTEE | 1 | 0 | 0 | 1710835 | YP_173236.1 | A0A140H1G9 |
| YSVLETFQNNVVNYQH | 1 | 0 | 0 | 1710838 | YP_009555238.1 | P0C6X6 |
| YSVLSVDSVSEESQGN | 1 | 0 | 0 | 1710839 | YP_173236.1 | A0A140H1G9 |
| YSVSSFEAKPSGSVVE | 1 | 0 | 0 | 1710848 | YP_009047204.1 | K9N5Q8 |
| YSYATHHDKFTDGVCL | 1 | 0 | 0 | 1710857 | NP_828849.2 | P0C6X7 |
| YSYATHSDKFKDGVCL | 1 | 0 | 0 | 1710858 | QIQ54047.1 | P0C6X7 |
| YSYATHSDKFTDGVCL | 1 | 0 | 0 | 1710859 | YP_009724389.1 | P0DTD1 |
| YSYLMPVLSMTDCLAA | 1 | 0 | 0 | 1710863 | YP_009047202.1 | K9N7C7 |
| YTAEQCIRGAMGDVGL | 1 | 0 | 0 | 1710872 | NP_073549.1 | A0A0P0LKV0 |
| YTANDVEVQSSGMIKP | 1 | 0 | 0 | 1710877 | YP_009047202.1 | K9N7C7 |
| YTARVVYTAASHAAVD | 1 | 0 | 0 | 1710878 | YP_173236.1 | A0A140H1G9 |
| YTCSITFEMSCDCGKK | 1 | 0 | 0 | 1710882 | YP_003766.2 | H9EJ42 |
| YTDFAEDVCTCYDNSI | 1 | 0 | 0 | 1710885 | NP_073549.1 | A0A0P0LKV0 |
| YTDQYLCGADGKPVMS | 1 | 0 | 0 | 1710892 | NP_073549.1 | A0A0P0LKV0 |
| YTEIEPKLDGYYKKDN | 1 | 0 | 0 | 1710896 | NP_828849.2 | P0C6X7 |
| YTELEPPCRFVTDTPK | 1 | 0 | 0 | 1710900 | YP_009724389.1 | P0DTD1 |
| YTERSEKSYELQTPFE | 1 | 0 | 0 | 1710903 | YP_009724389.1 | P0DTD1 |
| YTESAVNSTKALAKQD | 1 | 0 | 0 | 1710904 | YP_009047206.1 | K9N4V0 |
| YTGAIKLDDKDPSFKD | 1 | 0 | 0 | 1710916 | QIQ54056.1 | P59595 |
| YTGQAGEMPCAIINDK | 1 | 0 | 0 | 1710924 | YP_009555238.1 | P0C6X6 |
| YTGSLADGLEAYADKT | 1 | 0 | 0 | 1710926 | YP_009555238.1 | P0C6X6 |
| YTGSSFYAPEPITSLN | 1 | 0 | 0 | 1710930 | YP_009047204.1 | K9N5Q8 |
| YTIKFSGFKLDGFNHQ | 1 | 0 | 0 | 1710941 | YP_003766.2 | H9EJ42 |
| YTKVDGIDVEIFENKT | 1 | 0 | 0 | 1710957 | NP_828849.2 | P0C6X7 |
| YTKVDGVDVELFENKT | 1 | 0 | 0 | 1710958 | YP_009724389.1 | P0DTD1 |
| YTLANTTTEPLKLFAA | 1 | 0 | 0 | 1710960 | YP_009047202.1 | K9N7C7 |
| YTLEEAKTALKKCKSA | 1 | 0 | 0 | 1710962 | NP_828849.2 | P0C6X7 |
| YTMLDLCYALRHFDRN | 1 | 0 | 0 | 1710968 | YP_009555238.1 | P0C6X6 |
| YTMSLGADSSIAYSNN | 1 | 0 | 0 | 1710969 | NP_828851.1 | P59594 |
| YTMSLGAENSVAYSNN | 1 | 0 | 0 | 1710970 | YP_009724390.1 | P0DTC2 |
| YTNEIIGYKAIQDGVT | 1 | 0 | 0 | 1710973 | NP_828849.2 | P0C6X7 |
| YTNSCLVSSDGQPGGD | 1 | 0 | 0 | 1710978 | YP_009047202.1 | K9N7C7 |
| YTPFHYERDNTSCPEW | 1 | 0 | 0 | 1710982 | YP_009047202.1 | K9N7C7 |
| YTPGHYAGSRSSSGNR | 1 | 0 | 0 | 1710983 | YP_173242.1 | Q19U29 |
| YTPSFKKGAKLLHKPI | 1 | 0 | 0 | 1710991 | YP_009724389.1 | P0DTD1 |
| YTPSKLIEYSDFATSA | 1 | 0 | 0 | 1710992 | NP_828849.2 | P0C6X7 |
| YTPSKLIEYTDFATSA | 1 | 0 | 0 | 1710993 | YP_009724389.1 | P0DTD1 |
| YTQCAVIAVVAAVLNS | 1 | 0 | 0 | 1710994 | YP_009047202.1 | K9N7C7 |
| YTQGRNLQDFLPRSTM | 1 | 0 | 0 | 1710995 | NP_073549.1 | A0A0P0LKV0 |
| YTQGRNLSDFTPRSDM | 1 | 0 | 0 | 1710996 | YP_003766.2 | H9EJ42 |
| YTQLCQYLNSTTMCVP | 1 | 0 | 0 | 1710999 | YP_003766.2 | H9EJ42 |
| YTSACKTIEDALRNSA | 1 | 0 | 0 | 1711008 | NP_073551.1 | A0A0P0K6L9 |
| YTSKDTLDDILQQANK | 1 | 0 | 0 | 1711013 | YP_009047202.1 | K9N7C7 |
| YTSKTTVASLINTLND | 1 | 0 | 0 | 1711016 | YP_009724389.1 | P0DTD1 |
| YTSTSGNLLGFKDVTK | 1 | 0 | 0 | 1711025 | NP_073551.1 | A0A0P0K6L9 |
| YTTTIKPVSYKLDGVT | 1 | 0 | 0 | 1711038 | NP_828849.2 | P0C6X7 |
| YTVCDDCNGYTDNIFS | 1 | 0 | 0 | 1711044 | YP_003767.1 | A0A0P0G321 |
| YTVEEAKTVLKKCKSA | 1 | 0 | 0 | 1711046 | YP_009724389.1 | P0DTD1 |
| YTVEEARTVLKKCKSA | 1 | 0 | 0 | 1711047 | QHR63299.1 | P0C6W2 |
| YTVPDLVVEQYNQTIL | 1 | 0 | 0 | 1711052 | NP_073551.1 | A0A0P0K6L9 |
| YTVQKLQTLIDNINST | 1 | 0 | 0 | 1711053 | NP_073551.1 | A0A0P0K6L9 |
| YTVYDTAKKSMYDGDR | 1 | 0 | 0 | 1711059 | NP_073549.1 | A0A0P0LKV0 |
| YTYKMLVAKDNKNLPK | 1 | 0 | 0 | 1711063 | YP_003771.1 | H9EJ69 |
| YTYSFLPKEDGDVLLA | 1 | 0 | 0 | 1711065 | YP_009047202.1 | K9N7C7 |
| YVAGYKVLPPLMDVNM | 1 | 0 | 0 | 1711072 | YP_009047204.1 | K9N5Q8 |
| YVAKRVLGVQTQTVDS | 1 | 0 | 0 | 1711074 | YP_173236.1 | A0A140H1G9 |
| YVALRGRIDKDLSLSR | 1 | 0 | 0 | 1711077 | NP_073553.1 | A0A0P0KWE8 |
| YVAPQVTYQNISTNLP | 1 | 0 | 0 | 1711080 | YP_009047204.1 | K9N5Q8 |
| YVAPVNTVKPKPVINQ | 1 | 0 | 0 | 1711081 | NP_073549.1 | A0A0P0LKV0 |
| YVARLMKGDKGDAEDT | 1 | 0 | 0 | 1711083 | NP_073549.1 | A0A0P0LKV0 |
| YVASRFLGLQIQTVDS | 1 | 0 | 0 | 1711084 | YP_003766.2 | H9EJ42 |
| YVCKPHQVNISLNGNT | 1 | 0 | 0 | 1711090 | YP_003767.1 | A0A0P0G321 |
| YVCNAPGCDVNDVTKL | 1 | 0 | 0 | 1711093 | YP_009555238.1 | P0C6X6 |
| YVCNASGCGVSDVKKL | 1 | 0 | 0 | 1711097 | YP_003766.2 | H9EJ42 |
| YVCNGNVRCVELLKQY | 1 | 0 | 0 | 1711101 | NP_073551.1 | A0A0P0K6L9 |
| YVCNTSGCNVNDVTKL | 1 | 0 | 0 | 1711102 | NP_073549.1 | A0A0P0LKV0 |
| YVDFKPQSGGGKCFNC | 1 | 0 | 0 | 1711106 | NP_073551.1 | A0A0P0K6L9 |
| YVDGHRLASGVQVHNL | 1 | 0 | 0 | 1711107 | NP_073555.1 | A0A0P0KXN3 |
| YVDGLESKQVDYVPLR | 1 | 0 | 0 | 1711108 | YP_173236.1 | A0A140H1G9 |
| YVDKWDDFCRQYSNES | 1 | 0 | 0 | 1711109 | NP_073549.1 | A0A0P0LKV0 |
| YVDNNFCGPDGYPLDC | 1 | 0 | 0 | 1711112 | NP_828849.2 | P0C6X7 |
| YVDNNFCGPDGYPLEC | 1 | 0 | 0 | 1711113 | YP_009724389.1 | P0DTD1 |
| YVDNSSLTIKKPNELS | 1 | 0 | 0 | 1711114 | YP_009724389.1 | P0DTD1 |
| YVDNTSLTIKKPNELS | 1 | 0 | 0 | 1711115 | QIQ54046.1 | P0C6U8 |
| YVDQYGCDYTGSLADG | 1 | 0 | 0 | 1711117 | YP_009555238.1 | P0C6X6 |
| YVDTENNTEFTRVNAK | 1 | 0 | 0 | 1711119 | NP_828849.2 | P0C6X7 |

| | | | | | | |
|---|---|---|---|---|---|---|
| YVDTFSATFSVPMEKL | 1 | 0 | 0 | 1711122 | NP_828849.2 | P0C6X7 |
| YVDTPNNTDFSRVSAK | 1 | 0 | 0 | 1711124 | YP_009724389.1 | P0DTD1 |
| YVDVNKTLQEFAQNLP | 1 | 0 | 0 | 1711127 | YP_003767.1 | A0A0P0G321 |
| YVFSTVNALPEVNADI | 1 | 0 | 0 | 1711144 | NP_073549.1 | A0A0P0LKV0 |
| YVGKYDINHGDSGFIS | 1 | 0 | 0 | 1711163 | YP_173237.1 | A0A140H1H0 |
| YVGTGIIRSSNQSLAG | 1 | 0 | 0 | 1711168 | YP_003767.1 | A0A0P0G321 |
| YVGVQTVEDLRARMTY | 1 | 0 | 0 | 1711169 | YP_009047202.1 | K9N7C7 |
| YVHARLKGGLILKFDS | 1 | 0 | 0 | 1711173 | YP_009047202.1 | K9N7C7 |
| YVIDGALLSKTSEYKG | 1 | 0 | 0 | 1711185 | QIQ54046.1 | P0C6U8 |
| YVKDGKYFTSEPPVTY | 1 | 0 | 0 | 1711198 | YP_009047202.1 | K9N7C7 |
| YVKPGGTSSGDATTAF | 1 | 0 | 0 | 1711204 | YP_009555238.1 | P0C6X6 |
| YVKRRFSLLRHEDLSV | 1 | 0 | 0 | 1711207 | YP_009047207.1 | K9N643 |
| YVKSKVGNYRLPSTQK | 1 | 0 | 0 | 1711210 | YP_009555244.1 | Q01455 |
| YVLLDWVKHLNKNLNE | 1 | 0 | 0 | 1711226 | YP_003766.2 | H9EJ42 |
| YVLNNISLTTQPYFDS | 1 | 0 | 0 | 1711230 | YP_173238.1 | A0A140H1H1 |
| YVLPLTCNAISSNTDN | 1 | 0 | 0 | 1711231 | YP_173238.1 | A0A140H1H1 |
| YVLPSIIPNKKEEILG | 1 | 0 | 0 | 1711236 | QIQ54046.1 | P0C6U8 |
| YVLTGDSVKFVYMHQL | 1 | 0 | 0 | 1711243 | YP_173236.1 | A0A140H1G9 |
| YVMLNTSIPNLPDFKE | 1 | 0 | 0 | 1711251 | YP_009555241.1 | P36334 |
| YVNKHAFHTPAFDKSA | 1 | 0 | 0 | 1711259 | YP_009724389.1 | P0DTD1 |
| YVNSNQNTDSDSLSSK | 1 | 0 | 0 | 1711267 | YP_173242.1 | Q19U29 |
| YVNTFSSTFNVPMDKL | 1 | 0 | 0 | 1711268 | QIQ54046.1 | P0C6U8 |
| YVNTFSSTFNVPMEKL | 1 | 0 | 0 | 1711269 | YP_009724389.1 | P0DTD1 |
| YVNVYRSTSPDPKFVD | 1 | 0 | 0 | 1711271 | YP_009047202.1 | K9N7C7 |
| YVPSYVKPDSVSTSDL | 1 | 0 | 0 | 1711287 | YP_009047202.1 | K9N7C7 |
| YVQQGIFNKATVAQIK | 1 | 0 | 0 | 1711293 | YP_009555238.1 | P0C6X6 |
| YVRDYVDKWDDFCRQY | 1 | 0 | 0 | 1711296 | NP_073549.1 | A0A0P0LKV0 |
| YVRKNGQFQDHYDGFY | 1 | 0 | 0 | 1711301 | YP_003766.2 | H9EJ42 |
| YVRPIIEDYHTLTVTI | 1 | 0 | 0 | 1711303 | YP_009555244.1 | Q01455 |
| YVRVKHGDFSAVSSPM | 1 | 0 | 0 | 1711304 | NP_073555.1 | A0A0P0KXN3 |
| YVSNSGNLLGFKNVST | 1 | 0 | 0 | 1711315 | YP_003767.1 | A0A0P0G321 |
| YVTGGVVQMTTQWLTN | 1 | 0 | 0 | 1711326 | QIQ54046.1 | P0C6U8 |
| YVTHGFNLEEAARCMR | 1 | 0 | 0 | 1711327 | NP_828849.2 | P0C6X7 |
| YVTNEIGLNASVTLKI | 1 | 0 | 0 | 1711330 | YP_003767.1 | A0A0P0G321 |
| YVTTKPGMKYVKWEHD | 1 | 0 | 0 | 1711332 | NP_073549.1 | A0A0P0LKV0 |
| YVTYAGNVWQIQTIQD | 1 | 0 | 0 | 1711337 | YP_009555238.1 | P0C6X6 |
| YVTYAGSVWHIQTVQD | 1 | 0 | 0 | 1711338 | YP_173236.1 | A0A140H1G9 |
| YVVDSVAVKNGSLHLY | 1 | 0 | 0 | 1711340 | QIQ54046.1 | P0C6U8 |
| YVVKNSTAFPFETLSA | 1 | 0 | 0 | 1711344 | YP_009047202.1 | K9N7C7 |
| YVVVVESDADRANFLN | 1 | 0 | 0 | 1711351 | YP_173236.1 | A0A140H1G9 |
| YVVYDAANNAVYDGAR | 1 | 0 | 0 | 1711353 | YP_003766.2 | H9EJ42 |
| YVYDVHKDNFKDGLCM | 1 | 0 | 0 | 1711364 | YP_173236.1 | A0A140H1G9 |
| YVYSAGHATGTTPQKL | 1 | 0 | 0 | 1711378 | YP_009047204.1 | K9N5Q8 |
| YVYSRVKNLNSSEGVP | 1 | 0 | 0 | 1711379 | NP_828854.1 | P59637 |
| YWDQTYHANCVNCTDD | 1 | 0 | 0 | 1711382 | YP_009047202.1 | K9N7C7 |
| YWDQTYHPNCINCLDD | 1 | 0 | 0 | 1711383 | NP_828849.2 | P0C6X7 |
| YWDQTYHPNCVNCLDD | 1 | 0 | 0 | 1711384 | YP_009724389.1 | P0DTD1 |
| YWGQDYHPDCVDCHDE | 1 | 0 | 0 | 1711395 | NP_073549.1 | A0A0P0LKV0 |
| YWIGYGCNCDSLRQAA | 1 | 0 | 0 | 1711396 | YP_009047202.1 | K9N7C7 |
| YWRRQDRKINTGNGIK | 1 | 0 | 0 | 1711404 | YP_009047211.1 | K9N4V7 |
| YWSPQGNLGGNCAGNV | 1 | 0 | 0 | 1711406 | YP_173236.1 | A0A140H1G9 |
| YYCVTTKAKSVQQDLD | 1 | 0 | 0 | 1711434 | YP_009047202.1 | K9N7C7 |
| YYDDGSCEVVHDQVNY | 1 | 0 | 0 | 1711437 | NP_073549.1 | A0A0P0LKV0 |
| YYDDSQYYFNKDTGVI | 1 | 0 | 0 | 1711438 | YP_009555240.1 | P30215 |
| YYHTLDESFLGRYMSA | 1 | 0 | 0 | 1711465 | NP_828849.2 | P0C6X7 |
| YYIYVTNEIGLNASVT | 1 | 0 | 0 | 1711474 | YP_003767.1 | A0A0P0G321 |
| YYKGVTTHESSSAVNM | 1 | 0 | 0 | 1711475 | YP_009555238.1 | P0C6X6 |
| YYKKDNAYFTEQPIDL | 1 | 0 | 0 | 1711476 | QIQ54046.1 | P0C6U8 |
| YYKKDNAYYTEQPIDL | 1 | 0 | 0 | 1711477 | NP_828849.2 | P0C6X7 |
| YYKKDNSYFTEQPIDL | 1 | 0 | 0 | 1711478 | YP_009724389.1 | P0DTD1 |
| YYKKVDGVVQQLPETY | 1 | 0 | 0 | 1711479 | YP_009724389.1 | P0DTD1 |
| YYKKVNGVVQQLPETY | 1 | 0 | 0 | 1711480 | QHR63299.1 | P0C6W2 |
| YYKVNCCRFQRIDDDG | 1 | 0 | 0 | 1711485 | YP_173236.1 | A0A140H1G9 |
| YYKYDNGNFIKLPEVI | 1 | 0 | 0 | 1711487 | NP_073549.1 | A0A0P0LKV0 |
| YYLDDVKKIEYKPDLS | 1 | 0 | 0 | 1711490 | YP_009555238.1 | P0C6X6 |
| YYLGTGPYANASYGES | 1 | 0 | 0 | 1711498 | YP_173242.1 | Q19U29 |
| YYNNSNNGKIVYAILS | 1 | 0 | 0 | 1711513 | YP_009555238.1 | P0C6X6 |
| YYNSWQNLLYDSNGNI | 1 | 0 | 0 | 1711514 | YP_173238.1 | A0A140H1H1 |
| YYNSWQNLLYDSNGNL | 1 | 0 | 0 | 1711515 | YP_009555241.1 | P36334 |
| YYPEPISDKNVVFMNT | 1 | 0 | 0 | 1711523 | YP_173238.1 | A0A140H1H1 |
| YYPEPITENNVVVMST | 1 | 0 | 0 | 1711524 | YP_009555241.1 | P36334 |
| YYQLESTQITTDTGIE | 1 | 0 | 0 | 1711538 | NP_828852.2 | P59632 |
| YYQLYSTQLSTDTGVE | 1 | 0 | 0 | 1711541 | YP_009724391.1 | P0DTC3 |
| YYQNGVFMSTAKCWTE | 1 | 0 | 0 | 1711542 | NP_073549.1 | A0A0P0LKV0 |
| YYQNNVFMSEAKCWTE | 1 | 0 | 0 | 1711543 | YP_009724389.1 | P0DTD1 |
| YYQNNVFMSEAKCWVE | 1 | 0 | 0 | 1711544 | YP_009047202.1 | K9N7C7 |
| YYQNNVFMSESKCWVE | 1 | 0 | 0 | 1711545 | YP_009555238.1 | P0C6X6 |
| YYRSLPGVFCGVDAVN | 1 | 0 | 0 | 1711557 | YP_009724389.1 | P0DTD1 |
| YYRSLPGVFCGVDSVN | 1 | 0 | 0 | 1711558 | QIQ54046.1 | P0C6U8 |
| YYSDSPCESHGKQVVS | 1 | 0 | 0 | 1711563 | YP_009724389.1 | P0DTD1 |
| YYSGNANEADYRCACY | 1 | 0 | 0 | 1711568 | NP_073549.1 | A0A0P0LKV0 |
| YYSLPAINVTINNYNP | 1 | 0 | 0 | 1711573 | YP_173238.1 | A0A140H1H1 |
| YYSTTPCEVHGNGSMI | 1 | 0 | 0 | 1711578 | YP_009047202.1 | K9N7C7 |
| YYVKPGGTSSGDATTA | 1 | 0 | 0 | 1711600 | YP_009047202.1 | K9N7C7 |
| HSVPKLSDFESELSHWFKNQTSIAP | 1 | 0 | 0 | 1857699 | ADN03339.1 | A0A140H1H1 |
| TSIPNLPDFKEELDQWFKNQTSVAP | 1 | 0 | 0 | 1860153 | QEG03757.1 | P36334 |
| CIEAKISNTTTDSRCPT | 1 | 0 | 0 | 1860813 | QFS19562.1 | P17763 |
| CRRDVVDRGWGNGCGLF | 1 | 0 | 0 | 1860815 | P09866.2 | P17763 |
| CVTTMAKDKPTLDIELL | 1 | 0 | 0 | 1860816 | QFS19562.1 | P17763 |
| DANFVCRRTFVDRGWG | 1 | 0 | 0 | 1860818 | QFS19562.1 | P17763 |
| EIKITPQSSITEAFLTGY | 1 | 0 | 0 | 1860820 | SRC326855 | P17763 |

| | | | | | | |
|---|---|---|---|---|---|---|
| FDSYIVIGVGNSALTLH | 1 | 0 | 0 | 1860825 | SRC326857 | P17763 |
| FHLTTRGGEPHMIVSK | 1 | 0 | 0 | 1860827 | QFS19150.1 | P17763 |
| GCVTTMAKNKPTLDIEL | 1 | 0 | 0 | 1860831 | AAT69740.1 | P17763 |
| GCVTTMAQGKPTLDFEL | 1 | 0 | 0 | 1860832 | QTX92128.1 | P17763 |
| GLETRTETWMSSEGAWKQ | 1 | 0 | 0 | 1860834 | QFS19150.1 | P17763 |
| IGIGVLLTWIGLNSK | 1 | 0 | 0 | 1860837 | AAT69740.1 | P17763 |
| IGILLTWLGLNSRSTSL | 1 | 0 | 0 | 1860838 | P17763.2 | P17763 |
| KALKINWYKKGSSIGKMF | 1 | 0 | 0 | 1860842 | AAT69740.1 | P17763 |
| KKGSSIGQMIETTMRGAK | 1 | 0 | 0 | 1860845 | AAA42941.1 | P17763 |
| KRFVCKHSMVDRGWGNGCGL | 1 | 0 | 0 | 1860847 | Q9Q4T1 | P17763 |
| LEKEVAETQHGTVLVQI | 1 | 0 | 0 | 1860849 | ACJ04262.1 | P17763 |
| LVLWIGTNSRNTSMAM | 1 | 0 | 0 | 1860850 | QTX92128.1 | P17763 |
| NFVCRRTFVDRGWGNG | 1 | 0 | 0 | 1860854 | QFS19562.1 | P17763 |
| PPFGDSYIVIGAGEKAL | 1 | 0 | 0 | 1860861 | SRC265914 | P17763 |
| RCPTQGEPSLNEEQDKRF | 1 | 0 | 0 | 1860867 | AAB33854.1 | P17763 |
| RDKRSVALAPHVGLGLETRT | 1 | 0 | 0 | 1860868 | QFS19150.1 | P17763 |
| RITEAEPDDVDCWCNATDTW | 1 | 0 | 0 | 1860872 | QFS19150.1 | P17763 |
| SIGGVFTSVGKLVHQI | 1 | 0 | 0 | 1860875 | QFS19562.1 | P17763 |
| SIGGVFTSVGKLVHQVF | 1 | 0 | 0 | 1860876 | P33478.1 | P17763 |
| SISNITTATRCPTQQGEPY | 1 | 0 | 0 | 1860878 | SRC326857 | P17763 |
| SRCPTQGEAILPEEQDQNH | 1 | 0 | 0 | 1860880 | AAT69740.1 | P17763 |
| SRCPTQGEATLVEEQDA | 1 | 0 | 0 | 1860881 | QFS19562.1 | P17763 |
| TAWDFGSIGGVFTSVGK | 1 | 0 | 0 | 1860882 | QFS19562.1 | P17763 |
| TGTIADYNYKLPDDF | 1 | 0 | 0 | 1860884 | QWE88956.1 | P0DTC2 |
| TRCPTQGEPYLKEEQDQQY | 1 | 0 | 0 | 1860887 | QTX92128.1 | P17763 |
| TWADFGSIGGVFTSVGK | 1 | 0 | 0 | 1860888 | SRC265914 | P17763 |
| VLNSLGKMVHQIFGSAY | 1 | 0 | 0 | 1860890 | AAT69740.1 | P17763 |
| WALRHPGFTIALFLAHA | 1 | 0 | 0 | 1860893 | SRC326855 | P17763 |
| WDFGSVGGLFTSLGKAVH | 1 | 0 | 0 | 1860895 | P09866.2 | P17763 |
| WDFGSVGGVLNSLGKMVH | 1 | 0 | 0 | 1860896 | QTE05704.1 | P17763 |
| YIVIGAGEKALKLSWF | 1 | 0 | 0 | 1860898 | QFS19562.1 | P17763 |
| YIVVGAGEKALKQCWFK | 1 | 0 | 0 | 1860899 | P33478.1 | P17763 |
| ATYKPNTWCIRCLWS | 1 | 0 | 0 | 1861077 | P0DTD1.1 | P0DTD1 |
| DQFKHLIPLMYKGLP | 1 | 0 | 0 | 1861161 | P0DTD1.1 | P0DTD1 |
| DTTDAVRDPQTLEILDITPCSF | 1 | 0 | 0 | 1861181 | P0DTC2.1 | P0DTC2 |
| FCAFAVDAAKAYKDY | 1 | 0 | 0 | 1861302 | P0DTD1.1 | P0DTD1 |
| GWEIVKFISTCACEI | 1 | 0 | 0 | 1861511 | P0DTC1.1 | P0DTD1 |
| KDKSAQCFKMFYKGV | 1 | 0 | 0 | 1861688 | P0DTD1.1 | P0DTD1 |
| LGLAAIMQLFFSYFA | 1 | 0 | 0 | 1861852 | P0DTD1.1 | P0DTD1 |
| LKTLVATAEAELAKN | 1 | 0 | 0 | 1861870 | P0DTD1.1 | P0DTD1 |
| LNRGMVLGSLAATVR | 1 | 0 | 0 | 1861902 | P0DTD1.1 | P0DTD1 |
| MQTMLFTMLRKLDND | 1 | 0 | 0 | 1862021 | P0DTD1.1 | P0DTD1 |
| MYLKLRSDVLLPLTQ | 1 | 0 | 0 | 1862039 | P0DTD1.1 | P0DTD1 |
| NCYLATALLTLQQIE | 1 | 0 | 0 | 1862041 | P0DTC1.1 | P0DTD1 |
| NLDKSAGFPFNKWGK | 1 | 0 | 0 | 1862064 | P0DTD1.1 | P0DTD1 |
| PIHFYSKWYIRVGARKSAPLIE | 1 | 0 | 0 | 1862123 | P0DTC8.1 | P0DTC8 |
| SEVVLKKLKKSLNVA | 1 | 0 | 0 | 1862345 | P0DTD1.1 | P0DTD1 |
| SFCYMHHMELPTGVH | 1 | 0 | 0 | 1862346 | P0DTD1.1 | P0DTD1 |
| SFKWDLTAFGLVAEW | 1 | 0 | 0 | 1862347 | P0DTD1.1 | P0DTD1 |
| VLSTFISAARQGFVD | 1 | 0 | 0 | 1862657 | P0DTD1.1 | P0DTD1 |
| VPLNIIPLTTAAKLM | 1 | 0 | 0 | 1862664 | P0DTD1.1 | P0DTD1 |
| VVYRGTTTYKLNVGD | 1 | 0 | 0 | 1862715 | P0DTD1.1 | P0DTD1 |
| WADNNCYLATALLTL | 1 | 0 | 0 | 1862719 | P0DTC1.1 | P0DTD1 |
| YEDLLIRKSNHNFLV | 1 | 0 | 0 | 1862739 | P0DTD1.1 | P0DTD1 |
| YRRATRRIRGGDGKMKDLSPRW | 1 | 0 | 0 | 1862769 | P0DTC9.1 | P0DTC9 |
| GALNTPKDHIGTRNANNAA | 1 | 0 | 0 | 1863199 | SRC558394 | P0DTC9 |
| LPKGFYAEGSRGGSQASSRS | 1 | 0 | 0 | 1863428 | P0DTC9.1 | P0DTC9 |
| PTNGVGYQPYRVVVLSFELL | 1 | 0 | 0 | 1863584 | P0DTC2.1 | P0DTC2 |
| AAASAIQGNVTSIHSL | 0 | 1 | 0 | 60 | P0A564.2 | P9WNK7 |
| AAGTAAQAAVVRFQE | 0 | 1 | 0 | 187 | CAA17966.1 | P9WNK5 |
| AAIGLSMAGSSAMILAAYHP | 0 | 1 | 0 | 223 | P0C5B9.1 | P9WQP1 |
| AAVVLPGLVGLAGGAATAGA | 0 | 1 | 0 | 503 | P0C5B9.1 | P9WQP1 |
| ADEEQQQALSSQMGF | 0 | 1 | 0 | 655 | CAA17966.1 | P9WNK5 |
| AGGGHNGVFDFPDSG | 0 | 1 | 0 | 1522 | P0A4V2.1 | P9WQP3 |
| AGGYKAADMWGPSSDPAWER | 0 | 1 | 0 | 1545 | P0C5B9.1 | P9WQP1 |
| AGRRVGGRRRGRSNSG | 0 | 1 | 0 | 1670 | P24094.2 | P24094 |
| AIDREQSAAIREWA | 0 | 1 | 0 | 1880 | P24094.2 | P24094 |
| AKPRKISVDRGNNGH | 0 | 1 | 0 | 2261 | P12809.1 | P12809 |
| ALLDPSQGMGPSLIGLAMGD | 0 | 1 | 0 | 2695 | P0C5B9.1 | P9WQP1 |
| AMAAWSERAGEA | 0 | 1 | 0 | 3053 | P04977.1 | P04977 |
| AMEKLGQDSQALGQAIYEATQAASK | 0 | 1 | 0 | 3082 | AAA25362.1 | P19993 |
| AMKPDLQNTLMAVPR | 0 | 1 | 0 | 3127 | P31951.2 | P31951 |
| ANDVDNSNPVVQAEQLNWL | 0 | 1 | 0 | 3245 | BAA26110.1 | P13470 |
| ANLFSGEGAQTAQ | 0 | 1 | 0 | 3341 | P11089.1 | P11089 |
| ANRAVKPTGSAAIGLSMAGS | 0 | 1 | 0 | 3400 | P0C5B9.1 | P9WQP1 |
| ANRHVKPTGSAVVGL | 0 | 1 | 0 | 3402 | P0A4V2.1 | P9WQP3 |
| APSQGGVNSPVNV | 0 | 1 | 0 | 3820 | P11089.1 | P11089 |
| AQAAVVRFQEAANKQ | 0 | 1 | 0 | 3934 | CAA17966.1 | P9WNK5 |
| ASAFSRPGLPVEYLQ | 0 | 1 | 0 | 4308 | P31951.2 | P31951 |
| ASGAKEEAEKKAAEQRALL | 0 | 1 | 0 | 4412 | AAA27481.1 | P07643 |
| ASYVKKPKEDVD | 0 | 1 | 0 | 4856 | ACI16088.1 | P17835 |
| ATLVVNRIRGGF | 0 | 1 | 0 | 5054 | AAS19616.1 | P0C0Z7 |
| CGNGTPNELGGANIPAEFLE | 0 | 1 | 0 | 6323 | P0C5B9.1 | P9WQP1 |
| CSETFTNKLKEKHTDLGKEGV | 0 | 1 | 0 | 6984 | Q07337.1 | Q07337 |
| DAATAQTLQAFLHWAITDGN | 0 | 1 | 0 | 7472 | P15712.1 | P9WGU1 |
| DAWREGEEFVVEFDL | 0 | 1 | 0 | 7680 | P12809.1 | P12809 |
| DGDGAAAMRYTEARMTK | 0 | 1 | 0 | 8360 | Q5HJZ0.1 | Q2G2Q0 |
| DGKTLVSKKVTS | 0 | 1 | 0 | 8429 | P14013.1 | P0CL66 |
| DIKVQFQSGGNNSPAVYLLD | 0 | 1 | 0 | 8688 | P0C5B9.1 | P9WQP1 |
| DKILVQAGEAETMTP | 0 | 1 | 0 | 8898 | P24301.2 | P24301 |
| DKILVQANEAETTTA | 0 | 1 | 0 | 8900 | P09621.3 | P9WPE5 |

| | | | | | | |
|---|---|---|---|---|---|---|
| DKKAINLDKAQQKLD | 0 | 1 | 0 | 8906 | AAC67090.1 | Q45013 |
| DNIGNANIGFGN | 0 | 1 | 0 | 9474 | YP_177893.1 | P9WHZ5 |
| DQVHFQPLPPAVVKLSDALI | 0 | 1 | 0 | 9924 | P15712.1 | P9WGU1 |
| DREKLQERLAKLAGGVAVIHV | 0 | 1 | 0 | 9964 | NP_712836.1 | P61439 |
| DRNLGIGNTGNW | 0 | 1 | 0 | 10022 | YP_177893.1 | P9WHZ5 |
| DSDSESDSDSDSDSDSD | 0 | 1 | 0 | 10098 | Q5HCR7.1 | Q2FUY2 |
| DSLDIDIERNVVTVR | 0 | 1 | 0 | 10159 | P12809.1 | P12809 |
| DSSAATKKTAAW | 0 | 1 | 0 | 10226 | P14013.1 | P0CL66 |
| DVFQNGFTAWGNND | 0 | 1 | 0 | 10594 | P04977.1 | P04977 |
| DWYSPACGKAGCQTYKWETF | 0 | 1 | 0 | 10841 | P0C5B9.1 | P9WQP1 |
| DWYSPACGKAGCQTYKWETL | 0 | 1 | 0 | 10842 | CAA44268.1 | P9WQP1 |
| EFGLDGVTYEIDLTNKNA | 0 | 1 | 0 | 11999 | P24094.2 | P24094 |
| EFVVEFDLPGIKA | 0 | 1 | 0 | 12105 | P12809.1 | P12809 |
| EIAAKAIGKKIHQNNG | 0 | 1 | 0 | 12383 | Q07337.1 | Q07337 |
| ELGGTNVPAEFLENFVHGSNLKFQD | 0 | 1 | 0 | 13075 | P31951.2 | P31951 |
| EQAISALPDYASQPGKPPREDLK | 0 | 1 | 0 | 13799 | 1IKQ_A | P11439 |
| ERLAQHMTDFDVTHHR | 0 | 1 | 0 | 14016 | Q5HER3.1 | Q2G282 |
| ETMTPSGLVIPENAK | 0 | 1 | 0 | 14487 | P24301.2 | P24301 |
| ETTTASGLVIPDTAK | 0 | 1 | 0 | 14574 | P09621.3 | P9WPE5 |
| FEWYYQSGLSVVMPV | 0 | 1 | 0 | 15770 | P31951.2 | P31951 |
| FGLQLELTEGMRFDK | 0 | 1 | 0 | 16006 | P06806 | P9WPE7 |
| FSRPGLPVEYLQVPSPSMGR | 0 | 1 | 0 | 17838 | P0C5B9.1 | P9WQP1 |
| FVDSIGQSFQNTWEG | 0 | 1 | 0 | 18139 | O33084.3 | O33084 |
| GANIPAEFLENFVRSSNLKF | 0 | 1 | 0 | 18700 | P0C5B9.1 | P9WQP1 |
| GAQLNAMKPDLQNTL | 0 | 1 | 0 | 18743 | P31951.2 | P31951 |
| GCQTYKWETFLTSELPQWLS | 0 | 1 | 0 | 18898 | P0C5B9.1 | P9WQP1 |
| GFKVRPSFSFFAVGPDGMP | 0 | 1 | 0 | 19562 | CAE55588.1 | Q6MWX9 |
| GGRRRGRSNSGRGRG | 0 | 1 | 0 | 20005 | P24094.2 | P24094 |
| GIGNIGDRNLGI | 0 | 1 | 0 | 20281 | YP_177893.1 | P9WHZ5 |
| GKPTEKHIQIRSTN | 0 | 1 | 0 | 20662 | P0A5B7.2 | P9WMK1 |
| GLRAQDDYNGWDINTPAFEW | 0 | 1 | 0 | 21096 | P0C5B9.1 | P9WQP1 |
| GMTFRAQEGAFLTG | 0 | 1 | 0 | 21353 | Q45010.1 | Q45010 |
| GNFERISGDLKTQID | 0 | 1 | 0 | 21422 | CAA17966.1 | P9WNK5 |
| GNNGHQTINKTAHEIIDA | 0 | 1 | 0 | 21479 | P12809.1 | P12809 |
| GPSSDPAWERNDPTQQIPKL | 0 | 1 | 0 | 21797 | P0C5B9.1 | P9WQP1 |
| GRFDEAMQDQIRQLE | 0 | 1 | 0 | 22028 | O33084.3 | O33084 |
| GRGRGAIDREQSAA | 0 | 1 | 0 | 22055 | P24094.2 | P24094 |
| GRSNSGRGRGAIDREQSA | 0 | 1 | 0 | 22204 | P24094.2 | P24094 |
| GSINTGWFNTGNANTG | 0 | 1 | 0 | 22387 | CAE55588.1 | Q6MWX9 |
| GTVVAVGPGRWDEDG | 0 | 1 | 0 | 22882 | P24301.2 | P24301 |
| GVIFFFNPGELLPEA | 0 | 1 | 0 | 22997 | CAA98382.1 | P9WIN9 |
| GVTYEIDLTNKNAAKLRGD | 0 | 1 | 0 | 23196 | CAA37572.1 | P24094 |
| HHAERNGSQSTTSQSN | 0 | 1 | 0 | 23976 | Q5HJ48.2 | Q2G155 |
| HPTTTYKAFDWDQAYRKPIT | 0 | 1 | 0 | 24522 | P0A5Q4.1 | P9WIN9 |
| ISGDLKTQIDQVEST | 0 | 1 | 0 | 28505 | CAA17966.1 | P9WNK5 |
| ISTLIKQKLDGLKNE | 0 | 1 | 0 | 28749 | Q07337.1 | Q07337 |
| ITKGKSQKSLGD | 0 | 1 | 0 | 28953 | AAC67090.1 | Q45013 |
| ITNDGVSIAKEIELEDPYEKIGAEL | 0 | 1 | 0 | 29004 | AAA25354.1 | P09239 |
| ITVNSKKTKDLV | 0 | 1 | 0 | 29139 | P14013.1 | P0CL66 |
| IYEATQAASKVGGEASAPGGSNSTD | 0 | 1 | 0 | 29570 | AAA25362.1 | P19993 |
| KAASKETPPALNKC | 0 | 1 | 0 | 29759 | CAF34030.1 | G5IXI6 |
| KDLVFTKENTIT | 0 | 1 | 0 | 30212 | P14013.1 | P0CL66 |
| KEGTVTLSKNIS | 0 | 1 | 0 | 30415 | P14013.1 | P0CL66 |
| KLRGDLRQWVSAGRRVGG | 0 | 1 | 0 | 32128 | P24094.2 | P24094 |
| KNAAKLRGDLRQWVSAG | 0 | 1 | 0 | 32452 | P24094.2 | P24094 |
| KQELDEISTNIRQAG | 0 | 1 | 0 | 32971 | CAA17966.1 | P9WNK5 |
| KRGIEKAVDKVTETLLKDAKEVETK | 0 | 1 | 0 | 33137 | AAA25354.1 | P09239 |
| KTQIDQVESTAGSLQ | 0 | 1 | 0 | 33831 | CAA17966.1 | P9WNK5 |
| KVVQLPKISKNALKANG | 0 | 1 | 0 | 34243 | ACI16083.1 | P05788 |
| KVVQLPKISKNALRNDG | 0 | 1 | 0 | 34244 | ACI16087.1 | P17835 |
| LAGGAATAGAFSRPGLPVEY | 0 | 1 | 0 | 34776 | P0C5B9.1 | P9WQP1 |
| LAQEAGNFERISGDL | 0 | 1 | 0 | 34927 | CAA17966.1 | P9WNK5 |
| LDEGKQSLTKLAAAWG | 0 | 1 | 0 | 35139 | P0A564.2 | P9WNK7 |
| LDVAEGDTVIYSKYG | 0 | 1 | 0 | 35316 | P09621.3 | P9WPE5 |
| LFAKYAEEVGEAVIDDDFGY | 0 | 1 | 0 | 35794 | YP_186940.1 | Q2FWC4 |
| LKLSIPVAERAKPRK | 0 | 1 | 0 | 36950 | P12809.1 | P12809 |
| LKTQIDQVESTAG | 0 | 1 | 0 | 37040 | P0A566.2 | P9WNK5 |
| LNAMKGDLQSSLGAG | 0 | 1 | 0 | 38169 | P0C5B9.1 | P9WQP1 |
| LNRSGGNYTKTDDEA | 0 | 1 | 0 | 38330 | O33084.3 | O33084 |
| LPDWLAANRGLAPGG | 0 | 1 | 0 | 38439 | P0A4V6.1 | P9WQN7 |
| LQSLGAEIAVEQAAL | 0 | 1 | 0 | 38988 | P0A568.2 | P9WNK3 |
| LQVPSPSMGRDIKVQFQSGG | 0 | 1 | 0 | 39011 | P0C5B9.1 | P9WQP1 |
| LRRDLDASRDVKKHVGNALE | 0 | 1 | 0 | 39180 | SRC280368 | Q99XV0 |
| LRRDLEASREANKKVTSELT | 0 | 1 | 0 | 39184 | SRC280368 | Q99XV0 |
| LSRDLEASREAKKKVEAALE | 0 | 1 | 0 | 39623 | SRC280368 | Q99XV0 |
| LTSELPQWLSANRAVKPTGS | 0 | 1 | 0 | 40165 | P0C5B9.1 | P9WQP1 |
| LVRSDYKFYEDANGTRDHKKGRHT | 0 | 1 | 0 | 40566 | P45996.1 | P43840 |
| LVSKKVTSKDKS | 0 | 1 | 0 | 40589 | P14013.1 | P0CL66 |
| MAEMKTDAATLAQEA | 0 | 1 | 0 | 41028 | CAA17966.1 | P9WNK5 |
| MAKVKIKPLEDKILV | 0 | 1 | 0 | 41069 | P24301.2 | P24301 |
| MAKVNIKPLEDKILV | 0 | 1 | 0 | 41071 | P09621.3 | P9WPE5 |
| MGDAGGYKAADMWGPPNDPAWQRND | 0 | 1 | 0 | 41557 | P31951.2 | P31951 |
| MKKDDQIAAAIALRGMAKDGKFAVK | 0 | 1 | 0 | 41844 | ABK60035.1 | G5IXI6 |
| MKKDDQIAAAMVLRGMAKDGQFALK | 0 | 1 | 0 | 41846 | AAN87834.1 | B8F1F6 |
| MLMRTDPFRELDRFA | 0 | 1 | 0 | 42007 | P12809.1 | P12809 |
| MTDVSRKIRAWGRRLMIGTA | 0 | 1 | 0 | 42790 | P0C5B9.1 | P9WQP1 |
| MTEQQWNFAGIEAAAS | 0 | 1 | 0 | 42795 | P0A564.2 | P9WNK7 |
| NAVFNFPPNGTHSWEYWGAQ | 0 | 1 | 0 | 43332 | P0C5B9.1 | P9WQP1 |
| NDPTQQIPKLVANNTRLWVY | 0 | 1 | 0 | 43514 | P0C5B9.1 | P9WQP1 |
| NGEEYLILSARDVLA | 0 | 1 | 0 | 43977 | CAB01005.1 | P9WPE5 |
| NGTPSELGGTNVPAE | 0 | 1 | 0 | 44088 | P31951.2 | P31951 |

| | | | | | | |
|---|---|---|---|---|---|---|
| NHSQTPRYRRPKFPWFK | 0 | 1 | 0 | 44168 | AAW38306.1 | Q2FXJ6 |
| NIGFGNRGDANI | 0 | 1 | 0 | 44231 | YP_177893.1 | P9WHZ5 |
| NITSATYQSAIPPRGTQAVV | 0 | 1 | 0 | 44381 | P0A5Q4.1 | P9WIN9 |
| NNSPAVYLLDGLRAQDDYNG | 0 | 1 | 0 | 45250 | P0C5B9.1 | P9WQP1 |
| NVTSIHSLLDEGKQSL | 0 | 1 | 0 | 46500 | P0A564.2 | P9WNK7 |
| PAGGAYSMYTNWEQD | 0 | 1 | 0 | 46806 | P0A4V6.1 | P9WQN7 |
| PASCGRPSRALHPASVSNGGCRHPV | 0 | 1 | 0 | 46936 | AAA25354.1 | P09239 |
| PDTAKEKPQEGTVVA | 0 | 1 | 0 | 47209 | P09621.3 | P9WPE5 |
| PENAKEKPQEGTVVA | 0 | 1 | 0 | 47366 | P24301.2 | P24301 |
| PPATVYRYDSRPPE | 0 | 1 | 0 | 48751 | P04977.1 | P04977 |
| PQPGPQPPQPPQPQPEAPAPQPPAG | 0 | 1 | 0 | 49057 | P14283.3 | P14283 |
| PSDLGGNNLPAKFLE | 0 | 1 | 0 | 49333 | P0A4V2.1 | P9WQP3 |
| PSLIGLAMGDAGGYKAADMW | 0 | 1 | 0 | 49421 | P0C5B9.1 | P9WQP1 |
| PVGGQSSFYSDWYSPACGKA | 0 | 1 | 0 | 49872 | CAA44268.1 | P9WQP1 |
| QDAYNAAGGHNAVFNFPPNG | 0 | 1 | 0 | 50442 | P0C5B9.1 | P9WQP1 |
| QEAGNFERISGDLKT | 0 | 1 | 0 | 50549 | CAA17966.1 | P9WNK5 |
| QLVLGENLDTERILAS | 0 | 1 | 0 | 51572 | P12809.1 | P12809 |
| QPACRKAGCQTYKWE | 0 | 1 | 0 | 51790 | SRC280101 | P9WQP3 |
| QPAPATAPSQGGV | 0 | 1 | 0 | 51793 | P11089.1 | P11089 |
| QQEGAQQPAPATA | 0 | 1 | 0 | 52010 | P11089.1 | P11089 |
| QQFIYAGSLSALLDPSQGMG | 0 | 1 | 0 | 52026 | P0C5B9.1 | P9WQP1 |
| QQKWDATATELNN | 0 | 1 | 0 | 52048 | P0A564.2 | P9WNK7 |
| QSAAIREWARRNGHNV | 0 | 1 | 0 | 52280 | P24094.2 | P24094 |
| QSQDNSGDQRQVDLTPKK | 0 | 1 | 0 | 52411 | YP_187306.1 | P14738 |
| QTAQAAPVQEGVQ | 0 | 1 | 0 | 52474 | P11089.1 | P11089 |
| RAIVAIGCMFMITFVVLFYTNAT | 0 | 1 | 0 | 53123 | YP_185099.1 | Q2G1E2 |
| RALLGLAVCLFLMALCT | 0 | 1 | 0 | 53152 | YP_185007.1 | Q2G1N0 |
| RDGQLTIKAERTEQKDFDGR | 0 | 1 | 0 | 53355 | P0A5B7.2 | P9WMK1 |
| REWARRNGHNVSTRGRI | 0 | 1 | 0 | 53676 | P24094.2 | P24094 |
| RGDANIGIGNIG | 0 | 1 | 0 | 53833 | YP_177893.1 | P9WHZ5 |
| RILASYQEGVLKLSI | 0 | 1 | 0 | 54181 | P12809.1 | P12809 |
| RIPADVIDAFHAAT | 0 | 1 | 0 | 54206 | P24094.2 | P24094 |
| RKLFLVALVLFTIGSLICAI | 0 | 1 | 0 | 54382 | YP_187153.1 | Q2G2W1 |
| RLAKLSGGVAVIRVG | 0 | 1 | 0 | 54469 | AAS19616.1 | P0C0Z7 |
| RNGHNVSTRGRIPAD | 0 | 1 | 0 | 54992 | P24094.2 | P24094 |
| RPGLDELSFTLTGNPNRPDGGILT | 0 | 1 | 0 | 55158 | YP_177893.1 | P9WHZ5 |
| RPGLPVEYLQVPSEA | 0 | 1 | 0 | 55162 | P31951.2 | P31951 |
| RQWVSAGRRVGGRRRG | 0 | 1 | 0 | 55511 | P24094.2 | P24094 |
| RTEQKDFDGRSEFAYGSFVR | 0 | 1 | 0 | 56043 | P0A5B7.2 | P9WMK1 |
| SAMILAAYHPQQFIYAGSLS | 0 | 1 | 0 | 56895 | CAA44268.1 | P9WQP1 |
| SANNDAEIGNLI | 0 | 1 | 0 | 56918 | AAS19616.1 | P0C0Z7 |
| SGEGAQTAQAAPV | 0 | 1 | 0 | 58007 | P11089.1 | P11089 |
| SIVDKLNRSGGNYTK | 0 | 1 | 0 | 58733 | O33084.3 | O33084 |
| SKKTKDLVFTKE | 0 | 1 | 0 | 58875 | P14013.1 | P0CL66 |
| SQERNFVDSIGQSFQ | 0 | 1 | 0 | 60433 | O33084.3 | O33084 |
| SRKGLRRDLDASREAKKQVEK | 0 | 1 | 0 | 60712 | AIQ02470.1 | Q99XV0 |
| STEGNVTGMFA | 0 | 1 | 0 | 61578 | P0A564.2 | P9WNK7 |
| STLTITVNSKKT | 0 | 1 | 0 | 61760 | P14013.1 | P0CL66 |
| TAQAAPVQEGVQQEGAQQPAPA | 0 | 1 | 0 | 62968 | P11089.1 | P11089 |
| TEVELKERKHRIEDAVRNAK | 0 | 1 | 0 | 63573 | AAA25354.1 | P09239 |
| THSWEYWGAQLNAMKGDLQS | 0 | 1 | 0 | 64079 | P0C5B9.1 | P9WQP1 |
| TQNISGTQVYQDPAIVQPK | 0 | 1 | 0 | 65902 | Q5HH31.1 | Q2FZK7 |
| TRVGKYRPQDKPNTAINYNPWIG | 0 | 1 | 0 | 66118 | AAA24959.1 | P43840 |
| TSARPAVMPMDAWRE | 0 | 1 | 0 | 66163 | P12809.1 | P12809 |
| TVEESNTFGLQLELTEGMR | 0 | 1 | 0 | 66982 | AAA25354.1 | P09239 |
| TVSLPVGADEDDIKATYDKG | 0 | 1 | 0 | 67156 | P0A5B7.2 | P9WMK1 |
| VAKVNIKPLEDKILV | 0 | 1 | 0 | 67656 | SRC280164 | P9WPE5 |
| VANNTRLWVYCGNGTPNELG | 0 | 1 | 0 | 67697 | P0C5B9.1 | P9WQP1 |
| VDGVHAPDKEVKVHADKS | 0 | 1 | 0 | 67969 | YP_187447.1 | Q2FUX4 |
| VDPDREMLAAERPRG | 0 | 1 | 0 | 68035 | P12809.1 | P12809 |
| VDVSEGDIVIYSKYG | 0 | 1 | 0 | 68087 | P24301.2 | P24301 |
| VEFDLPGIKADSLDI | 0 | 1 | 0 | 68174 | P12809.1 | P12809 |
| VETSRSVFHQNGQVTEVTTATGI | 0 | 1 | 0 | 68387 | Q51271 | P30690 |
| VFNLNADGTHSWEYWGAQLNAMKPD | 0 | 1 | 0 | 68534 | P31951.2 | P31951 |
| VMPVGGQSSFYSDWY | 0 | 1 | 0 | 70010 | P0A4V2.1 | P9WQP3 |
| VQTGGTSRTVTMRYLAS | 0 | 1 | 0 | 70651 | ACI16083.1 | P05788 |
| VSILIIFILVSGLSMFVDQL | 0 | 1 | 0 | 70991 | YP_187238.1 | Q2FVI5 |
| VSRLTEKVLGWLCWVMLLVLTVITM | 0 | 1 | 0 | 71099 | YP_185976.1 | Q2G2G6 |
| VSTRGRIPADVIDA | 0 | 1 | 0 | 71165 | P24094.2 | P24094 |
| VVTVRAERPGVDPDR | 0 | 1 | 0 | 71862 | P12809.1 | P12809 |
| WDEDGAKRIPVDVSE | 0 | 1 | 0 | 72291 | P24301.2 | P24301 |
| WDEDGEKRIPLDVAE | 0 | 1 | 0 | 72294 | P09621.3 | P9WPE5 |
| WDINTPAFEWYYQSGLSIVM | 0 | 1 | 0 | 72314 | P0C5B9.1 | P9WQP1 |
| WGRRLMIGTAAAVVLPGLVG | 0 | 1 | 0 | 72515 | P0C5B9.1 | P9WQP1 |
| YDEEARRGLERGLNA | 0 | 1 | 0 | 73535 | YP_976577.1 | P9WPE7 |
| YNEEARKKIQKGVKT | 0 | 1 | 0 | 75140 | AAS19616.1 | P0C0Z7 |
| YNGEEYLILSARDVL | 0 | 1 | 0 | 75152 | P09621.3 | P9WPE5 |
| YYQSGLSIVMPVGGQSSFYS | 0 | 1 | 0 | 76584 | P0C5B9.1 | P9WQP1 |
| KLHASFEDGKKAFDKELQYSNNSRV | 0 | 1 | 0 | 78705 | P10641.1 | P43838 |
| AGTYDFAIAAPQANAKIWIA | 0 | 1 | 0 | 78925 | CAA57997.1 | Q7MTE2 |
| ALLEETITAKGVRSPEAIRG | 0 | 1 | 0 | 78927 | CAA57997.1 | Q7MTE2 |
| AVGQKVTLKWDAPNGTPNPN | 0 | 1 | 0 | 78934 | CAA57997.1 | Q7MTE2 |
| DADGDGHGWKPGNAPGIAGY | 0 | 1 | 0 | 78944 | CAA57997.1 | Q7MTE2 |
| DAPNGTPNPNPNPNPNPNPG | 0 | 1 | 0 | 78945 | CAA57997.1 | Q7MTE2 |
| FGLGGIGVLTPDNYLITPAL | 0 | 1 | 0 | 78973 | CAA57997.1 | Q7MTE2 |
| GIPASWKTIDADGDGHGWKP | 0 | 1 | 0 | 78979 | CAA57997.1 | Q7MTE2 |
| GQGPTKEDDYVFEAGKKYHF | 0 | 1 | 0 | 78986 | CAA57997.1 | Q7MTE2 |
| KDVTVEGSNEFAPVQNLTGS | 0 | 1 | 0 | 79001 | CAA57997.1 | Q7MTE2 |
| LMKKMGSGDGTELTISEGGG | 0 | 1 | 0 | 79016 | CAA57997.1 | Q7MTE2 |
| MDGTASVNIPAGTYDFAIAA | 0 | 1 | 0 | 79023 | CAA57997.1 | Q7MTE2 |
| NSNGCVYSESFGLGGIGVLT | 0 | 1 | 0 | 79031 | CAA57997.1 | Q7MTE2 |

| | | | | | | |
|---|---|---|---|---|---|---|
| PDNYLITPALDLPNGGKLTF | 0 | 1 | 0 | 79035 | CAA57997.1 | Q7MTE2 |
| PGNAPGIAGYNSNGCVYSES | 0 | 1 | 0 | 79039 | CAA57997.1 | Q7MTE2 |
| PNPNPNPNPGTTTLSESFEN | 0 | 1 | 0 | 79044 | CAA57997.1 | Q7MTE2 |
| PQANAKIWIAGQGPTKEDDY | 0 | 1 | 0 | 79045 | CAA57997.1 | Q7MTE2 |
| PSCSPTNMIMDGTASVNIPA | 0 | 1 | 0 | 79047 | CAA57997.1 | Q7MTE2 |
| RIQSTWRQKTVDLPAGTKYV | 0 | 1 | 0 | 79059 | CAA57997.1 | Q7MTE2 |
| SEHYAVYASSTGNDASNFTN | 0 | 1 | 0 | 79079 | CAA57997.1 | Q7MTE2 |
| TTTLSESFENGIPASWKTID | 0 | 1 | 0 | 79103 | CAA57997.1 | Q7MTE2 |
| VDLPAGTKYVAFRHFQSTDM | 0 | 1 | 0 | 79106 | CAA57997.1 | Q7MTE2 |
| AKLTDASNSHNS | 0 | 1 | 0 | 80318 | Q51271 | P30690 |
| ALNDNAGRHNSE | 0 | 1 | 0 | 80367 | Q51271 | P30690 |
| ALVSAGWLQEGK | 0 | 1 | 0 | 80394 | Q51271 | P30690 |
| ALYASVAVQQQD | 0 | 1 | 0 | 80395 | Q51271 | P30690 |
| ATAASVGLRHKF | 0 | 1 | 0 | 80506 | Q51271 | P30690 |
| ATGIVDLGSKIG | 0 | 1 | 0 | 80521 | Q51271 | P30690 |
| DADIGNEYDQVV | 0 | 1 | 0 | 80885 | Q51271 | P30690 |
| DVTLYGTIKAGV | 0 | 1 | 0 | 81337 | Q51271 | P30690 |
| EYDQVVVGAEYD | 0 | 1 | 0 | 81910 | Q51271 | P30690 |
| FFVQYGGAYKRH | 0 | 1 | 0 | 82034 | Q51271 | P30690 |
| FGKLRVGRLNSV | 0 | 1 | 0 | 82048 | Q51271 | P30690 |
| FKGLVDDADIGN | 0 | 1 | 0 | 82146 | Q51271 | P30690 |
| FSKRTSALVSAG | 0 | 1 | 0 | 82374 | Q51271 | P30690 |
| GAYKRHHQVQEG | 0 | 1 | 0 | 82573 | Q51271 | P30690 |
| GEKTREQAVLFGVDHKLHKQLL | 0 | 1 | 0 | 82603 | P20149.1 | P43839 |
| GENKFVATAASV | 0 | 1 | 0 | 82608 | Q51271 | P30690 |
| GNRQSFIGLKGG | 0 | 1 | 0 | 82809 | Q51271 | P30690 |
| GNVTPRVSYAHG | 0 | 1 | 0 | 82816 | Q51271 | P30690 |
| GRHNSESYHAGF | 0 | 1 | 0 | 82850 | Q51271 | P30690 |
| GTDSGWGNRQSF | 0 | 1 | 0 | 82909 | Q51271 | P30690 |
| HQVQEGLNIEKY | 0 | 1 | 0 | 83209 | Q51271 | P30690 |
| IGLKGGFGKLRV | 0 | 1 | 0 | 83458 | Q51271 | P30690 |
| INPWDSKSDYLG | 0 | 1 | 0 | 83781 | Q51271 | P30690 |
| KSDYLGVNKIAE | 0 | 1 | 0 | 84698 | Q51271 | P30690 |
| LGNGLKAIWQVE | 0 | 1 | 0 | 85207 | Q51271 | P30690 |
| LGSKIGFKGQED | 0 | 1 | 0 | 85221 | Q51271 | P30690 |
| LKDTGDINPWDS | 0 | 1 | 0 | 85372 | Q51271 | P30690 |
| LNIEKYQIHRLV | 0 | 1 | 0 | 85615 | Q51271 | P30690 |
| LSIIAEQSNSTVDNQK | 0 | 1 | 0 | 85833 | P20149.1 | P43839 |
| NYKNGGFFVQYG | 0 | 1 | 0 | 87165 | Q51271 | P30690 |
| PEARLISVRYDS | 0 | 1 | 0 | 87229 | Q51271 | P30690 |
| PEFAGLSGSVQY | 0 | 1 | 0 | 87232 | Q51271 | P30690 |
| QIHRLVSGYDND | 0 | 1 | 0 | 87580 | Q51271 | P30690 |
| QKASIAGTDSGW | 0 | 1 | 0 | 87595 | Q51271 | P30690 |
| QTEVAATLAYRF | 0 | 1 | 0 | 87725 | Q51271 | P30690 |
| SGSVQYALNDNA | 0 | 1 | 0 | 88585 | Q51271 | P30690 |
| SGYDNDALYASV | 0 | 1 | 0 | 88617 | Q51271 | P30690 |
| SNSHNSQTEVAA | 0 | 1 | 0 | 88944 | Q51271 | P30690 |
| SVRYDSPEFAGL | 0 | 1 | 0 | 89258 | Q51271 | P30690 |
| SYHAGFNYKNGG | 0 | 1 | 0 | 89292 | Q51271 | P30690 |
| TIKAGVETSRSV | 0 | 1 | 0 | 89474 | Q51271 | P30690 |
| TLAYRFGNVTPR | 0 | 1 | 0 | 89533 | Q51271 | P30690 |
| VGAEYDFSKRTS | 0 | 1 | 0 | 89966 | Q51271 | P30690 |
| VNKIAEPEARLI | 0 | 1 | 0 | 90217 | Q51271 | P30690 |
| VSYAHGFKGLVD | 0 | 1 | 0 | 90424 | Q51271 | P30690 |
| VTEVTTATGIVD | 0 | 1 | 0 | 90437 | Q51271 | P30690 |
| WLQEGKGENKFV | 0 | 1 | 0 | 90599 | Q51271 | P30690 |
| YAKTKNYKIKHEKRYFVSPGFQYEL | 0 | 1 | 0 | 90667 | P20149.1 | P43839 |
| AIWQVEQKASIA | 0 | 1 | 0 | 94775 | Q51271 | P30690 |
| FKGQEDLGNGLK | 0 | 1 | 0 | 94819 | Q51271 | P30690 |
| AGNKEGIASRITQIK | 0 | 1 | 0 | 97186 | BAA04222.1 | P42375 |
| GTTYIRAIAALEGLK | 0 | 1 | 0 | 97368 | BAA04222.1 | P42375 |
| IAEAMRKVKKEGVIT | 0 | 1 | 0 | 97391 | BAA04222.1 | P42375 |
| PGFGDRRKAMLEDIA | 0 | 1 | 0 | 97548 | BAA04222.1 | P42375 |
| QERLAKLAGGVAVLY | 0 | 1 | 0 | 97573 | BAA04222.1 | P42375 |
| ELSGGQMRRVAIAGV | 0 | 1 | 0 | 97997 | EAN09431.1 | Q3XYW0 |
| LKPIRKKVGIVFQFP | 0 | 1 | 0 | 98216 | EAN09431.1 | Q3XYW0 |
| MRRVAIAGVLAMRPE | 0 | 1 | 0 | 98255 | EAN09431.1 | Q3XYW0 |
| DFWATWCGPCKMVAPVLEEI | 0 | 1 | 0 | 98728 | NP_218431.1 | P9WG67 |
| GAKGKAALLRELSDVVPNLN | 0 | 1 | 0 | 98784 | NP_218431.1 | P9WG67 |
| GEQFMQWFLQEQIEEVALMA | 0 | 1 | 0 | 98793 | NP_218358.1 | P9WNE5 |
| KMVAPVLEEIATERATDLTV | 0 | 1 | 0 | 98852 | NP_218431.1 | P9WG67 |
| MLVQHLLDRDLRVEIPGVDT | 0 | 1 | 0 | 98906 | NP_218358.1 | P9WNE5 |
| AAAGDLVGPGCAEYA | 0 | 1 | 0 | 103000 | CAA98373.1 | P9WNF5 |
| AAASLAAIAIAFLAG | 0 | 1 | 0 | 103002 | CAA98350.1 | P9WNF3 |
| AAHKGLMNIALAISA | 0 | 1 | 0 | 103004 | YP_177770.1 | P9WGU1 |
| AAQAVQTAAQNGVRA | 0 | 1 | 0 | 103007 | CAE55371.1 | L7N675 |
| ACGGGTNSSSSGAGG | 0 | 1 | 0 | 103010 | YP_177769.1 | P9WGT9 |
| AEGDTVIYSKYGGTE | 0 | 1 | 0 | 103013 | CAB01005.1 | P9WPE5 |
| AFDLDRLLFKRNICH | 0 | 1 | 0 | 103017 | CAB05419.1 | P9WMX7 |
| AFLHWAITDGNKASF | 0 | 1 | 0 | 103018 | YP_177770.1 | P9WGU1 |
| AGAGTVATTPASSPV | 0 | 1 | 0 | 103021 | YP_177770.1 | P9WGU1 |
| AGFTAPATTLLSALG | 0 | 1 | 0 | 103022 | YP_177963.1 | Q6MWX9 |
| AGTVNIGASDAYLSE | 0 | 1 | 0 | 103029 | YP_177770.1 | P9WGU1 |
| AIANAYWSPQARRRF | 0 | 1 | 0 | 103031 | CAB05418.1 | P9WFR1 |
| AKVRRERMGHIELAA | 0 | 1 | 0 | 103037 | AAA21417.1 | P9WGY7 |
| ALMSGNFSNGILWRG | 0 | 1 | 0 | 103038 | YP_177963.1 | Q6MWX9 |
| ALTGLGDKFGESIVN | 0 | 1 | 0 | 103040 | YP_177701.1 | Q79FZ9 |
| AMEDLVRAYHAMSST | 0 | 1 | 0 | 103042 | CAA17363.1 | P9WNK3 |
| AMMARDTAEAAKWGG | 0 | 1 | 0 | 103043 | CAA17363.1 | P9WNK3 |
| AQAQHLTMANIVTSA | 0 | 1 | 0 | 103055 | YP_177768.1 | P9WGT7 |
| ASFLDQVHFQPLPPA | 0 | 1 | 0 | 103061 | YP_177770.1 | P9WGU1 |

| | | | | | | |
|---|---|---|---|---|---|---|
| ATIPQGEQSTGKIYF | 0 | 1 | 0 | 103067 | CAB06500.1 | P9WIP1 |
| AYVITSVDEEMRHNE | 0 | 1 | 0 | 103078 | AAA21417.1 | P9WGY7 |
| CGGVSTANATVYMID | 0 | 1 | 0 | 103082 | CAA98373.1 | P9WNF5 |
| CVAYIGISFLDQASQ | 0 | 1 | 0 | 103090 | YP_177770.1 | P9WGU1 |
| DEMTRVIWKLIKDML | 0 | 1 | 0 | 103102 | CAA17111.1 | P9WKL1 |
| DGYPIINYEYAIVNN | 0 | 1 | 0 | 103105 | YP_177770.1 | P9WGU1 |
| DPRYNQLLGPLPFRH | 0 | 1 | 0 | 103112 | CAB05415.1 | P9WN09 |
| DRFADFPALPLDPSA | 0 | 1 | 0 | 103113 | CAB09453.1 | O07175 |
| DSIYYVDANASIQEM | 0 | 1 | 0 | 103116 | NP_217142.1 | P9WJA3 |
| DTLNGGEYTVFAPTN | 0 | 1 | 0 | 103118 | CAA98350.1 | P9WNF3 |
| DVGSLNGTYVNREPV | 0 | 1 | 0 | 103121 | CAB01474.1 | P9WJA9 |
| DVVQHVARRPGESPP | 0 | 1 | 0 | 103125 | WP_009939718.1 | P9WQ59 |
| EAEHQAIVRDVLAAG | 0 | 1 | 0 | 103128 | YP_177838.1 | P9WNJ3 |
| EDEMKEGRYEVRAEL | 0 | 1 | 0 | 103130 | CAA17245.1 | P9WMK1 |
| EGWGKSPGFGTTVDF | 0 | 1 | 0 | 103137 | YP_177770.1 | P9WGU1 |
| EKVAKEAADEAKAKL | 0 | 1 | 0 | 103141 | CAB07109.1 | P9WHE3 |
| EMIEAVGYRSWPRYF | 0 | 1 | 0 | 103143 | YP_005911842.1 | O53732 |
| ENYIAQTRDKFLSAA | 0 | 1 | 0 | 103145 | CAA98382.1 | P9WIN9 |
| ESGTTDNFQRYLQAA | 0 | 1 | 0 | 103153 | YP_177768.1 | P9WGT7 |
| FCGRIHTRYSSAYEL | 0 | 1 | 0 | 103162 | CAA15575.1 | P9WQB1 |
| FEETFEVTAAAPVAV | 0 | 1 | 0 | 103165 | CAB07109.1 | P9WHE3 |
| FGDVDAHGAMIRAQA | 0 | 1 | 0 | 103170 | YP_177838.1 | P9WNJ3 |
| FLGGFAGLPSLGFGN | 0 | 1 | 0 | 103176 | CAB06237.1 | P9WJD3 |
| FPHHPIHTVPSEEVL | 0 | 1 | 0 | 103179 | CAB05418.1 | P9WFR1 |
| FPSFAGLRPTFDTRL | 0 | 1 | 0 | 103180 | CAA17245.1 | P9WMK1 |
| FRMELLSLPQDEWAG | 0 | 1 | 0 | 103185 | AAA25369.1 | I6Y231 |
| GASVTVTGQGNSLKV | 0 | 1 | 0 | 103195 | CAA98373.1 | P9WNF5 |
| GDDDRLHGMLTDRDI | 0 | 1 | 0 | 103197 | NP_217142.1 | P9WJA3 |
| GDLPTIGTAVSARNT | 0 | 1 | 0 | 103198 | CAE55335.1 | I6Y936 |
| GDLVGPGCAEYAAAN | 0 | 1 | 0 | 103199 | CAA98373.1 | P9WNF5 |
| GELDNRGSQFYLAMY | 0 | 1 | 0 | 103203 | CAA16247.1 | O53611 |
| GFLMPPSDGSGVTPG | 0 | 1 | 0 | 103208 | AAK31576.1 | P9WJD9 |
| GFSEIMRSTLEKDNT | 0 | 1 | 0 | 103209 | AAA21416.1 | P9WGY9 |
| GNKASFLDQVHFQPL | 0 | 1 | 0 | 103227 | YP_177770.1 | P9WGU1 |
| GPGQEGLDQYGSIPL | 0 | 1 | 0 | 103230 | YP_177769.1 | P9WGT9 |
| GVPSRLGYLLDLAPK | 0 | 1 | 0 | 103244 | AAA21417.1 | P9WGY7 |
| GVYKVCKGLEKIPLL | 0 | 1 | 0 | 103245 | CAB06237.1 | P9WJD3 |
| GYIPILGAPWLADLV | 0 | 1 | 0 | 103246 | AAA25369.1 | I6Y231 |
| GYPVHKPVTAGWNGY | 0 | 1 | 0 | 103247 | CAB08634.1 | P9WL63 |
| HGTCANQCPIVDAGD | 0 | 1 | 0 | 103255 | AAA21416.1 | P9WGY9 |
| HQGQGGAKSKGSQQE | 0 | 1 | 0 | 103258 | AAK31576.1 | P9WJD9 |
| HRSDGSGDTFLFTQY | 0 | 1 | 0 | 103260 | YP_177770.1 | P9WGU1 |
| HVIAGQASPSRIDGT | 0 | 1 | 0 | 103262 | CAA98350.1 | P9WNF3 |
| HWLRPITPTFRPSWP | 0 | 1 | 0 | 103264 | YP_005911842.1 | O53732 |
| IADIEAYPQWISEYK | 0 | 1 | 0 | 103266 | WP_019256904.1 | O53519 |
| IAFLAGCSSTKPVSQ | 0 | 1 | 0 | 103267 | CAA98350.1 | P9WNF3 |
| ILKDESYKVTGTAPI | 0 | 1 | 0 | 103274 | CAB08889.1 | P9WFC9 |
| IRGSVTPAVSQFNAR | 0 | 1 | 0 | 103283 | CAB06500.1 | P9WIP1 |
| IRLAKCGPWHRGGWD | 0 | 1 | 0 | 103284 | CAA16235.1 | P9WFL9 |
| ISMIDGPAPDGYPII | 0 | 1 | 0 | 103286 | YP_177770.1 | P9WGU1 |
| ITSAGRHPDSDIFLD | 0 | 1 | 0 | 103291 | CAB01474.1 | P9WJA9 |
| IVMSTNNDDPAGACR | 0 | 1 | 0 | 103296 | CAA94642.1 | P9WQD7 |
| IWKLIKDMLILPYLD | 0 | 1 | 0 | 103297 | CAA17111.1 | P9WKL1 |
| KAMLFDGRSGEPFPY | 0 | 1 | 0 | 103301 | AAA21416.1 | P9WGY9 |
| KGLMNIALAISAQQV | 0 | 1 | 0 | 103304 | YP_177770.1 | P9WGU1 |
| KTINLCAPDDPICTG | 0 | 1 | 0 | 103327 | NP_216500.1 | P9WP43 |
| KTLDAAIGKLLDNDK | 0 | 1 | 0 | 103328 | CAA16247.1 | O53611 |
| LAALPDFLGGFAGLP | 0 | 1 | 0 | 103331 | CAB06237.1 | P9WJD3 |
| LAPDTVPLAGVVLAG | 0 | 1 | 0 | 103338 | CAA16030.1 | P9WJQ9 |
| LASTLTDALKSHGPQ | 0 | 1 | 0 | 103339 | AAA25369.1 | I6Y231 |
| LCKTTSNFIWGQLLL | 0 | 1 | 0 | 103340 | CAB06237.1 | P9WJD3 |
| LEAAGDKKIGVIKVV | 0 | 1 | 0 | 103344 | CAB07109.1 | P9WHE3 |
| LEPGVVAEKVRNLPA | 0 | 1 | 0 | 103346 | P06806 | P9WPE7 |
| LGDVGEAFVDSLTSQ | 0 | 1 | 0 | 103348 | NP_216500.1 | P9WP43 |
| LGENGNGGMVTGCAE | 0 | 1 | 0 | 103349 | YP_177770.1 | P9WGU1 |
| LGGGVGAFLNALFAG | 0 | 1 | 0 | 103352 | YP_177963.1 | Q6MWX9 |
| LGWKVSDLKSSTAVI | 0 | 1 | 0 | 103355 | CAB06500.1 | P9WIP1 |
| LILTSYTSDEAMLDA | 0 | 1 | 0 | 103359 | NP_217649.1 | P9WMF9 |
| LLDIYRKLRPGEPPT | 0 | 1 | 0 | 103361 | AAA21416.1 | P9WGY9 |
| LMILIATNLLGQNTP | 0 | 1 | 0 | 103365 | CAE55371.1 | L7N675 |
| LPADTAAQLTSAGRE | 0 | 1 | 0 | 103371 | AAK31576.1 | P9WJD9 |
| LPGVSEHLKLNGKVL | 0 | 1 | 0 | 103373 | YP_177770.1 | P9WGU1 |
| LPLLEKVIGAGKPLL | 0 | 1 | 0 | 103378 | P06806 | P9WPE7 |
| LQAHPHAVNGTAMRE | 0 | 1 | 0 | 103384 | AAA25369.1 | I6Y231 |
| LRPASCRFVPTCSQY | 0 | 1 | 0 | 103385 | CAA16235.1 | P9WFL9 |
| LSEGDMAAHKGLMNI | 0 | 1 | 0 | 103386 | YP_177770.1 | P9WGU1 |
| LTTYILLPSQDLPLL | 0 | 1 | 0 | 103393 | CAE55504.1 | P9WHZ5 |
| MNIALAISAQQVNYN | 0 | 1 | 0 | 103415 | YP_177770.1 | P9WGU1 |
| MSFVTIQPVVLAAAT | 0 | 1 | 0 | 103420 | CAE55335.1 | I6Y936 |
| MTAHTHDGTRTWRTG | 0 | 1 | 0 | 103425 | CAB10947.1 | O33192 |
| MTLLELSDFVKKFEE | 0 | 1 | 0 | 103426 | CAB07109.1 | P9WHE3 |
| MWELYLAYSEAGFRS | 0 | 1 | 0 | 103431 | YP_005911842.1 | O53732 |
| MWLSPNGTIRNILGG | 0 | 1 | 0 | 103432 | CAA17111.1 | P9WKL1 |
| NAFLTALTNAGIAYD | 0 | 1 | 0 | 103434 | CAB10947.1 | O33192 |
| NTPAIAVNEAEYGEM | 0 | 1 | 0 | 103452 | CAE55371.1 | L7N675 |
| PDRSAERCGSPAWDL | 0 | 1 | 0 | 103462 | YP_177769.1 | P9WGT9 |
| PETINYRTLKPEKDG | 0 | 1 | 0 | 103464 | AAA21417.1 | P9WGY7 |
| PFNVNLKLQFLHDAF | 0 | 1 | 0 | 103469 | YP_177963.1 | Q6MWX9 |
| PGGTTTLVEHMVGIL | 0 | 1 | 0 | 103472 | CAA16030.1 | P9WJQ9 |
| PGQGAYAAANSWVDV | 0 | 1 | 0 | 103476 | AAA25369.1 | I6Y231 |
| PGYPVAGQVWEATAT | 0 | 1 | 0 | 103479 | CAB06500.1 | P9WIP1 |

| | | | | | | |
|---|---|---|---|---|---|---|
| PILGAPWLADLVRRS | 0 | 1 | 0 | 103480 | AAA25369.1 | I6Y231 |
| PLDTPPAPAPPPFRL | 0 | 1 | 0 | 103482 | YP_177963.1 | Q6MWX9 |
| PPGYDITALLQAHPH | 0 | 1 | 0 | 103490 | AAA25369.1 | I6Y231 |
| PPTPISVIFRSDKSG | 0 | 1 | 0 | 103493 | YP_177769.1 | P9WGT9 |
| PVAGQVWEATATVNA | 0 | 1 | 0 | 103501 | CAB06500.1 | P9WIP1 |
| PVIAENRAELMILIA | 0 | 1 | 0 | 103503 | CAE55371.1 | L7N675 |
| QAQLISSQAQQGGQQ | 0 | 1 | 0 | 103510 | CAB06237.1 | P9WJD3 |
| QAQYVAQLHVWARRE | 0 | 1 | 0 | 103511 | AAK31576.1 | P9WJD9 |
| QDPEGWGKSPGFGTT | 0 | 1 | 0 | 103516 | YP_177770.1 | P9WGU1 |
| QGQWRGAAGTAAQAA | 0 | 1 | 0 | 103519 | CAA17966.1 | P9WNK5 |
| QNAGGTHPTTTYKAF | 0 | 1 | 0 | 103525 | CAA98382.1 | P9WIN9 |
| QSGGNNSPAVYLLDG | 0 | 1 | 0 | 103530 | CAB10044.1 | P9WQP1 |
| QSGLGDNGYIPIPDE | 0 | 1 | 0 | 103531 | YP_177768.1 | P9WGT7 |
| QSVCPILAEPGGSFN | 0 | 1 | 0 | 103534 | CAA15739.1 | O50383 |
| QVGSQFGPYQLLRLL | 0 | 1 | 0 | 103536 | NP_215446.1 | P9WI79 |
| QVVLGWKVSDLKSST | 0 | 1 | 0 | 103537 | CAB06500.1 | P9WIP1 |
| RGEHRDEHTQDAGDK | 0 | 1 | 0 | 103544 | CAA17343.1 | O53673 |
| RGLGEAQLGNSSGNF | 0 | 1 | 0 | 103547 | YP_177770.1 | P9WGU1 |
| RGLRAAAEAGVRLAF | 0 | 1 | 0 | 103548 | CAA16030.1 | P9WJQ9 |
| RHGLSSLGWNLCRIF | 0 | 1 | 0 | 103549 | CAB05415.1 | P9WN09 |
| RIFYKDAFAKHQELF | 0 | 1 | 0 | 103550 | CAA16247.1 | O53611 |
| RKVVVTKDETTIVEG | 0 | 1 | 0 | 103554 | P06806 | P9WPE7 |
| RRFPLPDVPWTRFFG | 0 | 1 | 0 | 103566 | CAB05418.1 | P9WFR1 |
| RSCPGYTLDYNANGS | 0 | 1 | 0 | 103569 | YP_177769.1 | P9WGT9 |
| SAAAKNTPGSITYNE | 0 | 1 | 0 | 103577 | YP_177768.1 | P9WGT7 |
| SAAIGLSMAGSSAMI | 0 | 1 | 0 | 103578 | CAB10044.1 | P9WQP1 |
| SAERCGSPAWDLPTV | 0 | 1 | 0 | 103579 | YP_177769.1 | P9WGT9 |
| SAFAAKAGLMRHTIG | 0 | 1 | 0 | 103580 | O53692.1 | O53692 |
| SEFDVILEAAGDKKI | 0 | 1 | 0 | 103589 | CAB07109.1 | P9WHE3 |
| SKVSTVKDLLPLLEK | 0 | 1 | 0 | 103605 | P06806 | P9WPE7 |
| SPHPLLTHAVEQTGR | 0 | 1 | 0 | 103612 | AAA25369.1 | I6Y231 |
| SVPANVSRRAKVDVL | 0 | 1 | 0 | 103620 | CAB08889.1 | P9WFC9 |
| TAALRAAEDDCPPPG | 0 | 1 | 0 | 103629 | AAA25369.1 | I6Y231 |
| TAVVLLCCSGVATAA | 0 | 1 | 0 | 103635 | CAA98382.1 | P9WIN9 |
| TGGGGGSGFSNSGSG | 0 | 1 | 0 | 103645 | YP_177963.1 | Q6MWX9 |
| TGNMISNQAKYVSDT | 0 | 1 | 0 | 103649 | CAB06237.1 | P9WJD3 |
| TLQAFLHWAITDGNK | 0 | 1 | 0 | 103659 | YP_177770.1 | P9WGU1 |
| TNDGVIFFFNPGELL | 0 | 1 | 0 | 103661 | CAA98382.1 | P9WIN9 |
| TPPAPAPPPFRLPLL | 0 | 1 | 0 | 103664 | YP_177963.1 | Q6MWX9 |
| TSEQIVERFGFSEIM | 0 | 1 | 0 | 103669 | AAA21416.1 | P9WGY9 |
| TVIAATAGRNHLKNV | 0 | 1 | 0 | 103677 | CAB05418.1 | P9WFR1 |
| TWQTKSGGTRTGNVT | 0 | 1 | 0 | 103680 | CAB09453.1 | O07175 |
| VALFGEPSSGFSSML | 0 | 1 | 0 | 103686 | NP_216500.1 | P9WP43 |
| VCAVDMPYLTVELIE | 0 | 1 | 0 | 103687 | CAA16030.1 | P9WJQ9 |
| VFGGAASCAAPIQAD | 0 | 1 | 0 | 103693 | CAB10947.1 | O33192 |
| VGNVGLSTIAGRLLG | 0 | 1 | 0 | 103694 | CAB08889.1 | P9WFC9 |
| VGPGRWDEDGEKRIP | 0 | 1 | 0 | 103695 | CAB01005.1 | P9WPE5 |
| VGQVTATMGQLQQLV | 0 | 1 | 0 | 103697 | CAB06237.1 | P9WJD3 |
| VHNVAALPGAAYCEM | 0 | 1 | 0 | 103700 | AAA25369.1 | I6Y231 |
| VLCNGGSPTTQQALA | 0 | 1 | 0 | 103703 | CAB05418.1 | P9WFR1 |
| VRERMTTQDVEAITP | 0 | 1 | 0 | 103716 | AAA21416.1 | P9WGY9 |
| VRRRHGLSSLGWNLC | 0 | 1 | 0 | 103717 | CAB05415.1 | P9WN09 |
| VSVAVSEGKPTEKHI | 0 | 1 | 0 | 103718 | CAA17245.1 | P9WMK1 |
| VTVTGQGNSLKVGNA | 0 | 1 | 0 | 103720 | CAA98373.1 | P9WNF5 |
| WGPSSDPAWERNDPT | 0 | 1 | 0 | 103729 | CAB10044.1 | P9WQP1 |
| WISEYKEVEILEADD | 0 | 1 | 0 | 103731 | WP_019256904.1 | O53519 |
| WLSANRAVKPTGSAA | 0 | 1 | 0 | 103732 | CAB10044.1 | P9WQP1 |
| WRSLDVEMTAVQRSF | 0 | 1 | 0 | 103737 | CAE55489.1 | Q79FE1 |
| WSFAVGKQLNMAQII | 0 | 1 | 0 | 103738 | YP_177769.1 | P9WGT9 |
| YDREKLQERLAKLAG | 0 | 1 | 0 | 103744 | P06806 | P9WPE7 |
| YEEQMQDAFETGVMF | 0 | 1 | 0 | 103745 | CAA16247.1 | O53611 |
| YHVVAGQTSPANVVG | 0 | 1 | 0 | 103746 | CAA98373.1 | P9WNF5 |
| YIGISFLDQASQRGL | 0 | 1 | 0 | 103747 | YP_177770.1 | P9WGU1 |
| YNANGSGAGVTQFLN | 0 | 1 | 0 | 103751 | YP_177769.1 | P9WGT9 |
| YRADSVERLIAAHLM | 0 | 1 | 0 | 103754 | NP_215446.1 | P9WI79 |
| YRLERPLLFALQCMP | 0 | 1 | 0 | 103755 | CAB05415.1 | P9WN09 |
| DLQDVASHESGVSDQPAQVVTER | 0 | 1 | 0 | 108316 | AAQ75372.1 | Q2GI62 |
| ETEKESGITESHQKEDEIVSQ | 0 | 1 | 0 | 108352 | AAQ75372.1 | Q2GI62 |
| QVVTERESEIESHQGETEKESG | 0 | 1 | 0 | 108605 | AAQ75372.1 | Q2GI62 |
| SKVEQEETNPEVLIKDLQDVAS | 0 | 1 | 0 | 108635 | AAQ75372.1 | Q2GI62 |
| SSSEPFVAESEVSKVEQEETNP | 0 | 1 | 0 | 108642 | AAQ75372.1 | Q2GI62 |
| DIGAQAVSPSTSQGADVKKSSCQSK | 0 | 1 | 0 | 119556 | YP_507490.1 | Q2GGE3 |
| EDPSVEEEEIAPPLPP | 0 | 1 | 0 | 119564 | YP_507490.1 | Q2GGE3 |
| EPDLEEIVSILKNDKEGISE | 0 | 1 | 0 | 119567 | YP_507490.1 | Q2GGE3 |
| EPVQVDIPNNPVREGRNVMTLLHLA | 0 | 1 | 0 | 119568 | YP_507490.1 | Q2GGE3 |
| ETGETVEEGLYAVPLPKD | 0 | 1 | 0 | 119570 | YP_507490.1 | Q2GGE3 |
| INEPVQVDIPNNPVREGRNV | 0 | 1 | 0 | 119597 | YP_507490.1 | Q2GGE3 |
| QRPTPTQVLEEDPSVEEEE | 0 | 1 | 0 | 119631 | YP_507490.1 | Q2GGE3 |
| SEEQLQELSEEITDIVQGLPPITSE | 0 | 1 | 0 | 119640 | YP_507490.1 | Q2GGE3 |
| SGIFDKGEGQHKASEEQLQELSEEI | 0 | 1 | 0 | 119643 | YP_507490.1 | Q2GGE3 |
| SILKNDKEGISELINEPVQV | 0 | 1 | 0 | 119644 | YP_507490.1 | Q2GGE3 |
| TDIVQGLPPITSEDIGAQAVSPSTS | 0 | 1 | 0 | 119650 | YP_507490.1 | Q2GGE3 |
| VPLPKDQRPTPTQVLE | 0 | 1 | 0 | 119668 | YP_507490.1 | Q2GGE3 |
| AAVQAAWAVLVAYFNRLRCGTGDY | 0 | 1 | 0 | 120392 | NP_216382.1 | P95149 |
| AIWRRIRVVPFEVVIPADEQD | 0 | 1 | 0 | 120399 | NP_216098.1 | O06608 |
| APYNVRRLPPPVVEP | 0 | 1 | 0 | 120408 | NP_218400.1 | O05461 |
| ARLVVLASGTGSLLR | 0 | 1 | 0 | 120410 | NP_215471.1 | P9WHM5 |
| AVVRFQEAANKQK | 0 | 1 | 0 | 120421 | NP_218391.1 | P9WNK5 |
| DDLVGAGVIDAVA | 0 | 1 | 0 | 120428 | NP_218400.1 | O05461 |
| DKVLPLVAIVVSVGLAVSYD | 0 | 1 | 0 | 120441 | NP_216495.1 | P9WQM5 |
| DRYRHLVALSITDFGAAGPRSSWRA | 0 | 1 | 0 | 120452 | NP_216382.1 | P95149 |

| | | | | | | |
|---|---|---|---|---|---|---|
| EHLKLNGKVLAAMYQ | 0 | 1 | 0 | 120481 | YP_177770.1 | P9WGU1 |
| ETLHERIKVTERRLLVAAVAALATH | 0 | 1 | 0 | 120511 | NP_215471.1 | P9WHM5 |
| FPVVTHDEVLRLVGRRRLWGRG | 0 | 1 | 0 | 120537 | NP_215630.1 | P9WF73 |
| GDVQPAEVVAAAR | 0 | 1 | 0 | 120550 | NP_218395.1 | P9WJC3 |
| GSLIFAAVLVMLIAVLARLMMRGWR | 0 | 1 | 0 | 120569 | NP_215898.1 | P71810 |
| HSLLWIGVFLFLGGMAAGGCNS | 0 | 1 | 0 | 120587 | NP_217510.1 | P9WJW7 |
| IMIVSLGVTASSFLFINGVAFLIPR | 0 | 1 | 0 | 120630 | NP_217510.1 | P9WJW7 |
| IPVVSVTKSVGFQLRGQSGPTTVK | 0 | 1 | 0 | 120639 | NP_214932.1 | P96264 |
| LTFLGFIGVLLAINLFGNRAIKWAN | 0 | 1 | 0 | 120788 | NP_216495.1 | P9WQM5 |
| MRASPAERVDGAYAGAGPHTQSV | 0 | 1 | 0 | 120802 | NP_215950.1 | O06824 |
| MVNKSRMMPAV | 0 | 1 | 0 | 120810 | NP_214932.1 | P96264 |
| PHVKPAALAEQPG | 0 | 1 | 0 | 120870 | NP_218396.1 | P9WJC1 |
| PSWGLVVTMFAWGYLLDHVGERMV | 0 | 1 | 0 | 120878 | NP_217510.1 | P9WJW7 |
| QDVGNYDAFISIRAAIRAIESA | 0 | 1 | 0 | 120887 | NP_218253.1 | O69703 |
| QKVQAAGNNMAQT | 0 | 1 | 0 | 120892 | P95242.1 | P9WNI7 |
| RALKRRNARTKAR | 0 | 1 | 0 | 120904 | NP_217735.1 | P9WF43 |
| RGVAGKVVAGIGILAIRWRLPSGT | 0 | 1 | 0 | 120912 | NP_215898.1 | P71810 |
| SPYRGSSILWRIHAASALLMMP | 0 | 1 | 0 | 120961 | NP_217510.1 | P9WJW7 |
| TTIFPDHVVGRVGALLALAALIHRD | 0 | 1 | 0 | 121017 | NP_216382.1 | P95149 |
| VTVTFMLVWLINHHGWSVAQ | 0 | 1 | 0 | 121057 | NP_217510.1 | P9WJW7 |
| VVHPAVVQANRVRTWLLAVSNVFGQ | 0 | 1 | 0 | 121059 | YP_177753.1 | P9WI37 |
| FPGWQPGMPTIPTAPPTTPV | 0 | 1 | 0 | 122097 | NP_215052.1 | O06404 |
| LQDSGVHDVAVISEAQAATA | 0 | 1 | 0 | 122140 | WP_003911223.1 | O06404 |
| LSVVGDPDAPPTMVAVAPVA | 0 | 1 | 0 | 122143 | NP_215052.1 | O06404 |
| PIPVPIIIPPFPGWQPGMPT | 0 | 1 | 0 | 122162 | NP_215052.1 | O06404 |
| EIERDTERDRFLSADEAV | 0 | 1 | 0 | 125029 | YP_001007316.1 | A1JNN2 |
| AFSLPDRSLSAIPLFDVSFE | 0 | 1 | 0 | 127919 | AAO67531.1 | P38006 |
| DLVNCNLNTNCVAVAFSLPD | 0 | 1 | 0 | 127944 | AAO67531.1 | P38006 |
| FTNLKFKVRKITSSHRGN | 0 | 1 | 0 | 127984 | AAO67537.1 | P38006 |
| GMIEVQSNYGFVWDVSLKKV | 0 | 1 | 0 | 128004 | AAO67531.1 | P38006 |
| NSYDVLSALSGNLKL | 0 | 1 | 0 | 128066 | AAO67531.1 | P38006 |
| QKNACSFDLCNSYDVL | 0 | 1 | 0 | 128084 | AAO67531.1 | P38006 |
| VRMMLIPLDATLIKV | 0 | 1 | 0 | 130285 | AAT48493.1 | Q8F8E1 |
| AIGDRSKTDRENSVSIG | 0 | 1 | 0 | 131344 | CAA32086.1 | A1JUB7 |
| ASDVVKKMVGETVESA | 0 | 1 | 0 | 133518 | ABA10360.1 | G1UB25 |
| DALVAKAQEVS | 0 | 1 | 0 | 133540 | ABA10360.1 | G1UB25 |
| KKLLVRGLYRISFTTYK | 0 | 1 | 0 | 133618 | AAS73248.1 | O34094 |
| VKTLLPYGSVINYYGYVKPG | 0 | 1 | 0 | 133733 | AAS73248.1 | O34094 |
| GGGTGPADGTNATTI | 0 | 1 | 0 | 136315 | P69996.1 | P69996 |
| TLHDMGIFSITSSDS | 0 | 1 | 0 | 136686 | P69996.1 | P69996 |
| WMLRAAEEYSMNLGF | 0 | 1 | 0 | 136749 | P69996.1 | P69996 |
| ADYWAGELGQSR | 0 | 1 | 0 | 137797 | P29723.2 | P29723 |
| AFRQQFQYAVEV | 0 | 1 | 0 | 137798 | P29723.2 | P29723 |
| AGELGQSRDVLL | 0 | 1 | 0 | 137800 | P29723.2 | P29723 |
| DAGMFDAVSRAT | 0 | 1 | 0 | 137817 | P29723.2 | P29723 |
| DVLLAGNAEADR | 0 | 1 | 0 | 137828 | P29723.2 | P29723 |
| DYVMNFNTVRYD | 0 | 1 | 0 | 137830 | P29723.2 | P29723 |
| EDSRVTENTNGL | 0 | 1 | 0 | 137834 | P29723.2 | P29723 |
| EGAVALADRASS | 0 | 1 | 0 | 137837 | P29723.2 | P29723 |
| EYETDPSVTKMV | 0 | 1 | 0 | 137844 | P29723.2 | P29723 |
| FDRFKGSGPGYY | 0 | 1 | 0 | 137848 | P29723.2 | P29723 |
| FTPGTEYTVEVR | 0 | 1 | 0 | 137859 | P29723.2 | P29723 |
| GEIKFEAVEGAV | 0 | 1 | 0 | 137867 | P29723.2 | P29723 |
| GQNVLSADFTPG | 0 | 1 | 0 | 137883 | P29723.2 | P29723 |
| GQSRDVLLAGNA | 0 | 1 | 0 | 137884 | P29723.2 | P29723 |
| GSSHHETHYGYA | 0 | 1 | 0 | 137887 | P29723.2 | P29723 |
| GTGYHSRFGSDA | 0 | 1 | 0 | 137889 | P29723.2 | P29723 |
| HETHYGYATLSY | 0 | 1 | 0 | 137891 | P29723.2 | P29723 |
| HGHGAFRQQFQY | 0 | 1 | 0 | 137892 | P29723.2 | P29723 |
| IANGYRDVVADV | 0 | 1 | 0 | 137896 | P29723.2 | P29723 |
| IGGADAETLMDA | 0 | 1 | 0 | 137900 | P29723.2 | P29723 |
| KGSGPGYYRLTL | 0 | 1 | 0 | 137907 | P29723.2 | P29723 |
| KTMLTEDSFSAR | 0 | 1 | 0 | 137912 | P29723.2 | P29723 |
| LMDAAVDVFADG | 0 | 1 | 0 | 137923 | P29723.2 | P29723 |
| LQLLVVGCGSSH | 0 | 1 | 0 | 137924 | P29723.2 | P29723 |
| LSADFTPGTEYT | 0 | 1 | 0 | 137928 | P29723.2 | P29723 |
| MESPHDLVVDTV | 0 | 1 | 0 | 137930 | P29723.2 | P29723 |
| NFNTVRYDYYGD | 0 | 1 | 0 | 137935 | P29723.2 | P29723 |
| NIDIGLKGKVLT | 0 | 1 | 0 | 137936 | P29723.2 | P29723 |
| PGYYRLTLIANG | 0 | 1 | 0 | 137946 | P29723.2 | P29723 |
| RFLPKYEGNIDI | 0 | 1 | 0 | 137965 | P29723.2 | P29723 |
| RLTLIANGYRDV | 0 | 1 | 0 | 137967 | P29723.2 | P29723 |
| RVPRISCGINYG | 0 | 1 | 0 | 138025 | P29723.2 | P29723 |
| SAGALQLLVVGC | 0 | 1 | 0 | 138028 | P29723.2 | P29723 |
| TLSYADYWAGEL | 0 | 1 | 0 | 138054 | P29723.2 | P29723 |
| VDTVGTGYHSRF | 0 | 1 | 0 | 138063 | P29723.2 | P29723 |
| VSDQAVSLGQNV | 0 | 1 | 0 | 138079 | P29723.2 | P29723 |
| VVGCGSSHHETH | 0 | 1 | 0 | 138082 | P29723.2 | P29723 |
| YGYATLSYADYW | 0 | 1 | 0 | 138094 | P29723.2 | P29723 |
| YRDVVADVRFLP | 0 | 1 | 0 | 138098 | P29723.2 | P29723 |
| EAAAYIKGRLSPIVKTERIKVFVK | 0 | 1 | 0 | 141535 | AAT48493.1 | Q8F8E1 |
| KELLQEGYEEIVGETPSFKK | 0 | 1 | 0 | 141544 | AAT48493.1 | Q8F8E1 |
| LSPFPAYPIVVGGQIYRFGYKHEL | 0 | 1 | 0 | 141548 | AAT48511.1 | G1UB30 |
| SSIVIPAAVGIKLNVTEDA | 0 | 1 | 0 | 141566 | AAT48511.1 | G1UB30 |
| VFPKDKEGRALQKFLGTIR | 0 | 1 | 0 | 141568 | AAT48493.1 | Q8F8E1 |
| VRSSNTCTVGPSDPACFQNP | 0 | 1 | 0 | 141569 | AAT48511.1 | G1UB30 |
| YIGVAPRKAIPA | 0 | 1 | 0 | 141580 | AAT48511.1 | G1UB30 |
| AENSAIKVPLENGS | 0 | 1 | 0 | 141847 | NP_045581.1 | O50811 |
| CIKTLMTNVETYFK | 0 | 1 | 0 | 141862 | NP_045581.1 | O50811 |
| CKWHVDNPIDEATA | 0 | 1 | 0 | 141863 | NP_045581.1 | O50811 |
| EATAESKSALTSVD | 0 | 1 | 0 | 141873 | NP_045581.1 | O50811 |

| | | | | | | |
|---|---|---|---|---|---|---|
| EKSKKEIEDQKNTK | 0 | 1 | 0 | 141879 | NP_045581.1 | O50811 |
| ENGSKIEAKKCIKT | 0 | 1 | 0 | 141881 | NP_045581.1 | O50811 |
| ILAALSEAANKIEN | 0 | 1 | 0 | 141926 | NP_045581.1 | O50811 |
| KDKKDDKYTKILAA | 0 | 1 | 0 | 141932 | NP_045581.1 | O50811 |
| KIENAAMAIHLCFNN | 0 | 1 | 0 | 141933 | NP_045581.1 | O50811 |
| KNTKESKNIEVKDT | 0 | 1 | 0 | 141937 | NP_045581.1 | O50811 |
| NGDKSTQKYNELKT | 0 | 1 | 0 | 141967 | NP_045581.1 | O50811 |
| NLAKEAQDDSEKSK | 0 | 1 | 0 | 141973 | NP_045581.1 | O50811 |
| SEATGLSSEKITKL | 0 | 1 | 0 | 142008 | NP_045581.1 | O50811 |
| TSVDQVLDEISEAT | 0 | 1 | 0 | 142042 | NP_045581.1 | O50811 |
| TYFKGVSTELKDKK | 0 | 1 | 0 | 142047 | NP_045581.1 | O50811 |
| VKDTPRLIKLIKNS | 0 | 1 | 0 | 142055 | NP_045581.1 | O50811 |
| GLRRDLDASREAKKQVEKAL | 0 | 1 | 0 | 142167 | AAP32473.1 | Q99XV0 |
| DDSKLPIVKVEDKSKLQDTKDKKR | 0 | 1 | 0 | 148558 | YP_303107.1 | Q3YRZ6 |
| DDSKLPVIKVEDKSKLQDTKDKKR | 0 | 1 | 0 | 148560 | YP_303107.1 | Q3YRZ6 |
| DDSKLPVVKVEDKSKLQDTKDKKR | 0 | 1 | 0 | 148562 | YP_303107.1 | Q3YRZ6 |
| DKNKLQDVKDNKPSDVKLPVIKAE | 0 | 1 | 0 | 148566 | YP_507370.1 | Q2GGR3 |
| DKSKLQDVKDNKPSDVKLPVIKAE | 0 | 1 | 0 | 148568 | YP_507370.1 | Q2GGR3 |
| DKSKLRDVKDNKSTDVKLPVVKAE | 0 | 1 | 0 | 148570 | YP_507370.1 | Q2GGR3 |
| DSVKAASKETPPAL | 0 | 1 | 0 | 156262 | AAC45733.1 | G5IXI6 |
| FKSDPKKSDVKTYF | 0 | 1 | 0 | 156293 | AAC45733.1 | G5IXI6 |
| KDGKFAVKDGEKEK | 0 | 1 | 0 | 156341 | AAC45733.1 | G5IXI6 |
| KGAGKLFGKAGAAA | 0 | 1 | 0 | 156343 | AAC45733.1 | G5IXI6 |
| LRGMAKDGKFAVKD | 0 | 1 | 0 | 156370 | AAC45733.1 | G5IXI6 |
| LRKVGDSVKAASKE | 0 | 1 | 0 | 156371 | AAC45733.1 | G5IXI6 |
| QDEMKKDDQIAAAI | 0 | 1 | 0 | 156407 | AAC45733.1 | G5IXI6 |
| SDISSTTGKPDSTG | 0 | 1 | 0 | 156452 | AAC45733.1 | G5IXI6 |
| SQVADKDDPTNKFYQSVIQLGNGF | 0 | 1 | 0 | 156459 | AAC45733.1 | G5IXI6 |
| VKAASKETPPALNK | 0 | 1 | 0 | 156474 | AAC45733.1 | G5IXI6 |
| ARTISEAGQAMASTE | 0 | 1 | 0 | 161597 | P0A564.2 | P9WNK7 |
| LQGQWRGAAGTAAQA | 0 | 1 | 0 | 161652 | CAA17966.1 | P9WNK5 |
| AEANQKLDDGARAALLTQAHDLA | 0 | 1 | 0 | 178474 | Q63ID0 | Q63ID0 |
| ELRPGLQLATYYYYLK | 0 | 1 | 0 | 178752 | Q63ID0 | Q63ID0 |
| GVKGVQRPFTPDWA | 0 | 1 | 0 | 178952 | Q63ID0 | Q63ID0 |
| LAAAAWDALAAELY | 0 | 1 | 0 | 179111 | SGB09933.1 | P9WI05 |
| WSNGQPVTAADFVYAWQR | 0 | 1 | 0 | 179691 | Q63ID0 | Q63ID0 |
| AKKAILITDAAKDKG | 0 | 1 | 0 | 181187 | AAB86554.1 | Q07337 |
| FISCNNSGKDGNTSA | 0 | 1 | 0 | 181204 | AAB86554.1 | Q07337 |
| MTLFLFISCNNSGKD | 0 | 1 | 0 | 181231 | AAB86554.1 | Q07337 |
| TLLAGAYTISKLITQ | 0 | 1 | 0 | 181244 | AAB86554.1 | Q07337 |
| SYSVQDQYQALLQQHAQYLK | 0 | 1 | 0 | 189498 | WP_038772256.1 | Q63RA7 |
| APIYIYGNSYLFRND | 0 | 1 | 0 | 226464 | NP_212463.1 | H7C7K8 |
| IFFLTFLCCNNKERK | 0 | 1 | 0 | 226478 | NP_212463.1 | H7C7K8 |
| KKICEFIQNQWKKNLNIDVE | 0 | 1 | 0 | 226485 | NP_212463.1 | H7C7K8 |
| LAEAGYPNGNGFPILKLKYN | 0 | 1 | 0 | 226492 | NP_212463.1 | H7C7K8 |
| QKITDKVDNLNQAVSETKL | 0 | 1 | 0 | 230582 | GI:22335942 | P55980 |
| SETKLTGDFSKVEQALAEL | 0 | 1 | 0 | 230586 | GI:22335942 | P55980 |
| SIGVLGYQKTVDHTKVNSKLSLF | 0 | 1 | 0 | 236322 | AAV70486.1 | Q6NK15 |
| NATAMVAQINAVNKPQTVS | 0 | 1 | 0 | 429098 | YP_109915.1 | H7C7G3 |
| TRMQTQINGLNQGVSNAND | 0 | 1 | 0 | 429129 | YP_109915.1 | H7C7G3 |
| AAWGGSGSEAYQGVQQKWDATA | 0 | 1 | 0 | 429190 | P0A564.2 | P9WNK7 |
| AVVPVLAYAAARLLL | 0 | 1 | 0 | 429447 | Q73SP6 | Q73SP6 |
| NYPASDDYRASASNGSDDASAH | 0 | 1 | 0 | 431692 | NP_216500.1 | P9WP43 |
| PGRRPFTRKELQ | 0 | 1 | 0 | 431703 | Q745A5 | Q745A5 |
| PKTYCEELKGTDTGQACQ | 0 | 1 | 0 | 431708 | P0A5Q4.1 | P9WIN9 |
| RFQEAANKQKQELDEISTNIRQA | 0 | 1 | 0 | 432087 | CAA17966.1 | P9WNK5 |
| SAATSSTPREAPYELN | 0 | 1 | 0 | 432168 | P0A5Q4.1 | P9WIN9 |
| VQYSRADEEQQQAL | 0 | 1 | 0 | 432955 | CAA17966.1 | P9WNK5 |
| VTSIHSLLDEGKQSLTKL | 0 | 1 | 0 | 433149 | P0A564.2 | P9WNK7 |
| AVVAARAQGRYTTDDGYIFN | 0 | 1 | 0 | 489486 | COC35529.1 | Q8DQ08 |
| GDAYIVPHGDHYHYIPKNEL | 0 | 1 | 0 | 489548 | AAK06760.1 | Q8DQ08 |
| GVNHVIPYELFAGDGMLTRL | 0 | 1 | 0 | 489571 | WP_041179080.1 | Q8DRK2 |
| HYYNGKVPYDAIISEELLMK | 0 | 1 | 0 | 489580 | CIV64423.1 | Q8DQ08 |
| LPPLEGLTKGKYFYEVDLNG | 0 | 1 | 0 | 489651 | WP_044727941.1 | Q8DRK2 |
| RTPNGVAIPHGDHYHFIPYS | 0 | 1 | 0 | 489761 | CMX28928.1 | Q8DQ07 |
| TAGVNQVIPYELFAGDGMLT | 0 | 1 | 0 | 489800 | WP_044727941.1 | Q8DRK2 |
| VSPTDYSTASYNALGPVLET | 0 | 1 | 0 | 489842 | WP_044814256.1 | Q8DQN5 |
| YVTSHGDHYHYYNGKVPYDA | 0 | 1 | 0 | 489865 | CMX28928.1 | Q8DQ07 |
| YYPGEKVHYDQILEKDGYKW | 0 | 1 | 0 | 489867 | WP_041178981.1 | Q8DMZ4 |
| GRIFNQFTASYPLLR | 0 | 1 | 0 | 540209 | EGE26266.1 | D5VCH6 |
| RIFNQFTASYPLLRS | 0 | 1 | 0 | 540259 | EGE26266.1 | D5VCH6 |
| RLNIKITPNLVSKLL | 0 | 1 | 0 | 540260 | EGE27725.1 | D5V9Y8 |
| ALDITLPMPPR | 0 | 1 | 0 | 547962 | P9WIM9.1 | P9WIM9 |
| WTQVPDPNVPDAFVVIADR | 0 | 1 | 0 | 548667 | P9WIM9.1 | P9WIM9 |
| KGPNLVEISKKITDSN | 0 | 1 | 0 | 559957 | CAA57244.1 | Q07337 |
| GAIPGGWWLTFGQIL | 0 | 1 | 0 | 690384 | SGB09933.1 | P9WI05 |
| GGLFGIGGAGGGCGS | 0 | 1 | 0 | 690417 | WP_021036394.1 | Q79FE6 |
| IPIPGLPDIPGLPDF | 0 | 1 | 0 | 690848 | WP_003907147.1 | P9WJD3 |
| MFSGFDPWLPSLGNP | 0 | 1 | 0 | 691529 | SGA45816.1 | P9WI47 |
| NKPVLVDFWATWCGP | 0 | 1 | 0 | 691684 | WP_003907254.1 | P9WG67 |
| TRYSSAYELEGAVKR | 0 | 1 | 0 | 692695 | P30234.2 | P9WQB1 |
| VYVAQKRKISDGDKL | 0 | 1 | 0 | 693146 | AAA21416.1 | P9WGY9 |
| AGEHEAAAAGYVCAL | 0 | 1 | 0 | 693779 | WP_072466325.1 | P9WHY9 |
| AGPPQRWFVVWLGTA | 0 | 1 | 0 | 693795 | SGL61026.1 | P9WIR7 |
| CGSAVAIGGDGGAGG | 0 | 1 | 0 | 694015 | WP_075956638.1 | Q79FE6 |
| FEAAFAMTVPPAEVA | 0 | 1 | 0 | 694405 | SGO07857.1 | Q79FA9 |
| FFQELADLDRQLISL | 0 | 1 | 0 | 694418 | WP_075841292.1 | P9WJE1 |
| FGELLFTNPTGAFQF | 0 | 1 | 0 | 694421 | SGM78117.1 | P9WI23 |
| FLEETFAAYDQYLSA | 0 | 1 | 0 | 694439 | WP_072466325.1 | P9WHY9 |
| GYATGGMSTAALSSG | 0 | 1 | 0 | 694812 | SGI07003.1 | P9WI41 |
| IMQLTTEQWLGPASM | 0 | 1 | 0 | 694968 | SGO07857.1 | Q79FA9 |

| | | | | | | |
|---|---|---|---|---|---|---|
| LSTLTGEEWHGPASA | 0 | 1 | 0 | 695386 | SGN39511.1 | L0T7Y7 |
| NANLGDYNVGSGNVG | 0 | 1 | 0 | 695528 | SIP65580.1 | Q79FI9 |
| NSNTGGFNMGQYNTG | 0 | 1 | 0 | 695595 | WP_072532728.1 | I6Y7L4 |
| SAYASPRIGQPVGSE | 0 | 1 | 0 | 696332 | SGI07003.1 | P9WI41 |
| SSTAATFASGPSGLL | 0 | 1 | 0 | 696515 | CCP44573.1 | L0T7Y7 |
| STKIVIAGGFGSGKT | 0 | 1 | 0 | 696528 | WP_070900639.1 | O50391 |
| TVLVGGLRVLGANYK | 0 | 1 | 0 | 696742 | WP_075963877.1 | P9WIE5 |
| SYSVKDEYQPLMQQHAQYLK | 0 | 1 | 0 | 697122 | WP_077199646.1 | B1JWH2 |
| AAILKTNGTKDKG | 0 | 1 | 0 | 745097 | ACC94173.1 | Q07337 |
| DLENLEKQFDIKFD | 0 | 1 | 0 | 745099 | ACK75301.1 | Q9S070 |
| EMKEESPGLFDKGNSILE | 0 | 1 | 0 | 745104 | AAL84596.1 | O50835 |
| FDWNTPDPRIGFKDNML | 0 | 1 | 0 | 745105 | WP_044108210.1 | Q2GIP9 |
| GITKIKEEFDKKVAEIKA | 0 | 1 | 0 | 745108 | WP_012664925.1 | Q9RZX8 |
| KAILTTDAAKDKG | 0 | 1 | 0 | 745113 | CAA59253.1 | Q07337 |
| KFDKLENHPFLGYPYK | 0 | 1 | 0 | 745115 | ACL33989.1 | Q9S0D2 |
| KNPIAAAIGDKDG | 0 | 1 | 0 | 745116 | AAC45733.1 | G5IXI6 |
| LNVCYTDAIAALAKAKN | 0 | 1 | 0 | 745117 | WP_010890334.1 | O50850 |
| NSGKDGNTSANSA | 0 | 1 | 0 | 745120 | NP_047005.1 | Q07337 |
| NSVKELTSPVVVES | 0 | 1 | 0 | 745122 | AAM19715.1 | Q07337 |
| QTLSKEKAEELLQHAE | 0 | 1 | 0 | 745125 | WP_012672397.1 | O50889 |
| RKEKDGIETGLNAG | 0 | 1 | 0 | 745126 | AAF40177.1 | H7C7N5 |
| RKVLGAITGLIGDAV | 0 | 1 | 0 | 745127 | AAC45733.1 | G5IXI6 |
| SDIDIDSLVTDKVVAA | 0 | 1 | 0 | 745128 | CAA54797.1 | Q45013 |
| SEKITKLTPEELENLAK | 0 | 1 | 0 | 745129 | NP_045581.1 | O50811 |
| VAYNMYINGELDFL | 0 | 1 | 0 | 745132 | AAB97671.1 | O51307 |
| ARFNDAQSQPFTTEP | 0 | 1 | 0 | 746003 | CKM53108.1 | I6Y293 |
| AWALRMSPDGNVWGA | 0 | 1 | 0 | 746049 | COV25996.1 | I6Y293 |
| EPDVVRKDTHAHAWA | 0 | 1 | 0 | 746495 | COV25996.1 | I6Y293 |
| FPQGGGFPRNNDDKT | 0 | 1 | 0 | 746954 | COV25996.1 | I6Y293 |
| GAVEEISISAEPKVK | 0 | 1 | 0 | 747131 | CKM53108.1 | I6Y293 |
| HAHAWALRMSPDGNV | 0 | 1 | 0 | 747305 | COV25996.1 | I6Y293 |
| KDTHAHAWALRMSPD | 0 | 1 | 0 | 748225 | COV25996.1 | I6Y293 |
| LFPQGGGFPRNNDDK | 0 | 1 | 0 | 748529 | COV25996.1 | I6Y293 |
| PTTALSGIGSGGGLC | 0 | 1 | 0 | 750214 | CKM53108.1 | I6Y293 |
| PVDPAGDGGLMDIVL | 0 | 1 | 0 | 750219 | CKM53108.1 | I6Y293 |
| QSQPFTTEPELRPQP | 0 | 1 | 0 | 750740 | CKM53108.1 | I6Y293 |
| SGCARFNDAQSQPFT | 0 | 1 | 0 | 751051 | CKM53108.1 | I6Y293 |
| SSTPPPPPPLPPVPF | 0 | 1 | 0 | 751510 | CKM53108.1 | I6Y293 |
| TVAVRLAPSTGAVTG | 0 | 1 | 0 | 751960 | COV25996.1 | I6Y293 |
| VRLAPSTGAVTGEPD | 0 | 1 | 0 | 752785 | COV25996.1 | I6Y293 |
| YISTPTDNRVVRVAD | 0 | 1 | 0 | 752949 | CKM53108.1 | I6Y293 |
| FQMGAKPTTDTGNSAAPSTLTARE | 0 | 1 | 0 | 753350 | ANH58353.1 | Q46409 |
| PETISDPENRNKPSAE | 0 | 1 | 0 | 753408 | CRI42875.1 | Q9Z7F3 |
| MIRLVRHSIALVAAGLAAAL | 0 | 1 | 0 | 753510 | CNJ09075.1 | I6X7X3 |
| PPNDPAWQRNDPILQ | 0 | 1 | 0 | 767376 | CAA43269.1 | P31951 |
| AGTTLETTTTNNTDGS | 0 | 1 | 0 | 780258 | CRI45923.1 | Q9Z899 |
| APSLTEAESDQTDQTE | 0 | 1 | 0 | 780263 | WP_071804540.1 | O84419 |
| AQENSTAKRRRRRAAV | 0 | 1 | 0 | 780265 | CRI33036.1 | Q9Z827 |
| EEQNNNDASNQGESAN | 0 | 1 | 0 | 780277 | WP_071804540.1 | O84419 |
| EKVEEVEPAPEQKDNN | 0 | 1 | 0 | 780283 | AAQ73610.1 | O84818 |
| FGVKGTTVNANELPNV | 0 | 1 | 0 | 780289 | AAD33511.1 | P27455 |
| FKGKNGSADKVILVTQ | 0 | 1 | 0 | 780290 | WP_011324578.1 | P0DJI4 |
| FSKFSGDWDSLVEPMV | 0 | 1 | 0 | 780291 | WP_055347997.1 | O84462 |
| GNLKQNKPTEGTSKENGFMARL | 0 | 1 | 0 | 780293 | CAX10183.1 | O84623 |
| HGQVSYGRNHHNMTTK | 0 | 1 | 0 | 780298 | CRI45923.1 | Q9Z899 |
| HTKKTTPGSIPSKVFS | 0 | 1 | 0 | 780299 | WP_010883340.1 | Q9Z7K3 |
| IMDITEIPSINPEFVE | 0 | 1 | 0 | 780304 | CRH49556.1 | O84801 |
| INNTAKRSGGGIYAPK | 0 | 1 | 0 | 780305 | CRI45923.1 | Q9Z899 |
| KAFFTQPNNKMARVVN | 0 | 1 | 0 | 780307 | CRH44689.1 | O84534 |
| KRSPEEIEGAARPSDQ | 0 | 1 | 0 | 780312 | WP_009873573.1 | P0DPS6 |
| KTNPDGSFQLDPVSQQRTLLSP | 0 | 1 | 0 | 780313 | CAX10183.1 | O84623 |
| LLRGSNNYVYNSNCEL | 0 | 1 | 0 | 780319 | AAF38159.1 | O86164 |
| QALFASEDGDLSPESS | 0 | 1 | 0 | 780330 | WP_071804673.1 | O84818 |
| RSNPKLEQKDSGENIN | 0 | 1 | 0 | 780338 | WP_010895395.1 | Q9Z6U5 |
| SFNLVGLFGDNENQKT | 0 | 1 | 0 | 780342 | APQ41636.1 | Q46409 |
| SGAFDDINKDNSRSRS | 0 | 1 | 0 | 780343 | WP_009873573.1 | P0DPS6 |
| SHMKAASQKTQEGDEG | 0 | 1 | 0 | 780345 | CRH44689.1 | O84534 |
| TVSETQQQQLSTIETT | 0 | 1 | 0 | 780359 | CAX10183.1 | O84623 |
| TVVNYTNSASAPNVTV | 0 | 1 | 0 | 780361 | WP_010725202.1 | O84462 |
| VVESLSRRNSLVDQTQ | 0 | 1 | 0 | 780381 | CRH48743.1 | O84449 |
| AAGNIVPNTTGAAKAI | 0 | 1 | 0 | 840797 | WP_070112929.1 | Q181T9 |
| AAKAIGLVIPSLKGKL | 0 | 1 | 0 | 840798 | WP_070112929.1 | Q181T9 |
| FTSKDKAEAHIQAGAK | 0 | 1 | 0 | 840839 | WP_070112929.1 | Q181T9 |
| KGKLDGAAQRVPVVTG | 0 | 1 | 0 | 840863 | WP_102815282.1 | Q181T9 |
| TLDGPHPKGDLRRARA | 0 | 1 | 0 | 840950 | WP_070112929.1 | Q181T9 |
| TTNCLAPMAKVLNDKY | 0 | 1 | 0 | 840955 | WP_070112929.1 | Q181T9 |
| EEPIYGLTKPVTTFDPVYT | 0 | 1 | 0 | 871965 | WP_001162077.1 | Q8F2Z1 |
| GFISCDLFIRYEMKE | 0 | 1 | 0 | 910468 | AAL84596.1 | O50835 |
| KGKSKVSRKEPYIHS | 0 | 1 | 0 | 910479 | AAL84596.1 | O50835 |
| KIARKKGKSKVSRKE | 0 | 1 | 0 | 910480 | AAL84596.1 | O50835 |
| KKGKGKIARKKGKSK | 0 | 1 | 0 | 910481 | AAL84596.1 | O50835 |
| KKPMNKKGKGKIARK | 0 | 1 | 0 | 910482 | AAL84596.1 | O50835 |
| ESDVKTGNNNNVAL | 0 | 1 | 0 | 945065 | AAO59310.1 | F5GUL7 |
| ETDVQSGTKLKVAL | 0 | 1 | 0 | 945074 | AAQ16674.1 | F5GUP5 |
| GPTPVHNDNAKAVAK | 0 | 1 | 0 | 945174 | AAO59310.1 | F5GUL7 |
| GPVPVQNDNAEAVAK | 0 | 1 | 0 | 945175 | AAQ16674.1 | F5GUP5 |
| IVGAGDSSNAGTSL | 0 | 1 | 0 | 945263 | WP_044108210.1 | Q2GIP9 |
| KAVGVSHPTIDGKV | 0 | 1 | 0 | 945273 | AAO59310.1 | F5GUL7 |
| KTSGKDIVQFAKAVE | 0 | 1 | 0 | 945332 | WP_044108210.1 | Q2GIP9 |
| LVQELTPEEKTIVAG | 0 | 1 | 0 | 945413 | AAO59310.1 | F5GUL7 |
| TGSSSPQVLRDFINA | 0 | 1 | 0 | 945841 | AAO59310.1 | F5GUL7 |

| | | | | | | |
|---|---|---|---|---|---|---|
| TSKSNNDNAKAVAGD | 0 | 1 | 0 | 945871 | WP_044108210.1 | Q2GIP9 |
| VEKAQALHDFVSNTL | 0 | 1 | 0 | 945893 | WP_044108210.1 | Q2GIP9 |
| ATGKKGSETPTGKTKV | 0 | 1 | 0 | 952794 | WP_009890957 | Q183E7 |
| DGLSVTGWQEIDGNKY | 0 | 1 | 0 | 952801 | WP_009890957 | Q183E7 |
| EFRVATGKKGSETPTG | 0 | 1 | 0 | 952814 | WP_009890957 | Q183E7 |
| FDTAKKISSVGNWNAD | 0 | 1 | 0 | 952826 | WP_009890957 | Q183E7 |
| FVNNKLVKEFRVATGK | 0 | 1 | 0 | 952833 | WP_009890957 | Q183E7 |
| GGCIRMHNKDVRWLFD | 0 | 1 | 0 | 952838 | WP_009890957 | Q183E7 |
| GKKGSETPTGKTKVVN | 0 | 1 | 0 | 952841 | WP_009890957 | Q183E7 |
| GKSYYLDKDGKMLTGS | 0 | 1 | 0 | 952842 | WP_009890957 | Q183E7 |
| GSDVIIDYSNDSYVKI | 0 | 1 | 0 | 952846 | WP_009890957 | Q183E7 |
| GTYQKNSWLKVNGKMY | 0 | 1 | 0 | 952849 | WP_009890957 | Q183E7 |
| GYFVNNKLVKEFRVAT | 0 | 1 | 0 | 952850 | WP_009890957 | Q183E7 |
| KGSETPTGKTKVVNKI | 0 | 1 | 0 | 952865 | WP_009890957 | Q183E7 |
| KLVKEFRVATGKKGSE | 0 | 1 | 0 | 952867 | WP_009890957 | Q183E7 |
| KNTLGYFVNNKLVKEF | 0 | 1 | 0 | 952868 | WP_009890957 | Q183E7 |
| KTKVVNKIKNRPYYKG | 0 | 1 | 0 | 952872 | WP_009890957 | Q183E7 |
| KVVNKIKNRPYYKGNI | 0 | 1 | 0 | 952877 | WP_009890957 | Q183E7 |
| NKKYYLGTDGARVSGW | 0 | 1 | 0 | 952902 | WP_009890957 | Q183E7 |
| NNKLVKEFRVATGKKG | 0 | 1 | 0 | 952904 | WP_009890957 | Q183E7 |
| QTGWQEKNGKKYYLGS | 0 | 1 | 0 | 952924 | WP_009890957 | Q183E7 |
| RVATGKKGSETPTGKT | 0 | 1 | 0 | 952930 | WP_009890957 | Q183E7 |
| SETPTGKTKVVNKIKN | 0 | 1 | 0 | 952938 | WP_009890957 | Q183E7 |
| SRKNTLGYFVNNKLVK | 0 | 1 | 0 | 952947 | WP_009890957 | Q183E7 |
| TFNEDGTINPTWDTII | 0 | 1 | 0 | 952960 | WP_009890957 | Q183E7 |
| TGKTKVVNKIKNRPYY | 0 | 1 | 0 | 952961 | WP_009890957 | Q183E7 |
| TGWKTENGKKYYVKSD | 0 | 1 | 0 | 952962 | WP_009890957 | Q183E7 |
| TLGYFVNNKLVKEFRV | 0 | 1 | 0 | 952964 | WP_009890957 | Q183E7 |
| TPTGKTKVVNKIKNRP | 0 | 1 | 0 | 952966 | WP_009890957 | Q183E7 |
| VKEFRVATGKKGSETP | 0 | 1 | 0 | 952972 | WP_009890957 | Q183E7 |
| VNKIKNRPYYKGNIPG | 0 | 1 | 0 | 952977 | WP_009890957 | Q183E7 |
| AGQATSPEEPIYAQVAKKV | 0 | 1 | 0 | 956318 | GI:217315723 | P55980 |
| AGQATSPEEPIYAQVNKKK | 0 | 1 | 0 | 956320 | BAF45300.1 | P55980 |
| AGQATSPEEPIYATIDFDE | 0 | 1 | 0 | 956322 | BAF45300.1 | P55980 |
| ATSAINRKIDRINKIASAG | 0 | 1 | 0 | 956329 | GI:22335942 | P55980 |
| DYSDSFKFSTKLNNAVKDI | 0 | 1 | 0 | 956340 | GI:22335393 | P55980 |
| ESFRKFGDQRYQIFTSWVS | 0 | 1 | 0 | 956347 | GI:22335393 | P55980 |
| GMLTQKNPEWLKLVNDKIV | 0 | 1 | 0 | 956357 | GI:22335393 | P55980 |
| GRSASPEPIYAQVAKKV | 0 | 1 | 0 | 956359 | WP_000180527.1 | P55980 |
| IASAGKGVGGFSGAGRSAS | 0 | 1 | 0 | 956368 | GI:22335942 | P55980 |
| INSNVQSGTINEKATGMLTQ | 0 | 1 | 0 | 956374 | GI:446102890 | P55980 |
| IQGLPPEARDLLDERGNFF | 0 | 1 | 0 | 956375 | GI:22335393 | P55980 |
| IYAQVAKKVSAKIDQLNEA | 0 | 1 | 0 | 956379 | AAZ23952.1 | P55980 |
| IYAQVNKKKAGQATSPEEP | 0 | 1 | 0 | 956380 | GI:217315723 | P55980 |
| KANVEHGVKNTTKSGFQKS | 0 | 1 | 0 | 956381 | GI:12225004 | P55980 |
| KARQEAEKNAEPAGGDWLD | 0 | 1 | 0 | 956382 | GI:22335402 | P55980 |
| KQSSDLKETLNQEPRPDFE | 0 | 1 | 0 | 956394 | GI:22335393 | P55980 |
| KSPDKGVGATNGVSHLEAN | 0 | 1 | 0 | 956395 | GI:22335402 | P55980 |
| LDVEGVADKDPNYKFNQLL | 0 | 1 | 0 | 956396 | GI:22335393 | P55980 |
| LLYGDNGGPEARHDWNATV | 0 | 1 | 0 | 956399 | GI:22335393 | P55980 |
| LTKNFSDIRKELNEKLFGN | 0 | 1 | 0 | 956404 | GI:22335942 | P55980 |
| MFVNYSNFKYTNASKSPD | 0 | 1 | 0 | 956407 | GI:22335447 | P55980 |
| MTNETIDQTTTPDQTPNQT | 0 | 1 | 0 | 956410 | GI:5921294 | P55980 |
| NGLKNNTEPIYAQVNKKKA | 0 | 1 | 0 | 956412 | AAZ23952.1 | P55980 |
| NKKKAGQATSPEEPIYAQV | 0 | 1 | 0 | 956413 | GI:217315723 | P55980 |
| NNTEPIYAQVNKKKAGQAT | 0 | 1 | 0 | 956416 | GI:217315723 | P55980 |
| NSDLQKSVKNGVNGTLVGN | 0 | 1 | 0 | 956417 | GI:22335942 | P55980 |
| QAKLDNYATNSHTRINSN | 0 | 1 | 0 | 956428 | GI:22335393 | P55980 |
| QTPNQTDFVPQRFINNLQV | 0 | 1 | 0 | 956433 | GI:5921294 | P55980 |
| REEYANKAIKNPTKKNQYF | 0 | 1 | 0 | 956434 | GI:5921286 | P55980 |
| RGNFFKFTLGDVEMLDVEG | 0 | 1 | 0 | 956437 | GI:22335393 | P55980 |
| SAAVNDLSKVGLSREQELT | 0 | 1 | 0 | 956448 | GI:22335942 | P55980 |
| SEYDKIGFNQKNMKDYSDS | 0 | 1 | 0 | 956450 | GI:22335393 | P55980 |
| SNNNNNGLKNNTEPIYAQV | 0 | 1 | 0 | 956455 | GI:22335942 | P55980 |
| SPEEPIYAQVAKKVSAKID | 0 | 1 | 0 | 956456 | AAZ23952.1 | P55980 |
| TLVGNGLSKTEATTLTKNF | 0 | 1 | 0 | 956459 | GI:22335942 | P55980 |
| VAKKVSAKIDQLNEATSAI | 0 | 1 | 0 | 956464 | GI:22335942 | P55980 |
| AAGIALRAMAKD | 0 | 1 | 0 | 958538 | WP_025444236.1 | A0A1D8TF85 |
| AGIALRAMAKDG | 0 | 1 | 0 | 958543 | WP_025444236.1 | A0A1D8TF85 |
| AKDAASIAAAKV | 0 | 1 | 0 | 958547 | A0A481YLK4 | A0A1D8TF93 |
| AKDAASIASAKK | 0 | 1 | 0 | 958548 | A0A481YHP3 | A0A1D8TF92 |
| AKIETAKDAASI | 0 | 1 | 0 | 958550 | A0A481YLK4 | A0A1D8TF93 |
| AKKDAIIAAGIA | 0 | 1 | 0 | 958551 | WP_025444236.1 | A0A1D8TF85 |
| AKNAAEIAAAKA | 0 | 1 | 0 | 958552 | WP_025444482.1 | A0A1D8TFY1 |
| ALRAMAKDGKFI | 0 | 1 | 0 | 958554 | WP_025444236.1 | A0A1D8TF85 |
| AMAKDGKFIVKD | 0 | 1 | 0 | 958556 | WP_025444236.1 | A0A1D8TF85 |
| ASKAFIDKVKGENASLG | 0 | 1 | 0 | 958561 | ALZ45557.1 | A0A1D8TF77 |
| EAKDAASIASAK | 0 | 1 | 0 | 958583 | A0A481YHP3 | A0A1D8TF92 |
| EAKIETAKDAAS | 0 | 1 | 0 | 958584 | A0A481YLK4 | A0A1D8TF93 |
| EAKNAAEIAAAK | 0 | 1 | 0 | 958585 | WP_025444482.1 | A0A1D8TFY1 |
| EEAKDAASIASA | 0 | 1 | 0 | 958586 | A0A481YHP3 | A0A1D8TF92 |
| EEAKNAAEIAAA | 0 | 1 | 0 | 958587 | WP_025444482.1 | A0A1D8TFY1 |
| EIKIEEAKNAAE | 0 | 1 | 0 | 958588 | WP_025444482.1 | A0A1D8TFY1 |
| ETAKDAASIAAA | 0 | 1 | 0 | 958591 | A0A481YLK4 | A0A1D8TF93 |
| GIALRAMAKDGK | 0 | 1 | 0 | 958606 | WP_025444236.1 | A0A1D8TF85 |
| IAAGIALRAMAK | 0 | 1 | 0 | 958616 | WP_025444236.1 | A0A1D8TF85 |
| IALRAMAKDGKF | 0 | 1 | 0 | 958617 | WP_025444236.1 | A0A1D8TF85 |
| IEEAKDAASIAS | 0 | 1 | 0 | 958619 | A0A481YHP3 | A0A1D8TF92 |
| IEEAKNAAEIAA | 0 | 1 | 0 | 958620 | WP_025444482.1 | A0A1D8TFY1 |
| IETAKDAASIAA | 0 | 1 | 0 | 958622 | A0A481YLK4 | A0A1D8TF93 |
| IGKKIKNADELDTVAD | 0 | 1 | 0 | 958623 | A0A2D1YWX0 | A0A1D8TF91 |

| | | | | | | |
|---|---|---|---|---|---|---|
| IKDAVDFAASVK | 0 | 1 | 0 | 958627 | A0A2D1YWX0 | A0A1D8TF91 |
| IKIEEAKNAAEI | 0 | 1 | 0 | 958632 | WP_025444482.1 | A0A1D8TFY1 |
| KDAVDFAASVKE | 0 | 1 | 0 | 958639 | A0A2D1YWX0 | A0A1D8TF91 |
| KDSVEFAASVKE | 0 | 1 | 0 | 958644 | ALN43425.1 | A0A1D8TFC5 |
| KIEEAKNAAEIA | 0 | 1 | 0 | 958645 | WP_025444482.1 | A0A1D8TFY1 |
| KIETAKDAASIA | 0 | 1 | 0 | 958646 | A0A481YLK4 | A0A1D8TF93 |
| KIKDAVDFAASV | 0 | 1 | 0 | 958648 | A0A2D1YWX0 | A0A1D8TF91 |
| KKDAIIAAGIAL | 0 | 1 | 0 | 958653 | WP_025444236.1 | A0A1D8TF85 |
| KKIKDAVDFAAS | 0 | 1 | 0 | 958656 | A0A2D1YWX0 | A0A1D8TF91 |
| KNIEEAKDAASI | 0 | 1 | 0 | 958667 | A0A481YHP3 | A0A1D8TF92 |
| KTTTKKNDVGVYFNSLG | 0 | 1 | 0 | 958671 | AGS80213.1 | A0A1D8TF96 |
| LKFAKGGSDAHLSNSAN | 0 | 1 | 0 | 958678 | AGS80213.1 | A0A1D8TF96 |
| LRAMAKDGKFIV | 0 | 1 | 0 | 958682 | WP_025444236.1 | A0A1D8TF85 |
| NIEEAKDAASIA | 0 | 1 | 0 | 958690 | A0A481YHP3 | A0A1D8TF92 |
| RAMAKDGKFIVK | 0 | 1 | 0 | 958715 | WP_025444236.1 | A0A1D8TF85 |
| SKKIKDASELMESVKE | 0 | 1 | 0 | 958728 | BAA11304.1 | A0A1D8TF77 |
| SKKIKDAVDFAA | 0 | 1 | 0 | 958729 | A0A2D1YWX0 | A0A1D8TF91 |
| SKKIKDAVEFAANVKE | 0 | 1 | 0 | 958731 | AGS80212.1 | A0A1D8TFC5 |
| SKKIKDSVEFAA | 0 | 1 | 0 | 958733 | ALN43425.1 | A0A1D8TFC5 |
| SKNIEEAKDAAS | 0 | 1 | 0 | 958734 | A0A481YHP3 | A0A1D8TF92 |
| TAKDAASIAAAK | 0 | 1 | 0 | 958740 | A0A481YLK4 | A0A1D8TF93 |
| TKLEVLEKTAEISNELNGK | 0 | 1 | 0 | 958742 | ALU64345.1 | A0A1D8TF77 |
| YLPKNKIDSAD | 0 | 1 | 0 | 1066021 | P0A074 | Q2FVK3 |
| YLPKNKIESTN | 0 | 1 | 0 | 1066022 | SHD09795.1 | Q2FVK2 |
| YLPKNKIETTD | 0 | 1 | 0 | 1066023 | Q2FXB0 | Q2FXB0 |
| LREKMSEKGTSTHVIT | 0 | 1 | 0 | 1067693 | WP_009893549.1 | Q185D0 |
| REGATLAEKLSKKGIK | 0 | 1 | 0 | 1068156 | WP_009893549.1 | Q185D0 |
| AIKRSTHYGNSYLDG | 0 | 1 | 0 | 1074844 | WP_060578220.1 | Q2FVK2 |
| ATRRTTHYGNSYLEG | 0 | 1 | 0 | 1074861 | WP_086046373.1 | Q2FVK2 |
| KDSYTKKAKAFKKEA | 0 | 1 | 0 | 1125077 | WP_050151559.1 | Q8DZ80 |
| EVENPDPKVAQE | 0 | 1 | 0 | 1308568 | WP_000620910.1 | Q8F2W1 |
| ATGPKQDSCFGRMY | 0 | 1 | 0 | 1310293 | GI:40725 | P0CC04 |
| DPLGRRTPNYQSKNP | 0 | 1 | 0 | 1310334 | GI:809287703 | O84883 |
| EFTINKPKGYVGKEF | 0 | 1 | 0 | 1310353 | GI:129150 | Q46409 |
| FDTTTLNPTIAGAGDVK | 0 | 1 | 0 | 1310387 | GI:1450244218 | Q46409 |
| LEGSQGDTADTGT | 0 | 1 | 0 | 1310567 | GI:1279531111 | O84419 |
| LNQVLKNSKGENV | 0 | 1 | 0 | 1310608 | GI:497558025 | P18584 |
| MNSCSSSRGNQPADESIY | 0 | 1 | 0 | 1310646 | GI:497557232 | Q9S529 |
| NIYESIGGSRTSGPEN | 0 | 1 | 0 | 1310668 | GI:1279507774 | O84462 |
| QAYNCATHRNG | 0 | 1 | 0 | 1310731 | GI:815044328 | O84145 |
| RTTSSSGVSED | 0 | 1 | 0 | 1310779 | GI:809291693 | O84701 |
| VINDLPLGRSI | 0 | 1 | 0 | 1310890 | GI:1105030046 | O84608 |
| MTVKVGINGFGRIGRLAFRRIK | 0 | 1 | 0 | 1329775 | EAC2648702.1 | Q8Y4I1 |
| KAALTRTASNMNAAA | 0 | 1 | 0 | 1334921 | WP_010924727.1 | P9WJC3 |
| EYWGAQLNAMKGDLQSSL | 0 | 1 | 0 | 1380202 | P0C5B9.1 | P9WQP1 |
| KFQDAYNAAGGHNAVF | 0 | 1 | 0 | 1383366 | P0C5B9.1 | P9WQP1 |
| LLDGLRAQDDYNGW | 0 | 1 | 0 | 1384633 | P0C5B9.1 | P9WQP1 |
| SMAGSSAMILAAYHPQQF | 0 | 1 | 0 | 1388770 | P0C5B9.1 | P9WQP1 |
| VRSSNLKFQDAY | 0 | 1 | 0 | 1390855 | P0C5B9.1 | P9WQP1 |
| ASGNKVASVYGKNLEA | 0 | 1 | 0 | 1562434 | CBE01872.1 | Q18CX9 |
| ASMIRDKEKVDKAKQE | 0 | 1 | 0 | 1562457 | CBE01872.1 | Q18CX9 |
| ASVIYVPPAGAADAII | 0 | 1 | 0 | 1562509 | WP_035454724.1 | A1UQW0 |
| DTGTEGGGGTEGGGGT | 0 | 1 | 0 | 1566255 | ABA60112.1 | A1UU13 |
| EGILRNDDELRGFVED | 0 | 1 | 0 | 1567310 | CBE01872.1 | Q18CX9 |
| EGKEKTGANASVIYVP | 0 | 1 | 0 | 1567312 | WP_035454724.1 | A1UQW0 |
| EKTGANASVIYVPPAG | 0 | 1 | 0 | 1567656 | WP_035454724.1 | A1UQW0 |
| EQALAYHGTQMVGGIN | 0 | 1 | 0 | 1568631 | WP_035454724.1 | A1UQW0 |
| ETWTGAKGETLPIFAS | 0 | 1 | 0 | 1569211 | WP_035454724.1 | A1UQW0 |
| GADRIMPYVSGGVAYT | 0 | 1 | 0 | 1571376 | ABA60112.1 | A1UU13 |
| GGGSASKAVRSEALDV | 0 | 1 | 0 | 1571683 | ABA60112.1 | A1UU13 |
| GGGTEGGGGSASKAVR | 0 | 1 | 0 | 1571685 | ABA60112.1 | A1UU13 |
| GGGTEGGGGTEGGGGS | 0 | 1 | 0 | 1571686 | ABA60112.1 | A1UU13 |
| GGINPKKGGETWTGAK | 0 | 1 | 0 | 1571688 | WP_035454724.1 | A1UQW0 |
| GSAILKGTKDTGTEGG | 0 | 1 | 0 | 1572288 | ABA60112.1 | A1UU13 |
| GTKDTGTEGGGGTEGG | 0 | 1 | 0 | 1572457 | ABA60112.1 | A1UU13 |
| HGTQMVGGINPKKGGE | 0 | 1 | 0 | 1572826 | WP_035454724.1 | A1UQW0 |
| IFASVAEGKEKTGANA | 0 | 1 | 0 | 1574652 | WP_035454724.1 | A1UQW0 |
| IGFGADRIMPYVSGGV | 0 | 1 | 0 | 1574737 | ABA60112.1 | A1UU13 |
| ILKGTKDTGTEGGGGT | 0 | 1 | 0 | 1575006 | ABA60112.1 | A1UU13 |
| IQSAFFGDGKNIIALR | 0 | 1 | 0 | 1575432 | CBE01872.1 | Q18CX9 |
| ISRLVEKMNDKEKRLY | 0 | 1 | 0 | 1575565 | CBE01872.1 | Q18CX9 |
| KAVRSEALDVLASGTI | 0 | 1 | 0 | 1575897 | ABA60112.1 | A1UU13 |
| KGETLPIFASVAEGKE | 0 | 1 | 0 | 1576155 | WP_035454724.1 | A1UQW0 |
| KGGETWTGAKGETLPI | 0 | 1 | 0 | 1576161 | WP_035454724.1 | A1UQW0 |
| KLGINESENYNKKGQI | 0 | 1 | 0 | 1576348 | CBE01872.1 | Q18CX9 |
| KSMEKLSSGVRIKRA | 0 | 1 | 0 | 1576645 | CBE01876.1 | Q18CX7 |
| KVYKTLAGYSSNYDDK | 0 | 1 | 0 | 1576803 | CBE01872.1 | Q18CX9 |
| LAYHGTQMVGGINPKK | 0 | 1 | 0 | 1576984 | WP_035454724.1 | A1UQW0 |
| LIANNQMGRNVNAQS | 0 | 1 | 0 | 1577468 | CBE01876.1 | Q18CX7 |
| NILQRMRTLSVQSSN | 0 | 1 | 0 | 1581724 | CBE01876.1 | Q18CX7 |
| NPKKGGETWTGAKGET | 0 | 1 | 0 | 1582580 | WP_035454724.1 | A1UQW0 |
| NTSSIMSKAGITSST | 0 | 1 | 0 | 1582997 | CBE01876.1 | Q18CX7 |
| QAIGSAILKGTKDTGT | 0 | 1 | 0 | 1583736 | ABA60112.1 | A1UU13 |
| QMVGGINPKKGGETWT | 0 | 1 | 0 | 1584569 | WP_035454724.1 | A1UQW0 |
| QQIVKWKQEIYRNVIQ | 0 | 1 | 0 | 1584779 | CBE01872.1 | Q18CX9 |
| RIMPYVSGGVAYTQVQ | 0 | 1 | 0 | 1585615 | ABA60112.1 | A1UU13 |
| SASKAVRSEALDVLAS | 0 | 1 | 0 | 1586322 | ABA60112.1 | A1UU13 |
| SGGVAYTQVQAIGSAI | 0 | 1 | 0 | 1586652 | ABA60112.1 | A1UU13 |
| STSKLPKKAGIEKTAS | 0 | 1 | 0 | 1588672 | CBE01872.1 | Q18CX9 |
| SVAEGKEKTGANASVI | 0 | 1 | 0 | 1588689 | WP_035454724.1 | A1UQW0 |
| TEGGGGSASKAVRSEA | 0 | 1 | 0 | 1589272 | ABA60112.1 | A1UU13 |

| | | | | | | |
|---|---|---|---|---|---|---|
| TEGGGGTEGGGGSASK | 0 | 1 | 0 | 1589273 | ABA60112.1 | A1UU13 |
| TEGGGGTEGGGGTEGG | 0 | 1 | 0 | 1589274 | ABA60112.1 | A1UU13 |
| TGAKGETLPIFASVAE | 0 | 1 | 0 | 1589415 | WP_035454724.1 | A1UQW0 |
| TKEGKIMRVNTNVSA | 0 | 1 | 0 | 1589713 | CBE01876.1 | Q18CX7 |
| TLPIFASVAEGKEKTG | 0 | 1 | 0 | 1590179 | WP_035454724.1 | A1UQW0 |
| TQVQAIGSAILKGTKD | 0 | 1 | 0 | 1591058 | ABA60112.1 | A1UU13 |
| TSRAKLGAQQNRLES | 0 | 1 | 0 | 1591179 | CBE01876.1 | Q18CX7 |
| TTKKPKDYPPLTDAQK | 0 | 1 | 0 | 1591225 | CBE01872.1 | Q18CX9 |
| VAYTQVQAIGSAILKG | 0 | 1 | 0 | 1591762 | ABA60112.1 | A1UU13 |
| ADNIKAAVRLGSDMG | 0 | 1 | 0 | 1596592 | P46842.2 | P46842 |
| APSLDPNAPYPLAVD | 0 | 1 | 0 | 1596596 | P46842.2 | P46842 |
| AQLNAMKPDLQNTLM | 0 | 1 | 0 | 1596597 | CAA43269.1 | P31951 |
| DPNAPSLDPNAPYPL | 0 | 1 | 0 | 1596610 | P46842.2 | P46842 |
| EFYLPYPGTRINQET | 0 | 1 | 0 | 1596617 | P46842.2 | P46842 |
| EIITPLPGAPVSSEA | 0 | 1 | 0 | 1596618 | P46842.2 | P46842 |
| KPDLQNTLMAVPRSG | 0 | 1 | 0 | 1596679 | CAA43269.1 | P31951 |
| LGSDMGEFYLPYPGT | 0 | 1 | 0 | 1596692 | P46842.2 | P46842 |
| LPYPGTRINQETIPL | 0 | 1 | 0 | 1596705 | P46842.2 | P46842 |
| LYASAEADNIKAAVR | 0 | 1 | 0 | 1596712 | P46842.2 | P46842 |
| MWGPPNDPAWQRNDP | 0 | 1 | 0 | 1596717 | CAA43269.1 | P31951 |
| NAMKPDLQNTLMAVP | 0 | 1 | 0 | 1596719 | CAA43269.1 | P31951 |
| PAQEIITPLPGAPVS | 0 | 1 | 0 | 1596741 | P46842.2 | P46842 |
| PGAPVSSEAQPGDPN | 0 | 1 | 0 | 1596743 | P46842.2 | P46842 |
| PVSSEAQPGDPNAPS | 0 | 1 | 0 | 1596748 | P46842.2 | P46842 |
| QPGDPNAPSLDPNAP | 0 | 1 | 0 | 1596752 | P46842.2 | P46842 |
| SAEADNIKAAVRLGS | 0 | 1 | 0 | 1596776 | P46842.2 | P46842 |
| SAFSRPGLPVEYLQV | 0 | 1 | 0 | 1596777 | CAA43269.1 | P31951 |
| SEAQPGDPNAPSLDP | 0 | 1 | 0 | 1596779 | P46842.2 | P46842 |
| SRPGLPVEYLQVPSE | 0 | 1 | 0 | 1596795 | CAA43269.1 | P31951 |
| TPLPGAPVSSEAQPG | 0 | 1 | 0 | 1596820 | P46842.2 | P46842 |
| YWGAQLNAMKPDLQN | 0 | 1 | 0 | 1596839 | CAA43269.1 | P31951 |
| ALQKKTEAQQEEHAQQAIKE | 0 | 1 | 0 | 1712126 | WP_010958530.1 | H7C7D7 |
| EIQNDESTLRQQQQQF | 0 | 1 | 0 | 1713104 | WP_042525978.1 | Q83DT1 |
| KKDMDNPAIQKQLRDN | 0 | 1 | 0 | 1714195 | WP_010958530.1 | H7C7D7 |
| LSYDYALYRSKSN | 0 | 1 | 0 | 1714510 | AAP04426.1 | Q83EK8 |
| SATASAEGTAIG | 0 | 1 | 0 | 1715380 | AAP04426.1 | Q83EK8 |
| AEEEEDDDMGFGLFD | 0 | 0 | 1 | 876 | CAA49261.1 | Q4DWU6 |
| AEENVEENVEEVEENVEENV | 0 | 0 | 1 | 885 | CAA00077.1 | Q8I0U6 |
| AEIERMVREAAKYEA | 0 | 0 | 1 | 986 | P05456.1 | Q4DTM8 |
| AEPKSAEPKPAEPKS | 0 | 0 | 1 | 1116 | XP_820186.1 | Q4E0M7 |
| AGGGQGQNNEGANAPNEDSV | 0 | 0 | 1 | 1523 | SRC280381 | A5K1G9 |
| AHFTDEFKRKNKGKDLSTNL | 0 | 0 | 1 | 1758 | P05456.1 | Q4DTM8 |
| ALKTELEDTLDSTATQQELR | 0 | 0 | 1 | 2677 | P35749.3 | P35749 |
| ANDQGNRTTPSYVAFTDSERL | 0 | 0 | 1 | 3243 | P05456.1 | Q4DTM8 |
| ANGAGNQPPGEDGAGNQPG | 0 | 0 | 1 | 3295 | SRC280381 | A5K1G9 |
| ANNSANNGKKNNAEE | 0 | 0 | 1 | 3367 | P22620.1 | A0A144A026 |
| APMSTPSATSVR | 0 | 0 | 1 | 3656 | SRC269007 | Q26874 |
| ASGGSVASGGSVASG | 0 | 0 | 1 | 4417 | P04933.2 | Q8I0U8 |
| ASSHRALARLRTACERAKRT | 0 | 0 | 1 | 4774 | CAA59793.1 | E9AHH0 |
| ASVASGASGGSGNSK | 0 | 0 | 1 | 4815 | SRC279972 | Q8I0U8 |
| ATNGDTHLGGEDFDN | 0 | 0 | 1 | 5066 | P05456.1 | Q4DTM8 |
| AVQAFILTGGKSKQTEGLLL | 0 | 0 | 1 | 5519 | CAA59793.1 | E9AHH0 |
| CAPESIEFPVSEARVLED | 0 | 0 | 1 | 6015 | SRC300 | P05106 |
| CECTKPDSYPLFDGIFCSSSN | 0 | 0 | 1 | 6149 | P04933.2 | Q8I0U8 |
| CSVTCGVGVRVRRRVNAANK | 0 | 0 | 1 | 7099 | P08677.2 | A5KB63 |
| DATCTEEDSGSSRKKITCECTKPDS | 0 | 0 | 1 | 7650 | A26868 | Q8I0U8 |
| DDEHVEEPTVADDEHVEEPTVA | 0 | 0 | 1 | 7766 | CAA00077.1 | Q8I0U6 |
| DDIVGYIMHGISTINKEMK | 0 | 0 | 1 | 7803 | SRC7364 | Q8I0U6 |
| DEWWKVIKKDVWNVISWVF | 0 | 0 | 1 | 8216 | P19214.2 | Q8IBE8 |
| DICLGTDISVKQGDV | 0 | 0 | 1 | 8594 | AAK28130.1 | Q8I220 |
| DIGDIVRGKDLY | 0 | 0 | 1 | 8646 | AAB60251.1 | Q8IBD8 |
| DKDLDEPEQFRLPEE | 0 | 0 | 1 | 8851 | GI:124513464 | Q76NM2 |
| DKIDWRESGYVTELKDQGNC | 0 | 0 | 1 | 8887 | AAD23996.1 | A0A504Y6Y7 |
| DKIDWRESGYVTEVKDQGNC | 0 | 0 | 1 | 8888 | AAF44678.1 | A0A504Y6Y7 |
| DKNEKGQHEIVEVEEILPE | 0 | 0 | 1 | 8942 | AAG12326.1 | Q8IJ56 |
| DKVEDGKERNVLIFDL | 0 | 0 | 1 | 8997 | P05456.1 | Q4DTM8 |
| DLDANGILNVSAEEKGTGKR | 0 | 0 | 1 | 9038 | CAA59793.1 | E9AHH0 |
| DNEILQIVKELSEKITKYFMKL | 0 | 0 | 1 | 9450 | SRC280508 | A0A143ZZD7 |
| DRAAGQPAGDRAAGQPAGDRC | 0 | 0 | 1 | 9938 | SRC280381 | A5K1G9 |
| DRAAGQPAGDRADGQPAGDR | 0 | 0 | 1 | 9939 | P08677.2 | A5KB63 |
| DRAAGQPAGNGAGGQAAGGN | 0 | 0 | 1 | 9940 | P08677.2 | A5KB63 |
| DRADGQPAGDRAAGQPAGDR | 0 | 0 | 1 | 9943 | CAC82081.1 | A5KB63 |
| DRADGQPAGDRADGQPAGDR | 0 | 0 | 1 | 9944 | AAA29535.1 | A5KB63 |
| DSEPFPRQKHKKVD | 0 | 0 | 1 | 10106 | AAG12326.1 | Q8IJ56 |
| DSERLIGDAAKNQVAMNPHN | 0 | 0 | 1 | 10108 | CAA59793.1 | E9AHH0 |
| DTDKDGKVSLEEFCR | 0 | 0 | 1 | 10312 | AAC67308.1 | O01372 |
| DTHLGGEDFDNRLVTFFTEE | 0 | 0 | 1 | 10349 | CAA59793.1 | E9AHH0 |
| DTRTEATSQASDDATAGATTTQAPV | 0 | 0 | 1 | 10480 | CAA31690.1 | P21250 |
| DVENVDGLSLWEMYK | 0 | 0 | 1 | 10581 | AAA29485.1 | Q8I562 |
| EAAKYEAEDKDQVRQ | 0 | 0 | 1 | 10991 | P05456.1 | Q4DTM8 |
| EAFIKTAAAETFLAW | 0 | 0 | 1 | 11043 | AAK28130.1 | Q8I220 |
| ECKCLLNYKQEGDKCVENPNPTCNE | 0 | 0 | 1 | 11252 | P04933.2 | Q8I0U8 |
| EDGKERNVLIFDLGGGTFDVT | 0 | 0 | 1 | 11335 | P05456.1 | Q4DTM8 |
| EEADDDMGFGLFD | 0 | 0 | 1 | 11489 | XP_001563563.1 | A4H884 |
| EEEITTKGASAQGSS | 0 | 0 | 1 | 11589 | AAA29701.1 | Q8I0U8 |
| EEEITTKGASAQSGT | 0 | 0 | 1 | 11590 | P50495.1 | Q8I0U8 |
| EENVEENVEENVEENVEENV | 0 | 0 | 1 | 11816 | CAA00077.1 | Q8I0U6 |
| EENVEHDAEENVEHDAEENVEENV | 0 | 0 | 1 | 11822 | CAA00077.1 | Q8I0U6 |
| EENVEHDAEENVEHDAEENVEHDA | 0 | 0 | 1 | 11823 | CAA00077.1 | Q8I0U6 |
| EEPEESDEDDFGMGGLF | 0 | 0 | 1 | 11825 | P39096.1 | P39096 |
| EGLLLLDVTPLTLGIETAGG | 0 | 0 | 1 | 12218 | CAA59793.1 | E9AHH0 |

| | | | | | | |
|---|---|---|---|---|---|---|
| EGTSGTAVTTSTPGS | 0 | 0 | 1 | 12289 | P04933.2 | Q8I0U8 |
| EINHVQDHALPKENI | 0 | 0 | 1 | 12531 | AAG12326.1 | Q8IJ56 |
| EKVDNLGRSGGDIIK | 0 | 0 | 1 | 12888 | CAA00077.1 | Q8I0U6 |
| ELDQENEAALENGIKNEENT | 0 | 0 | 1 | 12994 | P43243.2 | P43243 |
| ELENLCTPIMTNMYQGMAGAG | 0 | 0 | 1 | 13033 | P05456.1 | Q4DTM8 |
| ENDVLNQETEEEMEK | 0 | 0 | 1 | 13460 | P22620.1 | A0A144A026 |
| ENLLASSPRLAKYLSNRPATPF | 0 | 0 | 1 | 13513 | P09792.1 | P09792 |
| EPLEPFPTQIHKDYK | 0 | 0 | 1 | 13688 | AAG12326.1 | Q8IJ56 |
| EPQVYTLPPSR | 0 | 0 | 1 | 13735 | P01857.1 | P01857 |
| EPTAAAIAYGLDKVEDGKER | 0 | 0 | 1 | 13770 | P05456.1 | Q4DTM8 |
| EQNKKFFADKPDESTL | 0 | 0 | 1 | 13888 | EAN87014.1 | Q4D7Y4 |
| ETAGGVMTALIKRNTTIPTK | 0 | 0 | 1 | 14331 | CAA59793.1 | E9AHH0 |
| FADKPDESTLSPEMKEHYDK | 0 | 0 | 1 | 15152 | AAC61835.1 | A0A088S0A8 |
| FADKPDESTLSPEMKEHYEK | 0 | 0 | 1 | 15153 | Q25297.1 | A4IBB3 |
| FEGERAMTKDCHLLGT | 0 | 0 | 1 | 15536 | P05456.1 | Q4DTM8 |
| FFTEEFKRKNKGKNLASSHR | 0 | 0 | 1 | 15911 | CAA59793.1 | E9AHH0 |
| FKHKFAELLEQQKAAQYPSK | 0 | 0 | 1 | 16375 | AAC61835.1 | A0A088S0A8 |
| FKYSNYVCLLWRTPS | 0 | 0 | 1 | 16487 | AAC67308.1 | O01372 |
| FNDSQRQATKDAGTIAGLEV | 0 | 0 | 1 | 17140 | CAA59793.1 | E9AHH0 |
| FRGETKTFNPEEVSS | 0 | 0 | 1 | 17589 | P05456.1 | Q4DTM8 |
| FRGKELKKSIQPDEA | 0 | 0 | 1 | 17590 | P05456.1 | Q4DTM8 |
| GAANPSDDSSDSDAK | 0 | 0 | 1 | 18517 | P19598.2 | Q8I0U8 |
| GAGGTAGGSAGGSAGGSAGG | 0 | 0 | 1 | 18591 | AAA29686.1 | A0A143ZX41 |
| GANAPNEKSVKEYLDKVRAT | 0 | 0 | 1 | 18694 | P08677.2 | A5KB63 |
| GASAQGSSGPSGTPS | 0 | 0 | 1 | 18758 | AAA29701.1 | Q8I0U8 |
| GASAQSGASAQSGAS | 0 | 0 | 1 | 18759 | CAA37136.1 | Q8I0U8 |
| GASAQSGTSGTSGTS | 0 | 0 | 1 | 18761 | P50495.1 | Q8I0U8 |
| GDAKKKKDGKKAEPKNPREN | 0 | 0 | 1 | 18933 | P08677.2 | A5KB63 |
| GDLLLKLNNYRYNKD | 0 | 0 | 1 | 19029 | AAA63423.1 | A5KDR7 |
| GDRAAGQAAGDRAAGQAA | 0 | 0 | 1 | 19091 | P13826.1 | A5KB63 |
| GDRADGQPAGDRADGQPA | 0 | 0 | 1 | 19098 | AAA29535.1 | A5KB63 |
| GFTTNEERYNVFAE | 0 | 0 | 1 | 19636 | SRC14833 | G4V6W4 |
| GFTTNEPRYNVFAE | 0 | 0 | 1 | 19637 | SRC14833 | G4V6W4 |
| GGMPGGMPGGMPGGMNFP | 0 | 0 | 1 | 19928 | B31238 | Q8IB24 |
| GGSGGSVASGGSVAS | 0 | 0 | 1 | 20012 | P04933.2 | Q8I0U8 |
| GGSGGSVASVASGGS | 0 | 0 | 1 | 20013 | ABF55313.1 | Q8I0U8 |
| GGSGNSRRTNPSDNS | 0 | 0 | 1 | 20014 | P04933.2 | Q8I0U8 |
| GGSVASVASGGSGGS | 0 | 0 | 1 | 20039 | AAA29707.1 | Q8I0U8 |
| GGSVTSGGSGASVAS | 0 | 0 | 1 | 20042 | SRC279972 | Q8I0U8 |
| GKAKTKAVSRSQRAGLQFPV | 0 | 0 | 1 | 20512 | P0C0S5.2 | P0C0S5 |
| GKDSADKMPLETQLA | 0 | 0 | 1 | 20538 | AAA29733.1 | Q6ZMA7 |
| GMPGGMPGGMPGGMPG | 0 | 0 | 1 | 21322 | P05456.1 | Q4DTM8 |
| GRRGQKAKSNPNRPL | 0 | 0 | 1 | 22185 | BAA00906.1 | A0A2I3RRI1 |
| GSTMMQTKTFSSSSSSKKMG | 0 | 0 | 1 | 22545 | P53814.7 | P53814 |
| GTGKRNQITITNDKGRLSKD | 0 | 0 | 1 | 22662 | CAA59793.1 | E9AHH0 |
| GTSAPSGSGTSPSSR | 0 | 0 | 1 | 22809 | SRC279972 | Q8I0U8 |
| GTSGPSGTGPSGTSP | 0 | 0 | 1 | 22815 | SRC279972 | Q8I0U8 |
| GTSGTSAQSGTSGTS | 0 | 0 | 1 | 22828 | P50495.1 | Q8I0U8 |
| GYYYPSDPNTFYAPPYS | 0 | 0 | 1 | 23511 | AAA74982.1 | Q26874 |
| HAVGPNQGGLSQGLS | 0 | 0 | 1 | 23576 | P17503.2 | Q6ZMA7 |
| HLGGEDFDNRLVAHF | 0 | 0 | 1 | 24211 | P05456.1 | Q4DTM8 |
| IAYGLDKGDDGKERNVLIFD | 0 | 0 | 1 | 25481 | CAA59793.1 | E9AHH0 |
| IDALFENVDFQATITRARFE | 0 | 0 | 1 | 25534 | CAA59793.1 | E9AHH0 |
| IEEADKKTITSAVEEALEWL | 0 | 0 | 1 | 25766 | P05456.1 | Q4DTM8 |
| IEEEKSLNMLDKKT | 0 | 0 | 1 | 25768 | SRC280011 | Q7K740 |
| IENEYEVLYLKPLAGVYRSLK | 0 | 0 | 1 | 25906 | 1202270A | Q8I0U8 |
| IFSTYADNQPGVHIQ | 0 | 0 | 1 | 26169 | P05456.1 | Q4DTM8 |
| IGDAAKNQVAMNPRN | 0 | 0 | 1 | 26204 | P05456.1 | Q4DTM8 |
| IKPGSADKPKDQLDYENDIE | 0 | 0 | 1 | 26865 | BAD73952.1 | Q7K740 |
| ILGWEFGGGAP | 0 | 0 | 1 | 27084 | AAS13447.1 | Q8IJ54 |
| ILTGGKSKQTEGLLLL | 0 | 0 | 1 | 27379 | P05456.1 | Q4DTM8 |
| IMLIILSGIVGFKVK | 0 | 0 | 1 | 27517 | P17503.2 | Q6ZMA7 |
| IPKVMQLVSDFFRGKELKKS | 0 | 0 | 1 | 27909 | P05456.1 | Q4DTM8 |
| IPPPPRGVPQIEVTF | 0 | 0 | 1 | 27979 | P05456.1 | Q4DTM8 |
| IQPVERVLQDAKMDKRSVHD | 0 | 0 | 1 | 28205 | CAA59793.1 | E9AHH0 |
| IQQLNLNDNAIHLYA | 0 | 0 | 1 | 28207 | GI:124513464 | Q76NM2 |
| IRQLDTSVERRALGEIQNV | 0 | 0 | 1 | 28374 | AAI08400.1 | Q32NY8 |
| ISTEPFPNQKHKDLQ | 0 | 0 | 1 | 28742 | AAG12326.1 | Q8IJ56 |
| IVGYIMHGISTINTEMK | 0 | 0 | 1 | 29294 | P13830.1 | Q8I0U6 |
| KCVENPNPTCNENNGGCDADATCTE | 0 | 0 | 1 | 30095 | P08569.3 | Q8I0U8 |
| KDAGTIAGLEVLRIINEPTAA | 0 | 0 | 1 | 30110 | P05456.1 | Q4DTM8 |
| KDNTDEGDEGDDS | 0 | 0 | 1 | 30222 | AAA29733.1 | Q6ZMA7 |
| KEALISSLTRGKLTYTQQ | 0 | 0 | 1 | 30302 | SRC248015 | P12883 |
| KEEKEEKEEKEEKEKEKE | 0 | 0 | 1 | 30349 | SRC280156 | Q8ID94 |
| KEKMILNEEEITTKG | 0 | 0 | 1 | 30476 | P50495.1 | Q8I0U8 |
| KEKMLLNEEEITTKG | 0 | 0 | 1 | 30477 | AAA29704.1 | Q8I0U8 |
| KEKMVLKDGANTQVV | 0 | 0 | 1 | 30478 | P19598.2 | Q8I0U8 |
| KEKMVLNEEEITTKG | 0 | 0 | 1 | 30479 | CAA37136.1 | Q8I0U8 |
| KEKMVLNEGTSGTAV | 0 | 0 | 1 | 30480 | P04933.2 | Q8I0U8 |
| KEYLDKVRATVGTEWTPCSV | 0 | 0 | 1 | 30677 | P08677.2 | A5KB63 |
| KFAELLEQQKNAQFPGK | 0 | 0 | 1 | 30696 | Q9U6Z1.2 | Q4D7Y4 |
| KFNKKMHEHSEHFKHKFAEL | 0 | 0 | 1 | 30794 | AAC61835.1 | A0A088S0A8 |
| KIENEPLVHENLSIPNDPIEQILNQ | 0 | 0 | 1 | 31224 | AAG12326.1 | Q8IJ56 |
| KKDGKKAEPKNPRENKLDQP | 0 | 0 | 1 | 31502 | SRC280381 | A5K1G9 |
| KLQPAEIETCMKTIT | 0 | 0 | 1 | 32113 | NP_704366.1 | Q8IB24 |
| KMKEIAESYLGKQVK | 0 | 0 | 1 | 32337 | P05456.1 | Q4DTM8 |
| KNGLENYAFSMKNAVNDPN | 0 | 0 | 1 | 32501 | P05456.1 | Q4DTM8 |
| KNGLENYAYSMKNTLSDSNV | 0 | 0 | 1 | 32502 | CAA59793.1 | E9AHH0 |
| KNQLAGKDEYEAKQKEAE | 0 | 0 | 1 | 32596 | NP_704366.1 | Q8IB24 |
| KNQVAMNPRNTVFDA | 0 | 0 | 1 | 32597 | P05456.1 | Q4DTM8 |
| KNTLTPLEELYPT | 0 | 0 | 1 | 32619 | NP_701990.1 | Q8ILZ1 |

| | | | | | | |
|---|---|---|---|---|---|---|
| KPKDQLDYENDIEKKICKMEKCS | 0 | 0 | 1 | 32761 | BAD73952.1 | Q7K740 |
| KPKHKKLKQPGDGNPD | 0 | 0 | 1 | 32770 | AAW78213.1 | Q7K740 |
| KPPPFGQAAAGDKPA | 0 | 0 | 1 | 32832 | XP_818927.1 | Q4DX15 |
| KPPPFGQAAAGDKPP | 0 | 0 | 1 | 32833 | XP_818927.1 | Q4DX15 |
| KPPPFGQAAAGDKPS | 0 | 0 | 1 | 32834 | XP_818927.1 | Q4DX15 |
| KPPPFGQAAEGDKPP | 0 | 0 | 1 | 32835 | XP_818927.1 | Q4DX15 |
| KPQEEKEKITKEILNGK | 0 | 0 | 1 | 32844 | 1304302A | P09792 |
| KPSPFGQAAAGDKPP | 0 | 0 | 1 | 32884 | XP_818927.1 | Q4DX15 |
| KRAVHDVVLVGGSTRIPKVM | 0 | 0 | 1 | 33111 | P05456.1 | Q4DTM8 |
| KRNQIVLTNDKGRLS | 0 | 0 | 1 | 33199 | P05456.1 | Q4DTM8 |
| KTKNADDDKGDKDKD | 0 | 0 | 1 | 33736 | P17503.2 | Q6ZMA7 |
| KTSKRQAVPVTL | 0 | 0 | 1 | 33860 | AAM21050.1 | P0DJ06 |
| KVITKGDDKPVIQVQ | 0 | 0 | 1 | 34057 | P05456.1 | Q4DTM8 |
| KYEEDDKAQRDRVEAKNGLE | 0 | 0 | 1 | 34391 | CAA59793.1 | E9AHH0 |
| LAQMYQANDNSLEDV | 0 | 0 | 1 | 34935 | AAA29485.1 | Q8I562 |
| LDENEDNIKKMKSKIDDMEKEIKYR | 0 | 0 | 1 | 35142 | AAC71819.1 | O96133 |
| LDNIKGNVGKMEDYIKKNNK | 0 | 0 | 1 | 35235 | SRC279972 | Q8I0U8 |
| LENNLIEIFKKIHENL | 0 | 0 | 1 | 35630 | AAB60251.1 | Q8IBD8 |
| LESVCNPIMTKMYQSMGGAA | 0 | 0 | 1 | 35713 | CAA59793.1 | E9AHH0 |
| LFDYNEKVDNLGRSG | 0 | 0 | 1 | 35817 | CAA00077.1 | Q8I0U6 |
| LGKQVKKAVVTVPAYFNDSQ | 0 | 0 | 1 | 36192 | P05456.1 | Q4DTM8 |
| LGRSGGDIIKKMQTL | 0 | 0 | 1 | 36309 | CAA00077.1 | Q8I0U6 |
| LIKRNTTIPTKKSQIF | 0 | 0 | 1 | 36580 | P05456.1 | Q4DTM8 |
| LIYDAAVEGDLLFKL | 0 | 0 | 1 | 36769 | AAY34061.1 | A5KDR7 |
| LIYDAAVEGDLLLKL | 0 | 0 | 1 | 36770 | AAA63423.1 | A5KDR7 |
| LKKDKEGKVSTLPLD | 0 | 0 | 1 | 36903 | AAC67308.1 | O01372 |
| LKNKIFPKKKEDNQAVDT | 0 | 0 | 1 | 36974 | P22620.1 | A0A144A026 |
| LKRDKEGKVSTLPLD | 0 | 0 | 1 | 37004 | AAA29940.1 | O01372 |
| LLSKDKEIEEKNKKIKELNNDIKKL | 0 | 0 | 1 | 37797 | AAC71819.1 | O96133 |
| LMIKEHILAIAIYESRILKR | 0 | 0 | 1 | 38053 | P19214.2 | Q8IBE8 |
| LNDITKEYEKLLNEI | 0 | 0 | 1 | 38186 | AAL88455.1 | Q8I0U8 |
| LNFKGRWLRDRL | 0 | 0 | 1 | 38215 | AAA66352.1 | Q4CZ79 |
| LNKSINPDEAVAYGAAVQAF | 0 | 0 | 1 | 38257 | CAA59793.1 | E9AHH0 |
| LPRSNTSSGAIPPAD | 0 | 0 | 1 | 38693 | CAA43932.1 | Q8I0U8 |
| LPRSNTSSGASPPAD | 0 | 0 | 1 | 38694 | P50495.1 | Q8I0U8 |
| LRQLDFVEEVLRKHPDKVE | 0 | 0 | 1 | 39174 | XP_806822.1 | Q4CXH3 |
| LRRLRTACERAKRTL | 0 | 0 | 1 | 39196 | P05456.1 | Q4DTM8 |
| LTIDGGIFEVKATNGDTHLG | 0 | 0 | 1 | 39950 | CAA59793.1 | E9AHH0 |
| LTMSNVKNVQTNFKSLLRNLGVS | 0 | 0 | 1 | 40041 | SRC280508 | A0A143ZZD7 |
| MGGAAGGMPGGMPGMSGMSG | 0 | 0 | 1 | 41575 | CAA59793.1 | E9AHH0 |
| MNPHNTVFDAKRLIGRKFND | 0 | 0 | 1 | 42227 | CAA59793.1 | E9AHH0 |
| MNPRNTVFDAKRLIGRKFSD | 0 | 0 | 1 | 42235 | P05456.1 | Q4DTM8 |
| MPPYPTGGPPPV | 0 | 0 | 1 | 42308 | AAA74982.1 | Q26874 |
| MQTLWDEIMDINKRK | 0 | 0 | 1 | 42416 | P13830.1 | Q8I0U6 |
| MRQLDTNVERRALGEIQNV | 0 | 0 | 1 | 42494 | Q9UL49.1 | F8W9A4 |
| MSKAKQYNICCKFKE | 0 | 0 | 1 | 42580 | AAC67308.1 | O01372 |
| MTFEGAIGIDLGTTYSCVGV | 0 | 0 | 1 | 42802 | CAA59793.1 | E9AHH0 |
| MTKDCHLLGTFDLSGIPPAP | 0 | 0 | 1 | 42818 | CAA59793.1 | E9AHH0 |
| NAAAAALDKKQRNFD | 0 | 0 | 1 | 43107 | P12883.5 | P12883 |
| NADMNEITERYFKLAENYY | 0 | 0 | 1 | 43134 | P13830.1 | Q8I0U6 |
| NAKNVNDMYRDGEMS | 0 | 0 | 1 | 43198 | AAA29485.1 | Q8I562 |
| NANPNANPNANPNANPNANP | 0 | 0 | 1 | 43248 | AAA29555.1 | Q7K740 |
| NDKGRLSRAEIERMV | 0 | 0 | 1 | 43462 | P05456.1 | Q4DTM8 |
| NGKQVESPAPSGHTHLKKPA | 0 | 0 | 1 | 44018 | AAF67727.1 | A4I4C9 |
| NIISCNKNDKNQ | 0 | 0 | 1 | 44251 | P22620.1 | A0A144A026 |
| NISGTGEKAQQHRKQ | 0 | 0 | 1 | 44342 | SRC280377 | A5KDR7 |
| NISQHCVKKQCPQNSGCFRHLDERE | 0 | 0 | 1 | 44355 | SRC279972 | Q8I0U8 |
| NKNLQEEISDLTEQLGSS | 0 | 0 | 1 | 44559 | AAA51837.1 | P12883 |
| NKYLKVQLKKGETVENTNS | 0 | 0 | 1 | 44634 | AAA29485.1 | Q8I562 |
| NNGGCDADAKCTEEDSGSNGKKITC | 0 | 0 | 1 | 45112 | P04933.2 | Q8I0U8 |
| NNRNRLTKLFRYKSSRSESE | 0 | 0 | 1 | 45225 | CAM69637.1 | A4I4C9 |
| NNYNNNAKNVNDMYRDGE | 0 | 0 | 1 | 45291 | AAA29485.1 | Q8I562 |
| NPPGATVPSGTASTK | 0 | 0 | 1 | 45450 | P19598.2 | Q8I0U8 |
| NPRENKLKQPGDRADGQPAG | 0 | 0 | 1 | 45464 | AAA29535.1 | A5KB63 |
| NQEAAKEEYEHKQKELESVC | 0 | 0 | 1 | 45532 | CAA59793.1 | E9AHH0 |
| NQPGVHIQVFEGERAMTKDC | 0 | 0 | 1 | 45603 | CAA59793.1 | E9AHH0 |
| NSGCFRHLDEREECKCLLNYKQEGD | 0 | 0 | 1 | 45819 | P04933.2 | Q8I0U8 |
| NTSSGASPPADASDS | 0 | 0 | 1 | 46271 | P50495.1 | Q8I0U8 |
| NVDPNANPNANPNANP | 0 | 0 | 1 | 46354 | P19597 | Q7K740 |
| NVMFRQNNSHLAQMY | 0 | 0 | 1 | 46432 | AAA29485.1 | Q8I562 |
| NVSINNNNNNNNSNT | 0 | 0 | 1 | 46479 | AAA29485.1 | Q8I562 |
| PKVQSLVSDFFGGKELNKSI | 0 | 0 | 1 | 48168 | CAA59793.1 | E9AHH0 |
| PLCNTRKSRQSGI | 0 | 0 | 1 | 48282 | SRC269007 | Q26874 |
| PNANPNANPNANPNANPNAN | 0 | 0 | 1 | 48600 | P02893.1 | Q7K740 |
| PPADASDSDAKSYAD | 0 | 0 | 1 | 48741 | P50495.1 | Q8I0U8 |
| PQTWWEKNAKDIWEG | 0 | 0 | 1 | 49140 | AAK28130.1 | Q8I220 |
| PSDKHIEQYLKKIQNSLSTE | 0 | 0 | 1 | 49325 | AAA29555.1 | Q7K740 |
| PSGTASTKGAIRSPG | 0 | 0 | 1 | 49391 | P19598.2 | Q8I0U8 |
| PSSRSNTLPRSN | 0 | 0 | 1 | 49518 | P50495.1 | Q8I0U8 |
| PSTLQTPGSSSGASL | 0 | 0 | 1 | 49529 | AAA29733.1 | Q6ZMA7 |
| QKYLSSTIQKQVD | 0 | 0 | 1 | 51271 | CAA31690.1 | P21250 |
| QSGASATSAQSGPSG | 0 | 0 | 1 | 52324 | SRC279972 | Q8I0U8 |
| RAKRTLSSATQATIEIDALF | 0 | 0 | 1 | 53135 | CAA59793.1 | E9AHH0 |
| RARFEELCGDLFRSTIQPVE | 0 | 0 | 1 | 53191 | CAA59793.1 | E9AHH0 |
| RGQKAKSNPNRPLPVPRN | 0 | 0 | 1 | 53955 | BAA00906.1 | A0A2I3RRI1 |
| RKFNDSVVQSDMKHWPFKVT | 0 | 0 | 1 | 54345 | CAA59793.1 | E9AHH0 |
| RKNKGKDLSTNLRAL | 0 | 0 | 1 | 54396 | P05456.1 | Q4DTM8 |
| RLAKLSDGVAVLKVG | 0 | 0 | 1 | 54468 | P10809.2 | A0A7I2V599 |
| RLIGRKFSDPVVQSD | 0 | 0 | 1 | 54589 | P05456.1 | Q4DTM8 |
| RLRSSVPGVRLLQDSVDFSL | 0 | 0 | 1 | 54747 | P08670.4 | P05783 |
| RLSKDEIERMVNDAMKYEED | 0 | 0 | 1 | 54765 | CAA59793.1 | E9AHH0 |

| | | | | | | |
|---|---|---|---|---|---|---|
| RNNHPQNTSDSQKECTDGNK | 0 | 0 | 1 | 55038 | AAA29686.1 | A0A143ZX41 |
| RSFQNKKSLVAFKIMPLEDM | 0 | 0 | 1 | 55838 | P15927.1 | P15927 |
| RSVHDVVLVGGSTRIPKVQS | 0 | 0 | 1 | 55987 | CAA59793.1 | E9AHH0 |
| RTNPSDNSSDSNIKT | 0 | 0 | 1 | 56142 | SRC279972 | Q8I0U8 |
| RVNAANKKPEDLTLNDLETD | 0 | 0 | 1 | 56366 | P08677.2 | A5KB63 |
| SAEEKGTGKRNQIVL | 0 | 0 | 1 | 56747 | P05456.1 | Q4DTM8 |
| SAQSGPSGTSAPSGS | 0 | 0 | 1 | 56956 | SRC279972 | Q8I0U8 |
| SAQSGTSGTSAQSGT | 0 | 0 | 1 | 56958 | P50495.1 | Q8I0U8 |
| SDANDKAKKPAGKGS | 0 | 0 | 1 | 57162 | AAA29733.1 | Q6ZMA7 |
| SDARQRVADRKRKEKYDSFF | 0 | 0 | 1 | 57166 | AAF67727.1 | A4I4C9 |
| SDSNVSGKLEDSDKATLNKE | 0 | 0 | 1 | 57286 | CAA59793.1 | E9AHH0 |
| SERVQLLHSQNTSLINQK | 0 | 0 | 1 | 57618 | AAA51837.1 | P12883 |
| SETNKNPTSHSNSTTTSLNNN | 0 | 0 | 1 | 57650 | AAN35543.1 | Q8IJ54 |
| SGDEDVDSDEL | 0 | 0 | 1 | 57998 | Q05866.1 | Q8I2X4 |
| SGGAGPAGGASSGPKVEEVD | 0 | 0 | 1 | 58044 | CAA59793.1 | E9AHH0 |
| SGGSVASGGSGNSRR | 0 | 0 | 1 | 58057 | P04933.2 | Q8I0U8 |
| SGGSVTSGGSGGSVA | 0 | 0 | 1 | 58058 | ABD85256.1 | Q8I0U8 |
| SGGSVTSGGSVTSGG | 0 | 0 | 1 | 58059 | ABD85256.1 | Q8I0U8 |
| SGPSGPSGTSPSSRS | 0 | 0 | 1 | 58195 | P04932.2 | Q8I0U8 |
| SGPSGTPSGTSGPSG | 0 | 0 | 1 | 58196 | AAA29701.1 | Q8I0U8 |
| SGPSGTSPSSRSNTL | 0 | 0 | 1 | 58197 | P04932.2 | Q8I0U8 |
| SGSNGKKITCECTKPDSYPLFDGIF | 0 | 0 | 1 | 58239 | P04933.2 | Q8I0U8 |
| SGTSGPSGTSGPSGT | 0 | 0 | 1 | 58266 | AAA29701.1 | Q8I0U8 |
| SIYKMNNEIVNSTID | 0 | 0 | 1 | 58759 | AAA29485.1 | Q8I562 |
| SKGSVASSGSVASGG | 0 | 0 | 1 | 58841 | AAA29674.2 | Q8I0U8 |
| SKIYQDAAGAAGGMPG | 0 | 0 | 1 | 58859 | NP_704366.1 | Q8IB24 |
| SLAQVKYTSASSI | 0 | 0 | 1 | 59016 | AAM21050.1 | P0DJ06 |
| SLRWIFKHVAKTHLK | 0 | 0 | 1 | 59446 | P13830.1 | Q8I0U6 |
| SLVTCPAQGSLQSSPSMEIK | 0 | 0 | 1 | 59571 | CAC24700.1 | F8W9A4 |
| SNTLPRSNTSSGASP | 0 | 0 | 1 | 59927 | P50495.1 | Q8I0U8 |
| SNTSSGAPPADASDS | 0 | 0 | 1 | 59932 | SRC279972 | Q8I0U8 |
| SSDSNTKTYADLKHR | 0 | 0 | 1 | 60956 | P08569.3 | Q8I0U8 |
| SSGSVASGGSVASVA | 0 | 0 | 1 | 61053 | AAA29707.1 | Q8I0U8 |
| SSMVLLKMKETAEAYLGKQV | 0 | 0 | 1 | 61219 | CAA59793.1 | E9AHH0 |
| SSVFNVVNSSIGLIM | 0 | 0 | 1 | 61443 | GI:124504759 | Q7K740 |
| SVASGGSGGSVASGG | 0 | 0 | 1 | 62051 | P04933.2 | Q8I0U8 |
| SVASVASVASGGSGN | 0 | 0 | 1 | 62054 | ABD85257.1 | Q8I0U8 |
| TAADTPTATESISPSPP | 0 | 0 | 1 | 62779 | CAA73082.1 | A0A143ZX41 |
| TCGVGVRVRRRVNAANKKPE | 0 | 0 | 1 | 63077 | P08677.2 | A5KB63 |
| TGYSLFHKEKMILNE | 0 | 0 | 1 | 64011 | P04932.2 | Q8I0U8 |
| TIPTKKSQIFSTYADNQPGV | 0 | 0 | 1 | 64428 | CAA59793.1 | E9AHH0 |
| TLNKEIDVVLEWLSSNQEAA | 0 | 0 | 1 | 64979 | CAA59793.1 | E9AHH0 |
| TLTKEYEDIVLKSHMNRESDD | 0 | 0 | 1 | 65097 | AAA29600.1 | Q8IBE8 |
| TNDEEDTNDEEDTNDEED | 0 | 0 | 1 | 65298 | SRC280156 | Q8ID94 |
| TPQRKTTEDRPQ | 0 | 0 | 1 | 65726 | XP_812850.1 | Q4DEP9 |
| TQGVVKERRWTLVNENRPIW | 0 | 0 | 1 | 65863 | AAF67727.1 | A4I4C9 |
| TSGTSGPSGPSGTSP | 0 | 0 | 1 | 66237 | P04932.2 | Q8I0U8 |
| TSGTSGPSGTSPSSR | 0 | 0 | 1 | 66238 | AAC69726.1 | Q8I0U8 |
| TSGTSGTSGPSGPSG | 0 | 0 | 1 | 66240 | P04932.2 | Q8I0U8 |
| TSGTSGTSGTSGTSG | 0 | 0 | 1 | 66241 | P50495.1 | Q8I0U8 |
| TSGTSGTSGTSPSSR | 0 | 0 | 1 | 66242 | P50495.1 | Q8I0U8 |
| TTQETVTTQETV | 0 | 0 | 1 | 66787 | NP_703360.1 | Q8I486 |
| TVAEEHVEEPTVAEE | 0 | 0 | 1 | 66933 | CAA00077.1 | Q8I0U6 |
| TVNRNLIISTAVTNM | 0 | 0 | 1 | 67098 | AAA29733.1 | Q6ZMA7 |
| VAKPADAVSTQSAKN | 0 | 0 | 1 | 67648 | P19598.2 | Q8I0U8 |
| VAKPVPAVSTQSAKN | 0 | 0 | 1 | 67650 | AAA29705.1 | Q8I0U8 |
| VAKQHVRDGRKDSLDGF | 0 | 0 | 1 | 67651 | BAA00906.1 | A0A2I3RRI1 |
| VASGGSGNSRRTNPS | 0 | 0 | 1 | 67745 | P04933.2 | Q8I0U8 |
| VASVASGGSGNSRRT | 0 | 0 | 1 | 67762 | ABF55315.1 | Q8I0U8 |
| VDREGRDNVRSKGV | 0 | 0 | 1 | 68054 | AAA63551.1 | A2E7I1 |
| VERVLQDAKMDKRAV | 0 | 0 | 1 | 68334 | P05456.1 | Q4DTM8 |
| VGTEWTPCSVTCGVGVRVRR | 0 | 0 | 1 | 68801 | AAA29535.1 | A5KB63 |
| VLIFDLGGGTFDVTLLTIDG | 0 | 0 | 1 | 69544 | CAA59793.1 | E9AHH0 |
| VNNTDTNFHRDITFR | 0 | 0 | 1 | 70138 | AAA63423.1 | A5KDR7 |
| VNNTDTNFHSDITFR | 0 | 0 | 1 | 70139 | ABA39300.1 | A5KDR7 |
| VPPTQSKKKNKNET | 0 | 0 | 1 | 70437 | P22620.1 | A0A144A026 |
| VQYRGEEKTFTPEEISSMVL | 0 | 0 | 1 | 70683 | CAA59793.1 | E9AHH0 |
| VRRRVNAANKKPEDLTLNDL | 0 | 0 | 1 | 70817 | P08677.2 | A5KB63 |
| VSTQSAKNPPGATVP | 0 | 0 | 1 | 71163 | P19598.2 | Q8I0U8 |
| VTHESYQELVKKLEALEDA | 0 | 0 | 1 | 71353 | AAA29611.1 | Q8I0U8 |
| VTTSTPGSGGSVASG | 0 | 0 | 1 | 71533 | SRC279972 | Q8I0U8 |
| VTTSTPGSSGSVASG | 0 | 0 | 1 | 71536 | AAA29707.1 | Q8I0U8 |
| VTTSTPGSSGSVTSG | 0 | 0 | 1 | 71537 | P08569.3 | Q8I0U8 |
| WEGMLCALTNGLTDA | 0 | 0 | 1 | 72351 | AAK28130.1 | Q8I220 |
| YDELLKRKENELF | 0 | 0 | 1 | 73540 | AAB60251.1 | Q8IBD8 |
| YHQKFKETVLGRNSGFGFVSYD | 0 | 0 | 1 | 74131 | AAA29485.1 | Q8I562 |
| YKAYVSYKKRKAQEK | 0 | 0 | 1 | 74430 | P22620.1 | A0A144A026 |
| YNNDDDNDDDNDDNNDDNN | 0 | 0 | 1 | 75197 | CAA70130.1 | Q8I586 |
| YVKELPTEVQKLKEK | 0 | 0 | 1 | 76307 | AAY34070.1 | A5KDR7 |
| YVSELPTEVQKLKEK | 0 | 0 | 1 | 76408 | P22290.2 | A5KDR7 |
| YYALDVYAYDVT | 0 | 0 | 1 | 76485 | P87020.1 | P87020 |
| PPPYAPNPGPPPPYTGA | 0 | 0 | 1 | 77968 | AAA74982.1 | Q26874 |
| CRVWNQTDRTQ | 0 | 0 | 1 | 78354 | P32022.1 | P32022 |
| EFMKEADKEAY | 0 | 0 | 1 | 78370 | P32022.1 | P32022 |
| CESLKGSTGKLAVG | 0 | 0 | 1 | 79167 | 1304302A | P09792 |
| ETDEEYYSVEKLIGQAEDVEHEYHK | 0 | 0 | 1 | 79204 | 1304302A | P09792 |
| LADLVLIAVIDHVTDLDK | 0 | 0 | 1 | 79279 | 1304302A | P09792 |
| LVAAGVDYEDERISFQDWPK | 0 | 0 | 1 | 79303 | 1304302A | P09792 |
| GRGWTLRDSSKTTIG | 0 | 0 | 1 | 101690 | AAB68960.1 | A0A044RRG7 |
| GQKAKSNPNRPLPVPRNPCR | 0 | 0 | 1 | 103946 | BAA00906.1 | A0A2I3RRI1 |
| NCALEGWGGHWRGAN | 0 | 0 | 1 | 106672 | P79017.2 | P79017 |

| | | | | | | |
|---|---|---|---|---|---|---|
| TIDVPSNCHTHEGGQLHCT | 0 | 0 | 1 | 106908 | P79017.2 | P79017 |
| ELWDKRYGGRSNI | 0 | 0 | 1 | 107824 | XP_001350415.1 | Q8I639 |
| KETELLYEYHDTGTAIISKN | 0 | 0 | 1 | 107935 | AAQ73926.1 | Q8I639 |
| ENPVVHFFKNIVTP | 0 | 0 | 1 | 114806 | P02686.3 | P02686 |
| RRREGPDRSPR | 0 | 0 | 1 | 117366 | P13985.1 | P13985 |
| APGSAAPAAGSAPAA | 0 | 0 | 1 | 118268 | NP_000995.1 | P05387 |
| ESEESDDDMGFGLFD | 0 | 0 | 1 | 118338 | NP_000995.1 | P05387 |
| KDIKKILDSVGIEAD | 0 | 0 | 1 | 118446 | NP_000995.1 | P05387 |
| KVISELNGKNIEDVI | 0 | 0 | 1 | 118478 | NP_000995.1 | P05387 |
| LNGKNIEDVIAQGIG | 0 | 0 | 1 | 118497 | NP_000995.1 | P05387 |
| PAGGAVAVSAAPGSA | 0 | 0 | 1 | 118546 | NP_000995.1 | P05387 |
| VAVSAAPGSAAPAAG | 0 | 0 | 1 | 118645 | NP_000995.1 | P05387 |
| GQYFAKPRNQGG | 0 | 0 | 1 | 119910 | NP_002127.1 | A0A7I2YQX9 |
| SSQRGRSGSGNF | 0 | 0 | 1 | 120220 | NP_002127.1 | A0A7I2YQX9 |
| MILHPDVQRRVQQEIDDVI | 0 | 0 | 1 | 122486 | P10635.2 | P10635 |
| GQVRRPEMGDQA | 0 | 0 | 1 | 122781 | P10635.2 | P10635 |
| EYDDPRFLRLLDLAQEGLKEESG | 0 | 0 | 1 | 123159 | P10635.2 | P10635 |
| LLTEHRMTWDPAQPPRDLTE | 0 | 0 | 1 | 123443 | P10635.2 | P10635 |
| ELLTEHRMTWDPAQPPRDLT | 0 | 0 | 1 | 125401 | P10635.2 | P10635 |
| FLPFSAGRRACLGEPLARME | 0 | 0 | 1 | 125408 | P10635.2 | P10635 |
| GKVLRFQKAFLTQLDELLTE | 0 | 0 | 1 | 125423 | P10635.2 | P10635 |
| HGVFAFLVSPSPYELCAVPR | 0 | 0 | 1 | 125438 | P10635.2 | P10635 |
| RRFEYDDPRFLRLLDLAQEG | 0 | 0 | 1 | 125552 | P10635.2 | P10635 |
| RRPEMGDQAHMPYTTAVIHE | 0 | 0 | 1 | 125553 | P10635.2 | P10635 |
| PTRAPSGPRPPSRSQAQTPPR | 0 | 0 | 1 | 132022 | P13985.1 | P13985 |
| SRKCPFYDIACML | 0 | 0 | 1 | 135296 | CAC00543.1 | W2TN28 |
| DKVSFFCKNKEKKC | 0 | 0 | 1 | 135650 | CAA37664.1 | P02749 |
| ALLRCIPALDSLTPANEDC | 0 | 0 | 1 | 135895 | SRC248227 | P10809 |
| KEQFEFALTAVAEEVC | 0 | 0 | 1 | 135944 | SRC244411 | Q92932 |
| SIQSIVPALEIA | 0 | 0 | 1 | 135995 | AAH02676.1 | A0A7I2V599 |
| SIVPALEIANAH | 0 | 0 | 1 | 135996 | AAH02676.1 | A0A7I2V599 |
| SEIEKLKNKYEVAGIP | 0 | 0 | 1 | 137290 | XP_001900803.1 | A0A0J9XTJ8 |
| EVVETDGKKKECSS | 0 | 0 | 1 | 138844 | AAC35355.1 | A0A3P7EGK8 |
| LAGIRPLGPKNKAPDFSGTAVV | 0 | 0 | 1 | 138871 | ADP24700.1 | A0A5S6PAI1 |
| NKLLIAFGLVILLVTLPCAS | 0 | 0 | 1 | 138884 | AAC35355.1 | A0A3P7EGK8 |
| SEGGDEYVTKGEVVETD | 0 | 0 | 1 | 138910 | AAC35355.1 | A0A3P7EGK8 |
| WTDKGCFCEDKLHSCVIERKN | 0 | 0 | 1 | 138933 | AAC35355.1 | A0A3P7EGK8 |
| YDQREPQAWCRPNENQSWT | 0 | 0 | 1 | 138935 | AAC35355.1 | A0A3P7EGK8 |
| SESDEEFDDGSNDETDDKEDEGNS | 0 | 0 | 1 | 139247 | AAC35355.1 | A0A3P7EGK8 |
| FLTLEEQCDWAFNQYWHLNR | 0 | 0 | 1 | 139501 | SRC313672 | A0A2D4C4Z9 |
| WTSIASTQPVGTTTFEHWPIR | 0 | 0 | 1 | 139639 | BAS04784.1 | A0A2D4BV21 |
| FSKWRKNHMRQKSNK | 0 | 0 | 1 | 141646 | AAC70783.1 | O61723 |
| IAESKQLTVKQKTEQIK | 0 | 0 | 1 | 141671 | AAC70783.1 | O61723 |
| RADESKTDPQTEA | 0 | 0 | 1 | 141734 | AAC70783.1 | O61723 |
| AEDAAEEAKEAAKK | 0 | 0 | 1 | 142382 | ABZ89526.1 | A5KAW0 |
| AEQIQAELQKVKTA | 0 | 0 | 1 | 142383 | ACM88500.1 | A5KAW0 |
| DAANIAHQKWLKAT | 0 | 0 | 1 | 142391 | ABZ89526.1 | A5KAW0 |
| DKTIAAAKKAKKARE | 0 | 0 | 1 | 142407 | AAD43006.1 | A5KAW0 |
| EAPNSSRHHLRNGF | 0 | 0 | 1 | 142410 | ADD39051.1 | A5KAW0 |
| EAQKACEKAKKAHA | 0 | 0 | 1 | 142411 | ACZ67789.1 | A5KAW0 |
| EIAAEVAKAKVAK | 0 | 0 | 1 | 142413 | ABZ89526.1 | A5KAW0 |
| EVAKAEVLNAEVK | 0 | 0 | 1 | 142415 | ACM88500.1 | A5KAW0 |
| GDEFDTYDDIKKVT | 0 | 0 | 1 | 142424 | ADD39051.1 | A5KAW0 |
| KAAYGLLKTKNQYVL | 0 | 0 | 1 | 142426 | ADD39051.1 | A5KAW0 |
| KAQKEATAAKLKA | 0 | 0 | 1 | 142428 | ABZ89526.1 | A5KAW0 |
| KKAEEAKKIVDKI | 0 | 0 | 1 | 142431 | ACM88500.1 | A5KAW0 |
| KNDSLPHEEPNNLE | 0 | 0 | 1 | 142433 | ADD39051.1 | A5KAW0 |
| KSTEDLSVAKDK | 0 | 0 | 1 | 142434 | ACZ67779.1 | A5KAW0 |
| LDISPESADNITSK | 0 | 0 | 1 | 142440 | ADD39051.1 | A5KAW0 |
| SATAAETAKNNAVS | 0 | 0 | 1 | 142472 | ABZ89526.1 | A5KAW0 |
| SASDQPTQYEEEMTDYQK | 0 | 0 | 1 | 145997 | ACY68791.1 | A5K4Z2 |
| ALDSKSYTSILYG | 0 | 0 | 1 | 146764 | P19111.2 | P19111 |
| DDASVNGVRKRKQ | 0 | 0 | 1 | 146816 | P19111.2 | P19111 |
| DVAVFARGPQAHL | 0 | 0 | 1 | 146852 | P19111.2 | P19111 |
| ETHGGEDVAVFAR | 0 | 0 | 1 | 146901 | P19111.2 | P19111 |
| EYPDDASVNGVRK | 0 | 0 | 1 | 146910 | P19111.2 | P19111 |
| FGGYTLRGTSIFG | 0 | 0 | 1 | 146920 | P19111.2 | P19111 |
| GAQYVWNRTALLQ | 0 | 0 | 1 | 146952 | AAA30571.1 | P19111 |
| GGEDVAVFARGPQ | 0 | 0 | 1 | 146963 | P19111.2 | P19111 |
| GLAPSKALDSKSY | 0 | 0 | 1 | 146979 | P19111.2 | P19111 |
| GNGPGYALGGGSR | 0 | 0 | 1 | 146986 | P19111.2 | P19111 |
| GVRKRKQNLVQAW | 0 | 0 | 1 | 147005 | P19111.2 | P19111 |
| HSHVFSFGGYTLR | 0 | 0 | 1 | 147019 | P19111.2 | P19111 |
| ILYGNGPGYALGG | 0 | 0 | 1 | 147038 | P19111.2 | P19111 |
| KHQGAQYVWNRTA | 0 | 0 | 1 | 147064 | P19111.2 | P19111 |
| KRKQNLVQAWQAK | 0 | 0 | 1 | 147074 | P19111.2 | P19111 |
| MTEVALRVVSRNP | 0 | 0 | 1 | 147142 | P19111.2 | P19111 |
| PSKALDSKSYTSI | 0 | 0 | 1 | 147206 | P19111.2 | P19111 |
| QNLVQAWQAKHQG | 0 | 0 | 1 | 147227 | P19111.2 | P19111 |
| RGTSIFGLAPSKA | 0 | 0 | 1 | 147252 | P19111.2 | P19111 |
| SIFGLAPSKALDS | 0 | 0 | 1 | 147284 | P19111.2 | P19111 |
| SKSYTSILYGNGP | 0 | 0 | 1 | 147290 | P19111.2 | P19111 |
| SVNGVRKRKQNLV | 0 | 0 | 1 | 147310 | P19111.2 | P19111 |
| VALRVVSRNPRGF | 0 | 0 | 1 | 147353 | P19111.2 | P19111 |
| VFARGPQAHLVHG | 0 | 0 | 1 | 147366 | P19111.2 | P19111 |
| VFSFGGYTLRGTS | 0 | 0 | 1 | 147368 | P19111.2 | P19111 |
| VQAWQAKHQGAQY | 0 | 0 | 1 | 147388 | P19111.2 | P19111 |
| WQAKHQGAQYVWN | 0 | 0 | 1 | 147410 | P19111.2 | P19111 |
| YTLRGTSIFGLAP | 0 | 0 | 1 | 147441 | P19111.2 | P19111 |
| YTSILYGNGPGYA | 0 | 0 | 1 | 147442 | P19111.2 | P19111 |
| SPPVSAPAKAAAPPAAARSAEPHVG | 0 | 0 | 1 | 149109 | AAC47657.1 | Q4D528 |

| | | | | | | |
|---|---|---|---|---|---|---|
| AFLPVGAFNSDNFKSKGRG | 0 | 0 | 1 | 156236 | CAA76546.1 | A5K4Z2 |
| EIPYVKEVEAEDLRECNRIVFGAS | 0 | 0 | 1 | 156276 | CAA76546.1 | A5K4Z2 |
| MTDYQKIQQGFRQNNREMIKSAF | 0 | 0 | 1 | 156379 | CAA76546.1 | A5K4Z2 |
| RIVFGASASDQPTQYEEEMTDYQ | 0 | 0 | 1 | 156437 | CAA76546.1 | A5K4Z2 |
| RKNLGNAKFGLWVDGNCEEIPYVKE | 0 | 0 | 1 | 156438 | CAA76546.1 | A5K4Z2 |
| NFLRGKLKLYTGEACRTGDR | 0 | 0 | 1 | 164545 | P01588.1 | P01588 |
| RTITADTFRKLFRVYSNFLR | 0 | 0 | 1 | 164548 | P01588.1 | P01588 |
| TVPDTKVNFYAWKRMEVGQQ | 0 | 0 | 1 | 164553 | P01588.1 | P01588 |
| APPRLICDSRVLERYLLEAK | 0 | 0 | 1 | 164596 | P01588.1 | P01588 |
| AISPPDAASAAPLRTITADT | 0 | 0 | 1 | 165582 | P01588.1 | P01588 |
| ALLVNSSQPWEPLQLHVDKA | 0 | 0 | 1 | 165586 | P01588.1 | P01588 |
| GLALLSEAVLRGQALLVNSS | 0 | 0 | 1 | 165602 | P01588.1 | P01588 |
| LTTLLRALGAQKEAISPPDA | 0 | 0 | 1 | 165637 | P01588.1 | P01588 |
| QLHVDKAVSGLRSLTTLLRA | 0 | 0 | 1 | 165642 | P01588.1 | P01588 |
| RMEVGQQAVEVWQGLALLSE | 0 | 0 | 1 | 165647 | P01588.1 | P01588 |
| RYLLEAKEAENITTGCAEHC | 0 | 0 | 1 | 165649 | P01588.1 | P01588 |
| TGCAEHCSLNENITVPDTKV | 0 | 0 | 1 | 165671 | P01588.1 | P01588 |
| EEEDDDDDFGMGALF | 0 | 0 | 1 | 168420 | XP_821118.1 | Q4E3A3 |
| AGNMMGKDIYEKG | 0 | 0 | 1 | 170250 | Q7Z1H1 | W2U161 |
| AKDGAGGNAPKAA | 0 | 0 | 1 | 170279 | Q7Z1H1 | W2U161 |
| ARQKFLELHNSLR | 0 | 0 | 1 | 170383 | Q7Z1H1 | W2U161 |
| EQASKAWFGELAE | 0 | 0 | 1 | 170905 | Q7Z1H1 | W2U161 |
| ETVKLGCYVEACS | 0 | 0 | 1 | 170964 | Q7Z1H1 | W2U161 |
| FSRGVGHYTQMVW | 0 | 0 | 1 | 171100 | Q7Z1H1 | W2U161 |
| MCYVVCQYGPAGN | 0 | 0 | 1 | 172338 | Q7Z1H1 | W2U161 |
| MKTMAYDCEVEKT | 0 | 0 | 1 | 172375 | Q7Z1H1 | W2U161 |
| MVWQETVKLGCYV | 0 | 0 | 1 | 172432 | Q7Z1H1 | W2U161 |
| NGMSEEARQKFLE | 0 | 0 | 1 | 172484 | Q7Z1H1 | W2U161 |
| NIFMSSDSGMDKA | 0 | 0 | 1 | 172491 | Q7Z1H1 | W2U161 |
| PCSKCENCDKEKG | 0 | 0 | 1 | 172627 | Q7Z1H1 | W2U161 |
| PDNGMSEEARQKF | 0 | 0 | 1 | 172632 | Q7Z1H1 | W2U161 |
| SNMCYVVCQYGPA | 0 | 0 | 1 | 173272 | Q7Z1H1 | W2U161 |
| WQETVKLGCYVEA | 0 | 0 | 1 | 173808 | Q7Z1H1 | W2U161 |
| EDVRLTQETFAPLKA | 0 | 0 | 1 | 174084 | Q9NJQ6 | A0A183FXU3 |
| EDYRFQEGDWPNLKP | 0 | 0 | 1 | 174085 | O18598.3 | O18598 |
| EIDGKQTHQSVAISRYLGKQ | 0 | 0 | 1 | 174094 | O18598.3 | O18598 |
| EVDGQQLAQSQAICRYLAKT | 0 | 0 | 1 | 174101 | Q9NJQ6 | A0A183FXU3 |
| TDDRINASDWPSMKS | 0 | 0 | 1 | 174290 | Q86LL8 | A0A3P7DNC0 |
| TKKFDEVVKANGGYLAAGKLTWADF | 0 | 0 | 1 | 174296 | O18598.3 | O18598 |
| YDGDHQIVQSGAILRHLARK | 0 | 0 | 1 | 174318 | Q86LL8 | A0A3P7DNC0 |
| YHYDADENSKQKKWDPLKKE | 0 | 0 | 1 | 174324 | O18598.3 | O18598 |
| APPTYLDYVPQFLRW | 0 | 0 | 1 | 181556 | ACJ68328.1 | Q8IAK2 |
| DVLEKIATGIYNQEK | 0 | 0 | 1 | 181590 | CAA73831.1 | Q8I218 |
| FECGNDKIIGNSDKY | 0 | 0 | 1 | 181614 | ABM88777.1 | Q8IIZ4 |
| KRFDEGQKFECGNDK | 0 | 0 | 1 | 181696 | ABM88777.1 | Q8IIZ4 |
| LSATDVLEKIATGIY | 0 | 0 | 1 | 181722 | CAA73831.1 | Q8I218 |
| NEIELSARDVLENIG | 0 | 0 | 1 | 181737 | ABM88777.1 | Q8IIZ4 |
| PRRRHICDQNLEFLD | 0 | 0 | 1 | 181764 | ABM88777.1 | Q8IIZ4 |
| RFSNEGEAKCGSDKI | 0 | 0 | 1 | 181787 | CAA73831.1 | Q8I218 |
| SVPTNLDYVPQFLRW | 0 | 0 | 1 | 181806 | ABM88777.1 | Q8IIZ4 |
| TNVPTNLDYVPQFLR | 0 | 0 | 1 | 181818 | CAA73831.1 | Q8I218 |
| TVNNLSATDVLEKIA | 0 | 0 | 1 | 181823 | CAA73831.1 | Q8I218 |
| WKALTCSAPYYADYF | 0 | 0 | 1 | 181840 | ACJ68328.1 | Q8IAK2 |
| ADRAFLDQKPERVP | 0 | 0 | 1 | 195003 | Q26921 | Q4CPQ6 |
| AQELAREKKLADRA | 0 | 0 | 1 | 195008 | Q26921 | Q4CPQ6 |
| ERVPLADVPLDDDS | 0 | 0 | 1 | 195021 | Q26921 | Q4CPQ6 |
| LADVPLDDDSDFVA | 0 | 0 | 1 | 195035 | Q26921 | Q4CPQ6 |
| LDQKPERVPLADVP | 0 | 0 | 1 | 195036 | Q26921 | Q4CPQ6 |
| REKKLADRAFLDQK | 0 | 0 | 1 | 195053 | Q26921 | Q4CPQ6 |
| ENSANPPPPDRSLPTP | 0 | 0 | 1 | 196427 | XP_804683.1 | Q4CRD0 |
| FFQPQPQPQPQPQPQQF | 0 | 0 | 1 | 196456 | XP_816009.1 | Q4DNS1 |
| MDDDDDETYRGG | 0 | 0 | 1 | 196664 | SRC269190 | Q4DH88 |
| QRMSNASGGGGGGMRQNE | 0 | 0 | 1 | 196744 | XP_819174.1 | Q4DXQ2 |
| ATAASSSSLEK | 0 | 0 | 1 | 202815 | P30162.1 | P30162 |
| ETSEAAATVDLFAFTLDGGK | 0 | 0 | 1 | 208014 | ABM55675.1 | Q5CXH4 |
| GTDKKTAAGEAPSPSG | 0 | 0 | 1 | 226599 | AAQ73324.1 | Q4CTF7 |
| GTDKKTAAGGTPSPSG | 0 | 0 | 1 | 226600 | AFF60282.1 | Q4D3H3 |
| GTEKKAAAGEAPSPSG | 0 | 0 | 1 | 226602 | AAQ73322.1 | Q4CTF7 |
| GTENKPAAGEAPSQPG | 0 | 0 | 1 | 226603 | SRC292632 | Q4D3H3 |
| GTENKPATGEAPSQPG | 0 | 0 | 1 | 226604 | ACY02872.1 | Q4D3H3 |
| DAENPGGEVFNDNKKGLSRV | 0 | 0 | 1 | 226835 | XP_820771.1 | Q4E2A4 |
| EEDESKKKSCGDEGEPKVE | 0 | 0 | 1 | 227910 | XP_001567804.1 | A4HL70 |
| EVAEAPPAEAAPA | 0 | 0 | 1 | 227916 | XP_001567804.1 | A4HL70 |
| AWWKAHRDQVWKAIT | 0 | 0 | 1 | 236156 | SRC280142 | Q8I639 |
| DFFADPAHSLLSHAITTSPSAISK | 0 | 0 | 1 | 236162 | Q4DQ97 | Q4DQ97 |
| DQVPTYFDYVPQYLR | 0 | 0 | 1 | 236163 | ABB71270.1 | Q8IFQ2 |
| EDWWNANRQEIWKAL | 0 | 0 | 1 | 236165 | AAL12845.1 | C6KTF7 |
| EFGQKVHDEVHGEAK | 0 | 0 | 1 | 236166 | ABM88775.1 | Q8IAS4 |
| EQTSLGAIDVYATK | 0 | 0 | 1 | 236169 | Q4DAD9 | Q4DAD9 |
| ETAFTVKSMQTESKY | 0 | 0 | 1 | 236171 | AAC05730.1 | Q8IAS4 |
| EWAEDFCTKRKHKLQ | 0 | 0 | 1 | 236172 | AAL12845.1 | C6KTF7 |
| IDNQQKEFEKQKKKY | 0 | 0 | 1 | 236221 | AAL12845.1 | C6KTF7 |
| IFSKNIRNGNTILFD | 0 | 0 | 1 | 236222 | CAJ40439.1 | C6KSK8 |
| KVDYFIKNSDGSRGF | 0 | 0 | 1 | 236250 | ACT21622.1 | Q8I218 |
| NPCELKYQWHTNATR | 0 | 0 | 1 | 236266 | AAL12845.1 | C6KTF7 |
| NTKLQTLTLHQVREY | 0 | 0 | 1 | 236273 | SRC280142 | Q8I639 |
| PFSAYRPNYGNPCEL | 0 | 0 | 1 | 236277 | AEI26313.1 | Q8I098 |
| PPTYFDYVPQYLRWF | 0 | 0 | 1 | 236279 | AAL12845.1 | C6KTF7 |
| PQFLRWFEEWAEEFC | 0 | 0 | 1 | 236280 | AEI26313.1 | Q8I098 |
| SFADIGDIIRGRDLY | 0 | 0 | 1 | 236321 | ABM88775.1 | Q8IAS4 |
| SKKLYCSHNGYDCTK | 0 | 0 | 1 | 236324 | AEI26313.1 | Q8I098 |

| | | | | | | |
|---|---|---|---|---|---|---|
| SSYFRATCSDTGQGP | 0 | 0 | 1 | 236346 | SRC280142 | Q8I639 |
| TFPGLAAGTEGWR | 0 | 0 | 1 | 236348 | Q4DQ97 | Q4DQ97 |
| VDTVAAFFDAGSVR | 0 | 0 | 1 | 236361 | Q4DAD9 | Q4DAD9 |
| VGIIGYGDIGQASAK | 0 | 0 | 1 | 236362 | Q4DAE1 | Q4DAE1 |
| VLTNLDYVPQFLRWF | 0 | 0 | 1 | 236363 | AEI26313.1 | Q8I098 |
| YCSHNGYDCTKTIRN | 0 | 0 | 1 | 236371 | AEI26313.1 | Q8I098 |
| YDFEEADQQIR | 0 | 0 | 1 | 236372 | Q8I6L9 | Q4E4V7 |
| YRPNYGNPCELDYRF | 0 | 0 | 1 | 236374 | AEI26313.1 | Q8I098 |
| YTGYGIYKSGICTSL | 0 | 0 | 1 | 236375 | AEI26313.1 | Q8I098 |
| QTSGSTTPGPTTTT | 0 | 0 | 1 | 237898 | CAM37171.1 | A4H5E6 |
| SGTENKPATGEAPSQ | 0 | 0 | 1 | 242071 | XP_808931.1 | Q4D3H3 |
| TANGGSTSSTPPSGT | 0 | 0 | 1 | 242105 | XP_808931.1 | Q4D3H3 |
| TSSTPPSGTENKPAT | 0 | 0 | 1 | 242138 | XP_808931.1 | Q4D3H3 |
| AAAADKPSLFGQAAA | 0 | 0 | 1 | 244401 | XP_818927.1 | Q4DX15 |
| AAAAEEEEDDDMGFG | 0 | 0 | 1 | 244411 | XP_818851.1 | Q4DWU6 |
| AAAAEPEEEDDDDDF | 0 | 0 | 1 | 244419 | XP_821118.1 | Q4E3A3 |
| AAAAPAAAAEPEEED | 0 | 0 | 1 | 244468 | XP_821118.1 | Q4E3A3 |
| AAADKPSLFGQAAAG | 0 | 0 | 1 | 244526 | XP_818927.1 | Q4DX15 |
| AAAEEEEDDDMGFGL | 0 | 0 | 1 | 244595 | XP_818851.1 | Q4DWU6 |
| AAAEPEEEDDDDDFG | 0 | 0 | 1 | 244616 | XP_821118.1 | Q4E3A3 |
| AAAGDKLSLFGQAAA | 0 | 0 | 1 | 244642 | XP_818927.1 | Q4DX15 |
| AAAGDKPAPFGQAAA | 0 | 0 | 1 | 244643 | XP_818927.1 | Q4DX15 |
| AAAGDKPPPFGQAAA | 0 | 0 | 1 | 244644 | XP_818927.1 | Q4DX15 |
| AAAGDKPPPFGQAAE | 0 | 0 | 1 | 244645 | XP_818927.1 | Q4DX15 |
| AAAGDKPSLFGQAAA | 0 | 0 | 1 | 244646 | XP_818927.1 | Q4DX15 |
| AAAGDKPSPFGQAAA | 0 | 0 | 1 | 244647 | XP_818927.1 | Q4DX15 |
| AAAGDKPSPFGQGTA | 0 | 0 | 1 | 244648 | XP_818927.1 | Q4DX15 |
| AAAGGKPSPFGQAAA | 0 | 0 | 1 | 244654 | XP_818927.1 | Q4DX15 |
| AAAMDKVFTTGFAQE | 0 | 0 | 1 | 244757 | XP_819176.1 | Q4DXP9 |
| AAAMKTTTATTGDSD | 0 | 0 | 1 | 244759 | XP_808183.1 | Q4D1D1 |
| AAAMKTTTATTGNSD | 0 | 0 | 1 | 244760 | XP_807739.1 | Q4D033 |
| AAAPAAAAEEEEDDD | 0 | 0 | 1 | 244776 | XP_818851.1 | Q4DWU6 |
| AAAPKKAAAPKKAAA | 0 | 0 | 1 | 244793 | XP_814064.1 | Q4DI49 |
| AAAPKKAAAPQKAAA | 0 | 0 | 1 | 244794 | XP_814064.1 | Q4DI49 |
| AAAPKKTAAPKKAAA | 0 | 0 | 1 | 244795 | XP_814064.1 | Q4DI49 |
| AAAPKNGTATTGDSD | 0 | 0 | 1 | 244798 | XP_809903.1 | Q4D6A2 |
| AAAPQKAAAPKKTAA | 0 | 0 | 1 | 244811 | XP_814064.1 | Q4DI49 |
| AAAPQKAAVAKKAVR | 0 | 0 | 1 | 244812 | XP_814064.1 | Q4DI49 |
| AADKPSLFGQAAAGD | 0 | 0 | 1 | 245075 | XP_818927.1 | Q4DX15 |
| AADVEAGDGPDKTTT | 0 | 0 | 1 | 245110 | XP_804663.1 | Q4CRB0 |
| AAEEEEDDDMGFGLF | 0 | 0 | 1 | 245271 | XP_818851.1 | Q4DWU6 |
| AAEGDKPPPFGQAAA | 0 | 0 | 1 | 245290 | XP_818927.1 | Q4DX15 |
| AAEPEEEDDDDDFGM | 0 | 0 | 1 | 245372 | XP_821118.1 | Q4E3A3 |
| AAGDKLSLFGQAAAG | 0 | 0 | 1 | 245500 | XP_818927.1 | Q4DX15 |
| AAGDKPAPFGQAAAA | 0 | 0 | 1 | 245501 | XP_818927.1 | Q4DX15 |
| AAGDKPAPFGQAAAG | 0 | 0 | 1 | 245502 | XP_818927.1 | Q4DX15 |
| AAGDKPPPFGQAAAG | 0 | 0 | 1 | 245503 | XP_818927.1 | Q4DX15 |
| AAGDKPPPFGQAAEG | 0 | 0 | 1 | 245504 | XP_818927.1 | Q4DX15 |
| AAGDKPSLFGQAAAG | 0 | 0 | 1 | 245505 | XP_818927.1 | Q4DX15 |
| AAGDKPSPFGQAAAG | 0 | 0 | 1 | 245506 | XP_818927.1 | Q4DX15 |
| AAGDKPSPFGQGTAF | 0 | 0 | 1 | 245507 | XP_818927.1 | Q4DX15 |
| AAGGKPSPFGQAAAG | 0 | 0 | 1 | 245542 | XP_818927.1 | Q4DX15 |
| AAKAPAPKAAAPAPK | 0 | 0 | 1 | 245808 | XP_819301.1 | Q4DY30 |
| AAKPAAKAPAPKAEK | 0 | 0 | 1 | 245880 | XP_819301.1 | Q4DY30 |
| AAKPAAKPAAKPAAK | 0 | 0 | 1 | 245881 | XP_821284.1 | Q4E3R2 |
| AAKPAAKPAAKTAAK | 0 | 0 | 1 | 245882 | XP_821284.1 | Q4E3R2 |
| AAKPAAKSAAKPAAK | 0 | 0 | 1 | 245883 | XP_821284.1 | Q4E3R2 |
| AAKPAAKTAAKPAKK | 0 | 0 | 1 | 245884 | XP_821284.1 | Q4E3R2 |
| AAKPAAKTAAKTAAK | 0 | 0 | 1 | 245885 | XP_821284.1 | Q4E3R2 |
| AAKPAKKPAVKPTVK | 0 | 0 | 1 | 245886 | XP_821284.1 | Q4E3R2 |
| AAKPPAESPFKNVFG | 0 | 0 | 1 | 245894 | XP_818928.1 | Q4DX14 |
| AAKPPAESPFKSVFG | 0 | 0 | 1 | 245895 | XP_818928.1 | Q4DX14 |
| AAKPPAESPFRSVFG | 0 | 0 | 1 | 245896 | XP_818928.1 | Q4DX14 |
| AAKSAAKPAAKPAAK | 0 | 0 | 1 | 245918 | XP_821284.1 | Q4E3R2 |
| AAKTAAKPAAKSAAK | 0 | 0 | 1 | 245927 | XP_821284.1 | Q4E3R2 |
| AAKTAAKPAKKPAVK | 0 | 0 | 1 | 245928 | XP_821284.1 | Q4E3R2 |
| AAKTAAKTAAKPAAK | 0 | 0 | 1 | 245929 | XP_821284.1 | Q4E3R2 |
| AAMKTTTATTGDSDG | 0 | 0 | 1 | 246125 | XP_810231.1 | Q4D780 |
| AAMKTTTATTGDSDS | 0 | 0 | 1 | 246126 | XP_808183.1 | Q4D1D1 |
| AAMKTTTATTGNSDG | 0 | 0 | 1 | 246127 | XP_807739.1 | Q4D033 |
| AAPAAAAEEEEDDDM | 0 | 0 | 1 | 246207 | XP_818851.1 | Q4DWU6 |
| AAPKKAAAPKKAAAP | 0 | 0 | 1 | 246279 | XP_814064.1 | Q4DI49 |
| AAPKKAAAPQKAAAP | 0 | 0 | 1 | 246280 | XP_814064.1 | Q4DI49 |
| AAPKKAAAPQKAAVA | 0 | 0 | 1 | 246281 | XP_814064.1 | Q4DI49 |
| AAPKKTAAPKKAAAP | 0 | 0 | 1 | 246284 | XP_814064.1 | Q4DI49 |
| AAPKNGTATTGDSDG | 0 | 0 | 1 | 246290 | XP_809903.1 | Q4D6A2 |
| AAPQKAAAPKKTAAP | 0 | 0 | 1 | 246333 | XP_814064.1 | Q4DI49 |
| AARSMEIDGGEWTNG | 0 | 0 | 1 | 246547 | XP_815234.1 | Q4DEP9 |
| AATVIKTTTAATGDS | 0 | 0 | 1 | 246973 | XP_820637.1 | Q4E1Z5 |
| AAVDADTAALAALLE | 0 | 0 | 1 | 247003 | XP_803309.1 | Q4CNL0 |
| ADDDDDDDDDDGETK | 0 | 0 | 1 | 247437 | XP_804979.1 | Q4CS76 |
| ADGAGGGDGSAGEDL | 0 | 0 | 1 | 247579 | XP_817817.1 | Q4DTW8 |
| ADGDEEDYDVVDDDD | 0 | 0 | 1 | 247601 | XP_821672.1 | Q4E4Z2 |
| ADGDEEDYDVVDDLE | 0 | 0 | 1 | 247602 | XP_822030.1 | Q4E5T2 |
| ADGVGGGDGSAGEDL | 0 | 0 | 1 | 247679 | XP_821138.1 | Q4E3E7 |
| ADKPSLFGQAAAGDK | 0 | 0 | 1 | 247742 | XP_818927.1 | Q4DX15 |
| ADQPTQFSVGTPDAA | 0 | 0 | 1 | 247873 | XP_820186.1 | Q4E0M7 |
| ADSENSKTGCGEEAS | 0 | 0 | 1 | 247931 | XP_813566.1 | Q4DGQ5 |
| ADTAALAALLEVLQH | 0 | 0 | 1 | 247969 | XP_803309.1 | Q4CNL0 |
| AEDVGKKKDEEQEPN | 0 | 0 | 1 | 248418 | XP_809910.1 | Q4D6A6 |
| AEEDDEEDKDDDVKD | 0 | 0 | 1 | 248459 | XP_821399.1 | Q4E444 |

| | | | | | | |
|---|---|---|---|---|---|---|
| AEEEEDEDDEEEEEE | 0 | 0 | 1 | 248477 | XP_815190.1 | Q4DLD3 |
| AEGDKPPPFGQAAAG | 0 | 0 | 1 | 248666 | XP_818927.1 | Q4DX15 |
| AEPEEEDDDDDFGMG | 0 | 0 | 1 | 249142 | XP_821118.1 | Q4E3A3 |
| AEPKPAEPKSAEPKP | 0 | 0 | 1 | 249160 | XP_820186.1 | Q4E0M7 |
| AEPKPAESKSAEPKP | 0 | 0 | 1 | 249165 | XP_820186.1 | Q4E0M7 |
| AEPKPAESKSAGPKP | 0 | 0 | 1 | 249166 | XP_820186.1 | Q4E0M7 |
| AEPKSAEPKPAEPNA | 0 | 0 | 1 | 249171 | XP_820186.1 | Q4E0M7 |
| AEPKSAEPKPAEPNS | 0 | 0 | 1 | 249172 | XP_820186.1 | Q4E0M7 |
| AEPKSAEPKPAESKS | 0 | 0 | 1 | 249173 | XP_820186.1 | Q4E0M7 |
| AEPNSAEPKPAEPKS | 0 | 0 | 1 | 249180 | XP_820186.1 | Q4E0M7 |
| AEQEEISDDDDEGED | 0 | 0 | 1 | 249216 | XP_803649.1 | Q4CNY5 |
| AESKSAEPKPAEPKS | 0 | 0 | 1 | 249381 | XP_820186.1 | Q4E0M7 |
| AESTVIKTTTATTGD | 0 | 0 | 1 | 249434 | XP_820022.1 | Q4E058 |
| AETEVSRDPTDEHSD | 0 | 0 | 1 | 249490 | XP_814355.1 | Q4DIZ9 |
| AEVDEDEDDDDDDND | 0 | 0 | 1 | 249578 | XP_818224.1 | Q4DV33 |
| AGDKLSLFGQAAAGD | 0 | 0 | 1 | 250191 | XP_818927.1 | Q4DX15 |
| AGDKPAPFGQAAAAD | 0 | 0 | 1 | 250192 | XP_818927.1 | Q4DX15 |
| AGDKPAPFGQAAAGD | 0 | 0 | 1 | 250193 | XP_818927.1 | Q4DX15 |
| AGDKPPPFGQAAAGD | 0 | 0 | 1 | 250194 | XP_818927.1 | Q4DX15 |
| AGDKPPPFGQAAEGD | 0 | 0 | 1 | 250195 | XP_818927.1 | Q4DX15 |
| AGDKPSLFGQAAAGD | 0 | 0 | 1 | 250196 | XP_818927.1 | Q4DX15 |
| AGDKPSPFGQAAAGD | 0 | 0 | 1 | 250197 | XP_818927.1 | Q4DX15 |
| AGDKPSPFGQAAAGG | 0 | 0 | 1 | 250198 | XP_818927.1 | Q4DX15 |
| AGDKPSPFGQGTAFD | 0 | 0 | 1 | 250199 | XP_818927.1 | Q4DX15 |
| AGGKPSPFGQAAAGD | 0 | 0 | 1 | 250499 | XP_818927.1 | Q4DX15 |
| AGGKPSPFGQAAAGG | 0 | 0 | 1 | 250500 | XP_818927.1 | Q4DX15 |
| AGIKTTTATTGDSDG | 0 | 0 | 1 | 250657 | XP_821134.1 | Q4E3F2 |
| AGKSSEEGEIDDDSD | 0 | 0 | 1 | 250737 | XP_813526.1 | Q4DGM5 |
| AGPKPAEPKSAEPKP | 0 | 0 | 1 | 250910 | XP_820186.1 | Q4E0M7 |
| AGSDEEDQDEDEDEG | 0 | 0 | 1 | 251068 | XP_821081.1 | Q4E380 |
| AKAPAPKAAAPAPKA | 0 | 0 | 1 | 252233 | XP_819301.1 | Q4DY30 |
| AKKPAVKPTVKPAAK | 0 | 0 | 1 | 252558 | XP_821284.1 | Q4E3R2 |
| AKLGKSVGLTAALSP | 0 | 0 | 1 | 252587 | XP_807065.1 | Q4CY60 |
| AKPAAKAPAPKAEKK | 0 | 0 | 1 | 252653 | XP_819301.1 | Q4DY30 |
| AKPAAKPAAKPAAKP | 0 | 0 | 1 | 252654 | XP_821284.1 | Q4E3R2 |
| AKPAAKPAAKPAAKT | 0 | 0 | 1 | 252655 | XP_821284.1 | Q4E3R2 |
| AKPAAKPAAKTAAKP | 0 | 0 | 1 | 252656 | XP_821284.1 | Q4E3R2 |
| AKPAAKSAAKPAAKP | 0 | 0 | 1 | 252657 | XP_821284.1 | Q4E3R2 |
| AKPAAKTAAKPAKKP | 0 | 0 | 1 | 252658 | XP_821284.1 | Q4E3R2 |
| AKPAAKTAAKTAAKP | 0 | 0 | 1 | 252659 | XP_821284.1 | Q4E3R2 |
| AKPAKKPAVKPTVKP | 0 | 0 | 1 | 252660 | XP_821284.1 | Q4E3R2 |
| AKPPAESPFKNVFGA | 0 | 0 | 1 | 252682 | XP_818928.1 | Q4DX14 |
| AKPPAESPFKSVFGA | 0 | 0 | 1 | 252684 | XP_813515.1 | Q4DGM1 |
| AKPPAESPFRSVFGA | 0 | 0 | 1 | 252685 | XP_818928.1 | Q4DX14 |
| AKQSEREDDEENDEE | 0 | 0 | 1 | 252728 | XP_821138.1 | Q4E3E7 |
| AKSAAKPAAKPAAKP | 0 | 0 | 1 | 252771 | XP_821284.1 | Q4E3R2 |
| AKSANKPASKPAAKP | 0 | 0 | 1 | 252772 | XP_819301.1 | Q4DY30 |
| AKTAAKPAAKSAAKP | 0 | 0 | 1 | 252823 | XP_821284.1 | Q4E3R2 |
| AKTAAKPAKKPAVKP | 0 | 0 | 1 | 252824 | XP_821284.1 | Q4E3R2 |
| AKTAAKTAAKPAAKS | 0 | 0 | 1 | 252825 | XP_821284.1 | Q4E3R2 |
| AKTAVSKAAAPKKAA | 0 | 0 | 1 | 252840 | XP_814064.1 | Q4DI49 |
| ALELMLRESDGECAP | 0 | 0 | 1 | 253304 | XP_812097.1 | Q4DCJ6 |
| ALHEVLQYGRLIDDF | 0 | 0 | 1 | 253432 | XP_808204.1 | Q4D1F2 |
| ALSSFLLKDKKKVAT | 0 | 0 | 1 | 253999 | XP_807065.1 | Q4CY60 |
| AMKNTTATGDSDGST | 0 | 0 | 1 | 254277 | XP_815190.1 | Q4DLD3 |
| AMKNTTATTGDSDSS | 0 | 0 | 1 | 254278 | XP_811361.1 | Q4DAF3 |
| AMKTTAATTGDSDGS | 0 | 0 | 1 | 254288 | XP_820410.1 | Q4E1A7 |
| AMKTTTATTGDSDGS | 0 | 0 | 1 | 254290 | XP_815852.1 | Q4DNA4 |
| AMKTTTATTGDSDSS | 0 | 0 | 1 | 254291 | XP_808183.1 | Q4D1D1 |
| AMKTTTATTGNSDGS | 0 | 0 | 1 | 254292 | XP_807739.1 | Q4D033 |
| AMKTTTTTTGDSDGS | 0 | 0 | 1 | 254293 | XP_820308.1 | Q4E0Z8 |
| ANGGSTSSTPPSGTE | 0 | 0 | 1 | 254633 | XP_808931.1 | Q4D3H3 |
| ANKPASKPAAKPAAK | 0 | 0 | 1 | 254701 | XP_819301.1 | Q4DY30 |
| ANTLLYSDDELFLSQ | 0 | 0 | 1 | 254949 | XP_814919.1 | Q4DKL5 |
| APAAAAEEEEDDDMG | 0 | 0 | 1 | 255019 | XP_818851.1 | Q4DWU6 |
| APAAAAEPEEEDDDD | 0 | 0 | 1 | 255020 | XP_821118.1 | Q4E3A3 |
| APAAPAAPAAAAEAT | 0 | 0 | 1 | 255044 | XP_805627.1 | Q4CU24 |
| APAPKAAAPAPKAAA | 0 | 0 | 1 | 255133 | XP_819301.1 | Q4DY30 |
| APFGQAAAADKPSLF | 0 | 0 | 1 | 255309 | XP_818927.1 | Q4DX15 |
| APFGQAAAGDKLSLF | 0 | 0 | 1 | 255310 | XP_818927.1 | Q4DX15 |
| APFGQAAAGDKPAPF | 0 | 0 | 1 | 255311 | XP_818927.1 | Q4DX15 |
| APIEATATKTTTATT | 0 | 0 | 1 | 255422 | XP_815165.1 | Q4DLB5 |
| APKAAAPAPKAAAAA | 0 | 0 | 1 | 255442 | XP_819301.1 | Q4DY30 |
| APKKAAAPKKAAAPQ | 0 | 0 | 1 | 255460 | XP_814064.1 | Q4DI49 |
| APKKAAAPQKAAAPK | 0 | 0 | 1 | 255461 | XP_814064.1 | Q4DI49 |
| APKKAAAPQKAAVAK | 0 | 0 | 1 | 255462 | XP_814064.1 | Q4DI49 |
| APKKTAAPKKAAAPQ | 0 | 0 | 1 | 255467 | XP_814064.1 | Q4DI49 |
| APKNGTATTGDSDGS | 0 | 0 | 1 | 255478 | XP_809903.1 | Q4D6A2 |
| APQARKQQGRDAPQA | 0 | 0 | 1 | 255700 | XP_821251.1 | Q4E3M0 |
| APQARKQQGRNAPRS | 0 | 0 | 1 | 255701 | XP_821251.1 | Q4E3M0 |
| APQARKQQGRNEPQA | 0 | 0 | 1 | 255702 | XP_821251.1 | Q4E3M0 |
| APQKAAAPKKTAAPK | 0 | 0 | 1 | 255717 | XP_814064.1 | Q4DI49 |
| APSRLREIDGSLSSS | 0 | 0 | 1 | 255886 | XP_821144.1 | Q4E3E1 |
| APTTTTTNAPSRLRE | 0 | 0 | 1 | 256029 | XP_821144.1 | Q4E3E1 |
| AQEEGQYDAADVEAG | 0 | 0 | 1 | 256163 | XP_804663.1 | Q4CRB0 |
| AQKGNVRKLQPQPQK | 0 | 0 | 1 | 256304 | XP_821251.1 | Q4E3M0 |
| AQSEADADDDDPQRP | 0 | 0 | 1 | 256567 | XP_803335.1 | Q4CNN7 |
| AQSEADADVDDPQRP | 0 | 0 | 1 | 256568 | XP_808315.1 | Q4D1R3 |
| ARKQQGRDAPQARKQ | 0 | 0 | 1 | 256968 | XP_821251.1 | Q4E3M0 |
| ARKQQGRNEPQARRQ | 0 | 0 | 1 | 256970 | XP_821251.1 | Q4E3M0 |
| ARRQQGRNEPQARKQ | 0 | 0 | 1 | 257192 | XP_821251.1 | Q4E3M0 |

| | | | | | | |
|---|---|---|---|---|---|---|
| ARSMEIDGGEWTNGT | 0 | 0 | 1 | 257237 | XP_815234.1 | Q4DEP9 |
| ASKPAAKPAAKAPAP | 0 | 0 | 1 | 257986 | XP_819301.1 | Q4DY30 |
| ASREKDGEDTTSEDE | 0 | 0 | 1 | 258347 | XP_818224.1 | Q4DV33 |
| ASSEDEEEDDADGDE | 0 | 0 | 1 | 258410 | XP_821672.1 | Q4E4Z2 |
| ASSEDKEEDDADGDE | 0 | 0 | 1 | 258411 | XP_822030.1 | Q4E5T2 |
| ASSSDDAQTVGTEGG | 0 | 0 | 1 | 258486 | XP_820186.1 | Q4E0M7 |
| ASTPKVNGRETIDPD | 0 | 0 | 1 | 258601 | XP_816182.1 | Q4DP76 |
| ATAMKNTTATGDSDG | 0 | 0 | 1 | 258820 | XP_815190.1 | Q4DLD3 |
| ATAMKNTTATTGDSD | 0 | 0 | 1 | 258821 | XP_811361.1 | Q4DAF3 |
| ATAMKTTAATTGDSD | 0 | 0 | 1 | 258822 | XP_820410.1 | Q4E1A7 |
| ATAMKTTTATTGDSD | 0 | 0 | 1 | 258824 | XP_804905.1 | Q4CS02 |
| ATAMKTTTTTTGDSD | 0 | 0 | 1 | 258825 | XP_820308.1 | Q4E0Z8 |
| ATAQEEGQYDAADVE | 0 | 0 | 1 | 258854 | XP_804663.1 | Q4CRB0 |
| ATATKNTTATTGDSD | 0 | 0 | 1 | 258897 | XP_809011.1 | Q4D3Q7 |
| ATATKTTTATTGDSD | 0 | 0 | 1 | 258898 | XP_815165.1 | Q4DLB5 |
| ATATKTTTATTGNSD | 0 | 0 | 1 | 258899 | XP_820650.1 | Q4E1Y0 |
| ATAVKATTATTGDSD | 0 | 0 | 1 | 258908 | XP_822030.1 | Q4E5T2 |
| ATAVKATTATTGESD | 0 | 0 | 1 | 258909 | XP_821672.1 | Q4E4Z2 |
| ATEKKKNPEAASKKA | 0 | 0 | 1 | 259023 | XP_821284.1 | Q4E3R2 |
| ATGEAPSQPGASSGE | 0 | 0 | 1 | 259108 | XP_808931.1 | Q4D3H3 |
| ATIEEEGEFDEEEQY | 0 | 0 | 1 | 259200 | XP_816690.1 | Q4DQP2 |
| ATKNTTATTGDSDGS | 0 | 0 | 1 | 259281 | XP_821138.1 | Q4E3E7 |
| ATKNTTATTGDSDSS | 0 | 0 | 1 | 259282 | XP_809011.1 | Q4D3Q7 |
| ATKTTTATTGDSDGG | 0 | 0 | 1 | 259297 | XP_821379.1 | Q4E442 |
| ATKTTTATTGDSDGS | 0 | 0 | 1 | 259298 | XP_815165.1 | Q4DLB5 |
| ATKTTTATTGNSDGG | 0 | 0 | 1 | 259299 | XP_820650.1 | Q4E1Y0 |
| ATSAKLGKSVGLTAA | 0 | 0 | 1 | 259659 | XP_807065.1 | Q4CY60 |
| ATTATTGDSDSSTAA | 0 | 0 | 1 | 259766 | XP_822030.1 | Q4E5T2 |
| ATTATTGESDGSTAV | 0 | 0 | 1 | 259767 | XP_821672.1 | Q4E4Z2 |
| ATVIKTTTAATGDSD | 0 | 0 | 1 | 259928 | XP_820637.1 | Q4E1Z5 |
| AVDADTAALAALLEV | 0 | 0 | 1 | 260127 | XP_803309.1 | Q4CNL0 |
| AVKATTATTGDSDSS | 0 | 0 | 1 | 260505 | XP_822030.1 | Q4E5T2 |
| AVKATTATTGESDGS | 0 | 0 | 1 | 260506 | XP_821672.1 | Q4E4Z2 |
| AVSKAAAPKKAAAPK | 0 | 0 | 1 | 260956 | XP_814064.1 | Q4DI49 |
| AVSSVVPSLPADSEN | 0 | 0 | 1 | 261003 | XP_813566.1 | Q4DGQ5 |
| CATAQEEGQYDAADV | 0 | 0 | 1 | 261862 | XP_804663.1 | Q4CRB0 |
| CRDEGGDEGNDEAEG | 0 | 0 | 1 | 263336 | XP_820569.1 | Q4E1T5 |
| CVSLAARSMEIDGGE | 0 | 0 | 1 | 264136 | XP_815234.1 | Q4DEP9 |
| DAADVEAGDGPDKTT | 0 | 0 | 1 | 264274 | XP_804663.1 | Q4CRB0 |
| DADEEDEGEDEEEVE | 0 | 0 | 1 | 264415 | XP_813432.1 | Q4DGC2 |
| DADGDEEDYDVVDDD | 0 | 0 | 1 | 264425 | XP_821672.1 | Q4E4Z2 |
| DADGDEEDYDVVDDL | 0 | 0 | 1 | 264426 | XP_822030.1 | Q4E5T2 |
| DADTAALAALLEVLQ | 0 | 0 | 1 | 264449 | XP_803309.1 | Q4CNL0 |
| DAEVDEDEDDDDDDN | 0 | 0 | 1 | 264548 | XP_818224.1 | Q4DV33 |
| DAPQARKQQGRDAPQ | 0 | 0 | 1 | 264906 | XP_821251.1 | Q4E3M0 |
| DAPQARKQQGRNAPR | 0 | 0 | 1 | 264907 | XP_821251.1 | Q4E3M0 |
| DAPQARKQQGRNEPQ | 0 | 0 | 1 | 264908 | XP_821251.1 | Q4E3M0 |
| DATIEEEGEFDEEEQ | 0 | 0 | 1 | 265121 | XP_816690.1 | Q4DQP2 |
| DDADGDEEDYDVVDD | 0 | 0 | 1 | 265381 | XP_821672.1 | Q4E4Z2 |
| DDDDDDDDDDGETKA | 0 | 0 | 1 | 265466 | XP_804979.1 | Q4CS76 |
| DDDDDDDDDGETKAE | 0 | 0 | 1 | 265467 | XP_804979.1 | Q4CS76 |
| DDDDDDDEPEEGSKE | 0 | 0 | 1 | 265469 | XP_809163.1 | Q4D466 |
| DDDDDDEPEEEKEKS | 0 | 0 | 1 | 265471 | XP_815012.1 | Q4DKW6 |
| DDDDDDEPEEGSKEE | 0 | 0 | 1 | 265472 | XP_809163.1 | Q4D466 |
| DDDDDDNDNDADGEE | 0 | 0 | 1 | 265474 | XP_818224.1 | Q4DV33 |
| DDDDDNDNDADGEEE | 0 | 0 | 1 | 265478 | XP_818224.1 | Q4DV33 |
| DDDDEEDDSDEAEAQ | 0 | 0 | 1 | 265484 | XP_812035.1 | Q4DCD8 |
| DDDDEGEDEEEGPSP | 0 | 0 | 1 | 265485 | XP_803649.1 | Q4CNY5 |
| DDDDNDNDADGEEED | 0 | 0 | 1 | 265495 | XP_818224.1 | Q4DV33 |
| DDDDPAADAAGTREE | 0 | 0 | 1 | 265497 | XP_821399.1 | Q4E444 |
| DDDDPAADAAGTRKE | 0 | 0 | 1 | 265498 | XP_809910.1 | Q4D6A6 |
| DDDDPAADGAGTREE | 0 | 0 | 1 | 265499 | XP_809011.1 | Q4D3Q7 |
| DDDEDEEENEEDAEM | 0 | 0 | 1 | 265507 | XP_820493.1 | Q4E1H5 |
| DDDEEDDSDEAEAQE | 0 | 0 | 1 | 265508 | XP_812035.1 | Q4DCD8 |
| DDDEGEDEEEGPSPP | 0 | 0 | 1 | 265511 | XP_803649.1 | Q4CNY5 |
| DDDGDNDADDVCRFS | 0 | 0 | 1 | 265522 | XP_806363.1 | Q4CW66 |
| DDDNDNDADGEEEDN | 0 | 0 | 1 | 265542 | XP_818224.1 | Q4DV33 |
| DDDPAADAAGTREEK | 0 | 0 | 1 | 265546 | XP_821399.1 | Q4E444 |
| DDDPAADAAGTRKEK | 0 | 0 | 1 | 265547 | XP_809910.1 | Q4D6A6 |
| DDDPAADGAGTREEK | 0 | 0 | 1 | 265550 | XP_809011.1 | Q4D3Q7 |
| DDDPAADVAGTAEGK | 0 | 0 | 1 | 265551 | XP_815852.1 | Q4DNA4 |
| DDDSDVQNEIKVLDD | 0 | 0 | 1 | 265560 | XP_813526.1 | Q4DGM5 |
| DDEDEEENEEDAEMS | 0 | 0 | 1 | 265585 | XP_820493.1 | Q4E1H5 |
| DDEEDDSDEAEAQEG | 0 | 0 | 1 | 265589 | XP_812035.1 | Q4DCD8 |
| DDEEDKDDDVKDEEE | 0 | 0 | 1 | 265590 | XP_821399.1 | Q4E444 |
| DDEGEDEEEGPSPPT | 0 | 0 | 1 | 265600 | XP_803649.1 | Q4CNY5 |
| DDEKEVEKIDSENSK | 0 | 0 | 1 | 265614 | XP_821826.1 | Q4E572 |
| DDNDNDADGEEEDNE | 0 | 0 | 1 | 265848 | XP_818224.1 | Q4DV33 |
| DDPAADAAGTREEKQ | 0 | 0 | 1 | 265881 | XP_821399.1 | Q4E444 |
| DDPAADAAGTRKEKQ | 0 | 0 | 1 | 265882 | XP_809910.1 | Q4D6A6 |
| DDPAADGAGTREEKQ | 0 | 0 | 1 | 265885 | XP_808645.1 | Q4D2N7 |
| DDPAADVAGTAEGKQ | 0 | 0 | 1 | 265887 | XP_815852.1 | Q4DNA4 |
| DDSDVQNEIKVLDDG | 0 | 0 | 1 | 265964 | XP_813526.1 | Q4DGM5 |
| DDVEDGDGEDEEEGQ | 0 | 0 | 1 | 266065 | XP_809061.1 | Q4D3W5 |
| DEAAAPKNGTATTGD | 0 | 0 | 1 | 266139 | XP_809903.1 | Q4D6A2 |
| DEAEEEEDEDDEEEE | 0 | 0 | 1 | 266160 | XP_815190.1 | Q4DLD3 |
| DEAQFQQHQQHEHPA | 0 | 0 | 1 | 266199 | XP_818475.1 | Q4DVR9 |
| DEAQVQQHQRHERPA | 0 | 0 | 1 | 266203 | XP_817818.1 | Q4DTW6 |
| DEDDDDDDNDNDADG | 0 | 0 | 1 | 266242 | XP_818224.1 | Q4DV33 |
| DEDDDGDNDADDVCR | 0 | 0 | 1 | 266243 | XP_806363.1 | Q4CW66 |
| DEDDEEEEEEEEEEK | 0 | 0 | 1 | 266244 | XP_815190.1 | Q4DLD3 |

| | | | | | | |
|---|---|---|---|---|---|---|
| DEDEDDDDDNDNDA | 0 | 0 | 1 | 266251 | XP_818224.1 | Q4DV33 |
| DEEDKDDDVKDEEEE | 0 | 0 | 1 | 266308 | XP_821399.1 | Q4E444 |
| DEEEDDADGDEEDYD | 0 | 0 | 1 | 266316 | XP_821672.1 | Q4E4Z2 |
| DEEEEEDEEEEEDEE | 0 | 0 | 1 | 266327 | XP_815015.1 | Q4DKW3 |
| DEEEEEDEEEEGEEV | 0 | 0 | 1 | 266328 | XP_815015.1 | Q4DKW3 |
| DEEKEEDEEEEEDEE | 0 | 0 | 1 | 266374 | XP_815015.1 | Q4DKW3 |
| DEELKDEEKEEDEEE | 0 | 0 | 1 | 266379 | XP_815015.1 | Q4DKW3 |
| DEEQEKKEEEEEEDG | 0 | 0 | 1 | 266397 | XP_807741.1 | Q4D040 |
| DEGEDEEEGPSPPTP | 0 | 0 | 1 | 266457 | XP_803649.1 | Q4CNY5 |
| DEGGDEGNDEAEGGN | 0 | 0 | 1 | 266469 | XP_820569.1 | Q4E1T5 |
| DEGLLLPVDNDGDES | 0 | 0 | 1 | 266492 | XP_818651.1 | Q4DW89 |
| DEHGVDPTGTYQGDS | 0 | 0 | 1 | 266523 | XP_816690.1 | Q4DQP2 |
| DEHSDFLTMGSKMIA | 0 | 0 | 1 | 266533 | XP_814355.1 | Q4DIZ9 |
| DEKEDNDDDDDDEPE | 0 | 0 | 1 | 266573 | XP_815012.1 | Q4DKW6 |
| DEKEVEKIDSENSKL | 0 | 0 | 1 | 266582 | XP_821826.1 | Q4E572 |
| DELSFSDFLLLMTRE | 0 | 0 | 1 | 266685 | XP_805245.1 | Q4CSZ0 |
| DETQVQQHQRHEHPA | 0 | 0 | 1 | 266942 | XP_815868.1 | Q4DNB3 |
| DEVEEDEKEDNDDDD | 0 | 0 | 1 | 266958 | XP_815012.1 | Q4DKW6 |
| DFLLLMTREVDDTMA | 0 | 0 | 1 | 267111 | XP_805245.1 | Q4CSZ0 |
| DGDAEVDEDEDDDDD | 0 | 0 | 1 | 267320 | XP_818224.1 | Q4DV33 |
| DGDEEDYDVVDDDDD | 0 | 0 | 1 | 267336 | XP_821672.1 | Q4E4Z2 |
| DGDEEDYDVVDDLEE | 0 | 0 | 1 | 267337 | XP_822030.1 | Q4E5T2 |
| DGDESIDEEEEKEVE | 0 | 0 | 1 | 267340 | XP_820569.1 | Q4E1T5 |
| DGDESSDKEDAEPTA | 0 | 0 | 1 | 267341 | XP_818651.1 | Q4DW89 |
| DGDNEDEEVEDEAKI | 0 | 0 | 1 | 267359 | XP_820304.1 | Q4E102 |
| DGDPAADGAGTAEGK | 0 | 0 | 1 | 267361 | XP_807739.1 | Q4D033 |
| DGEICRDEGGDEGND | 0 | 0 | 1 | 267441 | XP_820569.1 | Q4E1T5 |
| DGHDFIREHIQIRTA | 0 | 0 | 1 | 267599 | XP_816335.1 | Q4DPM2 |
| DGIEQVEEEEEEEDE | 0 | 0 | 1 | 267617 | XP_807741.1 | Q4D040 |
| DGVDSREQDGEVTTS | 0 | 0 | 1 | 268186 | XP_807741.1 | Q4D040 |
| DGVGGGDGSAGEDLL | 0 | 0 | 1 | 268199 | XP_821138.1 | Q4E3E7 |
| DHEKQHQSDETQVQQ | 0 | 0 | 1 | 268292 | XP_809166.1 | Q4D462 |
| DHEKQQQSDEAQFQQ | 0 | 0 | 1 | 268293 | XP_818475.1 | Q4DVR9 |
| DHEKQQQSDEAQVQQ | 0 | 0 | 1 | 268294 | XP_817818.1 | Q4DTW6 |
| DHEKQQQSDETQVQQ | 0 | 0 | 1 | 268295 | XP_815868.1 | Q4DNB3 |
| DKDEDDDGDNDADDV | 0 | 0 | 1 | 268712 | XP_806363.1 | Q4CW66 |
| DKDPAADGAGTQEEK | 0 | 0 | 1 | 268724 | XP_813432.1 | Q4DGC2 |
| DKEEDDADGDEEDYD | 0 | 0 | 1 | 268741 | XP_822030.1 | Q4E5T2 |
| DKEEEEDDDDDDDEP | 0 | 0 | 1 | 268743 | XP_809163.1 | Q4D466 |
| DKLSLFGQAAAGDKP | 0 | 0 | 1 | 268893 | XP_818927.1 | Q4DX15 |
| DKPAPFGQAAAADKP | 0 | 0 | 1 | 268945 | XP_818927.1 | Q4DX15 |
| DKPAPFGQAAAGDKL | 0 | 0 | 1 | 268946 | XP_818927.1 | Q4DX15 |
| DKPAPFGQAAAGDKP | 0 | 0 | 1 | 268947 | XP_818927.1 | Q4DX15 |
| DKPPPFGQAAAGDKL | 0 | 0 | 1 | 268951 | XP_818927.1 | Q4DX15 |
| DKPPPFGQAAAGDKP | 0 | 0 | 1 | 268952 | XP_818927.1 | Q4DX15 |
| DKPPPFGQAAEGDKP | 0 | 0 | 1 | 268953 | XP_818927.1 | Q4DX15 |
| DKPSPFGQAAAGDKP | 0 | 0 | 1 | 268956 | XP_818927.1 | Q4DX15 |
| DKPSPFGQAAAGGKP | 0 | 0 | 1 | 268957 | XP_818927.1 | Q4DX15 |
| DKPSPFGQGTAFDAS | 0 | 0 | 1 | 268958 | XP_818927.1 | Q4DX15 |
| DNDDDDDDEPEEEKE | 0 | 0 | 1 | 269797 | XP_815012.1 | Q4DKW6 |
| DNDDDDLEEAEEERS | 0 | 0 | 1 | 269798 | XP_821418.1 | Q4E455 |
| DNDGDESSDKEDAEP | 0 | 0 | 1 | 269802 | XP_818651.1 | Q4DW89 |
| DNEKQQQSDEAQVQQ | 0 | 0 | 1 | 269853 | XP_809948.1 | Q4D6E7 |
| DPAADAAGTREEKQN | 0 | 0 | 1 | 270140 | XP_821399.1 | Q4E444 |
| DPAADAAGTRKEKQN | 0 | 0 | 1 | 270141 | XP_809910.1 | Q4D6A6 |
| DPAADGAETREEKKN | 0 | 0 | 1 | 270142 | XP_814148.1 | Q4DIF3 |
| DPAADGAGTAEGKQN | 0 | 0 | 1 | 270144 | XP_815190.1 | Q4DLD3 |
| DPAADGAGTQEEKQN | 0 | 0 | 1 | 270146 | XP_813432.1 | Q4DGC2 |
| DPTDEHSDFLTMGSK | 0 | 0 | 1 | 270492 | XP_814355.1 | Q4DIZ9 |
| DPTGTYQGDSDLQLE | 0 | 0 | 1 | 270496 | XP_816690.1 | Q4DQP2 |
| DQPTQFSVGTPDAAN | 0 | 0 | 1 | 270684 | XP_820186.1 | Q4E0M7 |
| DQTSAAAAANDSSPA | 0 | 0 | 1 | 270777 | XP_815868.1 | Q4DNB3 |
| DREFDYEDEDYEGYK | 0 | 0 | 1 | 270855 | XP_815234.1 | Q4DEP9 |
| DRFLAAAMDKVFTTG | 0 | 0 | 1 | 270881 | XP_819176.1 | Q4DXP9 |
| DSDLQLERINVYFDE | 0 | 0 | 1 | 271314 | XP_816690.1 | Q4DQP2 |
| DSENSKTGCGEEASD | 0 | 0 | 1 | 271389 | XP_813566.1 | Q4DGQ5 |
| DSGGEGSSGPGASPP | 0 | 0 | 1 | 271438 | XP_809166.1 | Q4D462 |
| DSREQDGEDATSEGK | 0 | 0 | 1 | 271751 | XP_808131.1 | Q4D180 |
| DSREQDGEGTTSEGH | 0 | 0 | 1 | 271752 | XP_809849.1 | Q4D646 |
| DSREQDGEVTTSEGE | 0 | 0 | 1 | 271753 | XP_807741.1 | Q4D040 |
| DSTTGDDGCDGDDGD | 0 | 0 | 1 | 271966 | XP_805004.1 | Q4CSA0 |
| DSVASREKDGEDTTS | 0 | 0 | 1 | 271974 | XP_818224.1 | Q4DV33 |
| DSVDSREQDGEDATS | 0 | 0 | 1 | 271982 | XP_808131.1 | Q4D180 |
| DSVDSREQDGEGTTS | 0 | 0 | 1 | 271983 | XP_809849.1 | Q4D646 |
| DVATEKKKNPEAASK | 0 | 0 | 1 | 272725 | XP_821284.1 | Q4E3R2 |
| DVEDGDGEDEEEGQL | 0 | 0 | 1 | 272800 | XP_809061.1 | Q4D3W5 |
| DVGEKKDEEQEKKEE | 0 | 0 | 1 | 272890 | XP_807741.1 | Q4D040 |
| DVGKKKDEEQEPNEE | 0 | 0 | 1 | 272900 | XP_809910.1 | Q4D6A6 |
| DVNSGTDGDESIDEE | 0 | 0 | 1 | 273059 | XP_820569.1 | Q4E1T5 |
| EAAAPKNGTATTGDS | 0 | 0 | 1 | 273576 | XP_809903.1 | Q4D6A2 |
| EADADDDDPQRPNPE | 0 | 0 | 1 | 273853 | XP_810637.1 | Q4D8D6 |
| EADDDDDDDDDDGET | 0 | 0 | 1 | 273869 | XP_804979.1 | Q4CS76 |
| EAEEEEDEDDEEEEE | 0 | 0 | 1 | 273973 | XP_815190.1 | Q4DLD3 |
| EAGSDEEDQDEDEDE | 0 | 0 | 1 | 274199 | XP_821081.1 | Q4E380 |
| EALHEVLQYGRLIDD | 0 | 0 | 1 | 274383 | XP_808204.1 | Q4D1F2 |
| EAPIEATATKTTTAT | 0 | 0 | 1 | 274538 | XP_815165.1 | Q4DLB5 |
| EATAMKNTTATGDSD | 0 | 0 | 1 | 274825 | XP_815190.1 | Q4DLD3 |
| EATAMKNTTATTGDS | 0 | 0 | 1 | 274826 | XP_811361.1 | Q4DAF3 |
| EATAMKTTAATTGDS | 0 | 0 | 1 | 274827 | XP_820410.1 | Q4E1A7 |
| EATAMKTTTATTGDS | 0 | 0 | 1 | 274829 | XP_804905.1 | Q4CS02 |
| EATAMKTTTTTTGDS | 0 | 0 | 1 | 274830 | XP_820308.1 | Q4E0Z8 |

| | | | | | | |
|---|---|---|---|---|---|---|
| EATATKNTTATTGDS | 0 | 0 | 1 | 274835 | XP_809011.1 | Q4D3Q7 |
| EATATKTTTATTGDS | 0 | 0 | 1 | 274836 | XP_815165.1 | Q4DLB5 |
| EATATKTTTATTGNS | 0 | 0 | 1 | 274837 | XP_820650.1 | Q4E1Y0 |
| EATAVKATTATTGDS | 0 | 0 | 1 | 274838 | XP_822030.1 | Q4E5T2 |
| EATAVKATTATTGES | 0 | 0 | 1 | 274839 | XP_821672.1 | Q4E4Z2 |
| EDDADGDEEDYDVVD | 0 | 0 | 1 | 275273 | XP_821672.1 | Q4E4Z2 |
| EDDDDDDDEPEEGSK | 0 | 0 | 1 | 275282 | XP_809163.1 | Q4D466 |
| EDDDDDDNDNDADGE | 0 | 0 | 1 | 275283 | XP_818224.1 | Q4DV33 |
| EDDDDEEDDSDEAEA | 0 | 0 | 1 | 275284 | XP_812035.1 | Q4DCD8 |
| EDDDEDEEENEEDAE | 0 | 0 | 1 | 275288 | XP_820493.1 | Q4E1H5 |
| EDDDGDNDADDVCRF | 0 | 0 | 1 | 275289 | XP_806363.1 | Q4CW66 |
| EDDEEDKDDDVKDEE | 0 | 0 | 1 | 275292 | XP_821399.1 | Q4E444 |
| EDDEEEEEEEEEEKE | 0 | 0 | 1 | 275293 | XP_815190.1 | Q4DLD3 |
| EDDEENDEEEDGSEE | 0 | 0 | 1 | 275294 | XP_821138.1 | Q4E3E7 |
| EDEDDDDDDNDNDAD | 0 | 0 | 1 | 275366 | XP_818224.1 | Q4DV33 |
| EDEDDEEEEEEEEEE | 0 | 0 | 1 | 275367 | XP_815190.1 | Q4DLD3 |
| EDEEEDDADGDEEDY | 0 | 0 | 1 | 275383 | XP_821672.1 | Q4E4Z2 |
| EDEEEEEDEEEEEDE | 0 | 0 | 1 | 275388 | XP_815015.1 | Q4DKW3 |
| EDEEEEEDEEEEGEE | 0 | 0 | 1 | 275389 | XP_815015.1 | Q4DKW3 |
| EDEKEDNDDDDDDEP | 0 | 0 | 1 | 275432 | XP_815012.1 | Q4DKW6 |
| EDGDNEDEEVEDEAK | 0 | 0 | 1 | 275511 | XP_820304.1 | Q4E102 |
| EDGIEQVEEEEEEED | 0 | 0 | 1 | 275544 | XP_807741.1 | Q4D040 |
| EDGVDSREQDGEVTT | 0 | 0 | 1 | 275596 | XP_807741.1 | Q4D040 |
| EDHEKQHQSDETQVQ | 0 | 0 | 1 | 275602 | XP_809166.1 | Q4D462 |
| EDHEKQQQSDEAQFQ | 0 | 0 | 1 | 275603 | XP_818475.1 | Q4DVR9 |
| EDHEKQQQSDEAQVQ | 0 | 0 | 1 | 275604 | XP_817818.1 | Q4DTW6 |
| EDHEKQQQSDETQVQ | 0 | 0 | 1 | 275605 | XP_815868.1 | Q4DNB3 |
| EDKDDDVKDEEEEEE | 0 | 0 | 1 | 275651 | XP_821399.1 | Q4E444 |
| EDKEEDDADGDEEDY | 0 | 0 | 1 | 275657 | XP_822030.1 | Q4E5T2 |
| EDKEEEEDDDDDDDE | 0 | 0 | 1 | 275659 | XP_809163.1 | Q4D466 |
| EDNDDDDDDEPEEEK | 0 | 0 | 1 | 275780 | XP_815012.1 | Q4DKW6 |
| EDNDDDDLEEAEEER | 0 | 0 | 1 | 275781 | XP_821418.1 | Q4E455 |
| EDNEKQQQSDEAQVQ | 0 | 0 | 1 | 275786 | XP_809948.1 | Q4D6E7 |
| EDSVASREKDGEDTT | 0 | 0 | 1 | 275959 | XP_818224.1 | Q4DV33 |
| EDSVDSREQDGEDAT | 0 | 0 | 1 | 275963 | XP_808131.1 | Q4D180 |
| EDVGKKKDEEQEPNE | 0 | 0 | 1 | 276060 | XP_809910.1 | Q4D6A6 |
| EEADDDDDDDDDDGE | 0 | 0 | 1 | 276159 | XP_804979.1 | Q4CS76 |
| EEAEDEDSDEEEETA | 0 | 0 | 1 | 276169 | XP_807807.1 | Q4D0A4 |
| EEAGSDEEDQDEDED | 0 | 0 | 1 | 276202 | XP_821081.1 | Q4E380 |
| EEDDADGDEEDYDVV | 0 | 0 | 1 | 276352 | XP_821672.1 | Q4E4Z2 |
| EEDDDDDDDEPEEGS | 0 | 0 | 1 | 276354 | XP_809163.1 | Q4D466 |
| EEDDDDEEDDSDEAE | 0 | 0 | 1 | 276355 | XP_812035.1 | Q4DCD8 |
| EEDDEEDKDDDVKDE | 0 | 0 | 1 | 276358 | XP_821399.1 | Q4E444 |
| EEDEAFDEADDDELA | 0 | 0 | 1 | 276372 | XP_806363.1 | Q4CW66 |
| EEDEDDEEEEEEEEE | 0 | 0 | 1 | 276374 | XP_815190.1 | Q4DLD3 |
| EEDEEEEEDEEEEED | 0 | 0 | 1 | 276385 | XP_815015.1 | Q4DKW3 |
| EEDEEEEEDEEEEGE | 0 | 0 | 1 | 276386 | XP_815015.1 | Q4DKW3 |
| EEDEKEDNDDDDDDE | 0 | 0 | 1 | 276399 | XP_815012.1 | Q4DKW6 |
| EEDGIEQVEEEEEEE | 0 | 0 | 1 | 276416 | XP_807741.1 | Q4D040 |
| EEDHEKQHQSDETQV | 0 | 0 | 1 | 276423 | XP_809166.1 | Q4D462 |
| EEDHEKQQQSDEAQF | 0 | 0 | 1 | 276424 | XP_818475.1 | Q4DVR9 |
| EEDHEKQQQSDEAQV | 0 | 0 | 1 | 276425 | XP_817818.1 | Q4DTW6 |
| EEDHEKQQQSDETQV | 0 | 0 | 1 | 276426 | XP_815868.1 | Q4DNB3 |
| EEDKDDDVKDEEEEE | 0 | 0 | 1 | 276434 | XP_821399.1 | Q4E444 |
| EEDKEEEEDDDDDDD | 0 | 0 | 1 | 276438 | XP_809163.1 | Q4D466 |
| EEDNDDDDLEEAEEE | 0 | 0 | 1 | 276452 | XP_821418.1 | Q4E455 |
| EEDNEKQQQSDEAQV | 0 | 0 | 1 | 276453 | XP_809948.1 | Q4D6E7 |
| EEEAGSDEEDQDEDE | 0 | 0 | 1 | 276522 | XP_821081.1 | Q4E380 |
| EEEDDADGDEEDYDV | 0 | 0 | 1 | 276535 | XP_821672.1 | Q4E4Z2 |
| EEEDDDDDDDEPEEG | 0 | 0 | 1 | 276536 | XP_809163.1 | Q4D466 |
| EEEDDDDEEDDSDEA | 0 | 0 | 1 | 276537 | XP_812035.1 | Q4DCD8 |
| EEEDEDDEEEEEEEE | 0 | 0 | 1 | 276546 | XP_815190.1 | Q4DLD3 |
| EEEDEEEEEDEEEEE | 0 | 0 | 1 | 276553 | XP_815015.1 | Q4DKW3 |
| EEEDEEEEEDEEEEG | 0 | 0 | 1 | 276554 | XP_815015.1 | Q4DKW3 |
| EEEDGIEQVEEEEEE | 0 | 0 | 1 | 276564 | XP_807741.1 | Q4D040 |
| EEEDHEKQHQSDETQ | 0 | 0 | 1 | 276567 | XP_809166.1 | Q4D462 |
| EEEDHEKQQQSDEAQ | 0 | 0 | 1 | 276568 | XP_817818.1 | Q4DTW6 |
| EEEDHEKQQQSDETQ | 0 | 0 | 1 | 276569 | XP_815868.1 | Q4DNB3 |
| EEEDKEEEEDDDDDD | 0 | 0 | 1 | 276573 | XP_809163.1 | Q4D466 |
| EEEDNDDDDLEEAEE | 0 | 0 | 1 | 276576 | XP_821418.1 | Q4E455 |
| EEEDNEKQQQSDEAQ | 0 | 0 | 1 | 276577 | XP_809948.1 | Q4D6E7 |
| EEEEDDDDDDDEPEE | 0 | 0 | 1 | 276597 | XP_809163.1 | Q4D466 |
| EEEEDEDDEEEEEEE | 0 | 0 | 1 | 276599 | XP_815190.1 | Q4DLD3 |
| EEEEDEEEEEDEEEE | 0 | 0 | 1 | 276603 | XP_815015.1 | Q4DKW3 |
| EEEEDGIEQVEEEEE | 0 | 0 | 1 | 276609 | XP_807741.1 | Q4D040 |
| EEEEDHEKQHQSDET | 0 | 0 | 1 | 276611 | XP_809166.1 | Q4D462 |
| EEEEDHEKQQQSDEA | 0 | 0 | 1 | 276612 | XP_817818.1 | Q4DTW6 |
| EEEEDHEKQQQSDET | 0 | 0 | 1 | 276613 | XP_815868.1 | Q4DNB3 |
| EEEEDNEKQQQSDEA | 0 | 0 | 1 | 276616 | XP_809948.1 | Q4D6E7 |
| EEEEEDEEEEEDEEE | 0 | 0 | 1 | 276626 | XP_815015.1 | Q4DKW3 |
| EEEEEDEEEEGEEVE | 0 | 0 | 1 | 276627 | XP_815015.1 | Q4DKW3 |
| EEEEEDGIEQVEEEE | 0 | 0 | 1 | 276629 | XP_807741.1 | Q4D040 |
| EEEEEDHEKQHQSDE | 0 | 0 | 1 | 276630 | XP_809166.1 | Q4D462 |
| EEEEEDHEKQQQSDE | 0 | 0 | 1 | 276631 | XP_817818.1 | Q4DTW6 |
| EEEEEDNEKQQQSDE | 0 | 0 | 1 | 276632 | XP_809948.1 | Q4D6E7 |
| EEEEEEDGIEQVEEE | 0 | 0 | 1 | 276636 | XP_807741.1 | Q4D040 |
| EEEEEEDHEKQQQSD | 0 | 0 | 1 | 276638 | XP_817818.1 | Q4DTW6 |
| EEEEEEDNEKQQQSD | 0 | 0 | 1 | 276639 | XP_809948.1 | Q4D6E7 |
| EEEEEEEDHEKQQQS | 0 | 0 | 1 | 276643 | XP_817818.1 | Q4DTW6 |
| EEEEEEEEGSDEEDQ | 0 | 0 | 1 | 276675 | XP_820742.1 | Q4E281 |
| EEEEEEEGSDEEDQD | 0 | 0 | 1 | 276684 | XP_820742.1 | Q4E281 |

| | | | | | | |
|---|---|---|---|---|---|---|
| EEEEEEGSDEEDQDE | 0 | 0 | 1 | 276702 | XP_820742.1 | Q4E281 |
| EEEEEENERQQQSDE | 0 | 0 | 1 | 276717 | XP_819538.1 | Q4DYS6 |
| EEEEEGSDEEDQDER | 0 | 0 | 1 | 276731 | XP_820742.1 | Q4E281 |
| EEEEENERQQQSDET | 0 | 0 | 1 | 276751 | XP_819538.1 | Q4DYS6 |
| EEEENERQQQSDETQ | 0 | 0 | 1 | 276809 | XP_819538.1 | Q4DYS6 |
| EEEGEDHEKQQQSDE | 0 | 0 | 1 | 276849 | XP_812183.1 | Q4DCT1 |
| EEENERQQQSDETQV | 0 | 0 | 1 | 276975 | XP_819538.1 | Q4DYS6 |
| EEGEDHEKQQQSDEA | 0 | 0 | 1 | 277143 | XP_812183.1 | Q4DCT1 |
| EEGEEEDHEKQQQSD | 0 | 0 | 1 | 277146 | XP_815870.1 | Q4DNB0 |
| EEGEIDDDSDVQNEI | 0 | 0 | 1 | 277161 | XP_813526.1 | Q4DGM5 |
| EEGERMTEKEDGDNE | 0 | 0 | 1 | 277169 | XP_820304.1 | Q4E102 |
| EEGNQDADEEDEGED | 0 | 0 | 1 | 277227 | XP_813432.1 | Q4DGC2 |
| EEGQYDAADVEAGDG | 0 | 0 | 1 | 277245 | XP_804663.1 | Q4CRB0 |
| EEIRENDGDIWKTEA | 0 | 0 | 1 | 277361 | XP_814919.1 | Q4DKL5 |
| EEISDDDDEGEDEEE | 0 | 0 | 1 | 277367 | XP_803649.1 | Q4CNY5 |
| EEKEEDEEEEEDEEE | 0 | 0 | 1 | 277420 | XP_815015.1 | Q4DKW3 |
| EELKDEEKEEDEEEE | 0 | 0 | 1 | 277652 | XP_815015.1 | Q4DKW3 |
| EENERQQQSDETQVQ | 0 | 0 | 1 | 277812 | XP_819538.1 | Q4DYS6 |
| EEQEKKEEEEEEDGI | 0 | 0 | 1 | 277955 | XP_807741.1 | Q4D040 |
| EEVELYSFEDDEEDE | 0 | 0 | 1 | 278419 | XP_811579.1 | Q4DB20 |
| EFDYEDEDYEGYKSF | 0 | 0 | 1 | 278592 | XP_815234.1 | Q4DEP9 |
| EGDKPPPFGQAAAGD | 0 | 0 | 1 | 278906 | XP_818927.1 | Q4DX15 |
| EGEDEEEGPSPPTPP | 0 | 0 | 1 | 278949 | XP_803649.1 | Q4CNY5 |
| EGEDHEKQQQSDEAQ | 0 | 0 | 1 | 278951 | XP_812183.1 | Q4DCT1 |
| EGEEEDHEKQQQSDE | 0 | 0 | 1 | 278958 | XP_815870.1 | Q4DNB0 |
| EGEIDDDSDVQNEIK | 0 | 0 | 1 | 279007 | XP_813526.1 | Q4DGM5 |
| EGEIMTENEDDDEDE | 0 | 0 | 1 | 279010 | XP_820493.1 | Q4E1H5 |
| EGERMTEKEDGDNED | 0 | 0 | 1 | 279059 | XP_820304.1 | Q4E102 |
| EGESSSFYSREVDVS | 0 | 0 | 1 | 279071 | XP_807066.1 | Q4CY59 |
| EGGDEGNDEAEGGND | 0 | 0 | 1 | 279121 | XP_820569.1 | Q4E1T5 |
| EGGERMTEKEDGDNE | 0 | 0 | 1 | 279133 | XP_806129.1 | Q4CVH7 |
| EGLLLPVDNDGDESS | 0 | 0 | 1 | 279407 | XP_818651.1 | Q4DW89 |
| EGNQDADEEDEGEDE | 0 | 0 | 1 | 279494 | XP_813432.1 | Q4DGC2 |
| EGQYDAADVEAGDGP | 0 | 0 | 1 | 279628 | XP_804663.1 | Q4CRB0 |
| EGSQSTLAATDTANN | 0 | 0 | 1 | 279751 | XP_809167.1 | Q4D463 |
| EGTAMKTTTATTGDS | 0 | 0 | 1 | 279801 | XP_820304.1 | Q4E102 |
| EGTGGDDVEDGDGED | 0 | 0 | 1 | 279818 | XP_809061.1 | Q4D3W5 |
| EHGVDPTGTYQGDSD | 0 | 0 | 1 | 280047 | XP_816690.1 | Q4DQP2 |
| EHSDFLTMGSKMIAA | 0 | 0 | 1 | 280178 | XP_814355.1 | Q4DIZ9 |
| EICRDEGGDEGNDEA | 0 | 0 | 1 | 280260 | XP_820569.1 | Q4E1T5 |
| EIDDDSDVQNEIKVL | 0 | 0 | 1 | 280264 | XP_813526.1 | Q4DGM5 |
| EIDGSLSSSAWVCAP | 0 | 0 | 1 | 280272 | XP_815995.1 | Q4DNQ0 |
| EIDGSLSSSAWVCAS | 0 | 0 | 1 | 280273 | XP_821144.1 | Q4E3E1 |
| EIMTENEDDDEDEEE | 0 | 0 | 1 | 280422 | XP_820493.1 | Q4E1H5 |
| EIRENDGDIWKTEAW | 0 | 0 | 1 | 280502 | XP_814919.1 | Q4DKL5 |
| EISDDDDEGEDEEEG | 0 | 0 | 1 | 280528 | XP_803649.1 | Q4CNY5 |
| EKAAVDADTAALAAL | 0 | 0 | 1 | 280623 | XP_803309.1 | Q4CNL0 |
| EKDGEDTTSEDEKNV | 0 | 0 | 1 | 280764 | XP_818224.1 | Q4DV33 |
| EKEDGDNEDEEVEDE | 0 | 0 | 1 | 280819 | XP_820304.1 | Q4E102 |
| EKEDNDDDDDDEPEE | 0 | 0 | 1 | 280824 | XP_815012.1 | Q4DKW6 |
| EKEEDEEEEEDEEEE | 0 | 0 | 1 | 280829 | XP_815015.1 | Q4DKW3 |
| EKEVEKIDSENSKLQ | 0 | 0 | 1 | 281003 | XP_821826.1 | Q4E572 |
| EKIDSENSKLQQEMN | 0 | 0 | 1 | 281123 | XP_821826.1 | Q4E572 |
| EKKDEEQEKKEEEEE | 0 | 0 | 1 | 281150 | XP_807741.1 | Q4D040 |
| EKKEEEEEEDGIEQV | 0 | 0 | 1 | 281164 | XP_807741.1 | Q4D040 |
| EKKGAAKAPAPKAAA | 0 | 0 | 1 | 281176 | XP_819301.1 | Q4DY30 |
| EKKKNPEAASKKAPL | 0 | 0 | 1 | 281200 | XP_821284.1 | Q4E3R2 |
| EKQHQSDETQVQQHQ | 0 | 0 | 1 | 281530 | XP_809166.1 | Q4D462 |
| EKQQQSDEAQFQQHQ | 0 | 0 | 1 | 281574 | XP_818475.1 | Q4DVR9 |
| EKQQQSDEAQVQQHQ | 0 | 0 | 1 | 281575 | XP_809948.1 | Q4D6E7 |
| EKQQQSDETQVQQHQ | 0 | 0 | 1 | 281576 | XP_815868.1 | Q4DNB3 |
| EKTDEAEEEEDEDDE | 0 | 0 | 1 | 281757 | XP_815190.1 | Q4DLD3 |
| ELKDEEKEEDEEEEE | 0 | 0 | 1 | 282229 | XP_815015.1 | Q4DKW3 |
| ELMLRESDGECAPFT | 0 | 0 | 1 | 282376 | XP_812097.1 | Q4DCJ6 |
| ELSFSDFLLLMTREV | 0 | 0 | 1 | 282627 | XP_805245.1 | Q4CSZ0 |
| ENEDDDEDEEENEED | 0 | 0 | 1 | 283193 | XP_820493.1 | Q4E1H5 |
| ENERQQQSDETQVQQ | 0 | 0 | 1 | 283273 | XP_819538.1 | Q4DYS6 |
| ENKPATGEAPSQPGA | 0 | 0 | 1 | 283441 | XP_808931.1 | Q4D3H3 |
| ENLDGDEEEGDVSQD | 0 | 0 | 1 | 283468 | XP_813607.1 | Q4DGU9 |
| EPEEEDDDDDFGMGA | 0 | 0 | 1 | 283911 | XP_821118.1 | Q4E3A3 |
| EPKPAEPKSAEPKPA | 0 | 0 | 1 | 284041 | XP_820186.1 | Q4E0M7 |
| EPKPAESKSAEPKPA | 0 | 0 | 1 | 284046 | XP_820186.1 | Q4E0M7 |
| EPKPAESKSAGPKPA | 0 | 0 | 1 | 284047 | XP_820186.1 | Q4E0M7 |
| EPKSAEPKPAEPKSA | 0 | 0 | 1 | 284056 | XP_820186.1 | Q4E0M7 |
| EPKSAEPKPAEPNAA | 0 | 0 | 1 | 284057 | XP_820186.1 | Q4E0M7 |
| EPKSAEPKPAEPNSA | 0 | 0 | 1 | 284058 | XP_820186.1 | Q4E0M7 |
| EPKSAEPKPAESKSA | 0 | 0 | 1 | 284060 | XP_820186.1 | Q4E0M7 |
| EPNSAEPKPAEPKSA | 0 | 0 | 1 | 284136 | XP_820186.1 | Q4E0M7 |
| EPQARKQQGRDAPQA | 0 | 0 | 1 | 284197 | XP_821251.1 | Q4E3M0 |
| EPQARRQQGRNEPQA | 0 | 0 | 1 | 284198 | XP_821251.1 | Q4E3M0 |
| EPQTTIKQPGGIKTP | 0 | 0 | 1 | 284237 | XP_819902.1 | Q4DZS8 |
| EQDGEDTTSEGEKNV | 0 | 0 | 1 | 284450 | XP_812835.1 | Q4DEM8 |
| EQEEISDDDDEGEDE | 0 | 0 | 1 | 284466 | XP_803649.1 | Q4CNY5 |
| EQEKKEEEEEEDGIE | 0 | 0 | 1 | 284479 | XP_807741.1 | Q4D040 |
| EQKTAPFGQAAAGDK | 0 | 0 | 1 | 284627 | XP_818927.1 | Q4DX15 |
| EREDDEENDEEEDGS | 0 | 0 | 1 | 285122 | XP_821138.1 | Q4E3E7 |
| EREVDGHDFIREHIQ | 0 | 0 | 1 | 285203 | XP_816335.1 | Q4DPM2 |
| ERIEEADDDDDDDDD | 0 | 0 | 1 | 285302 | XP_804979.1 | Q4CS76 |
| ERMTEKEDGDNEDEE | 0 | 0 | 1 | 285453 | XP_820304.1 | Q4E102 |
| ERPVEATATKNTTAT | 0 | 0 | 1 | 285538 | XP_810952.1 | Q4D9A4 |
| ERQQQSDETQVQQQE | 0 | 0 | 1 | 285586 | XP_819538.1 | Q4DYS6 |

| | | | | | | |
|---|---|---|---|---|---|---|
| ERTLQVAGIKTTTAT | 0 | 0 | 1 | 285750 | XP_821134.1 | Q4E3F2 |
| ESKSAEPKPAEPKSA | 0 | 0 | 1 | 286273 | XP_820186.1 | Q4E0M7 |
| ESSSFYSREVDVSQC | 0 | 0 | 1 | 286656 | XP_807066.1 | Q4CY59 |
| ESTAMKTTTATTGDS | 0 | 0 | 1 | 286679 | XP_815852.1 | Q4DNA4 |
| ESTVIKTTTATTGDS | 0 | 0 | 1 | 286740 | XP_820022.1 | Q4E058 |
| ESTVMKTTTATTGDS | 0 | 0 | 1 | 286741 | XP_813432.1 | Q4DGC2 |
| ETEVSRDPTDEHSDF | 0 | 0 | 1 | 287109 | XP_814355.1 | Q4DIZ9 |
| ETPAESTVIKTTTAT | 0 | 0 | 1 | 287413 | XP_820022.1 | Q4E058 |
| ETPLEATATKNTTAT | 0 | 0 | 1 | 287457 | XP_821138.1 | Q4E3E7 |
| ETPSTTTTRAPSRLR | 0 | 0 | 1 | 287487 | XP_815995.1 | Q4DNQ0 |
| ETPVAATVIKTTTAA | 0 | 0 | 1 | 287510 | XP_820637.1 | Q4E1Z5 |
| ETPVEATAMKNTTAT | 0 | 0 | 1 | 287517 | XP_811361.1 | Q4DAF3 |
| ETPVEATAMKTTTAT | 0 | 0 | 1 | 287518 | XP_804905.1 | Q4CS02 |
| ETPVEATATKNTTAT | 0 | 0 | 1 | 287520 | XP_809011.1 | Q4D3Q7 |
| ETPVEATATKTTTAT | 0 | 0 | 1 | 287521 | XP_820650.1 | Q4E1Y0 |
| ETPVEATAVKATTAT | 0 | 0 | 1 | 287522 | XP_821672.1 | Q4E4Z2 |
| ETPVEGTAMKTTTAT | 0 | 0 | 1 | 287525 | XP_820304.1 | Q4E102 |
| ETPVESTAMKTTTAT | 0 | 0 | 1 | 287526 | XP_815852.1 | Q4DNA4 |
| ETPVESTVMKTTTAT | 0 | 0 | 1 | 287528 | XP_813432.1 | Q4DGC2 |
| ETPVETTTMKNTTAT | 0 | 0 | 1 | 287529 | XP_806129.1 | Q4CVH7 |
| ETPVTAAAMKTTTAT | 0 | 0 | 1 | 287537 | XP_808183.1 | Q4D1D1 |
| ETTTMKNTTATTGDS | 0 | 0 | 1 | 287756 | XP_806129.1 | Q4CVH7 |
| EVDDTMADELRSAFF | 0 | 0 | 1 | 287928 | XP_805245.1 | Q4CSZ0 |
| EVDEDEDDDDDDNDN | 0 | 0 | 1 | 287932 | XP_818224.1 | Q4DV33 |
| EVDGHDFIREHIQIR | 0 | 0 | 1 | 287942 | XP_816335.1 | Q4DPM2 |
| EVEEDEKEDNDDDDD | 0 | 0 | 1 | 287976 | XP_815012.1 | Q4DKW6 |
| EVEKIDSENSKLQQE | 0 | 0 | 1 | 288015 | XP_821826.1 | Q4E572 |
| EVELYSFEDDEEDEN | 0 | 0 | 1 | 288028 | XP_811579.1 | Q4DB20 |
| EVKKAAKPAAKTAAK | 0 | 0 | 1 | 288182 | XP_821284.1 | Q4E3R2 |
| EVLQYGRLIDDFIVV | 0 | 0 | 1 | 288261 | XP_808204.1 | Q4D1F2 |
| EVSRDPTDEHSDFLT | 0 | 0 | 1 | 288498 | XP_814355.1 | Q4DIZ9 |
| FANTLLYSDDELFLS | 0 | 0 | 1 | 289147 | XP_814919.1 | Q4DKL5 |
| FDDEKEVEKIDSENS | 0 | 0 | 1 | 289399 | XP_821826.1 | Q4E572 |
| FDVATEKKKNPEAAS | 0 | 0 | 1 | 289539 | XP_821284.1 | Q4E3R2 |
| FEATAMKTTTTTTGD | 0 | 0 | 1 | 289575 | XP_820308.1 | Q4E0Z8 |
| FEREVDGHDFIREHI | 0 | 0 | 1 | 289706 | XP_816335.1 | Q4DPM2 |
| FGQAAAADKPSLFGQ | 0 | 0 | 1 | 290260 | XP_818927.1 | Q4DX15 |
| FGQAAAGDKLSLFGQ | 0 | 0 | 1 | 290261 | XP_818927.1 | Q4DX15 |
| FGQAAAGDKPAPFGQ | 0 | 0 | 1 | 290262 | XP_818927.1 | Q4DX15 |
| FGQAAAGDKPPPFGQ | 0 | 0 | 1 | 290263 | XP_818927.1 | Q4DX15 |
| FGQAAAGDKPSLFGQ | 0 | 0 | 1 | 290264 | XP_818927.1 | Q4DX15 |
| FGQAAAGGKPSPFGQ | 0 | 0 | 1 | 290266 | XP_818927.1 | Q4DX15 |
| FGQAAEGDKPPPFGQ | 0 | 0 | 1 | 290267 | XP_818927.1 | Q4DX15 |
| FLAAAMDKVFTTGFA | 0 | 0 | 1 | 290815 | XP_819176.1 | Q4DXP9 |
| FLLKDKKKVATAKVA | 0 | 0 | 1 | 290997 | XP_807065.1 | Q4CY60 |
| FLLLMTREVDDTMAD | 0 | 0 | 1 | 291008 | XP_805245.1 | Q4CSZ0 |
| FSDFLLLMTREVDDT | 0 | 0 | 1 | 292071 | XP_805245.1 | Q4CSZ0 |
| FTAKPPAESPFKSVF | 0 | 0 | 1 | 292416 | XP_813515.1 | Q4DGM1 |
| FYSREVDVSQCNMQE | 0 | 0 | 1 | 293133 | XP_807066.1 | Q4CY59 |
| GAAAAPAAAAEPEEE | 0 | 0 | 1 | 293157 | XP_821118.1 | Q4E3A3 |
| GAAKAPAPKAAAPAP | 0 | 0 | 1 | 293247 | XP_819301.1 | Q4DY30 |
| GAQKGNVRKLQPQPQ | 0 | 0 | 1 | 294063 | XP_821251.1 | Q4E3M0 |
| GAQSEADADDDDPQR | 0 | 0 | 1 | 294081 | XP_810637.1 | Q4D8D6 |
| GAQSEADADVDDPQR | 0 | 0 | 1 | 294082 | XP_808315.1 | Q4D1R3 |
| GASSSDDAQTVGTEG | 0 | 0 | 1 | 294224 | XP_820186.1 | Q4E0M7 |
| GDAEVDEDEDDDDDD | 0 | 0 | 1 | 294680 | XP_818224.1 | Q4DV33 |
| GDDDPAADGAGTPEG | 0 | 0 | 1 | 294761 | XP_804905.1 | Q4CS02 |
| GDDDPAADVAGTAEG | 0 | 0 | 1 | 294762 | XP_815852.1 | Q4DNA4 |
| GDDVEDGDGEDEEEG | 0 | 0 | 1 | 294806 | XP_809061.1 | Q4D3W5 |
| GDEEDYDVVDDDDDV | 0 | 0 | 1 | 294826 | XP_821672.1 | Q4E4Z2 |
| GDEEDYDVVDDLEEE | 0 | 0 | 1 | 294827 | XP_822030.1 | Q4E5T2 |
| GDEGNDEAEGGNDAR | 0 | 0 | 1 | 294840 | XP_820569.1 | Q4E1T5 |
| GDESSDKEDAEPTAA | 0 | 0 | 1 | 294877 | XP_818651.1 | Q4DW89 |
| GDGDPAADGAGTAEG | 0 | 0 | 1 | 294901 | XP_807739.1 | Q4D033 |
| GDKDPAADGAGTQEE | 0 | 0 | 1 | 295019 | XP_813432.1 | Q4DGC2 |
| GDKLSLFGQAAAGDK | 0 | 0 | 1 | 295037 | XP_818927.1 | Q4DX15 |
| GDKPAPFGQAAAADK | 0 | 0 | 1 | 295041 | XP_818927.1 | Q4DX15 |
| GDKPAPFGQAAAGDK | 0 | 0 | 1 | 295042 | XP_818927.1 | Q4DX15 |
| GDKPPPFGQAAAGDK | 0 | 0 | 1 | 295043 | XP_818927.1 | Q4DX15 |
| GDKPPPFGQAAEGDK | 0 | 0 | 1 | 295044 | XP_818927.1 | Q4DX15 |
| GDKPSPFGQAAAGDK | 0 | 0 | 1 | 295047 | XP_818927.1 | Q4DX15 |
| GDKPSPFGQAAAGGK | 0 | 0 | 1 | 295048 | XP_818927.1 | Q4DX15 |
| GDKPSPFGQGTAFDA | 0 | 0 | 1 | 295049 | XP_818927.1 | Q4DX15 |
| GDPAADGAGTAEGKQ | 0 | 0 | 1 | 295129 | XP_807739.1 | Q4D033 |
| GDSDLQLERINVYFD | 0 | 0 | 1 | 295269 | XP_816690.1 | Q4DQP2 |
| GDSGGEGSSGPGASP | 0 | 0 | 1 | 295305 | XP_809166.1 | Q4D462 |
| GEDHEKQQQSDEAQV | 0 | 0 | 1 | 295659 | XP_812183.1 | Q4DCT1 |
| GEEEDHEKQQQSDET | 0 | 0 | 1 | 295718 | XP_815870.1 | Q4DNB0 |
| GEICRDEGGDEGNDE | 0 | 0 | 1 | 295996 | XP_820569.1 | Q4E1T5 |
| GEIDDDSDVQNEIKV | 0 | 0 | 1 | 295998 | XP_813526.1 | Q4DGM5 |
| GEIMTENEDDDEDEE | 0 | 0 | 1 | 296009 | XP_820493.1 | Q4E1H5 |
| GEKKDEEQEKKEEEE | 0 | 0 | 1 | 296053 | XP_807741.1 | Q4D040 |
| GENLDGDEEEGDVSQ | 0 | 0 | 1 | 296218 | XP_813607.1 | Q4DGU9 |
| GERMTEKEDGDNEDE | 0 | 0 | 1 | 296333 | XP_820304.1 | Q4E102 |
| GESSSFYSREVDVSQ | 0 | 0 | 1 | 296424 | XP_807066.1 | Q4CY59 |
| GGAQKGNVRKLQPQP | 0 | 0 | 1 | 296990 | XP_821251.1 | Q4E3M0 |
| GGAQSEADADDDDPQ | 0 | 0 | 1 | 296994 | XP_810637.1 | Q4D8D6 |
| GGAQSEADADVDDPQ | 0 | 0 | 1 | 296995 | XP_808315.1 | Q4D1R3 |
| GGDDVEDGDGEDEEE | 0 | 0 | 1 | 297106 | XP_809061.1 | Q4D3W5 |
| GGDEGNDEAEGGNDA | 0 | 0 | 1 | 297110 | XP_820569.1 | Q4E1T5 |
| GGDSGGEGSSGPGAS | 0 | 0 | 1 | 297157 | XP_809166.1 | Q4D462 |

| | | | | | | |
|---|---|---|---|---|---|---|
| GGERMTEKEDGDNED | 0 | 0 | 1 | 297257 | XP_806129.1 | Q4CVH7 |
| GGKPSPFGQAAAGDK | 0 | 0 | 1 | 297883 | XP_818927.1 | Q4DX15 |
| GGKPSPFGQAAAGGK | 0 | 0 | 1 | 297884 | XP_818927.1 | Q4DX15 |
| GGRDAPQARKQQGRN | 0 | 0 | 1 | 298253 | XP_821251.1 | Q4E3M0 |
| GGSTSSTPPSGTENK | 0 | 0 | 1 | 298511 | XP_808931.1 | Q4D3H3 |
| GIAADGVGGGDGSAG | 0 | 0 | 1 | 299043 | XP_821138.1 | Q4E3E7 |
| GIEQVEEEEEEEDEE | 0 | 0 | 1 | 299124 | XP_807741.1 | Q4D040 |
| GIKTTTATTGDSDGV | 0 | 0 | 1 | 299221 | XP_821134.1 | Q4E3F2 |
| GKEVKKAAKPAAKTA | 0 | 0 | 1 | 299702 | XP_821284.1 | Q4E3R2 |
| GKKEEEDNDDDDLEE | 0 | 0 | 1 | 299817 | XP_821418.1 | Q4E455 |
| GKKKDEEQEPNEEEE | 0 | 0 | 1 | 299836 | XP_809910.1 | Q4D6A6 |
| GKPSPFGQAAAGDKP | 0 | 0 | 1 | 299989 | XP_818927.1 | Q4DX15 |
| GKPSPFGQAAAGGKP | 0 | 0 | 1 | 299990 | XP_818927.1 | Q4DX15 |
| GKRIYDREFDYEDED | 0 | 0 | 1 | 300052 | XP_815234.1 | Q4DEP9 |
| GKSSEEGEIDDDSDV | 0 | 0 | 1 | 300114 | XP_813526.1 | Q4DGM5 |
| GKSVGLTAALSPRQR | 0 | 0 | 1 | 300123 | XP_807065.1 | Q4CY60 |
| GLLLPVDNDGDESSD | 0 | 0 | 1 | 300539 | XP_818651.1 | Q4DW89 |
| GLTAALSPRQRSLPR | 0 | 0 | 1 | 300810 | XP_807065.1 | Q4CY60 |
| GNGRPGRVKRVKRKA | 0 | 0 | 1 | 301345 | XP_817338.1 | Q4DSJ7 |
| GNQDADEEDEGEDEE | 0 | 0 | 1 | 301536 | XP_813432.1 | Q4DGC2 |
| GPKPAEPKSAEPKPA | 0 | 0 | 1 | 302066 | XP_820186.1 | Q4E0M7 |
| GQAAAADKPSLFGQA | 0 | 0 | 1 | 302403 | XP_818927.1 | Q4DX15 |
| GQAAAGDKLSLFGQA | 0 | 0 | 1 | 302404 | XP_818927.1 | Q4DX15 |
| GQAAAGDKPAPFGQA | 0 | 0 | 1 | 302405 | XP_818927.1 | Q4DX15 |
| GQAAAGDKPPPFGQA | 0 | 0 | 1 | 302406 | XP_818927.1 | Q4DX15 |
| GQAAAGDKPSLFGQA | 0 | 0 | 1 | 302407 | XP_818927.1 | Q4DX15 |
| GQAAAGDKPSPFGQG | 0 | 0 | 1 | 302409 | XP_818927.1 | Q4DX15 |
| GQAAAGGKPSPFGQA | 0 | 0 | 1 | 302410 | XP_818927.1 | Q4DX15 |
| GQAAEGDKPPPFGQA | 0 | 0 | 1 | 302412 | XP_818927.1 | Q4DX15 |
| GQHRHDTDTEDSTKN | 0 | 0 | 1 | 302592 | XP_811740.1 | Q4DBI1 |
| GQYDAADVEAGDGPD | 0 | 0 | 1 | 302888 | XP_804663.1 | Q4CRB0 |
| GRDAPQARKQQGRDA | 0 | 0 | 1 | 302974 | XP_821251.1 | Q4E3M0 |
| GRDAPQARKQQGRNA | 0 | 0 | 1 | 302975 | XP_821251.1 | Q4E3M0 |
| GRDAPQARKQQGRNE | 0 | 0 | 1 | 302976 | XP_821251.1 | Q4E3M0 |
| GRHSHDTDTEDSTKN | 0 | 0 | 1 | 303157 | XP_817854.1 | Q4DU19 |
| GRILPDLPGGRAGRI | 0 | 0 | 1 | 303169 | XP_811374.1 | Q4DAG2 |
| GRNEPQARKQQGRDA | 0 | 0 | 1 | 303264 | XP_821251.1 | Q4E3M0 |
| GRNEPQARRQQGRNE | 0 | 0 | 1 | 303265 | XP_821251.1 | Q4E3M0 |
| GRPGRVKRVKRKAAK | 0 | 0 | 1 | 303305 | XP_817338.1 | Q4DSJ7 |
| GRVKRVKRKAAKKGA | 0 | 0 | 1 | 303513 | XP_817338.1 | Q4DSJ7 |
| GSDEEDQDEDEDEGI | 0 | 0 | 1 | 303740 | XP_821081.1 | Q4E380 |
| GSGGAQSEADADDDD | 0 | 0 | 1 | 304001 | XP_810637.1 | Q4D8D6 |
| GSQSTLAATDTANNE | 0 | 0 | 1 | 304528 | XP_809167.1 | Q4D463 |
| GSTSSTPPSGTENKP | 0 | 0 | 1 | 304876 | XP_808931.1 | Q4D3H3 |
| GTAMKTTTATTGDSD | 0 | 0 | 1 | 305067 | XP_820304.1 | Q4E102 |
| GTDEGLLLPVDNDGD | 0 | 0 | 1 | 305142 | XP_818651.1 | Q4DW89 |
| GTDGDESIDEEEEKE | 0 | 0 | 1 | 305148 | XP_820569.1 | Q4E1T5 |
| GTGENLDGDEEEGDV | 0 | 0 | 1 | 305280 | XP_813607.1 | Q4DGU9 |
| GTGGDDVEDGDGEDE | 0 | 0 | 1 | 305289 | XP_809061.1 | Q4D3W5 |
| GTPNNDPAADGAETR | 0 | 0 | 1 | 305582 | XP_814148.1 | Q4DIF3 |
| GTYQGDSDLQLERIN | 0 | 0 | 1 | 305968 | XP_816690.1 | Q4DQP2 |
| GVDPTGTYQGDSDLQ | 0 | 0 | 1 | 306143 | XP_816690.1 | Q4DQP2 |
| GVDSREQDGEVTTSE | 0 | 0 | 1 | 306148 | XP_807741.1 | Q4D040 |
| GVGGGDGSAGEDLLL | 0 | 0 | 1 | 306280 | XP_821138.1 | Q4E3E7 |
| GVVEGESSSFYSREV | 0 | 0 | 1 | 306897 | XP_807066.1 | Q4CY59 |
| HALSSFLLKDKKKVA | 0 | 0 | 1 | 307398 | XP_807065.1 | Q4CY60 |
| HEKQHQSDETQVQQH | 0 | 0 | 1 | 307935 | XP_809166.1 | Q4D462 |
| HEKQQQSDEAQFQQH | 0 | 0 | 1 | 307938 | XP_818475.1 | Q4DVR9 |
| HEKQQQSDEAQVQQH | 0 | 0 | 1 | 307939 | XP_817818.1 | Q4DTW6 |
| HEKQQQSDETQVQQH | 0 | 0 | 1 | 307940 | XP_815868.1 | Q4DNB3 |
| HGVDPTGTYQGDSDL | 0 | 0 | 1 | 308416 | XP_816690.1 | Q4DQP2 |
| HPASSEDEEEDDADG | 0 | 0 | 1 | 309240 | XP_821672.1 | Q4E4Z2 |
| HPASSEDKEEDDADG | 0 | 0 | 1 | 309241 | XP_822030.1 | Q4E5T2 |
| HPEEDEAFDEADDDE | 0 | 0 | 1 | 309254 | XP_806363.1 | Q4CW66 |
| HQSDETQVQQHQQHE | 0 | 0 | 1 | 309512 | XP_809166.1 | Q4D462 |
| HRHDTDTEDSTKNAA | 0 | 0 | 1 | 309591 | XP_811740.1 | Q4DBI1 |
| HSHDTDTEDSTKNAA | 0 | 0 | 1 | 309801 | XP_817627.1 | Q4DTB1 |
| IAADGVGGGDGSAGE | 0 | 0 | 1 | 310702 | XP_821138.1 | Q4E3E7 |
| ICRDEGGDEGNDEAE | 0 | 0 | 1 | 311163 | XP_820569.1 | Q4E1T5 |
| IDDDSDVQNEIKVLD | 0 | 0 | 1 | 311218 | XP_813526.1 | Q4DGM5 |
| IDGDAEVDEDEDDDD | 0 | 0 | 1 | 311254 | XP_818224.1 | Q4DV33 |
| IDGSLSSSAWVCAPL | 0 | 0 | 1 | 311278 | XP_815995.1 | Q4DNQ0 |
| IDGSLSSSAWVCASL | 0 | 0 | 1 | 311279 | XP_821144.1 | Q4E3E1 |
| IEATATKTTTATTGD | 0 | 0 | 1 | 311450 | XP_815165.1 | Q4DLB5 |
| IEEADDDDDDDDDDG | 0 | 0 | 1 | 311468 | XP_804979.1 | Q4CS76 |
| IEQVEEEEEEEDEEE | 0 | 0 | 1 | 311661 | XP_807741.1 | Q4D040 |
| IKQPGGIKTPSSTTP | 0 | 0 | 1 | 312732 | XP_819902.1 | Q4DZS8 |
| IKTTTAATGDSDGST | 0 | 0 | 1 | 312784 | XP_820637.1 | Q4E1Z5 |
| IKTTTATTGDSDGST | 0 | 0 | 1 | 312786 | XP_820022.1 | Q4E058 |
| IKTTTATTGDSDGVK | 0 | 0 | 1 | 312787 | XP_821134.1 | Q4E3F2 |
| ILDRFLAAAMDKVFT | 0 | 0 | 1 | 312861 | XP_819176.1 | Q4DXP9 |
| IMTENEDDDEDEEEN | 0 | 0 | 1 | 313291 | XP_820493.1 | Q4E1H5 |
| INGTGENLDGDEEEG | 0 | 0 | 1 | 313374 | XP_813607.1 | Q4DGU9 |
| IPMLNLLKSHSKTVG | 0 | 0 | 1 | 313693 | XP_813297.1 | Q4DFZ7 |
| ISDDDDEGEDEEEGP | 0 | 0 | 1 | 314440 | XP_803649.1 | Q4CNY5 |
| ITDDDDPAADAAGTR | 0 | 0 | 1 | 314918 | XP_809910.1 | Q4D6A6 |
| IYDREFDYEDEDYEG | 0 | 0 | 1 | 315684 | XP_815234.1 | Q4DEP9 |
| KAAAPKKAAAPKKAA | 0 | 0 | 1 | 315851 | XP_814064.1 | Q4DI49 |
| KAAAPKKAAAPQKAA | 0 | 0 | 1 | 315852 | XP_814064.1 | Q4DI49 |
| KAAAPKKTAAPKKAA | 0 | 0 | 1 | 315853 | XP_814064.1 | Q4DI49 |
| KAAAPQKAAAPKKTA | 0 | 0 | 1 | 315860 | XP_814064.1 | Q4DI49 |

| | | | | | | |
|---|---|---|---|---|---|---|
| KAAAPQKAAVAKKAV | 0 | 0 | 1 | 315861 | XP_814064.1 | Q4DI49 |
| KAAKPAAKTAAKTAA | 0 | 0 | 1 | 315936 | XP_821284.1 | Q4E3R2 |
| KAAVDADTAALAALL | 0 | 0 | 1 | 315983 | XP_803309.1 | Q4CNL0 |
| KAPAAPAAPAAAAEA | 0 | 0 | 1 | 316381 | XP_805627.1 | Q4CU24 |
| KAPAPKAAAPAPKAA | 0 | 0 | 1 | 316384 | XP_819301.1 | Q4DY30 |
| KATTATTGDSDSSTA | 0 | 0 | 1 | 316612 | XP_822030.1 | Q4E5T2 |
| KATTATTGESDGSTA | 0 | 0 | 1 | 316613 | XP_821672.1 | Q4E4Z2 |
| KDEDDDGDNDADDVC | 0 | 0 | 1 | 316928 | XP_806363.1 | Q4CW66 |
| KDEEKEEDEEEEEDE | 0 | 0 | 1 | 316936 | XP_815015.1 | Q4DKW3 |
| KDEEQEPNEEEEEEE | 0 | 0 | 1 | 316939 | XP_809910.1 | Q4D6A6 |
| KDPAADGAGTQEEKQ | 0 | 0 | 1 | 317165 | XP_813432.1 | Q4DGC2 |
| KDSVDSREQDGEGTT | 0 | 0 | 1 | 317281 | XP_809849.1 | Q4D646 |
| KEAPIEATATKTTTA | 0 | 0 | 1 | 317429 | XP_815165.1 | Q4DLB5 |
| KEDGDNEDEEVEDEA | 0 | 0 | 1 | 317491 | XP_820304.1 | Q4E102 |
| KEDNDDDDDEPEEE | 0 | 0 | 1 | 317503 | XP_815012.1 | Q4DKW6 |
| KEEDDADGDEEDYDV | 0 | 0 | 1 | 317548 | XP_822030.1 | Q4E5T2 |
| KEEDEEEEEDEEEEE | 0 | 0 | 1 | 317553 | XP_815015.1 | Q4DKW3 |
| KEEEAGSDEEDQDED | 0 | 0 | 1 | 317563 | XP_821081.1 | Q4E380 |
| KEEEDNDDDDLEEAE | 0 | 0 | 1 | 317566 | XP_821418.1 | Q4E455 |
| KEEEEDDDDDDDEPE | 0 | 0 | 1 | 317569 | XP_809163.1 | Q4D466 |
| KEEEEEEDGIEQVEE | 0 | 0 | 1 | 317572 | XP_807741.1 | Q4D040 |
| KEEEEEEDHEKQQQS | 0 | 0 | 1 | 317573 | XP_815868.1 | Q4DNB3 |
| KEKTDEAEEEEDEDD | 0 | 0 | 1 | 318002 | XP_815190.1 | Q4DLD3 |
| KERTLQVAGIKTTTA | 0 | 0 | 1 | 318307 | XP_821134.1 | Q4E3F2 |
| KETPAESTVIKTTTA | 0 | 0 | 1 | 318434 | XP_820022.1 | Q4E058 |
| KETPLEATATKNTTA | 0 | 0 | 1 | 318443 | XP_821138.1 | Q4E3E7 |
| KETPVAATVIKTTTA | 0 | 0 | 1 | 318448 | XP_820637.1 | Q4E1Z5 |
| KETPVEATAMKNTTA | 0 | 0 | 1 | 318452 | XP_811361.1 | Q4DAF3 |
| KETPVEATAMKTTTA | 0 | 0 | 1 | 318453 | XP_804905.1 | Q4CS02 |
| KETPVEATATKTTTA | 0 | 0 | 1 | 318455 | XP_820650.1 | Q4E1Y0 |
| KETPVEGTAMKTTTA | 0 | 0 | 1 | 318459 | XP_820304.1 | Q4E102 |
| KETPVESTAMKTTTA | 0 | 0 | 1 | 318460 | XP_815852.1 | Q4DNA4 |
| KETPVESTVMKTTTA | 0 | 0 | 1 | 318462 | XP_813432.1 | Q4DGC2 |
| KETPVETTTMKNTTA | 0 | 0 | 1 | 318463 | XP_806129.1 | Q4CVH7 |
| KETPVTAAAMKTTTA | 0 | 0 | 1 | 318466 | XP_808183.1 | Q4D1D1 |
| KEVEKIDSENSKLQQ | 0 | 0 | 1 | 318506 | XP_821826.1 | Q4E572 |
| KEVKKAAKPAAKTAA | 0 | 0 | 1 | 318516 | XP_821284.1 | Q4E3R2 |
| KGAAKAPAPKAAAPA | 0 | 0 | 1 | 318702 | XP_819301.1 | Q4DY30 |
| KGNVRKLQPQPQKQR | 0 | 0 | 1 | 319047 | XP_821251.1 | Q4E3M0 |
| KKAAAPKKAAAPQKA | 0 | 0 | 1 | 319629 | XP_814064.1 | Q4DI49 |
| KKAAAPQKAAAPKKT | 0 | 0 | 1 | 319630 | XP_814064.1 | Q4DI49 |
| KKAAAPQKAAVAKKA | 0 | 0 | 1 | 319631 | XP_814064.1 | Q4DI49 |
| KKAAKPAAKTAAKTA | 0 | 0 | 1 | 319639 | XP_821284.1 | Q4E3R2 |
| KKDEEQEKKEEEEEE | 0 | 0 | 1 | 319713 | XP_807741.1 | Q4D040 |
| KKDEEQEPNEEEEEE | 0 | 0 | 1 | 319714 | XP_809910.1 | Q4D6A6 |
| KKEEEDNDDDDLEEA | 0 | 0 | 1 | 319768 | XP_821418.1 | Q4E455 |
| KKEEEEEEDGIEQVE | 0 | 0 | 1 | 319769 | XP_807741.1 | Q4D040 |
| KKGAAKAPAPKAAAP | 0 | 0 | 1 | 319871 | XP_819301.1 | Q4DY30 |
| KKKDEEQEPNEEEEE | 0 | 0 | 1 | 319983 | XP_809910.1 | Q4D6A6 |
| KKKNPEAASKKAPLS | 0 | 0 | 1 | 320020 | XP_821284.1 | Q4E3R2 |
| KKNPEAASKKAPLSV | 0 | 0 | 1 | 320154 | XP_821284.1 | Q4E3R2 |
| KKPAVKPTVKPAAKA | 0 | 0 | 1 | 320174 | XP_821284.1 | Q4E3R2 |
| KKTAAPKKAAAPQKA | 0 | 0 | 1 | 320313 | XP_814064.1 | Q4DI49 |
| KLGKSVGLTAALSPR | 0 | 0 | 1 | 320581 | XP_807065.1 | Q4CY60 |
| KLSLFGQAAAGDKPA | 0 | 0 | 1 | 320876 | XP_818927.1 | Q4DX15 |
| KLSLFGQAAAGDKPS | 0 | 0 | 1 | 320877 | XP_818927.1 | Q4DX15 |
| KNGTATTGDSDGSTA | 0 | 0 | 1 | 321356 | XP_809903.1 | Q4D6A2 |
| KNNDKDEDDDGDNDA | 0 | 0 | 1 | 321477 | XP_806363.1 | Q4CW66 |
| KNPEAASKKAPLSVV | 0 | 0 | 1 | 321517 | XP_821284.1 | Q4E3R2 |
| KNTTATGDSDGSTAV | 0 | 0 | 1 | 321632 | XP_815190.1 | Q4DLD3 |
| KNTTATTGDSDGSTA | 0 | 0 | 1 | 321633 | XP_806129.1 | Q4CVH7 |
| KNTTATTGDSDSSTA | 0 | 0 | 1 | 321634 | XP_809011.1 | Q4D3Q7 |
| KPAAKAPAPKAEKKG | 0 | 0 | 1 | 321697 | XP_819301.1 | Q4DY30 |
| KPAAKPAAKAPAPKA | 0 | 0 | 1 | 321698 | XP_819301.1 | Q4DY30 |
| KPAAKPAAKPAAKPA | 0 | 0 | 1 | 321699 | XP_821284.1 | Q4E3R2 |
| KPAAKPAAKPAAKTA | 0 | 0 | 1 | 321700 | XP_821284.1 | Q4E3R2 |
| KPAAKPAAKTAAKPA | 0 | 0 | 1 | 321701 | XP_821284.1 | Q4E3R2 |
| KPAAKSAAKPAAKPA | 0 | 0 | 1 | 321702 | XP_821284.1 | Q4E3R2 |
| KPAAKTAAKPAKKPA | 0 | 0 | 1 | 321703 | XP_821284.1 | Q4E3R2 |
| KPAAKTAAKTAAKPA | 0 | 0 | 1 | 321704 | XP_821284.1 | Q4E3R2 |
| KPAEPKSAEPKPAEP | 0 | 0 | 1 | 321717 | XP_820186.1 | Q4E0M7 |
| KPAEPKSAEPKPAES | 0 | 0 | 1 | 321718 | XP_820186.1 | Q4E0M7 |
| KPAEPNSAEPKPAEP | 0 | 0 | 1 | 321721 | XP_820186.1 | Q4E0M7 |
| KPAESKSAEPKPAEP | 0 | 0 | 1 | 321724 | XP_820186.1 | Q4E0M7 |
| KPAKKPAVKPTVKPA | 0 | 0 | 1 | 321732 | XP_821284.1 | Q4E3R2 |
| KPAPFGQAAAADKPS | 0 | 0 | 1 | 321736 | XP_818927.1 | Q4DX15 |
| KPAPFGQAAAGDKLS | 0 | 0 | 1 | 321737 | XP_818927.1 | Q4DX15 |
| KPAPFGQAAAGDKPA | 0 | 0 | 1 | 321738 | XP_818927.1 | Q4DX15 |
| KPAPFGQAAAGDKPS | 0 | 0 | 1 | 321739 | XP_818927.1 | Q4DX15 |
| KPASKPAAKPAAKAP | 0 | 0 | 1 | 321748 | XP_819301.1 | Q4DY30 |
| KPATGEAPSQPGASS | 0 | 0 | 1 | 321752 | XP_808931.1 | Q4D3H3 |
| KPAVKPTVKPAAKAA | 0 | 0 | 1 | 321755 | XP_821284.1 | Q4E3R2 |
| KPPAESPFRSVFGAP | 0 | 0 | 1 | 321984 | XP_818928.1 | Q4DX14 |
| KPPPFGQAAAGDKLS | 0 | 0 | 1 | 321998 | XP_818927.1 | Q4DX15 |
| KPSLFGQAAAGDKPP | 0 | 0 | 1 | 322093 | XP_818927.1 | Q4DX15 |
| KPSPFGQAAAGDKPS | 0 | 0 | 1 | 322098 | XP_818927.1 | Q4DX15 |
| KPSPFGQAAAGGKPS | 0 | 0 | 1 | 322099 | XP_818927.1 | Q4DX15 |
| KPSPFGQGTAFDASR | 0 | 0 | 1 | 322100 | XP_818927.1 | Q4DX15 |
| KQHQSDETQVQQHQQ | 0 | 0 | 1 | 322314 | XP_809166.1 | Q4D462 |
| KQPGGIKTPSSTTPP | 0 | 0 | 1 | 322461 | XP_819902.1 | Q4DZS8 |
| KQQGRDAPQARKQQG | 0 | 0 | 1 | 322490 | XP_821251.1 | Q4E3M0 |

| | | | | | | |
|---|---|---|---|---|---|---|
| KQQGRNEPQARRQQG | 0 | 0 | 1 | 322491 | XP_821251.1 | Q4E3M0 |
| KQQQSDEAQFQQHQQ | 0 | 0 | 1 | 322513 | XP_818475.1 | Q4DVR9 |
| KQQQSDEAQVQQHQQ | 0 | 0 | 1 | 322514 | XP_809948.1 | Q4D6E7 |
| KQQQSDEAQVQQHQR | 0 | 0 | 1 | 322515 | XP_817818.1 | Q4DTW6 |
| KQQQSDETQVQQHQQ | 0 | 0 | 1 | 322516 | XP_815870.1 | Q4DNB0 |
| KQQQSDETQVQQHQR | 0 | 0 | 1 | 322517 | XP_815868.1 | Q4DNB3 |
| KQSEREDDEENDEEE | 0 | 0 | 1 | 322571 | XP_821138.1 | Q4E3E7 |
| KRIYDREFDYEDEDY | 0 | 0 | 1 | 322805 | XP_815234.1 | Q4DEP9 |
| KSAAKPAAKPAAKPA | 0 | 0 | 1 | 323102 | XP_821284.1 | Q4E3R2 |
| KSAEPKPAEPKSAEP | 0 | 0 | 1 | 323117 | XP_820186.1 | Q4E0M7 |
| KSAEPKPAESKSAEP | 0 | 0 | 1 | 323122 | XP_820186.1 | Q4E0M7 |
| KSAEPKPAESKSAGP | 0 | 0 | 1 | 323123 | XP_820186.1 | Q4E0M7 |
| KSAGPKPAEPKSAEP | 0 | 0 | 1 | 323128 | XP_820186.1 | Q4E0M7 |
| KSANKPASKPAAKPA | 0 | 0 | 1 | 323146 | XP_819301.1 | Q4DY30 |
| KSPTLKAPAAPAAPA | 0 | 0 | 1 | 323507 | XP_805627.1 | Q4CU24 |
| KSSEEGEIDDDSDVQ | 0 | 0 | 1 | 323565 | XP_813526.1 | Q4DGM5 |
| KSTQTTGDDDPAADG | 0 | 0 | 1 | 323644 | XP_821138.1 | Q4E3E7 |
| KSVGLTAALSPRQRS | 0 | 0 | 1 | 323689 | XP_807065.1 | Q4CY60 |
| KTAAKPAAKSAAKPA | 0 | 0 | 1 | 323744 | XP_821284.1 | Q4E3R2 |
| KTAAKPAKKPAVKPT | 0 | 0 | 1 | 323745 | XP_821284.1 | Q4E3R2 |
| KTAAKTAAKPAAKSA | 0 | 0 | 1 | 323746 | XP_821284.1 | Q4E3R2 |
| KTAAPKKAAAPQKAA | 0 | 0 | 1 | 323749 | XP_814064.1 | Q4DI49 |
| KTAVSKAAAPKKAAA | 0 | 0 | 1 | 323818 | XP_814064.1 | Q4DI49 |
| KTDEAEEEEDEDDEE | 0 | 0 | 1 | 323833 | XP_815190.1 | Q4DLD3 |
| KTTTAATGDSDGSTA | 0 | 0 | 1 | 324305 | XP_820637.1 | Q4E1Z5 |
| KTTTATTGDSDGGTA | 0 | 0 | 1 | 324308 | XP_821379.1 | Q4E442 |
| KTTTATTGDSDGSTA | 0 | 0 | 1 | 324309 | XP_815852.1 | Q4DNA4 |
| KTTTATTGDSDGVKA | 0 | 0 | 1 | 324310 | XP_821134.1 | Q4E3F2 |
| KTTTATTGDSDSSTA | 0 | 0 | 1 | 324311 | XP_808183.1 | Q4D1D1 |
| KTTTATTGNSDGGTA | 0 | 0 | 1 | 324312 | XP_820650.1 | Q4E1Y0 |
| KTTTATTGNSDGSTA | 0 | 0 | 1 | 324313 | XP_807739.1 | Q4D033 |
| KTTTTTTGDSDGSTA | 0 | 0 | 1 | 324325 | XP_820308.1 | Q4E0Z8 |
| KVNGRETIDPDPQKQ | 0 | 0 | 1 | 324650 | XP_816182.1 | Q4DP76 |
| LAAAMDKVFTTGFAQ | 0 | 0 | 1 | 325034 | XP_819176.1 | Q4DXP9 |
| LAARSMEIDGGEWTN | 0 | 0 | 1 | 325120 | XP_815234.1 | Q4DEP9 |
| LAEQEEISDDDDEGE | 0 | 0 | 1 | 325265 | XP_803649.1 | Q4CNY5 |
| LDRFLAAAMDKVFTT | 0 | 0 | 1 | 326606 | XP_819176.1 | Q4DXP9 |
| LEATATKNTTATTGD | 0 | 0 | 1 | 326794 | XP_821138.1 | Q4E3E7 |
| LELMLRESDGECAPF | 0 | 0 | 1 | 327183 | XP_812097.1 | Q4DCJ6 |
| LFDVATEKKKNPEAA | 0 | 0 | 1 | 327619 | XP_821284.1 | Q4E3R2 |
| LFGQAAAGDKPAPFG | 0 | 0 | 1 | 327704 | XP_818927.1 | Q4DX15 |
| LFGQAAAGDKPPPFG | 0 | 0 | 1 | 327705 | XP_818927.1 | Q4DX15 |
| LFGQAAAGDKPSPFG | 0 | 0 | 1 | 327706 | XP_818927.1 | Q4DX15 |
| LGKSVGLTAALSPRQ | 0 | 0 | 1 | 328351 | XP_807065.1 | Q4CY60 |
| LHEVLQYGRLIDDFI | 0 | 0 | 1 | 328783 | XP_808204.1 | Q4D1F2 |
| LILDRFLAAAMDKVF | 0 | 0 | 1 | 329162 | XP_819176.1 | Q4DXP9 |
| LIPMLNLLKSHSKTV | 0 | 0 | 1 | 329224 | XP_813297.1 | Q4DFZ7 |
| LKAPAAPAAPAAAAE | 0 | 0 | 1 | 329407 | XP_805627.1 | Q4CU24 |
| LKDEEKEEDEEEEED | 0 | 0 | 1 | 329440 | XP_815015.1 | Q4DKW3 |
| LKEEEAGSDEEDQDE | 0 | 0 | 1 | 329485 | XP_821081.1 | Q4E380 |
| LLKDKKKVATAKVAV | 0 | 0 | 1 | 330399 | XP_807065.1 | Q4CY60 |
| LLLMTREVDDTMADE | 0 | 0 | 1 | 330731 | XP_805245.1 | Q4CSZ0 |
| LLLPVDNDGDESSDK | 0 | 0 | 1 | 330752 | XP_818651.1 | Q4DW89 |
| LLMTREVDDTMADEL | 0 | 0 | 1 | 330874 | XP_805245.1 | Q4CSZ0 |
| LLPVDNDGDESSDKE | 0 | 0 | 1 | 330986 | XP_818651.1 | Q4DW89 |
| LLYSDDELFLSQQRG | 0 | 0 | 1 | 331538 | XP_814919.1 | Q4DKL5 |
| LMLRESDGECAPFTT | 0 | 0 | 1 | 331672 | XP_812097.1 | Q4DCJ6 |
| LMTREVDDTMADELR | 0 | 0 | 1 | 331749 | XP_805245.1 | Q4CSZ0 |
| LPADSENSKTGCGEE | 0 | 0 | 1 | 332157 | XP_813566.1 | Q4DGQ5 |
| LPHPEEDEAFDEADD | 0 | 0 | 1 | 332414 | XP_806363.1 | Q4CW66 |
| LPIDGDAEVDEDEDD | 0 | 0 | 1 | 332426 | XP_818224.1 | Q4DV33 |
| LPSPPEEDVASREQD | 0 | 0 | 1 | 332817 | XP_812835.1 | Q4DEM8 |
| LPVDNDGDESSDKED | 0 | 0 | 1 | 332917 | XP_818651.1 | Q4DW89 |
| LREIDGSLSSSAWVC | 0 | 0 | 1 | 333829 | XP_821144.1 | Q4E3E1 |
| LSFSDFLLLMTREVD | 0 | 0 | 1 | 334833 | XP_805245.1 | Q4CSZ0 |
| LSLFGQAAAGDKPAP | 0 | 0 | 1 | 335019 | XP_818927.1 | Q4DX15 |
| LSLFGQAAAGDKPSP | 0 | 0 | 1 | 335020 | XP_818927.1 | Q4DX15 |
| LSSFLLKDKKKVATA | 0 | 0 | 1 | 335360 | XP_807065.1 | Q4CY60 |
| LTAAKPPAESPFKSV | 0 | 0 | 1 | 335605 | XP_813515.1 | Q4DGM1 |
| LVVRLWFEREVDGHD | 0 | 0 | 1 | 337020 | XP_816335.1 | Q4DPM2 |
| LWFEREVDGHDFIRE | 0 | 0 | 1 | 337253 | XP_816335.1 | Q4DPM2 |
| MEIDGGEWTNGTAIV | 0 | 0 | 1 | 338202 | XP_815234.1 | Q4DEP9 |
| MKNTTATGDSDGSTA | 0 | 0 | 1 | 338913 | XP_815190.1 | Q4DLD3 |
| MKNTTATTGDSDGST | 0 | 0 | 1 | 338914 | XP_806129.1 | Q4CVH7 |
| MKNTTATTGDSDSST | 0 | 0 | 1 | 338915 | XP_811361.1 | Q4DAF3 |
| MKTTAATTGDSDGST | 0 | 0 | 1 | 338967 | XP_820410.1 | Q4E1A7 |
| MKTTTATTGDSDGST | 0 | 0 | 1 | 338971 | XP_815852.1 | Q4DNA4 |
| MKTTTATTGDSDSST | 0 | 0 | 1 | 338972 | XP_808183.1 | Q4D1D1 |
| MKTTTATTGNSDGST | 0 | 0 | 1 | 338973 | XP_807739.1 | Q4D033 |
| MKTTTTTTGDSDGST | 0 | 0 | 1 | 338974 | XP_820308.1 | Q4E0Z8 |
| MLNLLKSHSKTVGIR | 0 | 0 | 1 | 339113 | XP_813297.1 | Q4DFZ7 |
| MLPHPEEDEAFDEAD | 0 | 0 | 1 | 339121 | XP_806363.1 | Q4CW66 |
| MLRESDGECAPFTTL | 0 | 0 | 1 | 339141 | XP_812097.1 | Q4DCJ6 |
| MPAKSANKPASKPAA | 0 | 0 | 1 | 339522 | XP_819301.1 | Q4DY30 |
| MPAKTAVSKAAAPKK | 0 | 0 | 1 | 339523 | XP_814064.1 | Q4DI49 |
| MPGKEVKKAAKPAAK | 0 | 0 | 1 | 339563 | XP_821284.1 | Q4E3R2 |
| MQADQPTQFSVGTPD | 0 | 0 | 1 | 339684 | XP_820186.1 | Q4E0M7 |
| MTEKEDGDNEDEEVE | 0 | 0 | 1 | 340337 | XP_820304.1 | Q4E102 |
| MTENEDDDEDEEENE | 0 | 0 | 1 | 340342 | XP_820493.1 | Q4E1H5 |
| MTFANTLLYSDDELF | 0 | 0 | 1 | 340348 | XP_814919.1 | Q4DKL5 |
| MTREVDDTMADELRS | 0 | 0 | 1 | 340438 | XP_805245.1 | Q4CSZ0 |

| | | | | | | |
|---|---|---|---|---|---|---|
| MVVNPLVVRLWFERE | 0 | 0 | 1 | 340638 | XP_816335.1 | Q4DPM2 |
| NAPSRLREIDGSLSS | 0 | 0 | 1 | 341176 | XP_821144.1 | Q4E3E1 |
| NDDDDDDEPEEEKEK | 0 | 0 | 1 | 341620 | XP_815012.1 | Q4DKW6 |
| NDEELKDEEKEEDEE | 0 | 0 | 1 | 341664 | XP_815015.1 | Q4DKW3 |
| NDGDESSDKEDAEPT | 0 | 0 | 1 | 341702 | XP_818651.1 | Q4DW89 |
| NDKDEDDDGDNDADD | 0 | 0 | 1 | 341757 | XP_806363.1 | Q4CW66 |
| NDPAADGAETREEKK | 0 | 0 | 1 | 341835 | XP_814148.1 | Q4DIF3 |
| NEDDDEDEEENEEDA | 0 | 0 | 1 | 342117 | XP_820493.1 | Q4E1H5 |
| NEEGNQDADEEDEGE | 0 | 0 | 1 | 342196 | XP_813432.1 | Q4DGC2 |
| NEKQQQSDEAQVQQH | 0 | 0 | 1 | 342404 | XP_809948.1 | Q4D6E7 |
| NEPQARKQQGRDAPQ | 0 | 0 | 1 | 342542 | XP_821251.1 | Q4E3M0 |
| NEPQARRQQGRNEPQ | 0 | 0 | 1 | 342543 | XP_821251.1 | Q4E3M0 |
| NERQQQSDETQVQQQ | 0 | 0 | 1 | 342609 | XP_819538.1 | Q4DYS6 |
| NGGSTSSTPPSGTEN | 0 | 0 | 1 | 343136 | XP_808931.1 | Q4D3H3 |
| NGKRIYDREFDYEDE | 0 | 0 | 1 | 343187 | XP_815234.1 | Q4DEP9 |
| NGRPGRVKRVKRKAA | 0 | 0 | 1 | 343331 | XP_817338.1 | Q4DSJ7 |
| NGTGENLDGDEEEGD | 0 | 0 | 1 | 343417 | XP_813607.1 | Q4DGU9 |
| NKNNDKDEDDDGDND | 0 | 0 | 1 | 344056 | XP_806363.1 | Q4CW66 |
| NKPASKPAAKPAAKA | 0 | 0 | 1 | 344076 | XP_819301.1 | Q4DY30 |
| NKPATGEAPSQPGAS | 0 | 0 | 1 | 344078 | XP_808931.1 | Q4D3H3 |
| NMTFANTLLYSDDEL | 0 | 0 | 1 | 344708 | XP_814919.1 | Q4DKL5 |
| NNDKDEDDDGDNDAD | 0 | 0 | 1 | 344779 | XP_806363.1 | Q4CW66 |
| NNDPAADGAETREEK | 0 | 0 | 1 | 344786 | XP_814148.1 | Q4DIF3 |
| NPEAASKKAPLSVVT | 0 | 0 | 1 | 345216 | XP_821284.1 | Q4E3R2 |
| NPKSTQTTGDDDPAA | 0 | 0 | 1 | 345312 | XP_821138.1 | Q4E3E7 |
| NPLVVRLWFEREVDG | 0 | 0 | 1 | 345346 | XP_816335.1 | Q4DPM2 |
| NPNSTQTTGDDDPAA | 0 | 0 | 1 | 345364 | XP_815190.1 | Q4DLD3 |
| NPTEGSQSTLAATDT | 0 | 0 | 1 | 345487 | XP_809167.1 | Q4D463 |
| NQDADEEDEGEDEEE | 0 | 0 | 1 | 345551 | XP_813432.1 | Q4DGC2 |
| NSAEPKPAEPKSAEP | 0 | 0 | 1 | 346068 | XP_820186.1 | Q4E0M7 |
| NSGTDGDESIDEEEE | 0 | 0 | 1 | 346241 | XP_820569.1 | Q4E1T5 |
| NSTPTTGDDDPAADG | 0 | 0 | 1 | 346581 | XP_820308.1 | Q4E0Z8 |
| NSTPTTGDDDPAADV | 0 | 0 | 1 | 346582 | XP_815852.1 | Q4DNA4 |
| NSTQTTGDDDPAADG | 0 | 0 | 1 | 346587 | XP_815190.1 | Q4DLD3 |
| NTLLYSDDELFLSQQ | 0 | 0 | 1 | 346967 | XP_814919.1 | Q4DKL5 |
| NTNSTPTTGDDDPAA | 0 | 0 | 1 | 347018 | XP_820308.1 | Q4E0Z8 |
| NTTATTGDSDGSTAV | 0 | 0 | 1 | 347171 | XP_806129.1 | Q4CVH7 |
| NTTATTGDSDSSTAV | 0 | 0 | 1 | 347172 | XP_809011.1 | Q4D3Q7 |
| PAAAAEEEEDDDMGF | 0 | 0 | 1 | 347915 | XP_818851.1 | Q4DWU6 |
| PAAAAEPEEEDDDDD | 0 | 0 | 1 | 347916 | XP_821118.1 | Q4E3A3 |
| PAADGAETREEKKNE | 0 | 0 | 1 | 347947 | XP_814148.1 | Q4DIF3 |
| PAADGAGTAEGKQNE | 0 | 0 | 1 | 347949 | XP_815190.1 | Q4DLD3 |
| PAADGAGTQEEKQNE | 0 | 0 | 1 | 347951 | XP_813432.1 | Q4DGC2 |
| PAAKPAAKAPAPKAE | 0 | 0 | 1 | 348014 | XP_819301.1 | Q4DY30 |
| PAAKPAAKPAAKPAA | 0 | 0 | 1 | 348015 | XP_821284.1 | Q4E3R2 |
| PAAKPAAKPAAKTAA | 0 | 0 | 1 | 348016 | XP_821284.1 | Q4E3R2 |
| PAAKPAAKTAAKPAK | 0 | 0 | 1 | 348017 | XP_821284.1 | Q4E3R2 |
| PAAKSAAKPAAKPAA | 0 | 0 | 1 | 348018 | XP_821284.1 | Q4E3R2 |
| PAAKTAAKPAKKPAV | 0 | 0 | 1 | 348019 | XP_821284.1 | Q4E3R2 |
| PAAKTAAKTAAKPAA | 0 | 0 | 1 | 348020 | XP_821284.1 | Q4E3R2 |
| PADSENSKTGCGEEA | 0 | 0 | 1 | 348167 | XP_813566.1 | Q4DGQ5 |
| PAEPKSAEPKPAEPK | 0 | 0 | 1 | 348235 | XP_820186.1 | Q4E0M7 |
| PAEPKSAEPKPAEPN | 0 | 0 | 1 | 348236 | XP_820186.1 | Q4E0M7 |
| PAEPKSAEPKPAESK | 0 | 0 | 1 | 348237 | XP_820186.1 | Q4E0M7 |
| PAEPNSAEPKPAEPK | 0 | 0 | 1 | 348240 | XP_820186.1 | Q4E0M7 |
| PAESKSAEPKPAEPK | 0 | 0 | 1 | 348252 | XP_820186.1 | Q4E0M7 |
| PAESTVIKTTTATTG | 0 | 0 | 1 | 348263 | XP_820022.1 | Q4E058 |
| PAKKPAVKPTVKPAA | 0 | 0 | 1 | 348480 | XP_821284.1 | Q4E3R2 |
| PAKSANKPASKPAAK | 0 | 0 | 1 | 348498 | XP_819301.1 | Q4DY30 |
| PAKTAVSKAAAPKKA | 0 | 0 | 1 | 348507 | XP_814064.1 | Q4DI49 |
| PAPAKAAAPPAAARS | 0 | 0 | 1 | 348613 | XP_809477.1 | Q4D528 |
| PAPFGQAAAADKPSL | 0 | 0 | 1 | 348626 | XP_818927.1 | Q4DX15 |
| PAPFGQAAAGDKLSL | 0 | 0 | 1 | 348627 | XP_818927.1 | Q4DX15 |
| PAPFGQAAAGDKPAP | 0 | 0 | 1 | 348628 | XP_818927.1 | Q4DX15 |
| PAPFGQAAAGDKPSP | 0 | 0 | 1 | 348629 | XP_818927.1 | Q4DX15 |
| PAPKAAAPAPKAAAA | 0 | 0 | 1 | 348636 | XP_819301.1 | Q4DY30 |
| PASKPAAKPAAKAPA | 0 | 0 | 1 | 348816 | XP_819301.1 | Q4DY30 |
| PASSEDEEEDDADGD | 0 | 0 | 1 | 348848 | XP_821672.1 | Q4E4Z2 |
| PASSEDKEEDDADGD | 0 | 0 | 1 | 348849 | XP_822030.1 | Q4E5T2 |
| PATGEAPSQPGASSG | 0 | 0 | 1 | 348898 | XP_808931.1 | Q4D3H3 |
| PAVKPTVKPAAKAAA | 0 | 0 | 1 | 348979 | XP_821284.1 | Q4E3R2 |
| PDDDDPAADAAGTRE | 0 | 0 | 1 | 349177 | XP_821399.1 | Q4E444 |
| PDDDDPAADGAGTRE | 0 | 0 | 1 | 349178 | XP_809011.1 | Q4D3Q7 |
| PEAASKKAPLSVVTG | 0 | 0 | 1 | 349511 | XP_821284.1 | Q4E3R2 |
| PEALHEVLQYGRLID | 0 | 0 | 1 | 349536 | XP_808204.1 | Q4D1F2 |
| PEDGVDSREQDGEVT | 0 | 0 | 1 | 349603 | XP_807741.1 | Q4D040 |
| PEDSVASREKDGEDT | 0 | 0 | 1 | 349624 | XP_818224.1 | Q4DV33 |
| PEEDEAFDEADDDEL | 0 | 0 | 1 | 349637 | XP_806363.1 | Q4CW66 |
| PEEEDDDDDFGMGAL | 0 | 0 | 1 | 349643 | XP_821118.1 | Q4E3A3 |
| PFEATAMKTTTTTTG | 0 | 0 | 1 | 350251 | XP_820308.1 | Q4E0Z8 |
| PFGQAAAADKPSLFG | 0 | 0 | 1 | 350280 | XP_818927.1 | Q4DX15 |
| PFGQAAAGDKLSLFG | 0 | 0 | 1 | 350281 | XP_818927.1 | Q4DX15 |
| PFGQAAAGDKPAPFG | 0 | 0 | 1 | 350282 | XP_818927.1 | Q4DX15 |
| PFGQAAAGDKPPPFG | 0 | 0 | 1 | 350283 | XP_818927.1 | Q4DX15 |
| PFGQAAAGDKPSLFG | 0 | 0 | 1 | 350284 | XP_818927.1 | Q4DX15 |
| PFGQAAAGGKPSPFG | 0 | 0 | 1 | 350286 | XP_818927.1 | Q4DX15 |
| PFGQAAEGDKPPPFG | 0 | 0 | 1 | 350287 | XP_818927.1 | Q4DX15 |
| PGKEVKKAAKPAAKT | 0 | 0 | 1 | 350816 | XP_821284.1 | Q4E3R2 |
| PGRVKRVKRKAAKKG | 0 | 0 | 1 | 351055 | XP_817338.1 | Q4DSJ7 |
| PHPEEDEAFDEADDD | 0 | 0 | 1 | 351388 | XP_806363.1 | Q4CW66 |
| PIDGDAEVDEDEDDD | 0 | 0 | 1 | 351486 | XP_818224.1 | Q4DV33 |

| | | | | | | |
|---|---|---|---|---|---|---|
| PIEATATKTTTATTG | 0 | 0 | 1 | 351500 | XP_815165.1 | Q4DLB5 |
| PKDSVDSREQDGEGT | 0 | 0 | 1 | 351823 | XP_809849.1 | Q4D646 |
| PKKAAAPKKAAAPQK | 0 | 0 | 1 | 351956 | XP_814064.1 | Q4DI49 |
| PKKAAAPQKAAAPKK | 0 | 0 | 1 | 351957 | XP_814064.1 | Q4DI49 |
| PKKAAAPQKAAVAKK | 0 | 0 | 1 | 351958 | XP_814064.1 | Q4DI49 |
| PKKTAAPKKAAAPQK | 0 | 0 | 1 | 351987 | XP_814064.1 | Q4DI49 |
| PKNGTATTGDSDGST | 0 | 0 | 1 | 352055 | XP_809903.1 | Q4D6A2 |
| PKPAEPKSAEPKPAE | 0 | 0 | 1 | 352076 | XP_820186.1 | Q4E0M7 |
| PKPAEPNSAEPKPAE | 0 | 0 | 1 | 352079 | XP_820186.1 | Q4E0M7 |
| PKPAESKSAEPKPAE | 0 | 0 | 1 | 352082 | XP_820186.1 | Q4E0M7 |
| PKSAEPKPAEPKSAE | 0 | 0 | 1 | 352169 | XP_820186.1 | Q4E0M7 |
| PKSAEPKPAEPNSAE | 0 | 0 | 1 | 352172 | XP_820186.1 | Q4E0M7 |
| PKSAEPKPAESKSAE | 0 | 0 | 1 | 352174 | XP_820186.1 | Q4E0M7 |
| PKSAEPKPAESKSAG | 0 | 0 | 1 | 352175 | XP_820186.1 | Q4E0M7 |
| PKSAGPKPAEPKSAE | 0 | 0 | 1 | 352176 | XP_820186.1 | Q4E0M7 |
| PKSTQTTGDDDPAAD | 0 | 0 | 1 | 352209 | XP_821138.1 | Q4E3E7 |
| PKVNGRETIDPDPQK | 0 | 0 | 1 | 352253 | XP_816182.1 | Q4DP76 |
| PLEATATKNTTATTG | 0 | 0 | 1 | 352377 | XP_821138.1 | Q4E3E7 |
| PLPIDGDAEVDEDED | 0 | 0 | 1 | 352823 | XP_818224.1 | Q4DV33 |
| PLVVRLWFEREVDGH | 0 | 0 | 1 | 353120 | XP_816335.1 | Q4DPM2 |
| PMLNLLKSHSKTVGI | 0 | 0 | 1 | 353205 | XP_813297.1 | Q4DFZ7 |
| PNNDPAADGAETREE | 0 | 0 | 1 | 353440 | XP_814148.1 | Q4DIF3 |
| PNSAEPKPAEPKSAE | 0 | 0 | 1 | 353516 | XP_820186.1 | Q4E0M7 |
| PNSTQTTGDDDPAAD | 0 | 0 | 1 | 353544 | XP_815190.1 | Q4DLD3 |
| PPAESPFRSVFGAPS | 0 | 0 | 1 | 353662 | XP_818928.1 | Q4DX14 |
| PPEDSVASREKDGED | 0 | 0 | 1 | 353817 | XP_818224.1 | Q4DV33 |
| PPEEDVASREQDGED | 0 | 0 | 1 | 353820 | XP_812835.1 | Q4DEM8 |
| PPFGQAAAGDKLSLF | 0 | 0 | 1 | 353873 | XP_818927.1 | Q4DX15 |
| PPFGQAAAGDKPAPF | 0 | 0 | 1 | 353874 | XP_818927.1 | Q4DX15 |
| PPFGQAAAGDKPPPF | 0 | 0 | 1 | 353875 | XP_818927.1 | Q4DX15 |
| PPFGQAAAGDKPSLF | 0 | 0 | 1 | 353876 | XP_818927.1 | Q4DX15 |
| PPFGQAAAGDKPSPF | 0 | 0 | 1 | 353877 | XP_818927.1 | Q4DX15 |
| PPFGQAAEGDKPPPF | 0 | 0 | 1 | 353878 | XP_818927.1 | Q4DX15 |
| PPKDSVDSREQDGEG | 0 | 0 | 1 | 354015 | XP_809849.1 | Q4D646 |
| PPKKSWKQISNLSGG | 0 | 0 | 1 | 354026 | XP_820569.1 | Q4E1T5 |
| PPPFGQAAAGDKLSL | 0 | 0 | 1 | 354197 | XP_818927.1 | Q4DX15 |
| PPPFGQAAAGDKPAP | 0 | 0 | 1 | 354198 | XP_818927.1 | Q4DX15 |
| PPPFGQAAAGDKPPP | 0 | 0 | 1 | 354199 | XP_818927.1 | Q4DX15 |
| PPPFGQAAAGDKPSL | 0 | 0 | 1 | 354200 | XP_818927.1 | Q4DX15 |
| PPPFGQAAAGDKPSP | 0 | 0 | 1 | 354201 | XP_818927.1 | Q4DX15 |
| PPPFGQAAEGDKPPP | 0 | 0 | 1 | 354202 | XP_818927.1 | Q4DX15 |
| PPPKDSVDSREQDGE | 0 | 0 | 1 | 354239 | XP_809849.1 | Q4D646 |
| PPPPKDSVDSREQDG | 0 | 0 | 1 | 354284 | XP_809849.1 | Q4D646 |
| PPPTEDSVDSREQDG | 0 | 0 | 1 | 354407 | XP_808131.1 | Q4D180 |
| PPSGTENKPATGEAP | 0 | 0 | 1 | 354548 | XP_808931.1 | Q4D3H3 |
| PPTEDSVDSREQDGE | 0 | 0 | 1 | 354652 | XP_808131.1 | Q4D180 |
| PPTEEAEDEDSDEEE | 0 | 0 | 1 | 354653 | XP_807807.1 | Q4D0A4 |
| PQARKQQGRDAPQAR | 0 | 0 | 1 | 354822 | XP_821251.1 | Q4E3M0 |
| PQARKQQGRNAPRSQ | 0 | 0 | 1 | 354823 | XP_821251.1 | Q4E3M0 |
| PQARKQQGRNEPQAR | 0 | 0 | 1 | 354824 | XP_821251.1 | Q4E3M0 |
| PQKAAAPKKTAAPKK | 0 | 0 | 1 | 355009 | XP_814064.1 | Q4DI49 |
| PQTTIKQPGGIKTPS | 0 | 0 | 1 | 355323 | XP_819902.1 | Q4DZS8 |
| PSGTENKPATGEAPS | 0 | 0 | 1 | 356104 | XP_808931.1 | Q4D3H3 |
| PSLFGQAAAGDKPPP | 0 | 0 | 1 | 356216 | XP_818927.1 | Q4DX15 |
| PSLFGQAAAGDKPSP | 0 | 0 | 1 | 356217 | XP_818927.1 | Q4DX15 |
| PSLPADSENSKTGCG | 0 | 0 | 1 | 356238 | XP_813566.1 | Q4DGQ5 |
| PSPFGQAAAGDKPPP | 0 | 0 | 1 | 356350 | XP_818927.1 | Q4DX15 |
| PSPFGQAAAGDKPSL | 0 | 0 | 1 | 356351 | XP_818927.1 | Q4DX15 |
| PSPFGQAAAGDKPSP | 0 | 0 | 1 | 356352 | XP_818927.1 | Q4DX15 |
| PSPFGQAAAGGKPSP | 0 | 0 | 1 | 356353 | XP_818927.1 | Q4DX15 |
| PSPFGQGTAFDASRS | 0 | 0 | 1 | 356354 | XP_818927.1 | Q4DX15 |
| PSPPEEDVASREQDG | 0 | 0 | 1 | 356402 | XP_812835.1 | Q4DEM8 |
| PSRLREIDGSLSSSA | 0 | 0 | 1 | 356530 | XP_821144.1 | Q4E3E1 |
| PSTTTTRAPSRLREI | 0 | 0 | 1 | 356766 | XP_815995.1 | Q4DNQ0 |
| PTDEHSDFLTMGSKM | 0 | 0 | 1 | 356977 | XP_814355.1 | Q4DIZ9 |
| PTEDSVDSREQDGED | 0 | 0 | 1 | 357020 | XP_808131.1 | Q4D180 |
| PTEEAEDEDSDEEEE | 0 | 0 | 1 | 357021 | XP_807807.1 | Q4D0A4 |
| PTEEEDDDDEEDDSD | 0 | 0 | 1 | 357022 | XP_812035.1 | Q4DCD8 |
| PTEGSQSTLAATDTA | 0 | 0 | 1 | 357031 | XP_809167.1 | Q4D463 |
| PTGTYQGDSDLQLER | 0 | 0 | 1 | 357150 | XP_816690.1 | Q4DQP2 |
| PTINGTGENLDGDEE | 0 | 0 | 1 | 357173 | XP_813607.1 | Q4DGU9 |
| PTLKAPAAPAAPAAA | 0 | 0 | 1 | 357238 | XP_805627.1 | Q4CU24 |
| PTQFSVGTPDAANAA | 0 | 0 | 1 | 357407 | XP_820186.1 | Q4E0M7 |
| PTTGDDDPAADGAGT | 0 | 0 | 1 | 357528 | XP_820308.1 | Q4E0Z8 |
| PTTGDDDPAADVAGT | 0 | 0 | 1 | 357529 | XP_815852.1 | Q4DNA4 |
| PTTTTTNAPSRLREI | 0 | 0 | 1 | 357593 | XP_821144.1 | Q4E3E1 |
| PVAATVIKTTTAATG | 0 | 0 | 1 | 357660 | XP_820637.1 | Q4E1Z5 |
| PVDNDGDESSDKEDA | 0 | 0 | 1 | 357733 | XP_818651.1 | Q4DW89 |
| PVEATAMKNTTATTG | 0 | 0 | 1 | 357758 | XP_811361.1 | Q4DAF3 |
| PVEATAMKTTTATTG | 0 | 0 | 1 | 357760 | XP_804905.1 | Q4CS02 |
| PVEATATKNTTATTG | 0 | 0 | 1 | 357762 | XP_809011.1 | Q4D3Q7 |
| PVEATATKTTTATTG | 0 | 0 | 1 | 357763 | XP_820650.1 | Q4E1Y0 |
| PVEATAVKATTATTG | 0 | 0 | 1 | 357764 | XP_821672.1 | Q4E4Z2 |
| PVEGTAMKTTTATTG | 0 | 0 | 1 | 357773 | XP_820304.1 | Q4E102 |
| PVESTAMKTTTATTG | 0 | 0 | 1 | 357796 | XP_815852.1 | Q4DNA4 |
| PVESTVMKTTTATTG | 0 | 0 | 1 | 357798 | XP_813432.1 | Q4DGC2 |
| PVETTTMKNTTATTG | 0 | 0 | 1 | 357807 | XP_806129.1 | Q4CVH7 |
| PVTAAAMKTTTATTG | 0 | 0 | 1 | 358123 | XP_808183.1 | Q4D1D1 |
| QAAAADKPSLFGQAA | 0 | 0 | 1 | 358422 | XP_818927.1 | Q4DX15 |
| QAAAGDKLSLFGQAA | 0 | 0 | 1 | 358427 | XP_818927.1 | Q4DX15 |
| QAAAGDKPAPFGQAA | 0 | 0 | 1 | 358428 | XP_818927.1 | Q4DX15 |

| | | | | | | |
|---|---|---|---|---|---|---|
| QAAAGDKPPPFGQAA | 0 | 0 | 1 | 358429 | XP_818927.1 | Q4DX15 |
| QAAAGDKPSLFGQAA | 0 | 0 | 1 | 358430 | XP_818927.1 | Q4DX15 |
| QAAAGDKPSPFGQAA | 0 | 0 | 1 | 358431 | XP_818927.1 | Q4DX15 |
| QAAAGDKPSPFGQGT | 0 | 0 | 1 | 358432 | XP_818927.1 | Q4DX15 |
| QAAAGGKPSPFGQAA | 0 | 0 | 1 | 358433 | XP_818927.1 | Q4DX15 |
| QAAEGDKPPPFGQAA | 0 | 0 | 1 | 358455 | XP_818927.1 | Q4DX15 |
| QADQPTQFSVGTPDA | 0 | 0 | 1 | 358510 | XP_820186.1 | Q4E0M7 |
| QARKQQGRDAPQARK | 0 | 0 | 1 | 358767 | XP_821251.1 | Q4E3M0 |
| QARKQQGRNAPRSQK | 0 | 0 | 1 | 358768 | XP_821251.1 | Q4E3M0 |
| QARKQQGRNEPQARR | 0 | 0 | 1 | 358769 | XP_821251.1 | Q4E3M0 |
| QARRQQGRNEPQARK | 0 | 0 | 1 | 358783 | XP_821251.1 | Q4E3M0 |
| QDADEEDEGEDEEEV | 0 | 0 | 1 | 358948 | XP_813432.1 | Q4DGC2 |
| QDATIEEEGEFDEEE | 0 | 0 | 1 | 358968 | XP_816690.1 | Q4DQP2 |
| QDGEDTTSEGEKNVP | 0 | 0 | 1 | 359036 | XP_812835.1 | Q4DEM8 |
| QEEGERMTEKEDGDN | 0 | 0 | 1 | 359403 | XP_820304.1 | Q4E102 |
| QEEGQYDAADVEAGD | 0 | 0 | 1 | 359410 | XP_804663.1 | Q4CRB0 |
| QEEISDDDDEGEDEE | 0 | 0 | 1 | 359420 | XP_803649.1 | Q4CNY5 |
| QEGEIMTENEDDDED | 0 | 0 | 1 | 359492 | XP_820493.1 | Q4E1H5 |
| QEGGERMTEKEDGDN | 0 | 0 | 1 | 359499 | XP_806129.1 | Q4CVH7 |
| QEKKEEEEEEDGIEQ | 0 | 0 | 1 | 359598 | XP_807741.1 | Q4D040 |
| QFSVGTPDAANAATH | 0 | 0 | 1 | 360099 | XP_820186.1 | Q4E0M7 |
| QGDSDLQLERINVYF | 0 | 0 | 1 | 360177 | XP_816690.1 | Q4DQP2 |
| QGQHRHDTDTEDSTK | 0 | 0 | 1 | 360384 | XP_811740.1 | Q4DBI1 |
| QGRDAPQARKQQGRD | 0 | 0 | 1 | 360400 | XP_821251.1 | Q4E3M0 |
| QGRDAPQARKQQGRN | 0 | 0 | 1 | 360401 | XP_821251.1 | Q4E3M0 |
| QGRHSHDTDTEDSTK | 0 | 0 | 1 | 360411 | XP_817854.1 | Q4DU19 |
| QGRNEPQARKQQGRD | 0 | 0 | 1 | 360416 | XP_821251.1 | Q4E3M0 |
| QGRNEPQARRQQGRN | 0 | 0 | 1 | 360417 | XP_821251.1 | Q4E3M0 |
| QHQSDETQVQQHQQH | 0 | 0 | 1 | 360717 | XP_809166.1 | Q4D462 |
| QHRHDTDTEDSTKNA | 0 | 0 | 1 | 360722 | XP_811740.1 | Q4DBI1 |
| QHSHDTDTEDSTKNA | 0 | 0 | 1 | 360734 | XP_817627.1 | Q4DTB1 |
| QITDDDDPAADAAGT | 0 | 0 | 1 | 360941 | XP_809910.1 | Q4D6A6 |
| QKAAAPKKTAAPKKA | 0 | 0 | 1 | 360980 | XP_814064.1 | Q4DI49 |
| QKEKTDEAEEEEDED | 0 | 0 | 1 | 361086 | XP_815190.1 | Q4DLD3 |
| QKGNVRKLQPQPQKQ | 0 | 0 | 1 | 361140 | XP_821251.1 | Q4E3M0 |
| QPTQFSVGTPDAANA | 0 | 0 | 1 | 362859 | XP_820186.1 | Q4E0M7 |
| QQGRDAPQARKQQGR | 0 | 0 | 1 | 363053 | XP_821251.1 | Q4E3M0 |
| QQGRNEPQARKQQGR | 0 | 0 | 1 | 363056 | XP_821251.1 | Q4E3M0 |
| QQGRNEPQARRQQGR | 0 | 0 | 1 | 363057 | XP_821251.1 | Q4E3M0 |
| QQQSDEAQFQQHQQH | 0 | 0 | 1 | 363386 | XP_818475.1 | Q4DVR9 |
| QQQSDEAQVQQHQQH | 0 | 0 | 1 | 363387 | XP_809948.1 | Q4D6E7 |
| QQQSDEAQVQQHQRH | 0 | 0 | 1 | 363388 | XP_817818.1 | Q4DTW6 |
| QQQSDETQVQQHQQH | 0 | 0 | 1 | 363389 | XP_815870.1 | Q4DNB0 |
| QQQSDETQVQQHQRH | 0 | 0 | 1 | 363390 | XP_815868.1 | Q4DNB3 |
| QQQSDETQVQQQEHE | 0 | 0 | 1 | 363391 | XP_819538.1 | Q4DYS6 |
| QQSDEAQFQQHQQHE | 0 | 0 | 1 | 363450 | XP_818475.1 | Q4DVR9 |
| QQSDEAQVQQHQQHE | 0 | 0 | 1 | 363451 | XP_809948.1 | Q4D6E7 |
| QQSDEAQVQQHQRHE | 0 | 0 | 1 | 363452 | XP_817818.1 | Q4DTW6 |
| QQSDETQVQQHQQHE | 0 | 0 | 1 | 363453 | XP_815870.1 | Q4DNB0 |
| QQSDETQVQQHQRHE | 0 | 0 | 1 | 363454 | XP_815868.1 | Q4DNB3 |
| QQSDETQVQQQEHEH | 0 | 0 | 1 | 363455 | XP_819538.1 | Q4DYS6 |
| QRGGAQKGNVRKLQP | 0 | 0 | 1 | 363733 | XP_821251.1 | Q4E3M0 |
| QRGGRDAPQARKQQG | 0 | 0 | 1 | 363735 | XP_821251.1 | Q4E3M0 |
| QSDEAQFQQHQQHEH | 0 | 0 | 1 | 364193 | XP_818475.1 | Q4DVR9 |
| QSDEAQVQQHQQHEH | 0 | 0 | 1 | 364194 | XP_809948.1 | Q4D6E7 |
| QSDEAQVQQHQRHER | 0 | 0 | 1 | 364195 | XP_817818.1 | Q4DTW6 |
| QSDETQVQQHQQHEH | 0 | 0 | 1 | 364196 | XP_809166.1 | Q4D462 |
| QSDETQVQQHQRHEH | 0 | 0 | 1 | 364197 | XP_815868.1 | Q4DNB3 |
| QSDETQVQQQEHEHP | 0 | 0 | 1 | 364198 | XP_819538.1 | Q4DYS6 |
| QSEADADDDDPQRPN | 0 | 0 | 1 | 364220 | XP_810637.1 | Q4D8D6 |
| QSEADADVDDPQRPN | 0 | 0 | 1 | 364221 | XP_808315.1 | Q4D1R3 |
| QSEREDDEENDEEED | 0 | 0 | 1 | 364261 | XP_821138.1 | Q4E3E7 |
| QTEQKTAPFGQAAAG | 0 | 0 | 1 | 364782 | XP_818927.1 | Q4DX15 |
| QTPDDDDPAADAAGT | 0 | 0 | 1 | 364975 | XP_821399.1 | Q4E444 |
| QTSAAAAANDSSPAE | 0 | 0 | 1 | 365060 | XP_815868.1 | Q4DNB3 |
| QTTGDGDPAADGAGT | 0 | 0 | 1 | 365113 | XP_807739.1 | Q4D033 |
| QTTIKQPGGIKTPSS | 0 | 0 | 1 | 365120 | XP_819902.1 | Q4DZS8 |
| QYDAADVEAGDGPDK | 0 | 0 | 1 | 365590 | XP_804663.1 | Q4CRB0 |
| RAPSRLREIDGSLSS | 0 | 0 | 1 | 366095 | XP_815995.1 | Q4DNQ0 |
| RDAPQARKQQGRDAP | 0 | 0 | 1 | 366497 | XP_821251.1 | Q4E3M0 |
| RDAPQARKQQGRNAP | 0 | 0 | 1 | 366498 | XP_821251.1 | Q4E3M0 |
| RDAPQARKQQGRNEP | 0 | 0 | 1 | 366499 | XP_821251.1 | Q4E3M0 |
| RDEGGDEGNDEAEGG | 0 | 0 | 1 | 366573 | XP_820569.1 | Q4E1T5 |
| RDPTDEHSDFLTMGS | 0 | 0 | 1 | 366777 | XP_814355.1 | Q4DIZ9 |
| REDDEENDEEEDGSE | 0 | 0 | 1 | 367053 | XP_821138.1 | Q4E3E7 |
| REEVELYSFEDDEED | 0 | 0 | 1 | 367191 | XP_811579.1 | Q4DB20 |
| REFDYEDEDYEGYKS | 0 | 0 | 1 | 367205 | XP_815234.1 | Q4DEP9 |
| REIDGSLSSSAWVCA | 0 | 0 | 1 | 367298 | XP_821144.1 | Q4E3E1 |
| REKDGEDTTSEDEKN | 0 | 0 | 1 | 367330 | XP_818224.1 | Q4DV33 |
| REPQTTIKQPGGIKT | 0 | 0 | 1 | 367537 | XP_819902.1 | Q4DZS8 |
| REQDGEDATSEGKKN | 0 | 0 | 1 | 367543 | XP_808131.1 | Q4D180 |
| REQDGEDTTSEGEKN | 0 | 0 | 1 | 367544 | XP_812835.1 | Q4DEM8 |
| REVDDTMADELRSAF | 0 | 0 | 1 | 367746 | XP_805245.1 | Q4CSZ0 |
| REVDGHDFIREHIQI | 0 | 0 | 1 | 367747 | XP_816335.1 | Q4DPM2 |
| RFLAAAMDKVFTTGF | 0 | 0 | 1 | 367898 | XP_819176.1 | Q4DXP9 |
| RGGAQKGNVRKLQPQ | 0 | 0 | 1 | 368264 | XP_821251.1 | Q4E3M0 |
| RGGRDAPQARKQQGR | 0 | 0 | 1 | 368310 | XP_821251.1 | Q4E3M0 |
| RHDTDTEDSTKNAAA | 0 | 0 | 1 | 368771 | XP_811740.1 | Q4DBI1 |
| RHSHDTDTEDSTKNA | 0 | 0 | 1 | 368893 | XP_817854.1 | Q4DU19 |
| RIEEADDDDDDDDDD | 0 | 0 | 1 | 368992 | XP_804979.1 | Q4CS76 |
| RILPDLPGGRAGRIF | 0 | 0 | 1 | 369092 | XP_811374.1 | Q4DAG2 |

| | | | | | | |
|---|---|---|---|---|---|---|
| RIYDREFDYEDEDYE | 0 | 0 | 1 | 369244 | XP_815234.1 | Q4DEP9 |
| RKQQGRDAPQARKQQ | 0 | 0 | 1 | 369567 | XP_821251.1 | Q4E3M0 |
| RKQQGRNEPQARRQQ | 0 | 0 | 1 | 369568 | XP_821251.1 | Q4E3M0 |
| RLREIDGSLSSSAWV | 0 | 0 | 1 | 370200 | XP_821144.1 | Q4E3E1 |
| RLWFEREVDGHDFIR | 0 | 0 | 1 | 370382 | XP_816335.1 | Q4DPM2 |
| RMTEKEDGDNEDEEV | 0 | 0 | 1 | 370574 | XP_820304.1 | Q4E102 |
| RNEPQARKQQGRDAP | 0 | 0 | 1 | 370697 | XP_821251.1 | Q4E3M0 |
| RNEPQARRQQGRNEP | 0 | 0 | 1 | 370698 | XP_821251.1 | Q4E3M0 |
| RPGRVKRVKRKAAKK | 0 | 0 | 1 | 371114 | XP_817338.1 | Q4DSJ7 |
| RPPKKSWKQISNLSG | 0 | 0 | 1 | 371245 | XP_820569.1 | Q4E1T5 |
| RPVEATATKNTTATT | 0 | 0 | 1 | 371387 | XP_810952.1 | Q4D9A4 |
| RQQGRNEPQARKQQG | 0 | 0 | 1 | 371763 | XP_821251.1 | Q4E3M0 |
| RQQQSDETQVQQQEH | 0 | 0 | 1 | 371779 | XP_819538.1 | Q4DYS6 |
| RRQQGRNEPQARKQQ | 0 | 0 | 1 | 372446 | XP_821251.1 | Q4E3M0 |
| RSMEIDGGEWTNGTA | 0 | 0 | 1 | 373123 | XP_815234.1 | Q4DEP9 |
| RTLQVAGIKTTTATT | 0 | 0 | 1 | 373701 | XP_821134.1 | Q4E3F2 |
| RVGKKEEEDNDDDDL | 0 | 0 | 1 | 374136 | XP_821418.1 | Q4E455 |
| RVKRVKRKAAKKGAA | 0 | 0 | 1 | 374205 | XP_817338.1 | Q4DSJ7 |
| SAAKPAAKPAAKPAA | 0 | 0 | 1 | 374969 | XP_821284.1 | Q4E3R2 |
| SAEPKPAEPKSAEPK | 0 | 0 | 1 | 375203 | XP_820186.1 | Q4E0M7 |
| SAEPKPAESKSAEPK | 0 | 0 | 1 | 375208 | XP_820186.1 | Q4E0M7 |
| SAEPKPAESKSAGPK | 0 | 0 | 1 | 375209 | XP_820186.1 | Q4E0M7 |
| SAGPKPAEPKSAEPK | 0 | 0 | 1 | 375376 | XP_820186.1 | Q4E0M7 |
| SAKLGKSVGLTAALS | 0 | 0 | 1 | 375523 | XP_807065.1 | Q4CY60 |
| SANKPASKPAAKPAA | 0 | 0 | 1 | 375662 | XP_819301.1 | Q4DY30 |
| SAVSSVVPSLPADSE | 0 | 0 | 1 | 376144 | XP_813566.1 | Q4DGQ5 |
| SCVSLAARSMEIDGG | 0 | 0 | 1 | 376353 | XP_815234.1 | Q4DEP9 |
| SDDDDEGEDEEEGPS | 0 | 0 | 1 | 376470 | XP_803649.1 | Q4CNY5 |
| SDEAQFQQHQQHEHP | 0 | 0 | 1 | 376527 | XP_818475.1 | Q4DVR9 |
| SDEAQVQQHQQHEHP | 0 | 0 | 1 | 376528 | XP_809948.1 | Q4D6E7 |
| SDEAQVQQHQRHERP | 0 | 0 | 1 | 376529 | XP_817818.1 | Q4DTW6 |
| SDEEDQDEDEDEGIT | 0 | 0 | 1 | 376535 | XP_821081.1 | Q4E380 |
| SDETQVQQHQQHEHP | 0 | 0 | 1 | 376581 | XP_809166.1 | Q4D462 |
| SDETQVQQHQRHEHP | 0 | 0 | 1 | 376582 | XP_815868.1 | Q4DNB3 |
| SDETQVQQQEHEHPA | 0 | 0 | 1 | 376583 | XP_819538.1 | Q4DYS6 |
| SDFLLLMTREVDDTM | 0 | 0 | 1 | 376602 | XP_805245.1 | Q4CSZ0 |
| SEADADDDDPQRPNP | 0 | 0 | 1 | 377238 | XP_810637.1 | Q4D8D6 |
| SEDEEEDDADGDEED | 0 | 0 | 1 | 377344 | XP_821672.1 | Q4E4Z2 |
| SEDKEEDDADGDEED | 0 | 0 | 1 | 377367 | XP_822030.1 | Q4E5T2 |
| SEEEDKEEEEDDDDD | 0 | 0 | 1 | 377424 | XP_809163.1 | Q4D466 |
| SEEGEIDDDSDVQNE | 0 | 0 | 1 | 377449 | XP_813526.1 | Q4DGM5 |
| SEREDDEENDEEEDG | 0 | 0 | 1 | 378008 | XP_821138.1 | Q4E3E7 |
| SFLLKDKKKVATAKV | 0 | 0 | 1 | 378432 | XP_807065.1 | Q4CY60 |
| SFSDFLLLMTREVDD | 0 | 0 | 1 | 378556 | XP_805245.1 | Q4CSZ0 |
| SFYSREVDVSQCNMQ | 0 | 0 | 1 | 378652 | XP_807066.1 | Q4CY59 |
| SGGAQSEADADDDDP | 0 | 0 | 1 | 379030 | XP_810637.1 | Q4D8D6 |
| SGGAQSEADADVDDP | 0 | 0 | 1 | 379031 | XP_808315.1 | Q4D1R3 |
| SGTDGDESIDEEEEK | 0 | 0 | 1 | 379862 | XP_820569.1 | Q4E1T5 |
| SKAAAPKKAAAPKKA | 0 | 0 | 1 | 380939 | XP_814064.1 | Q4DI49 |
| SKPAAKPAAKAPAPK | 0 | 0 | 1 | 381300 | XP_819301.1 | Q4DY30 |
| SKSAEPKPAEPKSAE | 0 | 0 | 1 | 381379 | XP_820186.1 | Q4E0M7 |
| SKSAGPKPAEPKSAE | 0 | 0 | 1 | 381380 | XP_820186.1 | Q4E0M7 |
| SLAARSMEIDGGEWT | 0 | 0 | 1 | 381512 | XP_815234.1 | Q4DEP9 |
| SLAEQEEISDDDDEG | 0 | 0 | 1 | 381519 | XP_803649.1 | Q4CNY5 |
| SLFGQAAAGDKPAPF | 0 | 0 | 1 | 381745 | XP_818927.1 | Q4DX15 |
| SLFGQAAAGDKPPPF | 0 | 0 | 1 | 381746 | XP_818927.1 | Q4DX15 |
| SLFGQAAAGDKPSPF | 0 | 0 | 1 | 381747 | XP_818927.1 | Q4DX15 |
| SLILDRFLAAAMDKV | 0 | 0 | 1 | 381881 | XP_819176.1 | Q4DXP9 |
| SLPADSENSKTGCGE | 0 | 0 | 1 | 382095 | XP_813566.1 | Q4DGQ5 |
| SMEIDGGEWTNGTAI | 0 | 0 | 1 | 382574 | XP_815234.1 | Q4DEP9 |
| SPFGQAAAGDKPPPF | 0 | 0 | 1 | 383580 | XP_818927.1 | Q4DX15 |
| SPFGQAAAGDKPSLF | 0 | 0 | 1 | 383581 | XP_818927.1 | Q4DX15 |
| SPFGQAAAGDKPSPF | 0 | 0 | 1 | 383582 | XP_818927.1 | Q4DX15 |
| SPFGQAAAGGKPSPF | 0 | 0 | 1 | 383583 | XP_818927.1 | Q4DX15 |
| SPPEEDVASREQDGE | 0 | 0 | 1 | 384028 | XP_812835.1 | Q4DEM8 |
| SPTLKAPAAPAAPAA | 0 | 0 | 1 | 384396 | XP_805627.1 | Q4CU24 |
| SRDPTDEHSDFLTMG | 0 | 0 | 1 | 385133 | XP_814355.1 | Q4DIZ9 |
| SREEVELYSFEDDEE | 0 | 0 | 1 | 385155 | XP_811579.1 | Q4DB20 |
| SREKDGEDTTSEDEK | 0 | 0 | 1 | 385168 | XP_818224.1 | Q4DV33 |
| SREQDGEDATSEGKK | 0 | 0 | 1 | 385184 | XP_808131.1 | Q4D180 |
| SREQDGEDTTSEGEK | 0 | 0 | 1 | 385185 | XP_812835.1 | Q4DEM8 |
| SREQDGEGTTSEGHK | 0 | 0 | 1 | 385186 | XP_809849.1 | Q4D646 |
| SRLREIDGSLSSSAW | 0 | 0 | 1 | 385377 | XP_821144.1 | Q4E3E1 |
| SSEDEEEDDADGDEE | 0 | 0 | 1 | 386031 | XP_821672.1 | Q4E4Z2 |
| SSEDKEEDDADGDEE | 0 | 0 | 1 | 386034 | XP_822030.1 | Q4E5T2 |
| SSEEGEIDDDSDVQN | 0 | 0 | 1 | 386045 | XP_813526.1 | Q4DGM5 |
| SSFLLKDKKKVATAK | 0 | 0 | 1 | 386157 | XP_807065.1 | Q4CY60 |
| SSFYSREVDVSQCNM | 0 | 0 | 1 | 386184 | XP_807066.1 | Q4CY59 |
| SSSDDAQTVGTEGGA | 0 | 0 | 1 | 386973 | XP_820186.1 | Q4E0M7 |
| SSSFYSREVDVSQCN | 0 | 0 | 1 | 387012 | XP_807066.1 | Q4CY59 |
| SSTPPSGTENKPATG | 0 | 0 | 1 | 387341 | XP_808931.1 | Q4D3H3 |
| SSVVPSLPADSENSK | 0 | 0 | 1 | 387477 | XP_813566.1 | Q4DGQ5 |
| STAAKPPAESPFKNA | 0 | 0 | 1 | 387523 | XP_813515.1 | Q4DGM1 |
| STAMKTTTATTGDSD | 0 | 0 | 1 | 387612 | XP_815852.1 | Q4DNA4 |
| STATRIQRTRPRVDG | 0 | 0 | 1 | 387654 | XP_802787.1 | Q4CM38 |
| STPKVNGRETIDPDP | 0 | 0 | 1 | 388229 | XP_816182.1 | Q4DP76 |
| STPPSGTENKPATGE | 0 | 0 | 1 | 388240 | XP_808931.1 | Q4D3H3 |
| STPTTGDDDPAADGA | 0 | 0 | 1 | 388282 | XP_820308.1 | Q4E0Z8 |
| STPTTGDDDPAADVA | 0 | 0 | 1 | 388283 | XP_815852.1 | Q4DNA4 |
| STQTTGDDDPAADGA | 0 | 0 | 1 | 388335 | XP_815190.1 | Q4DLD3 |
| STSSTPPSGTENKPA | 0 | 0 | 1 | 388493 | XP_808931.1 | Q4D3H3 |

| | | | | | | |
|---|---|---|---|---|---|---|
| STTGDDGCDGDDGDG | 0 | 0 | 1 | 388550 | XP_805004.1 | Q4CSA0 |
| STTTTRAPSRLREID | 0 | 0 | 1 | 388624 | XP_815995.1 | Q4DNQ0 |
| STVIKTTTATTGDSD | 0 | 0 | 1 | 388653 | XP_820022.1 | Q4E058 |
| STVMKTTTATTGDSD | 0 | 0 | 1 | 388665 | XP_813432.1 | Q4DGC2 |
| SVASREKDGEDTTSE | 0 | 0 | 1 | 388788 | XP_818224.1 | Q4DV33 |
| SVDSREQDGEDATSE | 0 | 0 | 1 | 388890 | XP_808131.1 | Q4D180 |
| SVDSREQDGEGTTSE | 0 | 0 | 1 | 388891 | XP_809849.1 | Q4D646 |
| SVGLTAALSPRQRSL | 0 | 0 | 1 | 389037 | XP_807065.1 | Q4CY60 |
| SVVPSLPADSENSKT | 0 | 0 | 1 | 389616 | XP_813566.1 | Q4DGQ5 |
| SYEEIRENDGDIWKT | 0 | 0 | 1 | 389796 | XP_814919.1 | Q4DKL5 |
| TAAAMKTTTATTGDS | 0 | 0 | 1 | 389983 | XP_808183.1 | Q4D1D1 |
| TAAAMKTTTATTGNS | 0 | 0 | 1 | 389984 | XP_807739.1 | Q4D033 |
| TAAKPAAKSAAKPAA | 0 | 0 | 1 | 390058 | XP_821284.1 | Q4E3R2 |
| TAAKPAKKPAVKPTV | 0 | 0 | 1 | 390059 | XP_821284.1 | Q4E3R2 |
| TAAKPPAESPFKNAF | 0 | 0 | 1 | 390064 | XP_813515.1 | Q4DGM1 |
| TAAKPPAESPFKNVF | 0 | 0 | 1 | 390065 | XP_818928.1 | Q4DX14 |
| TAAKPPAESPFKSVF | 0 | 0 | 1 | 390066 | XP_818928.1 | Q4DX14 |
| TAAKPPAESPFRSVF | 0 | 0 | 1 | 390067 | XP_818928.1 | Q4DX14 |
| TAAKTAAKPAAKSAA | 0 | 0 | 1 | 390076 | XP_821284.1 | Q4E3R2 |
| TAAPKKAAAPQKAAV | 0 | 0 | 1 | 390111 | XP_814064.1 | Q4DI49 |
| TAKPPAESPFKSVFG | 0 | 0 | 1 | 390582 | XP_813515.1 | Q4DGM1 |
| TAMKNTTATGDSDGS | 0 | 0 | 1 | 390677 | XP_815190.1 | Q4DLD3 |
| TAMKNTTATTGDSDS | 0 | 0 | 1 | 390678 | XP_811361.1 | Q4DAF3 |
| TAMKTTAATTGDSDG | 0 | 0 | 1 | 390679 | XP_820410.1 | Q4E1A7 |
| TAMKTTTATTGDSDG | 0 | 0 | 1 | 390681 | XP_815852.1 | Q4DNA4 |
| TAMKTTTTTTGDSDG | 0 | 0 | 1 | 390682 | XP_820308.1 | Q4E0Z8 |
| TAQEEGQYDAADVEA | 0 | 0 | 1 | 390884 | XP_804663.1 | Q4CRB0 |
| TATKNTTATTGDSDG | 0 | 0 | 1 | 391172 | XP_821138.1 | Q4E3E7 |
| TATKNTTATTGDSDS | 0 | 0 | 1 | 391173 | XP_809011.1 | Q4D3Q7 |
| TATKTTTATTGDSDG | 0 | 0 | 1 | 391174 | XP_815165.1 | Q4DLB5 |
| TATKTTTATTGNSDG | 0 | 0 | 1 | 391175 | XP_820650.1 | Q4E1Y0 |
| TATRIQRTRPRVDGG | 0 | 0 | 1 | 391231 | XP_802787.1 | Q4CM38 |
| TATTGDSDGGTAVSH | 0 | 0 | 1 | 391246 | XP_821379.1 | Q4E442 |
| TATTGDSDGSTAASH | 0 | 0 | 1 | 391247 | XP_820304.1 | Q4E102 |
| TATTGDSDGSTAISH | 0 | 0 | 1 | 391248 | XP_815852.1 | Q4DNA4 |
| TATTGDSDGSTAVPH | 0 | 0 | 1 | 391250 | XP_804905.1 | Q4CS02 |
| TATTGDSDGSTAVSH | 0 | 0 | 1 | 391251 | XP_813432.1 | Q4DGC2 |
| TATTGDSDGVKATSP | 0 | 0 | 1 | 391252 | XP_821134.1 | Q4E3F2 |
| TATTGNSDGGTAVSH | 0 | 0 | 1 | 391258 | XP_820650.1 | Q4E1Y0 |
| TAVKATTATTGDSDS | 0 | 0 | 1 | 391332 | XP_822030.1 | Q4E5T2 |
| TAVKATTATTGESDG | 0 | 0 | 1 | 391333 | XP_821672.1 | Q4E4Z2 |
| TAVSKAAAPKKAAAP | 0 | 0 | 1 | 391424 | XP_814064.1 | Q4DI49 |
| TDDDDPAADAAGTRK | 0 | 0 | 1 | 391631 | XP_809910.1 | Q4D6A6 |
| TDEAEEEEDEDDEEE | 0 | 0 | 1 | 391680 | XP_815190.1 | Q4DLD3 |
| TDEGLLLPVDNDGDE | 0 | 0 | 1 | 391698 | XP_818651.1 | Q4DW89 |
| TDEHSDFLTMGSKMI | 0 | 0 | 1 | 391704 | XP_814355.1 | Q4DIZ9 |
| TDGDESIDEEEEKEV | 0 | 0 | 1 | 391759 | XP_820569.1 | Q4E1T5 |
| TEDSVDSREQDGEDA | 0 | 0 | 1 | 392342 | XP_808131.1 | Q4D180 |
| TEEAEDEDSDEEEET | 0 | 0 | 1 | 392353 | XP_807807.1 | Q4D0A4 |
| TEEEDDDDEEDDSDE | 0 | 0 | 1 | 392370 | XP_812035.1 | Q4DCD8 |
| TEGSQSTLAATDTAN | 0 | 0 | 1 | 392567 | XP_809167.1 | Q4D463 |
| TEKEDGDNEDEEVED | 0 | 0 | 1 | 392642 | XP_820304.1 | Q4E102 |
| TEKKKNPEAASKKAP | 0 | 0 | 1 | 392654 | XP_821284.1 | Q4E3R2 |
| TENEDDDEDEEENEE | 0 | 0 | 1 | 392785 | XP_820493.1 | Q4E1H5 |
| TEQKTAPFGQAAAGD | 0 | 0 | 1 | 392865 | XP_818927.1 | Q4DX15 |
| TEVSRDPTDEHSDFL | 0 | 0 | 1 | 393095 | XP_814355.1 | Q4DIZ9 |
| TFANTLLYSDDELFL | 0 | 0 | 1 | 393144 | XP_814919.1 | Q4DKL5 |
| TGDDDPAADGAGTPE | 0 | 0 | 1 | 393453 | XP_804905.1 | Q4CS02 |
| TGDDDPAADVAGTAE | 0 | 0 | 1 | 393454 | XP_815852.1 | Q4DNA4 |
| TGDGDPAADGAGTAE | 0 | 0 | 1 | 393466 | XP_807739.1 | Q4D033 |
| TGDKDPAADGAGTQE | 0 | 0 | 1 | 393473 | XP_813432.1 | Q4DGC2 |
| TGEAPSQPGASSGEA | 0 | 0 | 1 | 393540 | XP_808931.1 | Q4D3H3 |
| TGENLDGDEEEGDVS | 0 | 0 | 1 | 393585 | XP_813607.1 | Q4DGU9 |
| TGGDDVEDGDGEDEE | 0 | 0 | 1 | 393630 | XP_809061.1 | Q4D3W5 |
| TGTYQGDSDLQLERI | 0 | 0 | 1 | 394184 | XP_816690.1 | Q4DQP2 |
| TIKQPGGIKTPSSTT | 0 | 0 | 1 | 394616 | XP_819902.1 | Q4DZS8 |
| TINGTGENLDGDEEE | 0 | 0 | 1 | 394652 | XP_813607.1 | Q4DGU9 |
| TKNTTATTGDSDGST | 0 | 0 | 1 | 395213 | XP_821138.1 | Q4E3E7 |
| TKNTTATTGDSDSST | 0 | 0 | 1 | 395214 | XP_809011.1 | Q4D3Q7 |
| TKTTTATTGDSDGGT | 0 | 0 | 1 | 395398 | XP_821379.1 | Q4E442 |
| TKTTTATTGDSDGST | 0 | 0 | 1 | 395399 | XP_815165.1 | Q4DLB5 |
| TKTTTATTGNSDGGT | 0 | 0 | 1 | 395400 | XP_820650.1 | Q4E1Y0 |
| TLKAPAAPAAPAAAA | 0 | 0 | 1 | 395723 | XP_805627.1 | Q4CU24 |
| TLLYSDDELFLSQQR | 0 | 0 | 1 | 395822 | XP_814919.1 | Q4DKL5 |
| TLQVAGIKTTTATTG | 0 | 0 | 1 | 396003 | XP_821134.1 | Q4E3F2 |
| TMKNTTATTGDSDGS | 0 | 0 | 1 | 396302 | XP_806129.1 | Q4CVH7 |
| TNAPSRLREIDGSLS | 0 | 0 | 1 | 396415 | XP_821144.1 | Q4E3E1 |
| TNSTPTTGDDDPAAD | 0 | 0 | 1 | 396787 | XP_820308.1 | Q4E0Z8 |
| TPAESTVIKTTTATT | 0 | 0 | 1 | 396934 | XP_820022.1 | Q4E058 |
| TPDDDDPAADAAGTR | 0 | 0 | 1 | 397003 | XP_821399.1 | Q4E444 |
| TPFEATAMKTTTTTT | 0 | 0 | 1 | 397109 | XP_820308.1 | Q4E0Z8 |
| TPKVNGRETIDPDPQ | 0 | 0 | 1 | 397327 | XP_816182.1 | Q4DP76 |
| TPLEATATKNTTATT | 0 | 0 | 1 | 397343 | XP_821138.1 | Q4E3E7 |
| TPNNDPAADGAETRE | 0 | 0 | 1 | 397415 | XP_814148.1 | Q4DIF3 |
| TPPSGTENKPATGEA | 0 | 0 | 1 | 397488 | XP_808931.1 | Q4D3H3 |
| TPSTTTTRAPSRLRE | 0 | 0 | 1 | 397716 | XP_815995.1 | Q4DNQ0 |
| TPTEEEDDDDEEDDS | 0 | 0 | 1 | 397741 | XP_812035.1 | Q4DCD8 |
| TPTTGDDDPAADGAG | 0 | 0 | 1 | 397798 | XP_820308.1 | Q4E0Z8 |
| TPTTGDDDPAADVAG | 0 | 0 | 1 | 397799 | XP_815852.1 | Q4DNA4 |
| TPVAATVIKTTTAAT | 0 | 0 | 1 | 397816 | XP_820637.1 | Q4E1Z5 |
| TPVEATAMKNTTATT | 0 | 0 | 1 | 397832 | XP_811361.1 | Q4DAF3 |

| | | | | | | |
|---|---|---|---|---|---|---|
| TPVEATAMKTTTATT | 0 | 0 | 1 | 397834 | XP_804905.1 | Q4CS02 |
| TPVEATATKNTTATT | 0 | 0 | 1 | 397836 | XP_809011.1 | Q4D3Q7 |
| TPVEATATKTTTATT | 0 | 0 | 1 | 397837 | XP_820650.1 | Q4E1Y0 |
| TPVEATAVKATTATT | 0 | 0 | 1 | 397838 | XP_821672.1 | Q4E4Z2 |
| TPVEGTAMKTTTATT | 0 | 0 | 1 | 397841 | XP_820304.1 | Q4E102 |
| TPVESTAMKTTTATT | 0 | 0 | 1 | 397842 | XP_815852.1 | Q4DNA4 |
| TPVESTVMKTTTATT | 0 | 0 | 1 | 397844 | XP_813432.1 | Q4DGC2 |
| TPVETTTMKNTTATT | 0 | 0 | 1 | 397848 | XP_806129.1 | Q4CVH7 |
| TPVTAAAMKTTTATT | 0 | 0 | 1 | 397876 | XP_808183.1 | Q4D1D1 |
| TQFSVGTPDAANAAT | 0 | 0 | 1 | 398000 | XP_820186.1 | Q4E0M7 |
| TQGRHSHDTDTEDST | 0 | 0 | 1 | 398025 | XP_817854.1 | Q4DU19 |
| TQITDDDDPAADAAG | 0 | 0 | 1 | 398048 | XP_809910.1 | Q4D6A6 |
| TQRGGRDAPQARKQQ | 0 | 0 | 1 | 398220 | XP_821251.1 | Q4E3M0 |
| TQTEQKTAPFGQAAA | 0 | 0 | 1 | 398298 | XP_818927.1 | Q4DX15 |
| TQTPDDDDPAADAAG | 0 | 0 | 1 | 398326 | XP_821399.1 | Q4E444 |
| TQTTGDDDPAADGAG | 0 | 0 | 1 | 398341 | XP_815190.1 | Q4DLD3 |
| TRAPSRLREIDGSLS | 0 | 0 | 1 | 398399 | XP_815995.1 | Q4DNQ0 |
| TREVDDTMADELRSA | 0 | 0 | 1 | 398505 | XP_805245.1 | Q4CSZ0 |
| TSAAAAANDSSPAEG | 0 | 0 | 1 | 398908 | XP_815868.1 | Q4DNB3 |
| TSAKLGKSVGLTAAL | 0 | 0 | 1 | 398965 | XP_807065.1 | Q4CY60 |
| TTAATGDSDGSTAVP | 0 | 0 | 1 | 400055 | XP_820637.1 | Q4E1Z5 |
| TTATTGDSDGGTAVS | 0 | 0 | 1 | 400154 | XP_821379.1 | Q4E442 |
| TTATTGDSDGSTAAS | 0 | 0 | 1 | 400155 | XP_820304.1 | Q4E102 |
| TTATTGDSDGSTAIS | 0 | 0 | 1 | 400156 | XP_815852.1 | Q4DNA4 |
| TTATTGDSDGSTAVF | 0 | 0 | 1 | 400157 | XP_808645.1 | Q4D2N7 |
| TTATTGDSDGSTAVP | 0 | 0 | 1 | 400158 | XP_804905.1 | Q4CS02 |
| TTATTGDSDGSTAVS | 0 | 0 | 1 | 400159 | XP_813432.1 | Q4DGC2 |
| TTATTGDSDGVKATS | 0 | 0 | 1 | 400160 | XP_821134.1 | Q4E3F2 |
| TTATTGDSDSSTAAS | 0 | 0 | 1 | 400161 | XP_822030.1 | Q4E5T2 |
| TTATTGDSDSSTAVF | 0 | 0 | 1 | 400162 | XP_811361.1 | Q4DAF3 |
| TTATTGDSDSSTAVS | 0 | 0 | 1 | 400163 | XP_809011.1 | Q4D3Q7 |
| TTATTGESDGSTAVS | 0 | 0 | 1 | 400164 | XP_821672.1 | Q4E4Z2 |
| TTATTGNSDGGTAVS | 0 | 0 | 1 | 400165 | XP_820650.1 | Q4E1Y0 |
| TTATTGNSDGSTAVS | 0 | 0 | 1 | 400166 | XP_807739.1 | Q4D033 |
| TTGDDDPAADGAGTP | 0 | 0 | 1 | 400359 | XP_804905.1 | Q4CS02 |
| TTGDDDPAADVAGTA | 0 | 0 | 1 | 400360 | XP_815852.1 | Q4DNA4 |
| TTGDDGCDGDDGDGA | 0 | 0 | 1 | 400361 | XP_805004.1 | Q4CSA0 |
| TTGDGDPAADGAGTA | 0 | 0 | 1 | 400362 | XP_807739.1 | Q4D033 |
| TTIKQPGGIKTPSST | 0 | 0 | 1 | 400481 | XP_819902.1 | Q4DZS8 |
| TTMKNTTATTGDSDG | 0 | 0 | 1 | 400628 | XP_806129.1 | Q4CVH7 |
| TTNAPSRLREIDGSL | 0 | 0 | 1 | 400646 | XP_821144.1 | Q4E3E1 |
| TTRAPSRLREIDGSL | 0 | 0 | 1 | 400854 | XP_815995.1 | Q4DNQ0 |
| TTTAATGDSDGSTAV | 0 | 0 | 1 | 401091 | XP_820637.1 | Q4E1Z5 |
| TTTATTGDSDGGTAV | 0 | 0 | 1 | 401105 | XP_821379.1 | Q4E442 |
| TTTATTGDSDGSTAA | 0 | 0 | 1 | 401106 | XP_820304.1 | Q4E102 |
| TTTATTGDSDGSTAI | 0 | 0 | 1 | 401107 | XP_815852.1 | Q4DNA4 |
| TTTATTGDSDGSTAV | 0 | 0 | 1 | 401108 | XP_813432.1 | Q4DGC2 |
| TTTATTGDSDGVKAT | 0 | 0 | 1 | 401109 | XP_821134.1 | Q4E3F2 |
| TTTATTGDSDSSTAV | 0 | 0 | 1 | 401110 | XP_808183.1 | Q4D1D1 |
| TTTATTGNSDGGTAV | 0 | 0 | 1 | 401111 | XP_820650.1 | Q4E1Y0 |
| TTTATTGNSDGSTAV | 0 | 0 | 1 | 401112 | XP_807739.1 | Q4D033 |
| TTTMKNTTATTGDSD | 0 | 0 | 1 | 401174 | XP_806129.1 | Q4CVH7 |
| TTTNAPSRLREIDGS | 0 | 0 | 1 | 401178 | XP_821144.1 | Q4E3E1 |
| TTTRAPSRLREIDGS | 0 | 0 | 1 | 401205 | XP_815995.1 | Q4DNQ0 |
| TTTTNAPSRLREIDG | 0 | 0 | 1 | 401241 | XP_821144.1 | Q4E3E1 |
| TTTTRAPSRLREIDG | 0 | 0 | 1 | 401248 | XP_815995.1 | Q4DNQ0 |
| TTTTTNAPSRLREID | 0 | 0 | 1 | 401265 | XP_821144.1 | Q4E3E1 |
| TTTTTTGDSDGSTAV | 0 | 0 | 1 | 401274 | XP_820308.1 | Q4E0Z8 |
| TVIKTTTAATGDSDG | 0 | 0 | 1 | 401686 | XP_820637.1 | Q4E1Z5 |
| TVIKTTTATTGDSDG | 0 | 0 | 1 | 401688 | XP_820022.1 | Q4E058 |
| TVMKTTTATTGDSDG | 0 | 0 | 1 | 401807 | XP_813432.1 | Q4DGC2 |
| TYQGDSDLQLERINV | 0 | 0 | 1 | 402369 | XP_816690.1 | Q4DQP2 |
| VAATVIKTTTAATGD | 0 | 0 | 1 | 402545 | XP_820637.1 | Q4E1Z5 |
| VADGAGGGDGSAGED | 0 | 0 | 1 | 402582 | XP_817817.1 | Q4DTW8 |
| VAGIKTTTATTGDSD | 0 | 0 | 1 | 402786 | XP_821134.1 | Q4E3F2 |
| VASREKDGEDTTSED | 0 | 0 | 1 | 403248 | XP_818224.1 | Q4DV33 |
| VATEKKKNPEAASKK | 0 | 0 | 1 | 403289 | XP_821284.1 | Q4E3R2 |
| VDADTAALAALLEVL | 0 | 0 | 1 | 403764 | XP_803309.1 | Q4CNL0 |
| VDEDEDDDDDNDND | 0 | 0 | 1 | 403879 | XP_818224.1 | Q4DV33 |
| VDGHDFIREHIQIRT | 0 | 0 | 1 | 403969 | XP_816335.1 | Q4DPM2 |
| VDNDGDESSDKEDAE | 0 | 0 | 1 | 404124 | XP_818651.1 | Q4DW89 |
| VDPTGTYQGDSDLQL | 0 | 0 | 1 | 404177 | XP_816690.1 | Q4DQP2 |
| VDSREQDGEDATSEG | 0 | 0 | 1 | 404270 | XP_808131.1 | Q4D180 |
| VDSREQDGEGTTSEG | 0 | 0 | 1 | 404271 | XP_809849.1 | Q4D646 |
| VDSREQDGEVTTSEG | 0 | 0 | 1 | 404272 | XP_807741.1 | Q4D040 |
| VEATAMKNTTATGDS | 0 | 0 | 1 | 404475 | XP_815190.1 | Q4DLD3 |
| VEATAMKNTTATTGD | 0 | 0 | 1 | 404476 | XP_811361.1 | Q4DAF3 |
| VEATAMKTTAATTGD | 0 | 0 | 1 | 404477 | XP_820410.1 | Q4E1A7 |
| VEATAMKTTTATTGD | 0 | 0 | 1 | 404479 | XP_804905.1 | Q4CS02 |
| VEATATKNTTATTGD | 0 | 0 | 1 | 404481 | XP_809011.1 | Q4D3Q7 |
| VEATATKTTTATTGD | 0 | 0 | 1 | 404482 | XP_821379.1 | Q4E442 |
| VEATATKTTTATTGN | 0 | 0 | 1 | 404483 | XP_820650.1 | Q4E1Y0 |
| VEATAVKATTATTGD | 0 | 0 | 1 | 404484 | XP_822030.1 | Q4E5T2 |
| VEATAVKATTATTGE | 0 | 0 | 1 | 404485 | XP_821672.1 | Q4E4Z2 |
| VEDGDGEDEEEGQLV | 0 | 0 | 1 | 404523 | XP_809061.1 | Q4D3W5 |
| VEEDEKEDNDDDDDD | 0 | 0 | 1 | 404591 | XP_815012.1 | Q4DKW6 |
| VEGESSSFYSREVDV | 0 | 0 | 1 | 404729 | XP_807066.1 | Q4CY59 |
| VEGTAMKTTTATTGD | 0 | 0 | 1 | 404773 | XP_820304.1 | Q4E102 |
| VEKIDSENSKLQQEM | 0 | 0 | 1 | 404849 | XP_821826.1 | Q4E572 |
| VELYSFEDDEEDENT | 0 | 0 | 1 | 404934 | XP_811579.1 | Q4DB20 |
| VESTAMKTTTATTGD | 0 | 0 | 1 | 405147 | XP_815852.1 | Q4DNA4 |

| | | | | | | |
|---|---|---|---|---|---|---|
| VESTVMKTTTATTGD | 0 | 0 | 1 | 405151 | XP_813432.1 | Q4DGC2 |
| VETTTMKNTTATTGD | 0 | 0 | 1 | 405201 | XP_806129.1 | Q4CVH7 |
| VGASSSDDAQTVGTE | 0 | 0 | 1 | 405694 | XP_820186.1 | Q4E0M7 |
| VGEKKDEEQEKKEEE | 0 | 0 | 1 | 405809 | XP_807741.1 | Q4D040 |
| VGKKEEEDNDDDDLE | 0 | 0 | 1 | 406019 | XP_821418.1 | Q4E455 |
| VGKKKDEEQEPNEEE | 0 | 0 | 1 | 406021 | XP_809910.1 | Q4D6A6 |
| VGLTAALSPRQRSLP | 0 | 0 | 1 | 406072 | XP_807065.1 | Q4CY60 |
| VIKTTTAATGDSDGS | 0 | 0 | 1 | 406705 | XP_820637.1 | Q4E1Z5 |
| VIKTTTATTGDSDGS | 0 | 0 | 1 | 406707 | XP_820022.1 | Q4E058 |
| VKATTATTGDSDSST | 0 | 0 | 1 | 406958 | XP_822030.1 | Q4E5T2 |
| VKATTATTGESDGST | 0 | 0 | 1 | 406959 | XP_821672.1 | Q4E4Z2 |
| VKKAAKPAAKTAAKT | 0 | 0 | 1 | 407130 | XP_821284.1 | Q4E3R2 |
| VKRVKRKAAKKGAAG | 0 | 0 | 1 | 407358 | XP_817338.1 | Q4DSJ7 |
| VLQYGRLIDDFIVVA | 0 | 0 | 1 | 408158 | XP_808204.1 | Q4D1F2 |
| VMKTTTATTGDSDGS | 0 | 0 | 1 | 408688 | XP_813432.1 | Q4DGC2 |
| VNGKRIYDREFDYED | 0 | 0 | 1 | 408930 | XP_815234.1 | Q4DEP9 |
| VNGRETIDPDPQKQL | 0 | 0 | 1 | 408941 | XP_816182.1 | Q4DP76 |
| VNPLVVRLWFEREVD | 0 | 0 | 1 | 409039 | XP_816335.1 | Q4DPM2 |
| VNSGTDGDESIDEEE | 0 | 0 | 1 | 409087 | XP_820569.1 | Q4E1T5 |
| VPSLPADSENSKTGC | 0 | 0 | 1 | 409756 | XP_813566.1 | Q4DGQ5 |
| VRLWFEREVDGHDFI | 0 | 0 | 1 | 410638 | XP_816335.1 | Q4DPM2 |
| VRPPKKSWKQISNLS | 0 | 0 | 1 | 410689 | XP_820569.1 | Q4E1T5 |
| VSKAAAPKKAAAPKK | 0 | 0 | 1 | 411333 | XP_814064.1 | Q4DI49 |
| VSLAARSMEIDGGEW | 0 | 0 | 1 | 411363 | XP_815234.1 | Q4DEP9 |
| VSLAEQEEISDDDDE | 0 | 0 | 1 | 411364 | XP_803649.1 | Q4CNY5 |
| VSLILDRFLAAAMDK | 0 | 0 | 1 | 411393 | XP_819176.1 | Q4DXP9 |
| VSRDPTDEHSDFLTM | 0 | 0 | 1 | 411598 | XP_814355.1 | Q4DIZ9 |
| VSSVVPSLPADSENS | 0 | 0 | 1 | 411746 | XP_813566.1 | Q4DGQ5 |
| VTAAAMKTTTATTGD | 0 | 0 | 1 | 411910 | XP_808183.1 | Q4D1D1 |
| VTAAAMKTTTATTGN | 0 | 0 | 1 | 411911 | XP_807739.1 | Q4D033 |
| VVEGESSSFYSREVD | 0 | 0 | 1 | 412847 | XP_807066.1 | Q4CY59 |
| VVNPLVVRLWFEREV | 0 | 0 | 1 | 413145 | XP_816335.1 | Q4DPM2 |
| VVPSLPADSENSKTG | 0 | 0 | 1 | 413199 | XP_813566.1 | Q4DGQ5 |
| VVRLWFEREVDGHDF | 0 | 0 | 1 | 413267 | XP_816335.1 | Q4DPM2 |
| WFEREVDGHDFIREH | 0 | 0 | 1 | 414277 | XP_816335.1 | Q4DPM2 |
| YDAADVEAGDGPDKT | 0 | 0 | 1 | 415533 | XP_804663.1 | Q4CRB0 |
| YDREFDYEDEDYEGY | 0 | 0 | 1 | 415627 | XP_815234.1 | Q4DEP9 |
| YEEIRENDGDIWKTE | 0 | 0 | 1 | 415727 | XP_814919.1 | Q4DKL5 |
| YQDATIEEEGEFDEE | 0 | 0 | 1 | 417041 | XP_816690.1 | Q4DQP2 |
| YQGDSDLQLERINVY | 0 | 0 | 1 | 417050 | XP_816690.1 | Q4DQP2 |
| YSREVDVSQCNMQEL | 0 | 0 | 1 | 417440 | XP_807066.1 | Q4CY59 |
| YVNGKRIYDREFDYE | 0 | 0 | 1 | 417767 | XP_815234.1 | Q4DEP9 |
| VVPVAARLLLE | 0 | 0 | 1 | 433186 | Q13568.2 | Q13568 |
| AQEKGLPEPTVTNEE | 0 | 0 | 1 | 463650 | AAA29462.1 | A0A144A026 |
| IVEVEEILPEDDKNE | 0 | 0 | 1 | 466930 | AAG12326.1 | Q8IJ56 |
| KGLPEPTVTNEEYVE | 0 | 0 | 1 | 467193 | AAA29462.1 | A0A144A026 |
| KNEKVEHEIVEVEEI | 0 | 0 | 1 | 467355 | AAG12326.1 | Q8IJ56 |
| KSTYLTEPILTEEHF | 0 | 0 | 1 | 467437 | AAO12051.1 | C0H571 |
| KVEHEIVEVEEILPE | 0 | 0 | 1 | 467462 | AAG12326.1 | Q8IJ56 |
| NWDDMEKIWHHTF | 0 | 0 | 1 | 469132 | P30162.1 | P30162 |
| PEPTVTNEEYVEELK | 0 | 0 | 1 | 469234 | AAA29462.1 | A0A144A026 |
| TYLTEPILTEEHFNLT | 0 | 0 | 1 | 472212 | AAO12051.1 | C0H571 |
| AEKYDKMDEPQDYGK | 0 | 0 | 1 | 509326 | GI:1373027 | Q7KQK5 |
| LKKNSRSLGENDDGN | 0 | 0 | 1 | 509806 | GI:124504759 | Q7K740 |
| MPNDPNRNVDENANA | 0 | 0 | 1 | 509997 | GI:124504759 | Q7K740 |
| NNNNEEPSDKHIKEY | 0 | 0 | 1 | 510032 | GI:124504759 | Q7K740 |
| RIKPGSANKPKDELD | 0 | 0 | 1 | 510113 | GI:124504759 | Q7K740 |
| RKKIYTHKTTRKKHKDNPDYEKALL | 0 | 0 | 1 | 538748 | CZT99494.1 | A0A144A1C7 |
| EKKICKMEKCSSVFN | 0 | 0 | 1 | 540750 | GI:124504759 | Q7K740 |
| ALPKVCHGLPPKSLD | 0 | 0 | 1 | 548925 | AAA30325.1 | A2DHT2 |
| AQPLYDEAIAFKEEV | 0 | 0 | 1 | 548927 | AAC72899.1 | A2DKH3 |
| DGRMVMTGTIRRLFV | 0 | 0 | 1 | 548932 | AAK73099.1 | A2F6T1 |
| DKVPKKFKLPSPFFN | 0 | 0 | 1 | 548934 | AAW78351.1 | Q5EFD8 |
| DYENWTKLNARLGQR | 0 | 0 | 1 | 548935 | AAW78351.1 | Q5EFD8 |
| EAHSRPDYVTVEGEL | 0 | 0 | 1 | 548936 | SRC337200 | A2F6T1 |
| EDDVKAEKHTYTRPE | 0 | 0 | 1 | 548937 | AAK73099.1 | A2F6T1 |
| EQLQAIITASVKTES | 0 | 0 | 1 | 548942 | AAK73099.1 | A2F6T1 |
| EWMYSCRCADIFHRL | 0 | 0 | 1 | 548945 | AAA30325.1 | A2DHT2 |
| FKDTFKYFDKDKSNS | 0 | 0 | 1 | 548946 | AAC72899.1 | A2DKH3 |
| HTYTRPEEVQDFVSK | 0 | 0 | 1 | 548955 | AAK73099.1 | A2F6T1 |
| KDAEGKIKKDDGYDGH | 0 | 0 | 1 | 548956 | SRC337199 | A2DHT2 |
| KKHPAIVSIED | 0 | 0 | 1 | 548958 | AAW78351.1 | Q5EFD8 |
| KLGGLLVKKYGLSAK | 0 | 0 | 1 | 548959 | AAW78351.1 | Q5EFD8 |
| LGEARTKLTEMYMRK | 0 | 0 | 1 | 548964 | AAK73099.1 | A2F6T1 |
| LVKKYGLSAKN | 0 | 0 | 1 | 548982 | AAW78351.1 | Q5EFD8 |
| LYTTNPITIKKGLEG | 0 | 0 | 1 | 548984 | AAW78351.1 | Q5EFD8 |
| NGGKHAGGNLK | 0 | 0 | 1 | 548988 | AAW78351.1 | Q5EFD8 |
| PIVLHLDHGDS | 0 | 0 | 1 | 548990 | AAK73099.1 | A2F6T1 |
| QEFTVGEGADKWVVK | 0 | 0 | 1 | 548991 | AAA30325.1 | A2DHT2 |
| QLRMVAEVYQK | 0 | 0 | 1 | 548993 | AAW78351.1 | Q5EFD8 |
| RACRKLYPKDIQVVA | 0 | 0 | 1 | 548995 | AAA30325.1 | A2DHT2 |
| RKYANQTMLRY | 0 | 0 | 1 | 548997 | AAK73099.1 | A2F6T1 |
| SLVDLTVNVNA | 0 | 0 | 1 | 549003 | AAA30325.1 | A2DHT2 |
| SSEFYDEEKKLYEVE | 0 | 0 | 1 | 549006 | AAW78351.1 | Q5EFD8 |
| SSSIPQEYVEMVNKY | 0 | 0 | 1 | 549007 | AAK73099.1 | A2F6T1 |
| SVNRHHSQLITYIKH | 0 | 0 | 1 | 549009 | AAC72899.1 | A2DKH3 |
| TDGTVLKKNIGGNAC | 0 | 0 | 1 | 549010 | AAW78351.1 | Q5EFD8 |
| VKYLGRVTLAARSSA | 0 | 0 | 1 | 549014 | AAW78351.1 | Q5EFD8 |
| VLSWYDNEWMY | 0 | 0 | 1 | 549015 | AAA30325.1 | A2DHT2 |
| VTLAARSSAPSGAST | 0 | 0 | 1 | 549018 | AAW78351.1 | Q5EFD8 |
| WVVKSIGGRLGPSQL | 0 | 0 | 1 | 549019 | AAA30325.1 | A2DHT2 |

| | | | | | | |
|---|---|---|---|---|---|---|
| YLLKYDTAHRA | 0 | 0 | 1 | 549021 | AAA30325.1 | A2DHT2 |
| YSSIVDALSTK | 0 | 0 | 1 | 549022 | AAA30325.1 | A2DHT2 |
| KKDVTPKTGKRKRSEQQKDE | 0 | 0 | 1 | 549546 | XP_001214344.1 | Q0CMB8 |
| AYFLLGPVVKTLFNK | 0 | 0 | 1 | 582208 | Q53UB7 | A5JZX5 |
| EGGSEFSERIGNSLS | 0 | 0 | 1 | 582469 | Q53UB7 | A5JZX5 |
| EVIGNELADNIANEIVSSLQK | 0 | 0 | 1 | 582616 | Q53UB7 | A5JZX5 |
| FDVKTQLKATAKKVL | 0 | 0 | 1 | 582645 | Q53UB7 | A5JZX5 |
| FNKVEDVLHKPIPDT | 0 | 0 | 1 | 582742 | Q53UB7 | A5JZX5 |
| KVLVEALKAALEPTE | 0 | 0 | 1 | 585603 | Q53UB7 | A5JZX5 |
| LALFCFVNVLSLRGK | 0 | 0 | 1 | 585702 | Q53UB7 | A5JZX5 |
| LEEEEAEDEFSDELLD | 0 | 0 | 1 | 585778 | Q53UB7 | A5JZX5 |
| LKAALEPTEKIVAST | 0 | 0 | 1 | 585971 | Q53UB7 | A5JZX5 |
| LKATAKKVLVEALKA | 0 | 0 | 1 | 585974 | Q53UB7 | A5JZX5 |
| LQKDSASFLQSGFDV | 0 | 0 | 1 | 586203 | Q53UB7 | A5JZX5 |
| MHLFNKPPKGKMNKV | 0 | 0 | 1 | 586451 | Q53UB7 | A5JZX5 |
| NEIVSSLQKDSASFL | 0 | 0 | 1 | 586539 | Q53UB7 | A5JZX5 |
| NKVNRVSIICAFLALFCFVNV | 0 | 0 | 1 | 586586 | Q53UB7 | A5JZX5 |
| PDTIWEYESKGSLEE | 0 | 0 | 1 | 586825 | Q53UB7 | A5JZX5 |
| PPKGKMNKVNRVSII | 0 | 0 | 1 | 587135 | Q53UB7 | A5JZX5 |
| PVVKTLFNKVEDVLH | 0 | 0 | 1 | 587634 | Q53UB7 | A5JZX5 |
| SERIGNSLSSFLSES | 0 | 0 | 1 | 589793 | Q53UB7 | A5JZX5 |
| SESASLEVIGNELAD | 0 | 0 | 1 | 589800 | Q53UB7 | A5JZX5 |
| SFLQSGFDVKTQLKA | 0 | 0 | 1 | 589814 | Q53UB7 | A5JZX5 |
| SLSSFLSESASLEVI | 0 | 0 | 1 | 589910 | Q53UB7 | A5JZX5 |
| TASSSLEGGSEFSER | 0 | 0 | 1 | 590217 | Q53UB7 | A5JZX5 |
| VLHKPIPDTIWEYES | 0 | 0 | 1 | 590783 | Q53UB7 | A5JZX5 |
| VNVLSLRGKSGSTAS | 0 | 0 | 1 | 590836 | Q53UB7 | A5JZX5 |
| YESKGSLEEEEAEDE | 0 | 0 | 1 | 591245 | Q53UB7 | A5JZX5 |
| IDERKDTECQTDFPLLDVSRNTNR | 0 | 0 | 1 | 644539 | CZT98473.1 | A0A143ZXM2 |
| NEDRGIYDELLENDMCDLYN | 0 | 0 | 1 | 647826 | CZT98473.1 | A0A143ZXM2 |
| NNNMNDIAIGENVIYENDIHENNID | 0 | 0 | 1 | 647940 | CZT98473.1 | A0A143ZXM2 |
| SYINNDENLQAFDLLDNFHMDDY | 0 | 0 | 1 | 650560 | CZT98473.1 | A0A143ZXM2 |
| HTLVKSRRLNGTIEFP | 0 | 0 | 1 | 689227 | XP_018649193.1 | A0A3Q0KSB0 |
| MGFATGYFVHDIYHNI | 0 | 0 | 1 | 689242 | XP_018648220.1 | G4V9U1 |
| QLCKDTSSRMKAAFQF | 0 | 0 | 1 | 689245 | XP_018652741.1 | A0A3Q0KQZ3 |
| QVFYYKDMYSLWLADT | 0 | 0 | 1 | 689246 | XP_018645745.1 | G4LWV6 |
| LQVAGIKTTTATTGDS | 0 | 0 | 1 | 697114 | XP_821134.1 | Q4E3F2 |
| PEEDVASREQDGEDTTSEGE | 0 | 0 | 1 | 697116 | XP_812835.1 | Q4DEM8 |
| QTTGDDDPAADGAGTAEGKQ | 0 | 0 | 1 | 697118 | XP_815190.1 | Q4DLD3 |
| FNTNSNFINDLVNLT | 0 | 0 | 1 | 738133 | XP_018651993.1 | G4VIY7 |
| RRVAVLVLLDRL | 0 | 0 | 1 | 738366 | XP_003861458.1 | A0A504WVT0 |
| FDLFLFSNGAVVWWG | 0 | 0 | 1 | 741400 | XP_003862935.1 | A0A504XDX0 |
| IMLYALIATQFSDDA | 0 | 0 | 1 | 741559 | XP_003857910.1 | A0A504X632 |
| LALLIMLYALIATQF | 0 | 0 | 1 | 741634 | XP_003857910.1 | A0A504X632 |
| LIMLYALIATQFSDD | 0 | 0 | 1 | 741660 | XP_003857910.1 | A0A504X632 |
| EKNKKFNENLVKIEKR | 0 | 0 | 1 | 745103 | XP_012648767.1 | A0A0K3AT66 |
| GENDIIQPPWEDTAP | 0 | 0 | 1 | 745106 | AAF68253.1 | I7J5Q4 |
| GTGWPSEAGGPSEAGG | 0 | 0 | 1 | 745109 | AAF68236.1 | I7I905 |
| HQEQNNANDSSNPTG | 0 | 0 | 1 | 745111 | XP_012648767.1 | A0A0K3AT66 |
| NKSEKAERKSHDTQT | 0 | 0 | 1 | 745119 | AAF68253.1 | I7J5Q4 |
| NANPNVDPNANPNVD | 0 | 0 | 1 | 756264 | GI:124504759 | Q7K740 |
| EKIRKQEEEEEERINNQRRA | 0 | 0 | 1 | 768141 | SCO74394.1 | A5K3N8 |
| KLQQEQNELNEEKERQRQEN | 0 | 0 | 1 | 768198 | SCO74394.1 | A5K3N8 |
| QEQEEEEDDNDPNGSKKNGK | 0 | 0 | 1 | 768247 | SCO74394.1 | A5K3N8 |
| YGRTRNKRYMHRNPGEKYKG | 0 | 0 | 1 | 768320 | SCO74394.1 | A5K3N8 |
| KQPADGNPDPNANPN | 0 | 0 | 1 | 768504 | GI:124504759 | Q7K740 |
| AHFEFATTPDDQNSVSEPRA | 0 | 0 | 1 | 840803 | AHF49837.1 | A5K1W7 |
| AKIRVRKRSGEEYDKEIFNL | 0 | 0 | 1 | 840804 | AHF49837.1 | A5K1W7 |
| DETVETSKLSESCRKE | 0 | 0 | 1 | 840816 | XP_002569898.1 | P32070 |
| EKFIQTYLTLLRDGDE | 0 | 0 | 1 | 840825 | XP_002569898.1 | P32070 |
| FNLSGKKEQLRTKIELTDLV | 0 | 0 | 1 | 840836 | AHF49837.1 | A5K1W7 |
| FQSIQFQYDNKIILAW | 0 | 0 | 1 | 840838 | XP_002569898.1 | P32070 |
| GVSIIASTMPKPKQVE | 0 | 0 | 1 | 840851 | XP_002569898.1 | P32070 |
| KDQKITFEEFCRGLGL | 0 | 0 | 1 | 840862 | XP_002569898.1 | P32070 |
| KSELERRYGRVWQVNA | 0 | 0 | 1 | 840872 | XP_002569898.1 | P32070 |
| NEMRIERNHIKTVQSG | 0 | 0 | 1 | 840906 | XP_002569898.1 | P32070 |
| PKQVEVTQLFKDIYNE | 0 | 0 | 1 | 840916 | XP_002569898.1 | P32070 |
| RGLGLKQNEMRIERNH | 0 | 0 | 1 | 840924 | XP_002569898.1 | P32070 |
| RNHIKTVQSGREQSLP | 0 | 0 | 1 | 840926 | XP_002569898.1 | P32070 |
| SESCRKEKLDMKQVNE | 0 | 0 | 1 | 840933 | XP_002569898.1 | P32070 |
| VKRKQKETNADEKKSYDDVS | 0 | 0 | 1 | 840963 | AEX92981.1 | A5K1W7 |
| VNAVTHSYWASFSHEP | 0 | 0 | 1 | 840964 | XP_002569898.1 | P32070 |
| VNEWIALFDVDKDQKI | 0 | 0 | 1 | 840965 | XP_002569898.1 | P32070 |
| YWASFSHEPFQSIQFQ | 0 | 0 | 1 | 840977 | XP_002569898.1 | P32070 |
| CVEEDMPEKVINDSQMTLRI | 0 | 0 | 1 | 842226 | CAA75924.1 | A0A044TPU2 |
| IRRLINEESEEQKAV | 0 | 0 | 1 | 845508 | A0A044TPU2 | A0A044TPU2 |
| QQIDPLDAEHSRGFV | 0 | 0 | 1 | 847913 | CAA75924.1 | A0A044TPU2 |
| AAAAIDNMNDSE | 0 | 0 | 1 | 850918 | A0A044U9P0 | A0A044U9P0 |
| AAAKEAEAHEEI | 0 | 0 | 1 | 850922 | A0A044UK00 | A0A044UK00 |
| AADESTGSMDK | 0 | 0 | 1 | 850926 | AAB52600.1 | A0A044UUD9 |
| AAEPEAPAAPASA | 0 | 0 | 1 | 850928 | A0A044R196 | A0A044R196 |
| AAESSAKFKENSN | 0 | 0 | 1 | 850929 | A0A044UVG8 | A0A044UVG8 |
| AAFMSENSTPNEP | 0 | 0 | 1 | 850930 | A0A044U0M2 | A0A044U0M2 |
| AAGPMKGILEYTEDQ | 0 | 0 | 1 | 850932 | AAB52599.1 | A0A158N7S3 |
| AAIGALEAYNKKK | 0 | 0 | 1 | 850934 | AAK16514.1 | A0A2K6W829 |
| AAKDADARMSAIA | 0 | 0 | 1 | 850936 | P36991.1 | P36991 |
| AAKKRHQEEEAA | 0 | 0 | 1 | 850937 | A0A044R196 | A0A044R196 |
| AAKQDGLNFYKYM | 0 | 0 | 1 | 850939 | A0A044UIG6 | A0A044UIG6 |
| AALADLQKNPDLI | 0 | 0 | 1 | 850940 | A0A044V420 | A0A044V420 |
| AALGHPLISISVA | 0 | 0 | 1 | 850942 | A0A044UMF0 | A0A044UMF0 |
| AANCEDLIEWDDN | 0 | 0 | 1 | 850947 | A0A044UK00 | A0A044UK00 |

| | | | | | | |
|---|---|---|---|---|---|---|
| AAPPAQFGMAP | 0 | 0 | 1 | 850950 | A0A2K6WJQ8 | A0A2K6WJQ8 |
| AAVKEAAAGPMKGI | 0 | 0 | 1 | 850955 | AAB52599.1 | A0A158N7S3 |
| ACEANCDSNEDSKQN | 0 | 0 | 1 | 850958 | A0A044UK00 | A0A044UK00 |
| ACEKITSIINEAL | 0 | 0 | 1 | 850959 | A0A044U9P0 | A0A044U9P0 |
| ACIIERKNRGELEYT | 0 | 0 | 1 | 850960 | A0A044QNT8 | A0A044QNT8 |
| ACKRFEELGVQF | 0 | 0 | 1 | 850961 | A0A044RAH9 | A0A044RAH9 |
| ACMPDNTKGCKSHP | 0 | 0 | 1 | 850962 | A0A044RJ74 | A0A044RJ74 |
| ADFSGNDVPLHIS | 0 | 0 | 1 | 850968 | AAA20541.1 | A0A044QNW3 |
| ADGPPGEPGQPGAPG | 0 | 0 | 1 | 850970 | A0A044SF60 | A0A044SF60 |
| ADISQGDPSGVA | 0 | 0 | 1 | 850972 | AAF64251.1 | A0A044V6Z2 |
| ADMSETGDSKRG | 0 | 0 | 1 | 850973 | A0A2K6WPE1 | A0A2K6WPE1 |
| ADNRNTFTIYEDGI | 0 | 0 | 1 | 850974 | A0A044UK00 | A0A044UK00 |
| ADRSQLFYQCQI | 0 | 0 | 1 | 850977 | A0A044QSA2 | A0A044QSA2 |
| ADTFPRRPVEE | 0 | 0 | 1 | 850980 | AAC48290.1 | A0A044VA40 |
| ADVDNGNVPYDYGSI | 0 | 0 | 1 | 850982 | A0A044VAG6 | A0A044VAG6 |
| AEAESDDEKKH | 0 | 0 | 1 | 850985 | A0A044T7P0 | A0A044T7P0 |
| AEANVDVKRPDALFD | 0 | 0 | 1 | 850987 | A0A044UK00 | A0A044UK00 |
| AEDEETKETTSQLR | 0 | 0 | 1 | 850990 | A0A044R196 | A0A044R196 |
| AEDKKKEAAAER | 0 | 0 | 1 | 850991 | A0A044SS43 | A0A044SS43 |
| AEEGSNGNVRE | 0 | 0 | 1 | 850992 | A0A044U9P0 | A0A044U9P0 |
| AEKAEAEEKKQEA | 0 | 0 | 1 | 850995 | A0A044TX11 | A0A044TX11 |
| AEMAMLAAKKRHQE | 0 | 0 | 1 | 850999 | A0A044R196 | A0A044R196 |
| AENFRQLCTGEKGFG | 0 | 0 | 1 | 851001 | A0A044U9P0 | A0A044U9P0 |
| AEREIVRDIKE | 0 | 0 | 1 | 851006 | P30162.1 | P30162 |
| AETSESNEERETR | 0 | 0 | 1 | 851009 | A0A044UK00 | A0A044UK00 |
| AEWFKDDKPLKT | 0 | 0 | 1 | 851012 | A0A044R9D0 | A0A044R9D0 |
| AFAERGEVHHKAMQ | 0 | 0 | 1 | 851013 | A0A044RF80 | A0A044RF80 |
| AFEQEGISAGDVN | 0 | 0 | 1 | 851015 | AAF64251.1 | A0A044V6Z2 |
| AFGGFSSTQDT | 0 | 0 | 1 | 851016 | A0A044RF80 | A0A044RF80 |
| AFGYVNETNSTL | 0 | 0 | 1 | 851018 | A0A044V420 | A0A044V420 |
| AGAPGEMGPPGKAG | 0 | 0 | 1 | 851029 | A0A044TIV1 | A0A044TIV1 |
| AGEWGNEEREGK | 0 | 0 | 1 | 851030 | AAA20541.1 | A0A044QNW3 |
| AGGKNFVIQKSE | 0 | 0 | 1 | 851035 | A0A044R196 | A0A044R196 |
| AGKEHSIGYIDVKKE | 0 | 0 | 1 | 851038 | A0A044UK00 | A0A044UK00 |
| AGNITTVLQTQVP | 0 | 0 | 1 | 851041 | A0A044UK00 | A0A044UK00 |
| AGPPGKDGERGKGER | 0 | 0 | 1 | 851043 | A0A044TIV1 | A0A044TIV1 |
| AGPQGPPGAAG | 0 | 0 | 1 | 851044 | A0A044UZ11 | A0A044UZ11 |
| AGSPGAQGPVGPQ | 0 | 0 | 1 | 851047 | A0A044SAV6 | A0A044SAV6 |
| AGSSGGSGMPGA | 0 | 0 | 1 | 851049 | A0A044SAV6 | A0A044SAV6 |
| AIAIAKDAEGMLKQD | 0 | 0 | 1 | 851061 | AAC48290.1 | A0A044VA40 |
| AIASSEIQSQENNAS | 0 | 0 | 1 | 851062 | A0A044UK00 | A0A044UK00 |
| AIDTSKKALKVVDR | 0 | 0 | 1 | 851064 | AAF64251.1 | A0A044V6Z2 |
| AIFESNELMTEA | 0 | 0 | 1 | 851065 | AAK16514.1 | A0A2K6W829 |
| AIFPRDANGRFI | 0 | 0 | 1 | 851066 | A0A044UVG8 | A0A044UVG8 |
| AIGGGVSAIGGG | 0 | 0 | 1 | 851067 | A0A044S628 | A0A044S628 |
| AIICEQKDVNKLN | 0 | 0 | 1 | 851068 | A0A044UK00 | A0A044UK00 |
| AIKSAAAQAPEA | 0 | 0 | 1 | 851071 | A0A044R521 | A0A044R521 |
| AILKTLCESQFD | 0 | 0 | 1 | 851072 | AAD00842.1 | A0A044SS43 |
| AILSIRASTIVERTN | 0 | 0 | 1 | 851074 | A0A044UK00 | A0A044UK00 |
| AIMQHETAHYETS | 0 | 0 | 1 | 851075 | A0A044UK00 | A0A044UK00 |
| AINHELEGEATATTR | 0 | 0 | 1 | 851076 | A0A044RRG7 | A0A044RRG7 |
| AIPEAEFRWMLNNFE | 0 | 0 | 1 | 851077 | A0A044UK00 | A0A044UK00 |
| AIQGSGWGWLAYDKE | 0 | 0 | 1 | 851080 | P41981.1 | A0A158N7R1 |
| AIQRGDELGLMHAVA | 0 | 0 | 1 | 851082 | AAK16514.1 | A0A2K6W829 |
| AIRDIRNEYDQ | 0 | 0 | 1 | 851083 | AAA74283.1 | A0A044V2F8 |
| AITMVRAVPEKQELK | 0 | 0 | 1 | 851087 | A0A044UK00 | A0A044UK00 |
| AIVFLVDCADVERIA | 0 | 0 | 1 | 851089 | A0A044TDC0 | A0A044TDC0 |
| AKCKEELENLVAE | 0 | 0 | 1 | 851093 | A0A044SS43 | A0A044SS43 |
| AKEENSYIKTHFE | 0 | 0 | 1 | 851094 | A0A044UK00 | A0A044UK00 |
| AKFQLIDTNHDGRI | 0 | 0 | 1 | 851095 | A0A044VDG0 | A0A044VDG0 |
| AKIERDKIEIKEEIA | 0 | 0 | 1 | 851097 | A0A044UK00 | A0A044UK00 |
| AKKGFLTPERKKKLR | 0 | 0 | 1 | 851099 | A0A044TX11 | A0A044TX11 |
| AKLHPTKGETSGTGI | 0 | 0 | 1 | 851102 | AAB68960.1 | A0A044RRG7 |
| AKPPIQPIPLNLPG | 0 | 0 | 1 | 851103 | A0A044R196 | A0A044R196 |
| AKQMRSLKDACPECW | 0 | 0 | 1 | 851105 | A0A044RF80 | A0A044RF80 |
| AKSLTQPQVPFVPQM | 0 | 0 | 1 | 851111 | AAC48290.1 | A0A044VA40 |
| AKTNQESETVVSA | 0 | 0 | 1 | 851113 | CAA04390.1 | A0A044R436 |
| AKVVAQETSEAWIEA | 0 | 0 | 1 | 851114 | A0A044UK00 | A0A044UK00 |
| ALANVSYEDNRITRD | 0 | 0 | 1 | 851118 | P46434.1 | P46434 |
| ALLDGVTTPATGFG | 0 | 0 | 1 | 851127 | A0A044U0M2 | A0A044U0M2 |
| ALLENGVTDEKAQ | 0 | 0 | 1 | 851128 | A0A044RF80 | A0A044RF80 |
| ALLKETQQGVE | 0 | 0 | 1 | 851130 | A0A044VBX6 | A0A044VBX6 |
| ALLNGTLRPEDLASG | 0 | 0 | 1 | 851132 | A0A044U9P0 | A0A044U9P0 |
| ALNVSLCCGSEAITE | 0 | 0 | 1 | 851135 | A0A044QU24 | A0A044QU24 |
| ALQIATFLESNPRIE | 0 | 0 | 1 | 851139 | A0A044R2U8 | A0A044R2U8 |
| AMARRQDDMEPKE | 0 | 0 | 1 | 851149 | AAF64251.1 | A0A044V6Z2 |
| AMEKHEGSVAEYRA | 0 | 0 | 1 | 851150 | Q9NJD8.1 | Q9NJD8 |
| AMESKYPYVGTEQ | 0 | 0 | 1 | 851151 | A0A2K6W829 | A0A2K6W829 |
| AMPIQIRDDSINRLD | 0 | 0 | 1 | 851156 | AAC48290.1 | A0A044VA40 |
| ANCKCKLGSAGPQ | 0 | 0 | 1 | 851159 | A0A044UZ11 | A0A044UZ11 |
| ANDDGQMIPQF | 0 | 0 | 1 | 851160 | AAC48290.1 | A0A044VA40 |
| ANDNSEIAHEEPD | 0 | 0 | 1 | 851162 | A0A044UVG8 | A0A044UVG8 |
| ANFTGITDVPLYINA | 0 | 0 | 1 | 851164 | A0A044V3Q0 | A0A044V3Q0 |
| ANGRFIFPSGPD | 0 | 0 | 1 | 851165 | A0A044UVG8 | A0A044UVG8 |
| ANKDGNSMPKGKNM | 0 | 0 | 1 | 851167 | A0A044UNG9 | A0A044UNG9 |
| ANNSNLDISMRE | 0 | 0 | 1 | 851175 | A0A044S628 | A0A044S628 |
| ANQAITSENSIKPKG | 0 | 0 | 1 | 851177 | P46434.1 | P46434 |
| ANSIESTDAGQAFDA | 0 | 0 | 1 | 851182 | A0A044UK00 | A0A044UK00 |
| APFAVHKTDADN | 0 | 0 | 1 | 851185 | A0A2K6WB53 | A0A2K6WB53 |
| APRASSSDAEDSSH | 0 | 0 | 1 | 851194 | A0A044SS43 | A0A044SS43 |
| AQANSLAQLGKYTG | 0 | 0 | 1 | 851204 | AAB52600.1 | A0A044UUD9 |

| | | | | | | |
|---|---|---|---|---|---|---|
| AQFRFQQEWFDDRLR | 0 | 0 | 1 | 851207 | A0A044QWA5 | A0A044QWA5 |
| AQLGKYTGGAGSGA | 0 | 0 | 1 | 851209 | AAB52600.1 | A0A044UUD9 |
| AQLYDRVNLQNTFEA | 0 | 0 | 1 | 851211 | A0A044UK00 | A0A044UK00 |
| AQMQQQMLQTAI | 0 | 0 | 1 | 851212 | AAC48290.1 | A0A044VA40 |
| AQRIQDELYFAKYD | 0 | 0 | 1 | 851214 | A0A044UH02 | A0A044UH02 |
| AQSAKAGDAVEQVL | 0 | 0 | 1 | 851215 | A0A044RF80 | A0A044RF80 |
| AQSNSYVNLDVPI | 0 | 0 | 1 | 851216 | A0A044UK00 | A0A044UK00 |
| AQVKLTGEAKKIKEE | 0 | 0 | 1 | 851218 | A0A044UK00 | A0A044UK00 |
| AQYIEDSNRYKEEVK | 0 | 0 | 1 | 851219 | AAA29419.1 | P21250 |
| ARCVLNNKENKEF | 0 | 0 | 1 | 851221 | A0A044U175 | A0A044U175 |
| ARDTTSENREFFKN | 0 | 0 | 1 | 851224 | AAA74283.1 | A0A044V2F8 |
| ARFDVIGDLLGSALA | 0 | 0 | 1 | 851227 | A0A044RF80 | A0A044RF80 |
| ARGVVSMAGNGPN | 0 | 0 | 1 | 851229 | A0A2K6VZ59 | A0A2K6VZ59 |
| ARLIMKNADKNDD | 0 | 0 | 1 | 851234 | AAF64251.1 | A0A044V6Z2 |
| ARLSANQEEAA | 0 | 0 | 1 | 851235 | A0A044RAH9 | A0A044RAH9 |
| ARNKGNMCHIATLA | 0 | 0 | 1 | 851238 | AAK16514.1 | A0A2K6W829 |
| ARSSPSVPMQTRQ | 0 | 0 | 1 | 851245 | A0A2K6WJQ8 | A0A2K6WJQ8 |
| ARTDRWDEQRVVA | 0 | 0 | 1 | 851246 | A0A044V9G2 | A0A044V9G2 |
| ARTIPPQPLPKPM | 0 | 0 | 1 | 851247 | A0A044UK00 | A0A044UK00 |
| ARVEIFIKISKENVS | 0 | 0 | 1 | 851248 | A0A044UK00 | A0A044UK00 |
| ASAAASGKAAA | 0 | 0 | 1 | 851250 | A0A044R196 | A0A044R196 |
| ASEIPNDPIEQKR | 0 | 0 | 1 | 851254 | A0A044RAH9 | A0A044RAH9 |
| ASGARCSNCGSD | 0 | 0 | 1 | 851255 | A0A044U9P0 | A0A044U9P0 |
| ASGRTTGIVLDSGDG | 0 | 0 | 1 | 851256 | P30162.1 | P30162 |
| ASHGLNGHVMSDG | 0 | 0 | 1 | 851258 | Q9NJD8.1 | Q9NJD8 |
| ASKEIDSNNKFKNPI | 0 | 0 | 1 | 851260 | AAD00842.1 | A0A044SS43 |
| ASNDTYEQKAKN | 0 | 0 | 1 | 851262 | A0A044UVG8 | A0A044UVG8 |
| ASNLEETPIMSSA | 0 | 0 | 1 | 851263 | AAC48290.1 | A0A044VA40 |
| ASNSSQSMAQSTA | 0 | 0 | 1 | 851264 | A0A044UVG8 | A0A044UVG8 |
| ASNVLASGFPTDF | 0 | 0 | 1 | 851265 | AAK66564.1 | Q963F6 |
| ASTIVERTNMVFS | 0 | 0 | 1 | 851269 | A0A044UK00 | A0A044UK00 |
| ASTSNTRISTKESL | 0 | 0 | 1 | 851270 | A0A044UK00 | A0A044UK00 |
| ASVPTEDRIKLGIGD | 0 | 0 | 1 | 851271 | A0A044R2U8 | A0A044R2U8 |
| ATAGATTTQAPVEA | 0 | 0 | 1 | 851272 | AAA29419.1 | P21250 |
| ATDHFKEFDMNKDKI | 0 | 0 | 1 | 851273 | AAF64251.1 | A0A044V6Z2 |
| ATFTNEDAALEALKN | 0 | 0 | 1 | 851277 | AAA65186.1 | Q25619 |
| ATKFGLAMESKYPY | 0 | 0 | 1 | 851281 | A0A2K6W829 | A0A2K6W829 |
| ATNSNVTSCVPPSCF | 0 | 0 | 1 | 851284 | A0A044SS43 | A0A044SS43 |
| ATQALEKYNIEKDIA | 0 | 0 | 1 | 851285 | AAF64249.1 | A0A044UWQ7 |
| ATRGNGRNLILGDKI | 0 | 0 | 1 | 851287 | P46427.1 | P46427 |
| ATRKLRLGDFMGNPD | 0 | 0 | 1 | 851288 | A0A044U9P0 | A0A044U9P0 |
| ATSFMEVSVMENVYD | 0 | 0 | 1 | 851290 | A0A044UK00 | A0A044UK00 |
| AVGSQSGKTSKDVV | 0 | 0 | 1 | 851300 | AAC47233.1 | Q25633 |
| AVHNSKDPAEIPWG | 0 | 0 | 1 | 851301 | AAB52599.1 | A0A158N7S3 |
| AVIQNNNRIPDNMIM | 0 | 0 | 1 | 851304 | A0A044VA40 | A0A044VA40 |
| AVKKYGDGKESQK | 0 | 0 | 1 | 851305 | AAK16514.1 | A0A2K6W829 |
| AVLPCNLCVTDIGEQ | 0 | 0 | 1 | 851306 | A0A044UK00 | A0A044UK00 |
| AVPEKQELKYQPVKR | 0 | 0 | 1 | 851316 | A0A044UK00 | A0A044UK00 |
| AVPFDEDERDKKT | 0 | 0 | 1 | 851317 | A0A044TU88 | A0A044TU88 |
| AVPGIRTQEACTSAK | 0 | 0 | 1 | 851318 | A0A044UVG8 | A0A044UVG8 |
| AVPSSETINKSAK | 0 | 0 | 1 | 851319 | A0A044UVG8 | A0A044UVG8 |
| AVWQESNRIKFPKGK | 0 | 0 | 1 | 851325 | A0A044RAH9 | A0A044RAH9 |
| AVWSETTVKPPSL | 0 | 0 | 1 | 851326 | A0A044UK00 | A0A044UK00 |
| AVYANTKEIAHAIVQ | 0 | 0 | 1 | 851327 | A0A044TDA1 | A0A044TDA1 |
| AVYKVPQIFDSTIKI | 0 | 0 | 1 | 851328 | A0A044UK00 | A0A044UK00 |
| AYDELIQEAVKNIDE | 0 | 0 | 1 | 851335 | A0A044SS43 | A0A044SS43 |
| AYRIQCFYMEADKT | 0 | 0 | 1 | 851340 | A0A044R521 | A0A044R521 |
| AYTTSDIIYKWDRTA | 0 | 0 | 1 | 851341 | A0A044QWA5 | A0A044QWA5 |
| CADNQASCRAETDTF | 0 | 0 | 1 | 851345 | A0A044R153 | A0A044R153 |
| CAGEMYSVDRDGYW | 0 | 0 | 1 | 851346 | A0A044UVG8 | A0A044UVG8 |
| CALRQVGIIDKEE | 0 | 0 | 1 | 851348 | A0A044UK00 | A0A044UK00 |
| CASLPGKSSIQQMD | 0 | 0 | 1 | 851350 | A0A044UW71 | A0A044UW71 |
| CASQEFDYEFDDVNE | 0 | 0 | 1 | 851352 | A0A044QNT8 | A0A044QNT8 |
| CATSDDEPKGPKVTD | 0 | 0 | 1 | 851353 | A0A2K6WB53 | A0A2K6WB53 |
| CCSFDAIYELTNM | 0 | 0 | 1 | 851359 | A0A044V1C1 | A0A044V1C1 |
| CDDVYGGKDSAII | 0 | 0 | 1 | 851362 | A0A044R2U8 | A0A044R2U8 |
| CDLCDEAWHTWCLHP | 0 | 0 | 1 | 851364 | A0A044SS43 | A0A044SS43 |
| CDLNENQTWVDNGC | 0 | 0 | 1 | 851365 | A0A044QNT8 | A0A044QNT8 |
| CDNTSGGLKNFVQI | 0 | 0 | 1 | 851366 | A0A2K6VRV9 | A0A2K6VRV9 |
| CDSAHPEYDPESSID | 0 | 0 | 1 | 851369 | AAA20215.1 | A0A044V456 |
| CEFEWKQFVECTETQ | 0 | 0 | 1 | 851375 | A0A2K6WJQ8 | A0A2K6WJQ8 |
| CENFIKHCKNGYYN | 0 | 0 | 1 | 851380 | CAA04390.1 | A0A044R436 |
| CENRMRQYGFPWPE | 0 | 0 | 1 | 851381 | A0A044TBR7 | A0A044TBR7 |
| CESQFDFNIKFKE | 0 | 0 | 1 | 851383 | AAD00842.1 | A0A044SS43 |
| CGEIAVDETCLHSG | 0 | 0 | 1 | 851390 | A0A044UVG8 | A0A044UVG8 |
| CGGVGSCVYCDICDN | 0 | 0 | 1 | 851393 | A0A2K6VRV9 | A0A2K6VRV9 |
| CIKPFLENCDVNND | 0 | 0 | 1 | 851398 | A0A044T7P0 | A0A044T7P0 |
| CKGGGAGVSAIGGA | 0 | 0 | 1 | 851405 | A0A044S628 | A0A044S628 |
| CKRPEIWEDCGGC | 0 | 0 | 1 | 851409 | AAY67786.1 | A0A158N7U2 |
| CKTSDSRNRSSSR | 0 | 0 | 1 | 851410 | AAF64250.1 | A0A044UVG8 |
| CKVIINEPKCCC | 0 | 0 | 1 | 851411 | A0A044UMF0 | A0A044UMF0 |
| CLGLKEGWPFLDDK | 0 | 0 | 1 | 851413 | A0A044T7P0 | A0A044T7P0 |
| CLPLSIEDNFDRIYV | 0 | 0 | 1 | 851415 | A0A044UMF0 | A0A044UMF0 |
| CNMEEYFDLQFSMQS | 0 | 0 | 1 | 851423 | A0A044UK00 | A0A044UK00 |
| CNSFTTPDIHPHPDL | 0 | 0 | 1 | 851424 | A0A044UK00 | A0A044UK00 |
| CPDAPNVTANIVCTA | 0 | 0 | 1 | 851428 | A0A044U9P0 | A0A044U9P0 |
| CPEPIRPWTKAVGLT | 0 | 0 | 1 | 851429 | AAF64254.1 | A0A044QLL7 |
| CPKYCAIDGGVFFED | 0 | 0 | 1 | 851431 | A0A044V9C9 | A0A044V9C9 |
| CPLGTYCEERDI | 0 | 0 | 1 | 851432 | A0A044RJ74 | A0A044RJ74 |
| CPPGPPGPPGEPG | 0 | 0 | 1 | 851434 | A0A044TIV1 | A0A044TIV1 |
| CRNDEGGRQVA | 0 | 0 | 1 | 851445 | A0A044QSA2 | A0A044QSA2 |

| | | | | | | |
|---|---|---|---|---|---|---|
| CRYEILDGGPSGQ | 0 | 0 | 1 | 851447 | A0A044QSA2 | A0A044QSA2 |
| CSDRSQKDVEIQLWE | 0 | 0 | 1 | 851449 | A0A044V9S3 | A0A044V9S3 |
| CSKSSNPEVLLLCD | 0 | 0 | 1 | 851450 | A0A044SS43 | A0A044SS43 |
| CSPDENWTCKYD | 0 | 0 | 1 | 851451 | AAD27587.1 | A0A158N7U8 |
| CSSCSQVSTYENI | 0 | 0 | 1 | 851453 | A0A044TBR7 | A0A044TBR7 |
| CSSSNNSENQM | 0 | 0 | 1 | 851454 | A0A044UK00 | A0A044UK00 |
| CTACGGAGHIA | 0 | 0 | 1 | 851455 | A0A044U9P0 | A0A044U9P0 |
| CVVKNREGQNEQSLN | 0 | 0 | 1 | 851463 | A0A044R9D0 | A0A044R9D0 |
| CVYCDICDNTSGGLK | 0 | 0 | 1 | 851464 | A0A2K6VRV9 | A0A2K6VRV9 |
| DAALEALKNKSDKL | 0 | 0 | 1 | 851467 | AAA65186.1 | Q25619 |
| DAEGMLKQDDTTPQ | 0 | 0 | 1 | 851470 | AAC48290.1 | A0A044VA40 |
| DAFTDDELASDSF | 0 | 0 | 1 | 851473 | A0A044SL50 | A0A044SL50 |
| DAFTSAANFTGI | 0 | 0 | 1 | 851474 | A0A044V3Q0 | A0A044V3Q0 |
| DAIDEAATKAKEAAC | 0 | 0 | 1 | 851477 | A0A044VBX6 | A0A044VBX6 |
| DAIVSIVQKERQFTN | 0 | 0 | 1 | 851479 | A0A044UK00 | A0A044UK00 |
| DAKLEHVVNGLKH | 0 | 0 | 1 | 851480 | A0A044UK00 | A0A044UK00 |
| DAKYRPVVEQKIQN | 0 | 0 | 1 | 851481 | A0A2K6VZ59 | A0A2K6VZ59 |
| DALKSINADAFGNLY | 0 | 0 | 1 | 851483 | A0A044UMF0 | A0A044UMF0 |
| DARGNFKIYGEENEV | 0 | 0 | 1 | 851484 | A0A044V9S3 | A0A044V9S3 |
| DCADVERIAESRHE | 0 | 0 | 1 | 851487 | A0A044TDC0 | A0A044TDC0 |
| DDAVRLAIAIAKDAE | 0 | 0 | 1 | 851496 | AAC48290.1 | A0A044VA40 |
| DDCRYCYESGQKDVC | 0 | 0 | 1 | 851497 | A0A044V1C1 | A0A044V1C1 |
| DDDFQEIITDKD | 0 | 0 | 1 | 851499 | A0A044QNT8 | A0A044QNT8 |
| DDDPLDQVCSNTN | 0 | 0 | 1 | 851500 | A0A044T7P0 | A0A044T7P0 |
| DDDVQVLGMPYA | 0 | 0 | 1 | 851503 | A0A044V3Q0 | A0A044V3Q0 |
| DDFILAETSTSSD | 0 | 0 | 1 | 851504 | A0A044SS43 | A0A044SS43 |
| DDGKGDTVEILN | 0 | 0 | 1 | 851507 | A0A044R521 | A0A044R521 |
| DDHGKLEAIYNQL | 0 | 0 | 1 | 851508 | A0A044TX11 | A0A044TX11 |
| DDIGYDFLQKDGT | 0 | 0 | 1 | 851509 | A0A044RF80 | A0A044RF80 |
| DDLENQPVVTRT | 0 | 0 | 1 | 851510 | AAF64254.1 | A0A044QLL7 |
| DDLLLGDENKQ | 0 | 0 | 1 | 851511 | A0A044U0M2 | A0A044U0M2 |
| DDMEPKETHNLFVMA | 0 | 0 | 1 | 851512 | AAF64251.1 | A0A044V6Z2 |
| DEDDFQQIITD | 0 | 0 | 1 | 851520 | A0A044QNT8 | A0A044QNT8 |
| DEEDELSRLRYEA | 0 | 0 | 1 | 851524 | A0A044UVG8 | A0A044UVG8 |
| DEEGGNDENED | 0 | 0 | 1 | 851526 | AAC79423.1 | A0A158N7U8 |
| DEGYRSNSLSRQN | 0 | 0 | 1 | 851534 | A0A044V9G2 | A0A044V9G2 |
| DEIAYRNQNQG | 0 | 0 | 1 | 851535 | A0A044UZ11 | A0A044UZ11 |
| DEIFRELQQRALG | 0 | 0 | 1 | 851536 | AAC48289.1 | A0A044VA40 |
| DEIISSLSHDQR | 0 | 0 | 1 | 851537 | A0A044R436 | A0A044R436 |
| DELEELIELKYK | 0 | 0 | 1 | 851541 | AAF64251.1 | A0A044V6Z2 |
| DELIDFNSLRMIDNN | 0 | 0 | 1 | 851542 | A0A044QWA5 | A0A044QWA5 |
| DENEDVPRGSFVNSM | 0 | 0 | 1 | 851545 | AAC79423.1 | A0A158N7U8 |
| DEQRVVAAKPKI | 0 | 0 | 1 | 851547 | A0A044V9G2 | A0A044V9G2 |
| DERDEKGNTLTAR | 0 | 0 | 1 | 851548 | AAA50214.2 | P52570 |
| DESYNQCISECL | 0 | 0 | 1 | 851550 | A0A044UK00 | A0A044UK00 |
| DETSASSIHTAPQFA | 0 | 0 | 1 | 851552 | A0A044UK00 | A0A044UK00 |
| DFDWEYPIGMAQEYA | 0 | 0 | 1 | 851555 | AAC47023.1 | A0A044RRG7 |
| DFEEEGEGEEM | 0 | 0 | 1 | 851556 | AAD27587.1 | A0A158N7U8 |
| DFILETDSVTYRN | 0 | 0 | 1 | 851559 | AAY67786.1 | A0A158N7U2 |
| DFIRRVQDQEIRDLQ | 0 | 0 | 1 | 851560 | AAA74283.1 | A0A044V2F8 |
| DFQAETNEGFIS | 0 | 0 | 1 | 851562 | P52570.1 | P52570 |
| DFRSAQHRMIAERD | 0 | 0 | 1 | 851566 | A0A044UK00 | A0A044UK00 |
| DFSEYGTLATKDCGE | 0 | 0 | 1 | 851567 | A0A044UVG8 | A0A044UVG8 |
| DFSSIKSQFQFGQ | 0 | 0 | 1 | 851569 | P46427.1 | P46427 |
| DFTRGNGTGGESI | 0 | 0 | 1 | 851570 | AAC47233.1 | Q25633 |
| DFVVSGSESEYEP | 0 | 0 | 1 | 851571 | A0A044SS43 | A0A044SS43 |
| DGAEEADQKNN | 0 | 0 | 1 | 851572 | A0A044SAZ1 | A0A044SAZ1 |
| DGCALDKYLLNNLE | 0 | 0 | 1 | 851574 | A0A044QSA2 | A0A044QSA2 |
| DGCMVQNDKIYVGRA | 0 | 0 | 1 | 851575 | AAA29419.1 | P21250 |
| DGDGYINNSEAVAMA | 0 | 0 | 1 | 851576 | AAF64251.1 | A0A044V6Z2 |
| DGEKIDEISGKTF | 0 | 0 | 1 | 851580 | A0A044SG03 | A0A044SG03 |
| DGERGKGERGPQGP | 0 | 0 | 1 | 851581 | A0A044TIV1 | A0A044TIV1 |
| DGGMRAVIGPNGTA | 0 | 0 | 1 | 851583 | A0A2K6WPE1 | A0A2K6WPE1 |
| DGKESQKSTVLQ | 0 | 0 | 1 | 851589 | AAK16514.1 | A0A2K6W829 |
| DGKPGPPGPVGDA | 0 | 0 | 1 | 851590 | A0A044TIV1 | A0A044TIV1 |
| DGLPGHPGMPG | 0 | 0 | 1 | 851592 | A0A044SAV6 | A0A044SAV6 |
| DGNGDKVELLNAD | 0 | 0 | 1 | 851593 | A0A044QSA2 | A0A044QSA2 |
| DGPPGRDGRDGA | 0 | 0 | 1 | 851595 | A0A044V9I3 | A0A044V9I3 |
| DGQLQQVNFTQEDAI | 0 | 0 | 1 | 851596 | A0A044V3Q0 | A0A044V3Q0 |
| DGRKGNDGRPGPPG | 0 | 0 | 1 | 851598 | A0A044V9I3 | A0A044V9I3 |
| DGRPGNDGRKGNDGR | 0 | 0 | 1 | 851599 | A0A044V9I3 | A0A044V9I3 |
| DGRPGPPGSPGLPG | 0 | 0 | 1 | 851600 | A0A044V9I3 | A0A044V9I3 |
| DHDNDNFTRAMFINC | 0 | 0 | 1 | 851605 | A0A044V9S3 | A0A044V9S3 |
| DHYGFDEKTLGPF | 0 | 0 | 1 | 851610 | AAF64251.1 | A0A044V6Z2 |
| DIAAYIKKEFDK | 0 | 0 | 1 | 851612 | AAF64249.1 | A0A044UWQ7 |
| DIANSFAVPFDEDER | 0 | 0 | 1 | 851615 | A0A044TU88 | A0A044TU88 |
| DIDDVCPSTSND | 0 | 0 | 1 | 851617 | A0A044SS43 | A0A044SS43 |
| DIEETKVSKTGICE | 0 | 0 | 1 | 851618 | A0A044UK00 | A0A044UK00 |
| DIEKNLTGKKLLH | 0 | 0 | 1 | 851619 | A0A044V3Q0 | A0A044V3Q0 |
| DIIERYIELLNAAE | 0 | 0 | 1 | 851624 | A0A044UK00 | A0A044UK00 |
| DIKFIDDRIAKDD | 0 | 0 | 1 | 851626 | P46427.1 | P46427 |
| DIKQFNKANVYDS | 0 | 0 | 1 | 851628 | A0A044UVG8 | A0A044UVG8 |
| DIPDEVKPFLDRRVL | 0 | 0 | 1 | 851632 | AAD00842.1 | A0A044SS43 |
| DISDLQRAELDKEDT | 0 | 0 | 1 | 851638 | A0A044RF80 | A0A044RF80 |
| DISKTVQDLLGG | 0 | 0 | 1 | 851639 | Q9NJD8.1 | Q9NJD8 |
| DISMREKNAVNAIEK | 0 | 0 | 1 | 851641 | A0A044S628 | A0A044S628 |
| DISSRPMEVFMCSVL | 0 | 0 | 1 | 851642 | A0A044TDC0 | A0A044TDC0 |
| DIVLDAEQAYQIYE | 0 | 0 | 1 | 851644 | AAV71152.1 | A0A044VAG6 |
| DIYQDESAVKKK | 0 | 0 | 1 | 851646 | AAD00842.1 | A0A044SS43 |
| DKANNHIISNA | 0 | 0 | 1 | 851648 | AAB52599.1 | A0A158N7S3 |
| DKINEYIAQFPVIDA | 0 | 0 | 1 | 851654 | A0A044SAZ1 | A0A044SAZ1 |

| | | | | | | |
|---|---|---|---|---|---|---|
| DKIRKDIEDEIKMK | 0 | 0 | 1 | 851655 | A0A044UK00 | A0A044UK00 |
| DKISYADYALFE | 0 | 0 | 1 | 851656 | P46427.1 | P46427 |
| DKIVTIDELEELIEL | 0 | 0 | 1 | 851657 | AAF64251.1 | A0A044V6Z2 |
| DKKQSDITTPTLE | 0 | 0 | 1 | 851658 | A0A044UK00 | A0A044UK00 |
| DKKQTEDIDDVCP | 0 | 0 | 1 | 851659 | A0A044SS43 | A0A044SS43 |
| DKMSMLQPLANDMQ | 0 | 0 | 1 | 851660 | AAA63412.2 | Q99099 |
| DKNTLPSDNAIEVIV | 0 | 0 | 1 | 851662 | A0A044SG03 | A0A044SG03 |
| DKPLKTTDRIK | 0 | 0 | 1 | 851665 | A0A044R9D0 | A0A044R9D0 |
| DKPNALGEDQLKWH | 0 | 0 | 1 | 851666 | A0A044TDC0 | A0A044TDC0 |
| DLFWGKDRKDQGQ | 0 | 0 | 1 | 851672 | A0A044UW71 | A0A044UW71 |
| DLIDEVLKVPSDNTN | 0 | 0 | 1 | 851675 | A0A044UVG8 | A0A044UVG8 |
| DLNTLDINEEFTPL | 0 | 0 | 1 | 851684 | A0A044QSA2 | A0A044QSA2 |
| DLPVGRSVDETLR | 0 | 0 | 1 | 851687 | AAC48312.1 | A0A044VFS3 |
| DLSIQNAVDCTWTLG | 0 | 0 | 1 | 851690 | AAK16514.1 | A0A2K6W829 |
| DLSSLLAPPAP | 0 | 0 | 1 | 851692 | A0A044U9P0 | A0A044U9P0 |
| DLTPDEVTQLKTFDA | 0 | 0 | 1 | 851693 | AAA29419.1 | P21250 |
| DMESWYKLKVQEIQ | 0 | 0 | 1 | 851700 | AAA21472.1 | A0A2K6WLA7 |
| DMTDLKVFSQAL | 0 | 0 | 1 | 851703 | A0A044R2U8 | A0A044R2U8 |
| DNAIFSCTAMNLFG | 0 | 0 | 1 | 851705 | A0A044UK00 | A0A044UK00 |
| DNDENEKDENDAD | 0 | 0 | 1 | 851706 | A0A044SS43 | A0A044SS43 |
| DNKKWVTENERIQLE | 0 | 0 | 1 | 851710 | A0A044UK00 | A0A044UK00 |
| DNKLSLQKNERDKEK | 0 | 0 | 1 | 851711 | A0A044TU88 | A0A044TU88 |
| DNLAFLDNLKSLSDN | 0 | 0 | 1 | 851712 | A0A044UK00 | A0A044UK00 |
| DNLKSLSDNTGGI | 0 | 0 | 1 | 851713 | A0A044UK00 | A0A044UK00 |
| DNNADPIWTFQEVSE | 0 | 0 | 1 | 851715 | A0A044QWA5 | A0A044QWA5 |
| DNYQSEESNSSGDRR | 0 | 0 | 1 | 851717 | A0A044SS43 | A0A044SS43 |
| DNYVTVRWENIDRDS | 0 | 0 | 1 | 851718 | AAV71152.1 | A0A044VAG6 |
| DPAEIPWGVEGA | 0 | 0 | 1 | 851719 | AAB52599.1 | A0A158N7S3 |
| DPIEQKRYALSTLA | 0 | 0 | 1 | 851724 | A0A044RAH9 | A0A044RAH9 |
| DPREINGLQIGGD | 0 | 0 | 1 | 851727 | AAA20541.1 | A0A044QNW3 |
| DPTGTGKGGDSIWG | 0 | 0 | 1 | 851732 | CAA04390.1 | A0A044R436 |
| DQAKYKCVVKNREGQ | 0 | 0 | 1 | 851734 | A0A044R9D0 | A0A044R9D0 |
| DQDDSEDNDENEKD | 0 | 0 | 1 | 851735 | A0A044SS43 | A0A044SS43 |
| DQENKFIKLRSDGK | 0 | 0 | 1 | 851737 | A0A044QWA5 | A0A044QWA5 |
| DQHILQNRQSVINA | 0 | 0 | 1 | 851738 | A0A044UK00 | A0A044UK00 |
| DQKNNNHEFDLIN | 0 | 0 | 1 | 851740 | A0A044SAZ1 | A0A044SAZ1 |
| DQLGQPLPSASNYMA | 0 | 0 | 1 | 851742 | A0A044SAZ1 | A0A044SAZ1 |
| DQQLCETFDQLD | 0 | 0 | 1 | 851745 | A0A044VDG0 | A0A044VDG0 |
| DQRKHKCKRPEIWED | 0 | 0 | 1 | 851746 | AAY67786.1 | A0A158N7U2 |
| DQTAGTLSQRITD | 0 | 0 | 1 | 851747 | A0A044TU88 | A0A044TU88 |
| DRALKNVLVELYEDD | 0 | 0 | 1 | 851748 | A0A044V9S3 | A0A044V9S3 |
| DRASNEDICDDVSYT | 0 | 0 | 1 | 851749 | A0A044QNT8 | A0A044QNT8 |
| DRDGSGTVTMDEAKR | 0 | 0 | 1 | 851752 | AAF64251.1 | A0A044V6Z2 |
| DRDTIFKELRSRAQM | 0 | 0 | 1 | 851754 | AAC48290.1 | A0A044VA40 |
| DRFQTSSQSGDYAQF | 0 | 0 | 1 | 851756 | A0A044QU24 | A0A044QU24 |
| DRKDQGQVVGFENI | 0 | 0 | 1 | 851761 | A0A044UW71 | A0A044UW71 |
| DRLEAEQRAEKEKT | 0 | 0 | 1 | 851762 | AAC48290.1 | A0A044VA40 |
| DRNMYDNNNSLATL | 0 | 0 | 1 | 851763 | A0A044SG03 | A0A044SG03 |
| DRRDEGTYTCRAEN | 0 | 0 | 1 | 851766 | A0A044UK00 | A0A044UK00 |
| DSFETIKYVETVQ | 0 | 0 | 1 | 851778 | A0A044UK00 | A0A044UK00 |
| DSHSKWADDILA | 0 | 0 | 1 | 851780 | AAA50214.2 | P52570 |
| DSKYTIYVDQKEVKE | 0 | 0 | 1 | 851781 | AAA20541.1 | A0A044QNW3 |
| DSNEDSKQNLDNPGN | 0 | 0 | 1 | 851782 | A0A044UK00 | A0A044UK00 |
| DSQKGMTGFGAPRD | 0 | 0 | 1 | 851784 | AAA20215.1 | A0A044V456 |
| DSTTVAPHEAEL | 0 | 0 | 1 | 851785 | A0A044V9R9 | A0A044V9R9 |
| DSVTYRNISEDSMK | 0 | 0 | 1 | 851786 | AAY67786.1 | A0A158N7U2 |
| DTFSGMKWASNGQSI | 0 | 0 | 1 | 851790 | A0A044R153 | A0A044R153 |
| DTKLMDVKLCDLRSW | 0 | 0 | 1 | 851792 | A0A044R077 | A0A044R077 |
| DTKQISEETTINVI | 0 | 0 | 1 | 851793 | A0A044UK00 | A0A044UK00 |
| DTNHDGRISFDEFM | 0 | 0 | 1 | 851796 | A0A044VDG0 | A0A044VDG0 |
| DTPCLISEDLLAEIQ | 0 | 0 | 1 | 851797 | A0A044RFT9 | A0A044RFT9 |
| DTSNRCPPGPPGRAG | 0 | 0 | 1 | 851799 | A0A044SAV6 | A0A044SAV6 |
| DVDIYIARKDNIQRQ | 0 | 0 | 1 | 851807 | A0A044UK00 | A0A044UK00 |
| DVERIWRLVLPNLN | 0 | 0 | 1 | 851809 | AAF64251.1 | A0A044V6Z2 |
| DVEVQLWEEDTGDPD | 0 | 0 | 1 | 851810 | A0A044V9S3 | A0A044V9S3 |
| DVMPFLGRYEKFLKE | 0 | 0 | 1 | 851814 | P46434.1 | P46434 |
| DVNEYNDDFQQN | 0 | 0 | 1 | 851815 | A0A044QNT8 | A0A044QNT8 |
| DVVKDLNVDTFRKFM | 0 | 0 | 1 | 851820 | AAF64254.1 | A0A044QLL7 |
| DVVQEQEVESGAAA | 0 | 0 | 1 | 851822 | A0A044SF60 | A0A044SF60 |
| DWEMMNYRRAMRE | 0 | 0 | 1 | 851825 | AAC48290.1 | A0A044VA40 |
| DWKDGDDCVVLP | 0 | 0 | 1 | 851826 | AAA50214.2 | P52570 |
| DWRKNLQPITQNERI | 0 | 0 | 1 | 851828 | A0A044UK00 | A0A044UK00 |
| DYEFDDVNEYND | 0 | 0 | 1 | 851830 | A0A044QNT8 | A0A044QNT8 |
| DYGALEPILSAEI | 0 | 0 | 1 | 851831 | P41981.1 | A0A158N7R1 |
| DYIEIRNSTDFAN | 0 | 0 | 1 | 851832 | AAV71152.1 | A0A044VAG6 |
| DYKEGDKEKEKEKGI | 0 | 0 | 1 | 851833 | A0A044TU88 | A0A044TU88 |
| DYLESMYGMFHKVA | 0 | 0 | 1 | 851834 | A0A044TU88 | A0A044TU88 |
| EAAERTLGRLNLDLK | 0 | 0 | 1 | 851836 | A0A044TDA1 | A0A044TDA1 |
| EAASSNANQAITSEN | 0 | 0 | 1 | 851837 | P46434.1 | P46434 |
| EAFLKRAQANSLAQL | 0 | 0 | 1 | 851844 | AAB52600.1 | A0A044UUD9 |
| EAGIKIYDPNRLNF | 0 | 0 | 1 | 851845 | AAF64254.1 | A0A044QLL7 |
| EAGKKKAKKGFL | 0 | 0 | 1 | 851846 | A0A044TX11 | A0A044TX11 |
| EAGKSHSVPPEI | 0 | 0 | 1 | 851847 | A0A044UK00 | A0A044UK00 |
| EAHNVQTNDQLMC | 0 | 0 | 1 | 851849 | A0A044TU88 | A0A044TU88 |
| EAIYNQLFQHML | 0 | 0 | 1 | 851851 | A0A044TX11 | A0A044TX11 |
| EAQNRKLNADLDMLR | 0 | 0 | 1 | 851857 | AAA74283.1 | A0A044V2F8 |
| EASENRNLSVSEMET | 0 | 0 | 1 | 851859 | A0A044UK00 | A0A044UK00 |
| EATNGIGSVQTKSF | 0 | 0 | 1 | 851863 | A0A044UK00 | A0A044UK00 |
| EAVHDSSETAETRSI | 0 | 0 | 1 | 851864 | A0A044UVG8 | A0A044UVG8 |
| EAVTDTPPKFIQQIP | 0 | 0 | 1 | 851867 | A0A044UK00 | A0A044UK00 |
| EAWIEAAKEENSYIK | 0 | 0 | 1 | 851869 | A0A044UK00 | A0A044UK00 |

| | | | | | | |
|---|---|---|---|---|---|---|
| ECVPTSPTSYLNSPE | 0 | 0 | 1 | 851874 | A0A044QWA5 | A0A044QWA5 |
| ECWQTEEPNEISA | 0 | 0 | 1 | 851875 | A0A044RF80 | A0A044RF80 |
| EDATLNLNAINQFPG | 0 | 0 | 1 | 851877 | AAB52600.1 | A0A044UUD9 |
| EDDGISSKLKDL | 0 | 0 | 1 | 851878 | A0A044UVG8 | A0A044UVG8 |
| EDDQRSYEAALNDR | 0 | 0 | 1 | 851879 | AAA21472.1 | A0A2K6WLA7 |
| EDDTIDPDDLLNATK | 0 | 0 | 1 | 851880 | A0A044V9S3 | A0A044V9S3 |
| EDEANSLHEIESIDD | 0 | 0 | 1 | 851881 | A0A044UK00 | A0A044UK00 |
| EDENFKEKHTGPGVL | 0 | 0 | 1 | 851884 | AAC47233.1 | Q25633 |
| EDGDEGSEECVERG | 0 | 0 | 1 | 851886 | A0A044SL50 | A0A044SL50 |
| EDIDGGDEEGGN | 0 | 0 | 1 | 851889 | AAD27587.1 | A0A158N7U8 |
| EDKGKGSLSNKKLNS | 0 | 0 | 1 | 851892 | A0A044U9P0 | A0A044U9P0 |
| EDNDGGDEEGGN | 0 | 0 | 1 | 851894 | AAC79423.1 | A0A158N7U8 |
| EDNGELRAMEKLETE | 0 | 0 | 1 | 851896 | AAK16514.1 | A0A2K6W829 |
| EDQVVSSDFVGDPH | 0 | 0 | 1 | 851898 | O01360.1 | O01360 |
| EDRPRYGGPLGEDDD | 0 | 0 | 1 | 851899 | A0A044TX11 | A0A044TX11 |
| EDSSSAPAVAEIGE | 0 | 0 | 1 | 851901 | AAD00842.1 | A0A044SS43 |
| EEAEIISNTKKQ | 0 | 0 | 1 | 851904 | A0A044UVG8 | A0A044UVG8 |
| EEDENAPPIQAEQDA | 0 | 0 | 1 | 851907 | A0A044R196 | A0A044R196 |
| EEDESTDETSAS | 0 | 0 | 1 | 851908 | A0A044UK00 | A0A044UK00 |
| EEEASKARIEAEKQR | 0 | 0 | 1 | 851910 | A0A044R196 | A0A044R196 |
| EEEIEERLKNVR | 0 | 0 | 1 | 851914 | A0A044SS43 | A0A044SS43 |
| EEESWSKKAIDSL | 0 | 0 | 1 | 851915 | A0A044V1C1 | A0A044V1C1 |
| EEETEEAAEPEA | 0 | 0 | 1 | 851916 | A0A044R196 | A0A044R196 |
| EEGCAVDKYLLNN | 0 | 0 | 1 | 851920 | A0A044R521 | A0A044R521 |
| EEGDEGDEEAE | 0 | 0 | 1 | 851921 | A0A044TX11 | A0A044TX11 |
| EEGEVPYLMLVKEAL | 0 | 0 | 1 | 851922 | A0A044SL50 | A0A044SL50 |
| EEGLEEKERKIVK | 0 | 0 | 1 | 851924 | A0A044SS43 | A0A044SS43 |
| EEIHKPKLLKS | 0 | 0 | 1 | 851925 | AAC48290.1 | A0A044VA40 |
| EEIPWGKVSMYI | 0 | 0 | 1 | 851927 | A0A044UK00 | A0A044UK00 |
| EEKKQEAPVEESA | 0 | 0 | 1 | 851929 | A0A044TX11 | A0A044TX11 |
| EEKRRQEEEASKA | 0 | 0 | 1 | 851930 | A0A044R196 | A0A044R196 |
| EENITQKNPTDE | 0 | 0 | 1 | 851936 | A0A044UVG8 | A0A044UVG8 |
| EEQNMQDEIAYRN | 0 | 0 | 1 | 851939 | A0A044UZ11 | A0A044UZ11 |
| EEQRELAQYIEDSNR | 0 | 0 | 1 | 851942 | AAA29419.1 | P21250 |
| EEREFAERQREIE | 0 | 0 | 1 | 851943 | A0A044R196 | A0A044R196 |
| EEREGKMILEKG | 0 | 0 | 1 | 851944 | AAA20541.1 | A0A044QNW3 |
| EEREGTLPIVKGEIA | 0 | 0 | 1 | 851945 | AAD00843.1 | A0A044VHF7 |
| EERRKGWQLARD | 0 | 0 | 1 | 851946 | AAA29419.1 | P21250 |
| EESEIVNAIVKLCS | 0 | 0 | 1 | 851947 | A0A044UK00 | A0A044UK00 |
| EESEKTDGEKIDE | 0 | 0 | 1 | 851948 | A0A044SG03 | A0A044SG03 |
| EESGEDDGERKSKK | 0 | 0 | 1 | 851949 | A0A044SS43 | A0A044SS43 |
| EESNLSTFKQKSE | 0 | 0 | 1 | 851951 | A0A044UK00 | A0A044UK00 |
| EETKEELKATFPN | 0 | 0 | 1 | 851953 | AAA65186.1 | Q25619 |
| EETTDTGTSKPTA | 0 | 0 | 1 | 851954 | A0A044U0M2 | A0A044U0M2 |
| EETTINVIYKSCS | 0 | 0 | 1 | 851955 | A0A044UK00 | A0A044UK00 |
| EEYFTISTAFNTQ | 0 | 0 | 1 | 851957 | A0A044UK00 | A0A044UK00 |
| EEYQYSLQWNDYYSR | 0 | 0 | 1 | 851959 | A0A044UVG8 | A0A044UVG8 |
| EFADFERIIRSRAID | 0 | 0 | 1 | 851961 | AAF64251.1 | A0A044V6Z2 |
| EFDMNKDKIVTIDEL | 0 | 0 | 1 | 851962 | AAF64251.1 | A0A044V6Z2 |
| EFEKIFQDDGLHEF | 0 | 0 | 1 | 851963 | A0A044UK00 | A0A044UK00 |
| EFHDKPHIVQRDG | 0 | 0 | 1 | 851967 | A0A044R9D0 | A0A044R9D0 |
| EFKMQAHCDIDYSNF | 0 | 0 | 1 | 851968 | A0A044QU24 | A0A044QU24 |
| EGEEMPEDIDGGD | 0 | 0 | 1 | 851978 | A0A044RUV7 | A0A044RUV7 |
| EGEEMPEDNDGGD | 0 | 0 | 1 | 851979 | A0A158N7U8 | A0A158N7U8 |
| EGEKYDLEKRHERQE | 0 | 0 | 1 | 851980 | A0A044R196 | A0A044R196 |
| EGGVPLTAYVYAA | 0 | 0 | 1 | 851984 | A0A044RF80 | A0A044RF80 |
| EGIGSFGITVND | 0 | 0 | 1 | 851985 | AAF64253.1 | A0A2K6VYS3 |
| EGIRGVPGASGEPGP | 0 | 0 | 1 | 851987 | A0A044SAV6 | A0A044SAV6 |
| EGLIPGEPGPPGDIG | 0 | 0 | 1 | 851989 | A0A044SF60 | A0A044SF60 |
| EGLYICEATNGIG | 0 | 0 | 1 | 851990 | A0A044UK00 | A0A044UK00 |
| EGPEFDFVSNIRTE | 0 | 0 | 1 | 851992 | A0A044R153 | A0A044R153 |
| EGQNEQSLNLCFDE | 0 | 0 | 1 | 851993 | A0A044R9D0 | A0A044R9D0 |
| EGRNSLLEKQIQELN | 0 | 0 | 1 | 851995 | AAA74283.1 | A0A044V2F8 |
| EGTTQGLDDLN | 0 | 0 | 1 | 851996 | AAB52600.1 | A0A044UUD9 |
| EGVGFKVGGNTGIQ | 0 | 0 | 1 | 851997 | A0A044U175 | A0A044U175 |
| EGYNIGANDNSEIA | 0 | 0 | 1 | 851998 | A0A044UVG8 | A0A044UVG8 |
| EHETVDEVIEQAA | 0 | 0 | 1 | 852000 | A0A044TBR7 | A0A044TBR7 |
| EHGIVTNWDDMEKI | 0 | 0 | 1 | 852001 | P30162.1 | P30162 |
| EHKTWECPDAPNVTA | 0 | 0 | 1 | 852002 | A0A044U9P0 | A0A044U9P0 |
| EHYIVGYKPFATMQK | 0 | 0 | 1 | 852006 | A0A044U175 | A0A044U175 |
| EIADITVINERYSFQ | 0 | 0 | 1 | 852008 | AAD00843.1 | A0A044VHF7 |
| EIESYGYTTADID | 0 | 0 | 1 | 852012 | A0A044UW71 | A0A044UW71 |
| EIFAIWKPIQQQD | 0 | 0 | 1 | 852014 | A0A044UK00 | A0A044UK00 |
| EIFYTESAKTEDVK | 0 | 0 | 1 | 852015 | A0A044SS43 | A0A044SS43 |
| EIIVKNGDVKVIPQ | 0 | 0 | 1 | 852019 | AAC48290.1 | A0A044VA40 |
| EIKNFDKELAGS | 0 | 0 | 1 | 852022 | A0A044R9D0 | A0A044R9D0 |
| EINSLHNNVMDEMSI | 0 | 0 | 1 | 852024 | A0A044V9I3 | A0A044V9I3 |
| EIPLQAGTNKFASQ | 0 | 0 | 1 | 852027 | AAA20215.1 | A0A044V456 |
| EIQFVADEKDLA | 0 | 0 | 1 | 852029 | A0A044TL01 | A0A044TL01 |
| EIRLKAGEKLVLESK | 0 | 0 | 1 | 852032 | A0A044UK00 | A0A044UK00 |
| EISKDVFLERKPATS | 0 | 0 | 1 | 852036 | A0A044UK00 | A0A044UK00 |
| EITEEEEDEEA | 0 | 0 | 1 | 852038 | A0A044R196 | A0A044R196 |
| EKDGRYHDDGRLITN | 0 | 0 | 1 | 852043 | A0A044V9G2 | A0A044V9G2 |
| EKEIDIHREPSYA | 0 | 0 | 1 | 852045 | A0A044UK00 | A0A044UK00 |
| EKFSEVLTVLAE | 0 | 0 | 1 | 852046 | A0A044SS43 | A0A044SS43 |
| EKGAFEVYGEDKEVT | 0 | 0 | 1 | 852047 | A0A044V9S3 | A0A044V9S3 |
| EKGEVSIYASDYSDP | 0 | 0 | 1 | 852048 | A0A044UK00 | A0A044UK00 |
| EKGVGKSGKPLHYKG | 0 | 0 | 1 | 852050 | AAC47233.1 | Q25633 |
| EKHDKITSDSYPVPV | 0 | 0 | 1 | 852051 | A0A044U0M2 | A0A044U0M2 |
| EKIHTIDLPSGKHYL | 0 | 0 | 1 | 852055 | P52570.1 | P52570 |
| EKLKADNITEAMNKL | 0 | 0 | 1 | 852062 | A0A044UVG8 | A0A044UVG8 |

| | | | | | | |
|---|---|---|---|---|---|---|
| EKQDQRRQQRAQEER | 0 | 0 | 1 | 852068 | A0A044R196 | A0A044R196 |
| EKTDWLDNKHVVFGH | 0 | 0 | 1 | 852073 | A0A044U9P0 | A0A044U9P0 |
| EKVRFLEAQNRKLNA | 0 | 0 | 1 | 852077 | AAA74283.1 | A0A044V2F8 |
| EKVSGWVVRENQ | 0 | 0 | 1 | 852078 | A0A044U175 | A0A044U175 |
| EKWTNRGCFCSSKGE | 0 | 0 | 1 | 852079 | AAC79423.1 | A0A158N7U8 |
| ELAKCNERNLEPGK | 0 | 0 | 1 | 852083 | A0A2K6VRV9 | A0A2K6VRV9 |
| ELASKHATFTNEDAA | 0 | 0 | 1 | 852085 | AAA65186.1 | Q25619 |
| ELENSDEVQELVWRK | 0 | 0 | 1 | 852088 | A0A044UK00 | A0A044UK00 |
| ELIGERNPLLPQMFE | 0 | 0 | 1 | 852093 | A0A044U175 | A0A044U175 |
| ELLSVTVTSEKIIE | 0 | 0 | 1 | 852102 | A0A044RXP7 | A0A044RXP7 |
| ELMTEATNRKYEQ | 0 | 0 | 1 | 852104 | AAK16514.1 | A0A2K6W829 |
| ELNFQLEDDQRSYEA | 0 | 0 | 1 | 852105 | AAA74283.1 | A0A044V2F8 |
| ELQRARTDLDQETLN | 0 | 0 | 1 | 852109 | AAA74283.1 | A0A044V2F8 |
| ELVDFRHEIRRF | 0 | 0 | 1 | 852113 | AAF64251.1 | A0A044V6Z2 |
| ELVNSLQEMGINDAF | 0 | 0 | 1 | 852114 | A0A044V3Q0 | A0A044V3Q0 |
| ELYMESFDFHADHNI | 0 | 0 | 1 | 852117 | A0A044UK00 | A0A044UK00 |
| EMFSTVNRDLH | 0 | 0 | 1 | 852120 | A0A044RXP7 | A0A044RXP7 |
| EMGDPGLDGEPGRPG | 0 | 0 | 1 | 852121 | A0A044V9I3 | A0A044V9I3 |
| EMLSPRVSYQQLAL | 0 | 0 | 1 | 852125 | A0A044SFY3 | A0A044SFY3 |
| EMNCYEDMATFKSYC | 0 | 0 | 1 | 852126 | A0A044SL50 | A0A044SL50 |
| EMSMEINDKENIEAV | 0 | 0 | 1 | 852127 | A0A044UK00 | A0A044UK00 |
| EMVEQDLFEEE | 0 | 0 | 1 | 852129 | A0A044UK00 | A0A044UK00 |
| EMYDADKLHSNEI | 0 | 0 | 1 | 852131 | A0A2K6VJ98 | A0A2K6VJ98 |
| ENALDFSDLLQWFDS | 0 | 0 | 1 | 852132 | A0A044UMF0 | A0A044UMF0 |
| ENCDVNNDGNISI | 0 | 0 | 1 | 852133 | A0A044T7P0 | A0A044T7P0 |
| ENDMKELNEEIK | 0 | 0 | 1 | 852135 | A0A044UK00 | A0A044UK00 |
| ENDPNFNYHNGNKEP | 0 | 0 | 1 | 852136 | A0A044RAH9 | A0A044RAH9 |
| ENEDGQLDPVEFAG | 0 | 0 | 1 | 852137 | AAF64251.1 | A0A044V6Z2 |
| ENEETSTIDDEKQR | 0 | 0 | 1 | 852138 | A0A044SS43 | A0A044SS43 |
| ENEYGKAVLPCNLCV | 0 | 0 | 1 | 852141 | A0A044UK00 | A0A044UK00 |
| ENFEDIQKSAQHLHA | 0 | 0 | 1 | 852143 | A0A044UK00 | A0A044UK00 |
| ENFGGEKFDVIEQDT | 0 | 0 | 1 | 852144 | A0A044UVG8 | A0A044UVG8 |
| ENGQPVFPTGVLGRP | 0 | 0 | 1 | 852146 | A0A044UVG8 | A0A044UVG8 |
| ENGRLVQTETKHLNP | 0 | 0 | 1 | 852147 | A0A044SAZ1 | A0A044SAZ1 |
| ENINISKKANSEQN | 0 | 0 | 1 | 852152 | A0A044UK00 | A0A044UK00 |
| ENISEEAKQAAKQAY | 0 | 0 | 1 | 852153 | A0A044R196 | A0A044R196 |
| ENKEAKDYKEGDKEK | 0 | 0 | 1 | 852155 | A0A044TU88 | A0A044TU88 |
| ENKTDQQTEADVEA | 0 | 0 | 1 | 852156 | P36991.1 | P36991 |
| ENREFFKNELSSA | 0 | 0 | 1 | 852160 | AAA74283.1 | A0A044V2F8 |
| ENSESSEESGEDDGE | 0 | 0 | 1 | 852161 | A0A044SS43 | A0A044SS43 |
| ENVDTSSEKINSDIH | 0 | 0 | 1 | 852164 | A0A044UK00 | A0A044UK00 |
| ENVEENRRLYRQL | 0 | 0 | 1 | 852165 | AAB52600.1 | A0A044UUD9 |
| ENVITISPDKESEII | 0 | 0 | 1 | 852166 | AAC48290.1 | A0A044VA40 |
| ENVYDISKRDVSKTP | 0 | 0 | 1 | 852167 | A0A044UK00 | A0A044UK00 |
| EPAAEKNDKITL | 0 | 0 | 1 | 852168 | A0A044UK00 | A0A044UK00 |
| EPETLPDGEHDVHRC | 0 | 0 | 1 | 852170 | AAB52600.1 | A0A044UUD9 |
| EPGVPGIAVPIESVK | 0 | 0 | 1 | 852178 | A0A044SAV6 | A0A044SAV6 |
| EPIFSKSLNEEDL | 0 | 0 | 1 | 852179 | A0A044UK00 | A0A044UK00 |
| EPITHQKAAILDAD | 0 | 0 | 1 | 852181 | A0A044R436 | A0A044R436 |
| EPKDLGLPTEAYVE | 0 | 0 | 1 | 852183 | A0A044TU88 | A0A044TU88 |
| EPQGFGAIKSAA | 0 | 0 | 1 | 852190 | A0A044R521 | A0A044R521 |
| EPTGESSDIKCPE | 0 | 0 | 1 | 852193 | AAB68960.1 | A0A044RRG7 |
| EPTTGLIPLFCID | 0 | 0 | 1 | 852194 | P41981.1 | A0A158N7R1 |
| EQAPPEIEQYVK | 0 | 0 | 1 | 852197 | A0A2K6VJ98 | A0A2K6VJ98 |
| EQAYQIYESTIQHGR | 0 | 0 | 1 | 852198 | AAV71152.1 | A0A044VAG6 |
| EQHNHSSKTTEDR | 0 | 0 | 1 | 852202 | A0A044UK00 | A0A044UK00 |
| EQNNQRSDGIRG | 0 | 0 | 1 | 852207 | A0A044UK00 | A0A044UK00 |
| ERAEKTAVPGIRTQE | 0 | 0 | 1 | 852212 | A0A044UVG8 | A0A044UVG8 |
| ERCLCKHKFKDCKN | 0 | 0 | 1 | 852214 | A0A044T7P0 | A0A044T7P0 |
| EREKKEMSDLNDR | 0 | 0 | 1 | 852217 | AAA74283.1 | A0A044V2F8 |
| ERIEKKDTADGFE | 0 | 0 | 1 | 852219 | A0A044UK00 | A0A044UK00 |
| ERMTFNKMNDVRT | 0 | 0 | 1 | 852221 | A0A2K6VYS3 | A0A2K6VYS3 |
| ERNLEPGKYKNISLR | 0 | 0 | 1 | 852223 | A0A2K6VRV9 | A0A2K6VRV9 |
| ERPDQSEIPLQAGTN | 0 | 0 | 1 | 852224 | AAA20215.1 | A0A044V456 |
| ERQFTNVQVPSLENI | 0 | 0 | 1 | 852228 | A0A044UK00 | A0A044UK00 |
| ERRDDEIQENRII | 0 | 0 | 1 | 852232 | A0A044VAG6 | A0A044VAG6 |
| ERSVSQCKTSDSRNR | 0 | 0 | 1 | 852235 | AAF64250.1 | A0A044UVG8 |
| ERYDPTLEKLPRLFD | 0 | 0 | 1 | 852237 | AAF64254.1 | A0A044QLL7 |
| ESAGIHESTYNSI | 0 | 0 | 1 | 852238 | P30162.1 | P30162 |
| ESDEKPKQLDGIA | 0 | 0 | 1 | 852243 | AAC48290.1 | A0A044VA40 |
| ESFIEVNEKGTEAAA | 0 | 0 | 1 | 852247 | A0A044V3Q0 | A0A044V3Q0 |
| ESKVDSSPESQFKW | 0 | 0 | 1 | 852250 | A0A044UK00 | A0A044UK00 |
| ESLAELPAAMTHA | 0 | 0 | 1 | 852251 | A0A044R2U8 | A0A044R2U8 |
| ESLGGFESLAELPA | 0 | 0 | 1 | 852252 | A0A044R2U8 | A0A044R2U8 |
| ESNLRMAIFESNE | 0 | 0 | 1 | 852255 | AAK16514.1 | A0A2K6W829 |
| ESSDSPSPFFNFLDK | 0 | 0 | 1 | 852260 | AAC48290.1 | A0A044VA40 |
| ESTYNSIMKCDIDIR | 0 | 0 | 1 | 852265 | P30162.1 | P30162 |
| ESYSYNSAEVRI | 0 | 0 | 1 | 852268 | A0A044SG03 | A0A044SG03 |
| ETAEGYHSVRL | 0 | 0 | 1 | 852269 | A0A044RFT9 | A0A044RFT9 |
| ETEWNDYVMALGKH | 0 | 0 | 1 | 852270 | AAK16514.1 | A0A2K6W829 |
| ETGKHRGFGFVEYE | 0 | 0 | 1 | 852272 | A0A044U9P0 | A0A044U9P0 |
| ETGRRGPPGERGRKG | 0 | 0 | 1 | 852274 | A0A044V9I3 | A0A044V9I3 |
| ETIDEQGVGACM | 0 | 0 | 1 | 852276 | AAB68960.1 | A0A044RRG7 |
| ETKEILKKLEKEYI | 0 | 0 | 1 | 852278 | CAA04390.1 | A0A044R436 |
| ETLCMKCSKSSNPEV | 0 | 0 | 1 | 852279 | A0A044SS43 | A0A044SS43 |
| ETMIDKLNAKTIAIQ | 0 | 0 | 1 | 852281 | P41981.1 | A0A158N7R1 |
| ETQHDLSLCQSFNE | 0 | 0 | 1 | 852286 | A0A2K6WJQ8 | A0A2K6WJQ8 |
| ETQLSSEEQEFR | 0 | 0 | 1 | 852287 | A0A044UK00 | A0A044UK00 |
| ETSEAWIEAAKEENS | 0 | 0 | 1 | 852288 | A0A044UK00 | A0A044UK00 |
| ETSQRNFTDSVTKVE | 0 | 0 | 1 | 852290 | A0A044UK00 | A0A044UK00 |
| ETSTSEAKIERDKIE | 0 | 0 | 1 | 852291 | A0A044UK00 | A0A044UK00 |

| | | | | | | |
|---|---|---|---|---|---|---|
| ETSTSSDDDFMPL | 0 | 0 | 1 | 852292 | A0A044SS43 | A0A044SS43 |
| EVEKEEENDMKELN | 0 | 0 | 1 | 852299 | A0A044UK00 | A0A044UK00 |
| EVESGAAAAAA | 0 | 0 | 1 | 852300 | A0A044SF60 | A0A044SF60 |
| EVFNTSAALFAIES | 0 | 0 | 1 | 852301 | AAB97282.1 | A0A044UNG9 |
| EVGVEHLLRDI | 0 | 0 | 1 | 852304 | A0A044TU88 | A0A044TU88 |
| EVHHKAMQGGAA | 0 | 0 | 1 | 852305 | A0A044RF80 | A0A044RF80 |
| EVIYQRGNPCKVDKD | 0 | 0 | 1 | 852313 | A0A044VD00 | A0A044VD00 |
| EVKESLEERRKGWQL | 0 | 0 | 1 | 852314 | AAA29419.1 | P21250 |
| EVKNFYKNLTQEDRQ | 0 | 0 | 1 | 852316 | AAA65186.1 | Q25619 |
| EVLDPSISWQNRKSQ | 0 | 0 | 1 | 852318 | AAA20215.1 | A0A044V456 |
| EVLGTFDQDYKRGNM | 0 | 0 | 1 | 852319 | AAK16514.1 | A0A2K6W829 |
| EVPYEERKFWATI | 0 | 0 | 1 | 852321 | A0A044V1C1 | A0A044V1C1 |
| EVQSDPPAIFEWFCN | 0 | 0 | 1 | 852323 | A0A044UK00 | A0A044UK00 |
| EWEELIAEAESDD | 0 | 0 | 1 | 852332 | A0A044T7P0 | A0A044T7P0 |
| EWGTESAITMVRAVP | 0 | 0 | 1 | 852333 | A0A044UK00 | A0A044UK00 |
| EWSKKGAEKAEAEE | 0 | 0 | 1 | 852336 | A0A044TX11 | A0A044TX11 |
| EYAKLVNEMKKAF | 0 | 0 | 1 | 852337 | A0A044RRG7 | A0A044RRG7 |
| EYILKEYGIGNRDVE | 0 | 0 | 1 | 852340 | AAF64251.1 | A0A044V6Z2 |
| EYLGDGQLRAVHANK | 0 | 0 | 1 | 852341 | A0A044UMF0 | A0A044UMF0 |
| EYTYCAPEHKVDD | 0 | 0 | 1 | 852345 | A0A044QNT8 | A0A044QNT8 |
| EYVVESTGVFT | 0 | 0 | 1 | 852347 | AAB52599.1 | A0A158N7S3 |
| FAAGTAAGGKN | 0 | 0 | 1 | 852348 | A0A044R196 | A0A044R196 |
| FANRSIYSKQKDSVK | 0 | 0 | 1 | 852358 | A0A044UK00 | A0A044UK00 |
| FAVNSKAAESSAK | 0 | 0 | 1 | 852362 | A0A044UVG8 | A0A044UVG8 |
| FCNDKPVQQNRQK | 0 | 0 | 1 | 852367 | A0A044UK00 | A0A044UK00 |
| FCSAGDTTQYYFDGC | 0 | 0 | 1 | 852368 | AAA29419.1 | P21250 |
| FCSTADIAGKRN | 0 | 0 | 1 | 852369 | A0A044UK00 | A0A044UK00 |
| FCVLMIIDPRNKIP | 0 | 0 | 1 | 852370 | A0A2K6WPE1 | A0A2K6WPE1 |
| FDDDDFQKIINYKDN | 0 | 0 | 1 | 852375 | A0A044QNT8 | A0A044QNT8 |
| FDDFKGTSCGKGPYP | 0 | 0 | 1 | 852376 | A0A044SXH1 | A0A044SXH1 |
| FDEKAIVRNSLIAGE | 0 | 0 | 1 | 852379 | AAA20541.1 | A0A044QNW3 |
| FDEQTYDKNTLP | 0 | 0 | 1 | 852380 | A0A044SG03 | A0A044SG03 |
| FDGTVAFTDGID | 0 | 0 | 1 | 852383 | AAF64254.1 | A0A044QLL7 |
| FDKDSDGTDYLLSIG | 0 | 0 | 1 | 852384 | A0A044R436 | A0A044R436 |
| FDSVNDGNITIEID | 0 | 0 | 1 | 852386 | A0A044UMF0 | A0A044UMF0 |
| FEANNNLQQSSL | 0 | 0 | 1 | 852389 | A0A044R153 | A0A044R153 |
| FEAQTKAGNITTVLQ | 0 | 0 | 1 | 852390 | A0A044UK00 | A0A044UK00 |
| FEDCNNTITFNRTPE | 0 | 0 | 1 | 852393 | A0A044UK00 | A0A044UK00 |
| FEDHLSDCIMHGERY | 0 | 0 | 1 | 852394 | AAF64254.1 | A0A044QLL7 |
| FEGAPETLEKRQMEI | 0 | 0 | 1 | 852396 | A0A044UK00 | A0A044UK00 |
| FESPKLTSNICKNLN | 0 | 0 | 1 | 852399 | A0A044UVG8 | A0A044UVG8 |
| FEVFSASANPRLQHN | 0 | 0 | 1 | 852401 | A0A044UK00 | A0A044UK00 |
| FFAQADENEDGQLD | 0 | 0 | 1 | 852403 | AAF64251.1 | A0A044V6Z2 |
| FFPDYITVDGSE | 0 | 0 | 1 | 852409 | A0A044TL01 | A0A044TL01 |
| FGFHRIIIDKSELED | 0 | 0 | 1 | 852418 | A0A044UK00 | A0A044UK00 |
| FGGGGGDTVAV | 0 | 0 | 1 | 852420 | A0A044S628 | A0A044S628 |
| FGGSQNIFDSYLTER | 0 | 0 | 1 | 852421 | A0A044S6B8 | A0A044S6B8 |
| FGMAPPSRGPGL | 0 | 0 | 1 | 852425 | A0A2K6WJQ8 | A0A2K6WJQ8 |
| FGTPTPQVRWYLNEQ | 0 | 0 | 1 | 852427 | A0A044UK00 | A0A044UK00 |
| FGTTKTESRMIVTES | 0 | 0 | 1 | 852428 | A0A044UK00 | A0A044UK00 |
| FHETFYQKGDDGTPF | 0 | 0 | 1 | 852429 | AAB52600.1 | A0A044UUD9 |
| FHLANSSKKEEALR | 0 | 0 | 1 | 852434 | A0A044RF80 | A0A044RF80 |
| FHPIQESTTPELDES | 0 | 0 | 1 | 852435 | A0A044UK00 | A0A044UK00 |
| FIWTLDFDDFKGTSC | 0 | 0 | 1 | 852446 | AAB68960.1 | A0A044RRG7 |
| FKGKTIMQIVPGKTD | 0 | 0 | 1 | 852453 | A0A044UK00 | A0A044UK00 |
| FKKMAGQKEDKQDFR | 0 | 0 | 1 | 852458 | A0A044TX11 | A0A044TX11 |
| FKTDLRFQSAAIGAL | 0 | 0 | 1 | 852462 | A0A044TMQ9 | A0A044TMQ9 |
| FKTEVATHDTSTPSE | 0 | 0 | 1 | 852463 | A0A044UMF0 | A0A044UMF0 |
| FKTSSILSEETDLN | 0 | 0 | 1 | 852464 | A0A044UVG8 | A0A044UVG8 |
| FLESNPRIERVLYPE | 0 | 0 | 1 | 852468 | A0A044R2U8 | A0A044R2U8 |
| FLIMADQNDRGDFIA | 0 | 0 | 1 | 852470 | A0A044TBR7 | A0A044TBR7 |
| FLKELPADVDNGNVP | 0 | 0 | 1 | 852471 | A0A044VAG6 | A0A044VAG6 |
| FLQKDGTTKKFISKN | 0 | 0 | 1 | 852474 | A0A044RF80 | A0A044RF80 |
| FMCQGGDFTRGNG | 0 | 0 | 1 | 852481 | AAC47233.1 | Q25633 |
| FNDIVPKTAENFRAL | 0 | 0 | 1 | 852484 | AAC47233.1 | Q25633 |
| FNEWKTDTLNIRSYG | 0 | 0 | 1 | 852486 | A0A044QU24 | A0A044QU24 |
| FNIKFKESLLNSCAA | 0 | 0 | 1 | 852487 | AAD00842.1 | A0A044SS43 |
| FNLFADTRTEATSQA | 0 | 0 | 1 | 852489 | AAA29419.1 | P21250 |
| FNSVTSNTDPIGVAL | 0 | 0 | 1 | 852491 | A0A044R153 | A0A044R153 |
| FPCSFSSGPSKENIR | 0 | 0 | 1 | 852494 | A0A2K6VRV9 | A0A2K6VRV9 |
| FPSGPDGRPTAPIN | 0 | 0 | 1 | 852505 | A0A044UVG8 | A0A044UVG8 |
| FPTGVLGRPIVPVDT | 0 | 0 | 1 | 852507 | A0A044UVG8 | A0A044UVG8 |
| FQDIITDEDDNGDRA | 0 | 0 | 1 | 852511 | A0A044QNT8 | A0A044QNT8 |
| FQNERDGWYHMLDQE | 0 | 0 | 1 | 852517 | A0A044QWA5 | A0A044QWA5 |
| FQQNIIDEDDFQQ | 0 | 0 | 1 | 852518 | A0A044QNT8 | A0A044QNT8 |
| FQSAAIGALQEASEA | 0 | 0 | 1 | 852519 | A0A044TMQ9 | A0A044TMQ9 |
| FRETMSNPKMDFTN | 0 | 0 | 1 | 852521 | AAA63412.2 | Q99099 |
| FRFDGDQLSGLMD | 0 | 0 | 1 | 852522 | A0A044RF80 | A0A044RF80 |
| FSAIEISEQNPNL | 0 | 0 | 1 | 852533 | A0A044R521 | A0A044R521 |
| FSDRSASWQNLSVP | 0 | 0 | 1 | 852536 | A0A044UK00 | A0A044UK00 |
| FSHNHIEYLGDGQLR | 0 | 0 | 1 | 852539 | A0A044UMF0 | A0A044UMF0 |
| FSQSIRRRNDD | 0 | 0 | 1 | 852541 | A0A044V9G2 | A0A044V9G2 |
| FTAIAKSAEKRKH | 0 | 0 | 1 | 852544 | AAC47023.1 | A0A044RRG7 |
| FTDSVTKVELDFWSK | 0 | 0 | 1 | 852549 | A0A044UK00 | A0A044UK00 |
| FTEEAKKSGSEQ | 0 | 0 | 1 | 852550 | A0A044RRG7 | A0A044RRG7 |
| FTIYEDGIVMLKITE | 0 | 0 | 1 | 852553 | A0A044UK00 | A0A044UK00 |
| FTTFDLGGHEQAR | 0 | 0 | 1 | 852559 | A0A044TDC0 | A0A044TDC0 |
| FTVGNTANNEMCN | 0 | 0 | 1 | 852561 | A0A044U175 | A0A044U175 |
| FTYVDADQRKHKC | 0 | 0 | 1 | 852562 | AAY67786.1 | A0A158N7U2 |
| FVGYKPASEIPND | 0 | 0 | 1 | 852565 | A0A044RAH9 | A0A044RAH9 |
| FVNKIGQMPGIKEWI | 0 | 0 | 1 | 852574 | P46434.1 | P46434 |

| | | | | | | |
|---|---|---|---|---|---|---|
| FVREQRVTTEASS | 0 | 0 | 1 | 852575 | AAC48290.1 | A0A044VA40 |
| FVSYAAVTFSTVA | 0 | 0 | 1 | 852579 | A0A044SF60 | A0A044SF60 |
| FYLVKREESEIVNA | 0 | 0 | 1 | 852588 | A0A044UK00 | A0A044UK00 |
| FYMEADKTVSAELE | 0 | 0 | 1 | 852589 | A0A044R521 | A0A044R521 |
| FYNELRVAPEEH | 0 | 0 | 1 | 852590 | P30162.1 | P30162 |
| GAAAASRQVAA | 0 | 0 | 1 | 852592 | A0A044QSA2 | A0A044QSA2 |
| GADVVKVGIGSGS | 0 | 0 | 1 | 852595 | Q9NJD8.1 | Q9NJD8 |
| GAEEAEEVGVEH | 0 | 0 | 1 | 852596 | A0A044TU88 | A0A044TU88 |
| GAGGISAIGGG | 0 | 0 | 1 | 852599 | A0A044S628 | A0A044S628 |
| GAKAIMVCFGAAVA | 0 | 0 | 1 | 852602 | AAF64254.1 | A0A044QLL7 |
| GALIIVKNADEKNEN | 0 | 0 | 1 | 852604 | A0A044SS43 | A0A044SS43 |
| GALQEASEAYLVGLF | 0 | 0 | 1 | 852605 | A0A044TMQ9 | A0A044TMQ9 |
| GAPLHIYSGPVPVTV | 0 | 0 | 1 | 852610 | A0A044SS43 | A0A044SS43 |
| GCFCSSKGECTIERQ | 0 | 0 | 1 | 852616 | AAC79423.1 | A0A158N7U8 |
| GCGIAMHCDDGEAFI | 0 | 0 | 1 | 852617 | AAB97282.1 | A0A044UNG9 |
| GDAGPDGTPGQPGGV | 0 | 0 | 1 | 852622 | A0A044SF60 | A0A044SF60 |
| GDCASSWAQST | 0 | 0 | 1 | 852624 | A0A044RFT9 | A0A044RFT9 |
| GDEVLNAPIKLLATS | 0 | 0 | 1 | 852626 | AAC48290.1 | A0A044VA40 |
| GDGVTHTVPIYEGYA | 0 | 0 | 1 | 852629 | P30162.1 | P30162 |
| GDIAENFESPKLTSN | 0 | 0 | 1 | 852630 | A0A044UVG8 | A0A044UVG8 |
| GDIIRDSDVYQTIYE | 0 | 0 | 1 | 852631 | A0A044UK00 | A0A044UK00 |
| GDIKQQQQKQLQNA | 0 | 0 | 1 | 852632 | A0A044UVG8 | A0A044UVG8 |
| GDKCESPKQEVNAHI | 0 | 0 | 1 | 852633 | AAD00842.1 | A0A044SS43 |
| GDLSPGELVDFRHEI | 0 | 0 | 1 | 852636 | AAF64251.1 | A0A044V6Z2 |
| GDMTSAGPNFQRAPV | 0 | 0 | 1 | 852637 | A0A044U175 | A0A044U175 |
| GDPNDRIKHIGDLGN | 0 | 0 | 1 | 852638 | AAB64227.1 | O18483 |
| GDTQAAVAGTKLMNF | 0 | 0 | 1 | 852639 | P41981.1 | A0A158N7R1 |
| GDVKVIPQHSAEDDA | 0 | 0 | 1 | 852641 | AAC48290.1 | A0A044VA40 |
| GEDNTVTYAPLTCPP | 0 | 0 | 1 | 852645 | A0A044TIV1 | A0A044TIV1 |
| GEGDGPGLKQLVE | 0 | 0 | 1 | 852647 | AAA74283.1 | A0A044V2F8 |
| GEGKNGNFGDNSPKQ | 0 | 0 | 1 | 852648 | A0A044U0M2 | A0A044U0M2 |
| GEHATTIVDTRKIVP | 0 | 0 | 1 | 852649 | A0A044UK00 | A0A044UK00 |
| GEIEETTTETKDKIQ | 0 | 0 | 1 | 852651 | A0A044UK00 | A0A044UK00 |
| GEIVSISIPLDYETG | 0 | 0 | 1 | 852652 | A0A044U9P0 | A0A044U9P0 |
| GEKFADENFKIKHYG | 0 | 0 | 1 | 852653 | A0A2K6WB53 | A0A2K6WB53 |
| GEKPTSRPYIAAG | 0 | 0 | 1 | 852655 | A0A044U9P0 | A0A044U9P0 |
| GEKSNLFNLFAD | 0 | 0 | 1 | 852656 | AAA29419.1 | P21250 |
| GELEYTYCAPEVCWR | 0 | 0 | 1 | 852657 | A0A044QNT8 | A0A044QNT8 |
| GELITEKDVSPKMRE | 0 | 0 | 1 | 852658 | A0A044UVG8 | A0A044UVG8 |
| GELNAPEFADFERII | 0 | 0 | 1 | 852659 | AAF64251.1 | A0A044V6Z2 |
| GENQDRPVLLGGWED | 0 | 0 | 1 | 852662 | AAA29418.1 | P22085 |
| GEQWSESTHKMMR | 0 | 0 | 1 | 852668 | A0A044UK00 | A0A044UK00 |
| GERGPQGPSGPPG | 0 | 0 | 1 | 852669 | A0A044TIV1 | A0A044TIV1 |
| GEVCPANWQPGSDTI | 0 | 0 | 1 | 852672 | A0A044VFS3 | A0A044VFS3 |
| GFEHTYCSPDENWTC | 0 | 0 | 1 | 852674 | AAC79423.1 | A0A158N7U8 |
| GFGADCGGRIEL | 0 | 0 | 1 | 852675 | AAV71152.1 | A0A044VAG6 |
| GFGEEVDEKVLQAG | 0 | 0 | 1 | 852676 | A0A044U9P0 | A0A044U9P0 |
| GFGFVEYELAEDAA | 0 | 0 | 1 | 852677 | A0A044U9P0 | A0A044U9P0 |
| GFGIGAGALTMDTGE | 0 | 0 | 1 | 852678 | A0A044V9R9 | A0A044V9R9 |
| GFTDPTSNSSKICLG | 0 | 0 | 1 | 852685 | A0A044V1C1 | A0A044V1C1 |
| GGCIECPPGQPGPPG | 0 | 0 | 1 | 852689 | A0A044V9I3 | A0A044V9I3 |
| GGCTNPGDVAKALGA | 0 | 0 | 1 | 852690 | Q9NJD8.1 | Q9NJD8 |
| GGKSIYGEKFAD | 0 | 0 | 1 | 852694 | A0A2K6WB53 | A0A2K6WB53 |
| GGMVDQKDLYSKE | 0 | 0 | 1 | 852695 | A0A044R153 | A0A044R153 |
| GGNMVGEVIYQRGNP | 0 | 0 | 1 | 852697 | A0A044VD00 | A0A044VD00 |
| GGPGLPGNDAAYC | 0 | 0 | 1 | 852698 | A0A044TIV1 | A0A044TIV1 |
| GGQQDFGIGDYPTFN | 0 | 0 | 1 | 852701 | AAB69626.2 | O16855 |
| GHVYVKGLYDRDECR | 0 | 0 | 1 | 852721 | A0A044R521 | A0A044R521 |
| GIGDYPTFNAGGSFA | 0 | 0 | 1 | 852724 | AAB69626.2 | O16855 |
| GIIDKEEIRRM | 0 | 0 | 1 | 852727 | A0A044UK00 | A0A044UK00 |
| GIIFNNGPTWKDIRRFSLTTL | 0 | 0 | 1 | 852729 | P05181.1 | P05181 |
| GINNQNTRVYQSQDK | 0 | 0 | 1 | 852735 | A0A044UK00 | A0A044UK00 |
| GISSGISDYDF | 0 | 0 | 1 | 852738 | Q9NJD8.1 | Q9NJD8 |
| GITVNDDMIPR | 0 | 0 | 1 | 852739 | AAF64253.1 | A0A2K6VYS3 |
| GIVLDSGDGVTHTVP | 0 | 0 | 1 | 852740 | P30162.1 | P30162 |
| GIVMLKITEGDKEG | 0 | 0 | 1 | 852741 | A0A044UK00 | A0A044UK00 |
| GKDSGRGSLTPPIS | 0 | 0 | 1 | 852745 | A0A044UMF0 | A0A044UMF0 |
| GKEDSKVLSALAEK | 0 | 0 | 1 | 852748 | AAA29419.1 | P21250 |
| GKEVSAKLNDFSGSK | 0 | 0 | 1 | 852749 | A0A044UVG8 | A0A044UVG8 |
| GKIGDLISQVNSFRE | 0 | 0 | 1 | 852752 | AAA63412.2 | Q99099 |
| GKIVFNTFSKGEWGK | 0 | 0 | 1 | 852754 | AAA20541.1 | A0A044QNW3 |
| GKLADLEGRNSLLE | 0 | 0 | 1 | 852755 | AAA21472.1 | A0A2K6WLA7 |
| GKLPPKKRETFETA | 0 | 0 | 1 | 852756 | A0A044U175 | A0A044U175 |
| GKLTNDNEDDEE | 0 | 0 | 1 | 852757 | AAD51087.1 | Q9U9A1 |
| GKPGLGSGSGGSG | 0 | 0 | 1 | 852760 | A0A044U9P0 | A0A044U9P0 |
| GKSIYGPKFADEN | 0 | 0 | 1 | 852767 | A0A044U9P0 | A0A044U9P0 |
| GKYYPVPYESGL | 0 | 0 | 1 | 852779 | AAA20541.1 | A0A044QNW3 |
| GLAGADGKPGL | 0 | 0 | 1 | 852780 | A0A044TIV1 | A0A044TIV1 |
| GLEESEEFLSSL | 0 | 0 | 1 | 852784 | A0A044R2U8 | A0A044R2U8 |
| GLGQMNIPILADTN | 0 | 0 | 1 | 852789 | AAC48312.1 | A0A044VFS3 |
| GLKGFKTCVDQFC | 0 | 0 | 1 | 852791 | A0A044RJ74 | A0A044RJ74 |
| GLKNFVQILEKD | 0 | 0 | 1 | 852792 | A0A2K6VRV9 | A0A2K6VRV9 |
| GLPKDVEVFNTSAAL | 0 | 0 | 1 | 852796 | AAB97282.1 | A0A044UNG9 |
| GLPRPEIFWTKDE | 0 | 0 | 1 | 852797 | A0A044UK00 | A0A044UK00 |
| GLQDNTGYVKKKYTD | 0 | 0 | 1 | 852799 | A0A044UK00 | A0A044UK00 |
| GLQIGGDLEVSGIQM | 0 | 0 | 1 | 852800 | AAA20541.1 | A0A044QNW3 |
| GLSRQSTKEWIS | 0 | 0 | 1 | 852801 | A0A044V420 | A0A044V420 |
| GLTNIRVPDTAPINP | 0 | 0 | 1 | 852802 | AAF64254.1 | A0A044QLL7 |
| GLYDRDECRSDSGGR | 0 | 0 | 1 | 852805 | A0A044R521 | A0A044R521 |
| GMAFRVPTPDVSVVD | 0 | 0 | 1 | 852807 | AAB52599.1 | A0A158N7S3 |
| GMEGYSSKGGAPP | 0 | 0 | 1 | 852808 | A0A044U9P0 | A0A044U9P0 |

| | | | | | | |
|---|---|---|---|---|---|---|
| GMQIEQDRLEAEQRA | 0 | 0 | 1 | 852813 | AAC48290.1 | A0A044VA40 |
| GMRLLKQMDYPDG | 0 | 0 | 1 | 852814 | A0A044RAH9 | A0A044RAH9 |
| GMTGFGTGRDVCRE | 0 | 0 | 1 | 852817 | AAA20215.1 | A0A044V456 |
| GMVTPVKNQGVCG | 0 | 0 | 1 | 852818 | AAK16514.1 | A0A2K6W829 |
| GNCTNVETVHRNE | 0 | 0 | 1 | 852820 | A0A044UK00 | A0A044UK00 |
| GNDWDELETLLENID | 0 | 0 | 1 | 852823 | A0A044T7P0 | A0A044T7P0 |
| GNFSSSRIASLNLN | 0 | 0 | 1 | 852824 | A0A044UK00 | A0A044UK00 |
| GNVIESLFNEVDKNG | 0 | 0 | 1 | 852831 | A0A044V6Z2 | A0A044V6Z2 |
| GNVRESYAPTVPE | 0 | 0 | 1 | 852832 | A0A044U9P0 | A0A044U9P0 |
| GNVSIQETSPDGKFI | 0 | 0 | 1 | 852833 | AAA74283.1 | A0A044V2F8 |
| GPAGPAGPPGKD | 0 | 0 | 1 | 852834 | A0A044TIV1 | A0A044TIV1 |
| GPDGEKGPDGQPGPA | 0 | 0 | 1 | 852835 | A0A044TIV1 | A0A044TIV1 |
| GPDGQPGPAGPAG | 0 | 0 | 1 | 852837 | A0A044TIV1 | A0A044TIV1 |
| GPGQNPNLSVTEAMM | 0 | 0 | 1 | 852839 | A0A044UW71 | A0A044UW71 |
| GPMPGENEPLHAYVT | 0 | 0 | 1 | 852843 | A0A044U9P0 | A0A044U9P0 |
| GPPGEPGSDGPPG | 0 | 0 | 1 | 852847 | A0A044TIV1 | A0A044TIV1 |
| GPPGKDGLAGAD | 0 | 0 | 1 | 852848 | A0A044TIV1 | A0A044TIV1 |
| GPPGPAGPDGEKG | 0 | 0 | 1 | 852850 | A0A044TIV1 | A0A044TIV1 |
| GPPGPPGEDNTVTYA | 0 | 0 | 1 | 852854 | A0A044TIV1 | A0A044TIV1 |
| GQAITKCFLQTITDA | 0 | 0 | 1 | 852869 | A0A044UK00 | A0A044UK00 |
| GQEAEQWDTNQII | 0 | 0 | 1 | 852872 | A0A044R2U8 | A0A044R2U8 |
| GQISRMDISSRPME | 0 | 0 | 1 | 852876 | A0A044TDC0 | A0A044TDC0 |
| GRPIVPVDTNGLAI | 0 | 0 | 1 | 852889 | A0A044UVG8 | A0A044UVG8 |
| GRVIDGFDVLDELEK | 0 | 0 | 1 | 852902 | A0A2K6VZ59 | A0A2K6VZ59 |
| GRYEKFLKESTTG | 0 | 0 | 1 | 852904 | P46434.1 | P46434 |
| GSDKTLKLSILYPA | 0 | 0 | 1 | 852909 | AAA50214.2 | P52570 |
| GSEIAANNDGE | 0 | 0 | 1 | 852910 | AAC48290.1 | A0A044VA40 |
| GSKDGKVQPELIL | 0 | 0 | 1 | 852914 | A0A044UVG8 | A0A044UVG8 |
| GSLLSNSVEHQMIN | 0 | 0 | 1 | 852919 | A0A044UK00 | A0A044UK00 |
| GSLLVQKGKLGDTIE | 0 | 0 | 1 | 852920 | A0A044UU92 | A0A044UU92 |
| GSREAYGYNSYSDE | 0 | 0 | 1 | 852924 | A0A044V9G2 | A0A044V9G2 |
| GSSNGNIQSGRIVA | 0 | 0 | 1 | 852927 | A0A044R153 | A0A044R153 |
| GSSPRKEETTDTG | 0 | 0 | 1 | 852928 | A0A044U0M2 | A0A044U0M2 |
| GSVAEYRASEGK | 0 | 0 | 1 | 852929 | Q9NJD8.1 | Q9NJD8 |
| GSVKEGKLGRREG | 0 | 0 | 1 | 852930 | A0A044U9P0 | A0A044U9P0 |
| GTETGEDDYREYLVP | 0 | 0 | 1 | 852938 | A0A044UK00 | A0A044UK00 |
| GTGGESIYGEKFEDE | 0 | 0 | 1 | 852940 | AAC47233.1 | Q25633 |
| GTGVISETGQPEKK | 0 | 0 | 1 | 852942 | A0A044U9P0 | A0A044U9P0 |
| GTPGQPGGVGAPGPV | 0 | 0 | 1 | 852949 | A0A044SF60 | A0A044SF60 |
| GTPGRPGKPGRPGAP | 0 | 0 | 1 | 852950 | A0A044SF60 | A0A044SF60 |
| GTPRNTTYEAEAGE | 0 | 0 | 1 | 852953 | AAA20215.1 | A0A044V456 |
| GTSGPRCADNQASCR | 0 | 0 | 1 | 852954 | A0A044R153 | A0A044R153 |
| GTSKPTALLDGVT | 0 | 0 | 1 | 852955 | A0A044U0M2 | A0A044U0M2 |
| GVGACMVKGNQWYG | 0 | 0 | 1 | 852960 | A0A044RRG7 | A0A044RRG7 |
| GVNNYDGDYHRSS | 0 | 0 | 1 | 852964 | A0A044V9G2 | A0A044V9G2 |
| GVQVFRVETDSAWTE | 0 | 0 | 1 | 852966 | A0A044TIV1 | A0A044TIV1 |
| GVSAIGGAGGISA | 0 | 0 | 1 | 852967 | A0A044S628 | A0A044S628 |
| GVYSNRDCGDLNHAV | 0 | 0 | 1 | 852970 | AAK16514.1 | A0A2K6W829 |
| GWGEDYSTGSPMKYW | 0 | 0 | 1 | 852972 | A0A044RFT9 | A0A044RFT9 |
| GWGRDAPDFYLPEG | 0 | 0 | 1 | 852973 | A0A044U175 | A0A044U175 |
| GWKQGDEEESWSK | 0 | 0 | 1 | 852974 | A0A044V1C1 | A0A044V1C1 |
| GYNSYSDEGYRSN | 0 | 0 | 1 | 852986 | A0A044V9G2 | A0A044V9G2 |
| GYSPENQREGVGENI | 0 | 0 | 1 | 852989 | AAB97282.1 | A0A044UNG9 |
| HCAAEEDFEEEGE | 0 | 0 | 1 | 853002 | AAD27587.1 | A0A158N7U8 |
| HCCLFKYASKEIS | 0 | 0 | 1 | 853003 | A0A044UMF0 | A0A044UMF0 |
| HCDIDYSNFPDDDKN | 0 | 0 | 1 | 853005 | A0A044QU24 | A0A044QU24 |
| HDDGRLITNGSLSN | 0 | 0 | 1 | 853011 | A0A044V9G2 | A0A044V9G2 |
| HDPSHAKEVNYKKFI | 0 | 0 | 1 | 853018 | A0A044VHF7 | A0A044VHF7 |
| HEKSDVNVPVTWSQF | 0 | 0 | 1 | 853025 | A0A044SAZ1 | A0A044SAZ1 |
| HELKAIDDCRYCYE | 0 | 0 | 1 | 853027 | A0A044V1C1 | A0A044V1C1 |
| HENPDDLGQGTGNMR | 0 | 0 | 1 | 853028 | AAB64227.1 | O18483 |
| HESYMRSVTTRRE | 0 | 0 | 1 | 853030 | A0A044V9G2 | A0A044V9G2 |
| HETAHYETSTSEAKI | 0 | 0 | 1 | 853031 | A0A044UK00 | A0A044UK00 |
| HETLEDGELITEKDV | 0 | 0 | 1 | 853032 | A0A044UVG8 | A0A044UVG8 |
| HETPTPHQTFHASMH | 0 | 0 | 1 | 853033 | A0A044UK00 | A0A044UK00 |
| HGEPKIIQDTKTNTV | 0 | 0 | 1 | 853040 | A0A044R9D0 | A0A044R9D0 |
| HGNQDGIALPEKQEM | 0 | 0 | 1 | 853042 | A0A2K6WJQ8 | A0A2K6WJQ8 |
| HGPVKEDQAKYKCVV | 0 | 0 | 1 | 853043 | A0A044R9D0 | A0A044R9D0 |
| HIDGSYNVRSLGENF | 0 | 0 | 1 | 853047 | A0A044RRG7 | A0A044RRG7 |
| HIMKDPAFGYVNE | 0 | 0 | 1 | 853050 | A0A044V420 | A0A044V420 |
| HKIGATTPSHMALVE | 0 | 0 | 1 | 853059 | AAB52600.1 | A0A044UUD9 |
| HKSKNTRKIKGKRKR | 0 | 0 | 1 | 853064 | A0A044SS43 | A0A044SS43 |
| HKVRNNVSSMIDLI | 0 | 0 | 1 | 853065 | A0A044UK00 | A0A044UK00 |
| HLNAPEDFILETDSV | 0 | 0 | 1 | 853074 | AAY67786.1 | A0A158N7U2 |
| HLPEIKDDPYA | 0 | 0 | 1 | 853076 | A0A044RF80 | A0A044RF80 |
| HPIQDQGDCASSWA | 0 | 0 | 1 | 853086 | A0A044RFT9 | A0A044RFT9 |
| HQTQLPPLPSDH | 0 | 0 | 1 | 853092 | A0A044UK00 | A0A044UK00 |
| HQYGDQTNGCTSAGG | 0 | 0 | 1 | 853093 | AAB64227.1 | O18483 |
| HRVIPNFMCQGGDF | 0 | 0 | 1 | 853105 | AAC47233.1 | Q25633 |
| HSCFVDDGNGDK | 0 | 0 | 1 | 853107 | A0A044QSA2 | A0A044QSA2 |
| HSTTPRSEKSS | 0 | 0 | 1 | 853117 | AAF64250.1 | A0A044UVG8 |
| HTIFGRVVGGTETLN | 0 | 0 | 1 | 853122 | CAA04390.1 | A0A044R436 |
| HTVGQKYTYDPNE | 0 | 0 | 1 | 853127 | AAK16514.1 | A0A2K6W829 |
| HYGAGWVSMANAGPD | 0 | 0 | 1 | 853141 | A0A2K6WB53 | A0A2K6WB53 |
| HYHWCQLKDVNC | 0 | 0 | 1 | 853142 | A0A044UW71 | A0A044UW71 |
| IAANSRDFFERLI | 0 | 0 | 1 | 853149 | A0A044R153 | A0A044R153 |
| IADDTWERMVIMH | 0 | 0 | 1 | 853151 | A0A044V9I3 | A0A044V9I3 |
| IADYKTSNEYYNYA | 0 | 0 | 1 | 853153 | A0A044UMF0 | A0A044UMF0 |
| IAEEEDDEPVP | 0 | 0 | 1 | 853154 | A0A044SAZ1 | A0A044SAZ1 |
| IAHEEPDFSEYGTLA | 0 | 0 | 1 | 853160 | A0A044UVG8 | A0A044UVG8 |
| IAHGTARPPPEWGR | 0 | 0 | 1 | 853161 | A0A044R196 | A0A044R196 |

| | | | | | | |
|---|---|---|---|---|---|---|
| IAKVQPGEGKNGNFG | 0 | 0 | 1 | 853164 | A0A044U0M2 | A0A044U0M2 |
| IALPEKQEMQQQQ | 0 | 0 | 1 | 853165 | A0A2K6WJQ8 | A0A2K6WJQ8 |
| IAQFPVIDASESNKD | 0 | 0 | 1 | 853170 | A0A044SAZ1 | A0A044SAZ1 |
| IAVEKDPFEMVESEE | 0 | 0 | 1 | 853174 | A0A044SS43 | A0A044SS43 |
| IAVEKERENIRLAKT | 0 | 0 | 1 | 853175 | CAA04390.1 | A0A044R436 |
| IAVPIESVKDYGSCR | 0 | 0 | 1 | 853177 | A0A044SAV6 | A0A044SAV6 |
| ICHYSPGGNMVGEVI | 0 | 0 | 1 | 853179 | AAB69625.2 | A0A044VD00 |
| IDASESNKDGVLHEK | 0 | 0 | 1 | 853184 | A0A044SAZ1 | A0A044SAZ1 |
| IDDEGEALLEPEFED | 0 | 0 | 1 | 853185 | A0A044UMF0 | A0A044UMF0 |
| IDESAARQMNRAN | 0 | 0 | 1 | 853188 | A0A044SS43 | A0A044SS43 |
| IDGGVFFEDGTRR | 0 | 0 | 1 | 853191 | A0A044SF60 | A0A044SF60 |
| IDGKREIVYTLPRD | 0 | 0 | 1 | 853193 | AAA74283.1 | A0A044V2F8 |
| IDLRNVFQLRRIIE | 0 | 0 | 1 | 853197 | A0A044UK00 | A0A044UK00 |
| IDLTQKNGSAEMK | 0 | 0 | 1 | 853198 | A0A044RF80 | A0A044RF80 |
| IDPRNKIPSVAH | 0 | 0 | 1 | 853200 | A0A2K6WPE1 | A0A2K6WPE1 |
| IDYAVEVSNAYGAVW | 0 | 0 | 1 | 853204 | A0A044UK00 | A0A044UK00 |
| IEAAKEENSYIKTH | 0 | 0 | 1 | 853205 | A0A044UK00 | A0A044UK00 |
| IEDEIKMKSKDNEK | 0 | 0 | 1 | 853207 | A0A044UK00 | A0A044UK00 |
| IEDELNAALKHDARG | 0 | 0 | 1 | 853208 | A0A2K6VZ59 | A0A2K6VZ59 |
| IELLNAAETSESNE | 0 | 0 | 1 | 853216 | A0A044UK00 | A0A044UK00 |
| IEQKSDICCRQI | 0 | 0 | 1 | 853221 | A0A044TDA1 | A0A044TDA1 |
| IERDKIEIKEEIAA | 0 | 0 | 1 | 853224 | A0A044UK00 | A0A044UK00 |
| IERKNRGELEYTYCA | 0 | 0 | 1 | 853225 | A0A044QNT8 | A0A044QNT8 |
| IESGLDFADKVYLDC | 0 | 0 | 1 | 853227 | A0A044UK00 | A0A044UK00 |
| IEYESLVEDEATLNV | 0 | 0 | 1 | 853230 | A0A044UK00 | A0A044UK00 |
| IFHRNIKDFMVQTGD | 0 | 0 | 1 | 853238 | A0A2K6VZ59 | A0A2K6VZ59 |
| IFKKILYSNGGSN | 0 | 0 | 1 | 853239 | A0A044UK00 | A0A044UK00 |
| IGAFSVDGAEEAD | 0 | 0 | 1 | 853254 | A0A044SAZ1 | A0A044SAZ1 |
| IGFDLEIKNEEYA | 0 | 0 | 1 | 853257 | AAA20541.1 | A0A044QNW3 |
| IGKTYDMHIVRSD | 0 | 0 | 1 | 853260 | A0A044V9S3 | A0A044V9S3 |
| IGQRDQLSFNDIRL | 0 | 0 | 1 | 853266 | AAV71152.1 | A0A044VAG6 |
| IGSAVGHAVGSMLTG | 0 | 0 | 1 | 853267 | A0A2K6WJQ8 | A0A2K6WJQ8 |
| IGYTWPHEGVEESIA | 0 | 0 | 1 | 853270 | A0A044U0M2 | A0A044U0M2 |
| IHFLDLNPEDIGCNG | 0 | 0 | 1 | 853273 | AAK66564.1 | Q963F6 |
| IIADEDRSLATELGM | 0 | 0 | 1 | 853275 | AAA50214.2 | P52570 |
| IIAFSDRISEFKKLD | 0 | 0 | 1 | 853276 | AAC48312.1 | A0A044VFS3 |
| IIDEDDFQQNIID | 0 | 0 | 1 | 853281 | A0A044QNT8 | A0A044QNT8 |
| IIDKSELEDQGT | 0 | 0 | 1 | 853282 | A0A044UK00 | A0A044UK00 |
| IIKDFMVQGGDFIN | 0 | 0 | 1 | 853290 | A0A2K6WB53 | A0A2K6WB53 |
| IISDTGKQREELE | 0 | 0 | 1 | 853297 | A0A044V2F8 | A0A044V2F8 |
| IISEMFSTVNR | 0 | 0 | 1 | 853298 | A0A044UK00 | A0A044UK00 |
| IITDKDDNGDKASD | 0 | 0 | 1 | 853301 | A0A044QNT8 | A0A044QNT8 |
| IKAPKIVEKTNRLF | 0 | 0 | 1 | 853306 | A0A044UK00 | A0A044UK00 |
| IKHIGDLGNIVAGAN | 0 | 0 | 1 | 853310 | AAB64227.1 | O18483 |
| IKRKNSNQLYLK | 0 | 0 | 1 | 853320 | A0A044UK00 | A0A044UK00 |
| ILDRNDDQDDSEDN | 0 | 0 | 1 | 853331 | A0A044SS43 | A0A044SS43 |
| ILDVSNNVVKTSGDI | 0 | 0 | 1 | 853332 | A0A044UVG8 | A0A044UVG8 |
| ILFSHPRDFTP | 0 | 0 | 1 | 853336 | P52570.1 | P52570 |
| ILICEDDTKQISEE | 0 | 0 | 1 | 853339 | A0A044UK00 | A0A044UK00 |
| ILMNTKLEIDVQNSV | 0 | 0 | 1 | 853346 | A0A044VBX6 | A0A044VBX6 |
| ILPTNACGEKAF | 0 | 0 | 1 | 853349 | A0A044UVG8 | A0A044UVG8 |
| ILQKSIAFLNAQQQ | 0 | 0 | 1 | 853350 | A0A044RF80 | A0A044RF80 |
| ILTERGYSFTTTA | 0 | 0 | 1 | 853354 | P30162.1 | P30162 |
| INENSKMIENPRNSS | 0 | 0 | 1 | 853372 | A0A044UK00 | A0A044UK00 |
| INGDGTGGKSIYGE | 0 | 0 | 1 | 853373 | A0A2K6WB53 | A0A2K6WB53 |
| INKDGNPTDFK | 0 | 0 | 1 | 853375 | A0A044V9G2 | A0A044V9G2 |
| INPRIDLTSDHIYN | 0 | 0 | 1 | 853382 | AAF64254.1 | A0A044QLL7 |
| INTSTSFEAQTKAGN | 0 | 0 | 1 | 853385 | A0A044UK00 | A0A044UK00 |
| IPGADGPPGPPG | 0 | 0 | 1 | 853388 | A0A044SF60 | A0A044SF60 |
| IPILADTNHAISKA | 0 | 0 | 1 | 853389 | AAC48312.1 | A0A044VFS3 |
| IPISMENNQENDGFD | 0 | 0 | 1 | 853390 | A0A044SS43 | A0A044SS43 |
| IPLPECFEDHLSDC | 0 | 0 | 1 | 853395 | AAF64254.1 | A0A044QLL7 |
| IPPISLSTTFKQPK | 0 | 0 | 1 | 853397 | A0A044R2U8 | A0A044R2U8 |
| IPRFMVQGGDFTKGD | 0 | 0 | 1 | 853400 | A0A044U9P0 | A0A044U9P0 |
| IPSQMGSNKYASQ | 0 | 0 | 1 | 853403 | AAA20215.1 | A0A044V456 |
| IPTNKIGSVYNMGI | 0 | 0 | 1 | 853405 | A0A044V9S3 | A0A044V9S3 |
| IQDEIEKIMATLPMR | 0 | 0 | 1 | 853408 | A0A044UK00 | A0A044UK00 |
| IQDLDQALKKAIPKI | 0 | 0 | 1 | 853409 | A0A044R2U8 | A0A044R2U8 |
| IQDTKTNTVRLEVIV | 0 | 0 | 1 | 853410 | A0A044R9D0 | A0A044R9D0 |
| IQENRIIFTDVDGCA | 0 | 0 | 1 | 853411 | A0A044VAG6 | A0A044VAG6 |
| IQEVAEKAMIQVNEK | 0 | 0 | 1 | 853413 | AAD51087.1 | Q9U9A1 |
| IQPIESNPAKPNAMP | 0 | 0 | 1 | 853417 | AAC48289.1 | A0A044VA40 |
| IQPSAGPVSQGWSP | 0 | 0 | 1 | 853418 | A0A044SS43 | A0A044SS43 |
| IQSQIERSLEVEKEL | 0 | 0 | 1 | 853423 | A0A044UK00 | A0A044UK00 |
| IQTDEEGDKCESPKQ | 0 | 0 | 1 | 853424 | AAD00842.1 | A0A044SS43 |
| IQTQSARQTMEQG | 0 | 0 | 1 | 853425 | AAA74283.1 | A0A044V2F8 |
| IRIEEAWDEIQHK | 0 | 0 | 1 | 853431 | A0A044UK00 | A0A044UK00 |
| IRTATTNDESV | 0 | 0 | 1 | 853441 | A0A044V3Q0 | A0A044V3Q0 |
| IRWYVDGDIIRD | 0 | 0 | 1 | 853442 | A0A044UK00 | A0A044UK00 |
| ISAFENKSCDSTTN | 0 | 0 | 1 | 853444 | A0A044QWA5 | A0A044QWA5 |
| ISAQNWADQCIFG | 0 | 0 | 1 | 853446 | AAB97282.1 | A0A044UNG9 |
| ISKEHDTWLQMQDKI | 0 | 0 | 1 | 853458 | A0A044SAZ1 | A0A044SAZ1 |
| ISKNPCEDHICGWG | 0 | 0 | 1 | 853460 | A0A044T7P0 | A0A044T7P0 |
| ISNNFELFDNLLN | 0 | 0 | 1 | 853464 | A0A044UIG6 | A0A044UIG6 |
| ISPAVEEAGIKIYD | 0 | 0 | 1 | 853465 | AAF64254.1 | A0A044QLL7 |
| ISPSSPSTTNPA | 0 | 0 | 1 | 853467 | AAB68960.1 | A0A044RRG7 |
| ISSYRFQCFQNEPIA | 0 | 0 | 1 | 853473 | A0A044UMF0 | A0A044UMF0 |
| ISTELTSEQWTHRI | 0 | 0 | 1 | 853475 | A0A044SAZ1 | A0A044SAZ1 |
| ISTYDDALFDD | 0 | 0 | 1 | 853477 | A0A044UK00 | A0A044UK00 |
| ISVIESQEQIATEDF | 0 | 0 | 1 | 853479 | A0A044UK00 | A0A044UK00 |
| ISVNIVARSMHDVA | 0 | 0 | 1 | 853480 | A0A044UK00 | A0A044UK00 |

| | | | | | | |
|---|---|---|---|---|---|---|
| ISYFELNTRVGEQYK | 0 | 0 | 1 | 853482 | A0A044V1C1 | A0A044V1C1 |
| ITCLNFERRDDE | 0 | 0 | 1 | 853487 | A0A044VAG6 | A0A044VAG6 |
| ITDDFPIPTNKIG | 0 | 0 | 1 | 853488 | A0A044V9S3 | A0A044V9S3 |
| ITEGDKEGLYICEA | 0 | 0 | 1 | 853490 | A0A044UK00 | A0A044UK00 |
| ITEMDGTSAKDYHQD | 0 | 0 | 1 | 853493 | A0A044QU24 | A0A044QU24 |
| ITEVTEEWGTESAI | 0 | 0 | 1 | 853494 | A0A044UK00 | A0A044UK00 |
| ITKEDFIRYMSTPPA | 0 | 0 | 1 | 853499 | A0A044VDG0 | A0A044VDG0 |
| ITNGSLSNGRYGS | 0 | 0 | 1 | 853504 | A0A044V9G2 | A0A044V9G2 |
| ITNTRPGCTAIGETF | 0 | 0 | 1 | 853505 | A0A044QS56 | A0A044QS56 |
| ITNTSAYIQIEKGEI | 0 | 0 | 1 | 853506 | A0A044UK00 | A0A044UK00 |
| IVDTRKIVPELFAIA | 0 | 0 | 1 | 853514 | A0A044UK00 | A0A044UK00 |
| IVERTNMVFSSDF | 0 | 0 | 1 | 853519 | A0A044UK00 | A0A044UK00 |
| IVIEDSDALKSINAD | 0 | 0 | 1 | 853521 | A0A044UMF0 | A0A044UMF0 |
| IVLAGANPSAEGED | 0 | 0 | 1 | 853523 | A0A044SL50 | A0A044SL50 |
| IVPELFAIAAAA | 0 | 0 | 1 | 853527 | A0A044UK00 | A0A044UK00 |
| IVPMPVCRYEILDG | 0 | 0 | 1 | 853529 | A0A044QSA2 | A0A044QSA2 |
| IVQTTAPTFIPNGA | 0 | 0 | 1 | 853531 | A0A044SS43 | A0A044SS43 |
| IVRNSNGDLGGI | 0 | 0 | 1 | 853532 | A0A044TDA1 | A0A044TDA1 |
| IVSEMFSTVNRD | 0 | 0 | 1 | 853533 | A0A044RXP7 | A0A044RXP7 |
| IWLETPSNPLLKVMD | 0 | 0 | 1 | 853539 | A0A044R2U8 | A0A044R2U8 |
| IWTFQEVSELIDVGQ | 0 | 0 | 1 | 853541 | A0A044QWA5 | A0A044QWA5 |
| IYASDYSDPVNIKD | 0 | 0 | 1 | 853542 | A0A044UK00 | A0A044UK00 |
| IYFQQDSGGSITTI | 0 | 0 | 1 | 853548 | AAB64227.1 | O18483 |
| IYGEKFEDENFKEKH | 0 | 0 | 1 | 853549 | AAC47233.1 | Q25633 |
| IYHHTSYNESEHR | 0 | 0 | 1 | 853550 | A0A044UK00 | A0A044UK00 |
| IYQDEDGSFYVPPE | 0 | 0 | 1 | 853553 | A0A044U0M2 | A0A044U0M2 |
| KAAILDADTVKYA | 0 | 0 | 1 | 853557 | CAA04390.1 | A0A044R436 |
| KAGNAPEFHDKPHIV | 0 | 0 | 1 | 853565 | A0A044R9D0 | A0A044R9D0 |
| KAKCLKVMNESGRVG | 0 | 0 | 1 | 853566 | A0A044UIG6 | A0A044UIG6 |
| KAMIQVNEKMKYDNL | 0 | 0 | 1 | 853568 | AAD51087.1 | Q9U9A1 |
| KANMVDEIFAI | 0 | 0 | 1 | 853569 | A0A044UK00 | A0A044UK00 |
| KAYGVLKEDEGIAYR | 0 | 0 | 1 | 853574 | AAC48312.1 | A0A044VFS3 |
| KCALCNKLLDSTNCC | 0 | 0 | 1 | 853575 | A0A044V9R9 | A0A044V9R9 |
| KCKKLEGHPEKVM | 0 | 0 | 1 | 853578 | P22085.2 | P22085 |
| KDDSNQLREEQ | 0 | 0 | 1 | 853582 | A0A044UK00 | A0A044UK00 |
| KDDTKSLTSQANSTL | 0 | 0 | 1 | 853583 | A0A044UK00 | A0A044UK00 |
| KDENDADDFILAE | 0 | 0 | 1 | 853585 | A0A044SS43 | A0A044SS43 |
| KDFGSLETMIDKLNA | 0 | 0 | 1 | 853586 | P41981.1 | A0A158N7R1 |
| KDFMVQTGDPTGTGK | 0 | 0 | 1 | 853587 | A0A2K6VZ59 | A0A2K6VZ59 |
| KDLYSKEGPEFDFVS | 0 | 0 | 1 | 853593 | A0A044R153 | A0A044R153 |
| KDMKEITFSSTKTSF | 0 | 0 | 1 | 853594 | A0A044SS43 | A0A044SS43 |
| KDNDSQEIINDEDD | 0 | 0 | 1 | 853595 | A0A044QNT8 | A0A044QNT8 |
| KDPKFYIRRMNNE | 0 | 0 | 1 | 853596 | CAA04390.1 | A0A044R436 |
| KDSKKVTIINEEDH | 0 | 0 | 1 | 853597 | A0A044UK00 | A0A044UK00 |
| KDSRGVIQTDEEGD | 0 | 0 | 1 | 853598 | AAD00842.1 | A0A044SS43 |
| KDTVEVVAVNDPFIN | 0 | 0 | 1 | 853599 | AAB52599.1 | A0A158N7S3 |
| KDVEIQLWERDTLDP | 0 | 0 | 1 | 853600 | A0A044V9S3 | A0A044V9S3 |
| KDVNKLNTTSNQIES | 0 | 0 | 1 | 853602 | A0A044UK00 | A0A044UK00 |
| KEALAKGDTQAAVAG | 0 | 0 | 1 | 853604 | P41981.1 | A0A158N7R1 |
| KEAQKIAFEQEGISA | 0 | 0 | 1 | 853605 | AAF64251.1 | A0A044V6Z2 |
| KEAVEKDIEETKVSK | 0 | 0 | 1 | 853606 | A0A044UK00 | A0A044UK00 |
| KEFIPEEVKNFYKNL | 0 | 0 | 1 | 853610 | Q25619.1 | Q25619 |
| KEGKIGEVKREISRS | 0 | 0 | 1 | 853612 | A0A044SD24 | A0A044SD24 |
| KEKIEGVEKKGESKE | 0 | 0 | 1 | 853615 | A0A044TU88 | A0A044TU88 |
| KEPRGFGHIGIAVKD | 0 | 0 | 1 | 853623 | A0A044RAH9 | A0A044RAH9 |
| KEQLQSLECMLEEVE | 0 | 0 | 1 | 853624 | A0A044UK00 | A0A044UK00 |
| KERRNTDTEKSDVMK | 0 | 0 | 1 | 853626 | A0A044UVG8 | A0A044UVG8 |
| KETTSQLRRPPPQE | 0 | 0 | 1 | 853628 | A0A044R196 | A0A044R196 |
| KETYGGHKDDTGR | 0 | 0 | 1 | 853629 | CAA04390.1 | A0A044R436 |
| KEVNYKKFIGERQYS | 0 | 0 | 1 | 853630 | A0A044VHF7 | A0A044VHF7 |
| KFCSLIPSLETEYLQ | 0 | 0 | 1 | 853633 | AAD00842.1 | A0A044SS43 |
| KFDVIEQDTKENQSS | 0 | 0 | 1 | 853634 | A0A044UVG8 | A0A044UVG8 |
| KFETTTTASEI | 0 | 0 | 1 | 853638 | AAB68960.1 | A0A044RRG7 |
| KFKENSNDADIRST | 0 | 0 | 1 | 853644 | A0A044UVG8 | A0A044UVG8 |
| KFTENLDVVKDLNVD | 0 | 0 | 1 | 853649 | AAF64254.1 | A0A044QLL7 |
| KFVKTLENMTAIVGQ | 0 | 0 | 1 | 853651 | A0A044UK00 | A0A044UK00 |
| KGAENFTPTDVQNSD | 0 | 0 | 1 | 853652 | A0A044UIG6 | A0A044UIG6 |
| KGDGTGGKSIYGPK | 0 | 0 | 1 | 853654 | A0A044U9P0 | A0A044U9P0 |
| KGECTIERQKGDGFE | 0 | 0 | 1 | 853656 | AAD27587.1 | A0A158N7U8 |
| KGETSGTGIFNTEFA | 0 | 0 | 1 | 853658 | AAB68960.1 | A0A044RRG7 |
| KGGDSIWGKPFKDEI | 0 | 0 | 1 | 853659 | A0A044R436 | A0A044R436 |
| KGKLGDTIELALGAK | 0 | 0 | 1 | 853660 | A0A044UU92 | A0A044UU92 |
| KGLKGLDGPPGRD | 0 | 0 | 1 | 853662 | A0A044V9I3 | A0A044V9I3 |
| KGPEEQNENQESQIN | 0 | 0 | 1 | 853663 | A0A044U9P0 | A0A044U9P0 |
| KGVGFGLGAGALTMD | 0 | 0 | 1 | 853666 | A0A044V9R9 | A0A044V9R9 |
| KHYDSNESNLRMAI | 0 | 0 | 1 | 853674 | AAK16514.1 | A0A2K6W829 |
| KIEIKEEIAAAKEA | 0 | 0 | 1 | 853676 | A0A044UK00 | A0A044UK00 |
| KIENRTELVNSLQEM | 0 | 0 | 1 | 853677 | A0A044V3Q0 | A0A044V3Q0 |
| KIERRQSLTEAQAEA | 0 | 0 | 1 | 853678 | A0A044UK00 | A0A044UK00 |
| KIESSDEFEPDEE | 0 | 0 | 1 | 853679 | A0A044SS43 | A0A044SS43 |
| KIETDESSRPIASFQ | 0 | 0 | 1 | 853680 | A0A044R436 | A0A044R436 |
| KIFVTEGLNKVIPL | 0 | 0 | 1 | 853682 | AAA63412.2 | Q99099 |
| KIGINGKPERCVNAD | 0 | 0 | 1 | 853684 | A0A2K6VRV9 | A0A2K6VRV9 |
| KIQDYEERRQIEREK | 0 | 0 | 1 | 853691 | A0A044R196 | A0A044R196 |
| KIYGEENEVNNIEP | 0 | 0 | 1 | 853698 | A0A044V9S3 | A0A044V9S3 |
| KKAGVGYPQLSAVLE | 0 | 0 | 1 | 853700 | Q9NJD8.1 | Q9NJD8 |
| KKANSEQNEEM | 0 | 0 | 1 | 853703 | A0A044UK00 | A0A044UK00 |
| KKCLSKSGLCRK | 0 | 0 | 1 | 853704 | AAB69625.2 | A0A044VD00 |
| KKEFDKKYNPT | 0 | 0 | 1 | 853705 | AAF64249.1 | A0A044UWQ7 |
| KKEPIVEQPISMTLE | 0 | 0 | 1 | 853707 | AAC48290.1 | A0A044VA40 |
| KKGENVQLKCVVSGN | 0 | 0 | 1 | 853709 | A0A044UK00 | A0A044UK00 |

| | | | | | | |
|---|---|---|---|---|---|---|
| KKRNNMSRSRD | 0 | 0 | 1 | 853727 | A0A044U9P0 | A0A044U9P0 |
| KLCGLKQHIDDD | 0 | 0 | 1 | 853737 | A0A044UVG8 | A0A044UVG8 |
| KLEEEKYDINQAVMA | 0 | 0 | 1 | 853739 | A0A044TX11 | A0A044TX11 |
| KLEQSEQINVKYVN | 0 | 0 | 1 | 853740 | A0A044UK00 | A0A044UK00 |
| KLHSNEIFKRRVRK | 0 | 0 | 1 | 853744 | A0A2K6VJ98 | A0A2K6VJ98 |
| KLIQGLFDEQTYD | 0 | 0 | 1 | 853747 | A0A044SG03 | A0A044SG03 |
| KLKENDPEMKILLS | 0 | 0 | 1 | 853749 | AAC47023.1 | A0A044RRG7 |
| KLLKSSTKLRDD | 0 | 0 | 1 | 853755 | AAC48290.1 | A0A044VA40 |
| KLNDFSGSKDGKVQ | 0 | 0 | 1 | 853759 | A0A044UVG8 | A0A044UVG8 |
| KMESKTGENQDRPVL | 0 | 0 | 1 | 853768 | P22085.2 | P22085 |
| KMPRLIISPAVEEA | 0 | 0 | 1 | 853770 | AAF64254.1 | A0A044QLL7 |
| KNADEKNENYSLNIK | 0 | 0 | 1 | 853774 | A0A044SS43 | A0A044SS43 |
| KNAFDILQEIDSDGD | 0 | 0 | 1 | 853775 | AAF64251.1 | A0A044V6Z2 |
| KNAVNAIEKQDLPRS | 0 | 0 | 1 | 853776 | A0A044S628 | A0A044S628 |
| KNGDGYLTSIEFNDI | 0 | 0 | 1 | 853779 | AAF64251.1 | A0A044V6Z2 |
| KNGDIALHFNPRFDE | 0 | 0 | 1 | 853780 | AAA20541.1 | A0A044QNW3 |
| KNIKSAQFNEKD | 0 | 0 | 1 | 853782 | A0A044UK00 | A0A044UK00 |
| KNITIRQGDVVAA | 0 | 0 | 1 | 853783 | A0A044U175 | A0A044U175 |
| KNLTQEDRQILRELA | 0 | 0 | 1 | 853787 | AAA65186.1 | Q25619 |
| KNNKIKTTTIAVE | 0 | 0 | 1 | 853788 | AAB68960.1 | A0A044RRG7 |
| KNPTDELEFITEE | 0 | 0 | 1 | 853789 | A0A044UVG8 | A0A044UVG8 |
| KNRGELEYTYCAPEH | 0 | 0 | 1 | 853792 | A0A044QNT8 | A0A044QNT8 |
| KNRNGTYMPRGKN | 0 | 0 | 1 | 853794 | AAB69625.2 | A0A044VD00 |
| KNSAPTTFATYLF | 0 | 0 | 1 | 853795 | AAA63412.2 | Q99099 |
| KPERCVNADADGCG | 0 | 0 | 1 | 853797 | A0A2K6VRV9 | A0A2K6VRV9 |
| KPFLDRRVLPRRSAR | 0 | 0 | 1 | 853798 | AAD00842.1 | A0A044SS43 |
| KPLIHYETSQRNFTD | 0 | 0 | 1 | 853807 | A0A044UK00 | A0A044UK00 |
| KPNSRSNMAVAEVDA | 0 | 0 | 1 | 853811 | A0A044RF80 | A0A044RF80 |
| KPQKQWGKMDTRI | 0 | 0 | 1 | 853813 | AAF64254.1 | A0A044QLL7 |
| KQCKEHPDCYDQRE | 0 | 0 | 1 | 853816 | AAD27587.1 | A0A158N7U8 |
| KQDDTTPQSQVNFVR | 0 | 0 | 1 | 853817 | AAC48290.1 | A0A044VA40 |
| KQEERNHVLEKYKSC | 0 | 0 | 1 | 853818 | A0A044SG03 | A0A044SG03 |
| KQGNSTISNNFEL | 0 | 0 | 1 | 853821 | A0A044UIG6 | A0A044UIG6 |
| KQIGCDIDYAVEVSN | 0 | 0 | 1 | 853822 | A0A044UK00 | A0A044UK00 |
| KQINSDKSAQSESI | 0 | 0 | 1 | 853823 | AAF64251.1 | A0A044V6Z2 |
| KQMEIENAKQLKILD | 0 | 0 | 1 | 853828 | A0A044SS43 | A0A044SS43 |
| KQNLDNPGNKKLHSK | 0 | 0 | 1 | 853829 | A0A044UK00 | A0A044UK00 |
| KQSGLVDIEKNLTG | 0 | 0 | 1 | 853834 | A0A044V3Q0 | A0A044V3Q0 |
| KQWDWPLNKDDGTVN | 0 | 0 | 1 | 853835 | A0A044SD24 | A0A044SD24 |
| KRAGTQIENKAKKYM | 0 | 0 | 1 | 853836 | A0A044RF80 | A0A044RF80 |
| KRDAAKVPVNGNG | 0 | 0 | 1 | 853838 | P46427.1 | P46427 |
| KRETFETACIVDED | 0 | 0 | 1 | 853839 | A0A044U175 | A0A044U175 |
| KRFAGFNVAGIGGNA | 0 | 0 | 1 | 853840 | CAA31690.1 | P21250 |
| KRGALLDIYQDESA | 0 | 0 | 1 | 853842 | AAD00842.1 | A0A044SS43 |
| KRKWEEDEEDEL | 0 | 0 | 1 | 853846 | A0A044UVG8 | A0A044UVG8 |
| KRNSSDDSEVKGQAT | 0 | 0 | 1 | 853847 | A0A044UVG8 | A0A044UVG8 |
| KRPSLTVDNDGDE | 0 | 0 | 1 | 853848 | A0A044SS43 | A0A044SS43 |
| KRQYSCGQCGGGG | 0 | 0 | 1 | 853850 | A0A044S628 | A0A044S628 |
| KSCIPVNAELRK | 0 | 0 | 1 | 853859 | A0A044SG03 | A0A044SG03 |
| KSDKDHCQDVVGL | 0 | 0 | 1 | 853861 | A0A044SFY3 | A0A044SFY3 |
| KSDRTLDIANSFAVP | 0 | 0 | 1 | 853862 | A0A044TU88 | A0A044TU88 |
| KSEQMKNLTAKREN | 0 | 0 | 1 | 853863 | A0A044R196 | A0A044R196 |
| KSEVEINEFKRKIQA | 0 | 0 | 1 | 853864 | A0A044SL50 | A0A044SL50 |
| KSGKLDKVELADFLR | 0 | 0 | 1 | 853866 | AAF64251.1 | A0A044V6Z2 |
| KSGSEQLLLTA | 0 | 0 | 1 | 853867 | A0A044RRG7 | A0A044RRG7 |
| KSNPYKKGDDIDIR | 0 | 0 | 1 | 853872 | AAA20541.1 | A0A044QNW3 |
| KSNTKMIWLETPSNP | 0 | 0 | 1 | 853873 | A0A044R2U8 | A0A044R2U8 |
| KSQQNKKQMEIENAK | 0 | 0 | 1 | 853875 | A0A044SS43 | A0A044SS43 |
| KSRADVDLVREF | 0 | 0 | 1 | 853876 | Q9NJD8.1 | Q9NJD8 |
| KSSIMSATEDMLVL | 0 | 0 | 1 | 853878 | AAV71152.1 | A0A044VAG6 |
| KSSIQQMDMLLGG | 0 | 0 | 1 | 853879 | A0A044UW71 | A0A044UW71 |
| KSYVMPQPPKFIQ | 0 | 0 | 1 | 853883 | A0A044UK00 | A0A044UK00 |
| KTAGTDAGKSW | 0 | 0 | 1 | 853884 | AAB69625.2 | A0A044VD00 |
| KTILSAEDIVKMN | 0 | 0 | 1 | 853891 | A0A044RJ74 | A0A044RJ74 |
| KTRISQPFQEGQTVH | 0 | 0 | 1 | 853893 | AAD00843.1 | A0A044VHF7 |
| KTTDPDVPVLMIAD | 0 | 0 | 1 | 853894 | AAC48290.1 | A0A044VA40 |
| KTTQPGDKPLKDI | 0 | 0 | 1 | 853896 | A0A2K6WB53 | A0A2K6WB53 |
| KTWLEQTHEVNEE | 0 | 0 | 1 | 853899 | A0A044UK00 | A0A044UK00 |
| KVAWAEMIAFANDR | 0 | 0 | 1 | 853900 | A0A044SAZ1 | A0A044SAZ1 |
| KVELDFWSKTERSLF | 0 | 0 | 1 | 853902 | A0A044UK00 | A0A044UK00 |
| KVENDMKDMKEITFS | 0 | 0 | 1 | 853903 | A0A044SS43 | A0A044SS43 |
| KVEWCKDNKKWVTEN | 0 | 0 | 1 | 853904 | A0A044UK00 | A0A044UK00 |
| KVLVGPTVSGSQEVY | 0 | 0 | 1 | 853909 | A0A044UK00 | A0A044UK00 |
| KVSVADLTVFNML | 0 | 0 | 1 | 853912 | P46434.1 | P46434 |
| KVTVSNSATETEDIK | 0 | 0 | 1 | 853913 | A0A044UVG8 | A0A044UVG8 |
| KWASNGQSILQSCC | 0 | 0 | 1 | 853918 | A0A044R153 | A0A044R153 |
| KWPNFIHPIQDQGDC | 0 | 0 | 1 | 853920 | A0A044RFT9 | A0A044RFT9 |
| KYAPLPQQPNDSPEN | 0 | 0 | 1 | 853922 | A0A044UVG8 | A0A044UVG8 |
| KYATEFNVAGDNLRI | 0 | 0 | 1 | 853923 | A0A044VFS3 | A0A044VFS3 |
| KYCAIDGGVFFEDG | 0 | 0 | 1 | 853924 | A0A044V9C9 | A0A044V9C9 |
| KYEALNEETKEEL | 0 | 0 | 1 | 853925 | AAA65186.1 | Q25619 |
| KYELSWQKFEDPE | 0 | 0 | 1 | 853926 | A0A044UK00 | A0A044UK00 |
| KYFRLYDNDGDGAI | 0 | 0 | 1 | 853927 | A0A044VDG0 | A0A044VDG0 |
| KYGSGVGKYINLPE | 0 | 0 | 1 | 853928 | CAA04390.1 | A0A044R436 |
| KYISIESTDDPQCFQ | 0 | 0 | 1 | 853931 | A0A044TDA1 | A0A044TDA1 |
| KYLLNNLEYPTDL | 0 | 0 | 1 | 853934 | A0A044RMM8 | A0A044RMM8 |
| KYMSANANDGIE | 0 | 0 | 1 | 853936 | A0A044SAZ1 | A0A044SAZ1 |
| KYNLNGENEMETT | 0 | 0 | 1 | 853938 | P46427.1 | P46427 |
| KYQIFANRVEAGTFD | 0 | 0 | 1 | 853941 | A0A044VHF7 | A0A044VHF7 |
| KYSTKFKEAVEKDIE | 0 | 0 | 1 | 853945 | A0A044UK00 | A0A044UK00 |
| KYVEQVKADGDAA | 0 | 0 | 1 | 853947 | A0A044R196 | A0A044R196 |

| | | | | | | |
|---|---|---|---|---|---|---|
| KYVETVQVPTEG | 0 | 0 | 1 | 853948 | A0A044UK00 | A0A044UK00 |
| LADIIGEQKLIGKQE | 0 | 0 | 1 | 853952 | A0A044SG03 | A0A044SG03 |
| LAEIPRKSVIKGEN | 0 | 0 | 1 | 853954 | A0A044SAZ1 | A0A044SAZ1 |
| LASSSATPVQQYD | 0 | 0 | 1 | 853970 | A0A044SS43 | A0A044SS43 |
| LATIELTHNWGTEND | 0 | 0 | 1 | 853972 | A0A044RAH9 | A0A044RAH9 |
| LCKFASNVTDVEELA | 0 | 0 | 1 | 853977 | A0A044QU24 | A0A044QU24 |
| LCPPGPTGPPGDKG | 0 | 0 | 1 | 853981 | A0A044V9I3 | A0A044V9I3 |
| LCTGEKGFGYEGSHF | 0 | 0 | 1 | 853984 | A0A044U9P0 | A0A044U9P0 |
| LCVTDIGEQWSESTH | 0 | 0 | 1 | 853987 | A0A044UK00 | A0A044UK00 |
| LDGEPGRPGPPGRDG | 0 | 0 | 1 | 853994 | A0A044V9I3 | A0A044V9I3 |
| LDGRHVVFGKVLEG | 0 | 0 | 1 | 853995 | A0A2K6WB53 | A0A2K6WB53 |
| LDKEINENVITISPD | 0 | 0 | 1 | 854000 | AAC48290.1 | A0A044VA40 |
| LDLEAQGNGIVIAQ | 0 | 0 | 1 | 854002 | A0A044RF80 | A0A044RF80 |
| LDLKLSNTVEVNL | 0 | 0 | 1 | 854003 | A0A044UK00 | A0A044UK00 |
| LDVHIESGEVCAMVF | 0 | 0 | 1 | 854007 | A0A044QLL7 | A0A044QLL7 |
| LECMLEEVEKEEEND | 0 | 0 | 1 | 854008 | A0A044UK00 | A0A044UK00 |
| LEDILPTIPSVTSL | 0 | 0 | 1 | 854009 | A0A044SAZ1 | A0A044SAZ1 |
| LEFITEEGYNIGAN | 0 | 0 | 1 | 854011 | A0A044UVG8 | A0A044UVG8 |
| LEGDYGETGRRGPPG | 0 | 0 | 1 | 854012 | A0A044V9I3 | A0A044V9I3 |
| LEIDVQNSVLPKYYP | 0 | 0 | 1 | 854016 | A0A044VBX6 | A0A044VBX6 |
| LEKQIQELNFQLEDD | 0 | 0 | 1 | 854020 | AAA74283.1 | A0A044V2F8 |
| LEPCTDELMIQFP | 0 | 0 | 1 | 854026 | A0A044T7P0 | A0A044T7P0 |
| LESPKEFPSPPPSPM | 0 | 0 | 1 | 854030 | A0A044UVG8 | A0A044UVG8 |
| LETLLENIDESVTEP | 0 | 0 | 1 | 854033 | A0A044T7P0 | A0A044T7P0 |
| LEVSEMTTAFQTQV | 0 | 0 | 1 | 854034 | A0A044R521 | A0A044R521 |
| LEYPTDLMAGQEAHV | 0 | 0 | 1 | 854036 | A0A044QSA2 | A0A044QSA2 |
| LFAGHDQCGGDT | 0 | 0 | 1 | 854037 | Q9NJD8.1 | Q9NJD8 |
| LFAKDEKQLQQS | 0 | 0 | 1 | 854040 | AAA63412.2 | Q99099 |
| LFAKELVEGIIN | 0 | 0 | 1 | 854041 | A0A044UK00 | A0A044UK00 |
| LFDNLLNYGINTNVE | 0 | 0 | 1 | 854043 | A0A044UIG6 | A0A044UIG6 |
| LFEEEITTDQD | 0 | 0 | 1 | 854045 | A0A044UK00 | A0A044UK00 |
| LFNEVDKNGDGYL | 0 | 0 | 1 | 854055 | AAF64251.1 | A0A044V6Z2 |
| LFQHMLKLEEEK | 0 | 0 | 1 | 854056 | A0A044TX11 | A0A044TX11 |
| LFVALPCASQEFDYE | 0 | 0 | 1 | 854062 | A0A044QNT8 | A0A044QNT8 |
| LFWEGLTAVKNSGE | 0 | 0 | 1 | 854064 | P41981.1 | A0A158N7R1 |
| LFYEEIPASAA | 0 | 0 | 1 | 854065 | A0A044R077 | A0A044R077 |
| LGAGALTMDDGSRFA | 0 | 0 | 1 | 854069 | A0A044V9R9 | A0A044V9R9 |
| LGDFMGNPDPALRSP | 0 | 0 | 1 | 854072 | A0A044U9P0 | A0A044U9P0 |
| LGDKFPDFQAETNE | 0 | 0 | 1 | 854073 | A0A044SIU8 | A0A044SIU8 |
| LGFWHEQSRPDRDNY | 0 | 0 | 1 | 854074 | AAV71152.1 | A0A044VAG6 |
| LGGRFGLLGTNDWE | 0 | 0 | 1 | 854075 | P46434.1 | P46434 |
| LGGSYKVRFTQFME | 0 | 0 | 1 | 854076 | P36991.1 | P36991 |
| LGLKPQNFAKMENM | 0 | 0 | 1 | 854078 | AAD00842.1 | A0A044SS43 |
| LGMPYAGDEVHL | 0 | 0 | 1 | 854081 | A0A044V3Q0 | A0A044V3Q0 |
| LGNIVAGANGVAE | 0 | 0 | 1 | 854082 | AAB64227.1 | O18483 |
| LGQGTGNMREESLK | 0 | 0 | 1 | 854083 | AAB64227.1 | O18483 |
| LGQSDFDDLENQPV | 0 | 0 | 1 | 854085 | AAF64254.1 | A0A044QLL7 |
| LGSAGPQGPPGAA | 0 | 0 | 1 | 854088 | A0A044UZ11 | A0A044UZ11 |
| LGSYYTGGMVDQ | 0 | 0 | 1 | 854089 | A0A044R153 | A0A044R153 |
| LHGFHVHQYGDQTNG | 0 | 0 | 1 | 854098 | AAB64227.1 | O18483 |
| LHLMLMRKLTFKSAR | 0 | 0 | 1 | 854101 | AAD00842.1 | A0A044SS43 |
| LHPTPSLTNEAFAQ | 0 | 0 | 1 | 854104 | A0A044UK00 | A0A044UK00 |
| LHQFYQSPAVEDIES | 0 | 0 | 1 | 854105 | A0A044U0M2 | A0A044U0M2 |
| LIHGKLKNRNGTYMP | 0 | 0 | 1 | 854118 | A0A044VD00 | A0A044VD00 |
| LILELQQRIGGSNEN | 0 | 0 | 1 | 854122 | A0A044UVG8 | A0A044UVG8 |
| LISQSDISTDTKLL | 0 | 0 | 1 | 854131 | A0A044V3Q0 | A0A044V3Q0 |
| LKATFPNTTKIITN | 0 | 0 | 1 | 854138 | AAA65186.1 | Q25619 |
| LKDACPECWQTEEPN | 0 | 0 | 1 | 854139 | A0A044RF80 | A0A044RF80 |
| LKEGGKETIDEQ | 0 | 0 | 1 | 854142 | AAB68960.1 | A0A044RRG7 |
| LKESTTGHIVGN | 0 | 0 | 1 | 854146 | P46434.1 | P46434 |
| LKNKSDKLYQKAVE | 0 | 0 | 1 | 854153 | AAA65186.1 | Q25619 |
| LKPIGLENVEENRRL | 0 | 0 | 1 | 854154 | AAB52600.1 | A0A044UUD9 |
| LKQPEPEIRLKAGE | 0 | 0 | 1 | 854155 | A0A044UK00 | A0A044UK00 |
| LKSMPGYKVSDFENG | 0 | 0 | 1 | 854158 | A0A044SAZ1 | A0A044SAZ1 |
| LKTEDDGTGVISE | 0 | 0 | 1 | 854159 | A0A044U9P0 | A0A044U9P0 |
| LKVPSDNTNTSKLEK | 0 | 0 | 1 | 854162 | A0A044UVG8 | A0A044UVG8 |
| LKYKLAVNMDEFMVM | 0 | 0 | 1 | 854163 | A0A044SAV6 | A0A044SAV6 |
| LKYTVFGRVIDGFDV | 0 | 0 | 1 | 854164 | A0A2K6VZ59 | A0A2K6VZ59 |
| LLDLQCYLHEVAEGK | 0 | 0 | 1 | 854167 | A0A044TU88 | A0A044TU88 |
| LLFTAGDEMSK | 0 | 0 | 1 | 854174 | AAB52600.1 | A0A044UUD9 |
| LLGTNDWEEAKIMA | 0 | 0 | 1 | 854178 | P46434.1 | P46434 |
| LLILSCTSDTQNSLS | 0 | 0 | 1 | 854180 | A0A044UK00 | A0A044UK00 |
| LLKEQERQKVLSQR | 0 | 0 | 1 | 854182 | A0A044TX11 | A0A044TX11 |
| LLPELSPTETEH | 0 | 0 | 1 | 854196 | A0A044SG03 | A0A044SG03 |
| LLRDIKDQTAG | 0 | 0 | 1 | 854197 | A0A044TU88 | A0A044TU88 |
| LLREKLTDLEETL | 0 | 0 | 1 | 854198 | A0A044UK00 | A0A044UK00 |
| LLSEAGDFFTKHF | 0 | 0 | 1 | 854200 | AAA63412.2 | Q99099 |
| LMAGQEAHVYKYADR | 0 | 0 | 1 | 854207 | A0A044R521 | A0A044R521 |
| LMDKDLQQNDEF | 0 | 0 | 1 | 854210 | A0A044UK00 | A0A044UK00 |
| LMKVNEQSNDE | 0 | 0 | 1 | 854214 | AAA29418.1 | P22085 |
| LMQKNGKSEVEIN | 0 | 0 | 1 | 854215 | A0A044SL50 | A0A044SL50 |
| LMTAIKKDFGSLE | 0 | 0 | 1 | 854218 | P41981.1 | A0A158N7R1 |
| LMTLDDEVKLEEYPQ | 0 | 0 | 1 | 854219 | P46434.1 | P46434 |
| LNADLDMLRGRWGKD | 0 | 0 | 1 | 854221 | A0A044V2F8 | A0A044V2F8 |
| LNAINQFPGKKP | 0 | 0 | 1 | 854222 | AAB52600.1 | A0A044UUD9 |
| LNEEIKVTIPVYSD | 0 | 0 | 1 | 854226 | A0A044UK00 | A0A044UK00 |
| LNKDDGTVNVINTD | 0 | 0 | 1 | 854233 | A0A044SD32 | A0A044SD32 |
| LNLNTSLSHLADEEA | 0 | 0 | 1 | 854234 | A0A044UVG8 | A0A044UVG8 |
| LNPEENGRYYEGDI | 0 | 0 | 1 | 854235 | AAV71152.1 | A0A044VAG6 |
| LNPTGKESDEKPKQ | 0 | 0 | 1 | 854239 | AAC48290.1 | A0A044VA40 |
| LNRTVSYDTEMLNEV | 0 | 0 | 1 | 854240 | A0A044TDA1 | A0A044TDA1 |

| | | | | | | |
|---|---|---|---|---|---|---|
| LNTTKTDARGNFKIY | 0 | 0 | 1 | 854245 | A0A044V9S3 | A0A044V9S3 |
| LNVTEEDNAIFSCTA | 0 | 0 | 1 | 854247 | A0A044UK00 | A0A044UK00 |
| LPDLPYDYGALEPI | 0 | 0 | 1 | 854249 | P41981.1 | A0A158N7R1 |
| LPFLDQNSERAEQLW | 0 | 0 | 1 | 854253 | A0A2K6VRV9 | A0A2K6VRV9 |
| LPHFIQPFALFEKDF | 0 | 0 | 1 | 854261 | A0A044UK00 | A0A044UK00 |
| LPQEAKRKAEEREMK | 0 | 0 | 1 | 854278 | A0A044TX11 | A0A044TX11 |
| LPQESRENALDFSDL | 0 | 0 | 1 | 854279 | A0A044UMF0 | A0A044UMF0 |
| LPTEAYVEVQEVHDD | 0 | 0 | 1 | 854285 | A0A044TU88 | A0A044TU88 |
| LQCKASKANMVDEI | 0 | 0 | 1 | 854290 | A0A044UK00 | A0A044UK00 |
| LQEIDSDGDGYINNS | 0 | 0 | 1 | 854291 | AAF64251.1 | A0A044V6Z2 |
| LQPLDEEKIRLSDG | 0 | 0 | 1 | 854301 | AAA20215.1 | A0A044V456 |
| LQQRALGMQIEQDRL | 0 | 0 | 1 | 854302 | AAC48290.1 | A0A044VA40 |
| LQQSSLSTKCLPN | 0 | 0 | 1 | 854303 | A0A044R153 | A0A044R153 |
| LQSGTNKFESQRGM | 0 | 0 | 1 | 854304 | AAA20215.1 | A0A044V456 |
| LQSYQLSNLDEKLD | 0 | 0 | 1 | 854305 | A0A044RF80 | A0A044RF80 |
| LQTTLGDIKIELYCD | 0 | 0 | 1 | 854307 | A0A2K6VZ59 | A0A2K6VZ59 |
| LQTVDKDHNGLL | 0 | 0 | 1 | 854308 | AAF64251.1 | A0A044V6Z2 |
| LQVDTGENEKIPFVL | 0 | 0 | 1 | 854309 | AAF64254.1 | A0A044QLL7 |
| LRAAVEKDTVEVVA | 0 | 0 | 1 | 854311 | O01360.1 | O01360 |
| LRENIYEEDESTD | 0 | 0 | 1 | 854320 | A0A044UK00 | A0A044UK00 |
| LRPEDLASGARCSNC | 0 | 0 | 1 | 854330 | A0A044U9P0 | A0A044U9P0 |
| LRRGELRYNPNGDIR | 0 | 0 | 1 | 854333 | A0A044V9G2 | A0A044V9G2 |
| LRVSAVPRMSDISEH | 0 | 0 | 1 | 854338 | A0A044UMF0 | A0A044UMF0 |
| LSCDSADSHSKWAD | 0 | 0 | 1 | 854346 | AAA50214.2 | P52570 |
| LSCNQHRQKGCEG | 0 | 0 | 1 | 854347 | A0A044RFT9 | A0A044RFT9 |
| LSEETDLNLNTSLSH | 0 | 0 | 1 | 854349 | A0A044UVG8 | A0A044UVG8 |
| LSEIIQADIFKLSR | 0 | 0 | 1 | 854350 | A0A044UK00 | A0A044UK00 |
| LSENCIITDDF | 0 | 0 | 1 | 854352 | A0A044V9S3 | A0A044V9S3 |
| LSIQTMEKEIDIHRE | 0 | 0 | 1 | 854360 | A0A044UK00 | A0A044UK00 |
| LSQMGPPPNPIS | 0 | 0 | 1 | 854372 | A0A044V1C1 | A0A044V1C1 |
| LSSITSTASQDGSL | 0 | 0 | 1 | 854376 | A0A044UK00 | A0A044UK00 |
| LSWAQQSNKFG | 0 | 0 | 1 | 854380 | AAC48290.1 | A0A044VA40 |
| LSYGGYNFGSS | 0 | 0 | 1 | 854382 | AAC47023.1 | A0A044RRG7 |
| LSYQVGYLTGGETD | 0 | 0 | 1 | 854383 | A0A044UK00 | A0A044UK00 |
| LTDAQKDELRKIAN | 0 | 0 | 1 | 854387 | AAB52600.1 | A0A044UUD9 |
| LTDEPFQRKDIIVLQ | 0 | 0 | 1 | 854388 | CAA04390.1 | A0A044R436 |
| LTDGLKRMIGLDQN | 0 | 0 | 1 | 854389 | AAC48290.1 | A0A044VA40 |
| LTGGSGHGNQDGIA | 0 | 0 | 1 | 854395 | A0A2K6WJQ8 | A0A2K6WJQ8 |
| LTICNIDRRDEGT | 0 | 0 | 1 | 854396 | A0A044UK00 | A0A044UK00 |
| LTINFNTRNTFEGH | 0 | 0 | 1 | 854399 | A0A044R521 | A0A044R521 |
| LTNEAFAQLYDRVN | 0 | 0 | 1 | 854404 | A0A044UK00 | A0A044UK00 |
| LTRDGSMDEAQAD | 0 | 0 | 1 | 854406 | A0A044V3Q0 | A0A044V3Q0 |
| LTSDHIYNDSV | 0 | 0 | 1 | 854408 | AAF64254.1 | A0A044QLL7 |
| LTSQANSTLPVKENI | 0 | 0 | 1 | 854410 | A0A044UK00 | A0A044UK00 |
| LTSSVASQDVLHFTQ | 0 | 0 | 1 | 854413 | AAB97282.1 | A0A044UNG9 |
| LTSTTMEPITHQKAA | 0 | 0 | 1 | 854414 | A0A044R436 | A0A044R436 |
| LTTTEWKETYGG | 0 | 0 | 1 | 854415 | CAA04390.1 | A0A044R436 |
| LTVFNMLMTLDDE | 0 | 0 | 1 | 854417 | P46434.1 | P46434 |
| LTYMIRDDTDKAL | 0 | 0 | 1 | 854420 | A0A044RF80 | A0A044RF80 |
| LVARFYIQDEIEKI | 0 | 0 | 1 | 854422 | A0A044UK00 | A0A044UK00 |
| LVATPVDWKDGDDC | 0 | 0 | 1 | 854423 | AAA50214.2 | P52570 |
| LVIKRQGNCVRVA | 0 | 0 | 1 | 854428 | A0A044TL01 | A0A044TL01 |
| LVKGNFANKDGNSMP | 0 | 0 | 1 | 854431 | A0A044UNG9 | A0A044UNG9 |
| LVKVAEIVASKLVA | 0 | 0 | 1 | 854432 | A0A044UK00 | A0A044UK00 |
| LVQNVTVSESNETEK | 0 | 0 | 1 | 854438 | A0A044UK00 | A0A044UK00 |
| LVTDQTFWPAPFMKP | 0 | 0 | 1 | 854440 | A0A044UW71 | A0A044UW71 |
| LVVSIVTSKKKKGND | 0 | 0 | 1 | 854444 | A0A044T7P0 | A0A044T7P0 |
| LWERDTLDPDDLLN | 0 | 0 | 1 | 854446 | A0A044V9S3 | A0A044V9S3 |
| LYAIMQHETAHYE | 0 | 0 | 1 | 854448 | A0A044UK00 | A0A044UK00 |
| LYDQAGCRNDEG | 0 | 0 | 1 | 854450 | A0A044RMM8 | A0A044RMM8 |
| LYKMKCVGCDSEKKL | 0 | 0 | 1 | 854454 | AAA50214.2 | P52570 |
| LYLNSDDRNMYDNN | 0 | 0 | 1 | 854455 | A0A044SG03 | A0A044SG03 |
| MAALKLPNTPEIVR | 0 | 0 | 1 | 854460 | A0A2K6WPE1 | A0A2K6WPE1 |
| MAGTVGEGTTQG | 0 | 0 | 1 | 854467 | AAB52600.1 | A0A044UUD9 |
| MAPYKEDEGKVNS | 0 | 0 | 1 | 854472 | A0A044TU88 | A0A044TU88 |
| MDDEYTALMAELGE | 0 | 0 | 1 | 854479 | A0A044U9P0 | A0A044U9P0 |
| MDEAQADFALDLIRT | 0 | 0 | 1 | 854480 | A0A044V3Q0 | A0A044V3Q0 |
| MDENKSYEVPECEDR | 0 | 0 | 1 | 854481 | A0A044TX11 | A0A044TX11 |
| MDITRSSQPHLEP | 0 | 0 | 1 | 854483 | A0A044UK00 | A0A044UK00 |
| MDMLLGGPGQNPNLS | 0 | 0 | 1 | 854484 | A0A044UW71 | A0A044UW71 |
| MDSLSESVRREI | 0 | 0 | 1 | 854488 | P36991.1 | P36991 |
| MEIREEQSGNLAEDN | 0 | 0 | 1 | 854495 | A0A044UK00 | A0A044UK00 |
| MEIVENGNCTNVET | 0 | 0 | 1 | 854496 | A0A044UK00 | A0A044UK00 |
| MELMASTGETLTKYS | 0 | 0 | 1 | 854497 | A0A044UK00 | A0A044UK00 |
| MFESVNKNITIRQ | 0 | 0 | 1 | 854505 | A0A044U175 | A0A044U175 |
| MGPPGKAGPPGKD | 0 | 0 | 1 | 854516 | A0A044TIV1 | A0A044TIV1 |
| MGQMAATAGGVAIG | 0 | 0 | 1 | 854517 | A0A2K6WJQ8 | A0A2K6WJQ8 |
| MGYPGPPGAPGLQGT | 0 | 0 | 1 | 854519 | A0A044SAV6 | A0A044SAV6 |
| MHLKASKEENISLVA | 0 | 0 | 1 | 854523 | A0A044UK00 | A0A044UK00 |
| MIAEMEREAAERRMA | 0 | 0 | 1 | 854526 | AAC48290.1 | A0A044VA40 |
| MIENPRNSSSEDENF | 0 | 0 | 1 | 854529 | A0A044UK00 | A0A044UK00 |
| MIKPDAKRIDINFHK | 0 | 0 | 1 | 854532 | AAD00843.1 | A0A044VHF7 |
| MIMTPESRTDEKASN | 0 | 0 | 1 | 854538 | AAC48290.1 | A0A044VA40 |
| MIQLFSTTTASML | 0 | 0 | 1 | 854544 | A0A044RJ74 | A0A044RJ74 |
| MKFENLLGQRGTAVA | 0 | 0 | 1 | 854549 | A0A044V1C1 | A0A044V1C1 |
| MKLLKKVRPEDIDL | 0 | 0 | 1 | 854556 | A0A044UVG8 | A0A044UVG8 |
| MKSKDNEKVFQIERA | 0 | 0 | 1 | 854559 | A0A044UK00 | A0A044UK00 |
| MKVGQEKPKDTQHYF | 0 | 0 | 1 | 854561 | A0A044RF80 | A0A044RF80 |
| MLNVSGNVLGESHAI | 0 | 0 | 1 | 854571 | P46434.1 | P46434 |
| MLPVMIANNSNLDI | 0 | 0 | 1 | 854572 | A0A044S628 | A0A044S628 |
| MLQSQLMITYEKGK | 0 | 0 | 1 | 854574 | A0A044S6B8 | A0A044S6B8 |

| | | | | | | |
|---|---|---|---|---|---|---|
| MLQTAIPIMSPLAPS | 0 | 0 | 1 | 854575 | AAC48290.1 | A0A044VA40 |
| MMPIKLMPPQMINR | 0 | 0 | 1 | 854587 | AAC48290.1 | A0A044VA40 |
| MMQVTIVECGELKQE | 0 | 0 | 1 | 854588 | A0A044U9P0 | A0A044U9P0 |
| MNQADSANLPSQA | 0 | 0 | 1 | 854596 | A0A044VBX6 | A0A044VBX6 |
| MPCNIDFSHYSDAVM | 0 | 0 | 1 | 854607 | A0A044V1C1 | A0A044V1C1 |
| MPLPNMSNNTGGN | 0 | 0 | 1 | 854621 | A0A044V9G2 | A0A044V9G2 |
| MPPQMINREEPIIQ | 0 | 0 | 1 | 854625 | AAC48290.1 | A0A044VA40 |
| MQEEVAEYRRKFD | 0 | 0 | 1 | 854632 | AAA74283.1 | A0A044V2F8 |
| MQGGAAEGGVPL | 0 | 0 | 1 | 854633 | A0A044RF80 | A0A044RF80 |
| MQIVPGKTDRISHDL | 0 | 0 | 1 | 854636 | A0A044UK00 | A0A044UK00 |
| MQKNDGNEMICIEEN | 0 | 0 | 1 | 854638 | A0A044UVG8 | A0A044UVG8 |
| MQKTTLGKIGDLISQ | 0 | 0 | 1 | 854639 | AAA63412.2 | Q99099 |
| MQKYAIFNSVTSNTD | 0 | 0 | 1 | 854640 | A0A044R153 | A0A044R153 |
| MREIMVNYATSESEA | 0 | 0 | 1 | 854649 | A0A044UVG8 | A0A044UVG8 |
| MRNTAPSMMYVE | 0 | 0 | 1 | 854653 | AAC48290.1 | A0A044VA40 |
| MRRARERSVDTSVD | 0 | 0 | 1 | 854655 | A0A044UVG8 | A0A044UVG8 |
| MSDLNDRLADYIEK | 0 | 0 | 1 | 854658 | AAA74283.1 | A0A044V2F8 |
| MSVQAYQIEKDNPNF | 0 | 0 | 1 | 854673 | A0A044SAZ1 | A0A044SAZ1 |
| MTFEQYRYIVKTDAI | 0 | 0 | 1 | 854678 | A0A044VBX6 | A0A044VBX6 |
| MTKIFALNPTGKESD | 0 | 0 | 1 | 854684 | AAC48290.1 | A0A044VA40 |
| MTLINDCASLPGKS | 0 | 0 | 1 | 854687 | A0A044UW71 | A0A044UW71 |
| MTNEYETNYPVPYR | 0 | 0 | 1 | 854688 | AAA20541.1 | A0A044QNW3 |
| MTPLTEEEYQYSLQW | 0 | 0 | 1 | 854689 | A0A044UVG8 | A0A044UVG8 |
| MTPVIHEPLRGQNVS | 0 | 0 | 1 | 854690 | A0A044UK00 | A0A044UK00 |
| MTTIDGNSDNLSES | 0 | 0 | 1 | 854692 | AAC48289.1 | A0A044VA40 |
| MTVDVTERAEKTAVP | 0 | 0 | 1 | 854694 | A0A044UVG8 | A0A044UVG8 |
| MVNHQEYFPHFGTNA | 0 | 0 | 1 | 854699 | A0A044R2U8 | A0A044R2U8 |
| MYSRVTKLKENDPEM | 0 | 0 | 1 | 854706 | AAC47023.1 | A0A044RRG7 |
| NADADGCGGVGSC | 0 | 0 | 1 | 854708 | A0A2K6VRV9 | A0A2K6VRV9 |
| NADKNDDGHID | 0 | 0 | 1 | 854710 | AAF64251.1 | A0A044V6Z2 |
| NAENKRIPISMENNQ | 0 | 0 | 1 | 854712 | A0A044SS43 | A0A044SS43 |
| NAGPDTNGSQFFLC | 0 | 0 | 1 | 854715 | AAC47233.1 | Q25633 |
| NAIVKLCSSSNNSEN | 0 | 0 | 1 | 854718 | A0A044UK00 | A0A044UK00 |
| NAKQLKILDRNDD | 0 | 0 | 1 | 854720 | A0A044SS43 | A0A044SS43 |
| NAMSNYNREQLESDA | 0 | 0 | 1 | 854721 | A0A044UVG8 | A0A044UVG8 |
| NATTTVVFEKP | 0 | 0 | 1 | 854726 | A0A044UK00 | A0A044UK00 |
| NCCEHQAELYC | 0 | 0 | 1 | 854728 | A0A044V9R9 | A0A044V9R9 |
| NCTSSAVSDFFASIE | 0 | 0 | 1 | 854730 | A0A044UMF0 | A0A044UMF0 |
| NDDINVWRDKAG | 0 | 0 | 1 | 854733 | A0A044TDA1 | A0A044TDA1 |
| NDDNKFKYELS | 0 | 0 | 1 | 854734 | A0A044UK00 | A0A044UK00 |
| NDEELQKDSRG | 0 | 0 | 1 | 854736 | AAD00842.1 | A0A044SS43 |
| NDGNISIMEWGKC | 0 | 0 | 1 | 854738 | A0A044T7P0 | A0A044T7P0 |
| NDIIIENNENYS | 0 | 0 | 1 | 854739 | A0A044UK00 | A0A044UK00 |
| NDKENIEAVTDTPPK | 0 | 0 | 1 | 854742 | A0A044UK00 | A0A044UK00 |
| NDKIKNLISQSDI | 0 | 0 | 1 | 854743 | A0A044V3Q0 | A0A044V3Q0 |
| NDKITLSNFGEPQSN | 0 | 0 | 1 | 854744 | A0A044UK00 | A0A044UK00 |
| NDLNVDEMTESD | 0 | 0 | 1 | 854747 | A0A044SS43 | A0A044SS43 |
| NDMCIGAVSEKATNL | 0 | 0 | 1 | 854748 | A0A044TBR7 | A0A044TBR7 |
| NEDDEEIQEVAE | 0 | 0 | 1 | 854753 | AAD51087.1 | Q9U9A1 |
| NEERETRFKLRQFE | 0 | 0 | 1 | 854754 | A0A044UK00 | A0A044UK00 |
| NEILGKHDDKDE | 0 | 0 | 1 | 854757 | A0A044UVG8 | A0A044UVG8 |
| NEITELRMPGAE | 0 | 0 | 1 | 854758 | A0A044U175 | A0A044U175 |
| NEIYEIEEFATH | 0 | 0 | 1 | 854761 | A0A044UK00 | A0A044UK00 |
| NEKGTEAAAASA | 0 | 0 | 1 | 854762 | A0A044V3Q0 | A0A044V3Q0 |
| NEKMKYDNLYKLVK | 0 | 0 | 1 | 854764 | AAD51087.1 | Q9U9A1 |
| NEMICIEENITQKN | 0 | 0 | 1 | 854765 | A0A044UVG8 | A0A044UVG8 |
| NEMIRHPSNNPHE | 0 | 0 | 1 | 854767 | A0A044U0M2 | A0A044U0M2 |
| NENQESQINANTVG | 0 | 0 | 1 | 854771 | A0A044U9P0 | A0A044U9P0 |
| NENQSWVDKGCYCDR | 0 | 0 | 1 | 854772 | A0A044QNT8 | A0A044QNT8 |
| NEPLHAYVTGTDYTV | 0 | 0 | 1 | 854774 | A0A044U9P0 | A0A044U9P0 |
| NERIHKSFSNRDVYD | 0 | 0 | 1 | 854778 | A0A044V9G2 | A0A044V9G2 |
| NESEVVGLQDNTGY | 0 | 0 | 1 | 854779 | A0A044UK00 | A0A044UK00 |
| NEVNNIEPYLII | 0 | 0 | 1 | 854783 | A0A044V9S3 | A0A044V9S3 |
| NEVNNIPESKFA | 0 | 0 | 1 | 854784 | A0A044TDA1 | A0A044TDA1 |
| NEYDQMTNANRN | 0 | 0 | 1 | 854785 | AAA74283.1 | A0A044V2F8 |
| NFAKMENMFTSFHI | 0 | 0 | 1 | 854787 | AAD00842.1 | A0A044SS43 |
| NFGDNSPKQTNADDL | 0 | 0 | 1 | 854792 | A0A044U0M2 | A0A044U0M2 |
| NGCPETNIQQQYPN | 0 | 0 | 1 | 854797 | A0A044S6B8 | A0A044S6B8 |
| NGEFDEEAVLNFMIT | 0 | 0 | 1 | 854798 | A0A044SAZ1 | A0A044SAZ1 |
| NGETGPVGKEGP | 0 | 0 | 1 | 854800 | A0A044SAV6 | A0A044SAV6 |
| NGKEIQADNRNTFTI | 0 | 0 | 1 | 854802 | A0A044UK00 | A0A044UK00 |
| NGKIESLTVGVDYLD | 0 | 0 | 1 | 854803 | A0A044UW71 | A0A044UW71 |
| NGSAEMKKQIDMQ | 0 | 0 | 1 | 854806 | A0A044RF80 | A0A044RF80 |
| NGYLPAHESYMRS | 0 | 0 | 1 | 854812 | A0A044V9G2 | A0A044V9G2 |
| NIPEEYKEFIPEE | 0 | 0 | 1 | 854826 | Q25619.1 | Q25619 |
| NIQQQYPNINEAMLQ | 0 | 0 | 1 | 854828 | A0A044S6B8 | A0A044S6B8 |
| NISEDSMKFRNITS | 0 | 0 | 1 | 854830 | AAY67786.1 | A0A158N7U2 |
| NISKVETIIGNATKR | 0 | 0 | 1 | 854831 | A0A044UK00 | A0A044UK00 |
| NIVIRESLNGDTSKS | 0 | 0 | 1 | 854837 | A0A044UVG8 | A0A044UVG8 |
| NKALQKGLDPEEAA | 0 | 0 | 1 | 854839 | A0A044R196 | A0A044R196 |
| NKCFKLSGGEVEVYP | 0 | 0 | 1 | 854840 | A0A2K6VRV9 | A0A2K6VRV9 |
| NKDGVLHEKSDVNVP | 0 | 0 | 1 | 854841 | A0A044SAZ1 | A0A044SAZ1 |
| NKENKEFTVGNTAN | 0 | 0 | 1 | 854842 | A0A044U175 | A0A044U175 |
| NKEVHVKKGENVQLK | 0 | 0 | 1 | 854843 | A0A044UK00 | A0A044UK00 |
| NKFDGFDFDWEYPIG | 0 | 0 | 1 | 854845 | AAC47023.1 | A0A044RRG7 |
| NKLKELRGYIKSK | 0 | 0 | 1 | 854848 | A0A044UVG8 | A0A044UVG8 |
| NKTRLNNVIENDL | 0 | 0 | 1 | 854851 | A0A044QWA5 | A0A044QWA5 |
| NLHNTSADFSGNDVP | 0 | 0 | 1 | 854862 | AAA20541.1 | A0A044QNW3 |
| NLLKQHEDSSSA | 0 | 0 | 1 | 854864 | AAD00842.1 | A0A044SS43 |
| NLLKTDENKTD | 0 | 0 | 1 | 854865 | P36991.1 | P36991 |
| NLNEFQVDNQVFGKE | 0 | 0 | 1 | 854868 | A0A044UVG8 | A0A044UVG8 |

| | | | | | | |
|---|---|---|---|---|---|---|
| NLNGVPLDIAGFVYN | 0 | 0 | 1 | 854869 | A0A044SAZ1 | A0A044SAZ1 |
| NLSDDQRQAYLLQ | 0 | 0 | 1 | 854872 | A0A044U9P0 | A0A044U9P0 |
| NMLIFKDAFTDDELA | 0 | 0 | 1 | 854881 | A0A044SL50 | A0A044SL50 |
| NMREESLKTGNAG | 0 | 0 | 1 | 854887 | AAB64227.1 | O18483 |
| NMTTINTSKFNSLM | 0 | 0 | 1 | 854888 | A0A044U175 | A0A044U175 |
| NMYERDHRHEQI | 0 | 0 | 1 | 854891 | A0A044V9G2 | A0A044V9G2 |
| NNDGETRQSDDN | 0 | 0 | 1 | 854893 | AAC48290.1 | A0A044VA40 |
| NNENYSTISDDG | 0 | 0 | 1 | 854894 | A0A044UK00 | A0A044UK00 |
| NNFLAPGSLIDWFKG | 0 | 0 | 1 | 854896 | A0A044UK00 | A0A044UK00 |
| NNIIDTVRDAK | 0 | 0 | 1 | 854897 | A0A044SS43 | A0A044SS43 |
| NNMTWKVAGQG | 0 | 0 | 1 | 854901 | AAB69625.2 | A0A044VD00 |
| NNNNNSNSNRI | 0 | 0 | 1 | 854902 | A0A044QSA2 | A0A044QSA2 |
| NNNSLATLWSPDYPS | 0 | 0 | 1 | 854903 | A0A044SG03 | A0A044SG03 |
| NNPSNKLTSSVA | 0 | 0 | 1 | 854904 | AAB97282.1 | A0A044UNG9 |
| NNQENDGFDGQKA | 0 | 0 | 1 | 854905 | A0A044SS43 | A0A044SS43 |
| NNSEAVAMARRQD | 0 | 0 | 1 | 854908 | AAF64251.1 | A0A044V6Z2 |
| NNVWGREEREGTL | 0 | 0 | 1 | 854910 | AAD00843.1 | A0A044VHF7 |
| NPAKPNAMPIQIRD | 0 | 0 | 1 | 854912 | AAC48290.1 | A0A044VA40 |
| NPEDIGCNGTTMKEL | 0 | 0 | 1 | 854914 | AAK66564.1 | Q963F6 |
| NPEICQIQKSEHN | 0 | 0 | 1 | 854915 | A0A044RFT9 | A0A044RFT9 |
| NPIENEQEDCRSNDE | 0 | 0 | 1 | 854918 | A0A044SS43 | A0A044SS43 |
| NPKCNCDTSNRCPPG | 0 | 0 | 1 | 854920 | A0A044SAV6 | A0A044SAV6 |
| NPKMDFTNKENKWE | 0 | 0 | 1 | 854921 | AAA63412.2 | Q99099 |
| NPMLQASSVEQAPAA | 0 | 0 | 1 | 854924 | A0A2K6VJ98 | A0A2K6VJ98 |
| NPSAEGEDGDEG | 0 | 0 | 1 | 854926 | A0A044SL50 | A0A044SL50 |
| NQGTDTNKSQF | 0 | 0 | 1 | 854929 | CAA04390.1 | A0A044R436 |
| NQNQGKANCKC | 0 | 0 | 1 | 854931 | A0A044UZ11 | A0A044UZ11 |
| NQSFELQEETTND | 0 | 0 | 1 | 854933 | A0A044QWA5 | A0A044QWA5 |
| NQSISDKYISIES | 0 | 0 | 1 | 854934 | A0A044TDA1 | A0A044TDA1 |
| NQWINNISTEL | 0 | 0 | 1 | 854937 | A0A044SAZ1 | A0A044SAZ1 |
| NRDLHTSVEAKVVA | 0 | 0 | 1 | 854939 | A0A044RXP7 | A0A044RXP7 |
| NREEPIIQPIESNPA | 0 | 0 | 1 | 854940 | AAC48290.1 | A0A044VA40 |
| NRIPDNMIMTPESRT | 0 | 0 | 1 | 854944 | AAC48290.1 | A0A044VA40 |
| NRQSVINATTTV | 0 | 0 | 1 | 854946 | A0A044UK00 | A0A044UK00 |
| NRRLSTGKDSGRG | 0 | 0 | 1 | 854948 | A0A044UMF0 | A0A044UMF0 |
| NRVEAGTFDQKSPLT | 0 | 0 | 1 | 854949 | AAD00843.1 | A0A044VHF7 |
| NSDNLSESSDSPSPF | 0 | 0 | 1 | 854950 | AAC48290.1 | A0A044VA40 |
| NSDRFAKQGNSTISN | 0 | 0 | 1 | 854952 | A0A044UIG6 | A0A044UIG6 |
| NSDRIISKEGDFYSC | 0 | 0 | 1 | 854953 | A0A044UK00 | A0A044UK00 |
| NSERAEQLWNLF | 0 | 0 | 1 | 854954 | A0A2K6VRV9 | A0A2K6VRV9 |
| NSLRMIDNNADPIWT | 0 | 0 | 1 | 854962 | A0A044QWA5 | A0A044QWA5 |
| NSLSRQNGYLPAHE | 0 | 0 | 1 | 854963 | A0A044V9G2 | A0A044V9G2 |
| NSSSEDENFEDIQK | 0 | 0 | 1 | 854967 | A0A044UK00 | A0A044UK00 |
| NSTDFANTGMRYC | 0 | 0 | 1 | 854968 | AAV71152.1 | A0A044VAG6 |
| NSTRSDEKGAFEVYG | 0 | 0 | 1 | 854970 | A0A044V9S3 | A0A044V9S3 |
| NSVSSIPIDNGVEGE | 0 | 0 | 1 | 854973 | A0A044R521 | A0A044R521 |
| NSWGTDWGENGTFRI | 0 | 0 | 1 | 854974 | A0A044RFT9 | A0A044RFT9 |
| NTAEQKASNSSQ | 0 | 0 | 1 | 854975 | A0A044UVG8 | A0A044UVG8 |
| NTFFNDKIKVCDH | 0 | 0 | 1 | 854978 | AAB68960.1 | A0A044RRG7 |
| NTIGELIYEQGSPCK | 0 | 0 | 1 | 854981 | AAB97282.1 | A0A044UNG9 |
| NTNNSSDNLAFLDNL | 0 | 0 | 1 | 854982 | A0A044UK00 | A0A044UK00 |
| NTNTSKLEKRETKIT | 0 | 0 | 1 | 854983 | A0A044UVG8 | A0A044UVG8 |
| NTPRPRPSYMPNAA | 0 | 0 | 1 | 854984 | A0A044U9P0 | A0A044U9P0 |
| NTRVGEQYKVSSPT | 0 | 0 | 1 | 854985 | A0A044V1C1 | A0A044V1C1 |
| NTTSNQIESGLDFA | 0 | 0 | 1 | 854988 | A0A044UK00 | A0A044UK00 |
| NVAGDNLRISREM | 0 | 0 | 1 | 854993 | A0A044VFS3 | A0A044VFS3 |
| NVEFWERFGSNASGK | 0 | 0 | 1 | 854998 | A0A044UIG6 | A0A044UIG6 |
| NVIDDIVNKQLAK | 0 | 0 | 1 | 855001 | A0A044TX11 | A0A044TX11 |
| NVIENDLLFSSGEC | 0 | 0 | 1 | 855002 | A0A044QWA5 | A0A044QWA5 |
| NVKLIGLSCDSADSH | 0 | 0 | 1 | 855004 | AAA50214.2 | P52570 |
| NVLGESHAIELLLGG | 0 | 0 | 1 | 855005 | P46434.1 | P46434 |
| NVNRNQQIESTR | 0 | 0 | 1 | 855006 | A0A044V1C1 | A0A044V1C1 |
| NVPVTWSQFEVQMMD | 0 | 0 | 1 | 855007 | A0A044SAZ1 | A0A044SAZ1 |
| NVRPKSGAGRGKDMA | 0 | 0 | 1 | 855009 | A0A044SS43 | A0A044SS43 |
| NVSSQNQPVKNIVLG | 0 | 0 | 1 | 855012 | A0A044UK00 | A0A044UK00 |
| NVTGRLEFKMQAHCD | 0 | 0 | 1 | 855015 | A0A044QU24 | A0A044QU24 |
| NVVKTSGDIAENFES | 0 | 0 | 1 | 855017 | A0A044UVG8 | A0A044UVG8 |
| NWQPGSDTIKPGVKE | 0 | 0 | 1 | 855019 | A0A044VFS3 | A0A044VFS3 |
| NYATSESEAEDNITV | 0 | 0 | 1 | 855021 | A0A044UVG8 | A0A044UVG8 |
| NYGINTNVEFWERFG | 0 | 0 | 1 | 855024 | A0A044UIG6 | A0A044UIG6 |
| NYYSNKRTQSDDD | 0 | 0 | 1 | 855032 | A0A044SS43 | A0A044SS43 |
| PAAPASAEDEETKE | 0 | 0 | 1 | 855036 | A0A044R196 | A0A044R196 |
| PAEQTEAEMAMLAA | 0 | 0 | 1 | 855037 | A0A044R196 | A0A044R196 |
| PAIFEWFCNDKPVQ | 0 | 0 | 1 | 855040 | A0A044UK00 | A0A044UK00 |
| PAISGKNEMIRHP | 0 | 0 | 1 | 855041 | A0A044U0M2 | A0A044U0M2 |
| PANWNLQQRDSPIAA | 0 | 0 | 1 | 855042 | A0A044R153 | A0A044R153 |
| PANWQPGSETIKPER | 0 | 0 | 1 | 855043 | A0A044VFS3 | A0A044VFS3 |
| PAQISSDEIFR | 0 | 0 | 1 | 855045 | AAC48290.1 | A0A044VA40 |
| PATTGRNFDEILR | 0 | 0 | 1 | 855048 | AAA50214.2 | P52570 |
| PCQPILACMPDNTKG | 0 | 0 | 1 | 855053 | A0A044RJ74 | A0A044RJ74 |
| PDCYDQREPGDWCI | 0 | 0 | 1 | 855057 | AAD27587.1 | A0A158N7U8 |
| PDDLLNATKSNSKGY | 0 | 0 | 1 | 855058 | A0A044V9S3 | A0A044V9S3 |
| PDFVLTHTENKNTTL | 0 | 0 | 1 | 855060 | A0A044SG03 | A0A044SG03 |
| PDFYLPEGVGFKVG | 0 | 0 | 1 | 855061 | A0A044U175 | A0A044U175 |
| PDIDIPTTSQDVQQ | 0 | 0 | 1 | 855063 | A0A044UK00 | A0A044UK00 |
| PDIHPHPDLSTRT | 0 | 0 | 1 | 855064 | A0A044UK00 | A0A044UK00 |
| PDLQSHHELKAIDD | 0 | 0 | 1 | 855065 | A0A044V1C1 | A0A044V1C1 |
| PDLSTRTYLLHSA | 0 | 0 | 1 | 855066 | A0A044UK00 | A0A044UK00 |
| PDVDTIDDHGKLEAI | 0 | 0 | 1 | 855070 | A0A044TX11 | A0A044TX11 |
| PEFKKRNVKLIGLSC | 0 | 0 | 1 | 855073 | P52570.1 | P52570 |
| PEHKVDDVNEYN | 0 | 0 | 1 | 855074 | A0A044QNT8 | A0A044QNT8 |

| | | | | | | |
|---|---|---|---|---|---|---|
| PENAGKLESPKEFPS | 0 | 0 | 1 | 855081 | A0A044UVG8 | A0A044UVG8 |
| PEQTAEEITEP | 0 | 0 | 1 | 855083 | AAC48290.1 | A0A044VA40 |
| PESKFAIVRNSNGD | 0 | 0 | 1 | 855085 | A0A044TDA1 | A0A044TDA1 |
| PEVTAATKRTANDIA | 0 | 0 | 1 | 855089 | CAA04390.1 | A0A044R436 |
| PEVVDKLKTEDDG | 0 | 0 | 1 | 855090 | A0A044U9P0 | A0A044U9P0 |
| PFAFRAHTHRHGEKV | 0 | 0 | 1 | 855091 | A0A044U175 | A0A044U175 |
| PFALFEKDFSTGLR | 0 | 0 | 1 | 855092 | A0A044UK00 | A0A044UK00 |
| PFEEAEIAVEKEREN | 0 | 0 | 1 | 855094 | CAA04390.1 | A0A044R436 |
| PFEMVESEEDEDSDD | 0 | 0 | 1 | 855095 | A0A044SS43 | A0A044SS43 |
| PFGHVPMLNVSGNVL | 0 | 0 | 1 | 855098 | P46434.1 | P46434 |
| PFIEFPPTLPRDRDA | 0 | 0 | 1 | 855099 | A0A044V9G2 | A0A044V9G2 |
| PFQEGQTVHAVGMIK | 0 | 0 | 1 | 855100 | AAD00843.1 | A0A044VHF7 |
| PFTVDSANQVHCV | 0 | 0 | 1 | 855103 | A0A044QS56 | A0A044QS56 |
| PGAAGPRGPPGAA | 0 | 0 | 1 | 855105 | A0A044UZ11 | A0A044UZ11 |
| PGASGEPGPRGPIGP | 0 | 0 | 1 | 855108 | A0A044SAV6 | A0A044SAV6 |
| PGEPGPDGRPGNDGR | 0 | 0 | 1 | 855112 | A0A044V9I3 | A0A044V9I3 |
| PGGYETYDREVKSN | 0 | 0 | 1 | 855116 | A0A044V9G2 | A0A044V9G2 |
| PGIVFLSGGQSEEDA | 0 | 0 | 1 | 855118 | AAB52600.1 | A0A044UUD9 |
| PGKGILAADES | 0 | 0 | 1 | 855120 | AAB52600.1 | A0A044UUD9 |
| PGMPGPVGQSGDNGE | 0 | 0 | 1 | 855123 | A0A044SAV6 | A0A044SAV6 |
| PGPAGSDGKPGP | 0 | 0 | 1 | 855126 | A0A044TIV1 | A0A044TIV1 |
| PGPNGERGADGEPGE | 0 | 0 | 1 | 855131 | A0A044V9I3 | A0A044V9I3 |
| PGPPGHMGYPGPPGA | 0 | 0 | 1 | 855132 | A0A044SAV6 | A0A044SAV6 |
| PGPVGDAGAPGEMGP | 0 | 0 | 1 | 855138 | A0A044TIV1 | A0A044TIV1 |
| PGSGVFNVGDGGMDD | 0 | 0 | 1 | 855144 | A0A044U9P0 | A0A044U9P0 |
| PGSPGLPGPDAHYCP | 0 | 0 | 1 | 855145 | A0A044V9I3 | A0A044V9I3 |
| PGVDEFEEDENAPPI | 0 | 0 | 1 | 855147 | A0A044R196 | A0A044R196 |
| PIDNGVEGEPEIECG | 0 | 0 | 1 | 855156 | A0A044R521 | A0A044R521 |
| PIGMAQEYAKLVNEM | 0 | 0 | 1 | 855158 | AAC47023.1 | A0A044RRG7 |
| PIKTFEHVPSEIGAE | 0 | 0 | 1 | 855161 | A0A044TU88 | A0A044TU88 |
| PISIPDPSSKNLPD | 0 | 0 | 1 | 855166 | A0A044SG03 | A0A044SG03 |
| PISNEVTVSGSTLS | 0 | 0 | 1 | 855167 | AAC48290.1 | A0A044VA40 |
| PIVASNMDTVGTFEV | 0 | 0 | 1 | 855168 | Q9NJD8.1 | Q9NJD8 |
| PKFADENFKLKHSMA | 0 | 0 | 1 | 855172 | A0A044U9P0 | A0A044U9P0 |
| PKGKLQPQMEKYTLT | 0 | 0 | 1 | 855173 | P46434.1 | P46434 |
| PKGPPGPPGQPGADG | 0 | 0 | 1 | 855175 | A0A044SF60 | A0A044SF60 |
| PKPGKPVKHDYSRAS | 0 | 0 | 1 | 855179 | A0A044R2U8 | A0A044R2U8 |
| PKQTNADDLLLGDEN | 0 | 0 | 1 | 855182 | A0A044U0M2 | A0A044U0M2 |
| PLLQKLKNSAPTTFA | 0 | 0 | 1 | 855189 | AAA63412.2 | Q99099 |
| PLPENPPPQLLNQ | 0 | 0 | 1 | 855191 | A0A044V9F8 | A0A044V9F8 |
| PLPSDHIYHHTS | 0 | 0 | 1 | 855192 | A0A044UK00 | A0A044UK00 |
| PLSNMNSPVKALQQN | 0 | 0 | 1 | 855194 | A0A044SG03 | A0A044SG03 |
| PLTSNIKEQLQSLEC | 0 | 0 | 1 | 855196 | A0A044UK00 | A0A044UK00 |
| PMEMIPTITVPIAQP | 0 | 0 | 1 | 855198 | AAC48290.1 | A0A044VA40 |
| PMFVMGVNNDKYD | 0 | 0 | 1 | 855199 | AAB52599.1 | A0A158N7S3 |
| PMPFVRKFGAAAVA | 0 | 0 | 1 | 855200 | A0A044U9P0 | A0A044U9P0 |
| PMQLLCEDSDYLPES | 0 | 0 | 1 | 855201 | A0A044UK00 | A0A044UK00 |
| PMYNQITSNADAKM | 0 | 0 | 1 | 855203 | A0A044UW71 | A0A044UW71 |
| PNFKPPADYRAPTIR | 0 | 0 | 1 | 855206 | A0A044U9P0 | A0A044U9P0 |
| PNFKTTAVVNGDFKE | 0 | 0 | 1 | 855207 | AAC48312.1 | A0A044VFS3 |
| PNFNANNDLIEPFHQ | 0 | 0 | 1 | 855208 | A0A044SAZ1 | A0A044SAZ1 |
| PNINEAMLQSQLMIT | 0 | 0 | 1 | 855210 | A0A044S6B8 | A0A044S6B8 |
| PNKTGSITKEDFIRY | 0 | 0 | 1 | 855211 | A0A044VDG0 | A0A044VDG0 |
| PNSECARPQCAEPQG | 0 | 0 | 1 | 855215 | A0A044QSA2 | A0A044QSA2 |
| PPASLCPTCRKQNGE | 0 | 0 | 1 | 855219 | A0A044SAZ1 | A0A044SAZ1 |
| PPETNDFYFQQTMLR | 0 | 0 | 1 | 855222 | A0A044RAH9 | A0A044RAH9 |
| PPGERGRKGSNGPSG | 0 | 0 | 1 | 855226 | A0A044V9I3 | A0A044V9I3 |
| PPGPDGLEGDYGETG | 0 | 0 | 1 | 855227 | A0A044V9I3 | A0A044V9I3 |
| PPGPPGIPGADG | 0 | 0 | 1 | 855229 | A0A044SF60 | A0A044SF60 |
| PPGQPGADGPPGEPG | 0 | 0 | 1 | 855236 | A0A044SF60 | A0A044SF60 |
| PPGTKIHNDLYWDVY | 0 | 0 | 1 | 855238 | A0A044R077 | A0A044R077 |
| PPGWVYHGESDGYRY | 0 | 0 | 1 | 855240 | A0A044UVG8 | A0A044UVG8 |
| PPILAFTSLKKVE | 0 | 0 | 1 | 855242 | A0A044UMF0 | A0A044UMF0 |
| PPIQAEQDAAEVNEA | 0 | 0 | 1 | 855243 | A0A044R196 | A0A044R196 |
| PPIRLYPAISGKNE | 0 | 0 | 1 | 855244 | A0A044U0M2 | A0A044U0M2 |
| PPKANERSQRPVWAD | 0 | 0 | 1 | 855245 | A0A044U9P0 | A0A044U9P0 |
| PPKVNVVSKFDRQID | 0 | 0 | 1 | 855248 | A0A044R196 | A0A044R196 |
| PPLGGSSVSGSTE | 0 | 0 | 1 | 855249 | A0A044U9P0 | A0A044U9P0 |
| PQHSAEDDAVRLAIA | 0 | 0 | 1 | 855265 | AAC48290.1 | A0A044VA40 |
| PQIPQFINPLPGETQ | 0 | 0 | 1 | 855267 | A0A044UK00 | A0A044UK00 |
| PQLIRQGSSNGNIQ | 0 | 0 | 1 | 855268 | A0A044R153 | A0A044R153 |
| PQPHVEWYFGEQKLE | 0 | 0 | 1 | 855271 | A0A044UK00 | A0A044UK00 |
| PQSQVNFVREQRVT | 0 | 0 | 1 | 855274 | AAC48290.1 | A0A044VA40 |
| PQVPFVPQMEAKDWE | 0 | 0 | 1 | 855278 | AAC48290.1 | A0A044VA40 |
| PRADFTVVVAVA | 0 | 0 | 1 | 855279 | A0A044SFY3 | A0A044SFY3 |
| PRELPISFDARQK | 0 | 0 | 1 | 855281 | A0A044RFT9 | A0A044RFT9 |
| PRFTVALMDKDLQ | 0 | 0 | 1 | 855282 | A0A044UK00 | A0A044UK00 |
| PRGSFVNSMGTKKQC | 0 | 0 | 1 | 855283 | AAD27587.1 | A0A158N7U8 |
| PRKLPCQRGGYTDPR | 0 | 0 | 1 | 855288 | AAV71152.1 | A0A044VAG6 |
| PRMSDISEHAISE | 0 | 0 | 1 | 855291 | A0A044UMF0 | A0A044UMF0 |
| PRPQCPEPQGFGAIK | 0 | 0 | 1 | 855294 | A0A044R521 | A0A044R521 |
| PRQPAIPSTPAE | 0 | 0 | 1 | 855295 | A0A044UH02 | A0A044UH02 |
| PRTGVADKIRKDIED | 0 | 0 | 1 | 855297 | A0A044UK00 | A0A044UK00 |
| PSDDPAFPKQQF | 0 | 0 | 1 | 855298 | A0A044SAZ1 | A0A044SAZ1 |
| PSGKMPNSDQKG | 0 | 0 | 1 | 855304 | A0A044QWA5 | A0A044QWA5 |
| PSKSFNFLSTNRVDN | 0 | 0 | 1 | 855306 | A0A044V9F8 | A0A044V9F8 |
| PSKTVKIWGRGQGG | 0 | 0 | 1 | 855307 | AAA74283.1 | A0A044V2F8 |
| PSNNPHEKHDKITSD | 0 | 0 | 1 | 855311 | A0A044U0M2 | A0A044U0M2 |
| PSRGEEVRKPP | 0 | 0 | 1 | 855313 | A0A2K6VJ98 | A0A2K6VJ98 |
| PSSASLSESSS | 0 | 0 | 1 | 855315 | A0A044UK00 | A0A044UK00 |
| PSSKNLPDFVLTH | 0 | 0 | 1 | 855316 | A0A044SG03 | A0A044SG03 |

| | | | | | | |
|---|---|---|---|---|---|---|
| PSSKSDGVKFTA | 0 | 0 | 1 | 855317 | A0A2K6VRV9 | A0A2K6VRV9 |
| PSTSNDVKLTDETA | 0 | 0 | 1 | 855321 | A0A044SS43 | A0A044SS43 |
| PTCRKQNGEFDEEAV | 0 | 0 | 1 | 855326 | A0A044SAZ1 | A0A044SAZ1 |
| PTDKSYQNDEI | 0 | 0 | 1 | 855328 | A0A044UK00 | A0A044UK00 |
| PTEIIEDIPDEVK | 0 | 0 | 1 | 855329 | AAD00842.1 | A0A044SS43 |
| PTETEHLADIIGE | 0 | 0 | 1 | 855330 | A0A044SG03 | A0A044SG03 |
| PTILPSNLSDD | 0 | 0 | 1 | 855335 | A0A044U9P0 | A0A044U9P0 |
| PTKREVQEENEEE | 0 | 0 | 1 | 855336 | A0A044UK00 | A0A044UK00 |
| PTKSELLPFLDQNSE | 0 | 0 | 1 | 855337 | A0A2K6VRV9 | A0A2K6VRV9 |
| PTLKKVSKYDNKFK | 0 | 0 | 1 | 855339 | A0A044TX11 | A0A044TX11 |
| PTPDVSVVDLTCRLQ | 0 | 0 | 1 | 855344 | AAB52599.1 | A0A158N7S3 |
| PTTVCKITNGI | 0 | 0 | 1 | 855348 | A0A044V1C1 | A0A044V1C1 |
| PTVAAGETLDAEI | 0 | 0 | 1 | 855349 | A0A044QU24 | A0A044QU24 |
| PTVEIDGFTDP | 0 | 0 | 1 | 855350 | A0A044V1C1 | A0A044V1C1 |
| PVLLGGWEDRDPKDE | 0 | 0 | 1 | 855362 | P22085.2 | P22085 |
| PVPKEFIGKTYDMHI | 0 | 0 | 1 | 855366 | A0A044V9S3 | A0A044V9S3 |
| PVSQGWSPYPPSNSF | 0 | 0 | 1 | 855369 | A0A044SS43 | A0A044SS43 |
| PVTLSTTTSENYIDS | 0 | 0 | 1 | 855370 | A0A044UK00 | A0A044UK00 |
| PWTAALEPAAEKN | 0 | 0 | 1 | 855374 | A0A044UK00 | A0A044UK00 |
| PWTKAVGLTNIRVPD | 0 | 0 | 1 | 855375 | AAF64254.1 | A0A044QLL7 |
| PYESGLSGEGLVPG | 0 | 0 | 1 | 855378 | AAA20541.1 | A0A044QNW3 |
| PYIKSILPGELAK | 0 | 0 | 1 | 855380 | P46427.1 | P46427 |
| PYPLLSAINHELEGE | 0 | 0 | 1 | 855382 | A0A044RRG7 | A0A044RRG7 |
| PYTEKRYEEITSGVS | 0 | 0 | 1 | 855385 | A0A044UK00 | A0A044UK00 |
| PYVGTEQKCKWQE | 0 | 0 | 1 | 855387 | A0A158N8I3 | A0A2K6W829 |
| QAGTNKYDSQKGM | 0 | 0 | 1 | 855391 | AAA20215.1 | A0A044V456 |
| QAYMEAIQRRGGNVA | 0 | 0 | 1 | 855394 | A0A044R196 | A0A044R196 |
| QCEAPDLCPPGPTG | 0 | 0 | 1 | 855395 | A0A044V9I3 | A0A044V9I3 |
| QCGGDTVEKDGQK | 0 | 0 | 1 | 855397 | Q9NJD8.1 | Q9NJD8 |
| QCNLIFESYSYNSAE | 0 | 0 | 1 | 855399 | A0A044SG03 | A0A044SG03 |
| QCPTRKHLNAPEDFI | 0 | 0 | 1 | 855400 | AAY67786.1 | A0A158N7U2 |
| QDAIDCATQALEKYN | 0 | 0 | 1 | 855403 | AAF64249.1 | A0A044UWQ7 |
| QDDGLHEFFEDKDI | 0 | 0 | 1 | 855404 | A0A044UK00 | A0A044UK00 |
| QDDHSGGSWNL | 0 | 0 | 1 | 855405 | AAD00842.1 | A0A044SS43 |
| QDFSGLPDYCHCE | 0 | 0 | 1 | 855406 | A0A044TIV1 | A0A044TIV1 |
| QDKQYKEAGKSHSVP | 0 | 0 | 1 | 855407 | A0A044UK00 | A0A044UK00 |
| QDNRDNRIVGNQGSE | 0 | 0 | 1 | 855409 | AAC48290.1 | A0A044VA40 |
| QDQEIRDLQAMAAR | 0 | 0 | 1 | 855410 | AAA74283.1 | A0A044V2F8 |
| QDTKENQSSDGNSTG | 0 | 0 | 1 | 855412 | A0A044UVG8 | A0A044UVG8 |
| QEAVKNIDESAARQ | 0 | 0 | 1 | 855414 | A0A044SS43 | A0A044SS43 |
| QEDCRSNDEELQKD | 0 | 0 | 1 | 855415 | A0A044SS43 | A0A044SS43 |
| QEETTNDELIDFN | 0 | 0 | 1 | 855420 | A0A044QWA5 | A0A044QWA5 |
| QEKEEAPAEQTEAE | 0 | 0 | 1 | 855423 | A0A044R196 | A0A044R196 |
| QELEQLRHEKIQALL | 0 | 0 | 1 | 855424 | A0A044U9P0 | A0A044U9P0 |
| QEMGINDAFTSAAN | 0 | 0 | 1 | 855425 | A0A044V3Q0 | A0A044V3Q0 |
| QEMQQQQQYRN | 0 | 0 | 1 | 855426 | A0A2K6WJQ8 | A0A2K6WJQ8 |
| QEVPEAYINLNVNVE | 0 | 0 | 1 | 855430 | A0A044UK00 | A0A044UK00 |
| QEWFDDRLRYTDHSD | 0 | 0 | 1 | 855431 | A0A044QWA5 | A0A044QWA5 |
| QEYDESGPSIVHRKC | 0 | 0 | 1 | 855432 | P30162.1 | P30162 |
| QEYNNKYINNNN | 0 | 0 | 1 | 855433 | A0A044QSA2 | A0A044QSA2 |
| QFAGNLNEIYEIE | 0 | 0 | 1 | 855434 | A0A044UK00 | A0A044UK00 |
| QFFICTEKTDWLDNK | 0 | 0 | 1 | 855436 | A0A044U9P0 | A0A044U9P0 |
| QFLLEIHGPVKEDQA | 0 | 0 | 1 | 855438 | A0A044R9D0 | A0A044R9D0 |
| QFNEKDIIVMIEER | 0 | 0 | 1 | 855439 | A0A044UK00 | A0A044UK00 |
| QFVDKHGEVCPANWQ | 0 | 0 | 1 | 855442 | A0A044VFS3 | A0A044VFS3 |
| QGDKHGSTWLTAF | 0 | 0 | 1 | 855446 | A0A044RF80 | A0A044RF80 |
| QGDKKKPAAID | 0 | 0 | 1 | 855447 | A0A044UU92 | A0A044UU92 |
| QGEPGQSGQRGYAGS | 0 | 0 | 1 | 855449 | A0A044SAV6 | A0A044SAV6 |
| QGGDFTKGDGTGG | 0 | 0 | 1 | 855451 | AAD09564.1 | A0A044S6Z0 |
| QGPPGAAGPQG | 0 | 0 | 1 | 855454 | A0A044UZ11 | A0A044UZ11 |
| QGPVGPQGEPGQSGQ | 0 | 0 | 1 | 855457 | A0A044SAV6 | A0A044SAV6 |
| QGTPGNDGLPGH | 0 | 0 | 1 | 855459 | A0A044SAV6 | A0A044SAV6 |
| QHIDDDGDIKQ | 0 | 0 | 1 | 855461 | A0A044UVG8 | A0A044UVG8 |
| QHNESFSTPPHP | 0 | 0 | 1 | 855463 | A0A044UK00 | A0A044UK00 |
| QIDRRSYGDRRDL | 0 | 0 | 1 | 855467 | A0A044R196 | A0A044R196 |
| QIEFSDDAKLEHV | 0 | 0 | 1 | 855468 | A0A044UK00 | A0A044UK00 |
| QIEQQGSKNGK | 0 | 0 | 1 | 855470 | A0A044U9P0 | A0A044U9P0 |
| QIGKEFSDECIKH | 0 | 0 | 1 | 855474 | A0A044UIG6 | A0A044UIG6 |
| QILEKDELAKCNER | 0 | 0 | 1 | 855476 | A0A2K6VRV9 | A0A2K6VRV9 |
| QINANTVGSAKLKA | 0 | 0 | 1 | 855477 | A0A044U9P0 | A0A044U9P0 |
| QIPKPKKYGSGVG | 0 | 0 | 1 | 855479 | CAA04390.1 | A0A044R436 |
| QIPNLTLRPGTEAVI | 0 | 0 | 1 | 855480 | A0A044UK00 | A0A044UK00 |
| QKDKLYLTTTEWKE | 0 | 0 | 1 | 855484 | CAA04390.1 | A0A044R436 |
| QKFEDPEINTNIL | 0 | 0 | 1 | 855486 | A0A044UK00 | A0A044UK00 |
| QKGDDGTPFVQILQ | 0 | 0 | 1 | 855487 | AAB52600.1 | A0A044UUD9 |
| QKGEYKVIAENRAGK | 0 | 0 | 1 | 855488 | A0A044UK00 | A0A044UK00 |
| QKGMTGFGQPRWEVL | 0 | 0 | 1 | 855489 | AAA20215.1 | A0A044V456 |
| QKIINYKDNDSQEII | 0 | 0 | 1 | 855490 | A0A044QNT8 | A0A044QNT8 |
| QKNERDKEKIEGVE | 0 | 0 | 1 | 855492 | A0A044TU88 | A0A044TU88 |
| QLCHETFANRSIYSK | 0 | 0 | 1 | 855498 | A0A044UK00 | A0A044UK00 |
| QLEVEDATRKLRLGD | 0 | 0 | 1 | 855501 | A0A044U9P0 | A0A044U9P0 |
| QLFPMDSQKCKLEIE | 0 | 0 | 1 | 855502 | A0A044UW71 | A0A044UW71 |
| QLPCLYDGDQQ | 0 | 0 | 1 | 855508 | P46427.1 | P46427 |
| QLSGLMDISDLQRA | 0 | 0 | 1 | 855513 | A0A044RF80 | A0A044RF80 |
| QLSKNTKGEGYIGSK | 0 | 0 | 1 | 855514 | A0A2K6WB53 | A0A2K6WB53 |
| QLSVEVYQHLDL | 0 | 0 | 1 | 855515 | A0A044UK00 | A0A044UK00 |
| QLYKSGKFTENLD | 0 | 0 | 1 | 855522 | AAF64254.1 | A0A044QLL7 |
| QMNACMANFRVSTC | 0 | 0 | 1 | 855524 | A0A044UK00 | A0A044UK00 |
| QMWISKQEYDESGP | 0 | 0 | 1 | 855526 | P30163.1 | P30163 |
| QNDEILQTSEIS | 0 | 0 | 1 | 855527 | A0A044UK00 | A0A044UK00 |
| QNEEMVRYKDEQE | 0 | 0 | 1 | 855530 | A0A044UK00 | A0A044UK00 |

| | | | | | | |
|---|---|---|---|---|---|---|
| QNFEVTADNSMVLQS | 0 | 0 | 1 | 855532 | A0A044RF80 | A0A044RF80 |
| QNGYIIQDNRDNRIV | 0 | 0 | 1 | 855534 | AAC48290.1 | A0A044VA40 |
| QNQKYPWTVEQLGA | 0 | 0 | 1 | 855537 | AAF64254.1 | A0A044QLL7 |
| QNSSPEVKNRTGNGC | 0 | 0 | 1 | 855538 | A0A044S6B8 | A0A044S6B8 |
| QPITQNERIEKKDTA | 0 | 0 | 1 | 855543 | A0A044UK00 | A0A044UK00 |
| QPLANDMQKTTLG | 0 | 0 | 1 | 855544 | AAA63412.2 | Q99099 |
| QPLPKPMTPVIHEPL | 0 | 0 | 1 | 855545 | A0A044UK00 | A0A044UK00 |
| QPQQVQKADLL | 0 | 0 | 1 | 855550 | A0A044UK00 | A0A044UK00 |
| QPSESSLPLFLSMQ | 0 | 0 | 1 | 855551 | AAC48290.1 | A0A044VA40 |
| QPSILYGWGRDAPD | 0 | 0 | 1 | 855552 | A0A044U175 | A0A044U175 |
| QPSVTNLTRDG | 0 | 0 | 1 | 855553 | A0A044V3Q0 | A0A044V3Q0 |
| QPVKNIVLGSDYLSH | 0 | 0 | 1 | 855556 | A0A044UK00 | A0A044UK00 |
| QQCITRNDDTLG | 0 | 0 | 1 | 855559 | A0A044V9F8 | A0A044V9F8 |
| QQIVVEDKKQSD | 0 | 0 | 1 | 855563 | A0A044UK00 | A0A044UK00 |
| QQKQLQNAMSNYNRE | 0 | 0 | 1 | 855565 | A0A044UVG8 | A0A044UVG8 |
| QQNGSFLYLNSDDRN | 0 | 0 | 1 | 855569 | A0A044SG03 | A0A044SG03 |
| QQPNDSPENAGKLES | 0 | 0 | 1 | 855570 | A0A044UVG8 | A0A044UVG8 |
| QQQENGAFAERGEVH | 0 | 0 | 1 | 855572 | A0A044RF80 | A0A044RF80 |
| QQQKQRQQQLQ | 0 | 0 | 1 | 855573 | AAC48290.1 | A0A044VA40 |
| QQRDSPIAANSRD | 0 | 0 | 1 | 855584 | A0A044R153 | A0A044R153 |
| QQYRNPCEFEWKQ | 0 | 0 | 1 | 855586 | A0A2K6WJQ8 | A0A2K6WJQ8 |
| QRAEKEKTTDPDVP | 0 | 0 | 1 | 855587 | AAC48290.1 | A0A044VA40 |
| QRAVHEVTCLYDDKA | 0 | 0 | 1 | 855588 | A0A044UK00 | A0A044UK00 |
| QREGVGENIYALGL | 0 | 0 | 1 | 855589 | AAB97282.1 | A0A044UNG9 |
| QRIGGSNENALALLE | 0 | 0 | 1 | 855592 | A0A044UVG8 | A0A044UVG8 |
| QRKDIIVLQDPNHLE | 0 | 0 | 1 | 855594 | CAA04390.1 | A0A044R436 |
| QRLKHVLPDLPYD | 0 | 0 | 1 | 855595 | P41981.1 | A0A158N7R1 |
| QRQFKASSVEDTFVS | 0 | 0 | 1 | 855600 | A0A044UK00 | A0A044UK00 |
| QRTIPLPDVDTIDDH | 0 | 0 | 1 | 855605 | A0A044TX11 | A0A044TX11 |
| QRWDCKLESSAQRWA | 0 | 0 | 1 | 855606 | SRC447106 | A0A044VD00 |
| QSGNLAEDNGVFTV | 0 | 0 | 1 | 855610 | A0A044UK00 | A0A044UK00 |
| QSNSFSLNSLQDSQE | 0 | 0 | 1 | 855616 | A0A044UK00 | A0A044UK00 |
| QSSVFREGESLELR | 0 | 0 | 1 | 855620 | A0A044UK00 | A0A044UK00 |
| QTEADVEAFINRLGG | 0 | 0 | 1 | 855623 | P36991.1 | P36991 |
| QTGNVYSLEAIEELN | 0 | 0 | 1 | 855625 | CAA04390.1 | A0A044R436 |
| QTLLVKGKTAEDS | 0 | 0 | 1 | 855628 | AAA20541.1 | A0A044QNW3 |
| QTSEISTSQFESEQN | 0 | 0 | 1 | 855629 | A0A044UK00 | A0A044UK00 |
| QTWQDPRLAFGDMF | 0 | 0 | 1 | 855631 | A0A044UW71 | A0A044UW71 |
| QTWVDNGCYCDRKEN | 0 | 0 | 1 | 855632 | A0A044QNT8 | A0A044QNT8 |
| QVCSNTNETFPSLC | 0 | 0 | 1 | 855635 | A0A044T7P0 | A0A044T7P0 |
| QVDLPVADISQ | 0 | 0 | 1 | 855636 | AAF64251.1 | A0A044V6Z2 |
| QVDSLFGEKSNLFN | 0 | 0 | 1 | 855637 | AAA29419.1 | P21250 |
| QVENITAVLPTRAP | 0 | 0 | 1 | 855639 | A0A044UK00 | A0A044UK00 |
| QVHYKKKLGPDYSGI | 0 | 0 | 1 | 855640 | A0A044U175 | A0A044U175 |
| QVITVGNERFRCPEA | 0 | 0 | 1 | 855642 | P30162.1 | P30162 |
| QVKRKAAEDKKKEA | 0 | 0 | 1 | 855643 | A0A044SS43 | A0A044SS43 |
| QVPTEGHETPTPHQT | 0 | 0 | 1 | 855644 | A0A044UK00 | A0A044UK00 |
| QVQQLREVLGTFDQD | 0 | 0 | 1 | 855645 | AAK16514.1 | A0A2K6W829 |
| QVVAGVKYKMDVQ | 0 | 0 | 1 | 855647 | AAA29418.1 | P22085 |
| QYGFPWPEMLSCNK | 0 | 0 | 1 | 855651 | A0A044TBR7 | A0A044TBR7 |
| QYICLDVANGYSEVF | 0 | 0 | 1 | 855652 | Q9NJD8.1 | Q9NJD8 |
| QYKVSSPTVEIDG | 0 | 0 | 1 | 855653 | A0A044V1C1 | A0A044V1C1 |
| RAELDKEDTRMNLYF | 0 | 0 | 1 | 855658 | A0A044RF80 | A0A044RF80 |
| RALCTGEKGVGKSGK | 0 | 0 | 1 | 855661 | AAC47233.1 | Q25633 |
| RAMEKLETEWNDYVM | 0 | 0 | 1 | 855662 | AAK16514.1 | A0A2K6W829 |
| RASEGKTITVPGDI | 0 | 0 | 1 | 855665 | A0A044SN49 | A0A044SN49 |
| RCAQYKKDGAQFA | 0 | 0 | 1 | 855671 | AAB52600.1 | A0A044UUD9 |
| RCLRESKEGKIGEVK | 0 | 0 | 1 | 855673 | A0A044SD24 | A0A044SD24 |
| RCRCPDGFTGKLC | 0 | 0 | 1 | 855675 | AAV71152.1 | A0A044VAG6 |
| RDDLPFYCTKEVR | 0 | 0 | 1 | 855678 | A0A044RF80 | A0A044RF80 |
| RDDSINRLDLEPKKE | 0 | 0 | 1 | 855679 | AAC48290.1 | A0A044VA40 |
| RDEFEASIEDSIKRQ | 0 | 0 | 1 | 855681 | A0A044UK00 | A0A044UK00 |
| RDFTPVCTTELA | 0 | 0 | 1 | 855684 | P52570.1 | P52570 |
| RDGRRGTGTPGPIG | 0 | 0 | 1 | 855686 | A0A044V9I3 | A0A044V9I3 |
| RDSSKTTIGAEG | 0 | 0 | 1 | 855691 | AAB68960.1 | A0A044RRG7 |
| RDTIGQIEVYYPMAN | 0 | 0 | 1 | 855692 | A0A044UK00 | A0A044UK00 |
| RDVSKTPQQETETSI | 0 | 0 | 1 | 855696 | A0A044UK00 | A0A044UK00 |
| REIAQDFKTDLRFQ | 0 | 0 | 1 | 855703 | A0A044UVZ9 | A0A044UVZ9 |
| RELHNLEWEELIAEA | 0 | 0 | 1 | 855708 | A0A044T7P0 | A0A044T7P0 |
| RELHSGNRLQLDLCC | 0 | 0 | 1 | 855709 | A0A044RF80 | A0A044RF80 |
| RENIRLAKTNQESE | 0 | 0 | 1 | 855712 | CAA04390.1 | A0A044R436 |
| RFGSNASGKIQNCDF | 0 | 0 | 1 | 855720 | A0A044UIG6 | A0A044UIG6 |
| RFKFVTFGGSQNIFD | 0 | 0 | 1 | 855721 | A0A044S6B8 | A0A044S6B8 |
| RFKGHVSAEGGKLI | 0 | 0 | 1 | 855722 | AAB52599.1 | A0A158N7S3 |
| RFSGSVEEYFTI | 0 | 0 | 1 | 855726 | A0A044UK00 | A0A044UK00 |
| RGADGEPGEPGPDGR | 0 | 0 | 1 | 855730 | A0A044V9I3 | A0A044V9I3 |
| RGYIKSKVTVSNSA | 0 | 0 | 1 | 855745 | A0A044UVG8 | A0A044UVG8 |
| RHKDEITKLDIETC | 0 | 0 | 1 | 855748 | A0A044RXP7 | A0A044RXP7 |
| RIDECTCKQEIFLVA | 0 | 0 | 1 | 855756 | A0A044UW71 | A0A044UW71 |
| RILINTYDDSETFKA | 0 | 0 | 1 | 855765 | A0A044UIG6 | A0A044UIG6 |
| RIPTEMPMYNQ | 0 | 0 | 1 | 855767 | A0A044UW71 | A0A044UW71 |
| RISVGIEDAEDLIQD | 0 | 0 | 1 | 855773 | A0A044R2U8 | A0A044R2U8 |
| RIVGNQGSEIAA | 0 | 0 | 1 | 855776 | AAC48290.1 | A0A044VA40 |
| RIYSAEQDDHSG | 0 | 0 | 1 | 855779 | AAD00842.1 | A0A044SS43 |
| RKGSNGPSGSPGEMG | 0 | 0 | 1 | 855789 | A0A044V9I3 | A0A044V9I3 |
| RKMENTKTTQPGDK | 0 | 0 | 1 | 855795 | A0A2K6WB53 | A0A2K6WB53 |
| RKTSNLMNQADSA | 0 | 0 | 1 | 855804 | A0A044VBX6 | A0A044VBX6 |
| RLDLEPKKEPIVEQ | 0 | 0 | 1 | 855810 | AAC48290.1 | A0A044VA40 |
| RLQKGASMDEIKAA | 0 | 0 | 1 | 855826 | AAB52599.1 | A0A158N7S3 |
| RLTDRDQIDEL | 0 | 0 | 1 | 855832 | AAA74283.1 | A0A044V2F8 |
| RLVLPNLNEEMDA | 0 | 0 | 1 | 855834 | AAF64251.1 | A0A044V6Z2 |

| | | | | | | |
|---|---|---|---|---|---|---|
| RMIGLDQNGYIIQDN | 0 | 0 | 1 | 855837 | AAC48290.1 | A0A044VA40 |
| RMIYMAYETEKDPYI | 0 | 0 | 1 | 855838 | P46427.1 | P46427 |
| RMPGAEPVMNDS | 0 | 0 | 1 | 855842 | A0A044U175 | A0A044U175 |
| RNFAEEHWSSKI | 0 | 0 | 1 | 855847 | A0A044UK00 | A0A044UK00 |
| RNLILGDKISYAD | 0 | 0 | 1 | 855848 | P46427.1 | P46427 |
| RNRSSSRRNSPE | 0 | 0 | 1 | 855854 | AAF64250.1 | A0A044UVG8 |
| RPPPEWGRRENE | 0 | 0 | 1 | 855866 | A0A044R196 | A0A044R196 |
| RPQCAEPQGFGAVRS | 0 | 0 | 1 | 855867 | A0A044QSA2 | A0A044QSA2 |
| RQEYDLKELHERERQ | 0 | 0 | 1 | 855874 | A0A044R196 | A0A044R196 |
| RQFFKKVDVDNSGD | 0 | 0 | 1 | 855875 | AAF64251.1 | A0A044V6Z2 |
| RQKGCEGGYLDRAW | 0 | 0 | 1 | 855877 | A0A044RFT9 | A0A044RFT9 |
| RQQRAQEEREFAER | 0 | 0 | 1 | 855883 | A0A044R196 | A0A044R196 |
| RQSDDNMIPDRFAA | 0 | 0 | 1 | 855886 | AAC48290.1 | A0A044VA40 |
| RRISAVEESEK | 0 | 0 | 1 | 855893 | A0A044SG03 | A0A044SG03 |
| RRNDDMARLEN | 0 | 0 | 1 | 855899 | A0A044V9G2 | A0A044V9G2 |
| RRNSPEYDISQI | 0 | 0 | 1 | 855900 | AAF64250.1 | A0A044UVG8 |
| RRRGRKIAVEKDPFE | 0 | 0 | 1 | 855902 | A0A044SS43 | A0A044SS43 |
| RRYSEKVQLPQQ | 0 | 0 | 1 | 855911 | A0A044UK00 | A0A044UK00 |
| RSEIVNASTSNTRI | 0 | 0 | 1 | 855914 | A0A044UK00 | A0A044UK00 |
| RSIYNGGKTHSQ | 0 | 0 | 1 | 855917 | A0A044UVG8 | A0A044UVG8 |
| RSLATELGMMDPDER | 0 | 0 | 1 | 855920 | AAA50214.2 | P52570 |
| RSLEHAEASENRNLS | 0 | 0 | 1 | 855921 | A0A044UK00 | A0A044UK00 |
| RSPSPEPIYDASGKR | 0 | 0 | 1 | 855923 | A0A044U9P0 | A0A044U9P0 |
| RSQNSEISTYDDA | 0 | 0 | 1 | 855925 | A0A044UK00 | A0A044UK00 |
| RSQRPVWADDEWL | 0 | 0 | 1 | 855926 | A0A044U9P0 | A0A044U9P0 |
| RSSSRQSFDWTEF | 0 | 0 | 1 | 855935 | A0A044UK00 | A0A044UK00 |
| RTCATHSDCYDQRE | 0 | 0 | 1 | 855940 | A0A044QNT8 | A0A044QNT8 |
| RTGSPPKGPEEQNEN | 0 | 0 | 1 | 855942 | A0A044U9P0 | A0A044U9P0 |
| RTHKVDTYKSTME | 0 | 0 | 1 | 855944 | A0A2K6VJ98 | A0A2K6VJ98 |
| RTLYVGGFGEEVD | 0 | 0 | 1 | 855949 | A0A044U9P0 | A0A044U9P0 |
| RTPASIIQPSAGP | 0 | 0 | 1 | 855951 | A0A044SS43 | A0A044SS43 |
| RTQSDDDFVVSG | 0 | 0 | 1 | 855952 | A0A044SS43 | A0A044SS43 |
| RVELDVIFQEMDPNK | 0 | 0 | 1 | 855955 | A0A044VDG0 | A0A044VDG0 |
| RVFEDHPTVIHGEAQ | 0 | 0 | 1 | 855957 | A0A044V9F9 | A0A044V9F9 |
| RVGNCDAAKQDGLNF | 0 | 0 | 1 | 855959 | A0A044UIG6 | A0A044UIG6 |
| RVNVTRATTSSDEYF | 0 | 0 | 1 | 855966 | A0A044UW71 | A0A044UW71 |
| RWENIDRDSKGQFLK | 0 | 0 | 1 | 855979 | AAV71152.1 | A0A044VAG6 |
| RYDDLFTLNTNI | 0 | 0 | 1 | 855984 | AAV71152.1 | A0A044VAG6 |
| RYIVKTDAIDEAATK | 0 | 0 | 1 | 855989 | A0A044VBX6 | A0A044VBX6 |
| RYKDEQEESQIPI | 0 | 0 | 1 | 855991 | A0A044UK00 | A0A044UK00 |
| RYLKITKRAGTQIEN | 0 | 0 | 1 | 855992 | A0A044RF80 | A0A044RF80 |
| RYNPNGDIREVHH | 0 | 0 | 1 | 855993 | A0A044V9G2 | A0A044V9G2 |
| RYTRAEEVIQTNDI | 0 | 0 | 1 | 855995 | A0A044UK00 | A0A044UK00 |
| RYYPDDDPLFP | 0 | 0 | 1 | 855996 | A0A158N7T4 | A0A158N7T4 |
| SAANINVHPSLEE | 0 | 0 | 1 | 855997 | A0A044UK00 | A0A044UK00 |
| SAEGGKLIVTNGK | 0 | 0 | 1 | 856002 | AAB52599.1 | A0A158N7S3 |
| SAIGGGFGGGG | 0 | 0 | 1 | 856009 | A0A044S628 | A0A044S628 |
| SAKTEDVKKLEVKVS | 0 | 0 | 1 | 856011 | A0A044SS43 | A0A044SS43 |
| SATETEDIKQFNKAN | 0 | 0 | 1 | 856022 | A0A044UVG8 | A0A044UVG8 |
| SATFTVKAGNAPEFH | 0 | 0 | 1 | 856023 | A0A044R9D0 | A0A044R9D0 |
| SATNATSDTAAVGAS | 0 | 0 | 1 | 856024 | A0A044UUD9 | A0A044UUD9 |
| SCQRNISKTSVDAA | 0 | 0 | 1 | 856032 | A0A044UK00 | A0A044UK00 |
| SCRAETDTFSGMKWA | 0 | 0 | 1 | 856033 | A0A044R153 | A0A044R153 |
| SDANFLPQLIRQGS | 0 | 0 | 1 | 856037 | A0A044R153 | A0A044R153 |
| SDCYDQREPSS | 0 | 0 | 1 | 856040 | A0A044QNT8 | A0A044QNT8 |
| SDDDQEKKAKKER | 0 | 0 | 1 | 856041 | A0A044SS43 | A0A044SS43 |
| SDEDISDDDSYT | 0 | 0 | 1 | 856044 | A0A044QNT8 | A0A044QNT8 |
| SDFVGDPHSSIFDAL | 0 | 0 | 1 | 856045 | O01360.1 | O01360 |
| SDGIRGIQINEAME | 0 | 0 | 1 | 856046 | A0A044UK00 | A0A044UK00 |
| SDGPTLSGEVEKKA | 0 | 0 | 1 | 856047 | A0A044UVG8 | A0A044UVG8 |
| SDHIVCCDDVYGG | 0 | 0 | 1 | 856048 | A0A044R2U8 | A0A044R2U8 |
| SDLLQWFDSVNDGN | 0 | 0 | 1 | 856051 | A0A044UMF0 | A0A044UMF0 |
| SDPSDDTDEYLP | 0 | 0 | 1 | 856056 | A0A044SS43 | A0A044SS43 |
| SDPVNIKDLEDAEE | 0 | 0 | 1 | 856058 | A0A044UK00 | A0A044UK00 |
| SDRISEFKKLDVAVM | 0 | 0 | 1 | 856059 | AAC48312.1 | A0A044VFS3 |
| SDRQFILKDGEITD | 0 | 0 | 1 | 856060 | A0A044UU92 | A0A044UU92 |
| SEASYSDATVLVVQK | 0 | 0 | 1 | 856068 | A0A044UK00 | A0A044UK00 |
| SEDLLAEIQHGRYTW | 0 | 0 | 1 | 856073 | A0A044RFT9 | A0A044RFT9 |
| SEHAISEKNVTEEAL | 0 | 0 | 1 | 856076 | A0A044UMF0 | A0A044UMF0 |
| SEKINSDIHFISPPQ | 0 | 0 | 1 | 856079 | A0A044UK00 | A0A044UK00 |
| SEKSSVREFEVK | 0 | 0 | 1 | 856080 | AAF64250.1 | A0A044UVG8 |
| SELPKLYENNPSNN | 0 | 0 | 1 | 856081 | A0A044VD00 | A0A044VD00 |
| SENQMSRSQNSEIS | 0 | 0 | 1 | 856082 | A0A044UK00 | A0A044UK00 |
| SENVESYGQKGVEQA | 0 | 0 | 1 | 856084 | A0A2K6VJ98 | A0A2K6VJ98 |
| SEQDVTVTLKDGDDI | 0 | 0 | 1 | 856086 | A0A044RF80 | A0A044RF80 |
| SEQNPNLSANRLLQ | 0 | 0 | 1 | 856087 | A0A044R521 | A0A044R521 |
| SEQRLKNYFKPNTEM | 0 | 0 | 1 | 856088 | AAD00843.1 | A0A044VHF7 |
| SESYDHYFSSIQPE | 0 | 0 | 1 | 856094 | A0A044UK00 | A0A044UK00 |
| SETAETRSIYNGG | 0 | 0 | 1 | 856095 | A0A044UVG8 | A0A044UVG8 |
| SETEELQLKCIVTGE | 0 | 0 | 1 | 856097 | A0A044UK00 | A0A044UK00 |
| SFDWTEFSDRSASWQ | 0 | 0 | 1 | 856105 | A0A044UK00 | A0A044UK00 |
| SFLFFEPTKREVQEE | 0 | 0 | 1 | 856114 | A0A044UK00 | A0A044UK00 |
| SFLKKMEDNGELRAM | 0 | 0 | 1 | 856115 | AAK16514.1 | A0A2K6W829 |
| SFPPRFETNIECNVV | 0 | 0 | 1 | 856117 | A0A044UK00 | A0A044UK00 |
| SFSADKNIKYSGSI | 0 | 0 | 1 | 856121 | AAK66564.1 | Q963F6 |
| SFSAFGQGDKHGSTW | 0 | 0 | 1 | 856122 | A0A044RF80 | A0A044RF80 |
| SFSNRDVYDSGSGVN | 0 | 0 | 1 | 856123 | A0A044V9G2 | A0A044V9G2 |
| SFVKGWGAEYQRQDV | 0 | 0 | 1 | 856126 | A0A044V1C1 | A0A044V1C1 |
| SFYLKGGLEESEE | 0 | 0 | 1 | 856132 | A0A044R2U8 | A0A044R2U8 |
| SGDSDSPRNRNRAAP | 0 | 0 | 1 | 856133 | A0A158N7T4 | A0A158N7T4 |
| SGFPTDFSVQF | 0 | 0 | 1 | 856137 | AAK66564.1 | Q963F6 |

| | | | | | | |
|---|---|---|---|---|---|---|
| SGGEVEVYPPGIDGS | 0 | 0 | 1 | 856138 | A0A2K6VRV9 | A0A2K6VRV9 |
| SGGQSEEDATLNLNA | 0 | 0 | 1 | 856140 | AAB52600.1 | A0A044UUD9 |
| SGIREGENFGGEKFD | 0 | 0 | 1 | 856143 | A0A044UVG8 | A0A044UVG8 |
| SGKDIYPRADFTV | 0 | 0 | 1 | 856145 | A0A044SFY3 | A0A044SFY3 |
| SGMSPFGQNAASTIR | 0 | 0 | 1 | 856151 | AAA74283.1 | A0A044V2F8 |
| SGPSKENIRIRFPS | 0 | 0 | 1 | 856155 | A0A2K6VRV9 | A0A2K6VRV9 |
| SGQRGYAGSSGGSG | 0 | 0 | 1 | 856156 | A0A044SAV6 | A0A044SAV6 |
| SGTGITELPTNG | 0 | 0 | 1 | 856160 | A0A044UMF0 | A0A044UMF0 |
| SGTRNSLVQNVTVSE | 0 | 0 | 1 | 856161 | A0A044UK00 | A0A044UK00 |
| SIDSSTIPSQMGSN | 0 | 0 | 1 | 856172 | AAA20215.1 | A0A044V456 |
| SIECSLFDGTVAFTD | 0 | 0 | 1 | 856173 | AAF64254.1 | A0A044QLL7 |
| SIEDSIKRQKYFPQS | 0 | 0 | 1 | 856175 | A0A044UK00 | A0A044UK00 |
| SIHTAPQFAGNLNEI | 0 | 0 | 1 | 856178 | A0A044UK00 | A0A044UK00 |
| SIINEALMIPDGQNE | 0 | 0 | 1 | 856181 | A0A044U9P0 | A0A044U9P0 |
| SIKNVTVKGQVACSD | 0 | 0 | 1 | 856184 | A0A044V9S3 | A0A044V9S3 |
| SILNQFVPTESPQVD | 0 | 0 | 1 | 856186 | AAF64251.1 | A0A044V6Z2 |
| SIPLDYETGKHRGFG | 0 | 0 | 1 | 856188 | A0A044U9P0 | A0A044U9P0 |
| SIRASTIVERTNMVF | 0 | 0 | 1 | 856189 | A0A044UK00 | A0A044UK00 |
| SISELSDNYQSEESN | 0 | 0 | 1 | 856192 | A0A044SS43 | A0A044SS43 |
| SIWGGPIEDELNAAL | 0 | 0 | 1 | 856197 | A0A2K6VZ59 | A0A2K6VZ59 |
| SKEEKKENKEAKDYK | 0 | 0 | 1 | 856200 | A0A044TU88 | A0A044TU88 |
| SKEGDFYSCSYTWPS | 0 | 0 | 1 | 856201 | A0A044UK00 | A0A044UK00 |
| SKGGAPPPLGGSS | 0 | 0 | 1 | 856204 | A0A044U9P0 | A0A044U9P0 |
| SKLKDLTSDFNEITA | 0 | 0 | 1 | 856209 | A0A044UVG8 | A0A044UVG8 |
| SKQESVWSIGKKYEK | 0 | 0 | 1 | 856216 | A0A044UK00 | A0A044UK00 |
| SKQIDQEIFYTESAK | 0 | 0 | 1 | 856217 | A0A044SS43 | A0A044SS43 |
| SKRDVSKTPQQETET | 0 | 0 | 1 | 856219 | A0A044UK00 | A0A044UK00 |
| SKSLDSKIMEKLKRN | 0 | 0 | 1 | 856223 | A0A044UVG8 | A0A044UVG8 |
| SKSNAQDEEQSEK | 0 | 0 | 1 | 856224 | AAF64251.1 | A0A044V6Z2 |
| SKYDNKFKKMAGQKE | 0 | 0 | 1 | 856229 | A0A044TX11 | A0A044TX11 |
| SLEAIEELNLKPGH | 0 | 0 | 1 | 856234 | CAA04390.1 | A0A044R436 |
| SLKEAQEYILKEYGI | 0 | 0 | 1 | 856241 | AAF64251.1 | A0A044V6Z2 |
| SLLAVIVENNPAKVA | 0 | 0 | 1 | 856245 | A0A044SAZ1 | A0A044SAZ1 |
| SLNEEDLYAVRQS | 0 | 0 | 1 | 856253 | A0A044UK00 | A0A044UK00 |
| SLNGDTSKSLDSKIM | 0 | 0 | 1 | 856254 | A0A044UVG8 | A0A044UVG8 |
| SLPGSDEVIWDCGD | 0 | 0 | 1 | 856258 | A0A044U175 | A0A044U175 |
| SLPSSTFKTEVATHD | 0 | 0 | 1 | 856259 | A0A044UMF0 | A0A044UMF0 |
| SLSGWRTYENNMEQ | 0 | 0 | 1 | 856265 | A0A044UW71 | A0A044UW71 |
| SLSPMHQNMTTNEKG | 0 | 0 | 1 | 856267 | A0A044UK00 | A0A044UK00 |
| SLTEAQAEAGEELL | 0 | 0 | 1 | 856268 | A0A044UK00 | A0A044UK00 |
| SMAAKKKKRNN | 0 | 0 | 1 | 856270 | A0A044U9P0 | A0A044U9P0 |
| SMANCGPDTNGSQ | 0 | 0 | 1 | 856272 | A0A044U9P0 | A0A044U9P0 |
| SMAQSTASNDTYE | 0 | 0 | 1 | 856273 | A0A044UVG8 | A0A044UVG8 |
| SMDEIKAAVKEAAA | 0 | 0 | 1 | 856274 | AAB52599.1 | A0A158N7S3 |
| SMHDEKITANYEART | 0 | 0 | 1 | 856275 | A0A044UK00 | A0A044UK00 |
| SMMLQETGINNAPWT | 0 | 0 | 1 | 856277 | A0A044UK00 | A0A044UK00 |
| SMMYVESANMQSPME | 0 | 0 | 1 | 856278 | AAC48290.1 | A0A044VA40 |
| SNEYYNYAVAWQTG | 0 | 0 | 1 | 856283 | A0A044UMF0 | A0A044UMF0 |
| SNFGEPQSNSFSLNS | 0 | 0 | 1 | 856284 | A0A044UK00 | A0A044UK00 |
| SNLDEKLDLEAQGNG | 0 | 0 | 1 | 856293 | A0A044RF80 | A0A044RF80 |
| SNNKFKNPIENEQED | 0 | 0 | 1 | 856295 | AAD00842.1 | A0A044SS43 |
| SNNTGGNMYERDHR | 0 | 0 | 1 | 856296 | A0A044V9G2 | A0A044V9G2 |
| SNRYKEEVKESLEER | 0 | 0 | 1 | 856299 | AAA29419.1 | P21250 |
| SNSGRRINKDGNPTD | 0 | 0 | 1 | 856300 | A0A044V9G2 | A0A044V9G2 |
| SNSKHLTDESNSSA | 0 | 0 | 1 | 856301 | A0A044UUD9 | A0A044UUD9 |
| SNTKKQTNDYKSDRT | 0 | 0 | 1 | 856304 | A0A044UVG8 | A0A044UVG8 |
| SPAINESRATFLKPV | 0 | 0 | 1 | 856306 | A0A044UK00 | A0A044UK00 |
| SPDKESEIIVKNGDV | 0 | 0 | 1 | 856311 | AAC48290.1 | A0A044VA40 |
| SPESQFKWYQNNFEV | 0 | 0 | 1 | 856313 | A0A044UK00 | A0A044UK00 |
| SPFFNFLDKEINENV | 0 | 0 | 1 | 856314 | AAC48290.1 | A0A044VA40 |
| SPFMAESEAEEKFKT | 0 | 0 | 1 | 856315 | A0A044UVG8 | A0A044UVG8 |
| SPYPPSNSFPPNVT | 0 | 0 | 1 | 856323 | A0A044SS43 | A0A044SS43 |
| SQAGWNKGDSQKLMT | 0 | 0 | 1 | 856324 | P37801.2 | P37801 |
| SQDANKMQGKANAI | 0 | 0 | 1 | 856326 | A0A044QU24 | A0A044QU24 |
| SQFQFGQLPCLYDGD | 0 | 0 | 1 | 856331 | P46427.1 | P46427 |
| SQLEMTAEWFKDDK | 0 | 0 | 1 | 856334 | A0A044R9D0 | A0A044R9D0 |
| SQQQPSQEFNLP | 0 | 0 | 1 | 856337 | A0A044V9I3 | A0A044V9I3 |
| SQQSGGPFIEFPP | 0 | 0 | 1 | 856338 | A0A044V9G2 | A0A044V9G2 |
| SQSGDYAQFYNLVA | 0 | 0 | 1 | 856339 | A0A044QU24 | A0A044QU24 |
| SRDRSPSADKSSR | 0 | 0 | 1 | 856343 | AAF64250.1 | A0A044UVG8 |
| SRFEIGVQDKEYNPR | 0 | 0 | 1 | 856344 | A0A044R077 | A0A044R077 |
| SRFFSYKTVDLIA | 0 | 0 | 1 | 856345 | A0A044UK00 | A0A044UK00 |
| SRIVKLEGEKYDLEK | 0 | 0 | 1 | 856350 | A0A044R196 | A0A044R196 |
| SRSNKDGVNYLDSYK | 0 | 0 | 1 | 856356 | A0A044UK00 | A0A044UK00 |
| SRSRDKTGGNTEPM | 0 | 0 | 1 | 856357 | A0A044U9P0 | A0A044U9P0 |
| SRSYADWDEGRG | 0 | 0 | 1 | 856362 | A0A044V9G2 | A0A044V9G2 |
| SRSYNLPPDVDVP | 0 | 0 | 1 | 856363 | A0A044SD24 | A0A044SD24 |
| SRTDEKASNLEETPI | 0 | 0 | 1 | 856365 | AAC48290.1 | A0A044VA40 |
| SSDFTKIVSEMFS | 0 | 0 | 1 | 856369 | A0A044RXP7 | A0A044RXP7 |
| SSDLAKIISEMF | 0 | 0 | 1 | 856370 | A0A044UK00 | A0A044UK00 |
| SSEVSLTTTMELEGA | 0 | 0 | 1 | 856371 | A0A044UK00 | A0A044UK00 |
| SSGRTTRNFAEE | 0 | 0 | 1 | 856374 | A0A044UK00 | A0A044UK00 |
| SSIEYSNMVRQVNMQ | 0 | 0 | 1 | 856376 | A0A044UZ11 | A0A044UZ11 |
| SSKEFVEMEKEQLHE | 0 | 0 | 1 | 856377 | A0A044UK00 | A0A044UK00 |
| SSLEKSYELPDGQ | 0 | 0 | 1 | 856379 | P30162.1 | P30162 |
| SSPTTDSETLGNVLA | 0 | 0 | 1 | 856386 | A0A044SS43 | A0A044SS43 |
| SSREKDIMSEI | 0 | 0 | 1 | 856390 | A0A044RFT9 | A0A044RFT9 |
| SSRGTSSNSKHLTD | 0 | 0 | 1 | 856391 | A0A044UUD9 | A0A044UUD9 |
| SSRKFSDTTAE | 0 | 0 | 1 | 856392 | A0A044UK00 | A0A044UK00 |
| SSSDKSEEIHK | 0 | 0 | 1 | 856394 | AAC48290.1 | A0A044VA40 |
| SSSIDSNNKSKS | 0 | 0 | 1 | 856396 | A0A044V9F8 | A0A044V9F8 |

| | | | | | | |
|---|---|---|---|---|---|---|
| SSTIQKQVDSLFGEK | 0 | 0 | 1 | 856398 | AAA29419.1 | P21250 |
| SSVEQAPAAAPL | 0 | 0 | 1 | 856401 | A0A2K6VJ98 | A0A2K6VJ98 |
| SSVFSQQQLPIE | 0 | 0 | 1 | 856402 | A0A044UMF0 | A0A044UMF0 |
| SSVGNSYESNIENE | 0 | 0 | 1 | 856403 | A0A044UVG8 | A0A044UVG8 |
| SSYKLNGDEVLNAPI | 0 | 0 | 1 | 856405 | A0A044VA40 | A0A044VA40 |
| STADERYTKNLEEIR | 0 | 0 | 1 | 856407 | A0A044UMF0 | A0A044UMF0 |
| STAFNTQSELNAALA | 0 | 0 | 1 | 856408 | A0A044UK00 | A0A044UK00 |
| STDGYESENVESYGQ | 0 | 0 | 1 | 856410 | A0A2K6VJ98 | A0A2K6VJ98 |
| STEINRKQEVATMQK | 0 | 0 | 1 | 856411 | A0A044R153 | A0A044R153 |
| STERDSNIVIRESLN | 0 | 0 | 1 | 856412 | A0A044UVG8 | A0A044UVG8 |
| STIGSTKFETTT | 0 | 0 | 1 | 856419 | AAB68960.1 | A0A044RRG7 |
| STPPHPSANKF | 0 | 0 | 1 | 856424 | A0A044UK00 | A0A044UK00 |
| STSEAKIERDKIEIK | 0 | 0 | 1 | 856426 | A0A044UK00 | A0A044UK00 |
| STTNPAGGTAA | 0 | 0 | 1 | 856430 | AAB68960.1 | A0A044RRG7 |
| STTPELDESYNQC | 0 | 0 | 1 | 856431 | A0A044UK00 | A0A044UK00 |
| STVLSFDLIDEVLK | 0 | 0 | 1 | 856433 | A0A044UVG8 | A0A044UVG8 |
| STVNRDLHTSVEAK | 0 | 0 | 1 | 856434 | A0A044UK00 | A0A044UK00 |
| SVDRDGYWYMQDNSG | 0 | 0 | 1 | 856440 | A0A044UVG8 | A0A044UVG8 |
| SVEAKVVAQETSEAW | 0 | 0 | 1 | 856441 | A0A044UK00 | A0A044UK00 |
| SVIASNKYGTAKSA | 0 | 0 | 1 | 856446 | A0A044UK00 | A0A044UK00 |
| SVLLDDDTCLM | 0 | 0 | 1 | 856451 | A0A044UZ11 | A0A044UZ11 |
| SVQWKRNDIIIEN | 0 | 0 | 1 | 856455 | A0A044UK00 | A0A044UK00 |
| SVRFDEGKIVFNTF | 0 | 0 | 1 | 856456 | AAA20541.1 | A0A044QNW3 |
| SVRFTINLHNTSADF | 0 | 0 | 1 | 856457 | AAA20541.1 | A0A044QNW3 |
| SVTKIGDIVKPHCAS | 0 | 0 | 1 | 856462 | A0A044TU88 | A0A044TU88 |
| SVTTRREKDGRYHD | 0 | 0 | 1 | 856463 | A0A044V9G2 | A0A044V9G2 |
| SWISYWIDWKSTAA | 0 | 0 | 1 | 856465 | A0A044QWA5 | A0A044QWA5 |
| SYCKSFMKKVVEL | 0 | 0 | 1 | 856468 | A0A044SL50 | A0A044SL50 |
| SYGRLWVPDINSDRF | 0 | 0 | 1 | 856469 | A0A044QU24 | A0A044QU24 |
| SYTTKGKFVESDG | 0 | 0 | 1 | 856478 | A0A044QNT8 | A0A044QNT8 |
| TACIVDEDIEIHPFA | 0 | 0 | 1 | 856482 | A0A044U175 | A0A044U175 |
| TAGGVAIGSAVGHA | 0 | 0 | 1 | 856487 | A0A2K6WJQ8 | A0A2K6WJQ8 |
| TAGIFSTYPQP | 0 | 0 | 1 | 856488 | A0A2K6WPE1 | A0A2K6WPE1 |
| TAHYETSTSEAKIER | 0 | 0 | 1 | 856492 | A0A044UK00 | A0A044UK00 |
| TALERKKIVGQNNKY | 0 | 0 | 1 | 856498 | AAB69625.2 | A0A044VD00 |
| TASQDGSLLEIDDVQ | 0 | 0 | 1 | 856504 | A0A044UK00 | A0A044UK00 |
| TATQKTVDGPSG | 0 | 0 | 1 | 856507 | AAB52599.1 | A0A158N7S3 |
| TAVKNSGEPNSELMT | 0 | 0 | 1 | 856508 | P41981.1 | A0A158N7R1 |
| TAYEPINPLQYRPAN | 0 | 0 | 1 | 856510 | A0A044R153 | A0A044R153 |
| TCPMKLQLFPMDSQK | 0 | 0 | 1 | 856515 | A0A044UW71 | A0A044UW71 |
| TDAGQAFDASEEFEV | 0 | 0 | 1 | 856519 | A0A044UK00 | A0A044UK00 |
| TDEERISATSDYEQ | 0 | 0 | 1 | 856521 | A0A044UK00 | A0A044UK00 |
| TDESNSSATNATSDT | 0 | 0 | 1 | 856522 | A0A044UUD9 | A0A044UUD9 |
| TDEYLPSDASCTRRA | 0 | 0 | 1 | 856523 | A0A044SS43 | A0A044SS43 |
| TDLDQETLNRIDYQN | 0 | 0 | 1 | 856528 | AAA74283.1 | A0A044V2F8 |
| TDVDQKINENSKMIE | 0 | 0 | 1 | 856534 | A0A044UK00 | A0A044UK00 |
| TEIIAFSDRISEFK | 0 | 0 | 1 | 856538 | AAC48312.1 | A0A044VFS3 |
| TEKAFQQIGKEFSD | 0 | 0 | 1 | 856539 | A0A044UIG6 | A0A044UIG6 |
| TEQESEKSSHQLVED | 0 | 0 | 1 | 856546 | AAF64250.1 | A0A044UVG8 |
| TEQKYEEVHTRSDR | 0 | 0 | 1 | 856548 | A0A044UK00 | A0A044UK00 |
| TETSITIEQPTFIEH | 0 | 0 | 1 | 856551 | A0A044UK00 | A0A044UK00 |
| TFAHRGSPDDIKTLE | 0 | 0 | 1 | 856554 | AAD00843.1 | A0A044VHF7 |
| TFCAVVHSCFVDDGN | 0 | 0 | 1 | 856556 | A0A044QSA2 | A0A044QSA2 |
| TFDQKSPLTEVDHVS | 0 | 0 | 1 | 856557 | AAD00843.1 | A0A044VHF7 |
| TFSKGEWGKEERKSN | 0 | 0 | 1 | 856568 | AAA20541.1 | A0A044QNW3 |
| TFSSTKTSFIDDDK | 0 | 0 | 1 | 856569 | A0A044SS43 | A0A044SS43 |
| TGDPTGTGKGGDSIW | 0 | 0 | 1 | 856572 | A0A2K6VZ59 | A0A2K6VZ59 |
| TGGAGSGAAGENLYI | 0 | 0 | 1 | 856579 | AAB52600.1 | A0A044UUD9 |
| TGIFNTEFAANYWL | 0 | 0 | 1 | 856581 | AAB68960.1 | A0A044RRG7 |
| TGINEVADTETMRV | 0 | 0 | 1 | 856583 | AAA74283.1 | A0A044V2F8 |
| TGINNAPWTAALEPA | 0 | 0 | 1 | 856584 | A0A044UK00 | A0A044UK00 |
| TGKGGDSIWGGPIED | 0 | 0 | 1 | 856585 | A0A2K6VZ59 | A0A2K6VZ59 |
| TGPPGDKGLKGLDG | 0 | 0 | 1 | 856588 | A0A044V9I3 | A0A044V9I3 |
| TGSSTPVCTTNV | 0 | 0 | 1 | 856591 | A0A044UK00 | A0A044UK00 |
| TGVFTTTEKASAH | 0 | 0 | 1 | 856593 | AAB52599.1 | A0A158N7S3 |
| THEVNEEMSMEIND | 0 | 0 | 1 | 856596 | A0A044UK00 | A0A044UK00 |
| THFEVQSEASYSDAI | 0 | 0 | 1 | 856597 | A0A044UK00 | A0A044UK00 |
| THFEVQSEASYSDAT | 0 | 0 | 1 | 856598 | A0A044RXP7 | A0A044RXP7 |
| THGEVRLQSGTNKFE | 0 | 0 | 1 | 856599 | AAA20215.1 | A0A044V456 |
| THNSFLRIDPDGTVF | 0 | 0 | 1 | 856601 | A0A044UW71 | A0A044UW71 |
| THNWGTENDPNFNYH | 0 | 0 | 1 | 856602 | A0A044RAH9 | A0A044RAH9 |
| TIDDEKQRQSDEE | 0 | 0 | 1 | 856604 | A0A044SS43 | A0A044SS43 |
| TIDDNEAKKLFGEKI | 0 | 0 | 1 | 856605 | AAA50214.2 | P52570 |
| TIDKNEFQKTLKT | 0 | 0 | 1 | 856606 | A0A044TDA1 | A0A044TDA1 |
| TIELALGAKVLDVQA | 0 | 0 | 1 | 856607 | A0A044UU92 | A0A044UU92 |
| TIGQPVYHKWTCDSE | 0 | 0 | 1 | 856610 | A0A044QSA2 | A0A044QSA2 |
| TIKILPDEDDRKTQ | 0 | 0 | 1 | 856614 | A0A044QU24 | A0A044QU24 |
| TILTAPTTCRKNSG | 0 | 0 | 1 | 856616 | AAD51087.1 | Q9U9A1 |
| TINERVEFEKIFQDD | 0 | 0 | 1 | 856617 | A0A044UK00 | A0A044UK00 |
| TIQSSQIDLRNVFQL | 0 | 0 | 1 | 856622 | A0A044UK00 | A0A044UK00 |
| TIRWSCNGKEIQADN | 0 | 0 | 1 | 856625 | A0A044UK00 | A0A044UK00 |
| TISGSVSGLTPGLHG | 0 | 0 | 1 | 856628 | AAB64227.1 | O18483 |
| TITDEYPVPKEFIGK | 0 | 0 | 1 | 856630 | A0A044V9S3 | A0A044V9S3 |
| TITFNRTPEREFCDA | 0 | 0 | 1 | 856631 | A0A044UK00 | A0A044UK00 |
| TITNEHGINNQNTR | 0 | 0 | 1 | 856632 | A0A044UK00 | A0A044UK00 |
| TITNPLEDEANSLHE | 0 | 0 | 1 | 856633 | A0A044UK00 | A0A044UK00 |
| TITVPGDISKTVQDL | 0 | 0 | 1 | 856635 | A0A044SN49 | A0A044SN49 |
| TKARESTITNPLEDE | 0 | 0 | 1 | 856639 | A0A044UK00 | A0A044UK00 |
| TKEKQEVDILIRNDD | 0 | 0 | 1 | 856640 | A0A044UK00 | A0A044UK00 |
| TKKGDVAHQALVIEY | 0 | 0 | 1 | 856643 | A0A044UK00 | A0A044UK00 |
| TKLRDDSSYKLNGDE | 0 | 0 | 1 | 856646 | A0A044VA40 | A0A044VA40 |

| | | | | | | |
|---|---|---|---|---|---|---|
| TKRTANDIAEFGT | 0 | 0 | 1 | 856650 | CAA04390.1 | A0A044R436 |
| TKTSLRDVENIVMLE | 0 | 0 | 1 | 856652 | A0A044TL01 | A0A044TL01 |
| TLEGTPDRDTIFKEL | 0 | 0 | 1 | 856656 | AAC48290.1 | A0A044VA40 |
| TLEKRQMEIREEQ | 0 | 0 | 1 | 856659 | A0A044UK00 | A0A044UK00 |
| TLKALEAETGAKIII | 0 | 0 | 1 | 856665 | A0A044U9P0 | A0A044U9P0 |
| TLNRIDYQNQVQTLQ | 0 | 0 | 1 | 856668 | AAA74283.1 | A0A044V2F8 |
| TLNTNIMDYQKTIGQ | 0 | 0 | 1 | 856669 | AAV71152.1 | A0A044VAG6 |
| TLQEEIDFIRRVQDQ | 0 | 0 | 1 | 856670 | AAA74283.1 | A0A044V2F8 |
| TLVVEERIDEREVTI | 0 | 0 | 1 | 856679 | A0A044UK00 | A0A044UK00 |
| TMDDGSRFANK | 0 | 0 | 1 | 856682 | A0A044V9R9 | A0A044V9R9 |
| TMNFVVEAASSNAN | 0 | 0 | 1 | 856686 | P46434.1 | P46434 |
| TNAIHAGQEAEQWDT | 0 | 0 | 1 | 856687 | A0A044R2U8 | A0A044R2U8 |
| TNANRNDMESWYKL | 0 | 0 | 1 | 856688 | AAA74283.1 | A0A044V2F8 |
| TNDYKSDRTKKKKE | 0 | 0 | 1 | 856690 | A0A044UVG8 | A0A044UVG8 |
| TNEKFKRIANSFLQK | 0 | 0 | 1 | 856691 | AAA65186.1 | Q25619 |
| TNGADKQNFEVTADN | 0 | 0 | 1 | 856692 | A0A044RF80 | A0A044RF80 |
| TNGCTSAGGHYNP | 0 | 0 | 1 | 856693 | AAB64227.1 | O18483 |
| TNGLNHLADLTD | 0 | 0 | 1 | 856694 | AAK16514.1 | A0A2K6W829 |
| TNKENKWEELLK | 0 | 0 | 1 | 856699 | AAA63412.2 | Q99099 |
| TNYPVPYRSKLTESF | 0 | 0 | 1 | 856705 | AAA20541.1 | A0A044QNW3 |
| TPIMSSAKSLTQPQ | 0 | 0 | 1 | 856712 | AAC48290.1 | A0A044VA40 |
| TPRNTQTRVKSENLA | 0 | 0 | 1 | 856715 | P37801.2 | P37801 |
| TPSHMALVEIAEVLA | 0 | 0 | 1 | 856717 | AAB52600.1 | A0A044UUD9 |
| TPTDVQNSDRFAKQG | 0 | 0 | 1 | 856719 | A0A044UIG6 | A0A044UIG6 |
| TPVQQYDPSYRN | 0 | 0 | 1 | 856720 | A0A044SS43 | A0A044SS43 |
| TQDENVEVEIFVKPL | 0 | 0 | 1 | 856721 | A0A044UK00 | A0A044UK00 |
| TQEACTSAKIDLVDR | 0 | 0 | 1 | 856722 | A0A044UVG8 | A0A044UVG8 |
| TQLLPPPLPENP | 0 | 0 | 1 | 856723 | A0A044V9F8 | A0A044V9F8 |
| TRSISDSGTRNSLVQ | 0 | 0 | 1 | 856733 | A0A044UK00 | A0A044UK00 |
| TRVKSENLAEIPEE | 0 | 0 | 1 | 856736 | P37801.2 | P37801 |
| TRVYQSQDKQYKEAG | 0 | 0 | 1 | 856737 | A0A044UK00 | A0A044UK00 |
| TSCVPPSCFPSTSA | 0 | 0 | 1 | 856740 | A0A044SS43 | A0A044SS43 |
| TSDFNEITAREAE | 0 | 0 | 1 | 856741 | A0A044UVG8 | A0A044UVG8 |
| TSDTQNSLSPMHQNM | 0 | 0 | 1 | 856743 | A0A044UK00 | A0A044UK00 |
| TSFIDDDKKQTEDID | 0 | 0 | 1 | 856744 | A0A044SS43 | A0A044SS43 |
| TSITIEQPTFIEHAE | 0 | 0 | 1 | 856745 | A0A044UK00 | A0A044UK00 |
| TSKKKKGNDWDELE | 0 | 0 | 1 | 856748 | A0A044T7P0 | A0A044T7P0 |
| TSLQRDIIVDEHNK | 0 | 0 | 1 | 856750 | AAB97282.1 | A0A044UNG9 |
| TSNADAKMSYQNMTL | 0 | 0 | 1 | 856751 | A0A044UW71 | A0A044UW71 |
| TSNICKNLNEFQVDN | 0 | 0 | 1 | 856752 | A0A044UVG8 | A0A044UVG8 |
| TSPQFVEAHNVQTND | 0 | 0 | 1 | 856754 | A0A044TU88 | A0A044TU88 |
| TSQFESEQNNQRSDG | 0 | 0 | 1 | 856755 | A0A044UK00 | A0A044UK00 |
| TSSEEYINTSTSFEA | 0 | 0 | 1 | 856756 | A0A044UK00 | A0A044UK00 |
| TSYSQFLTDAQKDEL | 0 | 0 | 1 | 856760 | AAD38403.1 | A0A044UUD9 |
| TTAFQTQVVPMPVCR | 0 | 0 | 1 | 856763 | A0A044QZV7 | A0A044QZV7 |
| TTCRKNSGMSPANCA | 0 | 0 | 1 | 856765 | AAD51087.1 | Q9U9A1 |
| TTEIWSQLCHETFAN | 0 | 0 | 1 | 856768 | A0A044UK00 | A0A044UK00 |
| TTHSDCYDQLEP | 0 | 0 | 1 | 856771 | A0A044QNT8 | A0A044QNT8 |
| TTITQSSTIGSTK | 0 | 0 | 1 | 856772 | AAB68960.1 | A0A044RRG7 |
| TTLLRIPGVQQVA | 0 | 0 | 1 | 856775 | AAC48290.1 | A0A044VA40 |
| TTQYYFDGCMVQND | 0 | 0 | 1 | 856778 | AAA29419.1 | P21250 |
| TTSQDVQQIVVEDK | 0 | 0 | 1 | 856782 | A0A044UK00 | A0A044UK00 |
| TTTDELPRTGVA | 0 | 0 | 1 | 856784 | A0A044UK00 | A0A044UK00 |
| TTTIAVEPTGESSD | 0 | 0 | 1 | 856785 | AAB68960.1 | A0A044RRG7 |
| TTVDEETEKAFQ | 0 | 0 | 1 | 856789 | A0A044UIG6 | A0A044UIG6 |
| TVDLTDNDDINVWR | 0 | 0 | 1 | 856794 | A0A044TDA1 | A0A044TDA1 |
| TVGNVKISKFDGSQM | 0 | 0 | 1 | 856797 | A0A044TDA1 | A0A044TDA1 |
| TVIPSQAGWNRGDSQ | 0 | 0 | 1 | 856800 | AAA20215.1 | A0A044V456 |
| TVNFNTRNPFEGH | 0 | 0 | 1 | 856806 | A0A044QSB0 | A0A044QSB0 |
| TVNVINTDDKFEVD | 0 | 0 | 1 | 856807 | A0A044SD32 | A0A044SD32 |
| TVNVPFSRSNKDGVN | 0 | 0 | 1 | 856808 | A0A044UK00 | A0A044UK00 |
| TVPIYEGYALPHAI | 0 | 0 | 1 | 856809 | P30162.1 | P30162 |
| TVSAQLEVSEMTTA | 0 | 0 | 1 | 856811 | A0A044QSA2 | A0A044QSA2 |
| TVSGSQEVYSHTAEI | 0 | 0 | 1 | 856812 | A0A044UK00 | A0A044UK00 |
| TVTMDEAKRIAFDHY | 0 | 0 | 1 | 856814 | AAF64251.1 | A0A044V6Z2 |
| TVVHCIANAMGYAA | 0 | 0 | 1 | 856817 | A0A044UK00 | A0A044UK00 |
| TWLQMQDKINEYIAQ | 0 | 0 | 1 | 856821 | A0A044SAZ1 | A0A044SAZ1 |
| TYAVYVDAINDFDG | 0 | 0 | 1 | 856824 | A0A2K6WPE1 | A0A2K6WPE1 |
| TYEAEAGELPYEDMK | 0 | 0 | 1 | 856825 | AAA20215.1 | A0A044V456 |
| TYENNMEQKAC | 0 | 0 | 1 | 856827 | A0A044UW71 | A0A044UW71 |
| TYKSTMEMYDADKLH | 0 | 0 | 1 | 856832 | A0A2K6VJ98 | A0A2K6VJ98 |
| TYLLHSANSIESTDA | 0 | 0 | 1 | 856833 | A0A044UK00 | A0A044UK00 |
| TYPQPYMTHFGTYLD | 0 | 0 | 1 | 856836 | A0A2K6WPE1 | A0A2K6WPE1 |
| TYTCRAENTYGSSET | 0 | 0 | 1 | 856837 | A0A044UK00 | A0A044UK00 |
| TYVFSQHDEGG | 0 | 0 | 1 | 856838 | A0A044R9D0 | A0A044R9D0 |
| TYVGAEGETKKQM | 0 | 0 | 1 | 856839 | A0A044V3Q0 | A0A044V3Q0 |
| TYYCLAENEYGKA | 0 | 0 | 1 | 856840 | A0A044UK00 | A0A044UK00 |
| VACLIVENVDTSSEK | 0 | 0 | 1 | 856843 | A0A044UK00 | A0A044UK00 |
| VAGTKLMNFNTGGH | 0 | 0 | 1 | 856847 | P41981.1 | A0A158N7R1 |
| VAKKDDKDKEGFQ | 0 | 0 | 1 | 856850 | A0A044R9D0 | A0A044R9D0 |
| VAKNNGNVTGRLEFK | 0 | 0 | 1 | 856851 | A0A044QU24 | A0A044QU24 |
| VANPFMSMSTNTVI | 0 | 0 | 1 | 856858 | A0A044UK00 | A0A044UK00 |
| VAQETSEAWIEAAKE | 0 | 0 | 1 | 856861 | A0A044UK00 | A0A044UK00 |
| VARFSPAAKDADAR | 0 | 0 | 1 | 856863 | P36991.1 | P36991 |
| VAVNDPFINIDYMVY | 0 | 0 | 1 | 856865 | AAB52599.1 | A0A158N7S3 |
| VCTTNVVFDGWEL | 0 | 0 | 1 | 856868 | A0A044UK00 | A0A044UK00 |
| VDDNNDGELNAPEFA | 0 | 0 | 1 | 856870 | AAF64251.1 | A0A044V6Z2 |
| VDGPSGKLWRDGR | 0 | 0 | 1 | 856873 | AAB52599.1 | A0A158N7S3 |
| VDILIRNDDNKFK | 0 | 0 | 1 | 856875 | A0A044UK00 | A0A044UK00 |
| VDKGCYCDRKENACI | 0 | 0 | 1 | 856877 | A0A044QNT8 | A0A044QNT8 |
| VDLEAVYFQPKEIEV | 0 | 0 | 1 | 856879 | A0A044SD32 | A0A044SD32 |

| | | | | | | |
|---|---|---|---|---|---|---|
| VDNDGDEEEEED | 0 | 0 | 1 | 856880 | A0A044SS43 | A0A044SS43 |
| VDNGVEGEPEIECGP | 0 | 0 | 1 | 856881 | A0A044RMM8 | A0A044RMM8 |
| VDNNKSNFAEQS | 0 | 0 | 1 | 856882 | A0A044V9F8 | A0A044V9F8 |
| VDNQVFGKEVSAKLN | 0 | 0 | 1 | 856883 | A0A044UVG8 | A0A044UVG8 |
| VDSYEESSFYERS | 0 | 0 | 1 | 856887 | A0A044UK00 | A0A044UK00 |
| VDTSDSSETDDSEDY | 0 | 0 | 1 | 856889 | A0A044UVG8 | A0A044UVG8 |
| VDVDNSGDLSPGELV | 0 | 0 | 1 | 856890 | AAF64251.1 | A0A044V6Z2 |
| VDYKFGAYDELIQEA | 0 | 0 | 1 | 856894 | A0A044SS43 | A0A044SS43 |
| VECGELKQERKSSMA | 0 | 0 | 1 | 856896 | A0A044U9P0 | A0A044U9P0 |
| VEDEATLNVSTLTAP | 0 | 0 | 1 | 856897 | A0A044UK00 | A0A044UK00 |
| VEDLTSIDGTVVMHK | 0 | 0 | 1 | 856898 | AAF64250.1 | A0A044UVG8 |
| VEEESKKENSAPP | 0 | 0 | 1 | 856899 | AAD00842.1 | A0A044SS43 |
| VEGIINITYEH | 0 | 0 | 1 | 856901 | A0A044UK00 | A0A044UK00 |
| VEIPKFKIENRTELV | 0 | 0 | 1 | 856903 | A0A044V3Q0 | A0A044V3Q0 |
| VEKDGQKYKLFYGM | 0 | 0 | 1 | 856904 | Q9NJD8.1 | Q9NJD8 |
| VEKKGESKEEKKENK | 0 | 0 | 1 | 856905 | A0A044TU88 | A0A044TU88 |
| VELLNADGCALDKYL | 0 | 0 | 1 | 856909 | A0A044QSA2 | A0A044QSA2 |
| VENNPAKVAWAEMIA | 0 | 0 | 1 | 856912 | A0A044SAZ1 | A0A044SAZ1 |
| VEQYPKNMTTINTSK | 0 | 0 | 1 | 856914 | A0A044U175 | A0A044U175 |
| VETDSAWTELMDVQI | 0 | 0 | 1 | 856916 | A0A044TIV1 | A0A044TIV1 |
| VFDGWELLDEHQRS | 0 | 0 | 1 | 856921 | A0A044UK00 | A0A044UK00 |
| VFIFASLIEYAIVNY | 0 | 0 | 1 | 856924 | A0A044QWA5 | A0A044QWA5 |
| VGAYCDLLSEAGDFF | 0 | 0 | 1 | 856937 | AAA63412.2 | Q99099 |
| VGCDSEKKLPYPIIA | 0 | 0 | 1 | 856938 | AAA50214.2 | P52570 |
| VGGQSPVFQLPLNKE | 0 | 0 | 1 | 856943 | A0A044UK00 | A0A044UK00 |
| VGKCPAGLAPNSGYG | 0 | 0 | 1 | 856947 | A0A044UZ11 | A0A044UZ11 |
| VGPPGREGIRGVPGA | 0 | 0 | 1 | 856951 | A0A044SAV6 | A0A044SAV6 |
| VGQSGDNGETGPVGK | 0 | 0 | 1 | 856955 | A0A044SAV6 | A0A044SAV6 |
| VGQSVTEIPKNLTYN | 0 | 0 | 1 | 856956 | A0A044UMF0 | A0A044UMF0 |
| VHSCTVDDGKGDT | 0 | 0 | 1 | 856964 | A0A044R521 | A0A044R521 |
| VHYFFMANLSAAD | 0 | 0 | 1 | 856966 | A0A044UMF0 | A0A044UMF0 |
| VIAENRAGKEHSIGY | 0 | 0 | 1 | 856967 | A0A044UK00 | A0A044UK00 |
| VICPNVYGISDGSTE | 0 | 0 | 1 | 856969 | A0A044R153 | A0A044R153 |
| VIELRSDIVPKTAEN | 0 | 0 | 1 | 856972 | A0A044U9P0 | A0A044U9P0 |
| VIGPNGTAGIFS | 0 | 0 | 1 | 856974 | A0A2K6WPE1 | A0A2K6WPE1 |
| VIKNADMSEDMQQDA | 0 | 0 | 1 | 856975 | A0A044UWQ7 | A0A044UWQ7 |
| VIKSIGSFSADKNI | 0 | 0 | 1 | 856976 | AAK66564.1 | Q963F6 |
| VINPACTITDEYPVP | 0 | 0 | 1 | 856981 | A0A044V9S3 | A0A044V9S3 |
| VIPSLYNAINEVHDA | 0 | 0 | 1 | 856983 | A0A044TIV1 | A0A044TIV1 |
| VIQKSEKSEQMKNL | 0 | 0 | 1 | 856984 | A0A044R196 | A0A044R196 |
| VITQRQWATTENSS | 0 | 0 | 1 | 856991 | A0A044V9F8 | A0A044V9F8 |
| VIVDYAEEQQVQ | 0 | 0 | 1 | 856992 | A0A044UK00 | A0A044UK00 |
| VIVEVTVQSITEISE | 0 | 0 | 1 | 856993 | A0A044SG03 | A0A044SG03 |
| VKAKIDSLKPDAKAF | 0 | 0 | 1 | 856997 | AAA65186.1 | Q25619 |
| VKGNQWYGYDNEETI | 0 | 0 | 1 | 857002 | A0A044RRG7 | A0A044RRG7 |
| VKGQVACSDRSQKDV | 0 | 0 | 1 | 857003 | A0A044V9S3 | A0A044V9S3 |
| VKKLEVKVSEKMVVS | 0 | 0 | 1 | 857005 | A0A044SS43 | A0A044SS43 |
| VKKPDDGRMKGLAFI | 0 | 0 | 1 | 857006 | A0A044RAH9 | A0A044RAH9 |
| VKNRTGNGCPETNIQ | 0 | 0 | 1 | 857017 | A0A044S6B8 | A0A044S6B8 |
| VKSFYRFEHYYFDH | 0 | 0 | 1 | 857024 | A0A044R077 | A0A044R077 |
| VLDKTATSVPLPQQQ | 0 | 0 | 1 | 857033 | AAF64251.1 | A0A044V6Z2 |
| VLECADASHGLNGHV | 0 | 0 | 1 | 857034 | Q9NJD8.1 | Q9NJD8 |
| VLKVEPEGYRVDRN | 0 | 0 | 1 | 857041 | A0A044RF80 | A0A044RF80 |
| VLLGSMKSDRTLDIA | 0 | 0 | 1 | 857044 | A0A044TU88 | A0A044TU88 |
| VLMTFITMNFVVEAA | 0 | 0 | 1 | 857045 | AAG44696.1 | A0A044UM60 |
| VLQDPNHLEKFNIEQ | 0 | 0 | 1 | 857047 | CAA04390.1 | A0A044R436 |
| VLVELYEDDTIDPDD | 0 | 0 | 1 | 857051 | A0A044V9S3 | A0A044V9S3 |
| VMADQDKSGKLDKVE | 0 | 0 | 1 | 857054 | AAF64251.1 | A0A044V6Z2 |
| VMDFQATALKSDSDR | 0 | 0 | 1 | 857056 | A0A044UU92 | A0A044UU92 |
| VMKKASHETLEDGEL | 0 | 0 | 1 | 857060 | A0A044UVG8 | A0A044UVG8 |
| VMLSVTAQSAKAGDA | 0 | 0 | 1 | 857061 | A0A044RF80 | A0A044RF80 |
| VMNESGRVGNCDAAK | 0 | 0 | 1 | 857062 | A0A044UIG6 | A0A044UIG6 |
| VMQPLMTRSISDSGT | 0 | 0 | 1 | 857063 | A0A044UK00 | A0A044UK00 |
| VMSAKVETTVRQ | 0 | 0 | 1 | 857066 | A0A044V9F8 | A0A044V9F8 |
| VMTNIKSLDENLK | 0 | 0 | 1 | 857067 | A0A044R2U8 | A0A044R2U8 |
| VNDLPVGRSVDE | 0 | 0 | 1 | 857070 | AAC48312.1 | A0A044VFS3 |
| VNEYNDDFQQI | 0 | 0 | 1 | 857071 | A0A044QNT8 | A0A044QNT8 |
| VNFTQEDAIDDIITE | 0 | 0 | 1 | 857072 | A0A044V3Q0 | A0A044V3Q0 |
| VNLDVPIKRKNSNQL | 0 | 0 | 1 | 857080 | A0A044UK00 | A0A044UK00 |
| VNLQNTFEAYQLCNM | 0 | 0 | 1 | 857082 | A0A044UK00 | A0A044UK00 |
| VNNDKYDKANN | 0 | 0 | 1 | 857084 | AAB52599.1 | A0A158N7S3 |
| VNPVTGKKMAAKE | 0 | 0 | 1 | 857085 | CAA04390.1 | A0A044R436 |
| VNYDNTVDVSAGFSA | 0 | 0 | 1 | 857088 | A0A044R521 | A0A044R521 |
| VNYVGIFDEHRE | 0 | 0 | 1 | 857089 | A0A044QWA5 | A0A044QWA5 |
| VPDINSDRFQTSSQS | 0 | 0 | 1 | 857104 | A0A044QU24 | A0A044QU24 |
| VPFSMSNIPEEYKEF | 0 | 0 | 1 | 857110 | Q25619.1 | Q25619 |
| VPMQTRQPTAPSAA | 0 | 0 | 1 | 857118 | A0A2K6WJQ8 | A0A2K6WJQ8 |
| VPTESPQVDLPVAD | 0 | 0 | 1 | 857131 | AAF64251.1 | A0A044V6Z2 |
| VPVLKEIAKVQPGEG | 0 | 0 | 1 | 857132 | A0A044U0M2 | A0A044U0M2 |
| VPVLMIADTFPRRP | 0 | 0 | 1 | 857133 | AAC48290.1 | A0A044VA40 |
| VQASFPQKNDSEVMD | 0 | 0 | 1 | 857142 | A0A044UU92 | A0A044UU92 |
| VQDKEYNPRVHGPYC | 0 | 0 | 1 | 857143 | A0A044R077 | A0A044R077 |
| VQGGDFINGDGTGG | 0 | 0 | 1 | 857145 | A0A2K6WB53 | A0A2K6WB53 |
| VQKERQFTNVQAPSL | 0 | 0 | 1 | 857148 | A0A044UK00 | A0A044UK00 |
| VQKERQFTNVQVPSL | 0 | 0 | 1 | 857149 | A0A044UK00 | A0A044UK00 |
| VQKTAEIQSQIERSL | 0 | 0 | 1 | 857150 | A0A044UK00 | A0A044UK00 |
| VQLPQQPIFIDRLP | 0 | 0 | 1 | 857152 | A0A044UK00 | A0A044UK00 |
| VQSITEISEITSSFK | 0 | 0 | 1 | 857157 | A0A044SG03 | A0A044SG03 |
| VRDAKVENSESSEE | 0 | 0 | 1 | 857161 | A0A044SS43 | A0A044SS43 |
| VRKRLEEAGKKKA | 0 | 0 | 1 | 857166 | A0A044TX11 | A0A044TX11 |
| VRNAMENNVWGREER | 0 | 0 | 1 | 857167 | AAD00843.1 | A0A044VHF7 |

| | | | | | | |
|---|---|---|---|---|---|---|
| VRNSLIAGEWGNEER | 0 | 0 | 1 | 857169 | AAA20541.1 | A0A044QNW3 |
| VRPEDIDLPPCVPP | 0 | 0 | 1 | 857170 | A0A044UVG8 | A0A044UVG8 |
| VRSRNGGAAAA | 0 | 0 | 1 | 857175 | A0A044QSA2 | A0A044QSA2 |
| VRSTREFWTLQDRTI | 0 | 0 | 1 | 857176 | A0A044UK00 | A0A044UK00 |
| VSANIAMTTIDGNSD | 0 | 0 | 1 | 857178 | AAC48289.1 | A0A044VA40 |
| VSESNETEKRSLHSH | 0 | 0 | 1 | 857181 | A0A044UK00 | A0A044UK00 |
| VSGKTRNPEICQIQ | 0 | 0 | 1 | 857183 | A0A044RFT9 | A0A044RFT9 |
| VSIVQKERQFTNVQV | 0 | 0 | 1 | 857186 | A0A044UK00 | A0A044UK00 |
| VSKTGICEEINKDVF | 0 | 0 | 1 | 857188 | A0A044UK00 | A0A044UK00 |
| VSMANAGPDSNGSQF | 0 | 0 | 1 | 857190 | A0A2K6WB53 | A0A2K6WB53 |
| VSNAYGAVWSETTVK | 0 | 0 | 1 | 857193 | A0A044UK00 | A0A044UK00 |
| VSSMTNFAVNSKAAE | 0 | 0 | 1 | 857198 | A0A044UVG8 | A0A044UVG8 |
| VSTINLQKVEIVDE | 0 | 0 | 1 | 857200 | A0A044UK00 | A0A044UK00 |
| VTATTTSSPTTDSE | 0 | 0 | 1 | 857209 | A0A044SS43 | A0A044SS43 |
| VTDKDDFQDIITDED | 0 | 0 | 1 | 857212 | A0A044QNT8 | A0A044QNT8 |
| VTERNSQRPHDSSKS | 0 | 0 | 1 | 857214 | AAF64251.1 | A0A044V6Z2 |
| VTIPVYSDDKRHEVH | 0 | 0 | 1 | 857222 | A0A044UK00 | A0A044UK00 |
| VTISALGTETGEDDY | 0 | 0 | 1 | 857223 | A0A044UK00 | A0A044UK00 |
| VTLKDGDDIGYDFLQ | 0 | 0 | 1 | 857226 | A0A044RF80 | A0A044RF80 |
| VTPADGVKWLWMTHN | 0 | 0 | 1 | 857230 | A0A044SAZ1 | A0A044SAZ1 |
| VTQLKTFDAKMTAYQ | 0 | 0 | 1 | 857232 | AAA29419.1 | P21250 |
| VTQQTNEQYATFEI | 0 | 0 | 1 | 857233 | Q9NJD8.1 | Q9NJD8 |
| VTTEASSSSDKSE | 0 | 0 | 1 | 857237 | AAC48290.1 | A0A044VA40 |
| VTTNARFEFTSTSVA | 0 | 0 | 1 | 857238 | A0A044UK00 | A0A044UK00 |
| VTVPSMVTATTTSSP | 0 | 0 | 1 | 857241 | A0A044SS43 | A0A044SS43 |
| VVDLTCRLQKGASMD | 0 | 0 | 1 | 857247 | AAB52599.1 | A0A158N7S3 |
| VVDNKLFANSFFLRE | 0 | 0 | 1 | 857248 | AAA29419.1 | P21250 |
| VVDVRTDLNTLDINE | 0 | 0 | 1 | 857249 | A0A044QSA2 | A0A044QSA2 |
| VVKRYNKRFAGFNVA | 0 | 0 | 1 | 857255 | CAA31690.1 | P21250 |
| VVNYEFDRTEPLSAE | 0 | 0 | 1 | 857259 | A0A044UK00 | A0A044UK00 |
| VVRENQYGQDIWELI | 0 | 0 | 1 | 857262 | A0A044U175 | A0A044U175 |
| VVSNMTATNSNVTSC | 0 | 0 | 1 | 857263 | A0A044SS43 | A0A044SS43 |
| VVTEIDSSIEYSNMV | 0 | 0 | 1 | 857265 | A0A044UZ11 | A0A044UZ11 |
| VWRDWEPISIPDPS | 0 | 0 | 1 | 857271 | A0A044SG03 | A0A044SG03 |
| VYDISKRDVSKTPQQ | 0 | 0 | 1 | 857272 | A0A044UK00 | A0A044UK00 |
| VYFKGNWKKSFNKKM | 0 | 0 | 1 | 857274 | A0A044V3Q0 | A0A044V3Q0 |
| VYGEDKEVTAIEPYL | 0 | 0 | 1 | 857276 | A0A044V9S0 | A0A044V9S0 |
| VYNDWVYQKPVEWP | 0 | 0 | 1 | 857281 | A0A044SAZ1 | A0A044SAZ1 |
| VYNTLVNGIWGSSE | 0 | 0 | 1 | 857282 | AAD00843.1 | A0A044VHF7 |
| VYPPGIDGSKKYYIH | 0 | 0 | 1 | 857283 | A0A2K6VRV9 | A0A2K6VRV9 |
| WAAGILLEQRSR | 0 | 0 | 1 | 857286 | A0A044U175 | A0A044U175 |
| WATTENSSSIDSNN | 0 | 0 | 1 | 857292 | A0A044V9F8 | A0A044V9F8 |
| WDCELEISAQNWA | 0 | 0 | 1 | 857293 | AAB97282.1 | A0A044UNG9 |
| WEDRDPKDEEILEL | 0 | 0 | 1 | 857299 | P22085.2 | P22085 |
| WEEDTGDPDDLLNS | 0 | 0 | 1 | 857301 | A0A044V9S3 | A0A044V9S3 |
| WEICKYLKEGGKE | 0 | 0 | 1 | 857303 | AAB68960.1 | A0A044RRG7 |
| WERKGNKWIPAKESK | 0 | 0 | 1 | 857305 | AAF64253.1 | A0A2K6VYS3 |
| WGGKPENIQAAKEAF | 0 | 0 | 1 | 857311 | AAB52600.1 | A0A044UUD9 |
| WGKECIVDKKGEPIC | 0 | 0 | 1 | 857312 | A0A044T7P0 | A0A044T7P0 |
| WGKEERKSNPYKKGD | 0 | 0 | 1 | 857313 | AAA20541.1 | A0A044QNW3 |
| WGYSYYGGQQDFGIG | 0 | 0 | 1 | 857317 | AAB69626.2 | O16855 |
| WIPDTFFQNERDGW | 0 | 0 | 1 | 857323 | A0A044QWA5 | A0A044QWA5 |
| WKKSFNKKMTKK | 0 | 0 | 1 | 857326 | A0A044V3Q0 | A0A044V3Q0 |
| WKLKRKIDGKREIVY | 0 | 0 | 1 | 857327 | AAA74283.1 | A0A044V2F8 |
| WLLGDKEIEPTETY | 0 | 0 | 1 | 857329 | A0A044R9D0 | A0A044R9D0 |
| WPSFQTSDNYVRLED | 0 | 0 | 1 | 857357 | A0A044SAZ1 | A0A044SAZ1 |
| WQLARDGKEDSKVL | 0 | 0 | 1 | 857361 | AAA29419.1 | P21250 |
| WRKNDIQIEFSDDA | 0 | 0 | 1 | 857364 | A0A044UK00 | A0A044UK00 |
| WSIGKKYEKAGEIEA | 0 | 0 | 1 | 857368 | A0A044UK00 | A0A044UK00 |
| WYFGEQKLEQSEQIN | 0 | 0 | 1 | 857378 | A0A044UK00 | A0A044UK00 |
| YAAEEMNAGGYK | 0 | 0 | 1 | 857381 | A0A044V9R9 | A0A044V9R9 |
| YAETYKQLANQNLQ | 0 | 0 | 1 | 857384 | A0A044SAZ1 | A0A044SAZ1 |
| YAPTVPEYLSPKQRP | 0 | 0 | 1 | 857386 | A0A044U9P0 | A0A044U9P0 |
| YASKEISTADERYTK | 0 | 0 | 1 | 857387 | A0A044UMF0 | A0A044UMF0 |
| YATVAYSKKKYEDE | 0 | 0 | 1 | 857388 | A0A044UW71 | A0A044UW71 |
| YDDSETFKAYARET | 0 | 0 | 1 | 857400 | A0A044UIG6 | A0A044UIG6 |
| YDISQITEQESEK | 0 | 0 | 1 | 857403 | AAF64250.1 | A0A044UVG8 |
| YDKEGLAYLYQQDAD | 0 | 0 | 1 | 857404 | AAD00842.1 | A0A044SS43 |
| YDPEQQVKTTYA | 0 | 0 | 1 | 857406 | A0A044RF80 | A0A044RF80 |
| YDPMKEETPIGVRK | 0 | 0 | 1 | 857408 | A0A044UH02 | A0A044UH02 |
| YDPNRLNFFTTEAR | 0 | 0 | 1 | 857409 | AAF64254.1 | A0A044QLL7 |
| YDREVKSNSGRRINK | 0 | 0 | 1 | 857411 | A0A044V9G2 | A0A044V9G2 |
| YDTEMLNEVNNIPE | 0 | 0 | 1 | 857415 | A0A044TDA1 | A0A044TDA1 |
| YEALPKVMQQKPLTD | 0 | 0 | 1 | 857417 | A0A044UK00 | A0A044UK00 |
| YEDNRITRDEWKYLK | 0 | 0 | 1 | 857418 | P46434.1 | P46434 |
| YEHLQEIVIEDSDAL | 0 | 0 | 1 | 857420 | A0A044UMF0 | A0A044UMF0 |
| YEKAGEIEADVVAA | 0 | 0 | 1 | 857421 | A0A044UK00 | A0A044UK00 |
| YELAEDAAAAIDNMN | 0 | 0 | 1 | 857423 | A0A044U9P0 | A0A044U9P0 |
| YELPDGQVITVGNER | 0 | 0 | 1 | 857424 | P30162.1 | P30162 |
| YENNPSNNMTWK | 0 | 0 | 1 | 857425 | AAB69625.2 | A0A044VD00 |
| YESGQKDVCINPYHY | 0 | 0 | 1 | 857426 | A0A044V1C1 | A0A044V1C1 |
| YESNIENEILGKHDD | 0 | 0 | 1 | 857427 | A0A044UVG8 | A0A044UVG8 |
| YETEKDPYIKSIL | 0 | 0 | 1 | 857429 | P46427.1 | P46427 |
| YEVPECEDRPRYGGP | 0 | 0 | 1 | 857430 | A0A044TX11 | A0A044TX11 |
| YFDVVGKDDTKSLTS | 0 | 0 | 1 | 857432 | A0A044UK00 | A0A044UK00 |
| YFPHFGTNAIHAGQE | 0 | 0 | 1 | 857436 | A0A044R2U8 | A0A044R2U8 |
| YFPYYEDDDVQVLG | 0 | 0 | 1 | 857437 | A0A044V3Q0 | A0A044V3Q0 |
| YGCGEQNMINFVPNI | 0 | 0 | 1 | 857441 | A0A044RF80 | A0A044RF80 |
| YGEECEVGGIEPYLR | 0 | 0 | 1 | 857443 | A0A044V9S3 | A0A044V9S3 |
| YGISDGSTEINRKQE | 0 | 0 | 1 | 857445 | A0A044R153 | A0A044R153 |
| YGQDIWELIGERNPL | 0 | 0 | 1 | 857448 | A0A044U175 | A0A044U175 |

| | | | | | | |
|---|---|---|---|---|---|---|
| YGQKGVEQAPPEIEQ | 0 | 0 | 1 | 857449 | A0A2K6VJ98 | A0A2K6VJ98 |
| YGVLKEDEGIAYRG | 0 | 0 | 1 | 857453 | AAC48312.1 | A0A044VFS3 |
| YGYDNEETIRMKMRW | 0 | 0 | 1 | 857454 | AAB68960.1 | A0A044RRG7 |
| YHHRGGGSREAYG | 0 | 0 | 1 | 857459 | A0A044V9G2 | A0A044V9G2 |
| YHRTTMRDDLPFYC | 0 | 0 | 1 | 857464 | A0A044RF80 | A0A044RF80 |
| YIHLETKIGINGKPE | 0 | 0 | 1 | 857470 | A0A2K6VRV9 | A0A2K6VRV9 |
| YINLNVNVERTPSLF | 0 | 0 | 1 | 857473 | A0A044UK00 | A0A044UK00 |
| YIPTKIEQIEEET | 0 | 0 | 1 | 857475 | CAA04390.1 | A0A044R436 |
| YIQEVLNLLPDVTSP | 0 | 0 | 1 | 857476 | A0A044TU88 | A0A044TU88 |
| YKAKFYSSGGENSAT | 0 | 0 | 1 | 857479 | A0A044R9D0 | A0A044R9D0 |
| YKPEEIGEHATTIVD | 0 | 0 | 1 | 857486 | A0A044UK00 | A0A044UK00 |
| YKPFATMQKAHHMLL | 0 | 0 | 1 | 857487 | A0A044U175 | A0A044U175 |
| YKVSDFENGRLVQTE | 0 | 0 | 1 | 857488 | A0A044SAZ1 | A0A044SAZ1 |
| YLGVLAIADYKTSNE | 0 | 0 | 1 | 857493 | A0A044UMF0 | A0A044UMF0 |
| YLHEVAEGKLPINHA | 0 | 0 | 1 | 857494 | A0A044TU88 | A0A044TU88 |
| YLKKYGVNPVTGK | 0 | 0 | 1 | 857498 | CAA04390.1 | A0A044R436 |
| YLTGGETDEERISA | 0 | 0 | 1 | 857505 | A0A044UK00 | A0A044UK00 |
| YMEADKTVSAQLEVS | 0 | 0 | 1 | 857508 | A0A044QSA2 | A0A044QSA2 |
| YNDSVVLPRAKENL | 0 | 0 | 1 | 857513 | AAF64254.1 | A0A044QLL7 |
| YNESEHREMPVQ | 0 | 0 | 1 | 857514 | A0A044UK00 | A0A044UK00 |
| YNFSKSTQDENVEVE | 0 | 0 | 1 | 857515 | A0A044UK00 | A0A044UK00 |
| YPDGRYELISPQCAP | 0 | 0 | 1 | 857522 | A0A044UK00 | A0A044UK00 |
| YPQLASFVNKIGQMP | 0 | 0 | 1 | 857530 | P46434.1 | P46434 |
| YQCHVSREFCADTIF | 0 | 0 | 1 | 857536 | A0A044UK00 | A0A044UK00 |
| YQHLDLTSPAQTSEC | 0 | 0 | 1 | 857537 | A0A044UK00 | A0A044UK00 |
| YQKPVEWPSFQTSDN | 0 | 0 | 1 | 857539 | A0A044SAZ1 | A0A044SAZ1 |
| YQNQVQTLQEEIDFI | 0 | 0 | 1 | 857542 | AAA74283.1 | A0A044V2F8 |
| YRIQCFYMEADKTVS | 0 | 0 | 1 | 857548 | A0A044QSA2 | A0A044QSA2 |
| YRKMLEGEGDGPGL | 0 | 0 | 1 | 857549 | AAA74283.1 | A0A044V2F8 |
| YRMQKLSDNTHTLTI | 0 | 0 | 1 | 857550 | A0A044UK00 | A0A044UK00 |
| YRRDDHSFSAFGQ | 0 | 0 | 1 | 857553 | A0A044RF80 | A0A044RF80 |
| YRSKLTESFEPGQTL | 0 | 0 | 1 | 857554 | AAA20541.1 | A0A044QNW3 |
| YSNGGSNTNNSSDN | 0 | 0 | 1 | 857564 | A0A044UK00 | A0A044UK00 |
| YSPISGRYEVIAMNS | 0 | 0 | 1 | 857566 | A0A044UK00 | A0A044UK00 |
| YSRYYKNTAEQKASN | 0 | 0 | 1 | 857567 | A0A044UVG8 | A0A044UVG8 |
| YTRSAKGNVSIQETS | 0 | 0 | 1 | 857574 | A0A044V2F8 | A0A044V2F8 |
| YTSTSVCLPLSIEDN | 0 | 0 | 1 | 857575 | A0A044UMF0 | A0A044UMF0 |
| YTYDPNEKLPVSVD | 0 | 0 | 1 | 857580 | AAK16514.1 | A0A2K6W829 |
| YVDELFDFDPVKA | 0 | 0 | 1 | 857582 | A0A044UK00 | A0A044UK00 |
| YVDQKEVKEYEHRVP | 0 | 0 | 1 | 857584 | AAA20541.1 | A0A044QNW3 |
| YVMALGKHYDSNESN | 0 | 0 | 1 | 857586 | AAK16514.1 | A0A2K6W829 |
| YVNALNQAEEKVKEA | 0 | 0 | 1 | 857587 | P41981.1 | A0A158N7R1 |
| YWYMQDNSGNFIPYT | 0 | 0 | 1 | 857593 | A0A044UVG8 | A0A044UVG8 |
| YYLAQGEEGLDL | 0 | 0 | 1 | 857597 | A0A044QU24 | A0A044QU24 |
| MQKDEESGEFKCEL | 0 | 0 | 1 | 886666 | XP_003873457.1 | E9APT0 |
| ACPKENLKMIRERVN | 0 | 0 | 1 | 912918 | GI:124511718 | Q8IC11 |
| AEPAPQEQNLFSSIE | 0 | 0 | 1 | 912920 | GI:1373027 | Q7KQK5 |
| AFFIALKYFNRCLGD | 0 | 0 | 1 | 912922 | GI:124506543 | A0A5K1K7X4 |
| AFNTEKTNVNNDALK | 0 | 0 | 1 | 912923 | GI:124804085 | Q8IIE8 |
| AHETVEHEETVSQES | 0 | 0 | 1 | 912925 | GI:124802897 | Q8IJ56 |
| AHKEDRQSSNSSVSF | 0 | 0 | 1 | 912926 | GI:124806619 | Q8I544 |
| AHQEEIVPEQNDEES | 0 | 0 | 1 | 912927 | GI:124802897 | Q8IJ56 |
| AHVMNTIDKYVELTV | 0 | 0 | 1 | 912928 | GI:124511708 | Q8IC16 |
| AIGISREINRKMPFV | 0 | 0 | 1 | 912929 | GI:124803531 | Q8IIU3 |
| AIQFMNLNSDNCNNN | 0 | 0 | 1 | 912931 | GI:124512538 | Q8IAV9 |
| AIYYPKNVDDDGAIT | 0 | 0 | 1 | 912932 | GI:258597905 | Q8IKI0 |
| AKINLGPNFTYNHLQ | 0 | 0 | 1 | 912934 | GI:124512258 | Q8IB96 |
| AKPGDILYLLGVKRK | 0 | 0 | 1 | 912935 | GI:124512538 | Q8IAV9 |
| AMKLIQQLNLNDNAI | 0 | 0 | 1 | 912937 | GI:124513464 | Q76NM2 |
| AMYSLYDSMNNDLQH | 0 | 0 | 1 | 912938 | GI:124507147 | Q8I0U8 |
| ANPDADSESNGEPNA | 0 | 0 | 1 | 912940 | GI:124804079 | Q8IIF0 |
| ANPNANPNVDPNANP | 0 | 0 | 1 | 912942 | GI:124504759 | Q7K740 |
| ANPNKNNQGNGQGHN | 0 | 0 | 1 | 912943 | GI:124504759 | Q7K740 |
| ANTIYVNVGQSGING | 0 | 0 | 1 | 912944 | GI:124802266 | Q8IJQ4 |
| AQNLISKNQNNNEKN | 0 | 0 | 1 | 912945 | GI:124802901 | Q8IJ55 |
| AREIIRLHSDASKNK | 0 | 0 | 1 | 912949 | GI:124513464 | Q76NM2 |
| ASCGVWDEWSPCSVT | 0 | 0 | 1 | 912952 | GI:124513464 | Q76NM2 |
| ASVDPEVKEKENVSE | 0 | 0 | 1 | 912954 | GI:124802897 | Q8IJ56 |
| ATKKVTENVITNQIL | 0 | 0 | 1 | 912955 | GI:124511650 | Q8IC44 |
| AVEKPNENIIDNNPQ | 0 | 0 | 1 | 912959 | GI:124513464 | Q76NM2 |
| AVPLAMKLIQQLNLN | 0 | 0 | 1 | 912961 | GI:124513464 | Q76NM2 |
| AYDLKIISSCNKYYV | 0 | 0 | 1 | 912963 | GI:124804085 | Q8IIE8 |
| AYTYYLLLCELIKNK | 0 | 0 | 1 | 912965 | GI:1371546548 | Q8IEB7 |
| CASYCMNITDVLELS | 0 | 0 | 1 | 912966 | GI:124802266 | Q8IJQ4 |
| CIKHTEIRQCDILSY | 0 | 0 | 1 | 912973 | GI:124504927 | O77338 |
| CKQLFDNINLILKTN | 0 | 0 | 1 | 912975 | GI:1371546465 | Q8I369 |
| CNIYNFNLNLFHPLN | 0 | 0 | 1 | 912977 | GI:296004981 | C0H4T3 |
| CNYSNINNALDDAEE | 0 | 0 | 1 | 912978 | GI:124511718 | Q8IC11 |
| DDDDNNNDHHNNDDD | 0 | 0 | 1 | 912982 | GI:124807012 | Q8I4T2 |
| DDDDNNNMNDIAIGE | 0 | 0 | 1 | 912983 | GI:1371546738 | A0A143ZXM2 |
| DDDNNDNHHNNDDNN | 0 | 0 | 1 | 912984 | GI:124807012 | Q8I4T2 |
| DDHHNDDDDNNNDHH | 0 | 0 | 1 | 912986 | GI:124807012 | Q8I4T2 |
| DDNEENNDDDNTYEL | 0 | 0 | 1 | 912987 | GI:124513450 | Q8IDX8 |
| DDSLVVLPIFGESED | 0 | 0 | 1 | 912988 | GI:124507147 | Q8I0U8 |
| DDTFNIEKKSNYKDS | 0 | 0 | 1 | 912989 | GI:124804085 | Q8IIE8 |
| DEEEMNNEEDHKESD | 0 | 0 | 1 | 912992 | GI:124511718 | Q8IC11 |
| DELENYNKQNENEFD | 0 | 0 | 1 | 912993 | GI:1371546738 | A0A143ZXM2 |
| DELSEDITKYFMKLG | 0 | 0 | 1 | 912994 | SRC7359 | A0A143ZZD7 |
| DENTLQHAYPIDHEG | 0 | 0 | 1 | 912995 | GI:1373027 | Q7KQK5 |
| DGDKYLINEKIYEGE | 0 | 0 | 1 | 912996 | GI:124804085 | Q8IIE8 |
| DGIPDSIQDSLKESR | 0 | 0 | 1 | 912997 | GI:124513464 | Q76NM2 |

| | | | | | | |
|---|---|---|---|---|---|---|
| DGSSYNNTINIQFYI | 0 | 0 | 1 | 912998 | GI:1371545989 | O97230 |
| DIDNELIDMKDGNVD | 0 | 0 | 1 | 913001 | GI:1371546738 | A0A143ZXM2 |
| DIIIILKSFCILKYD | 0 | 0 | 1 | 913003 | GI:124511718 | Q8IC11 |
| DILYHDHMKFLYNLM | 0 | 0 | 1 | 913004 | GI:1371546050 | A0A143ZV68 |
| DIQIDNDHTIEDIEN | 0 | 0 | 1 | 913005 | GI:258597476 | Q8I5R4 |
| DISNQMNVKNGNVEV | 0 | 0 | 1 | 913006 | GI:1371546738 | A0A143ZXM2 |
| DKDKDKDNTDEGDEG | 0 | 0 | 1 | 913007 | GI:124505267 | Q6ZMA7 |
| DKDSLKCPCDPEMVS | 0 | 0 | 1 | 913008 | GI:1373027 | Q7KQK5 |
| DKIVNTEIKYYLFFL | 0 | 0 | 1 | 913009 | GI:124807012 | Q8I4T2 |
| DKIYHGIYKKEHVDD | 0 | 0 | 1 | 913010 | GI:124511718 | Q8IC11 |
| DKKCHILYIAAQENN | 0 | 0 | 1 | 913011 | GI:1373027 | Q7KQK5 |
| DKKTTVNRNLIISTA | 0 | 0 | 1 | 913012 | GI:124505267 | Q6ZMA7 |
| DKLDYLNFDQTIKSF | 0 | 0 | 1 | 913013 | GI:258597736 | Q8ILH5 |
| DKNEKVEHEIVEVEE | 0 | 0 | 1 | 913014 | GI:124802897 | Q8IJ56 |
| DKSVTEPAEHEEVVS | 0 | 0 | 1 | 913015 | GI:124802897 | Q8IJ56 |
| DKTKIAYITIQINNG | 0 | 0 | 1 | 913016 | GI:308191562 | C0H5F4 |
| DKTYTDINKNVTKYA | 0 | 0 | 1 | 913017 | GI:124505325 | Q9U0J6 |
| DKYVELKGLIAEKII | 0 | 0 | 1 | 913019 | GI:124506239 | Q8I3U8 |
| DLEQERRAKEKLQEQ | 0 | 0 | 1 | 913020 | GI:1371546755 | A0A143ZZD7 |
| DLISDMIKKEEELVE | 0 | 0 | 1 | 913021 | GI:124804079 | Q8IIF0 |
| DLLKKSCKSSKAIEQ | 0 | 0 | 1 | 913022 | SRC7364 | Q8I0U6 |
| DLNNEEVYMDKHEMD | 0 | 0 | 1 | 913023 | GI:1371546738 | A0A143ZXM2 |
| DNDEDKKEEMYKDVY | 0 | 0 | 1 | 913025 | GI:1371546738 | A0A143ZXM2 |
| DNDEGGFEEAHHENF | 0 | 0 | 1 | 913026 | GI:124802897 | Q8IJ56 |
| DNDIKQSDDIYMNED | 0 | 0 | 1 | 913027 | GI:1371546738 | A0A143ZXM2 |
| DNDNADTILTDNDNN | 0 | 0 | 1 | 913028 | GI:124511650 | Q8IC44 |
| DNDNADTKATDNNNN | 0 | 0 | 1 | 913029 | GI:124511650 | Q8IC44 |
| DNDNNTDIILTDNNN | 0 | 0 | 1 | 913030 | GI:124511650 | Q8IC44 |
| DNDYNNNNNNNNNNK | 0 | 0 | 1 | 913031 | GI:124804085 | Q8IIE8 |
| DNHHNNDDNNDDHHN | 0 | 0 | 1 | 913032 | GI:124807012 | Q8I4T2 |
| DNIFALLHIAKTLNK | 0 | 0 | 1 | 913033 | GI:124802985 | Q8IJ31 |
| DNIKKDITINENNNN | 0 | 0 | 1 | 913034 | GI:124511718 | Q8IC11 |
| DNIQNENDHIIDNIQ | 0 | 0 | 1 | 913035 | GI:258597476 | Q8I5R4 |
| DNKLNVINEDNIKEE | 0 | 0 | 1 | 913036 | GI:1371546738 | A0A143ZXM2 |
| DNKNTDIKATDNNNI | 0 | 0 | 1 | 913037 | GI:124511650 | Q8IC44 |
| DNKYKIAGGIAGGLA | 0 | 0 | 1 | 913038 | GI:124513464 | Q76NM2 |
| DNLKNRLFKIEKVIG | 0 | 0 | 1 | 913039 | GI:1371547110 | A0A144A160 |
| DNNNTKTISNDNNNT | 0 | 0 | 1 | 913040 | GI:124511650 | Q8IC44 |
| DNQNDNMNIVLGKIH | 0 | 0 | 1 | 913041 | GI:124802266 | Q8IJQ4 |
| DNSKTNTISTDNDNA | 0 | 0 | 1 | 913042 | GI:124511650 | Q8IC44 |
| DNYISFNVIFGEFKK | 0 | 0 | 1 | 913043 | GI:124808066 | Q8IM47 |
| DPADNANPDADSESN | 0 | 0 | 1 | 913044 | GI:124804079 | Q8IIF0 |
| DQPRPRGDNFAVEKP | 0 | 0 | 1 | 913045 | GI:124513464 | Q76NM2 |
| DSEKEVPSDVPKNPE | 0 | 0 | 1 | 913047 | GI:124513464 | Q76NM2 |
| DSESNGEPNAGPQVT | 0 | 0 | 1 | 913049 | GI:124804079 | Q8IIF0 |
| DTILTDNDNNTDIIL | 0 | 0 | 1 | 913052 | GI:124511650 | Q8IC44 |
| DTSEDYHINTGAKKL | 0 | 0 | 1 | 913053 | GI:258597581 | Q8I4Y2 |
| DTTFFWINQKYQDVK | 0 | 0 | 1 | 913054 | GI:258597793 | Q8IL72 |
| DVLAEDLYGRLEIPA | 0 | 0 | 1 | 913055 | GI:1371546755 | A0A143ZZD7 |
| DYEVQNYLYLLIFQR | 0 | 0 | 1 | 913057 | GI:124511718 | Q8IC11 |
| DYKSEDIPLNTQPNT | 0 | 0 | 1 | 913058 | GI:124505545 | Q8IFQ5 |
| DYYHKDIFSIFEGVS | 0 | 0 | 1 | 913059 | GI:124809259 | Q8IL90 |
| EDDDDIDFEGLSRKD | 0 | 0 | 1 | 913062 | GI:124802897 | Q8IJ56 |
| EDDNNDDDNNDNHHN | 0 | 0 | 1 | 913063 | GI:124807012 | Q8I4T2 |
| EDDQPRPRGDNFAVE | 0 | 0 | 1 | 913064 | GI:124513464 | Q76NM2 |
| EDKNEKVQHEIVEVE | 0 | 0 | 1 | 913073 | GI:124802897 | Q8IJ56 |
| EDKNNENAYLNKTLR | 0 | 0 | 1 | 913074 | GI:124504821 | O77318 |
| EDLIEKNENLDDLDE | 0 | 0 | 1 | 913076 | GI:1371546755 | A0A143ZZD7 |
| EDNIKEEDDKESVVH | 0 | 0 | 1 | 913077 | GI:1371546738 | A0A143ZXM2 |
| EDNKNRYNITTNKNN | 0 | 0 | 1 | 913078 | GI:124804085 | Q8IIE8 |
| EDSGEDENTLQHAYP | 0 | 0 | 1 | 913079 | GI:1373027 | Q7KQK5 |
| EDVCHNNNVEDVCHN | 0 | 0 | 1 | 913080 | GI:1371546738 | A0A143ZXM2 |
| EDVDHFKQPSLRLEV | 0 | 0 | 1 | 913081 | GI:124504889 | O77365 |
| EDVLLIETITINNGT | 0 | 0 | 1 | 913082 | GI:124802266 | Q8IJQ4 |
| EEENKSKSLQISYNG | 0 | 0 | 1 | 913089 | GI:124804085 | Q8IIE8 |
| EEGQNKNQNVDHAHT | 0 | 0 | 1 | 913090 | GI:1371546548 | Q8IEB7 |
| EEKIRKFLLDKNINL | 0 | 0 | 1 | 913091 | GI:262527542 | Q8IBG1 |
| EEKQNEQKLKDQIKI | 0 | 0 | 1 | 913093 | GI:124802266 | Q8IJQ4 |
| EERREQYDNYGKNAT | 0 | 0 | 1 | 913096 | GI:124803007 | Q8IJ23 |
| EFIGKEKNTDIYNNI | 0 | 0 | 1 | 913098 | GI:258597476 | Q8I5R4 |
| EFIGKNKKYNNSNNN | 0 | 0 | 1 | 913099 | GI:124506535 | Q8I3F7 |
| EGDFEEPNHEEFEPD | 0 | 0 | 1 | 913103 | GI:124802897 | Q8IJ56 |
| EGGFLLDNDEDDNNN | 0 | 0 | 1 | 913105 | GI:1371546309 | A0A143ZWE7 |
| EGVSKIFNVITDMYK | 0 | 0 | 1 | 913108 | GI:124505699 | Q8I2A9 |
| EHFNDNYDPVKIETS | 0 | 0 | 1 | 913111 | GI:74864292 | Q8ILR9 |
| EHIFNNNDINNNVQY | 0 | 0 | 1 | 913112 | GI:124804085 | Q8IIE8 |
| EHNNNINTNTNSTEN | 0 | 0 | 1 | 913113 | GI:124511650 | Q8IC44 |
| EIDMHDPGQDLSNSI | 0 | 0 | 1 | 913115 | GI:124804085 | Q8IIE8 |
| EIDNEYKYEDTNMNN | 0 | 0 | 1 | 913116 | GI:1371546309 | A0A143ZWE7 |
| EIGMYKENDMETLNM | 0 | 0 | 1 | 913117 | GI:124506255 | Q8I3U0 |
| EILPEDDKNEKVEHE | 0 | 0 | 1 | 913118 | GI:124802897 | Q8IJ56 |
| EILPEDKNEKGQHEI | 0 | 0 | 1 | 913119 | GI:124802897 | Q8IJ56 |
| EIPGISNLLRHAIFY | 0 | 0 | 1 | 913121 | GI:124804085 | Q8IIE8 |
| EIVEVEEILPEDKNE | 0 | 0 | 1 | 913123 | GI:124802897 | Q8IJ56 |
| EIVSEKSVSEPAEHV | 0 | 0 | 1 | 913124 | GI:124802897 | Q8IJ56 |
| EKALIIIKSLLSTNL | 0 | 0 | 1 | 913126 | GI:124513464 | Q76NM2 |
| EKDGCKFEEINKEKK | 0 | 0 | 1 | 913127 | GI:1371546251 | C6KTC4 |
| EKEQSNENNDQKKDM | 0 | 0 | 1 | 913129 | GI:124802901 | Q8IJ55 |
| EKERAKIGDFGLAKF | 0 | 0 | 1 | 913130 | GI:296004592 | C0H4G2 |
| EKKRNEILQMLILKP | 0 | 0 | 1 | 913132 | GI:124808412 | Q8ILW0 |
| EKNENLDDLDEGIEK | 0 | 0 | 1 | 913133 | GI:1371546755 | A0A143ZZD7 |

| | | | | | | |
|---|---|---|---|---|---|---|
| EKNTDIYNNISSEKK | 0 | 0 | 1 | 913134 | GI:258597476 | Q8I5R4 |
| EKRICVSYNMKSHTF | 0 | 0 | 1 | 913136 | GI:124504821 | O77318 |
| EKSFNHSCSDIQFNQ | 0 | 0 | 1 | 913137 | GI:258597796 | Q8IL70 |
| ELAEISTSNLFYPKK | 0 | 0 | 1 | 913138 | GI:124802266 | Q8IJQ4 |
| ELEKRKKNLGKTNNT | 0 | 0 | 1 | 913140 | GI:124808513 | Q8ILT4 |
| ELKTDNILDNTSEQD | 0 | 0 | 1 | 913141 | GI:124804085 | Q8IIE8 |
| ELVNDYILLRKNGLT | 0 | 0 | 1 | 913142 | GI:124511718 | Q8IC11 |
| ENANANSAVKNNNNE | 0 | 0 | 1 | 913143 | GI:124504759 | Q7K740 |
| ENDDVLDEKEEEAEE | 0 | 0 | 1 | 913144 | GI:124802901 | Q8IJ55 |
| ENGDNTFNNNNNNMD | 0 | 0 | 1 | 913145 | GI:124802266 | Q8IJQ4 |
| ENIKEVKEQGSKNNN | 0 | 0 | 1 | 913146 | GI:124505377 | Q9U0H1 |
| ENKLNKEGKLIEHII | 0 | 0 | 1 | 913147 | GI:1371546755 | A0A143ZZD7 |
| ENKNTEDVLLIETIT | 0 | 0 | 1 | 913148 | GI:124802266 | Q8IJQ4 |
| ENKQSGDENKQSGDE | 0 | 0 | 1 | 913149 | GI:1371546738 | A0A143ZXM2 |
| ENMIPLYYNKNINYL | 0 | 0 | 1 | 913151 | GI:124506415 | Q8I3L6 |
| ENNIDDNDMYNNYVN | 0 | 0 | 1 | 913152 | GI:1371546738 | A0A143ZXM2 |
| ENNNKEEENKSKSLQ | 0 | 0 | 1 | 913153 | GI:124804085 | Q8IIE8 |
| ENSNTEQNDNNNNDN | 0 | 0 | 1 | 913155 | GI:124802266 | Q8IJQ4 |
| ENVETEEIIDDVPSP | 0 | 0 | 1 | 913157 | GI:124802897 | Q8IJ56 |
| ENVPSGLDIDDIPKE | 0 | 0 | 1 | 913158 | GI:124802897 | Q8IJ56 |
| ENWYSLKKNSRSLGE | 0 | 0 | 1 | 913159 | GI:124504759 | Q7K740 |
| EPAEHVESVSEQSNN | 0 | 0 | 1 | 913160 | GI:124802897 | Q8IJ56 |
| EPNFVDSQPNPQEPV | 0 | 0 | 1 | 913161 | GI:124802897 | Q8IJ56 |
| EPSDKHIKEYLNKIQ | 0 | 0 | 1 | 913166 | GI:124504759 | Q7K740 |
| EQGKWACPKENLKMI | 0 | 0 | 1 | 913169 | GI:124511718 | Q8IC11 |
| EQIKKYIVCSKNSQG | 0 | 0 | 1 | 913170 | GI:1371546548 | Q8IEB7 |
| EQMVVPCYNGECKPE | 0 | 0 | 1 | 913172 | GI:124512538 | Q8IAV9 |
| EQNDEESGESGLVDN | 0 | 0 | 1 | 913173 | GI:124802897 | Q8IJ56 |
| ERLAKEKLQEQQSDL | 0 | 0 | 1 | 913176 | GI:1371546755 | A0A143ZZD7 |
| ESVEEIPVNEDEFED | 0 | 0 | 1 | 913181 | GI:124802897 | Q8IJ56 |
| ESYIDEKISILKHGK | 0 | 0 | 1 | 913182 | GI:124810474 | Q8IK92 |
| ETNENISESNSGNPE | 0 | 0 | 1 | 913184 | GI:124802266 | Q8IJQ4 |
| ETQSEKPCNNNQTKQ | 0 | 0 | 1 | 913185 | SRC280142 | Q8I639 |
| EVEEILPEDKNEKVE | 0 | 0 | 1 | 913186 | GI:124802897 | Q8IJ56 |
| FDNNIKKKNETDIKY | 0 | 0 | 1 | 913192 | GI:124808154 | Q8IM20 |
| FENVDDDAVENEQED | 0 | 0 | 1 | 913195 | GI:124513450 | Q8IDX8 |
| FFNFFDFREKSIDSN | 0 | 0 | 1 | 913197 | GI:124802266 | Q8IJQ4 |
| FGDHGQLDTGDHGGY | 0 | 0 | 1 | 913198 | GI:258597476 | Q8I5R4 |
| FIPYIRIFPLKGVIN | 0 | 0 | 1 | 913200 | GI:124806515 | Q8I572 |
| FKEVRYLQSINLRVP | 0 | 0 | 1 | 913201 | GI:262527542 | Q8IBG1 |
| FKIKNFEYPNTHPVY | 0 | 0 | 1 | 913202 | GI:124806575 | Q8I556 |
| FKNIAIEINGPFHYK | 0 | 0 | 1 | 913203 | GI:124511718 | Q8IC11 |
| FKNNNDEQHICLEYD | 0 | 0 | 1 | 913204 | GI:1371546738 | A0A143ZXM2 |
| FLDFYIDIFLFNHFI | 0 | 0 | 1 | 913207 | GI:258597957 | Q8IK94 |
| FMTKDMRKEKESFIE | 0 | 0 | 1 | 913216 | GI:124511718 | Q8IC11 |
| FNDHVSGIIPNSKNK | 0 | 0 | 1 | 913217 | GI:124511896 | Q8IBS2 |
| FNINENIMTNKMSAP | 0 | 0 | 1 | 913218 | GI:124802266 | Q8IJQ4 |
| FPDVCKIGKKIALCY | 0 | 0 | 1 | 913219 | GI:1371546548 | Q8IEB7 |
| FPRQKHKKVDNHNEE | 0 | 0 | 1 | 913221 | GI:124802897 | Q8IJ56 |
| FPYTYYNPYYMFRLT | 0 | 0 | 1 | 913223 | GI:124802266 | Q8IJQ4 |
| FSDDNISEDDAPRRK | 0 | 0 | 1 | 913227 | GI:86170720 | C6KSS9 |
| FSFSKKYTLINYIIT | 0 | 0 | 1 | 913229 | GI:258597211 | Q8IIU1 |
| FSNNNGYFEKYVNKN | 0 | 0 | 1 | 913230 | GI:124800697 | Q8I669 |
| FSSIEIVERSNYMGN | 0 | 0 | 1 | 913231 | GI:1373027 | Q7KQK5 |
| FSVEAKSNMLAHKLN | 0 | 0 | 1 | 913234 | GI:124804529 | Q8II16 |
| FSVLINKNRCAVVVN | 0 | 0 | 1 | 913235 | GI:1371547014 | A0A144A1V5 |
| FTENVCASYCMNITD | 0 | 0 | 1 | 913236 | GI:124802266 | Q8IJQ4 |
| FVVLRHETYFPTEMG | 0 | 0 | 1 | 913239 | GI:296004524 | C0H4D2 |
| FVVYSKSKKDMQGNN | 0 | 0 | 1 | 913240 | GI:124802266 | Q8IJQ4 |
| FYDKMVFHFSKDGMM | 0 | 0 | 1 | 913241 | GI:124802266 | Q8IJQ4 |
| FYSSEEKKIACEMIK | 0 | 0 | 1 | 913242 | GI:124504821 | O77318 |
| GAAKGATRLIELIQS | 0 | 0 | 1 | 913244 | GI:124505929 | Q8I2C5 |
| GAIASTSEFYIEPEG | 0 | 0 | 1 | 913246 | GI:124512788 | Q8IEU1 |
| GEEKRASHTTPVLME | 0 | 0 | 1 | 913255 | GI:124804478 | Q7KQK5 |
| GEPNAGPQVTAQDVT | 0 | 0 | 1 | 913259 | GI:124804079 | Q8IIF0 |
| GETEVTEETEETEET | 0 | 0 | 1 | 913260 | GI:124507147 | Q8I0U8 |
| GFDLDENPENPPNPP | 0 | 0 | 1 | 913264 | GI:124513464 | Q76NM2 |
| GGCDADATCTEEDSG | 0 | 0 | 1 | 913267 | GI:124507147 | Q8I0U8 |
| GGDGHDCTENGELKH | 0 | 0 | 1 | 913268 | GI:124505159 | Q8I218 |
| GKFKLLGLYGSNSIN | 0 | 0 | 1 | 913274 | GI:6649938 | Q7KQL7 |
| GLFGKNVLSKVKAQS | 0 | 0 | 1 | 913276 | GI:124513450 | Q8IDX8 |
| GLTHSANGSLEVSDI | 0 | 0 | 1 | 913281 | GI:124507147 | Q8I0U8 |
| GNGGNEEMSYINNDE | 0 | 0 | 1 | 913284 | GI:1371546738 | A0A143ZXM2 |
| GNIVNDINLILPEIK | 0 | 0 | 1 | 913285 | GI:124804085 | Q8IIE8 |
| GNPDPNANPNVDPNA | 0 | 0 | 1 | 913286 | GI:124504759 | Q7K740 |
| GNSEEFSFDNELPEQ | 0 | 0 | 1 | 913288 | GI:124802266 | Q8IJQ4 |
| GNYNKYEIGKIKKNN | 0 | 0 | 1 | 913289 | GI:124804085 | Q8IIE8 |
| GPKHVEQNIQEDDND | 0 | 0 | 1 | 913290 | GI:124802897 | Q8IJ56 |
| GPRYCNKDESKRNSM | 0 | 0 | 1 | 913292 | GI:1373027 | Q7KQK5 |
| GQGHNMPNDPNRNVD | 0 | 0 | 1 | 913295 | GI:124504759 | Q7K740 |
| GQHEIVEVEEILPED | 0 | 0 | 1 | 913296 | GI:124802897 | Q8IJ56 |
| GSGHRRKISVKQKNT | 0 | 0 | 1 | 913305 | GI:124506525 | Q8I3G2 |
| GSGIRVDLGEDAEVA | 0 | 0 | 1 | 913306 | GI:1373027 | Q7KQK5 |
| GTGKTMLVKAVANET | 0 | 0 | 1 | 913311 | GI:124505405 | Q8I1V1 |
| GYNWGNYNTETQKCE | 0 | 0 | 1 | 913324 | GI:1373027 | Q7KQK5 |
| HDIIDKKKVHEQNTN | 0 | 0 | 1 | 913329 | GI:124802266 | Q8IJQ4 |
| HEIVEVEEILPEDKN | 0 | 0 | 1 | 913331 | GI:124802897 | Q8IJ56 |
| HFLHSISFILKKFNN | 0 | 0 | 1 | 913333 | GI:296004508 | C0H4C4 |
| HGNSKGGTGTEVGAC | 0 | 0 | 1 | 913338 | GI:124511794 | Q8IBX3 |
| HHLQRVLLVGEAEAE | 0 | 0 | 1 | 913341 | GI:124803377 | Q8IIZ4 |
| HICLEYDDTYNLKDT | 0 | 0 | 1 | 913342 | GI:1371546738 | A0A143ZXM2 |

| | | | | | | |
|---|---|---|---|---|---|---|
| HIFKSNILYNIPCLI | 0 | 0 | 1 | 913343 | GI:124804085 | Q8IIE8 |
| HIILDEFATLECKSK | 0 | 0 | 1 | 913344 | GI:124506797 | Q8I328 |
| HIPLSVNNKGYNKSS | 0 | 0 | 1 | 913345 | GI:124804631 | Q8IHY4 |
| HIVIKTITLLKLKFV | 0 | 0 | 1 | 913348 | GI:124513318 | Q8IE43 |
| HKESDSNDKNNNDYS | 0 | 0 | 1 | 913349 | GI:124511718 | Q8IC11 |
| HKNILTLLLSSIKNT | 0 | 0 | 1 | 913351 | GI:1371546029 | O97298 |
| HKNNTFIDNDCNNNH | 0 | 0 | 1 | 913352 | GI:124512230 | Q8IBB0 |
| HKTVDPEIVEVEEIP | 0 | 0 | 1 | 913353 | GI:124802897 | Q8IJ56 |
| HLSNDQKDSHVNNME | 0 | 0 | 1 | 913354 | GI:124512788 | Q8IEU1 |
| HNDDENMENIQSKRD | 0 | 0 | 1 | 913356 | GI:124807012 | Q8I4T2 |
| HNNHNNHNNHNNHSN | 0 | 0 | 1 | 913357 | GI:124801310 | O96234 |
| HNNNVEDVCHNNNVE | 0 | 0 | 1 | 913358 | GI:1371546738 | A0A143ZXM2 |
| HNQNQNNNQNDAYSN | 0 | 0 | 1 | 913359 | GI:124807012 | Q8I4T2 |
| HNTNYDDILQKKIKG | 0 | 0 | 1 | 913360 | GI:1371546724 | C6S3C6 |
| HPEIVEIEEVFPEPN | 0 | 0 | 1 | 913361 | GI:124802897 | Q8IJ56 |
| HSNHETFEEETSESE | 0 | 0 | 1 | 913367 | GI:124802897 | Q8IJ56 |
| HTEQLDLDHKTVDPE | 0 | 0 | 1 | 913368 | GI:124802897 | Q8IJ56 |
| HTIKNRPVVVNNEIV | 0 | 0 | 1 | 913369 | GI:124513182 | Q8IEA6 |
| HTYQQEDSGEDENTL | 0 | 0 | 1 | 913370 | GI:1373027 | Q7KQK5 |
| HVLSHNSYEKTKNNE | 0 | 0 | 1 | 913371 | GI:1371546755 | A0A143ZZD7 |
| ICDKNEDVDDLFVSK | 0 | 0 | 1 | 913378 | GI:1371546738 | A0A143ZXM2 |
| ICSNNNDNNISNSLM | 0 | 0 | 1 | 913379 | GI:124809320 | Q8IL77 |
| IDDIPKESIFIQEDQ | 0 | 0 | 1 | 913380 | GI:124802897 | Q8IJ56 |
| IDEIVDNMIVDKALD | 0 | 0 | 1 | 913381 | GI:1371546913 | A0A144A2Q6 |
| IDLFECNKLVFELSA | 0 | 0 | 1 | 913383 | GI:1373027 | Q7KQK5 |
| IDNIKLYDIYTVHSN | 0 | 0 | 1 | 913384 | GI:124809744 | Q8IKV7 |
| IDNNNTDIKSTDNYN | 0 | 0 | 1 | 913385 | GI:124511650 | Q8IC44 |
| IDPSPISSIRNLNIY | 0 | 0 | 1 | 913386 | GI:258597531 | Q8I5A6 |
| IDQLKEKINIIIIPN | 0 | 0 | 1 | 913388 | GI:1371546738 | A0A143ZXM2 |
| IDSEIILPENVETEE | 0 | 0 | 1 | 913389 | GI:124802897 | Q8IJ56 |
| IDYGDIFLTIHDTLK | 0 | 0 | 1 | 913390 | GI:1371546738 | A0A143ZXM2 |
| IEKSSEELSEEKIKK | 0 | 0 | 1 | 913392 | GI:1371546755 | A0A143ZZD7 |
| IEKVDVQPKIVDLQI | 0 | 0 | 1 | 913393 | GI:124802897 | Q8IJ56 |
| IELCKKYNINVNFLS | 0 | 0 | 1 | 913394 | GI:258597503 | Q8I5H5 |
| IENKNNVDVSIKRYN | 0 | 0 | 1 | 913395 | GI:124512788 | Q8IEU1 |
| IENTNRESITTNVEG | 0 | 0 | 1 | 913396 | GI:1371546755 | A0A143ZZD7 |
| IENVTYNLTEQVITN | 0 | 0 | 1 | 913397 | GI:296005025 | C0H4V1 |
| IENYFFLLPLINRNI | 0 | 0 | 1 | 913398 | GI:296004889 | C0H4P4 |
| IETITINNGTTSNTI | 0 | 0 | 1 | 913399 | GI:124802266 | Q8IJQ4 |
| IFFFLKKNKKSHWKK | 0 | 0 | 1 | 913400 | GI:124807012 | Q8I4T2 |
| IFGESEDNDEYLDQV | 0 | 0 | 1 | 913401 | GI:124507147 | Q8I0U8 |
| IFNPFVAKNIYTEIT | 0 | 0 | 1 | 913404 | GI:124506581 | Q8I3D4 |
| IGIIIPRYKEGNENI | 0 | 0 | 1 | 913407 | GI:124512538 | Q8IAV9 |
| IHIKTKFIENKINIL | 0 | 0 | 1 | 913409 | GI:124804314 | Q8II80 |
| IHPVWTVYMNRFSLF | 0 | 0 | 1 | 913410 | GI:258597569 | Q8I533 |
| IHSDISTCIEKKNNP | 0 | 0 | 1 | 913411 | GI:124511650 | Q8IC44 |
| IINEKETKITKFCNK | 0 | 0 | 1 | 913412 | GI:124505477 | Q8I1R5 |
| IINSKQLENIPSNNK | 0 | 0 | 1 | 913413 | GI:258597330 | Q8II86 |
| IISDLKGEENITKNN | 0 | 0 | 1 | 913414 | GI:124804085 | Q8IIE8 |
| IISVDKNNTDRDNNN | 0 | 0 | 1 | 913415 | GI:124804085 | Q8IIE8 |
| IITNSFQILRSNYFL | 0 | 0 | 1 | 913416 | GI:258597476 | Q8I5R4 |
| IKDDKLNNQYNNDDN | 0 | 0 | 1 | 913417 | GI:124512538 | Q8IAV9 |
| IKDNLLTVEKFVSLN | 0 | 0 | 1 | 913419 | GI:124512666 | Q8IAP6 |
| IKIDNKETVGVTNDD | 0 | 0 | 1 | 913421 | GI:124807012 | Q8I4T2 |
| IKKGILSFISSNINI | 0 | 0 | 1 | 913422 | GI:124803460 | Q8IIW6 |
| IKKNNEENKQNVTVE | 0 | 0 | 1 | 913423 | GI:124804085 | Q8IIE8 |
| IKNEQKNNYGFCNNH | 0 | 0 | 1 | 913424 | GI:74873275 | O97239 |
| IKRYNNFVDIARLSI | 0 | 0 | 1 | 913425 | GI:124512788 | Q8IEU1 |
| IKVIKIHNDLTHLKK | 0 | 0 | 1 | 913426 | GI:124505901 | Q8I2D9 |
| ILDIIQGKENMNNNS | 0 | 0 | 1 | 913427 | GI:124806698 | Q8I516 |
| ILHEVMEQQTVTIAK | 0 | 0 | 1 | 913428 | GI:124513100 | Q8IEE5 |
| ILHNNYNRVLNEENI | 0 | 0 | 1 | 913429 | GI:258597796 | Q8IL70 |
| ILKTLCTFDQIIIYM | 0 | 0 | 1 | 913430 | GI:296005516 | Q8IDI9 |
| ILKYDHANIYKYIYF | 0 | 0 | 1 | 913431 | GI:124511718 | Q8IC11 |
| ILKYYIINLCIHLYN | 0 | 0 | 1 | 913432 | GI:1371546374 | C0H4W8 |
| ILLNKDLREDVPNIN | 0 | 0 | 1 | 913433 | GI:1371546550 | A0A5K1K9E8 |
| ILNSLLTCKERIINN | 0 | 0 | 1 | 913435 | GI:258597011 | Q8IJT7 |
| ILPEDKNEKVEHEIV | 0 | 0 | 1 | 913436 | GI:124802897 | Q8IJ56 |
| ILTDNDNNTDIILTD | 0 | 0 | 1 | 913437 | GI:124511650 | Q8IC44 |
| ILTFFFHDVEDEHIE | 0 | 0 | 1 | 913438 | GI:296005351 | C0H5A6 |
| IMDHMIYKDNNNMSK | 0 | 0 | 1 | 913439 | GI:1371546738 | A0A143ZXM2 |
| IMEDICHNNNVEDMY | 0 | 0 | 1 | 913440 | GI:1371546738 | A0A143ZXM2 |
| INDNHNVSNASKKNR | 0 | 0 | 1 | 913442 | GI:258597215 | Q8IIT8 |
| INDNIYILNHQTHSE | 0 | 0 | 1 | 913443 | GI:124807012 | Q8I4T2 |
| INEDIYMKYYINTHN | 0 | 0 | 1 | 913444 | GI:124804685 | Q8IHW8 |
| INEEDSEPFPRQKHK | 0 | 0 | 1 | 913445 | GI:124802897 | Q8IJ56 |
| INIESHIVPSEKRIC | 0 | 0 | 1 | 913447 | GI:124504821 | O77318 |
| INITNNIKSIIRNLQN | 0 | 0 | 1 | 913448 | GI:296004458 | C0H496 |
| INKEYEKLLSTIKNT | 0 | 0 | 1 | 913449 | GI:124802515 | Q8IJI8 |
| INNDESNKMESIKTL | 0 | 0 | 1 | 913450 | GI:124511718 | Q8IC11 |
| INNGDKDSHAEEEMA | 0 | 0 | 1 | 913451 | GI:1371546738 | A0A143ZXM2 |
| INNGTTSNTIENNKD | 0 | 0 | 1 | 913452 | GI:124802266 | Q8IJQ4 |
| INNKTDNLYGIINNN | 0 | 0 | 1 | 913453 | GI:124512862 | Q8IEQ5 |
| INNNEDNQNDNMNIV | 0 | 0 | 1 | 913454 | GI:124802266 | Q8IJQ4 |
| INNTVNMNNMYNVDN | 0 | 0 | 1 | 913455 | GI:296005564 | A0A5K1K8Q4 |
| INTTQNLTTSTNTTT | 0 | 0 | 1 | 913456 | GI:124511650 | Q8IC44 |
| IPAHMLENITSSIRF | 0 | 0 | 1 | 913457 | GI:124809747 | Q8IKV6 |
| IPENSFFHENVVVPE | 0 | 0 | 1 | 913458 | GI:124506449 | Q8I3J9 |
| IPNNKITNVLTYDNS | 0 | 0 | 1 | 913459 | GI:124504821 | O77318 |
| IPQIVKGQDQFCNST | 0 | 0 | 1 | 913460 | GI:124505749 | Q8I284 |
| IQEDDNDEEDDDDID | 0 | 0 | 1 | 913462 | GI:124802897 | Q8IJ56 |

| | | | | | | |
|---|---|---|---|---|---|---|
| IQVTMLSAVEAPSAN | 0 | 0 | 1 | 913463 | GI:296005630 | C0H5N2 |
| IQYSKNDPITDIQNS | 0 | 0 | 1 | 913464 | GI:124804085 | Q8IIE8 |
| ISDLNKKTLEVPVNV | 0 | 0 | 1 | 913466 | GI:124513926 | Q76NL8 |
| ISEDEEEEEEEEKE | 0 | 0 | 1 | 913467 | GI:124802901 | Q8IJ55 |
| ISENEMFGGQSANQP | 0 | 0 | 1 | 913468 | GI:124809506 | Q8IL32 |
| ISGESNLSVHMIDSN | 0 | 0 | 1 | 913469 | GI:124806828 | Q8I4X5 |
| ISIKNRKLYYFNSIN | 0 | 0 | 1 | 913470 | GI:296005161 | C0H518 |
| ISNEEIIDKKINQPN | 0 | 0 | 1 | 913472 | GI:258597507 | Q8I5G7 |
| ISSEDYNNKNIIIIK | 0 | 0 | 1 | 913474 | GI:124505325 | Q9U0J6 |
| ISSETNISNESSVPN | 0 | 0 | 1 | 913475 | GI:124802919 | Q8IJ50 |
| ITFSRNPVLYQALVQ | 0 | 0 | 1 | 913477 | GI:86170640 | C6KSR3 |
| ITHLMKHFSKELVYE | 0 | 0 | 1 | 913478 | GI:258597476 | Q8I5R4 |
| ITSINMLMGKNDVSY | 0 | 0 | 1 | 913480 | GI:1371546366 | C0H4V8 |
| ITTNRNFQRIVGSIN | 0 | 0 | 1 | 913481 | GI:124511718 | Q8IC11 |
| IVEVEEILPEDKNEK | 0 | 0 | 1 | 913482 | GI:124802897 | Q8IJ56 |
| IVEVEEILPEIVEIE | 0 | 0 | 1 | 913483 | GI:124802897 | Q8IJ56 |
| IVQENKPNKKETPVV | 0 | 0 | 1 | 913484 | GI:124802897 | Q8IJ56 |
| IVSEKSTSEPAEHVE | 0 | 0 | 1 | 913485 | GI:124802897 | Q8IJ56 |
| IYIYQTLQLIYHPYF | 0 | 0 | 1 | 913487 | GI:124806463 | Q8I586 |
| IYKQKCSSKVKENQI | 0 | 0 | 1 | 913488 | GI:124807012 | Q8I4T2 |
| IYNLCQAVHLCIIKK | 0 | 0 | 1 | 913489 | GI:124511718 | Q8IC11 |
| KAIEQERRAKEKLQE | 0 | 0 | 1 | 913491 | SRC7359 | A0A143ZZD7 |
| KCKGLSKRQEPEIYK | 0 | 0 | 1 | 913492 | GI:124513586 | Q8IDR1 |
| KDILFIKKNVLPVNV | 0 | 0 | 1 | 913495 | GI:258597961 | Q8IK84 |
| KDTYSEDDDNNKSNN | 0 | 0 | 1 | 913497 | GI:124511788 | Q8IBX6 |
| KDYKEKNLINEEDSE | 0 | 0 | 1 | 913498 | GI:124802897 | Q8IJ56 |
| KEDINKMEGEYQYMA | 0 | 0 | 1 | 913499 | GI:1371547073 | A0A144A1R4 |
| KEGKLIEHIINDDDD | 0 | 0 | 1 | 913501 | GI:1371546755 | A0A143ZZD7 |
| KEHYIYDEEIVNPNM | 0 | 0 | 1 | 913502 | GI:124802266 | Q8IJQ4 |
| KENKDNNNDSSTECV | 0 | 0 | 1 | 913504 | GI:1371546738 | A0A143ZXM2 |
| KEYGSPLIINDGVTI | 0 | 0 | 1 | 913507 | GI:258597535 | Q8I0V3 |
| KEYPDNFIEHINNDE | 0 | 0 | 1 | 913508 | GI:124511718 | Q8IC11 |
| KFHFAFLVTNIIFSK | 0 | 0 | 1 | 913509 | GI:124513864 | A0A5K1K903 |
| KFTNGSFHNIINYNI | 0 | 0 | 1 | 913510 | GI:124809797 | Q8IKU2 |
| KGKGEVQEPNQANKE | 0 | 0 | 1 | 913511 | GI:124800995 | A0A143ZX41 |
| KGLSLINLIVHSNLI | 0 | 0 | 1 | 913512 | GI:124802266 | Q8IJQ4 |
| KHKKLKQPADGNPDP | 0 | 0 | 1 | 913514 | GI:124504759 | Q7K740 |
| KIEQDGHKYNDNNNK | 0 | 0 | 1 | 913517 | GI:296004474 | B9ZSI2 |
| KIIKNSEKDEIIKSN | 0 | 0 | 1 | 913518 | GI:1371546755 | A0A143ZZD7 |
| KINLFHKWLDRHRDM | 0 | 0 | 1 | 913519 | GI:1371546343 | A0A143ZXL1 |
| KIVDLQIIEPNFVDS | 0 | 0 | 1 | 913520 | GI:124802897 | Q8IJ56 |
| KKEKEQINNEKNILK | 0 | 0 | 1 | 913521 | GI:86170793 | C6KSU3 |
| KKEYEDFVLNSKEYD | 0 | 0 | 1 | 913522 | GI:124513450 | Q8IDX8 |
| KKKKHHNQNQNNNQN | 0 | 0 | 1 | 913523 | GI:124807012 | Q8I4T2 |
| KKKKNASNDPLSICD | 0 | 0 | 1 | 913524 | GI:124807012 | Q8I4T2 |
| KKKTKFPPDPYIPSE | 0 | 0 | 1 | 913525 | GI:124504821 | O77318 |
| KKLKKKKQKCMDKYT | 0 | 0 | 1 | 913526 | GI:1371546738 | A0A143ZXM2 |
| KKLKQPADGNPDPNA | 0 | 0 | 1 | 913527 | GI:124504759 | Q7K740 |
| KKNNPRNQITHLNNL | 0 | 0 | 1 | 913528 | GI:124511650 | Q8IC44 |
| KKPENKHDNQNNLPN | 0 | 0 | 1 | 913529 | GI:124513464 | Q76NM2 |
| KKTSEDMLIYMNFLN | 0 | 0 | 1 | 913530 | GI:124808549 | Q8ILS4 |
| KLKSFDEHLKDEKIE | 0 | 0 | 1 | 913531 | GI:124802897 | Q8IJ56 |
| KLLTCADDIGNTNQM | 0 | 0 | 1 | 913532 | GI:124801204 | O96205 |
| KLNNEEVKKLNHINN | 0 | 0 | 1 | 913533 | GI:124506347 | Q8I3P9 |
| KLSSSPRPEGPRANN | 0 | 0 | 1 | 913535 | GI:124802925 | Q8IJ48 |
| KMNNIINSASKYTFY | 0 | 0 | 1 | 913536 | GI:124505261 | Q8I1W9 |
| KNEDNKIPHIDPSNN | 0 | 0 | 1 | 913537 | GI:124804085 | Q8IIE8 |
| KNENTNIINVVDTFS | 0 | 0 | 1 | 913539 | GI:124804085 | Q8IIE8 |
| KNFVTYIDPTKKTFS | 0 | 0 | 1 | 913540 | GI:124801059 | P68874 |
| KNGNVEVTGNGGNEE | 0 | 0 | 1 | 913541 | GI:1371546738 | A0A143ZXM2 |
| KNHVEKEYPDNFIEH | 0 | 0 | 1 | 913542 | GI:124511718 | Q8IC11 |
| KNINLYLNNIEENII | 0 | 0 | 1 | 913543 | GI:124806489 | Q8I579 |
| KNIYFLLSYKSIIMK | 0 | 0 | 1 | 913544 | GI:296004512 | C0H4C6 |
| KNKNNNIYINKNNNI | 0 | 0 | 1 | 913545 | GI:124513254 | Q8IE74 |
| KNKVNIISVDKNNTD | 0 | 0 | 1 | 913546 | GI:124804085 | Q8IIE8 |
| KNLEPGPMAPYLTLI | 0 | 0 | 1 | 913547 | GI:124506897 | Q8I2X8 |
| KNNKVNRDVIKNESS | 0 | 0 | 1 | 913548 | GI:124804085 | Q8IIE8 |
| KNNTTEGNHMHNSVS | 0 | 0 | 1 | 913549 | GI:124807012 | Q8I4T2 |
| KNNYKKYDNNKYYNN | 0 | 0 | 1 | 913550 | GI:124809485 | Q8IL37 |
| KNRDNYNRNKDKNNT | 0 | 0 | 1 | 913552 | GI:124802266 | Q8IJQ4 |
| KNTFSIINAPYDKVV | 0 | 0 | 1 | 913553 | GI:258597476 | Q8I5R4 |
| KNTNQNLKISKNNTT | 0 | 0 | 1 | 913554 | GI:124807012 | Q8I4T2 |
| KPAQGGESTFQKDLD | 0 | 0 | 1 | 913555 | GI:124513450 | Q8IDX8 |
| KQDISVIESDIMTCI | 0 | 0 | 1 | 913558 | GI:74862587 | C0H5F4 |
| KQTDIKGQNKNDLNN | 0 | 0 | 1 | 913562 | GI:124804085 | Q8IIE8 |
| KRAPVAYNINCPFRK | 0 | 0 | 1 | 913563 | GI:124512842 | Q8IER5 |
| KRAQYNRTNVECITS | 0 | 0 | 1 | 913564 | GI:258597602 | Q8I4Q6 |
| KRISENLNRFFMESN | 0 | 0 | 1 | 913566 | GI:124802266 | Q8IJQ4 |
| KRNYKNLTEFLENTE | 0 | 0 | 1 | 913567 | GI:124802266 | Q8IJQ4 |
| KSSLHLHILYNLSIN | 0 | 0 | 1 | 913569 | GI:124809112 | Q8ILC4 |
| KSSYGYNIYFKRGLT | 0 | 0 | 1 | 913570 | GI:124804085 | Q8IIE8 |
| KSTTHLNNTDNCDVN | 0 | 0 | 1 | 913571 | GI:1371546738 | A0A143ZXM2 |
| KTNVEEKRSLTINKE | 0 | 0 | 1 | 913574 | GI:124809561 | Q8IL18 |
| KTNVNNDALKEERNN | 0 | 0 | 1 | 913575 | GI:124804085 | Q8IIE8 |
| KTYVQSFVSQAETTA | 0 | 0 | 1 | 913576 | GI:124505151 | O97311 |
| KVREVEKIRHVPGPI | 0 | 0 | 1 | 913579 | GI:124808542 | Q8ILS6 |
| KVTGDSVENINEQTN | 0 | 0 | 1 | 913582 | GI:124802266 | Q8IJQ4 |
| KYIYFFIVHFIHKFD | 0 | 0 | 1 | 913583 | GI:124511718 | Q8IC11 |
| KYNSENTNTNNNIPE | 0 | 0 | 1 | 913585 | GI:124511650 | Q8IC44 |
| KYVEDIINSSIIQRS | 0 | 0 | 1 | 913587 | GI:1371547134 | A0A144A0N6 |
| LAKYKDDLESIKKVI | 0 | 0 | 1 | 913588 | GI:124507147 | Q8I0U8 |

| | | | | | | |
|---|---|---|---|---|---|---|
| LALLKNSLVDNKDYN | 0 | 0 | 1 | 913589 | GI:124802266 | Q8IJQ4 |
| LDKNKKIEEHEEKIK | 0 | 0 | 1 | 913593 | GI:124507147 | Q8I0U8 |
| LDPEASFWGEEKRAS | 0 | 0 | 1 | 913594 | GI:124804478 | Q7KQK5 |
| LDVLESDLMQFKHIS | 0 | 0 | 1 | 913595 | GI:124507147 | Q8I0U8 |
| LEDLIEKNENLDDLD | 0 | 0 | 1 | 913597 | GI:1371546755 | A0A143ZZD7 |
| LEEGHHEEIVPEQNN | 0 | 0 | 1 | 913598 | GI:124802897 | Q8IJ56 |
| LEEHIKWEILMNKQE | 0 | 0 | 1 | 913599 | GI:86170764 | C6KST7 |
| LEIPAIELPSENERG | 0 | 0 | 1 | 913601 | GI:1371546755 | A0A143ZZD7 |
| LELFSRKVFGDEYVE | 0 | 0 | 1 | 913603 | GI:258597476 | Q8I5R4 |
| LENIQDDVGDNNIFK | 0 | 0 | 1 | 913604 | GI:124511718 | Q8IC11 |
| LENTERINTFVRNHM | 0 | 0 | 1 | 913605 | GI:124802266 | Q8IJQ4 |
| LFIITCPYIYENVTT | 0 | 0 | 1 | 913611 | GI:124512840 | Q8IER6 |
| LGKDDHDLEHSAKNF | 0 | 0 | 1 | 913614 | GI:124802266 | Q8IJQ4 |
| LIFQRLHLLNYRNND | 0 | 0 | 1 | 913616 | GI:124511718 | Q8IC11 |
| LIKFYSIKNETPKKS | 0 | 0 | 1 | 913617 | GI:124807012 | Q8I4T2 |
| LITDVNYNNDNKYIE | 0 | 0 | 1 | 913618 | GI:124804085 | Q8IIE8 |
| LKDEKIENEPLVHEN | 0 | 0 | 1 | 913619 | GI:124802897 | Q8IJ56 |
| LKDNVDKIYHGIYKK | 0 | 0 | 1 | 913620 | GI:124511718 | Q8IC11 |
| LKNTKDKSINMNLNI | 0 | 0 | 1 | 913622 | GI:124809435 | Q8IL49 |
| LKSIYNNKIKGNSEE | 0 | 0 | 1 | 913623 | GI:124802266 | Q8IJQ4 |
| LKVGLINFPPISYFL | 0 | 0 | 1 | 913624 | GI:124804685 | Q8IHW8 |
| LKVKKKDFDNCIGIH | 0 | 0 | 1 | 913625 | GI:1371546527 | A0A143ZVU1 |
| LKYFNRCLGDQLDMG | 0 | 0 | 1 | 913626 | GI:124506543 | A0A5K1K7X4 |
| LKYYKEDRNNIIGIT | 0 | 0 | 1 | 913627 | GI:124506531 | Q8I3F9 |
| LLNGNNHMYDEQMVV | 0 | 0 | 1 | 913629 | GI:124512538 | Q8IAV9 |
| LNIKKLLYNFHKKCY | 0 | 0 | 1 | 913631 | GI:124804085 | Q8IIE8 |
| LNINANTIALNCTCK | 0 | 0 | 1 | 913632 | GI:124802266 | Q8IJQ4 |
| LNINILQTLNIGINA | 0 | 0 | 1 | 913633 | GI:308191562 | C0H5F4 |
| LNKICSLVLKTIKNK | 0 | 0 | 1 | 913634 | GI:124802126 | Q8IJV3 |
| LNKYVDENYMKENNY | 0 | 0 | 1 | 913635 | GI:296004883 | C0H4P2 |
| LNNLNMPTFEKFINK | 0 | 0 | 1 | 913636 | GI:124506247 | Q8I3U4 |
| LNNMKISSYASSSLH | 0 | 0 | 1 | 913637 | GI:124511718 | Q8IC11 |
| LNNQYNNDDNKFLVE | 0 | 0 | 1 | 913638 | GI:124512538 | Q8IAV9 |
| LNTYDLSKYEKQLAL | 0 | 0 | 1 | 913639 | GI:1371546309 | A0A143ZWE7 |
| LPEDKNEKGQHEIVE | 0 | 0 | 1 | 913640 | GI:124802897 | Q8IJ56 |
| LPKREPLDVPDEPED | 0 | 0 | 1 | 913642 | GI:124513464 | Q76NM2 |
| LPLVSGYVGLYKYVW | 0 | 0 | 1 | 913643 | GI:258597476 | Q8I5R4 |
| LPQDNRGNSRDSKEI | 0 | 0 | 1 | 913644 | GI:1371546755 | A0A143ZZD7 |
| LQNHENELFFCNINK | 0 | 0 | 1 | 913648 | GI:124804543 | Q8II12 |
| LRIFIIKTSNKFKKE | 0 | 0 | 1 | 913650 | GI:296005009 | C0H4U6 |
| LRMKENVNNQGDIKN | 0 | 0 | 1 | 913651 | GI:124513066 | Q8IEG2 |
| LSGCDYSNDFHITGM | 0 | 0 | 1 | 913654 | GI:124512040 | Q8IBK1 |
| LSGKDSADKMPLETQ | 0 | 0 | 1 | 913655 | GI:124505267 | Q6ZMA7 |
| LSTNLPYGKTNLTDA | 0 | 0 | 1 | 913657 | GI:124513464 | Q76NM2 |
| LSVRVSGSSGSTKEE | 0 | 0 | 1 | 913659 | GI:124507147 | Q8I0U8 |
| LTEKMVNDTWITPYA | 0 | 0 | 1 | 913660 | GI:124802266 | Q8IJQ4 |
| LTELQQVVQLQNYDE | 0 | 0 | 1 | 913661 | GI:124507147 | Q8I0U8 |
| LTLCSRHAGNMIPDN | 0 | 0 | 1 | 913662 | GI:1373027 | Q7KQK5 |
| LTNNLDRVYDGTKIE | 0 | 0 | 1 | 913663 | GI:124511718 | Q8IC11 |
| LVFELSASDQPKQYE | 0 | 0 | 1 | 913665 | GI:1373027 | Q7KQK5 |
| MDKHEMDLNNEEVYM | 0 | 0 | 1 | 913668 | GI:1371546738 | A0A143ZXM2 |
| MDNILSGFENEYDVI | 0 | 0 | 1 | 913669 | GI:124507147 | Q8I0U8 |
| MDSNLKDKTIVKYNT | 0 | 0 | 1 | 913670 | GI:124809253 | Q8IL92 |
| MEVNENFSPTTSVED | 0 | 0 | 1 | 913672 | GI:258597476 | Q8I5R4 |
| MEYNNYMHKVKNTSL | 0 | 0 | 1 | 913673 | GI:296004580 | C0H4F8 |
| MFILFYLKYEYRLPL | 0 | 0 | 1 | 913674 | GI:124505449 | Q8I1S9 |
| MIYVISKMVLKNVNN | 0 | 0 | 1 | 913676 | GI:124511718 | Q8IC11 |
| MKCKGEKPFLLCINK | 0 | 0 | 1 | 913677 | GI:296005295 | C0H581 |
| MKEYFKKNYEKGLSL | 0 | 0 | 1 | 913678 | GI:124802266 | Q8IJQ4 |
| MKRNVINKNTHIKNS | 0 | 0 | 1 | 913679 | GI:296005544 | A0A5K1K8U0 |
| MLNLKMKMKIKKFSE | 0 | 0 | 1 | 913680 | GI:1371546574 | A0A5K1K8L6 |
| MLQSEQPKDVPNDYS | 0 | 0 | 1 | 913681 | GI:124805331 | Q8I643 |
| MNITDVLELSHYDMI | 0 | 0 | 1 | 913682 | GI:124802266 | Q8IJQ4 |
| MNNSTEKKINSISKG | 0 | 0 | 1 | 913683 | GI:124804085 | Q8IIE8 |
| MNTTSNNNNNNNNIS | 0 | 0 | 1 | 913684 | GI:258597361 | Q8II07 |
| MPKEMNEKFYNCNNL | 0 | 0 | 1 | 913685 | GI:124809688 | Q8IKX1 |
| MQFKHISSNEYIIED | 0 | 0 | 1 | 913686 | GI:124507147 | Q8I0U8 |
| MYFNKYVEDIINSSI | 0 | 0 | 1 | 913689 | GI:1371547134 | A0A144A0N6 |
| MYRNNNVEDVCHNNN | 0 | 0 | 1 | 913690 | GI:1371546738 | A0A143ZXM2 |
| NAEKYDKMDEPQDYG | 0 | 0 | 1 | 913691 | GI:1373027 | Q7KQK5 |
| NAILYNQNPRNICVT | 0 | 0 | 1 | 913692 | GI:51968317 | O97224 |
| NANPNANPNANPNVD | 0 | 0 | 1 | 913693 | GI:124504759 | Q7K740 |
| NANPNANPNKNNQGN | 0 | 0 | 1 | 913694 | GI:124504759 | Q7K740 |
| NATYTLQKEALETAN | 0 | 0 | 1 | 913697 | GI:124505659 | Q9NFA9 |
| NCVLINLPYRKCIIF | 0 | 0 | 1 | 913700 | GI:124808866 | Q8ILI8 |
| NDDNNDDHHNDDDDN | 0 | 0 | 1 | 913701 | GI:124807012 | Q8I4T2 |
| NDEGLKNENNINNNE | 0 | 0 | 1 | 913702 | GI:124802266 | Q8IJQ4 |
| NDFSGGEFLWPGYTE | 0 | 0 | 1 | 913703 | GI:124802901 | Q8IJ55 |
| NDHTIEDIENQSEQK | 0 | 0 | 1 | 913704 | GI:258597476 | Q8I5R4 |
| NDIKNMNNSTEKKIN | 0 | 0 | 1 | 913706 | GI:124804085 | Q8IIE8 |
| NDKEERLQFEKEKKQ | 0 | 0 | 1 | 913707 | GI:124805736 | Q8I5T5 |
| NDKNSNYKYPAVYDD | 0 | 0 | 1 | 913708 | GI:1373027 | Q7KQK5 |
| NDLYINNMQDDAMDD | 0 | 0 | 1 | 913709 | GI:1371546738 | A0A143ZXM2 |
| NDNKSRCNEKTNSYD | 0 | 0 | 1 | 913710 | GI:124511718 | Q8IC11 |
| NDNNTKQSDNVLKGN | 0 | 0 | 1 | 913711 | GI:124513326 | Q8IE40 |
| NDTKNNATLSEDNKN | 0 | 0 | 1 | 913712 | GI:124804085 | Q8IIE8 |
| NEEEKELYINRKPLN | 0 | 0 | 1 | 913713 | GI:124804106 | Q8IIE1 |
| NEEETESEISEDEEE | 0 | 0 | 1 | 913714 | GI:124802901 | Q8IJ55 |
| NEESSIEEAHQEEIV | 0 | 0 | 1 | 913715 | GI:124802897 | Q8IJ56 |
| NEIIENGEEKYVNEY | 0 | 0 | 1 | 913717 | GI:1371546738 | A0A143ZXM2 |
| NEIPNENIVEEQGEG | 0 | 0 | 1 | 913718 | GI:124801292 | O96230 |

| | | | | | | |
|---|---|---|---|---|---|---|
| NEKVEHEIVEVEEIL | 0 | 0 | 1 | 913719 | GI:124802897 | Q8IJ56 |
| NELKFDGKCLCREKV | 0 | 0 | 1 | 913720 | GI:296005088 | Q8IB97 |
| NELNYDNAGTNLYNE | 0 | 0 | 1 | 913721 | GI:124504759 | Q7K740 |
| NENENVTPKPSEGES | 0 | 0 | 1 | 913722 | GI:124802897 | Q8IJ56 |
| NENEYAFNTEKTNVN | 0 | 0 | 1 | 913723 | GI:124804085 | Q8IIE8 |
| NENIDNYKKYHSYLK | 0 | 0 | 1 | 913724 | GI:258597476 | Q8I5R4 |
| NEYIIEDSFKLLNSE | 0 | 0 | 1 | 913725 | GI:124507147 | Q8I0U8 |
| NFESYNNFFKNNNDE | 0 | 0 | 1 | 913726 | GI:1371546738 | A0A143ZXM2 |
| NFFDKIFFFLKKNKK | 0 | 0 | 1 | 913727 | GI:124807012 | Q8I4T2 |
| NFFNNICSKDFLIKE | 0 | 0 | 1 | 913728 | GI:124511718 | Q8IC11 |
| NFIEHINNDESNKME | 0 | 0 | 1 | 913729 | GI:124511718 | Q8IC11 |
| NFLGGGYSAALKLVN | 0 | 0 | 1 | 913730 | GI:124511650 | Q8IC44 |
| NFLLKFESTKELINI | 0 | 0 | 1 | 913731 | GI:124513344 | A0A5K1K8J1 |
| NFTDEEKRILHISGV | 0 | 0 | 1 | 913732 | GI:124802266 | Q8IJQ4 |
| NFYYNNNNLSTYVIK | 0 | 0 | 1 | 913733 | GI:124511650 | Q8IC44 |
| NGDLKYNNDLIKEGE | 0 | 0 | 1 | 913735 | GI:124804085 | Q8IIE8 |
| NGDTGTKTNIATNNT | 0 | 0 | 1 | 913736 | GI:296004378 | C0H484 |
| NGDVIENNNGDIIED | 0 | 0 | 1 | 913737 | GI:1371546738 | A0A143ZXM2 |
| NGEYNIKYQDGAIAS | 0 | 0 | 1 | 913738 | GI:124512788 | Q8IEU1 |
| NGLINSIHPNIIQRL | 0 | 0 | 1 | 913742 | GI:124506389 | A0A5K1K8Q3 |
| NGYNIINHIYLTVKV | 0 | 0 | 1 | 913747 | GI:124801292 | O96230 |
| NHNNEMDVSKFIFSM | 0 | 0 | 1 | 913748 | GI:124512818 | Q8IES7 |
| NHVVDISNKKNKINL | 0 | 0 | 1 | 913749 | GI:124513450 | Q8IDX8 |
| NIFNIKKKNQQVIWN | 0 | 0 | 1 | 913751 | GI:296005570 | A0A5K1K998 |
| NIFQGSNEFVDFYKK | 0 | 0 | 1 | 913752 | GI:124804685 | Q8IHW8 |
| NIINVVDTFSIRNQS | 0 | 0 | 1 | 913753 | GI:124804085 | Q8IIE8 |
| NIKINKKSMSSFIYL | 0 | 0 | 1 | 913754 | GI:296004450 | C0H4B8 |
| NIKNCSNILFGMNNN | 0 | 0 | 1 | 913755 | GI:1371546953 | A0A144A3L0 |
| NILKDFNINENIMTN | 0 | 0 | 1 | 913756 | GI:124802266 | Q8IJQ4 |
| NIQNDHMNDNMLNNA | 0 | 0 | 1 | 913757 | GI:86171446 | C6KT67 |
| NITQNFNQQINSNTT | 0 | 0 | 1 | 913758 | GI:124507217 | Q8I2H8 |
| NKDYNNSEELKYALE | 0 | 0 | 1 | 913760 | GI:124802266 | Q8IJQ4 |
| NKEKEKFIKSLFHIF | 0 | 0 | 1 | 913761 | GI:1371546755 | A0A143ZZD7 |
| NKETNGNNNNIINNN | 0 | 0 | 1 | 913762 | GI:124505007 | O77382 |
| NKFIIIRTNKNGVVN | 0 | 0 | 1 | 913763 | GI:1371546078 | C0H4B3 |
| NKIAETYEGDWVDGK | 0 | 0 | 1 | 913764 | GI:258597137 | Q8IJ93 |
| NKIEIEEEKKNKIEI | 0 | 0 | 1 | 913765 | GI:1371546000 | O77328 |
| NKIINFIANKKGFDR | 0 | 0 | 1 | 913766 | GI:124806238 | Q8I5E9 |
| NKKKNVNINNMNNNN | 0 | 0 | 1 | 913767 | GI:124504769 | O97236 |
| NKKLINNDVDNNSSL | 0 | 0 | 1 | 913768 | GI:124512538 | Q8IAV9 |
| NKMDIKKCILNNKGN | 0 | 0 | 1 | 913770 | GI:1371546299 | A0A143ZY62 |
| NKNEFIFSIERVIEN | 0 | 0 | 1 | 913771 | GI:1371546309 | A0A143ZWE7 |
| NKNNSETNENISESN | 0 | 0 | 1 | 913772 | GI:124802266 | Q8IJQ4 |
| NKRNNKLNNYKFNMN | 0 | 0 | 1 | 913773 | GI:124804085 | Q8IIE8 |
| NLAYITNDTNNMNNM | 0 | 0 | 1 | 913776 | GI:124513274 | Q8IE65 |
| NLESEGKGIVYDDVN | 0 | 0 | 1 | 913778 | GI:1371546738 | A0A143ZXM2 |
| NLKISKNNTTEGNHM | 0 | 0 | 1 | 913779 | GI:124807012 | Q8I4T2 |
| NLNDLINFKNIHLNN | 0 | 0 | 1 | 913781 | GI:124802266 | Q8IJQ4 |
| NLNIPRRNNCGVTSN | 0 | 0 | 1 | 913782 | GI:124513604 | Q8IDQ2 |
| NLNRFFMESNSNAPQ | 0 | 0 | 1 | 913783 | GI:124802266 | Q8IJQ4 |
| NLPINYYEDNIKENN | 0 | 0 | 1 | 913784 | GI:124505749 | Q8I284 |
| NLPPPPPNNLGMQPN | 0 | 0 | 1 | 913785 | GI:296005007 | C0H4U5 |
| NLTDALLQVRKHLND | 0 | 0 | 1 | 913788 | GI:124513464 | Q76NM2 |
| NLTEFLENTERINTF | 0 | 0 | 1 | 913789 | GI:124802266 | Q8IJQ4 |
| NMDNIYNVNNNDNNN | 0 | 0 | 1 | 913791 | GI:1371546029 | O97298 |
| NMENMNMDNSNMDNS | 0 | 0 | 1 | 913792 | GI:1371546142 | Q8I255 |
| NMKKSNENEYAFNTE | 0 | 0 | 1 | 913793 | GI:124804085 | Q8IIE8 |
| NMLEDFILIIINLYN | 0 | 0 | 1 | 913794 | GI:23499289 | C0H4L9 |
| NNDDDNNDDHHNDDE | 0 | 0 | 1 | 913795 | GI:124807012 | Q8I4T2 |
| NNDHHNNDDDNNDDH | 0 | 0 | 1 | 913796 | GI:124807012 | Q8I4T2 |
| NNDIFVNIPLPKIFK | 0 | 0 | 1 | 913797 | GI:124512800 | Q8IET6 |
| NNDNGDMVHKNIYNN | 0 | 0 | 1 | 913798 | GI:124804085 | Q8IIE8 |
| NNDSHFENVDDDAVE | 0 | 0 | 1 | 913799 | GI:124513450 | Q8IDX8 |
| NNENRSYNRKHNNTP | 0 | 0 | 1 | 913800 | GI:124513464 | Q76NM2 |
| NNFKENDEGLKNENN | 0 | 0 | 1 | 913801 | GI:124802266 | Q8IJQ4 |
| NNFPCSLYKDEIMKE | 0 | 0 | 1 | 913802 | GI:1373027 | Q7KQK5 |
| NNIFDNKKSMNYNEG | 0 | 0 | 1 | 913803 | GI:124804085 | Q8IIE8 |
| NNINNNNNNNNNNNN | 0 | 0 | 1 | 913804 | GI:296004981 | C0H4T3 |
| NNINPDATFFRFIHN | 0 | 0 | 1 | 913805 | GI:124804412 | Q8II50 |
| NNINTNTNSTDNSNT | 0 | 0 | 1 | 913806 | GI:124511650 | Q8IC44 |
| NNIPINNTLKKNDQN | 0 | 0 | 1 | 913807 | GI:124512030 | Q8IBK6 |
| NNKFFDKDKELTMSN | 0 | 0 | 1 | 913808 | GI:1371546755 | A0A143ZZD7 |
| NNLKNDDMDDDDNNN | 0 | 0 | 1 | 913809 | GI:1371546738 | A0A143ZXM2 |
| NNNIHNINENQNANV | 0 | 0 | 1 | 913810 | GI:124804085 | Q8IIE8 |
| NNNINNNNNNNDNKE | 0 | 0 | 1 | 913811 | GI:124802266 | Q8IJQ4 |
| NNNMLNINHVNNNRN | 0 | 0 | 1 | 913812 | GI:124512262 | Q8IB94 |
| NNNNNNDNKEEDMNE | 0 | 0 | 1 | 913813 | GI:124802266 | Q8IJQ4 |
| NNNNNNNNKRKKINF | 0 | 0 | 1 | 913814 | GI:124513174 | A0A5K1K8B7 |
| NNNNNNNNNKDGDKY | 0 | 0 | 1 | 913815 | GI:124804085 | Q8IIE8 |
| NNNNNNNNNNNDNNN | 0 | 0 | 1 | 913816 | GI:296004909 | Q8IBF8 |
| NNNNNNNYNINKYYI | 0 | 0 | 1 | 913817 | GI:124805979 | Q8I5L7 |
| NNNNSKVTGDSVENI | 0 | 0 | 1 | 913818 | GI:124802266 | Q8IJQ4 |
| NNNNYNNNNKGNINK | 0 | 0 | 1 | 913819 | GI:124505027 | O77372 |
| NNNSTSQKKKIISAQ | 0 | 0 | 1 | 913820 | GI:258597111 | Q8IJE8 |
| NNNTKTTSTDNNNNT | 0 | 0 | 1 | 913821 | GI:124511650 | Q8IC44 |
| NNNVEDMYRNNNVED | 0 | 0 | 1 | 913822 | GI:1371546738 | A0A143ZXM2 |
| NNNYYYYNNFGKIHN | 0 | 0 | 1 | 913823 | GI:124507036 | Q8I2R3 |
| NNQYPNTEYNTIQRS | 0 | 0 | 1 | 913824 | GI:124802266 | Q8IJQ4 |
| NNSSDLLENSNHIPM | 0 | 0 | 1 | 913825 | GI:124511838 | Q8IBV1 |
| NNSTTINTNSTENTS | 0 | 0 | 1 | 913826 | GI:124511650 | Q8IC44 |
| NNTDTISTDNDNADT | 0 | 0 | 1 | 913827 | GI:124511650 | Q8IC44 |

| | | | | | | |
|---|---|---|---|---|---|---|
| NNVEDMYHDNNIEDV | 0 | 0 | 1 | 913828 | GI:1371546738 | A0A143ZXM2 |
| NNVEDMYRNNNVEDM | 0 | 0 | 1 | 913829 | GI:1371546738 | A0A143ZXM2 |
| NNYNDNEEDGDGDGD | 0 | 0 | 1 | 913830 | GI:1371546738 | A0A143ZXM2 |
| NNYYYNNSDNNNYNE | 0 | 0 | 1 | 913831 | GI:124802266 | Q8IJQ4 |
| NPHTKEKINEKIITD | 0 | 0 | 1 | 913834 | GI:124507147 | Q8I0U8 |
| NPNVDPNANPNANPN | 0 | 0 | 1 | 913837 | GI:124504759 | Q7K740 |
| NPNVDPNANPNVDPN | 0 | 0 | 1 | 913838 | GI:124504759 | Q7K740 |
| NPQEPSPNPEEGKGE | 0 | 0 | 1 | 913839 | GI:124513464 | Q76NM2 |
| NPSMPNIVLTTTINV | 0 | 0 | 1 | 913840 | GI:124802266 | Q8IJQ4 |
| NQGNGQGHNMPNDPN | 0 | 0 | 1 | 913843 | GI:124504759 | Q7K740 |
| NQIENTNFDLFIQIK | 0 | 0 | 1 | 913845 | GI:86171608 | C6KT98 |
| NQKHKDLQQDLNDEP | 0 | 0 | 1 | 913846 | GI:124802897 | Q8IJ56 |
| NQSYYAYTKNNNINT | 0 | 0 | 1 | 913849 | GI:124511718 | Q8IC11 |
| NRITKKEDFLFLINA | 0 | 0 | 1 | 913851 | GI:124804296 | Q8II85 |
| NRNHLQEYKIFNMIQ | 0 | 0 | 1 | 913853 | GI:1371546544 | Q8IEI2 |
| NRNVDENANANSAVK | 0 | 0 | 1 | 913854 | GI:124504759 | Q7K740 |
| NSAVKNNNNEEPSDK | 0 | 0 | 1 | 913856 | GI:124504759 | Q7K740 |
| NSEELKYALEHIQNS | 0 | 0 | 1 | 913857 | GI:124802266 | Q8IJQ4 |
| NSHDNYNMNDIENFV | 0 | 0 | 1 | 913859 | GI:124804085 | Q8IIE8 |
| NSLVDNKDYNNSEEL | 0 | 0 | 1 | 913861 | GI:124802266 | Q8IJQ4 |
| NSNKNRDNSNKNRDN | 0 | 0 | 1 | 913862 | GI:124802266 | Q8IJQ4 |
| NSQIENEENVNTTIT | 0 | 0 | 1 | 913864 | GI:124802901 | Q8IJ55 |
| NSQLFGGISLKKTYS | 0 | 0 | 1 | 913865 | GI:124506447 | Q8I3K0 |
| NSRDSKEISIIENTN | 0 | 0 | 1 | 913866 | GI:1371546755 | A0A143ZZD7 |
| NSRSLGENDDGNNED | 0 | 0 | 1 | 913867 | GI:124504759 | Q7K740 |
| NSYEKTKNNENNKFF | 0 | 0 | 1 | 913868 | GI:1371546755 | A0A143ZZD7 |
| NSYVNVLYFRKLKLN | 0 | 0 | 1 | 913870 | GI:74815308 | Q8IC19 |
| NTDISAIQFMNLNSD | 0 | 0 | 1 | 913871 | GI:124512538 | Q8IAV9 |
| NTISTDNDNADTILT | 0 | 0 | 1 | 913872 | GI:124511650 | Q8IC44 |
| NTLTNTNNLDNSLKI | 0 | 0 | 1 | 913873 | GI:1371546282 | Q8IC35 |
| NTNSTENTSATKKVT | 0 | 0 | 1 | 913874 | GI:124511650 | Q8IC44 |
| NVDENANANSAVKNN | 0 | 0 | 1 | 913875 | GI:124504759 | Q7K740 |
| NVDGDDNTTIQNKGP | 0 | 0 | 1 | 913876 | GI:124513538 | A0A5K1K8A9 |
| NVDINNTDISAIQFM | 0 | 0 | 1 | 913877 | GI:124512538 | Q8IAV9 |
| NVEDVYHNNNVEDMY | 0 | 0 | 1 | 913878 | GI:1371546738 | A0A143ZXM2 |
| NVHTLKRPEGLYNIA | 0 | 0 | 1 | 913879 | XP_001348074.1 | Q8IHX2 |
| NVISTDNSKTNVIST | 0 | 0 | 1 | 913880 | GI:124511650 | Q8IC44 |
| NVKKDMLLNISHDNN | 0 | 0 | 1 | 913881 | GI:124511718 | Q8IC11 |
| NVKLNEDSDINNKMK | 0 | 0 | 1 | 913882 | GI:124512526 | A0A5K1K7Z9 |
| NVKYTKQVNLMSTLS | 0 | 0 | 1 | 913883 | GI:258597476 | Q8I5R4 |
| NVNRIMCLKHNNEKL | 0 | 0 | 1 | 913884 | GI:124804913 | Q8IHP8 |
| NVNTNDKCNVPNNNN | 0 | 0 | 1 | 913885 | GI:124504973 | O97273 |
| NYKDSRLVHNVTQNN | 0 | 0 | 1 | 913886 | GI:124804085 | Q8IIE8 |
| NYKKYHSYLKDRNKK | 0 | 0 | 1 | 913887 | GI:258597476 | Q8I5R4 |
| NYNYNCSNNYNYNCS | 0 | 0 | 1 | 913888 | GI:124504769 | O97236 |
| NYPNLEEQKCKNKNK | 0 | 0 | 1 | 913889 | GI:124513126 | Q8IED3 |
| NYYNNTNNNNCYPLK | 0 | 0 | 1 | 913890 | GI:124505219 | Q8I1Z0 |
| PAENEESSLEEGHHE | 0 | 0 | 1 | 913891 | GI:124802897 | Q8IJ56 |
| PEDKNEKVQHEIVEV | 0 | 0 | 1 | 913894 | GI:124802897 | Q8IJ56 |
| PEIVEIEEVPSQTNN | 0 | 0 | 1 | 913895 | GI:124802897 | Q8IJ56 |
| PETSEHSKDLNNNDS | 0 | 0 | 1 | 913896 | GI:124802897 | Q8IJ56 |
| PLDVPDEPEDDQPRP | 0 | 0 | 1 | 913902 | GI:124513464 | Q76NM2 |
| PLLDVSRNTNRTPRR | 0 | 0 | 1 | 913903 | GI:1371546738 | A0A143ZXM2 |
| PLSPKVLDNERKQSD | 0 | 0 | 1 | 913904 | GI:124513464 | Q76NM2 |
| PLSQQNYKILKEHIE | 0 | 0 | 1 | 913905 | GI:86171769 | C6KTC8 |
| PNANPNANPNANPNK | 0 | 0 | 1 | 913908 | GI:124504759 | Q7K740 |
| PNANPNANPNVDPNA | 0 | 0 | 1 | 913909 | GI:124504759 | Q7K740 |
| PNANPNVDPNANPNA | 0 | 0 | 1 | 913910 | GI:124504759 | Q7K740 |
| PNDKSLYDEHIKKYK | 0 | 0 | 1 | 913911 | GI:1371546755 | A0A143ZZD7 |
| PNINDTRAFPSLTSK | 0 | 0 | 1 | 913912 | GI:124809568 | Q8IL16 |
| PNNTSDKKTKIESIN | 0 | 0 | 1 | 913913 | GI:258597848 | Q8IKU9 |
| PNPQEPVEPSFVKIE | 0 | 0 | 1 | 913914 | GI:124802897 | Q8IJ56 |
| PNVDPNANPNANPNA | 0 | 0 | 1 | 913915 | GI:124504759 | Q7K740 |
| PSDVPKNPEDDREEN | 0 | 0 | 1 | 913927 | GI:124513464 | Q76NM2 |
| PVKLNEVNDLKTIII | 0 | 0 | 1 | 913931 | GI:124802266 | Q8IJQ4 |
| PYAGEPAPFDETLGE | 0 | 0 | 1 | 913932 | GI:124513464 | Q76NM2 |
| PYSNVNIINKIILHI | 0 | 0 | 1 | 913933 | GI:124511718 | Q8IC11 |
| QAVHLCIIKKIYIKP | 0 | 0 | 1 | 913934 | GI:124511718 | Q8IC11 |
| QCGKGVACICDKNED | 0 | 0 | 1 | 913935 | GI:1371546738 | A0A143ZXM2 |
| QDDAMDDIVYDEEEI | 0 | 0 | 1 | 913937 | GI:1371546738 | A0A143ZXM2 |
| QDNRGNSRDSKEISI | 0 | 0 | 1 | 913939 | GI:1371546755 | A0A143ZZD7 |
| QEEGQNKNQNVDHAH | 0 | 0 | 1 | 913942 | GI:1371546548 | Q8IEB7 |
| QEKLAKEKLQGQQSD | 0 | 0 | 1 | 913945 | GI:1371546755 | A0A143ZZD7 |
| QEPVVPTLNENENVT | 0 | 0 | 1 | 913946 | GI:124802897 | Q8IJ56 |
| QERRAKEKLQEQQSD | 0 | 0 | 1 | 913947 | GI:1371546755 | A0A143ZZD7 |
| QGGLSQGLSGKDSAD | 0 | 0 | 1 | 913952 | GI:124505267 | Q6ZMA7 |
| QGQHSDLERTKASKE | 0 | 0 | 1 | 913954 | GI:1371546755 | A0A143ZZD7 |
| QHEIVEVEEILPEDK | 0 | 0 | 1 | 913956 | GI:124802897 | Q8IJ56 |
| QHSDINSHVSFNNNL | 0 | 0 | 1 | 913958 | GI:1371546084 | Q8IFP4 |
| QKDSHVNNMEYMQKF | 0 | 0 | 1 | 913960 | GI:124512788 | Q8IEU1 |
| QKIYVKKKVTKNQLI | 0 | 0 | 1 | 913961 | GI:124512038 | Q8IBK2 |
| QKTAPFGQAAAGDKPSPFGQA | 0 | 0 | 1 | 913964 | EAN97076.1 | Q4DX15 |
| QLDMGSVHNNNSVVG | 0 | 0 | 1 | 913966 | GI:124506543 | A0A5K1K7X4 |
| QLQNYDEEDDSLVVL | 0 | 0 | 1 | 913967 | GI:124507147 | Q8I0U8 |
| QLQQQMNHQQINKHQ | 0 | 0 | 1 | 913968 | GI:74864292 | Q8ILR9 |
| QNENEFDENNNNKNN | 0 | 0 | 1 | 913971 | GI:1371546738 | A0A143ZXM2 |
| QNHFESLSDLELLEN | 0 | 0 | 1 | 913972 | GI:124802897 | Q8IJ56 |
| QNKSFKIKCTYDETL | 0 | 0 | 1 | 913973 | GI:124805948 | Q8I5M6 |
| QPADKVVSRSGASPD | 0 | 0 | 1 | 913977 | GI:124505933 | Q8I495 |
| QQLNLNDNAIHLYAS | 0 | 0 | 1 | 913990 | GI:124513464 | Q76NM2 |
| QQSDLEQDRLAKEKL | 0 | 0 | 1 | 913993 | GI:1371546755 | A0A143ZZD7 |

| | | | | | | |
|---|---|---|---|---|---|---|
| QSDLEQERRAKEKLQ | 0 | 0 | 1 | 914001 | GI:1371546755 | A0A143ZZD7 |
| QSSLPQDNRGNSRDS | 0 | 0 | 1 | 914003 | GI:1371546755 | A0A143ZZD7 |
| QTHSEIKIDNKETVG | 0 | 0 | 1 | 914006 | GI:124807012 | Q8I4T2 |
| QTPGSSSGASLHAVG | 0 | 0 | 1 | 914007 | GI:124505267 | Q6ZMA7 |
| QTVCINSTNDVLSIK | 0 | 0 | 1 | 914008 | GI:124512884 | Q8IEP4 |
| QTYYNENSKSTQLKR | 0 | 0 | 1 | 914009 | GI:124805343 | Q8I640 |
| QYNKTLTFDINYDTN | 0 | 0 | 1 | 914012 | GI:124802266 | Q8IJQ4 |
| RALDNLLNINEGKAT | 0 | 0 | 1 | 914015 | GI:124803966 | Q8III3 |
| RDNNNLYETNNGDLK | 0 | 0 | 1 | 914020 | GI:124804085 | Q8IIE8 |
| RDVKEINKIKKIKDQ | 0 | 0 | 1 | 914021 | GI:86171402 | C6KT58 |
| REAQLYEIGKGVKQS | 0 | 0 | 1 | 914022 | GI:124504919 | O77341 |
| REQQSDLEQEKLAKE | 0 | 0 | 1 | 914025 | GI:1371546755 | A0A143ZZD7 |
| RHFYKDNKYVKNLDE | 0 | 0 | 1 | 914029 | GI:1373027 | Q7KQK5 |
| RINTFVRNHMMLSNE | 0 | 0 | 1 | 914031 | GI:124802266 | Q8IJQ4 |
| RKYKNISFQNILNKF | 0 | 0 | 1 | 914033 | GI:124809891 | Q8IKR6 |
| RLHSDASKNKEKALI | 0 | 0 | 1 | 914034 | GI:124513464 | Q76NM2 |
| RNNENNYNKNHFVNL | 0 | 0 | 1 | 914037 | GI:296004730 | C6KTB5 |
| RRAKEKLQEQQSDLE | 0 | 0 | 1 | 914045 | GI:1371546755 | A0A143ZZD7 |
| RRHNWVNHAVPLAMK | 0 | 0 | 1 | 914047 | GI:124513464 | Q76NM2 |
| RSRKREILHEGCTSE | 0 | 0 | 1 | 914049 | GI:124513464 | Q76NM2 |
| RTTIKSRLLAQTQIH | 0 | 0 | 1 | 914054 | GI:1371546293 | A0A143ZXI1 |
| SAKNFYNISNENGDN | 0 | 0 | 1 | 914055 | GI:124802266 | Q8IJQ4 |
| SANKPKDELDYANDI | 0 | 0 | 1 | 914057 | GI:124504759 | Q7K740 |
| SCEEVDMTNTNYKNN | 0 | 0 | 1 | 914060 | GI:124511896 | Q8IBS2 |
| SDDELNKDSCDYSEE | 0 | 0 | 1 | 914062 | GI:124511650 | Q8IC44 |
| SDDSQNDEEKVSYNN | 0 | 0 | 1 | 914063 | GI:1371546309 | A0A143ZWE7 |
| SDINTSIEKRIKKGT | 0 | 0 | 1 | 914064 | GI:124807012 | Q8I4T2 |
| SEYQSNFLGGGYSAA | 0 | 0 | 1 | 914073 | GI:124511650 | Q8IC44 |
| SFNLMNCYFLKFISN | 0 | 0 | 1 | 914074 | GI:258597292 | Q8IIF4 |
| SFNVFDFYSLDIAAK | 0 | 0 | 1 | 914075 | GI:258597476 | Q8I5R4 |
| SGKICLLEKLLIRLK | 0 | 0 | 1 | 914077 | GI:124802575 | Q8IJG6 |
| SHFQNDNANVPDLIQ | 0 | 0 | 1 | 914078 | GI:86171687 | C6KTB3 |
| SIEKRIKKGTHKSNE | 0 | 0 | 1 | 914080 | GI:124807012 | Q8I4T2 |
| SIKLQKLINIKNIDF | 0 | 0 | 1 | 914081 | GI:124806110 | Q8I0X3 |
| SIKMHTQIVPIIYPN | 0 | 0 | 1 | 914082 | GI:1331227850 | A0A140KXX5 |
| SIPNDPIEQILNQPE | 0 | 0 | 1 | 914083 | GI:124802897 | Q8IJ56 |
| SKDKEEKIKFIEHIN | 0 | 0 | 1 | 914084 | GI:124804085 | Q8IIE8 |
| SKEYDIIKNLIIMFG | 0 | 0 | 1 | 914085 | GI:124513450 | Q8IDX8 |
| SKLNIVYKPNGSFSK | 0 | 0 | 1 | 914086 | GI:124511718 | Q8IC11 |
| SLSNKKNCDYKNHVE | 0 | 0 | 1 | 914088 | GI:124511718 | Q8IC11 |
| SLYKDEIMKEIERES | 0 | 0 | 1 | 914089 | GI:1373027 | Q7KQK5 |
| SNEDDMFINLSLIIS | 0 | 0 | 1 | 914091 | GI:124513704 | A0A5K1K9F8 |
| SNKEAENSNTEQNDN | 0 | 0 | 1 | 914095 | GI:124802266 | Q8IJQ4 |
| SNNIKSNNIKSNNIK | 0 | 0 | 1 | 914097 | GI:296005114 | C0H4Z5 |
| SNQNNKNEKMVENNN | 0 | 0 | 1 | 914098 | GI:124511760 | Q8IBZ0 |
| SPSLINDNKVGDQKK | 0 | 0 | 1 | 914103 | GI:124804755 | Q8IHU4 |
| SQEFSYDKYSMKDRM | 0 | 0 | 1 | 914104 | GI:1371546738 | A0A143ZXM2 |
| SRYTDNSNKNRDNSN | 0 | 0 | 1 | 914111 | GI:124802266 | Q8IJQ4 |
| SSLNYHMDKIISMVN | 0 | 0 | 1 | 914112 | GI:1371546239 | C6KSZ5 |
| SSNEKLSSYEKNKNV | 0 | 0 | 1 | 914113 | GI:124511718 | Q8IC11 |
| SSQEDTLALNEVQTR | 0 | 0 | 1 | 914114 | GI:124801080 | Q9TY94 |
| SSYDYILGWEFGGGV | 0 | 0 | 1 | 914116 | GI:124802901 | Q8IJ55 |
| STCIEKKNNPRNQIT | 0 | 0 | 1 | 914117 | GI:124511650 | Q8IC44 |
| STETLHEQQSDLEQE | 0 | 0 | 1 | 914118 | GI:1371546755 | A0A143ZZD7 |
| STHTLSQSGETEVTE | 0 | 0 | 1 | 914119 | GI:124507147 | Q8I0U8 |
| STLPQDNISIPVQNK | 0 | 0 | 1 | 914120 | GI:124801302 | O96232 |
| STMKNNTHDNITSNN | 0 | 0 | 1 | 914121 | GI:86170978 | C6KSX8 |
| STSNLFYPKKDIILR | 0 | 0 | 1 | 914122 | GI:124802266 | Q8IJQ4 |
| SVENINEQTNNNQYP | 0 | 0 | 1 | 914123 | GI:124802266 | Q8IJQ4 |
| SVHNNNSVVGNSSSH | 0 | 0 | 1 | 914124 | GI:124506543 | A0A5K1K7X4 |
| SYAKQNSHLSEKKIK | 0 | 0 | 1 | 914127 | GI:124513450 | Q8IDX8 |
| SYIATTALSHPIEVE | 0 | 0 | 1 | 914128 | GI:1373027 | Q7KQK5 |
| TAGGCGPGVASDNKK | 0 | 0 | 1 | 914134 | GI:124505387 | Q9U0G6 |
| TDIILTDNNNTDTIS | 0 | 0 | 1 | 914136 | GI:124511650 | Q8IC44 |
| TDIKATDNNNITTTT | 0 | 0 | 1 | 914137 | GI:124511650 | Q8IC44 |
| TDIKSTDNYNTGTKE | 0 | 0 | 1 | 914138 | GI:124511650 | Q8IC44 |
| TDITTYTNSSSSLYN | 0 | 0 | 1 | 914139 | GI:1371546084 | Q8IFP4 |
| TDNCDVNLSEQTNEI | 0 | 0 | 1 | 914140 | GI:1371546738 | A0A143ZXM2 |
| TDNDNADTILTDNDN | 0 | 0 | 1 | 914141 | GI:124511650 | Q8IC44 |
| TDNDNADTKATDNNN | 0 | 0 | 1 | 914142 | GI:124511650 | Q8IC44 |
| TDNKNTDIKATDNNN | 0 | 0 | 1 | 914143 | GI:124511650 | Q8IC44 |
| TDNNNTDTISTDNDN | 0 | 0 | 1 | 914144 | GI:124511650 | Q8IC44 |
| TDNTSTTKKLTDNIN | 0 | 0 | 1 | 914145 | GI:124511650 | Q8IC44 |
| TEIFIKLFKDKEYHE | 0 | 0 | 1 | 914147 | GI:124513450 | Q8IDX8 |
| TFLGQNNDSHFENVD | 0 | 0 | 1 | 914151 | GI:124513450 | Q8IDX8 |
| TFNKKNYHSSGNVNN | 0 | 0 | 1 | 914152 | GI:124809472 | Q8IL40 |
| TGTSSTSSPGNTTVN | 0 | 0 | 1 | 914153 | GI:124507147 | Q8I0U8 |
| TIDNIQSENDHTIDN | 0 | 0 | 1 | 914155 | GI:258597476 | Q8I5R4 |
| TIPNYMEVNENFSPT | 0 | 0 | 1 | 914156 | GI:258597476 | Q8I5R4 |
| TISTEPFPNQKHKDL | 0 | 0 | 1 | 914157 | GI:124802897 | Q8IJ56 |
| TIVQNSYNSFSDINK | 0 | 0 | 1 | 914158 | GI:308191562 | C0H5F4 |
| TKASTETLREQQSDL | 0 | 0 | 1 | 914159 | GI:1371546755 | A0A143ZZD7 |
| TKKINENFQKNKNEF | 0 | 0 | 1 | 914160 | GI:1371546309 | A0A143ZWE7 |
| TKNNENNKFFDKDKE | 0 | 0 | 1 | 914161 | GI:1371546755 | A0A143ZZD7 |
| TKPIDNNNTDIKSTD | 0 | 0 | 1 | 914162 | GI:124511650 | Q8IC44 |
| TLFIFFGDHGQLDTG | 0 | 0 | 1 | 914163 | GI:258597476 | Q8I5R4 |
| TLFKKIINELNNKLN | 0 | 0 | 1 | 914164 | GI:258597750 | Q8ILE1 |
| TNIRRTGHLNIKVND | 0 | 0 | 1 | 914165 | GI:124806029 | Q8I5K4 |
| TNKYTDNDYNNNNNN | 0 | 0 | 1 | 914166 | GI:124804085 | Q8IIE8 |
| TNNLGSTNFNVNNKV | 0 | 0 | 1 | 914167 | GI:124512374 | Q8IB40 |
| TNNVDKTNNIDKTNN | 0 | 0 | 1 | 914168 | GI:124513672 | A0A5K1K9A2 |

| | | | | | | |
|---|---|---|---|---|---|---|
| TNTLLQNKKENILLN | 0 | 0 | 1 | 914169 | GI:86170539 | C6KSP3 |
| TNVEGRRDIHKGHLE | 0 | 0 | 1 | 914170 | GI:1371546755 | A0A143ZZD7 |
| TPALKNHQENSINLI | 0 | 0 | 1 | 914171 | GI:1371546884 | A0A144A3K9 |
| TQLAIEEIKSLSNML | 0 | 0 | 1 | 914175 | GI:124505267 | Q6ZMA7 |
| TQPLVAASETTEDGG | 0 | 0 | 1 | 914176 | GI:124507147 | Q8I0U8 |
| TREQNAEKDIETLTN | 0 | 0 | 1 | 914179 | GI:1371546309 | A0A143ZWE7 |
| TRIHIIILQKKIERF | 0 | 0 | 1 | 914180 | GI:124804085 | Q8IIE8 |
| TSDEKKQTDIKGQNK | 0 | 0 | 1 | 914181 | GI:124804085 | Q8IIE8 |
| TSDILYKDIEENKNT | 0 | 0 | 1 | 914182 | GI:124802266 | Q8IJQ4 |
| TSEQDDINEKNNDNG | 0 | 0 | 1 | 914183 | GI:124804085 | Q8IIE8 |
| TTQNLTTSTNTTTVS | 0 | 0 | 1 | 914186 | GI:124511650 | Q8IC44 |
| TTSTNTTTVSTDNNN | 0 | 0 | 1 | 914187 | GI:124511650 | Q8IC44 |
| TTTDNTNTNVISTDN | 0 | 0 | 1 | 914188 | GI:124511650 | Q8IC44 |
| TVSYIIDSETLLNNV | 0 | 0 | 1 | 914191 | GI:124804085 | Q8IIE8 |
| TYDKNISDSTKAHQN | 0 | 0 | 1 | 914192 | GI:86171715 | C6KTB8 |
| TYTVKVNLEYLVFLL | 0 | 0 | 1 | 914194 | GI:124802266 | Q8IJQ4 |
| VAVSSGPAVVEESHD | 0 | 0 | 1 | 914195 | GI:124507147 | Q8I0U8 |
| VCNDEVDLYLLMDCS | 0 | 0 | 1 | 914197 | GI:124513464 | Q76NM2 |
| VCYSWWVLSSLSSLN | 0 | 0 | 1 | 914198 | GI:258549155 | C6S3K4 |
| VDHAHTYFLLNNIIQ | 0 | 0 | 1 | 914201 | GI:1371546548 | Q8IEB7 |
| VDIESFVTFLKNVYN | 0 | 0 | 1 | 914202 | GI:86171135 | C6KT08 |
| VDILNIAESFNPISL | 0 | 0 | 1 | 914203 | GI:124513206 | Q8IE96 |
| VDLKMDPLLEKLNKY | 0 | 0 | 1 | 914204 | GI:124505093 | O97292 |
| VDNHNEEKNVFHENG | 0 | 0 | 1 | 914205 | GI:124802897 | Q8IJ56 |
| VDPNANPNVDPNANP | 0 | 0 | 1 | 914206 | GI:124504759 | Q7K740 |
| VDTFSIRNQSKIPYS | 0 | 0 | 1 | 914207 | GI:124804085 | Q8IIE8 |
| VDTINSREKDVKNYD | 0 | 0 | 1 | 914208 | GI:1371546738 | A0A143ZXM2 |
| VEDVCHNNNVEDVYH | 0 | 0 | 1 | 914209 | GI:1371546738 | A0A143ZXM2 |
| VEEADVTEESAVETK | 0 | 0 | 1 | 914210 | GI:124506543 | A0A5K1K7X4 |
| VEEILPEDKNEKGQH | 0 | 0 | 1 | 914212 | GI:124802897 | Q8IJ56 |
| VEEKQNSQESVEEIP | 0 | 0 | 1 | 914213 | GI:124802897 | Q8IJ56 |
| VEVEEILPEDKNEKV | 0 | 0 | 1 | 914217 | GI:124802897 | Q8IJ56 |
| VGKLILAFALYMLIN | 0 | 0 | 1 | 914219 | GI:124511650 | Q8IC44 |
| VHHILQNEKNNTLLF | 0 | 0 | 1 | 914221 | GI:258597476 | Q8I5R4 |
| VHIRLPGFKEKINYN | 0 | 0 | 1 | 914222 | GI:124512490 | Q8IAY3 |
| VIIGATNRPDMIDDA | 0 | 0 | 1 | 914227 | GI:124505303 | Q9U0K7 |
| VIKKSESTYYTKKLI | 0 | 0 | 1 | 914228 | GI:124804085 | Q8IIE8 |
| VINFKETIIGKKKMA | 0 | 0 | 1 | 914231 | GI:258597814 | Q8IL30 |
| VIYENDIHENNIDDN | 0 | 0 | 1 | 914232 | GI:1371546738 | A0A143ZXM2 |
| VKEEYKDEYADIPEH | 0 | 0 | 1 | 914233 | GI:1373027 | Q7KQK5 |
| VKFINVPENMNQNYI | 0 | 0 | 1 | 914234 | GI:124809112 | Q8ILC4 |
| VKHNENKKLINNDVD | 0 | 0 | 1 | 914235 | GI:124512538 | Q8IAV9 |
| VKPTCLINNSSYIAT | 0 | 0 | 1 | 914236 | GI:1373027 | Q7KQK5 |
| VLEKRIDTLKKNENI | 0 | 0 | 1 | 914237 | GI:124507147 | Q8I0U8 |
| VLSHNSYEKTKNNEN | 0 | 0 | 1 | 914240 | GI:1371546755 | A0A143ZZD7 |
| VNEKHVQTIENQRIV | 0 | 0 | 1 | 914242 | GI:124806077 | Q8I5J1 |
| VNNMEYMQKFVQGLQ | 0 | 0 | 1 | 914243 | GI:124512788 | Q8IEU1 |
| VNNNYFLRSINNVDK | 0 | 0 | 1 | 914244 | GI:124506311 | Q8I3R5 |
| VNRNLIISTAVTNMI | 0 | 0 | 1 | 914245 | GI:124505267 | Q6ZMA7 |
| VNTKYNSENTNTNNN | 0 | 0 | 1 | 914246 | GI:124511650 | Q8IC44 |
| VNTLNEENNNNENID | 0 | 0 | 1 | 914247 | GI:258597476 | Q8I5R4 |
| VPSEENKHASVDPEV | 0 | 0 | 1 | 914249 | GI:124802897 | Q8IJ56 |
| VPSKPFEEIEKVDVQ | 0 | 0 | 1 | 914251 | GI:124802897 | Q8IJ56 |
| VPSQTNNNENIETIK | 0 | 0 | 1 | 914252 | GI:124802897 | Q8IJ56 |
| VQGLQENRNISLSKY | 0 | 0 | 1 | 914254 | GI:124512788 | Q8IEU1 |
| VRKHLNDRINRENAN | 0 | 0 | 1 | 914257 | GI:124513464 | Q76NM2 |
| VRNMEKLKIICEDNE | 0 | 0 | 1 | 914258 | GI:1371546309 | A0A143ZWE7 |
| VTENVITNQILTGNN | 0 | 0 | 1 | 914265 | GI:124511650 | Q8IC44 |
| VVEEKAIPQEPVVPT | 0 | 0 | 1 | 914269 | GI:124802897 | Q8IJ56 |
| VVEESHDPLTVLSIS | 0 | 0 | 1 | 914270 | GI:124507147 | Q8I0U8 |
| VVEVRTFFTYNGLGG | 0 | 0 | 1 | 914271 | GI:124804085 | Q8IIE8 |
| VVILTDGIPDSIQDS | 0 | 0 | 1 | 914273 | GI:124513464 | Q76NM2 |
| VYMDKNEMDLNNEEV | 0 | 0 | 1 | 914276 | GI:1371546738 | A0A143ZXM2 |
| VYNLINYNIWNNVVP | 0 | 0 | 1 | 914277 | GI:258597221 | Q8IIT2 |
| WEFGGGVPEHKKEEN | 0 | 0 | 1 | 914278 | GI:124802901 | Q8IJ55 |
| YADIPEHKPTYDKMK | 0 | 0 | 1 | 914281 | GI:1373027 | Q7KQK5 |
| YDEEIVNPNMDNINT | 0 | 0 | 1 | 914282 | GI:124802266 | Q8IJQ4 |
| YDKICVENQYAYDLK | 0 | 0 | 1 | 914283 | GI:124804085 | Q8IIE8 |
| YDKNNNNNNNYFKNM | 0 | 0 | 1 | 914284 | GI:124511718 | Q8IC11 |
| YDVIQYINKINSSVI | 0 | 0 | 1 | 914285 | GI:124807098 | Q8I4R0 |
| YEPIICFIGTPCVIK | 0 | 0 | 1 | 914286 | GI:258597961 | Q8IK84 |
| YICIKRIMNIINNKV | 0 | 0 | 1 | 914290 | GI:124804163 | Q8IIC5 |
| YIINKNLNKYFVFYL | 0 | 0 | 1 | 914291 | GI:124506307 | Q8I0W6 |
| YIKTNCFNEVSLESS | 0 | 0 | 1 | 914292 | GI:124512538 | Q8IAV9 |
| YILLRKNGLTIKELM | 0 | 0 | 1 | 914293 | GI:124511718 | Q8IC11 |
| YITKDDLIKSVNKLK | 0 | 0 | 1 | 914295 | GI:124804624 | Q8IHY7 |
| YKDIEENKNTEDVLL | 0 | 0 | 1 | 914296 | GI:124802266 | Q8IJQ4 |
| YKEYTIIIYNDNPSM | 0 | 0 | 1 | 914297 | GI:124802266 | Q8IJQ4 |
| YKKFLEYSFNLLNTM | 0 | 0 | 1 | 914298 | GI:124513450 | Q8IDX8 |
| YKSILKYYKNENKTN | 0 | 0 | 1 | 914299 | GI:124804469 | Q8II33 |
| YKYEDTNMNNNINNN | 0 | 0 | 1 | 914300 | GI:1371546309 | A0A143ZWE7 |
| YKYLNNQTDVNINNL | 0 | 0 | 1 | 914301 | GI:124804085 | Q8IIE8 |
| YKYVWPISQFYIFNH | 0 | 0 | 1 | 914302 | GI:258597476 | Q8I5R4 |
| YLEYRNGEYNIKYQD | 0 | 0 | 1 | 914303 | GI:124512788 | Q8IEU1 |
| YLFFLKANRNIIFSL | 0 | 0 | 1 | 914304 | GI:124807012 | Q8I4T2 |
| YLRKNRDENKINDNI | 0 | 0 | 1 | 914306 | GI:124807012 | Q8I4T2 |
| YNCTLYVDDDVSTNK | 0 | 0 | 1 | 914308 | GI:124512538 | Q8IAV9 |
| YNLKKHHTNVYEPND | 0 | 0 | 1 | 914310 | GI:124802266 | Q8IJQ4 |
| YNNDLIKEGENKRNN | 0 | 0 | 1 | 914311 | GI:124804085 | Q8IIE8 |
| YNNIENAMEGYSSNP | 0 | 0 | 1 | 914312 | GI:124511718 | Q8IC11 |
| YPCSYDNQIGRMNLI | 0 | 0 | 1 | 914313 | GI:124810339 | Q8IKD0 |

| | | | | | | |
|---|---|---|---|---|---|---|
| YREKALITDVNYNND | 0 | 0 | 1 | 914314 | GI:124804085 | Q8IIE8 |
| YRIFKNINLLIINSA | 0 | 0 | 1 | 914315 | GI:124512286 | Q8IB82 |
| YRNGHHIQNENDHII | 0 | 0 | 1 | 914318 | GI:258597476 | Q8I5R4 |
| YTEYISSILHIKKVK | 0 | 0 | 1 | 914321 | GI:124512538 | Q8IAV9 |
| YTLHRKLSGCYLICM | 0 | 0 | 1 | 914322 | GI:124512582 | Q8IAT7 |
| YVGLYKYVWPISQFY | 0 | 0 | 1 | 914323 | GI:258597476 | Q8I5R4 |
| YVTFQDDMNKSVASQ | 0 | 0 | 1 | 914324 | GI:124511718 | Q8IC11 |
| AAAAAPAGPTAAAAS | 0 | 0 | 1 | 918337 | SGX78861.1 | A5K7Y9 |
| AAALPQPASPVVSAP | 0 | 0 | 1 | 918339 | SGX78853.1 | A5K7Z7 |
| ADEKFQAEFDNFVHG | 0 | 0 | 1 | 918356 | SGX78860.1 | A5K7Z0 |
| ADEQDDDNEAVQWGP | 0 | 0 | 1 | 918357 | SGX78861.1 | A5K7Y9 |
| AEAEAEAVRGDTSHQ | 0 | 0 | 1 | 918364 | SGX78853.1 | A5K7Z7 |
| AFFKKDGNPSPADAV | 0 | 0 | 1 | 918370 | SGX78853.1 | A5K7Z7 |
| AKIKYLDKLYDEVLT | 0 | 0 | 1 | 918386 | SGX78860.1 | A5K7Z0 |
| ANMSESPADMSEEPF | 0 | 0 | 1 | 918401 | SGX78851.1 | A5K7Z9 |
| APAGPTAAAASPQVK | 0 | 0 | 1 | 918402 | SGX78861.1 | A5K7Y9 |
| ASEKGDTSAGGEGEL | 0 | 0 | 1 | 918412 | SGX78861.1 | A5K7Y9 |
| ASGSQGASDSSNDSA | 0 | 0 | 1 | 918414 | SGX78861.1 | A5K7Y9 |
| ASSDATPLVDVFKKA | 0 | 0 | 1 | 918416 | SGX78860.1 | A5K7Z0 |
| ATDNGEIFNKEIDAL | 0 | 0 | 1 | 918418 | SGX78860.1 | A5K7Z0 |
| ATDSPTSQEPSQTHT | 0 | 0 | 1 | 918419 | SGX78853.1 | A5K7Z7 |
| ATPLVDVFKKALADE | 0 | 0 | 1 | 918420 | SGX78860.1 | A5K7Z0 |
| ATPPKGTADSNDAKI | 0 | 0 | 1 | 918421 | SGX78860.1 | A5K7Z0 |
| AVASHGRTFITGTNY | 0 | 0 | 1 | 918422 | SGX78851.1 | A5K7Z9 |
| CDELLAGENKKNVLD | 0 | 0 | 1 | 918428 | SGX78861.1 | A5K7Y9 |
| CLLLLCAGPVLGDDD | 0 | 0 | 1 | 918431 | SGX78861.1 | A5K7Y9 |
| DADDSGKKTIFGVDE | 0 | 0 | 1 | 918437 | SGX78854.1 | A5K7Z6 |
| DAHKSLFVNTLSLLD | 0 | 0 | 1 | 918443 | SGX78853.1 | A5K7Z7 |
| DDAQGGEEGGEESGE | 0 | 0 | 1 | 918450 | SGX78851.1 | A5K7Z9 |
| DDDKKDKENEHKEDA | 0 | 0 | 1 | 918451 | SGX78861.1 | A5K7Y9 |
| DDDNEAVQWGPATEE | 0 | 0 | 1 | 918452 | SGX78861.1 | A5K7Y9 |
| DDEDEEAVEGDEAEA | 0 | 0 | 1 | 918453 | SGX78853.1 | A5K7Z7 |
| DEIHVPPFHSKYNDF | 0 | 0 | 1 | 918464 | SGX78853.1 | A5K7Z7 |
| DELKTLKGKLQKIRV | 0 | 0 | 1 | 918466 | SGX78861.1 | A5K7Y9 |
| DKTSSEGPANMSESP | 0 | 0 | 1 | 918480 | SGX78851.1 | A5K7Z9 |
| DMSEEPFRGQNITKY | 0 | 0 | 1 | 918486 | SGX78851.1 | A5K7Z9 |
| DNSEIFNKEIESLKK | 0 | 0 | 1 | 918490 | SGX78857.1 | A5K7Z3 |
| DRVPLPVERSDAPPQ | 0 | 0 | 1 | 918498 | SGX78855.1 | A5K7Z5 |
| DSAEPTSAAAAAAPA | 0 | 0 | 1 | 918500 | SGX78861.1 | A5K7Y9 |
| DSDSAQGSQPSSGTD | 0 | 0 | 1 | 918503 | SGX78860.1 | A5K7Z0 |
| DSGKKTIFGVDEDDL | 0 | 0 | 1 | 918504 | SGX78860.1 | A5K7Z0 |
| DSNDAKIKYLDKLYD | 0 | 0 | 1 | 918505 | SGX78860.1 | A5K7Z0 |
| DSSNDSAEPTSAAAA | 0 | 0 | 1 | 918507 | SGX78861.1 | A5K7Y9 |
| EAADDSGKKTIFGVD | 0 | 0 | 1 | 918512 | SGX78860.1 | A5K7Z0 |
| EAKTYLTLFENVINL | 0 | 0 | 1 | 918520 | SGX78851.1 | A5K7Z9 |
| EAVQWGPATEEVVAE | 0 | 0 | 1 | 918526 | SGX78861.1 | A5K7Y9 |
| EDADKKNNDELKTLK | 0 | 0 | 1 | 918527 | SGX78861.1 | A5K7Y9 |
| EEAVEGDEAEAEAEA | 0 | 0 | 1 | 918536 | SGX78853.1 | A5K7Z7 |
| EEESLGHLLESEDAD | 0 | 0 | 1 | 918543 | SGX78854.1 | A5K7Z6 |
| EEGTENNNKYNEFKK | 0 | 0 | 1 | 918545 | SGX78851.1 | A5K7Z9 |
| EEVPTSDEEDGDEGE | 0 | 0 | 1 | 918549 | SGX78851.1 | A5K7Z9 |
| EFVGQNVKANGDAAG | 0 | 0 | 1 | 918554 | SGX78861.1 | A5K7Y9 |
| EGENAPGEDDAQGGE | 0 | 0 | 1 | 918562 | SGX78851.1 | A5K7Z9 |
| EGTTVTQESALAATP | 0 | 0 | 1 | 918568 | SGX78860.1 | A5K7Z0 |
| EIIEKLVDAFSMYND | 0 | 0 | 1 | 918574 | SGX78851.1 | A5K7Z9 |
| ENNNKYNEFKKEYDM | 0 | 0 | 1 | 918589 | SGX78851.1 | A5K7Z9 |
| EPSQTHTTGSSRNVG | 0 | 0 | 1 | 918597 | SGX78853.1 | A5K7Z7 |
| EQDAMHALMKKLESL | 0 | 0 | 1 | 918599 | SGX78854.1 | A5K7Z6 |
| ERAPDRAPESPADRV | 0 | 0 | 1 | 918604 | SGX78855.1 | A5K7Z5 |
| EREKKNTLIKTFKKA | 0 | 0 | 1 | 918605 | SGX78861.1 | A5K7Y9 |
| ESLYKLSATDNSEIF | 0 | 0 | 1 | 918615 | SGX78854.1 | A5K7Z6 |
| ESPADMSEEPFRGQN | 0 | 0 | 1 | 918616 | SGX78851.1 | A5K7Z9 |
| ESPADRVPLPVERSD | 0 | 0 | 1 | 918617 | SGX78855.1 | A5K7Z5 |
| ESTEGPAVTPRPSST | 0 | 0 | 1 | 918619 | SGX78857.1 | A5K7Z3 |
| EYSMNPVEYEIVKNL | 0 | 0 | 1 | 918630 | SGX78855.1 | A5K7Z5 |
| FIDGDKVKEGDYSKL | 0 | 0 | 1 | 918647 | SGX78861.1 | A5K7Y9 |
| FQAEFDNFVHGLYGF | 0 | 0 | 1 | 918668 | SGX78860.1 | A5K7Z0 |
| GEPEKGRVPNVKYLD | 0 | 0 | 1 | 918706 | SGX78853.1 | A5K7Z7 |
| GSSRNVGQSSTSSIN | 0 | 0 | 1 | 918778 | SGX78853.1 | A5K7Z7 |
| HAGPHEGEAEATLPV | 0 | 0 | 1 | 918787 | SGX78851.1 | A5K7Z9 |
| HEGEAEATLPVQGEN | 0 | 0 | 1 | 918791 | SGX78851.1 | A5K7Z9 |
| HNYLSEARMKDADRY | 0 | 0 | 1 | 918797 | SGX78857.1 | A5K7Z3 |
| HQQPPTSPPANNEGT | 0 | 0 | 1 | 918799 | SGX78860.1 | A5K7Z0 |
| HSKYNDFRKKYEFTM | 0 | 0 | 1 | 918800 | SGX78853.1 | A5K7Z7 |
| IDEDADSVYEAIKKS | 0 | 0 | 1 | 918803 | SGX78851.1 | A5K7Z9 |
| IFRKQYDKMVLNKTE | 0 | 0 | 1 | 918815 | SGX78861.1 | A5K7Y9 |
| IKDDKLPQKISEEQI | 0 | 0 | 1 | 918823 | SGX78861.1 | A5K7Y9 |
| IRVQIKDDKLPQKIS | 0 | 0 | 1 | 918830 | SGX78861.1 | A5K7Y9 |
| KEEYEIIEKLVDAFS | 0 | 0 | 1 | 918845 | SGX78851.1 | A5K7Z9 |
| KEMEALKKQIDHLQQ | 0 | 0 | 1 | 918849 | SGX78854.1 | A5K7Z6 |
| KERMGDTTRYSDLLK | 0 | 0 | 1 | 918851 | SGX78855.1 | A5K7Z5 |
| KISEEQISVLKKKLE | 0 | 0 | 1 | 918866 | SGX78861.1 | A5K7Y9 |
| KKALADEKFQAEFDN | 0 | 0 | 1 | 918867 | SGX78860.1 | A5K7Z0 |
| KKALYDKQYSEKLRN | 0 | 0 | 1 | 918868 | SGX78861.1 | A5K7Y9 |
| KLEDFKNLKSEHEAK | 0 | 0 | 1 | 918885 | SGX78861.1 | A5K7Y9 |
| KLLDTMLTNGQVERE | 0 | 0 | 1 | 918889 | SGX78861.1 | A5K7Y9 |
| KLQKIRVQIKDDKLP | 0 | 0 | 1 | 918891 | SGX78861.1 | A5K7Y9 |
| KLVDAFSMYNDSIDE | 0 | 0 | 1 | 918893 | SGX78851.1 | A5K7Z9 |
| KNTLIKTFKKALYDK | 0 | 0 | 1 | 918903 | SGX78861.1 | A5K7Y9 |
| KRNNFIDGDKVKEGD | 0 | 0 | 1 | 918912 | SGX78861.1 | A5K7Y9 |
| KTEYNISLKLLDTML | 0 | 0 | 1 | 918918 | SGX78861.1 | A5K7Y9 |

| | | | | | | |
|---|---|---|---|---|---|---|
| KTIFGVDEDDLDNYD | 0 | 0 | 1 | 918919 | SGX78860.1 | A5K7Z0 |
| KTTDAKDEIHVPPFH | 0 | 0 | 1 | 918920 | SGX78854.1 | A5K7Z6 |
| KYNEFKKEYDMFISL | 0 | 0 | 1 | 918926 | SGX78851.1 | A5K7Z9 |
| LAGENKKNVLDEGED | 0 | 0 | 1 | 918929 | SGX78861.1 | A5K7Y9 |
| LCAGPVLGDDDKKDK | 0 | 0 | 1 | 918933 | SGX78861.1 | A5K7Y9 |
| LENGYESIHGEDEPQ | 0 | 0 | 1 | 918947 | SGX78860.1 | A5K7Z0 |
| LKYSDELYEDILTSL | 0 | 0 | 1 | 918974 | SGX78851.1 | A5K7Z9 |
| LLESEAADDSGKKTI | 0 | 0 | 1 | 918977 | SGX78860.1 | A5K7Z0 |
| LLSCSPASSEQLGIQ | 0 | 0 | 1 | 918981 | SGX78860.1 | A5K7Z0 |
| LYDEVLTTSDNTNWI | 0 | 0 | 1 | 919021 | SGX78860.1 | A5K7Z0 |
| MHALMKKLESLYKLS | 0 | 0 | 1 | 919033 | SGX78859.1 | A5K7Z1 |
| MLNDPNVQKEFDNIV | 0 | 0 | 1 | 919038 | SGX78853.1 | A5K7Z7 |
| MVLNKTEYNISLKLL | 0 | 0 | 1 | 919051 | SGX78861.1 | A5K7Y9 |
| NEHKEDADKKNNDEL | 0 | 0 | 1 | 919061 | SGX78861.1 | A5K7Y9 |
| NKIRQKYEYSMNPVE | 0 | 0 | 1 | 919077 | SGX78860.1 | A5K7Z0 |
| NKKNVLDEGEDHSQY | 0 | 0 | 1 | 919079 | SGX78861.1 | A5K7Y9 |
| NPTDDQAEEATPGPS | 0 | 0 | 1 | 919094 | SGX78861.1 | A5K7Y9 |
| NVGQSSTSSINPPPA | 0 | 0 | 1 | 919107 | SGX78853.1 | A5K7Z7 |
| PANNEGTTVTQESAL | 0 | 0 | 1 | 919117 | SGX78860.1 | A5K7Z0 |
| PATARGETATDSPTS | 0 | 0 | 1 | 919119 | SGX78853.1 | A5K7Z7 |
| PNVKYLDKLYDEVLK | 0 | 0 | 1 | 919143 | SGX78853.1 | A5K7Z7 |
| PPAAAAALPQPASPV | 0 | 0 | 1 | 919145 | SGX78853.1 | A5K7Z7 |
| PPQEQHAQREEVRNE | 0 | 0 | 1 | 919151 | SGX78855.1 | A5K7Z5 |
| PQPASPVVSAPEGEP | 0 | 0 | 1 | 919155 | SGX78853.1 | A5K7Z7 |
| PSSPADEQDDDNEAV | 0 | 0 | 1 | 919163 | SGX78861.1 | A5K7Y9 |
| PTAAAASPQVKHVDS | 0 | 0 | 1 | 919167 | SGX78861.1 | A5K7Y9 |
| PTSAAAAAAPAGPTA | 0 | 0 | 1 | 919168 | SGX78861.1 | A5K7Y9 |
| PTSPPANNEGTTVTQ | 0 | 0 | 1 | 919169 | SGX78860.1 | A5K7Z0 |
| PTSQEPSQTHTTGSS | 0 | 0 | 1 | 919170 | SGX78853.1 | A5K7Z7 |
| PVEYEIVKNLFNVGF | 0 | 0 | 1 | 919171 | SGX78860.1 | A5K7Z0 |
| PVLGDDDKKDKENEH | 0 | 0 | 1 | 919172 | SGX78861.1 | A5K7Y9 |
| QGASDSSNDSAEPTS | 0 | 0 | 1 | 919188 | SGX78861.1 | A5K7Y9 |
| QKYEYSMNPVEYEIV | 0 | 0 | 1 | 919202 | SGX78860.1 | A5K7Z0 |
| QSKGKIKGQADTTAV | 0 | 0 | 1 | 919216 | SGX78860.1 | A5K7Z0 |
| QYDKMVLNKTEYNIS | 0 | 0 | 1 | 919223 | SGX78861.1 | A5K7Y9 |
| RGETATDSPTSQEPS | 0 | 0 | 1 | 919231 | SGX78853.1 | A5K7Z7 |
| RKKIVFVCSIFVLLS | 0 | 0 | 1 | 919235 | SGX78860.1 | A5K7Z0 |
| RYSDLLKNAISLMNT | 0 | 0 | 1 | 919251 | SGX78855.1 | A5K7Z5 |
| SALAATPPKGTADSN | 0 | 0 | 1 | 919254 | SGX78860.1 | A5K7Z0 |
| SALGESAEVPKGDSD | 0 | 0 | 1 | 919255 | SGX78860.1 | A5K7Z0 |
| SAPEGEPEKGRVPNV | 0 | 0 | 1 | 919256 | SGX78853.1 | A5K7Z7 |
| SEARMKDADRYTNLL | 0 | 0 | 1 | 919265 | SGX78857.1 | A5K7Z3 |
| SEGPANMSESPADMS | 0 | 0 | 1 | 919266 | SGX78851.1 | A5K7Z9 |
| SGSNLKYSDELYEDI | 0 | 0 | 1 | 919294 | SGX78851.1 | A5K7Z9 |
| SIFVLLSCSPASSEQ | 0 | 0 | 1 | 919298 | SGX78860.1 | A5K7Z0 |
| SINPPPAAAAALPQP | 0 | 0 | 1 | 919300 | SGX78853.1 | A5K7Z7 |
| SMDPVEYEIVKNLFN | 0 | 0 | 1 | 919314 | SGX78857.1 | A5K7Z3 |
| SPASSEQLGIQKKKK | 0 | 0 | 1 | 919317 | SGX78860.1 | A5K7Z0 |
| SPVVSAPEGEPEKGR | 0 | 0 | 1 | 919318 | SGX78853.1 | A5K7Z7 |
| SQVSEEVPTSDEEDG | 0 | 0 | 1 | 919322 | SGX78851.1 | A5K7Z9 |
| SSTSSINPPPAAAAA | 0 | 0 | 1 | 919330 | SGX78853.1 | A5K7Z7 |
| SVATLENGYESIHGE | 0 | 0 | 1 | 919339 | SGX78860.1 | A5K7Z0 |
| TAVAKPPTGSGAGAH | 0 | 0 | 1 | 919350 | SGX78860.1 | A5K7Z0 |
| TDAKDEIHVPPFHSK | 0 | 0 | 1 | 919352 | SGX78853.1 | A5K7Z7 |
| TDPKFRKEFKDFMNC | 0 | 0 | 1 | 919355 | SGX78851.1 | A5K7Z9 |
| TEEVVAEAMSDEGPQ | 0 | 0 | 1 | 919357 | SGX78861.1 | A5K7Y9 |
| TGEQSDNQSDQQSGA | 0 | 0 | 1 | 919362 | SGX78851.1 | A5K7Z9 |
| THTTGSSRNVGQSST | 0 | 0 | 1 | 919367 | SGX78853.1 | A5K7Z7 |
| TLKGKLQKIRVQIKD | 0 | 0 | 1 | 919379 | SGX78861.1 | A5K7Y9 |
| TMLTNGQVEREKKNT | 0 | 0 | 1 | 919381 | SGX78861.1 | A5K7Y9 |
| TQTEEAKTYLTLFEN | 0 | 0 | 1 | 919390 | SGX78851.1 | A5K7Z9 |
| TSGQPESGRVPNVKY | 0 | 0 | 1 | 919394 | SGX78854.1 | A5K7Z6 |
| TVEDPREGDEGNEND | 0 | 0 | 1 | 919399 | SGX78855.1 | A5K7Z5 |
| VAEAMSDEGPQEQGA | 0 | 0 | 1 | 919402 | SGX78861.1 | A5K7Y9 |
| VEYEIVKNLFNVGFK | 0 | 0 | 1 | 919413 | SGX78857.1 | A5K7Z3 |
| VFVCSIFVLLSCSPA | 0 | 0 | 1 | 919416 | SGX78860.1 | A5K7Z0 |
| VGKNDKTSSEGPANM | 0 | 0 | 1 | 919419 | SGX78851.1 | A5K7Z9 |
| VKKLFDAFFKKDGNP | 0 | 0 | 1 | 919426 | SGX78854.1 | A5K7Z6 |
| VKYLDKLYDEVLKTT | 0 | 0 | 1 | 919427 | SGX78854.1 | A5K7Z6 |
| VLDEGEDHSQYNIFR | 0 | 0 | 1 | 919428 | SGX78861.1 | A5K7Y9 |
| VLKTTDAKDEIHVPP | 0 | 0 | 1 | 919430 | SGX78853.1 | A5K7Z7 |
| VLTTSDNTNWIHVPD | 0 | 0 | 1 | 919433 | SGX78860.1 | A5K7Z0 |
| VPDYHSKYNKIRQKY | 0 | 0 | 1 | 919440 | SGX78860.1 | A5K7Z0 |
| VPKGDSDSAQGSQPS | 0 | 0 | 1 | 919441 | SGX78860.1 | A5K7Z0 |
| VPPFHSKYNDFRKKY | 0 | 0 | 1 | 919442 | SGX78853.1 | A5K7Z7 |
| VPSLADRADAVPDGR | 0 | 0 | 1 | 919443 | SGX78860.1 | A5K7Z0 |
| WGPATEEVVAEAMSD | 0 | 0 | 1 | 919455 | SGX78861.1 | A5K7Y9 |
| WIHVPDYHSKYNTIR | 0 | 0 | 1 | 919456 | SGX78857.1 | A5K7Z3 |
| YESIHGEDEPQVVPS | 0 | 0 | 1 | 919472 | SGX78860.1 | A5K7Z0 |
| YIRGTQTEEAKTYLT | 0 | 0 | 1 | 919479 | SGX78851.1 | A5K7Z9 |
| YLDKLYDEVLKTTDA | 0 | 0 | 1 | 919482 | SGX78853.1 | A5K7Z7 |
| YLDKLYDEVLTTSDN | 0 | 0 | 1 | 919483 | SGX78860.1 | A5K7Z0 |
| YSMNPVEYEIVKNLF | 0 | 0 | 1 | 919493 | SGX78860.1 | A5K7Z0 |
| AFNMKQTIEDEKLKD | 0 | 0 | 1 | 944885 | AAA29417.1 | P11503 |
| FNMKQTIEDEKLKDK | 0 | 0 | 1 | 945108 | AAA29417.1 | P11503 |
| QPTKVSWDAEPGAL | 0 | 0 | 1 | 945625 | AAA29411.1 | P31729 |
| SGIDAKNQGEKLTYA | 0 | 0 | 1 | 945736 | AAA50363.1 | A0A044TBB6 |
| VMNCADNTGAKNLFV | 0 | 0 | 1 | 945914 | AAB30262.2 | A0A044V939 |
| VNEIINGNGNDF | 0 | 0 | 1 | 945915 | AAB96970.1 | Q9NFN6 |
| ECEEGECPVEPEPLPVPAPLPT | 0 | 0 | 1 | 947237 | AAC47463.1 | A5K806 |
| EQPEDKAKKSSNNGYK | 0 | 0 | 1 | 947260 | AAC47463.1 | A5K806 |

| | | | | | | |
|---|---|---|---|---|---|---|
| EWKNKCLDVTDNLPEKII | 0 | 0 | 1 | 956558 | CAX83041.1 | A0A4Z2DDJ1 |
| TDNLPEKIIQFAEHMNIL | 0 | 0 | 1 | 956721 | CAX83041.1 | A0A4Z2DDJ1 |
| ALLMNVFASRTLRLS | 0 | 0 | 1 | 985857 | AAA29259.1 | A0A504X2M8 |
| EKISRLMPSAHAIIR | 0 | 0 | 1 | 987702 | AAA29259.1 | A0A504X2M8 |
| PGRYFTAASHALLMN | 0 | 0 | 1 | 990770 | AAA29259.1 | A0A504X2M8 |
| SERIRMAPPASASKA | 0 | 0 | 1 | 991485 | AAA29259.1 | A0A504X2M8 |
| YNVVTRLPASPAALA | 0 | 0 | 1 | 992920 | AAA29259.1 | A0A504X2M8 |
| LPEHFRWIHVGS | 0 | 0 | 1 | 1075462 | BAF79609.1 | A0A068Y6Q2 |
| PGEDGADRAGG | 0 | 0 | 1 | 1075560 | BAF79609.1 | A0A068Y6Q2 |
| NREIAKYFVKKNDKHNKLFC | 0 | 0 | 1 | 1162255 | XP_002808883.1 | C0H4Y6 |
| NVYKNKIKSRILKENKEESL | 0 | 0 | 1 | 1162868 | XP_001350974.1 | Q8I298 |
| SCNYSTPSSVFNNDKENTAS | 0 | 0 | 1 | 1168108 | XP_001350974.1 | Q8I298 |
| YPTLILFNKKNKINYGGGRT | 0 | 0 | 1 | 1180873 | XP_002808883.1 | C0H4Y6 |
| VVLESTGIFRTK | 0 | 0 | 1 | 1300654 | AAA30325.1 | A2DHT2 |
| ADAKGSQKRQKKVLR | 0 | 0 | 1 | 1335434 | XP_001562231.1 | A4HNK4 |
| AYVEDIVRCSTAYTE | 0 | 0 | 1 | 1335480 | XP_001562231.1 | A4HNK4 |
| EDIVRCSTAYTEYAR | 0 | 0 | 1 | 1335518 | XP_001562231.1 | A4HNK4 |
| KPHRLTPRIMMTAVR | 0 | 0 | 1 | 1335632 | XP_001564819.1 | A4HBV1 |
| SLHKAITKKKGGKKS | 0 | 0 | 1 | 1335815 | XP_001564819.1 | A4HBV1 |
| YEEVRRVLKAYVEDI | 0 | 0 | 1 | 1335888 | XP_001562231.1 | A4HNK4 |
| ARAGLQGDLLGLNVLC | 0 | 0 | 1 | 1393692 | CDW54327.1 | A0A077Z1H0 |
| DKIVSEASRYKDMLQF | 0 | 0 | 1 | 1393723 | XP_013301487.1 | W2TB67 |
| ELELDPAKTNVNGGAI | 0 | 0 | 1 | 1393750 | CDW59723.1 | A0A077ZJA3 |
| FVVIENLSNDEKQIFT | 0 | 0 | 1 | 1393778 | CDW53379.1 | A0A077Z068 |
| GENYEQQLQQPKAYGI | 0 | 0 | 1 | 1393784 | XP_018646263.1 | A0A3Q0KQ41 |
| GIDEAVLRRTVDDLLT | 0 | 0 | 1 | 1393789 | KIH67114.1 | A0A0C2H6G3 |
| GIDERLAHVWQKTTNA | 0 | 0 | 1 | 1393790 | CDW54303.1 | A0A077Z3Q3 |
| GIDHVLDKMAVMLDRP | 0 | 0 | 1 | 1393791 | KIH59500.1 | A0A0C2GE87 |
| GIEAIKQNDESEFPNN | 0 | 0 | 1 | 1393793 | XP_018648222.1 | A0A3Q0KE79 |
| GIEQLRPRELCGDLIV | 0 | 0 | 1 | 1393796 | XP_013305594.1 | W2TQR8 |
| GIEQQVILQPAAPVAM | 0 | 0 | 1 | 1393797 | XP_018653281.1 | G4VMH7 |
| GPWHQKLTAKKVAKSM | 0 | 0 | 1 | 1393806 | KIH63297.1 | A0A0C2DL13 |
| GTVRKEMQKYSGESRG | 0 | 0 | 1 | 1393813 | KIH50814.1 | A0A0C2FVW4 |
| GTVYKLQEMKPSVVKE | 0 | 0 | 1 | 1393814 | CDW60598.1 | A0A077ZLH3 |
| GVEHLARVNCVLWQND | 0 | 0 | 1 | 1393816 | KIH47972.1 | A0A0C2CDQ5 |
| GVTANDLRTAEAMVRS | 0 | 0 | 1 | 1393819 | CDW60144.1 | A0A077ZK78 |
| IDREIVRILEEKLDHL | 0 | 0 | 1 | 1393833 | RCN38490.1 | A0A368G635 |
| IELDPSAATSNTLHDV | 0 | 0 | 1 | 1393835 | XP_013301076.1 | W2TB84 |
| ILKKHDKVLQNTAGNE | 0 | 0 | 1 | 1393847 | CDW59643.1 | A0A077ZJ59 |
| INESSMKQASLISNVM | 0 | 0 | 1 | 1393849 | XP_013306973.1 | W2TUJ0 |
| KNLISAQLNQTATGQP | 0 | 0 | 1 | 1393873 | XP_013304895.1 | W2TLM2 |
| KTESLISNSPSTKKSK | 0 | 0 | 1 | 1393878 | XP_018647234.1 | A0A5K4ESG5 |
| KTVPPRKKDSEISNSV | 0 | 0 | 1 | 1393883 | CDW59163.1 | A0A077ZHW5 |
| LIERQLKMESIIANIE | 0 | 0 | 1 | 1393898 | VDK49170.1 | A0A0M3JZ62 |
| LKKIIGGHLQMPTAIV | 0 | 0 | 1 | 1393900 | XP_013295903.1 | W2SXR1 |
| MAKEDGLISNLRSYPP | 0 | 0 | 1 | 1393919 | XP_013291651.1 | W2SIE5 |
| MEIANGKEWKSDVRET | 0 | 0 | 1 | 1393923 | KIH64126.1 | A0A0C2H3X2 |
| MTKPVKKTRWHQELSA | 0 | 0 | 1 | 1393932 | KIH45323.1 | A0A0C2BNB7 |
| QDILNNKALLGDEKAR | 0 | 0 | 1 | 1393999 | VDK44519.1 | A0A0M3JU71 |
| SKFDSVILNGYNIVGD | 0 | 0 | 1 | 1394059 | KIH51412.1 | A0A0C2CNU3 |
| SPLIGKEIVSLSLNGD | 0 | 0 | 1 | 1394067 | CDW53085.1 | A0A077Z310 |
| VATKKLIGDQMLNRLQ | 0 | 0 | 1 | 1394112 | XP_013300475.1 | W2T8R5 |
| VSKQTISAHLKPLGSN | 0 | 0 | 1 | 1394142 | CDW56559.1 | A0A077Z9V6 |
| YKHVIGIDLNKEAIEN | 0 | 0 | 1 | 1394166 | CDW59548.1 | A0A077ZIW4 |
| AAGFNRIHWAEGRR | 0 | 0 | 1 | 1596890 | EAN90963.1 | Q4DEM3 |
| AETPPVTSVNRSLV | 0 | 0 | 1 | 1596895 | EAN90963.1 | Q4DEM3 |
| ALRPASVAEAADVM | 0 | 0 | 1 | 1596912 | EAN90963.1 | Q4DEM3 |
| ANSDWTALMNAAGF | 0 | 0 | 1 | 1596916 | EAN90963.1 | Q4DEM3 |
| ATAERGFSGDIPPA | 0 | 0 | 1 | 1596918 | EAN90963.1 | Q4DEM3 |
| AVANKHTQGLMPPI | 0 | 0 | 1 | 1596922 | EAN90963.1 | Q4DEM3 |
| AVKSDDQSNNSVAY | 0 | 0 | 1 | 1596923 | EAN90963.1 | Q4DEM3 |
| CRFSGDRRSGEELL | 0 | 0 | 1 | 1596941 | EAN90963.1 | Q4DEM3 |
| CTALAYKEKLNFTT | 0 | 0 | 1 | 1596945 | EAN90963.1 | Q4DEM3 |
| DDSRDVYAFFFPGE | 0 | 0 | 1 | 1596948 | EAN90963.1 | Q4DEM3 |
| DFNAIDTFLVVDQV | 0 | 0 | 1 | 1596952 | EAN90963.1 | Q4DEM3 |
| DRRSGEELLLPREV | 0 | 0 | 1 | 1596968 | EAN90963.1 | Q4DEM3 |
| DSRVGKTGSAQQQK | 0 | 0 | 1 | 1596969 | EAN90963.1 | Q4DEM3 |
| DSRYELHVMSFWRG | 0 | 0 | 1 | 1596970 | EAN90963.1 | Q4DEM3 |
| DTFLVVDQVAYQDA | 0 | 0 | 1 | 1596971 | EAN90963.1 | Q4DEM3 |
| DTQAAIEAALHRIG | 0 | 0 | 1 | 1596972 | EAN90963.1 | Q4DEM3 |
| DTVALIFLMLSITA | 0 | 0 | 1 | 1596973 | EAN90963.1 | Q4DEM3 |
| EELLLPREVRSVYD | 0 | 0 | 1 | 1596978 | EAN90963.1 | Q4DEM3 |
| EGRRYPQNVLETRN | 0 | 0 | 1 | 1596980 | EAN90963.1 | Q4DEM3 |
| EKVFLTLTRYNTTF | 0 | 0 | 1 | 1596986 | EAN90963.1 | Q4DEM3 |
| ELEMEKSGVRRAST | 0 | 0 | 1 | 1596989 | EAN90963.1 | Q4DEM3 |
| ETRNGMSLAELLSS | 0 | 0 | 1 | 1596993 | EAN90963.1 | Q4DEM3 |
| FPGEHVGSVGSARF | 0 | 0 | 1 | 1597022 | EAN90963.1 | Q4DEM3 |
| FWRGNIGARSTMVG | 0 | 0 | 1 | 1597032 | EAN90963.1 | Q4DEM3 |
| GATWEQDRDWMRLH | 0 | 0 | 1 | 1597034 | EAN90963.1 | Q4DEM3 |
| GRENRVKTNTQYDM | 0 | 0 | 1 | 1597046 | EAN90963.1 | Q4DEM3 |
| HRIGWTDVARSPAV | 0 | 0 | 1 | 1597061 | EAN90963.1 | Q4DEM3 |
| IEAALHRIGWTDVA | 0 | 0 | 1 | 1597065 | EAN90963.1 | Q4DEM3 |
| IGARSTMVGSDTVA | 0 | 0 | 1 | 1597072 | EAN90963.1 | Q4DEM3 |
| IPPAPHDAAYAAYF | 0 | 0 | 1 | 1597093 | EAN90963.1 | Q4DEM3 |
| IRGLSIPDTVFSSG | 0 | 0 | 1 | 1597095 | EAN90963.1 | Q4DEM3 |
| KSLRIPHGDWGATW | 0 | 0 | 1 | 1597154 | EAN90963.1 | Q4DEM3 |
| KTGSAQQQKAELEM | 0 | 0 | 1 | 1597157 | EAN90963.1 | Q4DEM3 |
| LLVASTAVSLVQDA | 0 | 0 | 1 | 1597187 | EAN90963.1 | Q4DEM3 |
| LYYHVDSRVGKTGS | 0 | 0 | 1 | 1597200 | EAN90963.1 | Q4DEM3 |
| MPPITAVAFSDDNR | 0 | 0 | 1 | 1597214 | EAN90963.1 | Q4DEM3 |
| MRLHNDSRYELHVM | 0 | 0 | 1 | 1597215 | EAN90963.1 | Q4DEM3 |

| | | | | | | |
|---|---|---|---|---|---|---|
| NFTTVDFNAIDTFL | 0 | 0 | 1 | 1597229 | EAN90963.1 | Q4DEM3 |
| NPDVFTVVDKTAEH | 0 | 0 | 1 | 1597234 | EAN90963.1 | Q4DEM3 |
| NTTFANPDVFTVVD | 0 | 0 | 1 | 1597240 | EAN90963.1 | Q4DEM3 |
| PHGDWGATWEQDRD | 0 | 0 | 1 | 1597247 | EAN90963.1 | Q4DEM3 |
| PHGLLLLHATEVAQ | 0 | 0 | 1 | 1597248 | EAN90963.1 | Q4DEM3 |
| PITTLLDLLPAVMA | 0 | 0 | 1 | 1597249 | EAN90963.1 | Q4DEM3 |
| PLSGWTVWTSTADM | 0 | 0 | 1 | 1597250 | EAN90963.1 | Q4DEM3 |
| PQNVLETRNGMSLA | 0 | 0 | 1 | 1597251 | EAN90963.1 | Q4DEM3 |
| QCKFLSAPNDVAEF | 0 | 0 | 1 | 1597259 | EAN90963.1 | Q4DEM3 |
| QDRDWMRLHNDSRY | 0 | 0 | 1 | 1597260 | EAN90963.1 | Q4DEM3 |
| RASTTRLPYSPITT | 0 | 0 | 1 | 1597283 | EAN90963.1 | Q4DEM3 |
| RIHWAEGRRYPQNV | 0 | 0 | 1 | 1597289 | EAN90963.1 | Q4DEM3 |
| RLPYSPITTLLDLL | 0 | 0 | 1 | 1597302 | EAN90963.1 | Q4DEM3 |
| RRVGGDDSRDVYAF | 0 | 0 | 1 | 1597311 | EAN90963.1 | Q4DEM3 |
| RVLLPPTQNAAETP | 0 | 0 | 1 | 1597314 | EAN90963.1 | Q4DEM3 |
| SLQDIIRGLSIPDT | 0 | 0 | 1 | 1597334 | EAN90963.1 | Q4DEM3 |
| SPAVPKSLRIPHGD | 0 | 0 | 1 | 1597343 | EAN90963.1 | Q4DEM3 |
| SVAEAADVMLRVLL | 0 | 0 | 1 | 1597353 | EAN90963.1 | Q4DEM3 |
| SVYDFVILYFPSAA | 0 | 0 | 1 | 1597356 | EAN90963.1 | Q4DEM3 |
| TADMRWEWNETSLN | 0 | 0 | 1 | 1597358 | EAN90963.1 | Q4DEM3 |
| TAEHSALRPASVAE | 0 | 0 | 1 | 1597359 | EAN90963.1 | Q4DEM3 |
| TLTRYNTTFANPDV | 0 | 0 | 1 | 1597384 | EAN90963.1 | Q4DEM3 |
| TVWTSTADMRWEWN | 0 | 0 | 1 | 1597393 | EAN90963.1 | Q4DEM3 |
| VDQVAYQDAPLYYH | 0 | 0 | 1 | 1597396 | EAN90963.1 | Q4DEM3 |
| VGSVGSARFISDAT | 0 | 0 | 1 | 1597401 | EAN90963.1 | Q4DEM3 |
| VLEALRRVGGDDSR | 0 | 0 | 1 | 1597405 | EAN90963.1 | Q4DEM3 |
| VTSVNRSLVEQLWG | 0 | 0 | 1 | 1597419 | EAN90963.1 | Q4DEM3 |
| VYAFFFPGEHVGSV | 0 | 0 | 1 | 1597421 | EAN90963.1 | Q4DEM3 |
| WEWNETSLNFEKKT | 0 | 0 | 1 | 1597422 | EAN90963.1 | Q4DEM3 |
| YKEKLNFTTVDFNA | 0 | 0 | 1 | 1597428 | EAN90963.1 | Q4DEM3 |
| YQDAPLYYHVDSRV | 0 | 0 | 1 | 1597442 | EAN90963.1 | Q4DEM3 |
| EEFRDYYENGEEKSNKQM | 0 | 0 | 1 | 1711654 | ACY68791.1 | A5K4Z2 |
| GDQRLKDGGFAFPNADDH | 0 | 0 | 1 | 1711656 | ACB42437.1 | A5K4Z2 |
| SSGVRVDLGEDAEVENAK | 0 | 0 | 1 | 1711667 | ACY68791.1 | A5K4Z2 |
