## Supplementary Dataset 2 for "Analysis of virus-specific B cell epitopes reveals extensive antigen processing prior to recognition"

**Supplementary Dataset 2.** Virus-specific linear B cell epitopes with location

| Sequence | IEDB | Classification | Visible |
|---|---|---|---|
| AAAGAPFYLNVQ | 1406433 | Ectodomain | YES |
| AAAGVPLYINVQ | 1406437 | Ectodomain | YES |
| AAAGVPLYLNVQ | 1406438 | Ectodomain | YES |
| AAAINSVPWSKI | 1406440 | Cell/Virus Enclosed | NO |
| AAALEDEFTPFD | 1406448 | Cell/Virus Enclosed | NO |
| AAALKSLGFDKPQEKD | 1642919 | Capsid | UN |
| AAALKSLGFEKP | 1406450 | Cell/Virus Enclosed | NO |
| AAANATGTVIIS | 1406455 | Ectodomain | YES |
| AAANSTGTVIIR | 1406459 | Ectodomain | YES |
| AAAPFTKPFAVY | 1406460 | Cell/Virus Enclosed | NO |
| AAAQGFQACRFI | 1406461 | Cell/Virus Enclosed | NO |
| AAAVDEAFPLDEAEDV | 1642924 | Cell/Virus Enclosed | NO |
| AAAVDEAFPLNE | 1406465 | Cell/Virus Enclosed | NO |
| AAAVDEVFPLDE | 1406466 | Cell/Virus Enclosed | NO |
| AAAYMKLSDALK | 1406471 | Cell/Virus Enclosed | NO |
| AACAVDVNVCHS | 1406477 | Cell/Virus Enclosed | NO |
| AACHDLLSEYGT | 1406480 | Ectodomain | YES |
| AACHLAKALQTYSETG | 1642935 | Cell/Virus Enclosed | NO |
| AADAERMAMYTG | 1406490 | Ectodomain | YES |
| AADINIHDLPIC | 1406494 | Ectodomain | YES |
| AADINIPDLLIC | 1406495 | Ectodomain | YES |
| AADINIPHLPIC | 1406497 | Ectodomain | YES |
| AADINNPDLPIC | 1406498 | Ectodomain | YES |
| AADINTPDVPIC | 1406499 | Ectodomain | YES |
| AADLDEFSKQLQ | 1406502 | Cell/Virus Enclosed | NO |
| AADLGVLIQNGAKHVQ | 1642942 | Cell/Virus Enclosed | NO |
| AADLGVLIQNSAKHVQ | 1642943 | Cell/Virus Enclosed | NO |
| AADLPCTLHFSL | 1406505 | Cell/Virus Enclosed | NO |
| AADPALHVASASALYD | 1642946 | Cell/Virus Enclosed | NO |
| AADPAMHIASSNAFLD | 1642948 | Cell/Virus Enclosed | NO |
| AAEASKKPRQKRTATKAYNV | 1392030 | Capsid | UN |
| AAEELASPMKSD | 1406515 | Cell/Virus Enclosed | NO |
| AAEFKYMVANGLHAPH | 1642955 | Cell/Virus Enclosed | NO |
| AAEGVPFYLNVQ | 1406518 | Ectodomain | YES |
| AAEKLASPMKSD | 1406522 | Cell/Virus Enclosed | NO |
| AAEQDRNTREVFAQVKQM | 1087450 | Ectodomain | YES |
| AAESKAFVDTIVPPCP | 1642960 | Cell/Virus Enclosed | NO |
| AAETHRDCDFNKPLIE | 1642961 | Cell/Virus Enclosed | NO |
| AAETQKATEEAFKQSY | 1642965 | Cell/Virus Enclosed | NO |
| AAETVKAKEESVKSSY | 1642966 | Cell/Virus Enclosed | NO |
| AAFATHVCFKDC | 1406539 | Cell/Virus Enclosed | NO |
| AAFDCGDYAACK | 1406541 | Ectodomain | YES |
| AAFDFSLDVFKIGDVK | 1642969 | Cell/Virus Enclosed | NO |
| AAFDFSSDVFKI | 1406543 | Cell/Virus Enclosed | NO |
| AAFDLSKELKRP | 1406544 | Cell/Virus Enclosed | NO |
| AAFHANSSEPALLFRN | 1642974 | Ectodomain | YES |
| AAFHVNSSEPAL | 1406552 | Ectodomain | YES |
| AAFISPYNSQNF | 1406553 | Cell/Virus Enclosed | NO |
| AAFLELLPNVFK | 1406554 | Cell/Virus Enclosed | NO |
| AAFNAVCSVIDL | 1406555 | Cell/Virus Enclosed | NO |
| AAFYVYKLQPLT | 1406572 | Ectodomain | YES |
| AAGADTTITLNV | 1406574 | Cell/Virus Enclosed | NO |
| AAGGKVSFSDDVEVKD | 1642983 | Cell/Virus Enclosed | NO |
| AAGLEFTDENYNNLVP | 1642990 | Cell/Virus Enclosed | NO |
| AAGQQFVKETTD | 1406587 | Cell/Virus Enclosed | NO |
| AAGTDTTITLNV | 1406589 | Cell/Virus Enclosed | NO |
| AAGVGNTFICEE | 1406592 | Cell/Virus Enclosed | NO |
| AAGYGGFGCKDS | 1406597 | Cell/Virus Enclosed | NO |
| AAHPPPGEQFKH | 1406600 | Cell/Virus Enclosed | NO |
| AAIASNCYSSLI | 1406602 | Ectodomain | YES |
| AAIDFVQYEVDRRVLF | 1643004 | Cell/Virus Enclosed | NO |
| AAIELSKELKRPVNPT | 1643005 | Cell/Virus Enclosed | NO |
| AAIGDLKCPLDS | 1406607 | Ectodomain | YES |
| AAIGDLNCTLDP | 1406608 | Ectodomain | YES |
| AAIHADQLTPAW | 1406609 | Ectodomain | YES |
| AAIHYLNSKIVDLAQH | 1643007 | Cell/Virus Enclosed | NO |
| AAIKLVPTTTGG | 1406614 | Cell/Virus Enclosed | NO |
| AAIKLVPTTTGI | 1406615 | Cell/Virus Enclosed | NO |
| AAIKLVPTTTGVLVRF | 1643011 | Cell/Virus Enclosed | NO |
| AAIKLYNPKAIHDIGN | 1643012 | Cell/Virus Enclosed | NO |
| AAIKRFKNGFQG | 1406617 | Cell/Virus Enclosed | NO |
| AAIKRLKNGFQG | 1406618 | Cell/Virus Enclosed | NO |
| AAIKRLYMGFQG | 1406619 | Cell/Virus Enclosed | NO |
| AAILNGCAWFVKPNRT | 1643015 | Cell/Virus Enclosed | NO |

| | | | |
|---|---|---|---|
| AAINSVPWSKIF | 1406632 | Cell/Virus Enclosed | NO |
| AAIPDYNYYKYN | 1406634 | Cell/Virus Enclosed | NO |
| AAIQNCAWDYHI | 1406639 | Cell/Virus Enclosed | NO |
| AAIQNCAWDYHV | 1406640 | Cell/Virus Enclosed | NO |
| AAISSNCYSSLI | 1406647 | Ectodomain | YES |
| AAITAPYVVLTGLGES | 1643026 | Cell/Virus Enclosed | NO |
| AAITDYNYYKYNLPTM | 1643027 | Cell/Virus Enclosed | NO |
| AAITDYNYYSYNLPTM | 1643028 | Cell/Virus Enclosed | NO |
| AAITILDGISEQ | 1406650 | Cell/Virus Enclosed | NO |
| AAITSGIKFQTVKPGN | 1643030 | Cell/Virus Enclosed | NO |
| AAITSGVKFQTVKPGN | 1643031 | Cell/Virus Enclosed | NO |
| AAIVSNCYSSLI | 1406657 | Ectodomain | YES |
| AAIYGGDHYVVLGGET | 1643035 | Cell/Virus Enclosed | NO |
| AAIYYLNSKIVD | 1406660 | Cell/Virus Enclosed | NO |
| AAKGYIAGIQNFKETL | 1643037 | Cell/Virus Enclosed | NO |
| AAKICIVDNVYM | 1406667 | Cell/Virus Enclosed | NO |
| AAKLQNNEIKPS | 1406679 | Cell/Virus Enclosed | NO |
| AAKLQNNELSPV | 1406680 | Cell/Virus Enclosed | NO |
| AAKNKMRHKRIS | 1406682 | Cell/Virus Enclosed | NO |
| AAKRVLGLQTQTVDSA | 1643047 | Cell/Virus Enclosed | NO |
| AALAMEKVNECVKSQS | 1643051 | Ectodomain | YES |
| AALAVNACLIPL | 1406693 | Cell/Virus Enclosed | NO |
| AALAVNVCLIPLLLLL | 1643052 | Cell/Virus Enclosed | NO |
| AALDLSKELKRPIQPT | 1643054 | Cell/Virus Enclosed | NO |
| AALDYLATVKTVPAGN | 1643055 | Cell/Virus Enclosed | NO |
| AALFASLDVKPY | 1406700 | Cell/Virus Enclosed | NO |
| AALGLTVSDDDF | 1406707 | Cell/Virus Enclosed | NO |
| AALKSEKHQDHV | 1406719 | Cell/Virus Enclosed | NO |
| AALKSLGFDKPQEKDK | 1643064 | Capsid | UN |
| AALLAGSFDKVYDIGN | 1643066 | Cell/Virus Enclosed | NO |
| AALLAGSFDRIY | 1406724 | Cell/Virus Enclosed | NO |
| AALNAFVSQVLNKYTE | 1643068 | Ectodomain | YES |
| AALNSEKHQDHV | 1406730 | Cell/Virus Enclosed | NO |
| AALNVFVSHTLT | 1406732 | Ectodomain | YES |
| AALPQSKDSNFLNESG | 1643072 | Cell/Virus Enclosed | NO |
| AALPQSKDSNFLNRVR | 1643073 | Cell/Virus Enclosed | NO |
| AALPQSKESNFL | 1406738 | Cell/Virus Enclosed | NO |
| AALRAGSFDKVY | 1406743 | Cell/Virus Enclosed | NO |
| AALRAVSFADNN | 1406744 | Cell/Virus Enclosed | NO |
| AALSAQLAKDKVNECV | 1643077 | Ectodomain | YES |
| AALSFKENVDFI | 1406747 | Cell/Virus Enclosed | NO |
| AALSTGLTSQTVKPGH | 1643080 | Cell/Virus Enclosed | NO |
| AALTNNVAFQTV | 1406749 | Cell/Virus Enclosed | NO |
| AALTNSVAFQTV | 1406750 | Cell/Virus Enclosed | NO |
| AALVSGTATAGWTFGAG | 307 | Ectodomain | YES |
| AAMLFDSHIVSKESGN | 1643090 | Capsid | UN |
| AAMLSDSHIVSK | 1406765 | Cell/Virus Enclosed | NO |
| AAMSGQILLNCA | 1406767 | Cell/Virus Enclosed | NO |
| AAMTEQLQMGFG | 1406768 | Ectodomain | YES |
| AAMYKEARAVNR | 1406771 | Cell/Virus Enclosed | NO |
| AANATGTVIISP | 1406775 | Ectodomain | YES |
| AANGGPASRPPPVPSPATTK | 741180 | Ectodomain | YES |
| AANPTNCITPVN | 1406797 | Ectodomain | YES |
| AANSSEWAYCVDLHNK | 1643109 | Cell/Virus Enclosed | NO |
| AANSTGTVIINP | 1406801 | Ectodomain | YES |
| AANSTGTVIIRP | 1406802 | Ectodomain | YES |
| AANSTGTVIISP | 1406803 | Ectodomain | YES |
| AANTHLKHGGGIAGAI | 1643112 | Cell/Virus Enclosed | NO |
| AANTLNIIVPDKSVYD | 1643113 | Cell/Virus Enclosed | NO |
| AANVGRWSASIN | 1406812 | Ectodomain | YES |
| AANVSVSRFDPS | 1406814 | Ectodomain | YES |
| AANVSVSRFNPS | 1406815 | Ectodomain | YES |
| AAPAAPRAVSFA | 1406817 | Capsid | UN |
| AAPCAVSFADNN | 1406818 | Cell/Virus Enclosed | NO |
| AAPKSRVQQNKS | 1406823 | Cell/Virus Enclosed | NO |
| AAPNNTVSWYTG | 1406826 | Cell/Virus Enclosed | NO |
| AAPNTTVSWYTG | 1406827 | Cell/Virus Enclosed | NO |
| AAPRAISFADNN | 1406829 | Cell/Virus Enclosed | NO |
| AAPRAVSFADNNDITN | 1643125 | Cell/Virus Enclosed | NO |
| AAPRAVSFAYNN | 1406832 | Capsid | UN |
| AAPSGTWLTYHG | 1406834 | Capsid | UN |
| AAQADLQHEDSD | 1406835 | Cell/Virus Enclosed | NO |
| AAQAMEKVNERV | 1406837 | Ectodomain | YES |
| AAQCDCLCTPDP | 1406838 | Ectodomain | YES |
| AAQCNCLCTPDP | 1406839 | Ectodomain | YES |
| AAQGESSTQKVRPSHTNNPK | 387 | Cell/Virus Enclosed | NO |

| | | | |
|---|---|---|---|
| AAQKAYQEAMDSGDTS | 1643130 | Cell/Virus Enclosed | NO |
| AARGPAGVEFDV | 1406858 | Cell/Virus Enclosed | NO |
| AARLLGLQTQTVDSAQ | 1643139 | Cell/Virus Enclosed | NO |
| AARPPPGEHFKH | 1406871 | Cell/Virus Enclosed | NO |
| AASAVNVCLIPL | 1406885 | Cell/Virus Enclosed | NO |
| AASDITLKCCIV | 1406887 | Cell/Virus Enclosed | NO |
| AASDITLKCCTV | 1406888 | Cell/Virus Enclosed | NO |
| AASFNKAINNIVASFS | 1643153 | Ectodomain | YES |
| AASFNKAMTNIVDAFT | 1643154 | Ectodomain | YES |
| AASNGFRACRFV | 1406898 | Cell/Virus Enclosed | NO |
| AASNIFVINRDT | 1406901 | Cell/Virus Enclosed | NO |
| AASVNQTVLRNS | 1406910 | Cell/Virus Enclosed | NO |
| AASYNKYKYYSGSASE | 1643164 | Cell/Virus Enclosed | NO |
| AASYNRYKYYSGNANE | 1643165 | Cell/Virus Enclosed | NO |
| AATEKSNVIRGW | 1406917 | Ectodomain | YES |
| AATSANLFPPWS | 1406936 | Ectodomain | YES |
| AATSASLFPPWS | 1406938 | Ectodomain | YES |
| AATVAAMFPPWSAAAG | 1643176 | Ectodomain | YES |
| AATVRLQAGSNTEFAS | 1643179 | Cell/Virus Enclosed | NO |
| AAVDALCEKALK | 1406952 | Cell/Virus Enclosed | NO |
| AAVDEAFPLDEAEDVT | 1643183 | Cell/Virus Enclosed | NO |
| AAVDEAFPLNEA | 1406955 | Cell/Virus Enclosed | NO |
| AAVDLSMDKLCR | 1406956 | Cell/Virus Enclosed | NO |
| AAVDLSMDKLRRVDWK | 1643184 | Cell/Virus Enclosed | NO |
| AAVDLSMDRLRR | 1406958 | Cell/Virus Enclosed | NO |
| AAVDSLCAKAMT | 1406959 | Cell/Virus Enclosed | NO |
| AAVHEWKMVVCD | 1406968 | Cell/Virus Enclosed | NO |
| AAVHGWKMVVCDKVRS | 1643190 | Cell/Virus Enclosed | NO |
| AAVHKWKMVVCD | 1406970 | Cell/Virus Enclosed | NO |
| AAVHRWKMVVCD | 1406971 | Cell/Virus Enclosed | NO |
| AAVIAGDRWFLN | 1406972 | Cell/Virus Enclosed | NO |
| AAVINIPDLPIC | 1406975 | Ectodomain | YES |
| AAVKAAESKAFVDTIV | 1643195 | Cell/Virus Enclosed | NO |
| AAVKDFDFYRYNKPTI | 1643196 | Cell/Virus Enclosed | NO |
| AAVLNSLCICFV | 1406983 | Cell/Virus Enclosed | NO |
| AAVLNSLCICLV | 1406984 | Cell/Virus Enclosed | NO |
| AAVNLALKNLGF | 1406985 | Cell/Virus Enclosed | NO |
| AAVPVLAFDAARLRLLE | 429189 | Cell/Virus Enclosed | NO |
| AAVRSVTVEYNI | 1406992 | Cell/Virus Enclosed | NO |
| AAVSAMMRISYF | 1406994 | Transmembrane | NO |
| AAVSAMMWISYF | 1406995 | Transmembrane | NO |
| AAVSAMMWLSYF | 1406996 | Transmembrane | NO |
| AAVSRINWVTGG | 1406998 | Undetermined | UN |
| AAVTILDGIFEQ | 1406999 | Cell/Virus Enclosed | NO |
| AAVTLALKNLGFDNQS | 1643204 | Capsid | UN |
| AAVTTTNCIKRC | 1407003 | Cell/Virus Enclosed | NO |
| AAVYRINWATGG | 1407004 | Undetermined | UN |
| AAVYRINWVTGG | 1407006 | Undetermined | UN |
| AAYAEVHAEPKG | 1407011 | Cell/Virus Enclosed | NO |
| AAYANFNSQDVL | 1407012 | Cell/Virus Enclosed | NO |
| AAYENFNSQDTL | 1407017 | Cell/Virus Enclosed | NO |
| AAYFVGYLKPTTFMLKY | 525 | Ectodomain | YES |
| AAYHANSPEPAL | 1407026 | Ectodomain | YES |
| AAYHANSSDPAL | 1407027 | Ectodomain | YES |
| AAYHANSSEPAL | 1407028 | Ectodomain | YES |
| AAYHANSSEPTL | 1407029 | Ectodomain | YES |
| AAYINIPDLPIC | 1407031 | Ectodomain | YES |
| AAYINIPDLPVC | 1407032 | Ectodomain | YES |
| AAYLDAVKQGAKPVGN | 1643213 | Cell/Virus Enclosed | NO |
| AAYNGKPQGAFH | 1407041 | Cell/Virus Enclosed | NO |
| AAYNGPPQGAFH | 1407042 | Cell/Virus Enclosed | NO |
| AAYNNCTGGAEI | 1407043 | Ectodomain | YES |
| AAYREAAACHLA | 1407046 | Cell/Virus Enclosed | NO |
| AAYSRYRIGNYKLNTD | 1377315 | Underneath Membrane | NO |
| AAYTAALVSGIA | 1407052 | Ectodomain | YES |
| AAYTSSSLGSIA | 1407056 | Ectodomain | YES |
| AAYVEETSITIK | 1407059 | Cell/Virus Enclosed | NO |
| AAYVENTSITIK | 1407061 | Cell/Virus Enclosed | NO |
| ACADGTRHTYQL | 1407070 | Cell/Virus Enclosed | NO |
| ACAHAAVDSLCA | 1407072 | Cell/Virus Enclosed | NO |
| ACAQYYNGIMVLPGVA | 1643235 | Ectodomain | YES |
| ACAQYYNGIMVV | 1407076 | Ectodomain | YES |
| ACAVDVNVCHSV | 1407077 | Cell/Virus Enclosed | NO |
| ACCYVGVQTVED | 1407089 | Cell/Virus Enclosed | NO |
| ACDGLALAYRAN | 1407092 | Cell/Virus Enclosed | NO |
| ACDGLASAYRANSFDV | 1643242 | Cell/Virus Enclosed | NO |

| | | | |
|---|---|---|---|
| ACDYDRLPNEIT | 1407098 | Underneath Membrane | NO |
| ACDYDRLSNEVT | 1407100 | Underneath Membrane | NO |
| ACDYLYNPLIVD | 1407101 | Cell/Virus Enclosed | NO |
| ACEHISSTMFQY | 1407106 | Ectodomain | YES |
| ACEHISSTMSQH | 1407107 | Ectodomain | YES |
| ACEHISSTMSQYSRST | 1643247 | Ectodomain | YES |
| ACENSFMVNQLA | 1407114 | Cell/Virus Enclosed | NO |
| ACESQTPTSEEV | 1407117 | Cell/Virus Enclosed | NO |
| ACEVVVTNLNKSAGWP | 1643251 | Cell/Virus Enclosed | NO |
| ACEYAEEGICFN | 1407119 | Cell/Virus Enclosed | NO |
| ACFSKCHNCCDEYGGH | 1643254 | Underneath Membrane | NO |
| ACGCGASYTANDVEVQ | 1643259 | Cell/Virus Enclosed | NO |
| ACGCGTSYTAND | 1407138 | Cell/Virus Enclosed | NO |
| ACGNCTFGAPDD | 1407139 | Cell/Virus Enclosed | NO |
| ACGSPGYNLKNGEVEF | 1643260 | Cell/Virus Enclosed | NO |
| ACGSPGYNVRNDGTVE | 1643261 | Cell/Virus Enclosed | NO |
| ACGTNVPLQLGFSTGV | 1643262 | Cell/Virus Enclosed | NO |
| ACIIATIFLLWF | 1407157 | Cell/Virus Enclosed | NO |
| ACIKIAQADTAG | 1407158 | Cell/Virus Enclosed | NO |
| ACIKTAQADTAGLYTN | 1643274 | Cell/Virus Enclosed | NO |
| ACINGHAVPTLF | 1407165 | Cell/Virus Enclosed | NO |
| ACKCGVKQEQRTGLDA | 1643278 | Cell/Virus Enclosed | NO |
| ACKFAVCGDGFV | 1407178 | Cell/Virus Enclosed | NO |
| ACKGFYSNASSD | 1407179 | Ectodomain | YES |
| ACKGFYSNSSSD | 1407180 | Ectodomain | YES |
| ACKRGPGSGFFSRLN | 570 | Ectodomain | YES |
| ACKSQLGEYGSF | 1407184 | Ectodomain | YES |
| ACKSQLVEYGSF | 1407185 | Ectodomain | YES |
| ACKTIEDALRNSARLE | 1643282 | Ectodomain | YES |
| ACLCKHKKVSTN | 1407191 | Cell/Virus Enclosed | NO |
| ACLKGGLILKFD | 1407195 | Cell/Virus Enclosed | NO |
| ACLVGLMWLSYFIAS | 1312104 | Transmembrane | NO |
| ACNANRFNVAIT | 1407207 | Cell/Virus Enclosed | NO |
| ACNFELCDNPFF | 1407208 | Ectodomain | YES |
| ACNFELCDNPFI | 1407209 | Ectodomain | YES |
| ACNGHKIEDFSI | 1407210 | Cell/Virus Enclosed | NO |
| ACNGISVTQIMF | 1407212 | Cell/Virus Enclosed | NO |
| ACNGISVTQTMF | 1407213 | Cell/Virus Enclosed | NO |
| ACNIAKSAYERDRAVA | 1643294 | Cell/Virus Enclosed | NO |
| ACNIAKSVYERDKAVA | 1643295 | Cell/Virus Enclosed | NO |
| ACNKVSSRSAIE | 1407214 | Ectodomain | YES |
| ACNLGGAVCRKHATEY | 1643296 | Cell/Virus Enclosed | NO |
| ACNVNRFNVAIT | 1407220 | Cell/Virus Enclosed | NO |
| ACQDPEVGPEHS | 1407234 | Cell/Virus Enclosed | NO |
| ACQKHTPPAPKEDDPLKKYT | 111088 | Capsid | UN |
| ACQLPYCYFRNSSANY | 1643306 | Underneath Membrane | NO |
| ACQNLDCNCLGF | 1407239 | Cell/Virus Enclosed | NO |
| ACQPPYCYFRNSTTNY | 1643308 | Ectodomain | YES |
| ACQYTMCEFPHT | 1407242 | Ectodomain | YES |
| ACQYTMCEYPYT | 1407244 | Ectodomain | YES |
| ACRAYLDSCTTQ | 1407246 | Cell/Virus Enclosed | NO |
| ACRAYLNSRTTQ | 1407249 | Cell/Virus Enclosed | NO |
| ACRFIAFGLQDC | 1407250 | Cell/Virus Enclosed | NO |
| ACRFVAFGLQDC | 1407251 | Cell/Virus Enclosed | NO |
| ACSCGSCCKFDEDDSE | 1643318 | Underneath Membrane | NO |
| ACSSLTVLSDGR | 1407271 | Cell/Virus Enclosed | NO |
| ACTNGHALPTLF | 1407281 | Cell/Virus Enclosed | NO |
| ACTNGHAVPTLF | 1407282 | Cell/Virus Enclosed | NO |
| ACTPESKPLADD | 1407283 | Cell/Virus Enclosed | NO |
| ACTRLEGLGGNN | 1407286 | Cell/Virus Enclosed | NO |
| ACTYTMCEYPHT | 1407291 | Ectodomain | YES |
| ACVCGDYAACKS | 1407292 | Ectodomain | YES |
| ACVIAETVVKTL | 1407294 | Cell/Virus Enclosed | NO |
| ACVKDFDFKFYDAQPI | 1643339 | Cell/Virus Enclosed | NO |
| ACVKNYEFKFYD | 1407297 | Cell/Virus Enclosed | NO |
| ACVLCNSQTSLR | 1407299 | Cell/Virus Enclosed | NO |
| ACVSVPVSVIYDKETK | 1643346 | Ectodomain | YES |
| ACVVAEAVVKTL | 1407305 | Cell/Virus Enclosed | NO |
| ACVYYSQVLCKS | 1407314 | Cell/Virus Enclosed | NO |
| ACWNHSVGFDYV | 1407318 | Cell/Virus Enclosed | NO |
| ACYAYLAKAMLDFSRD | 1643360 | Cell/Virus Enclosed | NO |
| ACYDGCAQGVFGVNMR | 1643361 | Cell/Virus Enclosed | NO |
| ACYEGIASGVFGVNLR | 1643362 | Cell/Virus Enclosed | NO |
| ACYNGRPIGTFT | 1407325 | Cell/Virus Enclosed | NO |
| ACYNGRPTGTFT | 1407326 | Cell/Virus Enclosed | NO |
| ADAALSIKENVD | 1407330 | Cell/Virus Enclosed | NO |

| | | | |
|---|---|---|---|
| ADAEALNNLLQQ | 1407331 | Ectodomain | YES |
| ADAERMAMYTGS | 1407334 | Ectodomain | YES |
| ADAGFMKQYGECLGDI | 1643368 | Ectodomain | YES |
| ADAGFMKQYGQC | 1407338 | Ectodomain | YES |
| ADAPEPQRGRQG | 1407346 | Cell/Virus Enclosed | NO |
| ADASEPQRGRQS | 1407349 | Cell/Virus Enclosed | NO |
| ADASETQRGRHG | 1407350 | Cell/Virus Enclosed | NO |
| ADASETQRGRQG | 1407351 | Capsid | UN |
| ADASTFFKRVCG | 1407352 | Cell/Virus Enclosed | NO |
| ADASTFLNGFAV | 1407353 | Cell/Virus Enclosed | NO |
| ADASTFLNGVCG | 1407354 | Cell/Virus Enclosed | NO |
| ADAYFADTVWYV | 1407359 | Cell/Virus Enclosed | NO |
| ADAYLADTVWYV | 1407360 | Cell/Virus Enclosed | NO |
| ADCQTTNGLNTS | 1407364 | Ectodomain | YES |
| ADCVACSKSARL | 1407366 | Cell/Virus Enclosed | NO |
| ADDEGFITLKNN | 1407368 | Cell/Virus Enclosed | NO |
| ADDFVIASDTTL | 1407369 | Cell/Virus Enclosed | NO |
| ADDKPIVLFTDS | 1407371 | Cell/Virus Enclosed | NO |
| ADDLNQMTGFKK | 1407374 | Cell/Virus Enclosed | NO |
| ADDSGNPVVEYL | 1407378 | Cell/Virus Enclosed | NO |
| ADDSGTPVVECL | 1407379 | Cell/Virus Enclosed | NO |
| ADDSGTPVVEYL | 1407380 | Cell/Virus Enclosed | NO |
| ADEIANLVLAKLGKDS | 1643395 | Capsid | UN |
| ADEIKALKELYG | 1407385 | Cell/Virus Enclosed | NO |
| ADELQALPQRQK | 1407386 | Cell/Virus Enclosed | NO |
| ADEPDSRAPSRS | 1407387 | Cell/Virus Enclosed | NO |
| ADESEPQRGRQG | 1407389 | Cell/Virus Enclosed | NO |
| ADETQALPQRQKKQQTVTLLPAADL | 1407393 | Capsid | UN |
| ADFACKFAVCGD | 1407396 | Cell/Virus Enclosed | NO |
| ADFHLSGCDEYI | 1407400 | Ectodomain | YES |
| ADFIKHVRTNGG | 1407401 | Cell/Virus Enclosed | NO |
| ADFKSFTLIFGG | 1407402 | Cell/Virus Enclosed | NO |
| ADFLAGSSDVLV | 1407403 | Cell/Virus Enclosed | NO |
| ADFVQIENCDVT | 1407407 | Ectodomain | YES |
| ADFVQIENCNVT | 1407408 | Ectodomain | YES |
| ADFYLSGCDEYI | 1407410 | Ectodomain | YES |
| ADGAEEEVTVDT | 1407411 | Cell/Virus Enclosed | NO |
| ADGELPVAISGDKVFV | 1643407 | Cell/Virus Enclosed | NO |
| ADGFMPFLLDDL | 1407417 | Cell/Virus Enclosed | NO |
| ADGIIHPQGRTY | 1407418 | Ectodomain | YES |
| ADGIQEAKPSGS | 1407424 | Ectodomain | YES |
| ADGKPVISEDLW | 1407425 | Cell/Virus Enclosed | NO |
| ADGLEAYADKTLQEMK | 1643413 | Cell/Virus Enclosed | NO |
| ADGNLRQLLPRW | 1407428 | Cell/Virus Enclosed | NO |
| ADGQQKHLLPRW | 1407430 | Cell/Virus Enclosed | NO |
| ADGQQKPLLPRW | 1407431 | Cell/Virus Enclosed | NO |
| ADGSIIAVQPRN | 1407433 | Ectodomain | YES |
| ADGSLIPVRLRN | 1407434 | Ectodomain | YES |
| ADGSLIPVRPRNSSDN | 1643417 | Ectodomain | YES |
| ADGSLQPYCYTD | 1407435 | Cell/Virus Enclosed | NO |
| ADGSPQPYCYTD | 1407436 | Cell/Virus Enclosed | NO |
| ADGSPQPYCYTE | 1407437 | Cell/Virus Enclosed | NO |
| ADGSPQSYCYTD | 1407438 | Cell/Virus Enclosed | NO |
| ADGSQRQLLPRW | 1407439 | Cell/Virus Enclosed | NO |
| ADGTRHTYQLRA | 1407441 | Cell/Virus Enclosed | NO |
| ADGVECDFSPLL | 1407442 | Ectodomain | YES |
| ADGVQCYTPHSQ | 1407444 | Cell/Virus Enclosed | NO |
| ADHIASLVLAKL | 1407445 | Cell/Virus Enclosed | NO |
| ADIASRVDDEGF | 1407446 | Cell/Virus Enclosed | NO |
| ADICYKFVLWDYERSN | 1643423 | Cell/Virus Enclosed | NO |
| ADIDKLIDYAASGD | 677 | Capsid | UN |
| ADILDRDPSSDE | 1407457 | Cell/Virus Enclosed | NO |
| ADILVVDEVSMC | 1407458 | Cell/Virus Enclosed | NO |
| ADINIHDLPICV | 1407462 | Ectodomain | YES |
| ADINIPDLPICVYDPL | 1643428 | Ectodomain | YES |
| ADINIPDLPISV | 1407464 | Ectodomain | YES |
| ADINIPHLPICV | 1407465 | Ectodomain | YES |
| ADINNPDLPICV | 1407466 | Ectodomain | YES |
| ADINTPDVPICV | 1407467 | Ectodomain | YES |
| ADISAVQSSFGFDQQQ | 1643431 | Cell/Virus Enclosed | NO |
| ADITLFGDRVTV | 1407469 | Cell/Virus Enclosed | NO |
| ADITLLGDCVTV | 1407470 | Cell/Virus Enclosed | NO |
| ADIVDRDPSNDE | 1407471 | Cell/Virus Enclosed | NO |
| ADIVDRDPSSDEAIPT | 1643434 | Cell/Virus Enclosed | NO |
| ADIVDRDQSSEE | 1407473 | Capsid | UN |
| ADIVVVDEVSMCTNYD | 1643438 | Cell/Virus Enclosed | NO |

| | | | |
|---|---|---|---|
| ADKANFLNAVVF | 1407480 | Cell/Virus Enclosed | NO |
| ADKDKDLSGAVPSDLN | 1643440 | Cell/Virus Enclosed | NO |
| ADKFSVLHDIGN | 1407487 | Cell/Virus Enclosed | NO |
| ADKICIVDNVYM | 1407488 | Cell/Virus Enclosed | NO |
| ADKILYQYENLSLADI | 1643443 | Cell/Virus Enclosed | NO |
| ADKLCCAVVSKS | 1407489 | Cell/Virus Enclosed | NO |
| ADKLCHAVVSKS | 1407490 | Cell/Virus Enclosed | NO |
| ADKLCHAVVYKS | 1407491 | Cell/Virus Enclosed | NO |
| ADKLVEVGLVGV | 1407492 | Cell/Virus Enclosed | NO |
| ADKPNHFINFPLTHFQ | 1643446 | Cell/Virus Enclosed | NO |
| ADKPSQLKRPRW | 1407496 | Cell/Virus Enclosed | NO |
| ADKTLQEMKALL | 1407497 | Cell/Virus Enclosed | NO |
| ADKTLQEMKALY | 1407498 | Cell/Virus Enclosed | NO |
| ADLALTNMYKEARIND | 1643452 | Cell/Virus Enclosed | NO |
| ADLDEFSKQLQQ | 1407503 | Cell/Virus Enclosed | NO |
| ADLGVLIQNGAKHVQG | 1643453 | Cell/Virus Enclosed | NO |
| ADLGVLIQNSAKHVQG | 1643454 | Cell/Virus Enclosed | NO |
| ADLHSSQVCGLF | 1407506 | Cell/Virus Enclosed | NO |
| ADLIIERREGSDVCYPGKFVNEEAL | 1126060 | Ectodomain | YES |
| ADLNGVIVDKVGELKV | 1643459 | Cell/Virus Enclosed | NO |
| ADLPAYMSVAKV | 1407510 | Ectodomain | YES |
| ADLPAYMTVAKV | 1407511 | Ectodomain | YES |
| ADLPCTLHFSLF | 1407512 | Cell/Virus Enclosed | NO |
| ADLQHEDSDDDG | 1407514 | Cell/Virus Enclosed | NO |
| ADLSGAICELEL | 1407515 | Cell/Virus Enclosed | NO |
| ADNGIITVEELK | 1407526 | Ectodomain | YES |
| ADNGTITVEKLK | 1407528 | Ectodomain | YES |
| ADNKFALTCTST | 1407532 | Cell/Virus Enclosed | NO |
| ADNLNADETKAL | 1407536 | Cell/Virus Enclosed | NO |
| ADNLTADEIKAL | 1407539 | Cell/Virus Enclosed | NO |
| ADNLTDYSVSSC | 1407540 | Ectodomain | YES |
| ADNNCYLSSVLL | 1407543 | Cell/Virus Enclosed | NO |
| ADNNDITNTNLSRGRG | 1643481 | Cell/Virus Enclosed | NO |
| ADNNDITNTNMS | 1407544 | Cell/Virus Enclosed | NO |
| ADNNDTTTNQPR | 1407546 | Cell/Virus Enclosed | NO |
| ADNNQNGGRNGA | 1407548 | Capsid | UN |
| ADNNSYLSSVLL | 1407549 | Cell/Virus Enclosed | NO |
| ADNPNHFINFPL | 1407550 | Cell/Virus Enclosed | NO |
| ADNSIFRSFHFT | 1407551 | Ectodomain | YES |
| ADNSTASAVQTC | 1407552 | Ectodomain | YES |
| ADNSTITVEELK | 1407553 | Ectodomain | YES |
| ADNSYTSFATYH | 1407554 | Ectodomain | YES |
| ADPAMHIASSNAFLDL | 1643484 | Cell/Virus Enclosed | NO |
| ADPGYMQGYDDC | 1407556 | Ectodomain | YES |
| ADPKCKYAQNLL | 1407559 | Cell/Virus Enclosed | NO |
| ADPKSFTLIFGG | 1407562 | Cell/Virus Enclosed | NO |
| ADPNITNLCPFG | 1407565 | Ectodomain | YES |
| ADPSVVNSCVKS | 1407568 | Ectodomain | YES |
| ADPTFLHRFYSL | 1407569 | Cell/Virus Enclosed | NO |
| ADQAMTSMYKQARAED | 1643493 | Cell/Virus Enclosed | NO |
| ADQELGDAPFLDRLRRDQKS | 97170 | Cell/Virus Enclosed | NO |
| ADQLSDPNYDAL | 1407577 | Cell/Virus Enclosed | NO |
| ADQLTPAWRVYS | 1407580 | Ectodomain | YES |
| ADQSDQFRNFQT | 1407585 | Cell/Virus Enclosed | NO |
| ADQSDQFRNVQI | 1407586 | Cell/Virus Enclosed | NO |
| ADQSDQFRNVQTRGRRAQPK | 2060791 | Cell/Virus Enclosed | NO |
| ADQSDQFRSVQT | 1407588 | Cell/Virus Enclosed | NO |
| ADQSDQSRNVQT | 1407589 | Capsid | UN |
| ADQSDQVRNVQT | 1407590 | Cell/Virus Enclosed | NO |
| ADQSRKWTGHCT | 1407593 | Cell/Virus Enclosed | NO |
| ADRGQESLARSA | 1407595 | Cell/Virus Enclosed | NO |
| ADRRAFVDYTGV | 1407599 | Cell/Virus Enclosed | NO |
| ADRRVFVDYTGV | 1407600 | Cell/Virus Enclosed | NO |
| ADRSVVNSCVKS | 1407601 | Ectodomain | YES |
| ADRTLQEMKALF | 1407602 | Cell/Virus Enclosed | NO |
| ADSCLQGDKCNIFANF | 1377371 | Ectodomain | YES |
| ADSCLQGNKCNI | 1407605 | Ectodomain | YES |
| ADSFTCNIIDAA | 1407606 | Ectodomain | YES |
| ADSFTCNNIDAAKIYG | 1643506 | Ectodomain | YES |
| ADSFTCNNIDVA | 1407608 | Ectodomain | YES |
| ADSGDSSESGAK | 1407611 | Cell/Virus Enclosed | NO |
| ADSGMPTTFLFS | 1407612 | Ectodomain | YES |
| ADSKHVQLSEIN | 1407614 | Cell/Virus Enclosed | NO |
| ADSKIVQLSEIN | 1407615 | Cell/Virus Enclosed | NO |
| ADSKSFTLIFGG | 1407617 | Cell/Virus Enclosed | NO |
| ADSNGFIRFPDV | 1407620 | Cell/Virus Enclosed | NO |

| | | | |
|---|---|---|---|
| ADSPSKDTFELR | 1407623 | Cell/Virus Enclosed | NO |
| ADSPVKDVFELH | 1407624 | Cell/Virus Enclosed | NO |
| ADSPVKDVFELR | 1407625 | Cell/Virus Enclosed | NO |
| ADSSIAYSNNTIAIPT | 1643520 | Ectodomain | YES |
| ADSVLSAVSAGL | 1407630 | Cell/Virus Enclosed | NO |
| ADSVMSAVSAGL | 1407631 | Cell/Virus Enclosed | NO |
| ADSVMSAVSSGL | 1407632 | Cell/Virus Enclosed | NO |
| ADSVVEVVTTSLTPCG | 1377374 | Cell/Virus Enclosed | NO |
| ADSYFSYMMPML | 1407633 | Cell/Virus Enclosed | NO |
| ADSYQKYVEAYN | 1407634 | Cell/Virus Enclosed | NO |
| ADSYYSYIMPMLTMCH | 1377375 | Cell/Virus Enclosed | NO |
| ADSYYSYMMPMLTMCH | 1643523 | Cell/Virus Enclosed | NO |
| ADTAGLFRIDVP | 1407637 | Cell/Virus Enclosed | NO |
| ADTAGLFTNFRI | 1407638 | Cell/Virus Enclosed | NO |
| ADTAGLYTNFPI | 1407639 | Cell/Virus Enclosed | NO |
| ADTAGLYTNFRIDVPS | 1643524 | Cell/Virus Enclosed | NO |
| ADTAGLYTNSRI | 1407641 | Cell/Virus Enclosed | NO |
| ADTAHANNINRFNVAI | 1643525 | Cell/Virus Enclosed | NO |
| ADTEQTQNVTVKPKRL | 1643527 | Cell/Virus Enclosed | NO |
| ADTEQTQSAIAK | 1407644 | Cell/Virus Enclosed | NO |
| ADTEQTQSVIVK | 1407645 | Cell/Virus Enclosed | NO |
| ADTEQTQSVNVK | 1407646 | Cell/Virus Enclosed | NO |
| ADTEQTQSVTVK | 1407647 | Cell/Virus Enclosed | NO |
| ADTLMQGVTLSSNLNT | 1643528 | Ectodomain | YES |
| ADTLQEIPVVSD | 1407652 | Cell/Virus Enclosed | NO |
| ADTLQETPVVPDTVEV | 1643529 | Cell/Virus Enclosed | NO |
| ADTQHISNDFTP | 1407653 | Cell/Virus Enclosed | NO |
| ADTSIPSDVSAR | 1407654 | Cell/Virus Enclosed | NO |
| ADTSTPSDVSSRDPTT | 1643532 | Capsid | UN |
| ADTVMSAVSAGL | 1407658 | Cell/Virus Enclosed | NO |
| ADTVWYVGQIIF | 1407660 | Cell/Virus Enclosed | NO |
| ADVAARADDEGF | 1407662 | Cell/Virus Enclosed | NO |
| ADVAARADYEGF | 1407663 | Cell/Virus Enclosed | NO |
| ADVDDPKLMGWDYPKC | 1643536 | Cell/Virus Enclosed | NO |
| ADVEADIASRVD | 1407667 | Cell/Virus Enclosed | NO |
| ADVEADVAARAD | 1407668 | Cell/Virus Enclosed | NO |
| ADVGMPTTFLFS | 1407671 | Ectodomain | YES |
| ADVIVNPANGHMLHGG | 1643540 | Cell/Virus Enclosed | NO |
| ADVLVNVSQTAI | 1407672 | Ectodomain | YES |
| ADVMRYNFNFSS | 1407674 | Ectodomain | YES |
| ADVMRYNLNFSA | 1407675 | Ectodomain | YES |
| ADVMRYNLNFSS | 1407676 | Ectodomain | YES |
| ADVNIPADIVDR | 1407678 | Cell/Virus Enclosed | NO |
| ADVNTPADILDR | 1407680 | Cell/Virus Enclosed | NO |
| ADVNTSADIVDR | 1407683 | Cell/Virus Enclosed | NO |
| ADVPDAFQPVYR | 1407684 | Cell/Virus Enclosed | NO |
| ADVPDAFQSVYH | 1407685 | Cell/Virus Enclosed | NO |
| ADVSEMLTFDKKAFTL | 1643545 | Ectodomain | YES |
| ADVTKLYLGGMS | 1407692 | Cell/Virus Enclosed | NO |
| ADVVKEQVSDLLVKLL | 1643550 | Cell/Virus Enclosed | NO |
| ADVYRRIPNLPD | 1407695 | Ectodomain | YES |
| ADVYRRKLNLPN | 1407696 | Ectodomain | YES |
| ADVYRRKPDLPN | 1407697 | Ectodomain | YES |
| ADVYRRKPDPPN | 1407698 | Underneath Membrane | NO |
| ADVYRRKPNLPNCNIE | 1643551 | Ectodomain | YES |
| ADVYRRRPDLPN | 1407700 | Ectodomain | YES |
| ADWIELFPHNDR | 1407701 | Cell/Virus Enclosed | NO |
| ADWKPGHSMPSL | 1407703 | Cell/Virus Enclosed | NO |
| ADWKPGQAMPSL | 1407704 | Cell/Virus Enclosed | NO |
| ADWKPGYAMPSL | 1407705 | Cell/Virus Enclosed | NO |
| ADYDIAVTAGPF | 1407710 | Cell/Virus Enclosed | NO |
| ADYDIVVTAGPF | 1407711 | Cell/Virus Enclosed | NO |
| ADYDVAVTAGPFMDNA | 1643553 | Cell/Virus Enclosed | NO |
| ADYDVAVTVGPF | 1407713 | Cell/Virus Enclosed | NO |
| ADYDVVVTAGPF | 1407714 | Cell/Virus Enclosed | NO |
| ADYEGFITLKNN | 1407715 | Cell/Virus Enclosed | NO |
| ADYHNHSNVETR | 1407719 | Cell/Virus Enclosed | NO |
| ADYILPLVLSSL | 1407720 | Cell/Virus Enclosed | NO |
| ADYKKCTKGLSIADLA | 1643556 | Ectodomain | YES |
| ADYLAGSSDVLV | 1407724 | Cell/Virus Enclosed | NO |
| ADYNAKDEISVA | 1407726 | Cell/Virus Enclosed | NO |
| ADYRCACYAYLA | 1407730 | Cell/Virus Enclosed | NO |
| ADYSVLYNSTFF | 1407732 | Ectodomain | YES |
| ADYSVLYNSTSF | 1407733 | Ectodomain | YES |
| ADYTVLYNSTSF | 1407741 | Ectodomain | YES |
| AEAEWIELFPHN | 1407746 | Cell/Virus Enclosed | NO |

| | | | |
|---|---|---|---|
| AEAHNNLLQQLS | 1407749 | Ectodomain | YES |
| AEALNNLLQQLS | 1407753 | Ectodomain | YES |
| AEAQIDRLINGRLTAL | 1643572 | Ectodomain | YES |
| AEAQIIAIIKDW | 1407756 | Underneath Membrane | NO |
| AEAREHLKNGTC | 1407757 | Cell/Virus Enclosed | NO |
| AECKAALGLTVS | 1407762 | Cell/Virus Enclosed | NO |
| AECKATLGLTVS | 1407763 | Cell/Virus Enclosed | NO |
| AECKRALGLSISDHEF | 1643579 | Cell/Virus Enclosed | NO |
| AECTMFRDAEGR | 1407769 | Cell/Virus Enclosed | NO |
| AEDCMSDFMSEI | 1407770 | Ectodomain | YES |
| AEDGITKLADVE | 1407772 | Cell/Virus Enclosed | NO |
| AEDIFCPCWSAL | 1407773 | Cell/Virus Enclosed | NO |
| AEDIFGPCWSALASAL | 1643584 | Cell/Virus Enclosed | NO |
| AEDIFGPCWYAL | 1407775 | Cell/Virus Enclosed | NO |
| AEDISLLRKMDE | 1407777 | Capsid | UN |
| AEDTLTKLSKYLANEA | 1643588 | Cell/Virus Enclosed | NO |
| AEDVTEFVQEEA | 1407780 | Cell/Virus Enclosed | NO |
| AEDVTEFVQEEV | 1407781 | Cell/Virus Enclosed | NO |
| AEDVTESVQEKA | 1407783 | Cell/Virus Enclosed | NO |
| AEDVVDQVEEVNSIFD | 1643591 | Cell/Virus Enclosed | NO |
| AEEALSKLSEYLISDS | 1643592 | Cell/Virus Enclosed | NO |
| AEEARKLMPICM | 1407786 | Cell/Virus Enclosed | NO |
| AEEDGTGCFEIFHKCDDDCMASIRN | 1126261 | Ectodomain | YES |
| AEEEVTVDTTVE | 1407788 | Cell/Virus Enclosed | NO |
| AEEKLDNPNSSE | 1407791 | Cell/Virus Enclosed | NO |
| AEEKLKKKSSFYQS | 881 | Capsid | UN |
| AEELASPMKSDE | 1407792 | Cell/Virus Enclosed | NO |
| AEEPAKDIPEKV | 1407794 | Cell/Virus Enclosed | NO |
| AEEPDSRAPSRS | 1407795 | Cell/Virus Enclosed | NO |
| AEEPDSRVPSRS | 1407796 | Cell/Virus Enclosed | NO |
| AEEPVVGIVYSS | 1407797 | Cell/Virus Enclosed | NO |
| AEETRKLMLICM | 1407799 | Cell/Virus Enclosed | NO |
| AEEYCNYLESYNIVTT | 1643602 | Cell/Virus Enclosed | NO |
| AEEYREYLESYNTATT | 1643603 | Cell/Virus Enclosed | NO |
| AEFDRESSVQKKINRM | 1643604 | Cell/Virus Enclosed | NO |
| AEFDTYDPIYKNGAMY | 1643605 | Cell/Virus Enclosed | NO |
| AEFDTYDPTYKN | 1407806 | Cell/Virus Enclosed | NO |
| AEFKYMVANGLHAPHG | 1643606 | Cell/Virus Enclosed | NO |
| AEFKYMVANGLNAPNG | 1643607 | Cell/Virus Enclosed | NO |
| AEFSLWFKNHTS | 1407811 | Ectodomain | YES |
| AEFVEPGCGGILAVID | 1643608 | Cell/Virus Enclosed | NO |
| AEGAHATLDSIG | 1407815 | Cell/Virus Enclosed | NO |
| AEGAHATRDSIGTNFP | 1643609 | Cell/Virus Enclosed | NO |
| AEGDAFWSSTMI | 1407817 | Cell/Virus Enclosed | NO |
| AEGDASWSSTMI | 1407818 | Cell/Virus Enclosed | NO |
| AEGDASWYSTMI | 1407819 | Cell/Virus Enclosed | NO |
| AEGDGGVLGDGN | 1407820 | Cell/Virus Enclosed | NO |
| AEGEETDLTVTS | 1407821 | Cell/Virus Enclosed | NO |
| AEGFECDFSPLL | 1407822 | Ectodomain | YES |
| AEGGASWSSTMI | 1407823 | Cell/Virus Enclosed | NO |
| AEGISLLKKMDE | 1407824 | Cell/Virus Enclosed | NO |
| AEGQGVPIAPGV | 1407826 | Cell/Virus Enclosed | NO |
| AEGRITPYCHDP | 1407827 | Cell/Virus Enclosed | NO |
| AEGSHATLDSIG | 1407831 | Cell/Virus Enclosed | NO |
| AEGVECDFSLLL | 1407837 | Ectodomain | YES |
| AEGVECDFSPLL | 1407838 | Ectodomain | YES |
| AEGVPFYLNVQY | 1407840 | Ectodomain | YES |
| AEHDFFTFDIDGSRVP | 1643616 | Cell/Virus Enclosed | NO |
| AEHDFFTFDVEGSRVP | 1643617 | Cell/Virus Enclosed | NO |
| AEHDFFTWKDGRVIYG | 1643618 | Cell/Virus Enclosed | NO |
| AEHVDSSYECDI | 1407845 | Ectodomain | YES |
| AEHVDTSYDCDI | 1407846 | Ectodomain | YES |
| AEHVDTSYECDIPIGA | 971 | Ectodomain | YES |
| AEHVDTSYECDT | 1407848 | Ectodomain | YES |
| AEIKDLICVQSY | 1407853 | Ectodomain | YES |
| AEILIIIMRTFR | 1407855 | Cell/Virus Enclosed | NO |
| AEIVNTVSELVYENKF | 1643631 | Cell/Virus Enclosed | NO |
| AEKLASPMKSDE | 1407868 | Cell/Virus Enclosed | NO |
| AELAPPAGAFFF | 1407873 | Cell/Virus Enclosed | NO |
| AELAPTAGAFFF | 1407877 | Cell/Virus Enclosed | NO |
| AELAPTASAFMG | 1407878 | Cell/Virus Enclosed | NO |
| AELAPTPGAFFFGSKL | 1643638 | Capsid | UN |
| AELCCFARMVWR | 1407880 | Cell/Virus Enclosed | NO |
| AELCCSACMVWR | 1407881 | Cell/Virus Enclosed | NO |
| AELCCSARMIWR | 1407882 | Cell/Virus Enclosed | NO |
| AELCCSARMVWR | 1407883 | Cell/Virus Enclosed | NO |

| | | | |
|---|---|---|---|
| AELEAARKAYQE | 1407888 | Cell/Virus Enclosed | NO |
| AELEPPCRFVTD | 1407890 | Cell/Virus Enclosed | NO |
| AELGYIADISSF | 1407894 | Cell/Virus Enclosed | NO |
| AELIPNHAALFF | 1407896 | Cell/Virus Enclosed | NO |
| AELIPNQAALFFDSEV | 1643646 | Capsid | UN |
| AELKCSVKSFEIDKGI | 1643647 | Ectodomain | YES |
| AELNGVMVDKVG | 1407905 | Cell/Virus Enclosed | NO |
| AELNYTVQKLQS | 1407907 | Ectodomain | YES |
| AELTGTQSVSVD | 1407915 | Cell/Virus Enclosed | NO |
| AELVPSTAAMLFDSHI | 1643652 | Capsid | UN |
| AELVPSTAAMLS | 1407917 | Cell/Virus Enclosed | NO |
| AELVPTASAFMG | 1407918 | Cell/Virus Enclosed | NO |
| AEMLKLGTNDPQFPIL | 1643655 | Capsid | UN |
| AEMNRLQEAIKV | 1407921 | Ectodomain | YES |
| AEMTSNVRRQVASGVI | 1643656 | Cell/Virus Enclosed | NO |
| AENDTITVDQLK | 1407924 | Ectodomain | YES |
| AENKDGFYETIV | 1407926 | Cell/Virus Enclosed | NO |
| AENVTESVQEEA | 1407932 | Cell/Virus Enclosed | NO |
| AEPKGKYSQKAYALLR | 1643660 | Cell/Virus Enclosed | NO |
| AEPVRDEFSIET | 1407936 | Capsid | UN |
| AEPVRDEVSIETDIID | 1643661 | Capsid | UN |
| AEQAAAAMYKEARAVN | 1643662 | Cell/Virus Enclosed | NO |
| AEQAATQMYKEARSVN | 1643663 | Cell/Virus Enclosed | NO |
| AEQCIRGAIGDV | 1407938 | Cell/Virus Enclosed | NO |
| AEQDRNTREVFV | 1407941 | Ectodomain | YES |
| AEQLTPAWRIYS | 1407942 | Ectodomain | YES |
| AERAVEVDPDDVNK | 1146 | Capsid | UN |
| AERDCYIFKRTF | 1407943 | Cell/Virus Enclosed | NO |
| AERDCYTFKKTVAKAP | 1643667 | Cell/Virus Enclosed | NO |
| AERDCYTLKKTV | 1407946 | Cell/Virus Enclosed | NO |
| AERMAMYTGSLIGGIA | 1643668 | Ectodomain | YES |
| AESKAFVDTIVPPCPS | 1643675 | Cell/Virus Enclosed | NO |
| AESKPRFVRKAE | 1407954 | Cell/Virus Enclosed | NO |
| AESKPWFVRKAEDIFG | 1643676 | Cell/Virus Enclosed | NO |
| AESTGTQSVSVDLEST | 1643677 | Cell/Virus Enclosed | NO |
| AESVIVEDDVIENVKS | 1643678 | Cell/Virus Enclosed | NO |
| AETGTSDLDRCT | 1407959 | Ectodomain | YES |
| AETGTVDVGPDS | 1407960 | Ectodomain | YES |
| AETHRDCDFNKPLIEW | 1643681 | Cell/Virus Enclosed | NO |
| AETIKAKEESVKSSYA | 1643682 | Cell/Virus Enclosed | NO |
| AETLRATEEASKQSYA | 1643684 | Cell/Virus Enclosed | NO |
| AETLRATEEASR | 1407968 | Cell/Virus Enclosed | NO |
| AETPALQVSLTQ | 1407969 | Cell/Virus Enclosed | NO |
| AETQKATEEAFKQSYA | 1407970 | Cell/Virus Enclosed | NO |
| AETTDFCTVALA | 1407973 | Ectodomain | YES |
| AETTDFFTVALA | 1407974 | Ectodomain | YES |
| AETTDFLTVALA | 1407975 | Ectodomain | YES |
| AETTVCTVALAS | 1407976 | Ectodomain | YES |
| AETVKAKEESVKSSYA | 1643686 | Cell/Virus Enclosed | NO |
| AETVVKTLQPVS | 1407979 | Cell/Virus Enclosed | NO |
| AEVSRQVIQDVKDGTV | 1643692 | Cell/Virus Enclosed | NO |
| AEVSRQVIQEVK | 1407990 | Cell/Virus Enclosed | NO |
| AEVVKQMYGVNH | 1407992 | Cell/Virus Enclosed | NO |
| AEVVKQMYGVNLQSGK | 1643694 | Cell/Virus Enclosed | NO |
| AEVVVNPANGHMAHGG | 1643695 | Cell/Virus Enclosed | NO |
| AEVVVNSANGHM | 1407994 | Cell/Virus Enclosed | NO |
| AEWIELFPHNDRIKSF | 1643698 | Cell/Virus Enclosed | NO |
| AEWKCGYAMPQI | 1408002 | Cell/Virus Enclosed | NO |
| AEWKCGYSMPGI | 1408003 | Cell/Virus Enclosed | NO |
| AEYCRHGTCERS | 1408005 | Cell/Virus Enclosed | NO |
| AEYIFGPCWSAL | 1408008 | Cell/Virus Enclosed | NO |
| AEYITTYPGQGC | 1408009 | Cell/Virus Enclosed | NO |
| AEYKLQQGTFLC | 1408010 | Cell/Virus Enclosed | NO |
| AEYVDTSYECDI | 1408015 | Ectodomain | YES |
| AEYVTEFVQEEA | 1408016 | Cell/Virus Enclosed | NO |
| AFAAIGDLKCPL | 1408017 | Ectodomain | YES |
| AFAAIGDLNCTL | 1408018 | Ectodomain | YES |
| AFACADGTRHTY | 1408021 | Cell/Virus Enclosed | NO |
| AFADYTPCAVIA | 1408025 | Cell/Virus Enclosed | NO |
| AFADYTQCAVIA | 1408026 | Cell/Virus Enclosed | NO |
| AFALHSGYYDAN | 1408032 | Ectodomain | YES |
| AFALHTGYYDAN | 1408033 | Ectodomain | YES |
| AFALHTGYYDVN | 1408034 | Ectodomain | YES |
| AFANSVFNICQAVSAN | 1643715 | Cell/Virus Enclosed | NO |
| AFANWLAPTNAY | 1408036 | Cell/Virus Enclosed | NO |
| AFASDSVQCYTP | 1408037 | Cell/Virus Enclosed | NO |

| | | | |
|---|---|---|---|
| AFATLKEVVGPKELLL | 1643722 | Cell/Virus Enclosed | NO |
| AFAVDPAAAYLD | 1408042 | Cell/Virus Enclosed | NO |
| AFAVDPAKAYKD | 1408043 | Cell/Virus Enclosed | NO |
| AFAVIGDLKCTSDNIN | 1643726 | Ectodomain | YES |
| AFAVIGDLNCPL | 1408047 | Ectodomain | YES |
| AFAVVGDLNCPL | 1408049 | Ectodomain | YES |
| AFAYADKPNHFI | 1408051 | Cell/Virus Enclosed | NO |
| AFAYESAVVNGI | 1408052 | Cell/Virus Enclosed | NO |
| AFCDEMVLKGVV | 1408055 | Cell/Virus Enclosed | NO |
| AFCDYADKLCCA | 1408056 | Cell/Virus Enclosed | NO |
| AFCDYADKLCHA | 1408057 | Cell/Virus Enclosed | NO |
| AFCDYAGKFCHA | 1408058 | Cell/Virus Enclosed | NO |
| AFCDYTDKLCHA | 1408059 | Cell/Virus Enclosed | NO |
| AFCGDQVHEVAA | 1408060 | Cell/Virus Enclosed | NO |
| AFCKPGTLVLNN | 1408061 | Ectodomain | YES |
| AFCKSGSLVLNN | 1408063 | Ectodomain | YES |
| AFCVYKLQPLTF | 1408068 | Ectodomain | YES |
| AFDCGDYAACKS | 1408070 | Ectodomain | YES |
| AFDFSLDVFKIGDVKF | 1643734 | Cell/Virus Enclosed | NO |
| AFDFSSDVFKIG | 1408073 | Cell/Virus Enclosed | NO |
| AFDGTMYGAFMD | 1408076 | Cell/Virus Enclosed | NO |
| AFDIQQWGYVGS | 1408078 | Cell/Virus Enclosed | NO |
| AFDIYNERVAGF | 1408081 | Cell/Virus Enclosed | NO |
| AFDLSKELKRPI | 1408088 | Cell/Virus Enclosed | NO |
| AFDQDGIIFNAV | 1408089 | Ectodomain | YES |
| AFDVPDGFYSTS | 1408092 | Ectodomain | YES |
| AFDVYNKDASFIGKNL | 1643750 | Cell/Virus Enclosed | NO |
| AFEAFKRSNNGV | 1408095 | Cell/Virus Enclosed | NO |
| AFEFAEGQGVPI | 1408097 | Cell/Virus Enclosed | NO |
| AFELFAKRKMGLTPPL | 1643761 | Cell/Virus Enclosed | NO |
| AFELFAKRKVGLTPPL | 1643762 | Cell/Virus Enclosed | NO |
| AFFFGSKFELAK | 1408113 | Cell/Virus Enclosed | NO |
| AFFFGSKLDLVKRDSE | 1643770 | Capsid | UN |
| AFFFGSKLELAK | 1408115 | Cell/Virus Enclosed | NO |
| AFFSILQDMRNTIM | 1299 | Capsid | UN |
| AFGEYNHVVAAN | 1408131 | Cell/Virus Enclosed | NO |
| AFGGDQIHEVAA | 1408135 | Cell/Virus Enclosed | NO |
| AFGGDQVHEVAAVRSV | 1643783 | Cell/Virus Enclosed | NO |
| AFGGDQVHEVVA | 1408137 | Cell/Virus Enclosed | NO |
| AFGGVSVITPGTNASS | 1643784 | Ectodomain | YES |
| AFGIPPSEAKGY | 1408139 | Cell/Virus Enclosed | NO |
| AFGLQDCVTGINDDDY | 1643786 | Cell/Virus Enclosed | NO |
| AFGLQDCVTGVN | 1408141 | Cell/Virus Enclosed | NO |
| AFGLRGPGDLQGNFGD | 1643787 | Cell/Virus Enclosed | NO |
| AFGNAGVCYDIF | 1408145 | Cell/Virus Enclosed | NO |
| AFGNTGVCYDFD | 1408146 | Cell/Virus Enclosed | NO |
| AFGRRGPEQTQGNFGD | 1643793 | Cell/Virus Enclosed | NO |
| AFGSPFITNGMS | 1408152 | Cell/Virus Enclosed | NO |
| AFGVLLSNLGIP | 1408154 | Cell/Virus Enclosed | NO |
| AFGVPPSEAKGYWYRHSRRS | 2060793 | Capsid | UN |
| AFHANSSEPALI | 1408157 | Ectodomain | YES |
| AFHANSSEPALLFRNI | 1643797 | Ectodomain | YES |
| AFHKPAYDVSAF | 1408160 | Cell/Virus Enclosed | NO |
| AFHKVQDAVNNN | 1408161 | Ectodomain | YES |
| AFHRIESTDPIV | 1408168 | Cell/Virus Enclosed | NO |
| AFHTPAYDGSAF | 1408172 | Cell/Virus Enclosed | NO |
| AFHVNSSEPALL | 1408175 | Ectodomain | YES |
| AFIAFNHPIQVD | 1408178 | Ectodomain | YES |
| AFIDTIKSLDYK | 1408179 | Cell/Virus Enclosed | NO |
| AFIECGSCGNDS | 1408181 | Cell/Virus Enclosed | NO |
| AFIECGSCGNES | 1408182 | Cell/Virus Enclosed | NO |
| AFIECGSCGNNS | 1408183 | Cell/Virus Enclosed | NO |
| AFIGVLDITDAA | 1408186 | Cell/Virus Enclosed | NO |
| AFIKTAQADTAG | 1408190 | Cell/Virus Enclosed | NO |
| AFISVFDITDAA | 1408196 | Cell/Virus Enclosed | NO |
| AFISVLDITDAAVKAA | 1643813 | Cell/Virus Enclosed | NO |
| AFKAVCSVIDLK | 1408203 | Cell/Virus Enclosed | NO |
| AFKCNRFGSNDI | 1408207 | Ectodomain | YES |
| AFKQSYASATIQ | 1408216 | Cell/Virus Enclosed | NO |
| AFKRQIRIACRK | 1408218 | Cell/Virus Enclosed | NO |
| AFLCANEYTGSY | 1408224 | Cell/Virus Enclosed | NO |
| AFLDTVVSTVKIGGLT | 1643836 | Cell/Virus Enclosed | NO |
| AFLDVLNKALEM | 1408228 | Cell/Virus Enclosed | NO |
| AFLGASVTEDVKFAAS | 1643838 | Cell/Virus Enclosed | NO |
| AFLMAEDGITKL | 1408247 | Cell/Virus Enclosed | NO |
| AFLNGSCGSVGF | 1408249 | Cell/Virus Enclosed | NO |

| | | | |
|---|---|---|---|
| AFLPAQDTWGTS | 1408251 | Ectodomain | YES |
| AFLQKLEDNPLF | 1408257 | Cell/Virus Enclosed | NO |
| AFLQKLEDNSLF | 1408258 | Cell/Virus Enclosed | NO |
| AFLTLLFYYINK | 1408264 | Cell/Virus Enclosed | NO |
| AFLVHDNVAFYQGDVD | 1643859 | Cell/Virus Enclosed | NO |
| AFLVVLIILWFV | 1408267 | Cell/Virus Enclosed | NO |
| AFMDKQVHQVQLTDKY | 1643863 | Cell/Virus Enclosed | NO |
| AFMGMSQFKLTH | 1408271 | Cell/Virus Enclosed | NO |
| AFMKHKGGDLTD | 1408273 | Cell/Virus Enclosed | NO |
| AFMKHKGGDSTDFIAL | 1643865 | Cell/Virus Enclosed | NO |
| AFMLGSSVGNFSDGKM | 1643866 | Ectodomain | YES |
| AFMPVQSRKFIV | 1408283 | Cell/Virus Enclosed | NO |
| AFNHPIQIDQLN | 1408294 | Ectodomain | YES |
| AFNHPIQVDQFN | 1408296 | Ectodomain | YES |
| AFNHPIQVDQLNSSYF | 1643871 | Ectodomain | YES |
| AFNHPIQVGQLN | 1408298 | Ectodomain | YES |
| AFNNALDAIQEG | 1408301 | Ectodomain | YES |
| AFNNALGAIQEG | 1408302 | Ectodomain | YES |
| AFNNALHAIQQG | 1408303 | Ectodomain | YES |
| AFNNALHSIQQG | 1408304 | Ectodomain | YES |
| AFNNALNAIQQG | 1408305 | Ectodomain | YES |
| AFNNALYAIQQG | 1408307 | Ectodomain | YES |
| AFNNWFLLTNTSSVVD | 1643874 | Ectodomain | YES |
| AFNPETNAIISL | 1408311 | Underneath Membrane | NO |
| AFNQDGIIFNAE | 1408312 | Ectodomain | YES |
| AFNQDGIIFNAVDCMS | 1643876 | Ectodomain | YES |
| AFNQDGVIFNAV | 1408314 | Ectodomain | YES |
| AFNQLTADLQHK | 1408317 | Cell/Virus Enclosed | NO |
| AFNVNVMGTRPEKFPT | 1643883 | Cell/Virus Enclosed | NO |
| AFNVSDAYANLVPYYQ | 1643884 | Cell/Virus Enclosed | NO |
| AFNWLLLAGTLH | 1408337 | Cell/Virus Enclosed | NO |
| AFPFETFSACRA | 1408342 | Cell/Virus Enclosed | NO |
| AFPLDEAEDFTE | 1408346 | Cell/Virus Enclosed | NO |
| AFPLDEAEDVTESVQE | 1643892 | Cell/Virus Enclosed | NO |
| AFPLDEAENVTE | 1408348 | Cell/Virus Enclosed | NO |
| AFPLDEAEYVTE | 1408349 | Cell/Virus Enclosed | NO |
| AFPLNEAEDVTE | 1408352 | Cell/Virus Enclosed | NO |
| AFPPTEPKKDKK | 1408353 | Cell/Virus Enclosed | NO |
| AFPSAKVEGCMV | 1408354 | Cell/Virus Enclosed | NO |
| AFQKVQDAVNNI | 1408370 | Ectodomain | YES |
| AFQKVQDAVNNNAQAL | 1643899 | Ectodomain | YES |
| AFQPVYRTFTQA | 1408374 | Cell/Virus Enclosed | NO |
| AFQSVYRTFTQA | 1408381 | Cell/Virus Enclosed | NO |
| AFRDLKPLPFFY | 1408385 | Cell/Virus Enclosed | NO |
| AFRKARNGVFISTEKL | 1643908 | Cell/Virus Enclosed | NO |
| AFRNACGTNVPL | 1408394 | Cell/Virus Enclosed | NO |
| AFSASLFKPVQLVPVS | 1643910 | Cell/Virus Enclosed | NO |
| AFSLDVSEKSGNFKHL | 1643916 | Ectodomain | YES |
| AFSNALDAIQEG | 1408416 | Ectodomain | YES |
| AFSPAQDTWGTS | 1408419 | Ectodomain | YES |
| AFSPAQGTWGTS | 1408420 | Ectodomain | YES |
| AFSVDPKKTYLDFIQQ | 1643919 | Cell/Virus Enclosed | NO |
| AFSVDPKKTYLDYIQQ | 1643920 | Cell/Virus Enclosed | NO |
| AFTKAAGGKVSFSDDV | 1643924 | Cell/Virus Enclosed | NO |
| AFTKLAGGKISFSDDV | 1643925 | Cell/Virus Enclosed | NO |
| AFTREMQQHPLLNPSA | 1643931 | Capsid | UN |
| AFTREMQQQPLL | 1408438 | Cell/Virus Enclosed | NO |
| AFTRYVDNNFCG | 1408439 | Cell/Virus Enclosed | NO |
| AFVCGDCAACKS | 1408448 | Ectodomain | YES |
| AFVCGDFAACKS | 1408449 | Ectodomain | YES |
| AFVCGDYAACKL | 1408450 | Ectodomain | YES |
| AFVCGDYAACKS | 1408451 | Ectodomain | YES |
| AFVDTIVPPCPSILKV | 1643937 | Cell/Virus Enclosed | NO |
| AFVDTVKGLDYK | 1408454 | Cell/Virus Enclosed | NO |
| AFVECGSCGNDS | 1408455 | Cell/Virus Enclosed | NO |
| AFVKCNCGSENW | 1408460 | Cell/Virus Enclosed | NO |
| AFVKCNCGSESW | 1408461 | Cell/Virus Enclosed | NO |
| AFVNSVFNICQA | 1408469 | Cell/Virus Enclosed | NO |
| AFVPQPTGVFTN | 1408470 | Ectodomain | YES |
| AFVRGDYAACKL | 1408471 | Ectodomain | YES |
| AFVSQVLNKYTEVRGS | 1643950 | Ectodomain | YES |
| AFVVFLLVTLAILTA | 1419 | Undetermined | UN |
| AFWETKKNLTRKIRSEEL | 999356 | Secreted | YES |
| AFWVFSYCRKLG | 1408487 | Cell/Virus Enclosed | NO |
| AFYGVALKDCGF | 1408492 | Cell/Virus Enclosed | NO |
| AFYPACTNGHAV | 1408502 | Cell/Virus Enclosed | NO |

| | | | |
|---|---|---|---|
| AFYQGDVDTVVNGVDF | 1643966 | Cell/Virus Enclosed | NO |
| AFYRDLYSSPTT | 1408505 | Cell/Virus Enclosed | NO |
| AFYVKDGKYFTSEPPV | 1643968 | Cell/Virus Enclosed | NO |
| AFYVLPSETPNA | 1408509 | Cell/Virus Enclosed | NO |
| AGADTTITLNVL | 1408514 | Cell/Virus Enclosed | NO |
| AGAGWTAGLSSS | 1408523 | Ectodomain | YES |
| AGAINAASKGAVQKES | 1643977 | Cell/Virus Enclosed | NO |
| AGAMLSENFTSY | 1408540 | Ectodomain | YES |
| AGAPFYLNVQYR | 1408541 | Ectodomain | YES |
| AGASRAIEKVNE | 1408544 | Ectodomain | YES |
| AGCEQSYFTKKD | 1408550 | Cell/Virus Enclosed | NO |
| AGCFVNDLLKTD | 1408552 | Cell/Virus Enclosed | NO |
| AGCLIGAEHVDS | 1408555 | Ectodomain | YES |
| AGCLIGAEHVDTSYECD | 1460 | Ectodomain | YES |
| AGCLIGAEYVDT | 1408558 | Ectodomain | YES |
| AGCQTTNGMNTS | 1408561 | Ectodomain | YES |
| AGCQTTNGTNTI | 1408562 | Ectodomain | YES |
| AGDAANFCALIL | 1408563 | Cell/Virus Enclosed | NO |
| AGDAVHSLTWLW | 1408565 | Cell/Virus Enclosed | NO |
| AGDLQGNFGDLQ | 1408571 | Cell/Virus Enclosed | NO |
| AGDSGFAAYSRYRIGN | 1377468 | Underneath Membrane | NO |
| AGDSSWHIYLKS | 1408579 | Ectodomain | YES |
| AGEDIRLLKAAY | 1408587 | Cell/Virus Enclosed | NO |
| AGEDVQLLKAAY | 1408588 | Cell/Virus Enclosed | NO |
| AGEEKLSSRMYC | 1408589 | Cell/Virus Enclosed | NO |
| AGEENFSSRMYC | 1408590 | Cell/Virus Enclosed | NO |
| AGEEVFAPKLYC | 1408592 | Cell/Virus Enclosed | NO |
| AGEMPCAIINDKVVAK | 1644009 | Cell/Virus Enclosed | NO |
| AGEPCVASEQEE | 1408594 | Cell/Virus Enclosed | NO |
| AGFDSWFSQRGG | 1408601 | Cell/Virus Enclosed | NO |
| AGFGVDEEKCGV | 1408604 | Ectodomain | YES |
| AGFKRTYNFVWY | 1408606 | Cell/Virus Enclosed | NO |
| AGFMKQYGECLGDINA | 1644017 | Ectodomain | YES |
| AGFNIWVPHSFD | 1408611 | Cell/Virus Enclosed | NO |
| AGFNTEFASNSS | 1408612 | Cell/Virus Enclosed | NO |
| AGGITYVSNSGNLLGF | 1644032 | Ectodomain | YES |
| AGGQLFYSRPVVSANGEPTVKLYTS | 1533 | Undetermined | UN |
| AGGTTEMLSRAL | 1408640 | Cell/Virus Enclosed | NO |
| AGGVYTNSCLVS | 1408644 | Cell/Virus Enclosed | NO |
| AGHATGTTPQKLFVAN | 1644041 | Ectodomain | YES |
| AGHATLREHLRDIKAENTDA | 1547 | Ectodomain | YES |
| AGHHLGRCDIKDLPKE | 1377483 | Underneath Membrane | NO |
| AGHPLGRCDIKD | 1408657 | Underneath Membrane | NO |
| AGHYTGQAGEMP | 1408663 | Cell/Virus Enclosed | NO |
| AGICASYHTVSS | 1408670 | Ectodomain | YES |
| AGIQNFKETLYYQNNV | 1644067 | Cell/Virus Enclosed | NO |
| AGIRDLICVQSY | 1408698 | Ectodomain | YES |
| AGKSMCTLSEQL | 1408708 | Cell/Virus Enclosed | NO |
| AGKVKVGTGVMVECDS | 1644078 | Cell/Virus Enclosed | NO |
| AGLEFTDENYNNLVPT | 1644082 | Cell/Virus Enclosed | NO |
| AGLFRIDVPSAE | 1408716 | Cell/Virus Enclosed | NO |
| AGLFRIFLRKAY | 1408718 | Cell/Virus Enclosed | NO |
| AGLKLPSGIMFN | 1408728 | Cell/Virus Enclosed | NO |
| AGLPYGANKEGIIWVA | 1644090 | Cell/Virus Enclosed | NO |
| AGLSSSAAIPFA | 1408741 | Ectodomain | YES |
| AGLVGVLTLDNQDLNG | 1644094 | Cell/Virus Enclosed | NO |
| AGLWLDDTVYCP | 1408750 | Cell/Virus Enclosed | NO |
| AGLYTNFPIDFP | 1408751 | Cell/Virus Enclosed | NO |
| AGLYTNFRIDIP | 1408753 | Cell/Virus Enclosed | NO |
| AGLYTNSRIDVP | 1408755 | Cell/Virus Enclosed | NO |
| AGMEAQFLYLYA | 1408757 | Secreted | YES |
| AGMFGLYDDILTNNKP | 1644100 | Cell/Virus Enclosed | NO |
| AGMHFGACDYDRLPNE | 1644101 | Ectodomain | YES |
| AGMYRDWSYSGQ | 1408762 | Cell/Virus Enclosed | NO |
| AGNATEVPAHST | 1408763 | Cell/Virus Enclosed | NO |
| AGNATEVPANSA | 1408764 | Cell/Virus Enclosed | NO |
| AGNGGDTALALL | 1408768 | Cell/Virus Enclosed | NO |
| AGNIFGLYKNSATGSL | 1644105 | Cell/Virus Enclosed | NO |
| AGNKFGLYKNSA | 1408772 | Cell/Virus Enclosed | NO |
| AGNSYTSFATYHTPAT | 1644109 | Ectodomain | YES |
| AGNVIGNDALTRFTIF | 1644113 | Cell/Virus Enclosed | NO |
| AGNVQLRVIGHS | 1408785 | Cell/Virus Enclosed | NO |
| AGNVWQIQTIQDSDGT | 1644115 | Cell/Virus Enclosed | NO |
| AGPFMDDAINVG | 1408792 | Cell/Virus Enclosed | NO |
| AGPFMDNAINVGGTGL | 1644117 | Cell/Virus Enclosed | NO |
| AGPFMGNAINVG | 1408794 | Cell/Virus Enclosed | NO |

| | | | |
|---|---|---|---|
| AGPRVRQPARPLGSAWRDQAQRPAV | 1646 | Undetermined | UN |
| AGPSVETSDSQV | 1408800 | Cell/Virus Enclosed | NO |
| AGQEQTNCNISS | 1408806 | Cell/Virus Enclosed | NO |
| AGQEQTNCNTSA | 1408807 | Cell/Virus Enclosed | NO |
| AGQEQTNCNTSS | 1408808 | Cell/Virus Enclosed | NO |
| AGQPCVASEQDE | 1408815 | Cell/Virus Enclosed | NO |
| AGQPCVASEQEE | 1408816 | Cell/Virus Enclosed | NO |
| AGQPCVVSEQEE | 1408817 | Cell/Virus Enclosed | NO |
| AGQPYVASEQEE | 1408818 | Cell/Virus Enclosed | NO |
| AGQQFVKETTDM | 1408819 | Cell/Virus Enclosed | NO |
| AGQQFVKETTNM | 1408820 | Cell/Virus Enclosed | NO |
| AGQSCVVSEQEE | 1408821 | Cell/Virus Enclosed | NO |
| AGQVQTFYPRLQ | 1408825 | Cell/Virus Enclosed | NO |
| AGRKVNFNEKPVVMEI | 1644128 | Cell/Virus Enclosed | NO |
| AGRPCVASEQEE | 1408827 | Cell/Virus Enclosed | NO |
| AGRQTTNGMNTS | 1408828 | Ectodomain | YES |
| AGSAAAPFTKPF | 1408835 | Cell/Virus Enclosed | NO |
| AGSALQIPFAMQ | 1408836 | Ectodomain | YES |
| AGSDKGIAPGTSVLRQ | 1644135 | Cell/Virus Enclosed | NO |
| AGSDKGITPGTS | 1408839 | Cell/Virus Enclosed | NO |
| AGSDKGVAPGTTVLKR | 1644137 | Cell/Virus Enclosed | NO |
| AGSFDKVYDIGNPKGI | 1644140 | Cell/Virus Enclosed | NO |
| AGSFDRIYDIGN | 1408844 | Cell/Virus Enclosed | NO |
| AGSLELTTMRYF | 1408852 | Cell/Virus Enclosed | NO |
| AGSNDIWMQNKG | 1408853 | Ectodomain | YES |
| AGSNISAVIYSG | 1408855 | Cell/Virus Enclosed | NO |
| AGSNTEFAFNSS | 1408857 | Cell/Virus Enclosed | NO |
| AGSQTTNGTNTN | 1408861 | Ectodomain | YES |
| AGSRSRANSGNRTPTS | 1644147 | Cell/Virus Enclosed | NO |
| AGSTFISDEVAR | 1408867 | Cell/Virus Enclosed | NO |
| AGSVGFNIDYDC | 1408870 | Cell/Virus Enclosed | NO |
| AGSVGYTKEGSV | 1408871 | Cell/Virus Enclosed | NO |
| AGSVWHIQTVQDADGI | 1644155 | Cell/Virus Enclosed | NO |
| AGSVYTNSCLVS | 1408874 | Cell/Virus Enclosed | NO |
| AGSYRDWSYSGQ | 1408877 | Cell/Virus Enclosed | NO |
| AGTAVINLRADQINDM | 1644161 | Cell/Virus Enclosed | NO |
| AGTAVINLRSDQ | 1408879 | Cell/Virus Enclosed | NO |
| AGTAVINLRTDQ | 1408880 | Cell/Virus Enclosed | NO |
| AGTDTTITLNVL | 1408886 | Cell/Virus Enclosed | NO |
| AGTILVDNDLHP | 1408891 | Cell/Virus Enclosed | NO |
| AGTILVDNDLYPFVSD | 1644174 | Cell/Virus Enclosed | NO |
| AGTKLRSLNLGE | 1408895 | Cell/Virus Enclosed | NO |
| AGTKYRHCDLDT | 1408896 | Ectodomain | YES |
| AGTKYRHCDVDS | 1408897 | Ectodomain | YES |
| AGTKYRHCDVDT | 1408898 | Ectodomain | YES |
| AGTNHLTCDNLC | 1408901 | Ectodomain | YES |
| AGTNHLTCHNLC | 1408902 | Ectodomain | YES |
| AGTNSLTCDNLC | 1408903 | Ectodomain | YES |
| AGTNSLTCGNLC | 1408904 | Ectodomain | YES |
| AGTNYLNCDNLC | 1408905 | Ectodomain | YES |
| AGTNYLTCDNLCTPDP | 1644177 | Ectodomain | YES |
| AGTNYLTCHNAA | 1408907 | Ectodomain | YES |
| AGTNYLTCHNAV | 1408908 | Ectodomain | YES |
| AGTNYLTCHNLC | 1408909 | Ectodomain | YES |
| AGVCLGAGQFMG | 1408917 | Cell/Virus Enclosed | NO |
| AGVFTDHDVVYA | 1408925 | Ectodomain | YES |
| AGVFTNHDVVYA | 1408926 | Ectodomain | YES |
| AGVGVIRISNDT | 1408928 | Ectodomain | YES |
| AGVGWTTGLSSF | 1408931 | Ectodomain | YES |
| AGVIAQGARGTY | 1408932 | Cell/Virus Enclosed | NO |
| AGVKLRSLNLGE | 1408936 | Cell/Virus Enclosed | NO |
| AGVKYFCGMTLK | 1408937 | Cell/Virus Enclosed | NO |
| AGVLTNHDVVYA | 1408939 | Ectodomain | YES |
| AGVNITLANFNETKGP | 1644200 | Ectodomain | YES |
| AGVSICSTMTNR | 1408945 | Cell/Virus Enclosed | NO |
| AGVSILSTVTGR | 1408946 | Cell/Virus Enclosed | NO |
| AGVTAQGARGTY | 1408949 | Cell/Virus Enclosed | NO |
| AGVVANGVKAKGYPQF | 1644207 | Capsid | UN |
| AGVVANGVTAKG | 1408950 | Cell/Virus Enclosed | NO |
| AGVVTGNVVKQCCTAA | 1644208 | Cell/Virus Enclosed | NO |
| AGVVWTLNDVKDNDGR | 1644209 | Cell/Virus Enclosed | NO |
| AGVVWTLQEVKDNDGK | 1644210 | Cell/Virus Enclosed | NO |
| AGWAFGAGAALQ | 1408961 | Ectodomain | YES |
| AGWPLNKFGKAGLYYE | 1644211 | Cell/Virus Enclosed | NO |
| AGWTAGLSSSAA | 1408965 | Ectodomain | YES |
| AGYGGFGCKDSF | 1408970 | Cell/Virus Enclosed | NO |

| | | | |
|---|---|---|---|
| AGYKLQQGTFLC | 1408972 | Cell/Virus Enclosed | NO |
| AGYKVLPPLMDVNMEA | 1644216 | Ectodomain | YES |
| AGYTESAVNSTKALAK | 1644218 | Cell/Virus Enclosed | NO |
| AGYTLEEAKTAF | 1408984 | Cell/Virus Enclosed | NO |
| AGYTLEEARTAL | 1408986 | Cell/Virus Enclosed | NO |
| AGYWRRQDRKINTGNG | 1644220 | Cell/Virus Enclosed | NO |
| AHAPSFLAVDDKYKAT | 1644228 | Cell/Virus Enclosed | NO |
| AHAPSFLAVDDKYKVN | 1644229 | Cell/Virus Enclosed | NO |
| AHATLDSIGTNF | 1409006 | Cell/Virus Enclosed | NO |
| AHATRDSIGTNFPLQL | 1644232 | Cell/Virus Enclosed | NO |
| AHATRENIGTNFPLQI | 1644233 | Cell/Virus Enclosed | NO |
| AHGGGLAKALDVYTKG | 1644238 | Cell/Virus Enclosed | NO |
| AHIPSGTVLGNF | 1409031 | Ectodomain | YES |
| AHKHDIQITNESYNNY | 1644248 | Cell/Virus Enclosed | NO |
| AHKHDIQLTNES | 1409043 | Cell/Virus Enclosed | NO |
| AHLSLRGLPVCAFSS | 1799 | Cell/Virus Enclosed | NO |
| AHNSRSTSRTSS | 1409058 | Capsid | UN |
| AHSSIKQGTQIY | 1409078 | Cell/Virus Enclosed | NO |
| AHSTVLSFCAFA | 1409079 | Cell/Virus Enclosed | NO |
| AHTAGVCIKYYA | 1409081 | Cell/Virus Enclosed | NO |
| AHVASCDAIMTR | 1409088 | Cell/Virus Enclosed | NO |
| AHVCGDNIGTNVPLQV | 1644270 | Cell/Virus Enclosed | NO |
| AIAAKPEPEPLP | 1409104 | Cell/Virus Enclosed | NO |
| AIAFCKSGNLVL | 1409106 | Ectodomain | YES |
| AIAKIVANWLSV | 1409110 | Cell/Virus Enclosed | NO |
| AIAKPKRLRKKR | 1409111 | Cell/Virus Enclosed | NO |
| AIAMACLVGLMWLSY | 1312122 | Transmembrane | NO |
| AIANIIYCNSVI | 1409115 | Ectodomain | YES |
| AIASNCYSSLIV | 1409122 | Ectodomain | YES |
| AIATVKSEDGNIKNIN | 1644291 | Cell/Virus Enclosed | NO |
| AIAVAAGKKFSKETAA | 1644292 | Cell/Virus Enclosed | NO |
| AICAAFDFSSDV | 1409129 | Cell/Virus Enclosed | NO |
| AIDAYPLSKHPKPEYR | 1644304 | Cell/Virus Enclosed | NO |
| AIDCAQNPLSEL | 1409155 | Ectodomain | YES |
| AIDCSDGNYNRN | 1409159 | Ectodomain | YES |
| AIDGFGVFDIKNSSVN | 1644313 | Cell/Virus Enclosed | NO |
| AIDILTEGQLQSLSKD | 1644315 | Cell/Virus Enclosed | NO |
| AIDVEHLELKET | 1409166 | Cell/Virus Enclosed | NO |
| AIDVVQYEADRR | 1409168 | Cell/Virus Enclosed | NO |
| AIDYDHEYSIMRLHNF | 1644320 | Cell/Virus Enclosed | NO |
| AIEDILFSKLVTSGLG | 1644324 | Ectodomain | YES |
| AIEDLLFDKVKLSDVG | 1644325 | Ectodomain | YES |
| AIEDLLFDKVTIADPG | 1644326 | Ectodomain | YES |
| AIEDLLFSKVKL | 1409176 | Ectodomain | YES |
| AIEDLLFSKVRL | 1409177 | Ectodomain | YES |
| AIEDVEDVISSEAVED | 1644327 | Cell/Virus Enclosed | NO |
| AIEEIEYVHGDALHTL | 1644328 | Cell/Virus Enclosed | NO |
| AIEEKLSPCKEI | 1409180 | Cell/Virus Enclosed | NO |
| AIELSKELKRPVNPTD | 1644331 | Cell/Virus Enclosed | NO |
| AIENMWLSYKVG | 1409186 | Cell/Virus Enclosed | NO |
| AIETAAYREAAA | 1409188 | Cell/Virus Enclosed | NO |
| AIFAYSPQLTLV | 1409192 | Cell/Virus Enclosed | NO |
| AIFNDHHLNRPGVYCG | 1644340 | Cell/Virus Enclosed | NO |
| AIGDLKCPLDSR | 1409211 | Ectodomain | YES |
| AIGDLNCTLDPR | 1409212 | Ectodomain | YES |
| AIGDVGLCMYRL | 1409213 | Cell/Virus Enclosed | NO |
| AIGKKIKHEGDWDSFC | 1644347 | Cell/Virus Enclosed | NO |
| AIGSSLQEILSR | 1409223 | Ectodomain | YES |
| AIGTANLSIPSN | 1409225 | Ectodomain | YES |
| AIIAVELLLLDFKTAV | 1644363 | Cell/Virus Enclosed | NO |
| AIIIDNDINDYVSDAD | 1644365 | Cell/Virus Enclosed | NO |
| AIIIDNDLNEFVSDAD | 1644366 | Cell/Virus Enclosed | NO |
| AIIKDWNFAWSI | 1409240 | Undetermined | UN |
| AIINDKVLTKID | 1409249 | Cell/Virus Enclosed | NO |
| AIINDKVVAKIDKEDV | 1644373 | Cell/Virus Enclosed | NO |
| AIINDKVVTKID | 1409251 | Cell/Virus Enclosed | NO |
| AIIRDGDVVKQP | 1409252 | Cell/Virus Enclosed | NO |
| AIIRDSDFVKQP | 1409253 | Cell/Virus Enclosed | NO |
| AIIRNSDVVKQP | 1409258 | Cell/Virus Enclosed | NO |
| AIITANLSIPSNWTTS | 1644376 | Ectodomain | YES |
| AIKCVDTVKEAQ | 1409267 | Cell/Virus Enclosed | NO |
| AIKDCVDINKAIPSGN | 1644384 | Cell/Virus Enclosed | NO |
| AIKDDVLYSEFP | 1409271 | Ectodomain | YES |
| AIKDFGGFNFSQ | 1409272 | Ectodomain | YES |
| AIKITEFSWNADLYKL | 1644388 | Cell/Virus Enclosed | NO |
| AIKLDDKDPNFKDQVI | 1377559 | Cell/Virus Enclosed | NO |

| | | | |
|---|---|---|---|
| AIKLDDKDPSFKDNVI | 1644394 | Cell/Virus Enclosed | NO |
| AIKLDPKNPNYNKWLE | 1644395 | Cell/Virus Enclosed | NO |
| AIKLNFGMLNGNAIAT | 1644396 | Cell/Virus Enclosed | NO |
| AIKLVPTTTGAL | 1409287 | Cell/Virus Enclosed | NO |
| AIKNLTPIKLNF | 1409290 | Cell/Virus Enclosed | NO |
| AIKSDHCGGNPC | 1409298 | Ectodomain | YES |
| AIKSDYCGGNPC | 1409299 | Ectodomain | YES |
| AIKVTEYSWNKKLYEL | 1644404 | Cell/Virus Enclosed | NO |
| AIKVTEYSWNKY | 1409301 | Cell/Virus Enclosed | NO |
| AILCATCPFAKG | 1409304 | Cell/Virus Enclosed | NO |
| AILMSEDVKLNPSEDF | 1644410 | Cell/Virus Enclosed | NO |
| AILNNCNWFIQS | 1409316 | Cell/Virus Enclosed | NO |
| AILNSFKEYFNL | 1409318 | Ectodomain | YES |
| AILNTVKFCDHI | 1409320 | Cell/Virus Enclosed | NO |
| AILNTVKFCDHMVKAG | 1644415 | Cell/Virus Enclosed | NO |
| AILNTVTFCDHM | 1409321 | Cell/Virus Enclosed | NO |
| AILSGTTTNVDK | 1409328 | Cell/Virus Enclosed | NO |
| AILSGTTTNVEK | 1409329 | Cell/Virus Enclosed | NO |
| AILTAFLPAQDT | 1409332 | Ectodomain | YES |
| AILTAFSPAQDT | 1409334 | Ectodomain | YES |
| AIMNDKVVVKINNVDT | 1644428 | Cell/Virus Enclosed | NO |
| AIMSMAECKATL | 1409355 | Cell/Virus Enclosed | NO |
| AINAASKGAVQKESDE | 1644439 | Cell/Virus Enclosed | NO |
| AINDLNTTIPRI | 1409365 | Underneath Membrane | NO |
| AINDVDTGVPST | 1409366 | Ectodomain | YES |
| AINLPIGCMMNV | 1409372 | Cell/Virus Enclosed | NO |
| AINNIVASFSSV | 1409375 | Ectodomain | YES |
| AINSVPWSKIFA | 1409383 | Cell/Virus Enclosed | NO |
| AINVTINNYNPSSWNR | 1644448 | Ectodomain | YES |
| AIPDYNYYKYNL | 1409392 | Cell/Virus Enclosed | NO |
| AIPNGRKVDLHL | 1409396 | Ectodomain | YES |
| AIPNGRKVDLQLGNLG | 1644452 | Ectodomain | YES |
| AIPNGRRVDLQL | 1409398 | Ectodomain | YES |
| AIPNRRKFDLQL | 1409399 | Ectodomain | YES |
| AIPNRRKVDLQL | 1409400 | Ectodomain | YES |
| AIPNRRRDDLQL | 1409401 | Ectodomain | YES |
| AIPNRRRVDLQL | 1409402 | Ectodomain | YES |
| AIPNSRKVDLQL | 1409403 | Ectodomain | YES |
| AIPNSRRSDLQF | 1409404 | Ectodomain | YES |
| AIPSGNLIKFDIGSVV | 1644454 | Cell/Virus Enclosed | NO |
| AIPSLAANTLNIIVPD | 1644455 | Cell/Virus Enclosed | NO |
| AIPTNFSIGITT | 1409414 | Ectodomain | YES |
| AIPTNFSISITTEVMPV | 2053 | Ectodomain | YES |
| AIQDGVTRDIMA | 1409423 | Cell/Virus Enclosed | NO |
| AIQDGVTRDIVS | 1409424 | Cell/Virus Enclosed | NO |
| AIQEGFDATNSALVKI | 1644464 | Ectodomain | YES |
| AIQEVIDDMLWG | 1409426 | Cell/Virus Enclosed | NO |
| AIQGFACGCGAS | 1409427 | Cell/Virus Enclosed | NO |
| AIQNCAWDYHIT | 1409430 | Cell/Virus Enclosed | NO |
| AIQNCAWDYHVT | 1409431 | Cell/Virus Enclosed | NO |
| AIQQGFDATNFA | 1409432 | Ectodomain | YES |
| AIQQGSDATNSA | 1409433 | Ectodomain | YES |
| AIQQLYTGFQGK | 1409437 | Cell/Virus Enclosed | NO |
| AIQVAKCYGESV | 1409443 | Cell/Virus Enclosed | NO |
| AIRGSCEKNLQL | 1409449 | Cell/Virus Enclosed | NO |
| AIRNFDEQNCDV | 1409453 | Cell/Virus Enclosed | NO |
| AIRNVRGWLGMDVESA | 1644478 | Cell/Virus Enclosed | NO |
| AISAKNRARTVA | 1409460 | Cell/Virus Enclosed | NO |
| AISASIGDIIQRLDVLEQD | 1377571 | Ectodomain | YES |
| AISFADNNDITN | 1409469 | Cell/Virus Enclosed | NO |
| AISGDKVFVRDGNTDN | 1644483 | Cell/Virus Enclosed | NO |
| AISLSFNNLLGFDSSK | 1644485 | Cell/Virus Enclosed | NO |
| AISNELFLTVPT | 1409481 | Ectodomain | YES |
| AISNEQHYKSWV | 1409482 | Ectodomain | YES |
| AISNEWYIGAII | 1409483 | Cell/Virus Enclosed | NO |
| AISNSIQAIYDRLDSI | 1644491 | Ectodomain | YES |
| AISPKIDNETLE | 1409486 | Ectodomain | YES |
| AISPKTDKETLE | 1409487 | Ectodomain | YES |
| AISPNTDNETLQ | 1409489 | Ectodomain | YES |
| AISSNTDNETLE | 1409493 | Ectodomain | YES |
| AISSNTDNETLQYWVT | 1644495 | Ectodomain | YES |
| AISSTTDNETLE | 1409497 | Ectodomain | YES |
| AISSVLNDVLSR | 1409500 | Ectodomain | YES |
| AISVQTCDLTVGSGYC | 1644500 | Ectodomain | YES |
| AITDYNYYKYNLPTMV | 1644503 | Cell/Virus Enclosed | NO |
| AITDYNYYSYNLPTMC | 1644504 | Cell/Virus Enclosed | NO |

| | | | |
|---|---|---|---|
| AITIKPEANINQ | 1409515 | Cell/Virus Enclosed | NO |
| AITPYVCNASGC | 1409520 | Cell/Virus Enclosed | NO |
| AITPYVCNTSGCNVND | 1644510 | Cell/Virus Enclosed | NO |
| AITQTSPALQTV | 1409522 | Ectodomain | YES |
| AITQTSQALQTVATAL | 1644512 | Ectodomain | YES |
| AITSGIKFQTVKPGNF | 1644521 | Cell/Virus Enclosed | NO |
| AITSGVKFQTVKPGNF | 1644522 | Cell/Virus Enclosed | NO |
| AITTENRFTNFE | 1409527 | Ectodomain | YES |
| AITTGYRFTNFEPFTV | 1644523 | Ectodomain | YES |
| AIVCASVKRDGVQVGY | 1644529 | Cell/Virus Enclosed | NO |
| AIVGAMLSENFT | 1409549 | Ectodomain | YES |
| AIVGIPVCVNGL | 1409550 | Cell/Virus Enclosed | NO |
| AIVGTPVCVNGL | 1409551 | Cell/Virus Enclosed | NO |
| AIVIDNDLNEFV | 1409552 | Cell/Virus Enclosed | NO |
| AIVKETNLICKA | 1409557 | Cell/Virus Enclosed | NO |
| AIVLPQGGFVAD | 1409561 | Cell/Virus Enclosed | NO |
| AIVLVNGVHHPT | 1409565 | Cell/Virus Enclosed | NO |
| AIVLVNGVHYPT | 1409566 | Cell/Virus Enclosed | NO |
| AIVLVNGVRYPT | 1409567 | Cell/Virus Enclosed | NO |
| AIVMVTILLCCV | 1409570 | Undetermined | UN |
| AIVQASESKVLLERFL | 1644546 | Cell/Virus Enclosed | NO |
| AIVQATESKVLL | 1409575 | Cell/Virus Enclosed | NO |
| AIVTANLSIPSNWTTS | 1644548 | Ectodomain | YES |
| AIVTGLLPTHWI | 1409581 | Ectodomain | YES |
| AIVYGDFSHGQL | 1409587 | Cell/Virus Enclosed | NO |
| AIWDAFVAAIKLVPTT | 1644554 | Cell/Virus Enclosed | NO |
| AIYDRLDSIQADQQVD | 1644559 | Ectodomain | YES |
| AIYDRLDTIQADQQVD | 1644560 | Ectodomain | YES |
| AIYHRYKAGNYR | 1409606 | Ectodomain | YES |
| AIYPIDLASQII | 1409612 | Undetermined | UN |
| AIYPIDLSSQII | 1409613 | Undetermined | UN |
| AIYPIHLASQII | 1409614 | Undetermined | UN |
| AKAELCCFARMV | 1409622 | Cell/Virus Enclosed | NO |
| AKAELCCSACMV | 1409623 | Cell/Virus Enclosed | NO |
| AKAGDKYYPVVVDDHV | 1644569 | Cell/Virus Enclosed | NO |
| AKAKNTKVIKDD | 1409631 | Ectodomain | YES |
| AKAMDTFTNNNGSDVL | 1644578 | Cell/Virus Enclosed | NO |
| AKAMETFNHNNGNDVL | 1644579 | Cell/Virus Enclosed | NO |
| AKAMLDFSRDHNDILY | 1644580 | Cell/Virus Enclosed | NO |
| AKAMLDYAKDHNDMLY | 1644581 | Cell/Virus Enclosed | NO |
| AKAQEGCGIISR | 1409643 | Undetermined | UN |
| AKCWTEEDLSIGPHEF | 1644591 | Cell/Virus Enclosed | NO |
| AKCYGEPVLVNA | 1409652 | Cell/Virus Enclosed | NO |
| AKDDLYVKRYER | 1409655 | Cell/Virus Enclosed | NO |
| AKDNTVLQALQS | 1409667 | Cell/Virus Enclosed | NO |
| AKDQYGSDINGV | 1409668 | Cell/Virus Enclosed | NO |
| AKDQYGTDFDGV | 1409669 | Cell/Virus Enclosed | NO |
| AKDQYGTDIDGI | 1409670 | Cell/Virus Enclosed | NO |
| AKDQYGTDINGV | 1409672 | Cell/Virus Enclosed | NO |
| AKDYGFTVFVCT | 1409676 | Cell/Virus Enclosed | NO |
| AKEESVKSSYAYATLK | 1644607 | Cell/Virus Enclosed | NO |
| AKEGAKTVNTSLGNRK | 1644608 | Capsid | UN |
| AKEIRQKILNKP | 1409686 | Capsid | UN |
| AKERCTHWCTYL | 1409691 | Cell/Virus Enclosed | NO |
| AKFGFTPLNKQS | 1409702 | Cell/Virus Enclosed | NO |
| AKFKNSVASINSAIVC | 1644620 | Cell/Virus Enclosed | NO |
| AKFKNSVTSINS | 1409706 | Cell/Virus Enclosed | NO |
| AKFLWEWASVRFSWL | 1644622 | Undetermined | UN |
| AKFPLKLAGTAVINLR | 1644623 | Cell/Virus Enclosed | NO |
| AKGDSCSSNCKHSVAQ | 1644628 | Cell/Virus Enclosed | NO |
| AKGFPLLAELIP | 1409724 | Cell/Virus Enclosed | NO |
| AKGFPQFAELIP | 1409725 | Cell/Virus Enclosed | NO |
| AKGFPQLAELIPNQAA | 1644630 | Capsid | UN |
| AKGGFKFGDPADSRDF | 1644631 | Cell/Virus Enclosed | NO |
| AKGHFKFDEPSDATDF | 1644632 | Cell/Virus Enclosed | NO |
| AKGLLQVGDSVL | 1409729 | Cell/Virus Enclosed | NO |
| AKGYIAGIQNFKETLY | 1644639 | Cell/Virus Enclosed | NO |
| AKGYPQFAELVPSTAA | 1644640 | Capsid | UN |
| AKHGDFSGVASQ | 1409742 | Ectodomain | YES |
| AKHIVYVGDPAQLPAP | 1644647 | Cell/Virus Enclosed | NO |
| AKICIVDNVYMAKAGD | 1644652 | Cell/Virus Enclosed | NO |
| AKILAPSRSSET | 1409757 | Cell/Virus Enclosed | NO |
| AKIQSVVNSNAQALNS | 1644658 | Ectodomain | YES |
| AKIVGGAPIWFN | 1409769 | Cell/Virus Enclosed | NO |
| AKIYGMCFSSITIDKF | 1644661 | Ectodomain | YES |
| AKKLERMADLALTNMY | 1644674 | Cell/Virus Enclosed | NO |

| | | | |
|---|---|---|---|
| AKKNLDEARLSG | 1409792 | Cell/Virus Enclosed | NO |
| AKKNNIPFRLTC | 1409795 | Cell/Virus Enclosed | NO |
| AKKQVVLVSNNQEDFD | 1644682 | Cell/Virus Enclosed | NO |
| AKKSMYDGDRFVKHDL | 1644683 | Cell/Virus Enclosed | NO |
| AKLGFDSSKEFVEYKI | 1644688 | Cell/Virus Enclosed | NO |
| AKLGKESKPQQV | 1409819 | Cell/Virus Enclosed | NO |
| AKLKIQVVNSGPDQTC | 1644694 | Cell/Virus Enclosed | NO |
| AKLQNNELSPVA | 1409832 | Cell/Virus Enclosed | NO |
| AKMIERCYTAEQ | 1409840 | Cell/Virus Enclosed | NO |
| AKNAYEKDKAVARKLE | 1644706 | Cell/Virus Enclosed | NO |
| AKNILHVVGPDA | 1409850 | Cell/Virus Enclosed | NO |
| AKNKMRHKRIST | 1409852 | Cell/Virus Enclosed | NO |
| AKNLNESLIDLR | 1409857 | Ectodomain | YES |
| AKNSGSVNQQQI | 1409862 | Cell/Virus Enclosed | NO |
| AKPEEKLSAYDLACCM | 1644723 | Cell/Virus Enclosed | NO |
| AKPKRLRKKRNV | 1409874 | Cell/Virus Enclosed | NO |
| AKPLQSLEVCVQ | 1409878 | Cell/Virus Enclosed | NO |
| AKPNVLIALKMV | 1409881 | Ectodomain | YES |
| AKPNVVIALKMV | 1409882 | Ectodomain | YES |
| AKPTVIVNAANV | 1409892 | Cell/Virus Enclosed | NO |
| AKQDVSLLSKCYKAMN | 1644737 | Cell/Virus Enclosed | NO |
| AKQGAGDAGHSLTWLW | 1644738 | Cell/Virus Enclosed | NO |
| AKQKPIYLKGSGSDFS | 1644740 | Cell/Virus Enclosed | NO |
| AKQNVSLLSKCY | 1409913 | Cell/Virus Enclosed | NO |
| AKQVDYVPLKSATCIT | 1644741 | Cell/Virus Enclosed | NO |
| AKRALRSHPDFK | 1409917 | Cell/Virus Enclosed | NO |
| AKRAVRSHPDFKLLHN | 1644742 | Cell/Virus Enclosed | NO |
| AKRKMGLTPPLSILKN | 1644746 | Cell/Virus Enclosed | NO |
| AKRKVGLTPPLSILKN | 1644747 | Cell/Virus Enclosed | NO |
| AKRVLGLQTQTVDSAQ | 1644755 | Cell/Virus Enclosed | NO |
| AKRVLGVQTQTVDSAQ | 1644756 | Cell/Virus Enclosed | NO |
| AKRVVRSHPDFK | 1409937 | Cell/Virus Enclosed | NO |
| AKSIQQDLDVIL | 1409947 | Cell/Virus Enclosed | NO |
| AKTDLFNGYKIGCNCA | 1644775 | Cell/Virus Enclosed | NO |
| AKTELCCSARMV | 1409977 | Cell/Virus Enclosed | NO |
| AKTGTPKPSRNQSPAS | 1644777 | Capsid | UN |
| AKVEGCMVQVTC | 1409997 | Cell/Virus Enclosed | NO |
| AKVEISVTPIKNTVDT | 1644787 | Cell/Virus Enclosed | NO |
| AKVGDKYYPVVV | 1410001 | Cell/Virus Enclosed | NO |
| AKVHNLSGNPDE | 1410005 | Cell/Virus Enclosed | NO |
| AKVKNTKAIKDD | 1410008 | Ectodomain | YES |
| AKVKNTKVFIDG | 1410010 | Ectodomain | YES |
| AKVKNTKVFKHG | 1410012 | Ectodomain | YES |
| AKVKNTKVIKDD | 1410013 | Ectodomain | YES |
| AKVKNTKVIKDG | 1410014 | Ectodomain | YES |
| AKVKNTKVIKDRVMYS | 1644792 | Ectodomain | YES |
| AKVKNTKVVGKD | 1410018 | Ectodomain | YES |
| AKVRVDCYDKFKINDTTCKY | 1410027 | Cell/Virus Enclosed | NO |
| AKVRVECYDKFKINDTTRKYV | 1410028 | Cell/Virus Enclosed | NO |
| AKVSAEVSRQVIQDVK | 1644797 | Cell/Virus Enclosed | NO |
| AKVSPGLCISGD | 1410033 | Ectodomain | YES |
| AKVSTGLCIAGD | 1410034 | Ectodomain | YES |
| AKWAVPTTRTDDKLR | 2322 | Ectodomain | YES |
| AKWVFDSSKEFV | 1410040 | Cell/Virus Enclosed | NO |
| AKWYCYDKNPINSNVK | 1644804 | Cell/Virus Enclosed | NO |
| AKYIPLVGTPLH | 1410043 | Cell/Virus Enclosed | NO |
| AKYTPLVGTSLH | 1410048 | Cell/Virus Enclosed | NO |
| AKYTPLVGTSVH | 1410049 | Cell/Virus Enclosed | NO |
| ALAEAREHLKNG | 1410060 | Cell/Virus Enclosed | NO |
| ALAEHDFFTWKD | 1410061 | Cell/Virus Enclosed | NO |
| ALAFAWVLFHVL | 1410062 | Cell/Virus Enclosed | NO |
| ALAKQYAAQRIA | 1410072 | Cell/Virus Enclosed | NO |
| ALALLLLDRLNALESK | 1644822 | Cell/Virus Enclosed | NO |
| ALANQFTEFVGTQSVD | 1644825 | Cell/Virus Enclosed | NO |
| ALANQFTEFVST | 1410089 | Cell/Virus Enclosed | NO |
| ALASALKQLKVTTGEL | 1644827 | Cell/Virus Enclosed | NO |
| ALASDIFATPSV | 1410094 | Cell/Virus Enclosed | NO |
| ALASDIFDTPSVLQAT | 1644828 | Cell/Virus Enclosed | NO |
| ALASMTGVSIETLLAA | 1644829 | Cell/Virus Enclosed | NO |
| ALASMTGVSLETLLAA | 1644830 | Cell/Virus Enclosed | NO |
| ALAVNACLIPLL | 1410102 | Cell/Virus Enclosed | NO |
| ALAVNVCIIPLL | 1410103 | Cell/Virus Enclosed | NO |
| ALAVRLTANVGR | 1410106 | Cell/Virus Enclosed | NO |
| ALAYRANSFDVP | 1410109 | Cell/Virus Enclosed | NO |
| ALCFDKSKQFVPVQVV | 1644843 | Underneath Membrane | NO |
| ALCFNKPKQFVP | 1410121 | Underneath Membrane | NO |

| | | | |
|---|---|---|---|
| ALDAYVSQQLSD | 1410139 | Ectodomain | YES |
| ALDDATRNYIIT | 1410141 | Cell/Virus Enclosed | NO |
| ALDDDARTFIQS | 1410142 | Cell/Virus Enclosed | NO |
| ALDDDARTFVQS | 1410143 | Cell/Virus Enclosed | NO |
| ALDDVTLNYIIT | 1410144 | Cell/Virus Enclosed | NO |
| ALDDVTRNYIIT | 1410145 | Cell/Virus Enclosed | NO |
| ALDLSKELKRPIQPTD | 1644857 | Cell/Virus Enclosed | NO |
| ALDLSLDYINVT | 1410148 | Ectodomain | YES |
| ALDQAISMWALV | 1410153 | Cell/Virus Enclosed | NO |
| ALDQDVFIQKYD | 1410154 | Cell/Virus Enclosed | NO |
| ALDQDVFIQKYG | 1410155 | Cell/Virus Enclosed | NO |
| ALDSLGAAIHYL | 1410157 | Cell/Virus Enclosed | NO |
| ALDTANHSIPDL | 1410158 | Cell/Virus Enclosed | NO |
| ALDTILQEANKS | 1410159 | Cell/Virus Enclosed | NO |
| ALDTTIPFGPSS | 1410160 | Ectodomain | YES |
| ALDWELYVNNAY | 1410164 | Cell/Virus Enclosed | NO |
| ALEAQVQIDRLINGRL | 1644870 | Ectodomain | YES |
| ALEDEFTPFDVV | 1410171 | Cell/Virus Enclosed | NO |
| ALEDLLFSKVVTSGLG | 1644871 | Ectodomain | YES |
| ALEFNPSQTSPATAEP | 1644872 | Capsid | UN |
| ALEQFMSTPNAI | 1410187 | Cell/Virus Enclosed | NO |
| ALESKVLLERFL | 1410194 | Cell/Virus Enclosed | NO |
| ALESPEHCSPHHTALRQAIL | 96088 | Capsid | UN |
| ALFASLDVKPYGYPND | 1644881 | Cell/Virus Enclosed | NO |
| ALFASLNVKPYG | 1410204 | Cell/Virus Enclosed | NO |
| ALFDSLEFTELS | 1410208 | Cell/Virus Enclosed | NO |
| ALFFDSEVSTDEVGDN | 1644884 | Capsid | UN |
| ALFVSYGSSVETETYF | 1644896 | Cell/Virus Enclosed | NO |
| ALFVVEQKWCVI | 1410239 | Cell/Virus Enclosed | NO |
| ALGAIQEGFDAT | 1410242 | Ectodomain | YES |
| ALGAMQTGFTTTNEAF | 1644898 | Ectodomain | YES |
| ALGGSVAVKITE | 1410250 | Cell/Virus Enclosed | NO |
| ALGGWGLHQPPPQIFLKILP | 2558 | Capsid | UN |
| ALGKVVANAMLKCVAF | 1644910 | Cell/Virus Enclosed | NO |
| ALGSMDIDVFNKLSTS | 1644914 | Cell/Virus Enclosed | NO |
| ALGVINTLEWIPRFK | 2598 | Ectodomain | YES |
| ALHLVNPQIQLT | 1410276 | Cell/Virus Enclosed | NO |
| ALHLYFDKAGQK | 1410277 | Cell/Virus Enclosed | NO |
| ALHPYFDKAGQK | 1410280 | Cell/Virus Enclosed | NO |
| ALHSGYYDANQY | 1410283 | Ectodomain | YES |
| ALHSIQQGFDAT | 1410284 | Ectodomain | YES |
| ALHSKCLTSIEE | 1410285 | Cell/Virus Enclosed | NO |
| ALHTGYYDVNQY | 1410287 | Ectodomain | YES |
| ALIATAHSSIKQGTQI | 1644925 | Cell/Virus Enclosed | NO |
| ALIAVANWLAPTNAYM | 1644926 | Cell/Virus Enclosed | NO |
| ALIGANGNKIVD | 1410296 | Cell/Virus Enclosed | NO |
| ALIMACGNCTFG | 1410301 | Cell/Virus Enclosed | NO |
| ALIPDYAKILVD | 1410307 | Cell/Virus Enclosed | NO |
| ALISCNGNKIED | 1410312 | Cell/Virus Enclosed | NO |
| ALKELYGPADPT | 1410336 | Cell/Virus Enclosed | NO |
| ALKELYGPIDPT | 1410337 | Cell/Virus Enclosed | NO |
| ALKELYGPLDPT | 1410338 | Cell/Virus Enclosed | NO |
| ALKELYGSVDPT | 1410340 | Cell/Virus Enclosed | NO |
| ALKKVPIDEYIT | 1410350 | Cell/Virus Enclosed | NO |
| ALKNAFQTFRWS | 1410364 | Cell/Virus Enclosed | NO |
| ALKNLGFDNQSK | 1410366 | Cell/Virus Enclosed | NO |
| ALKSLGFDKPQEKDKK | 1644959 | Capsid | UN |
| ALKSLGFEKPQE | 1410374 | Cell/Virus Enclosed | NO |
| ALKTAGVVTGNVVKQC | 1644960 | Cell/Virus Enclosed | NO |
| ALLAGSFDKVYDIGNP | 1644966 | Cell/Virus Enclosed | NO |
| ALLDWVKHLNKTLNEG | 1644972 | Cell/Virus Enclosed | NO |
| ALLFSLAKAQEG | 1410403 | Undetermined | UN |
| ALLHIADCQTTN | 1410405 | Ectodomain | YES |
| ALLHIAGCQTTNGLNT | 1644977 | Ectodomain | YES |
| ALLHIAGSQTTN | 1410407 | Ectodomain | YES |
| ALLLLDRLNALESKMS | 1644983 | Cell/Virus Enclosed | NO |
| ALLLLDRLNQLESKMSG | 1410414 | Capsid | UN |
| ALLQNGMNGRTI | 1410424 | Cell/Virus Enclosed | NO |
| ALLRKTLVLNAS | 1410426 | Cell/Virus Enclosed | NO |
| ALLSDHQDLKWA | 1410428 | Cell/Virus Enclosed | NO |
| ALLVPKIFKARA | 1410443 | Cell/Virus Enclosed | NO |
| ALMGANGNKIVDKEVK | 1645005 | Cell/Virus Enclosed | NO |
| ALNAFVSQVLNKYTEV | 1645008 | Ectodomain | YES |
| ALNAIQQGFDAT | 1410454 | Ectodomain | YES |
| ALNAYVSQQLSDISLV | 1645009 | Ectodomain | YES |
| ALNAYVSQQLSDSTLV | 1645010 | Ectodomain | YES |

| | | | |
|---|---|---|---|
| ALNCYWPLKDYG | 1410456 | Ectodomain | YES |
| ALNCYWPLSDYG | 1410459 | Ectodomain | YES |
| ALNDFSNLGADV | 1410460 | Cell/Virus Enclosed | NO |
| ALNESYIDLKELGNYT | 1645014 | Ectodomain | YES |
| ALNGSCGSVGFN | 1410472 | Cell/Virus Enclosed | NO |
| ALNKIHDVVNQQ | 1410484 | Ectodomain | YES |
| ALNKIQDVVNQQGNSL | 1645025 | Ectodomain | YES |
| ALNKIQDVVNQQGSAL | 1645026 | Ectodomain | YES |
| ALNNVYFGLSIV | 1410488 | Undetermined | UN |
| ALNSEKHQDHVSLTVP | 1645035 | Cell/Virus Enclosed | NO |
| ALNSILDNAVKGCVPL | 1645036 | Cell/Virus Enclosed | NO |
| ALNTICIKDADYNAKV | 1645038 | Cell/Virus Enclosed | NO |
| ALNTPKDHIGTRNPAN | 1645040 | Cell/Virus Enclosed | NO |
| ALPASTVHATAT | 1410506 | Secreted | YES |
| ALPDTHSTLTPR | 1410507 | Ectodomain | YES |
| ALPDTPITLTPR | 1410508 | Ectodomain | YES |
| ALPDTPSPLTPR | 1410509 | Ectodomain | YES |
| ALPDTPSTFTPR | 1410510 | Ectodomain | YES |
| ALPDTPSTLAPR | 1410511 | Ectodomain | YES |
| ALPDTPSTLKPR | 1410512 | Ectodomain | YES |
| ALPDTPSTLTLR | 1410513 | Ectodomain | YES |
| ALPDTPSTLTPA | 1410514 | Ectodomain | YES |
| ALPDTPSTLTPC | 1410515 | Ectodomain | YES |
| ALPDTPSTLTPRSVRS | 1645043 | Ectodomain | YES |
| ALPGVFCGVDAM | 1410523 | Cell/Virus Enclosed | NO |
| ALPIYYQHTDINFTAT | 1645052 | Ectodomain | YES |
| ALPKGFYAEGSRGGSQ | 1645053 | Cell/Virus Enclosed | NO |
| ALPNMIRTISAM | 1410530 | Cell/Virus Enclosed | NO |
| ALPNRRKVDLQL | 1410532 | Ectodomain | YES |
| ALPQSKDSNFLK | 1410535 | Cell/Virus Enclosed | NO |
| ALPQSKDSNFLNESGV | 1645057 | Cell/Virus Enclosed | NO |
| ALPQSKDSNFLNRVRG | 1645058 | Cell/Virus Enclosed | NO |
| ALPQSKESNFLN | 1410537 | Cell/Virus Enclosed | NO |
| ALPVYYQHTDIN | 1410542 | Ectodomain | YES |
| ALQELQIQQWIQ | 1410551 | Cell/Virus Enclosed | NO |
| ALQEQCRFPNITNSHVPILQERP | 2838 | Ectodomain | YES |
| ALQHAFMKHKGGDSTD | 1645065 | Cell/Virus Enclosed | NO |
| ALQQIEVKFNAP | 1410570 | Cell/Virus Enclosed | NO |
| ALQTDVLQEDQK | 1410575 | Ectodomain | YES |
| ALQTDVLQENQKILAA | 1645076 | Ectodomain | YES |
| ALQTDVLQENQR | 1410577 | Ectodomain | YES |
| ALQTVATALNKIQDVV | 1645078 | Ectodomain | YES |
| ALRAGSFDKVYD | 1410583 | Cell/Virus Enclosed | NO |
| ALRHAFMKHKGG | 1410594 | Cell/Virus Enclosed | NO |
| ALRHFDQNIEVL | 1410598 | Cell/Virus Enclosed | NO |
| ALRHFDRNDCML | 1410601 | Cell/Virus Enclosed | NO |
| ALRMGSGFCVDYNSPS | 1645095 | Ectodomain | YES |
| ALRNFDEKDCEVFKEI | 1645096 | Cell/Virus Enclosed | NO |
| ALRNSLTNDAYSRFLG | 1645098 | Cell/Virus Enclosed | NO |
| ALSEAREHLKSG | 1410629 | Cell/Virus Enclosed | NO |
| ALSFEEQDEIYAYTKR | 1645106 | Cell/Virus Enclosed | NO |
| ALSFEEQNEIYAYTKR | 1645107 | Cell/Virus Enclosed | NO |
| ALSGIAVEQDRN | 1410637 | Ectodomain | YES |
| ALSIFDTWANWDSNWA | 1645113 | Undetermined | UN |
| ALSNGFSQVKSD | 1410659 | Cell/Virus Enclosed | NO |
| ALSPGLLATNNV | 1410660 | Cell/Virus Enclosed | NO |
| ALSTGLTSQTVKPGHF | 1645127 | Cell/Virus Enclosed | NO |
| ALSTNHCHKVVE | 1410667 | Cell/Virus Enclosed | NO |
| ALTGIAIEQDKN | 1410679 | Ectodomain | YES |
| ALTGTNQLCAKI | 1410681 | Cell/Virus Enclosed | NO |
| ALTNSVAFQTVK | 1410688 | Cell/Virus Enclosed | NO |
| ALTQHGKEDLKFPRG | 2000982 | Cell/Virus Enclosed | NO |
| ALTQHGKENLTF | 1410692 | Cell/Virus Enclosed | NO |
| ALVDQRTICFSV | 1410708 | Cell/Virus Enclosed | NO |
| ALVFIILCFTIK | 1410709 | Cell/Virus Enclosed | NO |
| ALVFMIVQFMPF | 1410711 | Cell/Virus Enclosed | NO |
| ALVKIQAVVNANAEAL | 1645154 | Ectodomain | YES |
| ALVKSLNLNCVS | 1410719 | Cell/Virus Enclosed | NO |
| ALVLAKGGFKFGDPAD | 1645157 | Cell/Virus Enclosed | NO |
| ALVQCNCGTKSW | 1410727 | Cell/Virus Enclosed | NO |
| ALVSAMERTEHKARKKGTSA | 741209 | Ectodomain | YES |
| ALVSGIATAGWT | 1410732 | Ectodomain | YES |
| ALVYDNKLKAKN | 1410739 | Cell/Virus Enclosed | NO |
| ALYAIQEGFDATNSAL | 1645179 | Ectodomain | YES |
| ALYAIQQGFDAT | 1410750 | Ectodomain | YES |
| ALYLDYVQCKCGNLEQ | 1645184 | Cell/Virus Enclosed | NO |

| | | | |
|---|---|---|---|
| ALYLDYVQHKCGNLHQ | 1645185 | Cell/Virus Enclosed | NO |
| ALYTPWFYPLLA | 1410777 | Cell/Virus Enclosed | NO |
| ALYVTWSEGNSITGVP | 1645195 | Ectodomain | YES |
| AMANGYTSVSSVECYS | 1645204 | Cell/Virus Enclosed | NO |
| AMCNWCLISQDS | 1410807 | Cell/Virus Enclosed | NO |
| AMCNWCLVSQDS | 1410808 | Cell/Virus Enclosed | NO |
| AMDAWNQPSGISSPPSPSLN | 3069 | Cell/Virus Enclosed | NO |
| AMDPIYDEPTMT | 1410813 | Secreted | YES |
| AMDTFTNNNGSDVLYQ | 1645212 | Cell/Virus Enclosed | NO |
| AMETFNHNNGNDVLYQ | 1645219 | Cell/Virus Enclosed | NO |
| AMETFTNNNGSD | 1410825 | Cell/Virus Enclosed | NO |
| AMFPPWSAAAGV | 1410833 | Ectodomain | YES |
| AMFQPRLPVLSD | 1410834 | Ectodomain | YES |
| AMFTKVVVGPTTEVKF | 1645223 | Cell/Virus Enclosed | NO |
| AMGDVGLCMYRL | 1410838 | Cell/Virus Enclosed | NO |
| AMGQGQNNADPK | 1410843 | Cell/Virus Enclosed | NO |
| AMHIASSNAFLDLRTS | 1645230 | Cell/Virus Enclosed | NO |
| AMIMPKTCFGPI | 1410855 | Cell/Virus Enclosed | NO |
| AMIPLQASLPFV | 1410857 | Secreted | YES |
| AMLDFSRDHNDILYTP | 1645239 | Cell/Virus Enclosed | NO |
| AMLFDSHIVSKESGNT | 1645241 | Capsid | UN |
| AMLFKKEPLLYV | 1410870 | Cell/Virus Enclosed | NO |
| AMLGAEMTSNVRRQVA | 1645243 | Cell/Virus Enclosed | NO |
| AMLSENFTSYGFSNVV | 1645249 | Ectodomain | YES |
| AMLTGFTTTNEA | 1410880 | Ectodomain | YES |
| AMMRISYFVQSI | 1410886 | Transmembrane | NO |
| AMMWLSYFVQSI | 1410888 | Transmembrane | NO |
| AMNGEDAFSILAAKTG | 1645257 | Cell/Virus Enclosed | NO |
| AMNGEDAFSILS | 1410893 | Cell/Virus Enclosed | NO |
| AMNVAKAEFDRESSVQ | 1645260 | Cell/Virus Enclosed | NO |
| AMNVIGQHIELP | 1410898 | Cell/Virus Enclosed | NO |
| AMNVIGQHIKLP | 1410899 | Cell/Virus Enclosed | NO |
| AMPIMANLANLN | 1410900 | Ectodomain | YES |
| AMPIMANLVNSK | 1410902 | Ectodomain | YES |
| AMPIVANLAYIN | 1410904 | Ectodomain | YES |
| AMPIVANLAYLN | 1410905 | Ectodomain | YES |
| AMPNLYKIQRML | 1410909 | Cell/Virus Enclosed | NO |
| AMQKESDGYIKL | 1410920 | Cell/Virus Enclosed | NO |
| AMQKESGDYIKL | 1410921 | Cell/Virus Enclosed | NO |
| AMQQESDDYIKK | 1410926 | Cell/Virus Enclosed | NO |
| AMQTGFTTTNEAFQKV | 1645279 | Ectodomain | YES |
| AMQTMLFTMLRK | 1410931 | Cell/Virus Enclosed | NO |
| AMRDAGIVGVLT | 1410934 | Cell/Virus Enclosed | NO |
| AMRHVRGWLGMDVEGA | 1645285 | Cell/Virus Enclosed | NO |
| AMRPNYTIKGSF | 1410945 | Cell/Virus Enclosed | NO |
| AMSGQILLNCAL | 1410951 | Cell/Virus Enclosed | NO |
| AMTAVNESRYGLQNLL | 1645295 | Ectodomain | YES |
| AMTSMYKQARAEDKKA | 1645303 | Cell/Virus Enclosed | NO |
| AMVAKGVVGVLT | 1410969 | Cell/Virus Enclosed | NO |
| AMVLGSKHVNCC | 1410973 | Cell/Virus Enclosed | NO |
| ANAEAHNNLLQQ | 1410985 | Ectodomain | YES |
| ANAFNNALHSIQ | 1410987 | Ectodomain | YES |
| ANAFSNALDAIQ | 1410991 | Ectodomain | YES |
| ANAKGFIRFPEVLREG | 1645324 | Cell/Virus Enclosed | NO |
| ANALPECNADIV | 1411001 | Cell/Virus Enclosed | NO |
| ANALSRAVNVPTIRKF | 1645326 | Cell/Virus Enclosed | NO |
| ANAQALNSLLQQ | 1411005 | Ectodomain | YES |
| ANASSSEGFLLGINYL | 1645329 | Cell/Virus Enclosed | NO |
| ANAYYKYDNGNFIKLP | 1645332 | Cell/Virus Enclosed | NO |
| ANCASILCKCYTTGT | 3236 | Ectodomain | YES |
| ANCVNCTDDRCV | 1411021 | Cell/Virus Enclosed | NO |
| ANDDLYVKRYER | 1411022 | Cell/Virus Enclosed | NO |
| ANDEKDTLRLFA | 1411023 | Cell/Virus Enclosed | NO |
| ANDFTTALRRPI | 1411024 | Cell/Virus Enclosed | NO |
| ANDKFYRLSNEL | 1411027 | Cell/Virus Enclosed | NO |
| ANDLKDTLRLFA | 1411029 | Cell/Virus Enclosed | NO |
| ANDLTTTLRRPI | 1411033 | Cell/Virus Enclosed | NO |
| ANDNILSVKFTA | 1411034 | Cell/Virus Enclosed | NO |
| ANDTKIASQLGM | 1411039 | Ectodomain | YES |
| ANDTKIVSQLGN | 1411041 | Ectodomain | YES |
| ANDVEVQSSGII | 1411045 | Cell/Virus Enclosed | NO |
| ANECAQILSEIV | 1411052 | Cell/Virus Enclosed | NO |
| ANECAQVLSETV | 1411055 | Cell/Virus Enclosed | NO |
| ANECAQVLSEYVLCGG | 1645349 | Cell/Virus Enclosed | NO |
| ANELTTALRRPI | 1411062 | Cell/Virus Enclosed | NO |
| ANENAINKGGRTVQSH | 1645354 | Cell/Virus Enclosed | NO |

| | | | |
|---|---|---|---|
| ANENLAHGGGLAKALD | 1645355 | Cell/Virus Enclosed | NO |
| ANEYTGNYQCGH | 1411066 | Cell/Virus Enclosed | NO |
| ANEYTGSYQCGH | 1411067 | Cell/Virus Enclosed | NO |
| ANFFDVPTFCAN | 1411071 | Cell/Virus Enclosed | NO |
| ANFILHDVNNGL | 1411074 | Ectodomain | YES |
| ANFILHDVNSGT | 1411076 | Ectodomain | YES |
| ANFLDDNVLPETYVAL | 1645364 | Ectodomain | YES |
| ANFNETKGPLCVDTSH | 1645367 | Ectodomain | YES |
| ANFNETNGPLCV | 1411081 | Ectodomain | YES |
| ANFNILFSMILP | 1411082 | Cell/Virus Enclosed | NO |
| ANFNSQDVLLAP | 1411083 | Cell/Virus Enclosed | NO |
| ANFNVLFSTVFP | 1411084 | Cell/Virus Enclosed | NO |
| ANFQHDNVLPET | 1411085 | Ectodomain | YES |
| ANFRDLALKGSK | 1411090 | Ectodomain | YES |
| ANFTDLALKGST | 1411093 | Ectodomain | YES |
| ANGDSEAVLKKL | 1411102 | Cell/Virus Enclosed | NO |
| ANGFFYIDVGKH | 1411105 | Ectodomain | YES |
| ANGFTAMNGEDA | 1411109 | Cell/Virus Enclosed | NO |
| ANGFTVMNGEDA | 1411110 | Cell/Virus Enclosed | NO |
| ANGFVLRIGAAA | 1411111 | Ectodomain | YES |
| ANGLFYIDVGNH | 1411122 | Ectodomain | YES |
| ANGLNAPNGPFDALFL | 1645391 | Cell/Virus Enclosed | NO |
| ANGNKIVDKEVKDMQF | 1645395 | Cell/Virus Enclosed | NO |
| ANGQVFGLYKNT | 1411127 | Cell/Virus Enclosed | NO |
| ANGRLVGTTLQG | 1411129 | Cell/Virus Enclosed | NO |
| ANGSLDGTTLQG | 1411131 | Cell/Virus Enclosed | NO |
| ANGSLVGTTLQG | 1411134 | Cell/Virus Enclosed | NO |
| ANGTHTGSAFDGTMYG | 1645401 | Cell/Virus Enclosed | NO |
| ANGVKAKGYPQFAELV | 1645402 | Capsid | UN |
| ANHQADTSTPSDVSSR | 1645405 | Capsid | UN |
| ANHYSPCVSIVP | 1411149 | Ectodomain | YES |
| ANICHSNILSLN | 1411150 | Ectodomain | YES |
| ANICNSNILSLN | 1411151 | Ectodomain | YES |
| ANIFIGDKVGHYVHVK | 1645409 | Cell/Virus Enclosed | NO |
| ANIFTPLVQPVG | 1411155 | Cell/Virus Enclosed | NO |
| ANIGASHTGITG | 1411156 | Cell/Virus Enclosed | NO |
| ANIIYCNSVINRLRCD | 1645414 | Ectodomain | YES |
| ANIKSDILSLNN | 1411164 | Ectodomain | YES |
| ANINQDSYGGAS | 1411166 | Cell/Virus Enclosed | NO |
| ANISMLQLGAPD | 1411171 | Ectodomain | YES |
| ANISMLQLGVPD | 1411172 | Ectodomain | YES |
| ANISTLQLGVPD | 1411173 | Ectodomain | YES |
| ANIVFPQGGYVA | 1411174 | Cell/Virus Enclosed | NO |
| ANIVHEPTTPIS | 1411175 | Cell/Virus Enclosed | NO |
| ANIVYEPTTPISSALI | 1645419 | Cell/Virus Enclosed | NO |
| ANKEGIIWVATEGALN | 1645421 | Cell/Virus Enclosed | NO |
| ANKEGIVWVATEGALN | 3325 | Capsid | UN |
| ANKFEQFDQWYESTFG | 1645423 | Cell/Virus Enclosed | NO |
| ANKFFQFDQWYESTFG | 1645424 | Cell/Virus Enclosed | NO |
| ANKFFQFNQWYE | 1411192 | Cell/Virus Enclosed | NO |
| ANKFNQALGAMQTGFT | 1645425 | Ectodomain | YES |
| ANKHAGFDAWFS | 1411199 | Cell/Virus Enclosed | NO |
| ANKHAGFDSWFS | 1411200 | Cell/Virus Enclosed | NO |
| ANKIVGGAHTWF | 1411203 | Cell/Virus Enclosed | NO |
| ANKIVGGAPTWFNALR | 1645430 | Cell/Virus Enclosed | NO |
| ANKLILINDVAFVSTF | 1645431 | Cell/Virus Enclosed | NO |
| ANKQGIVWVATE | 1411207 | Capsid | UN |
| ANKSVGIISMPLGYVS | 1645432 | Cell/Virus Enclosed | NO |
| ANLANLNSHNIG | 1411218 | Ectodomain | YES |
| ANLANLNSHTIG | 1411219 | Ectodomain | YES |
| ANLAYINSYTIG | 1411221 | Ectodomain | YES |
| ANLAYSNSYTIG | 1411223 | Ectodomain | YES |
| ANLESAKRVLNV | 1411226 | Cell/Virus Enclosed | NO |
| ANLFPPWSAAAG | 1411232 | Ectodomain | YES |
| ANLGERVRQALL | 1411237 | Cell/Virus Enclosed | NO |
| ANLGERVRQSLL | 1411238 | Cell/Virus Enclosed | NO |
| ANLILHDVNSGL | 1411239 | Ectodomain | YES |
| ANLKLPPHDVTVLRDN | 1645440 | Cell/Virus Enclosed | NO |
| ANLKLPSHDVTV | 1411241 | Cell/Virus Enclosed | NO |
| ANLNFVALIPDY | 1411247 | Cell/Virus Enclosed | NO |
| ANLNILFSMVLP | 1411248 | Cell/Virus Enclosed | NO |
| ANLRQDDFVRNL | 1411252 | Ectodomain | YES |
| ANLRQDDSVRNLFASV | 1645444 | Ectodomain | YES |
| ANLRVVGHAMQGTLLK | 1645445 | Cell/Virus Enclosed | NO |
| ANLSAPTLVPQENYSS | 1645446 | Cell/Virus Enclosed | NO |
| ANLSIPSNWTIS | 1411258 | Ectodomain | YES |

| | | | |
|---|---|---|---|
| ANLSMLQLSVPD | 1411261 | Ectodomain | YES |
| ANLVLAKLGKDSKPQQ | 1645449 | Capsid | UN |
| ANLVNHKSHNIG | 1411264 | Ectodomain | YES |
| ANLVNLNSHNIG | 1411265 | Ectodomain | YES |
| ANLVNYKSHNIG | 1411267 | Ectodomain | YES |
| ANMDQGSFGGAS | 1411270 | Cell/Virus Enclosed | NO |
| ANMSLAECKRALGLSI | 1645460 | Cell/Virus Enclosed | NO |
| ANMSMAECKAAL | 1411280 | Cell/Virus Enclosed | NO |
| ANNAAIVLQLPQGTTLPKGFY | 1334455 | Capsid | UN |
| ANNLMNGVTLST | 1411288 | Ectodomain | YES |
| ANNLTAPRNSWEAMAL | 1645468 | Cell/Virus Enclosed | NO |
| ANNYSKLFDEGH | 1411297 | Cell/Virus Enclosed | NO |
| ANNYSKLFDEGYTVVI | 1645473 | Cell/Virus Enclosed | NO |
| ANNYSKLSDEGY | 1411299 | Cell/Virus Enclosed | NO |
| ANPAAVDGKCLT | 1411301 | Cell/Virus Enclosed | NO |
| ANPAAVDSKCLA | 1411302 | Cell/Virus Enclosed | NO |
| ANPAAVDSKCLT | 1411303 | Cell/Virus Enclosed | NO |
| ANPATDPIYDEP | 1411305 | Secreted | YES |
| ANPLWHKAAFIS | 1411308 | Cell/Virus Enclosed | NO |
| ANPMVIVNAANI | 1411309 | Cell/Virus Enclosed | NO |
| ANPNSVLTTAVY | 1411310 | Cell/Virus Enclosed | NO |
| ANPSTPAYTFTTVKPG | 1645481 | Cell/Virus Enclosed | NO |
| ANPSTPVYTFTT | 1411313 | Cell/Virus Enclosed | NO |
| ANPTNCIAPVNGYFIK | 1645483 | Ectodomain | YES |
| ANPTNCITPVNG | 1411317 | Ectodomain | YES |
| ANPTVIVNAANI | 1411318 | Cell/Virus Enclosed | NO |
| ANQERYVKITGL | 1411322 | Cell/Virus Enclosed | NO |
| ANQFNKAISQIQESLT | 1645487 | Ectodomain | YES |
| ANQFTEFVGTQSVDML | 1645489 | Cell/Virus Enclosed | NO |
| ANQFTEFVSTQS | 1411330 | Cell/Virus Enclosed | NO |
| ANQNTSVYSANG | 1411332 | Ectodomain | YES |
| ANQQQLKQLEKACNIA | 1645491 | Cell/Virus Enclosed | NO |
| ANQSTSSYPANG | 1411333 | Ectodomain | YES |
| ANQSTSTYLANG | 1411334 | Ectodomain | YES |
| ANQSTSTYPANG | 1411335 | Ectodomain | YES |
| ANQSTSTYSANG | 1411336 | Ectodomain | YES |
| ANQSTSVYSAKG | 1411337 | Ectodomain | YES |
| ANQYSPCVPIVP | 1411339 | Ectodomain | YES |
| ANQYSPCVSIAP | 1411340 | Ectodomain | YES |
| ANQYSPCVSILP | 1411341 | Ectodomain | YES |
| ANQYSPCVSTVP | 1411343 | Ectodomain | YES |
| ANRCNEVSATVL | 1411349 | Cell/Virus Enclosed | NO |
| ANRNRFLYIIKLIFL | 2000984 | Undetermined | UN |
| ANRSAMCNWCLV | 1411356 | Cell/Virus Enclosed | NO |
| ANRYNEVANAVMQNNE | 1645499 | Cell/Virus Enclosed | NO |
| ANRYNEVPATVL | 1411358 | Cell/Virus Enclosed | NO |
| ANSAAKLQNNEL | 1411360 | Cell/Virus Enclosed | NO |
| ANSAVLSFCAFA | 1411362 | Cell/Virus Enclosed | NO |
| ANSDNRTPTSGV | 1411363 | Cell/Virus Enclosed | NO |
| ANSEALNNLLQQ | 1411364 | Ectodomain | YES |
| ANSFDAPTFCAN | 1411365 | Cell/Virus Enclosed | NO |
| ANSFVVKGDDVR | 1411369 | Ectodomain | YES |
| ANSGDYYHKVEA | 1411370 | Ectodomain | YES |
| ANSGDYYYKVEA | 1411371 | Ectodomain | YES |
| ANSGGYYYKVEA | 1411372 | Ectodomain | YES |
| ANSGNRAPTSGV | 1411373 | Cell/Virus Enclosed | NO |
| ANSGNRTPTPGV | 1411376 | Cell/Virus Enclosed | NO |
| ANSGNRTPTSGVTPDM | 1645505 | Cell/Virus Enclosed | NO |
| ANSIDVPTFCAN | 1411379 | Cell/Virus Enclosed | NO |
| ANSLDNLKSGVI | 1411387 | Ectodomain | YES |
| ANSNGFIRFPDVISEG | 1645511 | Cell/Virus Enclosed | NO |
| ANSPEPALLFRN | 1411393 | Ectodomain | YES |
| ANSSDPALLFRN | 1411396 | Ectodomain | YES |
| ANSSEPALLFRNIKCN | 1645513 | Ectodomain | YES |
| ANSSEPTLLFRN | 1411397 | Ectodomain | YES |
| ANSSEWAYCVDLHNKI | 1645514 | Cell/Virus Enclosed | NO |
| ANSTGTVIIRPS | 1411400 | Ectodomain | YES |
| ANSTGTVIISPSTSAT | 1645516 | Ectodomain | YES |
| ANSTPAQNAGYWRRQD | 1645517 | Cell/Virus Enclosed | NO |
| ANSTPSQNAGYW | 1411403 | Cell/Virus Enclosed | NO |
| ANSVFNICQAVSANVC | 1645521 | Cell/Virus Enclosed | NO |
| ANSVLNICQAVT | 1411410 | Cell/Virus Enclosed | NO |
| ANTDIILGVCVN | 1411412 | Ectodomain | YES |
| ANTDIKLDVCVN | 1411413 | Ectodomain | YES |
| ANTEIELGVCVN | 1411415 | Ectodomain | YES |
| ANTGSVCYTPIDEIPY | 1645528 | Cell/Virus Enclosed | NO |

| | | | |
|---|---|---|---|
| ANTHLKHGGGIAGAIN | 1645530 | Cell/Virus Enclosed | NO |
| ANTLNIIVPDKSVYDQ | 1645533 | Cell/Virus Enclosed | NO |
| ANTLTIIIPDKQ | 1411422 | Cell/Virus Enclosed | NO |
| ANTNQDSYGGASICLY | 1645535 | Cell/Virus Enclosed | NO |
| ANTTTAPLKLFA | 1411427 | Cell/Virus Enclosed | NO |
| ANVCALISCNGN | 1411434 | Cell/Virus Enclosed | NO |
| ANVCSPMACNGH | 1411436 | Cell/Virus Enclosed | NO |
| ANVGRVIKFETDAYKL | 1645544 | Cell/Virus Enclosed | NO |
| ANVGRWSASINTGNCP | 1645545 | Ectodomain | YES |
| ANVISFGDYNLS | 1411441 | Ectodomain | YES |
| ANVNWADDRAAR | 1411446 | Cell/Virus Enclosed | NO |
| ANVPAIDVLKKLLSSL | 1645551 | Cell/Virus Enclosed | NO |
| ANVRSLTACNGH | 1411450 | Cell/Virus Enclosed | NO |
| ANVSALIGANGN | 1411451 | Cell/Virus Enclosed | NO |
| ANVSALMGANGNKIVD | 1645553 | Cell/Virus Enclosed | NO |
| ANVSVSRFDPST | 1411452 | Ectodomain | YES |
| ANVSVSRFNPSI | 1411453 | Ectodomain | YES |
| ANVSVSRFNPST | 1411454 | Ectodomain | YES |
| ANVSYVYNGSAQ | 1411455 | Ectodomain | YES |
| ANVTQAFGRRGP | 1411456 | Capsid | UN |
| ANVTSFGDYNLS | 1411457 | Ectodomain | YES |
| ANVYRRKPDLPN | 1411460 | Ectodomain | YES |
| ANWLAPTNAYMRTTHT | 1645563 | Cell/Virus Enclosed | NO |
| ANWLAPTNAYTR | 1411467 | Cell/Virus Enclosed | NO |
| ANYFQDVKQFAN | 1411473 | Ectodomain | YES |
| ANYLGKLKEQID | 1411485 | Cell/Virus Enclosed | NO |
| ANYVGKYDINHGDSGF | 1645580 | Undetermined | UN |
| APAAPRAVSFAD | 1411496 | Capsid | UN |
| APAHDTPPVPDVDSRGAILRRQYNL | 3476 | Undetermined | UN |
| APAHVSTIGVCT | 1411499 | Cell/Virus Enclosed | NO |
| APCAVSFADNND | 1411520 | Cell/Virus Enclosed | NO |
| APCIKVAAMQSK | 1411522 | Cell/Virus Enclosed | NO |
| APDDASRLLHTV | 1411527 | Cell/Virus Enclosed | NO |
| APDFSLDYINVT | 1411530 | Ectodomain | YES |
| APDLSFDYINVT | 1411535 | Ectodomain | YES |
| APDLSLDYINAT | 1411536 | Ectodomain | YES |
| APDNPSAIVTGL | 1411538 | Ectodomain | YES |
| APDVMLNISTPN | 1411541 | Ectodomain | YES |
| APEDDDFLEGSD | 1411546 | Cell/Virus Enclosed | NO |
| APEPQRGRQGRI | 1411555 | Cell/Virus Enclosed | NO |
| APEYDDFLEESD | 1411558 | Cell/Virus Enclosed | NO |
| APFIYLNNSIPN | 1411560 | Ectodomain | YES |
| APGCDVNDVTKLYLGG | 1645606 | Cell/Virus Enclosed | NO |
| APGETGVIADYN | 1411577 | Ectodomain | YES |
| APGGDLLYLDLL | 1411578 | Cell/Virus Enclosed | NO |
| APGVPATQAKGY | 1411590 | Cell/Virus Enclosed | NO |
| APHDFFIFDVDK | 1411592 | Cell/Virus Enclosed | NO |
| APHGPFDALWLS | 1411596 | Cell/Virus Enclosed | NO |
| APIDIDAFPVKPICSS | 1645630 | Cell/Virus Enclosed | NO |
| APIDIDAFPVRP | 1411611 | Cell/Virus Enclosed | NO |
| APIELENKFTPLSVES | 1645631 | Cell/Virus Enclosed | NO |
| APKEESTDQMSEPPKE | 1645634 | Cell/Virus Enclosed | NO |
| APKNFCPCKLIS | 1411638 | Ectodomain | YES |
| APKNFCPCKLNS | 1411640 | Ectodomain | YES |
| APKNFCPCKSND | 1411641 | Ectodomain | YES |
| APKNFCPCKSNG | 1411642 | Ectodomain | YES |
| APKNFCPCSSCF | 1411644 | Ectodomain | YES |
| APKNFCPCSSCH | 1411645 | Ectodomain | YES |
| APKNFCPCSSCP | 1411646 | Ectodomain | YES |
| APKNFCPCSSCS | 1411647 | Ectodomain | YES |
| APKQEEPPKTEH | 1411649 | Cell/Virus Enclosed | NO |
| APKQGYFIKHND | 1411651 | Ectodomain | YES |
| APKRIRLPHIREDDQPSSHQ | 741216 | Underneath Membrane | NO |
| APKSGYFVNVNNTWMY | 1645644 | Ectodomain | YES |
| APKSRVQQNKIR | 1411655 | Cell/Virus Enclosed | NO |
| APLRAPTMANQEKYST | 1645657 | Cell/Virus Enclosed | NO |
| APLSPLLPLQDGTNTHIMATEASNY | 3648 | Undetermined | UN |
| APLTAVAPAHDTPPVPDVDSRGAIL | 3651 | Undetermined | UN |
| APLTNCVKMLTP | 1411676 | Cell/Virus Enclosed | NO |
| APLTRGSCRKRNRSPER | 3652 | Cell/Virus Enclosed | NO |
| APLVYLNHSVPKLSDF | 1645662 | Ectodomain | YES |
| APNAKEEILGAV | 1411681 | Cell/Virus Enclosed | NO |
| APNCYWPLNDYG | 1411684 | Ectodomain | YES |
| APNCYWPLNGYG | 1411685 | Ectodomain | YES |
| APNGLYFMHVGY | 1411691 | Ectodomain | YES |
| APNLTFNSHINA | 1411693 | Ectodomain | YES |

| | | | |
|---|---|---|---|
| APNLTLNLHTIN | 1411695 | Ectodomain | YES |
| APNNTVSWYTGL | 1411699 | Cell/Virus Enclosed | NO |
| APNSRSTSRTSSRASS | 1645671 | Cell/Virus Enclosed | NO |
| APNYTQHSSSMR | 1411705 | Ectodomain | YES |
| APPAEYKLQQGT | 1411708 | Cell/Virus Enclosed | NO |
| APPAGYKLQQGT | 1411710 | Cell/Virus Enclosed | NO |
| APPGEQFTHIVP | 1411715 | Cell/Virus Enclosed | NO |
| APPSEYTLQQGA | 1411719 | Cell/Virus Enclosed | NO |
| APPSQPFPWTHCYQPRL | 3702 | Ectodomain | YES |
| APPTGVYELHGY | 1411720 | Ectodomain | YES |
| APQANFGDYYYK | 1411726 | Ectodomain | YES |
| APQANSGGYYYK | 1411729 | Ectodomain | YES |
| APQATYQNISTN | 1411731 | Ectodomain | YES |
| APQTIQPEVKEVAPVY | 1645683 | Cell/Virus Enclosed | NO |
| APQVTYQNISIN | 1411738 | Ectodomain | YES |
| APQVTYQNISTNLPPP | 1645684 | Ectodomain | YES |
| APRAISFADNND | 1411740 | Cell/Virus Enclosed | NO |
| APRAVSFADNNDITNT | 1645685 | Cell/Virus Enclosed | NO |
| APRAVSFAYNND | 1411744 | Capsid | UN |
| APRITFGGPIDS | 1411750 | Capsid | UN |
| APRITFGGPSDSTDNN | 1645686 | Cell/Virus Enclosed | NO |
| APRITFGGPTDSTDNNQN | 3779 | Capsid | UN |
| APRLGPSSVYTPGASRPTPP | 741218 | Ectodomain | YES |
| APRSVRSVPGEM | 1411758 | Ectodomain | YES |
| APRVLISKGVMEPIDY | 1645692 | Cell/Virus Enclosed | NO |
| APRVMITKGVMEPVDY | 1645695 | Cell/Virus Enclosed | NO |
| APRWYFYYTGTGPEAA | 1645696 | Cell/Virus Enclosed | NO |
| APSDVYQQLAGVKLQS | 1645698 | Cell/Virus Enclosed | NO |
| APSFLAVDDKYKATGD | 1645699 | Cell/Virus Enclosed | NO |
| APSFLAVDDKYKVNEN | 1645700 | Cell/Virus Enclosed | NO |
| APSGTWLTYHGA | 1411769 | Capsid | UN |
| APSRSQSRSQSRGRGE | 1645701 | Capsid | UN |
| APSRSSETKEQK | 1411774 | Cell/Virus Enclosed | NO |
| APSTFGTRNPNNDSAI | 1645702 | Cell/Virus Enclosed | NO |
| APSTVWEDGDYY | 1411778 | Ectodomain | YES |
| APTIANQERYVK | 1411793 | Cell/Virus Enclosed | NO |
| APTLIPQENYSS | 1411798 | Cell/Virus Enclosed | NO |
| APTLVPQENYASIRFS | 1645715 | Cell/Virus Enclosed | NO |
| APTLVPQENYSSIRFA | 1645716 | Cell/Virus Enclosed | NO |
| APTNAYMRTTHTDIGV | 1645718 | Cell/Virus Enclosed | NO |
| APTNAYTRTTHT | 1411805 | Cell/Virus Enclosed | NO |
| APTNFCPCKLDG | 1411806 | Ectodomain | YES |
| APTNVYMRTTHT | 1411808 | Cell/Virus Enclosed | NO |
| APTPGAFFFGSKLDLV | 1645719 | Capsid | UN |
| APTSGVIPDMAD | 1411810 | Cell/Virus Enclosed | NO |
| APTYVSVDDKYKTSDE | 1645722 | Cell/Virus Enclosed | NO |
| APVEDIAQVVIA | 1411819 | Cell/Virus Enclosed | NO |
| APVEFYEDRILDFKVL | 1645723 | Cell/Virus Enclosed | NO |
| APVNTVKPKPVINQLD | 1645726 | Cell/Virus Enclosed | NO |
| APVSPAFHRIES | 1411835 | Cell/Virus Enclosed | NO |
| APVVNIQKEIDR | 1411839 | Ectodomain | YES |
| APVYEADTEQTQNVTV | 1645730 | Cell/Virus Enclosed | NO |
| APVYIWTADEAIKFLK | 1645731 | Underneath Membrane | NO |
| APVYTWTADEAI | 1411841 | Ectodomain | YES |
| APYIVGDVISSG | 1411847 | Cell/Virus Enclosed | NO |
| APYVMLNTSIPNLPDF | 1645739 | Ectodomain | YES |
| APYVMLNTSSPN | 1411857 | Ectodomain | YES |
| APYVMLNTSTSN | 1411858 | Ectodomain | YES |
| AQAAGTDTTITL | 1411863 | Cell/Virus Enclosed | NO |
| AQADTAGLFRID | 1411866 | Cell/Virus Enclosed | NO |
| AQADTAGLFTNF | 1411867 | Cell/Virus Enclosed | NO |
| AQADTAGLYTNFRIDV | 1645744 | Cell/Virus Enclosed | NO |
| AQADTAGLYTNS | 1411869 | Cell/Virus Enclosed | NO |
| AQAFGLRGPGDL | 1411874 | Cell/Virus Enclosed | NO |
| AQALAKLASELS | 1411879 | Ectodomain | YES |
| AQALSKLASELSNTFG | 1645747 | Ectodomain | YES |
| AQALTEVVYSNG | 1411883 | Cell/Virus Enclosed | NO |
| AQANGFTVMNGE | 1411886 | Cell/Virus Enclosed | NO |
| AQAQIDRLINGR | 1411887 | Ectodomain | YES |
| AQASVNGVTLIG | 1411888 | Cell/Virus Enclosed | NO |
| AQATSFCKSGNL | 1411891 | Ectodomain | YES |
| AQAVRRYSEAAS | 1411894 | Cell/Virus Enclosed | NO |
| AQCDCLCTPDPI | 1411896 | Ectodomain | YES |
| AQCNCLCTPDPI | 1411901 | Ectodomain | YES |
| AQDGCGHISTKP | 1411905 | Ectodomain | YES |
| AQDIWGTSTAAY | 1411910 | Ectodomain | YES |

| | | | |
|---|---|---|---|
| AQDTWGTSAAAY | 1411911 | Ectodomain | YES |
| AQEGICITTNGS | 1411917 | Cell/Virus Enclosed | NO |
| AQEGVCITTNGL | 1411918 | Cell/Virus Enclosed | NO |
| AQEGVCITTNGS | 1411919 | Cell/Virus Enclosed | NO |
| AQFKTFEKVDGVYTNF | 1645770 | Cell/Virus Enclosed | NO |
| AQFNYRRDGQPC | 1411928 | Cell/Virus Enclosed | NO |
| AQGAHLFSSRYV | 1411930 | Ectodomain | YES |
| AQGARGTYRAALNSEK | 1645773 | Cell/Virus Enclosed | NO |
| AQGFQACRFIAF | 1411935 | Cell/Virus Enclosed | NO |
| AQGFQACRFVAF | 1411936 | Cell/Virus Enclosed | NO |
| AQGTNHGYKVVE | 1411947 | Cell/Virus Enclosed | NO |
| AQGTWGTSAAAY | 1411948 | Ectodomain | YES |
| AQHAKPTVIVNA | 1411953 | Cell/Virus Enclosed | NO |
| AQHFSDFGTSFV | 1411961 | Cell/Virus Enclosed | NO |
| AQIDRLINGRLTALNA | 1645789 | Ectodomain | YES |
| AQIDRLINGRLTTLNA | 1645790 | Ectodomain | YES |
| AQIIAIIKDWNFAWSL | 1645791 | Undetermined | UN |
| AQIKALFLDKVDILLT | 1645792 | Cell/Virus Enclosed | NO |
| AQILSEIVMCGG | 1411970 | Cell/Virus Enclosed | NO |
| AQKFNGLTVLPPLLTDD | 3983 | Ectodomain | YES |
| AQKLETDGICPE | 1411979 | Cell/Virus Enclosed | NO |
| AQKLKTDGICPE | 1411982 | Cell/Virus Enclosed | NO |
| AQLNAKLGFNVD | 1411991 | Cell/Virus Enclosed | NO |
| AQLPAPRVLLSKGSLE | 1645805 | Cell/Virus Enclosed | NO |
| AQLPAPRVLLSKGTLE | 1645806 | Cell/Virus Enclosed | NO |
| AQNAGYWRRQDRKINT | 1645811 | Cell/Virus Enclosed | NO |
| AQNANPMVIVNA | 1412009 | Cell/Virus Enclosed | NO |
| AQNANPTVIVNA | 1412010 | Cell/Virus Enclosed | NO |
| AQNPLSELKCTI | 1412015 | Ectodomain | YES |
| AQNTVLNACTPE | 1412017 | Cell/Virus Enclosed | NO |
| AQNTVLNASIPESKPL | 1645817 | Capsid | UN |
| AQNTVLNASISE | 1412019 | Cell/Virus Enclosed | NO |
| AQNTVLNDSIPE | 1412020 | Capsid | UN |
| AQPEGCVLTNTG | 1412023 | Cell/Virus Enclosed | NO |
| AQPKQTATSQQPSGGN | 1645822 | Cell/Virus Enclosed | NO |
| AQPKQTSTSQQP | 1412027 | Cell/Virus Enclosed | NO |
| AQPLNKKVQQLF | 1412030 | Cell/Virus Enclosed | NO |
| AQPLTRKVQQLF | 1412035 | Cell/Virus Enclosed | NO |
| AQPNQTATSQQP | 1412036 | Cell/Virus Enclosed | NO |
| AQPVKIDNASPASTVH | 1645830 | Secreted | YES |
| AQQDQGQYYYRS | 1412047 | Ectodomain | YES |
| AQQEATHIAEDV | 1412048 | Cell/Virus Enclosed | NO |
| AQQEVTHIAEDV | 1412051 | Cell/Virus Enclosed | NO |
| AQQKVNECVESQ | 1412054 | Ectodomain | YES |
| AQQLVRSESAAI | 1412058 | Ectodomain | YES |
| AQRISWLLHKDG | 1412063 | Cell/Virus Enclosed | NO |
| AQRTRTRPSVQP | 1412065 | Capsid | UN |
| AQRVFNNYMPYV | 1412066 | Cell/Virus Enclosed | NO |
| AQSSETKEQKHE | 1412080 | Cell/Virus Enclosed | NO |
| AQTGSSKCGCSV | 1412086 | Cell/Virus Enclosed | NO |
| AQTGSSKCVRSV | 1412088 | Cell/Virus Enclosed | NO |
| AQTSDTAHACNANRFN | 1645851 | Cell/Virus Enclosed | NO |
| AQVAKSHNVSLI | 1412098 | Cell/Virus Enclosed | NO |
| AQVASAMQTMLF | 1412100 | Cell/Virus Enclosed | NO |
| AQVLSETVMCGG | 1412107 | Cell/Virus Enclosed | NO |
| AQVLTEVVYSNGGFYF | 1645862 | Cell/Virus Enclosed | NO |
| AQVQIDRLINGRLTAL | 1645863 | Ectodomain | YES |
| AQVQLEHYSSCV | 1412114 | Cell/Virus Enclosed | NO |
| AQVVIADTLQETPVVP | 1645865 | Cell/Virus Enclosed | NO |
| AQVVQLPIQDYI | 1412119 | Cell/Virus Enclosed | NO |
| AQVVQLPVKDYI | 1412120 | Cell/Virus Enclosed | NO |
| AQVVQLPVKDYVQTVN | 1645867 | Cell/Virus Enclosed | NO |
| AQYIFSTVNALPECNA | 1645871 | Cell/Virus Enclosed | NO |
| AQYVFSTVNALPEVNA | 1645874 | Cell/Virus Enclosed | NO |
| AQYYNGIMVLPGVADA | 1645875 | Ectodomain | YES |
| ARADDEGFITLK | 1412139 | Cell/Virus Enclosed | NO |
| ARADYEGFITLK | 1412140 | Cell/Virus Enclosed | NO |
| ARAEDKKAKIVSAMQT | 1645879 | Cell/Virus Enclosed | NO |
| ARAGDAANFCAL | 1412146 | Cell/Virus Enclosed | NO |
| ARAIDILTEGQLQSLS | 1645886 | Cell/Virus Enclosed | NO |
| ARALQICVALCD | 1412162 | Cell/Virus Enclosed | NO |
| ARAMLDYAKDHN | 1412164 | Cell/Virus Enclosed | NO |
| ARAMLKCVALCDAMVA | 1645890 | Cell/Virus Enclosed | NO |
| ARAQSSETKEQK | 1412166 | Cell/Virus Enclosed | NO |
| ARATIRYRPLVPNAVGGYAISISFW | 4121 | Undetermined | UN |
| ARDALDTILQEA | 1412176 | Cell/Virus Enclosed | NO |

| | | | |
|---|---|---|---|
| ARDGCVPLNIVP | 1412179 | Cell/Virus Enclosed | NO |
| ARDGIRRGDNFH | 1412180 | Cell/Virus Enclosed | NO |
| ARDGVRRGDNFHSVLT | 1645902 | Cell/Virus Enclosed | NO |
| ARDGVVPLSVIPATSA | 1645903 | Cell/Virus Enclosed | NO |
| ARDLICAQKFNA | 1412183 | Ectodomain | YES |
| ARDLICSQYVAG | 1412186 | Ectodomain | YES |
| ARDLSLDYINVT | 1412187 | Ectodomain | YES |
| ARDPIIQEAIPT | 1412191 | Cell/Virus Enclosed | NO |
| ARDPTIQEAIPT | 1412192 | Cell/Virus Enclosed | NO |
| AREANFGDYYYK | 1412194 | Ectodomain | YES |
| ARELICAQYVAG | 1412199 | Ectodomain | YES |
| ARESIFCSNTYG | 1412200 | Cell/Virus Enclosed | NO |
| ARGPLQVGESAL | 1412220 | Cell/Virus Enclosed | NO |
| ARGTYRAALNSEKHQD | 1645921 | Cell/Virus Enclosed | NO |
| ARGWVVGTISST | 1412224 | Cell/Virus Enclosed | NO |
| ARHSDNMTAVAC | 1412230 | Ectodomain | YES |
| ARIAGNGGDAALALLL | 1645927 | Cell/Virus Enclosed | NO |
| ARIRAKHYVYIGDPAQ | 1645934 | Cell/Virus Enclosed | NO |
| ARIVFTACSHAAVDSL | 1645935 | Cell/Virus Enclosed | NO |
| ARIVFVACAHAAVDSL | 1645936 | Cell/Virus Enclosed | NO |
| ARKHDTCCSQSD | 1412239 | Cell/Virus Enclosed | NO |
| ARKHETCCSQID | 1412240 | Cell/Virus Enclosed | NO |
| ARKHETCCSQSD | 1412243 | Cell/Virus Enclosed | NO |
| ARKLERMADLALTNMY | 1645944 | Cell/Virus Enclosed | NO |
| ARKRCHSPTKKL | 1412255 | Cell/Virus Enclosed | NO |
| ARKRKTKKPPKRPEATPPPD | 741230 | Ectodomain | YES |
| ARKRSYSPTKKL | 1412258 | Cell/Virus Enclosed | NO |
| ARKSCSYDSDVD | 1412262 | Cell/Virus Enclosed | NO |
| ARLEIVKLGAEKKDGF | 1645950 | Cell/Virus Enclosed | NO |
| ARLELVKLGAEK | 1412273 | Cell/Virus Enclosed | NO |
| ARLEPCNGTDIDKCVR | 1645951 | Cell/Virus Enclosed | NO |
| ARLKGGLILKFDSGTV | 1645956 | Cell/Virus Enclosed | NO |
| ARLKRFPVNTIVNGVQ | 1645957 | Cell/Virus Enclosed | NO |
| ARLKRVPLQTIINGMH | 1645958 | Cell/Virus Enclosed | NO |
| ARLLGLQTQTVDSAQG | 1645959 | Cell/Virus Enclosed | NO |
| ARLNCGLILKFD | 1412286 | Cell/Virus Enclosed | NO |
| ARLNYVALQTDV | 1412287 | Ectodomain | YES |
| ARLVKGDKGDAE | 1412298 | Cell/Virus Enclosed | NO |
| ARLYYEALSFEEQDEI | 1645969 | Cell/Virus Enclosed | NO |
| ARMAGNGGDAALALLL | 1377826 | Cell/Virus Enclosed | NO |
| ARMASGSGETAL | 1412306 | Cell/Virus Enclosed | NO |
| ARMTYTCQCGSE | 1412308 | Cell/Virus Enclosed | NO |
| ARNGVVPLSVIPATSA | 1645981 | Cell/Virus Enclosed | NO |
| ARNHGTCCTTRD | 1412313 | Cell/Virus Enclosed | NO |
| ARNPIDLPPSHATTYL | 1645983 | Cell/Virus Enclosed | NO |
| ARNPVDLPPSHA | 1412320 | Cell/Virus Enclosed | NO |
| ARPKQRRPQGLPNNTA | 1645984 | Cell/Virus Enclosed | NO |
| ARQAYEDAIANGSSSQ | 1645986 | Cell/Virus Enclosed | NO |
| ARQEYENAVANS | 1412328 | Cell/Virus Enclosed | NO |
| ARQFDNMTAVAC | 1412329 | Ectodomain | YES |
| ARQSDNMTAVACQPPY | 1645994 | Ectodomain | YES |
| ARRHETCCSQSD | 1412340 | Cell/Virus Enclosed | NO |
| ARRNIGFTLEYW | 1412342 | Ectodomain | YES |
| ARRYETCCSQSD | 1412347 | Cell/Virus Enclosed | NO |
| ARSEDKRAQVAS | 1412352 | Cell/Virus Enclosed | NO |
| ARSLDWTKVTLDGRPLSTIQQYSKT | 4260 | Undetermined | UN |
| ARSMLGLITQTV | 1412357 | Cell/Virus Enclosed | NO |
| ARSQSSETKEQK | 1412361 | Cell/Virus Enclosed | NO |
| ARSVVSRIGDSI | 1412367 | Cell/Virus Enclosed | NO |
| ARTFVQSNVDVL | 1412375 | Cell/Virus Enclosed | NO |
| ARTFWAWNPEVN | 1412378 | Underneath Membrane | NO |
| ARTRSMWSFNSE | 1412387 | Underneath Membrane | NO |
| ARTVYDDAARRV | 1412394 | Cell/Virus Enclosed | NO |
| ARVECYDRFKVNETNS | 1646025 | Cell/Virus Enclosed | NO |
| ARVECYSGFKPNNNSA | 1646026 | Cell/Virus Enclosed | NO |
| ARVECYSGFKPNNTSA | 1646027 | Cell/Virus Enclosed | NO |
| ARVEHPDVDGICKLRG | 1646029 | Cell/Virus Enclosed | NO |
| ARVSPGLCIAGDRGIA | 1646031 | Ectodomain | YES |
| ARVVYTAASHAAVDAL | 1646034 | Cell/Virus Enclosed | NO |
| ARVVYTACSHAAVDAL | 1646035 | Cell/Virus Enclosed | NO |
| ARWSASIDTGNC | 1412413 | Ectodomain | YES |
| ASADIGSGHTGI | 1412426 | Cell/Virus Enclosed | NO |
| ASADWKPGQAMP | 1412427 | Cell/Virus Enclosed | NO |
| ASADWKPGYAMP | 1412428 | Cell/Virus Enclosed | NO |
| ASAFFGMSRIGMEVTPSGTW | 1392055 | Capsid | UN |
| ASALKQLKVTTGELVR | 1646049 | Cell/Virus Enclosed | NO |

| | | | |
|---|---|---|---|
| ASAMQTMLFTML | 1412440 | Cell/Virus Enclosed | NO |
| ASANIGASHTGI | 1412441 | Cell/Virus Enclosed | NO |
| ASAPATGWLLYQ | 1412444 | Cell/Virus Enclosed | NO |
| ASAPVEPSTVDV | 1412445 | Cell/Virus Enclosed | NO |
| ASARDLICSQYV | 1412449 | Ectodomain | YES |
| ASARELICAQYV | 1412450 | Ectodomain | YES |
| ASARNLICAQYV | 1412452 | Underneath Membrane | NO |
| ASATDFWTVAFA | 1412454 | Ectodomain | YES |
| ASATGPTYIRVADSELYADW | 741238 | Underneath Membrane | NO |
| ASATIQEIVSDREVIL | 1646061 | Cell/Virus Enclosed | NO |
| ASATIQEIVSERELIL | 1646062 | Cell/Virus Enclosed | NO |
| ASAVAQAFQSVA | 1412457 | Cell/Virus Enclosed | NO |
| ASAVAQAFQSVV | 1412458 | Cell/Virus Enclosed | NO |
| ASAVQTCDLTVG | 1412460 | Ectodomain | YES |
| ASAYGLCDAANP | 1412462 | Ectodomain | YES |
| ASAYRANFFDVP | 1412463 | Cell/Virus Enclosed | NO |
| ASAYRANSFDAP | 1412464 | Cell/Virus Enclosed | NO |
| ASAYRANSFDVPTFCA | 1646065 | Cell/Virus Enclosed | NO |
| ASAYRANSIDVP | 1412466 | Cell/Virus Enclosed | NO |
| ASCDAIMTRCLA | 1412468 | Cell/Virus Enclosed | NO |
| ASCELYHYQECV | 1412469 | Cell/Virus Enclosed | NO |
| ASCFFTCNSNAN | 1412470 | Ectodomain | YES |
| ASCFSTCNSNAN | 1412472 | Ectodomain | YES |
| ASCPIGTNYRSCESTT | 1646072 | Ectodomain | YES |
| ASDDLYVKRYFKGCET | 1646074 | Cell/Virus Enclosed | NO |
| ASDGYFRLMANP | 1412487 | Cell/Virus Enclosed | NO |
| ASDIFATPSVLQ | 1412488 | Cell/Virus Enclosed | NO |
| ASDISLSVACQL | 1412490 | Underneath Membrane | NO |
| ASDKFYRLSNELAQVL | 1646078 | Cell/Virus Enclosed | NO |
| ASDLSTLAVTAI | 1412496 | Cell/Virus Enclosed | NO |
| ASDNIKDCVKCF | 1412498 | Cell/Virus Enclosed | NO |
| ASDNLKECVKAF | 1412499 | Cell/Virus Enclosed | NO |
| ASDSVQCYTPHI | 1412505 | Cell/Virus Enclosed | NO |
| ASDTSLSDACQL | 1412506 | Underneath Membrane | NO |
| ASDVMFEDAQGL | 1412509 | Cell/Virus Enclosed | NO |
| ASDWKSGYSMPV | 1412512 | Cell/Virus Enclosed | NO |
| ASECFMKSDIFGQDFK | 1646091 | Cell/Virus Enclosed | NO |
| ASECFVKSDIFGSDFK | 1646092 | Cell/Virus Enclosed | NO |
| ASEIYCANIHRF | 1412522 | Cell/Virus Enclosed | NO |
| ASEIYCANIHSF | 1412523 | Cell/Virus Enclosed | NO |
| ASEIYCANLHRF | 1412524 | Cell/Virus Enclosed | NO |
| ASEPQRGRQSRI | 1412528 | Cell/Virus Enclosed | NO |
| ASESKVLLERFLPKCP | 1646100 | Cell/Virus Enclosed | NO |
| ASETQRGRHGRI | 1412531 | Cell/Virus Enclosed | NO |
| ASETQRGRQGRI | 1412532 | Capsid | UN |
| ASEVVVNNLDKSAGHP | 1646101 | Cell/Virus Enclosed | NO |
| ASFGNRSGNGIL | 1412542 | Cell/Virus Enclosed | NO |
| ASFIKLFSLFRH | 1412547 | Cell/Virus Enclosed | NO |
| ASFKKGAKILHK | 1412548 | Cell/Virus Enclosed | NO |
| ASFKKGAKLLHK | 1412549 | Cell/Virus Enclosed | NO |
| ASFKVIDNGVVRDISV | 1646112 | Cell/Virus Enclosed | NO |
| ASFSASTSAFID | 1412557 | Cell/Virus Enclosed | NO |
| ASFSSVNDAITQ | 1412559 | Ectodomain | YES |
| ASFVEYEVAKKNLADA | 1646125 | Cell/Virus Enclosed | NO |
| ASFVEYEVAKKNLDEA | 1646126 | Cell/Virus Enclosed | NO |
| ASGPTVAMTEQL | 1412582 | Ectodomain | YES |
| ASGSGETALALL | 1412585 | Cell/Virus Enclosed | NO |
| ASGTHTGSAFDG | 1412591 | Cell/Virus Enclosed | NO |
| ASGYGGFGCKDS | 1412596 | Cell/Virus Enclosed | NO |
| ASGYVAASAVNV | 1412599 | Cell/Virus Enclosed | NO |
| ASHTGITGDNVE | 1412607 | Cell/Virus Enclosed | NO |
| ASHYVVTDIKQV | 1412609 | Cell/Virus Enclosed | NO |
| ASIAFNHPIQID | 1412610 | Ectodomain | YES |
| ASIAFNHPIQVD | 1412611 | Ectodomain | YES |
| ASIEEVCDDYAK | 1412615 | Cell/Virus Enclosed | NO |
| ASIGDIIQRLDF | 1412617 | Ectodomain | YES |
| ASIGDIIQRLDG | 1412618 | Ectodomain | YES |
| ASIGDIIQRLDP | 1412619 | Ectodomain | YES |
| ASIGDIIRRLDV | 1412621 | Ectodomain | YES |
| ASIPESKPLADDDSAI | 1646167 | Capsid | UN |
| ASIPLCILPYTA | 1412635 | Cell/Virus Enclosed | NO |
| ASIPLCIVPYTV | 1412637 | Cell/Virus Enclosed | NO |
| ASIQADVNTPAD | 1412638 | Cell/Virus Enclosed | NO |
| ASIQHLHEGFGGKNIL | 1646168 | Cell/Virus Enclosed | NO |
| ASISESKPLADD | 1412642 | Cell/Virus Enclosed | NO |
| ASIVDIIQRLDV | 1412647 | Ectodomain | YES |

| | | | |
|---|---|---|---|
| ASKGAVQKESDEYILA | 1646175 | Cell/Virus Enclosed | NO |
| ASKLTIVSPDLESYSK | 1646184 | Cell/Virus Enclosed | NO |
| ASLDVKPYGYPNDFVG | 1646189 | Cell/Virus Enclosed | NO |
| ASLFKPVQLAPV | 1412678 | Cell/Virus Enclosed | NO |
| ASLFKPVQLVPVSPAF | 1646191 | Cell/Virus Enclosed | NO |
| ASLFPLWTAAAG | 1412680 | Ectodomain | YES |
| ASLFPPWSAAAG | 1412681 | Ectodomain | YES |
| ASLFPPWTAAEG | 1412683 | Ectodomain | YES |
| ASLNSLKEYFNL | 1412699 | Ectodomain | YES |
| ASLNSLTYFNKP | 1412700 | Cell/Virus Enclosed | NO |
| ASLNSLTYFNRS | 1412702 | Cell/Virus Enclosed | NO |
| ASLNVKPYGYPN | 1412703 | Cell/Virus Enclosed | NO |
| ASLPTTIAKNSV | 1412708 | Cell/Virus Enclosed | NO |
| ASLPYGANKQGI | 1412711 | Capsid | UN |
| ASLQEIISRLDA | 1412712 | Ectodomain | YES |
| ASLQEILSRLDALEAE | 1646211 | Ectodomain | YES |
| ASLRQIFDVAPIELEN | 1646213 | Cell/Virus Enclosed | NO |
| ASLRQIFDVPPI | 1412722 | Cell/Virus Enclosed | NO |
| ASLSAPTLVPQENYAS | 1646214 | Cell/Virus Enclosed | NO |
| ASLVLAKLGKDATKPQ | 1646215 | Cell/Virus Enclosed | NO |
| ASMEVKTIKVFT | 1412734 | Cell/Virus Enclosed | NO |
| ASMYENYTILQA | 1412742 | Cell/Virus Enclosed | NO |
| ASNAAPSASPRNASAPRTTP | 741243 | Ectodomain | YES |
| ASNDAIMTRCLA | 1412745 | Cell/Virus Enclosed | NO |
| ASNPTNCIAPVN | 1412760 | Ectodomain | YES |
| ASNQADVNIPAD | 1412761 | Cell/Virus Enclosed | NO |
| ASNQADVNTPADIVDR | 1646233 | Cell/Virus Enclosed | NO |
| ASNQADVNTSAD | 1412763 | Cell/Virus Enclosed | NO |
| ASNSKLWQYCST | 1412765 | Cell/Virus Enclosed | NO |
| ASNSRPGSRSQSRGPN | 1646238 | Capsid | UN |
| ASNSRSTSRTSS | 1412767 | Cell/Virus Enclosed | NO |
| ASNSSVLSLVNF | 1412768 | Cell/Virus Enclosed | NO |
| ASNVKKVTDVTGKLSD | 1646243 | Cell/Virus Enclosed | NO |
| ASNVPSIVSEKI | 1412769 | Cell/Virus Enclosed | NO |
| ASPAAPCAVSFA | 1412772 | Cell/Virus Enclosed | NO |
| ASPAAPRAISFA | 1412773 | Cell/Virus Enclosed | NO |
| ASPAAPRAVFFA | 1412774 | Cell/Virus Enclosed | NO |
| ASPAAPRAVSFA | 1412775 | Cell/Virus Enclosed | NO |
| ASPASTAHATAT | 1412776 | Secreted | YES |
| ASPASTVHAIAT | 1412777 | Secreted | YES |
| ASPASTVHATAM | 1412778 | Secreted | YES |
| ASPGPSGVGAPG | 1412782 | Cell/Virus Enclosed | NO |
| ASPMKSDEGGLCPSTG | 1646246 | Cell/Virus Enclosed | NO |
| ASPQECNNMHLS | 1412786 | Cell/Virus Enclosed | NO |
| ASPQLEGFSAESRKL | 504100 | Cell/Virus Enclosed | NO |
| ASPYDLEVLLQDALQS | 1646251 | Cell/Virus Enclosed | NO |
| ASQEGVLSEREKLLHLI | 1412797 | Ectodomain | YES |
| ASQIISGTVAAV | 1412804 | Undetermined | UN |
| ASQLGNCVEYHH | 1412807 | Ectodomain | YES |
| ASQLGNCVEYSL | 1412808 | Ectodomain | YES |
| ASQRVAGDSGFAAYSR | 1377882 | Underneath Membrane | NO |
| ASQRVGNDSGFA | 1412812 | Underneath Membrane | NO |
| ASQVIVNNYDKSAGYP | 1646258 | Cell/Virus Enclosed | NO |
| ASQVIVNNYEKS | 1412815 | Cell/Virus Enclosed | NO |
| ASRAIEKVNECV | 1412816 | Ectodomain | YES |
| ASRELSVTFFPD | 1412821 | Cell/Virus Enclosed | NO |
| ASRELVVSYVNVNMG | 1646263 | Capsid | UN |
| ASRFLGLQIQTVDSSQ | 1646265 | Cell/Virus Enclosed | NO |
| ASRFPNVYAWER | 1412823 | Ectodomain | YES |
| ASRMYCTFAIEDVEDV | 1646267 | Cell/Virus Enclosed | NO |
| ASRNSSRSSSQG | 1412834 | Cell/Virus Enclosed | NO |
| ASRNSSRSSSRG | 1412835 | Cell/Virus Enclosed | NO |
| ASSAAPRAVSFA | 1412847 | Cell/Virus Enclosed | NO |
| ASSAGSRSKANS | 1412849 | Cell/Virus Enclosed | NO |
| ASSDVIHYNINF | 1412859 | Ectodomain | YES |
| ASSDVIRYNINF | 1412860 | Ectodomain | YES |
| ASSEFAVLYQDV | 1412861 | Ectodomain | YES |
| ASSELYHYQECV | 1412862 | Cell/Virus Enclosed | NO |
| ASSEVAVLYQGV | 1412864 | Ectodomain | YES |
| ASSFVGMPSFVAYETA | 1646278 | Cell/Virus Enclosed | NO |
| ASSGNRAPTSGV | 1412867 | Cell/Virus Enclosed | NO |
| ASSGNRASTSGV | 1412868 | Cell/Virus Enclosed | NO |
| ASSGNRPGNGIL | 1412869 | Capsid | UN |
| ASSGNRSGNGILKWAD | 1646280 | Cell/Virus Enclosed | NO |
| ASSGNRSSNGIL | 1412871 | Cell/Virus Enclosed | NO |
| ASSGNRSVNGIL | 1412872 | Cell/Virus Enclosed | NO |

| | | | |
|---|---|---|---|
| ASSGYRSGNGIL | 1412875 | Cell/Virus Enclosed | NO |
| ASSIRGCCESTKLPYY | 1646283 | Underneath Membrane | NO |
| ASSKPLYVPEHCQNYS | 1646284 | Cell/Virus Enclosed | NO |
| ASSLSRNSSRSSSQGS | 1646286 | Cell/Virus Enclosed | NO |
| ASSNQSPQGNLG | 1412885 | Cell/Virus Enclosed | NO |
| ASSPALVDKRTVCFSV | 1646288 | Cell/Virus Enclosed | NO |
| ASSQADVNTPAD | 1412889 | Capsid | UN |
| ASSQSAAKILAP | 1412890 | Cell/Virus Enclosed | NO |
| ASSQSAAKILAR | 1412891 | Cell/Virus Enclosed | NO |
| ASSQSSAKSLAR | 1412894 | Cell/Virus Enclosed | NO |
| ASSQSSAKSLSR | 1412895 | Cell/Virus Enclosed | NO |
| ASSQTSAKSLARSQSS | 1646291 | Capsid | UN |
| ASSRSSSRSRGNSRNST | 4782 | Capsid | UN |
| ASSRSSTRNNSRDSSR | 1646295 | Capsid | UN |
| ASSSEAFLIGVN | 1412904 | Cell/Virus Enclosed | NO |
| ASSSLRTFVVDK | 1412906 | Cell/Virus Enclosed | NO |
| ASSSYDALLHRNSLDP | 1646303 | Cell/Virus Enclosed | NO |
| ASSTFVEEEHVN | 1412907 | Cell/Virus Enclosed | NO |
| ASSTGSRSRANS | 1412908 | Cell/Virus Enclosed | NO |
| ASTAHATATIPL | 1412910 | Secreted | YES |
| ASTCAFVDTVKG | 1412912 | Cell/Virus Enclosed | NO |
| ASTDFHQLECIV | 1412913 | Cell/Virus Enclosed | NO |
| ASTDFHQLEFVR | 1412914 | Cell/Virus Enclosed | NO |
| ASTDFHQLEYDS | 1412915 | Cell/Virus Enclosed | NO |
| ASTDSIVFTYIP | 1412916 | Cell/Virus Enclosed | NO |
| ASTEELPDEFVV | 1412918 | Cell/Virus Enclosed | NO |
| ASTFLFSCIITA | 1412924 | Cell/Virus Enclosed | NO |
| ASTFLNGVCGVS | 1412926 | Cell/Virus Enclosed | NO |
| ASTGVIDISAGM | 1412928 | Cell/Virus Enclosed | NO |
| ASTNRQSGRQPTPISPPLRDSHPQ | 4802 | Underneath Membrane | NO |
| ASTPESKPLADD | 1412933 | Cell/Virus Enclosed | NO |
| ASTPVEPPTVDV | 1412934 | Cell/Virus Enclosed | NO |
| ASTPVEPSTVDV | 1412935 | Cell/Virus Enclosed | NO |
| ASTPVESPTVDV | 1412936 | Cell/Virus Enclosed | NO |
| ASTSAFIDTIKS | 1412941 | Cell/Virus Enclosed | NO |
| ASTSAVKLQNNEIKPS | 1646315 | Cell/Virus Enclosed | NO |
| ASTSGVTPDMAD | 1412945 | Cell/Virus Enclosed | NO |
| ASTSQKSIVAYT | 1412948 | Ectodomain | YES |
| ASTVHAIATIPL | 1412949 | Secreted | YES |
| ASTVHATAMIPL | 1412950 | Secreted | YES |
| ASTWFNALRDFT | 1412957 | Cell/Virus Enclosed | NO |
| ASVDARLVPCAN | 1412963 | Cell/Virus Enclosed | NO |
| ASVDARLVPCAS | 1412964 | Cell/Virus Enclosed | NO |
| ASVFKDGCDVGF | 1412966 | Cell/Virus Enclosed | NO |
| ASVGSVPGEMRL | 1412968 | Ectodomain | YES |
| ASVIQDYENVCF | 1412971 | Cell/Virus Enclosed | NO |
| ASVKIYQSSPII | 1412973 | Ectodomain | YES |
| ASVKLPKGVGSA | 1412974 | Cell/Virus Enclosed | NO |
| ASVKNSQSSPII | 1412976 | Ectodomain | YES |
| ASVKRDGVQVGYCVHG | 1646325 | Cell/Virus Enclosed | NO |
| ASVKSSQSSPIIPGFG | 1646326 | Ectodomain | YES |
| ASVKSYQSSPII | 1412979 | Ectodomain | YES |
| ASVLKDGCDVGF | 1412981 | Cell/Virus Enclosed | NO |
| ASVLRSTGQKSI | 1412983 | Ectodomain | YES |
| ASVNGVTLIGES | 1412984 | Cell/Virus Enclosed | NO |
| ASVPEKFLNAFDVFVT | 1646331 | Cell/Virus Enclosed | NO |
| ASVRDLICAQYV | 1412991 | Ectodomain | YES |
| ASVTEDVKFAASTGVI | 1646334 | Cell/Virus Enclosed | NO |
| ASVVAGGIIAIL | 1413003 | Cell/Virus Enclosed | NO |
| ASVYSVLETFQNNVVN | 1646338 | Cell/Virus Enclosed | NO |
| ASWFTALTQHGKEDL | 1310292 | Capsid | UN |
| ASWFTALTQHGKEELRFPRGQ | 4838 | Capsid | UN |
| ASYADVLVNVSQ | 1413021 | Ectodomain | YES |
| ASYDINLNKFNR | 1413024 | Cell/Virus Enclosed | NO |
| ASYDTNLNKFNG | 1413025 | Cell/Virus Enclosed | NO |
| ASYGESLEGVFW | 1413031 | Cell/Virus Enclosed | NO |
| ASYNKYKYYSGSASEA | 1646353 | Cell/Virus Enclosed | NO |
| ASYNRYKYYSGNANEA | 1646354 | Cell/Virus Enclosed | NO |
| ASYRFVTFEPFNVSFV | 1646357 | Ectodomain | YES |
| ASYTANDVEVQSSGMI | 1646358 | Cell/Virus Enclosed | NO |
| ATAEPVRDEFSI | 1413053 | Capsid | UN |
| ATAEPVRDEVSIETDI | 1646361 | Capsid | UN |
| ATAFNNALLSIQNGFS | 1646363 | Ectodomain | YES |
| ATAGWAFGAGAA | 1413056 | Ectodomain | YES |
| ATAGWTFGAGSA | 1413058 | Ectodomain | YES |
| ATAHSSIKQGTL | 1413062 | Cell/Virus Enclosed | NO |

| | | | |
|---|---|---|---|
| ATAKVSPGLCIA | 1413065 | Ectodomain | YES |
| ATALNKIHDVVN | 1413068 | Ectodomain | YES |
| ATALNKIQDVVNQQGN | 1646370 | Ectodomain | YES |
| ATAMIPLQASLP | 1413069 | Secreted | YES |
| ATAVKTGGKSLPAIKL | 1646376 | Cell/Virus Enclosed | NO |
| ATCANDPVGFTL | 1413079 | Cell/Virus Enclosed | NO |
| ATCDWTESGDYTLANT | 1646377 | Cell/Virus Enclosed | NO |
| ATCEHCGTENLV | 1413085 | Cell/Virus Enclosed | NO |
| ATCEQCGTENLV | 1413088 | Cell/Virus Enclosed | NO |
| ATCPFAKGDSCSSNCK | 1646391 | Cell/Virus Enclosed | NO |
| ATCPFVKGDSCS | 1413096 | Cell/Virus Enclosed | NO |
| ATCYNSVSNEYC | 1413098 | Cell/Virus Enclosed | NO |
| ATDAPSTFGTRN | 1413100 | Cell/Virus Enclosed | NO |
| ATDCSDGNYNCN | 1413101 | Ectodomain | YES |
| ATDCSDGNYNRNASLN | 1646398 | Ectodomain | YES |
| ATDCSDGNYNRY | 1413103 | Ectodomain | YES |
| ATDDCFANKHAG | 1413104 | Cell/Virus Enclosed | NO |
| ATDHAQAIKVHTAAMKVG | 558229 | Undetermined | UN |
| ATDPIYDEPTTT | 1413109 | Secreted | YES |
| ATDPSEAFEKFI | 1413110 | Cell/Virus Enclosed | NO |
| ATDRFCRLGNEL | 1413111 | Cell/Virus Enclosed | NO |
| ATDYREYMEAYN | 1413114 | Cell/Virus Enclosed | NO |
| ATEAKGYWYRHN | 1413117 | Cell/Virus Enclosed | NO |
| ATEATFNYLLDL | 1413119 | Cell/Virus Enclosed | NO |
| ATEEAFKQSYAS | 1413121 | Cell/Virus Enclosed | NO |
| ATEFADKLVEVG | 1413123 | Cell/Virus Enclosed | NO |
| ATEFANKLVEVG | 1413124 | Cell/Virus Enclosed | NO |
| ATEKSNVIRGWI | 1413129 | Ectodomain | YES |
| ATESKGYWYRHN | 1413135 | Capsid | UN |
| ATEVPAHSTVLS | 1413137 | Cell/Virus Enclosed | NO |
| ATEVPANSAVLS | 1413138 | Cell/Virus Enclosed | NO |
| ATFAELEAAQKAYQEA | 1646420 | Cell/Virus Enclosed | NO |
| ATFAELEAARKA | 1413143 | Cell/Virus Enclosed | NO |
| ATFLDLYYEMNLIQES | 1646427 | Ectodomain | YES |
| ATFNFLLDLTGL | 1413157 | Cell/Virus Enclosed | NO |
| ATFNYLLDLAGL | 1413158 | Cell/Virus Enclosed | NO |
| ATFSVPMEELKA | 1413161 | Cell/Virus Enclosed | NO |
| ATFVDVLVNVSATNIQ | 1646431 | Ectodomain | YES |
| ATFYPQLQSAEWKCGY | 1646435 | Cell/Virus Enclosed | NO |
| ATGDLAVCLGIGDSAV | 1646438 | Cell/Virus Enclosed | NO |
| ATGDLAVCLGIS | 1413171 | Cell/Virus Enclosed | NO |
| ATGDLAVCLGVG | 1413172 | Cell/Virus Enclosed | NO |
| ATGDVVLASDDLYVKR | 1646439 | Cell/Virus Enclosed | NO |
| ATGDVVLATDDLYVKR | 1646440 | Cell/Virus Enclosed | NO |
| ATGGIAIAMACL | 1413176 | Undetermined | UN |
| ATGITPQKLFVA | 1413178 | Ectodomain | YES |
| ATGMKNVPEIPKGRGLFGAIAGFIE | 1129863 | Ectodomain | YES |
| ATGNLVVPRYAFAMERNAGSGIIIS | 1129864 | Ectodomain | YES |
| ATGSDVNCNGYQHNSV | 1646443 | Ectodomain | YES |
| ATGSLDVEVFNRLATS | 1646444 | Cell/Virus Enclosed | NO |
| ATGSTPQKLFVA | 1413186 | Ectodomain | YES |
| ATGTTLQKLFVA | 1413189 | Ectodomain | YES |
| ATGTTPQKLFVANYSQ | 1646445 | Ectodomain | YES |
| ATGTTPRKLFVA | 1413191 | Ectodomain | YES |
| ATGTYKCPQTKS | 1413193 | Ectodomain | YES |
| ATHIAEDVVDQV | 1413207 | Cell/Virus Enclosed | NO |
| ATHVCFKDCYNI | 1413211 | Cell/Virus Enclosed | NO |
| ATHVCFKDCYNV | 1413212 | Cell/Virus Enclosed | NO |
| ATHVCFKDCYSI | 1413213 | Cell/Virus Enclosed | NO |
| ATIGVCKYTDIDVNSA | 1646456 | Cell/Virus Enclosed | NO |
| ATIKEIVGERQL | 1413224 | Cell/Virus Enclosed | NO |
| ATIPLQASLPFG | 1413242 | Secreted | YES |
| ATIPYKVCNSVKDTLA | 1646464 | Cell/Virus Enclosed | NO |
| ATIQRKYKGVKI | 1413250 | Cell/Virus Enclosed | NO |
| ATIRLQAGKQTELAVN | 1646468 | Cell/Virus Enclosed | NO |
| ATKFTLWNYLFE | 1413260 | Cell/Virus Enclosed | NO |
| ATKGEGDGGIIS | 1413263 | Cell/Virus Enclosed | NO |
| ATKGEGDGGITSEGNA | 1646472 | Cell/Virus Enclosed | NO |
| ATKIQNLLYCDS | 1413266 | Ectodomain | YES |
| ATKMFDSFVNSFVSLY | 1646475 | Cell/Virus Enclosed | NO |
| ATKMSECVPGQS | 1413270 | Ectodomain | YES |
| ATKQYNVTQAFGRRGP | 1646478 | Cell/Virus Enclosed | NO |
| ATKSYNVTQAFG | 1413279 | Cell/Virus Enclosed | NO |
| ATKVYFDNPTLP | 1413283 | Ectodomain | YES |
| ATKYCRVGECVD | 1413284 | Cell/Virus Enclosed | NO |
| ATLDKCTKYNIY | 1413289 | Ectodomain | YES |

| | | | |
|---|---|---|---|
| ATLINNISYVVT | 1413294 | Cell/Virus Enclosed | NO |
| ATLLCVLAALFC | 1413296 | Cell/Virus Enclosed | NO |
| ATLMAPTIVNQE | 1413298 | Cell/Virus Enclosed | NO |
| ATLMTFSGNVDLDALA | 1646490 | Cell/Virus Enclosed | NO |
| ATLPDIAEDVVD | 1413301 | Cell/Virus Enclosed | NO |
| ATLPVYDTIKYY | 1413307 | Ectodomain | YES |
| ATLQAENVTGLF | 1413308 | Cell/Virus Enclosed | NO |
| ATLQEILSRLDA | 1413311 | Ectodomain | YES |
| ATLTAPTIVNQERYVK | 1646498 | Cell/Virus Enclosed | NO |
| ATLTASTIVNQE | 1413321 | Cell/Virus Enclosed | NO |
| ATLYYQNGVFMSTAKC | 1646504 | Cell/Virus Enclosed | NO |
| ATNCSDGNYNRN | 1413331 | Ectodomain | YES |
| ATNGAMQKESDG | 1413336 | Cell/Virus Enclosed | NO |
| ATNGAMQKESGD | 1413337 | Cell/Virus Enclosed | NO |
| ATNHDRYCSVHQ | 1413342 | Cell/Virus Enclosed | NO |
| ATNNVFRLKGGA | 1413352 | Cell/Virus Enclosed | NO |
| ATNNVFSLKGGA | 1413354 | Cell/Virus Enclosed | NO |
| ATNSALVKIQAVVNAN | 1646523 | Ectodomain | YES |
| ATPAAPRAVSFA | 1413362 | Cell/Virus Enclosed | NO |
| ATPFDVVRQCSG | 1413364 | Cell/Virus Enclosed | NO |
| ATPHKMVLSVSP | 1413366 | Cell/Virus Enclosed | NO |
| ATPQECNDMHLS | 1413369 | Cell/Virus Enclosed | NO |
| ATPSTTIVCDRV | 1413374 | Ectodomain | YES |
| ATPSVLQATLSE | 1413375 | Cell/Virus Enclosed | NO |
| ATQAKSGIITKV | 1413379 | Ectodomain | YES |
| ATQAMEKVNECV | 1413380 | Ectodomain | YES |
| ATQDGVNKLQGL | 1413381 | Ectodomain | YES |
| ATQGFQACRFVA | 1413383 | Cell/Virus Enclosed | NO |
| ATRDSIGTNFPLQLGF | 1646536 | Cell/Virus Enclosed | NO |
| ATRENIGTNFPLQIGF | 1646540 | Cell/Virus Enclosed | NO |
| ATRGVTCVIGTT | 1413400 | Cell/Virus Enclosed | NO |
| ATRLCVQCITGF | 1413402 | Cell/Virus Enclosed | NO |
| ATRLCVQCMTGF | 1413404 | Cell/Virus Enclosed | NO |
| ATRLCVQCMTGL | 1413405 | Cell/Virus Enclosed | NO |
| ATRNNCWLRSVL | 1413410 | Cell/Virus Enclosed | NO |
| ATRNYIITMPCF | 1413411 | Cell/Virus Enclosed | NO |
| ATRRVRGGDGKMKDLS | 1646551 | Cell/Virus Enclosed | NO |
| ATRTFTATTVVG | 1413421 | Cell/Virus Enclosed | NO |
| ATRVQVGQLPKY | 1413425 | Ectodomain | YES |
| ATRYCRVGECRD | 1413427 | Cell/Virus Enclosed | NO |
| ATSANLFPPWSA | 1413431 | Ectodomain | YES |
| ATSASKLTIVSPDLES | 1646562 | Cell/Virus Enclosed | NO |
| ATSASLFPLWTA | 1413432 | Ectodomain | YES |
| ATSASLFPPWSA | 1413433 | Ectodomain | YES |
| ATSASLFPPWTAAAGV | 1646563 | Ectodomain | YES |
| ATSCQLYYNLPAANVS | 1646564 | Ectodomain | YES |
| ATSCQLYYNLPS | 1413437 | Ectodomain | YES |
| ATSCQLYYSLPA | 1413438 | Ectodomain | YES |
| ATSDRTDVRDYK | 1413441 | Cell/Virus Enclosed | NO |
| ATSDWTDVRDYKLAND | 1646566 | Cell/Virus Enclosed | NO |
| ATSFCKSGNLVL | 1413444 | Ectodomain | YES |
| ATSIVAKQGAGDAGHS | 1646567 | Cell/Virus Enclosed | NO |
| ATSLSFNNLLGF | 1413455 | Cell/Virus Enclosed | NO |
| ATSNVTQCFGPR | 1413456 | Cell/Virus Enclosed | NO |
| ATSNYSGVVTTI | 1413457 | Cell/Virus Enclosed | NO |
| ATSQQPSGGNVI | 1413458 | Cell/Virus Enclosed | NO |
| ATSQQPSGGNVVPYYS | 1646569 | Cell/Virus Enclosed | NO |
| ATSQQPSGRNVV | 1413460 | Capsid | UN |
| ATSSYDALLHRN | 1413464 | Cell/Virus Enclosed | NO |
| ATSTGNYNYKHR | 1413468 | Ectodomain | YES |
| ATSYGMCFSSIT | 1413473 | Ectodomain | YES |
| ATTANVSALIGA | 1413475 | Cell/Virus Enclosed | NO |
| ATTANVSALMGANGNK | 1646575 | Cell/Virus Enclosed | NO |
| ATTAYANSVFNI | 1413482 | Cell/Virus Enclosed | NO |
| ATTAYANSVLNI | 1413483 | Cell/Virus Enclosed | NO |
| ATTQNDRMPGPELSG | 227054 | Cell/Virus Enclosed | NO |
| ATTSQDSYGGAS | 1413496 | Cell/Virus Enclosed | NO |
| ATTYCRVGECVD | 1413506 | Cell/Virus Enclosed | NO |
| ATTYLEAVKHGAKPVS | 1646592 | Cell/Virus Enclosed | NO |
| ATVAAMFPPWSAAAGIPF | 2060799 | Ectodomain | YES |
| ATVCGPKLSTDLIKNQ | 1646597 | Ectodomain | YES |
| ATVCGPKLSTGL | 1413512 | Ectodomain | YES |
| ATVDHSAFAYES | 1413515 | Cell/Virus Enclosed | NO |
| ATVDVTNGLGTY | 1413517 | Ectodomain | YES |
| ATVEPVRDEVSI | 1413518 | Cell/Virus Enclosed | NO |
| ATVEPVRDEVST | 1413519 | Cell/Virus Enclosed | NO |

| | | | |
|---|---|---|---|
| ATVIRGHLYIQGVKLG | 1646603 | Underneath Membrane | NO |
| ATVKSEDGNIKNINWF | 1646604 | Cell/Virus Enclosed | NO |
| ATVKWADAPEPQ | 1413528 | Cell/Virus Enclosed | NO |
| ATVKWADESEPQ | 1413530 | Cell/Virus Enclosed | NO |
| ATVLQNNELMPA | 1413532 | Cell/Virus Enclosed | NO |
| ATVLQNNELMPV | 1413533 | Cell/Virus Enclosed | NO |
| ATVNLRLVDCAP | 1413537 | Cell/Virus Enclosed | NO |
| ATVNLRLVDCVP | 1413538 | Cell/Virus Enclosed | NO |
| ATVNQVCVLLAKKIDV | 1646610 | Cell/Virus Enclosed | NO |
| ATVQSKMSDVRC | 1413546 | Cell/Virus Enclosed | NO |
| ATVRLQAGKQTEFVSN | 1646616 | Cell/Virus Enclosed | NO |
| ATVTANLSIPSN | 1413555 | Ectodomain | YES |
| ATVVVTLKDSDVNDMV | 1646627 | Cell/Virus Enclosed | NO |
| ATVYRRIPDLPD | 1413561 | Ectodomain | YES |
| ATYEGNSPFHPL | 1413568 | Ectodomain | YES |
| ATYHTPAIDCSD | 1413570 | Ectodomain | YES |
| ATYHTPATDCSDGNYN | 1646633 | Ectodomain | YES |
| ATYHTPATNCSD | 1413572 | Ectodomain | YES |
| ATYHTPSTDCSD | 1413573 | Ectodomain | YES |
| ATYQNISTNLPP | 1413582 | Ectodomain | YES |
| ATYYNSWQNLLYDSNG | 1646641 | Ectodomain | YES |
| ATYYTPATDCSD | 1413586 | Ectodomain | YES |
| AVAAALKSLGFDKPQE | 1646642 | Capsid | UN |
| AVAAGLEFTDENYNNL | 1646644 | Cell/Virus Enclosed | NO |
| AVADSYFSYMMP | 1413592 | Cell/Virus Enclosed | NO |
| AVADSYYSYMMPMLTM | 1646648 | Cell/Virus Enclosed | NO |
| AVAKAPPGEQFK | 1413595 | Cell/Virus Enclosed | NO |
| AVAKCNEKHDEEFCD | 540184 | Ectodomain | YES |
| AVAKHDFFTWHEGRTI | 1646651 | Cell/Virus Enclosed | NO |
| AVANAHRYDVLLSDLS | 1646654 | Cell/Virus Enclosed | NO |
| AVANGDSEAVLK | 1413601 | Cell/Virus Enclosed | NO |
| AVANSTGTVIIS | 1413604 | Ectodomain | YES |
| AVANWLAPTNAYMRTT | 1646657 | Cell/Virus Enclosed | NO |
| AVARKLERMADLALTN | 1646660 | Cell/Virus Enclosed | NO |
| AVASRRRPTTAGAAPLTAVAPAHDT | 5275 | Undetermined | UN |
| AVATFYPQLQSAEWKC | 1646665 | Cell/Virus Enclosed | NO |
| AVAYADKPNHFINFPL | 1646668 | Cell/Virus Enclosed | NO |
| AVAYADNPNHFI | 1413621 | Cell/Virus Enclosed | NO |
| AVAYSSPLQSYA | 1413622 | Cell/Virus Enclosed | NO |
| AVAYVDKPNHFI | 1413624 | Cell/Virus Enclosed | NO |
| AVCITTCNLGGA | 1413630 | Cell/Virus Enclosed | NO |
| AVCLGISDSAVT | 1413632 | Cell/Virus Enclosed | NO |
| AVCLKHAEEYREYLES | 1646674 | Cell/Virus Enclosed | NO |
| AVCNGCVGHSEN | 1413635 | Ectodomain | YES |
| AVCRKHATDYRE | 1413641 | Cell/Virus Enclosed | NO |
| AVCSKHAEEYCN | 1413649 | Cell/Virus Enclosed | NO |
| AVCSKHANLYQKYVEA | 1646679 | Cell/Virus Enclosed | NO |
| AVCSVIDLKLDD | 1413653 | Cell/Virus Enclosed | NO |
| AVDCKSDFMSEI | 1413663 | Ectodomain | YES |
| AVDCMSDFMSGI | 1413665 | Ectodomain | YES |
| AVDDKYKVNENLAVNL | 1646694 | Cell/Virus Enclosed | NO |
| AVDEAFPLDEAEDVTE | 1646695 | Cell/Virus Enclosed | NO |
| AVDEAFPLNEAE | 1413673 | Cell/Virus Enclosed | NO |
| AVDGAKTEPTGH | 1413676 | Capsid | UN |
| AVDINRLCEEVL | 1413681 | Cell/Virus Enclosed | NO |
| AVDIVSIRSSNQ | 1413682 | Cell/Virus Enclosed | NO |
| AVDLSMDKLCRV | 1413684 | Cell/Virus Enclosed | NO |
| AVDLSMDRLRRV | 1413687 | Cell/Virus Enclosed | NO |
| AVDPAAAYLDAVKQGA | 1646701 | Cell/Virus Enclosed | NO |
| AVDSAARIHDFRYSQLAKLG | 5298 | Capsid | UN |
| AVDSLCAKAVTS | 1413691 | Cell/Virus Enclosed | NO |
| AVDTGSGGGGQPHDTAPRGARKKQ | 5303 | Capsid | UN |
| AVDVVSIRTSNQ | 1413694 | Cell/Virus Enclosed | NO |
| AVEDTIDGVVEDTIND | 1646710 | Cell/Virus Enclosed | NO |
| AVEGTKKLAERL | 1413700 | Cell/Virus Enclosed | NO |
| AVESGGYIPPNF | 1413721 | Ectodomain | YES |
| AVESGGYIPSDFAFNN | 1646718 | Ectodomain | YES |
| AVESGGYIPSNF | 1413722 | Ectodomain | YES |
| AVESNVMDVNDC | 1413723 | Cell/Virus Enclosed | NO |
| AVEVDPDDVNKNTL | 5312 | Capsid | UN |
| AVFISPYNSQNFAAKR | 1646727 | Cell/Virus Enclosed | NO |
| AVFISPYNSQNYVAKR | 1646729 | Cell/Virus Enclosed | NO |
| AVGIMTVTMTFKLGPRKATG | 5346 | Capsid | UN |
| AVGVRQQRFVYDAYQN | 1646747 | Ectodomain | YES |
| AVHEWKMVVCDK | 1413788 | Cell/Virus Enclosed | NO |
| AVHSDGMFVASS | 1413796 | Cell/Virus Enclosed | NO |

| | | | |
|---|---|---|---|
| AVHYYQNNVFMS | 1413799 | Cell/Virus Enclosed | NO |
| AVIAGVAGARIP | 1413803 | Cell/Virus Enclosed | NO |
| AVIDEHVFYKKD | 1413809 | Cell/Virus Enclosed | NO |
| AVIDQDFGSTVFNVPT | 1646758 | Cell/Virus Enclosed | NO |
| AVIGDFNCTNFAINDL | 1646760 | Ectodomain | YES |
| AVIGDFNCTNSF | 1413811 | Ectodomain | YES |
| AVIGDLKCPLDS | 1413812 | Ectodomain | YES |
| AVIGDLKCPLES | 1413814 | Ectodomain | YES |
| AVIGDLKCTSDNINDK | 1646761 | Ectodomain | YES |
| AVIGDLKCTSDT | 1413815 | Ectodomain | YES |
| AVIGDLKCTTVA | 1413816 | Ectodomain | YES |
| AVIGDLKCTTVS | 1413817 | Ectodomain | YES |
| AVIGDLNCPLDP | 1413818 | Ectodomain | YES |
| AVIGDLNCPLDS | 1413819 | Ectodomain | YES |
| AVIGDLNCPLDT | 1413820 | Ectodomain | YES |
| AVIGDLNCTLDP | 1413821 | Ectodomain | YES |
| AVIGDLNCTLHP | 1413822 | Ectodomain | YES |
| AVIGTPVCVNGL | 1413823 | Cell/Virus Enclosed | NO |
| AVILRGHLRIAGHHL | 2000989 | Underneath Membrane | NO |
| AVIMTLDLLKDIKFVI | 1646768 | Cell/Virus Enclosed | NO |
| AVINIPDLPICV | 1413833 | Ectodomain | YES |
| AVINLRADQINDMVYS | 1646771 | Cell/Virus Enclosed | NO |
| AVIRRANNVLKNEMKRYKGL | 5387 | Capsid | UN |
| AVITTQQAGAGIKYFC | 1646775 | Cell/Virus Enclosed | NO |
| AVIYSGRESLIFPSGT | 1646778 | Cell/Virus Enclosed | NO |
| AVKAAESKAFVDTIVP | 1646779 | Cell/Virus Enclosed | NO |
| AVKDFDFYRYNKPTIL | 1646780 | Cell/Virus Enclosed | NO |
| AVKGCVPLNAIPSLAA | 1646782 | Cell/Virus Enclosed | NO |
| AVKGCVPLSAIPALAA | 1646783 | Cell/Virus Enclosed | NO |
| AVKHGAKPVSNC | 1413860 | Cell/Virus Enclosed | NO |
| AVKNGALHLYFD | 1413870 | Cell/Virus Enclosed | NO |
| AVKNGALHPYFD | 1413871 | Cell/Virus Enclosed | NO |
| AVKQGAKPVGNCVKML | 1646792 | Cell/Virus Enclosed | NO |
| AVKRVRAWVGFDAEGA | 1646794 | Cell/Virus Enclosed | NO |
| AVKSDYCGGNLC | 1413875 | Ectodomain | YES |
| AVKSDYCGGNSCTCRP | 1646795 | Ectodomain | YES |
| AVLDTLLASSSF | 1413894 | Cell/Virus Enclosed | NO |
| AVLFCTPIKKAF | 1413896 | Cell/Virus Enclosed | NO |
| AVLFSATAVKTGGKSL | 1646804 | Cell/Virus Enclosed | NO |
| AVLFSTVVIKNLTPIK | 1646805 | Cell/Virus Enclosed | NO |
| AVLKKLKKSLNV | 1413904 | Cell/Virus Enclosed | NO |
| AVLNSLCICFVA | 1413916 | Cell/Virus Enclosed | NO |
| AVLSDVDGLKYTKIMK | 1646819 | Cell/Virus Enclosed | NO |
| AVLTIANNYSKLFDEG | 1646820 | Cell/Virus Enclosed | NO |
| AVMFEDAQGLKK | 1413945 | Cell/Virus Enclosed | NO |
| AVNDQDLYDHVV | 1413965 | Cell/Virus Enclosed | NO |
| AVNESRYGLHNL | 1413967 | Ectodomain | YES |
| AVNESRYGLQNLLQLP | 1646843 | Ectodomain | YES |
| AVNFNVTNAATI | 1413969 | Ectodomain | YES |
| AVNFNVTTAETTDFCT | 1646844 | Ectodomain | YES |
| AVNFNVTTASAT | 1413970 | Ectodomain | YES |
| AVNIAKNAYEKDKAVA | 1646847 | Cell/Virus Enclosed | NO |
| AVNLALKNLGFD | 1413978 | Cell/Virus Enclosed | NO |
| AVNLNICEPVLT | 1413980 | Cell/Virus Enclosed | NO |
| AVNNIAQALSKL | 1413984 | Ectodomain | YES |
| AVNNNAHALSKL | 1413985 | Ectodomain | YES |
| AVNNNAQALAKL | 1413986 | Ectodomain | YES |
| AVNNNAQALSKLASEL | 1646853 | Ectodomain | YES |
| AVNPTNCIAPVN | 1413990 | Ectodomain | YES |
| AVNQSLSNAKVSAEVS | 1646854 | Cell/Virus Enclosed | NO |
| AVNSTKALAKQY | 1413999 | Cell/Virus Enclosed | NO |
| AVNYTKAPDVML | 1414013 | Ectodomain | YES |
| AVPGGCNFPLEA | 1414020 | Ectodomain | YES |
| AVPGSCNFPLEA | 1414021 | Ectodomain | YES |
| AVQIGNNNVCTYEHVI | 1646877 | Cell/Virus Enclosed | NO |
| AVQIGSNNVCTYEHVI | 1646878 | Cell/Virus Enclosed | NO |
| AVQKESDEYILAKGPL | 1646879 | Cell/Virus Enclosed | NO |
| AVQMGSNNVCTY | 1414047 | Cell/Virus Enclosed | NO |
| AVQNTVFNASIS | 1414048 | Cell/Virus Enclosed | NO |
| AVQQESDEYILA | 1414053 | Cell/Virus Enclosed | NO |
| AVQSKDLNFLNRVRGT | 1646882 | Cell/Virus Enclosed | NO |
| AVQTCDLTVGSG | 1414057 | Ectodomain | YES |
| AVRKEGQDVIFSQFDS | 1646885 | Cell/Virus Enclosed | NO |
| AVRKFLDGLDLP | 1414063 | Cell/Virus Enclosed | NO |
| AVRNKISACFNF | 1414066 | Cell/Virus Enclosed | NO |
| AVRNKTFACFNF | 1414068 | Cell/Virus Enclosed | NO |

| | | | |
|---|---|---|---|
| AVRNKTSACFNF | 1414069 | Cell/Virus Enclosed | NO |
| AVRQVRSWIGFDVEGA | 1646888 | Cell/Virus Enclosed | NO |
| AVRSHPDFKLLHNLQA | 1646890 | Cell/Virus Enclosed | NO |
| AVRSSFNFDQKELLAY | 1646891 | Cell/Virus Enclosed | NO |
| AVRTYSEAAAQG | 1414080 | Cell/Virus Enclosed | NO |
| AVSAGLEFTDES | 1414083 | Cell/Virus Enclosed | NO |
| AVSANVCALISC | 1414087 | Cell/Virus Enclosed | NO |
| AVSANVSALMSC | 1414088 | Cell/Virus Enclosed | NO |
| AVSCGHYQTNIYSQNL | 1646897 | Cell/Virus Enclosed | NO |
| AVSDNVRALMSC | 1414091 | Cell/Virus Enclosed | NO |
| AVSFADNNDITNTNLS | 1646898 | Cell/Virus Enclosed | NO |
| AVSFADNNDTTN | 1414094 | Cell/Virus Enclosed | NO |
| AVSFADNNDTTT | 1414095 | Cell/Virus Enclosed | NO |
| AVSFAYNNDITN | 1414096 | Capsid | UN |
| AVSFDYLIREAE | 1414098 | Cell/Virus Enclosed | NO |
| AVSFDYLIREAKTRVL | 1646899 | Cell/Virus Enclosed | NO |
| AVSFDYLIREAR | 1414100 | Cell/Virus Enclosed | NO |
| AVSKVVKVPIDL | 1414114 | Cell/Virus Enclosed | NO |
| AVSNGTKVNTLTERGVEVVNATETV | 1130264 | Ectodomain | YES |
| AVSRINWVTGGI | 1414118 | Undetermined | UN |
| AVSRSMLGLTTQTVDS | 1646907 | Cell/Virus Enclosed | NO |
| AVSSNINCVLSVNSSN | 1646909 | Cell/Virus Enclosed | NO |
| AVSSNINMLLSV | 1414124 | Cell/Virus Enclosed | NO |
| AVSSNINRILGV | 1414125 | Cell/Virus Enclosed | NO |
| AVSSNINRLFSV | 1414126 | Cell/Virus Enclosed | NO |
| AVSSNINRLLSVPSDS | 1646910 | Cell/Virus Enclosed | NO |
| AVSSNINRVLSV | 1414127 | Cell/Virus Enclosed | NO |
| AVSSPMSNMTENERLL | 1646911 | Underneath Membrane | NO |
| AVSWNLREMLAH | 1414131 | Cell/Virus Enclosed | NO |
| AVTANVCSLMACNGHK | 1646916 | Cell/Virus Enclosed | NO |
| AVTANVCSPMAC | 1414136 | Cell/Virus Enclosed | NO |
| AVTANVRSLTAC | 1414138 | Cell/Virus Enclosed | NO |
| AVTDAKWYCYDKNPIN | 1646920 | Cell/Virus Enclosed | NO |
| AVTDAKWYCYDKQPVN | 1646921 | Cell/Virus Enclosed | NO |
| AVVCASVFKDGC | 1414161 | Cell/Virus Enclosed | NO |
| AVVCDAIENALNSCKE | 1646932 | Cell/Virus Enclosed | NO |
| AVVDGKQIDDYR | 1414166 | Cell/Virus Enclosed | NO |
| AVVGAMLSENFTSYGF | 1646938 | Ectodomain | YES |
| AVVGDLNCPLDP | 1414173 | Ectodomain | YES |
| AVVGTPVCINGL | 1414175 | Cell/Virus Enclosed | NO |
| AVVGTPVCVNGL | 1414176 | Cell/Virus Enclosed | NO |
| AVVLLERYVSLAIDAY | 1646947 | Cell/Virus Enclosed | NO |
| AVVNADAEALNN | 1414190 | Ectodomain | YES |
| AVVNANAEAHNN | 1414191 | Ectodomain | YES |
| AVVNANAEALNNLLQQ | 1646949 | Ectodomain | YES |
| AVVNDKIFTKSGVNDI | 1646950 | Cell/Virus Enclosed | NO |
| AVVNDKVFTKSG | 1414193 | Cell/Virus Enclosed | NO |
| AVVNGVRVLKTS | 1414197 | Cell/Virus Enclosed | NO |
| AVVNLRPDQLNDLVYS | 1646953 | Cell/Virus Enclosed | NO |
| AVVNSQDVYKSL | 1414199 | Cell/Virus Enclosed | NO |
| AVVYFGGCVYWS | 1414210 | Cell/Virus Enclosed | NO |
| AVYANVGRWSASINTG | 1646959 | Ectodomain | YES |
| AVYQQSIIGAMTAVNE | 1646966 | Ectodomain | YES |
| AVYRINWITGGIAIA | 1312246 | Undetermined | UN |
| AVYVKSKVGNYR | 77840 | Ectodomain | YES |
| AWDFMLKTLYKD | 1414261 | Cell/Virus Enclosed | NO |
| AWDVIRKDLSYEQQNV | 1646979 | Cell/Virus Enclosed | NO |
| AWDYHITKVLAE | 1414265 | Cell/Virus Enclosed | NO |
| AWEDGDYYRKQL | 1414267 | Ectodomain | YES |
| AWFVQNDVCSTE | 1414289 | Cell/Virus Enclosed | NO |
| AWHVVRDPRYVMRLQS | 1646990 | Cell/Virus Enclosed | NO |
| AWHVVRDSRFVMRLQT | 1646991 | Cell/Virus Enclosed | NO |
| AWLLHKDGGIPD | 1414304 | Cell/Virus Enclosed | NO |
| AWLLHKDGGVPD | 1414305 | Cell/Virus Enclosed | NO |
| AWLNDKAVPSPL | 1414308 | Ectodomain | YES |
| AWLNDKLVPSPL | 1414309 | Ectodomain | YES |
| AWLNDKSVPSPLNWER | 1647000 | Ectodomain | YES |
| AWLNGKSVPSPL | 1414311 | Ectodomain | YES |
| AWLSYYVPSVEY | 1414313 | Cell/Virus Enclosed | NO |
| AWLTKRKPLDYK | 1414314 | Cell/Virus Enclosed | NO |
| AWNIGQEKSILT | 1414323 | Cell/Virus Enclosed | NO |
| AWNIGQQRSVLT | 1414325 | Cell/Virus Enclosed | NO |
| AWNPEVNAITVT | 1414330 | Underneath Membrane | NO |
| AWNTAKQDQGQY | 1414332 | Ectodomain | YES |
| AWNTAQQDQGQY | 1414336 | Ectodomain | YES |
| AWNTRNIDATSTGNYN | 1647016 | Ectodomain | YES |

| | | | |
|---|---|---|---|
| AWRIYSTGNNVFQTQA | 1647021 | Ectodomain | YES |
| AWSGLCVDGRNG | 1414355 | Ectodomain | YES |
| AWSGLCVDGTNGYVLR | 1647026 | Ectodomain | YES |
| AWVGFDAEGAHATRDS | 1647029 | Cell/Virus Enclosed | NO |
| AYADKPNHFINFPLTH | 1647039 | Cell/Virus Enclosed | NO |
| AYADNPNHFINF | 1414382 | Cell/Virus Enclosed | NO |
| AYADRTLQEMKA | 1414383 | Cell/Virus Enclosed | NO |
| AYALIRGYRGVK | 1414392 | Cell/Virus Enclosed | NO |
| AYANFNSQDVLL | 1414397 | Cell/Virus Enclosed | NO |
| AYANSVFNILQATTAN | 1647053 | Cell/Virus Enclosed | NO |
| AYAPTYVSVDDKYKTS | 1647055 | Cell/Virus Enclosed | NO |
| AYCVDLHNKINLCDDP | 1647064 | Cell/Virus Enclosed | NO |
| AYDGSAFRDLKP | 1414424 | Cell/Virus Enclosed | NO |
| AYDKRAMAKLKPAPFF | 1647068 | Cell/Virus Enclosed | NO |
| AYDQVHEEPKGK | 1414433 | Cell/Virus Enclosed | NO |
| AYDVSAFRDLKPLPFF | 1647072 | Cell/Virus Enclosed | NO |
| AYEDAIANGSSSQLIK | 1647073 | Cell/Virus Enclosed | NO |
| AYEKDKAVARKLERMA | 1647074 | Cell/Virus Enclosed | NO |
| AYENARQAYEDAIANG | 1647075 | Cell/Virus Enclosed | NO |
| AYENFNSQDTLL | 1414443 | Cell/Virus Enclosed | NO |
| AYETARQEYENAVANG | 1647084 | Cell/Virus Enclosed | NO |
| AYFNGAIIRDGD | 1414459 | Cell/Virus Enclosed | NO |
| AYFNGAIIRDSDVVKQ | 1647088 | Cell/Virus Enclosed | NO |
| AYFNGAIIRNSD | 1414461 | Cell/Virus Enclosed | NO |
| AYFPREGVFVFNGTSWF | 5773 | Ectodomain | YES |
| AYFSDSKFGTTE | 1414470 | Cell/Virus Enclosed | NO |
| AYFSKKHVEVFTDGKL | 1647092 | Cell/Virus Enclosed | NO |
| AYFTDNGVVNKF | 1414477 | Ectodomain | YES |
| AYFTDTGFVTKF | 1414478 | Ectodomain | YES |
| AYGIPLSEAKGY | 1414491 | Cell/Virus Enclosed | NO |
| AYGIPPSEAKGY | 1414492 | Cell/Virus Enclosed | NO |
| AYGLCDASNPTN | 1414494 | Ectodomain | YES |
| AYGLCDLANPTN | 1414496 | Ectodomain | YES |
| AYGLCDVANPTN | 1414498 | Ectodomain | YES |
| AYGNCTFGAPDDASRL | 1647100 | Cell/Virus Enclosed | NO |
| AYGNCTFGDPDD | 1414503 | Cell/Virus Enclosed | NO |
| AYGNCTFGSPDD | 1414504 | Cell/Virus Enclosed | NO |
| AYHANSPEPALL | 1414509 | Ectodomain | YES |
| AYHANSSDPALL | 1414510 | Ectodomain | YES |
| AYHANSSEPTLL | 1414511 | Ectodomain | YES |
| AYHNLVGYYSDD | 1414513 | Ectodomain | YES |
| AYHTVIDVKQVG | 1414514 | Cell/Virus Enclosed | NO |
| AYIAPQAKSGII | 1414517 | Ectodomain | YES |
| AYIAPQANFGDY | 1414519 | Ectodomain | YES |
| AYIAPQANSGGY | 1414520 | Ectodomain | YES |
| AYIAPQANSGII | 1414521 | Ectodomain | YES |
| AYIAREANFGDY | 1414523 | Ectodomain | YES |
| AYIDVLRNSFGKDLNA | 1647103 | Cell/Virus Enclosed | NO |
| AYIKKLYNDLGNQILD | 1647105 | Cell/Virus Enclosed | NO |
| AYINILDLPICV | 1414538 | Ectodomain | YES |
| AYINIPDLPICV | 1414539 | Ectodomain | YES |
| AYINIPDLPVCV | 1414540 | Ectodomain | YES |
| AYISNKADLKFVKWEY | 1647108 | Cell/Virus Enclosed | NO |
| AYITFSGSFDNG | 1414546 | Cell/Virus Enclosed | NO |
| AYKAFPPTEPKK | 1414553 | Cell/Virus Enclosed | NO |
| AYKFLNINDCTRIVPA | 1647115 | Cell/Virus Enclosed | NO |
| AYKHLNKYDCCL | 1414556 | Cell/Virus Enclosed | NO |
| AYLAGTVWYVGQ | 1414571 | Cell/Virus Enclosed | NO |
| AYLAKAMLDFSRDHND | 1647124 | Cell/Virus Enclosed | NO |
| AYLDAVKQGAKPVGNC | 1647125 | Cell/Virus Enclosed | NO |
| AYLDFVNAGGAPLTNC | 1647126 | Cell/Virus Enclosed | NO |
| AYLDLRTTQQLT | 1414575 | Cell/Virus Enclosed | NO |
| AYLDSCTTQQLT | 1414576 | Cell/Virus Enclosed | NO |
| AYLDSRATQQLT | 1414577 | Cell/Virus Enclosed | NO |
| AYLDSRTTQQLT | 1414578 | Cell/Virus Enclosed | NO |
| AYLNSRTTQQLT | 1414595 | Cell/Virus Enclosed | NO |
| AYLQSSNYRIDT | 1414597 | Ectodomain | YES |
| AYNCTYDRVEQS | 1414620 | Ectodomain | YES |
| AYNGKPQGAFHV | 1414623 | Cell/Virus Enclosed | NO |
| AYNGPPQGAFHV | 1414624 | Cell/Virus Enclosed | NO |
| AYNKDHGCGGFG | 1414629 | Cell/Virus Enclosed | NO |
| AYNKDHKRVGFG | 1414632 | Cell/Virus Enclosed | NO |
| AYNNCTGGAEIK | 1414636 | Ectodomain | YES |
| AYNNCTGGAGIR | 1414638 | Ectodomain | YES |
| AYNNDITNTNLS | 1414639 | Capsid | UN |
| AYNNTAISVQTC | 1414640 | Ectodomain | YES |

| | | | |
|---|---|---|---|
| AYNPDLSQYYCDNGKY | 1647151 | Cell/Virus Enclosed | NO |
| AYNSILFENGIPLMPL | 1647154 | Cell/Virus Enclosed | NO |
| AYNSILVENGIP | 1414644 | Cell/Virus Enclosed | NO |
| AYNSTAISVKTC | 1414647 | Ectodomain | YES |
| AYNTFTQAGFNI | 1414648 | Cell/Virus Enclosed | NO |
| AYNTINNEQGTPLTPI | 1647157 | Cell/Virus Enclosed | NO |
| AYPLTKHEDIEYQNVF | 1647163 | Cell/Virus Enclosed | NO |
| AYPLVHHENEEYQKVF | 1647165 | Cell/Virus Enclosed | NO |
| AYPLVYHENEEYQKVF | 1647167 | Cell/Virus Enclosed | NO |
| AYQDYMQIAPVPAEVL | 1647168 | Cell/Virus Enclosed | NO |
| AYQLYMIDPAGV | 1414666 | Cell/Virus Enclosed | NO |
| AYQNLLGYYSDD | 1414669 | Ectodomain | YES |
| AYQNLVGYYSDDGNYY | 1647171 | Ectodomain | YES |
| AYQQQDGMMNIS | 1414672 | Cell/Virus Enclosed | NO |
| AYQQQDGMMNMS | 1414673 | Cell/Virus Enclosed | NO |
| AYQQQDGMNMSP | 1414674 | Cell/Virus Enclosed | NO |
| AYQQQDGMVNMS | 1414675 | Cell/Virus Enclosed | NO |
| AYQVAARYFDCYEGGC | 1647173 | Cell/Virus Enclosed | NO |
| AYRANFFDVPTF | 1414678 | Cell/Virus Enclosed | NO |
| AYRANSFDAPTF | 1414679 | Cell/Virus Enclosed | NO |
| AYRANSIDVPTF | 1414681 | Cell/Virus Enclosed | NO |
| AYSKYYTIGTLY | 1414698 | Ectodomain | YES |
| AYSLFDMAKFPLKLAG | 1647187 | Cell/Virus Enclosed | NO |
| AYSNNTIAIPTNFSISI | 5927 | Ectodomain | YES |
| AYSNRTVGELGD | 1414707 | Cell/Virus Enclosed | NO |
| AYSPIDIPRGLP | 1414708 | Ectodomain | YES |
| AYSPQLTLVFLE | 1414710 | Cell/Virus Enclosed | NO |
| AYSPQLTLVFPEVKMI | 1647193 | Cell/Virus Enclosed | NO |
| AYSQMRPHVRYDLYDG | 1647194 | Cell/Virus Enclosed | NO |
| AYSRYRIGNYKLNTDH | 1378028 | Underneath Membrane | NO |
| AYSSPLQSYACWTHSC | 1647196 | Cell/Virus Enclosed | NO |
| AYSSSANGILVG | 1414722 | Cell/Virus Enclosed | NO |
| AYSSSANGRLVG | 1414723 | Cell/Virus Enclosed | NO |
| AYSSSANGSLDG | 1414724 | Cell/Virus Enclosed | NO |
| AYSSSANGSLVC | 1414725 | Cell/Virus Enclosed | NO |
| AYSSSANGSLVGTTLQ | 1647197 | Cell/Virus Enclosed | NO |
| AYSTMRLHNFSVSHNG | 1647198 | Cell/Virus Enclosed | NO |
| AYSVDKCTRIIPARAR | 1647199 | Cell/Virus Enclosed | NO |
| AYTAFSGPVDKGHYTV | 1647201 | Cell/Virus Enclosed | NO |
| AYTFTTVKPSAA | 1414740 | Cell/Virus Enclosed | NO |
| AYTMSLGADSSIAYSN | 1647207 | Ectodomain | YES |
| AYTQGRSCSDFE | 1414749 | Cell/Virus Enclosed | NO |
| AYTTFSGSFDNGHYVV | 1647212 | Cell/Virus Enclosed | NO |
| AYVANPTEDLCDGSVV | 1647215 | Cell/Virus Enclosed | NO |
| AYVANPTEDLCN | 1414761 | Cell/Virus Enclosed | NO |
| AYVDKPNHFINF | 1414762 | Cell/Virus Enclosed | NO |
| AYVDVLCNSFFKELTA | 1647219 | Cell/Virus Enclosed | NO |
| AYVGCYNKRAYW | 1414772 | Cell/Virus Enclosed | NO |
| AYVIIDKVDFVNGFYR | 1647223 | Cell/Virus Enclosed | NO |
| AYVMIDKVEFENGFYR | 1647225 | Cell/Virus Enclosed | NO |
| AYVNSNQNTDSDSLSS | 1647226 | Capsid | UN |
| AYVSQQLSDISLVKFG | 1647229 | Ectodomain | YES |
| AYVSQQLSDSTLVKFS | 1647230 | Ectodomain | YES |
| AYVTGGLVQQIT | 1414785 | Cell/Virus Enclosed | NO |
| AYVTGGLVQQTS | 1414786 | Cell/Virus Enclosed | NO |
| AYWFLNQFDINA | 1414794 | Cell/Virus Enclosed | NO |
| AYYADSGMPTTF | 1414800 | Ectodomain | YES |
| AYYADVGMPTTF | 1414801 | Ectodomain | YES |
| AYYEPVQGRKMLMALL | 1647245 | Cell/Virus Enclosed | NO |
| AYYKYDAKNYVRFPEI | 1647248 | Cell/Virus Enclosed | NO |
| AYYKYDNGNFIKLPEV | 1647249 | Cell/Virus Enclosed | NO |
| AYYNDWQNLLYD | 1414814 | Ectodomain | YES |
| AYYNNWQNLLYD | 1414819 | Ectodomain | YES |
| CAACVCGDYAAC | 1414831 | Ectodomain | YES |
| CAAFDCGDYAAC | 1414832 | Ectodomain | YES |
| CAFAVDPAKAYK | 1414867 | Cell/Virus Enclosed | NO |
| CAFGGVSVITPGTNAS | 1647285 | Ectodomain | YES |
| CAFSVDPKKTYLDYIQ | 1647290 | Cell/Virus Enclosed | NO |
| CAFVDTVKGLDY | 1414882 | Cell/Virus Enclosed | NO |
| CAGNVIGNDALTRFTI | 1647294 | Cell/Virus Enclosed | NO |
| CAGRKVNFNEKPVVME | 1647296 | Cell/Virus Enclosed | NO |
| CAGRRVTFKEQPTVKE | 1647297 | Cell/Virus Enclosed | NO |
| CAGYTLEEARTA | 1414892 | Cell/Virus Enclosed | NO |
| CAIINDKVVAKIDKED | 1647302 | Cell/Virus Enclosed | NO |
| CAIVQASESKVF | 1414906 | Cell/Virus Enclosed | NO |
| CAIVQASESKVLLERF | 1647307 | Cell/Virus Enclosed | NO |

| | | | |
|---|---|---|---|
| CAIVQATESKVL | 1414908 | Cell/Virus Enclosed | NO |
| CAKILPFSDRPL | 1414914 | Cell/Virus Enclosed | NO |
| CAKSPPGEQFRR | 1414924 | Cell/Virus Enclosed | NO |
| CALILAYSNKTV | 1414935 | Cell/Virus Enclosed | NO |
| CALISCNGNKIE | 1414937 | Cell/Virus Enclosed | NO |
| CALPDTPSTFTP | 1414944 | Ectodomain | YES |
| CALPDTPSTLAP | 1414945 | Ectodomain | YES |
| CALPDTPSTLTL | 1414947 | Ectodomain | YES |
| CALPDTPSTLTPRSVR | 1647325 | Ectodomain | YES |
| CALSPGLLATNN | 1414953 | Cell/Virus Enclosed | NO |
| CAMLLVKHKHAF | 1414959 | Cell/Virus Enclosed | NO |
| CAMPIMANLANL | 1414960 | Ectodomain | YES |
| CAMRPNYTIKGS | 1414973 | Cell/Virus Enclosed | NO |
| CANEYTGNYQCG | 1414980 | Cell/Virus Enclosed | NO |
| CANEYTGSYQCG | 1414981 | Cell/Virus Enclosed | NO |
| CANFNVLFAMTM | 1414983 | Cell/Virus Enclosed | NO |
| CANFTILFSMVL | 1414985 | Cell/Virus Enclosed | NO |
| CANLLLQYGSFCTQLNRALSGIA | 6011 | Ectodomain | YES |
| CANQSTSTYLAN | 1414996 | Ectodomain | YES |
| CANQSTSVYSAK | 1414999 | Ectodomain | YES |
| CAPVVCPKGKIVVIAG | 1647349 | Cell/Virus Enclosed | NO |
| CAQVLSEYVLCGGGYY | 1647357 | Cell/Virus Enclosed | NO |
| CARKSCSYDSDV | 1415029 | Cell/Virus Enclosed | NO |
| CARKSFSIDSDV | 1415030 | Cell/Virus Enclosed | NO |
| CARNPIDLPPSHATTY | 1647361 | Cell/Virus Enclosed | NO |
| CARNPVDLPPSH | 1415033 | Cell/Virus Enclosed | NO |
| CASVFKDGCDVG | 1415040 | Cell/Virus Enclosed | NO |
| CASVKRDGVQVGYCVH | 1647368 | Cell/Virus Enclosed | NO |
| CASYHTVSLLRSTSQKS | 6040 | Ectodomain | YES |
| CATCPFAKGDSCSSNC | 1647374 | Cell/Virus Enclosed | NO |
| CATTRQVVNVIT | 1415060 | Cell/Virus Enclosed | NO |
| CAVSFADNNDIT | 1415084 | Cell/Virus Enclosed | NO |
| CAWDYHVTKVLAEKFD | 1647390 | Cell/Virus Enclosed | NO |
| CAYCCNIVNVSLVKPS | 1647395 | Ectodomain | YES |
| CAYTQGRSCSDF | 1415099 | Cell/Virus Enclosed | NO |
| CCCTGCGPSCFK | 1415105 | Undetermined | UN |
| CCCTGCGTSCFK | 1415107 | Undetermined | UN |
| CCDATYFENKLW | 1415113 | Cell/Virus Enclosed | NO |
| CCDCGSTKLPYYEFEK | 1647406 | Underneath Membrane | NO |
| CCDDYTGHQELV | 1415118 | Underneath Membrane | NO |
| CCDDYTGYQEFV | 1415119 | Underneath Membrane | NO |
| CCDDYTGYQELVIKTS | 1647408 | Underneath Membrane | NO |
| CCDDYTGYQESV | 1415121 | Underneath Membrane | NO |
| CCDEYGGHHDFVIKTS | 1647409 | Underneath Membrane | NO |
| CCDEYGGHNDFV | 1415123 | Underneath Membrane | NO |
| CCDRYEEYDLQP | 1415131 | Underneath Membrane | NO |
| CCEDYTGYQELV | 1415134 | Underneath Membrane | NO |
| CCESTKLPYYDVEKIH | 1647413 | Underneath Membrane | NO |
| CCFSVAAITSGVKFQT | 1647417 | Cell/Virus Enclosed | NO |
| CCGTPAKKLCVV | 1415146 | Cell/Virus Enclosed | NO |
| CCLDVVRKDLEE | 1415164 | Cell/Virus Enclosed | NO |
| CCLDVVRKELEE | 1415165 | Cell/Virus Enclosed | NO |
| CCLKCGFSFDLNGLDA | 1647436 | Cell/Virus Enclosed | NO |
| CCNIVNVSLVKPSFY | 1539674 | Ectodomain | YES |
| CCREHEHEIVWF | 1415194 | Cell/Virus Enclosed | NO |
| CCRFQRIDDDGNKLDK | 1647456 | Cell/Virus Enclosed | NO |
| CCRFQRVDENGDKLDQ | 1647457 | Cell/Virus Enclosed | NO |
| CCSACMVWREWC | 1415208 | Cell/Virus Enclosed | NO |
| CCSQIDRFYRLA | 1415219 | Cell/Virus Enclosed | NO |
| CCTASDKFYRLSNELA | 1647469 | Cell/Virus Enclosed | NO |
| CCTVYDSSEVKAQSSG | 1647479 | Cell/Virus Enclosed | NO |
| CCVTSCCSCLKG | 1415251 | Undetermined | UN |
| CCVVEASTDFHQ | 1415252 | Cell/Virus Enclosed | NO |
| CCYDHVIATPHKMVLS | 1647481 | Cell/Virus Enclosed | NO |
| CCYDHVMATDHKYVLS | 1647483 | Cell/Virus Enclosed | NO |
| CCYDHVMATNHKYVLS | 1647484 | Cell/Virus Enclosed | NO |
| CCYVGVQTVEDLRARM | 1647486 | Cell/Virus Enclosed | NO |
| CDAIENALNSCKEHPV | 1647488 | Cell/Virus Enclosed | NO |
| CDAKFKNSVASINSAI | 1647490 | Cell/Virus Enclosed | NO |
| CDAKFKNSVTSI | 1415270 | Cell/Virus Enclosed | NO |
| CDAMRDAGIVGV | 1415271 | Cell/Virus Enclosed | NO |
| CDATYFENKLWF | 1415275 | Cell/Virus Enclosed | NO |
| CDCGIKQESRVG | 1415284 | Cell/Virus Enclosed | NO |
| CDCGKKFDEQVG | 1415286 | Cell/Virus Enclosed | NO |
| CDCGSTKLPYYEFEKVHVQ | 1415288 | Underneath Membrane | NO |
| CDCLCTPDPITS | 1415289 | Ectodomain | YES |

| | | | |
|---|---|---|---|
| CDDCNGYTDNIFSVQQ | 1647499 | Ectodomain | YES |
| CDDYTGHQELVI | 1415296 | Underneath Membrane | NO |
| CDDYTGYQELVIKTSH | 1647504 | Underneath Membrane | NO |
| CDDYTGYQESVI | 1415299 | Underneath Membrane | NO |
| CDEECSEVEASDLEEG | 1647505 | Cell/Virus Enclosed | NO |
| CDEFTLAEVVKQMYGV | 1647506 | Cell/Virus Enclosed | NO |
| CDEYGGHHDFVIKTSH | 1647512 | Underneath Membrane | NO |
| CDEYGGHNDFVI | 1415315 | Underneath Membrane | NO |
| CDFEIACKCGVKQEQR | 1647515 | Cell/Virus Enclosed | NO |
| CDFKGWIPGNMI | 1415321 | Cell/Virus Enclosed | NO |
| CDFNKPLIEWPLTEYD | 1647518 | Cell/Virus Enclosed | NO |
| CDFYGWVSGNMMDGFS | 1647520 | Cell/Virus Enclosed | NO |
| CDGGKYYTQRII | 1415338 | Cell/Virus Enclosed | NO |
| CDGLASAYRANF | 1415346 | Cell/Virus Enclosed | NO |
| CDGLASAYRANSFDVP | 1647526 | Cell/Virus Enclosed | NO |
| CDGLKFTEESFYKNMY | 1647527 | Cell/Virus Enclosed | NO |
| CDGSLGCKMCLFGYQE | 1647531 | Cell/Virus Enclosed | NO |
| CDGVNVTKHKCDINYK | 1647535 | Cell/Virus Enclosed | NO |
| CDGVRESYVNSS | 1415357 | Cell/Virus Enclosed | NO |
| CDHMVKDGLVGV | 1415361 | Cell/Virus Enclosed | NO |
| CDICKSTVVEVKSAIV | 1647540 | Cell/Virus Enclosed | NO |
| CDIDKWLNNFNVPSPL | 1647541 | Ectodomain | YES |
| CDIDKWLNNFSV | 1415366 | Ectodomain | YES |
| CDIDNWLNKVSV | 1415367 | Ectodomain | YES |
| CDIENNPGLCYASQM | 227058 | Cell/Virus Enclosed | NO |
| CDINKPLQVFVY | 1415373 | Cell/Virus Enclosed | NO |
| CDINYKGKVFFQFDNL | 1647545 | Cell/Virus Enclosed | NO |
| CDIPIGAGICASYHTVS | 6110 | Ectodomain | YES |
| CDIPVGAGICAS | 1415377 | Ectodomain | YES |
| CDISYKGKVFFQ | 1415379 | Cell/Virus Enclosed | NO |
| CDKVRSLKLSDNNCYL | 1647556 | Cell/Virus Enclosed | NO |
| CDLALKLKGLDA | 1415395 | Cell/Virus Enclosed | NO |
| CDLRMGSGFCID | 1415408 | Ectodomain | YES |
| CDLYSIHDVGFK | 1415413 | Cell/Virus Enclosed | NO |
| CDLYSMQDVGFK | 1415414 | Cell/Virus Enclosed | NO |
| CDMCKSTVVEVK | 1415415 | Cell/Virus Enclosed | NO |
| CDNDRCIIHCAN | 1415417 | Cell/Virus Enclosed | NO |
| CDNINSILDEVNGLLD | 1647571 | Ectodomain | YES |
| CDNLCTLDPIAL | 1415423 | Transmembrane | NO |
| CDNPFFAVSKPMGTQT | 1647573 | Ectodomain | YES |
| CDNPFIAVSKPM | 1415431 | Ectodomain | YES |
| CDNSYFDSKGWYDPVE | 1647574 | Cell/Virus Enclosed | NO |
| CDQLAFDVPDGF | 1415435 | Ectodomain | YES |
| CDQLSFDVPDGFYSTS | 1647578 | Ectodomain | YES |
| CDRTAIQSFDNSYLNR | 1647587 | Cell/Virus Enclosed | NO |
| CDRTAIQSFDSS | 1415453 | Cell/Virus Enclosed | NO |
| CDRYEEYDLQPH | 1415459 | Underneath Membrane | NO |
| CDSFGPGNTFINGDIA | 1647591 | Cell/Virus Enclosed | NO |
| CDSLNAKLGFDSSKEF | 1647593 | Cell/Virus Enclosed | NO |
| CDSLNAKWIFDS | 1415464 | Cell/Virus Enclosed | NO |
| CDSLNAKWVFDS | 1415465 | Cell/Virus Enclosed | NO |
| CDSLNFKLGFDS | 1415466 | Cell/Virus Enclosed | NO |
| CDSLRQAALPQS | 1415469 | Cell/Virus Enclosed | NO |
| CDSLRQVALPQS | 1415471 | Cell/Virus Enclosed | NO |
| CDSPFEKLQCEHLQFG | 1647596 | Ectodomain | YES |
| CDTPIGAGICAS | 1415489 | Ectodomain | YES |
| CDVADVTKLYLG | 1415490 | Cell/Virus Enclosed | NO |
| CDVDTTTLYVNN | 1415494 | Ectodomain | YES |
| CDVNDVTKLYLGGMSY | 1647610 | Cell/Virus Enclosed | NO |
| CDVSDVTKLYLGGMSY | 1647611 | Cell/Virus Enclosed | NO |
| CDVVQEQVNYVPLRAS | 1647616 | Cell/Virus Enclosed | NO |
| CDWTESGDYTLANTTT | 1647618 | Cell/Virus Enclosed | NO |
| CDYAGKFCHAVV | 1415512 | Cell/Virus Enclosed | NO |
| CDYLADYDVAVT | 1415517 | Cell/Virus Enclosed | NO |
| CDYTGSLADGLE | 1415522 | Cell/Virus Enclosed | NO |
| CDYTGSLADGLG | 1415523 | Cell/Virus Enclosed | NO |
| CEDFYASMYENS | 1415530 | Cell/Virus Enclosed | NO |
| CEDFYASMYENY | 1415531 | Cell/Virus Enclosed | NO |
| CEEADEGICFNL | 1415536 | Cell/Virus Enclosed | NO |
| CEEGTMYFVPRAKSVV | 1647637 | Cell/Virus Enclosed | NO |
| CEEVANDFTTAL | 1415549 | Cell/Virus Enclosed | NO |
| CEEVANDLTTALRRPI | 1647640 | Cell/Virus Enclosed | NO |
| CEHLQFGLQDGF | 1415577 | Ectodomain | YES |
| CEILCEYADCKESYFS | 1647654 | Cell/Virus Enclosed | NO |
| CEKAIGKKIKHEGDWD | 1647657 | Cell/Virus Enclosed | NO |
| CEKNLQLMRKVDLYNG | 1647662 | Cell/Virus Enclosed | NO |

| | | | |
|---|---|---|---|
| CELCHSGFDMLDTYAA | 1647665 | Cell/Virus Enclosed | NO |
| CELELICDCGIK | 1415600 | Cell/Virus Enclosed | NO |
| CELFVNDSYRQFDLVQ | 1647667 | Cell/Virus Enclosed | NO |
| CELHGNGSMIED | 1415601 | Cell/Virus Enclosed | NO |
| CELVNYKQSIPM | 1415603 | Cell/Virus Enclosed | NO |
| CELYVNNAYRIF | 1415605 | Cell/Virus Enclosed | NO |
| CELYVNNAYRLFDLVQ | 1647670 | Cell/Virus Enclosed | NO |
| CENLESAVLFVNDKIT | 1647672 | Cell/Virus Enclosed | NO |
| CEPVLTYSRLIS | 1415619 | Cell/Virus Enclosed | NO |
| CEQAYQLYDASN | 1415620 | Cell/Virus Enclosed | NO |
| CEQDLCDFKGWV | 1415623 | Cell/Virus Enclosed | NO |
| CEQFYQLYDASN | 1415625 | Cell/Virus Enclosed | NO |
| CEQSYQLYDACN | 1415628 | Cell/Virus Enclosed | NO |
| CEQSYQLYDASNVKKV | 1647680 | Cell/Virus Enclosed | NO |
| CEQTPCGFKGGI | 1415630 | Cell/Virus Enclosed | NO |
| CESHGKQVVSDV | 1415641 | Cell/Virus Enclosed | NO |
| CESIYEFDTYKA | 1415642 | Cell/Virus Enclosed | NO |
| CESTKLPYYDVEKIHI | 1647687 | Underneath Membrane | NO |
| CESTTVLDHTDW | 1415647 | Transmembrane | NO |
| CESTTVLDRTDW | 1415648 | Ectodomain | YES |
| CETFWRYNFDITESKY | 1647690 | Cell/Virus Enclosed | NO |
| CETSNVPPIVSE | 1415651 | Cell/Virus Enclosed | NO |
| CEVLKAILVKYG | 1415656 | Cell/Virus Enclosed | NO |
| CEVVHDQVNYVPLRAT | 1647693 | Cell/Virus Enclosed | NO |
| CEVVVTNLNKSAGWPL | 1647694 | Cell/Virus Enclosed | NO |
| CEYPQTICHPNL | 1415673 | Ectodomain | YES |
| CFALRNFDEKDCEVFK | 1647701 | Cell/Virus Enclosed | NO |
| CFALRNFDERDC | 1415679 | Cell/Virus Enclosed | NO |
| CFANKFEQFDQWYEST | 1647702 | Cell/Virus Enclosed | NO |
| CFCPRDFNHNMG | 1415698 | Cell/Virus Enclosed | NO |
| CFDIRDNGSLEK | 1415704 | Cell/Virus Enclosed | NO |
| CFDIRDNVSLEK | 1415706 | Cell/Virus Enclosed | NO |
| CFDKSKQFVPVQVVDS | 1647715 | Underneath Membrane | NO |
| CFDNSIQGSYER | 1415711 | Cell/Virus Enclosed | NO |
| CFEATEATFNYL | 1415721 | Cell/Virus Enclosed | NO |
| CFFTCNSNANIS | 1415732 | Ectodomain | YES |
| CFFTCNSNANLS | 1415733 | Ectodomain | YES |
| CFGKNNGIGTCP | 1415741 | Ectodomain | YES |
| CFGKRGPNQNFGGGEM | 1647726 | Cell/Virus Enclosed | NO |
| CFGKRGPSQNFGNAEM | 1647727 | Capsid | UN |
| CFGPRDFNHNMG | 1415750 | Cell/Virus Enclosed | NO |
| CFIDVVNKALEM | 1415758 | Cell/Virus Enclosed | NO |
| CFILLAIVMGLVFICVKNGNMRCTI | 1130442 | Undetermined | UN |
| CFINSTINTTHV | 1415762 | Ectodomain | YES |
| CFINSTINTTPV | 1415763 | Transmembrane | NO |
| CFKCCRRTRGLFKYRHLKD | 244173 | Underneath Membrane | NO |
| CFKKDHVVLKIT | 1415790 | Cell/Virus Enclosed | NO |
| CFKNDNVVLKITEDGI | 1647749 | Cell/Virus Enclosed | NO |
| CFKVFFKGNVQVDNGS | 1647751 | Cell/Virus Enclosed | NO |
| CFKVYFKGQTTHESSS | 1647752 | Cell/Virus Enclosed | NO |
| CFLNISNTHVSV | 6241 | Ectodomain | YES |
| CFMKSDIFGQDFKTFD | 1647762 | Cell/Virus Enclosed | NO |
| CFNAEGDAFWSS | 1415815 | Cell/Virus Enclosed | NO |
| CFNKRKYFTPVQ | 1415829 | Ectodomain | YES |
| CFRIYPQQMFVT | 1415846 | Cell/Virus Enclosed | NO |
| CFRTYPQQMFVT | 1415850 | Cell/Virus Enclosed | NO |
| CFSFVKMLFMLS | 1415856 | Cell/Virus Enclosed | NO |
| CFSKCHNCCDEYGGHH | 1647778 | Underneath Membrane | NO |
| CFSTCNSNANIS | 1415872 | Ectodomain | YES |
| CFSTCNSNASIS | 1415873 | Transmembrane | NO |
| CFSTVAVPGGCN | 1415878 | Transmembrane | NO |
| CFSTVEVPGSCN | 1415879 | Underneath Membrane | NO |
| CFSVAAITSGVKFQTV | 1647788 | Cell/Virus Enclosed | NO |
| CFSVAALSTGLTSQTV | 1647790 | Cell/Virus Enclosed | NO |
| CFTGVDGELPVAVVND | 1647796 | Cell/Virus Enclosed | NO |
| CFVDDIVKTDGT | 1415895 | Cell/Virus Enclosed | NO |
| CFVDDLLKTDSI | 1415896 | Cell/Virus Enclosed | NO |
| CFVKNVDWSITYPMIA | 1647808 | Cell/Virus Enclosed | NO |
| CFVKSDIFGSDFKTFD | 1647812 | Cell/Virus Enclosed | NO |
| CFVLAAVYRINWITG | 2000994 | Undetermined | UN |
| CFVLSLVCFLGL | 1415912 | Cell/Virus Enclosed | NO |
| CFVNDLLKTDSV | 1415915 | Cell/Virus Enclosed | NO |
| CFVNFNVDWIFF | 1415916 | Undetermined | UN |
| CFVNTTIDNETT | 1415918 | Ectodomain | YES |
| CFVNTTIGNETTSAFV | 1647818 | Ectodomain | YES |
| CGACIRRPFLCC | 1415939 | Cell/Virus Enclosed | NO |

| | | | |
|---|---|---|---|
| CGADGKPVMSEDLWQF | 1647829 | Cell/Virus Enclosed | NO |
| CGASYTANDVEVQSSG | 1647830 | Cell/Virus Enclosed | NO |
| CGCGASYTANDVEVQS | 1647832 | Cell/Virus Enclosed | NO |
| CGCGTSYTANDV | 1415951 | Cell/Virus Enclosed | NO |
| CGDCAACKSQLV | 1415954 | Ectodomain | YES |
| CGDIVSGYEKSTFNKD | 1647839 | Cell/Virus Enclosed | NO |
| CGDKFFPIVMNDKNIC | 1647840 | Cell/Virus Enclosed | NO |
| CGDNIGTNVPLQVGFS | 1647841 | Cell/Virus Enclosed | NO |
| CGDNSAKFWEEA | 1415964 | Cell/Virus Enclosed | NO |
| CGDQVHEVAAVR | 1415966 | Cell/Virus Enclosed | NO |
| CGDYAACKSQLVEYGS | 1647846 | Ectodomain | YES |
| CGEPWFLTDYNAILQS | 1647847 | Cell/Virus Enclosed | NO |
| CGFDGKPMLPKN | 1415983 | Cell/Virus Enclosed | NO |
| CGFDGKPVLPKN | 1415984 | Cell/Virus Enclosed | NO |
| CGFVMQFGYIDC | 1415997 | Cell/Virus Enclosed | NO |
| CGFVMQYGYVDC | 1415998 | Cell/Virus Enclosed | NO |
| CGGCCDDYTGHQ | 1416000 | Underneath Membrane | NO |
| CGGCCDDYTGYQELVI | 1647862 | Underneath Membrane | NO |
| CGGCCEDYTGYQ | 1416002 | Underneath Membrane | NO |
| CGGCYYVKPGGTSSGD | 1647863 | Cell/Virus Enclosed | NO |
| CGGCYYVTPGGT | 1416005 | Cell/Virus Enclosed | NO |
| CGGERHQQLVEH | 1416006 | Cell/Virus Enclosed | NO |
| CGGFCDDYTGYQ | 1416009 | Underneath Membrane | NO |
| CGGGYYVKPGGTSSGD | 1647865 | Cell/Virus Enclosed | NO |
| CGGILAVIDEHV | 1416010 | Cell/Virus Enclosed | NO |
| CGHVYEVSDLIA | 1416032 | Cell/Virus Enclosed | NO |
| CGHYHTNSYSQN | 1416033 | Cell/Virus Enclosed | NO |
| CGHYQTNIYSQNLCVD | 1647874 | Cell/Virus Enclosed | NO |
| CGHYTHITAKET | 1416036 | Cell/Virus Enclosed | NO |
| CGIKDVVLQGLR | 1416049 | Cell/Virus Enclosed | NO |
| CGIKQESRVGVD | 1416050 | Cell/Virus Enclosed | NO |
| CGIVELEKGVLP | 1416053 | Cell/Virus Enclosed | NO |
| CGKGYHLMSFPQAAPH | 1647882 | Ectodomain | YES |
| CGLFKDCARNPIDLPP | 1647888 | Cell/Virus Enclosed | NO |
| CGLILKFDSGTV | 1416063 | Cell/Virus Enclosed | NO |
| CGNCTFGAPDDA | 1416071 | Cell/Virus Enclosed | NO |
| CGNDSWLTGHAI | 1416072 | Cell/Virus Enclosed | NO |
| CGNDSWLTGNAIQGFA | 1647896 | Cell/Virus Enclosed | NO |
| CGNESWLTGNAI | 1416074 | Cell/Virus Enclosed | NO |
| CGNLCTLDPITL | 1416080 | Ectodomain | YES |
| CGNLHQRELLGVSDVW | 1647905 | Cell/Virus Enclosed | NO |
| CGNYKVTKGKPV | 1416087 | Cell/Virus Enclosed | NO |
| CGPDGYPLDCIK | 1416089 | Cell/Virus Enclosed | NO |
| CGPKLSTDLIKNQCVNF | 6332 | Ectodomain | YES |
| CGQKSTTLTGVE | 1416100 | Cell/Virus Enclosed | NO |
| CGRDATQYLVQQ | 1416104 | Cell/Virus Enclosed | NO |
| CGSACFSKCHNC | 1416108 | Underneath Membrane | NO |
| CGSCGNDSWLTGNAIQ | 1647923 | Cell/Virus Enclosed | NO |
| CGSCGSVGYTKEGSVI | 1647924 | Cell/Virus Enclosed | NO |
| CGSENWSVGAWD | 1416120 | Cell/Virus Enclosed | NO |
| CGSGNLVEKLSPWFMD | 1647930 | Cell/Virus Enclosed | NO |
| CGSPGYNLKNGEVEFV | 1647932 | Cell/Virus Enclosed | NO |
| CGSPGYNVRNDGTVEF | 1647933 | Cell/Virus Enclosed | NO |
| CGSSSRSFFEDLLFDK | 1647938 | Ectodomain | YES |
| CGSVGYTKDGSV | 1416133 | Cell/Virus Enclosed | NO |
| CGSVGYVIMGDCVKFV | 1647942 | Cell/Virus Enclosed | NO |
| CGSVGYVLTGDSVKFV | 1647943 | Cell/Virus Enclosed | NO |
| CGTQLNHSSCSC | 1416157 | Ectodomain | YES |
| CGTQLNHTSCSC | 1416158 | Ectodomain | YES |
| CGTSYTANDVEV | 1416168 | Cell/Virus Enclosed | NO |
| CGVHGNGSMIED | 1416181 | Cell/Virus Enclosed | NO |
| CGVLDGSYNASC | 1416187 | Ectodomain | YES |
| CGVLDGSYNVSC | 1416188 | Underneath Membrane | NO |
| CGVSDVTKLYLGGMSY | 1647973 | Cell/Virus Enclosed | NO |
| CGYTDFAEDVCTCYDN | 1647984 | Cell/Virus Enclosed | NO |
| CHALDCELYVNNAYRL | 1647985 | Cell/Virus Enclosed | NO |
| CHDPTVLPGAFA | 1416223 | Cell/Virus Enclosed | NO |
| CHGKNNGIGTCP | 1416227 | Underneath Membrane | NO |
| CHILGPDCCIEPHDW | 6391 | Secreted | YES |
| CHMFCNRTVYGPIKNV | 1648004 | Cell/Virus Enclosed | NO |
| CHNAAQCDCLCT | 1416239 | Ectodomain | YES |
| CHNCCDEYGGHHDFVI | 1648006 | Underneath Membrane | NO |
| CHNLCNPDPITF | 1416251 | Ectodomain | YES |
| CHPKLGNHFKEL | 1416256 | Ectodomain | YES |
| CHPKLGNHFKES | 1416257 | Ectodomain | YES |
| CHPNLGNHRKEL | 1416259 | Ectodomain | YES |

| | | | |
|---|---|---|---|
| CHPNLGNRRVEL | 1416261 | Ectodomain | YES |
| CHSNILSLNNPI | 1416267 | Ectodomain | YES |
| CHSQTSLRCGTC | 1416269 | Cell/Virus Enclosed | NO |
| CHSVAVIGDEQIDGKF | 1648017 | Cell/Virus Enclosed | NO |
| CHTGTAFNGDFY | 1416272 | Cell/Virus Enclosed | NO |
| CHTGTDFNGDFY | 1416273 | Cell/Virus Enclosed | NO |
| CHTGTDFTGNFY | 1416274 | Cell/Virus Enclosed | NO |
| CHVLDCELFVNDSYRQ | 1648020 | Cell/Virus Enclosed | NO |
| CHYLVLPSGNYL | 1416279 | Ectodomain | YES |
| CIAGNRGIAPKS | 1416285 | Ectodomain | YES |
| CIALQYSKPHFI | 1416288 | Cell/Virus Enclosed | NO |
| CIAPVNGYFIKT | 1416292 | Ectodomain | YES |
| CICLVASIPLCI | 1416300 | Cell/Virus Enclosed | NO |
| CIDMKGKMFVRPVIED | 1648036 | Underneath Membrane | NO |
| CIDMKGTMYVRPIIED | 1648037 | Ectodomain | YES |
| CIDQVTIAGAKL | 1416319 | Cell/Virus Enclosed | NO |
| CIERLKLFAAET | 1416326 | Cell/Virus Enclosed | NO |
| CIESTDSIVFTY | 1416327 | Cell/Virus Enclosed | NO |
| CIFNGKFLSNTKYYDD | 1648044 | Ectodomain | YES |
| CIFNGKSFSNTK | 1416336 | Ectodomain | YES |
| CIGRGNIGFTLE | 1416343 | Underneath Membrane | NO |
| CIGRRNIGFTLE | 1416345 | Underneath Membrane | NO |
| CIKACEVVVTNLNKSA | 1648054 | Cell/Virus Enclosed | NO |
| CIKDADYNAKDE | 1416364 | Cell/Virus Enclosed | NO |
| CIKDADYNAKVEISVT | 1648056 | Cell/Virus Enclosed | NO |
| CIKDLLARAGKS | 1416368 | Cell/Virus Enclosed | NO |
| CIKIAQADTAGL | 1416371 | Cell/Virus Enclosed | NO |
| CIKISTVQSKLTDLKC | 1648060 | Cell/Virus Enclosed | NO |
| CIKTAQADTAGLYTNF | 1648064 | Cell/Virus Enclosed | NO |
| CILEPRPGNHCP | 1416393 | Ectodomain | YES |
| CILEPRSGNYCP | 1416395 | Ectodomain | YES |
| CILRQLRHLPSRVRYPHYS | 6429 | Undetermined | UN |
| CILSVNSSNCNN | 1416409 | Cell/Virus Enclosed | NO |
| CIPASQVIVNNYDKSA | 1648085 | Cell/Virus Enclosed | NO |
| CIPLSVIPLCAS | 1416425 | Cell/Virus Enclosed | NO |
| CIPYNSITDTIV | 1416428 | Secreted | YES |
| CIRGAIGDVGLC | 1416437 | Cell/Virus Enclosed | NO |
| CIRRAKDGFSPE | 1416440 | Ectodomain | YES |
| CIRRPKDGFSPE | 1416442 | Ectodomain | YES |
| CISRLKDGFSLE | 1416454 | Ectodomain | YES |
| CITIIKLTKLCF | 1416470 | Cell/Virus Enclosed | NO |
| CITIKKPNELSL | 1416471 | Cell/Virus Enclosed | NO |
| CITPNVCFVKGD | 1416473 | Cell/Virus Enclosed | NO |
| CITSREVVVTNLNKSA | 1416476 | Cell/Virus Enclosed | NO |
| CITTNGSWAIFNDHHL | 1648115 | Cell/Virus Enclosed | NO |
| CIVDNVYMAKAGDKYY | 1648117 | Cell/Virus Enclosed | NO |
| CIVFDVKGVEEL | 1416485 | Cell/Virus Enclosed | NO |
| CIVGNKFYNVSTGLIP | 1648120 | Cell/Virus Enclosed | NO |
| CIVNLSFTEQLG | 1416493 | Underneath Membrane | NO |
| CIWNAAVYMKLS | 1416504 | Cell/Virus Enclosed | NO |
| CKAGQVQTFYPR | 1416511 | Cell/Virus Enclosed | NO |
| CKAISSNTDNET | 1416513 | Ectodomain | YES |
| CKCCYDHVISTS | 1416521 | Cell/Virus Enclosed | NO |
| CKCGNLEQNHIL | 1416525 | Cell/Virus Enclosed | NO |
| CKCGVKQEQCTG | 1416526 | Cell/Virus Enclosed | NO |
| CKCGVKQEQRIG | 1416527 | Cell/Virus Enclosed | NO |
| CKCGVKQEQRTGLDAV | 1648146 | Cell/Virus Enclosed | NO |
| CKDGKVQTFYPR | 1416530 | Cell/Virus Enclosed | NO |
| CKDGQVQTFYPRLQAS | 1648150 | Cell/Virus Enclosed | NO |
| CKDNAVATFYPQLQSA | 1648151 | Cell/Virus Enclosed | NO |
| CKEPMYAISNEQ | 1416545 | Ectodomain | YES |
| CKFAFYVLPSEA | 1416547 | Cell/Virus Enclosed | NO |
| CKFAVCGDGFVP | 1416548 | Cell/Virus Enclosed | NO |
| CKFGNTTFDFLS | 1416555 | Underneath Membrane | NO |
| CKFKGKCVQVPIGCLD | 1648161 | Cell/Virus Enclosed | NO |
| CKFSSNTTFDFL | 1416564 | Ectodomain | YES |
| CKGFSSNVSSDV | 1416571 | Ectodomain | YES |
| CKGFYSNASSDV | 1416572 | Ectodomain | YES |
| CKGFYSNSSSDV | 1416573 | Ectodomain | YES |
| CKHEKVSTNLCT | 1416579 | Cell/Virus Enclosed | NO |
| CKLNGSCPDKNN | 1416613 | Ectodomain | YES |
| CKLNGSCPGKNN | 1416614 | Ectodomain | YES |
| CKLPLGQSLCALPDTP | 1648185 | Ectodomain | YES |
| CKMCLFRYQEFN | 1416627 | Cell/Virus Enclosed | NO |
| CKPHQVNISLNGNTSV | 1648193 | Ectodomain | YES |
| CKPHQVNLSLNG | 1416637 | Underneath Membrane | NO |

| | | | |
|---|---|---|---|
| CKPKVEIDGNVMHANY | 1648194 | Cell/Virus Enclosed | NO |
| CKQYICNGFQKC | 1416640 | Ectodomain | YES |
| CKRALGLSISDHEFTS | 1648196 | Cell/Virus Enclosed | NO |
| CKSALYVLPSET | 1416652 | Cell/Virus Enclosed | NO |
| CKSGSKFKNLVL | 1416658 | Ectodomain | YES |
| CKSGSLVLNNPA | 1416660 | Ectodomain | YES |
| CKSGSLVLNNPV | 1416662 | Ectodomain | YES |
| CKSHKPPSASCPIGTN | 1648203 | Ectodomain | YES |
| CKSIVKETALIT | 1416670 | Cell/Virus Enclosed | NO |
| CKSKGSFRNESW | 1416671 | Ectodomain | YES |
| CKSKGSSRNESWHFDK | 1648206 | Ectodomain | YES |
| CKSKNPLLHDAN | 1416673 | Secreted | YES |
| CKSNDSCPGKNN | 1416677 | Ectodomain | YES |
| CKSNGSCPGKNN | 1416678 | Ectodomain | YES |
| CKSNSSLCVGSG | 1416679 | Underneath Membrane | NO |
| CKSQLGEYGSFC | 1416680 | Underneath Membrane | NO |
| CKSQLVEYGSFCDNIN | 1648208 | Ectodomain | YES |
| CKSQNPLLYDAN | 1416683 | Secreted | YES |
| CKSVNWNLEYPIISNE | 1648210 | Cell/Virus Enclosed | NO |
| CKTAVTTGDVKI | 1416691 | Cell/Virus Enclosed | NO |
| CKTIEDALRNSARLES | 1648217 | Ectodomain | YES |
| CKTILNIVGPDARQDG | 1648218 | Cell/Virus Enclosed | NO |
| CKTKSLLPNTGVYDLS | 1648220 | Ectodomain | YES |
| CKTQFIAPPTGV | 1416699 | Underneath Membrane | NO |
| CKTVLNVVGPDARTQG | 1648223 | Cell/Virus Enclosed | NO |
| CKVTDVLFPGQKYSSD | 1648227 | Cell/Virus Enclosed | NO |
| CKVTDVLFSGQK | 1416716 | Cell/Virus Enclosed | NO |
| CLARSACVYYSQ | 1416746 | Cell/Virus Enclosed | NO |
| CLASECFMKSDIFGQD | 1648246 | Cell/Virus Enclosed | NO |
| CLASECFVKSDIFGSD | 1648247 | Cell/Virus Enclosed | NO |
| CLASIAFNHPIQ | 1416748 | Ectodomain | YES |
| CLASIEEVCDDY | 1416749 | Cell/Virus Enclosed | NO |
| CLDAGINYVKSP | 1416765 | Cell/Virus Enclosed | NO |
| CLDVVRKELEEG | 1416773 | Cell/Virus Enclosed | NO |
| CLENHVMVDCRR | 1416778 | Cell/Virus Enclosed | NO |
| CLFKKNFTYNVS | 1416786 | Ectodomain | YES |
| CLFKKNFTYNVY | 1416787 | Ectodomain | YES |
| CLFKSNFNQYYY | 1416788 | Ectodomain | YES |
| CLFSYQEFNDLD | 1416796 | Cell/Virus Enclosed | NO |
| CLFSYQEFNDLV | 1416797 | Cell/Virus Enclosed | NO |
| CLGDINARDLIC | 1416814 | Ectodomain | YES |
| CLGGINARDLIC | 1416821 | Ectodomain | YES |
| CLGPHCGSSSRSFFED | 1648295 | Ectodomain | YES |
| CLGSECNKVSSR | 1416835 | Ectodomain | YES |
| CLGSECSKASSR | 1416836 | Ectodomain | YES |
| CLGSLIYVTAAL | 1416841 | Cell/Virus Enclosed | NO |
| CLGVGDSAVTYS | 1416842 | Cell/Virus Enclosed | NO |
| CLIGAEYVDTSY | 1416855 | Ectodomain | YES |
| CLILATVSHNLT | 1416859 | Ectodomain | YES |
| CLKCGFSFDLNGLDAL | 1648315 | Cell/Virus Enclosed | NO |
| CLKGACSCGSCC | 1416877 | Underneath Membrane | NO |
| CLKHAEEYREYLESYN | 1648318 | Cell/Virus Enclosed | NO |
| CLKSIAATRGVP | 1416890 | Cell/Virus Enclosed | NO |
| CLLQFAYANRNRFLY | 1312263 | Undetermined | UN |
| CLMPTYTVHKSD | 1416938 | Cell/Virus Enclosed | NO |
| CLNASEVVVNNLDKSA | 1648345 | Cell/Virus Enclosed | NO |
| CLNGLDSLDSYP | 1416945 | Cell/Virus Enclosed | NO |
| CLPDPISTYSPN | 1416953 | Ectodomain | YES |
| CLPDPITAYDPRSCSQ | 1648351 | Ectodomain | YES |
| CLPTFSMAPVEFYEDR | 1648353 | Cell/Virus Enclosed | NO |
| CLQGDKCNIFSN | 1416962 | Underneath Membrane | NO |
| CLQHLHVASNSKLWHY | 1648356 | Cell/Virus Enclosed | NO |
| CLQHLHVASNSKLWQY | 1648357 | Cell/Virus Enclosed | NO |
| CLSGRKIYEVAR | 1416985 | Cell/Virus Enclosed | NO |
| CLSNNPCNVCQY | 1416992 | Cell/Virus Enclosed | NO |
| CLSNVSTSHHCI | 1416994 | Ectodomain | YES |
| CLSSMNIAANSN | 1416996 | Cell/Virus Enclosed | NO |
| CLSSMNIAANSSEWAY | 1648371 | Cell/Virus Enclosed | NO |
| CLSTGCCGCCNC | 1416999 | Undetermined | UN |
| CLTDCLYLKNLTQTFT | 1648376 | Cell/Virus Enclosed | NO |
| CLTFVVKHKVLF | 1417007 | Cell/Virus Enclosed | NO |
| CLTSSMRGCCDC | 1417016 | Underneath Membrane | NO |
| CLTYANIVHEPT | 1417017 | Cell/Virus Enclosed | NO |
| CLVSSDGQPGGDAISL | 1648386 | Cell/Virus Enclosed | NO |
| CLVSSNGQPGGD | 1417032 | Cell/Virus Enclosed | NO |
| CLYCRAHIEHPD | 1417052 | Cell/Virus Enclosed | NO |

| | | | |
|---|---|---|---|
| CLYKRNFTYDVN | 1417058 | Ectodomain | YES |
| CLYLKNLTQTFTSMLT | 1648403 | Cell/Virus Enclosed | NO |
| CLYRNRDVDQEF | 1417066 | Cell/Virus Enclosed | NO |
| CMFENKTTLPTNIAFE | 1648412 | Cell/Virus Enclosed | NO |
| CMGLNKPFVKDN | 1417089 | Cell/Virus Enclosed | NO |
| CMHVVGPNLNAG | 1417092 | Cell/Virus Enclosed | NO |
| CMKQGPASARDL | 1417095 | Ectodomain | YES |
| CMLRACIKTAQADTAG | 1648417 | Cell/Virus Enclosed | NO |
| CMLRAFIKTAQA | 1417103 | Cell/Virus Enclosed | NO |
| CMQQGPASAREL | 1417114 | Underneath Membrane | NO |
| CMQQGPASARNL | 1417115 | Underneath Membrane | NO |
| CMRAGSKVINHN | 1417119 | Cell/Virus Enclosed | NO |
| CMYTDLNFIDKL | 1417139 | Cell/Virus Enclosed | NO |
| CNAASVNQIVLRNSNG | 1648438 | Cell/Virus Enclosed | NO |
| CNADIVVVDEVSMCTN | 1648439 | Cell/Virus Enclosed | NO |
| CNAISPKTDNET | 1417147 | Ectodomain | YES |
| CNAISSNTDNETLQYW | 1648440 | Ectodomain | YES |
| CNAPGCDVNDVTKLYL | 1648442 | Cell/Virus Enclosed | NO |
| CNASGCGVSDVKKLYL | 1648449 | Cell/Virus Enclosed | NO |
| CNAVAKHDFFTWHEGR | 1648451 | Cell/Virus Enclosed | NO |
| CNCDSLRQAALP | 1417165 | Cell/Virus Enclosed | NO |
| CNCGSENWSVGA | 1417169 | Cell/Virus Enclosed | NO |
| CNCLCTPDPITS | 1417172 | Ectodomain | YES |
| CNCLTSSMRGCC | 1417174 | Underneath Membrane | NO |
| CNCTFGAPDDAS | 1417175 | Cell/Virus Enclosed | NO |
| CNDEKVMTFYPRLQAA | 1648457 | Cell/Virus Enclosed | NO |
| CNESIACQFCLA | 1417185 | Cell/Virus Enclosed | NO |
| CNFELCDNPFIA | 1417190 | Ectodomain | YES |
| CNGCVGHSENVF | 1417201 | Ectodomain | YES |
| CNGCVGYSENVF | 1417203 | Ectodomain | YES |
| CNGHKIEDLSIRNLQK | 1648471 | Cell/Virus Enclosed | NO |
| CNGNKIEDFSIR | 1417215 | Cell/Virus Enclosed | NO |
| CNGNKIEDLSIR | 1417216 | Cell/Virus Enclosed | NO |
| CNGNKIEDLSMR | 1417217 | Cell/Virus Enclosed | NO |
| CNGNPRCKNLLKQYTS | 1648474 | Ectodomain | YES |
| CNGNVRCVELLKQYTS | 1648475 | Ectodomain | YES |
| CNGSIACQFCLA | 1417225 | Cell/Virus Enclosed | NO |
| CNGTDIDYCVRA | 1417228 | Cell/Virus Enclosed | NO |
| CNGTDIDYCVRV | 1417229 | Cell/Virus Enclosed | NO |
| CNGYQHHSVADV | 1417235 | Ectodomain | YES |
| CNGYQHNSFADV | 1417236 | Ectodomain | YES |
| CNGYQHNSVVDV | 1417238 | Ectodomain | YES |
| CNGYTDNIFSVQQDGR | 1648486 | Ectodomain | YES |
| CNHIGNIKPHNY | 1417243 | Ectodomain | YES |
| CNHINNLKIKNFDYLD | 1648488 | Ectodomain | YES |
| CNHINNLKVKNY | 1417247 | Ectodomain | YES |
| CNIAKSAYERDRAVAK | 1648492 | Cell/Virus Enclosed | NO |
| CNIEAWLNDKAV | 1417254 | Ectodomain | YES |
| CNIEAWLNDKLV | 1417255 | Ectodomain | YES |
| CNIEAWLNDKSVPSPL | 1648494 | Ectodomain | YES |
| CNIEAWLNGKSV | 1417257 | Ectodomain | YES |
| CNIESWLNDKSV | 1417258 | Ectodomain | YES |
| CNIGAWLNDKSV | 1417264 | Underneath Membrane | NO |
| CNIGGAVCSKHANLYQ | 1648497 | Cell/Virus Enclosed | NO |
| CNIGGAVCSKHANLYR | 1648498 | Cell/Virus Enclosed | NO |
| CNLGGAVCRKHATEYR | 1648511 | Cell/Virus Enclosed | NO |
| CNLLLLFVTIYS | 1417285 | Secreted | YES |
| CNLYNYGAGIKLPSGI | 1648518 | Cell/Virus Enclosed | NO |
| CNNIDAAKIYGMCFSS | 1648526 | Ectodomain | YES |
| CNNIDVAKIYGM | 1417300 | Underneath Membrane | NO |
| CNNINWNVEYPIISNE | 1648527 | Cell/Virus Enclosed | NO |
| CNNLVPTYLKSDNIVA | 1648528 | Cell/Virus Enclosed | NO |
| CNNLVQTYLKSD | 1417304 | Cell/Virus Enclosed | NO |
| CNNMHLSTLMKC | 1417305 | Cell/Virus Enclosed | NO |
| CNNSDYASKGYI | 1417306 | Cell/Virus Enclosed | NO |
| CNPDPIRFTGPY | 1417309 | Ectodomain | YES |
| CNQPDQVAVYQQSIIG | 1648536 | Ectodomain | YES |
| CNRAGSNDIWMQ | 1417319 | Ectodomain | YES |
| CNRCCDRYEEYDLEPH | 1648537 | Underneath Membrane | NO |
| CNRSGSNDIWMQNKGL | 1648539 | Ectodomain | YES |
| CNSFFKDLTANM | 1417335 | Cell/Virus Enclosed | NO |
| CNSFFKELTAIM | 1417336 | Cell/Virus Enclosed | NO |
| CNSFFKELTANMSMAE | 1648545 | Cell/Virus Enclosed | NO |
| CNSKVKNGFTLE | 1417343 | Ectodomain | YES |
| CNSNANISMLQL | 1417346 | Ectodomain | YES |
| CNSNANISTLQL | 1417347 | Underneath Membrane | NO |

| | | | |
|---|---|---|---|
| CNSNANLSMLQLGVPD | 1648547 | Ectodomain | YES |
| CNSVINRLRCDQLSFD | 1648550 | Ectodomain | YES |
| CNSVKDTLTYYA | 1417357 | Cell/Virus Enclosed | NO |
| CNTGSCTVNKHV | 1417361 | Cell/Virus Enclosed | NO |
| CNTKCQTPMGAINSS | 148521 | Ectodomain | YES |
| CNTPTQCYYNNSNNGK | 1648559 | Cell/Virus Enclosed | NO |
| CNTSGCNVNDVTKLYL | 1648560 | Cell/Virus Enclosed | NO |
| CNVDRYPANAIV | 1417380 | Cell/Virus Enclosed | NO |
| CNVHRNEHVASGDAIM | 1648573 | Cell/Virus Enclosed | NO |
| CNVIADFKSFTL | 1417382 | Cell/Virus Enclosed | NO |
| CNVIADPKSFTL | 1417383 | Cell/Virus Enclosed | NO |
| CNVLDDFIVFNNNGTN | 1648577 | Cell/Virus Enclosed | NO |
| CNVSFVGELKAAEAKV | 1648585 | Cell/Virus Enclosed | NO |
| CNWFIQSDKCSI | 1417407 | Cell/Virus Enclosed | NO |
| CNWFIQSDKCSV | 1417408 | Cell/Virus Enclosed | NO |
| CNWFVQSDKCSV | 1417410 | Cell/Virus Enclosed | NO |
| CNYLESYNIVTTAGFT | 1648593 | Cell/Virus Enclosed | NO |
| CNYSKFWYLEHAK | 6742 | Ectodomain | YES |
| CNYVFNNSLTRQ | 1417422 | Ectodomain | YES |
| CPADNSYTSFAT | 1417430 | Ectodomain | YES |
| CPADQLSDPNYD | 1417431 | Cell/Virus Enclosed | NO |
| CPAGNSHTSFAT | 1417435 | Ectodomain | YES |
| CPAGNSYTSFAT | 1417437 | Ectodomain | YES |
| CPAGTNSLTCGN | 1417444 | Ectodomain | YES |
| CPAGTNYLNCDN | 1417445 | Ectodomain | YES |
| CPCADPSVVNSC | 1417451 | Ectodomain | YES |
| CPCADRSVVNSC | 1417452 | Ectodomain | YES |
| CPCKLNGSCVGSGPGK | 1648609 | Ectodomain | YES |
| CPCSSCFGKNNG | 1417466 | Ectodomain | YES |
| CPCSSCHGKNNG | 1417467 | Underneath Membrane | NO |
| CPCSSCPGKNNG | 1417468 | Ectodomain | YES |
| CPCSSCSCPGKN | 1417470 | Ectodomain | YES |
| CPCSSCSGKNNG | 1417471 | Ectodomain | YES |
| CPDKNNGIGICP | 1417475 | Ectodomain | YES |
| CPDKNNGIGTCP | 1417476 | Ectodomain | YES |
| CPECRPLGLQGCAFQSTVAELQRLK | 6757 | Undetermined | UN |
| CPEWMDDFEADPKGKY | 1648614 | Cell/Virus Enclosed | NO |
| CPEWMDEFEADP | 1417481 | Cell/Virus Enclosed | NO |
| CPEWMDYFEADP | 1417482 | Cell/Virus Enclosed | NO |
| CPFAKGDSCSSNCKHS | 1648615 | Cell/Virus Enclosed | NO |
| CPFVKGDSCSSN | 1417496 | Cell/Virus Enclosed | NO |
| CPGKNNGIGTCP | 1417497 | Ectodomain | YES |
| CPGKNNGIGTRP | 1417498 | Underneath Membrane | NO |
| CPGKNNGISTCP | 1417499 | Underneath Membrane | NO |
| CPGKNNGLGTCP | 1417500 | Ectodomain | YES |
| CPIGTNYRSCESTTVL | 1648624 | Ectodomain | YES |
| CPKEIVDTVSALVYEN | 1648632 | Cell/Virus Enclosed | NO |
| CPKEIVETVSALVYDN | 1648633 | Cell/Virus Enclosed | NO |
| CPKEIVSTVSALVYNN | 1648634 | Cell/Virus Enclosed | NO |
| CPKFEFANDTKI | 1417516 | Ectodomain | YES |
| CPKLEFANDTKIASQL | 1648637 | Ectodomain | YES |
| CPLCCTVYDSSEVKAQ | 1648638 | Cell/Virus Enclosed | NO |
| CPLDPKLKGSFN | 1417526 | Ectodomain | YES |
| CPLDPRLKGSFN | 1417527 | Underneath Membrane | NO |
| CPLDSRTGSFNN | 1417529 | Ectodomain | YES |
| CPPGMGIPYCTS | 1417550 | Cell/Virus Enclosed | NO |
| CPQSKSLVGIGE | 1417555 | Ectodomain | YES |
| CPRHVICPADQL | 1417561 | Cell/Virus Enclosed | NO |
| CPRHVMCPADQL | 1417566 | Cell/Virus Enclosed | NO |
| CPSATYEGNSPF | 1417568 | Ectodomain | YES |
| CPSKNNGIGTCP | 1417572 | Ectodomain | YES |
| CPTAADINIHDL | 1417575 | Ectodomain | YES |
| CPTAADINIPDLPICV | 1648657 | Ectodomain | YES |
| CPTAADINIPHL | 1417577 | Ectodomain | YES |
| CPTAADINNPDL | 1417578 | Ectodomain | YES |
| CPTAADINTPDV | 1417579 | Ectodomain | YES |
| CPTGTKYRHCDL | 1417583 | Ectodomain | YES |
| CPWLLRKFYQHV | 1417594 | Cell/Virus Enclosed | NO |
| CQAVSANVCALI | 1417597 | Cell/Virus Enclosed | NO |
| CQAVSANVCALMSCNG | 1648661 | Cell/Virus Enclosed | NO |
| CQAVTANVCSPM | 1417599 | Cell/Virus Enclosed | NO |
| CQAVTANVRSLT | 1417601 | Cell/Virus Enclosed | NO |
| CQDKILTHYENDSTD | 111167 | Cell/Virus Enclosed | NO |
| CQFKDKNLQGLW | 1417618 | Cell/Virus Enclosed | NO |
| CQFNMCEYPHTI | 1417619 | Ectodomain | YES |
| CQITSVVGTKAL | 1417626 | Cell/Virus Enclosed | NO |

| | | | |
|---|---|---|---|
| CQLPYCYFRNSSANYV | 1648675 | Underneath Membrane | NO |
| CQLSFAGYTESAVNST | 1648676 | Cell/Virus Enclosed | NO |
| CQLYYNLPSANV | 1417637 | Underneath Membrane | NO |
| CQNLDCNCLGFY | 1417644 | Cell/Virus Enclosed | NO |
| CQPVSEYHITLA | 1417666 | Cell/Virus Enclosed | NO |
| CQTADTAHANNINRFN | 1648686 | Cell/Virus Enclosed | NO |
| CQTTNGLNTSYSVCNG | 1648688 | Ectodomain | YES |
| CQTTNGMNTSQS | 1417681 | Ectodomain | YES |
| CQTTNGTNTIHS | 1417682 | Ectodomain | YES |
| CQTTNGTNTSNS | 1417684 | Ectodomain | YES |
| CQVHGNAHVASC | 1417686 | Cell/Virus Enclosed | NO |
| CQVHGNGSMIED | 1417687 | Cell/Virus Enclosed | NO |
| CQYKGKWVQVPIGTND | 1648694 | Cell/Virus Enclosed | NO |
| CQYNMCDYPHTI | 1417691 | Ectodomain | YES |
| CQYNMCEHPHTI | 1417692 | Underneath Membrane | NO |
| CQYNMCEHPHTS | 1417693 | Ectodomain | YES |
| CQYNMCEYPHTT | 1417696 | Underneath Membrane | NO |
| CQYNMCEYPQTI | 1417697 | Ectodomain | YES |
| CQYNMCEYPYTI | 1417698 | Ectodomain | YES |
| CQYTMCEFPHTV | 1417699 | Ectodomain | YES |
| CQYTMCEYPHTV | 1417701 | Ectodomain | YES |
| CQYTMCEYPNTI | 1417702 | Ectodomain | YES |
| CQYWIGYGCNCD | 1417704 | Cell/Virus Enclosed | NO |
| CRAYLDSCTTQQ | 1417711 | Cell/Virus Enclosed | NO |
| CRFDTRVLNNLNLPGC | 1648720 | Cell/Virus Enclosed | NO |
| CRFPNITNSHVPILQERPPLENR | 6919 | Ectodomain | YES |
| CRFQRIDDDGNKLDKF | 1648728 | Cell/Virus Enclosed | NO |
| CRFQRVDENGDKLDQF | 1648729 | Cell/Virus Enclosed | NO |
| CRIPNGFPFNNW | 1417773 | Underneath Membrane | NO |
| CRKCNLAFRLTS | 1417776 | Cell/Virus Enclosed | NO |
| CRKHATEYREYMEAYN | 1648746 | Cell/Virus Enclosed | NO |
| CRKIGVNVCSDSTFEE | 1648747 | Cell/Virus Enclosed | NO |
| CRKLGSSVRSDG | 1417789 | Cell/Virus Enclosed | NO |
| CRKLGTAVRSDG | 1417790 | Cell/Virus Enclosed | NO |
| CRLAMLGAEMTSNVRR | 1648753 | Cell/Virus Enclosed | NO |
| CRLKNSVSDVAYNRYL | 1648754 | Cell/Virus Enclosed | NO |
| CRLYIEESDLIISNFK | 1648757 | Cell/Virus Enclosed | NO |
| CRPIKLVDSELLSTLS | 1648760 | Cell/Virus Enclosed | NO |
| CRQYSNESWFEDDYRA | 1648763 | Cell/Virus Enclosed | NO |
| CRRLLKQECCVQ | 1417827 | Cell/Virus Enclosed | NO |
| CRTDMEKDFIALDQDV | 1648769 | Cell/Virus Enclosed | NO |
| CRVGECRDSHKG | 1417842 | Cell/Virus Enclosed | NO |
| CRVGECVESNAGVCFG | 1648773 | Cell/Virus Enclosed | NO |
| CSAFVCGDCAAC | 1417855 | Ectodomain | YES |
| CSASDMTNPDYTNLLC | 1648784 | Cell/Virus Enclosed | NO |
| CSCDQLREPMMQ | 1417868 | Cell/Virus Enclosed | NO |
| CSCPGKNNGIGT | 1417881 | Ectodomain | YES |
| CSCSPHAFLGWS | 1417884 | Ectodomain | YES |
| CSCVGSSVAVQSKDLN | 1648796 | Cell/Virus Enclosed | NO |
| CSCVSNDTTVQS | 1417888 | Cell/Virus Enclosed | NO |
| CSCVSTDTTVQSKDTN | 1648797 | Cell/Virus Enclosed | NO |
| CSDCYDDMCVIH | 1417890 | Cell/Virus Enclosed | NO |
| CSDFEPMSDMEN | 1417893 | Cell/Virus Enclosed | NO |
| CSDFIPLSDMEK | 1417896 | Cell/Virus Enclosed | NO |
| CSDFLPLSDMEKDFLS | 1648799 | Cell/Virus Enclosed | NO |
| CSDFYADKILYQ | 1417900 | Cell/Virus Enclosed | NO |
| CSDGNYNCNASL | 1417901 | Underneath Membrane | NO |
| CSDGNYNRNAIL | 1417902 | Ectodomain | YES |
| CSDGNYNRNASLNSFK | 1648801 | Ectodomain | YES |
| CSDGNYNRYASL | 1417904 | Ectodomain | YES |
| CSETVNGYAKSNFVKD | 1648806 | Cell/Virus Enclosed | NO |
| CSEVEASGLEEG | 1417915 | Cell/Virus Enclosed | NO |
| CSFYPPDEEEDC | 1417931 | Cell/Virus Enclosed | NO |
| CSGKNNGIGTCP | 1417935 | Ectodomain | YES |
| CSGLAIKSDYCG | 1417936 | Ectodomain | YES |
| CSGLAVKSDYCGGNSC | 1648819 | Ectodomain | YES |
| CSGVMWLYTYSIGEAS | 1648820 | Cell/Virus Enclosed | NO |
| CSGVTFQGKFKK | 1417942 | Cell/Virus Enclosed | NO |
| CSIEDFNVWALS | 1417951 | Cell/Virus Enclosed | NO |
| CSIGIGVYYPGA | 1417954 | Cell/Virus Enclosed | NO |
| CSKEEDGFAFYP | 1417966 | Cell/Virus Enclosed | NO |
| CSKHAEEYCNYLESYN | 1648837 | Cell/Virus Enclosed | NO |
| CSKHANLYQKYVEAYN | 1648838 | Cell/Virus Enclosed | NO |
| CSKIINGLHPTQ | 1417971 | Cell/Virus Enclosed | NO |
| CSKSCLGYHPAHAPSF | 1648845 | Cell/Virus Enclosed | NO |
| CSKSYSGYHPAHAPSF | 1648847 | Cell/Virus Enclosed | NO |

| | | | |
|---|---|---|---|
| CSNDLLQPNTEVFTDV | 1648866 | Ectodomain | YES |
| CSNDLLYSNTDV | 1418000 | Ectodomain | YES |
| CSNDLLYSNTEI | 1418001 | Underneath Membrane | NO |
| CSNSSSGALDTT | 1418007 | Ectodomain | YES |
| CSNSSSGVLDTTIPFG | 1648870 | Ectodomain | YES |
| CSNSYSGYHPAH | 1418009 | Cell/Virus Enclosed | NO |
| CSPMACNGHKIE | 1418016 | Cell/Virus Enclosed | NO |
| CSQHTHYIKDGD | 1418021 | Cell/Virus Enclosed | NO |
| CSQHTLYIKDGDDGYF | 1648881 | Cell/Virus Enclosed | NO |
| CSQHTMLVKMDGDDVY | 1648884 | Cell/Virus Enclosed | NO |
| CSQHTMQIVDENGKYY | 1648886 | Cell/Virus Enclosed | NO |
| CSQHTMQIVDKDGTYY | 1648887 | Cell/Virus Enclosed | NO |
| CSQISPAAIASN | 1418026 | Ectodomain | YES |
| CSQISPAAISSN | 1418027 | Ectodomain | YES |
| CSQNPLAELKCSVKSFE | 7064 | Ectodomain | YES |
| CSRETSGLSPAYAPTY | 1648895 | Cell/Virus Enclosed | NO |
| CSRLLSDDRTEVPQLV | 1648898 | Ectodomain | YES |
| CSSCFGKNNGIG | 1418066 | Ectodomain | YES |
| CSSCHGKNNGIG | 1418067 | Underneath Membrane | NO |
| CSSCPGKNNGIG | 1418068 | Ectodomain | YES |
| CSSCPGKNNGIS | 1418069 | Underneath Membrane | NO |
| CSSCPGKNNGLG | 1418070 | Ectodomain | YES |
| CSSCPSKNNGIG | 1418071 | Ectodomain | YES |
| CSSCSCPGKNNG | 1418072 | Ectodomain | YES |
| CSSCSGKNNGIG | 1418073 | Ectodomain | YES |
| CSSCSGKNNGVG | 1418074 | Ectodomain | YES |
| CSSHYWSPQGNLGGNC | 1648901 | Cell/Virus Enclosed | NO |
| CSSLTVLSDGRV | 1418082 | Cell/Virus Enclosed | NO |
| CSSSFLSEIQCK | 1418089 | Ectodomain | YES |
| CSTDLQKANIEI | 1418098 | Underneath Membrane | NO |
| CSTDLQKANTEI | 1418100 | Ectodomain | YES |
| CSTLHNEILATS | 1418109 | Cell/Virus Enclosed | NO |
| CSTNLFKDCSKSYSGY | 1648920 | Cell/Virus Enclosed | NO |
| CSTNLFKDCSNS | 1418113 | Cell/Virus Enclosed | NO |
| CSTRVNVDGFNEWAMA | 1648921 | Cell/Virus Enclosed | NO |
| CSVEDFNVWALSNGFS | 1378250 | Cell/Virus Enclosed | NO |
| CSVIDLLLNDFV | 1418131 | Cell/Virus Enclosed | NO |
| CSVKSFEIDKGIYQTS | 1648934 | Ectodomain | YES |
| CSYESFDVESGV | 1418153 | Ectodomain | YES |
| CTAEDMLNPNYE | 1418167 | Cell/Virus Enclosed | NO |
| CTANDKFYRLSN | 1418168 | Cell/Virus Enclosed | NO |
| CTARVVYTAASHAAVD | 1648944 | Cell/Virus Enclosed | NO |
| CTASDKFYRLSNELAQ | 1648945 | Cell/Virus Enclosed | NO |
| CTATDRFCRLGN | 1418169 | Cell/Virus Enclosed | NO |
| CTCDRTAIQSFDNSYL | 1648948 | Cell/Virus Enclosed | NO |
| CTCYDNSIQGSY | 1418186 | Cell/Virus Enclosed | NO |
| CTDNSANTKVLRNKGV | 1648962 | Cell/Virus Enclosed | NO |
| CTDVPTAIRADQ | 1418195 | Ectodomain | YES |
| CTDVPTSIHADQ | 1418197 | Ectodomain | YES |
| CTFEDELAPSDVYQQL | 1648972 | Cell/Virus Enclosed | NO |
| CTFGSPDDASRL | 1418221 | Cell/Virus Enclosed | NO |
| CTGGIDICSQDK | 1418246 | Cell/Virus Enclosed | NO |
| CTGLDAVMHFGT | 1418247 | Cell/Virus Enclosed | NO |
| CTGSPYIDDFNKIASC | 1648994 | Cell/Virus Enclosed | NO |
| CTGSWNKVNQIA | 1418255 | Cell/Virus Enclosed | NO |
| CTGSWNKVTQIA | 1418256 | Cell/Virus Enclosed | NO |
| CTHWCTYLFSLI | 1418262 | Cell/Virus Enclosed | NO |
| CTIANASSSEGF | 1418264 | Cell/Virus Enclosed | NO |
| CTIAQLTDAALSIKEN | 1378263 | Cell/Virus Enclosed | NO |
| CTIDSNTTQDTYGGAS | 1649003 | Cell/Virus Enclosed | NO |
| CTIKNFNVSKGI | 1418274 | Ectodomain | YES |
| CTIPAQGTSMFPSCCCTKPSDGNC | 7161 | Underneath Membrane | NO |
| CTIPAQGTSVFPSCCCTKPSDGNC | 7162 | Underneath Membrane | NO |
| CTKGLSIADLACAQYY | 1649011 | Ectodomain | YES |
| CTKHTMLVKMDG | 1418284 | Cell/Virus Enclosed | NO |
| CTKKFLTVDGVQ | 1418285 | Cell/Virus Enclosed | NO |
| CTKPSDGNCTCIPIP | 1649013 | Underneath Membrane | NO |
| CTKYNIYDVAGV | 1418293 | Ectodomain | YES |
| CTKYNIYDVSGVGVIR | 1649016 | Ectodomain | YES |
| CTKYNIYDVVGV | 1418295 | Ectodomain | YES |
| CTKYNIYDYVGTGIIR | 1649017 | Ectodomain | YES |
| CTLDPIALKAPD | 1418300 | Underneath Membrane | NO |
| CTLDPISFKAPD | 1418301 | Ectodomain | YES |
| CTLDPITLKAPDTYKCPQSK | 2060800 | Ectodomain | YES |
| CTLDPITLKTPD | 1418302 | Ectodomain | YES |
| CTLDSITFKAPG | 1418305 | Ectodomain | YES |

| | | | |
|---|---|---|---|
| CTMFMDAEGRMT | 1418327 | Cell/Virus Enclosed | NO |
| CTMFRDAEGRIT | 1418329 | Cell/Virus Enclosed | NO |
| CTMFRDAEGRMT | 1418330 | Cell/Virus Enclosed | NO |
| CTMFTMADGSLQ | 1418332 | Cell/Virus Enclosed | NO |
| CTMFTTADGSPQ | 1418334 | Cell/Virus Enclosed | NO |
| CTMYICGDSQEC | 1418339 | Ectodomain | YES |
| CTNFAINDLNTTVPRI | 1649037 | Ectodomain | YES |
| CTNGHALPTLFR | 1418344 | Cell/Virus Enclosed | NO |
| CTNNTLVSGDAHIPFG | 1649040 | Ectodomain | YES |
| CTPESKPLADDD | 1418360 | Cell/Virus Enclosed | NO |
| CTPPALNCYWPLNDYGF | 7193 | Ectodomain | YES |
| CTPTKKAFPYGTCLNC | 1649052 | Cell/Virus Enclosed | NO |
| CTQLNRALSGIAAEQD | 1649054 | Ectodomain | YES |
| CTQVYIAVNDKA | 1418377 | Cell/Virus Enclosed | NO |
| CTRASTSAVKLQNNEI | 1649056 | Cell/Virus Enclosed | NO |
| CTRKLNRLMCDI | 1418387 | Cell/Virus Enclosed | NO |
| CTRLEGLGGNNVYCYN | 1649064 | Cell/Virus Enclosed | NO |
| CTRTPLNLPPTH | 1418391 | Cell/Virus Enclosed | NO |
| CTSDNINDKDTGPPPI | 1649066 | Ectodomain | YES |
| CTSPGPSGIGAV | 1418402 | Cell/Virus Enclosed | NO |
| CTSYYAYMMPVM | 1418408 | Cell/Virus Enclosed | NO |
| CTSYYSYMMPIM | 1418409 | Cell/Virus Enclosed | NO |
| CTSYYSYMMPVM | 1418410 | Cell/Virus Enclosed | NO |
| CTTAVMTYSNFG | 1418411 | Ectodomain | YES |
| CTTCGHVYEVGD | 1418412 | Cell/Virus Enclosed | NO |
| CTTFDDVQAPNYTQHT | 1649082 | Ectodomain | YES |
| CTTIVNGMKRSF | 1418418 | Cell/Virus Enclosed | NO |
| CTTPAQGNSMFPSCCCTKPTDGNC | 7233 | Underneath Membrane | NO |
| CTVQGPPGTGKSHLAI | 1649095 | Cell/Virus Enclosed | NO |
| CTVQQCFGKRGPNQNF | 1649096 | Cell/Virus Enclosed | NO |
| CTVYDSSEVKAQSSGV | 1649101 | Cell/Virus Enclosed | NO |
| CTYDRVEKSFQL | 1418447 | Ectodomain | YES |
| CTYDRVEQSFQL | 1418448 | Ectodomain | YES |
| CTYISRFSKISL | 1418455 | Cell/Virus Enclosed | NO |
| CTYKRAFLDKLDVNSG | 1649107 | Underneath Membrane | NO |
| CTYMDLLLDDFVSVLK | 1649110 | Cell/Virus Enclosed | NO |
| CTYTMCEYPHTI | 1418466 | Underneath Membrane | NO |
| CVADQSRKWTGH | 1418472 | Cell/Virus Enclosed | NO |
| CVAETGTSDLDR | 1418478 | Underneath Membrane | NO |
| CVALCDAMVAKG | 1418482 | Cell/Virus Enclosed | NO |
| CVCFDNSIQGSY | 1418489 | Cell/Virus Enclosed | NO |
| CVDCDAAGVGNT | 1418496 | Cell/Virus Enclosed | NO |
| CVDCDIAGVGNT | 1418497 | Cell/Virus Enclosed | NO |
| CVDCDTAGVGNT | 1418500 | Cell/Virus Enclosed | NO |
| CVDGFGLNKIQP | 1418505 | Cell/Virus Enclosed | NO |
| CVDGFGVNKIQPWTND | 1649132 | Cell/Virus Enclosed | NO |
| CVDGIYGYVLRQ | 1418508 | Ectodomain | YES |
| CVDGRNGYVLRQ | 1418509 | Ectodomain | YES |
| CVDHRPVCSFPL | 1418510 | Cell/Virus Enclosed | NO |
| CVDINKAIPSGNLIKF | 1649137 | Cell/Virus Enclosed | NO |
| CVDLHNKINLCDDPEK | 1649140 | Cell/Virus Enclosed | NO |
| CVDLSVASTYFL | 1418517 | Cell/Virus Enclosed | NO |
| CVDTVKEAQSAN | 1418522 | Cell/Virus Enclosed | NO |
| CVDYSKNGGSGG | 1418528 | Underneath Membrane | NO |
| CVDYSKNRRSRK | 1418531 | Ectodomain | YES |
| CVECDAKFKNSVASIN | 1649147 | Cell/Virus Enclosed | NO |
| CVEMHNKINLCDDPET | 1649150 | Cell/Virus Enclosed | NO |
| CVESQSKRYGFG | 1418540 | Ectodomain | YES |
| CVETVRTQVYFA | 1418544 | Cell/Virus Enclosed | NO |
| CVFFNGVDISDT | 1418562 | Cell/Virus Enclosed | NO |
| CVFNADNLTDYS | 1418564 | Ectodomain | YES |
| CVFSVVNTTVTV | 1418565 | Ectodomain | YES |
| CVFYFAVRKEGQDVIF | 1649167 | Cell/Virus Enclosed | NO |
| CVGHSENVFVVE | 1418575 | Ectodomain | YES |
| CVGNGPGIDAGY | 1418577 | Ectodomain | YES |
| CVGSGPGKNNGM | 1418581 | Ectodomain | YES |
| CVGSSVAVQSKDLNFL | 1649171 | Cell/Virus Enclosed | NO |
| CVIANVPAIDVLKKLL | 1649175 | Cell/Virus Enclosed | NO |
| CVIGTTKFYGGW | 1418599 | Cell/Virus Enclosed | NO |
| CVKAFLDVLNKA | 1418605 | Cell/Virus Enclosed | NO |
| CVKCFIDVVNKA | 1418606 | Cell/Virus Enclosed | NO |
| CVKDFDFKFYDAQPIV | 1649187 | Cell/Virus Enclosed | NO |
| CVKDYEFKFYDAFPVA | 1649188 | Cell/Virus Enclosed | NO |
| CVKSQSPRINFCGNGN | 1649198 | Ectodomain | YES |
| CVLADSESGGHITHS | 7279 | Ectodomain | YES |
| CVLAWNTRNIDATSTGN | 7280 | Ectodomain | YES |

| | | | |
|---|---|---|---|
| CVLGLVNSSLSV | 1418635 | Ectodomain | YES |
| CVLSVNSSNCNNFNVK | 1649222 | Cell/Virus Enclosed | NO |
| CVMPTYTVHKSD | 1418649 | Cell/Virus Enclosed | NO |
| CVMSNMQLFESLNFIT | 1649226 | Cell/Virus Enclosed | NO |
| CVNCDSFGPGNTFING | 1649230 | Cell/Virus Enclosed | NO |
| CVPHNMRVLHYGAGSD | 1649242 | Cell/Virus Enclosed | NO |
| CVPLSAIPALAANTLT | 1649245 | Cell/Virus Enclosed | NO |
| CVPLYVRDYVDKWDDF | 1649246 | Cell/Virus Enclosed | NO |
| CVRAFDGYNKDA | 1418714 | Cell/Virus Enclosed | NO |
| CVSDFMSEIKCK | 1418732 | Ectodomain | YES |
| CVSIVPPTVWED | 1418737 | Ectodomain | YES |
| CVSIVPSTAWED | 1418738 | Ectodomain | YES |
| CVSKVVNLNVDF | 1418742 | Cell/Virus Enclosed | NO |
| CVSNDTTVQSKD | 1418744 | Cell/Virus Enclosed | NO |
| CVSTDTTVQSKDTNFL | 1649280 | Cell/Virus Enclosed | NO |
| CVSVPVSVIYDKETKT | 1649283 | Ectodomain | YES |
| CVTCPFAKGDSC | 1418752 | Cell/Virus Enclosed | NO |
| CVTSNVPPIISE | 1418763 | Cell/Virus Enclosed | NO |
| CVTSNVPPIVSE | 1418764 | Cell/Virus Enclosed | NO |
| CVTSNVSPIVNE | 1418768 | Cell/Virus Enclosed | NO |
| CVTSNVSPIVSE | 1418769 | Cell/Virus Enclosed | NO |
| CVTTKAKSIQQD | 1418770 | Cell/Virus Enclosed | NO |
| CVVDQSRKWIGH | 1418783 | Cell/Virus Enclosed | NO |
| CVVDQSRKWISH | 1418784 | Cell/Virus Enclosed | NO |
| CVVDYSVLYNST | 1418787 | Ectodomain | YES |
| CVVNAYNNTAIS | 1418797 | Ectodomain | YES |
| CVVPGNVKPGDAVITT | 1649311 | Cell/Virus Enclosed | NO |
| CVVPSINDLSFDKNLG | 1649314 | Cell/Virus Enclosed | NO |
| CVYELNGYTVQP | 1418818 | Ectodomain | YES |
| CVYTVAMCFRHF | 1418823 | Cell/Virus Enclosed | NO |
| CVYVDGLESKQVDYVP | 1649328 | Cell/Virus Enclosed | NO |
| CVYYSQVLSKSI | 1418830 | Cell/Virus Enclosed | NO |
| CWETGKVKPPLNKNYVFTG | 1418834 | Cell/Virus Enclosed | NO |
| CWGELMTLATWVGVNLEDPASRDLV | 7332 | Capsid | UN |
| CWHTNCFDYCIP | 1418842 | Secreted | YES |
| CWKCKSKNPLLYDANY | 1649341 | Secreted | YES |
| CWVEEDLTKGPHEFCS | 1649358 | Cell/Virus Enclosed | NO |
| CYALRHFDRNDC | 1418883 | Cell/Virus Enclosed | NO |
| CYCGNSATCYNSVSNE | 1649373 | Cell/Virus Enclosed | NO |
| CYDFDGVTTSDKCIFN | 1649375 | Cell/Virus Enclosed | NO |
| CYDGCAQGVFGVNMRT | 1649376 | Cell/Virus Enclosed | NO |
| CYDHVISTSHKL | 1418896 | Cell/Virus Enclosed | NO |
| CYDIFGVTTPEK | 1418900 | Cell/Virus Enclosed | NO |
| CYDKFKINDTTCKYVF | 1649385 | Cell/Virus Enclosed | NO |
| CYDKFKINDTTRKYVF | 1649386 | Cell/Virus Enclosed | NO |
| CYDKNPINSNVKTLEY | 1649387 | Cell/Virus Enclosed | NO |
| CYDKQPVNSNVKLLDY | 1649388 | Cell/Virus Enclosed | NO |
| CYDRFKVNETNSQYLF | 1649391 | Cell/Virus Enclosed | NO |
| CYEGIASGVFGVNLRT | 1649398 | Cell/Virus Enclosed | NO |
| CYFNLCKEPMYA | 1418939 | Ectodomain | YES |
| CYFRNSSANYVGKYDI | 1649406 | Underneath Membrane | NO |
| CYGEPVLVNAAN | 1418952 | Cell/Virus Enclosed | NO |
| CYGFNEPLNVVS | 1418954 | Ectodomain | YES |
| CYGVSATKLNDLCFSNV | 7383 | Ectodomain | YES |
| CYLFIGYVISCY | 1418987 | Cell/Virus Enclosed | NO |
| CYMHQMELANGT | 1418995 | Cell/Virus Enclosed | NO |
| CYNCRPAVVNIT | 1418999 | Ectodomain | YES |
| CYNGSPSGVYQC | 1419006 | Cell/Virus Enclosed | NO |
| CYNSDYASKGYIANIS | 1649439 | Cell/Virus Enclosed | NO |
| CYNTALMEGSLPYSSI | 1649444 | Cell/Virus Enclosed | NO |
| CYNTDLIEGSKPYSTL | 1649445 | Cell/Virus Enclosed | NO |
| CYPAGVNITLANFNET | 1649447 | Ectodomain | YES |
| CYPGKFVNEEALRQILRESGGIDKE | 1130598 | Ectodomain | YES |
| CYRCPKEIVDTVSALV | 1649450 | Cell/Virus Enclosed | NO |
| CYRCPKEIVETVSALV | 1649451 | Cell/Virus Enclosed | NO |
| CYRCPKEIVSTVSALV | 1649452 | Cell/Virus Enclosed | NO |
| CYRKSNVDESFV | 1419027 | Cell/Virus Enclosed | NO |
| CYRNSNVDEAFV | 1419029 | Cell/Virus Enclosed | NO |
| CYRNSNVGESFV | 1419030 | Cell/Virus Enclosed | NO |
| CYSDGVMKNASLYTSL | 1649456 | Cell/Virus Enclosed | NO |
| CYSGRVSAAFHANSSE | 1649459 | Ectodomain | YES |
| CYSILAAKTGVSVEQL | 1649460 | Cell/Virus Enclosed | NO |
| CYSSLILDYFSY | 1419047 | Ectodomain | YES |
| CYTAEQCIRGAI | 1419052 | Cell/Virus Enclosed | NO |
| CYTAEQCIRGAM | 1419053 | Cell/Virus Enclosed | NO |
| CYTARDALDTIL | 1419054 | Cell/Virus Enclosed | NO |

| | | | |
|---|---|---|---|
| CYTDGLMQNAFL | 1419055 | Cell/Virus Enclosed | NO |
| CYTEGLMQNASL | 1419058 | Cell/Virus Enclosed | NO |
| CYVCKPHQVNISLNGN | 1649471 | Ectodomain | YES |
| CYVCKPHQVNLS | 1419068 | Underneath Membrane | NO |
| CYVGVQTLEDLH | 1419073 | Cell/Virus Enclosed | NO |
| CYVGVQTVEDLC | 1419074 | Cell/Virus Enclosed | NO |
| CYVGVQTVEDLRARMT | 1649472 | Cell/Virus Enclosed | NO |
| CYWPLKDYGFYT | 1419081 | Underneath Membrane | NO |
| CYWPLNDYGFYTTTGIG | 7452 | Ectodomain | YES |
| CYWPLSDYGFYT | 1419086 | Ectodomain | YES |
| CYYVKPGGTGSG | 1419090 | Cell/Virus Enclosed | NO |
| CYYVKPGGTSSGDATT | 1649487 | Cell/Virus Enclosed | NO |
| CYYVTPGGTSSG | 1419092 | Cell/Virus Enclosed | NO |
| DAAEELASPMKS | 1419095 | Cell/Virus Enclosed | NO |
| DAAIKDFDYYRYNRPT | 1649491 | Cell/Virus Enclosed | NO |
| DAAKIYGMGFSS | 1419102 | Ectodomain | YES |
| DAALALLLLDRLNALE | 1649494 | Cell/Virus Enclosed | NO |
| DAALALLLLDRLNQLE | 1378299 | Cell/Virus Enclosed | NO |
| DAANPTNCITPV | 1419110 | Ectodomain | YES |
| DAASNIFVINKDTYAA | 1649506 | Cell/Virus Enclosed | NO |
| DAAVKAAESKAFVDTI | 1649507 | Cell/Virus Enclosed | NO |
| DAAVKDFDFYRYNKPT | 1649508 | Cell/Virus Enclosed | NO |
| DACNVKKVTDVT | 1419132 | Cell/Virus Enclosed | NO |
| DACQLPYCYFRN | 1419133 | Underneath Membrane | NO |
| DADGINKQLTDISVDS | 1649514 | Cell/Virus Enclosed | NO |
| DADKANFLNAAVFYAQ | 1649516 | Cell/Virus Enclosed | NO |
| DADSKHVQLSEI | 1419142 | Cell/Virus Enclosed | NO |
| DADYKMCTKGLS | 1419148 | Ectodomain | YES |
| DADYNAKVEISVTPIK | 1649525 | Cell/Virus Enclosed | NO |
| DAEDTLTKLSKYLANE | 1649527 | Cell/Virus Enclosed | NO |
| DAEEALSKLSEYLISD | 1649528 | Cell/Virus Enclosed | NO |
| DAEGAHATRDSIGTNF | 1649531 | Cell/Virus Enclosed | NO |
| DAERMAMYTGSLIGGI | 1649533 | Ectodomain | YES |
| DAERMAMYTGSLIGGM | 1649534 | Ectodomain | YES |
| DAFNQLTADLQH | 1419178 | Cell/Virus Enclosed | NO |
| DAFQSVYHTFTQ | 1419183 | Cell/Virus Enclosed | NO |
| DAFSLDVSEKSGNFKH | 1649546 | Ectodomain | YES |
| DAFVAAIKLVPTTTGV | 1649548 | Cell/Virus Enclosed | NO |
| DAGCGVKYYAGL | 1419188 | Cell/Virus Enclosed | NO |
| DAGEEKLSSRMY | 1419189 | Cell/Virus Enclosed | NO |
| DAGESSGTGAEVVPFN | 7517 | Cell/Virus Enclosed | NO |
| DAGFMKQYGECLGDIN | 1649551 | Ectodomain | YES |
| DAGNHRSAFALH | 1419200 | Underneath Membrane | NO |
| DAIENALNSCKEHPVV | 1649559 | Cell/Virus Enclosed | NO |
| DAIFSLTKRNILPTMT | 1649560 | Cell/Virus Enclosed | NO |
| DAIIIDNDINDYVSDA | 1649562 | Cell/Virus Enclosed | NO |
| DAIIIDNDLNEFVSDA | 1649563 | Cell/Virus Enclosed | NO |
| DAINVGGTGLQY | 1419226 | Cell/Virus Enclosed | NO |
| DAIQVAKCYGEP | 1419230 | Cell/Virus Enclosed | NO |
| DAITQTSPALQT | 1419235 | Ectodomain | YES |
| DAITQTSQALQTVATA | 1649573 | Ectodomain | YES |
| DAITRTAEAIHT | 1419236 | Ectodomain | YES |
| DAIVIDNDLNEF | 1419238 | Cell/Virus Enclosed | NO |
| DAIVVDNDVVDYVSDA | 1649575 | Cell/Virus Enclosed | NO |
| DAKETKEINIIK | 1419241 | Cell/Virus Enclosed | NO |
| DAKFKNSVASINSAIV | 1649579 | Cell/Virus Enclosed | NO |
| DAKFKNSVTSIN | 1419243 | Cell/Virus Enclosed | NO |
| DAKGFPQFAELI | 1419246 | Cell/Virus Enclosed | NO |
| DAKGFPQLAELIPNQA | 1649581 | Capsid | UN |
| DAKQVDYVPLKSATCI | 1649585 | Cell/Virus Enclosed | NO |
| DAKWYCYDKNPINSNV | 1649586 | Cell/Virus Enclosed | NO |
| DAKWYCYDKQPVNSNV | 1649587 | Cell/Virus Enclosed | NO |
| DALASMTGVSIETLLA | 1649589 | Cell/Virus Enclosed | NO |
| DALDTILQEANK | 1419261 | Cell/Virus Enclosed | NO |
| DALEAEAQIDRLINGR | 1649595 | Ectodomain | YES |
| DALEAQVQIDRLINGR | 1649596 | Ectodomain | YES |
| DALFVSYGSSVETETY | 1649602 | Cell/Virus Enclosed | NO |
| DALLHRNSLDPFCFDV | 1649606 | Cell/Virus Enclosed | NO |
| DALNTICIKDADYNAK | 1649609 | Cell/Virus Enclosed | NO |
| DALQSREAVLVT | 1419281 | Cell/Virus Enclosed | NO |
| DALRLSAHLETNDVSS | 1649611 | Ectodomain | YES |
| DALRNSAMLESA | 1419284 | Ectodomain | YES |
| DANKLILINDVAFVST | 1649629 | Cell/Virus Enclosed | NO |
| DANYKKCTKGLS | 1419321 | Underneath Membrane | NO |
| DAPCYCFNAEGG | 1419325 | Cell/Virus Enclosed | NO |
| DAPSTFGTRNPN | 1419327 | Cell/Virus Enclosed | NO |

| | | | |
|---|---|---|---|
| DAQGTNHGYKVV | 1419339 | Cell/Virus Enclosed | NO |
| DAQIDRLINGRLTTLN | 1649642 | Ectodomain | YES |
| DAQTGSSKCGCS | 1419346 | Cell/Virus Enclosed | NO |
| DAQVVQLPIQDY | 1419350 | Cell/Virus Enclosed | NO |
| DAQVVQLPVKDYVQTV | 1649649 | Cell/Virus Enclosed | NO |
| DARAKQDVSLLG | 1419352 | Cell/Virus Enclosed | NO |
| DARAKQNVSLLS | 1419354 | Cell/Virus Enclosed | NO |
| DASAPATGWLLY | 1419361 | Cell/Virus Enclosed | NO |
| DASEPQRGRQSR | 1419368 | Cell/Virus Enclosed | NO |
| DASFIGKNLKSNCVRF | 1649660 | Cell/Virus Enclosed | NO |
| DASHYVVTDIKQ | 1419375 | Cell/Virus Enclosed | NO |
| DASNVKKVIDVT | 1419379 | Cell/Virus Enclosed | NO |
| DASNVKNVTDVT | 1419381 | Cell/Virus Enclosed | NO |
| DASTFLNGVCGV | 1419394 | Cell/Virus Enclosed | NO |
| DATKPKQVTKHT | 1419409 | Cell/Virus Enclosed | NO |
| DATKPQQVTKHTAKEV | 1649674 | Cell/Virus Enclosed | NO |
| DATSTGNHNYKY | 1419424 | Ectodomain | YES |
| DATSTGNYNYKF | 1419426 | Ectodomain | YES |
| DATTAYANSVFN | 1419431 | Cell/Virus Enclosed | NO |
| DATTAYANSVLN | 1419432 | Cell/Virus Enclosed | NO |
| DAVATTHSWIPKRNRSIL | 97236 | Cell/Virus Enclosed | NO |
| DAVFKPQPAGVL | 1419447 | Ectodomain | YES |
| DAVILLERYVSLAIDA | 1649694 | Cell/Virus Enclosed | NO |
| DAVKQGAKPVGNCVKM | 1649696 | Cell/Virus Enclosed | NO |
| DAVMHFGTLAKTDLFN | 1649698 | Cell/Virus Enclosed | NO |
| DAVNNNAHALSK | 1419461 | Ectodomain | YES |
| DAVNNNAQALSKLASE | 1649703 | Ectodomain | YES |
| DAVNPTNCIAPV | 1419463 | Ectodomain | YES |
| DAVSRKKMDTKPTD | 7671 | Capsid | UN |
| DAVVLLERYVSLAIDA | 1649710 | Cell/Virus Enclosed | NO |
| DAYAEVHAEPKGKYSQ | 1649715 | Cell/Virus Enclosed | NO |
| DAYDLVHDEPKG | 1419489 | Cell/Virus Enclosed | NO |
| DAYDQVHDEPKGKFSK | 1649717 | Cell/Virus Enclosed | NO |
| DAYKAFPPTEPK | 1419494 | Cell/Virus Enclosed | NO |
| DAYKLFLSGDDC | 1419495 | Cell/Virus Enclosed | NO |
| DAYKTFPKKENK | 1419498 | Cell/Virus Enclosed | NO |
| DAYKTFPPTEPKKDKKKKADETQA | 1311663 | Cell/Virus Enclosed | NO |
| DAYPLSKHPKPEYRKV | 1649725 | Cell/Virus Enclosed | NO |
| DAYPLSKHPNSEYRKV | 1649726 | Cell/Virus Enclosed | NO |
| DAYPLVHHENEEYQKV | 1649729 | Cell/Virus Enclosed | NO |
| DAYPLVYHENEEYQKV | 1649730 | Cell/Virus Enclosed | NO |
| DAYQNLVGYYSDDGNY | 1649731 | Ectodomain | YES |
| DAYVSQQLSDST | 1419518 | Ectodomain | YES |
| DCAACVCGDYAA | 1419522 | Underneath Membrane | NO |
| DCAAFVCGDFAA | 1419524 | Ectodomain | YES |
| DCARNPIDLPPSHATT | 1649740 | Cell/Virus Enclosed | NO |
| DCDAVAPHDFFV | 1419541 | Cell/Virus Enclosed | NO |
| DCDFNKPLIEWPLTEY | 1649747 | Cell/Virus Enclosed | NO |
| DCDFNKPLIVWP | 1419545 | Cell/Virus Enclosed | NO |
| DCDIDKWLNNFNVPSP | 1649748 | Ectodomain | YES |
| DCDIDKWLNNFS | 1419549 | Underneath Membrane | NO |
| DCDIDNWLNNVS | 1419551 | Ectodomain | YES |
| DCDVVQEQVNYVPLRA | 1649753 | Cell/Virus Enclosed | NO |
| DCEQTPCGFKGG | 1419567 | Cell/Virus Enclosed | NO |
| DCFANKHAGFDS | 1419570 | Cell/Virus Enclosed | NO |
| DCFKNDNVVLKITEDG | 1649763 | Cell/Virus Enclosed | NO |
| DCGIKQESRVGV | 1419591 | Cell/Virus Enclosed | NO |
| DCGKKFDEQVGC | 1419593 | Cell/Virus Enclosed | NO |
| DCGVNALDGDIY | 1419596 | Cell/Virus Enclosed | NO |
| DCGVNALDGDTY | 1419597 | Cell/Virus Enclosed | NO |
| DCHAPTYLPAEVDGD | 7704 | Ectodomain | YES |
| DCKLPLGQSLCALPDT | 1649790 | Ectodomain | YES |
| DCKQYICNGFQK | 1419614 | Ectodomain | YES |
| DCLAAETHRDCDFNKP | 1649792 | Cell/Virus Enclosed | NO |
| DCLYLKNLTQTFTSML | 1649798 | Cell/Virus Enclosed | NO |
| DCMHQGPASARD | 1419635 | Ectodomain | YES |
| DCMKQGPASARD | 1419636 | Ectodomain | YES |
| DCMQQGPASARE | 1419642 | Underneath Membrane | NO |
| DCMQQGPASARN | 1419643 | Underneath Membrane | NO |
| DCMSDFMSGIKC | 1419647 | Ectodomain | YES |
| DCNGYTDNIFSVQQDG | 1649806 | Ectodomain | YES |
| DCNHIGNINPHN | 1419658 | Ectodomain | YES |
| DCNHINNLKIKNFDYL | 1649807 | Ectodomain | YES |
| DCNHINNLKVKN | 1419659 | Ectodomain | YES |
| DCNHVVNTNPRN | 1419661 | Ectodomain | YES |
| DCNIEAWLNDKS | 1419662 | Ectodomain | YES |

| | | | |
|---|---|---|---|
| DCNLPLGQSLCA | 1419664 | Ectodomain | YES |
| DCNMYICGDSTECANL | 1649809 | Ectodomain | YES |
| DCNVVRYSLDGN | 1419669 | Cell/Virus Enclosed | NO |
| DCRIPNGFPFNN | 1419684 | Underneath Membrane | NO |
| DCSAFVCGDCAA | 1419687 | Ectodomain | YES |
| DCSAFVCGDYAA | 1419688 | Ectodomain | YES |
| DCSDGNYNCNAS | 1419693 | Underneath Membrane | NO |
| DCSDGNYNRNAI | 1419694 | Ectodomain | YES |
| DCSDGNYNRNASLNSF | 1649821 | Ectodomain | YES |
| DCSDGNYNRYAS | 1419696 | Ectodomain | YES |
| DCSILCEILCEYADCK | 1649822 | Cell/Virus Enclosed | NO |
| DCSKIITGLHPT | 1419699 | Cell/Virus Enclosed | NO |
| DCSKSCLGYHPAHAPS | 1649824 | Cell/Virus Enclosed | NO |
| DCSKSYSGYHPAHAPS | 1649825 | Cell/Virus Enclosed | NO |
| DCSNSYSGYHPA | 1419704 | Cell/Virus Enclosed | NO |
| DCSRETSGLSPAYAPT | 1649830 | Cell/Virus Enclosed | NO |
| DCSSSFLSEIQC | 1419712 | Ectodomain | YES |
| DCVDINKAIPSGNLIK | 1649839 | Cell/Virus Enclosed | NO |
| DCVFYFAVRKEGQDVI | 1649841 | Cell/Virus Enclosed | NO |
| DCVGLSVASTYF | 1419738 | Cell/Virus Enclosed | NO |
| DCVKCFIDVVNK | 1419740 | Cell/Virus Enclosed | NO |
| DCVPVVCPKGKI | 1419744 | Cell/Virus Enclosed | NO |
| DCWTATDRFYRL | 1419756 | Cell/Virus Enclosed | NO |
| DCYDKFKINDTTCKYV | 1649855 | Cell/Virus Enclosed | NO |
| DCYIFKRTFAKA | 1419761 | Cell/Virus Enclosed | NO |
| DCYTLKKTVAKA | 1419768 | Cell/Virus Enclosed | NO |
| DDAINVGGTGLQ | 1419777 | Cell/Virus Enclosed | NO |
| DDARTFVQSNVDVVPE | 1649866 | Cell/Virus Enclosed | NO |
| DDASRLLHTVLAKAEL | 1649867 | Cell/Virus Enclosed | NO |
| DDATRNYIITMP | 1419788 | Cell/Virus Enclosed | NO |
| DDATSNVTQCFG | 1419791 | Cell/Virus Enclosed | NO |
| DDCMHQGPASAR | 1419799 | Ectodomain | YES |
| DDCMKQGPASAR | 1419800 | Ectodomain | YES |
| DDCMQQGPASVR | 1419804 | Ectodomain | YES |
| DDCMQQGPSSAR | 1419805 | Ectodomain | YES |
| DDCNGYTDNIFSVQQD | 1649874 | Ectodomain | YES |
| DDDDSQPRADKP | 1419816 | Cell/Virus Enclosed | NO |
| DDDFVSAVANAHRYDV | 1649879 | Cell/Virus Enclosed | NO |
| DDDSAIIEIVNEVLH | 1649881 | Capsid | UN |
| DDEDECDEYEEE | 1419834 | Cell/Virus Enclosed | NO |
| DDEDVVTGDNDDEDVV | 1649886 | Cell/Virus Enclosed | NO |
| DDEDVVTGDNDQ | 1419837 | Cell/Virus Enclosed | NO |
| DDEDVVTGDNNDEEIV | 1649887 | Cell/Virus Enclosed | NO |
| DDEGFITLKNNL | 1419841 | Cell/Virus Enclosed | NO |
| DDEWAAAVDEAFPLDE | 1649889 | Cell/Virus Enclosed | NO |
| DDEWAAAVDEVF | 1419845 | Cell/Virus Enclosed | NO |
| DDEWTAAVDEAF | 1419846 | Cell/Virus Enclosed | NO |
| DDEWVAAVDEAF | 1419847 | Cell/Virus Enclosed | NO |
| DDFCRQYSNESWFEDD | 1649890 | Cell/Virus Enclosed | NO |
| DDFFKNVSTSIP | 1419853 | Ectodomain | YES |
| DDFHTDEDKLDTNSVYEPYY | 7771 | Ectodomain | YES |
| DDFIVFNNNGTNVTQV | 1649893 | Cell/Virus Enclosed | NO |
| DDFNEHVTSVNL | 1419859 | Cell/Virus Enclosed | NO |
| DDFQGYLQKHFS | 1419862 | Cell/Virus Enclosed | NO |
| DDFSKQLQQSMSSADSTQA | 1864755 | Capsid | UN |
| DDFTEHVTSVNL | 1419868 | Cell/Virus Enclosed | NO |
| DDGGDNSESDAK | 1419881 | Cell/Virus Enclosed | NO |
| DDGNKLDKFFVVKRTN | 1649920 | Cell/Virus Enclosed | NO |
| DDGVVCYNSDYAAKGY | 1649925 | Cell/Virus Enclosed | NO |
| DDGVVCYNSDYASKGY | 1649926 | Cell/Virus Enclosed | NO |
| DDHETDMELKPAKVATRTSR | 741286 | Ectodomain | YES |
| DDHIPSGTVLGN | 1419904 | Ectodomain | YES |
| DDIDTINCESIY | 1419911 | Cell/Virus Enclosed | NO |
| DDIEQQADNMITEML | 80922 | Ectodomain | YES |
| DDIFAESKPRFV | 1419914 | Cell/Virus Enclosed | NO |
| DDIFAESKPWFVRKAE | 1649934 | Cell/Virus Enclosed | NO |
| DDILQQANKSVGIISM | 1649936 | Cell/Virus Enclosed | NO |
| DDINFAPVLGCL | 1419923 | Ectodomain | YES |
| DDIPFPGPINDDDNP | 227076 | Capsid | UN |
| DDITKTDAVILLERYV | 1649940 | Cell/Virus Enclosed | NO |
| DDIYYPASCNGVLPVA | 1649943 | Cell/Virus Enclosed | NO |
| DDKASVNWADDR | 1419934 | Cell/Virus Enclosed | NO |
| DDKDPQFKDNVILLNKHIDA | 7808 | Capsid | UN |
| DDKDPSFKDNVILLNKH | 1378399 | Cell/Virus Enclosed | NO |
| DDKLRMETCFQQACK | 7812 | Ectodomain | YES |
| DDKYKTIDELCV | 1419942 | Cell/Virus Enclosed | NO |

| | | | |
|---|---|---|---|
| DDKYKTSDELCV | 1419943 | Cell/Virus Enclosed | NO |
| DDLLKTDSVLLIERFV | 1649953 | Cell/Virus Enclosed | NO |
| DDLNQMTGFKKP | 1419954 | Cell/Virus Enclosed | NO |
| DDLPTTMFQPVRAPCIQTAW | 7822 | Cell/Virus Enclosed | NO |
| DDLQLGSSGFLQ | 1419956 | Ectodomain | YES |
| DDLVNYIRDKGCVADQ | 1649957 | Cell/Virus Enclosed | NO |
| DDMHWGDGFLIK | 1419968 | Cell/Virus Enclosed | NO |
| DDMLRHLIKDVDNPVL | 1649966 | Cell/Virus Enclosed | NO |
| DDMLRRLIKDVDNPVL | 1649967 | Cell/Virus Enclosed | NO |
| DDMLWGDGFQIK | 1419974 | Cell/Virus Enclosed | NO |
| DDNGDRPPEPAAQGESSTQK | 7833 | Cell/Virus Enclosed | NO |
| DDNKFDGLKVDD | 1419982 | Cell/Virus Enclosed | NO |
| DDNKFDVLKLDD | 1419983 | Cell/Virus Enclosed | NO |
| DDNQALLVPKIF | 1419985 | Cell/Virus Enclosed | NO |
| DDNQTLLVPQIF | 1419987 | Cell/Virus Enclosed | NO |
| DDNQTLLVSKIF | 1419988 | Cell/Virus Enclosed | NO |
| DDNTDSIVIDGV | 1419989 | Cell/Virus Enclosed | NO |
| DDNVLPDTYVAL | 1419991 | Underneath Membrane | NO |
| DDNVLPETYVALPIYY | 1649976 | Ectodomain | YES |
| DDPKLMGWDYPKCDRA | 1649980 | Cell/Virus Enclosed | NO |
| DDQIGYYRRATRRIRGGDGK | 1392100 | Capsid | UN |
| DDQIVVTGDDVDDIES | 1649985 | Cell/Virus Enclosed | NO |
| DDQPSLQVESANLMLS | 1649986 | Cell/Virus Enclosed | NO |
| DDRGVITNAVDC | 1420022 | Ectodomain | YES |
| DDSGDSSEGGAK | 1420034 | Cell/Virus Enclosed | NO |
| DDSGNPVVEYLS | 1420037 | Cell/Virus Enclosed | NO |
| DDSQPRADKPSQ | 1420043 | Cell/Virus Enclosed | NO |
| DDSQYFFNKDTG | 1420045 | Ectodomain | YES |
| DDSQYYFNKDTGVIYG | 1650002 | Ectodomain | YES |
| DDSQYYLNKDTG | 1420047 | Ectodomain | YES |
| DDSVREEVDIIE | 1420051 | Cell/Virus Enclosed | NO |
| DDTSETKYVKSLSIVD | 1650011 | Cell/Virus Enclosed | NO |
| DDVDDGGDSSES | 1420073 | Cell/Virus Enclosed | NO |
| DDVDDIDPINCE | 1420074 | Cell/Virus Enclosed | NO |
| DDVDDIESIYEF | 1420075 | Cell/Virus Enclosed | NO |
| DDVDVVKPFRVEGNFS | 1650021 | Cell/Virus Enclosed | NO |
| DDVITSLCKKSF | 1420083 | Cell/Virus Enclosed | NO |
| DDVIVHDVEPTH | 1420084 | Cell/Virus Enclosed | NO |
| DDVNASLFVDYSNLLH | 1650029 | Cell/Virus Enclosed | NO |
| DDVNFSPVLGCL | 1420089 | Ectodomain | YES |
| DDVQAPNYTQHS | 1420095 | Underneath Membrane | NO |
| DDVQAPNYTQHTSSMR | 1650030 | Ectodomain | YES |
| DDVVKTDAVVLLERYV | 1650035 | Cell/Virus Enclosed | NO |
| DDYCDGSLGCKMCLFG | 1650041 | Cell/Virus Enclosed | NO |
| DDYKAFISVFDI | 1420123 | Cell/Virus Enclosed | NO |
| DDYRITSITSDK | 1420132 | Cell/Virus Enclosed | NO |
| DDYTGYQELVIKTSHD | 1650052 | Underneath Membrane | NO |
| DEAFPLDEAEDVTESV | 1650059 | Cell/Virus Enclosed | NO |
| DEAFPLDEAENV | 1420148 | Cell/Virus Enclosed | NO |
| DEAIKRVKGWVG | 1420159 | Cell/Virus Enclosed | NO |
| DEAIPTRFPPGTVLPQ | 1650064 | Cell/Virus Enclosed | NO |
| DEAMASRMYCTF | 1420162 | Cell/Virus Enclosed | NO |
| DEAQPLLQRQKK | 1420163 | Capsid | UN |
| DEAQPLPQRKKQ | 1420164 | Cell/Virus Enclosed | NO |
| DEARFSGTANQQ | 1420167 | Cell/Virus Enclosed | NO |
| DEATPFDVVRQC | 1420170 | Cell/Virus Enclosed | NO |
| DECRLAMLGAEMTSNV | 1650069 | Cell/Virus Enclosed | NO |
| DEDVVTGDNDDEDVVT | 1650077 | Cell/Virus Enclosed | NO |
| DEDVVTGDNDQI | 1420193 | Cell/Virus Enclosed | NO |
| DEDVVTGDNNDEEIVT | 1650078 | Cell/Virus Enclosed | NO |
| DEEDLYDVSKPV | 1420197 | Cell/Virus Enclosed | NO |
| DEEGDNVQITYT | 1420207 | Cell/Virus Enclosed | NO |
| DEEIVTGDNDDQIVVT | 1650087 | Cell/Virus Enclosed | NO |
| DEFFKNVSTSIPNFGS | 1650089 | Ectodomain | YES |
| DEFSINEVVKQMFGVN | 1650092 | Cell/Virus Enclosed | NO |
| DEFTPFDVVAQC | 1420228 | Cell/Virus Enclosed | NO |
| DEGSELTLKHFL | 1420244 | Cell/Virus Enclosed | NO |
| DEHEFKFWDESF | 1420249 | Cell/Virus Enclosed | NO |
| DEIANLVLAKLGKDSK | 1650110 | Capsid | UN |
| DEIKALKELYGP | 1420255 | Cell/Virus Enclosed | NO |
| DEIPYKSFFDSG | 1420258 | Cell/Virus Enclosed | NO |
| DEKDCEVFKEILVLTG | 1650119 | Cell/Virus Enclosed | NO |
| DEKQNGHSLYLADNLT | 1650121 | Cell/Virus Enclosed | NO |
| DELAPSDVYQQLAGVK | 1650125 | Cell/Virus Enclosed | NO |
| DELDDFFKNVST | 1420270 | Ectodomain | YES |
| DELDEFFKNFST | 1420271 | Ectodomain | YES |

| | | | |
|---|---|---|---|
| DELDEFFKNVSTSIPN | 1650127 | Ectodomain | YES |
| DELFAMTKRNVIPTMT | 1650128 | Cell/Virus Enclosed | NO |
| DELKINSACRKV | 1420282 | Cell/Virus Enclosed | NO |
| DELKSLLSLREV | 1420283 | Cell/Virus Enclosed | NO |
| DEMNRLQEAIKV | 1420292 | Ectodomain | YES |
| DENGTITDAVDCSQNP | 1650142 | Ectodomain | YES |
| DENYNNLVPTYLKSDN | 1650144 | Cell/Virus Enclosed | NO |
| DEPDSRAPSRSQ | 1420299 | Cell/Virus Enclosed | NO |
| DEPQKDVYELRYNGAI | 1650148 | Cell/Virus Enclosed | NO |
| DEQEQQEEQEQQEEQEQ | 8131 | Cell/Virus Enclosed | NO |
| DEQIDGKFVTKFSGDK | 1650150 | Cell/Virus Enclosed | NO |
| DEQNCDVLKEVLVLTG | 1650152 | Cell/Virus Enclosed | NO |
| DESAAKSASVYY | 1420315 | Cell/Virus Enclosed | NO |
| DESCNNLVPTYLKSDN | 1650158 | Cell/Virus Enclosed | NO |
| DESCNNLVQTYL | 1420318 | Cell/Virus Enclosed | NO |
| DESFVDDFQGYL | 1420323 | Cell/Virus Enclosed | NO |
| DESQPLPQRQKKQQTVT | 1378496 | Cell/Virus Enclosed | NO |
| DETKALKELYGS | 1420340 | Cell/Virus Enclosed | NO |
| DETVPQSVREAFNNL | 227084 | Cell/Virus Enclosed | NO |
| DEVGDNVQITHT | 1420346 | Cell/Virus Enclosed | NO |
| DEVGLRTPQRFTDLV | 8195 | Ectodomain | YES |
| DEVNGLLDTTQLHVAD | 1650175 | Ectodomain | YES |
| DEWAAAVDEAFPLDEA | 1650184 | Cell/Virus Enclosed | NO |
| DEWSYTGSSFYAPEPI | 1650189 | Ectodomain | YES |
| DEWVAAVDEAFP | 1420373 | Cell/Virus Enclosed | NO |
| DEYGGHHDFVIKTSHD | 1650190 | Underneath Membrane | NO |
| DEYILAKGPLQVGDSV | 1650191 | Cell/Virus Enclosed | NO |
| DEYYKKDNAYYT | 1420383 | Cell/Virus Enclosed | NO |
| DFAFNNWFLLTNTSSV | 1650197 | Ectodomain | YES |
| DFALELEFRNLTPGNTNTRVSRYSS | 8233 | Undetermined | UN |
| DFAQGPFIDGNIIHAN | 1650200 | Cell/Virus Enclosed | NO |
| DFCGKGYHLMSFPQAAP | 8239 | Ectodomain | YES |
| DFCRQYSNESWFEDDY | 1650208 | Cell/Virus Enclosed | NO |
| DFDGVTTSDKCIFNSA | 1650214 | Cell/Virus Enclosed | NO |
| DFDIIQKCQITSVVGT | 1650215 | Cell/Virus Enclosed | NO |
| DFDITESKYSCKEVLK | 1650216 | Cell/Virus Enclosed | NO |
| DFDIVVAWHVVRDSRF | 1650217 | Cell/Virus Enclosed | NO |
| DFDLISKCQITAVEGT | 1650219 | Cell/Virus Enclosed | NO |
| DFEADPKGKYAQNLLK | 1650224 | Cell/Virus Enclosed | NO |
| DFEAELSLWFKNHTS | 1594868 | Ectodomain | YES |
| DFEEDDVEGEET | 1420427 | Cell/Virus Enclosed | NO |
| DFEIACKCGVKQEQRT | 1650225 | Cell/Virus Enclosed | NO |
| DFFIFDVDKFKT | 1420436 | Cell/Virus Enclosed | NO |
| DFFTWHEGRTIYGNVS | 1650234 | Cell/Virus Enclosed | NO |
| DFFVFDVDKTKT | 1420444 | Cell/Virus Enclosed | NO |
| DFGDFEQVAPGC | 1420445 | Cell/Virus Enclosed | NO |
| DFGDFVVTQPGS | 1420454 | Cell/Virus Enclosed | NO |
| DFGDISGINASV | 1420456 | Ectodomain | YES |
| DFGLGDLVDSYFENDS | 1650247 | Cell/Virus Enclosed | NO |
| DFGYVCDGSFYK | 1420463 | Cell/Virus Enclosed | NO |
| DFHSVEIEAGEVKPFA | 1650253 | Cell/Virus Enclosed | NO |
| DFHSVEIEVGEV | 1420472 | Cell/Virus Enclosed | NO |
| DFINGIFAKVKNTKVI | 1650262 | Ectodomain | YES |
| DFIPTDSTVKNY | 1420509 | Cell/Virus Enclosed | NO |
| DFIVNAANENLAHGGG | 1650275 | Cell/Virus Enclosed | NO |
| DFKDELEEFFKNVTS | 1594870 | Ectodomain | YES |
| DFKDFQFMLWGN | 1420521 | Cell/Virus Enclosed | NO |
| DFKEELDQWLKN | 1420524 | Ectodomain | YES |
| DFKEELEEFFKNVSS | 1594872 | Ectodomain | YES |
| DFKEELGQWFKN | 1420526 | Ectodomain | YES |
| DFKFLHNLQADI | 1420528 | Cell/Virus Enclosed | NO |
| DFKFSDGQGVPL | 1420529 | Cell/Virus Enclosed | NO |
| DFKGELDQWFKN | 1420530 | Ectodomain | YES |
| DFKPQSGGGKCFNCYP | 1650285 | Ectodomain | YES |
| DFKVEYAYDQVH | 1420546 | Cell/Virus Enclosed | NO |
| DFLEASDVEEDD | 1420555 | Cell/Virus Enclosed | NO |
| DFLNIDMGVFIN | 1420574 | Cell/Virus Enclosed | NO |
| DFLSFDSDVFVK | 1420580 | Cell/Virus Enclosed | NO |
| DFLSNSSSSFDC | 1420584 | Ectodomain | YES |
| DFLSSDSDVFIK | 1420585 | Cell/Virus Enclosed | NO |
| DFMLKTLYKDVDNPHL | 1650317 | Cell/Virus Enclosed | NO |
| DFNCTNFAINDLNTTV | 1650322 | Ectodomain | YES |
| DFNCTNSFINDY | 1420612 | Ectodomain | YES |
| DFNEDVCVCFDN | 1420614 | Cell/Virus Enclosed | NO |
| DFNHNMGDSDLV | 1420617 | Cell/Virus Enclosed | NO |
| DFNKPLIEWPLTEYDF | 1650330 | Cell/Virus Enclosed | NO |

| | | | |
|---|---|---|---|
| DFNTILNTACGVFEVD | 1650337 | Cell/Virus Enclosed | NO |
| DFNVILPGEFSK | 1420632 | Cell/Virus Enclosed | NO |
| DFNVNVIGTRPE | 1420633 | Cell/Virus Enclosed | NO |
| DFNVNVMGTRPE | 1420634 | Cell/Virus Enclosed | NO |
| DFPFAESTGTQS | 1420637 | Cell/Virus Enclosed | NO |
| DFQGYLQKHFSM | 1420649 | Cell/Virus Enclosed | NO |
| DFSAVSSPMSNMTENE | 1650350 | Underneath Membrane | NO |
| DFSLADSVVEVVTTSL | 1650360 | Cell/Virus Enclosed | NO |
| DFSLDVFKIGDVKFKR | 1650361 | Cell/Virus Enclosed | NO |
| DFSNLGADVLYQ | 1420679 | Cell/Virus Enclosed | NO |
| DFSRDHNDILYTPPTV | 1650367 | Cell/Virus Enclosed | NO |
| DFSSDVFKIGDV | 1420690 | Cell/Virus Enclosed | NO |
| DFTDYKLELFNKYFKH | 1650375 | Cell/Virus Enclosed | NO |
| DFTDYKLELFNKYFKY | 1650376 | Cell/Virus Enclosed | NO |
| DFTGSVYGNFDDQPSL | 1650387 | Cell/Virus Enclosed | NO |
| DFTIGNMEEFIQ | 1420719 | Underneath Membrane | NO |
| DFTIGNMGEFIQ | 1420720 | Underneath Membrane | NO |
| DFTPRSDMEYDFLNMD | 1650391 | Cell/Virus Enclosed | NO |
| DFTPVQVVDSRWNNAR | 1650392 | Ectodomain | YES |
| DFTVLFDRLSLT | 1420732 | Cell/Virus Enclosed | NO |
| DFVDFESVIQDY | 1420739 | Cell/Virus Enclosed | NO |
| DFVDLASVIQDY | 1420740 | Cell/Virus Enclosed | NO |
| DFVDLESVIQDYENVC | 1650399 | Cell/Virus Enclosed | NO |
| DFVENPDIINIYKKLG | 1650403 | Cell/Virus Enclosed | NO |
| DFVENPDIINVYKKLG | 1650404 | Cell/Virus Enclosed | NO |
| DFVTASDTTLRCCTVT | 1650427 | Cell/Virus Enclosed | NO |
| DFVVNAANENLLHGGG | 1650432 | Cell/Virus Enclosed | NO |
| DFYASMYENSTILQAA | 1650438 | Cell/Virus Enclosed | NO |
| DFYEFVKSKGLFKEGS | 1650442 | Cell/Virus Enclosed | NO |
| DFYGWVSGNMMDGFSC | 1650444 | Cell/Virus Enclosed | NO |
| DFYSKLGVGFTSHFCN | 1650454 | Cell/Virus Enclosed | NO |
| DFYTTEPEFVKV | 1420798 | Cell/Virus Enclosed | NO |
| DGAHLTKMSEYK | 1420805 | Cell/Virus Enclosed | NO |
| DGAKTEPTGHGA | 1420806 | Capsid | UN |
| DGARLFSSDLST | 1420811 | Cell/Virus Enclosed | NO |
| DGARLSASDLST | 1420812 | Cell/Virus Enclosed | NO |
| DGCAQGVFGVNMRTNW | 1650465 | Cell/Virus Enclosed | NO |
| DGCGPLHPSWVSLMPPKQVP | 741291 | Ectodomain | YES |
| DGCGTLLGAFYC | 1420821 | Underneath Membrane | NO |
| DGDRFVKHDLSLLSVT | 1650480 | Cell/Virus Enclosed | NO |
| DGDVLLAEFDTYDPIY | 1650481 | Cell/Virus Enclosed | NO |
| DGDVVKQPVKFY | 1420857 | Cell/Virus Enclosed | NO |
| DGDYYRKQLSPLEGGG | 1650485 | Ectodomain | YES |
| DGELPVAISGDKVFVR | 1650486 | Cell/Virus Enclosed | NO |
| DGELPVAVVNDKVFVR | 1650487 | Cell/Virus Enclosed | NO |
| DGEVLPLDELKS | 1420873 | Cell/Virus Enclosed | NO |
| DGFGLNKIQPWT | 1420885 | Cell/Virus Enclosed | NO |
| DGFGVFDIKNSSVNTV | 1650502 | Cell/Virus Enclosed | NO |
| DGFGVNKIQPWTNDAL | 1650503 | Cell/Virus Enclosed | NO |
| DGFLDIWTYNAELLVLLENERTLDY | 1131855 | Ectodomain | YES |
| DGFLSVYNAYSP | 1420890 | Ectodomain | YES |
| DGFLSVYQTYTA | 1420891 | Ectodomain | YES |
| DGFNEWAIANGY | 1420896 | Cell/Virus Enclosed | NO |
| DGFNFSQILPDP | 1420898 | Ectodomain | YES |
| DGFNHQFVNASS | 1420900 | Cell/Virus Enclosed | NO |
| DGFNLEYWVTPL | 1420902 | Ectodomain | YES |
| DGFTEHVTSVNL | 1420911 | Cell/Virus Enclosed | NO |
| DGFYSANFLDDNVLPE | 1650515 | Ectodomain | YES |
| DGFYTQGRNLSD | 1420927 | Cell/Virus Enclosed | NO |
| DGGCLNASEVVVNNLD | 1650522 | Cell/Virus Enclosed | NO |
| DGGDSSESGAKETKEI | 1650523 | Cell/Virus Enclosed | NO |
| DGGIPDGCSLYL | 1420932 | Cell/Virus Enclosed | NO |
| DGGITSEGNALYNNEG | 1650525 | Cell/Virus Enclosed | NO |
| DGGKIWNGVIKNVNSV | 1650526 | Cell/Virus Enclosed | NO |
| DGGKYYTHRIIK | 1420935 | Cell/Virus Enclosed | NO |
| DGGRSGARPKQR | 1420937 | Cell/Virus Enclosed | NO |
| DGHRLASGVQVHNLPE | 1650535 | Underneath Membrane | NO |
| DGHVGLLDQAWR | 1420947 | Cell/Virus Enclosed | NO |
| DGICSENHVMVD | 1420952 | Cell/Virus Enclosed | NO |
| DGIFKTEVDPNL | 1420957 | Cell/Virus Enclosed | NO |
| DGIIELFHYPLGNHCPRVVH | 741294 | Ectodomain | YES |
| DGIIWVATEGALNTPK | 1650544 | Cell/Virus Enclosed | NO |
| DGINVKDVFVES | 1420977 | Cell/Virus Enclosed | NO |
| DGIRRGDNFHSV | 1420982 | Cell/Virus Enclosed | NO |
| DGISEQSLRLID | 1420985 | Cell/Virus Enclosed | NO |
| DGISTPKLKEDYQIGG | 1650553 | Secreted | YES |

| | | | |
|---|---|---|---|
| DGIYFAATEKSNVVRGW | 8413 | Ectodomain | YES |
| DGKCLTSIEEVC | 1421004 | Cell/Virus Enclosed | NO |
| DGKFVTKFSGDKFDFI | 1650561 | Cell/Virus Enclosed | NO |
| DGKHVDHYDGFY | 1421006 | Cell/Virus Enclosed | NO |
| DGKLNCSFQDAASNIF | 1650562 | Cell/Virus Enclosed | NO |
| DGKMGLFFNHTL | 1421009 | Ectodomain | YES |
| DGKMGRFFYHTL | 1421012 | Ectodomain | YES |
| DGKMKDLSPRWYFYYL | 1650564 | Cell/Virus Enclosed | NO |
| DGKMKKLSPRWY | 1421015 | Cell/Virus Enclosed | NO |
| DGKPCTPPAFNC | 1421017 | Ectodomain | YES |
| DGKPCTPPAPNC | 1421019 | Ectodomain | YES |
| DGKPMLPKNMWE | 1421024 | Cell/Virus Enclosed | NO |
| DGKPVISEDLWQ | 1421025 | Cell/Virus Enclosed | NO |
| DGKPVLPKNMWE | 1421026 | Cell/Virus Enclosed | NO |
| DGKQIDDHRITS | 1421027 | Cell/Virus Enclosed | NO |
| DGKYFTSAPPVT | 1421038 | Cell/Virus Enclosed | NO |
| DGLASAYRANSFDVPT | 1650576 | Cell/Virus Enclosed | NO |
| DGLCMFWNCNVDKYPS | 1650581 | Cell/Virus Enclosed | NO |
| DGLEAYADKTLQEMKA | 1650583 | Cell/Virus Enclosed | NO |
| DGLEQDAQIDRL | 1421054 | Underneath Membrane | NO |
| DGLESKQVDYVPLRSA | 1650584 | Cell/Virus Enclosed | NO |
| DGLKFTEESFYKNMYL | 1650588 | Cell/Virus Enclosed | NO |
| DGLKYTKILKDDGNFV | 1650590 | Cell/Virus Enclosed | NO |
| DGLKYTKIMKDDGNCV | 1650591 | Cell/Virus Enclosed | NO |
| DGLMQNAFLYSS | 1421065 | Cell/Virus Enclosed | NO |
| DGLMQNASLYSS | 1421066 | Cell/Virus Enclosed | NO |
| DGLRGAIRKAKDYGFT | 1650593 | Cell/Virus Enclosed | NO |
| DGLTEHVTSVNL | 1421073 | Cell/Virus Enclosed | NO |
| DGMDAKQVDYVPLKSA | 1650595 | Cell/Virus Enclosed | NO |
| DGMMNMSPKPQL | 1421085 | Cell/Virus Enclosed | NO |
| DGMMNMSPKPQRQRGH | 1650597 | Cell/Virus Enclosed | NO |
| DGMNMSPKPQRQ | 1421087 | Cell/Virus Enclosed | NO |
| DGNAAIPDYNYY | 1421089 | Cell/Virus Enclosed | NO |
| DGNAAITDYNYYKYNL | 1650599 | Cell/Virus Enclosed | NO |
| DGNDVEIFENKT | 1421092 | Cell/Virus Enclosed | NO |
| DGNKLDKFFVVKRTNL | 1650610 | Cell/Virus Enclosed | NO |
| DGNYNCNASLNS | 1421107 | Underneath Membrane | NO |
| DGNYNRNAILNS | 1421108 | Ectodomain | YES |
| DGNYNRNASLNSFKEY | 1650617 | Ectodomain | YES |
| DGPCVETSDSQF | 1421116 | Cell/Virus Enclosed | NO |
| DGPNEHVTSVNL | 1421120 | Cell/Virus Enclosed | NO |
| DGPSVDTSDSQV | 1421121 | Cell/Virus Enclosed | NO |
| DGQGVPIAFGIP | 1421132 | Cell/Virus Enclosed | NO |
| DGQGVPIAYGIP | 1421133 | Cell/Virus Enclosed | NO |
| DGQGVPLAFGVP | 1421134 | Cell/Virus Enclosed | NO |
| DGQIGEAPVSII | 1421135 | Cell/Virus Enclosed | NO |
| DGQKFTDESFYKNMYL | 1650622 | Cell/Virus Enclosed | NO |
| DGQRVYDDVDASLFVD | 1650629 | Cell/Virus Enclosed | NO |
| DGQVQTFYPRFQ | 1421155 | Cell/Virus Enclosed | NO |
| DGRDNGAFEAFK | 1421156 | Cell/Virus Enclosed | NO |
| DGRDNGALEAFKRSNN | 1650632 | Cell/Virus Enclosed | NO |
| DGRDTGALEAFRKARN | 1650633 | Cell/Virus Enclosed | NO |
| DGRIPNDFSFNN | 1421160 | Ectodomain | YES |
| DGRPVHVKEITR | 1421168 | Cell/Virus Enclosed | NO |
| DGSAFRDLKPLP | 1421179 | Cell/Virus Enclosed | NO |
| DGSCEVVHDQVNYVPL | 1650646 | Cell/Virus Enclosed | NO |
| DGSCSCVSTDTTVQSK | 1650648 | Cell/Virus Enclosed | NO |
| DGSGVGVIRISN | 1421184 | Ectodomain | YES |
| DGSLGCKMCLFGYQEL | 1650653 | Cell/Virus Enclosed | NO |
| DGSLIPVRPRNSSDNG | 1650654 | Ectodomain | YES |
| DGSPQPYCYTDG | 1421193 | Cell/Virus Enclosed | NO |
| DGSPQSYCYTDG | 1421195 | Cell/Virus Enclosed | NO |
| DGSSGVANPAMDPIYD | 1650657 | Secreted | YES |
| DGTMYGAFMDKQ | 1421220 | Cell/Virus Enclosed | NO |
| DGTNKQLNEISDDCNW | 1650673 | Cell/Virus Enclosed | NO |
| DGTRDTLPQSPGP | 8503 | Ectodomain | YES |
| DGTTQPKLKEDY | 1421231 | Secreted | YES |
| DGVDCYNWDSAE | 1421237 | Cell/Virus Enclosed | NO |
| DGVDSFEGVLPINTDT | 1650683 | Cell/Virus Enclosed | NO |
| DGVENPMLMGWDYPKC | 1650686 | Cell/Virus Enclosed | NO |
| DGVMYGGFEDQPNLQV | 1650694 | Cell/Virus Enclosed | NO |
| DGVNFNIDDINF | 1421258 | Underneath Membrane | NO |
| DGVNFNVDDINFSPVL | 1650696 | Ectodomain | YES |
| DGVNFNVDDVNF | 1421259 | Ectodomain | YES |
| DGVNFTNRFVPVGESF | 1650698 | Cell/Virus Enclosed | NO |
| DGVNKLQGLLEI | 1421263 | Ectodomain | YES |

| | | | |
|---|---|---|---|
| DGVNKLQGLLEV | 1421264 | Ectodomain | YES |
| DGVNVTKHKCDINYKG | 1650700 | Cell/Virus Enclosed | NO |
| DGVNVTKHRCDI | 1421266 | Cell/Virus Enclosed | NO |
| DGVQCYTPHSQI | 1421272 | Cell/Virus Enclosed | NO |
| DGVRESYVNSSN | 1421276 | Cell/Virus Enclosed | NO |
| DGVTRDIVSTDD | 1421287 | Cell/Virus Enclosed | NO |
| DGVTTSDKCIFNSACT | 1650714 | Cell/Virus Enclosed | NO |
| DGVVCYNKDYAELGYI | 1650718 | Cell/Virus Enclosed | NO |
| DGVVCYNSDYAAKGYI | 1650719 | Cell/Virus Enclosed | NO |
| DGVVCYNSDYASKGYI | 1650720 | Cell/Virus Enclosed | NO |
| DGVVPLSVIPATSASK | 1650722 | Cell/Virus Enclosed | NO |
| DGVVWVAKEGAKTVNT | 1650725 | Capsid | UN |
| DGVYTNFKLIGH | 1421300 | Cell/Virus Enclosed | NO |
| DGVYWVASNQAD | 1421303 | Cell/Virus Enclosed | NO |
| DGVYYPSNGTNILPVA | 1650730 | Cell/Virus Enclosed | NO |
| DGYCKLFITKEEAVKR | 1650734 | Cell/Virus Enclosed | NO |
| DGYSKLFITKDEAIKR | 1650747 | Cell/Virus Enclosed | NO |
| DHAYSTMRLHNF | 1421345 | Cell/Virus Enclosed | NO |
| DHCPVVEVNGVTIQV | 8530 | Ectodomain | YES |
| DHEISVQKKINR | 1421356 | Cell/Virus Enclosed | NO |
| DHFGENEEIVIN | 1421361 | Cell/Virus Enclosed | NO |
| DHFNYNVSVLQM | 1421363 | Cell/Virus Enclosed | NO |
| DHGDMYVYSAGHATGT | 1650770 | Ectodomain | YES |
| DHIGTRNPNNNAAIVL | 1650775 | Cell/Virus Enclosed | NO |
| DHIGTRNPNNNV | 1421380 | Capsid | UN |
| DHILEPSIPWKSKLLTLVQL | 8544 | Underneath Membrane | NO |
| DHNDILYTPPTVSYGS | 1650785 | Cell/Virus Enclosed | NO |
| DHNDMLYSPPTI | 1421396 | Cell/Virus Enclosed | NO |
| DHPHLGKFLEEL | 1421399 | Cell/Virus Enclosed | NO |
| DHSGSNDNIALL | 1421414 | Underneath Membrane | NO |
| DHSLLATLDDPYVEDS | 1650795 | Capsid | UN |
| DHSQFYFNKDTG | 1421415 | Ectodomain | YES |
| DHSQYYFNKDTG | 1421416 | Ectodomain | YES |
| DHSQYYLNKDTG | 1421417 | Ectodomain | YES |
| DHTDWCRCSCLPDPITAYDP | 2060803 | Ectodomain | YES |
| DHTPPMPSIGLE | 741300 | Ectodomain | YES |
| DHTSLVWKHIRD | 1421422 | Cell/Virus Enclosed | NO |
| DHYDGFYTQGRN | 1421443 | Cell/Virus Enclosed | NO |
| DHYIVLGGETIT | 1421445 | Cell/Virus Enclosed | NO |
| DHYVVLGGETIT | 1421446 | Cell/Virus Enclosed | NO |
| DIAEDVVDQVEEVNSI | 1650818 | Cell/Virus Enclosed | NO |
| DIAQVVIADTLQETPV | 1650824 | Cell/Virus Enclosed | NO |
| DICAQLNAKLGF | 1421470 | Cell/Virus Enclosed | NO |
| DICMFENKTTLPTNIA | 1650829 | Cell/Virus Enclosed | NO |
| DIDKWLNNFNVPSPLN | 1650841 | Ectodomain | YES |
| DIDKWLNNFSVP | 1421500 | Underneath Membrane | NO |
| DIDVFNKLSTSDWSDI | 1650842 | Cell/Virus Enclosed | NO |
| DIDYCVRAFDVYNKDA | 1650844 | Cell/Virus Enclosed | NO |
| DIEEVELLDAEFVEPG | 1650847 | Cell/Virus Enclosed | NO |
| DIESIYEFDTYK | 1421524 | Cell/Virus Enclosed | NO |
| DIEYPYISHEKKLNSC | 1650853 | Cell/Virus Enclosed | NO |
| DIFAESKPRFVR | 1421533 | Cell/Virus Enclosed | NO |
| DIFAESKPWFVRKAED | 1650855 | Cell/Virus Enclosed | NO |
| DIFATPSVLQAT | 1421535 | Cell/Virus Enclosed | NO |
| DIFGPCWSALASALKQ | 1650857 | Cell/Virus Enclosed | NO |
| DIFGQDFKTFDLLKYD | 1650858 | Cell/Virus Enclosed | NO |
| DIFGSDFKTFDLLKYD | 1650859 | Cell/Virus Enclosed | NO |
| DIFRSDVLHLTQ | 1421547 | Ectodomain | YES |
| DIFVHEDLHVVEVLTK | 1650865 | Cell/Virus Enclosed | NO |
| DIGNPKGIRCAV | 1421564 | Cell/Virus Enclosed | NO |
| DIGNPKGVRCAV | 1421566 | Cell/Virus Enclosed | NO |
| DIHLPAYASFKVIDNG | 1650888 | Cell/Virus Enclosed | NO |
| DIIEQPFGEVEH | 1421580 | Cell/Virus Enclosed | NO |
| DIIQADQQVDRL | 1421586 | Ectodomain | YES |
| DIIQKCQITSVVGTKA | 1650902 | Cell/Virus Enclosed | NO |
| DIIQRLDGLEQD | 1421588 | Underneath Membrane | NO |
| DIIQRLDPPEQD | 1421589 | Underneath Membrane | NO |
| DIKNSSVNTVCFVDVD | 1650909 | Cell/Virus Enclosed | NO |
| DILQHANKSVGI | 1421624 | Cell/Virus Enclosed | NO |
| DILQQANKSVGIISMP | 1650924 | Cell/Virus Enclosed | NO |
| DILSLNNPIFINYSKE | 1650928 | Ectodomain | YES |
| DINAYANWCCLKCGFS | 1650938 | Cell/Virus Enclosed | NO |
| DINDYVSDADFSITGD | 1650939 | Cell/Virus Enclosed | NO |
| DINHGDNGFTSI | 1421662 | Undetermined | UN |
| DINHGDSGFISI | 1421663 | Undetermined | UN |
| DINKAIPSGNLIKFDI | 1650950 | Cell/Virus Enclosed | NO |

| | | | |
|---|---|---|---|
| DINNGPHEFCTK | 1421674 | Cell/Virus Enclosed | NO |
| DINRLCEEVLDN | 1421679 | Cell/Virus Enclosed | NO |
| DINTPDVPICVY | 1421681 | Ectodomain | YES |
| DINYKGKVFFQFDNLS | 1650957 | Cell/Virus Enclosed | NO |
| DIPGGCAMPIVANWAY | 1650958 | Ectodomain | YES |
| DIPVGAGICASY | 1421697 | Ectodomain | YES |
| DIPYNVSLFKFN | 1421698 | Ectodomain | YES |
| DIPYNVSLSKFNSCKS | 1650965 | Ectodomain | YES |
| DIQITNESYNNYVPSY | 1650967 | Cell/Virus Enclosed | NO |
| DIQLTNESYNNY | 1421704 | Cell/Virus Enclosed | NO |
| DIQQTFFDKTWPRPID | 1650974 | Ectodomain | YES |
| DIQQWGYVGSLS | 1421710 | Cell/Virus Enclosed | NO |
| DIRDNGSLEKFM | 1421714 | Cell/Virus Enclosed | NO |
| DIRDNVSLEKFM | 1421716 | Cell/Virus Enclosed | NO |
| DIRDYKLANDAKESLR | 1650980 | Cell/Virus Enclosed | NO |
| DISAVQSSFGFDQQQL | 1650987 | Cell/Virus Enclosed | NO |
| DISEATQVEQHHRRT | 187296 | Secreted | YES |
| DISFIDYTTTVT | 1421733 | Cell/Virus Enclosed | NO |
| DISGINAPVVNI | 1421735 | Ectodomain | YES |
| DISNVPFSPNGK | 1421750 | Ectodomain | YES |
| DISNVPFSSDGK | 1421751 | Ectodomain | YES |
| DISYLKRARGSS | 1421763 | Cell/Virus Enclosed | NO |
| DITDAAVKAAESKAFV | 1651007 | Cell/Virus Enclosed | NO |
| DITESKYSCKEVFKNC | 1651008 | Cell/Virus Enclosed | NO |
| DITESKYSCKEVLKNC | 1651009 | Cell/Virus Enclosed | NO |
| DITKTDAVILLERYVS | 1651011 | Cell/Virus Enclosed | NO |
| DITNINLSRGRG | 1421779 | Cell/Virus Enclosed | NO |
| DITNTNLSRDRG | 1421780 | Cell/Virus Enclosed | NO |
| DITNTNLSRGRGRNPK | 1651015 | Cell/Virus Enclosed | NO |
| DITNTNMSRGRG | 1421783 | Cell/Virus Enclosed | NO |
| DIVCKVENKAEA | 1421788 | Cell/Virus Enclosed | NO |
| DIVCKVENKTEAEWIE | 1651018 | Cell/Virus Enclosed | NO |
| DIVDRDPSNDEA | 1421792 | Cell/Virus Enclosed | NO |
| DIVDVPPQVVEL | 1421796 | Cell/Virus Enclosed | NO |
| DIVEVPPQVVEL | 1421801 | Cell/Virus Enclosed | NO |
| DIVKTDGTLMVERFVS | 1651027 | Cell/Virus Enclosed | NO |
| DIVSGYEKSTFNKDIY | 1651031 | Cell/Virus Enclosed | NO |
| DIVSSFEGYDFKYIEN | 1651033 | Cell/Virus Enclosed | NO |
| DIVVAWHVVRDSRFVM | 1651035 | Cell/Virus Enclosed | NO |
| DKALYEQVVMDY | 1421860 | Cell/Virus Enclosed | NO |
| DKANFLNAVVFY | 1421863 | Cell/Virus Enclosed | NO |
| DKAVPSPLNWER | 1421867 | Ectodomain | YES |
| DKCIFNSACTRLEGLG | 1651064 | Cell/Virus Enclosed | NO |
| DKCTKYNIYDVSGVGV | 1651069 | Ectodomain | YES |
| DKDKDLSGAVPSDLNT | 1651075 | Cell/Virus Enclosed | NO |
| DKDPNFKDQVILLNKHIDAY | 2116174 | Capsid | UN |
| DKDTGPPPISTD | 1421892 | Ectodomain | YES |
| DKEDVVIFINNTTYPT | 1651082 | Cell/Virus Enclosed | NO |
| DKEEDLQQAQQE | 1421894 | Cell/Virus Enclosed | NO |
| DKETLEYWVTPL | 1421901 | Ectodomain | YES |
| DKFAIPNGRKVDLQLG | 1651087 | Ectodomain | YES |
| DKFAIPNRRRVD | 1421911 | Ectodomain | YES |
| DKFDFIIGHGMSFSMT | 1651089 | Cell/Virus Enclosed | NO |
| DKFDLLISDMYDGRIK | 1651091 | Cell/Virus Enclosed | NO |
| DKFDLLISDMYDGRTK | 1651092 | Cell/Virus Enclosed | NO |
| DKFKINDTTCKYVFTT | 1651097 | Cell/Virus Enclosed | NO |
| DKFKINDTTRKYVFTT | 1651098 | Cell/Virus Enclosed | NO |
| DKFSVLHDIGNP | 1421925 | Cell/Virus Enclosed | NO |
| DKFYRLSNELAQVLTE | 1651104 | Cell/Virus Enclosed | NO |
| DKGCVVDQSRKW | 1421934 | Cell/Virus Enclosed | NO |
| DKGDAEDTLNKL | 1421935 | Cell/Virus Enclosed | NO |
| DKGDAEDTLTKLSKYL | 1651106 | Cell/Virus Enclosed | NO |
| DKGIAIPHDIDLGESRVVIQDYDNQ | 8875 | Undetermined | UN |
| DKGIAPGTSVLRQWLP | 1651108 | Cell/Virus Enclosed | NO |
| DKGIYQTSNFRVVPSG | 1651109 | Ectodomain | YES |
| DKGVAPGTTVLKRWLP | 1651111 | Cell/Virus Enclosed | NO |
| DKIEEIESKQKK | 1421949 | Underneath Membrane | NO |
| DKIFTKSGFNDI | 1421950 | Cell/Virus Enclosed | NO |
| DKIFTKSGTNDI | 1421951 | Cell/Virus Enclosed | NO |
| DKIHHHHHHNST | 1421956 | Cell/Virus Enclosed | NO |
| DKIHHHHHHNTA | 1421957 | Capsid | UN |
| DKIIEFQLDYSI | 1421958 | Cell/Virus Enclosed | NO |
| DKIRHLLKGVTL | 1421963 | Cell/Virus Enclosed | NO |
| DKISDTSGFAVY | 1421966 | Underneath Membrane | NO |
| DKITEFQLDYSIDVID | 1651119 | Cell/Virus Enclosed | NO |
| DKKAFTLANVSSFGDY | 1651121 | Ectodomain | YES |

| | | | |
|---|---|---|---|
| DKKKKADETQALPQRQKKQQ | 1392108 | Capsid | UN |
| DKLCHAVVSKSK | 1421982 | Cell/Virus Enclosed | NO |
| DKLDTGAQKFFQFGDF | 1651138 | Cell/Virus Enclosed | NO |
| DKLEKLIFTARD | 1421986 | Cell/Virus Enclosed | NO |
| DKLEKLISTARDGVRR | 1651140 | Cell/Virus Enclosed | NO |
| DKLGFNVDLPFVEYKV | 1651142 | Cell/Virus Enclosed | NO |
| DKLKPVFTWVEA | 1421999 | Cell/Virus Enclosed | NO |
| DKLRDYLADYDVAVTA | 1651149 | Cell/Virus Enclosed | NO |
| DKLSVYDIACCMRAGS | 1651153 | Cell/Virus Enclosed | NO |
| DKLVPSPLNWER | 1422018 | Ectodomain | YES |
| DKLYIWGVHHPSTN | 8939 | Ectodomain | YES |
| DKNGTITDAVDC | 1422021 | Underneath Membrane | NO |
| DKNICLLDQAWR | 1422023 | Cell/Virus Enclosed | NO |
| DKNKLIGYWNIQ | 1422024 | Cell/Virus Enclosed | NO |
| DKNKLIGYWNVQ | 1422025 | Cell/Virus Enclosed | NO |
| DKNLITTANTGTSVTE | 1651160 | Cell/Virus Enclosed | NO |
| DKNLQDLWLLYK | 1422028 | Cell/Virus Enclosed | NO |
| DKNNGIGICPAG | 1422032 | Ectodomain | YES |
| DKNNGIGTCPAG | 1422033 | Ectodomain | YES |
| DKNPINSNVKTLEYDY | 1651163 | Cell/Virus Enclosed | NO |
| DKPNHFINFPLS | 1422038 | Cell/Virus Enclosed | NO |
| DKPNHFINFPLTHFQG | 1651166 | Cell/Virus Enclosed | NO |
| DKPQEKDKKSAKTGTP | 1651167 | Capsid | UN |
| DKQGQLLDTKLNGILQ | 1651170 | Cell/Virus Enclosed | NO |
| DKQPVNSNVKLLDYDY | 1651171 | Cell/Virus Enclosed | NO |
| DKQVHQVQLIDK | 1422049 | Cell/Virus Enclosed | NO |
| DKSAGHPFNKFGKARV | 1651187 | Cell/Virus Enclosed | NO |
| DKSAIPNGRKVD | 1422064 | Underneath Membrane | NO |
| DKSCEHQTPFEI | 1422066 | Cell/Virus Enclosed | NO |
| DKSELTNGVYYR | 1422068 | Cell/Virus Enclosed | NO |
| DKSEPLCLFKKNFTYN | 1651192 | Ectodomain | YES |
| DKSFEQQVGVIADKDK | 1651193 | Cell/Virus Enclosed | NO |
| DKSKIFGSCFNS | 1422072 | Ectodomain | YES |
| DKTGDFMSTIII | 1422091 | Cell/Virus Enclosed | NO |
| DKTLQEMKALFL | 1422096 | Cell/Virus Enclosed | NO |
| DKTWPRPIDVSK | 1422105 | Ectodomain | YES |
| DKVDFVNGFYRLYSGD | 1651210 | Cell/Virus Enclosed | NO |
| DKVDILLTVDGVNFTN | 1651211 | Cell/Virus Enclosed | NO |
| DKVDSTFVTEYYEFLN | 1651212 | Cell/Virus Enclosed | NO |
| DKVFGYVQQGIFNKAT | 1651216 | Cell/Virus Enclosed | NO |
| DKVFNASRFPNV | 1422116 | Ectodomain | YES |
| DKVFVRDGNTDNLVFV | 1651220 | Cell/Virus Enclosed | NO |
| DKVGDTDCVFYF | 1422121 | Cell/Virus Enclosed | NO |
| DKVGELKVEFWFAMRK | 1651223 | Cell/Virus Enclosed | NO |
| DKVGHYVHVKCEQSYQ | 1651224 | Cell/Virus Enclosed | NO |
| DKVGNTDCVFYF | 1422123 | Cell/Virus Enclosed | NO |
| DKVIIADPGYMQ | 1422124 | Ectodomain | YES |
| DKVKLSDVGFVEAYNN | 1651225 | Ectodomain | YES |
| DKVKTPHIVRQR | 1422126 | Cell/Virus Enclosed | NO |
| DKVMTFYPRLQA | 1422132 | Cell/Virus Enclosed | NO |
| DKVPQAVETRVQ | 1422136 | Cell/Virus Enclosed | NO |
| DKVPQSVETKVQ | 1422137 | Cell/Virus Enclosed | NO |
| DKVRSLKLSDNNCYLN | 1651233 | Cell/Virus Enclosed | NO |
| DKVYCPRHVICSASDM | 1651239 | Cell/Virus Enclosed | NO |
| DKVYCPRHVICSSSNM | 1651240 | Cell/Virus Enclosed | NO |
| DKWFVNDGRVANGYVC | 1651243 | Cell/Virus Enclosed | NO |
| DKWLNNFNVPSPLNWE | 1651244 | Ectodomain | YES |
| DKWLNNFSVPSP | 1422155 | Underneath Membrane | NO |
| DKYKATGDLAVC | 1422171 | Cell/Virus Enclosed | NO |
| DKYKTIDELCVN | 1422172 | Cell/Virus Enclosed | NO |
| DKYMCGFDGKPM | 1422174 | Cell/Virus Enclosed | NO |
| DLAGLFRIFLRK | 1422184 | Cell/Virus Enclosed | NO |
| DLALTNMYKEARINDK | 1651266 | Cell/Virus Enclosed | NO |
| DLAVCLGIGDSAVTYS | 1651271 | Cell/Virus Enclosed | NO |
| DLAVQIGNNNVCTYEH | 1651272 | Cell/Virus Enclosed | NO |
| DLAVQIGSNNVCTYEH | 1651273 | Cell/Virus Enclosed | NO |
| DLCTHSFRKKQV | 1422235 | Cell/Virus Enclosed | NO |
| DLDCTYISLKPG | 1422239 | Underneath Membrane | NO |
| DLDEFSKQLQQS | 1422244 | Cell/Virus Enclosed | NO |
| DLDHNFGSAGVVANGV | 1651292 | Capsid | UN |
| DLDVILPGEFSKKQLG | 1651296 | Cell/Virus Enclosed | NO |
| DLDVMLPGEFSK | 1422257 | Cell/Virus Enclosed | NO |
| DLEADPKGKYAQ | 1422258 | Cell/Virus Enclosed | NO |
| DLEIGYTVDCSCGKKL | 1651303 | Cell/Virus Enclosed | NO |
| DLERDFIDMDDNLFIA | 1651304 | Cell/Virus Enclosed | NO |
| DLESVIQDYENVCFEF | 1651306 | Cell/Virus Enclosed | NO |

| | | | |
|---|---|---|---|
| DLEVLLQDALQSREAV | 1651308 | Cell/Virus Enclosed | NO |
| DLEVTGDSCYNF | 1422278 | Cell/Virus Enclosed | NO |
| DLEVTSDSCNNF | 1422279 | Cell/Virus Enclosed | NO |
| DLGAVSKVVKVP | 1422307 | Cell/Virus Enclosed | NO |
| DLGNQILDSYSV | 1422318 | Cell/Virus Enclosed | NO |
| DLGSLIDCNAAS | 1422321 | Cell/Virus Enclosed | NO |
| DLGVISKVHEVIIDNK | 1651337 | Cell/Virus Enclosed | NO |
| DLGVLIQNGAKHVQGN | 1651338 | Cell/Virus Enclosed | NO |
| DLGVLIQNSAKHVQGN | 1651339 | Cell/Virus Enclosed | NO |
| DLHNKINLCDDPEKAQ | 1651341 | Cell/Virus Enclosed | NO |
| DLHTLYVDSKYQ | 1422341 | Cell/Virus Enclosed | NO |
| DLHVVEVLTKTAVKSG | 1651346 | Cell/Virus Enclosed | NO |
| DLIAQSSGVLPV | 1422342 | Cell/Virus Enclosed | NO |
| DLICAQKFNALT | 1422343 | Ectodomain | YES |
| DLICVQSYNGIK | 1422348 | Ectodomain | YES |
| DLIEGSKPYSTLQPNA | 1651350 | Cell/Virus Enclosed | NO |
| DLIRQGTDYKYW | 1422361 | Cell/Virus Enclosed | NO |
| DLISKCQITAVEGTKK | 1651359 | Cell/Virus Enclosed | NO |
| DLISRCQITAVE | 1422363 | Cell/Virus Enclosed | NO |
| DLKCPLDTSLKG | 1422369 | Ectodomain | YES |
| DLKCTSDNINDKDTGP | 1651365 | Ectodomain | YES |
| DLKELGNYTYYK | 1422379 | Ectodomain | YES |
| DLKFIDKLNVLFDGRD | 1651369 | Cell/Virus Enclosed | NO |
| DLKFVKWEYEGG | 1422386 | Cell/Virus Enclosed | NO |
| DLKGKFVQIPTT | 1422387 | Cell/Virus Enclosed | NO |
| DLKSYDLGDELG | 1422399 | Cell/Virus Enclosed | NO |
| DLLALSADQALQPPCPNLVSYSS | 9157 | Ectodomain | YES |
| DLLAQSSGVLPV | 1422405 | Cell/Virus Enclosed | NO |
| DLLCLRDDNQTLLVPK | 1651387 | Cell/Virus Enclosed | NO |
| DLLCVADDKPIV | 1422411 | Cell/Virus Enclosed | NO |
| DLLCVANDKPIV | 1422412 | Cell/Virus Enclosed | NO |
| DLLDKGMQLLHT | 1422414 | Cell/Virus Enclosed | NO |
| DLLFDKVTIADPGYMQ | 1651392 | Ectodomain | YES |
| DLLIKAYNTINNEQGT | 1651396 | Cell/Virus Enclosed | NO |
| DLLISDMYDGRTKAID | 1651400 | Cell/Virus Enclosed | NO |
| DLLKDIKFVIPALQHA | 1651401 | Cell/Virus Enclosed | NO |
| DLLKTDSVLLIERFVS | 1651402 | Cell/Virus Enclosed | NO |
| DLLKYDFTEHKENLFN | 1651406 | Cell/Virus Enclosed | NO |
| DLLLGDFVEIIK | 1422441 | Cell/Virus Enclosed | NO |
| DLLNRLQSLESG | 1422447 | Cell/Virus Enclosed | NO |
| DLMAAYVEETSI | 1422475 | Cell/Virus Enclosed | NO |
| DLMAQSSGVLPV | 1422478 | Cell/Virus Enclosed | NO |
| DLMEILNRDRVN | 1422479 | Cell/Virus Enclosed | NO |
| DLNANMSLAECKRALG | 1651432 | Cell/Virus Enclosed | NO |
| DLNCHYLVLPFG | 1422488 | Ectodomain | YES |
| DLNCPLDSRLKG | 1422491 | Ectodomain | YES |
| DLNCPLDTRLKG | 1422492 | Ectodomain | YES |
| DLNEFVSDADITLFGD | 1651435 | Cell/Virus Enclosed | NO |
| DLNGVIVDKVGELKVE | 1651447 | Cell/Virus Enclosed | NO |
| DLNPALCDSGKI | 1422508 | Ectodomain | YES |
| DLNQMTGFKKPA | 1422511 | Cell/Virus Enclosed | NO |
| DLNWLNRVETYI | 1422518 | Ectodomain | YES |
| DLPDDVVAANMFMGVG | 1651459 | Cell/Virus Enclosed | NO |
| DLPENDDDAWFK | 1422522 | Cell/Virus Enclosed | NO |
| DLPICVYDRLPV | 1422528 | Undetermined | UN |
| DLPKEITVATSRTLSY | 1651464 | Underneath Membrane | NO |
| DLPNCNIEAWLN | 1422535 | Ectodomain | YES |
| DLPNCNIGAWLN | 1422536 | Underneath Membrane | NO |
| DLPSGFNTLKPIFKLPL | 9242 | Ectodomain | YES |
| DLQFNKLGTEDP | 1422552 | Cell/Virus Enclosed | NO |
| DLQGNFGDLKFN | 1422553 | Cell/Virus Enclosed | NO |
| DLQGNFGDLQFN | 1422554 | Cell/Virus Enclosed | NO |
| DLQGNFGDLQLNKLGT | 1651475 | Cell/Virus Enclosed | NO |
| DLQGNFGDLQLT | 1422556 | Cell/Virus Enclosed | NO |
| DLQKANIEIELG | 1422561 | Underneath Membrane | NO |
| DLQKANTDIILG | 1422562 | Ectodomain | YES |
| DLQKANTDIKLD | 1422563 | Underneath Membrane | NO |
| DLQKANTDIKLG | 1422564 | Ectodomain | YES |
| DLQKSNTDIILG | 1422566 | Ectodomain | YES |
| DLQLGNLAYLQS | 1422567 | Ectodomain | YES |
| DLQLGNLGNLQS | 1422568 | Ectodomain | YES |
| DLQLGSSGFLQSSNYK | 1651479 | Ectodomain | YES |
| DLQLNKLGTDDP | 1422571 | Cell/Virus Enclosed | NO |
| DLQLNKLGTEDPRWPQ | 1651480 | Cell/Virus Enclosed | NO |
| DLQLTKLGIEDP | 1422573 | Cell/Virus Enclosed | NO |
| DLQLTKLGTEDP | 1422574 | Cell/Virus Enclosed | NO |

| | | | |
|---|---|---|---|
| DLQQAQQEATHI | 1422580 | Cell/Virus Enclosed | NO |
| DLQRAAVTILDG | 1422583 | Cell/Virus Enclosed | NO |
| DLQVEMNRLQDA | 1422587 | Ectodomain | YES |
| DLREYVFKNKDG | 1422591 | Ectodomain | YES |
| DLRFPRGQGVPINTNS | 1651490 | Cell/Virus Enclosed | NO |
| DLSAFYVKDGKYFTSE | 1651495 | Cell/Virus Enclosed | NO |
| DLSKELKRPIQPTDVA | 1651508 | Cell/Virus Enclosed | NO |
| DLSKFQLKLKGTPVLQ | 1651511 | Cell/Virus Enclosed | NO |
| DLSNCMVALGELKLA | 9281 | Ectodomain | YES |
| DLSQYYCDNGKYYTKP | 1651524 | Cell/Virus Enclosed | NO |
| DLSVFYVKDGKY | 1422641 | Cell/Virus Enclosed | NO |
| DLSVSSAGPIFQ | 1422647 | Ectodomain | YES |
| DLTGAICDFEIA | 1422667 | Cell/Virus Enclosed | NO |
| DLTLDGYSKLFITKDE | 1651552 | Cell/Virus Enclosed | NO |
| DLTSDENGVRTL | 1422688 | Ectodomain | YES |
| DLTSLGLAAEWF | 1422689 | Cell/Virus Enclosed | NO |
| DLTVGSGYCVDC | 1422695 | Underneath Membrane | NO |
| DLTVVSKVHEVIIDNK | 1651560 | Cell/Virus Enclosed | NO |
| DLTYEMLSLQQV | 1422698 | Ectodomain | YES |
| DLVDSYFENDSILQSV | 1651563 | Cell/Virus Enclosed | NO |
| DLVHDEPKGKFS | 1422705 | Cell/Virus Enclosed | NO |
| DLVKFISDKIKFLNP | 9319 | Ectodomain | YES |
| DLVKRDSEADSP | 1422709 | Cell/Virus Enclosed | NO |
| DLVQNGVDAKGFPQLA | 1651573 | Capsid | UN |
| DLWQFVDHFGENEEII | 1651583 | Cell/Virus Enclosed | NO |
| DLYSMQDVGFKN | 1422760 | Cell/Virus Enclosed | NO |
| DLYVNVYRSTSPDPKF | 1651606 | Cell/Virus Enclosed | NO |
| DMADQIASLVLA | 1422771 | Cell/Virus Enclosed | NO |
| DMAKFPLKLAGTAVIN | 1651610 | Cell/Virus Enclosed | NO |
| DMDDFSRQLQNSMSGAS | 1378859 | Capsid | UN |
| DMDDNLFIAKYG | 1422782 | Cell/Virus Enclosed | NO |
| DMDSGGSSFSEL | 1422784 | Cell/Virus Enclosed | NO |
| DMDYVPLKSAVC | 1422788 | Cell/Virus Enclosed | NO |
| DMEKDFLSFDSDVFIK | 1651622 | Cell/Virus Enclosed | NO |
| DMEYDFLNMDMGVFIN | 1651623 | Cell/Virus Enclosed | NO |
| DMGVFINKYGLEDFNF | 1651625 | Cell/Virus Enclosed | NO |
| DMHWGDGFQIKF | 1422812 | Cell/Virus Enclosed | NO |
| DMIAAYTAALVSGTATA | 9365 | Ectodomain | YES |
| DMICSTTAQKLE | 1422816 | Cell/Virus Enclosed | NO |
| DMKGKMFVRPVIEDYH | 1651631 | Underneath Membrane | NO |
| DMKGTMYVRPIIEDYH | 1651632 | Ectodomain | YES |
| DMLRHLIKDVDNPVLM | 1651637 | Cell/Virus Enclosed | NO |
| DMLRRLIKDVDNPVLM | 1651638 | Cell/Virus Enclosed | NO |
| DMLYSPPTISYNSTLQ | 1651639 | Cell/Virus Enclosed | NO |
| DMSSVDNILNMA | 1422858 | Cell/Virus Enclosed | NO |
| DMSSVDTILNMARNGV | 1651650 | Cell/Virus Enclosed | NO |
| DMTKFSLKLAGTAVVN | 1651653 | Cell/Virus Enclosed | NO |
| DMTNPDYTNLLCRVTS | 1651654 | Cell/Virus Enclosed | NO |
| DMYDGRTKAIDGENVS | 1651664 | Cell/Virus Enclosed | NO |
| DMYDPITKNIGEYNVS | 1651665 | Cell/Virus Enclosed | NO |
| DMYDPLTKNIGDYNVS | 1651668 | Cell/Virus Enclosed | NO |
| DMYDPTTKNLTG | 1422877 | Cell/Virus Enclosed | NO |
| DMYDPTTKNVTGSNES | 1651670 | Cell/Virus Enclosed | NO |
| DMYVYSAGYATG | 1422884 | Ectodomain | YES |
| DMYVYSSGHATG | 1422885 | Ectodomain | YES |
| DNALSGVLSAAIKDCV | 1651681 | Cell/Virus Enclosed | NO |
| DNAQALNTLVKQ | 1422892 | Ectodomain | YES |
| DNASHASTVRAT | 1422893 | Secreted | YES |
| DNASPASTAHAT | 1422894 | Secreted | YES |
| DNASPASTVHAI | 1422895 | Secreted | YES |
| DNASPASTVHATATIP | 1651683 | Secreted | YES |
| DNAVATFYPQLQSAEW | 1651684 | Cell/Virus Enclosed | NO |
| DNAVKGCVPLNAIPSL | 1651685 | Cell/Virus Enclosed | NO |
| DNAVKGCVPLSAIPAL | 1651686 | Cell/Virus Enclosed | NO |
| DNAYLKWARVEGKDGF | 1651688 | Cell/Virus Enclosed | NO |
| DNCDFYGWVSGNMMDG | 1651690 | Cell/Virus Enclosed | NO |
| DNCSLEKFMSTPNAIF | 1651693 | Cell/Virus Enclosed | NO |
| DNCYRKSNVDES | 1422908 | Cell/Virus Enclosed | NO |
| DNCYRNSNVDEA | 1422910 | Cell/Virus Enclosed | NO |
| DNCYRNSNVGES | 1422911 | Cell/Virus Enclosed | NO |
| DNCYRNSNVNKS | 1422912 | Cell/Virus Enclosed | NO |
| DNCYRSTSVEES | 1422914 | Cell/Virus Enclosed | NO |
| DNDDDNDDQIVV | 1422916 | Cell/Virus Enclosed | NO |
| DNDDEDVVTGDNDDED | 1651698 | Cell/Virus Enclosed | NO |
| DNDDEDVVTGDNNDEE | 1651699 | Cell/Virus Enclosed | NO |
| DNDDEHVVTGDN | 1422921 | Cell/Virus Enclosed | NO |

| | | | |
|---|---|---|---|
| DNDDQIVVTGDDVDDI | 1651700 | Cell/Virus Enclosed | NO |
| DNDFNTILNTACGVFE | 1651701 | Cell/Virus Enclosed | NO |
| DNDGRPVHVKEITKEN | 1651703 | Cell/Virus Enclosed | NO |
| DNDINDYVSDADFSIT | 1651704 | Cell/Virus Enclosed | NO |
| DNDLYPFVSDSA | 1422931 | Cell/Virus Enclosed | NO |
| DNDVFIKKYGLE | 1422936 | Cell/Virus Enclosed | NO |
| DNETLEYWVTPL | 1422943 | Ectodomain | YES |
| DNFCAFCTPKKV | 1422948 | Cell/Virus Enclosed | NO |
| DNFNQWHEAKFG | 1422957 | Cell/Virus Enclosed | NO |
| DNFTSIWPYYSF | 1422963 | Ectodomain | YES |
| DNGALEAFKRSNNGVY | 1651726 | Cell/Virus Enclosed | NO |
| DNGIITVEELKQ | 1422975 | Ectodomain | YES |
| DNGKYYTKPIIKAQFK | 1651731 | Cell/Virus Enclosed | NO |
| DNGPQNQSSAPR | 1422982 | Cell/Virus Enclosed | NO |
| DNGPQQNQRSAP | 1422984 | Capsid | UN |
| DNGSLEKFMSTP | 1422987 | Cell/Virus Enclosed | NO |
| DNGSSINRKQLEIVKL | 1651736 | Cell/Virus Enclosed | NO |
| DNGSSINRRQLDVVKR | 1651737 | Cell/Virus Enclosed | NO |
| DNGVIRDVSVEDVCFA | 1651741 | Cell/Virus Enclosed | NO |
| DNHLSTFYPQLQSAEW | 1651743 | Cell/Virus Enclosed | NO |
| DNIFSVQEDGRI | 1423002 | Ectodomain | YES |
| DNIFSVQQDCRI | 1423003 | Underneath Membrane | NO |
| DNIGTNVPLQVGFSNG | 1651747 | Cell/Virus Enclosed | NO |
| DNINAILTEVNELLDT | 1651752 | Ectodomain | YES |
| DNINSILDEVNGLLDT | 1651756 | Ectodomain | YES |
| DNINSTLVDLKWLNRV | 1651757 | Ectodomain | YES |
| DNINSTLVDLNW | 1423020 | Ectodomain | YES |
| DNISVASPKSRV | 1423025 | Cell/Virus Enclosed | NO |
| DNISVATPKSRV | 1423026 | Cell/Virus Enclosed | NO |
| DNISVAVPKSRVQQNK | 1651758 | Cell/Virus Enclosed | NO |
| DNIVAADLGVLIQNSA | 1651760 | Cell/Virus Enclosed | NO |
| DNKFALTCTSTH | 1423034 | Cell/Virus Enclosed | NO |
| DNKFDVLKVDDVDDGG | 1651764 | Cell/Virus Enclosed | NO |
| DNKIMTFYPKMQ | 1423038 | Cell/Virus Enclosed | NO |
| DNKLKAHKEKSA | 1423041 | Cell/Virus Enclosed | NO |
| DNKPYRWMLWCKDNHL | 1651772 | Cell/Virus Enclosed | NO |
| DNLCTLDPITLK | 1423057 | Ectodomain | YES |
| DNLEPVSGLYEI | 1423060 | Ectodomain | YES |
| DNLSSEDLKAVRSSFN | 1651790 | Cell/Virus Enclosed | NO |
| DNLTADEIKALK | 1423080 | Cell/Virus Enclosed | NO |
| DNLTDYSVSSCA | 1423081 | Ectodomain | YES |
| DNLVFTNKTTLPTNVA | 1651793 | Cell/Virus Enclosed | NO |
| DNNDEEIVTGDNDDQI | 1651805 | Cell/Virus Enclosed | NO |
| DNNDITNTNLSRGRGR | 1651806 | Cell/Virus Enclosed | NO |
| DNNDITNTNMSR | 1423093 | Cell/Virus Enclosed | NO |
| DNNDTTNTNLSR | 1423097 | Cell/Virus Enclosed | NO |
| DNNDTTTNQPRG | 1423098 | Cell/Virus Enclosed | NO |
| DNNGEDVVTGDN | 1423100 | Cell/Virus Enclosed | NO |
| DNNQDGGRSGAR | 1423104 | Cell/Virus Enclosed | NO |
| DNNQNGDRSGARPKQR | 1651809 | Cell/Virus Enclosed | NO |
| DNNSTLQSVASS | 1423109 | Cell/Virus Enclosed | NO |
| DNNTDSIVIDGV | 1423111 | Cell/Virus Enclosed | NO |
| DNNVQRCTRKLN | 1423113 | Cell/Virus Enclosed | NO |
| DNPFFVVSKPMG | 1423119 | Ectodomain | YES |
| DNPHLMGWDYPKCDRA | 1651813 | Cell/Virus Enclosed | NO |
| DNPNHFINFPLT | 1423125 | Cell/Virus Enclosed | NO |
| DNPNSSEVDMIC | 1423127 | Cell/Virus Enclosed | NO |
| DNPSAIVTGLLP | 1423130 | Ectodomain | YES |
| DNPSSSEVDMICSTTA | 1651815 | Cell/Virus Enclosed | NO |
| DNPTLPFGDGIY | 1423133 | Ectodomain | YES |
| DNPVIPFKDGIY | 1423134 | Underneath Membrane | NO |
| DNPVLMGWDYPKCDRA | 1651816 | Cell/Virus Enclosed | NO |
| DNQALNSILDNAVKGC | 1651820 | Cell/Virus Enclosed | NO |
| DNQDLNGKWYDFGDFV | 1651821 | Cell/Virus Enclosed | NO |
| DNQDLNGKWYDFGDYV | 1651822 | Cell/Virus Enclosed | NO |
| DNQSKSPSSSGTSTPK | 1651828 | Capsid | UN |
| DNSAKFWEEAFYRDLY | 1651833 | Cell/Virus Enclosed | NO |
| DNSANTKVLKNK | 1423162 | Cell/Virus Enclosed | NO |
| DNSANTKVLRNKGVDY | 1651834 | Cell/Virus Enclosed | NO |
| DNSESDAKEPKE | 1423164 | Cell/Virus Enclosed | NO |
| DNSSTLQSVASSFVSM | 1651843 | Cell/Virus Enclosed | NO |
| DNSTITVEELKQ | 1423178 | Ectodomain | YES |
| DNSTSSVVQTCD | 1423179 | Ectodomain | YES |
| DNSTTVSVTVSSNMVE | 1651844 | Cell/Virus Enclosed | NO |
| DNSYFDSKGWYDPVEN | 1651845 | Cell/Virus Enclosed | NO |
| DNTDSIVIDGVT | 1423185 | Cell/Virus Enclosed | NO |

| | | | |
|---|---|---|---|
| DNTSCPEWMDDFEADP | 1651854 | Cell/Virus Enclosed | NO |
| DNVAFYQGDVDTVVNG | 1651859 | Cell/Virus Enclosed | NO |
| DNVAIKCVDIVK | 1423209 | Cell/Virus Enclosed | NO |
| DNVIMLNKHIDA | 1423218 | Capsid | UN |
| DNVLPETYVALPIYYQ | 1651864 | Ectodomain | YES |
| DNVSSVWPFYPY | 1423233 | Ectodomain | YES |
| DNVSSVWPLYPYGRCP | 1651870 | Ectodomain | YES |
| DNVYCYNTDLIEGSKP | 1651874 | Cell/Virus Enclosed | NO |
| DNVYCYNTGLIE | 1423239 | Cell/Virus Enclosed | NO |
| DNVYMAKAGDKYYPVV | 1651876 | Cell/Virus Enclosed | NO |
| DNVYMAKSGDKY | 1423243 | Cell/Virus Enclosed | NO |
| DNVYMAKVGDKY | 1423244 | Cell/Virus Enclosed | NO |
| DNWLNKVSVPSP | 1423245 | Underneath Membrane | NO |
| DPAAAYLDAVKQGAKP | 1651883 | Cell/Virus Enclosed | NO |
| DPAGVCLGAGQF | 1423259 | Cell/Virus Enclosed | NO |
| DPAMHIASSNAFLDLR | 1651890 | Cell/Virus Enclosed | NO |
| DPAPIDIDAFPVKPIC | 1651892 | Cell/Virus Enclosed | NO |
| DPAQLPAPRVLLSKGS | 1651895 | Cell/Virus Enclosed | NO |
| DPAQLPAPRVLLSKGT | 1651896 | Cell/Virus Enclosed | NO |
| DPAQLPTPRTLL | 1423271 | Cell/Virus Enclosed | NO |
| DPDDASRLLHTV | 1423278 | Cell/Virus Enclosed | NO |
| DPDLSAFYVKDGKYFT | 1651899 | Cell/Virus Enclosed | NO |
| DPGTNTTTEDHKIMA | 227099 | Secreted | YES |
| DPGYMQGYDDCMQQGP | 1651906 | Ectodomain | YES |
| DPIALKAPDTYK | 1423297 | Underneath Membrane | NO |
| DPINCESIYEFD | 1423304 | Cell/Virus Enclosed | NO |
| DPIRFCLENNVCNVCG | 1651913 | Cell/Virus Enclosed | NO |
| DPIRFCLENTVCKVCG | 1651914 | Cell/Virus Enclosed | NO |
| DPIRSCLENTVC | 1423305 | Cell/Virus Enclosed | NO |
| DPISFKAPDTYK | 1423306 | Ectodomain | YES |
| DPITAYDPRSCSQKKS | 1651915 | Ectodomain | YES |
| DPITFAGPYKCP | 1423308 | Ectodomain | YES |
| DPITFKAPDTYK | 1423309 | Ectodomain | YES |
| DPITLKTPDTYK | 1423313 | Ectodomain | YES |
| DPIYDEPTMTTS | 1423316 | Secreted | YES |
| DPKFVDKYYAFLNKHF | 1651919 | Cell/Virus Enclosed | NO |
| DPKKTYLDFIQQGGTP | 1651921 | Cell/Virus Enclosed | NO |
| DPKKTYLDYIQQGGVP | 1651922 | Cell/Virus Enclosed | NO |
| DPKLKGSFNDRD | 1423330 | Ectodomain | YES |
| DPKLMGWDYPKCDRAM | 1651925 | Cell/Virus Enclosed | NO |
| DPKNPNYNKWLELLEQ | 1651926 | Cell/Virus Enclosed | NO |
| DPLFSNIQPFIV | 1423351 | Cell/Virus Enclosed | NO |
| DPLKPTKRSFIEDLLF | 1651934 | Ectodomain | YES |
| DPLNLSADTQHI | 1423358 | Cell/Virus Enclosed | NO |
| DPLTKNIGDYNVSKDG | 1651940 | Cell/Virus Enclosed | NO |
| DPNGNLYGFRDY | 1423376 | Underneath Membrane | NO |
| DPNITNLCPFGE | 1423378 | Underneath Membrane | NO |
| DPNKTNVVPPAL | 1423380 | Cell/Virus Enclosed | NO |
| DPNLSGFYVKDG | 1423381 | Cell/Virus Enclosed | NO |
| DPNQTSVVPPAL | 1423391 | Cell/Virus Enclosed | NO |
| DPPCKFTVQDAKGLKI | 1651946 | Cell/Virus Enclosed | NO |
| DPPNCNIEAWLN | 1423398 | Underneath Membrane | NO |
| DPPNVQIHTIDGSSGV | 1651947 | Secreted | YES |
| DPPQADQQVDRL | 1423400 | Underneath Membrane | NO |
| DPQIQLTTTRME | 1423410 | Cell/Virus Enclosed | NO |
| DPQKAYLDFVNAGGAP | 1651952 | Cell/Virus Enclosed | NO |
| DPQQLPAPRVLI | 1423416 | Cell/Virus Enclosed | NO |
| DPQTLEILDISP | 1423418 | Ectodomain | YES |
| DPRLKGSFNDRD | 1423422 | Ectodomain | YES |
| DPRLRGSFNNRD | 1423424 | Underneath Membrane | NO |
| DPRSCSQKKSLVGVGE | 1651955 | Ectodomain | YES |
| DPRVRGLYFPAGGSS | 1651957 | Underneath Membrane | NO |
| DPRWPQIAELAPTASA | 1651958 | Cell/Virus Enclosed | NO |
| DPSFKDNVILLNKHID | 1651961 | Cell/Virus Enclosed | NO |
| DPSLKMADPNRFRGKNLPVL | 101555 | Ectodomain | YES |
| DPSSDEAIPTRFPPGT | 1651969 | Cell/Virus Enclosed | NO |
| DPTPAPGDTGTPAPASGERA | 741315 | Ectodomain | YES |
| DPTTKNLTGSNE | 1423453 | Cell/Virus Enclosed | NO |
| DPTTKNVTGSNESKAL | 1651977 | Cell/Virus Enclosed | NO |
| DPVGFCLSNTPCNVCQ | 1651983 | Cell/Virus Enclosed | NO |
| DPVIDRLYLSSHRGV | 9804 | Ectodomain | YES |
| DPVVDWHYFDAQPLTR | 1651988 | Cell/Virus Enclosed | NO |
| DPYKEFGASVELLSF | 1651990 | Capsid | UN |
| DQAMTSMYKQARAEDK | 1651997 | Cell/Virus Enclosed | NO |
| DQDFGSTVFNVPTKVL | 1652002 | Cell/Virus Enclosed | NO |
| DQDLIRQGIDYK | 1423502 | Capsid | UN |

| | | | |
|---|---|---|---|
| DQFKTTGDLAVQIGSN | 1652012 | Cell/Virus Enclosed | NO |
| DQFRSVQTRGRR | 1423524 | Cell/Virus Enclosed | NO |
| DQGLKKFYKEVR | 1423528 | Cell/Virus Enclosed | NO |
| DQGQYYYRSSRK | 1423530 | Ectodomain | YES |
| DQGQYYYRSYRK | 1423531 | Ectodomain | YES |
| DQIEQEADNMITEML | 540192 | Ectodomain | YES |
| DQIVVTGDDVDDIESI | 1652024 | Cell/Virus Enclosed | NO |
| DQLNDLVYSLIERGKL | 1652026 | Cell/Virus Enclosed | NO |
| DQLNSSYFKLSIPTNF | 1652027 | Ectodomain | YES |
| DQLNSSYKLSIP | 1423556 | Ectodomain | YES |
| DQLNSTYFKLSI | 1423557 | Ectodomain | YES |
| DQLREPMMQSAD | 1423560 | Cell/Virus Enclosed | NO |
| DQLSFDVPDGFYSTSP | 1652031 | Ectodomain | YES |
| DQMSELPKEQRV | 1423573 | Cell/Virus Enclosed | NO |
| DQMSEPPKEQRVQGSI | 1652034 | Cell/Virus Enclosed | NO |
| DQMSEPPNEQRV | 1423577 | Capsid | UN |
| DQPSLQVESANLMLSD | 1652037 | Cell/Virus Enclosed | NO |
| DQQAQQEATHIA | 1423588 | Cell/Virus Enclosed | NO |
| DQSDQVRNVQTR | 1423604 | Cell/Virus Enclosed | NO |
| DQSRKWISHCTI | 1423614 | Cell/Virus Enclosed | NO |
| DQSRKWTGHCTI | 1423615 | Cell/Virus Enclosed | NO |
| DQTPIVWHAKDFNSLS | 1652051 | Cell/Virus Enclosed | NO |
| DQVAVYQQSIIGAMTA | 1652055 | Ectodomain | YES |
| DQVEEVNSIFDIETVD | 1652056 | Cell/Virus Enclosed | NO |
| DQWFKNQTLVAP | 1423658 | Ectodomain | YES |
| DQWFKNQTSVAR | 1423661 | Ectodomain | YES |
| DQWLKNQTSVAP | 1423662 | Ectodomain | YES |
| DQYCADVAAEELMNA | 9929 | Ectodomain | YES |
| DQYGCDYSGKLADCLQ | 1652069 | Cell/Virus Enclosed | NO |
| DQYGTDINGVYW | 1423672 | Cell/Virus Enclosed | NO |
| DQYLCGADGKPVMSED | 1652072 | Cell/Virus Enclosed | NO |
| DQYMCGVDGKPISAYA | 1652073 | Cell/Virus Enclosed | NO |
| DRAAFVCGDYAA | 1423678 | Ectodomain | YES |
| DRCIKISTVQSKLTDL | 1652084 | Cell/Virus Enclosed | NO |
| DRCTTFDDVQAPNYTQH | 9956 | Ectodomain | YES |
| DRDGQRTYDDVNASLF | 1652089 | Cell/Virus Enclosed | NO |
| DRDGQRVYDDVDASLF | 1652090 | Cell/Virus Enclosed | NO |
| DRDPSNDEAIPT | 1423707 | Cell/Virus Enclosed | NO |
| DRDSTSTHDGPT | 1423712 | Cell/Virus Enclosed | NO |
| DRDTGPPSISTD | 1423713 | Ectodomain | YES |
| DRESTSTHDGFN | 1423724 | Cell/Virus Enclosed | NO |
| DRESTSTHDGLN | 1423727 | Cell/Virus Enclosed | NO |
| DRESTSTHDGPN | 1423730 | Cell/Virus Enclosed | NO |
| DRESTSTHDGPT | 1423731 | Cell/Virus Enclosed | NO |
| DRESTSTHDVPT | 1423733 | Cell/Virus Enclosed | NO |
| DREVILCWETGKVKPP | 1652098 | Cell/Virus Enclosed | NO |
| DRFKTSGDLAVQIGNN | 1652099 | Cell/Virus Enclosed | NO |
| DRFKVNETNSQYLFST | 1652100 | Cell/Virus Enclosed | NO |
| DRFKVNETTSQY | 1423735 | Cell/Virus Enclosed | NO |
| DRGIAPKSGYFVNVNN | 1652104 | Ectodomain | YES |
| DRGQESLARSAC | 1423748 | Cell/Virus Enclosed | NO |
| DRGRNPKPRAAP | 1423749 | Cell/Virus Enclosed | NO |
| DRIFARRYNATHIKVGQPQY | 741318 | Ectodomain | YES |
| DRIRFCLMLNSY | 1423763 | Cell/Virus Enclosed | NO |
| DRKIKKYPCMVGPDYA | 1652114 | Cell/Virus Enclosed | NO |
| DRLDIIQADQQV | 1423785 | Ectodomain | YES |
| DRLNALESKMSGKGSQ | 1652124 | Cell/Virus Enclosed | NO |
| DRLNQLENKISG | 1423797 | Cell/Virus Enclosed | NO |
| DRLNQLENKVSG | 1423798 | Cell/Virus Enclosed | NO |
| DRLNQLESKVSGKGQQQQ | 10010 | Capsid | UN |
| DRLNRLESKVSG | 1423801 | Capsid | UN |
| DRNHNSQDDIMKAVAA | 1652132 | Capsid | UN |
| DRNTREVFAQVKQMYK | 1652133 | Ectodomain | YES |
| DRSGARPKQRRPQGLP | 1652142 | Cell/Virus Enclosed | NO |
| DRSHYYVDSVTV | 1423837 | Cell/Virus Enclosed | NO |
| DRSNNSSRASSRSSTR | 1652144 | Capsid | UN |
| DRTAIQSFDSSY | 1423842 | Cell/Virus Enclosed | NO |
| DRTELPQLVNAN | 1423848 | Ectodomain | YES |
| DRVEKSFQLDTA | 1423858 | Ectodomain | YES |
| DRVLPNMIRMIS | 1423861 | Cell/Virus Enclosed | NO |
| DRWSASINTGNC | 1423880 | Ectodomain | YES |
| DRYEEYDLQPHK | 1423885 | Underneath Membrane | NO |
| DRYHNSQDDIMK | 1423888 | Capsid | UN |
| DRYPANAIVCRF | 1423889 | Cell/Virus Enclosed | NO |
| DRYVYSAGHATG | 1423891 | Ectodomain | YES |
| DSADNNQNGGRN | 1423894 | Capsid | UN |

| | | | |
|---|---|---|---|
| DSAFVPQPTGVF | 1423898 | Ectodomain | YES |
| DSAKSACIEVDI | 1423901 | Ectodomain | YES |
| DSAKSVCIEVDI | 1423902 | Ectodomain | YES |
| DSANPTNCIAPV | 1423903 | Ectodomain | YES |
| DSATKVYFDNPT | 1423905 | Ectodomain | YES |
| DSCISNNRCNIF | 1423913 | Ectodomain | YES |
| DSCNNVNVRDLQRRLY | 1652180 | Cell/Virus Enclosed | NO |
| DSCPGKNNGIGT | 1423921 | Ectodomain | YES |
| DSDCDVVQEQVNYVPL | 1652183 | Cell/Virus Enclosed | NO |
| DSDFVKQPVKFY | 1423931 | Cell/Virus Enclosed | NO |
| DSDGTNKQLNEISDDC | 1652184 | Cell/Virus Enclosed | NO |
| DSDINEMVLSLVRSGK | 1652185 | Cell/Virus Enclosed | NO |
| DSDQPWKVIPRN | 1423936 | Cell/Virus Enclosed | NO |
| DSDSLSSKPQRKRGVK | 1652188 | Capsid | UN |
| DSDVETRFITKSMISA | 1652192 | Cell/Virus Enclosed | NO |
| DSDVFIKKYGLENYAF | 1652193 | Cell/Virus Enclosed | NO |
| DSDVVKQPVKFY | 1423947 | Cell/Virus Enclosed | NO |
| DSDVVKQPVNFY | 1423948 | Cell/Virus Enclosed | NO |
| DSEAVLKKLKKS | 1423952 | Cell/Virus Enclosed | NO |
| DSEISANGCSTIAQAV | 1652197 | Cell/Virus Enclosed | NO |
| DSEISGFGFAIPSVAV | 1652198 | Cell/Virus Enclosed | NO |
| DSELLSTLSVDFNGVL | 1652199 | Cell/Virus Enclosed | NO |
| DSFEGVLPINTDTVLS | 1652207 | Cell/Virus Enclosed | NO |
| DSFGPGNTFINGDIAR | 1652208 | Cell/Virus Enclosed | NO |
| DSFNILACYEGIASGV | 1652212 | Cell/Virus Enclosed | NO |
| DSFRSTSRQQSR | 1423978 | Cell/Virus Enclosed | NO |
| DSFSCNNFDESKIYGS | 1652214 | Ectodomain | YES |
| DSFSCNNLDKSK | 1423980 | Ectodomain | YES |
| DSFTCNNIDAAKIYGM | 1652215 | Ectodomain | YES |
| DSFTCNNIDVAK | 1423982 | Underneath Membrane | NO |
| DSFTNGAVIGTP | 1423987 | Cell/Virus Enclosed | NO |
| DSFVKMMVDGFVHYAG | 1652219 | Cell/Virus Enclosed | NO |
| DSFVNSFVSLYNVTRD | 1652220 | Cell/Virus Enclosed | NO |
| DSFVVKGDDVRQIAPG | 1652221 | Ectodomain | YES |
| DSGFAAYSRYRIGNYK | 1379081 | Underneath Membrane | NO |
| DSGTVSKTSDWKCKVT | 1652232 | Cell/Virus Enclosed | NO |
| DSHIVSKESGNTVVLT | 1652236 | Capsid | UN |
| DSIEMQLFKVGKVDSI | 1652240 | Cell/Virus Enclosed | NO |
| DSIESVGGLYEIKIPT | 1652241 | Ectodomain | YES |
| DSIGTNFPLQLGFSTG | 1652244 | Cell/Virus Enclosed | NO |
| DSIKSACIEVDI | 1424050 | Ectodomain | YES |
| DSKCLTSIEEVC | 1424069 | Cell/Virus Enclosed | NO |
| DSKFQVGEFVFEKVDY | 1652264 | Cell/Virus Enclosed | NO |
| DSKGWYDPVENEDIHR | 1652265 | Cell/Virus Enclosed | NO |
| DSKIVQLSEINM | 1424077 | Cell/Virus Enclosed | NO |
| DSKPPLPPDEWF | 1424083 | Cell/Virus Enclosed | NO |
| DSKPQQVTKQNA | 1424087 | Cell/Virus Enclosed | NO |
| DSKTEEKSVEQK | 1424094 | Cell/Virus Enclosed | NO |
| DSKTEEKSVVQK | 1424095 | Cell/Virus Enclosed | NO |
| DSLCKTIQSALS | 1424100 | Cell/Virus Enclosed | NO |
| DSLCSSHYWSPQGNLG | 1652278 | Cell/Virus Enclosed | NO |
| DSLECSNLLLQY | 1424106 | Ectodomain | YES |
| DSLGASSNQSPQ | 1424112 | Cell/Virus Enclosed | NO |
| DSLKPVGGLYEI | 1424119 | Ectodomain | YES |
| DSLNAKLGFDSSKEFV | 1652289 | Cell/Virus Enclosed | NO |
| DSLNAKWIFDSS | 1424125 | Cell/Virus Enclosed | NO |
| DSLNAKWVFDSS | 1424126 | Cell/Virus Enclosed | NO |
| DSLNLSAGTQHISNDF | 1652290 | Capsid | UN |
| DSLNSKLGFDSS | 1424131 | Cell/Virus Enclosed | NO |
| DSLRQAALPQSKDSNF | 1652292 | Cell/Virus Enclosed | NO |
| DSLRQATLPQSK | 1424140 | Cell/Virus Enclosed | NO |
| DSLRVTFIDGLS | 1424145 | Cell/Virus Enclosed | NO |
| DSLTHYPALKMV | 1424150 | Cell/Virus Enclosed | NO |
| DSNAFSLANVIS | 1424159 | Ectodomain | YES |
| DSNAFSLANVTS | 1424160 | Ectodomain | YES |
| DSNCCVVEASTD | 1424161 | Cell/Virus Enclosed | NO |
| DSNFRTEVDPDL | 1424169 | Cell/Virus Enclosed | NO |
| DSNGCVVEASTDFHQL | 1652302 | Cell/Virus Enclosed | NO |
| DSNGFIRFPDVI | 1424171 | Cell/Virus Enclosed | NO |
| DSNGNIIGFKDFVTNK | 1652303 | Ectodomain | YES |
| DSNGNLYGFRDYITNR | 1652304 | Ectodomain | YES |
| DSNGNLYGLRDY | 1424173 | Ectodomain | YES |
| DSNKGVTAACPHAGAKSFYKNLIWL | 1134662 | Ectodomain | YES |
| DSNKYTYFDNPI | 1424176 | Ectodomain | YES |
| DSNTTQDTYGGASVCI | 1652306 | Cell/Virus Enclosed | NO |
| DSNVKNLYEKVRSQLKNNAKEIGNG | 1134664 | Ectodomain | YES |

| | | | |
|---|---|---|---|
| DSNWAFVAFSFL | 1424185 | Undetermined | UN |
| DSPCFSQQGVFR | 1424190 | Ectodomain | YES |
| DSPFEKLQCEHLQFGL | 1652311 | Ectodomain | YES |
| DSPLKLEDFIPT | 1424195 | Cell/Virus Enclosed | NO |
| DSPSTNGSGFEILSPTAPSV | 741322 | Underneath Membrane | NO |
| DSPVKDVFELHY | 1424198 | Cell/Virus Enclosed | NO |
| DSQFEEDVEMSD | 1424203 | Cell/Virus Enclosed | NO |
| DSQNMLRGDDMS | 1424206 | Cell/Virus Enclosed | NO |
| DSQNMLRGEDMF | 1424207 | Cell/Virus Enclosed | NO |
| DSQNMLRGEDMS | 1424208 | Cell/Virus Enclosed | NO |
| DSQPRADKPSQL | 1424210 | Cell/Virus Enclosed | NO |
| DSQYFFNKDTGV | 1424216 | Ectodomain | YES |
| DSRGAILRRQYNLSTSPLTSSVATG | 10221 | Undetermined | UN |
| DSRLKGSFNNRD | 1424224 | Ectodomain | YES |
| DSRSDCNHIGNI | 1424225 | Ectodomain | YES |
| DSRSDCNHINNLKIKN | 1652323 | Ectodomain | YES |
| DSRTGSFNNRDT | 1424229 | Underneath Membrane | NO |
| DSRTGSLNNVDT | 1424230 | Ectodomain | YES |
| DSRWNNARHSDN | 1424234 | Ectodomain | YES |
| DSRWNNARLSDN | 1424235 | Ectodomain | YES |
| DSRWNNARQSDNMTAV | 1652326 | Ectodomain | YES |
| DSSDHGQESLAR | 1424240 | Cell/Virus Enclosed | NO |
| DSSDNSQNGERN | 1424242 | Cell/Virus Enclosed | NO |
| DSSEGDAKETKE | 1424246 | Cell/Virus Enclosed | NO |
| DSSEPLCLFKKN | 1424248 | Ectodomain | YES |
| DSSESGAKETKEINII | 1652330 | Cell/Virus Enclosed | NO |
| DSSEVKAQSSGVIPEN | 1652331 | Cell/Virus Enclosed | NO |
| DSSFEKLHCEHL | 1424251 | Underneath Membrane | NO |
| DSSKSVTKKYTY | 1424264 | Cell/Virus Enclosed | NO |
| DSSRSTSRQQSRTRSD | 1652341 | Capsid | UN |
| DSTDNNQDGGRS | 1424279 | Cell/Virus Enclosed | NO |
| DSTDNNQNGDRSGARP | 1652345 | Cell/Virus Enclosed | NO |
| DSTDNNQSGGRN | 1424281 | Capsid | UN |
| DSTFVTEYYEFLNKHF | 1652353 | Cell/Virus Enclosed | NO |
| DSTGSNQNGERSGARP | 1652354 | Cell/Virus Enclosed | NO |
| DSTLVKFSAAQAMEKV | 1652359 | Ectodomain | YES |
| DSTQDGVNKLQG | 1424304 | Ectodomain | YES |
| DSTYGPLQTPVGCVLG | 1652365 | Ectodomain | YES |
| DSTYGPLQTPVS | 1424314 | Ectodomain | YES |
| DSTYGPLQTSVG | 1424315 | Ectodomain | YES |
| DSVAVKNGALHP | 1424318 | Cell/Virus Enclosed | NO |
| DSVFKPQPAGVL | 1424327 | Underneath Membrane | NO |
| DSVFYVVKNSTAFPFE | 1652374 | Cell/Virus Enclosed | NO |
| DSVISAVSAGIE | 1424334 | Cell/Virus Enclosed | NO |
| DSVIVNDDTSES | 1424337 | Cell/Virus Enclosed | NO |
| DSVIVNDDTSETKYVK | 1652375 | Cell/Virus Enclosed | NO |
| DSVIVNNDTSET | 1424338 | Cell/Virus Enclosed | NO |
| DSVKSACIKVDI | 1424344 | Ectodomain | YES |
| DSVLLIERFVSLAIDA | 1652379 | Cell/Virus Enclosed | NO |
| DSVQYAHKHDIQITNE | 1652385 | Cell/Virus Enclosed | NO |
| DSVRDPKTSEILDISP | 1652387 | Ectodomain | YES |
| DSVRDPQTLQIL | 1424360 | Ectodomain | YES |
| DSVREEVDIIEQPFEE | 1652388 | Cell/Virus Enclosed | NO |
| DSVRNLFESVKS | 1424363 | Ectodomain | YES |
| DSVSAIGTANLS | 1424366 | Ectodomain | YES |
| DSVTVKETVAQF | 1424374 | Cell/Virus Enclosed | NO |
| DSVVEVVTTSLTPCGY | 1652396 | Cell/Virus Enclosed | NO |
| DSYCKLKNSLSDVAFN | 1652400 | Cell/Virus Enclosed | NO |
| DSYCRLKNSVSDVAYN | 1652401 | Cell/Virus Enclosed | NO |
| DSYFDNSSTLQSVASS | 1652402 | Cell/Virus Enclosed | NO |
| DSYLGCVFNADN | 1424394 | Ectodomain | YES |
| DSYPALETIQVT | 1424397 | Cell/Virus Enclosed | NO |
| DTAAYREAACSQLAKA | 1652423 | Cell/Virus Enclosed | NO |
| DTAHANNINRFNVAIT | 1652430 | Cell/Virus Enclosed | NO |
| DTAKKSMYDGDRFVKH | 1652431 | Cell/Virus Enclosed | NO |
| DTALALLLLDRL | 1424417 | Cell/Virus Enclosed | NO |
| DTANHSIPDLQR | 1424418 | Cell/Virus Enclosed | NO |
| DTAPKTGNFKDL | 1424420 | Ectodomain | YES |
| DTASALYREALESPE | 1652434 | Capsid | UN |
| DTCFGVSKPNAI | 1424430 | Cell/Virus Enclosed | NO |
| DTCVSNNLCNIF | 1424438 | Ectodomain | YES |
| DTCVSNNRCNIF | 1424439 | Ectodomain | YES |
| DTEQTQNVTVKPKRLR | 1652448 | Cell/Virus Enclosed | NO |
| DTEQTQSAIAKP | 1424452 | Cell/Virus Enclosed | NO |
| DTEQTQSVIVKP | 1424453 | Cell/Virus Enclosed | NO |
| DTEQTQSVNVKL | 1424454 | Cell/Virus Enclosed | NO |

| | | | |
|---|---|---|---|
| DTEQTQSVNVKP | 1424455 | Cell/Virus Enclosed | NO |
| DTEWGNMLTGIS | 1424459 | Cell/Virus Enclosed | NO |
| DTFLSMFDVDKKSLNA | 1652457 | Cell/Virus Enclosed | NO |
| DTFTNNNGSDVLYQPP | 1652461 | Cell/Virus Enclosed | NO |
| DTGHPSISTDTV | 1424491 | Ectodomain | YES |
| DTGLPSISTDTV | 1424496 | Ectodomain | YES |
| DTGPPPISTDTVDVTN | 1652467 | Ectodomain | YES |
| DTGRPSISTDTV | 1424504 | Ectodomain | YES |
| DTGSFYGFSNVV | 1424505 | Underneath Membrane | NO |
| DTGVIYGFNSTE | 1424511 | Ectodomain | YES |
| DTGVPSTSTDIV | 1424513 | Ectodomain | YES |
| DTINDDEDVVTGDNDD | 1652481 | Cell/Virus Enclosed | NO |
| DTITVDQLKHLL | 1424539 | Undetermined | UN |
| DTIVVTEGDGISTPKL | 1652486 | Secreted | YES |
| DTKCLADTVMSA | 1424549 | Cell/Virus Enclosed | NO |
| DTKIASQLGMGG | 1424554 | Ectodomain | YES |
| DTKIVSQLGNCV | 1424556 | Ectodomain | YES |
| DTKLNGILQQAALDYL | 1652494 | Cell/Virus Enclosed | NO |
| DTLDDGPSVDTS | 1424560 | Cell/Virus Enclosed | NO |
| DTLDDILQQANKSVGI | 1652499 | Cell/Virus Enclosed | NO |
| DTLDKLSDYCTF | 1424563 | Cell/Virus Enclosed | NO |
| DTLFASSSLRTF | 1424566 | Cell/Virus Enclosed | NO |
| DTLKGLSDRVVF | 1424570 | Cell/Virus Enclosed | NO |
| DTLLNAILRCGSSGRSKR | 558237 | Undetermined | UN |
| DTLMQGVTLSSNLNTN | 1652506 | Ectodomain | YES |
| DTLNDGPSVETS | 1424577 | Cell/Virus Enclosed | NO |
| DTLNKLSKYLVN | 1424578 | Cell/Virus Enclosed | NO |
| DTLPQSPGPAFPLAEDVEKD | 741329 | Ectodomain | YES |
| DTLQETPVVPDTVEVP | 1652507 | Cell/Virus Enclosed | NO |
| DTLQETSVVSDT | 1424584 | Cell/Virus Enclosed | NO |
| DTLTKLSKYLANEAQV | 1652515 | Cell/Virus Enclosed | NO |
| DTNHGDAGFTSI | 1424607 | Ectodomain | YES |
| DTNLNKFNGASL | 1424610 | Cell/Virus Enclosed | NO |
| DTNSACPKLEFA | 1424613 | Ectodomain | YES |
| DTNSVCPKFEFA | 1424614 | Ectodomain | YES |
| DTNSVCPKLEFANDTK | 1652523 | Ectodomain | YES |
| DTNSVCPKLEHH | 1424618 | Underneath Membrane | NO |
| DTNSVCPKLHHH | 1424619 | Underneath Membrane | NO |
| DTNSVSPKLEFA | 1424622 | Ectodomain | YES |
| DTPIGAGICASY | 1424628 | Ectodomain | YES |
| DTPSPLTPRSVR | 1424637 | Ectodomain | YES |
| DTPSTLTPCSVR | 1424644 | Ectodomain | YES |
| DTPTEHVTSVNL | 1424648 | Cell/Virus Enclosed | NO |
| DTRTRSTLNLEGVNGG | 1652539 | Cell/Virus Enclosed | NO |
| DTRTRSVFNLEGVNGG | 1652540 | Cell/Virus Enclosed | NO |
| DTRVLNKLNLPGCNGG | 1652542 | Cell/Virus Enclosed | NO |
| DTRVLNNLNLPGCNGG | 1652543 | Cell/Virus Enclosed | NO |
| DTSDFTDSVRDP | 1424664 | Ectodomain | YES |
| DTSGFAVYVKSK | 1424670 | Underneath Membrane | NO |
| DTSLSDACQLPY | 1424681 | Underneath Membrane | NO |
| DTSPQVLKALQKAVNI | 1652554 | Cell/Virus Enclosed | NO |
| DTSRKGSFNNRD | 1424691 | Ectodomain | YES |
| DTSTPSDVSSRDPTTQ | 1652555 | Capsid | UN |
| DTSYDCDIPIGA | 1424695 | Ectodomain | YES |
| DTSYECDIPIGAGICA | 1379190 | Ectodomain | YES |
| DTSYINNKDTGP | 1424701 | Ectodomain | YES |
| DTTATSCQLYYNLPAA | 1652556 | Ectodomain | YES |
| DTTLYVNNWCRC | 1424725 | Ectodomain | YES |
| DTTNTNLSRGRG | 1424726 | Cell/Virus Enclosed | NO |
| DTTPKEEFVVKEKLNA | 1652565 | Cell/Virus Enclosed | NO |
| DTTQLHVADTLMQGVT | 1652566 | Ectodomain | YES |
| DTVDVTNGLGTY | 1424738 | Ectodomain | YES |
| DTVFEVQGYKNV | 1424749 | Cell/Virus Enclosed | NO |
| DTVFTSEEVVLK | 1424750 | Cell/Virus Enclosed | NO |
| DTVNRDSFNLYG | 1424762 | Cell/Virus Enclosed | NO |
| DTVSALVYENKLKAKN | 1652585 | Cell/Virus Enclosed | NO |
| DTVVSTVKIGGLTFKT | 1652591 | Cell/Virus Enclosed | NO |
| DTVYCPRHVICT | 1424777 | Cell/Virus Enclosed | NO |
| DTYAALRNSLTN | 1424784 | Cell/Virus Enclosed | NO |
| DTYDPTYKNGAM | 1424789 | Cell/Virus Enclosed | NO |
| DVAARADYEGFI | 1424805 | Cell/Virus Enclosed | NO |
| DVAFNRYLSLYN | 1424808 | Cell/Virus Enclosed | NO |
| DVAIQEVIDDML | 1424812 | Cell/Virus Enclosed | NO |
| DVALDSLGAAIH | 1424814 | Cell/Virus Enclosed | NO |
| DVANNAVYDGAR | 1424816 | Cell/Virus Enclosed | NO |
| DVANPTNCIAPV | 1424818 | Ectodomain | YES |

| | | | |
|---|---|---|---|
| DVAVTAGPFMDNAINV | 1652614 | Cell/Virus Enclosed | NO |
| DVDDIDPINCES | 1424841 | Cell/Virus Enclosed | NO |
| DVDDIESIYDFDTYKA | 1652630 | Cell/Virus Enclosed | NO |
| DVDDPKLMGWDYPKCD | 1652631 | Cell/Virus Enclosed | NO |
| DVDDYILANECI | 1424847 | Cell/Virus Enclosed | NO |
| DVDFHSVEIEAGEVKP | 1652633 | Cell/Virus Enclosed | NO |
| DVDGLKYTKILKDDGN | 1652636 | Cell/Virus Enclosed | NO |
| DVDGLKYTKIMKDDGN | 1652637 | Cell/Virus Enclosed | NO |
| DVDHEFVGEFYA | 1424856 | Cell/Virus Enclosed | NO |
| DVDKFKTPHIVR | 1424858 | Cell/Virus Enclosed | NO |
| DVDNIDFKSLLG | 1424864 | Ectodomain | YES |
| DVDNINFKSLVGCLGP | 1652642 | Ectodomain | YES |
| DVDNLVFTNKTTLPTN | 1652643 | Cell/Virus Enclosed | NO |
| DVDNPHLMGWDYPKCD | 1652644 | Cell/Virus Enclosed | NO |
| DVDNPVLMGWDYPKCD | 1652645 | Cell/Virus Enclosed | NO |
| DVDQEFVDEFYA | 1424868 | Cell/Virus Enclosed | NO |
| DVDTGVPSTSTD | 1424875 | Ectodomain | YES |
| DVDTHRYCLSLK | 1424877 | Cell/Virus Enclosed | NO |
| DVDTNCLADSVM | 1424882 | Cell/Virus Enclosed | NO |
| DVDTTTLYVNNW | 1424884 | Ectodomain | YES |
| DVDTVVNGVDFDFIVN | 1652652 | Cell/Virus Enclosed | NO |
| DVEEDDAEGEET | 1424900 | Cell/Virus Enclosed | NO |
| DVEGAHATRENIGTNF | 1652663 | Cell/Virus Enclosed | NO |
| DVESAHVCGDNIGTNV | 1652680 | Cell/Virus Enclosed | NO |
| DVESGVYSVSSFEAKP | 1652681 | Ectodomain | YES |
| DVFDLIYQLFKG | 1424927 | Cell/Virus Enclosed | NO |
| DVFDLSYQLFKG | 1424929 | Cell/Virus Enclosed | NO |
| DVFHETSDDEWT | 1424933 | Cell/Virus Enclosed | NO |
| DVFKIGDVKFKRLGDY | 1652695 | Cell/Virus Enclosed | NO |
| DVGFKNQYCNGSIACQ | 1652708 | Cell/Virus Enclosed | NO |
| DVGFVEAYNNCTGGAE | 1652709 | Ectodomain | YES |
| DVGFVEAYNNCTGGSE | 1652710 | Ectodomain | YES |
| DVGLDSAKSACI | 1424963 | Ectodomain | YES |
| DVGNHRGAFALH | 1424965 | Underneath Membrane | NO |
| DVGNHRSAFALH | 1424966 | Ectodomain | YES |
| DVGNRRSAFALH | 1424967 | Underneath Membrane | NO |
| DVHKDNFKDGLCMFWN | 1652717 | Cell/Virus Enclosed | NO |
| DVIDNEIIVKPNISLCVPL | 1424988 | Cell/Virus Enclosed | NO |
| DVIFSQFDSLGVSSNQ | 1652722 | Cell/Virus Enclosed | NO |
| DVIRKDLSYEQQNVLA | 1652730 | Cell/Virus Enclosed | NO |
| DVIRYNLNFEENLRRG | 1652731 | Ectodomain | YES |
| DVISSEAVEDTIDGVV | 1652733 | Cell/Virus Enclosed | NO |
| DVIVAWHVVRDPRYVM | 1652735 | Cell/Virus Enclosed | NO |
| DVIVHDVEPTHK | 1425018 | Cell/Virus Enclosed | NO |
| DVIVNPANGHMLHGGG | 1652738 | Cell/Virus Enclosed | NO |
| DVKGLEELHYDL | 1425033 | Cell/Virus Enclosed | NO |
| DVKPYGYPNDFVGGFR | 1652757 | Cell/Virus Enclosed | NO |
| DVKTIEVVGNVI | 1425052 | Cell/Virus Enclosed | NO |
| DVLCNSFFKELTANMS | 1652763 | Cell/Virus Enclosed | NO |
| DVLFSGQKYSSD | 1425064 | Cell/Virus Enclosed | NO |
| DVLFYCSNSSSGVLDT | 1652767 | Ectodomain | YES |
| DVLLAPLLSAGI | 1425079 | Cell/Virus Enclosed | NO |
| DVLNKNQKLIAN | 1425084 | Ectodomain | YES |
| DVLPEGWHVVNK | 1425086 | Cell/Virus Enclosed | NO |
| DVLQENQKILAASFNK | 1652783 | Ectodomain | YES |
| DVLRNSFGKDLNANMS | 1652785 | Cell/Virus Enclosed | NO |
| DVLSDVIRYNLNFEEN | 1652786 | Ectodomain | YES |
| DVLSQNQKLIANAFNN | 1652788 | Ectodomain | YES |
| DVLSRLDKVEAE | 1425096 | Ectodomain | YES |
| DVMFEDDQGLKK | 1425112 | Cell/Virus Enclosed | NO |
| DVMRYNFNFSSN | 1425120 | Ectodomain | YES |
| DVMRYNLNFSANSLDN | 1652797 | Ectodomain | YES |
| DVNAIDIEEPSGQT | 10677 | Capsid | UN |
| DVNCNGYQHHSF | 1425125 | Ectodomain | YES |
| DVNCNGYQHHSI | 1425126 | Underneath Membrane | NO |
| DVNCNGYQHNSF | 1425128 | Ectodomain | YES |
| DVNCTDVPTSIH | 1425132 | Ectodomain | YES |
| DVNDSHSMPVVD | 1425144 | Cell/Virus Enclosed | NO |
| DVNDVTKLYLGGMSYY | 1652806 | Cell/Virus Enclosed | NO |
| DVNFSPVLGCLG | 1425146 | Ectodomain | YES |
| DVNKTLQEFAKN | 1425152 | Ectodomain | YES |
| DVNKTLQEFAQNLPKY | 1652808 | Ectodomain | YES |
| DVNKTLQELSYKLPNY | 1652809 | Ectodomain | YES |
| DVNMEAAYTSSL | 1425155 | Ectodomain | YES |
| DVNMQIMGVVMQSGVR | 1652812 | Cell/Virus Enclosed | NO |
| DVNMQLMGVVMQ | 1425160 | Cell/Virus Enclosed | NO |

| | | | |
|---|---|---|---|
| DVNSVLNICFDI | 1425168 | Cell/Virus Enclosed | NO |
| DVNTLLSNQLRL | 1425172 | Cell/Virus Enclosed | NO |
| DVPDGFYSTSPIQSVE | 1652823 | Ectodomain | YES |
| DVPFLICSNTPA | 1425187 | Cell/Virus Enclosed | NO |
| DVPSISGTQVYS | 1425198 | Cell/Virus Enclosed | NO |
| DVPSTYGTQVYS | 1425202 | Cell/Virus Enclosed | NO |
| DVPTAIRADQLT | 1425203 | Ectodomain | YES |
| DVPTEHVTSVNL | 1425204 | Cell/Virus Enclosed | NO |
| DVPTTLAWLNNQ | 1425209 | Cell/Virus Enclosed | NO |
| DVPTTLAWVNNH | 1425210 | Cell/Virus Enclosed | NO |
| DVPYPKQSIFTINSVV | 1652831 | Cell/Virus Enclosed | NO |
| DVQAPNYTQHSS | 1425217 | Underneath Membrane | NO |
| DVQLRAFDICNTNRAG | 1652833 | Cell/Virus Enclosed | NO |
| DVQQAQQEATHI | 1425228 | Cell/Virus Enclosed | NO |
| DVQQAQQEVTHI | 1425229 | Cell/Virus Enclosed | NO |
| DVQQWGYVGNLA | 1425232 | Cell/Virus Enclosed | NO |
| DVRDYKLANDVKDTLR | 1652838 | Cell/Virus Enclosed | NO |
| DVRDYLLSKMTSLPKD | 1652839 | Cell/Virus Enclosed | NO |
| DVSARDPTIQEA | 1425255 | Cell/Virus Enclosed | NO |
| DVSEMLTFDKKAFTLA | 1652850 | Ectodomain | YES |
| DVSGVGVIRVSN | 1425277 | Underneath Membrane | NO |
| DVSHETSDDDWT | 1425280 | Cell/Virus Enclosed | NO |
| DVSKADGIIHPQ | 1425288 | Ectodomain | YES |
| DVSKADGIIYPQGRTY | 1652857 | Ectodomain | YES |
| DVSKPNAIDVEH | 1425290 | Cell/Virus Enclosed | NO |
| DVSLTYLLGTAKKQVV | 1652863 | Cell/Virus Enclosed | NO |
| DVSPFEMPFEELNGLK | 1652866 | Cell/Virus Enclosed | NO |
| DVSVEDVCFANKFEQF | 1652871 | Cell/Virus Enclosed | NO |
| DVTGKLSDCLYL | 1425319 | Cell/Virus Enclosed | NO |
| DVTLEKLRDYLA | 1425328 | Cell/Virus Enclosed | NO |
| DVTQLAEEPAKD | 1425333 | Cell/Virus Enclosed | NO |
| DVTVLRDNLKPVTTLS | 1652894 | Cell/Virus Enclosed | NO |
| DVTVTTDKSFEQQVGV | 1652895 | Cell/Virus Enclosed | NO |
| DVVAANMFMGVGVGHY | 1652897 | Cell/Virus Enclosed | NO |
| DVVDQVEEVNSIFDIE | 1652902 | Cell/Virus Enclosed | NO |
| DVVIAILSGTTT | 1425361 | Cell/Virus Enclosed | NO |
| DVVKEQVSDLLVKLLR | 1652915 | Cell/Virus Enclosed | NO |
| DVVKQPVKFYMY | 1425370 | Cell/Virus Enclosed | NO |
| DVVKTDAVVLLERYVS | 1652919 | Cell/Virus Enclosed | NO |
| DVVLASDDLYVKRYFK | 1652921 | Cell/Virus Enclosed | NO |
| DVVLATDDLYVKRYER | 1652922 | Cell/Virus Enclosed | NO |
| DVVNDNAQALNT | 1425376 | Ectodomain | YES |
| DVVNQQGNSLNHLTSQ | 1652926 | Ectodomain | YES |
| DVVNQQGNSLNY | 1425380 | Ectodomain | YES |
| DVVNQQGNSLSH | 1425381 | Ectodomain | YES |
| DVVNQQGSALNHLTSQ | 1652927 | Ectodomain | YES |
| DVVQYEADRRAFVDYT | 1652931 | Cell/Virus Enclosed | NO |
| DVVQYEADRRVF | 1425384 | Cell/Virus Enclosed | NO |
| DVVRDLPSGFNTLKPI | 10779 | Ectodomain | YES |
| DVVSIRSSNRGN | 1425396 | Cell/Virus Enclosed | NO |
| DVVSIRTSNQGN | 1425397 | Cell/Virus Enclosed | NO |
| DVVSSEAVEDTI | 1425398 | Cell/Virus Enclosed | NO |
| DVVTGDNDDDND | 1425400 | Cell/Virus Enclosed | NO |
| DVVTGDNDDEDVVTGD | 1652940 | Cell/Virus Enclosed | NO |
| DVVTGDNNDEDV | 1425402 | Cell/Virus Enclosed | NO |
| DVVTGDNNDEEIVTGD | 1652941 | Cell/Virus Enclosed | NO |
| DVVTGDNNGEDV | 1425405 | Cell/Virus Enclosed | NO |
| DVVWKHITDPLFSNMQ | 1652944 | Cell/Virus Enclosed | NO |
| DVVYAQHCFKVP | 1425412 | Ectodomain | YES |
| DVYDDIFAESKPWFVR | 1652954 | Cell/Virus Enclosed | NO |
| DVYDMWLTGCKY | 1425425 | Cell/Virus Enclosed | NO |
| DVYNDALFVSYGSSVE | 1652960 | Cell/Virus Enclosed | NO |
| DVYNKDASFIGKNLKS | 1652962 | Cell/Virus Enclosed | NO |
| DVYNLTMPSQVQ | 1425434 | Cell/Virus Enclosed | NO |
| DVYRRIPNLPDC | 1425436 | Ectodomain | YES |
| DVYRRKPDLPNCNIEAW | 2060808 | Ectodomain | YES |
| DVYRRKPDPPNC | 1425439 | Underneath Membrane | NO |
| DVYRRKPNLPNCNIEA | 1652966 | Ectodomain | YES |
| DVYRRRPDLPNC | 1425441 | Underneath Membrane | NO |
| DWDIEYPYISHEKKLN | 1652971 | Cell/Virus Enclosed | NO |
| DWDSFCKTIQSA | 1425445 | Cell/Virus Enclosed | NO |
| DWDSFGKTIQSA | 1425446 | Cell/Virus Enclosed | NO |
| DWHYFDAQPLNK | 1425459 | Cell/Virus Enclosed | NO |
| DWNSFCKTIQSA | 1425505 | Cell/Virus Enclosed | NO |
| DWQNLLYDSNGN | 1425509 | Ectodomain | YES |
| DWRLINKFDVINGVKT | 1652996 | Cell/Virus Enclosed | NO |

| | | | |
|---|---|---|---|
| DWRSCNYAVSSA | 1425511 | Cell/Virus Enclosed | NO |
| DWRSYNYAVSSA | 1425512 | Cell/Virus Enclosed | NO |
| DWSDIRDYKLANDAKE | 1652998 | Cell/Virus Enclosed | NO |
| DWSITYPMIANENAIN | 1652999 | Cell/Virus Enclosed | NO |
| DWTDVRDYKLANDVKD | 1653004 | Cell/Virus Enclosed | NO |
| DWTESGDYTLANTTTE | 1653005 | Cell/Virus Enclosed | NO |
| DWTVTYPFIANEKFIN | 1653009 | Cell/Virus Enclosed | NO |
| DWVKHLNKNLNEGVLE | 1653013 | Cell/Virus Enclosed | NO |
| DWVKHLNKTLNEGVLE | 1653014 | Cell/Virus Enclosed | NO |
| DWYDFVENPDIINIYK | 1653015 | Cell/Virus Enclosed | NO |
| DWYDFVENPDIINVYK | 1653016 | Cell/Virus Enclosed | NO |
| DYAAKGYIAGIQNFKE | 1653020 | Cell/Virus Enclosed | NO |
| DYADKLCHAVVS | 1425550 | Cell/Virus Enclosed | NO |
| DYADKLCHAVVY | 1425551 | Cell/Virus Enclosed | NO |
| DYAELGYIADISAFKA | 1653024 | Cell/Virus Enclosed | NO |
| DYAFEHIVYGNFNQKI | 1653026 | Cell/Virus Enclosed | NO |
| DYAGKFCHAVVS | 1425555 | Cell/Virus Enclosed | NO |
| DYAKDNTVLQAL | 1425557 | Cell/Virus Enclosed | NO |
| DYANKGYIANIS | 1425562 | Cell/Virus Enclosed | NO |
| DYASKGYIVNIS | 1425565 | Cell/Virus Enclosed | NO |
| DYAYFNGAIIRDSDVV | 1653035 | Cell/Virus Enclosed | NO |
| DYCDGSLGCKMCLFGY | 1653037 | Cell/Virus Enclosed | NO |
| DYCGGNLCTCQP | 1425576 | Ectodomain | YES |
| DYCGGNPCICQP | 1425577 | Underneath Membrane | NO |
| DYCGGNPCTCQP | 1425578 | Ectodomain | YES |
| DYCGGNSCTCQP | 1425580 | Ectodomain | YES |
| DYCGNNSCTCQP | 1425582 | Ectodomain | YES |
| DYCRGNFCTCQP | 1425587 | Ectodomain | YES |
| DYCRGNSCICQP | 1425588 | Ectodomain | YES |
| DYCRGNSCTCQP | 1425589 | Ectodomain | YES |
| DYCVRAFDGYNK | 1425591 | Cell/Virus Enclosed | NO |
| DYCVRAFDVYNKDASF | 1653043 | Cell/Virus Enclosed | NO |
| DYDIAVTAGPFM | 1425597 | Cell/Virus Enclosed | NO |
| DYDVAVTAGPFMDNAI | 1653050 | Cell/Virus Enclosed | NO |
| DYDVAVTVGPFM | 1425603 | Cell/Virus Enclosed | NO |
| DYDVVVTAGPFM | 1425604 | Cell/Virus Enclosed | NO |
| DYEAERPFTSYT | 1425606 | Cell/Virus Enclosed | NO |
| DYEFKFYDAFPVAKSV | 1653056 | Cell/Virus Enclosed | NO |
| DYERSNIYGTAT | 1425618 | Cell/Virus Enclosed | NO |
| DYFEMKNWFDPI | 1425624 | Cell/Virus Enclosed | NO |
| DYFSYPFSMKSD | 1425632 | Ectodomain | YES |
| DYFSYPLSMKSD | 1425633 | Ectodomain | YES |
| DYGSDTVTYKSTATTK | 1653071 | Cell/Virus Enclosed | NO |
| DYHNHSNVETRL | 1425655 | Cell/Virus Enclosed | NO |
| DYHVTKVLAEKFDYNV | 1653082 | Cell/Virus Enclosed | NO |
| DYINVTFLDLQVEMNR | 1653092 | Ectodomain | YES |
| DYISSNGPLKVGAGVM | 1653095 | Cell/Virus Enclosed | NO |
| DYKHWPQIAQFAPSAS | 1653100 | Cell/Virus Enclosed | NO |
| DYKKCTKGLSIADLAC | 1653102 | Ectodomain | YES |
| DYKLANDAKESLRLFA | 1653103 | Cell/Virus Enclosed | NO |
| DYKLANDVKDTLRLFA | 1653104 | Cell/Virus Enclosed | NO |
| DYKLELFNKYFKHWSM | 1653105 | Cell/Virus Enclosed | NO |
| DYKNVKAWSGIC | 1425701 | Ectodomain | YES |
| DYKQWPQIAQFA | 1425704 | Capsid | UN |
| DYKRQNNLAIEEIEYV | 1653110 | Cell/Virus Enclosed | NO |
| DYKSCTKGLSIADLAC | 1653111 | Ectodomain | YES |
| DYLADYDVAVTV | 1425722 | Cell/Virus Enclosed | NO |
| DYLDIHPSLCNNGKIS | 1653120 | Ectodomain | YES |
| DYLWYVVDDIYY | 1425740 | Cell/Virus Enclosed | NO |
| DYMTHGQMEGLC | 1425751 | Cell/Virus Enclosed | NO |
| DYNAILQSDNPQ | 1425752 | Cell/Virus Enclosed | NO |
| DYNAILQSNNPQCAIV | 1653134 | Cell/Virus Enclosed | NO |
| DYNAKVEISVTPIKNT | 1653135 | Cell/Virus Enclosed | NO |
| DYNKTIPRISED | 1425757 | Ectodomain | YES |
| DYNLSSVIPSLPTSGS | 1653136 | Ectodomain | YES |
| DYNLSSVLPQRNIRSS | 1653137 | Ectodomain | YES |
| DYNSPSSSSSRRKRRS | 1653139 | Ectodomain | YES |
| DYNYYKYNLPTMVDIK | 1653147 | Cell/Virus Enclosed | NO |
| DYPIISNEVSIN | 1425769 | Cell/Virus Enclosed | NO |
| DYQIGGYSEDRHSGVK | 1653159 | Secreted | YES |
| DYRITSITSDKF | 1425792 | Cell/Virus Enclosed | NO |
| DYSDAVSYKSSTTYKL | 1653166 | Cell/Virus Enclosed | NO |
| DYSIDVIDNEIIVKPN | 1653168 | Cell/Virus Enclosed | NO |
| DYSNLLHSKVKSVPNM | 1653170 | Cell/Virus Enclosed | NO |
| DYTGSLADGLEAYADK | 1653177 | Cell/Virus Enclosed | NO |
| DYTGSLADGLGA | 1425827 | Cell/Virus Enclosed | NO |

| | | | |
|---|---|---|---|
| DYTGYQEFVIKT | 1425829 | Underneath Membrane | NO |
| DYTGYQELVIKTSHDD | 1653179 | Underneath Membrane | NO |
| DYTGYQESVIKT | 1425831 | Underneath Membrane | NO |
| DYTKEGSVINFC | 1425833 | Cell/Virus Enclosed | NO |
| DYTLANTTTAPL | 1425836 | Cell/Virus Enclosed | NO |
| DYTLANTTTEPLKLFA | 1653182 | Cell/Virus Enclosed | NO |
| DYTQYYFNKDTG | 1425846 | Ectodomain | YES |
| DYVDKWDDFCRQYSNE | 1653188 | Cell/Virus Enclosed | NO |
| DYVDVNKTLQEFAQNL | 1653189 | Ectodomain | YES |
| DYVGTGIIRSSNQSLA | 1653191 | Ectodomain | YES |
| DYVHLPYPDPSR | 1425857 | Cell/Virus Enclosed | NO |
| DYVSLLNQIWQK | 1425879 | Cell/Virus Enclosed | NO |
| DYYNMLVNCFKW | 1425894 | Cell/Virus Enclosed | NO |
| EAAAQGFQACRF | 1425905 | Cell/Virus Enclosed | NO |
| EAAIELSKELKRPVNP | 1653239 | Cell/Virus Enclosed | NO |
| EAALDLSKELKRPIQP | 1653241 | Cell/Virus Enclosed | NO |
| EAARKAYQEAMD | 1425927 | Cell/Virus Enclosed | NO |
| EAASNGFRACRF | 1425931 | Cell/Virus Enclosed | NO |
| EAATQGFQACRF | 1425933 | Cell/Virus Enclosed | NO |
| EAAYAEVHAEPK | 1425937 | Cell/Virus Enclosed | NO |
| EAAYTSSSLGSI | 1425939 | Ectodomain | YES |
| EACYERGCNPKG | 1425946 | Cell/Virus Enclosed | NO |
| EADEGICFNFNGSWVL | 1653253 | Cell/Virus Enclosed | NO |
| EADPKCKYAQNL | 1425957 | Cell/Virus Enclosed | NO |
| EADTEQTQNVTVKPKR | 1653258 | Cell/Virus Enclosed | NO |
| EADTEQTQSVNV | 1425966 | Cell/Virus Enclosed | NO |
| EADWIELFPHND | 1425971 | Cell/Virus Enclosed | NO |
| EAEAQIDRLINGRLTA | 1653263 | Ectodomain | YES |
| EAFHKVQDAVNN | 1425997 | Ectodomain | YES |
| EAFKRFNNGVYI | 1425998 | Cell/Virus Enclosed | NO |
| EAFKRSNNGVYISTTK | 1653279 | Cell/Virus Enclosed | NO |
| EAFLIGANYLGK | 1426000 | Cell/Virus Enclosed | NO |
| EAFPLDEAEDVTESVQ | 1653283 | Cell/Virus Enclosed | NO |
| EAFPLDEAENVT | 1426005 | Cell/Virus Enclosed | NO |
| EAFQKVQDAVNNNAQA | 1653284 | Ectodomain | YES |
| EAFRKARNGVFISTEK | 1653285 | Cell/Virus Enclosed | NO |
| EAFTKPVFISAL | 1426012 | Cell/Virus Enclosed | NO |
| EAGLPYGANKEGIIWV | 1653294 | Cell/Virus Enclosed | NO |
| EAHNNLLQQLSN | 1426031 | Underneath Membrane | NO |
| EAIKRVKGWVGF | 1426044 | Cell/Virus Enclosed | NO |
| EAILMSEDVKLNPSED | 1653304 | Cell/Virus Enclosed | NO |
| EAIPTRFPPGTILPQG | 1653305 | Capsid | UN |
| EAIPTRFPPGTVLPQG | 1653306 | Cell/Virus Enclosed | NO |
| EAKFKNSVTSIN | 1426061 | Cell/Virus Enclosed | NO |
| EAKGYWYRHNRR | 1426066 | Cell/Virus Enclosed | NO |
| EAKLSAGIEFLR | 1426070 | Cell/Virus Enclosed | NO |
| EAKTASLFKTTV | 1426077 | Ectodomain | YES |
| EAKTASLFQTTV | 1426078 | Ectodomain | YES |
| EAKVITIKVTEDGVNV | 1653324 | Cell/Virus Enclosed | NO |
| EALAEAREHLKN | 1426083 | Cell/Virus Enclosed | NO |
| EALNNLLQQLSNRFGA | 1653328 | Ectodomain | YES |
| EALQFTTLTLDK | 1426095 | Cell/Virus Enclosed | NO |
| EALQFTTLTVDK | 1426096 | Cell/Virus Enclosed | NO |
| EALSFEEQDEIYAYTK | 1653333 | Cell/Virus Enclosed | NO |
| EALSFEEQNEIYAYTK | 1653334 | Cell/Virus Enclosed | NO |
| EAPLKCVRNVFENFED | 1653347 | Cell/Virus Enclosed | NO |
| EAPNAKEEILGA | 1426144 | Cell/Virus Enclosed | NO |
| EAPNAKEEILGT | 1426145 | Cell/Virus Enclosed | NO |
| EAPRDGQAYVRK | 1426146 | Underneath Membrane | NO |
| EAQAQIDRLING | 1426154 | Ectodomain | YES |
| EAQHAKPTVIVN | 1426157 | Cell/Virus Enclosed | NO |
| EAQIDRLINGRLTALN | 1653353 | Ectodomain | YES |
| EAQLVEVPVEDI | 1426164 | Cell/Virus Enclosed | NO |
| EAQNANPMVIVN | 1426165 | Cell/Virus Enclosed | NO |
| EAQNANPTVIVN | 1426166 | Cell/Virus Enclosed | NO |
| EAQPVEVHVEDI | 1426172 | Cell/Virus Enclosed | NO |
| EAQVQIDRLINGRLTA | 1653358 | Ectodomain | YES |
| EARFSGSANQQQLKQL | 1653363 | Cell/Virus Enclosed | NO |
| EARFSGTANQQQ | 1426182 | Cell/Virus Enclosed | NO |
| EARLSGSANQQQ | 1426184 | Cell/Virus Enclosed | NO |
| EARSVNRKSKVISAMH | 1653367 | Cell/Virus Enclosed | NO |
| EASLNSLTYFNK | 1426194 | Cell/Virus Enclosed | NO |
| EASLPYGANKEGIVWVA | 11189 | Capsid | UN |
| EASLPYGANKQG | 1426196 | Capsid | UN |
| EASTDFHQLECI | 1426200 | Cell/Virus Enclosed | NO |
| EASTVVCGGKRT | 1426204 | Cell/Virus Enclosed | NO |

| | | | |
|---|---|---|---|
| EATEATFNYLLD | 1426211 | Cell/Virus Enclosed | NO |
| EATGLFADRDGYSFKK | 1653385 | Cell/Virus Enclosed | NO |
| EATGLFAERDCYTFKK | 1653386 | Cell/Virus Enclosed | NO |
| EATSGLVMLCIHGSPVNSYTNTPYT | 11204 | Undetermined | UN |
| EAVEDTIDGVVEDTIN | 1653391 | Cell/Virus Enclosed | NO |
| EAVKHGAKPISN | 1426232 | Cell/Virus Enclosed | NO |
| EAWLEFRAGRPA | 1426252 | Cell/Virus Enclosed | NO |
| EAWLNDKAVPSP | 1426254 | Ectodomain | YES |
| EAWLNDKSVPSPLNWE | 1653409 | Ectodomain | YES |
| EAWLNGKSVPSP | 1426257 | Ectodomain | YES |
| EAWSGLCVDGRN | 1426259 | Ectodomain | YES |
| EAWSGLCVDGTN | 1426260 | Underneath Membrane | NO |
| EAWYEFRAGRPH | 1426261 | Cell/Virus Enclosed | NO |
| EAYADKTLQEMKALFP | 1653412 | Cell/Virus Enclosed | NO |
| EAYADRTLQEMK | 1426263 | Cell/Virus Enclosed | NO |
| EAYNSILFENGIPLMP | 1653418 | Cell/Virus Enclosed | NO |
| ECAQVLSEYVLCGGGY | 1653424 | Cell/Virus Enclosed | NO |
| ECDEECSEVEASDLEE | 1653428 | Cell/Virus Enclosed | NO |
| ECDTPIGAGICA | 1426305 | Ectodomain | YES |
| ECFMKSDIFGQDFKTF | 1653438 | Cell/Virus Enclosed | NO |
| ECFVKSDIFGSDFKTF | 1653441 | Cell/Virus Enclosed | NO |
| ECGSCGNDSWLTGNAI | 1653442 | Cell/Virus Enclosed | NO |
| ECKRALGLSISDHEFT | 1653452 | Cell/Virus Enclosed | NO |
| ECNADIVVVDEVSMCT | 1653459 | Cell/Virus Enclosed | NO |
| ECPNSIYICSQG | 1426362 | Cell/Virus Enclosed | NO |
| ECRDSRKGVCFG | 1426366 | Cell/Virus Enclosed | NO |
| ECRLAMLGAEMTSNVR | 1653469 | Cell/Virus Enclosed | NO |
| ECSKASSRSAIEDLLFDKVKLSDV | 1379358 | Ectodomain | YES |
| ECTERLKLFAEE | 1426376 | Cell/Virus Enclosed | NO |
| ECTMFKDAEGRM | 1426386 | Cell/Virus Enclosed | NO |
| ECTMFRDAEGRI | 1426389 | Cell/Virus Enclosed | NO |
| ECTMFRDAEGRM | 1426390 | Cell/Virus Enclosed | NO |
| ECTRASTSAVKLQNNE | 1653482 | Cell/Virus Enclosed | NO |
| ECTYIGRFSKIS | 1426396 | Cell/Virus Enclosed | NO |
| ECVKAFLDVLNK | 1426403 | Cell/Virus Enclosed | NO |
| ECYDRFKVNETNSQYL | 1653500 | Cell/Virus Enclosed | NO |
| ECYSGFKPNNNSAQYV | 1653502 | Cell/Virus Enclosed | NO |
| ECYSGFKPNNTSAQYI | 1653503 | Cell/Virus Enclosed | NO |
| ECYSILAAKTGVSVEQ | 1653504 | Cell/Virus Enclosed | NO |
| EDAEEKLDNPSSSEVD | 1653506 | Cell/Virus Enclosed | NO |
| EDALRNSAMLES | 1426436 | Ectodomain | YES |
| EDAVVHGQSNAD | 1426446 | Cell/Virus Enclosed | NO |
| EDAVVHGQSNAV | 1426447 | Cell/Virus Enclosed | NO |
| EDAYAEVHAEPKGKYS | 1653517 | Cell/Virus Enclosed | NO |
| EDAYDLVHDEPK | 1426450 | Cell/Virus Enclosed | NO |
| EDCEEYEDEEEI | 1426453 | Cell/Virus Enclosed | NO |
| EDCKLPLGQSLCALPD | 1653519 | Ectodomain | YES |
| EDCKLPLGRSLC | 1426455 | Ectodomain | YES |
| EDCNLPLGQSLC | 1426457 | Ectodomain | YES |
| EDDDFLEENDVE | 1426462 | Cell/Virus Enclosed | NO |
| EDDQGLKKFYKE | 1426467 | Cell/Virus Enclosed | NO |
| EDDYKGLPLEFG | 1426474 | Cell/Virus Enclosed | NO |
| EDECDEYEEEEE | 1426481 | Cell/Virus Enclosed | NO |
| EDEFTPEDVNMQIMGV | 1653533 | Cell/Virus Enclosed | NO |
| EDEILEWFGITQTAQG | 1653535 | Ectodomain | YES |
| EDEKLVDVCEKAIGKK | 1653536 | Cell/Virus Enclosed | NO |
| EDELAPSDVYQQLAGV | 1653537 | Cell/Virus Enclosed | NO |
| EDELTPSDVYQQLAGI | 1653538 | Cell/Virus Enclosed | NO |
| EDFDLISKCQITAVEG | 1653540 | Cell/Virus Enclosed | NO |
| EDFIKHIRTNGG | 1426503 | Cell/Virus Enclosed | NO |
| EDFIKHVRTNGD | 1426506 | Cell/Virus Enclosed | NO |
| EDFKPRSKMETD | 1426510 | Cell/Virus Enclosed | NO |
| EDFYASMYENSTILQA | 1653553 | Cell/Virus Enclosed | NO |
| EDGDVLLAEFDTYDPI | 1653555 | Cell/Virus Enclosed | NO |
| EDGFAFYPACIN | 1426532 | Cell/Virus Enclosed | NO |
| EDGINVKDVFVE | 1426535 | Cell/Virus Enclosed | NO |
| EDGRIPNDFSFN | 1426542 | Ectodomain | YES |
| EDHKIMASENSSAMV | 187314 | Secreted | YES |
| EDHSLLATLDDPYVED | 1653565 | Capsid | UN |
| EDIAQVVIADTLQETP | 1653567 | Cell/Virus Enclosed | NO |
| EDIFGPCWSALASALK | 1653570 | Cell/Virus Enclosed | NO |
| EDIHRVYAALGKVVAN | 1653572 | Cell/Virus Enclosed | NO |
| EDILFSKLVTSGLGTV | 1653574 | Ectodomain | YES |
| EDINGIRRPKHLYV | 11354 | Capsid | UN |
| EDIRLFKAAYEN | 1426560 | Cell/Virus Enclosed | NO |
| EDIRLLKAAYEN | 1426561 | Cell/Virus Enclosed | NO |

| | | | |
|---|---|---|---|
| EDISLLRKMDEP | 1426562 | Capsid | UN |
| EDKFDLLISDMYDGRI | 1653579 | Cell/Virus Enclosed | NO |
| EDKFDLLISDMYDGRT | 1653580 | Cell/Virus Enclosed | NO |
| EDKFWCEDFYAS | 1426566 | Cell/Virus Enclosed | NO |
| EDKLSVYDIACCMRAG | 1653583 | Cell/Virus Enclosed | NO |
| EDLEIGYTVDCSCGKK | 1653588 | Cell/Virus Enclosed | NO |
| EDLHTLYVDSKY | 1426589 | Cell/Virus Enclosed | NO |
| EDLKAVRSSFNFDQKE | 1653591 | Cell/Virus Enclosed | NO |
| EDLLFDKVKLSDVGFV | 1653596 | Ectodomain | YES |
| EDLLFDKVTIADPGYM | 1653597 | Ectodomain | YES |
| EDLLFNKVTLADAGFM | 1653599 | Ectodomain | YES |
| EDLLFNKVTLSD | 1426599 | Ectodomain | YES |
| EDLLFSKVVTSGLGTV | 1653600 | Ectodomain | YES |
| EDLMAAYVENTC | 1426608 | Cell/Virus Enclosed | NO |
| EDLMEILNRDRV | 1426610 | Cell/Virus Enclosed | NO |
| EDLQQAQQEATH | 1426614 | Cell/Virus Enclosed | NO |
| EDLRARMTYVCQCGGE | 1653605 | Cell/Virus Enclosed | NO |
| EDLRFPRGQGVPINTN | 1653606 | Cell/Virus Enclosed | NO |
| EDLSIRALQKHL | 1426618 | Cell/Virus Enclosed | NO |
| EDLSIRNLQKRLYSNV | 1653609 | Cell/Virus Enclosed | NO |
| EDLWQFVDHFGENEEI | 1653612 | Cell/Virus Enclosed | NO |
| EDLYDVSKPVMI | 1426633 | Cell/Virus Enclosed | NO |
| EDMFFLEKDAPY | 1426637 | Cell/Virus Enclosed | NO |
| EDNPLFLFDEAG | 1426642 | Cell/Virus Enclosed | NO |
| EDPPWPQIAELV | 1426646 | Cell/Virus Enclosed | NO |
| EDPRWPQIAELV | 1426649 | Cell/Virus Enclosed | NO |
| EDRHSGVKDYVV | 1426656 | Secreted | YES |
| EDRSNNSSRASSRSST | 1653624 | Capsid | UN |
| EDSITIPYQGSGKGV | 11441 | Ectodomain | YES |
| EDSVFKPQPAGV | 1426670 | Underneath Membrane | NO |
| EDSVFVPQPTGV | 1426673 | Ectodomain | YES |
| EDSVIINDDTSE | 1426674 | Cell/Virus Enclosed | NO |
| EDSVIVNDDTGE | 1426675 | Cell/Virus Enclosed | NO |
| EDSVIVNDDTSD | 1426676 | Cell/Virus Enclosed | NO |
| EDTINDDEDVVTGDND | 1653630 | Cell/Virus Enclosed | NO |
| EDTLTKLSKYLANEAQ | 1653633 | Cell/Virus Enclosed | NO |
| EDTQVVIADTLQ | 1426683 | Cell/Virus Enclosed | NO |
| EDTYRFVTSQAIACQKHTPPA | 110825 | Capsid | UN |
| EDVDYVPLKSAV | 1426689 | Cell/Virus Enclosed | NO |
| EDVEDVISSEAVEDTI | 1653641 | Cell/Virus Enclosed | NO |
| EDVKFAASTGVIDISA | 1653644 | Cell/Virus Enclosed | NO |
| EDVKLNPSEEFI | 1426697 | Cell/Virus Enclosed | NO |
| EDVQLLKAAYAN | 1426702 | Cell/Virus Enclosed | NO |
| EDVQQAQQEATH | 1426703 | Cell/Virus Enclosed | NO |
| EDVQQAQQEVTH | 1426704 | Cell/Virus Enclosed | NO |
| EDVTEFVQEEVQ | 1426707 | Cell/Virus Enclosed | NO |
| EDVTESVQEEAQPVEV | 1653648 | Cell/Virus Enclosed | NO |
| EDVVDQVEEVNSIFDI | 1653650 | Cell/Virus Enclosed | NO |
| EDVVIFINNTTYPTNV | 1653651 | Cell/Virus Enclosed | NO |
| EDVVTGDNDDEDVVTG | 1653652 | Cell/Virus Enclosed | NO |
| EDVVTGDNNDEEIVTG | 1653653 | Cell/Virus Enclosed | NO |
| EDWHYAKFGFTP | 1426721 | Cell/Virus Enclosed | NO |
| EDWLDDATELSE | 1426722 | Cell/Virus Enclosed | NO |
| EDWLEEAIAAKP | 1426726 | Cell/Virus Enclosed | NO |
| EDYAFDHIVYGSFNHK | 1653659 | Cell/Virus Enclosed | NO |
| EDYAFEHIVYGNFNQK | 1653660 | Cell/Virus Enclosed | NO |
| EDYQGLPLEFGA | 1426736 | Cell/Virus Enclosed | NO |
| EDYTGYQELVIK | 1426739 | Underneath Membrane | NO |
| EEAARCMRSLRA | 1426742 | Cell/Virus Enclosed | NO |
| EEADEGICFNFNGSWV | 1653670 | Cell/Virus Enclosed | NO |
| EEADGAEEEVTV | 1426747 | Cell/Virus Enclosed | NO |
| EEAFYRDLYSSP | 1426749 | Cell/Virus Enclosed | NO |
| EEAKLNREEIDGVKLESTRIYQILA | 1136292 | Undetermined | UN |
| EEAKTAFKKCKF | 1426757 | Cell/Virus Enclosed | NO |
| EEALSKLSEYLISDSI | 1653684 | Cell/Virus Enclosed | NO |
| EEASVTVVEGQVDYY | 111233 | Cell/Virus Enclosed | NO |
| EEDGFAFYPACTNGHA | 1653696 | Cell/Virus Enclosed | NO |
| EEDLTKGPYEFC | 1426788 | Cell/Virus Enclosed | NO |
| EEDWLEEAIAAK | 1426796 | Cell/Virus Enclosed | NO |
| EEDYQGLPLEFG | 1426798 | Cell/Virus Enclosed | NO |
| EEEDWLDDATEL | 1426803 | Cell/Virus Enclosed | NO |
| EEEEDWLDDATE | 1426808 | Cell/Virus Enclosed | NO |
| EEEEEEEGAGDGE | 11576 | Ectodomain | YES |
| EEEEEEGAGDGEHL | 741352 | Ectodomain | YES |
| EEEEGAGDGEHLE | 11579 | Ectodomain | YES |
| EEFASHVANFQKSGYS | 1653726 | Cell/Virus Enclosed | NO |

| | | | |
|---|---|---|---|
| EEFIKHVRTNGG | 1426842 | Cell/Virus Enclosed | NO |
| EEFIQTNSPKVTIDCS | 1653729 | Ectodomain | YES |
| EEFVAASDITLK | 1426851 | Cell/Virus Enclosed | NO |
| EEFVVKEKLNAFLVHD | 1653731 | Cell/Virus Enclosed | NO |
| EEFYNFLRLRGFFDEG | 1653733 | Cell/Virus Enclosed | NO |
| EEGESECISETS | 1426860 | Cell/Virus Enclosed | NO |
| EEIEYVHGDALHTLRN | 1653749 | Cell/Virus Enclosed | NO |
| EEIVINGRTYVC | 1426889 | Cell/Virus Enclosed | NO |
| EEIVTGDNDDQIVVTG | 1653754 | Cell/Virus Enclosed | NO |
| EELASPMKSDEG | 1426908 | Cell/Virus Enclosed | NO |
| EELDQWFKNQIS | 1426910 | Ectodomain | YES |
| EELDQWFKNQTL | 1426911 | Ectodomain | YES |
| EELDQWFKNQTSVAPD | 1653764 | Ectodomain | YES |
| EELGEEIRLKVFVLG | 11748 | Cell/Virus Enclosed | NO |
| EELHDDLVNYIRDKGC | 1653769 | Cell/Virus Enclosed | NO |
| EELPDEFVVVTA | 1426935 | Cell/Virus Enclosed | NO |
| EEMLKRSSTIHN | 1426950 | Cell/Virus Enclosed | NO |
| EEMNRLQEAIKV | 1426952 | Ectodomain | YES |
| EENFSSRMYCSF | 1426955 | Cell/Virus Enclosed | NO |
| EENISRGTILFK | 1426956 | Underneath Membrane | NO |
| EENLNAYVNSNQNTDS | 1653782 | Capsid | UN |
| EENVIQCFCPRD | 1426965 | Cell/Virus Enclosed | NO |
| EEQDAIFSLTKRNILP | 1653792 | Cell/Virus Enclosed | NO |
| EEQDEIYAYTKRNVLP | 1653794 | Cell/Virus Enclosed | NO |
| EEQNEIYAYTKRNVLP | 1653797 | Cell/Virus Enclosed | NO |
| EESDLIISNFKFDDYD | 1653803 | Cell/Virus Enclosed | NO |
| EESFYKNMYLKSAVMQ | 1653806 | Cell/Virus Enclosed | NO |
| EESGAVLFCTPTKKAF | 1653807 | Cell/Virus Enclosed | NO |
| EESLMDVPSTYG | 1426998 | Cell/Virus Enclosed | NO |
| EESLTDVPSISG | 1426999 | Cell/Virus Enclosed | NO |
| EESNDQEMTTPP | 1427000 | Cell/Virus Enclosed | NO |
| EESQGNVVTSVMESQI | 1653809 | Cell/Virus Enclosed | NO |
| EESQPVEVPVED | 1427002 | Cell/Virus Enclosed | NO |
| EESTDQMSEPPM | 1427008 | Cell/Virus Enclosed | NO |
| EESVKSSYAFATLKEV | 1653813 | Cell/Virus Enclosed | NO |
| EESVKSSYAYATLKEI | 1653814 | Cell/Virus Enclosed | NO |
| EETDLTVTSAGQPCVA | 1653815 | Cell/Virus Enclosed | NO |
| EETGTLIVNSVLLFL | 1540053 | Undetermined | UN |
| EETKFLTSRLLL | 1427030 | Cell/Virus Enclosed | NO |
| EETLHSTSSEGN | 11894 | Secreted | YES |
| EETRKLMLICMD | 1427036 | Cell/Virus Enclosed | NO |
| EETRKLMPVCMD | 1427038 | Cell/Virus Enclosed | NO |
| EETSITIKKPNE | 1427041 | Cell/Virus Enclosed | NO |
| EEVDIIEQPFEEVEHV | 1653832 | Cell/Virus Enclosed | NO |
| EEVLDNRATLQA | 1427059 | Cell/Virus Enclosed | NO |
| EEVNSIFDIETVDVKH | 1653843 | Cell/Virus Enclosed | NO |
| EEVSEESCAHEY | 1427068 | Cell/Virus Enclosed | NO |
| EEYCNYLESYNIVTTA | 1653855 | Cell/Virus Enclosed | NO |
| EEYREYLESYNTATTA | 1653859 | Cell/Virus Enclosed | NO |
| EFADKLVEVGLV | 1427100 | Cell/Virus Enclosed | NO |
| EFAKNLPKYVKP | 1427103 | Ectodomain | YES |
| EFANDTKIASQLGNCV | 1653866 | Ectodomain | YES |
| EFANDTKIVSQL | 1427106 | Ectodomain | YES |
| EFANHVANFQKS | 1427107 | Cell/Virus Enclosed | NO |
| EFATRTFTATTAVGSC | 1653870 | Cell/Virus Enclosed | NO |
| EFAVLYQDVNCT | 1427118 | Ectodomain | YES |
| EFAVNSTKALAK | 1427119 | Cell/Virus Enclosed | NO |
| EFCSQHTHYIKD | 1427125 | Cell/Virus Enclosed | NO |
| EFCSQHTMLVKMDGDD | 1653876 | Cell/Virus Enclosed | NO |
| EFCSQHTMQIVDENGK | 1653878 | Cell/Virus Enclosed | NO |
| EFCSQHTMQIVDKDGT | 1653879 | Cell/Virus Enclosed | NO |
| EFCTKHTMLVKM | 1427129 | Cell/Virus Enclosed | NO |
| EFDTYDPIYKNGAMYK | 1653884 | Cell/Virus Enclosed | NO |
| EFDTYDPTYKNG | 1427138 | Cell/Virus Enclosed | NO |
| EFDVETNEIVDS | 1427140 | Cell/Virus Enclosed | NO |
| EFEDRSNNSSRASSRS | 1653888 | Capsid | UN |
| EFEFAEGQGVPI | 1427147 | Cell/Virus Enclosed | NO |
| EFEFVEGQGPPI | 1427148 | Cell/Virus Enclosed | NO |
| EFEFVEGQGVPIAPGV | 1653889 | Cell/Virus Enclosed | NO |
| EFFKNFSTSIPN | 1427152 | Ectodomain | YES |
| EFGASTEMQVEE | 1427159 | Cell/Virus Enclosed | NO |
| EFGASVELLSFLPSD | 1653896 | Capsid | UN |
| EFIFEKVEYGSD | 1427176 | Cell/Virus Enclosed | NO |
| EFIKHVRTNGGY | 1427179 | Cell/Virus Enclosed | NO |
| EFIQTNSPKVTIDCSL | 1653905 | Ectodomain | YES |
| EFIQTSSPKVTIDCAA | 1653906 | Ectodomain | YES |

| | | | |
|---|---|---|---|
| EFKFWDESFYAS | 1427184 | Cell/Virus Enclosed | NO |
| EFKYMVANGLHAPHGP | 1653912 | Cell/Virus Enclosed | NO |
| EFKYMVANGLNAPNGP | 1653913 | Cell/Virus Enclosed | NO |
| EFLATFIVCKIV | 1427191 | Cell/Virus Enclosed | NO |
| EFLKRGEKIVYH | 1427202 | Cell/Virus Enclosed | NO |
| EFLRDAWEILKF | 1427209 | Cell/Virus Enclosed | NO |
| EFMLWCKDGKVQ | 1427213 | Cell/Virus Enclosed | NO |
| EFMLWCKDGQVQTFYP | 1653927 | Cell/Virus Enclosed | NO |
| EFNDLAHTSLVW | 1427216 | Cell/Virus Enclosed | NO |
| EFNPSQTSPATAEPVR | 1653930 | Capsid | UN |
| EFNRLATCDWTESGDY | 1653931 | Cell/Virus Enclosed | NO |
| EFPAITIGSNFV | 1427228 | Underneath Membrane | NO |
| EFQLDYSIDVIDNEII | 1653936 | Cell/Virus Enclosed | NO |
| EFRDYFDNNTDS | 1427244 | Cell/Virus Enclosed | NO |
| EFRDYFNDNTDS | 1427245 | Cell/Virus Enclosed | NO |
| EFRFMTANNLTAPRNS | 1653939 | Cell/Virus Enclosed | NO |
| EFRYMNSQGPLP | 1427258 | Cell/Virus Enclosed | NO |
| EFSHLATFAELEAAQK | 1653946 | Cell/Virus Enclosed | NO |
| EFSKKQLGLLQPTDNS | 1653950 | Cell/Virus Enclosed | NO |
| EFSKKQLGLLQS | 1427268 | Cell/Virus Enclosed | NO |
| EFSKQLQQSMSG | 1427269 | Cell/Virus Enclosed | NO |
| EFSWNADLYKLMSCFA | 1653958 | Cell/Virus Enclosed | NO |
| EFTDENYNNLVPTYLK | 1653960 | Cell/Virus Enclosed | NO |
| EFTIGNMEEFIQ | 1427286 | Ectodomain | YES |
| EFTPEDVNMQLM | 1427291 | Cell/Virus Enclosed | NO |
| EFTTGFVYFNGN | 1427298 | Ectodomain | YES |
| EFVDEFYAYLRK | 1427302 | Cell/Virus Enclosed | NO |
| EFVEGQGVPIAPGVPA | 1653970 | Cell/Virus Enclosed | NO |
| EFVEYKITQWPT | 1427311 | Cell/Virus Enclosed | NO |
| EFVFEKVDYGSD | 1427313 | Cell/Virus Enclosed | NO |
| EFVFKNKDGFPY | 1427316 | Ectodomain | YES |
| EFVGEFYAYLRK | 1427320 | Cell/Virus Enclosed | NO |
| EFVIARTGHFYI | 1427323 | Ectodomain | YES |
| EFVQEEAQPVEA | 1427328 | Cell/Virus Enclosed | NO |
| EFVQEEVQPVEV | 1427330 | Cell/Virus Enclosed | NO |
| EFVREEVDIIEQ | 1427331 | Cell/Virus Enclosed | NO |
| EFVSDADITLLG | 1427332 | Cell/Virus Enclosed | NO |
| EFVSDADSTLIG | 1427333 | Cell/Virus Enclosed | NO |
| EFVVKEKLNAFLVHDN | 1653984 | Cell/Virus Enclosed | NO |
| EFVVNDVVIAIL | 1427337 | Cell/Virus Enclosed | NO |
| EFWFAMRKDGDDVIFS | 1653987 | Cell/Virus Enclosed | NO |
| EFYDDRILEFKV | 1427342 | Cell/Virus Enclosed | NO |
| EFYEDRILDFKVLDNG | 1653992 | Cell/Virus Enclosed | NO |
| EFYEKVTGTDLELIQILKDH | 12111 | Capsid | UN |
| EFYGFKDAVTFA | 1427346 | Cell/Virus Enclosed | NO |
| EFYNDIKPPVLD | 1427349 | Cell/Virus Enclosed | NO |
| EFYNDVKPPVLDVDDV | 1653995 | Cell/Virus Enclosed | NO |
| EFYTTEPEFVKV | 1427351 | Cell/Virus Enclosed | NO |
| EGAGDGEHLEGGD | 12118 | Ectodomain | YES |
| EGAHASRNACGTNVPL | 1653998 | Cell/Virus Enclosed | NO |
| EGAHATRDSIGTNFPL | 1653999 | Cell/Virus Enclosed | NO |
| EGAHATRENIGTNFPL | 1654000 | Cell/Virus Enclosed | NO |
| EGAKTVNTSLGNRKRNQKPL | 2060812 | Capsid | UN |
| EGCHATRDAVGT | 1427366 | Cell/Virus Enclosed | NO |
| EGDASWYSTMIF | 1427376 | Cell/Virus Enclosed | NO |
| EGDCYHSGGTIISNLPFQNIDSRAV | 1136912 | Ectodomain | YES |
| EGDGGITSEGNALYNN | 1654014 | Cell/Virus Enclosed | NO |
| EGDGISTPKLKE | 1427381 | Secreted | YES |
| EGDLPENDDDAW | 1427384 | Cell/Virus Enclosed | NO |
| EGDMFPFLDGLINAPIDFDPVPN | 1204102 | Ectodomain | YES |
| EGDQIIFYEGVNFNPY | 1654020 | Ectodomain | YES |
| EGDWDSFGKTIQ | 1427394 | Cell/Virus Enclosed | NO |
| EGDYYRKQLSPL | 1427398 | Ectodomain | YES |
| EGEDVIFSQFDS | 1427401 | Cell/Virus Enclosed | NO |
| EGEETDLTVTSAGQPC | 1654023 | Cell/Virus Enclosed | NO |
| EGELPVAVVNDKIFTK | 1654025 | Cell/Virus Enclosed | NO |
| EGFGGKNILGYA | 1427413 | Cell/Virus Enclosed | NO |
| EGFGGKNILGYS | 1427414 | Cell/Virus Enclosed | NO |
| EGFLLGINYLGTIKEN | 1654036 | Cell/Virus Enclosed | NO |
| EGGAKETKEINI | 1427430 | Cell/Virus Enclosed | NO |
| EGGASWSSTMIF | 1427431 | Cell/Virus Enclosed | NO |
| EGGCNTIELDSPCRFM | 1654043 | Cell/Virus Enclosed | NO |
| EGGDGTRDTLPQS | 12169 | Ectodomain | YES |
| EGGKTFMYAYISNKAD | 1654045 | Cell/Virus Enclosed | NO |
| EGGLCPSTGQAMESVG | 1654046 | Cell/Virus Enclosed | NO |
| EGGRAFMYAYVTTKPG | 1654049 | Cell/Virus Enclosed | NO |

| | | | |
|---|---|---|---|
| EGHIIMEEMLKGSSTI | 1654052 | Cell/Virus Enclosed | NO |
| EGHIIMEEMLRG | 1427456 | Cell/Virus Enclosed | NO |
| EGIIWVATEGALNTPK | 1379562 | Cell/Virus Enclosed | NO |
| EGILSEREKLLH | 1427471 | Ectodomain | YES |
| EGIVWVATEGALNTPKD | 1379563 | Capsid | UN |
| EGKAYFPREGVF | 1427481 | Ectodomain | YES |
| EGKDGFVSVELQ | 1427482 | Cell/Virus Enclosed | NO |
| EGKTFFVLPSDD | 1427491 | Cell/Virus Enclosed | NO |
| EGLDCKIQTAPH | 1427499 | Cell/Virus Enclosed | NO |
| EGLVRIVRTRSM | 1427516 | Cell/Virus Enclosed | NO |
| EGNALYNNEGGRAFMY | 1654080 | Cell/Virus Enclosed | NO |
| EGPTTCGYLPAN | 1427537 | Cell/Virus Enclosed | NO |
| EGQDVIFSQFDSLGVS | 1654092 | Cell/Virus Enclosed | NO |
| EGQLGENNELRLTRDALEPC | 741366 | Ectodomain | YES |
| EGQLQSLSKDYISSNG | 1654095 | Cell/Virus Enclosed | NO |
| EGQLQSLSRDYI | 1427550 | Cell/Virus Enclosed | NO |
| EGQQIIFYEGVN | 1427552 | Ectodomain | YES |
| EGRITPYCHDPT | 1427554 | Cell/Virus Enclosed | NO |
| EGRTIYGNVSRQDLTK | 1654100 | Cell/Virus Enclosed | NO |
| EGSGKSAPNSRS | 1427564 | Cell/Virus Enclosed | NO |
| EGSGRPAPNSRS | 1427566 | Capsid | UN |
| EGSGRSASNSRPGSRS | 1654107 | Capsid | UN |
| EGSKPYSTLQPNAYYK | 1654109 | Cell/Virus Enclosed | NO |
| EGSLPYSSIQANAYYK | 1654110 | Cell/Virus Enclosed | NO |
| EGSRVPHIVRKDLTKY | 1654112 | Cell/Virus Enclosed | NO |
| EGTGGNIQSSSR | 1427588 | Capsid | UN |
| EGTGGNSQSSSRASSL | 1654126 | Cell/Virus Enclosed | NO |
| EGTKKLAARLSFNVGR | 1654127 | Cell/Virus Enclosed | NO |
| EGVCITTNGLWA | 1427594 | Cell/Virus Enclosed | NO |
| EGVCITTNGSWAIFND | 1654130 | Cell/Virus Enclosed | NO |
| EGVCLGAGQFLG | 1427596 | Cell/Virus Enclosed | NO |
| EGVDCKIQTAPH | 1427597 | Cell/Virus Enclosed | NO |
| EGVFVSNGTSWF | 1427604 | Ectodomain | YES |
| EGVFWVANHQADTSTP | 1654137 | Capsid | UN |
| EGVGAKVSAFLQKLED | 1654138 | Cell/Virus Enclosed | NO |
| EGVGARVSAFLQ | 1427608 | Cell/Virus Enclosed | NO |
| EGVLPINTDTVLSVAP | 1654142 | Cell/Virus Enclosed | NO |
| EGVNFNPHHRFK | 1427615 | Ectodomain | YES |
| EGVNGGSLYVNKHAFH | 1654147 | Cell/Virus Enclosed | NO |
| EGVNGGSLYVNNHAFH | 1654148 | Cell/Virus Enclosed | NO |
| EGVSSFMNVTLN | 1427623 | Ectodomain | YES |
| EGWHVVNKFYQI | 1427627 | Cell/Virus Enclosed | NO |
| EGYDFKYIENGQLKNF | 1654159 | Cell/Virus Enclosed | NO |
| EHALSIKQPFSF | 1427638 | Cell/Virus Enclosed | NO |
| EHCQNYSGSMLR | 1427646 | Cell/Virus Enclosed | NO |
| EHCSGLAIKSDH | 1427650 | Ectodomain | YES |
| EHCSGLAIKSDY | 1427651 | Ectodomain | YES |
| EHESKFWDESFYASMY | 1654181 | Cell/Virus Enclosed | NO |
| EHEYGTEDDYKG | 1427675 | Cell/Virus Enclosed | NO |
| EHGATDAPSTFG | 1427683 | Cell/Virus Enclosed | NO |
| EHGQPPPGPSLYEES | 227128 | Cell/Virus Enclosed | NO |
| EHLEGGDGTRDTL | 12346 | Ectodomain | YES |
| EHPDVSGVCKYK | 1427722 | Cell/Virus Enclosed | NO |
| EHPHTICHPNLG | 1427724 | Underneath Membrane | NO |
| EHRRGPGKCYNC | 1427732 | Ectodomain | YES |
| EHSLLVEAYNSILFEN | 1654208 | Cell/Virus Enclosed | NO |
| EHVLSIKQPFSF | 1427763 | Cell/Virus Enclosed | NO |
| EHVTSGNLFDVG | 1427773 | Cell/Virus Enclosed | NO |
| EHVTSVNLFDIG | 1427775 | Cell/Virus Enclosed | NO |
| EHVTSVNLFDVGYSVN | 1654227 | Cell/Virus Enclosed | NO |
| EHVTSVNLLDVG | 1427777 | Cell/Virus Enclosed | NO |
| EHVVTGDNDDEH | 1427778 | Cell/Virus Enclosed | NO |
| EHYNEWAQANGFTAMN | 1654230 | Cell/Virus Enclosed | NO |
| EHYRALPGVFCG | 1427783 | Cell/Virus Enclosed | NO |
| EIACKCGVKQEQRTGL | 1654235 | Cell/Virus Enclosed | NO |
| EIANLVLAKLGKDSKP | 1654238 | Capsid | UN |
| EIAPVYEADTEQ | 1427796 | Cell/Virus Enclosed | NO |
| EIATKMFDSFVNSFVS | 1654242 | Cell/Virus Enclosed | NO |
| EIDGNVMHANYLFWRN | 1654247 | Cell/Virus Enclosed | NO |
| EIDKGIYQTSNFRVVPS | 12417 | Ectodomain | YES |
| EIEAGEVKPFAVYKNV | 1654251 | Cell/Virus Enclosed | NO |
| EIEGVGAKVSAF | 1427821 | Cell/Virus Enclosed | NO |
| EIEKPRWKRQPN | 1427822 | Capsid | UN |
| EIEYVHGDALHTLRNG | 1654254 | Cell/Virus Enclosed | NO |
| EIGTYEMYVKWP | 1427842 | Ectodomain | YES |
| EIIVKPNISLCVPLYV | 1654272 | Cell/Virus Enclosed | NO |

| | | | |
|---|---|---|---|
| EIKALKELYGPV | 1427865 | Cell/Virus Enclosed | NO |
| EIKCKTQFIAPP | 1427868 | Underneath Membrane | NO |
| EIKPGKMKVKAT | 1427881 | Cell/Virus Enclosed | NO |
| EIKPSGLKTMIV | 1427882 | Cell/Virus Enclosed | NO |
| EILCEYADCKESYFSK | 1654289 | Cell/Virus Enclosed | NO |
| EILDISPCSFGG | 1427889 | Ectodomain | YES |
| EILSPTAPSVYPHSEGRKSR | 741370 | Underneath Membrane | NO |
| EINMDNSQNLAW | 1427930 | Cell/Virus Enclosed | NO |
| EINRLQEAIKVL | 1427931 | Ectodomain | YES |
| EIPEETCEHEYG | 1427932 | Cell/Virus Enclosed | NO |
| EIPHFNQKLPNGVTVV | 1654317 | Capsid | UN |
| EIPSLMTVNVMF | 1427938 | Cell/Virus Enclosed | NO |
| EIPVVSDTVEVP | 1427940 | Cell/Virus Enclosed | NO |
| EIPYKSFSDSGCILPS | 1654323 | Cell/Virus Enclosed | NO |
| EIQIPSDFTIGN | 1427946 | Underneath Membrane | NO |
| EIQQVDEEEEED | 1427951 | Cell/Virus Enclosed | NO |
| EIQQVVDADSKH | 1427952 | Cell/Virus Enclosed | NO |
| EIRDLICVQSYN | 1427958 | Ectodomain | YES |
| EIRQKILNKPRQ | 1427964 | Capsid | UN |
| EISCDCGKKFDE | 1427969 | Cell/Virus Enclosed | NO |
| EISCNVCTKRATVYNS | 1654333 | Cell/Virus Enclosed | NO |
| EISVQKKINRMAEQAA | 1654342 | Cell/Virus Enclosed | NO |
| EISVTPIKNTVDTTPK | 1654343 | Cell/Virus Enclosed | NO |
| EISYTTTIKPVS | 1427987 | Cell/Virus Enclosed | NO |
| EITACQYTMCEF | 1427988 | Ectodomain | YES |
| EITKENVETLTW | 1427990 | Cell/Virus Enclosed | NO |
| EITNKVNSVIEKMNTQFTAVGKEFN | 1137428 | Ectodomain | YES |
| EITVMSSEVLPSTNQEYI | 558247 | Undetermined | UN |
| EIVDTVSALVYENKLK | 1654355 | Cell/Virus Enclosed | NO |
| EIVETVSALVYDNKLK | 1654356 | Cell/Virus Enclosed | NO |
| EIVGERQLLLLW | 1428002 | Cell/Virus Enclosed | NO |
| EIVGPKELLLLWESGK | 1654359 | Cell/Virus Enclosed | NO |
| EIVKLGAEKKDGFYET | 1654363 | Cell/Virus Enclosed | NO |
| EIVKSSDVVDSNENLT | 1654364 | Cell/Virus Enclosed | NO |
| EIVSDREVILCWETGK | 1654371 | Cell/Virus Enclosed | NO |
| EIVSERELILSWEIGK | 1654372 | Cell/Virus Enclosed | NO |
| EIVTGDNDDEDV | 1428019 | Cell/Virus Enclosed | NO |
| EIVTGDNDDQIVVTGD | 1654374 | Cell/Virus Enclosed | NO |
| EKACNIAKSAYERDRA | 1654385 | Cell/Virus Enclosed | NO |
| EKACNIAKSVYERDKA | 1654386 | Cell/Virus Enclosed | NO |
| EKAIGKKIKHEGDWDS | 1654388 | Cell/Virus Enclosed | NO |
| EKANPVNDLCYPGDFNDYEELKH | 97267 | Ectodomain | YES |
| EKASLNSLTYFNRPSL | 1654392 | Cell/Virus Enclosed | NO |
| EKCGVLDGSYNA | 1428043 | Ectodomain | YES |
| EKDAPYIVGDVI | 1428054 | Cell/Virus Enclosed | NO |
| EKDAPYMVGDVI | 1428055 | Cell/Virus Enclosed | NO |
| EKDFIALDQDVFIQKY | 1654409 | Cell/Virus Enclosed | NO |
| EKDFLFFDSDVF | 1428060 | Cell/Virus Enclosed | NO |
| EKDFLSFDNDVF | 1428061 | Cell/Virus Enclosed | NO |
| EKDFLSFDSAVF | 1428062 | Cell/Virus Enclosed | NO |
| EKDFLSFDSDVFIKKY | 1654410 | Cell/Virus Enclosed | NO |
| EKDFLSSDSDVF | 1428067 | Cell/Virus Enclosed | NO |
| EKFINGCGRNVQGHVV | 1654415 | Cell/Virus Enclosed | NO |
| EKGAYNKDHGRGGFGH | 1654425 | Cell/Virus Enclosed | NO |
| EKGKLLIRDTNKEVFV | 1654430 | Cell/Virus Enclosed | NO |
| EKHQDHVSLTVPLCGS | 1654437 | Cell/Virus Enclosed | NO |
| EKICIIDNMYMGKCGD | 1654438 | Cell/Virus Enclosed | NO |
| EKIVYHTLESPV | 1428112 | Cell/Virus Enclosed | NO |
| EKLASPMKSDEG | 1428130 | Cell/Virus Enclosed | NO |
| EKLDNPSGSEVD | 1428133 | Cell/Virus Enclosed | NO |
| EKLFVEHYNEWAQANG | 1654453 | Cell/Virus Enclosed | NO |
| EKLHCEHLQFGL | 1428139 | Underneath Membrane | NO |
| EKLISTARDGVRRGDN | 1654456 | Cell/Virus Enclosed | NO |
| EKLKQLLEQWNL | 1428157 | Undetermined | UN |
| EKLNAFLVHGNV | 1428165 | Cell/Virus Enclosed | NO |
| EKLNNLLEGNDSDNDLSLEDF | 12799 | Cell/Virus Enclosed | NO |
| EKLSAYDLACCMRAGA | 1654467 | Cell/Virus Enclosed | NO |
| EKLSPCKEIEGV | 1428176 | Cell/Virus Enclosed | NO |
| EKLSPWFMDGENAYEV | 1654469 | Cell/Virus Enclosed | NO |
| EKLSRLSMIKGPQRAD | 1654470 | Cell/Virus Enclosed | NO |
| EKLYKDLTGYML | 1428191 | Cell/Virus Enclosed | NO |
| EKNEQELLELDKWASLW | 150180 | Underneath Membrane | NO |
| EKNVTVTHSVNLLED | 150978 | Ectodomain | YES |
| EKNYYELTKSCG | 1428200 | Cell/Virus Enclosed | NO |
| EKPQEKDKKSAK | 1428203 | Cell/Virus Enclosed | NO |
| EKPTINITKPNI | 12847 | Secreted | YES |

| | | | |
|---|---|---|---|
| EKPTINTTRTNI | 12848 | Secreted | YES |
| EKQNGHSLYLADNLTA | 1654485 | Cell/Virus Enclosed | NO |
| EKSAGYPFNQFG | 1428222 | Cell/Virus Enclosed | NO |
| EKSFQLDTAPKT | 1428224 | Ectodomain | YES |
| EKSGNFKHLREFVFKN | 1654490 | Ectodomain | YES |
| EKSTFNKDIYCGNSMV | 1654493 | Cell/Virus Enclosed | NO |
| EKSVEQKPVDVK | 1428236 | Cell/Virus Enclosed | NO |
| EKSYERQTPFEI | 1428241 | Cell/Virus Enclosed | NO |
| EKTGILTVTYHSETQRTK | 110571 | Cell/Virus Enclosed | NO |
| EKVDYGSDTVTYKSTA | 1654501 | Cell/Virus Enclosed | NO |
| EKVMTFYPRLQAASDW | 1654504 | Cell/Virus Enclosed | NO |
| EKVSSSRRGVYY | 1428257 | Ectodomain | YES |
| EKVTGTDLELIQILKDHYNI | 12901 | Capsid | UN |
| EKYFKYWDQTYHANCV | 1654513 | Cell/Virus Enclosed | NO |
| EKYFKYWDQTYY | 1428266 | Cell/Virus Enclosed | NO |
| EKYSTIYKLHPSFNVS | 1654515 | Cell/Virus Enclosed | NO |
| ELANGTHTGSAF | 1428281 | Cell/Virus Enclosed | NO |
| ELAPPAGAFFFG | 1428284 | Cell/Virus Enclosed | NO |
| ELAPSDVYQQLAGVKL | 1654526 | Cell/Virus Enclosed | NO |
| ELAPTAGAFFFG | 1428288 | Cell/Virus Enclosed | NO |
| ELAPTAGAFFFR | 1428289 | Cell/Virus Enclosed | NO |
| ELAPTPGAFFFGSKLD | 1654529 | Capsid | UN |
| ELAQALTEVVYS | 1428291 | Cell/Virus Enclosed | NO |
| ELAQVLTEVVYSNGGF | 1654530 | Cell/Virus Enclosed | NO |
| ELASPMKSDEGG | 1428298 | Cell/Virus Enclosed | NO |
| ELCDNPFFAVSKPMGT | 1654535 | Ectodomain | YES |
| ELCDNPFFVVSK | 1428307 | Ectodomain | YES |
| ELDDEELMELDY | 1428315 | Cell/Virus Enclosed | NO |
| ELDEFFKNVSTSIPNF | 1654543 | Ectodomain | YES |
| ELDGIQYGRSGT | 1428326 | Cell/Virus Enclosed | NO |
| ELDKWASLWNWFDITNWLWYIR | 688897 | Ectodomain | YES |
| ELDKWASLWNWFNITNWLWYIK | 236082 | Undetermined | UN |
| ELDNDFNTILNTACGV | 1654550 | Cell/Virus Enclosed | NO |
| ELDPPCKFSIQDVKGL | 1654552 | Cell/Virus Enclosed | NO |
| ELDQWFKNQTSVAPDL | 1654554 | Ectodomain | YES |
| ELEAARKAYQEA | 1428341 | Cell/Virus Enclosed | NO |
| ELEELPVIEKPV | 1428348 | Cell/Virus Enclosed | NO |
| ELELICDCGIKQ | 1428359 | Cell/Virus Enclosed | NO |
| ELFAKRKMGLTPPLSI | 1654571 | Cell/Virus Enclosed | NO |
| ELFAKRKVGLTPPLSI | 1654572 | Cell/Virus Enclosed | NO |
| ELFDSLNLSADT | 1428375 | Cell/Virus Enclosed | NO |
| ELFTKRSIRHHP | 1428389 | Cell/Virus Enclosed | NO |
| ELGHEDLMAAYM | 1428400 | Cell/Virus Enclosed | NO |
| ELGHEDLMAAYV | 1428401 | Cell/Virus Enclosed | NO |
| ELGKYEQYIKGS | 1428406 | Underneath Membrane | NO |
| ELGKYEQYIKIK | 1428407 | Underneath Membrane | NO |
| ELGNYTYYNKEF | 1428416 | Underneath Membrane | NO |
| ELGNYTYYNKGS | 1428417 | Underneath Membrane | NO |
| ELGSGSHVGSSFDGVM | 1654602 | Cell/Virus Enclosed | NO |
| ELGVVMNFDVDT | 1428431 | Cell/Virus Enclosed | NO |
| ELGYIADISSFK | 1428433 | Cell/Virus Enclosed | NO |
| ELHDDLVNYIRDKGCV | 1654614 | Cell/Virus Enclosed | NO |
| ELHGNGSMIEDI | 1428443 | Cell/Virus Enclosed | NO |
| ELHGYTVQPIAD | 1428444 | Ectodomain | YES |
| ELHTVIPDYVDVNKTL | 1654617 | Ectodomain | YES |
| ELHWAPEFPWMFEDAE | 1654618 | Cell/Virus Enclosed | NO |
| ELICDCGIKQES | 1428456 | Cell/Virus Enclosed | NO |
| ELILSWEIGKVKPPLN | 1654621 | Cell/Virus Enclosed | NO |
| ELIPNHAALFFD | 1428460 | Cell/Virus Enclosed | NO |
| ELIRQGTDYKHWPQIAQFAP | 2060814 | Capsid | UN |
| ELKAAEAKVITIKVTE | 1654628 | Cell/Virus Enclosed | NO |
| ELKETVFVEPKN | 1428479 | Cell/Virus Enclosed | NO |
| ELKLAALCHGEDSIT | 13126 | Ectodomain | YES |
| ELKQLEAKTASLFQTT | 1654642 | Ectodomain | YES |
| ELLDAEFVEPGCGGIL | 1654649 | Cell/Virus Enclosed | NO |
| ELLDTTQLQVANSLMN | 1654650 | Ectodomain | YES |
| ELLEQNIDAYKTFPKK | 1654652 | Cell/Virus Enclosed | NO |
| ELLLLWESGKAKPPLN | 1654663 | Cell/Virus Enclosed | NO |
| ELLLSWESGKVKPPLN | 1654664 | Cell/Virus Enclosed | NO |
| ELLNAPATVCGPKLST | 1654667 | Ectodomain | YES |
| ELLNAPATVSGL | 1428532 | Underneath Membrane | NO |
| ELLQLRTQTASL | 1428535 | Ectodomain | YES |
| ELLSELLKDKYT | 1428539 | Cell/Virus Enclosed | NO |
| ELLTKAIDVDWV | 1428546 | Cell/Virus Enclosed | NO |
| ELLVQDLCLNKL | 1428550 | Cell/Virus Enclosed | NO |
| ELMNLATWVGSNLED | 1654684 | Capsid | UN |

| | | | |
|---|---|---|---|
| ELNAFTREMQQQ | 1428563 | Cell/Virus Enclosed | NO |
| ELNGLKILKQLDNNCW | 1654693 | Cell/Virus Enclosed | NO |
| ELNGVMVDKVGD | 1428572 | Cell/Virus Enclosed | NO |
| ELNGVVVDKVGDTDCV | 1654694 | Cell/Virus Enclosed | NO |
| ELNGYPVQPIAD | 1428573 | Underneath Membrane | NO |
| ELNGYTVQPIAD | 1428574 | Ectodomain | YES |
| ELPFEPQTIQPE | 1428589 | Cell/Virus Enclosed | NO |
| ELPKFFYASNGTYNCT | 1654698 | Ectodomain | YES |
| ELPVAISGDKVFVRDG | 1654704 | Cell/Virus Enclosed | NO |
| ELPVAVVNDKIFTKSG | 1654705 | Cell/Virus Enclosed | NO |
| ELPVAVVNDKVFVRYG | 1654706 | Cell/Virus Enclosed | NO |
| ELPYEPRTIQPE | 1428609 | Cell/Virus Enclosed | NO |
| ELQALPQRQKKQ | 1428610 | Cell/Virus Enclosed | NO |
| ELQTIVPEYIDVNKTL | 1654712 | Ectodomain | YES |
| ELRRGPGRCYNC | 1428627 | Ectodomain | YES |
| ELRRGPGTCYNC | 1428628 | Ectodomain | YES |
| ELRYMNANGLRPPKNS | 1654717 | Cell/Virus Enclosed | NO |
| ELRYMNANGLRPPRNS | 1654718 | Cell/Virus Enclosed | NO |
| ELSFTNYKLQSQ | 1428637 | Cell/Virus Enclosed | NO |
| ELSKELKRPVNPTDAS | 1654725 | Cell/Virus Enclosed | NO |
| ELSNTFGAISASIGDI | 1654728 | Ectodomain | YES |
| ELSTGCHTGTDFNGDF | 1654737 | Cell/Virus Enclosed | NO |
| ELSTGCHTGTDFTGNF | 1654738 | Cell/Virus Enclosed | NO |
| ELSYKLPNYTVPDLVV | 1654744 | Ectodomain | YES |
| ELTAEDISLLKKMDEP | 1654745 | Cell/Virus Enclosed | NO |
| ELTAEDISLLRK | 1428668 | Capsid | UN |
| ELTASMSMAECK | 1428672 | Cell/Virus Enclosed | NO |
| ELTLKHFFFTQKGDAA | 1654754 | Cell/Virus Enclosed | NO |
| ELTPSDVYQQLAGIKL | 1654756 | Cell/Virus Enclosed | NO |
| ELTSASYFCKIGKEQK | 1654759 | Cell/Virus Enclosed | NO |
| ELVIGAVILRGHLRI | 1312367 | Underneath Membrane | NO |
| ELVTDIVVVDEVSMLT | 1654776 | Cell/Virus Enclosed | NO |
| ELVYENKFVPVKEASK | 1654778 | Cell/Virus Enclosed | NO |
| ELVYENKFVPVKPASK | 1654779 | Cell/Virus Enclosed | NO |
| ELYAKRAVRSHPDFKL | 1654783 | Cell/Virus Enclosed | NO |
| ELYGPLDPTFLH | 1428740 | Cell/Virus Enclosed | NO |
| ELYLSSSNVTTM | 1428749 | Cell/Virus Enclosed | NO |
| ELYVNNAYRLFDLVQY | 1654795 | Cell/Virus Enclosed | NO |
| EMAKEVKTSSKVVLSD | 1654796 | Cell/Virus Enclosed | NO |
| EMCILHCSNFNT | 1428765 | Cell/Virus Enclosed | NO |
| EMCLASIAFNHP | 1428766 | Ectodomain | YES |
| EMHNKINLCDDPETAQ | 1654804 | Cell/Virus Enclosed | NO |
| EMKALFPIWSQE | 1428783 | Cell/Virus Enclosed | NO |
| EMKALLPTWSQE | 1428785 | Cell/Virus Enclosed | NO |
| EMKWLLSNTDNAAFPQMTKSYKNTR | 1138304 | Ectodomain | YES |
| EMLKGSSTIHNY | 1428807 | Cell/Virus Enclosed | NO |
| EMLKLGTNDPQFPILA | 1654817 | Capsid | UN |
| EMLKLGTSDSQF | 1428810 | Cell/Virus Enclosed | NO |
| EMLKRSSTIHNY | 1428811 | Cell/Virus Enclosed | NO |
| EMLSLQQVVKALNESY | 1654819 | Ectodomain | YES |
| EMLTFDKKAFTLANVS | 1654821 | Ectodomain | YES |
| EMNRLQEAIKVL | 1428820 | Ectodomain | YES |
| EMPCAIINDKVL | 1428821 | Cell/Virus Enclosed | NO |
| EMPCAIINDKVVAKID | 1654826 | Cell/Virus Enclosed | NO |
| EMPKFFYASNGT | 1428825 | Ectodomain | YES |
| EMQKPRWKRQPNDDVT | 1654828 | Capsid | UN |
| EMQQHPLLNPGA | 1428829 | Cell/Virus Enclosed | NO |
| EMQQHPLLNPSALEFN | 1654830 | Capsid | UN |
| EMQQQPLLNPSA | 1428831 | Cell/Virus Enclosed | NO |
| EMQSQPSHAVQN | 1428833 | Cell/Virus Enclosed | NO |
| EMQSQPSHVAQN | 1428834 | Cell/Virus Enclosed | NO |
| EMRLASIAFNHP | 1428841 | Ectodomain | YES |
| EMSCDCGKKFDDQVGC | 1654833 | Cell/Virus Enclosed | NO |
| EMSCDCGKKFDE | 1428843 | Cell/Virus Enclosed | NO |
| EMSDFVDFESVI | 1428844 | Cell/Virus Enclosed | NO |
| EMSDFVDLESVI | 1428846 | Cell/Virus Enclosed | NO |
| EMSPFEMPFEEL | 1428847 | Cell/Virus Enclosed | NO |
| EMTSNVRRQVASGVID | 1654836 | Cell/Virus Enclosed | NO |
| EMYLKLRSETLL | 1428862 | Cell/Virus Enclosed | NO |
| ENALPASTVHAT | 1428871 | Secreted | YES |
| ENAVANGSSPQIIKQL | 1654853 | Cell/Virus Enclosed | NO |
| ENAYEVVKAMLF | 1428881 | Cell/Virus Enclosed | NO |
| ENDDDAWFKPSY | 1428889 | Cell/Virus Enclosed | NO |
| ENDFLNFDSDVF | 1428890 | Cell/Virus Enclosed | NO |
| ENDITNGPHEFCSQHT | 1654859 | Cell/Virus Enclosed | NO |
| ENDNISVAAPKS | 1428891 | Cell/Virus Enclosed | NO |

| | | | |
|---|---|---|---|
| ENDTITVDQLKH | 1428899 | Ectodomain | YES |
| ENEEIIINGHTYVCAW | 1654866 | Cell/Virus Enclosed | NO |
| ENEEIVINGHTY | 1428903 | Cell/Virus Enclosed | NO |
| ENEEIVINGRTY | 1428904 | Cell/Virus Enclosed | NO |
| ENEIARIKKIKL | 1428905 | Underneath Membrane | NO |
| ENFIVYNNSGSN | 1428909 | Cell/Virus Enclosed | NO |
| ENFNSQDTLLAP | 1428914 | Cell/Virus Enclosed | NO |
| ENFNSVTRLMCN | 1428916 | Cell/Virus Enclosed | NO |
| ENFSSRMYCSFY | 1428917 | Cell/Virus Enclosed | NO |
| ENGTITDAIDCA | 1428925 | Ectodomain | YES |
| ENIAFNVFKQGH | 1428934 | Cell/Virus Enclosed | NO |
| ENIAPYKFKATMYYKDVTVS | 741381 | Ectodomain | YES |
| ENIDGDAMHANY | 1428937 | Cell/Virus Enclosed | NO |
| ENIDYFNMRRAKFSLE | 1654894 | Cell/Virus Enclosed | NO |
| ENIGTNFPLQIGFSTG | 1654895 | Cell/Virus Enclosed | NO |
| ENKFTPLSVASA | 1428948 | Cell/Virus Enclosed | NO |
| ENKISGKGQQQQ | 1428954 | Cell/Virus Enclosed | NO |
| ENKLNSCCRIVE | 1428956 | Cell/Virus Enclosed | NO |
| ENKLWFDFIENP | 1428957 | Cell/Virus Enclosed | NO |
| ENKLWFDFVENP | 1428958 | Cell/Virus Enclosed | NO |
| ENKSAELNYTVQKLQT | 1654902 | Ectodomain | YES |
| ENKTEAEWIELF | 1428962 | Cell/Virus Enclosed | NO |
| ENKTTLPTNIAFELYA | 1654905 | Cell/Virus Enclosed | NO |
| ENLELKEQPAVE | 1428969 | Cell/Virus Enclosed | NO |
| ENLFVRSKEEDG | 1428971 | Cell/Virus Enclosed | NO |
| ENLIHGGGVARA | 1428972 | Cell/Virus Enclosed | NO |
| ENLIKRMKQIED | 1428973 | Underneath Membrane | NO |
| ENLLHGGGVARA | 1428976 | Cell/Virus Enclosed | NO |
| ENLMHGGGVARA | 1428978 | Cell/Virus Enclosed | NO |
| ENLNAYVNSNQNTDSD | 1654917 | Capsid | UN |
| ENLNILFDGRDT | 1428981 | Cell/Virus Enclosed | NO |
| ENLPWPLVLEFA | 1428982 | Cell/Virus Enclosed | NO |
| ENLSSEDLKAVR | 1428986 | Cell/Virus Enclosed | NO |
| ENLTFPRGQGVP | 1428989 | Cell/Virus Enclosed | NO |
| ENLYFQGGGGSG | 1428998 | Underneath Membrane | NO |
| ENNRVVVSSDVL | 1429010 | Cell/Virus Enclosed | NO |
| ENPDIINIYKKLGPIF | 1654934 | Cell/Virus Enclosed | NO |
| ENPDIINVYKKLGPIF | 1654935 | Cell/Virus Enclosed | NO |
| ENPFMVNQLAYSSSAN | 1654937 | Cell/Virus Enclosed | NO |
| ENPMLMGWDYPKCDRA | 1654940 | Cell/Virus Enclosed | NO |
| ENQINGYTLAAT | 1429042 | Ectodomain | YES |
| ENQISGYTLAATSASL | 1654950 | Ectodomain | YES |
| ENQKILAASFNKAINN | 1654951 | Ectodomain | YES |
| ENQKILAASFNKAMTN | 1654952 | Ectodomain | YES |
| ENQKLIANKFNQALGA | 1654953 | Ectodomain | YES |
| ENQKQIANQFNKAISQ | 1654954 | Ectodomain | YES |
| ENRYSVLSVDSVSEES | 1654957 | Cell/Virus Enclosed | NO |
| ENSESAVLFVND | 1429055 | Cell/Virus Enclosed | NO |
| ENSFMVNQLAYS | 1429057 | Cell/Virus Enclosed | NO |
| ENSSAMVQVHSQGRE | 227141 | Secreted | YES |
| ENTIPHYSWFSG | 1429068 | Cell/Virus Enclosed | NO |
| ENVFAVESGGYIPSDF | 1654974 | Ectodomain | YES |
| ENVIQGFGPRDF | 1429086 | Cell/Virus Enclosed | NO |
| ENYAFEHVVYGD | 1429095 | Cell/Virus Enclosed | NO |
| ENYNNLVPTYLKSDNI | 1654990 | Cell/Virus Enclosed | NO |
| EPAMDEFIQRYK | 1429109 | Cell/Virus Enclosed | NO |
| EPCNLWHYGAGI | 1429113 | Cell/Virus Enclosed | NO |
| EPCNLYNYGAGIKLPS | 1654999 | Cell/Virus Enclosed | NO |
| EPCPSGTYEGNS | 1429117 | Cell/Virus Enclosed | NO |
| EPEIPHFNQKLPNGVT | 1655011 | Capsid | UN |
| EPFNVSFVNDSIESVG | 1655016 | Ectodomain | YES |
| EPFNVSFVNDSV | 1429145 | Ectodomain | YES |
| EPFTVNSINDSL | 1429146 | Ectodomain | YES |
| EPFTVNSKNDNL | 1429147 | Ectodomain | YES |
| EPFTVNSVNDSLEPVG | 1655017 | Ectodomain | YES |
| EPFTVNSVSDSL | 1429149 | Underneath Membrane | NO |
| EPHDWTKNITDKIDQ | 13658 | Secreted | YES |
| EPIISQSSKLLN | 1429159 | Cell/Virus Enclosed | NO |
| EPITSFNTKYVA | 1429170 | Ectodomain | YES |
| EPITSLNTKYVAPQVT | 1655024 | Ectodomain | YES |
| EPITYLNTKYVA | 1429173 | Ectodomain | YES |
| EPKDGGHFFVSD | 1429177 | Cell/Virus Enclosed | NO |
| EPKDGGHFFVSG | 1429178 | Cell/Virus Enclosed | NO |
| EPKDGGQFFVSD | 1429179 | Cell/Virus Enclosed | NO |
| EPKDGGQFFVSG | 1429180 | Cell/Virus Enclosed | NO |
| EPKGKYSQKAYALLRQ | 1655028 | Cell/Virus Enclosed | NO |

| | | | |
|---|---|---|---|
| EPKKDKKKKTDEAQPL | 13680 | Capsid | UN |
| EPLMQSADASTF | 1429200 | Cell/Virus Enclosed | NO |
| EPMYAISNEQHY | 1429217 | Ectodomain | YES |
| EPPKEHRVQGTQ | 1429224 | Cell/Virus Enclosed | NO |
| EPPKEQRVQGII | 1429226 | Cell/Virus Enclosed | NO |
| EPPKEQRVQGSITQRT | 1655050 | Cell/Virus Enclosed | NO |
| EPPKEQRVQSSI | 1429229 | Cell/Virus Enclosed | NO |
| EPPKTEHPKIEE | 1429231 | Cell/Virus Enclosed | NO |
| EPPKVAAKICIV | 1429232 | Cell/Virus Enclosed | NO |
| EPPLYVPIRLAG | 1429234 | Cell/Virus Enclosed | NO |
| EPPNEQRVQGSI | 1429236 | Capsid | UN |
| EPQKDVYELRYNGAIR | 1655057 | Cell/Virus Enclosed | NO |
| EPQRGRQSRIPY | 1429246 | Cell/Virus Enclosed | NO |
| EPQTIKPEVKEV | 1429249 | Cell/Virus Enclosed | NO |
| EPQTIQPEVKEV | 1429251 | Cell/Virus Enclosed | NO |
| EPRPGNHCPAGN | 1429258 | Ectodomain | YES |
| EPRSGNHCPADN | 1429259 | Ectodomain | YES |
| EPSITYSPATIL | 1429266 | Cell/Virus Enclosed | NO |
| EPTDSTDNNQNG | 1429271 | Capsid | UN |
| EPTGYGVRRKNSEPEI | 1655066 | Capsid | UN |
| EPTISQPSKLLN | 1429275 | Cell/Virus Enclosed | NO |
| EPVHVYADDSII | 1429287 | Cell/Virus Enclosed | NO |
| EPVHVYANDSII | 1429288 | Cell/Virus Enclosed | NO |
| EPVLVNAANTHL | 1429292 | Cell/Virus Enclosed | NO |
| EPVRDEVSIETDIIDE | 1655080 | Capsid | UN |
| EPVSGLYEIQIP | 1429302 | Ectodomain | YES |
| EPVSISTGSLSA | 1429303 | Ectodomain | YES |
| EPVSISTGSPSA | 1429304 | Underneath Membrane | NO |
| EPVSISTGSRSA | 1429305 | Ectodomain | YES |
| EQAATQMYKEARSVNR | 1655086 | Cell/Virus Enclosed | NO |
| EQAEGFECDFSP | 1429312 | Ectodomain | YES |
| EQAEGVECDFSL | 1429313 | Ectodomain | YES |
| EQCIRGAIGDVG | 1429322 | Cell/Virus Enclosed | NO |
| EQCTGLDSVMHF | 1429325 | Cell/Virus Enclosed | NO |
| EQDAIFSLTKRNILPT | 1655091 | Cell/Virus Enclosed | NO |
| EQDAQIDRLINGRLTT | 1655093 | Ectodomain | YES |
| EQDEIYAYTKRNVLPT | 1655094 | Cell/Virus Enclosed | NO |
| EQDELFAMTKRNVIPT | 1655095 | Cell/Virus Enclosed | NO |
| EQDLCDFKGWVP | 1429330 | Cell/Virus Enclosed | NO |
| EQEAILMSEDVKLNPS | 1655099 | Cell/Virus Enclosed | NO |
| EQFDSLNLSAGTQHIS | 1655107 | Capsid | UN |
| EQGSGYIPEAPR | 1429361 | Undetermined | UN |
| EQIDGKFVTKFSGDKF | 1655115 | Cell/Virus Enclosed | NO |
| EQISAFTKPSSIKEMQ | 1655119 | Capsid | UN |
| EQISAFTKPSSV | 1429373 | Cell/Virus Enclosed | NO |
| EQKCCMCNRRAA | 1429377 | Cell/Virus Enclosed | NO |
| EQKCCMCNRRAV | 1429379 | Cell/Virus Enclosed | NO |
| EQKCCVCNRRAA | 1429381 | Cell/Virus Enclosed | NO |
| EQKHEIEKPRWK | 1429382 | Capsid | UN |
| EQLDTTQLQVAN | 1429397 | Undetermined | UN |
| EQLEPCAQSRLLSGVA | 1655127 | Cell/Virus Enclosed | NO |
| EQLGAPLGITIS | 1429402 | Ectodomain | YES |
| EQLTPAWRIYST | 1429422 | Ectodomain | YES |
| EQNEIYAYTKRNVLPT | 1655142 | Cell/Virus Enclosed | NO |
| EQNHILGVNNSW | 1429423 | Cell/Virus Enclosed | NO |
| EQNIDAYKTFPKKEKK | 1655144 | Cell/Virus Enclosed | NO |
| EQPFGEVEYSLP | 1429436 | Cell/Virus Enclosed | NO |
| EQPKIEPKPVSVIKVA | 1655151 | Cell/Virus Enclosed | NO |
| EQPTVKEIISKP | 1429446 | Cell/Virus Enclosed | NO |
| EQPTVKEIISMH | 1429447 | Cell/Virus Enclosed | NO |
| EQPWKVIPRNLVPINK | 1655155 | Capsid | UN |
| EQQNVLAIESIH | 1429460 | Cell/Virus Enclosed | NO |
| EQQVGVIADKDKDLSG | 1655161 | Cell/Virus Enclosed | NO |
| EQRVQGSITQHT | 1429475 | Cell/Virus Enclosed | NO |
| EQRVQGSITQRA | 1429476 | Capsid | UN |
| EQRVQGSITQRTRTRP | 1655165 | Cell/Virus Enclosed | NO |
| EQRVRGSITQRT | 1429483 | Cell/Virus Enclosed | NO |
| EQSFGEVEHSLP | 1429486 | Cell/Virus Enclosed | NO |
| EQSFYENRFAAI | 1429488 | Cell/Virus Enclosed | NO |
| EQSLRLIDAMVY | 1429491 | Cell/Virus Enclosed | NO |
| EQSMYNLLKGCNAVAK | 1655170 | Cell/Virus Enclosed | NO |
| EQSYFTKKDWYDFVEN | 1655172 | Cell/Virus Enclosed | NO |
| EQTNCNTSSLAYYEPV | 1655174 | Cell/Virus Enclosed | NO |
| EQTNMHSHDVIV | 1429508 | Cell/Virus Enclosed | NO |
| EQTQHSLNFVGE | 1429516 | Cell/Virus Enclosed | NO |
| EQTQNVTVKPKRLRKK | 1655179 | Cell/Virus Enclosed | NO |

| | | | |
|---|---|---|---|
| EQTQSAIAKPKR | 1429520 | Cell/Virus Enclosed | NO |
| EQTQSVNVKLKR | 1429522 | Cell/Virus Enclosed | NO |
| EQTQSVNVKPKR | 1429523 | Cell/Virus Enclosed | NO |
| EQTQSVTVKPKR | 1429524 | Cell/Virus Enclosed | NO |
| EQVAPGCGVPIV | 1429527 | Cell/Virus Enclosed | NO |
| EQVIPPTSDYLI | 1429536 | Cell/Virus Enclosed | NO |
| EQVSDLLVKLLRGMPI | 1655183 | Cell/Virus Enclosed | NO |
| EQYNQTILNLTSEIST | 1655190 | Ectodomain | YES |
| EQYSCDFKGWIP | 1429562 | Cell/Virus Enclosed | NO |
| ERARTVGGVPLL | 1429571 | Cell/Virus Enclosed | NO |
| ERCELANYKQSIPMPR | 1655198 | Cell/Virus Enclosed | NO |
| ERCNVIADFKSF | 1429580 | Cell/Virus Enclosed | NO |
| ERCVVRVCYGST | 1429585 | Cell/Virus Enclosed | NO |
| ERCYTAEQCIRG | 1429586 | Cell/Virus Enclosed | NO |
| ERDFIDMDDNLFIAKY | 1655204 | Cell/Virus Enclosed | NO |
| ERDISNVPFSPN | 1429595 | Ectodomain | YES |
| ERDLLIKAYNTINNEQ | 1655209 | Cell/Virus Enclosed | NO |
| ERDLTSDENGVR | 1429600 | Ectodomain | YES |
| ERELILSWEIGKVKPP | 1655212 | Cell/Virus Enclosed | NO |
| ERGKLLVRDTRKEIFV | 1655222 | Cell/Virus Enclosed | NO |
| ERGLLHSSQLPL | 1429616 | Cell/Virus Enclosed | NO |
| ERGLLHSSRLPL | 1429617 | Cell/Virus Enclosed | NO |
| ERGSVQVDNGSSINRR | 1655225 | Cell/Virus Enclosed | NO |
| ERIDYSDAVSYKSSTT | 1655232 | Cell/Virus Enclosed | NO |
| ERINVVPGVSTN | 1429636 | Cell/Virus Enclosed | NO |
| ERIRQAVLNTVK | 1429637 | Cell/Virus Enclosed | NO |
| ERKDVPYSKQSI | 1429639 | Cell/Virus Enclosed | NO |
| ERKKISNCVADY | 1429641 | Ectodomain | YES |
| ERKSMYDGDRFV | 1429645 | Cell/Virus Enclosed | NO |
| ERKTFSNCNFNM | 1429646 | Ectodomain | YES |
| ERLIACENPFMVNQLA | 1655240 | Cell/Virus Enclosed | NO |
| ERLINTISPEKL | 1429653 | Cell/Virus Enclosed | NO |
| ERLKLFAAETQK | 1429656 | Cell/Virus Enclosed | NO |
| ERMADLALTNMYKEAR | 1655247 | Cell/Virus Enclosed | NO |
| ERNGARPKQRRP | 1429672 | Cell/Virus Enclosed | NO |
| ERNVVRASLLAG | 1429680 | Cell/Virus Enclosed | NO |
| ERQLLLVWEAGKSKPP | 1655256 | Cell/Virus Enclosed | NO |
| ERQLLLVWESGK | 1429686 | Cell/Virus Enclosed | NO |
| ERQTPFEIKSAK | 1429688 | Cell/Virus Enclosed | NO |
| ERSDKSCEHQTP | 1429689 | Cell/Virus Enclosed | NO |
| ERSEKSYERQTP | 1429698 | Cell/Virus Enclosed | NO |
| ERSGARPKQRRPQGLP | 1655263 | Cell/Virus Enclosed | NO |
| ERSGARSKQRRPQGLPNNTA | 1392143 | Capsid | UN |
| ERSNIYGTATIG | 1429705 | Cell/Virus Enclosed | NO |
| ERSVSLAIDAYP | 1429708 | Cell/Virus Enclosed | NO |
| ERVDWDIEYPYISHEK | 1655271 | Cell/Virus Enclosed | NO |
| ERVNVVPGVPTN | 1429725 | Cell/Virus Enclosed | NO |
| ERVRKAILNTVK | 1429727 | Cell/Virus Enclosed | NO |
| ERVRQAILNTVT | 1429728 | Cell/Virus Enclosed | NO |
| ERVYKHFNNYDC | 1429738 | Cell/Virus Enclosed | NO |
| ERVYNHFNNYDC | 1429739 | Cell/Virus Enclosed | NO |
| ERYVKMFWVVPT | 1429750 | Cell/Virus Enclosed | NO |
| ESAALSAQLAKDKVNE | 1655291 | Ectodomain | YES |
| ESAAQADLQHED | 1429754 | Cell/Virus Enclosed | NO |
| ESADVSEMLTFDKKAF | 1655293 | Ectodomain | YES |
| ESAHVCGDNIGTNVPL | 1655294 | Cell/Virus Enclosed | NO |
| ESASAVNMQQIY | 1429765 | Cell/Virus Enclosed | NO |
| ESAVNSTKALAKQDAA | 1655302 | Cell/Virus Enclosed | NO |
| ESAYMPIADPTHFDIE | 1655304 | Cell/Virus Enclosed | NO |
| ESCLQGDRCNIF | 1429777 | Ectodomain | YES |
| ESCNNLVPTYLKSDNI | 1655308 | Cell/Virus Enclosed | NO |
| ESDADRANFLNAVVFY | 1655310 | Cell/Virus Enclosed | NO |
| ESDDYIKKNGPL | 1429783 | Cell/Virus Enclosed | NO |
| ESDEYILAKGPLQVGD | 1655314 | Cell/Virus Enclosed | NO |
| ESDIYFVKNIPRYASA | 1655316 | Cell/Virus Enclosed | NO |
| ESDKEEDLQQAQ | 1429792 | Cell/Virus Enclosed | NO |
| ESDKEEDVQQAQ | 1429793 | Cell/Virus Enclosed | NO |
| ESDLIISNFKFDDYDF | 1655317 | Cell/Virus Enclosed | NO |
| ESDQWFKNQTSV | 1429797 | Ectodomain | YES |
| ESELVIGAVILRGHL | 1540144 | Underneath Membrane | NO |
| ESEPQRGRQGRI | 1429809 | Cell/Virus Enclosed | NO |
| ESFDVESGVYSV | 1429815 | Ectodomain | YES |
| ESFENPTYIYHS | 1429817 | Cell/Virus Enclosed | NO |
| ESFIDDYYGYLM | 1429821 | Cell/Virus Enclosed | NO |
| ESFIDDYYGYLRKHFS | 1655329 | Cell/Virus Enclosed | NO |
| ESFKKVATIPYKVCNS | 1655330 | Cell/Virus Enclosed | NO |

| | | | |
|---|---|---|---|
| ESFLGRYMSAFN | 1429829 | Cell/Virus Enclosed | NO |
| ESFSVTLLDNQEDKFW | 1655333 | Cell/Virus Enclosed | NO |
| ESFVDDFQGYLQ | 1429832 | Cell/Virus Enclosed | NO |
| ESFVDDFYGYLQKHFS | 1655334 | Cell/Virus Enclosed | NO |
| ESFYKNMYLKSAVMQS | 1655336 | Cell/Virus Enclosed | NO |
| ESFYKNMYLSSA | 1429835 | Cell/Virus Enclosed | NO |
| ESGAKETKEINIIKLS | 1655338 | Cell/Virus Enclosed | NO |
| ESGDYTLANTTTEPLK | 1655341 | Cell/Virus Enclosed | NO |
| ESGGYIPSDFAFNNWF | 1655344 | Ectodomain | YES |
| ESGGYIPSNFAF | 1429852 | Ectodomain | YES |
| ESGGYIPSNFSF | 1429854 | Ectodomain | YES |
| ESGKAKPPLNRNSVFT | 1655346 | Cell/Virus Enclosed | NO |
| ESGKLKQSQPKV | 1429855 | Cell/Virus Enclosed | NO |
| ESGKVKPPLNRNSVFT | 1655347 | Cell/Virus Enclosed | NO |
| ESGVYSVSSFEAKPSG | 1655355 | Ectodomain | YES |
| ESHMDADLAKPL | 1429873 | Cell/Virus Enclosed | NO |
| ESIHYLGTTGHTLKSG | 1655362 | Cell/Virus Enclosed | NO |
| ESIKSLNNSYINLKDI | 1655363 | Ectodomain | YES |
| ESIPIVWGAKDF | 1429883 | Cell/Virus Enclosed | NO |
| ESIPIVWGVKDFNTLS | 1655364 | Cell/Virus Enclosed | NO |
| ESIPLADDDSAI | 1429885 | Capsid | UN |
| ESITAQPVKIDN | 1429886 | Secreted | YES |
| ESIVSLPVYHKH | 1429890 | Ectodomain | YES |
| ESIYEFDTYKAL | 1429892 | Cell/Virus Enclosed | NO |
| ESKAFVDTIVPPCPSI | 1655369 | Cell/Virus Enclosed | NO |
| ESKFWDESFYASMYEK | 1655373 | Cell/Virus Enclosed | NO |
| ESKIYGSCFKSIVLDK | 1655374 | Ectodomain | YES |
| ESKPQQVTKQNA | 1429907 | Cell/Virus Enclosed | NO |
| ESKPQSRNPSSDRNHN | 1655379 | Capsid | UN |
| ESKPWFVRKAEDIFGP | 1655381 | Cell/Virus Enclosed | NO |
| ESKQKKIENEIA | 1429913 | Undetermined | UN |
| ESKQVDYVPLRSATCI | 1655382 | Cell/Virus Enclosed | NO |
| ESKVFLERFLPK | 1429921 | Cell/Virus Enclosed | NO |
| ESKVLLERFLPKCPEI | 1655384 | Cell/Virus Enclosed | NO |
| ESKYSCKEVFKNCNVL | 1655386 | Cell/Virus Enclosed | NO |
| ESKYSCKEVLKNCNVL | 1655387 | Cell/Virus Enclosed | NO |
| ESKYVKSLSIVD | 1429927 | Cell/Virus Enclosed | NO |
| ESLALGVSEKTH | 1429929 | Cell/Virus Enclosed | NO |
| ESLIDLRELGKY | 1429936 | Ectodomain | YES |
| ESLIFPSGTYYC | 1429939 | Cell/Virus Enclosed | NO |
| ESLRLFAAETVKAKEE | 1655397 | Cell/Virus Enclosed | NO |
| ESLTTTSTALGKLQDVV | 14208 | Ectodomain | YES |
| ESMSYQEQDELFAMTK | 1655402 | Cell/Virus Enclosed | NO |
| ESNVVDVNDCFK | 1429974 | Cell/Virus Enclosed | NO |
| ESPVEFHLDGEV | 1429984 | Cell/Virus Enclosed | NO |
| ESQGNVVTSVMESQIS | 1655410 | Cell/Virus Enclosed | NO |
| ESQRGVYCCRDH | 1429993 | Cell/Virus Enclosed | NO |
| ESRYGLQNLLQLPNFY | 1655418 | Ectodomain | YES |
| ESSCGLFKDCARNPID | 1655421 | Cell/Virus Enclosed | NO |
| ESSEVLEDTLDA | 1430007 | Cell/Virus Enclosed | NO |
| ESSKSLGKQLGVVSDG | 1655424 | Cell/Virus Enclosed | NO |
| ESSVQKKINRMAEQAA | 1655430 | Cell/Virus Enclosed | NO |
| ESTADQETYGGASVCL | 1655431 | Cell/Virus Enclosed | NO |
| ESTDQMSELPKE | 1430022 | Cell/Virus Enclosed | NO |
| ESTDQMSEPPKEQRVQ | 1655432 | Cell/Virus Enclosed | NO |
| ESTDSIVFTYIPASGY | 1655433 | Cell/Virus Enclosed | NO |
| ESTDSTVLTYIP | 1430029 | Cell/Virus Enclosed | NO |
| ESTFGLSYYSNSMACP | 1655434 | Cell/Virus Enclosed | NO |
| ESTFGSVYYHNSMDCP | 1655435 | Cell/Virus Enclosed | NO |
| ESTGTQSVSVDLESTS | 1655436 | Cell/Virus Enclosed | NO |
| ESTGTQSVSVDR | 1430037 | Cell/Virus Enclosed | NO |
| ESTINKSVANLKLPPH | 1655437 | Cell/Virus Enclosed | NO |
| ESTKLPYYDVEKIHIQ | 1655438 | Underneath Membrane | NO |
| ESTLRITRTLST | 1430041 | Cell/Virus Enclosed | NO |
| ESTLTHYGLTEH | 1430044 | Cell/Virus Enclosed | NO |
| ESTPVELPTVDV | 1430045 | Cell/Virus Enclosed | NO |
| ESTPVEPSTVDV | 1430047 | Cell/Virus Enclosed | NO |
| ESTQITTDTGIENATF | 1655441 | Secreted | YES |
| ESTSTHDGFNEH | 1430057 | Cell/Virus Enclosed | NO |
| ESTSTHDGFTEH | 1430058 | Cell/Virus Enclosed | NO |
| ESTSTHDGPNEH | 1430063 | Cell/Virus Enclosed | NO |
| ESTSTHDGPTEHVTSV | 1655442 | Cell/Virus Enclosed | NO |
| ESTSTHDTPTEH | 1430066 | Cell/Virus Enclosed | NO |
| ESTSVEPPTVDV | 1430068 | Cell/Virus Enclosed | NO |
| ESTTVLDHTDWCRCSC | 1655443 | Ectodomain | YES |
| ESVKSSQSSPII | 1430080 | Ectodomain | YES |

| | | | |
|---|---|---|---|
| ESVLVNAVNTHL | 1430083 | Cell/Virus Enclosed | NO |
| ESWLNDKSVPSP | 1430100 | Ectodomain | YES |
| ESYIDLKELGNYTYYN | 1655459 | Ectodomain | YES |
| ESYNNYVPSYVKPDSV | 1655462 | Cell/Virus Enclosed | NO |
| ESYVDVGLDSVK | 1430109 | Ectodomain | YES |
| ESYVDVGPDSAK | 1430110 | Ectodomain | YES |
| ESYVDVGPDSLK | 1430113 | Ectodomain | YES |
| ESYVNSSNVTTM | 1430116 | Cell/Virus Enclosed | NO |
| ETALDLSKELKR | 1430125 | Cell/Virus Enclosed | NO |
| ETARQEYENAVANGSS | 1655477 | Cell/Virus Enclosed | NO |
| ETCTIAGKAFDKVFDY | 1655481 | Cell/Virus Enclosed | NO |
| ETDFLEPAMDEF | 1430144 | Cell/Virus Enclosed | NO |
| ETDGICLENHVM | 1430145 | Cell/Virus Enclosed | NO |
| ETDGICPENHVMVDCR | 1655485 | Cell/Virus Enclosed | NO |
| ETDLKKGPHEFCSQHT | 1655486 | Cell/Virus Enclosed | NO |
| ETETYFKVNGLWSPTI | 1655491 | Cell/Virus Enclosed | NO |
| ETFNHNNGNDVLYQPP | 1655494 | Cell/Virus Enclosed | NO |
| ETFSACRAYLDS | 1430162 | Cell/Virus Enclosed | NO |
| ETFTNNNGSDVL | 1430163 | Cell/Virus Enclosed | NO |
| ETFTVCADGFMP | 1430165 | Cell/Virus Enclosed | NO |
| ETGSDLLYQPPNCSIT | 1655507 | Cell/Virus Enclosed | NO |
| ETGTSDLDRCTT | 1430189 | Undetermined | UN |
| ETHRDCDFNKPLIEWP | 1655509 | Cell/Virus Enclosed | NO |
| ETIKAKEESVKSSYAF | 1655511 | Cell/Virus Enclosed | NO |
| ETIMKVLNENLNAYQQ | 1655513 | Cell/Virus Enclosed | NO |
| ETIMKVLTENLN | 1430199 | Cell/Virus Enclosed | NO |
| ETKALKELYGPVDPTF | 1655526 | Cell/Virus Enclosed | NO |
| ETKEINIIKLSGVKKP | 1655530 | Cell/Virus Enclosed | NO |
| ETKEQKHEMQKPRWKR | 1655531 | Capsid | UN |
| ETKGPLCVDTSHFTTK | 1655535 | Ectodomain | YES |
| ETKVQCSTNLFKDCSK | 1655538 | Cell/Virus Enclosed | NO |
| ETLNEDLMEILN | 1430260 | Cell/Virus Enclosed | NO |
| ETLSACRAYLNS | 1430272 | Cell/Virus Enclosed | NO |
| ETMFDVYVDTFL | 1430286 | Cell/Virus Enclosed | NO |
| ETNCLLNVPIGG | 1430297 | Ectodomain | YES |
| ETNILLNVPLHGTIL | 1540162 | Underneath Membrane | NO |
| ETNNLMCIDMKGTMYV | 1655577 | Ectodomain | YES |
| ETNSQYLFSTINALPE | 1655578 | Cell/Virus Enclosed | NO |
| ETNTAAFNAVCS | 1430318 | Cell/Virus Enclosed | NO |
| ETPNAKEEILGT | 1430331 | Cell/Virus Enclosed | NO |
| ETPVAYRNVLLR | 1430336 | Cell/Virus Enclosed | NO |
| ETPVDSFTNGAV | 1430339 | Cell/Virus Enclosed | NO |
| ETPVVPDTVEVPPQVV | 1655588 | Cell/Virus Enclosed | NO |
| ETPVVSDTVEVP | 1430347 | Cell/Virus Enclosed | NO |
| ETRKLMPVCMDV | 1430360 | Cell/Virus Enclosed | NO |
| ETRTTNQFLAVSKGN | 14536 | Ectodomain | YES |
| ETSITIKKPNEL | 1430377 | Cell/Virus Enclosed | NO |
| ETSTEQVDVSHE | 1430391 | Cell/Virus Enclosed | NO |
| ETTDFCTVALASYADV | 1655612 | Ectodomain | YES |
| ETTDMVKSKGVCATGD | 1655613 | Cell/Virus Enclosed | NO |
| ETTDNNCWVNAT | 1430405 | Cell/Virus Enclosed | NO |
| ETTSAFVGALPKTVRE | 1655614 | Ectodomain | YES |
| ETVFVEPKDGGH | 1430424 | Cell/Virus Enclosed | NO |
| ETVFVEPKNGGQ | 1430426 | Cell/Virus Enclosed | NO |
| ETVKAKEESVKSSYAY | 1655618 | Cell/Virus Enclosed | NO |
| ETVNGYAKSNFVKDDY | 1655623 | Cell/Virus Enclosed | NO |
| ETVRLHLYNATR | 1430436 | Undetermined | UN |
| ETYIKSGGYIPE | 1430457 | Undetermined | UN |
| ETYIKWPWWVWLCISV | 1655637 | Undetermined | UN |
| ETYVALPIYYQHTDIN | 1655638 | Ectodomain | YES |
| EVAAVRSVTVEY | 1430463 | Cell/Virus Enclosed | NO |
| EVDGDVKLSSNLVIL | 14641 | Ectodomain | YES |
| EVDIIEQPFEEVEHVL | 1655656 | Cell/Virus Enclosed | NO |
| EVDIQQTFFDKTWPRP | 1655657 | Ectodomain | YES |
| EVDPDLSAFYVKDGKY | 1655659 | Cell/Virus Enclosed | NO |
| EVDWVAFYGFEK | 1430518 | Cell/Virus Enclosed | NO |
| EVEAGEVKPFAV | 1430519 | Cell/Virus Enclosed | NO |
| EVFDDEWAAAVD | 1430549 | Cell/Virus Enclosed | NO |
| EVFIAQSKGLYR | 1430554 | Cell/Virus Enclosed | NO |
| EVFKNCNVLDDFIVFN | 1655676 | Cell/Virus Enclosed | NO |
| EVFNATKFPSVYAWER | 1655678 | Ectodomain | YES |
| EVFNRLATSDWTDVRD | 1655681 | Cell/Virus Enclosed | NO |
| EVFTDGKLNCSFQDAA | 1655682 | Cell/Virus Enclosed | NO |
| EVFVAFVYYVAR | 1430560 | Cell/Virus Enclosed | NO |
| EVGDNVQITHTY | 1430566 | Cell/Virus Enclosed | NO |
| EVGEVKPFAVYK | 1430568 | Cell/Virus Enclosed | NO |

| | | | |
|---|---|---|---|
| EVGPEHSVADYH | 1430577 | Cell/Virus Enclosed | NO |
| EVHGNGSMIEDIDYVP | 1655698 | Cell/Virus Enclosed | NO |
| EVHGNGSMIEDM | 1430583 | Cell/Virus Enclosed | NO |
| EVHGNGSMIEDV | 1430584 | Cell/Virus Enclosed | NO |
| EVHIQQTFFDKT | 1430587 | Ectodomain | YES |
| EVHQELYSPLFL | 1430589 | Cell/Virus Enclosed | NO |
| EVILCWETGKVKPPLN | 1655705 | Cell/Virus Enclosed | NO |
| EVKAQSSGVIPENPVL | 1655711 | Cell/Virus Enclosed | NO |
| EVKDNDGKNVHLKDVT | 1655714 | Cell/Virus Enclosed | NO |
| EVKFSVIELATG | 1430620 | Cell/Virus Enclosed | NO |
| EVKPGTPAQEIEML | 14751 | Capsid | UN |
| EVKTASLFQTTV | 1430633 | Ectodomain | YES |
| EVKVSALKTAGV | 1430638 | Cell/Virus Enclosed | NO |
| EVLDNRATLQAI | 1430642 | Cell/Virus Enclosed | NO |
| EVLEDTLDAGPS | 1430643 | Cell/Virus Enclosed | NO |
| EVLEDTLNDGPS | 1430645 | Cell/Virus Enclosed | NO |
| EVLEVEDTQGMD | 1430646 | Cell/Virus Enclosed | NO |
| EVLLDVCNTKEF | 1430656 | Cell/Virus Enclosed | NO |
| EVLLDVCNTKEVKVFV | 1655738 | Cell/Virus Enclosed | NO |
| EVLLKDALQSRE | 1430658 | Cell/Virus Enclosed | NO |
| EVLVTVDGVNFR | 1430681 | Cell/Virus Enclosed | NO |
| EVMPVSMAKTSV | 1430683 | Ectodomain | YES |
| EVNAPYYNSWQN | 1430685 | Ectodomain | YES |
| EVNELLDTTQLQVANS | 1655757 | Ectodomain | YES |
| EVNGLLDTTQLHVADT | 1655758 | Ectodomain | YES |
| EVNSIFDIETVDVKHD | 1655762 | Cell/Virus Enclosed | NO |
| EVPGSCNFPLEA | 1430700 | Undetermined | UN |
| EVPPQLVELPSE | 1430704 | Cell/Virus Enclosed | NO |
| EVPPQVVELPYE | 1430708 | Cell/Virus Enclosed | NO |
| EVPPQVVKLPSAPQTI | 1655767 | Cell/Virus Enclosed | NO |
| EVPQLVNSNQYS | 1430717 | Ectodomain | YES |
| EVQPVEVPVEDI | 1430742 | Cell/Virus Enclosed | NO |
| EVRASRQLAQQKVNEC | 1655783 | Ectodomain | YES |
| EVRGSRRLAQQKINEC | 1655785 | Ectodomain | YES |
| EVRQKILNKPRQKRSP | 1655787 | Cell/Virus Enclosed | NO |
| EVRQVAPGETGV | 1430758 | Ectodomain | YES |
| EVSDDEWVAAVD | 1430768 | Cell/Virus Enclosed | NO |
| EVSDLIAQSSGV | 1430769 | Cell/Virus Enclosed | NO |
| EVSMCTNYDLSIINAR | 1655795 | Cell/Virus Enclosed | NO |
| EVSQFQSKLTDVKCAN | 1655798 | Cell/Virus Enclosed | NO |
| EVSQIQSKLTDVKCAN | 1655799 | Cell/Virus Enclosed | NO |
| EVSRELGNITKTNVQP | 1655800 | Cell/Virus Enclosed | NO |
| EVSRQVIQDVKDGTVT | 1655801 | Cell/Virus Enclosed | NO |
| EVSRQVIQEVKD | 1430780 | Cell/Virus Enclosed | NO |
| EVSTDEVGDNVQITYTYKM | 2060821 | Capsid | UN |
| EVSVKMVDAYVD | 1430791 | Cell/Virus Enclosed | NO |
| EVSVQKKINRMA | 1430792 | Cell/Virus Enclosed | NO |
| EVTSDSCNNFML | 1430814 | Cell/Virus Enclosed | NO |
| EVTTALEETKFL | 1430815 | Cell/Virus Enclosed | NO |
| EVTVDTTVESAA | 1430821 | Cell/Virus Enclosed | NO |
| EVVAVRSVTVEY | 1430823 | Cell/Virus Enclosed | NO |
| EVVGNVILKPSD | 1430837 | Cell/Virus Enclosed | NO |
| EVVGNVILKPSE | 1430838 | Cell/Virus Enclosed | NO |
| EVVGPKELLLSWESGK | 1655827 | Cell/Virus Enclosed | NO |
| EVVHDQVNYVPLRATN | 1655828 | Cell/Virus Enclosed | NO |
| EVVKAMLFKKEP | 1430844 | Cell/Virus Enclosed | NO |
| EVVRFPNITNLC | 1430858 | Ectodomain | YES |
| EVVSAYGLCDSA | 1430864 | Ectodomain | YES |
| EVVSAYGLCDVA | 1430865 | Ectodomain | YES |
| EVVVNNLDKSAGHPFN | 1655839 | Cell/Virus Enclosed | NO |
| EVVVNPANGHMAHGGG | 1655840 | Cell/Virus Enclosed | NO |
| EVVVTNLNKSAGWPLN | 1655841 | Cell/Virus Enclosed | NO |
| EVVYSNGGFYFKPGGT | 1655843 | Cell/Virus Enclosed | NO |
| EVYLPYPNPSRI | 1430876 | Cell/Virus Enclosed | NO |
| EVYVHLRNWNFSWNLI | 1655847 | Undetermined | UN |
| EVYYQLESTQIT | 1430883 | Secreted | YES |
| EWAAAVDEAFPLDEAE | 1655849 | Cell/Virus Enclosed | NO |
| EWAMANGYTSVSSVEC | 1655851 | Cell/Virus Enclosed | NO |
| EWAYCVDLHNKINLCD | 1655853 | Cell/Virus Enclosed | NO |
| EWFGITQTAQGA | 1430903 | Ectodomain | YES |
| EWFGITQTAQGV | 1430904 | Ectodomain | YES |
| EWIELFPHNDRIKSFS | 1655864 | Cell/Virus Enclosed | NO |
| EWKMVVCDKVRS | 1430925 | Cell/Virus Enclosed | NO |
| EWMDDFEADPKGKYAQ | 1655868 | Cell/Virus Enclosed | NO |
| EWPLFDQQAQQE | 1430945 | Cell/Virus Enclosed | NO |
| EWPLSVQQAQQEATLP | 1655872 | Cell/Virus Enclosed | NO |

| | | | |
|---|---|---|---|
| EWPLTEYDFTDYKVQL | 1655873 | Cell/Virus Enclosed | NO |
| EWSYTGSSFYAPEPIT | 1655877 | Ectodomain | YES |
| EWVAAVDEAFPL | 1430958 | Cell/Virus Enclosed | NO |
| EYAYDQVHDEPK | 1430972 | Cell/Virus Enclosed | NO |
| EYDFLNMDMGVFINKY | 1655895 | Cell/Virus Enclosed | NO |
| EYDFTDYKVQLFEKYF | 1655896 | Cell/Virus Enclosed | NO |
| EYDLEPHKVHVH | 1430981 | Underneath Membrane | NO |
| EYDLQPHKVHVH | 1430982 | Underneath Membrane | NO |
| EYDNISVAVPKS | 1430983 | Cell/Virus Enclosed | NO |
| EYDYVIYSQTAETAHS | 1655901 | Cell/Virus Enclosed | NO |
| EYEEEEEVSEES | 1430988 | Cell/Virus Enclosed | NO |
| EYENAVANGSSPQIIK | 1655903 | Cell/Virus Enclosed | NO |
| EYEVAKKNLADAKNSG | 1655904 | Cell/Virus Enclosed | NO |
| EYGGHHDFVIKTSHDD | 1655908 | Underneath Membrane | NO |
| EYGGHNDFVIKA | 1431001 | Underneath Membrane | NO |
| EYGQFCSKINQA | 1431004 | Ectodomain | YES |
| EYGSDTVTYKSTVTTK | 1655910 | Cell/Virus Enclosed | NO |
| EYGTEDDYQGLP | 1431012 | Cell/Virus Enclosed | NO |
| EYGTFCDNINSILDEV | 1655916 | Ectodomain | YES |
| EYIDVNKTLQELSYKL | 1655921 | Ectodomain | YES |
| EYIFERIDYSDAVSYK | 1655922 | Cell/Virus Enclosed | NO |
| EYIKKLYNDLGTQILD | 1655923 | Cell/Virus Enclosed | NO |
| EYIQTTIQKVTVDCKQ | 1655925 | Ectodomain | YES |
| EYIRYGIKPITI | 1431029 | Cell/Virus Enclosed | NO |
| EYISDASSLDVS | 1431031 | Undetermined | UN |
| EYKHWPQIAQFAPSAS | 1655935 | Cell/Virus Enclosed | NO |
| EYKLQQGTFLCA | 1431041 | Cell/Virus Enclosed | NO |
| EYLESYNTATTAGFTF | 1655941 | Cell/Virus Enclosed | NO |
| EYLQITSIPIVV | 1431051 | Ectodomain | YES |
| EYNFIIYDSSDRGQES | 1655948 | Cell/Virus Enclosed | NO |
| EYNHVVAANALL | 1431065 | Cell/Virus Enclosed | NO |
| EYPIISNELSINTSCR | 1655955 | Cell/Virus Enclosed | NO |
| EYPIISNEVSINTSCR | 1655956 | Cell/Virus Enclosed | NO |
| EYPNTICHPNLG | 1431089 | Ectodomain | YES |
| EYPYISHENKLN | 1431093 | Cell/Virus Enclosed | NO |
| EYPYISHETKLN | 1431094 | Cell/Virus Enclosed | NO |
| EYREYLESYNTATTAG | 1655964 | Cell/Virus Enclosed | NO |
| EYREYMEAYNFV | 1431101 | Cell/Virus Enclosed | NO |
| EYSWNKKLYELVQRFS | 1655975 | Cell/Virus Enclosed | NO |
| EYTYTDEVIYGM | 1431134 | Cell/Virus Enclosed | NO |
| EYTYTDEVIYSM | 1431135 | Cell/Virus Enclosed | NO |
| EYVFKNKDGFLS | 1431139 | Ectodomain | YES |
| EYVSYFDGVKTV | 1431144 | Cell/Virus Enclosed | NO |
| EYVTEFVQEEAQ | 1431145 | Cell/Virus Enclosed | NO |
| EYYEFLNKHFNM | 1431161 | Cell/Virus Enclosed | NO |
| FAAETIKAKEESVKSS | 1655999 | Cell/Virus Enclosed | NO |
| FAAETVKAKEESVKSS | 1656003 | Cell/Virus Enclosed | NO |
| FAAIGDLKCPLD | 1431176 | Ectodomain | YES |
| FAAIGDLNCTLD | 1431177 | Undetermined | UN |
| FAAKRVLGLQTQTVDS | 1656008 | Cell/Virus Enclosed | NO |
| FAAYSRYRIGNYKLNT | 1380209 | Underneath Membrane | NO |
| FACADGTRHTYQ | 1431192 | Cell/Virus Enclosed | NO |
| FACGCGASYTANDVEV | 1656014 | Cell/Virus Enclosed | NO |
| FACGCGTSYTAN | 1431198 | Cell/Virus Enclosed | NO |
| FACKFAVCGDGF | 1431199 | Cell/Virus Enclosed | NO |
| FACTTCGHVYEV | 1431204 | Cell/Virus Enclosed | NO |
| FADDLNQMTGFK | 1431213 | Cell/Virus Enclosed | NO |
| FADNNDITNTNLSRGR | 1656030 | Cell/Virus Enclosed | NO |
| FADNNDTTNTNL | 1431226 | Cell/Virus Enclosed | NO |
| FADNNDTTTNQP | 1431227 | Cell/Virus Enclosed | NO |
| FADVVKGQVSDL | 1431232 | Cell/Virus Enclosed | NO |
| FAEDVCTCYDNSIQGS | 1656035 | Cell/Virus Enclosed | NO |
| FAELEAAQKAYQEAMD | 1656037 | Cell/Virus Enclosed | NO |
| FAELEAARKAYQ | 1431242 | Cell/Virus Enclosed | NO |
| FAELIPNQAALF | 1431244 | Cell/Virus Enclosed | NO |
| FAFNSSVLSLVN | 1431264 | Cell/Virus Enclosed | NO |
| FAGTHITMTTVY | 1431278 | Cell/Virus Enclosed | NO |
| FAGYTESAVNFT | 1431281 | Cell/Virus Enclosed | NO |
| FAGYTESAVNSTKALA | 1656054 | Cell/Virus Enclosed | NO |
| FAINDLNTTIPR | 1431287 | Underneath Membrane | NO |
| FAINDLNTTVPRISEY | 1656060 | Ectodomain | YES |
| FAIPNGRKVDLH | 1431290 | Ectodomain | YES |
| FAIPNGRKVDLQLGNL | 1656061 | Ectodomain | YES |
| FAIPNRRKFDLQ | 1431293 | Undetermined | UN |
| FAIPNRRKVDLQ | 1431294 | Ectodomain | YES |
| FAIPNRRRDDLQ | 1431295 | Ectodomain | YES |

| | | | |
|---|---|---|---|
| FAIPNRRRVDLQ | 1431296 | Ectodomain | YES |
| FAIPSVAVRTYSEAAA | 1656063 | Cell/Virus Enclosed | NO |
| FAKGDSCSSNCKHSVA | 1656067 | Cell/Virus Enclosed | NO |
| FAKRKMGLTPPLSILK | 1656070 | Cell/Virus Enclosed | NO |
| FAKRKVGLTPPLSILK | 1656071 | Cell/Virus Enclosed | NO |
| FAKVKNTKVFGK | 1431316 | Ectodomain | YES |
| FAKVKNTKVIKD | 1431319 | Ectodomain | YES |
| FAKVKNTKVVGK | 1431322 | Ectodomain | YES |
| FALRNFDEKDCEVFKE | 1656085 | Cell/Virus Enclosed | NO |
| FAMRHVRGWLGMDVEG | 1656095 | Cell/Virus Enclosed | NO |
| FANDTKIASQLGNCVE | 1656099 | Ectodomain | YES |
| FANFILHDVNNG | 1431371 | Ectodomain | YES |
| FANFILHDVNSGLTCS | 1656100 | Ectodomain | YES |
| FANGFVVRIGAAANST | 1656101 | Ectodomain | YES |
| FANGFVVRIGAV | 1431375 | Undetermined | UN |
| FANHVANFQKSG | 1431376 | Cell/Virus Enclosed | NO |
| FANKFEQFDQWYESTF | 1656102 | Cell/Virus Enclosed | NO |
| FANKFFQFDQWYESTF | 1656103 | Cell/Virus Enclosed | NO |
| FANKHAGFDAWF | 1431381 | Cell/Virus Enclosed | NO |
| FANPAALHSKCL | 1431388 | Cell/Virus Enclosed | NO |
| FANPAAVDSKCL | 1431389 | Cell/Virus Enclosed | NO |
| FANSVFNICQAVSANV | 1656111 | Cell/Virus Enclosed | NO |
| FANSVFNICQAVTANV | 1656112 | Cell/Virus Enclosed | NO |
| FANTGSVCYTPIDEIP | 1656113 | Cell/Virus Enclosed | NO |
| FANVSYVYNGSA | 1431397 | Ectodomain | YES |
| FANWLAPTNAYM | 1431398 | Cell/Virus Enclosed | NO |
| FAQSEEEVPFSN | 1431418 | Cell/Virus Enclosed | NO |
| FAQSEKEESFSN | 1431421 | Cell/Virus Enclosed | NO |
| FAQTSDTAHACNANRF | 1656124 | Cell/Virus Enclosed | NO |
| FARTRSMWSFNPETN | 1540201 | Underneath Membrane | NO |
| FASDSVQCYTPH | 1431443 | Cell/Virus Enclosed | NO |
| FASHFISNSWLM | 1431449 | Cell/Virus Enclosed | NO |
| FASLDVKPYGYPNDFV | 1656139 | Cell/Virus Enclosed | NO |
| FASPQECNNMHL | 1431458 | Cell/Virus Enclosed | NO |
| FASSCKSHKPSS | 1431464 | Undetermined | UN |
| FASVKSSQSSPIIPGF | 1656145 | Ectodomain | YES |
| FASVKSYQSSPI | 1431473 | Ectodomain | YES |
| FATLMTFSGNVDLDAL | 1656152 | Cell/Virus Enclosed | NO |
| FATTIPNTAFGP | 1431499 | Cell/Virus Enclosed | NO |
| FATYHTPATDCSDGNY | 1656161 | Ectodomain | YES |
| FATYHTPSTDCS | 1431506 | Ectodomain | YES |
| FAVDPAAAYLDAVKQG | 1656165 | Cell/Virus Enclosed | NO |
| FAVESGGYIPSDFAFN | 1656167 | Ectodomain | YES |
| FAVIGDLKCPLD | 1431517 | Ectodomain | YES |
| FAVIGDLKCPLE | 1431518 | Ectodomain | YES |
| FAVIGDLKCTSDNIND | 1656170 | Ectodomain | YES |
| FAVIGDLKCTTV | 1431520 | Ectodomain | YES |
| FAVIGDLNCPLD | 1431521 | Ectodomain | YES |
| FAVIGDLNCTLD | 1431522 | Ectodomain | YES |
| FAVLYQDVNCTD | 1431526 | Ectodomain | YES |
| FAVNDQDLYDHV | 1431527 | Cell/Virus Enclosed | NO |
| FAVRKEGQDVIFSQFD | 1380243 | Cell/Virus Enclosed | NO |
| FAVVGDLNCPLD | 1431535 | Ectodomain | YES |
| FAVYHRYRIGNY | 1431536 | Underneath Membrane | NO |
| FAVYKNVKFYLG | 1431537 | Cell/Virus Enclosed | NO |
| FAWWTVFCTIAN | 1431550 | Cell/Virus Enclosed | NO |
| FAYADKPNHFIN | 1431551 | Cell/Virus Enclosed | NO |
| FAYANRNRFLYIIKL | 1312401 | Undetermined | UN |
| FAYESAVVNGIRVLKT | 1656180 | Cell/Virus Enclosed | NO |
| FAYNNDITNTNL | 1431557 | Capsid | UN |
| FAYVKPFLGQAA | 1431570 | Cell/Virus Enclosed | NO |
| FCAFAVDPAKAY | 1431577 | Cell/Virus Enclosed | NO |
| FCAFCTPKKVFI | 1431580 | Cell/Virus Enclosed | NO |
| FCALCVISVAML | 1431585 | Cell/Virus Enclosed | NO |
| FCCFLVTKFRRM | 1431599 | Cell/Virus Enclosed | NO |
| FCDAMRDAGIVG | 1431604 | Cell/Virus Enclosed | NO |
| FCDGVNVTKHKCDINY | 1656208 | Cell/Virus Enclosed | NO |
| FCDHMVKDGLVG | 1431611 | Cell/Virus Enclosed | NO |
| FCDNINSILDEVNGLL | 1656212 | Ectodomain | YES |
| FCDYADKLCHAV | 1431619 | Cell/Virus Enclosed | NO |
| FCDYAGKFCHAV | 1431620 | Cell/Virus Enclosed | NO |
| FCDYTDKLCHAV | 1431621 | Cell/Virus Enclosed | NO |
| FCGDQVHEVAAV | 1431630 | Cell/Virus Enclosed | NO |
| FCGRGYHLMSFP | 1431642 | Ectodomain | YES |
| FCHRHNWNCVDC | 1431651 | Cell/Virus Enclosed | NO |
| FCIDYALPSSRR | 1431654 | Ectodomain | YES |

| | | | |
|---|---|---|---|
| FCKIGKEQKCCV | 1431660 | Cell/Virus Enclosed | NO |
| FCKIGREQKCCM | 1431661 | Cell/Virus Enclosed | NO |
| FCKSGSKFKNLV | 1431666 | Ectodomain | YES |
| FCPCADRSVVNS | 1431701 | Ectodomain | YES |
| FCPCKSNGSCPG | 1431712 | Ectodomain | YES |
| FCPCSSCFGKNN | 1431714 | Ectodomain | YES |
| FCPCSSCHGKNN | 1431715 | Undetermined | UN |
| FCPCSSCPGKNN | 1431716 | Ectodomain | YES |
| FCPCSSCPSKNN | 1431717 | Ectodomain | YES |
| FCPCSSCSGKNN | 1431719 | Ectodomain | YES |
| FCQTADTAHANNINRF | 1656273 | Cell/Virus Enclosed | NO |
| FCQYKGKWVQVPIGTN | 1656274 | Cell/Virus Enclosed | NO |
| FCRLYIEESDLI | 1431728 | Cell/Virus Enclosed | NO |
| FCRQYSNESWFEDDYR | 1656276 | Cell/Virus Enclosed | NO |
| FCSETVNGYAKSNFVK | 1656278 | Cell/Virus Enclosed | NO |
| FCSQHTLYIKDGDDGY | 1656282 | Cell/Virus Enclosed | NO |
| FCSQHTMQIVDENGKY | 1656286 | Cell/Virus Enclosed | NO |
| FCSQHTMQIVDKDGTY | 1656287 | Cell/Virus Enclosed | NO |
| FCTKHTMLVKMD | 1431749 | Cell/Virus Enclosed | NO |
| FCTPTKKAFPYGTCLN | 1656298 | Cell/Virus Enclosed | NO |
| FCTQLNRALSGIAAEQ | 1656299 | Ectodomain | YES |
| FCVNCDSFGPGN | 1431769 | Cell/Virus Enclosed | NO |
| FDAEGAHATRDSIGTN | 1656324 | Cell/Virus Enclosed | NO |
| FDAQPLNKKVQQ | 1431795 | Cell/Virus Enclosed | NO |
| FDAQPLTRKVQQ | 1431796 | Cell/Virus Enclosed | NO |
| FDATNSALVKIQAVVN | 1656331 | Ectodomain | YES |
| FDCGVNALDGDT | 1431813 | Cell/Virus Enclosed | NO |
| FDCYNLWQTFTE | 1431824 | Cell/Virus Enclosed | NO |
| FDDNFYAFCTPK | 1431832 | Cell/Virus Enclosed | NO |
| FDDNTDSIVIDG | 1431833 | Cell/Virus Enclosed | NO |
| FDDPVIPFKDGI | 1431834 | Ectodomain | YES |
| FDDQPSLQVESANLML | 1656349 | Cell/Virus Enclosed | NO |
| FDDSDCDVVQEQVNYV | 1656351 | Cell/Virus Enclosed | NO |
| FDDVQAPNYTQHTSSM | 1656352 | Ectodomain | YES |
| FDEAGEEVLAPKLYCA | 1656355 | Cell/Virus Enclosed | NO |
| FDEHGVITKAVD | 1431848 | Ectodomain | YES |
| FDENGDKLDQFF | 1431851 | Cell/Virus Enclosed | NO |
| FDEQNCDVLKEVLVLT | 1656364 | Cell/Virus Enclosed | NO |
| FDFIIGHGMSYS | 1431860 | Cell/Virus Enclosed | NO |
| FDFLSNVSTSHH | 1431864 | Undetermined | UN |
| FDFSLDVFKIGDVKFK | 1656374 | Cell/Virus Enclosed | NO |
| FDFSSDVFKIGD | 1431867 | Cell/Virus Enclosed | NO |
| FDFYTTEPEFVK | 1431869 | Cell/Virus Enclosed | NO |
| FDGFNFSQILPD | 1431872 | Ectodomain | YES |
| FDGHAGEAPVSI | 1431873 | Cell/Virus Enclosed | NO |
| FDGKPMLPKNMW | 1431874 | Cell/Virus Enclosed | NO |
| FDGKPVLPKNMW | 1431875 | Cell/Virus Enclosed | NO |
| FDGKSKCDESAA | 1431876 | Cell/Virus Enclosed | NO |
| FDGKSKCEESAA | 1431878 | Cell/Virus Enclosed | NO |
| FDGQIGEAPVSI | 1431883 | Cell/Virus Enclosed | NO |
| FDGRDNGALEAFKRSN | 1656387 | Cell/Virus Enclosed | NO |
| FDGRVEGQVDLF | 1431889 | Cell/Virus Enclosed | NO |
| FDGTMYGAFMDK | 1431891 | Cell/Virus Enclosed | NO |
| FDGVMYGGFEDQPNLQ | 1656394 | Cell/Virus Enclosed | NO |
| FDGVTTSDKCIFNSAC | 1656395 | Cell/Virus Enclosed | NO |
| FDHSQYYFNKDT | 1431901 | Ectodomain | YES |
| FDIETVDVKHDVSPFE | 1656402 | Cell/Virus Enclosed | NO |
| FDIIQKCQITSVVGTK | 1656404 | Cell/Virus Enclosed | NO |
| FDIKNSSVNTVCFVDV | 1656405 | Cell/Virus Enclosed | NO |
| FDIRDNGSLEKF | 1431913 | Cell/Virus Enclosed | NO |
| FDIRDNVSLEKF | 1431915 | Cell/Virus Enclosed | NO |
| FDITDAAVKAAE | 1431920 | Cell/Virus Enclosed | NO |
| FDIVVAWHVVRDSRFV | 1656413 | Cell/Virus Enclosed | NO |
| FDKPQEKDKKSAKTGT | 1656428 | Capsid | UN |
| FDKSELTNGVYY | 1431945 | Cell/Virus Enclosed | NO |
| FDKSEPLCLFKK | 1431946 | Ectodomain | YES |
| FDKTWPRPIDVF | 1431950 | Ectodomain | YES |
| FDKTWPRPIDVS | 1431951 | Ectodomain | YES |
| FDKVKLSDVGFVEAYN | 1656437 | Ectodomain | YES |
| FDKVVDNVYVTYAGSV | 1656439 | Cell/Virus Enclosed | NO |
| FDKVYDIGNPKGIPIV | 1656440 | Cell/Virus Enclosed | NO |
| FDKWFVNDGRVG | 1431955 | Cell/Virus Enclosed | NO |
| FDLISKCQITAVEGTK | 1656448 | Cell/Virus Enclosed | NO |
| FDLNGLDAVFFY | 1431977 | Cell/Virus Enclosed | NO |
| FDLQLGNLGYLQ | 1431978 | Undetermined | UN |
| FDLSKELKRPIQ | 1431980 | Cell/Virus Enclosed | NO |

| | | | |
|---|---|---|---|
| FDLSKFQLKLKGTPVL | 1656456 | Cell/Virus Enclosed | NO |
| FDLTPFNLTYFN | 1431981 | Ectodomain | YES |
| FDLTPFNLTYLN | 1431982 | Ectodomain | YES |
| FDLVQYDFTDYKLEL | 1431984 | Cell/Virus Enclosed | NO |
| FDLYVNVYRSTSPDPK | 1656459 | Cell/Virus Enclosed | NO |
| FDMAKFPLKLAGTAVI | 1656460 | Cell/Virus Enclosed | NO |
| FDMSQFPLKLRG | 1431995 | Cell/Virus Enclosed | NO |
| FDNLSSEDLTAV | 1432012 | Cell/Virus Enclosed | NO |
| FDNMTAVACQPP | 1432015 | Ectodomain | YES |
| FDNNTDSIVIDG | 1432017 | Cell/Virus Enclosed | NO |
| FDNPILDFGDGV | 1432018 | Ectodomain | YES |
| FDNPPTNVVSHV | 1432020 | Ectodomain | YES |
| FDNPVIPFKDGI | 1432022 | Undetermined | UN |
| FDNQSKLPSSSG | 1432024 | Cell/Virus Enclosed | NO |
| FDNQSKPPSSSG | 1432025 | Cell/Virus Enclosed | NO |
| FDNQSKSLSSSG | 1432026 | Cell/Virus Enclosed | NO |
| FDNQSKSPSSSGTSTP | 1656479 | Capsid | UN |
| FDNSIQGSYERF | 1432028 | Cell/Virus Enclosed | NO |
| FDNTTQSAIIVN | 1432030 | Ectodomain | YES |
| FDPIENEDIHRVYAAL | 1656485 | Cell/Virus Enclosed | NO |
| FDPITLKAPDTY | 1432033 | Ectodomain | YES |
| FDQNNCEVLKAI | 1432036 | Cell/Virus Enclosed | NO |
| FDQQAQQEATHI | 1432038 | Cell/Virus Enclosed | NO |
| FDQQQLLAYYNF | 1432040 | Cell/Virus Enclosed | NO |
| FDQRCSCSRCFY | 1432043 | Cell/Virus Enclosed | NO |
| FDRNDCMLLCDI | 1432051 | Cell/Virus Enclosed | NO |
| FDSCISNNRCNI | 1432056 | Ectodomain | YES |
| FDSEVSTDEEGD | 1432060 | Cell/Virus Enclosed | NO |
| FDSGTVSKTSDWKCKV | 1656503 | Cell/Virus Enclosed | NO |
| FDSGTVTKISDW | 1432070 | Cell/Virus Enclosed | NO |
| FDSHIVSKESGNTVVL | 1656504 | Capsid | UN |
| FDSKGWYDPVENEDIH | 1656505 | Cell/Virus Enclosed | NO |
| FDSLNLSAGTQHISND | 1656508 | Capsid | UN |
| FDSNAFSLANVI | 1432078 | Ectodomain | YES |
| FDSNAFSLANVT | 1432079 | Ectodomain | YES |
| FDTFKGECPKFV | 1432093 | Cell/Virus Enclosed | NO |
| FDTRVHSEFNLP | 1432099 | Cell/Virus Enclosed | NO |
| FDTRVLNKLNLPGCNG | 1656523 | Cell/Virus Enclosed | NO |
| FDTRVLNNLNLPGCNG | 1656524 | Cell/Virus Enclosed | NO |
| FDTVEVPPQVVK | 1432103 | Cell/Virus Enclosed | NO |
| FDVAPIELENKFTPLS | 1656531 | Cell/Virus Enclosed | NO |
| FDVDKFKTPHIV | 1432113 | Cell/Virus Enclosed | NO |
| FDVDKKSLNALIATAH | 1656532 | Cell/Virus Enclosed | NO |
| FDVDKVKTPHIV | 1432117 | Cell/Virus Enclosed | NO |
| FDVEGAHATRENIGTN | 1656537 | Cell/Virus Enclosed | NO |
| FDVESGVYSVSSFEAK | 1656542 | Ectodomain | YES |
| FDVINGVKTVKYFECP | 1656545 | Cell/Virus Enclosed | NO |
| FDVIVAWHVVRDPRYV | 1656546 | Cell/Virus Enclosed | NO |
| FDVKGVEELHDD | 1432130 | Cell/Virus Enclosed | NO |
| FDVPDGFYSTSPIQSV | 1656553 | Ectodomain | YES |
| FDVVFGHGAGSV | 1432142 | Cell/Virus Enclosed | NO |
| FDVVRQCSGVTF | 1432148 | Cell/Virus Enclosed | NO |
| FDVYNKDASFIGKNLK | 1656564 | Cell/Virus Enclosed | NO |
| FDVYVDTFLSMFDVDK | 1656566 | Cell/Virus Enclosed | NO |
| FDVYVDTFMSHFDVDR | 1656567 | Cell/Virus Enclosed | NO |
| FDWIKISADNKV | 1432152 | Cell/Virus Enclosed | NO |
| FDYCIPYNSITD | 1432153 | Secreted | YES |
| FDYHPNCSDCYDDMCV | 1656568 | Cell/Virus Enclosed | NO |
| FDYLDIHPSLCNNGKI | 1656569 | Ectodomain | YES |
| FDYLIREAETRV | 1432157 | Cell/Virus Enclosed | NO |
| FDYYRYNRPTIL | 1432165 | Cell/Virus Enclosed | NO |
| FEAHKDFFKDGL | 1432170 | Cell/Virus Enclosed | NO |
| FEAIPSGSVVEQ | 1432173 | Ectodomain | YES |
| FEDAQGLKEFYK | 1432196 | Cell/Virus Enclosed | NO |
| FEDDQGLKKFYK | 1432199 | Cell/Virus Enclosed | NO |
| FEDDVITSLCKK | 1432200 | Cell/Virus Enclosed | NO |
| FEDDVVTSLCKKSFGK | 1656589 | Cell/Virus Enclosed | NO |
| FEDEKLVDVCEK | 1432204 | Cell/Virus Enclosed | NO |
| FEDELAPSDVYQQLAG | 1656592 | Cell/Virus Enclosed | NO |
| FEDELTPSDVYQQLAG | 1656593 | Cell/Virus Enclosed | NO |
| FEDLLFDKVKLSDVGF | 1656594 | Ectodomain | YES |
| FEDRSNNSSRASSRSS | 1656596 | Capsid | UN |
| FEEQDEIYAYTKRNVL | 1656602 | Cell/Virus Enclosed | NO |
| FEFEDDVITSLC | 1432228 | Cell/Virus Enclosed | NO |
| FEFVEGQGPPIA | 1432234 | Cell/Virus Enclosed | NO |
| FEFVEGQGVPIAPGVP | 1656608 | Cell/Virus Enclosed | NO |

| | | | |
|---|---|---|---|
| FEFYHKCDNTCMESVKNGTYDYPKY | 1140522 | Ectodomain | YES |
| FEFYTTEPEFVK | 1432236 | Cell/Virus Enclosed | NO |
| FEGAQGLKKFYK | 1432237 | Cell/Virus Enclosed | NO |
| FEGVLPINTDTVLSVA | 1656613 | Cell/Virus Enclosed | NO |
| FEGYDFKYIENGQLKN | 1656614 | Cell/Virus Enclosed | NO |
| FEIACKCGVKQEQRTG | 1656623 | Cell/Virus Enclosed | NO |
| FEIAQLYGSCITPNVC | 1656624 | Cell/Virus Enclosed | NO |
| FEIDGNVMHANYLFWR | 1656625 | Cell/Virus Enclosed | NO |
| FEIDKGIYQTSNFRVV | 1656626 | Ectodomain | YES |
| FEIFPKTSSWPNHDSNKGVTAACPH | 1140533 | Ectodomain | YES |
| FEIKHADLHSSQVCGL | 1656627 | Cell/Virus Enclosed | NO |
| FEISCDCGKKFD | 1432268 | Cell/Virus Enclosed | NO |
| FEIYDGGCIPASQVIV | 1656632 | Cell/Virus Enclosed | NO |
| FEIYDGGCLNASEVVV | 1656633 | Cell/Virus Enclosed | NO |
| FEKLHCEHLQFG | 1432274 | Undetermined | UN |
| FEKPQEKDKKSA | 1432278 | Cell/Virus Enclosed | NO |
| FEKVDYGSDTVTYKST | 1656641 | Cell/Virus Enclosed | NO |
| FEKVEYGSDTVTYKST | 1656642 | Cell/Virus Enclosed | NO |
| FEKYFKYWDQTYHANC | 1656644 | Cell/Virus Enclosed | NO |
| FELAKVQNLSGN | 1432283 | Cell/Virus Enclosed | NO |
| FELFAKRKMGLTPPLS | 1656649 | Cell/Virus Enclosed | NO |
| FELFAKRKVGLTPPLS | 1380313 | Cell/Virus Enclosed | NO |
| FELLNAPATVCGPKLS | 1656654 | Ectodomain | YES |
| FELLNAPATVSG | 1432300 | Undetermined | UN |
| FELTCLRYFAKL | 1432310 | Cell/Virus Enclosed | NO |
| FELYAKRAVRSHPDFK | 1656662 | Cell/Virus Enclosed | NO |
| FEMKNWFDPIEN | 1432321 | Cell/Virus Enclosed | NO |
| FEMLKLGTNDPQ | 1432322 | Cell/Virus Enclosed | NO |
| FEMNRLQEAIKV | 1432323 | Ectodomain | YES |
| FEMPFEELNGLKILKQ | 1656664 | Cell/Virus Enclosed | NO |
| FENKTTLPTNIAFELY | 1656672 | Cell/Virus Enclosed | NO |
| FENYIKWPWWVWLIIS | 1656676 | Undetermined | UN |
| FEPFNVSFVNDSIESV | 1656677 | Ectodomain | YES |
| FEPFTVNSINDS | 1432341 | Ectodomain | YES |
| FEPFTVNSKNDN | 1432342 | Ectodomain | YES |
| FEPFTVNSVNDSLEPV | 1656678 | Ectodomain | YES |
| FEPFTVNSVSDS | 1432344 | Undetermined | UN |
| FEPRIPTIADFV | 1432350 | Ectodomain | YES |
| FEQQVGVIADKDKDLS | 1656684 | Cell/Virus Enclosed | NO |
| FERDISNVPFSS | 1432365 | Ectodomain | YES |
| FERDLTSDENGV | 1432368 | Ectodomain | YES |
| FERGSVQVDNGSSINR | 1656689 | Cell/Virus Enclosed | NO |
| FERIDYSDAVSYKSST | 1656690 | Cell/Virus Enclosed | NO |
| FERNLRRGTILF | 1432372 | Ectodomain | YES |
| FESKTTLPVNVA | 1432381 | Cell/Virus Enclosed | NO |
| FESTINKSVANL | 1432384 | Cell/Virus Enclosed | NO |
| FESVKSSQSSPI | 1432388 | Ectodomain | YES |
| FETIMKVLEENLNAYV | 1656700 | Capsid | UN |
| FETIMKVLNENLNAYQ | 1656701 | Cell/Virus Enclosed | NO |
| FEVLEVEDTQGM | 1432415 | Cell/Virus Enclosed | NO |
| FEYISDASSLDV | 1432427 | Undetermined | UN |
| FEYTSPTDKLAMIMGY | 1656710 | Cell/Virus Enclosed | NO |
| FFADNNDITNTN | 1432437 | Cell/Virus Enclosed | NO |
| FFAFCYYVWKSY | 1432438 | Cell/Virus Enclosed | NO |
| FFAVSKPMGTQTHTMIF | 15783 | Ectodomain | YES |
| FFDKTWPRPIDV | 1432459 | Ectodomain | YES |
| FFEDLLFDKVKLSDVG | 1656734 | Ectodomain | YES |
| FFEIKHADLHSSQVCG | 1656736 | Cell/Virus Enclosed | NO |
| FFEIKHADYTLA | 1432468 | Cell/Virus Enclosed | NO |
| FFEITMTDLQSESSCG | 1656737 | Cell/Virus Enclosed | NO |
| FFFLTHINTTLL | 1432479 | Ectodomain | YES |
| FFFTQDGNAAIP | 1432484 | Cell/Virus Enclosed | NO |
| FFFTQKGDAAIKDFDY | 1656745 | Cell/Virus Enclosed | NO |
| FFGSKLELVKRE | 1432499 | Cell/Virus Enclosed | NO |
| FFICCCTGCGPS | 1432508 | Undetermined | UN |
| FFIFNKLVKDPPNVQI | 1656761 | Secreted | YES |
| FFILCCTGCGTN | 1432519 | Undetermined | UN |
| FFKCTMGVYDFKVSAA | 1656769 | Cell/Virus Enclosed | NO |
| FFKELTANMSMAECKA | 1656772 | Cell/Virus Enclosed | NO |
| FFKGNVQVDNGSSINR | 1656775 | Cell/Virus Enclosed | NO |
| FFKNVSTSISNF | 1432542 | Ectodomain | YES |
| FFKTFTTSTALA | 1432545 | Cell/Virus Enclosed | NO |
| FFLSKHSDFGLGDLVD | 1656782 | Cell/Virus Enclosed | NO |
| FFPQFTSDQAIT | 1432585 | Underneath Membrane | NO |
| FFPYDIEFVTGK | 1432587 | Cell/Virus Enclosed | NO |
| FFQFGDFVMNNI | 1432592 | Cell/Virus Enclosed | NO |

| | | | |
|---|---|---|---|
| FFSNTKYYDHSQ | 1432607 | Ectodomain | YES |
| FFSPESVFSVFGDVKL | 1656810 | Cell/Virus Enclosed | NO |
| FFSPQIITTDNTFVSG | 1656811 | Ectodomain | YES |
| FFSTFKCYGVSATKLND | 15903 | Ectodomain | YES |
| FFTCNSNANISM | 1432619 | Ectodomain | YES |
| FFTCNSNANLSMLQLG | 1656816 | Ectodomain | YES |
| FFTPYFMYDIVS | 1432635 | Cell/Virus Enclosed | NO |
| FFTQDGNAAITDYNYY | 1656823 | Cell/Virus Enclosed | NO |
| FFTQKGDAAIKDFDYY | 1656824 | Cell/Virus Enclosed | NO |
| FFVNDFTCSQIS | 1432661 | Ectodomain | YES |
| FFVVSKPMGTQT | 1432669 | Ectodomain | YES |
| FFVVSKPMGTRT | 1432670 | Ectodomain | YES |
| FFYGDVVSHICK | 1432688 | Cell/Virus Enclosed | NO |
| FFYIDVGNHRGA | 1432694 | Undetermined | UN |
| FFYTSKEPVASI | 1432702 | Cell/Virus Enclosed | NO |
| FFYYSDSPCESH | 1432706 | Cell/Virus Enclosed | NO |
| FFYYSTTPCGVH | 1432710 | Cell/Virus Enclosed | NO |
| FGAALAMEKVNECVKS | 1380354 | Ectodomain | YES |
| FGACDYDRLPNEVTVA | 1656866 | Ectodomain | YES |
| FGACDYDRLSNE | 1432713 | Underneath Membrane | NO |
| FGAGSDKGIAPGTSVL | 1656870 | Cell/Virus Enclosed | NO |
| FGAISASIGDIIQRLD | 1656873 | Ectodomain | YES |
| FGASTEIQQVDE | 1432742 | Cell/Virus Enclosed | NO |
| FGCENPDCIACS | 1432751 | Cell/Virus Enclosed | NO |
| FGCKDSFTLYGQ | 1432755 | Cell/Virus Enclosed | NO |
| FGDFIQTTPGFG | 1432763 | Cell/Virus Enclosed | NO |
| FGDFVVTQPGSG | 1432771 | Cell/Virus Enclosed | NO |
| FGDGIYFAATEK | 1432773 | Ectodomain | YES |
| FGDGVYFAATEK | 1432774 | Ectodomain | YES |
| FGDISGINASVV | 1432775 | Ectodomain | YES |
| FGDLQFNKLGTE | 1432776 | Cell/Virus Enclosed | NO |
| FGDLQLNKLGTEDPRW | 1656903 | Cell/Virus Enclosed | NO |
| FGDPDDASRLLH | 1432784 | Cell/Virus Enclosed | NO |
| FGDQDLIRQGID | 1432785 | Capsid | UN |
| FGDQELIRQGTEYKHW | 1656910 | Cell/Virus Enclosed | NO |
| FGDSRSDCNHINNLKI | 1656911 | Ectodomain | YES |
| FGDSVEEALAEA | 1432794 | Cell/Virus Enclosed | NO |
| FGDVNKPVCAKS | 1432801 | Cell/Virus Enclosed | NO |
| FGDYNLSSVLPQRNIR | 1656919 | Ectodomain | YES |
| FGEDTVLEVQGY | 1432807 | Cell/Virus Enclosed | NO |
| FGEPTDSTDNNQ | 1432816 | Capsid | UN |
| FGEVEYSLPIKQ | 1432820 | Cell/Virus Enclosed | NO |
| FGEVFNATKFPSVYAW | 1656927 | Ectodomain | YES |
| FGFIEDAVFKPQ | 1432831 | Ectodomain | YES |
| FGFIENSVFKPQ | 1432836 | Ectodomain | YES |
| FGFSLEIQDLEE | 1432842 | Cell/Virus Enclosed | NO |
| FGGCVFSYVGCY | 1432849 | Cell/Virus Enclosed | NO |
| FGGDQVHEVAAVRSVT | 1656947 | Cell/Virus Enclosed | NO |
| FGGFNFSQILPDPLKP | 1656949 | Ectodomain | YES |
| FGGKQILGYSSLNDEF | 1656953 | Cell/Virus Enclosed | NO |
| FGGPSDSSDNSK | 1432865 | Cell/Virus Enclosed | NO |
| FGGPSDSTDNNQNGDR | 1656955 | Cell/Virus Enclosed | NO |
| FGGPSDSTGSNQNGER | 1656956 | Cell/Virus Enclosed | NO |
| FGGPTDSADNNQ | 1432869 | Capsid | UN |
| FGGQPVSEYHIT | 1432871 | Cell/Virus Enclosed | NO |
| FGGTTVVRPLVEDSTS | 1656959 | Ectodomain | YES |
| FGGVSVITPGTNASSE | 1656962 | Ectodomain | YES |
| FGHVYDFKVEDAYDQV | 1656965 | Cell/Virus Enclosed | NO |
| FGINLQSGKTTS | 1432894 | Cell/Virus Enclosed | NO |
| FGIPPSEAKGYW | 1432895 | Cell/Virus Enclosed | NO |
| FGITFQYGTDTN | 1432896 | Ectodomain | YES |
| FGKDGVMYSEFP | 1432907 | Ectodomain | YES |
| FGKDLNANMSLAECKR | 1656977 | Cell/Virus Enclosed | NO |
| FGKNNGIGTCPA | 1432910 | Ectodomain | YES |
| FGKRGGGNVTYT | 1432913 | Cell/Virus Enclosed | NO |
| FGKRGPNQNFGGGEML | 1656983 | Cell/Virus Enclosed | NO |
| FGKRGPSQNFGNAEML | 1656984 | Capsid | UN |
| FGKSIIYTGDWE | 1432921 | Cell/Virus Enclosed | NO |
| FGLGDLVDSYFENDSI | 1656990 | Cell/Virus Enclosed | NO |
| FGLQDCVTGINDDDYV | 1656993 | Cell/Virus Enclosed | NO |
| FGLQDCVTGVND | 1432938 | Cell/Virus Enclosed | NO |
| FGLRGPGDLQGNFGDL | 1656995 | Cell/Virus Enclosed | NO |
| FGLSYYSNSMDC | 1432946 | Cell/Virus Enclosed | NO |
| FGLVTICDFYSKLGVG | 1656999 | Cell/Virus Enclosed | NO |
| FGLYKNSATGSLDVEV | 1657002 | Cell/Virus Enclosed | NO |
| FGLYKQSCTGSPYIDD | 1657004 | Cell/Virus Enclosed | NO |

| | | | |
|---|---|---|---|
| FGMIKKLDNDVLNGII | 1657007 | Cell/Virus Enclosed | NO |
| FGMLNGNAIATVKSED | 1657009 | Cell/Virus Enclosed | NO |
| FGMSRIGMEVTPSGTWLTYTGAIKL | 1597682 | Cell/Virus Enclosed | NO |
| FGPFKFCSETVN | 1432983 | Cell/Virus Enclosed | NO |
| FGPFNFCSETVNGYAK | 1657022 | Cell/Virus Enclosed | NO |
| FGPGNTFINGDIAREL | 1657023 | Cell/Virus Enclosed | NO |
| FGPRDLDHNFGSAGVV | 1657031 | Capsid | UN |
| FGRDTSDFTDSV | 1433000 | Ectodomain | YES |
| FGRRGPEQTQGNFGDQ | 1657039 | Cell/Virus Enclosed | NO |
| FGSAGVVANGVKAKGY | 1657040 | Capsid | UN |
| FGSAGVVANGVT | 1433009 | Cell/Virus Enclosed | NO |
| FGSGSPGRRHSQASYPSVLW | 741410 | Underneath Membrane | NO |
| FGSKLELAKVQN | 1433020 | Cell/Virus Enclosed | NO |
| FGSKLELVKRES | 1433023 | Cell/Virus Enclosed | NO |
| FGSNDIWMQNKG | 1433032 | Ectodomain | YES |
| FGSPDDASRLLH | 1433033 | Cell/Virus Enclosed | NO |
| FGSRLELAKVQNLSGN | 1657051 | Cell/Virus Enclosed | NO |
| FGSSGFLQSSNY | 1433039 | Undetermined | UN |
| FGSTMDNTTQSA | 1433042 | Ectodomain | YES |
| FGSTMNNKSQTV | 1433044 | Ectodomain | YES |
| FGSVACEHISSTMSQY | 1657054 | Ectodomain | YES |
| FGTDHRFILAIT | 1433052 | Cell/Virus Enclosed | NO |
| FGTNETEYLFEVDNL | 227170 | Secreted | YES |
| FGTRNPNNDLAI | 1433057 | Cell/Virus Enclosed | NO |
| FGVFDIKNSSVNTVCF | 1657073 | Cell/Virus Enclosed | NO |
| FGVNKIQPWTNDALNT | 1657078 | Cell/Virus Enclosed | NO |
| FGVPLSEAKGYW | 1433097 | Cell/Virus Enclosed | NO |
| FGVSKPNAIDVE | 1433104 | Cell/Virus Enclosed | NO |
| FGVTQEYIQTTIQKVT | 1657086 | Ectodomain | YES |
| FGWIKISADNKVIYTT | 1657091 | Cell/Virus Enclosed | NO |
| FGWVKISADNKV | 1433123 | Cell/Virus Enclosed | NO |
| FGYIDCEQDSCDFKGW | 1657098 | Cell/Virus Enclosed | NO |
| FGYVCDGSFYKA | 1433132 | Cell/Virus Enclosed | NO |
| FGYVQQGIFNKATVAQ | 1657103 | Cell/Virus Enclosed | NO |
| FGYVQQGVFNKA | 1433134 | Cell/Virus Enclosed | NO |
| FHANSSEPALIF | 1433137 | Ectodomain | YES |
| FHANSSEPALLFRNIK | 1657106 | Ectodomain | YES |
| FHFRKACQNLDC | 1433149 | Cell/Virus Enclosed | NO |
| FHFRKSCQNLDC | 1433150 | Cell/Virus Enclosed | NO |
| FHIEGTGGNSQSSSRA | 1657113 | Cell/Virus Enclosed | NO |
| FHNYTGEGQQII | 1433188 | Ectodomain | YES |
| FHPPSSNQCESTPPIAMDAW | 16132 | Cell/Virus Enclosed | NO |
| FHRIESTDSTVL | 1433207 | Cell/Virus Enclosed | NO |
| FHRYGTTVNCIVEEVDARSV | 741412 | Ectodomain | YES |
| FHSSQLPLDVYD | 1433212 | Cell/Virus Enclosed | NO |
| FHSVEIEAGEVKPFAV | 1657137 | Cell/Virus Enclosed | NO |
| FHSVEIEVGEVK | 1433214 | Cell/Virus Enclosed | NO |
| FHSVLTTFIDAARGPA | 1657138 | Cell/Virus Enclosed | NO |
| FHTINHTFGNPI | 1433221 | Ectodomain | YES |
| FHTINHTFGNPV | 1433222 | Ectodomain | YES |
| FHTPAYDVSAFR | 1433230 | Cell/Virus Enclosed | NO |
| FHTVLLPTQYKD | 1433233 | Ectodomain | YES |
| FHYERDNTSCPEWMDD | 1657151 | Cell/Virus Enclosed | NO |
| FHYERDNTSYPE | 1433245 | Cell/Virus Enclosed | NO |
| FIAFNHPIQVDQ | 1433251 | Ectodomain | YES |
| FIANEKFINGCGRNVQ | 1657161 | Cell/Virus Enclosed | NO |
| FIAVSKPMGTQT | 1433270 | Ectodomain | YES |
| FICCCTGCGPSC | 1433272 | Undetermined | UN |
| FICCCTGCGTSC | 1433273 | Undetermined | UN |
| FICEEVANDLTTALRR | 1657169 | Cell/Virus Enclosed | NO |
| FICSQISPAAIA | 1433282 | Ectodomain | YES |
| FIDAARGPAGVESDVE | 1657175 | Cell/Virus Enclosed | NO |
| FIDDYYGYLMKR | 1433286 | Cell/Virus Enclosed | NO |
| FIDGVMYSEFPA | 1433290 | Ectodomain | YES |
| FIDVGNHRSAFA | 1433307 | Ectodomain | YES |
| FIECGSCGNNSW | 1433314 | Cell/Virus Enclosed | NO |
| FIEDAVFKPQPA | 1433315 | Ectodomain | YES |
| FIEDLLFDKVKL | 1433316 | Ectodomain | YES |
| FIEDLLYNKVTL | 1433318 | Ectodomain | YES |
| FIEDSVFVPQPT | 1433323 | Ectodomain | YES |
| FIENSVFKPQPA | 1433327 | Ectodomain | YES |
| FIFDVDKFKTPH | 1433340 | Cell/Virus Enclosed | NO |
| FIFDVDKVKTPH | 1433341 | Cell/Virus Enclosed | NO |
| FIFEKVEYGSDTVTYK | 1657201 | Cell/Virus Enclosed | NO |
| FIFNKLVKDPPNVQIH | 1657203 | Secreted | YES |
| FIGATIRLQAGKQTEL | 1657205 | Cell/Virus Enclosed | NO |

| | | | |
|---|---|---|---|
| FIGDNVGHYVHV | 1433358 | Cell/Virus Enclosed | NO |
| FIGVEGELPVAVVNDK | 1657210 | Cell/Virus Enclosed | NO |
| FIHRCQRRYRRPRGQIYNPG | 741413 | Undetermined | UN |
| FIIYDSSDRGQESLAR | 1657222 | Cell/Virus Enclosed | NO |
| FIKRSDAQGTNH | 1433436 | Cell/Virus Enclosed | NO |
| FIKTAQADTAGL | 1433439 | Cell/Virus Enclosed | NO |
| FIKTNNIRIVDE | 1433440 | Ectodomain | YES |
| FIKTNNTMIVDE | 1433441 | Ectodomain | YES |
| FIKTNNTRIVDEWSYT | 1657239 | Ectodomain | YES |
| FIKTNNTSIVDE | 1433445 | Ectodomain | YES |
| FIKTNNTTIVDE | 1433446 | Ectodomain | YES |
| FILHDVNNGLTC | 1433457 | Ectodomain | YES |
| FILHDVNSGTTC | 1433459 | Ectodomain | YES |
| FILLFGWVKISA | 1433462 | Cell/Virus Enclosed | NO |
| FILPLVYFSDSK | 1433466 | Cell/Virus Enclosed | NO |
| FILVSCIIGSLG | 1433471 | Ectodomain | YES |
| FIMVLGSKHVNC | 1433481 | Cell/Virus Enclosed | NO |
| FINGACGSPGYNLKNG | 1657253 | Cell/Virus Enclosed | NO |
| FINGACGSPGYNVRND | 1657254 | Cell/Virus Enclosed | NO |
| FINGIFAKVKNTKVIK | 1657257 | Ectodomain | YES |
| FINKYGLEDFNFEHVV | 1657258 | Cell/Virus Enclosed | NO |
| FINNTTYPTNVAVELF | 1657260 | Cell/Virus Enclosed | NO |
| FIQTAPGFGVAVADSY | 1657273 | Cell/Virus Enclosed | NO |
| FISPYNSQNAVS | 1433559 | Cell/Virus Enclosed | NO |
| FISPYNSQNFAAKRVL | 1657300 | Cell/Virus Enclosed | NO |
| FISPYNSQNYVAKRVL | 1657302 | Cell/Virus Enclosed | NO |
| FISVLDITDAAVKAAE | 1657308 | Cell/Virus Enclosed | NO |
| FITETNTAAFNA | 1433580 | Cell/Virus Enclosed | NO |
| FITKDEAIKRVRGWVG | 1657321 | Cell/Virus Enclosed | NO |
| FITLEEAVRQVR | 1433591 | Cell/Virus Enclosed | NO |
| FITMHSINHDIF | 1433595 | Cell/Virus Enclosed | NO |
| FITMSSKGQKGDAEEA | 1657329 | Cell/Virus Enclosed | NO |
| FITNGMSLLDII | 1433598 | Cell/Virus Enclosed | NO |
| FITPEVSRELGNITKT | 1657331 | Cell/Virus Enclosed | NO |
| FITVEAAFDLSK | 1433610 | Cell/Virus Enclosed | NO |
| FITVEAAIELSKELKR | 1657340 | Cell/Virus Enclosed | NO |
| FITVEAALDLSKELKR | 1657341 | Cell/Virus Enclosed | NO |
| FITVETAIELSK | 1433611 | Cell/Virus Enclosed | NO |
| FITVETALDLSK | 1433612 | Cell/Virus Enclosed | NO |
| FIVNAANENLAHGGGL | 1657354 | Cell/Virus Enclosed | NO |
| FIVVLQYGHYKF | 1433651 | Undetermined | UN |
| FIVYNNSGSNVT | 1433656 | Cell/Virus Enclosed | NO |
| FIYFITMSSKGQ | 1433659 | Cell/Virus Enclosed | NO |
| FIYVFSLLHFTN | 1433667 | Cell/Virus Enclosed | NO |
| FKADGIIYPQGR | 1433671 | Ectodomain | YES |
| FKAPKNFCPCKL | 1433678 | Ectodomain | YES |
| FKATGTYKCPQT | 1433688 | Ectodomain | YES |
| FKAVHYYQNNVF | 1433689 | Cell/Virus Enclosed | NO |
| FKCSSNGSNDVW | 1433698 | Ectodomain | YES |
| FKCTMGVYDFKVSAAE | 1657377 | Cell/Virus Enclosed | NO |
| FKCYGVSATKLNDLCF | 1657378 | Ectodomain | YES |
| FKDAEGRMTPYC | 1433703 | Cell/Virus Enclosed | NO |
| FKDCARNPIDLPPSHA | 1657384 | Cell/Virus Enclosed | NO |
| FKDCSKSCLGYHPAHA | 1657386 | Cell/Virus Enclosed | NO |
| FKDCSKSYSGYHPAHA | 1657387 | Cell/Virus Enclosed | NO |
| FKDCSRETSGLSPAYA | 1657389 | Cell/Virus Enclosed | NO |
| FKDFQFMLWCNDNKIM | 1657392 | Cell/Virus Enclosed | NO |
| FKDFQFMLWGND | 1433723 | Cell/Virus Enclosed | NO |
| FKDGIYFAATEKSNVV | 1657394 | Ectodomain | YES |
| FKDGVMYSEFPA | 1433730 | Ectodomain | YES |
| FKDGVMYSEFPT | 1433731 | Ectodomain | YES |
| FKDLTANMSMAE | 1433737 | Cell/Virus Enclosed | NO |
| FKDQVILLNKHIDAYK | 1331955 | Cell/Virus Enclosed | NO |
| FKEELDQWLKNQ | 1433743 | Ectodomain | YES |
| FKEESDQWFKNQ | 1433745 | Ectodomain | YES |
| FKELTANMSMAECKAT | 1657410 | Cell/Virus Enclosed | NO |
| FKELWHLDTGAV | 1433758 | Ectodomain | YES |
| FKELWHMDTGVV | 1433759 | Ectodomain | YES |
| FKENVDFINSMQ | 1433760 | Cell/Virus Enclosed | NO |
| FKEVSAAYYNDW | 1433767 | Ectodomain | YES |
| FKFCSETVNGYA | 1433774 | Cell/Virus Enclosed | NO |
| FKFDDYDFSVEA | 1433775 | Cell/Virus Enclosed | NO |
| FKFPDGQGVPIA | 1433785 | Cell/Virus Enclosed | NO |
| FKFRDLAIENMW | 1433786 | Cell/Virus Enclosed | NO |
| FKFSDGQGVPIAFGVP | 1657425 | Capsid | UN |
| FKGCETFGKPVI | 1433797 | Cell/Virus Enclosed | NO |

| | | | |
|---|---|---|---|
| FKGGIPGNMIDG | 1433800 | Cell/Virus Enclosed | NO |
| FKGNVQVDNGSSINRK | 1657435 | Cell/Virus Enclosed | NO |
| FKGWIPGNMIDGFACT | 1657438 | Cell/Virus Enclosed | NO |
| FKGWVPGNMIDG | 1433809 | Cell/Virus Enclosed | NO |
| FKHGVMYSEFPA | 1433811 | Ectodomain | YES |
| FKHLREFVFKNKDGFLY | 16376 | Ectodomain | YES |
| FKHWSFDYHPNC | 1433828 | Cell/Virus Enclosed | NO |
| FKHWSMPYHPNTVDCQ | 1657448 | Cell/Virus Enclosed | NO |
| FKIDKGIYQTSN | 1433831 | Ectodomain | YES |
| FKIGDVKFKRLGDYVL | 1657450 | Cell/Virus Enclosed | NO |
| FKIGRRRICLSG | 1433835 | Cell/Virus Enclosed | NO |
| FKKCGGCCDDYTGYQE | 1657462 | Underneath Membrane | NO |
| FKKCGGFCDDYT | 1433849 | Underneath Membrane | NO |
| FKKNFTYNVSTD | 1433863 | Ectodomain | YES |
| FKKNFTYNVYAD | 1433864 | Ectodomain | YES |
| FKLDAIVPGYPK | 1433876 | Cell/Virus Enclosed | NO |
| FKLGFDSSKEFV | 1433885 | Cell/Virus Enclosed | NO |
| FKLKISTQLFEGDKFI | 1657482 | Cell/Virus Enclosed | NO |
| FKLLGIGGVPIIEVSQ | 1657484 | Cell/Virus Enclosed | NO |
| FKLSDNNCYLNA | 1433908 | Cell/Virus Enclosed | NO |
| FKMFVLWLLWPS | 1433922 | Undetermined | UN |
| FKMNYQVNGYPH | 1433926 | Cell/Virus Enclosed | NO |
| FKNADLKDGYFV | 1433928 | Cell/Virus Enclosed | NO |
| FKNDNVVLKITEDGIN | 1657515 | Cell/Virus Enclosed | NO |
| FKNFSTSIPNFG | 1433931 | Ectodomain | YES |
| FKNKDGFLYVYKGYQPI | 16417 | Ectodomain | YES |
| FKNQHCNGSIAC | 1433948 | Cell/Virus Enclosed | NO |
| FKNQTLVAPDLS | 1433950 | Ectodomain | YES |
| FKNQTSVAPDFS | 1433951 | Ectodomain | YES |
| FKNSVTSINSAI | 1433960 | Cell/Virus Enclosed | NO |
| FKNVDKDDAFYIVKRC | 1657526 | Cell/Virus Enclosed | NO |
| FKNVSTSIPNFF | 1433966 | Ectodomain | YES |
| FKNVSTSISNFG | 1433969 | Ectodomain | YES |
| FKPGGTTSGDASTAYA | 1657530 | Cell/Virus Enclosed | NO |
| FKPGGTTSGEAI | 1433972 | Cell/Virus Enclosed | NO |
| FKPQPAGVFTDH | 1433985 | Ectodomain | YES |
| FKPQPAGVFTNH | 1433986 | Ectodomain | YES |
| FKPQPVGVFTHH | 1433989 | Ectodomain | YES |
| FKPQSGGGKCFNCYPA | 1657540 | Ectodomain | YES |
| FKPQSGGGKCYN | 1433992 | Ectodomain | YES |
| FKPQSGGGRCYN | 1433993 | Ectodomain | YES |
| FKPVHLVPVSPA | 1434000 | Cell/Virus Enclosed | NO |
| FKPVHLVPVSPV | 1434001 | Cell/Virus Enclosed | NO |
| FKPVQLAPVSPA | 1434003 | Cell/Virus Enclosed | NO |
| FKPVQLVPVSPAFHRI | 1657544 | Cell/Virus Enclosed | NO |
| FKQANNNCFVNVSCLM | 1657546 | Cell/Virus Enclosed | NO |
| FKQELDKYFKNH | 1434010 | Ectodomain | YES |
| FKQGHFIGVEGE | 1434011 | Cell/Virus Enclosed | NO |
| FKRDSDADSPSK | 1434015 | Cell/Virus Enclosed | NO |
| FKRGDGTPHPYCYSDG | 1657551 | Cell/Virus Enclosed | NO |
| FKRSINPTDQSS | 1434025 | Cell/Virus Enclosed | NO |
| FKSENRYSVLSVDSVS | 1657560 | Cell/Virus Enclosed | NO |
| FKSISLTVGEVFGKIL | 1657563 | Cell/Virus Enclosed | NO |
| FKSIVNTRNATV | 1434041 | Cell/Virus Enclosed | NO |
| FKSNFSQYYYNIDTGS | 1657567 | Undetermined | UN |
| FKTADGNQRQLLPRWY | 1657571 | Cell/Virus Enclosed | NO |
| FKTADGQQKHLL | 1434053 | Cell/Virus Enclosed | NO |
| FKTADGQQKPLL | 1434054 | Cell/Virus Enclosed | NO |
| FKTADGQQKQLLPRWY | 1657572 | Capsid | UN |
| FKTAVFVYTCVV | 1434056 | Cell/Virus Enclosed | NO |
| FKTEVDPNLSGF | 1434059 | Cell/Virus Enclosed | NO |
| FKTFDLLKYDFTEHKE | 1657575 | Cell/Virus Enclosed | NO |
| FKTFEKVDGVYTNFKL | 1657576 | Cell/Virus Enclosed | NO |
| FKTLKPGDSFNILACY | 1657580 | Cell/Virus Enclosed | NO |
| FKTSGDLAVQIGNNNV | 1657582 | Cell/Virus Enclosed | NO |
| FKTTGDLAVQIGSNNV | 1657584 | Cell/Virus Enclosed | NO |
| FKVEDAYDQVHDEPKG | 1657587 | Cell/Virus Enclosed | NO |
| FKVEDAYDQVHE | 1434071 | Cell/Virus Enclosed | NO |
| FKVEDSVIINDD | 1434072 | Cell/Virus Enclosed | NO |
| FKVEDSVIVNDA | 1434073 | Cell/Virus Enclosed | NO |
| FKVEDSVIVNND | 1434075 | Cell/Virus Enclosed | NO |
| FKVFFKGNVQVDNGSS | 1657589 | Cell/Virus Enclosed | NO |
| FKVGKVDSIVQKCYEL | 1657590 | Cell/Virus Enclosed | NO |
| FKVLDNGIIRDVNPDD | 1657592 | Cell/Virus Enclosed | NO |
| FKVNETNSQYLFSTIN | 1657593 | Cell/Virus Enclosed | NO |
| FKVNETTSQYLF | 1434086 | Cell/Virus Enclosed | NO |

| | | | |
|---|---|---|---|
| FKVQNVNLERCELANY | 1657596 | Cell/Virus Enclosed | NO |
| FKVTRGRAKKGA | 1434100 | Cell/Virus Enclosed | NO |
| FKVYFKGQTTHESSSA | 1657603 | Cell/Virus Enclosed | NO |
| FKWAPEFRWLLPDAAE | 1657605 | Cell/Virus Enclosed | NO |
| FKYMVANGLNAPNGPF | 1657616 | Cell/Virus Enclosed | NO |
| FKYWDQTYHANCVNCT | 1657618 | Cell/Virus Enclosed | NO |
| FKYWGMKYHPNTVDCD | 1657621 | Cell/Virus Enclosed | NO |
| FKYWGQDYHPDCVDCH | 1657622 | Cell/Virus Enclosed | NO |
| FLAFVVFLLAIL | 1434127 | Transmembrane | NO |
| FLAVDDKYKATGDLAV | 1657643 | Cell/Virus Enclosed | NO |
| FLAVDDKYKVNENLAV | 1657644 | Cell/Virus Enclosed | NO |
| FLCANEYTGSYQ | 1434155 | Cell/Virus Enclosed | NO |
| FLDDNVLPETYVALPI | 1657665 | Ectodomain | YES |
| FLDDVEMVAYNPDLSQ | 1657666 | Cell/Virus Enclosed | NO |
| FLDGNFRTEVDP | 1434177 | Cell/Virus Enclosed | NO |
| FLDKVDILLTVDGVNF | 1657670 | Cell/Virus Enclosed | NO |
| FLDLQAEMNRLQ | 1434183 | Ectodomain | YES |
| FLDLQDEMNRLQ | 1434184 | Ectodomain | YES |
| FLDLQEEMNRLQ | 1434185 | Ectodomain | YES |
| FLDLYYEMNLIQESIK | 1657673 | Ectodomain | YES |
| FLDTCFDVSKPN | 1434192 | Cell/Virus Enclosed | NO |
| FLDTVVSTVKIGGLTF | 1657676 | Cell/Virus Enclosed | NO |
| FLDVLNKALEMC | 1434194 | Cell/Virus Enclosed | NO |
| FLEELNAFTREMQQHP | 1657677 | Capsid | UN |
| FLEENDVEEDDV | 1434197 | Cell/Virus Enclosed | NO |
| FLEQDAQIDRLI | 1434211 | Ectodomain | YES |
| FLFDEAGEEVLAPKLY | 1657693 | Cell/Virus Enclosed | NO |
| FLFFDSDVFIKK | 1434227 | Cell/Virus Enclosed | NO |
| FLFGDSRSDCNHINNL | 1657695 | Ectodomain | YES |
| FLFGDSRSDCNHIVNI | 1657696 | Ectodomain | YES |
| FLFILLLCLIFLLVL | 1657697 | Transmembrane | NO |
| FLGASVTEDVKFAAST | 1657709 | Cell/Virus Enclosed | NO |
| FLGLQIQTVDSSQGSE | 1657715 | Cell/Virus Enclosed | NO |
| FLGLWCLMPTYT | 1434251 | Cell/Virus Enclosed | NO |
| FLHTVLLPTQYKDVEA | 1657738 | Ectodomain | YES |
| FLIAGPKGPAIR | 1434293 | Cell/Virus Enclosed | NO |
| FLIAGPKGPEIR | 1434294 | Cell/Virus Enclosed | NO |
| FLIHNFVIFFTW | 1434308 | Cell/Virus Enclosed | NO |
| FLITGVFDIVKG | 1434321 | Cell/Virus Enclosed | NO |
| FLKLVDDHALIV | 1434343 | Cell/Virus Enclosed | NO |
| FLKPLYVPEHCQ | 1434346 | Cell/Virus Enclosed | NO |
| FLLDFSVDGYIR | 1434367 | Ectodomain | YES |
| FLLGINYLGTIKENID | 1657788 | Cell/Virus Enclosed | NO |
| FLLNGYYPTSGS | 1434394 | Ectodomain | YES |
| FLLTNTSSVVDGVVRS | 1657807 | Ectodomain | YES |
| FLMAEDGITKLA | 1434425 | Cell/Virus Enclosed | NO |
| FLMLFVFQVYPI | 1434431 | Cell/Virus Enclosed | NO |
| FLNGITYTSTSGNLLG | 1657820 | Ectodomain | YES |
| FLNGSAGSVGFN | 1434445 | Cell/Virus Enclosed | NO |
| FLNGVCGVSAAR | 1434447 | Cell/Virus Enclosed | NO |
| FLNRVCGVSAAR | 1434460 | Cell/Virus Enclosed | NO |
| FLNRVRGTSVDARLVP | 1657834 | Cell/Virus Enclosed | NO |
| FLPAQDTWGTSA | 1434463 | Ectodomain | YES |
| FLPESAAYYVGN | 1434464 | Ectodomain | YES |
| FLPEVAAYFVGN | 1434465 | Ectodomain | YES |
| FLPFDSNLTQYF | 1434468 | Ectodomain | YES |
| FLPGVYSIFYLY | 1434477 | Cell/Virus Enclosed | NO |
| FLPKEDGDVLLAEFDT | 1657848 | Cell/Virus Enclosed | NO |
| FLPLLPIFFCLWVYI | 16811 | Undetermined | UN |
| FLPQFTSDQAVTFLKE | 1657850 | Undetermined | UN |
| FLPTFIGKIAQW | 1434494 | Cell/Virus Enclosed | NO |
| FLQSSNYKIDTTSSSC | 1657862 | Ectodomain | YES |
| FLRLIDDNGIVLNSIL | 1657868 | Cell/Virus Enclosed | NO |
| FLSDFINAIFAK | 1434542 | Ectodomain | YES |
| FLSDFINGIFAKVKNT | 1657876 | Ectodomain | YES |
| FLSFDSDVFIKKYGLE | 1657878 | Cell/Virus Enclosed | NO |
| FLSGDDCFVSNSSVIQ | 1657880 | Cell/Virus Enclosed | NO |
| FLSKHSDFGLDG | 1434554 | Cell/Virus Enclosed | NO |
| FLSKHSDFGLGDLVDS | 1657882 | Cell/Virus Enclosed | NO |
| FLSLSDQFKTTGDLAV | 1657885 | Cell/Virus Enclosed | NO |
| FLSMFDVDKKSLNALI | 1657888 | Cell/Virus Enclosed | NO |
| FLSNASSSFDCI | 1434560 | Ectodomain | YES |
| FLSNTKYYDDSQYYFN | 1657890 | Ectodomain | YES |
| FLSNTKYYDHSQ | 1434565 | Ectodomain | YES |
| FLSNTKYYDYSQ | 1434566 | Ectodomain | YES |
| FLTWICLLQFAYANR | 2001056 | Undetermined | UN |

| | | | |
|---|---|---|---|
| FLVHDNVAFYQGDVDT | 1657913 | Cell/Virus Enclosed | NO |
| FLVKNPSWSNAV | 1434631 | Cell/Virus Enclosed | NO |
| FLVVDDKYKVNE | 1434655 | Cell/Virus Enclosed | NO |
| FLYIIKLIFLWLLWP | 1312469 | Undetermined | UN |
| FMDGENAYEMVK | 1434695 | Cell/Virus Enclosed | NO |
| FMDNAINVGGTGLQYA | 1657951 | Cell/Virus Enclosed | NO |
| FMFGWRILFCFKCCRRTRG | 244201 | Undetermined | UN |
| FMIRSCYSGHVS | 1434728 | Ectodomain | YES |
| FMKHKGGDLTDF | 1434736 | Cell/Virus Enclosed | NO |
| FMKHKGGDSTDFIALI | 1657968 | Cell/Virus Enclosed | NO |
| FMKQYGQCLGDI | 1434743 | Ectodomain | YES |
| FMKSDIFGQDFKTFDL | 1657970 | Cell/Virus Enclosed | NO |
| FMLGSSVGNFSDGKMG | 1657972 | Ectodomain | YES |
| FMLKTLYKDVDNPHLM | 1657974 | Cell/Virus Enclosed | NO |
| FMLKYDENGTITDAVD | 1657975 | Ectodomain | YES |
| FMLNFNENGTIT | 1434760 | Ectodomain | YES |
| FMLSFNENGTIT | 1434766 | Ectodomain | YES |
| FMLSTLHWSFRL | 1434768 | Cell/Virus Enclosed | NO |
| FMLTYNKVENMT | 1434770 | Cell/Virus Enclosed | NO |
| FMLWCKDGQVQTFYPR | 1657981 | Cell/Virus Enclosed | NO |
| FMNSCSVNFTKA | 1434783 | Ectodomain | YES |
| FMNVTLDKCTKYNIYD | 1657988 | Ectodomain | YES |
| FMPVQSRKFIVP | 1434794 | Cell/Virus Enclosed | NO |
| FMPYFMYDVVSS | 1434799 | Cell/Virus Enclosed | NO |
| FMSHFDVDRKSFNNFV | 1658005 | Cell/Virus Enclosed | NO |
| FMSTSKCWVEEDLTKG | 1658010 | Cell/Virus Enclosed | NO |
| FMVDNGTRLHDA | 1434840 | Undetermined | UN |
| FMVDPTTVLNDP | 1434841 | Cell/Virus Enclosed | NO |
| FMVDPTTVLNGP | 1434842 | Cell/Virus Enclosed | NO |
| FMVGNVTRLHDA | 1434844 | Undetermined | UN |
| FMVNQLAYSSSANGSL | 1658016 | Cell/Virus Enclosed | NO |
| FMYAYISNKADL | 1434850 | Cell/Virus Enclosed | NO |
| FNADNLTDYSVS | 1434863 | Ectodomain | YES |
| FNAEDCMSDFMS | 1434864 | Ectodomain | YES |
| FNAEGDAFWSST | 1434865 | Cell/Virus Enclosed | NO |
| FNAEGDASWSST | 1434866 | Cell/Virus Enclosed | NO |
| FNAFVKCNCGSE | 1434869 | Cell/Virus Enclosed | NO |
| FNASISESKPLA | 1434876 | Cell/Virus Enclosed | NO |
| FNATGSDVNCNGYQHN | 1658030 | Ectodomain | YES |
| FNAVDCKSDFMS | 1434885 | Ectodomain | YES |
| FNAVDCMSDFMSEIKC | 1658034 | Ectodomain | YES |
| FNAVDCVSDFMS | 1434887 | Ectodomain | YES |
| FNCDTFCTGSTF | 1434890 | Cell/Virus Enclosed | NO |
| FNCFYALNNAFLAFSI | 1658035 | Undetermined | UN |
| FNCHSFKPGNTF | 1434894 | Cell/Virus Enclosed | NO |
| FNCTNFAINDKN | 1434900 | Ectodomain | YES |
| FNCTNFAINDLNTTVP | 1658039 | Ectodomain | YES |
| FNCTYDRVEKSF | 1434902 | Ectodomain | YES |
| FNDLSQLYCSYE | 1434914 | Ectodomain | YES |
| FNEDVCVCFDNSIQGS | 1658051 | Cell/Virus Enclosed | NO |
| FNEHVTSVNLFD | 1434922 | Cell/Virus Enclosed | NO |
| FNEHVTSVNLFS | 1434923 | Cell/Virus Enclosed | NO |
| FNENGTITDAVD | 1434929 | Ectodomain | YES |
| FNETKGPLCVDTSHFT | 1658056 | Ectodomain | YES |
| FNFCKSDILSLN | 1434941 | Ectodomain | YES |
| FNFCYGFNEPLN | 1434943 | Ectodomain | YES |
| FNFDQKELLAYYNMLV | 1658060 | Cell/Virus Enclosed | NO |
| FNFDQKELLDYY | 1434945 | Cell/Virus Enclosed | NO |
| FNFNGLTGTGVLTPSS | 1658067 | Ectodomain | YES |
| FNGAIIRDSDVVKQPV | 1658076 | Cell/Virus Enclosed | NO |
| FNGKFLSNTKYYDDSQ | 1658082 | Ectodomain | YES |
| FNGLTGTGVLTS | 1434987 | Ectodomain | YES |
| FNGSNDVWIFNK | 1434991 | Ectodomain | YES |
| FNGTGRGACKGF | 1434993 | Ectodomain | YES |
| FNGTGRGDCKGFSSDV | 1658089 | Ectodomain | YES |
| FNGTGRGVCKGF | 1434996 | Ectodomain | YES |
| FNHNMGDSDLVQ | 1435003 | Cell/Virus Enclosed | NO |
| FNHNNGNDVLYQPPTA | 1658100 | Cell/Virus Enclosed | NO |
| FNHPIQIDQLNS | 1435005 | Ectodomain | YES |
| FNHPIQVDQFNS | 1435007 | Ectodomain | YES |
| FNHPIQVDQLNSSYFK | 1658101 | Ectodomain | YES |
| FNHPIQVGQLNS | 1435009 | Ectodomain | YES |
| FNICQAVSANVCALMS | 1658104 | Cell/Virus Enclosed | NO |
| FNILQATTANVS | 1435032 | Cell/Virus Enclosed | NO |
| FNIQDMDIVASEHLLHQSLV | 1948343 | Capsid | UN |
| FNKAIDGFGVFDIKNS | 1658117 | Cell/Virus Enclosed | NO |

| | | | |
|---|---|---|---|
| FNKDTGVIYGLNSTET | 1658124 | Ectodomain | YES |
| FNKEFYDFLRSQGFFD | 1658125 | Cell/Virus Enclosed | NO |
| FNKLVKDPPNVQIHTI | 1658132 | Secreted | YES |
| FNKPLIEWPLIE | 1435058 | Cell/Virus Enclosed | NO |
| FNKPLIEWPLTEYDFT | 1658133 | Cell/Virus Enclosed | NO |
| FNKSKQFVPVQV | 1435066 | Underneath Membrane | NO |
| FNKVTLSDAGFM | 1435070 | Ectodomain | YES |
| FNKWGKARLYYD | 1435071 | Cell/Virus Enclosed | NO |
| FNKWQWQEAWYE | 1435072 | Cell/Virus Enclosed | NO |
| FNKYFKHWSMPYHPNT | 1658141 | Cell/Virus Enclosed | NO |
| FNKYKYFSGAMETAAY | 1658144 | Cell/Virus Enclosed | NO |
| FNMAQAFGLRGP | 1435113 | Cell/Virus Enclosed | NO |
| FNNALDAIQEGF | 1435132 | Ectodomain | YES |
| FNNALGAIQEGF | 1435133 | Ectodomain | YES |
| FNNALHSIQQGF | 1435136 | Ectodomain | YES |
| FNNALNAIQQGF | 1435137 | Ectodomain | YES |
| FNNALYAIQEGFDATN | 1658164 | Ectodomain | YES |
| FNNFFISYAKPEDKLS | 1658165 | Cell/Virus Enclosed | NO |
| FNNFVSSYAKPEEKLS | 1658168 | Cell/Virus Enclosed | NO |
| FNNGVYISTTKV | 1435146 | Cell/Virus Enclosed | NO |
| FNNLLGFDFSKP | 1435149 | Cell/Virus Enclosed | NO |
| FNNLLGFDSSKPVTKK | 1658170 | Cell/Virus Enclosed | NO |
| FNNLLGFDSSKS | 1435151 | Cell/Virus Enclosed | NO |
| FNNNGTNVTQVKNASV | 1658171 | Cell/Virus Enclosed | NO |
| FNNNTDSIIIDG | 1435154 | Cell/Virus Enclosed | NO |
| FNNRDTGHPSIS | 1435157 | Ectodomain | YES |
| FNNRDTGPLSIS | 1435159 | Ectodomain | YES |
| FNNRDTGPPSIS | 1435160 | Ectodomain | YES |
| FNNSLTRQLQPINYFD | 1658173 | Ectodomain | YES |
| FNNWFLLTNDST | 1435167 | Ectodomain | YES |
| FNNWFLLTNTSSVVDG | 1658175 | Ectodomain | YES |
| FNNYDCVITTLI | 1435170 | Cell/Virus Enclosed | NO |
| FNPETNCLLNVPFGGT | 1658179 | Ectodomain | YES |
| FNPETNILLNVPLHGT | 1658180 | Underneath Membrane | NO |
| FNPHHRFKCSSN | 1435184 | Ectodomain | YES |
| FNPNKDTWPDANKVG | 153257 | Underneath Membrane | NO |
| FNPSTWNKRFGFIEDS | 1658186 | Ectodomain | YES |
| FNQALEAMQTGF | 1435198 | Ectodomain | YES |
| FNQALGAMQTGFTTTN | 1658188 | Ectodomain | YES |
| FNQDGIIFNAVDCMSD | 1658192 | Ectodomain | YES |
| FNQVLSSDVMFE | 1435212 | Cell/Virus Enclosed | NO |
| FNQYYYNMATGF | 1435217 | Ectodomain | YES |
| FNRALVSATEFADKLV | 1658202 | Cell/Virus Enclosed | NO |
| FNRCCDRYEEYD | 1435222 | Underneath Membrane | NO |
| FNRGRQFYEFYNDVKP | 1658206 | Cell/Virus Enclosed | NO |
| FNRLATCDWTESGDYT | 1658208 | Cell/Virus Enclosed | NO |
| FNRPSLVDDNKFDVLK | 1658210 | Cell/Virus Enclosed | NO |
| FNRSSFVDDNKF | 1435236 | Cell/Virus Enclosed | NO |
| FNSCKSDILSLNNPIF | 1658214 | Ectodomain | YES |
| FNSNFLPESAAY | 1435249 | Ectodomain | YES |
| FNTEFASNSSVL | 1435263 | Cell/Virus Enclosed | NO |
| FNTILNTACGVFEVDD | 1658226 | Cell/Virus Enclosed | NO |
| FNTLFSTTIPNT | 1435266 | Cell/Virus Enclosed | NO |
| FNTMLDASAPATGWLL | 1658233 | Cell/Virus Enclosed | NO |
| FNVDDINFSPVLGCLG | 1658241 | Ectodomain | YES |
| FNVDWVFFGFSI | 1435286 | Undetermined | UN |
| FNVKKLQKQLYD | 1435301 | Cell/Virus Enclosed | NO |
| FNVPIKEAGEDCHAP | 17243 | Ectodomain | YES |
| FNVSDAYANLVPYYQL | 1658259 | Cell/Virus Enclosed | NO |
| FNVSFVNDSIESVGGL | 1658261 | Ectodomain | YES |
| FNVSKGIYQTSN | 1435316 | Ectodomain | YES |
| FNVVGPRTGKHE | 1435330 | Cell/Virus Enclosed | NO |
| FNVVPYYSWFSG | 1435334 | Cell/Virus Enclosed | NO |
| FNYNVSVLQMDVQGLV | 1658284 | Cell/Virus Enclosed | NO |
| FPCAGRKVNFNEKPVV | 1658293 | Cell/Virus Enclosed | NO |
| FPCAGRRVTFKE | 1435379 | Cell/Virus Enclosed | NO |
| FPDGQGVPIAYG | 1435385 | Cell/Virus Enclosed | NO |
| FPDHQLDPAFGANSN | 1658297 | Underneath Membrane | NO |
| FPEWIDDLEADP | 1435396 | Cell/Virus Enclosed | NO |
| FPFAESTGTQSV | 1435397 | Cell/Virus Enclosed | NO |
| FPGPSGIGAVGG | 1435403 | Cell/Virus Enclosed | NO |
| FPHNDRIKSFSTFESA | 1658310 | Cell/Virus Enclosed | NO |
| FPIDFPSAESTG | 1435407 | Cell/Virus Enclosed | NO |
| FPIVMNDKNICL | 1435413 | Cell/Virus Enclosed | NO |
| FPKSDGTGTIYA | 1435422 | Cell/Virus Enclosed | NO |
| FPKSGANFRDLA | 1435424 | Ectodomain | YES |

| | | | |
|---|---|---|---|
| FPKSGANFTDLA | 1435425 | Ectodomain | YES |
| FPLAEDVEKDKPNRPVVPSP | 741427 | Ectodomain | YES |
| FPLKLAGTAVINLRAD | 1435437 | Cell/Virus Enclosed | NO |
| FPNSNYVLQDFD | 1435469 | Cell/Virus Enclosed | NO |
| FPPGQGVPLNANSTPA | 1658342 | Cell/Virus Enclosed | NO |
| FPPGQGVPLNSN | 1435474 | Cell/Virus Enclosed | NO |
| FPPGTILPQGYY | 1435475 | Capsid | UN |
| FPPGTVLPQGYY | 1435477 | Cell/Virus Enclosed | NO |
| FPPWTAAAGAPF | 1435488 | Ectodomain | YES |
| FPPWTAAEGVPF | 1435491 | Ectodomain | YES |
| FPQFAELIPNQA | 1435494 | Cell/Virus Enclosed | NO |
| FPQFTSDQAITF | 1435495 | Underneath Membrane | NO |
| FPQIGGLTSIKW | 1435499 | Cell/Virus Enclosed | NO |
| FPQMTKSYKNTRKSPALIVWGIHHS | 1141159 | Ectodomain | YES |
| FPRGQGVPINTNSSPD | 1658360 | Cell/Virus Enclosed | NO |
| FPRGQGVPINTNSTKD | 1658361 | Cell/Virus Enclosed | NO |
| FPSAKVEGCMVQ | 1435515 | Cell/Virus Enclosed | NO |
| FPSGQGVPLNAN | 1435520 | Cell/Virus Enclosed | NO |
| FPTTVTCENLES | 1435537 | Cell/Virus Enclosed | NO |
| FPTWSQELPFDV | 1435541 | Cell/Virus Enclosed | NO |
| FPVAKSVKQLFY | 1435543 | Cell/Virus Enclosed | NO |
| FPYDIEFVTGKQ | 1435553 | Cell/Virus Enclosed | NO |
| FPYDMDFGVSSF | 1435556 | Cell/Virus Enclosed | NO |
| FPYDMDSGGSSF | 1435557 | Cell/Virus Enclosed | NO |
| FPYQGDHGDMYV | 1435568 | Ectodomain | YES |
| FPYQGDHGDRYV | 1435569 | Ectodomain | YES |
| FPYVYKGYQPID | 1435570 | Ectodomain | YES |
| FQASADWKPGHA | 1435575 | Cell/Virus Enclosed | NO |
| FQAVSSNINCIL | 1435576 | Cell/Virus Enclosed | NO |
| FQDAASNIFVINKDTY | 1658386 | Cell/Virus Enclosed | NO |
| FQDELDDFFKNV | 1435584 | Ectodomain | YES |
| FQDELDEFFKNF | 1435585 | Ectodomain | YES |
| FQFATFPVYDTI | 1435606 | Ectodomain | YES |
| FQFMLWCNDEKV | 1435618 | Cell/Virus Enclosed | NO |
| FQFMLWGNDEKV | 1435621 | Cell/Virus Enclosed | NO |
| FQGIETAVGHYV | 1435628 | Cell/Virus Enclosed | NO |
| FQGRQLMGSCSF | 1435637 | Cell/Virus Enclosed | NO |
| FQGYLQKHFSMM | 1435639 | Cell/Virus Enclosed | NO |
| FQHKLKKACCKT | 1435642 | Cell/Virus Enclosed | NO |
| FQILDEGLDCKI | 1435647 | Cell/Virus Enclosed | NO |
| FQKCEQLLREYGQFCS | 1658419 | Ectodomain | YES |
| FQKGKEFEFAEG | 1435660 | Cell/Virus Enclosed | NO |
| FQKGKEFEFVEGQGVP | 1658422 | Cell/Virus Enclosed | NO |
| FQKGRDFKFSDGQGVP | 1658423 | Capsid | UN |
| FQKVQDAVNNNAQALS | 1658425 | Ectodomain | YES |
| FQLDTAPQTGNF | 1435675 | Ectodomain | YES |
| FQLKLKGTPVLQLKES | 1658428 | Cell/Virus Enclosed | NO |
| FQLTFESTINKS | 1435686 | Cell/Virus Enclosed | NO |
| FQLTLESTINKS | 1435687 | Cell/Virus Enclosed | NO |
| FQNCTAVGVPQQ | 1435698 | Ectodomain | YES |
| FQNCTAVGVRQQ | 1435699 | Ectodomain | YES |
| FQQFGRDTSDFT | 1435716 | Ectodomain | YES |
| FQQFGRDVSDFTDSVRDPKT | 17524 | Ectodomain | YES |
| FQRIDDDGNKLDKFFV | 1658445 | Cell/Virus Enclosed | NO |
| FQRVDENGDKLDQFFV | 1658446 | Cell/Virus Enclosed | NO |
| FQSMRKHTPYRNGCLEPTLG | 17535 | Cell/Virus Enclosed | NO |
| FQTTVELQGLIDQINS | 1658462 | Ectodomain | YES |
| FQTVKPGNFNQDFYEF | 1658465 | Cell/Virus Enclosed | NO |
| FQTVRSGNFNQD | 1435752 | Cell/Virus Enclosed | NO |
| FRDAEGRMTPYCHDPT | 1658480 | Cell/Virus Enclosed | NO |
| FRDELGVRVLDQSDNN | 1658481 | Cell/Virus Enclosed | NO |
| FRDLALKGSKYL | 1435780 | Underneath Membrane | NO |
| FRDLTWCINPPERIK | 17572 | Ectodomain | YES |
| FRDYFDDNTDSI | 1435789 | Cell/Virus Enclosed | NO |
| FRDYFNNNTDSIVIGG | 1658485 | Cell/Virus Enclosed | NO |
| FREELDQWFKNQ | 1435797 | Ectodomain | YES |
| FRHSDSIVKPDM | 1435828 | Cell/Virus Enclosed | NO |
| FRIDFPFAESTG | 1435836 | Cell/Virus Enclosed | NO |
| FRIDFPSAELTG | 1435837 | Cell/Virus Enclosed | NO |
| FRIDFPSAESTG | 1435838 | Cell/Virus Enclosed | NO |
| FRIDVPSAESTGTQSV | 1658508 | Cell/Virus Enclosed | NO |
| FRKACQNLDCNC | 1435852 | Cell/Virus Enclosed | NO |
| FRKARNGVFISTEKLS | 1658512 | Cell/Virus Enclosed | NO |
| FRKMAFPSAKVE | 1435853 | Cell/Virus Enclosed | NO |
| FRKSCQNLDCNC | 1435856 | Cell/Virus Enclosed | NO |
| FRKVQDAVNNNA | 1435858 | Ectodomain | YES |

| | | | |
|---|---|---|---|
| FRLFARTRSMWSFNP | 2001061 | Undetermined | UN |
| FRLKGGAPIKGV | 1435865 | Cell/Virus Enclosed | NO |
| FRNSSANYVGKYDINH | 1658537 | Underneath Membrane | NO |
| FRNSTTNYVGVYDINH | 1658538 | Ectodomain | YES |
| FRPFNSLLCGDI | 1435904 | Cell/Virus Enclosed | NO |
| FRRAFGEYNHVV | 1435914 | Cell/Virus Enclosed | NO |
| FRRRSVCSTGRC | 1435922 | Cell/Virus Enclosed | NO |
| FRSDTLYLTQDLFLPFY | 17644 | Ectodomain | YES |
| FRSDVLHLTQDY | 1435930 | Ectodomain | YES |
| FRTFPQQMFVTP | 1435948 | Cell/Virus Enclosed | NO |
| FRTGVDPDLSAF | 1435950 | Cell/Virus Enclosed | NO |
| FRTRKGKRVDLS | 1435953 | Cell/Virus Enclosed | NO |
| FRVTSRVMFQPR | 1435977 | Ectodomain | YES |
| FRYDNVSSVWPL | 1435987 | Ectodomain | YES |
| FSAAQAMEKVNE | 1436000 | Ectodomain | YES |
| FSAFYPIDLASQ | 1436003 | Undetermined | UN |
| FSAGSRSRASSG | 1436004 | Cell/Virus Enclosed | NO |
| FSASLFKPVQLA | 1436013 | Cell/Virus Enclosed | NO |
| FSASLFKPVQLVPVSP | 1658588 | Cell/Virus Enclosed | NO |
| FSASTSAFIDTI | 1436016 | Cell/Virus Enclosed | NO |
| FSATFSVPMEEL | 1436019 | Cell/Virus Enclosed | NO |
| FSAVSSPMSNMTENER | 1658595 | Underneath Membrane | NO |
| FSCNNFDESKIYGSCF | 1658601 | Ectodomain | YES |
| FSCPLCCTVYDSSEVK | 1658602 | Cell/Virus Enclosed | NO |
| FSDDVIVHDVEP | 1436039 | Cell/Virus Enclosed | NO |
| FSDDVIVYDVEP | 1436040 | Cell/Virus Enclosed | NO |
| FSDGKMGRFFNY | 1436044 | Ectodomain | YES |
| FSDGQGVPIAFGVPPS | 1658609 | Capsid | UN |
| FSDSKFGTTEVK | 1436062 | Cell/Virus Enclosed | NO |
| FSFEQSHNNCYVNVAC | 1658620 | Cell/Virus Enclosed | NO |
| FSFFDCGVNALA | 1436082 | Cell/Virus Enclosed | NO |
| FSFGTVNNFVKF | 1436089 | Ectodomain | YES |
| FSFGVTQEYIQTTIQK | 1658624 | Ectodomain | YES |
| FSFIMFVGHLIS | 1436093 | Cell/Virus Enclosed | NO |
| FSFLNLKLRAPM | 1436096 | Cell/Virus Enclosed | NO |
| FSFRDELGVRVLDQSD | 1658627 | Cell/Virus Enclosed | NO |
| FSGFKLDGFNHQFVTA | 1658637 | Cell/Virus Enclosed | NO |
| FSGITQFQKGKE | 1436120 | Cell/Virus Enclosed | NO |
| FSGNVDLDALASDIFD | 1658640 | Cell/Virus Enclosed | NO |
| FSGVASQEGVLSEREK | 1658644 | Ectodomain | YES |
| FSIETDIIDEVN | 1436149 | Capsid | UN |
| FSIKANIKSDIL | 1436158 | Ectodomain | YES |
| FSIVNSAPDGLLFLHT | 1658668 | Ectodomain | YES |
| FSKCHNCCDEYGGHHD | 1658669 | Underneath Membrane | NO |
| FSKKDWYDFVENPDII | 1658674 | Cell/Virus Enclosed | NO |
| FSKKHVEVFTDGKLNC | 1658675 | Cell/Virus Enclosed | NO |
| FSKKQLGLLQPTDNST | 1658676 | Cell/Virus Enclosed | NO |
| FSKKQLGLLQST | 1436178 | Cell/Virus Enclosed | NO |
| FSKLLYGLKMSI | 1436184 | Undetermined | UN |
| FSKPFYVPEHCQ | 1436188 | Cell/Virus Enclosed | NO |
| FSKPIYVPEHCQ | 1436189 | Cell/Virus Enclosed | NO |
| FSLADSVVEVVTTSLT | 1658684 | Cell/Virus Enclosed | NO |
| FSLGDVEAVNFN | 1436228 | Ectodomain | YES |
| FSLGNVEAVNFN | 1436230 | Ectodomain | YES |
| FSLHPVECDEEC | 1436231 | Cell/Virus Enclosed | NO |
| FSLHPVECDEGC | 1436232 | Cell/Virus Enclosed | NO |
| FSLKDIPGGCAMPIVA | 1658698 | Ectodomain | YES |
| FSLKLAGTAVVNLRPD | 1658701 | Cell/Virus Enclosed | NO |
| FSMAPVEFYEDRILDF | 1658720 | Cell/Virus Enclosed | NO |
| FSMVRKLDNQALNSIL | 1658729 | Cell/Virus Enclosed | NO |
| FSNALDAIQEGF | 1436298 | Ectodomain | YES |
| FSNCNFNMSSLMSFIQ | 1658731 | Ectodomain | YES |
| FSNGVDFVAQPEGCVL | 1658733 | Cell/Virus Enclosed | NO |
| FSNLVLNNCTKYNIYD | 1658735 | Ectodomain | YES |
| FSNMVETVVGQL | 1436312 | Cell/Virus Enclosed | NO |
| FSNSNYVLQDFDVVFG | 1658741 | Cell/Virus Enclosed | NO |
| FSNVVELPKFFYASNG | 1658744 | Ectodomain | YES |
| FSNVYADSFVVKGDDV | 1658746 | Ectodomain | YES |
| FSPESVFNVFGD | 1436335 | Cell/Virus Enclosed | NO |
| FSPGQGVPLNAN | 1436339 | Cell/Virus Enclosed | NO |
| FSPGTILPQGYY | 1436340 | Cell/Virus Enclosed | NO |
| FSPLLSGTPPQV | 1436350 | Ectodomain | YES |
| FSPQIITTDNTFVSGN | 1658758 | Ectodomain | YES |
| FSQFDSLGASSN | 1436359 | Cell/Virus Enclosed | NO |
| FSQFDSLGVSSNQSPQ | 1658761 | Cell/Virus Enclosed | NO |
| FSQFNSNFLPES | 1436360 | Ectodomain | YES |

| | | | |
|---|---|---|---|
| FSQILPDPLKPI | 1436362 | Ectodomain | YES |
| FSQILPDPLKST | 1436364 | Ectodomain | YES |
| FSQILPDPSKPT | 1436365 | Ectodomain | YES |
| FSQQGVFRYDNV | 1436370 | Ectodomain | YES |
| FSQRGGSYKNDK | 1436374 | Cell/Virus Enclosed | NO |
| FSQSDSLGVSSN | 1436377 | Cell/Virus Enclosed | NO |
| FSQVDLTEAICD | 1436379 | Cell/Virus Enclosed | NO |
| FSQVKSDLVIDA | 1436381 | Cell/Virus Enclosed | NO |
| FSRDHNDILYTPPTVS | 1658774 | Cell/Virus Enclosed | NO |
| FSRNTTFDFLSNASSS | 1658775 | Ectodomain | YES |
| FSRTGDPAPNMENIT | 1658778 | Underneath Membrane | NO |
| FSRTLYQPVYKI | 1436395 | Cell/Virus Enclosed | NO |
| FSSDGKPCTPPA | 1436403 | Ectodomain | YES |
| FSSDVFKIGDVK | 1436405 | Cell/Virus Enclosed | NO |
| FSSSRRGVYYND | 1436422 | Ectodomain | YES |
| FSSVNDAITQTAEAIH | 1658800 | Ectodomain | YES |
| FSTCNSNANISM | 1436436 | Ectodomain | YES |
| FSTFESAYMPIADPTH | 1658804 | Cell/Virus Enclosed | NO |
| FSTINALPEISA | 1436458 | Cell/Virus Enclosed | NO |
| FSTMSQYSRSTR | 1436464 | Ectodomain | YES |
| FSTTIPNTAFGP | 1436471 | Cell/Virus Enclosed | NO |
| FSTVNALPEVNADIVV | 1658820 | Cell/Virus Enclosed | NO |
| FSTVVIKNLTPIKLNF | 1658821 | Cell/Virus Enclosed | NO |
| FSTVYPIDLASQ | 1436482 | Undetermined | UN |
| FSVAAITSGVKFQTVK | 1658823 | Cell/Virus Enclosed | NO |
| FSVAALSTGLTSQTVK | 1658825 | Cell/Virus Enclosed | NO |
| FSVDPATTYLEAVKHG | 1658830 | Cell/Virus Enclosed | NO |
| FSVDPKKTYLDFIQQG | 1658831 | Cell/Virus Enclosed | NO |
| FSVDPKKTYLDYIQQG | 1658832 | Cell/Virus Enclosed | NO |
| FSVLHDIGNPKA | 1436507 | Cell/Virus Enclosed | NO |
| FSVNDFICSQIS | 1436512 | Ectodomain | YES |
| FSVNDFTCRQIS | 1436513 | Ectodomain | YES |
| FSVNDFTCSQISPAAI | 1658839 | Ectodomain | YES |
| FSVNNTFCPCAKPSFA | 1658840 | Ectodomain | YES |
| FSVNSDFCPCAD | 1436516 | Ectodomain | YES |
| FSVPMEELKALV | 1436517 | Cell/Virus Enclosed | NO |
| FSVQEDGRIPND | 1436521 | Ectodomain | YES |
| FSVQKHIGAPANLRVV | 1658844 | Cell/Virus Enclosed | NO |
| FSWNADLYKLMSCFAF | 1658852 | Cell/Virus Enclosed | NO |
| FSWNAELYNLMG | 1436545 | Cell/Virus Enclosed | NO |
| FSWNLILTVFIV | 1436547 | Transmembrane | NO |
| FSYCRKIGVNVCSDST | 1658856 | Cell/Virus Enclosed | NO |
| FSYDGLRGAIRKAKDY | 1658858 | Cell/Virus Enclosed | NO |
| FSYPLSMKSDLG | 1436564 | Ectodomain | YES |
| FSYQEFNDLAHT | 1436567 | Cell/Virus Enclosed | NO |
| FSYQEFNDLVHT | 1436569 | Cell/Virus Enclosed | NO |
| FTACSHAAVDSLCAKA | 1658868 | Cell/Virus Enclosed | NO |
| FTANKIVGGAPTWFNA | 1658875 | Cell/Virus Enclosed | NO |
| FTARDGVRRGDN | 1436589 | Cell/Virus Enclosed | NO |
| FTCNNIDAAKIYGMCF | 1658883 | Ectodomain | YES |
| FTCNSNANISML | 1436603 | Ectodomain | YES |
| FTCNSNANLSMLQLGV | 1658884 | Ectodomain | YES |
| FTCRQISPAAIA | 1436607 | Ectodomain | YES |
| FTCRTDMEKDFIALDQ | 1658885 | Cell/Virus Enclosed | NO |
| FTCSQISPAAIS | 1436610 | Ectodomain | YES |
| FTCSQISPPAIA | 1436612 | Ectodomain | YES |
| FTDFHNYTGEGQ | 1436618 | Ectodomain | YES |
| FTDGKLNCSFQDAASN | 1658890 | Cell/Virus Enclosed | NO |
| FTDSVRDPKTSEILDIS | 17929 | Ectodomain | YES |
| FTDSVRDPQTLE | 1436638 | Ectodomain | YES |
| FTDYKLELFNKYFKHW | 1658898 | Cell/Virus Enclosed | NO |
| FTECAYTQGRSC | 1436649 | Cell/Virus Enclosed | NO |
| FTEDLLCVANDK | 1436650 | Cell/Virus Enclosed | NO |
| FTEESFYKNMYLKSAV | 1658905 | Cell/Virus Enclosed | NO |
| FTEFVGTQSIDM | 1436657 | Cell/Virus Enclosed | NO |
| FTEHVTSVNLFD | 1436659 | Cell/Virus Enclosed | NO |
| FTEQLGAPLGIT | 1436665 | Ectodomain | YES |
| FTEQSVFKPQPA | 1436669 | Ectodomain | YES |
| FTEQSVFKPQPV | 1436670 | Ectodomain | YES |
| FTETITTGFDLN | 1436675 | Ectodomain | YES |
| FTFDIDGSRVPHIVRR | 1658916 | Cell/Virus Enclosed | NO |
| FTFDVEGSRVPHIVRK | 1658917 | Cell/Virus Enclosed | NO |
| FTFKQANNNCFVNVSC | 1658918 | Cell/Virus Enclosed | NO |
| FTGDLSYTIKFS | 1436700 | Cell/Virus Enclosed | NO |
| FTGSSYYYPEPISDKN | 1658932 | Ectodomain | YES |
| FTGSVYGNFDDQPSLQ | 1658933 | Cell/Virus Enclosed | NO |

| | | | |
|---|---|---|---|
| FTGTYKCPQTKS | 1436713 | Ectodomain | YES |
| FTGVDGELPVAVVNDK | 1658935 | Cell/Virus Enclosed | NO |
| FTGYFPKAGANF | 1436715 | Ectodomain | YES |
| FTGYHFTKNGKTVLGE | 1658938 | Cell/Virus Enclosed | NO |
| FTGYHFTSTGKTVLGE | 1658939 | Cell/Virus Enclosed | NO |
| FTGYHITKNSKVQLGE | 1658940 | Cell/Virus Enclosed | NO |
| FTHHDVVYAQHC | 1436719 | Ectodomain | YES |
| FTIAGHEEFIQT | 1436727 | Ectodomain | YES |
| FTIGNIEEFIQT | 1436730 | Ectodomain | YES |
| FTIGNMEEFIQT | 1436731 | Ectodomain | YES |
| FTIGNMVEFIQT | 1436733 | Ectodomain | YES |
| FTIVGQEEFIQTNSPK | 1658961 | Ectodomain | YES |
| FTKAAGGKVSFSDDVE | 1658963 | Cell/Virus Enclosed | NO |
| FTKKDWYDFVENPDII | 1658966 | Cell/Virus Enclosed | NO |
| FTKLAGGKISFSDDVI | 1658967 | Cell/Virus Enclosed | NO |
| FTKNGKTVLGEYVFDK | 1658970 | Cell/Virus Enclosed | NO |
| FTKPASRELSVT | 1436765 | Cell/Virus Enclosed | NO |
| FTKPSSVKEMQS | 1436768 | Cell/Virus Enclosed | NO |
| FTKSGVNDICMFENKT | 1658975 | Cell/Virus Enclosed | NO |
| FTKSVCGYTDFAEDVC | 1658977 | Cell/Virus Enclosed | NO |
| FTKVQGLENIAF | 1436776 | Cell/Virus Enclosed | NO |
| FTKVQGLENIVF | 1436778 | Cell/Virus Enclosed | NO |
| FTKVVVGPTTEVKFSV | 1658979 | Cell/Virus Enclosed | NO |
| FTKWTWETKGDL | 17998 | Ectodomain | YES |
| FTLAEVVKQMYGVNLQ | 1658980 | Cell/Virus Enclosed | NO |
| FTLANVSSFGDYNLSS | 1658981 | Ectodomain | YES |
| FTLCCTGCGTNC | 1436782 | Undetermined | UN |
| FTLESITAQPVK | 1436784 | Secreted | YES |
| FTLGIMAIAACA | 1436793 | Cell/Virus Enclosed | NO |
| FTLKNTVCTVCG | 1436810 | Cell/Virus Enclosed | NO |
| FTLQAAFGNTGV | 1436820 | Cell/Virus Enclosed | NO |
| FTLQLVSAVNQSLSNA | 1658998 | Cell/Virus Enclosed | NO |
| FTLRSITAQPVK | 1436825 | Secreted | YES |
| FTLYGQTVVHFG | 1436839 | Cell/Virus Enclosed | NO |
| FTMADGSLQPYC | 1436842 | Cell/Virus Enclosed | NO |
| FTNDKTIFYTTEIASK | 1659011 | Cell/Virus Enclosed | NO |
| FTNEPAFIMHVSWYIM | 1659013 | Cell/Virus Enclosed | NO |
| FTNFEPFTVNSK | 1436856 | Ectodomain | YES |
| FTNFEPFTVNSVNDSL | 1659014 | Ectodomain | YES |
| FTNGAVIGTPVC | 1436861 | Cell/Virus Enclosed | NO |
| FTNKTTLPTNVAFELF | 1659016 | Cell/Virus Enclosed | NO |
| FTNNNGSDVLYQPPTA | 1659019 | Cell/Virus Enclosed | NO |
| FTNRFVPVGDSF | 1436871 | Cell/Virus Enclosed | NO |
| FTNSILMLDKQGQLLD | 1659021 | Cell/Virus Enclosed | NO |
| FTNSTDTVNHDSFNLY | 1659022 | Cell/Virus Enclosed | NO |
| FTPEDVNMQIMGVVMQ | 1659028 | Cell/Virus Enclosed | NO |
| FTPEDVNMQLMG | 1436879 | Cell/Virus Enclosed | NO |
| FTPFDVVAQCSG | 1436880 | Cell/Virus Enclosed | NO |
| FTPGKQSSSRASSGNR | 1659030 | Cell/Virus Enclosed | NO |
| FTPKKPNKPLSQ | 1436885 | Cell/Virus Enclosed | NO |
| FTPLSVEPTPIE | 1436892 | Cell/Virus Enclosed | NO |
| FTPLVQPVGALD | 1436896 | Cell/Virus Enclosed | NO |
| FTPRSDMEYDFLNMDM | 1659037 | Cell/Virus Enclosed | NO |
| FTPRSVRSVPGE | 1436898 | Ectodomain | YES |
| FTPYHAFKCNRSGSND | 1659039 | Ectodomain | YES |
| FTQDGNAAIPDY | 1436912 | Cell/Virus Enclosed | NO |
| FTQDGNAAITDYNYYK | 1659043 | Cell/Virus Enclosed | NO |
| FTQKGDAAIKDFDYYR | 1659044 | Cell/Virus Enclosed | NO |
| FTQWYHEHIGGV | 1436921 | Cell/Virus Enclosed | NO |
| FTREMQQHPLLNPSAL | 1659052 | Capsid | UN |
| FTREMQQQPLLN | 1436927 | Cell/Virus Enclosed | NO |
| FTRNYIITMPCF | 1436935 | Cell/Virus Enclosed | NO |
| FTRQEEVHQELY | 1436937 | Cell/Virus Enclosed | NO |
| FTRVSAKPPPGD | 1436945 | Cell/Virus Enclosed | NO |
| FTSACTRLEGLG | 1436947 | Cell/Virus Enclosed | NO |
| FTSDQAITFLKE | 1436955 | Underneath Membrane | NO |
| FTSDQAVTFLKEWNFS | 1659068 | Undetermined | UN |
| FTSEPPITYSPA | 1436958 | Cell/Virus Enclosed | NO |
| FTSEPPVTYSPATILA | 1659072 | Cell/Virus Enclosed | NO |
| FTSEPPVTYSTA | 1436960 | Cell/Virus Enclosed | NO |
| FTSTGKTVLGEYVFDK | 1659080 | Cell/Virus Enclosed | NO |
| FTSYGFSNVVEM | 1436975 | Ectodomain | YES |
| FTTADGSPQPYC | 1436977 | Cell/Virus Enclosed | NO |
| FTTQFVDNVKLA | 1436996 | Ectodomain | YES |
| FTTTNEAFQKVQDAVN | 1659097 | Ectodomain | YES |
| FTVCADGFMPFL | 1437014 | Cell/Virus Enclosed | NO |

| | | | |
|---|---|---|---|
| FTVDGVQYYCYT | 1437015 | Cell/Virus Enclosed | NO |
| FTVDPQKAYLDFVNAG | 1659104 | Cell/Virus Enclosed | NO |
| FTVLAAYNGPPQ | 1437028 | Cell/Virus Enclosed | NO |
| FTVMNGEDAFSI | 1437030 | Cell/Virus Enclosed | NO |
| FTVNSVNDSLEH | 1437033 | Ectodomain | YES |
| FTVNSVNDSLEPVGGL | 1659111 | Ectodomain | YES |
| FTVNSVNDSLKP | 1437034 | Ectodomain | YES |
| FTVSTQPQENTI | 1437039 | Cell/Virus Enclosed | NO |
| FTVSTQPQGNII | 1437040 | Cell/Virus Enclosed | NO |
| FTVSTQPQGNTIPHYS | 1659113 | Capsid | UN |
| FTYICYMIRDKL | 1437056 | Cell/Virus Enclosed | NO |
| FVAAYNNCTGGA | 1437071 | Ectodomain | YES |
| FVAFNVFQGIET | 1437080 | Cell/Virus Enclosed | NO |
| FVAGNCDVVIGI | 1437085 | Ectodomain | YES |
| FVANYFQDVKQF | 1437090 | Ectodomain | YES |
| FVANYSQDVKQFANGF | 1659154 | Ectodomain | YES |
| FVAYETARQEYENAVA | 1659164 | Cell/Virus Enclosed | NO |
| FVCTDNSANTKVLRNK | 1659172 | Cell/Virus Enclosed | NO |
| FVDDFQGYLQKH | 1437132 | Cell/Virus Enclosed | NO |
| FVDDIVKTDGTL | 1437134 | Cell/Virus Enclosed | NO |
| FVDGVPFVVSYG | 1437143 | Cell/Virus Enclosed | NO |
| FVDHFGENEEIIINGH | 1659186 | Cell/Virus Enclosed | NO |
| FVDHFGENEEIV | 1437144 | Cell/Virus Enclosed | NO |
| FVDQYGCDYSGKLADC | 1659192 | Cell/Virus Enclosed | NO |
| FVDYSNLLHSKVKSVP | 1659206 | Cell/Virus Enclosed | NO |
| FVEAYNNCTGGAEIRD | 1659208 | Ectodomain | YES |
| FVEAYNNCTGGSEIRD | 1659209 | Ectodomain | YES |
| FVEDCKLPLGQSLCAL | 1659210 | Ectodomain | YES |
| FVEGQGVPIAPGVPATEAKG | 2060823 | Cell/Virus Enclosed | NO |
| FVEHYNEWAQANGFTA | 1659213 | Cell/Virus Enclosed | NO |
| FVELDDQGHHLD | 1437187 | Cell/Virus Enclosed | NO |
| FVENPDIINIYKKLGP | 1659219 | Cell/Virus Enclosed | NO |
| FVEPKDGGHFFV | 1437195 | Cell/Virus Enclosed | NO |
| FVEPKNGGQFFV | 1437197 | Cell/Virus Enclosed | NO |
| FVEYEVAKKNLADAKN | 1659231 | Cell/Virus Enclosed | NO |
| FVFDVDKTKTPH | 1437214 | Cell/Virus Enclosed | NO |
| FVFEKVDYGSDTVTYK | 1659235 | Cell/Virus Enclosed | NO |
| FVFNGTSWFITQRNFFS | 18161 | Ectodomain | YES |
| FVGADGELPVAISGDK | 1659248 | Cell/Virus Enclosed | NO |
| FVGGFRVLRTTD | 1437246 | Cell/Virus Enclosed | NO |
| FVGMPSFVAYETARQE | 1659257 | Cell/Virus Enclosed | NO |
| FVGTQSVDMLAVKTGV | 1659261 | Cell/Virus Enclosed | NO |
| FVHEDLHVVEVLTKTA | 1659264 | Cell/Virus Enclosed | NO |
| FVHYAGVVWTLQEVKD | 1659265 | Cell/Virus Enclosed | NO |
| FVIDTPTGPQIK | 1437275 | Cell/Virus Enclosed | NO |
| FVKCNCGSENWS | 1437293 | Cell/Virus Enclosed | NO |
| FVKDDYCDGSLGCKMC | 1659280 | Cell/Virus Enclosed | NO |
| FVKDNVSFVVDD | 1437299 | Cell/Virus Enclosed | NO |
| FVKNIPCYASAV | 1437321 | Cell/Virus Enclosed | NO |
| FVKNLNNLHRGQ | 1437323 | Cell/Virus Enclosed | NO |
| FVKQPVKFYLYK | 1437331 | Cell/Virus Enclosed | NO |
| FVKRRADFACKF | 1437332 | Cell/Virus Enclosed | NO |
| FVKSDIFGSDFKTFDL | 1659307 | Cell/Virus Enclosed | NO |
| FVKSKVGNYRLPSSKP | 1659311 | Underneath Membrane | NO |
| FVKWEYEGGCNT | 1437340 | Cell/Virus Enclosed | NO |
| FVLLFFICCCTG | 1437362 | Undetermined | UN |
| FVLLRFYIVVTA | 1437364 | Cell/Virus Enclosed | NO |
| FVLTSHTVMPLT | 1437384 | Cell/Virus Enclosed | NO |
| FVMQFGYIDCEQDSCD | 1659356 | Cell/Virus Enclosed | NO |
| FVMQYGYVDCFNDNCD | 1659357 | Cell/Virus Enclosed | NO |
| FVNAGGAPLTNCVKML | 1659359 | Cell/Virus Enclosed | NO |
| FVNAGGVPLTNC | 1437415 | Cell/Virus Enclosed | NO |
| FVNDGRVANGYVCGTG | 1659360 | Cell/Virus Enclosed | NO |
| FVNDKITEFQLDYSID | 1659361 | Cell/Virus Enclosed | NO |
| FVNDSIESVGGLYEIK | 1659362 | Ectodomain | YES |
| FVNDSYRQFDLVQYDF | 1659363 | Cell/Virus Enclosed | NO |
| FVNEYYEFLCKHFSMM | 1659365 | Cell/Virus Enclosed | NO |
| FVNFNVDWIFFG | 1437423 | Undetermined | UN |
| FVNGFYRLYSGDTFWR | 1659367 | Cell/Virus Enclosed | NO |
| FVNIAHASLREGVQLE | 1659369 | Cell/Virus Enclosed | NO |
| FVNISRSELQTIVPEY | 1659371 | Ectodomain | YES |
| FVNISRVELHTVIPDY | 1659372 | Ectodomain | YES |
| FVNKTSLPTNIAFELF | 1659373 | Cell/Virus Enclosed | NO |
| FVNMASFVEYEVAKKN | 1659378 | Cell/Virus Enclosed | NO |
| FVNSFVSLYNVTRDKL | 1659380 | Cell/Virus Enclosed | NO |
| FVNTTIGNETTF | 1437442 | Ectodomain | YES |

| | | | |
|---|---|---|---|
| FVNTTIGNETTSAFVG | 1659384 | Ectodomain | YES |
| FVNVNNTWMYTGSGYY | 1659386 | Ectodomain | YES |
| FVNVPYVYNGSAQATA | 1659387 | Ectodomain | YES |
| FVNVPYVYNGSV | 1437450 | Ectodomain | YES |
| FVNVSYFYNGSA | 1437452 | Ectodomain | YES |
| FVNVSYVYNGSA | 1437453 | Ectodomain | YES |
| FVPQPTGVFTNH | 1437465 | Ectodomain | YES |
| FVPSPGARPQVNGLSGRIDFHWLML | 1141600 | Ectodomain | YES |
| FVPVQVVDSRWN | 1437471 | Underneath Membrane | NO |
| FVQDPAPIDIDAFPVK | 1659398 | Cell/Virus Enclosed | NO |
| FVQDPSPIDIDA | 1437473 | Cell/Virus Enclosed | NO |
| FVQERVGLFIVN | 1437481 | Cell/Virus Enclosed | NO |
| FVQIENCDVTFV | 1437482 | Ectodomain | YES |
| FVQIPAQCVRDPVGFC | 1659401 | Cell/Virus Enclosed | NO |
| FVQIPTTCANDP | 1437486 | Cell/Virus Enclosed | NO |
| FVQNDVCSTEDFNVWA | 1659403 | Cell/Virus Enclosed | NO |
| FVQQKPEPVNPVSFVV | 1659404 | Cell/Virus Enclosed | NO |
| FVQSDKCSVEDFNVWA | 1659405 | Cell/Virus Enclosed | NO |
| FVQSNVDVVPEGWRVV | 1659407 | Cell/Virus Enclosed | NO |
| FVQTCDLTVGSG | 1437494 | Ectodomain | YES |
| FVRHIIVGCNNADCVA | 1659415 | Cell/Virus Enclosed | NO |
| FVRKAEDIFCPC | 1437519 | Cell/Virus Enclosed | NO |
| FVRKASGLFDAI | 1437521 | Cell/Virus Enclosed | NO |
| FVRPIIEDYHTL | 1437523 | Underneath Membrane | NO |
| FVRPVIEDYHTLTATV | 1659421 | Underneath Membrane | NO |
| FVRYGDVDNLVFTNKT | 1659423 | Cell/Virus Enclosed | NO |
| FVSAYNNCTGGA | 1437536 | Ectodomain | YES |
| FVSDADITLLGD | 1437538 | Cell/Virus Enclosed | NO |
| FVSEETGTLIVNSVL | 2001065 | Undetermined | UN |
| FVSQVLNKYTEVRGSR | 1659460 | Ectodomain | YES |
| FVSSYAKPEEKLSAYD | 1659463 | Cell/Virus Enclosed | NO |
| FVSYGSSVETETYFKV | 1659468 | Cell/Virus Enclosed | NO |
| FVTDTPKGPKMK | 1437585 | Cell/Virus Enclosed | NO |
| FVTDTPRGPKVK | 1437587 | Cell/Virus Enclosed | NO |
| FVTEYYEFLNKHFSMM | 1659477 | Cell/Virus Enclosed | NO |
| FVTFEPFNVSFVNDSI | 1659478 | Ectodomain | YES |
| FVVAFLATIKLC | 1437618 | Cell/Virus Enclosed | NO |
| FVVDKSLSIEEFADVV | 1659499 | Cell/Virus Enclosed | NO |
| FVVESGGYIPSN | 1437626 | Ectodomain | YES |
| FVVKEKLNAFLVHDNV | 1659508 | Cell/Virus Enclosed | NO |
| FVVNAANENLLHGGGV | 1659520 | Cell/Virus Enclosed | NO |
| FVVQPVGVVDTEWGNM | 1659525 | Cell/Virus Enclosed | NO |
| FVVQTEGCVSTN | 1437650 | Cell/Virus Enclosed | NO |
| FVVRIGAAANAT | 1437653 | Ectodomain | YES |
| FVYDAYHNLVGY | 1437684 | Ectodomain | YES |
| FVYDAYQNLVGYYSDD | 1659538 | Ectodomain | YES |
| FVYFNATGSDVNCNGY | 1659543 | Ectodomain | YES |
| FVYFNGTGRGAC | 1437696 | Ectodomain | YES |
| FVYSSNEEQAVLKFLD | 1659548 | Cell/Virus Enclosed | NO |
| FVYTDTEVCKVK | 1437701 | Cell/Virus Enclosed | NO |
| FVYVIKMIILWL | 1437707 | Transmembrane | NO |
| FVYVRKDGKHVD | 1437711 | Cell/Virus Enclosed | NO |
| FVYYVARLMKGD | 1437712 | Cell/Virus Enclosed | NO |
| FWAWNPEVNAIT | 1437717 | Underneath Membrane | NO |
| FWFAMRKDGDDVIFSR | 1659566 | Cell/Virus Enclosed | NO |
| FWNCNVDRYPAN | 1437743 | Cell/Virus Enclosed | NO |
| FWRYDFDITESK | 1437757 | Cell/Virus Enclosed | NO |
| FWSFNPETNNLMCIDM | 1659593 | Ectodomain | YES |
| FWSSTMIFSLHP | 1437759 | Cell/Virus Enclosed | NO |
| FWTLFCTSVNTSSSEA | 1659594 | Cell/Virus Enclosed | NO |
| FWTMFCTSVNTSSSEA | 1659595 | Cell/Virus Enclosed | NO |
| FWTVFCTNANASSSEG | 1659597 | Cell/Virus Enclosed | NO |
| FWTVFCTNVNASSSEG | 1659598 | Cell/Virus Enclosed | NO |
| FWVANHQADTSTPSDV | 1659599 | Capsid | UN |
| FWVASNQADVNI | 1437763 | Cell/Virus Enclosed | NO |
| FWVASNQADVNT | 1437764 | Cell/Virus Enclosed | NO |
| FWVRRRAQMAPKRLRLPHIR | 741438 | Undetermined | UN |
| FYAEGSRGGSQASSRSSSRS | 1392162 | Capsid | UN |
| FYAPEPITSFNT | 1437783 | Ectodomain | YES |
| FYAPEPITSLNTKYVA | 1659609 | Ectodomain | YES |
| FYASNGTYNCTDAVLT | 1659616 | Ectodomain | YES |
| FYCFVNTTIGNETTSA | 1659622 | Ectodomain | YES |
| FYCSNSSSGVLDTTIP | 1659624 | Ectodomain | YES |
| FYCTNNTLVSGDAHIP | 1659626 | Ectodomain | YES |
| FYCTNTTLVSGD | 1437812 | Ectodomain | YES |
| FYDRDGQRIYDD | 1437822 | Cell/Virus Enclosed | NO |

| | | | |
|---|---|---|---|
| FYDRDGQRTYDDVNAS | 1659636 | Cell/Virus Enclosed | NO |
| FYDRDGQRVYDDVDAS | 1659637 | Cell/Virus Enclosed | NO |
| FYEFYNDIKPPV | 1437835 | Cell/Virus Enclosed | NO |
| FYEFYNDVKPPVLDVD | 1659644 | Cell/Virus Enclosed | NO |
| FYEGVNFSPHHR | 1437838 | Ectodomain | YES |
| FYELDKDFNTIL | 1437840 | Cell/Virus Enclosed | NO |
| FYELDNDFNTILNTAC | 1659647 | Cell/Virus Enclosed | NO |
| FYENRFAAIYGG | 1437842 | Cell/Virus Enclosed | NO |
| FYEPEPITSLNT | 1437843 | Ectodomain | YES |
| FYFAVRKEGEDV | 1437850 | Cell/Virus Enclosed | NO |
| FYFAVRKEGQDVIFSQ | 1659652 | Cell/Virus Enclosed | NO |
| FYFDDSDCDVVQEQVN | 1659654 | Cell/Virus Enclosed | NO |
| FYFKPGGTTSGE | 1437861 | Cell/Virus Enclosed | NO |
| FYGGWDNMLKNLMADV | 1659675 | Cell/Virus Enclosed | NO |
| FYGKESFENPTY | 1437881 | Cell/Virus Enclosed | NO |
| FYGKESLENPTY | 1437883 | Cell/Virus Enclosed | NO |
| FYGVALKDCGFV | 1437892 | Cell/Virus Enclosed | NO |
| FYGWVSGNMMDGFSCP | 1659685 | Cell/Virus Enclosed | NO |
| FYIEDMASRFAD | 1437904 | Cell/Virus Enclosed | NO |
| FYINGYRYFSLG | 1437917 | Ectodomain | YES |
| FYIYDEEGGYDVSKPV | 1659696 | Cell/Virus Enclosed | NO |
| FYKKDGVYYPSNGTNI | 1659704 | Cell/Virus Enclosed | NO |
| FYKNMYLKSAVMQSVG | 1659705 | Cell/Virus Enclosed | NO |
| FYLGDISHLINC | 1437948 | Cell/Virus Enclosed | NO |
| FYLGDISHLVNC | 1437949 | Cell/Virus Enclosed | NO |
| FYLLGCDEYIVP | 1437954 | Ectodomain | YES |
| FYLTMYIIPVLC | 1437964 | Cell/Virus Enclosed | NO |
| FYMEADTVVNAFYGVA | 1659724 | Cell/Virus Enclosed | NO |
| FYNDIKPPVLDV | 1437976 | Cell/Virus Enclosed | NO |
| FYNLWSTFTKLQ | 1437986 | Cell/Virus Enclosed | NO |
| FYPACTNGHALP | 1437997 | Cell/Virus Enclosed | NO |
| FYPACTNGHAVP | 1437998 | Cell/Virus Enclosed | NO |
| FYPNVPIEYQAT | 1438010 | Ectodomain | YES |
| FYPPDDEDECDE | 1438011 | Cell/Virus Enclosed | NO |
| FYPPDEEEDCEE | 1438013 | Cell/Virus Enclosed | NO |
| FYPRLQAASDWKPGYS | 1659747 | Cell/Virus Enclosed | NO |
| FYQGDVDTVVNGVDFD | 1659753 | Cell/Virus Enclosed | NO |
| FYQLYDASNVKK | 1438034 | Cell/Virus Enclosed | NO |
| FYRLANECAQVLSEYV | 1659761 | Cell/Virus Enclosed | NO |
| FYRLANGCAQVL | 1438050 | Cell/Virus Enclosed | NO |
| FYRLSNELAQVLTEVV | 1659764 | Cell/Virus Enclosed | NO |
| FYRPANECAQVL | 1438056 | Cell/Virus Enclosed | NO |
| FYSANFLDDNVLPETY | 1659768 | Ectodomain | YES |
| FYSPEPITSLNT | 1438080 | Ectodomain | YES |
| FYSPQIITTDNT | 1438081 | Ectodomain | YES |
| FYTTSGIGYQPY | 1438106 | Ectodomain | YES |
| FYVKDGKYFISE | 1438114 | Cell/Virus Enclosed | NO |
| FYVKDGKYFTSA | 1438115 | Cell/Virus Enclosed | NO |
| FYVKDGKYFTSEPPVT | 1659791 | Cell/Virus Enclosed | NO |
| FYVRVKHGDFSAVSSP | 1659804 | Underneath Membrane | NO |
| FYVVKNSTAFPFETLS | 1659806 | Cell/Virus Enclosed | NO |
| FYVYDVHKDNFKDGLC | 1659809 | Cell/Virus Enclosed | NO |
| FYYLGTGPYANA | 1438162 | Cell/Virus Enclosed | NO |
| GAAAINSVPWSK | 1438175 | Cell/Virus Enclosed | NO |
| GAAALEDEATPF | 1438177 | Cell/Virus Enclosed | NO |
| GAAIHYLNSKIVDLAQ | 1659836 | Cell/Virus Enclosed | NO |
| GAALAMEKVNECVKSQ | 1659837 | Ectodomain | YES |
| GACDYDRLPNEVTVAK | 1659839 | Ectodomain | YES |
| GACGSPGYNLKNGEVE | 1659841 | Cell/Virus Enclosed | NO |
| GACGSPGYNVRNDGTV | 1659842 | Cell/Virus Enclosed | NO |
| GACIRRPFLCCK | 1438197 | Cell/Virus Enclosed | NO |
| GACIWNAAVYMK | 1438200 | Cell/Virus Enclosed | NO |
| GACKGFYSNSSS | 1438202 | Ectodomain | YES |
| GACSCGSCCKFDEDDS | 1659848 | Underneath Membrane | NO |
| GADGELPVAISGDKVF | 1659851 | Cell/Virus Enclosed | NO |
| GADISDTIPDEKQNGH | 1659853 | Cell/Virus Enclosed | NO |
| GADSSIAYSNNTIAIP | 1659857 | Ectodomain | YES |
| GAEHVDTSYDCD | 1438227 | Ectodomain | YES |
| GAEIKDLICVQS | 1438232 | Ectodomain | YES |
| GAEIRDLICVQS | 1438233 | Ectodomain | YES |
| GAEKEDGFYETI | 1438235 | Cell/Virus Enclosed | NO |
| GAEMTSNVRRQVASGV | 1659867 | Cell/Virus Enclosed | NO |
| GAENKDGFYETI | 1438239 | Cell/Virus Enclosed | NO |
| GAEYVDTSYECD | 1438244 | Ectodomain | YES |
| GAFFFGSKLDLVKRDS | 1659871 | Capsid | UN |
| GAFFLYDRLAST | 852598 | Secreted | YES |

| | | | |
|---|---|---|---|
| GAFGPGFTPPHGGLL | 1659873 | Underneath Membrane | NO |
| GAFHVTMRSSYTIKGS | 1659874 | Cell/Virus Enclosed | NO |
| GAFLNGSCGSVG | 1438260 | Cell/Virus Enclosed | NO |
| GAFTRYVDNNFC | 1438266 | Cell/Virus Enclosed | NO |
| GAGCFVNDLLKT | 1438271 | Cell/Virus Enclosed | NO |
| GAGIRDLICVQS | 1438291 | Ectodomain | YES |
| GAGKALTGLSTGTSQNTRIS | 18595 | Capsid | UN |
| GAGLKLPSGIMF | 1438292 | Cell/Virus Enclosed | NO |
| GAGNGQAITTSVDANT | 1659891 | Cell/Virus Enclosed | NO |
| GAGSDKGIAPGTSVLR | 1659894 | Cell/Virus Enclosed | NO |
| GAGSDKGVAPGTTVLK | 1659896 | Cell/Virus Enclosed | NO |
| GAGSEKGVAPGSAVLR | 1659898 | Cell/Virus Enclosed | NO |
| GAGVMLECEKFNVFNV | 1659900 | Cell/Virus Enclosed | NO |
| GAGWTAGLSSSA | 1438307 | Ectodomain | YES |
| GAHAFRNACGTN | 1438309 | Cell/Virus Enclosed | NO |
| GAHATRDSIGTNFPLQ | 1659902 | Cell/Virus Enclosed | NO |
| GAHATRENIGTNFPLQ | 1659903 | Cell/Virus Enclosed | NO |
| GAIGDVGLCMYR | 1438325 | Cell/Virus Enclosed | NO |
| GAIGSSLQEILS | 1438326 | Ectodomain | YES |
| GAIIRDGDVVKQ | 1438327 | Cell/Virus Enclosed | NO |
| GAIIRDSDVVKQPVKF | 1659913 | Cell/Virus Enclosed | NO |
| GAIIRNSDVVKQ | 1438332 | Cell/Virus Enclosed | NO |
| GAIKLDDKDPSFKDNV | 1659916 | Cell/Virus Enclosed | NO |
| GAINAASKGAVQKESD | 1659921 | Cell/Virus Enclosed | NO |
| GAIRSDSTLSGF | 1438349 | Capsid | UN |
| GAISASIGDIIQRLDV | 1659924 | Ectodomain | YES |
| GAISSSLQEILS | 1438357 | Ectodomain | YES |
| GAITTGYRFTNFEPFT | 1659928 | Ectodomain | YES |
| GAKETKEINIIKLSGV | 1659931 | Cell/Virus Enclosed | NO |
| GAKFGRWSASIN | 1438375 | Ectodomain | YES |
| GAKSFYKNLIWLVKKGNSYPKLSKS | 1141843 | Ectodomain | YES |
| GALAEHDFFTWK | 1438399 | Cell/Virus Enclosed | NO |
| GALDTTIPFGPS | 1438403 | Ectodomain | YES |
| GALDVSASVVAG | 1438406 | Cell/Virus Enclosed | NO |
| GALHPYFDKAGQ | 1438413 | Cell/Virus Enclosed | NO |
| GALLVDSDLNDF | 1438417 | Cell/Virus Enclosed | NO |
| GALNTPKDHIGTRNPNNNA | 18671 | Capsid | UN |
| GAMLSENFTSYGFSNV | 1659970 | Ectodomain | YES |
| GAMQTGFTTTNEAFQK | 1659972 | Ectodomain | YES |
| GAMTAVNESRYGLQNL | 1659973 | Ectodomain | YES |
| GANFRDLSLKGTTYLS | 1659975 | Ectodomain | YES |
| GANGNKIVDKEVKDMQ | 1659976 | Cell/Virus Enclosed | NO |
| GANLRQDDFVRN | 1438462 | Ectodomain | YES |
| GANLRQDDSVRNLFAS | 1659979 | Ectodomain | YES |
| GAPDDASRLLHT | 1438473 | Cell/Virus Enclosed | NO |
| GAPDNPSAIVTG | 1438475 | Ectodomain | YES |
| GAPLGITISGET | 1438482 | Ectodomain | YES |
| GAPTKGVTFGED | 1438493 | Cell/Virus Enclosed | NO |
| GAPVKKVAFCGD | 1438499 | Cell/Virus Enclosed | NO |
| GAPVKKVAFGGDQVHE | 1659990 | Cell/Virus Enclosed | NO |
| GAQAVARSLDWTKVTLDGR | 18736 | Undetermined | UN |
| GAQGLKKFYKEV | 1438503 | Cell/Virus Enclosed | NO |
| GARGTYRAALNSEKHQ | 1659994 | Cell/Virus Enclosed | NO |
| GARIPDVPTTLAWVNN | 1659995 | Cell/Virus Enclosed | NO |
| GARLFASDLSTL | 1438515 | Cell/Virus Enclosed | NO |
| GARLFSSDLSTL | 1438516 | Cell/Virus Enclosed | NO |
| GARLSASDLSTL | 1438517 | Cell/Virus Enclosed | NO |
| GARPKQRRPQGLPNNT | 1660001 | Cell/Virus Enclosed | NO |
| GASLRQIFDVAP | 1438536 | Cell/Virus Enclosed | NO |
| GASPGPSGVGAP | 1438537 | Cell/Virus Enclosed | NO |
| GASQRVAGDSGFAAYS | 1380881 | Underneath Membrane | NO |
| GASRAIEKVNEC | 1438541 | Ectodomain | YES |
| GASSNQSPQGNL | 1438543 | Cell/Virus Enclosed | NO |
| GASTEIQQVDEE | 1438544 | Cell/Virus Enclosed | NO |
| GASVTEDVKFAASTGV | 1660011 | Cell/Virus Enclosed | NO |
| GATIRLQAGKQTELAV | 1660015 | Cell/Virus Enclosed | NO |
| GATIRLQTGKQT | 1438559 | Cell/Virus Enclosed | NO |
| GATVRLQAGKQTEFVS | 1660020 | Cell/Virus Enclosed | NO |
| GAVCLKHAEEYREYLE | 1660023 | Cell/Virus Enclosed | NO |
| GAVCRPHANEYR | 1438573 | Cell/Virus Enclosed | NO |
| GAVCSKHANLYQKYVE | 1660028 | Cell/Virus Enclosed | NO |
| GAVIGTPVCVNG | 1438592 | Cell/Virus Enclosed | NO |
| GAVILRGHLRIAGHH | 1312531 | Underneath Membrane | NO |
| GAVLFCTPTKKAFPYG | 1660038 | Cell/Virus Enclosed | NO |
| GAVQKESDEYILAKGP | 1660042 | Cell/Virus Enclosed | NO |
| GAVSCGHYQTNIYSQN | 1660043 | Cell/Virus Enclosed | NO |

| | | | |
|---|---|---|---|
| GAVSCLYKRNFT | 1438607 | Ectodomain | YES |
| GAVVGTPVCING | 1438616 | Cell/Virus Enclosed | NO |
| GAVVGTPVCVNG | 1438617 | Cell/Virus Enclosed | NO |
| GAWDGYLSSCCGTPAK | 1660045 | Cell/Virus Enclosed | NO |
| GAYNKDHGCGGF | 1438633 | Cell/Virus Enclosed | NO |
| GAYSLFDMAKFPLKLA | 1660050 | Cell/Virus Enclosed | NO |
| GCACDRTTIQSVDISY | 1660053 | Cell/Virus Enclosed | NO |
| GCAGYTLEEART | 1438650 | Cell/Virus Enclosed | NO |
| GCAMPIMANLVN | 1438655 | Ectodomain | YES |
| GCAQGVFGVNMRTNWT | 1660056 | Cell/Virus Enclosed | NO |
| GCCDATYFENKL | 1438662 | Cell/Virus Enclosed | NO |
| GCCDCGSTKLPYYEFE | 1660058 | Underneath Membrane | NO |
| GCCDDYTGYQELVIKT | 1660059 | Underneath Membrane | NO |
| GCCDDYTGYQES | 1438666 | Underneath Membrane | NO |
| GCCDNSYFDSKGWYDP | 1660060 | Cell/Virus Enclosed | NO |
| GCCDVTYFENKL | 1438668 | Cell/Virus Enclosed | NO |
| GCCESTKLPYYDVEKI | 1660062 | Underneath Membrane | NO |
| GCCNCLTSSMRG | 1438673 | Underneath Membrane | NO |
| GCCNTDYFEMKN | 1438674 | Cell/Virus Enclosed | NO |
| GCCSTDYFEMKN | 1438677 | Cell/Virus Enclosed | NO |
| GCDVADVTKLYL | 1438683 | Cell/Virus Enclosed | NO |
| GCDVNDVTKLYLGGMS | 1660071 | Cell/Virus Enclosed | NO |
| GCDYSGKLADCLQAYG | 1660073 | Cell/Virus Enclosed | NO |
| GCEEGTMYFVPRAKSV | 1660078 | Cell/Virus Enclosed | NO |
| GCEQSYFTKKDW | 1438694 | Cell/Virus Enclosed | NO |
| GCETSNVPPIVS | 1438696 | Cell/Virus Enclosed | NO |
| GCFVNDLLKTDS | 1438706 | Cell/Virus Enclosed | NO |
| GCGASYTANDVEVQSS | 1660087 | Cell/Virus Enclosed | NO |
| GCGTNCMGKLKCNRCC | 1660092 | Underneath Membrane | NO |
| GCGVSDVTKLYLGGMS | 1660098 | Cell/Virus Enclosed | NO |
| GCHATRDAVGTN | 1438737 | Cell/Virus Enclosed | NO |
| GCHTGTAFNGDF | 1438742 | Cell/Virus Enclosed | NO |
| GCIKACEVVVTNLNKS | 1660106 | Cell/Virus Enclosed | NO |
| GCILPSECTMFK | 1438748 | Cell/Virus Enclosed | NO |
| GCIPASQVIVNNYDKS | 1660110 | Cell/Virus Enclosed | NO |
| GCITSREVVVTNLNKS | 1660113 | Cell/Virus Enclosed | NO |
| GCKAIVKETNFI | 1438755 | Cell/Virus Enclosed | NO |
| GCKYVVRTANAL | 1438767 | Cell/Virus Enclosed | NO |
| GCLFCYKRNCSV | 1438770 | Cell/Virus Enclosed | NO |
| GCLGSDCNRVSS | 1438776 | Ectodomain | YES |
| GCLGSECNKVSS | 1438777 | Ectodomain | YES |
| GCLIGAEYVDTS | 1438786 | Ectodomain | YES |
| GCLNASEVVVNNLDKS | 1660130 | Cell/Virus Enclosed | NO |
| GCLSSMNIAANSSEWA | 1660131 | Cell/Virus Enclosed | NO |
| GCMLRACIKTAQADTA | 1660134 | Cell/Virus Enclosed | NO |
| GCMLVLTVTLQN | 1438796 | Cell/Virus Enclosed | NO |
| GCNNADCVACSK | 1438805 | Cell/Virus Enclosed | NO |
| GCNPNGWTMGLF | 1438807 | Cell/Virus Enclosed | NO |
| GCNVNDVTKLYLGGLN | 1660152 | Cell/Virus Enclosed | NO |
| GCQSWTCPYTGAVSSPYWKFQHDVN | 18895 | Ectodomain | YES |
| GCQTTNGLNTSH | 1438816 | Ectodomain | YES |
| GCQTTNGLNTSYSVCN | 1660155 | Ectodomain | YES |
| GCQTTNGTNTIH | 1438820 | Ectodomain | YES |
| GCQTTNGTNTSN | 1438822 | Ectodomain | YES |
| GCRWWLCSTRVN | 1438824 | Cell/Virus Enclosed | NO |
| GCSTIAQAVRRYSEAA | 1660163 | Cell/Virus Enclosed | NO |
| GCSVGNFSDGKM | 1438841 | Ectodomain | YES |
| GCSVIDLLLGDF | 1438842 | Cell/Virus Enclosed | NO |
| GCTCDRTAIQSFDNSY | 1660165 | Cell/Virus Enclosed | NO |
| GCTGSWNKVTQI | 1438848 | Cell/Virus Enclosed | NO |
| GCVADQSRKWTG | 1438855 | Cell/Virus Enclosed | NO |
| GCVFSYVGCYNK | 1438864 | Cell/Virus Enclosed | NO |
| GCVGHSENVFVV | 1438867 | Ectodomain | YES |
| GCVIAWNTAQQD | 1438874 | Ectodomain | YES |
| GCVLNAVNLTSY | 1438884 | Ectodomain | YES |
| GCVPLSAIPALAANTL | 1660189 | Cell/Virus Enclosed | NO |
| GCVSTNFGDVNK | 1438894 | Cell/Virus Enclosed | NO |
| GCVTSNVPPIIS | 1438896 | Cell/Virus Enclosed | NO |
| GCVTSNVPPIVS | 1438897 | Cell/Virus Enclosed | NO |
| GCVTSNVPSLVS | 1438898 | Cell/Virus Enclosed | NO |
| GCVTSNVPTIVS | 1438899 | Cell/Virus Enclosed | NO |
| GCVTSNVSPIVN | 1438900 | Cell/Virus Enclosed | NO |
| GCVTSNVSPIVS | 1438901 | Cell/Virus Enclosed | NO |
| GCVVDQSRKWIG | 1438902 | Cell/Virus Enclosed | NO |
| GCVVNADNSTAS | 1438904 | Ectodomain | YES |
| GCVVNAYNNTAI | 1438905 | Ectodomain | YES |

| | | | |
|---|---|---|---|
| GCVVNAYNSTAI | 1438906 | Ectodomain | YES |
| GCVVNAYNSTSI | 1438907 | Ectodomain | YES |
| GCWRHSVACDYL | 1438911 | Cell/Virus Enclosed | NO |
| GCYYVKPGGTSSGDAT | 1660202 | Cell/Virus Enclosed | NO |
| GDAAIKDFDYYRYNRP | 1660203 | Cell/Virus Enclosed | NO |
| GDAALALLLLDRLNAL | 1660204 | Cell/Virus Enclosed | NO |
| GDAALALLLLDRLNQL | 1074903 | Cell/Virus Enclosed | NO |
| GDAAVKDFDFYRYNKP | 1660207 | Cell/Virus Enclosed | NO |
| GDAEDTLTKLSKYLAN | 1660208 | Cell/Virus Enclosed | NO |
| GDAEEALSKLSEYLIS | 1660209 | Cell/Virus Enclosed | NO |
| GDAFWSSTMIFS | 1438927 | Cell/Virus Enclosed | NO |
| GDAGFTSILIGL | 1438929 | Ectodomain | YES |
| GDAHEGIFWVAS | 1438932 | Cell/Virus Enclosed | NO |
| GDAHIPSGTVLG | 1438935 | Ectodomain | YES |
| GDAMHANYIFWR | 1438945 | Cell/Virus Enclosed | NO |
| GDASWSSTMILS | 1438953 | Cell/Virus Enclosed | NO |
| GDASWYSTMIFS | 1438954 | Cell/Virus Enclosed | NO |
| GDATTAYANSVL | 1438957 | Cell/Virus Enclosed | NO |
| GDCKGFSSNVSS | 1438968 | Ectodomain | YES |
| GDCKGFYSNASS | 1438969 | Ectodomain | YES |
| GDCQGFYSNASS | 1438976 | Ectodomain | YES |
| GDCVVEASTDFH | 1438983 | Cell/Virus Enclosed | NO |
| GDDCFVSNSSVIQEVL | 1660239 | Cell/Virus Enclosed | NO |
| GDDDDSQPRADK | 1438986 | Cell/Virus Enclosed | NO |
| GDDVDDIDTINC | 1438993 | Cell/Virus Enclosed | NO |
| GDDVRQIAPGQTGVIA | 1660245 | Ectodomain | YES |
| GDEVYLPYPNPS | 1439012 | Cell/Virus Enclosed | NO |
| GDFAACKSQLVE | 1439013 | Ectodomain | YES |
| GDFEQVAPGCGV | 1439015 | Cell/Virus Enclosed | NO |
| GDFHLNEEVAII | 1439016 | Cell/Virus Enclosed | NO |
| GDFMQGPFIAGN | 1439029 | Cell/Virus Enclosed | NO |
| GDFNCTNFAINDLNTT | 1660270 | Ectodomain | YES |
| GDFNCTNSFIND | 1439034 | Ectodomain | YES |
| GDFNCTNSFINH | 1439035 | Ectodomain | YES |
| GDFQLNEEVAII | 1439041 | Cell/Virus Enclosed | NO |
| GDFQLTEEVAII | 1439042 | Cell/Virus Enclosed | NO |
| GDFSAVSSPMSNMTEN | 1660272 | Underneath Membrane | NO |
| GDFSHGQLGGLH | 1439044 | Cell/Virus Enclosed | NO |
| GDFTSTIITSCQ | 1439050 | Cell/Virus Enclosed | NO |
| GDFVVTQPGSGV | 1439057 | Cell/Virus Enclosed | NO |
| GDGEHLEGGDGTR | 18985 | Ectodomain | YES |
| GDGFVPFLLDGLIPRS | 1660288 | Cell/Virus Enclosed | NO |
| GDGGITSEGNALYNNE | 1660289 | Cell/Virus Enclosed | NO |
| GDGISTPKLKED | 1439065 | Secreted | YES |
| GDGITGVPKPVE | 1439066 | Ectodomain | YES |
| GDGIYFAATEKS | 1439069 | Ectodomain | YES |
| GDGKMKKLSPRW | 1439072 | Cell/Virus Enclosed | NO |
| GDGLIDLLINVL | 1439074 | Cell/Virus Enclosed | NO |
| GDGLIDLLVNVL | 1439075 | Cell/Virus Enclosed | NO |
| GDGNALYNTEGGKTFM | 1660295 | Cell/Virus Enclosed | NO |
| GDGVYFAATEKS | 1439084 | Ectodomain | YES |
| GDHGDMYVYSSG | 1439088 | Ectodomain | YES |
| GDHGDRYVYSAG | 1439089 | Ectodomain | YES |
| GDIARELGNVVK | 1439093 | Cell/Virus Enclosed | NO |
| GDICNTMHYTNWTHIYICEE | 110589 | Cell/Virus Enclosed | NO |
| GDIGIAPKQGYF | 1439095 | Ectodomain | YES |
| GDIIQRLDVLEQDAQI | 1660307 | Ectodomain | YES |
| GDINARDLICAQKFNG | 1660309 | Ectodomain | YES |
| GDINVIAASGSL | 1439105 | Cell/Virus Enclosed | NO |
| GDINVVAASGSL | 1439106 | Cell/Virus Enclosed | NO |
| GDIVSGYEKSTFNKDI | 1660314 | Cell/Virus Enclosed | NO |
| GDIVTHLKNWNF | 1439121 | Ectodomain | YES |
| GDKGDAEDTLTKLSKY | 1660324 | Cell/Virus Enclosed | NO |
| GDKLSVEHYNEW | 1439131 | Cell/Virus Enclosed | NO |
| GDKVFVRDGNTDNLVF | 1660330 | Cell/Virus Enclosed | NO |
| GDKVVYHTVGSP | 1439136 | Cell/Virus Enclosed | NO |
| GDLAVCLGIGDSAVTY | 1660335 | Cell/Virus Enclosed | NO |
| GDLAVCLGVGDS | 1439141 | Cell/Virus Enclosed | NO |
| GDLAVQIGNNNVCTYE | 1660336 | Cell/Virus Enclosed | NO |
| GDLAVQIGSNNVCTYE | 1660337 | Cell/Virus Enclosed | NO |
| GDLAVQMGSNNV | 1439143 | Cell/Virus Enclosed | NO |
| GDLFYTIKFSGFKLDG | 1660338 | Cell/Virus Enclosed | NO |
| GDLKCPLDSRTG | 1439149 | Ectodomain | YES |
| GDLKCPLDTSLK | 1439151 | Ectodomain | YES |
| GDLKCPLESRTG | 1439155 | Ectodomain | YES |
| GDLKCTSDNINDKDTG | 1660339 | Ectodomain | YES |

| | | | |
|---|---|---|---|
| GDLKCTTVAIND | 1439158 | Ectodomain | YES |
| GDLNCPLDPKLK | 1439162 | Ectodomain | YES |
| GDLNCPLDSRLK | 1439164 | Ectodomain | YES |
| GDLNCPLDTRLK | 1439165 | Ectodomain | YES |
| GDLNCTLDPRLK | 1439166 | Ectodomain | YES |
| GDLNCTLHPRLK | 1439168 | Ectodomain | YES |
| GDLPENDDDAWF | 1439170 | Cell/Virus Enclosed | NO |
| GDLQFNKLGTED | 1439171 | Cell/Virus Enclosed | NO |
| GDLQGNFGDLKF | 1439172 | Cell/Virus Enclosed | NO |
| GDLQGNFGDLQF | 1439173 | Cell/Virus Enclosed | NO |
| GDLQGNFGDLQLNKLG | 1660343 | Cell/Virus Enclosed | NO |
| GDLQLNKLGTEDPRWP | 1660344 | Cell/Virus Enclosed | NO |
| GDLQLTKLGIED | 1439176 | Cell/Virus Enclosed | NO |
| GDLTMVITQDCDTCH | 227198 | Secreted | YES |
| GDLVDSYFENDSILQS | 1660348 | Cell/Virus Enclosed | NO |
| GDNDDEDNKDEE | 1439194 | Cell/Virus Enclosed | NO |
| GDNDDEDVVTGDNDDE | 1660351 | Cell/Virus Enclosed | NO |
| GDNDDEDVVTGDNNDE | 1660352 | Cell/Virus Enclosed | NO |
| GDNDDEEIVTGD | 1439197 | Cell/Virus Enclosed | NO |
| GDNDDQIVVTGDDVDD | 1660353 | Cell/Virus Enclosed | NO |
| GDNGTITVEELK | 1439203 | Ectodomain | YES |
| GDNIGTNVPLQVGFSN | 1660355 | Cell/Virus Enclosed | NO |
| GDNNDEEIVTGDNDDQ | 1660357 | Cell/Virus Enclosed | NO |
| GDNNGEDVVTGD | 1439211 | Cell/Virus Enclosed | NO |
| GDNSAKFWEEAFYRDL | 1660358 | Cell/Virus Enclosed | NO |
| GDNSWRYDFDIT | 1439216 | Cell/Virus Enclosed | NO |
| GDNVGHYVHVKC | 1439219 | Cell/Virus Enclosed | NO |
| GDNVGTNVPLQVGFSN | 1660360 | Cell/Virus Enclosed | NO |
| GDNVQITHTYKM | 1439220 | Cell/Virus Enclosed | NO |
| GDNVYCYNTDLIEGSK | 1660362 | Cell/Virus Enclosed | NO |
| GDNYLRCFLLYF | 1439223 | Cell/Virus Enclosed | NO |
| GDPAQLPAPRVLLSKG | 1660368 | Cell/Virus Enclosed | NO |
| GDPAQLPTPRTL | 1439227 | Cell/Virus Enclosed | NO |
| GDPVQLPAPRTL | 1439231 | Cell/Virus Enclosed | NO |
| GDQDLIRQGIDY | 1439235 | Capsid | UN |
| GDQELIRQGTDYKHWP | 1660372 | Cell/Virus Enclosed | NO |
| GDQELIRQGTEYKHWP | 1660373 | Cell/Virus Enclosed | NO |
| GDQFKHLIPLMY | 1439239 | Cell/Virus Enclosed | NO |
| GDQIHEVAAVRS | 1439241 | Cell/Virus Enclosed | NO |
| GDRCIKISTVQSKLTD | 1660378 | Cell/Virus Enclosed | NO |
| GDRGIAPKSGYFVNVN | 1660381 | Ectodomain | YES |
| GDRSGARPKQRRPQGL | 1660383 | Cell/Virus Enclosed | NO |
| GDSCYNFMLTYN | 1439273 | Cell/Virus Enclosed | NO |
| GDSEAVLKKLKK | 1439275 | Cell/Virus Enclosed | NO |
| GDSGELLSELLK | 1439279 | Cell/Virus Enclosed | NO |
| GDSGFAAYSRYRIGNY | 1380962 | Underneath Membrane | NO |
| GDSGKLLSELLKEKYT | 1660398 | Cell/Virus Enclosed | NO |
| GDSIFKSYHFTR | 1439289 | Ectodomain | YES |
| GDSIFTGCTGSW | 1439290 | Cell/Virus Enclosed | NO |
| GDSLECSNLLLQ | 1439292 | Ectodomain | YES |
| GDSQECSNLLLQ | 1439293 | Ectodomain | YES |
| GDSSESGAKETKEINI | 1660407 | Cell/Virus Enclosed | NO |
| GDSSNTGAGKALTGLSTGTS | 19119 | Capsid | UN |
| GDSSWHIYLKSGTCPF | 1660409 | Ectodomain | YES |
| GDSVEEALAEAR | 1439310 | Cell/Virus Enclosed | NO |
| GDSVFTGCTGSW | 1439313 | Cell/Virus Enclosed | NO |
| GDTDCVFYFAVR | 1439318 | Cell/Virus Enclosed | NO |
| GDTVGHYVHVKC | 1439324 | Cell/Virus Enclosed | NO |
| GDTYLLFTNSIL | 1439327 | Cell/Virus Enclosed | NO |
| GDVDNLVFTNKTTLPT | 1660426 | Cell/Virus Enclosed | NO |
| GDVDTVVNGVEF | 1439333 | Cell/Virus Enclosed | NO |
| GDVEACMVQVTCGSMT | 1660428 | Cell/Virus Enclosed | NO |
| GDVGIAPKQGYFIKHN | 1660432 | Ectodomain | YES |
| GDVGLCMYRLLK | 1439337 | Cell/Virus Enclosed | NO |
| GDVIKPVCAKSPPGEQ | 1660436 | Cell/Virus Enclosed | NO |
| GDVKFKRLGDYVLTEN | 1660439 | Cell/Virus Enclosed | NO |
| GDVLLAEFDTYDPIYK | 1660442 | Cell/Virus Enclosed | NO |
| GDVNKPVCAKSP | 1439350 | Cell/Virus Enclosed | NO |
| GDVNVVAASGSL | 1439351 | Cell/Virus Enclosed | NO |
| GDVVLASDDLYVKRYF | 1660452 | Cell/Virus Enclosed | NO |
| GDVVLATDDLYVKRYE | 1660453 | Cell/Virus Enclosed | NO |
| GDVVSHICKCGE | 1439368 | Cell/Virus Enclosed | NO |
| GDWDSFGKTIQS | 1439371 | Cell/Virus Enclosed | NO |
| GDWNSFCKTIQS | 1439375 | Cell/Virus Enclosed | NO |
| GDWTGFKSSCCN | 1439376 | Cell/Virus Enclosed | NO |
| GDYIKLNGPLTV | 1439386 | Cell/Virus Enclosed | NO |

| | | | |
|---|---|---|---|
| GDYNLSSVLPQRNIRS | 1660468 | Ectodomain | YES |
| GDYSPSCCTLTIGVSSYHSKP | 19192 | Ectodomain | YES |
| GDYTLANTTTAP | 1439391 | Cell/Virus Enclosed | NO |
| GDYYRKQLSPLEGGGW | 1660478 | Ectodomain | YES |
| GEDAFSILAVKT | 1439425 | Cell/Virus Enclosed | NO |
| GEDFLGWIVPFGFMPS | 1660495 | Cell/Virus Enclosed | NO |
| GEDTVFEVQGYK | 1439440 | Cell/Virus Enclosed | NO |
| GEDVDDIESIYD | 1439444 | Cell/Virus Enclosed | NO |
| GEDVQLLKAAYA | 1439445 | Cell/Virus Enclosed | NO |
| GEDVVTGDNNDE | 1439447 | Cell/Virus Enclosed | NO |
| GEENFSSRMYCS | 1439453 | Cell/Virus Enclosed | NO |
| GEFASHVANFQK | 1439458 | Cell/Virus Enclosed | NO |
| GEFIFEKVEYGS | 1439461 | Cell/Virus Enclosed | NO |
| GEFSKKQLGLLQPTDN | 1660506 | Cell/Virus Enclosed | NO |
| GEFVFEKVDYGS | 1439469 | Cell/Virus Enclosed | NO |
| GEGAPPAKRARTDQME | 19269 | Cell/Virus Enclosed | NO |
| GEGDGGITSEGNALYN | 1660509 | Cell/Virus Enclosed | NO |
| GEGDQIIFYEGV | 1439478 | Ectodomain | YES |
| GEHCSGLAIKSD | 1439485 | Ectodomain | YES |
| GEHCSGLAVKSDYCGNNSCT | 2060824 | Ectodomain | YES |
| GEHFKHLVPLMH | 1439487 | Cell/Virus Enclosed | NO |
| GEIRPSAPPLPPVA | 19311 | Cell/Virus Enclosed | NO |
| GEKGAYNKDHGC | 1439497 | Cell/Virus Enclosed | NO |
| GEKGAYNKDHGRGGFG | 1660521 | Cell/Virus Enclosed | NO |
| GEKLFVEHYNEWAQAN | 1660522 | Cell/Virus Enclosed | NO |
| GELDQWFKNQTS | 1439504 | Ectodomain | YES |
| GELKVEFWFAMRKDGD | 1660529 | Cell/Virus Enclosed | NO |
| GELLVQDLRFNK | 1439519 | Cell/Virus Enclosed | NO |
| GELPVAISGDKVFVRD | 1660532 | Cell/Virus Enclosed | NO |
| GELPVAVVNDKIFTKS | 1660533 | Cell/Virus Enclosed | NO |
| GELPVAVVNDKVFVRY | 1660534 | Cell/Virus Enclosed | NO |
| GEMPCAIINDKVVAKI | 1660538 | Cell/Virus Enclosed | NO |
| GENDNISVAAPK | 1439540 | Cell/Virus Enclosed | NO |
| GENDNISVATPK | 1439542 | Cell/Virus Enclosed | NO |
| GENDNISVAVPKSRVQ | 1660542 | Cell/Virus Enclosed | NO |
| GENDNISVAVSK | 1439544 | Cell/Virus Enclosed | NO |
| GENDNTSVAVPK | 1439545 | Capsid | UN |
| GERHQQLVEHTT | 1439570 | Cell/Virus Enclosed | NO |
| GERNGARPKQRR | 1439574 | Cell/Virus Enclosed | NO |
| GERQLLLLWEAG | 1439575 | Cell/Virus Enclosed | NO |
| GERQLLLVWEAGKSKP | 1660556 | Cell/Virus Enclosed | NO |
| GERSGARPKQRRPQGL | 1660557 | Cell/Virus Enclosed | NO |
| GERSGARSKQRRPQGL | 1381027 | Cell/Virus Enclosed | NO |
| GERVRQAILNTVKFCD | 1660558 | Cell/Virus Enclosed | NO |
| GESKSQSRNSSS | 1439596 | Cell/Virus Enclosed | NO |
| GETALALLLLDRLNQLESKVSGKG | 1381034 | Capsid | UN |
| GETGVIADYNYK | 1439606 | Ectodomain | YES |
| GETKALKELYGP | 1439609 | Cell/Virus Enclosed | NO |
| GEVFNATKFPSVYAWE | 1660591 | Ectodomain | YES |
| GEWNAVYKAFGS | 1439642 | Cell/Virus Enclosed | NO |
| GEWNAVYRAFGS | 1439643 | Cell/Virus Enclosed | NO |
| GEYDNISVAVPK | 1439647 | Cell/Virus Enclosed | NO |
| GEYIFERIDYSDAVSY | 1660601 | Cell/Virus Enclosed | NO |
| GEYNHVVAANAL | 1439650 | Cell/Virus Enclosed | NO |
| GFAAYSRYRIGNYKLN | 1381044 | Underneath Membrane | NO |
| GFACGCGASYTANDVE | 1660610 | Cell/Virus Enclosed | NO |
| GFACGCGTSYTA | 1439664 | Cell/Virus Enclosed | NO |
| GFAIPSVAVRTYSEAA | 1660613 | Cell/Virus Enclosed | NO |
| GFAVYHRYRIGN | 1439676 | Underneath Membrane | NO |
| GFCDLKGKFVQI | 1439683 | Cell/Virus Enclosed | NO |
| GFCEEADEGICF | 1439685 | Cell/Virus Enclosed | NO |
| GFCKTHNWNCLN | 1439696 | Cell/Virus Enclosed | NO |
| GFCLSNNPCNVC | 1439697 | Cell/Virus Enclosed | NO |
| GFCVKHQWNCFI | 1439703 | Cell/Virus Enclosed | NO |
| GFDKPQEKDKKSAKTG | 1660634 | Capsid | UN |
| GFDKPQEKKKSA | 1439710 | Cell/Virus Enclosed | NO |
| GFDKWYVNDGRV | 1439712 | Cell/Virus Enclosed | NO |
| GFDLNCYYLVLP | 1439714 | Ectodomain | YES |
| GFDMLDNYKAIDVVQY | 1660638 | Cell/Virus Enclosed | NO |
| GFDNQSKLPSSS | 1439719 | Cell/Virus Enclosed | NO |
| GFDNQSKPPSSS | 1439720 | Cell/Virus Enclosed | NO |
| GFDNQSKSPSSSGTST | 1660640 | Capsid | UN |
| GFDSSKEFVEYKITEW | 1660642 | Cell/Virus Enclosed | NO |
| GFDSSKPVTKKYTYSF | 1660643 | Cell/Virus Enclosed | NO |
| GFDSSKSVTKKY | 1439726 | Cell/Virus Enclosed | NO |
| GFECDFSPLLSG | 1439733 | Ectodomain | YES |

| | | | |
|---|---|---|---|
| GFEDQPNLQVESANQM | 1660651 | Cell/Virus Enclosed | NO |
| GFEKAALFASLN | 1439737 | Cell/Virus Enclosed | NO |
| GFEKPQEKDKKS | 1439739 | Cell/Virus Enclosed | NO |
| GFELTSMKYFVK | 1439740 | Cell/Virus Enclosed | NO |
| GFENPPTNVVSH | 1439741 | Ectodomain | YES |
| GFETIMKVLEEN | 1439742 | Cell/Virus Enclosed | NO |
| GFETIMKVLNENLNAY | 1660656 | Cell/Virus Enclosed | NO |
| GFGGKNILGYAS | 1439777 | Cell/Virus Enclosed | NO |
| GFGGKNILGYSS | 1439778 | Cell/Virus Enclosed | NO |
| GFGGKQILGYSSLNDE | 1660679 | Cell/Virus Enclosed | NO |
| GFGITVQYGTDTNSVC | 1660681 | Ectodomain | YES |
| GFGPRDFNHNMG | 1439786 | Cell/Virus Enclosed | NO |
| GFGVAVADSYYSYMMP | 1660683 | Cell/Virus Enclosed | NO |
| GFGVDEEKCGVLDGSY | 1660684 | Ectodomain | YES |
| GFGVFDIKNSSVNTVC | 1660685 | Cell/Virus Enclosed | NO |
| GFGVNKIQPWTNDALN | 1660686 | Cell/Virus Enclosed | NO |
| GFHNPPTNVVSH | 1439796 | Ectodomain | YES |
| GFIEAVNFNVTT | 1439814 | Ectodomain | YES |
| GFIEDAVFKPQP | 1439815 | Ectodomain | YES |
| GFIEDSAFVPQP | 1439816 | Ectodomain | YES |
| GFIEDSVFKPRPAGVL | 1660696 | Ectodomain | YES |
| GFIEDSVFVPQP | 1439819 | Ectodomain | YES |
| GFIEGGWTGMVDGWYGYHHQNEQGS | 1142694 | Ectodomain | YES |
| GFIENSVFKPQP | 1439820 | Ectodomain | YES |
| GFITLKNNLYRL | 1439835 | Cell/Virus Enclosed | NO |
| GFKDAVTFATVDHSAF | 1660707 | Cell/Virus Enclosed | NO |
| GFKDFVTNKTYNIFPC | 1660708 | Ectodomain | YES |
| GFKDVTNGTIYS | 1439840 | Ectodomain | YES |
| GFKEELDQWFKN | 1439841 | Ectodomain | YES |
| GFKFGDPADSRDFLRV | 1660710 | Cell/Virus Enclosed | NO |
| GFKLDAIVPGYP | 1439851 | Cell/Virus Enclosed | NO |
| GFKLDANVPGYP | 1439854 | Cell/Virus Enclosed | NO |
| GFKNQHCNGSIA | 1439861 | Cell/Virus Enclosed | NO |
| GFKNQYCNESIA | 1439862 | Cell/Virus Enclosed | NO |
| GFKNVSTGNIFI | 1439864 | Ectodomain | YES |
| GFLETVCSVAHT | 1439879 | Cell/Virus Enclosed | NO |
| GFLFLTWICLLQFAY | 1312541 | Transmembrane | NO |
| GFLPTFVGKIAQ | 1439903 | Cell/Virus Enclosed | NO |
| GFLQSSNYKIDTTSSS | 1660736 | Ectodomain | YES |
| GFMKQYGECLGDINARD | 19585 | Ectodomain | YES |
| GFNDICMFENKT | 1439932 | Cell/Virus Enclosed | NO |
| GFNDLSQLHCSYESFD | 1660748 | Ectodomain | YES |
| GFNDLSQLYCSY | 1439934 | Ectodomain | YES |
| GFNEWAMANGYTSVSS | 1660751 | Cell/Virus Enclosed | NO |
| GFNFSQILPDPLKPTK | 1660753 | Ectodomain | YES |
| GFNLEDAARCMR | 1439949 | Cell/Virus Enclosed | NO |
| GFNTEFASNSSV | 1439956 | Cell/Virus Enclosed | NO |
| GFPQLAELIPNQAALF | 1660768 | Capsid | UN |
| GFQGKQILGSTM | 1439980 | Cell/Virus Enclosed | NO |
| GFQGRQILGSCTFEDE | 1660773 | Cell/Virus Enclosed | NO |
| GFQGRQIMGSCSFEDE | 1660774 | Cell/Virus Enclosed | NO |
| GFQIKFENPHIL | 1439985 | Cell/Virus Enclosed | NO |
| GFQKCEQLLREYGQFC | 1660776 | Ectodomain | YES |
| GFRDYITNRTFMIRSC | 1660779 | Ectodomain | YES |
| GFSCPLCCTVYDSSEV | 1660790 | Cell/Virus Enclosed | NO |
| GFSLEIQDLEEP | 1440014 | Cell/Virus Enclosed | NO |
| GFSNGVDFVAQPEGCV | 1660796 | Cell/Virus Enclosed | NO |
| GFSNVVELPKFFYASN | 1660798 | Ectodomain | YES |
| GFSNVVEMPKFF | 1440020 | Ectodomain | YES |
| GFSQVKSDLVID | 1440024 | Cell/Virus Enclosed | NO |
| GFSTGVNFVVQP | 1440030 | Cell/Virus Enclosed | NO |
| GFSVLRPILKLP | 1440038 | Ectodomain | YES |
| GFTAMNGEDAFSILAA | 1660808 | Cell/Virus Enclosed | NO |
| GFTEQSVFKPQP | 1440044 | Ectodomain | YES |
| GFTGNLQSNHDQ | 1440047 | Cell/Virus Enclosed | NO |
| GFTKPASRELSV | 1440050 | Cell/Virus Enclosed | NO |
| GFTLKNTVCTVC | 1440054 | Cell/Virus Enclosed | NO |
| GFTTTNEAFQKVQDAV | 1660821 | Ectodomain | YES |
| GFTVKPVATVHRRIPD | 1660823 | Ectodomain | YES |
| GFTVMNGEDAFS | 1440074 | Cell/Virus Enclosed | NO |
| GFTWTGVTQNG | 19639 | Ectodomain | YES |
| GFVLNYKGLQFQ | 1440085 | Cell/Virus Enclosed | NO |
| GFVLSIPTKALC | 1440090 | Underneath Membrane | NO |
| GFVVRIGAAANSTGTV | 1660843 | Ectodomain | YES |
| GFVYDNHVKIDC | 1440116 | Cell/Virus Enclosed | NO |
| GFVYFNATGSDVNCNG | 1660846 | Ectodomain | YES |

| | | | |
|---|---|---|---|
| GFWRDGSCSCVGSSVA | 1660848 | Cell/Virus Enclosed | NO |
| GFYFKPGGTTSGDAST | 1660852 | Cell/Virus Enclosed | NO |
| GFYNPLTNVVSH | 1440128 | Ectodomain | YES |
| GFYRLYSGDNSW | 1440134 | Cell/Virus Enclosed | NO |
| GFYSANFLDDNVLPET | 1660859 | Ectodomain | YES |
| GFYSNSSSDVIR | 1440141 | Ectodomain | YES |
| GFYSTSPIQPVE | 1440143 | Ectodomain | YES |
| GFYTTSGIGYQP | 1440149 | Ectodomain | YES |
| GFYTTTGIGYQPYRVVV | 19657 | Ectodomain | YES |
| GGAEIRDLICVS | 1440156 | Ectodomain | YES |
| GGAFTRYVDNNF | 1440157 | Cell/Virus Enclosed | NO |
| GGAKTPSDAVQNILQKIEKIKNTEE | 19678 | Cell/Virus Enclosed | NO |
| GGAPIWFNALRD | 1440168 | Cell/Virus Enclosed | NO |
| GGAPVKKVAFGGDQVH | 1660873 | Cell/Virus Enclosed | NO |
| GGAVCLKHAEEYREYL | 1660879 | Cell/Virus Enclosed | NO |
| GGAVCSKHANLYQKYV | 1660884 | Cell/Virus Enclosed | NO |
| GGAYSLFDMAKFPLKL | 1660888 | Cell/Virus Enclosed | NO |
| GGAYSLFDMTKFSLKL | 1660889 | Cell/Virus Enclosed | NO |
| GGCAMPIMANLV | 1440198 | Ectodomain | YES |
| GGCCDDYTGYQELVIK | 1660892 | Underneath Membrane | NO |
| GGCCEDYTGYQE | 1440203 | Underneath Membrane | NO |
| GGCIKACEVVVTNLNK | 1660893 | Cell/Virus Enclosed | NO |
| GGCIPASQVIVNNYDK | 1660895 | Cell/Virus Enclosed | NO |
| GGCITSREVVVTNLNK | 1660896 | Cell/Virus Enclosed | NO |
| GGCLNASEVVVNNLDK | 1660897 | Cell/Virus Enclosed | NO |
| GGCYYVKPGGTSSGDA | 1660905 | Cell/Virus Enclosed | NO |
| GGDAALALLLLDRLNA | 1660906 | Cell/Virus Enclosed | NO |
| GGDAALALLLLDRLNQ | 1381105 | Cell/Virus Enclosed | NO |
| GGDAISLSFNNLLGFD | 1660907 | Cell/Virus Enclosed | NO |
| GGDATSLSFNNL | 1440223 | Cell/Virus Enclosed | NO |
| GGDFNLTLLEPVSIST | 1660908 | Ectodomain | YES |
| GGDGKMKDLSPRWYFYY | 1381108 | Cell/Virus Enclosed | NO |
| GGDGKMKELSPRWYFY | 1381109 | Capsid | UN |
| GGDLLYLDLLNRLQAL | 1660910 | Cell/Virus Enclosed | NO |
| GGDNVYCYNTDLIEGS | 1660911 | Cell/Virus Enclosed | NO |
| GGDNVYCYNTGL | 1440236 | Cell/Virus Enclosed | NO |
| GGDQIHEVAAVR | 1440238 | Cell/Virus Enclosed | NO |
| GGDQVHEVVAVR | 1440240 | Cell/Virus Enclosed | NO |
| GGDRCIKISTVQSKLT | 1660914 | Cell/Virus Enclosed | NO |
| GGDSSESGAKETKEIN | 1660916 | Cell/Virus Enclosed | NO |
| GGEMLKLGTSDPQFPI | 1660919 | Cell/Virus Enclosed | NO |
| GGERHRQLVEHTTPWL | 1660920 | Cell/Virus Enclosed | NO |
| GGESSEELSESSFFNLIT | 19723 | Cell/Virus Enclosed | NO |
| GGFNFSQILPDPLKPT | 1660928 | Ectodomain | YES |
| GGFWGDRVDSQPFAIPYIHPTNPFA | 19736 | Undetermined | UN |
| GGFYRFMVDPTTVLND | 1660934 | Cell/Virus Enclosed | NO |
| GGGIAGAINAASKGAV | 1660938 | Cell/Virus Enclosed | NO |
| GGGKCFNCAPAV | 1440299 | Ectodomain | YES |
| GGGKCFNCYPAGVNIT | 1660941 | Ectodomain | YES |
| GGGLAKALDVYTKGKL | 1660942 | Cell/Virus Enclosed | NO |
| GGGWLVASGPTV | 1440314 | Ectodomain | YES |
| GGGYYVKPGGTSSGDA | 1660951 | Cell/Virus Enclosed | NO |
| GGIAGAINAASKGAVQ | 1660956 | Cell/Virus Enclosed | NO |
| GGINARDLICAQ | 1440341 | Ectodomain | YES |
| GGITGSMSLASGKSLLHEVDKD | 19891 | Ectodomain | YES |
| GGITQRTRTRPS | 1440347 | Cell/Virus Enclosed | NO |
| GGITSEGNALYNNEGG | 1660966 | Cell/Virus Enclosed | NO |
| GGITYVSNSGNLLGFK | 1660967 | Ectodomain | YES |
| GGKCFNCYPAGVNITL | 1660969 | Ectodomain | YES |
| GGKNILGYASLC | 1440360 | Cell/Virus Enclosed | NO |
| GGKNILGYSSLC | 1440361 | Cell/Virus Enclosed | NO |
| GGKQILGYSSLNDEFS | 1660980 | Cell/Virus Enclosed | NO |
| GGLAKALDVYTKGKLQ | 1660986 | Cell/Virus Enclosed | NO |
| GGLCPSTGQAMESVGF | 1660987 | Cell/Virus Enclosed | NO |
| GGLFQLTFESTI | 1440383 | Cell/Virus Enclosed | NO |
| GGLTSIKWADNN | 1440395 | Cell/Virus Enclosed | NO |
| GGMIRYYDVMANGGTG | 1661010 | Cell/Virus Enclosed | NO |
| GGMSYYCEDHKPQYSF | 1661012 | Cell/Virus Enclosed | NO |
| GGNCAGNVIGNDALTR | 1661017 | Cell/Virus Enclosed | NO |
| GGNEALSISTIF | 1440416 | Cell/Virus Enclosed | NO |
| GGNGTHIFSLVN | 1440420 | Ectodomain | YES |
| GGNLCTCQPQAF | 1440424 | Ectodomain | YES |
| GGNMFQFATFPV | 1440427 | Ectodomain | YES |
| GGNSCTCQPQAF | 1440435 | Ectodomain | YES |
| GGNSQSSSRASSLSRN | 1661026 | Cell/Virus Enclosed | NO |
| GGNVIPHYSWFS | 1440438 | Cell/Virus Enclosed | NO |

| | | | |
|---|---|---|---|
| GGNVVPHYSWFS | 1440440 | Cell/Virus Enclosed | NO |
| GGPIDSTDNNQN | 1440444 | Capsid | UN |
| GGPRTIKVSTVQSKLT | 1661031 | Cell/Virus Enclosed | NO |
| GGPSDSSDNSQN | 1440446 | Cell/Virus Enclosed | NO |
| GGPSDSTDNNQNGDRS | 1661032 | Cell/Virus Enclosed | NO |
| GGPSDSTGSNQNGERSGARSKQ | 1892918 | Cell/Virus Enclosed | NO |
| GGPTDSTDNNQD | 1440450 | Cell/Virus Enclosed | NO |
| GGPTDSTDNNQS | 1440452 | Capsid | UN |
| GGQPITNCVKML | 1440457 | Cell/Virus Enclosed | NO |
| GGSIAIKVTEYSWNKK | 1661054 | Cell/Virus Enclosed | NO |
| GGTSSGDATTAFANSV | 1661083 | Cell/Virus Enclosed | NO |
| GGTTSGDASTAYANSI | 1661089 | Cell/Virus Enclosed | NO |
| GGTTSGDATTAYANSV | 1661090 | Cell/Virus Enclosed | NO |
| GGTTVVRPLVEDSTSV | 1661091 | Ectodomain | YES |
| GGVLGDGNALYNTEGG | 1661099 | Cell/Virus Enclosed | NO |
| GGVPVIEVSQIQSKLT | 1661102 | Cell/Virus Enclosed | NO |
| GGVSVITPGTNASSEV | 1661106 | Ectodomain | YES |
| GGWDFMLKTLYKDVDN | 1661116 | Cell/Virus Enclosed | NO |
| GGWDNMLKNLMADVDD | 1661117 | Cell/Virus Enclosed | NO |
| GGWGNMLKNLMA | 1440583 | Cell/Virus Enclosed | NO |
| GGWNNMLRTLID | 1440590 | Cell/Virus Enclosed | NO |
| GGYHYTPFHYERDNTS | 1661124 | Cell/Virus Enclosed | NO |
| GGYIPPNFAFNN | 1440598 | Ectodomain | YES |
| GGYIPSNFAFNN | 1440600 | Ectodomain | YES |
| GGYNSWHLVEGG | 1440604 | Cell/Virus Enclosed | NO |
| GGYYVKPGGTSSGDAT | 1661134 | Cell/Virus Enclosed | NO |
| GGYYYKVEADFY | 1440610 | Ectodomain | YES |
| GHAGEAPVSIIN | 1440613 | Cell/Virus Enclosed | NO |
| GHAMPSLFKVQNVNLE | 1661136 | Cell/Virus Enclosed | NO |
| GHATGTTPKKLF | 1440621 | Ectodomain | YES |
| GHCTIAQLTDAA | 1440630 | Cell/Virus Enclosed | NO |
| GHCTIVQLTDAA | 1440632 | Cell/Virus Enclosed | NO |
| GHDICAQLNAKL | 1440637 | Cell/Virus Enclosed | NO |
| GHDICAQLNDKL | 1440638 | Cell/Virus Enclosed | NO |
| GHEEFIQTSSPK | 1440643 | Ectodomain | YES |
| GHFLAKNILHVV | 1440657 | Cell/Virus Enclosed | NO |
| GHFNKEFYDFLRSQGF | 1661151 | Cell/Virus Enclosed | NO |
| GHFTGVEGELPV | 1440661 | Cell/Virus Enclosed | NO |
| GHHLGRCDIKDLPKEI | 1381236 | Underneath Membrane | NO |
| GHIIMEEMLRGS | 1440691 | Cell/Virus Enclosed | NO |
| GHIVCDSLNSKL | 1440695 | Cell/Virus Enclosed | NO |
| GHKNGQGENDNISVAV | 1661164 | Cell/Virus Enclosed | NO |
| GHKNGQGEYDNI | 1440704 | Cell/Virus Enclosed | NO |
| GHKTGQGENDNI | 1440708 | Capsid | UN |
| GHLKMAGMHFGA | 1440717 | Ectodomain | YES |
| GHLRIAGHHLGRCDI | 1540477 | Underneath Membrane | NO |
| GHLYIQGIKLGTGYSL | 1661170 | Ectodomain | YES |
| GHLYIQGVKLGTGYTL | 1661171 | Underneath Membrane | NO |
| GHMAHDGGVAKA | 1440729 | Cell/Virus Enclosed | NO |
| GHMVHGGGVAKA | 1440736 | Cell/Virus Enclosed | NO |
| GHNLAKKCMHVV | 1440740 | Cell/Virus Enclosed | NO |
| GHPLGRCDIKDL | 1440751 | Underneath Membrane | NO |
| GHRLASGVQVHNLPEY | 1661187 | Underneath Membrane | NO |
| GHSEIVFDVESG | 1440763 | Ectodomain | YES |
| GHSENVFAVESG | 1440764 | Ectodomain | YES |
| GHSLGRCDIKDLPKEITVATSRTLS | 1381252 | Underneath Membrane | NO |
| GHTLKSGCKLIN | 1440785 | Cell/Virus Enclosed | NO |
| GHTVCDSLNAKLGFDS | 1661203 | Cell/Virus Enclosed | NO |
| GHTVCDSLNAKW | 1440789 | Cell/Virus Enclosed | NO |
| GHTVCDSLNSKL | 1440791 | Cell/Virus Enclosed | NO |
| GHVYEVSDLIAQ | 1440806 | Cell/Virus Enclosed | NO |
| GHYAGSRSSSGNRSGI | 1661213 | Capsid | UN |
| GHYQTNIYSQNLCVDG | 1661219 | Cell/Virus Enclosed | NO |
| GHYTGQAGEMPCAIIN | 1661222 | Cell/Virus Enclosed | NO |
| GHYTVYDTAKKSMYDG | 1661225 | Cell/Virus Enclosed | NO |
| GIAAEQDRNTREVFAQ | 1661229 | Ectodomain | YES |
| GIACVKDFDFKF | 1440841 | Cell/Virus Enclosed | NO |
| GIADDTYVMGLHGNQT | 1661232 | Cell/Virus Enclosed | NO |
| GIAIAMACLVGLMWL | 1540483 | Transmembrane | NO |
| GIAISVKPESTA | 1440850 | Cell/Virus Enclosed | NO |
| GIAISVKPESTV | 1440851 | Cell/Virus Enclosed | NO |
| GIALGGLTSAAS | 1440853 | Ectodomain | YES |
| GIALTGTNQLCA | 1440854 | Cell/Virus Enclosed | NO |
| GIASGVFGVNLRTNFT | 1661242 | Cell/Virus Enclosed | NO |
| GIATAGWTFGAG | 1440858 | Ectodomain | YES |
| GIAVLDMCAALK | 1440862 | Cell/Virus Enclosed | NO |

| | | | |
|---|---|---|---|
| GIAWFTERSDKS | 1440864 | Cell/Virus Enclosed | NO |
| GICADGSLIPVRPRNS | 1661248 | Ectodomain | YES |
| GICASYHTASVL | 1440872 | Ectodomain | YES |
| GICFEYTSPTDK | 1440876 | Cell/Virus Enclosed | NO |
| GICPAGTNYLTC | 1440884 | Ectodomain | YES |
| GICPENHVMVDCRRLL | 1661257 | Cell/Virus Enclosed | NO |
| GICSENHVMVDC | 1440886 | Cell/Virus Enclosed | NO |
| GIDAGYKNSGIG | 1440890 | Ectodomain | YES |
| GIDICSQDKVFGYVQQ | 1661262 | Cell/Virus Enclosed | NO |
| GIDLKSYDLGVE | 1440898 | Cell/Virus Enclosed | NO |
| GIDVTKLKCSDFYADK | 1661270 | Cell/Virus Enclosed | NO |
| GIEDPRWPQIAE | 1440905 | Cell/Virus Enclosed | NO |
| GIENATFFIFNKLVKD | 1661274 | Secreted | YES |
| GIETAVGHYVHARLKG | 1661275 | Cell/Virus Enclosed | NO |
| GIFAKVKNTKVT | 1440922 | Ectodomain | YES |
| GIFAKVKNTKVV | 1440923 | Ectodomain | YES |
| GIFEQSLRLVDA | 1440928 | Cell/Virus Enclosed | NO |
| GIFGAKPLQSLQ | 1440936 | Cell/Virus Enclosed | NO |
| GIFKEVSAVYYN | 1440942 | Ectodomain | YES |
| GIFKTEVDPNLS | 1440943 | Cell/Virus Enclosed | NO |
| GIFVASSSYDAL | 1440949 | Cell/Virus Enclosed | NO |
| GIFVEVNAPYYN | 1440950 | Ectodomain | YES |
| GIGAVGGDLLYLDLLN | 1661294 | Cell/Virus Enclosed | NO |
| GIGDSAVTYSRLISLM | 1661295 | Cell/Virus Enclosed | NO |
| GIGEHCSGLAIK | 1440960 | Ectodomain | YES |
| GIGGDRCIKISTVQSK | 1661297 | Cell/Virus Enclosed | NO |
| GIGGVPIIEVSQFQSK | 1661302 | Cell/Virus Enclosed | NO |
| GIGGVPVIEVSQIQSK | 1661303 | Cell/Virus Enclosed | NO |
| GIGICPAGTNYL | 1440976 | Ectodomain | YES |
| GIGLCAFLTLLF | 1440980 | Cell/Virus Enclosed | NO |
| GIGLHLKVNCCR | 1440983 | Cell/Virus Enclosed | NO |
| GIGTCPAGTNSL | 1440988 | Ectodomain | YES |
| GIGTCPAGTNYLTCDN | 1661308 | Ectodomain | YES |
| GIGVTQNVLYENQKQIA | 20288 | Ectodomain | YES |
| GIGYSLADLPAY | 1440999 | Ectodomain | YES |
| GIIRDVNPDDKCFANK | 1661322 | Cell/Virus Enclosed | NO |
| GIIRSSNQSLAGGITY | 1661323 | Ectodomain | YES |
| GIISNARNGCIPLSVI | 1661325 | Cell/Virus Enclosed | NO |
| GIITVEELKQLL | 1441033 | Undetermined | UN |
| GIIWVATEGALNTPKD | 1661327 | Cell/Virus Enclosed | NO |
| GIKDVVLQGLKACCYV | 1661331 | Cell/Virus Enclosed | NO |
| GIKDVVLQGLRA | 1441044 | Cell/Virus Enclosed | NO |
| GIKFQTVKPGNFNQDF | 1661332 | Cell/Virus Enclosed | NO |
| GIKLGIGYSLAD | 1441048 | Ectodomain | YES |
| GIKLGIGYSWAD | 1441049 | Underneath Membrane | NO |
| GIKLGTGYSLADLPAY | 1661337 | Ectodomain | YES |
| GIKLQSKRTRFF | 1441054 | Cell/Virus Enclosed | NO |
| GIKLQSKRTRLF | 1441055 | Cell/Virus Enclosed | NO |
| GIKPFTIPIDLL | 1441057 | Cell/Virus Enclosed | NO |
| GIKQESRVGVDA | 1441059 | Cell/Virus Enclosed | NO |
| GIKVLPPLLSDN | 1441061 | Ectodomain | YES |
| GIKVLPPLLSENQISG | 1661347 | Ectodomain | YES |
| GIKYVAQPTEDVVDGD | 1661349 | Cell/Virus Enclosed | NO |
| GIKYYSRVKSVR | 1441066 | Cell/Virus Enclosed | NO |
| GILDGDYAMQFFKMGR | 1661355 | Cell/Virus Enclosed | NO |
| GILGQGIFVEVN | 1441077 | Ectodomain | YES |
| GILKKTSWVDQS | 1441081 | Cell/Virus Enclosed | NO |
| GILKWADQSDQFRNVQ | 1661359 | Cell/Virus Enclosed | NO |
| GILLCLALLFVV | 1441085 | Transmembrane | NO |
| GILLEAYNAFLDTVVS | 1661361 | Cell/Virus Enclosed | NO |
| GILQQAALDYLATVKT | 1661365 | Cell/Virus Enclosed | NO |
| GILSNMNIASNSKEWA | 1661366 | Cell/Virus Enclosed | NO |
| GIMVLPGAADAE | 1441113 | Ectodomain | YES |
| GINDDDYVIALTGTNQ | 1661377 | Cell/Virus Enclosed | NO |
| GINEKTHVQLSL | 1441122 | Cell/Virus Enclosed | NO |
| GINSGTTCSNDLLQPN | 1661381 | Ectodomain | YES |
| GINYLGDFAQGPFIDG | 1661384 | Cell/Virus Enclosed | NO |
| GINYLGDFIQGPFIAG | 1661385 | Cell/Virus Enclosed | NO |
| GINYLGDFMQGP | 1441139 | Cell/Virus Enclosed | NO |
| GINYLGTIKENIDGGA | 1661387 | Cell/Virus Enclosed | NO |
| GIPFEYTSPTDK | 1441148 | Cell/Virus Enclosed | NO |
| GIPGNMIDGFAC | 1441150 | Cell/Virus Enclosed | NO |
| GIPLSEAKGYWY | 1441156 | Cell/Virus Enclosed | NO |
| GIPPSEAKGYWY | 1441157 | Cell/Virus Enclosed | NO |
| GIPVVEYLSKED | 1441160 | Cell/Virus Enclosed | NO |
| GIREFSNLVLNNCTKY | 1661402 | Ectodomain | YES |

| | | | |
|---|---|---|---|
| GIRFEYISPTDK | 1441179 | Cell/Virus Enclosed | NO |
| GIRFEYTSPTDKLAMI | 1661403 | Cell/Virus Enclosed | NO |
| GISFCRRHNWNCVDCD | 1661407 | Cell/Virus Enclosed | NO |
| GISGRGVFQNCT | 1441198 | Ectodomain | YES |
| GISLLKKMDEPF | 1441202 | Cell/Virus Enclosed | NO |
| GISRQGIFVEVN | 1441207 | Ectodomain | YES |
| GISSACDGLALA | 1441208 | Cell/Virus Enclosed | NO |
| GISTPKLKEDYQIGGY | 1661412 | Secreted | YES |
| GITFQYGTDTNS | 1441220 | Ectodomain | YES |
| GITGVPKPVEGV | 1441224 | Ectodomain | YES |
| GITQFQKGKAFE | 1441234 | Cell/Virus Enclosed | NO |
| GITQFQKGKEFEFVEG | 1661421 | Cell/Virus Enclosed | NO |
| GITQFQKGKEFK | 1441236 | Cell/Virus Enclosed | NO |
| GITQFQKGKEFQ | 1441237 | Capsid | UN |
| GITQFQKGRDFKFSDG | 1661422 | Capsid | UN |
| GITQQVLSDNQK | 1441241 | Ectodomain | YES |
| GITQQVLSKNQK | 1441243 | Ectodomain | YES |
| GITSEGNALYNNEGGR | 1661425 | Cell/Virus Enclosed | NO |
| GITVQYGTDTNSVCPK | 1661426 | Ectodomain | YES |
| GITYTSTSGNLLGFKD | 1661428 | Ectodomain | YES |
| GITYVSNSGNLLGFKN | 1661429 | Ectodomain | YES |
| GIVAAVSAMMRI | 1441254 | Transmembrane | NO |
| GIVEDTINDDED | 1441263 | Cell/Virus Enclosed | NO |
| GIVELEKGVLPQ | 1441264 | Cell/Virus Enclosed | NO |
| GIVKMVSPTSKI | 1441270 | Cell/Virus Enclosed | NO |
| GIVYSSQLYEDVPSIN | 1661444 | Cell/Virus Enclosed | NO |
| GIYGYVLRQSNL | 1441292 | Ectodomain | YES |
| GIYKPLLENIDY | 1441293 | Cell/Virus Enclosed | NO |
| GIYQTSNFRVSP | 1441297 | Ectodomain | YES |
| GKAFEFAEGQGV | 1441301 | Cell/Virus Enclosed | NO |
| GKAGLYYESISYEEQD | 1661453 | Cell/Virus Enclosed | NO |
| GKASSRSAIEDL | 1441310 | Ectodomain | YES |
| GKCGDKFFPIVMNDKN | 1661465 | Cell/Virus Enclosed | NO |
| GKCLKMNCVRFKNADL | 1661466 | Cell/Virus Enclosed | NO |
| GKCYNCRPAVVN | 1441324 | Ectodomain | YES |
| GKDGVMYSEFPA | 1441338 | Ectodomain | YES |
| GKDLNANMSLAD | 1441339 | Cell/Virus Enclosed | NO |
| GKDLNANMSLAECKRA | 1661474 | Cell/Virus Enclosed | NO |
| GKEDLKFPRGQGVPIN | 1661476 | Cell/Virus Enclosed | NO |
| GKEDLRFPRGQGVPIN | 1661477 | Cell/Virus Enclosed | NO |
| GKEFEFAEGQGV | 1441344 | Cell/Virus Enclosed | NO |
| GKEFEFVEGQGVPIAP | 1661479 | Cell/Virus Enclosed | NO |
| GKEFKFAEGQGV | 1441347 | Cell/Virus Enclosed | NO |
| GKEFQFAEGQGV | 1441348 | Capsid | UN |
| GKEQKCCVCNRR | 1441351 | Cell/Virus Enclosed | NO |
| GKERARTVGGVP | 1441354 | Cell/Virus Enclosed | NO |
| GKESLENPTYIY | 1441358 | Cell/Virus Enclosed | NO |
| GKEVEFVEGQGV | 1441361 | Cell/Virus Enclosed | NO |
| GKFLEELNAFTREMQQ | 1661490 | Capsid | UN |
| GKFLSNTKYYDDSQYY | 1661491 | Ectodomain | YES |
| GKGNKDEQIGYWNVQE | 1661501 | Capsid | UN |
| GKGSKDEQIGYW | 1441399 | Cell/Virus Enclosed | NO |
| GKGYHLMSFPQAAPHG | 1661505 | Ectodomain | YES |
| GKHEHSLLVEAYNSIL | 1661507 | Cell/Virus Enclosed | NO |
| GKHERDLLIKAYNTIN | 1661508 | Cell/Virus Enclosed | NO |
| GKIAHWIKNTFS | 1441408 | Cell/Virus Enclosed | NO |
| GKIWNGVIKNVNSVRD | 1661522 | Cell/Virus Enclosed | NO |
| GKKIKHEGDWDSFCKT | 1661525 | Cell/Virus Enclosed | NO |
| GKLADCLQAYGH | 1441434 | Cell/Virus Enclosed | NO |
| GKLKCNRCCDRYEEYD | 1661532 | Underneath Membrane | NO |
| GKLKQKPMKAEG | 1441444 | Cell/Virus Enclosed | NO |
| GKLLIRDNDVLS | 1441450 | Cell/Virus Enclosed | NO |
| GKLLIRDNGTLS | 1441454 | Cell/Virus Enclosed | NO |
| GKLLIRDTNKEVFVGD | 1661536 | Cell/Virus Enclosed | NO |
| GKLLVRDTNKEV | 1441457 | Cell/Virus Enclosed | NO |
| GKLNCSFQDAASNIFV | 1661540 | Cell/Virus Enclosed | NO |
| GKLQDVVNDNAQ | 1441464 | Ectodomain | YES |
| GKLRPFERDISNVPFS | 1661543 | Ectodomain | YES |
| GKMDTAAYREAACSQL | 1661546 | Cell/Virus Enclosed | NO |
| GKMFVRPVIEDYHTLT | 1661547 | Underneath Membrane | NO |
| GKMGLFFNHTLV | 1441478 | Ectodomain | YES |
| GKNILGYASLCD | 1441492 | Cell/Virus Enclosed | NO |
| GKNNGIGTCPVG | 1441497 | Ectodomain | YES |
| GKNNGLGTCPAG | 1441500 | Ectodomain | YES |
| GKNNGVGTCPAG | 1441502 | Ectodomain | YES |
| GKPGNVGGNDALSIST | 1661558 | Cell/Virus Enclosed | NO |

| | | | |
|---|---|---|---|
| GKPGNVGGNDDL | 1441510 | Cell/Virus Enclosed | NO |
| GKPILWVNKASYDTNL | 1661560 | Cell/Virus Enclosed | NO |
| GKPNAIDVEHLE | 1441525 | Cell/Virus Enclosed | NO |
| GKPVISEDLWQF | 1441532 | Cell/Virus Enclosed | NO |
| GKPVKGAWNIGQ | 1441534 | Cell/Virus Enclosed | NO |
| GKQIDDHRITSITSDK | 1661580 | Cell/Virus Enclosed | NO |
| GKQNILLRKYGR | 1441553 | Cell/Virus Enclosed | NO |
| GKQSSSRASFGN | 1441554 | Cell/Virus Enclosed | NO |
| GKQSSSRASSGNRSGN | 1661587 | Cell/Virus Enclosed | NO |
| GKQTEFVSNSHL | 1441556 | Cell/Virus Enclosed | NO |
| GKRGGGSVTYTD | 1441562 | Cell/Virus Enclosed | NO |
| GKRGHHKNFGGG | 1441563 | Cell/Virus Enclosed | NO |
| GKRGPNHKFGGG | 1441566 | Cell/Virus Enclosed | NO |
| GKRGPNQNFGGGEMLK | 1661593 | Cell/Virus Enclosed | NO |
| GKSAPNSRSTSR | 1441577 | Cell/Virus Enclosed | NO |
| GKSIIYTGDWEGLHEV | 1661603 | Cell/Virus Enclosed | NO |
| GKSLGNVFCDGVNVTK | 1661610 | Cell/Virus Enclosed | NO |
| GKTFFVLPSDDT | 1441603 | Cell/Virus Enclosed | NO |
| GKTVLGEYVFDKSELT | 1661623 | Cell/Virus Enclosed | NO |
| GKVDSIVQKCYELSHL | 1661625 | Cell/Virus Enclosed | NO |
| GKVFFQFDNLSSEDLK | 1661627 | Cell/Virus Enclosed | NO |
| GKVKQSQPKVIT | 1441629 | Cell/Virus Enclosed | NO |
| GKVKVGTGVMVECDSL | 1661632 | Cell/Virus Enclosed | NO |
| GKVNNFVKFGSV | 1441631 | Ectodomain | YES |
| GKVPLIFPPGQG | 1441632 | Cell/Virus Enclosed | NO |
| GKVSTLVKQVLD | 1441638 | Cell/Virus Enclosed | NO |
| GKWVQVPIGTNDPIRF | 1661639 | Cell/Virus Enclosed | NO |
| GKYAQNLLKKLIGGDV | 1661642 | Cell/Virus Enclosed | NO |
| GKYDINHGDNGF | 1441644 | Undetermined | UN |
| GKYFTFKQANNNCFVN | 1661646 | Cell/Virus Enclosed | NO |
| GKYSQKAYALLRQYRG | 1661648 | Cell/Virus Enclosed | NO |
| GKYYTKPIIKAQFKPF | 1661653 | Cell/Virus Enclosed | NO |
| GLAAEWVLAYML | 1441671 | Cell/Virus Enclosed | NO |
| GLADGLEAYADK | 1441678 | Cell/Virus Enclosed | NO |
| GLAGKVKVGTGVMVEC | 1661661 | Cell/Virus Enclosed | NO |
| GLAIKSDHCGGN | 1441683 | Ectodomain | YES |
| GLAIKSDYCGGN | 1441684 | Ectodomain | YES |
| GLALDNALSGVLSAAI | 1661668 | Cell/Virus Enclosed | NO |
| GLASAYRANSFDVPTF | 1661671 | Cell/Virus Enclosed | NO |
| GLASAYRANSID | 1441700 | Cell/Virus Enclosed | NO |
| GLAVKSDYCRGN | 1441707 | Ectodomain | YES |
| GLCDAANPTNCIAPVN | 1661677 | Ectodomain | YES |
| GLCDASNPTNCI | 1441712 | Ectodomain | YES |
| GLCDVANPTNCI | 1441716 | Ectodomain | YES |
| GLCVDIPGIPKD | 1441734 | Cell/Virus Enclosed | NO |
| GLDAAWNKFVLGDVEI | 1661692 | Cell/Virus Enclosed | NO |
| GLDAMFFYGDVV | 1441737 | Cell/Virus Enclosed | NO |
| GLDARTQAKQSY | 1441738 | Cell/Virus Enclosed | NO |
| GLDGLIDSYFDNSSTL | 1661696 | Cell/Virus Enclosed | NO |
| GLDLIQAGSVVR | 1441742 | Cell/Virus Enclosed | NO |
| GLDLLFWEQGGLCKALQEQCRFPN | 94530 | Ectodomain | YES |
| GLDLMQAGTIVR | 1441744 | Cell/Virus Enclosed | NO |
| GLDLTPVIDDVD | 1441747 | Cell/Virus Enclosed | NO |
| GLDYKTFKDIVE | 1441757 | Cell/Virus Enclosed | NO |
| GLEAYADKTLQEMKAL | 1661707 | Cell/Virus Enclosed | NO |
| GLEAYADRTLQE | 1441767 | Cell/Virus Enclosed | NO |
| GLEDFNFEHVVYGDVS | 1661708 | Cell/Virus Enclosed | NO |
| GLEDYAFDHIVYGSFN | 1661709 | Cell/Virus Enclosed | NO |
| GLEDYAFEHIVYGNFN | 1661710 | Cell/Virus Enclosed | NO |
| GLEFTDENYNNLVPTY | 1661711 | Cell/Virus Enclosed | NO |
| GLELHWAPEFPWMFED | 1661712 | Cell/Virus Enclosed | NO |
| GLELTDESCNNLVPTY | 1661713 | Cell/Virus Enclosed | NO |
| GLENHAFEHVVY | 1441777 | Cell/Virus Enclosed | NO |
| GLENIAFNVVKQGHFI | 1661715 | Cell/Virus Enclosed | NO |
| GLENIAFNVVNKGSFV | 1661716 | Cell/Virus Enclosed | NO |
| GLENIVFNVVKQ | 1441783 | Cell/Virus Enclosed | NO |
| GLENYAFEHVVY | 1441785 | Cell/Virus Enclosed | NO |
| GLESKQVDYVPLRSAT | 1661718 | Cell/Virus Enclosed | NO |
| GLFDAIWDAFVAAIKL | 1661722 | Cell/Virus Enclosed | NO |
| GLFHSSQLPLDV | 1441798 | Cell/Virus Enclosed | NO |
| GLFKDCARNPIDLPPS | 1661724 | Cell/Virus Enclosed | NO |
| GLFKDCSKIING | 1441802 | Cell/Virus Enclosed | NO |
| GLFKDCSKIITG | 1441803 | Cell/Virus Enclosed | NO |
| GLFKDCSRETSGLSPA | 1661727 | Cell/Virus Enclosed | NO |
| GLFPYQGDHGDR | 1441818 | Ectodomain | YES |
| GLFRIDVPSAES | 1441821 | Cell/Virus Enclosed | NO |

| | | | |
|---|---|---|---|
| GLFRIFFRNAYV | 1441822 | Cell/Virus Enclosed | NO |
| GLFRILLRNAYV | 1441825 | Cell/Virus Enclosed | NO |
| GLFRRRSVGNTG | 1441828 | Cell/Virus Enclosed | NO |
| GLFTLQLVSAVNQSLS | 1661738 | Cell/Virus Enclosed | NO |
| GLGDLVDSYFENDSIL | 1661741 | Cell/Virus Enclosed | NO |
| GLGLIQAGSVVR | 1441847 | Cell/Virus Enclosed | NO |
| GLGTVDADYKMC | 1441850 | Ectodomain | YES |
| GLGTVDVDYKSCTKGL | 1661748 | Ectodomain | YES |
| GLGVTMDVLNKNQKLI | 1661751 | Ectodomain | YES |
| GLGVTMDVLSQNQKLI | 1661752 | Ectodomain | YES |
| GLGYIADISAFKATLY | 1661753 | Cell/Virus Enclosed | NO |
| GLHLLIGLYRRQQTSN | 1661763 | Cell/Virus Enclosed | NO |
| GLHYPTGSCLIA | 1441875 | Cell/Virus Enclosed | NO |
| GLIDLIVNVLSI | 1441882 | Cell/Virus Enclosed | NO |
| GLIDLLVNVLSIFSSS | 1661773 | Cell/Virus Enclosed | NO |
| GLIDQINSTYVDLKLL | 1661774 | Ectodomain | YES |
| GLIDSYFDNSSTLQSV | 1661775 | Cell/Virus Enclosed | NO |
| GLILKFDSGTVT | 1441894 | Cell/Virus Enclosed | NO |
| GLINSSLFVEDC | 1441896 | Ectodomain | YES |
| GLKAKPKFTLGKRKATPTTS | 110863 | Capsid | UN |
| GLKAKPKLKRAAPTSTRTSS | 110864 | Capsid | UN |
| GLKFTEESFYKNMYLK | 1661780 | Cell/Virus Enclosed | NO |
| GLKNGQGENDNI | 1441919 | Cell/Virus Enclosed | NO |
| GLKSSTGFVYFNATGS | 1661793 | Ectodomain | YES |
| GLKYTKILKDDGNFVV | 1661802 | Cell/Virus Enclosed | NO |
| GLKYTKIMKDDGNCVV | 1661803 | Cell/Virus Enclosed | NO |
| GLLATNNVFSLK | 1441939 | Cell/Virus Enclosed | NO |
| GLLDLAWRVPCA | 1441940 | Cell/Virus Enclosed | NO |
| GLLDQAWRFPCA | 1441941 | Cell/Virus Enclosed | NO |
| GLLDTTQLHVADTLMQ | 1661806 | Ectodomain | YES |
| GLLEVSVCQYNMCEYP | 1661807 | Ectodomain | YES |
| GLLGWSPQAQGILTT | 1661812 | Underneath Membrane | NO |
| GLLHSSQLPLDI | 1441957 | Cell/Virus Enclosed | NO |
| GLLHSSQLPLDVYDLT | 1661813 | Cell/Virus Enclosed | NO |
| GLLNVQKVAQPK | 1441971 | Cell/Virus Enclosed | NO |
| GLLQPTANSTTV | 1441977 | Cell/Virus Enclosed | NO |
| GLLRPTDNSTTV | 1441985 | Cell/Virus Enclosed | NO |
| GLLRYHTYARFG | 21026 | Capsid | UN |
| GLLTACAFSVDPATTY | 1661826 | Cell/Virus Enclosed | NO |
| GLLYDSPCFSQQ | 1441992 | Ectodomain | YES |
| GLLYNSPCFSQQ | 1441995 | Ectodomain | YES |
| GLLYNSSCFSHQ | 1441996 | Ectodomain | YES |
| GLLYNSSCFSQQ | 1441997 | Ectodomain | YES |
| GLMQNAFLYSSL | 1442006 | Cell/Virus Enclosed | NO |
| GLNFTETITTGF | 1442015 | Ectodomain | YES |
| GLNKPFVKDNVSFVAD | 1661843 | Cell/Virus Enclosed | NO |
| GLNLEEAARCMR | 1442024 | Cell/Virus Enclosed | NO |
| GLNNLNRGMVLE | 1442027 | Cell/Virus Enclosed | NO |
| GLNTLLVQPALY | 1442032 | Cell/Virus Enclosed | NO |
| GLPLEFGASTEM | 1442042 | Cell/Virus Enclosed | NO |
| GLPNNTASWFTALTQHGKED | 2060826 | Capsid | UN |
| GLPNTCFGPLVR | 1442051 | Cell/Virus Enclosed | NO |
| GLPYGANKEGIIWVAT | 1661862 | Cell/Virus Enclosed | NO |
| GLQDCVTGINDDDYVI | 1661863 | Cell/Virus Enclosed | NO |
| GLQFQILDEGLD | 1442066 | Cell/Virus Enclosed | NO |
| GLQFQILDEGVDCKIQ | 1661865 | Cell/Virus Enclosed | NO |
| GLQIQTVDSSQGSEYD | 1661866 | Cell/Virus Enclosed | NO |
| GLQKCEQLLREY | 1442068 | Underneath Membrane | NO |
| GLQLFTTWLPDL | 1442069 | Cell/Virus Enclosed | NO |
| GLQTQTVDSAQGSEYD | 1661870 | Cell/Virus Enclosed | NO |
| GLQYAAITAPYV | 1442072 | Cell/Virus Enclosed | NO |
| GLRDYITNRTFM | 1442075 | Ectodomain | YES |
| GLRGAGDLQGNF | 1442078 | Cell/Virus Enclosed | NO |
| GLRGAIRKAKDYGFTV | 1661873 | Cell/Virus Enclosed | NO |
| GLRGPGDLQGNFGDLQ | 1661874 | Cell/Virus Enclosed | NO |
| GLRGTGVLTTSS | 1442083 | Ectodomain | YES |
| GLSGIGAVGGDL | 1442097 | Cell/Virus Enclosed | NO |
| GLSPAYAPTYVSVDDK | 1661885 | Cell/Virus Enclosed | NO |
| GLSSFAAIPFAQ | 1442109 | Ectodomain | YES |
| GLSSSAAIPFAQ | 1442111 | Ectodomain | YES |
| GLTCSTDLQKAN | 1442116 | Ectodomain | YES |
| GLTCVKDYEFKF | 1442118 | Cell/Virus Enclosed | NO |
| GLTFQTVRSGNF | 1442127 | Cell/Virus Enclosed | NO |
| GLTNCLASECFVKSDI | 1661900 | Cell/Virus Enclosed | NO |
| GLTNQVVKPGHFNEEF | 1661902 | Cell/Virus Enclosed | NO |
| GLTSQTVKPGHFNKEF | 1661910 | Cell/Virus Enclosed | NO |

| | | | |
|---|---|---|---|
| GLTTQTVDSSQGSEYQ | 1661911 | Cell/Virus Enclosed | NO |
| GLVGVLTLDNQDLNGK | 1661927 | Cell/Virus Enclosed | NO |
| GLVNSSLLVEDC | 1442176 | Ectodomain | YES |
| GLVNSSLSVEDC | 1442177 | Ectodomain | YES |
| GLVNVQKFAQPK | 1442180 | Cell/Virus Enclosed | NO |
| GLVNVQKVAQPK | 1442181 | Cell/Virus Enclosed | NO |
| GLVNVQKVEQPKIEPK | 1661933 | Cell/Virus Enclosed | NO |
| GLVQQITQWLSN | 1442186 | Cell/Virus Enclosed | NO |
| GLVRMYNLLACP | 1442195 | Cell/Virus Enclosed | NO |
| GLVTICDFYSKLGVGF | 1661937 | Cell/Virus Enclosed | NO |
| GLWSPTITHTNC | 1442214 | Cell/Virus Enclosed | NO |
| GLYDDILTNNKPWFVR | 1661950 | Cell/Virus Enclosed | NO |
| GLYEIKIPTNFT | 1442217 | Ectodomain | YES |
| GLYEIQIPSEFTIGNM | 1661952 | Ectodomain | YES |
| GLYKNMCTGSPP | 1442231 | Cell/Virus Enclosed | NO |
| GLYKNSATGSLDVEVF | 1661958 | Cell/Virus Enclosed | NO |
| GLYKQSCTGSPYIDDF | 1661960 | Cell/Virus Enclosed | NO |
| GLYRRQQTSNLVVQEF | 1661968 | Cell/Virus Enclosed | NO |
| GLYVPIATRNNC | 1442254 | Cell/Virus Enclosed | NO |
| GMAITIKPEANI | 1442262 | Cell/Virus Enclosed | NO |
| GMDAKQVDYVPLKSAT | 1661980 | Cell/Virus Enclosed | NO |
| GMDNLACESQTP | 1442273 | Cell/Virus Enclosed | NO |
| GMEAAPSGTWLT | 1442276 | Capsid | UN |
| GMEVTPLGTWLT | 1442284 | Cell/Virus Enclosed | NO |
| GMFGLYDDILTNNKPW | 1661992 | Cell/Virus Enclosed | NO |
| GMGTCPAGTNYL | 1442298 | Ectodomain | YES |
| GMGVSCTVTREDGTNRR | 21277 | Ectodomain | YES |
| GMHFGACDYDRLPNEV | 1661995 | Ectodomain | YES |
| GMIKKLDNDVLNGIIS | 1661998 | Cell/Virus Enclosed | NO |
| GMIRYYDVMANGGTGF | 1662000 | Cell/Virus Enclosed | NO |
| GMKYVKWEHDSG | 1442316 | Cell/Virus Enclosed | NO |
| GMLNGMPVSSIK | 1442318 | Cell/Virus Enclosed | NO |
| GMLNGNAIATVKSEDG | 1662008 | Cell/Virus Enclosed | NO |
| GMMNISPKPQRQ | 1442324 | Cell/Virus Enclosed | NO |
| GMMNMSPKPQLQ | 1442325 | Cell/Virus Enclosed | NO |
| GMNMSPKPQRQR | 1442333 | Cell/Virus Enclosed | NO |
| GMNTSHSVCNGC | 1442334 | Ectodomain | YES |
| GMNTSQSVCNGC | 1442335 | Ectodomain | YES |
| GMPIPDFDLDDFIDAP | 1662014 | Cell/Virus Enclosed | NO |
| GMPSFVAYETARQEYE | 1662015 | Cell/Virus Enclosed | NO |
| GMPVSSIKSDKG | 1442346 | Cell/Virus Enclosed | NO |
| GMQTVEDLRARM | 1442350 | Cell/Virus Enclosed | NO |
| GMSFYCEDHKPQ | 1442354 | Cell/Virus Enclosed | NO |
| GMSYSMTTFEIA | 1442368 | Cell/Virus Enclosed | NO |
| GMSYYCEDHKPQYSFK | 1662031 | Cell/Virus Enclosed | NO |
| GMTNCLASECFM | 1442376 | Cell/Virus Enclosed | NO |
| GMVNMSPKPQRQ | 1442381 | Cell/Virus Enclosed | NO |
| GNAAIPDYNYYK | 1442390 | Cell/Virus Enclosed | NO |
| GNAAITDYNYYKYNLP | 1662046 | Cell/Virus Enclosed | NO |
| GNAIATVKSEDGNIKN | 1662051 | Cell/Virus Enclosed | NO |
| GNAINVGGRGLQ | 1442394 | Cell/Virus Enclosed | NO |
| GNALYNNECGRA | 1442397 | Cell/Virus Enclosed | NO |
| GNCAGNVIGNDALTRF | 1662059 | Cell/Virus Enclosed | NO |
| GNCAKRFRRVLV | 1442409 | Cell/Virus Enclosed | NO |
| GNCTFGSPDDAS | 1442421 | Cell/Virus Enclosed | NO |
| GNCYRCPKEIVETVSA | 1662070 | Cell/Virus Enclosed | NO |
| GNDDLSISTIFT | 1442431 | Cell/Virus Enclosed | NO |
| GNDEKVMTFYPR | 1442432 | Cell/Virus Enclosed | NO |
| GNDLSLPVMFSE | 1442433 | Cell/Virus Enclosed | NO |
| GNDSGFAAYNRY | 1442434 | Underneath Membrane | NO |
| GNDVLYQPPTASVSTS | 1662075 | Cell/Virus Enclosed | NO |
| GNEALSISTIFT | 1442440 | Cell/Virus Enclosed | NO |
| GNELAQVLTEVVYSNG | 1662076 | Cell/Virus Enclosed | NO |
| GNESWLTGNAIQ | 1442442 | Cell/Virus Enclosed | NO |
| GNETTFAFVGAL | 1442443 | Ectodomain | YES |
| GNETTSSFVGAL | 1442445 | Ectodomain | YES |
| GNFDDQPSLQVESANL | 1662078 | Cell/Virus Enclosed | NO |
| GNFGDLQLNKLGTEDP | 1662079 | Cell/Virus Enclosed | NO |
| GNFGDLQLTKLG | 1442455 | Cell/Virus Enclosed | NO |
| GNFGDQDLIRQGTDYKHW | 21424 | Capsid | UN |
| GNFGDQELIRQGTEYK | 1662082 | Cell/Virus Enclosed | NO |
| GNFNQDFYDFIL | 1442470 | Cell/Virus Enclosed | NO |
| GNFNQDFYEFVKSKGL | 1662092 | Cell/Virus Enclosed | NO |
| GNFRTGVDPDLS | 1442473 | Cell/Virus Enclosed | NO |
| GNFYCFVNTTIGNETT | 1662100 | Ectodomain | YES |
| GNGGDAALALLLLDRL | 1381497 | Cell/Virus Enclosed | NO |

| | | | |
|---|---|---|---|
| GNGGDTALALLL | 1442490 | Cell/Virus Enclosed | NO |
| GNGILKWADQSDQFRN | 1662106 | Cell/Virus Enclosed | NO |
| GNGILKWSDQSD | 1442493 | Cell/Virus Enclosed | NO |
| GNGNHIISLVQN | 1442495 | Ectodomain | YES |
| GNGNHILSLVQN | 1442496 | Ectodomain | YES |
| GNGPGIDAGYKN | 1442497 | Ectodomain | YES |
| GNGSMIEDIDYVPLKS | 1662112 | Cell/Virus Enclosed | NO |
| GNGSMIEDMDYV | 1442502 | Cell/Virus Enclosed | NO |
| GNGSMIEDVDYV | 1442503 | Cell/Virus Enclosed | NO |
| GNHCPAGNSHTS | 1442513 | Ectodomain | YES |
| GNHCPAGNSYTS | 1442514 | Ectodomain | YES |
| GNHFKELWHLDT | 1442517 | Ectodomain | YES |
| GNHFKESWHMDT | 1442520 | Ectodomain | YES |
| GNHRSAFALHTG | 1442528 | Ectodomain | YES |
| GNIFGLYKNSATGSLD | 1662122 | Cell/Virus Enclosed | NO |
| GNIFIVTPCNQPDQVA | 1662123 | Ectodomain | YES |
| GNIIGFKDFVTNKTYN | 1662124 | Ectodomain | YES |
| GNIIPHYSWFSG | 1442540 | Cell/Virus Enclosed | NO |
| GNIKPHNYSYMD | 1442542 | Ectodomain | YES |
| GNITKTNVQPTG | 1442546 | Cell/Virus Enclosed | NO |
| GNKDEQIGYWNVQERW | 1662128 | Capsid | UN |
| GNKFYNVSTGLIPGFV | 1662129 | Cell/Virus Enclosed | NO |
| GNKIVDKEVKDMQFDL | 1662137 | Cell/Virus Enclosed | NO |
| GNLGNLQSFNYR | 1442579 | Ectodomain | YES |
| GNLGNLQSSNYR | 1442580 | Ectodomain | YES |
| GNLGSNGKPGNVGGND | 1662143 | Cell/Virus Enclosed | NO |
| GNLHQRELLGVSDVWH | 1662147 | Cell/Virus Enclosed | NO |
| GNLLDSYFVVKR | 1442593 | Cell/Virus Enclosed | NO |
| GNLQSNHDQHCQ | 1442600 | Cell/Virus Enclosed | NO |
| GNLQSSNYRIDT | 1442601 | Ectodomain | YES |
| GNLVEKLSPWFMDGEN | 1662159 | Cell/Virus Enclosed | NO |
| GNMEEFIQTSSP | 1442615 | Ectodomain | YES |
| GNMFQFATFPVY | 1442616 | Ectodomain | YES |
| GNMFQFATLPVYDTIK | 1662164 | Ectodomain | YES |
| GNMLTGIAAHPP | 1442625 | Cell/Virus Enclosed | NO |
| GNMLTGIAARPP | 1442626 | Cell/Virus Enclosed | NO |
| GNMLTGISARPP | 1442627 | Cell/Virus Enclosed | NO |
| GNMVEFIQTSSPKVTI | 1662169 | Ectodomain | YES |
| GNNSCTCQPLAF | 1442637 | Ectodomain | YES |
| GNNSCTCQPQAF | 1442638 | Ectodomain | YES |
| GNNSWLTGNAIQ | 1442639 | Cell/Virus Enclosed | NO |
| GNNVVVMSTCAV | 1442642 | Ectodomain | YES |
| GNNVYCYNTELM | 1442644 | Cell/Virus Enclosed | NO |
| GNPDEPHKDVYE | 1442648 | Cell/Virus Enclosed | NO |
| GNPDEPQKDVYELRYN | 1662175 | Cell/Virus Enclosed | NO |
| GNPIIPFKDGIY | 1442652 | Ectodomain | YES |
| GNPKGIPIFDDP | 1442657 | Cell/Virus Enclosed | NO |
| GNPKGIPIVDDPVVDW | 1662180 | Cell/Virus Enclosed | NO |
| GNPVVEYLSKED | 1442667 | Cell/Virus Enclosed | NO |
| GNPVYFLRYSGA | 1442668 | Cell/Virus Enclosed | NO |
| GNQHVDDRPSPAPAPKRALGTL | 21502 | Capsid | UN |
| GNRKRNQKPLEPKFSI | 1662190 | Capsid | UN |
| GNRPGNGILKWA | 1442681 | Capsid | UN |
| GNRRVELWHWDT | 1442683 | Ectodomain | YES |
| GNRSGNGILKWADQSD | 1662192 | Cell/Virus Enclosed | NO |
| GNRSGNGILKWS | 1442686 | Cell/Virus Enclosed | NO |
| GNRTPTPGVTPD | 1442689 | Cell/Virus Enclosed | NO |
| GNSLNHLTFQLR | 1442706 | Ectodomain | YES |
| GNSLNYLTSQLR | 1442708 | Ectodomain | YES |
| GNSQSSSRASSLSRNS | 1662203 | Cell/Virus Enclosed | NO |
| GNSTRGTFPGPS | 1442731 | Cell/Virus Enclosed | NO |
| GNSTRGTSPCPS | 1442732 | Cell/Virus Enclosed | NO |
| GNSTRGTSPGPSGIGA | 1662206 | Cell/Virus Enclosed | NO |
| GNSTRGTSPVPS | 1442735 | Capsid | UN |
| GNSTRSTSPGPS | 1442737 | Capsid | UN |
| GNSYTSFAIYHT | 1442739 | Ectodomain | YES |
| GNSYTSFATYHTPATD | 1662207 | Ectodomain | YES |
| GNTSFACLSNVS | 1442760 | Ectodomain | YES |
| GNVDLDALASDIFDTP | 1662225 | Cell/Virus Enclosed | NO |
| GNVEAVNFNVTN | 1442774 | Ectodomain | YES |
| GNVEAVNFNVTTAETT | 1662226 | Ectodomain | YES |
| GNVFCDGVNVTKHKCD | 1662228 | Cell/Virus Enclosed | NO |
| GNVGGNDALSTS | 1442779 | Cell/Virus Enclosed | NO |
| GNVGGNEALSIS | 1442780 | Cell/Virus Enclosed | NO |
| GNVILKPSEEGV | 1442785 | Cell/Virus Enclosed | NO |
| GNVINWTRDSITEVWSYNAELLVAM | 1144095 | Ectodomain | YES |

| | | | |
|---|---|---|---|
| GNVKPGDAVITTQQAG | 1662235 | Cell/Virus Enclosed | NO |
| GNVQVDNGSSINRKQL | 1662241 | Cell/Virus Enclosed | NO |
| GNVSRHNLTKYTMMDL | 1662243 | Cell/Virus Enclosed | NO |
| GNVSRQDLTKYTMMDL | 1662244 | Cell/Virus Enclosed | NO |
| GNVSTVFVPATRDSVP | 1662245 | Cell/Virus Enclosed | NO |
| GNVVKTAVQPTAPAYV | 1662249 | Cell/Virus Enclosed | NO |
| GNVVQQCCTAAV | 1442808 | Cell/Virus Enclosed | NO |
| GNVVTSVMESQISTKE | 1662252 | Cell/Virus Enclosed | NO |
| GNVWQIQTIQDSDGTN | 1662253 | Cell/Virus Enclosed | NO |
| GNYFTFKQANNN | 1442816 | Cell/Virus Enclosed | NO |
| GNYKLNTDHSNSSDNI | 1662258 | Underneath Membrane | NO |
| GNYNRNASLNSFKEYF | 1662262 | Ectodomain | YES |
| GNYNYKHRYLRH | 1442830 | Ectodomain | YES |
| GNYRLPSSKPSGMDTA | 1662266 | Underneath Membrane | NO |
| GNYRLPSTQKGSGMDT | 1662267 | Ectodomain | YES |
| GNYYLPYPDPSR | 1442864 | Cell/Virus Enclosed | NO |
| GPAGVESDVETNEIVD | 1662274 | Cell/Virus Enclosed | NO |
| GPASARDLICAQYVAG | 1662276 | Ectodomain | YES |
| GPASARNLICAQ | 1442882 | Underneath Membrane | NO |
| GPASVRDLICAQ | 1442883 | Ectodomain | YES |
| GPAYVMIDKVEFENGF | 1662278 | Cell/Virus Enclosed | NO |
| GPCKTCTTPAQGTSM | 1662279 | Underneath Membrane | NO |
| GPCWSALVSALK | 1442887 | Cell/Virus Enclosed | NO |
| GPDDQIGFYRRA | 1442895 | Capsid | UN |
| GPDMFLGTCRCC | 1442906 | Cell/Virus Enclosed | NO |
| GPDQICNTPTQC | 1442908 | Cell/Virus Enclosed | NO |
| GPDVFLHKCYRCPAEI | 1662293 | Cell/Virus Enclosed | NO |
| GPDVFLHKCYRY | 1442921 | Cell/Virus Enclosed | NO |
| GPEAALPFRAVKDGIV | 1662295 | Cell/Virus Enclosed | NO |
| GPEAGLPYGANKDGII | 1662296 | Cell/Virus Enclosed | NO |
| GPEAGLPYGANKEGII | 1662297 | Cell/Virus Enclosed | NO |
| GPEEFEGAGDGEPP | 741481 | Ectodomain | YES |
| GPFMDNAINVGGTGLQ | 1662312 | Cell/Virus Enclosed | NO |
| GPFMGNAINVGG | 1442947 | Cell/Virus Enclosed | NO |
| GPFNFCSETVNGYAKS | 1662313 | Cell/Virus Enclosed | NO |
| GPGDLQGNFGDLQLNK | 1662316 | Cell/Virus Enclosed | NO |
| GPGESKPQSRNP | 1442953 | Capsid | UN |
| GPGIDAGYKNSG | 1442955 | Ectodomain | YES |
| GPGKNNGIGTCP | 1442958 | Ectodomain | YES |
| GPGKNNGMGTCP | 1442959 | Ectodomain | YES |
| GPGNTFINGDIARELG | 1662318 | Cell/Virus Enclosed | NO |
| GPHAKDQYGSDI | 1442964 | Cell/Virus Enclosed | NO |
| GPHEFCSQHTML | 1442972 | Cell/Virus Enclosed | NO |
| GPHEFCSQHTMQIVDK | 1662326 | Cell/Virus Enclosed | NO |
| GPHKAAKFRERV | 1442974 | Cell/Virus Enclosed | NO |
| GPHKDAKFRERV | 1442975 | Cell/Virus Enclosed | NO |
| GPHKDLKFRQRS | 1442978 | Cell/Virus Enclosed | NO |
| GPIEHVTSVNLF | 1442984 | Cell/Virus Enclosed | NO |
| GPISQFNYKQSFSNPT | 1662333 | Ectodomain | YES |
| GPKELLLLWESGKAKP | 1662338 | Cell/Virus Enclosed | NO |
| GPKELLLSWESGKVKP | 1662339 | Cell/Virus Enclosed | NO |
| GPLGSSGLKKMA | 1443020 | Cell/Virus Enclosed | NO |
| GPLKVGAGVMLECEKF | 1662349 | Cell/Virus Enclosed | NO |
| GPLQNFGNSEML | 1443028 | Cell/Virus Enclosed | NO |
| GPLQVGDSVLLQ | 1443036 | Cell/Virus Enclosed | NO |
| GPNEHVTSVNLL | 1443049 | Cell/Virus Enclosed | NO |
| GPNKYCYCGNFA | 1443053 | Cell/Virus Enclosed | NO |
| GPNLNAGEDIRL | 1443055 | Cell/Virus Enclosed | NO |
| GPNLNAGEDVQL | 1443056 | Cell/Virus Enclosed | NO |
| GPNNRSLSRSNSNFRH | 1662362 | Capsid | UN |
| GPNTRSLSRSNS | 1443060 | Cell/Virus Enclosed | NO |
| GPPIAPGVPATE | 1443067 | Cell/Virus Enclosed | NO |
| GPPPISTDTVDVTNGL | 1662371 | Ectodomain | YES |
| GPPQGAFHVTMR | 1443070 | Cell/Virus Enclosed | NO |
| GPPRAELNGVVVDKVG | 1662372 | Cell/Virus Enclosed | NO |
| GPPSISTATVDV | 1443074 | Ectodomain | YES |
| GPPSISTDTVDV | 1443075 | Ectodomain | YES |
| GPQNQRSAPRIT | 1443078 | Cell/Virus Enclosed | NO |
| GPQNQSSAPRIT | 1443079 | Cell/Virus Enclosed | NO |
| GPQRADLNGVIVDKVG | 1662375 | Cell/Virus Enclosed | NO |
| GPRDLDHNFGSAGVVA | 1662380 | Capsid | UN |
| GPRTIKVSTVQSKLTD | 1662383 | Cell/Virus Enclosed | NO |
| GPSDSTDNNQDG | 1443100 | Cell/Virus Enclosed | NO |
| GPSDSTDNNQNGDRSG | 1662385 | Cell/Virus Enclosed | NO |
| GPSGIGTVGGDL | 1443106 | Cell/Virus Enclosed | NO |
| GPSQNFGNAEMLKLGT | 1662388 | Capsid | UN |

| | | | |
|---|---|---|---|
| GPSSARDLICAQ | 1443110 | Ectodomain | YES |
| GPSSQPYYCFINSTIN | 1662389 | Ectodomain | YES |
| GPSVETSDSQFE | 1443112 | Cell/Virus Enclosed | NO |
| GPTDSADNNQNG | 1443114 | Capsid | UN |
| GPTEHVTSVNLFDVGY | 1662392 | Cell/Virus Enclosed | NO |
| GPTTCGYLPTNA | 1443125 | Cell/Virus Enclosed | NO |
| GPVDKGHYTVYDTAKK | 1662397 | Cell/Virus Enclosed | NO |
| GPVDPTFLHRFYSLKA | 1662398 | Cell/Virus Enclosed | NO |
| GPVFTGDLFYTI | 1443134 | Cell/Virus Enclosed | NO |
| GPVVLLSVSPRPRSGPTAPQ | 741485 | Underneath Membrane | NO |
| GPVYVSDSVTLVNVATGAQAVARSL | 21834 | Undetermined | UN |
| GPYAKDQYGTDI | 1443137 | Cell/Virus Enclosed | NO |
| GPYANASYGQSL | 1443139 | Cell/Virus Enclosed | NO |
| GPYEFCSQHTMQ | 1443142 | Cell/Virus Enclosed | NO |
| GPYKDAQVVQLLIQDY | 1662402 | Cell/Virus Enclosed | NO |
| GPYKDAQVVQLP | 1443149 | Cell/Virus Enclosed | NO |
| GPYRDAQVVQLPVKDY | 1662403 | Cell/Virus Enclosed | NO |
| GQAAVTTTNCIK | 1443154 | Cell/Virus Enclosed | NO |
| GQAEPDRAHYNIVTFCCKCD | 110869 | Cell/Virus Enclosed | NO |
| GQAFFYSGGFYR | 1443158 | Cell/Virus Enclosed | NO |
| GQAGEMPCAIINDKVV | 1662407 | Cell/Virus Enclosed | NO |
| GQAMESVGFVYD | 1443162 | Cell/Virus Enclosed | NO |
| GQARVAYQVASR | 1443166 | Cell/Virus Enclosed | NO |
| GQDFKTFDLLKYDFTE | 1662415 | Cell/Virus Enclosed | NO |
| GQDVIFSQFDSLGVSS | 1662416 | Cell/Virus Enclosed | NO |
| GQECLARSACVY | 1443183 | Cell/Virus Enclosed | NO |
| GQEEFIQTNSPKVTID | 1662418 | Ectodomain | YES |
| GQEENDNISVAV | 1443185 | Capsid | UN |
| GQFVGWVIPLAF | 1443205 | Cell/Virus Enclosed | NO |
| GQFYINGFKYFD | 1443206 | Ectodomain | YES |
| GQGENDNISVAVPKSR | 1662429 | Cell/Virus Enclosed | NO |
| GQGENDNTSVAV | 1443211 | Capsid | UN |
| GQGEYDNISVAV | 1443212 | Cell/Virus Enclosed | NO |
| GQGIFFEVNATY | 1443214 | Ectodomain | YES |
| GQGQNNADPKVY | 1443220 | Cell/Virus Enclosed | NO |
| GQGVPIAYGIPP | 1443230 | Cell/Virus Enclosed | NO |
| GQGVPINTNSSPDDQI | 1662438 | Cell/Virus Enclosed | NO |
| GQGVPLAFGVPL | 1443235 | Cell/Virus Enclosed | NO |
| GQGVPLNANSTPAQNA | 1662440 | Cell/Virus Enclosed | NO |
| GQKGDAEEALSKLSEY | 1662449 | Cell/Virus Enclosed | NO |
| GQKNGQGENDNI | 1443253 | Capsid | UN |
| GQKSTTLTGVEA | 1443256 | Cell/Virus Enclosed | NO |
| GQLKNFEAPLKC | 1443272 | Cell/Virus Enclosed | NO |
| GQLLDTKLNGILQQAA | 1662461 | Cell/Virus Enclosed | NO |
| GQLQSLSKDYISSNGP | 1662463 | Cell/Virus Enclosed | NO |
| GQLSDHPHALSSSSSHAEPR | 21912 | Capsid | UN |
| GQNNADPKVYPI | 1443285 | Cell/Virus Enclosed | NO |
| GQPCVVSEQEES | 1443290 | Cell/Virus Enclosed | NO |
| GQQFVKETTDMV | 1443302 | Cell/Virus Enclosed | NO |
| GQQKHLLPRWYF | 1443307 | Cell/Virus Enclosed | NO |
| GQQKQLLPRWYF | 1443309 | Cell/Virus Enclosed | NO |
| GQQQQSQTVTKKSAAE | 1662482 | Cell/Virus Enclosed | NO |
| GQQVDLVISDMYDPTT | 1662485 | Cell/Virus Enclosed | NO |
| GQRIYDDVNASL | 1443315 | Cell/Virus Enclosed | NO |
| GQRVYDDVDASLFVDI | 1662489 | Cell/Virus Enclosed | NO |
| GQSLCALPDTHS | 1443327 | Ectodomain | YES |
| GQSLCALPDTPI | 1443328 | Ectodomain | YES |
| GQSLCALPDTPSTLTP | 1662491 | Ectodomain | YES |
| GQSLEGVFWVAN | 1443334 | Cell/Virus Enclosed | NO |
| GQSNADPKVYPI | 1443336 | Cell/Virus Enclosed | NO |
| GQSNAVPKVYPI | 1443337 | Cell/Virus Enclosed | NO |
| GQTGVIADYNYKLPDD | 1662496 | Ectodomain | YES |
| GQWFKNQTSVAP | 1443362 | Ectodomain | YES |
| GRAIIVSVEQLEPCAQ | 1662509 | Cell/Virus Enclosed | NO |
| GRCFNCRPAVVN | 1443374 | Ectodomain | YES |
| GRCIVFDVKGVE | 1443375 | Cell/Virus Enclosed | NO |
| GRCIVLDVKGVEELHD | 1662510 | Cell/Virus Enclosed | NO |
| GRCPTAADINIH | 1443379 | Ectodomain | YES |
| GRCPTAADINIPDLPI | 1662511 | Ectodomain | YES |
| GRCPTAADINNP | 1443381 | Ectodomain | YES |
| GRCPTAADINTP | 1443382 | Ectodomain | YES |
| GRCYNCRPAVIN | 1443389 | Ectodomain | YES |
| GRCYNCRPAVVN | 1443390 | Ectodomain | YES |
| GRDFKFPDGQGV | 1443394 | Cell/Virus Enclosed | NO |
| GRDFKFSDGQGVPIAF | 1662516 | Capsid | UN |
| GRDNGALEAFKRSNNG | 1662520 | Cell/Virus Enclosed | NO |

| | | | |
|---|---|---|---|
| GRDRIMAWTVVNSIC | 504102 | Cell/Virus Enclosed | NO |
| GRDTGALEAFRKARNG | 1662521 | Cell/Virus Enclosed | NO |
| GRDTSDFTDSVR | 1443402 | Ectodomain | YES |
| GREISCNVCTKRATVY | 1662524 | Cell/Virus Enclosed | NO |
| GRELNCNVCANR | 1443411 | Cell/Virus Enclosed | NO |
| GRESLIFPLGTY | 1443415 | Cell/Virus Enclosed | NO |
| GRGDCKGFSSDVLSDV | 1662533 | Ectodomain | YES |
| GRGESKPQSRNPSSDR | 1662534 | Capsid | UN |
| GRGFCKAHNWNC | 1443439 | Cell/Virus Enclosed | NO |
| GRGGYHYTPFHYERDN | 1662537 | Cell/Virus Enclosed | NO |
| GRGPGKCYNCRP | 1443458 | Ectodomain | YES |
| GRGYNVSSIVTMTSQ | 22063 | Ectodomain | YES |
| GRITPYCHDPTV | 1443487 | Cell/Virus Enclosed | NO |
| GRIVNYTVCDDC | 1443490 | Ectodomain | YES |
| GRKMLMALLSDNAYLK | 1662552 | Cell/Virus Enclosed | NO |
| GRKSLIFPSGTY | 1443503 | Cell/Virus Enclosed | NO |
| GRKVDLQLGNLGYLQS | 1662554 | Ectodomain | YES |
| GRKVNFNEKPVVMEIP | 1662555 | Cell/Virus Enclosed | NO |
| GRLAALNVFVSH | 1443509 | Ectodomain | YES |
| GRLNCSFQDAAS | 1443518 | Cell/Virus Enclosed | NO |
| GRLVGTTLQGKP | 1443526 | Cell/Virus Enclosed | NO |
| GRMTPYCHDPTV | 1443533 | Cell/Virus Enclosed | NO |
| GRNLSDFTPRSDMEYD | 1662573 | Cell/Virus Enclosed | NO |
| GRPQGAFHVTMR | 1443552 | Cell/Virus Enclosed | NO |
| GRPVHVKEITRE | 1443557 | Cell/Virus Enclosed | NO |
| GRQFYEFYNDIK | 1443559 | Cell/Virus Enclosed | NO |
| GRQFYEFYNDVKPPVL | 1662582 | Cell/Virus Enclosed | NO |
| GRQIMGSCSFEDELTP | 1662585 | Cell/Virus Enclosed | NO |
| GRQLYKSYSEHV | 1443564 | Cell/Virus Enclosed | NO |
| GRRAQPKQTSTS | 1443568 | Cell/Virus Enclosed | NO |
| GRRAQPKQTVTS | 1443569 | Cell/Virus Enclosed | NO |
| GRRGPEQTQGNFGDQE | 1662589 | Cell/Virus Enclosed | NO |
| GRSVYVKFQDSKPPLP | 1662602 | Cell/Virus Enclosed | NO |
| GRTEVPQLVNAN | 1443602 | Ectodomain | YES |
| GRTGELPCAIIN | 1443603 | Cell/Virus Enclosed | NO |
| GRTGELPCAIMNDKVV | 1662603 | Cell/Virus Enclosed | NO |
| GRTPTSGVTPDM | 1443616 | Cell/Virus Enclosed | NO |
| GRTVVRPLVEDS | 1443619 | Ectodomain | YES |
| GRVDDDYICGDG | 1443625 | Cell/Virus Enclosed | NO |
| GRVGNGYVCGTG | 1443632 | Cell/Virus Enclosed | NO |
| GRVIYGNVSRHN | 1443634 | Cell/Virus Enclosed | NO |
| GRVLRPYHANSS | 1443638 | Ectodomain | YES |
| GRVSAAFHANSSEPAL | 1662623 | Ectodomain | YES |
| GRVVNYTVCDDC | 1443649 | Ectodomain | YES |
| GRWNPQPGVYPPHAA | 22253 | Capsid | UN |
| GSAFRDLKPLPF | 1443667 | Cell/Virus Enclosed | NO |
| GSAGNATEVPAN | 1443668 | Cell/Virus Enclosed | NO |
| GSAGVVANGVKAKGYP | 1662635 | Capsid | UN |
| GSAIASEFSSLP | 1443675 | Cell/Virus Enclosed | NO |
| GSALNGSCGSVG | 1443678 | Cell/Virus Enclosed | NO |
| GSALQIPFAMQM | 1443680 | Ectodomain | YES |
| GSANIFIGDNVG | 1443683 | Cell/Virus Enclosed | NO |
| GSANIFIGDTVG | 1443684 | Cell/Virus Enclosed | NO |
| GSANQQQLKQLE | 1443685 | Cell/Virus Enclosed | NO |
| GSAQVTSFCKSG | 1443690 | Ectodomain | YES |
| GSAVLWANGVKA | 1443698 | Cell/Virus Enclosed | NO |
| GSCEYAQEGVCI | 1443706 | Cell/Virus Enclosed | NO |
| GSCFNSITVDKF | 1443707 | Ectodomain | YES |
| GSCGSVGYTKDG | 1443711 | Cell/Virus Enclosed | NO |
| GSCGSVGYTKEGSVIN | 1662652 | Cell/Virus Enclosed | NO |
| GSCNFPLEATWH | 1443729 | Ectodomain | YES |
| GSCPDKNNGIGI | 1443730 | Ectodomain | YES |
| GSCPDKNNGIGT | 1443731 | Ectodomain | YES |
| GSCPGKNNGIGT | 1443732 | Ectodomain | YES |
| GSCSFEDELTPSDVYQ | 1662665 | Cell/Virus Enclosed | NO |
| GSCTFEDELAPSDVYQ | 1662666 | Cell/Virus Enclosed | NO |
| GSCVGSGPGKNNGIGT | 1662667 | Ectodomain | YES |
| GSCYVCKPHQVNISLN | 1662670 | Ectodomain | YES |
| GSDAALALLLLDRLNQ | 1662671 | Cell/Virus Enclosed | NO |
| GSDDYVWLSRLLDLAPNYRQ | 125428 | Cell/Virus Enclosed | NO |
| GSDFSLADSVVEVVTT | 1662674 | Cell/Virus Enclosed | NO |
| GSDFTGSVYGNFDDQP | 1662675 | Cell/Virus Enclosed | NO |
| GSDINCNIPTQC | 1443746 | Cell/Virus Enclosed | NO |
| GSDKGIAPGTSVLRQW | 1662677 | Cell/Virus Enclosed | NO |
| GSDKGVAPGTTVLKRW | 1662679 | Cell/Virus Enclosed | NO |
| GSDLLYQLPNCS | 1443755 | Cell/Virus Enclosed | NO |

| | | | |
|---|---|---|---|
| GSDVLYQPPTASVSTS | 1662687 | Cell/Virus Enclosed | NO |
| GSECSKASSRSAIEDL | 1662690 | Ectodomain | YES |
| GSEKGVAPGSAVLRQW | 1662694 | Cell/Virus Enclosed | NO |
| GSENWSVGAWDG | 1443779 | Cell/Virus Enclosed | NO |
| GSFCTQLNRALSGIAA | 1662705 | Ectodomain | YES |
| GSFDNGHYVVYDAANN | 1662708 | Cell/Virus Enclosed | NO |
| GSFINGACGSPGYNLK | 1662710 | Cell/Virus Enclosed | NO |
| GSFKTADGNQRQ | 1443802 | Cell/Virus Enclosed | NO |
| GSFNHKVIGGLH | 1443810 | Cell/Virus Enclosed | NO |
| GSFNNIDTGPPS | 1443811 | Ectodomain | YES |
| GSFNNRDTGHPS | 1443814 | Ectodomain | YES |
| GSFNNRDTGLPS | 1443816 | Ectodomain | YES |
| GSFNNRDTGPLS | 1443817 | Ectodomain | YES |
| GSFNNRDTGPPS | 1443818 | Ectodomain | YES |
| GSFNNRDTGRPS | 1443819 | Ectodomain | YES |
| GSFNNRDTGSPP | 1443820 | Ectodomain | YES |
| GSFNNRDTGSPS | 1443821 | Ectodomain | YES |
| GSFYKATVNQVC | 1443833 | Cell/Virus Enclosed | NO |
| GSGDATTAFANS | 1443839 | Cell/Virus Enclosed | NO |
| GSGFCIDYALPS | 1443842 | Ectodomain | YES |
| GSGFCVDYNSPSSSSS | 1662724 | Ectodomain | YES |
| GSGKSAPNSRST | 1443852 | Cell/Virus Enclosed | NO |
| GSGNLVEKLSPWFMDG | 1662730 | Cell/Virus Enclosed | NO |
| GSGRPAPNSRST | 1443866 | Capsid | UN |
| GSGRSASNSRPGSRSQ | 1662734 | Capsid | UN |
| GSGVAIVDSYYSYLMP | 1662738 | Cell/Virus Enclosed | NO |
| GSGYYYPEPIIE | 1443891 | Ectodomain | YES |
| GSGYYYPEPITENNVV | 1662743 | Ectodomain | YES |
| GSGYYYPEPITG | 1443894 | Ectodomain | YES |
| GSHMDPLSNFEH | 1443896 | Cell/Virus Enclosed | NO |
| GSHMVLPSEAPN | 1443899 | Cell/Virus Enclosed | NO |
| GSIAIKVTEYSWNKKL | 1662749 | Cell/Virus Enclosed | NO |
| GSILVDNDLNPF | 1443916 | Cell/Virus Enclosed | NO |
| GSIRFDSTLPGFETIM | 1662760 | Capsid | UN |
| GSITAQPVKTDN | 1443923 | Secreted | YES |
| GSITQRTRSRPS | 1443933 | Cell/Virus Enclosed | NO |
| GSITQRTRTRPSVQPG | 1662762 | Cell/Virus Enclosed | NO |
| GSKDEQIGYWNV | 1443944 | Cell/Virus Enclosed | NO |
| GSKHVTCCTAND | 1443954 | Cell/Virus Enclosed | NO |
| GSKLELFKRDSD | 1443963 | Cell/Virus Enclosed | NO |
| GSLADGLEAYADKTLQ | 1662775 | Cell/Virus Enclosed | NO |
| GSLCVGSGPGKN | 1443982 | Ectodomain | YES |
| GSLDVEVFNRLATSDW | 1662776 | Cell/Virus Enclosed | NO |
| GSLEKFMSTPNA | 1443984 | Cell/Virus Enclosed | NO |
| GSLGDSVKLLSE | 1443993 | Cell/Virus Enclosed | NO |
| GSLGFHNPPTNV | 1443994 | Ectodomain | YES |
| GSLIGGMVLGGLTSAA | 1662786 | Ectodomain | YES |
| GSLIPVRPRNSSDNGI | 1662787 | Ectodomain | YES |
| GSLNNIDTGPPS | 1444007 | Ectodomain | YES |
| GSLNNVDTGPPS | 1444008 | Ectodomain | YES |
| GSLTHINTTLLD | 1444019 | Ectodomain | YES |
| GSMDIDVFNKLSTSDW | 1662810 | Cell/Virus Enclosed | NO |
| GSMIEDIDYVPLKSAV | 1662811 | Cell/Virus Enclosed | NO |
| GSMIEDVDYVPL | 1444044 | Cell/Virus Enclosed | NO |
| GSNDIWMQNKGLFYTQ | 1662815 | Ectodomain | YES |
| GSNDVWIFNKVR | 1444056 | Ectodomain | YES |
| GSNEPGNVGGND | 1444058 | Cell/Virus Enclosed | NO |
| GSNGKPGNVGGNDALS | 1662818 | Cell/Virus Enclosed | NO |
| GSNQNGERSGARPKQR | 1662824 | Cell/Virus Enclosed | NO |
| GSPGYNLKNGEVEFVY | 1662830 | Cell/Virus Enclosed | NO |
| GSPGYNVRNDGTVEFC | 1662831 | Cell/Virus Enclosed | NO |
| GSPQSYCYTDGL | 1444090 | Cell/Virus Enclosed | NO |
| GSPYIDDFNKIASCKW | 1662837 | Cell/Virus Enclosed | NO |
| GSQLELAKVQNL | 1444100 | Capsid | UN |
| GSQQQSQTVTKKSAAE | 1662843 | Cell/Virus Enclosed | NO |
| GSQTTNGTNTNH | 1444107 | Ectodomain | YES |
| GSQTVLRCGDCL | 1444108 | Cell/Virus Enclosed | NO |
| GSRGGSQASSRSSSRSRNSS | 2060829 | Capsid | UN |
| GSRSARSAIEDLLFDKVTIAD | 1381810 | Ectodomain | YES |
| GSRSGNSTRGTF | 1444124 | Cell/Virus Enclosed | NO |
| GSRSGNSTRGTSPGPS | 1662850 | Cell/Virus Enclosed | NO |
| GSRSGNSTRSTS | 1444125 | Capsid | UN |
| GSRSRANSGNRA | 1444130 | Cell/Virus Enclosed | NO |
| GSRSRANSGNRTPTSG | 1662852 | Cell/Virus Enclosed | NO |
| GSRVPHIVRKDLTKYT | 1662855 | Cell/Virus Enclosed | NO |
| GSSFDGVMYGGFEDQP | 1662859 | Cell/Virus Enclosed | NO |

| | | | |
|---|---|---|---|
| GSSFSELLFDCV | 1444150 | Cell/Virus Enclosed | NO |
| GSSFYAPEPITSLNTK | 1662860 | Ectodomain | YES |
| GSSFYAPEPITY | 1444153 | Ectodomain | YES |
| GSSGFLQSSNYKIDTT | 1662861 | Ectodomain | YES |
| GSSGVANPAMDPIYDE | 1662864 | Secreted | YES |
| GSSGVANPATDP | 1444159 | Secreted | YES |
| GSSLQEILSRLD | 1444170 | Ectodomain | YES |
| GSSMVCKMCLFS | 1444173 | Cell/Virus Enclosed | NO |
| GSSRGTSPARIAGNGG | 1662874 | Cell/Virus Enclosed | NO |
| GSSRGTSPARMAGNGG | 1662875 | Cell/Virus Enclosed | NO |
| GSSRGTSPARMAGNGS | 1662876 | Cell/Virus Enclosed | NO |
| GSSRNESWHFDKSEPL | 1662877 | Ectodomain | YES |
| GSSSRSFFEDLLFDKV | 1662880 | Ectodomain | YES |
| GSSSRSFIEDLL | 1444185 | Ectodomain | YES |
| GSSSRSLLEDLLFNK | 2135475 | Ectodomain | YES |
| GSSVETETYFKVNGLW | 1662885 | Cell/Virus Enclosed | NO |
| GSSVGNFSDGKMGRFF | 1662886 | Ectodomain | YES |
| GSSYYYPEPISDKNVV | 1662888 | Ectodomain | YES |
| GSTLVDGVSRRY | 1444208 | Ectodomain | YES |
| GSTMDNKSQSVI | 1444209 | Ectodomain | YES |
| GSTMNNKSQSVIIINN | 1662900 | Ectodomain | YES |
| GSTMNNKSQTVI | 1444214 | Ectodomain | YES |
| GSVACEHISSTMSQYS | 1662908 | Ectodomain | YES |
| GSVCFSLKYIPG | 1444234 | Ectodomain | YES |
| GSVDPTFLHRFY | 1444236 | Cell/Virus Enclosed | NO |
| GSVGYVLTGDSVKFVY | 1662921 | Cell/Virus Enclosed | NO |
| GSVLEMAKEVKTSSKV | 1662926 | Cell/Virus Enclosed | NO |
| GSVQVDNGSSINRRQL | 1662931 | Cell/Virus Enclosed | NO |
| GSVVEQADGVEC | 1444267 | Ectodomain | YES |
| GSVVEQAEGVECDFSP | 1662935 | Ectodomain | YES |
| GSVVFVDKYMCGFDGK | 1662936 | Cell/Virus Enclosed | NO |
| GSVVIKEPVHVYADDS | 1662937 | Cell/Virus Enclosed | NO |
| GSVVKPVRARAPPGEQ | 1662938 | Cell/Virus Enclosed | NO |
| GSVWHIQTVQDADGIN | 1662941 | Cell/Virus Enclosed | NO |
| GSVYGNFDDQPSLQVE | 1662943 | Cell/Virus Enclosed | NO |
| GSVYTNSCLVSSDGQP | 1662944 | Cell/Virus Enclosed | NO |
| GSVYYHNSMDCPIVVA | 1662945 | Cell/Virus Enclosed | NO |
| GSWAIFNDHHLNRPGV | 1662946 | Cell/Virus Enclosed | NO |
| GSWNKVTQIANM | 1444286 | Cell/Virus Enclosed | NO |
| GSWWSFNPETNC | 1444289 | Ectodomain | YES |
| GTAKKQVVLVSNNQED | 1662965 | Cell/Virus Enclosed | NO |
| GTALPKGFYAEGSRGG | 1662966 | Cell/Virus Enclosed | NO |
| GTANLSIPSNWT | 1444318 | Ectodomain | YES |
| GTANQQQLKQLE | 1444319 | Cell/Virus Enclosed | NO |
| GTATAGWAFGAG | 1444320 | Ectodomain | YES |
| GTAVINLRADQINDMV | 1662970 | Cell/Virus Enclosed | NO |
| GTAVLRQWLPTG | 1444326 | Cell/Virus Enclosed | NO |
| GTAVVNLRPDQLNDLV | 1662975 | Cell/Virus Enclosed | NO |
| GTCAKRFKRWLG | 1444332 | Cell/Virus Enclosed | NO |
| GTCCTTSYKFYR | 1444335 | Cell/Virus Enclosed | NO |
| GTCGIVELEKGV | 1444345 | Cell/Virus Enclosed | NO |
| GTCPFSFSKLNNFQKF | 1662988 | Ectodomain | YES |
| GTCYNCRPAVVN | 1444360 | Ectodomain | YES |
| GTCYPGDFIDYEELREQLSSVSSFE | 1145632 | Ectodomain | YES |
| GTCYRCPKEIVDTVSA | 1662990 | Cell/Virus Enclosed | NO |
| GTDDPRWPQIAE | 1444362 | Cell/Virus Enclosed | NO |
| GTDFNGDFYGPYKDAQ | 1662991 | Cell/Virus Enclosed | NO |
| GTDFTGNFYGPYRDAQ | 1662992 | Cell/Virus Enclosed | NO |
| GTDLLSTLWFKP | 1444376 | Ectodomain | YES |
| GTDTNSVCPKLEFAND | 1663004 | Ectodomain | YES |
| GTDYKYWPQIAQ | 1444399 | Cell/Virus Enclosed | NO |
| GTEDPPWPQIAE | 1444405 | Cell/Virus Enclosed | NO |
| GTEDPRWPQIAELAPT | 1663012 | Cell/Virus Enclosed | NO |
| GTENLVIEGPTT | 1444412 | Cell/Virus Enclosed | NO |
| GTFCDNINSILDEVNG | 1663020 | Ectodomain | YES |
| GTFEEMALTTFM | 1444422 | Cell/Virus Enclosed | NO |
| GTFPGPSGIGAV | 1444427 | Cell/Virus Enclosed | NO |
| GTFTPKKPNKPL | 1444430 | Cell/Virus Enclosed | NO |
| GTFTVVMRPNYTIKGS | 1663030 | Cell/Virus Enclosed | NO |
| GTGGNIQSSSRA | 1444453 | Capsid | UN |
| GTGGNSQSSSRASSLS | 1663038 | Cell/Virus Enclosed | NO |
| GTGIAISVKPES | 1444456 | Cell/Virus Enclosed | NO |
| GTGIIRSSNQSLAGGI | 1663040 | Ectodomain | YES |
| GTGLQYAAITAP | 1444458 | Cell/Virus Enclosed | NO |
| GTGMAITVKPDATTSQ | 1663048 | Cell/Virus Enclosed | NO |
| GTGPEAALPFRAVKDG | 1663049 | Cell/Virus Enclosed | NO |

| | | | |
|---|---|---|---|
| GTGPEAGLPYGANKEG | 1663051 | Cell/Virus Enclosed | NO |
| GTGPHKDLKFRQ | 1444471 | Cell/Virus Enclosed | NO |
| GTGPHSKDQYGT | 1444473 | Cell/Virus Enclosed | NO |
| GTGPYANASYGESLEG | 1663056 | Capsid | UN |
| GTGPYANASYGQ | 1444477 | Cell/Virus Enclosed | NO |
| GTGRGDCKGFSSDVLS | 1663058 | Ectodomain | YES |
| GTGRGDCKGFYS | 1444482 | Ectodomain | YES |
| GTGRGDCQGFYS | 1444483 | Ectodomain | YES |
| GTGSGDATTAFA | 1444486 | Cell/Virus Enclosed | NO |
| GTGTIYAELEPP | 1444490 | Cell/Virus Enclosed | NO |
| GTGVFVQDPSPI | 1444495 | Cell/Virus Enclosed | NO |
| GTGVLTTSSKRF | 1444500 | Ectodomain | YES |
| GTGYSLADLPAYMTVA | 1663066 | Ectodomain | YES |
| GTGYSLSDLPAY | 1444502 | Underneath Membrane | NO |
| GTIKENIDGGAMHANY | 1663078 | Cell/Virus Enclosed | NO |
| GTILQGKPIGMF | 1444526 | Cell/Virus Enclosed | NO |
| GTITDAIDCAQN | 1444534 | Ectodomain | YES |
| GTIYAELEPPCR | 1444546 | Cell/Virus Enclosed | NO |
| GTLIVNSVLLFLAFV | 22741 | Undetermined | UN |
| GTLLVDSDLNEF | 1444583 | Cell/Virus Enclosed | NO |
| GTLLVDSNLNDF | 1444585 | Cell/Virus Enclosed | NO |
| GTLMVERSVSLA | 1444589 | Cell/Virus Enclosed | NO |
| GTLVKTITDDQIEV | 22755 | Ectodomain | YES |
| GTMYVRPIIEDYHTLT | 1663123 | Ectodomain | YES |
| GTNCMGKLKRNR | 1444621 | Underneath Membrane | NO |
| GTNDICVFENKT | 1444622 | Cell/Virus Enclosed | NO |
| GTNKQLNEISDDCNWP | 1663136 | Cell/Virus Enclosed | NO |
| GTNSGVAIYHRY | 1444640 | Ectodomain | YES |
| GTNTSNSVCNGC | 1444648 | Ectodomain | YES |
| GTNYLTCDNLCTPDPI | 1663146 | Ectodomain | YES |
| GTNYLTCHNAVQ | 1444656 | Ectodomain | YES |
| GTPAKKLCVVPGNVVP | 1663148 | Cell/Virus Enclosed | NO |
| GTPKPSRNHSPA | 1444664 | Cell/Virus Enclosed | NO |
| GTPKPSRNQSPASSQT | 1663152 | Capsid | UN |
| GTPKPSRNQSPV | 1444667 | Cell/Virus Enclosed | NO |
| GTPLHPVLWNCQ | 1444669 | Cell/Virus Enclosed | NO |
| GTPNEKLVTTST | 1444671 | Cell/Virus Enclosed | NO |
| GTPSMDALSALLSNTLSLAS | 22786 | Cell/Virus Enclosed | NO |
| GTPVVEYLSKEDLHTL | 1663160 | Cell/Virus Enclosed | NO |
| GTQHISNDFTPEDHSL | 1663161 | Capsid | UN |
| GTQLNHTSCSCS | 1444691 | Ectodomain | YES |
| GTQSVSVDRELT | 1444700 | Cell/Virus Enclosed | NO |
| GTRIRFKDDTSFED | 22799 | Capsid | UN |
| GTRNPANNAAIVLQLP | 1663170 | Cell/Virus Enclosed | NO |
| GTRNPNNDLAIV | 1444725 | Cell/Virus Enclosed | NO |
| GTRNPNNNAAIVLQLP | 1663172 | Cell/Virus Enclosed | NO |
| GTSDPQFPILAE | 1444742 | Cell/Virus Enclosed | NO |
| GTSDSQFPILAE | 1444743 | Cell/Virus Enclosed | NO |
| GTSLHPVLWNCQ | 1444750 | Cell/Virus Enclosed | NO |
| GTSPARIAGNGGDAAL | 1663183 | Cell/Virus Enclosed | NO |
| GTSPCPSGIGAV | 1444755 | Cell/Virus Enclosed | NO |
| GTSPGLSGIGAV | 1444756 | Cell/Virus Enclosed | NO |
| GTSPGPSGIGAVGGDL | 1663186 | Cell/Virus Enclosed | NO |
| GTSPGPSGIGTV | 1444758 | Cell/Virus Enclosed | NO |
| GTSTPKKPNKPLSQPR | 1663191 | Capsid | UN |
| GTSVLRQWLPTD | 1444772 | Cell/Virus Enclosed | NO |
| GTSVTEDVKFAA | 1444774 | Cell/Virus Enclosed | NO |
| GTSYTANDVEVQ | 1444778 | Cell/Virus Enclosed | NO |
| GTTCSNDLLQPNTEVF | 1663199 | Ectodomain | YES |
| GTTCSNDLLYSN | 1444780 | Ectodomain | YES |
| GTTEVKVSALKT | 1444787 | Cell/Virus Enclosed | NO |
| GTTKFYGGWDDMLRRL | 1663208 | Cell/Virus Enclosed | NO |
| GTTKFYGGWGFM | 1444792 | Cell/Virus Enclosed | NO |
| GTTLGVLVPHVG | 1444797 | Cell/Virus Enclosed | NO |
| GTTQTACTDDNA | 1444809 | Cell/Virus Enclosed | NO |
| GTTVVRPLVEDSTSVT | 1663232 | Ectodomain | YES |
| GTVDADYKMCTK | 1444830 | Ectodomain | YES |
| GTVIINPSTSAT | 1444838 | Ectodomain | YES |
| GTVIISPFTSAT | 1444840 | Ectodomain | YES |
| GTVLRAINGDFL | 1444848 | Cell/Virus Enclosed | NO |
| GTVNNFVKFGSV | 1444852 | Ectodomain | YES |
| GTVSKTSDWKCKVTDV | 1663244 | Cell/Virus Enclosed | NO |
| GTVWYVGQIIFI | 1444861 | Cell/Virus Enclosed | NO |
| GTWLTYTGAIKLDDKD | 1663253 | Cell/Virus Enclosed | NO |
| GTYHCVTTKAKS | 1444875 | Cell/Virus Enclosed | NO |
| GTYTWTTVLSMA | 1444886 | Cell/Virus Enclosed | NO |

| | | | |
|---|---|---|---|
| GTYYCVTTKAKSVQQD | 1663265 | Cell/Virus Enclosed | NO |
| GVAFLAVFQSATKIIA | 1663272 | Secreted | YES |
| GVALEQDKNTQE | 1444908 | Ectodomain | YES |
| GVALKDCGFVMQFGYI | 1663286 | Cell/Virus Enclosed | NO |
| GVANPATDPIYD | 1444915 | Secreted | YES |
| GVARAIDILTEGQLQS | 1663295 | Cell/Virus Enclosed | NO |
| GVAVADSYYSYMMPML | 1663300 | Cell/Virus Enclosed | NO |
| GVAYTAFSGPVDKGHY | 1663302 | Cell/Virus Enclosed | NO |
| GVCITTNGSWAIFNDH | 1663308 | Cell/Virus Enclosed | NO |
| GVCKGFYSNASS | 1444941 | Ectodomain | YES |
| GVCKYTDIDVNSALNI | 1663310 | Cell/Virus Enclosed | NO |
| GVCMYTDLNFID | 1444956 | Cell/Virus Enclosed | NO |
| GVCYDFDGVTTSDKCI | 1663325 | Cell/Virus Enclosed | NO |
| GVDAKGFPLLAE | 1444968 | Cell/Virus Enclosed | NO |
| GVDAKGFPQFAE | 1444969 | Cell/Virus Enclosed | NO |
| GVDAKGFPQLAELIPN | 1663327 | Capsid | UN |
| GVDCYNWDSAEL | 1444977 | Cell/Virus Enclosed | NO |
| GVDFDFIVNAANENLA | 1663333 | Cell/Virus Enclosed | NO |
| GVDFVAQPEGCVLTNT | 1663334 | Cell/Virus Enclosed | NO |
| GVDGELPVAVVNDKVF | 1663336 | Cell/Virus Enclosed | NO |
| GVDSFEGVLPINTDTV | 1663340 | Cell/Virus Enclosed | NO |
| GVDYTKKFLTVDGVQY | 1663344 | Cell/Virus Enclosed | NO |
| GVDYTKKSLNVD | 1445003 | Cell/Virus Enclosed | NO |
| GVECDFSPLLSGTPPQ | 1663348 | Ectodomain | YES |
| GVEDTPPHYFTL | 1445009 | Cell/Virus Enclosed | NO |
| GVEELHDDLVNYIRDK | 1663349 | Cell/Virus Enclosed | NO |
| GVEFLKRGDKVV | 1445014 | Cell/Virus Enclosed | NO |
| GVEGELPVAVVNDKIF | 1663353 | Cell/Virus Enclosed | NO |
| GVELGTDPIEDY | 1445023 | Cell/Virus Enclosed | NO |
| GVENPMLMGWDYPKCD | 1663358 | Cell/Virus Enclosed | NO |
| GVENTPPHFFTL | 1445024 | Cell/Virus Enclosed | NO |
| GVENTPPHYFTL | 1445025 | Cell/Virus Enclosed | NO |
| GVESDVETNEIVDSVQ | 1663360 | Cell/Virus Enclosed | NO |
| GVFDIKNSSVNTVCFV | 1663364 | Cell/Virus Enclosed | NO |
| GVFLGVVGVTMH | 1445040 | Cell/Virus Enclosed | NO |
| GVFMSTAKCWTEEDLS | 1663375 | Cell/Virus Enclosed | NO |
| GVFQNCTAVGVP | 1445047 | Ectodomain | YES |
| GVFRVTSRAMFQ | 1445049 | Ectodomain | YES |
| GVFVQDPAPIDIDAFP | 1663384 | Cell/Virus Enclosed | NO |
| GVFWVANHQADTSTPS | 1663387 | Capsid | UN |
| GVGAPGGDLLYL | 1445080 | Cell/Virus Enclosed | NO |
| GVGITQQVLSKN | 1445089 | Ectodomain | YES |
| GVGNTFICEEVANDLT | 1663395 | Cell/Virus Enclosed | NO |
| GVGSANIFIGDKVGHY | 1663396 | Cell/Virus Enclosed | NO |
| GVGSANIFIGDN | 1445097 | Cell/Virus Enclosed | NO |
| GVHGNGSMIEDI | 1445115 | Cell/Virus Enclosed | NO |
| GVHKGAHVASSD | 1445117 | Cell/Virus Enclosed | NO |
| GVHMNIAKYMQL | 1445121 | Cell/Virus Enclosed | NO |
| GVIADKDKDLSGAVPS | 1663409 | Cell/Virus Enclosed | NO |
| GVIADYNYKLPDDFMG | 1663410 | Ectodomain | YES |
| GVIAQGARGTYR | 1445130 | Cell/Virus Enclosed | NO |
| GVIDISAGMFGLYDDI | 1663412 | Cell/Virus Enclosed | NO |
| GVIFNAVDCKSD | 1445133 | Ectodomain | YES |
| GVIIMCADVPDAFQSV | 1663414 | Cell/Virus Enclosed | NO |
| GVIKNVNSVRDWLKSL | 1663415 | Cell/Virus Enclosed | NO |
| GVIKNVNYVRDW | 1445136 | Cell/Virus Enclosed | NO |
| GVILTRSFKYNI | 1445137 | Cell/Virus Enclosed | NO |
| GVIPDMADQIAS | 1445143 | Cell/Virus Enclosed | NO |
| GVIRDVSVEDVCFANK | 1663420 | Cell/Virus Enclosed | NO |
| GVIRVSNDTFLNGITY | 1663423 | Ectodomain | YES |
| GVIRYYDITANGGTGF | 1663424 | Cell/Virus Enclosed | NO |
| GVISKVHEVIIDNKPY | 1663425 | Cell/Virus Enclosed | NO |
| GVISNARNGCIP | 1445146 | Cell/Virus Enclosed | NO |
| GVIYGFNSTETI | 1445151 | Ectodomain | YES |
| GVIYGLNFTETI | 1445152 | Ectodomain | YES |
| GVIYGLNSTETI | 1445153 | Ectodomain | YES |
| GVIYTEIQPKLD | 1445156 | Cell/Virus Enclosed | NO |
| GVKAKGYPQFAELVPS | 1663431 | Capsid | UN |
| GVKDFNTLSQEGKKYL | 1663434 | Cell/Virus Enclosed | NO |
| GVKDYVVVHGYFTEVY | 1663435 | Secreted | YES |
| GVKFQTVKPGNFNQDF | 1663436 | Cell/Virus Enclosed | NO |
| GVKIQEGIVDYG | 1445169 | Cell/Virus Enclosed | NO |
| GVKKTFKVEDSV | 1445171 | Cell/Virus Enclosed | NO |
| GVKLGTGYTLSDLPVY | 1663441 | Underneath Membrane | NO |
| GVKLRSLNLGEV | 1445173 | Cell/Virus Enclosed | NO |
| GVKPAVSFDYLI | 1445174 | Cell/Virus Enclosed | NO |

| | | | |
|---|---|---|---|
| GVKPAVSFDYLT | 1445175 | Cell/Virus Enclosed | NO |
| GVKQEQCTGLDS | 1445181 | Cell/Virus Enclosed | NO |
| GVKQEQRTGLDAVMHF | 1663445 | Cell/Virus Enclosed | NO |
| GVKQLPEQFDSLNLSA | 1663446 | Capsid | UN |
| GVKQSPELLDPL | 1445188 | Cell/Virus Enclosed | NO |
| GVKQSPESFDSL | 1445189 | Cell/Virus Enclosed | NO |
| GVLDGSYNASCL | 1445198 | Ectodomain | YES |
| GVLDGSYNVSCLCSTD | 1663451 | Ectodomain | YES |
| GVLDGYNVSCLC | 1445200 | Ectodomain | YES |
| GVLDTTIPFGPSSQPY | 1663452 | Ectodomain | YES |
| GVLDTTSYREAA | 1445204 | Cell/Virus Enclosed | NO |
| GVLESFSVTLLDNQED | 1663455 | Cell/Virus Enclosed | NO |
| GVLGCKAIVKETNLIC | 1663456 | Cell/Virus Enclosed | NO |
| GVLGDGNALYNTEGGK | 1663457 | Cell/Virus Enclosed | NO |
| GVLIQNGAKHVQGNVA | 1663460 | Cell/Virus Enclosed | NO |
| GVLKADDFVIAS | 1445214 | Cell/Virus Enclosed | NO |
| GVLLSNFGAPSY | 1445217 | Cell/Virus Enclosed | NO |
| GVLLVDGHKIAT | 1445219 | Undetermined | UN |
| GVLPINTDTVLSVAPE | 1663471 | Cell/Virus Enclosed | NO |
| GVLQQRFVYDAY | 1445225 | Ectodomain | YES |
| GVLQSGLVKMSHPSGD | 1663477 | Cell/Virus Enclosed | NO |
| GVLRELIRELNG | 1445227 | Cell/Virus Enclosed | NO |
| GVLSAAIKDCVDINKA | 1663478 | Cell/Virus Enclosed | NO |
| GVLSVDLSLTVELKI | 23048 | Ectodomain | YES |
| GVLTLDNQDLNGKWYD | 1663482 | Cell/Virus Enclosed | NO |
| GVLTPSSKRFQPFQQFG | 23051 | Ectodomain | YES |
| GVLTTSSKRFQS | 1445240 | Ectodomain | YES |
| GVLVPHVGETPI | 1445244 | Cell/Virus Enclosed | NO |
| GVLVPHVGETPV | 1445245 | Cell/Virus Enclosed | NO |
| GVMKNASLYTSLVPHT | 1663497 | Cell/Virus Enclosed | NO |
| GVMWLYTYSIGEASSP | 1663500 | Cell/Virus Enclosed | NO |
| GVMYGGFEDQPNLQVE | 1663501 | Cell/Virus Enclosed | NO |
| GVMYSEFPAITI | 1445261 | Ectodomain | YES |
| GVMYSEFPAIVI | 1445262 | Ectodomain | YES |
| GVMYSEFPTITI | 1445263 | Ectodomain | YES |
| GVNDAITQTSPA | 1445265 | Ectodomain | YES |
| GVNDAITQTSQALQTV | 1663503 | Ectodomain | YES |
| GVNDDDYVIALT | 1445266 | Cell/Virus Enclosed | NO |
| GVNDICIFENKT | 1445267 | Cell/Virus Enclosed | NO |
| GVNDICMFENKTTLPT | 1663504 | Cell/Virus Enclosed | NO |
| GVNFKSISLTVGEVFG | 1663506 | Cell/Virus Enclosed | NO |
| GVNFNPYHRFKCFPNG | 1663507 | Ectodomain | YES |
| GVNFNVDDINFA | 1445275 | Ectodomain | YES |
| GVNFNVDDINFSPVLG | 1663508 | Ectodomain | YES |
| GVNFNVDDVNFS | 1445276 | Ectodomain | YES |
| GVNFRTVVLNNKNTYR | 1663509 | Cell/Virus Enclosed | NO |
| GVNFTNRFVPVGESFG | 1663510 | Cell/Virus Enclosed | NO |
| GVNFTPYHAFKCNRSG | 1663511 | Ectodomain | YES |
| GVNFVVQPVGVVDTEW | 1663512 | Cell/Virus Enclosed | NO |
| GVNITLANFNETKGPL | 1663517 | Ectodomain | YES |
| GVNKIQPWTNDALNTI | 1663518 | Cell/Virus Enclosed | NO |
| GVNKLQGLLEIS | 1445284 | Ectodomain | YES |
| GVNLRTNFTIKGSFIN | 1663521 | Cell/Virus Enclosed | NO |
| GVNSSNCNNLNV | 1445288 | Cell/Virus Enclosed | NO |
| GVNVTKHKCDINYKGK | 1663528 | Cell/Virus Enclosed | NO |
| GVNVTKHRCDIN | 1445295 | Cell/Virus Enclosed | NO |
| GVNYLGDFIQGP | 1445296 | Cell/Virus Enclosed | NO |
| GVNYLGKLREQI | 1445297 | Cell/Virus Enclosed | NO |
| GVPATEAKGYWY | 1445301 | Cell/Virus Enclosed | NO |
| GVPATQAKGYWY | 1445303 | Cell/Virus Enclosed | NO |
| GVPDNSSTIVTGLLPT | 1663532 | Ectodomain | YES |
| GVPIAFGVPPSEAKGY | 1663537 | Capsid | UN |
| GVPIAYGIPLSE | 1445319 | Cell/Virus Enclosed | NO |
| GVPIELQVECFTWDQ | 23096 | Ectodomain | YES |
| GVPIINCVKMLCDHAG | 1663540 | Cell/Virus Enclosed | NO |
| GVPINTNSSKDD | 1445324 | Cell/Virus Enclosed | NO |
| GVPINTNSSPDDQIGYYRRA | 1392199 | Capsid | UN |
| GVPINTNSTKDDQIGY | 1663543 | Cell/Virus Enclosed | NO |
| GVPLLSTMTTRQ | 1445330 | Cell/Virus Enclosed | NO |
| GVPLNANSTPAQNAGY | 1663545 | Cell/Virus Enclosed | NO |
| GVPLNANSTPSQ | 1445332 | Cell/Virus Enclosed | NO |
| GVPLSEAKGYWY | 1445334 | Cell/Virus Enclosed | NO |
| GVPLTNCVKMLT | 1445335 | Cell/Virus Enclosed | NO |
| GVPVIEVSQIQSKLTD | 1663550 | Cell/Virus Enclosed | NO |
| GVQCYTPHSQIS | 1445350 | Cell/Virus Enclosed | NO |
| GVQIVEDLRARM | 1445354 | Cell/Virus Enclosed | NO |

| | | | |
|---|---|---|---|
| GVQTIEDLRARM | 1445358 | Cell/Virus Enclosed | NO |
| GVQTLEDLHARM | 1445359 | Cell/Virus Enclosed | NO |
| GVQTVEDLCARM | 1445360 | Cell/Virus Enclosed | NO |
| GVQTVEDLRARMTYVC | 1663561 | Cell/Virus Enclosed | NO |
| GVQVHNLPEYMA | 1445366 | Underneath Membrane | NO |
| GVRDARIKKAMFTKVV | 1663567 | Cell/Virus Enclosed | NO |
| GVRELYLSSSNV | 1445374 | Cell/Virus Enclosed | NO |
| GVRGASVDARLV | 1445377 | Cell/Virus Enclosed | NO |
| GVRGSGHYQTNL | 1445378 | Cell/Virus Enclosed | NO |
| GVRIENSLKALF | 1445381 | Cell/Virus Enclosed | NO |
| GVRQQRFVYDAYQNLV | 1663575 | Ectodomain | YES |
| GVRRKNSEPEIPHFNQ | 1663577 | Capsid | UN |
| GVRYPTGSCLIA | 1445398 | Cell/Virus Enclosed | NO |
| GVSATKLNDLCFSNVY | 1663582 | Ectodomain | YES |
| GVSCIWSVDAFN | 1445402 | Cell/Virus Enclosed | NO |
| GVSDLSLKQKLLYTFY | 1663584 | Cell/Virus Enclosed | NO |
| GVSDVKKLYLGGLNYY | 1663585 | Cell/Virus Enclosed | NO |
| GVSDVWHKQLLLNRGV | 1663587 | Cell/Virus Enclosed | NO |
| GVSEIVNTVAGIPSNV | 1663588 | Cell/Virus Enclosed | NO |
| GVSERINVVPGVPTNV | 1663589 | Cell/Virus Enclosed | NO |
| GVSERVNVVPGV | 1445408 | Cell/Virus Enclosed | NO |
| GVSFVVQPVGVV | 1445413 | Cell/Virus Enclosed | NO |
| GVSGCLTDCLYLKNLT | 1663592 | Cell/Virus Enclosed | NO |
| GVSGRGVFQNCTAVGV | 1663593 | Ectodomain | YES |
| GVSGSSITTLSSGPHSLSPA | 23142 | Capsid | UN |
| GVSICSTMTNRQ | 1445419 | Cell/Virus Enclosed | NO |
| GVSILSTMTGRV | 1445425 | Cell/Virus Enclosed | NO |
| GVSILSTVTGRM | 1445426 | Cell/Virus Enclosed | NO |
| GVSIPCVCGRDA | 1445427 | Cell/Virus Enclosed | NO |
| GVSKPNAIDVEHLELK | 1663599 | Cell/Virus Enclosed | NO |
| GVSNLSLKQKLL | 1445432 | Cell/Virus Enclosed | NO |
| GVSSAGPISQFN | 1445438 | Ectodomain | YES |
| GVSSFMNVTLNK | 1445440 | Ectodomain | YES |
| GVSSNQSPQGNLGSNG | 1663608 | Cell/Virus Enclosed | NO |
| GVSVITPGTNASSEVA | 1663611 | Ectodomain | YES |
| GVTAKGYPQFAE | 1445452 | Cell/Virus Enclosed | NO |
| GVTHEYIQTTIQ | 1445464 | Ectodomain | YES |
| GVTIDKLCDYLA | 1445465 | Cell/Virus Enclosed | NO |
| GVTIDKLRDYLA | 1445466 | Cell/Virus Enclosed | NO |
| GVTLDKLRDYLADYDV | 1663622 | Cell/Virus Enclosed | NO |
| GVTLKIKVSQTN | 1445474 | Cell/Virus Enclosed | NO |
| GVTLSSNLNTNLHFDV | 1663627 | Ectodomain | YES |
| GVTLSTKLKDGVNFNV | 1663628 | Ectodomain | YES |
| GVTMDVLNKNQKLIAT | 1663630 | Ectodomain | YES |
| GVTPDMADQIAS | 1445480 | Cell/Virus Enclosed | NO |
| GVTQNVLYDNQK | 1445483 | Ectodomain | YES |
| GVTQNVLYENQKQIAN | 1663637 | Ectodomain | YES |
| GVTRDIMATDDC | 1445489 | Cell/Virus Enclosed | NO |
| GVTRDMADQIAS | 1445491 | Capsid | UN |
| GVTRPSAPPLPHVV | 23176 | Cell/Virus Enclosed | NO |
| GVTTSDKCIFNSACTR | 1663645 | Cell/Virus Enclosed | NO |
| GVTVAEEPDSRV | 1445500 | Cell/Virus Enclosed | NO |
| GVTYQLAWDVIRKDLS | 1663649 | Cell/Virus Enclosed | NO |
| GVVANGVKAKGYPQFA | 1663652 | Capsid | UN |
| GVVCNNSDYASK | 1445510 | Cell/Virus Enclosed | NO |
| GVVCYNKDYAELGYIA | 1663656 | Cell/Virus Enclosed | NO |
| GVVCYNSDYAAKGYIA | 1663657 | Cell/Virus Enclosed | NO |
| GVVCYNSDYASKGYIA | 1663658 | Cell/Virus Enclosed | NO |
| GVVDKQVILVSNNKED | 1663659 | Cell/Virus Enclosed | NO |
| GVVDTDVDAKDV | 1445515 | Cell/Virus Enclosed | NO |
| GVVDTEWGNMLTGIAA | 1663661 | Cell/Virus Enclosed | NO |
| GVVEDTINDDEDVVTG | 1663664 | Cell/Virus Enclosed | NO |
| GVVLKRSLKYNLKASA | 1663676 | Cell/Virus Enclosed | NO |
| GVVMNFDVDTHR | 1445532 | Cell/Virus Enclosed | NO |
| GVVPLSVIPATSAARL | 1663682 | Cell/Virus Enclosed | NO |
| GVVPLSVIPATSASKL | 1663683 | Cell/Virus Enclosed | NO |
| GVVRDISVNDLCFANK | 1663686 | Cell/Virus Enclosed | NO |
| GVVTCNVVKQCC | 1445556 | Cell/Virus Enclosed | NO |
| GVVTGNVVKQCCTAAV | 1663693 | Cell/Virus Enclosed | NO |
| GVVTGNVVQQCC | 1445559 | Cell/Virus Enclosed | NO |
| GVVWTLNDVKDNDGRP | 1663702 | Cell/Virus Enclosed | NO |
| GVVWVAKEGAKTVNTS | 1663704 | Capsid | UN |
| GVVYVNATGSDV | 1445575 | Ectodomain | YES |
| GVYCCRDHEHGI | 1445581 | Cell/Virus Enclosed | NO |
| GVYDFCVSPAEFKYMV | 1663711 | Cell/Virus Enclosed | NO |
| GVYDFKVSAAEFKYMV | 1663712 | Cell/Virus Enclosed | NO |

| | | | |
|---|---|---|---|
| GVYDFKVSTQEFRFMT | 1663713 | Cell/Virus Enclosed | NO |
| GVYDTNHGDAGF | 1445588 | Ectodomain | YES |
| GVYELHGYTVQP | 1445591 | Ectodomain | YES |
| GVYHKLGERIRQ | 1445597 | Cell/Virus Enclosed | NO |
| GVYHKLGERLRQ | 1445598 | Cell/Virus Enclosed | NO |
| GVYKPLLENIDYFNMR | 1663723 | Cell/Virus Enclosed | NO |
| GVYSVSSFEAKPSGSV | 1663730 | Ectodomain | YES |
| GVYWVASNQADVNTPA | 1663733 | Cell/Virus Enclosed | NO |
| GVYYRATTTYKL | 1445636 | Cell/Virus Enclosed | NO |
| GWALYVRAKHGD | 1445640 | Ectodomain | YES |
| GWDDMLRHLIKDVDNP | 1663744 | Cell/Virus Enclosed | NO |
| GWDDMLRRLIKDVDNP | 1663745 | Cell/Virus Enclosed | NO |
| GWDFMLKTLYKDVDNP | 1663746 | Cell/Virus Enclosed | NO |
| GWDNMLKNLMADVDDP | 1663747 | Cell/Virus Enclosed | NO |
| GWDYPKCDRALPNMIR | 1663748 | Cell/Virus Enclosed | NO |
| GWDYPKCDRAMPNLLR | 1663750 | Cell/Virus Enclosed | NO |
| GWDYPKCDRAMPNMCR | 1663751 | Cell/Virus Enclosed | NO |
| GWDYPKCDRAMPSMIR | 1663753 | Cell/Virus Enclosed | NO |
| GWEGLIDGWYGFRHQNAQGEGTAAD | 1146121 | Ectodomain | YES |
| GWFDVYDDIFAESKPW | 1663756 | Cell/Virus Enclosed | NO |
| GWGLHQPPPQIFLKILPQSG | 23246 | Capsid | UN |
| GWGNMLKNLMAD | 1445649 | Cell/Virus Enclosed | NO |
| GWIFGSSFDNTT | 1445652 | Ectodomain | YES |
| GWIKISADNKVIYTTE | 1663762 | Cell/Virus Enclosed | NO |
| GWLGMDVEGAHVTGDN | 1663766 | Cell/Virus Enclosed | NO |
| GWLGMDVESAHVCGDN | 1663767 | Cell/Virus Enclosed | NO |
| GWVFGSTMNNKSQSVII | 23296 | Ectodomain | YES |
| GWVFYVRVKHGDFSAV | 1663790 | Underneath Membrane | NO |
| GWVGFDVEGAHATREN | 1663791 | Cell/Virus Enclosed | NO |
| GWVSGNMMDGFSCPLC | 1663793 | Cell/Virus Enclosed | NO |
| GWYDPVENEDIHRVYA | 1663796 | Cell/Virus Enclosed | NO |
| GYCCLDVVRKEL | 1445722 | Cell/Virus Enclosed | NO |
| GYCVDYSKIRRS | 1445730 | Ectodomain | YES |
| GYCVDYSKNRRSRGAI | 1663810 | Ectodomain | YES |
| GYDDCMHQGPAS | 1445740 | Ectodomain | YES |
| GYDDCMQQGPASARDL | 1663812 | Ectodomain | YES |
| GYDDCMQQGPSS | 1445744 | Ectodomain | YES |
| GYDFKYIENGQLKNFE | 1663813 | Cell/Virus Enclosed | NO |
| GYEKSTFNKDIYCGNS | 1663817 | Cell/Virus Enclosed | NO |
| GYFIKQNDSWMF | 1445759 | Ectodomain | YES |
| GYFIKTNNTRIVDEWS | 1663823 | Ectodomain | YES |
| GYFPKAGANFRD | 1445768 | Ectodomain | YES |
| GYFVNVNNTWMYTGSG | 1663832 | Ectodomain | YES |
| GYGCNCDSLRQV | 1445783 | Cell/Virus Enclosed | NO |
| GYGKSTFNKDIY | 1445789 | Cell/Virus Enclosed | NO |
| GYGSTFITPEVSRELG | 1663838 | Cell/Virus Enclosed | NO |
| GYGVRRKNSEPEIPHF | 1663839 | Capsid | UN |
| GYHPAHAPSFLAVDDK | 1663847 | Cell/Virus Enclosed | NO |
| GYHPAHAPSFLV | 1445799 | Cell/Virus Enclosed | NO |
| GYHYKELGLVMN | 1445804 | Cell/Virus Enclosed | NO |
| GYHYNELGIEMN | 1445806 | Cell/Virus Enclosed | NO |
| GYHYTPFHYERDNTSC | 1663852 | Cell/Virus Enclosed | NO |
| GYIDCEQDSCDFKGWI | 1663858 | Cell/Virus Enclosed | NO |
| GYIGATVRLQAGKQTE | 1663859 | Cell/Virus Enclosed | NO |
| GYIRTAIDCGFN | 1445839 | Ectodomain | YES |
| GYKIGCNCAGRI | 1445850 | Cell/Virus Enclosed | NO |
| GYKVLPPLMDVNMEAA | 1663878 | Ectodomain | YES |
| GYLDIHPSLCNN | 1445862 | Ectodomain | YES |
| GYNKDASFIGKN | 1445902 | Cell/Virus Enclosed | NO |
| GYNLKNGEVEFVYMHQ | 1663900 | Cell/Virus Enclosed | NO |
| GYNMTLIAADLP | 1445903 | Cell/Virus Enclosed | NO |
| GYNVRNDGTVEFCYLH | 1663903 | Cell/Virus Enclosed | NO |
| GYNVSSIVTMTSQGM | 23422 | Ectodomain | YES |
| GYPNDFVGGFRV | 1445924 | Cell/Virus Enclosed | NO |
| GYPQFAELVPSTAAML | 1663912 | Capsid | UN |
| GYQELSQFSHLD | 1445933 | Cell/Virus Enclosed | NO |
| GYQESVIKTSHD | 1445935 | Underneath Membrane | NO |
| GYQHHSVADVMR | 1445938 | Ectodomain | YES |
| GYQHNSFADVMR | 1445939 | Ectodomain | YES |
| GYQHNSVVDVMRYNLN | 1663916 | Ectodomain | YES |
| GYQPIDVVRDLPSGFN | 1663917 | Ectodomain | YES |
| GYQPYRVVVLSFELLNA | 23437 | Ectodomain | YES |
| GYQVRAFLNKLNENVV | 1663920 | Cell/Virus Enclosed | NO |
| GYRFTNFEPFTVNSVN | 1663923 | Ectodomain | YES |
| GYRGRSSIRTGVKRPAVSKA | 110880 | Capsid | UN |
| GYSEPPKVADKI | 1445966 | Cell/Virus Enclosed | NO |

| | | | |
|---|---|---|---|
| GYSKLFITKDEAIKRV | 1663933 | Cell/Virus Enclosed | NO |
| GYSLADLPAYMS | 1445969 | Ectodomain | YES |
| GYSNFVSSDLDC | 1445974 | Underneath Membrane | NO |
| GYSSLCDEFTLAEVVK | 1663940 | Cell/Virus Enclosed | NO |
| GYTDFAEDVCTCYDNS | 1663943 | Cell/Virus Enclosed | NO |
| GYTDNIFSVQED | 1445982 | Ectodomain | YES |
| GYTDNIFSVQQDGRIPNGFP | 2060831 | Ectodomain | YES |
| GYTESAVNSTKALAKQ | 1663947 | Cell/Virus Enclosed | NO |
| GYTLAATSASLFPPWT | 1663951 | Ectodomain | YES |
| GYTLEEARTALK | 1446004 | Cell/Virus Enclosed | NO |
| GYTVQPVADVYR | 1446016 | Ectodomain | YES |
| GYVDCFNDNCDFYGWV | 1663971 | Cell/Virus Enclosed | NO |
| GYVLTGDSVKFVYMHQ | 1663982 | Cell/Virus Enclosed | NO |
| GYVQQGIFNKATVAQI | 1663985 | Cell/Virus Enclosed | NO |
| GYVQQGVFNKAT | 1446046 | Cell/Virus Enclosed | NO |
| GYVTHGFNLEDA | 1446052 | Cell/Virus Enclosed | NO |
| GYWRRQDRKINTGNGI | 1663992 | Cell/Virus Enclosed | NO |
| GYYIEGSGRSAS | 1446083 | Cell/Virus Enclosed | NO |
| GYYIEGSGRSVP | 1446084 | Cell/Virus Enclosed | NO |
| GYYPSNHIEVAS | 1446091 | Ectodomain | YES |
| GYYVEGSGRSAS | 1446104 | Cell/Virus Enclosed | NO |
| GYYVKPGGTSSGDATT | 1664009 | Cell/Virus Enclosed | NO |
| GYYYPEPIIENN | 1446109 | Ectodomain | YES |
| GYYYPEPITGNN | 1446112 | Ectodomain | YES |
| HAAVTNHKKWQYNSPLVP | 2134434 | Undetermined | UN |
| HACLKGGLILKF | 1446124 | Cell/Virus Enclosed | NO |
| HAEEYREYLESYNTAT | 1664027 | Cell/Virus Enclosed | NO |
| HAENVTGLFKDC | 1446143 | Cell/Virus Enclosed | NO |
| HAEPKGKYSQKAYALL | 1664028 | Cell/Virus Enclosed | NO |
| HAEQLTPAWRIY | 1446144 | Ectodomain | YES |
| HAFKCTTSGSND | 1446157 | Ectodomain | YES |
| HAFMKHKGGDLT | 1446160 | Cell/Virus Enclosed | NO |
| HAFMKHKGGDSTDFIA | 1664039 | Cell/Virus Enclosed | NO |
| HAGEAPVSIINN | 1446168 | Cell/Virus Enclosed | NO |
| HAGFDAWFSQRG | 1446170 | Cell/Virus Enclosed | NO |
| HAIATIPLQASL | 1446184 | Secreted | YES |
| HAIQQGFDATNF | 1446190 | Ectodomain | YES |
| HAKDQYGTDFDG | 1446197 | Cell/Virus Enclosed | NO |
| HAKDQYGTDING | 1446199 | Cell/Virus Enclosed | NO |
| HAKPTVIVNAAN | 1446201 | Cell/Virus Enclosed | NO |
| HALDCELYVNNAYRLF | 1664056 | Cell/Virus Enclosed | NO |
| HALDDATRNYII | 1446203 | Cell/Virus Enclosed | NO |
| HALDDFTRNYII | 1446204 | Cell/Virus Enclosed | NO |
| HALDDVTLNYII | 1446205 | Cell/Virus Enclosed | NO |
| HALDDVTRNYII | 1446206 | Cell/Virus Enclosed | NO |
| HANSSEPALLFRNIKC | 1664073 | Ectodomain | YES |
| HANSSEPAMLFR | 1446236 | Ectodomain | YES |
| HANSYQKYVEAY | 1446238 | Cell/Virus Enclosed | NO |
| HANYIFWRNSTP | 1446241 | Cell/Virus Enclosed | NO |
| HAPSFLAVDDKYKATG | 1664082 | Cell/Virus Enclosed | NO |
| HAPSFLAVDDKYKVNE | 1664083 | Cell/Virus Enclosed | NO |
| HARLKGGLILKFDSGT | 1664084 | Cell/Virus Enclosed | NO |
| HATGTTPKKLFV | 1446276 | Ectodomain | YES |
| HATGTTPQKLFVANYS | 1664090 | Ectodomain | YES |
| HATRDSIGTNFPLQLG | 1664093 | Cell/Virus Enclosed | NO |
| HATRENIGTNFPLQIG | 1664096 | Cell/Virus Enclosed | NO |
| HATRLTITELLQ | 1446287 | Cell/Virus Enclosed | NO |
| HAYPLVYHENEE | 1446306 | Cell/Virus Enclosed | NO |
| HCAFAVDPAAAY | 1446308 | Cell/Virus Enclosed | NO |
| HCGSSSRSFFEDLLFD | 1664112 | Ectodomain | YES |
| HCGSSSRSFIED | 1446339 | Ectodomain | YES |
| HCHWFFNNYLRK | 1446342 | Cell/Virus Enclosed | NO |
| HCPADNSYTSFA | 1446352 | Ectodomain | YES |
| HCPAGNFYTSFA | 1446353 | Ectodomain | YES |
| HCPAGNSHTSFA | 1446354 | Ectodomain | YES |
| HCPAGNSYTSFA | 1446355 | Ectodomain | YES |
| HCQVHGNAHVAS | 1446370 | Cell/Virus Enclosed | NO |
| HCSFAVDPAAAY | 1446372 | Cell/Virus Enclosed | NO |
| HCSFAVDPAVAY | 1446373 | Cell/Virus Enclosed | NO |
| HCSGLAIKSDYC | 1446374 | Ectodomain | YES |
| HCSNFNTLFSTT | 1446378 | Cell/Virus Enclosed | NO |
| HCTIAQLADAAL | 1446383 | Cell/Virus Enclosed | NO |
| HCTIAQLTDAAL | 1446384 | Cell/Virus Enclosed | NO |
| HDAIVVDNDVVDYVSD | 1664136 | Cell/Virus Enclosed | NO |
| HDASSAINRPQLTFVK | 1664138 | Cell/Virus Enclosed | NO |
| HDDLVNYIRDKGCVAD | 1664140 | Cell/Virus Enclosed | NO |

| | | | |
|---|---|---|---|
| HDEEFCDMLRLFDFN | 540214 | Ectodomain | YES |
| HDFFIFDVDKFK | 1446418 | Cell/Virus Enclosed | NO |
| HDFFVFDVDKTK | 1446426 | Cell/Virus Enclosed | NO |
| HDFKEELDQWFK | 1446427 | Ectodomain | YES |
| HDGDWDSFCKTI | 1446429 | Cell/Virus Enclosed | NO |
| HDGLNEHVTSVN | 1446436 | Cell/Virus Enclosed | NO |
| HDGPIEHVTSVN | 1446438 | Cell/Virus Enclosed | NO |
| HDGPNEHVTSVN | 1446440 | Cell/Virus Enclosed | NO |
| HDICAQLNDKLGFNVD | 1664156 | Cell/Virus Enclosed | NO |
| HDIGNPKGVRCA | 1446454 | Cell/Virus Enclosed | NO |
| HDINNGPHEFCT | 1446457 | Cell/Virus Enclosed | NO |
| HDIQITNESYNNYVPS | 1664165 | Cell/Virus Enclosed | NO |
| HDIQLTNENYNN | 1446460 | Cell/Virus Enclosed | NO |
| HDIQLTNESYNN | 1446461 | Cell/Virus Enclosed | NO |
| HDITLDDDARTF | 1446465 | Cell/Virus Enclosed | NO |
| HDNVAFYQGDVDTVVN | 1664174 | Cell/Virus Enclosed | NO |
| HDPTVLPGAFAY | 1446483 | Cell/Virus Enclosed | NO |
| HDSQNMLRGDDM | 1446492 | Cell/Virus Enclosed | NO |
| HDSQNMLRGEDM | 1446493 | Cell/Virus Enclosed | NO |
| HDVIVLRDNLKP | 1446504 | Cell/Virus Enclosed | NO |
| HDVNNGLTCSTD | 1446506 | Ectodomain | YES |
| HDVPTEHVTSVN | 1446509 | Cell/Virus Enclosed | NO |
| HDVSPFEMPFEELNGL | 1664192 | Cell/Virus Enclosed | NO |
| HDVTVLRDNLKPVTTL | 1664194 | Cell/Virus Enclosed | NO |
| HDVTVTTDKSFEQQVG | 1664195 | Cell/Virus Enclosed | NO |
| HDVVSSFEGYDF | 1446515 | Cell/Virus Enclosed | NO |
| HEAKFGVVTTNSDKCP | 1664199 | Cell/Virus Enclosed | NO |
| HEDLHVVEVLTKTAVK | 1664204 | Cell/Virus Enclosed | NO |
| HEDLMAAYMENT | 1446534 | Cell/Virus Enclosed | NO |
| HEDLMAAYVEET | 1446535 | Cell/Virus Enclosed | NO |
| HEETIYNLIKDC | 1446547 | Cell/Virus Enclosed | NO |
| HEFAFLRKTLVL | 1446552 | Cell/Virus Enclosed | NO |
| HEFCSQHTMQIVDKDG | 1664218 | Cell/Virus Enclosed | NO |
| HEGDWDSFGKTI | 1446567 | Cell/Virus Enclosed | NO |
| HEGDWNSFCKTI | 1446569 | Cell/Virus Enclosed | NO |
| HEGFGGKNILGY | 1446570 | Cell/Virus Enclosed | NO |
| HEGKAYFPREGV | 23685 | Ectodomain | YES |
| HEGRTIYGNVSRQDLT | 1664228 | Cell/Virus Enclosed | NO |
| HEHIGGVYDNSI | 1446586 | Cell/Virus Enclosed | NO |
| HEHSLLVEAYNSILFE | 1664232 | Cell/Virus Enclosed | NO |
| HEIEKPRWKRQP | 1446596 | Capsid | UN |
| HEISDDEWAAAV | 1446598 | Cell/Virus Enclosed | NO |
| HEIVWFTERSEK | 1446600 | Cell/Virus Enclosed | NO |
| HELEPEPTSEEP | 1446607 | Cell/Virus Enclosed | NO |
| HEMQKPRWKRQPNDDV | 1664241 | Capsid | UN |
| HEMSPFEMPFEE | 1446610 | Cell/Virus Enclosed | NO |
| HENKLNSCCRIV | 1446611 | Cell/Virus Enclosed | NO |
| HEPTTPISSALI | 1446613 | Cell/Virus Enclosed | NO |
| HEQSMYNLLKGCNAVA | 1664244 | Cell/Virus Enclosed | NO |
| HERDLLIKAYNTINNE | 1664246 | Cell/Virus Enclosed | NO |
| HETSDDDWTAAV | 1446623 | Cell/Virus Enclosed | NO |
| HETSDDEWTAAV | 1446625 | Cell/Virus Enclosed | NO |
| HEVFDDEWAAAV | 1446630 | Cell/Virus Enclosed | NO |
| HEWKMVVCDKVR | 1446642 | Cell/Virus Enclosed | NO |
| HEYGATDAPSTF | 1446643 | Cell/Virus Enclosed | NO |
| HEYGTEDDYKGL | 1446644 | Cell/Virus Enclosed | NO |
| HEYGTEDDYQGL | 1446645 | Cell/Virus Enclosed | NO |
| HEYMLDLQPETTDLYCYE | 110884 | Cell/Virus Enclosed | NO |
| HFCPCKLNGSCV | 1446659 | Ectodomain | YES |
| HFDVDRKSFNNFVNIA | 1664278 | Cell/Virus Enclosed | NO |
| HFFFTQKGDAAIKDFD | 1664284 | Cell/Virus Enclosed | NO |
| HFGACDYDRLPNEVTV | 1664287 | Ectodomain | YES |
| HFGAGSDKGIAPGTSV | 1664288 | Cell/Virus Enclosed | NO |
| HFGENEEIVING | 1446696 | Cell/Virus Enclosed | NO |
| HFIGVEGELPVAVVND | 1664295 | Cell/Virus Enclosed | NO |
| HFISQGLDAAWN | 1446716 | Cell/Virus Enclosed | NO |
| HFKELWHLDTGA | 1446719 | Ectodomain | YES |
| HFKESWHMDTGV | 1446723 | Ectodomain | YES |
| HFMVGNVTRLHD | 1446736 | Undetermined | UN |
| HFNKEFYDFLRSQGFF | 1664308 | Cell/Virus Enclosed | NO |
| HFNQKLPNGVNV | 1446742 | Cell/Virus Enclosed | NO |
| HFNQKLPNGVTVVEEP | 1664309 | Capsid | UN |
| HFNYVPTKYVTA | 1446747 | Ectodomain | YES |
| HFRKACQNLDCN | 1446756 | Cell/Virus Enclosed | NO |
| HFRKSCQNLDCN | 1446757 | Cell/Virus Enclosed | NO |
| HFSLFDDNFYAF | 1446760 | Cell/Virus Enclosed | NO |

| | | | |
|---|---|---|---|
| HFSMMILSDEGG | 1446765 | Cell/Virus Enclosed | NO |
| HFSYKPTSFKTV | 1446770 | Ectodomain | YES |
| HFTNDKTIFYTT | 1446780 | Cell/Virus Enclosed | NO |
| HFTTQFVDNVKL | 1446786 | Ectodomain | YES |
| HFTTQFVGNVKF | 1446789 | Ectodomain | YES |
| HFTTQFVGVKFD | 1446791 | Ectodomain | YES |
| HFYYLGTGPHKA | 1446818 | Cell/Virus Enclosed | NO |
| HGAAAINSVPWS | 1446822 | Cell/Virus Enclosed | NO |
| HGAKPISNCIKM | 1446825 | Cell/Virus Enclosed | NO |
| HGCACDRTTIQSVDIS | 1664353 | Cell/Virus Enclosed | NO |
| HGCTCDRTAIQSFDNS | 1664354 | Cell/Virus Enclosed | NO |
| HGDFSGVASQEG | 1446843 | Ectodomain | YES |
| HGGGLAKALDVYTKGK | 1664368 | Cell/Virus Enclosed | NO |
| HGKEDLKFPRGQGVPI | 1664380 | Cell/Virus Enclosed | NO |
| HGKEELRFPRGQGVPI | 23917 | Capsid | UN |
| HGKENLTFPRGQ | 1446878 | Cell/Virus Enclosed | NO |
| HGKLLIRDNDTL | 1446883 | Cell/Virus Enclosed | NO |
| HGKLRPFERDISNVPF | 1664382 | Ectodomain | YES |
| HGLGLIQAGSVV | 1446899 | Cell/Virus Enclosed | NO |
| HGLLHSSQLPLD | 1446901 | Cell/Virus Enclosed | NO |
| HGLNLEEAARCM | 1446902 | Cell/Virus Enclosed | NO |
| HGMSFSMTTFEI | 1446907 | Cell/Virus Enclosed | NO |
| HGNGSMIEDIDYVPLK | 1664392 | Cell/Virus Enclosed | NO |
| HGNGSMIEDMDY | 1446913 | Cell/Virus Enclosed | NO |
| HGNGSMIEDVDY | 1446914 | Cell/Virus Enclosed | NO |
| HGNVAFYQGDVD | 1446919 | Cell/Virus Enclosed | NO |
| HGQSNADPKVYP | 1446928 | Cell/Virus Enclosed | NO |
| HGTCERSEAGIC | 1446940 | Cell/Virus Enclosed | NO |
| HGTCERSEVGIC | 1446942 | Cell/Virus Enclosed | NO |
| HGTILTRPLLESELV | 1312642 | Underneath Membrane | NO |
| HGTKKDAAAAKN | 1446947 | Cell/Virus Enclosed | NO |
| HGVITKAVDCSS | 1446954 | Ectodomain | YES |
| HGVITNAVDCSS | 1446955 | Ectodomain | YES |
| HGVRSKYALVDASLKMADPN | 741519 | Ectodomain | YES |
| HHAENVTGLFKD | 1446971 | Cell/Virus Enclosed | NO |
| HHFQSHGQLSDHPHALSSSS | 23985 | Capsid | UN |
| HHHAGNATEVPA | 1446990 | Cell/Virus Enclosed | NO |
| HHHAWSHPQFEK | 1446992 | Cell/Virus Enclosed | NO |
| HHHEEPVNQFTG | 1446993 | Cell/Virus Enclosed | NO |
| HHHHAGNATEVP | 1446995 | Cell/Virus Enclosed | NO |
| HHHHEEPVNQFT | 1446998 | Cell/Virus Enclosed | NO |
| HHHHHAENVTGL | 1446999 | Cell/Virus Enclosed | NO |
| HHHHHAGNATEV | 1447000 | Cell/Virus Enclosed | NO |
| HHHHHHAGNATE | 1447005 | Cell/Virus Enclosed | NO |
| HHHHHHEEPVNQ | 1447008 | Cell/Virus Enclosed | NO |
| HHHHHHMPVNQF | 1447012 | Cell/Virus Enclosed | NO |
| HHHMPVNQFTGY | 1447041 | Cell/Virus Enclosed | NO |
| HHHSGDDDDKAS | 1447048 | Cell/Virus Enclosed | NO |
| HHLGRCDIKDLPKEIT | 1382079 | Underneath Membrane | NO |
| HHLHSVYNWLKP | 1447062 | Cell/Virus Enclosed | NO |
| HHMPVNQFTGYL | 1447066 | Cell/Virus Enclosed | NO |
| HHPELKLFRNLNIDVC | 1664433 | Cell/Virus Enclosed | NO |
| HHRFKCSSNGSN | 1447076 | Ectodomain | YES |
| HHVINGCKDTAC | 1447094 | Cell/Virus Enclosed | NO |
| HIADCQTTNGLN | 1447104 | Ectodomain | YES |
| HIAGCQTTNGLNTSYS | 1664441 | Ectodomain | YES |
| HIAGCQTTNGTN | 1447108 | Ectodomain | YES |
| HIASSNAFLDLRTSCF | 1664442 | Cell/Virus Enclosed | NO |
| HIDPFPKRVIDL | 1447123 | Cell/Virus Enclosed | NO |
| HIDSSEPLCLFK | 1447125 | Ectodomain | YES |
| HIEVASAYGLCD | 1447133 | Ectodomain | YES |
| HIFYRRFLGKDGRRDPK | 24046 | Cell/Virus Enclosed | NO |
| HIGAPANLRVVGHAMQ | 1664453 | Cell/Virus Enclosed | NO |
| HIGLAGKVKVGTGVMV | 1664454 | Cell/Virus Enclosed | NO |
| HIGNINPHNYSY | 1447147 | Ectodomain | YES |
| HIGPPANLRVVG | 1447148 | Cell/Virus Enclosed | NO |
| HIGTRNPNNNAAIVLQ | 1664457 | Cell/Virus Enclosed | NO |
| HIGTRNPNNNVA | 1447152 | Capsid | UN |
| HILGRCIVLDVKGVEE | 1664464 | Cell/Virus Enclosed | NO |
| HINNLKIKNYGY | 1447192 | Ectodomain | YES |
| HINNLKVKNYGY | 1447193 | Ectodomain | YES |
| HIQTVQDADGINKQLT | 1664478 | Cell/Virus Enclosed | NO |
| HISSTMSQHSRS | 1447221 | Ectodomain | YES |
| HISTKPQPLMDK | 1447224 | Ectodomain | YES |
| HITAKETLYRID | 1447226 | Cell/Virus Enclosed | NO |
| HITDPLFSNMQP | 1447229 | Cell/Virus Enclosed | NO |

| | | | |
|---|---|---|---|
| HITHSGMEGMGVSCT | 24104 | Ectodomain | YES |
| HITKVLAEKFDY | 1447232 | Cell/Virus Enclosed | NO |
| HIVCDSLNSKLG | 1447243 | Cell/Virus Enclosed | NO |
| HIVGPDARAKQD | 1447244 | Cell/Virus Enclosed | NO |
| HIVSKESGNTVVLTFT | 1664502 | Capsid | UN |
| HIVYGDFSHGQL | 1447252 | Cell/Virus Enclosed | NO |
| HIVYGNFNQKII | 1447255 | Cell/Virus Enclosed | NO |
| HKAVFISPYNSQNFAA | 1664515 | Cell/Virus Enclosed | NO |
| HKAYIDVLRNSFGKDL | 1664516 | Cell/Virus Enclosed | NO |
| HKAYVDVLCNSF | 1447268 | Cell/Virus Enclosed | NO |
| HKCGNLHQRELL | 1447270 | Cell/Virus Enclosed | NO |
| HKDFFKDGLCMF | 1447275 | Cell/Virus Enclosed | NO |
| HKDITECVTIVL | 1447278 | Cell/Virus Enclosed | NO |
| HKELKFRQRSDG | 1447290 | Cell/Virus Enclosed | NO |
| HKGAAWPIVRRR | 1447292 | Cell/Virus Enclosed | NO |
| HKHAIVTVTYDSEEQRQQ | 110625 | Cell/Virus Enclosed | NO |
| HKHDIQITNESYNNYV | 1664540 | Cell/Virus Enclosed | NO |
| HKHFYLTMYIIP | 1447308 | Cell/Virus Enclosed | NO |
| HKIFKMEKGKVVKSVELDAPNYHY | 97383 | Ectodomain | YES |
| HKMQTYKLVNMK | 1447342 | Cell/Virus Enclosed | NO |
| HKNGQGEYDNIS | 1447348 | Cell/Virus Enclosed | NO |
| HKNVQGENDNIS | 1447353 | Cell/Virus Enclosed | NO |
| HKPPSASCPIGTNYRS | 1664569 | Ectodomain | YES |
| HKSAIVTLTYDSEWQRDQ | 110284 | Cell/Virus Enclosed | NO |
| HKSDFQLPVYAS | 1447378 | Cell/Virus Enclosed | NO |
| HLATFAELEAAQKAYQ | 1664591 | Cell/Virus Enclosed | NO |
| HLDGEVLPLDEL | 1447426 | Cell/Virus Enclosed | NO |
| HLDGEVLPLDKL | 1447427 | Cell/Virus Enclosed | NO |
| HLGKFLEELNAF | 1447454 | Cell/Virus Enclosed | NO |
| HLGLDDQEKKILMNF | 227244 | Capsid | UN |
| HLGRCDIKDLPKEITV | 1382130 | Underneath Membrane | NO |
| HLHVASNSKLWQYCSV | 1664614 | Cell/Virus Enclosed | NO |
| HLIKDVDNPVLMGWDY | 1664616 | Cell/Virus Enclosed | NO |
| HLKHGGGIAGAINAAS | 1664627 | Cell/Virus Enclosed | NO |
| HLKMAGMHFGAC | 1447487 | Ectodomain | YES |
| HLKVNCCRFQRVDENG | 1664634 | Cell/Virus Enclosed | NO |
| HLNKTLNEGVLESFSV | 1664657 | Cell/Virus Enclosed | NO |
| HLPAYASFKVIDNGVV | 1664662 | Cell/Virus Enclosed | NO |
| HLRIAGHHLGRCDIKD | 1664669 | Underneath Membrane | NO |
| HLSMTTSTTNFGEDFL | 1664675 | Cell/Virus Enclosed | NO |
| HLSTFYPQLQSAEWKC | 1664676 | Cell/Virus Enclosed | NO |
| HLSVDIKFKTEG | 1447586 | Cell/Virus Enclosed | NO |
| HLVNPQIQLTIT | 1447625 | Cell/Virus Enclosed | NO |
| HLYIQGIKLGTGYSLA | 1664697 | Ectodomain | YES |
| HLYIQGVKLGTGYTLS | 1664698 | Underneath Membrane | NO |
| HMPLSNFEHKVI | 1447678 | Cell/Virus Enclosed | NO |
| HMPVNQFTGYLK | 1447682 | Cell/Virus Enclosed | NO |
| HMTNYLEQPVSNDLS | 24352 | Ectodomain | YES |
| HMVKDGLVGVLT | 1447690 | Cell/Virus Enclosed | NO |
| HMVLPSEAPNAK | 1447692 | Cell/Virus Enclosed | NO |
| HMVVKSALLADK | 1447695 | Cell/Virus Enclosed | NO |
| HNAAQCDCLCTP | 1447701 | Ectodomain | YES |
| HNASGKEAKVCTISPIMGYS | 24362 | Capsid | UN |
| HNAVQCNCLCTP | 1447703 | Ectodomain | YES |
| HNCCDEYGGHHDFVIK | 1664723 | Underneath Membrane | NO |
| HNDMLYSPPTISYNST | 1664728 | Cell/Virus Enclosed | NO |
| HNGVLDITACQY | 1447731 | Ectodomain | YES |
| HNLCTLDPITFK | 1447753 | Ectodomain | YES |
| HNLPEYMAVAVP | 1447759 | Underneath Membrane | NO |
| HNLSGNPDEPQK | 1447762 | Cell/Virus Enclosed | NO |
| HNMGDSDLVQNG | 1447769 | Cell/Virus Enclosed | NO |
| HNMRVLHYGAGSDKGV | 1664765 | Cell/Virus Enclosed | NO |
| HNNGNDVLYQPPTASV | 1664768 | Cell/Virus Enclosed | NO |
| HNRRSFKTADGN | 1447783 | Cell/Virus Enclosed | NO |
| HNRRSFKTADGS | 1447784 | Cell/Virus Enclosed | NO |
| HNTASWFTALTQ | 1447800 | Capsid | UN |
| HNTPVYKLDISEATQ | 227248 | Secreted | YES |
| HNWLNGGQPVSE | 1447814 | Cell/Virus Enclosed | NO |
| HNYSYMDLNSAL | 1447833 | Ectodomain | YES |
| HPAFNVSDAYANLVPY | 1664794 | Cell/Virus Enclosed | NO |
| HPAHAPSFLAVDDKYK | 1664795 | Cell/Virus Enclosed | NO |
| HPDCVDCHDEMC | 1447846 | Cell/Virus Enclosed | NO |
| HPDFKLLHNLQADICY | 1664798 | Cell/Virus Enclosed | NO |
| HPHLGKFLEELN | 1447859 | Cell/Virus Enclosed | NO |
| HPIQIDQLNSSY | 1447866 | Ectodomain | YES |
| HPIQVDQFNSSY | 1447868 | Ectodomain | YES |

| | | | |
|---|---|---|---|
| HPIQVDQLNSSYFKLS | 1664809 | Ectodomain | YES |
| HPIQVDQLNSTY | 1447870 | Ectodomain | YES |
| HPIQVGQLNSSY | 1447871 | Ectodomain | YES |
| HPITIGKCPKYVKSTKLRLATGLRN | 1146809 | Ectodomain | YES |
| HPKLGNHFKESW | 1447876 | Ectodomain | YES |
| HPLLNPGALEFN | 1447884 | Cell/Virus Enclosed | NO |
| HPLLNPSALEFNPSQT | 1664814 | Capsid | UN |
| HPLSHFVNLDNL | 1447887 | Cell/Virus Enclosed | NO |
| HPNLGNHFKELW | 1447898 | Ectodomain | YES |
| HPNTPPKPRRRSSSSTTMPS | 741530 | Underneath Membrane | NO |
| HPPPGEQFKHLV | 1447907 | Cell/Virus Enclosed | NO |
| HPSFNVSDAYANLVPY | 1664827 | Cell/Virus Enclosed | NO |
| HPSISTDTVDVT | 1447917 | Ectodomain | YES |
| HPSLCNNGKISSSAGD | 1664829 | Ectodomain | YES |
| HPTNPFAPDVTAAAGAGPRVRQPAR | 24518 | Undetermined | UN |
| HPVECDEGCSEV | 1447926 | Cell/Virus Enclosed | NO |
| HPYCYSDGVMKNASLY | 1664839 | Cell/Virus Enclosed | NO |
| HQADTSTPSDVSSRDP | 1664840 | Capsid | UN |
| HQFVNASSATDA | 1447965 | Cell/Virus Enclosed | NO |
| HQGVFRYDNISS | 1447971 | Ectodomain | YES |
| HQKMLKSMAATRGATC | 1664854 | Cell/Virus Enclosed | NO |
| HQLECIVDDSVR | 1447982 | Cell/Virus Enclosed | NO |
| HQMELASGTHTG | 1447996 | Cell/Virus Enclosed | NO |
| HQNAQGEGTAADYKSTQSAIDQITG | 1146898 | Ectodomain | YES |
| HQTYNRGRKPQP | 1448017 | Cell/Virus Enclosed | NO |
| HQVNISLNGNTSVCVR | 1664869 | Ectodomain | YES |
| HQVQLIDKYCSV | 1448021 | Cell/Virus Enclosed | NO |
| HQVQLTDKYCSV | 1448022 | Cell/Virus Enclosed | NO |
| HQWNCIDCDSYK | 1448027 | Cell/Virus Enclosed | NO |
| HRCDINYKGKVF | 1448029 | Cell/Virus Enclosed | NO |
| HRDCDFNKPLIEWPLT | 1664874 | Cell/Virus Enclosed | NO |
| HRFKCFFNGSND | 1448038 | Ectodomain | YES |
| HRFYSLKAAVHK | 1448045 | Cell/Virus Enclosed | NO |
| HRIESIDSIVFT | 1448062 | Cell/Virus Enclosed | NO |
| HRIIKAQFKTFE | 1448069 | Cell/Virus Enclosed | NO |
| HRITSITSDKFD | 1448070 | Cell/Virus Enclosed | NO |
| HRITSVTSDKFD | 1448071 | Cell/Virus Enclosed | NO |
| HRLASGVQVHNLPEYM | 1664887 | Underneath Membrane | NO |
| HRNLGNHFKELW | 1448079 | Ectodomain | YES |
| HRSFTQWYHEHI | 1448096 | Cell/Virus Enclosed | NO |
| HRWKMVVCDKVR | 1448112 | Cell/Virus Enclosed | NO |
| HRYKAGNYRSPPITAD | 1664910 | Ectodomain | YES |
| HSAFVECGSCGN | 1448127 | Cell/Virus Enclosed | NO |
| HSDFGLDGLIDS | 1448141 | Cell/Virus Enclosed | NO |
| HSDHAESSWVNRGESSRKAY | 24730 | Ectodomain | YES |
| HSDVDNIDFKSL | 1448157 | Ectodomain | YES |
| HSEIVFDVESGG | 1448160 | Ectodomain | YES |
| HSFRTLKSGEGF | 1448170 | Cell/Virus Enclosed | NO |
| HSFSVQKHIGAPANLR | 1664931 | Cell/Virus Enclosed | NO |
| HSGDDDDSQPRA | 1448178 | Cell/Virus Enclosed | NO |
| HSGSNDNIALLV | 1448189 | Underneath Membrane | NO |
| HSIPDLQRAAIT | 1448205 | Cell/Virus Enclosed | NO |
| HSIQQGFDATNS | 1448207 | Ectodomain | YES |
| HSIRSFQSDRKA | 1448209 | Ectodomain | YES |
| HSIRSIKSDRKA | 1448210 | Ectodomain | YES |
| HSIRSIQSGRKA | 1448212 | Ectodomain | YES |
| HSKCLTSIEEVC | 1448213 | Cell/Virus Enclosed | NO |
| HSKEQYGTDIDG | 1448215 | Cell/Virus Enclosed | NO |
| HSKMSDVKCTSV | 1448221 | Cell/Virus Enclosed | NO |
| HSLLATLDDPYVEDSV | 1664953 | Capsid | UN |
| HSLNFVGEFVVN | 1448231 | Cell/Virus Enclosed | NO |
| HSLSVHDGYTNE | 1448238 | Cell/Virus Enclosed | NO |
| HSMAVVDGKQIDDHRI | 1664963 | Cell/Virus Enclosed | NO |
| HSMPVVDGKQID | 1448249 | Cell/Virus Enclosed | NO |
| HSQFYFNKDTGV | 1448269 | Ectodomain | YES |
| HSQTSLRCGTCI | 1448272 | Cell/Virus Enclosed | NO |
| HSRVKNLNSSEG | 1448280 | Ectodomain | YES |
| HSSIKEGTQIYK | 1448289 | Cell/Virus Enclosed | NO |
| HSSIKQGTQIYK | 1448292 | Cell/Virus Enclosed | NO |
| HSSLFAQSEEEE | 1448295 | Cell/Virus Enclosed | NO |
| HSSQLPLDIYDL | 1448307 | Cell/Virus Enclosed | NO |
| HSSQLPLDVYDLTMPS | 1664982 | Cell/Virus Enclosed | NO |
| HSSQLPLDVYNL | 1448309 | Cell/Virus Enclosed | NO |
| HSTRSIQSDRKA | 1448329 | Ectodomain | YES |
| HSVADVMRYNFN | 1448334 | Ectodomain | YES |
| HSVADYHNHSNI | 1448337 | Cell/Virus Enclosed | NO |

| | | | |
|---|---|---|---|
| HSVASLSAPTLVPQEN | 1664993 | Cell/Virus Enclosed | NO |
| HSVAVIGDEQIDGKFV | 1664995 | Cell/Virus Enclosed | NO |
| HSVCKGCVGHSE | 1448344 | Ectodomain | YES |
| HSVCNGCVGHSE | 1448345 | Ectodomain | YES |
| HSVEIEAGEVKPFAVY | 1664996 | Cell/Virus Enclosed | NO |
| HSVHNRLKPYGG | 1448353 | Cell/Virus Enclosed | NO |
| HSVHNWLKPFGG | 1448354 | Cell/Virus Enclosed | NO |
| HSVHNWLNGGQP | 1448357 | Cell/Virus Enclosed | NO |
| HSVLTTFIDAARGPAG | 1665000 | Cell/Virus Enclosed | NO |
| HSVPKLSDFESELSHWFKNQTSIAP | 1857699 | Ectodomain | YES |
| HTASTLRSVGQK | 1448396 | Ectodomain | YES |
| HTFDDFCPECRPLGLQGCAFQSTVA | 24910 | Undetermined | UN |
| HTFDDPVIPFKD | 1448413 | Ectodomain | YES |
| HTFGNPIIPFKD | 1448415 | Ectodomain | YES |
| HTFGNPVIPFKDGIYF | 1665027 | Ectodomain | YES |
| HTGFQGKQILGS | 1448430 | Cell/Virus Enclosed | NO |
| HTICKSIGSSRN | 1448456 | Ectodomain | YES |
| HTIDGSSGVANPAMDP | 1665047 | Secreted | YES |
| HTIKGSFLNGSA | 1448465 | Cell/Virus Enclosed | NO |
| HTINHTFDDPVI | 1448468 | Ectodomain | YES |
| HTINHTFGNPII | 1448470 | Ectodomain | YES |
| HTINHTFGNPVIPFKDG | 24923 | Ectodomain | YES |
| HTKKWKFPQIGG | 1448480 | Cell/Virus Enclosed | NO |
| HTKKWKIPQVGG | 1448482 | Cell/Virus Enclosed | NO |
| HTKSAAGYGGFG | 1448487 | Cell/Virus Enclosed | NO |
| HTLDDVTRNYII | 1448495 | Cell/Virus Enclosed | NO |
| HTLGNPVIPFKD | 1448500 | Ectodomain | YES |
| HTLRNGSVLEMAKEVK | 1665064 | Cell/Virus Enclosed | NO |
| HTLSIHDGYTNE | 1448509 | Cell/Virus Enclosed | NO |
| HTLTKHTEVRAS | 1448513 | Ectodomain | YES |
| HTLTKYTEVRAS | 1448514 | Ectodomain | YES |
| HTLYIKDGDYGY | 1448519 | Cell/Virus Enclosed | NO |
| HTMQIVDENGKYYLPY | 1665077 | Cell/Virus Enclosed | NO |
| HTMQIVDKDGTYYLPY | 1665078 | Cell/Virus Enclosed | NO |
| HTNPFTRTVFEN | 1448542 | Cell/Virus Enclosed | NO |
| HTPAFDKSAFTY | 1448548 | Cell/Virus Enclosed | NO |
| HTPAIDCSDGNY | 1448550 | Ectodomain | YES |
| HTPATDCSDGNYNRNA | 1665087 | Ectodomain | YES |
| HTPATNCSDGNY | 1448552 | Ectodomain | YES |
| HTPAYDGSAFRD | 1448553 | Cell/Virus Enclosed | NO |
| HTPKHVFKTLKPGDSF | 1665094 | Cell/Virus Enclosed | NO |
| HTPLFSFLTASPA | 24960 | Ectodomain | YES |
| HTPSTDCSDGNY | 1448562 | Ectodomain | YES |
| HTRYSLADSNGF | 1448572 | Cell/Virus Enclosed | NO |
| HTSCSCSPDAFL | 1448575 | Ectodomain | YES |
| HTSFATYHTPAT | 1448576 | Ectodomain | YES |
| HTSIAPNLTFNS | 1448578 | Ectodomain | YES |
| HTSSMRGVYYPDEIFRS | 24978 | Ectodomain | YES |
| HTTLGGLHLFIG | 1448594 | Cell/Virus Enclosed | NO |
| HTVCDSLNAKLGFDSS | 1665109 | Cell/Virus Enclosed | NO |
| HTVCDSLNAKWI | 1448600 | Cell/Virus Enclosed | NO |
| HTVCDSLNAKWV | 1448601 | Cell/Virus Enclosed | NO |
| HTVCDSLNFKLG | 1448602 | Cell/Virus Enclosed | NO |
| HTVIDVKQVGCS | 1448608 | Cell/Virus Enclosed | NO |
| HTVTDVKQVGCF | 1448624 | Cell/Virus Enclosed | NO |
| HTVTDVKQVGCS | 1448625 | Cell/Virus Enclosed | NO |
| HTVYPDYVDVNK | 1448629 | Ectodomain | YES |
| HVADTLMQGVTLSSNL | 1665124 | Ectodomain | YES |
| HVAQNTVLNACT | 1448645 | Cell/Virus Enclosed | NO |
| HVAQNTVLNASIPESK | 1665127 | Capsid | UN |
| HVCFKDCYNIWE | 1448653 | Cell/Virus Enclosed | NO |
| HVCGDNIGTNVPLQVG | 1665140 | Cell/Virus Enclosed | NO |
| HVDHYDGFYTQG | 1448661 | Cell/Virus Enclosed | NO |
| HVDTSYDCDIPI | 1448666 | Ectodomain | YES |
| HVDTSYECDIPIGAGIC | 25031 | Ectodomain | YES |
| HVEVFTDGKLNCSFQD | 1665148 | Cell/Virus Enclosed | NO |
| HVFKTLKPGDSFNILA | 1665150 | Cell/Virus Enclosed | NO |
| HVGLLDQAWRFP | 1448700 | Cell/Virus Enclosed | NO |
| HVGYYPSNHIDV | 1448707 | Ectodomain | YES |
| HVGYYPSNHIEV | 1448708 | Ectodomain | YES |
| HVHVGGVYDNSI | 1448710 | Cell/Virus Enclosed | NO |
| HVIASNTTSAIDYDHE | 1665164 | Cell/Virus Enclosed | NO |
| HVICPADQLSDP | 1448711 | Cell/Virus Enclosed | NO |
| HVIWDYVRESIF | 1448720 | Cell/Virus Enclosed | NO |
| HVKEITRENVET | 1448725 | Cell/Virus Enclosed | NO |
| HVKGTIDHPVLSKLKFTK | 558264 | Undetermined | UN |

| | | | |
|---|---|---|---|
| HVLDCELFVNDSYRQF | 1665178 | Cell/Virus Enclosed | NO |
| HVNSSEPALLFR | 1448773 | Ectodomain | YES |
| HVPHPSMDGYCKFKGK | 1665203 | Cell/Virus Enclosed | NO |
| HVRNNGGYNSWH | 1448788 | Cell/Virus Enclosed | NO |
| HVTCCTANDKFY | 1448814 | Cell/Virus Enclosed | NO |
| HVTKVLAEKFDYNVSV | 1665227 | Cell/Virus Enclosed | NO |
| HVTSVNLLDVGY | 1448824 | Cell/Virus Enclosed | NO |
| HVTYVPSQEKNF | 1448831 | Ectodomain | YES |
| HVVGPDARAKQDVSLL | 1665238 | Cell/Virus Enclosed | NO |
| HVVPLENDADKA | 1448858 | Cell/Virus Enclosed | NO |
| HVVRDPRYVMRLQSAA | 1665244 | Cell/Virus Enclosed | NO |
| HVVRDSRFVMRLQTIA | 1665245 | Cell/Virus Enclosed | NO |
| HVYDFKVEDAYDQVHD | 1665250 | Cell/Virus Enclosed | NO |
| HVYDFKVEYAYD | 1448871 | Cell/Virus Enclosed | NO |
| HVYEVGDLIAQS | 1448872 | Cell/Virus Enclosed | NO |
| HVYEVSDLIAQS | 1448874 | Cell/Virus Enclosed | NO |
| HWAPEFPWMFEDAEEK | 1665255 | Cell/Virus Enclosed | NO |
| HWMGIRFEYTSPTDKL | 1665269 | Cell/Virus Enclosed | NO |
| HWSFDYHPNCSDCYDD | 1665273 | Cell/Virus Enclosed | NO |
| HWSMPYHPNPVD | 1448937 | Cell/Virus Enclosed | NO |
| HWSMPYHPNTVDCQDD | 1665275 | Cell/Virus Enclosed | NO |
| HYAGSRSSSGNRSGIL | 1665276 | Capsid | UN |
| HYAGVVWTLNDVKDND | 1665277 | Cell/Virus Enclosed | NO |
| HYCSTLHNEILA | 1448943 | Cell/Virus Enclosed | NO |
| HYDGRTGELPCAIMND | 1665284 | Cell/Virus Enclosed | NO |
| HYDLVNYIRDKG | 1448951 | Cell/Virus Enclosed | NO |
| HYERDNTSYPEW | 1448957 | Cell/Virus Enclosed | NO |
| HYERNNTSCPEW | 1448958 | Cell/Virus Enclosed | NO |
| HYFDAQPLTRKV | 1448960 | Cell/Virus Enclosed | NO |
| HYGAGSDKGVAPGTTV | 1665292 | Cell/Virus Enclosed | NO |
| HYIKDGDDGYFL | 1448971 | Cell/Virus Enclosed | NO |
| HYKELGIVMNMD | 1448977 | Cell/Virus Enclosed | NO |
| HYKELGVVMNFD | 1448979 | Cell/Virus Enclosed | NO |
| HYKELGVVMNLDVDTH | 1665297 | Cell/Virus Enclosed | NO |
| HYKYSRFLYGLK | 1448985 | Undetermined | UN |
| HYMYSRFLYGLK | 1448999 | Undetermined | UN |
| HYNEWAQANGFTAMNG | 1665304 | Cell/Virus Enclosed | NO |
| HYQTNIYSQNLCVDGF | 1665309 | Cell/Virus Enclosed | NO |
| HYQTNLYSFNKAIDGF | 1665310 | Cell/Virus Enclosed | NO |
| HYTGQAGEMPCAIIND | 1665318 | Cell/Virus Enclosed | NO |
| HYTPFHYERDNTSCPE | 1665321 | Cell/Virus Enclosed | NO |
| HYTPVHYERDNT | 1449044 | Cell/Virus Enclosed | NO |
| HYTPVRYERDNT | 1449045 | Cell/Virus Enclosed | NO |
| HYTVYDTAKKSMYDGD | 1665324 | Cell/Virus Enclosed | NO |
| HYVHARLKCGLI | 1449050 | Cell/Virus Enclosed | NO |
| HYVHARLKGGLILKFD | 1665325 | Cell/Virus Enclosed | NO |
| HYVNRLVTQFKL | 1449065 | Cell/Virus Enclosed | NO |
| HYVRVTFSDPNL | 1449072 | Cell/Virus Enclosed | NO |
| HYVRVTFSDPNT | 1449073 | Cell/Virus Enclosed | NO |
| HYVVYDAANNAV | 1449077 | Cell/Virus Enclosed | NO |
| HYWSPQGNLGGNCAGN | 1665336 | Cell/Virus Enclosed | NO |
| IAADLPCTLHFS | 1449106 | Cell/Virus Enclosed | NO |
| IAAEQDRNTREVFAQVK | 25219 | Ectodomain | YES |
| IAAHPPPGEQFK | 1449109 | Cell/Virus Enclosed | NO |
| IAAKPEPEPLPE | 1449112 | Cell/Virus Enclosed | NO |
| IAARPPPGEHFK | 1449121 | Cell/Virus Enclosed | NO |
| IAARPPPGEQFKHLVP | 1665351 | Cell/Virus Enclosed | NO |
| IAAYTAALVSGTATAG | 1665356 | Ectodomain | YES |
| IACKCGVKQEQRTGLD | 1665359 | Cell/Virus Enclosed | NO |
| IACVKDFDFKFYDAQP | 1665364 | Cell/Virus Enclosed | NO |
| IADKDKDLSGAVPSDL | 1665369 | Cell/Virus Enclosed | NO |
| IADPGYMQGYDD | 1449165 | Ectodomain | YES |
| IADTLQEIPVVS | 1449173 | Cell/Virus Enclosed | NO |
| IADTLQETPVVPDTVE | 1665377 | Cell/Virus Enclosed | NO |
| IADTLQETSVVS | 1449177 | Cell/Virus Enclosed | NO |
| IADVYRRKLNLP | 1449181 | Ectodomain | YES |
| IADVYRRKPDLP | 1449182 | Ectodomain | YES |
| IADVYRRKPNLP | 1449184 | Ectodomain | YES |
| IADYNYKLPDDFMGCVL | 25250 | Ectodomain | YES |
| IAEDVVDQVEEVNSIF | 1665381 | Cell/Virus Enclosed | NO |
| IAFANWLAPTNA | 1449202 | Cell/Virus Enclosed | NO |
| IAFCKSGNLVLN | 1449203 | Ectodomain | YES |
| IAFELFAKRKVGLTPP | 1665389 | Cell/Virus Enclosed | NO |
| IAFELYAKRSVR | 1449209 | Cell/Virus Enclosed | NO |
| IAFGIPPSEAKG | 1449211 | Cell/Virus Enclosed | NO |
| IAFGLQDCVTGI | 1449212 | Cell/Virus Enclosed | NO |

| | | | |
|---|---|---|---|
| IAFGVPLSEAKG | 1449213 | Cell/Virus Enclosed | NO |
| IAFNHPIQIDQL | 1449218 | Ectodomain | YES |
| IAFNHPIQVDQF | 1449220 | Ectodomain | YES |
| IAFNHPIQVDQLNSSY | 1665396 | Ectodomain | YES |
| IAFNVVKQGHFT | 1449225 | Cell/Virus Enclosed | NO |
| IAGCQTTNGLNTSYSV | 1665403 | Ectodomain | YES |
| IAGHEEFIQTSS | 1449241 | Ectodomain | YES |
| IAGHHLGRCDIKDLPK | 1665405 | Underneath Membrane | NO |
| IAGIQNFKETLYYQNN | 1665406 | Cell/Virus Enclosed | NO |
| IAGLIAIVMVTILLCCM | 25293 | Undetermined | UN |
| IAGNGGDAALALLLLD | 1665412 | Cell/Virus Enclosed | NO |
| IAGPKGPEIRYL | 1449256 | Cell/Virus Enclosed | NO |
| IAGQAFFYSDGF | 1449257 | Cell/Virus Enclosed | NO |
| IAGVAGARIPDVPTTL | 1665417 | Cell/Virus Enclosed | NO |
| IAHASLREGVQLEKVL | 1665420 | Cell/Virus Enclosed | NO |
| IAHWIKNTFSLV | 1449274 | Cell/Virus Enclosed | NO |
| IAIEDLLFDKVT | 1449277 | Ectodomain | YES |
| IAIKVTEYSWNKKLYE | 1665427 | Cell/Virus Enclosed | NO |
| IAKPKRLRKKRN | 1449321 | Cell/Virus Enclosed | NO |
| IALDDDARTFIQ | 1449330 | Cell/Virus Enclosed | NO |
| IALDDDARTFVQ | 1449331 | Cell/Virus Enclosed | NO |
| IALNKIQDVVNQQGSA | 1665460 | Ectodomain | YES |
| IAMACLVGLMWLSYF | 2001106 | Transmembrane | NO |
| IANAFNNALDAI | 1449372 | Ectodomain | YES |
| IANAFNNALHAI | 1449373 | Ectodomain | YES |
| IANAFSNALDAI | 1449375 | Ectodomain | YES |
| IANENAINKGGRTVQS | 1665475 | Cell/Virus Enclosed | NO |
| IANKFNQALGAMQTGF | 1665482 | Ectodomain | YES |
| IANLVLAKLGKDSKPQ | 1665483 | Capsid | UN |
| IANMFLEQTQHSLNFV | 1665484 | Cell/Virus Enclosed | NO |
| IANNYSKLFDEGYTVV | 1665485 | Cell/Virus Enclosed | NO |
| IANQERYVKISG | 1449394 | Cell/Virus Enclosed | NO |
| IANQERYVKITG | 1449395 | Cell/Virus Enclosed | NO |
| IANQERYVKMFW | 1449396 | Cell/Virus Enclosed | NO |
| IANRYNEVANAVMQNN | 1665489 | Cell/Virus Enclosed | NO |
| IANRYNEVSATVLQNN | 1665490 | Cell/Virus Enclosed | NO |
| IANVPAIDVLKKLLSS | 1665491 | Cell/Virus Enclosed | NO |
| IAPDRASFLRGKSMGIQSGVQVDAN | 1147379 | Ectodomain | YES |
| IAPGQTGVIADYNYKLP | 25378 | Ectodomain | YES |
| IAPKSGYFVNVNNTWM | 1665496 | Ectodomain | YES |
| IAPNLTLNLHTI | 1449424 | Ectodomain | YES |
| IAPQANFGDYYY | 1449430 | Ectodomain | YES |
| IAPQANSGDYYH | 1449431 | Ectodomain | YES |
| IAPQANSGDYYY | 1449432 | Ectodomain | YES |
| IAPQANSGGYYY | 1449433 | Ectodomain | YES |
| IAPQANSGIITK | 1449434 | Ectodomain | YES |
| IAPSTVWEDGDY | 1449436 | Ectodomain | YES |
| IAPVNGYFIKTNNTRI | 1665501 | Ectodomain | YES |
| IAQRTRTRPSVQ | 1449453 | Capsid | UN |
| IAQVVIADTLQETPVV | 1665508 | Cell/Virus Enclosed | NO |
| IASFRLFARTRSMWS | 1312686 | Undetermined | UN |
| IASLVLAKLGKDATKP | 1665523 | Cell/Virus Enclosed | NO |
| IASNCYSSLILD | 1449484 | Ectodomain | YES |
| IASNSKEWAYCVEMHN | 1665525 | Cell/Virus Enclosed | NO |
| IASNTTSAIDYDHEYS | 1665526 | Cell/Virus Enclosed | NO |
| IASQLGNCVEYS | 1449489 | Ectodomain | YES |
| IATAFNNALLSIQNGF | 1665533 | Ectodomain | YES |
| IATAGWTFGAGA | 1449496 | Ectodomain | YES |
| IATKMFDSFVNSFVSL | 1665541 | Cell/Virus Enclosed | NO |
| IATPHKMVLSVS | 1449511 | Cell/Virus Enclosed | NO |
| IATRNNCWLRSV | 1449513 | Cell/Virus Enclosed | NO |
| IATSHKMVLSVS | 1449516 | Cell/Virus Enclosed | NO |
| IATVKSEDGNIKNINW | 1665548 | Cell/Virus Enclosed | NO |
| IAVAAGKKFSKETAAM | 1665552 | Cell/Virus Enclosed | NO |
| IAVANWLAPTNAYMRT | 1665554 | Cell/Virus Enclosed | NO |
| IAVANWLAPTNV | 1449523 | Cell/Virus Enclosed | NO |
| IAVDIQQTFFDK | 1449528 | Ectodomain | YES |
| IAWNTAQQDQGQ | 1449571 | Ectodomain | YES |
| IAYAILYFFATR | 1449575 | Cell/Virus Enclosed | NO |
| IAYENARQAYEDAIAN | 1665577 | Cell/Virus Enclosed | NO |
| IAYGIPLSEAKG | 1449580 | Cell/Virus Enclosed | NO |
| ICAAFDFSLDVFKIGD | 1665585 | Cell/Virus Enclosed | NO |
| ICAQKFNALTVL | 1449614 | Ectodomain | YES |
| ICCCTGCGTSCS | 1449629 | Undetermined | UN |
| ICDCGIKQESRV | 1449630 | Cell/Virus Enclosed | NO |
| ICEKLAIGGSIA | 1449643 | Cell/Virus Enclosed | NO |

| | | | |
|---|---|---|---|
| ICFDIRDNGSLE | 1449650 | Cell/Virus Enclosed | NO |
| ICFEYTSPTDKL | 1449654 | Cell/Virus Enclosed | NO |
| ICFIGLWCLMPT | 1449655 | Cell/Virus Enclosed | NO |
| ICGGCCDDYTGH | 1449674 | Underneath Membrane | NO |
| ICHEGKAYFPRE | 1449680 | Ectodomain | YES |
| ICHPKLGNHFKE | 1449685 | Ectodomain | YES |
| ICHPNLGNRRVE | 1449689 | Ectodomain | YES |
| ICIKDADYNAKD | 1449697 | Cell/Virus Enclosed | NO |
| ICIKDADYNAKVEISV | 1665631 | Cell/Virus Enclosed | NO |
| ICIVDNVYMAKAGDKY | 1665634 | Cell/Virus Enclosed | NO |
| ICKTTTGIPEYN | 1449724 | Cell/Virus Enclosed | NO |
| ICLSGRKIYEIE | 1449739 | Cell/Virus Enclosed | NO |
| ICMFENKTTLPTNIAF | 1665655 | Cell/Virus Enclosed | NO |
| ICNSNILSLNNP | 1449758 | Ectodomain | YES |
| ICNTNRAGIGLYYKVN | 1665659 | Cell/Virus Enclosed | NO |
| ICPAGTKYRHCD | 1449763 | Ectodomain | YES |
| ICPTGTKYRHCD | 1449766 | Ectodomain | YES |
| ICQAVSANVCALMSCN | 1665665 | Cell/Virus Enclosed | NO |
| ICQAVSDNVRAL | 1449767 | Cell/Virus Enclosed | NO |
| ICQAVTANVRSL | 1449770 | Cell/Virus Enclosed | NO |
| ICSASDMTNPDYTNLL | 1665669 | Cell/Virus Enclosed | NO |
| ICSNTPLSKDLPDDVV | 1665671 | Cell/Virus Enclosed | NO |
| ICSQISPAAIAS | 1449785 | Ectodomain | YES |
| ICSSSNMNEPDYSALL | 1665674 | Cell/Virus Enclosed | NO |
| ICSVAYVANPTE | 1449790 | Cell/Virus Enclosed | NO |
| ICVALCDAMVAK | 1449799 | Cell/Virus Enclosed | NO |
| ICVKNGNMRCTICI | 1147484 | Undetermined | UN |
| IDAAKIYGMGFS | 1449833 | Ectodomain | YES |
| IDAMVYTSDLLT | 1449848 | Cell/Virus Enclosed | NO |
| IDAYKTFPKKEN | 1449859 | Cell/Virus Enclosed | NO |
| IDAYPLSKHPKPEYRK | 1665711 | Cell/Virus Enclosed | NO |
| IDAYPLSKHPNSEYRK | 1665712 | Cell/Virus Enclosed | NO |
| IDAYPLTKHEDIEYQN | 1665713 | Cell/Virus Enclosed | NO |
| IDCAAFVRGDYA | 1449871 | Ectodomain | YES |
| IDCEQDLCDFKG | 1449873 | Cell/Virus Enclosed | NO |
| IDCEQTPCGFKG | 1449875 | Cell/Virus Enclosed | NO |
| IDCGFNDLSQLY | 1449878 | Ectodomain | YES |
| IDCSAFVCGDCA | 1449887 | Ectodomain | YES |
| IDCSAFVCGDYA | 1449888 | Ectodomain | YES |
| IDCSDGNYNRNA | 1449891 | Ectodomain | YES |
| IDDCYGYLRKHF | 1449893 | Cell/Virus Enclosed | NO |
| IDDGKIWNGVIK | 1449899 | Cell/Virus Enclosed | NO |
| IDEHVFYKKDGVYYPS | 1665736 | Cell/Virus Enclosed | NO |
| IDEIPYKSFSDS | 1449918 | Cell/Virus Enclosed | NO |
| IDEYITTYPGQG | 1449925 | Cell/Virus Enclosed | NO |
| IDFPFAESTGTQ | 1449931 | Cell/Virus Enclosed | NO |
| IDFPSAELTGTQ | 1449932 | Cell/Virus Enclosed | NO |
| IDFQDELDDFFK | 1449934 | Ectodomain | YES |
| IDGFGVFDIKNSSVNT | 1665755 | Cell/Virus Enclosed | NO |
| IDGGKIWNGVIKNVNS | 1665757 | Cell/Virus Enclosed | NO |
| IDGKFVTKFSGDKFDF | 1665759 | Cell/Virus Enclosed | NO |
| IDGSSGVANPAMDPIY | 1665764 | Secreted | YES |
| IDGVENPMLMGWDYPK | 1665768 | Cell/Virus Enclosed | NO |
| IDGVYWVASSQA | 1449976 | Capsid | UN |
| IDIDAFPVKPICSSVY | 1665779 | Cell/Virus Enclosed | NO |
| IDILTEGQLQSLSKDY | 1665783 | Cell/Virus Enclosed | NO |
| IDKEDVVIFINNTTYP | 1665795 | Cell/Virus Enclosed | NO |
| IDKFAIPNGRKVDLQL | 1665796 | Ectodomain | YES |
| IDKLNVLFDGRDNGAL | 1665798 | Cell/Virus Enclosed | NO |
| IDKLRDYLADYD | 1450020 | Cell/Virus Enclosed | NO |
| IDKWLNNFNVPSPLNW | 1665803 | Ectodomain | YES |
| IDLASQIISGII | 1450032 | Undetermined | UN |
| IDLKSYDLGDEL | 1450048 | Cell/Virus Enclosed | NO |
| IDLLCLRDDNQTLLVP | 1665813 | Cell/Virus Enclosed | NO |
| IDLLNLREIKPAVNVV | 1665816 | Cell/Virus Enclosed | NO |
| IDLTECIYTQSR | 1450067 | Cell/Virus Enclosed | NO |
| IDLTMIEFMLWC | 1450070 | Cell/Virus Enclosed | NO |
| IDMKGKMFVRPVIEDY | 1665827 | Underneath Membrane | NO |
| IDMKGRMYVRPI | 1450079 | Underneath Membrane | NO |
| IDMKGTMYVRPIIEDY | 1665828 | Ectodomain | YES |
| IDMPSWPPVQSTSSPGTPSM | 25631 | Cell/Virus Enclosed | NO |
| IDNASHASTVRA | 1450085 | Secreted | YES |
| IDNDINDYVSDADFSI | 1665832 | Cell/Virus Enclosed | NO |
| IDNINSTLVDLKWLNR | 1665836 | Ectodomain | YES |
| IDNMYMGKCGDK | 1450095 | Cell/Virus Enclosed | NO |
| IDPAGVCFGAGQ | 1450101 | Cell/Virus Enclosed | NO |

| | | | |
|---|---|---|---|
| IDPAGVCLGAGQ | 1450102 | Cell/Virus Enclosed | NO |
| IDQDFGSTVFNVPTKV | 1665844 | Cell/Virus Enclosed | NO |
| IDQINSTYVDLKLLNR | 1665845 | Ectodomain | YES |
| IDQKVKEISNQEPL | 25653 | Capsid | UN |
| IDSDVDTKCLADSVMS | 1665851 | Cell/Virus Enclosed | NO |
| IDSDVETRFITKSMIS | 1665852 | Cell/Virus Enclosed | NO |
| IDSNKYTYFDNP | 1450131 | Ectodomain | YES |
| IDSNTTQDTYGGASVC | 1665854 | Cell/Virus Enclosed | NO |
| IDSSEPLCLFKK | 1450133 | Ectodomain | YES |
| IDSTDNNQNGGR | 1450134 | Capsid | UN |
| IDSTQDGVNKLQ | 1450135 | Ectodomain | YES |
| IDSTVLTYIPAS | 1450137 | Cell/Virus Enclosed | NO |
| IDTINCESIYEF | 1450150 | Cell/Virus Enclosed | NO |
| IDTTLLDLTYEM | 1450157 | Ectodomain | YES |
| IDTVDVTNGLGT | 1450158 | Ectodomain | YES |
| IDVFNKLSTSDWSDIR | 1665871 | Cell/Virus Enclosed | NO |
| IDVGNHRSAFAL | 1450172 | Ectodomain | YES |
| IDVIDNEIIVKPNISL | 1665873 | Cell/Virus Enclosed | NO |
| IDVLLTVDGVNFKSIS | 1665875 | Cell/Virus Enclosed | NO |
| IDVLRNSFGKDLNANM | 1665876 | Cell/Virus Enclosed | NO |
| IDVNKTLQELSYKLPN | 1665877 | Ectodomain | YES |
| IDVNPALNICFD | 1450176 | Cell/Virus Enclosed | NO |
| IDVNSALNICFDIRDN | 1665878 | Cell/Virus Enclosed | NO |
| IDVPFAELTGTQ | 1450180 | Cell/Virus Enclosed | NO |
| IDVPSAESTGTQSVSV | 1665879 | Cell/Virus Enclosed | NO |
| IDVSKADGIIYPQGRT | 1665881 | Ectodomain | YES |
| IDVTGKLSDCLY | 1450189 | Cell/Virus Enclosed | NO |
| IDVVQYEADRRAFVDY | 1665884 | Cell/Virus Enclosed | NO |
| IDVVRDLPSGFNTLKP | 1665885 | Ectodomain | YES |
| IDVVSAYGLCDA | 1450194 | Ectodomain | YES |
| IDYCVRAFDVYNKDAS | 1665889 | Cell/Virus Enclosed | NO |
| IEAAAQGFQACR | 1450219 | Cell/Virus Enclosed | NO |
| IEAGEVKPFAVYKNVK | 1665903 | Cell/Virus Enclosed | NO |
| IECDVKTIEVVG | 1450235 | Cell/Virus Enclosed | NO |
| IECGSCGNDSWLTGNA | 1665908 | Cell/Virus Enclosed | NO |
| IECGSCGNESWL | 1450238 | Cell/Virus Enclosed | NO |
| IECGSCGNNSWL | 1450239 | Cell/Virus Enclosed | NO |
| IEDAETPALQVS | 1450250 | Cell/Virus Enclosed | NO |
| IEDAIIVNDENSSIKV | 1665912 | Cell/Virus Enclosed | NO |
| IEDALRLSAHLETNDV | 1665913 | Ectodomain | YES |
| IEDIDYVPLKSAVCIT | 1665917 | Cell/Virus Enclosed | NO |
| IEDILFSKIVTS | 1450259 | Ectodomain | YES |
| IEDILFSKLVTSGLGT | 1665918 | Ectodomain | YES |
| IEDLLFDKVKLSDVGF | 1665919 | Ectodomain | YES |
| IEDLLFDKVTIADPGY | 1665920 | Ectodomain | YES |
| IEDLLFSKVRLS | 1450266 | Ectodomain | YES |
| IEDLRARMTYVC | 1450269 | Cell/Virus Enclosed | NO |
| IEDLSIRAFQKR | 1450270 | Cell/Virus Enclosed | NO |
| IEDLSIRALQKR | 1450272 | Cell/Virus Enclosed | NO |
| IEDLSIRNLQKRLYSN | 1665923 | Cell/Virus Enclosed | NO |
| IEDSVFKPRPAGVLTN | 1665924 | Ectodomain | YES |
| IEDVEDVISSEV | 1450288 | Cell/Virus Enclosed | NO |
| IEEFADVVKEQVSDLL | 1665930 | Cell/Virus Enclosed | NO |
| IEEFADVVKGQV | 1450295 | Cell/Virus Enclosed | NO |
| IEEFIQTSSPKV | 1450297 | Ectodomain | YES |
| IEEKSVVQKPVD | 1450300 | Cell/Virus Enclosed | NO |
| IEESDLIISNFKFDDY | 1665936 | Cell/Virus Enclosed | NO |
| IEEVELLDAEFVEPGC | 1665938 | Cell/Virus Enclosed | NO |
| IEFANDTKIASQ | 1450312 | Ectodomain | YES |
| IEFLRDAWEILK | 1450316 | Cell/Virus Enclosed | NO |
| IEFMLWCKDGQVQTFY | 1665943 | Cell/Virus Enclosed | NO |
| IEGSGRPAPNSR | 1450330 | Capsid | UN |
| IEGSGRSAPNSRSTSR | 1665947 | Cell/Virus Enclosed | NO |
| IEGSGRSASNSR | 1450333 | Cell/Virus Enclosed | NO |
| IEGSGRSVPNSR | 1450334 | Cell/Virus Enclosed | NO |
| IEGSKPYSTLQPNAYY | 1665948 | Cell/Virus Enclosed | NO |
| IEGTGGNIQSSS | 1450337 | Capsid | UN |
| IEGTGGNSQSSSRASS | 1665949 | Cell/Virus Enclosed | NO |
| IEHVTSVNLFDV | 1450344 | Cell/Virus Enclosed | NO |
| IEKDFLSFDSDV | 1450349 | Cell/Virus Enclosed | NO |
| IEKLYKDLTGHMLDSY | 1665953 | Cell/Virus Enclosed | NO |
| IELGSGAHVGSD | 1450364 | Cell/Virus Enclosed | NO |
| IELGSGSHVGSSFDGV | 1665962 | Cell/Virus Enclosed | NO |
| IELGVCVNYDLY | 1450366 | Ectodomain | YES |
| IELSKELKRPVNPTDA | 1665965 | Cell/Virus Enclosed | NO |
| IELTAEDISLLR | 1450373 | Capsid | UN |

| | | | |
|---|---|---|---|
| IEMQLFKVGKVDSIVQ | 1665968 | Cell/Virus Enclosed | NO |
| IENALPASTVHA | 1450381 | Secreted | YES |
| IENSLNALFSCD | 1450393 | Cell/Virus Enclosed | NO |
| IEPVYRVKLCFEFEDE | 1665985 | Cell/Virus Enclosed | NO |
| IERGKLLVRDTRKEIF | 1665993 | Cell/Virus Enclosed | NO |
| IERGLLHSSRLP | 1450422 | Cell/Virus Enclosed | NO |
| IERVDWDIEYPYISHE | 1665995 | Cell/Virus Enclosed | NO |
| IESIHYLGTTGHTLKS | 1665998 | Cell/Virus Enclosed | NO |
| IESIYEFDTYKA | 1450435 | Cell/Virus Enclosed | NO |
| IESQRGVYCCRD | 1450440 | Cell/Virus Enclosed | NO |
| IESRDETVPQSVREA | 227263 | Cell/Virus Enclosed | NO |
| IESWLNDKSVPS | 1450443 | Ectodomain | YES |
| IETAVGHYFHAR | 1450446 | Cell/Virus Enclosed | NO |
| IETAVGHYVHAC | 1450447 | Cell/Virus Enclosed | NO |
| IETVDVKHEMSP | 1450460 | Cell/Virus Enclosed | NO |
| IEVDIQQTFFDKTWPR | 1666010 | Ectodomain | YES |
| IEVGEVKPFAVY | 1450468 | Cell/Virus Enclosed | NO |
| IEVKFNAPALQE | 1450470 | Cell/Virus Enclosed | NO |
| IEVNELLDTTQL | 1450472 | Ectodomain | YES |
| IEVPETLVHKLTGKK | 227264 | Secreted | YES |
| IEVSQFQSKLTDVKCA | 1666015 | Cell/Virus Enclosed | NO |
| IEVSQIQSKLTDVKCA | 1666016 | Cell/Virus Enclosed | NO |
| IEVVSAYGLCDA | 1450481 | Ectodomain | YES |
| IEVVSAYGLCDS | 1450483 | Ectodomain | YES |
| IEWPLTEYDFTDYKVQ | 1666019 | Cell/Virus Enclosed | NO |
| IEYPYISHENKL | 1450492 | Cell/Virus Enclosed | NO |
| IEYPYISHETKL | 1450493 | Cell/Virus Enclosed | NO |
| IFADTLVEVGLVGVLT | 1666027 | Cell/Virus Enclosed | NO |
| IFAEEPVVGIVY | 1450499 | Cell/Virus Enclosed | NO |
| IFAKVKNNKGFK | 1450503 | Ectodomain | YES |
| IFAKVKNTKVVG | 1450507 | Ectodomain | YES |
| IFANFILHDVNN | 1450511 | Ectodomain | YES |
| IFANFILHDVNS | 1450512 | Ectodomain | YES |
| IFAQTSDTAHACNANR | 1666032 | Cell/Virus Enclosed | NO |
| IFARTLDAANHS | 1450515 | Cell/Virus Enclosed | NO |
| IFASVPEKFLNA | 1450520 | Cell/Virus Enclosed | NO |
| IFATPSVLQATL | 1450523 | Cell/Virus Enclosed | NO |
| IFCQTADTAHANNINR | 1666040 | Cell/Virus Enclosed | NO |
| IFDIETVDVKHDVSPF | 1666045 | Cell/Virus Enclosed | NO |
| IFDSSKEFVEYK | 1450551 | Cell/Virus Enclosed | NO |
| IFDVAPIELENKFTPL | 1666049 | Cell/Virus Enclosed | NO |
| IFDVDKFKTPHI | 1450557 | Cell/Virus Enclosed | NO |
| IFDVDKVKTPHI | 1450558 | Cell/Virus Enclosed | NO |
| IFERGSVQVDNGSSIN | 1666055 | Cell/Virus Enclosed | NO |
| IFERIDYSDAVSYKSS | 1666056 | Cell/Virus Enclosed | NO |
| IFEWIEAKLSAG | 1450570 | Cell/Virus Enclosed | NO |
| IFGLYKNSATGSLDVE | 1666075 | Cell/Virus Enclosed | NO |
| IFGQDFKTFDLLKYDF | 1666078 | Cell/Virus Enclosed | NO |
| IFICLFVALLLT | 1450622 | Cell/Virus Enclosed | NO |
| IFIETNLQSLENIAFN | 1666090 | Cell/Virus Enclosed | NO |
| IFIGDNVGHYVH | 1450624 | Cell/Virus Enclosed | NO |
| IFIGDTVGHYVH | 1450625 | Cell/Virus Enclosed | NO |
| IFINNTTYPTNVAVEL | 1666094 | Cell/Virus Enclosed | NO |
| IFKDDGNFVVLE | 1450643 | Cell/Virus Enclosed | NO |
| IFKNNTSFPTNIAVEL | 1666107 | Cell/Virus Enclosed | NO |
| IFKTEVDPNLSG | 1450654 | Cell/Virus Enclosed | NO |
| IFLAWIMLLQFA | 1450655 | Transmembrane | NO |
| IFLWLLWPVTLACFV | 2001114 | Transmembrane | NO |
| IFNAVDCKSDFM | 1450685 | Ectodomain | YES |
| IFNAVDCMSDFMSEIK | 1666125 | Ectodomain | YES |
| IFNCFYALNNAFLAFS | 1666126 | Undetermined | UN |
| IFNGINSGTTCS | 1450698 | Ectodomain | YES |
| IFNGKFFSNTKY | 1450699 | Ectodomain | YES |
| IFNGKFLSNTKYYDDS | 1666129 | Ectodomain | YES |
| IFNIFQGVSSNI | 1450703 | Cell/Virus Enclosed | NO |
| IFNKATVAQIKALFLD | 1666131 | Cell/Virus Enclosed | NO |
| IFNKLVKDPPNVQIHT | 1666132 | Secreted | YES |
| IFNSACTRLEGLGGDN | 1666136 | Cell/Virus Enclosed | NO |
| IFPPGQGVPLNA | 1450714 | Cell/Virus Enclosed | NO |
| IFPSGTYYCVTT | 1450717 | Cell/Virus Enclosed | NO |
| IFQAVSSNINCI | 1450720 | Cell/Virus Enclosed | NO |
| IFQGVSSNINRL | 1450726 | Cell/Virus Enclosed | NO |
| IFRSDVLHLTQD | 1450736 | Ectodomain | YES |
| IFSAIYPIDLAS | 1450741 | Undetermined | UN |
| IFSAVYPIDLAS | 1450744 | Undetermined | UN |
| IFSIVNSAPDGL | 1450751 | Ectodomain | YES |

| | | | |
|---|---|---|---|
| IFSLHPVECDEE | 1450753 | Cell/Virus Enclosed | NO |
| IFSLHPVECDEG | 1450754 | Cell/Virus Enclosed | NO |
| IFSLVNAAPEGL | 1450757 | Ectodomain | YES |
| IFSQFDSLGVSSNQSP | 1666158 | Cell/Virus Enclosed | NO |
| IFSQSDSLGVSS | 1450766 | Cell/Virus Enclosed | NO |
| IFSTANALPECN | 1450775 | Cell/Virus Enclosed | NO |
| IFSTVNALPECNADIV | 1666166 | Cell/Virus Enclosed | NO |
| IFSTVYPIDLAS | 1450782 | Undetermined | UN |
| IFSVQQDGRIPNGFPF | 1666169 | Ectodomain | YES |
| IFTARDGVRRGD | 1450793 | Cell/Virus Enclosed | NO |
| IFTINSVVQKDGVENT | 1666172 | Cell/Virus Enclosed | NO |
| IFTKSGFNDICM | 1450804 | Cell/Virus Enclosed | NO |
| IFTLGSITAQPV | 1450809 | Secreted | YES |
| IFVDGVPFVVST | 1450833 | Cell/Virus Enclosed | NO |
| IFVDGVPFVVSY | 1450834 | Cell/Virus Enclosed | NO |
| IFVEVNAPYYNS | 1450837 | Ectodomain | YES |
| IFVHEDLHVVEVLTKT | 1666190 | Cell/Virus Enclosed | NO |
| IFVQQKPEPVNPVSFV | 1666194 | Cell/Virus Enclosed | NO |
| IFYTTEIASKFT | 1450873 | Cell/Virus Enclosed | NO |
| IGAAANATGTVI | 1450875 | Ectodomain | YES |
| IGAEHVDSSYEC | 1450879 | Ectodomain | YES |
| IGAMTAVNESRYGLQN | 1666221 | Ectodomain | YES |
| IGANGNKIVDKE | 1450893 | Cell/Virus Enclosed | NO |
| IGAPANLRVVGHAMQG | 1666223 | Cell/Virus Enclosed | NO |
| IGASHTGITGDN | 1450899 | Cell/Virus Enclosed | NO |
| IGATIRLQAGKQTELA | 1666224 | Cell/Virus Enclosed | NO |
| IGATVRLQAGKQTEFV | 1666225 | Cell/Virus Enclosed | NO |
| IGAVGGDLLYLDLLNR | 1666226 | Cell/Virus Enclosed | NO |
| IGDDVDDIESIY | 1450924 | Cell/Virus Enclosed | NO |
| IGDFMSTIITSC | 1450929 | Cell/Virus Enclosed | NO |
| IGDFNCTNFAINDLNT | 1666238 | Ectodomain | YES |
| IGDIIQRLDFLE | 1450931 | Ectodomain | YES |
| IGDIIQRLDVLEQDAQ | 1666239 | Ectodomain | YES |
| IGDLKCTSDNINDKDT | 1666242 | Ectodomain | YES |
| IGDLKCTTVAIN | 1450945 | Ectodomain | YES |
| IGDLNCPLDPKL | 1450947 | Ectodomain | YES |
| IGDLNCPLDSRL | 1450949 | Ectodomain | YES |
| IGDLNCPLDTRL | 1450950 | Ectodomain | YES |
| IGDLNCTLDPRL | 1450951 | Ectodomain | YES |
| IGDLNCTLHPRL | 1450952 | Ectodomain | YES |
| IGDTVGHYVHVK | 1450963 | Cell/Virus Enclosed | NO |
| IGDVKFKRLGDYVLTE | 1666250 | Cell/Virus Enclosed | NO |
| IGEASSPIAYLV | 1450969 | Cell/Virus Enclosed | NO |
| IGEHCSGLAVKSDYCG | 1666256 | Ectodomain | YES |
| IGFDVEGAHAFR | 1450983 | Cell/Virus Enclosed | NO |
| IGFKDFVTNKTYNIFP | 1666268 | Ectodomain | YES |
| IGGAVCSKHANLYQKY | 1666274 | Cell/Virus Enclosed | NO |
| IGGDRCIKISTVQSKL | 1666276 | Cell/Virus Enclosed | NO |
| IGGGDLLYLDLL | 1451017 | Cell/Virus Enclosed | NO |
| IGGLTSIKWADN | 1451020 | Cell/Virus Enclosed | NO |
| IGGTIVVRPLVE | 1451026 | Ectodomain | YES |
| IGGVPIIEVSQFQSKL | 1666290 | Cell/Virus Enclosed | NO |
| IGGVPVIEVSQIQSKL | 1666291 | Cell/Virus Enclosed | NO |
| IGHGMSYSMTTF | 1451043 | Cell/Virus Enclosed | NO |
| IGHTVCDSLNAKLGFD | 1666300 | Cell/Virus Enclosed | NO |
| IGHTVCDSLNFK | 1451050 | Cell/Virus Enclosed | NO |
| IGHTVCDSLNSK | 1451051 | Cell/Virus Enclosed | NO |
| IGICPAGTNYLT | 1451054 | Ectodomain | YES |
| IGIDFQDELDEF | 1451055 | Ectodomain | YES |
| IGINYLCKPKVEIDGN | 1666307 | Cell/Virus Enclosed | NO |
| IGINYLGKSSFEIDGN | 1666309 | Cell/Virus Enclosed | NO |
| IGKGNKDEQIGYWNVQ | 1666313 | Capsid | UN |
| IGKGSKDEQIGY | 1451077 | Cell/Virus Enclosed | NO |
| IGKIAQWIKNTF | 1451079 | Cell/Virus Enclosed | NO |
| IGLAGKVKVGTGVMVE | 1666321 | Cell/Virus Enclosed | NO |
| IGLDPVAPGDLTMVI | 227266 | Secreted | YES |
| IGLYKKQQEGHI | 1451122 | Cell/Virus Enclosed | NO |
| IGLYRRQQTSNLVVQE | 1666339 | Cell/Virus Enclosed | NO |
| IGNIEEFIQTSS | 1451142 | Ectodomain | YES |
| IGNMVEFIQTSSPKVT | 1666347 | Ectodomain | YES |
| IGNNNVCTYEHVISYM | 1666348 | Cell/Virus Enclosed | NO |
| IGNPKGIPIVDDPVVD | 1666353 | Cell/Virus Enclosed | NO |
| IGNPKGIRCAVTDAKW | 1666354 | Cell/Virus Enclosed | NO |
| IGNYKLNTDHSNSSDN | 1666358 | Underneath Membrane | NO |
| IGPDMFLGTCRC | 1451168 | Cell/Virus Enclosed | NO |
| IGPDVFLHKCYRCPAE | 1666363 | Cell/Virus Enclosed | NO |

| | | | |
|---|---|---|---|
| IGPHEFCSQHTM | 1451174 | Cell/Virus Enclosed | NO |
| IGPIKYCYCGNF | 1451175 | Cell/Virus Enclosed | NO |
| IGPNKYCYCGNF | 1451177 | Cell/Virus Enclosed | NO |
| IGREQKCCMCNR | 1451191 | Cell/Virus Enclosed | NO |
| IGSLGFHNPPTN | 1451206 | Ectodomain | YES |
| IGSNNVCTYEHVISFM | 1666378 | Cell/Virus Enclosed | NO |
| IGSSLQEILSRL | 1451214 | Ectodomain | YES |
| IGTANLSIPSNW | 1451222 | Ectodomain | YES |
| IGTCPAGTNHLT | 1451223 | Ectodomain | YES |
| IGTCPAGTNYLN | 1451225 | Ectodomain | YES |
| IGTCPVGTNYLT | 1451227 | Ectodomain | YES |
| IGTNDPIRFCLENTVC | 1666385 | Cell/Virus Enclosed | NO |
| IGTNFPLQIGFSTGVD | 1666386 | Cell/Virus Enclosed | NO |
| IGTNFPLQLGFSTGID | 1666387 | Cell/Virus Enclosed | NO |
| IGTNVPLQVGFSNGVN | 1666389 | Cell/Virus Enclosed | NO |
| IGTNYRSCESTTVLDH | 1666390 | Ectodomain | YES |
| IGTRNPANNAAIVLQL | 1666392 | Cell/Virus Enclosed | NO |
| IGTRNPNNNAATVLQL | 26329 | Capsid | UN |
| IGTTKFCGGWNN | 1451257 | Cell/Virus Enclosed | NO |
| IGTTKFYGGWGN | 1451259 | Cell/Virus Enclosed | NO |
| IGTYEMYVKWPW | 1451262 | Undetermined | UN |
| IGVCKYTDIDVNSALN | 1666405 | Cell/Virus Enclosed | NO |
| IGVEGELPVAVVNDKI | 1666410 | Cell/Virus Enclosed | NO |
| IGVFVQKYGLED | 1451274 | Cell/Virus Enclosed | NO |
| IGVGGAPCIKVA | 1451275 | Cell/Virus Enclosed | NO |
| IGVNVCSDSTFEETSL | 1666412 | Cell/Virus Enclosed | NO |
| IGVTGVIIAVIALFC | 26343 | Secreted | YES |
| IGVTQNVLYDNQ | 1451287 | Ectodomain | YES |
| IGVVREFLTRNP | 1451290 | Cell/Virus Enclosed | NO |
| IGVYHKLGERVR | 1451294 | Cell/Virus Enclosed | NO |
| IGYHYKELGVVMNLDV | 1666424 | Cell/Virus Enclosed | NO |
| IGYSLADLPAYM | 1451308 | Ectodomain | YES |
| IGYVTHGFNLED | 1451312 | Cell/Virus Enclosed | NO |
| IHADQLTPAWRIYSTGNNV | 1382573 | Ectodomain | YES |
| IHAEQLTPAWRI | 1451329 | Ectodomain | YES |
| IHDVVNQQGNSL | 1451346 | Ectodomain | YES |
| IHFGAGSDKGIAPGTS | 1666448 | Cell/Virus Enclosed | NO |
| IHHPSTSADQQSLYQNADTYVFVGS | 1148302 | Ectodomain | YES |
| IHIVSFVVNAPN | 1451366 | Ectodomain | YES |
| IHLPAYASFKVIDNGV | 1666457 | Cell/Virus Enclosed | NO |
| IHSVCNGCVGHS | 1451405 | Ectodomain | YES |
| IHSVYLLMFLLT | 1451407 | Ectodomain | YES |
| IHTIDGSSGVANPAMD | 1666472 | Secreted | YES |
| IHTKPLFIRNFD | 1451413 | Cell/Virus Enclosed | NO |
| IHYNINFEENLR | 1451423 | Ectodomain | YES |
| IIANENAINKGG | 1451443 | Cell/Virus Enclosed | NO |
| IIANRCNEVSAT | 1451445 | Cell/Virus Enclosed | NO |
| IIANRYNEVANAVMQN | 1666491 | Cell/Virus Enclosed | NO |
| IIAVIYSGRESLIFPS | 1666497 | Cell/Virus Enclosed | NO |
| IIDAATSYGMCF | 1451461 | Ectodomain | YES |
| IIDNDINDYVSDADFS | 1666502 | Cell/Virus Enclosed | NO |
| IIEQPFGEVEHS | 1451477 | Cell/Virus Enclosed | NO |
| IIEQPFGEVEYS | 1451478 | Cell/Virus Enclosed | NO |
| IIEQSFGEVEHS | 1451479 | Cell/Virus Enclosed | NO |
| IIEVSQFQSKLTDVKC | 1666510 | Cell/Virus Enclosed | NO |
| IIFNAEDCMSDF | 1451495 | Ectodomain | YES |
| IIFNAVDCMSDFMSEI | 1666517 | Ectodomain | YES |
| IIFVQQKPEPVNPVSF | 1666520 | Cell/Virus Enclosed | NO |
| IIGAMTAVNESRYGLQ | 1666525 | Ectodomain | YES |
| IIGDELKINSAC | 1451513 | Cell/Virus Enclosed | NO |
| IIGSLGFYNPPTNVVS | 1666533 | Ectodomain | YES |
| IIGVPYPVSGIR | 1451529 | Ectodomain | YES |
| IIHDFVDKTLPDQGD | 26510 | Secreted | YES |
| IIHDIGNPKGVR | 1451537 | Cell/Virus Enclosed | NO |
| IIIDKVDFVNGF | 1451543 | Cell/Virus Enclosed | NO |
| IIIDNDINDYVSDADF | 1666540 | Cell/Virus Enclosed | NO |
| IIIDNDLNEFVSDADI | 1666541 | Cell/Virus Enclosed | NO |
| IIINNSTNVVIRACNF | 1666546 | Ectodomain | YES |
| IIISCQTAVSKF | 1451562 | Cell/Virus Enclosed | NO |
| IIKAQFKTFEKVDGVY | 1666549 | Cell/Virus Enclosed | NO |
| IIKPNALLCATC | 1451582 | Cell/Virus Enclosed | NO |
| IIKSSVKSYDDLVYSG | 1666561 | Cell/Virus Enclosed | NO |
| IILASFSASTCA | 1451594 | Cell/Virus Enclosed | NO |
| IINARIKAKHIVYVGD | 1666597 | Cell/Virus Enclosed | NO |
| IINDKVLTKIDK | 1451659 | Cell/Virus Enclosed | NO |
| IINDKVVAKIDKEDVV | 1666599 | Cell/Virus Enclosed | NO |

| | | | |
|---|---|---|---|
| IINNSTNVVIRACNFEL | 26610 | Ectodomain | YES |
| IIPHSTRSIQSD | 1451690 | Ectodomain | YES |
| IIQADQQVDRLI | 1451698 | Ectodomain | YES |
| IIRDGDVVKQPV | 1451713 | Cell/Virus Enclosed | NO |
| IIRDVNPDDKCFANKH | 1666629 | Cell/Virus Enclosed | NO |
| IIRGHLYIQGIKLGTG | 1666634 | Ectodomain | YES |
| IIRGHLYMQGIK | 1451726 | Underneath Membrane | NO |
| IIRSSNQSLAGGITYV | 1666637 | Ectodomain | YES |
| IISEKISVMDKL | 1451750 | Cell/Virus Enclosed | NO |
| IISIVQMAPVSA | 1451765 | Cell/Virus Enclosed | NO |
| IISNFKFDDYDFSVED | 1666660 | Cell/Virus Enclosed | NO |
| IISPSTRATIRK | 1451780 | Ectodomain | YES |
| IISRKPQPKMEK | 1451784 | Ectodomain | YES |
| IISTQDGVNKLQ | 1451789 | Ectodomain | YES |
| IITKLNLLNEPL | 1451799 | Cell/Virus Enclosed | NO |
| IITMPCFRTFPQ | 1451807 | Cell/Virus Enclosed | NO |
| IITSCQTAVSKL | 1451817 | Cell/Virus Enclosed | NO |
| IITSTDAGCGVK | 1451818 | Cell/Virus Enclosed | NO |
| IITSTSAGCGVKYYAG | 1666680 | Cell/Virus Enclosed | NO |
| IITTDNTFVAGN | 1451821 | Ectodomain | YES |
| IITVEELKQLLE | 1451825 | Undetermined | UN |
| IIVFLCAVLMYL | 1451828 | Cell/Virus Enclosed | NO |
| IIVKPNISLCVPLYVR | 1666687 | Cell/Virus Enclosed | NO |
| IIVPDKSVYDQI | 1451836 | Cell/Virus Enclosed | NO |
| IIVPDKSVYDQVVDNV | 1666690 | Cell/Virus Enclosed | NO |
| IIWHINQTTNKT | 1451842 | Cell/Virus Enclosed | NO |
| IIWVATEGALNTPKDH | 1666695 | Cell/Virus Enclosed | NO |
| IIYDSADRGQES | 1451848 | Cell/Virus Enclosed | NO |
| IIYDSSDRGQESLARS | 1666697 | Cell/Virus Enclosed | NO |
| IIYTGDWEGLHEVLTS | 1666700 | Cell/Virus Enclosed | NO |
| IKACEVVVTNLNKSAG | 1666701 | Cell/Virus Enclosed | NO |
| IKANIKSDILSL | 1451872 | Ectodomain | YES |
| IKAQFKTFEKVDGVYT | 1666709 | Cell/Virus Enclosed | NO |
| IKAYNTINNEQGTPLT | 1666712 | Cell/Virus Enclosed | NO |
| IKCNYVFNNTLS | 1451888 | Ectodomain | YES |
| IKCVDTVKEAQS | 1451895 | Cell/Virus Enclosed | NO |
| IKDADYNAKVEISVTP | 1666720 | Cell/Virus Enclosed | NO |
| IKDCVDINKAIPSGNL | 1666723 | Cell/Virus Enclosed | NO |
| IKDFDYYRYNRPTMLD | 1666725 | Cell/Virus Enclosed | NO |
| IKDGDDGYFLPYPDPS | 1666729 | Cell/Virus Enclosed | NO |
| IKDGDYGYFLPY | 1451911 | Cell/Virus Enclosed | NO |
| IKDGVMYSEFPA | 1451912 | Ectodomain | YES |
| IKDRVMYSEFPAITIG | 1666735 | Ectodomain | YES |
| IKDVDNPVLMGWDYPK | 1666737 | Cell/Virus Enclosed | NO |
| IKDWNFVWSLIF | 1451934 | Undetermined | UN |
| IKENVDFINSKQ | 1451950 | Cell/Virus Enclosed | NO |
| IKENVDFINSMQFNYK | 1666744 | Cell/Virus Enclosed | NO |
| IKEPVHVYANDS | 1451952 | Cell/Virus Enclosed | NO |
| IKESIPIVWGVKDFNT | 1666747 | Cell/Virus Enclosed | NO |
| IKFCDGENVSKDGFFT | 1666751 | Cell/Virus Enclosed | NO |
| IKFETDAYKLFL | 1451968 | Cell/Virus Enclosed | NO |
| IKFPEVVFESTL | 1451974 | Cell/Virus Enclosed | NO |
| IKFQTVKPGNFNQDFY | 1666759 | Cell/Virus Enclosed | NO |
| IKFSGFKLDGFNHQFV | 1666760 | Cell/Virus Enclosed | NO |
| IKGCCESTKLPY | 1451982 | Underneath Membrane | NO |
| IKGPQRADLNGVIVDK | 1666766 | Cell/Virus Enclosed | NO |
| IKHEGDWNSFCK | 1452003 | Cell/Virus Enclosed | NO |
| IKHGVMYSEFPA | 1452004 | Ectodomain | YES |
| IKHPNQEYADVF | 1452011 | Cell/Virus Enclosed | NO |
| IKHVRNNGGYNS | 1452015 | Cell/Virus Enclosed | NO |
| IKHVRTNGDYNS | 1452016 | Cell/Virus Enclosed | NO |
| IKIAQADTAGLY | 1452018 | Cell/Virus Enclosed | NO |
| IKIPTNFTIVGQEEFI | 1666780 | Ectodomain | YES |
| IKISADNKVIYTTEIA | 1666784 | Cell/Virus Enclosed | NO |
| IKISTVQSKLTDLKCT | 1666785 | Cell/Virus Enclosed | NO |
| IKITEFSWNADLYKLM | 1666786 | Cell/Virus Enclosed | NO |
| IKITEFSWNAELYKLM | 1666787 | Cell/Virus Enclosed | NO |
| IKKAMFTKVVVG | 1452035 | Cell/Virus Enclosed | NO |
| IKKGVMYSEFPA | 1452037 | Ectodomain | YES |
| IKKLDNDVLNGIISNA | 1666795 | Cell/Virus Enclosed | NO |
| IKKLYNDLGNQILDSY | 1666796 | Cell/Virus Enclosed | NO |
| IKKYPCIVGPDY | 1452055 | Cell/Virus Enclosed | NO |
| IKLDDKDPNFKDQVIL | 1452063 | Cell/Virus Enclosed | NO |
| IKLDDKDPSFKDNVIL | 1666809 | Cell/Virus Enclosed | NO |
| IKLDPKNPNYNKWLEL | 1666810 | Cell/Virus Enclosed | NO |
| IKLGTGYSLADLPAYM | 1666813 | Ectodomain | YES |

| | | | |
|---|---|---|---|
| IKLIFLWLLWPVTLA | 1312710 | Undetermined | UN |
| IKLKRAFGDYTS | 1452082 | Cell/Virus Enclosed | NO |
| IKLNFGMLNGMPVSSI | 1666822 | Cell/Virus Enclosed | NO |
| IKLNFGMLNGNAIATV | 1666823 | Cell/Virus Enclosed | NO |
| IKLPQFYIYDEEGGYD | 1666826 | Cell/Virus Enclosed | NO |
| IKLPSGIMLNVVKYTQ | 1666827 | Cell/Virus Enclosed | NO |
| IKLQSKRTRLFK | 1452092 | Cell/Virus Enclosed | NO |
| IKNLTPIKLNFGMLNG | 1666846 | Cell/Virus Enclosed | NO |
| IKNTVDTIPKEK | 1452130 | Cell/Virus Enclosed | NO |
| IKNVNSVRDWLKSLKL | 1666852 | Cell/Virus Enclosed | NO |
| IKPEVKEVAPVY | 1452139 | Cell/Virus Enclosed | NO |
| IKPNAILCATCP | 1452149 | Cell/Virus Enclosed | NO |
| IKPNALLCATCP | 1452151 | Cell/Virus Enclosed | NO |
| IKPNARLCATCP | 1452153 | Cell/Virus Enclosed | NO |
| IKPQQVTKHTAK | 1452156 | Capsid | UN |
| IKPVSYKLDGVI | 1452165 | Cell/Virus Enclosed | NO |
| IKQGTLIYKVLD | 1452172 | Cell/Virus Enclosed | NO |
| IKQLKKAMNVAKAEFD | 1666876 | Cell/Virus Enclosed | NO |
| IKQLKRAMNIAKSEFD | 1666877 | Cell/Virus Enclosed | NO |
| IKQPFSFSFRDELGVR | 1666880 | Cell/Virus Enclosed | NO |
| IKRSFIEDLLFN | 1452201 | Ectodomain | YES |
| IKRVKGWVGFDV | 1452202 | Cell/Virus Enclosed | NO |
| IKRVRGWVGFDVEGAH | 1666890 | Cell/Virus Enclosed | NO |
| IKSDHCGGNPCT | 1452206 | Ectodomain | YES |
| IKSFSTFESAYMPIAD | 1666892 | Cell/Virus Enclosed | NO |
| IKSGRLLLRNNG | 1452212 | Cell/Virus Enclosed | NO |
| IKSLNNSYINLKDIGT | 1666896 | Ectodomain | YES |
| IKSSVKSYDDLVYSGV | 1666900 | Cell/Virus Enclosed | NO |
| IKSTQDGDNKLQ | 1452225 | Ectodomain | YES |
| IKSVMDHEQSMYNLLK | 1666901 | Cell/Virus Enclosed | NO |
| IKTAQADTAGLF | 1452233 | Cell/Virus Enclosed | NO |
| IKTAQADTAGLYTNFR | 1666903 | Cell/Virus Enclosed | NO |
| IKTNNIRIVDEW | 1452243 | Ectodomain | YES |
| IKTNNTMIVDEW | 1452244 | Ectodomain | YES |
| IKTNNTRIVDEWSYTG | 1666908 | Ectodomain | YES |
| IKTNNTRVVDEW | 1452247 | Ectodomain | YES |
| IKTNNTSIVDEW | 1452248 | Ectodomain | YES |
| IKTSKYGLGFKW | 1452252 | Cell/Virus Enclosed | NO |
| IKVFYELDKDFN | 1452263 | Cell/Virus Enclosed | NO |
| IKVFYELDNDFNTILN | 1666918 | Cell/Virus Enclosed | NO |
| IKVLPPLLSDNQ | 1452266 | Ectodomain | YES |
| IKVLPPLLSENQISGY | 1666923 | Ectodomain | YES |
| IKVQEGIVDYGV | 1452269 | Cell/Virus Enclosed | NO |
| IKVSQTNMHTPR | 1452272 | Cell/Virus Enclosed | NO |
| IKVTEDGVNVHDVTVT | 1666928 | Cell/Virus Enclosed | NO |
| IKVTEYSWNKKLYELV | 1666929 | Cell/Virus Enclosed | NO |
| IKWADNNCYLSS | 1452281 | Cell/Virus Enclosed | NO |
| IKWLFVLLFDWI | 1452287 | Cell/Virus Enclosed | NO |
| ILAASFNKAINNIVAS | 1666953 | Ectodomain | YES |
| ILAASFNKAMTNIVDA | 1666954 | Ectodomain | YES |
| ILARSQSSETKE | 1452322 | Cell/Virus Enclosed | NO |
| ILASVPEKFLNAFDVF | 1666971 | Cell/Virus Enclosed | NO |
| ILCATCPFAKGD | 1452336 | Cell/Virus Enclosed | NO |
| ILCEYADCKESYFSKK | 1666981 | Cell/Virus Enclosed | NO |
| ILCWETGKVKPPLNKN | 1666987 | Cell/Virus Enclosed | NO |
| ILDEVNGLLDTTQLHV | 1666989 | Ectodomain | YES |
| ILDFKVLDNGIIRDVN | 1666990 | Cell/Virus Enclosed | NO |
| ILDNAVKGCVPLNAIP | 1666997 | Cell/Virus Enclosed | NO |
| ILDNAVKGCVPLSAIP | 1666998 | Cell/Virus Enclosed | NO |
| ILFANPAAVDSK | 1452396 | Cell/Virus Enclosed | NO |
| ILFTGYFPKAGA | 1452417 | Ectodomain | YES |
| ILGGCIVLDVKG | 1452427 | Cell/Virus Enclosed | NO |
| ILGNPECESLSTASS | 150998 | Ectodomain | YES |
| ILGRCIVLDVKGVEEL | 1667034 | Cell/Virus Enclosed | NO |
| ILGSCTFEDELAPSDV | 1667037 | Cell/Virus Enclosed | NO |
| ILGYSSLNDEFSINEV | 1667046 | Cell/Virus Enclosed | NO |
| ILHDVNNGLTCS | 1452452 | Ectodomain | YES |
| ILHDVNSGTTCS | 1452454 | Ectodomain | YES |
| ILHHPGFLNPSS | 1452456 | Cell/Virus Enclosed | NO |
| ILIIIMRTFRIA | 1452472 | Cell/Virus Enclosed | NO |
| ILKFDSGTVSKTSDWK | 1667063 | Cell/Virus Enclosed | NO |
| ILKSQDLGAVSK | 1452509 | Cell/Virus Enclosed | NO |
| ILKWADQSDQFRNVQT | 1667077 | Cell/Virus Enclosed | NO |
| ILKWSDQSDQFR | 1452514 | Cell/Virus Enclosed | NO |
| ILLDDFVALVKSLNLNC | 1382727 | Cell/Virus Enclosed | NO |
| ILLEAYNAFLNT | 1452525 | Cell/Virus Enclosed | NO |

| | | | |
|---|---|---|---|
| ILLERYVSLAIDAYPL | 1667085 | Cell/Virus Enclosed | NO |
| ILLLDQSLVSDV | 1452548 | Cell/Virus Enclosed | NO |
| ILLNKHIDAYKTFPPTEPKK | 1392224 | Capsid | UN |
| ILLNVPLHGTILTRP | 2001119 | Underneath Membrane | NO |
| ILLVDKKLITTACNGI | 1667114 | Cell/Virus Enclosed | NO |
| ILMLDKQGQLLDTKLN | 1667118 | Cell/Virus Enclosed | NO |
| ILMSEDVKLNPSEDFI | 1667119 | Cell/Virus Enclosed | NO |
| ILMTARTVYDDA | 1452585 | Cell/Virus Enclosed | NO |
| ILMVDKNLITTANTGT | 1667123 | Cell/Virus Enclosed | NO |
| ILNGCTWWLKGE | 1452595 | Cell/Virus Enclosed | NO |
| ILNGINSGTTCS | 1452596 | Ectodomain | YES |
| ILNIVGPDARQDGRQS | 1667128 | Cell/Virus Enclosed | NO |
| ILNKPRQKRSPN | 1452601 | Cell/Virus Enclosed | NO |
| ILNKPRQKSSPN | 1452602 | Cell/Virus Enclosed | NO |
| ILNLTSEISTLENKSA | 1667130 | Ectodomain | YES |
| ILNRDRVNINIV | 1452613 | Cell/Virus Enclosed | NO |
| ILNTVKFCDHMVKAGL | 1667136 | Cell/Virus Enclosed | NO |
| ILPDPLKPTKRSFIED | 1667137 | Ectodomain | YES |
| ILPFSDRPLNLR | 1452635 | Cell/Virus Enclosed | NO |
| ILPGEFSKKQLGLLQP | 1667139 | Cell/Virus Enclosed | NO |
| ILPLAYFSDSKL | 1452637 | Cell/Virus Enclosed | NO |
| ILPQGYYIEGSG | 1452648 | Capsid | UN |
| ILPTMTQLNLKYAISG | 1667146 | Cell/Virus Enclosed | NO |
| ILPVAFTKAAGGKVSF | 1667148 | Cell/Virus Enclosed | NO |
| ILQDKTGDFMST | 1452662 | Cell/Virus Enclosed | NO |
| ILQEVKDNDGKN | 1452664 | Cell/Virus Enclosed | NO |
| ILQGKPIGMFFP | 1452667 | Cell/Virus Enclosed | NO |
| ILQQANKSVGIISMPL | 1667161 | Cell/Virus Enclosed | NO |
| ILQSNNPQCAIVQASE | 1667162 | Cell/Virus Enclosed | NO |
| ILQVVSFCRFIM | 1452680 | Secreted | YES |
| ILRIGTQVLKTM | 1452688 | Cell/Virus Enclosed | NO |
| ILRQHNLVDIMS | 1452697 | Cell/Virus Enclosed | NO |
| ILSCGIFGIKLETSLE | 1667178 | Cell/Virus Enclosed | NO |
| ILSCNFNMSSLM | 1452710 | Ectodomain | YES |
| ILSDDGVVCYNKDYAE | 1667183 | Cell/Virus Enclosed | NO |
| ILSDDGVVCYNSDYAA | 1667184 | Cell/Virus Enclosed | NO |
| ILSDDGVVCYNSDYAS | 1667185 | Cell/Virus Enclosed | NO |
| ILSDDSVVCYNKTYAG | 1667186 | Cell/Virus Enclosed | NO |
| ILSLNNPIFINYSKEV | 1667197 | Undetermined | UN |
| ILSNMNIASNSKEWAY | 1667200 | Cell/Virus Enclosed | NO |
| ILSRLDVLEAQV | 1452754 | Ectodomain | YES |
| ILSTSDLSVAFDKLAQ | 1667211 | Cell/Virus Enclosed | NO |
| ILSWEIGKVKPPLNKN | 1667213 | Cell/Virus Enclosed | NO |
| ILSWELGKVKPP | 1452766 | Cell/Virus Enclosed | NO |
| ILTALRLCAYCCNIV | 1312738 | Undetermined | UN |
| ILTEVNELLDTTQLQV | 1667218 | Ectodomain | YES |
| ILTLTRALAAES | 1452790 | Cell/Virus Enclosed | NO |
| ILTNNKPWFVRKASGL | 1667229 | Cell/Virus Enclosed | NO |
| ILVDNDLYPFVSDSVA | 1667240 | Cell/Virus Enclosed | NO |
| ILVGTTLQGKPI | 1452813 | Cell/Virus Enclosed | NO |
| ILVSNNKEDFDIIQKC | 1667250 | Cell/Virus Enclosed | NO |
| ILWREGDMFPFLDGLINAP | 1233642 | Ectodomain | YES |
| ILWVNKASYDTNLNKF | 1667260 | Cell/Virus Enclosed | NO |
| IMACGNCTFGAP | 1452868 | Cell/Virus Enclosed | NO |
| IMANLANLNSHT | 1452871 | Ectodomain | YES |
| IMASENSSAMVQVHS | 27448 | Secreted | YES |
| IMASKTVGTAEEKL | 27451 | Capsid | UN |
| IMCADVPDAFQSVYRT | 1667278 | Cell/Virus Enclosed | NO |
| IMFQPRLPVLSD | 1452908 | Ectodomain | YES |
| IMGSCSFEDELTPSDV | 1667289 | Cell/Virus Enclosed | NO |
| IMGYSTPWRYLDFNALNLFF | 27498 | Capsid | UN |
| IMKDDGNCVVLELDPP | 1667296 | Cell/Virus Enclosed | NO |
| IMKHDIKVIAKA | 1452933 | Cell/Virus Enclosed | NO |
| IMKNHLRDIMG | 189683 | Ectodomain | YES |
| IMKVLNENLNAYQQQD | 1667300 | Cell/Virus Enclosed | NO |
| IMLNKHIDAYKT | 1452950 | Capsid | UN |
| IMNDKVVVKINNVDTV | 1667311 | Cell/Virus Enclosed | NO |
| IMRTFRIAIWNL | 1452983 | Cell/Virus Enclosed | NO |
| IMSCFYMEADAV | 1452984 | Cell/Virus Enclosed | NO |
| IMSMAECKATLG | 1452989 | Cell/Virus Enclosed | NO |
| IMTVTMTFKLGPRKATGRWN | 27573 | Capsid | UN |
| INAASKGAVQKESDEY | 1667349 | Cell/Virus Enclosed | NO |
| INAQVAKSHNVS | 1453036 | Cell/Virus Enclosed | NO |
| INARIRAKHYVYIGDP | 1667362 | Cell/Virus Enclosed | NO |
| INCILSVNSSNC | 1453049 | Cell/Virus Enclosed | NO |
| INCVLSVNSSNCNNFN | 1667371 | Cell/Virus Enclosed | NO |

| | | | |
|---|---|---|---|
| INDDEDVVTGDNDDED | 1667375 | Cell/Virus Enclosed | NO |
| INDKGKEVLVLWGIHHPSTSADQQS | 1148935 | Ectodomain | YES |
| INDKVLTKIDKE | 1453069 | Cell/Virus Enclosed | NO |
| INDKVVAKIDKEDVVI | 1667378 | Cell/Virus Enclosed | NO |
| INDYNKTIPRIS | 1453080 | Ectodomain | YES |
| INECVKSQSNRY | 1453082 | Ectodomain | YES |
| INEVVKQMFGVNLQSG | 1667392 | Cell/Virus Enclosed | NO |
| INFVKFGSVCFS | 1453110 | Ectodomain | YES |
| INGACGSPGYNLKNGE | 1667402 | Cell/Virus Enclosed | NO |
| INGACGSPGYNVRNDG | 1667403 | Cell/Virus Enclosed | NO |
| INGDFLHFLPRV | 1453114 | Cell/Virus Enclosed | NO |
| INGLGVTMDVLNKNQK | 1667417 | Ectodomain | YES |
| INGMHKSFYVNANGGT | 1667421 | Cell/Virus Enclosed | NO |
| INGYTLAATSAS | 1453152 | Ectodomain | YES |
| INHTFDDPVIPF | 1453160 | Ectodomain | YES |
| INHTFGNPVIPFKDGI | 1667434 | Ectodomain | YES |
| INHTLGNPVIPF | 1453164 | Ectodomain | YES |
| INKAIPSGNLIKFDIG | 1667445 | Cell/Virus Enclosed | NO |
| INKCSRLLSDDRTEVP | 1667447 | Ectodomain | YES |
| INKFLKANPLWHKAVF | 1667450 | Cell/Virus Enclosed | NO |
| INKQLTDISVDSNWPL | 1667456 | Cell/Virus Enclosed | NO |
| INKSVANLKLPPHDVT | 1667457 | Cell/Virus Enclosed | NO |
| INKYGLEDFNFEHVVY | 1667459 | Cell/Virus Enclosed | NO |
| INKYGLELHWAPEFPW | 1667460 | Cell/Virus Enclosed | NO |
| INLCDDPEKAQG | 1453218 | Cell/Virus Enclosed | NO |
| INLKDIGTYEMY | 1453226 | Ectodomain | YES |
| INLKDIGTYEYY | 1453227 | Ectodomain | YES |
| INLKFNKWQWQE | 1453228 | Cell/Virus Enclosed | NO |
| INLPPPLLGNST | 1453232 | Ectodomain | YES |
| INLRADQINDMVYSLL | 1667470 | Cell/Virus Enclosed | NO |
| INMDNSQNLAWP | 1453243 | Cell/Virus Enclosed | NO |
| INNFNPSSWNRR | 1453251 | Ectodomain | YES |
| INNGPHEFCTKH | 1453253 | Cell/Virus Enclosed | NO |
| INNKDTGPPFIS | 1453255 | Ectodomain | YES |
| INNLKIKNFDYLDIHP | 1667481 | Ectodomain | YES |
| INNTVYDPLQPELDSF | 1667487 | Ectodomain | YES |
| INNVDNEIVKSS | 1453265 | Cell/Virus Enclosed | NO |
| INPNALLCATCP | 1453275 | Cell/Virus Enclosed | NO |
| INQLDEKAQKFFDFGD | 1667501 | Cell/Virus Enclosed | NO |
| INRGIYKPLLEN | 1453295 | Cell/Virus Enclosed | NO |
| INRLFSVPSDSC | 1453300 | Cell/Virus Enclosed | NO |
| INRMAEQAAAAM | 1453305 | Cell/Virus Enclosed | NO |
| INRVLSVPSDSC | 1453312 | Cell/Virus Enclosed | NO |
| INSAIVCASVKRDGVQ | 1667515 | Cell/Virus Enclosed | NO |
| INSILDEVNGLLDTTQ | 1667518 | Ectodomain | YES |
| INSILNEVNDLL | 1453319 | Ectodomain | YES |
| INSNVKTLEYDYMTHG | 1667520 | Cell/Virus Enclosed | NO |
| INSQDGVNKLQG | 1453325 | Ectodomain | YES |
| INSTLVDLKWLNRVET | 1667522 | Ectodomain | YES |
| INSTQDGDNKLQGLLE | 1667523 | Ectodomain | YES |
| INSTQDGVNKLQ | 1453333 | Ectodomain | YES |
| INSVVQKDGVED | 1453343 | Cell/Virus Enclosed | NO |
| INTGNGIKQLAP | 1453351 | Cell/Virus Enclosed | NO |
| INTISPEKLKNYAASY | 1667531 | Cell/Virus Enclosed | NO |
| INTNSSKDDQIG | 1453360 | Cell/Virus Enclosed | NO |
| INTNSTKDDQIGYYRR | 1667535 | Cell/Virus Enclosed | NO |
| INVKDVFVESSK | 1453384 | Cell/Virus Enclosed | NO |
| INVLSIFSSSFS | 1453385 | Cell/Virus Enclosed | NO |
| INVTINNYNPSF | 1453390 | Ectodomain | YES |
| INVVAASGSLTR | 1453393 | Cell/Virus Enclosed | NO |
| INWITGGIAIAMACL | 1312747 | Undetermined | UN |
| INWNVEYPIISNELSI | 1667560 | Cell/Virus Enclosed | NO |
| INYKGKVFFQFDNLSS | 1667563 | Cell/Virus Enclosed | NO |
| INYLGTIKENIDGGAM | 1667568 | Cell/Virus Enclosed | NO |
| IPAHCVRDPVGF | 1453419 | Cell/Virus Enclosed | NO |
| IPAKARVECYDR | 1453420 | Cell/Virus Enclosed | NO |
| IPAKKAGGTTEM | 1453421 | Cell/Virus Enclosed | NO |
| IPAKVRVDCYDKFKIN | 1667573 | Cell/Virus Enclosed | NO |
| IPASGYVAALAV | 1453430 | Cell/Virus Enclosed | NO |
| IPASQVIVNNYDKSAG | 1667581 | Cell/Virus Enclosed | NO |
| IPATCANDPVGF | 1453433 | Cell/Virus Enclosed | NO |
| IPATSASKLTIVSPDL | 1667583 | Cell/Virus Enclosed | NO |
| IPATSIVAKQGAGDAG | 1667584 | Cell/Virus Enclosed | NO |
| IPCVCGRDATQY | 1453439 | Cell/Virus Enclosed | NO |
| IPDFDLDDFIDAPCYC | 1667592 | Cell/Virus Enclosed | NO |
| IPDKQVFDKVVDNVYV | 1667595 | Cell/Virus Enclosed | NO |

| | | | |
|---|---|---|---|
| IPDLPICVYDSL | 1453461 | Ectodomain | YES |
| IPDMADQIASLV | 1453468 | Cell/Virus Enclosed | NO |
| IPDYNYYKYNLP | 1453474 | Cell/Virus Enclosed | NO |
| IPDYVDVNKTLQEFAQ | 1667602 | Ectodomain | YES |
| IPESKPLADDDSAIIE | 1667604 | Capsid | UN |
| IPFKDGIYFAATEKSN | 1667612 | Ectodomain | YES |
| IPFNWTTSVQVE | 1453503 | Ectodomain | YES |
| IPGGCAMPIMAN | 1453510 | Ectodomain | YES |
| IPHDIDLGESRVVIQDYDNQHEQDR | 27861 | Undetermined | UN |
| IPHFNQKLPNGVTVVE | 1667626 | Capsid | UN |
| IPHYSWFSGITQ | 1453532 | Cell/Virus Enclosed | NO |
| IPIDLLCLRDDN | 1453535 | Cell/Virus Enclosed | NO |
| IPIDLLNLREIKPAVN | 1667630 | Cell/Virus Enclosed | NO |
| IPIFDDPVVDWH | 1453538 | Cell/Virus Enclosed | NO |
| IPIVDDLVVDWH | 1453549 | Cell/Virus Enclosed | NO |
| IPIVWGVKDFNTLSQE | 1667638 | Cell/Virus Enclosed | NO |
| IPKPSPWAPKKHRRLSSDQD | 110293 | Cell/Virus Enclosed | NO |
| IPLADDDSAIIE | 1453567 | Capsid | UN |
| IPLGVIHNSTLQVSDVD | 1021615 | Secreted | YES |
| IPLLVDSEQPWK | 1453582 | Cell/Virus Enclosed | NO |
| IPLSDMEKDFLS | 1453593 | Cell/Virus Enclosed | NO |
| IPNFGSLTQINI | 1453616 | Ectodomain | YES |
| IPNFGSLTQINTTLLD | 1667664 | Ectodomain | YES |
| IPNGRKVDLQLGNLGY | 1667666 | Ectodomain | YES |
| IPNLPDFKEELG | 1453629 | Ectodomain | YES |
| IPNLPDFKEESD | 1453630 | Ectodomain | YES |
| IPNLPDFKGELD | 1453631 | Ectodomain | YES |
| IPNLSDFEAEFS | 1453634 | Ectodomain | YES |
| IPNRRKVDLQLG | 1453638 | Ectodomain | YES |
| IPNRRRVDLQLG | 1453640 | Ectodomain | YES |
| IPQVGGLTSIKW | 1453663 | Cell/Virus Enclosed | NO |
| IPRFKVSPYLFNVPI | 28041 | Ectodomain | YES |
| IPRISEDGVDVS | 1453665 | Ectodomain | YES |
| IPRYASAVAQAF | 1453679 | Cell/Virus Enclosed | NO |
| IPSDVSARDPII | 1453684 | Cell/Virus Enclosed | NO |
| IPSDVSARDPTI | 1453685 | Cell/Virus Enclosed | NO |
| IPSEFTIGNIEE | 1453686 | Ectodomain | YES |
| IPSEFTIGNMEE | 1453687 | Ectodomain | YES |
| IPSGNLIKFDIGSVVV | 1667683 | Cell/Virus Enclosed | NO |
| IPSIQDAATPYHPPATPNNM | 741563 | Ectodomain | YES |
| IPSIQSRGLFGAIAG | 150999 | Ectodomain | YES |
| IPSLAANTLNIIVPDK | 1667685 | Cell/Virus Enclosed | NO |
| IPSNWTTSVQVE | 1453702 | Ectodomain | YES |
| IPTDSTVKNYFI | 1453710 | Cell/Virus Enclosed | NO |
| IPTIEDWLLVQG | 1453712 | Cell/Virus Enclosed | NO |
| IPTIKDWLLVQG | 1453713 | Cell/Virus Enclosed | NO |
| IPTMTHMNLKYA | 1453721 | Cell/Virus Enclosed | NO |
| IPTMTQMNLKYAISAK | 1667696 | Cell/Virus Enclosed | NO |
| IPTNADKICLGHHAVSNGTKVNTLT | 1149354 | Ectodomain | YES |
| IPTNFTIVGQEEFIQT | 1667700 | Ectodomain | YES |
| IPTQCYYNNGSSGRIV | 1667701 | Cell/Virus Enclosed | NO |
| IPTQCYYNNVSS | 1453731 | Cell/Virus Enclosed | NO |
| IPVRPRNSSDNGISAI | 1667710 | Ectodomain | YES |
| IPYCNYSKYWYLNHT | 83829 | Ectodomain | YES |
| IPYKSFSDSGCILPSE | 1667713 | Cell/Virus Enclosed | NO |
| IPYNSITDTIVL | 1453764 | Secreted | YES |
| IPYNVSLSKFNSCKSD | 1667718 | Ectodomain | YES |
| IQADSFTCNNIDAAKI | 1667721 | Ectodomain | YES |
| IQADVNTPADIV | 1453778 | Cell/Virus Enclosed | NO |
| IQAGSIVRRVNV | 1453779 | Cell/Virus Enclosed | NO |
| IQAIVSSPCHNSLILPPFSLSPV | 78425 | Ectodomain | YES |
| IQAIYDRLDTIQADQQ | 1667723 | Ectodomain | YES |
| IQANAYYKYDNGNFIK | 1667724 | Cell/Virus Enclosed | NO |
| IQCKTKSLLPNTGVYD | 1667730 | Ectodomain | YES |
| IQDSDGTNKQLNEISD | 1667732 | Cell/Virus Enclosed | NO |
| IQDVGFKNQYCNGSIA | 1667733 | Cell/Virus Enclosed | NO |
| IQDVKDGTVTFNLLAY | 1667734 | Cell/Virus Enclosed | NO |
| IQDVKGLKIKYLYFIK | 1667735 | Cell/Virus Enclosed | NO |
| IQDVVNQQGNSLNHLT | 1667736 | Ectodomain | YES |
| IQDVVNQQGSALNHLT | 1667737 | Ectodomain | YES |
| IQDYIQSVNFVA | 1453812 | Cell/Virus Enclosed | NO |
| IQEAIPTRFSPG | 1453813 | Cell/Virus Enclosed | NO |
| IQEGFDATNSALVKIQ | 1667741 | Ectodomain | YES |
| IQGFACGCGASYTAND | 1667754 | Cell/Virus Enclosed | NO |
| IQGFGPRDFNHN | 1453847 | Cell/Virus Enclosed | NO |
| IQGIKLGTGYSLADLP | 1667756 | Ectodomain | YES |

| | | | |
|---|---|---|---|
| IQGVKLGTGYTLSDLP | 1667762 | Underneath Membrane | NO |
| IQIDQLNSSYFK | 1453858 | Ectodomain | YES |
| IQILASVPEKFLNAFD | 1667764 | Cell/Virus Enclosed | NO |
| IQIPSKFTIGNM | 1453864 | Ectodomain | YES |
| IQISYNDFYASGCVLS | 1667767 | Cell/Virus Enclosed | NO |
| IQITNESYNNYVPSYV | 1667769 | Cell/Virus Enclosed | NO |
| IQKVTVDCKQYVCNGF | 1667776 | Ectodomain | YES |
| IQKYGLEDFNFEHVVY | 1667777 | Cell/Virus Enclosed | NO |
| IQLNNDVRDYLLSKMT | 1667786 | Cell/Virus Enclosed | NO |
| IQLTTTRMEDAM | 1453913 | Cell/Virus Enclosed | NO |
| IQNGAKHVQGNVAKAA | 1667793 | Cell/Virus Enclosed | NO |
| IQNLLYCDSPFEKLQC | 1667795 | Ectodomain | YES |
| IQNQTLPRLHCT | 1453928 | Cell/Virus Enclosed | NO |
| IQNSAKHVQGNVAKIA | 1667797 | Cell/Virus Enclosed | NO |
| IQPVELPVSIVS | 1453954 | Ectodomain | YES |
| IQPWTNDALNTICIKD | 1667806 | Cell/Virus Enclosed | NO |
| IQQDLDVILPGE | 1453959 | Cell/Virus Enclosed | NO |
| IQQGFDATNFAL | 1453960 | Ectodomain | YES |
| IQQGFDATNSAL | 1453961 | Ectodomain | YES |
| IQQGGIPIANCV | 1453962 | Cell/Virus Enclosed | NO |
| IQQLPETYFTQS | 1453973 | Cell/Virus Enclosed | NO |
| IQQVVDADSKHV | 1453980 | Cell/Virus Enclosed | NO |
| IQQWGYVGSLSQ | 1453982 | Cell/Virus Enclosed | NO |
| IQQWGYVGSLST | 1453983 | Cell/Virus Enclosed | NO |
| IQRMLLEKCDLQ | 1453996 | Cell/Virus Enclosed | NO |
| IQSFDSSYLNES | 1454006 | Cell/Virus Enclosed | NO |
| IQSKLTDLKCTN | 1454012 | Cell/Virus Enclosed | NO |
| IQSNVDVVPEGW | 1454013 | Cell/Virus Enclosed | NO |
| IQSVELPVSIVA | 1454023 | Ectodomain | YES |
| IQSVVNSNAQALNSLL | 1667836 | Ectodomain | YES |
| IQTIQDSDGTNKQLNE | 1667839 | Cell/Virus Enclosed | NO |
| IQTNSPKVTIDCSLFV | 1667841 | Ectodomain | YES |
| IQTTIQKVTVDCKQYV | 1667843 | Ectodomain | YES |
| IQTTPGFGVAVA | 1454039 | Cell/Virus Enclosed | NO |
| IQTVFDCAVETC | 1454042 | Cell/Virus Enclosed | NO |
| IQTVQDADGINKQLTD | 1667848 | Cell/Virus Enclosed | NO |
| IQVAKCYGESVLVNAA | 1667849 | Cell/Virus Enclosed | NO |
| IQVDQFNSSYFK | 1454051 | Ectodomain | YES |
| IQVDQLNSSYFKLSIP | 1667851 | Ectodomain | YES |
| IQVDQLNSSYKL | 1454053 | Ectodomain | YES |
| IQVDQLNSTYFK | 1454054 | Ectodomain | YES |
| IQVSLQIGIKTDDSHDWT | 2134442 | Undetermined | UN |
| IQVVNSGPDQIC | 1454061 | Cell/Virus Enclosed | NO |
| IQVVNSGPDQTCNTPT | 1667855 | Cell/Virus Enclosed | NO |
| IQVVNSGSDINC | 1454063 | Cell/Virus Enclosed | NO |
| IQVVNSGSDMNCNIPT | 1667856 | Cell/Virus Enclosed | NO |
| IRDIVCKVENKTEAEW | 1667873 | Cell/Virus Enclosed | NO |
| IRDKGCVADQSRKWIG | 1667874 | Cell/Virus Enclosed | NO |
| IRDNGSLEKFMS | 1454113 | Cell/Virus Enclosed | NO |
| IRDNVSLEKFMS | 1454115 | Cell/Virus Enclosed | NO |
| IRDVSVEDVCFANKFE | 1667885 | Cell/Virus Enclosed | NO |
| IRDYKLANDAKESLRL | 1667886 | Cell/Virus Enclosed | NO |
| IREIVMNASPYD | 1454137 | Cell/Virus Enclosed | NO |
| IRFEYTSPTDKLAMIM | 1667902 | Cell/Virus Enclosed | NO |
| IRFKDDTSFEDING | 28291 | Capsid | UN |
| IRGAIGDVGLCM | 1454169 | Cell/Virus Enclosed | NO |
| IRGCCESTKLPYYDVE | 1667908 | Underneath Membrane | NO |
| IRGHLYIQGIKLGTGY | 1667911 | Ectodomain | YES |
| IRGHLYIQGVKLGTGY | 1667912 | Underneath Membrane | NO |
| IRGKEQLQLLMP | 1454181 | Cell/Virus Enclosed | NO |
| IRGPPRAELNGVVVDK | 1667914 | Cell/Virus Enclosed | NO |
| IRHKILTKPRQK | 1454199 | Cell/Virus Enclosed | NO |
| IRIVDEWSYTGS | 1454205 | Ectodomain | YES |
| IRKDLSYEQQNVLAIE | 1667927 | Cell/Virus Enclosed | NO |
| IRLQAGVATEYAANSS | 1667940 | Cell/Virus Enclosed | NO |
| IRLTITELLQFV | 1454236 | Cell/Virus Enclosed | NO |
| IRNESWHIDSSE | 1454244 | Ectodomain | YES |
| IRNLQKRLYFNV | 1454250 | Cell/Virus Enclosed | NO |
| IRNSDVVKQPVK | 1454255 | Cell/Virus Enclosed | NO |
| IRPIDFFSLTAS | 1454258 | Cell/Virus Enclosed | NO |
| IRQGTDYKQWPQ | 1454268 | Capsid | UN |
| IRQKILNKPRQK | 1454270 | Capsid | UN |
| IRRAIDCGFNDLSQLH | 1667957 | Ectodomain | YES |
| IRRLDVLEQDAQ | 1454286 | Ectodomain | YES |
| IRRPKDGFNLEY | 1454290 | Ectodomain | YES |
| IRSDSTLSGFET | 1454306 | Capsid | UN |

| | | | |
|---|---|---|---|
| IRSEELSFTVVSNGA | 28398 | Secreted | YES |
| IRSSNQSLAGGITYVS | 1667967 | Ectodomain | YES |
| IRSVYPFASPQE | 1454322 | Cell/Virus Enclosed | NO |
| IRSYEKLANSIS | 1454326 | Cell/Virus Enclosed | NO |
| IRTPPAYRPPNAPILSTLPETTV | 79577 | Capsid | UN |
| IRYGIKPITIPI | 1454359 | Cell/Virus Enclosed | NO |
| ISAGIFSVPADVSLTY | 1667998 | Cell/Virus Enclosed | NO |
| ISAGIFSVPSDVSLTY | 1667999 | Cell/Virus Enclosed | NO |
| ISAMQTMLFGMI | 1454392 | Cell/Virus Enclosed | NO |
| ISAQLAKDKVNE | 1454395 | Ectodomain | YES |
| ISASYRFVTFEPFNVS | 1454404 | Ectodomain | YES |
| ISATSNYSGVVT | 1454406 | Cell/Virus Enclosed | NO |
| ISAYGCSTIAQA | 1454411 | Cell/Virus Enclosed | NO |
| ISCDCGKKFDEQ | 1454412 | Cell/Virus Enclosed | NO |
| ISCIWFIDAFNQ | 1454415 | Cell/Virus Enclosed | NO |
| ISCNGNKIEDLS | 1454417 | Cell/Virus Enclosed | NO |
| ISDKNVVFMNSC | 1454439 | Ectodomain | YES |
| ISDMYDGRIKFC | 1454443 | Cell/Virus Enclosed | NO |
| ISDMYDGRTKAIDGEN | 1668027 | Cell/Virus Enclosed | NO |
| ISDMYDPKTKHV | 1454446 | Cell/Virus Enclosed | NO |
| ISDMYDPLTKNIGDYN | 1668031 | Cell/Virus Enclosed | NO |
| ISDMYDPTTKNL | 1454449 | Cell/Virus Enclosed | NO |
| ISDMYDPTTKNVTGSN | 1668033 | Cell/Virus Enclosed | NO |
| ISDRKIKKYPCI | 1454453 | Cell/Virus Enclosed | NO |
| ISDTSGFAVYVK | 1454460 | Ectodomain | YES |
| ISEDLWQFVDHF | 1454464 | Cell/Virus Enclosed | NO |
| ISEEFSSNVANY | 1454465 | Cell/Virus Enclosed | NO |
| ISEGNALYNNEG | 1454467 | Cell/Virus Enclosed | NO |
| ISEQSLRLIDAM | 1454478 | Cell/Virus Enclosed | NO |
| ISEWPLFDQQAQ | 1454484 | Cell/Virus Enclosed | NO |
| ISEWPLSESDKE | 1454486 | Cell/Virus Enclosed | NO |
| ISEWPLSVQQAQQEAT | 1668046 | Cell/Virus Enclosed | NO |
| ISFADNNDITNT | 1454490 | Cell/Virus Enclosed | NO |
| ISFCCRHNWNCV | 1454492 | Cell/Virus Enclosed | NO |
| ISFCKSGSLVLN | 1454494 | Ectodomain | YES |
| ISFSDDVIVHDV | 1454508 | Cell/Virus Enclosed | NO |
| ISGDKVFVRDGNTDNL | 1668057 | Cell/Virus Enclosed | NO |
| ISGDRGIAPKSG | 1454512 | Ectodomain | YES |
| ISGFLSRVIRRE | 1454515 | Cell/Virus Enclosed | NO |
| ISGIVSAVSAMM | 1454524 | Transmembrane | NO |
| ISGKELARTVGG | 1454526 | Cell/Virus Enclosed | NO |
| ISGKGQQQQGQT | 1454527 | Cell/Virus Enclosed | NO |
| ISGYTLAATSASLFPP | 1668066 | Ectodomain | YES |
| ISIMKHDIKVIA | 1454559 | Cell/Virus Enclosed | NO |
| ISINLPPPLLGN | 1454561 | Ectodomain | YES |
| ISKDSKFQIGEFIFEK | 1668081 | Cell/Virus Enclosed | NO |
| ISLICFILLWAL | 1454605 | Cell/Virus Enclosed | NO |
| ISLNAPVTLKIC | 1454618 | Ectodomain | YES |
| ISLPTAFAAIGD | 1454624 | Ectodomain | YES |
| ISLPTAFAVVGD | 1454626 | Ectodomain | YES |
| ISLSFNNLLGFDSSKP | 1668106 | Cell/Virus Enclosed | NO |
| ISLTMLDIQPEDYKSV | 1668108 | Cell/Virus Enclosed | NO |
| ISLTVGEVFGKILGNV | 1668110 | Cell/Virus Enclosed | NO |
| ISLVSVTDFSLNGNVS | 1668113 | Cell/Virus Enclosed | NO |
| ISMLQLGAPDNP | 1454644 | Ectodomain | YES |
| ISMSLVLVIVVK | 1454653 | Cell/Virus Enclosed | NO |
| ISMTNHSFSVQK | 1454654 | Cell/Virus Enclosed | NO |
| ISNDSNGCVVEASTDF | 1668130 | Cell/Virus Enclosed | NO |
| ISNDTVDVTNGL | 1454667 | Ectodomain | YES |
| ISNFKFDDYDFSVEDA | 1668136 | Cell/Virus Enclosed | NO |
| ISNKADLKFVKWEYEG | 1668137 | Cell/Virus Enclosed | NO |
| ISNKLCVVPGNVKPGD | 1668138 | Cell/Virus Enclosed | NO |
| ISNSIQAIYDRLDSIQADQQ | 2060836 | Ectodomain | YES |
| ISNVPFSPDGKP | 1454685 | Ectodomain | YES |
| ISNVPFSPNGKP | 1454686 | Ectodomain | YES |
| ISNVPFSSDGKP | 1454687 | Ectodomain | YES |
| ISPAAISSNCYS | 1454691 | Ectodomain | YES |
| ISPCAFGGVSVITPGTN | 28643 | Ectodomain | YES |
| ISPKTDKETLEY | 1454706 | Ectodomain | YES |
| ISPSTSATIQKI | 1454716 | Ectodomain | YES |
| ISPYNSQNAVSRSMLG | 1668152 | Cell/Virus Enclosed | NO |
| ISPYNSQNFAAKRVLG | 1668153 | Cell/Virus Enclosed | NO |
| ISPYNSQNYVAA | 1454729 | Cell/Virus Enclosed | NO |
| ISPYNSQNYVAKRVLGVQ | 1454730 | Cell/Virus Enclosed | NO |
| ISQDSITHYPALKMVQ | 1668157 | Cell/Virus Enclosed | NO |
| ISQDSITHYSAL | 1454734 | Cell/Virus Enclosed | NO |

| | | | |
|---|---|---|---|
| ISQFNYKQSSSN | 1454737 | Ectodomain | YES |
| ISQGLDAAWNKF | 1454739 | Cell/Virus Enclosed | NO |
| ISQWPISNDSNC | 1454750 | Cell/Virus Enclosed | NO |
| ISQWPISNDSNGCVVE | 1668165 | Cell/Virus Enclosed | NO |
| ISRCQITAVEGT | 1454755 | Cell/Virus Enclosed | NO |
| ISRMGFKLDANV | 1454768 | Cell/Virus Enclosed | NO |
| ISRMGFKLDATVPGYP | 1668169 | Cell/Virus Enclosed | NO |
| ISRRNIGFTLEY | 1454776 | Ectodomain | YES |
| ISRSELQTIVPEYIDV | 1668171 | Ectodomain | YES |
| ISRVELHTVYPD | 1454781 | Ectodomain | YES |
| ISSGIFSVPSDV | 1454792 | Cell/Virus Enclosed | NO |
| ISSKVGNSIFRS | 1454797 | Ectodomain | YES |
| ISSLAYYEPVQG | 1454798 | Cell/Virus Enclosed | NO |
| ISSNCYSSLILD | 1454802 | Ectodomain | YES |
| ISSNGPLKVGAGVMLE | 1668184 | Cell/Virus Enclosed | NO |
| ISSNGPLKVGTG | 1454804 | Cell/Virus Enclosed | NO |
| ISSNTDNETLQYWVTP | 1668185 | Ectodomain | YES |
| ISSVLNDVLSRL | 1454821 | Ectodomain | YES |
| ISSYKLDLTILG | 1454826 | Cell/Virus Enclosed | NO |
| ISTARDGVRRGDNFHS | 1668193 | Cell/Virus Enclosed | NO |
| ISTDIVDVTNGL | 1454835 | Ectodomain | YES |
| ISTEKLSRLSMIKGPQ | 1668200 | Cell/Virus Enclosed | NO |
| ISTGSRSARSAIEDLL | 1668203 | Ectodomain | YES |
| ISTLENKSAELNYTVQ | 1668211 | Ectodomain | YES |
| ISTNLPPPLLSN | 1454859 | Ectodomain | YES |
| ISTNYSKDVYFT | 1454860 | Undetermined | UN |
| ISTPKLKEDYQIGGYS | 1668213 | Secreted | YES |
| ISTPNLPDFKEE | 1454864 | Ectodomain | YES |
| ISTQDGVNKLQG | 1454865 | Ectodomain | YES |
| ISTQLFEGDKFIGTFE | 1668214 | Cell/Virus Enclosed | NO |
| ISTRENKSAELN | 1454867 | Ectodomain | YES |
| ISTSKVKSLSMI | 1454869 | Cell/Virus Enclosed | NO |
| ISTTLIQLQLTK | 1454873 | Cell/Virus Enclosed | NO |
| ISTVQSKLTDLKCTNV | 1668219 | Cell/Virus Enclosed | NO |
| ISVAPIKNTVDT | 1454883 | Cell/Virus Enclosed | NO |
| ISVLDITDAAVKAAES | 1668229 | Cell/Virus Enclosed | NO |
| ISVMDKLDTGAQKFFQ | 1668230 | Cell/Virus Enclosed | NO |
| ISVNDLCFANKFFQFD | 1668231 | Cell/Virus Enclosed | NO |
| ISVQELRYMNANGLRP | 1383031 | Cell/Virus Enclosed | NO |
| ISVTPIKNTVDTTPKE | 1668235 | Cell/Virus Enclosed | NO |
| ISWLLHKDGGIP | 1454927 | Cell/Virus Enclosed | NO |
| ISYEEQDAIFLL | 1454932 | Cell/Virus Enclosed | NO |
| ISYEEQDALFALTKRN | 1383032 | Cell/Virus Enclosed | NO |
| ISYLNEQGVLVQ | 1454937 | Cell/Virus Enclosed | NO |
| ISYLNRARGSSAARLEPC | 1454938 | Cell/Virus Enclosed | NO |
| ISYNSTLQSGLK | 1454940 | Cell/Virus Enclosed | NO |
| ITACQYTMCEFP | 1454945 | Ectodomain | YES |
| ITADIELALLRA | 1383036 | Ectodomain | YES |
| ITADSELALLRA | 1454948 | Underneath Membrane | NO |
| ITANGGISFCCR | 1454955 | Cell/Virus Enclosed | NO |
| ITANGGISFCRR | 1454957 | Cell/Virus Enclosed | NO |
| ITAPYVVLTGLGESFK | 1668264 | Cell/Virus Enclosed | NO |
| ITAQPVKIDNASPAST | 1668265 | Secreted | YES |
| ITAQPVKTDNAS | 1454967 | Secreted | YES |
| ITCEEAVRQVRS | 1454972 | Cell/Virus Enclosed | NO |
| ITDAAVKAAESKAFVD | 1668272 | Cell/Virus Enclosed | NO |
| ITDPLFSNIQPF | 1454994 | Cell/Virus Enclosed | NO |
| ITDYNYYKYNLPTMVD | 1668283 | Cell/Virus Enclosed | NO |
| ITEDEILEWFGI | 1455005 | Ectodomain | YES |
| ITEDGINVKDVF | 1455008 | Cell/Virus Enclosed | NO |
| ITEDGINVKDVV | 1455009 | Cell/Virus Enclosed | NO |
| ITEFQLDYSIDVIDNE | 1668290 | Cell/Virus Enclosed | NO |
| ITEFSWNADLYKLMSC | 1668291 | Cell/Virus Enclosed | NO |
| ITEGPTTCGYLP | 1455014 | Cell/Virus Enclosed | NO |
| ITENNVVVTSTC | 1455023 | Ectodomain | YES |
| ITETNTAAFKAV | 1455027 | Cell/Virus Enclosed | NO |
| ITFGGPSDSTDNNQNG | 1668315 | Cell/Virus Enclosed | NO |
| ITGGIAIAMACLVGL | 1319003 | Undetermined | UN |
| ITGLYPTITVPG | 1455081 | Cell/Virus Enclosed | NO |
| ITGLYPTLNISE | 1455083 | Cell/Virus Enclosed | NO |
| ITGNTLQCVMLV | 1455085 | Cell/Virus Enclosed | NO |
| ITHALSADGVQC | 1455097 | Cell/Virus Enclosed | NO |
| ITIDKFAIPNGRKVDL | 1383054 | Ectodomain | YES |
| ITIDKFAIPNRR | 1455111 | Ectodomain | YES |
| ITIDKFAIPNSR | 1455112 | Ectodomain | YES |
| ITIDKFALPNRR | 1455113 | Ectodomain | YES |

| | | | |
|---|---|---|---|
| ITIEAALDLSKE | 1455115 | Cell/Virus Enclosed | NO |
| ITIKVTEDGVNVHDVT | 1668356 | Cell/Virus Enclosed | NO |
| ITILDGISEQSL | 1455124 | Cell/Virus Enclosed | NO |
| ITINPSSPARLE | 1455127 | Cell/Virus Enclosed | NO |
| ITITYQGLFPYQGDHG | 1668362 | Ectodomain | YES |
| ITKDEAIKRVRGWVGF | 1383061 | Cell/Virus Enclosed | NO |
| ITKDSKFQVGEFVFEK | 1668367 | Cell/Virus Enclosed | NO |
| ITKENVETLTWPLILN | 1668370 | Cell/Virus Enclosed | NO |
| ITKKHAAAAKNK | 1455145 | Capsid | UN |
| ITKLADVEADIA | 1455146 | Cell/Virus Enclosed | NO |
| ITKLEADFYLSG | 1455148 | Ectodomain | YES |
| ITKPLKYSYIDK | 1455157 | Ectodomain | YES |
| ITKTDAVILLERYVSL | 1668383 | Cell/Virus Enclosed | NO |
| ITKVEADFYLSG | 1455165 | Ectodomain | YES |
| ITLANFNETKGPLCVD | 1668386 | Ectodomain | YES |
| ITLDDDARTFVQ | 1455177 | Cell/Virus Enclosed | NO |
| ITLEEAVRQVRS | 1455178 | Cell/Virus Enclosed | NO |
| ITLFGDCVTVRV | 1455179 | Cell/Virus Enclosed | NO |
| ITLKCCTVTYLNDPSS | 1668392 | Cell/Virus Enclosed | NO |
| ITLKTPDTYKCP | 1455191 | Ectodomain | YES |
| ITLMNLKYAISA | 1455196 | Cell/Virus Enclosed | NO |
| ITLVYKVYYGNA | 1455206 | Cell/Virus Enclosed | NO |
| ITMPCFITYPQQ | 1455212 | Cell/Virus Enclosed | NO |
| ITMTDLQSESSCGLFK | 1668407 | Cell/Virus Enclosed | NO |
| ITNESYNNYVPSYVKP | 1668411 | Cell/Virus Enclosed | NO |
| ITNINLSRGRGR | 1455228 | Cell/Virus Enclosed | NO |
| ITNLCPFGEVFNATKF | 1668417 | Ectodomain | YES |
| ITNTNLSRGRGRNPKP | 1668421 | Cell/Virus Enclosed | NO |
| ITNVGEPTISQP | 1455243 | Cell/Virus Enclosed | NO |
| ITPEVSRELGNITKTN | 1668426 | Cell/Virus Enclosed | NO |
| ITPGTNASSEFA | 1455248 | Ectodomain | YES |
| ITPNVCFVKGDIINVA | 1668431 | Cell/Virus Enclosed | NO |
| ITPYCHDPTVLP | 1455259 | Cell/Virus Enclosed | NO |
| ITQFQKGKEFEFVEGQ | 1668437 | Cell/Virus Enclosed | NO |
| ITQFQKGKEFQF | 1455267 | Capsid | UN |
| ITQHTRTRPSVQ | 1455269 | Cell/Virus Enclosed | NO |
| ITQQVLSDNQKL | 1455276 | Ectodomain | YES |
| ITQQVLSENQKLIANK | 1668442 | Ectodomain | YES |
| ITQQVLSKNQKL | 1455279 | Ectodomain | YES |
| ITQRNFYSPQII | 1455283 | Ectodomain | YES |
| ITQRTRTRPSVQPGPM | 1668445 | Cell/Virus Enclosed | NO |
| ITQTSPALQTVA | 1455295 | Ectodomain | YES |
| ITQTSQALQTVATALN | 1668449 | Ectodomain | YES |
| ITQTSQALQTVS | 1455296 | Ectodomain | YES |
| ITQWPTATGDVV | 1455300 | Cell/Virus Enclosed | NO |
| ITREEAVRQVRSWIGF | 1668462 | Cell/Virus Enclosed | NO |
| ITSAGQPCVASE | 1455331 | Cell/Virus Enclosed | NO |
| ITSEGNALYNNEGGRA | 1668472 | Cell/Virus Enclosed | NO |
| ITSFKVVMSMFS | 1455342 | Ectodomain | YES |
| ITSFNTKYVAPQ | 1455343 | Ectodomain | YES |
| ITSGIKFQTVKPGNFN | 1668476 | Cell/Virus Enclosed | NO |
| ITSGVKFQTVKPGNFN | 1668478 | Cell/Virus Enclosed | NO |
| ITSLNTKYVAPQVTYQ | 1668484 | Ectodomain | YES |
| ITSREVVVTNLNKSAG | 1668485 | Cell/Virus Enclosed | NO |
| ITSVTSDKFDFI | 1455368 | Cell/Virus Enclosed | NO |
| ITTGYRFTNFEPFTVN | 1668500 | Ectodomain | YES |
| ITTKISLKGGKI | 1455390 | Cell/Virus Enclosed | NO |
| ITTNGSWAIFNDHHLN | 1668504 | Cell/Virus Enclosed | NO |
| ITTNTGIENATF | 1455401 | Secreted | YES |
| ITTSVDANTNQDSYGG | 1668507 | Cell/Virus Enclosed | NO |
| ITTYPGQGCAGY | 1455407 | Cell/Virus Enclosed | NO |
| ITVDKFAIPNRR | 1455413 | Ectodomain | YES |
| ITVDQLKHLLEQ | 1455414 | Undetermined | UN |
| ITVDVLKDMPVL | 1455415 | Cell/Virus Enclosed | NO |
| ITVEAAFDLSKE | 1455416 | Cell/Virus Enclosed | NO |
| ITVEAAIELSKELKRP | 1383099 | Cell/Virus Enclosed | NO |
| ITVEAALDLSKELKRP | 1383100 | Cell/Virus Enclosed | NO |
| ITVEAALDLSRE | 1455417 | Cell/Virus Enclosed | NO |
| ITVEKLKQLLEQ | 1455420 | Undetermined | UN |
| ITVETAIELSKE | 1455421 | Cell/Virus Enclosed | NO |
| ITVETALDLSKE | 1455422 | Cell/Virus Enclosed | NO |
| ITVPEEFASHVANFQK | 1383102 | Cell/Virus Enclosed | NO |
| ITVPGEFASHVA | 1455436 | Cell/Virus Enclosed | NO |
| ITVQYGTDTISV | 1455437 | Ectodomain | YES |
| ITVQYGTDTNSA | 1455438 | Ectodomain | YES |
| ITVSQIQLSLLKVTAF | 1383103 | Cell/Virus Enclosed | NO |

| | | | |
|---|---|---|---|
| ITVSQIQLSLLQ | 1455442 | Cell/Virus Enclosed | NO |
| ITYPMIANENAINKGG | 1668525 | Cell/Virus Enclosed | NO |
| ITYSPATILAGS | 1455456 | Cell/Virus Enclosed | NO |
| ITYTSTSGNLLGFKDV | 1668527 | Ectodomain | YES |
| ITYTYKMLVAKDNKNL | 1668528 | Capsid | UN |
| ITYVSNSGNLLGFKNV | 1668529 | Ectodomain | YES |
| IVAAVSAMMRIS | 1455465 | Transmembrane | NO |
| IVAICLLVTIVL | 1455471 | Cell/Virus Enclosed | NO |
| IVAKQGAGDAGHSLTW | 1668542 | Cell/Virus Enclosed | NO |
| IVAWHVVRDPRYVMRL | 1668551 | Cell/Virus Enclosed | NO |
| IVCASVKRDGVQVGYC | 1668554 | Cell/Virus Enclosed | NO |
| IVCRLDTRVHSE | 1455513 | Cell/Virus Enclosed | NO |
| IVDENGKYYLPYPDPS | 1668569 | Cell/Virus Enclosed | NO |
| IVDEWSYTGSSFYAPE | 1668571 | Ectodomain | YES |
| IVDIIQRLDVLE | 1455525 | Ectodomain | YES |
| IVDKDGTYYLPYPDPS | 1668575 | Cell/Virus Enclosed | NO |
| IVDKEVKDMQFDLYVN | 1668576 | Cell/Virus Enclosed | NO |
| IVDNVYMAKAGDKYYP | 1668579 | Cell/Virus Enclosed | NO |
| IVDNVYMAKSGD | 1455537 | Cell/Virus Enclosed | NO |
| IVDNVYMAKVGD | 1455538 | Cell/Virus Enclosed | NO |
| IVDRDQSSEEAI | 1455543 | Capsid | UN |
| IVDTLQETPVVS | 1455549 | Cell/Virus Enclosed | NO |
| IVDTVSALVYENKLKA | 1668589 | Cell/Virus Enclosed | NO |
| IVDVTNGLGTYY | 1455554 | Ectodomain | YES |
| IVEDDVIENVKSSLSS | 1668593 | Cell/Virus Enclosed | NO |
| IVEDTINDDEDV | 1455560 | Cell/Virus Enclosed | NO |
| IVETVSALVYDNKLKA | 1668603 | Cell/Virus Enclosed | NO |
| IVFNNNGTNVTQVKNA | 1668615 | Cell/Virus Enclosed | NO |
| IVFNVVKQGHFI | 1455599 | Cell/Virus Enclosed | NO |
| IVFTACSHAAVDSLCA | 1668617 | Cell/Virus Enclosed | NO |
| IVGCNNADCVAC | 1455613 | Cell/Virus Enclosed | NO |
| IVGDFHLNEEVA | 1455615 | Cell/Virus Enclosed | NO |
| IVGERQLLLLWE | 1455625 | Cell/Virus Enclosed | NO |
| IVGERQLLLVWEAGKS | 1668633 | Cell/Virus Enclosed | NO |
| IVGNKFYNVSTGLIPG | 1668641 | Cell/Virus Enclosed | NO |
| IVGPDARQDGRQSYVL | 1668642 | Cell/Virus Enclosed | NO |
| IVGPKELLLLWESGKA | 1668643 | Cell/Virus Enclosed | NO |
| IVGQEEFIQTNSPKVT | 1668644 | Ectodomain | YES |
| IVHSHPILHHPG | 1455663 | Cell/Virus Enclosed | NO |
| IVHSYPILHHSG | 1455667 | Cell/Virus Enclosed | NO |
| IVIDNDLNEFVS | 1455672 | Cell/Virus Enclosed | NO |
| IVIPDKQVFDKVVDNV | 1668663 | Cell/Virus Enclosed | NO |
| IVIRDIVCKVEN | 1455687 | Cell/Virus Enclosed | NO |
| IVKCKGLNKPFVKDNV | 1668668 | Cell/Virus Enclosed | NO |
| IVKEAQNANPTV | 1455705 | Cell/Virus Enclosed | NO |
| IVKLGAEKEDGF | 1455715 | Cell/Virus Enclosed | NO |
| IVKLGAEKKDGFYETI | 1668678 | Cell/Virus Enclosed | NO |
| IVKMVSPTSKIE | 1455718 | Cell/Virus Enclosed | NO |
| IVKSSDVVDSNENLTW | 1668687 | Cell/Virus Enclosed | NO |
| IVLDKFAIPNSRRSDL | 1668692 | Ectodomain | YES |
| IVLDVKGVEELHDDLV | 1668693 | Cell/Virus Enclosed | NO |
| IVLPQGGYVADF | 1455760 | Cell/Virus Enclosed | NO |
| IVLVCYLFIGYI | 1455776 | Cell/Virus Enclosed | NO |
| IVMNASPYDLEV | 1455797 | Cell/Virus Enclosed | NO |
| IVNAANENLAHGGGLA | 1668721 | Cell/Virus Enclosed | NO |
| IVNARIEPCSSGLSTD | 1668724 | Cell/Virus Enclosed | NO |
| IVNDDTSETKYVKSLS | 1668725 | Cell/Virus Enclosed | NO |
| IVNGYAKSNFVK | 1455827 | Cell/Virus Enclosed | NO |
| IVNNYDKSAGYPFNKF | 1668738 | Cell/Virus Enclosed | NO |
| IVNNYEKSAGYP | 1455843 | Cell/Virus Enclosed | NO |
| IVNPANGHMLHGGGVA | 1668739 | Cell/Virus Enclosed | NO |
| IVNSVLLFLAFVVFL | 1312788 | Transmembrane | NO |
| IVNTVSELVYENKFVP | 1668745 | Cell/Virus Enclosed | NO |
| IVNVSLVKPSFYVYS | 1597717 | Ectodomain | YES |
| IVPDKSVYDQVVDNVY | 1668750 | Cell/Virus Enclosed | NO |
| IVPEYIDVNKTLQELS | 1668751 | Ectodomain | YES |
| IVPLCIFNGKFLSNTK | 1668756 | Ectodomain | YES |
| IVPPCPSILKVI | 1455876 | Cell/Virus Enclosed | NO |
| IVPSTAWEDGDY | 1455878 | Ectodomain | YES |
| IVPSTVWEDGDYYRKQ | 1668760 | Ectodomain | YES |
| IVPSTVWEEGDY | 1455882 | Ectodomain | YES |
| IVQASESKVFLE | 1455890 | Cell/Virus Enclosed | NO |
| IVQASESKVLLERFLP | 1668763 | Cell/Virus Enclosed | NO |
| IVQMISDYLSNLSDIL | 1668773 | Cell/Virus Enclosed | NO |
| IVQMLSDTLKGL | 1455904 | Cell/Virus Enclosed | NO |
| IVQMLSDYLLDLSDSV | 1668777 | Cell/Virus Enclosed | NO |

| | | | |
|---|---|---|---|
| IVRIVRTRSMTY | 1455929 | Cell/Virus Enclosed | NO |
| IVRRDAAFATHV | 1455934 | Cell/Virus Enclosed | NO |
| IVSDREVILCWETGKV | 1668796 | Cell/Virus Enclosed | NO |
| IVSEKISVMDKLDTGA | 1668798 | Cell/Virus Enclosed | NO |
| IVSERELILSWEIGKV | 1668799 | Cell/Virus Enclosed | NO |
| IVSGYEKSTFNKDIYC | 1668801 | Cell/Virus Enclosed | NO |
| IVSIRSSNQGNK | 1455958 | Cell/Virus Enclosed | NO |
| IVSKESGNTVVLTFTT | 1668804 | Capsid | UN |
| IVSLPVYHKHTF | 1455961 | Ectodomain | YES |
| IVSSFEGYDFKYIENG | 1668808 | Cell/Virus Enclosed | NO |
| IVTANLSIPSNWTTSV | 1668819 | Ectodomain | YES |
| IVTGDNDDQIVVTGDD | 1668821 | Cell/Virus Enclosed | NO |
| IVTGLFKDCSRETSGL | 1668822 | Cell/Virus Enclosed | NO |
| IVTLEQYSTCDICKST | 1668825 | Cell/Virus Enclosed | NO |
| IVVAVMDEDIGSTMFN | 1668837 | Cell/Virus Enclosed | NO |
| IVVAWHVVRDSRFVMR | 1668838 | Cell/Virus Enclosed | NO |
| IVVDCATYVCNG | 1456015 | Ectodomain | YES |
| IVVQPHNGVLEI | 1456036 | Ectodomain | YES |
| IVVTGDDVDDIESIYD | 1668859 | Cell/Virus Enclosed | NO |
| IVWGAKDFNTLS | 1456050 | Cell/Virus Enclosed | NO |
| IVWGVKDFNTLSQEGK | 1668867 | Cell/Virus Enclosed | NO |
| IVWHAKDFNSLSAEGR | 1668868 | Cell/Virus Enclosed | NO |
| IVWPLTEYDFTD | 1456054 | Cell/Virus Enclosed | NO |
| IVWVATEGALNTPKDHI | 29504 | Capsid | UN |
| IVYAVLSDVDGLKYTK | 1668876 | Cell/Virus Enclosed | NO |
| IVYGDFSHGQLG | 1456071 | Cell/Virus Enclosed | NO |
| IVYKAACVVDVNDSHS | 1668888 | Cell/Virus Enclosed | NO |
| IVYNNSGSNITQIKNA | 1668890 | Cell/Virus Enclosed | NO |
| IVYVGDPAQLPT | 1456083 | Cell/Virus Enclosed | NO |
| IWHIAYIVAYFL | 1456107 | Cell/Virus Enclosed | NO |
| IWHINQTTNKTT | 1456108 | Cell/Virus Enclosed | NO |
| IWLSHEKASLSS | 1456123 | Cell/Virus Enclosed | NO |
| IWMMWYWGPSLYNIL | 1668917 | Undetermined | UN |
| IWNGVIKNVNSVRDWL | 1668921 | Cell/Virus Enclosed | NO |
| IWNKRFGFIEDS | 1456132 | Ectodomain | YES |
| IWSVDAFNQFSS | 1456154 | Cell/Virus Enclosed | NO |
| IWTADEAIKFLKEWNF | 1668931 | Undetermined | UN |
| IWVATEGALNTPKDHI | 1668932 | Cell/Virus Enclosed | NO |
| IYAQTSDTAHACNVNR | 1668940 | Cell/Virus Enclosed | NO |
| IYCNSVINRLRCDQLS | 1668945 | Ectodomain | YES |
| IYCRAHVAHPTM | 1456185 | Cell/Virus Enclosed | NO |
| IYDEEDLYDVSK | 1456190 | Cell/Virus Enclosed | NO |
| IYDEEGGYDVSKPVMI | 1668954 | Cell/Virus Enclosed | NO |
| IYDEPTMTTSVP | 1456193 | Secreted | YES |
| IYDKKTKTHATL | 1456201 | Ectodomain | YES |
| IYDRLDIIQADQ | 1456207 | Ectodomain | YES |
| IYDRLDSIQADQQVDR | 1668960 | Ectodomain | YES |
| IYDRLDTIQADQQVDR | 1668961 | Ectodomain | YES |
| IYDSSDRGQECL | 1456214 | Cell/Virus Enclosed | NO |
| IYDSSDRGQESLARSA | 1668962 | Cell/Virus Enclosed | NO |
| IYEACLYHPQLPECLSPADA | 741576 | Ectodomain | YES |
| IYFAATEKSNVVRGWV | 1668967 | Ectodomain | YES |
| IYFITMSSKGQKGDAE | 1668969 | Cell/Virus Enclosed | NO |
| IYFLVRLLQDKT | 1456240 | Cell/Virus Enclosed | NO |
| IYGFNSTETITT | 1456247 | Ectodomain | YES |
| IYGLNSTETIST | 1456252 | Ectodomain | YES |
| IYGLNSTETITT | 1456253 | Ectodomain | YES |
| IYLINKFLKANP | 1456311 | Cell/Virus Enclosed | NO |
| IYLKGSGSDFSLADSV | 1668998 | Cell/Virus Enclosed | NO |
| IYMCVVPSINDLSFDK | 1669002 | Cell/Virus Enclosed | NO |
| IYNLFNVGHYDG | 1456344 | Cell/Virus Enclosed | NO |
| IYNRVKSGSPGDSSWH | 1669015 | Ectodomain | YES |
| IYPQGRTYSNIT | 1456370 | Ectodomain | YES |
| IYQLDENAQKFF | 1456377 | Cell/Virus Enclosed | NO |
| IYQSYMHIDPFP | 1456380 | Cell/Virus Enclosed | NO |
| IYSQNLCVDGFGVNKI | 1669036 | Cell/Virus Enclosed | NO |
| IYSQTAETAHSVNVNR | 1669037 | Cell/Virus Enclosed | NO |
| IYSTGNNVFQTQAGCL | 1669042 | Ectodomain | YES |
| IYTEIQPKLDEY | 1456412 | Cell/Virus Enclosed | NO |
| IYTGDWEGLHEVLTSA | 1669044 | Cell/Virus Enclosed | NO |
| IYVPEHCQNYSG | 1456428 | Cell/Virus Enclosed | NO |
| IYVRKNGQFQDHYDGF | 1669052 | Cell/Virus Enclosed | NO |
| IYVTNEIGLNASVTLK | 1669053 | Ectodomain | YES |
| KAACVVDVNDSHSMAV | 1669059 | Cell/Virus Enclosed | NO |
| KAAESKAFVDTIVPPC | 1669061 | Cell/Virus Enclosed | NO |
| KAAGGKVSFSDDVEVK | 1669062 | Cell/Virus Enclosed | NO |

| | | | |
|---|---|---|---|
| KAAYANFNSQDV | 1456469 | Cell/Virus Enclosed | NO |
| KAAYENFNSQDT | 1456471 | Cell/Virus Enclosed | NO |
| KACCYVGMQTVE | 1456474 | Cell/Virus Enclosed | NO |
| KACEVVVTNLNKSAGW | 1669073 | Cell/Virus Enclosed | NO |
| KACIDEVTTALE | 1456480 | Cell/Virus Enclosed | NO |
| KACNIAKSAYERDRAV | 1669075 | Cell/Virus Enclosed | NO |
| KACNIAKSVYERDKAV | 1669076 | Cell/Virus Enclosed | NO |
| KACQNLDCNCLG | 1456490 | Cell/Virus Enclosed | NO |
| KADDFVIASDTT | 1456495 | Cell/Virus Enclosed | NO |
| KADDFVTASDTTLRCC | 1669081 | Cell/Virus Enclosed | NO |
| KADELQALPQRQ | 1456497 | Cell/Virus Enclosed | NO |
| KADGIIYPQGRTYSNI | 1669083 | Ectodomain | YES |
| KADLVLDALASM | 1456506 | Cell/Virus Enclosed | NO |
| KAEDIFGPCWSA | 1456511 | Cell/Virus Enclosed | NO |
| KAEFDRESSVQK | 1456513 | Cell/Virus Enclosed | NO |
| KAEVVVNPANGHMAHG | 1669092 | Cell/Virus Enclosed | NO |
| KAFEFAEGQGVP | 1456527 | Cell/Virus Enclosed | NO |
| KAFGSPFITNGI | 1456528 | Cell/Virus Enclosed | NO |
| KAFLDVLNKALE | 1456532 | Cell/Virus Enclosed | NO |
| KAFPPTEPKKDK | 1456534 | Cell/Virus Enclosed | NO |
| KAGDKYYPVVVDDHVG | 1669101 | Cell/Virus Enclosed | NO |
| KAGHYTGQAGEM | 1456548 | Cell/Virus Enclosed | NO |
| KAGLLVSDGGPNLY | 97407 | Cell/Virus Enclosed | NO |
| KAIACVKDFDFKFYDA | 1669114 | Cell/Virus Enclosed | NO |
| KAIAVAAGKKFSKETA | 1669115 | Cell/Virus Enclosed | NO |
| KAIDVDWVEFYGFKDA | 1669121 | Cell/Virus Enclosed | NO |
| KAIGKKIKHEGDWDSF | 1669124 | Cell/Virus Enclosed | NO |
| KAILNTVKFCDH | 1456578 | Cell/Virus Enclosed | NO |
| KAIPSGNLIKFDIGSV | 1669133 | Cell/Virus Enclosed | NO |
| KAIQDGVTRDII | 1456586 | Cell/Virus Enclosed | NO |
| KAIQDGVTRDIM | 1456588 | Cell/Virus Enclosed | NO |
| KAIQDGVTRDIV | 1456589 | Cell/Virus Enclosed | NO |
| KAISQIQESLTTTSTAL | 29832 | Ectodomain | YES |
| KAKEESVKSSYAFATL | 1669140 | Cell/Virus Enclosed | NO |
| KAKGYPQFAELVPSTA | 1669144 | Capsid | UN |
| KAKPPLNRNYVF | 1456609 | Cell/Virus Enclosed | NO |
| KAKQKPIYLKGSGSDF | 1669152 | Cell/Virus Enclosed | NO |
| KALCFDKSKQFVPVQV | 1669157 | Underneath Membrane | NO |
| KALCFNKPKQFV | 1456626 | Underneath Membrane | NO |
| KALENKKKQLSAGGKNLS | 29852 | Cell/Virus Enclosed | NO |
| KALKELYGPIDP | 1456646 | Cell/Virus Enclosed | NO |
| KALNDFSNLGAD | 1456658 | Cell/Virus Enclosed | NO |
| KALNESYIDLKELGNY | 1669170 | Ectodomain | YES |
| KALTGLSTGTSQNTRISLRP | 29879 | Capsid | UN |
| KAMDTFTNNNGSDVLY | 1669180 | Cell/Virus Enclosed | NO |
| KAMETFNHNNGNDVLY | 1669181 | Cell/Virus Enclosed | NO |
| KAMETFTNNNGS | 1456677 | Cell/Virus Enclosed | NO |
| KAMFTKVVVGPTTEVK | 1669182 | Cell/Virus Enclosed | NO |
| KAMLDFSRDHNDILYT | 1669183 | Cell/Virus Enclosed | NO |
| KAMLDYAKDHNDMLYS | 1669184 | Cell/Virus Enclosed | NO |
| KAMNVAKAEFDRESSV | 1669187 | Cell/Virus Enclosed | NO |
| KANGNGSGDNGDTTW | 29899 | Capsid | UN |
| KANPLWHKAVFISPYN | 1669192 | Cell/Virus Enclosed | NO |
| KANSGNRTPTSG | 1456704 | Cell/Virus Enclosed | NO |
| KANTDIKLGVCV | 1456708 | Ectodomain | YES |
| KAPKEESTDQIS | 1456718 | Cell/Virus Enclosed | NO |
| KAPKNFCPCKLNGSCV | 1669200 | Ectodomain | YES |
| KAPLVYLNHSVPKLSD | 1669202 | Ectodomain | YES |
| KAPYVMLNTSIPNLPD | 1669207 | Ectodomain | YES |
| KAQEGCGIISRK | 1456739 | Ectodomain | YES |
| KAQFKPFAKVDGVYTN | 1669209 | Cell/Virus Enclosed | NO |
| KAQFKTFEKVDGVYTN | 1669210 | Cell/Virus Enclosed | NO |
| KAQSSGVIPENPVLFT | 1669214 | Cell/Virus Enclosed | NO |
| KARIEEVNTSLE | 1456749 | Cell/Virus Enclosed | NO |
| KARLYYEALSFEEQDE | 1669219 | Cell/Virus Enclosed | NO |
| KARNGVFISTEKLSRL | 1669221 | Cell/Virus Enclosed | NO |
| KARVECYDRFKVNETN | 1669223 | Cell/Virus Enclosed | NO |
| KARVYYESMSYQEQDE | 1669224 | Cell/Virus Enclosed | NO |
| KASYDTNLNKFNRASL | 1669232 | Cell/Virus Enclosed | NO |
| KATGDLAVCLGIGDSA | 1669240 | Cell/Virus Enclosed | NO |
| KATGDLAVCLGV | 1456789 | Cell/Virus Enclosed | NO |
| KAVAAALKSLGFDKPQ | 1669258 | Capsid | UN |
| KAVARKLERMADQAMT | 1669261 | Cell/Virus Enclosed | NO |
| KAVDCSSSFLSE | 1456811 | Ectodomain | YES |
| KAVFISPYNSQNFAAK | 1669266 | Cell/Virus Enclosed | NO |
| KAVNIAKNAYEKDKAV | 1669270 | Cell/Virus Enclosed | NO |

| | | | |
|---|---|---|---|
| KAVPSPLNWERK | 1456818 | Ectodomain | YES |
| KAVRNKISVCFN | 1456820 | Cell/Virus Enclosed | NO |
| KAVRSSFNFDQKELLA | 1669272 | Cell/Virus Enclosed | NO |
| KAYIDVLRNSFGKDLN | 1669279 | Cell/Virus Enclosed | NO |
| KAYLDFVNAGGAPLTN | 1669285 | Cell/Virus Enclosed | NO |
| KAYNTINNEQGTPLTP | 1669286 | Cell/Virus Enclosed | NO |
| KAYNVTQAFGRRGPEQ | 1669287 | Cell/Virus Enclosed | NO |
| KCCSIDSDVETRFITK | 1669297 | Cell/Virus Enclosed | NO |
| KCDDDCMASIRNNTYDHSKYREEAM | 1150868 | Ectodomain | YES |
| KCDINYKGKVFFQFDN | 1669306 | Cell/Virus Enclosed | NO |
| KCDSTLRLCVQSTHVDIRTLE | 110640 | Cell/Virus Enclosed | NO |
| KCEQFYQLYDAS | 1456901 | Cell/Virus Enclosed | NO |
| KCFFNGSNDVWI | 1456912 | Ectodomain | YES |
| KCFNCYPAGVNITLAN | 1669323 | Ectodomain | YES |
| KCFPNGSNDVWL | 1456917 | Ectodomain | YES |
| KCGDKFFPIVMNDKNI | 1669325 | Cell/Virus Enclosed | NO |
| KCGFSFDLNGLDALFF | 1669327 | Cell/Virus Enclosed | NO |
| KCGGCCDDYTGH | 1456922 | Underneath Membrane | NO |
| KCGGCCDDYTGYQELV | 1669328 | Underneath Membrane | NO |
| KCGNLEQNHILGVNNS | 1669330 | Cell/Virus Enclosed | NO |
| KCGNLHQRELLGVSDV | 1669331 | Cell/Virus Enclosed | NO |
| KCGVKQEQCTGL | 1456932 | Cell/Virus Enclosed | NO |
| KCGVKQEQRTGLDAVM | 1669333 | Cell/Virus Enclosed | NO |
| KCGVKQEQRTGV | 1456935 | Cell/Virus Enclosed | NO |
| KCGVLDGSYNAS | 1456936 | Ectodomain | YES |
| KCHNCCDEYGGHHDFV | 1669337 | Underneath Membrane | NO |
| KCHNDILAATDP | 1456941 | Cell/Virus Enclosed | NO |
| KCKGLNKPFVKDNVSF | 1669342 | Cell/Virus Enclosed | NO |
| KCKTQSIAPPTGVYEL | 1669346 | Ectodomain | YES |
| KCKVTDVIFPGQ | 1456969 | Cell/Virus Enclosed | NO |
| KCKVTDVLFPGQKYSS | 1669347 | Cell/Virus Enclosed | NO |
| KCKVTDVLFSGQ | 1456971 | Cell/Virus Enclosed | NO |
| KCNCGSESWSVGAWDG | 1669356 | Cell/Virus Enclosed | NO |
| KCNEKHDEEFCDMLRL | 189691 | Ectodomain | YES |
| KCNIGGAVCSKHANLY | 1669359 | Cell/Virus Enclosed | NO |
| KCNRAGSNDIWM | 1457002 | Ectodomain | YES |
| KCNRCCDRYEEYDLEP | 1669361 | Underneath Membrane | NO |
| KCNRSGSNDIWMQNKG | 1669362 | Ectodomain | YES |
| KCNYVFNNTLSR | 1457011 | Ectodomain | YES |
| KCPLDSRTGSFN | 1457015 | Ectodomain | YES |
| KCPRYVKQRSLLLATGMKNVPEIPK | 1150881 | Ectodomain | YES |
| KCSRLLSDDRTEVPQL | 1669376 | Ectodomain | YES |
| KCSSNGSNDVWI | 1457043 | Ectodomain | YES |
| KCSVKSFKIDKG | 1457050 | Ectodomain | YES |
| KCTIKNFNVSKG | 1457055 | Ectodomain | YES |
| KCTKGLSIADLACAQY | 1669384 | Ectodomain | YES |
| KCTKYNIYDVAG | 1457058 | Ectodomain | YES |
| KCTKYNIYDVSGVGVI | 1669385 | Ectodomain | YES |
| KCTKYNIYDVVG | 1457060 | Ectodomain | YES |
| KCTSDNINDKDTGPPP | 1669393 | Ectodomain | YES |
| KCTSDTSYINDV | 1457069 | Ectodomain | YES |
| KCVALCDAMVAK | 1457076 | Cell/Virus Enclosed | NO |
| KCVDTVKEAQSA | 1457083 | Cell/Virus Enclosed | NO |
| KCWTEEDLSIGPHEFC | 1669411 | Cell/Virus Enclosed | NO |
| KCWVEEDLTKGPHEFC | 1669413 | Cell/Virus Enclosed | NO |
| KCYGVSATKLNDLCFS | 1669422 | Ectodomain | YES |
| KDADYNAKVEISVTPI | 1669428 | Cell/Virus Enclosed | NO |
| KDAIKPQQVTKH | 1457124 | Capsid | UN |
| KDALQSREAVLV | 1457126 | Cell/Virus Enclosed | NO |
| KDAPYIVGDVIS | 1457130 | Cell/Virus Enclosed | NO |
| KDAPYMVGDVIT | 1457132 | Cell/Virus Enclosed | NO |
| KDAQVVQLPIQD | 1457133 | Cell/Virus Enclosed | NO |
| KDASFIGKNLKSNCVR | 1669435 | Cell/Virus Enclosed | NO |
| KDATKPQQVTKHTAKE | 1669437 | Cell/Virus Enclosed | NO |
| KDATKPQQVTKQ | 1457139 | Capsid | UN |
| KDCARNPIDLPPSHAT | 1669441 | Cell/Virus Enclosed | NO |
| KDCSKIINGLHP | 1457151 | Cell/Virus Enclosed | NO |
| KDCSKIITGLHP | 1457152 | Cell/Virus Enclosed | NO |
| KDCSKSYSGYHPAHAP | 1669450 | Cell/Virus Enclosed | NO |
| KDCSNSYSGYHP | 1457156 | Cell/Virus Enclosed | NO |
| KDCVDINKAIPSGNLI | 1669453 | Cell/Virus Enclosed | NO |
| KDDGNCVVLELDPPCK | 1669461 | Cell/Virus Enclosed | NO |
| KDDGNFVVLELDPPCK | 1669462 | Cell/Virus Enclosed | NO |
| KDDQIGYYRRAT | 1457177 | Cell/Virus Enclosed | NO |
| KDDYCDGSLGCKMCLF | 1669464 | Cell/Virus Enclosed | NO |
| KDEAIKRVKGWV | 1457181 | Cell/Virus Enclosed | NO |

| | | | |
|---|---|---|---|
| KDEISVTPIKNT | 1457185 | Cell/Virus Enclosed | NO |
| KDFDFKFYDAQPIVKS | 1669469 | Cell/Virus Enclosed | NO |
| KDFDFYRYNKPTILDI | 1669470 | Cell/Virus Enclosed | NO |
| KDFGYVCDGSFY | 1457193 | Cell/Virus Enclosed | NO |
| KDFKPQQVTKQN | 1457194 | Cell/Virus Enclosed | NO |
| KDFLSFDNDVFI | 1457197 | Cell/Virus Enclosed | NO |
| KDFNTILNTACG | 1457205 | Cell/Virus Enclosed | NO |
| KDFQFMLWCNDD | 1457207 | Cell/Virus Enclosed | NO |
| KDFQFMLWCNDE | 1457208 | Cell/Virus Enclosed | NO |
| KDFQFMLWCNDH | 1457209 | Cell/Virus Enclosed | NO |
| KDFQFMLWGNDE | 1457211 | Cell/Virus Enclosed | NO |
| KDGDDGYFLPYPDPSR | 1669487 | Cell/Virus Enclosed | NO |
| KDGDYYRKQLSP | 1457219 | Ectodomain | YES |
| KDGEGAPPAKKL | 30152 | Cell/Virus Enclosed | NO |
| KDGFLYVYKGYQPIDV | 1669492 | Ectodomain | YES |
| KDGFNLEYWVTP | 1457224 | Ectodomain | YES |
| KDGFSPEYWVTP | 1457229 | Ectodomain | YES |
| KDGFVSVELQPP | 1457231 | Cell/Virus Enclosed | NO |
| KDGFYETIVSYW | 1457234 | Cell/Virus Enclosed | NO |
| KDGGHFFVSGDY | 1457236 | Cell/Virus Enclosed | NO |
| KDGITKLADVEA | 1457243 | Cell/Virus Enclosed | NO |
| KDGIVWVHEHGA | 1457246 | Cell/Virus Enclosed | NO |
| KDGIVWVHEYGA | 1457247 | Cell/Virus Enclosed | NO |
| KDGIYFAATEKSNVVR | 1669501 | Ectodomain | YES |
| KDGKVQTFYPRL | 1457252 | Cell/Virus Enclosed | NO |
| KDGKYFTSEPPI | 1457255 | Cell/Virus Enclosed | NO |
| KDGKYFTSEPPVTYSP | 1669503 | Cell/Virus Enclosed | NO |
| KDGQVQTFYPRF | 1457267 | Cell/Virus Enclosed | NO |
| KDGQVQTFYPRLQASA | 1669505 | Cell/Virus Enclosed | NO |
| KDGVEDTPPHYF | 1457273 | Cell/Virus Enclosed | NO |
| KDGVENTPPHFF | 1457274 | Cell/Virus Enclosed | NO |
| KDGVENTPPHYF | 1457275 | Cell/Virus Enclosed | NO |
| KDGVMYSEFPAI | 1457277 | Ectodomain | YES |
| KDGVMYSEFPTI | 1457278 | Ectodomain | YES |
| KDGVNFNVDDINFSPV | 1669512 | Ectodomain | YES |
| KDGVNFNVDDVN | 1457281 | Ectodomain | YES |
| KDGVYYPSNGTNILPV | 1669513 | Cell/Virus Enclosed | NO |
| KDHIGTRNPNNNAAIV | 1669518 | Cell/Virus Enclosed | NO |
| KDIGTYEMYVKW | 1457299 | Ectodomain | YES |
| KDIPEKVSLVSV | 1457308 | Cell/Virus Enclosed | NO |
| KDIPGGCAMPIM | 1457309 | Ectodomain | YES |
| KDISQLTQAIVKNHKNLLKIAQY | 30178 | Ectodomain | YES |
| KDKDLSGAVPSDLNTS | 1669533 | Cell/Virus Enclosed | NO |
| KDKKKKTDESQPLPQR | 1669537 | Cell/Virus Enclosed | NO |
| KDKNKLIGYWNI | 1457328 | Cell/Virus Enclosed | NO |
| KDLKFRQRSDGV | 1457344 | Cell/Virus Enclosed | NO |
| KDLNANMSLADC | 1457349 | Cell/Virus Enclosed | NO |
| KDLNANMSLAECKRAL | 1669553 | Cell/Virus Enclosed | NO |
| KDLNFLNRVRGTSVNA | 1669554 | Cell/Virus Enclosed | NO |
| KDLREYVFKNKD | 1457353 | Ectodomain | YES |
| KDMQFDLYVNVY | 1457363 | Cell/Virus Enclosed | NO |
| KDNAVATFYPQLQSAE | 1669566 | Cell/Virus Enclosed | NO |
| KDNDGKNVHLKDVTKE | 1669569 | Cell/Virus Enclosed | NO |
| KDNFKDGLCMFWNCNV | 1669571 | Cell/Virus Enclosed | NO |
| KDNHLSTFYPQLQSAE | 1669572 | Cell/Virus Enclosed | NO |
| KDNKNLPKFIEQISAF | 1669573 | Capsid | UN |
| KDNRIPSYGVLSVDL | 30220 | Ectodomain | YES |
| KDNTVLQALQSEFVNM | 1669575 | Cell/Virus Enclosed | NO |
| KDNVIFVADDSG | 1457376 | Cell/Virus Enclosed | NO |
| KDPPNVQIHTIDGSSG | 1669581 | Secreted | YES |
| KDPSFKDNVILLNKHI | 1669583 | Cell/Virus Enclosed | NO |
| KDQTPIVWHAKD | 1457394 | Cell/Virus Enclosed | NO |
| KDQVILLNKHIDAYKT | 1669586 | Cell/Virus Enclosed | NO |
| KDQYGSDINGVY | 1457396 | Cell/Virus Enclosed | NO |
| KDQYGTDIDGIY | 1457398 | Cell/Virus Enclosed | NO |
| KDQYGTDINGVY | 1457401 | Cell/Virus Enclosed | NO |
| KDSFKDGLCMFWNCNV | 1669591 | Cell/Virus Enclosed | NO |
| KDSFTLYDQTVV | 1457406 | Cell/Virus Enclosed | NO |
| KDSNFLNRVRGSIVNA | 1669598 | Cell/Virus Enclosed | NO |
| KDSYCRLKNSVSDVAY | 1669603 | Cell/Virus Enclosed | NO |
| KDTGPPPISTDT | 1457425 | Ectodomain | YES |
| KDTGVIYGFNST | 1457427 | Ectodomain | YES |
| KDTGVIYGLNFT | 1457428 | Ectodomain | YES |
| KDTLDDILQQANKSVG | 1669610 | Cell/Virus Enclosed | NO |
| KDTNFLNGFGVR | 1457441 | Cell/Virus Enclosed | NO |
| KDTNFLNGVRGA | 1457442 | Cell/Virus Enclosed | NO |

| | | | |
|---|---|---|---|
| KDTNFLNRVRGA | 1457443 | Cell/Virus Enclosed | NO |
| KDTNFLNRVRGTSVDA | 1669612 | Cell/Virus Enclosed | NO |
| KDVDNPHLMGWDYPKC | 1669617 | Cell/Virus Enclosed | NO |
| KDVDNPVLMGWDYPKC | 1669618 | Cell/Virus Enclosed | NO |
| KDVNTHATRLTI | 1457469 | Cell/Virus Enclosed | NO |
| KDWRLINKFDVINGVK | 1669635 | Cell/Virus Enclosed | NO |
| KDWYDFVENPDIINIY | 1669638 | Cell/Virus Enclosed | NO |
| KDWYDFVENPDIINVY | 1669639 | Cell/Virus Enclosed | NO |
| KDYEFKFYDAFPVAKS | 1669642 | Cell/Virus Enclosed | NO |
| KDYISSNGPLKVGAGV | 1669644 | Cell/Virus Enclosed | NO |
| KEAPTTKPTEKP | 30304 | Secreted | YES |
| KEDDVDDSGDSS | 1457529 | Cell/Virus Enclosed | NO |
| KEDGDVLLAEFDTYDP | 1669655 | Cell/Virus Enclosed | NO |
| KEDGDVVLAEFD | 1457532 | Cell/Virus Enclosed | NO |
| KEDLHTLYVDSK | 1457535 | Cell/Virus Enclosed | NO |
| KEDLKFPRGQGVPINTNSSP | 1392232 | Capsid | UN |
| KEDLRFPRGQGVPINT | 1669658 | Cell/Virus Enclosed | NO |
| KEDVVIFINNTTYPTN | 1669659 | Cell/Virus Enclosed | NO |
| KEDYQIGGYSEDRHSG | 1669660 | Secreted | YES |
| KEEDVQQAQQEA | 1457542 | Cell/Virus Enclosed | NO |
| KEEFVVKEKLNAFLVH | 1669663 | Cell/Virus Enclosed | NO |
| KEEGDVLLAEFA | 1457545 | Cell/Virus Enclosed | NO |
| KEELDQWFKNQI | 1457549 | Ectodomain | YES |
| KEELTDQMSEPP | 1457553 | Cell/Virus Enclosed | NO |
| KEESTDQISEPP | 1457558 | Cell/Virus Enclosed | NO |
| KEESTDQVSEPP | 1457562 | Cell/Virus Enclosed | NO |
| KEESVKSSYAFATLKE | 1669669 | Cell/Virus Enclosed | NO |
| KEESVKSSYAYATLKE | 1669670 | Cell/Virus Enclosed | NO |
| KEFEFAEGQGVP | 1457573 | Cell/Virus Enclosed | NO |
| KEFEFVEGQGVPIAPG | 1669675 | Cell/Virus Enclosed | NO |
| KEFQFAEGQGVP | 1457581 | Capsid | UN |
| KEFVEYKITEWP | 1457587 | Cell/Virus Enclosed | NO |
| KEGAKTVNTSLGNRKR | 1669680 | Capsid | UN |
| KEGIIWVHEDGA | 1457602 | Cell/Virus Enclosed | NO |
| KEGVMYSEFPAI | 1457621 | Ectodomain | YES |
| KEHRVQGTQRTR | 1457626 | Cell/Virus Enclosed | NO |
| KEIAPVYEADTE | 1457628 | Cell/Virus Enclosed | NO |
| KEIGTYEMYVKW | 1457633 | Ectodomain | YES |
| KEIISIPNIIKV | 1457638 | Cell/Virus Enclosed | NO |
| KEITRENVETLT | 1457656 | Cell/Virus Enclosed | NO |
| KEIVDTVSALVYENKL | 1669715 | Cell/Virus Enclosed | NO |
| KEIVETVSALVYDNKL | 1669716 | Cell/Virus Enclosed | NO |
| KEKKQKAPKEESTDQM | 1669720 | Cell/Virus Enclosed | NO |
| KEKTYYELTKSCGVVA | 1669722 | Cell/Virus Enclosed | NO |
| KELGIVMNMDVDTHRY | 1669728 | Cell/Virus Enclosed | NO |
| KELGLVMNMDVS | 1457688 | Cell/Virus Enclosed | NO |
| KELGVVMNFDVD | 1457695 | Cell/Virus Enclosed | NO |
| KELGVVMNLDVDTHRY | 1669731 | Cell/Virus Enclosed | NO |
| KELKFRQRSDGV | 1457696 | Cell/Virus Enclosed | NO |
| KELLDYYNMLVN | 1457700 | Cell/Virus Enclosed | NO |
| KELLLLWESGKAKPPL | 1669736 | Cell/Virus Enclosed | NO |
| KELLLSWESGKVKPPL | 1669737 | Cell/Virus Enclosed | NO |
| KELWHLDTGAVS | 1457714 | Ectodomain | YES |
| KELWHLDTGVVS | 1457716 | Ectodomain | YES |
| KELYGPADPTFL | 1457721 | Cell/Virus Enclosed | NO |
| KELYGSVDPTFL | 1457726 | Cell/Virus Enclosed | NO |
| KENIDGDAMHAN | 1457737 | Cell/Virus Enclosed | NO |
| KENVDFINSMQFNYKI | 1669759 | Cell/Virus Enclosed | NO |
| KEPVHVYANDSI | 1457757 | Cell/Virus Enclosed | NO |
| KEQLQLLMPLRA | 1457772 | Cell/Virus Enclosed | NO |
| KEQRVQGGITQR | 1457779 | Cell/Virus Enclosed | NO |
| KEQRVQGSITQRTRTR | 1669772 | Cell/Virus Enclosed | NO |
| KEQRVQGSTTQR | 1457784 | Capsid | UN |
| KEQRVQSSITQR | 1457786 | Cell/Virus Enclosed | NO |
| KEQYGTDIDGVY | 1457793 | Cell/Virus Enclosed | NO |
| KERSSLRYGNVLDV | 30597 | Capsid | UN |
| KESDEYILAKGPLQVG | 1669780 | Cell/Virus Enclosed | NO |
| KESIPIVWGAKD | 1457811 | Cell/Virus Enclosed | NO |
| KESWHMDTGVVP | 1457825 | Ectodomain | YES |
| KESYFSKKDWYDFVEN | 1669789 | Cell/Virus Enclosed | NO |
| KETLHSTSPEGN | 30629 | Secreted | YES |
| KETLYRIDGAYL | 1457835 | Cell/Virus Enclosed | NO |
| KETSYTTTIKPL | 1457840 | Cell/Virus Enclosed | NO |
| KETVFVEPKDGG | 1457846 | Cell/Virus Enclosed | NO |
| KETVFVEPKNGG | 1457847 | Cell/Virus Enclosed | NO |
| KEVCKTTTGIPEYNFI | 1669807 | Cell/Virus Enclosed | NO |

| | | | |
|---|---|---|---|
| KEVLVLTGCCDN | 1457869 | Cell/Virus Enclosed | NO |
| KEVRQKILNKPR | 1457872 | Cell/Virus Enclosed | NO |
| KEVSAAYYNDWQ | 1457873 | Ectodomain | YES |
| KEVSAVYYNSWQ | 1457875 | Ectodomain | YES |
| KEVTFFEGDSHD | 1457876 | Cell/Virus Enclosed | NO |
| KFAEGQGVPIAP | 1457896 | Cell/Virus Enclosed | NO |
| KFAIPNGRKVDLQLGN | 1669834 | Ectodomain | YES |
| KFAVESNVMDVN | 1457911 | Cell/Virus Enclosed | NO |
| KFCDHMVKDGLV | 1457920 | Cell/Virus Enclosed | NO |
| KFCSETVNGYAK | 1457928 | Cell/Virus Enclosed | NO |
| KFDNRGVITNAVDCSS | 1669857 | Ectodomain | YES |
| KFDSGTVSKTSDWKCK | 1669858 | Cell/Virus Enclosed | NO |
| KFEFANDTKIAS | 1457954 | Ectodomain | YES |
| KFEHGRGPGKCY | 1457955 | Ectodomain | YES |
| KFEHQRGPGKCY | 1457956 | Ectodomain | YES |
| KFELAKVQNLSG | 1457957 | Cell/Virus Enclosed | NO |
| KFFEIKHADYTL | 1457968 | Cell/Virus Enclosed | NO |
| KFFYASNGTYNCTDAV | 1669876 | Ectodomain | YES |
| KFGFIEDSVFVP | 1457987 | Ectodomain | YES |
| KFGFTPLNKQRC | 1457988 | Cell/Virus Enclosed | NO |
| KFGIITKVEADF | 1457990 | Ectodomain | YES |
| KFGNTSFACLSN | 1458000 | Ectodomain | YES |
| KFGSVCFSLKDI | 1458005 | Ectodomain | YES |
| KFIDKLNVLFDGRDNG | 1669890 | Cell/Virus Enclosed | NO |
| KFIGTFESAAAG | 1458011 | Cell/Virus Enclosed | NO |
| KFKEVCKTTTGIPEYN | 1669905 | Cell/Virus Enclosed | NO |
| KFKGKCVQVPIGCLDP | 1669906 | Cell/Virus Enclosed | NO |
| KFKNSVASINSAIVCA | 1669912 | Cell/Virus Enclosed | NO |
| KFKTPHIVRQRL | 1458041 | Cell/Virus Enclosed | NO |
| KFLADHFNYNVS | 1458045 | Cell/Virus Enclosed | NO |
| KFLDGLDLTPVI | 1458048 | Cell/Virus Enclosed | NO |
| KFLEELNAFTREMQQH | 1669922 | Capsid | UN |
| KFLNPDREYDFRDLT | 30774 | Ectodomain | YES |
| KFLSNTKYFDHS | 1458072 | Ectodomain | YES |
| KFLSNTKYYDDSQYYF | 1669936 | Ectodomain | YES |
| KFNKWQWQEAWY | 1458098 | Cell/Virus Enclosed | NO |
| KFNQALGAMLTG | 1458102 | Ectodomain | YES |
| KFNQALGAMQTGFTTT | 1669951 | Ectodomain | YES |
| KFNSCKSDILSLNNPI | 1669953 | Ectodomain | YES |
| KFPLKLAGTAVINLRA | 1669958 | Cell/Virus Enclosed | NO |
| KFPPPSFYMPLL | 1458124 | Cell/Virus Enclosed | NO |
| KFQDSKPPLPPH | 1458134 | Cell/Virus Enclosed | NO |
| KFQTVKPGNFNQDFYD | 1669970 | Cell/Virus Enclosed | NO |
| KFQTVKPGNFNQDFYE | 1669971 | Cell/Virus Enclosed | NO |
| KFRNTTFDFLSN | 1458146 | Ectodomain | YES |
| KFRQLLWFHISCLTFGRETVIEY | 79583 | Capsid | UN |
| KFSGFKLDGFNHQFVT | 1669983 | Cell/Virus Enclosed | NO |
| KFSKLFTIAMWL | 1458166 | Cell/Virus Enclosed | NO |
| KFTEESFYKNMYLKSA | 1669996 | Cell/Virus Enclosed | NO |
| KFTIGNMEEFIQ | 1458182 | Ectodomain | YES |
| KFTPLSVASTLV | 1458189 | Cell/Virus Enclosed | NO |
| KFTPLSVESTPV | 1458193 | Cell/Virus Enclosed | NO |
| KFTPLSVESTSV | 1458194 | Cell/Virus Enclosed | NO |
| KFTVSTQPQENT | 1458197 | Cell/Virus Enclosed | NO |
| KFTVSTQPQGNS | 1458200 | Cell/Virus Enclosed | NO |
| KFVGTFESAAAG | 1458212 | Cell/Virus Enclosed | NO |
| KFVVESNVMDVNDCFK | 1670029 | Cell/Virus Enclosed | NO |
| KFVYMHQLELST | 1458232 | Cell/Virus Enclosed | NO |
| KFYDAFPVAKSVKQLF | 1670034 | Cell/Virus Enclosed | NO |
| KFYGGWDDMLRHLIKD | 1670037 | Cell/Virus Enclosed | NO |
| KFYGGWDNMLKNLMAD | 1670040 | Cell/Virus Enclosed | NO |
| KFYLYKKVNNEFIDPT | 1670047 | Cell/Virus Enclosed | NO |
| KFYQHVINGCKDTACL | 1670049 | Cell/Virus Enclosed | NO |
| KGAFLNGSCGSV | 1458270 | Cell/Virus Enclosed | NO |
| KGAHVASADAIMTRCL | 1670058 | Cell/Virus Enclosed | NO |
| KGAHVASSDAIMTRCL | 1670059 | Cell/Virus Enclosed | NO |
| KGAINTSLPFQNIHPITIGKCPKYV | 1151775 | Ectodomain | YES |
| KGAVQKESDEYILAKG | 1670062 | Cell/Virus Enclosed | NO |
| KGAYNKDHGRGGFGHV | 1670065 | Cell/Virus Enclosed | NO |
| KGAYNKDHKRVG | 1458290 | Cell/Virus Enclosed | NO |
| KGCFTGVDGELPVAVV | 1670068 | Cell/Virus Enclosed | NO |
| KGCNAVAKHDFFTWHE | 1670069 | Cell/Virus Enclosed | NO |
| KGCVADQSRKWIGHCT | 1670072 | Cell/Virus Enclosed | NO |
| KGCVPLNAIPSLAANT | 1670073 | Cell/Virus Enclosed | NO |
| KGCVPLSAIPALAANT | 1670074 | Cell/Virus Enclosed | NO |
| KGCVVDQSRKWI | 1458303 | Cell/Virus Enclosed | NO |

| | | | |
|---|---|---|---|
| KGDAAIKDFDYYRYNR | 1670075 | Cell/Virus Enclosed | NO |
| KGDAEDTLTKLSKYLA | 1670076 | Cell/Virus Enclosed | NO |
| KGDAEEALSKLSEYLI | 1670077 | Cell/Virus Enclosed | NO |
| KGDKLFVEHYNE | 1458313 | Cell/Virus Enclosed | NO |
| KGDKLSVEHYNE | 1458314 | Cell/Virus Enclosed | NO |
| KGDSCSSNCKHSVAQL | 1670084 | Cell/Virus Enclosed | NO |
| KGECCICHKMQT | 1458325 | Cell/Virus Enclosed | NO |
| KGEGDGGIISEG | 1458330 | Cell/Virus Enclosed | NO |
| KGEGDGGITSEGNALY | 1670091 | Cell/Virus Enclosed | NO |
| KGFIRFPEVLREGLVR | 1670097 | Cell/Virus Enclosed | NO |
| KGFKKGVMYSEF | 1458344 | Ectodomain | YES |
| KGFPQFAELIPN | 1458346 | Cell/Virus Enclosed | NO |
| KGFPQLAELIPNQAAL | 1670099 | Capsid | UN |
| KGGAPVKKVAFGGDQV | 1670105 | Cell/Virus Enclosed | NO |
| KGGFKFGDPADSRDFL | 1670108 | Cell/Virus Enclosed | NO |
| KGGLILKFDSGTVSKT | 1670112 | Cell/Virus Enclosed | NO |
| KGGLILRFDSGT | 1458379 | Cell/Virus Enclosed | NO |
| KGHFDGQIGEAP | 1458392 | Cell/Virus Enclosed | NO |
| KGHFKFDEPSDATDFI | 1670123 | Cell/Virus Enclosed | NO |
| KGHYTVYDTAKKSMYD | 1670124 | Cell/Virus Enclosed | NO |
| KGIAPGTSVLRQWLPT | 1670125 | Cell/Virus Enclosed | NO |
| KGIFANCFSFVK | 1458402 | Cell/Virus Enclosed | NO |
| KGIFCVMCDKTLFDSL | 1670127 | Cell/Virus Enclosed | NO |
| KGIKVLPPLLSENQIS | 1670133 | Ectodomain | YES |
| KGILTDCFSFVK | 1458411 | Cell/Virus Enclosed | NO |
| KGIPIVDDLVVD | 1458416 | Cell/Virus Enclosed | NO |
| KGIYQTSNFRVVPSGDVVRF | 30988 | Ectodomain | YES |
| KGKEFEFVEGQGVPIA | 1670145 | Cell/Virus Enclosed | NO |
| KGKEFQFAEGQG | 1458435 | Capsid | UN |
| KGKFVQIPAQCVRDPV | 1670148 | Cell/Virus Enclosed | NO |
| KGKHERDLLIKAYNTI | 1670149 | Cell/Virus Enclosed | NO |
| KGKLLIRDTNKEVFVG | 1670151 | Cell/Virus Enclosed | NO |
| KGKLQRLSKEHIGLAG | 1670152 | Cell/Virus Enclosed | NO |
| KGKMFVRPVIEDYHTL | 1670153 | Underneath Membrane | NO |
| KGKPILWVNKASYDTN | 1670154 | Cell/Virus Enclosed | NO |
| KGKVFFQFDNLSSEDL | 1670157 | Cell/Virus Enclosed | NO |
| KGKVFFQFENLS | 1458456 | Cell/Virus Enclosed | NO |
| KGKWVQVPIGTNDPIR | 1670158 | Cell/Virus Enclosed | NO |
| KGKYAQNLLKKLIGGD | 1670159 | Cell/Virus Enclosed | NO |
| KGLACVKDYEFKFYDA | 1670162 | Cell/Virus Enclosed | NO |
| KGLDYKTFKDIV | 1458464 | Cell/Virus Enclosed | NO |
| KGLEELHDDLVN | 1458466 | Cell/Virus Enclosed | NO |
| KGLEELHYDLVN | 1458467 | Cell/Virus Enclosed | NO |
| KGLFKEGSTVDLKHFF | 1670168 | Cell/Virus Enclosed | NO |
| KGLLKEGSSVDLKHFF | 1670172 | Cell/Virus Enclosed | NO |
| KGLNVLWNKFVTGDVG | 1670175 | Cell/Virus Enclosed | NO |
| KGLQFQILDEGL | 1458476 | Cell/Virus Enclosed | NO |
| KGLSIADLACAQYYNG | 1670182 | Ectodomain | YES |
| KGLSKPFVKDNV | 1458483 | Cell/Virus Enclosed | NO |
| KGLWSPTITHTN | 1458491 | Cell/Virus Enclosed | NO |
| KGNKDEQIGYWNVQER | 1670190 | Capsid | UN |
| KGNVTHDASSAINRPQ | 1670192 | Cell/Virus Enclosed | NO |
| KGPKVKYLYFIK | 1458510 | Cell/Virus Enclosed | NO |
| KGPLCVDTSHFTTKYV | 1670201 | Ectodomain | YES |
| KGPQRADLNGVIVDKV | 1670203 | Cell/Virus Enclosed | NO |
| KGPVTDVFYKEI | 1458512 | Cell/Virus Enclosed | NO |
| KGPYEFCSQHTM | 1458514 | Cell/Virus Enclosed | NO |
| KGQKGDAEEALSKLSE | 1670207 | Cell/Virus Enclosed | NO |
| KGQQQQGQTVTKKSAAEASKKPRQ | 1311721 | Cell/Virus Enclosed | NO |
| KGQTTHESSSAVNIQQ | 1670213 | Cell/Virus Enclosed | NO |
| KGQVDLFRNARN | 1458519 | Cell/Virus Enclosed | NO |
| KGQVSDLLVKLL | 1458520 | Cell/Virus Enclosed | NO |
| KGRDFKFSDGQGVPIA | 1670214 | Capsid | UN |
| KGSALNGSCGSV | 1458528 | Cell/Virus Enclosed | NO |
| KGSFINGACGSPGYNV | 1670218 | Cell/Virus Enclosed | NO |
| KGSFLCGSCGSVGYTK | 1670219 | Cell/Virus Enclosed | NO |
| KGSFNDRDTGPP | 1458532 | Ectodomain | YES |
| KGSFNNRDTGPL | 1458537 | Ectodomain | YES |
| KGSFVGADGELPVAIS | 1670223 | Cell/Virus Enclosed | NO |
| KGSGSDFSLADSVVEV | 1670226 | Cell/Virus Enclosed | NO |
| KGSKDEQIGYWN | 1458551 | Cell/Virus Enclosed | NO |
| KGSNLEEELLLM | 1458556 | Cell/Virus Enclosed | NO |
| KGSPCTNVAVNPGDCPPLEL | 111450 | Capsid | UN |
| KGSSRNESWHFDKSEP | 1670233 | Ectodomain | YES |
| KGTDKLSTLWFK | 1458564 | Ectodomain | YES |
| KGTGVFVQDPAPIDID | 1670236 | Cell/Virus Enclosed | NO |

| | | | |
|---|---|---|---|
| KGTGVFVQDPSP | 1458570 | Cell/Virus Enclosed | NO |
| KGTIYSITPCNPPDQL | 1670239 | Ectodomain | YES |
| KGTMYVRPIIEDYHTL | 1670242 | Ectodomain | YES |
| KGTPVLQLKESQINEL | 1670243 | Cell/Virus Enclosed | NO |
| KGTQQSSWVYQS | 1458580 | Ectodomain | YES |
| KGTVCWIMASTF | 1458582 | Cell/Virus Enclosed | NO |
| KGVAPGTAVLRQ | 1458586 | Cell/Virus Enclosed | NO |
| KGVAPGTTVLKRWLPP | 1670249 | Cell/Virus Enclosed | NO |
| KGVCATGDCYVSTGGK | 1670250 | Cell/Virus Enclosed | NO |
| KGVCFGFDKWYVNDGR | 1670251 | Cell/Virus Enclosed | NO |
| KGVCQVGDCYVSTGGK | 1670252 | Cell/Virus Enclosed | NO |
| KGVDCTKKFLTV | 1458592 | Cell/Virus Enclosed | NO |
| KGVDYTKKFLIV | 1458594 | Cell/Virus Enclosed | NO |
| KGVDYTKKFLTVDGVQ | 1670253 | Cell/Virus Enclosed | NO |
| KGVDYTKKSLNV | 1458597 | Cell/Virus Enclosed | NO |
| KGVEELHDDLVNYIRD | 1670255 | Cell/Virus Enclosed | NO |
| KGVEKLVSWYIY | 1458601 | Cell/Virus Enclosed | NO |
| KGVGSANIFIGDKVGH | 1670257 | Cell/Virus Enclosed | NO |
| KGVKIQEGIVDY | 1458605 | Cell/Virus Enclosed | NO |
| KGVMEPVDYNVVTQRM | 1670264 | Cell/Virus Enclosed | NO |
| KGVMYSEFPAIT | 1458612 | Ectodomain | YES |
| KGVRCAVTDAKW | 1458615 | Cell/Virus Enclosed | NO |
| KGVREGGLNKVK | 1458616 | Cell/Virus Enclosed | NO |
| KGVRKTVKIEDA | 1458618 | Cell/Virus Enclosed | NO |
| KGVTIDKLRDYL | 1458625 | Cell/Virus Enclosed | NO |
| KGVTLDKLRDYLADYD | 1670271 | Cell/Virus Enclosed | NO |
| KGVTTHESSSAVNMQQ | 1670272 | Cell/Virus Enclosed | NO |
| KGVVGVLTLDNQ | 1458627 | Cell/Virus Enclosed | NO |
| KGWIPGNMIDGFACTT | 1670275 | Cell/Virus Enclosed | NO |
| KGWVGFDVEGAH | 1458630 | Cell/Virus Enclosed | NO |
| KGWYDPVENEDIHRVY | 1670277 | Cell/Virus Enclosed | NO |
| KGYHLMSFPQAAPHGV | 1670281 | Ectodomain | YES |
| KGYIAGIQNFKETLYY | 1670283 | Cell/Virus Enclosed | NO |
| KGYPQFAELVPSTAAM | 1670286 | Capsid | UN |
| KGYQPIDVVRDLPSGF | 1670287 | Ectodomain | YES |
| KHDIQITNESYNNYVP | 1670319 | Cell/Virus Enclosed | NO |
| KHDVSPFEMPFEELNG | 1670321 | Cell/Virus Enclosed | NO |
| KHEGDWDSFGKT | 1458705 | Cell/Virus Enclosed | NO |
| KHEHSLLVEAYNSILF | 1670325 | Cell/Virus Enclosed | NO |
| KHEMSPFEMPFE | 1458712 | Cell/Virus Enclosed | NO |
| KHERDLLIKAYNTINN | 1670327 | Cell/Virus Enclosed | NO |
| KHFFFTQKGDAAIKDF | 1670333 | Cell/Virus Enclosed | NO |
| KHGDFSAVSSPMSNMT | 1670342 | Underneath Membrane | NO |
| KHIDAYKTFPPTEPKKDKKKK | 1397350 | Capsid | UN |
| KHIGAPANLRVVGHAM | 1670355 | Cell/Virus Enclosed | NO |
| KHKATVVVQLKDSDIN | 1670363 | Cell/Virus Enclosed | NO |
| KHKFLFLQVFLL | 1458774 | Cell/Virus Enclosed | NO |
| KHKGGDSIDFIA | 1458777 | Cell/Virus Enclosed | NO |
| KHKKVSTNLCTHSFRKKQVR | 31124 | Cell/Virus Enclosed | NO |
| KHLIPLMSKGQK | 1458790 | Cell/Virus Enclosed | NO |
| KHLLEQWNLVIG | 1458794 | Undetermined | UN |
| KHLLPRWYFYYL | 1458795 | Cell/Virus Enclosed | NO |
| KHLNKTLNEGVLESFS | 1670381 | Cell/Virus Enclosed | NO |
| KHLNKYDCCLST | 1458797 | Cell/Virus Enclosed | NO |
| KHLVPLIHKGAA | 1458804 | Cell/Virus Enclosed | NO |
| KHLVPLMHKGAA | 1458805 | Cell/Virus Enclosed | NO |
| KHLYVSMPTAQSTM | 31133 | Capsid | UN |
| KHNGKLCKLRGVAPLHLGKCNIAGW | 1152132 | Ectodomain | YES |
| KHPCMVGPDYAY | 1458817 | Cell/Virus Enclosed | NO |
| KHQWNCFNCHSF | 1458826 | Cell/Virus Enclosed | NO |
| KHRCDINYKGKV | 1458829 | Cell/Virus Enclosed | NO |
| KHRRLSSDQDQSQTPETPAT | 110303 | Cell/Virus Enclosed | NO |
| KHSDFGLGDLVDSYFEND | 1458838 | Cell/Virus Enclosed | NO |
| KHVAYGCSKSGC | 1458865 | Cell/Virus Enclosed | NO |
| KHVDHYDGFYTQ | 1458868 | Cell/Virus Enclosed | NO |
| KHVNCCNATDRF | 1458878 | Cell/Virus Enclosed | NO |
| KHVRTNGDYNSW | 1458890 | Cell/Virus Enclosed | NO |
| KHWPQIAQFAPSASAFFGMS | 2060837 | Capsid | UN |
| KHWSMPYHPNPV | 1458903 | Cell/Virus Enclosed | NO |
| KHWSMPYHPNTVDCQD | 1670432 | Cell/Virus Enclosed | NO |
| KHYVYIGDPVQL | 1458909 | Cell/Virus Enclosed | NO |
| KIAQADTAGLYT | 1458921 | Cell/Virus Enclosed | NO |
| KICGGCCDDYTG | 1458929 | Underneath Membrane | NO |
| KICIVDNVYMAKAGDK | 1670448 | Cell/Virus Enclosed | NO |
| KIDGDYVYLPYPDPSR | 1670451 | Cell/Virus Enclosed | NO |
| KIDNASPASTVHATAT | 1670453 | Secreted | YES |

| | | | |
|---|---|---|---|
| KIDTTSSSCQLYYSLP | 1670454 | Ectodomain | YES |
| KIDVLLTVDGVNFKSI | 1670455 | Cell/Virus Enclosed | NO |
| KIEDAIIVNDENSSIK | 1670456 | Cell/Virus Enclosed | NO |
| KIEDLSIRAFQK | 1458953 | Cell/Virus Enclosed | NO |
| KIEDLSMRALQK | 1458956 | Cell/Virus Enclosed | NO |
| KIEYKPDLSQYY | 1458968 | Cell/Virus Enclosed | NO |
| KIFAYVKPFLGQ | 1458969 | Cell/Virus Enclosed | NO |
| KIFERGSVQVDNGSSI | 1670466 | Cell/Virus Enclosed | NO |
| KIFKARAIECYG | 1458973 | Cell/Virus Enclosed | NO |
| KIFKDDGNFVVL | 1458975 | Cell/Virus Enclosed | NO |
| KIFPTGFVLSIP | 1458977 | Underneath Membrane | NO |
| KIGDVKFKRLGDYVLT | 1670477 | Cell/Virus Enclosed | NO |
| KIGGLTFKTYAFDKPY | 1670479 | Cell/Virus Enclosed | NO |
| KIGTTEVKVSAL | 1459011 | Cell/Virus Enclosed | NO |
| KIHDVVNQQGNS | 1459013 | Ectodomain | YES |
| KIHHHHHHNTAS | 1459015 | Capsid | UN |
| KIKHDGDWDSFC | 1459042 | Cell/Virus Enclosed | NO |
| KIKHEGDWDSLC | 1459045 | Cell/Virus Enclosed | NO |
| KIKKYPCMVGPDYAYF | 1670502 | Cell/Virus Enclosed | NO |
| KIKNYGYLDIHP | 1459053 | Ectodomain | YES |
| KILAASFNKAINNIVA | 1670510 | Ectodomain | YES |
| KILAASFNKAMTNIVD | 1670511 | Ectodomain | YES |
| KILARAQSSETK | 1459066 | Cell/Virus Enclosed | NO |
| KILARSQSSETK | 1459067 | Cell/Virus Enclosed | NO |
| KILGNVFCDGID | 1459073 | Cell/Virus Enclosed | NO |
| KILKDDGNFVVLELDP | 1670516 | Cell/Virus Enclosed | NO |
| KILPFSDRPLNL | 1459085 | Cell/Virus Enclosed | NO |
| KILSMIRGPPRA | 1459089 | Cell/Virus Enclosed | NO |
| KILTLTPRNKWN | 1459092 | Cell/Virus Enclosed | NO |
| KINKYGLELHWAPEFP | 1670538 | Cell/Virus Enclosed | NO |
| KINRALHGANLR | 1459113 | Ectodomain | YES |
| KINSACRKVQHM | 1459115 | Cell/Virus Enclosed | NO |
| KINTGNGIKQLA | 1459116 | Cell/Virus Enclosed | NO |
| KIPANIVLPQGGYVAD | 1670546 | Cell/Virus Enclosed | NO |
| KIPANIVLSQGG | 1459124 | Cell/Virus Enclosed | NO |
| KIPLLGKMAIYICRMSNHP | 244229 | Ectodomain | YES |
| KIPTNFTIVGQEEFIQ | 1670547 | Ectodomain | YES |
| KIQAVVNADAEA | 1459131 | Ectodomain | YES |
| KIQAVVNANAEALNNL | 1670548 | Ectodomain | YES |
| KIQDVVNQQGNSLNHL | 1670550 | Ectodomain | YES |
| KIQDVVNQQGSALNHL | 1670551 | Ectodomain | YES |
| KIQPWTNDALNTICIK | 1670556 | Cell/Virus Enclosed | NO |
| KIQTAPHISLTM | 1459148 | Cell/Virus Enclosed | NO |
| KIQVVNSGPDQI | 1459150 | Cell/Virus Enclosed | NO |
| KIQVVNSGPDQTCNTP | 1670559 | Cell/Virus Enclosed | NO |
| KIQVVNSGSDIN | 1459151 | Cell/Virus Enclosed | NO |
| KIRELTAEDISL | 1459154 | Cell/Virus Enclosed | NO |
| KISADNKVIYTTEIAS | 1670562 | Cell/Virus Enclosed | NO |
| KISGKGQQQQGQ | 1459170 | Cell/Virus Enclosed | NO |
| KISSKAGNSIFR | 1459183 | Ectodomain | YES |
| KISTQLFEGDKFIGTF | 1670571 | Cell/Virus Enclosed | NO |
| KISTVQSKLTDLKCTN | 1670572 | Cell/Virus Enclosed | NO |
| KITEDGINVKDV | 1459192 | Cell/Virus Enclosed | NO |
| KITEFQLDYSIDVIDN | 1670579 | Cell/Virus Enclosed | NO |
| KITEFSWNADLYKLMS | 1670580 | Cell/Virus Enclosed | NO |
| KITNFRAILTAF | 1459203 | Ectodomain | YES |
| KITQWPTATGDV | 1459205 | Cell/Virus Enclosed | NO |
| KIVARAMLKCVA | 1459208 | Cell/Virus Enclosed | NO |
| KIVDKEVKDMQFDLYV | 1670593 | Cell/Virus Enclosed | NO |
| KIVGGAHTWFNA | 1459218 | Cell/Virus Enclosed | NO |
| KIVGGAPTWFNALRDF | 1670595 | Cell/Virus Enclosed | NO |
| KIVQWIKNTFSL | 1459235 | Cell/Virus Enclosed | NO |
| KIVYAILSDVDGLKYT | 1670612 | Cell/Virus Enclosed | NO |
| KIWNGVIKNVNSVRDW | 1670614 | Cell/Virus Enclosed | NO |
| KIYEIERGLLHS | 1459252 | Cell/Virus Enclosed | NO |
| KKAKIVSAMQTMLFGM | 1670632 | Cell/Virus Enclosed | NO |
| KKAMFTKVVVGPTTEV | 1670633 | Cell/Virus Enclosed | NO |
| KKAMNVAKAEFDRESS | 1670634 | Cell/Virus Enclosed | NO |
| KKCGCCCDDYTG | 1459284 | Underneath Membrane | NO |
| KKCGGCCDDYTGYQEL | 1670637 | Underneath Membrane | NO |
| KKCTKGLSIADLACAQ | 1670641 | Ectodomain | YES |
| KKDGFYETIVSHWMGI | 1670646 | Cell/Virus Enclosed | NO |
| KKDGVYYPSNGTNILP | 1670647 | Cell/Virus Enclosed | NO |
| KKDKKKKADELQ | 1459303 | Cell/Virus Enclosed | NO |
| KKDKKKKADETQALPQRQ | 1087574 | Cell/Virus Enclosed | NO |
| KKDKKKKTDESQPLPQ | 1670651 | Cell/Virus Enclosed | NO |

| | | | |
|---|---|---|---|
| KKDWYDFVENPDIINI | 1670656 | Cell/Virus Enclosed | NO |
| KKDWYDFVENPDIINV | 1670657 | Cell/Virus Enclosed | NO |
| KKEAAAAKNKMR | 1459315 | Cell/Virus Enclosed | NO |
| KKEFSGQCFKIL | 1459316 | Cell/Virus Enclosed | NO |
| KKEKKQKAPKEESTDQ | 1670660 | Cell/Virus Enclosed | NO |
| KKENKQKAPKEE | 1459325 | Cell/Virus Enclosed | NO |
| KKEPLLYVPIRL | 1459326 | Cell/Virus Enclosed | NO |
| KKGAKILHKPIV | 1459348 | Cell/Virus Enclosed | NO |
| KKGAKLLHKPIL | 1459350 | Cell/Virus Enclosed | NO |
| KKGAWNIGQEKS | 1459354 | Cell/Virus Enclosed | NO |
| KKGCFTGVDGELPVAV | 1670676 | Cell/Virus Enclosed | NO |
| KKGNSYPKLSKSYINDKGKEVLVLW | 1152423 | Ectodomain | YES |
| KKGVMYSEFPAI | 1459357 | Ectodomain | YES |
| KKHPCMVGPDYA | 1459361 | Cell/Virus Enclosed | NO |
| KKHVEVFTDGRL | 1459365 | Cell/Virus Enclosed | NO |
| KKIDVLLTVDGVNFKS | 1670686 | Cell/Virus Enclosed | NO |
| KKIKHEGDWDSFCKTI | 1670690 | Cell/Virus Enclosed | NO |
| KKISNCVADYSVLYNST | 31581 | Ectodomain | YES |
| KKKKADELQALP | 1459391 | Cell/Virus Enclosed | NO |
| KKKKTDESQPLPQRQK | 1670698 | Cell/Virus Enclosed | NO |
| KKKTDESQPLPQRQKK | 1670702 | Cell/Virus Enclosed | NO |
| KKLCVVPGNVVPGDVI | 1670705 | Cell/Virus Enclosed | NO |
| KKLDNDVLNGIISNAR | 1670707 | Cell/Virus Enclosed | NO |
| KKLERMADLALTNMYK | 1670709 | Cell/Virus Enclosed | NO |
| KKLFVANYSQDV | 1459411 | Ectodomain | YES |
| KKLGPIFNRALVSATE | 1670711 | Cell/Virus Enclosed | NO |
| KKLIGGDVTPVDQYMC | 1670712 | Cell/Virus Enclosed | NO |
| KKLQRQLYDNCYRNSN | 1670720 | Cell/Virus Enclosed | NO |
| KKLYNDLGNQILDSYS | 1670725 | Cell/Virus Enclosed | NO |
| KKLYNDLGTQILDSYS | 1670726 | Cell/Virus Enclosed | NO |
| KKMAQPSGCVERCVVR | 1670727 | Cell/Virus Enclosed | NO |
| KKMDEPFTEDTS | 1459442 | Cell/Virus Enclosed | NO |
| KKNFTYNVSTDF | 1459448 | Ectodomain | YES |
| KKNFTYNVSTDG | 1459449 | Ectodomain | YES |
| KKNFTYNVSTDW | 1459450 | Ectodomain | YES |
| KKNLADAKNSGSVNQQ | 1670731 | Cell/Virus Enclosed | NO |
| KKNLDEARFSGSANQQ | 1670732 | Cell/Virus Enclosed | NO |
| KKNLDEARFSGT | 1459454 | Cell/Virus Enclosed | NO |
| KKNYSELDDEEL | 1459462 | Cell/Virus Enclosed | NO |
| KKNYSKLDDEEP | 1459464 | Cell/Virus Enclosed | NO |
| KKPFKVEDSVII | 1459469 | Cell/Virus Enclosed | NO |
| KKPNKPLSQPRADKPS | 1670744 | Capsid | UN |
| KKPNRQGLGYYSPSYNDP | 31664 | Ectodomain | YES |
| KKPRQKRTATKAYNVT | 1670745 | Cell/Virus Enclosed | NO |
| KKQPTVTLLPAADMDDFSRQLQNSM | 1383528 | Capsid | UN |
| KKQVVLVSNNQEDFDL | 1670756 | Cell/Virus Enclosed | NO |
| KKRNVDLLSNFE | 1459510 | Cell/Virus Enclosed | NO |
| KKRNVDPLSNFEHKVI | 1670757 | Cell/Virus Enclosed | NO |
| KKRVEVFTDGKL | 1459513 | Cell/Virus Enclosed | NO |
| KKSAAEASKKPRQKRTATKQYNVTQ | 31693 | Capsid | UN |
| KKSAKTGTPKPSRNQS | 1670762 | Capsid | UN |
| KKSFGKSIIYTS | 1459521 | Cell/Virus Enclosed | NO |
| KKSKVVSALQTMLFSM | 1670765 | Cell/Virus Enclosed | NO |
| KKSMYDGDRFVKHDLS | 1670769 | Cell/Virus Enclosed | NO |
| KKSNLLIQEFLQYDSS | 1670770 | Cell/Virus Enclosed | NO |
| KKTDESQPLPQRQKKQ | 1670774 | Cell/Virus Enclosed | NO |
| KKTSWADQSERNYQTF | 1670776 | Capsid | UN |
| KKTYLDYIQQGGVPII | 1670780 | Cell/Virus Enclosed | NO |
| KKVAFCGDQVHE | 1459552 | Cell/Virus Enclosed | NO |
| KKVNNEFIDPTECIYT | 1670790 | Cell/Virus Enclosed | NO |
| KKVPIDEYITTY | 1459570 | Cell/Virus Enclosed | NO |
| KKWKIPQVGGLT | 1459586 | Cell/Virus Enclosed | NO |
| KKYGLEKYAFEH | 1459589 | Cell/Virus Enclosed | NO |
| KKYPCIVGPDYA | 1459598 | Cell/Virus Enclosed | NO |
| KKYPCMVGPDYAYFNG | 1670800 | Cell/Virus Enclosed | NO |
| KKYSELDDEEPM | 1459600 | Cell/Virus Enclosed | NO |
| KKYTYSFLPKEDGDVL | 1670802 | Cell/Virus Enclosed | NO |
| KLADCLQAYGHYSLQD | 1670804 | Cell/Virus Enclosed | NO |
| KLADVEADIASR | 1459611 | Cell/Virus Enclosed | NO |
| KLAGGKISFSDDVIVH | 1670807 | Cell/Virus Enclosed | NO |
| KLAGTAVINLRADQIN | 1670808 | Cell/Virus Enclosed | NO |
| KLAGTAVVNLRPDQLN | 1670809 | Cell/Virus Enclosed | NO |
| KLAGVSCIWSVD | 1459614 | Cell/Virus Enclosed | NO |
| KLANDAKESLRLFAAE | 1670818 | Cell/Virus Enclosed | NO |
| KLANDVKDTLRLFAAE | 1670819 | Cell/Virus Enclosed | NO |
| KLARWSASIDTG | 1459625 | Ectodomain | YES |

| | | | |
|---|---|---|---|
| KLASELSNTFGAISAS | 1670822 | Ectodomain | YES |
| KLASPMKSDEGG | 1459628 | Cell/Virus Enclosed | NO |
| KLASTEELPDEF | 1459629 | Cell/Virus Enclosed | NO |
| KLCHAVVSKSKE | 1459642 | Cell/Virus Enclosed | NO |
| KLCHAVVYKSKE | 1459643 | Cell/Virus Enclosed | NO |
| KLCMQLCGFCNF | 1459650 | Cell/Virus Enclosed | NO |
| KLCVVPGNVKPGDAVI | 1670835 | Cell/Virus Enclosed | NO |
| KLDANVPGYPKL | 1459655 | Cell/Virus Enclosed | NO |
| KLDDKDPNFKDQVILLNKH | 1383548 | Cell/Virus Enclosed | NO |
| KLDDKDPQFKDNVILL | 31780 | Capsid | UN |
| KLDDKDPSFKDNVILL | 1383549 | Cell/Virus Enclosed | NO |
| KLDEYYKKDNAY | 1459663 | Cell/Virus Enclosed | NO |
| KLDGFNHQFVTASSAT | 1670840 | Cell/Virus Enclosed | NO |
| KLDGSLCVGNGP | 1459668 | Ectodomain | YES |
| KLDNDVLNGIISNARN | 1670851 | Cell/Virus Enclosed | NO |
| KLDNPNSSEVDM | 1459686 | Cell/Virus Enclosed | NO |
| KLDNPSGSEVDM | 1459687 | Cell/Virus Enclosed | NO |
| KLDNPSSSEVDM | 1459688 | Cell/Virus Enclosed | NO |
| KLDNQALNSILDNAVK | 1670853 | Cell/Virus Enclosed | NO |
| KLDTGAQKFFQFGDFV | 1670857 | Cell/Virus Enclosed | NO |
| KLEDFIPTDSTV | 1459701 | Cell/Virus Enclosed | NO |
| KLEFANDTKIASQLGN | 1670862 | Ectodomain | YES |
| KLEKLISTARDGVRRG | 1670865 | Cell/Virus Enclosed | NO |
| KLERMADLALTNMYKE | 1670869 | Cell/Virus Enclosed | NO |
| KLERMADQAMTSMYKQ | 1670870 | Cell/Virus Enclosed | NO |
| KLESTRIYQILAIYSTVASSLVLVV | 1152749 | Undetermined | UN |
| KLETDGICLENH | 1459727 | Cell/Virus Enclosed | NO |
| KLETDRICPENH | 1459730 | Cell/Virus Enclosed | NO |
| KLFAAESQKATE | 1459732 | Cell/Virus Enclosed | NO |
| KLFCAFITKRIV | 1459737 | Cell/Virus Enclosed | NO |
| KLFITKDEAIKRVRGW | 1670882 | Cell/Virus Enclosed | NO |
| KLFLSGDDCFVSNSSV | 1670886 | Cell/Virus Enclosed | NO |
| KLFVESCTIYMCVVPS | 1670894 | Cell/Virus Enclosed | NO |
| KLGAEKEDGFYE | 1459777 | Cell/Virus Enclosed | NO |
| KLGAEKKDGFYETIVS | 1670896 | Cell/Virus Enclosed | NO |
| KLGAENKDGFYE | 1459779 | Cell/Virus Enclosed | NO |
| KLGASQRVAGDSGFAA | 1670897 | Underneath Membrane | NO |
| KLGERVRKAILN | 1459789 | Cell/Virus Enclosed | NO |
| KLGFDSSKEFVEYKIT | 1670900 | Cell/Virus Enclosed | NO |
| KLGNHFKESWHM | 1459805 | Ectodomain | YES |
| KLGPIFNRALVSATEF | 1670905 | Cell/Virus Enclosed | NO |
| KLGRELNCNVCA | 1459809 | Cell/Virus Enclosed | NO |
| KLGTGYSLADLPAYMT | 1670910 | Ectodomain | YES |
| KLGTSDSQFPIL | 1459824 | Cell/Virus Enclosed | NO |
| KLGVWKSPTDMQSWV | 31894 | Ectodomain | YES |
| KLHALDDATRNY | 1459828 | Cell/Virus Enclosed | NO |
| KLHPAFNVSDAYANLV | 1670921 | Cell/Virus Enclosed | NO |
| KLHPSFNVSDAYANLV | 1670922 | Cell/Virus Enclosed | NO |
| KLIASAFNNALH | 1459849 | Ectodomain | YES |
| KLIFEFEDDVIT | 1459854 | Cell/Virus Enclosed | NO |
| KLIQLCFTCHYF | 1459880 | Cell/Virus Enclosed | NO |
| KLISTARDGVRRGDNF | 1670942 | Cell/Virus Enclosed | NO |
| KLKAKNDNSSLCFKVY | 1670947 | Cell/Virus Enclosed | NO |
| KLKAKNESSSLCFKVY | 1670948 | Cell/Virus Enclosed | NO |
| KLKCNRCCDRYEEYDL | 1670952 | Underneath Membrane | NO |
| KLKCSDFYADKI | 1459898 | Cell/Virus Enclosed | NO |
| KLKDGVNFNVDDINFS | 1670956 | Ectodomain | YES |
| KLKEDYQIGGYSEDRH | 1670957 | Secreted | YES |
| KLKIQVVNSGSDMNCN | 1670965 | Cell/Virus Enclosed | NO |
| KLKISTQLFEGDKFIG | 1670966 | Cell/Virus Enclosed | NO |
| KLKLTYNKQMAN | 1459920 | Cell/Virus Enclosed | NO |
| KLKPMPFFYFDDSDCD | 1670976 | Cell/Virus Enclosed | NO |
| KLKVSTNLFEGDKFVG | 1670988 | Cell/Virus Enclosed | NO |
| KLLDICFEATEA | 1459945 | Cell/Virus Enclosed | NO |
| KLLDVYNGFLET | 1459948 | Cell/Virus Enclosed | NO |
| KLLHWSDQTICYKDSV | 1671008 | Cell/Virus Enclosed | NO |
| KLLIRDTNKEVFVGDS | 1671010 | Cell/Virus Enclosed | NO |
| KLLRGMPIPDFD | 1459994 | Cell/Virus Enclosed | NO |
| KLMGWDYPKCDRAMPS | 1671032 | Cell/Virus Enclosed | NO |
| KLNDLCFSNVYADSFVV | 32037 | Ectodomain | YES |
| KLNFGMLNGNAIATVK | 1671050 | Cell/Virus Enclosed | NO |
| KLNGILQQAALDYLAT | 1671051 | Cell/Virus Enclosed | NO |
| KLNGSCPGKNNG | 1460044 | Ectodomain | YES |
| KLNLPNCNIEAW | 1460052 | Ectodomain | YES |
| KLNPSEDFIKHI | 1460055 | Cell/Virus Enclosed | NO |
| KLNPSEEFIKHV | 1460057 | Cell/Virus Enclosed | NO |

| | | | |
|---|---|---|---|
| KLNSSLCVGSGP | 1460063 | Ectodomain | YES |
| KLNTDHAGSNDNIALLVQ | 32059 | Underneath Membrane | NO |
| KLNTDHSNSSDNIALL | 1671065 | Underneath Membrane | NO |
| KLNTDHSSSSDNIALLVQ | 1383583 | Underneath Membrane | NO |
| KLPCQFKDKNLQDLWV | 1671070 | Cell/Virus Enclosed | NO |
| KLPDDFMGCVLAWNTRN | 32070 | Ectodomain | YES |
| KLPFKFKDLAIENMWL | 1671076 | Cell/Virus Enclosed | NO |
| KLPKGVGSANIFIGDK | 1671077 | Cell/Virus Enclosed | NO |
| KLPLGQSLCALPDTPS | 1671080 | Ectodomain | YES |
| KLPNGVTVADEP | 1460090 | Cell/Virus Enclosed | NO |
| KLPNYTVPDLVVEQYN | 1671082 | Ectodomain | YES |
| KLPPHDVIVLRD | 1460092 | Cell/Virus Enclosed | NO |
| KLPPHDVTVLRDNLKP | 1671083 | Cell/Virus Enclosed | NO |
| KLPQFYIYDEEGGYDV | 1671084 | Cell/Virus Enclosed | NO |
| KLPSAPQTIQPEVKEV | 1671085 | Cell/Virus Enclosed | NO |
| KLPSEPRTIQPE | 1460101 | Cell/Virus Enclosed | NO |
| KLPSSSGTSTPK | 1460102 | Cell/Virus Enclosed | NO |
| KLQNNEIMPGKM | 1460128 | Cell/Virus Enclosed | NO |
| KLQSGVGRCFNC | 1460137 | Ectodomain | YES |
| KLQSKRTRLFKS | 1460140 | Cell/Virus Enclosed | NO |
| KLQTLIDNINSTLVDL | 1671111 | Ectodomain | YES |
| KLRANDNILSVR | 1460150 | Cell/Virus Enclosed | NO |
| KLRDYLADYDIA | 1460153 | Cell/Virus Enclosed | NO |
| KLRDYLADYDVAVTAG | 1671115 | Cell/Virus Enclosed | NO |
| KLRDYLADYDVV | 1460156 | Cell/Virus Enclosed | NO |
| KLREQIDGYTMH | 1460157 | Cell/Virus Enclosed | NO |
| KLRPFERDISNVPFSP | 1671120 | Ectodomain | YES |
| KLRSETLLPLTQ | 1460174 | Cell/Virus Enclosed | NO |
| KLSAGIEFLRDA | 1460189 | Cell/Virus Enclosed | NO |
| KLSDALKRQIRI | 1460193 | Cell/Virus Enclosed | NO |
| KLSDCLYFKNLK | 1460195 | Cell/Virus Enclosed | NO |
| KLSDNNCYFNAV | 1460198 | Cell/Virus Enclosed | NO |
| KLSDNNCYLNAVIMTL | 1671134 | Cell/Virus Enclosed | NO |
| KLSDVGFVEAYNNCTG | 1671135 | Ectodomain | YES |
| KLSPWFMDGENAYEVV | 1671147 | Cell/Virus Enclosed | NO |
| KLSRLSMIKGPQRADL | 1671148 | Cell/Virus Enclosed | NO |
| KLSTDLIKNQCVNFNF | 1671149 | Ectodomain | YES |
| KLTDLKCTNVVLMGIL | 1671162 | Cell/Virus Enclosed | NO |
| KLTDVKCANVVLLNCL | 1671167 | Cell/Virus Enclosed | NO |
| KLTFNKQEASVPILTT | 1671169 | Cell/Virus Enclosed | NO |
| KLTYNKQMANVS | 1460273 | Cell/Virus Enclosed | NO |
| KLVGHTVCDSLN | 1460287 | Cell/Virus Enclosed | NO |
| KLVKDPPNVQIHTIDG | 1671190 | Secreted | YES |
| KLVKDVDHVQIH | 1460292 | Secreted | YES |
| KLVSMKGTGVFVQDPA | 1671202 | Cell/Virus Enclosed | NO |
| KLVTVGSSNYQQSFVPSPGARPQVN | 1152983 | Ectodomain | YES |
| KLWFDFIENPSV | 1460317 | Cell/Virus Enclosed | NO |
| KLYCAFTAPEDDDFLE | 1671213 | Cell/Virus Enclosed | NO |
| KLYERVKRQLRENAEEDGTGCFEIF | 1152987 | Ectodomain | YES |
| KLYLGGMSYYCEDHKP | 1671222 | Cell/Virus Enclosed | NO |
| KLYRLVWYVERK | 1460342 | Cell/Virus Enclosed | NO |
| KMCLFRYQEFND | 1460360 | Cell/Virus Enclosed | NO |
| KMCLFSYQEFND | 1460361 | Cell/Virus Enclosed | NO |
| KMDEPFTEDTSD | 1460365 | Cell/Virus Enclosed | NO |
| KMDEPFTEHTSE | 1460367 | Capsid | UN |
| KMDEPFTVDTSE | 1460368 | Cell/Virus Enclosed | NO |
| KMDGDDVYLPYPNPSR | 1671239 | Cell/Virus Enclosed | NO |
| KMDGDEVYLPYP | 1460370 | Cell/Virus Enclosed | NO |
| KMDTAAYREAACSQLA | 1671240 | Cell/Virus Enclosed | NO |
| KMDTKPTDPTGIEP | 32296 | Capsid | UN |
| KMFDSFVNSFVL | 1460381 | Cell/Virus Enclosed | NO |
| KMFDSFVNSFVSLYNV | 1671243 | Cell/Virus Enclosed | NO |
| KMFVRPVIEDYHTLTA | 1671245 | Underneath Membrane | NO |
| KMGLTPPLSILKNLGV | 1671248 | Cell/Virus Enclosed | NO |
| KMGVLKADDFVI | 1460397 | Cell/Virus Enclosed | NO |
| KMGVLKADDFVTASDT | 1671251 | Cell/Virus Enclosed | NO |
| KMKDLSPRWYFYYLG | 1541073 | Cell/Virus Enclosed | NO |
| KMLEYKGPVTDV | 1460417 | Cell/Virus Enclosed | NO |
| KMLKSMAATRGATCVI | 1671266 | Cell/Virus Enclosed | NO |
| KMLTPKTGTGIA | 1460440 | Cell/Virus Enclosed | NO |
| KMLVAKDNKNLPKFIE | 1671271 | Capsid | UN |
| KMNCVRFKNADLKDGY | 1671275 | Cell/Virus Enclosed | NO |
| KMQTYKLVSMKG | 1460456 | Cell/Virus Enclosed | NO |
| KMTSLPKDWRLINKFD | 1671290 | Cell/Virus Enclosed | NO |
| KMVDAYVDTFSA | 1460477 | Cell/Virus Enclosed | NO |
| KMVKRQSYGTNSGVAI | 1671291 | Ectodomain | YES |

| | | | |
|---|---|---|---|
| KMVNWFNTMLDASAPA | 1671294 | Cell/Virus Enclosed | NO |
| KMVSPTSKIEPC | 1460492 | Cell/Virus Enclosed | NO |
| KMVVCDKVRSLK | 1460498 | Cell/Virus Enclosed | NO |
| KNAFQTFRWSIF | 1460506 | Cell/Virus Enclosed | NO |
| KNASLYTSLVPHTRYS | 1671305 | Cell/Virus Enclosed | NO |
| KNAYEKDKAVARKLER | 1671308 | Cell/Virus Enclosed | NO |
| KNFCPCSSCPSK | 1460534 | Ectodomain | YES |
| KNFHIEGTGGNI | 1460541 | Capsid | UN |
| KNFHIEGTGGNSQSSS | 1671322 | Cell/Virus Enclosed | NO |
| KNFIAACAVDVN | 1460543 | Cell/Virus Enclosed | NO |
| KNFSTSIPNFGS | 1460551 | Ectodomain | YES |
| KNFTYNVSTDFL | 1460555 | Ectodomain | YES |
| KNGFQGRQIMGSCSFE | 1671334 | Cell/Virus Enclosed | NO |
| KNGFQGRQLMGS | 1460572 | Cell/Virus Enclosed | NO |
| KNGKTVLGEYVFDKSE | 1671335 | Cell/Virus Enclosed | NO |
| KNGNKRAGGHSY | 1460583 | Cell/Virus Enclosed | NO |
| KNGQEENDNISV | 1460586 | Capsid | UN |
| KNGQFQDHYDGFYTQG | 1671339 | Cell/Virus Enclosed | NO |
| KNGQGENDNTSV | 1460589 | Capsid | UN |
| KNGQGEYDNISV | 1460590 | Cell/Virus Enclosed | NO |
| KNHKNLLKIAQYAAQNRRGLDLL | 32506 | Ectodomain | YES |
| KNHTSPDVDFGD | 1460602 | Ectodomain | YES |
| KNIDLFVKRRAD | 1460606 | Cell/Virus Enclosed | NO |
| KNIGDYNVSKDGFFTY | 1671348 | Cell/Virus Enclosed | NO |
| KNIGEYNVSKDGFFTY | 1671349 | Cell/Virus Enclosed | NO |
| KNIPCYASAVAQ | 1460614 | Cell/Virus Enclosed | NO |
| KNIPRYASAVAQ | 1460615 | Cell/Virus Enclosed | NO |
| KNKDGFLSVYNA | 1460620 | Ectodomain | YES |
| KNKMRHKRTSTK | 1460630 | Cell/Virus Enclosed | NO |
| KNLDEARFSGSANQQQ | 1671359 | Cell/Virus Enclosed | NO |
| KNLDEARFSGTA | 1460637 | Cell/Virus Enclosed | NO |
| KNLGFDNQSKPP | 1460643 | Cell/Virus Enclosed | NO |
| KNLGFDNQSKSPSSSG | 1671360 | Capsid | UN |
| KNLKSNCVRFKNVDKD | 1671365 | Cell/Virus Enclosed | NO |
| KNLNEGVLEYFY | 1460653 | Cell/Virus Enclosed | NO |
| KNLNESLIDLRE | 1460655 | Ectodomain | YES |
| KNLNTLRRGAVP | 1460661 | Cell/Virus Enclosed | NO |
| KNLPKYVKPNFD | 1460663 | Ectodomain | YES |
| KNLTPIKLNFGMLNGM | 1671385 | Cell/Virus Enclosed | NO |
| KNMCTGSPSIVEFNRL | 1671389 | Cell/Virus Enclosed | NO |
| KNMWEFRDYFNNNTDS | 1671390 | Cell/Virus Enclosed | NO |
| KNMYLKSAVMQSVGAC | 1671391 | Cell/Virus Enclosed | NO |
| KNNGIGTCPVGT | 1460692 | Ectodomain | YES |
| KNNTSFPTNIAVELFT | 1671398 | Cell/Virus Enclosed | NO |
| KNPINSNVKTLEYDYM | 1671399 | Cell/Virus Enclosed | NO |
| KNQCVNFNFNGLTGTG | 1671404 | Ectodomain | YES |
| KNQISVAPDLSL | 1460724 | Ectodomain | YES |
| KNQKLIANAFNK | 1460725 | Ectodomain | YES |
| KNQKLIATAFNNALLS | 1671405 | Ectodomain | YES |
| KNQTLVAPDLSL | 1460728 | Ectodomain | YES |
| KNRVRTVAGVSI | 1460752 | Cell/Virus Enclosed | NO |
| KNSATGSLDVEVFNRL | 1671412 | Cell/Virus Enclosed | NO |
| KNSGIGTCPAGT | 1460757 | Ectodomain | YES |
| KNSGSVNQQQIK | 1460758 | Cell/Virus Enclosed | NO |
| KNSLSDVAFNRH | 1460761 | Cell/Virus Enclosed | NO |
| KNSQSSPIIPGF | 1460763 | Ectodomain | YES |
| KNSSVNTVCFVDVDFH | 1671422 | Cell/Virus Enclosed | NO |
| KNSVSDVAYNRYLSLY | 1671427 | Cell/Virus Enclosed | NO |
| KNTKLYVNKTLYSEFS | 1671433 | Ectodomain | YES |
| KNTVDTIPKEEF | 1460801 | Cell/Virus Enclosed | NO |
| KNTYISQLGCVF | 1460805 | Cell/Virus Enclosed | NO |
| KNVDWSITYPII | 1460808 | Cell/Virus Enclosed | NO |
| KNVDWSITYPMIANEN | 1671443 | Cell/Virus Enclosed | NO |
| KNVDWTVTYPFIANEK | 1671444 | Cell/Virus Enclosed | NO |
| KNVHLKDVTKENQEIL | 1671445 | Cell/Virus Enclosed | NO |
| KNVKAWSGICVD | 1460812 | Ectodomain | YES |
| KNVSFLGKCLKM | 1460824 | Cell/Virus Enclosed | NO |
| KNVSTSIPNFSS | 1460832 | Ectodomain | YES |
| KNVTDVTGNLSD | 1460833 | Cell/Virus Enclosed | NO |
| KNVTGSNESKALFFTY | 1671456 | Cell/Virus Enclosed | NO |
| KNWFDPIENEDIHRVY | 1671460 | Cell/Virus Enclosed | NO |
| KNYEFKFYDAFP | 1460843 | Cell/Virus Enclosed | NO |
| KNYVFTGYHFTK | 1460853 | Cell/Virus Enclosed | NO |
| KPAFNVVKAVRN | 1460859 | Cell/Virus Enclosed | NO |
| KPAMADEIASLV | 1460864 | Cell/Virus Enclosed | NO |
| KPCTPPALNCYWPLND | 1671481 | Ectodomain | YES |

| | | | |
|---|---|---|---|
| KPDLSQYYCDGGKYYT | 1671483 | Cell/Virus Enclosed | NO |
| KPEANINQDSYG | 1460893 | Cell/Virus Enclosed | NO |
| KPEEKLSAYDLACCMR | 1671488 | Cell/Virus Enclosed | NO |
| KPEYRKVFYALL | 1460903 | Cell/Virus Enclosed | NO |
| KPFARAAFEHLKPMPF | 1671493 | Cell/Virus Enclosed | NO |
| KPFKVEDSVIVNDDTS | 1671497 | Cell/Virus Enclosed | NO |
| KPFLSDFNNGIF | 1460920 | Ectodomain | YES |
| KPFSQPRADKPS | 1460922 | Cell/Virus Enclosed | NO |
| KPFYVPEHCQNY | 1460932 | Cell/Virus Enclosed | NO |
| KPGDAVITTQQAGAGI | 1671506 | Cell/Virus Enclosed | NO |
| KPGDSFNILACYEGIA | 1671507 | Cell/Virus Enclosed | NO |
| KPGGTSSGDATTAFAN | 1671510 | Cell/Virus Enclosed | NO |
| KPGHFNEEFYNFLRLR | 1671515 | Cell/Virus Enclosed | NO |
| KPGHFNKEFYDFLRSQ | 1671516 | Cell/Virus Enclosed | NO |
| KPGHSMPSLFKV | 1460945 | Cell/Virus Enclosed | NO |
| KPGKRQRMALKLEADRLF | 558280 | Undetermined | UN |
| KPGNFNQDFYDFVLSK | 1671520 | Cell/Virus Enclosed | NO |
| KPGNFNQDFYEFVKSK | 1671521 | Cell/Virus Enclosed | NO |
| KPGNTFITIEAA | 1460951 | Cell/Virus Enclosed | NO |
| KPGNVGGNDDLS | 1460953 | Cell/Virus Enclosed | NO |
| KPGNVGGNEALS | 1460954 | Cell/Virus Enclosed | NO |
| KPGPDNSTHNTPVYK | 227307 | Secreted | YES |
| KPHQVNISLNGNTSVC | 1671532 | Ectodomain | YES |
| KPICSSVYLGVR | 1460981 | Cell/Virus Enclosed | NO |
| KPIGMFFPYDIE | 1460986 | Cell/Virus Enclosed | NO |
| KPIIKAQFKPFAKVDG | 1671539 | Cell/Virus Enclosed | NO |
| KPILWVNKASYDTNLN | 1671542 | Cell/Virus Enclosed | NO |
| KPIVWHINQATT | 1461010 | Cell/Virus Enclosed | NO |
| KPIYLKGSGFDF | 1461012 | Cell/Virus Enclosed | NO |
| KPIYLKGSGSDFSLAD | 1671553 | Cell/Virus Enclosed | NO |
| KPIYVPEHCQNY | 1461014 | Cell/Virus Enclosed | NO |
| KPKDQIDGYTMH | 1461015 | Cell/Virus Enclosed | NO |
| KPKPVINQLDEKAQKF | 1671557 | Cell/Virus Enclosed | NO |
| KPKPVISQLDEN | 1461023 | Cell/Virus Enclosed | NO |
| KPKRLRKKRNVDPLSN | 1671558 | Cell/Virus Enclosed | NO |
| KPKVEIDGNVMHANYL | 1671559 | Cell/Virus Enclosed | NO |
| KPLEPKFSIALPPELS | 1671563 | Capsid | UN |
| KPLEPKFSTALP | 1461038 | Cell/Virus Enclosed | NO |
| KPLFAFNVNVMGTRPE | 1671564 | Cell/Virus Enclosed | NO |
| KPLIEWPLTEYDFTDY | 1671567 | Cell/Virus Enclosed | NO |
| KPLLENIDYFNMRRAK | 1671571 | Cell/Virus Enclosed | NO |
| KPLLVDDNKFDV | 1461052 | Cell/Virus Enclosed | NO |
| KPLQSLEVCVQT | 1461055 | Cell/Virus Enclosed | NO |
| KPLQVFVYSSNEEQAV | 1671575 | Cell/Virus Enclosed | NO |
| KPLSAICPAGTK | 1461060 | Ectodomain | YES |
| KPLSQPRSDKPS | 1461063 | Cell/Virus Enclosed | NO |
| KPLVDFNVNVMG | 1461071 | Cell/Virus Enclosed | NO |
| KPNFDLTPFNLT | 1461094 | Ectodomain | YES |
| KPNKPLSQPRSD | 1461102 | Cell/Virus Enclosed | NO |
| KPNLPNCNIESW | 1461105 | Ectodomain | YES |
| KPNRPVVPSPDPN | 32812 | Ectodomain | YES |
| KPNRTSVVSFNEWALA | 1671601 | Cell/Virus Enclosed | NO |
| KPPISFPLCANG | 1461115 | Cell/Virus Enclosed | NO |
| KPPSASCPIGTNYRSC | 1671614 | Ectodomain | YES |
| KPQEKDKKSAKTGTPK | 1671615 | Capsid | UN |
| KPQPAGVFTDHD | 1461129 | Ectodomain | YES |
| KPQPAGVFTNHD | 1461130 | Ectodomain | YES |
| KPQPAGVLTNHD | 1461132 | Ectodomain | YES |
| KPQPKFTVSTQP | 1461133 | Cell/Virus Enclosed | NO |
| KPQPVGVFTHHD | 1461136 | Ectodomain | YES |
| KPQRKRGVKQLPEQFD | 1671620 | Capsid | UN |
| KPQRQRGHKNGQGEND | 1671621 | Cell/Virus Enclosed | NO |
| KPQSGGGKCFNCYPAG | 1671622 | Ectodomain | YES |
| KPRAAPNNTVSW | 1461156 | Cell/Virus Enclosed | NO |
| KPRQKRSPNTQR | 1461167 | Cell/Virus Enclosed | NO |
| KPRQKRTATKAYNVTQ | 1671630 | Cell/Virus Enclosed | NO |
| KPRWKRQPNDDVTSNV | 1671636 | Capsid | UN |
| KPSFYVYSRVKNLNSSRVPDLLV | 1625395 | Ectodomain | YES |
| KPSGMDTALLRA | 1461199 | Underneath Membrane | NO |
| KPSGSFVEQAEG | 1461200 | Ectodomain | YES |
| KPSIIHDIGNPK | 1461203 | Cell/Virus Enclosed | NO |
| KPSSIKEMQSQSSHVA | 1671647 | Capsid | UN |
| KPSSVKEMQSQS | 1461216 | Cell/Virus Enclosed | NO |
| KPTAKYTPLVGT | 1461230 | Cell/Virus Enclosed | NO |
| KPTAKYTPLVRT | 1461231 | Cell/Virus Enclosed | NO |
| KPTEKPTINITK | 32891 | Secreted | YES |

| | | | |
|---|---|---|---|
| KPTKRSFIEDLLFNKVTL | 1070805 | Ectodomain | YES |
| KPTTFMLKYDENGTITD | 32912 | Ectodomain | YES |
| KPTVIVNAANVH | 1461254 | Cell/Virus Enclosed | NO |
| KPTVYVYSRVKN | 1461258 | Ectodomain | YES |
| KPVATVHRRIPDLPDC | 1671663 | Ectodomain | YES |
| KPVDHYDGFYTQ | 1461261 | Cell/Virus Enclosed | NO |
| KPVFFNAFVKCN | 1461268 | Cell/Virus Enclosed | NO |
| KPVISQLDENAQ | 1461281 | Cell/Virus Enclosed | NO |
| KPVIWLSHEQAS | 1461285 | Cell/Virus Enclosed | NO |
| KPVKGAWNIGQQ | 1461287 | Cell/Virus Enclosed | NO |
| KPVMISQWPISD | 1461292 | Cell/Virus Enclosed | NO |
| KPVMISQWPISNDSNG | 1671682 | Cell/Virus Enclosed | NO |
| KPVRARAPPGEQFTHI | 1671690 | Cell/Virus Enclosed | NO |
| KPVYRVNTAIPT | 1461324 | Cell/Virus Enclosed | NO |
| KPYSTLQPNAYYKYDA | 1671710 | Cell/Virus Enclosed | NO |
| KQADLSGAICEL | 1461339 | Cell/Virus Enclosed | NO |
| KQANDLKDTLRL | 1461340 | Cell/Virus Enclosed | NO |
| KQANNNCFVNVSCLML | 1671713 | Cell/Virus Enclosed | NO |
| KQARAEDKKAKIVSAM | 1671714 | Cell/Virus Enclosed | NO |
| KQCCTAAVDLSM | 1461352 | Cell/Virus Enclosed | NO |
| KQCTVQQCFGKRGPNQ | 1671723 | Cell/Virus Enclosed | NO |
| KQDQGQYYYRSS | 1461368 | Ectodomain | YES |
| KQDVSLLGKCYK | 1461373 | Cell/Virus Enclosed | NO |
| KQDVSLLSKCYKAMNA | 1671727 | Cell/Virus Enclosed | NO |
| KQEASVPILTTPFSLK | 1671728 | Cell/Virus Enclosed | NO |
| KQELDKYFKNHT | 1461382 | Ectodomain | YES |
| KQEQRIGLDAVM | 1461385 | Cell/Virus Enclosed | NO |
| KQFVPVQVVDSR | 1461395 | Underneath Membrane | NO |
| KQGAGDAGHSLTWLWL | 1671739 | Cell/Virus Enclosed | NO |
| KQGAKPVGNCVKMLTN | 1671741 | Cell/Virus Enclosed | NO |
| KQGDDYVHLPYP | 1461397 | Cell/Virus Enclosed | NO |
| KQGHFIGVDGEL | 1461404 | Cell/Virus Enclosed | NO |
| KQGPASARDLIC | 1461412 | Ectodomain | YES |
| KQGQLLDTKLNGILQQ | 1671747 | Cell/Virus Enclosed | NO |
| KQGTLIYKVLDT | 1461416 | Cell/Virus Enclosed | NO |
| KQGTQIYKVLDT | 1461418 | Cell/Virus Enclosed | NO |
| KQIANQFNKAISQIQE | 1671751 | Ectodomain | YES |
| KQIDDHRITSITSDKF | 1671752 | Cell/Virus Enclosed | NO |
| KQILGSTMLEDE | 1461435 | Cell/Virus Enclosed | NO |
| KQKEGHIIMEEM | 1461460 | Cell/Virus Enclosed | NO |
| KQKPIYLKGSGSDFSL | 1671764 | Cell/Virus Enclosed | NO |
| KQLEAKTASLFK | 1461476 | Ectodomain | YES |
| KQLEVKTASLFQ | 1461479 | Ectodomain | YES |
| KQLFYVYDVHKDNFKD | 1671773 | Cell/Virus Enclosed | NO |
| KQLGLVWNKDVN | 1461487 | Cell/Virus Enclosed | NO |
| KQLKKAMNVAKAEFDR | 1671779 | Cell/Virus Enclosed | NO |
| KQLKRAMNIAKSEFDH | 1671780 | Cell/Virus Enclosed | NO |
| KQLLINRGVYKP | 1461494 | Cell/Virus Enclosed | NO |
| KQLNEISDDCNWPLVI | 1671787 | Cell/Virus Enclosed | NO |
| KQLSPLEGGGRL | 1461512 | Ectodomain | YES |
| KQNDSWMFTGSS | 1461525 | Ectodomain | YES |
| KQNGCSLYLADN | 1461526 | Cell/Virus Enclosed | NO |
| KQNGHSLYLADNLTAD | 1671799 | Cell/Virus Enclosed | NO |
| KQPFSFSFRDELGVRV | 1671801 | Cell/Virus Enclosed | NO |
| KQPTVTPLPAAD | 1461542 | Capsid | UN |
| KQPVNSNVKLLDYDYA | 1671804 | Cell/Virus Enclosed | NO |
| KQQYSQLFVDTL | 1461551 | Cell/Virus Enclosed | NO |
| KQSCTGSPYIDDFNKI | 1671809 | Cell/Virus Enclosed | NO |
| KQSCTGSPYIDDFNRI | 1671810 | Cell/Virus Enclosed | NO |
| KQSIFTINSVVQKDGV | 1671812 | Cell/Virus Enclosed | NO |
| KQSSSRASSGNRSGNG | 1671815 | Cell/Virus Enclosed | NO |
| KQTATSQQPSGGNVVP | 1671821 | Cell/Virus Enclosed | NO |
| KQTATSQQPSRG | 1461583 | Cell/Virus Enclosed | NO |
| KQTSTSQQPLGG | 1461589 | Cell/Virus Enclosed | NO |
| KQTVTSQQPSGG | 1461592 | Cell/Virus Enclosed | NO |
| KQVGCMMRLFYDRDGQ | 1671830 | Cell/Virus Enclosed | NO |
| KQVGCSMRLFYDRDGQ | 1671831 | Cell/Virus Enclosed | NO |
| KQVGCYMRLFYD | 1461599 | Cell/Virus Enclosed | NO |
| KQVLDLLDKGMQ | 1461604 | Cell/Virus Enclosed | NO |
| KQVLDLLNKGMQLLHT | 1671834 | Cell/Virus Enclosed | NO |
| KQVVLVSNNQEDFDLI | 1671835 | Cell/Virus Enclosed | NO |
| KQVVSDVDYVPL | 1461609 | Cell/Virus Enclosed | NO |
| KQYGECLGDINARDLI | 1671839 | Ectodomain | YES |
| KQYLLAFNQDGV | 1461626 | Ectodomain | YES |
| KQYNVTQAFGRRGPEQ | 1671840 | Cell/Virus Enclosed | NO |
| KRAFLDKLDVNSGFAV | 1671847 | Underneath Membrane | NO |

| | | | |
|---|---|---|---|
| KRAGGHSYGIDL | 1461638 | Cell/Virus Enclosed | NO |
| KRALGLSISDHEFTSA | 1671850 | Cell/Virus Enclosed | NO |
| KRAMAKLKPAPFFYYD | 1671851 | Cell/Virus Enclosed | NO |
| KRAMNIAKSEFDHEIS | 1671852 | Cell/Virus Enclosed | NO |
| KRARGSSAARLE | 1461646 | Cell/Virus Enclosed | NO |
| KRAVRSHPDFKLLHNL | 1671857 | Cell/Virus Enclosed | NO |
| KRCIKSVMDHEQSMYN | 1671859 | Cell/Virus Enclosed | NO |
| KRCTKSVMEHEQSMYN | 1671860 | Cell/Virus Enclosed | NO |
| KRDAAAAKNKMR | 1461659 | Cell/Virus Enclosed | NO |
| KRDGVQVGYCVHGIKY | 1671862 | Cell/Virus Enclosed | NO |
| KRDSEADSPVKDVFEL | 1671863 | Capsid | UN |
| KRESEADSPVKD | 1461671 | Cell/Virus Enclosed | NO |
| KRFGFIEDSVFKPRPA | 1671866 | Ectodomain | YES |
| KRFIHKNSTWSK | 1461679 | Cell/Virus Enclosed | NO |
| KRFQSFQQFGRD | 1461685 | Ectodomain | YES |
| KRFRRVLVKLLD | 1461687 | Cell/Virus Enclosed | NO |
| KRGEKGAYNKDHGRGG | 1671881 | Cell/Virus Enclosed | NO |
| KRGFLDKIGDTS | 1461699 | Underneath Membrane | NO |
| KRGGGNVTYTDQYLCG | 1671883 | Cell/Virus Enclosed | NO |
| KRGMQLYKSYSEQVIP | 1671884 | Cell/Virus Enclosed | NO |
| KRGPNQNFGGGEMLKL | 1671885 | Cell/Virus Enclosed | NO |
| KRGVKQLPEQFDSLNL | 1671887 | Capsid | UN |
| KRIVQMIADFLAGSSD | 1671898 | Cell/Virus Enclosed | NO |
| KRIVYKAACVVDVNDS | 1671899 | Cell/Virus Enclosed | NO |
| KRKMGLTPPLSILKNL | 1671901 | Cell/Virus Enclosed | NO |
| KRKPLDYKRQNNLAIE | 1671902 | Cell/Virus Enclosed | NO |
| KRKVGLTPPLSILKNL | 1671903 | Cell/Virus Enclosed | NO |
| KRLGDYVLTENALVRL | 1671905 | Cell/Virus Enclosed | NO |
| KRLKNGFQGRQIMGSC | 1671906 | Cell/Virus Enclosed | NO |
| KRLKNGFQGRQL | 1461744 | Cell/Virus Enclosed | NO |
| KRLRKKRNVDPLSNFE | 1671907 | Cell/Virus Enclosed | NO |
| KRLVFTNCNYNLTKLL | 1671908 | Ectodomain | YES |
| KRLYPNVYRTDY | 1461759 | Cell/Virus Enclosed | NO |
| KRLYSHVYRSDKVDST | 1671911 | Cell/Virus Enclosed | NO |
| KRNFTYDVNADYLYFH | 1671914 | Ectodomain | YES |
| KRNILPTMTQLNLKYA | 1671917 | Cell/Virus Enclosed | NO |
| KRNQKPLEPKFSIALP | 1671918 | Capsid | UN |
| KRNRSLRVKCSTIVGG | 1671921 | Cell/Virus Enclosed | NO |
| KRNVDPLSNFEHKVIT | 1671922 | Cell/Virus Enclosed | NO |
| KRNVIPTMTQMNLKYA | 1671924 | Cell/Virus Enclosed | NO |
| KRNVLPTLTQMNLKYA | 1671925 | Cell/Virus Enclosed | NO |
| KRNVLPTMTQLNLKYA | 1671926 | Cell/Virus Enclosed | NO |
| KRPIQPTDVAYHTVTD | 1671930 | Cell/Virus Enclosed | NO |
| KRPRWKRVPTRE | 1461786 | Cell/Virus Enclosed | NO |
| KRPVNPTDASHYVVTD | 1671931 | Cell/Virus Enclosed | NO |
| KRQIRIACRKCNLAFR | 1671932 | Cell/Virus Enclosed | NO |
| KRQIRIACRKFN | 1461792 | Cell/Virus Enclosed | NO |
| KRQNNLAIEEIEYVHG | 1671933 | Cell/Virus Enclosed | NO |
| KRQPNDDATSNV | 1461795 | Cell/Virus Enclosed | NO |
| KRQPNDDVTSNVTQCF | 1671934 | Capsid | UN |
| KRQSYGTNSGVAIYHR | 1671935 | Ectodomain | YES |
| KRQYLLKFDNRGVITN | 1671936 | Ectodomain | YES |
| KRRDSTYGPLQTPVGC | 1671938 | Ectodomain | YES |
| KRRFSLLRPEDL | 1461808 | Cell/Virus Enclosed | NO |
| KRRSISASYRFVTFEP | 1671941 | Ectodomain | YES |
| KRSDAQGTNHGY | 1461827 | Cell/Virus Enclosed | NO |
| KRSIRHHPELKILRNL | 1671948 | Cell/Virus Enclosed | NO |
| KRSVRHHPELKLFRNL | 1671955 | Cell/Virus Enclosed | NO |
| KRSVRSHPDFKL | 1461850 | Cell/Virus Enclosed | NO |
| KRTATKAYNVTQAFGRRGPE | 1392246 | Capsid | UN |
| KRTATKQYNVTQAFGRR | 33220 | Capsid | UN |
| KRTDLTIYNREMKCYE | 1671958 | Cell/Virus Enclosed | NO |
| KRTHSPTKKLRY | 1461864 | Cell/Virus Enclosed | NO |
| KRTNLEVYNKEKTYYE | 1671962 | Cell/Virus Enclosed | NO |
| KRTRLFKSTVCW | 1461873 | Cell/Virus Enclosed | NO |
| KRTVCFSVAALSTGLT | 1671968 | Cell/Virus Enclosed | NO |
| KRVLGLQTQTVDSAQG | 1671976 | Cell/Virus Enclosed | NO |
| KRVLGVQTQTVDSAQG | 1671977 | Cell/Virus Enclosed | NO |
| KRVPTIEENVIQ | 1461902 | Cell/Virus Enclosed | NO |
| KRVPTREENVIQCFGP | 1671982 | Capsid | UN |
| KRVRAWVGFDAEGAHA | 1671983 | Cell/Virus Enclosed | NO |
| KRVRGWVGFDVEGAHA | 1671984 | Cell/Virus Enclosed | NO |
| KRWLGILLEAYNAFLD | 1671985 | Cell/Virus Enclosed | NO |
| KRWLPHDAIVVDNDVV | 1671986 | Cell/Virus Enclosed | NO |
| KRWLPPDAIIIDNDIN | 1671987 | Cell/Virus Enclosed | NO |
| KRYCTVQGPPGTGKSH | 1671992 | Cell/Virus Enclosed | NO |

| | | | |
|---|---|---|---|
| KRYVVHHLSMTTSTTN | 1671996 | Cell/Virus Enclosed | NO |
| KSAAGYGGFGCK | 1461925 | Cell/Virus Enclosed | NO |
| KSAELNYTVQKLQTLI | 1671999 | Ectodomain | YES |
| KSAGWPLNKFGKAGLY | 1672006 | Cell/Virus Enclosed | NO |
| KSAGYPFNQFGK | 1461940 | Cell/Virus Enclosed | NO |
| KSAKTGTPKPSRNQSP | 1672011 | Capsid | UN |
| KSALGLSISDHE | 1461946 | Cell/Virus Enclosed | NO |
| KSALLADKFSVL | 1461948 | Cell/Virus Enclosed | NO |
| KSARLKRFPVNTIVNG | 1672015 | Cell/Virus Enclosed | NO |
| KSAYERDRAVAKKLER | 1672023 | Cell/Virus Enclosed | NO |
| KSCLGYHPAHAPSFLA | 1672026 | Cell/Virus Enclosed | NO |
| KSCQNLDCNCLG | 1461975 | Cell/Virus Enclosed | NO |
| KSCTKGLSIADLACAQ | 1672029 | Ectodomain | YES |
| KSDEGGLCPSTGQAME | 1672030 | Cell/Virus Enclosed | NO |
| KSDIFGQDFKTFDLLK | 1672033 | Cell/Virus Enclosed | NO |
| KSDIFGSDFKTFDLLK | 1672034 | Cell/Virus Enclosed | NO |
| KSDIHLPAYASFKVID | 1672035 | Cell/Virus Enclosed | NO |
| KSDILSLNNPIFINYS | 1672036 | Ectodomain | YES |
| KSDYCRGNSCIC | 1462004 | Ectodomain | YES |
| KSEFDHEISVQK | 1462008 | Cell/Virus Enclosed | NO |
| KSENRYSVLSVDSVSE | 1672050 | Cell/Virus Enclosed | NO |
| KSFEIDKGIYQTSNFR | 1672054 | Ectodomain | YES |
| KSFEQQVGVIADKDKD | 1672055 | Cell/Virus Enclosed | NO |
| KSFFPQFTSDQA | 1462021 | Ectodomain | YES |
| KSFFSQFTSDQA | 1462022 | Underneath Membrane | NO |
| KSFLPQFTSDQAVTFL | 1672061 | Ectodomain | YES |
| KSFQLDTAPKTG | 1462047 | Ectodomain | YES |
| KSFSTFESAYMPIADP | 1672065 | Cell/Virus Enclosed | NO |
| KSGANFRDLSLK | 1462061 | Ectodomain | YES |
| KSGCKLINAKPP | 1462063 | Cell/Virus Enclosed | NO |
| KSGCKLTNAKPPKYSS | 1672070 | Cell/Virus Enclosed | NO |
| KSGKKGNSADLTSPE | 227312 | Secreted | YES |
| KSGSKFKNLVLN | 1462081 | Ectodomain | YES |
| KSGSLVLNNPAYIAPQ | 1672076 | Ectodomain | YES |
| KSGSPGDSSWHIYLKS | 1672077 | Ectodomain | YES |
| KSGTAILIKSPLHSLG | 1672078 | Cell/Virus Enclosed | NO |
| KSGTCGIVELEK | 1462089 | Cell/Virus Enclosed | NO |
| KSGTITKVEADF | 1462090 | Ectodomain | YES |
| KSGTLVLNNPAY | 1462091 | Ectodomain | YES |
| KSGVNDICMFENKTTL | 1672081 | Cell/Virus Enclosed | NO |
| KSGYFVNVNNTWMYTG | 1672082 | Ectodomain | YES |
| KSHKPPSASCPIGTNY | 1672089 | Ectodomain | YES |
| KSHNVSLIWNVK | 1462109 | Cell/Virus Enclosed | NO |
| KSIELTAEDISL | 1462116 | Cell/Virus Enclosed | NO |
| KSIIYTGDWEGLHEVL | 1672099 | Cell/Virus Enclosed | NO |
| KSILNLRTNTETNNPCAKTI | 1950312 | Capsid | UN |
| KSISLTVGEVFGKILG | 1672104 | Cell/Virus Enclosed | NO |
| KSIYNVDLCDCG | 1462140 | Cell/Virus Enclosed | NO |
| KSKELLDVALDS | 1462145 | Cell/Virus Enclosed | NO |
| KSKELLDVSLDSLGAA | 1672114 | Cell/Virus Enclosed | NO |
| KSKGLFKEGSTVDLKH | 1672115 | Cell/Virus Enclosed | NO |
| KSKGLNVLWNKFVTGD | 1672116 | Cell/Virus Enclosed | NO |
| KSKGSFRNESWH | 1462148 | Ectodomain | YES |
| KSKGSSRNESWHFDKS | 1672117 | Ectodomain | YES |
| KSKGVCQVGDCYVSTG | 1672119 | Cell/Virus Enclosed | NO |
| KSKNPLLHDANY | 1462155 | Secreted | YES |
| KSKNPLLYEANY | 1462157 | Secreted | YES |
| KSKVGNYRLPSSKPSG | 1672126 | Underneath Membrane | NO |
| KSKVGNYRLPSTQKGS | 1672127 | Ectodomain | YES |
| KSLGFDKPQEKDKKSA | 1672137 | Capsid | UN |
| KSLGFDKPQEKK | 1462175 | Cell/Virus Enclosed | NO |
| KSLKLNLTQQGLLGTC | 1672140 | Cell/Virus Enclosed | NO |
| KSLLPNTGVYDLSGFT | 1672141 | Ectodomain | YES |
| KSLNVDGVQYYC | 1462185 | Cell/Virus Enclosed | NO |
| KSLSRSQSSETK | 1462194 | Cell/Virus Enclosed | NO |
| KSLSSTEFIPN | 144712 | Cell/Virus Enclosed | NO |
| KSLTIVDIPLSL | 1462200 | Cell/Virus Enclosed | NO |
| KSLTTVSIPQGL | 1462202 | Cell/Virus Enclosed | NO |
| KSMAATRGATCVIGTT | 1672159 | Cell/Virus Enclosed | NO |
| KSMYDGDRFVKHDLSL | 1672162 | Cell/Virus Enclosed | NO |
| KSNCVRFKNVDKDDAF | 1672163 | Cell/Virus Enclosed | NO |
| KSNDSCPGKNNG | 1462214 | Ectodomain | YES |
| KSNFSQYYYNIDTGSV | 1672164 | Underneath Membrane | NO |
| KSNLLIQEFLQYDSSI | 1672172 | Cell/Virus Enclosed | NO |
| KSNVDESFVDDF | 1462230 | Cell/Virus Enclosed | NO |
| KSPSSSGTSTPKKPNK | 1672180 | Capsid | UN |

| | | | |
|---|---|---|---|
| KSQLVEYGSFCDNINA | 1672185 | Ectodomain | YES |
| KSQSNRYGFCGN | 1462259 | Ectodomain | YES |
| KSQSPRINFCGNGNHI | 1672189 | Ectodomain | YES |
| KSQSSRINFCGNGNHI | 1672190 | Ectodomain | YES |
| KSQSVIIINNSTNVVI | 1672191 | Ectodomain | YES |
| KSRVQQNKIREL | 1462275 | Cell/Virus Enclosed | NO |
| KSRVQQNKSRELTAED | 1672195 | Cell/Virus Enclosed | NO |
| KSSALGSMDIDVFNKL | 1672196 | Cell/Virus Enclosed | NO |
| KSSCCNLISNKL | 1462279 | Cell/Virus Enclosed | NO |
| KSSDVVDSNENLTWPL | 1672200 | Cell/Virus Enclosed | NO |
| KSSFEIDGNVMHANYL | 1672203 | Cell/Virus Enclosed | NO |
| KSSTGFVYFNATGSDV | 1672209 | Ectodomain | YES |
| KSTFNKDIYCGNSMVC | 1672217 | Cell/Virus Enclosed | NO |
| KSTQSAIDQITGKLNRLIEKTNQQF | 1154324 | Ectodomain | YES |
| KSVANLKLPPHDVTVL | 1672225 | Cell/Virus Enclosed | NO |
| KSVDVAIQEVID | 1462330 | Cell/Virus Enclosed | NO |
| KSVEQKPVDVKP | 1462333 | Cell/Virus Enclosed | NO |
| KSVMDHEQSMYNLLKG | 1672239 | Cell/Virus Enclosed | NO |
| KSVMEHEQSMYNLLNF | 1672240 | Cell/Virus Enclosed | NO |
| KSVNWNLEYPIISNEV | 1672242 | Cell/Virus Enclosed | NO |
| KSVVSKIGDSIF | 1462366 | Cell/Virus Enclosed | NO |
| KSVVSKIGNSIF | 1462367 | Cell/Virus Enclosed | NO |
| KSVYDQIVDNVY | 1462370 | Cell/Virus Enclosed | NO |
| KSVYERDKAVARKLER | 1672250 | Cell/Virus Enclosed | NO |
| KSWYDPVENEDI | 1462380 | Cell/Virus Enclosed | NO |
| KSYAASYNRYKYYSGN | 1672255 | Cell/Virus Enclosed | NO |
| KSYSEHVIPSTL | 1462400 | Cell/Virus Enclosed | NO |
| KSYSGYHPAHAPSFLA | 1672265 | Cell/Virus Enclosed | NO |
| KTADGNQRQLLPRWYF | 1672269 | Cell/Virus Enclosed | NO |
| KTADGQQKPLLP | 1462413 | Cell/Virus Enclosed | NO |
| KTAGVVTCNVVK | 1462417 | Cell/Virus Enclosed | NO |
| KTAGVVTGNVVKQCCT | 1672272 | Cell/Virus Enclosed | NO |
| KTALRKCKSVFY | 1462420 | Cell/Virus Enclosed | NO |
| KTAQADTAGLFT | 1462422 | Cell/Virus Enclosed | NO |
| KTAQAYTAGLYT | 1462425 | Cell/Virus Enclosed | NO |
| KTCVIANVPAIDVLKK | 1672284 | Cell/Virus Enclosed | NO |
| KTDAVILLERYVSLAI | 1672285 | Cell/Virus Enclosed | NO |
| KTDAVVLLERYVSLAI | 1672286 | Cell/Virus Enclosed | NO |
| KTDESQPLPQRQKKQQ | 1672288 | Cell/Virus Enclosed | NO |
| KTDLFNGYKIGCNCAG | 1672291 | Cell/Virus Enclosed | NO |
| KTDSVLLIERFVSLAI | 1672292 | Cell/Virus Enclosed | NO |
| KTEADWIELFPH | 1462456 | Cell/Virus Enclosed | NO |
| KTEAEWIELFPHNDRI | 1672293 | Cell/Virus Enclosed | NO |
| KTEPTGYGVRRK | 1462470 | Cell/Virus Enclosed | NO |
| KTEVDPNLSGFY | 1462472 | Cell/Virus Enclosed | NO |
| KTFDLLKYDFTEHKEN | 1672300 | Cell/Virus Enclosed | NO |
| KTFDLLKYDFTEHKEV | 1672301 | Cell/Virus Enclosed | NO |
| KTFKDIVESCGN | 1462484 | Cell/Virus Enclosed | NO |
| KTFPPTEPKKDKKKKTDE | 1087598 | Capsid | UN |
| KTFSNCNFNMSS | 1462493 | Ectodomain | YES |
| KTGHYTGQAGEMPCAI | 1672319 | Cell/Virus Enclosed | NO |
| KTGNFKDLREYV | 1462507 | Ectodomain | YES |
| KTGQGENDNISV | 1462508 | Capsid | UN |
| KTGTPKPSRNQSPASS | 1672323 | Capsid | UN |
| KTIATHGIAAIN | 1462531 | Cell/Virus Enclosed | NO |
| KTIATHGVAAIN | 1462534 | Cell/Virus Enclosed | NO |
| KTIDELCVNLNL | 1462538 | Cell/Virus Enclosed | NO |
| KTIEDALRNSAM | 1462540 | Ectodomain | YES |
| KTIEDALRNSARLESA | 1383944 | Ectodomain | YES |
| KTILNIVGPDARQDGR | 1672339 | Cell/Virus Enclosed | NO |
| KTIWAFNPETNA | 1462556 | Underneath Membrane | NO |
| KTKSLLPNTGVYDLSG | 1672351 | Ectodomain | YES |
| KTLFDSLKFFEI | 1462572 | Cell/Virus Enclosed | NO |
| KTLKPGDSFNILACYE | 1672359 | Cell/Virus Enclosed | NO |
| KTLLYSFEAHKDSFKD | 1672362 | Cell/Virus Enclosed | NO |
| KTLQEFAQNLPKYVKP | 1672364 | Ectodomain | YES |
| KTLQELSYKLPNYTVP | 1672365 | Ectodomain | YES |
| KTLQEMKALFLT | 1462584 | Cell/Virus Enclosed | NO |
| KTLQEMKALFPTWSQE | 1672366 | Cell/Virus Enclosed | NO |
| KTLYKDVDNPHLMGWD | 1672375 | Cell/Virus Enclosed | NO |
| KTNNIRIVDEWS | 1462618 | Ectodomain | YES |
| KTNNTRIVDEWSYTGS | 1672383 | Ectodomain | YES |
| KTNNTSIVDEWS | 1462623 | Ectodomain | YES |
| KTPDTYKCPQSK | 1462629 | Ectodomain | YES |
| KTPTLKDFGGFN | 1462640 | Ectodomain | YES |
| KTPTLKYFDGFN | 1462641 | Ectodomain | YES |

| | | | |
|---|---|---|---|
| KTPTLKYFGGFNFSQI | 1672391 | Ectodomain | YES |
| KTRVLAVVNSQD | 1462663 | Cell/Virus Enclosed | NO |
| KTSEEHFIETVS | 1462671 | Cell/Virus Enclosed | NO |
| KTSEILDISPRS | 1462676 | Ectodomain | YES |
| KTSGDLAVQIGNNNVC | 1672408 | Cell/Virus Enclosed | NO |
| KTSQADTAGLYT | 1462685 | Cell/Virus Enclosed | NO |
| KTSVDCNMYICGDSTEC | 33874 | Ectodomain | YES |
| KTSWADQSERNYQTFN | 1672417 | Capsid | UN |
| KTSWVDQFERSH | 1462698 | Cell/Virus Enclosed | NO |
| KTSWVDQSERSH | 1462699 | Cell/Virus Enclosed | NO |
| KTTGDLAVQIGSNNVC | 1672424 | Cell/Virus Enclosed | NO |
| KTTPAPVYIWTADEAI | 1672433 | Underneath Membrane | NO |
| KTVGELGDVRETMTHL | 1383978 | Cell/Virus Enclosed | NO |
| KTVKIEDAIIVNDENS | 1672441 | Cell/Virus Enclosed | NO |
| KTVKYFECPNSI | 1462720 | Cell/Virus Enclosed | NO |
| KTVLGEYVFDKSELTN | 1383981 | Cell/Virus Enclosed | NO |
| KTVQADTAGLYT | 1462731 | Cell/Virus Enclosed | NO |
| KTVQFCDAMRDA | 1462732 | Cell/Virus Enclosed | NO |
| KTVREFVIARTG | 1462735 | Ectodomain | YES |
| KTVYSDVENPHL | 1462737 | Cell/Virus Enclosed | NO |
| KTVYSDVETPHL | 1462739 | Cell/Virus Enclosed | NO |
| KTWPRPIDVSKADGII | 1672455 | Ectodomain | YES |
| KTYAELGYIADI | 1462746 | Cell/Virus Enclosed | NO |
| KTYLDFIQQGGI | 1462754 | Cell/Virus Enclosed | NO |
| KTYLDFIQQGGTPIAN | 1672461 | Cell/Virus Enclosed | NO |
| KTYLDYIQQGGVPIIN | 1672462 | Cell/Virus Enclosed | NO |
| KVAFGGDQVHEVAAVR | 1672468 | Cell/Virus Enclosed | NO |
| KVAQPKIEQKPL | 1462776 | Cell/Virus Enclosed | NO |
| KVDIQQTFFDKT | 1462809 | Ectodomain | YES |
| KVDLQLGNLGYLQSFN | 1672493 | Ectodomain | YES |
| KVDSTFVTEYYEFLNK | 1672496 | Cell/Virus Enclosed | NO |
| KVDYGSDTVTYKSTAT | 1672500 | Cell/Virus Enclosed | NO |
| KVEDAYDQVHEE | 1462825 | Cell/Virus Enclosed | NO |
| KVEDSVIVNDAT | 1462827 | Cell/Virus Enclosed | NO |
| KVEDSVIVNNDT | 1462828 | Cell/Virus Enclosed | NO |
| KVEFWFAMRKDGDDVI | 1672506 | Cell/Virus Enclosed | NO |
| KVEISVTPIKNTVDTT | 1672509 | Cell/Virus Enclosed | NO |
| KVEQPKIEPKPVSVIK | 1672515 | Cell/Virus Enclosed | NO |
| KVFFKGNVQVDNGSSI | 1672520 | Cell/Virus Enclosed | NO |
| KVFFQFDNLSSEDLKA | 1672521 | Cell/Virus Enclosed | NO |
| KVFFQFENLSSE | 1462851 | Cell/Virus Enclosed | NO |
| KVFGKDGVMYSE | 1462852 | Ectodomain | YES |
| KVFGYVQQGIFNKATV | 1672523 | Cell/Virus Enclosed | NO |
| KVFGYVQQGVFN | 1462856 | Cell/Virus Enclosed | NO |
| KVFKEGVMYSEF | 1462861 | Ectodomain | YES |
| KVFNASRFPNVY | 1462866 | Ectodomain | YES |
| KVGAGVMLECEKFNVF | 1672540 | Cell/Virus Enclosed | NO |
| KVGELKVEFWFAMRKD | 1672542 | Cell/Virus Enclosed | NO |
| KVGKVDSIVQKCYELS | 1672547 | Cell/Virus Enclosed | NO |
| KVGNYRLPSSKPSGMD | 1672551 | Underneath Membrane | NO |
| KVGNYRLPSTQKGSGM | 1672552 | Ectodomain | YES |
| KVGTGVMVECDSLRIF | 1672554 | Cell/Virus Enclosed | NO |
| KVHNLSGNPDEP | 1462909 | Cell/Virus Enclosed | NO |
| KVIDGGKIWNGV | 1462916 | Cell/Virus Enclosed | NO |
| KVIKDGVMYSEF | 1462921 | Ectodomain | YES |
| KVIKHGVMYSEF | 1462923 | Ectodomain | YES |
| KVINHNVLIKES | 1462927 | Cell/Virus Enclosed | NO |
| KVITKKDAAAAK | 1462939 | Cell/Virus Enclosed | NO |
| KVITKRDAAAAK | 1462942 | Cell/Virus Enclosed | NO |
| KVKATKGEGDGGITSE | 1672578 | Cell/Virus Enclosed | NO |
| KVKDFVSGLLNV | 1462945 | Cell/Virus Enclosed | NO |
| KVKLIFEFEDDVVTSL | 1672580 | Cell/Virus Enclosed | NO |
| KVKLSDVGFVEAYNNC | 1672582 | Ectodomain | YES |
| KVKNTKVFKDGV | 1462957 | Ectodomain | YES |
| KVKNTKVIKDDV | 1462960 | Ectodomain | YES |
| KVKNTKVIKDGV | 1462961 | Ectodomain | YES |
| KVKNTKVIKHGV | 1462963 | Ectodomain | YES |
| KVKNTKVTKNGL | 1462965 | Ectodomain | YES |
| KVKVGTGVMVECDSLR | 1672594 | Cell/Virus Enclosed | NO |
| KVLAEKFDYNVSVMQM | 1672599 | Cell/Virus Enclosed | NO |
| KVLDNGIIRDVNPDDK | 1672600 | Cell/Virus Enclosed | NO |
| KVLEENLNAYVNSNQN | 1672604 | Capsid | UN |
| KVLHKRTLGLSAMSTT | 34103 | Cell/Virus Enclosed | NO |
| KVLLERFLPKCPEILL | 1672607 | Cell/Virus Enclosed | NO |
| KVLNENLNAYQQQDGM | 1672613 | Cell/Virus Enclosed | NO |
| KVLPPILSETQI | 1463009 | Ectodomain | YES |

| | | | |
|---|---|---|---|
| KVLPPLLSDNQI | 1463010 | Ectodomain | YES |
| KVLPPLLSENQISGYT | 1672617 | Ectodomain | YES |
| KVLSENLNAYQQ | 1463021 | Cell/Virus Enclosed | NO |
| KVLTENLNAYQQ | 1463023 | Cell/Virus Enclosed | NO |
| KVNCCRFQRIDDDGNK | 1672626 | Cell/Virus Enclosed | NO |
| KVNCCRFQRVDENGDK | 1672627 | Cell/Virus Enclosed | NO |
| KVNECVKSQSPRINFC | 1672630 | Ectodomain | YES |
| KVNETNSQYLFSTINA | 1672633 | Cell/Virus Enclosed | NO |
| KVNFNEKPVVMEIPSL | 1672634 | Cell/Virus Enclosed | NO |
| KVNNEFIDPTECIYTQ | 1672639 | Cell/Virus Enclosed | NO |
| KVNNEFVDFTEC | 1463048 | Cell/Virus Enclosed | NO |
| KVQCSTNLFKDCSKSY | 1672649 | Cell/Virus Enclosed | NO |
| KVQDAVNNNAQALSKL | 1672650 | Ectodomain | YES |
| KVQNLFGNPDEP | 1463102 | Capsid | UN |
| KVQNLSGNPDEPQKDV | 1672658 | Cell/Virus Enclosed | NO |
| KVQNVNLERCELANYK | 1672659 | Cell/Virus Enclosed | NO |
| KVQQLFYTEDMA | 1463109 | Cell/Virus Enclosed | NO |
| KVQTFYPRLQAS | 1463112 | Cell/Virus Enclosed | NO |
| KVRNLSGNPDEP | 1463116 | Cell/Virus Enclosed | NO |
| KVRSLKLSDNNCYLNA | 1672663 | Cell/Virus Enclosed | NO |
| KVSAEVSRQVIQDVKD | 1672667 | Cell/Virus Enclosed | NO |
| KVSALKTAGVVTGNVV | 1672669 | Cell/Virus Enclosed | NO |
| KVSPDLCTHSFR | 1463138 | Cell/Virus Enclosed | NO |
| KVSTNLFEGDKFVGTF | 1672678 | Cell/Virus Enclosed | NO |
| KVSTQEFRFMTANNLT | 1672679 | Cell/Virus Enclosed | NO |
| KVSTVQSKLTDLKCTN | 1672681 | Cell/Virus Enclosed | NO |
| KVTAFQHQNSKKTTK | 34210 | Cell/Virus Enclosed | NO |
| KVTDVLFPGQKYSSDC | 1672684 | Cell/Virus Enclosed | NO |
| KVTDVLFSGQKY | 1463162 | Cell/Virus Enclosed | NO |
| KVTEDGVNVHDVTVTT | 1672686 | Cell/Virus Enclosed | NO |
| KVTEYSWNKKLYELVQ | 1672687 | Cell/Virus Enclosed | NO |
| KVTLADAGFMKQYGEC | 1672701 | Ectodomain | YES |
| KVTSAMQTMLFT | 1463192 | Cell/Virus Enclosed | NO |
| KVVAKIDKEDVVIFIN | 1672711 | Cell/Virus Enclosed | NO |
| KVVGKDGVMYSE | 1463208 | Ectodomain | YES |
| KVVMSMFSRTTS | 1463220 | Ectodomain | YES |
| KVVNANVLTKDQTPIV | 1672726 | Cell/Virus Enclosed | NO |
| KVVNINVDFKDFQFML | 1672728 | Cell/Virus Enclosed | NO |
| KVVNVNVDFKDFQFML | 1672729 | Cell/Virus Enclosed | NO |
| KVVVGPTTEVKFSVIE | 1672735 | Cell/Virus Enclosed | NO |
| KVVVGPTTEVKS | 1463230 | Cell/Virus Enclosed | NO |
| KVYFDNPTLPFG | 1463236 | Ectodomain | YES |
| KVYFKGQTTHESSSAV | 1672741 | Cell/Virus Enclosed | NO |
| KWADASEPQRGRQGRI | 1672745 | Capsid | UN |
| KWADASETQRGR | 1463249 | Capsid | UN |
| KWADESEPQRGR | 1463250 | Cell/Virus Enclosed | NO |
| KWADNNCYLSSV | 1463253 | Cell/Virus Enclosed | NO |
| KWADNNSYLSSV | 1463254 | Cell/Virus Enclosed | NO |
| KWADQSDQFRNVQTRG | 1672748 | Cell/Virus Enclosed | NO |
| KWADQSDQFRSV | 1463257 | Cell/Virus Enclosed | NO |
| KWADQSDQVRNV | 1463258 | Cell/Virus Enclosed | NO |
| KWDDFCRQYSNESWFE | 1672753 | Cell/Virus Enclosed | NO |
| KWEYEGGCNTIE | 1463276 | Cell/Virus Enclosed | NO |
| KWIFDSSKEFVE | 1463280 | Cell/Virus Enclosed | NO |
| KWKFPQIGGLTS | 1463285 | Cell/Virus Enclosed | NO |
| KWLELLEQNIDAYKTF | 1672774 | Cell/Virus Enclosed | NO |
| KWLNNFNVPSPLNWER | 1672777 | Ectodomain | YES |
| KWNGVSDLSLKQ | 1463309 | Cell/Virus Enclosed | NO |
| KWSGISDLSLKQ | 1463323 | Cell/Virus Enclosed | NO |
| KWSGVSDLSSNK | 1463325 | Cell/Virus Enclosed | NO |
| KWTDVDDYILAN | 1463329 | Cell/Virus Enclosed | NO |
| KWVQVPIGTNDPIRFC | 1672796 | Cell/Virus Enclosed | NO |
| KWYCYDKNPINSNVKT | 1672798 | Cell/Virus Enclosed | NO |
| KYATVVVGSTEEVKSS | 1672809 | Cell/Virus Enclosed | NO |
| KYDAKNYVRFPEILAR | 1672814 | Cell/Virus Enclosed | NO |
| KYDENGTITDAVDCSQ | 1672815 | Ectodomain | YES |
| KYDFTEHKENLFNKYF | 1672819 | Cell/Virus Enclosed | NO |
| KYDNGNFIKLPEVIAQ | 1672822 | Cell/Virus Enclosed | NO |
| KYFDLGFIEAVN | 1463397 | Ectodomain | YES |
| KYFEIYDGGCIPASQV | 1672830 | Cell/Virus Enclosed | NO |
| KYFEIYDGGCLNASEV | 1672831 | Cell/Virus Enclosed | NO |
| KYFISEPPVTYS | 1463404 | Cell/Virus Enclosed | NO |
| KYFKHWSMPYHPNTVD | 1672835 | Cell/Virus Enclosed | NO |
| KYFNTVTKLMCC | 1463413 | Cell/Virus Enclosed | NO |
| KYFSGAMETAAYREAA | 1672843 | Cell/Virus Enclosed | NO |
| KYGCCDATYFEN | 1463431 | Cell/Virus Enclosed | NO |

| | | | |
|---|---|---|---|
| KYIENGQLKNFEAPLK | 1672857 | Cell/Virus Enclosed | NO |
| KYKATGDLAVCLGIGD | 1672861 | Cell/Virus Enclosed | NO |
| KYKTSDELCVNLNLPA | 1672867 | Cell/Virus Enclosed | NO |
| KYLANEAQVQLEHYSS | 1672873 | Cell/Virus Enclosed | NO |
| KYMCGFDGKPML | 1463489 | Cell/Virus Enclosed | NO |
| KYMVANGLHAPHGPFD | 1672891 | Cell/Virus Enclosed | NO |
| KYMVANGLNAPNGPFD | 1672892 | Cell/Virus Enclosed | NO |
| KYNIYDVAGVGV | 1463501 | Ectodomain | YES |
| KYNIYDVSGVGVIRVS | 1672895 | Ectodomain | YES |
| KYNIYDYVGTGIIRSS | 1672896 | Ectodomain | YES |
| KYPCMVGPDYAYFNGA | 1672901 | Cell/Virus Enclosed | NO |
| KYRHCDVDTTTL | 1463530 | Ectodomain | YES |
| KYSELDDEEPME | 1463537 | Cell/Virus Enclosed | NO |
| KYSTFMLNFNEN | 1463553 | Ectodomain | YES |
| KYSYINKCSRLLSDDR | 1672919 | Ectodomain | YES |
| KYTDFNEDVCVC | 1463564 | Cell/Virus Enclosed | NO |
| KYTEVRASRQLAQQKV | 1672924 | Ectodomain | YES |
| KYTFGNVKPGETFTVL | 1672926 | Cell/Virus Enclosed | NO |
| KYTGVSGCLTDC | 1463578 | Cell/Virus Enclosed | NO |
| KYTKIFKDDGNF | 1463579 | Cell/Virus Enclosed | NO |
| KYTKILKDDGNFVVLE | 1672929 | Cell/Virus Enclosed | NO |
| KYTKIMKDDGNCVVLE | 1672930 | Cell/Virus Enclosed | NO |
| KYTPLVGTSVHP | 1463592 | Cell/Virus Enclosed | NO |
| KYTQSTALQELQ | 1463596 | Cell/Virus Enclosed | NO |
| KYVIVATPSTTI | 1463617 | Ectodomain | YES |
| KYVKWEHDSGVV | 1463623 | Cell/Virus Enclosed | NO |
| KYWDQTYHANCVNCTD | 1672960 | Cell/Virus Enclosed | NO |
| KYWGQDYHPDCVDCHD | 1672964 | Cell/Virus Enclosed | NO |
| KYYDDSQYFFNK | 1463646 | Ectodomain | YES |
| KYYDDSQYYFNKDTGV | 1672968 | Ectodomain | YES |
| KYYDDSQYYLNK | 1463648 | Ectodomain | YES |
| KYYSGNANEADYRCAC | 1672972 | Cell/Virus Enclosed | NO |
| LAANTLNIIVPDKSVY | 1672995 | Cell/Virus Enclosed | NO |
| LAASFNKAINNIVASF | 1672998 | Ectodomain | YES |
| LAASFNKAMTNIVDAF | 1672999 | Ectodomain | YES |
| LAATSANLFPPW | 1463708 | Ectodomain | YES |
| LAATSASLFPPWTAAA | 1673002 | Ectodomain | YES |
| LAAVYRINWITGGIA | 1541186 | Undetermined | UN |
| LAAYNGPPQGAF | 1463723 | Cell/Virus Enclosed | NO |
| LACYNGRPTGTF | 1463756 | Cell/Virus Enclosed | NO |
| LADAGFMKQYGECLGD | 1673027 | Ectodomain | YES |
| LADDDSAIIEIVNEVL | 1673030 | Capsid | UN |
| LADGLEAYADKTLQEM | 1673032 | Cell/Virus Enclosed | NO |
| LADGLEAYADRT | 1463766 | Cell/Virus Enclosed | NO |
| LADKFSVLHDIG | 1463773 | Cell/Virus Enclosed | NO |
| LADNLTADEIKA | 1463787 | Cell/Virus Enclosed | NO |
| LADNLTADETKA | 1463788 | Cell/Virus Enclosed | NO |
| LADSNGFIRFPD | 1463791 | Cell/Virus Enclosed | NO |
| LADSVMSAVSSG | 1463793 | Cell/Virus Enclosed | NO |
| LADSVVEVVTTSLTPC | 1673042 | Cell/Virus Enclosed | NO |
| LADTVMSAVSAG | 1463797 | Cell/Virus Enclosed | NO |
| LADVEADVAARA | 1463801 | Cell/Virus Enclosed | NO |
| LADYDIAVTAGP | 1463803 | Cell/Virus Enclosed | NO |
| LADYDVAVTAGPFMDN | 1673046 | Cell/Virus Enclosed | NO |
| LAECKRALGLSISDHE | 1673047 | Cell/Virus Enclosed | NO |
| LAEFDTYDPIYKNGAM | 1673048 | Cell/Virus Enclosed | NO |
| LAEFDTYDPTYK | 1463819 | Cell/Virus Enclosed | NO |
| LAELAPPAGAFF | 1463823 | Cell/Virus Enclosed | NO |
| LAELIPNHAALF | 1463824 | Cell/Virus Enclosed | NO |
| LAELKCSVKSFEIDKG | 1673054 | Ectodomain | YES |
| LAERLSFNVGRS | 1463831 | Cell/Virus Enclosed | NO |
| LAFGVPLSEAKG | 1463844 | Cell/Virus Enclosed | NO |
| LAFLGASVTEDVKFAA | 1673063 | Cell/Virus Enclosed | NO |
| LAFMPVQSRRFI | 1463851 | Cell/Virus Enclosed | NO |
| LAFNQDGIIFNAVDCM | 1673067 | Ectodomain | YES |
| LAGFDMLDNYKAIDVV | 1673072 | Cell/Virus Enclosed | NO |
| LAGGITYVSNSGNLLG | 1673073 | Ectodomain | YES |
| LAGHTGHLPGPR | 1463868 | Cell/Virus Enclosed | NO |
| LAGKVKVGTGVMVECD | 1673077 | Cell/Virus Enclosed | NO |
| LAHGGGLAKALDVYTK | 1673094 | Cell/Virus Enclosed | NO |
| LAIDAYPLIKHP | 1463901 | Cell/Virus Enclosed | NO |
| LAIDAYPLSKHPKPEY | 1673099 | Cell/Virus Enclosed | NO |
| LAIDAYPLTKHEDIEY | 1673101 | Cell/Virus Enclosed | NO |
| LAIEEIEYVHGDALHT | 1673105 | Cell/Virus Enclosed | NO |
| LAIKSDHCGGNP | 1463912 | Ectodomain | YES |
| LAIKSDYCGGNP | 1463913 | Ectodomain | YES |

| | | | |
|---|---|---|---|
| LAILTALRLCAYCCN | 532641 | Undetermined | UN |
| LAKAELCCSARM | 1463933 | Cell/Virus Enclosed | NO |
| LAKALNDFSNLG | 1463936 | Cell/Virus Enclosed | NO |
| LAKAMDTFTNNNGSDV | 1673130 | Cell/Virus Enclosed | NO |
| LAKAMETFNHNNGNDV | 1673131 | Cell/Virus Enclosed | NO |
| LAKAMLDFSRDHNDIL | 1673132 | Cell/Virus Enclosed | NO |
| LAKDDLYVKRYE | 1463943 | Cell/Virus Enclosed | NO |
| LAKGEKANVLIGQGD | 504103 | Cell/Virus Enclosed | NO |
| LAKGGFKFGDPADSRD | 1673138 | Cell/Virus Enclosed | NO |
| LAKGHFKFDEPSDATD | 1673139 | Cell/Virus Enclosed | NO |
| LAKKIDVLLTVDGVNF | 1673148 | Cell/Virus Enclosed | NO |
| LAKLGKDATKPQQVTK | 1673149 | Cell/Virus Enclosed | NO |
| LAKLGKDFKPQQ | 1463960 | Cell/Virus Enclosed | NO |
| LAKLGRDATKPQ | 1463963 | Cell/Virus Enclosed | NO |
| LAKQDAAQRISW | 1463972 | Cell/Virus Enclosed | NO |
| LAKTELCCSARM | 1463982 | Cell/Virus Enclosed | NO |
| LAKVHNLSGNPD | 1463985 | Cell/Virus Enclosed | NO |
| LAKVQNLFGNPD | 1463986 | Capsid | UN |
| LAKVRNLSGNPD | 1463988 | Cell/Virus Enclosed | NO |
| LALLLLDRLNALESKM | 1673181 | Cell/Virus Enclosed | NO |
| LALLLLDRLNQLESKM | 1384139 | Cell/Virus Enclosed | NO |
| LALLLLDRLNRL | 1464021 | Capsid | UN |
| LALLNLTDEDLA | 1464022 | Cell/Virus Enclosed | NO |
| LALPAPHLTLPFNWTHCFDPQIQ | 34858 | Ectodomain | YES |
| LALSADQALQPPCPNLVSYSSYH | 34862 | Ectodomain | YES |
| LALTNMYKEARINDKK | 1673195 | Cell/Virus Enclosed | NO |
| LAMLGAEMTSNVRRQV | 1673206 | Cell/Virus Enclosed | NO |
| LANAKGFIRFPE | 1464066 | Cell/Virus Enclosed | NO |
| LANDDLYVKRYE | 1464070 | Cell/Virus Enclosed | NO |
| LANECAQVLSEYVLCG | 1673217 | Cell/Virus Enclosed | NO |
| LANFNETKGPLCVDTS | 1673220 | Ectodomain | YES |
| LANGTHTGSAFD | 1464082 | Cell/Virus Enclosed | NO |
| LANKEQEAILMSEDVK | 1673222 | Cell/Virus Enclosed | NO |
| LANLNSHTIGSL | 1464093 | Ectodomain | YES |
| LANPTNCIAPVN | 1464096 | Ectodomain | YES |
| LANQFTEFVGTQSVDM | 1673224 | Cell/Virus Enclosed | NO |
| LANQFTEFVSTQ | 1464098 | Cell/Virus Enclosed | NO |
| LANSNGFIRFPDVISE | 1673226 | Cell/Virus Enclosed | NO |
| LANTTTAPLKLF | 1464102 | Cell/Virus Enclosed | NO |
| LANTTTEPLKLFAAET | 1673228 | Cell/Virus Enclosed | NO |
| LANVISFGDYNL | 1464103 | Ectodomain | YES |
| LAPHSALALLEDTMDYPARAHTFDD | 34904 | Undetermined | UN |
| LAPQAKSGIITK | 1464121 | Ectodomain | YES |
| LAPRSVRSVPGE | 1464122 | Ectodomain | YES |
| LAPSDVYQQLAGVKLQ | 1673239 | Cell/Virus Enclosed | NO |
| LAPSTLRSLRKRRLSSP | 34914 | Cell/Virus Enclosed | NO |
| LAPTNAYMRTTHTDIG | 1673242 | Cell/Virus Enclosed | NO |
| LAPTPGAFFFGSKLDL | 1673243 | Capsid | UN |
| LAPVPPPATRRRRAVPI | 34925 | Ectodomain | YES |
| LAQPVDFLALTASSIA | 1673246 | Cell/Virus Enclosed | NO |
| LARDGVVPLSVIPATS | 1673257 | Cell/Virus Enclosed | NO |
| LARKHDTCCSQS | 1464172 | Cell/Virus Enclosed | NO |
| LARKHSTCCNLS | 1464176 | Cell/Virus Enclosed | NO |
| LARRYETCCSQS | 1464178 | Cell/Virus Enclosed | NO |
| LASALKQLKVTTGELV | 1673270 | Cell/Virus Enclosed | NO |
| LASAYRANSFDVPTFC | 1673271 | Cell/Virus Enclosed | NO |
| LASCFFTCNSNA | 1464187 | Ectodomain | YES |
| LASDDLYVKRYFKGCE | 1673273 | Cell/Virus Enclosed | NO |
| LASECFMKSDIFGQDF | 1673275 | Cell/Virus Enclosed | NO |
| LASECFVKSDIFGSDF | 1673276 | Cell/Virus Enclosed | NO |
| LASIAFNHPIQI | 1464209 | Ectodomain | YES |
| LASIAFNHPIQV | 1464210 | Ectodomain | YES |
| LASMTGVSIETLLAAI | 1673287 | Cell/Virus Enclosed | NO |
| LASPMKSDEGGL | 1464217 | Cell/Virus Enclosed | NO |
| LASSSLRTFVVD | 1464228 | Cell/Virus Enclosed | NO |
| LASVIQDYENVC | 1464231 | Cell/Virus Enclosed | NO |
| LASVPEKFLNAFDVFV | 1673292 | Cell/Virus Enclosed | NO |
| LATCPEHQLTWDPIDGRVVS | 34981 | Cell/Virus Enclosed | NO |
| LATFAELEAAQKAYQE | 1673298 | Cell/Virus Enclosed | NO |
| LATFAELEAARK | 1464241 | Cell/Virus Enclosed | NO |
| LATWVGSNLEDPASR | 1673317 | Capsid | UN |
| LAVDDKYKVNENLAVN | 1673322 | Cell/Virus Enclosed | NO |
| LAVIGDFNCTNFAIND | 1384168 | Ectodomain | YES |
| LAVIGDLKCTTV | 1464292 | Ectodomain | YES |
| LAVQIGNNNVCTYEHV | 1673345 | Cell/Virus Enclosed | NO |
| LAVQIGSNNVCTYEHV | 1673346 | Cell/Virus Enclosed | NO |

| | | | |
|---|---|---|---|
| LAVSGQAFCDYADKLC | 1673348 | Cell/Virus Enclosed | NO |
| LAVSGQAFCDYT | 1464312 | Cell/Virus Enclosed | NO |
| LAVTAIVVVSGC | 1464315 | Cell/Virus Enclosed | NO |
| LAVVNSQDVYKS | 1464316 | Cell/Virus Enclosed | NO |
| LAWDVIRKDLSYEQQN | 1673354 | Cell/Virus Enclosed | NO |
| LAWNTRNIDATSTGNY | 1673357 | Ectodomain | YES |
| LAYFSDSKFGTT | 1464338 | Cell/Virus Enclosed | NO |
| LAYFSKKHVEVFTDGK | 1673362 | Cell/Virus Enclosed | NO |
| LAYQDYMQIAPVPAEV | 1673367 | Cell/Virus Enclosed | NO |
| LAYSSSANGSLD | 1464361 | Cell/Virus Enclosed | NO |
| LAYYNNSKGGRF | 1464372 | Cell/Virus Enclosed | NO |
| LCAAVINGDRWF | 1464376 | Cell/Virus Enclosed | NO |
| LCAKILPFSDRP | 1464385 | Cell/Virus Enclosed | NO |
| LCALPDTPSTLA | 1464391 | Ectodomain | YES |
| LCALPDTPSTLK | 1464392 | Ectodomain | YES |
| LCALPDTPSTLTPRSV | 1673392 | Ectodomain | YES |
| LCANEYTGNYQC | 1464394 | Cell/Virus Enclosed | NO |
| LCATCPFAKGDSCSSN | 1673399 | Cell/Virus Enclosed | NO |
| LCATCPFVKGDS | 1464400 | Cell/Virus Enclosed | NO |
| LCCKCCYDHVMA | 1464417 | Cell/Virus Enclosed | NO |
| LCCTVYDSSEVKAQSS | 1673412 | Cell/Virus Enclosed | NO |
| LCDDPEKAQGML | 1464436 | Cell/Virus Enclosed | NO |
| LCDEFTLAEVVKQMYG | 1673418 | Cell/Virus Enclosed | NO |
| LCDFKGWVPGNM | 1464441 | Cell/Virus Enclosed | NO |
| LCDLANPTNCIA | 1464447 | Ectodomain | YES |
| LCDNPFFAVSKPMGTQ | 1673426 | Ectodomain | YES |
| LCDNPFIAVSKP | 1464450 | Ectodomain | YES |
| LCDSGKISSKAGNSIF | 1673428 | Ectodomain | YES |
| LCDVANPTNCIA | 1464458 | Ectodomain | YES |
| LCEILCEYADCKESYF | 1673432 | Cell/Virus Enclosed | NO |
| LCEKASKYLPID | 1464470 | Cell/Virus Enclosed | NO |
| LCENPEWAPLKDNRI | 35050 | Ectodomain | YES |
| LCFDKSKQFVPVQVVD | 1673440 | Underneath Membrane | NO |
| LCFKVYFKGQTTHESS | 1673443 | Cell/Virus Enclosed | NO |
| LCGADGKPVMSEDLWQ | 1673456 | Cell/Virus Enclosed | NO |
| LCGDIVSGYEKSTFNK | 1673457 | Cell/Virus Enclosed | NO |
| LCGDIVSGYGKS | 1464503 | Cell/Virus Enclosed | NO |
| LCGDNSAKFWEE | 1464504 | Cell/Virus Enclosed | NO |
| LCIFNGKFLSNTKYYD | 1673473 | Ectodomain | YES |
| LCIFNGKSFSNT | 1464528 | Ectodomain | YES |
| LCIFNGKSLSNT | 1464529 | Ectodomain | YES |
| LCKEPMYAISNE | 1464540 | Ectodomain | YES |
| LCKTVLNVVGPDARTQ | 1673489 | Cell/Virus Enclosed | NO |
| LCLIGCKPPIGEHWGKGSPC | 110652 | Capsid | UN |
| LCLRDDNQALLV | 1464585 | Cell/Virus Enclosed | NO |
| LCLRDDNQTLLVPKIF | 1673501 | Cell/Virus Enclosed | NO |
| LCMFWNCNVDKYPSNS | 1673507 | Cell/Virus Enclosed | NO |
| LCNLINNNLALGGSVA | 1673511 | Cell/Virus Enclosed | NO |
| LCNNGKISSSAGDSIF | 1673512 | Ectodomain | YES |
| LCNPDPIRFTGP | 1464603 | Ectodomain | YES |
| LCNPDPITFAGP | 1464604 | Ectodomain | YES |
| LCNPDPITFTGS | 1464606 | Ectodomain | YES |
| LCPFGEVFNATKFPSVY | 35083 | Ectodomain | YES |
| LCPPKNSFEALM | 1464619 | Cell/Virus Enclosed | NO |
| LCRVTLGDFTIM | 1464626 | Cell/Virus Enclosed | NO |
| LCSSHYWSPQGNLGGN | 1673538 | Cell/Virus Enclosed | NO |
| LCSTRVNVDGFNEWAM | 1673540 | Cell/Virus Enclosed | NO |
| LCTPDPITFTGT | 1464657 | Ectodomain | YES |
| LCVADDKPIVLF | 1464665 | Cell/Virus Enclosed | NO |
| LCVDGFGVNKIQPWTN | 1673555 | Cell/Virus Enclosed | NO |
| LCVDGTNGYVLRQPNL | 1673556 | Ectodomain | YES |
| LCVGNGPGIDAG | 1464678 | Ectodomain | YES |
| LCVGSGPGKNNG | 1464679 | Ectodomain | YES |
| LCVTKGILTGCF | 1464702 | Cell/Virus Enclosed | NO |
| LCVVPGNVKPGDAVIT | 1673573 | Cell/Virus Enclosed | NO |
| LCWETGKVKPPLNKNY | 1673575 | Cell/Virus Enclosed | NO |
| LCWGELMTLATWVGVNLEDPASRDL | 79594 | Capsid | UN |
| LCYDIGNPKAIA | 1464715 | Cell/Virus Enclosed | NO |
| LDALASMTGVSIETLL | 1673591 | Cell/Virus Enclosed | NO |
| LDALEAEAQIDRLING | 1673592 | Ectodomain | YES |
| LDALEAQVQIDRLING | 1673593 | Ectodomain | YES |
| LDAVKQGAKPVGNCVK | 1673599 | Cell/Virus Enclosed | NO |
| LDDATRNYIITM | 1464766 | Cell/Virus Enclosed | NO |
| LDDEELMELDYP | 1464771 | Cell/Virus Enclosed | NO |
| LDDFIVFNNNGTNVTQ | 1673613 | Cell/Virus Enclosed | NO |
| LDDILQQANKSVGIIS | 1673622 | Cell/Virus Enclosed | NO |

| | | | |
|---|---|---|---|
| LDDKDPSFKDNVILLN | 1673624 | Cell/Virus Enclosed | NO |
| LDDNVLPETYVALPIY | 1673628 | Ectodomain | YES |
| LDDSIEMQLFKV | 1464794 | Cell/Virus Enclosed | NO |
| LDDVKKIEYKPDLSQY | 1673636 | Cell/Virus Enclosed | NO |
| LDEAEDVTEFVQ | 1464806 | Cell/Virus Enclosed | NO |
| LDEFFKNFSTSI | 1464815 | Ectodomain | YES |
| LDEGVDVKIQTA | 1464821 | Cell/Virus Enclosed | NO |
| LDEQESKFWDEN | 1464829 | Cell/Virus Enclosed | NO |
| LDEQESKFWDES | 1464830 | Cell/Virus Enclosed | NO |
| LDEYYKKDNAYY | 1464840 | Cell/Virus Enclosed | NO |
| LDFSRDHNDILYTPPT | 1673660 | Cell/Virus Enclosed | NO |
| LDFVNAGGAPLTNCVK | 1673662 | Cell/Virus Enclosed | NO |
| LDFVNAGGVPLT | 1464855 | Cell/Virus Enclosed | NO |
| LDGDYAMQFFKMGRVA | 1673667 | Cell/Virus Enclosed | NO |
| LDGEVLLLDKLK | 1464865 | Cell/Virus Enclosed | NO |
| LDGEVLPLDELK | 1464866 | Cell/Virus Enclosed | NO |
| LDGIFKTEVDPN | 1464871 | Cell/Virus Enclosed | NO |
| LDGNFRTGVDPD | 1464885 | Cell/Virus Enclosed | NO |
| LDGSLCVGNGPG | 1464886 | Ectodomain | YES |
| LDGSYNASCLCS | 1464887 | Ectodomain | YES |
| LDGVIYTEIQPK | 1464891 | Cell/Virus Enclosed | NO |
| LDGYSKLFITKDEAIK | 1673689 | Cell/Virus Enclosed | NO |
| LDHTSLVWKHIR | 1464907 | Cell/Virus Enclosed | NO |
| LDICFEATEATF | 1464908 | Cell/Virus Enclosed | NO |
| LDIHPSLCNNGKISSS | 1673697 | Ectodomain | YES |
| LDITDAAVKAAESKAF | 1673703 | Cell/Virus Enclosed | NO |
| LDIYDLTMPSQV | 1464937 | Cell/Virus Enclosed | NO |
| LDKCTKYNIYDVSGVG | 1673706 | Ectodomain | YES |
| LDKDFNTILNTA | 1464939 | Cell/Virus Enclosed | NO |
| LDKIQNQTLPRL | 1464943 | Cell/Virus Enclosed | NO |
| LDKNHVADIDKLID | 35199 | Capsid | UN |
| LDKQGQLLDTKLNGIL | 1673716 | Cell/Virus Enclosed | NO |
| LDKVDILLTVDGVNFT | 1673720 | Cell/Virus Enclosed | NO |
| LDLLKDIKLVIP | 1464978 | Cell/Virus Enclosed | NO |
| LDLLNRLQALESGKVK | 1673731 | Cell/Virus Enclosed | NO |
| LDLQAEMNRLQE | 1464986 | Ectodomain | YES |
| LDLQDEMNRLQE | 1464988 | Ectodomain | YES |
| LDLQEEMNRLQE | 1464989 | Ectodomain | YES |
| LDLQVEINRLQE | 1464990 | Ectodomain | YES |
| LDLQVEMNRLQD | 1464991 | Ectodomain | YES |
| LDLRTSCFSVAA | 1464993 | Cell/Virus Enclosed | NO |
| LDLSKELKRPIQPTDV | 1673738 | Cell/Virus Enclosed | NO |
| LDLTLDGYSKLFITKD | 1673742 | Cell/Virus Enclosed | NO |
| LDLTPVIDDVDV | 1465006 | Cell/Virus Enclosed | NO |
| LDLVKRDSEADS | 1465012 | Cell/Virus Enclosed | NO |
| LDNAVKGCVPLNAIPS | 1673760 | Cell/Virus Enclosed | NO |
| LDNDFNTILNTACGVF | 1673763 | Cell/Virus Enclosed | NO |
| LDNDVLNGIISNARNG | 1673764 | Cell/Virus Enclosed | NO |
| LDNGVIRDVSVEDVCF | 1673767 | Cell/Virus Enclosed | NO |
| LDNPLENPSSLFDLVARIKN | 35244 | Capsid | UN |
| LDNQDLNGKWYDFGDF | 1673777 | Cell/Virus Enclosed | NO |
| LDNQDLNGKWYDFGDY | 1673778 | Cell/Virus Enclosed | NO |
| LDPFCFDVNTLLSNQL | 1673788 | Cell/Virus Enclosed | NO |
| LDPISFKAPDTY | 1465067 | Ectodomain | YES |
| LDPITFKAPDTY | 1465068 | Ectodomain | YES |
| LDPKLKGSFNDR | 1465073 | Ectodomain | YES |
| LDPKNPNYNKWLELLE | 1673790 | Cell/Virus Enclosed | NO |
| LDPPCKFSIQDVKGLK | 1673794 | Cell/Virus Enclosed | NO |
| LDPRLKGSFNDR | 1465084 | Ectodomain | YES |
| LDQDVFIQKYGLEDYA | 1673803 | Cell/Virus Enclosed | NO |
| LDQPPTASVSTS | 1465100 | Cell/Virus Enclosed | NO |
| LDQWLKNQTSVA | 1465109 | Ectodomain | YES |
| LDRLNALESKMSGKGS | 1673809 | Cell/Virus Enclosed | NO |
| LDRLNQLENKIS | 1465114 | Cell/Virus Enclosed | NO |
| LDRLNQLENKVS | 1465115 | Cell/Virus Enclosed | NO |
| LDSIQADQQVDRLITG | 1673816 | Ectodomain | YES |
| LDSNFRTEVDPD | 1465158 | Cell/Virus Enclosed | NO |
| LDSRTGSFNNID | 1465162 | Ectodomain | YES |
| LDSTYGPLQTPV | 1465164 | Ectodomain | YES |
| LDTANHSIPDLQ | 1465170 | Cell/Virus Enclosed | NO |
| LDTAPQTGNFKD | 1465172 | Ectodomain | YES |
| LDTCFEAIEATF | 1465173 | Cell/Virus Enclosed | NO |
| LDTCFEATEATL | 1465176 | Cell/Virus Enclosed | NO |
| LDTCFGVSKPNA | 1465178 | Cell/Virus Enclosed | NO |
| LDTCFSVSKPNA | 1465179 | Cell/Virus Enclosed | NO |
| LDTKLNGILQQAALDY | 1673840 | Cell/Virus Enclosed | NO |

| | | | |
|---|---|---|---|
| LDTLLASSSFRT | 1465189 | Cell/Virus Enclosed | NO |
| LDTTLYVNNWCR | 1465203 | Ectodomain | YES |
| LDTTQLHVADTLMQGV | 1673843 | Ectodomain | YES |
| LDTTQLQVANNL | 1465204 | Ectodomain | YES |
| LDTVVSTVKIGGLTFK | 1673846 | Cell/Virus Enclosed | NO |
| LDVCINYVKSPK | 1465210 | Cell/Virus Enclosed | NO |
| LDVCNTKEFKVF | 1465211 | Cell/Virus Enclosed | NO |
| LDVFKIGDVKFKRLGD | 1673852 | Cell/Virus Enclosed | NO |
| LDVKGLEELHDD | 1465218 | Cell/Virus Enclosed | NO |
| LDVKGLEELHYD | 1465219 | Cell/Virus Enclosed | NO |
| LDVKGVEELHDDLVNY | 1673855 | Cell/Virus Enclosed | NO |
| LDVKPYGYPNDFVGGF | 1673856 | Cell/Virus Enclosed | NO |
| LDVLNKALEMCI | 1465227 | Cell/Virus Enclosed | NO |
| LDVMLPGEFSKK | 1465228 | Cell/Virus Enclosed | NO |
| LDVVRKDLEEGD | 1465237 | Cell/Virus Enclosed | NO |
| LDVVRKELEEGD | 1465238 | Cell/Virus Enclosed | NO |
| LDVYNLTMPSQV | 1465242 | Cell/Virus Enclosed | NO |
| LDWVKHLNKNLNEGVL | 1673871 | Cell/Virus Enclosed | NO |
| LDWVKHLNKTLNEGVL | 1673872 | Cell/Virus Enclosed | NO |
| LDYAKDHNDMLYSPPT | 1673873 | Cell/Virus Enclosed | NO |
| LDYKRQNNLAIEEIEY | 1673882 | Cell/Virus Enclosed | NO |
| LDYPKCDRAMPN | 1465268 | Cell/Virus Enclosed | NO |
| LDYPKCDRAMPS | 1465269 | Cell/Virus Enclosed | NO |
| LDYSIDVIDNEIIVKP | 1673887 | Cell/Virus Enclosed | NO |
| LEACYERGCNPK | 1465282 | Cell/Virus Enclosed | NO |
| LEACYVRGCNPN | 1465285 | Cell/Virus Enclosed | NO |
| LEAMQTGFTTTN | 1465302 | Ectodomain | YES |
| LEAQVQIDRLINGRLT | 1673904 | Ectodomain | YES |
| LEAYADKTLQEMKALF | 1673910 | Cell/Virus Enclosed | NO |
| LEAYADRTLQEM | 1465327 | Cell/Virus Enclosed | NO |
| LECTRASTSAVK | 1465335 | Cell/Virus Enclosed | NO |
| LEDAARCMRSLK | 1465338 | Cell/Virus Enclosed | NO |
| LEDEFTPEDVNMQIMG | 1673921 | Cell/Virus Enclosed | NO |
| LEDEFTPFDVVA | 1465344 | Cell/Virus Enclosed | NO |
| LEDFIPTDSTVK | 1465349 | Cell/Virus Enclosed | NO |
| LEDFKPRSKMET | 1465350 | Cell/Virus Enclosed | NO |
| LEDFNFEHVVYGDVSK | 1673925 | Cell/Virus Enclosed | NO |
| LEDKFDLLISDMYDGR | 1673926 | Cell/Virus Enclosed | NO |
| LEDLLFSKVVTSGLGT | 1384293 | Ectodomain | YES |
| LEDTLDDGPSVETSDS | 1673929 | Cell/Virus Enclosed | NO |
| LEEAARCMRSLR | 1465373 | Cell/Virus Enclosed | NO |
| LEEAKTALRKCK | 1465379 | Cell/Virus Enclosed | NO |
| LEEARTALKKCK | 1465380 | Cell/Virus Enclosed | NO |
| LEEAVRQVRSWI | 1465381 | Cell/Virus Enclosed | NO |
| LEEFDVVFGKRGGGNV | 1673938 | Cell/Virus Enclosed | NO |
| LEELHDDLVNYI | 1465390 | Cell/Virus Enclosed | NO |
| LEENLNAYVNSNQNTD | 1673944 | Capsid | UN |
| LEESGAVLFCTP | 1465398 | Cell/Virus Enclosed | NO |
| LEETKFLTSRLL | 1465402 | Cell/Virus Enclosed | NO |
| LEFANDTKIASQLGNC | 1673951 | Ectodomain | YES |
| LEFGASTEIQQV | 1465410 | Cell/Virus Enclosed | NO |
| LEFGASTEMQVE | 1465411 | Cell/Virus Enclosed | NO |
| LEFNPSQTSPATAEPV | 1673956 | Capsid | UN |
| LEFRAGRPARFV | 1465417 | Cell/Virus Enclosed | NO |
| LEFTDENYNNLVPTYL | 1673958 | Cell/Virus Enclosed | NO |
| LEFVREEVDIIE | 1465422 | Cell/Virus Enclosed | NO |
| LEGGGWLVASGP | 1465429 | Ectodomain | YES |
| LEGGSRGAPGGGFVPNLQ | 35484 | Cell/Virus Enclosed | NO |
| LEGLGGNNVYCYNTAL | 1673968 | Cell/Virus Enclosed | NO |
| LEIGYTVDCSCGKKLI | 1673982 | Cell/Virus Enclosed | NO |
| LEISVCQYNMCE | 1465466 | Ectodomain | YES |
| LEITACQYTMCEYPHT | 1673991 | Ectodomain | YES |
| LEIVKLGAEKED | 1465469 | Cell/Virus Enclosed | NO |
| LEIVKLGAEKKDGFYE | 1673993 | Cell/Virus Enclosed | NO |
| LEKACNIAKSAYERDR | 1673995 | Cell/Virus Enclosed | NO |
| LEKCDLQNYGEN | 1465475 | Cell/Virus Enclosed | NO |
| LEKFMSTPNAVF | 1465479 | Cell/Virus Enclosed | NO |
| LEKGKLLIRDTNKEVF | 1674001 | Cell/Virus Enclosed | NO |
| LEKHCYELLRDC | 1465486 | Cell/Virus Enclosed | NO |
| LEKLISTARDGVRRGD | 1674005 | Cell/Virus Enclosed | NO |
| LEKRIENLNKKVDDGFLDIWTYNAE | 1156056 | Ectodomain | YES |
| LELAKVRNLSGN | 1465502 | Cell/Virus Enclosed | NO |
| LELDPPCKFSIQ | 1465507 | Cell/Virus Enclosed | NO |
| LELHWAPEFPWMFEDA | 1674019 | Cell/Virus Enclosed | NO |
| LELICDCGIKQE | 1465517 | Cell/Virus Enclosed | NO |
| LELLEQNIDAYKTFPK | 1674022 | Cell/Virus Enclosed | NO |

| | | | |
|---|---|---|---|
| LELLPNVFKLKISTQL | 1674023 | Cell/Virus Enclosed | NO |
| LELSTGCHTGTDFTGN | 1674026 | Cell/Virus Enclosed | NO |
| LELTTMRYFVRI | 1465537 | Cell/Virus Enclosed | NO |
| LEMAKEVKTSSKVVLS | 1674030 | Cell/Virus Enclosed | NO |
| LENGRLIIRENN | 1465554 | Cell/Virus Enclosed | NO |
| LENHVMVDCRRL | 1465556 | Cell/Virus Enclosed | NO |
| LENIAFNVFKQG | 1465557 | Cell/Virus Enclosed | NO |
| LENIDYFNMRRAKFSL | 1674041 | Cell/Virus Enclosed | NO |
| LENKISGKGQQQ | 1465564 | Cell/Virus Enclosed | NO |
| LENKSAELNYTVQKLQ | 1674044 | Ectodomain | YES |
| LENPTYIYHSAF | 1465567 | Cell/Virus Enclosed | NO |
| LENVIYNLFNVG | 1465571 | Cell/Virus Enclosed | NO |
| LEPCNLWHYGAG | 1465576 | Cell/Virus Enclosed | NO |
| LEPCNLYNYGAGIKLP | 1674057 | Cell/Virus Enclosed | NO |
| LEPKFSIALPPELSVV | 1674061 | Capsid | UN |
| LEPPCRFVIDTPTGPQ | 1674064 | Cell/Virus Enclosed | NO |
| LEPRSGNHCPAD | 1465595 | Ectodomain | YES |
| LEPRSGNYCPAG | 1465597 | Ectodomain | YES |
| LEQDAQIDRLINGRLT | 1674071 | Ectodomain | YES |
| LEQDKNTQEVFA | 1465605 | Ectodomain | YES |
| LEQFMSTPNAIL | 1465610 | Cell/Virus Enclosed | NO |
| LEQNIDAYKTFPKKEK | 1674073 | Cell/Virus Enclosed | NO |
| LEQTNMHSHDVI | 1465627 | Cell/Virus Enclosed | NO |
| LEQTQHSLNFVR | 1465633 | Cell/Virus Enclosed | NO |
| LEQWNLVIGFLC | 1465636 | Transmembrane | NO |
| LEQWNLVIGFLFLTW | 1312964 | Undetermined | UN |
| LERCELANYKQSIPMP | 1674086 | Cell/Virus Enclosed | NO |
| LERDFIDMDDNLFIAK | 1674087 | Cell/Virus Enclosed | NO |
| LERVSLWNYGKPINLP | 1674091 | Cell/Virus Enclosed | NO |
| LERVYKHFNNYD | 1465655 | Cell/Virus Enclosed | NO |
| LERVYKHLNNYDCVVT | 1674092 | Cell/Virus Enclosed | NO |
| LERVYNHFNNYD | 1465657 | Cell/Virus Enclosed | NO |
| LESADVSEMLTFDKKA | 1674095 | Ectodomain | YES |
| LESARRVLNVVC | 1465660 | Cell/Virus Enclosed | NO |
| LESFSVTLLDNQEDKF | 1674100 | Cell/Virus Enclosed | NO |
| LESGKTKQSQPK | 1465672 | Cell/Virus Enclosed | NO |
| LESKMSGKGQQQQSQTVTKK | 1384356 | Cell/Virus Enclosed | NO |
| LESKQVDYVPLRSATC | 1674105 | Cell/Virus Enclosed | NO |
| LESKVLLERFLP | 1465676 | Cell/Virus Enclosed | NO |
| LESTQITTNTGI | 1465688 | Secreted | YES |
| LESTQLSTDTGA | 1465689 | Secreted | YES |
| LESTSTHDGPTEHVTS | 1674110 | Cell/Virus Enclosed | NO |
| LETDGICLENHV | 1465700 | Cell/Virus Enclosed | NO |
| LETDGICPENHVMVDC | 1674118 | Cell/Virus Enclosed | NO |
| LETFQNNVVNYQHIGM | 1674120 | Cell/Virus Enclosed | NO |
| LETPVGCVLGLV | 1465718 | Ectodomain | YES |
| LETVCSVAHTAG | 1465720 | Cell/Virus Enclosed | NO |
| LETVCSVAYTAG | 1465721 | Cell/Virus Enclosed | NO |
| LETVCSVVHTAG | 1465722 | Cell/Virus Enclosed | NO |
| LETVCSVVRTAG | 1465723 | Cell/Virus Enclosed | NO |
| LETVCSVVYTAG | 1465724 | Cell/Virus Enclosed | NO |
| LEVCVQTVRTQV | 1465729 | Cell/Virus Enclosed | NO |
| LEVEDTQGMDNL | 1465731 | Cell/Virus Enclosed | NO |
| LEVKTASLFQTT | 1465736 | Ectodomain | YES |
| LEVLLDVCNTKEVKVF | 1674133 | Cell/Virus Enclosed | NO |
| LEVLLKDALQSR | 1465742 | Cell/Virus Enclosed | NO |
| LEVLLQDALQSREAVL | 1674134 | Cell/Virus Enclosed | NO |
| LEVQGYKNVKIT | 1465750 | Cell/Virus Enclosed | NO |
| LEVQGYKNVRIT | 1465751 | Cell/Virus Enclosed | NO |
| LEVSVCQFNMCE | 1465754 | Ectodomain | YES |
| LEVTGDSCYNFM | 1465758 | Cell/Virus Enclosed | NO |
| LEVYNKEKTYYELTKS | 1674139 | Cell/Virus Enclosed | NO |
| LEVYVHLRNWNFSWNL | 1674140 | Undetermined | UN |
| LEYIKKLYNDLGTQIL | 1674144 | Cell/Virus Enclosed | NO |
| LEYLVSFGVWIRTPP | 956863 | Capsid | UN |
| LEYPIISNEVSINTSC | 1674145 | Cell/Virus Enclosed | NO |
| LEYWVTPLTPRQ | 1465786 | Ectodomain | YES |
| LFAAETIKAKEESVKS | 1674147 | Cell/Virus Enclosed | NO |
| LFAAETVKAKEESVKS | 1674151 | Cell/Virus Enclosed | NO |
| LFAKRKMGLTPPLSIL | 1674159 | Cell/Virus Enclosed | NO |
| LFAKRKVGLTPPLSIL | 1674160 | Cell/Virus Enclosed | NO |
| LFANPAAVDSKC | 1465814 | Cell/Virus Enclosed | NO |
| LFAQSEEQEPFS | 1465818 | Cell/Virus Enclosed | NO |
| LFAQSEEQESFS | 1465819 | Cell/Virus Enclosed | NO |
| LFAQSEKEESFS | 1465820 | Cell/Virus Enclosed | NO |
| LFAQSQEEESFS | 1465821 | Cell/Virus Enclosed | NO |

| | | | |
|---|---|---|---|
| LFASVKNSQSSP | 1465827 | Ectodomain | YES |
| LFASVKSSQSSPIIPG | 1674171 | Ectodomain | YES |
| LFASVKSYQSSP | 1465830 | Ectodomain | YES |
| LFATLMTFSGNVDLDA | 1674172 | Cell/Virus Enclosed | NO |
| LFCNIMKFSDRP | 1465845 | Cell/Virus Enclosed | NO |
| LFCSSTYGVCKY | 1465846 | Cell/Virus Enclosed | NO |
| LFDCVGLSVAST | 1465856 | Cell/Virus Enclosed | NO |
| LFDEAGEEVFAP | 1465862 | Cell/Virus Enclosed | NO |
| LFDEAGEEVLAPKLYC | 1674198 | Cell/Virus Enclosed | NO |
| LFDGRDNGALEAFKRS | 1674202 | Cell/Virus Enclosed | NO |
| LFDGRVEGQVDL | 1465869 | Cell/Virus Enclosed | NO |
| LFDLDEDDEDTKPVP | 227330 | Capsid | UN |
| LFDLSKFQLKLKGTPV | 1674208 | Cell/Virus Enclosed | NO |
| LFDQQAQQEATH | 1465878 | Cell/Virus Enclosed | NO |
| LFDSHIVSKESGNTVV | 1674216 | Capsid | UN |
| LFDSLNLSADTQ | 1465884 | Cell/Virus Enclosed | NO |
| LFDVIVAWHVVRDPRY | 1674218 | Cell/Virus Enclosed | NO |
| LFDWIKISADNK | 1465888 | Cell/Virus Enclosed | NO |
| LFEGDKFIGTFESAAA | 1674221 | Cell/Virus Enclosed | NO |
| LFEGDKFVGTFESAAA | 1674222 | Cell/Virus Enclosed | NO |
| LFELGKQPRGTKKMKKALLS | 1950813 | Capsid | UN |
| LFENGIPLMPLLSCGI | 1674224 | Cell/Virus Enclosed | NO |
| LFFDSEVSTDEE | 1465908 | Cell/Virus Enclosed | NO |
| LFFDSEVSTEEV | 1465909 | Cell/Virus Enclosed | NO |
| LFFICCCTGCGP | 1465914 | Undetermined | UN |
| LFFNDTAWYIGQ | 1465923 | Cell/Virus Enclosed | NO |
| LFGAIAGFIEGGWQGMVDGWYG | 97441 | Ectodomain | YES |
| LFGCENPDCIAC | 1465941 | Cell/Virus Enclosed | NO |
| LFGDSRSDCNHINNLK | 1674247 | Ectodomain | YES |
| LFGMIKKLDNDVLNGI | 1674249 | Cell/Virus Enclosed | NO |
| LFGNPDEPQKDV | 1465950 | Capsid | UN |
| LFGSVACEHIFS | 1465951 | Ectodomain | YES |
| LFGWIKISADNK | 1465957 | Cell/Virus Enclosed | NO |
| LFGWIKISADNN | 1465958 | Cell/Virus Enclosed | NO |
| LFITKDEAIKRVRGWV | 1674272 | Cell/Virus Enclosed | NO |
| LFIVDVYDMWLT | 1465988 | Cell/Virus Enclosed | NO |
| LFIYTNTIWINP | 1465993 | Cell/Virus Enclosed | NO |
| LFKDCARNPIDLPPSH | 1674280 | Cell/Virus Enclosed | NO |
| LFKDCARNPVDL | 1465999 | Cell/Virus Enclosed | NO |
| LFKDCSKSCLGYHPAH | 1674282 | Cell/Virus Enclosed | NO |
| LFKDCSKSYSGYHPAH | 1674283 | Cell/Virus Enclosed | NO |
| LFKDCSNSYSGY | 1466004 | Cell/Virus Enclosed | NO |
| LFKKNFTYNVSA | 1466018 | Ectodomain | YES |
| LFKPLTARGRVV | 1466026 | Cell/Virus Enclosed | NO |
| LFKPVQLVPVSPAFHR | 1674294 | Cell/Virus Enclosed | NO |
| LFKRDSDADSPS | 1466036 | Cell/Virus Enclosed | NO |
| LFKVGKVDSIVQKCYE | 1674297 | Cell/Virus Enclosed | NO |
| LFLAFVVFLLVTLAI | 1541267 | Transmembrane | NO |
| LFLFDEAGEEVF | 1466056 | Cell/Virus Enclosed | NO |
| LFLFDEAGEEVLAPKL | 1674303 | Cell/Virus Enclosed | NO |
| LFLSGDDCFVSNSSVI | 1674320 | Cell/Virus Enclosed | NO |
| LFLTLTSGSDLDRCTT | 1674323 | Ectodomain | YES |
| LFMLPLVYFSDS | 1466116 | Cell/Virus Enclosed | NO |
| LFNKVTLSDAGF | 1466137 | Ectodomain | YES |
| LFNKYFKHWSFD | 1466138 | Cell/Virus Enclosed | NO |
| LFNKYKYFSGAMETAA | 1674349 | Cell/Virus Enclosed | NO |
| LFNVTKLCFISG | 1466147 | Cell/Virus Enclosed | NO |
| LFNVTKLCFVLG | 1466148 | Cell/Virus Enclosed | NO |
| LFPHNDRIKSFSTFES | 1674354 | Cell/Virus Enclosed | NO |
| LFPPWSAAAGVP | 1466167 | Ectodomain | YES |
| LFPPWTAAAGAP | 1466168 | Ectodomain | YES |
| LFPTQIQVRNIMSP | 35947 | Capsid | UN |
| LFPTWSQELPFD | 1466175 | Cell/Virus Enclosed | NO |
| LFPYQGDHGDRY | 1466178 | Ectodomain | YES |
| LFQLCTFTKSTN | 1466188 | Cell/Virus Enclosed | NO |
| LFQTTVELQGLIDQIN | 1384395 | Ectodomain | YES |
| LFRIDVPSAEST | 1466206 | Cell/Virus Enclosed | NO |
| LFSCDINKLLQV | 1466232 | Cell/Virus Enclosed | NO |
| LFSDRPLNLRGW | 1466234 | Cell/Virus Enclosed | NO |
| LFSMVRKLDNQALNSI | 1674394 | Cell/Virus Enclosed | NO |
| LFSSDLSTLAVT | 1466255 | Cell/Virus Enclosed | NO |
| LFSTINALPEIS | 1466257 | Cell/Virus Enclosed | NO |
| LFSTVAIKNLTP | 1466260 | Cell/Virus Enclosed | NO |
| LFSTVVIKNLTPIKLN | 1674400 | Cell/Virus Enclosed | NO |
| LFSVNDFTCSQISPAA | 1674402 | Ectodomain | YES |
| LFSVPSDSCNNV | 1466270 | Cell/Virus Enclosed | NO |

| | | | |
|---|---|---|---|
| LFSYQEFNDLVH | 1466274 | Cell/Virus Enclosed | NO |
| LFTGYFPKAGAN | 1466280 | Ectodomain | YES |
| LFTLQLVSAVNQSLSN | 1674413 | Cell/Virus Enclosed | NO |
| LFTNSILMLDKQGQLL | 1674415 | Cell/Virus Enclosed | NO |
| LFTNSTDTVNHDSFNL | 1674416 | Cell/Virus Enclosed | NO |
| LFTRQEEVHQEL | 1466303 | Cell/Virus Enclosed | NO |
| LFVANYSQDVKQFANG | 1674422 | Ectodomain | YES |
| LFVCSNYAACHD | 1466313 | Ectodomain | YES |
| LFVDTLVNKIPANIVL | 1674426 | Cell/Virus Enclosed | NO |
| LFVEDCKLPLGQSLCA | 1674428 | Ectodomain | YES |
| LFVEDCKLPLGR | 1466319 | Ectodomain | YES |
| LFVEDCNLPLGQ | 1466320 | Ectodomain | YES |
| LFVEHYNEWAQANGFT | 1674429 | Cell/Virus Enclosed | NO |
| LFVNDKIIEFQL | 1466335 | Cell/Virus Enclosed | NO |
| LFVRSKEEDGFA | 1466343 | Cell/Virus Enclosed | NO |
| LFVSYGSSVETETYFK | 1674443 | Cell/Virus Enclosed | NO |
| LFWNCNVPKYPN | 1466367 | Cell/Virus Enclosed | NO |
| LFYCSNSSSGVLDTTI | 1674465 | Ectodomain | YES |
| LFYDRDGQRTYDDVNA | 1674466 | Cell/Virus Enclosed | NO |
| LFYVNANGGSKF | 1466397 | Cell/Virus Enclosed | NO |
| LFYVYDVHKDNFKDGL | 1674478 | Cell/Virus Enclosed | NO |
| LGAAALEDEATP | 1466400 | Cell/Virus Enclosed | NO |
| LGAAIHYLNSKIVDLA | 1674480 | Cell/Virus Enclosed | NO |
| LGADVLYQPPQT | 1466406 | Cell/Virus Enclosed | NO |
| LGAEMTSNVRRQVASG | 1674485 | Cell/Virus Enclosed | NO |
| LGAISFWMCSNGSLQCRICI | 1156351 | Undetermined | UN |
| LGAMLTGFTTTN | 1466420 | Ectodomain | YES |
| LGAMQTGFTTTNEAFQ | 1674494 | Ectodomain | YES |
| LGAPDNPSAIVT | 1466422 | Ectodomain | YES |
| LGASQRVAGDSGFAAY | 1384412 | Underneath Membrane | NO |
| LGASQRVGNDSG | 1466425 | Underneath Membrane | NO |
| LGASSNQSPQGN | 1466427 | Cell/Virus Enclosed | NO |
| LGASVTEDVKFAASTG | 1674497 | Cell/Virus Enclosed | NO |
| LGAVSKVVKVPI | 1466428 | Cell/Virus Enclosed | NO |
| LGAYADKTLQEM | 1466430 | Cell/Virus Enclosed | NO |
| LGCLGSECSKAS | 1466445 | Ectodomain | YES |
| LGCLGSQCGSSS | 1466447 | Ectodomain | YES |
| LGCLSSMNIAANSSEW | 1674504 | Cell/Virus Enclosed | NO |
| LGCVVNADNSTS | 1466455 | Ectodomain | YES |
| LGCVVNAYNNTA | 1466456 | Ectodomain | YES |
| LGDAIQVAKCYG | 1466463 | Cell/Virus Enclosed | NO |
| LGDFIQGPFIAGNTVH | 1674514 | Cell/Virus Enclosed | NO |
| LGDINARDLICA | 1466477 | Ectodomain | YES |
| LGDISGINAPVV | 1466479 | Ectodomain | YES |
| LGDSVKLLSELL | 1466484 | Cell/Virus Enclosed | NO |
| LGESFKKVATIP | 1466502 | Cell/Virus Enclosed | NO |
| LGEYIFERIDYSDAVS | 1674539 | Cell/Virus Enclosed | NO |
| LGEYVFDKSELTNGVY | 1674540 | Cell/Virus Enclosed | NO |
| LGFDKPQEKDKKSAKT | 1674541 | Capsid | UN |
| LGFDKPQEKKKS | 1466514 | Cell/Virus Enclosed | NO |
| LGFDNPPTNVVS | 1466515 | Ectodomain | YES |
| LGFDNQSKLPSS | 1466516 | Cell/Virus Enclosed | NO |
| LGFDNQSKPPSS | 1466517 | Cell/Virus Enclosed | NO |
| LGFDNQSKSLSS | 1466518 | Cell/Virus Enclosed | NO |
| LGFDNQSKSPSSSGTS | 1674542 | Capsid | UN |
| LGFDSSKEFVEYKITE | 1674543 | Cell/Virus Enclosed | NO |
| LGFDSSKPVAKK | 1466521 | Cell/Virus Enclosed | NO |
| LGFDSSKPVIKK | 1466522 | Cell/Virus Enclosed | NO |
| LGFDSSKPVTKKYTYS | 1674544 | Cell/Virus Enclosed | NO |
| LGFEKPQEKDKK | 1466525 | Cell/Virus Enclosed | NO |
| LGFENPPTNVVS | 1466526 | Ectodomain | YES |
| LGFHNPPTNVVS | 1466528 | Ectodomain | YES |
| LGFYNPLTNVVS | 1466552 | Ectodomain | YES |
| LGGMSYFCVDHR | 1466580 | Cell/Virus Enclosed | NO |
| LGHEDLMAAYVE | 1466594 | Cell/Virus Enclosed | NO |
| LGHGCACDRTTIQSVD | 1674594 | Cell/Virus Enclosed | NO |
| LGIGDSAVTYSR | 1466602 | Cell/Virus Enclosed | NO |
| LGIGGDRCIKISTVQS | 1674602 | Cell/Virus Enclosed | NO |
| LGIGYSLADLPA | 1466613 | Ectodomain | YES |
| LGINEEKCGTQL | 1466623 | Ectodomain | YES |
| LGINEKTHVQLS | 1466624 | Cell/Virus Enclosed | NO |
| LGINITNFRAILTAFSP | 36167 | Ectodomain | YES |
| LGINYLGTIKENIDGG | 1674618 | Cell/Virus Enclosed | NO |
| LGISDSAVTYSR | 1466629 | Cell/Virus Enclosed | NO |
| LGISSACDGLAS | 1466630 | Cell/Virus Enclosed | NO |
| LGKCYKAMNAYP | 1466633 | Cell/Virus Enclosed | NO |

| | | | |
|---|---|---|---|
| LGKDAIKPQQVT | 1466634 | Capsid | UN |
| LGKFLEELNAFTREMQ | 1674625 | Capsid | UN |
| LGKLREQIDGYT | 1466646 | Cell/Virus Enclosed | NO |
| LGKPKDQIDGYT | 1466648 | Cell/Virus Enclosed | NO |
| LGKVKPPLNKNY | 1466653 | Cell/Virus Enclosed | NO |
| LGLLQPTDNSTT | 1466677 | Cell/Virus Enclosed | NO |
| LGLQIQTVDSSQGSEY | 1674648 | Cell/Virus Enclosed | NO |
| LGLSAIMQVFFG | 1466688 | Cell/Virus Enclosed | NO |
| LGLTTQTVDSSQGSEY | 1674654 | Cell/Virus Enclosed | NO |
| LGLVNSSLFVED | 1466695 | Ectodomain | YES |
| LGLVNSSLLVED | 1466696 | Ectodomain | YES |
| LGMSLEACYVRGCNPK | 1674667 | Cell/Virus Enclosed | NO |
| LGNCYRCPKEIVETVS | 1674670 | Cell/Virus Enclosed | NO |
| LGNELAQVLTEVVYSN | 1674671 | Cell/Virus Enclosed | NO |
| LGNHRKELWHLD | 1466732 | Ectodomain | YES |
| LGNLGNLQSSNY | 1466736 | Ectodomain | YES |
| LGNPECESLSTASSWSYIVETPSSD | 1156513 | Ectodomain | YES |
| LGNPVIPFKDGI | 1466741 | Ectodomain | YES |
| LGNRKRNQKPLEPKFS | 1674677 | Capsid | UN |
| LGNSIGIDFQDE | 1466744 | Ectodomain | YES |
| LGNSTGIDFQDELDEFFKNVSTSIP | 1384455 | Ectodomain | YES |
| LGNVEAVNFNVTTAET | 1674679 | Ectodomain | YES |
| LGNVFCDGVNVTKHKC | 1674681 | Cell/Virus Enclosed | NO |
| LGNVVKTAVQPTAPAY | 1674682 | Cell/Virus Enclosed | NO |
| LGPDIFLGNCYRCPKE | 1674684 | Cell/Virus Enclosed | NO |
| LGPDIFLGTCYRCPKE | 1674685 | Cell/Virus Enclosed | NO |
| LGPDIFLSMCYRCPKE | 1674686 | Cell/Virus Enclosed | NO |
| LGPHCGSSSRSFFEDL | 1674687 | Ectodomain | YES |
| LGQSKRVGFCGK | 1466771 | Ectodomain | YES |
| LGQSLCALPDTPSTLT | 1674695 | Ectodomain | YES |
| LGRCIVLDVKGVEELH | 1674697 | Cell/Virus Enclosed | NO |
| LGRELNCNVCAN | 1466783 | Cell/Virus Enclosed | NO |
| LGRELNCNVCSNRATC | 1674699 | Cell/Virus Enclosed | NO |
| LGSAGNATEVPA | 1466798 | Cell/Virus Enclosed | NO |
| LGSCSFEDELTP | 1466804 | Cell/Virus Enclosed | NO |
| LGSCTFEDELAPSDVY | 1674706 | Cell/Virus Enclosed | NO |
| LGSECSKASSRSAIED | 1674707 | Ectodomain | YES |
| LGSGAHVGSDFT | 1466809 | Cell/Virus Enclosed | NO |
| LGSGCGKASSRS | 1466810 | Ectodomain | YES |
| LGSITAQPVKID | 1466816 | Secreted | YES |
| LGSLIDCNAASVNQIV | 1674715 | Cell/Virus Enclosed | NO |
| LGSMDIDVFNKLSTSD | 1674720 | Cell/Virus Enclosed | NO |
| LGSNEPGNVGGN | 1466838 | Cell/Virus Enclosed | NO |
| LGSNGKPGNVGGNDAL | 1674721 | Cell/Virus Enclosed | NO |
| LGSQLSLSTARR | 1466845 | Cell/Virus Enclosed | NO |
| LGSSGFLQSSNYKIDT | 1674723 | Ectodomain | YES |
| LGSSVGNFSDGKMGRF | 1674724 | Ectodomain | YES |
| LGTAKKQVVLVSNNQE | 1674728 | Cell/Virus Enclosed | NO |
| LGTCYRCPKEIVDTVS | 1674731 | Cell/Virus Enclosed | NO |
| LGTEDPRCPQIA | 1466874 | Cell/Virus Enclosed | NO |
| LGTGPEAGLPYGANKE | 1674738 | Cell/Virus Enclosed | NO |
| LGTGPHAKDQYGTDID | 1674740 | Cell/Virus Enclosed | NO |
| LGTGPHSKDQYG | 1466888 | Cell/Virus Enclosed | NO |
| LGTGPYASSSYG | 1466892 | Cell/Virus Enclosed | NO |
| LGTGYSLADLPAYMTV | 1674744 | Ectodomain | YES |
| LGTGYSLSDLPA | 1466894 | Underneath Membrane | NO |
| LGTIKENIDGGAMHAN | 1674746 | Cell/Virus Enclosed | NO |
| LGTNDPQFPILAELAP | 1674748 | Capsid | UN |
| LGTSDPQFPIFA | 1466907 | Cell/Virus Enclosed | NO |
| LGTSDSQFPILA | 1466909 | Cell/Virus Enclosed | NO |
| LGTVDADYKKCTKGLSIADL | 2060846 | Ectodomain | YES |
| LGTVDADYKMCT | 1466918 | Ectodomain | YES |
| LGVEFLKRGDKV | 1466940 | Cell/Virus Enclosed | NO |
| LGVGDSAVTYSR | 1466944 | Cell/Virus Enclosed | NO |
| LGVNSSNCNNLN | 1466960 | Cell/Virus Enclosed | NO |
| LGVRGSGHYQTN | 1466966 | Cell/Virus Enclosed | NO |
| LGVSDVWHKQLL | 1466968 | Cell/Virus Enclosed | NO |
| LGVSSNQSPQGNLGSN | 1674789 | Cell/Virus Enclosed | NO |
| LGVTMDVLNKNQKLIA | 1674791 | Ectodomain | YES |
| LGVTMDVLSQNQKLIA | 1674792 | Ectodomain | YES |
| LGWSADSCLQGN | 1466987 | Ectodomain | YES |
| LGWSVDSCLQGD | 1466989 | Ectodomain | YES |
| LGWSVESCLQGD | 1466990 | Ectodomain | YES |
| LGWSYDTCVSNN | 1466991 | Ectodomain | YES |
| LGYHPAHAPSFLAVDD | 1674812 | Cell/Virus Enclosed | NO |
| LHAIQQGFDATN | 1467015 | Ectodomain | YES |

| | | | |
|---|---|---|---|
| LHALDDATRNYI | 1467017 | Cell/Virus Enclosed | NO |
| LHALDDVTRNYI | 1467020 | Cell/Virus Enclosed | NO |
| LHARMTYTCQCG | 1467027 | Cell/Virus Enclosed | NO |
| LHCANFNVLFAM | 1467029 | Cell/Virus Enclosed | NO |
| LHCTTNLFKDCS | 1467038 | Cell/Virus Enclosed | NO |
| LHCVRNVFDNFN | 1467039 | Cell/Virus Enclosed | NO |
| LHDDLVNYIRDKGCVA | 1674831 | Cell/Virus Enclosed | NO |
| LHDVNNGLTCST | 1467049 | Ectodomain | YES |
| LHDVNSGLTCST | 1467050 | Ectodomain | YES |
| LHFSLFDDNFCA | 1467066 | Cell/Virus Enclosed | NO |
| LHFSTYIECTYI | 1467068 | Cell/Virus Enclosed | NO |
| LHFTNDKTIFYT | 1467069 | Cell/Virus Enclosed | NO |
| LHGANLRQDDFV | 1467074 | Ectodomain | YES |
| LHGNGSMIEDID | 1467080 | Cell/Virus Enclosed | NO |
| LHGTILTRPLLESEL | 2001178 | Underneath Membrane | NO |
| LHIAGCQTTNGLNTSY | 1674856 | Ectodomain | YES |
| LHIAGCQTTNGT | 1467092 | Ectodomain | YES |
| LHIAVCQTTNGT | 1467095 | Ectodomain | YES |
| LHKAYIDVLRNSFGKD | 1674857 | Cell/Virus Enclosed | NO |
| LHKEWPHSRNWRCNPTWCW | 244240 | Ectodomain | YES |
| LHKLLQTLVLKM | 1467106 | Cell/Virus Enclosed | NO |
| LHLEANGRAWAF | 1467115 | Cell/Virus Enclosed | NO |
| LHLKVNCCRFQRVDEN | 1674869 | Cell/Virus Enclosed | NO |
| LHLTQDYFLPFD | 1467141 | Ectodomain | YES |
| LHLVDPQIQLTT | 1467144 | Cell/Virus Enclosed | NO |
| LHPAFNVSDAYANLVP | 1674895 | Cell/Virus Enclosed | NO |
| LHPRLKGSFNNR | 1467176 | Ectodomain | YES |
| LHPSFNVSDAYANLVP | 1674898 | Cell/Virus Enclosed | NO |
| LHQPPPQIFLKILPQSGPIG | 36413 | Capsid | UN |
| LHQRELLGVSDVWHKQ | 1674905 | Cell/Virus Enclosed | NO |
| LHRFKPVYRVNT | 1467201 | Cell/Virus Enclosed | NO |
| LHSSQLPLDVYDLTMP | 1674915 | Cell/Virus Enclosed | NO |
| LHSSQLPLDVYN | 1467220 | Cell/Virus Enclosed | NO |
| LHSTSPEGNPSP | 36426 | Secreted | YES |
| LHSTSSEGNTSP | 36429 | Secreted | YES |
| LHTGFQGKQILG | 1467235 | Cell/Virus Enclosed | NO |
| LHTGYYDANQYY | 1467236 | Ectodomain | YES |
| LHTLDDVTRNYI | 1467247 | Cell/Virus Enclosed | NO |
| LHTLSIHDGYTN | 1467249 | Cell/Virus Enclosed | NO |
| LHTVLLPTDYKN | 1467260 | Ectodomain | YES |
| LHVADTLMQGVTLSSN | 1674939 | Ectodomain | YES |
| LHVTYVPSQEKN | 1467277 | Ectodomain | YES |
| LHVVGPDARAKQDVSL | 1674946 | Cell/Virus Enclosed | NO |
| LHYGAGSDKGVAPGTT | 1674955 | Cell/Virus Enclosed | NO |
| LHYRNQGWRSVETSGVAEEEATSGL | 36443 | Undetermined | UN |
| LIAGPKGPEIRY | 1467307 | Cell/Virus Enclosed | NO |
| LIANAFNNALDA | 1467322 | Ectodomain | YES |
| LIANAFNNALNA | 1467326 | Ectodomain | YES |
| LIANAFNNALYAIQEG | 1674966 | Ectodomain | YES |
| LIANKFNQALEA | 1467330 | Ectodomain | YES |
| LIANKFNQALGAMQTG | 1674968 | Ectodomain | YES |
| LIAVANWLAPTNAYMR | 1674975 | Cell/Virus Enclosed | NO |
| LICDCGIKQESR | 1467347 | Cell/Virus Enclosed | NO |
| LICMDVKALMST | 1467352 | Cell/Virus Enclosed | NO |
| LICSNTPLSKDL | 1467354 | Cell/Virus Enclosed | NO |
| LIDCNAASVNQT | 1467366 | Cell/Virus Enclosed | NO |
| LIDGVENPMLMGWDYP | 1674997 | Cell/Virus Enclosed | NO |
| LIDNEFNEVEKQIGNVINWTRDSIT | 1156755 | Ectodomain | YES |
| LIDNINSTLVDLKWLN | 1675001 | Ectodomain | YES |
| LIDQINSTYVDLKLLN | 1675002 | Ectodomain | YES |
| LIDSYFDNSSTLQSVA | 1675003 | Cell/Virus Enclosed | NO |
| LIEGSKPYSTLQPNAY | 1675009 | Cell/Virus Enclosed | NO |
| LIEHVTSVNLFD | 1467390 | Cell/Virus Enclosed | NO |
| LIELENKFTPLS | 1467394 | Cell/Virus Enclosed | NO |
| LIERFVSLAIDAYPLV | 1675013 | Cell/Virus Enclosed | NO |
| LIERGKLLVRDTRKEI | 1675014 | Cell/Virus Enclosed | NO |
| LIEWPLTEYDFTDYKV | 1675015 | Cell/Virus Enclosed | NO |
| LIEYDFTDYKVQ | 1467398 | Cell/Virus Enclosed | NO |
| LIFEFEDDVITS | 1467402 | Cell/Virus Enclosed | NO |
| LIFLLVLLDYQGMLP | 956865 | Undetermined | UN |
| LIFPNSNYVLQD | 1467414 | Cell/Virus Enclosed | NO |
| LIFPPGQGVPLN | 1467416 | Cell/Virus Enclosed | NO |
| LIFRNIKCNYVF | 1467420 | Ectodomain | YES |
| LIGAEHVDSSYE | 1467432 | Ectodomain | YES |
| LIGANGNKIVDK | 1467438 | Cell/Virus Enclosed | NO |
| LIGANYLGKLKE | 1467439 | Cell/Virus Enclosed | NO |

| | | | |
|---|---|---|---|
| LIGINYLCKPKVEIDG | 1675042 | Cell/Virus Enclosed | NO |
| LIGLLAIAGIRLHRAAIYTAEIHK | 36538 | Undetermined | UN |
| LIGLYRRQQTSNLVVQ | 1675053 | Cell/Virus Enclosed | NO |
| LIHAEQLTPAWR | 1467482 | Ectodomain | YES |
| LIIGALAGSTLAVLVIGGIA | 741658 | Transmembrane | NO |
| LIISDMYDPITKNIGE | 1675074 | Cell/Virus Enclosed | NO |
| LIKAYNTINNEQGTPL | 1675082 | Cell/Virus Enclosed | NO |
| LIKDVDNPVLMGWDYP | 1675083 | Cell/Virus Enclosed | NO |
| LIKESIPIVWGVKDFN | 1675085 | Cell/Virus Enclosed | NO |
| LIKNQCVNFNFNGLTGT | 36579 | Ectodomain | YES |
| LIKQLKRAMNIAKSEF | 1675092 | Cell/Virus Enclosed | NO |
| LILAYSNKTVGE | 1467552 | Cell/Virus Enclosed | NO |
| LILDYFSYPFSM | 1467556 | Ectodomain | YES |
| LILSWEIGKVKPPLNK | 1675111 | Cell/Virus Enclosed | NO |
| LIMAYGNCTFGAPDDA | 1675115 | Cell/Virus Enclosed | NO |
| LIMAYGNCTFGD | 1467600 | Cell/Virus Enclosed | NO |
| LINKFDVINGVKTVKY | 1675123 | Cell/Virus Enclosed | NO |
| LINKFLKANPLWHKAV | 1675124 | Cell/Virus Enclosed | NO |
| LINNISYVVTQN | 1467617 | Cell/Virus Enclosed | NO |
| LINRGVYKPLLE | 1467620 | Cell/Virus Enclosed | NO |
| LINSSLFVEDCK | 1467621 | Ectodomain | YES |
| LINTISPEKLKN | 1467624 | Cell/Virus Enclosed | NO |
| LIPGFVLPCNAQEQQI | 1675130 | Cell/Virus Enclosed | NO |
| LIPNHAALFFDS | 1467644 | Cell/Virus Enclosed | NO |
| LIPVRPRNSSDNGISA | 1675141 | Ectodomain | YES |
| LIQESIKSLNNSYINL | 1675145 | Ectodomain | YES |
| LIQNGAKHVQGNVAKA | 1675148 | Cell/Virus Enclosed | NO |
| LIQQWIPFMMSR | 1467673 | Cell/Virus Enclosed | NO |
| LIRDNDVLSVST | 1467683 | Cell/Virus Enclosed | NO |
| LIRGYRGIKPLL | 1467697 | Cell/Virus Enclosed | NO |
| LIRQGTDYKHWPQIAQFAPS | 1392265 | Capsid | UN |
| LIRQGTDYKYWP | 1467707 | Cell/Virus Enclosed | NO |
| LIRQLQPINYFD | 1467708 | Ectodomain | YES |
| LISAGIFSVPADVSLT | 1675168 | Cell/Virus Enclosed | NO |
| LISAGIFSVPSDVSLT | 1675169 | Cell/Virus Enclosed | NO |
| LISCNGNKIEDL | 1467712 | Cell/Virus Enclosed | NO |
| LISDMYDGRTKAIDGE | 1675171 | Cell/Virus Enclosed | NO |
| LISGSLGDSGEL | 1467719 | Cell/Virus Enclosed | NO |
| LISQDSITHYPALKMV | 1675187 | Cell/Virus Enclosed | NO |
| LISQFRLSKMGV | 1467744 | Cell/Virus Enclosed | NO |
| LISQVRLSKMGI | 1467745 | Cell/Virus Enclosed | NO |
| LISSGIFSVPSD | 1467748 | Cell/Virus Enclosed | NO |
| LISTARDGVRRGDNFH | 1675190 | Cell/Virus Enclosed | NO |
| LITNTIAGVAGLITGGRR | 1029633 | Secreted | YES |
| LIVNVLSIFSSS | 1467814 | Cell/Virus Enclosed | NO |
| LKAAVHEWKMVV | 1467843 | Cell/Virus Enclosed | NO |
| LKAAVHGWKMVVCDKV | 1675239 | Cell/Virus Enclosed | NO |
| LKAAYANFNSQD | 1467848 | Cell/Virus Enclosed | NO |
| LKADDFVTASDTTLRC | 1675242 | Cell/Virus Enclosed | NO |
| LKAVRSSFNFDQKELL | 1675267 | Cell/Virus Enclosed | NO |
| LKCGFSFDLNGLDALF | 1675269 | Cell/Virus Enclosed | NO |
| LKCGSPYQHYDACSVK | 1675270 | Cell/Virus Enclosed | NO |
| LKCNRCCDRYEEYDLE | 1675271 | Underneath Membrane | NO |
| LKCSVKSFEIDKGIYQT | 36815 | Ectodomain | YES |
| LKCSVKSFKIDK | 1467909 | Ectodomain | YES |
| LKCTSDNINDKDTGPP | 1675277 | Ectodomain | YES |
| LKCTTVAINDVD | 1467917 | Ectodomain | YES |
| LKDAERAVEVDPDD | 36816 | Capsid | UN |
| LKDALQSREAVL | 1467923 | Cell/Virus Enclosed | NO |
| LKDDGNFVVLELDPPC | 1675289 | Cell/Virus Enclosed | NO |
| LKDGVNFNVDDINFSP | 1675292 | Ectodomain | YES |
| LKDGVNFNVDDV | 1467937 | Ectodomain | YES |
| LKDIKFVIPALQHAFM | 1675297 | Cell/Virus Enclosed | NO |
| LKDLLLYAADPALHVA | 1675301 | Cell/Virus Enclosed | NO |
| LKDLLLYAADPAMHVA | 1675302 | Cell/Virus Enclosed | NO |
| LKDNVLSFMLRL | 1467953 | Cell/Virus Enclosed | NO |
| LKDSDINEMVLSLVRS | 1675305 | Cell/Virus Enclosed | NO |
| LKDSDVNDMVLSLIKS | 1675306 | Cell/Virus Enclosed | NO |
| LKEDDVDDSGDS | 1467964 | Cell/Virus Enclosed | NO |
| LKEDYQIGGYSEDRHS | 1675309 | Secreted | YES |
| LKEIVGPKELLLLWES | 1675315 | Cell/Virus Enclosed | NO |
| LKELYGPADPTF | 1467979 | Cell/Virus Enclosed | NO |
| LKELYGPIDPTF | 1467980 | Cell/Virus Enclosed | NO |
| LKELYGPLDPTF | 1467981 | Cell/Virus Enclosed | NO |
| LKELYGPVDPTFLHRF | 1675321 | Cell/Virus Enclosed | NO |
| LKETVFVEPKNG | 1467994 | Cell/Virus Enclosed | NO |

| | | | |
|---|---|---|---|
| LKEVVGPKELLLSWES | 1675329 | Cell/Virus Enclosed | NO |
| LKEWNFSLGIILLFIT | 1675330 | Undetermined | UN |
| LKFDNRGVITNAVDCS | 1675332 | Ectodomain | YES |
| LKFDSGTVSKTSDWKC | 1675333 | Cell/Virus Enclosed | NO |
| LKFFEIKHADYT | 1468004 | Cell/Virus Enclosed | NO |
| LKFIDKLNVLFDGRDN | 1675336 | Cell/Virus Enclosed | NO |
| LKFTEESFYKNMYLKS | 1675345 | Cell/Virus Enclosed | NO |
| LKGACSCGSCCKFDEDD | 36856 | Underneath Membrane | NO |
| LKGGAPVKKVAFGGDQ | 1675354 | Cell/Virus Enclosed | NO |
| LKGGKIVSTWFK | 1468040 | Cell/Virus Enclosed | NO |
| LKGGLILKFDSGTVSK | 1675359 | Cell/Virus Enclosed | NO |
| LKGGLILRFDSG | 1468044 | Cell/Virus Enclosed | NO |
| LKGLDAMFFYGD | 1468049 | Cell/Virus Enclosed | NO |
| LKGSFNNRDTGH | 1468052 | Ectodomain | YES |
| LKGSFNNRDTGL | 1468053 | Ectodomain | YES |
| LKGSFNNRDTGP | 1468054 | Ectodomain | YES |
| LKGSGSDFSLADSVVE | 1675364 | Cell/Virus Enclosed | NO |
| LKGVTLDKLRDYLADY | 1675372 | Cell/Virus Enclosed | NO |
| LKHAEEYREYLESYNT | 1675375 | Cell/Virus Enclosed | NO |
| LKHFFFTQKGDAAIKD | 1675381 | Cell/Virus Enclosed | NO |
| LKICKFGNTSFA | 1468096 | Ectodomain | YES |
| LKICKFSSNTTF | 1468100 | Ectodomain | YES |
| LKIKIASGFGPLITH | 36890 | Ectodomain | YES |
| LKILPQSGPIGGIKSMGITT | 36893 | Capsid | UN |
| LKISTQLFEGDKFIGT | 1675402 | Cell/Virus Enclosed | NO |
| LKKAMNVAKAEFDRES | 1675409 | Cell/Virus Enclosed | NO |
| LKKEPLLYVPIR | 1468132 | Cell/Virus Enclosed | NO |
| LKKLIGGDVTPVDQYM | 1675418 | Cell/Virus Enclosed | NO |
| LKKLLEQWNLVIGFL | 1312987 | Undetermined | UN |
| LKKMDEPFTEDT | 1468147 | Cell/Virus Enclosed | NO |
| LKKPRWKRVPTR | 1468152 | Cell/Virus Enclosed | NO |
| LKKTSWVDQSER | 1468159 | Cell/Virus Enclosed | NO |
| LKLAGTAVINLRADQI | 1675433 | Cell/Virus Enclosed | NO |
| LKLAGTAVVNLRPDQL | 1675434 | Cell/Virus Enclosed | NO |
| LKLGTNDPQFPILAEL | 1675442 | Capsid | UN |
| LKLKVSTNLFEGDKFV | 1675451 | Cell/Virus Enclosed | NO |
| LKLLGIGGDRCI | 1468187 | Cell/Virus Enclosed | NO |
| LKLPPHDVIVLR | 1468194 | Cell/Virus Enclosed | NO |
| LKLPPHDVTVLRDNLK | 1675456 | Cell/Virus Enclosed | NO |
| LKLRSETLLPLT | 1468203 | Cell/Virus Enclosed | NO |
| LKLSDNNCYLNAVIMT | 1675464 | Cell/Virus Enclosed | NO |
| LKLTFNKQEASVPILT | 1675470 | Cell/Virus Enclosed | NO |
| LKLTLTSNGLLG | 1468220 | Cell/Virus Enclosed | NO |
| LKLTYNKQKANV | 1468221 | Cell/Virus Enclosed | NO |
| LKLVDDHALIVN | 1468222 | Cell/Virus Enclosed | NO |
| LKMAGMHFGACD | 1468227 | Ectodomain | YES |
| LKMVKRQSYGTNSGVA | 1675482 | Ectodomain | YES |
| LKMVQTHLSHYV | 1468241 | Cell/Virus Enclosed | NO |
| LKNLGFDNQSKSPSSS | 1675492 | Capsid | UN |
| LKNLTQTFTSMLTNYF | 1675497 | Cell/Virus Enclosed | NO |
| LKNNLYRLVWHVERKD | 1675498 | Cell/Virus Enclosed | NO |
| LKNSVSDVAYNRYLSL | 1675500 | Cell/Virus Enclosed | NO |
| LKNYAASYNKYKYYSG | 1675503 | Cell/Virus Enclosed | NO |
| LKPFERDLTSDE | 1468279 | Ectodomain | YES |
| LKPGDSFNILACYEGI | 1675508 | Cell/Virus Enclosed | NO |
| LKPIFKLPLGINITNFR | 36991 | Ectodomain | YES |
| LKPIKRSFIEDL | 1468286 | Ectodomain | YES |
| LKPLPFLYYSTT | 1468290 | Cell/Virus Enclosed | NO |
| LKPLYVPEHCQN | 1468293 | Cell/Virus Enclosed | NO |
| LKPMPFFYFDDSDCDV | 1675514 | Cell/Virus Enclosed | NO |
| LKPTFKSKGLNV | 1468303 | Cell/Virus Enclosed | NO |
| LKQADLSGAICE | 1468318 | Cell/Virus Enclosed | NO |
| LKQRDSTYGPLQ | 1468343 | Ectodomain | YES |
| LKRAMNIAKSEFDHEI | 1675544 | Cell/Virus Enclosed | NO |
| LKRGEKIVYHTL | 1468356 | Cell/Virus Enclosed | NO |
| LKRQIRIACHKC | 1468367 | Cell/Virus Enclosed | NO |
| LKRRDSTYGPLE | 1468372 | Ectodomain | YES |
| LKRRDSTYGPLQTPVG | 1675554 | Ectodomain | YES |
| LKSIVNTRNATVVIGT | 1675576 | Cell/Virus Enclosed | NO |
| LKSLGFDKPQEKDKKS | 1675580 | Capsid | UN |
| LKSLKLNLTQQG | 1468404 | Cell/Virus Enclosed | NO |
| LKSMAATRGATCVIGT | 1675583 | Cell/Virus Enclosed | NO |
| LKSNRCCDRYEE | 1468412 | Underneath Membrane | NO |
| LKSSTGFVYFNATGSD | 1675589 | Ectodomain | YES |
| LKSTKRSFIEDL | 1468419 | Ectodomain | YES |
| LKTAGVVTGNVVKQCC | 1675594 | Cell/Virus Enclosed | NO |

| | | | |
|---|---|---|---|
| LKTCVIANVPAIDVLK | 1675595 | Cell/Virus Enclosed | NO |
| LKTDSVLLIERFVSLA | 1675596 | Cell/Virus Enclosed | NO |
| LKTGVTIPCICG | 1468438 | Cell/Virus Enclosed | NO |
| LKTLKLTLTSNG | 1468449 | Cell/Virus Enclosed | NO |
| LKTMSLYMAISP | 1468457 | Cell/Virus Enclosed | NO |
| LKTPDTYKCPQS | 1468463 | Ectodomain | YES |
| LKVEFWFAMRKDGDDV | 1675633 | Cell/Virus Enclosed | NO |
| LKVGAGVMLECEKFNV | 1675635 | Cell/Virus Enclosed | NO |
| LKVNCCRFQRVDENGD | 1675639 | Cell/Virus Enclosed | NO |
| LKVNCCRFQRVG | 1468498 | Cell/Virus Enclosed | NO |
| LKVSTNLFEGDKFVGT | 1675641 | Cell/Virus Enclosed | NO |
| LKVTAFQHQNSK | 1468502 | Cell/Virus Enclosed | NO |
| LKWADQSDQFRNVQTR | 1675645 | Cell/Virus Enclosed | NO |
| LKWADQSDQFRS | 1468505 | Cell/Virus Enclosed | NO |
| LKWADQSDQVRN | 1468506 | Cell/Virus Enclosed | NO |
| LKWRKDIGFRVNANTAALSN | 1951006 | Capsid | UN |
| LKWSDQSDQFRN | 1468515 | Cell/Virus Enclosed | NO |
| LKYFGGFNFSQILPDPL | 1384627 | Ectodomain | YES |
| LKYNPSRVEAFHRYGTTVNC | 226063 | Ectodomain | YES |
| LKYSYINKCSRLLSDD | 1675663 | Ectodomain | YES |
| LKYTKILKDDGNFVVL | 1675664 | Cell/Virus Enclosed | NO |
| LKYTKIMKDDGNCVVL | 1675665 | Cell/Virus Enclosed | NO |
| LLAFNQDGIIFNAVDC | 1675674 | Ectodomain | YES |
| LLAGSFDKVYDIGNPK | 1675675 | Cell/Virus Enclosed | NO |
| LLCATCPFAKGDSCSS | 1675707 | Cell/Virus Enclosed | NO |
| LLCATCPFVKGD | 1468639 | Cell/Virus Enclosed | NO |
| LLCEPIKLVNSE | 1468650 | Cell/Virus Enclosed | NO |
| LLCLRDDNQTLLVPKI | 1675717 | Cell/Virus Enclosed | NO |
| LLCVANDKPIVL | 1468662 | Cell/Virus Enclosed | NO |
| LLCVTCPFAKGD | 1468667 | Cell/Virus Enclosed | NO |
| LLDLAGLFRILL | 1468693 | Cell/Virus Enclosed | NO |
| LLDNQEDKFWCEDFYA | 1675749 | Cell/Virus Enclosed | NO |
| LLDQAWRFPCAGRKVN | 1675752 | Cell/Virus Enclosed | NO |
| LLDQAWRVPCAGRRVT | 1675753 | Cell/Virus Enclosed | NO |
| LLDRLNALESKMSGKG | 1675754 | Cell/Virus Enclosed | NO |
| LLDRLNRLESKV | 1468719 | Capsid | UN |
| LLDTKLNGILQQAALD | 1675758 | Cell/Virus Enclosed | NO |
| LLDTTQLHVADTLMQG | 1675759 | Ectodomain | YES |
| LLDVCNTKEFKV | 1468727 | Cell/Virus Enclosed | NO |
| LLDWVKHLNKNLNEGV | 1675764 | Cell/Virus Enclosed | NO |
| LLDWVKHLNKTLNEGV | 1675765 | Cell/Virus Enclosed | NO |
| LLEAYNAFLNTV | 1468732 | Cell/Virus Enclosed | NO |
| LLEKGKLLIRDTNKEV | 1675779 | Cell/Virus Enclosed | NO |
| LLENGRLIIREN | 1468754 | Cell/Virus Enclosed | NO |
| LLENIDYFNMRRAKFS | 1675781 | Cell/Virus Enclosed | NO |
| LLEQNIDAYKTFPKKE | 1675783 | Cell/Virus Enclosed | NO |
| LLEQWNLVIGFLFLT | 1320697 | Undetermined | UN |
| LLESELVIGAVILRG | 1312993 | Underneath Membrane | NO |
| LLEYDFTEHKEN | 1468780 | Cell/Virus Enclosed | NO |
| LLFDWIKISADN | 1468789 | Cell/Virus Enclosed | NO |
| LLFQLCTFTRST | 1468824 | Cell/Virus Enclosed | NO |
| LLGCLSSMNIAANSSE | 1675831 | Cell/Virus Enclosed | NO |
| LLGFDSSKPVAK | 1468855 | Cell/Virus Enclosed | NO |
| LLGFDSSKPVIK | 1468856 | Cell/Virus Enclosed | NO |
| LLGFDSSKPVTKKYTY | 1675833 | Cell/Virus Enclosed | NO |
| LLGINYLGTIKENIDG | 1675844 | Cell/Virus Enclosed | NO |
| LLGNSIGIDFQD | 1468875 | Ectodomain | YES |
| LLGTAKKQVVLVSNNQ | 1675853 | Cell/Virus Enclosed | NO |
| LLGTTVEKLRPI | 1468894 | Cell/Virus Enclosed | NO |
| LLGVSDVWHKQL | 1468897 | Cell/Virus Enclosed | NO |
| LLHFTNDKTIFY | 1468905 | Cell/Virus Enclosed | NO |
| LLHIAGCQTTNGLNTS | 1675865 | Ectodomain | YES |
| LLHKPIIWHINQ | 1468917 | Cell/Virus Enclosed | NO |
| LLHNLQADICYN | 1468923 | Cell/Virus Enclosed | NO |
| LLHSKVKVVPNLYVVV | 1468930 | Cell/Virus Enclosed | NO |
| LLHSSQLPLDVYDLTM | 1675874 | Cell/Virus Enclosed | NO |
| LLHTKVSWAGSK | 1468937 | Cell/Virus Enclosed | NO |
| LLICVHDPLPVI | 1468951 | Undetermined | UN |
| LLIERFVSLAIDAYPL | 1675880 | Cell/Virus Enclosed | NO |
| LLIKAYNTINNEQGTP | 1675891 | Cell/Virus Enclosed | NO |
| LLIKRFVSLAIH | 1468970 | Cell/Virus Enclosed | NO |
| LLIQEFLQYDSSIHSY | 1675899 | Cell/Virus Enclosed | NO |
| LLISDMYDGRTKAIDG | 1675906 | Cell/Virus Enclosed | NO |
| LLKDALQSREAV | 1469025 | Cell/Virus Enclosed | NO |
| LLKDIKFVIPALQHAF | 1675924 | Cell/Virus Enclosed | NO |
| LLKDIKLVIPAL | 1469029 | Cell/Virus Enclosed | NO |

| | | | |
|---|---|---|---|
| LLKFDNRGVITNAVDC | 1675930 | Ectodomain | YES |
| LLKIAQYAAQNRRGLDLLFWEQG | 37441 | Ectodomain | YES |
| LLKKMDEPFTED | 1469051 | Cell/Virus Enclosed | NO |
| LLKKMDEPFTEH | 1469052 | Capsid | UN |
| LLKKMDEPFTVD | 1469053 | Cell/Virus Enclosed | NO |
| LLKLKVSTNLFEGDKF | 1675937 | Cell/Virus Enclosed | NO |
| LLKTDSVLLIERFVSL | 1675948 | Cell/Virus Enclosed | NO |
| LLKVTAFQHQNS | 1469084 | Cell/Virus Enclosed | NO |
| LLKYDFTEHKENLFNK | 1675957 | Cell/Virus Enclosed | NO |
| LLKYDFTEHKEVLFNK | 1675958 | Cell/Virus Enclosed | NO |
| LLLCLIFLLVLLDYQ | 1675964 | Undetermined | UN |
| LLLDDFVSVLKSLDLT | 1675968 | Cell/Virus Enclosed | NO |
| LLLDQVLVSDVG | 1469109 | Cell/Virus Enclosed | NO |
| LLLDRLNALESKMSGK | 1675974 | Cell/Virus Enclosed | NO |
| LLLECITVDVLK | 1469113 | Cell/Virus Enclosed | NO |
| LLLEYGQFCSKI | 1469116 | Ectodomain | YES |
| LLLGDFVEIIKS | 1469123 | Cell/Virus Enclosed | NO |
| LLLLDRLNALESKMSG | 1675987 | Cell/Virus Enclosed | NO |
| LLLLDRLNQLEN | 1469148 | Cell/Virus Enclosed | NO |
| LLLLDRLNQLESKMSGKGQQQQGQ | 1311743 | Cell/Virus Enclosed | NO |
| LLLLDRLNRLES | 1469150 | Capsid | UN |
| LLLLWESGKAKPPLNR | 1675995 | Cell/Virus Enclosed | NO |
| LLLMLGTTMVLL | 1469163 | Cell/Virus Enclosed | NO |
| LLLNDFVEIIKS | 1469168 | Cell/Virus Enclosed | NO |
| LLLNRGVYKPLLENID | 1676000 | Cell/Virus Enclosed | NO |
| LLLRHDIQLNNDVRDY | 1676005 | Cell/Virus Enclosed | NO |
| LLLSRLWFKPPF | 1469194 | Ectodomain | YES |
| LLLSWESGKVKPPLNR | 1676012 | Cell/Virus Enclosed | NO |
| LLLWESGKAKPPLNRN | 1676021 | Cell/Virus Enclosed | NO |
| LLMPLRAPKEVT | 1469242 | Cell/Virus Enclosed | NO |
| LLNCLQHLHVASNSKL | 1676040 | Cell/Virus Enclosed | NO |
| LLNGIAYTTFSGSFDN | 1676045 | Cell/Virus Enclosed | NO |
| LLNKHIDAYKAF | 1469275 | Cell/Virus Enclosed | NO |
| LLNPSALEFNPSQTSP | 1676060 | Capsid | UN |
| LLNRGDYNMLLKNIDL | 1676063 | Cell/Virus Enclosed | NO |
| LLNRGVYKPLLENIDY | 1676064 | Cell/Virus Enclosed | NO |
| LLNVPLHGTILTRPL | 1541374 | Underneath Membrane | NO |
| LLNVPLRGTILT | 1469307 | Underneath Membrane | NO |
| LLNVPLRGTIVT | 1469308 | Underneath Membrane | NO |
| LLPHSNLDHILEPSIPWKSKLLT | 37661 | Ectodomain | YES |
| LLPLLFVTMAFV | 1469330 | Cell/Virus Enclosed | NO |
| LLPNKSGKKGNSADL | 227347 | Secreted | YES |
| LLQDKAGDFMST | 1469362 | Cell/Virus Enclosed | NO |
| LLQDKIGDFMST | 1469363 | Cell/Virus Enclosed | NO |
| LLQDKTGDFMSI | 1469364 | Cell/Virus Enclosed | NO |
| LLQFAYANRNRFLYI | 2001189 | Undetermined | UN |
| LLQGKTGDFMST | 1469370 | Cell/Virus Enclosed | NO |
| LLQHANLESAKR | 1469371 | Cell/Virus Enclosed | NO |
| LLQHANLESARR | 1469372 | Cell/Virus Enclosed | NO |
| LLQLPNFYYVSNGGNN | 1676106 | Ectodomain | YES |
| LLQPTANSTTVS | 1469385 | Cell/Virus Enclosed | NO |
| LLQVGDSVLLQG | 1469397 | Cell/Virus Enclosed | NO |
| LLQVTAFQHQNS | 1469400 | Cell/Virus Enclosed | NO |
| LLRGMPIPDFDL | 1469411 | Cell/Virus Enclosed | NO |
| LLRKMDEPFTED | 1469420 | Capsid | UN |
| LLRNTSQKSIVA | 1469436 | Ectodomain | YES |
| LLRSTSQKSIVAYTMSL | 37758 | Ectodomain | YES |
| LLRTSIMKRDIK | 1469449 | Cell/Virus Enclosed | NO |
| LLRVRTATRDYAGLYVLRVW | 741670 | Ectodomain | YES |
| LLSDDRTEVPQLVNAN | 1676147 | Ectodomain | YES |
| LLSDHQDLKWAR | 1469464 | Cell/Virus Enclosed | NO |
| LLSENQFSGYTL | 1469470 | Ectodomain | YES |
| LLSGDRTEVPQL | 1469482 | Ectodomain | YES |
| LLSKGSLEPRHFNSIT | 1676170 | Cell/Virus Enclosed | NO |
| LLSKLWFKPPFL | 1469506 | Ectodomain | YES |
| LLSNQLRLAFLGASVT | 1676184 | Cell/Virus Enclosed | NO |
| LLSQGKLLIRDNDTLS | 1676185 | Cell/Virus Enclosed | NO |
| LLSSLTLTVKFA | 1469540 | Cell/Virus Enclosed | NO |
| LLSWESGKVKPPLNRN | 1676203 | Cell/Virus Enclosed | NO |
| LLTFNDNQAITQVPAT | 1676210 | Cell/Virus Enclosed | NO |
| LLTHCAFAVDPA | 1469573 | Cell/Virus Enclosed | NO |
| LLTHCSFAVDPA | 1469574 | Cell/Virus Enclosed | NO |
| LLTLVQLTLQSTNYTCIVCIDRA | 94590 | Ectodomain | YES |
| LLTNTSSVVDGVVRSF | 1676224 | Ectodomain | YES |
| LLTNTSSVVDSV | 1469596 | Ectodomain | YES |
| LLTPMGIDLDEW | 1469600 | Cell/Virus Enclosed | NO |

| | | | |
|---|---|---|---|
| LLTRGTLEPENFNSVT | 1676228 | Cell/Virus Enclosed | NO |
| LLTTNTTGNPEY | 37876 | Secreted | YES |
| LLVAAGMEAQFL | 1469617 | Secreted | YES |
| LLVDKKLITTACNGIS | 1676243 | Cell/Virus Enclosed | NO |
| LLVDSEQPWKVV | 1469633 | Cell/Virus Enclosed | NO |
| LLVDSNLNDFVS | 1469635 | Cell/Virus Enclosed | NO |
| LLVILAGPCILRQLRHLPSRVRY | 37899 | Undetermined | UN |
| LLVQDLCLNKLL | 1469675 | Cell/Virus Enclosed | NO |
| LLVQDLRLNKLL | 1469677 | Cell/Virus Enclosed | NO |
| LLVQNQSFNKLL | 1469679 | Cell/Virus Enclosed | NO |
| LLVSSDKAPHRV | 1469686 | Cell/Virus Enclosed | NO |
| LLVTASSKPLYVPEHC | 1676276 | Cell/Virus Enclosed | NO |
| LLWESGKAKPPLNRNS | 1676284 | Cell/Virus Enclosed | NO |
| LLYCDSPFEKLQCEHL | 1676295 | Ectodomain | YES |
| LLYDSNGNIIGFKDFV | 1676298 | Ectodomain | YES |
| LLYDSNGNLYGFRDYI | 1676299 | Ectodomain | YES |
| LLYDSNGNLYGL | 1469730 | Ectodomain | YES |
| LLYDSPCFSQQG | 1469731 | Ectodomain | YES |
| LLYFMVDNGTRL | 1469734 | Undetermined | UN |
| LLYGLKMLVLWL | 1469736 | Transmembrane | NO |
| LLYLDLLNRLQALESG | 1676307 | Cell/Virus Enclosed | NO |
| LLYNSPCFSQQG | 1469751 | Ectodomain | YES |
| LLYNTPCFSQQG | 1469755 | Ectodomain | YES |
| LLYQPPNCSITSGVLQ | 1676312 | Cell/Virus Enclosed | NO |
| LLYSFEAHKDSFKDGL | 1676314 | Cell/Virus Enclosed | NO |
| LLYTFYGKESFE | 1469766 | Cell/Virus Enclosed | NO |
| LMAAYVEETSIT | 1469782 | Cell/Virus Enclosed | NO |
| LMACNGHKIEDF | 1469785 | Cell/Virus Enclosed | NO |
| LMADVDDPKLMGWDYP | 1676325 | Cell/Virus Enclosed | NO |
| LMAQSSGVLPVN | 1469794 | Cell/Virus Enclosed | NO |
| LMEGSLPYSSIQANAY | 1676340 | Cell/Virus Enclosed | NO |
| LMEILNRDRVNI | 1469826 | Cell/Virus Enclosed | NO |
| LMESELVIGAVIIRGHLR | 38014 | Underneath Membrane | NO |
| LMGANGNKIVDKEVKD | 1676344 | Cell/Virus Enclosed | NO |
| LMGFKLDLTLDGYSKL | 1676345 | Cell/Virus Enclosed | NO |
| LMGILSNMNIASNSKE | 1676350 | Cell/Virus Enclosed | NO |
| LMGWDYPKCDRALPNM | 1676352 | Cell/Virus Enclosed | NO |
| LMGWDYPKCDRAMPNI | 1676353 | Cell/Virus Enclosed | NO |
| LMGWDYPKCDRAMPNL | 1676354 | Cell/Virus Enclosed | NO |
| LMGWDYPKCDRAMPSM | 1676356 | Cell/Virus Enclosed | NO |
| LMKPRQKRTPNK | 1469865 | Cell/Virus Enclosed | NO |
| LMLDKQGQLLDI | 1469873 | Cell/Virus Enclosed | NO |
| LMLDKQGQLLDTKLNG | 1676370 | Cell/Virus Enclosed | NO |
| LMMYAADPAMHI | 1469893 | Cell/Virus Enclosed | NO |
| LMNVITLVYKVY | 1469895 | Cell/Virus Enclosed | NO |
| LMPHKLKIQVVNSGSD | 1676388 | Cell/Virus Enclosed | NO |
| LMPTYTVHKSDF | 1469912 | Cell/Virus Enclosed | NO |
| LMPTYTVHKSGF | 1469913 | Cell/Virus Enclosed | NO |
| LMQGVTLSSNLNTNLH | 1676402 | Ectodomain | YES |
| LMSEDVKLNPSEDFIK | 1676410 | Cell/Virus Enclosed | NO |
| LMSFPQAAPHGVVFLHV | 38118 | Ectodomain | YES |
| LMTFSGNVDLDALASD | 1676424 | Cell/Virus Enclosed | NO |
| LMTGRIPVSDIFCDI | 227350 | Cell/Virus Enclosed | NO |
| LMVDKNLITTANTGTS | 1676427 | Cell/Virus Enclosed | NO |
| LMYLCLPTFSMA | 1469983 | Cell/Virus Enclosed | NO |
| LNADETKALKEL | 1469992 | Cell/Virus Enclosed | NO |
| LNAFVAQQLVRSESAA | 1676446 | Ectodomain | YES |
| LNAKLGFDSSKEFVEY | 1676450 | Cell/Virus Enclosed | NO |
| LNALESKMSGKGSQQQSQTVTK | 1384735 | Cell/Virus Enclosed | NO |
| LNALIATAHSSIKQGT | 1676452 | Cell/Virus Enclosed | NO |
| LNANMSLAECKRALGL | 1676453 | Cell/Virus Enclosed | NO |
| LNASEVVVNNLDKSAG | 1676459 | Cell/Virus Enclosed | NO |
| LNASTPESKPLA | 1470021 | Cell/Virus Enclosed | NO |
| LNASVTLKICKFSRNT | 1676461 | Ectodomain | YES |
| LNAYQQQDGMNM | 1470028 | Cell/Virus Enclosed | NO |
| LNAYQQQDGMVN | 1470029 | Cell/Virus Enclosed | NO |
| LNAYVNSNQNTDSDSL | 1676465 | Capsid | UN |
| LNAYVSQQLSDSTLVK | 1676467 | Ectodomain | YES |
| LNCERVVKLQNNEIMP | 1676470 | Cell/Virus Enclosed | NO |
| LNCLQHLHVASNSKLW | 1676471 | Cell/Virus Enclosed | NO |
| LNCPLDSRLKGS | 1470049 | Ectodomain | YES |
| LNCSFQDAASNIFVIN | 1676475 | Cell/Virus Enclosed | NO |
| LNCTLHPRLKGS | 1470056 | Ectodomain | YES |
| LNDKLGFNVDLP | 1470076 | Cell/Virus Enclosed | NO |
| LNDKLVPSPLNW | 1470077 | Ectodomain | YES |
| LNDPSSKTVCTYMDLL | 1676497 | Cell/Virus Enclosed | NO |

| | | | |
|---|---|---|---|
| LNDVKDNDGRPVHVKE | 1676499 | Cell/Virus Enclosed | NO |
| LNDVLSRLDKVE | 1470094 | Ectodomain | YES |
| LNEFVSDADSTL | 1470106 | Cell/Virus Enclosed | NO |
| LNEHVTSVNLFD | 1470110 | Cell/Virus Enclosed | NO |
| LNEIAKNLNESL | 1470111 | Ectodomain | YES |
| LNENLNAYQQQDGMMN | 1676513 | Cell/Virus Enclosed | NO |
| LNESYIDLKELGNYTY | 1676517 | Ectodomain | YES |
| LNEVNDLLDITQ | 1470130 | Ectodomain | YES |
| LNFGMLNGNAIATVKS | 1676522 | Cell/Virus Enclosed | NO |
| LNFLGGTPVCLGQNS | 1676527 | Ectodomain | YES |
| LNFNENGTITDA | 1470147 | Ectodomain | YES |
| LNFSGALAEHDF | 1470151 | Cell/Virus Enclosed | NO |
| LNFVGEFVINDV | 1470155 | Cell/Virus Enclosed | NO |
| LNGCTWWLKGEK | 1470165 | Cell/Virus Enclosed | NO |
| LNGDFLHFLPRV | 1470166 | Cell/Virus Enclosed | NO |
| LNGIAYITFSGS | 1470181 | Cell/Virus Enclosed | NO |
| LNGIISNARNGCIPLS | 1676543 | Cell/Virus Enclosed | NO |
| LNGITYTSTSGNLLGF | 1676547 | Ectodomain | YES |
| LNGLDSLDSYPA | 1470191 | Cell/Virus Enclosed | NO |
| LNGMPVSSIKGD | 1470197 | Cell/Virus Enclosed | NO |
| LNGMPVSSIKSD | 1470198 | Cell/Virus Enclosed | NO |
| LNGNAIATVKSEDGNI | 1676559 | Cell/Virus Enclosed | NO |
| LNGNVSTVFVPATRDS | 1676563 | Cell/Virus Enclosed | NO |
| LNGSAGSVGFNI | 1470208 | Cell/Virus Enclosed | NO |
| LNGSCPDKNNGI | 1470211 | Ectodomain | YES |
| LNGSCPGKNNGI | 1470212 | Ectodomain | YES |
| LNGTVSTVFVPA | 1470216 | Cell/Virus Enclosed | NO |
| LNGVCGVSAARL | 1470217 | Cell/Virus Enclosed | NO |
| LNGVIREKLAIG | 1470220 | Cell/Virus Enclosed | NO |
| LNGVIVDKVGELKVEF | 1676572 | Cell/Virus Enclosed | NO |
| LNGVMVDKVGDT | 1470221 | Cell/Virus Enclosed | NO |
| LNHGCTCDRTAIQSFD | 1676582 | Cell/Virus Enclosed | NO |
| LNHSVPKLSDFESELS | 1676586 | Ectodomain | YES |
| LNICFDIRDNCSLEKF | 1676592 | Cell/Virus Enclosed | NO |
| LNIIVPDKSVYDQVVD | 1676598 | Cell/Virus Enclosed | NO |
| LNILFDGRDTGALEAF | 1676601 | Cell/Virus Enclosed | NO |
| LNILQATTANVS | 1470263 | Cell/Virus Enclosed | NO |
| LNISEEFSSNVA | 1470268 | Cell/Virus Enclosed | NO |
| LNIVGPDARQDGRQSY | 1676605 | Cell/Virus Enclosed | NO |
| LNKALEMCIDQV | 1470276 | Cell/Virus Enclosed | NO |
| LNKATNGAMQQE | 1470278 | Cell/Virus Enclosed | NO |
| LNKEMYLKLRSE | 1470285 | Cell/Virus Enclosed | NO |
| LNKFGKASLYYESISY | 1676613 | Cell/Virus Enclosed | NO |
| LNKFNRASLRQIFDVA | 1676614 | Cell/Virus Enclosed | NO |
| LNKHFNMMILSD | 1470291 | Cell/Virus Enclosed | NO |
| LNKHFSMMILSDDGVV | 1676617 | Cell/Virus Enclosed | NO |
| LNKIHDVVNQQG | 1470295 | Ectodomain | YES |
| LNKIQDVVNQQGNSLN | 1676619 | Ectodomain | YES |
| LNKIQDVVNQQGSALN | 1676620 | Ectodomain | YES |
| LNKLGTEDPPWP | 1470301 | Cell/Virus Enclosed | NO |
| LNKLGTEDPRWPQIAE | 1676621 | Cell/Virus Enclosed | NO |
| LNKLLHWSDQTICYKD | 1676622 | Cell/Virus Enclosed | NO |
| LNKLNLPGCNGGSLYV | 1676624 | Cell/Virus Enclosed | NO |
| LNKNLNEGVLESFSVT | 1676625 | Cell/Virus Enclosed | NO |
| LNKNQKLIATAFNNAL | 1676626 | Ectodomain | YES |
| LNKRWQLALYKG | 1470320 | Secreted | YES |
| LNKSAGWPLNKFGKAG | 1676634 | Cell/Virus Enclosed | NO |
| LNKSAGWPLNKFGKAS | 1676635 | Cell/Virus Enclosed | NO |
| LNKTLNEGVLESFSVT | 1676636 | Cell/Virus Enclosed | NO |
| LNLGEIFIAQSK | 1470328 | Cell/Virus Enclosed | NO |
| LNLGEVFIAQSK | 1470330 | Cell/Virus Enclosed | NO |
| LNLQFEMNRLQE | 1470342 | Ectodomain | YES |
| LNLSADTQHISN | 1470347 | Cell/Virus Enclosed | NO |
| LNLSAGTQHISNDFTP | 1676660 | Capsid | UN |
| LNLSSELKQLEAKTAS | 1676662 | Ectodomain | YES |
| LNMDMGVFINKYGLED | 1676668 | Cell/Virus Enclosed | NO |
| LNNCTKYNIYDYVGTG | 1676672 | Ectodomain | YES |
| LNNDYYRSVPGT | 1470371 | Cell/Virus Enclosed | NO |
| LNNGQGENDNIS | 1470378 | Cell/Virus Enclosed | NO |
| LNNKNIYRSQLG | 1470382 | Cell/Virus Enclosed | NO |
| LNNKSTYRSQLG | 1470386 | Cell/Virus Enclosed | NO |
| LNNPAYIAPQAN | 1470394 | Ectodomain | YES |
| LNNPAYIAPRAN | 1470395 | Ectodomain | YES |
| LNNPAYIAREAN | 1470396 | Ectodomain | YES |
| LNNPAYIASQAK | 1470397 | Ectodomain | YES |
| LNNSIPNLSDFE | 1470409 | Ectodomain | YES |

| | | | |
|---|---|---|---|
| LNNSYINLKDIGTYEM | 1676692 | Ectodomain | YES |
| LNNVSVPSPFNW | 1470414 | Ectodomain | YES |
| LNPGALEFNPSQ | 1470420 | Cell/Virus Enclosed | NO |
| LNPSALEFNPSQTSPA | 1676701 | Capsid | UN |
| LNPSEDFIKHVRTNGG | 1676702 | Cell/Virus Enclosed | NO |
| LNPSEEFIKHVR | 1470426 | Cell/Virus Enclosed | NO |
| LNQIWQKYLNSS | 1470435 | Cell/Virus Enclosed | NO |
| LNQLESKMSGKGQQQQS | 1384789 | Cell/Virus Enclosed | NO |
| LNQMTGFKKPAS | 1470444 | Cell/Virus Enclosed | NO |
| LNQSYINLKDIGTYEY | 1676710 | Ectodomain | YES |
| LNQSYINLKYIG | 1470446 | Ectodomain | YES |
| LNRARGSSAARLEPCN | 1676713 | Cell/Virus Enclosed | NO |
| LNRGDYNMLLKNIDLF | 1676716 | Cell/Virus Enclosed | NO |
| LNRGGDENPRGPVSHEHVDW | 1951138 | Capsid | UN |
| LNRGVYKPLLENIDYF | 1676718 | Cell/Virus Enclosed | NO |
| LNRLIEKTNQQFELIDNEFNEVEKQ | 1157672 | Ectodomain | YES |
| LNRLQALESGKT | 1470466 | Cell/Virus Enclosed | NO |
| LNRNYVFTGYRV | 1470468 | Cell/Virus Enclosed | NO |
| LNSCCRIVERNVVRAA | 1676737 | Cell/Virus Enclosed | NO |
| LNSEKHQDHVSLTVPL | 1676739 | Cell/Virus Enclosed | NO |
| LNSILDNAVKGCVPLN | 1676742 | Cell/Virus Enclosed | NO |
| LNSILDNAVKGCVPLS | 1676743 | Cell/Virus Enclosed | NO |
| LNSLTYFNKPSFKSEN | 1676752 | Cell/Virus Enclosed | NO |
| LNSLTYFNRPSLVDDN | 1676753 | Cell/Virus Enclosed | NO |
| LNSNSTPAQNAG | 1470512 | Cell/Virus Enclosed | NO |
| LNSSQGIPDLLV | 1470525 | Ectodomain | YES |
| LNSSSCVGSGPG | 1470526 | Ectodomain | YES |
| LNTDHSGSNDNI | 1470542 | Underneath Membrane | NO |
| LNTDHSNSSDNIALLV | 1676769 | Underneath Membrane | NO |
| LNTICIKDADYNAKVE | 1676770 | Cell/Virus Enclosed | NO |
| LNTNLHFDVDNINFKS | 1676774 | Ectodomain | YES |
| LNTNLHSDVDNI | 1470559 | Ectodomain | YES |
| LNTPIPNLPDFR | 1470560 | Ectodomain | YES |
| LNTPKDHIGTRNPANN | 1676775 | Cell/Virus Enclosed | NO |
| LNTSIPNLHDFK | 1470565 | Ectodomain | YES |
| LNTSIPNLPDFKEELD | 1676778 | Ectodomain | YES |
| LNTSYSVCNGCV | 1470570 | Ectodomain | YES |
| LNTTFLLNGYYP | 1470571 | Ectodomain | YES |
| LNTTIPRISEYV | 1470573 | Underneath Membrane | NO |
| LNTTLFLNGYYPTSGS | 1676781 | Ectodomain | YES |
| LNVKPYGYPNDF | 1470593 | Cell/Virus Enclosed | NO |
| LNVQKVAQPKIE | 1470609 | Cell/Virus Enclosed | NO |
| LNVQYRINGLGVTMDV | 1676799 | Ectodomain | YES |
| LNVTINNFNPSS | 1470613 | Ectodomain | YES |
| LNVVGLDARTQA | 1470619 | Cell/Virus Enclosed | NO |
| LNVVGPDARTQGKQSY | 1676802 | Cell/Virus Enclosed | NO |
| LNWLNRVETYIK | 1470626 | Ectodomain | YES |
| LNYAGALWDITV | 1470629 | Cell/Virus Enclosed | NO |
| LNYTVQKLQTLIDNIN | 1676811 | Ectodomain | YES |
| LPAADLDEFSKQ | 1470640 | Cell/Virus Enclosed | NO |
| LPAIKLNFGMLNGNAI | 1676820 | Cell/Virus Enclosed | NO |
| LPAINVTINNYNPSSW | 1676821 | Ectodomain | YES |
| LPAPRVLLSKGSLEPR | 1676826 | Cell/Virus Enclosed | NO |
| LPAPRVLLSKGTLEPK | 1676827 | Cell/Virus Enclosed | NO |
| LPAQDTWGTSAA | 1470652 | Ectodomain | YES |
| LPAYASFKVIDNGVVR | 1676829 | Cell/Virus Enclosed | NO |
| LPCQFKDKNLQDLWVL | 1676833 | Cell/Virus Enclosed | NO |
| LPDAAEKLASPM | 1470668 | Cell/Virus Enclosed | NO |
| LPDCDIDKWLNNFNVP | 1676836 | Ectodomain | YES |
| LPDCDIDNWLNN | 1470671 | Ectodomain | YES |
| LPDFKEELDQWL | 1470679 | Ectodomain | YES |
| LPDFKEELGQWF | 1470680 | Ectodomain | YES |
| LPDFKEESDQWF | 1470681 | Ectodomain | YES |
| LPDIAEDVVDQV | 1470691 | Cell/Virus Enclosed | NO |
| LPDPLKPIKRSF | 1470696 | Ectodomain | YES |
| LPDPLKPTKRSFIEDL | 1676847 | Ectodomain | YES |
| LPDTPSTLKPRS | 1470706 | Ectodomain | YES |
| LPENDDDAWFKP | 1470725 | Cell/Virus Enclosed | NO |
| LPEQFDSLNLSAGTQH | 1676856 | Capsid | UN |
| LPETYVALPIYYQHTD | 1676862 | Ectodomain | YES |
| LPEYMAVAVPST | 1470735 | Underneath Membrane | NO |
| LPEYMTVAVPST | 1470736 | Underneath Membrane | NO |
| LPFDSNLTQYFS | 1470740 | Ectodomain | YES |
| LPFKFKDLAIENMWLS | 1676878 | Cell/Virus Enclosed | NO |
| LPGEFSKKQLGLLQPT | 1676892 | Cell/Virus Enclosed | NO |
| LPHDAIVVDNDVVDYV | 1676906 | Cell/Virus Enclosed | NO |

| | | | |
|---|---|---|---|
| LPHSNLDHILEPSIPWKSKLLTLV | 38524 | Ectodomain | YES |
| LPHVVDLPQLGPRR | 38528 | Cell/Virus Enclosed | NO |
| LPICVYHPLPVI | 1470834 | Undetermined | UN |
| LPINTDTVLSVAPEVD | 1676911 | Cell/Virus Enclosed | NO |
| LPIQDYIQSVNF | 1470847 | Cell/Virus Enclosed | NO |
| LPIYYQHTDINFTATA | 1676915 | Ectodomain | YES |
| LPKDWRLINKFDVING | 1676917 | Cell/Virus Enclosed | NO |
| LPKEDGDVLLAEFDTY | 1676918 | Cell/Virus Enclosed | NO |
| LPKEEGDVLLAE | 1470857 | Cell/Virus Enclosed | NO |
| LPKFFYASNGTYNCTD | 1676919 | Ectodomain | YES |
| LPKFIEQISAFTKPSS | 1676920 | Capsid | UN |
| LPKGFYAEGSRGGSQASSRS | 1863428 | Capsid | UN |
| LPKGVGSANIFIGDKV | 1676923 | Cell/Virus Enclosed | NO |
| LPKGVGSANIFK | 1470868 | Cell/Virus Enclosed | NO |
| LPKYIGLDPVAPGDL | 227357 | Secreted | YES |
| LPLAYFSDSKFG | 1470883 | Cell/Virus Enclosed | NO |
| LPLAYFSDSKLGTTEV | 1676931 | Cell/Virus Enclosed | NO |
| LPLDELKSLLSL | 1470887 | Cell/Virus Enclosed | NO |
| LPLDIYDLTMPS | 1470888 | Cell/Virus Enclosed | NO |
| LPLDKIQNQTLP | 1470889 | Cell/Virus Enclosed | NO |
| LPLDVYNLTMPS | 1470892 | Cell/Virus Enclosed | NO |
| LPLEFGASTEIQ | 1470894 | Cell/Virus Enclosed | NO |
| LPLEFGASTEMQ | 1470895 | Cell/Virus Enclosed | NO |
| LPLRGKLSFWEAGTTKAGYPYNYNT | 38586 | Undetermined | UN |
| LPLSDIEKDFLS | 1470924 | Cell/Virus Enclosed | NO |
| LPLTCNAISSNTDNET | 1676942 | Ectodomain | YES |
| LPMSDMEKDFLS | 1470942 | Cell/Virus Enclosed | NO |
| LPNCNIEAWLNDKSVP | 1676948 | Ectodomain | YES |
| LPNCSITSGVLQ | 1470950 | Cell/Virus Enclosed | NO |
| LPNFYYVSNGGNNCTT | 1676953 | Ectodomain | YES |
| LPNRRKVDLQLG | 1470969 | Ectodomain | YES |
| LPNYTVPDLVVEQYNQ | 1676962 | Ectodomain | YES |
| LPPDAIIIDNDINDYV | 1676965 | Cell/Virus Enclosed | NO |
| LPPELSVVEFEDRSNN | 1676966 | Capsid | UN |
| LPPHDVTVLRDNLKPV | 1676968 | Cell/Virus Enclosed | NO |
| LPPLLSDNQISG | 1470989 | Ectodomain | YES |
| LPPLLSENQFSG | 1470990 | Ectodomain | YES |
| LPPLLSENQING | 1470991 | Ectodomain | YES |
| LPPLLSENQISGYTLA | 1676974 | Ectodomain | YES |
| LPPPLLGNSIGI | 1471001 | Ectodomain | YES |
| LPPPLLSNSTGI | 1471006 | Ectodomain | YES |
| LPPTAPPLLPHSNLDHILEPSIPW | 78454 | Ectodomain | YES |
| LPPVADLPQPGLRR | 38632 | Cell/Virus Enclosed | NO |
| LPQFTSDQAVTFLKEW | 1676983 | Undetermined | UN |
| LPQFYIYDEEGGYDVS | 1676984 | Cell/Virus Enclosed | NO |
| LPQGGFVADFAY | 1471014 | Cell/Virus Enclosed | NO |
| LPQGYYIEGSGK | 1471018 | Cell/Virus Enclosed | NO |
| LPQLVNANQYSP | 1471024 | Ectodomain | YES |
| LPQRKKQPTVTL | 1471027 | Cell/Virus Enclosed | NO |
| LPQRNIRSSRIAGRSA | 1676995 | Ectodomain | YES |
| LPQRQKKQQTVTLLPAADLDDFSK | 1311750 | Cell/Virus Enclosed | NO |
| LPQSGPIGGIKSMGITTLVQ | 38659 | Capsid | UN |
| LPQSKDSNFLNESGVL | 1676997 | Cell/Virus Enclosed | NO |
| LPQSKDSNFLNRVRGS | 1676998 | Cell/Virus Enclosed | NO |
| LPQSKESNFLNE | 1471038 | Cell/Virus Enclosed | NO |
| LPQSKESNFLNR | 1471039 | Cell/Virus Enclosed | NO |
| LPSAPQTIQPEVKEVA | 1677006 | Cell/Virus Enclosed | NO |
| LPSGNYLAISNE | 1471066 | Ectodomain | YES |
| LPSGNYLSISNE | 1471067 | Ectodomain | YES |
| LPSSKSSGMDTA | 1471082 | Ectodomain | YES |
| LPSTQKGSGMDTALLR | 1677022 | Ectodomain | YES |
| LPTAFAAIGDLK | 1471093 | Ectodomain | YES |
| LPTAFAVIGDLKCTSD | 1677025 | Ectodomain | YES |
| LPTAFAVVGDLN | 1471097 | Ectodomain | YES |
| LPTDAIIIDNDLNEFV | 1677026 | Cell/Virus Enclosed | NO |
| LPTDYKNVKAWS | 1471101 | Ectodomain | YES |
| LPTFSMAPVEFYEDRI | 1677030 | Cell/Virus Enclosed | NO |
| LPTHWICANQST | 1471123 | Ectodomain | YES |
| LPTKALCFDKSKQFVP | 1677038 | Underneath Membrane | NO |
| LPTLTQMNLKYAISAK | 1677039 | Cell/Virus Enclosed | NO |
| LPTMCDIKQMLF | 1471128 | Cell/Virus Enclosed | NO |
| LPTSGSRVAGRSAIED | 1677050 | Ectodomain | YES |
| LPTWSQELPFDV | 1471140 | Cell/Virus Enclosed | NO |
| LPVAFTKAAGGKVSFS | 1677056 | Cell/Virus Enclosed | NO |
| LPVAVVNDKVFT | 1471146 | Cell/Virus Enclosed | NO |
| LPVGFSVLKPIL | 1471150 | Ectodomain | YES |

| | | | |
|---|---|---|---|
| LPVKDYIQTVNV | 1471155 | Cell/Virus Enclosed | NO |
| LPVMISEWPLFE | 1471164 | Cell/Virus Enclosed | NO |
| LPVMISEWPLSE | 1471165 | Cell/Virus Enclosed | NO |
| LPVYASYKVLDN | 1471174 | Cell/Virus Enclosed | NO |
| LPVYTSYKVLDN | 1471180 | Cell/Virus Enclosed | NO |
| LPYCYFRNSSANYVGK | 1677080 | Underneath Membrane | NO |
| LPYGANKDGIIWVATEGALN | 1392281 | Capsid | UN |
| LPYGANKQGIVW | 1471191 | Capsid | UN |
| LQAAFGNTGVCY | 1471199 | Cell/Virus Enclosed | NO |
| LQAASDWKPGYSMPVL | 1677094 | Cell/Virus Enclosed | NO |
| LQAGLRKMAQPSGFVE | 1677101 | Cell/Virus Enclosed | NO |
| LQAGNATEVPAH | 1471208 | Cell/Virus Enclosed | NO |
| LQAGSNTEFASNSSVL | 1677103 | Cell/Virus Enclosed | NO |
| LQAGTATEYASNSSIL | 1677104 | Cell/Virus Enclosed | NO |
| LQALESGKTKQS | 1471215 | Cell/Virus Enclosed | NO |
| LQALESGKVKQS | 1471216 | Cell/Virus Enclosed | NO |
| LQALPQRQKKQQ | 1471217 | Cell/Virus Enclosed | NO |
| LQALQSEFVNMASFVE | 1677108 | Cell/Virus Enclosed | NO |
| LQAMRKYSPFRNGYMEPTLG | 38810 | Cell/Virus Enclosed | NO |
| LQASADWKPGHAMPSL | 1677110 | Cell/Virus Enclosed | NO |
| LQDCIVGIADDTYVMG | 1677126 | Cell/Virus Enclosed | NO |
| LQDCVTGINDDDYVIA | 1677127 | Cell/Virus Enclosed | NO |
| LQDFDVVFGHGA | 1471245 | Cell/Virus Enclosed | NO |
| LQDFLPRSTMEEDFLN | 1677129 | Cell/Virus Enclosed | NO |
| LQDKIGDFMSTI | 1471251 | Cell/Virus Enclosed | NO |
| LQDKTGDFMSII | 1471253 | Cell/Virus Enclosed | NO |
| LQDKTGDFTSTI | 1471255 | Cell/Virus Enclosed | NO |
| LQEIISRLDALE | 1471277 | Ectodomain | YES |
| LQELSYKLPNYTVPDL | 1677145 | Ectodomain | YES |
| LQENQKILAASFNKAI | 1677147 | Ectodomain | YES |
| LQENQKILAASFNKAM | 1677148 | Ectodomain | YES |
| LQENQRILAASF | 1471298 | Ectodomain | YES |
| LQETPVVPDTVEVPPQ | 1677149 | Cell/Virus Enclosed | NO |
| LQEVKDNDGKNVHLKD | 1677150 | Cell/Virus Enclosed | NO |
| LQFGLQDGFYSANFLD | 1677155 | Ectodomain | YES |
| LQFTTLTLDKVP | 1471333 | Cell/Virus Enclosed | NO |
| LQGDKCNIFANL | 1471337 | Ectodomain | YES |
| LQGDRCNIFANF | 1471340 | Ectodomain | YES |
| LQGHFLAKNILH | 1471341 | Cell/Virus Enclosed | NO |
| LQGLENIAFNVVNKGS | 1677170 | Cell/Virus Enclosed | NO |
| LQGLIDQINSTYVDLK | 1677171 | Ectodomain | YES |
| LQGLKACCYVGM | 1471351 | Cell/Virus Enclosed | NO |
| LQGNFGDLQFNK | 1471360 | Cell/Virus Enclosed | NO |
| LQGNFGDLQLNKLGTE | 1677174 | Cell/Virus Enclosed | NO |
| LQGNFGDLQLTK | 1471362 | Cell/Virus Enclosed | NO |
| LQGTIIFVQQKP | 1471366 | Cell/Virus Enclosed | NO |
| LQHAFMKHKGGDSTDF | 1677177 | Cell/Virus Enclosed | NO |
| LQHAFMKHKGGG | 1471369 | Cell/Virus Enclosed | NO |
| LQHANLESAKRV | 1471371 | Cell/Virus Enclosed | NO |
| LQHANLESARRV | 1471372 | Cell/Virus Enclosed | NO |
| LQHLHVASNSKLWQYC | 1677182 | Cell/Virus Enclosed | NO |
| LQIQLTTTRMED | 1471382 | Cell/Virus Enclosed | NO |
| LQIQTVDSSQGSEYDY | 1677189 | Cell/Virus Enclosed | NO |
| LQKHLYSHVYRS | 1471402 | Cell/Virus Enclosed | NO |
| LQKLFVANYSQD | 1471403 | Ectodomain | YES |
| LQKRLYSHVYRSDKVD | 1677198 | Cell/Virus Enclosed | NO |
| LQLFLRATTELRTFSILN | 1031292 | Secreted | YES |
| LQLGAPDNPSAI | 1471419 | Ectodomain | YES |
| LQLGFSTGVSFV | 1471422 | Cell/Virus Enclosed | NO |
| LQLGSSGFLQSSNYKI | 1677206 | Ectodomain | YES |
| LQLKESQINELV | 1471427 | Cell/Virus Enclosed | NO |
| LQLNKLGTDDPR | 1471436 | Cell/Virus Enclosed | NO |
| LQLNKLGTEDPP | 1471437 | Cell/Virus Enclosed | NO |
| LQLNKLGTEDPRWPQI | 1677212 | Cell/Virus Enclosed | NO |
| LQLPNFYYVSNGGNNC | 1677213 | Ectodomain | YES |
| LQLPQGTTLPKGFYAEGSRGGS | 1870797 | Capsid | UN |
| LQLVNANQYSPC | 1471449 | Ectodomain | YES |
| LQLVSAVNQSLSNAKV | 1677216 | Cell/Virus Enclosed | NO |
| LQMDVQGLVNVL | 1471452 | Cell/Virus Enclosed | NO |
| LQMGFGITVQYGTDTN | 1677218 | Ectodomain | YES |
| LQNFGNFEMLKL | 1471459 | Cell/Virus Enclosed | NO |
| LQNNEIKPSGLKTMVV | 1677223 | Cell/Virus Enclosed | NO |
| LQNNEIMPGKMK | 1471465 | Cell/Virus Enclosed | NO |
| LQPNAYYKYDVK | 1471488 | Cell/Virus Enclosed | NO |
| LQPNTEVYTDVC | 1471489 | Ectodomain | YES |
| LQPVSDLLTNMG | 1471499 | Cell/Virus Enclosed | NO |

| | | | |
|---|---|---|---|
| LQQAQQEATHIA | 1471506 | Cell/Virus Enclosed | NO |
| LQQEMTIVKCKGLNKP | 1677250 | Cell/Virus Enclosed | NO |
| LQQGAFLCANEY | 1471512 | Cell/Virus Enclosed | NO |
| LQQIEVKFNAPA | 1471518 | Cell/Virus Enclosed | NO |
| LQQMFVTPLAVD | 1471528 | Cell/Virus Enclosed | NO |
| LQQVVKALNESYIDLK | 1677259 | Ectodomain | YES |
| LQRAAVTILDGI | 1471534 | Cell/Virus Enclosed | NO |
| LQSFNYRIDTTATSCQ | 1677278 | Ectodomain | YES |
| LQSGVGRCFNCR | 1471565 | Ectodomain | YES |
| LQSKRTRLFKST | 1471570 | Cell/Virus Enclosed | NO |
| LQSKTKRFIKET | 1471571 | Cell/Virus Enclosed | NO |
| LQSLSKDYISSNGPLK | 1677298 | Cell/Virus Enclosed | NO |
| LQSNNPQCAIVQASES | 1677303 | Cell/Virus Enclosed | NO |
| LQSSNYKIDTTSSSCQ | 1677306 | Ectodomain | YES |
| LQSYACWTHSCG | 1471597 | Cell/Virus Enclosed | NO |
| LQTDVLQEDQKI | 1471601 | Ectodomain | YES |
| LQTGKQTELAVN | 1471605 | Cell/Virus Enclosed | NO |
| LQTHVGCVLGLV | 1471606 | Ectodomain | YES |
| LQTIINGMHKSF | 1471607 | Cell/Virus Enclosed | NO |
| LQTIVPEYIDVNKTLQ | 1677318 | Ectodomain | YES |
| LQTLIDNINSTLVDLK | 1677319 | Ectodomain | YES |
| LQTTGRPSHEAPN | 39001 | Ectodomain | YES |
| LQTVATALNKIQDVVN | 1677325 | Ectodomain | YES |
| LQTYSETGSDLL | 1471624 | Cell/Virus Enclosed | NO |
| LQVANNLMNGVT | 1471628 | Ectodomain | YES |
| LQVESANLMLSDNVVA | 1677331 | Cell/Virus Enclosed | NO |
| LQVESANQMLTVNVVA | 1677332 | Cell/Virus Enclosed | NO |
| LQVFVYSSNEEQAVLK | 1677335 | Cell/Virus Enclosed | NO |
| LQVGFSNGVDFVAQPE | 1677337 | Cell/Virus Enclosed | NO |
| LQYDSSIHSYFITDQE | 1677345 | Cell/Virus Enclosed | NO |
| LQYDVLFYCSNSSSGV | 1677346 | Ectodomain | YES |
| LQYSKPHFISQGLDAA | 1677354 | Cell/Virus Enclosed | NO |
| LRACCYVGVQTV | 1471674 | Cell/Virus Enclosed | NO |
| LRACIKIAQADT | 1471676 | Cell/Virus Enclosed | NO |
| LRACIKTAQADTAGLY | 1677356 | Cell/Virus Enclosed | NO |
| LRACIKTVQADT | 1471681 | Cell/Virus Enclosed | NO |
| LRAFDIYNANVA | 1471688 | Cell/Virus Enclosed | NO |
| LRAFDIYNASVAGIGL | 1677361 | Cell/Virus Enclosed | NO |
| LRAFIKTAQADT | 1471690 | Cell/Virus Enclosed | NO |
| LRAPKEVTFFEG | 1471705 | Cell/Virus Enclosed | NO |
| LRAPTMANQEKYSTIY | 1677372 | Cell/Virus Enclosed | NO |
| LRATEEASKQSYAIAT | 1677377 | Cell/Virus Enclosed | NO |
| LRAVSFADNNDI | 1471712 | Cell/Virus Enclosed | NO |
| LRDDNQALLVPK | 1471729 | Cell/Virus Enclosed | NO |
| LRDFMLKGYVLA | 1471733 | Cell/Virus Enclosed | NO |
| LRDNLKPVTTLSTITA | 1677393 | Cell/Virus Enclosed | NO |
| LRDYLADYDVAVTAGP | 1677394 | Cell/Virus Enclosed | NO |
| LRELGKYEQYIK | 1471764 | Ectodomain | YES |
| LREQLSSVSSFERFEIFPKTSSWPN | 1158864 | Ectodomain | YES |
| LRFPRGQGVPINTNST | 1677413 | Cell/Virus Enclosed | NO |
| LRFTSARRMETTVN | 39089 | Cell/Virus Enclosed | NO |
| LRGDDMSFLEKD | 1471796 | Cell/Virus Enclosed | NO |
| LRGHLRIAGHHLGRC | 1313076 | Underneath Membrane | NO |
| LRGMPIPDFDLDDFID | 1677424 | Cell/Virus Enclosed | NO |
| LRGPGDLQGNFGDLQL | 1677425 | Cell/Virus Enclosed | NO |
| LRGYGFYTTSGI | 1471829 | Ectodomain | YES |
| LRHDITLDDDAR | 1471833 | Cell/Virus Enclosed | NO |
| LRHFDRNDCMLL | 1471839 | Cell/Virus Enclosed | NO |
| LRHGKLRPFERDISNV | 1677440 | Ectodomain | YES |
| LRHLIKDVDNPVLMGW | 1677441 | Cell/Virus Enclosed | NO |
| LRIAGHHLGRCDIKDL | 1677444 | Underneath Membrane | NO |
| LRIFNVVGPRKG | 1471860 | Cell/Virus Enclosed | NO |
| LRIKVSQSNVHT | 1471866 | Cell/Virus Enclosed | NO |
| LRKKRNVDPLSNFEHK | 1677462 | Cell/Virus Enclosed | NO |
| LRKMDEPFTEDT | 1471894 | Capsid | UN |
| LRKYGRGGYHYT | 1471914 | Cell/Virus Enclosed | NO |
| LRLAFLGASVTEDVKF | 1677479 | Cell/Virus Enclosed | NO |
| LRLCAYCCNIVNVSL | 1313078 | Ectodomain | YES |
| LRLGSQLSLSMA | 1471924 | Cell/Virus Enclosed | NO |
| LRLHTVESGDINVVAA | 1677484 | Cell/Virus Enclosed | NO |
| LRLHTVESGDVN | 1471928 | Cell/Virus Enclosed | NO |
| LRLIDDNGIVLNSILW | 1677488 | Cell/Virus Enclosed | NO |
| LRLLLMLCTTMV | 1471936 | Cell/Virus Enclosed | NO |
| LRLPHIRDDDAPPSHQPLFY | 741691 | Underneath Membrane | NO |
| LRLVDCAPVVCPKGKI | 1677496 | Cell/Virus Enclosed | NO |
| LRMAGHSLGRCDIKDLPK | 39141 | Underneath Membrane | NO |

| | | | |
|---|---|---|---|
| LRMGSGFCVDYNSPSS | 1677500 | Ectodomain | YES |
| LRNCTFMYTYNITEDE | 1677502 | Ectodomain | YES |
| LRNFDEKDCEVFKEIL | 1677503 | Cell/Virus Enclosed | NO |
| LRNGSVLEMAKEVKTS | 1677504 | Cell/Virus Enclosed | NO |
| LRNKGVDCTKKF | 1471979 | Cell/Virus Enclosed | NO |
| LRNLQADICYKF | 1471983 | Cell/Virus Enclosed | NO |
| LRNNGVDYTKKF | 1471987 | Cell/Virus Enclosed | NO |
| LRNSFGKDLNANMSLA | 1677509 | Cell/Virus Enclosed | NO |
| LRNSGRFGGFSN | 1471988 | Cell/Virus Enclosed | NO |
| LRNVTFELHEFA | 1472003 | Cell/Virus Enclosed | NO |
| LRPFERDISNVPFSPD | 1677517 | Ectodomain | YES |
| LRPGPVSQPYHHWDTDK | 39164 | Capsid | UN |
| LRPILKLPFGIN | 1472018 | Ectodomain | YES |
| LRPILLPNKSGKKGN | 227365 | Secreted | YES |
| LRPPKNSFEALMLNFK | 1677519 | Cell/Virus Enclosed | NO |
| LRPPKNSFEALV | 1472022 | Cell/Virus Enclosed | NO |
| LRPPRNSFEAIL | 1472023 | Cell/Virus Enclosed | NO |
| LRQAALPQSKDSNFLN | 1677521 | Cell/Virus Enclosed | NO |
| LRQIFDVAPIELENKF | 1677524 | Cell/Virus Enclosed | NO |
| LRRLDMSSVDTILNMA | 1677539 | Cell/Virus Enclosed | NO |
| LRRLDMSSVETVLNLA | 1677540 | Cell/Virus Enclosed | NO |
| LRRLIKDVDNPVLMGW | 1677541 | Cell/Virus Enclosed | NO |
| LRSGHHLHSVHN | 1472099 | Cell/Virus Enclosed | NO |
| LRSINHDLFSFI | 1472108 | Cell/Virus Enclosed | NO |
| LRSITAQPVKIDNASP | 1677555 | Secreted | YES |
| LRSQGFFDEGSELTLK | 1677557 | Cell/Virus Enclosed | NO |
| LRSTRVNVDGFN | 1472125 | Cell/Virus Enclosed | NO |
| LRTDQINDMVYS | 1472136 | Cell/Virus Enclosed | NO |
| LRTNFTIKGSFINGAC | 1677567 | Cell/Virus Enclosed | NO |
| LRTSIMKRDIKV | 1472156 | Cell/Virus Enclosed | NO |
| LRVSSNQSPQGN | 1472187 | Cell/Virus Enclosed | NO |
| LRYMNANGLCPP | 1472211 | Cell/Virus Enclosed | NO |
| LSAHLETNDVSSMLTF | 1677608 | Ectodomain | YES |
| LSAMISGSKHVT | 1472246 | Cell/Virus Enclosed | NO |
| LSAPTLVPQENYASIR | 1677613 | Cell/Virus Enclosed | NO |
| LSAPTLVPQENYSSIR | 1677614 | Cell/Virus Enclosed | NO |
| LSARSAIEDLLF | 1472253 | Ectodomain | YES |
| LSCARKSCSIDS | 1472258 | Cell/Virus Enclosed | NO |
| LSCGIFGIKLETSLEV | 1677621 | Cell/Virus Enclosed | NO |
| LSCNFNMSSLMS | 1472268 | Ectodomain | YES |
| LSDDGVVCYNKDYAEL | 1677632 | Cell/Virus Enclosed | NO |
| LSDDGVVCYNSDYAAK | 1677633 | Cell/Virus Enclosed | NO |
| LSDDGVVCYNSDYASK | 1677634 | Cell/Virus Enclosed | NO |
| LSDFINGIFAKVKNTK | 1677638 | Ectodomain | YES |
| LSDFTPRSDMEYDFLN | 1677640 | Cell/Virus Enclosed | NO |
| LSDIEKDFLSFD | 1472303 | Cell/Virus Enclosed | NO |
| LSDMEKDFLSFDSDVF | 1677649 | Cell/Virus Enclosed | NO |
| LSDMEKDFLSSD | 1472313 | Cell/Virus Enclosed | NO |
| LSDNAYLKWARVEGKD | 1677650 | Cell/Virus Enclosed | NO |
| LSDNQKLIANKF | 1472318 | Ectodomain | YES |
| LSDQFKTTGDLAVQIG | 1677654 | Cell/Virus Enclosed | NO |
| LSDRFKTSGDLAVQIG | 1677657 | Cell/Virus Enclosed | NO |
| LSDTLDKLSDYC | 1472333 | Cell/Virus Enclosed | NO |
| LSDTLKGLSDRV | 1472334 | Cell/Virus Enclosed | NO |
| LSDVAGLKYTKI | 1472338 | Cell/Virus Enclosed | NO |
| LSDVDGLKYTKILKDD | 1677666 | Cell/Virus Enclosed | NO |
| LSDVDGLKYTKIMKDD | 1677667 | Cell/Virus Enclosed | NO |
| LSDVGFVEAYNNCTGG | 1677668 | Ectodomain | YES |
| LSDVIRYNLNFEENLR | 1677669 | Ectodomain | YES |
| LSDYLLDLSDSVVFIT | 1677671 | Cell/Virus Enclosed | NO |
| LSEFSHLATFAELEAA | 1677676 | Cell/Virus Enclosed | NO |
| LSEINMDNSQNL | 1472359 | Cell/Virus Enclosed | NO |
| LSEIQCKTQSFA | 1472360 | Ectodomain | YES |
| LSENLNAYQQQD | 1472376 | Cell/Virus Enclosed | NO |
| LSENQKLIANKFNQAL | 1677686 | Ectodomain | YES |
| LSERCNVIADSKSFTL | 1677690 | Cell/Virus Enclosed | NO |
| LSESQISGYTTAATVA | 1677691 | Ectodomain | YES |
| LSEYGTFCDNINSILD | 1677694 | Ectodomain | YES |
| LSFAGYTESAVNSTKA | 1677698 | Cell/Virus Enclosed | NO |
| LSFCAFAVDPAK | 1472408 | Cell/Virus Enclosed | NO |
| LSFDNDVFIKKY | 1472411 | Cell/Virus Enclosed | NO |
| LSFDSDVFIKKYGLEN | 1677703 | Cell/Virus Enclosed | NO |
| LSFDVPDGFYSTSPIQ | 1677704 | Ectodomain | YES |
| LSFEEQDEIYAYTKRN | 1677706 | Cell/Virus Enclosed | NO |
| LSFEEQNEIYAYTKRN | 1677707 | Cell/Virus Enclosed | NO |
| LSFKENVDFINS | 1472424 | Cell/Virus Enclosed | NO |

| | | | |
|---|---|---|---|
| LSFLPSDFFPSIRDL | 956866 | Capsid | UN |
| LSFNENGTITDA | 1472437 | Ectodomain | YES |
| LSFNNFVSSYAKPEEK | 1677716 | Cell/Virus Enclosed | NO |
| LSFNNLLGFDSSKPVT | 1677717 | Cell/Virus Enclosed | NO |
| LSFTNYKLQSQI | 1472449 | Cell/Virus Enclosed | NO |
| LSGAVPSDLNTSELLT | 1677723 | Cell/Virus Enclosed | NO |
| LSGDDCFVSNSSVIQE | 1677725 | Cell/Virus Enclosed | NO |
| LSGDRTEVPQLV | 1472457 | Ectodomain | YES |
| LSGEWNAVYKAF | 1472459 | Cell/Virus Enclosed | NO |
| LSGFETIMKVLNENLN | 1677728 | Cell/Virus Enclosed | NO |
| LSGIAVEQDRNT | 1472475 | Ectodomain | YES |
| LSGIGAVGGDLL | 1472476 | Cell/Virus Enclosed | NO |
| LSGNLDEPQKDV | 1472483 | Capsid | UN |
| LSGNPDEPQKDVYELR | 1677740 | Cell/Virus Enclosed | NO |
| LSGRIDFHWLMLNPNDTVTFSFNGA | 1159143 | Ectodomain | YES |
| LSGRKIYEVERG | 1472493 | Cell/Virus Enclosed | NO |
| LSGRKIYEVVRG | 1472494 | Cell/Virus Enclosed | NO |
| LSGTPNEKLVTT | 1472499 | Cell/Virus Enclosed | NO |
| LSGTPPQVYNFK | 1472501 | Ectodomain | YES |
| LSGVAYTAFSGP | 1472507 | Cell/Virus Enclosed | NO |
| LSHEKASLNSLT | 1472516 | Cell/Virus Enclosed | NO |
| LSHEKASLSSLT | 1472517 | Cell/Virus Enclosed | NO |
| LSIADLACAQYYNGIM | 1385058 | Ectodomain | YES |
| LSIDDGHLWNLFVEKF | 1677767 | Cell/Virus Enclosed | NO |
| LSIDDGRLWNLF | 1472549 | Cell/Virus Enclosed | NO |
| LSIDEFADVVKE | 1472550 | Cell/Virus Enclosed | NO |
| LSIEEFADVVKEQVSD | 1385059 | Cell/Virus Enclosed | NO |
| LSIEEFADVVKG | 1472553 | Cell/Virus Enclosed | NO |
| LSIEEFAVVVKE | 1472554 | Cell/Virus Enclosed | NO |
| LSINTSCRVLQR | 1472574 | Cell/Virus Enclosed | NO |
| LSISNDTVDVTN | 1472591 | Ectodomain | YES |
| LSKDLPDDVVAANMFM | 1677791 | Cell/Virus Enclosed | NO |
| LSKDYISSNGPLKVGA | 1677792 | Cell/Virus Enclosed | NO |
| LSKELKRPIQPTDVAY | 1677795 | Cell/Virus Enclosed | NO |
| LSKELKRPVNPTDASH | 1677796 | Cell/Virus Enclosed | NO |
| LSKFQLKLKGTPVLQL | 1677800 | Cell/Virus Enclosed | NO |
| LSKGSLEPRHFNSITK | 1677805 | Cell/Virus Enclosed | NO |
| LSKHSDFGLGDLVDSY | 1677812 | Cell/Virus Enclosed | NO |
| LSKLASELSNTFGAIS | 1677813 | Ectodomain | YES |
| LSKLLCVSPMCMMIEPPKLR | 111521 | Cell/Virus Enclosed | NO |
| LSKMGVLKADDFVTAS | 1677816 | Cell/Virus Enclosed | NO |
| LSKPFVKDNVSF | 1472641 | Cell/Virus Enclosed | NO |
| LSLASPPSPPREPPRPSRSL | 39473 | Cell/Virus Enclosed | NO |
| LSLCAFSVDPKKTYLD | 1677827 | Cell/Virus Enclosed | NO |
| LSLFAQSEEEEP | 1472663 | Cell/Virus Enclosed | NO |
| LSLHPVECDEEC | 1472677 | Cell/Virus Enclosed | NO |
| LSLKELMMYAADPAMH | 1677838 | Cell/Virus Enclosed | NO |
| LSLLDLYLGRGYNVS | 39483 | Ectodomain | YES |
| LSLQQVVKALNESYID | 1677855 | Ectodomain | YES |
| LSLSDRFKTSGDLAVQ | 1677860 | Cell/Virus Enclosed | NO |
| LSLSMVRRNLDS | 1472721 | Cell/Virus Enclosed | NO |
| LSLVNFTVDPQKAYLD | 1677867 | Cell/Virus Enclosed | NO |
| LSLYNKYRYYSGKMDT | 1677870 | Cell/Virus Enclosed | NO |
| LSMFDVDKKSLNALIA | 1677875 | Cell/Virus Enclosed | NO |
| LSMIKGPQRADLNGVI | 1677879 | Cell/Virus Enclosed | NO |
| LSMIRGPPRAELNGVV | 1677880 | Cell/Virus Enclosed | NO |
| LSMKSDLSVSSS | 1472754 | Ectodomain | YES |
| LSMLQLGVPDNSSTIV | 1677882 | Ectodomain | YES |
| LSMTTSTTNFGEDFLG | 1677886 | Cell/Virus Enclosed | NO |
| LSMVRRNLDSLE | 1472764 | Cell/Virus Enclosed | NO |
| LSNAKVSAEVSRQVIQ | 1677887 | Cell/Virus Enclosed | NO |
| LSNELAQVLTEVVYSN | 1677889 | Cell/Virus Enclosed | NO |
| LSNGFSQVKSDL | 1472777 | Cell/Virus Enclosed | NO |
| LSNMNIASNSKEWAYC | 1677899 | Cell/Virus Enclosed | NO |
| LSNQLRLAFLGASVTE | 1677900 | Cell/Virus Enclosed | NO |
| LSNTKYYDDSQYYFNK | 1677903 | Ectodomain | YES |
| LSNTKYYDHSQY | 1472806 | Ectodomain | YES |
| LSPAYAPTYVSVDDKY | 1677906 | Cell/Virus Enclosed | NO |
| LSPLEGGGRLVA | 1472824 | Ectodomain | YES |
| LSPWFMDGENAYEVVK | 1677916 | Cell/Virus Enclosed | NO |
| LSQPRADKPSQLKKPR | 1677923 | Capsid | UN |
| LSQPRSDKPSQL | 1472867 | Cell/Virus Enclosed | NO |
| LSQPRVDKPSQL | 1472868 | Cell/Virus Enclosed | NO |
| LSRDYISSNGPL | 1472875 | Cell/Virus Enclosed | NO |
| LSRLDALEAEAQIDRL | 1677932 | Ectodomain | YES |
| LSRLDVLEAQVQ | 1472884 | Ectodomain | YES |

| | | | |
|---|---|---|---|
| LSRLIIFFSPES | 1472887 | Cell/Virus Enclosed | NO |
| LSRNSSRSSSQGSRSG | 1677939 | Cell/Virus Enclosed | NO |
| LSRSNSNFRHSDSIVK | 1677940 | Capsid | UN |
| LSRSQSSETKEQ | 1472899 | Cell/Virus Enclosed | NO |
| LSSACTMFTMADGSPQ | 1677943 | Cell/Virus Enclosed | NO |
| LSSACTMFTTAD | 1472906 | Cell/Virus Enclosed | NO |
| LSSDSDVFIKKY | 1472911 | Cell/Virus Enclosed | NO |
| LSSDVMFEDAQG | 1472912 | Cell/Virus Enclosed | NO |
| LSSDVMFEDDQG | 1472914 | Cell/Virus Enclosed | NO |
| LSSDVMFEGAQG | 1472915 | Cell/Virus Enclosed | NO |
| LSSNLNTNLHFDVDNI | 1677960 | Ectodomain | YES |
| LSSSNVTTMDFC | 1472942 | Cell/Virus Enclosed | NO |
| LSSVIPSLPTSGSRVA | 1677964 | Ectodomain | YES |
| LSSVLPQRNIRSSRIA | 1677966 | Ectodomain | YES |
| LSSYEYCQPPKSVEKI | 1677968 | Cell/Virus Enclosed | NO |
| LSTCDGLKFTEESFYK | 1677969 | Cell/Virus Enclosed | NO |
| LSTCDGQKFTDESFYKNMY | 1472958 | Cell/Virus Enclosed | NO |
| LSTDTGAENATF | 1472962 | Secreted | YES |
| LSTFVSAARQGV | 1472971 | Cell/Virus Enclosed | NO |
| LSTGCCGCCNCL | 1472972 | Undetermined | UN |
| LSTGLTSQTVKPGHFN | 1677987 | Cell/Virus Enclosed | NO |
| LSTKLKDGVNFNVDDI | 1677989 | Ectodomain | YES |
| LSTLPETTVVRRRGRSPRRRTPSPR | 79609 | Capsid | UN |
| LSTWHVLYSPNVSVPSSSSTPLL | 39726 | Ectodomain | YES |
| LSVDLSLTVELKIKI | 39732 | Ectodomain | YES |
| LSVDSVSEESQGNVVT | 1678021 | Cell/Virus Enclosed | NO |
| LSVEDCKLPLGQ | 1473017 | Ectodomain | YES |
| LSVNSSNCNNFNVKKL | 1678045 | Cell/Virus Enclosed | NO |
| LSVNSSNCNNLN | 1473047 | Cell/Virus Enclosed | NO |
| LSVQQAQQEATH | 1473053 | Cell/Virus Enclosed | NO |
| LSVQQAQQEATLPDIA | 1678047 | Cell/Virus Enclosed | NO |
| LSVSPYVCNAPGCGVS | 1678050 | Cell/Virus Enclosed | NO |
| LSVVEFEDRSNNSSRA | 1678056 | Capsid | UN |
| LSWEIGKVKPPLNKNY | 1678062 | Cell/Virus Enclosed | NO |
| LSWESGKVKPPLNRNS | 1678063 | Cell/Virus Enclosed | NO |
| LSYEELKTGVTI | 1473089 | Cell/Virus Enclosed | NO |
| LSYHYENNSWFFRHVY | 1678076 | Cell/Virus Enclosed | NO |
| LSYKLPNYTVPDLVVE | 1678078 | Ectodomain | YES |
| LSYYRLGASQRV | 1473111 | Underneath Membrane | NO |
| LSYYVPSVEYTYTDEV | 1678086 | Cell/Virus Enclosed | NO |
| LTAATIVNQERY | 1473118 | Cell/Virus Enclosed | NO |
| LTACNGHKIEDL | 1473120 | Cell/Virus Enclosed | NO |
| LTADEIKALKEL | 1473121 | Cell/Virus Enclosed | NO |
| LTADETKALKELYGPV | 1678088 | Cell/Virus Enclosed | NO |
| LTAEDISLLKKMDEPY | 1678090 | Cell/Virus Enclosed | NO |
| LTAEDISLLRKM | 1473124 | Capsid | UN |
| LTAFSPAQDIWGTSAA | 1678094 | Ectodomain | YES |
| LTAFSPAQGTWG | 1473131 | Ectodomain | YES |
| LTAGETKALKEL | 1473133 | Cell/Virus Enclosed | NO |
| LTALDAYVSQQL | 1473136 | Ectodomain | YES |
| LTALNAYVSQQLSDIS | 1678095 | Ectodomain | YES |
| LTALNAYVSQQLSDST | 1678096 | Ectodomain | YES |
| LTALRLCAYCCNIVN | 2001195 | Undetermined | UN |
| LTANMSMAECKATLGL | 1678098 | Cell/Virus Enclosed | NO |
| LTASMSMAECKA | 1473150 | Cell/Virus Enclosed | NO |
| LTASTIVNQERY | 1473151 | Cell/Virus Enclosed | NO |
| LTCHNAAQCDCL | 1473165 | Ectodomain | YES |
| LTCIRRPKDGFN | 1473176 | Ectodomain | YES |
| LTCKAISSNTDN | 1473187 | Ectodomain | YES |
| LTCNAISPKTDK | 1473197 | Ectodomain | YES |
| LTCNAISSNTDNETLQ | 1678118 | Ectodomain | YES |
| LTCSTDLQKANT | 1473206 | Ectodomain | YES |
| LTCVKDYEFKFY | 1473210 | Cell/Virus Enclosed | NO |
| LTDAALSIKENVDFIN | 1678124 | Cell/Virus Enclosed | NO |
| LTDCLYLKNLTQTFTS | 1678126 | Cell/Virus Enclosed | NO |
| LTDESCNNLVPTYLKS | 1678130 | Cell/Virus Enclosed | NO |
| LTDESCNNLVQT | 1473220 | Cell/Virus Enclosed | NO |
| LTDHDVVYAQHC | 1473222 | Ectodomain | YES |
| LTDIQEDITRHEQQ | 39849 | Capsid | UN |
| LTDLKCTNVVLLGCLS | 1678133 | Cell/Virus Enclosed | NO |
| LTDLKCTNVVLMGILS | 1678134 | Cell/Virus Enclosed | NO |
| LTDQMSEPPKEQ | 1473237 | Cell/Virus Enclosed | NO |
| LTDVKCANVVLLNCLQ | 1678139 | Cell/Virus Enclosed | NO |
| LTDYNAILQSDN | 1473247 | Cell/Virus Enclosed | NO |
| LTDYNAILQSNNPQCA | 1678140 | Cell/Virus Enclosed | NO |
| LTEGQLQSLSRD | 1473258 | Cell/Virus Enclosed | NO |

| | | | |
|---|---|---|---|
| LTEHVTSVNFFD | 1473259 | Cell/Virus Enclosed | NO |
| LTEHVTSVNLFD | 1473260 | Cell/Virus Enclosed | NO |
| LTEVNELLDTTQLQVA | 1678151 | Ectodomain | YES |
| LTFDSNAFSLAN | 1473287 | Ectodomain | YES |
| LTFNDNQAITQVPATS | 1678164 | Cell/Virus Enclosed | NO |
| LTFNKQEASVPILTTP | 1678165 | Cell/Virus Enclosed | NO |
| LTFPPGQGVPLNANST | 1678166 | Cell/Virus Enclosed | NO |
| LTFSYDGLRGAIRKAK | 1678168 | Cell/Virus Enclosed | NO |
| LTFVNVPYVYNGSAQA | 1678172 | Ectodomain | YES |
| LTGCCDNSYFDSKGWY | 1678178 | Cell/Virus Enclosed | NO |
| LTGDSVKFVYMHQLEL | 1678183 | Cell/Virus Enclosed | NO |
| LTGHAIQGFACG | 1473344 | Cell/Virus Enclosed | NO |
| LTGIAIEQDKNT | 1473347 | Ectodomain | YES |
| LTGILDGDYAMQFFKM | 1678189 | Cell/Virus Enclosed | NO |
| LTGNAIQGFACG | 1473352 | Cell/Virus Enclosed | NO |
| LTGTNQLCAKILPFSD | 1678196 | Cell/Virus Enclosed | NO |
| LTGTQSVSVDRE | 1473362 | Cell/Virus Enclosed | NO |
| LTHCAFAVDPAA | 1473367 | Cell/Virus Enclosed | NO |
| LTHCSFAVDPAA | 1473368 | Cell/Virus Enclosed | NO |
| LTHCSFAVDPAV | 1473369 | Cell/Virus Enclosed | NO |
| LTHFQGFVLNYK | 1473370 | Cell/Virus Enclosed | NO |
| LTIANNYSKLFDEGYT | 1678206 | Cell/Virus Enclosed | NO |
| LTITELLQFVTDPSLI | 1385165 | Cell/Virus Enclosed | NO |
| LTITELLQFVTDPTLI | 1385166 | Cell/Virus Enclosed | NO |
| LTITRMEDAMGQ | 1473405 | Cell/Virus Enclosed | NO |
| LTITSAGQPCVA | 1473408 | Cell/Virus Enclosed | NO |
| LTIVDIPQSLTFSYDG | 1385167 | Cell/Virus Enclosed | NO |
| LTIVIPDKQVFDKVVD | 1678220 | Cell/Virus Enclosed | NO |
| LTKDQTPIVWHA | 1473424 | Cell/Virus Enclosed | NO |
| LTKKKYSELDDE | 1473434 | Cell/Virus Enclosed | NO |
| LTKLGTEDPRWP | 1473441 | Cell/Virus Enclosed | NO |
| LTKLSKYLANEAQVQL | 1678237 | Cell/Virus Enclosed | NO |
| LTKLSVKCYFNY | 1473450 | Ectodomain | YES |
| LTKSGSTYPVLNVT | 39998 | Ectodomain | YES |
| LTLDDRGLALDNALSG | 1678258 | Cell/Virus Enclosed | NO |
| LTLDGYSKLFITKDEA | 1678259 | Cell/Virus Enclosed | NO |
| LTLDNQDLNGKWYDFG | 1678261 | Cell/Virus Enclosed | NO |
| LTLTKALAAESH | 1473511 | Cell/Virus Enclosed | NO |
| LTLTPRNKWSGV | 1473517 | Cell/Virus Enclosed | NO |
| LTLTSGSDLDRCTTFD | 1678286 | Ectodomain | YES |
| LTLTVKFAVESN | 1473522 | Cell/Virus Enclosed | NO |
| LTMIEFMLWCKDGQVQ | 1678295 | Cell/Virus Enclosed | NO |
| LTMLDIQPEDYKSVDV | 1678297 | Cell/Virus Enclosed | NO |
| LTMVEFMLWCKD | 1473551 | Cell/Virus Enclosed | NO |
| LTNHDVVYAQHCFKAP | 1678312 | Ectodomain | YES |
| LTNKISLNAPVT | 1473572 | Ectodomain | YES |
| LTNQVVKPGHFNEEFY | 1678323 | Cell/Virus Enclosed | NO |
| LTNTGSVVKPVRARAP | 1678325 | Cell/Virus Enclosed | NO |
| LTNTSSVVDGVVRSFQ | 1678326 | Ectodomain | YES |
| LTPAWRIYSTGNNVFQ | 40075 | Ectodomain | YES |
| LTPCGYSEPPKVAAKI | 1678332 | Cell/Virus Enclosed | NO |
| LTPIKLNFGMLNGMPV | 1678338 | Cell/Virus Enclosed | NO |
| LTPILSCGIFGIKLET | 1678339 | Cell/Virus Enclosed | NO |
| LTPKTGTGIAIS | 1473612 | Cell/Virus Enclosed | NO |
| LTPLLTDDMIAA | 1473617 | Ectodomain | YES |
| LTPLYGFPSQAA | 1473619 | Cell/Virus Enclosed | NO |
| LTPMGIDLDEWS | 1473620 | Cell/Virus Enclosed | NO |
| LTPSDVYQQLAGIKLQ | 1678347 | Cell/Virus Enclosed | NO |
| LTPTESYVDVGL | 1473637 | Ectodomain | YES |
| LTPTESYVDVGPDSVK | 1678350 | Ectodomain | YES |
| LTQDHVEILGPL | 1473649 | Cell/Virus Enclosed | NO |
| LTQDLFLPFYSN | 1473650 | Ectodomain | YES |
| LTQDYFLPFDSN | 1473651 | Ectodomain | YES |
| LTQDYFLPFDTN | 1473652 | Ectodomain | YES |
| LTQHGKEDLKFPRGQ | 1321406 | Capsid | UN |
| LTQMNLKYAISAKNRA | 1678363 | Cell/Virus Enclosed | NO |
| LTQTFTSMLTNYFLDDVEM | 1385188 | Cell/Virus Enclosed | NO |
| LTRALAAESHMD | 1473677 | Cell/Virus Enclosed | NO |
| LTRGSDLDRCTT | 1473684 | Ectodomain | YES |
| LTRGTLEPENFNSVTR | 1678372 | Cell/Virus Enclosed | NO |
| LTRPLLESELVIGAV | 1313109 | Underneath Membrane | NO |
| LTRVEASTVVCG | 1473698 | Cell/Virus Enclosed | NO |
| LTSCFFTCNSNA | 1473704 | Ectodomain | YES |
| LTSDENGVRTLS | 1473705 | Ectodomain | YES |
| LTSDYMITVFVT | 1473706 | Cell/Virus Enclosed | NO |
| LTSEISILENKS | 1473715 | Ectodomain | YES |

| | | | |
|---|---|---|---|
| LTSEISTLENKSAELN | 1678387 | Ectodomain | YES |
| LTSGVTPDMADQ | 1473723 | Cell/Virus Enclosed | NO |
| LTSHSVANLSAPTLVP | 1678392 | Cell/Virus Enclosed | NO |
| LTSHSVASLSAPTLVP | 1678393 | Cell/Virus Enclosed | NO |
| LTSIAEESEPGPVVLLSVSP | 741706 | Underneath Membrane | NO |
| LTSQTVKPGHFNKEFY | 1678407 | Cell/Virus Enclosed | NO |
| LTSSMRGCCDCG | 1473756 | Underneath Membrane | NO |
| LTSTHDGPTEHV | 1473764 | Cell/Virus Enclosed | NO |
| LTTCAFSVDPAT | 1473773 | Cell/Virus Enclosed | NO |
| LTTFIDAARGPAGVES | 1678422 | Cell/Virus Enclosed | NO |
| LTTLNAFVAQQL | 1473789 | Ectodomain | YES |
| LTTQTVDSSQGSEYQY | 1678435 | Cell/Virus Enclosed | NO |
| LTTVPAAPPPASTNR | 1678444 | Underneath Membrane | NO |
| LTTYYLDDVKKIEYKP | 1678448 | Cell/Virus Enclosed | NO |
| LTVDGVNFTNRFVPVG | 1678452 | Cell/Virus Enclosed | NO |
| LTVDGVQYYCYT | 1473817 | Cell/Virus Enclosed | NO |
| LTVDMIAAYTAA | 1473818 | Ectodomain | YES |
| LTVDVANPSTPAYTFT | 1678454 | Cell/Virus Enclosed | NO |
| LTVGDIFVLTSHSVAT | 1385202 | Cell/Virus Enclosed | NO |
| LTVKFAVESNVM | 1473836 | Cell/Virus Enclosed | NO |
| LTVMSYQMQGCM | 1473848 | Cell/Virus Enclosed | NO |
| LTVMSYQMRGCM | 1473849 | Cell/Virus Enclosed | NO |
| LTVPFCGSGNLV | 1473851 | Cell/Virus Enclosed | NO |
| LTVSDDDFVSAVANAH | 1385206 | Cell/Virus Enclosed | NO |
| LTVTLQNSCTPK | 1473863 | Cell/Virus Enclosed | NO |
| LTVTSAGQPCVV | 1473866 | Cell/Virus Enclosed | NO |
| LTVTSAGQPYVA | 1473867 | Cell/Virus Enclosed | NO |
| LTVTSDGQPYVA | 1473868 | Cell/Virus Enclosed | NO |
| LTVVSKVHEVIIDNKP | 1678473 | Cell/Virus Enclosed | NO |
| LTWPLGLECTRA | 1473878 | Cell/Virus Enclosed | NO |
| LTWSQELPFDVI | 1473882 | Cell/Virus Enclosed | NO |
| LTYANIVHEPTT | 1473885 | Cell/Virus Enclosed | NO |
| LTYEMLSLQQVV | 1473888 | Ectodomain | YES |
| LTYFNKPSFKSENRYS | 1678484 | Cell/Virus Enclosed | NO |
| LTYFNRPSLVDDNKFD | 1678485 | Cell/Virus Enclosed | NO |
| LTYLNLSSELKQ | 1473904 | Ectodomain | YES |
| LTYNKGENMTPR | 1473907 | Cell/Virus Enclosed | NO |
| LTYNKQMANVSVLTTP | 1678490 | Cell/Virus Enclosed | NO |
| LTYQNISTNLPP | 1473910 | Ectodomain | YES |
| LTYTGAIKLDDKDPNFKDQV | 2116225 | Capsid | UN |
| LTYTGAIKLDDKDPSF | 1678495 | Cell/Virus Enclosed | NO |
| LTYYLHSVLYRV | 1473916 | Cell/Virus Enclosed | NO |
| LVAKCYGESVLV | 1473932 | Cell/Virus Enclosed | NO |
| LVALANMLPAHVFMRF | 1678510 | Cell/Virus Enclosed | NO |
| LVAQEFVSYDSS | 1473947 | Cell/Virus Enclosed | NO |
| LVAQNTVLNDSI | 1473948 | Capsid | UN |
| LVASGPTVAMTE | 1473951 | Ectodomain | YES |
| LVASIKNFKAVH | 1473955 | Cell/Virus Enclosed | NO |
| LVCAPAPCNFFTSA | 40283 | Cell/Virus Enclosed | NO |
| LVCASRELERFAVNPGLLE | 165638 | Undetermined | UN |
| LVCITIIKLTKL | 1473982 | Cell/Virus Enclosed | NO |
| LVCTTLQGKPIG | 1473992 | Cell/Virus Enclosed | NO |
| LVCYIVMPLHTL | 1473995 | Cell/Virus Enclosed | NO |
| LVDDNKFDGLKV | 1474004 | Cell/Virus Enclosed | NO |
| LVDGVSRLYQPLRLTC | 1678541 | Ectodomain | YES |
| LVDLKWLNRVETYIKW | 1678545 | Ectodomain | YES |
| LVDNDLYPFVSDSVAT | 1678549 | Cell/Virus Enclosed | NO |
| LVDQRTICFSVAALST | 1678551 | Cell/Virus Enclosed | NO |
| LVDSDLNEFVSD | 1474031 | Cell/Virus Enclosed | NO |
| LVDSEQPCLVIP | 1474034 | Cell/Virus Enclosed | NO |
| LVDSEQPWKVIP | 1474035 | Cell/Virus Enclosed | NO |
| LVDSEQSWKVIP | 1474037 | Capsid | UN |
| LVDSSLVTSVGDSSEI | 1678555 | Cell/Virus Enclosed | NO |
| LVDSYFENDSILQSVA | 1678556 | Cell/Virus Enclosed | NO |
| LVEAYNSILVEN | 1474044 | Cell/Virus Enclosed | NO |
| LVEDSTSVTAVV | 1474046 | Ectodomain | YES |
| LVEKLSPWFMDGENAY | 1678563 | Cell/Virus Enclosed | NO |
| LVEKPNLSSKRSELS | 40319 | Ectodomain | YES |
| LVEVPVEDIAQV | 1474068 | Cell/Virus Enclosed | NO |
| LVFQNNVANYQHIGMK | 1678586 | Cell/Virus Enclosed | NO |
| LVFSNSNYLLEEFDVV | 1678588 | Cell/Virus Enclosed | NO |
| LVFTNKTTLPTNVAFE | 1678591 | Cell/Virus Enclosed | NO |
| LVFVNKTSLPTNIAFE | 1678594 | Cell/Virus Enclosed | NO |
| LVFVTTLTSDYT | 1474114 | Cell/Virus Enclosed | NO |
| LVGCLGPHCGSS | 1474117 | Ectodomain | YES |
| LVGHDICAQLNA | 1474123 | Cell/Virus Enclosed | NO |

| | | | |
|---|---|---|---|
| LVGHDICAQLND | 1474124 | Cell/Virus Enclosed | NO |
| LVGQLEQTNMHS | 1474130 | Cell/Virus Enclosed | NO |
| LVGTSVHPVLWN | 1474141 | Cell/Virus Enclosed | NO |
| LVGVLTLDNQDLNGKW | 1678610 | Cell/Virus Enclosed | NO |
| LVHDNVAFYQGDVDTV | 1678613 | Cell/Virus Enclosed | NO |
| LVHHENEEYQKVFRVY | 1678614 | Cell/Virus Enclosed | NO |
| LVIGAVIIRGHLRMAGHSLGR | 40389 | Underneath Membrane | NO |
| LVIIANRYNEVANAVM | 1678625 | Cell/Virus Enclosed | NO |
| LVILLIFGNMYL | 1474183 | Cell/Virus Enclosed | NO |
| LVISATSNYSGV | 1474193 | Cell/Virus Enclosed | NO |
| LVKDPPNVQIHTIDGS | 1678638 | Secreted | YES |
| LVKDVDHVQIHT | 1474207 | Secreted | YES |
| LVKFSATQAMEK | 1474212 | Ectodomain | YES |
| LVKIQAVVNADA | 1474218 | Ectodomain | YES |
| LVKIQAVVNANAEALN | 1678646 | Ectodomain | YES |
| LVKKGNSYPKLSKSY | 151031 | Ectodomain | YES |
| LVKMDGDDVYLPYPNP | 1678652 | Cell/Virus Enclosed | NO |
| LVKPSFYVYSRVKNLNSSRVPDL | 1597756 | Ectodomain | YES |
| LVKTGHYIGQAG | 1474247 | Cell/Virus Enclosed | NO |
| LVKTGHYTGQAGEMPC | 1678665 | Cell/Virus Enclosed | NO |
| LVLAKGGFKFGDPADS | 1678668 | Cell/Virus Enclosed | NO |
| LVLAKGHFKFDEPSDA | 1678669 | Cell/Virus Enclosed | NO |
| LVLAKLGKDATKPQQV | 1678670 | Cell/Virus Enclosed | NO |
| LVLAKLGKDSKPQQVT | 1678671 | Capsid | UN |
| LVLAKLGRDATK | 1474258 | Cell/Virus Enclosed | NO |
| LVLGINEKTHVQ | 1474274 | Cell/Virus Enclosed | NO |
| LVLGSKHVNCCT | 1474275 | Cell/Virus Enclosed | NO |
| LVLLDYQGMLPVCPL | 40448 | Undetermined | UN |
| LVLLENERTLDYHDSNVKNLYEKVR | 1159730 | Ectodomain | YES |
| LVLNNCTKYNIYDYVG | 1678695 | Ectodomain | YES |
| LVLNNPAYIATQ | 1474304 | Ectodomain | YES |
| LVLNNPAYLAPQ | 1474305 | Ectodomain | YES |
| LVLNNPSYIAPQ | 1474306 | Ectodomain | YES |
| LVLNNPVYIAPQ | 1474307 | Ectodomain | YES |
| LVLPSGNYLSIS | 1474313 | Ectodomain | YES |
| LVMQKLPFKFKDLAIE | 1678722 | Cell/Virus Enclosed | NO |
| LVNAANTHLKHGGGIA | 1678725 | Cell/Virus Enclosed | NO |
| LVNANHYSPCVS | 1474351 | Ectodomain | YES |
| LVNANQYSPCVP | 1474352 | Ectodomain | YES |
| LVNANQYSPCVSIVPS | 1678726 | Ectodomain | YES |
| LVNFTVDPQKAYLDFV | 1678730 | Cell/Virus Enclosed | NO |
| LVNGLHYPTGSC | 1474363 | Cell/Virus Enclosed | NO |
| LVNGVHHPTGSC | 1474366 | Cell/Virus Enclosed | NO |
| LVNGVHYPTGGC | 1474367 | Cell/Virus Enclosed | NO |
| LVNGVHYPTGIC | 1474368 | Cell/Virus Enclosed | NO |
| LVNGVHYPTGSC | 1474369 | Cell/Virus Enclosed | NO |
| LVNKMVNWFNTMLDAS | 1678737 | Cell/Virus Enclosed | NO |
| LVNPTDKSHYYV | 1474378 | Cell/Virus Enclosed | NO |
| LVNSNQYSPCVS | 1474379 | Ectodomain | YES |
| LVNSSLFVEDCKLPLG | 1678739 | Ectodomain | YES |
| LVNSVSTKEGDSSNTGAGKA | 40495 | Capsid | UN |
| LVNVGHYDGRTGELPC | 1678740 | Cell/Virus Enclosed | NO |
| LVNVQKVEQPKIEPKP | 1678743 | Cell/Virus Enclosed | NO |
| LVNWYIYVRKNG | 1474392 | Cell/Virus Enclosed | NO |
| LVNYKQSIPMPR | 1474395 | Cell/Virus Enclosed | NO |
| LVPCASGLSTDVQLRA | 1678749 | Cell/Virus Enclosed | NO |
| LVPIGKGNKDEQIGYW | 1678760 | Capsid | UN |
| LVPIGKGSKDEQ | 1474416 | Cell/Virus Enclosed | NO |
| LVPTTTGGLGRF | 1474447 | Cell/Virus Enclosed | NO |
| LVPTTTGGLVRF | 1474448 | Cell/Virus Enclosed | NO |
| LVPTYLKSDNIVAADL | 1678774 | Cell/Virus Enclosed | NO |
| LVPVSPALHRIE | 1474455 | Cell/Virus Enclosed | NO |
| LVQAGNVHLRVI | 1474460 | Cell/Virus Enclosed | NO |
| LVQAGNVQLRVI | 1474461 | Cell/Virus Enclosed | NO |
| LVQCTCGTKSWS | 1474464 | Cell/Virus Enclosed | NO |
| LVQLTLQSTNYTCIVCIDRASLST | 40541 | Ectodomain | YES |
| LVQNGVDAKGFPQLAE | 1678788 | Capsid | UN |
| LVQNQSFNKLLH | 1474481 | Cell/Virus Enclosed | NO |
| LVQPVGALDASA | 1474486 | Cell/Virus Enclosed | NO |
| LVQQITQWLSNM | 1474491 | Cell/Virus Enclosed | NO |
| LVQTYLKSDNIV | 1474498 | Cell/Virus Enclosed | NO |
| LVQYDFTDYKLELFNK | 1678797 | Cell/Virus Enclosed | NO |
| LVRGFGDSVEEA | 1474508 | Cell/Virus Enclosed | NO |
| LVRLLQDKTGDF | 1474520 | Cell/Virus Enclosed | NO |
| LVRLLQGKTGDF | 1474522 | Cell/Virus Enclosed | NO |
| LVSAVNQSLSNAKVSA | 1678820 | Cell/Virus Enclosed | NO |

| | | | |
|---|---|---|---|
| LVSGDAHIPFGTVLGN | 1678826 | Ectodomain | YES |
| LVSGTATAGWTFGAGA | 1678827 | Ectodomain | YES |
| LVSKIFKARAIE | 1474562 | Cell/Virus Enclosed | NO |
| LVSNNKEDFDIIQKCQ | 1678831 | Cell/Virus Enclosed | NO |
| LVSNNQEDFDLISKCQ | 1678832 | Cell/Virus Enclosed | NO |
| LVSSDGQPGGDAISLS | 1678834 | Cell/Virus Enclosed | NO |
| LVSSDKAPHRVI | 1474578 | Cell/Virus Enclosed | NO |
| LVSSNGQPGGDA | 1474583 | Cell/Virus Enclosed | NO |
| LVSVTDFSLNGNVSTV | 1678841 | Cell/Virus Enclosed | NO |
| LVTAQDGCGHIS | 1474601 | Ectodomain | YES |
| LVTASSKPLYVPEHCQ | 1678843 | Cell/Virus Enclosed | NO |
| LVTDIVVVDEVSMLTN | 1678845 | Cell/Virus Enclosed | NO |
| LVTICDFYSKLGVGFT | 1678850 | Cell/Virus Enclosed | NO |
| LVTLAILTALRLCAY | 40612 | Undetermined | UN |
| LVTQFKLLGTTL | 1474631 | Cell/Virus Enclosed | NO |
| LVTTEVKVSALK | 1474637 | Cell/Virus Enclosed | NO |
| LVTTSMAPDFVA | 1474642 | Cell/Virus Enclosed | NO |
| LVTTSTAPDFVA | 1474643 | Cell/Virus Enclosed | NO |
| LVVDDKYKVNEN | 1474659 | Cell/Virus Enclosed | NO |
| LVVDWHYFDAQP | 1474661 | Cell/Virus Enclosed | NO |
| LVVEQYNQTILNLTSE | 1678876 | Ectodomain | YES |
| LVVVVPDHDSFVKMMV | 1678886 | Cell/Virus Enclosed | NO |
| LVVYQQAVVGAMLSEN | 1678890 | Ectodomain | YES |
| LVWEAGKSKPPLNRNY | 1678891 | Cell/Virus Enclosed | NO |
| LVWHVERKDVPYPKQS | 1678892 | Cell/Virus Enclosed | NO |
| LVYALRHFDQNSEVLK | 1678899 | Cell/Virus Enclosed | NO |
| LVYDNKLKAKNDNSSL | 1678904 | Cell/Virus Enclosed | NO |
| LVYENKFVPVKPASKQ | 1678907 | Cell/Virus Enclosed | NO |
| LVYHENEEYQKVFRVY | 1678911 | Cell/Virus Enclosed | NO |
| LVYKVYYGNSLD | 1474716 | Cell/Virus Enclosed | NO |
| LVYLNHSVPKLSDFES | 1678913 | Ectodomain | YES |
| LWCKDGKVQTFY | 1474738 | Cell/Virus Enclosed | NO |
| LWCKDGQVQTFYPRLQ | 1678927 | Cell/Virus Enclosed | NO |
| LWDITVINNVDNEIVK | 1678936 | Cell/Virus Enclosed | NO |
| LWDYEAERPFTS | 1474748 | Cell/Virus Enclosed | NO |
| LWESGKAKPPLNRNSV | 1678942 | Cell/Virus Enclosed | NO |
| LWFHISCLTFGRETVIEYLV | 96630 | Capsid | UN |
| LWIPIIKSSVKSYDDL | 1678952 | Cell/Virus Enclosed | NO |
| LWLDDTVYCPRH | 1474798 | Cell/Virus Enclosed | NO |
| LWNCQLSFAGYTESAV | 1678971 | Cell/Virus Enclosed | NO |
| LWNKFVTGDVGP | 1474824 | Cell/Virus Enclosed | NO |
| LWNYLFETIPTQ | 1474832 | Cell/Virus Enclosed | NO |
| LWPLVLALSIFD | 1474836 | Undetermined | UN |
| LWQFVDHFGENEEIII | 1678988 | Cell/Virus Enclosed | NO |
| LWQTFTDVNLQG | 1474848 | Cell/Virus Enclosed | NO |
| LWSVSGLRFTTG | 1474864 | Ectodomain | YES |
| LWSVSGSRFTTG | 1474869 | Ectodomain | YES |
| LWVKKASYDTNL | 1474876 | Cell/Virus Enclosed | NO |
| LWVNKASYDINL | 1474880 | Cell/Virus Enclosed | NO |
| LWVNKASYDTNLNKFN | 1679002 | Cell/Virus Enclosed | NO |
| LWYKGSNLEEEI | 1474890 | Cell/Virus Enclosed | NO |
| LWYVVDDIYYPA | 1474898 | Cell/Virus Enclosed | NO |
| LYAIQEGFDATNSALV | 1679021 | Ectodomain | YES |
| LYAIQQGFDATN | 1474913 | Ectodomain | YES |
| LYAKRALRSHPD | 1474920 | Cell/Virus Enclosed | NO |
| LYAKRAVRSHPDFKLL | 1679024 | Cell/Virus Enclosed | NO |
| LYCAFTAPEDDDFLEE | 1679028 | Cell/Virus Enclosed | NO |
| LYCDSPFEKLQCEHLQ | 1679029 | Ectodomain | YES |
| LYCGVHKGAHVA | 1474933 | Cell/Virus Enclosed | NO |
| LYCRAHIEHPDVSGVC | 1679032 | Cell/Virus Enclosed | NO |
| LYDACNVKKVTD | 1474946 | Cell/Virus Enclosed | NO |
| LYDASNVKKVTDVTGK | 1679039 | Cell/Virus Enclosed | NO |
| LYDASNVKNVTD | 1474951 | Cell/Virus Enclosed | NO |
| LYDFNGNIIGFK | 1474953 | Ectodomain | YES |
| LYDGNMFIKFPEVVFE | 1679041 | Cell/Virus Enclosed | NO |
| LYDNCYRLTSVEESFI | 1679047 | Cell/Virus Enclosed | NO |
| LYDNCYRNSNVG | 1474962 | Cell/Virus Enclosed | NO |
| LYDNCYRNSNVN | 1474963 | Cell/Virus Enclosed | NO |
| LYDSNGNIIGFKDFVT | 1679050 | Ectodomain | YES |
| LYDSNGNLYGFRDYIT | 1679051 | Ectodomain | YES |
| LYDSNGNLYGLR | 1474975 | Ectodomain | YES |
| LYDSPCFSQQGV | 1474976 | Ectodomain | YES |
| LYFIKGLNNLNR | 1475014 | Cell/Virus Enclosed | NO |
| LYGISGQGIFVE | 1475048 | Ectodomain | YES |
| LYGQTVFYFGGC | 1475066 | Cell/Virus Enclosed | NO |
| LYHDSQNMLRGD | 1475080 | Cell/Virus Enclosed | NO |

| | | | |
|---|---|---|---|
| LYIIKLIFLWLLWPV | 1321579 | Undetermined | UN |
| LYIKDGDDGYFLPYPD | 1679110 | Cell/Virus Enclosed | NO |
| LYIKGSGSTANLASSNYFPT | 110938 | Capsid | UN |
| LYIPKPTAKYTP | 1475103 | Cell/Virus Enclosed | NO |
| LYIQGIKLGTGYSLAD | 1679114 | Ectodomain | YES |
| LYIQGVKLGTGYTLSD | 1679115 | Underneath Membrane | NO |
| LYIYALIYILQI | 1475110 | Secreted | YES |
| LYKDLTGYMLDS | 1475115 | Cell/Virus Enclosed | NO |
| LYKDVDNPHLMGWDYP | 1679117 | Cell/Virus Enclosed | NO |
| LYKGFQFISNLL | 1475120 | Secreted | YES |
| LYKGNVTHDASS | 1475123 | Cell/Virus Enclosed | NO |
| LYKKQKEGHIIM | 1475128 | Cell/Virus Enclosed | NO |
| LYKKVNNEFIDPTECI | 1679123 | Cell/Virus Enclosed | NO |
| LYKNMCTGSPPI | 1475138 | Cell/Virus Enclosed | NO |
| LYKNMCTGSPSIVEFN | 1679131 | Cell/Virus Enclosed | NO |
| LYKNSATGSLDVEVFN | 1679132 | Cell/Virus Enclosed | NO |
| LYKQSCTGSPYIDDFN | 1679136 | Cell/Virus Enclosed | NO |
| LYKRNFTYDVNA | 1475145 | Ectodomain | YES |
| LYKSNHNNVYWLTIP | 40826 | Ectodomain | YES |
| LYKSSALGSMDIDVFN | 1679138 | Cell/Virus Enclosed | NO |
| LYKSYSEQVIPPTSDY | 1679139 | Cell/Virus Enclosed | NO |
| LYKYLNSPMERVSLWN | 1679143 | Cell/Virus Enclosed | NO |
| LYLADNLNADET | 1475155 | Cell/Virus Enclosed | NO |
| LYLADNLTAGET | 1475160 | Cell/Virus Enclosed | NO |
| LYLDLLNRLQALESGK | 1679147 | Cell/Virus Enclosed | NO |
| LYLDYVQCKCGNLEQN | 1679148 | Cell/Virus Enclosed | NO |
| LYLGGMRYFCVD | 1475170 | Cell/Virus Enclosed | NO |
| LYLGVMSYFCVD | 1475179 | Cell/Virus Enclosed | NO |
| LYLKNLKQTLKS | 1475181 | Cell/Virus Enclosed | NO |
| LYLKNLTQTFTSMLTN | 1679159 | Cell/Virus Enclosed | NO |
| LYLSSSNVTTMD | 1475194 | Cell/Virus Enclosed | NO |
| LYMIDPAGVCLG | 1475215 | Cell/Virus Enclosed | NO |
| LYNAPCFSQQGV | 1475219 | Ectodomain | YES |
| LYNCVLDYYIVK | 1475221 | Cell/Virus Enclosed | NO |
| LYNKYRYYSGKMDTAA | 1679184 | Cell/Virus Enclosed | NO |
| LYNSPCFSQQGV | 1475245 | Ectodomain | YES |
| LYNSSCFSHQGV | 1475248 | Ectodomain | YES |
| LYNSSCFSQQGV | 1475249 | Ectodomain | YES |
| LYNTEGGKTFMY | 1475254 | Cell/Virus Enclosed | NO |
| LYNTGRSVYVKFQDSK | 1679195 | Cell/Virus Enclosed | NO |
| LYNTPCFSQQGV | 1475257 | Ectodomain | YES |
| LYNVTRDKLEKL | 1475260 | Cell/Virus Enclosed | NO |
| LYNYGAGIKLPSGIML | 1679199 | Cell/Virus Enclosed | NO |
| LYPNVYRTDYVD | 1475268 | Cell/Virus Enclosed | NO |
| LYPTITVPEEFASHVA | 1679203 | Cell/Virus Enclosed | NO |
| LYPYGRCPTAADINIP | 1679205 | Ectodomain | YES |
| LYQDVNCTDVSTAIHAD | 40905 | Ectodomain | YES |
| LYQNADTYVFVGSSR | 151032 | Ectodomain | YES |
| LYQPPTASVSTSFLQS | 1679213 | Cell/Virus Enclosed | NO |
| LYREALESPEHCSPHHTALRQA | 79613 | Capsid | UN |
| LYRRQQTSNLVVQEFV | 1679229 | Cell/Virus Enclosed | NO |
| LYSFDKAIDGFG | 1475321 | Cell/Virus Enclosed | NO |
| LYSFEAHKDSFKDGLC | 1679235 | Cell/Virus Enclosed | NO |
| LYSGETFWRYNF | 1475325 | Cell/Virus Enclosed | NO |
| LYSHHFQSHGQLSDHPHALS | 40948 | Capsid | UN |
| LYSIQDVGFKNQ | 1475327 | Cell/Virus Enclosed | NO |
| LYSNVYRTDYVDYTFV | 1679241 | Cell/Virus Enclosed | NO |
| LYSPLLVDSDQP | 1475335 | Cell/Virus Enclosed | NO |
| LYSPLLVDSEQP | 1475336 | Cell/Virus Enclosed | NO |
| LYSPPTISYNSTLQSG | 1679247 | Cell/Virus Enclosed | NO |
| LYVDFKPQSGGGKCFN | 1679267 | Ectodomain | YES |
| LYVNFEHQRGPG | 1475407 | Ectodomain | YES |
| LYVNVYRSTNPD | 1475419 | Cell/Virus Enclosed | NO |
| LYVNVYRSTSPDPKFV | 1679285 | Cell/Virus Enclosed | NO |
| LYVPIRLAGHTR | 1475425 | Cell/Virus Enclosed | NO |
| LYVSWSDGDGITGVPQ | 1679290 | Ectodomain | YES |
| LYVVVVESDADRANFL | 1679292 | Cell/Virus Enclosed | NO |
| LYVYKGYQPIDVVRDL | 1309876 | Ectodomain | YES |
| LYYKVNCCRFQRIDDD | 1679302 | Cell/Virus Enclosed | NO |
| MAARLCCQLDPARDV | 41004 | Cell/Virus Enclosed | NO |
| MAATRGATCVIGTTKF | 1679314 | Cell/Virus Enclosed | NO |
| MAAYVEETSITI | 1475471 | Cell/Virus Enclosed | NO |
| MAAYVENTSITI | 1475473 | Cell/Virus Enclosed | NO |
| MACNGHKIEDFS | 1475477 | Cell/Virus Enclosed | NO |
| MADAYFADTVWY | 1475483 | Cell/Virus Enclosed | NO |
| MADGSPQSYCYT | 1475490 | Cell/Virus Enclosed | NO |

| | | | |
|---|---|---|---|
| MADLALTNMYKEARIN | 1679327 | Cell/Virus Enclosed | NO |
| MADNGTITVEELKKLLEQWNLVI | 1385326 | Undetermined | UN |
| MADNGTITVEELKQLLEQWNLVIG | 1475496 | Undetermined | UN |
| MADSNGTITVEELKKLLEQWNLVI | 1385329 | Undetermined | UN |
| MADVDDPKLMGWDYPK | 1679333 | Cell/Virus Enclosed | NO |
| MAEQAAAAMYKEARAV | 1679335 | Cell/Virus Enclosed | NO |
| MAFAYADKPNHF | 1475516 | Cell/Virus Enclosed | NO |
| MAFNPPWGDINKSGRSGIAL | 1951482 | Capsid | UN |
| MAFPSAKVEGCM | 1475521 | Cell/Virus Enclosed | NO |
| MAFSASLFKPVH | 1475525 | Cell/Virus Enclosed | NO |
| MAFSASLFKPVQLVPV | 1679340 | Cell/Virus Enclosed | NO |
| MAGNGGDAALALLLLDR | 1475537 | Capsid | UN |
| MAGRSALEDLLF | 1475539 | Ectodomain | YES |
| MAHFPGFGQSLLYGYPVYVF | 41048 | Cell/Virus Enclosed | NO |
| MAITIKPEANIN | 1475551 | Cell/Virus Enclosed | NO |
| MAKAGDKYYPVVVDDH | 1679352 | Cell/Virus Enclosed | NO |
| MAKEVKTSSKVVLSDA | 1679355 | Cell/Virus Enclosed | NO |
| MAKFPLKLAGTAVINL | 1679356 | Cell/Virus Enclosed | NO |
| MALNFKLIGVGG | 1475581 | Cell/Virus Enclosed | NO |
| MALSIFSAFYPI | 1475587 | Transmembrane | NO |
| MANQEKYSTIYKLHPS | 1679372 | Cell/Virus Enclosed | NO |
| MAPPAAPRAVSF | 1475617 | Cell/Virus Enclosed | NO |
| MAPVEFYEDRILDFKV | 1679376 | Cell/Virus Enclosed | NO |
| MAQAFGLRGPGD | 1475622 | Cell/Virus Enclosed | NO |
| MAQSSGVLPVNP | 1475623 | Cell/Virus Enclosed | NO |
| MARLGAELRSHPNTPPKPRR | 741723 | Underneath Membrane | NO |
| MARNGVVPLSVIPATS | 1679381 | Cell/Virus Enclosed | NO |
| MASFVEYEVAKKNLAD | 1679384 | Cell/Virus Enclosed | NO |
| MASGGGETALALLLLDRL | 41120 | Capsid | UN |
| MASPAAPRAVSF | 1475648 | Cell/Virus Enclosed | NO |
| MASQGTKRSYEQMETGGERQNATE | 97491 | Capsid | UN |
| MASSAAPRAVSF | 1475651 | Cell/Virus Enclosed | NO |
| MASVNWADDRAARKKF | 1679394 | Capsid | UN |
| MATDDCFANKHA | 1475658 | Cell/Virus Enclosed | NO |
| MATVKWADASEPQRGR | 1679399 | Capsid | UN |
| MATVKWADESEP | 1475676 | Cell/Virus Enclosed | NO |
| MAVAYADKPNHFINFP | 1679402 | Cell/Virus Enclosed | NO |
| MAVAYADNPNHF | 1475681 | Cell/Virus Enclosed | NO |
| MAVAYVDKPNHF | 1475682 | Cell/Virus Enclosed | NO |
| MAVVDGKQIDDY | 1475690 | Cell/Virus Enclosed | NO |
| MAYGNCTFGDPD | 1475698 | Cell/Virus Enclosed | NO |
| MAYGNCTFGSPD | 1475699 | Cell/Virus Enclosed | NO |
| MCDIKQMLFCMEVVNK | 1679417 | Cell/Virus Enclosed | NO |
| MCDYPHTICHPN | 1475723 | Ectodomain | YES |
| MCEHPHTSCHPN | 1475728 | Ectodomain | YES |
| MCEYPHTICHPN | 1475730 | Ectodomain | YES |
| MCEYPNTICHPN | 1475738 | Ectodomain | YES |
| MCEYPQTICHPN | 1475740 | Ectodomain | YES |
| MCEYPYTICHPN | 1475741 | Ectodomain | YES |
| MCGFDGKPMLPK | 1475749 | Cell/Virus Enclosed | NO |
| MCIDMKGTMYVRPIIE | 1679432 | Ectodomain | YES |
| MCIDQVTIAGVK | 1475771 | Cell/Virus Enclosed | NO |
| MCLASIAFNHPI | 1475777 | Ectodomain | YES |
| MCLEPCNLYNYGAGIK | 1679437 | Cell/Virus Enclosed | NO |
| MCLEPCNLYNYGAGLK | 1679438 | Cell/Virus Enclosed | NO |
| MCPADQLSDPNY | 1475797 | Cell/Virus Enclosed | NO |
| MCTGSPSIVEFNRLAT | 1679453 | Cell/Virus Enclosed | NO |
| MCVVPSINDLSFDKNL | 1679464 | Cell/Virus Enclosed | NO |
| MCVVSSINDLSF | 1475825 | Cell/Virus Enclosed | NO |
| MCYRCPKEIVSTVSAL | 1679466 | Cell/Virus Enclosed | NO |
| MDAEGRMTPYCH | 1475832 | Cell/Virus Enclosed | NO |
| MDAKQVDYVPLKSATC | 1679468 | Cell/Virus Enclosed | NO |
| MDDAINVGGTGL | 1475837 | Cell/Virus Enclosed | NO |
| MDFCEGSFPCSV | 1475856 | Cell/Virus Enclosed | NO |
| MDFVYALRHFDQ | 1475860 | Cell/Virus Enclosed | NO |
| MDGCTYSTCMMC | 1475863 | Cell/Virus Enclosed | NO |
| MDGFSCPLCCTVYDSS | 1679483 | Cell/Virus Enclosed | NO |
| MDGMDAKQVDYVPLKS | 1679486 | Cell/Virus Enclosed | NO |
| MDIDPYKEFGATVELLSFLP | 41238 | Capsid | UN |
| MDIDVFNKLSTSDWSD | 1679490 | Cell/Virus Enclosed | NO |
| MDKLDTGAQKFFQFGD | 1679494 | Cell/Virus Enclosed | NO |
| MDKQVHQVQLID | 1475893 | Cell/Virus Enclosed | NO |
| MDLNPVLCDSGKISSK | 1679506 | Ectodomain | YES |
| MDLVYALRHFDQNSEV | 1679507 | Cell/Virus Enclosed | NO |
| MDMGVFINKYGLEDFN | 1679509 | Cell/Virus Enclosed | NO |
| MDNAINVGGTGLQYAA | 1679510 | Cell/Virus Enclosed | NO |

| | | | |
|---|---|---|---|
| MDNKSQSVIIIN | 1475918 | Ectodomain | YES |
| MDNLACESQQPT | 1475920 | Cell/Virus Enclosed | NO |
| MDNTTQSAIIVN | 1475925 | Ectodomain | YES |
| MDPIYDEPTMTT | 1475927 | Secreted | YES |
| MDPLSNFEHKVI | 1475932 | Cell/Virus Enclosed | NO |
| MDPNKTNVVPPA | 1475933 | Cell/Virus Enclosed | NO |
| MDPNQTSVVPPA | 1475935 | Cell/Virus Enclosed | NO |
| MDQGSFGGASCC | 1475937 | Cell/Virus Enclosed | NO |
| MDSGGSSFSELL | 1475944 | Cell/Virus Enclosed | NO |
| MDTFTNNNGSDVLYQP | 1679525 | Cell/Virus Enclosed | NO |
| MDVLSQNQKLIANAFN | 1679532 | Ectodomain | YES |
| MDVPSTYGTQVY | 1475971 | Cell/Virus Enclosed | NO |
| MDVQGLVNVLAC | 1475972 | Cell/Virus Enclosed | NO |
| MDVSVHRHRLSL | 1475977 | Cell/Virus Enclosed | NO |
| MDYKCSCTSGRL | 1475980 | Cell/Virus Enclosed | NO |
| MDYPARAHTFDDFCPECRPLGLQGC | 41293 | Undetermined | UN |
| MECDFLNMDMGV | 1476001 | Cell/Virus Enclosed | NO |
| MEDAMGQGQNNA | 1476002 | Cell/Virus Enclosed | NO |
| MEEDFLNMDMGV | 1476007 | Cell/Virus Enclosed | NO |
| MEEFIQTNSPKV | 1476008 | Ectodomain | YES |
| MEELKALVATAH | 1476010 | Cell/Virus Enclosed | NO |
| MEESLTDVPSTS | 1476019 | Cell/Virus Enclosed | NO |
| MEESPTDVPSTS | 1476021 | Cell/Virus Enclosed | NO |
| MEILNRDRVNIN | 1476026 | Cell/Virus Enclosed | NO |
| MEKDFIALDQDVFIQK | 1679556 | Cell/Virus Enclosed | NO |
| MEKDFLSFDSDVFIKK | 1679557 | Cell/Virus Enclosed | NO |
| MEKVNECVKSQSSRIN | 1679561 | Ectodomain | YES |
| MEKVSSSRRGVY | 1476038 | Ectodomain | YES |
| MENFELEKHCYD | 1476050 | Cell/Virus Enclosed | NO |
| MERNAGSGIIISDTPVHDCNTTCQT | 1160155 | Ectodomain | YES |
| MESQISTKEVKLKGVR | 1679578 | Cell/Virus Enclosed | NO |
| METAAYREAAAC | 1476073 | Cell/Virus Enclosed | NO |
| METDFLEPAMDE | 1476076 | Cell/Virus Enclosed | NO |
| METFNHNNGNDVLYQP | 1679582 | Cell/Virus Enclosed | NO |
| METFTNNNGSDV | 1476078 | Cell/Virus Enclosed | NO |
| MEVTPSGTWLTYHGAIK | 41461 | Capsid | UN |
| MEYDFLNIDMGV | 1476095 | Cell/Virus Enclosed | NO |
| MEYDFLNMDMGVFINK | 1679588 | Cell/Virus Enclosed | NO |
| MFADHLIDLSDC | 1476099 | Cell/Virus Enclosed | NO |
| MFADHLTDLSDC | 1476100 | Cell/Virus Enclosed | NO |
| MFDLDSTFDDILGKVC | 1679594 | Cell/Virus Enclosed | NO |
| MFDSFVNSFVSLYNVT | 1679595 | Cell/Virus Enclosed | NO |
| MFDVDKKILNAL | 1476113 | Cell/Virus Enclosed | NO |
| MFDVDKKSLNALIATA | 1679596 | Cell/Virus Enclosed | NO |
| MFDVDKKSLNVL | 1476115 | Cell/Virus Enclosed | NO |
| MFDVYVDTFLSMFDVD | 1679597 | Cell/Virus Enclosed | NO |
| MFEDAQGLKEFY | 1476118 | Cell/Virus Enclosed | NO |
| MFEDAQSLKKFY | 1476120 | Cell/Virus Enclosed | NO |
| MFEDDQGLKKFY | 1476121 | Cell/Virus Enclosed | NO |
| MFENKTTLPTNIAFEL | 1679601 | Cell/Virus Enclosed | NO |
| MFEPRIPTMADFVQIE | 1679602 | Ectodomain | YES |
| MFFLEKDAPYIV | 1476131 | Cell/Virus Enclosed | NO |
| MFFPYDIEFVTG | 1476134 | Cell/Virus Enclosed | NO |
| MFGHIVPIWMTC | 1476139 | Cell/Virus Enclosed | NO |
| MFGLYDDILTNNKPWF | 1679606 | Cell/Virus Enclosed | NO |
| MFHQKCLKSIAA | 1476146 | Cell/Virus Enclosed | NO |
| MFIFLLFLTLTSGSDLD | 41504 | Undetermined | UN |
| MFKDAEGRMTPY | 1476162 | Cell/Virus Enclosed | NO |
| MFKRGDGTPHPYCYSD | 1679617 | Cell/Virus Enclosed | NO |
| MFLLPRFVLVSC | 1476189 | Ectodomain | YES |
| MFLLTPTESYVD | 1476193 | Ectodomain | YES |
| MFLPMLPAPPPGQPSGRRRGRRSGG | 41513 | Undetermined | UN |
| MFLRLIDDNGIVLNSI | 1679630 | Cell/Virus Enclosed | NO |
| MFMADAYFADTV | 1476198 | Cell/Virus Enclosed | NO |
| MFMADAYLADTV | 1476199 | Cell/Virus Enclosed | NO |
| MFMADAYLAGTV | 1476200 | Cell/Virus Enclosed | NO |
| MFMDAEGRMTPY | 1476202 | Cell/Virus Enclosed | NO |
| MFQPRLPVLSDF | 1476218 | Ectodomain | YES |
| MFRDAEGRITPY | 1476222 | Cell/Virus Enclosed | NO |
| MFRDAEGRMTPY | 1476223 | Cell/Virus Enclosed | NO |
| MFSQFNSNFLPE | 1476240 | Ectodomain | YES |
| MFTGSGYYYPEP | 1476245 | Ectodomain | YES |
| MFTGSRYYYPEP | 1476246 | Ectodomain | YES |
| MFTGSSYYYPEPISDK | 1679647 | Ectodomain | YES |
| MFTKVVVGPTTEVKFS | 1679648 | Cell/Virus Enclosed | NO |
| MFVATSSYDALL | 1476259 | Cell/Virus Enclosed | NO |

| | | | |
|---|---|---|---|
| MFVRPVIEDYHTLTAT | 1679662 | Underneath Membrane | NO |
| MGANGNKIVDKEVKDM | 1679672 | Cell/Virus Enclosed | NO |
| MGDCVKFVYMHQ | 1476296 | Cell/Virus Enclosed | NO |
| MGFKLDANVPGY | 1476306 | Cell/Virus Enclosed | NO |
| MGFRFDISIPGS | 1476313 | Cell/Virus Enclosed | NO |
| MGFTYSGIRTNGATSACRRSGSSFY | 1160199 | Ectodomain | YES |
| MGGWSSKPRKGMGTNLSVPNP | 41584 | Underneath Membrane | NO |
| MGHFSWWTAFVT | 1476318 | Cell/Virus Enclosed | NO |
| MGILSNMNIASNSKEW | 1679692 | Cell/Virus Enclosed | NO |
| MGIQSGVQVDANCEGDCYHSGGTII | 1160208 | Ectodomain | YES |
| MGIRFEYTSPTDKLAM | 1679694 | Cell/Virus Enclosed | NO |
| MGKCGDKFFPIVMNDK | 1679695 | Cell/Virus Enclosed | NO |
| MGLHGNQTLFCNIMKF | 1679701 | Cell/Virus Enclosed | NO |
| MGLNKPFVKDNV | 1476354 | Cell/Virus Enclosed | NO |
| MGLTICLASECF | 1476357 | Cell/Virus Enclosed | NO |
| MGLTNCLASECF | 1476358 | Cell/Virus Enclosed | NO |
| MGNAINVGGRGL | 1476363 | Cell/Virus Enclosed | NO |
| MGQGQNNADPKV | 1476366 | Cell/Virus Enclosed | NO |
| MGRFFNYTLVLL | 1476372 | Ectodomain | YES |
| MGRIRSVYPVAT | 1476377 | Cell/Virus Enclosed | NO |
| MGSCSFEDELTPSDVY | 1679712 | Cell/Virus Enclosed | NO |
| MGSGFCVDYNSPSSSS | 1679713 | Ectodomain | YES |
| MGSNNVCTYEHV | 1476387 | Cell/Virus Enclosed | NO |
| MGTKKSAAEASK | 1476390 | Capsid | UN |
| MGTNLSVPNPLGFFPDHQLDP | 41631 | Underneath Membrane | NO |
| MGTVNKPVVGVLMGFGIITG | 41636 | Undetermined | UN |
| MGVFINKYGLEDFNFE | 1679720 | Cell/Virus Enclosed | NO |
| MGVLKADDFVIA | 1476406 | Cell/Virus Enclosed | NO |
| MGVYDFKVSAAEFKYM | 1679725 | Cell/Virus Enclosed | NO |
| MGWDYPKCDRALPNMI | 1679728 | Cell/Virus Enclosed | NO |
| MGWDYPKCDRAMPNIL | 1679729 | Cell/Virus Enclosed | NO |
| MGWDYPKCDRAMPNLL | 1679730 | Cell/Virus Enclosed | NO |
| MGWDYPKCDRAMPNMC | 1679731 | Cell/Virus Enclosed | NO |
| MGWDYPKCDRAMPSMI | 1679733 | Cell/Virus Enclosed | NO |
| MHFGACDYDRLPNEVT | 1679741 | Ectodomain | YES |
| MHFGTLAKTDLFNGYK | 1679742 | Cell/Virus Enclosed | NO |
| MHGDTPTLHEYMLDLQPETT | 41660 | Cell/Virus Enclosed | NO |
| MHGSVLRIKVSQ | 1476437 | Cell/Virus Enclosed | NO |
| MHHMELPTGVHA | 1476444 | Cell/Virus Enclosed | NO |
| MHIASSNAFLDLRTSC | 1679748 | Cell/Virus Enclosed | NO |
| MHIDPFPKRVIDF | 1679749 | Cell/Virus Enclosed | NO |
| MHKIIKVFYELD | 1476449 | Cell/Virus Enclosed | NO |
| MHQKRTAMFQDPQERPRKLPQLC | 110669 | Cell/Virus Enclosed | NO |
| MHSINHDIFSLM | 1476465 | Cell/Virus Enclosed | NO |
| MHSPDVIVGDYV | 1476471 | Cell/Virus Enclosed | NO |
| MHVGYYPSNHID | 1476476 | Ectodomain | YES |
| MHVGYYPSNHIE | 1476477 | Ectodomain | YES |
| MHVGYYPSNQIE | 1476478 | Ectodomain | YES |
| MIANENAINKGGRTVQ | 1679773 | Cell/Virus Enclosed | NO |
| MICSTTAQKLKT | 1476498 | Cell/Virus Enclosed | NO |
| MIDPEGVCLGAG | 1476503 | Cell/Virus Enclosed | NO |
| MIDRPYVLLAVLFVM | 41693 | Undetermined | UN |
| MIEDMDYVPLKS | 1476509 | Cell/Virus Enclosed | NO |
| MIEDVDYVPLKS | 1476510 | Cell/Virus Enclosed | NO |
| MIEFMLWCKDGQVQTF | 1679782 | Cell/Virus Enclosed | NO |
| MIKGPQRADLNGVIVD | 1679796 | Cell/Virus Enclosed | NO |
| MIKKLDNDVLNGIISN | 1679797 | Cell/Virus Enclosed | NO |
| MIKPNAILCATC | 1476539 | Cell/Virus Enclosed | NO |
| MIKPNALLCATC | 1476541 | Cell/Virus Enclosed | NO |
| MIKPNARLCATC | 1476543 | Cell/Virus Enclosed | NO |
| MILKNQDLGVVS | 1476549 | Cell/Virus Enclosed | NO |
| MILKSQDLGAVS | 1476550 | Cell/Virus Enclosed | NO |
| MILSDDGVVCNN | 1476562 | Cell/Virus Enclosed | NO |
| MILSDDGVVCYNKDYA | 1679807 | Cell/Virus Enclosed | NO |
| MILSDDGVVCYNSDYA | 1679808 | Cell/Virus Enclosed | NO |
| MILSLHPVECDE | 1476563 | Cell/Virus Enclosed | NO |
| MINPNALLCATC | 1476570 | Cell/Virus Enclosed | NO |
| MIPLQASLPFVW | 1476572 | Secreted | YES |
| MIRGPPRAELNGVVVD | 1679814 | Cell/Virus Enclosed | NO |
| MIRMLSAMISGS | 1476584 | Cell/Virus Enclosed | NO |
| MIRSCYSGHVSA | 1476585 | Ectodomain | YES |
| MISEWPLSESDK | 1476595 | Cell/Virus Enclosed | NO |
| MISEWPLSVQQAQQEA | 1679824 | Cell/Virus Enclosed | NO |
| MISGPCPKAGQPSLVVQSSL | 41775 | Cell/Virus Enclosed | NO |
| MISQWPISNDSNGCVV | 1679825 | Cell/Virus Enclosed | NO |
| MISTVGVFLGYK | 1476602 | Cell/Virus Enclosed | NO |

| | | | |
|---|---|---|---|
| MIVDEWSYTGSS | 1476607 | Ectodomain | YES |
| MKAILVVLLYTFATANADTLCIGYH | 1160252 | Ectodomain | YES |
| MKALFPIWSQEL | 1476621 | Cell/Virus Enclosed | NO |
| MKALYPTWSQEL | 1476625 | Cell/Virus Enclosed | NO |
| MKAVAAALKSLGFDKP | 1679834 | Capsid | UN |
| MKDDGNCVVLELDPPC | 1679839 | Cell/Virus Enclosed | NO |
| MKDLSPRWYFYYLGTGPEAG | 1160254 | Capsid | UN |
| MKGKMFVRPVIEDYHT | 1679848 | Underneath Membrane | NO |
| MKGRMYVRPIIE | 1476645 | Underneath Membrane | NO |
| MKGTGVFVQDPAPIDI | 1679849 | Cell/Virus Enclosed | NO |
| MKGTGVFVQDPS | 1476648 | Cell/Virus Enclosed | NO |
| MKGTMYVRPIIEDYHT | 1679850 | Ectodomain | YES |
| MKHKGGDSIDFI | 1476654 | Cell/Virus Enclosed | NO |
| MKQYGECLGDINARDL | 1679864 | Ectodomain | YES |
| MKQYGQCLGDIN | 1476700 | Ectodomain | YES |
| MKSDEGGLCPSTGQAM | 1679867 | Cell/Virus Enclosed | NO |
| MKSDIFGQDFKTFDLL | 1679868 | Cell/Virus Enclosed | NO |
| MKTVAFDLSSPQKSGTGPQPGSAGM | 41884 | Cell/Virus Enclosed | NO |
| MKVKATKGEGDGGITS | 1679874 | Cell/Virus Enclosed | NO |
| MKVLNENLNAYQQQDG | 1679876 | Cell/Virus Enclosed | NO |
| MKVLTENLNAYQ | 1476728 | Cell/Virus Enclosed | NO |
| MLAKKELSGQCF | 1476743 | Cell/Virus Enclosed | NO |
| MLDFSRDHNDILYTPP | 1679900 | Cell/Virus Enclosed | NO |
| MLDKQGQLLDTKLNGI | 1679903 | Cell/Virus Enclosed | NO |
| MLDYAKDHNDMLYSPP | 1679911 | Cell/Virus Enclosed | NO |
| MLEYKGPVTDVF | 1476777 | Cell/Virus Enclosed | NO |
| MLFDSHIVSKESGNTV | 1679917 | Capsid | UN |
| MLFGMIKKLDNDVLNG | 1679918 | Cell/Virus Enclosed | NO |
| MLFSMVRKLDNQALNS | 1679922 | Cell/Virus Enclosed | NO |
| MLGAEMTSNVRRQVAS | 1679928 | Cell/Virus Enclosed | NO |
| MLGCSVGNFSDG | 1476796 | Ectodomain | YES |
| MLGLTTQTVDSSQGSE | 1679931 | Cell/Virus Enclosed | NO |
| MLGSSVGNFSDGKMGR | 1679932 | Ectodomain | YES |
| MLGTTVDKLKPV | 1476815 | Cell/Virus Enclosed | NO |
| MLKDSSTIHNYF | 1476838 | Cell/Virus Enclosed | NO |
| MLKHFFFAQDGN | 1476843 | Cell/Virus Enclosed | NO |
| MLKRRDSTYGPLQTPV | 1679948 | Ectodomain | YES |
| MLKRRDSTYSPL | 1476852 | Ectodomain | YES |
| MLKSMAATRGATCVIG | 1679949 | Cell/Virus Enclosed | NO |
| MLKTVYSDVENP | 1476858 | Cell/Virus Enclosed | NO |
| MLKYDENGTITDAVDCS | 41984 | Ectodomain | YES |
| MLLALLAFFLSK | 1476865 | Cell/Virus Enclosed | NO |
| MLLSVPSDSCNN | 1476886 | Cell/Virus Enclosed | NO |
| MLMGWDYPKCDRALPN | 1679978 | Cell/Virus Enclosed | NO |
| MLNGMPVSSIKS | 1476899 | Cell/Virus Enclosed | NO |
| MLNGNAIATVKSEDGN | 1679982 | Cell/Virus Enclosed | NO |
| MLNISTPNLHDF | 1476902 | Ectodomain | YES |
| MLNKHIDAYKTF | 1476903 | Capsid | UN |
| MLNTPIPNLPDF | 1476910 | Ectodomain | YES |
| MLNTSIPNLPDFKEEL | 1679985 | Ectodomain | YES |
| MLNTSSPNLPDF | 1476914 | Ectodomain | YES |
| MLNTSTPNLPDF | 1476915 | Ectodomain | YES |
| MLPAPPPGQPSGRRRGRRSGGSGGG | 42026 | Undetermined | UN |
| MLPGEFSKKQLG | 1476923 | Cell/Virus Enclosed | NO |
| MLPKNMWEFRDY | 1476924 | Cell/Virus Enclosed | NO |
| MLPLAYFSDSKLGTTE | 1679990 | Cell/Virus Enclosed | NO |
| MLQIQLTGILDG | 1476930 | Cell/Virus Enclosed | NO |
| MLRACIKIAQAD | 1476941 | Cell/Virus Enclosed | NO |
| MLRACIKTAQADTAGL | 1679998 | Cell/Virus Enclosed | NO |
| MLRACIKTVQAD | 1476946 | Cell/Virus Enclosed | NO |
| MLRGDDMSFLEK | 1476949 | Cell/Virus Enclosed | NO |
| MLRHLIKDVDNPVLMG | 1680000 | Cell/Virus Enclosed | NO |
| MLRLHTVESGDINVVA | 1680005 | Cell/Virus Enclosed | NO |
| MLRRLIKDVDNPVLMG | 1680008 | Cell/Virus Enclosed | NO |
| MLSDTLKGLSDR | 1476966 | Cell/Virus Enclosed | NO |
| MLSENFTSYGFSNVVE | 1680016 | Ectodomain | YES |
| MLSLQQVVKALNESYI | 1680017 | Ectodomain | YES |
| MLTFDKKAFTLANVSS | 1680021 | Ectodomain | YES |
| MLTFDSNAFSLA | 1476984 | Ectodomain | YES |
| MLTGIAARPPPG | 1476986 | Cell/Virus Enclosed | NO |
| MLTGISARPPPG | 1476987 | Cell/Virus Enclosed | NO |
| MLTLDDRGLALDNALS | 1680027 | Cell/Virus Enclosed | NO |
| MLVKMDGDDVYLPYPN | 1680041 | Cell/Virus Enclosed | NO |
| MLVKMDGDEVYL | 1477002 | Cell/Virus Enclosed | NO |
| MLVNCFKWQVVV | 1477012 | Cell/Virus Enclosed | NO |
| MLWCKDGKVQTF | 1477023 | Cell/Virus Enclosed | NO |

| | | | |
|---|---|---|---|
| MLWCKDGQVQTFYPRL | 1680050 | Cell/Virus Enclosed | NO |
| MLWGDGFQIKFE | 1477025 | Cell/Virus Enclosed | NO |
| MLYSPPTISYNSTLQS | 1680055 | Cell/Virus Enclosed | NO |
| MMILSDDGVVCYNKDY | 1680068 | Cell/Virus Enclosed | NO |
| MMNISPKPQRQR | 1477056 | Cell/Virus Enclosed | NO |
| MMQSADASTFLN | 1477065 | Cell/Virus Enclosed | NO |
| MMRLFYDRDGQRVYDD | 1680081 | Cell/Virus Enclosed | NO |
| MMSAPPAEYKLQ | 1477070 | Cell/Virus Enclosed | NO |
| MMYAADPAMHIA | 1477085 | Cell/Virus Enclosed | NO |
| MNALVNSTLLETRTT | 42187 | Ectodomain | YES |
| MNASPYDLEVLLQDAL | 1680092 | Cell/Virus Enclosed | NO |
| MNAYPFVVTPLV | 1477091 | Cell/Virus Enclosed | NO |
| MNDKNICLLDHA | 1477098 | Cell/Virus Enclosed | NO |
| MNGEDAFSILSA | 1477108 | Cell/Virus Enclosed | NO |
| MNIAKSEFDHEV | 1477114 | Cell/Virus Enclosed | NO |
| MNIAKYMQLCQY | 1477115 | Cell/Virus Enclosed | NO |
| MNKSFFPQFTSD | 1477118 | Ectodomain | YES |
| MNKSFFSQFTSD | 1477119 | Underneath Membrane | NO |
| MNKSFLPQFTSDQAVT | 1680110 | Ectodomain | YES |
| MNLSTYSLFDLSKFQL | 1680115 | Cell/Virus Enclosed | NO |
| MNMDVDTHRYCL | 1477127 | Cell/Virus Enclosed | NO |
| MNMDVSLHRHRL | 1477128 | Cell/Virus Enclosed | NO |
| MNNKSQSVIIINNSTN | 1680120 | Ectodomain | YES |
| MNRGFFNMLGRRPFPAAT | 558294 | Undetermined | UN |
| MNTQILVFALIAIIPTNADKICLGH | 1160366 | Ectodomain | YES |
| MNTSHSVCNGCV | 1477153 | Ectodomain | YES |
| MNVIGQHIELPQ | 1477159 | Cell/Virus Enclosed | NO |
| MNVIGQHIKLPQ | 1477160 | Cell/Virus Enclosed | NO |
| MNVTLDKCTKYNIYDV | 1680133 | Ectodomain | YES |
| MNVTLNKCTKYN | 1477164 | Ectodomain | YES |
| MPCAIINDKVVAKIDK | 1680139 | Cell/Virus Enclosed | NO |
| MPFEELNGLKILKQLD | 1680142 | Cell/Virus Enclosed | NO |
| MPFFYFDDSDCDVVQE | 1680143 | Cell/Virus Enclosed | NO |
| MPHKLKIQVVNSGSDM | 1680154 | Cell/Virus Enclosed | NO |
| MPIADPTHFDIEEVEL | 1680155 | Cell/Virus Enclosed | NO |
| MPILTLTRALAA | 1477207 | Cell/Virus Enclosed | NO |
| MPIMANLVNLNS | 1477210 | Ectodomain | YES |
| MPIPDFDLDDFIDAPC | 1680164 | Cell/Virus Enclosed | NO |
| MPKFFYASNGTY | 1477229 | Ectodomain | YES |
| MPKIIKVFYELDNDFN | 1680166 | Cell/Virus Enclosed | NO |
| MPKTRRRPRRSQRKRPPTPW | 78749 | Cell/Virus Enclosed | NO |
| MPLTRIARRNIG | 1477272 | Undetermined | UN |
| MPMLTMCHALDC | 1477277 | Cell/Virus Enclosed | NO |
| MPMLTMCHVLDC | 1477279 | Cell/Virus Enclosed | NO |
| MPSFVAYETARQEYEN | 1680191 | Cell/Virus Enclosed | NO |
| MPSLFKVQNVNLERCE | 1680192 | Cell/Virus Enclosed | NO |
| MPTYTVHKSDFQ | 1477314 | Cell/Virus Enclosed | NO |
| MPVHTLSIHDGY | 1477321 | Cell/Virus Enclosed | NO |
| MPVSIAKTSVDC | 1477332 | Ectodomain | YES |
| MPVSSIKSDKGV | 1477335 | Cell/Virus Enclosed | NO |
| MPVVDGKQIDDH | 1477336 | Cell/Virus Enclosed | NO |
| MPYFMYDVVSSF | 1477340 | Cell/Virus Enclosed | NO |
| MPYHPNPVDCQD | 1477341 | Cell/Virus Enclosed | NO |
| MPYHPNTVDCQDDRCI | 1680215 | Cell/Virus Enclosed | NO |
| MPYTKLFQKGECCICH | 1680216 | Cell/Virus Enclosed | NO |
| MQCFGPRDFNHN | 1477357 | Cell/Virus Enclosed | NO |
| MQFGYIDCEQDL | 1477364 | Cell/Virus Enclosed | NO |
| MQFNYKITINPS | 1477369 | Cell/Virus Enclosed | NO |
| MQGPFIAGNTVH | 1477375 | Cell/Virus Enclosed | NO |
| MQGVTLSSNLNTNLHF | 1680230 | Ectodomain | YES |
| MQIVDENGKYYLPYPD | 1680235 | Cell/Virus Enclosed | NO |
| MQIVDKDGNYYL | 1477391 | Cell/Virus Enclosed | NO |
| MQIVDKDGTYYLPYPD | 1680236 | Cell/Virus Enclosed | NO |
| MQKESDGYIKLN | 1477395 | Cell/Virus Enclosed | NO |
| MQKLPCQFKDKNLQDL | 1680238 | Cell/Virus Enclosed | NO |
| MQKLPFKFKDLAIENM | 1680239 | Cell/Virus Enclosed | NO |
| MQKPRWKRQPNDDVTS | 1680240 | Capsid | UN |
| MQLLHTKVSWAG | 1477410 | Cell/Virus Enclosed | NO |
| MQNKGLFYTQVY | 1477424 | Ectodomain | YES |
| MQQGPASARDLI | 1477433 | Ectodomain | YES |
| MQQHPLLNPGAL | 1477439 | Cell/Virus Enclosed | NO |
| MQQHPLLNPSALEFNP | 1680263 | Capsid | UN |
| MQQQPLLNPSAL | 1477442 | Cell/Virus Enclosed | NO |
| MQSQSSHAVQNT | 1477455 | Cell/Virus Enclosed | NO |
| MQSWVPLSTDDPVID | 42413 | Ectodomain | YES |
| MQTGFTTTNEAFQKVQ | 1680275 | Ectodomain | YES |

| | | | |
|---|---|---|---|
| MQTVEDLRARMT | 1477467 | Cell/Virus Enclosed | NO |
| MQTYKLVNMKGT | 1477469 | Cell/Virus Enclosed | NO |
| MRAGSKVVNHNVLIKE | 1680286 | Cell/Virus Enclosed | NO |
| MRGCCDCGSTKLPYYE | 1680293 | Underneath Membrane | NO |
| MRHVRGWLGMDVEGAH | 1680297 | Cell/Virus Enclosed | NO |
| MRISYFVQSIRL | 1477513 | Transmembrane | NO |
| MRKVDLYNGAVIYIFA | 1680304 | Cell/Virus Enclosed | NO |
| MRLFYDRDGQRTYDDV | 1680306 | Cell/Virus Enclosed | NO |
| MRLFYDRDGQRVYDDV | 1680307 | Cell/Virus Enclosed | NO |
| MRLFYNRDGQRT | 1477525 | Cell/Virus Enclosed | NO |
| MRNFDEQNCDVLKEVL | 1680314 | Cell/Virus Enclosed | NO |
| MRPNYTIKGSFL | 1477543 | Cell/Virus Enclosed | NO |
| MRRGQRVDLPPKVHFY | 1680321 | Capsid | UN |
| MRVLHLGAGSDKEVAP | 1680336 | Cell/Virus Enclosed | NO |
| MRVLHYGAGSDKGVAP | 1680339 | Cell/Virus Enclosed | NO |
| MRYNLNFSANSLDNLK | 1680343 | Ectodomain | YES |
| MRYNLNFSSNSV | 1477583 | Ectodomain | YES |
| MSANNGTITVEELKKLLEQWNLVI | 1385614 | Undetermined | UN |
| MSAPPSEYTLQQ | 1477597 | Cell/Virus Enclosed | NO |
| MSCDCGKKFDDQVGCL | 1680356 | Cell/Virus Enclosed | NO |
| MSCDCGKKFDEQ | 1477604 | Cell/Virus Enclosed | NO |
| MSCNGNKIEDFS | 1477607 | Cell/Virus Enclosed | NO |
| MSDFMSEIKCET | 1477610 | Ectodomain | YES |
| MSDFMSEIKCKTQSIA | 1680359 | Ectodomain | YES |
| MSDLTDIQEDITRH | 42532 | Capsid | UN |
| MSDNGPQNQRNAPRITFGGP | 1392311 | Capsid | UN |
| MSDNGPQNQRSA | 1477620 | Cell/Virus Enclosed | NO |
| MSDNGPQNQSSA | 1477621 | Cell/Virus Enclosed | NO |
| MSDNGPQPNQRS | 1477622 | Capsid | UN |
| MSDNGPQQNQRS | 1477623 | Capsid | UN |
| MSDNGPQSNQRSAPRIT | 42541 | Capsid | UN |
| MSEDLWQFVDHFGENE | 1680369 | Cell/Virus Enclosed | NO |
| MSEDVKLNPSEDFIKH | 1680370 | Cell/Virus Enclosed | NO |
| MSEIKCKTLSIA | 1477640 | Ectodomain | YES |
| MSELPKEQRVQG | 1477642 | Cell/Virus Enclosed | NO |
| MSEPPKEHRVQG | 1477643 | Cell/Virus Enclosed | NO |
| MSEPPKEQRVQGSITQ | 1680374 | Cell/Virus Enclosed | NO |
| MSEPPNEQRVQG | 1477648 | Capsid | UN |
| MSESKCWVEHDINNGP | 1680375 | Cell/Virus Enclosed | NO |
| MSESKCWVENDITNGP | 1680376 | Cell/Virus Enclosed | NO |
| MSFTPGKQSSSRASSG | 1680385 | Cell/Virus Enclosed | NO |
| MSHFDVDRKSFNNFVN | 1680394 | Cell/Virus Enclosed | NO |
| MSISKLHALDDV | 1477693 | Cell/Virus Enclosed | NO |
| MSKINKYGLELHWAPE | 1680403 | Cell/Virus Enclosed | NO |
| MSLAECKRALGLSISD | 1680404 | Cell/Virus Enclosed | NO |
| MSLEACYERGCN | 1477702 | Cell/Virus Enclosed | NO |
| MSLGADSSIAYSNNTI | 1680407 | Ectodomain | YES |
| MSLSLLDLYLGRGYN | 42603 | Ectodomain | YES |
| MSMAECKAALGL | 1477729 | Cell/Virus Enclosed | NO |
| MSMFSRTTSNFL | 1477731 | Ectodomain | YES |
| MSNDNCTGDIVT | 1477738 | Ectodomain | YES |
| MSNFKVRDPVIQERLDHDYA | 42614 | Capsid | UN |
| MSNITQLTEAQI | 1477740 | Underneath Membrane | NO |
| MSNMQLFESLNFITLP | 1680424 | Cell/Virus Enclosed | NO |
| MSNMTHLTEAQI | 1477744 | Ectodomain | YES |
| MSNSSVPLLEVYVHLR | 1680427 | Ectodomain | YES |
| MSPFEMPFEELN | 1477756 | Cell/Virus Enclosed | NO |
| MSPKPQRQRGLN | 1477761 | Cell/Virus Enclosed | NO |
| MSQFKLTHQNNDDHGN | 1680430 | Cell/Virus Enclosed | NO |
| MSQFSQEVSDMC | 1477769 | Cell/Virus Enclosed | NO |
| MSQYSRSTRSML | 1477775 | Ectodomain | YES |
| MSQYSRSTRSTL | 1477776 | Ectodomain | YES |
| MSRSESRKNRGGREEILE | 244254 | Cell/Virus Enclosed | NO |
| MSSVDNILNMAR | 1477794 | Cell/Virus Enclosed | NO |
| MSSVDTILNMARNGVV | 1680440 | Cell/Virus Enclosed | NO |
| MSSVETVLNLARDGVV | 1680441 | Cell/Virus Enclosed | NO |
| MSTPNAIFISDR | 1477803 | Cell/Virus Enclosed | NO |
| MSTPNAVFISDR | 1477805 | Cell/Virus Enclosed | NO |
| MSTSKCWVEEDLTKGP | 1680446 | Cell/Virus Enclosed | NO |
| MSYEDQDVLFAY | 1477818 | Cell/Virus Enclosed | NO |
| MSYTPGHYAGSRSSSG | 1680455 | Capsid | UN |
| MSYYCENHKPHY | 1477827 | Cell/Virus Enclosed | NO |
| MTAVNESRYGLH | 1477838 | Ectodomain | YES |
| MTAVNESRYGLQNLLQ | 1680464 | Ectodomain | YES |
| MTEQLQMGFGIT | 1477860 | Ectodomain | YES |
| MTFSGNVDLDALASDI | 1680476 | Cell/Virus Enclosed | NO |

| | | | |
|---|---|---|---|
| MTFYPKMQATND | 1477869 | Cell/Virus Enclosed | NO |
| MTFYPRLQAASDWKPG | 1680478 | Cell/Virus Enclosed | NO |
| MTGRVFHQKCLK | 1477878 | Cell/Virus Enclosed | NO |
| MTHMDDFSIKSI | 1477882 | Cell/Virus Enclosed | NO |
| MTIVKCKGLNKPFVKD | 1680487 | Cell/Virus Enclosed | NO |
| MTKLASTEELPD | 1477891 | Cell/Virus Enclosed | NO |
| MTKRNVIPTMTQ | 1477898 | Cell/Virus Enclosed | NO |
| MTKTSVDCNMYI | 1477899 | Ectodomain | YES |
| MTLDLLKDIKFVIPAL | 1385637 | Cell/Virus Enclosed | NO |
| MTLDLLKDIKLV | 1477902 | Cell/Virus Enclosed | NO |
| MTLNGLWLDNTV | 1477907 | Cell/Virus Enclosed | NO |
| MTNHSFSVQKHIGAPA | 1680501 | Cell/Virus Enclosed | NO |
| MTNIVDAFTGVN | 1477921 | Ectodomain | YES |
| MTPYCHDPTVLP | 1477925 | Cell/Virus Enclosed | NO |
| MTQLNLKYAISGKERA | 1680509 | Cell/Virus Enclosed | NO |
| MTQMNLKYAISAKNRA | 1680511 | Cell/Virus Enclosed | NO |
| MTSLPKDWRLINKFDV | 1680522 | Cell/Virus Enclosed | NO |
| MTSMYKQARAEDKKAK | 1680523 | Cell/Virus Enclosed | NO |
| MTSQALFDSLEF | 1477950 | Cell/Virus Enclosed | NO |
| MTTSTTNFGEEF | 1477959 | Cell/Virus Enclosed | NO |
| MTYVCQCGGERH | 1477980 | Cell/Virus Enclosed | NO |
| MVANGLHAPHGPFDAL | 1680546 | Cell/Virus Enclosed | NO |
| MVANGLNAPNGPFDAL | 1680547 | Cell/Virus Enclosed | NO |
| MVCKMCLFRYQE | 1477985 | Cell/Virus Enclosed | NO |
| MVDCRRLLKQDC | 1477988 | Cell/Virus Enclosed | NO |
| MVDPTTVLNDPV | 1477993 | Cell/Virus Enclosed | NO |
| MVDPTTVLNGPV | 1477994 | Cell/Virus Enclosed | NO |
| MVDVFFTDTAWY | 1477998 | Cell/Virus Enclosed | NO |
| MVERSVSLAIDA | 1478008 | Cell/Virus Enclosed | NO |
| MVETLVGQLEQT | 1478009 | Cell/Virus Enclosed | NO |
| MVETPNGPQVKY | 1478010 | Cell/Virus Enclosed | NO |
| MVITQDCDTCHSPAS | 227398 | Secreted | YES |
| MVIVNAANIHLK | 1478026 | Cell/Virus Enclosed | NO |
| MVKSKGVCATGDCYVS | 1680578 | Cell/Virus Enclosed | NO |
| MVKSKGVCQVGDCYVS | 1680579 | Cell/Virus Enclosed | NO |
| MVLESLAATVRL | 1478039 | Cell/Virus Enclosed | NO |
| MVLPGVADAERM | 1478049 | Ectodomain | YES |
| MVLTDVDVPFTA | 1478054 | Cell/Virus Enclosed | NO |
| MVNMSPKPQRQR | 1478066 | Cell/Virus Enclosed | NO |
| MVNWFNTMLDASAPAT | 1680601 | Cell/Virus Enclosed | NO |
| MVPVEFYEDRIL | 1478077 | Cell/Virus Enclosed | NO |
| MVQAFGLRGAGD | 1478083 | Cell/Virus Enclosed | NO |
| MVQTHLSHYVFN | 1478091 | Cell/Virus Enclosed | NO |
| MVRKLDNQALNSILDN | 1680610 | Cell/Virus Enclosed | NO |
| MVRRNLDSLEAR | 1478100 | Cell/Virus Enclosed | NO |
| MVTDIVVVDEVSMLTN | 1680615 | Cell/Virus Enclosed | NO |
| MVTILLCCMTSCCSCLK | 42999 | Undetermined | UN |
| MVVCDKVRSLKLSDNN | 1680619 | Cell/Virus Enclosed | NO |
| MVVKSALLADKF | 1478122 | Cell/Virus Enclosed | NO |
| MVVSAGQEQTNC | 1478126 | Cell/Virus Enclosed | NO |
| MVYTSDLLTNSV | 1478132 | Cell/Virus Enclosed | NO |
| MWEFRDYFNNNTDSIV | 1680632 | Cell/Virus Enclosed | NO |
| MWLSYFIASFRLFAR | 1310651 | Undetermined | UN |
| MWLYTYSIGEASSPIA | 1680646 | Cell/Virus Enclosed | NO |
| MYAADPAMHIASSNAF | 1680653 | Cell/Virus Enclosed | NO |
| MYAISNEQHYKS | 1478178 | Ectodomain | YES |
| MYCTFAIEDVEDVISS | 1680662 | Cell/Virus Enclosed | NO |
| MYDGDRFVKHDLSLLS | 1680663 | Cell/Virus Enclosed | NO |
| MYDGRTKAIDGENVSK | 1680665 | Cell/Virus Enclosed | NO |
| MYDPITKNIGEYNVSK | 1680667 | Cell/Virus Enclosed | NO |
| MYDPLTKNIGDYNVSK | 1680670 | Cell/Virus Enclosed | NO |
| MYDPTTKNLTGS | 1478201 | Cell/Virus Enclosed | NO |
| MYDPTTKNVTGSNESK | 1680672 | Cell/Virus Enclosed | NO |
| MYGAFMDKQVHQ | 1478212 | Cell/Virus Enclosed | NO |
| MYGGFEDQPNLQVESA | 1680680 | Cell/Virus Enclosed | NO |
| MYGVNLQSGKVI | 1478219 | Cell/Virus Enclosed | NO |
| MYKEARSVNRKSKVIS | 1680695 | Cell/Virus Enclosed | NO |
| MYKQARAEDKKAKIVS | 1680699 | Cell/Virus Enclosed | NO |
| MYKTPTLKDFGG | 1478248 | Ectodomain | YES |
| MYKVVGFIRHIV | 1478251 | Cell/Virus Enclosed | NO |
| MYLRKRGEKGAYNKDH | 1680709 | Cell/Virus Enclosed | NO |
| MYMGKCGDKFFP | 1478262 | Cell/Virus Enclosed | NO |
| MYNLLKGCNAVAKHDF | 1680719 | Cell/Virus Enclosed | NO |
| MYRVFEVSVIRNPGL | 43084 | Ectodomain | YES |
| MYSFVSEETGTLIVNSV | 1309883 | Undetermined | UN |
| MYTVSKGTQQSS | 1478301 | Ectodomain | YES |

| | | | |
|---|---|---|---|
| MYTYNITEDEIL | 1478302 | Ectodomain | YES |
| MYVRPIIEDYHTLTVT | 1680734 | Ectodomain | YES |
| MYVYSAGHATGS | 1478315 | Ectodomain | YES |
| MYVYSAGHATGTTPQK | 1680736 | Ectodomain | YES |
| NAAAYMKLSDAF | 1478320 | Cell/Virus Enclosed | NO |
| NAAIPDYNYYKY | 1478322 | Cell/Virus Enclosed | NO |
| NAAITDYNYYKYNLPT | 1680741 | Cell/Virus Enclosed | NO |
| NAAIVLQLPQGTALPKGFYAEGS | 1385685 | Cell/Virus Enclosed | NO |
| NAANTHLKHGGGIAGA | 1680747 | Cell/Virus Enclosed | NO |
| NAAQCDCLCTPD | 1478332 | Ectodomain | YES |
| NAAQCNCLCTPD | 1478333 | Ectodomain | YES |
| NAASVNQIVLRNSNGA | 1680752 | Cell/Virus Enclosed | NO |
| NAASVNQTVLRN | 1478338 | Cell/Virus Enclosed | NO |
| NACTPESKPLAD | 1478350 | Cell/Virus Enclosed | NO |
| NADAEALNNLLQ | 1478353 | Ectodomain | YES |
| NADIVVVDEVSMCTNY | 1680763 | Cell/Virus Enclosed | NO |
| NADYLYFHFYQE | 1478372 | Ectodomain | YES |
| NAFLVHDNVAFYQGDV | 1680782 | Cell/Virus Enclosed | NO |
| NAFNCTFEYISDAFSLD | 43145 | Ectodomain | YES |
| NAFTREMQQHPLLNPS | 1680787 | Capsid | UN |
| NAFVSQVLNKYTEVRG | 1680790 | Ectodomain | YES |
| NAFYGVALKDCG | 1478410 | Cell/Virus Enclosed | NO |
| NAGEDIRLLKAA | 1478414 | Cell/Virus Enclosed | NO |
| NAGYWRRQDRKINTGN | 1680798 | Cell/Virus Enclosed | NO |
| NAGYWRRQYRKI | 1478426 | Cell/Virus Enclosed | NO |
| NAHRYDALLSDL | 1478428 | Cell/Virus Enclosed | NO |
| NAIATVKSEDGNIKNI | 1680803 | Cell/Virus Enclosed | NO |
| NAILISDRKIKK | 1478440 | Cell/Virus Enclosed | NO |
| NAILTEVNELLDTTQL | 1680808 | Ectodomain | YES |
| NAIPSLAANTLNIIVP | 1680811 | Cell/Virus Enclosed | NO |
| NAISPKTDNETL | 1478455 | Ectodomain | YES |
| NAISSNTDNETLQYWV | 1680813 | Ectodomain | YES |
| NAKEEILGAVSW | 1478465 | Cell/Virus Enclosed | NO |
| NAKEIRHKILMK | 1478467 | Cell/Virus Enclosed | NO |
| NAKLGFDSSKEFVEYK | 1680820 | Cell/Virus Enclosed | NO |
| NAKLGFNVDLPF | 1478473 | Cell/Virus Enclosed | NO |
| NAKVEISVTPIKNTVD | 1680823 | Cell/Virus Enclosed | NO |
| NAKVSAEVSRQVIQDV | 1680824 | Cell/Virus Enclosed | NO |
| NAKWIFDSSKEF | 1478477 | Cell/Virus Enclosed | NO |
| NALFCATCPFAK | 1478484 | Cell/Virus Enclosed | NO |
| NALIATAHSSIKQGTQ | 1680832 | Cell/Virus Enclosed | NO |
| NALLCVTCPFAK | 1478493 | Cell/Virus Enclosed | NO |
| NALLSIQNGFSATNSA | 1680835 | Ectodomain | YES |
| NALMQGVTLSSN | 1478499 | Ectodomain | YES |
| NALSRAVNVPTIRKFI | 1680849 | Cell/Virus Enclosed | NO |
| NALYAIQEGFDATNSA | 1680852 | Ectodomain | YES |
| NANMSLAECKRALGLS | 1680868 | Cell/Virus Enclosed | NO |
| NANSEALNNLLQ | 1478542 | Ectodomain | YES |
| NAPATVCGPKLSTDLIK | 43264 | Ectodomain | YES |
| NAPGCDVNDVTKLYLG | 1680877 | Cell/Virus Enclosed | NO |
| NAPILSTLPETTVVR | 956879 | Capsid | UN |
| NAPRITFGGPSDSTGSNQNG | 1392317 | Capsid | UN |
| NAPYYNSWQNLL | 1478566 | Ectodomain | YES |
| NAQALSKLASELSNTF | 1680887 | Ectodomain | YES |
| NAQVAKSHNVSL | 1478577 | Cell/Virus Enclosed | NO |
| NARIRAKHYVYIGDPA | 1680898 | Cell/Virus Enclosed | NO |
| NARQAYEDAIANGSSS | 1680904 | Cell/Virus Enclosed | NO |
| NARQFDNMTAVA | 1478591 | Ectodomain | YES |
| NASEVVVNNLDKSAGH | 1680907 | Cell/Virus Enclosed | NO |
| NASGCGVSDVKKLYLG | 1680911 | Cell/Virus Enclosed | NO |
| NASISESKPLAD | 1478608 | Cell/Virus Enclosed | NO |
| NASLNSFKEYFNLRNC | 1680915 | Ectodomain | YES |
| NASLYTSLVPHTRYSL | 1680917 | Cell/Virus Enclosed | NO |
| NASPASTAHATA | 1478614 | Secreted | YES |
| NASPYDLEVLLQDALQ | 1680920 | Cell/Virus Enclosed | NO |
| NASRFPNVYAWE | 1478623 | Ectodomain | YES |
| NASSEVAVLYQDVNCTD | 43313 | Ectodomain | YES |
| NATFLDLQVEMN | 1478654 | Ectodomain | YES |
| NATFLDLYYEMN | 1478655 | Ectodomain | YES |
| NATGSDVNCNGYQHNS | 1680940 | Ectodomain | YES |
| NATKFPSVYAWERKKI | 1680941 | Ectodomain | YES |
| NAVAKHDFFTWHEGRT | 1680952 | Cell/Virus Enclosed | NO |
| NAVATFYPQLQSAEWK | 1680955 | Cell/Virus Enclosed | NO |
| NAVCSVIDLKLD | 1478685 | Cell/Virus Enclosed | NO |
| NAVDCMSDFMSEIKCK | 1680957 | Ectodomain | YES |
| NAVDCMSDFMSG | 1478688 | Ectodomain | YES |

| | | | |
|---|---|---|---|
| NAVDCSSSFFSEIQCK | 1680958 | Ectodomain | YES |
| NAVKGCVPLNAIPSLA | 1680961 | Cell/Virus Enclosed | NO |
| NAVLTIANNYSK | 1478700 | Cell/Virus Enclosed | NO |
| NAVMQNNELMPH | 1478702 | Cell/Virus Enclosed | NO |
| NAVSRSMLGLTTQTVD | 1680966 | Cell/Virus Enclosed | NO |
| NAVYDGVRLFAS | 1478725 | Cell/Virus Enclosed | NO |
| NAVYTKVDGNDV | 1478729 | Cell/Virus Enclosed | NO |
| NAYANWCCLKCGFSFD | 1680977 | Cell/Virus Enclosed | NO |
| NAYEKDKAVARKLERM | 1680978 | Cell/Virus Enclosed | NO |
| NAYNNTAISVQT | 1478744 | Ectodomain | YES |
| NAYNSTSISVQT | 1478749 | Ectodomain | YES |
| NAYQQQDGMMNI | 1478752 | Cell/Virus Enclosed | NO |
| NAYSPIDIPRGL | 1478758 | Ectodomain | YES |
| NAYVNSNQNTDSDSLS | 1680987 | Capsid | UN |
| NAYVSQQLSDSTLVKF | 1680989 | Ectodomain | YES |
| NAYYKYDAKNYVRFPE | 1680991 | Cell/Virus Enclosed | NO |
| NAYYKYDNGNFIKLPE | 1680992 | Cell/Virus Enclosed | NO |
| NCAGNVIGNDALTRFT | 1680994 | Cell/Virus Enclosed | NO |
| NCCDEYGGHHDFVIKT | 1681002 | Underneath Membrane | NO |
| NCCRFQRIDDDGNKLD | 1681006 | Cell/Virus Enclosed | NO |
| NCCRFQRVDENGDKLD | 1681007 | Cell/Virus Enclosed | NO |
| NCDFYGWVSGNMMDGF | 1681011 | Cell/Virus Enclosed | NO |
| NCDSFGPGNTFINGDI | 1681012 | Cell/Virus Enclosed | NO |
| NCDVSDVTKLYLGGMS | 1681020 | Cell/Virus Enclosed | NO |
| NCESIYEFDTYK | 1478819 | Cell/Virus Enclosed | NO |
| NCFICHSFKPGN | 1478824 | Cell/Virus Enclosed | NO |
| NCIDCDSYKPGNTFIT | 1681033 | Cell/Virus Enclosed | NO |
| NCLASECFMKSD | 1478859 | Cell/Virus Enclosed | NO |
| NCLQHLHVASNSKLWH | 1681048 | Cell/Virus Enclosed | NO |
| NCLQHLHVASNSKLWQ | 1681049 | Cell/Virus Enclosed | NO |
| NCNGYQHNSFAD | 1478903 | Ectodomain | YES |
| NCNIEAWLNGKS | 1478910 | Ectodomain | YES |
| NCNIESWLNDKS | 1478911 | Ectodomain | YES |
| NCNIPTQCYYNNGSSG | 1681058 | Cell/Virus Enclosed | NO |
| NCNVDRYPANAI | 1478922 | Cell/Virus Enclosed | NO |
| NCNWFVQSDKCS | 1478929 | Cell/Virus Enclosed | NO |
| NCQLSFAGYTESAVNS | 1681077 | Cell/Virus Enclosed | NO |
| NCSDCYDDMCVI | 1478954 | Cell/Virus Enclosed | NO |
| NCSDGNYNRNAS | 1478956 | Ectodomain | YES |
| NCSFQDAASNIFVINK | 1681079 | Cell/Virus Enclosed | NO |
| NCTDVPTSIHAD | 1478978 | Ectodomain | YES |
| NCTFGDPDDASR | 1478991 | Cell/Virus Enclosed | NO |
| NCTFGSPDDASR | 1478992 | Cell/Virus Enclosed | NO |
| NCTFMYIYNITE | 1478993 | Ectodomain | YES |
| NCTFMYTFNITE | 1478994 | Ectodomain | YES |
| NCTKYNIYDYVGTGII | 1681096 | Ectodomain | YES |
| NCTNFAINDLNTTVPR | 1681097 | Ectodomain | YES |
| NCVDCDIAGVGN | 1479022 | Cell/Virus Enclosed | NO |
| NCVDCDTAGVGN | 1479025 | Cell/Virus Enclosed | NO |
| NCVKMLTNGSGSGQAI | 1681107 | Cell/Virus Enclosed | NO |
| NCVLSVNSSNCNNFNV | 1681113 | Cell/Virus Enclosed | NO |
| NCYRCPKEIVETVSAL | 1681136 | Cell/Virus Enclosed | NO |
| NCYRNSNVDEAF | 1479057 | Cell/Virus Enclosed | NO |
| NCYRNSNVGESF | 1479058 | Cell/Virus Enclosed | NO |
| NDADKANFLNAV | 1479074 | Cell/Virus Enclosed | NO |
| NDAITQTSQALQTVAT | 1681148 | Ectodomain | YES |
| NDAITRTAEAIH | 1479081 | Ectodomain | YES |
| NDDDAWFKLSYHYENN | 1681161 | Cell/Virus Enclosed | NO |
| NDDDYGIALTGT | 1479097 | Cell/Virus Enclosed | NO |
| NDDEDVVTGDNDDEDV | 1681163 | Cell/Virus Enclosed | NO |
| NDDEDVVTGDNNDEEI | 1681164 | Cell/Virus Enclosed | NO |
| NDDLYVKRYERG | 1479108 | Cell/Virus Enclosed | NO |
| NDDQIVVTGDDVDDIE | 1681166 | Cell/Virus Enclosed | NO |
| NDDTSETKYVKSLSIV | 1681170 | Cell/Virus Enclosed | NO |
| NDDVTSNVTQCFGPRD | 1681171 | Capsid | UN |
| NDEEIVTGDNDDQIVV | 1681172 | Cell/Virus Enclosed | NO |
| NDEESVTGDNDD | 1479120 | Cell/Virus Enclosed | NO |
| NDEKVMTFYPRLQAAS | 1681174 | Cell/Virus Enclosed | NO |
| NDFNTILNTACGVFEV | 1681178 | Cell/Virus Enclosed | NO |
| NDGRVDDDYICG | 1479148 | Cell/Virus Enclosed | NO |
| NDHEPQISPGGLEPPSEKHFRETEV | 43449 | Cell/Virus Enclosed | NO |
| NDICMFENKTTLPTNI | 1681194 | Cell/Virus Enclosed | NO |
| NDINDYVSDADFSITG | 1681201 | Cell/Virus Enclosed | NO |
| NDITNTNLSRGRGRNP | 1681204 | Cell/Virus Enclosed | NO |
| NDKALYEQVVMD | 1479178 | Cell/Virus Enclosed | NO |
| NDKHSNGTIKDRSPY | 215815 | Ectodomain | YES |

| | | | |
|---|---|---|---|
| NDKIFTKSGTND | 1479186 | Cell/Virus Enclosed | NO |
| NDKLGFNVDLPF | 1479189 | Cell/Virus Enclosed | NO |
| NDKNICLLDHAW | 1479191 | Cell/Virus Enclosed | NO |
| NDKNTTVPRISE | 1479193 | Ectodomain | YES |
| NDLLDITQLQVA | 1479211 | Ectodomain | YES |
| NDLLQPNTEVYT | 1479214 | Ectodomain | YES |
| NDLNTTIPRISE | 1479219 | Underneath Membrane | NO |
| NDLYPFVSDSAA | 1479228 | Cell/Virus Enclosed | NO |
| NDMLYSPPTISYNSTL | 1681245 | Cell/Virus Enclosed | NO |
| NDNAQALNTLVK | 1479234 | Ectodomain | YES |
| NDNCDFYGWVSGNMMD | 1681248 | Cell/Virus Enclosed | NO |
| NDNKIMTFYPKM | 1479245 | Cell/Virus Enclosed | NO |
| NDPIRSCLENTV | 1479260 | Cell/Virus Enclosed | NO |
| NDQEMTTPPPKE | 1479269 | Cell/Virus Enclosed | NO |
| NDSCPGKNNGIG | 1479278 | Ectodomain | YES |
| NDSNGCVVEASTDFHQ | 1681276 | Cell/Virus Enclosed | NO |
| NDTFLNGITYTSTSGN | 1681281 | Ectodomain | YES |
| NDTITVDQLKHL | 1479297 | Undetermined | UN |
| NDTKIASQLGMG | 1479298 | Ectodomain | YES |
| NDTKIVSQLGNC | 1479300 | Ectodomain | YES |
| NDTSETKYVKSL | 1479303 | Cell/Virus Enclosed | NO |
| NDTTNTNLSRGR | 1479305 | Cell/Virus Enclosed | NO |
| NDTVDVTNGLGT | 1479308 | Ectodomain | YES |
| NDVEIFENKTTL | 1479317 | Cell/Virus Enclosed | NO |
| NDVSSMLTFDSN | 1479333 | Ectodomain | YES |
| NDWQNLLYDSNG | 1479345 | Ectodomain | YES |
| NDYNKTIPRISE | 1479350 | Ectodomain | YES |
| NEAFHKVQDAVN | 1479357 | Ectodomain | YES |
| NEAFQKVQDAVNNNAQ | 1681314 | Ectodomain | YES |
| NECAQVLSEYVLCGGG | 1681318 | Cell/Virus Enclosed | NO |
| NECTERLKLFAE | 1479375 | Cell/Virus Enclosed | NO |
| NECVESQSKRYG | 1479376 | Ectodomain | YES |
| NEDLMEILNRDR | 1479391 | Cell/Virus Enclosed | NO |
| NEEKCGTQLNHT | 1479396 | Ectodomain | YES |
| NEHVTSVNLLDV | 1479422 | Cell/Virus Enclosed | NO |
| NEIAKNLNESLI | 1479425 | Ectodomain | YES |
| NEIIVKPNISLCVPLY | 1681354 | Cell/Virus Enclosed | NO |
| NEIKPSGLKTMVVSAG | 1681355 | Cell/Virus Enclosed | NO |
| NEKLVTTSTAPD | 1479457 | Cell/Virus Enclosed | NO |
| NEKLVTTSTEPD | 1479458 | Cell/Virus Enclosed | NO |
| NELAQVLTEVVYSNGG | 1681374 | Cell/Virus Enclosed | NO |
| NELLDTTQLQVANSLM | 1681375 | Ectodomain | YES |
| NELMPVKLKIQV | 1479475 | Cell/Virus Enclosed | NO |
| NENGTITDAIDC | 1479487 | Ectodomain | YES |
| NENLNAYQQQDGMMNM | 1681395 | Cell/Virus Enclosed | NO |
| NENLTWPLGLEC | 1479492 | Cell/Virus Enclosed | NO |
| NEQHYKSWVYQN | 1479503 | Ectodomain | YES |
| NESIACQFCLAG | 1479511 | Cell/Virus Enclosed | NO |
| NESRYGLQNLLQLPNF | 1681412 | Ectodomain | YES |
| NESWFEDDYKAF | 1479519 | Cell/Virus Enclosed | NO |
| NESWHFDKSEPLCLFK | 1681415 | Ectodomain | YES |
| NESWLTGNAIQG | 1479523 | Cell/Virus Enclosed | NO |
| NESYNNYVPSYVKPDS | 1681417 | Cell/Virus Enclosed | NO |
| NETLQYWVTPLSKRQY | 1681419 | Ectodomain | YES |
| NETNSQYLFSTINALP | 1681421 | Cell/Virus Enclosed | NO |
| NEVNDLLDITQL | 1479546 | Ectodomain | YES |
| NEVVKQMFGVNLQSGK | 1681429 | Cell/Virus Enclosed | NO |
| NEWAIANGYTSV | 1479552 | Cell/Virus Enclosed | NO |
| NEWAQANGFTVM | 1479556 | Cell/Virus Enclosed | NO |
| NEYTGNYQCGHY | 1479565 | Cell/Virus Enclosed | NO |
| NFAAKRVLGLQTQTVD | 1681439 | Cell/Virus Enclosed | NO |
| NFAINDLNTTVPRISE | 1681440 | Ectodomain | YES |
| NFCPCKSNGSCP | 1479602 | Ectodomain | YES |
| NFCPCSSCFGKN | 1479604 | Ectodomain | YES |
| NFCPCSSCPGKN | 1479606 | Ectodomain | YES |
| NFCPCSSCPSKN | 1479607 | Ectodomain | YES |
| NFCPCSSCSGKN | 1479609 | Ectodomain | YES |
| NFCYMHQMELAS | 1479613 | Cell/Virus Enclosed | NO |
| NFDDQPSLQVESANLM | 1681453 | Cell/Virus Enclosed | NO |
| NFDEQNCDVLKEVLVL | 1681455 | Cell/Virus Enclosed | NO |
| NFDQKELLAYYNMLVN | 1681461 | Cell/Virus Enclosed | NO |
| NFDQKELLDYYN | 1479624 | Cell/Virus Enclosed | NO |
| NFDSDVFIKKYG | 1479629 | Cell/Virus Enclosed | NO |
| NFDYLDIHPSLCNNGK | 1681463 | Ectodomain | YES |
| NFEHKDITECVT | 1479639 | Cell/Virus Enclosed | NO |
| NFELCDNPFFAVSKPMG | 43825 | Ectodomain | YES |

| | | | |
|---|---|---|---|
| NFELCDNPFIAV | 1479649 | Ectodomain | YES |
| NFEPFTVNSIND | 1479655 | Ectodomain | YES |
| NFEPFTVNSKND | 1479656 | Ectodomain | YES |
| NFEPFTVNSVNDSLEP | 1681470 | Ectodomain | YES |
| NFFFLTHINTTL | 1479663 | Ectodomain | YES |
| NFFISYAKPEDKLSVY | 1681474 | Cell/Virus Enclosed | NO |
| NFFSPQIITTDNTFVS | 1681475 | Ectodomain | YES |
| NFGDLQLNKLGTEDPR | 1681479 | Cell/Virus Enclosed | NO |
| NFGDLQLTKLGI | 1479680 | Cell/Virus Enclosed | NO |
| NFGDQDLIRQGI | 1479683 | Capsid | UN |
| NFGDQELIRQGTEYKH | 1681482 | Cell/Virus Enclosed | NO |
| NFGDVNKPVCAK | 1479690 | Cell/Virus Enclosed | NO |
| NFGMLNGNAIATVKSE | 1681488 | Cell/Virus Enclosed | NO |
| NFIDKLNVLFDG | 1479712 | Cell/Virus Enclosed | NO |
| NFIIYDSSDRGQESLA | 1681496 | Cell/Virus Enclosed | NO |
| NFILHDVNNGLT | 1479723 | Ectodomain | YES |
| NFILHDVNSGTT | 1479725 | Ectodomain | YES |
| NFKDLREYVFKN | 1479737 | Ectodomain | YES |
| NFKSISLTVGEVFGKI | 1681523 | Cell/Virus Enclosed | NO |
| NFLDDNVLPETYVALP | 1681528 | Ectodomain | YES |
| NFNEKPVVMEIPSLMT | 1681547 | Cell/Virus Enclosed | NO |
| NFNENGTITDAI | 1479797 | Ectodomain | YES |
| NFNETKGPLCVDTSHF | 1681548 | Ectodomain | YES |
| NFNFSSNSVDNL | 1479802 | Ectodomain | YES |
| NFNGLTGTGVLTPSSKR | 43893 | Ectodomain | YES |
| NFNMSCLMSFIQ | 1479811 | Ectodomain | YES |
| NFNPSSWNRRYG | 1479817 | Ectodomain | YES |
| NFNQYYYNMATG | 1479824 | Ectodomain | YES |
| NFNTLFATTIPNTAFG | 1681575 | Cell/Virus Enclosed | NO |
| NFNTLFSTTIPN | 1479830 | Cell/Virus Enclosed | NO |
| NFNVDDINFAPV | 1479831 | Ectodomain | YES |
| NFNVDDINFSPVLGCL | 1681576 | Ectodomain | YES |
| NFNVTNAETTDF | 1479842 | Ectodomain | YES |
| NFPKKEKKQKAP | 1479844 | Cell/Virus Enclosed | NO |
| NFRDLALKGSTY | 1479870 | Ectodomain | YES |
| NFRIDIPSAELT | 1479877 | Cell/Virus Enclosed | NO |
| NFRIDVPSAESTGTQS | 1681597 | Cell/Virus Enclosed | NO |
| NFRTEVDPDLSA | 1479886 | Cell/Virus Enclosed | NO |
| NFSDGKMGLFFN | 1479905 | Ectodomain | YES |
| NFSDGKMGRFFNHTLV | 1681605 | Ectodomain | YES |
| NFSGALAEHDFF | 1479915 | Cell/Virus Enclosed | NO |
| NFSQILPDPLKPTKRS | 1681617 | Ectodomain | YES |
| NFSQYYYNIDTGSVYG | 1681618 | Underneath Membrane | NO |
| NFSSNSVDNLKS | 1479943 | Ectodomain | YES |
| NFSSVLGCLGSG | 1479946 | Ectodomain | YES |
| NFTIVGQEEFIQTNSP | 1681628 | Ectodomain | YES |
| NFTKAPFIYLNN | 1479971 | Ectodomain | YES |
| NFTVDPQKAYLDFVNA | 1681637 | Cell/Virus Enclosed | NO |
| NFVGEFVINDVV | 1480006 | Cell/Virus Enclosed | NO |
| NFVKDDYCDGSLGCKM | 1681647 | Cell/Virus Enclosed | NO |
| NFVKQGHFIGVE | 1480009 | Cell/Virus Enclosed | NO |
| NFVSSYAKPEEKLSAY | 1681651 | Cell/Virus Enclosed | NO |
| NFVVQPVGVVDTEWGN | 1681654 | Cell/Virus Enclosed | NO |
| NFYCFVNTTIGNETTS | 1681657 | Ectodomain | YES |
| NFYQHVINGCKD | 1480037 | Cell/Virus Enclosed | NO |
| NFYSPQIITTDN | 1480038 | Ectodomain | YES |
| NGACGSPGYNLKNGEV | 1681664 | Cell/Virus Enclosed | NO |
| NGACGSPGYNVRNDGT | 1681665 | Cell/Virus Enclosed | NO |
| NGAIIRDSDVVKQPVK | 1681669 | Cell/Virus Enclosed | NO |
| NGALEAFKRSNNGVYI | 1681673 | Cell/Virus Enclosed | NO |
| NGCVGHSENVFA | 1480102 | Ectodomain | YES |
| NGDAAVKDFDFYRYNK | 1681688 | Cell/Virus Enclosed | NO |
| NGDRSGARPKQRRPQG | 1681693 | Cell/Virus Enclosed | NO |
| NGDSEAVLKKLK | 1480121 | Cell/Virus Enclosed | NO |
| NGELEALETPVD | 1480131 | Cell/Virus Enclosed | NO |
| NGERSGARPKQRRPQG | 1681705 | Cell/Virus Enclosed | NO |
| NGFICEKLAIGG | 1480146 | Cell/Virus Enclosed | NO |
| NGFQGRQLLGSC | 1480159 | Cell/Virus Enclosed | NO |
| NGFQKCEQLLLE | 1480161 | Ectodomain | YES |
| NGFTAMNGEDAF | 1480168 | Cell/Virus Enclosed | NO |
| NGFVYFNGTGRG | 1480178 | Ectodomain | YES |
| NGGAYSLFDMAKFPLK | 1681727 | Cell/Virus Enclosed | NO |
| NGGAYSLFDMTKFSLK | 1681728 | Cell/Virus Enclosed | NO |
| NGGDAALALLLLDRLN | 1385897 | Cell/Virus Enclosed | NO |
| NGGDTALALLLL | 1480187 | Cell/Virus Enclosed | NO |
| NGHKIEDFSIRN | 1480211 | Cell/Virus Enclosed | NO |

| | | | |
|---|---|---|---|
| NGHKIEDLSIRNLQKR | 1681746 | Cell/Virus Enclosed | NO |
| NGIAYITFSGSF | 1480218 | Cell/Virus Enclosed | NO |
| NGIAYTTFSGSFDNGH | 1681754 | Cell/Virus Enclosed | NO |
| NGICWGNQLFVTVVDTTRST | 111581 | Capsid | UN |
| NGIGTCPAGTNY | 1480231 | Ectodomain | YES |
| NGIGVTQNVLYENQKQ | 1681759 | Ectodomain | YES |
| NGIIRDVNPDDKCFAN | 1681760 | Cell/Virus Enclosed | NO |
| NGIISNARNGCIPLSV | 1681761 | Cell/Virus Enclosed | NO |
| NGIITVEELKQL | 1480240 | Undetermined | UN |
| NGIRRPKHLYVSMP | 44002 | Capsid | UN |
| NGITYTSTSGNLLGFK | 1681774 | Ectodomain | YES |
| NGKFLSNTKYYDDSQY | 1681777 | Ectodomain | YES |
| NGKIVYAILSDV | 1480266 | Cell/Virus Enclosed | NO |
| NGKPCTPPALNC | 1480269 | Ectodomain | YES |
| NGKSVPSPLNWE | 1480278 | Ectodomain | YES |
| NGKTVLGEYVFDKSEL | 1385911 | Cell/Virus Enclosed | NO |
| NGKYYTKPIIKAQFKP | 1681788 | Cell/Virus Enclosed | NO |
| NGLDSLDSYPAL | 1480288 | Cell/Virus Enclosed | NO |
| NGLGVTMDVLNKNQKL | 1681792 | Ectodomain | YES |
| NGLLDTTQLHVADTLM | 1681796 | Ectodomain | YES |
| NGLTCSTDLQKA | 1480318 | Ectodomain | YES |
| NGMGTCPAGTNY | 1480337 | Ectodomain | YES |
| NGMNGRTILGAA | 1480341 | Cell/Virus Enclosed | NO |
| NGMNTSQSVCNG | 1480345 | Ectodomain | YES |
| NGMPVSSIKSDK | 1480348 | Cell/Virus Enclosed | NO |
| NGNAIATVKSEDGNIK | 1681827 | Cell/Virus Enclosed | NO |
| NGNKIVDKEVKDMQFD | 1681839 | Cell/Virus Enclosed | NO |
| NGNVRCVELLKQYTSA | 1681846 | Ectodomain | YES |
| NGNVSTVFVPAM | 1480371 | Cell/Virus Enclosed | NO |
| NGNVSTVFVPATRDSV | 1681847 | Cell/Virus Enclosed | NO |
| NGNWYDFGDFEQ | 1480373 | Cell/Virus Enclosed | NO |
| NGPGIDAGYKNS | 1480379 | Ectodomain | YES |
| NGPHEFCTKHTM | 1480381 | Cell/Virus Enclosed | NO |
| NGPLKVGAGVMLECEK | 1681855 | Cell/Virus Enclosed | NO |
| NGPQNQSSAPRI | 1480393 | Cell/Virus Enclosed | NO |
| NGQAITTSVDANTNQD | 1681864 | Cell/Virus Enclosed | NO |
| NGRKVDLQLGNLGYLQ | 1681875 | Ectodomain | YES |
| NGSAQAISFCKS | 1480441 | Ectodomain | YES |
| NGSAQATALCKS | 1480443 | Ectodomain | YES |
| NGSAQVTSFCKS | 1480446 | Ectodomain | YES |
| NGSCPDKNNGIG | 1480449 | Ectodomain | YES |
| NGSCPGKNNGIG | 1480450 | Ectodomain | YES |
| NGSDAALALLLLDRLN | 1681888 | Cell/Virus Enclosed | NO |
| NGSDVLDQPPTA | 1480452 | Cell/Virus Enclosed | NO |
| NGSDVLYQPPTASVST | 1681889 | Cell/Virus Enclosed | NO |
| NGSGSGQAITCTIDSN | 1681891 | Cell/Virus Enclosed | NO |
| NGSMIEDIDYVPLKSA | 1681896 | Cell/Virus Enclosed | NO |
| NGSMIEDVDYVP | 1480463 | Cell/Virus Enclosed | NO |
| NGSVLEMAKEVKTSSK | 1681907 | Cell/Virus Enclosed | NO |
| NGSWAIFNDHHLNRPG | 1681908 | Cell/Virus Enclosed | NO |
| NGTDIDKCVRAFDIYN | 1681911 | Cell/Virus Enclosed | NO |
| NGTGRGDCKGFSSDVL | 1681913 | Ectodomain | YES |
| NGTITDAIDCAQ | 1480502 | Ectodomain | YES |
| NGTITVEKLKQL | 1480507 | Undetermined | UN |
| NGTNTSHSVCKG | 1480518 | Ectodomain | YES |
| NGTNTSHSVCNG | 1480519 | Ectodomain | YES |
| NGVCGVSAARLT | 1480530 | Cell/Virus Enclosed | NO |
| NGVDAKGFPLLA | 1480531 | Cell/Virus Enclosed | NO |
| NGVDAKGFPQFA | 1480532 | Cell/Virus Enclosed | NO |
| NGVDAKGFPQLAELIP | 1681929 | Capsid | UN |
| NGVDFDFIVNAANENL | 1681930 | Cell/Virus Enclosed | NO |
| NGVDYTKKFLTV | 1480537 | Cell/Virus Enclosed | NO |
| NGVFMSTAKCWTEEDL | 1681935 | Cell/Virus Enclosed | NO |
| NGVGITQQVLSK | 1480546 | Ectodomain | YES |
| NGVIKNVNSVRDWLKS | 1681942 | Cell/Virus Enclosed | NO |
| NGVIRDVSVEDVCFAN | 1681943 | Cell/Virus Enclosed | NO |
| NGVIVDKVGELKVEFW | 1681945 | Cell/Virus Enclosed | NO |
| NGVKAKGYPQFAELVP | 1681946 | Capsid | UN |
| NGVSDLSLKQKL | 1480575 | Cell/Virus Enclosed | NO |
| NGVTLSTKLKDGVNFN | 1681967 | Ectodomain | YES |
| NGVVPLSVIPATSAAR | 1681969 | Cell/Virus Enclosed | NO |
| NGVYWVASNQAD | 1480590 | Cell/Virus Enclosed | NO |
| NGYFIKTNNTMI | 1480598 | Ectodomain | YES |
| NGYFIKTNNTRIVDEW | 1681978 | Ectodomain | YES |
| NGYFIKTNNTRV | 1480600 | Ectodomain | YES |
| NGYPHMFITREE | 1480613 | Cell/Virus Enclosed | NO |

| | | | |
|---|---|---|---|
| NGYQHNSFADVM | 1480620 | Ectodomain | YES |
| NGYQHNSVVDVM | 1480622 | Ectodomain | YES |
| NGYTDNIFSVQQDGRI | 1681987 | Ectodomain | YES |
| NGYYPTSGSTYL | 1480635 | Ectodomain | YES |
| NHFKELWHMDTG | 1480676 | Ectodomain | YES |
| NHFKESWHMDTG | 1480679 | Ectodomain | YES |
| NHIEVVSAYGLC | 1480698 | Ectodomain | YES |
| NHINNLKVKNYG | 1480707 | Ectodomain | YES |
| NHLHSVHNWLKP | 1480716 | Cell/Virus Enclosed | NO |
| NHLSTFYPQLQSAEWK | 1682031 | Cell/Virus Enclosed | NO |
| NHNMGDSDLVQN | 1480732 | Cell/Virus Enclosed | NO |
| NHNNGNDVLYQPPTAS | 1682037 | Cell/Virus Enclosed | NO |
| NHPIQIDQLNSS | 1480737 | Ectodomain | YES |
| NHPIQVDQFNSS | 1480739 | Ectodomain | YES |
| NHPIQVDQLNSSYFKL | 1682040 | Ectodomain | YES |
| NHPIQVDQLNST | 1480741 | Ectodomain | YES |
| NHQADTSTPSDVSSRD | 1682041 | Capsid | UN |
| NHQFVTASSATD | 1480749 | Cell/Virus Enclosed | NO |
| NHSFSVQKHIGAPANL | 1682046 | Cell/Virus Enclosed | NO |
| NHSGSNDIWMQN | 1480763 | Ectodomain | YES |
| NHSIPDLQRAAI | 1480765 | Cell/Virus Enclosed | NO |
| NHSVPKLSDFESELSH | 1682051 | Ectodomain | YES |
| NHTFGNPIIPFK | 1480784 | Ectodomain | YES |
| NHTIKGAFLNGS | 1480790 | Cell/Virus Enclosed | NO |
| NHTLGNPVIPFK | 1480798 | Ectodomain | YES |
| NHVADIDKLIDYAA | 44172 | Capsid | UN |
| NHVMVDCRRLLK | 1480816 | Cell/Virus Enclosed | NO |
| NIAFELFAKRKVGLTP | 1682065 | Cell/Virus Enclosed | NO |
| NICFDIRDNGAL | 1480839 | Cell/Virus Enclosed | NO |
| NICQAVSANVCALMSC | 1682085 | Cell/Virus Enclosed | NO |
| NICQAVSDNVRA | 1480848 | Cell/Virus Enclosed | NO |
| NICQAVTANVRS | 1480850 | Cell/Virus Enclosed | NO |
| NIDAAKIYGMGF | 1480855 | Ectodomain | YES |
| NIDATSTGNYNYKYRYL | 44198 | Ectodomain | YES |
| NIDMGVFINKYG | 1480869 | Cell/Virus Enclosed | NO |
| NIDTGSFYGFSN | 1480872 | Underneath Membrane | NO |
| NIEAWLNDKSVPSPLN | 1682103 | Ectodomain | YES |
| NIEEFIQTSSPK | 1480883 | Ectodomain | YES |
| NIESWLNDKSVP | 1480886 | Ectodomain | YES |
| NIFGLYKNSATGSLDV | 1682109 | Cell/Virus Enclosed | NO |
| NIFQAVSSNINM | 1480901 | Cell/Virus Enclosed | NO |
| NIFSVQEDGRIP | 1480907 | Ectodomain | YES |
| NIFVINKDTYAALRNS | 1682123 | Cell/Virus Enclosed | NO |
| NIGGAVCSKHANLYQK | 1682130 | Cell/Virus Enclosed | NO |
| NIGGAVCSKHANLYRA | 1682131 | Cell/Virus Enclosed | NO |
| NIGQQRSVLTPL | 1480928 | Cell/Virus Enclosed | NO |
| NIGTNFPLQIGFSTGV | 1682133 | Cell/Virus Enclosed | NO |
| NIGTNVPLQVGFSNGV | 1682134 | Cell/Virus Enclosed | NO |
| NIHAVLDTLLAS | 1480938 | Cell/Virus Enclosed | NO |
| NIHLKHGGGVAG | 1480940 | Cell/Virus Enclosed | NO |
| NIHSSRIAGRSA | 1480945 | Ectodomain | YES |
| NIIDAATSYGMC | 1480951 | Ectodomain | YES |
| NIIVPDKSVYDQVVDN | 1682149 | Cell/Virus Enclosed | NO |
| NILACYDGCAQG | 1480981 | Cell/Virus Enclosed | NO |
| NILGYSSLCDEF | 1480989 | Cell/Virus Enclosed | NO |
| NILHVVGPDARAKQDV | 1682167 | Cell/Virus Enclosed | NO |
| NILSPFLPLLPIFFC | 1682176 | Transmembrane | NO |
| NINAILTEVNELLDTT | 1682179 | Ectodomain | YES |
| NINCILSVNSSN | 1481014 | Cell/Virus Enclosed | NO |
| NINFERNLRRGT | 1481022 | Ectodomain | YES |
| NINPHNYSYMDLNPVL | 1682189 | Ectodomain | YES |
| NINRFNVAITRT | 1481037 | Cell/Virus Enclosed | NO |
| NINRILGVNSSN | 1481038 | Cell/Virus Enclosed | NO |
| NINSTLVDLKWLNRVE | 1682193 | Ectodomain | YES |
| NINWNVEYPIISNELS | 1682196 | Cell/Virus Enclosed | NO |
| NIPADIVDRDPS | 1481048 | Cell/Virus Enclosed | NO |
| NIPCYASAVAQA | 1481050 | Cell/Virus Enclosed | NO |
| NIPDLPICVYDR | 1481055 | Ectodomain | YES |
| NIPDLPISVYDP | 1481058 | Ectodomain | YES |
| NIPISIDIPNAAFIRTSD | 558299 | Undetermined | UN |
| NIPTQCYYNNGSSGRI | 1682200 | Cell/Virus Enclosed | NO |
| NIPYSRVISRMG | 1481070 | Cell/Virus Enclosed | NO |
| NIQNLLYCDSPFEKLQ | 1682201 | Ectodomain | YES |
| NIRIVDEWSYTG | 1481079 | Ectodomain | YES |
| NISEEFSSNVAN | 1481090 | Cell/Virus Enclosed | NO |
| NISGQSPARTSSDPG | 227409 | Secreted | YES |

| | | | |
|---|---|---|---|
| NISLDNPLENPSSLFDLVAR | 44350 | Capsid | UN |
| NISLNGNTSVCVRTSH | 1682210 | Ectodomain | YES |
| NISMLQLGVPDN | 1481103 | Ectodomain | YES |
| NISTPNLHDFKE | 1481117 | Ectodomain | YES |
| NISTPNLPDFKE | 1481118 | Ectodomain | YES |
| NISYVVTQNLLF | 1481125 | Cell/Virus Enclosed | NO |
| NITNLCPFGEVFNATK | 1682228 | Ectodomain | YES |
| NIVAADLGVLIQNSAK | 1682235 | Cell/Virus Enclosed | NO |
| NIVGDFQLTEEV | 1481162 | Cell/Virus Enclosed | NO |
| NIVGPDARQDGRQSYV | 1682241 | Cell/Virus Enclosed | NO |
| NIVNVSLVKPTV | 1481171 | Ectodomain | YES |
| NIYDYVGTGIIRSSNQ | 1682254 | Ectodomain | YES |
| NIYGTATIGVCK | 1481193 | Cell/Virus Enclosed | NO |
| NIYSQNLCVDGFGVNK | 1682257 | Cell/Virus Enclosed | NO |
| NKAIDGFGVFDIKNSS | 1682260 | Cell/Virus Enclosed | NO |
| NKAIPSGNLIKFDIGS | 1682262 | Cell/Virus Enclosed | NO |
| NKASYDTNLNKFNRAS | 1682267 | Cell/Virus Enclosed | NO |
| NKATNGAMQQES | 1481217 | Cell/Virus Enclosed | NO |
| NKCSRLLSDDRTEVPQ | 1682278 | Ectodomain | YES |
| NKCTKYNIYDGS | 1481232 | Ectodomain | YES |
| NKCTKYNIYDVS | 1481234 | Ectodomain | YES |
| NKCTKYNIYDVV | 1481235 | Ectodomain | YES |
| NKCTKYNIYGVS | 1481236 | Ectodomain | YES |
| NKDASFIGKNLKSNCV | 1682279 | Cell/Virus Enclosed | NO |
| NKDGFLYVYKGYQPID | 1682282 | Ectodomain | YES |
| NKDGIIWVATEGALNTP | 1386017 | Cell/Virus Enclosed | NO |
| NKDIAVFGPVIDWNV | 922915 | Ectodomain | YES |
| NKDIAVLGPVIDWNV | 922916 | Ectodomain | YES |
| NKDIQVLGPVIDWNV | 922921 | Ectodomain | YES |
| NKDQEAILMSED | 1481255 | Cell/Virus Enclosed | NO |
| NKEDFDIIQKCQITSV | 1682291 | Cell/Virus Enclosed | NO |
| NKEQEAILMSEDVKLN | 1682298 | Cell/Virus Enclosed | NO |
| NKEVFVGDSLVN | 1481275 | Cell/Virus Enclosed | NO |
| NKFCNVQQCFGK | 1481281 | Cell/Virus Enclosed | NO |
| NKFGAISSSLQE | 1481285 | Ectodomain | YES |
| NKFGFPDTSFYNPDTQRLVW | 111583 | Capsid | UN |
| NKFNQALEAMQT | 1481292 | Ectodomain | YES |
| NKFNQALGAMQTGFTT | 1682315 | Ectodomain | YES |
| NKFTPLSVASAP | 1481300 | Cell/Virus Enclosed | NO |
| NKFVPVKPASKQCFKV | 1682320 | Cell/Virus Enclosed | NO |
| NKGHFDGQIGEA | 1481322 | Cell/Virus Enclosed | NO |
| NKGVDCTKKFLT | 1481333 | Cell/Virus Enclosed | NO |
| NKGVDYTKKFLT | 1481336 | Cell/Virus Enclosed | NO |
| NKHAFHTPAYDG | 1481345 | Cell/Virus Enclosed | NO |
| NKHCNVQQCFGKRGPS | 1682345 | Capsid | UN |
| NKHIDAYKAFPP | 1481356 | Cell/Virus Enclosed | NO |
| NKHRSFTQWYHE | 1481359 | Cell/Virus Enclosed | NO |
| NKIEDLSMRALQ | 1481364 | Cell/Virus Enclosed | NO |
| NKIHDVVNQQGN | 1481366 | Ectodomain | YES |
| NKINLCDDPEKAQSML | 1682358 | Cell/Virus Enclosed | NO |
| NKINLCDDPETAQELL | 1682359 | Cell/Virus Enclosed | NO |
| NKIPANIVLPQG | 1481375 | Cell/Virus Enclosed | NO |
| NKIQDVVNQQGNSLNH | 1682361 | Ectodomain | YES |
| NKIQDVVNQQGSALNH | 1682362 | Ectodomain | YES |
| NKIQPWTNDALNTICI | 1682363 | Cell/Virus Enclosed | NO |
| NKISGKGQQQQG | 1481383 | Cell/Virus Enclosed | NO |
| NKISLNAPVTLK | 1481384 | Ectodomain | YES |
| NKIVDKEVKDMQFDLY | 1682366 | Cell/Virus Enclosed | NO |
| NKIVGGAPTWFNALRD | 1682367 | Cell/Virus Enclosed | NO |
| NKKFGFIEDSVF | 1481397 | Ectodomain | YES |
| NKLCVVPGNVKPGDAV | 1682373 | Cell/Virus Enclosed | NO |
| NKLLHWSDQTICYKDS | 1682382 | Cell/Virus Enclosed | NO |
| NKLNLPGCNGGSLYVN | 1682385 | Cell/Virus Enclosed | NO |
| NKLSTSDWSDIRDYKL | 1682389 | Cell/Virus Enclosed | NO |
| NKLVKDPPNVQIHTID | 1682391 | Secreted | YES |
| NKLWFDFIENPS | 1481440 | Cell/Virus Enclosed | NO |
| NKMVNWFNTMLDASAP | 1682394 | Cell/Virus Enclosed | NO |
| NKNTYRSQLGCV | 1481461 | Cell/Virus Enclosed | NO |
| NKPLIEWPLTEYDFTD | 1682404 | Cell/Virus Enclosed | NO |
| NKPLSQPRADKPSQLK | 1682406 | Capsid | UN |
| NKPLSQPRVDKP | 1481477 | Cell/Virus Enclosed | NO |
| NKPRQKRSPNKQCTVQ | 1682407 | Cell/Virus Enclosed | NO |
| NKPWRWMLWCKDNAVA | 1682411 | Cell/Virus Enclosed | NO |
| NKQEASVPILTTPFSL | 1682414 | Cell/Virus Enclosed | NO |
| NKQLNEISDDCNWPLV | 1682416 | Cell/Virus Enclosed | NO |
| NKQMANVSVLTTPFSL | 1682418 | Cell/Virus Enclosed | NO |

| | | | |
|---|---|---|---|
| NKRFGFIEDSAF | 1481506 | Ectodomain | YES |
| NKRFGFIEDSVFKPRP | 1682421 | Ectodomain | YES |
| NKSAGWPLNKFGKAGL | 1682425 | Cell/Virus Enclosed | NO |
| NKSEVPLDICTSICKYPDYI | 111585 | Capsid | UN |
| NKSFFPQFTSDQ | 1481511 | Ectodomain | YES |
| NKSFFSQFTSDQ | 1481512 | Underneath Membrane | NO |
| NKSFLPQFTSDQAVTF | 1682427 | Ectodomain | YES |
| NKSVANLKLPPHDVTV | 1682431 | Cell/Virus Enclosed | NO |
| NKTEADWIELFP | 1481536 | Cell/Virus Enclosed | NO |
| NKTLQEFAKNLP | 1481547 | Ectodomain | YES |
| NKTLQEFAQNLPKYVK | 1682437 | Ectodomain | YES |
| NKTTYKPNTWCL | 1481553 | Cell/Virus Enclosed | NO |
| NKVIYTTEIASKLTCK | 1682452 | Cell/Virus Enclosed | NO |
| NKVTLADAGFMKQYGE | 1682457 | Ectodomain | YES |
| NKWQWQEAWYEF | 1481585 | Cell/Virus Enclosed | NO |
| NKWSGVSDLSLK | 1481588 | Cell/Virus Enclosed | NO |
| NKWSGVSDLSSN | 1481589 | Cell/Virus Enclosed | NO |
| NKWSGVSNLSLK | 1481590 | Cell/Virus Enclosed | NO |
| NKYFKHWSMPYHPNTV | 1682471 | Cell/Virus Enclosed | NO |
| NKYKYYSGSASEADYR | 1682480 | Cell/Virus Enclosed | NO |
| NKYLYELIQKFA | 1481603 | Cell/Virus Enclosed | NO |
| NKYTYFDNPILD | 1481611 | Ectodomain | YES |
| NLAIEEIEYVHGDALH | 1682491 | Cell/Virus Enclosed | NO |
| NLARDGVVPLSVIPAT | 1682500 | Cell/Virus Enclosed | NO |
| NLAYSNSYTIGS | 1481647 | Ectodomain | YES |
| NLCDDPEKAQGM | 1481648 | Cell/Virus Enclosed | NO |
| NLCNPDPIRFTG | 1481655 | Ectodomain | YES |
| NLCPFGEVFNATKFPS | 1682507 | Ectodomain | YES |
| NLCVDGFGLNKI | 1481673 | Cell/Virus Enclosed | NO |
| NLCVDGFGVNKIQPWT | 1682512 | Cell/Virus Enclosed | NO |
| NLDAYVNSNQNT | 1481675 | Cell/Virus Enclosed | NO |
| NLDEARFSGSANQQQL | 1682515 | Cell/Virus Enclosed | NO |
| NLDEARFSGTAN | 1481679 | Cell/Virus Enclosed | NO |
| NLDEARLSGSAN | 1481680 | Cell/Virus Enclosed | NO |
| NLDEPQKDVYEL | 1481681 | Capsid | UN |
| NLEAQVQIDRLI | 1481701 | Ectodomain | YES |
| NLERCELVNYKQ | 1481716 | Cell/Virus Enclosed | NO |
| NLEYPIISNEVSINTS | 1682542 | Cell/Virus Enclosed | NO |
| NLFASVKNSQSS | 1481725 | Ectodomain | YES |
| NLFEGDKFVGTFESAA | 1682545 | Cell/Virus Enclosed | NO |
| NLFGNPDEPQKD | 1481734 | Capsid | UN |
| NLFIAKYGLEDY | 1481736 | Cell/Virus Enclosed | NO |
| NLFKDCSKSCLGYHPA | 1682548 | Cell/Virus Enclosed | NO |
| NLFKDCSKSYSGYHPA | 1682549 | Cell/Virus Enclosed | NO |
| NLFKDCSNSYSG | 1481738 | Cell/Virus Enclosed | NO |
| NLGEVFIAQSKG | 1481762 | Cell/Virus Enclosed | NO |
| NLGFDNQSKLPS | 1481764 | Cell/Virus Enclosed | NO |
| NLGFDNQSKSPSSSGT | 1682558 | Capsid | UN |
| NLGIPSYCDGVR | 1481775 | Cell/Virus Enclosed | NO |
| NLGNHRKELWHL | 1481782 | Ectodomain | YES |
| NLGNLQSFNYRI | 1481784 | Ectodomain | YES |
| NLGSNEPGNVGG | 1481789 | Cell/Virus Enclosed | NO |
| NLGSNGKPGNVGGNDA | 1682570 | Cell/Virus Enclosed | NO |
| NLHFDVDNINFKSLVG | 1682575 | Ectodomain | YES |
| NLICAQYVAGYK | 1481809 | Underneath Membrane | NO |
| NLILHDVNSGLT | 1481819 | Ectodomain | YES |
| NLKLPPHDVTVLRDNL | 1682605 | Cell/Virus Enclosed | NO |
| NLKSNCVRFKNVDKDD | 1682614 | Cell/Virus Enclosed | NO |
| NLLFTEQLGAPL | 1481869 | Ectodomain | YES |
| NLLGFDSSKPVI | 1481871 | Cell/Virus Enclosed | NO |
| NLLGFDSSKPVTKKYT | 1682627 | Cell/Virus Enclosed | NO |
| NLLIQEFLQYDSSIHS | 1682634 | Cell/Virus Enclosed | NO |
| NLLYCDSPFEKLQCEH | 1682651 | Ectodomain | YES |
| NLLYDFNGNIIG | 1481902 | Ectodomain | YES |
| NLLYDSNGNIIGFKDF | 1682652 | Ectodomain | YES |
| NLMCIDMKGKMY | 1481910 | Ectodomain | YES |
| NLNAGEDVQLLK | 1481921 | Cell/Virus Enclosed | NO |
| NLNAYQQQDGMN | 1481923 | Cell/Virus Enclosed | NO |
| NLNAYVNSNQNTDSDS | 1682661 | Capsid | UN |
| NLNFSANSLDNLKSGV | 1682666 | Ectodomain | YES |
| NLNSSEGVSDLL | 1481947 | Underneath Membrane | NO |
| NLNTLRRGAALG | 1481949 | Cell/Virus Enclosed | NO |
| NLNTNLHFDVDNINFK | 1682683 | Ectodomain | YES |
| NLPDFKEELGQW | 1481959 | Ectodomain | YES |
| NLPEYMAVAVPS | 1481964 | Underneath Membrane | NO |
| NLPFQNIDSRAVGKCPRYVKQRSLL | 1161651 | Ectodomain | YES |

| | | | |
|---|---|---|---|
| NLPKFIEQISAFTKPS | 1682691 | Capsid | UN |
| NLPPPLLGNSTR | 1481982 | Ectodomain | YES |
| NLPPPLLSNSTG | 1481984 | Ectodomain | YES |
| NLPTYYIYDEEGGNDL | 1682701 | Cell/Virus Enclosed | NO |
| NLQFEMNRLQEA | 1482000 | Ectodomain | YES |
| NLQVESANQMLTVNVV | 1682717 | Cell/Virus Enclosed | NO |
| NLRQDDSVRNLFASVK | 1682731 | Ectodomain | YES |
| NLRVVGHAMQGTLLKL | 1682735 | Cell/Virus Enclosed | NO |
| NLSAGTQHISNDFTPE | 1682736 | Capsid | UN |
| NLSAPTLIPQEN | 1482052 | Cell/Virus Enclosed | NO |
| NLSAPTLVPQENYSSI | 1682737 | Cell/Virus Enclosed | NO |
| NLSGNLDEPQKD | 1482061 | Capsid | UN |
| NLSGNPDEPHKD | 1482062 | Cell/Virus Enclosed | NO |
| NLSGNPDEPQKDVYEL | 1682744 | Cell/Virus Enclosed | NO |
| NLSMLQLGVPDNSSTI | 1682751 | Ectodomain | YES |
| NLSSVLPQRNIRSSRI | 1682757 | Ectodomain | YES |
| NLSSVVPSLPTS | 1482090 | Ectodomain | YES |
| NLTADEIKALKE | 1482097 | Cell/Virus Enclosed | NO |
| NLTPIKLNFGMLNGMP | 1682769 | Cell/Virus Enclosed | NO |
| NLTYLNFSFELL | 1482144 | Ectodomain | YES |
| NLVEKLSPWFMDGENA | 1682786 | Cell/Virus Enclosed | NO |
| NLVFTNKTTLPTNVAF | 1682788 | Cell/Virus Enclosed | NO |
| NLVFVNKTSLPTNIAF | 1682789 | Cell/Virus Enclosed | NO |
| NLVILPGQDLQYVLA | 44915 | Ectodomain | YES |
| NLVKTGHYTGQAGEMP | 1682797 | Cell/Virus Enclosed | NO |
| NLVLAKLGKDSKPQQV | 1682798 | Capsid | UN |
| NLVLNNCTKYNIYDYV | 1682799 | Ectodomain | YES |
| NLVNSKSHNIGS | 1482174 | Ectodomain | YES |
| NLVNVGHYDGRTGELP | 1682801 | Cell/Virus Enclosed | NO |
| NLVNYKSHNIGS | 1482175 | Ectodomain | YES |
| NLVPIGKGNKDEQIGY | 1682802 | Capsid | UN |
| NLVPINKKDKNK | 1482180 | Cell/Virus Enclosed | NO |
| NLVPTYLKSDNIVAAD | 1682804 | Cell/Virus Enclosed | NO |
| NLYGFRDYIINR | 1482209 | Ectodomain | YES |
| NLYNYGAGIKLPSGIM | 1682825 | Cell/Virus Enclosed | NO |
| NLYRLVWHVERKDVPY | 1682830 | Cell/Virus Enclosed | NO |
| NLYSFDKAIDGF | 1482222 | Cell/Virus Enclosed | NO |
| NLYVVVVESDADRANF | 1682832 | Cell/Virus Enclosed | NO |
| NMALKGTDLLSK | 1482228 | Ectodomain | YES |
| NMARNGVVPLSVIPAT | 1682835 | Cell/Virus Enclosed | NO |
| NMASFVEYEVAKKNLA | 1682836 | Cell/Virus Enclosed | NO |
| NMCEHPHTSCHP | 1482247 | Ectodomain | YES |
| NMCEYPQTICHP | 1482252 | Ectodomain | YES |
| NMCEYPYTICHP | 1482253 | Ectodomain | YES |
| NMDMGVFINKYGLEDF | 1682843 | Cell/Virus Enclosed | NO |
| NMDQGSFGGASC | 1482266 | Cell/Virus Enclosed | NO |
| NMDTGFAYGYSN | 1482268 | Underneath Membrane | NO |
| NMEAAYTSSSLG | 1482274 | Ectodomain | YES |
| NMEEFIQTNSPK | 1482276 | Ectodomain | YES |
| NMFQFATLPVYDTIKY | 1682855 | Ectodomain | YES |
| NMHSPDVVGDYV | 1482304 | Cell/Virus Enclosed | NO |
| NMNIASNSKEWAYCVE | 1682884 | Cell/Virus Enclosed | NO |
| NMQLFEALQFTT | 1482347 | Cell/Virus Enclosed | NO |
| NMRVIHFGAGSDKGIA | 1682892 | Cell/Virus Enclosed | NO |
| NMRVLHLGAGSDKEVA | 1682894 | Cell/Virus Enclosed | NO |
| NMRVLHLGAGSDYGVA | 1682895 | Cell/Virus Enclosed | NO |
| NMRVLHLGAGSEKGVA | 1682896 | Cell/Virus Enclosed | NO |
| NMRVLHYGAGSDKGVA | 1682897 | Cell/Virus Enclosed | NO |
| NMSLAECKRALGLSIS | 1682898 | Cell/Virus Enclosed | NO |
| NMSPKPQRQRGHKNGQ | 1682900 | Cell/Virus Enclosed | NO |
| NMSPKPQRQRGQ | 1482368 | Capsid | UN |
| NMSSLMSFIQAD | 1482371 | Ectodomain | YES |
| NMVEFIQTSSPKVTID | 1682908 | Ectodomain | YES |
| NMVETVVGQLEQTNMH | 1682909 | Cell/Virus Enclosed | NO |
| NMWEFRDYFNDN | 1482400 | Cell/Virus Enclosed | NO |
| NMWEFRDYFNNNTDSI | 1682912 | Cell/Virus Enclosed | NO |
| NMYMGKCGDKFF | 1482412 | Cell/Virus Enclosed | NO |
| NNAAIVLQLPQGTALP | 1682921 | Cell/Virus Enclosed | NO |
| NNAAIVLQLPQGTTLPKGFYAEGS | 1311774 | Cell/Virus Enclosed | NO |
| NNAHALSKLASE | 1482423 | Ectodomain | YES |
| NNALDAIQEGFD | 1482425 | Ectodomain | YES |
| NNALHAIQQGFD | 1482426 | Ectodomain | YES |
| NNALHSIQQGFD | 1482428 | Ectodomain | YES |
| NNALNAIQQGFD | 1482429 | Ectodomain | YES |
| NNALYAIQEGFDATNS | 1682927 | Ectodomain | YES |
| NNALYAIQQGFD | 1482430 | Ectodomain | YES |

| | | | |
|---|---|---|---|
| NNAQALAKLASE | 1482434 | Ectodomain | YES |
| NNARQSDNKTAV | 1482439 | Ectodomain | YES |
| NNARQSDNMTAVACQP | 1682933 | Ectodomain | YES |
| NNAVYTKVDGND | 1482453 | Cell/Virus Enclosed | NO |
| NNCTKYNIYDYVGTGI | 1682946 | Ectodomain | YES |
| NNCTTAVMTYSN | 1482467 | Ectodomain | YES |
| NNDDAGNPVYFL | 1482478 | Cell/Virus Enclosed | NO |
| NNDEEIVTGDNDDQIV | 1682958 | Cell/Virus Enclosed | NO |
| NNDITNTNLSRGRGRN | 1682959 | Cell/Virus Enclosed | NO |
| NNDTTNTNLSRG | 1482493 | Cell/Virus Enclosed | NO |
| NNDTTTNQPRGR | 1482494 | Cell/Virus Enclosed | NO |
| NNDYCCFKHALG | 1482495 | Cell/Virus Enclosed | NO |
| NNECGRAFMYAY | 1482499 | Cell/Virus Enclosed | NO |
| NNEFIDLTECIY | 1482500 | Cell/Virus Enclosed | NO |
| NNEFIDPTECIYTQSR | 1682965 | Cell/Virus Enclosed | NO |
| NNEHYRALPGVF | 1482506 | Cell/Virus Enclosed | NO |
| NNEIKPSGLKTMVVSA | 1682968 | Cell/Virus Enclosed | NO |
| NNELMPAKLKIQVVNS | 1682971 | Cell/Virus Enclosed | NO |
| NNELMPVKLKIQ | 1482514 | Cell/Virus Enclosed | NO |
| NNEQGTPLTPIL | 1482517 | Cell/Virus Enclosed | NO |
| NNFMLTYNKVEN | 1482529 | Cell/Virus Enclosed | NO |
| NNFVSSYAKPEEKLSA | 1682988 | Cell/Virus Enclosed | NO |
| NNGIGICPAGTN | 1482548 | Ectodomain | YES |
| NNGIGTCPAGTN | 1482549 | Ectodomain | YES |
| NNGIGTCPVGTN | 1482550 | Ectodomain | YES |
| NNGLGTCPAGTN | 1482555 | Ectodomain | YES |
| NNGLTCSTDLQK | 1482556 | Ectodomain | YES |
| NNGMGTCPAGTN | 1482557 | Ectodomain | YES |
| NNGNDVLYQPPTASVS | 1682994 | Cell/Virus Enclosed | NO |
| NNGQGENDNISV | 1482560 | Cell/Virus Enclosed | NO |
| NNGRFGGFSNHL | 1482561 | Cell/Virus Enclosed | NO |
| NNGVGTCPAGTN | 1482566 | Ectodomain | YES |
| NNHAFHTPAYDKRAMA | 1683002 | Cell/Virus Enclosed | NO |
| NNIDAAKIYGMG | 1482575 | Ectodomain | YES |
| NNISLTTQPYFDSYLG | 1683008 | Ectodomain | YES |
| NNIVASFSSVND | 1482590 | Ectodomain | YES |
| NNKEDFDIIQKCQITS | 1683012 | Cell/Virus Enclosed | NO |
| NNKNTYRSQLGC | 1482604 | Cell/Virus Enclosed | NO |
| NNKPWFVRKASGLFDA | 1683015 | Cell/Virus Enclosed | NO |
| NNKSQSVIIINNSTNVV | 45151 | Ectodomain | YES |
| NNKSTYRSQLGC | 1482610 | Cell/Virus Enclosed | NO |
| NNLAIEEIEYVHGDAL | 1683020 | Cell/Virus Enclosed | NO |
| NNLKIKNFDYLDIHPS | 1683027 | Ectodomain | YES |
| NNLKIKNYGYLD | 1482627 | Ectodomain | YES |
| NNLLGFDSSKPVTKKY | 1683028 | Cell/Virus Enclosed | NO |
| NNLMCIDMKGTMYVRP | 1683032 | Ectodomain | YES |
| NNLMNGVTLSTK | 1482637 | Ectodomain | YES |
| NNLTAPRNSWEAMALN | 1683037 | Cell/Virus Enclosed | NO |
| NNLVPTYLKSDNIVAA | 1683038 | Cell/Virus Enclosed | NO |
| NNNGSDVLDQPP | 1482659 | Cell/Virus Enclosed | NO |
| NNNGSDVLYQPPTASV | 1683048 | Cell/Virus Enclosed | NO |
| NNNGTNVTQVKNASVY | 1683049 | Cell/Virus Enclosed | NO |
| NNNLALGGSVAI | 1482662 | Cell/Virus Enclosed | NO |
| NNNTDSIIIDGV | 1482664 | Cell/Virus Enclosed | NO |
| NNNTDSIVIDGV | 1482665 | Cell/Virus Enclosed | NO |
| NNNTRKSINIGPGRALYTT | 187169 | Ectodomain | YES |
| NNPAYIAPQANF | 1482669 | Ectodomain | YES |
| NNPAYIAPQANS | 1482670 | Ectodomain | YES |
| NNPAYIASQAKF | 1482673 | Ectodomain | YES |
| NNPAYIATQAKS | 1482674 | Ectodomain | YES |
| NNPGLCYASQMQQTK | 227416 | Cell/Virus Enclosed | NO |
| NNPQCAIVQATE | 1482683 | Cell/Virus Enclosed | NO |
| NNQEDFDLISRC | 1482689 | Cell/Virus Enclosed | NO |
| NNQNGDRSGARPKQRR | 1683060 | Cell/Virus Enclosed | NO |
| NNRDTGHPSISI | 1482702 | Ectodomain | YES |
| NNRDTGLPSIST | 1482705 | Ectodomain | YES |
| NNRDTGPLSISN | 1482706 | Ectodomain | YES |
| NNRDTGRPSIST | 1482709 | Ectodomain | YES |
| NNRSLSRSNSNFRHSD | 1683063 | Capsid | UN |
| NNSAQYVFSTVNALPE | 1683066 | Cell/Virus Enclosed | NO |
| NNSCTCQPQAFL | 1482719 | Ectodomain | YES |
| NNSGSNITQIKNACVY | 1683067 | Cell/Virus Enclosed | NO |
| NNSIPNLSDFEA | 1482726 | Ectodomain | YES |
| NNSLIRQLQPIN | 1482731 | Ectodomain | YES |
| NNSLTRQLQPINYFDS | 1683071 | Ectodomain | YES |
| NNSLTRQLRPIN | 1482734 | Ectodomain | YES |

| | | | |
|---|---|---|---|
| NNSPARPETSRPK | 45247 | Ectodomain | YES |
| NNSRDSSRSTSRQQSR | 1683073 | Capsid | UN |
| NNSSRASSRSSTRNNS | 1683074 | Capsid | UN |
| NNSTDTVDTVLEKNVTVTHSVNLLE | 1161800 | Ectodomain | YES |
| NNSTHIIIRVCN | 1482740 | Ectodomain | YES |
| NNSTNVVIRACNFELC | 1683075 | Ectodomain | YES |
| NNSWLTGNAIQG | 1482746 | Cell/Virus Enclosed | NO |
| NNSYTIVVQPHN | 1482750 | Ectodomain | YES |
| NNTAISVQTCDL | 1482751 | Ectodomain | YES |
| NNTDSIVIDGVT | 1482755 | Cell/Virus Enclosed | NO |
| NNTEFTRVSAKP | 1482759 | Cell/Virus Enclosed | NO |
| NNTLVSGDAHIPFGTV | 1683090 | Ectodomain | YES |
| NNTLVSGDDHIP | 1482769 | Ectodomain | YES |
| NNTRIVDEWSYTGSSF | 1683091 | Ectodomain | YES |
| NNTRVVDEWSYT | 1482774 | Ectodomain | YES |
| NNTSAQYIFSTVNALP | 1683092 | Cell/Virus Enclosed | NO |
| NNTSCPEWMDDF | 1482776 | Cell/Virus Enclosed | NO |
| NNTTIVDEWSYT | 1482779 | Ectodomain | YES |
| NNTVSWYTGLTQ | 1482782 | Cell/Virus Enclosed | NO |
| NNVDTVIFKNNTSFPT | 1683105 | Cell/Virus Enclosed | NO |
| NNVFMSEAKCWL | 1482797 | Cell/Virus Enclosed | NO |
| NNVFMSEAKCWVETDL | 1683107 | Cell/Virus Enclosed | NO |
| NNVFMSESKCWVEHDI | 1683108 | Cell/Virus Enclosed | NO |
| NNVFMSESKCWVENDI | 1683109 | Cell/Virus Enclosed | NO |
| NNVSVPSPFNWE | 1482809 | Ectodomain | YES |
| NNVYCYNTALMEGSLP | 1683118 | Cell/Virus Enclosed | NO |
| NNWFLLTNTSSVVDGV | 1683121 | Ectodomain | YES |
| NNWQNLLYDSNG | 1482827 | Ectodomain | YES |
| NNYDKSAGYPFNKFGK | 1683125 | Cell/Virus Enclosed | NO |
| NNYEKSAGYPFN | 1482830 | Cell/Virus Enclosed | NO |
| NNYMPYVITLLF | 1482836 | Cell/Virus Enclosed | NO |
| NNYVPSYVKPDSVSTS | 1683132 | Cell/Virus Enclosed | NO |
| NPANNAAIVLQLPQGT | 1683139 | Cell/Virus Enclosed | NO |
| NPATDPIYDEPT | 1482860 | Secreted | YES |
| NPAYIAPQANFG | 1482865 | Ectodomain | YES |
| NPAYIAPRANSG | 1482867 | Ectodomain | YES |
| NPAYIATQAKSG | 1482870 | Ectodomain | YES |
| NPCTCRPQAFLG | 1482877 | Ectodomain | YES |
| NPDDKCFANKHRSFTQ | 1683144 | Cell/Virus Enclosed | NO |
| NPDEPHKDVYEL | 1482883 | Cell/Virus Enclosed | NO |
| NPDEPQKDVYELRYNG | 1683145 | Cell/Virus Enclosed | NO |
| NPDIINIYKKLGPIFN | 1683146 | Cell/Virus Enclosed | NO |
| NPDIINVYKKLGPIFN | 1683147 | Cell/Virus Enclosed | NO |
| NPDPITFAGPYK | 1482892 | Ectodomain | YES |
| NPDPKFVDRYYA | 1482896 | Cell/Virus Enclosed | NO |
| NPETNCLLNVPFGGTT | 1683153 | Ectodomain | YES |
| NPETNCLLNVPL | 1482908 | Underneath Membrane | NO |
| NPETNILLNVPLHGTI | 1683154 | Underneath Membrane | NO |
| NPEVNAITVTTVLGQT | 1683159 | Underneath Membrane | NO |
| NPFMVNQLAYSSSANG | 1683163 | Cell/Virus Enclosed | NO |
| NPFTRTVFENLKPMPF | 1683164 | Cell/Virus Enclosed | NO |
| NPHHRFKCSSNG | 1482930 | Ectodomain | YES |
| NPHNYSYMDLNS | 1482939 | Ectodomain | YES |
| NPIDLPPSHATTYLSL | 1683170 | Cell/Virus Enclosed | NO |
| NPINSNVKTLEYDYMT | 1683172 | Cell/Virus Enclosed | NO |
| NPITKNIGEYNV | 1482952 | Cell/Virus Enclosed | NO |
| NPKGIPIFDDPV | 1482962 | Cell/Virus Enclosed | NO |
| NPKGIPIVDDPVVDWH | 1683181 | Cell/Virus Enclosed | NO |
| NPKPRAAPNNTV | 1482972 | Cell/Virus Enclosed | NO |
| NPLYGSRLGVPDTLGGDPKF | 45398 | Capsid | UN |
| NPMLMGWDYPKCDRAL | 1683197 | Cell/Virus Enclosed | NO |
| NPNNNAAIVLQLPQGT | 1683200 | Cell/Virus Enclosed | NO |
| NPNSSEVDMICS | 1483008 | Cell/Virus Enclosed | NO |
| NPPDQLVVYQQA | 1483019 | Ectodomain | YES |
| NPPTNVVSHLNG | 1483021 | Ectodomain | YES |
| NPPTNVVSHVNG | 1483022 | Ectodomain | YES |
| NPSALEFNPSQTSPAT | 1683211 | Capsid | UN |
| NPSEDFIKHVRN | 1483035 | Cell/Virus Enclosed | NO |
| NPSEDFIKHVRTNGGY | 1683212 | Cell/Virus Enclosed | NO |
| NPSEEFIKHVRT | 1483037 | Cell/Virus Enclosed | NO |
| NPSSLFDLVARIKNNLKNSP | 45480 | Capsid | UN |
| NPSSPARLEIVK | 1483052 | Cell/Virus Enclosed | NO |
| NPSSSEVDMICSTTAQ | 1683219 | Cell/Virus Enclosed | NO |
| NPSTPAYTFTTVKPGA | 1683221 | Cell/Virus Enclosed | NO |
| NPSTWNKKFGFI | 1483058 | Ectodomain | YES |
| NPSTWNKRFGFIEDSV | 1683222 | Ectodomain | YES |

| | | | |
|---|---|---|---|
| NPTEDLCDGSVVIKEP | 1683230 | Cell/Virus Enclosed | NO |
| NPTSKVESCVVS | 1483085 | Cell/Virus Enclosed | NO |
| NPVDCQDDRCII | 1483092 | Cell/Virus Enclosed | NO |
| NPVDLPPSHATT | 1483093 | Cell/Virus Enclosed | NO |
| NPVIPFKDGIYFAATEK | 45506 | Ectodomain | YES |
| NPVLFTNSTDTVNHDS | 1683242 | Cell/Virus Enclosed | NO |
| NPVLMGWDYPKCDRAM | 1683244 | Cell/Virus Enclosed | NO |
| NPVRASVLRYDDFHTDEDKL | 45512 | Ectodomain | YES |
| NPVSFVVKPVCSSIFR | 1683247 | Cell/Virus Enclosed | NO |
| NPYHRFKCFPNGSNDV | 1683251 | Ectodomain | YES |
| NPYTPKYTFGNVKPGE | 1683252 | Cell/Virus Enclosed | NO |
| NQADVNIPADIV | 1483115 | Cell/Virus Enclosed | NO |
| NQALGAMQTGFTTTNE | 1683259 | Ectodomain | YES |
| NQALHGANLRQD | 1483121 | Ectodomain | YES |
| NQALNSILDNAVKGCV | 1683261 | Cell/Virus Enclosed | NO |
| NQEKYSTIYKLHPSFN | 1683284 | Cell/Virus Enclosed | NO |
| NQETLVWPLILT | 1483155 | Cell/Virus Enclosed | NO |
| NQFTEFVGTQSVDMLA | 1683291 | Cell/Virus Enclosed | NO |
| NQFTEFVSTQSV | 1483169 | Cell/Virus Enclosed | NO |
| NQGNKQIVHSCP | 1483174 | Cell/Virus Enclosed | NO |
| NQHCNGSIACQF | 1483180 | Cell/Virus Enclosed | NO |
| NQHTIDLADSEMDKLYERVKRQLRE | 1162178 | Ectodomain | YES |
| NQISVAPDLSLD | 1483194 | Ectodomain | YES |
| NQIVLRNSNGAC | 1483196 | Cell/Virus Enclosed | NO |
| NQKILAASFNKAINNI | 1683304 | Ectodomain | YES |
| NQKILAASFNKAMTNI | 1683305 | Ectodomain | YES |
| NQKLIANAFSNA | 1483204 | Ectodomain | YES |
| NQKLIANKFNQALGAM | 1683307 | Ectodomain | YES |
| NQKPLEPKFSIALPPE | 1683311 | Capsid | UN |
| NQKQIANQFNKAISQI | 1683312 | Ectodomain | YES |
| NQLAYSSSANGI | 1483215 | Cell/Virus Enclosed | NO |
| NQLAYSSSANGSLVGT | 1683313 | Cell/Virus Enclosed | NO |
| NQLESKMSGKGQQQQGQTVTKKSA | 1483226 | Capsid | UN |
| NQLTADLQHKLKKACV | 1683319 | Cell/Virus Enclosed | NO |
| NQMTGFKKPASR | 1483241 | Cell/Virus Enclosed | NO |
| NQNGDRSGARPKQRRP | 1683328 | Cell/Virus Enclosed | NO |
| NQNGERSGARPKQRRP | 1683329 | Cell/Virus Enclosed | NO |
| NQPDQVAVYQQSIIGA | 1683334 | Ectodomain | YES |
| NQQGNSLNHLTSQLRQ | 1683336 | Ectodomain | YES |
| NQQGSALNHLTSQLRH | 1683337 | Ectodomain | YES |
| NQRSAPRITFGGPTDSTDNN | 1386304 | Capsid | UN |
| NQSKPPSSSGTS | 1483279 | Cell/Virus Enclosed | NO |
| NQSKSPSSSGTSTPKK | 1683346 | Capsid | UN |
| NQSLAGGITYVSNSGN | 1683347 | Ectodomain | YES |
| NQSLSNAKVSAEVSRQ | 1683348 | Cell/Virus Enclosed | NO |
| NQSPASSQSAAT | 1483289 | Cell/Virus Enclosed | NO |
| NQSPASSQSVAK | 1483290 | Cell/Virus Enclosed | NO |
| NQSPASSQTSAKSLAR | 1683349 | Capsid | UN |
| NQSPQGNLGSNGKPGN | 1683350 | Cell/Virus Enclosed | NO |
| NQSSAPRITFGG | 1483293 | Cell/Virus Enclosed | NO |
| NQTSVAPDFSLD | 1483320 | Ectodomain | YES |
| NQTSVAPDLALD | 1483321 | Ectodomain | YES |
| NQTSVAPDLSLG | 1483324 | Ectodomain | YES |
| NQTSVAPHLSLD | 1483325 | Ectodomain | YES |
| NQVLSSDVMFEG | 1483343 | Cell/Virus Enclosed | NO |
| NQVLSSDVMFEN | 1483344 | Cell/Virus Enclosed | NO |
| NQVVKPGHFNEEFYNF | 1683369 | Cell/Virus Enclosed | NO |
| NQWHEAKFGVVTTNSD | 1683371 | Cell/Virus Enclosed | NO |
| NQYCNGSIACQF | 1483354 | Cell/Virus Enclosed | NO |
| NQYYIYVTNEIGLNAS | 1683374 | Ectodomain | YES |
| NQYYYNMATGFA | 1483367 | Ectodomain | YES |
| NRAGSNDIWMQN | 1483369 | Ectodomain | YES |
| NRALSGIAAEQDRNTR | 1683377 | Ectodomain | YES |
| NRALTGIAIEQD | 1483374 | Ectodomain | YES |
| NRALVSATEFADKLVE | 1683379 | Cell/Virus Enclosed | NO |
| NRASLRQIFDVP | 1483385 | Cell/Virus Enclosed | NO |
| NRATCYNSRTGY | 1483388 | Cell/Virus Enclosed | NO |
| NRCNEVSATVLQ | 1483398 | Cell/Virus Enclosed | NO |
| NRCPFDKVFNAS | 1483401 | Ectodomain | YES |
| NRDINVFGPVIDRNV | 922929 | Ectodomain | YES |
| NRDIQVLGPVIDWNV | 922932 | Ectodomain | YES |
| NRDTGRPSISTD | 1483417 | Ectodomain | YES |
| NRDVDQEFVDEF | 1483422 | Cell/Virus Enclosed | NO |
| NRFGSNDIWMQN | 1483437 | Ectodomain | YES |
| NRFLYIIKLIFLWLL | 1541814 | Undetermined | UN |
| NRGGDENPRGPVSR | 189777 | Capsid | UN |

| | | | |
|---|---|---|---|
| NRGRQFYEFYNDVKPP | 1683409 | Cell/Virus Enclosed | NO |
| NRGVYKPLLENIDYFN | 1683411 | Cell/Virus Enclosed | NO |
| NRILGVNSSNCN | 1483467 | Cell/Virus Enclosed | NO |
| NRIQIDPVKLSSGYKDVILWFSFGA | 1162286 | Undetermined | UN |
| NRLLSVPSDSCNNVNV | 1683423 | Cell/Virus Enclosed | NO |
| NRLSRRAITTGY | 1483490 | Ectodomain | YES |
| NRMAEQAAAAMY | 1483494 | Cell/Virus Enclosed | NO |
| NRNASLNSFKEYFNLR | 1683433 | Ectodomain | YES |
| NRNRFLYIIKLIFLW | 1310685 | Undetermined | UN |
| NRNYVFTGYRVT | 1483504 | Cell/Virus Enclosed | NO |
| NRRAAAYSSPLQSYAC | 1683447 | Cell/Virus Enclosed | NO |
| NRRGLDLLFWEQGGLCKALQEQCRF | 45728 | Ectodomain | YES |
| NRRKVDLQLGNL | 1483523 | Ectodomain | YES |
| NRRQLDVVKRFIHKNS | 1683448 | Cell/Virus Enclosed | NO |
| NRRSFKTADGNQRQLL | 1683449 | Cell/Virus Enclosed | NO |
| NRRSFKTADGSQ | 1483531 | Cell/Virus Enclosed | NO |
| NRRSRGAITTGYRFTN | 1683450 | Ectodomain | YES |
| NRRSRRAITTEN | 1483535 | Ectodomain | YES |
| NRRYGFNNFILS | 1483540 | Ectodomain | YES |
| NRSGSNDIWMQNKGLF | 1683455 | Ectodomain | YES |
| NRSLSRSNSNFRHSDS | 1683457 | Capsid | UN |
| NRSSFVDDNKFD | 1483552 | Cell/Virus Enclosed | NO |
| NRTLTSGVTPDM | 1483561 | Cell/Virus Enclosed | NO |
| NRTPTSGVTRDM | 1483564 | Capsid | UN |
| NRVKSGSPGDSSWHIY | 1683465 | Ectodomain | YES |
| NRVLSVPSDSCN | 1483581 | Cell/Virus Enclosed | NO |
| NRYGFCGNGTHI | 1483597 | Ectodomain | YES |
| NRYKYYSGNANEADYR | 1683476 | Cell/Virus Enclosed | NO |
| NRYNEVANAVMQNNEL | 1683479 | Cell/Virus Enclosed | NO |
| NRYSVLSVDSVSEESQ | 1683483 | Cell/Virus Enclosed | NO |
| NSACPKLEFAND | 1483605 | Ectodomain | YES |
| NSAIVCASVKRDGVQV | 1683490 | Cell/Virus Enclosed | NO |
| NSALAKIQSVVNSNAQ | 1683493 | Ectodomain | YES |
| NSALVKIQAVVNANAE | 1683495 | Ectodomain | YES |
| NSANTKVLKNKG | 1483624 | Cell/Virus Enclosed | NO |
| NSAQYVFSTVNALPEV | 1683498 | Cell/Virus Enclosed | NO |
| NSATGSLDVEVFNRLA | 1683501 | Cell/Virus Enclosed | NO |
| NSAVFISPYNSQNYVA | 1683503 | Cell/Virus Enclosed | NO |
| NSCLVSSDGQPD | 1483651 | Cell/Virus Enclosed | NO |
| NSCLVSSDGQPGGDAI | 1683508 | Cell/Virus Enclosed | NO |
| NSCLVSSNGQPG | 1483653 | Cell/Virus Enclosed | NO |
| NSCVKSKPPSAI | 1483659 | Ectodomain | YES |
| NSDFCPCADPSV | 1483660 | Ectodomain | YES |
| NSDVVKQPVKFY | 1483666 | Cell/Virus Enclosed | NO |
| NSDYANKGYIAN | 1483669 | Cell/Virus Enclosed | NO |
| NSEKHQDHVSLTVPLC | 1683515 | Cell/Virus Enclosed | NO |
| NSEMLKLGTSDP | 1483677 | Cell/Virus Enclosed | NO |
| NSFDVPTFCANRSAMC | 1683523 | Cell/Virus Enclosed | NO |
| NSFFKELTANMSMAEC | 1683527 | Cell/Virus Enclosed | NO |
| NSFGKDLNANMSLAEC | 1683528 | Cell/Virus Enclosed | NO |
| NSFKEYFNLRNCTFMY | 1683529 | Ectodomain | YES |
| NSGIGTCPAGTN | 1483723 | Ectodomain | YES |
| NSGLTCSTDLQK | 1483728 | Ectodomain | YES |
| NSGNLLGFKNVSTGNI | 1683540 | Ectodomain | YES |
| NSGNRTPTSGVTPDMA | 1683541 | Cell/Virus Enclosed | NO |
| NSGNRTSTSGVT | 1483736 | Cell/Virus Enclosed | NO |
| NSGSDMNCNIPTQCYY | 1683544 | Cell/Virus Enclosed | NO |
| NSGSNITQIKNACVYF | 1683546 | Cell/Virus Enclosed | NO |
| NSGTTCSTDLQK | 1483751 | Ectodomain | YES |
| NSHNIGSLYVSW | 1483760 | Ectodomain | YES |
| NSIFDIETVDVKHDVS | 1683555 | Cell/Virus Enclosed | NO |
| NSILDEVNGLLDTTQL | 1683559 | Ectodomain | YES |
| NSILDNAVKGCVPLNA | 1683560 | Cell/Virus Enclosed | NO |
| NSILDNAVKGCVPLSA | 1683561 | Cell/Virus Enclosed | NO |
| NSILMLDKQGQLLDTK | 1683563 | Cell/Virus Enclosed | NO |
| NSISPEKLKSYAASYN | 1683570 | Cell/Virus Enclosed | NO |
| NSITDTIVLTSG | 1483794 | Secreted | YES |
| NSKIVDLAQHFSDFGT | 1683581 | Cell/Virus Enclosed | NO |
| NSKQFNYKITIN | 1483820 | Cell/Virus Enclosed | NO |
| NSKRKNGFTLEY | 1483823 | Ectodomain | YES |
| NSKRKYGSAIEDLLFDKV | 1868229 | Ectodomain | YES |
| NSKRKYRSAIEDLLFDKV | 1868230 | Ectodomain | YES |
| NSLILPPFSLSPVPTLGSRSRRA | 45894 | Ectodomain | YES |
| NSLTYFNRPSLVDDNK | 1683609 | Cell/Virus Enclosed | NO |
| NSNANISMLQLG | 1483879 | Ectodomain | YES |
| NSNANLSMLQLGVPDN | 1683615 | Ectodomain | YES |

| | | | |
|---|---|---|---|
| NSNQNTDSDSLSSKPQ | 1683624 | Capsid | UN |
| NSNVGESFVDDF | 1483901 | Cell/Virus Enclosed | NO |
| NSNYAAQGLVAS | 1483902 | Cell/Virus Enclosed | NO |
| NSPKVTIDCAAF | 1483912 | Ectodomain | YES |
| NSQDDIMKAVAAALKS | 1683643 | Capsid | UN |
| NSQDILLAPLLS | 1483928 | Cell/Virus Enclosed | NO |
| NSQLRQNFQAIS | 1483937 | Ectodomain | YES |
| NSQNAVSRSMLGLTTQ | 1683650 | Cell/Virus Enclosed | NO |
| NSQNFAAKRVLGLQTQ | 1683651 | Cell/Virus Enclosed | NO |
| NSQNYVAKRVLGVQTQ | 1683654 | Cell/Virus Enclosed | NO |
| NSQSSSRASSLSRNSS | 1683656 | Cell/Virus Enclosed | NO |
| NSQYLFSTINALPETS | 1683658 | Cell/Virus Enclosed | NO |
| NSRIDVPSAEST | 1483958 | Cell/Virus Enclosed | NO |
| NSRKVDLQLGNL | 1483962 | Ectodomain | YES |
| NSRSTSRTSSRASSAG | 1683668 | Cell/Virus Enclosed | NO |
| NSSANYVGKYDINHGD | 1683673 | Underneath Membrane | NO |
| NSSDNGISAIITANLS | 1683674 | Ectodomain | YES |
| NSSEWAYCVDLHNKIN | 1683676 | Cell/Virus Enclosed | NO |
| NSSFINLKEIGT | 1483994 | Ectodomain | YES |
| NSSKDDQIGYYR | 1484000 | Cell/Virus Enclosed | NO |
| NSSLFVEDCNLP | 1484003 | Ectodomain | YES |
| NSSLLVEDCKLP | 1484004 | Ectodomain | YES |
| NSSLSVEDCKLP | 1484005 | Ectodomain | YES |
| NSSPDDQIGYYRRAT | 1313250 | Cell/Virus Enclosed | NO |
| NSSRASSRSSTRNNSR | 1683685 | Capsid | UN |
| NSSRNSTPGSSRGTSP | 1683686 | Cell/Virus Enclosed | NO |
| NSSRSSSQGSRSGNST | 1683687 | Cell/Virus Enclosed | NO |
| NSSSDRNHNSQD | 1484025 | Cell/Virus Enclosed | NO |
| NSSSGALDTTIP | 1484028 | Ectodomain | YES |
| NSSSGVLDTTIPFGPS | 1683688 | Ectodomain | YES |
| NSSSSFDCIVNL | 1484030 | Ectodomain | YES |
| NSSVPLLEVYVHLRNW | 1683696 | Ectodomain | YES |
| NSSVPLSEVYVH | 1484035 | Ectodomain | YES |
| NSTASAVQTCDL | 1484044 | Ectodomain | YES |
| NSTDTVNHDSFNLYGY | 1683701 | Cell/Virus Enclosed | NO |
| NSTHNTPVYKLDISE | 929450 | Secreted | YES |
| NSTITVEELKQL | 1484063 | Undetermined | UN |
| NSTPGSSRGTSPARIA | 1683722 | Cell/Virus Enclosed | NO |
| NSTPGSSRGTSPARMA | 1683723 | Cell/Virus Enclosed | NO |
| NSTQDGDNKLQGLLEV | 1683725 | Ectodomain | YES |
| NSTQDGYNKLQG | 1484078 | Ectodomain | YES |
| NSTRGTFPGPSG | 1484082 | Cell/Virus Enclosed | NO |
| NSTRGTSPGPSGIGAV | 1683726 | Cell/Virus Enclosed | NO |
| NSTTFHQALLDPRVRGLYFPAGG | 840115 | Ectodomain | YES |
| NSTTMCVPHNMRVLHY | 1683729 | Cell/Virus Enclosed | NO |
| NSTTNYVGVYDINHGD | 1683730 | Ectodomain | YES |
| NSTTVSVTVSSNMVET | 1683731 | Cell/Virus Enclosed | NO |
| NSVADVMRYNLN | 1484103 | Ectodomain | YES |
| NSVAFQTVKPGN | 1484104 | Cell/Virus Enclosed | NO |
| NSVCNGCVGHSE | 1484106 | Ectodomain | YES |
| NSVCPKFEFAND | 1484107 | Ectodomain | YES |
| NSVCPKLEFANDTKIA | 1683743 | Ectodomain | YES |
| NSVFKPQPAGVL | 1484119 | Ectodomain | YES |
| NSVFNICQAVSANVCA | 1683745 | Cell/Virus Enclosed | NO |
| NSVINRLRCDQLSFDV | 1683754 | Ectodomain | YES |
| NSVNDSLEPVGGLYEI | 1683763 | Ectodomain | YES |
| NSVPVNKKDKNK | 1484147 | Cell/Virus Enclosed | NO |
| NSVPWSKIFAYV | 1484151 | Cell/Virus Enclosed | NO |
| NSVSPKLEFAND | 1484157 | Ectodomain | YES |
| NSVTDTIVVTEG | 1484162 | Secreted | YES |
| NSWLTGNAIQGF | 1484180 | Cell/Virus Enclosed | NO |
| NSWQNLLYDSNGNIIG | 1683786 | Ectodomain | YES |
| NSWQNLLYDSNGNLYG | 1683787 | Ectodomain | YES |
| NSWQNLLYDSSG | 1484184 | Ectodomain | YES |
| NSWRYDFDITES | 1484185 | Cell/Virus Enclosed | NO |
| NSYFDSKGWYDPVENE | 1683790 | Cell/Virus Enclosed | NO |
| NSYTSFATYHTPATDC | 1683798 | Ectodomain | YES |
| NSYTSFATYYTP | 1484206 | Ectodomain | YES |
| NTAKQDQGQYYY | 1484216 | Ectodomain | YES |
| NTAQQDQGQYYY | 1484221 | Ectodomain | YES |
| NTCITIKKPNEL | 1484231 | Cell/Virus Enclosed | NO |
| NTDHSNSSDNIALLVQ | 1683820 | Underneath Membrane | NO |
| NTDLIEGSKPYSTLQP | 1683822 | Cell/Virus Enclosed | NO |
| NTEFTRVSAKPP | 1484265 | Cell/Virus Enclosed | NO |
| NTEIELGVCVNY | 1484267 | Ectodomain | YES |
| NTFICEEVANDF | 1484279 | Cell/Virus Enclosed | NO |

| | | | |
|---|---|---|---|
| NTGRCNVNKHVA | 1484310 | Cell/Virus Enclosed | NO |
| NTGRSVYVKFQDSKPP | 1683855 | Cell/Virus Enclosed | NO |
| NTGSVCYTPIDEIPYK | 1683856 | Cell/Virus Enclosed | NO |
| NTGSVVKPVRARAPPG | 1683857 | Cell/Virus Enclosed | NO |
| NTGVCYDFDGVTTSDK | 1683859 | Cell/Virus Enclosed | NO |
| NTGVYDLSGFTVKPVA | 1683860 | Ectodomain | YES |
| NTHLKHGGGIAGAINA | 1683861 | Cell/Virus Enclosed | NO |
| NTICIKDADYNAKVEI | 1683866 | Cell/Virus Enclosed | NO |
| NTIHSVCNGCVG | 1484336 | Ectodomain | YES |
| NTILNTACGVFEVDDT | 1683869 | Cell/Virus Enclosed | NO |
| NTISPEKLKNYAASYN | 1683873 | Cell/Virus Enclosed | NO |
| NTKHGGGVLREL | 1484352 | Cell/Virus Enclosed | NO |
| NTKLYVDNTLYS | 1484358 | Undetermined | UN |
| NTKVFKDGVMYS | 1484364 | Ectodomain | YES |
| NTKVIKDGVMYS | 1484368 | Ectodomain | YES |
| NTKVIKDRVMYSEFPA | 1683888 | Ectodomain | YES |
| NTKVTKNGLMYS | 1484376 | Ectodomain | YES |
| NTKYVAPQVTYQNIST | 1683890 | Ectodomain | YES |
| NTKYYDDSQYYFNKDT | 1683891 | Ectodomain | YES |
| NTLNIIVPDKSVYDQV | 1683901 | Cell/Virus Enclosed | NO |
| NTLTIIIPDKQV | 1484415 | Cell/Virus Enclosed | NO |
| NTLVSGDAHIPFGTVL | 1683910 | Ectodomain | YES |
| NTMIVDEWSYTG | 1484425 | Ectodomain | YES |
| NTNLHFDVDNINFKSL | 1683912 | Ectodomain | YES |
| NTNLSRGRGRNPKPRA | 1683914 | Cell/Virus Enclosed | NO |
| NTNSGPDDQIGF | 1484448 | Capsid | UN |
| NTNSGPDDQIGYYRRATR | 46198 | Capsid | UN |
| NTNSSKDDQIGY | 1484450 | Cell/Virus Enclosed | NO |
| NTNSTKDDQIGYYRRA | 1683920 | Cell/Virus Enclosed | NO |
| NTPADILDRDPS | 1484456 | Cell/Virus Enclosed | NO |
| NTPADIVDRDQS | 1484457 | Capsid | UN |
| NTPCNVCQYWIG | 1484461 | Cell/Virus Enclosed | NO |
| NTPDVPICVYDP | 1484463 | Ectodomain | YES |
| NTPIPNLPDFRE | 1484464 | Ectodomain | YES |
| NTPKDHIGTRNPANNA | 1683925 | Cell/Virus Enclosed | NO |
| NTPTQCYYNNSNNGKI | 1683929 | Cell/Virus Enclosed | NO |
| NTPVSVKLPKGV | 1484471 | Cell/Virus Enclosed | NO |
| NTQEVFAQVKQM | 1484475 | Ectodomain | YES |
| NTQFTAVGKEFNHLEKRIENLNKKV | 1162644 | Ectodomain | YES |
| NTREVFAQVKQMYKTP | 1683932 | Ectodomain | YES |
| NTRNIDATSTGNYNYK | 1683936 | Ectodomain | YES |
| NTSADIVDRDPS | 1484494 | Cell/Virus Enclosed | NO |
| NTSAQYIFSTVNALPE | 1683937 | Cell/Virus Enclosed | NO |
| NTSCPEWMDDFEADPK | 1683938 | Cell/Virus Enclosed | NO |
| NTSCPEWMDEFE | 1484498 | Cell/Virus Enclosed | NO |
| NTSFPEWIDDLE | 1484505 | Cell/Virus Enclosed | NO |
| NTSGCNVNDVTKLYLG | 1683943 | Cell/Virus Enclosed | NO |
| NTSGPEWMDDFE | 1484506 | Cell/Virus Enclosed | NO |
| NTSHAVCNGCVG | 1484507 | Ectodomain | YES |
| NTSHSVCNGCVG | 1484509 | Ectodomain | YES |
| NTSIPNLPDFKEELDQ | 1683944 | Ectodomain | YES |
| NTSIPNLPDFKG | 1484513 | Ectodomain | YES |
| NTSIPNLPDFRE | 1484514 | Ectodomain | YES |
| NTSIPNLPGFKE | 1484515 | Ectodomain | YES |
| NTSLGNRKRNQKPLEP | 1683947 | Capsid | UN |
| NTSSPNLPDFRE | 1484533 | Ectodomain | YES |
| NTSTPNLPDFKE | 1484536 | Ectodomain | YES |
| NTSTPNLPDFRE | 1484537 | Ectodomain | YES |
| NTSTSNLPDFRE | 1484538 | Ectodomain | YES |
| NTSYSVCNGCVG | 1484545 | Ectodomain | YES |
| NTTASDQLLVENAAGHRVA | 46276 | Undetermined | UN |
| NTTQLTEAQIIA | 1484571 | Underneath Membrane | NO |
| NTTTEDHKIMASENS | 227422 | Secreted | YES |
| NTTVPRISEYVV | 1484579 | Ectodomain | YES |
| NTTVTVNVTTHN | 1484581 | Ectodomain | YES |
| NTVDCQDDRCFI | 1484587 | Cell/Virus Enclosed | NO |
| NTVDTTPKEEFVVKEK | 1683977 | Cell/Virus Enclosed | NO |
| NTVKFCDHIVKA | 1484599 | Cell/Virus Enclosed | NO |
| NTVKPKPVINQLDEKA | 1683984 | Cell/Virus Enclosed | NO |
| NTVLNASIPESKPLAD | 1683986 | Capsid | UN |
| NTVLNASISESK | 1484610 | Cell/Virus Enclosed | NO |
| NTVLNASTPESK | 1484611 | Cell/Virus Enclosed | NO |
| NTVSELVYENKFVPVKE | 1386438 | Cell/Virus Enclosed | NO |
| NTVVLTFTTRVTVPKD | 1683992 | Capsid | UN |
| NVAFELFAKRKMGLTP | 1684009 | Cell/Virus Enclosed | NO |
| NVAIKCVDTVKE | 1484655 | Cell/Virus Enclosed | NO |

| | | | |
|---|---|---|---|
| NVCALMSCNGNKIEDL | 1684037 | Cell/Virus Enclosed | NO |
| NVCGIKDVFIQG | 1484683 | Cell/Virus Enclosed | NO |
| NVCHSVAVIGDEQIDG | 1684043 | Cell/Virus Enclosed | NO |
| NVDDINFAPVLG | 1484703 | Ectodomain | YES |
| NVDDINFSPVLGCLGS | 1684053 | Ectodomain | YES |
| NVDESFVDDFYGYLQK | 1684054 | Cell/Virus Enclosed | NO |
| NVDGFNEWAMANGYTS | 1684058 | Cell/Virus Enclosed | NO |
| NVDKIRQLLKDV | 1484714 | Cell/Virus Enclosed | NO |
| NVDLDALASDIFDTPS | 1684066 | Cell/Virus Enclosed | NO |
| NVDPLSNFEHKVITEC | 1684070 | Cell/Virus Enclosed | NO |
| NVDWSITYPMIANENA | 1684075 | Cell/Virus Enclosed | NO |
| NVEAVNFNVTTAETTD | 1684078 | Ectodomain | YES |
| NVETLNEDLLEI | 1484744 | Cell/Virus Enclosed | NO |
| NVEYPIISNELSINTS | 1684081 | Cell/Virus Enclosed | NO |
| NVFAVESGGYIPSDFA | 1684083 | Ectodomain | YES |
| NVFCDGVNVTKHKCDI | 1684085 | Cell/Virus Enclosed | NO |
| NVFFLLLFSLTHFPLAQ | 46375 | Undetermined | UN |
| NVFKLKISTQLFEGDK | 1684089 | Cell/Virus Enclosed | NO |
| NVFMSEAKCWVETDLK | 1684091 | Cell/Virus Enclosed | NO |
| NVFMSESKCWVEHDIN | 1684092 | Cell/Virus Enclosed | NO |
| NVFMSESKCWVENDIT | 1684093 | Cell/Virus Enclosed | NO |
| NVFQTQAGCLIGAEHVD | 46379 | Ectodomain | YES |
| NVFVSHTLTKYT | 1484775 | Ectodomain | YES |
| NVGEPIISQPSKLLNG | 1484780 | Cell/Virus Enclosed | NO |
| NVGHYDGRTGELPCAI | 1684107 | Cell/Virus Enclosed | NO |
| NVGRWSASINTGNCPF | 1684110 | Ectodomain | YES |
| NVHDVTVTTDKSFEQQ | 1684112 | Cell/Virus Enclosed | NO |
| NVHRNEHVASGDAIMT | 1684117 | Cell/Virus Enclosed | NO |
| NVHTPKHVFKTLKPGD | 1684118 | Cell/Virus Enclosed | NO |
| NVIADSKSFTLI | 1484802 | Cell/Virus Enclosed | NO |
| NVIYTTEIASKL | 1484837 | Cell/Virus Enclosed | NO |
| NVKAWSGICVDD | 1484838 | Ectodomain | YES |
| NVKLLDYDYATHGQLD | 1684148 | Cell/Virus Enclosed | NO |
| NVKLPRWSASID | 1484854 | Ectodomain | YES |
| NVKNVTDVTGNL | 1484856 | Cell/Virus Enclosed | NO |
| NVKPGDAVITTQQAGA | 1684150 | Cell/Virus Enclosed | NO |
| NVKSSLSSYEYCQPPK | 1684153 | Cell/Virus Enclosed | NO |
| NVLDDFIVFNNNGTNV | 1684158 | Cell/Virus Enclosed | NO |
| NVLDVNAIDIEEPS | 46417 | Capsid | UN |
| NVLFDGRDNGALEAFK | 1684163 | Cell/Virus Enclosed | NO |
| NVLIKESIPIVWGVKD | 1684166 | Cell/Virus Enclosed | NO |
| NVLLRKNGNKRA | 1484880 | Cell/Virus Enclosed | NO |
| NVLPETYVALPIYYQH | 1684169 | Ectodomain | YES |
| NVLQDVLSLRHDIALD | 1684172 | Cell/Virus Enclosed | NO |
| NVLSSMLRLHTV | 1484886 | Cell/Virus Enclosed | NO |
| NVLYENQKQIANQFNK | 1684181 | Ectodomain | YES |
| NVMDVNDCFKKD | 1484900 | Cell/Virus Enclosed | NO |
| NVMDVNDCFKNDNVVL | 1684183 | Cell/Virus Enclosed | NO |
| NVMFDLDSTFDD | 1484902 | Cell/Virus Enclosed | NO |
| NVMQCFGPRDFN | 1484905 | Cell/Virus Enclosed | NO |
| NVNNTWMFTGSG | 1484911 | Ectodomain | YES |
| NVPAIDVLKKLLSSLT | 1684205 | Cell/Virus Enclosed | NO |
| NVPFSSDGKPCT | 1484929 | Ectodomain | YES |
| NVPLGGTTVVRP | 1484938 | Underneath Membrane | NO |
| NVPLHGTILTRPLLE | 1313260 | Underneath Membrane | NO |
| NVPLQVGFSNGVDFVA | 1684213 | Cell/Virus Enclosed | NO |
| NVPYSRVISRMG | 1484956 | Cell/Virus Enclosed | NO |
| NVPYVYNGSVQA | 1484960 | Ectodomain | YES |
| NVQIHTIDGSSGVANP | 1684229 | Secreted | YES |
| NVQKVAQPKIEQ | 1484969 | Cell/Virus Enclosed | NO |
| NVQKVEQPKIEPKPVS | 1684232 | Cell/Virus Enclosed | NO |
| NVQVDNGSSINRKQLE | 1684240 | Cell/Virus Enclosed | NO |
| NVQYRINGLGVTMDVL | 1684241 | Ectodomain | YES |
| NVSALIGANGNK | 1484995 | Cell/Virus Enclosed | NO |
| NVSALMGANGNKIVDK | 1684250 | Cell/Virus Enclosed | NO |
| NVSDAYANLVPYYQLI | 1684255 | Cell/Virus Enclosed | NO |
| NVSFVGELKAAEAKVI | 1684259 | Cell/Virus Enclosed | NO |
| NVSFVNDSIESVGGLY | 1684260 | Ectodomain | YES |
| NVSFVVDDSGTP | 1485004 | Cell/Virus Enclosed | NO |
| NVSHVYNGSAQA | 1485005 | Ectodomain | YES |
| NVSLEKFMSTPN | 1485011 | Cell/Virus Enclosed | NO |
| NVSSDVIRYNIN | 1485026 | Ectodomain | YES |
| NVSSFGDYNLSSVIPS | 1684274 | Ectodomain | YES |
| NVSSVWPHYPYG | 1485031 | Ectodomain | YES |
| NVSTDGLYFHFY | 1485036 | Ectodomain | YES |
| NVSTSIPNFGSL | 1485041 | Ectodomain | YES |

| | | | |
|---|---|---|---|
| NVSTSIPNFSSL | 1485042 | Ectodomain | YES |
| NVSTSISNFGSL | 1485043 | Ectodomain | YES |
| NVSTVFVPVTRD | 1485046 | Cell/Virus Enclosed | NO |
| NVSVPSPFNWER | 1485050 | Ectodomain | YES |
| NVSYDSVSAIGT | 1485055 | Ectodomain | YES |
| NVSYDSVSATVT | 1485056 | Ectodomain | YES |
| NVTGSNDSKALF | 1485076 | Cell/Virus Enclosed | NO |
| NVTGSNESKALFFTYL | 1684294 | Cell/Virus Enclosed | NO |
| NVTINNYNPSFW | 1485082 | Ectodomain | YES |
| NVTKHKCDINYKGKVF | 1684300 | Cell/Virus Enclosed | NO |
| NVTKHKCDISYK | 1485085 | Cell/Virus Enclosed | NO |
| NVTLDKCTKYNIYDVS | 1684302 | Ectodomain | YES |
| NVTQCFGPRDLD | 1485096 | Cell/Virus Enclosed | NO |
| NVTRDKLEKLIF | 1485097 | Cell/Virus Enclosed | NO |
| NVTREKLEKLIS | 1485102 | Cell/Virus Enclosed | NO |
| NVTSFGDYNLSS | 1485103 | Ectodomain | YES |
| NVTTASATDFWT | 1485106 | Ectodomain | YES |
| NVTVTHSVNLLEDKHNGKLCKLRGV | 1162859 | Ectodomain | YES |
| NVVDVNDCFKND | 1485124 | Cell/Virus Enclosed | NO |
| NVVELPKFFYASNGTY | 1684326 | Ectodomain | YES |
| NVVEMPKFFYAS | 1485127 | Ectodomain | YES |
| NVVGPDARTQGKQSYV | 1684328 | Cell/Virus Enclosed | NO |
| NVVKAVRNKISA | 1485141 | Cell/Virus Enclosed | NO |
| NVVPGDVIITST | 1485158 | Cell/Virus Enclosed | NO |
| NVVPGVSTNVYL | 1485160 | Cell/Virus Enclosed | NO |
| NVVRYSLDGNFRTEVD | 1684356 | Cell/Virus Enclosed | NO |
| NVVSHVNGDWFLFGDS | 1684358 | Ectodomain | YES |
| NVWQIQTIQDSDGTNK | 1684372 | Cell/Virus Enclosed | NO |
| NVYADSFVVKGDDVRQI | 46522 | Ectodomain | YES |
| NVYAWERTKISD | 1485199 | Ectodomain | YES |
| NVYCYNTDLIEGSKPY | 1684378 | Cell/Virus Enclosed | NO |
| NVYMAKAGDKYYPVVV | 1684388 | Cell/Virus Enclosed | NO |
| NVYMAKVGDKYY | 1485218 | Cell/Virus Enclosed | NO |
| NVYMRTTHTDIG | 1485220 | Cell/Virus Enclosed | NO |
| NVYRSTSPDPKFVDKY | 1684389 | Cell/Virus Enclosed | NO |
| NVYRTDYVDYTFVNEY | 1684390 | Cell/Virus Enclosed | NO |
| NVYVDSFVVKGD | 1485226 | Ectodomain | YES |
| NWADDRAARKKFPPPSFYMP | 2060873 | Cell/Virus Enclosed | NO |
| NWFDISKWLWYI | 150874 | Ectodomain | YES |
| NWFDPIENEDIHRVYA | 1684401 | Cell/Virus Enclosed | NO |
| NWFIQSDKCSVE | 1485248 | Cell/Virus Enclosed | NO |
| NWFLLTNTSSVVDGVV | 1684405 | Ectodomain | YES |
| NWFNTMLDASAPATGW | 1684406 | Cell/Virus Enclosed | NO |
| NWLAPTNAYMRTTHTD | 1684411 | Cell/Virus Enclosed | NO |
| NWLAPTNAYTRT | 1485262 | Cell/Virus Enclosed | NO |
| NWNLEYPIISNEVSIN | 1684422 | Cell/Virus Enclosed | NO |
| NWNVEYPIISNELSIN | 1684427 | Cell/Virus Enclosed | NO |
| NWQNLLYDSNGN | 1485309 | Ectodomain | YES |
| NWVTGGIAIAMA | 1485313 | Undetermined | UN |
| NYEKSAGYPFNQ | 1485348 | Cell/Virus Enclosed | NO |
| NYFTFKQANNNC | 1485371 | Cell/Virus Enclosed | NO |
| NYGAGIKLPSGIMLNV | 1684469 | Cell/Virus Enclosed | NO |
| NYKGKVFFQFEN | 1485406 | Cell/Virus Enclosed | NO |
| NYKIDTTSSSCQLYYS | 1684489 | Ectodomain | YES |
| NYKLNTDHSNSSDNIA | 1684493 | Underneath Membrane | NO |
| NYKYRYLKHGKL | 1485431 | Ectodomain | YES |
| NYLGTIKENIDGGAMH | 1684514 | Cell/Virus Enclosed | NO |
| NYLTCHNAAQCD | 1485462 | Ectodomain | YES |
| NYLTCHNAAQCN | 1485463 | Ectodomain | YES |
| NYLVVYKHTFRG | 1485468 | Cell/Virus Enclosed | NO |
| NYNNLVPTYLKSDNIV | 1684533 | Cell/Virus Enclosed | NO |
| NYNRNASLNSFKEYFN | 1684535 | Ectodomain | YES |
| NYNVSVLQMDVQGLVN | 1684536 | Cell/Virus Enclosed | NO |
| NYNYKYRYLRHGKLRPF | 46681 | Ectodomain | YES |
| NYRLPSTQKGSGMDTA | 1684553 | Ectodomain | YES |
| NYSGSMLRACIK | 1485547 | Cell/Virus Enclosed | NO |
| NYSKDVYFTLSG | 1485551 | Undetermined | UN |
| NYSKLFDEGHTV | 1485554 | Cell/Virus Enclosed | NO |
| NYSKLSDEGYTV | 1485556 | Cell/Virus Enclosed | NO |
| NYSYMDLNPALC | 1485561 | Ectodomain | YES |
| NYTVCDDCNGYTDNIF | 1684572 | Ectodomain | YES |
| NYTVPDLVVEQYNQTI | 1684574 | Ectodomain | YES |
| NYTVQKLQSLID | 1485587 | Ectodomain | YES |
| NYTVQKLQTLIDNINS | 1684575 | Ectodomain | YES |
| NYVAKRVLGVQTQTVD | 1684580 | Cell/Virus Enclosed | NO |
| NYVGKYDINHGDSGFI | 1684590 | Undetermined | UN |

| | | | |
|---|---|---|---|
| NYVGVYDINHGDAGFT | 1684591 | Ectodomain | YES |
| NYVPSYVKPDSVSTSD | 1684596 | Cell/Virus Enclosed | NO |
| PAAAYLDAVKQGAKPV | 1684608 | Cell/Virus Enclosed | NO |
| PAADLDEFSKQL | 1485637 | Cell/Virus Enclosed | NO |
| PAAISSNCYSSL | 1485641 | Ectodomain | YES |
| PAANQVGVGAFGPRLTPPHGGI | 46745 | Underneath Membrane | NO |
| PAANVSVSRFDP | 1485646 | Ectodomain | YES |
| PAANVSVSRFNP | 1485647 | Ectodomain | YES |
| PAAPPPASTNRQSGR | 1684612 | Underneath Membrane | NO |
| PAAPRAISFADN | 1485649 | Cell/Virus Enclosed | NO |
| PAAPRAVSFADNNDIT | 1684613 | Cell/Virus Enclosed | NO |
| PAASSCHNASGKEAKVCTIS | 46753 | Capsid | UN |
| PAAVDTKCLASIDEVS | 1684615 | Cell/Virus Enclosed | NO |
| PACIEVDIQQTF | 1485657 | Ectodomain | YES |
| PACINGHAVPTL | 1485658 | Cell/Virus Enclosed | NO |
| PACTNGHALPTL | 1485663 | Cell/Virus Enclosed | NO |
| PACTNGHAVPTL | 1485664 | Cell/Virus Enclosed | NO |
| PADILDRDPSSD | 1485667 | Cell/Virus Enclosed | NO |
| PADIVDRDPSND | 1485668 | Cell/Virus Enclosed | NO |
| PADIVDRDPSSD | 1485669 | Cell/Virus Enclosed | NO |
| PADIVDRDQSSE | 1485670 | Capsid | UN |
| PADNSYTSFATY | 1485672 | Ectodomain | YES |
| PADQLSDPNYDA | 1485676 | Cell/Virus Enclosed | NO |
| PADSRDFLRVVFSQVD | 1684624 | Cell/Virus Enclosed | NO |
| PADYILPLVLSS | 1485679 | Cell/Virus Enclosed | NO |
| PAECTMFRDAEG | 1485682 | Cell/Virus Enclosed | NO |
| PAEFKYMVANGLNAPN | 1684627 | Cell/Virus Enclosed | NO |
| PAEYKLQQGTFL | 1485690 | Cell/Virus Enclosed | NO |
| PAFGANSNNPDWDFNPVKDDWP | 46793 | Underneath Membrane | NO |
| PAFHRIESPDSI | 1485702 | Cell/Virus Enclosed | NO |
| PAFHRIESTDPI | 1485703 | Cell/Virus Enclosed | NO |
| PAFMLGSSVGNFSDGK | 1684636 | Ectodomain | YES |
| PAFNVSDAYANLVPYY | 1684637 | Cell/Virus Enclosed | NO |
| PAFPLAEDVEKDK | 46796 | Ectodomain | YES |
| PAGNSHTSFATY | 1485725 | Ectodomain | YES |
| PAGNSYTSFATYHTPA | 1684641 | Ectodomain | YES |
| PAGTILVDNDLH | 1485730 | Cell/Virus Enclosed | NO |
| PAGTILVDNDLYPFVS | 1684642 | Cell/Virus Enclosed | NO |
| PAGTKYRHCDLD | 1485732 | Ectodomain | YES |
| PAGTKYRHCDVD | 1485733 | Ectodomain | YES |
| PAGTNHLTCDNL | 1485734 | Ectodomain | YES |
| PAGTNSLTCDNL | 1485736 | Ectodomain | YES |
| PAGTNSLTCGNL | 1485737 | Ectodomain | YES |
| PAGTNYLNCDNL | 1485738 | Ectodomain | YES |
| PAGTNYLTCHNA | 1485740 | Ectodomain | YES |
| PAGVCLGAGQFM | 1485743 | Cell/Virus Enclosed | NO |
| PAGVCLGAGQFV | 1485744 | Cell/Virus Enclosed | NO |
| PAGVNITLANFNETKG | 1684647 | Ectodomain | YES |
| PAGYKLQQGTFL | 1485752 | Cell/Virus Enclosed | NO |
| PAHAPSFLAVDDKYKA | 1684648 | Cell/Virus Enclosed | NO |
| PAHAPSFLAVDDKYKV | 1684649 | Cell/Virus Enclosed | NO |
| PAICHEGKAYFP | 1485768 | Ectodomain | YES |
| PAIDCSDGNYNR | 1485770 | Ectodomain | YES |
| PAIKDFGGFNFS | 1485772 | Ectodomain | YES |
| PAIKLNFGMLNGNAIA | 1684658 | Cell/Virus Enclosed | NO |
| PAINVTINNYNPSSWN | 1684659 | Ectodomain | YES |
| PAKDIPEKVSLV | 1485781 | Cell/Virus Enclosed | NO |
| PAKLKIQVVNSGPDQT | 1684664 | Cell/Virus Enclosed | NO |
| PAKVRVDCYDKFKIND | 1684665 | Cell/Virus Enclosed | NO |
| PAKVRVECYDKFKIND | 1684666 | Cell/Virus Enclosed | NO |
| PALAANTLTIVIPDKQ | 1684667 | Cell/Virus Enclosed | NO |
| PALHLVNPQIQL | 1485795 | Cell/Virus Enclosed | NO |
| PALLFRNIKCNY | 1485802 | Ectodomain | YES |
| PALQHAFMKHKGGDST | 1684679 | Cell/Virus Enclosed | NO |
| PALQTVATALNK | 1485820 | Ectodomain | YES |
| PAMDEFIQRYKL | 1485829 | Cell/Virus Enclosed | NO |
| PAMDPIYDEPTM | 1485830 | Secreted | YES |
| PAMHIASSNAFLDLRT | 1684686 | Cell/Virus Enclosed | NO |
| PAMRDSVPLHII | 1485835 | Cell/Virus Enclosed | NO |
| PANAVVKMPCPA | 1485837 | Cell/Virus Enclosed | NO |
| PANECAQVLSEI | 1485838 | Cell/Virus Enclosed | NO |
| PANGHMAHDGGV | 1485841 | Cell/Virus Enclosed | NO |
| PANGHMVHGGGV | 1485844 | Cell/Virus Enclosed | NO |
| PANNAAIVLQLPQGTTLPKG | 1392345 | Capsid | UN |
| PANSTVLSFCAF | 1485859 | Cell/Virus Enclosed | NO |
| PAPIDIDAFPVKPICS | 1684702 | Cell/Virus Enclosed | NO |

| | | | |
|---|---|---|---|
| PAPIDIDAFPVR | 1485864 | Cell/Virus Enclosed | NO |
| PAPNCYWPLNGY | 1485867 | Ectodomain | YES |
| PAPQAPYQGYQEPPAPQAPY | 46895 | Cell/Virus Enclosed | NO |
| PAPRTLLTRGTLEPEN | 1684704 | Cell/Virus Enclosed | NO |
| PAPRVLISKGVMEPID | 1684705 | Cell/Virus Enclosed | NO |
| PAPRVLLSKGSLEPRH | 1684706 | Cell/Virus Enclosed | NO |
| PAPRVLLSKGTLEPKY | 1684707 | Cell/Virus Enclosed | NO |
| PAPVPTPTPDDYDEEDDAGV | 741786 | Ectodomain | YES |
| PAPVYIWTADEAIKFL | 1684709 | Underneath Membrane | NO |
| PAPVYTWTADEA | 1485877 | Ectodomain | YES |
| PAQASVNGVTLI | 1485878 | Cell/Virus Enclosed | NO |
| PAQCIRDPVGFC | 1485879 | Cell/Virus Enclosed | NO |
| PAQDTWGTSAAA | 1485884 | Ectodomain | YES |
| PAQGTWGTSAAA | 1485886 | Ectodomain | YES |
| PAQNAGYWRRQDRKIN | 1684720 | Cell/Virus Enclosed | NO |
| PARDVLCLRPVGAES | 46927 | Cell/Virus Enclosed | NO |
| PARFVIDTPTGP | 1485895 | Cell/Virus Enclosed | NO |
| PARIAGNGGDAALALL | 1684726 | Cell/Virus Enclosed | NO |
| PARLEIVKLGAEKKDG | 1684727 | Cell/Virus Enclosed | NO |
| PARMAGNGGDTA | 1485899 | Cell/Virus Enclosed | NO |
| PARPETSRPKTPP | 46933 | Ectodomain | YES |
| PASARNLICAQY | 1485909 | Underneath Membrane | NO |
| PASCNGVLPVAF | 1485910 | Cell/Virus Enclosed | NO |
| PASGYVAASAVN | 1485913 | Cell/Virus Enclosed | NO |
| PASKQCFKVFFK | 1485915 | Cell/Virus Enclosed | NO |
| PASQVIVNNYDKSAGY | 1485918 | Cell/Virus Enclosed | NO |
| PASQVIVNNYEK | 1485919 | Cell/Virus Enclosed | NO |
| PASSQSAAKILA | 1485925 | Cell/Virus Enclosed | NO |
| PASSQSAATILA | 1485926 | Cell/Virus Enclosed | NO |
| PASSQSSAKILA | 1485927 | Cell/Virus Enclosed | NO |
| PASSQTSAKSLARSQS | 1684740 | Capsid | UN |
| PASTAHATATIP | 1485933 | Secreted | YES |
| PASTNRQSGRQPTPISPPLRDSHP | 46957 | Underneath Membrane | NO |
| PASTVHATAMIP | 1485935 | Secreted | YES |
| PASTVHATATIPLQAS | 1684741 | Secreted | YES |
| PASVKLPKGVGS | 1485939 | Cell/Virus Enclosed | NO |
| PASVRDLICAQY | 1485941 | Ectodomain | YES |
| PATAEPVRDEFS | 1485944 | Capsid | UN |
| PATAEPVRDEVS | 1485945 | Capsid | UN |
| PATCANDPVGFT | 1485946 | Cell/Virus Enclosed | NO |
| PATDCSDGNYNRNASL | 1684746 | Ectodomain | YES |
| PATDPIYDEPTT | 1485949 | Secreted | YES |
| PATNCSDGNYNR | 1485957 | Ectodomain | YES |
| PATSASKLTIVSPDLE | 1684752 | Cell/Virus Enclosed | NO |
| PATSIVAKQGAGDAGH | 1684753 | Cell/Virus Enclosed | NO |
| PATVCGPKLSTDLIKN | 1684757 | Ectodomain | YES |
| PAVAFTLCRSTHHAHSPAYP | 741790 | Ectodomain | YES |
| PAVETLTQENLE | 1485976 | Cell/Virus Enclosed | NO |
| PAVNVSVSRFNP | 1485980 | Ectodomain | YES |
| PAVSFDYLTREA | 1485984 | Cell/Virus Enclosed | NO |
| PAYAPTYVSVDDKYKT | 1684773 | Cell/Virus Enclosed | NO |
| PAYDGSAFRDLK | 1485994 | Cell/Virus Enclosed | NO |
| PAYIAPQANSGD | 1486000 | Ectodomain | YES |
| PAYIAPQANSGG | 1486001 | Ectodomain | YES |
| PAYIAPQANSGI | 1486002 | Ectodomain | YES |
| PCADPSVVNSCA | 1486015 | Ectodomain | YES |
| PCADPSVVNSCV | 1486016 | Ectodomain | YES |
| PCADRSVVNSCV | 1486017 | Ectodomain | YES |
| PCAFGGVSVITPGTNA | 1684786 | Ectodomain | YES |
| PCAIINDKVLTK | 1486021 | Cell/Virus Enclosed | NO |
| PCAIINDKVVAKIDKE | 1684789 | Cell/Virus Enclosed | NO |
| PCAVSFADNNDI | 1486027 | Cell/Virus Enclosed | NO |
| PCCTSYYSYMMP | 1486031 | Cell/Virus Enclosed | NO |
| PCELHGNGSMIE | 1486032 | Cell/Virus Enclosed | NO |
| PCEVHGNGSMIEDIDY | 1684796 | Cell/Virus Enclosed | NO |
| PCFITYPQQMFV | 1486038 | Cell/Virus Enclosed | NO |
| PCFRIYPQQMFV | 1486040 | Cell/Virus Enclosed | NO |
| PCFRTFPQQMFV | 1486041 | Cell/Virus Enclosed | NO |
| PCGFKGGIPGNM | 1486046 | Cell/Virus Enclosed | NO |
| PCGVHGNGSMIE | 1486050 | Cell/Virus Enclosed | NO |
| PCGYSEPPKVAAKICI | 1684801 | Cell/Virus Enclosed | NO |
| PCGYSEPPKVAD | 1486052 | Cell/Virus Enclosed | NO |
| PCICGRDATQYL | 1486053 | Cell/Virus Enclosed | NO |
| PCKEIEGVGAKV | 1486058 | Cell/Virus Enclosed | NO |
| PCKELEGVGARV | 1486060 | Cell/Virus Enclosed | NO |
| PCKFTVQDAKGLKIKY | 1684809 | Cell/Virus Enclosed | NO |

| | | | |
|---|---|---|---|
| PCKLDGSLCVGN | 1486063 | Ectodomain | YES |
| PCKLNSSLCVGS | 1486068 | Ectodomain | YES |
| PCKLNSSSCVGS | 1486069 | Ectodomain | YES |
| PCKSNDSCPGKN | 1486070 | Ectodomain | YES |
| PCKSNGSCPGKN | 1486071 | Ectodomain | YES |
| PCNAQEQQIYFFEGVA | 1684812 | Cell/Virus Enclosed | NO |
| PCNAQKQQIYFF | 1486081 | Cell/Virus Enclosed | NO |
| PCNGTDIDYCVR | 1486083 | Cell/Virus Enclosed | NO |
| PCNLYNYGAGIKLPSG | 1684818 | Cell/Virus Enclosed | NO |
| PCNPPDQLVVYQ | 1486086 | Ectodomain | YES |
| PCPSATYEGNSP | 1486096 | Ectodomain | YES |
| PCPSGIGAVGGD | 1486097 | Cell/Virus Enclosed | NO |
| PCQFKDKNLQGL | 1486103 | Cell/Virus Enclosed | NO |
| PCSLKCPYLGCQSWTCPYTGAVS | 47047 | Ectodomain | YES |
| PCSSCFGKNNGI | 1486112 | Ectodomain | YES |
| PCSSCPGKNNGI | 1486114 | Ectodomain | YES |
| PCSSCPGKNNGL | 1486115 | Ectodomain | YES |
| PCSSCPSKNNGI | 1486116 | Ectodomain | YES |
| PCSSCSCPGKNN | 1486117 | Ectodomain | YES |
| PCSSCSGKNNGI | 1486118 | Ectodomain | YES |
| PCSSCSGKNNGV | 1486119 | Ectodomain | YES |
| PCSVRSVPGEMR | 1486123 | Ectodomain | YES |
| PCVCGRDATQYL | 1486137 | Cell/Virus Enclosed | NO |
| PCVETSDSQVEE | 1486140 | Cell/Virus Enclosed | NO |
| PCVSIVPSTVWEDGDY | 1684846 | Ectodomain | YES |
| PCVYVDGLESKQVDYV | 1684849 | Cell/Virus Enclosed | NO |
| PCYCFNAEGGAS | 1486165 | Cell/Virus Enclosed | NO |
| PDAAEKLASPMK | 1486174 | Cell/Virus Enclosed | NO |
| PDAFQSVYHTFT | 1486176 | Cell/Virus Enclosed | NO |
| PDAIIIDNDINDYVSD | 1684855 | Cell/Virus Enclosed | NO |
| PDARAKQDFSLL | 1486180 | Cell/Virus Enclosed | NO |
| PDARAKQDVSLLSKCY | 1684857 | Cell/Virus Enclosed | NO |
| PDCCIEPHDWTKNIT | 47064 | Secreted | YES |
| PDCDIDNWLNNV | 1486192 | Ectodomain | YES |
| PDCNIEAWLNDK | 1486195 | Ectodomain | YES |
| PDDASRLLHTVLAKAE | 1684866 | Cell/Virus Enclosed | NO |
| PDDDPQPGPSREYRYVL | 47068 | Cell/Virus Enclosed | NO |
| PDDEDECDEYEE | 1486201 | Cell/Virus Enclosed | NO |
| PDDQIGFYRRAT | 1486208 | Capsid | UN |
| PDDQIGYYRRATRRI | 1541910 | Capsid | UN |
| PDEKQNGCSLYL | 1486221 | Cell/Virus Enclosed | NO |
| PDEKQNGHSLYL | 1486222 | Cell/Virus Enclosed | NO |
| PDEPHKDVYELR | 1486224 | Cell/Virus Enclosed | NO |
| PDEPQKDVYELRYNGA | 1684881 | Cell/Virus Enclosed | NO |
| PDFKEELDQWLK | 1486232 | Ectodomain | YES |
| PDFKEESDQWFK | 1486233 | Ectodomain | YES |
| PDFKGELDQWFK | 1486235 | Ectodomain | YES |
| PDFKLLRNLQAD | 1486237 | Cell/Virus Enclosed | NO |
| PDFREELDQWFK | 1486239 | Ectodomain | YES |
| PDGFYSTSPIQP | 1486256 | Ectodomain | YES |
| PDGFYSTSPTQP | 1486258 | Ectodomain | YES |
| PDGKPCTPPAFN | 1486260 | Ectodomain | YES |
| PDGKPCTPPAPN | 1486262 | Ectodomain | YES |
| PDGQGVPIAYGI | 1486267 | Cell/Virus Enclosed | NO |
| PDGYPLDCIKDF | 1486272 | Cell/Virus Enclosed | NO |
| PDHTCNTPTQCY | 1486278 | Cell/Virus Enclosed | NO |
| PDIINIYKKLGPIFNR | 1684901 | Cell/Virus Enclosed | NO |
| PDIINVYKKLGPIFNR | 1684902 | Cell/Virus Enclosed | NO |
| PDIVDVPPQVVE | 1486289 | Cell/Virus Enclosed | NO |
| PDKNNGIGICPA | 1486293 | Ectodomain | YES |
| PDKNNGIGTCPA | 1486294 | Ectodomain | YES |
| PDKQVFDKVVDNVYVT | 1684904 | Cell/Virus Enclosed | NO |
| PDKSVYDQIVDN | 1486296 | Cell/Virus Enclosed | NO |
| PDKSVYDQVVDNVYVT | 1684905 | Cell/Virus Enclosed | NO |
| PDLALDYINVTF | 1486299 | Ectodomain | YES |
| PDLESYSKIVCD | 1486303 | Cell/Virus Enclosed | NO |
| PDLPDCDIDKWLNNFN | 1684914 | Ectodomain | YES |
| PDLPICVHDPLP | 1486315 | Ectodomain | YES |
| PDLPICVYDPLP | 1486317 | Ectodomain | YES |
| PDLPICVYDRLP | 1486318 | Ectodomain | YES |
| PDLPICVYDSLP | 1486320 | Ectodomain | YES |
| PDLPICVYHPLP | 1486321 | Ectodomain | YES |
| PDLPISVYDPLP | 1486322 | Ectodomain | YES |
| PDLPVCVYDPLP | 1486325 | Ectodomain | YES |
| PDLQRAAITILD | 1486327 | Cell/Virus Enclosed | NO |
| PDLQRAAVTILD | 1486328 | Cell/Virus Enclosed | NO |

| | | | |
|---|---|---|---|
| PDLSFDYINVTF | 1486331 | Ectodomain | YES |
| PDLSLDYINATF | 1486332 | Ectodomain | YES |
| PDLSQYYCDNGKYYTK | 1684920 | Cell/Virus Enclosed | NO |
| PDMADEIANLVL | 1486348 | Cell/Virus Enclosed | NO |
| PDMADHIASLVL | 1486349 | Cell/Virus Enclosed | NO |
| PDMADQIASLVL | 1486350 | Cell/Virus Enclosed | NO |
| PDMFLGTCRCCP | 1486351 | Cell/Virus Enclosed | NO |
| PDNSSTIVTGLLPTHW | 1684925 | Ectodomain | YES |
| PDPITFAGPYKC | 1486358 | Ectodomain | YES |
| PDPITSKSTGPY | 1486363 | Ectodomain | YES |
| PDPKFVDKYYAFLNKH | 1684928 | Cell/Virus Enclosed | NO |
| PDPKFVDRYYAF | 1486365 | Cell/Virus Enclosed | NO |
| PDPLKPTKRSFIEDLL | 1684929 | Ectodomain | YES |
| PDPLKSTKRSFI | 1486370 | Ectodomain | YES |
| PDPNNSPARPETS | 47159 | Ectodomain | YES |
| PDPPNCNIEAWL | 1486372 | Underneath Membrane | NO |
| PDPSKPTKRSFI | 1486375 | Ectodomain | YES |
| PDQICNTPTQCY | 1486377 | Cell/Virus Enclosed | NO |
| PDQLNDLVYSLI | 1486378 | Cell/Virus Enclosed | NO |
| PDQLVVYQQAIV | 1486379 | Ectodomain | YES |
| PDQLVVYQQAVV | 1486381 | Ectodomain | YES |
| PDQTCNTPTQCYYNNS | 1684937 | Cell/Virus Enclosed | NO |
| PDQVAVYQQSIIGAMT | 1684938 | Ectodomain | YES |
| PDSAKSACIEVD | 1486386 | Ectodomain | YES |
| PDSAKSVCIEVD | 1486387 | Ectodomain | YES |
| PDSVKPACIEVD | 1486402 | Ectodomain | YES |
| PDSVKSACIAVD | 1486403 | Ectodomain | YES |
| PDSVKSACIEVH | 1486405 | Ectodomain | YES |
| PDSVKSACIKVD | 1486406 | Ectodomain | YES |
| PDSVSTSDLGSL | 1486408 | Cell/Virus Enclosed | NO |
| PDTHSTLTPRSV | 1486410 | Ectodomain | YES |
| PDTPITLTPRSV | 1486412 | Ectodomain | YES |
| PDTPSPLTPRSV | 1486413 | Ectodomain | YES |
| PDTPSTFTPRSV | 1486414 | Ectodomain | YES |
| PDTPSTLAPRSV | 1486415 | Ectodomain | YES |
| PDTPSTLKPRSV | 1486416 | Ectodomain | YES |
| PDTPSTLTLRSV | 1486417 | Ectodomain | YES |
| PDTPSTLTPCSV | 1486419 | Ectodomain | YES |
| PDTVGVPPQVVE | 1486426 | Cell/Virus Enclosed | NO |
| PDTYKCPQTKSL | 1486428 | Ectodomain | YES |
| PDVDFGDISGIN | 1486431 | Ectodomain | YES |
| PDVFLHKCYRYP | 1486434 | Cell/Virus Enclosed | NO |
| PDVMLNISTPNL | 1486437 | Ectodomain | YES |
| PDVPICVYDPLP | 1486438 | Ectodomain | YES |
| PDVTAAAGAGPRVRQPARPLGSAWR | 47224 | Undetermined | UN |
| PDYAYFNGAIIRDSDV | 1684957 | Cell/Virus Enclosed | NO |
| PDYNYYKYNLPT | 1486451 | Cell/Virus Enclosed | NO |
| PDYTNLLCRVTSSDFT | 1684962 | Cell/Virus Enclosed | NO |
| PDYVDVNKTLQEFAQN | 1684963 | Ectodomain | YES |
| PEAALPFRAVKDGIVW | 1684964 | Cell/Virus Enclosed | NO |
| PEAALPFRAVKE | 1486459 | Cell/Virus Enclosed | NO |
| PEAGLPYGANKDGIIWVAT | 1386574 | Cell/Virus Enclosed | NO |
| PEAGLPYGANKEGIIW | 1684965 | Cell/Virus Enclosed | NO |
| PEANINQDSYGG | 1486467 | Cell/Virus Enclosed | NO |
| PEANMDQGSFGG | 1486471 | Cell/Virus Enclosed | NO |
| PEASLPYGANKQ | 1486474 | Capsid | UN |
| PEDDDFLEASDV | 1486476 | Cell/Virus Enclosed | NO |
| PEDDDFLEENDV | 1486477 | Cell/Virus Enclosed | NO |
| PEDDDFLEGSDV | 1486480 | Cell/Virus Enclosed | NO |
| PEDFSVIVQPTH | 1486482 | Cell/Virus Enclosed | NO |
| PEDHSLLATLDDPYVE | 1684972 | Capsid | UN |
| PEDLSVIIQPTH | 1486486 | Cell/Virus Enclosed | NO |
| PEDLSVIQPTHY | 1486487 | Cell/Virus Enclosed | NO |
| PEDLSVMVQPTH | 1486488 | Cell/Virus Enclosed | NO |
| PEDVNMQLMGVV | 1486493 | Cell/Virus Enclosed | NO |
| PEEFASHVANFQKSGY | 1684979 | Cell/Virus Enclosed | NO |
| PEELSVIVQPTH | 1486503 | Cell/Virus Enclosed | NO |
| PEETCEHEYGTE | 1486505 | Cell/Virus Enclosed | NO |
| PEFSIVCRFDTR | 1486509 | Cell/Virus Enclosed | NO |
| PEGCVLTNTGSV | 1486513 | Cell/Virus Enclosed | NO |
| PEGLVFFHTVLL | 1486514 | Ectodomain | YES |
| PEGVCLGAGQFL | 1486515 | Cell/Virus Enclosed | NO |
| PEGVCLGAGQFV | 1486516 | Cell/Virus Enclosed | NO |
| PEGWHVVNKFYQ | 1486517 | Cell/Virus Enclosed | NO |
| PEHCQNYSGCML | 1486519 | Cell/Virus Enclosed | NO |
| PEHCQNYSGFML | 1486520 | Cell/Virus Enclosed | NO |

| | | | |
|---|---|---|---|
| PEHCQNYSGSML | 1486522 | Cell/Virus Enclosed | NO |
| PEHSVADYHNHS | 1486529 | Cell/Virus Enclosed | NO |
| PEIPHFNQKLPNGVTV | 1685003 | Capsid | UN |
| PEKAQGMLLALL | 1486536 | Cell/Virus Enclosed | NO |
| PEKAQSMLLALL | 1486537 | Cell/Virus Enclosed | NO |
| PEKCIFTSACTR | 1486538 | Cell/Virus Enclosed | NO |
| PEKFPTTVTCEN | 1486540 | Cell/Virus Enclosed | NO |
| PELLDPLNLSAD | 1486556 | Cell/Virus Enclosed | NO |
| PELVPEDPEDSALLEDPAGT | 101960 | Ectodomain | YES |
| PENDDDAWFKLS | 1486563 | Cell/Virus Enclosed | NO |
| PENDDDAWFKPS | 1486564 | Cell/Virus Enclosed | NO |
| PEPALLFRNIKC | 1486566 | Ectodomain | YES |
| PEPEPLPEEPVN | 1486567 | Cell/Virus Enclosed | NO |
| PEPIIENNVVVM | 1486571 | Ectodomain | YES |
| PEPINENNVVVM | 1486572 | Ectodomain | YES |
| PEPISDKNVVFMNTCS | 1685024 | Ectodomain | YES |
| PEPITENNVVVT | 1486575 | Ectodomain | YES |
| PEPITGNNVVVM | 1486576 | Ectodomain | YES |
| PEPITSFNTKYV | 1486579 | Ectodomain | YES |
| PEPITSLNIKYV | 1486580 | Ectodomain | YES |
| PEPQRGRQGRIP | 1486585 | Cell/Virus Enclosed | NO |
| PEPTSEEPVNQF | 1486588 | Cell/Virus Enclosed | NO |
| PEQFDSLNLSAGTQHI | 1685029 | Capsid | UN |
| PERIKLDYDQYCADV | 47398 | Ectodomain | YES |
| PESAAYYVGNLK | 1486597 | Ectodomain | YES |
| PESFDSLNLSAD | 1486598 | Cell/Virus Enclosed | NO |
| PESIPLADDDSA | 1486599 | Capsid | UN |
| PESKPLADDDSAIIEI | 1685036 | Capsid | UN |
| PETNAIISLQVY | 1486607 | Underneath Membrane | NO |
| PETNCLLNVPIG | 1486609 | Ectodomain | YES |
| PETNCLLNVPLG | 1486610 | Underneath Membrane | NO |
| PETNILPNVPLR | 1486614 | Underneath Membrane | NO |
| PETNNLMCIDMKGKMF | 1685044 | Underneath Membrane | NO |
| PETNNLMCIDMKGTMY | 1685045 | Ectodomain | YES |
| PETTDLYCYEQLNDSSEEEDE | 110960 | Cell/Virus Enclosed | NO |
| PEVGPEHSVADY | 1486626 | Cell/Virus Enclosed | NO |
| PEVKEIAPVYEA | 1486627 | Cell/Virus Enclosed | NO |
| PEVKEVAPVYEA | 1486628 | Cell/Virus Enclosed | NO |
| PEVSRELGNITKTNVQ | 1685062 | Cell/Virus Enclosed | NO |
| PEWIDDLEADPK | 1486641 | Cell/Virus Enclosed | NO |
| PEWMDDFEADPKGKYA | 1685064 | Cell/Virus Enclosed | NO |
| PEWMDEFEADPK | 1486643 | Cell/Virus Enclosed | NO |
| PEWMDYFEADPK | 1486644 | Cell/Virus Enclosed | NO |
| PEYDDFLEESDV | 1486648 | Cell/Virus Enclosed | NO |
| PEYIDVNKTLQELSYK | 1685066 | Ectodomain | YES |
| PEYMAVAVPSTT | 1486650 | Underneath Membrane | NO |
| PEYMTVAVPSTT | 1486651 | Underneath Membrane | NO |
| PEYNFIIYDSSD | 1486653 | Cell/Virus Enclosed | NO |
| PFAELTGTQSVS | 1486656 | Cell/Virus Enclosed | NO |
| PFASPQECNNMH | 1486669 | Cell/Virus Enclosed | NO |
| PFAVYKNVKFYL | 1486671 | Cell/Virus Enclosed | NO |
| PFCGSGNLVEKL | 1486672 | Cell/Virus Enclosed | NO |
| PFDSNLTQYFSL | 1486682 | Ectodomain | YES |
| PFDVVAQCSGVT | 1486685 | Cell/Virus Enclosed | NO |
| PFDVVRQCSGVT | 1486686 | Cell/Virus Enclosed | NO |
| PFEEVEHVLSIK | 1486691 | Cell/Virus Enclosed | NO |
| PFEEVEHVLSIR | 1486692 | Cell/Virus Enclosed | NO |
| PFEPQTIQPEVK | 1486699 | Cell/Virus Enclosed | NO |
| PFERDLTSDENG | 1486703 | Ectodomain | YES |
| PFEYTSPTDKLA | 1486709 | Cell/Virus Enclosed | NO |
| PFFSFEQSHNNCYVNV | 1685101 | Cell/Virus Enclosed | NO |
| PFFVDVQQWGYV | 1486715 | Cell/Virus Enclosed | NO |
| PFFYYSDTSCVY | 1486720 | Cell/Virus Enclosed | NO |
| PFGEVEHALSIK | 1486727 | Cell/Virus Enclosed | NO |
| PFGEVEHSLPIK | 1486728 | Cell/Virus Enclosed | NO |
| PFGEVEHSLTIK | 1486729 | Cell/Virus Enclosed | NO |
| PFGEVEYSLPIK | 1486730 | Cell/Virus Enclosed | NO |
| PFGEVFNATKFPSVYA | 1685112 | Ectodomain | YES |
| PFGGQPVSEYHI | 1486735 | Cell/Virus Enclosed | NO |
| PFGGTTVVRPLVEDST | 1685117 | Ectodomain | YES |
| PFGVVDTEWGNM | 1486746 | Cell/Virus Enclosed | NO |
| PFHYERDNTSCPEWMD | 1685128 | Cell/Virus Enclosed | NO |
| PFHYERDNTSYP | 1486756 | Cell/Virus Enclosed | NO |
| PFHYERNNTSCP | 1486757 | Cell/Virus Enclosed | NO |
| PFIANEKFINGCGRNV | 1685130 | Cell/Virus Enclosed | NO |
| PFIAVSKPMGTQ | 1486759 | Ectodomain | YES |

| | | | |
|---|---|---|---|
| PFKDGIYFAATEKSNV | 1685135 | Ectodomain | YES |
| PFKFCSETVNGY | 1486769 | Cell/Virus Enclosed | NO |
| PFLGQAAVTTTN | 1486782 | Cell/Virus Enclosed | NO |
| PFLSDFTNGIFA | 1486797 | Ectodomain | YES |
| PFMDDAINVGGT | 1486805 | Cell/Virus Enclosed | NO |
| PFMDNAINVGGTGLQY | 1685155 | Cell/Virus Enclosed | NO |
| PFMGNAINVGGR | 1486807 | Cell/Virus Enclosed | NO |
| PFMIDVQQWGFT | 1486808 | Cell/Virus Enclosed | NO |
| PFMVNQLAYSSSANGS | 1685159 | Cell/Virus Enclosed | NO |
| PFNVSFVNDSIESVGG | 1685168 | Ectodomain | YES |
| PFSDRPLNLRGW | 1486842 | Cell/Virus Enclosed | NO |
| PFSPNGKPCTPP | 1486848 | Ectodomain | YES |
| PFSSDGKPCTPP | 1486850 | Ectodomain | YES |
| PFTVNSINDSLE | 1486864 | Ectodomain | YES |
| PFTVNSKNDNLE | 1486865 | Ectodomain | YES |
| PFTVNSVNDSLEPVGG | 1685191 | Ectodomain | YES |
| PFTVNSVNDSLK | 1486866 | Ectodomain | YES |
| PFVDRQTAQAAG | 1486870 | Cell/Virus Enclosed | NO |
| PFVKGDSCSSNC | 1486880 | Cell/Virus Enclosed | NO |
| PFVSDSVATYFG | 1486888 | Cell/Virus Enclosed | NO |
| PFVSFIYFITMS | 1486889 | Cell/Virus Enclosed | NO |
| PFYLNVQYRING | 1486905 | Ectodomain | YES |
| PFYVPEHCQNYS | 1486910 | Cell/Virus Enclosed | NO |
| PFYYERDNTSCP | 1486911 | Cell/Virus Enclosed | NO |
| PGAADAERMAMY | 1486912 | Ectodomain | YES |
| PGAFAYSQMRPH | 1486915 | Cell/Virus Enclosed | NO |
| PGAFFFGSKLDLVKRD | 1685215 | Capsid | UN |
| PGALEFNPSQTS | 1486922 | Cell/Virus Enclosed | NO |
| PGCDGGSLYVNK | 1486925 | Cell/Virus Enclosed | NO |
| PGCDVADVTKLY | 1486926 | Cell/Virus Enclosed | NO |
| PGCDVNDVTKLYLGGM | 1685219 | Cell/Virus Enclosed | NO |
| PGCGGILSVIDD | 1486928 | Cell/Virus Enclosed | NO |
| PGDLQGNFGDLK | 1486931 | Cell/Virus Enclosed | NO |
| PGDLQGNFGDLQLNKL | 1685228 | Cell/Virus Enclosed | NO |
| PGDQFKHLIPLM | 1486933 | Cell/Virus Enclosed | NO |
| PGDVIITSTDAG | 1486935 | Cell/Virus Enclosed | NO |
| PGDVIITSTGAG | 1486936 | Cell/Virus Enclosed | NO |
| PGEFASHVANFQ | 1486938 | Cell/Virus Enclosed | NO |
| PGEFSKKQLGLLQPTD | 1685234 | Cell/Virus Enclosed | NO |
| PGEHFKHLVPLM | 1486940 | Cell/Virus Enclosed | NO |
| PGEMRLASIAFN | 1486945 | Ectodomain | YES |
| PGEQFKHLIPLV | 1486947 | Cell/Virus Enclosed | NO |
| PGEQFKHLVPLI | 1486948 | Cell/Virus Enclosed | NO |
| PGESKPQSRNPS | 1486950 | Capsid | UN |
| PGETGVIADYNY | 1486951 | Ectodomain | YES |
| PGFGVAVADSYF | 1486959 | Cell/Virus Enclosed | NO |
| PGFGVAVADSYYSYMM | 1685245 | Cell/Virus Enclosed | NO |
| PGFKEELDQWFK | 1486961 | Ectodomain | YES |
| PGFLTPFMILLV | 1486962 | Cell/Virus Enclosed | NO |
| PGGCAMPIMANL | 1486969 | Ectodomain | YES |
| PGGCAMPIVANL | 1486970 | Ectodomain | YES |
| PGGDAISLSFNNLLGF | 1685252 | Cell/Virus Enclosed | NO |
| PGGDATSLSFNN | 1486975 | Cell/Virus Enclosed | NO |
| PGGDLLYLDLLN | 1486976 | Cell/Virus Enclosed | NO |
| PGGIDICSQDKV | 1486978 | Cell/Virus Enclosed | NO |
| PGGTGSGDATTA | 1486984 | Cell/Virus Enclosed | NO |
| PGGTSSGDATTAFANS | 1685253 | Cell/Virus Enclosed | NO |
| PGHFNKEFYDFLRSQG | 1685260 | Cell/Virus Enclosed | NO |
| PGHYAGSRSSSGNRSG | 1685261 | Capsid | UN |
| PGIDAGYKNSGI | 1486996 | Ectodomain | YES |
| PGKCYNCRPAVI | 1486999 | Ectodomain | YES |
| PGKCYNCRPSVI | 1487000 | Ectodomain | YES |
| PGKNNGIGTCPA | 1487004 | Ectodomain | YES |
| PGKNNGIGTCPV | 1487005 | Ectodomain | YES |
| PGKNNGLGTCPA | 1487008 | Ectodomain | YES |
| PGKNNGMGTCPA | 1487009 | Ectodomain | YES |
| PGKQSSSRASSGNRSG | 1685269 | Cell/Virus Enclosed | NO |
| PGLCIAGNRGIA | 1487015 | Ectodomain | YES |
| PGLCISGDRGIA | 1487017 | Ectodomain | YES |
| PGLCLSGDIGIA | 1487019 | Ectodomain | YES |
| PGLCLSGDRGIA | 1487020 | Ectodomain | YES |
| PGLGAPVFHMTNYLE | 47694 | Ectodomain | YES |
| PGLGINEEKCGT | 1487022 | Ectodomain | YES |
| PGLSGIGAVGGD | 1487031 | Cell/Virus Enclosed | NO |
| PGMIFVLTSHNV | 1487036 | Cell/Virus Enclosed | NO |
| PGMKYVKWEHDS | 1487037 | Cell/Virus Enclosed | NO |

| | | | |
|---|---|---|---|
| PGNFNQDFYDFVLSKG | 1685284 | Cell/Virus Enclosed | NO |
| PGNFNQDFYDFVVSKG | 1685285 | Cell/Virus Enclosed | NO |
| PGNFNQDFYEFVKSKG | 1685286 | Cell/Virus Enclosed | NO |
| PGNGILKWADQS | 1487042 | Capsid | UN |
| PGNHCPAGNSYT | 1487043 | Ectodomain | YES |
| PGNMIDGFACTS | 1487044 | Cell/Virus Enclosed | NO |
| PGNTFINGDIARELGN | 1685288 | Cell/Virus Enclosed | NO |
| PGNVGGNDALST | 1487053 | Cell/Virus Enclosed | NO |
| PGNVGGNDDLSI | 1487054 | Cell/Virus Enclosed | NO |
| PGNVGGNEALSI | 1487055 | Cell/Virus Enclosed | NO |
| PGNVKPGDAVITTQQA | 1685292 | Cell/Virus Enclosed | NO |
| PGNVVPGDVIIT | 1487056 | Cell/Virus Enclosed | NO |
| PGPSGIGGGDLL | 1487064 | Cell/Virus Enclosed | NO |
| PGPSGIGTVGGD | 1487065 | Cell/Virus Enclosed | NO |
| PGPSGVGAPGGD | 1487066 | Cell/Virus Enclosed | NO |
| PGQAMPSLFKVQ | 1487067 | Cell/Virus Enclosed | NO |
| PGQGVPLNANSTPAQN | 1685300 | Cell/Virus Enclosed | NO |
| PGQGVPLNSNST | 1487072 | Cell/Virus Enclosed | NO |
| PGQKYSSDCNVV | 1487073 | Cell/Virus Enclosed | NO |
| PGQTGVIADYNYKLPD | 1685304 | Ectodomain | YES |
| PGRCYNCRPAVI | 1487081 | Ectodomain | YES |
| PGRCYNCRPAVV | 1487082 | Ectodomain | YES |
| PGRTSSGDATTA | 1487083 | Cell/Virus Enclosed | NO |
| PGSCNFPLEATW | 1487091 | Ectodomain | YES |
| PGSSRGTSPARIAGNG | 1685319 | Cell/Virus Enclosed | NO |
| PGSSRGTSPARMAGNG | 1685320 | Cell/Virus Enclosed | NO |
| PGTAVLRPWLPT | 1487106 | Cell/Virus Enclosed | NO |
| PGTCYNCRPAVV | 1487108 | Ectodomain | YES |
| PGTGKSHFAIGL | 1487110 | Cell/Virus Enclosed | NO |
| PGTGPHKDLKFR | 1487111 | Cell/Virus Enclosed | NO |
| PGTPAQEIEMLKRN | 47765 | Capsid | UN |
| PGTVLRAINGDF | 1487132 | Cell/Virus Enclosed | NO |
| PGVADAERMAMY | 1487134 | Ectodomain | YES |
| PGVAMPNLYKIQ | 1487135 | Cell/Virus Enclosed | NO |
| PGVAMPNLYKMQ | 1487136 | Cell/Virus Enclosed | NO |
| PGVPATEAKGYW | 1487144 | Cell/Virus Enclosed | NO |
| PGVPATESKGYW | 1487145 | Capsid | UN |
| PGVSTNVYLVGK | 1487152 | Cell/Virus Enclosed | NO |
| PGVTPDMADQIA | 1487153 | Cell/Virus Enclosed | NO |
| PGYMQGYDDCMH | 1487163 | Ectodomain | YES |
| PGYMQGYDDCMQQGPA | 1685353 | Ectodomain | YES |
| PGYNLKNGEVEFVYMH | 1685354 | Cell/Virus Enclosed | NO |
| PHAKDQYGSDIN | 1487174 | Cell/Virus Enclosed | NO |
| PHAKDQYGTDFD | 1487175 | Cell/Virus Enclosed | NO |
| PHAKDQYGTDIN | 1487177 | Cell/Virus Enclosed | NO |
| PHANEYRQYLDA | 1487180 | Cell/Virus Enclosed | NO |
| PHCGSSSRSFFEDLLF | 1685363 | Ectodomain | YES |
| PHDAIVVDNDVVDYVS | 1685364 | Cell/Virus Enclosed | NO |
| PHDFFIFDVDKV | 1487184 | Cell/Virus Enclosed | NO |
| PHDFFVFDVDKT | 1487185 | Cell/Virus Enclosed | NO |
| PHDVIVLRDNLK | 1487186 | Cell/Virus Enclosed | NO |
| PHDVTVLRDNLKPVTT | 1685366 | Cell/Virus Enclosed | NO |
| PHEADTEQTQSV | 1487188 | Cell/Virus Enclosed | NO |
| PHEFCSQHTHYI | 1487189 | Cell/Virus Enclosed | NO |
| PHEFCSQHTLYIKDGD | 1685367 | Cell/Virus Enclosed | NO |
| PHEFCSQHTMLV | 1487192 | Cell/Virus Enclosed | NO |
| PHEFCSQHTMQIVDKD | 1685372 | Cell/Virus Enclosed | NO |
| PHEFCTKHTMLV | 1487194 | Cell/Virus Enclosed | NO |
| PHFNQKLPNGVN | 1487198 | Cell/Virus Enclosed | NO |
| PHGVVFLHVTYVPSQER | 47823 | Ectodomain | YES |
| PHKDLKFRQRSD | 1487224 | Cell/Virus Enclosed | NO |
| PHKDVYELRYNG | 1487225 | Cell/Virus Enclosed | NO |
| PHKLKIQVVNSGSDMN | 1685390 | Cell/Virus Enclosed | NO |
| PHLMGWDYPKFD | 1487232 | Cell/Virus Enclosed | NO |
| PHMFITREEAIR | 1487236 | Cell/Virus Enclosed | NO |
| PHNGVLDITACQ | 1487240 | Ectodomain | YES |
| PHNYFYMDLNPA | 1487248 | Ectodomain | YES |
| PHNYSYMDLNPA | 1487249 | Ectodomain | YES |
| PHNYSYMDLNSA | 1487251 | Ectodomain | YES |
| PHPYCYSDGVMKNASL | 1685403 | Cell/Virus Enclosed | NO |
| PHQVNISLNGNTSVCV | 1685404 | Ectodomain | YES |
| PHSIRSIKSDRK | 1487266 | Ectodomain | YES |
| PHSIRSIQSDRK | 1487267 | Ectodomain | YES |
| PHSIRSIQSGRK | 1487268 | Ectodomain | YES |
| PHSKDQYGTDIN | 1487269 | Cell/Virus Enclosed | NO |
| PHSKEQYGTDID | 1487270 | Cell/Virus Enclosed | NO |

| | | | |
|---|---|---|---|
| PHTRYSLADSNG | 1487282 | Cell/Virus Enclosed | NO |
| PHTSCHPNLGNH | 1487284 | Ectodomain | YES |
| PHTTNLDNKLQG | 1487287 | Ectodomain | YES |
| PHTVLLVPRGSG | 1487289 | Cell/Virus Enclosed | NO |
| PHVGETPVAYRN | 1487296 | Cell/Virus Enclosed | NO |
| PHVNHEGKTFFV | 1487301 | Cell/Virus Enclosed | NO |
| PHWTKKPNRNGGGYYSASYSDP | 47861 | Ectodomain | YES |
| PIADPTHFDIEEVELL | 1685422 | Cell/Virus Enclosed | NO |
| PIADVYRRKPDL | 1487321 | Ectodomain | YES |
| PIAFGIPPSEAK | 1487326 | Cell/Virus Enclosed | NO |
| PIAFGVPPSEAKGYWY | 1685424 | Capsid | UN |
| PIAYGIPLSEAK | 1487344 | Cell/Virus Enclosed | NO |
| PIAYGIPPSEAK | 1487345 | Cell/Virus Enclosed | NO |
| PIDAIIIDNDLN | 1487363 | Cell/Virus Enclosed | NO |
| PIDEIPYKSFFD | 1487365 | Cell/Virus Enclosed | NO |
| PIDEIPYKSFSD | 1487366 | Cell/Virus Enclosed | NO |
| PIDEYITTYPGQ | 1487367 | Cell/Virus Enclosed | NO |
| PIDFPSAESTGT | 1487369 | Cell/Virus Enclosed | NO |
| PIDIDAFPVKPICSSV | 1685439 | Cell/Virus Enclosed | NO |
| PIDIDAFPVRPI | 1487373 | Cell/Virus Enclosed | NO |
| PIDIPRGLPVGF | 1487374 | Ectodomain | YES |
| PIDLASQIISGT | 1487378 | Undetermined | UN |
| PIDLLCLRDDNQ | 1487381 | Cell/Virus Enclosed | NO |
| PIDLSSQIISGI | 1487382 | Undetermined | UN |
| PIDSTDNNQNGG | 1487392 | Capsid | UN |
| PIDVFKADGIIY | 1487395 | Ectodomain | YES |
| PIDYNVVTQRMCAIGP | 1685452 | Cell/Virus Enclosed | NO |
| PIEDAETPALQV | 1487398 | Cell/Virus Enclosed | NO |
| PIEHVTSVNLFD | 1487404 | Cell/Virus Enclosed | NO |
| PIELQVECFTWDQKL | 47890 | Ectodomain | YES |
| PIFDDPVVDWHY | 1487413 | Cell/Virus Enclosed | NO |
| PIGKGNKDEQIGYWNV | 1685472 | Capsid | UN |
| PIGKGSKDEQIG | 1487439 | Cell/Virus Enclosed | NO |
| PIGMFFPYDIELVTGK | 1685473 | Cell/Virus Enclosed | NO |
| PIGTKYRHCDLD | 1487444 | Ectodomain | YES |
| PIGTNDPIRFCLENTV | 1685474 | Cell/Virus Enclosed | NO |
| PIGTNDPIRSCL | 1487446 | Cell/Virus Enclosed | NO |
| PIGTNYRSCESTTVLD | 1685475 | Ectodomain | YES |
| PIGVPSQVVYVPPAYIDMPS | 47910 | Cell/Virus Enclosed | NO |
| PIIEVSQFQSKLTDVK | 1685480 | Cell/Virus Enclosed | NO |
| PIIGDELKINSA | 1487463 | Cell/Virus Enclosed | NO |
| PIIKAQFKPFAKVDGV | 1685483 | Cell/Virus Enclosed | NO |
| PIIKSSVKSYDDLVYS | 1685484 | Cell/Virus Enclosed | NO |
| PIIQEAIPTRFS | 1487476 | Cell/Virus Enclosed | NO |
| PIISEKISVMDK | 1487477 | Cell/Virus Enclosed | NO |
| PIKKAFPYGTCL | 1487485 | Cell/Virus Enclosed | NO |
| PIKNTVDTIPKE | 1487488 | Cell/Virus Enclosed | NO |
| PILAELAPPAGA | 1487497 | Cell/Virus Enclosed | NO |
| PILAELAPTAGA | 1487498 | Cell/Virus Enclosed | NO |
| PILHHPGFLNPS | 1487503 | Cell/Virus Enclosed | NO |
| PILKLPFGINIT | 1487506 | Ectodomain | YES |
| PILKSSVKSYDSLVYT | 1685506 | Cell/Virus Enclosed | NO |
| PILLLDQSLVSD | 1487513 | Cell/Virus Enclosed | NO |
| PILMVDKNLITTANTG | 1685509 | Cell/Virus Enclosed | NO |
| PILSCGIFGIKLETSL | 1685512 | Cell/Virus Enclosed | NO |
| PILTLTKALAAE | 1487525 | Cell/Virus Enclosed | NO |
| PILWVKKASYDT | 1487530 | Cell/Virus Enclosed | NO |
| PILWVNKASYDTNLNK | 1685519 | Cell/Virus Enclosed | NO |
| PIMANLVNHKSH | 1487535 | Ectodomain | YES |
| PIMANLVNLNSH | 1487536 | Ectodomain | YES |
| PIMANLVNSKSH | 1487537 | Ectodomain | YES |
| PINATDRSHYYVDSVT | 1685522 | Cell/Virus Enclosed | NO |
| PINKKDKNKLIG | 1487543 | Cell/Virus Enclosed | NO |
| PINKKEKNKLIG | 1487544 | Capsid | UN |
| PINSNVKTLEYDYMTH | 1685530 | Cell/Virus Enclosed | NO |
| PINTDTVLSVAPEVDW | 1685531 | Cell/Virus Enclosed | NO |
| PINTNSGKDDQI | 1487552 | Cell/Virus Enclosed | NO |
| PINTNSSKDDQI | 1487554 | Cell/Virus Enclosed | NO |
| PINTNSTKDDQIGYYR | 1685534 | Cell/Virus Enclosed | NO |
| PIPDFDLDDFIDAPCY | 1685538 | Cell/Virus Enclosed | NO |
| PIPNLPDFREEL | 1487562 | Ectodomain | YES |
| PIQDYIQSVNFL | 1487568 | Cell/Virus Enclosed | NO |
| PIQIDQLNSSYF | 1487570 | Ectodomain | YES |
| PIQPTDVAYHTVTDVK | 1685542 | Cell/Virus Enclosed | NO |
| PIQPVELPESIV | 1487578 | Ectodomain | YES |
| PIQPVELPMSIV | 1487579 | Ectodomain | YES |

| | | | |
|---|---|---|---|
| PIQPVELPVSIV | 1487580 | Ectodomain | YES |
| PIQSAELPVSIV | 1487582 | Ectodomain | YES |
| PIQVDQFNSSYF | 1487585 | Ectodomain | YES |
| PIQVDQLNSSYK | 1487587 | Ectodomain | YES |
| PIQVDQLNSTYF | 1487588 | Ectodomain | YES |
| PIQVGQLNSSYF | 1487589 | Ectodomain | YES |
| PISASFTPFDHKVVIHRG | 558303 | Undetermined | UN |
| PISAYAFLMAED | 1487601 | Cell/Virus Enclosed | NO |
| PISLLFNEKEADDNDHEPQI | 47984 | Cell/Virus Enclosed | NO |
| PISNCIKMLSNG | 1487613 | Cell/Virus Enclosed | NO |
| PISNDSNCCVVE | 1487614 | Cell/Virus Enclosed | NO |
| PISNDSNGCVVEASTD | 1685558 | Cell/Virus Enclosed | NO |
| PISTYSPNTCPQ | 1487624 | Ectodomain | YES |
| PITENNVVVMSTCAVN | 1685568 | Ectodomain | YES |
| PITENNVVVTST | 1487631 | Ectodomain | YES |
| PITFAGPYKCPQ | 1487632 | Ectodomain | YES |
| PITFKAPDTYKC | 1487633 | Ectodomain | YES |
| PITGNNVVVMST | 1487639 | Ectodomain | YES |
| PITKNIGEYNVSKDGF | 1685573 | Cell/Virus Enclosed | NO |
| PITLKTPDTYKC | 1487650 | Ectodomain | YES |
| PITLTPRSVRSV | 1487652 | Ectodomain | YES |
| PITSFNTKYVAP | 1487657 | Ectodomain | YES |
| PITSLNTKYVAPQVTY | 1685575 | Ectodomain | YES |
| PIVDDPVVDWHY | 1487671 | Cell/Virus Enclosed | NO |
| PIVMNDKNICLLDQAW | 1685584 | Cell/Virus Enclosed | NO |
| PIVNEKISVMDK | 1487683 | Cell/Virus Enclosed | NO |
| PIVSEKISVMDK | 1487692 | Cell/Virus Enclosed | NO |
| PIVWGAKDFNTL | 1487700 | Cell/Virus Enclosed | NO |
| PIVWGVKDFNTLSQEG | 1685596 | Cell/Virus Enclosed | NO |
| PIVWHAKDFNSLSAEG | 1685597 | Cell/Virus Enclosed | NO |
| PIVYETDANKLI | 1487703 | Cell/Virus Enclosed | NO |
| PIWFLNTEPSQLPPTAPPLLPHS | 48016 | Ectodomain | YES |
| PIYDEPTMTTSV | 1487711 | Secreted | YES |
| PIYKNGAMYKGKPILW | 1685603 | Cell/Virus Enclosed | NO |
| PIYLKGSGFDFS | 1487719 | Cell/Virus Enclosed | NO |
| PIYLKGSGSDFSLADS | 1685604 | Cell/Virus Enclosed | NO |
| PKAGANFRDLSL | 1487723 | Ectodomain | YES |
| PKAIACVKDFDF | 1487724 | Cell/Virus Enclosed | NO |
| PKDGFNLEYWVT | 1487741 | Ectodomain | YES |
| PKDGFSPEYWVT | 1487743 | Ectodomain | YES |
| PKDGGHFFVSDD | 1487744 | Cell/Virus Enclosed | NO |
| PKDGGHFFVSGD | 1487745 | Cell/Virus Enclosed | NO |
| PKDGGQFFVSDD | 1487746 | Cell/Virus Enclosed | NO |
| PKDGGQFFVSGD | 1487747 | Cell/Virus Enclosed | NO |
| PKDHIGTRNPNNNAAI | 1685622 | Cell/Virus Enclosed | NO |
| PKDQIDGYTMHA | 1487752 | Cell/Virus Enclosed | NO |
| PKDWRLINKFDVINGV | 1685627 | Cell/Virus Enclosed | NO |
| PKEDGDVLLAEFDTYD | 1685628 | Cell/Virus Enclosed | NO |
| PKEDGDVVLAEF | 1487755 | Cell/Virus Enclosed | NO |
| PKEEFVVKEKLNAFLV | 1685629 | Cell/Virus Enclosed | NO |
| PKEEGDVLLAEF | 1487756 | Cell/Virus Enclosed | NO |
| PKEELTDQMSEP | 1487757 | Cell/Virus Enclosed | NO |
| PKEESNDQEMTT | 1487759 | Cell/Virus Enclosed | NO |
| PKEESTDQISEP | 1487760 | Cell/Virus Enclosed | NO |
| PKEESTDQMSEL | 1487762 | Cell/Virus Enclosed | NO |
| PKEHRVQGTQRT | 1487766 | Cell/Virus Enclosed | NO |
| PKEITVATSRML | 1487770 | Underneath Membrane | NO |
| PKEITVATSRTLSYYKL | 48052 | Underneath Membrane | NO |
| PKEIVDTVSALVYENK | 1685635 | Cell/Virus Enclosed | NO |
| PKEIVETVSALVYDNK | 1685636 | Cell/Virus Enclosed | NO |
| PKELLLLWESGKAKPP | 1685638 | Cell/Virus Enclosed | NO |
| PKELLLSWESGKVKPP | 1685639 | Cell/Virus Enclosed | NO |
| PKEQRVQGGITQ | 1487775 | Cell/Virus Enclosed | NO |
| PKEQRVQGIITQ | 1487776 | Cell/Virus Enclosed | NO |
| PKEQRVQGSIAQ | 1487777 | Capsid | UN |
| PKEQRVQGSITH | 1487778 | Cell/Virus Enclosed | NO |
| PKEQRVQGSITQRTRT | 1685641 | Cell/Virus Enclosed | NO |
| PKEQRVQGSTTQ | 1487780 | Capsid | UN |
| PKEVTFFEGDSH | 1487783 | Cell/Virus Enclosed | NO |
| PKFEFANDTKIA | 1487786 | Ectodomain | YES |
| PKFFYASNGTYNCTDA | 1685644 | Ectodomain | YES |
| PKFIEQISAFTKPSSI | 1685645 | Capsid | UN |
| PKFTVSTQPQEN | 1487794 | Cell/Virus Enclosed | NO |
| PKGIPIFDDPVV | 1487804 | Cell/Virus Enclosed | NO |
| PKGKYSQKAYALLRQY | 1685660 | Cell/Virus Enclosed | NO |
| PKGVGSANIFIGDKVG | 1685664 | Cell/Virus Enclosed | NO |

| | | | |
|---|---|---|---|
| PKHVFKTLKPGDSFNI | 1685669 | Cell/Virus Enclosed | NO |
| PKIEFANDTKIA | 1487827 | Ectodomain | YES |
| PKIFKARAIECY | 1487828 | Cell/Virus Enclosed | NO |
| PKIFKARAIEFY | 1487829 | Cell/Virus Enclosed | NO |
| PKIIKVFYELDNDFNT | 1685673 | Cell/Virus Enclosed | NO |
| PKKDKKKKADEL | 1487840 | Cell/Virus Enclosed | NO |
| PKKDKKKKTDESQPLP | 1685677 | Cell/Virus Enclosed | NO |
| PKKEKKQKAPKEESTD | 1685678 | Cell/Virus Enclosed | NO |
| PKKPNKPLSQPRADKP | 1685680 | Capsid | UN |
| PKKTYLDYIQQGGVPI | 1685683 | Cell/Virus Enclosed | NO |
| PKLEFANDTKIASQLG | 1685688 | Ectodomain | YES |
| PKLEFANDTKIV | 1487858 | Ectodomain | YES |
| PKLGNHFKESWH | 1487867 | Ectodomain | YES |
| PKLKEDYQIGGYSEDR | 1685694 | Secreted | YES |
| PKLKGSFNNRDT | 1487874 | Ectodomain | YES |
| PKLMGWDYPKCDRAMP | 1685695 | Cell/Virus Enclosed | NO |
| PKLSTDLIKNQCVNFN | 1685699 | Ectodomain | YES |
| PKLSTGLVKNQC | 1487880 | Ectodomain | YES |
| PKLYCAFTAPEY | 1487883 | Cell/Virus Enclosed | NO |
| PKNFCPCKLNGSCVGS | 1685703 | Ectodomain | YES |
| PKNFCPCSSCFG | 1487897 | Ectodomain | YES |
| PKNFCPCSSCPG | 1487899 | Ectodomain | YES |
| PKNFCPCSSCPS | 1487900 | Ectodomain | YES |
| PKNFCPCSSCSC | 1487901 | Ectodomain | YES |
| PKNFCPCSSCSG | 1487902 | Ectodomain | YES |
| PKNFHIEGTGGN | 1487903 | Cell/Virus Enclosed | NO |
| PKNGGQFFVSGD | 1487905 | Cell/Virus Enclosed | NO |
| PKNGQGENDNIS | 1487906 | Cell/Virus Enclosed | NO |
| PKNMWEFRDYFD | 1487907 | Cell/Virus Enclosed | NO |
| PKNSFEALMLNFKLLG | 1685707 | Cell/Virus Enclosed | NO |
| PKPQRQRGHRNG | 1487918 | Cell/Virus Enclosed | NO |
| PKPQRQRGQKNG | 1487922 | Capsid | UN |
| PKPSRNQSPVSS | 1487931 | Cell/Virus Enclosed | NO |
| PKPVEGVSSFMN | 1487937 | Ectodomain | YES |
| PKPVIYQLDENA | 1487940 | Cell/Virus Enclosed | NO |
| PKQEEPPKTEHP | 1487944 | Cell/Virus Enclosed | NO |
| PKQFVPVQVVDS | 1487947 | Underneath Membrane | NO |
| PKQGYFIKHNDH | 1487950 | Ectodomain | YES |
| PKQGYFIKQNDS | 1487951 | Ectodomain | YES |
| PKQRRPQGLPNNTASWFT | 48131 | Capsid | UN |
| PKQSIFTINSVVQKDG | 1685725 | Cell/Virus Enclosed | NO |
| PKQTSTSQQPLG | 1487958 | Cell/Virus Enclosed | NO |
| PKQTSTSQQPSG | 1487959 | Cell/Virus Enclosed | NO |
| PKQTSTSQQPSR | 1487960 | Cell/Virus Enclosed | NO |
| PKQTVTSQQPSG | 1487961 | Cell/Virus Enclosed | NO |
| PKRLRIKRNVDP | 1487962 | Cell/Virus Enclosed | NO |
| PKRLRKKRNVDPLSNF | 1685727 | Cell/Virus Enclosed | NO |
| PKRLRLPHIRDDDAPPSHQP | 110364 | Underneath Membrane | NO |
| PKRTGVILTRSL | 1487967 | Cell/Virus Enclosed | NO |
| PKSDGTGTIYAE | 1487968 | Cell/Virus Enclosed | NO |
| PKSGYFVNVNNTWMYT | 1685731 | Ectodomain | YES |
| PKSRVQQNKSIE | 1487974 | Cell/Virus Enclosed | NO |
| PKSRVQQNKSRELTAE | 1685733 | Cell/Virus Enclosed | NO |
| PKTDNETLEYWV | 1487979 | Ectodomain | YES |
| PKTEHPKIEEKS | 1487980 | Cell/Virus Enclosed | NO |
| PKTGTGIAISVK | 1487985 | Cell/Virus Enclosed | NO |
| PKTKHVTKENDS | 1487987 | Cell/Virus Enclosed | NO |
| PKTSEILDISPCAFGGV | 48155 | Ectodomain | YES |
| PKTSEILDISPR | 1487996 | Ectodomain | YES |
| PKTSVVPPALHL | 1488000 | Cell/Virus Enclosed | NO |
| PKVADKICIVDN | 1488004 | Cell/Virus Enclosed | NO |
| PKVEIDGNVMHANYLF | 1685744 | Cell/Virus Enclosed | NO |
| PKVHFYYPGTGP | 1488008 | Cell/Virus Enclosed | NO |
| PKVITKRDAAAA | 1488012 | Cell/Virus Enclosed | NO |
| PKVRDQEGRMNYYWTLVEPGDKITF | 1163235 | Ectodomain | YES |
| PKYKFVRIQPGQ | 1488025 | Cell/Virus Enclosed | NO |
| PKYTFGNVKPGETFTV | 1685757 | Cell/Virus Enclosed | NO |
| PLAELKCSVKSFEIDK | 1685765 | Ectodomain | YES |
| PLAFGVPLSEAK | 1488042 | Cell/Virus Enclosed | NO |
| PLAFMPVQSRRF | 1488045 | Cell/Virus Enclosed | NO |
| PLARYGSRVQIRCRFPNSTR | 741815 | Ectodomain | YES |
| PLASCFFTCNSN | 1488053 | Ectodomain | YES |
| PLAYFSDSKIGT | 1488061 | Cell/Virus Enclosed | NO |
| PLCASNKLRVVIPDFT | 1685774 | Cell/Virus Enclosed | NO |
| PLCGFPSQAAGV | 1488068 | Cell/Virus Enclosed | NO |
| PLCGSGNLVEKLSPWF | 1685777 | Cell/Virus Enclosed | NO |

| | | | |
|---|---|---|---|
| PLCIFNGKFLSNTKYY | 1685778 | Ectodomain | YES |
| PLCLFKSNFNQY | 1488077 | Ectodomain | YES |
| PLCSAGNIFGLYKNSA | 1685784 | Cell/Virus Enclosed | NO |
| PLDCIKDFLARA | 1488084 | Cell/Virus Enclosed | NO |
| PLDEAEDFTESV | 1488086 | Cell/Virus Enclosed | NO |
| PLDEAEDVTEFV | 1488087 | Cell/Virus Enclosed | NO |
| PLDEAENVTESV | 1488088 | Cell/Virus Enclosed | NO |
| PLDEAEYVTEFV | 1488089 | Cell/Virus Enclosed | NO |
| PLDELKSLLSLR | 1488091 | Cell/Virus Enclosed | NO |
| PLDIYDLTMPSQ | 1488095 | Cell/Virus Enclosed | NO |
| PLDPKLKGSFND | 1488099 | Ectodomain | YES |
| PLDPRLKGSFND | 1488100 | Ectodomain | YES |
| PLDSRLKGSFNN | 1488103 | Ectodomain | YES |
| PLDSRTGSFNNI | 1488104 | Ectodomain | YES |
| PLDTRLKGSFNN | 1488106 | Ectodomain | YES |
| PLDTSLKGSFNN | 1488108 | Ectodomain | YES |
| PLDTSYINNKDT | 1488110 | Ectodomain | YES |
| PLDVYNLTMPSQ | 1488113 | Cell/Virus Enclosed | NO |
| PLDYKRQNNLAIEEIE | 1685791 | Cell/Virus Enclosed | NO |
| PLEFGASAETVR | 1488122 | Cell/Virus Enclosed | NO |
| PLEFGASTEIQQ | 1488123 | Cell/Virus Enclosed | NO |
| PLEFGASTEMQV | 1488124 | Cell/Virus Enclosed | NO |
| PLEGGGRLVASG | 1488129 | Ectodomain | YES |
| PLEGGGWLVASGSTVA | 1685799 | Ectodomain | YES |
| PLEGGGWLVVSG | 1488131 | Ectodomain | YES |
| PLENDADKANFL | 1488133 | Cell/Virus Enclosed | NO |
| PLENPSSLFDLVARIKNNLK | 48307 | Capsid | UN |
| PLENRVLTGWGLNWDLGLSQWAR | 48309 | Ectodomain | YES |
| PLEPKFSIALPPELSV | 1685802 | Capsid | UN |
| PLESRTGSLNNI | 1488136 | Ectodomain | YES |
| PLETPVGCVLGL | 1488137 | Ectodomain | YES |
| PLFALISIQILT | 1488140 | Cell/Virus Enclosed | NO |
| PLFDQQAQQEAT | 1488143 | Cell/Virus Enclosed | NO |
| PLFESDKEEDVQ | 1488145 | Cell/Virus Enclosed | NO |
| PLFIRNFDQRCS | 1488146 | Cell/Virus Enclosed | NO |
| PLFLFDEAGEEVLAPK | 1685811 | Cell/Virus Enclosed | NO |
| PLGFFPDHQLDPAFG | 48332 | Underneath Membrane | NO |
| PLGFLPDHQLDPAFGANSTNP | 48339 | Ectodomain | YES |
| PLGGTTVVRPLV | 1488153 | Underneath Membrane | NO |
| PLGIKDPILYVL | 1488155 | Cell/Virus Enclosed | NO |
| PLGMSLEACYVRGCNP | 1685820 | Cell/Virus Enclosed | NO |
| PLGRCDIKDLPK | 1488166 | Underneath Membrane | NO |
| PLGRSLCALPDT | 1488167 | Ectodomain | YES |
| PLGSAGNATEVP | 1488168 | Cell/Virus Enclosed | NO |
| PLGSAIASEFSS | 1488169 | Cell/Virus Enclosed | NO |
| PLGSAWRDQAQRPAVASRRRPTTAG | 48350 | Undetermined | UN |
| PLGSMYKQARSE | 1488172 | Cell/Virus Enclosed | NO |
| PLGSSKMSDVKC | 1488175 | Cell/Virus Enclosed | NO |
| PLHGTILTRPLLESE | 1541971 | Underneath Membrane | NO |
| PLHLGKCNIAGWILGNPECESLSTA | 1163247 | Ectodomain | YES |
| PLHMSISKLHAL | 1488195 | Cell/Virus Enclosed | NO |
| PLHPYGRCPTAA | 1488200 | Ectodomain | YES |
| PLHSLGGFPKGYVMGL | 1685828 | Cell/Virus Enclosed | NO |
| PLHTLSIHDGYT | 1488203 | Cell/Virus Enclosed | NO |
| PLIEWPLTEYDFTDYK | 1685834 | Cell/Virus Enclosed | NO |
| PLIKHPNQEYAD | 1488218 | Cell/Virus Enclosed | NO |
| PLITHGSGMDLYKSN | 48370 | Ectodomain | YES |
| PLKLAGTAVINLRADQ | 1685846 | Cell/Virus Enclosed | NO |
| PLKPIKRSFIED | 1488241 | Ectodomain | YES |
| PLKPTKRSFIEDLLFNKV | 1309917 | Ectodomain | YES |
| PLKVGAGVMLECEKFN | 1685854 | Cell/Virus Enclosed | NO |
| PLLAELIPNHAA | 1488261 | Cell/Virus Enclosed | NO |
| PLLEVYVHLRNWNFSW | 1685862 | Undetermined | UN |
| PLLGNSTGIDFQ | 1488275 | Ectodomain | YES |
| PLLNPGALEFNP | 1488290 | Cell/Virus Enclosed | NO |
| PLLNPSALEFNPSQTS | 1685869 | Capsid | UN |
| PLLQRQKKQPTV | 1488294 | Capsid | UN |
| PLLSDNQISGYT | 1488299 | Ectodomain | YES |
| PLLSENQFSGYT | 1488300 | Ectodomain | YES |
| PLLSENQINGYT | 1488301 | Ectodomain | YES |
| PLLSENQISGYTLAAT | 1685877 | Ectodomain | YES |
| PLLSGTPPQVYN | 1488305 | Ectodomain | YES |
| PLLSNSTGIDFQ | 1488307 | Ectodomain | YES |
| PLLVDDNKFDVL | 1488316 | Cell/Virus Enclosed | NO |
| PLLVDSDQPWKV | 1488317 | Cell/Virus Enclosed | NO |
| PLLVDSEQPCLV | 1488318 | Cell/Virus Enclosed | NO |

| | | | |
|---|---|---|---|
| PLLVDSEQPWKV | 1488319 | Cell/Virus Enclosed | NO |
| PLLVDSEQSWKV | 1488320 | Capsid | UN |
| PLLVLFGMYLDS | 1488322 | Cell/Virus Enclosed | NO |
| PLLVSSDKAPHR | 1488323 | Cell/Virus Enclosed | NO |
| PLMESELVIGAV | 1488335 | Underneath Membrane | NO |
| PLMKRGHRWDVV | 1488339 | Cell/Virus Enclosed | NO |
| PLMQSADASTFF | 1488345 | Cell/Virus Enclosed | NO |
| PLMQSADASTFL | 1488346 | Cell/Virus Enclosed | NO |
| PLNAIPSLAANTLNII | 1685901 | Cell/Virus Enclosed | NO |
| PLNANSTPAQNAGYWR | 1685902 | Cell/Virus Enclosed | NO |
| PLNANSTPSQNA | 1488353 | Cell/Virus Enclosed | NO |
| PLNDYGFYTTSG | 1488354 | Ectodomain | YES |
| PLNLPPTHAHTF | 1488362 | Cell/Virus Enclosed | NO |
| PLNLPPTQAHTF | 1488363 | Cell/Virus Enclosed | NO |
| PLNSNSTPAQNA | 1488375 | Cell/Virus Enclosed | NO |
| PLNWERKTFSNC | 1488379 | Ectodomain | YES |
| PLPAADMDDFSR | 1488381 | Capsid | UN |
| PLPEEPVNQFTG | 1488382 | Cell/Virus Enclosed | NO |
| PLQASLPFGWLVIGVA | 1685929 | Secreted | YES |
| PLQSLKMCVETV | 1488405 | Cell/Virus Enclosed | NO |
| PLQSLQVCVQTV | 1488406 | Cell/Virus Enclosed | NO |
| PLRAPKEVTFFE | 1488418 | Cell/Virus Enclosed | NO |
| PLRAPTIANQEKYSSI | 1685943 | Cell/Virus Enclosed | NO |
| PLRAPTMANQEKYSTI | 1685944 | Cell/Virus Enclosed | NO |
| PLRASSCVTRCNIGGA | 1685945 | Cell/Virus Enclosed | NO |
| PLRATNCITKCNIGGA | 1685946 | Cell/Virus Enclosed | NO |
| PLSAICPTGTKY | 1488441 | Ectodomain | YES |
| PLSDIEKDFLSF | 1488447 | Cell/Virus Enclosed | NO |
| PLSDMEKDFLFF | 1488448 | Cell/Virus Enclosed | NO |
| PLSDMEKDFLSFDSDV | 1685955 | Cell/Virus Enclosed | NO |
| PLSDMEKDFLSS | 1488450 | Cell/Virus Enclosed | NO |
| PLSDYGFYTTTG | 1488452 | Ectodomain | YES |
| PLSEVYVHLRNW | 1488459 | Ectodomain | YES |
| PLSEVYVQLRNW | 1488460 | Undetermined | UN |
| PLSHFQGFVLNY | 1488462 | Cell/Virus Enclosed | NO |
| PLSISNDTVDVT | 1488464 | Ectodomain | YES |
| PLSKDLPDDVVAANMF | 1685960 | Cell/Virus Enclosed | NO |
| PLSMKSDLGVSS | 1488470 | Ectodomain | YES |
| PLSMKSDLSVSS | 1488472 | Ectodomain | YES |
| PLSMKSDLSVTS | 1488473 | Ectodomain | YES |
| PLSNFEHKDITE | 1488476 | Cell/Virus Enclosed | NO |
| PLSQPRADKPSQLKKP | 1685966 | Capsid | UN |
| PLSQPRVDKPSQ | 1488484 | Cell/Virus Enclosed | NO |
| PLSVASAPVEPS | 1488485 | Cell/Virus Enclosed | NO |
| PLSVASTLVEPP | 1488486 | Cell/Virus Enclosed | NO |
| PLSVASTPVEPP | 1488487 | Cell/Virus Enclosed | NO |
| PLSVASTPVEPS | 1488488 | Cell/Virus Enclosed | NO |
| PLSVASTPVESP | 1488489 | Cell/Virus Enclosed | NO |
| PLSVEPTPIEDA | 1488491 | Cell/Virus Enclosed | NO |
| PLSVESTPVELP | 1488492 | Cell/Virus Enclosed | NO |
| PLSVESTPVEPP | 1488493 | Cell/Virus Enclosed | NO |
| PLSVESTPVEPS | 1488494 | Cell/Virus Enclosed | NO |
| PLSVQQAQQEATLPDI | 1685971 | Cell/Virus Enclosed | NO |
| PLTAPTLVPQEH | 1488503 | Cell/Virus Enclosed | NO |
| PLTCISRLKDGF | 1488516 | Ectodomain | YES |
| PLTCKAISSNTD | 1488520 | Ectodomain | YES |
| PLTCNAISPKTD | 1488522 | Ectodomain | YES |
| PLTCNAISSNTDNETL | 1685973 | Ectodomain | YES |
| PLTCNSAMTLEY | 1488524 | Ectodomain | YES |
| PLTCNSKVKNGF | 1488528 | Ectodomain | YES |
| PLTFPPGQGVPL | 1488533 | Cell/Virus Enclosed | NO |
| PLTFPSGQGVPL | 1488534 | Cell/Virus Enclosed | NO |
| PLTFSPGQGVPL | 1488535 | Cell/Virus Enclosed | NO |
| PLTKKKYSELDD | 1488543 | Cell/Virus Enclosed | NO |
| PLTKNIGDYNVSKDGF | 1685986 | Cell/Virus Enclosed | NO |
| PLTNVVSHVNGD | 1488549 | Ectodomain | YES |
| PLTPILSCGIFGIKLE | 1685988 | Cell/Virus Enclosed | NO |
| PLTQDHVEILGP | 1488557 | Cell/Virus Enclosed | NO |
| PLTVGGSCLLSG | 1488571 | Cell/Virus Enclosed | NO |
| PLVDFNVNVMGT | 1488576 | Cell/Virus Enclosed | NO |
| PLVEDSTSVTAV | 1488579 | Ectodomain | YES |
| PLVQPVGALDAS | 1488600 | Cell/Virus Enclosed | NO |
| PLVQPVGALDVS | 1488601 | Cell/Virus Enclosed | NO |
| PLVYFSDSKPST | 1488611 | Cell/Virus Enclosed | NO |
| PLVYHENEEYQKVFRV | 1686024 | Cell/Virus Enclosed | NO |
| PLVYLNHSVPKLSDFE | 1686025 | Ectodomain | YES |

| | | | |
|---|---|---|---|
| PLWFCLLYIAVV | 1488613 | Cell/Virus Enclosed | NO |
| PLYGFPSQAAGV | 1488625 | Cell/Virus Enclosed | NO |
| PLYPYGRCPTAADINI | 1686030 | Ectodomain | YES |
| PMACNGHKIEDL | 1488635 | Cell/Virus Enclosed | NO |
| PMDTFIVSTNPNTVTSSTPI | 110685 | Capsid | UN |
| PMEELKALVATA | 1488650 | Cell/Virus Enclosed | NO |
| PMEKLKALVATA | 1488651 | Cell/Virus Enclosed | NO |
| PMEQRVQGSITQ | 1488654 | Cell/Virus Enclosed | NO |
| PMERVSLWNYGK | 1488655 | Cell/Virus Enclosed | NO |
| PMEYVSYFDGVK | 1488656 | Cell/Virus Enclosed | NO |
| PMGTQTHTMIFDNAFNC | 48554 | Ectodomain | YES |
| PMGTRTHTMIFD | 1488663 | Ectodomain | YES |
| PMGVYDFKVSTQ | 1488664 | Cell/Virus Enclosed | NO |
| PMIANENAINKGGRTV | 1686047 | Cell/Virus Enclosed | NO |
| PMKAEGDGGVLG | 1488667 | Cell/Virus Enclosed | NO |
| PMLMGWDYPKCDRALP | 1686051 | Cell/Virus Enclosed | NO |
| PMLPKNMWEFRD | 1488672 | Cell/Virus Enclosed | NO |
| PMMQSADASTFL | 1488680 | Cell/Virus Enclosed | NO |
| PMSDMEKDFLSF | 1488696 | Cell/Virus Enclosed | NO |
| PMSDMENDFLNF | 1488697 | Cell/Virus Enclosed | NO |
| PMYTVSKGTQQS | 1488714 | Ectodomain | YES |
| PNAKEEILGTVS | 1488722 | Cell/Virus Enclosed | NO |
| PNALLCATCPFAKGDS | 1686067 | Cell/Virus Enclosed | NO |
| PNARLCATCPFA | 1488727 | Cell/Virus Enclosed | NO |
| PNCNIEAWLNGK | 1488737 | Ectodomain | YES |
| PNCNIESWLNDK | 1488738 | Ectodomain | YES |
| PNCYWPLNDYGF | 1488743 | Ectodomain | YES |
| PNCYWPLNGYGF | 1488744 | Ectodomain | YES |
| PNDDATSNVTQC | 1488746 | Cell/Virus Enclosed | NO |
| PNDDVTSNVTQCFGPR | 1686082 | Capsid | UN |
| PNDFSFNNWFLL | 1488750 | Ectodomain | YES |
| PNDFVGGFRVLG | 1488752 | Cell/Virus Enclosed | NO |
| PNDKEEILGTVS | 1488755 | Cell/Virus Enclosed | NO |
| PNDTVTFSFNGAFIAPDRASFLRGK | 1163297 | Ectodomain | YES |
| PNEHVTSVNLLD | 1488761 | Cell/Virus Enclosed | NO |
| PNEITVAKPNVL | 1488762 | Underneath Membrane | NO |
| PNEKLVTTFTAP | 1488764 | Cell/Virus Enclosed | NO |
| PNEKLVTTSMAP | 1488765 | Cell/Virus Enclosed | NO |
| PNEKLVTTSTAP | 1488766 | Cell/Virus Enclosed | NO |
| PNEQRVQGSITQ | 1488773 | Capsid | UN |
| PNEVTVAKPNVV | 1488776 | Ectodomain | YES |
| PNFDLTPFNLTY | 1488779 | Ectodomain | YES |
| PNFGSLTHINTT | 1488782 | Ectodomain | YES |
| PNGRRVDLQLGN | 1488810 | Ectodomain | YES |
| PNGVNVAEEPDS | 1488813 | Cell/Virus Enclosed | NO |
| PNGWTMGLFRRR | 1488817 | Cell/Virus Enclosed | NO |
| PNHAALFFDSEV | 1488818 | Cell/Virus Enclosed | NO |
| PNHFINFPLSHF | 1488819 | Cell/Virus Enclosed | NO |
| PNHTIKGAFLNG | 1488823 | Cell/Virus Enclosed | NO |
| PNHTIKGSALNG | 1488824 | Cell/Virus Enclosed | NO |
| PNIIKVFYELDN | 1488826 | Cell/Virus Enclosed | NO |
| PNITNRCPFDKV | 1488835 | Ectodomain | YES |
| PNKFCNVQQCFG | 1488840 | Cell/Virus Enclosed | NO |
| PNKHCNVQQCFGKRGP | 1686117 | Capsid | UN |
| PNKPLSQPRADKPSQL | 1686119 | Capsid | UN |
| PNKYCYCGNFAT | 1488849 | Cell/Virus Enclosed | NO |
| PNLGNHRTELWH | 1488853 | Ectodomain | YES |
| PNLGNRRVELWH | 1488854 | Ectodomain | YES |
| PNLHDFKEELDQ | 1488855 | Ectodomain | YES |
| PNLNAGEDIRLF | 1488858 | Cell/Virus Enclosed | NO |
| PNLNAGEDIRLL | 1488859 | Cell/Virus Enclosed | NO |
| PNLNAGEDVQLL | 1488860 | Cell/Virus Enclosed | NO |
| PNLPDFKEELGQ | 1488864 | Ectodomain | YES |
| PNLPDFKEESDQ | 1488865 | Ectodomain | YES |
| PNLPDFKGELDQ | 1488866 | Ectodomain | YES |
| PNLPDFREELDQ | 1488867 | Ectodomain | YES |
| PNLPGFKEELDQ | 1488868 | Ectodomain | YES |
| PNLYVVVVESDADRAN | 1686135 | Cell/Virus Enclosed | NO |
| PNMENITSGFLGPLL | 1686137 | Ectodomain | YES |
| PNMGVPCCTSYY | 1488891 | Cell/Virus Enclosed | NO |
| PNMHVVPLENDA | 1488892 | Cell/Virus Enclosed | NO |
| PNMWCLRSGHHL | 1488900 | Cell/Virus Enclosed | NO |
| PNMWYLRSGYHL | 1488903 | Cell/Virus Enclosed | NO |
| PNNDSAIVTQFAPGTK | 1686148 | Cell/Virus Enclosed | NO |
| PNNNKILVPKVSGLQYRVFR | 110965 | Capsid | UN |
| PNNNSAQYVFSTVNAL | 1686151 | Cell/Virus Enclosed | NO |

| | | | |
|---|---|---|---|
| PNNRSLSRSNSNFRHS | 1686152 | Capsid | UN |
| PNQAALFFDSEN | 1488931 | Cell/Virus Enclosed | NO |
| PNQTATSQQPSG | 1488939 | Cell/Virus Enclosed | NO |
| PNQTSVVPPALH | 1488941 | Cell/Virus Enclosed | NO |
| PNRRKVDLQLGN | 1488944 | Ectodomain | YES |
| PNRRRVDLQLGN | 1488946 | Ectodomain | YES |
| PNRTIKGSFLNG | 1488947 | Cell/Virus Enclosed | NO |
| PNSIYICSQGKD | 1488951 | Cell/Virus Enclosed | NO |
| PNSNYVLQDFDV | 1488953 | Cell/Virus Enclosed | NO |
| PNSRRSDLQLAS | 1488956 | Ectodomain | YES |
| PNSRSTSRTSSRASSA | 1686170 | Cell/Virus Enclosed | NO |
| PNSSEVDMICST | 1488959 | Cell/Virus Enclosed | NO |
| PNTRSLSRSNSN | 1488973 | Cell/Virus Enclosed | NO |
| PNTVDCDNDRCIIHCA | 1686177 | Cell/Virus Enclosed | NO |
| PNTVDCQDDRCF | 1488976 | Cell/Virus Enclosed | NO |
| PNTVDCQDDRCIIHCA | 1686178 | Cell/Virus Enclosed | NO |
| PNTWYLRSGHHL | 1488980 | Cell/Virus Enclosed | NO |
| PNVPIEYQATRV | 1488988 | Ectodomain | YES |
| PNVQIHTIDGSSGVAN | 1686190 | Secreted | YES |
| PNYEDLLIRKAN | 1489000 | Cell/Virus Enclosed | NO |
| PNYTVPDLVVEQYNQT | 1686199 | Ectodomain | YES |
| PPAAPRAVSFAD | 1489011 | Cell/Virus Enclosed | NO |
| PPAEYKLQQGTF | 1489014 | Cell/Virus Enclosed | NO |
| PPAIASNCYSSL | 1489018 | Ectodomain | YES |
| PPALHLVNPQIQ | 1489021 | Cell/Virus Enclosed | NO |
| PPANLRVVGHAM | 1489027 | Cell/Virus Enclosed | NO |
| PPAPNCYWPLND | 1489029 | Ectodomain | YES |
| PPAPNCYWPLNG | 1489030 | Ectodomain | YES |
| PPARFVIDTPTG | 1489033 | Cell/Virus Enclosed | NO |
| PPCKFTVQDAKGLKIK | 1686211 | Cell/Virus Enclosed | NO |
| PPDAIIIDNDINDYVS | 1686215 | Cell/Virus Enclosed | NO |
| PPDDEDECDEYE | 1489042 | Cell/Virus Enclosed | NO |
| PPDEEEDCEEYE | 1489044 | Cell/Virus Enclosed | NO |
| PPDQLVVYQQAI | 1489047 | Ectodomain | YES |
| PPDQLVVYQQAV | 1489048 | Ectodomain | YES |
| PPFLSDFINGIF | 1489054 | Ectodomain | YES |
| PPGDQFKHLIPL | 1489056 | Cell/Virus Enclosed | NO |
| PPGEHFKHLVPL | 1489057 | Cell/Virus Enclosed | NO |
| PPGMGIPYCTSY | 1489060 | Cell/Virus Enclosed | NO |
| PPGQGVPLNANSTPAQ | 1686231 | Cell/Virus Enclosed | NO |
| PPGQGVPLNSNS | 1489062 | Cell/Virus Enclosed | NO |
| PPGTGKSHFAIG | 1489063 | Cell/Virus Enclosed | NO |
| PPHDVIVLRDNL | 1489066 | Cell/Virus Enclosed | NO |
| PPHDVTVLRDNLKPVT | 1686241 | Cell/Virus Enclosed | NO |
| PPHGGLLGWSPQAQG | 1686242 | Underneath Membrane | NO |
| PPHRPGVRMRRVPVTHP | 48799 | Cell/Virus Enclosed | NO |
| PPIAPGVPATEA | 1489076 | Cell/Virus Enclosed | NO |
| PPIELENKFTPL | 1489078 | Cell/Virus Enclosed | NO |
| PPILSESQISGYTTAA | 1686245 | Ectodomain | YES |
| PPILSETQISGY | 1489082 | Ectodomain | YES |
| PPKEQRVQGIIT | 1489093 | Cell/Virus Enclosed | NO |
| PPKEQRVQGSITQRTR | 1686249 | Cell/Virus Enclosed | NO |
| PPKEQRVQSSIT | 1489096 | Cell/Virus Enclosed | NO |
| PPKSSIDAFKLT | 1489101 | Cell/Virus Enclosed | NO |
| PPKTEHPKIEEK | 1489103 | Cell/Virus Enclosed | NO |
| PPKTGPPKTSSEPVRCNRHD | 741823 | Ectodomain | YES |
| PPLLGNSIGIDF | 1489110 | Ectodomain | YES |
| PPLLGNSTRIDF | 1489111 | Ectodomain | YES |
| PPLLSDNQISGY | 1489113 | Ectodomain | YES |
| PPLLSENQFSGY | 1489114 | Ectodomain | YES |
| PPLLSENQINGY | 1489115 | Ectodomain | YES |
| PPLLSENQISGYTLAA | 1686259 | Ectodomain | YES |
| PPLLSNSTGIDF | 1489118 | Ectodomain | YES |
| PPLLSVNQISGY | 1489120 | Ectodomain | YES |
| PPLLTDDMIAAYTAAL | 1686260 | Ectodomain | YES |
| PPLYVPIRLAGH | 1489134 | Cell/Virus Enclosed | NO |
| PPMEQRVQGSIT | 1489135 | Cell/Virus Enclosed | NO |
| PPMPSIGLEEEEEEEGAGDG | 741824 | Ectodomain | YES |
| PPNCNIEAWLND | 1489140 | Underneath Membrane | NO |
| PPNEQRVQGSIT | 1489141 | Capsid | UN |
| PPNSTRSASESRHRLTVAQV | 741825 | Undetermined | UN |
| PPNVQIHTIDGSSGVA | 1686274 | Secreted | YES |
| PPPGEHFKHLVP | 1489150 | Cell/Virus Enclosed | NO |
| PPPGEQFKHLVPLMHK | 1686276 | Cell/Virus Enclosed | NO |
| PPPLLGNSIGID | 1489155 | Ectodomain | YES |
| PPPLLGNSTRID | 1489157 | Ectodomain | YES |

| | | | |
|---|---|---|---|
| PPPLLSNSTGID | 1489161 | Ectodomain | YES |
| PPPQIFLKILPQSGPIGGIK | 48879 | Capsid | UN |
| PPPSFYMPLLVS | 1489163 | Cell/Virus Enclosed | NO |
| PPQLVELPSEPQ | 1489168 | Cell/Virus Enclosed | NO |
| PPQVMKLPSEPQ | 1489172 | Cell/Virus Enclosed | NO |
| PPQVVELPSEPQ | 1489174 | Cell/Virus Enclosed | NO |
| PPQVVELPYEPR | 1489175 | Cell/Virus Enclosed | NO |
| PPQVVKLPFEPQ | 1489176 | Cell/Virus Enclosed | NO |
| PPQVVKLPSAPQTIQP | 1686281 | Cell/Virus Enclosed | NO |
| PPQVVKLPSEPQ | 1489178 | Cell/Virus Enclosed | NO |
| PPQVVKLPSEPR | 1489179 | Cell/Virus Enclosed | NO |
| PPQVVKLQFEPQ | 1489180 | Cell/Virus Enclosed | NO |
| PPRAELNGVMVD | 1489185 | Cell/Virus Enclosed | NO |
| PPRAELNGVVVDKVGD | 1686283 | Cell/Virus Enclosed | NO |
| PPSASCPIGTKY | 1489190 | Ectodomain | YES |
| PPSASCPIGTNYRSCE | 1686285 | Ectodomain | YES |
| PPSEYTLQQGAF | 1489195 | Cell/Virus Enclosed | NO |
| PPSSSGTSTPKK | 1489200 | Cell/Virus Enclosed | NO |
| PPTCVYELNGYT | 1489202 | Ectodomain | YES |
| PPTEPKKDKKKKADETQALPQR | 1870805 | Capsid | UN |
| PPTEPKKDKKKKTDES | 1686292 | Cell/Virus Enclosed | NO |
| PPTGVYELHGYT | 1489206 | Ectodomain | YES |
| PPTGVYELNGYT | 1489208 | Ectodomain | YES |
| PPTNVVSHLNGD | 1489221 | Ectodomain | YES |
| PPTNVVSHVNGD | 1489222 | Ectodomain | YES |
| PPVIDDGDGVKP | 1489232 | Cell/Virus Enclosed | NO |
| PPVIDDVDVVKP | 1489233 | Cell/Virus Enclosed | NO |
| PPVPSPATTKARKRKTKKPP | 741832 | Ectodomain | YES |
| PPVTYLPATILA | 1489236 | Cell/Virus Enclosed | NO |
| PPVTYSTATILA | 1489239 | Cell/Virus Enclosed | NO |
| PPWPQIAELVPT | 1489243 | Cell/Virus Enclosed | NO |
| PPWSAAAGVPFS | 1489245 | Ectodomain | YES |
| PPWTAAAGVPFYLNVQ | 1686306 | Ectodomain | YES |
| PQAKSGIITKLE | 1489264 | Ectodomain | YES |
| PQAKSLVGIGEH | 1489267 | Ectodomain | YES |
| PQANSGDYYHKV | 1489269 | Ectodomain | YES |
| PQANSGDYYYKV | 1489270 | Ectodomain | YES |
| PQANSGIITKVE | 1489272 | Ectodomain | YES |
| PQASTTPFPTGDPKPQPPGV | 741833 | Ectodomain | YES |
| PQATYQNISTNL | 1489273 | Ectodomain | YES |
| PQAVETKVQCST | 1489274 | Cell/Virus Enclosed | NO |
| PQAVETRVQCST | 1489275 | Cell/Virus Enclosed | NO |
| PQCAIVQALESK | 1489276 | Cell/Virus Enclosed | NO |
| PQCAIVQASESK | 1489277 | Cell/Virus Enclosed | NO |
| PQCAIVQATESK | 1489278 | Cell/Virus Enclosed | NO |
| PQECNDMHLSTL | 1489279 | Cell/Virus Enclosed | NO |
| PQECNNMHLSTL | 1489280 | Cell/Virus Enclosed | NO |
| PQEKDKKSAKTGTPKP | 1686319 | Capsid | UN |
| PQENTIPHYSWF | 1489285 | Cell/Virus Enclosed | NO |
| PQFAELIPNQAA | 1489288 | Cell/Virus Enclosed | NO |
| PQFPILAELAPP | 1489292 | Cell/Virus Enclosed | NO |
| PQFTSDQAITFL | 1489295 | Underneath Membrane | NO |
| PQFTSDQAVTFLKEWN | 1686326 | Undetermined | UN |
| PQFYIYDEEGGYDVSK | 1686327 | Cell/Virus Enclosed | NO |
| PQGAFHVTMRSS | 1489299 | Cell/Virus Enclosed | NO |
| PQGGYVADFAYW | 1489302 | Cell/Virus Enclosed | NO |
| PQGNLGGNCAGNVIGN | 1686333 | Cell/Virus Enclosed | NO |
| PQGNLGSNGKPGNVGG | 1686334 | Cell/Virus Enclosed | NO |
| PQGNSIPHYSWF | 1489312 | Cell/Virus Enclosed | NO |
| PQGRTYSNITIT | 1489316 | Ectodomain | YES |
| PQGTALPKGFYAEGSR | 1686337 | Cell/Virus Enclosed | NO |
| PQGYYIEGSGKS | 1489320 | Cell/Virus Enclosed | NO |
| PQGYYIEGSGRP | 1489322 | Capsid | UN |
| PQGYYIEGSGRSAPNS | 1686338 | Cell/Virus Enclosed | NO |
| PQGYYVEGSGRS | 1489325 | Cell/Virus Enclosed | NO |
| PQIAELVPTASA | 1489329 | Cell/Virus Enclosed | NO |
| PQIITTDNTFVA | 1489334 | Ectodomain | YES |
| PQIQLTTTRMED | 1489342 | Cell/Virus Enclosed | NO |
| PQKAYLDFVNAGGAPL | 1686351 | Cell/Virus Enclosed | NO |
| PQLSFPLCSAGN | 1489365 | Cell/Virus Enclosed | NO |
| PQLTYQNISTNL | 1489371 | Ectodomain | YES |
| PQLVELPSEPQT | 1489372 | Cell/Virus Enclosed | NO |
| PQLVNANHYSPC | 1489373 | Ectodomain | YES |
| PQLVNSNQYSPC | 1489375 | Ectodomain | YES |
| PQNQRNAPRITFGGPS | 1686370 | Cell/Virus Enclosed | NO |
| PQNQSSAPRITF | 1489381 | Cell/Virus Enclosed | NO |

| | | | |
|---|---|---|---|
| PQPAGVFTDHDV | 1489382 | Ectodomain | YES |
| PQPAGVFTNHDV | 1489383 | Ectodomain | YES |
| PQPAGVLTNHDV | 1489384 | Ectodomain | YES |
| PQPKFTVSTQPQ | 1489385 | Cell/Virus Enclosed | NO |
| PQPLMDKFSSSR | 1489387 | Ectodomain | YES |
| PQPTGVFTNHSV | 1489391 | Ectodomain | YES |
| PQPVEGVSSFMN | 1489392 | Ectodomain | YES |
| PQPVGVFTHHDV | 1489393 | Ectodomain | YES |
| PQPYCYTDGLMQ | 1489394 | Cell/Virus Enclosed | NO |
| PQPYCYTEGLMQ | 1489395 | Cell/Virus Enclosed | NO |
| PQQMFVTPLAVD | 1489398 | Cell/Virus Enclosed | NO |
| PQQMFVTPLSVD | 1489399 | Cell/Virus Enclosed | NO |
| PQQMFVTPLTVD | 1489400 | Cell/Virus Enclosed | NO |
| PQQNQRSAPRIT | 1489402 | Capsid | UN |
| PQQRFVYDAYQN | 1489403 | Ectodomain | YES |
| PQQVTKQNAKEI | 1489406 | Cell/Virus Enclosed | NO |
| PQRADLNGVIVDKVGE | 1686379 | Cell/Virus Enclosed | NO |
| PQRKRGVKQLPEQFDS | 1686381 | Capsid | UN |
| PQRKRGVKQSPE | 1489414 | Cell/Virus Enclosed | NO |
| PQRQKKQHTVTL | 1489418 | Capsid | UN |
| PQRQRGHKNGQE | 1489423 | Capsid | UN |
| PQRQRGHKNGQGENDN | 1686385 | Cell/Virus Enclosed | NO |
| PQRQRGHKNGQV | 1489425 | Cell/Virus Enclosed | NO |
| PQRQRGHKNVQG | 1489426 | Cell/Virus Enclosed | NO |
| PQRQRGHKYGQG | 1489428 | Cell/Virus Enclosed | NO |
| PQRQRGHRNGQG | 1489429 | Cell/Virus Enclosed | NO |
| PQRQRGQKNGQG | 1489432 | Capsid | UN |
| PQSAVDSAARIHDFRYSQLA | 49120 | Capsid | UN |
| PQSGGGKCFNCA | 1489436 | Ectodomain | YES |
| PQSGGGKCFNCYPAGV | 1686388 | Ectodomain | YES |
| PQSGGGKCYNCR | 1489438 | Ectodomain | YES |
| PQSGGGRCYNCR | 1489439 | Ectodomain | YES |
| PQSKDSNFLNESGVLL | 1686389 | Cell/Virus Enclosed | NO |
| PQSKDSNFLNRVRGSI | 1686390 | Cell/Virus Enclosed | NO |
| PQSKESNFLNES | 1489444 | Cell/Virus Enclosed | NO |
| PQSVETKVQCST | 1489452 | Cell/Virus Enclosed | NO |
| PQSVREAFNNLNSTT | 227458 | Cell/Virus Enclosed | NO |
| PQSYCYTDGLMQ | 1489453 | Cell/Virus Enclosed | NO |
| PQTGNFKDLREY | 1489454 | Ectodomain | YES |
| PQTICHPKLGNH | 1489457 | Ectodomain | YES |
| PQTICHPNLGNH | 1489458 | Ectodomain | YES |
| PQTIKPEVKEVA | 1489459 | Cell/Virus Enclosed | NO |
| PQTIQPEVKEIA | 1489460 | Cell/Virus Enclosed | NO |
| PQTKYLVGIGEH | 1489464 | Ectodomain | YES |
| PQVMKLPSEPQT | 1489481 | Cell/Virus Enclosed | NO |
| PQVTYQNISTNLPPPL | 1686403 | Ectodomain | YES |
| PQVVELLSEPQT | 1489489 | Cell/Virus Enclosed | NO |
| PQVVELPSEPQT | 1489490 | Cell/Virus Enclosed | NO |
| PQVVELPYEPRT | 1489491 | Cell/Virus Enclosed | NO |
| PQVVKLPFEPQT | 1489492 | Cell/Virus Enclosed | NO |
| PQVVKLPSAPQTIQPE | 1686404 | Cell/Virus Enclosed | NO |
| PQVVKLPSEPQT | 1489494 | Cell/Virus Enclosed | NO |
| PQVVKLPSEPRT | 1489495 | Cell/Virus Enclosed | NO |
| PQVVKLQFEPQT | 1489496 | Cell/Virus Enclosed | NO |
| PQVYNFKRLVFT | 1489499 | Ectodomain | YES |
| PQXRAPRITFGGPSDS | 1686406 | Cell/Virus Enclosed | NO |
| PRAAPNNTVSWY | 1489502 | Cell/Virus Enclosed | NO |
| PRAAPNTTVSWY | 1489503 | Cell/Virus Enclosed | NO |
| PRAELNGVMVDK | 1489507 | Cell/Virus Enclosed | NO |
| PRAELNGVVVDKVGDT | 1686411 | Cell/Virus Enclosed | NO |
| PRAGKHEHSLLV | 1489508 | Cell/Virus Enclosed | NO |
| PRAISFADNNDI | 1489509 | Cell/Virus Enclosed | NO |
| PRANSGIITKVE | 1489517 | Ectodomain | YES |
| PRASADIGSGHT | 1489520 | Cell/Virus Enclosed | NO |
| PRASANIGASHT | 1489521 | Cell/Virus Enclosed | NO |
| PRAVSFADNNDITNTN | 1686415 | Cell/Virus Enclosed | NO |
| PRAVSFADNNDT | 1489527 | Cell/Virus Enclosed | NO |
| PRCPQIAELVPT | 1489535 | Cell/Virus Enclosed | NO |
| PRDFNHNMGDSD | 1489536 | Cell/Virus Enclosed | NO |
| PRDLDHNFGSAV | 1489538 | Cell/Virus Enclosed | NO |
| PREGVFVSNGTS | 1489546 | Ectodomain | YES |
| PRGIACVKDFDF | 1489555 | Cell/Virus Enclosed | NO |
| PRGQGVPINTNSSPDD | 1686429 | Cell/Virus Enclosed | NO |
| PRGQGVPINTNSTKDD | 1686430 | Cell/Virus Enclosed | NO |
| PRGRGRNPKPRA | 1489559 | Cell/Virus Enclosed | NO |
| PRGVGSANIFIG | 1489568 | Cell/Virus Enclosed | NO |

| | | | |
|---|---|---|---|
| PRHVICSSSNMNEPDY | 1686437 | Cell/Virus Enclosed | NO |
| PRHVMCPADQLS | 1489579 | Cell/Virus Enclosed | NO |
| PRITFGEPTDST | 1489587 | Capsid | UN |
| PRITFGGPSDSS | 1489589 | Cell/Virus Enclosed | NO |
| PRITFGGPSDSTDNNQ | 1686444 | Cell/Virus Enclosed | NO |
| PRITFGGPTDSA | 1489591 | Capsid | UN |
| PRIVQMFADHLT | 1489595 | Cell/Virus Enclosed | NO |
| PRLKGSFNNRDT | 1489605 | Ectodomain | YES |
| PRLPVLSDFVQI | 1489606 | Ectodomain | YES |
| PRLQAASDWKPGYSMP | 1686451 | Cell/Virus Enclosed | NO |
| PRLQASADWKPGHAMP | 1686452 | Cell/Virus Enclosed | NO |
| PRNKWNGVSDLS | 1489621 | Cell/Virus Enclosed | NO |
| PRNKWSGISDLS | 1489623 | Cell/Virus Enclosed | NO |
| PRNKWSGVSDLS | 1489624 | Cell/Virus Enclosed | NO |
| PRNLVPIGKGSK | 1489629 | Cell/Virus Enclosed | NO |
| PRNLVPINKKDK | 1489630 | Cell/Virus Enclosed | NO |
| PRNSSDNGISAIITAN | 1686458 | Ectodomain | YES |
| PRNSWEAMALNFKLIG | 1686459 | Cell/Virus Enclosed | NO |
| PRNVLYDSVSAI | 1489639 | Ectodomain | YES |
| PRNVSYDSVSAI | 1489640 | Ectodomain | YES |
| PRNVSYDSVSAT | 1489641 | Ectodomain | YES |
| PRNYSYMDLNPA | 1489642 | Ectodomain | YES |
| PRPAGVLTNHDV | 1489643 | Ectodomain | YES |
| PRPGNHCPAGNS | 1489648 | Ectodomain | YES |
| PRPIDVFKADGI | 1489651 | Ectodomain | YES |
| PRPIDVSKADGIIYPQ | 1686463 | Ectodomain | YES |
| PRQKRTATKAYNVTQA | 1686466 | Cell/Virus Enclosed | NO |
| PRQKRTATKQYNVTQA | 1686467 | Cell/Virus Enclosed | NO |
| PRQKRTPNKHCN | 1489664 | Cell/Virus Enclosed | NO |
| PRSDMECDFLNM | 1489674 | Cell/Virus Enclosed | NO |
| PRSDMEYDFLNI | 1489675 | Cell/Virus Enclosed | NO |
| PRSDMEYDFLNMDMGV | 1686473 | Cell/Virus Enclosed | NO |
| PRSGNHCPADNS | 1489678 | Ectodomain | YES |
| PRSGNHCPAGNF | 1489679 | Ectodomain | YES |
| PRSGNHCPAGNS | 1489680 | Ectodomain | YES |
| PRSGNHCPAGNY | 1489681 | Ectodomain | YES |
| PRSGNYCPAGNS | 1489683 | Ectodomain | YES |
| PRSQMETDFLEP | 1489691 | Cell/Virus Enclosed | NO |
| PRSVRSVPGEMC | 1489694 | Ectodomain | YES |
| PRTGKHEHSLLVEAYN | 1686481 | Cell/Virus Enclosed | NO |
| PRTIDSTQDGVN | 1489702 | Ectodomain | YES |
| PRTIKVSTVQSKLTDL | 1686482 | Cell/Virus Enclosed | NO |
| PRTINATQDGVN | 1489704 | Ectodomain | YES |
| PRTINSQDGVNK | 1489707 | Ectodomain | YES |
| PRTINSTQDGDNKLQG | 1686483 | Ectodomain | YES |
| PRTINSTQDGVN | 1489709 | Ectodomain | YES |
| PRTINSTQDGYN | 1489710 | Ectodomain | YES |
| PRTIQPEVKEVA | 1489713 | Cell/Virus Enclosed | NO |
| PRVFSAVGNICY | 1489725 | Cell/Virus Enclosed | NO |
| PRWKRQPNDDAT | 1489734 | Cell/Virus Enclosed | NO |
| PRWKRQPNDDVTSNVT | 1686497 | Capsid | UN |
| PRWPQIAELVPT | 1489740 | Cell/Virus Enclosed | NO |
| PSAELTGTQSVS | 1489753 | Cell/Virus Enclosed | NO |
| PSAKVEGCMVQV | 1489757 | Cell/Virus Enclosed | NO |
| PSALEFNPSQTSPATA | 1686508 | Capsid | UN |
| PSASAFFGMSRIGMEV | 1686511 | Cell/Virus Enclosed | NO |
| PSASCPIGTKYR | 1489768 | Ectodomain | YES |
| PSASCPIGTNYRSCES | 1686512 | Ectodomain | YES |
| PSASVVEQAEGV | 1489770 | Ectodomain | YES |
| PSATYEGNSPFH | 1489771 | Ectodomain | YES |
| PSCCCTKPSDGNCTC | 1686514 | Underneath Membrane | NO |
| PSCFKKCGGCCD | 1489775 | Underneath Membrane | NO |
| PSDDRTEVPQLV | 1489780 | Ectodomain | YES |
| PSDDTLRSEAFE | 1489781 | Cell/Virus Enclosed | NO |
| PSDEGVKVTQEL | 1489783 | Cell/Virus Enclosed | NO |
| PSDFAFNNWFLLTNTS | 1686518 | Ectodomain | YES |
| PSDFFPSIRDLLDTA | 1686519 | Capsid | UN |
| PSDLNTSELLTK | 1489789 | Cell/Virus Enclosed | NO |
| PSDSCNNVNVRD | 1489791 | Cell/Virus Enclosed | NO |
| PSDSSDNSKNGE | 1489793 | Cell/Virus Enclosed | NO |
| PSDSSDNSQNGE | 1489794 | Cell/Virus Enclosed | NO |
| PSDSTDNNQDGG | 1489795 | Cell/Virus Enclosed | NO |
| PSDSTDNNQNGDRSGA | 1686524 | Cell/Virus Enclosed | NO |
| PSDVSARDPIIQ | 1489797 | Cell/Virus Enclosed | NO |
| PSDVSARDPTIQ | 1489798 | Cell/Virus Enclosed | NO |
| PSDVSLTYLLGTAKKQ | 1686527 | Cell/Virus Enclosed | NO |

| | | | |
|---|---|---|---|
| PSEAKGYWYKHN | 1489802 | Cell/Virus Enclosed | NO |
| PSEAKGYWYRHN | 1489803 | Cell/Virus Enclosed | NO |
| PSEAPNAKEEIL | 1489805 | Cell/Virus Enclosed | NO |
| PSECTMFMDAEG | 1489809 | Cell/Virus Enclosed | NO |
| PSECTMFRDAEG | 1489810 | Cell/Virus Enclosed | NO |
| PSEDFIKHIRTN | 1489811 | Cell/Virus Enclosed | NO |
| PSEDFIKHVRNN | 1489812 | Cell/Virus Enclosed | NO |
| PSEDFIKHVRTNGGYN | 1686536 | Cell/Virus Enclosed | NO |
| PSEEFIKHVRTN | 1489814 | Cell/Virus Enclosed | NO |
| PSEEGVKVTQEL | 1489815 | Cell/Virus Enclosed | NO |
| PSEFTIGNIEEF | 1489816 | Ectodomain | YES |
| PSEFTIGNMEEF | 1489817 | Ectodomain | YES |
| PSEFTIGNMVEF | 1489818 | Ectodomain | YES |
| PSEKDKHLDNNVQRCT | 1686540 | Cell/Virus Enclosed | NO |
| PSEPQTIQPEVK | 1489821 | Cell/Virus Enclosed | NO |
| PSEPRTIQPEVK | 1489822 | Cell/Virus Enclosed | NO |
| PSETPNAKEEIL | 1489823 | Cell/Virus Enclosed | NO |
| PSEYTLQQGAFL | 1489824 | Cell/Virus Enclosed | NO |
| PSFASSCKSHKPPSAS | 1686541 | Ectodomain | YES |
| PSFKDNVILLNKHIDA | 1686542 | Cell/Virus Enclosed | NO |
| PSFLAVDDKYKATGDL | 1686545 | Cell/Virus Enclosed | NO |
| PSFLAVDDKYKVNENL | 1686546 | Cell/Virus Enclosed | NO |
| PSFLVVDDKYKV | 1489829 | Cell/Virus Enclosed | NO |
| PSFNVSDAYANLVPYY | 1686548 | Cell/Virus Enclosed | NO |
| PSFPTQRTSKTLKVLTPPTT | 49366 | Cell/Virus Enclosed | NO |
| PSFSSSRRKRRS | 1489830 | Ectodomain | YES |
| PSFVAYETARQEYENA | 1686549 | Cell/Virus Enclosed | NO |
| PSGDVEACMVQV | 1489838 | Cell/Virus Enclosed | NO |
| PSGGNVIPHYSW | 1489841 | Cell/Virus Enclosed | NO |
| PSGGNVVPHYSW | 1489842 | Cell/Virus Enclosed | NO |
| PSGIGAVGGDLLYLDL | 1686558 | Cell/Virus Enclosed | NO |
| PSGIGGGDLLYL | 1489848 | Cell/Virus Enclosed | NO |
| PSGIGTVGGDLL | 1489849 | Cell/Virus Enclosed | NO |
| PSGLKTMIVSAG | 1489852 | Cell/Virus Enclosed | NO |
| PSGNLIKFDIDS | 1489856 | Cell/Virus Enclosed | NO |
| PSGNYLAISNEL | 1489858 | Ectodomain | YES |
| PSGNYLSISNEL | 1489860 | Ectodomain | YES |
| PSGQGVPLNANS | 1489862 | Cell/Virus Enclosed | NO |
| PSGSFVEQAEGV | 1489864 | Ectodomain | YES |
| PSGSILVDNDLN | 1489865 | Cell/Virus Enclosed | NO |
| PSGSVVEQADGV | 1489866 | Ectodomain | YES |
| PSGSVVEQAEGF | 1489867 | Ectodomain | YES |
| PSGSVVEQAEGV | 1489868 | Ectodomain | YES |
| PSGTWLTYTGAIKLD | 1310726 | Capsid | UN |
| PSGVGAPGGDLL | 1489876 | Cell/Virus Enclosed | NO |
| PSHAVQNTVLNA | 1489879 | Cell/Virus Enclosed | NO |
| PSHDVTVLRDNL | 1489883 | Cell/Virus Enclosed | NO |
| PSHEAPNMTQTGT | 49395 | Ectodomain | YES |
| PSHVAQNTVLNA | 1489885 | Cell/Virus Enclosed | NO |
| PSIIHDIGNPKG | 1489888 | Cell/Virus Enclosed | NO |
| PSILKVIDDGKI | 1489891 | Cell/Virus Enclosed | NO |
| PSINDLSFDKNLGRCV | 1686579 | Cell/Virus Enclosed | NO |
| PSIPVEYQATRV | 1489896 | Ectodomain | YES |
| PSIQSRGLFGAIAGFIEGGWTGMVD | 1163468 | Ectodomain | YES |
| PSKDTFELRYSG | 1489907 | Cell/Virus Enclosed | NO |
| PSKIENALPAST | 1489913 | Secreted | YES |
| PSKLLNGIAYIT | 1489919 | Cell/Virus Enclosed | NO |
| PSKLLNGIAYTTFSGS | 1686587 | Cell/Virus Enclosed | NO |
| PSKNNGIGTCPA | 1489921 | Ectodomain | YES |
| PSKPTKRSFIED | 1489924 | Ectodomain | YES |
| PSKTPNDKEEIL | 1489928 | Cell/Virus Enclosed | NO |
| PSLAANTLNIIVPDKS | 1686588 | Cell/Virus Enclosed | NO |
| PSLCNNGKISSSAGDS | 1686590 | Ectodomain | YES |
| PSLVDDNKFDGL | 1489944 | Cell/Virus Enclosed | NO |
| PSLYNILSPFLPLLP | 1686602 | Transmembrane | NO |
| PSMDGYCKFKGKCVQV | 1686604 | Cell/Virus Enclosed | NO |
| PSNDEAIPTRFP | 1489953 | Cell/Virus Enclosed | NO |
| PSNFSFNNWFLL | 1489957 | Ectodomain | YES |
| PSNGTNILPVAF | 1489958 | Cell/Virus Enclosed | NO |
| PSNHIDVVSAYG | 1489959 | Ectodomain | YES |
| PSNHIEVASAYG | 1489960 | Ectodomain | YES |
| PSNWTTSVQVEY | 1489967 | Ectodomain | YES |
| PSPFNWERRIFS | 1489968 | Ectodomain | YES |
| PSPIDIDAFPVR | 1489970 | Cell/Virus Enclosed | NO |
| PSQLKKPRWKRV | 1489985 | Cell/Virus Enclosed | NO |
| PSQNAGYWRRQD | 1489987 | Cell/Virus Enclosed | NO |

| | | | |
|---|---|---|---|
| PSQNFGNSEMLK | 1489989 | Cell/Virus Enclosed | NO |
| PSQTSPATVEPV | 1489992 | Cell/Virus Enclosed | NO |
| PSRDVVRFPNIT | 1489993 | Ectodomain | YES |
| PSRNQSPASSQS | 1489999 | Cell/Virus Enclosed | NO |
| PSRNQSPSSFQS | 1490001 | Cell/Virus Enclosed | NO |
| PSRNQSPSSSQS | 1490002 | Cell/Virus Enclosed | NO |
| PSRNQSPVSSQS | 1490003 | Cell/Virus Enclosed | NO |
| PSRSLPLPPLLSPPRFHPPS | 49498 | Cell/Virus Enclosed | NO |
| PSRSQSRSQSRGRGES | 1686627 | Capsid | UN |
| PSRSQSRSQSRS | 1490010 | Cell/Virus Enclosed | NO |
| PSSARDLICAQY | 1490013 | Ectodomain | YES |
| PSSDEAIPTRFPPGTV | 1686628 | Cell/Virus Enclosed | NO |
| PSSDEAISTRFP | 1490017 | Cell/Virus Enclosed | NO |
| PSSDRYHNSQDD | 1490019 | Capsid | UN |
| PSSIKEMQSQLS | 1490022 | Capsid | UN |
| PSSIKEMQSQPS | 1490023 | Cell/Virus Enclosed | NO |
| PSSIKEMQSQSSHVAQ | 1686630 | Capsid | UN |
| PSSKSSGMDTAL | 1490026 | Ectodomain | YES |
| PSSPARLEIVKL | 1490032 | Cell/Virus Enclosed | NO |
| PSSPARLELVKL | 1490033 | Cell/Virus Enclosed | NO |
| PSSSGTSTPKKPNKPL | 1686638 | Capsid | UN |
| PSSVKEMQSQSS | 1490043 | Cell/Virus Enclosed | NO |
| PSTAAMLSDSHI | 1490050 | Cell/Virus Enclosed | NO |
| PSTAWEDGDYYR | 1490051 | Ectodomain | YES |
| PSTDCSDGNYNR | 1490052 | Ectodomain | YES |
| PSTFGTRNPNNDSAIV | 1686642 | Cell/Virus Enclosed | NO |
| PSTGVYELNGYT | 1490058 | Ectodomain | YES |
| PSTLTPRSVRSVPGEM | 1686644 | Ectodomain | YES |
| PSTPAYTFTTVKPGAA | 1686645 | Cell/Virus Enclosed | NO |
| PSTQKGSGMDTALLRN | 1686646 | Ectodomain | YES |
| PSTQKVSGMDTA | 1490072 | Underneath Membrane | NO |
| PSTTVLIDYDHAYSTM | 1686652 | Cell/Virus Enclosed | NO |
| PSTVDVVALQQE | 1490091 | Cell/Virus Enclosed | NO |
| PSTVWEDGDYYRKQLS | 1686653 | Ectodomain | YES |
| PSTVWEDGNYYR | 1490093 | Ectodomain | YES |
| PSTVWKDGDYYR | 1490095 | Ectodomain | YES |
| PSTWNKKFGFIE | 1490096 | Ectodomain | YES |
| PSTWNKRFGFIEDSVF | 1686654 | Ectodomain | YES |
| PSTWNRRFGFTE | 1490098 | Ectodomain | YES |
| PSVAPGPMTDVN | 1490100 | Cell/Virus Enclosed | NO |
| PSVLQATLSEFS | 1490113 | Cell/Virus Enclosed | NO |
| PSVQPGPMIDVN | 1490115 | Cell/Virus Enclosed | NO |
| PSVYAWERKKISNCVAD | 49557 | Ectodomain | YES |
| PSYCDGVRESYV | 1490130 | Cell/Virus Enclosed | NO |
| PSYIAYENARQAYEDA | 1686671 | Cell/Virus Enclosed | NO |
| PTAADINIHDLP | 1490147 | Ectodomain | YES |
| PTAADINIPDLL | 1490148 | Ectodomain | YES |
| PTAADINIPDLPICVY | 1686675 | Ectodomain | YES |
| PTAADINIPHLP | 1490150 | Ectodomain | YES |
| PTAADINNPDLP | 1490151 | Ectodomain | YES |
| PTAAYINIPDLP | 1490153 | Ectodomain | YES |
| PTADSPLTASPPATAPGPSA | 523862 | Ectodomain | YES |
| PTAFAAIGDLKC | 1490154 | Ectodomain | YES |
| PTAFAVVGDLNC | 1490158 | Ectodomain | YES |
| PTAGSTYRNMAL | 1490165 | Ectodomain | YES |
| PTAIRADQLTPA | 1490168 | Ectodomain | YES |
| PTAKYTPLVGTS | 1490173 | Cell/Virus Enclosed | NO |
| PTARVVYTACSHAAVD | 1686681 | Cell/Virus Enclosed | NO |
| PTASAFMGMSQF | 1490180 | Cell/Virus Enclosed | NO |
| PTATGDVVLAKD | 1490181 | Cell/Virus Enclosed | NO |
| PTDAIVIDNDLN | 1490192 | Cell/Virus Enclosed | NO |
| PTDKLAMIMGYC | 1490194 | Cell/Virus Enclosed | NO |
| PTDNSTTVSVTVSSNM | 1686691 | Cell/Virus Enclosed | NO |
| PTDSADNNQNGG | 1490202 | Capsid | UN |
| PTDSTDNNQDGG | 1490203 | Cell/Virus Enclosed | NO |
| PTDSTDNNQNGGRNGARPKQRRPQ | 49608 | Capsid | UN |
| PTDSTDNNQSGG | 1490205 | Capsid | UN |
| PTDSTVKNYFIT | 1490206 | Cell/Virus Enclosed | NO |
| PTDVAYHTVIDV | 1490208 | Cell/Virus Enclosed | NO |
| PTDVPSTSGTQV | 1490210 | Cell/Virus Enclosed | NO |
| PTDYKNVKAWSG | 1490211 | Ectodomain | YES |
| PTECIYTQSRSC | 1490212 | Cell/Virus Enclosed | NO |
| PTEDLCNGSVVI | 1490214 | Cell/Virus Enclosed | NO |
| PTEHVTSGNLFD | 1490218 | Cell/Virus Enclosed | NO |
| PTEHVTSVNLFDVGYS | 1686703 | Cell/Virus Enclosed | NO |
| PTESACSSLTVL | 1490227 | Cell/Virus Enclosed | NO |

| | | | |
|---|---|---|---|
| PTESYVDVGSDS | 1490230 | Ectodomain | YES |
| PTFSMAPVEFYEDRIL | 1686718 | Cell/Virus Enclosed | NO |
| PTGALLVDSDLN | 1490249 | Cell/Virus Enclosed | NO |
| PTGPAYVMIDKV | 1490263 | Cell/Virus Enclosed | NO |
| PTGTFTVVMRPNYTIK | 1686725 | Cell/Virus Enclosed | NO |
| PTGTKYRHCDLD | 1490270 | Ectodomain | YES |
| PTGTLLVDSNLN | 1490272 | Cell/Virus Enclosed | NO |
| PTGVCLGAGQFV | 1490276 | Cell/Virus Enclosed | NO |
| PTGYGVRRKNSEPEIP | 1686732 | Capsid | UN |
| PTHEVIRFPNIT | 1490293 | Ectodomain | YES |
| PTIADFVQIENC | 1490310 | Ectodomain | YES |
| PTIEDWLLVQGF | 1490314 | Cell/Virus Enclosed | NO |
| PTIEENVIQCFG | 1490315 | Cell/Virus Enclosed | NO |
| PTISQPSKLLNG | 1490325 | Cell/Virus Enclosed | NO |
| PTIVSEKISVMDKLDT | 1686757 | Cell/Virus Enclosed | NO |
| PTKALCFDKSKQFVPV | 1686761 | Underneath Membrane | NO |
| PTKVLRYGYHVL | 1490349 | Cell/Virus Enclosed | NO |
| PTLIPQENYSSI | 1490359 | Cell/Virus Enclosed | NO |
| PTLKDFGGFNFS | 1490360 | Ectodomain | YES |
| PTLKYFDGFNFS | 1490361 | Ectodomain | YES |
| PTLTQMNLKYAISAKN | 1686778 | Cell/Virus Enclosed | NO |
| PTLVPQENYASIRFSS | 1686780 | Cell/Virus Enclosed | NO |
| PTLVPQENYSSIRFAS | 1686781 | Cell/Virus Enclosed | NO |
| PTMCDIKQMLFC | 1490383 | Cell/Virus Enclosed | NO |
| PTMLDIGQARVAYQVA | 1686787 | Cell/Virus Enclosed | NO |
| PTMTHMNLKYAI | 1490387 | Cell/Virus Enclosed | NO |
| PTMTQLNLKYAISGKE | 1686788 | Cell/Virus Enclosed | NO |
| PTMTQMNLKYAISAKN | 1686789 | Cell/Virus Enclosed | NO |
| PTNAVVKMPCPA | 1490389 | Cell/Virus Enclosed | NO |
| PTNAYMRTTHTDIGVY | 1686792 | Cell/Virus Enclosed | NO |
| PTNAYTRTTHTD | 1490391 | Cell/Virus Enclosed | NO |
| PTNCITPVNGYF | 1490393 | Ectodomain | YES |
| PTNFCPCKLDGS | 1490396 | Ectodomain | YES |
| PTNFTIVGQEEFIQTN | 1686797 | Ectodomain | YES |
| PTNIAFELFAKRKVGL | 1686799 | Cell/Virus Enclosed | NO |
| PTNVAFELFAKRKMGL | 1686802 | Cell/Virus Enclosed | NO |
| PTNVVSHVNGDW | 1490415 | Ectodomain | YES |
| PTNVYMRTTHTD | 1490417 | Cell/Virus Enclosed | NO |
| PTPGAFFFGSKLDLVK | 1686807 | Capsid | UN |
| PTPIEDAETPAL | 1490420 | Cell/Virus Enclosed | NO |
| PTPSDNPFSKLYKETIETFD | 49749 | Cell/Virus Enclosed | NO |
| PTPSPAPSRPFSVLRANDVL | 49750 | Undetermined | UN |
| PTQAPTHLSVDI | 1490423 | Cell/Virus Enclosed | NO |
| PTQCYYNNSNNGKIVY | 1686814 | Cell/Virus Enclosed | NO |
| PTQENLELKEQP | 1490426 | Cell/Virus Enclosed | NO |
| PTQEVVRFPNIT | 1490427 | Ectodomain | YES |
| PTQYKDVEAWSGLCVD | 1686821 | Ectodomain | YES |
| PTREENVMQCFG | 1490444 | Cell/Virus Enclosed | NO |
| PTSEEPVNQFTG | 1490451 | Cell/Virus Enclosed | NO |
| PTSGSRVAGRSAIEDI | 1686828 | Ectodomain | YES |
| PTSGSTYLNMAL | 1490456 | Ectodomain | YES |
| PTSGSTYRNMAL | 1490457 | Ectodomain | YES |
| PTSGVIPDMADQ | 1490458 | Cell/Virus Enclosed | NO |
| PTSGVTPDMADE | 1490459 | Cell/Virus Enclosed | NO |
| PTSGVTPDMADH | 1490460 | Cell/Virus Enclosed | NO |
| PTSIHADQLTPA | 1490463 | Ectodomain | YES |
| PTSKIEPCIVSV | 1490464 | Cell/Virus Enclosed | NO |
| PTSKIEPCLVSV | 1490465 | Cell/Virus Enclosed | NO |
| PTSSSTYRNMAL | 1490471 | Ectodomain | YES |
| PTSTFLLHTDFEQVMC | 49778 | Cell/Virus Enclosed | NO |
| PTTCGYLPANAV | 1490478 | Cell/Virus Enclosed | NO |
| PTTCGYLPTNAV | 1490480 | Cell/Virus Enclosed | NO |
| PTTDCSDGNYNR | 1490483 | Ectodomain | YES |
| PTTKNLTGSNES | 1490497 | Cell/Virus Enclosed | NO |
| PTTKNVTGSNESKALF | 1686846 | Cell/Virus Enclosed | NO |
| PTTPISSALIAF | 1490506 | Cell/Virus Enclosed | NO |
| PTTVLNGPVFTG | 1490515 | Cell/Virus Enclosed | NO |
| PTVAMTEQLQMG | 1490518 | Ectodomain | YES |
| PTVDVVALQQEM | 1490519 | Cell/Virus Enclosed | NO |
| PTVKEIISIPNI | 1490530 | Cell/Virus Enclosed | NO |
| PTVKEIISKPKI | 1490531 | Cell/Virus Enclosed | NO |
| PTVKEIISMPKIIKVF | 1686858 | Cell/Virus Enclosed | NO |
| PTVKLYTSVENAQQDKGIAIPHDID | 49810 | Undetermined | UN |
| PTVNEIASTPKT | 1490539 | Cell/Virus Enclosed | NO |
| PTVTPLPAADMD | 1490545 | Capsid | UN |
| PTVWEDGDYYRK | 1490548 | Ectodomain | YES |

| | | | |
|---|---|---|---|
| PTYKNGAMYKGK | 1490562 | Cell/Virus Enclosed | NO |
| PTYLKSDNIVAADLGV | 1686873 | Cell/Virus Enclosed | NO |
| PTYTVHKSDFQLPVYA | 1686876 | Cell/Virus Enclosed | NO |
| PTYVSVDDKYKTSDEL | 1686877 | Cell/Virus Enclosed | NO |
| PVAEYITTYPGQ | 1490573 | Cell/Virus Enclosed | NO |
| PVAFTKAAGGKVSFSD | 1686879 | Cell/Virus Enclosed | NO |
| PVAFTKLAGGKISFSD | 1686880 | Cell/Virus Enclosed | NO |
| PVATGDVVLASDDLYV | 1686889 | Cell/Virus Enclosed | NO |
| PVATPQECNDMH | 1490583 | Cell/Virus Enclosed | NO |
| PVAVVNDKIFTKSGVN | 1686892 | Cell/Virus Enclosed | NO |
| PVCSFPLCANGL | 1490594 | Cell/Virus Enclosed | NO |
| PVDCQDDRCIIH | 1490602 | Cell/Virus Enclosed | NO |
| PVDHYDGFYTQG | 1490608 | Cell/Virus Enclosed | NO |
| PVDLKHFFFTQD | 1490610 | Cell/Virus Enclosed | NO |
| PVDLPPKVYFYY | 1490611 | Cell/Virus Enclosed | NO |
| PVDQYMCGVDGKPISA | 1686908 | Cell/Virus Enclosed | NO |
| PVDSFTNGAIVG | 1490615 | Cell/Virus Enclosed | NO |
| PVDSFTNGAVIG | 1490616 | Cell/Virus Enclosed | NO |
| PVDSFTNGAVVG | 1490617 | Cell/Virus Enclosed | NO |
| PVDVKFKARIEE | 1490620 | Cell/Virus Enclosed | NO |
| PVEAPVEDIAQV | 1490623 | Cell/Virus Enclosed | NO |
| PVECDEGCSEVE | 1490624 | Cell/Virus Enclosed | NO |
| PVEDTQVVIADT | 1490629 | Cell/Virus Enclosed | NO |
| PVEFHLDGEVLP | 1490631 | Cell/Virus Enclosed | NO |
| PVEFYEDRILDFKVLD | 1686916 | Cell/Virus Enclosed | NO |
| PVEGVSSFMNVT | 1490635 | Ectodomain | YES |
| PVELPTVDVVAL | 1490638 | Cell/Virus Enclosed | NO |
| PVEPPTVDVVAL | 1490643 | Cell/Virus Enclosed | NO |
| PVEPSTVDVVAL | 1490644 | Cell/Virus Enclosed | NO |
| PVESPTVDVVAL | 1490645 | Cell/Virus Enclosed | NO |
| PVEVHVEDIVQV | 1490651 | Cell/Virus Enclosed | NO |
| PVEVPVEDIAQV | 1490652 | Cell/Virus Enclosed | NO |
| PVEVPVEDIVQV | 1490653 | Cell/Virus Enclosed | NO |
| PVEVPVEDTQVV | 1490654 | Cell/Virus Enclosed | NO |
| PVEYVSYFDGVK | 1490656 | Cell/Virus Enclosed | NO |
| PVFPAFHRIEST | 1490667 | Cell/Virus Enclosed | NO |
| PVGALDASASVV | 1490671 | Cell/Virus Enclosed | NO |
| PVGALDVSASAV | 1490672 | Cell/Virus Enclosed | NO |
| PVGALDVSASVV | 1490673 | Cell/Virus Enclosed | NO |
| PVGCVLGLINSS | 1490676 | Ectodomain | YES |
| PVGCVLGLVNSS | 1490677 | Ectodomain | YES |
| PVGDSFGKSLGN | 1490678 | Cell/Virus Enclosed | NO |
| PVGESFGKSLGN | 1490679 | Cell/Virus Enclosed | NO |
| PVGFCLSNNPCN | 1490680 | Cell/Virus Enclosed | NO |
| PVGGLYEIQIPS | 1490691 | Ectodomain | YES |
| PVGIRDPVSYVL | 1490694 | Cell/Virus Enclosed | NO |
| PVGNCVKMLTNGSGSG | 1686937 | Cell/Virus Enclosed | NO |
| PVGTNYLTCHNL | 1490699 | Ectodomain | YES |
| PVGVVDTEWGNMLTGI | 1686939 | Cell/Virus Enclosed | NO |
| PVHVKEITKENVETLT | 1686943 | Cell/Virus Enclosed | NO |
| PVHVYADDSIIL | 1490718 | Cell/Virus Enclosed | NO |
| PVHYERDNTSCP | 1490721 | Cell/Virus Enclosed | NO |
| PVHYERDNTSGP | 1490722 | Cell/Virus Enclosed | NO |
| PVIDDGDGVKPF | 1490723 | Cell/Virus Enclosed | NO |
| PVIEVSQIQSKLTDVK | 1686949 | Cell/Virus Enclosed | NO |
| PVISQLDENAQK | 1490735 | Cell/Virus Enclosed | NO |
| PVIWLSHEQASL | 1490739 | Cell/Virus Enclosed | NO |
| PVIYQLDENAQK | 1490740 | Cell/Virus Enclosed | NO |
| PVKDVFELRYSG | 1490742 | Cell/Virus Enclosed | NO |
| PVKDYIQTVNVI | 1490743 | Cell/Virus Enclosed | NO |
| PVKDYVQTVNVIAWLY | 1686959 | Cell/Virus Enclosed | NO |
| PVKEASKQCFKIFERG | 1686960 | Cell/Virus Enclosed | NO |
| PVKGAWNIGQQR | 1490752 | Cell/Virus Enclosed | NO |
| PVKIDNASPASTVHAT | 1686963 | Secreted | YES |
| PVKKVAFGGDQI | 1490758 | Cell/Virus Enclosed | NO |
| PVKKVAFGGDQVHEVA | 1686964 | Cell/Virus Enclosed | NO |
| PVKMFDAYVDTF | 1490761 | Cell/Virus Enclosed | NO |
| PVKPICSSVYLGVKGS | 1686966 | Cell/Virus Enclosed | NO |
| PVKTDNASPAST | 1490764 | Secreted | YES |
| PVLKTINGLVCIVGNK | 1686976 | Cell/Virus Enclosed | NO |
| PVLMGWDYPKCDRAMP | 1686977 | Cell/Virus Enclosed | NO |
| PVLPKNMWEFRDYFNN | 1686979 | Cell/Virus Enclosed | NO |
| PVLVNAANTHLK | 1490789 | Cell/Virus Enclosed | NO |
| PVLYKYLNSPME | 1490795 | Cell/Virus Enclosed | NO |
| PVMAAPTGITLT | 1490796 | Transmembrane | NO |
| PVMAAPTGVTLT | 1490797 | Transmembrane | NO |

| | | | |
|---|---|---|---|
| PVMISEWPLFDQ | 1490802 | Cell/Virus Enclosed | NO |
| PVMISEWPLSES | 1490804 | Cell/Virus Enclosed | NO |
| PVMISEWPLSVQQAQQ | 1686992 | Cell/Virus Enclosed | NO |
| PVMISQWPISDD | 1490806 | Cell/Virus Enclosed | NO |
| PVMISQWPISNDSNGC | 1686993 | Cell/Virus Enclosed | NO |
| PVMSEDLWQFVD | 1490808 | Cell/Virus Enclosed | NO |
| PVMTNGSNILEAFTKP | 1686995 | Cell/Virus Enclosed | NO |
| PVNGYFIKTNNI | 1490812 | Ectodomain | YES |
| PVNPVLHTKSAA | 1490818 | Cell/Virus Enclosed | NO |
| PVNSNVKLLDYDYATH | 1687003 | Cell/Virus Enclosed | NO |
| PVNTVKPKPVINQLDE | 1687005 | Cell/Virus Enclosed | NO |
| PVQLAPVSPAFH | 1490841 | Cell/Virus Enclosed | NO |
| PVQSRKFIVPWV | 1490849 | Cell/Virus Enclosed | NO |
| PVRDEVSIETDIIDEV | 1687021 | Capsid | UN |
| PVRDEVSTETDI | 1490860 | Cell/Virus Enclosed | NO |
| PVRLRNSSDNGI | 1490863 | Ectodomain | YES |
| PVRNIWIPILKS | 1490864 | Cell/Virus Enclosed | NO |
| PVRPRNSSDNGISAII | 1687022 | Ectodomain | YES |
| PVRYERDNTSCP | 1490868 | Cell/Virus Enclosed | NO |
| PVSEYITTYPGQ | 1490878 | Cell/Virus Enclosed | NO |
| PVSGIREFSNLV | 1490880 | Ectodomain | YES |
| PVSGLYEIQIPS | 1490882 | Ectodomain | YES |
| PVSMAKTSVDCNMYICGDS | 49968 | Ectodomain | YES |
| PVSMTKTSVDCN | 1490899 | Ectodomain | YES |
| PVSNCIKMLSNG | 1490902 | Cell/Virus Enclosed | NO |
| PVSPAFHCIEST | 1490903 | Cell/Virus Enclosed | NO |
| PVSPAFHRIESP | 1490906 | Cell/Virus Enclosed | NO |
| PVSPVFHRIESI | 1490910 | Cell/Virus Enclosed | NO |
| PVSSIKGDKGVE | 1490913 | Cell/Virus Enclosed | NO |
| PVSVIYDKETKTHATL | 1687043 | Ectodomain | YES |
| PVTKKYTYSFLPKEDG | 1687048 | Cell/Virus Enclosed | NO |
| PVTRDSVPLHII | 1490941 | Cell/Virus Enclosed | NO |
| PVVDGKQIDDHR | 1490953 | Cell/Virus Enclosed | NO |
| PVVEYLSKEDLHTLYV | 1687059 | Cell/Virus Enclosed | NO |
| PVVGIVYSSQLYEDVP | 1687060 | Cell/Virus Enclosed | NO |
| PVVMEIPSLMTV | 1490968 | Cell/Virus Enclosed | NO |
| PVVNIQKEIDRL | 1490969 | Ectodomain | YES |
| PVVPDIVDVPPQ | 1490971 | Cell/Virus Enclosed | NO |
| PVVPDTVEVPPQVVKL | 1687064 | Cell/Virus Enclosed | NO |
| PVVPDTVGVPPQ | 1490974 | Cell/Virus Enclosed | NO |
| PVVSDTVDVPPQ | 1490976 | Cell/Virus Enclosed | NO |
| PVVSDTVEVPPH | 1490977 | Cell/Virus Enclosed | NO |
| PVVSDTVEVPPQ | 1490978 | Cell/Virus Enclosed | NO |
| PVVSDTVEVSPQ | 1490979 | Cell/Virus Enclosed | NO |
| PVVSDTVEVTPQ | 1490980 | Cell/Virus Enclosed | NO |
| PVVVDDHVGLLDQAWR | 1687067 | Cell/Virus Enclosed | NO |
| PVYASYKVLDNGVIRD | 1687070 | Cell/Virus Enclosed | NO |
| PVYEADTEQTQNVTVK | 1687072 | Cell/Virus Enclosed | NO |
| PVYFLWYSGAIK | 1490993 | Capsid | UN |
| PVYIAPQANSGI | 1490995 | Ectodomain | YES |
| PVYIWTADEAIKFLKE | 1687075 | Underneath Membrane | NO |
| PVYVFGDCVQADWCPVSGGL | 50022 | Cell/Virus Enclosed | NO |
| PWFMDGENAYEVVKAM | 1687085 | Cell/Virus Enclosed | NO |
| PWMFEDAEEKLDNPSS | 1687091 | Cell/Virus Enclosed | NO |
| PWPQIAELVPTA | 1491033 | Cell/Virus Enclosed | NO |
| PWRYLDFNALNLFFSPLEFQ | 50051 | Capsid | UN |
| PWSAAAGVPFSL | 1491035 | Ectodomain | YES |
| PWTAAEGVPFYL | 1491042 | Ectodomain | YES |
| PWTNDALNTICIKDAD | 1687099 | Cell/Virus Enclosed | NO |
| PWYVWLGFIAGLIAIVM | 50058 | Undetermined | UN |
| PWYVWLLIGFAG | 1491058 | Undetermined | UN |
| PYAKDQYGTDID | 1491062 | Cell/Virus Enclosed | NO |
| PYANASYGQSLE | 1491065 | Cell/Virus Enclosed | NO |
| PYCHDPTVLPGA | 1491069 | Cell/Virus Enclosed | NO |
| PYCYDTNLLEGS | 1491071 | Cell/Virus Enclosed | NO |
| PYCYFRNSSANYVGKY | 1687119 | Underneath Membrane | NO |
| PYCYSDGVMKNASLYT | 1687121 | Cell/Virus Enclosed | NO |
| PYCYTEGLMQNA | 1491074 | Cell/Virus Enclosed | NO |
| PYDIEFVTGKQN | 1491077 | Cell/Virus Enclosed | NO |
| PYDIELVTGKQN | 1491078 | Cell/Virus Enclosed | NO |
| PYDLEVLLQDALQSRE | 1687124 | Cell/Virus Enclosed | NO |
| PYDMDFGVSSFS | 1491081 | Cell/Virus Enclosed | NO |
| PYDMDSGGSSFS | 1491082 | Cell/Virus Enclosed | NO |
| PYDMDSGVSSIS | 1491084 | Cell/Virus Enclosed | NO |
| PYEFCSQHTMQI | 1491090 | Cell/Virus Enclosed | NO |
| PYEPRTIQPEVK | 1491093 | Cell/Virus Enclosed | NO |

| | | | |
|---|---|---|---|
| PYFDKAGQKTYE | 1491097 | Cell/Virus Enclosed | NO |
| PYFMHDVVSSFE | 1491102 | Cell/Virus Enclosed | NO |
| PYFMYDVVSSFE | 1491104 | Cell/Virus Enclosed | NO |
| PYGCQPVSEYHI | 1491111 | Cell/Virus Enclosed | NO |
| PYGRCPTAADINIPDL | 1687136 | Ectodomain | YES |
| PYGRCPTAAVIN | 1491118 | Ectodomain | YES |
| PYHANSSEPAML | 1491129 | Ectodomain | YES |
| PYHPNTVDCQDDRCII | 1687140 | Cell/Virus Enclosed | NO |
| PYHRFKCFPNGSNDVW | 1687141 | Ectodomain | YES |
| PYIDDFNRIASCKWTD | 1687143 | Cell/Virus Enclosed | NO |
| PYISHENKLNSC | 1491141 | Cell/Virus Enclosed | NO |
| PYISHETKLNSC | 1491142 | Cell/Virus Enclosed | NO |
| PYKDAQVVQLPI | 1491156 | Cell/Virus Enclosed | NO |
| PYKVCNSVKDTLAYYA | 1687152 | Cell/Virus Enclosed | NO |
| PYLYNGSAQATA | 1491174 | Ectodomain | YES |
| PYMVGDVITSGD | 1491176 | Cell/Virus Enclosed | NO |
| PYNSQNAVARSM | 1491181 | Cell/Virus Enclosed | NO |
| PYNSQNAVSRSMLGLT | 1687158 | Cell/Virus Enclosed | NO |
| PYNSQNFAAKRVLGLQ | 1687159 | Cell/Virus Enclosed | NO |
| PYNSQNYVAARLLGLQ | 1687160 | Cell/Virus Enclosed | NO |
| PYNSVTDTIVVT | 1491186 | Secreted | YES |
| PYNVSLSKFNSCKSDI | 1687165 | Ectodomain | YES |
| PYNVSLSNFNSC | 1491191 | Ectodomain | YES |
| PYPVSGIREFSN | 1491198 | Ectodomain | YES |
| PYQGDHGDIYVY | 1491199 | Ectodomain | YES |
| PYQGDHGDMYVY | 1491200 | Ectodomain | YES |
| PYQGDHGDRYVY | 1491201 | Ectodomain | YES |
| PYQHYDACSVKKYTGV | 1687176 | Cell/Virus Enclosed | NO |
| PYQRNNTSFPEW | 1491204 | Cell/Virus Enclosed | NO |
| PYTALYYYATFY | 1491222 | Cell/Virus Enclosed | NO |
| PYTKLFQKGECY | 1491229 | Cell/Virus Enclosed | NO |
| PYTPKYTFGNVKPGET | 1687189 | Cell/Virus Enclosed | NO |
| PYVASEQEESSD | 1491234 | Cell/Virus Enclosed | NO |
| PYVASEQEESSE | 1491235 | Cell/Virus Enclosed | NO |
| PYVCNAPGCDVNDVTK | 1687192 | Cell/Virus Enclosed | NO |
| PYVCNAPGCGAS | 1491244 | Cell/Virus Enclosed | NO |
| PYVCNASGCGVSDVKK | 1687196 | Cell/Virus Enclosed | NO |
| PYVCNTPGCDVN | 1491245 | Cell/Virus Enclosed | NO |
| PYVFIKRSDAQG | 1491249 | Cell/Virus Enclosed | NO |
| PYVMLNTPIPNL | 1491256 | Ectodomain | YES |
| PYVMLNTSIPNLPDFK | 1687207 | Ectodomain | YES |
| PYVMLNTSSPNL | 1491258 | Ectodomain | YES |
| PYVMLNTSTSNL | 1491260 | Ectodomain | YES |
| PYVYKGYQPIDV | 1491263 | Ectodomain | YES |
| PYVYNGSAHATA | 1491264 | Ectodomain | YES |
| PYVYNGSAQAIA | 1491265 | Ectodomain | YES |
| PYVYNGSAQSTA | 1491267 | Ectodomain | YES |
| QAALDYLATVKTVPAG | 1687226 | Cell/Virus Enclosed | NO |
| QAALPQSKDSNFLNES | 1687228 | Cell/Virus Enclosed | NO |
| QAALPQSKDSNFLNRV | 1687229 | Cell/Virus Enclosed | NO |
| QAASDWKPGYSMPVLY | 1687231 | Cell/Virus Enclosed | NO |
| QADQQVDRLITGRLAA | 1687236 | Ectodomain | YES |
| QADSFTCNNIDAAKIY | 1687237 | Ectodomain | YES |
| QADTAGLYTNFRIDVP | 1687238 | Cell/Virus Enclosed | NO |
| QADTSIPSDVSA | 1491318 | Cell/Virus Enclosed | NO |
| QADTSTPSDVSSRDPT | 1687240 | Capsid | UN |
| QADVNIPADIVD | 1491321 | Cell/Virus Enclosed | NO |
| QAEMNRLQEAIK | 1491328 | Ectodomain | YES |
| QAENVTGLFKDC | 1491329 | Cell/Virus Enclosed | NO |
| QAFCDYADKLCC | 1491335 | Cell/Virus Enclosed | NO |
| QAFCDYADKLCH | 1491336 | Cell/Virus Enclosed | NO |
| QAFCDYTDKLCH | 1491338 | Cell/Virus Enclosed | NO |
| QAFGLRGAGDLQ | 1491343 | Cell/Virus Enclosed | NO |
| QAFGLRGPGDLQGNFG | 1687249 | Cell/Virus Enclosed | NO |
| QAGCLIGAEYVD | 1491364 | Ectodomain | YES |
| QAGEMPCAIINDKVVA | 1687256 | Cell/Virus Enclosed | NO |
| QAGNATEVPAHS | 1491373 | Cell/Virus Enclosed | NO |
| QAGSIVRRVNVP | 1491379 | Cell/Virus Enclosed | NO |
| QAILNTVKFCDH | 1491393 | Cell/Virus Enclosed | NO |
| QAIYDRLDSIQADQQV | 1687284 | Ectodomain | YES |
| QAIYDRLDTIQADQQV | 1687285 | Ectodomain | YES |
| QAKFGIITKVEA | 1491407 | Ectodomain | YES |
| QAKSGIITKLEA | 1491409 | Ectodomain | YES |
| QAKSGIITKVEA | 1491410 | Ectodomain | YES |
| QALAKLASELSN | 1491411 | Ectodomain | YES |
| QALFDSLEFTEL | 1491418 | Cell/Virus Enclosed | NO |

| | | | |
|---|---|---|---|
| QALGAMQTGFTTTNEA | 1687288 | Ectodomain | YES |
| QALHGANLRQDDSVRN | 1687289 | Ectodomain | YES |
| QALIDQKVKEISNQ | 50307 | Capsid | UN |
| QALNSILDNAVKGCVP | 1687292 | Cell/Virus Enclosed | NO |
| QALNSILDNAVR | 1491431 | Cell/Virus Enclosed | NO |
| QALQTVATALNKIQDV | 1687296 | Ectodomain | YES |
| QALQTVSTALNK | 1491440 | Ectodomain | YES |
| QALSKLASELSNTFGA | 1687297 | Ectodomain | YES |
| QAMESVGFVYDN | 1491449 | Cell/Virus Enclosed | NO |
| QAMPSLFKVQNV | 1491451 | Cell/Virus Enclosed | NO |
| QAMTSMYKQARAEDKK | 1687304 | Cell/Virus Enclosed | NO |
| QANAYYKYDNGNFIKL | 1687305 | Cell/Virus Enclosed | NO |
| QANLNFVALIPD | 1491460 | Cell/Virus Enclosed | NO |
| QANNNCFVNVSCLMLQ | 1687309 | Cell/Virus Enclosed | NO |
| QANSGGYYYKVE | 1491464 | Ectodomain | YES |
| QANSGIITKVEA | 1491465 | Ectodomain | YES |
| QAPTHLSVDTKF | 1491471 | Cell/Virus Enclosed | NO |
| QAQGLLTTVPAAPP | 180917 | Ectodomain | YES |
| QAQRPAVASRRRPTTAGAAPLTAVA | 50343 | Undetermined | UN |
| QARSEDKRAQVA | 1491486 | Cell/Virus Enclosed | NO |
| QASADWKPGHAMPSLF | 1687322 | Cell/Virus Enclosed | NO |
| QASADWKPGQAM | 1491489 | Cell/Virus Enclosed | NO |
| QASADWKPGYAM | 1491490 | Cell/Virus Enclosed | NO |
| QASESKVLLERFLPKC | 1687323 | Cell/Virus Enclosed | NO |
| QASSRSSSRSRNSSRNSTPG | 1392366 | Capsid | UN |
| QATLPQSKDSNF | 1491514 | Cell/Virus Enclosed | NO |
| QATNDWKPGYSMPVLY | 1687339 | Cell/Virus Enclosed | NO |
| QATSFCKSGSKF | 1491520 | Ectodomain | YES |
| QATTANVSALMGANGN | 1687342 | Cell/Virus Enclosed | NO |
| QAVGVCVLCNSQ | 1491531 | Cell/Virus Enclosed | NO |
| QAVRRYSEAASN | 1491539 | Cell/Virus Enclosed | NO |
| QAVSDNVRALMS | 1491541 | Cell/Virus Enclosed | NO |
| QAVSSNINMLLS | 1491542 | Cell/Virus Enclosed | NO |
| QAVSSNINRILG | 1491543 | Cell/Virus Enclosed | NO |
| QAVSSNINRLLSVPSD | 1687353 | Cell/Virus Enclosed | NO |
| QAVVGAMLSENFTSYG | 1687358 | Ectodomain | YES |
| QAWRFPCAGRKVNFNE | 1687361 | Cell/Virus Enclosed | NO |
| QAYEDAIANGSSSQLI | 1687363 | Cell/Virus Enclosed | NO |
| QCAIVQALESKV | 1491573 | Cell/Virus Enclosed | NO |
| QCAIVQATESKV | 1491575 | Cell/Virus Enclosed | NO |
| QCDCLCTPDPIT | 1491592 | Ectodomain | YES |
| QCFGKRGPNQNFGGGE | 1687372 | Cell/Virus Enclosed | NO |
| QCFGKRGPSQNFGNAE | 1687373 | Capsid | UN |
| QCIRGAIGDVGL | 1491633 | Cell/Virus Enclosed | NO |
| QCKCGNLEQNHI | 1491639 | Cell/Virus Enclosed | NO |
| QCKTKSLLPNTGVYDL | 1687391 | Ectodomain | YES |
| QCLGDINARDLI | 1491644 | Ectodomain | YES |
| QCNCGTKSWSVG | 1491650 | Cell/Virus Enclosed | NO |
| QCSTNLFKDCSKSYSG | 1687396 | Cell/Virus Enclosed | NO |
| QCSTNLFKDCSN | 1491660 | Cell/Virus Enclosed | NO |
| QCTCGTKSWSVG | 1491662 | Cell/Virus Enclosed | NO |
| QCTVQQCFGKRGPNQN | 1687398 | Cell/Virus Enclosed | NO |
| QCYYNNSNNGKIVYAI | 1687408 | Cell/Virus Enclosed | NO |
| QDADGINKQLTDISVD | 1687411 | Cell/Virus Enclosed | NO |
| QDAQIDRLINGRLTTL | 1687418 | Ectodomain | YES |
| QDAVNNNAHALS | 1491703 | Ectodomain | YES |
| QDAVNNNAQALA | 1491704 | Ectodomain | YES |
| QDAVNNNAQALSKLAS | 1687419 | Ectodomain | YES |
| QDDFVRNLFASV | 1491712 | Ectodomain | YES |
| QDDIMKAVAAALKSLG | 1687424 | Capsid | UN |
| QDDRCIIHCANF | 1491718 | Cell/Virus Enclosed | NO |
| QDDRCIIHCANL | 1491719 | Cell/Virus Enclosed | NO |
| QDELDDFFKNVS | 1491722 | Ectodomain | YES |
| QDELDEFFKNFS | 1491723 | Ectodomain | YES |
| QDELFAMTKRNVIPTM | 1687430 | Cell/Virus Enclosed | NO |
| QDEMNRLQEAIK | 1491725 | Ectodomain | YES |
| QDFDVVFGHGAG | 1491730 | Cell/Virus Enclosed | NO |
| QDFGSTVFNVPTKVLR | 1687432 | Cell/Virus Enclosed | NO |
| QDFNVILPGEFS | 1491734 | Cell/Virus Enclosed | NO |
| QDGFYSANFLDDNVLP | 1687439 | Ectodomain | YES |
| QDGMMNISPKPQ | 1491753 | Cell/Virus Enclosed | NO |
| QDGNAAIPDYNY | 1491757 | Cell/Virus Enclosed | NO |
| QDGNAAITDYNYYKYN | 1687443 | Cell/Virus Enclosed | NO |
| QDGTNTHIMATEASNYAQYRVARAT | 50464 | Undetermined | UN |
| QDGVIFNAVDCK | 1491766 | Ectodomain | YES |
| QDGYNKLQGLLE | 1491773 | Ectodomain | YES |

| | | | |
|---|---|---|---|
| QDHVEILGPLSA | 1491777 | Cell/Virus Enclosed | NO |
| QDKGIAIPHDIDLGESRVVI | 50481 | Undetermined | UN |
| QDKTGDFISTII | 1491791 | Cell/Virus Enclosed | NO |
| QDKVFGYVQQGIFNKA | 1687458 | Cell/Virus Enclosed | NO |
| QDKVFGYVQQGV | 1491798 | Cell/Virus Enclosed | NO |
| QDLCDFKGWIPG | 1491799 | Cell/Virus Enclosed | NO |
| QDLCDFKGWVPG | 1491800 | Cell/Virus Enclosed | NO |
| QDLDVILPGEFS | 1491802 | Cell/Virus Enclosed | NO |
| QDLDVMLPGEFS | 1491803 | Cell/Virus Enclosed | NO |
| QDLIRQGIDYKQ | 1491810 | Capsid | UN |
| QDLNGKWYDFGDFVIT | 1687466 | Cell/Virus Enclosed | NO |
| QDLNGKWYDFGDYVIA | 1687467 | Cell/Virus Enclosed | NO |
| QDLTKYTMMDLC | 1491824 | Cell/Virus Enclosed | NO |
| QDMRNTIMASKTVG | 50494 | Capsid | UN |
| QDPAPIDIDAFPVKPI | 1687480 | Cell/Virus Enclosed | NO |
| QDPEVGPEHSVA | 1491834 | Cell/Virus Enclosed | NO |
| QDPSPIDIDAFP | 1491838 | Cell/Virus Enclosed | NO |
| QDRNTREVFAQVKQMY | 1687483 | Ectodomain | YES |
| QDRPTPSPAPSRPFSVLRA | 50514 | Undetermined | UN |
| QDSKPPLPPHEW | 1491870 | Cell/Virus Enclosed | NO |
| QDSKPPLPSDEW | 1491871 | Cell/Virus Enclosed | NO |
| QDSQNMLRGEDM | 1491879 | Cell/Virus Enclosed | NO |
| QDSRVRALYLPA | 180919 | Underneath Membrane | NO |
| QDSTVLQALQSE | 1491883 | Cell/Virus Enclosed | NO |
| QDVFIQKYDLED | 1491894 | Cell/Virus Enclosed | NO |
| QDVFIQKYGLEDYAFE | 1687495 | Cell/Virus Enclosed | NO |
| QDVFIQKYGLEY | 1491896 | Cell/Virus Enclosed | NO |
| QDVGFKNQHCNG | 1491897 | Cell/Virus Enclosed | NO |
| QDVGFKNQYCNGSIAC | 1687496 | Cell/Virus Enclosed | NO |
| QDVIFSQFDSLGVSSN | 1687497 | Cell/Virus Enclosed | NO |
| QDVKQFANGFVL | 1491907 | Ectodomain | YES |
| QDVLDKNHVADIDK | 50528 | Capsid | UN |
| QDVNCTDVPTAI | 1491913 | Ectodomain | YES |
| QDVNCTDVPTSI | 1491915 | Ectodomain | YES |
| QDVNCTDVSTAIHADQLTPAWRIYS | 1387042 | Ectodomain | YES |
| QDVSLLGKCYKA | 1491927 | Cell/Virus Enclosed | NO |
| QDVVNDNAQALN | 1491930 | Ectodomain | YES |
| QDVVNQQGNSLNHLTS | 1687507 | Ectodomain | YES |
| QDVVNQQGSALNHLTS | 1687508 | Ectodomain | YES |
| QDVYKSLTIVDIPQSL | 1687509 | Cell/Virus Enclosed | NO |
| QDYDNQHEQDRPTPSPAPSRPFSVL | 50539 | Undetermined | UN |
| QDYFLPFDTNLT | 1491940 | Ectodomain | YES |
| QDYIQSVNFLAW | 1491942 | Cell/Virus Enclosed | NO |
| QDYMQIAPVPAEVLNV | 1687513 | Cell/Virus Enclosed | NO |
| QEAILMSEDVKLNPSE | 1687515 | Cell/Virus Enclosed | NO |
| QEAMDSGDTSPQ | 1491954 | Cell/Virus Enclosed | NO |
| QEAWLEFRAGRP | 1491962 | Cell/Virus Enclosed | NO |
| QECNNMHLSTLM | 1491975 | Cell/Virus Enclosed | NO |
| QEDFDLISKCQITAVE | 1687534 | Cell/Virus Enclosed | NO |
| QEDFDLISRCQI | 1491984 | Cell/Virus Enclosed | NO |
| QEDQKILAASFN | 1491987 | Ectodomain | YES |
| QEEFIQTNSPKVTIDC | 1687540 | Ectodomain | YES |
| QEEQEQQEEQEQELEEQ | 50578 | Cell/Virus Enclosed | NO |
| QEFAQNLPKYVKPNFD | 1687548 | Ectodomain | YES |
| QEFNDLAHTSLV | 1492013 | Cell/Virus Enclosed | NO |
| QEFRYMNSQGPL | 1492018 | Cell/Virus Enclosed | NO |
| QEGCGIISRKPQ | 1492024 | Ectodomain | YES |
| QEGFDATNSALVKIQA | 1687556 | Ectodomain | YES |
| QEIISRLDALEA | 1492047 | Ectodomain | YES |
| QEILSRLDVLEA | 1492050 | Ectodomain | YES |
| QEKDKKSAKTGTPKPS | 1687577 | Capsid | UN |
| QELDKYFKNHTS | 1492069 | Ectodomain | YES |
| QELIRQGTDYKHWPQ | 1313331 | Cell/Virus Enclosed | NO |
| QELPFDVIVAWH | 1492085 | Cell/Virus Enclosed | NO |
| QELSYKLPNYTVPDLV | 1687593 | Ectodomain | YES |
| QEMKALYPTWSQ | 1492101 | Cell/Virus Enclosed | NO |
| QEMTTPPPKEQR | 1492105 | Cell/Virus Enclosed | NO |
| QENQKILAASFNKAIN | 1687600 | Ectodomain | YES |
| QENQKILAASFNKAMT | 1687601 | Ectodomain | YES |
| QENQRILAASFN | 1492109 | Ectodomain | YES |
| QEQDELFAMTKR | 1492121 | Cell/Virus Enclosed | NO |
| QEQEEQELEEVEEQEQE | 50674 | Cell/Virus Enclosed | NO |
| QEQRTGLDAVMY | 1492127 | Cell/Virus Enclosed | NO |
| QEQTNCNTSALA | 1492129 | Cell/Virus Enclosed | NO |
| QERWRMRRGQRVDLPP | 1687615 | Capsid | UN |
| QESIKSLNNSYINLKD | 1687618 | Ectodomain | YES |

| | | | |
|---|---|---|---|
| QESKFWDENFYA | 1492150 | Cell/Virus Enclosed | NO |
| QESLARSACVYYSQVL | 1687619 | Cell/Virus Enclosed | NO |
| QESLTTTSTALG | 1492156 | Ectodomain | YES |
| QETPVVFDTVEV | 1492171 | Cell/Virus Enclosed | NO |
| QETPVVPDTVEVPPQV | 1687625 | Cell/Virus Enclosed | NO |
| QEVFAQVKQMYK | 1492174 | Ectodomain | YES |
| QEVIDDMHWGDGFQIK | 1687627 | Cell/Virus Enclosed | NO |
| QEVKDNDGKNVHLKDV | 1687628 | Cell/Virus Enclosed | NO |
| QEVLLLRHDIQLNNDV | 1687629 | Cell/Virus Enclosed | NO |
| QEYIQTTIQKVTVDCK | 1687635 | Ectodomain | YES |
| QFAALGGWGLHQPPPQIFLK | 50735 | Capsid | UN |
| QFANGFVLRIGA | 1492200 | Ectodomain | YES |
| QFAPSASAFFGMSRIGM | 50741 | Capsid | UN |
| QFATFPVYDTIK | 1492205 | Ectodomain | YES |
| QFCIDNCGEPWF | 1492214 | Cell/Virus Enclosed | NO |
| QFCLAGFDMLDN | 1492215 | Cell/Virus Enclosed | NO |
| QFDINAYANWCCLKCG | 1687656 | Cell/Virus Enclosed | NO |
| QFDNLSSEDLKAVRSS | 1687659 | Cell/Virus Enclosed | NO |
| QFDQWYESTFGS | 1492228 | Cell/Virus Enclosed | NO |
| QFDSLGVSSNQSPQGN | 1687662 | Cell/Virus Enclosed | NO |
| QFDSLNLSAGTQHISN | 1687663 | Capsid | UN |
| QFEMNRLQEAIK | 1492237 | Ectodomain | YES |
| QFGLQDGFYSANFLDDNVL | 1492255 | Ectodomain | YES |
| QFGYIDCEQDLC | 1492264 | Cell/Virus Enclosed | NO |
| QFISTFGSFLGF | 1492277 | Cell/Virus Enclosed | NO |
| QFKDKNLQDLWL | 1492278 | Cell/Virus Enclosed | NO |
| QFKDKNLQGLWV | 1492280 | Cell/Virus Enclosed | NO |
| QFKDNVILLNKHIDAYK | 50781 | Capsid | UN |
| QFKTFEKVDGVYTNFK | 1687694 | Cell/Virus Enclosed | NO |
| QFKTTGDLAVQIGSNN | 1687695 | Cell/Virus Enclosed | NO |
| QFKTTGDLAVQM | 1492296 | Cell/Virus Enclosed | NO |
| QFMLWGNDEKVM | 1492314 | Cell/Virus Enclosed | NO |
| QFNKLGTEDPRW | 1492321 | Cell/Virus Enclosed | NO |
| QFNSNFLPESAA | 1492327 | Ectodomain | YES |
| QFNYKQSSSNPT | 1492336 | Ectodomain | YES |
| QFPNIYLEGSVR | 1492347 | Cell/Virus Enclosed | NO |
| QFQILDEGLDCK | 1492352 | Cell/Virus Enclosed | NO |
| QFQILDEGVDCKIQTA | 1687717 | Cell/Virus Enclosed | NO |
| QFQKGKAFEFAE | 1492356 | Cell/Virus Enclosed | NO |
| QFQKGKEFEFAE | 1492357 | Cell/Virus Enclosed | NO |
| QFQKGKEFEFVEGQGV | 1687718 | Cell/Virus Enclosed | NO |
| QFQKGKEFQFAE | 1492360 | Capsid | UN |
| QFTEFVGTQSID | 1492381 | Cell/Virus Enclosed | NO |
| QFTEFVGTQSVDMLAV | 1687728 | Cell/Virus Enclosed | NO |
| QFTSDQAVTFLKEWNF | 1687731 | Undetermined | UN |
| QFTTLTLDKVPQ | 1492391 | Cell/Virus Enclosed | NO |
| QFTTLTVDKVPQ | 1492392 | Cell/Virus Enclosed | NO |
| QFVDHFGENEEIIING | 1687735 | Cell/Virus Enclosed | NO |
| QFVEFYEKVTGTDLELIQIL | 50820 | Capsid | UN |
| QFVKETTDMVKS | 1492401 | Cell/Virus Enclosed | NO |
| QFVKETTNMVKS | 1492402 | Cell/Virus Enclosed | NO |
| QFYEFYNDIKPP | 1492410 | Cell/Virus Enclosed | NO |
| QFYEFYNDVKPPVLDV | 1687742 | Cell/Virus Enclosed | NO |
| QFYIYDEEGGYDVSKP | 1687744 | Cell/Virus Enclosed | NO |
| QGAHLFSSRYVD | 1492423 | Ectodomain | YES |
| QGAKPVGNCVKMLTNG | 1687750 | Cell/Virus Enclosed | NO |
| QGALLNDKHSNGTIK | 504114 | Ectodomain | YES |
| QGARGTYRAALNSEKH | 1687751 | Cell/Virus Enclosed | NO |
| QGCAFQSTVAELQRLKMKVGKTREL | 50837 | Undetermined | UN |
| QGDDYVHLPYPD | 1492441 | Cell/Virus Enclosed | NO |
| QGDHGDMYVYSS | 1492445 | Ectodomain | YES |
| QGDKCNIFANFILHDV | 1687759 | Ectodomain | YES |
| QGDRCNIFANFI | 1492453 | Ectodomain | YES |
| QGDVDTVVNGVE | 1492454 | Cell/Virus Enclosed | NO |
| QGENDNISVAAP | 1492458 | Cell/Virus Enclosed | NO |
| QGENDNTSVAVP | 1492463 | Capsid | UN |
| QGFACGCGASYTANDV | 1687766 | Cell/Virus Enclosed | NO |
| QGFDATNSALVK | 1492472 | Ectodomain | YES |
| QGFGPRDFNHNM | 1492474 | Cell/Virus Enclosed | NO |
| QGFVVVNGKVST | 1492487 | Cell/Virus Enclosed | NO |
| QGGITQRTRTRP | 1492493 | Cell/Virus Enclosed | NO |
| QGGLCKALQEQCRFPNITNSHVP | 78492 | Ectodomain | YES |
| QGHFIGVDGELP | 1492495 | Cell/Virus Enclosed | NO |
| QGIDYKQWPQIA | 1492507 | Capsid | UN |
| QGIETAVGHYFH | 1492508 | Cell/Virus Enclosed | NO |
| QGIETAVGHYVH | 1492509 | Cell/Virus Enclosed | NO |

| | | | |
|---|---|---|---|
| QGIFVEVNAPYY | 1492512 | Ectodomain | YES |
| QGIKLGTGYSLADLPA | 1687790 | Ectodomain | YES |
| QGKFKKIVKGTH | 1492527 | Cell/Virus Enclosed | NO |
| QGKLLIRDNDVL | 1492530 | Cell/Virus Enclosed | NO |
| QGKPIGMFFPYD | 1492536 | Cell/Virus Enclosed | NO |
| QGKTGDFMSTII | 1492544 | Cell/Virus Enclosed | NO |
| QGLIDQINSTYVDLKL | 1687806 | Ectodomain | YES |
| QGLKEFYKEVRA | 1492562 | Cell/Virus Enclosed | NO |
| QGLLEVSVCQYN | 1492570 | Ectodomain | YES |
| QGLPLEFGASAE | 1492576 | Cell/Virus Enclosed | NO |
| QGLPLEFGASTE | 1492577 | Cell/Virus Enclosed | NO |
| QGMDNLACESQQ | 1492589 | Cell/Virus Enclosed | NO |
| QGMDNLACESQT | 1492590 | Cell/Virus Enclosed | NO |
| QGNFGDLQFNKL | 1492596 | Cell/Virus Enclosed | NO |
| QGNFGDLQLNKLGTED | 1687823 | Cell/Virus Enclosed | NO |
| QGNFGDLQLTKL | 1492598 | Cell/Virus Enclosed | NO |
| QGNFGDQELIRQGTEY | 1687826 | Cell/Virus Enclosed | NO |
| QGNKQIVHSCPI | 1492606 | Cell/Virus Enclosed | NO |
| QGNLGGNCAGNVIGND | 1687830 | Cell/Virus Enclosed | NO |
| QGNLGSNGKPGNVGGN | 1687831 | Cell/Virus Enclosed | NO |
| QGNSLNYLTSQL | 1492614 | Ectodomain | YES |
| QGNVVTSVMESQISTK | 1687836 | Cell/Virus Enclosed | NO |
| QGPASARDLICAQYVA | 1687837 | Ectodomain | YES |
| QGPASVRDLICA | 1492624 | Ectodomain | YES |
| QGPPIAPGVPAT | 1492635 | Cell/Virus Enclosed | NO |
| QGPSSARDLICA | 1492639 | Ectodomain | YES |
| QGQLLDTKLNGILQQA | 1687844 | Cell/Virus Enclosed | NO |
| QGQNNADPKVYP | 1492641 | Cell/Virus Enclosed | NO |
| QGRSCSDFEPMS | 1492657 | Cell/Virus Enclosed | NO |
| QGSGKGVSFQLVKLG | 50957 | Ectodomain | YES |
| QGSIAQRTRTRP | 1492669 | Capsid | UN |
| QGSITQRARTRP | 1492672 | Capsid | UN |
| QGSITQRTRTRPSVQP | 1687860 | Cell/Virus Enclosed | NO |
| QGSRSGNSTRGTSPGP | 1687861 | Cell/Virus Enclosed | NO |
| QGSRSRNSTRGT | 1492680 | Cell/Virus Enclosed | NO |
| QGTALPKGFYAEGSRG | 1387119 | Cell/Virus Enclosed | NO |
| QGTDYKQWPQIA | 1492685 | Capsid | UN |
| QGTPLTPILSCG | 1492693 | Cell/Virus Enclosed | NO |
| QGVFNKATVAQI | 1492701 | Cell/Virus Enclosed | NO |
| QGVFRYDSVSSV | 1492704 | Ectodomain | YES |
| QGVFRYNNVSSV | 1492705 | Ectodomain | YES |
| QGVKLGTGYTLSDLPV | 1687875 | Underneath Membrane | NO |
| QGVPESPFSRTGEGLD | 50977 | Cell/Virus Enclosed | NO |
| QGVPIAFGIPPS | 1492714 | Cell/Virus Enclosed | NO |
| QGVPIAFGVPPSEAKG | 1687876 | Capsid | UN |
| QGVPINTNSSKD | 1492722 | Cell/Virus Enclosed | NO |
| QGVPINTNSSPDDQIG | 1687879 | Cell/Virus Enclosed | NO |
| QGVPINTNSTKDDQIG | 1687880 | Cell/Virus Enclosed | NO |
| QGVPLNANSTPAQNAG | 1687881 | Cell/Virus Enclosed | NO |
| QGVTLSSNLNTNLHFD | 1687882 | Ectodomain | YES |
| QGYDDCMHQGPA | 1492731 | Ectodomain | YES |
| QGYDDCMKQGPA | 1492732 | Ectodomain | YES |
| QGYDDCMQQGPS | 1492736 | Ectodomain | YES |
| QGYYIEGSGKSA | 1492745 | Cell/Virus Enclosed | NO |
| QGYYIEGSGRSV | 1492749 | Cell/Virus Enclosed | NO |
| QGYYVEGSGRSA | 1492751 | Cell/Virus Enclosed | NO |
| QHAFMKHKGGDSTDFI | 1687891 | Cell/Virus Enclosed | NO |
| QHAKPTVIVNAA | 1492757 | Cell/Virus Enclosed | NO |
| QHANLESARRVL | 1492760 | Cell/Virus Enclosed | NO |
| QHCFKAPKNFCP | 1492762 | Ectodomain | YES |
| QHCNGSIACQFC | 1492767 | Cell/Virus Enclosed | NO |
| QHDVNFTQEVSRLNINLHFSKCG | 50995 | Ectodomain | YES |
| QHEFCCSHGDRF | 1492782 | Cell/Virus Enclosed | NO |
| QHGKEDLKFPRGQGVP | 1687902 | Cell/Virus Enclosed | NO |
| QHGKEDLRFPRGQGVP | 1687903 | Cell/Virus Enclosed | NO |
| QHGKVPLTFPPGQGVP | 1687905 | Cell/Virus Enclosed | NO |
| QHIGMKRYCTVQGPPG | 1687906 | Cell/Virus Enclosed | NO |
| QHKCGNLHQREL | 1492809 | Cell/Virus Enclosed | NO |
| QHLHVASNSKLWQYCS | 1687915 | Cell/Virus Enclosed | NO |
| QHPLLNPGALEF | 1492825 | Cell/Virus Enclosed | NO |
| QHPLLNPSALEFNPSQ | 1687921 | Capsid | UN |
| QHTDINFTATAS | 1492842 | Ectodomain | YES |
| QHTLYIKDGDDGYFLP | 1687930 | Cell/Virus Enclosed | NO |
| QHTMLVKMDGDE | 1492847 | Cell/Virus Enclosed | NO |
| QHTMQIVDENGKYYLP | 1687934 | Cell/Virus Enclosed | NO |
| QHTMQIVDKDGTYYLP | 1687935 | Cell/Virus Enclosed | NO |

| | | | |
|---|---|---|---|
| QHVIKGCKDTAC | 1492854 | Cell/Virus Enclosed | NO |
| QIANMFLEQTQH | 1492868 | Cell/Virus Enclosed | NO |
| QIANQFNKAISQIQESL | 51043 | Ectodomain | YES |
| QIAPGQTGVIADYNYK | 1687943 | Ectodomain | YES |
| QIDDYRITSITS | 1492886 | Cell/Virus Enclosed | NO |
| QIDGKFVTKFSGDKFD | 1687949 | Cell/Virus Enclosed | NO |
| QIDQLNSSYFKL | 1492892 | Ectodomain | YES |
| QIDRFYRLANEC | 1492893 | Cell/Virus Enclosed | NO |
| QIELGSGAHVGS | 1492902 | Cell/Virus Enclosed | NO |
| QIEVKFNAPALQ | 1492907 | Cell/Virus Enclosed | NO |
| QIFASVPEKFLN | 1492908 | Cell/Virus Enclosed | NO |
| QIFDVAPIELENKFTP | 1687960 | Cell/Virus Enclosed | NO |
| QIFLKILPQSGPIGGIKSMG | 51063 | Capsid | UN |
| QIGSNNVCTYEHVISF | 1687969 | Cell/Virus Enclosed | NO |
| QIGYYRRATRRVRGGDGK | 51074 | Capsid | UN |
| QIHTIDGSSGVANPAM | 1687975 | Secreted | YES |
| QIIAIIKDWSFA | 1492944 | Undetermined | UN |
| QIILNLTSEIST | 1492955 | Ectodomain | YES |
| QIKFENPHILGT | 1492969 | Cell/Virus Enclosed | NO |
| QILASVPEKFLNAFDV | 1687993 | Cell/Virus Enclosed | NO |
| QILDEGLDCKIQ | 1492974 | Cell/Virus Enclosed | NO |
| QILDITPCSFGG | 1492976 | Ectodomain | YES |
| QILGSCTFEDELAPSD | 1687997 | Cell/Virus Enclosed | NO |
| QILGYSSLNDEFSINE | 1687999 | Cell/Virus Enclosed | NO |
| QILIDNINSTLV | 1492980 | Ectodomain | YES |
| QILLNCALGALA | 1492982 | Cell/Virus Enclosed | NO |
| QILPDPLKPIKR | 1492985 | Ectodomain | YES |
| QILPDPLKPTKRSFIE | 1688001 | Ectodomain | YES |
| QILPDPLKSTKR | 1492987 | Ectodomain | YES |
| QILPDPSKPTKR | 1492988 | Ectodomain | YES |
| QILRESGGIDKEAMGFTYSGIRTNG | 1164116 | Ectodomain | YES |
| QIMGSCSFEDELTPSD | 1688004 | Cell/Virus Enclosed | NO |
| QINDMIYSLLEN | 1493002 | Cell/Virus Enclosed | NO |
| QINDMVYSLLEKGKLL | 1688008 | Cell/Virus Enclosed | NO |
| QINGYTLAATSA | 1493006 | Ectodomain | YES |
| QINSTYVDLKLLNRFE | 1688011 | Ectodomain | YES |
| QIPAHCVRDPVG | 1493012 | Cell/Virus Enclosed | NO |
| QIPATSIVAKQGAGDA | 1688014 | Cell/Virus Enclosed | NO |
| QIPSEFTIGNMV | 1493024 | Ectodomain | YES |
| QIPSKFTIGNME | 1493025 | Ectodomain | YES |
| QIQSKLTDVKCANVVL | 1688026 | Cell/Virus Enclosed | NO |
| QIQTIQDSDGTNKQLN | 1688027 | Cell/Virus Enclosed | NO |
| QIQTVDSSQGSEYDYV | 1688028 | Cell/Virus Enclosed | NO |
| QITKDSKFQVGEFVFE | 1688043 | Cell/Virus Enclosed | NO |
| QITNESYNNYVPSYVK | 1688044 | Cell/Virus Enclosed | NO |
| QITQWLSNMLGT | 1493079 | Cell/Virus Enclosed | NO |
| QITYTYKMLVAKDNKN | 1688049 | Capsid | UN |
| QIVDENGKYYLPYPDP | 1688050 | Cell/Virus Enclosed | NO |
| QIVDKDGNYYLP | 1493091 | Cell/Virus Enclosed | NO |
| QIVDKDGTYYLPYPDP | 1688051 | Cell/Virus Enclosed | NO |
| QIVHSHPILHHP | 1493099 | Cell/Virus Enclosed | NO |
| QIVNMFLEQTQH | 1493110 | Cell/Virus Enclosed | NO |
| QIVVTGDDVDDIESIY | 1688058 | Cell/Virus Enclosed | NO |
| QKAKQKPIYLKGSGSD | 1688067 | Cell/Virus Enclosed | NO |
| QKANTEIELGVC | 1493144 | Ectodomain | YES |
| QKAYLDFVNAGGAPLT | 1688073 | Cell/Virus Enclosed | NO |
| QKELLAYYNMLVNCFK | 1688089 | Cell/Virus Enclosed | NO |
| QKESDEYILAKGPLQV | 1688091 | Cell/Virus Enclosed | NO |
| QKESDGYIKLNG | 1493190 | Cell/Virus Enclosed | NO |
| QKESGDYIKLNG | 1493192 | Cell/Virus Enclosed | NO |
| QKFKTICFSTVA | 1493201 | Ectodomain | YES |
| QKGDAAIKDFDYYRYN | 1688099 | Cell/Virus Enclosed | NO |
| QKGDAEEALSKLSEYL | 1688100 | Cell/Virus Enclosed | NO |
| QKGILCVMTSQA | 1493213 | Cell/Virus Enclosed | NO |
| QKGKAFEFAEGQ | 1493216 | Cell/Virus Enclosed | NO |
| QKGKEFEFAEGQ | 1493217 | Cell/Virus Enclosed | NO |
| QKGKEFEFVEGQGVPI | 1688103 | Cell/Virus Enclosed | NO |
| QKGRDFKFPDGQ | 1493222 | Cell/Virus Enclosed | NO |
| QKHIGAPANLRVVGHA | 1688108 | Cell/Virus Enclosed | NO |
| QKHLKSIVNTRN | 1493234 | Cell/Virus Enclosed | NO |
| QKILAASFNKAINNIV | 1688112 | Ectodomain | YES |
| QKILAASFNKAMTNIV | 1688113 | Ectodomain | YES |
| QKITTIQGPPGS | 1493246 | Cell/Virus Enclosed | NO |
| QKKNEISFQQTNAMV | 227489 | Capsid | UN |
| QKKQPTVTPLPA | 1493253 | Capsid | UN |
| QKLALGGSVAVK | 1493259 | Cell/Virus Enclosed | NO |

| | | | |
|---|---|---|---|
| QKLEDNSLFLFD | 1493261 | Cell/Virus Enclosed | NO |
| QKLETDGICPENHVMV | 1688124 | Cell/Virus Enclosed | NO |
| QKLIANAFNNAL | 1493269 | Ectodomain | YES |
| QKLIANKFNQALGAMQ | 1688127 | Ectodomain | YES |
| QKLPCQFKDKNLQDLW | 1688132 | Cell/Virus Enclosed | NO |
| QKLPFKFKDLAIENMW | 1688133 | Cell/Virus Enclosed | NO |
| QKLPNGVTVVEEPDSR | 1688134 | Capsid | UN |
| QKLQSLIDNINS | 1493285 | Ectodomain | YES |
| QKLQTLIDNINSTLVD | 1688135 | Ectodomain | YES |
| QKMLKSMAATRGATCV | 1688136 | Cell/Virus Enclosed | NO |
| QKNGQGENDNIS | 1493291 | Capsid | UN |
| QKPIYLKGSGSDFSLA | 1688139 | Cell/Virus Enclosed | NO |
| QKPLEPKFSIALPPEL | 1688140 | Capsid | UN |
| QKPRWKRQPNDDVTSN | 1688142 | Capsid | UN |
| QKQIANQFNKAISQIQ | 1688144 | Ectodomain | YES |
| QKRLYSHVYRSDKVDS | 1688149 | Cell/Virus Enclosed | NO |
| QKRSPNTQRTVQ | 1493326 | Cell/Virus Enclosed | NO |
| QKRTATKAYNVTQAFG | 1688153 | Cell/Virus Enclosed | NO |
| QKRTATKQYNVTQAFG | 1688154 | Cell/Virus Enclosed | NO |
| QKRTPNKFCNVQ | 1493330 | Cell/Virus Enclosed | NO |
| QKRTPNKHCNVQQCFG | 1688155 | Capsid | UN |
| QKSGYSKYVTVQ | 1493332 | Cell/Virus Enclosed | NO |
| QKVEQPKIEPKPVSVI | 1688163 | Cell/Virus Enclosed | NO |
| QKVQDAVNNNAQALSK | 1688169 | Ectodomain | YES |
| QKVTVDCKQYVCNGFQ | 1688172 | Ectodomain | YES |
| QKYDLEDYAFEH | 1493371 | Cell/Virus Enclosed | NO |
| QKYLNSSYTTCL | 1493377 | Cell/Virus Enclosed | NO |
| QKYVEAYNTFTQAGFN | 1688180 | Cell/Virus Enclosed | NO |
| QLADAALSIKEN | 1493385 | Cell/Virus Enclosed | NO |
| QLAEEPAKDIPE | 1493388 | Cell/Virus Enclosed | NO |
| QLAFDVPDGFYS | 1493390 | Ectodomain | YES |
| QLAGIKLQSKHT | 1493392 | Cell/Virus Enclosed | NO |
| QLAGVKLQSKTK | 1493394 | Cell/Virus Enclosed | NO |
| QLCLAGFDMLDN | 1493428 | Cell/Virus Enclosed | NO |
| QLCQYLNTCTLA | 1493431 | Cell/Virus Enclosed | NO |
| QLCQYLNTLTLA | 1493433 | Cell/Virus Enclosed | NO |
| QLDEKAQKFFDFGDFL | 1688205 | Cell/Virus Enclosed | NO |
| QLDENAQKFFDF | 1493439 | Cell/Virus Enclosed | NO |
| QLDPAFGANSNNPDW | 1688209 | Underneath Membrane | NO |
| QLDTAPKTGNFK | 1493445 | Ectodomain | YES |
| QLDYSIDVIDNEIIVK | 1688212 | Cell/Virus Enclosed | NO |
| QLEAKTASLFKT | 1493450 | Ectodomain | YES |
| QLECIVDDSREE | 1493452 | Cell/Virus Enclosed | NO |
| QLEFVREEVDII | 1493457 | Cell/Virus Enclosed | NO |
| QLEHYSSCVECE | 1493461 | Cell/Virus Enclosed | NO |
| QLEKACNIAKSAYERD | 1688217 | Cell/Virus Enclosed | NO |
| QLELAKVQNLSG | 1493465 | Capsid | UN |
| QLESKMSGKGQQQQSQ | 1688230 | Cell/Virus Enclosed | NO |
| QLESTQITTNTG | 1493485 | Secreted | YES |
| QLEVKTASLFQT | 1493488 | Ectodomain | YES |
| QLFEGDKFIGTFESAA | 1688235 | Cell/Virus Enclosed | NO |
| QLFYTEDMASRFADGL | 1688247 | Cell/Virus Enclosed | NO |
| QLFYVYDVHKDNFKDG | 1688248 | Cell/Virus Enclosed | NO |
| QLGAPDNPSAIV | 1493508 | Ectodomain | YES |
| QLGEYIFERIDYSDAV | 1688251 | Cell/Virus Enclosed | NO |
| QLGGLHLMIGLA | 1493519 | Cell/Virus Enclosed | NO |
| QLGLVWNKDVNTHSTR | 1688260 | Cell/Virus Enclosed | NO |
| QLGNLGNLQSFN | 1493530 | Ectodomain | YES |
| QLGSSGFLQSSNYKID | 1688264 | Ectodomain | YES |
| QLHTGFQGKQIL | 1493548 | Cell/Virus Enclosed | NO |
| QLHVADTLMQGVTLSS | 1688270 | Ectodomain | YES |
| QLIDKYCSVNVV | 1493552 | Cell/Virus Enclosed | NO |
| QLIGKQKITTIQGPPG | 1688271 | Cell/Virus Enclosed | NO |
| QLIGKQRITTIQGPPG | 1688272 | Cell/Virus Enclosed | NO |
| QLKKAMNVAKAEFDRE | 1688278 | Cell/Virus Enclosed | NO |
| QLKNNAKEIGNGCFEFYHKCDNTCM | 1164264 | Ectodomain | YES |
| QLKRAMNIAKSEFDHE | 1688285 | Cell/Virus Enclosed | NO |
| QLLDTKLNGILQQAAL | 1688293 | Cell/Virus Enclosed | NO |
| QLLHTKISWAGS | 1493599 | Cell/Virus Enclosed | NO |
| QLLHTKVSWAGF | 1493601 | Cell/Virus Enclosed | NO |
| QLLIQDYIQSVN | 1493606 | Cell/Virus Enclosed | NO |
| QLLKDVTLEKLR | 1493610 | Cell/Virus Enclosed | NO |
| QLLLEYGQFCSK | 1493615 | Ectodomain | YES |
| QLLLVWEAGKSKPPLN | 1688311 | Cell/Virus Enclosed | NO |
| QLLMPLKAPKEI | 1493626 | Cell/Virus Enclosed | NO |
| QLLMPLRAPKEV | 1493628 | Cell/Virus Enclosed | NO |

| | | | |
|---|---|---|---|
| QLLYAIQQLHTG | 1493641 | Cell/Virus Enclosed | NO |
| QLNHSSCFCSPN | 1493659 | Ectodomain | YES |
| QLNHTSCSCSPD | 1493662 | Ectodomain | YES |
| QLNKLGTEDPRWPQIA | 1688324 | Cell/Virus Enclosed | NO |
| QLNRALSGIAAEQDRNT | 51456 | Ectodomain | YES |
| QLNSSYKLSIPT | 1493677 | Ectodomain | YES |
| QLPAPRVLLSKGSLEP | 1688333 | Cell/Virus Enclosed | NO |
| QLPAPRVLLSKGTLEP | 1688334 | Cell/Virus Enclosed | NO |
| QLPIQDYIQSVN | 1493690 | Cell/Virus Enclosed | NO |
| QLPLDVYDLTMPSQVQ | 1688342 | Cell/Virus Enclosed | NO |
| QLPLDVYNLTMP | 1493696 | Cell/Virus Enclosed | NO |
| QLPNFYYVSNGGNNCT | 1688343 | Ectodomain | YES |
| QLPQGTTLPKGFYAEGSRGGSQ | 51485 | Capsid | UN |
| QLPVKDYIQTVN | 1493705 | Cell/Virus Enclosed | NO |
| QLPVKDYVQTVNVIAW | 1688347 | Cell/Virus Enclosed | NO |
| QLPVYTSYKVLD | 1493707 | Cell/Virus Enclosed | NO |
| QLPYCYFRNSSANYVG | 1688349 | Underneath Membrane | NO |
| QLPYCYFRNSSG | 1493709 | Underneath Membrane | NO |
| QLQLLMPLKAPK | 1493716 | Cell/Virus Enclosed | NO |
| QLQLLMPLRAPK | 1493717 | Cell/Virus Enclosed | NO |
| QLQPINYSFDSY | 1493722 | Ectodomain | YES |
| QLQSAEWKCGYA | 1493728 | Cell/Virus Enclosed | NO |
| QLQSLSKDYISSNGPL | 1688359 | Cell/Virus Enclosed | NO |
| QLQVANALMQGV | 1493732 | Ectodomain | YES |
| QLQVANSLMNGV | 1493734 | Ectodomain | YES |
| QLRAFDIYNANV | 1493735 | Cell/Virus Enclosed | NO |
| QLREPLMQSADA | 1493740 | Cell/Virus Enclosed | NO |
| QLRLAFLGASVTEDVK | 1688369 | Cell/Virus Enclosed | NO |
| QLSDPNYDALLI | 1493760 | Cell/Virus Enclosed | NO |
| QLSEINMDNSQN | 1493765 | Cell/Virus Enclosed | NO |
| QLSFAGYTESAVNSTK | 1688380 | Cell/Virus Enclosed | NO |
| QLSFDVPDGFYSTSPI | 1688381 | Ectodomain | YES |
| QLSPLEGGGRLV | 1493789 | Ectodomain | YES |
| QLTDKYCSVNVA | 1493804 | Cell/Virus Enclosed | NO |
| QLTGILDGDYAMQFFK | 1688398 | Cell/Virus Enclosed | NO |
| QLTLESTINKSVANLK | 1688404 | Cell/Virus Enclosed | NO |
| QLTPAWRVYSTG | 1493838 | Ectodomain | YES |
| QLTTTRMEDAMG | 1493848 | Cell/Virus Enclosed | NO |
| QLTYQNISTNLP | 1493854 | Ectodomain | YES |
| QLVEHTTPWLLL | 1493858 | Cell/Virus Enclosed | NO |
| QLVNANHYSPCV | 1493864 | Ectodomain | YES |
| QLVNANQYSPCV | 1493865 | Ectodomain | YES |
| QLVNSNQYSPCV | 1493866 | Ectodomain | YES |
| QLVPVSPAFHRIESTD | 1688416 | Cell/Virus Enclosed | NO |
| QLVSAVNQSLSNAKVS | 1688418 | Cell/Virus Enclosed | NO |
| QLVTYPSLNYAG | 1493885 | Cell/Virus Enclosed | NO |
| QLYDASNVKNVT | 1493894 | Cell/Virus Enclosed | NO |
| QLYDNCYRKSNV | 1493895 | Cell/Virus Enclosed | NO |
| QLYYNLPAVNVS | 1493913 | Ectodomain | YES |
| QMAPVSAMVRMY | 1493922 | Cell/Virus Enclosed | NO |
| QMDIQGFVNIFI | 1493926 | Cell/Virus Enclosed | NO |
| QMETDFLEPAMD | 1493935 | Cell/Virus Enclosed | NO |
| QMFVTPLSVDVV | 1493942 | Cell/Virus Enclosed | NO |
| QMGSNNVCTYEH | 1493948 | Cell/Virus Enclosed | NO |
| QMRPHVRYDLYDGNMF | 1688461 | Cell/Virus Enclosed | NO |
| QMSELPKEQRVQ | 1493963 | Cell/Virus Enclosed | NO |
| QMYKEARSVNRKSKVI | 1688471 | Cell/Virus Enclosed | NO |
| QMYKTPTLKDFG | 1493982 | Ectodomain | YES |
| QMYKTPTLKYFGGFNFSQ | 1087693 | Ectodomain | YES |
| QNAGYWRRQDRKINTG | 1688474 | Cell/Virus Enclosed | NO |
| QNAGYWRRQYRK | 1493987 | Cell/Virus Enclosed | NO |
| QNAKEIRHKILTKPRQ | 1688475 | Capsid | UN |
| QNAVSRSMLGLTTQTV | 1688483 | Cell/Virus Enclosed | NO |
| QNCTAVGVLQQR | 1494020 | Ectodomain | YES |
| QNCTAVGVPQQR | 1494021 | Ectodomain | YES |
| QNDVCSTEDFNVWAMA | 1688494 | Cell/Virus Enclosed | NO |
| QNFAAKRVLGLQTQTV | 1688496 | Cell/Virus Enclosed | NO |
| QNFGNFEMLKLG | 1494033 | Cell/Virus Enclosed | NO |
| QNFGNSEMLKLG | 1494034 | Cell/Virus Enclosed | NO |
| QNGAKHVQGNVAKAAN | 1688501 | Cell/Virus Enclosed | NO |
| QNGCSLYLADNL | 1494039 | Cell/Virus Enclosed | NO |
| QNGDAAVKDFDFYRYN | 1688502 | Cell/Virus Enclosed | NO |
| QNGDRSGARPKQRRPQ | 1688503 | Cell/Virus Enclosed | NO |
| QNGERSGARPKQRRPQ | 1387258 | Cell/Virus Enclosed | NO |
| QNGMNGRTILGA | 1494050 | Cell/Virus Enclosed | NO |
| QNGVDAKGFPQLAELI | 1688510 | Capsid | UN |

| | | | |
|---|---|---|---|
| QNGVFMSTAKCWTEED | 1688511 | Cell/Virus Enclosed | NO |
| QNIDAYKTFPKKEKKQ | 1688515 | Cell/Virus Enclosed | NO |
| QNILLRKYGRGG | 1494065 | Cell/Virus Enclosed | NO |
| QNISINLPPPLL | 1494067 | Ectodomain | YES |
| QNISTNLPPPLLGNST | 1688517 | Ectodomain | YES |
| QNKGLFYTQVYKNMAV | 1688518 | Ectodomain | YES |
| QNKSIELTAEGI | 1494076 | Cell/Virus Enclosed | NO |
| QNKSRELTAEDISLLK | 1688519 | Cell/Virus Enclosed | NO |
| QNKSRELTSEDI | 1494079 | Cell/Virus Enclosed | NO |
| QNLCVDGFGVNKIQPW | 1688520 | Cell/Virus Enclosed | NO |
| QNLDCNCLGFYE | 1494084 | Cell/Virus Enclosed | NO |
| QNLLYCDSPFEKLQCE | 1688525 | Ectodomain | YES |
| QNLLYDSNGNIIGFKD | 1688526 | Ectodomain | YES |
| QNLLYDSNGNLYGFRD | 1688527 | Ectodomain | YES |
| QNLSGNPDEPQKDVYE | 1688529 | Cell/Virus Enclosed | NO |
| QNNADPKVYPII | 1494108 | Cell/Virus Enclosed | NO |
| QNNDDHGNPVYF | 1494111 | Cell/Virus Enclosed | NO |
| QNNEIKPSGLKTMVVS | 1688534 | Cell/Virus Enclosed | NO |
| QNNEIMPGKLKQ | 1494112 | Cell/Virus Enclosed | NO |
| QNNEIMPGKMKV | 1494113 | Cell/Virus Enclosed | NO |
| QNNELMPVKLKI | 1494114 | Cell/Virus Enclosed | NO |
| QNNLAIEEIEYVHGDA | 1688540 | Cell/Virus Enclosed | NO |
| QNNVFMSEAKCWVETD | 1688543 | Cell/Virus Enclosed | NO |
| QNNVFMSESKCWVEHD | 1688544 | Cell/Virus Enclosed | NO |
| QNNVFMSESKCWVEND | 1688545 | Cell/Virus Enclosed | NO |
| QNPYTPKYTFGNVKPG | 1688549 | Cell/Virus Enclosed | NO |
| QNQKLIASAFNN | 1494131 | Ectodomain | YES |
| QNQRNAPRITFGGPSDSTGSN | 1387279 | Cell/Virus Enclosed | NO |
| QNTVLNASIPESKPLA | 1688562 | Capsid | UN |
| QNTVLNDSIPES | 1494161 | Capsid | UN |
| QNVLYDNQKQIA | 1494165 | Ectodomain | YES |
| QNVLYENQKQIANQFN | 1688565 | Ectodomain | YES |
| QNVNLERCELVN | 1494170 | Cell/Virus Enclosed | NO |
| QNVTVKPKRLRKKRNV | 1688567 | Cell/Virus Enclosed | NO |
| QNYVAKRVLGVQTQTV | 1688574 | Cell/Virus Enclosed | NO |
| QPAGVFTDHDVV | 1494192 | Ectodomain | YES |
| QPALYLYNTGRL | 1494197 | Cell/Virus Enclosed | NO |
| QPCLVIPRNSVP | 1494200 | Cell/Virus Enclosed | NO |
| QPCVASEQDESS | 1494203 | Cell/Virus Enclosed | NO |
| QPDQVAVYQQSIIGAM | 1688579 | Ectodomain | YES |
| QPEDYKSVDVAI | 1494213 | Cell/Virus Enclosed | NO |
| QPFAIPYIHPTNPFAPDVTAAAGAG | 51811 | Undetermined | UN |
| QPFEEVEHVLSI | 1494226 | Cell/Virus Enclosed | NO |
| QPFGEVEHSLPI | 1494228 | Cell/Virus Enclosed | NO |
| QPFGEVEYSLPI | 1494230 | Cell/Virus Enclosed | NO |
| QPGGDATSLSFN | 1494242 | Cell/Virus Enclosed | NO |
| QPGPSREYRYVLRTSPP | 51844 | Cell/Virus Enclosed | NO |
| QPIADVYRRKPD | 1494256 | Ectodomain | YES |
| QPKIEPKPVSVIKVAP | 1688614 | Cell/Virus Enclosed | NO |
| QPKIEQKPLSLI | 1494276 | Cell/Virus Enclosed | NO |
| QPKLDEYYKKDN | 1494277 | Cell/Virus Enclosed | NO |
| QPKQTSTSQQPS | 1494283 | Cell/Virus Enclosed | NO |
| QPKQTVTSQQPS | 1494284 | Cell/Virus Enclosed | NO |
| QPKVITKKEAAA | 1494286 | Cell/Virus Enclosed | NO |
| QPKVITKRDAAA | 1494288 | Cell/Virus Enclosed | NO |
| QPLGGNVVPYYS | 1494294 | Cell/Virus Enclosed | NO |
| QPLPQRQKKQQTVTLLPA | 1387305 | Cell/Virus Enclosed | NO |
| QPLRLTCLWPVS | 1494313 | Ectodomain | YES |
| QPNDDATSNVTQ | 1494326 | Cell/Virus Enclosed | NO |
| QPNDDVTSNVTQCFGP | 1688627 | Capsid | UN |
| QPNLALYKEGNY | 1494328 | Ectodomain | YES |
| QPNQTATSQQPS | 1494331 | Cell/Virus Enclosed | NO |
| QPNTEVFTDVCV | 1494333 | Ectodomain | YES |
| QPPKSVEKICIIDNMY | 1688633 | Cell/Virus Enclosed | NO |
| QPPPVPSPATTKARKRKTKK | 741875 | Ectodomain | YES |
| QPPTASVSTSFLQSGI | 1688636 | Cell/Virus Enclosed | NO |
| QPRADKPSQLKKPRWK | 1688641 | Capsid | UN |
| QPRGRGRNPKPR | 1494359 | Cell/Virus Enclosed | NO |
| QPRLPVLSDFVQ | 1494360 | Ectodomain | YES |
| QPRNVLYDSVSA | 1494361 | Ectodomain | YES |
| QPRNVSYDSVSA | 1494362 | Ectodomain | YES |
| QPRSDKPSQLKK | 1494363 | Cell/Virus Enclosed | NO |
| QPRTIDSTQDGV | 1494365 | Ectodomain | YES |
| QPRTIISTQDGV | 1494366 | Ectodomain | YES |
| QPRTINATQDGV | 1494368 | Ectodomain | YES |
| QPRTINSTQDGDNKLQ | 1688645 | Ectodomain | YES |

| | | | |
|---|---|---|---|
| QPRTINSTQDGV | 1494374 | Ectodomain | YES |
| QPRTINSTQDGY | 1494375 | Ectodomain | YES |
| QPSGGNVIPHYS | 1494383 | Cell/Virus Enclosed | NO |
| QPSGGNVVPHYS | 1494384 | Cell/Virus Enclosed | NO |
| QPSGRRRGRRSGGSGGGFWGDRVDS | 51965 | Undetermined | UN |
| QPSHAVQNTVLN | 1494387 | Cell/Virus Enclosed | NO |
| QPSKLLNGIAYT | 1494391 | Cell/Virus Enclosed | NO |
| QPSLQVESANLMLSDN | 1688653 | Cell/Virus Enclosed | NO |
| QPSRGNVVPHYS | 1494395 | Cell/Virus Enclosed | NO |
| QPSRGNVVPYYS | 1494396 | Cell/Virus Enclosed | NO |
| QPTANSTTVSVT | 1494401 | Cell/Virus Enclosed | NO |
| QPTDNSTTVSVTVSSN | 1688656 | Cell/Virus Enclosed | NO |
| QPTGPAYVMIDK | 1494412 | Cell/Virus Enclosed | NO |
| QPTRNSTTVSVT | 1494423 | Cell/Virus Enclosed | NO |
| QPTVKEIISIPK | 1494428 | Cell/Virus Enclosed | NO |
| QPTVKEIISIPN | 1494429 | Cell/Virus Enclosed | NO |
| QPVDLPPKVYFY | 1494437 | Cell/Virus Enclosed | NO |
| QPVEVHVEDIVQ | 1494448 | Cell/Virus Enclosed | NO |
| QPVEVPVEDTQV | 1494451 | Cell/Virus Enclosed | NO |
| QPVGALDVSASA | 1494453 | Cell/Virus Enclosed | NO |
| QPVGVVDTEWGNMLTG | 1688674 | Cell/Virus Enclosed | NO |
| QPVKIDNASPASTVHA | 1688676 | Secreted | YES |
| QPVKTDNASPAS | 1494463 | Secreted | YES |
| QPVNSNVKLLDYDYAT | 1688677 | Cell/Virus Enclosed | NO |
| QPVYRTFTQAIC | 1494473 | Cell/Virus Enclosed | NO |
| QPYCYTEGLMQN | 1494479 | Cell/Virus Enclosed | NO |
| QPYVFIKRSDAQ | 1494491 | Cell/Virus Enclosed | NO |
| QQAALDYLATVKTVPA | 1688698 | Cell/Virus Enclosed | NO |
| QQACKGKIQALCENP | 52000 | Ectodomain | YES |
| QQAGAGIKYFCG | 1494499 | Cell/Virus Enclosed | NO |
| QQAQQEATHIAE | 1494504 | Cell/Virus Enclosed | NO |
| QQAVVGAMLSENFTSY | 1688703 | Ectodomain | YES |
| QQCFGKRGPNHK | 1494512 | Cell/Virus Enclosed | NO |
| QQCFGKRGPNQNFGGG | 1688704 | Cell/Virus Enclosed | NO |
| QQCFGKRGPSQNFGNA | 1688705 | Capsid | UN |
| QQDGMNMSPKPQ | 1494529 | Cell/Virus Enclosed | NO |
| QQDGRIPNGFPFNNWF | 1688708 | Ectodomain | YES |
| QQFGRDTSDFTD | 1494558 | Ectodomain | YES |
| QQFVKETTNMVK | 1494564 | Cell/Virus Enclosed | NO |
| QQGAFLCANEYT | 1494566 | Cell/Virus Enclosed | NO |
| QQGFDATNFALV | 1494568 | Ectodomain | YES |
| QQGGIPIANCVK | 1494569 | Cell/Virus Enclosed | NO |
| QQGIFNKATVAQIKAL | 1688727 | Cell/Virus Enclosed | NO |
| QQGLLGTCAKRF | 1494572 | Cell/Virus Enclosed | NO |
| QQGNLLNHLNSQ | 1494573 | Ectodomain | YES |
| QQGNSLNHLTFQ | 1494574 | Ectodomain | YES |
| QQGNSLNHLTSQLRQN | 1688729 | Ectodomain | YES |
| QQGNSLNYLTSQ | 1494576 | Ectodomain | YES |
| QQGNSLSHLTSQ | 1494577 | Ectodomain | YES |
| QQGPASARDLIC | 1494578 | Ectodomain | YES |
| QQGPASVRDLIC | 1494581 | Ectodomain | YES |
| QQGSFNKATVAQ | 1494588 | Cell/Virus Enclosed | NO |
| QQGVFNKATVAQ | 1494591 | Cell/Virus Enclosed | NO |
| QQGVFRYDSVSS | 1494594 | Ectodomain | YES |
| QQGVFRYNNVSS | 1494595 | Ectodomain | YES |
| QQHFVYDAYQNL | 1494596 | Ectodomain | YES |
| QQHPLLNPGALE | 1494597 | Cell/Virus Enclosed | NO |
| QQHPLLNPSALEFNPS | 1688735 | Capsid | UN |
| QQIEVKFNAPAL | 1494602 | Cell/Virus Enclosed | NO |
| QQITQWLSNMLG | 1494608 | Cell/Virus Enclosed | NO |
| QQKQLLPRWYFY | 1494619 | Cell/Virus Enclosed | NO |
| QQLAGIKLQSKH | 1494626 | Cell/Virus Enclosed | NO |
| QQLFYTEDMASRFADG | 1688751 | Cell/Virus Enclosed | NO |
| QQLHLEANGRAW | 1494637 | Cell/Virus Enclosed | NO |
| QQLHTGFQGKQI | 1494640 | Cell/Virus Enclosed | NO |
| QQLSNRFGAIGS | 1494654 | Ectodomain | YES |
| QQLVEHTTPWLL | 1494659 | Cell/Virus Enclosed | NO |
| QQMFVTPLSVDV | 1494668 | Cell/Virus Enclosed | NO |
| QQNKSIELTAEG | 1494674 | Cell/Virus Enclosed | NO |
| QQNQRSAPRITF | 1494678 | Capsid | UN |
| QQPLGGNVVPYY | 1494681 | Cell/Virus Enclosed | NO |
| QQPSGGNVIPHY | 1494684 | Cell/Virus Enclosed | NO |
| QQPSRGNVVPHY | 1494688 | Cell/Virus Enclosed | NO |
| QQQDGMMNISPK | 1494693 | Cell/Virus Enclosed | NO |
| QQQDGMMNMSPK | 1494694 | Cell/Virus Enclosed | NO |
| QQQQSQTVTKKSAAEA | 1688779 | Cell/Virus Enclosed | NO |

| | | | |
|---|---|---|---|
| QQQSQTVTKKSAAEAS | 1688780 | Cell/Virus Enclosed | NO |
| QQRFVYDAYQNLVGYY | 1688782 | Ectodomain | YES |
| QQRIQFMMSRRR | 1494715 | Cell/Virus Enclosed | NO |
| QQSIIGAMTAVNESRY | 1688785 | Ectodomain | YES |
| QQSRTRSDSNQSSSDL | 1688787 | Capsid | UN |
| QQTSNLVVQEFVSYDS | 1688789 | Cell/Virus Enclosed | NO |
| QQTVTLLPAADLDDFSKQ | 2060880 | Capsid | UN |
| QQVDLVISDMYDPTTK | 1688795 | Cell/Virus Enclosed | NO |
| QQVGVIADKDKDLSGA | 1688798 | Cell/Virus Enclosed | NO |
| QQVLSENQKLIANKFN | 1688799 | Ectodomain | YES |
| QQVLYYQNNVFMSESK | 1688800 | Cell/Virus Enclosed | NO |
| QQVVDADSKHVQ | 1494747 | Cell/Virus Enclosed | NO |
| QQVVKALNESYIDLKE | 1688804 | Ectodomain | YES |
| QQWGYVGSLSQN | 1494750 | Cell/Virus Enclosed | NO |
| QRADLNGVIVDKVGEL | 1688817 | Cell/Virus Enclosed | NO |
| QRFVYDAYQNLVGYYS | 1688825 | Ectodomain | YES |
| QRGHKNGQGENDNISV | 1688829 | Cell/Virus Enclosed | NO |
| QRGLNNGQGEND | 1494805 | Cell/Virus Enclosed | NO |
| QRIDDDGNKLDKFFVV | 1688832 | Cell/Virus Enclosed | NO |
| QRILAASFNKAM | 1494820 | Ectodomain | YES |
| QRKKQPTVTLLP | 1494828 | Cell/Virus Enclosed | NO |
| QRKRGVKQLPEQFDSL | 1688838 | Capsid | UN |
| QRLDVLEQDAQIDRLI | 1688841 | Ectodomain | YES |
| QRNNTSFPEWID | 1494876 | Cell/Virus Enclosed | NO |
| QRQKKQPTVTLLPAADMDDFSRQ | 52244 | Capsid | UN |
| QRQKKQPTVTPL | 1494883 | Capsid | UN |
| QRQRGHKNGQEE | 1494891 | Capsid | UN |
| QRQRGHKNGQGENDNI | 1688859 | Cell/Virus Enclosed | NO |
| QRQRGHRNGQGE | 1494897 | Cell/Virus Enclosed | NO |
| QRQRGLKNGQGE | 1494898 | Cell/Virus Enclosed | NO |
| QRQRGLNNGQGE | 1494899 | Cell/Virus Enclosed | NO |
| QRSAPRITFGEP | 1494908 | Capsid | UN |
| QRVAGDSGFAAYSRYR | 1387385 | Underneath Membrane | NO |
| QRVDENGDKLDQFFVV | 1688873 | Cell/Virus Enclosed | NO |
| QRVFNNYMPYVL | 1494933 | Cell/Virus Enclosed | NO |
| QRVQGGITQRTR | 1494942 | Cell/Virus Enclosed | NO |
| QRVQGIITQRTR | 1494943 | Cell/Virus Enclosed | NO |
| QRVQGSITQHTR | 1494946 | Cell/Virus Enclosed | NO |
| QRVQGSITQRSK | 1494949 | Cell/Virus Enclosed | NO |
| QRVQGSITQRTRTRPS | 1688879 | Cell/Virus Enclosed | NO |
| QRVYDDVDASLFVDIN | 1688880 | Cell/Virus Enclosed | NO |
| QSAIAKPKRLRK | 1494974 | Cell/Virus Enclosed | NO |
| QSATKIIALNKRWQLA | 1688895 | Secreted | YES |
| QSCPIVVGVSEI | 1494991 | Cell/Virus Enclosed | NO |
| QSCTGSPYIDDFNKIA | 1688900 | Cell/Virus Enclosed | NO |
| QSCTGSPYIDDFNRIA | 1688901 | Cell/Virus Enclosed | NO |
| QSDQVRNVQTRG | 1495006 | Cell/Virus Enclosed | NO |
| QSDRFYRLANEC | 1495007 | Cell/Virus Enclosed | NO |
| QSDRFYRPANEC | 1495009 | Cell/Virus Enclosed | NO |
| QSDSLGVSSNQS | 1495013 | Cell/Virus Enclosed | NO |
| QSERYGFCGNGT | 1495021 | Ectodomain | YES |
| QSESSCGLFKDCARNP | 1688914 | Cell/Virus Enclosed | NO |
| QSFQQFGRDTSD | 1495039 | Ectodomain | YES |
| QSGGGKCFNCAP | 1495050 | Ectodomain | YES |
| QSGGGKCFNCYPAGVN | 1688924 | Ectodomain | YES |
| QSGGRNGARPKQ | 1495056 | Capsid | UN |
| QSGLVKMSHPSGDVEA | 1688931 | Cell/Virus Enclosed | NO |
| QSGVRKVTYGTV | 1495065 | Cell/Virus Enclosed | NO |
| QSHGQLSDHPHALSSSSSHA | 52333 | Capsid | UN |
| QSIFTINSVVQKDGVE | 1688937 | Cell/Virus Enclosed | NO |
| QSIIGAMTAVNESRYG | 1688939 | Ectodomain | YES |
| QSIPMPRGVHMN | 1495082 | Cell/Virus Enclosed | NO |
| QSKDSNFLKRVR | 1495088 | Cell/Virus Enclosed | NO |
| QSKDSNFLNRVRGSIV | 1688945 | Cell/Virus Enclosed | NO |
| QSKDTNFLNAFG | 1495092 | Cell/Virus Enclosed | NO |
| QSKDTNFLNGFG | 1495093 | Cell/Virus Enclosed | NO |
| QSKESNFLNRVR | 1495096 | Cell/Virus Enclosed | NO |
| QSKLPSSSGTST | 1495100 | Cell/Virus Enclosed | NO |
| QSKMSDVRCTSV | 1495104 | Cell/Virus Enclosed | NO |
| QSKRSGFCGQGI | 1495108 | Ectodomain | YES |
| QSKRTRFFKGTV | 1495110 | Cell/Virus Enclosed | NO |
| QSKRTRLFKGTV | 1495112 | Cell/Virus Enclosed | NO |
| QSKRVDFCGRGY | 1495115 | Ectodomain | YES |
| QSKRVGFCGKGY | 1495116 | Ectodomain | YES |
| QSKRYGFGGNGT | 1495117 | Ectodomain | YES |
| QSKSPSSSGTSTPKKP | 1688957 | Capsid | UN |

| | | | |
|---|---|---|---|
| QSKTKRFIKETI | 1495124 | Cell/Virus Enclosed | NO |
| QSLCALPDTHST | 1495128 | Ectodomain | YES |
| QSLCALPDTPSP | 1495130 | Ectodomain | YES |
| QSLCALPDTPSTLTPR | 1688961 | Ectodomain | YES |
| QSLENVIYNLFN | 1495141 | Cell/Virus Enclosed | NO |
| QSLENVIYNLVNVGHY | 1688966 | Cell/Virus Enclosed | NO |
| QSLENVVYNLVN | 1495142 | Cell/Virus Enclosed | NO |
| QSLIDNINSTLV | 1495148 | Ectodomain | YES |
| QSLLKTVQFCDA | 1495154 | Cell/Virus Enclosed | NO |
| QSLSKDYISSNGPLKV | 1688976 | Cell/Virus Enclosed | NO |
| QSLSNAKVSAEVSRQV | 1688977 | Cell/Virus Enclosed | NO |
| QSLTFSYDGLRGAIRK | 1688978 | Cell/Virus Enclosed | NO |
| QSLVSDVGDSTE | 1495171 | Cell/Virus Enclosed | NO |
| QSNADPKVYPII | 1495178 | Cell/Virus Enclosed | NO |
| QSNAVPKVYPII | 1495179 | Cell/Virus Enclosed | NO |
| QSNQRSAPRITFGGPTDS | 52389 | Capsid | UN |
| QSNRYGFCGNGT | 1495187 | Ectodomain | YES |
| QSNVDVLPEGWR | 1495188 | Cell/Virus Enclosed | NO |
| QSNVDVVPEGWRVVNK | 1688991 | Cell/Virus Enclosed | NO |
| QSNVEVLPEGWR | 1495190 | Cell/Virus Enclosed | NO |
| QSPARTSSDPGTNTT | 52394 | Secreted | YES |
| QSPASSQSAAKI | 1495192 | Cell/Virus Enclosed | NO |
| QSPASSQSAATI | 1495193 | Cell/Virus Enclosed | NO |
| QSPASSQSSAKI | 1495194 | Cell/Virus Enclosed | NO |
| QSPASSQSSAKS | 1495195 | Cell/Virus Enclosed | NO |
| QSPASSQSVAKI | 1495196 | Cell/Virus Enclosed | NO |
| QSPASSQTSAKSLARS | 1688993 | Capsid | UN |
| QSPELFDSLNLS | 1495198 | Cell/Virus Enclosed | NO |
| QSPQGNLGSNEP | 1495200 | Cell/Virus Enclosed | NO |
| QSPQGNLGSNGKPGNV | 1688994 | Cell/Virus Enclosed | NO |
| QSPVSSQSAAKI | 1495206 | Cell/Virus Enclosed | NO |
| QSQAITCQKPTPEKEKPDPYK | 110696 | Capsid | UN |
| QSQPKVITKRDA | 1495214 | Cell/Virus Enclosed | NO |
| QSQSSHAVQNTV | 1495218 | Cell/Virus Enclosed | NO |
| QSQSSHVAQNTVLNAS | 1688999 | Capsid | UN |
| QSQSSHVVQNTV | 1495220 | Cell/Virus Enclosed | NO |
| QSQSSLVAQNTV | 1495221 | Capsid | UN |
| QSQTPETPATPLSCCTETQW | 110388 | Cell/Virus Enclosed | NO |
| QSQTVTKKSAAEASKKPRQ | 1387411 | Cell/Virus Enclosed | NO |
| QSRGPNNRSLSRSNSN | 1689003 | Capsid | UN |
| QSRGRGESKPQSRNPS | 1689004 | Capsid | UN |
| QSRKPPNPTPPPPGASANAS | 741879 | Ectodomain | YES |
| QSRNPSSDRYHN | 1495244 | Capsid | UN |
| QSRRFIVPWVMY | 1495247 | Cell/Virus Enclosed | NO |
| QSRSCSDFIPLS | 1495248 | Cell/Virus Enclosed | NO |
| QSRSQSRSRGES | 1495255 | Cell/Virus Enclosed | NO |
| QSRTRSDSNQSSSDLV | 1689013 | Capsid | UN |
| QSRVISSFTCRT | 1495259 | Cell/Virus Enclosed | NO |
| QSSAPRITFGGP | 1495263 | Cell/Virus Enclosed | NO |
| QSSETKEQKHEI | 1495266 | Capsid | UN |
| QSSGMIKPNALF | 1495274 | Cell/Virus Enclosed | NO |
| QSSGMINPNALL | 1495277 | Cell/Virus Enclosed | NO |
| QSSHAVQNTVFN | 1495281 | Cell/Virus Enclosed | NO |
| QSSHVAQNTVLNASIP | 1689021 | Capsid | UN |
| QSSITQRTRTRP | 1495289 | Cell/Virus Enclosed | NO |
| QSSLVAQNTVLN | 1495293 | Capsid | UN |
| QSSNYKIDTTSSSCQL | 1689023 | Ectodomain | YES |
| QSSNYRIDTTAT | 1495295 | Ectodomain | YES |
| QSSPIIPGFGGDFNLT | 1689024 | Ectodomain | YES |
| QSSSDLVAAVNL | 1495300 | Cell/Virus Enclosed | NO |
| QSSSRASSGNRSGNGI | 1689028 | Cell/Virus Enclosed | NO |
| QSSSRASSLSRNSSRS | 1689029 | Cell/Virus Enclosed | NO |
| QSTALCKSGSLV | 1495313 | Ectodomain | YES |
| QSTALQELQIQQ | 1495315 | Cell/Virus Enclosed | NO |
| QSVIIINNSTNVVIRA | 1689047 | Ectodomain | YES |
| QSVNVKPKRLRK | 1495355 | Cell/Virus Enclosed | NO |
| QSVSVDRDSTST | 1495361 | Cell/Virus Enclosed | NO |
| QSVSVDRESTLT | 1495363 | Cell/Virus Enclosed | NO |
| QSVTVKPKRLRK | 1495368 | Cell/Virus Enclosed | NO |
| QSVVNSNAQALNSLLQ | 1689051 | Ectodomain | YES |
| QSYQLYDASNVKKVTD | 1689064 | Cell/Virus Enclosed | NO |
| QTADTAHANNINRFNV | 1689071 | Cell/Virus Enclosed | NO |
| QTAQAAGADTTI | 1495402 | Cell/Virus Enclosed | NO |
| QTAQGAHLFSSR | 1495405 | Ectodomain | YES |
| QTATSQQPSGGNVVPY | 1689080 | Cell/Virus Enclosed | NO |
| QTCNTPTQCYYNNSNN | 1689084 | Cell/Virus Enclosed | NO |

| | | | |
|---|---|---|---|
| QTFKSVLTTYYLDDVK | 1689094 | Cell/Virus Enclosed | NO |
| QTFQAISSSIQA | 1495436 | Ectodomain | YES |
| QTFYPRLQASADWKPG | 1689100 | Cell/Virus Enclosed | NO |
| QTGFTTTNEAFQKVQD | 1689103 | Ectodomain | YES |
| QTGNFKDLREYV | 1495455 | Ectodomain | YES |
| QTGSSKCGCSVI | 1495456 | Cell/Virus Enclosed | NO |
| QTGSSKCVRSVI | 1495458 | Cell/Virus Enclosed | NO |
| QTGVIADYNYKLPDDF | 1689109 | Ectodomain | YES |
| QTIEDLRARMTY | 1495477 | Cell/Virus Enclosed | NO |
| QTIINGMHKSFY | 1495483 | Cell/Virus Enclosed | NO |
| QTIKPEVKEVAP | 1495484 | Cell/Virus Enclosed | NO |
| QTIQDSDGTNKQLNEI | 1689119 | Cell/Virus Enclosed | NO |
| QTIQPEVKEIAP | 1495490 | Cell/Virus Enclosed | NO |
| QTIVPEYIDVNKTLQE | 1689122 | Ectodomain | YES |
| QTKWVFNSPDLIRHADHT | 558309 | Undetermined | UN |
| QTKYLVGIGEHC | 1495498 | Ectodomain | YES |
| QTLAGCLIGAEH | 1495499 | Ectodomain | YES |
| QTLEDLHARMTY | 1495502 | Cell/Virus Enclosed | NO |
| QTLEILDISPCS | 1495503 | Ectodomain | YES |
| QTLIDNINSTLVDLKW | 1689126 | Ectodomain | YES |
| QTLVLKMLHSSS | 1495520 | Cell/Virus Enclosed | NO |
| QTNIYSQNLCVDGFGV | 1689138 | Cell/Virus Enclosed | NO |
| QTNLYSFDKAID | 1495528 | Cell/Virus Enclosed | NO |
| QTNLYSFNKAIDGFGV | 1689139 | Cell/Virus Enclosed | NO |
| QTNSPKVTIDCA | 1495541 | Ectodomain | YES |
| QTPIVWHAKDFNSLSA | 1689148 | Cell/Virus Enclosed | NO |
| QTQGNFGDQELIRQGTDYKHW | 1333233 | Cell/Virus Enclosed | NO |
| QTQHSLNFVGEF | 1495559 | Cell/Virus Enclosed | NO |
| QTQSAIAKPKRL | 1495563 | Cell/Virus Enclosed | NO |
| QTQSVIVKPKRL | 1495564 | Cell/Virus Enclosed | NO |
| QTQSVTVKPKRL | 1495567 | Cell/Virus Enclosed | NO |
| QTRGRRAQPKQTATSQ | 1689159 | Cell/Virus Enclosed | NO |
| QTSDTAHACNANRFNV | 1689161 | Cell/Virus Enclosed | NO |
| QTSNFRVSPTHE | 1495590 | Ectodomain | YES |
| QTSNFRVVPSRD | 1495593 | Ectodomain | YES |
| QTSNLVVQEFVSYDSS | 1689174 | Cell/Virus Enclosed | NO |
| QTSTSQQPSGGN | 1495607 | Cell/Virus Enclosed | NO |
| QTSVAPDLSLGY | 1495611 | Ectodomain | YES |
| QTTNGMNTSHSV | 1495626 | Ectodomain | YES |
| QTTNGTNTIHSV | 1495627 | Ectodomain | YES |
| QTTVELQGLIDQINSTY | 1387449 | Ectodomain | YES |
| QTVATALNKIHD | 1495641 | Ectodomain | YES |
| QTVATALNKIQDVVNQ | 1689191 | Ectodomain | YES |
| QTVDSSQGSEYQYVIF | 1689196 | Cell/Virus Enclosed | NO |
| QTVEDLCARMTY | 1495649 | Cell/Virus Enclosed | NO |
| QTVEDLRVRMTY | 1495651 | Cell/Virus Enclosed | NO |
| QTVIIINNSTNV | 1495657 | Ectodomain | YES |
| QTVKPGHFNKEFYDFL | 1689200 | Cell/Virus Enclosed | NO |
| QTVKPGNFNQDFYDFV | 1689202 | Cell/Virus Enclosed | NO |
| QTVKPGNFNQDFYEFV | 1689203 | Cell/Virus Enclosed | NO |
| QTVQDADGINKQLTDI | 1689207 | Cell/Virus Enclosed | NO |
| QTVRPGNFNQDFYDFV | 1689208 | Cell/Virus Enclosed | NO |
| QTVRSGNFNQDF | 1495663 | Cell/Virus Enclosed | NO |
| QTVRTQVYIAVN | 1495665 | Cell/Virus Enclosed | NO |
| QTVSTALNKIQD | 1495668 | Ectodomain | YES |
| QTVTKKSAAEASKKPRQK | 1495671 | Capsid | UN |
| QTVTSQQPSGGN | 1495673 | Cell/Virus Enclosed | NO |
| QTYLKSDNIVAA | 1495683 | Cell/Virus Enclosed | NO |
| QTYYQPIQQAPT | 1495692 | Underneath Membrane | NO |
| QVAKSHNVSLIW | 1495698 | Cell/Virus Enclosed | NO |
| QVALPQSKDSNF | 1495700 | Cell/Virus Enclosed | NO |
| QVANNLMNGVTL | 1495703 | Ectodomain | YES |
| QVAPGETGVIAD | 1495706 | Ectodomain | YES |
| QVASDNLKECVK | 1495710 | Cell/Virus Enclosed | NO |
| QVASVVLVVEPAPVPTPTPD | 741882 | Ectodomain | YES |
| QVAVYQQSIIGAMTAV | 1689231 | Ectodomain | YES |
| QVCAFLNKLNEN | 1495719 | Cell/Virus Enclosed | NO |
| QVCVQTVCTQVY | 1495723 | Cell/Virus Enclosed | NO |
| QVDLTEAICDFE | 1495732 | Cell/Virus Enclosed | NO |
| QVDLVISDMYDPTTKN | 1689239 | Cell/Virus Enclosed | NO |
| QVDNGSSINRKQLEIV | 1689240 | Cell/Virus Enclosed | NO |
| QVDNGSSINRRQLDVV | 1689241 | Cell/Virus Enclosed | NO |
| QVDQLNSSYFKLSIPT | 1689242 | Ectodomain | YES |
| QVDQLNSSYKLS | 1495737 | Ectodomain | YES |
| QVDQLNSTYFKL | 1495738 | Ectodomain | YES |
| QVDSFTCNNIDA | 1495740 | Ectodomain | YES |

| | | | |
|---|---|---|---|
| QVDVSHEVFDDE | 1495746 | Cell/Virus Enclosed | NO |
| QVEEDVEMSDFV | 1495748 | Cell/Virus Enclosed | NO |
| QVEEVNSIFDIETVDV | 1689250 | Cell/Virus Enclosed | NO |
| QVEINRLQEAIK | 1495750 | Ectodomain | YES |
| QVEMNRLQEAIK | 1495753 | Ectodomain | YES |
| QVESANLMLSDNVVAF | 1689254 | Cell/Virus Enclosed | NO |
| QVESANQMLTVNVVAF | 1689255 | Cell/Virus Enclosed | NO |
| QVEYLQITSKPI | 1495764 | Ectodomain | YES |
| QVFDKVVDNVYVTYAG | 1689259 | Cell/Virus Enclosed | NO |
| QVGDSVLLQGHF | 1495783 | Cell/Virus Enclosed | NO |
| QVGDSVLLQGHS | 1495784 | Cell/Virus Enclosed | NO |
| QVGFSNGVDFVAQPEG | 1689273 | Cell/Virus Enclosed | NO |
| QVGGLTSIKWAD | 1495786 | Cell/Virus Enclosed | NO |
| QVGVIADKDKDLSGAV | 1689278 | Cell/Virus Enclosed | NO |
| QVHGNGSMIEDI | 1495799 | Cell/Virus Enclosed | NO |
| QVHNLPEYMAVA | 1495801 | Underneath Membrane | NO |
| QVILCPTSVFSSNEV | 111693 | Cell/Virus Enclosed | NO |
| QVILLNKHIDAYKTF | 1313394 | Cell/Virus Enclosed | NO |
| QVIQDVKDGTVF | 1495814 | Cell/Virus Enclosed | NO |
| QVIVNNYDKSAGYPFN | 1689291 | Cell/Virus Enclosed | NO |
| QVKNASVYFSQL | 1495825 | Cell/Virus Enclosed | NO |
| QVKQMYKTPAIK | 1495827 | Ectodomain | YES |
| QVLSENQKLIANKFNQ | 1689309 | Ectodomain | YES |
| QVLSSDVMFEGA | 1495860 | Cell/Virus Enclosed | NO |
| QVLSSDVMFENA | 1495861 | Cell/Virus Enclosed | NO |
| QVLVSDVGDSTE | 1495866 | Cell/Virus Enclosed | NO |
| QVMKLPSEPQTI | 1495868 | Cell/Virus Enclosed | NO |
| QVNISLNGNTSVCVRT | 1689317 | Ectodomain | YES |
| QVPATSIVAKQGAGFK | 1689321 | Cell/Virus Enclosed | NO |
| QVPIGTNDPIRFCLEN | 1689323 | Cell/Virus Enclosed | NO |
| QVPIGTNDPIRS | 1495881 | Cell/Virus Enclosed | NO |
| QVPVGIKDPVSYVLTH | 1689325 | Cell/Virus Enclosed | NO |
| QVQLIDKYCSVN | 1495892 | Cell/Virus Enclosed | NO |
| QVQLTDKYCSVN | 1495893 | Cell/Virus Enclosed | NO |
| QVRNIMSPVMGVIG | 52778 | Capsid | UN |
| QVRSWIGFDVEGAHAS | 1689334 | Cell/Virus Enclosed | NO |
| QVSEPPKEQRVQ | 1495906 | Cell/Virus Enclosed | NO |
| QVTAFQHQNSKK | 1495914 | Cell/Virus Enclosed | NO |
| QVTYQNISFNLP | 1495936 | Ectodomain | YES |
| QVTYQNISTNLPPPLL | 1689347 | Ectodomain | YES |
| QVVDSRWNNARL | 1495946 | Ectodomain | YES |
| QVVDSRWNNARQSDNM | 1689353 | Ectodomain | YES |
| QVVELLSEPQTI | 1495951 | Cell/Virus Enclosed | NO |
| QVVIADTLQETPVVPD | 1689355 | Cell/Virus Enclosed | NO |
| QVVIADTLQETS | 1495956 | Cell/Virus Enclosed | NO |
| QVVKALNESYIDLKEL | 1689356 | Ectodomain | YES |
| QVVKLPSAPQTIQPEV | 1689357 | Cell/Virus Enclosed | NO |
| QVVKLPSEPRTI | 1495962 | Cell/Virus Enclosed | NO |
| QVVKPGHFNEEFYNFL | 1689358 | Cell/Virus Enclosed | NO |
| QVVLVSNNQEDFDLIS | 1689359 | Cell/Virus Enclosed | NO |
| QVVMDYLDSLKP | 1495965 | Cell/Virus Enclosed | NO |
| QVVNAITTKISL | 1495967 | Cell/Virus Enclosed | NO |
| QVVNSGPDHTCN | 1495969 | Cell/Virus Enclosed | NO |
| QVVNSGSDINCN | 1495972 | Cell/Virus Enclosed | NO |
| QVVNVITTKISL | 1495973 | Cell/Virus Enclosed | NO |
| QVVQLPVKDYIQ | 1495978 | Cell/Virus Enclosed | NO |
| QVVQLPVKDYVQTVNV | 1689366 | Cell/Virus Enclosed | NO |
| QVVSDVDYVPLK | 1495981 | Cell/Virus Enclosed | NO |
| QVVTYPSLNYAGALWD | 1689367 | Cell/Virus Enclosed | NO |
| QVYFAVNDQDLY | 1495992 | Cell/Virus Enclosed | NO |
| QWDLIISDMYDPITKN | 1689376 | Cell/Virus Enclosed | NO |
| QWDLIISDMYNP | 1496019 | Cell/Virus Enclosed | NO |
| QWFKNQISVAPD | 1496024 | Ectodomain | YES |
| QWFKNQTSVAPV | 1496030 | Ectodomain | YES |
| QWFKNQTSVARD | 1496031 | Ectodomain | YES |
| QWIQFMMSRRRL | 1496040 | Cell/Virus Enclosed | NO |
| QWLKNQTSVAPD | 1496042 | Ectodomain | YES |
| QWLPAGTILVDNDLYP | 1689393 | Cell/Virus Enclosed | NO |
| QWLPTDAIIIDNDLNE | 1689395 | Cell/Virus Enclosed | NO |
| QWNCIDCDSYKP | 1496058 | Cell/Virus Enclosed | NO |
| QWNLVIGFLFLTWIC | 1542219 | Undetermined | UN |
| QWPISNDSNGCVVEAS | 1689407 | Cell/Virus Enclosed | NO |
| QWYDFGDFIQTT | 1496082 | Cell/Virus Enclosed | NO |
| QWYHEHVGGVYDNSIT | 1689415 | Cell/Virus Enclosed | NO |
| QXRAPRITFGGPSDST | 1689416 | Cell/Virus Enclosed | NO |
| QYAHKHDIQITNESYN | 1689419 | Cell/Virus Enclosed | NO |

| | | | |
|---|---|---|---|
| QYAHKHDIQLTN | 1496095 | Cell/Virus Enclosed | NO |
| QYCNGSIACQFC | 1496103 | Cell/Virus Enclosed | NO |
| QYCNGSIACQLC | 1496104 | Cell/Virus Enclosed | NO |
| QYCSTLHNEILA | 1496107 | Cell/Virus Enclosed | NO |
| QYDFTDYKLELFNKYF | 1689425 | Cell/Virus Enclosed | NO |
| QYDVLFYCSNSSSGVL | 1689427 | Ectodomain | YES |
| QYENLSLADISAVQSS | 1689431 | Cell/Virus Enclosed | NO |
| QYGCDYSGKLAA | 1496124 | Cell/Virus Enclosed | NO |
| QYGCDYSGKLADCLQA | 1689437 | Cell/Virus Enclosed | NO |
| QYGQCLGDINAR | 1496138 | Ectodomain | YES |
| QYGTDTISVCPK | 1496151 | Ectodomain | YES |
| QYGTDTNSVCPKLEFA | 1689448 | Ectodomain | YES |
| QYIFSTVNALPECNAD | 1689457 | Cell/Virus Enclosed | NO |
| QYKGKWVQVPIGTNDP | 1689462 | Cell/Virus Enclosed | NO |
| QYLCGADGKPVMSEDL | 1689463 | Cell/Virus Enclosed | NO |
| QYLFSTINALPETSAD | 1689465 | Cell/Virus Enclosed | NO |
| QYLLAFNQDGIIFNAV | 1689467 | Ectodomain | YES |
| QYLLKFDDRGVI | 1496188 | Ectodomain | YES |
| QYLLKFDNRGVITNAV | 1689468 | Ectodomain | YES |
| QYMCGVDGKPISAYAF | 1689475 | Cell/Virus Enclosed | NO |
| QYNMCEYPQTIC | 1496210 | Ectodomain | YES |
| QYRINGLGVTMDVLNK | 1689482 | Ectodomain | YES |
| QYRINGLGVTMDVLSQ | 1689483 | Ectodomain | YES |
| QYRKINTGNGIK | 1496221 | Cell/Virus Enclosed | NO |
| QYSCDFKGWIPG | 1496222 | Cell/Virus Enclosed | NO |
| QYSPCVSIAPST | 1496232 | Ectodomain | YES |
| QYSPCVSIVPSTVWED | 1689489 | Ectodomain | YES |
| QYSPCVSTVPST | 1496236 | Ectodomain | YES |
| QYSTCDICKSTVVEVK | 1689492 | Cell/Virus Enclosed | NO |
| QYTMCEYPNTIC | 1496250 | Ectodomain | YES |
| QYTSACKTIEDALRNS | 1689495 | Ectodomain | YES |
| QYVAGYKVLPPLMDVN | 1689497 | Ectodomain | YES |
| QYVFSTVNALPEVNAD | 1689501 | Cell/Virus Enclosed | NO |
| QYVLATYDTSRVEHA | 53046 | Ectodomain | YES |
| QYYCDGGKYYTQ | 1496276 | Cell/Virus Enclosed | NO |
| QYYFNKDTGVIYGLNS | 1689508 | Ectodomain | YES |
| QYYIYVTNEIGLNASV | 1689509 | Ectodomain | YES |
| QYYNGIMVVPGV | 1496290 | Ectodomain | YES |
| QYYYNMATGFAY | 1496294 | Ectodomain | YES |
| QYYYNMDTGFAY | 1496295 | Underneath Membrane | NO |
| RAAFEYLKPMPF | 1496303 | Cell/Virus Enclosed | NO |
| RAAPNNTVSWYT | 1496322 | Cell/Virus Enclosed | NO |
| RAAPNTTVSWYT | 1496323 | Cell/Virus Enclosed | NO |
| RACIKTAQADTAGLYT | 1689524 | Cell/Virus Enclosed | NO |
| RACIKTAQAGLY | 1496337 | Cell/Virus Enclosed | NO |
| RACIKTVQADTA | 1496340 | Cell/Virus Enclosed | NO |
| RADLNGVIVDKVGELK | 1689531 | Cell/Virus Enclosed | NO |
| RADQINDMVYSLLEKG | 1689532 | Cell/Virus Enclosed | NO |
| RAELNGVMVDKV | 1496359 | Cell/Virus Enclosed | NO |
| RAELNGVVVDKVGDTD | 1689535 | Cell/Virus Enclosed | NO |
| RAFADYTPCAVI | 1496361 | Cell/Virus Enclosed | NO |
| RAFDIYNERVAG | 1496369 | Cell/Virus Enclosed | NO |
| RAFDVYNKDASFIGKN | 1689543 | Cell/Virus Enclosed | NO |
| RAFISVLDITDAAVKA | 1689548 | Cell/Virus Enclosed | NO |
| RAGKHEHSLLVE | 1496394 | Cell/Virus Enclosed | NO |
| RAIDCGFNDLSQLHCS | 1689572 | Ectodomain | YES |
| RAIIVSVEQLEPCAQS | 1689575 | Cell/Virus Enclosed | NO |
| RAILTAFSPAQDIWGTS | 53116 | Ectodomain | YES |
| RAILTAFSPAQG | 1496418 | Ectodomain | YES |
| RAIMATIQRKYK | 1496419 | Cell/Virus Enclosed | NO |
| RAITTENRFTNF | 1496423 | Ectodomain | YES |
| RAITTGYRFTNF | 1496425 | Ectodomain | YES |
| RAIVFVCVEYYP | 1496427 | Cell/Virus Enclosed | NO |
| RAKSVVSKIGNS | 1496442 | Cell/Virus Enclosed | NO |
| RALMSCNGNKIE | 1496456 | Cell/Virus Enclosed | NO |
| RALSGIAAEQDRNTRE | 1689600 | Ectodomain | YES |
| RALSGIAVEQDR | 1496474 | Ectodomain | YES |
| RALTGIAIEQDK | 1496477 | Ectodomain | YES |
| RAMFQPRLPVLS | 1496486 | Ectodomain | YES |
| RAMLDYAKDHND | 1496487 | Cell/Virus Enclosed | NO |
| RAMNIAKSEFDHEISV | 1689608 | Cell/Virus Enclosed | NO |
| RANFFDVPTFCA | 1496499 | Cell/Virus Enclosed | NO |
| RANSGNRTPTPG | 1496509 | Cell/Virus Enclosed | NO |
| RANSGNRTPTSGVTPD | 1689620 | Cell/Virus Enclosed | NO |
| RAPHPAPSARGPLRLGAVLG | 741889 | Undetermined | UN |
| RAPRITFGGPSDSTDN | 1689623 | Cell/Virus Enclosed | NO |

| | | | |
|---|---|---|---|
| RAPTMANQEKYSTIYK | 1689626 | Cell/Virus Enclosed | NO |
| RAQPKQTVTSQQ | 1496527 | Cell/Virus Enclosed | NO |
| RARGGDGKMKEL | 1496534 | Capsid | UN |
| RARTRPSVQPGP | 1496541 | Capsid | UN |
| RARVEHPDVDGLCKLR | 1689647 | Cell/Virus Enclosed | NO |
| RASDTSLSDACQ | 1496552 | Underneath Membrane | NO |
| RASGGVAATVAANGGPASRP | 741891 | Ectodomain | YES |
| RASLRQIFDVAPIELE | 1689653 | Cell/Virus Enclosed | NO |
| RASLSTWHVLYSPNVSVPSS | 78500 | Ectodomain | YES |
| RASSAGSRSRANSGNR | 1689656 | Cell/Virus Enclosed | NO |
| RASSGNRPGNGI | 1496578 | Capsid | UN |
| RASSGNRSGNGILKWA | 1689658 | Cell/Virus Enclosed | NO |
| RASSGNRSVNGI | 1496580 | Cell/Virus Enclosed | NO |
| RASSLSRNSSRSSSQG | 1689659 | Cell/Virus Enclosed | NO |
| RASSRSSTRNNSRDSS | 1689660 | Capsid | UN |
| RATRRVRGGDGKMKDL | 1689671 | Cell/Virus Enclosed | NO |
| RAVKDGIVWVHEDGATD | 1387520 | Cell/Virus Enclosed | NO |
| RAVRSHPDFKFL | 1496623 | Cell/Virus Enclosed | NO |
| RAVRSHPDFKLLHNLQ | 1689682 | Cell/Virus Enclosed | NO |
| RAVSFADNNDITNTNL | 1689683 | Cell/Virus Enclosed | NO |
| RAVSFAYNNDIT | 1496627 | Capsid | UN |
| RAWAFCVKCHNDILAA | 1689684 | Cell/Virus Enclosed | NO |
| RAWVGFDAEGAHATRD | 1689687 | Cell/Virus Enclosed | NO |
| RAYIEAAAQGFQ | 1496631 | Cell/Virus Enclosed | NO |
| RAYLDLRTTQQL | 1496635 | Cell/Virus Enclosed | NO |
| RAYLDSCTTQQL | 1496636 | Cell/Virus Enclosed | NO |
| RAYSEAAAQGFQ | 1496646 | Cell/Virus Enclosed | NO |
| RAYSEAATQGFQ | 1496647 | Cell/Virus Enclosed | NO |
| RAYWVPRASADI | 1496648 | Cell/Virus Enclosed | NO |
| RCDINYKGKVFF | 1496670 | Cell/Virus Enclosed | NO |
| RCDQLAFDVPDG | 1496672 | Ectodomain | YES |
| RCDQLSFDVPDG | 1496673 | Ectodomain | YES |
| RCIKISTVQSKLTDLK | 1689717 | Cell/Virus Enclosed | NO |
| RCIVFDVKGVEE | 1496701 | Cell/Virus Enclosed | NO |
| RCIVLDVKGLEE | 1496702 | Cell/Virus Enclosed | NO |
| RCIVLDVKGVEELHDD | 1689721 | Cell/Virus Enclosed | NO |
| RCNIGGAVCSKHANLY | 1689738 | Cell/Virus Enclosed | NO |
| RCNVIADFKSFT | 1496727 | Cell/Virus Enclosed | NO |
| RCNVIADPKSFT | 1496728 | Cell/Virus Enclosed | NO |
| RCPKEIVDTVSALVYE | 1689745 | Cell/Virus Enclosed | NO |
| RCPTAADINIHD | 1496741 | Ectodomain | YES |
| RCPTAADINIPDLPIC | 1689748 | Ectodomain | YES |
| RCPTAADINIPH | 1496743 | Ectodomain | YES |
| RCPTAADINNPD | 1496744 | Ectodomain | YES |
| RCPTAADINTPD | 1496745 | Ectodomain | YES |
| RCPTAAYINILD | 1496747 | Ectodomain | YES |
| RCYTAEQCIRGA | 1496795 | Cell/Virus Enclosed | NO |
| RDAQVVQLPVKDYVQT | 1689770 | Cell/Virus Enclosed | NO |
| RDATKPQQVTKH | 1496812 | Cell/Virus Enclosed | NO |
| RDCDFNKPLIEWPLTE | 1689776 | Cell/Virus Enclosed | NO |
| RDDLQLGSSGFL | 1496827 | Ectodomain | YES |
| RDDNQALLVPKI | 1496829 | Cell/Virus Enclosed | NO |
| RDDNQTLLVPQI | 1496831 | Cell/Virus Enclosed | NO |
| RDEFSIETDIID | 1496834 | Capsid | UN |
| RDELGVRVLDQSDNNC | 1689779 | Cell/Virus Enclosed | NO |
| RDFKFPDGQGVP | 1496840 | Cell/Virus Enclosed | NO |
| RDFNHNMGDSDL | 1496843 | Cell/Virus Enclosed | NO |
| RDGDVVKQPVKF | 1496855 | Cell/Virus Enclosed | NO |
| RDGQPCYEQFPL | 1496862 | Cell/Virus Enclosed | NO |
| RDGQPCYERFPL | 1496863 | Cell/Virus Enclosed | NO |
| RDGQPFYEPFPL | 1496864 | Cell/Virus Enclosed | NO |
| RDGQPFYERFPL | 1496865 | Cell/Virus Enclosed | NO |
| RDGQRIYDDVNA | 1496866 | Cell/Virus Enclosed | NO |
| RDGQRVYDDVDASLFV | 1689794 | Cell/Virus Enclosed | NO |
| RDGVRRGDNFHN | 1496876 | Cell/Virus Enclosed | NO |
| RDGVRRGDNFHSVLTT | 1689798 | Cell/Virus Enclosed | NO |
| RDGVVPLSVIPATSAS | 1689799 | Cell/Virus Enclosed | NO |
| RDHNDILYTPPTVSYG | 1689804 | Cell/Virus Enclosed | NO |
| RDISNVPFSSDG | 1496903 | Ectodomain | YES |
| RDKAVARKLERMADLA | 1689816 | Cell/Virus Enclosed | NO |
| RDKLEKLISTARDGVR | 1689819 | Cell/Virus Enclosed | NO |
| RDLEDFKPRSKM | 1496919 | Cell/Virus Enclosed | NO |
| RDLLDTASALYREAL | 1689831 | Capsid | UN |
| RDLLIKAYNTINNEQG | 1689832 | Cell/Virus Enclosed | NO |
| RDLQRRLYDNCY | 1496940 | Cell/Virus Enclosed | NO |
| RDLSLQFKRSIN | 1496943 | Cell/Virus Enclosed | NO |

| | | | |
|---|---|---|---|
| RDLTSDENGVRT | 1496947 | Ectodomain | YES |
| RDMADQIASLVL | 1496952 | Capsid | UN |
| RDNGALEAFKRSNNGV | 1689846 | Cell/Virus Enclosed | NO |
| RDNGSLEKFMST | 1496963 | Cell/Virus Enclosed | NO |
| RDNLKPVTTLSTITAY | 1689847 | Cell/Virus Enclosed | NO |
| RDPILISLQPFV | 1496975 | Cell/Virus Enclosed | NO |
| RDPQTLEILDIS | 1496980 | Ectodomain | YES |
| RDPRYVMRLQSAATIR | 1689851 | Cell/Virus Enclosed | NO |
| RDPSNDEAIPTR | 1496988 | Cell/Virus Enclosed | NO |
| RDPSSDEAIPTRFPPG | 1689852 | Cell/Virus Enclosed | NO |
| RDPTIQEAIPTR | 1496989 | Cell/Virus Enclosed | NO |
| RDPVGFCLSNTP | 1496991 | Cell/Virus Enclosed | NO |
| RDRAVAKKLERMADLA | 1689855 | Cell/Virus Enclosed | NO |
| RDSDFVKQPVKF | 1497004 | Cell/Virus Enclosed | NO |
| RDSDVVKQPVNF | 1497006 | Cell/Virus Enclosed | NO |
| RDSHKGVCFGFDKWYV | 1689859 | Cell/Virus Enclosed | NO |
| RDSIGTNFPLQLGFST | 1689860 | Cell/Virus Enclosed | NO |
| RDSRFVMRLQTIATIC | 1689861 | Cell/Virus Enclosed | NO |
| RDSSRSTSRQHS | 1497017 | Cell/Virus Enclosed | NO |
| RDSSRSTSRQQSRTRS | 1689862 | Capsid | UN |
| RDSTSTHDGPTE | 1497019 | Cell/Virus Enclosed | NO |
| RDSTYGPLQTPVGCVL | 1689863 | Ectodomain | YES |
| RDSTYSPLQTPV | 1497024 | Ectodomain | YES |
| RDSVPLHIIAPSSLIV | 1689864 | Cell/Virus Enclosed | NO |
| RDTGHPSISTDT | 1497026 | Ectodomain | YES |
| RDTHPQAMQWNSTTF | 1689866 | Ectodomain | YES |
| RDTLPQSPGPAFP | 53429 | Ectodomain | YES |
| RDVSDFTDSVRDPKTSEI | 53435 | Ectodomain | YES |
| RDVSVEDVCFANKFEQ | 1689877 | Cell/Virus Enclosed | NO |
| RDYFDDNTDSIV | 1497065 | Cell/Virus Enclosed | NO |
| RDYFDNNTDSIV | 1497066 | Cell/Virus Enclosed | NO |
| RDYKLANDAKESLRLF | 1689882 | Cell/Virus Enclosed | NO |
| RDYKLANDVKDTLRLF | 1689883 | Cell/Virus Enclosed | NO |
| RDYLADYDIAVT | 1497080 | Cell/Virus Enclosed | NO |
| RDYLADYDIVVT | 1497081 | Cell/Virus Enclosed | NO |
| RDYLADYDVVVT | 1497083 | Cell/Virus Enclosed | NO |
| REAPAHVSTIGV | 1497097 | Cell/Virus Enclosed | NO |
| REELDQWFKNQT | 1497109 | Ectodomain | YES |
| REENVMQCFGPR | 1497113 | Cell/Virus Enclosed | NO |
| REEVDIIEQPFEEVEH | 1689904 | Cell/Virus Enclosed | NO |
| REEVDIIEQPFG | 1497117 | Cell/Virus Enclosed | NO |
| REFVFKNKDGFP | 1497126 | Ectodomain | YES |
| REGGLNKVKYAT | 1497131 | Cell/Virus Enclosed | NO |
| REHLKSGTCGIV | 1497150 | Cell/Virus Enclosed | NO |
| REIKPAFNVVKA | 1497155 | Cell/Virus Enclosed | NO |
| REISCNVCTKRATVYN | 1689926 | Cell/Virus Enclosed | NO |
| REIVMNASPYDL | 1497159 | Cell/Virus Enclosed | NO |
| REIVMNASPYHL | 1497160 | Cell/Virus Enclosed | NO |
| REIVMNASPYYL | 1497161 | Cell/Virus Enclosed | NO |
| REKLEKLISTAR | 1497164 | Cell/Virus Enclosed | NO |
| RELILSWEIGKVKPPL | 1689935 | Cell/Virus Enclosed | NO |
| RELIRELNGGAF | 1497171 | Cell/Virus Enclosed | NO |
| RELTAEDISLLKKMDE | 1689944 | Cell/Virus Enclosed | NO |
| RELTSTHDGPTE | 1497187 | Cell/Virus Enclosed | NO |
| RELYLSSSNVTT | 1497191 | Cell/Virus Enclosed | NO |
| REMLAHAEEARK | 1497195 | Cell/Virus Enclosed | NO |
| REMQQHPLLNPSALEF | 1689952 | Capsid | UN |
| REMQQQPLLNPS | 1497200 | Cell/Virus Enclosed | NO |
| RENIGTNFPLQIGFST | 1689954 | Cell/Virus Enclosed | NO |
| RENKSAELNYTV | 1497201 | Ectodomain | YES |
| REPMMQSADAST | 1497211 | Cell/Virus Enclosed | NO |
| REPVHLLSADAI | 1497213 | Cell/Virus Enclosed | NO |
| REQIDGYTMHAN | 1497214 | Cell/Virus Enclosed | NO |
| RERGLNKVKYAT | 1497220 | Cell/Virus Enclosed | NO |
| RESTSTHDDFTE | 1497238 | Cell/Virus Enclosed | NO |
| RESTSTHDGFNE | 1497239 | Cell/Virus Enclosed | NO |
| RESTSTHDVPTE | 1497248 | Cell/Virus Enclosed | NO |
| RESYVNSSNVTT | 1497249 | Cell/Virus Enclosed | NO |
| RETMTHLLQHAN | 1497256 | Cell/Virus Enclosed | NO |
| RETVLEYLVSFGVWI | 1689976 | Capsid | UN |
| REVFAQVKQMYKTPTL | 1689977 | Ectodomain | YES |
| REVFVQVKQMYK | 1497263 | Ectodomain | YES |
| REVILCWETGKVKPPL | 1689979 | Cell/Virus Enclosed | NO |
| REVVVTNLNKSAGWPL | 1689985 | Cell/Virus Enclosed | NO |
| REWCNVCGIKDV | 1497277 | Cell/Virus Enclosed | NO |
| REYLESYNTATTAGFT | 1689989 | Cell/Virus Enclosed | NO |

| | | | |
|---|---|---|---|
| RFDENGDKLDQF | 1497308 | Cell/Virus Enclosed | NO |
| RFDTRVLNNLNLPGCN | 1690010 | Cell/Virus Enclosed | NO |
| RFEYISPTDKLA | 1497326 | Cell/Virus Enclosed | NO |
| RFEYTSPTDKLAMIMG | 1690016 | Cell/Virus Enclosed | NO |
| RFGAISASLQEI | 1497344 | Ectodomain | YES |
| RFGFIEDAVFKP | 1497347 | Ectodomain | YES |
| RFGFIEDSAFVP | 1497348 | Ectodomain | YES |
| RFGFIEDSVFKPRPAG | 1690025 | Ectodomain | YES |
| RFGFIEDSVFVP | 1497350 | Ectodomain | YES |
| RFGPFEFCSEIV | 1497355 | Cell/Virus Enclosed | NO |
| RFKCSSNGSNDV | 1497375 | Ectodomain | YES |
| RFKTSGDLAVQIGNNN | 1690044 | Cell/Virus Enclosed | NO |
| RFKVNETNSQYLFSTI | 1690045 | Cell/Virus Enclosed | NO |
| RFLGLQIQTVDSSQGS | 1690047 | Cell/Virus Enclosed | NO |
| RFMTANNLTAPRNSWE | 1690051 | Cell/Virus Enclosed | NO |
| RFMVDPTTVLNG | 1497398 | Cell/Virus Enclosed | NO |
| RFNPSTWNKRFGFIED | 1690055 | Ectodomain | YES |
| RFPCAGRKVNFNEKPV | 1690062 | Cell/Virus Enclosed | NO |
| RFPNITNRCPFD | 1497423 | Ectodomain | YES |
| RFPPGTVLPQGY | 1497428 | Cell/Virus Enclosed | NO |
| RFPRGQGVPINTNSGPDDQI | 53758 | Capsid | UN |
| RFPRGQGVPINTNSTK | 1690072 | Cell/Virus Enclosed | NO |
| RFQRFDENGDKL | 1497436 | Cell/Virus Enclosed | NO |
| RFQRIDDDGNKLDKFF | 1690078 | Cell/Virus Enclosed | NO |
| RFQRVDENGDKLDQFF | 1690079 | Cell/Virus Enclosed | NO |
| RFQRVGENGDKL | 1497438 | Cell/Virus Enclosed | NO |
| RFTGPYKCPQTK | 1497477 | Ectodomain | YES |
| RFTNFEPFTVNSVNDS | 1690095 | Ectodomain | YES |
| RFVELDDQGHHL | 1497490 | Cell/Virus Enclosed | NO |
| RFVTFEPFNVSFVNDS | 1690114 | Ectodomain | YES |
| RFVYDAYQNLVGYYSD | 1690115 | Ectodomain | YES |
| RFYRLANECAQI | 1497528 | Cell/Virus Enclosed | NO |
| RFYRLANECAQVLSEY | 1690124 | Cell/Virus Enclosed | NO |
| RGADGTAELTTTAATRFMKDLYFTS | 53816 | Undetermined | UN |
| RGAGDLQGNFGD | 1497546 | Cell/Virus Enclosed | NO |
| RGAIGDVGLCMY | 1497547 | Cell/Virus Enclosed | NO |
| RGAITTGYRFTNFEPF | 1690129 | Ectodomain | YES |
| RGAVSCGHYHTN | 1497556 | Cell/Virus Enclosed | NO |
| RGCCDCGSTKLPYYEF | 1690136 | Underneath Membrane | NO |
| RGCCESTKLPYYDVEK | 1690137 | Underneath Membrane | NO |
| RGDDMSFLEKDA | 1497573 | Cell/Virus Enclosed | NO |
| RGDYNMLLKNIDLFVK | 1690150 | Cell/Virus Enclosed | NO |
| RGEKGAYNKDHGRGGF | 1690153 | Cell/Virus Enclosed | NO |
| RGFCKTHNWNCF | 1497602 | Cell/Virus Enclosed | NO |
| RGFFDEGSELTL | 1497604 | Cell/Virus Enclosed | NO |
| RGFLDKIGDTSG | 1497609 | Underneath Membrane | NO |
| RGGDGKMKKLSP | 1497619 | Cell/Virus Enclosed | NO |
| RGGGSVTYTDQY | 1497623 | Cell/Virus Enclosed | NO |
| RGGYHYTPFHYERDNT | 1690173 | Cell/Virus Enclosed | NO |
| RGGYHYTPVRYE | 1497645 | Cell/Virus Enclosed | NO |
| RGHKNGQGENDNISVA | 1690174 | Cell/Virus Enclosed | NO |
| RGHLYIQGIKLGTGYS | 1690177 | Ectodomain | YES |
| RGHLYIQGVKLGTGYT | 1690178 | Underneath Membrane | NO |
| RGIACVKDFDFK | 1497662 | Cell/Virus Enclosed | NO |
| RGIAPKSGYFVNVNNT | 1690180 | Ectodomain | YES |
| RGKFVQVPLGIKDPIL | 1690185 | Cell/Virus Enclosed | NO |
| RGKFVQVPVGIKDPVS | 1690186 | Cell/Virus Enclosed | NO |
| RGKLLVRDTRKEIFVG | 1690187 | Cell/Virus Enclosed | NO |
| RGLALDNALSGVLSAA | 1690188 | Cell/Virus Enclosed | NO |
| RGLFGAIAGFIENGWEGLIDGWYGF | 1165813 | Ectodomain | YES |
| RGLFHSSQLPLD | 1497684 | Cell/Virus Enclosed | NO |
| RGLKNGQGENDN | 1497685 | Cell/Virus Enclosed | NO |
| RGLLHSSQLPLDVYDL | 1690190 | Cell/Virus Enclosed | NO |
| RGMPIPDFDLDDFIDA | 1690194 | Cell/Virus Enclosed | NO |
| RGNGKCLSFSNHLVST | 1690197 | Cell/Virus Enclosed | NO |
| RGPGDLQGNFGDLQLN | 1690203 | Cell/Virus Enclosed | NO |
| RGPGKCYNCRPA | 1497726 | Ectodomain | YES |
| RGPGKCYNCRPS | 1497727 | Ectodomain | YES |
| RGPGRCYNCRPA | 1497728 | Ectodomain | YES |
| RGPKNGQGENDN | 1497730 | Cell/Virus Enclosed | NO |
| RGPLQNFGNFEM | 1497732 | Cell/Virus Enclosed | NO |
| RGPNQNFGGGEMLKLG | 1690205 | Cell/Virus Enclosed | NO |
| RGPPRAELNGVVVDKV | 1690206 | Cell/Virus Enclosed | NO |
| RGQGVPINTNSSPDDQ | 1690210 | Cell/Virus Enclosed | NO |
| RGQGVPINTNSTKDDQ | 1690211 | Cell/Virus Enclosed | NO |
| RGRAIIVSVEQLEPCA | 1690215 | Cell/Virus Enclosed | NO |

| | | | |
|---|---|---|---|
| RGRESKSQSRNP | 1497755 | Cell/Virus Enclosed | NO |
| RGRKTQPKFTVS | 1497765 | Cell/Virus Enclosed | NO |
| RGRQFYEFYNDVKPPV | 1690221 | Cell/Virus Enclosed | NO |
| RGSGHYQTNLYS | 1497791 | Cell/Virus Enclosed | NO |
| RGSIVNARIEPCSSGL | 1690227 | Cell/Virus Enclosed | NO |
| RGSRSGNSSRGA | 1497803 | Cell/Virus Enclosed | NO |
| RGSVQVDNGSSINRRQ | 1690230 | Cell/Virus Enclosed | NO |
| RGTFPGPSGIGA | 1497818 | Cell/Virus Enclosed | NO |
| RGTGVLTTSSKR | 1497823 | Ectodomain | YES |
| RGTILFKTSYGA | 1497825 | Ectodomain | YES |
| RGTLEPENFNSVTRLM | 1690236 | Cell/Virus Enclosed | NO |
| RGTSPARIAGNGGDAA | 1690238 | Cell/Virus Enclosed | NO |
| RGTSPARMAGNGSDAA | 1690240 | Cell/Virus Enclosed | NO |
| RGTSPGPSGIGAVGGD | 1690241 | Cell/Virus Enclosed | NO |
| RGTYRAALNSEKHQDH | 1690248 | Cell/Virus Enclosed | NO |
| RGVEVVNATETVERTNIPRICSKGK | 1165884 | Ectodomain | YES |
| RGVKQLPEQFDSLNLS | 1690253 | Capsid | UN |
| RGVKQSPELFDS | 1497867 | Cell/Virus Enclosed | NO |
| RGVKQSPELLDP | 1497868 | Cell/Virus Enclosed | NO |
| RGVKQSPESFDS | 1497869 | Cell/Virus Enclosed | NO |
| RGVYKPLLENIDYFNM | 1690260 | Cell/Virus Enclosed | NO |
| RGWLGMDVEGAHVTGD | 1690265 | Cell/Virus Enclosed | NO |
| RGWLGMDVESAHVCGD | 1690266 | Cell/Virus Enclosed | NO |
| RGWVFGSTMDNK | 1497886 | Ectodomain | YES |
| RGWVFGSTMNNKSQSV | 1690268 | Ectodomain | YES |
| RGWVGFDVEGAHATRE | 1690269 | Cell/Virus Enclosed | NO |
| RHANEYRQYLDA | 1497898 | Cell/Virus Enclosed | NO |
| RHFFWVLAYFSK | 1497922 | Cell/Virus Enclosed | NO |
| RHGKLRPFERDISNVP | 1690285 | Ectodomain | YES |
| RHKRTSTKSFNM | 1497947 | Cell/Virus Enclosed | NO |
| RHLIKDVDNPVLMGWD | 1690302 | Cell/Virus Enclosed | NO |
| RHRLRRGADGTAELTTTAATRFMKD | 54090 | Undetermined | UN |
| RHSFRTLKSGEG | 1497988 | Cell/Virus Enclosed | NO |
| RHSSLSAQSEEE | 1498004 | Cell/Virus Enclosed | NO |
| RHVICPADQLSD | 1498019 | Cell/Virus Enclosed | NO |
| RHVMCPADQLSD | 1498025 | Cell/Virus Enclosed | NO |
| RIAGHHLGRCDIKDLP | 1690340 | Underneath Membrane | NO |
| RIAIEDLLFDKV | 1498040 | Ectodomain | YES |
| RIASCKWTDVDD | 1498045 | Cell/Virus Enclosed | NO |
| RIDDDGNKLDKFFVVK | 1690350 | Cell/Virus Enclosed | NO |
| RIDFQDELDEFF | 1498063 | Ectodomain | YES |
| RIDGAHLTKMSE | 1498065 | Cell/Virus Enclosed | NO |
| RIDTTATSCQLY | 1498074 | Ectodomain | YES |
| RIDVPSAESTGTQSVS | 1690357 | Cell/Virus Enclosed | NO |
| RIEDPPFNSLL | 144763 | Cell/Virus Enclosed | NO |
| RIGMEAAPSGTW | 1498119 | Capsid | UN |
| RIGNYKLNTDHSNSSD | 1690374 | Underneath Membrane | NO |
| RIGPIKYCYCGN | 1498125 | Cell/Virus Enclosed | NO |
| RIGTQVLKTMSL | 1498129 | Cell/Virus Enclosed | NO |
| RIHDFRYSQLAKLGINPYTH | 54158 | Capsid | UN |
| RIKFCDGENVSK | 1498138 | Cell/Virus Enclosed | NO |
| RILGAGCFVNDL | 1498151 | Cell/Virus Enclosed | NO |
| RILGVNSSNCNN | 1498152 | Cell/Virus Enclosed | NO |
| RINGLGVTMDVLNKNQ | 1690409 | Ectodomain | YES |
| RINWATGGIAIA | 1498172 | Undetermined | UN |
| RINWVTGGIAIA | 1498174 | Undetermined | UN |
| RIPNDFSFNNWF | 1498180 | Ectodomain | YES |
| RIPNLPDCDIDN | 1498183 | Ectodomain | YES |
| RIPTIADFVQIE | 1498185 | Ectodomain | YES |
| RIPVSDIFCDIENNP | 227512 | Cell/Virus Enclosed | NO |
| RIQSRNPSSDRN | 1498190 | Cell/Virus Enclosed | NO |
| RIRAKHYVYIGDPAQL | 1690421 | Cell/Virus Enclosed | NO |
| RIRSVYPFASPQ | 1498204 | Cell/Virus Enclosed | NO |
| RIRTRPSVQPGP | 1498208 | Cell/Virus Enclosed | NO |
| RISEYVVDVSHG | 1498215 | Ectodomain | YES |
| RITFGEPTDSTD | 1498224 | Capsid | UN |
| RITFGGPSDSTDNNQN | 1690435 | Cell/Virus Enclosed | NO |
| RITFGGPTDSAD | 1498229 | Capsid | UN |
| RITFGGPTDSTDNNQNGGR | 54296 | Capsid | UN |
| RITPYCHDPTVL | 1498234 | Cell/Virus Enclosed | NO |
| RITRTLSTQYCR | 1498235 | Cell/Virus Enclosed | NO |
| RIVDEWSYTGSSFYAP | 1690443 | Ectodomain | YES |
| RIVFTACSHAAVDSLC | 1690445 | Cell/Virus Enclosed | NO |
| RIVHCTKLNVPF | 1498249 | Cell/Virus Enclosed | NO |
| RIVQMLSDTLDKLSDY | 1690453 | Cell/Virus Enclosed | NO |
| RIVYAVLSDVDGLKYT | 1690461 | Cell/Virus Enclosed | NO |

| | | | |
|---|---|---|---|
| RIYPQQMFVTPL | 1498273 | Cell/Virus Enclosed | NO |
| RKACQNLDCNCL | 1498275 | Cell/Virus Enclosed | NO |
| RKAEDIFCPCWS | 1498276 | Cell/Virus Enclosed | NO |
| RKARNGVFISTEKLSR | 1690470 | Cell/Virus Enclosed | NO |
| RKARNRSHSPTK | 1498294 | Cell/Virus Enclosed | NO |
| RKASGLFDAIWDAFVAAI | 1498296 | Cell/Virus Enclosed | NO |
| RKATKRSHSPTK | 1498297 | Cell/Virus Enclosed | NO |
| RKDFTPVQVVDS | 1498311 | Ectodomain | YES |
| RKDLSYEQQNVLAIES | 1690481 | Cell/Virus Enclosed | NO |
| RKEIFVGDSLVN | 1498322 | Cell/Virus Enclosed | NO |
| RKELWHLDTGVV | 1498325 | Ectodomain | YES |
| RKFIKFGMTFVS | 1498330 | Cell/Virus Enclosed | NO |
| RKFYPAFMLGSS | 1498337 | Ectodomain | YES |
| RKGKHERDLLIKAYNT | 1690495 | Cell/Virus Enclosed | NO |
| RKGKRVDLSPKL | 1498352 | Cell/Virus Enclosed | NO |
| RKGMGTNLSVPNPLG | 1690497 | Ectodomain | YES |
| RKHATDYREYME | 1498361 | Cell/Virus Enclosed | NO |
| RKIKKYPCMVGPDYAY | 1690512 | Cell/Virus Enclosed | NO |
| RKINTGNGIKQL | 1498386 | Cell/Virus Enclosed | NO |
| RKKISNCVADYS | 1498399 | Ectodomain | YES |
| RKKKSNLLIQEFLQYD | 1690519 | Cell/Virus Enclosed | NO |
| RKKRNVDPLSNFEHKV | 1690520 | Cell/Virus Enclosed | NO |
| RKLDNQALNSILDNAV | 1690522 | Cell/Virus Enclosed | NO |
| RKLERMADLALTNMYK | 1690524 | Cell/Virus Enclosed | NO |
| RKLERMADQAMTSMYK | 1690525 | Cell/Virus Enclosed | NO |
| RKLNLPNCNIEA | 1498427 | Ectodomain | YES |
| RKMAFPSAKVEG | 1498430 | Cell/Virus Enclosed | NO |
| RKMDEPFTEDTS | 1498435 | Capsid | UN |
| RKNGQFQDHYDGFYTQ | 1690543 | Cell/Virus Enclosed | NO |
| RKPLDYKRQNNLAIEE | 1690545 | Cell/Virus Enclosed | NO |
| RKPLVSFKEDPTPSDNPFSK | 54408 | Cell/Virus Enclosed | NO |
| RKPNLPNCNIEAWLND | 1690548 | Ectodomain | YES |
| RKRGEKGAYNKDHGRG | 1690554 | Cell/Virus Enclosed | NO |
| RKRGVKQLPEQFDSLN | 1690555 | Capsid | UN |
| RKRGVKQSPESF | 1498476 | Cell/Virus Enclosed | NO |
| RKRNQKPLEPKFSIAL | 1690559 | Capsid | UN |
| RKRRSISASYRFVTFE | 1690560 | Ectodomain | YES |
| RKRTHSPTKKLR | 1498489 | Cell/Virus Enclosed | NO |
| RKSCQNLDCNCL | 1498497 | Cell/Virus Enclosed | NO |
| RKSCSYDSDVDT | 1498499 | Cell/Virus Enclosed | NO |
| RKSNVDESFVDD | 1498508 | Cell/Virus Enclosed | NO |
| RKTKLKPFERDL | 1498515 | Ectodomain | YES |
| RKTLVLNASEIY | 1498518 | Cell/Virus Enclosed | NO |
| RKTQPKFTVSTQ | 1498522 | Cell/Virus Enclosed | NO |
| RKVNTGNGVKQL | 1498537 | Cell/Virus Enclosed | NO |
| RKVQDAVNNNAH | 1498540 | Ectodomain | YES |
| RKVQQLFYIEDM | 1498545 | Cell/Virus Enclosed | NO |
| RLAFLGASVTEDVKFA | 1690604 | Cell/Virus Enclosed | NO |
| RLAMLGAEMTSNVRRQ | 1690606 | Cell/Virus Enclosed | NO |
| RLANECAQVLSEYVLC | 1690609 | Cell/Virus Enclosed | NO |
| RLANELAQVLTE | 1498584 | Cell/Virus Enclosed | NO |
| RLASGVQVHNLPEYMT | 1690612 | Underneath Membrane | NO |
| RLASIAFNHPIQ | 1498592 | Ectodomain | YES |
| RLCATCPFAKGD | 1498596 | Cell/Virus Enclosed | NO |
| RLDALEAQVQIDRLIN | 1690623 | Ectodomain | YES |
| RLDMSSVDNILN | 1498624 | Cell/Virus Enclosed | NO |
| RLDMSSVDTILNMARN | 1690625 | Cell/Virus Enclosed | NO |
| RLDSTYGPLQTP | 1498630 | Ectodomain | YES |
| RLDVLEAQVQID | 1498633 | Ectodomain | YES |
| RLEESGAVLFCT | 1498638 | Cell/Virus Enclosed | NO |
| RLEGLGGDNVYCYNTD | 1690631 | Cell/Virus Enclosed | NO |
| RLEGLGGNNVYCYNTA | 1690632 | Cell/Virus Enclosed | NO |
| RLEIVKLGAEKE | 1498641 | Cell/Virus Enclosed | NO |
| RLEIVKLGAEKKDGFY | 1690633 | Cell/Virus Enclosed | NO |
| RLEIVKLGAENK | 1498643 | Cell/Virus Enclosed | NO |
| RLELAKVQNLSGNPDEPQKD | 2060885 | Cell/Virus Enclosed | NO |
| RLELVKLGAEKK | 1498646 | Cell/Virus Enclosed | NO |
| RLEPCNGTDIDKCVRA | 1690635 | Cell/Virus Enclosed | NO |
| RLEPCNGTDIDY | 1498648 | Cell/Virus Enclosed | NO |
| RLESADVSEMLTFDKK | 1690637 | Ectodomain | YES |
| RLESKVSGKGQQ | 1498650 | Capsid | UN |
| RLFARTRSMWSFNPET | 1690640 | Underneath Membrane | NO |
| RLFASDLSTLAV | 1498653 | Cell/Virus Enclosed | NO |
| RLFSVPSDSCNN | 1498669 | Cell/Virus Enclosed | NO |
| RLFYDRDGQRTYDDVN | 1690652 | Cell/Virus Enclosed | NO |
| RLFYDRDGQRVYDDVD | 1690653 | Cell/Virus Enclosed | NO |

| | | | |
|---|---|---|---|
| RLGASQRVGTDS | 1498674 | Underneath Membrane | NO |
| RLGSQLSLSMVR | 1498681 | Cell/Virus Enclosed | NO |
| RLGVPDTLGGDPKFRSLTHE | 54566 | Capsid | UN |
| RLHCTTNLFKDC | 1498685 | Cell/Virus Enclosed | NO |
| RLHTVESGDVNV | 1498691 | Cell/Virus Enclosed | NO |
| RLIKDVDNPVLMGWDY | 1690671 | Cell/Virus Enclosed | NO |
| RLINGRLTALDA | 1498709 | Ectodomain | YES |
| RLINKFDVINGVKTVK | 1690674 | Cell/Virus Enclosed | NO |
| RLKGGLILKFDSGTVS | 1690688 | Cell/Virus Enclosed | NO |
| RLKGSFNDRDTG | 1498732 | Ectodomain | YES |
| RLKGSFNNRDTG | 1498733 | Ectodomain | YES |
| RLKNSVSDVAYNRYLS | 1690694 | Cell/Virus Enclosed | NO |
| RLLGLQTQTVDSAQGS | 1690700 | Cell/Virus Enclosed | NO |
| RLLLRNSGRFGG | 1498771 | Cell/Virus Enclosed | NO |
| RLLQDKAGDFMS | 1498776 | Cell/Virus Enclosed | NO |
| RLLQDKIGDFMS | 1498777 | Cell/Virus Enclosed | NO |
| RLLQDKTGDFTS | 1498780 | Cell/Virus Enclosed | NO |
| RLLSDDRTEVPQLVNA | 1690708 | Ectodomain | YES |
| RLLSDGRTEVPQ | 1498784 | Ectodomain | YES |
| RLNALESKMSGKGSQQ | 1690719 | Cell/Virus Enclosed | NO |
| RLNQLENKVSGK | 1498810 | Cell/Virus Enclosed | NO |
| RLPSSKSSGMDT | 1498819 | Ectodomain | YES |
| RLPSTQKGSGMDTALL | 1690729 | Ectodomain | YES |
| RLQAASDWKPGYSMPV | 1690731 | Cell/Virus Enclosed | NO |
| RLQAGFNTEFAS | 1498825 | Cell/Virus Enclosed | NO |
| RLQAGSNTEFASNSSV | 1690735 | Cell/Virus Enclosed | NO |
| RLQAGTATEYASNSSI | 1690736 | Cell/Virus Enclosed | NO |
| RLQAGVATEYAANSSI | 1690737 | Cell/Virus Enclosed | NO |
| RLQASADWKPGHAMPS | 1690739 | Cell/Virus Enclosed | NO |
| RLQASADWKPGY | 1498833 | Cell/Virus Enclosed | NO |
| RLQEAIKVLNQS | 1498836 | Ectodomain | YES |
| RLQSLENVAYNV | 1498841 | Cell/Virus Enclosed | NO |
| RLQTGKQTELAV | 1498843 | Cell/Virus Enclosed | NO |
| RLRKKRNVDPLSNFEH | 1690751 | Cell/Virus Enclosed | NO |
| RLSAHLETNDVSSMLT | 1690752 | Ectodomain | YES |
| RLSDVGFVEAYN | 1498870 | Ectodomain | YES |
| RLSKMGILKAEEFVAA | 1690756 | Cell/Virus Enclosed | NO |
| RLSLKELMMYAADPAM | 1690760 | Cell/Virus Enclosed | NO |
| RLTANVGRVIKFETDA | 1690768 | Cell/Virus Enclosed | NO |
| RLTSKGRPLVPTPQHTPLFS | 741919 | Ectodomain | YES |
| RLTTLNAFVAQQ | 1498918 | Ectodomain | YES |
| RLTTSKLRANDN | 1498922 | Cell/Virus Enclosed | NO |
| RLVASGSTVAMT | 1498925 | Ectodomain | YES |
| RLYDNCYRLTSVEESF | 1690799 | Cell/Virus Enclosed | NO |
| RLYSHVYRSDKVDSTF | 1690810 | Cell/Virus Enclosed | NO |
| RMADAVSRKKMDTK | 54848 | Capsid | UN |
| RMADLALTNMYKEARI | 1690815 | Cell/Virus Enclosed | NO |
| RMAGNGGDTALA | 1498998 | Cell/Virus Enclosed | NO |
| RMEDAMGQGQNN | 1499016 | Cell/Virus Enclosed | NO |
| RMGFKLDAIVPG | 1499022 | Cell/Virus Enclosed | NO |
| RMGFKLDATIPG | 1499027 | Cell/Virus Enclosed | NO |
| RMGFKLDATVPGYPKL | 1690835 | Cell/Virus Enclosed | NO |
| RMGSGFCVDYNSPSSS | 1690836 | Ectodomain | YES |
| RMRRGQRVDLPPKVHF | 1690846 | Capsid | UN |
| RMTPYCHDPTVL | 1499061 | Cell/Virus Enclosed | NO |
| RMVWREWCNVCG | 1499066 | Cell/Virus Enclosed | NO |
| RMYCSFYPPDDE | 1499067 | Cell/Virus Enclosed | NO |
| RMYCTFAIEDVEDVIS | 1690853 | Cell/Virus Enclosed | NO |
| RNASAPRTTPTPPQPRKATK | 741924 | Ectodomain | YES |
| RNASLNSFKEYFNLRN | 1690861 | Ectodomain | YES |
| RNAYVYTSQGFV | 1499090 | Cell/Virus Enclosed | NO |
| RNDCMLLCDILSIYAG | 1690866 | Cell/Virus Enclosed | NO |
| RNDCSILCEILCEYAD | 1690867 | Cell/Virus Enclosed | NO |
| RNEKWREQLGQKFEE | 54980 | Cell/Virus Enclosed | NO |
| RNESWHFDKSEPLCLF | 1690871 | Ectodomain | YES |
| RNFDEQNCDVLKEVLV | 1690873 | Cell/Virus Enclosed | NO |
| RNFFSPQIITTD | 1499107 | Ectodomain | YES |
| RNFTTAPAICHEGKAYF | 54989 | Ectodomain | YES |
| RNFTYDVNATYL | 1499114 | Ectodomain | YES |
| RNFYSPQIITTD | 1499119 | Ectodomain | YES |
| RNGQGENDNISV | 1499126 | Cell/Virus Enclosed | NO |
| RNGSVLEMAKEVKTSS | 1690882 | Cell/Virus Enclosed | NO |
| RNGVVPLSVIPATSAA | 1690887 | Cell/Virus Enclosed | NO |
| RNHNSQDDIMKAVAAA | 1690888 | Capsid | UN |
| RNIDATSTGNYNYKYR | 1690889 | Ectodomain | YES |
| RNKWSGVSDLSL | 1499162 | Cell/Virus Enclosed | NO |

| | | | |
|---|---|---|---|
| RNKWSGVSDLSS | 1499163 | Cell/Virus Enclosed | NO |
| RNKWSGVSNLSL | 1499164 | Cell/Virus Enclosed | NO |
| RNLQKRLYPNVY | 1499180 | Cell/Virus Enclosed | NO |
| RNLSGNPDEPQK | 1499185 | Cell/Virus Enclosed | NO |
| RNLTPGNTNTRVSRYSSTARHRLRR | 55026 | Undetermined | UN |
| RNNGCFGGFSNH | 1499202 | Cell/Virus Enclosed | NO |
| RNNSRDSFRSTS | 1499206 | Cell/Virus Enclosed | NO |
| RNNTSCPEWMDD | 1499208 | Cell/Virus Enclosed | NO |
| RNPANNAAIVLQLPQG | 1690917 | Cell/Virus Enclosed | NO |
| RNPNNDLAIVTQ | 1499220 | Cell/Virus Enclosed | NO |
| RNPSSDRNHNSQDDIM | 1690924 | Capsid | UN |
| RNPVDLPPSHAT | 1499231 | Cell/Virus Enclosed | NO |
| RNQSPASSQSAA | 1499236 | Cell/Virus Enclosed | NO |
| RNQSPASSQSVA | 1499237 | Cell/Virus Enclosed | NO |
| RNQSPASSQTSA | 1499238 | Capsid | UN |
| RNQSPSSSQSAA | 1499240 | Cell/Virus Enclosed | NO |
| RNQSPVSSQSAA | 1499241 | Cell/Virus Enclosed | NO |
| RNRFLYIIKLVF | 1499249 | Undetermined | UN |
| RNRSPERCDLGDDLHLQ | 55055 | Cell/Virus Enclosed | NO |
| RNSDVVKQPVKF | 1499253 | Cell/Virus Enclosed | NO |
| RNSFGKDLNANMSLAE | 1690939 | Cell/Virus Enclosed | NO |
| RNSLDPFCFDVN | 1499259 | Cell/Virus Enclosed | NO |
| RNSNGACIWNAA | 1499262 | Cell/Virus Enclosed | NO |
| RNSNGVCIWNAA | 1499263 | Cell/Virus Enclosed | NO |
| RNSSANYVGKYDINHG | 1690944 | Underneath Membrane | NO |
| RNSSDNGISAIITANL | 1690945 | Ectodomain | YES |
| RNSSRNSTPGSSRGTS | 1690946 | Cell/Virus Enclosed | NO |
| RNSSRSSSQGSRSGNS | 1690947 | Cell/Virus Enclosed | NO |
| RNSTPGSSRGTSPARM | 1690951 | Cell/Virus Enclosed | NO |
| RNSTTNYVGVYDINHG | 1690953 | Ectodomain | YES |
| RNSTTVSVTVSS | 1499282 | Cell/Virus Enclosed | NO |
| RNSTVWNGGAYS | 1499284 | Cell/Virus Enclosed | NO |
| RNSWEAMALNFT | 1499291 | Cell/Virus Enclosed | NO |
| RNTIMASKTVGTAE | 55066 | Capsid | UN |
| RNTREVFAQVKQMYKT | 1690962 | Ectodomain | YES |
| RNVFENFEDWHYAKFG | 1690968 | Cell/Virus Enclosed | NO |
| RNVSYDSVSAIV | 1499322 | Ectodomain | YES |
| RNYIITMPCFRT | 1499338 | Cell/Virus Enclosed | NO |
| RPAVINITLANF | 1499351 | Ectodomain | YES |
| RPEDLSVIQPTH | 1499367 | Cell/Virus Enclosed | NO |
| RPFERDISNVPFSPDGK | 55149 | Ectodomain | YES |
| RPGNFNQDFYDFVVSK | 1690998 | Cell/Virus Enclosed | NO |
| RPGNGILKWADQ | 1499385 | Capsid | UN |
| RPIDVFKADGII | 1499398 | Ectodomain | YES |
| RPIDVSKADGIIYPQG | 1691004 | Ectodomain | YES |
| RPIIEDYHTLTVTIIR | 1691006 | Ectodomain | YES |
| RPIVYETDANKL | 1499416 | Cell/Virus Enclosed | NO |
| RPKQRRPQGLPNNTASWF | 55196 | Capsid | UN |
| RPLGLQGCAFQSTVAELQRLKMKVG | 55207 | Undetermined | UN |
| RPLLESELVIGAVIL | 1324008 | Underneath Membrane | NO |
| RPLLVDDNKFDV | 1499427 | Cell/Virus Enclosed | NO |
| RPLMESELVIGA | 1499428 | Underneath Membrane | NO |
| RPLTETRGDLFSGDEDSDSSD | 55227 | Cell/Virus Enclosed | NO |
| RPLTTFGSGSPGRRHSQASY | 741930 | Underneath Membrane | NO |
| RPNRTIKGSFLN | 1499440 | Cell/Virus Enclosed | NO |
| RPNYTIKGSFLN | 1499441 | Cell/Virus Enclosed | NO |
| RPPKNSFEALIL | 1499442 | Cell/Virus Enclosed | NO |
| RPPPGEHFKHLV | 1499447 | Cell/Virus Enclosed | NO |
| RPPTPWPISQDSDRASYMDT | 55275 | Cell/Virus Enclosed | NO |
| RPQGLPNNTASWFTALTQHGK | 1397407 | Capsid | UN |
| RPRGQIYNPGGVSCAVNEAA | 741931 | Underneath Membrane | NO |
| RPRNSSDNGISAIITA | 1691038 | Ectodomain | YES |
| RPSLVDDNKFDG | 1499470 | Cell/Virus Enclosed | NO |
| RPSVAPGPMTDV | 1499473 | Cell/Virus Enclosed | NO |
| RPSVQPGPMIDV | 1499475 | Cell/Virus Enclosed | NO |
| RPTDNSTTVSVT | 1499478 | Cell/Virus Enclosed | NO |
| RPTTAGAAPLTAVAPAHDTPPVPDV | 55362 | Undetermined | UN |
| RPVIEDYHTLTATVIR | 1691046 | Underneath Membrane | NO |
| RPVIWLSHEKAS | 1499491 | Cell/Virus Enclosed | NO |
| RPYDLEVLLQDA | 1499498 | Cell/Virus Enclosed | NO |
| RPYHANSSEPAM | 1499499 | Ectodomain | YES |
| RQAALPQSKDSNFLNE | 1691049 | Cell/Virus Enclosed | NO |
| RQAALPQSKDSNFLNR | 1691050 | Cell/Virus Enclosed | NO |
| RQAALPQSKESN | 1499502 | Cell/Virus Enclosed | NO |
| RQAILCWGELMNLAT | 956912 | Capsid | UN |
| RQAYEDAIANGSSSQL | 1691055 | Cell/Virus Enclosed | NO |

| | | | |
|---|---|---|---|
| RQFDNMTAVACQ | 1499533 | Ectodomain | YES |
| RQFYEFYNDVKPPVLD | 1691073 | Cell/Virus Enclosed | NO |
| RQGIDYKQWPQI | 1499536 | Capsid | UN |
| RQGIFVEVNATY | 1499537 | Ectodomain | YES |
| RQIAPGQTGVIADYNY | 1691084 | Ectodomain | YES |
| RQIFDVAPIELENKFT | 1691085 | Cell/Virus Enclosed | NO |
| RQIFDVPPIELE | 1499565 | Cell/Virus Enclosed | NO |
| RQILGSCTFEDELAPS | 1691087 | Cell/Virus Enclosed | NO |
| RQIMGSCSFEDELTPS | 1691088 | Cell/Virus Enclosed | NO |
| RQKKQHTVTLLP | 1499579 | Capsid | UN |
| RQKLKDAERAVEVD | 55436 | Capsid | UN |
| RQKQSVWLANCDFDIV | 1691092 | Cell/Virus Enclosed | NO |
| RQKRTATKAYNVTQAF | 1691095 | Cell/Virus Enclosed | NO |
| RQKRTATKQYNVTQAF | 1691096 | Cell/Virus Enclosed | NO |
| RQKRTPNKFCNV | 1499589 | Cell/Virus Enclosed | NO |
| RQLMGSCSFEDE | 1499604 | Cell/Virus Enclosed | NO |
| RQLRPINYSFDS | 1499610 | Ectodomain | YES |
| RQLYDNCYRNSNVDES | 1691111 | Cell/Virus Enclosed | NO |
| RQNKPPSKPNNDFHFEVFNFVP | 125242 | Secreted | YES |
| RQNNLAIEEIEYVHGD | 1691115 | Cell/Virus Enclosed | NO |
| RQPFSFSFRDEL | 1499622 | Cell/Virus Enclosed | NO |
| RQPNDDVTSNVTQCFG | 1691116 | Capsid | UN |
| RQQEGHIIMEEM | 1499627 | Cell/Virus Enclosed | NO |
| RQQRFVYDAYQNLVGY | 1691119 | Ectodomain | YES |
| RQQSRTRSDSNQSSSD | 1691120 | Capsid | UN |
| RQQTSNLVAQEF | 1499635 | Cell/Virus Enclosed | NO |
| RQQTSNLVVQEFVSYD | 1691121 | Cell/Virus Enclosed | NO |
| RQRGHKNGQGENDNIS | 1691123 | Cell/Virus Enclosed | NO |
| RQRGLKNGQGEN | 1499646 | Cell/Virus Enclosed | NO |
| RQRGPKNGQGEN | 1499648 | Cell/Virus Enclosed | NO |
| RQRLTEYTMMDL | 1499652 | Cell/Virus Enclosed | NO |
| RQSDNMTAVACQPPYC | 1691127 | Ectodomain | YES |
| RQTPFEIKSAKK | 1499668 | Cell/Virus Enclosed | NO |
| RQVAPGETGVIA | 1499674 | Ectodomain | YES |
| RQVIQDVKDGTVTFNL | 1691135 | Cell/Virus Enclosed | NO |
| RQVIQEVKDGTV | 1499680 | Cell/Virus Enclosed | NO |
| RQVRSWIGFDVEGAHA | 1691136 | Cell/Virus Enclosed | NO |
| RQVVNVITTKIS | 1499687 | Cell/Virus Enclosed | NO |
| RQWLPTDAIIIDNDLN | 1691141 | Cell/Virus Enclosed | NO |
| RQYLLAFNQDGIIFNA | 1691146 | Ectodomain | YES |
| RQYNLSTSPLTSSVATGTNLVLYAA | 55519 | Undetermined | UN |
| RQYSNESWFEDDYRAF | 1691150 | Cell/Virus Enclosed | NO |
| RRAIDCGFNDLSQLHC | 1691155 | Ectodomain | YES |
| RRAITTENRFTN | 1499721 | Ectodomain | YES |
| RRAQPKQTVTSQ | 1499729 | Cell/Virus Enclosed | NO |
| RRAQPNQTATSQ | 1499730 | Cell/Virus Enclosed | NO |
| RRARGGDGKMKE | 1499731 | Capsid | UN |
| RRDGQPCYEQFP | 1499750 | Cell/Virus Enclosed | NO |
| RRDGQPFYEPFP | 1499752 | Cell/Virus Enclosed | NO |
| RRDSTYGPLQTPVGCV | 1691167 | Ectodomain | YES |
| RRFLVKLLDVYN | 1499767 | Cell/Virus Enclosed | NO |
| RRGPEQTQGNFGDQELIRQGTDYK | 1928137 | Cell/Virus Enclosed | NO |
| RRGQPVDLPPKV | 1499795 | Cell/Virus Enclosed | NO |
| RRGQRVDLPPKVHFYY | 1691176 | Capsid | UN |
| RRGRRSGGSGGGFWGDRVDSQPFAI | 55587 | Undetermined | UN |
| RRGVYYNDDIFR | 1499801 | Ectodomain | YES |
| RRIVQMLSDTLDKLSD | 1691184 | Cell/Virus Enclosed | NO |
| RRKKKSNLLIQEFLQY | 1691185 | Cell/Virus Enclosed | NO |
| RRKNSEPEIPHFNQKL | 1691186 | Capsid | UN |
| RRKPDPPNCNIE | 1499822 | Underneath Membrane | NO |
| RRKPNLPNCNIEAWLN | 1691187 | Ectodomain | YES |
| RRLIKDVDNPVLMGWD | 1691193 | Cell/Virus Enclosed | NO |
| RRLSKERFTHWF | 1499849 | Cell/Virus Enclosed | NO |
| RRLVNPTDKSHY | 1499853 | Cell/Virus Enclosed | NO |
| RRLYDNCYRLTSVEES | 1691199 | Cell/Virus Enclosed | NO |
| RRPKHLYVSMPTAQ | 55675 | Capsid | UN |
| RRPQGLPNNTASWFTA | 1387894 | Cell/Virus Enclosed | NO |
| RRQQTSNLVVQEFVSY | 1691210 | Cell/Virus Enclosed | NO |
| RRQYLLNFDEHG | 1499879 | Ectodomain | YES |
| RRSFKTADGQQKQLLP | 1691220 | Capsid | UN |
| RRSIDCGFNDLS | 1499906 | Ectodomain | YES |
| RRSISASYRFVTFEPF | 1691223 | Ectodomain | YES |
| RRSRGAITTGYRFTNF | 1691224 | Ectodomain | YES |
| RRTRREVIVNAQPKCNPN | 1045294 | Secreted | YES |
| RRVRGGDGKMKDLSPR | 1691233 | Cell/Virus Enclosed | NO |
| RRYSEAASNGFR | 1499955 | Cell/Virus Enclosed | NO |

| | | | |
|---|---|---|---|
| RSAIEDILFSKI | 1499966 | Ectodomain | YES |
| RSAPNSRSTSRTSSRA | 1691252 | Cell/Virus Enclosed | NO |
| RSAPRITFGEPT | 1499981 | Capsid | UN |
| RSAPRITFGGPS | 1499983 | Cell/Virus Enclosed | NO |
| RSCNYAVSSAFW | 1499995 | Cell/Virus Enclosed | NO |
| RSCSDFLPLSDI | 1500000 | Cell/Virus Enclosed | NO |
| RSDCNHIGNIKP | 1500014 | Ectodomain | YES |
| RSDCNHIGNINP | 1500015 | Ectodomain | YES |
| RSDCNHINNLKIKNFD | 1691264 | Ectodomain | YES |
| RSDGVVWVAKEGAKTV | 1691267 | Capsid | UN |
| RSDLQLGSSGFLQSSN | 1691272 | Ectodomain | YES |
| RSDMEYDFLNMDMGVF | 1691273 | Cell/Virus Enclosed | NO |
| RSDSNQSSSDLVAAVT | 1691274 | Capsid | UN |
| RSDVLHLTQDYF | 1500035 | Ectodomain | YES |
| RSELQTIVPEYIDVNK | 1691282 | Ectodomain | YES |
| RSELSQLSMYRVFEV | 55826 | Ectodomain | YES |
| RSEVGICLSTSG | 1500060 | Cell/Virus Enclosed | NO |
| RSFFEDLLFDKVKLSD | 1691286 | Ectodomain | YES |
| RSFHFTDFYNYTGEGQ | 1691287 | Ectodomain | YES |
| RSFKTADGNQRQLLPR | 1691288 | Cell/Virus Enclosed | NO |
| RSFKTADGQQKH | 1500072 | Cell/Virus Enclosed | NO |
| RSGARPKQRRPQGLPN | 1691296 | Cell/Virus Enclosed | NO |
| RSGILKKTSWADQSERNYQT | 2060889 | Capsid | UN |
| RSGILKKTSWVD | 1500100 | Cell/Virus Enclosed | NO |
| RSGNGILKWADQSDQF | 1691305 | Cell/Virus Enclosed | NO |
| RSGNSSRGASPG | 1500112 | Cell/Virus Enclosed | NO |
| RSGNSTRCTSPG | 1500114 | Cell/Virus Enclosed | NO |
| RSGNSTRGTSPGPSGI | 1691307 | Cell/Virus Enclosed | NO |
| RSGNSTRSTSPG | 1500119 | Capsid | UN |
| RSHFIRVSTVSS | 1500132 | Cell/Virus Enclosed | NO |
| RSHPDFKLLHNLQADI | 1691312 | Cell/Virus Enclosed | NO |
| RSHYYVDSVTVK | 1500145 | Cell/Virus Enclosed | NO |
| RSIDCGFNDLSQ | 1500149 | Ectodomain | YES |
| RSIKSDRKAWAA | 1500159 | Ectodomain | YES |
| RSINPTDQSSYI | 1500163 | Cell/Virus Enclosed | NO |
| RSIQSGRKAWAA | 1500167 | Ectodomain | YES |
| RSIRHHPELKILRNLN | 1691323 | Cell/Virus Enclosed | NO |
| RSISASYRFVTFEPFN | 1691324 | Ectodomain | YES |
| RSKEEDGFAFYP | 1500178 | Cell/Virus Enclosed | NO |
| RSKQRRPQGLPNNTAS | 1387924 | Cell/Virus Enclosed | NO |
| RSLSRSNSNFRHSDSI | 1691342 | Capsid | UN |
| RSMLGLITQTVD | 1500231 | Cell/Virus Enclosed | NO |
| RSMLKQRDSTYG | 1500234 | Ectodomain | YES |
| RSMLKRRDSTYGPLQT | 1691347 | Ectodomain | YES |
| RSMWSFNPETNILLNVPLHG | 1597799 | Underneath Membrane | NO |
| RSNIYGTATIGV | 1500250 | Cell/Virus Enclosed | NO |
| RSNSNFRHSASI | 1500252 | Cell/Virus Enclosed | NO |
| RSNSNFRHSDSIVKPD | 1691355 | Capsid | UN |
| RSPNKQCTVQQC | 1500256 | Cell/Virus Enclosed | NO |
| RSPPITADIELA | 1500258 | Ectodomain | YES |
| RSQMETDFLEPA | 1500267 | Cell/Virus Enclosed | NO |
| RSQSRGPGESKP | 1500269 | Capsid | UN |
| RSRASSGNRAPT | 1500287 | Cell/Virus Enclosed | NO |
| RSRCNSRNSTPG | 1500290 | Capsid | UN |
| RSRGAITTGYRFTNFE | 1691368 | Ectodomain | YES |
| RSRGNSRNSTPGSSR | 55940 | Capsid | UN |
| RSRNSSRNSTPGSSRG | 1691371 | Cell/Virus Enclosed | NO |
| RSRRAITTENRF | 1500305 | Ectodomain | YES |
| RSRRAVPVAVWLVSALAMGAGVA | 55950 | Ectodomain | YES |
| RSSFNFDQKELLAYYN | 1691373 | Cell/Virus Enclosed | NO |
| RSSFVDDNKFDV | 1500319 | Cell/Virus Enclosed | NO |
| RSSRIAGRSALEDLLFSKVVTSGL | 1387943 | Ectodomain | YES |
| RSSRIAIEDLLF | 1500333 | Ectodomain | YES |
| RSSSQGSRSGNSTRGT | 1691379 | Cell/Virus Enclosed | NO |
| RSSSQGSRSRNS | 1500337 | Cell/Virus Enclosed | NO |
| RSSTRNNSRDSF | 1500346 | Cell/Virus Enclosed | NO |
| RSSTRNNSRDSSRSTS | 1691382 | Capsid | UN |
| RSTKGGQQKNSQKGQ | 227532 | Capsid | UN |
| RSTRSMLKRRDSTYGP | 1691390 | Ectodomain | YES |
| RSTSPDPKFVDKYYAF | 1691391 | Cell/Virus Enclosed | NO |
| RSTSQKSIVAYTMSLG | 1691392 | Ectodomain | YES |
| RSTSRQQSRTRSDSNQ | 1691393 | Capsid | UN |
| RSTSRTSSRASSAGSR | 1691394 | Cell/Virus Enclosed | NO |
| RSVAYVANPTED | 1500381 | Cell/Virus Enclosed | NO |
| RSVFNLEGVNGGSLYV | 1691397 | Cell/Virus Enclosed | NO |
| RSVRGRAIIVSVEQLE | 1691406 | Cell/Virus Enclosed | NO |

| | | | |
|---|---|---|---|
| RSVRHHPELKLFRNLN | 1691407 | Cell/Virus Enclosed | NO |
| RSVRSHPDFKLL | 1500411 | Cell/Virus Enclosed | NO |
| RSVRSVPGEMCL | 1500412 | Ectodomain | YES |
| RSVRSVPGEMRLASIA | 1691408 | Ectodomain | YES |
| RSVRSVPGQMRL | 1500414 | Ectodomain | YES |
| RSVTVEYNIHAV | 1500419 | Cell/Virus Enclosed | NO |
| RSVYPFASPQEC | 1500424 | Cell/Virus Enclosed | NO |
| RSVYPVATPQEC | 1500427 | Cell/Virus Enclosed | NO |
| RSVYVKFQDSKPPLPP | 1691415 | Cell/Virus Enclosed | NO |
| RSWIGFDVEGAHASRN | 1691416 | Cell/Virus Enclosed | NO |
| RSYERLINTISP | 1500430 | Cell/Virus Enclosed | NO |
| RTAEAIHTVTIA | 1500451 | Ectodomain | YES |
| RTAIDCGFNDLS | 1500456 | Ectodomain | YES |
| RTDMEKDFIALDQDVF | 1691437 | Cell/Virus Enclosed | NO |
| RTDQINDMVYSL | 1500477 | Cell/Virus Enclosed | NO |
| RTDSLCSSHYWSPQGN | 1691438 | Cell/Virus Enclosed | NO |
| RTDVRDYKQAND | 1500479 | Cell/Virus Enclosed | NO |
| RTEVDPDLSAFY | 1500494 | Cell/Virus Enclosed | NO |
| RTEVDPDLSVFY | 1500495 | Cell/Virus Enclosed | NO |
| RTEVLQLVNANQ | 1500496 | Ectodomain | YES |
| RTEVPQLVNANQYSPC | 1691443 | Ectodomain | YES |
| RTFAKAPPGDNF | 1500500 | Cell/Virus Enclosed | NO |
| RTFPQQMFVTPL | 1500510 | Cell/Virus Enclosed | NO |
| RTFVVDKSLSID | 1500519 | Cell/Virus Enclosed | NO |
| RTFVVDKSLSIEEFAD | 1691453 | Cell/Virus Enclosed | NO |
| RTGKHEHSLLVEAYNS | 1691460 | Cell/Virus Enclosed | NO |
| RTGSFNNIDTGP | 1500537 | Ectodomain | YES |
| RTGVDPDLSAFY | 1500544 | Cell/Virus Enclosed | NO |
| RTGYIPLGTRPPT | 111013 | Capsid | UN |
| RTIDSTQDGVNK | 1500554 | Ectodomain | YES |
| RTIKGSFLNGSC | 1500556 | Cell/Virus Enclosed | NO |
| RTIKSTQDGDNK | 1500558 | Ectodomain | YES |
| RTILGAAALEDE | 1500561 | Cell/Virus Enclosed | NO |
| RTINSTQDGDNKLQGL | 1691480 | Ectodomain | YES |
| RTIQPEVKEVAP | 1500574 | Cell/Virus Enclosed | NO |
| RTITVGPRGNASNAAPSASP | 741947 | Ectodomain | YES |
| RTLDAANHSIPD | 1500592 | Cell/Virus Enclosed | NO |
| RTLFDALKFFEI | 1500596 | Cell/Virus Enclosed | NO |
| RTLFDALKFFEV | 1500597 | Cell/Virus Enclosed | NO |
| RTLIDGVENPML | 1500598 | Cell/Virus Enclosed | NO |
| RTLQEMKALFPT | 1500605 | Cell/Virus Enclosed | NO |
| RTLYQPVYKIFL | 1500618 | Cell/Virus Enclosed | NO |
| RTNGGYNSWHLVEGEL | 1691502 | Cell/Virus Enclosed | NO |
| RTNIPRICSKGKRTVDLGQCGLLGT | 1167503 | Ectodomain | YES |
| RTNLEVYNKEKTYYEL | 1691503 | Cell/Virus Enclosed | NO |
| RTPLNLPPTHAH | 1500635 | Cell/Virus Enclosed | NO |
| RTPNKHCNVQQCFGKR | 1691508 | Capsid | UN |
| RTPTPGVTPDMA | 1500641 | Cell/Virus Enclosed | NO |
| RTPTSGVTPDMADQIA | 1691509 | Cell/Virus Enclosed | NO |
| RTPTSGVTRDMA | 1500643 | Capsid | UN |
| RTRKGKRVDLSP | 1500657 | Cell/Virus Enclosed | NO |
| RTRSDSNQSSSDLVAA | 1691520 | Capsid | UN |
| RTRSMWSFNPETNIL | 1313501 | Underneath Membrane | NO |
| RTSLHPVLWNCQ | 1500683 | Cell/Virus Enclosed | NO |
| RTSNQGNKQIVH | 1500684 | Cell/Virus Enclosed | NO |
| RTSSDPGTNTTTEDH | 187509 | Secreted | YES |
| RTSSRASSAGSRSRAN | 1691531 | Cell/Virus Enclosed | NO |
| RTSSRASSTGSR | 1500691 | Cell/Virus Enclosed | NO |
| RTSSRTSSAGSR | 1500692 | Capsid | UN |
| RTSTSGVTPDMA | 1500693 | Cell/Virus Enclosed | NO |
| RTTSNFLPEVAA | 1500716 | Ectodomain | YES |
| RTTTSPSSPRDPTPAPGDTG | 741952 | Ectodomain | YES |
| RTVCFSVAALSTGLTS | 1691542 | Cell/Virus Enclosed | NO |
| RTVFENLKPMPFFYYS | 1691543 | Cell/Virus Enclosed | NO |
| RTVGELGDVRET | 1500725 | Cell/Virus Enclosed | NO |
| RTVGGVPLLSTM | 1500726 | Cell/Virus Enclosed | NO |
| RTVVRPLVEDST | 1500741 | Ectodomain | YES |
| RTYDDVNASLFVDYSN | 1691556 | Cell/Virus Enclosed | NO |
| RTYSEAAAQGFQ | 1500757 | Cell/Virus Enclosed | NO |
| RVAGDSGFAAYSRYRI | 1387989 | Underneath Membrane | NO |
| RVCIKTAQADTA | 1500776 | Cell/Virus Enclosed | NO |
| RVCYFNLCKEPM | 1500781 | Ectodomain | YES |
| RVDENGDKLDQFFVVK | 1691576 | Cell/Virus Enclosed | NO |
| RVDLNTTILFTG | 1500796 | Underneath Membrane | NO |
| RVDLQLGNLGYL | 1500799 | Ectodomain | YES |
| RVDWDIEYPYISHEKK | 1691585 | Cell/Virus Enclosed | NO |

| | | | |
|---|---|---|---|
| RVECYDRFKVNETNSQ | 1691596 | Cell/Virus Enclosed | NO |
| RVECYSGFKPNNNSAQ | 1691597 | Cell/Virus Enclosed | NO |
| RVECYSGFKPNNTSAQ | 1691598 | Cell/Virus Enclosed | NO |
| RVEHAVVYYVYSPSR | 56275 | Ectodomain | YES |
| RVEVFTDGKLNC | 1500838 | Cell/Virus Enclosed | NO |
| RVFEVGVIRNPGLGA | 56283 | Ectodomain | YES |
| RVFPYDMDSGVSSFSE | 1691617 | Cell/Virus Enclosed | NO |
| RVGLCEEADEGICFNF | 1691623 | Cell/Virus Enclosed | NO |
| RVGRSVNETSQTGWAF | 1691625 | Ectodomain | YES |
| RVGRSVNSQNSTGWVF | 1691626 | Underneath Membrane | NO |
| RVHSEFNLPGCDGGSL | 1691628 | Cell/Virus Enclosed | NO |
| RVIPRNLVPIGKGNKDEQI | 2060890 | Capsid | UN |
| RVIRRERCDMTFPCVS | 1691637 | Cell/Virus Enclosed | NO |
| RVISRMGFKLDATVPG | 1691638 | Cell/Virus Enclosed | NO |
| RVISSFTCRTDM | 1500893 | Cell/Virus Enclosed | NO |
| RVIYGNVSRHNL | 1500894 | Cell/Virus Enclosed | NO |
| RVKGQVDLFRNA | 1500900 | Cell/Virus Enclosed | NO |
| RVKGWVGFDVEG | 1500901 | Cell/Virus Enclosed | NO |
| RVKHGDFSAVSSPMSN | 1691645 | Underneath Membrane | NO |
| RVKLHTVIPDYV | 1500905 | Ectodomain | YES |
| RVKNLNSSEGVS | 1500907 | Underneath Membrane | NO |
| RVKNLNSSQGIP | 1500908 | Ectodomain | YES |
| RVKNTKLYVNKTLYSE | 1691649 | Ectodomain | YES |
| RVKSGSPGDSSWHIYL | 1691650 | Ectodomain | YES |
| RVKSVNGRVVIT | 1500914 | Cell/Virus Enclosed | NO |
| RVKSVSPGDSSW | 1500917 | Ectodomain | YES |
| RVLDQSDNNCWISTTL | 1691652 | Cell/Virus Enclosed | NO |
| RVLDQSENNCWI | 1500922 | Cell/Virus Enclosed | NO |
| RVLGLQTQTVDSAQGS | 1691656 | Cell/Virus Enclosed | NO |
| RVLGVQTQTVDSAQGS | 1691658 | Cell/Virus Enclosed | NO |
| RVLHLGAGSDKEVAPG | 1691659 | Cell/Virus Enclosed | NO |
| RVLNNLNLPGCNGGSL | 1691668 | Cell/Virus Enclosed | NO |
| RVMITKGVMEPI | 1500954 | Cell/Virus Enclosed | NO |
| RVNSACRKVQHM | 1500965 | Cell/Virus Enclosed | NO |
| RVNTAIPTIKDW | 1500967 | Cell/Virus Enclosed | NO |
| RVNVDGFNEWAMANGY | 1691687 | Cell/Virus Enclosed | NO |
| RVNVPYVCLLANKEQE | 1691688 | Cell/Virus Enclosed | NO |
| RVPHIVRKDLTKYTML | 1691690 | Cell/Virus Enclosed | NO |
| RVPSRSQSRSQS | 1500982 | Cell/Virus Enclosed | NO |
| RVPTIEENVIQC | 1500983 | Cell/Virus Enclosed | NO |
| RVPYQRNNTSFP | 1500988 | Cell/Virus Enclosed | NO |
| RVQGGITQRTRT | 1500990 | Cell/Virus Enclosed | NO |
| RVQGSITQHTRT | 1500994 | Cell/Virus Enclosed | NO |
| RVQGSITQRIRT | 1500996 | Cell/Virus Enclosed | NO |
| RVQGSITQRTRS | 1500998 | Cell/Virus Enclosed | NO |
| RVQGSITQRTRTRPSV | 1691695 | Cell/Virus Enclosed | NO |
| RVQQNKIRELTA | 1501007 | Cell/Virus Enclosed | NO |
| RVQQNKSRELTAEDIS | 1691699 | Cell/Virus Enclosed | NO |
| RVRAWVGFDAEGAHAT | 1691702 | Cell/Virus Enclosed | NO |
| RVRGGDGKMKDLSPRW | 1691703 | Cell/Virus Enclosed | NO |
| RVRGSIVNARIEPCSS | 1691705 | Cell/Virus Enclosed | NO |
| RVRGSSAARLEPCNGT | 1691706 | Cell/Virus Enclosed | NO |
| RVRGWVGFDVEGAHAT | 1691709 | Cell/Virus Enclosed | NO |
| RVRNPKPRAAPN | 1501028 | Cell/Virus Enclosed | NO |
| RVRSVRGRAIIVSVEQ | 1691714 | Cell/Virus Enclosed | NO |
| RVSAAFHANSSEPALL | 1691715 | Ectodomain | YES |
| RVSSNQSPQGNL | 1501055 | Cell/Virus Enclosed | NO |
| RVTLGDFTIMSG | 1501081 | Cell/Virus Enclosed | NO |
| RVTVPKDHPHLGKFLE | 1691734 | Capsid | UN |
| RVVDEWSYTGSS | 1501093 | Ectodomain | YES |
| RVVIPDFTVWNQ | 1501109 | Cell/Virus Enclosed | NO |
| RVVITNVGEPTI | 1501112 | Cell/Virus Enclosed | NO |
| RVVPSGDVVRFPNITN | 1691749 | Ectodomain | YES |
| RVVYTAASHAAVDALC | 1691757 | Cell/Virus Enclosed | NO |
| RVYANLGERVRQ | 1501139 | Cell/Virus Enclosed | NO |
| RVYDDVDASLFVDINN | 1691765 | Cell/Virus Enclosed | NO |
| RVYHIQPSLEDPFQPPSIPI | 741959 | Ectodomain | YES |
| RVYLAYIKKLYN | 1501145 | Cell/Virus Enclosed | NO |
| RVYNHFNNYDCV | 1501157 | Cell/Virus Enclosed | NO |
| RVYYESMSYQEQDELF | 1691776 | Cell/Virus Enclosed | NO |
| RWKRQPNDDATS | 1501171 | Cell/Virus Enclosed | NO |
| RWKRQPNDDVTSNVTQ | 1691781 | Capsid | UN |
| RWLLPDAAEELASPMK | 1691784 | Cell/Virus Enclosed | NO |
| RWLPHDAIVVDNDVVD | 1691785 | Cell/Virus Enclosed | NO |
| RWLPPDAIIIDNDIND | 1691786 | Cell/Virus Enclosed | NO |
| RWNNARHSDNMT | 1501183 | Ectodomain | YES |

| | | | |
|---|---|---|---|
| RWNNARQSDNKT | 1501186 | Ectodomain | YES |
| RWNNERASDISLSVAC | 1691790 | Underneath Membrane | NO |
| RWPQIAELAPTASAFM | 1691791 | Cell/Virus Enclosed | NO |
| RWRMRRGQRVDLPPKV | 1691796 | Capsid | UN |
| RWTGRCMSCCRSSRTRRETQL | 111750 | Cell/Virus Enclosed | NO |
| RYASAVAQAFQS | 1501218 | Cell/Virus Enclosed | NO |
| RYDALLSDLSFN | 1501229 | Cell/Virus Enclosed | NO |
| RYDNISSVWPLH | 1501234 | Ectodomain | YES |
| RYDNVSSVWPFY | 1501236 | Ectodomain | YES |
| RYGDVDNLVFTNKTTL | 1691832 | Cell/Virus Enclosed | NO |
| RYGIKPFTIPID | 1501269 | Cell/Virus Enclosed | NO |
| RYGNVLDVNAIDIE | 56537 | Capsid | UN |
| RYIYNRVKSGSPGDSS | 1691842 | Ectodomain | YES |
| RYIYNRVKSVSP | 1501281 | Ectodomain | YES |
| RYKAGNYRSPPITADI | 1691843 | Ectodomain | YES |
| RYKYYSGNANEADYRC | 1691845 | Cell/Virus Enclosed | NO |
| RYLRHGKLRPFERDIS | 1691848 | Ectodomain | YES |
| RYMNANGLRPPKNSFE | 1691852 | Cell/Virus Enclosed | NO |
| RYMNANGLRPPRNSFE | 1691853 | Cell/Virus Enclosed | NO |
| RYMSAFNHTKKW | 1501310 | Cell/Virus Enclosed | NO |
| RYNEVANAVMQNNELM | 1691859 | Cell/Virus Enclosed | NO |
| RYNFDITESKYSCKEV | 1691861 | Cell/Virus Enclosed | NO |
| RYNLNFSANSLDNLKS | 1691867 | Ectodomain | YES |
| RYNRPTMLDIGQARVA | 1691869 | Cell/Virus Enclosed | NO |
| RYQEFNDLDHTS | 1501343 | Cell/Virus Enclosed | NO |
| RYRIGNYKLNTDHSNS | 1691874 | Underneath Membrane | NO |
| RYRIGNYKLNTDHSSSSDNIA | 1625429 | Underneath Membrane | NO |
| RYSEAASNGFRA | 1501349 | Cell/Virus Enclosed | NO |
| RYSKKFKPEIAIRPKVRDQEGRMNY | 1167778 | Ectodomain | YES |
| RYSLADSNGFIR | 1501354 | Cell/Virus Enclosed | NO |
| RYSSTARHRLRRGADGTAELTTTAA | 56660 | Undetermined | UN |
| RYSVLSVDSVSEESQG | 1691881 | Cell/Virus Enclosed | NO |
| RYYSGKMDTAAYREAA | 1691896 | Cell/Virus Enclosed | NO |
| SAAEFKYMVANGLHAP | 1691902 | Cell/Virus Enclosed | NO |
| SAAFHANSSEPALLFR | 1691903 | Ectodomain | YES |
| SAAFHQNASSLA | 1501401 | Ectodomain | YES |
| SAAFHQSASSLA | 1501402 | Ectodomain | YES |
| SAAGYGGFGCKD | 1501405 | Cell/Virus Enclosed | NO |
| SAAIKDCVDINKAIPS | 1691906 | Cell/Virus Enclosed | NO |
| SAAKLQNNELSP | 1501420 | Cell/Virus Enclosed | NO |
| SAALSAQLAKDKVNEC | 1691911 | Ectodomain | YES |
| SAAPAAPAAPRASGGVAATV | 741963 | Undetermined | UN |
| SAAQADLQHEDS | 1501430 | Cell/Virus Enclosed | NO |
| SAAQAMEKVNECVKSQ | 1691912 | Ectodomain | YES |
| SAARIHDFRYSQLAKLGINP | 56712 | Capsid | UN |
| SAARLEPCNGTDIDKC | 1691913 | Cell/Virus Enclosed | NO |
| SAARLVVVVPDHDSFV | 1691916 | Cell/Virus Enclosed | NO |
| SAAVLKSKWWLLAKFT | 1691921 | Cell/Virus Enclosed | NO |
| SAAYHANSPEPA | 1501446 | Ectodomain | YES |
| SAAYHANSSDPA | 1501447 | Ectodomain | YES |
| SACDGLASAYRANSFD | 1691922 | Cell/Virus Enclosed | NO |
| SACKTIEDALRLSAHLETND | 1388052 | Ectodomain | YES |
| SACKTIEDALRNSARL | 1691926 | Ectodomain | YES |
| SACPKLEFANDT | 1501467 | Ectodomain | YES |
| SACTRLEGLGGN | 1501480 | Cell/Virus Enclosed | NO |
| SADAIVLKLPSLMKVM | 1691938 | Cell/Virus Enclosed | NO |
| SADFIKHVRTNG | 1501490 | Cell/Virus Enclosed | NO |
| SADGVQCYTPHS | 1501492 | Cell/Virus Enclosed | NO |
| SADIVDRDPSSD | 1501495 | Cell/Virus Enclosed | NO |
| SADSCLQGDKCNIFAN | 1691948 | Ectodomain | YES |
| SADVSEMLTFDKKAFT | 1691949 | Ectodomain | YES |
| SADWKPGHAMPSLFKV | 1691950 | Cell/Virus Enclosed | NO |
| SADWKPGHSMPS | 1501511 | Cell/Virus Enclosed | NO |
| SAEGRKYIVKTS | 1501517 | Cell/Virus Enclosed | NO |
| SAEVSRQVIQDVKDGT | 1691956 | Cell/Virus Enclosed | NO |
| SAEWKCGYAMPQ | 1501526 | Cell/Virus Enclosed | NO |
| SAEWKCGYSMPG | 1501527 | Cell/Virus Enclosed | NO |
| SAFFGMSRIGMEVTPSGTWL | 2060891 | Capsid | UN |
| SAFIDTIKSLDY | 1501537 | Cell/Virus Enclosed | NO |
| SAFIECGSCGND | 1501539 | Cell/Virus Enclosed | NO |
| SAFIECGSCGNN | 1501541 | Cell/Virus Enclosed | NO |
| SAFVECGSCGND | 1501573 | Cell/Virus Enclosed | NO |
| SAFYVKDGKYFTSEPP | 1691983 | Cell/Virus Enclosed | NO |
| SAGDSIFKSFHF | 1501586 | Ectodomain | YES |
| SAGGQLFYSRPVVSANGE | 56796 | Undetermined | UN |
| SAGHATGITPQK | 1501596 | Ectodomain | YES |

| | | | |
|---|---|---|---|
| SAGHATGTTLQK | 1501598 | Ectodomain | YES |
| SAGHATGTTPQKLFVA | 1691992 | Ectodomain | YES |
| SAGIEFLRDAWE | 1501605 | Cell/Virus Enclosed | NO |
| SAGLELTDESCNNLVP | 1692001 | Cell/Virus Enclosed | NO |
| SAGMFGLYDDILTNNK | 1692002 | Cell/Virus Enclosed | NO |
| SAGNATEVPANS | 1501620 | Cell/Virus Enclosed | NO |
| SAGNIFGLYKNSATGS | 1692003 | Cell/Virus Enclosed | NO |
| SAGQEQTNCNTS | 1501627 | Cell/Virus Enclosed | NO |
| SAGSRSRANSGNRTPT | 1692008 | Cell/Virus Enclosed | NO |
| SAGSVGFNIDYD | 1501639 | Cell/Virus Enclosed | NO |
| SAGTQHISNDFTPEDH | 1692009 | Capsid | UN |
| SAGVVANGVKAKGYPQ | 1692013 | Capsid | UN |
| SAGWPLNKFGKAGLYY | 1692014 | Cell/Virus Enclosed | NO |
| SAGYPFNQFGKA | 1501648 | Cell/Virus Enclosed | NO |
| SAHLETNDVSSMLTFD | 1692017 | Ectodomain | YES |
| SAHVCGDNIGTNVPLQ | 1692018 | Cell/Virus Enclosed | NO |
| SAIAKPKRLRKK | 1501654 | Cell/Virus Enclosed | NO |
| SAIDYDHEYSIMRLHN | 1692019 | Cell/Virus Enclosed | NO |
| SAIEDLLFDKVTIADP | 1692022 | Ectodomain | YES |
| SAIEDLLFSKVR | 1501660 | Ectodomain | YES |
| SAIITANLSIPSNWTT | 1692025 | Ectodomain | YES |
| SAIVCASVKRDGVQVG | 1692033 | Cell/Virus Enclosed | NO |
| SAIVCASVLKDGCDVG | 1692034 | Cell/Virus Enclosed | NO |
| SAIVTANLSIPSNWTT | 1692035 | Ectodomain | YES |
| SAIVTQFAPGTKLPKN | 1692036 | Cell/Virus Enclosed | NO |
| SAIYPIDLSSQI | 1501679 | Undetermined | UN |
| SAKSLARSQSSETKEQ | 1692045 | Capsid | UN |
| SAKTGTPKPSRNQSPA | 1692046 | Capsid | UN |
| SALASALKQLKVTTGE | 1692048 | Cell/Virus Enclosed | NO |
| SALEFNPSQTSPATAE | 1692050 | Capsid | UN |
| SALEWIDLDSSL | 56856 | Ectodomain | YES |
| SALFVVKQKWCV | 1501722 | Cell/Virus Enclosed | NO |
| SALGSMDIDVFNKLST | 1692054 | Cell/Virus Enclosed | NO |
| SALIAVANWLAPTNAY | 1692055 | Cell/Virus Enclosed | NO |
| SALIGANGNKIV | 1501730 | Cell/Virus Enclosed | NO |
| SALKQLKVTTGELVRF | 1692056 | Cell/Virus Enclosed | NO |
| SALKTAGVVTGNVVKQ | 1692057 | Cell/Virus Enclosed | NO |
| SALMGANGNKIVDKEV | 1692063 | Cell/Virus Enclosed | NO |
| SALMSCNGNKIE | 1501743 | Cell/Virus Enclosed | NO |
| SALNGSCGSVGF | 1501744 | Cell/Virus Enclosed | NO |
| SALVKIQAVVNANAEA | 1692071 | Ectodomain | YES |
| SAMNVIGQHIEL | 1501780 | Cell/Virus Enclosed | NO |
| SAMNVIGQHIKL | 1501781 | Cell/Virus Enclosed | NO |
| SAMSTTDLEAYFKDC | 56900 | Cell/Virus Enclosed | NO |
| SANFLDDNVLPETYVA | 1692091 | Ectodomain | YES |
| SANGEPTVKLYTSVENAQQDKGIAI | 56908 | Undetermined | UN |
| SANIFIGDNVGH | 1501806 | Cell/Virus Enclosed | NO |
| SANLFPPWSAAA | 1501811 | Ectodomain | YES |
| SANVCALISCNG | 1501826 | Cell/Virus Enclosed | NO |
| SANVSALMSCNG | 1501828 | Cell/Virus Enclosed | NO |
| SANYVGKYDINHGDSG | 1692108 | Underneath Membrane | NO |
| SAPNSRSTSRTSSRAS | 1692115 | Cell/Virus Enclosed | NO |
| SAPPSEYTLQQG | 1501843 | Cell/Virus Enclosed | NO |
| SAPPVTYSPATI | 1501844 | Cell/Virus Enclosed | NO |
| SAPQTIQPEVKEVAPV | 1692120 | Cell/Virus Enclosed | NO |
| SAPRITFGGPID | 1501847 | Capsid | UN |
| SAPTLIPQENYS | 1501850 | Cell/Virus Enclosed | NO |
| SAPTLVPQENYASIRF | 1692123 | Cell/Virus Enclosed | NO |
| SAPTLVPQENYSSIRF | 1692124 | Cell/Virus Enclosed | NO |
| SAQAIAFCKSGS | 1501857 | Ectodomain | YES |
| SAQSEEEESFSN | 1501869 | Cell/Virus Enclosed | NO |
| SAQYIFSTANAL | 1501874 | Cell/Virus Enclosed | NO |
| SAQYIFSTVNALPECN | 1692132 | Cell/Virus Enclosed | NO |
| SAQYVFSTVNALPEVN | 1692133 | Cell/Virus Enclosed | NO |
| SARDGIRDAMIGLREEMIEK | 56962 | Cell/Virus Enclosed | NO |
| SARDPIIQEAIP | 1501880 | Cell/Virus Enclosed | NO |
| SARDPTIQEAIP | 1501881 | Cell/Virus Enclosed | NO |
| SARNLICAQYVA | 1501894 | Underneath Membrane | NO |
| SASAFFGMSRIGMEVT | 1692144 | Cell/Virus Enclosed | NO |
| SASAVNMQQIYL | 1501904 | Cell/Virus Enclosed | NO |
| SASCPIGTNYRSCEST | 1692147 | Ectodomain | YES |
| SASDMTNPDYTNLLCR | 1692148 | Cell/Virus Enclosed | NO |
| SASEADYRCACYAHLA | 1692149 | Cell/Virus Enclosed | NO |
| SASGYGGFGCKD | 1501922 | Cell/Virus Enclosed | NO |
| SASIGDIIRRLD | 1501924 | Ectodomain | YES |
| SASINTGNCPFSFGKV | 1692155 | Ectodomain | YES |

| | | | |
|---|---|---|---|
| SASIVKPDMADE | 1501930 | Cell/Virus Enclosed | NO |
| SASKLTIVSPDLESYS | 1692160 | Cell/Virus Enclosed | NO |
| SASLFKPVHLVP | 1501934 | Cell/Virus Enclosed | NO |
| SASLFKPVHLVS | 1501935 | Cell/Virus Enclosed | NO |
| SASLFKPVQLAP | 1501936 | Cell/Virus Enclosed | NO |
| SASLFKPVQLVPVSPA | 1692162 | Cell/Virus Enclosed | NO |
| SASLQEILSRLDALEA | 1692164 | Ectodomain | YES |
| SASNSRPGSRSQSRGP | 1692165 | Capsid | UN |
| SASNSRSTSRTS | 1501943 | Cell/Virus Enclosed | NO |
| SASTCAFVDTVK | 1501949 | Cell/Virus Enclosed | NO |
| SASYFCKIGREQ | 1501960 | Cell/Virus Enclosed | NO |
| SASYRFVTFEPFNVSF | 1692172 | Ectodomain | YES |
| SASYVLPSEAPN | 1501962 | Cell/Virus Enclosed | NO |
| SATEFADKLVEV | 1501967 | Cell/Virus Enclosed | NO |
| SATFSVPMEELK | 1501969 | Cell/Virus Enclosed | NO |
| SATGSLDVEVFNRLAT | 1692182 | Cell/Virus Enclosed | NO |
| SATIRKFYPAFM | 1501976 | Ectodomain | YES |
| SATKIIALNKRWQLAL | 1692186 | Secreted | YES |
| SATKLNDLCFSNVYAD | 1692187 | Ectodomain | YES |
| SATKVYFDNPTL | 1501982 | Ectodomain | YES |
| SATNSALAKIQS | 1501989 | Ectodomain | YES |
| SATQPYFDSYLG | 1501991 | Ectodomain | YES |
| SATSNYSGVVTT | 1501993 | Cell/Virus Enclosed | NO |
| SATVTANLSIPS | 1501996 | Ectodomain | YES |
| SATYEGNSPFHP | 1501997 | Ectodomain | YES |
| SAVANAHRYDVL | 1502002 | Cell/Virus Enclosed | NO |
| SAVAVVGGTIQILASV | 1692197 | Cell/Virus Enclosed | NO |
| SAVFISPYNSQNYVAK | 1692199 | Cell/Virus Enclosed | NO |
| SAVGVRQQRFVY | 1502014 | Ectodomain | YES |
| SAVLWANGVKAK | 1502025 | Cell/Virus Enclosed | NO |
| SAVNFTKALAKQ | 1502029 | Cell/Virus Enclosed | NO |
| SAVNQSLSNAKVSAEV | 1692211 | Cell/Virus Enclosed | NO |
| SAVNSTKALAKQDAAQ | 1692212 | Cell/Virus Enclosed | NO |
| SAVSFDYLIREA | 1502041 | Cell/Virus Enclosed | NO |
| SAVSSPMSNMTENERL | 1692217 | Underneath Membrane | NO |
| SAVVNGIRVLKTSDNN | 1692220 | Cell/Virus Enclosed | NO |
| SAYGCSTIAQAV | 1502052 | Cell/Virus Enclosed | NO |
| SAYGLCDAANPTNCIA | 1692228 | Ectodomain | YES |
| SAYGLCDLANPT | 1502056 | Ectodomain | YES |
| SAYIKLNCFYDP | 1502059 | Ectodomain | YES |
| SAYRANSFDAPT | 1502066 | Cell/Virus Enclosed | NO |
| SAYRANSFDVPTFCAN | 1692230 | Cell/Virus Enclosed | NO |
| SCAHEYGTEEDY | 1502078 | Cell/Virus Enclosed | NO |
| SCAKSKPLSAIC | 1502079 | Ectodomain | YES |
| SCARKSFSIDSD | 1502087 | Cell/Virus Enclosed | NO |
| SCDCGKKFDDQVGCLF | 1692246 | Cell/Virus Enclosed | NO |
| SCDCGKKFDEQV | 1502105 | Cell/Virus Enclosed | NO |
| SCDFKGWIPGNM | 1502106 | Cell/Virus Enclosed | NO |
| SCDINKPLQVFV | 1502111 | Cell/Virus Enclosed | NO |
| SCDLRMGSGFCI | 1502112 | Ectodomain | YES |
| SCDPAQLPFSAPRLGPSSVY | 741972 | Ectodomain | YES |
| SCDQLREPMMQS | 1502115 | Cell/Virus Enclosed | NO |
| SCEKNLQLMRKVDLYN | 1692253 | Cell/Virus Enclosed | NO |
| SCELYHYQECVR | 1502120 | Cell/Virus Enclosed | NO |
| SCESTTVLDRTD | 1502124 | Ectodomain | YES |
| SCEVVHDQVNYVPLRA | 1692257 | Cell/Virus Enclosed | NO |
| SCFFTCNSNANI | 1502133 | Ectodomain | YES |
| SCFFTCNSNANL | 1502134 | Ectodomain | YES |
| SCFGKNNGIGTC | 1502136 | Ectodomain | YES |
| SCGHYQTNIYSQNLCV | 1692267 | Cell/Virus Enclosed | NO |
| SCGLFKDCARNPIDLP | 1692271 | Cell/Virus Enclosed | NO |
| SCGNDSWLTGNAIQGF | 1692272 | Cell/Virus Enclosed | NO |
| SCGNESWLTGNA | 1502168 | Cell/Virus Enclosed | NO |
| SCGNNSWLTGNA | 1502171 | Cell/Virus Enclosed | NO |
| SCGYDYVYSPFF | 1502185 | Cell/Virus Enclosed | NO |
| SCGYHYKELGLV | 1502186 | Cell/Virus Enclosed | NO |
| SCIIGSLGFYNPPTNV | 1692284 | Ectodomain | YES |
| SCIWSVDAFNQF | 1502214 | Cell/Virus Enclosed | NO |
| SCIYWSPRPGLW | 1502216 | Cell/Virus Enclosed | NO |
| SCLVSSDGQPGGDAIS | 1692313 | Cell/Virus Enclosed | NO |
| SCLVSSNGQPGG | 1502264 | Cell/Virus Enclosed | NO |
| SCNFNMSSLMSF | 1502268 | Ectodomain | YES |
| SCNFPLEATWHY | 1502269 | Ectodomain | YES |
| SCNGNKIEDFSI | 1502271 | Cell/Virus Enclosed | NO |
| SCNGNKIEDLSI | 1502272 | Cell/Virus Enclosed | NO |
| SCNGNKIEDLSM | 1502273 | Cell/Virus Enclosed | NO |

| | | | |
|---|---|---|---|
| SCNNLVPTYLKSDNIV | 1692321 | Cell/Virus Enclosed | NO |
| SCNNVNVRDLQRRLYD | 1692322 | Cell/Virus Enclosed | NO |
| SCPDKNNGIGIC | 1502284 | Ectodomain | YES |
| SCPDKNNGIGTC | 1502285 | Ectodomain | YES |
| SCPEWMDDFEADPKGK | 1692327 | Cell/Virus Enclosed | NO |
| SCPEWMDEFEAD | 1502287 | Cell/Virus Enclosed | NO |
| SCPGKNNGIGTC | 1502289 | Ectodomain | YES |
| SCPGKNNGLGTC | 1502292 | Ectodomain | YES |
| SCPIGTNYRSCESTTV | 1692328 | Ectodomain | YES |
| SCPLCCTVYDSSEVKA | 1692330 | Cell/Virus Enclosed | NO |
| SCQLYYSLPLLN | 1502307 | Ectodomain | YES |
| SCQNLDCNCLGF | 1502310 | Cell/Virus Enclosed | NO |
| SCRLLQRVMLKA | 1502317 | Cell/Virus Enclosed | NO |
| SCRVLQRVMLKA | 1502320 | Cell/Virus Enclosed | NO |
| SCSCPGKNNGIG | 1502322 | Ectodomain | YES |
| SCSCVSTDTTVQSKDT | 1692339 | Cell/Virus Enclosed | NO |
| SCSDFIPLSDME | 1502327 | Cell/Virus Enclosed | NO |
| SCSDFLPLSDMEKDFL | 1692340 | Cell/Virus Enclosed | NO |
| SCSDFLPMSDME | 1502330 | Cell/Virus Enclosed | NO |
| SCSFEDELTPSDVYQQ | 1692341 | Cell/Virus Enclosed | NO |
| SCSGKNNGIGTC | 1502332 | Ectodomain | YES |
| SCSGKNNGVGTC | 1502333 | Ectodomain | YES |
| SCSIDSDVDTKC | 1502334 | Cell/Virus Enclosed | NO |
| SCSSNCKHSVAQLVSY | 1692347 | Cell/Virus Enclosed | NO |
| SCTCQPQAFLGW | 1502349 | Ectodomain | YES |
| SCTCRPQAFLGW | 1502351 | Ectodomain | YES |
| SCTFEDELAPSDVYQQ | 1692349 | Cell/Virus Enclosed | NO |
| SCTGSPYIDDFNKIAS | 1692350 | Cell/Virus Enclosed | NO |
| SCTGSPYIDDFNRIAS | 1692351 | Cell/Virus Enclosed | NO |
| SCTKGLSIADLACAQY | 1692353 | Ectodomain | YES |
| SCTTQQLTIEVL | 1502361 | Cell/Virus Enclosed | NO |
| SCVGSSVAVQSKDLNF | 1692360 | Cell/Virus Enclosed | NO |
| SCVSTDTTVQSKDTNF | 1692363 | Cell/Virus Enclosed | NO |
| SCVVCHSQTSLR | 1502382 | Cell/Virus Enclosed | NO |
| SCVYAICYFYAT | 1502383 | Cell/Virus Enclosed | NO |
| SCYSGRVSAAFHANSS | 1692369 | Ectodomain | YES |
| SDAALALLLLDRLNQL | 1692374 | Cell/Virus Enclosed | NO |
| SDAFSLDVSEKSGNFK | 1692380 | Ectodomain | YES |
| SDAHKKNLYDHAL | 189839 | Ectodomain | YES |
| SDALSTNHCHKV | 1502416 | Cell/Virus Enclosed | NO |
| SDAQGTNHGYKV | 1502422 | Cell/Virus Enclosed | NO |
| SDCDVVQEQVNYVPLR | 1692389 | Cell/Virus Enclosed | NO |
| SDCNHIGNIKPH | 1502437 | Ectodomain | YES |
| SDCNHIGNINPH | 1502438 | Ectodomain | YES |
| SDCNHINNLKIKNFDY | 1692391 | Ectodomain | YES |
| SDCNHVVNTNPR | 1502441 | Ectodomain | YES |
| SDCNRVSSRSAI | 1502443 | Ectodomain | YES |
| SDCNVVRYFLDG | 1502444 | Cell/Virus Enclosed | NO |
| SDCNVVRYSLDG | 1502445 | Cell/Virus Enclosed | NO |
| SDDDFVSAVANAHRYD | 1692399 | Cell/Virus Enclosed | NO |
| SDDEWAAAVDEAFPLD | 1692400 | Cell/Virus Enclosed | NO |
| SDDEWVAAVDEA | 1502462 | Cell/Virus Enclosed | NO |
| SDDGVVCYNKDYAELG | 1692402 | Cell/Virus Enclosed | NO |
| SDDGVVCYNSDYASKG | 1692404 | Cell/Virus Enclosed | NO |
| SDDVIVHDVEPT | 1502479 | Cell/Virus Enclosed | NO |
| SDDVIVYDVEPT | 1502480 | Cell/Virus Enclosed | NO |
| SDEAIPTRFPPGTVLP | 1692415 | Cell/Virus Enclosed | NO |
| SDEAISTRFPPG | 1502487 | Cell/Virus Enclosed | NO |
| SDEYILAKGLLQ | 1502501 | Cell/Virus Enclosed | NO |
| SDEYILAKGPLQVGDS | 1692421 | Cell/Virus Enclosed | NO |
| SDFAFNNWFLLTNTSS | 1692422 | Ectodomain | YES |
| SDFFPSVRDLLDTASALYRE | 96862 | Capsid | UN |
| SDFINAIFAKVK | 1502518 | Ectodomain | YES |
| SDFINGIFAKVKNTKV | 1692429 | Ectodomain | YES |
| SDFLPLSDIEKD | 1502526 | Cell/Virus Enclosed | NO |
| SDFLPLSDMEKDFLSF | 1692431 | Cell/Virus Enclosed | NO |
| SDFNNGIFSRVKNTKL | 1692433 | Ectodomain | YES |
| SDFQLPVYASYK | 1502533 | Cell/Virus Enclosed | NO |
| SDFQLPVYTSYK | 1502534 | Cell/Virus Enclosed | NO |
| SDFSLADSVVEVVTTS | 1692436 | Cell/Virus Enclosed | NO |
| SDFTGSVYGNFDDQPS | 1692438 | Cell/Virus Enclosed | NO |
| SDFTPRSDMEYDFLNM | 1692439 | Cell/Virus Enclosed | NO |
| SDFVDFESVIQD | 1502543 | Cell/Virus Enclosed | NO |
| SDFVDLESVIQDYENV | 1692441 | Cell/Virus Enclosed | NO |
| SDGMFVATSSYD | 1502570 | Cell/Virus Enclosed | NO |
| SDGNCTCIPIPSSWA | 1692450 | Underneath Membrane | NO |

| | | | |
|---|---|---|---|
| SDGNYNRNAILN | 1502573 | Ectodomain | YES |
| SDGNYNRNASLNSFKE | 1692451 | Ectodomain | YES |
| SDGNYNRYASLN | 1502575 | Ectodomain | YES |
| SDGQGVPIAFGI | 1502576 | Cell/Virus Enclosed | NO |
| SDGQGVPLAFGV | 1502577 | Cell/Virus Enclosed | NO |
| SDGQPYVASEQE | 1502580 | Cell/Virus Enclosed | NO |
| SDGRVEGQVDLF | 1502582 | Cell/Virus Enclosed | NO |
| SDGTNKQLNEISDDCN | 1692456 | Cell/Virus Enclosed | NO |
| SDGVDSFEGVLPINTD | 1692457 | Cell/Virus Enclosed | NO |
| SDGVMKNASLYTSLVP | 1692458 | Cell/Virus Enclosed | NO |
| SDGVVWVAKEGAKTVN | 1692459 | Capsid | UN |
| SDHQDLKWARFP | 1502598 | Cell/Virus Enclosed | NO |
| SDIFATPSVLQA | 1502604 | Cell/Virus Enclosed | NO |
| SDIFCDIENNPGLCY | 227547 | Cell/Virus Enclosed | NO |
| SDIFGQDFKTFDLLKY | 1692468 | Cell/Virus Enclosed | NO |
| SDIFGSDFKTFDLLKY | 1692469 | Cell/Virus Enclosed | NO |
| SDIHLPAYASFKVIDN | 1692470 | Cell/Virus Enclosed | NO |
| SDILSLNNPIFINYSK | 1692472 | Ectodomain | YES |
| SDINCNIPTQCY | 1502611 | Cell/Virus Enclosed | NO |
| SDINGVYWVASN | 1502612 | Cell/Virus Enclosed | NO |
| SDIRDYKLANDAKESL | 1692475 | Cell/Virus Enclosed | NO |
| SDISLSVACQLPYCYF | 1692476 | Underneath Membrane | NO |
| SDISLVKFGAALAMEK | 1692477 | Ectodomain | YES |
| SDITLKCCTVTYLNDP | 1692478 | Cell/Virus Enclosed | NO |
| SDIYFVKNIPCY | 1502618 | Cell/Virus Enclosed | NO |
| SDIYFVKNIPRYASAV | 1692479 | Cell/Virus Enclosed | NO |
| SDKCSIEDFNVW | 1502627 | Cell/Virus Enclosed | NO |
| SDKFDFIIGHGMSFSM | 1692486 | Cell/Virus Enclosed | NO |
| SDKFYRLSNELAQVLT | 1692489 | Cell/Virus Enclosed | NO |
| SDKIHHHHHHNT | 1502638 | Capsid | UN |
| SDKNVVFMNSCS | 1502641 | Ectodomain | YES |
| SDLEEGESECIS | 1502653 | Cell/Virus Enclosed | NO |
| SDLEESESECIS | 1502654 | Cell/Virus Enclosed | NO |
| SDLEVTSDSCNN | 1502658 | Cell/Virus Enclosed | NO |
| SDLIISNFKFDDYDFS | 1692503 | Cell/Virus Enclosed | NO |
| SDLNTSELLTKAIDVD | 1692509 | Cell/Virus Enclosed | NO |
| SDLQLGSSGFLQSSNY | 1692512 | Ectodomain | YES |
| SDLVIDALASMT | 1502687 | Cell/Virus Enclosed | NO |
| SDLVQNGVDAKGFPQL | 1692522 | Capsid | UN |
| SDMEKDFLSFDSDVFI | 1692525 | Cell/Virus Enclosed | NO |
| SDMEYDFLNMDMGVFI | 1692526 | Cell/Virus Enclosed | NO |
| SDMVETVVGQLE | 1502696 | Cell/Virus Enclosed | NO |
| SDMYDGRTKAIDGENV | 1692530 | Cell/Virus Enclosed | NO |
| SDMYDPKTKHVT | 1502700 | Cell/Virus Enclosed | NO |
| SDMYDPLTKNIGDYNV | 1692534 | Cell/Virus Enclosed | NO |
| SDMYDPTTKNLT | 1502703 | Cell/Virus Enclosed | NO |
| SDMYDPTTKNVTGSNE | 1692536 | Cell/Virus Enclosed | NO |
| SDNGPQNQRSAP | 1502710 | Cell/Virus Enclosed | NO |
| SDNGPQPNQRSA | 1502712 | Capsid | UN |
| SDNINDKDTGPPPIST | 1692544 | Ectodomain | YES |
| SDNKTAVACQPP | 1502719 | Ectodomain | YES |
| SDNNCYFNAVIM | 1502722 | Cell/Virus Enclosed | NO |
| SDNPQCAIVQAS | 1502725 | Cell/Virus Enclosed | NO |
| SDNQISGYTLAA | 1502726 | Ectodomain | YES |
| SDNQKLIANKFN | 1502727 | Ectodomain | YES |
| SDPNLWYLRSGH | 1502743 | Cell/Virus Enclosed | NO |
| SDPNMWCLRSGH | 1502745 | Cell/Virus Enclosed | NO |
| SDPNMWYLRSGH | 1502746 | Cell/Virus Enclosed | NO |
| SDPNMWYLRSGN | 1502747 | Cell/Virus Enclosed | NO |
| SDPNTWYLRSGH | 1502749 | Cell/Virus Enclosed | NO |
| SDPNYDALLISM | 1502750 | Cell/Virus Enclosed | NO |
| SDPQFPILAELAPTAG | 1692555 | Cell/Virus Enclosed | NO |
| SDQFKTTGDLAVQIGS | 1692557 | Cell/Virus Enclosed | NO |
| SDQSDQFRNVQT | 1502764 | Cell/Virus Enclosed | NO |
| SDREVILCWETGKVKP | 1692564 | Cell/Virus Enclosed | NO |
| SDRFKTSGDLAVQIGN | 1692565 | Cell/Virus Enclosed | NO |
| SDRFYRLANGCA | 1502779 | Cell/Virus Enclosed | NO |
| SDRGQECLARSA | 1502781 | Cell/Virus Enclosed | NO |
| SDRKIKKHPCMV | 1502787 | Cell/Virus Enclosed | NO |
| SDRYHNSQDDIM | 1502798 | Capsid | UN |
| SDSCNNVNVRDLQRRL | 1692576 | Cell/Virus Enclosed | NO |
| SDSGCILPSECTMFRD | 1692579 | Cell/Virus Enclosed | NO |
| SDSLGVSSNQSP | 1502823 | Cell/Virus Enclosed | NO |
| SDSNMWYLRSGH | 1502829 | Cell/Virus Enclosed | NO |
| SDSNQSSSDLVAAVTL | 1692586 | Capsid | UN |
| SDSQFEEDVEMS | 1502835 | Cell/Virus Enclosed | NO |

| | | | |
|---|---|---|---|
| SDSQFPILAELA | 1502836 | Cell/Virus Enclosed | NO |
| SDSTDNNQNGDRSGAR | 1692589 | Cell/Virus Enclosed | NO |
| SDSTGSNQNGERSGARSKQR | 1392416 | Capsid | UN |
| SDSTLVKFSAAQAMEK | 1692592 | Ectodomain | YES |
| SDSVATYFGDCITLPF | 1692593 | Cell/Virus Enclosed | NO |
| SDSVKSACIEVD | 1502844 | Ectodomain | YES |
| SDSVQCYTPHIQISYN | 1692594 | Cell/Virus Enclosed | NO |
| SDTAHACNANRFNVAI | 1692597 | Cell/Virus Enclosed | NO |
| SDTLKGLSDRVV | 1502854 | Cell/Virus Enclosed | NO |
| SDTPCVYMDGMD | 1502859 | Cell/Virus Enclosed | NO |
| SDTSLSDACQLP | 1502863 | Underneath Membrane | NO |
| SDTSYINDKDTG | 1502864 | Ectodomain | YES |
| SDTVEVPPHEAD | 1502870 | Cell/Virus Enclosed | NO |
| SDTVEVSPQVVK | 1502875 | Cell/Virus Enclosed | NO |
| SDTVTYKSTATTKLVP | 1692609 | Cell/Virus Enclosed | NO |
| SDVDGLKYTKILKDDG | 1692616 | Cell/Virus Enclosed | NO |
| SDVDGLKYTKIMKDDG | 1692617 | Cell/Virus Enclosed | NO |
| SDVDYVPLKSAT | 1502886 | Cell/Virus Enclosed | NO |
| SDVFIKKYGLENYAFE | 1692626 | Cell/Virus Enclosed | NO |
| SDVFVKKYGLEN | 1502897 | Cell/Virus Enclosed | NO |
| SDVGFVEAYNNCTGGA | 1692630 | Ectodomain | YES |
| SDVGFVEAYNNCTGGS | 1692631 | Ectodomain | YES |
| SDVGFVSAYNNC | 1502902 | Ectodomain | YES |
| SDVIHYNINFEE | 1502904 | Ectodomain | YES |
| SDVIRYNLNFEENLRR | 1692632 | Ectodomain | YES |
| SDVLSDVIRYNLNFEE | 1692636 | Ectodomain | YES |
| SDVMFEDAQSLK | 1502919 | Cell/Virus Enclosed | NO |
| SDVMFEGAQGLK | 1502921 | Cell/Virus Enclosed | NO |
| SDVNCNGYQHHS | 1502924 | Ectodomain | YES |
| SDVSARDPIIQE | 1502928 | Cell/Virus Enclosed | NO |
| SDVSLTYLLGTAKKQV | 1692643 | Cell/Virus Enclosed | NO |
| SDVSSRDPTTQEAIPT | 1692644 | Capsid | UN |
| SDVVKQPVKFYM | 1502940 | Cell/Virus Enclosed | NO |
| SDWSDIRDYKLANDAK | 1692654 | Cell/Virus Enclosed | NO |
| SDWTDVRDYKLANDVK | 1692655 | Cell/Virus Enclosed | NO |
| SDYANKGYIANI | 1502957 | Cell/Virus Enclosed | NO |
| SDYCGGNPCTCR | 1502963 | Ectodomain | YES |
| SDYCGGNSCTCQ | 1502964 | Ectodomain | YES |
| SEAAAQGFQACR | 1502985 | Cell/Virus Enclosed | NO |
| SEAASNGFRACR | 1502988 | Cell/Virus Enclosed | NO |
| SEADYRCACYAHLAKA | 1692673 | Cell/Virus Enclosed | NO |
| SEAFEKFISLFA | 1502992 | Cell/Virus Enclosed | NO |
| SEAFEKFVSLFA | 1502994 | Cell/Virus Enclosed | NO |
| SEASKKPRQKRT | 1503017 | Cell/Virus Enclosed | NO |
| SEAVEDTIDGVVEDTI | 1692692 | Cell/Virus Enclosed | NO |
| SEAVLKKLKKSL | 1503022 | Cell/Virus Enclosed | NO |
| SECFMKSDIFGQDFKT | 1692694 | Cell/Virus Enclosed | NO |
| SECFVKSDIFGSDFKT | 1692695 | Cell/Virus Enclosed | NO |
| SECSKASSRSAIEDLL | 1692697 | Ectodomain | YES |
| SECTMFKDAEGR | 1503031 | Cell/Virus Enclosed | NO |
| SEDFIKHVRTNGGYNS | 1692702 | Cell/Virus Enclosed | NO |
| SEDKRAQVASAM | 1503048 | Cell/Virus Enclosed | NO |
| SEDLKAVRSSFNFDQK | 1692705 | Cell/Virus Enclosed | NO |
| SEDLWQFVDHFGENEE | 1692706 | Cell/Virus Enclosed | NO |
| SEDVKLNPSEDFIKHV | 1692712 | Cell/Virus Enclosed | NO |
| SEEAIPTRFPPG | 1503058 | Capsid | UN |
| SEEFSSNVANYQ | 1503059 | Cell/Virus Enclosed | NO |
| SEESQGNVVTSVMESQ | 1692715 | Cell/Virus Enclosed | NO |
| SEFNLPGCDGGSLYVN | 1692726 | Cell/Virus Enclosed | NO |
| SEFTIGNMEEFI | 1503103 | Ectodomain | YES |
| SEGFLLGINYLGTIKE | 1692738 | Cell/Virus Enclosed | NO |
| SEGLNVLWNKFV | 1503111 | Cell/Virus Enclosed | NO |
| SEGNALYNNECG | 1503113 | Cell/Virus Enclosed | NO |
| SEGNALYNNEGGRAFM | 1692741 | Cell/Virus Enclosed | NO |
| SEGNSIIGVPYP | 1503115 | Ectodomain | YES |
| SEHLLHQSLVGKQSPFQNAY | 1952656 | Capsid | UN |
| SEIATKMFDSFVNSFV | 1692744 | Cell/Virus Enclosed | NO |
| SEIKCKTQSIAPPTGV | 1692747 | Ectodomain | YES |
| SEILDISPCAFGGVSV | 1692748 | Ectodomain | YES |
| SEILDISPCSFG | 1503131 | Ectodomain | YES |
| SEINMDNSQNLA | 1503135 | Cell/Virus Enclosed | NO |
| SEIRDLLCVQSFNGIK | 1692751 | Ectodomain | YES |
| SEISANGCSTIAQAVR | 1692752 | Cell/Virus Enclosed | NO |
| SEISGFGFAIPSVAVR | 1692753 | Cell/Virus Enclosed | NO |
| SEISVIMHLQWIVMYG | 1692756 | Cell/Virus Enclosed | NO |
| SEISVIMHLQWLVMYG | 1692757 | Cell/Virus Enclosed | NO |

| | | | |
|---|---|---|---|
| SEIVMCGGCYYVKPGG | 1692761 | Cell/Virus Enclosed | NO |
| SEIVNTVAGIPSNVYL | 1692762 | Cell/Virus Enclosed | NO |
| SEKHQDHVSLTVPLCG | 1692766 | Cell/Virus Enclosed | NO |
| SEKISVMDKLDTGAQK | 1692767 | Cell/Virus Enclosed | NO |
| SEKLFVRSREED | 1503160 | Cell/Virus Enclosed | NO |
| SEKSGNFKHLREFVFK | 1692772 | Ectodomain | YES |
| SEKSYERQTPFE | 1503170 | Cell/Virus Enclosed | NO |
| SELKQLEAKTASLFQT | 1692777 | Ectodomain | YES |
| SELLKEKYTCSITFEM | 1692780 | Cell/Virus Enclosed | NO |
| SELPKEQRVQGS | 1503189 | Cell/Virus Enclosed | NO |
| SELQTIVPEYIDVNKT | 1692785 | Ectodomain | YES |
| SELVIGAVILRGHLR | 1310793 | Underneath Membrane | NO |
| SELVYENKFVPVKEAS | 1692796 | Cell/Virus Enclosed | NO |
| SELVYENKFVPVKPAS | 1692797 | Cell/Virus Enclosed | NO |
| SEMLKLGTSDPQ | 1503205 | Cell/Virus Enclosed | NO |
| SEMLTFDKKAFTLANV | 1692798 | Ectodomain | YES |
| SENLFVRSKEED | 1503215 | Cell/Virus Enclosed | NO |
| SENQISGYTLAATSAS | 1692801 | Ectodomain | YES |
| SENQKLIANKFNQALG | 1692802 | Ectodomain | YES |
| SENRYSVLSVDSVSEE | 1692803 | Cell/Virus Enclosed | NO |
| SEPALLSRNIKC | 1503232 | Ectodomain | YES |
| SEPPKEQRVQGI | 1503242 | Cell/Virus Enclosed | NO |
| SEPPKEQRVQGSITQR | 1692808 | Cell/Virus Enclosed | NO |
| SEPQRGRQSRIP | 1503255 | Cell/Virus Enclosed | NO |
| SEPVRCNRHDPLARYGSRVQ | 741979 | Ectodomain | YES |
| SEQPWKVVPRNL | 1503276 | Cell/Virus Enclosed | NO |
| SEQSWKVIPRNL | 1503279 | Capsid | UN |
| SEQVIPPTSDYL | 1503280 | Cell/Virus Enclosed | NO |
| SERCNVIADFKS | 1503281 | Cell/Virus Enclosed | NO |
| SERELILSWEIGKVKP | 1692820 | Cell/Virus Enclosed | NO |
| SERINVVPGVST | 1503287 | Cell/Virus Enclosed | NO |
| SERVNVVPGVPT | 1503291 | Cell/Virus Enclosed | NO |
| SESAAISAQLAK | 1503294 | Ectodomain | YES |
| SESAALSAQLAK | 1503295 | Ectodomain | YES |
| SESGAKETKEINIIKL | 1692824 | Cell/Virus Enclosed | NO |
| SESGVKETKEIN | 1503302 | Cell/Virus Enclosed | NO |
| SESKVLLERFLPKCPE | 1692827 | Cell/Virus Enclosed | NO |
| SESQISGYTTAATVAA | 1692829 | Ectodomain | YES |
| SESSCGLFKDCARNPI | 1692830 | Cell/Virus Enclosed | NO |
| SETKEQKHEIEK | 1503322 | Capsid | UN |
| SETPNAKEEILG | 1503328 | Cell/Virus Enclosed | NO |
| SETQRGRQGRIP | 1503331 | Capsid | UN |
| SETVNGYAKSNFVKDD | 1692841 | Cell/Virus Enclosed | NO |
| SEVKAQSSGVIPENPV | 1692847 | Cell/Virus Enclosed | NO |
| SEVSTDEVGDNV | 1503353 | Cell/Virus Enclosed | NO |
| SEVVVNNLDKSAGHPF | 1692853 | Cell/Virus Enclosed | NO |
| SEWAYCVDLHNKINLC | 1692854 | Cell/Virus Enclosed | NO |
| SEWPLSESDKEE | 1503361 | Cell/Virus Enclosed | NO |
| SEYGTFCDNINSILDE | 1692862 | Ectodomain | YES |
| SEYLVGDSIVTL | 1503373 | Cell/Virus Enclosed | NO |
| SEYPSQPPSPSN | 57707 | Secreted | YES |
| SFADNNDITNIN | 1503388 | Cell/Virus Enclosed | NO |
| SFADNNDITNTNLSRG | 1692877 | Cell/Virus Enclosed | NO |
| SFADNNDTTNTN | 1503391 | Cell/Virus Enclosed | NO |
| SFADVMRYNLNF | 1503393 | Ectodomain | YES |
| SFAGYTESAVNF | 1503398 | Cell/Virus Enclosed | NO |
| SFAGYTESAVNSTKAL | 1692878 | Cell/Virus Enclosed | NO |
| SFAIYHTPATDC | 1503400 | Ectodomain | YES |
| SFASSCKSHKPPSASC | 1692881 | Ectodomain | YES |
| SFATYHTPAIDC | 1503409 | Ectodomain | YES |
| SFATYHTPATDCSDGN | 1692882 | Ectodomain | YES |
| SFAVDPAAAYLDAVKQ | 1692883 | Cell/Virus Enclosed | NO |
| SFAYNNDITNTN | 1503417 | Capsid | UN |
| SFCAFAVDPAKA | 1503419 | Cell/Virus Enclosed | NO |
| SFCDNINAILTEVNEL | 1692887 | Ectodomain | YES |
| SFCKSGNLVLNN | 1503425 | Ectodomain | YES |
| SFCKTIQSALSV | 1503428 | Cell/Virus Enclosed | NO |
| SFCRQLNRALSG | 1503432 | Ectodomain | YES |
| SFCTQLNRALSGIAAE | 1692890 | Ectodomain | YES |
| SFDDLYVAIRGSCEKN | 1692894 | Cell/Virus Enclosed | NO |
| SFDGVMYGGFEDQPNL | 1692896 | Cell/Virus Enclosed | NO |
| SFDNSYLNRVRGSSAA | 1692906 | Cell/Virus Enclosed | NO |
| SFDSDVFIKKYGLENY | 1692907 | Cell/Virus Enclosed | NO |
| SFDSSYLNESGA | 1503469 | Cell/Virus Enclosed | NO |
| SFDVESGVYSVS | 1503472 | Ectodomain | YES |
| SFDVPDGFYSTSPIQS | 1692911 | Ectodomain | YES |

| | | | |
|---|---|---|---|
| SFDVPTFCANRSAMCN | 1692912 | Cell/Virus Enclosed | NO |
| SFDYHPNCSDCYDDMC | 1692914 | Cell/Virus Enclosed | NO |
| SFDYLTREAKTR | 1503483 | Cell/Virus Enclosed | NO |
| SFEAHKDFFKDG | 1503484 | Cell/Virus Enclosed | NO |
| SFEAHKDSFKDGLCMF | 1692916 | Cell/Virus Enclosed | NO |
| SFEDELTPSDVYQQLA | 1692920 | Cell/Virus Enclosed | NO |
| SFEEQDEIYAYTKRNV | 1692921 | Cell/Virus Enclosed | NO |
| SFEEQNEIYAYTKRNV | 1692922 | Cell/Virus Enclosed | NO |
| SFEGVLPINTDTVLSV | 1692923 | Cell/Virus Enclosed | NO |
| SFEGYDFKYIENGQLK | 1692924 | Cell/Virus Enclosed | NO |
| SFEIDKGIYQTSNFRV | 1692926 | Ectodomain | YES |
| SFEQQVGVIADKDKDL | 1692933 | Cell/Virus Enclosed | NO |
| SFEVLEDTLDDG | 1503512 | Cell/Virus Enclosed | NO |
| SFEVLEVEDTQG | 1503513 | Cell/Virus Enclosed | NO |
| SFFDCGVNALAG | 1503516 | Cell/Virus Enclosed | NO |
| SFFDCGVNALDG | 1503517 | Cell/Virus Enclosed | NO |
| SFFEDLLFDKVKLSDV | 1692937 | Ectodomain | YES |
| SFFKELTANMSMAECK | 1692938 | Cell/Virus Enclosed | NO |
| SFFPQFTSDQAI | 1503524 | Underneath Membrane | NO |
| SFGDYNLSSVIPSLPT | 1692941 | Ectodomain | YES |
| SFGDYNLSSVLPQRNI | 1692942 | Ectodomain | YES |
| SFGKDLNANMSLAECK | 1692949 | Cell/Virus Enclosed | NO |
| SFGKSIIYTGDW | 1503536 | Cell/Virus Enclosed | NO |
| SFGNRSGNGILK | 1503542 | Cell/Virus Enclosed | NO |
| SFGPGNTFINGDIARE | 1692953 | Cell/Virus Enclosed | NO |
| SFGVTQEYIQTTIQKV | 1692958 | Ectodomain | YES |
| SFHFTRFYNYTG | 1503558 | Ectodomain | YES |
| SFHNLHLLFDEYTNIPVSIL | 57789 | Cell/Virus Enclosed | NO |
| SFIDDYYGYLMK | 1503564 | Cell/Virus Enclosed | NO |
| SFIDYTTTVTSFHGYD | 1692965 | Cell/Virus Enclosed | NO |
| SFIEDLLFDKVK | 1503567 | Ectodomain | YES |
| SFIEDLLYNKVT | 1503568 | Ectodomain | YES |
| SFINGACGSPGYNLKN | 1692971 | Cell/Virus Enclosed | NO |
| SFIQADSFTCNNIDAA | 1692973 | Ectodomain | YES |
| SFISQPFYFDSY | 1503597 | Ectodomain | YES |
| SFKAPDTYKCPQ | 1503604 | Ectodomain | YES |
| SFKDNVILLNKHIDAY | 1692976 | Cell/Virus Enclosed | NO |
| SFKENVDFINSM | 1503608 | Cell/Virus Enclosed | NO |
| SFKEYFNLRNCTFMYT | 1692978 | Ectodomain | YES |
| SFKSENRYSVLSVDSV | 1692988 | Cell/Virus Enclosed | NO |
| SFKTADGQQKPL | 1503638 | Cell/Virus Enclosed | NO |
| SFKTADGQQKQLLPRW | 1692990 | Capsid | UN |
| SFLAVDDKYKATGDLA | 1693000 | Cell/Virus Enclosed | NO |
| SFLAVDDKYKVNENLA | 1693001 | Cell/Virus Enclosed | NO |
| SFLEKDAPYMVG | 1503652 | Cell/Virus Enclosed | NO |
| SFLPKEDGDVLLAEFD | 1693017 | Cell/Virus Enclosed | NO |
| SFLPKEEGDVLL | 1503677 | Cell/Virus Enclosed | NO |
| SFLPQFTSDQAVTFLK | 1693018 | Ectodomain | YES |
| SFLQSGIVKMVS | 1503681 | Cell/Virus Enclosed | NO |
| SFNENGTITDAV | 1503704 | Ectodomain | YES |
| SFNEWALANQFTEFVG | 1693026 | Cell/Virus Enclosed | NO |
| SFNFDQKELLAYYNML | 1693027 | Cell/Virus Enclosed | NO |
| SFNFDQKELLDY | 1503710 | Cell/Virus Enclosed | NO |
| SFNIGRSIVYET | 1503713 | Cell/Virus Enclosed | NO |
| SFNKAIDGFGVF | 1503716 | Cell/Virus Enclosed | NO |
| SFNNFVSSYAKPEEKL | 1693038 | Cell/Virus Enclosed | NO |
| SFNNIDTGPPSI | 1503726 | Ectodomain | YES |
| SFNNLLGFDSSKPVTK | 1693039 | Cell/Virus Enclosed | NO |
| SFNNRDTGPLSI | 1503729 | Ectodomain | YES |
| SFNNRDTGRPSI | 1503730 | Ectodomain | YES |
| SFNPETNCLLNVPFGG | 1693040 | Ectodomain | YES |
| SFNPETNILPNV | 1503733 | Underneath Membrane | NO |
| SFNVSDAYANLVPYYQ | 1693045 | Cell/Virus Enclosed | NO |
| SFQDAASNIFVINKDT | 1693059 | Cell/Virus Enclosed | NO |
| SFRDELGVRVLDQSDN | 1693061 | Cell/Virus Enclosed | NO |
| SFRLFARTRSMWSFN | 1542448 | Undetermined | UN |
| SFRTLKSGEGFN | 1503776 | Cell/Virus Enclosed | NO |
| SFSASTSAFIDT | 1503779 | Cell/Virus Enclosed | NO |
| SFSCNNFDESKIYGSC | 1693071 | Ectodomain | YES |
| SFSFRDELGVRVLDQS | 1693077 | Cell/Virus Enclosed | NO |
| SFSIKANICNSN | 1503796 | Ectodomain | YES |
| SFSNTKYYDHSQ | 1503802 | Ectodomain | YES |
| SFSSVNDAITQT | 1503808 | Ectodomain | YES |
| SFSTFESAYMPIADPT | 1693085 | Cell/Virus Enclosed | NO |
| SFSVQKHIGAPANLRV | 1693088 | Cell/Virus Enclosed | NO |
| SFSVQKHIGPPA | 1503816 | Cell/Virus Enclosed | NO |

| | | | |
|---|---|---|---|
| SFTCNNIDAAKIYGMC | 1693092 | Ectodomain | YES |
| SFTCRTDMEKDFIALD | 1693093 | Cell/Virus Enclosed | NO |
| SFTPGKQSSSRASSGN | 1693100 | Cell/Virus Enclosed | NO |
| SFTQWYHEHIGG | 1503849 | Cell/Virus Enclosed | NO |
| SFVAYETARQEYENAV | 1693108 | Cell/Virus Enclosed | NO |
| SFVDDFQGYLQK | 1503861 | Cell/Virus Enclosed | NO |
| SFVDDFYGYLQKHFSM | 1693109 | Cell/Virus Enclosed | NO |
| SFVDDNKFDVLK | 1503863 | Cell/Virus Enclosed | NO |
| SFVDYLLTTIPKAIVL | 1693110 | Cell/Virus Enclosed | NO |
| SFVEYEVAKKNLADAK | 1693111 | Cell/Virus Enclosed | NO |
| SFVEYEVAKKNLDEAR | 1693112 | Cell/Virus Enclosed | NO |
| SFVGELKAAEAKVITI | 1693114 | Cell/Virus Enclosed | NO |
| SFVGMPSFVAYETARQ | 1693115 | Cell/Virus Enclosed | NO |
| SFVMMSAPPAGY | 1503886 | Cell/Virus Enclosed | NO |
| SFVMMSAPPSEY | 1503887 | Cell/Virus Enclosed | NO |
| SFVNDSIESVGGLYEI | 1693122 | Ectodomain | YES |
| SFVSKIIHFFKT | 1503901 | Cell/Virus Enclosed | NO |
| SFVSKIVHFFET | 1503902 | Cell/Virus Enclosed | NO |
| SFVSMPSYIAYENARQ | 1693128 | Cell/Virus Enclosed | NO |
| SFVVKGDDVRQIAPGQ | 1693129 | Ectodomain | YES |
| SFWEAGTTKAGYPYNYNTTA | 57943 | Undetermined | UN |
| SFYASMYEKSTVLQAA | 1693136 | Cell/Virus Enclosed | NO |
| SFYCEDHKPQYS | 1503923 | Cell/Virus Enclosed | NO |
| SFYGFSNVVYPD | 1503929 | Underneath Membrane | NO |
| SFYMPLLVSSDKAPYR | 1693144 | Capsid | UN |
| SFYPPDDEDECD | 1503935 | Cell/Virus Enclosed | NO |
| SFYYIWKSYVHI | 1503952 | Cell/Virus Enclosed | NO |
| SGAIKLDPKNPNYNKW | 1693160 | Cell/Virus Enclosed | NO |
| SGAKETKEINIIKLSG | 1693161 | Cell/Virus Enclosed | NO |
| SGALAEHDFFTW | 1503964 | Cell/Virus Enclosed | NO |
| SGARPKQRRPQGLPNN | 1693168 | Cell/Virus Enclosed | NO |
| SGAVLFCTPTKKAFPY | 1693169 | Cell/Virus Enclosed | NO |
| SGCETSNVPPIV | 1503978 | Cell/Virus Enclosed | NO |
| SGCMLRACIKTAQADT | 1693177 | Cell/Virus Enclosed | NO |
| SGCMLRVCIKTA | 1503992 | Cell/Virus Enclosed | NO |
| SGCNVNDVTKLYLGGL | 1693178 | Cell/Virus Enclosed | NO |
| SGDAHIPFGTVLGNFY | 1693182 | Ectodomain | YES |
| SGDCVVEASTDF | 1504004 | Cell/Virus Enclosed | NO |
| SGDDCFVSNSSVIQEV | 1693190 | Cell/Virus Enclosed | NO |
| SGDKVFVRDGNTDNLV | 1693198 | Cell/Virus Enclosed | NO |
| SGDKYYPVVVDD | 1504017 | Cell/Virus Enclosed | NO |
| SGDLAVQIGNNNVCTY | 1693199 | Cell/Virus Enclosed | NO |
| SGDNGTITVEEL | 1504021 | Ectodomain | YES |
| SGDRTEVPQLVN | 1504025 | Ectodomain | YES |
| SGDSSEGGAKET | 1504026 | Cell/Virus Enclosed | NO |
| SGDVNVVAASGS | 1504034 | Cell/Virus Enclosed | NO |
| SGDVVRFPNITNLCPFG | 58001 | Ectodomain | YES |
| SGDYIKLNGPLT | 1504037 | Cell/Virus Enclosed | NO |
| SGDYTLANTTTA | 1504039 | Cell/Virus Enclosed | NO |
| SGDYTLANTTTEPLKL | 1693206 | Cell/Virus Enclosed | NO |
| SGEIATKMFDSF | 1504044 | Cell/Virus Enclosed | NO |
| SGFAAYSRYRIGNYKL | 1388327 | Underneath Membrane | NO |
| SGFCGQGIHIVS | 1504063 | Ectodomain | YES |
| SGFETIMKVLNENLNA | 1693222 | Cell/Virus Enclosed | NO |
| SGFKLDGFNHQFVTAS | 1693225 | Cell/Virus Enclosed | NO |
| SGFKPNNTSAQYIFST | 1693228 | Cell/Virus Enclosed | NO |
| SGFLQSSNYKIDTTSS | 1693229 | Ectodomain | YES |
| SGGFYRFMVDST | 1504099 | Cell/Virus Enclosed | NO |
| SGGGKCFNCAPA | 1504101 | Ectodomain | YES |
| SGGGKCFNCYPAGVNI | 1693238 | Ectodomain | YES |
| SGGGKCYNCRPA | 1504103 | Ectodomain | YES |
| SGGNVVPHYSWF | 1504108 | Cell/Virus Enclosed | NO |
| SGGQPITNCVKM | 1504110 | Cell/Virus Enclosed | NO |
| SGGYIPSDFAFNNWFL | 1693242 | Ectodomain | YES |
| SGHYQTNLYSFNKAID | 1693249 | Cell/Virus Enclosed | NO |
| SGIAVEQDRNTR | 1504150 | Ectodomain | YES |
| SGIGAVGGDLLYLDLL | 1693253 | Cell/Virus Enclosed | NO |
| SGIGGGDLLYLD | 1504157 | Cell/Virus Enclosed | NO |
| SGIGTCPAGTNY | 1504158 | Ectodomain | YES |
| SGIKFQTVKPGNFNQD | 1693254 | Cell/Virus Enclosed | NO |
| SGISDLSLKQKL | 1504176 | Cell/Virus Enclosed | NO |
| SGITQFQKGKEFEFVE | 1693262 | Cell/Virus Enclosed | NO |
| SGKDDQIGYYRR | 1504192 | Cell/Virus Enclosed | NO |
| SGKGVSFQLVKLGVW | 58103 | Ectodomain | YES |
| SGKISSKADNSI | 1504195 | Ectodomain | YES |
| SGKNNGIGTCPA | 1504202 | Ectodomain | YES |

| | | | |
|---|---|---|---|
| SGKNNGVGTCPA | 1504203 | Ectodomain | YES |
| SGKSLLHEVDKDISQLTQAIVK | 58115 | Ectodomain | YES |
| SGLAIKSDYCGG | 1504220 | Ectodomain | YES |
| SGLCVDGRNGYV | 1504226 | Ectodomain | YES |
| SGLEEGESECIS | 1504231 | Cell/Virus Enclosed | NO |
| SGLGTVDADYKK | 1504233 | Ectodomain | YES |
| SGLKTMIVSAGQ | 1504241 | Cell/Virus Enclosed | NO |
| SGLLNVQKVAQP | 1504245 | Cell/Virus Enclosed | NO |
| SGLSPAYAPTYVSVDD | 1693305 | Cell/Virus Enclosed | NO |
| SGLTCSTDLQKA | 1504265 | Ectodomain | YES |
| SGLVNVQKVEQPKIEP | 1693311 | Cell/Virus Enclosed | NO |
| SGLYPTITVPEE | 1504275 | Cell/Virus Enclosed | NO |
| SGMIRYYDVMAN | 1504283 | Cell/Virus Enclosed | NO |
| SGNCDVVIGIINNTVYD | 58143 | Ectodomain | YES |
| SGNGILKWADQSDQFR | 1693319 | Cell/Virus Enclosed | NO |
| SGNHCPADNSYT | 1504299 | Ectodomain | YES |
| SGNHCPAGNFYT | 1504300 | Ectodomain | YES |
| SGNHCPAGNSYT | 1504302 | Ectodomain | YES |
| SGNHCPAGNYYT | 1504303 | Ectodomain | YES |
| SGNHLHSVHNWL | 1504304 | Cell/Virus Enclosed | NO |
| SGNLDEPQKDVY | 1504305 | Capsid | UN |
| SGNLLGFKDVTKGTIY | 1693322 | Ectodomain | YES |
| SGNLLGFKNVSTGNIF | 1693323 | Ectodomain | YES |
| SGNLVEKLSPWFMDGE | 1693326 | Cell/Virus Enclosed | NO |
| SGNLYGFRDYIT | 1504313 | Ectodomain | YES |
| SGNPDEPHKDVY | 1504316 | Cell/Virus Enclosed | NO |
| SGNPDEPQKDVYELRY | 1693328 | Cell/Virus Enclosed | NO |
| SGNPVVEYLSKE | 1504319 | Cell/Virus Enclosed | NO |
| SGNRAPTSGVIP | 1504320 | Cell/Virus Enclosed | NO |
| SGNRPGNGILKW | 1504324 | Capsid | UN |
| SGNRSGILKKTS | 1504325 | Cell/Virus Enclosed | NO |
| SGNRSGNGILKWADQS | 1693330 | Cell/Virus Enclosed | NO |
| SGNSSRGASPGP | 1504333 | Cell/Virus Enclosed | NO |
| SGNSTRGTSPGPSGIG | 1693332 | Cell/Virus Enclosed | NO |
| SGNTVVLTFTTK | 1504345 | Cell/Virus Enclosed | NO |
| SGNVDLDALASDIFDT | 1693334 | Cell/Virus Enclosed | NO |
| SGNYCPAGNSYT | 1504350 | Ectodomain | YES |
| SGNYLSISNELL | 1504352 | Ectodomain | YES |
| SGPDQICNTPTQ | 1504356 | Cell/Virus Enclosed | NO |
| SGPDQTCNTPTQ | 1504357 | Cell/Virus Enclosed | NO |
| SGPEWMDDFEAD | 1504358 | Cell/Virus Enclosed | NO |
| SGPGKNNGIGTC | 1504359 | Ectodomain | YES |
| SGPGKNNGMGTC | 1504360 | Ectodomain | YES |
| SGPISQFNYKQS | 1504365 | Ectodomain | YES |
| SGPVDKGHYTVYDTAK | 1693339 | Cell/Virus Enclosed | NO |
| SGQAFCDYTDKL | 1504376 | Cell/Virus Enclosed | NO |
| SGQGIFFEVNAT | 1504378 | Ectodomain | YES |
| SGQGVPLNANST | 1504379 | Cell/Virus Enclosed | NO |
| SGRKIYEIERGL | 1504396 | Cell/Virus Enclosed | NO |
| SGRKIYEVERGL | 1504399 | Cell/Virus Enclosed | NO |
| SGRQPTPLSPPLRDT | 1693355 | Ectodomain | YES |
| SGRSASNSRPGSRSQS | 1693357 | Capsid | UN |
| SGRTPTSGVTPD | 1504413 | Cell/Virus Enclosed | NO |
| SGRVSAAFHANSSEPA | 1693358 | Ectodomain | YES |
| SGSDINCNIPTQ | 1504424 | Cell/Virus Enclosed | NO |
| SGSHVGSSFDGVMYGG | 1693370 | Cell/Virus Enclosed | NO |
| SGSIRFDSTLPG | 1504434 | Cell/Virus Enclosed | NO |
| SGSLGTLSSPSPSAV | 58234 | Cell/Virus Enclosed | NO |
| SGSLVLNNPAYIAPQA | 1693375 | Ectodomain | YES |
| SGSNDIWMQNKGLFYT | 1693376 | Ectodomain | YES |
| SGSRVAGRSAIEDILFSKLVTSGL | 1388480 | Ectodomain | YES |
| SGTATAGWAFGA | 1504477 | Ectodomain | YES |
| SGTCGIVELEKG | 1504480 | Cell/Virus Enclosed | NO |
| SGTHTGSAFDGT | 1504488 | Cell/Virus Enclosed | NO |
| SGTNDICVFENK | 1504491 | Cell/Virus Enclosed | NO |
| SGTPVVEYLSKEDLHT | 1693393 | Cell/Virus Enclosed | NO |
| SGTSTPKKPNKPLSQP | 1693395 | Capsid | UN |
| SGTTCSNDLLQPNTEV | 1693396 | Ectodomain | YES |
| SGTTCSTDLQKS | 1504511 | Ectodomain | YES |
| SGTTTNVEKIRQ | 1504516 | Cell/Virus Enclosed | NO |
| SGTVSKTSDWKCKVTD | 1693398 | Cell/Virus Enclosed | NO |
| SGTWLTYTGAIKLDDK | 1693400 | Cell/Virus Enclosed | NO |
| SGTYHCVTTKAK | 1504526 | Cell/Virus Enclosed | NO |
| SGVANPATDPIY | 1504531 | Secreted | YES |
| SGVFGVNLRTNFTIKG | 1693411 | Cell/Virus Enclosed | NO |
| SGVIPDMADQIA | 1504550 | Cell/Virus Enclosed | NO |

| | | | |
|---|---|---|---|
| SGVKETKEINII | 1504555 | Cell/Virus Enclosed | NO |
| SGVKFQTVKPGNFNQD | 1693419 | Cell/Virus Enclosed | NO |
| SGVKKPFKVEDS | 1504556 | Cell/Virus Enclosed | NO |
| SGVKKTFKVEDS | 1504557 | Cell/Virus Enclosed | NO |
| SGVLDTTIPFGPSSQP | 1693421 | Ectodomain | YES |
| SGVLDTTSYREA | 1504561 | Cell/Virus Enclosed | NO |
| SGVLPVNPVLHT | 1504562 | Cell/Virus Enclosed | NO |
| SGVLQSGLVKMSHPSG | 1693424 | Cell/Virus Enclosed | NO |
| SGVMWLYTYSIGEASS | 1693427 | Cell/Virus Enclosed | NO |
| SGVNDICMFENKTTLP | 1693428 | Cell/Virus Enclosed | NO |
| SGVRELYLSSSN | 1504573 | Cell/Virus Enclosed | NO |
| SGVSDLSLKQKL | 1504575 | Cell/Virus Enclosed | NO |
| SGVSNLSLKQKL | 1504576 | Cell/Virus Enclosed | NO |
| SGVTPDMADEIA | 1504584 | Cell/Virus Enclosed | NO |
| SGVTPDMADHIA | 1504585 | Cell/Virus Enclosed | NO |
| SGVTRDMADQIA | 1504588 | Capsid | UN |
| SGVVGCKSIVKE | 1504591 | Cell/Virus Enclosed | NO |
| SGVYSVSSFEAKPSGS | 1693449 | Ectodomain | YES |
| SGYEKSTFNKDIYCGN | 1693454 | Cell/Virus Enclosed | NO |
| SGYGKSTFNKDI | 1504627 | Cell/Virus Enclosed | NO |
| SGYHPAHAPSFLAVDD | 1693456 | Cell/Virus Enclosed | NO |
| SGYIPANTTIPF | 112837 | Capsid | UN |
| SGYYYPEPIIEN | 1504648 | Ectodomain | YES |
| SHATTYLSLSDR | 1504657 | Cell/Virus Enclosed | NO |
| SHAVQNTVFNAS | 1504658 | Cell/Virus Enclosed | NO |
| SHAVQNTVLNAS | 1504659 | Cell/Virus Enclosed | NO |
| SHDVSDDEWAAA | 1504671 | Cell/Virus Enclosed | NO |
| SHEISDDEWAAA | 1504675 | Cell/Virus Enclosed | NO |
| SHENKLNSCCRI | 1504678 | Cell/Virus Enclosed | NO |
| SHETSDDDWTAA | 1504680 | Cell/Virus Enclosed | NO |
| SHEVSDDEWAAA | 1504685 | Cell/Virus Enclosed | NO |
| SHFTTQFVDNVK | 1504704 | Ectodomain | YES |
| SHFTTQFVGNVK | 1504706 | Ectodomain | YES |
| SHFTTQFVSNVK | 1504708 | Ectodomain | YES |
| SHGKQVVSDVDY | 1504718 | Cell/Virus Enclosed | NO |
| SHIVSKESGNTVVLTF | 1693503 | Capsid | UN |
| SHKPPSASCPIGTNYR | 1693508 | Ectodomain | YES |
| SHLATFAELEAAQKAY | 1693511 | Cell/Virus Enclosed | NO |
| SHMDADLAKPLI | 1504790 | Cell/Virus Enclosed | NO |
| SHMPLSNFEHKV | 1504793 | Cell/Virus Enclosed | NO |
| SHPDFKLLHNLQADIC | 1693531 | Cell/Virus Enclosed | NO |
| SHRGVIADNQAKWAV | 58403 | Ectodomain | YES |
| SHSMAVVDGKQI | 1504829 | Cell/Virus Enclosed | NO |
| SHSVCNGCVGYS | 1504836 | Ectodomain | YES |
| SHSVVYSRYCFS | 1504838 | Ectodomain | YES |
| SHTGITGDNVET | 1504839 | Cell/Virus Enclosed | NO |
| SHTLTKHTEVRA | 1504842 | Ectodomain | YES |
| SHVAQNTVLNAC | 1504850 | Cell/Virus Enclosed | NO |
| SHVNGDWFLFGDSRSD | 1693556 | Ectodomain | YES |
| SHWMGIRFEYTSPTDK | 1693560 | Cell/Virus Enclosed | NO |
| SHYVLNIDWLWF | 1504872 | Cell/Virus Enclosed | NO |
| SHYVVTDIKQVG | 1504874 | Cell/Virus Enclosed | NO |
| SHYWSPQGNLGGNCAG | 1693563 | Cell/Virus Enclosed | NO |
| SIADLACAQYYNGIMV | 1693571 | Ectodomain | YES |
| SIAFNHPIQIDQ | 1504892 | Ectodomain | YES |
| SIAFNHPIQVDQLNSS | 1693573 | Ectodomain | YES |
| SIAFNHPIQVGQ | 1504895 | Ectodomain | YES |
| SIAGVGWTAGLSSFAA | 1693575 | Ectodomain | YES |
| SIAIKVTEYSWNKKLY | 1693576 | Cell/Virus Enclosed | NO |
| SIAPSTVWEDGD | 1504913 | Ectodomain | YES |
| SICGNNQLCKSICKT | 58458 | Secreted | YES |
| SIDEFADVVKEQ | 1504938 | Cell/Virus Enclosed | NO |
| SIDEVSDDYVQDSTVL | 1693593 | Cell/Virus Enclosed | NO |
| SIDSTVLTYIPA | 1504946 | Cell/Virus Enclosed | NO |
| SIDVIDNEIIVKPNIS | 1693597 | Cell/Virus Enclosed | NO |
| SIEDFNVWALSN | 1504952 | Cell/Virus Enclosed | NO |
| SIEEFADVVKGQ | 1504955 | Cell/Virus Enclosed | NO |
| SIEHQVKDVLTPLFK | 58485 | Ectodomain | YES |
| SIESVGGLYEIKIPTN | 1693602 | Ectodomain | YES |
| SIFDIETVDVKHDVSP | 1693607 | Cell/Virus Enclosed | NO |
| SIFNIFQGVSSN | 1504980 | Cell/Virus Enclosed | NO |
| SIFPHWNSDQAI | 1504981 | Underneath Membrane | NO |
| SIFSTVYPIDLA | 1504994 | Undetermined | UN |
| SIFTINSVVQKDGVEN | 1693621 | Cell/Virus Enclosed | NO |
| SIGDIIQRLDFL | 1505010 | Ectodomain | YES |
| SIGDIIRRLDVL | 1505014 | Ectodomain | YES |

| | | | |
|---|---|---|---|
| SIGTNFPLQLGFSTGI | 1693631 | Cell/Virus Enclosed | NO |
| SIGVLGYQKTVDHTKVNSKLSLF | 236322 | Secreted | YES |
| SIHADQLTPAWR | 1505031 | Ectodomain | YES |
| SIHSYFITDEKSGGSK | 1693636 | Cell/Virus Enclosed | NO |
| SIIAVQPRNVSYDSVS | 1693639 | Ectodomain | YES |
| SIIGAMTAVNESRYGL | 1693640 | Ectodomain | YES |
| SIIYTGDWEGLHEVLT | 1693657 | Cell/Virus Enclosed | NO |
| SIKEMQSQPSHA | 1505087 | Cell/Virus Enclosed | NO |
| SIKENVDFINSMQFNY | 1693659 | Cell/Virus Enclosed | NO |
| SIKGDKGVEKLVNWYI | 1693660 | Cell/Virus Enclosed | NO |
| SIKSDKGVEKLV | 1505105 | Cell/Virus Enclosed | NO |
| SIKSDRKAWAAF | 1505106 | Ectodomain | YES |
| SIKSLNNSYINLKDIG | 1693667 | Ectodomain | YES |
| SILDEVNGLLDTTQLH | 1693674 | Ectodomain | YES |
| SILDNAVKGCVPLNAI | 1693675 | Cell/Virus Enclosed | NO |
| SILDNAVKGCVPLSAI | 1693676 | Cell/Virus Enclosed | NO |
| SILMLDKQGQLLDTKL | 1693682 | Cell/Virus Enclosed | NO |
| SILNEVNDLLDI | 1505127 | Ectodomain | YES |
| SILQDMRNTIMASK | 58614 | Capsid | UN |
| SILSGLLYNSPC | 1505135 | Ectodomain | YES |
| SILVENGIPLMP | 1505144 | Cell/Virus Enclosed | NO |
| SILWCLWPLVLA | 1505146 | Transmembrane | NO |
| SINDIDTGAPSI | 1505160 | Ectodomain | YES |
| SINDLSFDKNLGRCVR | 1693696 | Cell/Virus Enclosed | NO |
| SINDSLEPVGGL | 1505162 | Ectodomain | YES |
| SINSAIVCASVKRDGV | 1693703 | Cell/Virus Enclosed | NO |
| SINTGNCPFSFGKVNN | 1693704 | Ectodomain | YES |
| SIPCVCGRDATQ | 1505180 | Cell/Virus Enclosed | NO |
| SIPESKPLADDDSAII | 1693712 | Capsid | UN |
| SIPIDLLNLREIKPAV | 1693715 | Cell/Virus Enclosed | NO |
| SIPIVWGAKDFN | 1505196 | Cell/Virus Enclosed | NO |
| SIPIVWGVKDFNTLSQ | 1693717 | Cell/Virus Enclosed | NO |
| SIPLADDDSAII | 1505201 | Capsid | UN |
| SIPLCIVPYTAL | 1505203 | Cell/Virus Enclosed | NO |
| SIPNFGSLTHIN | 1505210 | Ectodomain | YES |
| SIPNFGSLTQID | 1505211 | Ectodomain | YES |
| SIPNFGSLTQINTTLL | 1693720 | Ectodomain | YES |
| SIPNLPDFKEES | 1505216 | Ectodomain | YES |
| SIPNLPDFKGEL | 1505217 | Ectodomain | YES |
| SIPNLPDFREEL | 1505218 | Ectodomain | YES |
| SIPNLPGFKEEL | 1505219 | Ectodomain | YES |
| SIPNLSDFEAEF | 1505220 | Ectodomain | YES |
| SIQADVNTPADI | 1505235 | Cell/Virus Enclosed | NO |
| SIQAIYDRLDTIQADQ | 1693726 | Ectodomain | YES |
| SIQANAYYKYDNGNFI | 1693727 | Cell/Virus Enclosed | NO |
| SIQDVGFKNQYCNGSI | 1693728 | Cell/Virus Enclosed | NO |
| SIQNGFTATNSA | 1505247 | Ectodomain | YES |
| SIQQGFDATNSA | 1505250 | Ectodomain | YES |
| SIQSGRKAWAAF | 1505253 | Ectodomain | YES |
| SIRFDSTLPGFETIMK | 1693739 | Capsid | UN |
| SIRGCCESTKLPYYDV | 1693741 | Underneath Membrane | NO |
| SIRHHPELKILRNLNI | 1693742 | Cell/Virus Enclosed | NO |
| SIRNESWHIDSS | 1505274 | Ectodomain | YES |
| SIRNGTYDHDVYRDE | 58691 | Ectodomain | YES |
| SIRNLQKRLYPN | 1505276 | Cell/Virus Enclosed | NO |
| SIRSIKSDRKAW | 1505282 | Ectodomain | YES |
| SIRSIQSDRKAW | 1505283 | Ectodomain | YES |
| SISASYRFVTFEPFNV | 1693750 | Ectodomain | YES |
| SISFWPQTTTTPTSVDMNSITSTDV | 58700 | Undetermined | UN |
| SISKLHALDDAT | 1505305 | Cell/Virus Enclosed | NO |
| SISLTVGEVFGKILGN | 1693755 | Cell/Virus Enclosed | NO |
| SISNDTVDVTNG | 1505313 | Ectodomain | YES |
| SISNELLLTVPT | 1505314 | Ectodomain | YES |
| SISNFGSLTQIN | 1505315 | Ectodomain | YES |
| SISTDIVDVTNG | 1505319 | Ectodomain | YES |
| SISTGSLSARSA | 1505323 | Ectodomain | YES |
| SISTGSRSARSAIEDL | 1693757 | Ectodomain | YES |
| SITAQPVKIDNASPAS | 1693762 | Secreted | YES |
| SITAQPVKTDNA | 1505332 | Secreted | YES |
| SITFCALCVISV | 1505339 | Cell/Virus Enclosed | NO |
| SITFEMSCDCGKKFDD | 1693766 | Cell/Virus Enclosed | NO |
| SITKIMCCLGPDIFLG | 1693772 | Cell/Virus Enclosed | NO |
| SITQRTRSRPSV | 1505364 | Cell/Virus Enclosed | NO |
| SITQRTRTRPSVQPGP | 1693774 | Cell/Virus Enclosed | NO |
| SITSDKFDFIIGHGMS | 1693776 | Cell/Virus Enclosed | NO |
| SITVDKFAIPNR | 1505377 | Ectodomain | YES |

| | | | |
|---|---|---|---|
| SITYPMIANENAINKG | 1693780 | Cell/Virus Enclosed | NO |
| SIVAKQGAGDAGHSLT | 1693783 | Cell/Virus Enclosed | NO |
| SIVDEWSYTGSS | 1505392 | Ectodomain | YES |
| SIVDIIQRLDVL | 1505393 | Ectodomain | YES |
| SIVDVYDMWLTG | 1505394 | Cell/Virus Enclosed | NO |
| SIVFLLITIVLQ | 1505396 | Transmembrane | NO |
| SIVKPDMADEIA | 1505406 | Cell/Virus Enclosed | NO |
| SIVPSTAWEDGD | 1505414 | Ectodomain | YES |
| SIVPSTLWEDGD | 1505415 | Ectodomain | YES |
| SIVPSTVWEDGDYYRK | 1693810 | Ectodomain | YES |
| SIVPSTVWEEGD | 1505418 | Ectodomain | YES |
| SIVQMAPVSAMV | 1505422 | Cell/Virus Enclosed | NO |
| SIVSEKISVMDK | 1505428 | Cell/Virus Enclosed | NO |
| SIVYETDANKLILIND | 1693818 | Cell/Virus Enclosed | NO |
| SIYICSQGKDFG | 1505450 | Cell/Virus Enclosed | NO |
| SKADGIIYPQGRTYSN | 1693827 | Ectodomain | YES |
| SKAFVDTIVPPCPSIL | 1693828 | Cell/Virus Enclosed | NO |
| SKAGNSIFRSFH | 1505466 | Ectodomain | YES |
| SKANSGNRTPTS | 1505475 | Cell/Virus Enclosed | NO |
| SKASTAKPAPPPKTGPPKTS | 741984 | Ectodomain | YES |
| SKCHNCCDEYGGHHDF | 1693839 | Underneath Membrane | NO |
| SKCLTSIEEVCD | 1505491 | Cell/Virus Enclosed | NO |
| SKCQITAVEGTKKLAA | 1693841 | Cell/Virus Enclosed | NO |
| SKCWVEEDLTKGPHEF | 1693843 | Cell/Virus Enclosed | NO |
| SKDQYGTDINGV | 1505512 | Cell/Virus Enclosed | NO |
| SKDSNFLKRVRG | 1505513 | Cell/Virus Enclosed | NO |
| SKDSNFLNRVRGSIVN | 1693854 | Cell/Virus Enclosed | NO |
| SKDTLDDILQQANKSV | 1693856 | Cell/Virus Enclosed | NO |
| SKDTNFLNRVRGTSVD | 1693857 | Cell/Virus Enclosed | NO |
| SKDYISSNGPLKVGAG | 1693858 | Cell/Virus Enclosed | NO |
| SKEGFFTYLCGF | 1505532 | Cell/Virus Enclosed | NO |
| SKELKRPIQPTDVAYH | 1693866 | Cell/Virus Enclosed | NO |
| SKELKRPVNPTDASHY | 1693867 | Cell/Virus Enclosed | NO |
| SKESNFLNESGV | 1505541 | Cell/Virus Enclosed | NO |
| SKESNFLNRVRG | 1505542 | Cell/Virus Enclosed | NO |
| SKETAAMVKSKG | 1505543 | Cell/Virus Enclosed | NO |
| SKFNSCKSDILSLNNP | 1693883 | Ectodomain | YES |
| SKGAVQKESDEYILAK | 1693892 | Cell/Virus Enclosed | NO |
| SKGIYQTSNFRV | 1505570 | Ectodomain | YES |
| SKGSFRNESWHF | 1505583 | Ectodomain | YES |
| SKGSLEPRHFNSITKI | 1693909 | Cell/Virus Enclosed | NO |
| SKGSSRNESWHFDKSE | 1693911 | Ectodomain | YES |
| SKGTLEPKYFNT | 1505588 | Cell/Virus Enclosed | NO |
| SKGWYDPVENEDIHRV | 1693918 | Cell/Virus Enclosed | NO |
| SKHAEEYCNYLESYNI | 1693921 | Cell/Virus Enclosed | NO |
| SKHANLYQKYVEAYNT | 1693922 | Cell/Virus Enclosed | NO |
| SKHSDFGLDGLI | 1505604 | Cell/Virus Enclosed | NO |
| SKHSDFGLGDLVDSYF | 1693929 | Cell/Virus Enclosed | NO |
| SKIGNSIFTGCT | 1505631 | Cell/Virus Enclosed | NO |
| SKIGTTEVKVSA | 1505632 | Cell/Virus Enclosed | NO |
| SKINKYGLELHWAPEF | 1693944 | Cell/Virus Enclosed | NO |
| SKISLVSVTDFSLNGN | 1693946 | Cell/Virus Enclosed | NO |
| SKIVCDGSVYYA | 1505653 | Cell/Virus Enclosed | NO |
| SKIVDLAQHFSDFGTS | 1693948 | Cell/Virus Enclosed | NO |
| SKIVTSGLGTVD | 1505660 | Ectodomain | YES |
| SKIYGSCFKSIVLDKF | 1693953 | Ectodomain | YES |
| SKKDWYDFVENPDIIN | 1693956 | Cell/Virus Enclosed | NO |
| SKKHVEVFTDGKLNCS | 1693957 | Cell/Virus Enclosed | NO |
| SKLASELSNTFGAISA | 1693963 | Ectodomain | YES |
| SKLDDEEPMELD | 1505690 | Cell/Virus Enclosed | NO |
| SKLELAKVHNLS | 1505693 | Cell/Virus Enclosed | NO |
| SKLELAKVRNLS | 1505696 | Cell/Virus Enclosed | NO |
| SKLELVKRESEA | 1505698 | Cell/Virus Enclosed | NO |
| SKLFITKDEAIKRVRG | 1693966 | Cell/Virus Enclosed | NO |
| SKLHALDDATRN | 1505705 | Cell/Virus Enclosed | NO |
| SKLHALDDVTLN | 1505707 | Cell/Virus Enclosed | NO |
| SKLLDTCFEATEATFN | 1693976 | Cell/Virus Enclosed | NO |
| SKLLTLVQLTLQSTNYTCIVCID | 58884 | Ectodomain | YES |
| SKLNNFQKFKTI | 1505727 | Ectodomain | YES |
| SKLPSSSGTSTP | 1505729 | Cell/Virus Enclosed | NO |
| SKLRANDNILSV | 1505732 | Cell/Virus Enclosed | NO |
| SKLTIVSPDLESYSKI | 1693987 | Cell/Virus Enclosed | NO |
| SKLVTSGLGTVD | 1505743 | Ectodomain | YES |
| SKNDNLEPVSGL | 1505761 | Ectodomain | YES |
| SKNPLLHDANYF | 1505767 | Secreted | YES |
| SKPFYVPEHCQN | 1505786 | Cell/Virus Enclosed | NO |

| | | | |
|---|---|---|---|
| SKPIYVPEHCQN | 1505791 | Cell/Virus Enclosed | NO |
| SKPQSRNPSSDRNHNS | 1694014 | Capsid | UN |
| SKPTKRSFIEDL | 1505815 | Ectodomain | YES |
| SKPVMISQWPISNDSN | 1694017 | Cell/Virus Enclosed | NO |
| SKPYSTLQPNAYYKYD | 1694020 | Cell/Virus Enclosed | NO |
| SKQRRPQGLPNNTASW | 1388658 | Cell/Virus Enclosed | NO |
| SKRFQSFQQFGR | 1505842 | Ectodomain | YES |
| SKRQYLLKFDDR | 1505849 | Ectodomain | YES |
| SKSCLGYHPAHAPSFL | 1694038 | Cell/Virus Enclosed | NO |
| SKSHNIGSLYVS | 1505873 | Ectodomain | YES |
| SKSPSSSGTSTPKKPN | 1694044 | Capsid | UN |
| SKSQSRNPSSDR | 1505889 | Cell/Virus Enclosed | NO |
| SKSSGMDTALLR | 1505895 | Ectodomain | YES |
| SKSYSGYHPAHAPSFL | 1694047 | Cell/Virus Enclosed | NO |
| SKTEEKSVEQKP | 1505902 | Cell/Virus Enclosed | NO |
| SKTRRSRRAITT | 1505912 | Ectodomain | YES |
| SKTSDWKCKVTDVLFP | 1694056 | Cell/Virus Enclosed | NO |
| SKTTLPVNVAFE | 1505917 | Cell/Virus Enclosed | NO |
| SKTVGTAEEKLKKK | 58954 | Capsid | UN |
| SKVEPCVVSVTY | 1505922 | Cell/Virus Enclosed | NO |
| SKVGNYRLPSSKPSGM | 1694066 | Underneath Membrane | NO |
| SKVGNYRLPSTQKGSG | 1694067 | Ectodomain | YES |
| SKVKNGFTLEYW | 1505936 | Ectodomain | YES |
| SKVKVVPNLYVVVVES | 1694075 | Cell/Virus Enclosed | NO |
| SKVLLERFLPKCPEIL | 1694076 | Cell/Virus Enclosed | NO |
| SKVRLSDVGFVE | 1505948 | Ectodomain | YES |
| SKVSGKGQQQQGQTVTKKS | 1388672 | Capsid | UN |
| SKVVKFPIDLTM | 1505952 | Cell/Virus Enclosed | NO |
| SKVVKVPIDLTI | 1505955 | Cell/Virus Enclosed | NO |
| SKVVKVPIDLTM | 1505956 | Cell/Virus Enclosed | NO |
| SKVVKVTIDYAE | 1505959 | Cell/Virus Enclosed | NO |
| SKVVLSDALDKLYKVF | 1694085 | Cell/Virus Enclosed | NO |
| SKVVLSGEWNAV | 1505961 | Cell/Virus Enclosed | NO |
| SKVVNINVDFKDFQFM | 1694088 | Cell/Virus Enclosed | NO |
| SKVVNVNVDFKDFQFM | 1694089 | Cell/Virus Enclosed | NO |
| SKYLANEAQVQLEHYS | 1694096 | Cell/Virus Enclosed | NO |
| SKYQVIVLKDNV | 1505972 | Cell/Virus Enclosed | NO |
| SKYTMLDLCYAL | 1505976 | Cell/Virus Enclosed | NO |
| SKYVTVQGPPGT | 1505977 | Cell/Virus Enclosed | NO |
| SLAANTLNIIVPDKSV | 1694102 | Cell/Virus Enclosed | NO |
| SLACEHISSTMS | 1505986 | Ectodomain | YES |
| SLADGLEAYADKTLQE | 1694104 | Cell/Virus Enclosed | NO |
| SLADGLEAYADR | 1505988 | Cell/Virus Enclosed | NO |
| SLADGLGAYADK | 1505989 | Cell/Virus Enclosed | NO |
| SLADISAVQSSFGFDQ | 1694105 | Cell/Virus Enclosed | NO |
| SLADLPAYMSVA | 1505990 | Ectodomain | YES |
| SLADSNGFIRFP | 1505992 | Cell/Virus Enclosed | NO |
| SLADSVVEVVTTSLTP | 1694107 | Cell/Virus Enclosed | NO |
| SLAECKRALGLSISDH | 1694108 | Cell/Virus Enclosed | NO |
| SLAIDAYPLIKH | 1506004 | Cell/Virus Enclosed | NO |
| SLAIDAYPLTKHEDIE | 1694119 | Cell/Virus Enclosed | NO |
| SLALGVSEKTHV | 1506014 | Cell/Virus Enclosed | NO |
| SLANSNGFIRFPDVIS | 1694129 | Cell/Virus Enclosed | NO |
| SLANVISFGDYN | 1506021 | Ectodomain | YES |
| SLAYYEPVQGRK | 1506031 | Cell/Virus Enclosed | NO |
| SLCAFSVDPKKTYLDF | 1694135 | Cell/Virus Enclosed | NO |
| SLCAFSVDPKKTYLDY | 1694136 | Cell/Virus Enclosed | NO |
| SLCALPDTPSTLTPRS | 1694139 | Ectodomain | YES |
| SLCDEFTLAEVVKQMY | 1694140 | Cell/Virus Enclosed | NO |
| SLCFKVYFKGQTTHES | 1694141 | Cell/Virus Enclosed | NO |
| SLCKTIQSALSV | 1506050 | Cell/Virus Enclosed | NO |
| SLCNNGKISSSAGDSI | 1694147 | Ectodomain | YES |
| SLCSSHYWSPQGNLGG | 1694149 | Cell/Virus Enclosed | NO |
| SLCVGSGPGKNN | 1506064 | Ectodomain | YES |
| SLDGNFRTGVDP | 1506072 | Cell/Virus Enclosed | NO |
| SLDGTTLQGKPI | 1506074 | Cell/Virus Enclosed | NO |
| SLDLQDCIVGIADDTY | 1694156 | Cell/Virus Enclosed | NO |
| SLDPFCFDVNTLLSNQ | 1694160 | Cell/Virus Enclosed | NO |
| SLDSNFRTEVDP | 1506082 | Cell/Virus Enclosed | NO |
| SLDVEVFNRLATSDWT | 1694165 | Cell/Virus Enclosed | NO |
| SLDVFKIGDVKFKRLG | 1694166 | Cell/Virus Enclosed | NO |
| SLDVKPYGYPNDFVGG | 1694167 | Cell/Virus Enclosed | NO |
| SLDYINVTFLDL | 1506090 | Ectodomain | YES |
| SLEACYLRGCNP | 1506096 | Cell/Virus Enclosed | NO |
| SLEGVFWVANHQADTS | 1694175 | Capsid | UN |
| SLENPTYIYHSA | 1506125 | Cell/Virus Enclosed | NO |

| | | | |
|---|---|---|---|
| SLEVLLDVCNTKEVKV | 1694193 | Cell/Virus Enclosed | NO |
| SLEYWVTPLTSK | 1506151 | Ectodomain | YES |
| SLFAQSDEEESF | 1506153 | Cell/Virus Enclosed | NO |
| SLFAQSEEEEPFSN | 1694196 | Cell/Virus Enclosed | NO |
| SLFAQSEEQESF | 1506160 | Cell/Virus Enclosed | NO |
| SLFATLMTFSGNVDLD | 1694197 | Cell/Virus Enclosed | NO |
| SLFDMSQFPLKL | 1506171 | Cell/Virus Enclosed | NO |
| SLFKPVQLVPVF | 1506188 | Cell/Virus Enclosed | NO |
| SLFKPVQLVPVSPAFH | 1694212 | Cell/Virus Enclosed | NO |
| SLFLFDEAGEEV | 1506192 | Cell/Virus Enclosed | NO |
| SLFPLWTAAAGV | 1506198 | Ectodomain | YES |
| SLFSVNDFTCRQ | 1506211 | Ectodomain | YES |
| SLFVEDCKLPLGQSLC | 1694224 | Ectodomain | YES |
| SLGADSSIAYSNNTIAI | 59120 | Ectodomain | YES |
| SLGASSNQSPQG | 1506226 | Cell/Virus Enclosed | NO |
| SLGDSVKLLSEL | 1506232 | Cell/Virus Enclosed | NO |
| SLGFDKPQEKDKKSAK | 1694233 | Capsid | UN |
| SLGFDKPQEKKK | 1506235 | Cell/Virus Enclosed | NO |
| SLGFDNPPTNVV | 1506236 | Ectodomain | YES |
| SLGNVFCDGVNVTKHK | 1694241 | Cell/Virus Enclosed | NO |
| SLGVSSNQSPQGNLGS | 1694244 | Cell/Virus Enclosed | NO |
| SLHPVECDEECS | 1506266 | Cell/Virus Enclosed | NO |
| SLHSVLWNCQLS | 1506272 | Cell/Virus Enclosed | NO |
| SLIDCNAASVNQIVLR | 1694255 | Cell/Virus Enclosed | NO |
| SLIDLRELGKYE | 1506279 | Ectodomain | YES |
| SLIERGKLLVRDTRKE | 1694258 | Cell/Virus Enclosed | NO |
| SLIGGMVLGGLTSAAA | 1694263 | Ectodomain | YES |
| SLIIASSPALVDQRTI | 1694264 | Cell/Virus Enclosed | NO |
| SLIPVRLRNSSD | 1506293 | Ectodomain | YES |
| SLIPVRPRNSSDNGIS | 1694269 | Ectodomain | YES |
| SLKDLLLYAADPALHV | 1694281 | Cell/Virus Enclosed | NO |
| SLKDLLLYAADPAMHV | 1694282 | Cell/Virus Enclosed | NO |
| SLKFFEIKHADY | 1506327 | Cell/Virus Enclosed | NO |
| SLKGGKIVSTCF | 1506329 | Cell/Virus Enclosed | NO |
| SLKLAGTAVVNLRPDQ | 1694296 | Cell/Virus Enclosed | NO |
| SLKLESSCNFDL | 59222 | Ectodomain | YES |
| SLKLSDNNCYLNAVIM | 1694298 | Cell/Virus Enclosed | NO |
| SLKMCVETVRTQ | 1506341 | Cell/Virus Enclosed | NO |
| SLKYNLKASQAV | 1506358 | Cell/Virus Enclosed | NO |
| SLLATLDDPYVEDSVA | 1506363 | Cell/Virus Enclosed | NO |
| SLLGCLGSQCGS | 1506372 | Ectodomain | YES |
| SLLHEVDKDISQLTQAIVKNHKN | 94690 | Ectodomain | YES |
| SLLKIHRMCKYT | 1506384 | Cell/Virus Enclosed | NO |
| SLLKKMDEPYTEDTSE | 1694317 | Cell/Virus Enclosed | NO |
| SLLKLKVSTNLFEGDK | 1694318 | Cell/Virus Enclosed | NO |
| SLLLEVDKDISHLTQAIVKNHQN | 59270 | Ectodomain | YES |
| SLLNQIWQKYLNSPYT | 1694327 | Cell/Virus Enclosed | NO |
| SLLTEVETPTRNEWECRCSDSSDP | 97651 | Ectodomain | YES |
| SLLVEAYNSILFENGI | 1694343 | Cell/Virus Enclosed | NO |
| SLLVEDCKLPLG | 1506439 | Ectodomain | YES |
| SLLVTASSKPLY | 1506451 | Cell/Virus Enclosed | NO |
| SLMKVMTHMDDFSIKS | 1694354 | Cell/Virus Enclosed | NO |
| SLMNSLFRMPLG | 1506459 | Cell/Virus Enclosed | NO |
| SLNAKLGFDSSKEFVE | 1694360 | Cell/Virus Enclosed | NO |
| SLNALIATAHSSIKQG | 1694361 | Cell/Virus Enclosed | NO |
| SLNDEFSINEVVKQMF | 1694362 | Cell/Virus Enclosed | NO |
| SLNFKLGFDSSK | 1506472 | Cell/Virus Enclosed | NO |
| SLNFVGEFVIND | 1506474 | Cell/Virus Enclosed | NO |
| SLNFVGEFVVND | 1506476 | Cell/Virus Enclosed | NO |
| SLNGNVSTVFVPATRD | 1694369 | Cell/Virus Enclosed | NO |
| SLNGTVSTVFVP | 1506478 | Cell/Virus Enclosed | NO |
| SLNLSCVSKVVNINVD | 1694376 | Cell/Virus Enclosed | NO |
| SLNNPIFINYSKEVYF | 1694377 | Undetermined | UN |
| SLNNPIFISTNY | 1506493 | Ectodomain | YES |
| SLNNSYINLKDIGTYE | 1694378 | Ectodomain | YES |
| SLNVQYRINGLGVTMD | 1694384 | Ectodomain | YES |
| SLPAIKLNFGMLNGNA | 1694386 | Cell/Virus Enclosed | NO |
| SLPAINVTINNYNPSS | 1694387 | Ectodomain | YES |
| SLPIKQPFSFSF | 1506516 | Cell/Virus Enclosed | NO |
| SLPKDWRLINKFDVIN | 1694396 | Cell/Virus Enclosed | NO |
| SLPNMGVPCCTS | 1506525 | Cell/Virus Enclosed | NO |
| SLPRSGSRVAGRSAIEDILF | 2060893 | Ectodomain | YES |
| SLPTAFAAIGDL | 1506530 | Ectodomain | YES |
| SLPVMISEWPLS | 1506542 | Cell/Virus Enclosed | NO |
| SLPYGANKQGIV | 1506549 | Capsid | UN |
| SLQDMRQKQSVW | 1506551 | Cell/Virus Enclosed | NO |

| | | | |
|---|---|---|---|
| SLQEILSRLDVL | 1506554 | Ectodomain | YES |
| SLQQVVKALNESYIDL | 1694417 | Ectodomain | YES |
| SLQVESANLMLSDNVV | 1694423 | Cell/Virus Enclosed | NO |
| SLQVYGHNYYLPVMAA | 1694424 | Undetermined | UN |
| SLRQIFDVAPIELENK | 1694440 | Cell/Virus Enclosed | NO |
| SLRYAWIWCAAY | 1506627 | Cell/Virus Enclosed | NO |
| SLSAPTLVPQENYASI | 1694447 | Cell/Virus Enclosed | NO |
| SLSDRFKTSGDLAVQI | 1694449 | Cell/Virus Enclosed | NO |
| SLSFNNLLGFDSSKPV | 1694454 | Cell/Virus Enclosed | NO |
| SLSHLTSQLRQN | 1506645 | Ectodomain | YES |
| SLSIEEFADVVKEQVS | 1694462 | Cell/Virus Enclosed | NO |
| SLSKDYISSNGPLKVG | 1694464 | Cell/Virus Enclosed | NO |
| SLSKFNSCKSDILSLN | 1694465 | Ectodomain | YES |
| SLSLLVTASSKP | 1506660 | Cell/Virus Enclosed | NO |
| SLSMIRGPPRAELNGV | 1694470 | Cell/Virus Enclosed | NO |
| SLSMVRRNLDSL | 1506665 | Cell/Virus Enclosed | NO |
| SLSNAKVSAEVSRQVI | 1694471 | Cell/Virus Enclosed | NO |
| SLSNFALALCSG | 1506666 | Cell/Virus Enclosed | NO |
| SLSNTKYYDHSQ | 1506670 | Ectodomain | YES |
| SLSPVPTLGSRSRRAVPVAV | 59467 | Ectodomain | YES |
| SLSRNSSRSSSQGSRS | 1694475 | Cell/Virus Enclosed | NO |
| SLSRSNSNFRHSDSIV | 1694476 | Capsid | UN |
| SLSRSQSSETKE | 1506677 | Cell/Virus Enclosed | NO |
| SLSSKPQRKRGVKQLP | 1694477 | Capsid | UN |
| SLSSNHDLYCSVHKGA | 1694478 | Cell/Virus Enclosed | NO |
| SLSSSGTSTPKK | 1506681 | Cell/Virus Enclosed | NO |
| SLSSYEYCQPPKSVEK | 1694480 | Cell/Virus Enclosed | NO |
| SLSTARRNLDSL | 1506683 | Cell/Virus Enclosed | NO |
| SLSTNLDVTNSIEHQ | 59483 | Ectodomain | YES |
| SLSVESTPVEPP | 1506688 | Cell/Virus Enclosed | NO |
| SLSVHDGYTNEI | 1506689 | Cell/Virus Enclosed | NO |
| SLTASSIFGAIL | 1506695 | Cell/Virus Enclosed | NO |
| SLTDVRSTSGTQ | 1506703 | Cell/Virus Enclosed | NO |
| SLTFSYDGLRGAIRKA | 1694488 | Cell/Virus Enclosed | NO |
| SLTFVNVPYVYNGSAQ | 1694489 | Ectodomain | YES |
| SLTHINTTLLDL | 1506712 | Ectodomain | YES |
| SLTMLDIQPEDYKSVD | 1694495 | Cell/Virus Enclosed | NO |
| SLTPCGYSEPPKVAAK | 1694497 | Cell/Virus Enclosed | NO |
| SLTQIDTTLLDL | 1506728 | Ectodomain | YES |
| SLTQINTTLLDLTYEM | 1694498 | Ectodomain | YES |
| SLTSLLKIHRMC | 1506739 | Cell/Virus Enclosed | NO |
| SLTTVSIPQGLT | 1506746 | Cell/Virus Enclosed | NO |
| SLTWLWLLCGLV | 1506760 | Cell/Virus Enclosed | NO |
| SLVAQNTVLNDS | 1506767 | Capsid | UN |
| SLVCTTLQGKPI | 1506771 | Cell/Virus Enclosed | NO |
| SLVGTTLQGKPIGMFF | 1694523 | Cell/Virus Enclosed | NO |
| SLVLAKLGKDATKPQQ | 1694531 | Cell/Virus Enclosed | NO |
| SLVLAKLGKDFK | 1506794 | Cell/Virus Enclosed | NO |
| SLVLAKLGKDSK | 1506795 | Cell/Virus Enclosed | NO |
| SLVLGVNEKTHV | 1506801 | Cell/Virus Enclosed | NO |
| SLVLNNPAYIAS | 1506803 | Ectodomain | YES |
| SLVLNNPSYIAP | 1506805 | Ectodomain | YES |
| SLVNFTVDPQKAYLDF | 1694541 | Cell/Virus Enclosed | NO |
| SLVSVQELVQNT | 1506828 | Cell/Virus Enclosed | NO |
| SLVSVTDFSLNGNVST | 1694548 | Cell/Virus Enclosed | NO |
| SLVTFVGDSSEI | 1506830 | Cell/Virus Enclosed | NO |
| SLWNWFDITKWLWYIK | 745647 | Undetermined | UN |
| SLWNYGKPVTLP | 1506852 | Cell/Virus Enclosed | NO |
| SLWQDPPERPDSPSTNGSGF | 741988 | Underneath Membrane | NO |
| SLWRVTAVHSDGMFVA | 1694559 | Cell/Virus Enclosed | NO |
| SLWYKGSNLEEE | 1506857 | Cell/Virus Enclosed | NO |
| SLYLADNLNADE | 1506866 | Cell/Virus Enclosed | NO |
| SLYLADNLTAGE | 1506870 | Cell/Virus Enclosed | NO |
| SLYNKYRYYSGKMDTA | 1694570 | Cell/Virus Enclosed | NO |
| SLYVNNHAFHTPAYDK | 1694584 | Cell/Virus Enclosed | NO |
| SLYVSWSDGDGITGVP | 1694585 | Ectodomain | YES |
| SMAATRGATCVIGTTK | 1694587 | Cell/Virus Enclosed | NO |
| SMALSIFSAFYP | 1506910 | Transmembrane | NO |
| SMALSIFSAIYP | 1506911 | Transmembrane | NO |
| SMALSIFSAVYP | 1506912 | Transmembrane | NO |
| SMAPVEFYEDRILDFK | 1694592 | Cell/Virus Enclosed | NO |
| SMAVAKVIAKWV | 1506923 | Cell/Virus Enclosed | NO |
| SMCYRCPKEIVSTVSA | 1694602 | Cell/Virus Enclosed | NO |
| SMDIDVFNKLSTSDWS | 1694605 | Cell/Virus Enclosed | NO |
| SMDVLSQNQKLI | 1506943 | Ectodomain | YES |
| SMEVKTIKVFTT | 1506946 | Cell/Virus Enclosed | NO |

| | | | |
|---|---|---|---|
| SMFDVDKKILNA | 1506948 | Cell/Virus Enclosed | NO |
| SMFDVDKKSLNALIAT | 1694608 | Cell/Virus Enclosed | NO |
| SMFSRTTSNFLP | 1506956 | Ectodomain | YES |
| SMGLPNTCFGPL | 1506963 | Cell/Virus Enclosed | NO |
| SMIEDMDYVPLK | 1506967 | Cell/Virus Enclosed | NO |
| SMIKGPQRADLNGVIV | 1694616 | Cell/Virus Enclosed | NO |
| SMIRGPPRAELNGVVV | 1694617 | Cell/Virus Enclosed | NO |
| SMKGTGVFVQDPAPID | 1694620 | Cell/Virus Enclosed | NO |
| SMKSDLSVSSAGPISQ | 1694622 | Ectodomain | YES |
| SMKSDLSVSSSG | 1506983 | Ectodomain | YES |
| SMLGLITQTVDS | 1506989 | Cell/Virus Enclosed | NO |
| SMLGLTTQTVDSSQGS | 1694624 | Cell/Virus Enclosed | NO |
| SMLKQRDSTYGP | 1506992 | Ectodomain | YES |
| SMLKRRDSTYGPLQTP | 1694625 | Ectodomain | YES |
| SMLTFDSNAFSL | 1507005 | Ectodomain | YES |
| SMNIAANSNEWA | 1507014 | Cell/Virus Enclosed | NO |
| SMPGIYKTQRMCLEPC | 1694642 | Cell/Virus Enclosed | NO |
| SMPKIIKVFYELDNDF | 1694645 | Cell/Virus Enclosed | NO |
| SMPSYIAYENARQAYE | 1694647 | Cell/Virus Enclosed | NO |
| SMPVVDGKQIDD | 1507028 | Cell/Virus Enclosed | NO |
| SMPYHPNPVDCQ | 1507029 | Cell/Virus Enclosed | NO |
| SMPYHPNTVDCQDDRC | 1694651 | Cell/Virus Enclosed | NO |
| SMQDVGFKNQYC | 1507031 | Cell/Virus Enclosed | NO |
| SMRLFYDRDGQRTYDD | 1694661 | Cell/Virus Enclosed | NO |
| SMSSFELPQLYGLCIT | 1694663 | Cell/Virus Enclosed | NO |
| SMSYEDQDVLFA | 1507058 | Cell/Virus Enclosed | NO |
| SMTTSTTNFGEE | 1507079 | Cell/Virus Enclosed | NO |
| SMVCKMCLFRYQ | 1507086 | Cell/Virus Enclosed | NO |
| SMVLIDVDVPFTAHFA | 1694684 | Cell/Virus Enclosed | NO |
| SMVPVEFYEDRI | 1507092 | Cell/Virus Enclosed | NO |
| SMVRKLDNQALNSILD | 1694686 | Cell/Virus Enclosed | NO |
| SMWSFNSETNIL | 1507102 | Underneath Membrane | NO |
| SMYDGDRFVKHDLSLL | 1694691 | Cell/Virus Enclosed | NO |
| SMYKQARAEDKKAKIV | 1694694 | Cell/Virus Enclosed | NO |
| SMYKQARSEDKR | 1507107 | Cell/Virus Enclosed | NO |
| SNAFSLANVISF | 1507115 | Ectodomain | YES |
| SNAHRCDVLLSD | 1507117 | Cell/Virus Enclosed | NO |
| SNAKVSAEVSRQVIQD | 1694701 | Cell/Virus Enclosed | NO |
| SNANISMLQLGA | 1507124 | Ectodomain | YES |
| SNANISMLQLGV | 1507125 | Ectodomain | YES |
| SNANLSMLQLGVPDNS | 1694702 | Ectodomain | YES |
| SNASSDVIHYNI | 1507132 | Ectodomain | YES |
| SNASSSFDCIVNLLFT | 1388793 | Ectodomain | YES |
| SNCNFNMSCLMS | 1507147 | Ectodomain | YES |
| SNCVRFKNVDKDDAFY | 1694714 | Cell/Virus Enclosed | NO |
| SNDFTPEDHSLLATLDDPYV | 2060894 | Capsid | UN |
| SNDIWMQNKGLFYTQV | 1694718 | Ectodomain | YES |
| SNDLLQPNTEVFTDVC | 1694719 | Ectodomain | YES |
| SNDLLQPNTEVY | 1507170 | Ectodomain | YES |
| SNDNCTGDIVTHLKNW | 1694721 | Ectodomain | YES |
| SNDQEMTTPPPK | 1507176 | Cell/Virus Enclosed | NO |
| SNDSCPGKNNGI | 1507177 | Ectodomain | YES |
| SNDSKALFFTYL | 1507178 | Cell/Virus Enclosed | NO |
| SNDSNGCVVEASTDFH | 1694722 | Cell/Virus Enclosed | NO |
| SNDYCCFKHALGCDYV | 1694725 | Cell/Virus Enclosed | NO |
| SNEEQAVLKFLDGLDL | 1694726 | Cell/Virus Enclosed | NO |
| SNELAQVLTEVVYSNG | 1694728 | Cell/Virus Enclosed | NO |
| SNENLTWPLGLE | 1507196 | Cell/Virus Enclosed | NO |
| SNENLTWPLVLE | 1507197 | Cell/Virus Enclosed | NO |
| SNEPGNVGGNDA | 1507198 | Cell/Virus Enclosed | NO |
| SNESWFEDDYKA | 1507202 | Cell/Virus Enclosed | NO |
| SNESWFEDDYRAFISV | 1694733 | Cell/Virus Enclosed | NO |
| SNEVTVAKPNVL | 1507205 | Underneath Membrane | NO |
| SNFGICADGSLIPVRP | 1694742 | Ectodomain | YES |
| SNFNQYYYNMAT | 1507229 | Ectodomain | YES |
| SNFNSCKSDILS | 1507230 | Ectodomain | YES |
| SNFSQYYYNIDTGSVY | 1694752 | Underneath Membrane | NO |
| SNFVKDDYCDGSLGCK | 1694753 | Cell/Virus Enclosed | NO |
| SNFYASGCVLSSACTM | 1694754 | Cell/Virus Enclosed | NO |
| SNGCVVEASTDFHQLE | 1694757 | Cell/Virus Enclosed | NO |
| SNGILKWADQSD | 1507264 | Cell/Virus Enclosed | NO |
| SNGKPGNVGGNE | 1507266 | Cell/Virus Enclosed | NO |
| SNGNIIGFKDFVTNKT | 1694765 | Ectodomain | YES |
| SNGPLKVGAGVMLECE | 1694767 | Cell/Virus Enclosed | NO |
| SNGPLKVGTGVM | 1507273 | Cell/Virus Enclosed | NO |
| SNGSCPGKNNGI | 1507276 | Ectodomain | YES |

| | | | |
|---|---|---|---|
| SNGVIIMCADVP | 1507286 | Cell/Virus Enclosed | NO |
| SNHDIICNVHKGAHVA | 1694776 | Cell/Virus Enclosed | NO |
| SNHDLYCSVHKGAHVA | 1694778 | Cell/Virus Enclosed | NO |
| SNHIEVASAYGL | 1507300 | Ectodomain | YES |
| SNIFVINKDTYAALRN | 1694788 | Cell/Virus Enclosed | NO |
| SNINCVLSVNSSNCNN | 1694792 | Cell/Virus Enclosed | NO |
| SNINRVLSVPSD | 1507325 | Cell/Virus Enclosed | NO |
| SNITITYQGLFPYQGD | 1694794 | Ectodomain | YES |
| SNKLCVVPGNVKPGDA | 1694799 | Cell/Virus Enclosed | NO |
| SNLNTNLHFDVDNINF | 1694814 | Ectodomain | YES |
| SNLVLNNCTKYNIYDY | 1694817 | Ectodomain | YES |
| SNMNIASNSKEWAYCV | 1694821 | Cell/Virus Enclosed | NO |
| SNMTQLAEAQII | 1507398 | Underneath Membrane | NO |
| SNMTQLTEAQII | 1507401 | Underneath Membrane | NO |
| SNMVETVVGQLEQTNM | 1694827 | Cell/Virus Enclosed | NO |
| SNNGVYISTTKVKSLS | 1694830 | Cell/Virus Enclosed | NO |
| SNNKEDFDIIQKCQIT | 1694831 | Cell/Virus Enclosed | NO |
| SNNPQCAIVQAL | 1507420 | Cell/Virus Enclosed | NO |
| SNNQEDFDLISR | 1507424 | Cell/Virus Enclosed | NO |
| SNNSSRASSRSSTRNN | 1694837 | Capsid | UN |
| SNPAAVDTKCLA | 1507430 | Cell/Virus Enclosed | NO |
| SNPTNCIAPVNG | 1507442 | Ectodomain | YES |
| SNQADVNIPADI | 1507445 | Cell/Virus Enclosed | NO |
| SNQADVNTPADIVDRD | 1694844 | Cell/Virus Enclosed | NO |
| SNQGNKQIVHSC | 1507447 | Cell/Virus Enclosed | NO |
| SNQGNKQIVHSH | 1507448 | Cell/Virus Enclosed | NO |
| SNQNGERSGARPKQRR | 1694847 | Cell/Virus Enclosed | NO |
| SNQNTDSDSLSSKPQR | 1694849 | Capsid | UN |
| SNQSPQGNLGSNGKPG | 1694852 | Cell/Virus Enclosed | NO |
| SNSGNLLGFKNVSTGN | 1694863 | Ectodomain | YES |
| SNSNFRHSASIV | 1507492 | Cell/Virus Enclosed | NO |
| SNSRPGSRSQSRGPNN | 1694875 | Capsid | UN |
| SNSSILSLCAFSVDPK | 1694877 | Cell/Virus Enclosed | NO |
| SNSSSGVLDTTIPFGP | 1694878 | Ectodomain | YES |
| SNSSVIQEVLLLRHDI | 1694879 | Cell/Virus Enclosed | NO |
| SNSSVLSLVNFTVDPQ | 1694880 | Cell/Virus Enclosed | NO |
| SNSSVPLLEVYVHLRN | 1694881 | Ectodomain | YES |
| SNSSVPLSEVYV | 1507507 | Ectodomain | YES |
| SNSTPAQNAGYW | 1507511 | Cell/Virus Enclosed | NO |
| SNSVCNGCVGHS | 1507512 | Ectodomain | YES |
| SNTDNETLQYWVTPLS | 1694884 | Ectodomain | YES |
| SNTKYYDDSQYYFNKD | 1694889 | Ectodomain | YES |
| SNTTFDFLSNAS | 1507536 | Ectodomain | YES |
| SNVCALMSCNGN | 1507546 | Cell/Virus Enclosed | NO |
| SNVDESFVDDFYGYLQ | 1694898 | Cell/Virus Enclosed | NO |
| SNVDVLPEGWHV | 1507548 | Cell/Virus Enclosed | NO |
| SNVDVLPEGWRV | 1507549 | Cell/Virus Enclosed | NO |
| SNVDVVPEGWRVVNKF | 1694899 | Cell/Virus Enclosed | NO |
| SNVEVLPEGWRV | 1507553 | Cell/Virus Enclosed | NO |
| SNVKKVTDVTGKLSDC | 1694902 | Cell/Virus Enclosed | NO |
| SNVKLLDYDYATHGQL | 1694903 | Cell/Virus Enclosed | NO |
| SNVKTLEYDYMTHGQM | 1694904 | Cell/Virus Enclosed | NO |
| SNVMDVNDCFKK | 1507566 | Cell/Virus Enclosed | NO |
| SNVMDVNDCFKN | 1507567 | Cell/Virus Enclosed | NO |
| SNVPFSPDGKPCTPPAL | 59944 | Ectodomain | YES |
| SNVPFSSDGKPC | 1507570 | Ectodomain | YES |
| SNVSPIVNEKIS | 1507577 | Cell/Virus Enclosed | NO |
| SNVTGFHTINHT | 1507581 | Ectodomain | YES |
| SNVTQIKNACVY | 1507583 | Cell/Virus Enclosed | NO |
| SNVVDVNDCFKN | 1507586 | Cell/Virus Enclosed | NO |
| SNVVELPKFFYASNGT | 1694914 | Ectodomain | YES |
| SNVVRGWVFGST | 1507589 | Ectodomain | YES |
| SNVVYPDLDCIYISLK | 1694916 | Underneath Membrane | NO |
| SNVYRTDYVDYTFVNE | 1694919 | Cell/Virus Enclosed | NO |
| SNVYVDSFVVKG | 1507594 | Ectodomain | YES |
| SNYKIDTTSSSCQLYY | 1694926 | Ectodomain | YES |
| SNYLLEEFDVVFGKRG | 1694928 | Cell/Virus Enclosed | NO |
| SPAAISSNCYSS | 1507623 | Ectodomain | YES |
| SPAAPRAISFAD | 1507626 | Cell/Virus Enclosed | NO |
| SPAAPRAVSFADNNDI | 1694938 | Cell/Virus Enclosed | NO |
| SPAEFKYMVANGLNAP | 1694939 | Cell/Virus Enclosed | NO |
| SPALHRIESTDS | 1507636 | Cell/Virus Enclosed | NO |
| SPALIVWGIHHSVSTAEQTKLYGSG | 1170135 | Ectodomain | YES |
| SPAPSRPFSVLRANDVLWLS | 59984 | Undetermined | UN |
| SPAQDIWGTSAAAYFVG | 59985 | Ectodomain | YES |
| SPAQGTWGTSAA | 1507643 | Ectodomain | YES |

| | | | |
|---|---|---|---|
| SPARIAGNGGDAALAL | 1694944 | Cell/Virus Enclosed | NO |
| SPARLEIVKLGAEKKD | 1694945 | Cell/Virus Enclosed | NO |
| SPARMAGNGGDAALAL | 1694946 | Cell/Virus Enclosed | NO |
| SPARMAGNGGDT | 1507645 | Cell/Virus Enclosed | NO |
| SPARMAGNGSDAALAL | 1694947 | Cell/Virus Enclosed | NO |
| SPASSQSAATIL | 1507651 | Cell/Virus Enclosed | NO |
| SPASSQSSAKSL | 1507652 | Cell/Virus Enclosed | NO |
| SPASSQTSAKSLARSQ | 1388859 | Capsid | UN |
| SPASTVHATAMI | 1507657 | Secreted | YES |
| SPATAEPVRDEF | 1507660 | Capsid | UN |
| SPAYAPTYVSVDDKYK | 1694952 | Cell/Virus Enclosed | NO |
| SPCFSQQGVFRY | 1507671 | Ectodomain | YES |
| SPCKELEGVGAKVSAF | 1694956 | Cell/Virus Enclosed | NO |
| SPCKELEGVGAR | 1507675 | Cell/Virus Enclosed | NO |
| SPCPSGIGAVGG | 1507676 | Cell/Virus Enclosed | NO |
| SPCVSILPSTVW | 1507681 | Ectodomain | YES |
| SPCVSIVPSTVWEDGD | 1694958 | Ectodomain | YES |
| SPDDQIGYYRRATRR | 2001314 | Cell/Virus Enclosed | NO |
| SPDGKPCTPPAF | 1507697 | Ectodomain | YES |
| SPDGKPCTPPALNCYWP | 60014 | Ectodomain | YES |
| SPDPKFVDKYYAFLNK | 1694964 | Cell/Virus Enclosed | NO |
| SPELLDPLNLSA | 1507716 | Cell/Virus Enclosed | NO |
| SPEPALLFRNIK | 1507717 | Ectodomain | YES |
| SPFEKLQCEHLQFGLQ | 1694969 | Ectodomain | YES |
| SPFEMPFEELNGLKIL | 1694971 | Cell/Virus Enclosed | NO |
| SPGLSGIGAVGG | 1507744 | Cell/Virus Enclosed | NO |
| SPGPSGIGGGDL | 1507749 | Cell/Virus Enclosed | NO |
| SPGPSGIGTVGG | 1507750 | Cell/Virus Enclosed | NO |
| SPGPSGVGAPGG | 1507751 | Cell/Virus Enclosed | NO |
| SPGQGVPLNANS | 1507752 | Cell/Virus Enclosed | NO |
| SPGYNLKNGEVEFVYM | 1694985 | Cell/Virus Enclosed | NO |
| SPGYNVRNDGTVEFCY | 1694986 | Cell/Virus Enclosed | NO |
| SPIDIPRGLPVG | 1507771 | Ectodomain | YES |
| SPIIPGFGGDFNLTLL | 1694991 | Ectodomain | YES |
| SPIQSVELPVSIVSLP | 1694992 | Ectodomain | YES |
| SPKLFTRQEEVH | 1507798 | Cell/Virus Enclosed | NO |
| SPKPQRKRGVKQ | 1507803 | Cell/Virus Enclosed | NO |
| SPKPQRQRGHKT | 1507805 | Capsid | UN |
| SPKPQRQRGLKN | 1507808 | Cell/Virus Enclosed | NO |
| SPKSGYFVNVNN | 1507812 | Ectodomain | YES |
| SPKTDKETLEYW | 1507814 | Ectodomain | YES |
| SPKVTIDCAAFV | 1507818 | Ectodomain | YES |
| SPKVTIDCVAFV | 1507821 | Ectodomain | YES |
| SPKVTIDRAAFV | 1507822 | Ectodomain | YES |
| SPLEGGGRLVAS | 1507823 | Ectodomain | YES |
| SPLLVDSEQPWK | 1507839 | Cell/Virus Enclosed | NO |
| SPLLVDSEQSWK | 1507840 | Capsid | UN |
| SPLNWERKTILS | 1507844 | Ectodomain | YES |
| SPMACNGHKIED | 1507854 | Cell/Virus Enclosed | NO |
| SPMKSDEGGLCPSTGQ | 1695021 | Cell/Virus Enclosed | NO |
| SPMSNMTENERLLHFF | 1695022 | Underneath Membrane | NO |
| SPNGKPCTPPAL | 1507864 | Ectodomain | YES |
| SPNKQCTVQQCFGKRG | 1695029 | Cell/Virus Enclosed | NO |
| SPNLPDFREELD | 1507867 | Ectodomain | YES |
| SPNTQRTVQHWF | 1507871 | Cell/Virus Enclosed | NO |
| SPPAIASNCYSS | 1507875 | Ectodomain | YES |
| SPPGEQFKHLVP | 1507876 | Cell/Virus Enclosed | NO |
| SPPISTDTVDVT | 1507881 | Ectodomain | YES |
| SPPLRDTHPQAMQWN | 1695034 | Underneath Membrane | NO |
| SPPLVHDSDLEHVLTPS | 60186 | Ectodomain | YES |
| SPQGNLGGNCAGNVIG | 1695037 | Cell/Virus Enclosed | NO |
| SPQGNLGSNGKPGNVG | 1695038 | Cell/Virus Enclosed | NO |
| SPQIITTDNTFVSGNC | 1695040 | Ectodomain | YES |
| SPRRRSFIEDLLFTS | 2135567 | Ectodomain | YES |
| SPSAVPTDHGAHLSL | 60249 | Cell/Virus Enclosed | NO |
| SPSKDTFELRYS | 1507928 | Cell/Virus Enclosed | NO |
| SPSLNLASVPKTSTPPGEKP | 60259 | Cell/Virus Enclosed | NO |
| SPSSFQSAAKIL | 1507933 | Cell/Virus Enclosed | NO |
| SPSSSGTFTPKK | 1507935 | Cell/Virus Enclosed | NO |
| SPSSSGTSTPKKPNKP | 1695054 | Capsid | UN |
| SPTSKIEPCLVS | 1507959 | Cell/Virus Enclosed | NO |
| SPTVWLSVIWMMWYW | 1695065 | Undetermined | UN |
| SPVFHRIESIDS | 1507973 | Cell/Virus Enclosed | NO |
| SPVFHRIESTDS | 1507974 | Cell/Virus Enclosed | NO |
| SPVKDVFELHYSGSIR | 1695068 | Capsid | UN |
| SPVMTNGSNILEAFTK | 1695070 | Cell/Virus Enclosed | NO |

| | | | |
|---|---|---|---|
| SPYDLEVLLQDALQSR | 1695072 | Cell/Virus Enclosed | NO |
| SPYIDDFNKIASCKWT | 1695074 | Cell/Virus Enclosed | NO |
| SPYIDDFNRIASCKWT | 1695075 | Cell/Virus Enclosed | NO |
| SPYNSQNFAAKRVLGL | 1695079 | Cell/Virus Enclosed | NO |
| SPYRFVTFEPFN | 1507997 | Ectodomain | YES |
| SPYVCNAPGCDVNDVT | 1695085 | Cell/Virus Enclosed | NO |
| SPYVCNAPGCGA | 1508000 | Cell/Virus Enclosed | NO |
| SQAKFGIITKVE | 1508008 | Ectodomain | YES |
| SQALQTVSTALN | 1508012 | Ectodomain | YES |
| SQASSRSSSRSRGNSR | 1388946 | Capsid | UN |
| SQASSRSSSRSRNSSR | 1695093 | Cell/Virus Enclosed | NO |
| SQDDIMKAVAAALKSL | 1695098 | Capsid | UN |
| SQDKVFGYVQQGIFNK | 1695100 | Cell/Virus Enclosed | NO |
| SQDSITHYPALKMVQT | 1695104 | Cell/Virus Enclosed | NO |
| SQDSITHYSALK | 1508045 | Cell/Virus Enclosed | NO |
| SQDTLLAPLLSA | 1508051 | Cell/Virus Enclosed | NO |
| SQDVKQFANGFVVRIG | 1695107 | Ectodomain | YES |
| SQDVYKSLTIVDIPQS | 1695108 | Cell/Virus Enclosed | NO |
| SQDVYKSLTTVS | 1508057 | Cell/Virus Enclosed | NO |
| SQECSNLLLQYG | 1508059 | Ectodomain | YES |
| SQEGILSEREKL | 1508061 | Ectodomain | YES |
| SQEGKKYLVKTTKAKG | 1695109 | Cell/Virus Enclosed | NO |
| SQERNFTTAPAICHEG | 1695112 | Ectodomain | YES |
| SQFDSLGASSNQ | 1508070 | Cell/Virus Enclosed | NO |
| SQFDSLGVSSNQSPQG | 1695114 | Cell/Virus Enclosed | NO |
| SQGFVVVNGKFS | 1508090 | Cell/Virus Enclosed | NO |
| SQGKLLIRDNDV | 1508094 | Cell/Virus Enclosed | NO |
| SQGNVVTSVMESQIST | 1695129 | Cell/Virus Enclosed | NO |
| SQGSRSGNSTRC | 1508106 | Cell/Virus Enclosed | NO |
| SQGSRSGNSTRGTSPG | 1695133 | Cell/Virus Enclosed | NO |
| SQHTHYIKDGDD | 1508111 | Cell/Virus Enclosed | NO |
| SQHTLYIKDGDDGYFL | 1695134 | Cell/Virus Enclosed | NO |
| SQHTMQIVDEKG | 1508115 | Cell/Virus Enclosed | NO |
| SQHTMQIVDENGKYYL | 1695138 | Cell/Virus Enclosed | NO |
| SQHTMQIVDKDGTYYL | 1695139 | Cell/Virus Enclosed | NO |
| SQIISGIVAAVS | 1508120 | Undetermined | UN |
| SQILPDPLKPIK | 1508124 | Ectodomain | YES |
| SQILPDPLKPTKRSFI | 1695141 | Ectodomain | YES |
| SQILPDPLKSTK | 1508126 | Ectodomain | YES |
| SQILPDPSKPTK | 1508127 | Ectodomain | YES |
| SQIQSKLTDVKCANVV | 1695146 | Cell/Virus Enclosed | NO |
| SQISGYTTAATVAAMF | 1695147 | Ectodomain | YES |
| SQISPAAIASNCYSSL | 1695148 | Ectodomain | YES |
| SQISPAAISSNC | 1508136 | Ectodomain | YES |
| SQISSHSPTCCPPICPGYRW | 180928 | Ectodomain | YES |
| SQISYSNFYASGCVLS | 1695150 | Cell/Virus Enclosed | NO |
| SQIVTGLFKDCSRETS | 1695151 | Cell/Virus Enclosed | NO |
| SQLAKAMDTFTN | 1508151 | Cell/Virus Enclosed | NO |
| SQLFVDTLVNKIPANI | 1695157 | Cell/Virus Enclosed | NO |
| SQLGCVFFNGAD | 1508161 | Cell/Virus Enclosed | NO |
| SQLKNNAKEIGNGCF | 151056 | Ectodomain | YES |
| SQLPLDIYDLTM | 1508179 | Cell/Virus Enclosed | NO |
| SQLPLDVYDLTMPSQV | 1695167 | Cell/Virus Enclosed | NO |
| SQLPLDVYNLTM | 1508181 | Cell/Virus Enclosed | NO |
| SQMETDFLEPAM | 1508200 | Cell/Virus Enclosed | NO |
| SQNAVSRSMLGLTTQT | 1695177 | Cell/Virus Enclosed | NO |
| SQNFAAKRVLGLQTQT | 1695178 | Cell/Virus Enclosed | NO |
| SQNFGNAEMLKLGTND | 1695179 | Capsid | UN |
| SQNMLRGDDMSF | 1508214 | Cell/Virus Enclosed | NO |
| SQNMLRGEDMFF | 1508215 | Cell/Virus Enclosed | NO |
| SQNMLRGEDMSF | 1508216 | Cell/Virus Enclosed | NO |
| SQNVKQFANGFV | 1508227 | Ectodomain | YES |
| SQNYVAKRVLGVQTQT | 1695187 | Cell/Virus Enclosed | NO |
| SQPKVITKRDAA | 1508233 | Cell/Virus Enclosed | NO |
| SQPLPQRQKKQQTVTL | 1695190 | Cell/Virus Enclosed | NO |
| SQPRADKPSQLKKPRW | 1695191 | Capsid | UN |
| SQPRVDKPSQLK | 1508238 | Cell/Virus Enclosed | NO |
| SQPSHAVQNTVL | 1508239 | Cell/Virus Enclosed | NO |
| SQQGVFRYDNIS | 1508246 | Ectodomain | YES |
| SQQGVFRYDNVSSVWP | 1695196 | Ectodomain | YES |
| SQQGVFRYDSVS | 1508248 | Ectodomain | YES |
| SQQPLGGNVVPY | 1508252 | Cell/Virus Enclosed | NO |
| SQQPSGGNVVPH | 1508254 | Cell/Virus Enclosed | NO |
| SQQPSGRNVVPH | 1508256 | Capsid | UN |
| SQQPSRGNVVPH | 1508257 | Cell/Virus Enclosed | NO |
| SQQPSRGNVVPY | 1508258 | Cell/Virus Enclosed | NO |

| | | | |
|---|---|---|---|
| SQQQSQTVTKKSAAEA | 1695201 | Cell/Virus Enclosed | NO |
| SQRGGSYKNDKS | 1508266 | Cell/Virus Enclosed | NO |
| SQRGVYCCRDHE | 1508269 | Cell/Virus Enclosed | NO |
| SQRVAGDSGFAAYSRY | 1388977 | Underneath Membrane | NO |
| SQSAAKILAPSR | 1508278 | Cell/Virus Enclosed | NO |
| SQSAAKILARSQ | 1508280 | Cell/Virus Enclosed | NO |
| SQSDRFYRPANE | 1508284 | Cell/Virus Enclosed | NO |
| SQSDSLGVSSNQ | 1508285 | Cell/Virus Enclosed | NO |
| SQSERYGFCGNG | 1508286 | Ectodomain | YES |
| SQSNRYGFCGNG | 1508295 | Ectodomain | YES |
| SQSRGPGESKPQ | 1508298 | Capsid | UN |
| SQSRGRGESKPQSRNP | 1695214 | Capsid | UN |
| SQSRGRGESKSQ | 1508301 | Cell/Virus Enclosed | NO |
| SQSRNPSSDRNH | 1508303 | Cell/Virus Enclosed | NO |
| SQSRNSSSDRNH | 1508304 | Cell/Virus Enclosed | NO |
| SQSRSQSRGRES | 1508306 | Cell/Virus Enclosed | NO |
| SQSRSQSRGRGESKPQ | 1695215 | Capsid | UN |
| SQSRSQSRIQSR | 1508308 | Cell/Virus Enclosed | NO |
| SQSSAKSLARSQ | 1508314 | Cell/Virus Enclosed | NO |
| SQSSAKSPARSQ | 1508316 | Cell/Virus Enclosed | NO |
| SQSSETKEQKHEMQKP | 1695216 | Capsid | UN |
| SQSSHAVQNTVF | 1508318 | Cell/Virus Enclosed | NO |
| SQSSHVAQNTVLNASI | 1695217 | Capsid | UN |
| SQSSHVVQNTVL | 1508322 | Cell/Virus Enclosed | NO |
| SQSSPIIPGFGGDFNL | 1695218 | Ectodomain | YES |
| SQSSRINFCGNGNHII | 1695219 | Ectodomain | YES |
| SQSSSRASSLSRNSSR | 1695220 | Cell/Virus Enclosed | NO |
| SQSVAKILARSQ | 1508329 | Cell/Virus Enclosed | NO |
| SQSVCNGCVGHS | 1508330 | Ectodomain | YES |
| SQSVIIINNSTNVVIR | 1695221 | Ectodomain | YES |
| SQTSAKSLARSQSSET | 1695225 | Capsid | UN |
| SQTVKPGHFNKEFYDF | 1695231 | Cell/Virus Enclosed | NO |
| SQVIVNNYDKSAGYPF | 1508356 | Cell/Virus Enclosed | NO |
| SQVIVNNYEKSA | 1508357 | Cell/Virus Enclosed | NO |
| SQVKADLVLDAL | 1508358 | Cell/Virus Enclosed | NO |
| SQVYTTSEYPSQ | 60589 | Secreted | YES |
| SQWAREALQTGITLVALLLLVIL | 60591 | Undetermined | UN |
| SQWLSNLLGTTV | 1508365 | Cell/Virus Enclosed | NO |
| SQWPISDDSDGC | 1508368 | Cell/Virus Enclosed | NO |
| SQWPISDDSGDC | 1508369 | Cell/Virus Enclosed | NO |
| SQWPISNDSNCC | 1508370 | Cell/Virus Enclosed | NO |
| SQWPISNDSNGCVVEA | 1695247 | Cell/Virus Enclosed | NO |
| SQYLFSTINALPETSA | 1695248 | Cell/Virus Enclosed | NO |
| SQYSRSTRSILK | 1508377 | Ectodomain | YES |
| SQYSRSTRSMLK | 1508379 | Ectodomain | YES |
| SQYYFNKDTGVIYGLN | 1695254 | Ectodomain | YES |
| SRALKKVPIDEY | 1508392 | Cell/Virus Enclosed | NO |
| SRALKKVPVAEY | 1508393 | Cell/Virus Enclosed | NO |
| SRANSGNRAPTS | 1508397 | Cell/Virus Enclosed | NO |
| SRANSGNRTPTSGVTP | 1695257 | Cell/Virus Enclosed | NO |
| SRANSGNRTSTS | 1508402 | Cell/Virus Enclosed | NO |
| SRASSAGSRSKA | 1508406 | Cell/Virus Enclosed | NO |
| SRASSAGSRSRANSGN | 1695259 | Cell/Virus Enclosed | NO |
| SRASSGNRSGNGILKW | 1388989 | Cell/Virus Enclosed | NO |
| SRASSGNRSSNG | 1508412 | Cell/Virus Enclosed | NO |
| SRASSLSRNSSRSSSQ | 1388990 | Cell/Virus Enclosed | NO |
| SRASSRSSTRNNSRDS | 1695260 | Capsid | UN |
| SRAWAFCVKCHN | 1508421 | Cell/Virus Enclosed | NO |
| SRDHNDILYTPPTVSY | 1695265 | Cell/Virus Enclosed | NO |
| SRDLEDFKPRSK | 1508433 | Cell/Virus Enclosed | NO |
| SRDPTTQEAIPTRFPP | 1695267 | Capsid | UN |
| SRDSSRSTSRQQSRTR | 1695268 | Capsid | UN |
| SRDVVRFPNITN | 1508447 | Ectodomain | YES |
| SRELSVTFFPDL | 1508459 | Cell/Virus Enclosed | NO |
| SRELTAEDISLLKKMD | 1695277 | Cell/Virus Enclosed | NO |
| SRETFGLSPAYA | 1508462 | Cell/Virus Enclosed | NO |
| SREVVVTNLNKSAGWP | 1695280 | Cell/Virus Enclosed | NO |
| SRFLGLQIQTVDSSQG | 1695283 | Cell/Virus Enclosed | NO |
| SRFNPSTWNKKF | 1508473 | Ectodomain | YES |
| SRGAITTGYRFTNFEP | 1695287 | Ectodomain | YES |
| SRGAPGGGFVPSLQGVP | 60663 | Cell/Virus Enclosed | NO |
| SRGAPGGGFVPSMQ | 60664 | Cell/Virus Enclosed | NO |
| SRGESKSQSRNP | 1508486 | Cell/Virus Enclosed | NO |
| SRGNSPARMASGGGETAL | 60681 | Capsid | UN |
| SRGNVVPHYSWF | 1508495 | Cell/Virus Enclosed | NO |
| SRGPGESKPQSR | 1508497 | Capsid | UN |

| | | | |
|---|---|---|---|
| SRGPNNRSLSRSNSNF | 1695292 | Capsid | UN |
| SRGTSPARIAGNGGDA | 1695295 | Cell/Virus Enclosed | NO |
| SRILGAGCFVND | 1508527 | Cell/Virus Enclosed | NO |
| SRIMFEPRIPTMADFV | 1695310 | Ectodomain | YES |
| SRINFCGNGNHIISLV | 1695311 | Ectodomain | YES |
| SRKAMKRSHSPT | 1508539 | Cell/Virus Enclosed | NO |
| SRKARNRSHSPT | 1508546 | Cell/Virus Enclosed | NO |
| SRKAYDHNSPYIWPRNDYDG | 60710 | Ectodomain | YES |
| SRKFIVPWVMYL | 1508548 | Cell/Virus Enclosed | NO |
| SRKKMDTKPTDPTG | 60714 | Capsid | UN |
| SRKSYKKMLKEMGEVAPEYR | 60721 | Capsid | UN |
| SRLDALEAEAQIDRLI | 1695318 | Ectodomain | YES |
| SRLDALEAQVQIDRLI | 1695319 | Ectodomain | YES |
| SRLELAKVQNLSGNPD | 1695321 | Cell/Virus Enclosed | NO |
| SRLLSDDRTEVPQLVN | 1695327 | Ectodomain | YES |
| SRLPLDVYDLTM | 1508597 | Cell/Virus Enclosed | NO |
| SRLRSDDRTEVP | 1508600 | Ectodomain | YES |
| SRMGFKLDAIVP | 1508616 | Cell/Virus Enclosed | NO |
| SRMGFKLDATVPGYPK | 1695336 | Cell/Virus Enclosed | NO |
| SRNESWHFDKSEPLCL | 1695341 | Ectodomain | YES |
| SRNPSSDRNHNSQDDI | 1695344 | Capsid | UN |
| SRNQSPASSQSV | 1508638 | Cell/Virus Enclosed | NO |
| SRNQSPSSSQSA | 1508640 | Cell/Virus Enclosed | NO |
| SRNQSPVSSQSA | 1508641 | Cell/Virus Enclosed | NO |
| SRNSSRNSTPGSSRGT | 1695346 | Cell/Virus Enclosed | NO |
| SRNSSRSSSQGSRSGN | 1695347 | Cell/Virus Enclosed | NO |
| SRNSTPGSSRGNSPARMA | 60749 | Capsid | UN |
| SRNSTPGSSRGTSPAR | 1695349 | Cell/Virus Enclosed | NO |
| SRNSTRGTSPGP | 1508649 | Cell/Virus Enclosed | NO |
| SRNTTFDFLSNASSSF | 1695350 | Ectodomain | YES |
| SRQLQPINYFDS | 1508667 | Ectodomain | YES |
| SRQVIQDVKDGTVTFN | 1695359 | Cell/Virus Enclosed | NO |
| SRQVIQEVKDGT | 1508671 | Cell/Virus Enclosed | NO |
| SRRAITTENRFT | 1508677 | Ectodomain | YES |
| SRRFIVPWVMYL | 1508682 | Cell/Virus Enclosed | NO |
| SRRGVYYNDDIF | 1508684 | Ectodomain | YES |
| SRRKRRSISASYRFVT | 1695361 | Ectodomain | YES |
| SRRRLLACLCKH | 1508690 | Cell/Virus Enclosed | NO |
| SRSCSDFIPLSD | 1508697 | Cell/Virus Enclosed | NO |
| SRSCSDFLPLSD | 1508698 | Cell/Virus Enclosed | NO |
| SRSDCNHIGNIN | 1508701 | Ectodomain | YES |
| SRSERRKDRGGREDIL | 60796 | Cell/Virus Enclosed | NO |
| SRSFFEDLLFDKVKLS | 1508706 | Ectodomain | YES |
| SRSGNSTRCTSP | 1508710 | Cell/Virus Enclosed | NO |
| SRSGNSTRGTSPGPSG | 1695372 | Cell/Virus Enclosed | NO |
| SRSNSNFRHSAS | 1508721 | Cell/Virus Enclosed | NO |
| SRSQSRIQSRNP | 1508726 | Cell/Virus Enclosed | NO |
| SRSQSRSQSRGP | 1508727 | Capsid | UN |
| SRSQSSETKEQK | 1508731 | Cell/Virus Enclosed | NO |
| SRSRANSGNRTL | 1508735 | Cell/Virus Enclosed | NO |
| SRSRANSGNRTPTSGV | 1695381 | Cell/Virus Enclosed | NO |
| SRSRASSGNRAP | 1508738 | Cell/Virus Enclosed | NO |
| SRSRNSSRNSTPGSSR | 1695383 | Cell/Virus Enclosed | NO |
| SRSRNSTRGTSP | 1508744 | Cell/Virus Enclosed | NO |
| SRSRVSSGNRAP | 1508745 | Cell/Virus Enclosed | NO |
| SRSSSQGSRSGNSTRG | 1695385 | Cell/Virus Enclosed | NO |
| SRSSSQGSRSRN | 1508749 | Cell/Virus Enclosed | NO |
| SRSSTRNNSRDSSRST | 1695388 | Capsid | UN |
| SRSTRSILKRRD | 1508756 | Ectodomain | YES |
| SRSTRSMLKRRDSTYG | 1695389 | Ectodomain | YES |
| SRSTSRTSSRASSAGS | 1695391 | Cell/Virus Enclosed | NO |
| SRTGHFYINGYR | 1508773 | Ectodomain | YES |
| SRTLSYYRLGAS | 1508784 | Underneath Membrane | NO |
| SRTPKYTFGVVK | 1508785 | Cell/Virus Enclosed | NO |
| SRTRSDSNQSSSDLVA | 1695404 | Capsid | UN |
| SRTSSRASSTGS | 1508790 | Cell/Virus Enclosed | NO |
| SRVESKSQSRNP | 1508804 | Cell/Virus Enclosed | NO |
| SRVGRSVNSQNC | 1508810 | Underneath Membrane | NO |
| SRVGRSVNSQNSTGWV | 1695411 | Underneath Membrane | NO |
| SRVIRRERCDMT | 1508812 | Cell/Virus Enclosed | NO |
| SRVISRMGFKLDATVP | 1695415 | Cell/Virus Enclosed | NO |
| SRVKNLNSSEGVPDLLV | 60827 | Ectodomain | YES |
| SRVLSSFEPRSDLERD | 1695423 | Cell/Virus Enclosed | NO |
| SRVPHIVRKDLTKYTM | 1695425 | Cell/Virus Enclosed | NO |
| SRVSSGNRAPTS | 1508832 | Cell/Virus Enclosed | NO |
| SRVVIQDYDNQHEQDRPTPSPAPSR | 60833 | Undetermined | UN |

| | | | |
|---|---|---|---|
| SRWNNARQSDNMTAVA | 1695430 | Ectodomain | YES |
| SRWNNERASDISLSVA | 1695431 | Underneath Membrane | NO |
| SRYRIGNYKLNTDHSN | 1695435 | Underneath Membrane | NO |
| SRYRIGNYKLNTDHSS | 1389032 | Underneath Membrane | NO |
| SRYSKKFKPEIAARP | 504115 | Ectodomain | YES |
| SRYVDLYGGNMF | 1508851 | Ectodomain | YES |
| SSAGSRSRANSGNRTP | 1695446 | Cell/Virus Enclosed | NO |
| SSALGSMDIDVFNKLS | 1695450 | Cell/Virus Enclosed | NO |
| SSANGSLDGTTL | 1508887 | Cell/Virus Enclosed | NO |
| SSANGSLVCTTL | 1508888 | Cell/Virus Enclosed | NO |
| SSANGSLVGTIL | 1508889 | Cell/Virus Enclosed | NO |
| SSANYVGKYDINHGDS | 1695455 | Underneath Membrane | NO |
| SSAPRITFGGPS | 1508893 | Cell/Virus Enclosed | NO |
| SSATQPYFDSYL | 1508903 | Ectodomain | YES |
| SSCFGKNNGIGT | 1508916 | Ectodomain | YES |
| SSCGLFKDCARNPIDL | 1695463 | Cell/Virus Enclosed | NO |
| SSCHNASGKEAKVCTISPIM | 60923 | Capsid | UN |
| SSCPGKNNGIGT | 1508925 | Ectodomain | YES |
| SSCPGKNNGLGT | 1508927 | Ectodomain | YES |
| SSCSCPGKNNGI | 1508932 | Ectodomain | YES |
| SSCSGKNNGIGT | 1508933 | Ectodomain | YES |
| SSCSGKNNGVGT | 1508934 | Ectodomain | YES |
| SSCVECDAKFKNSVAS | 1695467 | Cell/Virus Enclosed | NO |
| SSCVGSGPGKNN | 1508937 | Ectodomain | YES |
| SSDEAIPTRFPPGTVL | 1695472 | Cell/Virus Enclosed | NO |
| SSDGQPGGDAISLSFN | 1695475 | Cell/Virus Enclosed | NO |
| SSDKAPYRVIPR | 1508959 | Cell/Virus Enclosed | NO |
| SSDMVETVVGQL | 1508963 | Cell/Virus Enclosed | NO |
| SSDRYHNSQDDI | 1508972 | Capsid | UN |
| SSDSDVFIKKYG | 1508973 | Cell/Virus Enclosed | NO |
| SSDVFKIGDVKF | 1508975 | Cell/Virus Enclosed | NO |
| SSDVIHYNINFE | 1508976 | Ectodomain | YES |
| SSEGFLLGINYLGTIK | 1695497 | Cell/Virus Enclosed | NO |
| SSELKQLEAKTASLFQ | 1695501 | Ectodomain | YES |
| SSEPALIFRNIK | 1509012 | Ectodomain | YES |
| SSEPTLLFRNIK | 1509017 | Ectodomain | YES |
| SSESGAKETKEINIIK | 1695503 | Cell/Virus Enclosed | NO |
| SSETKEQKHEMQKPRW | 1695504 | Capsid | UN |
| SSEVAVLYQDVNCTDV | 60978 | Ectodomain | YES |
| SSEVKAQSSGVIPENP | 1695506 | Cell/Virus Enclosed | NO |
| SSEWAYCVDLHNKINL | 1695508 | Cell/Virus Enclosed | NO |
| SSFDGVMYGGFEDQPN | 1695513 | Cell/Virus Enclosed | NO |
| SSFEAKPSGSVVEQAE | 1695514 | Ectodomain | YES |
| SSFEGYDFKYIENGQL | 1695515 | Cell/Virus Enclosed | NO |
| SSFGDYNLSSVIPSLP | 1695520 | Ectodomain | YES |
| SSFGEYNLSSVV | 1509056 | Ectodomain | YES |
| SSFGFDQQQLLA | 1509057 | Cell/Virus Enclosed | NO |
| SSFIELLFDCVG | 1509059 | Cell/Virus Enclosed | NO |
| SSFKATLYYQNG | 1509062 | Cell/Virus Enclosed | NO |
| SSFMNVTLNKCT | 1509071 | Ectodomain | YES |
| SSFNFDQKELLAYYNM | 1695527 | Cell/Virus Enclosed | NO |
| SSFNFDQKELLD | 1509074 | Cell/Virus Enclosed | NO |
| SSFQSAAKILAR | 1509075 | Cell/Virus Enclosed | NO |
| SSFSEILFDCVD | 1509077 | Cell/Virus Enclosed | NO |
| SSFSELLFDCVD | 1509078 | Cell/Virus Enclosed | NO |
| SSFVGMPSFVAYETAR | 1695533 | Cell/Virus Enclosed | NO |
| SSFVMMSAPPSE | 1509092 | Cell/Virus Enclosed | NO |
| SSFYAPEPITSF | 1509093 | Ectodomain | YES |
| SSFYSPEPITSL | 1509098 | Ectodomain | YES |
| SSGFLQSSNYKIDTTS | 1695540 | Ectodomain | YES |
| SSGFRKMAFPSG | 1509108 | Cell/Virus Enclosed | NO |
| SSGIFSVPSDVS | 1509114 | Cell/Virus Enclosed | NO |
| SSGMINPNALLC | 1509128 | Cell/Virus Enclosed | NO |
| SSGNRPGNGILK | 1509132 | Capsid | UN |
| SSGNRSGNGILKWADQ | 1695545 | Cell/Virus Enclosed | NO |
| SSGNRSVNGILK | 1509134 | Cell/Virus Enclosed | NO |
| SSGNYVGKYDIN | 1509136 | Underneath Membrane | NO |
| SSGPISQFNYKQ | 1509141 | Ectodomain | YES |
| SSGTSTPKKPNKPLSQ | 1695547 | Capsid | UN |
| SSGVANPAMDPIYDEPTTTTSVPL | 1509151 | Secreted | YES |
| SSGVLDTTIPFGPSSQ | 1695551 | Ectodomain | YES |
| SSGYRSGNGILK | 1509164 | Cell/Virus Enclosed | NO |
| SSHAVQNTVFNA | 1509165 | Cell/Virus Enclosed | NO |
| SSHYWSPQGNLGGNCA | 1695561 | Cell/Virus Enclosed | NO |
| SSIAYSNNTIAIPTNF | 1695563 | Ectodomain | YES |
| SSIDAFKLTIKL | 1509186 | Cell/Virus Enclosed | NO |

| | | | |
|---|---|---|---|
| SSIKEMQSQSSHVAQN | 1695570 | Capsid | UN |
| SSIKGDKGVEKLVNWY | 1695571 | Cell/Virus Enclosed | NO |
| SSIKSDKGVEKL | 1509202 | Cell/Virus Enclosed | NO |
| SSINDLSFDKNL | 1509204 | Cell/Virus Enclosed | NO |
| SSIQAIYDRLDI | 1509206 | Ectodomain | YES |
| SSIQAIYDRLDTIQAD | 1695578 | Ectodomain | YES |
| SSIQANAYYKYDNGNF | 1695579 | Cell/Virus Enclosed | NO |
| SSIRGCCESTKLPYYD | 1695581 | Underneath Membrane | NO |
| SSITIDKFAIPNGRKV | 1695583 | Ectodomain | YES |
| SSIVRQLFKPLT | 1509223 | Cell/Virus Enclosed | NO |
| SSIYKLHPAFNVSDAY | 1695587 | Cell/Virus Enclosed | NO |
| SSKADNSIFRSF | 1509228 | Ectodomain | YES |
| SSKAGNSIFRSF | 1509229 | Ectodomain | YES |
| SSKEFVEYKITQ | 1509236 | Cell/Virus Enclosed | NO |
| SSKMSDVKCTSV | 1509243 | Cell/Virus Enclosed | NO |
| SSKPLYVPEHCQNYSG | 1695593 | Cell/Virus Enclosed | NO |
| SSKPVAKKYTYS | 1509248 | Cell/Virus Enclosed | NO |
| SSKPVIKKYTYS | 1509249 | Cell/Virus Enclosed | NO |
| SSKSLGKQLGVVSDGV | 1695599 | Cell/Virus Enclosed | NO |
| SSKVVLSDALDKLYKV | 1695606 | Cell/Virus Enclosed | NO |
| SSLCDEFTLAEVVKQM | 1695610 | Cell/Virus Enclosed | NO |
| SSLCFKVYFKGQTTHE | 1695611 | Cell/Virus Enclosed | NO |
| SSLCTMFKRGDGTPHP | 1695613 | Cell/Virus Enclosed | NO |
| SSLCVGSGPGKN | 1509268 | Ectodomain | YES |
| SSLFAQSEEDES | 1509274 | Cell/Virus Enclosed | NO |
| SSLFAQSEEEEL | 1509275 | Cell/Virus Enclosed | NO |
| SSLFAQSEEQES | 1509280 | Cell/Virus Enclosed | NO |
| SSLFVEDCKLPLGQSL | 1695615 | Ectodomain | YES |
| SSLILDYFSYPF | 1509291 | Ectodomain | YES |
| SSLIREIVMNAS | 1509293 | Cell/Virus Enclosed | NO |
| SSLNDEFSINEVVKQM | 1695621 | Cell/Virus Enclosed | NO |
| SSLQEILSRLDALEAQ | 1695624 | Ectodomain | YES |
| SSLQEILSRLDV | 1509312 | Ectodomain | YES |
| SSLQQVVKALNE | 1509313 | Ectodomain | YES |
| SSLRDHINVIVS | 61186 | Ectodomain | YES |
| SSLRYGNVLDVNAI | 61192 | Capsid | UN |
| SSLSAQSEEEES | 1509317 | Cell/Virus Enclosed | NO |
| SSLSRNSSRSSSQGSR | 1695626 | Cell/Virus Enclosed | NO |
| SSLSSYEYCQPPKSVE | 1695627 | Cell/Virus Enclosed | NO |
| SSLTCNVSFVGELKAA | 1695628 | Cell/Virus Enclosed | NO |
| SSMLRLHTVESG | 1509345 | Cell/Virus Enclosed | NO |
| SSMNIAANSNEW | 1509348 | Cell/Virus Enclosed | NO |
| SSMRGCCDCGST | 1509350 | Underneath Membrane | NO |
| SSNEEQAVLKFLDGLD | 1695647 | Cell/Virus Enclosed | NO |
| SSNGILKWADQS | 1509363 | Cell/Virus Enclosed | NO |
| SSNGPLKVGAGVMLEC | 1695649 | Cell/Virus Enclosed | NO |
| SSNGPLKVGTGV | 1509366 | Cell/Virus Enclosed | NO |
| SSNGQPGGDAIS | 1509367 | Cell/Virus Enclosed | NO |
| SSNGSNDVWIFN | 1509368 | Ectodomain | YES |
| SSNINCVLSVNSSNCN | 1695651 | Cell/Virus Enclosed | NO |
| SSNINRILGVNS | 1509373 | Cell/Virus Enclosed | NO |
| SSNINRLLSVPSDSCN | 1695652 | Cell/Virus Enclosed | NO |
| SSNINRVLSVPS | 1509374 | Cell/Virus Enclosed | NO |
| SSNLNTNLHFDVDNIN | 1695653 | Ectodomain | YES |
| SSNMVETLVGQL | 1509378 | Cell/Virus Enclosed | NO |
| SSNMVETVVGQLEQTN | 1695655 | Cell/Virus Enclosed | NO |
| SSNQSPQGNLGSNGKP | 1695658 | Cell/Virus Enclosed | NO |
| SSNRGNKQIVHS | 1509385 | Cell/Virus Enclosed | NO |
| SSNTDNETLQYWVTPL | 1695659 | Ectodomain | YES |
| SSNYKIDTTSSSCQLY | 1695663 | Ectodomain | YES |
| SSPARLEIVKLGAEKK | 1695666 | Cell/Virus Enclosed | NO |
| SSPCHNSLILPPFSLSPVPTLGSRS | 61260 | Ectodomain | YES |
| SSPKVTIDCVAF | 1509416 | Ectodomain | YES |
| SSPMSNMTENERLLHF | 1695674 | Underneath Membrane | NO |
| SSPNKQCTVQQC | 1509421 | Cell/Virus Enclosed | NO |
| SSPNLPDFREEL | 1509422 | Ectodomain | YES |
| SSPQIIKQLKKA | 1509423 | Cell/Virus Enclosed | NO |
| SSPTTLQAVGSC | 1509424 | Cell/Virus Enclosed | NO |
| SSPYRFVTFEPF | 1509425 | Ectodomain | YES |
| SSQGSRSGNSTRGTSP | 1695683 | Cell/Virus Enclosed | NO |
| SSQGSRSRNSTR | 1509436 | Cell/Virus Enclosed | NO |
| SSQLPLDIYDLT | 1509439 | Cell/Virus Enclosed | NO |
| SSQLPLDVYDLTMPSQ | 1695685 | Cell/Virus Enclosed | NO |
| SSQLPLDVYNLT | 1509440 | Cell/Virus Enclosed | NO |
| SSQPYYCFINSTINTT | 1695686 | Ectodomain | YES |
| SSQSSAKILARS | 1509450 | Cell/Virus Enclosed | NO |

| | | | |
|---|---|---|---|
| SSQSSPIIPGFGGDFN | 1695687 | Ectodomain | YES |
| SSQTSAKSLARSQSSE | 1695688 | Capsid | UN |
| SSRASSAGSRSRANSG | 1695691 | Cell/Virus Enclosed | NO |
| SSRASSGNRPGN | 1509461 | Capsid | UN |
| SSRASSGNRSGNGILK | 1695692 | Cell/Virus Enclosed | NO |
| SSRASSGNRSSN | 1509462 | Cell/Virus Enclosed | NO |
| SSRASSGNRSVN | 1509463 | Cell/Virus Enclosed | NO |
| SSRASSLSRNSSRSSS | 1695693 | Cell/Virus Enclosed | NO |
| SSRASSRSSTRNNSRD | 1695694 | Capsid | UN |
| SSRGTSPARIAGNGGD | 1695697 | Cell/Virus Enclosed | NO |
| SSRGTSPARMAGNGGDAALA | 1392436 | Capsid | UN |
| SSRNESWHFDKSEPLC | 1695706 | Ectodomain | YES |
| SSRNSTPGSSRGTSPA | 1695708 | Cell/Virus Enclosed | NO |
| SSRRKRRGISSP | 1509490 | Ectodomain | YES |
| SSRRKRRSISASYRFV | 1695709 | Ectodomain | YES |
| SSRRLAQQKVNE | 1509493 | Ectodomain | YES |
| SSRSFFEDLLFDKVKL | 1695711 | Ectodomain | YES |
| SSRSRNSSRNSTPGSS | 1695714 | Cell/Virus Enclosed | NO |
| SSRSSSQGSRSGNSTR | 1695715 | Cell/Virus Enclosed | NO |
| SSRSSTRNNSRDSSRS | 1695719 | Capsid | UN |
| SSRVERSNAVLT | 1509505 | Cell/Virus Enclosed | NO |
| SSSAAIPFAQSI | 1509508 | Ectodomain | YES |
| SSSANGRLVGTT | 1509514 | Cell/Virus Enclosed | NO |
| SSSANGSLDGTT | 1509515 | Cell/Virus Enclosed | NO |
| SSSANGSLVCTT | 1509516 | Cell/Virus Enclosed | NO |
| SSSANGSLVGTI | 1509517 | Cell/Virus Enclosed | NO |
| SSSAVNIQQIYLISKF | 1695726 | Cell/Virus Enclosed | NO |
| SSSAVNMQQIYLINKF | 1695727 | Cell/Virus Enclosed | NO |
| SSSCVGSGPGKN | 1509526 | Ectodomain | YES |
| SSSEAFLIGVNY | 1509534 | Cell/Virus Enclosed | NO |
| SSSEVDMICSTTAQKL | 1695737 | Cell/Virus Enclosed | NO |
| SSSFSVVAMSGH | 1509543 | Cell/Virus Enclosed | NO |
| SSSGPISQFNYK | 1509550 | Ectodomain | YES |
| SSSGTSTPKKPNKPLS | 1695743 | Capsid | UN |
| SSSGVLDTTIPFGPSS | 1695744 | Ectodomain | YES |
| SSSLQEILSRLDALEA | 1695751 | Ectodomain | YES |
| SSSLRTFVVDKS | 1509566 | Cell/Virus Enclosed | NO |
| SSSNPTCLILAT | 1509571 | Ectodomain | YES |
| SSSNVTTMDFCE | 1509572 | Cell/Virus Enclosed | NO |
| SSSPALVDQRTI | 1509573 | Cell/Virus Enclosed | NO |
| SSSQGSRSGNSTRGTS | 1695755 | Cell/Virus Enclosed | NO |
| SSSRASFGNRSG | 1509579 | Cell/Virus Enclosed | NO |
| SSSRASSGNRPG | 1509583 | Capsid | UN |
| SSSRASSGNRSGNGIL | 1695757 | Cell/Virus Enclosed | NO |
| SSSRASSLSRNSSRSS | 1695758 | Cell/Virus Enclosed | NO |
| SSSRRKRRSISASYRF | 1695759 | Ectodomain | YES |
| SSSRSFFEDLLFDKVK | 1695760 | Ectodomain | YES |
| SSSRSRNSSRNSTPGS | 1695762 | Cell/Virus Enclosed | NO |
| SSSSRRKRRSISASYR | 1695763 | Ectodomain | YES |
| SSSTYRNMALKG | 1509600 | Ectodomain | YES |
| SSSYDALLHRNSLDPF | 1695765 | Cell/Virus Enclosed | NO |
| SSSYGDAHEGIF | 1509605 | Cell/Virus Enclosed | NO |
| SSTFVEEEHVNRMDTF | 1695770 | Cell/Virus Enclosed | NO |
| SSTGFVYFNATGSDVN | 1695771 | Ectodomain | YES |
| SSTIHNYFITETNTAA | 1695772 | Cell/Virus Enclosed | NO |
| SSTIRLQAGVATEYAA | 1695773 | Cell/Virus Enclosed | NO |
| SSTMIFSLHPVE | 1509625 | Cell/Virus Enclosed | NO |
| SSTRGGSRVEGIFAGLFMNA | 61410 | Capsid | UN |
| SSTRNNSRDSSRSTSR | 1695779 | Capsid | UN |
| SSTVRLQAGTATEYAS | 1695781 | Cell/Virus Enclosed | NO |
| SSVAVQSKDLNFLNRV | 1695784 | Cell/Virus Enclosed | NO |
| SSVDLKHFFFTQDGNA | 1695785 | Cell/Virus Enclosed | NO |
| SSVDNILNMARN | 1509649 | Cell/Virus Enclosed | NO |
| SSVDTILNMARNGVVP | 1695786 | Cell/Virus Enclosed | NO |
| SSVETETYFKVNGLWS | 1695789 | Cell/Virus Enclosed | NO |
| SSVETVLNLARDGVVP | 1695790 | Cell/Virus Enclosed | NO |
| SSVKSYDDLVYSGVVG | 1695794 | Cell/Virus Enclosed | NO |
| SSVLPQRNIRSSRIAG | 1695800 | Ectodomain | YES |
| SSVNDAITQTAEAIHT | 1695802 | Ectodomain | YES |
| SSVNDAITRTAE | 1509674 | Ectodomain | YES |
| SSVPLLEVYVHLRNWN | 1695804 | Ectodomain | YES |
| SSVPLSEVYVHL | 1509677 | Ectodomain | YES |
| SSVRSDGTFEEM | 1509680 | Cell/Virus Enclosed | NO |
| SSVSNFSYGKMG | 1509682 | Ectodomain | YES |
| SSVVPSLPTSGS | 1509687 | Ectodomain | YES |
| SSVWPLYPYGRCPTAA | 1695809 | Ectodomain | YES |

| | | | |
|---|---|---|---|
| SSVYSVPLLFQT | 1509698 | Cell/Virus Enclosed | NO |
| SSWVLNNDYYRS | 1509705 | Cell/Virus Enclosed | NO |
| SSWVYQSAFNCT | 1509706 | Ectodomain | YES |
| SSYAKPEEKLSAYDLA | 1695817 | Cell/Virus Enclosed | NO |
| SSYDALLHRNSLDPFC | 1695819 | Cell/Virus Enclosed | NO |
| SSYECDIPIGAG | 1509712 | Ectodomain | YES |
| SSYKLDLTSLGL | 1509725 | Cell/Virus Enclosed | NO |
| SSYYYPEPISDKNVVF | 1695828 | Ectodomain | YES |
| STAEQTKLYGSGNKLVTVGSSNYQQ | 1171846 | Ectodomain | YES |
| STAHATATIPLQ | 1509751 | Secreted | YES |
| STALCKSGSLVL | 1509758 | Ectodomain | YES |
| STALGKLQDVVNQNAQ | 1695836 | Ectodomain | YES |
| STALQELQIQQW | 1509763 | Cell/Virus Enclosed | NO |
| STANALPECNAD | 1509764 | Cell/Virus Enclosed | NO |
| STARDGIRRGDN | 1509768 | Cell/Virus Enclosed | NO |
| STARDGVRRGDNFHSV | 1695839 | Cell/Virus Enclosed | NO |
| STASAVQTCDLT | 1509774 | Ectodomain | YES |
| STAVLTIANNYSKLFD | 1695842 | Cell/Virus Enclosed | NO |
| STCAFVDTVKGL | 1509789 | Cell/Virus Enclosed | NO |
| STCDGLKFTEESFYKN | 1695847 | Cell/Virus Enclosed | NO |
| STCNSNANISML | 1509800 | Ectodomain | YES |
| STDCSDGNYNRN | 1509810 | Ectodomain | YES |
| STDEEGDNVQIT | 1509813 | Cell/Virus Enclosed | NO |
| STDFHQLECIVD | 1509816 | Cell/Virus Enclosed | NO |
| STDFHQLEYDSV | 1509817 | Cell/Virus Enclosed | NO |
| STDGLYFHFYQE | 1509823 | Ectodomain | YES |
| STDLQKANTDII | 1509827 | Ectodomain | YES |
| STDLQKANTDIK | 1509828 | Ectodomain | YES |
| STDLQKSNTDII | 1509830 | Ectodomain | YES |
| STDNNQDGGRSG | 1509831 | Cell/Virus Enclosed | NO |
| STDNNQNGDRSGARPK | 1695862 | Cell/Virus Enclosed | NO |
| STDNNQSGGRNG | 1509833 | Capsid | UN |
| STDQMSELPKEQ | 1509838 | Cell/Virus Enclosed | NO |
| STDQMSEPPKEH | 1509839 | Cell/Virus Enclosed | NO |
| STDQMSEPPMEQ | 1509841 | Cell/Virus Enclosed | NO |
| STDSTVFTYIPA | 1509848 | Cell/Virus Enclosed | NO |
| STDTVNHDSFNLYGYS | 1695870 | Cell/Virus Enclosed | NO |
| STEDFNVWAMANGFSQ | 1695881 | Cell/Virus Enclosed | NO |
| STEEVGDNVQIT | 1509867 | Cell/Virus Enclosed | NO |
| STEKLSRLSMIKGPQR | 1695886 | Cell/Virus Enclosed | NO |
| STEQVDVFHETS | 1509885 | Cell/Virus Enclosed | NO |
| STFDDILGKVCS | 1509898 | Cell/Virus Enclosed | NO |
| STFGTRNPNNDL | 1509909 | Cell/Virus Enclosed | NO |
| STFITPEVSRELGNIT | 1695904 | Cell/Virus Enclosed | NO |
| STFKCYGVSATKLNDL | 1695905 | Ectodomain | YES |
| STFNKDIYCGNSMVCK | 1695910 | Cell/Virus Enclosed | NO |
| STFTPRSVRSVP | 1509933 | Ectodomain | YES |
| STFVGVLPPTVR | 1509936 | Ectodomain | YES |
| STFVTEYYEFLNKHFS | 1695920 | Cell/Virus Enclosed | NO |
| STFYPQLQSAEWKCGY | 1695921 | Cell/Virus Enclosed | NO |
| STGFDFNCHYLV | 1509953 | Ectodomain | YES |
| STGFVYFNATGSDVNC | 1695926 | Ectodomain | YES |
| STGKTVLGEYVFDKSE | 1695931 | Cell/Virus Enclosed | NO |
| STGLTSQTVKPGHFNK | 1695934 | Cell/Virus Enclosed | NO |
| STGLVKNQCVNF | 1509962 | Ectodomain | YES |
| STGNHNYKYRYL | 1509963 | Ectodomain | YES |
| STGNIFIVTPCN | 1509964 | Ectodomain | YES |
| STGNNVFQTLAG | 1509966 | Ectodomain | YES |
| STGNYNYKHRYL | 1509970 | Ectodomain | YES |
| STGQKSIVAYTM | 1509978 | Ectodomain | YES |
| STGRCTVNKHVA | 1509979 | Cell/Virus Enclosed | NO |
| STGSNQNGERSGARPK | 1695938 | Cell/Virus Enclosed | NO |
| STGSRSARSAIEDLLF | 1389156 | Ectodomain | YES |
| STGSRSRANSGN | 1509987 | Cell/Virus Enclosed | NO |
| STGTQSVSVDLESTST | 1695940 | Cell/Virus Enclosed | NO |
| STGTQSVSVDRE | 1509997 | Cell/Virus Enclosed | NO |
| STGTSQNTRISLRPGPVSQP | 61638 | Capsid | UN |
| STGTVIIRPSTS | 1509999 | Ectodomain | YES |
| STHDDFNEHVTS | 1510014 | Cell/Virus Enclosed | NO |
| STHDDFTEHVTS | 1510015 | Cell/Virus Enclosed | NO |
| STHDGFNEHVTS | 1510017 | Cell/Virus Enclosed | NO |
| STHDGLIEHVTS | 1510019 | Cell/Virus Enclosed | NO |
| STHDGPTEYVTI | 1510027 | Cell/Virus Enclosed | NO |
| STHDTPTEHVTS | 1510029 | Cell/Virus Enclosed | NO |
| STHNYEEIPMDTFIVST | 111808 | Capsid | UN |
| STIDSNTTQDTY | 1510041 | Cell/Virus Enclosed | NO |

| | | | |
|---|---|---|---|
| STIGDSIFTGCT | 1510045 | Cell/Virus Enclosed | NO |
| STINALPEISAD | 1510062 | Cell/Virus Enclosed | NO |
| STINTTHVSTFVGILP | 1695965 | Ectodomain | YES |
| STIRLQAGVATEYAAN | 1695967 | Cell/Virus Enclosed | NO |
| STITVEELKQLL | 1510072 | Undetermined | UN |
| STIVTGLLPTHWFCAN | 1695973 | Ectodomain | YES |
| STIYKLHPSFNVSDAY | 1695974 | Cell/Virus Enclosed | NO |
| STKALAKQDAAQRIAW | 1695975 | Cell/Virus Enclosed | NO |
| STKLKDGVNFNVDDIN | 1695980 | Ectodomain | YES |
| STKLPYYDVKKI | 1510084 | Underneath Membrane | NO |
| STKLRLATGLRNIPSIQSRGLFGAI | 1171943 | Ectodomain | YES |
| STLENKSAELNYTVQK | 1695988 | Ectodomain | YES |
| STLLGSIAGVGW | 1510121 | Ectodomain | YES |
| STLPGFETIMKV | 1510127 | Cell/Virus Enclosed | NO |
| STLQPNAYYKYDAKNY | 1696003 | Cell/Virus Enclosed | NO |
| STLQVSDVDKLVCRD | 61752 | Secreted | YES |
| STLSGFETIMKV | 1510148 | Cell/Virus Enclosed | NO |
| STLVDLNWLNRV | 1510157 | Ectodomain | YES |
| STMDNKSQSVII | 1510169 | Ectodomain | YES |
| STMDNTTQSAII | 1510170 | Ectodomain | YES |
| STMEEDFLNMDI | 1510171 | Cell/Virus Enclosed | NO |
| STMNNKSQTVII | 1510179 | Ectodomain | YES |
| STMTGRVFHQKC | 1510184 | Cell/Virus Enclosed | NO |
| STMTNRQFHQKL | 1510185 | Cell/Virus Enclosed | NO |
| STMTNRQYHQKM | 1510186 | Cell/Virus Enclosed | NO |
| STNLFEGDKFVGTFES | 1696035 | Cell/Virus Enclosed | NO |
| STNLFKDCSNSY | 1510200 | Cell/Virus Enclosed | NO |
| STNLPPPLLGNSTGID | 1696037 | Ectodomain | YES |
| STNLPPPLLSNS | 1510202 | Ectodomain | YES |
| STNPDPKFVDKY | 1510206 | Cell/Virus Enclosed | NO |
| STNPDPKFVDRY | 1510207 | Cell/Virus Enclosed | NO |
| STPAQNAGYWRRQDRK | 1696045 | Cell/Virus Enclosed | NO |
| STPGSSRGTSPARIAG | 1696050 | Cell/Virus Enclosed | NO |
| STPGSSRGTSPARMAG | 1696051 | Cell/Virus Enclosed | NO |
| STPKKPNKPLSQPRAD | 1696055 | Capsid | UN |
| STPKLKEDYQIGGYSE | 1696056 | Secreted | YES |
| STPNAIFISDRK | 1510235 | Cell/Virus Enclosed | NO |
| STPNLPDFKEEL | 1510239 | Ectodomain | YES |
| STPNLPDFREEL | 1510240 | Ectodomain | YES |
| STPSDVSSRDPTTQEA | 1696059 | Capsid | UN |
| STPSQNAGYWRR | 1510242 | Cell/Virus Enclosed | NO |
| STPVELPTVDVV | 1510245 | Cell/Virus Enclosed | NO |
| STQDGDNKLQGLLEVS | 1696061 | Ectodomain | YES |
| STQDGVNKLQGL | 1510254 | Ectodomain | YES |
| STQDGYNKLQGL | 1510256 | Ectodomain | YES |
| STQKGSGMDTALLRNN | 1696067 | Ectodomain | YES |
| STQPQGNIIPHY | 1510278 | Cell/Virus Enclosed | NO |
| STQWSLFFFVYE | 1510288 | Cell/Virus Enclosed | NO |
| STRATIRKIYPA | 1510291 | Ectodomain | YES |
| STRCTSPGPSGI | 1510292 | Cell/Virus Enclosed | NO |
| STRGSSPGPSGI | 1510294 | Cell/Virus Enclosed | NO |
| STRGTFPGPSGI | 1510295 | Cell/Virus Enclosed | NO |
| STRGTSPCPSGI | 1510296 | Cell/Virus Enclosed | NO |
| STRGTSPGLSGI | 1510297 | Cell/Virus Enclosed | NO |
| STRGTSPGPSGIGAVG | 1696077 | Cell/Virus Enclosed | NO |
| STRGTSPVPSGI | 1510299 | Capsid | UN |
| STRNNSRDSFRS | 1510304 | Cell/Virus Enclosed | NO |
| STRSMLKRLDST | 1510312 | Ectodomain | YES |
| STRSMLKRRDSTYGPL | 1696082 | Ectodomain | YES |
| STRSTSPGPSGI | 1510315 | Capsid | UN |
| STRVNVDGFNEWAMAN | 1696083 | Cell/Virus Enclosed | NO |
| STSAVKLQNNEIKPSG | 1696090 | Cell/Virus Enclosed | NO |
| STSDLSVAFDKLAQLL | 1696092 | Cell/Virus Enclosed | NO |
| STSDWSDIRDYKLAND | 1696093 | Cell/Virus Enclosed | NO |
| STSFLQSGIVKMVNPT | 1696094 | Cell/Virus Enclosed | NO |
| STSGNLLGFKDVTKGT | 1696097 | Ectodomain | YES |
| STSGTQVYPRKA | 1510339 | Cell/Virus Enclosed | NO |
| STSGVTPDMADQ | 1510343 | Cell/Virus Enclosed | NO |
| STSIPNFGSLTQINTT | 1696102 | Ectodomain | YES |
| STSISNFGSLTQ | 1510351 | Ectodomain | YES |
| STSISVQTCDLT | 1510352 | Ectodomain | YES |
| STSKCWVEEDLTKGPH | 1696103 | Cell/Virus Enclosed | NO |
| STSNLPDFREEL | 1510360 | Ectodomain | YES |
| STSPDPKFVDKYYAFL | 1696105 | Cell/Virus Enclosed | NO |
| STSPGPSGIGAV | 1510362 | Capsid | UN |
| STSPGPSGIGGG | 1510363 | Cell/Virus Enclosed | NO |

| | | | |
|---|---|---|---|
| STSQKSIVAYTMSLGADSSIAY | 1389187 | Ectodomain | YES |
| STSQQPLGGNVV | 1510371 | Cell/Virus Enclosed | NO |
| STSRQQSRTRSDSNQS | 1696108 | Capsid | UN |
| STSRTSSRASSAGSRS | 1696109 | Cell/Virus Enclosed | NO |
| STSTHDDFNEHV | 1510386 | Cell/Virus Enclosed | NO |
| STSTHDGLIEHV | 1510390 | Cell/Virus Enclosed | NO |
| STSTHDGLNEHV | 1510391 | Cell/Virus Enclosed | NO |
| STSTHDGPIEHV | 1510393 | Cell/Virus Enclosed | NO |
| STSTHDGPNEHV | 1510394 | Cell/Virus Enclosed | NO |
| STSTHDGPTEHVTSVN | 1696112 | Cell/Virus Enclosed | NO |
| STSVEESFIDDY | 1510403 | Cell/Virus Enclosed | NO |
| STTAQKLETDRI | 1510411 | Cell/Virus Enclosed | NO |
| STTDNETLEYWV | 1510413 | Ectodomain | YES |
| STTNYVGVYDINHGDA | 1696126 | Ectodomain | YES |
| STTNYVGVYDTN | 1510440 | Ectodomain | YES |
| STTPCELHGNGS | 1510441 | Cell/Virus Enclosed | NO |
| STTPCEVHGNGSMIED | 1696127 | Cell/Virus Enclosed | NO |
| STTPCQVHGNGS | 1510444 | Cell/Virus Enclosed | NO |
| STTSLTEENFGKPDI | 227589 | Cell/Virus Enclosed | NO |
| STTVSVTVSSNMVETV | 1696131 | Cell/Virus Enclosed | NO |
| STTWNGGAYSLFDMTK | 1696132 | Cell/Virus Enclosed | NO |
| STVAMTEQLQMG | 1510457 | Ectodomain | YES |
| STVDVVALQQEM | 1510463 | Cell/Virus Enclosed | NO |
| STVKNYFITDAK | 1510480 | Cell/Virus Enclosed | NO |
| STVLNGVWLGDTVTCP | 1696144 | Cell/Virus Enclosed | NO |
| STVPSTVWEDGD | 1510487 | Ectodomain | YES |
| STVSALVYNNKM | 1510495 | Cell/Virus Enclosed | NO |
| STVTGRMFHQKC | 1510501 | Cell/Virus Enclosed | NO |
| STVVIKNLTPIKLNFG | 1696162 | Cell/Virus Enclosed | NO |
| STVWEDGDYYRKQLSP | 1696163 | Ectodomain | YES |
| STVWEDGNYYRK | 1510509 | Ectodomain | YES |
| STVWNGGAYSLFDMAK | 1696164 | Cell/Virus Enclosed | NO |
| STWNKKFGFIED | 1510519 | Ectodomain | YES |
| STWNKRFGFIEDSVFK | 1696165 | Ectodomain | YES |
| STWPTTPPRPIPKPSPWAPK | 110427 | Cell/Virus Enclosed | NO |
| STYDFYPNVPIE | 1510528 | Ectodomain | YES |
| STYDFYPSIPVE | 1510529 | Ectodomain | YES |
| STYFLVRILQDK | 1510534 | Cell/Virus Enclosed | NO |
| STYGVCKYTDLKFIEN | 1696170 | Cell/Virus Enclosed | NO |
| STYRNMALKGTD | 1510553 | Ectodomain | YES |
| STYVDLKLLNRFENYI | 1696176 | Ectodomain | YES |
| SVAAITSGVKFQTVKP | 1696179 | Cell/Virus Enclosed | NO |
| SVAALSTGLTSQTVKP | 1696181 | Cell/Virus Enclosed | NO |
| SVAAPKSRVQQN | 1510567 | Cell/Virus Enclosed | NO |
| SVACEHIFSTMS | 1510569 | Ectodomain | YES |
| SVACEHISSTIS | 1510570 | Ectodomain | YES |
| SVACEHISSTMSQYSR | 1696187 | Ectodomain | YES |
| SVADVMRYNLNF | 1510580 | Ectodomain | YES |
| SVAFQTVKPGNF | 1510584 | Cell/Virus Enclosed | NO |
| SVANLSAPTLVPQENY | 1696201 | Cell/Virus Enclosed | NO |
| SVASLSAPTLVPQENY | 1389203 | Cell/Virus Enclosed | NO |
| SVASPKSRVQQN | 1510622 | Cell/Virus Enclosed | NO |
| SVASSFVGMPSFVAYE | 1389205 | Cell/Virus Enclosed | NO |
| SVASSFVSMPSYIAYE | 1389206 | Cell/Virus Enclosed | NO |
| SVASTPVEPSTV | 1510624 | Cell/Virus Enclosed | NO |
| SVASTYFIIRLL | 1510626 | Cell/Virus Enclosed | NO |
| SVAVIGDEQIDGKFVT | 1696210 | Cell/Virus Enclosed | NO |
| SVAYTAGVCIKY | 1510647 | Cell/Virus Enclosed | NO |
| SVAYVANPTEDL | 1510648 | Cell/Virus Enclosed | NO |
| SVCLSGLDSLDS | 1510659 | Cell/Virus Enclosed | NO |
| SVCNGCVGHSEN | 1510662 | Ectodomain | YES |
| SVCNGCVGHSKN | 1510663 | Ectodomain | YES |
| SVCNTGSCTVNK | 1510668 | Cell/Virus Enclosed | NO |
| SVCPKFEFANDT | 1510669 | Ectodomain | YES |
| SVCPKLEFANDTKIAS | 1696233 | Ectodomain | YES |
| SVCQYNMCDYPH | 1510679 | Ectodomain | YES |
| SVCQYNMCEHPH | 1510680 | Ectodomain | YES |
| SVCQYNMCEYPH | 1510681 | Ectodomain | YES |
| SVCQYNMCEYPQ | 1510682 | Ectodomain | YES |
| SVCTMTKWQEVDEMLRSEYG | 742012 | Ectodomain | YES |
| SVCVRTSHFSIR | 1510689 | Ectodomain | YES |
| SVDAFNQFSSDFQHKL | 1696240 | Cell/Virus Enclosed | NO |
| SVDARLVPCASG | 1510693 | Cell/Virus Enclosed | NO |
| SVDDKYKTIDEL | 1510696 | Cell/Virus Enclosed | NO |
| SVDGYIRRAIDC | 1510702 | Ectodomain | YES |
| SVDGYIRRSIDC | 1510704 | Ectodomain | YES |

| | | | |
|---|---|---|---|
| SVDISYLKRARG | 1510708 | Cell/Virus Enclosed | NO |
| SVDISYLNRARGSSAA | 1696252 | Cell/Virus Enclosed | NO |
| SVDLESTSTHDG | 1510711 | Cell/Virus Enclosed | NO |
| SVDLKHFFFTQDGNAA | 1696255 | Cell/Virus Enclosed | NO |
| SVDPATTYLEAVKHGA | 1696257 | Cell/Virus Enclosed | NO |
| SVDPKKTYLDFIQQGG | 1696258 | Cell/Virus Enclosed | NO |
| SVDPKKTYLDYIQQGG | 1696259 | Cell/Virus Enclosed | NO |
| SVDRESTSTHDD | 1510723 | Cell/Virus Enclosed | NO |
| SVDRESTSTHDG | 1510724 | Cell/Virus Enclosed | NO |
| SVDSVSEESQGNVVTS | 1696261 | Cell/Virus Enclosed | NO |
| SVDVIDNEIIVK | 1510733 | Cell/Virus Enclosed | NO |
| SVEAAYAEVHAE | 1510738 | Cell/Virus Enclosed | NO |
| SVECYSILAAKTGVSV | 1696265 | Cell/Virus Enclosed | NO |
| SVEDFNVWALSNGFSQ | 1696267 | Cell/Virus Enclosed | NO |
| SVEDVCFANKFEQFDQ | 1696268 | Cell/Virus Enclosed | NO |
| SVEIEAGEVKPFAVYK | 1696274 | Cell/Virus Enclosed | NO |
| SVELLSFLPSDFFPS | 1696277 | Capsid | UN |
| SVEQLEPCVQSR | 1510767 | Cell/Virus Enclosed | NO |
| SVETETYFKVNGLWSP | 1696286 | Cell/Virus Enclosed | NO |
| SVETVLNLARDGVVPL | 1696288 | Cell/Virus Enclosed | NO |
| SVFDITDAAVKA | 1510782 | Cell/Virus Enclosed | NO |
| SVFKPQPAGVFT | 1510788 | Ectodomain | YES |
| SVFKPQPAGVLT | 1510789 | Ectodomain | YES |
| SVFNICQAVSANVCAL | 1696296 | Cell/Virus Enclosed | NO |
| SVFNLEGVNGGSLYVN | 1696301 | Cell/Virus Enclosed | NO |
| SVFPIFVGRIVM | 1510802 | Cell/Virus Enclosed | NO |
| SVGESFGKSLGN | 1510828 | Cell/Virus Enclosed | NO |
| SVGQKSIVAYTM | 1510851 | Ectodomain | YES |
| SVGYTKGGSVIN | 1510858 | Cell/Virus Enclosed | NO |
| SVGYVLTGDSVKFVYM | 1696328 | Cell/Virus Enclosed | NO |
| SVHQGAHVASNDAIMT | 1696331 | Cell/Virus Enclosed | NO |
| SVIDLKLDDFVMILKS | 1696333 | Cell/Virus Enclosed | NO |
| SVIDLLLNDFVE | 1510877 | Cell/Virus Enclosed | NO |
| SVIELATVNLRLVDCA | 1696335 | Cell/Virus Enclosed | NO |
| SVIKVAPKPYRVDGKF | 1696340 | Cell/Virus Enclosed | NO |
| SVILSTCDGLKFTEES | 1696341 | Cell/Virus Enclosed | NO |
| SVINRLRCDQLSFDVP | 1696350 | Ectodomain | YES |
| SVIPATSASKLTIVSP | 1696352 | Cell/Virus Enclosed | NO |
| SVIQDYENVCFD | 1510904 | Cell/Virus Enclosed | NO |
| SVIQNYENVCFE | 1510907 | Cell/Virus Enclosed | NO |
| SVIVEDDVIENVKSSL | 1696361 | Cell/Virus Enclosed | NO |
| SVIVNDATSDTK | 1510924 | Cell/Virus Enclosed | NO |
| SVIVNDDTSDTK | 1510926 | Cell/Virus Enclosed | NO |
| SVIVNDDTSETKYVKS | 1696362 | Cell/Virus Enclosed | NO |
| SVIYDKETKTHATLFG | 1696365 | Ectodomain | YES |
| SVKLPKGVGSAN | 1510952 | Cell/Virus Enclosed | NO |
| SVKMVDAYVDTF | 1510955 | Cell/Virus Enclosed | NO |
| SVKNGTYDYPKYSEEAKLNREEIDG | 1172218 | Ectodomain | YES |
| SVKRDGVQVGYCVHGI | 1696379 | Cell/Virus Enclosed | NO |
| SVKSFEIDKGIYQTSN | 1696381 | Ectodomain | YES |
| SVKSFKIDKGIY | 1510969 | Ectodomain | YES |
| SVKSSQSSPIIPGFGG | 1696382 | Ectodomain | YES |
| SVKTCDLTVGSG | 1510977 | Ectodomain | YES |
| SVLAVMQKLPCQ | 1510985 | Cell/Virus Enclosed | NO |
| SVLDITDAAVKAAESK | 1696393 | Cell/Virus Enclosed | NO |
| SVLDLSKFECKH | 1510990 | Cell/Virus Enclosed | NO |
| SVLEMAKEVKTSSKVV | 1696396 | Cell/Virus Enclosed | NO |
| SVLETFQNNVVNYQHI | 1696397 | Cell/Virus Enclosed | NO |
| SVLGCLGSECSK | 1510996 | Ectodomain | YES |
| SVLKPILKLPIS | 1511005 | Ectodomain | YES |
| SVLLALQQIEVK | 1511008 | Cell/Virus Enclosed | NO |
| SVLLALQQLEVK | 1511009 | Cell/Virus Enclosed | NO |
| SVLLFLAFVVFLLVT | 62216 | Transmembrane | NO |
| SVLLIERFVSLAIDAY | 1696405 | Cell/Virus Enclosed | NO |
| SVLNDVLSRLDK | 1511024 | Ectodomain | YES |
| SVLPQRNIRSSRIAGR | 1696412 | Ectodomain | YES |
| SVLQMDVRGLVN | 1511031 | Cell/Virus Enclosed | NO |
| SVLQQLHLEANG | 1511033 | Cell/Virus Enclosed | NO |
| SVLRQWLPTDAI | 1511039 | Cell/Virus Enclosed | NO |
| SVLSVDSVSEESQGNV | 1696421 | Cell/Virus Enclosed | NO |
| SVLTTAVYKPLV | 1511046 | Cell/Virus Enclosed | NO |
| SVLTTFIDAARGPAGV | 1696424 | Cell/Virus Enclosed | NO |
| SVLTTYYLDDVKKIEY | 1696426 | Cell/Virus Enclosed | NO |
| SVMESQISTKEVKLKG | 1696436 | Cell/Virus Enclosed | NO |
| SVMHFGTLSRED | 1511062 | Cell/Virus Enclosed | NO |
| SVMLCGDNSAKF | 1511064 | Cell/Virus Enclosed | NO |

| | | | |
|---|---|---|---|
| SVMQMDIQGFVN | 1511067 | Cell/Virus Enclosed | NO |
| SVNDAITQTAEAIHTV | 1696446 | Ectodomain | YES |
| SVNDFTCSQISPAAIA | 1696447 | Ectodomain | YES |
| SVNDLCFANKFFQFDQ | 1696448 | Cell/Virus Enclosed | NO |
| SVNDSLEHVGGL | 1511079 | Ectodomain | YES |
| SVNDSLEPVGGLYEIQ | 1696449 | Ectodomain | YES |
| SVNDSLKPVGGL | 1511080 | Ectodomain | YES |
| SVNETSQTGWAL | 1511081 | Ectodomain | YES |
| SVNLNICEPVLT | 1511094 | Cell/Virus Enclosed | NO |
| SVNQISGYTLAA | 1511100 | Ectodomain | YES |
| SVPEKFLNAFDVFVTA | 1696475 | Cell/Virus Enclosed | NO |
| SVPLLEVYVHLRNWNF | 1696479 | Undetermined | UN |
| SVPLVFQNNVANYQHI | 1696480 | Cell/Virus Enclosed | NO |
| SVPMEELKALVA | 1511133 | Cell/Virus Enclosed | NO |
| SVPNMHVVPLEN | 1511135 | Cell/Virus Enclosed | NO |
| SVPNSRSTSRTS | 1511137 | Cell/Virus Enclosed | NO |
| SVPSPFNWERRI | 1511139 | Ectodomain | YES |
| SVPVSVIYDKETKTHA | 1696487 | Ectodomain | YES |
| SVQKHIGAPANLRVVG | 1696495 | Cell/Virus Enclosed | NO |
| SVQQAQQEATHI | 1511174 | Cell/Virus Enclosed | NO |
| SVQQAQQEATLPDIAE | 1696498 | Cell/Virus Enclosed | NO |
| SVQQDLDVICPG | 1511181 | Cell/Virus Enclosed | NO |
| SVQVDNGSSINRRQLD | 1696504 | Cell/Virus Enclosed | NO |
| SVQYAHKHDIQITNES | 1696506 | Cell/Virus Enclosed | NO |
| SVRDPKTSEILDISPC | 1696509 | Ectodomain | YES |
| SVRDPQTLQILD | 1511201 | Ectodomain | YES |
| SVRGRAIIVSVEQLEP | 1696513 | Cell/Virus Enclosed | NO |
| SVRHHPELKLFRNLNI | 1696514 | Cell/Virus Enclosed | NO |
| SVRNLFESVKSS | 1511214 | Ectodomain | YES |
| SVRSDGTFEEMA | 1511216 | Cell/Virus Enclosed | NO |
| SVRSHPDFKLLH | 1511217 | Cell/Virus Enclosed | NO |
| SVSAIVTANLSIPSNW | 1696523 | Ectodomain | YES |
| SVSFDYLIREIE | 1511238 | Cell/Virus Enclosed | NO |
| SVSKPNAIDVEH | 1511249 | Cell/Virus Enclosed | NO |
| SVSLAIDAYPLT | 1511250 | Cell/Virus Enclosed | NO |
| SVSNDYCCFKHALGCD | 1696528 | Cell/Virus Enclosed | NO |
| SVSNFSYGKMGR | 1511256 | Ectodomain | YES |
| SVSPKLEFANDT | 1511260 | Ectodomain | YES |
| SVSRFNPSTWNKRFGF | 1696535 | Ectodomain | YES |
| SVSSAGPISQFNYKQS | 1696536 | Ectodomain | YES |
| SVSSCALRIGSG | 1511271 | Ectodomain | YES |
| SVSSCALRMGSGFCVD | 1696537 | Ectodomain | YES |
| SVSSCDLRMGSG | 1511273 | Ectodomain | YES |
| SVSSFEAKPSGSVVEQ | 1696538 | Ectodomain | YES |
| SVSSVECYSILAAKTG | 1696541 | Cell/Virus Enclosed | NO |
| SVSSVWPLYPYG | 1511285 | Ectodomain | YES |
| SVSTKEGDSSNTGAGKALTG | 62336 | Capsid | UN |
| SVSTSDLGSLIDCNAA | 1696543 | Cell/Virus Enclosed | NO |
| SVTAVVTNGHLKMAGM | 1696547 | Ectodomain | YES |
| SVTDDFTLTSPYLGTCSY | 558330 | Undetermined | UN |
| SVTDECRLAMLG | 1511306 | Cell/Virus Enclosed | NO |
| SVTDFSLNGTVS | 1511309 | Cell/Virus Enclosed | NO |
| SVTDTIVVTEGDGIST | 1696551 | Secreted | YES |
| SVTEDVKFAASTGVID | 1696552 | Cell/Virus Enclosed | NO |
| SVTLKICKFGIN | 1511326 | Ectodomain | YES |
| SVTPIKNTVDTI | 1511339 | Cell/Virus Enclosed | NO |
| SVTPIKNTVDTTPKEE | 1696562 | Cell/Virus Enclosed | NO |
| SVTRLMCNLGPDIFLS | 1696565 | Cell/Virus Enclosed | NO |
| SVTSAGPISQFN | 1511344 | Ectodomain | YES |
| SVTVKRKRLRKK | 1511364 | Cell/Virus Enclosed | NO |
| SVTVSSNMVETVVGQL | 1696574 | Cell/Virus Enclosed | NO |
| SVTYTDQYLCGA | 1511366 | Cell/Virus Enclosed | NO |
| SVVDVMRYNLNFSANS | 1696581 | Ectodomain | YES |
| SVVEFEDRSNNSSRAS | 1696582 | Capsid | UN |
| SVVEVVTTSLTPCGYS | 1696584 | Cell/Virus Enclosed | NO |
| SVVHTAGVCIKY | 1511391 | Cell/Virus Enclosed | NO |
| SVVIQPRTINST | 1511395 | Ectodomain | YES |
| SVVKPVRARAPPGEQF | 1696591 | Cell/Virus Enclosed | NO |
| SVVMVGDYVAPV | 1511403 | Cell/Virus Enclosed | NO |
| SVVNSNAQALNSLLQQ | 1696600 | Ectodomain | YES |
| SVVNTTVTVNVT | 1511410 | Ectodomain | YES |
| SVVQKDGVEDTP | 1511416 | Cell/Virus Enclosed | NO |
| SVVSDTVEVPPQ | 1511430 | Cell/Virus Enclosed | NO |
| SVVSFNEWALANQFTE | 1696606 | Cell/Virus Enclosed | NO |
| SVVVQPRTIDST | 1511442 | Ectodomain | YES |
| SVVVQPRTIIST | 1511443 | Ectodomain | YES |

| | | | |
|---|---|---|---|
| SVVVQPRTINSTQDGD | 1696612 | Ectodomain | YES |
| SVVVYMCVVPSEKDKH | 1696613 | Cell/Virus Enclosed | NO |
| SVWHIQTVQDADGINK | 1696615 | Cell/Virus Enclosed | NO |
| SVYDIACCMRAGSKVV | 1696621 | Cell/Virus Enclosed | NO |
| SVYDIACCMRAS | 1511479 | Cell/Virus Enclosed | NO |
| SVYDQIVDNIYV | 1511481 | Cell/Virus Enclosed | NO |
| SVYDQVVDNVYVTYAG | 1696622 | Cell/Virus Enclosed | NO |
| SVYERDKAVARKLERM | 1696623 | Cell/Virus Enclosed | NO |
| SVYGFSNVVYPD | 1511485 | Underneath Membrane | NO |
| SVYGNFDDQPSLQVES | 1696626 | Cell/Virus Enclosed | NO |
| SVYKSDIHLPAYASFK | 1696628 | Cell/Virus Enclosed | NO |
| SVYLGVKGSGHYQTNL | 1696629 | Cell/Virus Enclosed | NO |
| SVYNAYSPIDIP | 1511493 | Ectodomain | YES |
| SVYSVLETFQNNVVNY | 1696635 | Cell/Virus Enclosed | NO |
| SVYSVPLLFQTN | 1511507 | Cell/Virus Enclosed | NO |
| SVYSVPLVFQNN | 1511508 | Cell/Virus Enclosed | NO |
| SVYTNSCLVSSDGQPG | 1696637 | Cell/Virus Enclosed | NO |
| SVYTNSCLVSSN | 1511511 | Cell/Virus Enclosed | NO |
| SWADLPAYMTVA | 1511520 | Ectodomain | YES |
| SWADQSERNYQTFNRG | 1696644 | Capsid | UN |
| SWAFAKFLWEWASVR | 1696645 | Underneath Membrane | NO |
| SWAGSKINAVIY | 1511527 | Cell/Virus Enclosed | NO |
| SWAIFNDHHLNRPGVY | 1696647 | Cell/Virus Enclosed | NO |
| SWEIGKVKPPLNKNYVFTG | 1511542 | Cell/Virus Enclosed | NO |
| SWESGKVKPPLNRNSV | 1696651 | Cell/Virus Enclosed | NO |
| SWFEDDYKAFIG | 1511544 | Cell/Virus Enclosed | NO |
| SWFTALTQHGKEDLKF | 1696659 | Cell/Virus Enclosed | NO |
| SWFTALTQHGKEDLRF | 1696660 | Cell/Virus Enclosed | NO |
| SWHMDTGVVPCL | 1511566 | Ectodomain | YES |
| SWIGFDVEGAHASRNA | 1696665 | Cell/Virus Enclosed | NO |
| SWLLHKDGGIPD | 1511571 | Cell/Virus Enclosed | NO |
| SWLNDKSVPSPL | 1511575 | Ectodomain | YES |
| SWNKVTQIANMF | 1511587 | Cell/Virus Enclosed | NO |
| SWNKYLYELIQK | 1511589 | Cell/Virus Enclosed | NO |
| SWNRRYGFGSFN | 1511594 | Ectodomain | YES |
| SWQNLLYDFNGN | 1511599 | Ectodomain | YES |
| SWQNLLYDSNGNIIGF | 1696681 | Ectodomain | YES |
| SWQNLLYDSNGNLYGF | 1696682 | Ectodomain | YES |
| SWQNLLYDSSGN | 1511602 | Ectodomain | YES |
| SWSDGDVITGVP | 1511613 | Ectodomain | YES |
| SWSSTMIFSLHP | 1511618 | Cell/Virus Enclosed | NO |
| SWSSTMILSLHP | 1511620 | Cell/Virus Enclosed | NO |
| SWSYIVETPSSDNGTCYPGDFIDYE | 1172375 | Ectodomain | YES |
| SWVDQFERSHQT | 1511627 | Cell/Virus Enclosed | NO |
| SWVDQSERSHQT | 1511628 | Cell/Virus Enclosed | NO |
| SWWSFNPETNNL | 1511643 | Ectodomain | YES |
| SWYDPVENEDIH | 1511645 | Cell/Virus Enclosed | NO |
| SYAASYNRYKYYSGNA | 1696703 | Cell/Virus Enclosed | NO |
| SYAIHHDKFTDG | 1511662 | Cell/Virus Enclosed | NO |
| SYAKPEEKLSAYDLAC | 1696710 | Cell/Virus Enclosed | NO |
| SYALYTAWFYPL | 1511666 | Cell/Virus Enclosed | NO |
| SYALYTPWFYPL | 1511667 | Cell/Virus Enclosed | NO |
| SYCRKLGSSVRS | 1511682 | Cell/Virus Enclosed | NO |
| SYCRLKNSVSDVAYNR | 1696722 | Cell/Virus Enclosed | NO |
| SYDGLRGAIRKAKDYG | 1696728 | Cell/Virus Enclosed | NO |
| SYDINLNKFNRA | 1511697 | Cell/Virus Enclosed | NO |
| SYDLGDELGTDP | 1511700 | Cell/Virus Enclosed | NO |
| SYDTNLNKFNGA | 1511708 | Cell/Virus Enclosed | NO |
| SYEEQDAIFSLTKRNI | 1696741 | Cell/Virus Enclosed | NO |
| SYELLNAPATVC | 1511719 | Ectodomain | YES |
| SYERQTPFEIKS | 1511727 | Cell/Virus Enclosed | NO |
| SYFCKIGKEQKCCMCN | 1696757 | Cell/Virus Enclosed | NO |
| SYFDSKGWYDPVENED | 1696761 | Cell/Virus Enclosed | NO |
| SYFIASFRLFARTRS | 2001328 | Undetermined | UN |
| SYFTEDLLCVADDKPI | 1696770 | Cell/Virus Enclosed | NO |
| SYFVQSIRLFMR | 1511755 | Undetermined | UN |
| SYFYNGSAQATS | 1511761 | Ectodomain | YES |
| SYGFSNVVEMPK | 1511773 | Ectodomain | YES |
| SYGSSVETETYFKVNG | 1696790 | Cell/Virus Enclosed | NO |
| SYHSKPCNPAQPVCSWTLDLLAL | 62567 | Ectodomain | YES |
| SYHTVSLLRSTSQKSI | 1696797 | Ectodomain | YES |
| SYIAYENARQAYEDAI | 1696799 | Cell/Virus Enclosed | NO |
| SYIDLKELGNYTYYNK | 1696800 | Ectodomain | YES |
| SYINDVDTGVPP | 1511816 | Ectodomain | YES |
| SYINNKDTGPPF | 1511820 | Ectodomain | YES |
| SYKIFSTGFVLS | 1511840 | Underneath Membrane | NO |

| | | | |
|---|---|---|---|
| SYKLPNYTVPDLVVEQ | 1696817 | Ectodomain | YES |
| SYLGCVLNAVNL | 1511864 | Ectodomain | YES |
| SYLGCVVNAYNS | 1511866 | Ectodomain | YES |
| SYLSERCNVIADSKSF | 1696834 | Cell/Virus Enclosed | NO |
| SYMDTPSTCLAIVYRPIGVP | 62650 | Cell/Virus Enclosed | NO |
| SYMMPVMGMTNC | 1511900 | Cell/Virus Enclosed | NO |
| SYNASCLCSTDA | 1511904 | Ectodomain | YES |
| SYNDFYASGCVLSSLC | 1696843 | Cell/Virus Enclosed | NO |
| SYNNYVPSYVKPDSVS | 1696848 | Cell/Virus Enclosed | NO |
| SYNRYKYYSGNANEAD | 1696850 | Cell/Virus Enclosed | NO |
| SYPLSMKSDLGV | 1511932 | Ectodomain | YES |
| SYPLSMKSDLSV | 1511933 | Ectodomain | YES |
| SYQEFNDLDHTS | 1511942 | Cell/Virus Enclosed | NO |
| SYQLYDASNVKKVTDV | 1696861 | Cell/Virus Enclosed | NO |
| SYQLYDASNVKN | 1511951 | Cell/Virus Enclosed | NO |
| SYRFVTFEPFNVSFVN | 1696868 | Ectodomain | YES |
| SYSDPCSLKCPYLGCQSWTCPYT | 62700 | Ectodomain | YES |
| SYSGYHPAHAPSFLAV | 1696875 | Cell/Virus Enclosed | NO |
| SYSKIVCDGSVHYAGV | 1696876 | Cell/Virus Enclosed | NO |
| SYSNFYASGCVLSSAC | 1696878 | Cell/Virus Enclosed | NO |
| SYSQNLCVDGFG | 1511989 | Cell/Virus Enclosed | NO |
| SYSVVVQPRTID | 1511995 | Ectodomain | YES |
| SYTGSSFYAPEPITSL | 1696886 | Ectodomain | YES |
| SYTGSSFYEPEP | 1512004 | Ectodomain | YES |
| SYTKSVCKYTDFNEDV | 1696891 | Cell/Virus Enclosed | NO |
| SYTPGHYAGSRSSSGN | 1696894 | Capsid | UN |
| SYTTTIKPLEHH | 1512018 | Cell/Virus Enclosed | NO |
| SYTTTILEHHHH | 1512022 | Cell/Virus Enclosed | NO |
| SYVLLERVYKHF | 1512053 | Cell/Virus Enclosed | NO |
| SYVLLERVYNHF | 1512055 | Cell/Virus Enclosed | NO |
| SYVLNNISLTTQPYFD | 1696909 | Ectodomain | YES |
| SYVNSSNVTTMD | 1512059 | Cell/Virus Enclosed | NO |
| SYVPTKYVTAKV | 1512060 | Ectodomain | YES |
| SYVYNGSAQVTS | 1512067 | Ectodomain | YES |
| SYYAYMMPVMGM | 1512069 | Cell/Virus Enclosed | NO |
| SYYLIQSGIFFT | 1512083 | Cell/Virus Enclosed | NO |
| SYYRLGASQRVG | 1512085 | Underneath Membrane | NO |
| TAADINIHDLPI | 1512097 | Ectodomain | YES |
| TAADINIPHLPI | 1512100 | Ectodomain | YES |
| TAADINNPDLPI | 1512101 | Ectodomain | YES |
| TAADINTPDVPI | 1512102 | Ectodomain | YES |
| TAAMLFDSHIVSKESG | 1696940 | Capsid | UN |
| TAATIVNQERYV | 1512128 | Cell/Virus Enclosed | NO |
| TAATVAAMFPPWSAAA | 1696943 | Ectodomain | YES |
| TAAVDLSMDKLC | 1512133 | Cell/Virus Enclosed | NO |
| TAAVINIPDLPI | 1512136 | Ectodomain | YES |
| TAAYINIPDLPI | 1512143 | Ectodomain | YES |
| TACAFSVDPATTYLEA | 1696947 | Cell/Virus Enclosed | NO |
| TACNGHKIEDLS | 1512151 | Cell/Virus Enclosed | NO |
| TACNGISVTQIM | 1512152 | Cell/Virus Enclosed | NO |
| TADEAIKFLKEWNFSL | 1696960 | Undetermined | UN |
| TADEIKALKELY | 1512161 | Cell/Virus Enclosed | NO |
| TADGSPQPYCYT | 1512169 | Cell/Virus Enclosed | NO |
| TADTAHANNINRFNVA | 1696967 | Cell/Virus Enclosed | NO |
| TAEDISLLKKMDEPYT | 1696970 | Cell/Virus Enclosed | NO |
| TAEDMLNPNYED | 1512182 | Cell/Virus Enclosed | NO |
| TAEELEAIKHQLNPKDNDVEL | 62828 | Capsid | UN |
| TAEPVRDEFSIE | 1512184 | Capsid | UN |
| TAEPVRDEVSIETDII | 1696973 | Capsid | UN |
| TAEQCIRGAIGD | 1512186 | Cell/Virus Enclosed | NO |
| TAETAHSVNVNRFNVA | 1696976 | Cell/Virus Enclosed | NO |
| TAFAAIGDLKCP | 1512192 | Ectodomain | YES |
| TAFAVIGDLKCP | 1512193 | Ectodomain | YES |
| TAFAVIGDLNCT | 1512196 | Ectodomain | YES |
| TAFNNALLSIQNGFSA | 1696989 | Ectodomain | YES |
| TAFSKPLYVPEH | 1512215 | Cell/Virus Enclosed | NO |
| TAFSPAQDIWGTSAAA | 1696993 | Ectodomain | YES |
| TAGLFTNFRIDV | 1512238 | Cell/Virus Enclosed | NO |
| TAGLYTNSRIDV | 1512248 | Cell/Virus Enclosed | NO |
| TAGVVTCNVVKQ | 1512256 | Cell/Virus Enclosed | NO |
| TAGVVTGNVVKQCCTA | 1697006 | Cell/Virus Enclosed | NO |
| TAGWTFGAGAALQIPFA | 62872 | Ectodomain | YES |
| TAGWTFGAGSAL | 1512262 | Ectodomain | YES |
| TAIDCGFNDLSQ | 1512280 | Ectodomain | YES |
| TAIPTIKDWLLV | 1512291 | Cell/Virus Enclosed | NO |
| TAIQTVFDCAVE | 1512292 | Cell/Virus Enclosed | NO |

| | | | |
|---|---|---|---|
| TAIRADQLTPAW | 1512293 | Ectodomain | YES |
| TAKEVRQKILNK | 1512304 | Cell/Virus Enclosed | NO |
| TAKKQVVLVSNNQEDF | 1697031 | Cell/Virus Enclosed | NO |
| TAKKSMYDGDRFVKHD | 1697032 | Cell/Virus Enclosed | NO |
| TAKYIPLVGTPL | 1512320 | Cell/Virus Enclosed | NO |
| TALDAYVSQQLS | 1512331 | Ectodomain | YES |
| TALGKLQDVVNQNAQAL | 62908 | Ectodomain | YES |
| TALNAYVSQQLSDISL | 1697043 | Ectodomain | YES |
| TALNAYVSQQLSDSTL | 1697044 | Ectodomain | YES |
| TALNKIQDVVNQQGNS | 1697045 | Ectodomain | YES |
| TALPKGFYAEGSRGGS | 1697046 | Cell/Virus Enclosed | NO |
| TALQELLIQQRI | 1512348 | Cell/Virus Enclosed | NO |
| TALQELQIQQWI | 1512350 | Cell/Virus Enclosed | NO |
| TALRLCAYCCNIVNV | 1542667 | Ectodomain | YES |
| TALRRPINATDR | 1512357 | Cell/Virus Enclosed | NO |
| TALTQHGKEDLKFPR | 1512360 | Capsid | UN |
| TAMIPLQASLPF | 1512370 | Secreted | YES |
| TANADTLCIGYHANNSTDTVDTVLE | 1172621 | Ectodomain | YES |
| TANHSIPDLQRA | 1512386 | Cell/Virus Enclosed | NO |
| TANKIVGGAPTWFNAL | 1697063 | Cell/Virus Enclosed | NO |
| TANVCSPMACNG | 1512396 | Cell/Virus Enclosed | NO |
| TANVGRVIKFETDAYK | 1697074 | Cell/Virus Enclosed | NO |
| TANVSALMGANGNKIV | 1697077 | Cell/Virus Enclosed | NO |
| TAPEDDDFLEEN | 1512411 | Cell/Virus Enclosed | NO |
| TAPEYDDFLEES | 1512412 | Cell/Virus Enclosed | NO |
| TAPKTGNFKDLR | 1512423 | Ectodomain | YES |
| TAPQTGNFKDLR | 1512425 | Ectodomain | YES |
| TAPRNSWEAMALNFKL | 1697089 | Cell/Virus Enclosed | NO |
| TAPTIVNQERYVKITG | 1697090 | Cell/Virus Enclosed | NO |
| TAPYVVLTGLGESFKK | 1697091 | Cell/Virus Enclosed | NO |
| TAQAAGADTTIT | 1512432 | Cell/Virus Enclosed | NO |
| TAQADTAGLFTN | 1512436 | Cell/Virus Enclosed | NO |
| TAQADTAGLYTNFRID | 1697095 | Cell/Virus Enclosed | NO |
| TAQELLLALLAF | 1512441 | Cell/Virus Enclosed | NO |
| TAQPVKIDNASPASTV | 1697106 | Secreted | YES |
| TARDALDTILQE | 1512461 | Cell/Virus Enclosed | NO |
| TARDGVRRGDNFHSVL | 1697107 | Cell/Virus Enclosed | NO |
| TARVVYTAASHAAVDA | 1697113 | Cell/Virus Enclosed | NO |
| TARVVYTACSHAAVDA | 1697114 | Cell/Virus Enclosed | NO |
| TARYFDCYEGGC | 1512480 | Cell/Virus Enclosed | NO |
| TASDKFYRLSNELAQV | 1697118 | Cell/Virus Enclosed | NO |
| TASLFQTTVKLQ | 1512492 | Ectodomain | YES |
| TASMSMAECKAT | 1512494 | Cell/Virus Enclosed | NO |
| TATGDVVLATDDLYVK | 1389324 | Cell/Virus Enclosed | NO |
| TATKQYNVTQAFGRRG | 1697140 | Cell/Virus Enclosed | NO |
| TATKSYNVTQAF | 1512530 | Cell/Virus Enclosed | NO |
| TATSQQPSGGNVVPYY | 1697143 | Cell/Virus Enclosed | NO |
| TATTVVGSCIKG | 1512542 | Cell/Virus Enclosed | NO |
| TATTVVGSCIKS | 1512543 | Cell/Virus Enclosed | NO |
| TAVGVPQQRFVY | 1512563 | Ectodomain | YES |
| TAVHSDGMFVAT | 1512569 | Cell/Virus Enclosed | NO |
| TAVINLRADQINDMVY | 1697155 | Cell/Virus Enclosed | NO |
| TAVKAGGKSLPA | 1512570 | Cell/Virus Enclosed | NO |
| TAVLRQWLPTGT | 1512577 | Cell/Virus Enclosed | NO |
| TAVLTIANNYSKLFDE | 1697160 | Cell/Virus Enclosed | NO |
| TAVNESRYGLHN | 1512584 | Ectodomain | YES |
| TAVNESRYGLQNLLQL | 1697164 | Ectodomain | YES |
| TAVNLPRGFPAG | 1512586 | Ectodomain | YES |
| TAVRSDGTFEEM | 1512588 | Cell/Virus Enclosed | NO |
| TAVRSSFNFDQK | 1512589 | Cell/Virus Enclosed | NO |
| TAVTTCDVKIMA | 1512596 | Cell/Virus Enclosed | NO |
| TAVTTGDVKIMA | 1512597 | Cell/Virus Enclosed | NO |
| TAVVNLRPDQLNDLVY | 1697169 | Cell/Virus Enclosed | NO |
| TAVVTNGYLKMA | 1512603 | Ectodomain | YES |
| TAVYKPLVDFNV | 1512606 | Cell/Virus Enclosed | NO |
| TAWYVGQIFFLV | 1512614 | Cell/Virus Enclosed | NO |
| TCAFVDTVKGLD | 1512630 | Cell/Virus Enclosed | NO |
| TCAKRFKRWLCI | 1512632 | Cell/Virus Enclosed | NO |
| TCATTRQVVNAI | 1512640 | Cell/Virus Enclosed | NO |
| TCATTRQVVNVI | 1512641 | Cell/Virus Enclosed | NO |
| TCAVNYTKAPDV | 1512643 | Ectodomain | YES |
| TCDGLKFTEESFYKNM | 1697207 | Cell/Virus Enclosed | NO |
| TCDICKSTVVEVKSAI | 1697211 | Cell/Virus Enclosed | NO |
| TCDNLCTLDSIT | 1512666 | Ectodomain | YES |
| TCDRTAIQSFDNSYLN | 1697214 | Cell/Virus Enclosed | NO |
| TCDVKIMAKAPQ | 1512671 | Cell/Virus Enclosed | NO |

| | | | |
|---|---|---|---|
| TCEHEYGTEDDY | 1512679 | Cell/Virus Enclosed | NO |
| TCFDVSKPNAID | 1512693 | Cell/Virus Enclosed | NO |
| TCFGVGKPNAID | 1512702 | Cell/Virus Enclosed | NO |
| TCFGVSKPNAID | 1512703 | Cell/Virus Enclosed | NO |
| TCFRCPKEIVDT | 1512707 | Cell/Virus Enclosed | NO |
| TCFSVSKPNAID | 1512713 | Cell/Virus Enclosed | NO |
| TCGIVELEKGVL | 1512720 | Cell/Virus Enclosed | NO |
| TCGTTTLAGLWL | 1512727 | Cell/Virus Enclosed | NO |
| TCHNAAQCDCLC | 1512736 | Ectodomain | YES |
| TCHNAAQCNCLC | 1512737 | Ectodomain | YES |
| TCIVLDVKGVEE | 1512769 | Cell/Virus Enclosed | NO |
| TCNAISPKTDKE | 1512792 | Ectodomain | YES |
| TCNAISSNTDNETLQY | 1697279 | Ectodomain | YES |
| TCNNIDAAKIYGMCFS | 1697280 | Ectodomain | YES |
| TCNSKVKNGFTL | 1512806 | Ectodomain | YES |
| TCNSNANISMLQ | 1512807 | Ectodomain | YES |
| TCNSNANLSMLQLGVP | 1697282 | Ectodomain | YES |
| TCNTPTQCYYNNSNNG | 1697283 | Cell/Virus Enclosed | NO |
| TCNVSFVGELKAAEAK | 1697284 | Cell/Virus Enclosed | NO |
| TCNVVKQCCTAA | 1512814 | Cell/Virus Enclosed | NO |
| TCPAGTNYLNCD | 1512819 | Ectodomain | YES |
| TCRFVELDDQGH | 1512842 | Cell/Virus Enclosed | NO |
| TCRQISPAAIAS | 1512848 | Ectodomain | YES |
| TCRTDMEKDFIALDQD | 1697295 | Cell/Virus Enclosed | NO |
| TCSNDLLQPNTEVFTD | 1697298 | Ectodomain | YES |
| TCTGCYACSSGISCKVRIH | 244287 | Ectodomain | YES |
| TCTIAGKAFDKVFDYV | 1697304 | Cell/Virus Enclosed | NO |
| TCTIDSNTTQDT | 1512871 | Cell/Virus Enclosed | NO |
| TCTSTHFAFACA | 1512876 | Cell/Virus Enclosed | NO |
| TCTTPAQGTSMFPSC | 1697309 | Underneath Membrane | NO |
| TCVIANVPAIDVLKKL | 1697311 | Cell/Virus Enclosed | NO |
| TCYRCPKEIVDTVSAL | 1697323 | Cell/Virus Enclosed | NO |
| TDAALSFKENVD | 1512911 | Cell/Virus Enclosed | NO |
| TDAAVKAAESKAFVDT | 1697325 | Cell/Virus Enclosed | NO |
| TDAFLGWSYDTC | 1512912 | Ectodomain | YES |
| TDAIIAVELLLS | 1512916 | Cell/Virus Enclosed | NO |
| TDAIIIDNDLNEFVSD | 1697328 | Cell/Virus Enclosed | NO |
| TDAKWYCYDKNPINSN | 1697329 | Cell/Virus Enclosed | NO |
| TDAKWYCYDKQPVNSN | 1697330 | Cell/Virus Enclosed | NO |
| TDANKLILINDVAFVS | 1697331 | Cell/Virus Enclosed | NO |
| TDAPSTFGTRNP | 1512921 | Cell/Virus Enclosed | NO |
| TDAQTGSSKCGC | 1512922 | Cell/Virus Enclosed | NO |
| TDAQTGSSKCVR | 1512924 | Cell/Virus Enclosed | NO |
| TDAVILLERYVSLAID | 1697338 | Cell/Virus Enclosed | NO |
| TDAVVLLERYVSLAID | 1697340 | Cell/Virus Enclosed | NO |
| TDAYKLFLSGDD | 1512932 | Cell/Virus Enclosed | NO |
| TDCLAAETHRDCDFNK | 1697342 | Cell/Virus Enclosed | NO |
| TDCLYLKNLTQTFTSM | 1697343 | Cell/Virus Enclosed | NO |
| TDCSDGNYNRNASLNS | 1697344 | Ectodomain | YES |
| TDCSDGNYNRYA | 1512941 | Ectodomain | YES |
| TDCVFYFAVRKEGQDV | 1697345 | Cell/Virus Enclosed | NO |
| TDDPRWPQIAEL | 1512954 | Cell/Virus Enclosed | NO |
| TDEAQPLPQRKK | 1512957 | Cell/Virus Enclosed | NO |
| TDEAQPLPQRQKKQPT | 1389360 | Capsid | UN |
| TDECRLAMLGAEMTSN | 1697353 | Cell/Virus Enclosed | NO |
| TDENYNNLVPTYLKSD | 1697356 | Cell/Virus Enclosed | NO |
| TDESFYKNMYLS | 1512966 | Cell/Virus Enclosed | NO |
| TDESQPLPQRQKKQQT | 1697360 | Cell/Virus Enclosed | NO |
| TDFHNYTGEGQQ | 1512976 | Ectodomain | YES |
| TDFNGNFYGPYR | 1512992 | Cell/Virus Enclosed | NO |
| TDGICSENHVMV | 1513005 | Cell/Virus Enclosed | NO |
| TDGKLNCSFQDAASNI | 1697381 | Cell/Virus Enclosed | NO |
| TDHDVVYAQHCF | 1513020 | Ectodomain | YES |
| TDHSGSNDNIAL | 1513024 | Underneath Membrane | NO |
| TDIDVNPALNIC | 1513036 | Cell/Virus Enclosed | NO |
| TDIDYCVRAFDVYNKD | 1697402 | Cell/Virus Enclosed | NO |
| TDINFTATASFG | 1513053 | Ectodomain | YES |
| TDINGVYWVASN | 1513054 | Cell/Virus Enclosed | NO |
| TDIVDVTNGLGT | 1513059 | Ectodomain | YES |
| TDIVVVDEVSMLTNYE | 1697410 | Cell/Virus Enclosed | NO |
| TDKETLEYWVTP | 1513063 | Ectodomain | YES |
| TDKINQIIHDFIDNPL | 769763 | Secreted | YES |
| TDKLCHAVVSKS | 1513065 | Cell/Virus Enclosed | NO |
| TDKSFEQQVGVIADKD | 1697412 | Cell/Virus Enclosed | NO |
| TDLLSTLWFKPP | 1513092 | Ectodomain | YES |
| TDLQKANTDIIL | 1513097 | Ectodomain | YES |

| | | | |
|---|---|---|---|
| TDLQKANTDIKL | 1513098 | Ectodomain | YES |
| TDLQKANTEIEL | 1513099 | Ectodomain | YES |
| TDLQKSNTDIIL | 1513100 | Ectodomain | YES |
| TDLQSESSCGLFKDCA | 1697428 | Cell/Virus Enclosed | NO |
| TDLTVTSDGQPY | 1513108 | Cell/Virus Enclosed | NO |
| TDMVKSKGVCATGDCY | 1697434 | Cell/Virus Enclosed | NO |
| TDNETLEYWVTP | 1513116 | Ectodomain | YES |
| TDNGTITVEELK | 1513120 | Ectodomain | YES |
| TDNIFSVQEDGR | 1513123 | Ectodomain | YES |
| TDNLVFVNKTSLPTNI | 1697437 | Cell/Virus Enclosed | NO |
| TDNNQDGGRSGA | 1513131 | Cell/Virus Enclosed | NO |
| TDNNQNGDRSGARPKQ | 1697439 | Cell/Virus Enclosed | NO |
| TDNNQSGGRNGA | 1513133 | Capsid | UN |
| TDNSANTKVLRNKGVD | 1697441 | Cell/Virus Enclosed | NO |
| TDNSTTVSVTVSSNMV | 1697442 | Cell/Virus Enclosed | NO |
| TDNTFVAGNCDV | 1513139 | Ectodomain | YES |
| TDNTFVSGKCDV | 1513140 | Ectodomain | YES |
| TDPIVFTYIPAS | 1513153 | Cell/Virus Enclosed | NO |
| TDPIYDEPTTTT | 1513154 | Secreted | YES |
| TDQINDMVYSLL | 1513162 | Cell/Virus Enclosed | NO |
| TDQMEVDSGPRKRPLRGG | 63217 | Cell/Virus Enclosed | NO |
| TDQYLCGADGKPVMSE | 1697463 | Cell/Virus Enclosed | NO |
| TDSADNNQNGGR | 1513185 | Capsid | UN |
| TDSDSLSSKPQRKRGV | 1697466 | Capsid | UN |
| TDSFSCNNFDESKIYG | 1697467 | Ectodomain | YES |
| TDSGFAVYHRYR | 1513190 | Underneath Membrane | NO |
| TDSIVFTYIPASGYVA | 1697469 | Cell/Virus Enclosed | NO |
| TDSIVIDGVTYQ | 1513195 | Cell/Virus Enclosed | NO |
| TDSLCSSHYWSPQGNL | 1697473 | Cell/Virus Enclosed | NO |
| TDSMLTLDDRGLALDN | 1697474 | Cell/Virus Enclosed | NO |
| TDSTDNNQDGGR | 1513202 | Cell/Virus Enclosed | NO |
| TDSTVFTYIPAS | 1513206 | Cell/Virus Enclosed | NO |
| TDSTVLTYIPAS | 1513208 | Cell/Virus Enclosed | NO |
| TDSVLLIERFVSLAID | 1697476 | Cell/Virus Enclosed | NO |
| TDSVRDPQTLEI | 1513216 | Ectodomain | YES |
| TDTIVVTEGDGISTPK | 1697484 | Secreted | YES |
| TDTNSVCPKFEF | 1513238 | Ectodomain | YES |
| TDTNSVCPKIEF | 1513239 | Ectodomain | YES |
| TDTNSVCPKLEFANDT | 1697485 | Ectodomain | YES |
| TDTNSVSPKLEF | 1513246 | Ectodomain | YES |
| TDTVDVTNGLGT | 1513258 | Ectodomain | YES |
| TDTVNHDSFNLYGYSV | 1697493 | Cell/Virus Enclosed | NO |
| TDTVNRDSFNLY | 1513263 | Cell/Virus Enclosed | NO |
| TDVAYHTVTDVKQVGC | 1697494 | Cell/Virus Enclosed | NO |
| TDVKQVGCFMRL | 1513280 | Cell/Virus Enclosed | NO |
| TDVKQVGCSMRL | 1513281 | Cell/Virus Enclosed | NO |
| TDVLFPGQKYSSDCNV | 1697502 | Cell/Virus Enclosed | NO |
| TDVLFSGQKYSS | 1513283 | Cell/Virus Enclosed | NO |
| TDVLQEDQKILA | 1513289 | Ectodomain | YES |
| TDVLQENQKILAASFN | 1697503 | Ectodomain | YES |
| TDVPSISGTQVY | 1513299 | Cell/Virus Enclosed | NO |
| TDVPTAIRADQL | 1513301 | Ectodomain | YES |
| TDVPTSIHADQL | 1513303 | Ectodomain | YES |
| TDVQLRAFDIYNASVA | 1697506 | Cell/Virus Enclosed | NO |
| TDVRDYKLANDVKDTL | 1697507 | Cell/Virus Enclosed | NO |
| TDVSTAIHADQLTPAWRIYSTG | 558510 | Ectodomain | YES |
| TDVSTLIHAEQL | 1513312 | Ectodomain | YES |
| TDYKLELFNKYFKHWS | 1697520 | Cell/Virus Enclosed | NO |
| TDYKLELFNKYFKYWG | 1697521 | Cell/Virus Enclosed | NO |
| TDYNAILQSDNP | 1513335 | Cell/Virus Enclosed | NO |
| TDYNAILQSNNPQCAI | 1697524 | Cell/Virus Enclosed | NO |
| TDYNYYKYNLPTMVDI | 1697525 | Cell/Virus Enclosed | NO |
| TEADWIELFPHN | 1513342 | Cell/Virus Enclosed | NO |
| TEAEWIELFPHNDRIK | 1697530 | Cell/Virus Enclosed | NO |
| TEASNYAQYRVARATIRYRPLVPNA | 63307 | Undetermined | UN |
| TECANLLLQYGSFCTQL | 63309 | Ectodomain | YES |
| TEDDYKGLPLEF | 1513365 | Cell/Virus Enclosed | NO |
| TEDPPWPQIAEL | 1513380 | Cell/Virus Enclosed | NO |
| TEDPRWPQIAELAPTA | 1697550 | Cell/Virus Enclosed | NO |
| TEDVKFAASTGVIDIS | 1697552 | Cell/Virus Enclosed | NO |
| TEESFYKNMYLKSAVM | 1697563 | Cell/Virus Enclosed | NO |
| TEETQTTIQRPRSEP | 111821 | Cell/Virus Enclosed | NO |
| TEEVKSSRVERSTAVL | 1697567 | Cell/Virus Enclosed | NO |
| TEFAVNSTKALA | 1513407 | Cell/Virus Enclosed | NO |
| TEFQLDYSIDVIDNEI | 1697573 | Cell/Virus Enclosed | NO |
| TEFSWNADLYKLMSCF | 1697575 | Cell/Virus Enclosed | NO |

| | | | |
|---|---|---|---|
| TEFVGTQSIDML | 1513418 | Cell/Virus Enclosed | NO |
| TEGCVSTNFGDVIKPV | 1697581 | Cell/Virus Enclosed | NO |
| TEGGKTFMYAYI | 1513431 | Cell/Virus Enclosed | NO |
| TEGLMQNASLYSSLVP | 1697586 | Cell/Virus Enclosed | NO |
| TEGQLQSLSKDYISSN | 1697588 | Cell/Virus Enclosed | NO |
| TEGQLQSLSRDY | 1513438 | Cell/Virus Enclosed | NO |
| TEHVTSVNLFDVGYSV | 1697595 | Cell/Virus Enclosed | NO |
| TEIELGVCVNYD | 1513457 | Ectodomain | YES |
| TEKLSRLSMIKGPQRA | 1697608 | Cell/Virus Enclosed | NO |
| TEKSNVVRGWVF | 1513480 | Ectodomain | YES |
| TELAVNSGLLTA | 1513484 | Cell/Virus Enclosed | NO |
| TELPQLVNANQY | 1513496 | Ectodomain | YES |
| TENLNAYQQQDG | 1513515 | Cell/Virus Enclosed | NO |
| TENLVIEGPTTC | 1513518 | Cell/Virus Enclosed | NO |
| TENRFTNFEPFT | 1513527 | Ectodomain | YES |
| TEQLGAPLGITI | 1513537 | Ectodomain | YES |
| TEQTQNVTVKPKRLRK | 1697650 | Cell/Virus Enclosed | NO |
| TEQTQSAIAKPK | 1513551 | Cell/Virus Enclosed | NO |
| TEQTQSVNVKLK | 1513552 | Cell/Virus Enclosed | NO |
| TEQTQSVNVKPK | 1513553 | Cell/Virus Enclosed | NO |
| TERLKLFAEETQ | 1513564 | Cell/Virus Enclosed | NO |
| TESAVNSTKALAKQDA | 1697658 | Cell/Virus Enclosed | NO |
| TESYVDVGPDSF | 1513597 | Ectodomain | YES |
| TETAAPAVADARKPPSGKKK | 63545 | Capsid | UN |
| TETITTGFDFNC | 1513605 | Ectodomain | YES |
| TETITTGFDLNC | 1513606 | Ectodomain | YES |
| TETTSAFVGALP | 1513614 | Ectodomain | YES |
| TEVDDYVLANECIERL | 1697678 | Cell/Virus Enclosed | NO |
| TEVKVSALKTAG | 1513631 | Cell/Virus Enclosed | NO |
| TEVNELLDTTQLQVAN | 1697687 | Ectodomain | YES |
| TEVPQLVNANHY | 1513642 | Ectodomain | YES |
| TEVRGSRRLAQQKINE | 1697694 | Ectodomain | YES |
| TEVSVKMVDAYV | 1513652 | Cell/Virus Enclosed | NO |
| TEVVGNVILKPA | 1513656 | Cell/Virus Enclosed | NO |
| TEVVRFPNITNL | 1513659 | Ectodomain | YES |
| TEVVYSNGGFYFKPGG | 1697702 | Cell/Virus Enclosed | NO |
| TEYDFTDYKVQLFEKY | 1697708 | Cell/Virus Enclosed | NO |
| TEYREYMEAYNF | 1513674 | Cell/Virus Enclosed | NO |
| TEYSWNKKLYELVQRF | 1697711 | Cell/Virus Enclosed | NO |
| TEYYEFLNKHFN | 1513684 | Cell/Virus Enclosed | NO |
| TEYYEFLNKHFSMMIL | 1697714 | Cell/Virus Enclosed | NO |
| TFAELEAAQKAYQEAM | 1697715 | Cell/Virus Enclosed | NO |
| TFAELEAARKAY | 1513687 | Cell/Virus Enclosed | NO |
| TFCDNINSILDEVNGL | 1697724 | Ectodomain | YES |
| TFDDILGKVCSE | 1513718 | Cell/Virus Enclosed | NO |
| TFDDPVIPFKDG | 1513719 | Ectodomain | YES |
| TFDDVQAPNYTQHTSS | 1697734 | Ectodomain | YES |
| TFDFYNLWSTFT | 1513724 | Cell/Virus Enclosed | NO |
| TFDPITLKAPDT | 1513733 | Ectodomain | YES |
| TFEDELAPSDVYQQLA | 1697748 | Cell/Virus Enclosed | NO |
| TFEMSCDCGKKFDDQV | 1697757 | Cell/Virus Enclosed | NO |
| TFEPFNVSFVNDSIES | 1697758 | Ectodomain | YES |
| TFERGSVQVDNG | 1513752 | Cell/Virus Enclosed | NO |
| TFFEGDSHDTVF | 1513763 | Cell/Virus Enclosed | NO |
| TFFIFNKLVKDPPNVQ | 1697765 | Secreted | YES |
| TFFKRVCGVSAA | 1513769 | Cell/Virus Enclosed | NO |
| TFGAISASIGDIIQRL | 1697776 | Ectodomain | YES |
| TFGEPTDSTDNN | 1513796 | Capsid | UN |
| TFGGPIDSTDNN | 1513797 | Capsid | UN |
| TFGGPSDSSDNS | 1513798 | Cell/Virus Enclosed | NO |
| TFGGPSDSTDNNQNGD | 1697781 | Cell/Virus Enclosed | NO |
| TFGGPSDSTGSNQNGE | 1697782 | Cell/Virus Enclosed | NO |
| TFGNPIIPFKDG | 1513809 | Ectodomain | YES |
| TFGNVKPGETFTVLAA | 1697789 | Cell/Virus Enclosed | NO |
| TFGTRNPNNDLA | 1513817 | Cell/Virus Enclosed | NO |
| TFGVVKPGDTFT | 1513824 | Cell/Virus Enclosed | NO |
| TFGVVKPGETFTVLAA | 1697793 | Cell/Virus Enclosed | NO |
| TFIDAARGPAGVESDV | 1697800 | Cell/Virus Enclosed | NO |
| TFIQSNVDVVPE | 1513837 | Cell/Virus Enclosed | NO |
| TFITPEVSRELGNITK | 1697807 | Cell/Virus Enclosed | NO |
| TFITVEAAIELSKELK | 1697808 | Cell/Virus Enclosed | NO |
| TFITVEAALDLSKELK | 1697809 | Cell/Virus Enclosed | NO |
| TFKAPDTYKCPQ | 1513857 | Ectodomain | YES |
| TFKATGTYKCPQ | 1513859 | Ectodomain | YES |
| TFKCYGVSATKLNDLC | 1697812 | Ectodomain | YES |
| TFKIVQWQEAWF | 1513868 | Cell/Virus Enclosed | NO |

| | | | |
|---|---|---|---|
| TFKQANNNCFVNVSCL | 1697823 | Cell/Virus Enclosed | NO |
| TFKQTNNNCFVN | 1513877 | Cell/Virus Enclosed | NO |
| TFLDLQAEMNRL | 1513889 | Ectodomain | YES |
| TFLDLQDEMNRL | 1513890 | Ectodomain | YES |
| TFLDLQVEMNRLQEAI | 1697833 | Ectodomain | YES |
| TFLLTFNDNQAITQVP | 1697848 | Cell/Virus Enclosed | NO |
| TFLNGITYTSTSGNLL | 1697850 | Ectodomain | YES |
| TFLNGVCGVSAA | 1513917 | Cell/Virus Enclosed | NO |
| TFLNRVCGVSAA | 1513920 | Cell/Virus Enclosed | NO |
| TFLSCARKSFSI | 1513926 | Cell/Virus Enclosed | NO |
| TFLSLSDQFKTTGDLA | 1697853 | Cell/Virus Enclosed | NO |
| TFLSMFDVDKKI | 1513929 | Cell/Virus Enclosed | NO |
| TFLSMFDVDKKSLNAL | 1697854 | Cell/Virus Enclosed | NO |
| TFMLKYDENGTITDAV | 1697860 | Ectodomain | YES |
| TFNDNQAITQVPATSI | 1697864 | Cell/Virus Enclosed | NO |
| TFNHNNGNDVLYQPPT | 1697868 | Cell/Virus Enclosed | NO |
| TFNITEDEILEW | 1513956 | Ectodomain | YES |
| TFNKDIYCGNSMVCKM | 1697869 | Cell/Virus Enclosed | NO |
| TFNKQEASVPILTTPF | 1697870 | Cell/Virus Enclosed | NO |
| TFNRGRKTQPKF | 1513965 | Cell/Virus Enclosed | NO |
| TFPGPSGIGAVG | 1513975 | Cell/Virus Enclosed | NO |
| TFPPGQGVPLNANSTP | 1697879 | Cell/Virus Enclosed | NO |
| TFPQQMFVTPLA | 1513980 | Cell/Virus Enclosed | NO |
| TFPSGQGVPLNA | 1513982 | Cell/Virus Enclosed | NO |
| TFQTVRPGNFNQ | 1513997 | Cell/Virus Enclosed | NO |
| TFQTVRSGNFNQ | 1513998 | Cell/Virus Enclosed | NO |
| TFSATFSVPMEE | 1514011 | Cell/Virus Enclosed | NO |
| TFSGNVDLDALASDIF | 1697893 | Cell/Virus Enclosed | NO |
| TFSGSFDNGHYVVYDA | 1697894 | Cell/Virus Enclosed | NO |
| TFSMAPVEFYEDRILD | 1697896 | Cell/Virus Enclosed | NO |
| TFSPGQGVPLNA | 1514037 | Cell/Virus Enclosed | NO |
| TFSVPMEELKAL | 1514043 | Cell/Virus Enclosed | NO |
| TFSYDGLRGAIRKAKD | 1697905 | Cell/Virus Enclosed | NO |
| TFTATTVVGSCI | 1514050 | Cell/Virus Enclosed | NO |
| TFTDVNLQGLEN | 1514053 | Cell/Virus Enclosed | NO |
| TFTEVNLQGLEN | 1514054 | Cell/Virus Enclosed | NO |
| TFTGSYKCPQTK | 1514060 | Ectodomain | YES |
| TFTNNNGSDVLYQPPT | 1697915 | Cell/Virus Enclosed | NO |
| TFTQAGFNIWVP | 1514071 | Cell/Virus Enclosed | NO |
| TFTTKVTVPKDH | 1514077 | Cell/Virus Enclosed | NO |
| TFVIDMRSYEKLANSI | 1697940 | Cell/Virus Enclosed | NO |
| TFVNEYYEFLCKHFSM | 1697943 | Cell/Virus Enclosed | NO |
| TFVNISRVELHT | 1514110 | Ectodomain | YES |
| TFVQSNVEVLPE | 1514120 | Cell/Virus Enclosed | NO |
| TFVVDKSLSIDE | 1514129 | Cell/Virus Enclosed | NO |
| TFVVDKSLSIEEFADV | 1697957 | Cell/Virus Enclosed | NO |
| TFYAYFTDTGVV | 1514144 | Ectodomain | YES |
| TFYGKESFENPT | 1514150 | Cell/Virus Enclosed | NO |
| TFYGKESLENPT | 1514152 | Cell/Virus Enclosed | NO |
| TFYPRLQAASDWKPGY | 1697981 | Cell/Virus Enclosed | NO |
| TGAICDFEIACKCGVK | 1697988 | Cell/Virus Enclosed | NO |
| TGAIKLDDKDPSFKDN | 1697990 | Cell/Virus Enclosed | NO |
| TGALLVDSDLND | 1514183 | Cell/Virus Enclosed | NO |
| TGAPSLGDYVRPAYIVTPYW | 78850 | Cell/Virus Enclosed | NO |
| TGCCDNSYFDSKGWYD | 1697993 | Cell/Virus Enclosed | NO |
| TGCFSFVKMLFM | 1514198 | Cell/Virus Enclosed | NO |
| TGCTGSWNKVNQ | 1514209 | Cell/Virus Enclosed | NO |
| TGCVIAWNTAQQ | 1514215 | Ectodomain | YES |
| TGDCATVYLEDK | 1514218 | Cell/Virus Enclosed | NO |
| TGDLAVCLGIGDSAVT | 1698019 | Cell/Virus Enclosed | NO |
| TGDLAVQIGSNNVCTY | 1698020 | Cell/Virus Enclosed | NO |
| TGDLFYTIKFSGFKLD | 1698021 | Cell/Virus Enclosed | NO |
| TGDNDDEDVVTGDNDD | 1698023 | Cell/Virus Enclosed | NO |
| TGDNDDEDVVTGDNND | 1698024 | Cell/Virus Enclosed | NO |
| TGDNNDEEIVTGDNDD | 1698026 | Cell/Virus Enclosed | NO |
| TGDSVKFVYMHQLELS | 1698031 | Cell/Virus Enclosed | NO |
| TGDVKIMAKAPQ | 1514257 | Cell/Virus Enclosed | NO |
| TGDVVLASDDLYVKRY | 1698034 | Cell/Virus Enclosed | NO |
| TGDVVLATDDLYVKRY | 1698035 | Cell/Virus Enclosed | NO |
| TGEESASSGKLGLIT | 187541 | Secreted | YES |
| TGEGDQIIFYEG | 1514262 | Ectodomain | YES |
| TGFCSKHQWNCI | 1514268 | Cell/Virus Enclosed | NO |
| TGFTKPASRELS | 1514289 | Cell/Virus Enclosed | NO |
| TGFTTTNEAFQKVQDA | 1698052 | Ectodomain | YES |
| TGFTTTNEAFRK | 1514291 | Ectodomain | YES |
| TGFVYFNATGSDVNCN | 1698053 | Ectodomain | YES |

| | | | |
|---|---|---|---|
| TGGFMSTIITSC | 1514307 | Cell/Virus Enclosed | NO |
| TGGIDICSQDKV | 1514311 | Cell/Virus Enclosed | NO |
| TGGNIQSSSRAS | 1514317 | Capsid | UN |
| TGGNSQSSSRASSLSR | 1698064 | Cell/Virus Enclosed | NO |
| TGHFYINGYRYF | 1514331 | Ectodomain | YES |
| TGHYIGQAGEMP | 1514341 | Cell/Virus Enclosed | NO |
| TGHYTGQAGEMPCAII | 1698074 | Cell/Virus Enclosed | NO |
| TGIAIEQDKNTQ | 1514342 | Ectodomain | YES |
| TGIDFQDELDDF | 1514348 | Ectodomain | YES |
| TGIIRSSNQSLAGGIT | 1698083 | Ectodomain | YES |
| TGILDGDYAMQFFKMG | 1698084 | Cell/Virus Enclosed | NO |
| TGKLSDCLYFKN | 1514369 | Cell/Virus Enclosed | NO |
| TGKTVLGEYVFDKSEL | 1698098 | Cell/Virus Enclosed | NO |
| TGLCIAGDRGIA | 1514380 | Ectodomain | YES |
| TGLFKDCSRETSGLSP | 1698106 | Cell/Virus Enclosed | NO |
| TGLSTGTSQNTRISLRPGPV | 63887 | Capsid | UN |
| TGLTFQTVRSGN | 1514414 | Cell/Virus Enclosed | NO |
| TGLVKNQCVNFN | 1514420 | Ectodomain | YES |
| TGLYPTLNISEE | 1514422 | Cell/Virus Enclosed | NO |
| TGMAITIKPEAN | 1514423 | Cell/Virus Enclosed | NO |
| TGMQSWVPLSTDDPV | 63895 | Ectodomain | YES |
| TGNVVQQCCTAA | 1514455 | Cell/Virus Enclosed | NO |
| TGNYNYKFRYLR | 1514457 | Ectodomain | YES |
| TGNYNYKHRYLR | 1514458 | Ectodomain | YES |
| TGNYNYKYRYLK | 1514462 | Ectodomain | YES |
| TGPEAALPFRAVKDGI | 1698146 | Cell/Virus Enclosed | NO |
| TGPEAGLPYGANKEGI | 1698148 | Cell/Virus Enclosed | NO |
| TGPHKDAKFRER | 1514478 | Cell/Virus Enclosed | NO |
| TGPPPISTDTVDVTNG | 1698153 | Ectodomain | YES |
| TGPPQCDQFLEFSADLIIERREGSD | 1173378 | Ectodomain | YES |
| TGPYAKDQYGTD | 1514489 | Cell/Virus Enclosed | NO |
| TGPYANASYGQS | 1514491 | Cell/Virus Enclosed | NO |
| TGPYASSSYGEA | 1514493 | Cell/Virus Enclosed | NO |
| TGPYKCPQSKSL | 1514496 | Ectodomain | YES |
| TGQAGEMPCAIINDKV | 1698156 | Cell/Virus Enclosed | NO |
| TGQGIFKEVSAV | 1514502 | Ectodomain | YES |
| TGRCTVNKHVAY | 1514508 | Cell/Virus Enclosed | NO |
| TGRGDCKGFSSDVLSD | 1698162 | Ectodomain | YES |
| TGRPSHEAPNMTQ | 63953 | Ectodomain | YES |
| TGRSVYVKFQDSKPPL | 1698167 | Cell/Virus Enclosed | NO |
| TGSAFDGTMYGA | 1514529 | Cell/Virus Enclosed | NO |
| TGSFNNIDTGPP | 1514540 | Ectodomain | YES |
| TGSLADGLEAYADKTL | 1698177 | Cell/Virus Enclosed | NO |
| TGSLADGLGAYA | 1514551 | Cell/Virus Enclosed | NO |
| TGSNDSKALFFT | 1514560 | Cell/Virus Enclosed | NO |
| TGSNQNGERSGARPKQ | 1698183 | Cell/Virus Enclosed | NO |
| TGSNQNGERSGARSKQRRPQG | 1870816 | Capsid | UN |
| TGSPYIDDFNKIASCK | 1698186 | Cell/Virus Enclosed | NO |
| TGSPYIDDFNRIASCK | 1698187 | Cell/Virus Enclosed | NO |
| TGSRSARSAIEDLLFDKV | 1869859 | Ectodomain | YES |
| TGSRYYYPEPIT | 1514573 | Ectodomain | YES |
| TGSSFYAPEPITSLNT | 1698189 | Ectodomain | YES |
| TGSSFYEPEPIT | 1514578 | Ectodomain | YES |
| TGSSFYSPEPIT | 1514579 | Ectodomain | YES |
| TGSTPQKLFVAN | 1514587 | Ectodomain | YES |
| TGSVVKPVRARAPPGE | 1698194 | Cell/Virus Enclosed | NO |
| TGSVYGNFDDQPSLQV | 1698196 | Cell/Virus Enclosed | NO |
| TGSWNKVTQIAN | 1514596 | Cell/Virus Enclosed | NO |
| TGSWWSFNPETN | 1514598 | Ectodomain | YES |
| TGTAFNGDFYGP | 1514603 | Cell/Virus Enclosed | NO |
| TGTDFNGDFYGPYKDA | 1698200 | Cell/Virus Enclosed | NO |
| TGTDFNGNFYGP | 1514606 | Cell/Virus Enclosed | NO |
| TGTDLELIQILKDHYNISLD | 63973 | Capsid | UN |
| TGTFTVVMRPNYTIKG | 1698202 | Cell/Virus Enclosed | NO |
| TGTGPEAALPFRAVKD | 1698204 | Cell/Virus Enclosed | NO |
| TGTIYAELEPPC | 1514624 | Cell/Virus Enclosed | NO |
| TGTLIVNSVLPF | 1514632 | Undetermined | UN |
| TGTPKPSRNQSPASSQ | 1698213 | Capsid | UN |
| TGTQSVSVDREL | 1514645 | Cell/Virus Enclosed | NO |
| TGTVIIRPSTSA | 1514657 | Ectodomain | YES |
| TGTVIISPFTSA | 1514658 | Ectodomain | YES |
| TGTYKCPQTKSL | 1514662 | Ectodomain | YES |
| TGVAIEQLLYAI | 1514665 | Cell/Virus Enclosed | NO |
| TGVCYDFDGVTTSDKC | 1698222 | Cell/Virus Enclosed | NO |
| TGVDGELPVAVVNDKV | 1698224 | Cell/Virus Enclosed | NO |
| TGVIADYNYKLPDDFM | 1698233 | Ectodomain | YES |

| | | | |
|---|---|---|---|
| TGVIYGFNSTET | 1514689 | Ectodomain | YES |
| TGVIYGLNFTET | 1514690 | Ectodomain | YES |
| TGVLTTSSKRFQ | 1514701 | Ectodomain | YES |
| TGVNDDDYVIAL | 1514706 | Cell/Virus Enclosed | NO |
| TGVNFVVQPVGV | 1514708 | Cell/Virus Enclosed | NO |
| TGVSIEILLAAI | 1514720 | Cell/Virus Enclosed | NO |
| TGVVLKRSLKYNLKAS | 1698260 | Cell/Virus Enclosed | NO |
| TGVVYVNATGSD | 1514730 | Ectodomain | YES |
| TGWLLYQLLNGF | 1514739 | Cell/Virus Enclosed | NO |
| TGYFPKAGANFR | 1514743 | Ectodomain | YES |
| TGYGVRRKNSEPEIPH | 1698270 | Capsid | UN |
| TGYHFTKNGKTVLGEY | 1698272 | Cell/Virus Enclosed | NO |
| TGYHFTSTGKTVLGEY | 1698273 | Cell/Virus Enclosed | NO |
| TGYHITKNSKVQLGEY | 1698274 | Cell/Virus Enclosed | NO |
| TGYLKLTDNVAI | 1514749 | Cell/Virus Enclosed | NO |
| TGYRFTNFEPFTVNSV | 1698282 | Ectodomain | YES |
| TGYSLADLPAYMTVAK | 1698285 | Ectodomain | YES |
| TGYSWADLPAYM | 1514762 | Ectodomain | YES |
| TGYTLADLPVYV | 1514763 | Ectodomain | YES |
| THCAFAVDPAAA | 1514783 | Cell/Virus Enclosed | NO |
| THCSFAVDPAAA | 1514785 | Cell/Virus Enclosed | NO |
| THCSFAVDPAVA | 1514786 | Cell/Virus Enclosed | NO |
| THDGPNEHVTSV | 1514798 | Cell/Virus Enclosed | NO |
| THGAAAINSVPW | 1514818 | Cell/Virus Enclosed | NO |
| THGFNLEDAARC | 1514819 | Cell/Virus Enclosed | NO |
| THGLNLEEAARC | 1514825 | Cell/Virus Enclosed | NO |
| THGVAAINSVPW | 1514830 | Cell/Virus Enclosed | NO |
| THHAHSPAYPTLELGLARQP | 742040 | Ectodomain | YES |
| THHWMLLTFLTS | 1514837 | Cell/Virus Enclosed | NO |
| THIAEDVVDQVE | 1514838 | Cell/Virus Enclosed | NO |
| THLKHGGGIAGAINAA | 1698328 | Cell/Virus Enclosed | NO |
| THLKHGGGVAGA | 1514860 | Cell/Virus Enclosed | NO |
| THLSHYVFNIDW | 1514866 | Cell/Virus Enclosed | NO |
| THMNLKYAISAK | 1514875 | Cell/Virus Enclosed | NO |
| THNGRVVNYTVC | 1514876 | Ectodomain | YES |
| THQNNDDAGNPV | 1514884 | Cell/Virus Enclosed | NO |
| THQNNDDHGNPV | 1514885 | Cell/Virus Enclosed | NO |
| THRDCDFNKPLIEWPL | 1698340 | Cell/Virus Enclosed | NO |
| THWFCANQSTSVYSAN | 1698360 | Ectodomain | YES |
| THYIKDGDDGYF | 1514940 | Cell/Virus Enclosed | NO |
| THYVRVTFSDSN | 1514954 | Cell/Virus Enclosed | NO |
| TIADFVQIENCD | 1514955 | Ectodomain | YES |
| TIADFVQIENCN | 1514956 | Ectodomain | YES |
| TIALNKIQDVVNQQGS | 1698377 | Ectodomain | YES |
| TIANNYSKLFDEGYTV | 1698379 | Cell/Virus Enclosed | NO |
| TIATHGIAAINS | 1514989 | Cell/Virus Enclosed | NO |
| TICDFYSKLGVGFTSH | 1698390 | Cell/Virus Enclosed | NO |
| TICDLYSIHDVG | 1515002 | Cell/Virus Enclosed | NO |
| TICDLYSIQDVG | 1515003 | Cell/Virus Enclosed | NO |
| TICDLYSMQDVG | 1515004 | Cell/Virus Enclosed | NO |
| TICGIKYVAQPTEDVV | 1698394 | Cell/Virus Enclosed | NO |
| TICHPNLGNHRK | 1515012 | Ectodomain | YES |
| TICIKDADYNAKVEIS | 1698396 | Cell/Virus Enclosed | NO |
| TIDCSAFVCGDC | 1515034 | Ectodomain | YES |
| TIDELCVNLNLP | 1515038 | Cell/Virus Enclosed | NO |
| TIDGIVEDTIND | 1515039 | Cell/Virus Enclosed | NO |
| TIDGSSGVANPAMDPI | 1698401 | Secreted | YES |
| TIDKFAIPNGRKVDLQ | 1698406 | Ectodomain | YES |
| TIDKFAIPNRRK | 1515048 | Ectodomain | YES |
| TIDKLCDYLADY | 1515052 | Cell/Virus Enclosed | NO |
| TIDKLRDYLADY | 1515053 | Cell/Virus Enclosed | NO |
| TIDSNTTQDTYGGASV | 1698407 | Cell/Virus Enclosed | NO |
| TIDSTQDGVNKL | 1515060 | Ectodomain | YES |
| TIEDALRNSAML | 1515067 | Ectodomain | YES |
| TIEDALRNSARLESAD | 1698412 | Ectodomain | YES |
| TIEDLRARMTYV | 1515068 | Cell/Virus Enclosed | NO |
| TIEDWLLVQGFS | 1515069 | Cell/Virus Enclosed | NO |
| TIEENVIQCFGP | 1515072 | Cell/Virus Enclosed | NO |
| TIFNCFYALNNAFLAF | 1698424 | Undetermined | UN |
| TIFYTTEIASKF | 1515096 | Cell/Virus Enclosed | NO |
| TIGDSIFTGCTG | 1515097 | Cell/Virus Enclosed | NO |
| TIGNETTSAFVGALPK | 1698430 | Ectodomain | YES |
| TIGNIEEFIQTS | 1515107 | Ectodomain | YES |
| TIGNMEEFIQTN | 1515108 | Ectodomain | YES |
| TIGNMEEFIQTS | 1515109 | Ectodomain | YES |
| TIHSVCNGCVGH | 1515130 | Ectodomain | YES |

| | | | |
|---|---|---|---|
| TIIFVQQKPEPVNPVS | 1698441 | Cell/Virus Enclosed | NO |
| TIIIPDKQVFDK | 1515133 | Cell/Virus Enclosed | NO |
| TIILTTFNCVYA | 1515138 | Undetermined | UN |
| TIISTQDGVNKL | 1515143 | Ectodomain | YES |
| TIKAKEESVKSSYAFA | 1698452 | Cell/Virus Enclosed | NO |
| TIKENIDGGAMHANYI | 1698455 | Cell/Virus Enclosed | NO |
| TIKFSGFKLDGFNHQF | 1698456 | Cell/Virus Enclosed | NO |
| TIKPEANINQDS | 1515169 | Cell/Virus Enclosed | NO |
| TIKPLEHHHHHH | 1515170 | Cell/Virus Enclosed | NO |
| TIKPVSLEHHHH | 1515171 | Cell/Virus Enclosed | NO |
| TIKVSTVQSKLTDLKC | 1698475 | Cell/Virus Enclosed | NO |
| TIKVTEDGVNVHDVTV | 1698476 | Cell/Virus Enclosed | NO |
| TILNIVGPDARQDGRQ | 1698493 | Cell/Virus Enclosed | NO |
| TILNTACGVFEVDDTV | 1698496 | Cell/Virus Enclosed | NO |
| TILPCYSGRVSA | 1515225 | Ectodomain | YES |
| TILTRPLLESELVIG | 1542785 | Underneath Membrane | NO |
| TIMKVLNENLNAYQQQ | 1698508 | Cell/Virus Enclosed | NO |
| TINCESIYEFDT | 1515250 | Cell/Virus Enclosed | NO |
| TINDDEDVVTGDNDDE | 1698519 | Cell/Virus Enclosed | NO |
| TINNFNPSSWNR | 1515265 | Ectodomain | YES |
| TINPSSPARLEI | 1515269 | Cell/Virus Enclosed | NO |
| TINSQDGVNKLQ | 1515272 | Ectodomain | YES |
| TINSTQDGVNKL | 1515273 | Ectodomain | YES |
| TINSTQDGYNKL | 1515274 | Ectodomain | YES |
| TINSVVQKDGVENTPP | 1698530 | Cell/Virus Enclosed | NO |
| TIPCICGRDATQ | 1515285 | Cell/Virus Enclosed | NO |
| TIPDEKQNGHSL | 1515287 | Cell/Virus Enclosed | NO |
| TIQDSDGTNKQLNEIS | 1698546 | Cell/Virus Enclosed | NO |
| TIQTVFDCAVET | 1515338 | Cell/Virus Enclosed | NO |
| TIRGQFSNMSLSLLD | 64444 | Ectodomain | YES |
| TIRLQAGKQTELAVNS | 1698570 | Cell/Virus Enclosed | NO |
| TIRLQAGVATEYAANS | 1698571 | Cell/Virus Enclosed | NO |
| TIRYQATGFGTNETE | 187549 | Secreted | YES |
| TISSYKLDLTSL | 1515368 | Cell/Virus Enclosed | NO |
| TISTGFDFNCHY | 1515370 | Ectodomain | YES |
| TITDAIDCAQNP | 1515380 | Ectodomain | YES |
| TITDAVDCSQNPLAELK | 64477 | Ectodomain | YES |
| TITIGSTFVNTS | 1515389 | Ectodomain | YES |
| TITSAGRPCVAS | 1515408 | Cell/Virus Enclosed | NO |
| TITTGFDFNCHY | 1515412 | Ectodomain | YES |
| TITVDQLKHLLE | 1515415 | Undetermined | UN |
| TITVEKLKQLLE | 1515418 | Undetermined | UN |
| TIVDIPQSLTFSYDGL | 1698608 | Cell/Virus Enclosed | NO |
| TIVGQEEFIQTNSPKV | 1698615 | Ectodomain | YES |
| TIVIPDKQVFDKVVDN | 1698617 | Cell/Virus Enclosed | NO |
| TIVKCKGLNKPFVKDN | 1698618 | Cell/Virus Enclosed | NO |
| TIVPEYIDVNKTLQEL | 1698625 | Ectodomain | YES |
| TIVSEKISVMDKLDTG | 1698627 | Cell/Virus Enclosed | NO |
| TIYAELEPPCRF | 1515486 | Cell/Virus Enclosed | NO |
| TIYGNVSRQDLTKYTM | 1698643 | Cell/Virus Enclosed | NO |
| TIYMCVVPSINDLSFD | 1698645 | Cell/Virus Enclosed | NO |
| TIYNLVKECPAV | 1515495 | Cell/Virus Enclosed | NO |
| TKAAGGKVSFSDDVEV | 1698654 | Cell/Virus Enclosed | NO |
| TKAIDGENVSKEGFFT | 1698657 | Cell/Virus Enclosed | NO |
| TKALCFDKSKQFVPVQ | 1698665 | Underneath Membrane | NO |
| TKALKELYGPID | 1515528 | Cell/Virus Enclosed | NO |
| TKALKELYGPVDPTFL | 1698666 | Cell/Virus Enclosed | NO |
| TKALKELYGSVD | 1515531 | Cell/Virus Enclosed | NO |
| TKAPLVYLNHSVPKLS | 1698668 | Ectodomain | YES |
| TKAPYVMLNTSIPNLP | 1698669 | Ectodomain | YES |
| TKARATAPTTRNLLTTPKFT | 742045 | Ectodomain | YES |
| TKCNIGGAVCSKHANL | 1698673 | Cell/Virus Enclosed | NO |
| TKDEAIKRVRGWVGFD | 1698677 | Cell/Virus Enclosed | NO |
| TKDSKFQVGEFVFEKV | 1698679 | Cell/Virus Enclosed | NO |
| TKDSYCRLKNSVSDVA | 1698681 | Cell/Virus Enclosed | NO |
| TKEGDSSNTGAGKALTGLST | 64547 | Capsid | UN |
| TKEQKHEMQKPRWKRQPNDD | 2060898 | Capsid | UN |
| TKFQGLENIAFN | 1515592 | Cell/Virus Enclosed | NO |
| TKFVGANFGRWS | 1515598 | Ectodomain | YES |
| TKFYGGWDNMLKNLMA | 1698715 | Cell/Virus Enclosed | NO |
| TKGEGDGGITSEGNAL | 1698717 | Cell/Virus Enclosed | NO |
| TKGKPVKGAWNI | 1515619 | Cell/Virus Enclosed | NO |
| TKGLSIADLACAQYYN | 1698724 | Ectodomain | YES |
| TKGPYEFCSQHT | 1515624 | Cell/Virus Enclosed | NO |
| TKGVMEPIDYNV | 1515631 | Cell/Virus Enclosed | NO |
| TKGVMEPVDYNV | 1515632 | Cell/Virus Enclosed | NO |

| | | | |
|---|---|---|---|
| TKGVTFGEDTVL | 1515633 | Cell/Virus Enclosed | NO |
| TKHKCDINYKGKVFFQ | 1698739 | Cell/Virus Enclosed | NO |
| TKIFKDDGNFVV | 1515661 | Cell/Virus Enclosed | NO |
| TKKAFPYGTCLNCNAP | 1698758 | Cell/Virus Enclosed | NO |
| TKKDWYDFVENPDIIN | 1698762 | Cell/Virus Enclosed | NO |
| TKKGIAIGPGRTLY | 163621 | Ectodomain | YES |
| TKKKYSELDDEE | 1515694 | Cell/Virus Enclosed | NO |
| TKKMKKALLSTPMKWGKKLY | 1954169 | Capsid | UN |
| TKKPNRNGGGYYSASYSDPCSL | 64602 | Ectodomain | YES |
| TKKSLNVDGVQY | 1515704 | Cell/Virus Enclosed | NO |
| TKKWKFPQIGGL | 1515707 | Cell/Virus Enclosed | NO |
| TKKYTYSFLPKEDGDV | 1698771 | Cell/Virus Enclosed | NO |
| TKLAGGKISFSDDVIV | 1698773 | Cell/Virus Enclosed | NO |
| TKLASTEELPDE | 1515716 | Cell/Virus Enclosed | NO |
| TKLKDGVNFNVDDINF | 1698778 | Ectodomain | YES |
| TKLLFNVYLGMA | 1515740 | Ectodomain | YES |
| TKLMCCIGPDIF | 1515746 | Cell/Virus Enclosed | NO |
| TKLNDLCFSNVYADSF | 1698784 | Ectodomain | YES |
| TKLNGILQQAALDYLA | 1698786 | Cell/Virus Enclosed | NO |
| TKLPYYDVKKIH | 1515756 | Underneath Membrane | NO |
| TKLSKYLANEAQVQLE | 1698793 | Cell/Virus Enclosed | NO |
| TKMFDSFVNSFVSLYN | 1698805 | Cell/Virus Enclosed | NO |
| TKNGKTVLGEYVFDKS | 1698808 | Cell/Virus Enclosed | NO |
| TKNGLMYSEFPA | 1515782 | Ectodomain | YES |
| TKNIGDYNVSKDGFFT | 1698809 | Cell/Virus Enclosed | NO |
| TKNIGEYNVSKDGFFT | 1698810 | Cell/Virus Enclosed | NO |
| TKNITDKIDQIIHDFVDKTL | 931784 | Secreted | YES |
| TKNVTGSNDSKA | 1515790 | Cell/Virus Enclosed | NO |
| TKNVTGSNESKALFFT | 1698814 | Cell/Virus Enclosed | NO |
| TKPFVRAAFEHL | 1515799 | Cell/Virus Enclosed | NO |
| TKPIIKAQFKPFAKVD | 1698819 | Cell/Virus Enclosed | NO |
| TKPQPLMDKFSS | 1515811 | Ectodomain | YES |
| TKPTDPTGIEPDDH | 64645 | Capsid | UN |
| TKQYNVTQAFGRRGPE | 1698833 | Cell/Virus Enclosed | NO |
| TKRNILPTMTQLNLKY | 1698841 | Cell/Virus Enclosed | NO |
| TKRNVIPTMTQMNLKY | 1698843 | Cell/Virus Enclosed | NO |
| TKRNVLPTLTQMNLKY | 1698844 | Cell/Virus Enclosed | NO |
| TKRNVLPTMTQLNLKY | 1698845 | Cell/Virus Enclosed | NO |
| TKSAAGYGGFGC | 1515852 | Cell/Virus Enclosed | NO |
| TKSCGVVAEHDF | 1515854 | Cell/Virus Enclosed | NO |
| TKSGTNDICVFE | 1515860 | Cell/Virus Enclosed | NO |
| TKSGVNDICIFE | 1515861 | Cell/Virus Enclosed | NO |
| TKSGVNDICMFENKTT | 1698852 | Cell/Virus Enclosed | NO |
| TKSLLPNTGVYDLSGF | 1698853 | Ectodomain | YES |
| TKSWSVGDWTGF | 1515878 | Cell/Virus Enclosed | NO |
| TKTAVKSGTAIL | 1515880 | Cell/Virus Enclosed | NO |
| TKTDAVILLERYVSLA | 1698864 | Cell/Virus Enclosed | NO |
| TKTHATLFGSVA | 1515884 | Ectodomain | YES |
| TKTPHIVRQRLT | 1515888 | Cell/Virus Enclosed | NO |
| TKVDGNDVEIFE | 1515896 | Cell/Virus Enclosed | NO |
| TKVFKDGVMYSE | 1515907 | Ectodomain | YES |
| TKVFKEGVMYSE | 1515908 | Ectodomain | YES |
| TKVIKDGVMYSE | 1515911 | Ectodomain | YES |
| TKVLAEKFDYNVSVMQ | 1698877 | Cell/Virus Enclosed | NO |
| TKVLRYGYHVLY | 1515920 | Cell/Virus Enclosed | NO |
| TKVTKNGLMYSE | 1515933 | Ectodomain | YES |
| TKVTVPKDHPHL | 1515934 | Cell/Virus Enclosed | NO |
| TKVVVGPTTEVKFSVI | 1698886 | Cell/Virus Enclosed | NO |
| TKYNIYDGSGVG | 1515945 | Ectodomain | YES |
| TKYNIYDVAGVG | 1515946 | Ectodomain | YES |
| TKYNIYDVSGVGVIRV | 1698889 | Ectodomain | YES |
| TKYNIYDYVGTGIIRS | 1698890 | Ectodomain | YES |
| TKYPLLKLLGSTWPTTPPRP | 110434 | Cell/Virus Enclosed | NO |
| TKYTEVRASRQLAQQK | 1698891 | Ectodomain | YES |
| TKYVAPHVTYQN | 1515959 | Ectodomain | YES |
| TKYVAPQATYQN | 1515960 | Ectodomain | YES |
| TKYVAPQVTYQNISTN | 1698896 | Ectodomain | YES |
| TKYYDDSQYYFNKDTG | 1698900 | Ectodomain | YES |
| TKYYDHSQYYLN | 1515971 | Ectodomain | YES |
| TLAATSASLFPPWTAA | 1698901 | Ectodomain | YES |
| TLACFVLAAVYRINW | 1310849 | Undetermined | UN |
| TLACFVLAVVYR | 1515981 | Transmembrane | NO |
| TLADAGFMKQYGECLG | 1698906 | Ectodomain | YES |
| TLAKTDLFNGYKIGCN | 1698909 | Cell/Virus Enclosed | NO |
| TLALLNLTDEDL | 1515993 | Cell/Virus Enclosed | NO |
| TLANFNETKGPLCVDT | 1698912 | Ectodomain | YES |

| | | | |
|---|---|---|---|
| TLANTTTAPLKL | 1515999 | Cell/Virus Enclosed | NO |
| TLAVIGDFNCTN | 1516008 | Ectodomain | YES |
| TLDDILQHANKS | 1516036 | Cell/Virus Enclosed | NO |
| TLDDILQQANKSVGII | 1698938 | Cell/Virus Enclosed | NO |
| TLDDRGLALDNALSGV | 1698940 | Cell/Virus Enclosed | NO |
| TLDDVTRNYIIT | 1516040 | Cell/Virus Enclosed | NO |
| TLDGYSKLFITKDEAI | 1698943 | Cell/Virus Enclosed | NO |
| TLDKCTKYNIYDVSGV | 1698944 | Ectodomain | YES |
| TLDKLSDYCTFV | 1516049 | Cell/Virus Enclosed | NO |
| TLDNQDLNGKWYDFGD | 1698949 | Cell/Virus Enclosed | NO |
| TLEDLHARMTYT | 1516067 | Cell/Virus Enclosed | NO |
| TLEEAKTALRKC | 1516070 | Cell/Virus Enclosed | NO |
| TLEEAVRQVRSW | 1516072 | Cell/Virus Enclosed | NO |
| TLELGLARQPLLRVRTATRD | 742048 | Ectodomain | YES |
| TLENKSAELNYTVQKL | 1698960 | Ectodomain | YES |
| TLEQYSTCDMCK | 1516090 | Cell/Virus Enclosed | NO |
| TLESGKVKQSQP | 1516093 | Cell/Virus Enclosed | NO |
| TLESITAQPVKI | 1516094 | Secreted | YES |
| TLESTINKSVANLKLP | 1698968 | Cell/Virus Enclosed | NO |
| TLFAGTHITMTT | 1516114 | Cell/Virus Enclosed | NO |
| TLFGDCVTVRVGQQVD | 1698981 | Cell/Virus Enclosed | NO |
| TLGGDPKFRSLTHEDHAIQP | 64815 | Capsid | UN |
| TLGNVEAVNFNVTTAE | 1698997 | Ectodomain | YES |
| TLGSITAQPVKI | 1516164 | Secreted | YES |
| TLGVLVASAAPSSPGTPGVA | 742050 | Undetermined | UN |
| TLIDNINSTLVDLKWL | 1699016 | Ectodomain | YES |
| TLIPQENYSSIR | 1516214 | Cell/Virus Enclosed | NO |
| TLIVFTSCELYH | 1516220 | Ectodomain | YES |
| TLKEVVGPKELLLSWE | 1699038 | Cell/Virus Enclosed | NO |
| TLKHFFFAQDGN | 1516243 | Cell/Virus Enclosed | NO |
| TLKIKVSQTNMY | 1516255 | Cell/Virus Enclosed | NO |
| TLKNTVCTVCGT | 1516263 | Cell/Virus Enclosed | NO |
| TLKPGDSFNILACYEG | 1699057 | Cell/Virus Enclosed | NO |
| TLKYFDGFNFSQ | 1516278 | Ectodomain | YES |
| TLKYFGGFNFSQILPDP | 64888 | Ectodomain | YES |
| TLLDLQVEMNRL | 1516291 | Ectodomain | YES |
| TLLDNQEDKFWC | 1516297 | Cell/Virus Enclosed | NO |
| TLLPAADLDDFSKQLQQSMSSADS | 1516334 | Capsid | UN |
| TLLVPQIFKARA | 1516369 | Cell/Virus Enclosed | NO |
| TLLYSFEAHKDF | 1516374 | Cell/Virus Enclosed | NO |
| TLMNLKYAISAK | 1516383 | Cell/Virus Enclosed | NO |
| TLMQGVTLSSNLNTNL | 1699109 | Ectodomain | YES |
| TLMTFSGNVDLDALAS | 1699111 | Cell/Virus Enclosed | NO |
| TLNDVKDNDGRPVHVK | 1699117 | Cell/Virus Enclosed | NO |
| TLNEDLMEILNR | 1516397 | Cell/Virus Enclosed | NO |
| TLNIIVPDKSVYDQVV | 1699125 | Cell/Virus Enclosed | NO |
| TLNKDQLLSSSKYTIQRSTG | 64978 | Capsid | UN |
| TLNLEGVNGGSLYVNN | 1699127 | Cell/Virus Enclosed | NO |
| TLPDIAEDVVDQ | 1516424 | Cell/Virus Enclosed | NO |
| TLPFNWTHCFDPQIQAIVS | 65003 | Ectodomain | YES |
| TLPQSKDSNFLN | 1516439 | Cell/Virus Enclosed | NO |
| TLQAAFGNTGVC | 1516446 | Cell/Virus Enclosed | NO |
| TLQEFAQNLPKYVKPN | 1699154 | Ectodomain | YES |
| TLQELSYKLPNYTVPD | 1699155 | Ectodomain | YES |
| TLQEMKALFPTWSQEL | 1699156 | Cell/Virus Enclosed | NO |
| TLQETPVVPDTVEVPP | 1699157 | Cell/Virus Enclosed | NO |
| TLQLVSAVNQSLSNAK | 1699161 | Cell/Virus Enclosed | NO |
| TLQPNAYYKYDAKNYV | 1699163 | Cell/Virus Enclosed | NO |
| TLQQGAFLCANE | 1516485 | Cell/Virus Enclosed | NO |
| TLQSGLKKMAQP | 1516488 | Cell/Virus Enclosed | NO |
| TLQSLENVIYNLVNVG | 1699169 | Cell/Virus Enclosed | NO |
| TLRATEEASRQS | 1516498 | Cell/Virus Enclosed | NO |
| TLRCCTVTYLNELSSK | 1699175 | Cell/Virus Enclosed | NO |
| TLRNGSVLEMAKEVKT | 1699182 | Cell/Virus Enclosed | NO |
| TLRSITAQPVKIDNAS | 1699189 | Secreted | YES |
| TLSIHDGYTNEI | 1516563 | Cell/Virus Enclosed | NO |
| TLSLLDQYLLAF | 1516567 | Ectodomain | YES |
| TLSSNLNTNLHFDVDN | 1699201 | Ectodomain | YES |
| TLSTKLKDGVNFNVDD | 1699205 | Ectodomain | YES |
| TLSTYDFYPNVP | 1516578 | Ectodomain | YES |
| TLSVDFNGVLHK | 1516581 | Cell/Virus Enclosed | NO |
| TLSYYKLGASQRVAGDSGFA | 1389743 | Underneath Membrane | NO |
| TLSYYKLGASQRVGTDSGFA | 1389744 | Underneath Membrane | NO |
| TLTAPTIANQER | 1516593 | Cell/Virus Enclosed | NO |
| TLTAPTIVNQERYVKI | 1699216 | Cell/Virus Enclosed | NO |
| TLTKLSKYLANEAQVQ | 1699221 | Cell/Virus Enclosed | NO |

| | | | |
|---|---|---|---|
| TLTLDKVPQAVE | 1516613 | Cell/Virus Enclosed | NO |
| TLTPCSVRSVPG | 1516627 | Ectodomain | YES |
| TLTPRSVRSVPGEMRL | 1699228 | Ectodomain | YES |
| TLTQMNLKYAISAKNR | 1699229 | Cell/Virus Enclosed | NO |
| TLTRGSDLDRCT | 1516637 | Ectodomain | YES |
| TLTSGSDLDRCTTFDDV | 65110 | Undetermined | UN |
| TLTTPTIVNQER | 1516645 | Cell/Virus Enclosed | NO |
| TLVAPDLSLDYI | 1516656 | Ectodomain | YES |
| TLVDLKWLNRVETYIK | 1699242 | Ectodomain | YES |
| TLVDLNWLNRVE | 1516664 | Ectodomain | YES |
| TLVGQLEQTNMH | 1516671 | Cell/Virus Enclosed | NO |
| TLVKQLSFNFGA | 1516680 | Ectodomain | YES |
| TLVNKMVNWFNTMLDA | 1699258 | Cell/Virus Enclosed | NO |
| TLVSGDAHIPFGTVLG | 1699264 | Ectodomain | YES |
| TLVSGDDHIPSG | 1516710 | Ectodomain | YES |
| TLWNYLFETIPT | 1516725 | Cell/Virus Enclosed | NO |
| TLWYQKPFLSDF | 1516729 | Ectodomain | YES |
| TLYFNNEPAFIM | 1516732 | Cell/Virus Enclosed | NO |
| TLYGQAVVYFGG | 1516738 | Cell/Virus Enclosed | NO |
| TLYIKDGDDGYFLPYP | 1699277 | Cell/Virus Enclosed | NO |
| TLYVKNWCRCSC | 1516755 | Ectodomain | YES |
| TMCDIKQMLFCM | 1516771 | Cell/Virus Enclosed | NO |
| TMCEYPHTICKS | 1516776 | Ectodomain | YES |
| TMCEYPNTICHP | 1516778 | Ectodomain | YES |
| TMDGFCQYKGKW | 1516788 | Cell/Virus Enclosed | NO |
| TMDNKSQSVIII | 1516789 | Ectodomain | YES |
| TMENFELEKHCY | 1516794 | Cell/Virus Enclosed | NO |
| TMFDVYVDTFLS | 1516798 | Cell/Virus Enclosed | NO |
| TMFMDAEGRMTP | 1516804 | Cell/Virus Enclosed | NO |
| TMFRDAEGRITP | 1516811 | Cell/Virus Enclosed | NO |
| TMFTMADGSPQPYCYT | 1699317 | Cell/Virus Enclosed | NO |
| TMFTMADGSPQS | 1516815 | Cell/Virus Enclosed | NO |
| TMFTTADGSPQP | 1516816 | Cell/Virus Enclosed | NO |
| TMGVYDFKVSAAEFKY | 1699320 | Cell/Virus Enclosed | NO |
| TMHSINHDIFSL | 1516827 | Cell/Virus Enclosed | NO |
| TMIEFMLWCKDGQVQT | 1699327 | Cell/Virus Enclosed | NO |
| TMILSLHPVECD | 1516840 | Cell/Virus Enclosed | NO |
| TMIVDEWSYTGS | 1516847 | Ectodomain | YES |
| TMLDIGQARVAYQVAA | 1699332 | Cell/Virus Enclosed | NO |
| TMLDLCYALRHFDRND | 1699334 | Cell/Virus Enclosed | NO |
| TMLFGMIKKLDNDVLN | 1699336 | Cell/Virus Enclosed | NO |
| TMLFSMVRKLDNQALN | 1699337 | Cell/Virus Enclosed | NO |
| TMLVKMDGDDVYLPYP | 1699341 | Cell/Virus Enclosed | NO |
| TMMDLCFALRNFDEKD | 1699343 | Cell/Virus Enclosed | NO |
| TMPIGYVTHGLN | 1516879 | Cell/Virus Enclosed | NO |
| TMQIVDENGKYYLPYP | 1699353 | Cell/Virus Enclosed | NO |
| TMQIVDKDGNYY | 1516883 | Cell/Virus Enclosed | NO |
| TMQIVDKDGTYYLPYP | 1699354 | Cell/Virus Enclosed | NO |
| TMRYFVKIGPNK | 1516890 | Cell/Virus Enclosed | NO |
| TMSLGADSSIAYSNNT | 1699360 | Ectodomain | YES |
| TMSQHSRSTRSM | 1516901 | Ectodomain | YES |
| TMTGRVFHQKCL | 1516913 | Cell/Virus Enclosed | NO |
| TMTNRQFHQKLL | 1516916 | Cell/Virus Enclosed | NO |
| TMTQLNLKYAISGKER | 1699374 | Cell/Virus Enclosed | NO |
| TMTQMNLKYAISAKNR | 1699375 | Cell/Virus Enclosed | NO |
| TMTSQGMYGGTYPVE | 65269 | Ectodomain | YES |
| TMYGAFMDKQVH | 1516937 | Cell/Virus Enclosed | NO |
| TMYVRPIIEDYHTLTV | 1699390 | Ectodomain | YES |
| TNAETTDFFTVA | 1516949 | Ectodomain | YES |
| TNAKPPKYSSKVVLSG | 1699393 | Cell/Virus Enclosed | NO |
| TNANASSSEGFLIGIN | 1699394 | Cell/Virus Enclosed | NO |
| TNANASSSEGFLLGIN | 1699395 | Cell/Virus Enclosed | NO |
| TNCSDGNYNRNA | 1516994 | Ectodomain | YES |
| TNDICVFENKTT | 1517004 | Cell/Virus Enclosed | NO |
| TNDPIRSCLENT | 1517010 | Cell/Virus Enclosed | NO |
| TNEAFQKVQDAVNNNA | 1699434 | Ectodomain | YES |
| TNEIGLNASVTLKICK | 1699435 | Ectodomain | YES |
| TNESYNNYVPSYVKPD | 1699439 | Cell/Virus Enclosed | NO |
| TNFAINDKNTTV | 1517033 | Ectodomain | YES |
| TNFAINDLNTTI | 1517034 | Underneath Membrane | NO |
| TNFAINDLNTTVPRIS | 1699440 | Ectodomain | YES |
| TNFEPFTVNSIN | 1517038 | Ectodomain | YES |
| TNFEPFTVNSVNDSLE | 1699441 | Ectodomain | YES |
| TNFGDVIKPVCA | 1517043 | Cell/Virus Enclosed | NO |
| TNFGDVNKPVCA | 1517044 | Cell/Virus Enclosed | NO |
| TNFGEDFLGWIV | 1517047 | Cell/Virus Enclosed | NO |

| | | | |
|---|---|---|---|
| TNFPIDFPSAES | 1517056 | Cell/Virus Enclosed | NO |
| TNFRIDIPSAEL | 1517062 | Cell/Virus Enclosed | NO |
| TNFRIDVPFAEL | 1517063 | Cell/Virus Enclosed | NO |
| TNFRIDVPSAEL | 1517065 | Cell/Virus Enclosed | NO |
| TNFSFGVTQEYI | 1517069 | Ectodomain | YES |
| TNFTIVGQEEFIQTNS | 1699455 | Ectodomain | YES |
| TNGDYNSWHLVE | 1517090 | Cell/Virus Enclosed | NO |
| TNGHAVPTLFKL | 1517096 | Cell/Virus Enclosed | NO |
| TNGMNTSHSVCN | 1517104 | Ectodomain | YES |
| TNGMNTSQSVCN | 1517105 | Ectodomain | YES |
| TNGSGSGQAITCTIDS | 1699469 | Cell/Virus Enclosed | NO |
| TNGSWAIFNDHHLNRP | 1699472 | Cell/Virus Enclosed | NO |
| TNGTNTSHAVCN | 1517130 | Ectodomain | YES |
| TNGTNTSHSVCK | 1517131 | Ectodomain | YES |
| TNHDRYCSVHQGAHVA | 1699478 | Cell/Virus Enclosed | NO |
| TNHHAICNVHRNEHVA | 1699481 | Cell/Virus Enclosed | NO |
| TNHSFSVQKHIGAPAN | 1699484 | Cell/Virus Enclosed | NO |
| TNHSVVYAQHCF | 1517156 | Ectodomain | YES |
| TNIAFELFAKRKVGLT | 1699485 | Cell/Virus Enclosed | NO |
| TNILLNVPLRGS | 1517165 | Underneath Membrane | NO |
| TNIQNLLYCDSPFEKL | 1699495 | Ectodomain | YES |
| TNIYSQNLCVDGFGVN | 1699499 | Cell/Virus Enclosed | NO |
| TNKLLLFADING | 1517186 | Cell/Virus Enclosed | NO |
| TNKQLNEISDDCNWPL | 1699504 | Cell/Virus Enclosed | NO |
| TNKTTYKPNTWC | 1517189 | Cell/Virus Enclosed | NO |
| TNLCPFGEVFNATKFP | 1699508 | Ectodomain | YES |
| TNLEDPASRDLVVSY | 65360 | Capsid | UN |
| TNLHFDVDNINFKSLV | 1699518 | Ectodomain | YES |
| TNLPPPLLGNSI | 1517218 | Ectodomain | YES |
| TNLSRGRGRNPKPRAA | 1699530 | Cell/Virus Enclosed | NO |
| TNMSRGRGRNPK | 1517250 | Cell/Virus Enclosed | NO |
| TNNNGSDVLDQP | 1517264 | Cell/Virus Enclosed | NO |
| TNNTLVSGDAHIPFGT | 1699548 | Ectodomain | YES |
| TNNTLVSGDDHI | 1517269 | Ectodomain | YES |
| TNNTRIVDDWSY | 1517271 | Ectodomain | YES |
| TNNTRIVDEWSYTGSS | 1699549 | Ectodomain | YES |
| TNNTSIVDEWSY | 1517274 | Ectodomain | YES |
| TNNVFSLKGGAP | 1517279 | Cell/Virus Enclosed | NO |
| TNPDPKFVDKYY | 1517282 | Cell/Virus Enclosed | NO |
| TNPDPKFVDRYY | 1517283 | Cell/Virus Enclosed | NO |
| TNPFTRTVFENLKPMP | 1699553 | Cell/Virus Enclosed | NO |
| TNPRNYSYMDLN | 1517290 | Ectodomain | YES |
| TNQEDLLVLWGIHHPNDAAEQT | 97700 | Ectodomain | YES |
| TNQPRGRGRNPK | 1517298 | Cell/Virus Enclosed | NO |
| TNQVVKPGHFNEEFYN | 1699557 | Cell/Virus Enclosed | NO |
| TNRCPFDKVFNA | 1517302 | Ectodomain | YES |
| TNRFVSVGESFG | 1517307 | Cell/Virus Enclosed | NO |
| TNSALAKIQSVVNSNA | 1699564 | Ectodomain | YES |
| TNSALVKIQAVVNANA | 1699565 | Ectodomain | YES |
| TNSCLVSSDGQPGGDA | 1699566 | Cell/Virus Enclosed | NO |
| TNSCLVSSNGQP | 1517325 | Cell/Virus Enclosed | NO |
| TNSGPDDQIGFY | 1517331 | Capsid | UN |
| TNSGVAIYHRYK | 1517333 | Ectodomain | YES |
| TNSILMLDKQGQLLDT | 1699571 | Cell/Virus Enclosed | NO |
| TNSQYLFSTINALPET | 1699574 | Cell/Virus Enclosed | NO |
| TNSSKDDQIGYY | 1517347 | Cell/Virus Enclosed | NO |
| TNSTDTVNHDSFNLYG | 1699580 | Cell/Virus Enclosed | NO |
| TNSVCPKLEFANDTKI | 1699582 | Ectodomain | YES |
| TNSYSQNLCVDG | 1517363 | Cell/Virus Enclosed | NO |
| TNTAAFNAVCSV | 1517364 | Cell/Virus Enclosed | NO |
| TNTGSVVKPVRARAPP | 1699586 | Cell/Virus Enclosed | NO |
| TNTIHSVCNGCV | 1517368 | Ectodomain | YES |
| TNTNLSRDRGRN | 1517371 | Cell/Virus Enclosed | NO |
| TNTNLSRGRGRNPKPR | 1699588 | Cell/Virus Enclosed | NO |
| TNTRVSRYSSTARHRLRRGADGTAE | 65464 | Undetermined | UN |
| TNTSHAVCNGCV | 1517377 | Ectodomain | YES |
| TNTSHSVCKGCV | 1517378 | Ectodomain | YES |
| TNTSNSVCNGCV | 1517383 | Ectodomain | YES |
| TNTSSVVDGVVRSFQP | 1699590 | Ectodomain | YES |
| TNVAFELFAKRKMGLT | 1699591 | Cell/Virus Enclosed | NO |
| TNVDKIRHLLKG | 1517393 | Cell/Virus Enclosed | NO |
| TNVEKIRQLLKG | 1517397 | Cell/Virus Enclosed | NO |
| TNVGEPIISQSS | 1517401 | Cell/Virus Enclosed | NO |
| TNVVIRACIFEL | 1517414 | Ectodomain | YES |
| TNVVIRACNFELCDNPF | 65479 | Ectodomain | YES |
| TNVVLLGCLSSMNIAA | 1699611 | Cell/Virus Enclosed | NO |

| | | | |
|---|---|---|---|
| TNVVPPALHLVN | 1517417 | Cell/Virus Enclosed | NO |
| TNVYMRTTHTDI | 1517422 | Cell/Virus Enclosed | NO |
| TNYLEQPVSNDLSNC | 65486 | Ectodomain | YES |
| TNYLTCHNAAQC | 1517432 | Ectodomain | YES |
| TNYVGVYDINHGDAGF | 1699634 | Ectodomain | YES |
| TPAAPRAVSFAD | 1517443 | Cell/Virus Enclosed | NO |
| TPADILDRDPSS | 1517446 | Cell/Virus Enclosed | NO |
| TPADIVDRDPSN | 1517447 | Cell/Virus Enclosed | NO |
| TPADIVDRDPSS | 1517448 | Cell/Virus Enclosed | NO |
| TPADIVDRDQSS | 1517449 | Capsid | UN |
| TPAIDCSDGNYN | 1517458 | Ectodomain | YES |
| TPAIKDFGGFNF | 1517459 | Ectodomain | YES |
| TPAKKLCVVPGNVVPG | 1699638 | Cell/Virus Enclosed | NO |
| TPAPASGERAPPNSTRSASE | 742067 | Ectodomain | YES |
| TPAPVYIWTADEAIKF | 1699639 | Underneath Membrane | NO |
| TPAPVYTWTADE | 1517465 | Ectodomain | YES |
| TPAQNAGYWRRQDRKI | 1699640 | Cell/Virus Enclosed | NO |
| TPATDCSDGNYNRNAS | 1699642 | Ectodomain | YES |
| TPATNCSDGNYN | 1517471 | Ectodomain | YES |
| TPAWRIYSTGNNVFQT | 1699643 | Ectodomain | YES |
| TPAYDGSAFRDL | 1517476 | Cell/Virus Enclosed | NO |
| TPAYTFTTVKPS | 1517480 | Cell/Virus Enclosed | NO |
| TPCELHGNGSMI | 1517482 | Cell/Virus Enclosed | NO |
| TPCEVHGNGSMIEDID | 1699649 | Cell/Virus Enclosed | NO |
| TPCGFKGGIPGN | 1517487 | Cell/Virus Enclosed | NO |
| TPCGYSEPPKVAAKIC | 1699651 | Cell/Virus Enclosed | NO |
| TPCNPPDQLVVY | 1517497 | Ectodomain | YES |
| TPCNQPDQVAVYQQSI | 1699656 | Ectodomain | YES |
| TPCVYVDGLESKQVDY | 1699660 | Cell/Virus Enclosed | NO |
| TPDMADHIASLV | 1517506 | Cell/Virus Enclosed | NO |
| TPDMADQIASLV | 1517507 | Cell/Virus Enclosed | NO |
| TPDPITFKATGT | 1517508 | Ectodomain | YES |
| TPEDVNMQLMGV | 1517520 | Cell/Virus Enclosed | NO |
| TPESKPLADDDS | 1517526 | Cell/Virus Enclosed | NO |
| TPEVSRELGNITKTNV | 1699670 | Cell/Virus Enclosed | NO |
| TPFGSCIYWSPR | 1517535 | Cell/Virus Enclosed | NO |
| TPFHYERDNTSCPEWM | 1699676 | Cell/Virus Enclosed | NO |
| TPFHYERDNTSY | 1517537 | Cell/Virus Enclosed | NO |
| TPFYYERDNTSC | 1517546 | Cell/Virus Enclosed | NO |
| TPGAFFFGSKLDLVKR | 1699681 | Capsid | UN |
| TPGASRPTPPRTTTSPSSPR | 742069 | Ectodomain | YES |
| TPGFGVAVADSY | 1517550 | Cell/Virus Enclosed | NO |
| TPGHYAGSRSSSGNRS | 1699683 | Capsid | UN |
| TPGKQSSSRASF | 1517553 | Cell/Virus Enclosed | NO |
| TPGKQSSSRASSGNRS | 1699685 | Cell/Virus Enclosed | NO |
| TPGSSRGTSPARMAGN | 1699690 | Cell/Virus Enclosed | NO |
| TPGTNASSEFAV | 1517558 | Ectodomain | YES |
| TPHPYCYSDGVMKNAS | 1699698 | Cell/Virus Enclosed | NO |
| TPIDEIPYKSFF | 1517581 | Cell/Virus Enclosed | NO |
| TPIKLNFGMLNGMPVS | 1699706 | Cell/Virus Enclosed | NO |
| TPIKNTVDTTPKEEFV | 1699707 | Cell/Virus Enclosed | NO |
| TPILSCGIFGIKLETS | 1699709 | Cell/Virus Enclosed | NO |
| TPIVVDCSTYVCNGNV | 1699714 | Ectodomain | YES |
| TPIVWHAKDFNSLSAE | 1699716 | Cell/Virus Enclosed | NO |
| TPKDHIGTRNPANNAAIVLQ | 1392466 | Capsid | UN |
| TPKEEFVVKEKLNAFL | 1699719 | Cell/Virus Enclosed | NO |
| TPKGAINTSLPFQNI | 151064 | Ectodomain | YES |
| TPKHVFKTLKPGDSFN | 1699721 | Cell/Virus Enclosed | NO |
| TPKKPNKPFSQP | 1517619 | Cell/Virus Enclosed | NO |
| TPKKPNKPLSQPRADK | 1699723 | Capsid | UN |
| TPKLKEDYQIGGYSED | 1699724 | Secreted | YES |
| TPKPSRNQSPVS | 1517626 | Cell/Virus Enclosed | NO |
| TPKYTFGNVKPGETFT | 1699728 | Cell/Virus Enclosed | NO |
| TPLCGFPSQAAG | 1517636 | Cell/Virus Enclosed | NO |
| TPLFKIIGDEVGLRT | 65635 | Ectodomain | YES |
| TPLLYPSLALPAPHLTLPFNWTH | 65644 | Ectodomain | YES |
| TPLNLPPTQAHT | 1517657 | Cell/Virus Enclosed | NO |
| TPLPAADMDDFS | 1517659 | Capsid | UN |
| TPLSKRQYLLKF | 1517660 | Ectodomain | YES |
| TPLSVASAPVEP | 1517662 | Cell/Virus Enclosed | NO |
| TPLSVASTLVEP | 1517663 | Cell/Virus Enclosed | NO |
| TPLSVESTPVEL | 1517669 | Cell/Virus Enclosed | NO |
| TPLTSRQYLLAFNQDG | 1699742 | Ectodomain | YES |
| TPLVGTSLHPVL | 1517681 | Cell/Virus Enclosed | NO |
| TPLVGTSVHPVL | 1517682 | Cell/Virus Enclosed | NO |
| TPLYGFPSQAAG | 1517689 | Cell/Virus Enclosed | NO |

| | | | |
|---|---|---|---|
| TPNAVFISDRKI | 1517697 | Cell/Virus Enclosed | NO |
| TPNEKLVTTSMA | 1517699 | Cell/Virus Enclosed | NO |
| TPNKFCNVQQCF | 1517704 | Cell/Virus Enclosed | NO |
| TPNKHCNVQQCFGKRG | 1699753 | Capsid | UN |
| TPNLPDFKEELD | 1517708 | Ectodomain | YES |
| TPPAPNCYWPLN | 1517721 | Ectodomain | YES |
| TPPMPSIGLEEEE | 65691 | Ectodomain | YES |
| TPPPKEQRVQGS | 1517729 | Cell/Virus Enclosed | NO |
| TPPQPRKATKSKASTAKPAP | 742072 | Ectodomain | YES |
| TPPTTPVSPKVPPAFFQSMR | 65699 | Cell/Virus Enclosed | NO |
| TPQECNDMHLST | 1517736 | Cell/Virus Enclosed | NO |
| TPQKLFVANYSQDVKQ | 1699765 | Ectodomain | YES |
| TPRNKWNGVSDL | 1517747 | Cell/Virus Enclosed | NO |
| TPRNKWSGISDL | 1517749 | Cell/Virus Enclosed | NO |
| TPRQYLLAFNQD | 1517753 | Ectodomain | YES |
| TPRQYLLSFNQD | 1517754 | Ectodomain | YES |
| TPRSDMEYDFLNMDMG | 1699771 | Cell/Virus Enclosed | NO |
| TPRSVRSVPGEMRLAS | 1699772 | Ectodomain | YES |
| TPSDVSSRDPTTQEAI | 1699776 | Capsid | UN |
| TPSDVYQQLAGIKLQS | 1699777 | Cell/Virus Enclosed | NO |
| TPSMDALSAQLYSSLSLDSP | 78863 | Cell/Virus Enclosed | NO |
| TPSPLTPRSVRS | 1517774 | Ectodomain | YES |
| TPSSKRFQPFQQFGRDVSDFTDSVR | 1389873 | Ectodomain | YES |
| TPSTDCSDGNYN | 1517779 | Ectodomain | YES |
| TPSTLTLRSVRS | 1517784 | Ectodomain | YES |
| TPSTLTPRSVRSVPGE | 1699786 | Ectodomain | YES |
| TPTEHVTSVNLF | 1517790 | Cell/Virus Enclosed | NO |
| TPTESYVDVGLD | 1517792 | Ectodomain | YES |
| TPTESYVDVGPDSVKS | 1699790 | Ectodomain | YES |
| TPTESYVDVGSD | 1517794 | Ectodomain | YES |
| TPTKKAFPYGTCLNCN | 1699793 | Cell/Virus Enclosed | NO |
| TPTPGVTPDMAD | 1517806 | Cell/Virus Enclosed | NO |
| TPTQCYYNNSNNGKIV | 1699795 | Cell/Virus Enclosed | NO |
| TPTQENLELKEQ | 1517808 | Cell/Virus Enclosed | NO |
| TPTTDCSDGNYN | 1517813 | Ectodomain | YES |
| TPVAYRNVLLRK | 1517815 | Cell/Virus Enclosed | NO |
| TPVDSFTNGAIV | 1517819 | Cell/Virus Enclosed | NO |
| TPVDSFTNGAVV | 1517821 | Cell/Virus Enclosed | NO |
| TPVHDCNTTCQTPKGAINTSLPFQN | 1174631 | Ectodomain | YES |
| TPVHYERDNTSC | 1517828 | Cell/Virus Enclosed | NO |
| TPVHYERDNTSG | 1517829 | Cell/Virus Enclosed | NO |
| TPVIDDVDVVKP | 1517830 | Cell/Virus Enclosed | NO |
| TPVQVVDSRWNNARQS | 1699810 | Ectodomain | YES |
| TPVSVKLPKGVG | 1517842 | Cell/Virus Enclosed | NO |
| TPVVEYLSKEDLHTLY | 1699812 | Cell/Virus Enclosed | NO |
| TPVVPDTVEVPPQVVK | 1699813 | Cell/Virus Enclosed | NO |
| TPVVSDTVEVSP | 1517850 | Cell/Virus Enclosed | NO |
| TPYCHDPTVLPG | 1517861 | Cell/Virus Enclosed | NO |
| TPYHAFKCNRSGSNDI | 1699821 | Ectodomain | YES |
| TPYVCNASGCGVSDVK | 1699822 | Cell/Virus Enclosed | NO |
| TPYVCNTSGCNVNDVT | 1699823 | Cell/Virus Enclosed | NO |
| TQCFGPRDLDHN | 1517897 | Cell/Virus Enclosed | NO |
| TQCYYNNSNNGKIVYA | 1699834 | Cell/Virus Enclosed | NO |
| TQDGDNKLQGLLEVSV | 1699835 | Ectodomain | YES |
| TQDGNAAIPDYN | 1517903 | Cell/Virus Enclosed | NO |
| TQDGNAAITDYNYYKY | 1699836 | Cell/Virus Enclosed | NO |
| TQDLFLPFYSNVTGFHT | 65832 | Ectodomain | YES |
| TQDYFLPFDTNL | 1517916 | Ectodomain | YES |
| TQEFRFMTANNLTAPR | 1699843 | Cell/Virus Enclosed | NO |
| TQEVFAQVKQMY | 1517928 | Ectodomain | YES |
| TQEYIQTTIQKVTVDC | 1699846 | Ectodomain | YES |
| TQFQGFVLNYKG | 1517949 | Cell/Virus Enclosed | NO |
| TQFQKGKEFEFA | 1517951 | Cell/Virus Enclosed | NO |
| TQFQKGKEFEFVEGQG | 1699853 | Cell/Virus Enclosed | NO |
| TQFQKGKEFKFA | 1517953 | Cell/Virus Enclosed | NO |
| TQFQKGKEFQFA | 1517954 | Capsid | UN |
| TQFVGNVKFDRW | 1517962 | Ectodomain | YES |
| TQFVGNVKLDRW | 1517963 | Ectodomain | YES |
| TQFVGVKFDRWS | 1517964 | Ectodomain | YES |
| TQFVSNVKLDRW | 1517965 | Ectodomain | YES |
| TQGNFGDQELIRQGTE | 1699861 | Cell/Virus Enclosed | NO |
| TQGRNLQDFLPRSTME | 1699862 | Cell/Virus Enclosed | NO |
| TQHGKVPLTFPS | 1517986 | Cell/Virus Enclosed | NO |
| TQIPATSIVAKQGAGD | 1699877 | Cell/Virus Enclosed | NO |
| TQIQVRNIMSPVMG | 65881 | Capsid | UN |
| TQITTNTGIENA | 1518007 | Secreted | YES |

| | | | |
|---|---|---|---|
| TQKGDAAIKDFDYYRY | 1699881 | Cell/Virus Enclosed | NO |
| TQKGSGMDTALLRNNI | 1699882 | Ectodomain | YES |
| TQLAEEPAKDIP | 1518020 | Cell/Virus Enclosed | NO |
| TQLFPLLFVTMA | 1518026 | Cell/Virus Enclosed | NO |
| TQLHVADTLMQGVTLS | 1699890 | Ectodomain | YES |
| TQLNRALSGIAAEQDR | 1699892 | Ectodomain | YES |
| TQMYKEARSVNRKSKV | 1699903 | Cell/Virus Enclosed | NO |
| TQNLFFMLLYAI | 1518074 | Cell/Virus Enclosed | NO |
| TQNVLYDNQKQI | 1518078 | Ectodomain | YES |
| TQNVLYENQKQIANQF | 1699908 | Ectodomain | YES |
| TQNVTVKPKRLRKKRN | 1699909 | Cell/Virus Enclosed | NO |
| TQPKFTVSTQPQGNTI | 1699911 | Capsid | UN |
| TQPKLKEDYQIG | 1518088 | Secreted | YES |
| TQPQGNSIPHYS | 1518095 | Cell/Virus Enclosed | NO |
| TQQVLSENQKLIANKF | 1699920 | Ectodomain | YES |
| TQRIIKAQFKTF | 1518124 | Cell/Virus Enclosed | NO |
| TQRMCAIGPDVFLHKC | 1699922 | Cell/Virus Enclosed | NO |
| TQRNFYSPQIIT | 1518130 | Ectodomain | YES |
| TQSAIAKPKRLR | 1518141 | Cell/Virus Enclosed | NO |
| TQSFAPNTGVYD | 1518147 | Ectodomain | YES |
| TQSTALQELQIQ | 1518164 | Cell/Virus Enclosed | NO |
| TQSVNVKLKRLR | 1518166 | Cell/Virus Enclosed | NO |
| TQSVTVKPKRLR | 1518173 | Cell/Virus Enclosed | NO |
| TQTAQGAHLFSS | 1518177 | Ectodomain | YES |
| TQTSQALQTVATALNK | 1699949 | Ectodomain | YES |
| TQTSQALQTVST | 1518186 | Ectodomain | YES |
| TQTVDSSQGSEYQYVI | 1699953 | Cell/Virus Enclosed | NO |
| TQVYSRKARNRS | 1518215 | Cell/Virus Enclosed | NO |
| TQVYSRKATKRS | 1518216 | Cell/Virus Enclosed | NO |
| TQWLSNMLGTTV | 1518218 | Cell/Virus Enclosed | NO |
| TQWYHEHVGGVYDNSI | 1699965 | Cell/Virus Enclosed | NO |
| TQYNRYLALYNK | 1518234 | Cell/Virus Enclosed | NO |
| TQYYFNKDTGVI | 1518237 | Ectodomain | YES |
| TRCNIGGAVCSKHANL | 1699995 | Cell/Virus Enclosed | NO |
| TRCTSPGPSGIG | 1518267 | Cell/Virus Enclosed | NO |
| TRDFAIRNVRGWLGMD | 1700000 | Cell/Virus Enclosed | NO |
| TRDFAMRHVRGWLGMD | 1700001 | Cell/Virus Enclosed | NO |
| TRDIMATDDCFA | 1518279 | Cell/Virus Enclosed | NO |
| TRDIVSTDDCFA | 1518280 | Cell/Virus Enclosed | NO |
| TRDKLEKLISTARDGV | 1700005 | Cell/Virus Enclosed | NO |
| TRDMADQIASLV | 1518284 | Capsid | UN |
| TRDRFYRLANECAQVL | 1700007 | Cell/Virus Enclosed | NO |
| TRDSIGTNFPLQLGFS | 1700008 | Cell/Virus Enclosed | NO |
| TRDSVPLHIIAPSSLI | 1700009 | Cell/Virus Enclosed | NO |
| TREEAVRQVRSWIGFD | 1700016 | Cell/Virus Enclosed | NO |
| TREENVIQCFCP | 1518301 | Cell/Virus Enclosed | NO |
| TREENVIQCFGPRDFN | 1700017 | Capsid | UN |
| TREKLEKLISTA | 1518307 | Cell/Virus Enclosed | NO |
| TREMQQHPLLNPSALE | 1700021 | Capsid | UN |
| TRENIGTNFPLQIGFS | 1700023 | Cell/Virus Enclosed | NO |
| TRENKSAELNYT | 1518314 | Ectodomain | YES |
| TRENVETLTWPL | 1518315 | Cell/Virus Enclosed | NO |
| TREVFAQVKQMYKTPT | 1700024 | Ectodomain | YES |
| TRFITKSMISAV | 1518326 | Cell/Virus Enclosed | NO |
| TRFQSLENVAYN | 1518332 | Cell/Virus Enclosed | NO |
| TRGRRAQPKQTATSQQ | 1700037 | Cell/Virus Enclosed | NO |
| TRGTFPGPSGIG | 1518355 | Cell/Virus Enclosed | NO |
| TRGTSPGLSGIG | 1518357 | Cell/Virus Enclosed | NO |
| TRLEGLGGDNVYCYNT | 1700060 | Cell/Virus Enclosed | NO |
| TRLEGLGGNNVYCYNT | 1700061 | Cell/Virus Enclosed | NO |
| TRLKGSFNNRDT | 1518395 | Ectodomain | YES |
| TRLTSKGRPLVPT | 66066 | Ectodomain | YES |
| TRNIDATSTGNYNYKY | 1700071 | Ectodomain | YES |
| TRNNSRDSFRST | 1518415 | Cell/Virus Enclosed | NO |
| TRNPANNAAIVLQLPQ | 1700074 | Cell/Virus Enclosed | NO |
| TRNPNNDSAIVTQFAP | 1700076 | Cell/Virus Enclosed | NO |
| TRNPNNNAAIVLQLPQ | 1700077 | Cell/Virus Enclosed | NO |
| TRPLLESELVIGAVI | 2001342 | Underneath Membrane | NO |
| TRPLMESELVIG | 1518435 | Underneath Membrane | NO |
| TRPYLAHCPDCGEGHSC | 2134495 | Undetermined | UN |
| TRRARGGDGKMK | 1518452 | Capsid | UN |
| TRRIRGGDGKMKDLSPRWYF | 1392478 | Capsid | UN |
| TRRVRGGDGKMKDLSP | 1700091 | Cell/Virus Enclosed | NO |
| TRRVRGGDGKMKELSPRW | 66094 | Capsid | UN |
| TRSDSNQSSSDLVAAV | 1700093 | Capsid | UN |
| TRSKLKRRDSTY | 1518467 | Ectodomain | YES |

| | | | |
|---|---|---|---|
| TRSLKYNIRSAL | 1518469 | Cell/Virus Enclosed | NO |
| TRSMLKRRDSTYGPLQ | 1700096 | Ectodomain | YES |
| TRSMWSFNPETNILL | 1325865 | Underneath Membrane | NO |
| TRTAEAIHTVTI | 1518491 | Ectodomain | YES |
| TRTFTATTVVGS | 1518494 | Cell/Virus Enclosed | NO |
| TRTHTMIFDNAF | 1518496 | Ectodomain | YES |
| TRTRSVFNLEGVNGGS | 1700112 | Cell/Virus Enclosed | NO |
| TRTTHTDIGVYI | 1518510 | Cell/Virus Enclosed | NO |
| TRVECTTIVNGM | 1518513 | Cell/Virus Enclosed | NO |
| TRVLNNLNLPGCNGGS | 1700120 | Cell/Virus Enclosed | NO |
| TRVNVDGFNEWAMANG | 1700124 | Cell/Virus Enclosed | NO |
| TRVTVPKDHPHLGKFL | 1700126 | Capsid | UN |
| TRYSLADSNGFI | 1518529 | Cell/Virus Enclosed | NO |
| TRYSLANSNGFI | 1518530 | Cell/Virus Enclosed | NO |
| TSAAAIPFSLAL | 1518535 | Ectodomain | YES |
| TSACKTIEDALRNSAR | 1700136 | Ectodomain | YES |
| TSACRRSGSSFYAEMKWLLSNTDNA | 1174935 | Ectodomain | YES |
| TSADIVDRDPSS | 1518543 | Cell/Virus Enclosed | NO |
| TSAGQPCVVSEQ | 1518554 | Cell/Virus Enclosed | NO |
| TSAGQPYVASEQ | 1518555 | Cell/Virus Enclosed | NO |
| TSAGRPCVASEQ | 1518557 | Cell/Virus Enclosed | NO |
| TSAKSLARSQSSETKE | 1700150 | Capsid | UN |
| TSANLFPPWSAA | 1518568 | Ectodomain | YES |
| TSASKLTIVSPDLESY | 1700155 | Cell/Virus Enclosed | NO |
| TSASLFPLWTAA | 1518574 | Ectodomain | YES |
| TSASLFPPWSAA | 1518575 | Ectodomain | YES |
| TSASLFPPWTAAAGVP | 1700156 | Ectodomain | YES |
| TSATSCQLYYNL | 1518581 | Ectodomain | YES |
| TSCELYHYQECV | 1518588 | Ectodomain | YES |
| TSCFFTCNSNAN | 1518589 | Ectodomain | YES |
| TSCPEWMDDFEADPKG | 1700167 | Cell/Virus Enclosed | NO |
| TSCPPICPGYRWMCL | 1700168 | Ectodomain | YES |
| TSCQIAVSKFLD | 1518606 | Cell/Virus Enclosed | NO |
| TSCQTAVSKFLD | 1518613 | Cell/Virus Enclosed | NO |
| TSDDDWTAAVDE | 1518624 | Cell/Virus Enclosed | NO |
| TSDDEWAAAVDE | 1518625 | Cell/Virus Enclosed | NO |
| TSDELCVNLNLPANVP | 1700174 | Cell/Virus Enclosed | NO |
| TSDFTDSVRDPQ | 1518631 | Ectodomain | YES |
| TSDKCIFNSACTRLEG | 1700175 | Cell/Virus Enclosed | NO |
| TSDNINDKDTGPPPIS | 1700183 | Ectodomain | YES |
| TSDNNCWVNAVC | 1518643 | Cell/Virus Enclosed | NO |
| TSDPQFPILAEL | 1518644 | Cell/Virus Enclosed | NO |
| TSDQAVTFLKEWNFSL | 1700186 | Undetermined | UN |
| TSDTAHACNANRFNVA | 1700188 | Cell/Virus Enclosed | NO |
| TSDTAHACNVNRFNVA | 1700189 | Cell/Virus Enclosed | NO |
| TSDTSYINDKDT | 1518655 | Ectodomain | YES |
| TSEEPVNQFTGY | 1518672 | Cell/Virus Enclosed | NO |
| TSEGNALYNNEGGRAF | 1700201 | Cell/Virus Enclosed | NO |
| TSEILDISPRSF | 1518682 | Ectodomain | YES |
| TSEPPVTYLPAT | 1518688 | Cell/Virus Enclosed | NO |
| TSFACLSNVSTS | 1518698 | Ectodomain | YES |
| TSFAIYHTPATD | 1518699 | Ectodomain | YES |
| TSFATYHTPATDCSDG | 1700209 | Ectodomain | YES |
| TSFATYHTPATN | 1518701 | Ectodomain | YES |
| TSFATYHTPSTD | 1518702 | Ectodomain | YES |
| TSFATYHTPTTD | 1518703 | Ectodomain | YES |
| TSFATYYTPATD | 1518704 | Ectodomain | YES |
| TSGCNVNDVTKLYLGG | 1700221 | Cell/Virus Enclosed | NO |
| TSGDGTTQPKLK | 1518729 | Secreted | YES |
| TSGDLAVQIGNNNVCT | 1700227 | Cell/Virus Enclosed | NO |
| TSGHYDDDDDIPFPG | 227631 | Capsid | UN |
| TSGIKFQTVKPGNFNQ | 1700229 | Cell/Virus Enclosed | NO |
| TSGLGTVDVDYKSCTK | 1700231 | Ectodomain | YES |
| TSGLSPAYAPTYVSVD | 1700232 | Cell/Virus Enclosed | NO |
| TSGNLLGFKDVTKGTI | 1700233 | Ectodomain | YES |
| TSGSDLDRCTTFDDVQ | 1700238 | Ectodomain | YES |
| TSGSRVAGRSAIEDIL | 1700239 | Ectodomain | YES |
| TSGSTYLNMALK | 1518754 | Ectodomain | YES |
| TSGSTYRNMALK | 1518755 | Ectodomain | YES |
| TSGVKFQTVKPGNFNQ | 1700242 | Cell/Virus Enclosed | NO |
| TSGVTRDMADQI | 1518766 | Capsid | UN |
| TSHFTTQFVGVK | 1518777 | Ectodomain | YES |
| TSHPIEEIPLDTFVI | 111851 | Capsid | UN |
| TSHPIEEIPLDTFVV | 111852 | Capsid | UN |
| TSHTVMPLTAPT | 1518790 | Cell/Virus Enclosed | NO |
| TSIEEVCDDYAK | 1518796 | Cell/Virus Enclosed | NO |

| | | | |
|---|---|---|---|
| TSIFRNPVFTEPSVTQPQPPVE | 111853 | Capsid | UN |
| TSIHADQLTPAW | 1518800 | Ectodomain | YES |
| TSIKWADNNSYL | 1518802 | Cell/Virus Enclosed | NO |
| TSIPNFGSLTQINTTL | 1700265 | Ectodomain | YES |
| TSIPNLHDFKEE | 1518820 | Ectodomain | YES |
| TSIPNLPDFKEELDQWFKNQTSVAP | 1860153 | Ectodomain | YES |
| TSIPNLPDFKGE | 1518822 | Ectodomain | YES |
| TSIVAKQGAGDAGHSL | 1700270 | Cell/Virus Enclosed | NO |
| TSKCWVEEDLTKGPHE | 1700274 | Cell/Virus Enclosed | NO |
| TSKDTLDDILQQANKS | 1700275 | Cell/Virus Enclosed | NO |
| TSKVKSLSMIRG | 1518861 | Cell/Virus Enclosed | NO |
| TSLGAGPVSISAVAVLAPHSALALL | 66296 | Undetermined | UN |
| TSLGNRKRNQKPLEPK | 1700294 | Capsid | UN |
| TSLNDRQGNATK | 66304 | Capsid | UN |
| TSLNTKYVAPHV | 1518887 | Ectodomain | YES |
| TSLPKDWRLINKFDVI | 1700301 | Cell/Virus Enclosed | NO |
| TSLRRLVNPTDK | 1518895 | Cell/Virus Enclosed | NO |
| TSLSDACQLPYC | 1518896 | Underneath Membrane | NO |
| TSLSFNNLLGFD | 1518898 | Cell/Virus Enclosed | NO |
| TSLTPCGYSEPPKVAA | 1700311 | Cell/Virus Enclosed | NO |
| TSMLTNYFLDDVEMVA | 1700319 | Cell/Virus Enclosed | NO |
| TSMYKQARAEDKKAKI | 1700320 | Cell/Virus Enclosed | NO |
| TSNFRVSPTHEV | 1518934 | Ectodomain | YES |
| TSNFRVVPSGDVVRFPN | 66337 | Ectodomain | YES |
| TSNGLLGNCAKR | 1518938 | Cell/Virus Enclosed | NO |
| TSNLPDFREELD | 1518940 | Ectodomain | YES |
| TSNSFEVLEVED | 1518951 | Cell/Virus Enclosed | NO |
| TSNVPPIVSEKI | 1518957 | Cell/Virus Enclosed | NO |
| TSNVPSIVSEKI | 1518958 | Cell/Virus Enclosed | NO |
| TSNVPSLVSEKI | 1518959 | Cell/Virus Enclosed | NO |
| TSNVPTIVSEKI | 1518960 | Cell/Virus Enclosed | NO |
| TSNVSPIVNEKI | 1518961 | Cell/Virus Enclosed | NO |
| TSNYSGVVTTIM | 1518965 | Cell/Virus Enclosed | NO |
| TSPARIAGNGGDAALA | 1700342 | Cell/Virus Enclosed | NO |
| TSPARMAGNGSDAALA | 1700344 | Cell/Virus Enclosed | NO |
| TSPATVEPVRDE | 1518970 | Cell/Virus Enclosed | NO |
| TSPCPSGIGAVG | 1518971 | Cell/Virus Enclosed | NO |
| TSPDPKFVDKYYAFLN | 1700346 | Cell/Virus Enclosed | NO |
| TSPDVDFGDISG | 1518974 | Ectodomain | YES |
| TSPGLSGIGAVG | 1518976 | Cell/Virus Enclosed | NO |
| TSPGPSGIGAVGGDLL | 1700348 | Cell/Virus Enclosed | NO |
| TSPGPSGIGTVG | 1518979 | Cell/Virus Enclosed | NO |
| TSPIQPVELPAS | 1518981 | Ectodomain | YES |
| TSQALQTVATALNKIQ | 1700354 | Ectodomain | YES |
| TSQGFVVVNGKF | 1519003 | Cell/Virus Enclosed | NO |
| TSQGFVVVNGKV | 1519004 | Cell/Virus Enclosed | NO |
| TSQGMYGGTYLVEKP | 66385 | Ectodomain | YES |
| TSQKETLHSTSP | 66387 | Secreted | YES |
| TSQLRHNFQAISNSIQ | 1700358 | Ectodomain | YES |
| TSQNTRISLRPGPVSQPYHH | 66389 | Capsid | UN |
| TSQQPLGGNVVP | 1519015 | Cell/Virus Enclosed | NO |
| TSQQPSGGNVIP | 1519016 | Cell/Virus Enclosed | NO |
| TSQSVCNGCVGH | 1519020 | Ectodomain | YES |
| TSQTVKPGHFNKEFYD | 1700363 | Cell/Virus Enclosed | NO |
| TSREVVVTNLNKSAGW | 1700367 | Cell/Virus Enclosed | NO |
| TSRMLSYYKLGA | 1519042 | Underneath Membrane | NO |
| TSRTLSYYKLGASQRV | 66409 | Underneath Membrane | NO |
| TSRVEHAVVYYVYSP | 66410 | Ectodomain | YES |
| TSSKVVLSDALDKLYK | 1700383 | Cell/Virus Enclosed | NO |
| TSSLAYYEPVQG | 1519082 | Cell/Virus Enclosed | NO |
| TSSMRGCCDCGS | 1519086 | Underneath Membrane | NO |
| TSSPKVTIDCVA | 1519092 | Ectodomain | YES |
| TSSPKVTIDRAA | 1519093 | Ectodomain | YES |
| TSSRASSAGSRSRANS | 1700389 | Cell/Virus Enclosed | NO |
| TSSRTSSAGSRS | 1519099 | Capsid | UN |
| TSSSLGSIAGAG | 1519106 | Ectodomain | YES |
| TSSSTYRNMALK | 1519107 | Ectodomain | YES |
| TSTALGKLQDVVNQNA | 1700398 | Ectodomain | YES |
| TSTDRFYRLGNE | 1519131 | Cell/Virus Enclosed | NO |
| TSTEQVDVSHDV | 1519135 | Cell/Virus Enclosed | NO |
| TSTGKTVLGEYVFDKS | 1700403 | Cell/Virus Enclosed | NO |
| TSTGNHNYKYRY | 1519142 | Ectodomain | YES |
| TSTGNYNYKFRY | 1519144 | Ectodomain | YES |
| TSTGNYNYKYRC | 1519146 | Ectodomain | YES |
| TSTGNYNYKYRS | 1519148 | Ectodomain | YES |
| TSTHDGFNEHVT | 1519153 | Cell/Virus Enclosed | NO |

| | | | |
|---|---|---|---|
| TSTHDGLIEHVT | 1519155 | Cell/Virus Enclosed | NO |
| TSTHDGLNEHVT | 1519156 | Cell/Virus Enclosed | NO |
| TSTHDGLTEHVT | 1519157 | Cell/Virus Enclosed | NO |
| TSTHDGPNEHVT | 1519159 | Cell/Virus Enclosed | NO |
| TSTHDGPTEHVTSVNL | 1700405 | Cell/Virus Enclosed | NO |
| TSTPIVVDCATY | 1519179 | Ectodomain | YES |
| TSTPIVVDCSTY | 1519180 | Ectodomain | YES |
| TSTPKKPNKPLSQPRA | 1700411 | Capsid | UN |
| TSTPNLPDFKEE | 1519182 | Ectodomain | YES |
| TSTPSDVSSRDPTTQE | 1700412 | Capsid | UN |
| TSTSGNLLGFKDVTKG | 1700414 | Ectodomain | YES |
| TSTSNLPDFREE | 1519191 | Ectodomain | YES |
| TSTTNFGEEFLG | 1519200 | Cell/Virus Enclosed | NO |
| TSVAPDLSFDYI | 1519209 | Ectodomain | YES |
| TSVDANTNQDSYGGAS | 1700420 | Cell/Virus Enclosed | NO |
| TSVEESFIDDYY | 1519221 | Cell/Virus Enclosed | NO |
| TSVENAQQDKGIAIPHDIDLGESRV | 66481 | Undetermined | UN |
| TSVLRQWLPTDA | 1519229 | Cell/Virus Enclosed | NO |
| TSVMESQISTKEVKLK | 1700428 | Cell/Virus Enclosed | NO |
| TSVTEDVKFAAS | 1519246 | Cell/Virus Enclosed | NO |
| TSWADQSERNYQTFNR | 1700447 | Capsid | UN |
| TSWVDQSERSHQ | 1519265 | Cell/Virus Enclosed | NO |
| TSYDCDIPIGAG | 1519266 | Ectodomain | YES |
| TSYECDIPIGAGICAS | 1700451 | Ectodomain | YES |
| TSYECDIPVGAG | 1519268 | Ectodomain | YES |
| TSYECDTPIGAG | 1519269 | Ectodomain | YES |
| TSYGFSNVVEMP | 1519272 | Ectodomain | YES |
| TSYINDKDTGPP | 1519277 | Ectodomain | YES |
| TSYINDVDTGVP | 1519278 | Ectodomain | YES |
| TSYINNKDTGPP | 1519279 | Ectodomain | YES |
| TSYSVCNGCVGYSENV | 1700456 | Ectodomain | YES |
| TSYSVVIQPRTI | 1519291 | Ectodomain | YES |
| TSYTANDVEVQS | 1519293 | Cell/Virus Enclosed | NO |
| TSYTTTILEHHH | 1519299 | Cell/Virus Enclosed | NO |
| TTAATRFMKDLYFTSTNGVGEIGRG | 66511 | Undetermined | UN |
| TTAATVAAMFPPWSAA | 1700466 | Ectodomain | YES |
| TTANVSALMGANGNKI | 1700478 | Cell/Virus Enclosed | NO |
| TTAPLKLFAAET | 1519321 | Cell/Virus Enclosed | NO |
| TTAQKLETDRIC | 1519323 | Cell/Virus Enclosed | NO |
| TTAVSALVIGQATRPQNP | 558338 | Undetermined | UN |
| TTAVYKPLVDFN | 1519340 | Cell/Virus Enclosed | NO |
| TTCAFSVDPATT | 1519344 | Cell/Virus Enclosed | NO |
| TTCGYLPANAVV | 1519354 | Cell/Virus Enclosed | NO |
| TTCLYIPKPTAK | 1519362 | Cell/Virus Enclosed | NO |
| TTCSNDLLQPNTEVFT | 1700504 | Ectodomain | YES |
| TTCSTDLQKSNT | 1519367 | Ectodomain | YES |
| TTDCSDGNYNRN | 1519373 | Ectodomain | YES |
| TTDKSFEQQVGVIADK | 1700507 | Cell/Virus Enclosed | NO |
| TTDMVKSKGVCATGDC | 1700508 | Cell/Virus Enclosed | NO |
| TTDNTFVAGNCD | 1519383 | Ectodomain | YES |
| TTENRFTNFEPF | 1519405 | Ectodomain | YES |
| TTEVMPVSIAKT | 1519412 | Ectodomain | YES |
| TTEVMPVSMAKT | 1519413 | Ectodomain | YES |
| TTFDFLSNSSSS | 1519420 | Ectodomain | YES |
| TTFIDAARGPAGVESD | 1700547 | Cell/Virus Enclosed | NO |
| TTFLLNGYYPTS | 1519429 | Ectodomain | YES |
| TTFMLKYDENGTITDA | 1700552 | Ectodomain | YES |
| TTFTAPDFVAFN | 1519437 | Cell/Virus Enclosed | NO |
| TTGDLAVQIGSNNVCT | 1700556 | Cell/Virus Enclosed | NO |
| TTGYRFTNFEPFTVNS | 1700567 | Ectodomain | YES |
| TTIGNETTSAFVGALP | 1700575 | Ectodomain | YES |
| TTIQSVDISYLK | 1519500 | Cell/Virus Enclosed | NO |
| TTIQSVDISYLN | 1519501 | Cell/Virus Enclosed | NO |
| TTIQTVFDCAVE | 1519503 | Cell/Virus Enclosed | NO |
| TTIVNGMKRSFY | 1519514 | Cell/Virus Enclosed | NO |
| TTKAGYPYNYNTTASDQLLVENAAG | 66667 | Undetermined | UN |
| TTKFYGGWDNMLKNLM | 1700608 | Cell/Virus Enclosed | NO |
| TTKNLTGSNESK | 1519537 | Cell/Virus Enclosed | NO |
| TTKNVTGSNESKALFF | 1700618 | Cell/Virus Enclosed | NO |
| TTKVTVPKDHPH | 1519542 | Cell/Virus Enclosed | NO |
| TTLAGLWLDDTV | 1519546 | Cell/Virus Enclosed | NO |
| TTLAVIGDFNCT | 1519547 | Ectodomain | YES |
| TTLEETKFLTNKLLLF | 1390041 | Cell/Virus Enclosed | NO |
| TTLGGLHLLISR | 1519566 | Cell/Virus Enclosed | NO |
| TTLLDLTYEMLS | 1519576 | Ectodomain | YES |
| TTLLDLTYEMSS | 1519577 | Ectodomain | YES |

| | | | |
|---|---|---|---|
| TTLNAFVAQQLVRSES | 1700638 | Ectodomain | YES |
| TTLNGLWLDDIV | 1519580 | Cell/Virus Enclosed | NO |
| TTLNGLWLDDTV | 1519581 | Cell/Virus Enclosed | NO |
| TTLQSLENVIYNLVNV | 1700649 | Cell/Virus Enclosed | NO |
| TTLQSLENVVYN | 1519595 | Cell/Virus Enclosed | NO |
| TTLSTITAYLLV | 1519599 | Cell/Virus Enclosed | NO |
| TTLTSDYTITVF | 1519603 | Cell/Virus Enclosed | NO |
| TTLVQYAVGIMTVTMTFKLG | 66719 | Capsid | UN |
| TTMDFCEGSFPC | 1519612 | Cell/Virus Enclosed | NO |
| TTNEAFHKVQDA | 1519624 | Ectodomain | YES |
| TTNEAFQKVQDAVNNN | 1700665 | Ectodomain | YES |
| TTNEAFRKVQDA | 1519626 | Ectodomain | YES |
| TTNFGEDFLGWI | 1519628 | Cell/Virus Enclosed | NO |
| TTNGSWAIFNDHHLNR | 1700669 | Cell/Virus Enclosed | NO |
| TTNLDNKLQGLL | 1519644 | Ectodomain | YES |
| TTNQPRGRGRNP | 1519649 | Cell/Virus Enclosed | NO |
| TTNTGIENATFF | 1519652 | Secreted | YES |
| TTNYVGVYDINHGDAG | 1700676 | Ectodomain | YES |
| TTPAPVYIWTADEAIK | 1700677 | Underneath Membrane | NO |
| TTPCELHGNGSM | 1519665 | Cell/Virus Enclosed | NO |
| TTPCEVHGNGSMIEDI | 1700678 | Cell/Virus Enclosed | NO |
| TTPCGVHGNGSM | 1519668 | Cell/Virus Enclosed | NO |
| TTPCQVHGNGSM | 1519669 | Cell/Virus Enclosed | NO |
| TTPKEEFVVKEKLNAF | 1700685 | Cell/Virus Enclosed | NO |
| TTPLGMSLEACY | 1519677 | Cell/Virus Enclosed | NO |
| TTPLGMSLGACY | 1519678 | Cell/Virus Enclosed | NO |
| TTQFVDNVKLAR | 1519690 | Ectodomain | YES |
| TTQLHVADTLMQGVTL | 1700691 | Ectodomain | YES |
| TTQLQVANNLMN | 1519698 | Ectodomain | YES |
| TTQPKLKEDYQI | 1519700 | Secreted | YES |
| TTQSAIIVNNST | 1519705 | Ectodomain | YES |
| TTQTVDSSQGSEYQYV | 1700698 | Cell/Virus Enclosed | NO |
| TTRVTVPKDHPHLGKF | 1700710 | Capsid | UN |
| TTSDKCIFNSACTRLE | 1700713 | Cell/Virus Enclosed | NO |
| TTSGIGYQPYRV | 1519735 | Ectodomain | YES |
| TTSGSNDIWMQN | 1519736 | Ectodomain | YES |
| TTSLTPCGYSEPPKVA | 1700720 | Cell/Virus Enclosed | NO |
| TTSMAPDFVAFN | 1519745 | Cell/Virus Enclosed | NO |
| TTSNFLPEVAAY | 1519749 | Ectodomain | YES |
| TTSTALGKLQDVVNQN | 1700728 | Ectodomain | YES |
| TTSTGPCKTCTTPAQ | 66833 | Underneath Membrane | NO |
| TTTAPLKLFAAE | 1519766 | Cell/Virus Enclosed | NO |
| TTTGIPEYNFII | 1519770 | Cell/Virus Enclosed | NO |
| TTTIKPLEHHHH | 1519772 | Cell/Virus Enclosed | NO |
| TTTIKPVSLEHH | 1519773 | Cell/Virus Enclosed | NO |
| TTTLNGLWLDDI | 1519784 | Cell/Virus Enclosed | NO |
| TTTLNGLWLDDT | 1519785 | Cell/Virus Enclosed | NO |
| TTTNEAFHKVQD | 1519793 | Ectodomain | YES |
| TTTNEAFRKVQD | 1519795 | Ectodomain | YES |
| TTTNVDKIRHLL | 1519801 | Cell/Virus Enclosed | NO |
| TTTVTSFHGYDF | 1519811 | Cell/Virus Enclosed | NO |
| TTVAINDVDTGV | 1519815 | Ectodomain | YES |
| TTVDKLKPVFTW | 1519819 | Cell/Virus Enclosed | NO |
| TTVEKLRPIFEW | 1519827 | Cell/Virus Enclosed | NO |
| TTVESAAQADLQ | 1519831 | Cell/Virus Enclosed | NO |
| TTVLDRTDWCRC | 1519835 | Ectodomain | YES |
| TTVLGQTYYQPIQQAP | 1700767 | Underneath Membrane | NO |
| TTVLNDPVFTGE | 1519843 | Cell/Virus Enclosed | NO |
| TTVLNGPVFTGD | 1519844 | Cell/Virus Enclosed | NO |
| TTVNAHQILPKVLHK | 66895 | Cell/Virus Enclosed | NO |
| TTVRSDGTFEEM | 1519848 | Cell/Virus Enclosed | NO |
| TTVSINDIDTGA | 1519849 | Ectodomain | YES |
| TTVSVTVSSNMVETVV | 1700778 | Cell/Virus Enclosed | NO |
| TTVTVNVTTHNG | 1519857 | Ectodomain | YES |
| TTVVRPLVEDSTSVTA | 1700783 | Ectodomain | YES |
| TTWNGGAYSLFDMTKF | 1700787 | Cell/Virus Enclosed | NO |
| TTYLEAVKHGAKPVSN | 1700793 | Cell/Virus Enclosed | NO |
| TTYPGQGCAGYT | 1519870 | Cell/Virus Enclosed | NO |
| TTYYLDDVKKIEYKPD | 1700802 | Cell/Virus Enclosed | NO |
| TVAEEPDSRVPS | 1519881 | Cell/Virus Enclosed | NO |
| TVAKAPPGEKFK | 1519889 | Cell/Virus Enclosed | NO |
| TVAQFNYRRDGQ | 1519902 | Cell/Virus Enclosed | NO |
| TVATALNKIHDV | 1519907 | Ectodomain | YES |
| TVATALNKIQDVVNQQ | 1700818 | Ectodomain | YES |
| TVATSRMLSYYK | 1519908 | Underneath Membrane | NO |
| TVATSRTLSYYKLGA | 1313810 | Underneath Membrane | NO |

| | | | |
|---|---|---|---|
| TVAWHVVRDPRY | 1519913 | Cell/Virus Enclosed | NO |
| TVCDSLNAKLGFDSSK | 1700826 | Cell/Virus Enclosed | NO |
| TVCDSLNSKLGF | 1519922 | Cell/Virus Enclosed | NO |
| TVCFSVAALSTGLTSQ | 1700827 | Cell/Virus Enclosed | NO |
| TVCGPKLSTDLIKNQC | 1700831 | Ectodomain | YES |
| TVDCKQYVCNGFQKCE | 1700841 | Ectodomain | YES |
| TVDGVNFKSISLTVGE | 1700845 | Cell/Virus Enclosed | NO |
| TVDGVNFRTVVLNNKN | 1700846 | Cell/Virus Enclosed | NO |
| TVDGVNFTNRFVPVGE | 1700847 | Cell/Virus Enclosed | NO |
| TVDLGQCGLLGTITGPPQCDQFLEF | 1175353 | Ectodomain | YES |
| TVDLKHFFFTQDGNAA | 1700850 | Cell/Virus Enclosed | NO |
| TVDPQKAYLDFVNAGG | 1700854 | Cell/Virus Enclosed | NO |
| TVDQLKHLLEQW | 1519975 | Undetermined | UN |
| TVDSAQGSEYDYVIFA | 1700855 | Cell/Virus Enclosed | NO |
| TVDSSQGSEYDYVIYA | 1700858 | Cell/Virus Enclosed | NO |
| TVDSSQGSEYQYVIFC | 1700859 | Cell/Virus Enclosed | NO |
| TVDTTPKEEFVVKEKL | 1700860 | Cell/Virus Enclosed | NO |
| TVDTTVESAAQA | 1519982 | Cell/Virus Enclosed | NO |
| TVDVANPSTPAYTFTT | 1700861 | Cell/Virus Enclosed | NO |
| TVDVDYKSCTKGLSIA | 1700862 | Ectodomain | YES |
| TVEAAFDLSKEL | 1519993 | Cell/Virus Enclosed | NO |
| TVEAAIELSKELKRPV | 1700867 | Cell/Virus Enclosed | NO |
| TVEAALDLSKELKRPI | 1700868 | Cell/Virus Enclosed | NO |
| TVEDLRARMTYVCQCG | 1700870 | Cell/Virus Enclosed | NO |
| TVEPVRDEVSTE | 1520016 | Cell/Virus Enclosed | NO |
| TVESAAQADLQH | 1520018 | Cell/Virus Enclosed | NO |
| TVETALDLSKEL | 1520022 | Cell/Virus Enclosed | NO |
| TVEVSPQVVKLP | 1520036 | Cell/Virus Enclosed | NO |
| TVFSNMVETVVG | 1520059 | Cell/Virus Enclosed | NO |
| TVFVCTDNSANT | 1520064 | Cell/Virus Enclosed | NO |
| TVFVEPKDGGHF | 1520065 | Cell/Virus Enclosed | NO |
| TVFVEPKDGGQF | 1520066 | Cell/Virus Enclosed | NO |
| TVFVEPKNGGQF | 1520067 | Cell/Virus Enclosed | NO |
| TVGEVFGKILGNVFCD | 1700903 | Cell/Virus Enclosed | NO |
| TVGGVPLLSTMT | 1520088 | Cell/Virus Enclosed | NO |
| TVGPFMDNAINV | 1520091 | Cell/Virus Enclosed | NO |
| TVGRRVSARMLGDVMAVSTC | 742098 | Ectodomain | YES |
| TVGSGYCLDYSK | 1520095 | Ectodomain | YES |
| TVGSPIQFHLDG | 1520100 | Cell/Virus Enclosed | NO |
| TVGVFLGYKETD | 1520101 | Cell/Virus Enclosed | NO |
| TVHKSGFQLPVY | 1520115 | Cell/Virus Enclosed | NO |
| TVIDILLDDFVALVKS | 1700917 | Cell/Virus Enclosed | NO |
| TVIIINNSTNVV | 1520135 | Ectodomain | YES |
| TVIISPSTRATI | 1520139 | Ectodomain | YES |
| TVINNVDNEIIK | 1520144 | Cell/Virus Enclosed | NO |
| TVIPDYVDVNKTLQEF | 1700925 | Ectodomain | YES |
| TVIRGHLYIQGVKLGT | 1700926 | Underneath Membrane | NO |
| TVKAKEESVKSSYAYA | 1700931 | Cell/Virus Enclosed | NO |
| TVKEIISIPNII | 1520167 | Cell/Virus Enclosed | NO |
| TVKFCDHIVKAG | 1520174 | Cell/Virus Enclosed | NO |
| TVKIEDAIIVNDENSS | 1700940 | Cell/Virus Enclosed | NO |
| TVKIGGLTFKTYAFDK | 1700941 | Cell/Virus Enclosed | NO |
| TVKPGHFNKEFYDFLR | 1700946 | Cell/Virus Enclosed | NO |
| TVKPGNFNQDFYDFVL | 1700948 | Cell/Virus Enclosed | NO |
| TVKPGNFNQDFYEFVK | 1700949 | Cell/Virus Enclosed | NO |
| TVKPKPVINQLDEKAQ | 1700950 | Cell/Virus Enclosed | NO |
| TVKPVATVHRRIPDLP | 1700952 | Ectodomain | YES |
| TVKRKRLRKKRN | 1520194 | Cell/Virus Enclosed | NO |
| TVKSEDGNIKNINWFV | 1700953 | Cell/Virus Enclosed | NO |
| TVKWADASEPQRGRQG | 1700957 | Capsid | UN |
| TVKWADASETQR | 1520202 | Capsid | UN |
| TVKWADESEPQR | 1520203 | Cell/Virus Enclosed | NO |
| TVKYFECTGGID | 1520206 | Cell/Virus Enclosed | NO |
| TVLAAYNGPPQG | 1520208 | Cell/Virus Enclosed | NO |
| TVLEVQGYKNVR | 1520216 | Cell/Virus Enclosed | NO |
| TVLGEYVFDKSELTNG | 1700967 | Cell/Virus Enclosed | NO |
| TVLLPTQYKDVEAWSG | 1700978 | Ectodomain | YES |
| TVLNASIPESKPLADDDSA | 1520236 | Cell/Virus Enclosed | NO |
| TVLNASTPESKP | 1520238 | Cell/Virus Enclosed | NO |
| TVLNDPVFTGDLFYTI | 1700980 | Cell/Virus Enclosed | NO |
| TVLNDPVFTGEL | 1520240 | Cell/Virus Enclosed | NO |
| TVLNDPVFTGNL | 1520241 | Cell/Virus Enclosed | NO |
| TVLNVVGPDARTQGKQ | 1700984 | Cell/Virus Enclosed | NO |
| TVLPGAFAYSQM | 1520244 | Cell/Virus Enclosed | NO |
| TVLPPLLTDDMI | 1520249 | Ectodomain | YES |
| TVLQNNELMPVK | 1520256 | Cell/Virus Enclosed | NO |

| | | | |
|---|---|---|---|
| TVLRAINGDFLH | 1520257 | Cell/Virus Enclosed | NO |
| TVLRDNLKPVTTLSTI | 1700997 | Cell/Virus Enclosed | NO |
| TVLRNSNGACIW | 1520260 | Cell/Virus Enclosed | NO |
| TVLSVAPEVDWV | 1520268 | Cell/Virus Enclosed | NO |
| TVLSYYYVMPLT | 1520269 | Ectodomain | YES |
| TVMPLTAPTLVP | 1520285 | Cell/Virus Enclosed | NO |
| TVNGYAKSNFVKDDYC | 1701014 | Cell/Virus Enclosed | NO |
| TVNLAKVCTYAV | 1520296 | Cell/Virus Enclosed | NO |
| TVNNFVKFGSVC | 1520299 | Ectodomain | YES |
| TVNSINDSLEPV | 1520300 | Ectodomain | YES |
| TVNSVNDSLEHV | 1520302 | Ectodomain | YES |
| TVNSVNDSLEPVGGLY | 1701020 | Ectodomain | YES |
| TVNSVNDSLKPV | 1520303 | Ectodomain | YES |
| TVNVTTLNGRLV | 1520309 | Ectodomain | YES |
| TVPDLVVEQYNQTILN | 1701028 | Ectodomain | YES |
| TVPEEFASHVANFQKS | 1701029 | Cell/Virus Enclosed | NO |
| TVPSTVWEDGDY | 1520332 | Ectodomain | YES |
| TVQADTAGLYTN | 1520337 | Cell/Virus Enclosed | NO |
| TVQDADGINKQLTDIS | 1701036 | Cell/Virus Enclosed | NO |
| TVQDAKGLKIKYLYFV | 1701037 | Cell/Virus Enclosed | NO |
| TVQFCDAMRNAG | 1520342 | Cell/Virus Enclosed | NO |
| TVQGPPGTGKSHLAIG | 1701041 | Cell/Virus Enclosed | NO |
| TVQKLQSLIDNI | 1520350 | Ectodomain | YES |
| TVQKLQTLIDNINSTL | 1701042 | Ectodomain | YES |
| TVQQCFGKRGPNQNFG | 1701044 | Cell/Virus Enclosed | NO |
| TVQSKDTNFLNA | 1520360 | Cell/Virus Enclosed | NO |
| TVQSKMSDVRCT | 1520363 | Cell/Virus Enclosed | NO |
| TVQYGTDTNSVCPKLE | 1701050 | Ectodomain | YES |
| TVREYIRYGIKP | 1520373 | Cell/Virus Enclosed | NO |
| TVRLQAGKQTEFVSNS | 1701057 | Cell/Virus Enclosed | NO |
| TVRSGNFNQDFY | 1520380 | Cell/Virus Enclosed | NO |
| TVSALVYENKLKAKNE | 1701072 | Cell/Virus Enclosed | NO |
| TVSALVYNNKLLAKKE | 1701073 | Cell/Virus Enclosed | NO |
| TVSDDDFVSAVANAHR | 1701076 | Cell/Virus Enclosed | NO |
| TVSINDIDTGAP | 1520403 | Ectodomain | YES |
| TVSKTSDWKCKVTDVL | 1701080 | Cell/Virus Enclosed | NO |
| TVSLLRNTSQKS | 1520414 | Ectodomain | YES |
| TVSSHGMVPVIH | 1520423 | Cell/Virus Enclosed | NO |
| TVSSNMVETVVGQLEQ | 1701087 | Cell/Virus Enclosed | NO |
| TVSTALNKIQDV | 1520430 | Ectodomain | YES |
| TVSVTVSSNMVETVVG | 1701090 | Cell/Virus Enclosed | NO |
| TVTANLSIPSNW | 1520446 | Ectodomain | YES |
| TVTDVKQVGCFM | 1520449 | Cell/Virus Enclosed | NO |
| TVTIALNKIQDVVNQQ | 1701100 | Ectodomain | YES |
| TVTKTSDWKCKV | 1520457 | Cell/Virus Enclosed | NO |
| TVTLLPAADLDDFSKQLQQS | 1392482 | Capsid | UN |
| TVTSAGEPCVAS | 1520468 | Cell/Virus Enclosed | NO |
| TVTSDGQPYVAS | 1520471 | Cell/Virus Enclosed | NO |
| TVTTDKSFEQQVGVIA | 1701110 | Cell/Virus Enclosed | NO |
| TVTYLNDPSSKT | 1520479 | Cell/Virus Enclosed | NO |
| TVTYPFIANEKFINGC | 1701118 | Cell/Virus Enclosed | NO |
| TVVEEPDSRAPS | 1520487 | Cell/Virus Enclosed | NO |
| TVVFTVKIGGLT | 1520489 | Cell/Virus Enclosed | NO |
| TVVGQLEQTNMHSPDVI | 1390125 | Cell/Virus Enclosed | NO |
| TVVIGTTKFCGG | 1520496 | Cell/Virus Enclosed | NO |
| TVVIKNLTPIKLNFGM | 1701130 | Cell/Virus Enclosed | NO |
| TVVLNNKNTYRS | 1520502 | Cell/Virus Enclosed | NO |
| TVVLNNKSTYRS | 1520503 | Cell/Virus Enclosed | NO |
| TVVSKVHEVIIDNKPW | 1701141 | Cell/Virus Enclosed | NO |
| TVVSNGAKNISGQSP | 227646 | Secreted | YES |
| TVVSTVKIGGLTFKTY | 1701142 | Cell/Virus Enclosed | NO |
| TVVVPGSCNFPL | 1520525 | Ectodomain | YES |
| TVVVTLKDSDVNDMVL | 1701148 | Cell/Virus Enclosed | NO |
| TVWEDGDYYRKQLSPL | 1701151 | Ectodomain | YES |
| TVWKDGDYYRKQ | 1520534 | Ectodomain | YES |
| TVWNGGAYSLFDMAKF | 1701154 | Cell/Virus Enclosed | NO |
| TVYCNGVKKVIL | 67217 | Ectodomain | YES |
| TVYDSSEVKAQSSGVI | 1701162 | Cell/Virus Enclosed | NO |
| TVYDTAKKSMYDGDRF | 1701163 | Cell/Virus Enclosed | NO |
| TVYPDYVDVNKT | 1520553 | Ectodomain | YES |
| TVYRRIPDLPDC | 1520558 | Ectodomain | YES |
| TVYRRIPNLPDC | 1520559 | Ectodomain | YES |
| TVYSDVETPHLM | 1520562 | Cell/Virus Enclosed | NO |
| TWHEGRTIYGNVSRQD | 1701192 | Cell/Virus Enclosed | NO |
| TWLPELFMLNTL | 1520606 | Cell/Virus Enclosed | NO |
| TWLTYTGAIKLDDKDP | 1701205 | Cell/Virus Enclosed | NO |

| | | | |
|---|---|---|---|
| TWNGGAYSLFDMTKFS | 1701208 | Cell/Virus Enclosed | NO |
| TWNKKFGFIEDS | 1520616 | Ectodomain | YES |
| TWNKRFGFIEDA | 1520617 | Ectodomain | YES |
| TWNKRFGFIEDSVFKP | 1701209 | Ectodomain | YES |
| TWNLDLNSLTTDQRLH | 67283 | Ectodomain | YES |
| TWNSKGKTIKTTPEGTEE | 558342 | Undetermined | UN |
| TWPLGLECTRAS | 1520621 | Cell/Virus Enclosed | NO |
| TWPLVLECTRAS | 1520624 | Cell/Virus Enclosed | NO |
| TWPRPIDVFKAD | 1520625 | Ectodomain | YES |
| TWPRPIDVSKADGIIY | 1701212 | Ectodomain | YES |
| TWSEGNSITGVPYPVS | 1701218 | Ectodomain | YES |
| TWTADEAIKFLK | 1520638 | Ectodomain | YES |
| TWVEAKLSAGIE | 1520642 | Cell/Virus Enclosed | NO |
| TWVTGGIAIAMA | 1520643 | Undetermined | UN |
| TWWLKGDKLFVE | 1520644 | Cell/Virus Enclosed | NO |
| TWWLKGDKLSVE | 1520645 | Cell/Virus Enclosed | NO |
| TYAALRNSLTNDAYSR | 1701226 | Cell/Virus Enclosed | NO |
| TYAGNVWQIQTIQDSD | 1701233 | Cell/Virus Enclosed | NO |
| TYAGSVWHIQTVQDAD | 1701234 | Cell/Virus Enclosed | NO |
| TYANIVHEPTTP | 1520662 | Cell/Virus Enclosed | NO |
| TYANIVYEPTTP | 1520664 | Cell/Virus Enclosed | NO |
| TYDHSKYREEAMQNRIQIDPVKLSS | 1175592 | Ectodomain | YES |
| TYDPTYKNGAMY | 1520681 | Cell/Virus Enclosed | NO |
| TYERHPLSHFVN | 1520698 | Cell/Virus Enclosed | NO |
| TYFDNPILDFGD | 1520707 | Ectodomain | YES |
| TYFKVNGLWSPI | 1520714 | Cell/Virus Enclosed | NO |
| TYFLVRILQDKT | 1520716 | Cell/Virus Enclosed | NO |
| TYFNKPLLVDDN | 1520721 | Cell/Virus Enclosed | NO |
| TYFNLSSELKQL | 1520723 | Ectodomain | YES |
| TYFNRPSLVDDNKFDV | 1701260 | Cell/Virus Enclosed | NO |
| TYGNMTLNGLWLDDKV | 1701268 | Cell/Virus Enclosed | NO |
| TYGVCMYTDLNF | 1520756 | Cell/Virus Enclosed | NO |
| TYHTPATDCSDGNYNR | 1701281 | Ectodomain | YES |
| TYINGFICEKLA | 1520776 | Cell/Virus Enclosed | NO |
| TYIPASGYVAAS | 1520779 | Cell/Virus Enclosed | NO |
| TYKCPQSKSLVG | 1520794 | Ectodomain | YES |
| TYKLAMIMGYCC | 1520797 | Cell/Virus Enclosed | NO |
| TYKLVSMKGTGV | 1520801 | Cell/Virus Enclosed | NO |
| TYKMLVAKDNKNLPKF | 1701302 | Capsid | UN |
| TYKNTCDGNTFT | 1520803 | Cell/Virus Enclosed | NO |
| TYKSTATTKLVPGMLF | 1701310 | Cell/Virus Enclosed | NO |
| TYKSTVTTKLVPGMIF | 1701311 | Cell/Virus Enclosed | NO |
| TYLDFIQQGGIP | 1520819 | Cell/Virus Enclosed | NO |
| TYLDFIQQGGTPIANC | 1701314 | Cell/Virus Enclosed | NO |
| TYLDYIQQGGVPIINC | 1701318 | Cell/Virus Enclosed | NO |
| TYLKSDNIVAADLGVL | 1701323 | Cell/Virus Enclosed | NO |
| TYLNDPSSKTVCTYMD | 1701326 | Cell/Virus Enclosed | NO |
| TYLNLSSELKQLEAKT | 1701329 | Ectodomain | YES |
| TYLPATILAGSV | 1520837 | Cell/Virus Enclosed | NO |
| TYNKGENMTPRD | 1520863 | Cell/Virus Enclosed | NO |
| TYPFIANEKFINGCGR | 1701346 | Cell/Virus Enclosed | NO |
| TYPMIANENAINKGGR | 1701351 | Cell/Virus Enclosed | NO |
| TYPQQMFVTPLS | 1520882 | Cell/Virus Enclosed | NO |
| TYPQQMFVTPLT | 1520883 | Cell/Virus Enclosed | NO |
| TYQNISFNLPPP | 1520891 | Ectodomain | YES |
| TYQNISINLPPP | 1520892 | Ectodomain | YES |
| TYQNISTNLPPPLLGN | 1701359 | Ectodomain | YES |
| TYRNMALKGTDL | 1520899 | Ectodomain | YES |
| TYSEAAAQGFQA | 1520904 | Cell/Virus Enclosed | NO |
| TYSFLPKEDGDVLLAE | 1701366 | Cell/Virus Enclosed | NO |
| TYSPATILAGGV | 1520922 | Cell/Virus Enclosed | NO |
| TYTDQYLCGADGKPVM | 1701378 | Cell/Virus Enclosed | NO |
| TYTGAIKLDDKDPSFK | 1701381 | Cell/Virus Enclosed | NO |
| TYTSTSGNLLGFKDVT | 1701382 | Ectodomain | YES |
| TYTWTTVLSMAV | 1520949 | Cell/Virus Enclosed | NO |
| TYTYKMLVAKDNKNLP | 1701387 | Capsid | UN |
| TYVCNGNPRCKN | 1520956 | Ectodomain | YES |
| TYVCNGNVRCVELLKQ | 1701392 | Ectodomain | YES |
| TYVCQCGGERHQ | 1520958 | Cell/Virus Enclosed | NO |
| TYVCSLIAVAYT | 1520961 | Cell/Virus Enclosed | NO |
| TYVPSQEKNFTT | 1520971 | Ectodomain | YES |
| TYVSNSGNLLGFKNVS | 1701400 | Ectodomain | YES |
| TYVSVDDKYKTI | 1520974 | Cell/Virus Enclosed | NO |
| TYYAHSVLYRVF | 1520986 | Cell/Virus Enclosed | NO |
| TYYCVTTKAKSI | 1520988 | Cell/Virus Enclosed | NO |
| TYYCVTTKAKSVQQDL | 1701406 | Cell/Virus Enclosed | NO |

| | | | |
|---|---|---|---|
| TYYIYDEEGGNDLSLP | 1701409 | Cell/Virus Enclosed | NO |
| TYYLDDVKKIEYKPDL | 1701410 | Cell/Virus Enclosed | NO |
| TYYNSWQNLLYDSNGN | 1701415 | Ectodomain | YES |
| TYYQPIQQAPTG | 1521003 | Underneath Membrane | NO |
| VAAALKSLGFDKPQEK | 1701418 | Capsid | UN |
| VAADLGVLIQNSAKHV | 1701420 | Cell/Virus Enclosed | NO |
| VAAGQQFVKETT | 1521016 | Cell/Virus Enclosed | NO |
| VAAITSGVKFQTVKPG | 1701433 | Cell/Virus Enclosed | NO |
| VAALSTGLTNQV | 1521030 | Cell/Virus Enclosed | NO |
| VAALSTGLTSQTVKPG | 1701439 | Cell/Virus Enclosed | NO |
| VAALTNNVAFQT | 1521031 | Cell/Virus Enclosed | NO |
| VAANVGRWSASI | 1521041 | Ectodomain | YES |
| VAARADYEGFIT | 1521044 | Cell/Virus Enclosed | NO |
| VAASDITLKCCT | 1521046 | Cell/Virus Enclosed | NO |
| VAAVDEAFPLDE | 1521050 | Cell/Virus Enclosed | NO |
| VAAYNNCTGGAE | 1521066 | Ectodomain | YES |
| VACEHISSTMSQYSRS | 1701460 | Ectodomain | YES |
| VACQLPYCYFRNSSAN | 1701463 | Underneath Membrane | NO |
| VACQPPYCYFRNSTTN | 1701464 | Ectodomain | YES |
| VADDSGTPVVEC | 1521087 | Cell/Virus Enclosed | NO |
| VADDSGTPVVEY | 1521088 | Cell/Virus Enclosed | NO |
| VADQSRKWIGHCTIAQ | 1701472 | Cell/Virus Enclosed | NO |
| VADQSRKWTGHC | 1521093 | Cell/Virus Enclosed | NO |
| VADSELYADWSSDSEGERDG | 742103 | Underneath Membrane | NO |
| VADSYFSYMMPM | 1521094 | Cell/Virus Enclosed | NO |
| VADTLMQGVTLSSNLN | 1701474 | Ectodomain | YES |
| VADVMRYNFNFS | 1521097 | Ectodomain | YES |
| VADVMRYNLNFS | 1521098 | Ectodomain | YES |
| VADYHNHSNVET | 1521104 | Cell/Virus Enclosed | NO |
| VAEYITTYPGQG | 1521128 | Cell/Virus Enclosed | NO |
| VAFCGDQVHEVA | 1521131 | Cell/Virus Enclosed | NO |
| VAFELFAKRKMGLTPP | 1390173 | Cell/Virus Enclosed | NO |
| VAFGGDQIHEVA | 1521135 | Cell/Virus Enclosed | NO |
| VAFGGDQVHEVAAVRS | 1701494 | Cell/Virus Enclosed | NO |
| VAFGGDQVHEVV | 1521137 | Cell/Virus Enclosed | NO |
| VAFGLQDCVTGINDDD | 1701495 | Cell/Virus Enclosed | NO |
| VAFTKAAGGKVSFSDD | 1701510 | Cell/Virus Enclosed | NO |
| VAFTKLAGGKISFSDD | 1701511 | Cell/Virus Enclosed | NO |
| VAFYQGDVDTVVNGVD | 1701516 | Cell/Virus Enclosed | NO |
| VAGDSGFAAYSRYRIG | 1390174 | Underneath Membrane | NO |
| VAGRKKLEELERDLRKTK | 67615 | Cell/Virus Enclosed | NO |
| VAGYKVLPPLMDVNME | 1701533 | Ectodomain | YES |
| VAIADSYYSYIMPMLT | 1701536 | Cell/Virus Enclosed | NO |
| VAICLTYANIVH | 1521195 | Cell/Virus Enclosed | NO |
| VAIEQLLYAIQQLYTG | 1390179 | Cell/Virus Enclosed | NO |
| VAINDVDTGVPS | 1521217 | Ectodomain | YES |
| VAIQEVIDDMHWGDGF | 1701550 | Cell/Virus Enclosed | NO |
| VAIQEVIDDMLW | 1521220 | Cell/Virus Enclosed | NO |
| VAITRAKIGILC | 1521223 | Cell/Virus Enclosed | NO |
| VAITRAQKGILC | 1521225 | Cell/Virus Enclosed | NO |
| VAKAIAVAAGKKFSKE | 1701566 | Cell/Virus Enclosed | NO |
| VAKAIAVAAGQQ | 1521235 | Cell/Virus Enclosed | NO |
| VAKAPPGEQFKH | 1521236 | Cell/Virus Enclosed | NO |
| VAKEGAKTVNTSLGNR | 1701572 | Capsid | UN |
| VAKILARSQSSE | 1521248 | Cell/Virus Enclosed | NO |
| VAKLCLDAGINY | 1521254 | Cell/Virus Enclosed | NO |
| VAKPNVLIALKM | 1521266 | Ectodomain | YES |
| VAKQGAGDAGHSLTWL | 1701585 | Cell/Virus Enclosed | NO |
| VAKRVLGVQTQTVDSA | 1701587 | Cell/Virus Enclosed | NO |
| VALEQDKNTQEV | 1521291 | Ectodomain | YES |
| VALPQSKDSNFL | 1521303 | Cell/Virus Enclosed | NO |
| VALQTDVLQEDQ | 1521306 | Ectodomain | YES |
| VALQTDVLQENQ | 1521307 | Ectodomain | YES |
| VANDFTTALRRP | 1521328 | Cell/Virus Enclosed | NO |
| VANGDSEAVLKK | 1521335 | Cell/Virus Enclosed | NO |
| VANGLNAPNGPFDALF | 1701636 | Cell/Virus Enclosed | NO |
| VANGVKAKGYPQFAEL | 1701638 | Capsid | UN |
| VANGVTAKGYPQ | 1521340 | Cell/Virus Enclosed | NO |
| VANHQADTSTPSDVSS | 1701640 | Capsid | UN |
| VANLKLPPHDVI | 1521346 | Cell/Virus Enclosed | NO |
| VANLKLPPHDVTVLRD | 1701642 | Cell/Virus Enclosed | NO |
| VANLSAPTLVPQENYS | 1701643 | Cell/Virus Enclosed | NO |
| VANNLMNGVTLS | 1521349 | Ectodomain | YES |
| VANPSTPAYTFTTVKP | 1701646 | Cell/Virus Enclosed | NO |
| VANPTEDLCNGS | 1521355 | Cell/Virus Enclosed | NO |
| VANPTNCIAPVN | 1521356 | Ectodomain | YES |

| | | | |
|---|---|---|---|
| VANWLAPTNAYMRTTH | 1701650 | Cell/Virus Enclosed | NO |
| VANWLAPTNAYT | 1521361 | Cell/Virus Enclosed | NO |
| VANWLAPTNVYM | 1521362 | Cell/Virus Enclosed | NO |
| VANYFQDVKQFA | 1521364 | Ectodomain | YES |
| VANYSQNVKQFA | 1521368 | Ectodomain | YES |
| VAPDLSFDYINV | 1521373 | Ectodomain | YES |
| VAPGETGVIADY | 1521378 | Ectodomain | YES |
| VAPGTAVLKRWLPHDA | 1701661 | Cell/Virus Enclosed | NO |
| VAPHLSLDYINV | 1521386 | Ectodomain | YES |
| VAPIELENKFTPLSVE | 1701665 | Cell/Virus Enclosed | NO |
| VAPLLSLDYINV | 1521390 | Ectodomain | YES |
| VAPQATYQNIST | 1521392 | Ectodomain | YES |
| VAPQVTYQNISTNLPP | 1701668 | Ectodomain | YES |
| VAPVNTVKPKPVINQL | 1701669 | Cell/Virus Enclosed | NO |
| VAPVYEADTEQTQNVT | 1701670 | Cell/Virus Enclosed | NO |
| VAQNTVLNACTP | 1521414 | Cell/Virus Enclosed | NO |
| VAQNTVLNASIPESKP | 1701676 | Capsid | UN |
| VAQNTVLNDSIP | 1521417 | Capsid | UN |
| VARDLSLQFKRS | 1521430 | Cell/Virus Enclosed | NO |
| VARLMKGDKGDAEDTL | 1701687 | Cell/Virus Enclosed | NO |
| VASDLTTSLRRL | 1521449 | Cell/Virus Enclosed | NO |
| VASDSEISANGCSTIA | 1701695 | Cell/Virus Enclosed | NO |
| VASDSEISGFGFAIPS | 1701696 | Cell/Virus Enclosed | NO |
| VASIITKLNLLN | 1521463 | Cell/Virus Enclosed | NO |
| VASINSAIVCASVKRD | 1701707 | Cell/Virus Enclosed | NO |
| VASIPLCILPYT | 1521468 | Cell/Virus Enclosed | NO |
| VASIQADVNTPA | 1521470 | Cell/Virus Enclosed | NO |
| VASLSAPTLVPQENYA | 1701710 | Cell/Virus Enclosed | NO |
| VASNDAIMTRCL | 1521477 | Cell/Virus Enclosed | NO |
| VASNQADVNIPA | 1521478 | Cell/Virus Enclosed | NO |
| VASNQADVNTPADIVD | 1701712 | Cell/Virus Enclosed | NO |
| VASNSKLWHYCSTLHN | 1701713 | Cell/Virus Enclosed | NO |
| VASNSKLWQYCSVLHN | 1701714 | Cell/Virus Enclosed | NO |
| VASPKSRVQQNK | 1521480 | Cell/Virus Enclosed | NO |
| VASRFLGLQIQTVDSS | 1701719 | Cell/Virus Enclosed | NO |
| VASSFVGMPSFVAYET | 1701721 | Cell/Virus Enclosed | NO |
| VASSSYDALLHRNSLD | 1701724 | Cell/Virus Enclosed | NO |
| VASTLVEPPTVD | 1521490 | Cell/Virus Enclosed | NO |
| VATALNKIHDVV | 1521501 | Ectodomain | YES |
| VATALNKIQDVVNQQG | 1701730 | Ectodomain | YES |
| VATCYNSVNNDY | 1521502 | Cell/Virus Enclosed | NO |
| VATEGALNTPKDHIGTRNPA | 1334138 | Cell/Virus Enclosed | NO |
| VATFYPQLQSAEWKCG | 1701736 | Cell/Virus Enclosed | NO |
| VATGDVVLASDDLYVK | 1390205 | Cell/Virus Enclosed | NO |
| VATIPYKVCNSVKDTL | 1701737 | Cell/Virus Enclosed | NO |
| VATLMAPTIVNQ | 1521515 | Cell/Virus Enclosed | NO |
| VATLTAPTIANQ | 1521518 | Cell/Virus Enclosed | NO |
| VATRNATVVIGTTKFY | 1701742 | Cell/Virus Enclosed | NO |
| VATSRTLSYYRL | 1521531 | Underneath Membrane | NO |
| VATSSYDALLHR | 1521532 | Cell/Virus Enclosed | NO |
| VATVFLFWFNFL | 1521533 | Cell/Virus Enclosed | NO |
| VAVADSYFSYMM | 1521541 | Cell/Virus Enclosed | NO |
| VAVADSYYSYMMPMLT | 1701750 | Cell/Virus Enclosed | NO |
| VAVDGAKTEPTGYGVR | 1701751 | Capsid | UN |
| VAVIGDEQIDGKFVTK | 1701756 | Cell/Virus Enclosed | NO |
| VAVKNGALHPYF | 1521555 | Cell/Virus Enclosed | NO |
| VAVSKSRVQQNK | 1521581 | Cell/Virus Enclosed | NO |
| VAVTAGPFMGNA | 1521584 | Cell/Virus Enclosed | NO |
| VAVVNDKIFTKSGVND | 1701776 | Cell/Virus Enclosed | NO |
| VAVVNDKVFVRYGDVD | 1701777 | Cell/Virus Enclosed | NO |
| VAVYANVGRWSASINT | 1701778 | Ectodomain | YES |
| VAVYQQSIIGAMTAVN | 1701779 | Ectodomain | YES |
| VAWDWVPKRPSVCTMTKWQE | 742106 | Ectodomain | YES |
| VAWHVVRDPRYVMRLQ | 1701780 | Cell/Virus Enclosed | NO |
| VAWHVVRDSRFVMRLQ | 1701781 | Cell/Virus Enclosed | NO |
| VAYADKPNHFINFPLT | 1701784 | Cell/Virus Enclosed | NO |
| VAYADNPNHFIN | 1521598 | Cell/Virus Enclosed | NO |
| VAYHTVIDVKQV | 1521615 | Cell/Virus Enclosed | NO |
| VAYQLYMIDPAG | 1521621 | Cell/Virus Enclosed | NO |
| VAYTMSLGADSSIAYS | 1701805 | Ectodomain | YES |
| VAYVANPTEDLC | 1521636 | Cell/Virus Enclosed | NO |
| VCAFLNKLNENV | 1521641 | Cell/Virus Enclosed | NO |
| VCALMSCNGNKIEDLS | 1701813 | Cell/Virus Enclosed | NO |
| VCASVKRDGVQVGYCV | 1701814 | Cell/Virus Enclosed | NO |
| VCAWLTKRKPLD | 1521655 | Cell/Virus Enclosed | NO |
| VCDAIENALNSCKEHP | 1701819 | Cell/Virus Enclosed | NO |

| | | | |
|---|---|---|---|
| VCDDCNGYTDNIFSVQ | 1701820 | Ectodomain | YES |
| VCDKVCSLKLSD | 1521666 | Cell/Virus Enclosed | NO |
| VCDSLNAKLGFDSSKE | 1701829 | Cell/Virus Enclosed | NO |
| VCDSLNAKWIFD | 1521673 | Cell/Virus Enclosed | NO |
| VCDSLNAKWVFD | 1521674 | Cell/Virus Enclosed | NO |
| VCDSLNFKLGFD | 1521675 | Cell/Virus Enclosed | NO |
| VCEDVTQLAEEP | 1521677 | Cell/Virus Enclosed | NO |
| VCFDNSIQGSYE | 1521684 | Cell/Virus Enclosed | NO |
| VCFGFDKWFVNDGRVA | 1701836 | Cell/Virus Enclosed | NO |
| VCFSVAALSTGLTSQT | 1701843 | Cell/Virus Enclosed | NO |
| VCFVDVDFHSVK | 1521702 | Cell/Virus Enclosed | NO |
| VCGDNIGTNVPLQVGF | 1701852 | Cell/Virus Enclosed | NO |
| VCGDYAACKSQL | 1521712 | Ectodomain | YES |
| VCGPKLSTDLIKNQCV | 67844 | Ectodomain | YES |
| VCGYTDFAEDVCTCYD | 1701868 | Cell/Virus Enclosed | NO |
| VCHSVAVIGDEQIDGK | 1701870 | Cell/Virus Enclosed | NO |
| VCITTNGSWAIFNDHH | 1701877 | Cell/Virus Enclosed | NO |
| VCKGCVGHSENV | 1521756 | Ectodomain | YES |
| VCKPHQVNISLNGNTS | 1701885 | Ectodomain | YES |
| VCKVCGCWLNHG | 1521773 | Cell/Virus Enclosed | NO |
| VCKVENKAEAEW | 1521774 | Cell/Virus Enclosed | NO |
| VCKYKGKFVQIP | 1521779 | Cell/Virus Enclosed | NO |
| VCKYTDIDVNSALNIC | 1701894 | Cell/Virus Enclosed | NO |
| VCLGISDSAVTY | 1521795 | Cell/Virus Enclosed | NO |
| VCLKHAEEYREYLESY | 1701907 | Cell/Virus Enclosed | NO |
| VCLNGLDSLDSY | 1521808 | Cell/Virus Enclosed | NO |
| VCLSGLDSLDSY | 1521810 | Cell/Virus Enclosed | NO |
| VCMAFLTATRLC | 1521817 | Cell/Virus Enclosed | NO |
| VCNAPGCDVNDVTKLY | 1701915 | Cell/Virus Enclosed | NO |
| VCNASGCGVSDVKKLY | 1701919 | Cell/Virus Enclosed | NO |
| VCNFNLCKEPMY | 1521827 | Ectodomain | YES |
| VCNGCVGHSEIV | 1521829 | Ectodomain | YES |
| VCNGCVGHSKNV | 1521831 | Ectodomain | YES |
| VCNGFQKCEQLL | 1521836 | Ectodomain | YES |
| VCNGLQKCEQLL | 1521837 | Underneath Membrane | NO |
| VCNGNPRCKNLL | 1521838 | Ectodomain | YES |
| VCNGNVRCVELLKQYT | 1701924 | Ectodomain | YES |
| VCNNSDYASKGY | 1521845 | Cell/Virus Enclosed | NO |
| VCNTSGCNVNDVTKLY | 1701929 | Cell/Virus Enclosed | NO |
| VCPKLEFANDTKIASQ | 1701932 | Ectodomain | YES |
| VCQTTNGTNTSH | 1521871 | Ectodomain | YES |
| VCQYNMCEHPHT | 1521873 | Ectodomain | YES |
| VCQYNMCEYPQT | 1521875 | Ectodomain | YES |
| VCRKHATDYREY | 1521890 | Cell/Virus Enclosed | NO |
| VCRKHATEYREY | 1521892 | Cell/Virus Enclosed | NO |
| VCRSTAKYVRNNMETTAFHR | 742108 | Ectodomain | YES |
| VCSKHAEEYCNYLESY | 1701952 | Cell/Virus Enclosed | NO |
| VCSKHANLYQKYVEAY | 1701953 | Cell/Virus Enclosed | NO |
| VCSLKLSDNNCY | 1521905 | Cell/Virus Enclosed | NO |
| VCSSVKDTLTYY | 1521911 | Cell/Virus Enclosed | NO |
| VCTDNSANTKVLRNKG | 1701968 | Cell/Virus Enclosed | NO |
| VCVFQLQDGKTLGLK | 227651 | Cell/Virus Enclosed | NO |
| VCVNGLMLLELK | 1521945 | Cell/Virus Enclosed | NO |
| VCVYDPLPVILL | 1521952 | Undetermined | UN |
| VCYNSDYASKGYIANI | 1702011 | Cell/Virus Enclosed | NO |
| VDADSKHVQLSE | 1521980 | Cell/Virus Enclosed | NO |
| VDAFNQFSSDFQHKLK | 1702020 | Cell/Virus Enclosed | NO |
| VDAKDVIECLKL | 1521984 | Cell/Virus Enclosed | NO |
| VDAKGFPLLAEL | 1521985 | Cell/Virus Enclosed | NO |
| VDAKGFPQFAEL | 1521986 | Cell/Virus Enclosed | NO |
| VDAKGFPQLAELIPNQ | 1702022 | Capsid | UN |
| VDALCEKAYKFLNIND | 1702025 | Cell/Virus Enclosed | NO |
| VDANTNQDSYGGASIC | 1702028 | Cell/Virus Enclosed | NO |
| VDAYVDTFSATF | 1522004 | Cell/Virus Enclosed | NO |
| VDCATYVCNGNP | 1522006 | Ectodomain | YES |
| VDCDAAGVGNTF | 1522007 | Cell/Virus Enclosed | NO |
| VDCDIAGVGNTF | 1522008 | Cell/Virus Enclosed | NO |
| VDCDTAGVGNTF | 1522011 | Cell/Virus Enclosed | NO |
| VDCKIQTAPHIS | 1522014 | Cell/Virus Enclosed | NO |
| VDCKQYICNGFQ | 1522015 | Ectodomain | YES |
| VDCKQYVCNGFH | 1522018 | Ectodomain | YES |
| VDCKQYVCNGLQ | 1522021 | Underneath Membrane | NO |
| VDCKSDFMSEIK | 1522022 | Ectodomain | YES |
| VDCMSDFMSGIK | 1522024 | Ectodomain | YES |
| VDCQDDRCFIHC | 1522026 | Cell/Virus Enclosed | NO |
| VDCRRLLKQDCC | 1522027 | Cell/Virus Enclosed | NO |

| | | | |
|---|---|---|---|
| VDCSTYVCNGNVRCVE | 1702052 | Ectodomain | YES |
| VDCTMYICGDSL | 1522037 | Ectodomain | YES |
| VDCTMYICGDSQ | 1522038 | Ectodomain | YES |
| VDCVSDFMSEIK | 1522041 | Ectodomain | YES |
| VDCWTATDRFYR | 1522042 | Cell/Virus Enclosed | NO |
| VDDDYICGDGLI | 1522044 | Cell/Virus Enclosed | NO |
| VDDFQGYLQKHF | 1522046 | Cell/Virus Enclosed | NO |
| VDDFYGYLQRHF | 1522047 | Cell/Virus Enclosed | NO |
| VDDGFLDIWTYNAEL | 151067 | Ectodomain | YES |
| VDDGGDNSESDA | 1522048 | Cell/Virus Enclosed | NO |
| VDDGYICGDGLI | 1522050 | Cell/Virus Enclosed | NO |
| VDDHALIVNVLL | 1522052 | Cell/Virus Enclosed | NO |
| VDDHVGLLDLAW | 1522054 | Cell/Virus Enclosed | NO |
| VDDHVRLLDQAW | 1522056 | Cell/Virus Enclosed | NO |
| VDDIDPINCESI | 1522057 | Cell/Virus Enclosed | NO |
| VDDIDTINCESI | 1522058 | Cell/Virus Enclosed | NO |
| VDDIESIYDFDTYKAL | 1702062 | Cell/Virus Enclosed | NO |
| VDDIESIYEFDT | 1522060 | Cell/Virus Enclosed | NO |
| VDDINFSPVLGCLGSE | 1702063 | Ectodomain | YES |
| VDDIYGYVLRQP | 1522063 | Ectodomain | YES |
| VDDIYYPASCNGVLPV | 1702067 | Cell/Virus Enclosed | NO |
| VDDKYKTIDELC | 1522064 | Cell/Virus Enclosed | NO |
| VDDKYKTSDELC | 1522065 | Cell/Virus Enclosed | NO |
| VDDLLKTDSILL | 1522067 | Cell/Virus Enclosed | NO |
| VDDLLKTDSVLLIERF | 1702071 | Cell/Virus Enclosed | NO |
| VDDLVVDWHYFD | 1522068 | Cell/Virus Enclosed | NO |
| VDDPKLMGWDYPKCDR | 1702074 | Cell/Virus Enclosed | NO |
| VDDPVVDWHYFD | 1522072 | Cell/Virus Enclosed | NO |
| VDDSGDSSEGGA | 1522076 | Cell/Virus Enclosed | NO |
| VDDSGDSSESGA | 1522077 | Cell/Virus Enclosed | NO |
| VDDSGTPVVEYL | 1522078 | Cell/Virus Enclosed | NO |
| VDDSIILRQYNL | 1522079 | Cell/Virus Enclosed | NO |
| VDDVNFSPVLGC | 1522082 | Ectodomain | YES |
| VDDVVKTDAVVLLERY | 1702079 | Cell/Virus Enclosed | NO |
| VDDWSYTGSSFY | 1522084 | Ectodomain | YES |
| VDEAFPLDEAEDVTES | 1702082 | Cell/Virus Enclosed | NO |
| VDEAFPLDEAEN | 1522086 | Cell/Virus Enclosed | NO |
| VDEAFPLNEAED | 1522089 | Cell/Virus Enclosed | NO |
| VDENGDKLDQFFVVKR | 1702085 | Cell/Virus Enclosed | NO |
| VDENGKYYLPYPDPSR | 1702086 | Cell/Virus Enclosed | NO |
| VDESFVDDFQGY | 1522101 | Cell/Virus Enclosed | NO |
| VDEVSMCTNYDLSIIN | 1702092 | Cell/Virus Enclosed | NO |
| VDEVSMCTNYDLSVIN | 1702093 | Cell/Virus Enclosed | NO |
| VDEWSYTGSSFYAPEP | 1702095 | Ectodomain | YES |
| VDFCGRGYHLMS | 1522109 | Ectodomain | YES |
| VDFESVIQDYEN | 1522110 | Cell/Virus Enclosed | NO |
| VDFHSVEIEAGEVKPF | 1702100 | Cell/Virus Enclosed | NO |
| VDFINSMQFNYKITIN | 1702101 | Cell/Virus Enclosed | NO |
| VDFKDFQFMLWG | 1522117 | Cell/Virus Enclosed | NO |
| VDFKPQSGGGKCFNCY | 1702105 | Ectodomain | YES |
| VDFNVNVIGTRP | 1522125 | Cell/Virus Enclosed | NO |
| VDFNVNVMGTRP | 1522126 | Cell/Virus Enclosed | NO |
| VDFTECAYTQGR | 1522130 | Cell/Virus Enclosed | NO |
| VDGAKTEPTGHG | 1522132 | Capsid | UN |
| VDGDMVPHISRQ | 1522138 | Cell/Virus Enclosed | NO |
| VDGDVVIREPVH | 1522139 | Cell/Virus Enclosed | NO |
| VDGELPVAVVNDKVFV | 1702120 | Cell/Virus Enclosed | NO |
| VDGFGLNKIQPW | 1522141 | Cell/Virus Enclosed | NO |
| VDGFGVNKIQPWTNDA | 1702121 | Cell/Virus Enclosed | NO |
| VDGFNEWAIANG | 1522144 | Cell/Virus Enclosed | NO |
| VDGHRLASGVQVHNLP | 1702125 | Underneath Membrane | NO |
| VDGHVGLLDQAW | 1522148 | Cell/Virus Enclosed | NO |
| VDGIDVEIFESK | 1522150 | Cell/Virus Enclosed | NO |
| VDGKPISAYAFL | 1522154 | Cell/Virus Enclosed | NO |
| VDGKQIDDHRIT | 1522156 | Cell/Virus Enclosed | NO |
| VDGLESKQVDYVPLRS | 1702134 | Cell/Virus Enclosed | NO |
| VDGLKYTKILKDDGNF | 1702135 | Cell/Virus Enclosed | NO |
| VDGLKYTKIMKDDGNC | 1702136 | Cell/Virus Enclosed | NO |
| VDGNDVEIFENK | 1522159 | Cell/Virus Enclosed | NO |
| VDGVNFKSISLTVGEV | 1702141 | Cell/Virus Enclosed | NO |
| VDGVNFRTVVLNNKNT | 1702142 | Cell/Virus Enclosed | NO |
| VDGVNFTNRFVPVGES | 1702143 | Cell/Virus Enclosed | NO |
| VDGVPFVVSYGY | 1522168 | Cell/Virus Enclosed | NO |
| VDGVSRLYQPLRLTCL | 1702148 | Ectodomain | YES |
| VDGVSRRYQPLR | 1522171 | Ectodomain | YES |
| VDGYIRRAIDCG | 1522175 | Ectodomain | YES |

| | | | |
|---|---|---|---|
| VDGYIRRSIDCG | 1522177 | Ectodomain | YES |
| VDGYIRTAIDCG | 1522178 | Ectodomain | YES |
| VDHFGENEEIIINGHT | 1702156 | Cell/Virus Enclosed | NO |
| VDHKPHLSFPLC | 1522186 | Cell/Virus Enclosed | NO |
| VDHRPVCSFPLC | 1522187 | Cell/Virus Enclosed | NO |
| VDHSAFAYESAV | 1522188 | Cell/Virus Enclosed | NO |
| VDHVQIHTIDGS | 1522190 | Secreted | YES |
| VDIIEQPFGEVE | 1522196 | Cell/Virus Enclosed | NO |
| VDIIEQPFGEVK | 1522197 | Cell/Virus Enclosed | NO |
| VDIIEQSFGEVE | 1522198 | Cell/Virus Enclosed | NO |
| VDIIQRLDVLEQ | 1522199 | Ectodomain | YES |
| VDIMSCFYMEAD | 1522204 | Cell/Virus Enclosed | NO |
| VDINKAIPSGNLIKFD | 1702169 | Cell/Virus Enclosed | NO |
| VDINRLCEEVLD | 1522209 | Cell/Virus Enclosed | NO |
| VDIQQTFFDKTWPRPI | 1702176 | Ectodomain | YES |
| VDIQQWGYIGSL | 1522216 | Cell/Virus Enclosed | NO |
| VDIVKEAQHAKP | 1522221 | Cell/Virus Enclosed | NO |
| VDKDGNYYLPYP | 1522226 | Cell/Virus Enclosed | NO |
| VDKEVKDMQFDLYVNV | 1702185 | Cell/Virus Enclosed | NO |
| VDKFAIPNRRRD | 1522231 | Ectodomain | YES |
| VDKFKTPHIVRQ | 1522232 | Cell/Virus Enclosed | NO |
| VDKIRHLLKGVT | 1522234 | Cell/Virus Enclosed | NO |
| VDKLKPVFTWVE | 1522237 | Cell/Virus Enclosed | NO |
| VDKPNHFINFPL | 1522238 | Cell/Virus Enclosed | NO |
| VDKQVILVSNNKEDFDIIQ | 1522241 | Cell/Virus Enclosed | NO |
| VDKTKTPHIVRQ | 1522246 | Cell/Virus Enclosed | NO |
| VDKTLPDQGDNDNWWTGWRQWIPAG | 932264 | Secreted | YES |
| VDKYMCGFDGKPVLPK | 1702202 | Cell/Virus Enclosed | NO |
| VDLDALASDIFA | 1522253 | Cell/Virus Enclosed | NO |
| VDLHNKINLCDDPEKA | 1702213 | Cell/Virus Enclosed | NO |
| VDLLSNFEHKVI | 1522268 | Cell/Virus Enclosed | NO |
| VDLNTTILFTGY | 1522272 | Underneath Membrane | NO |
| VDLNWLNRVETY | 1522273 | Ectodomain | YES |
| VDLPFVEYKVTV | 1522274 | Cell/Virus Enclosed | NO |
| VDLPPKVHFYYP | 1522276 | Cell/Virus Enclosed | NO |
| VDLPPSHATTYL | 1522278 | Cell/Virus Enclosed | NO |
| VDLQIQLTTTRM | 1522279 | Cell/Virus Enclosed | NO |
| VDLQLGNLAYLQ | 1522280 | Ectodomain | YES |
| VDLSMDKLRRAD | 1522286 | Cell/Virus Enclosed | NO |
| VDLTEAICDFEI | 1522293 | Cell/Virus Enclosed | NO |
| VDLTGAICDFEI | 1522294 | Cell/Virus Enclosed | NO |
| VDLYGGNMFQFATLPV | 1702226 | Ectodomain | YES |
| VDMIAAYTAALV | 1522302 | Ectodomain | YES |
| VDMNSITSTDVRILVQPGIASELVI | 68025 | Undetermined | UN |
| VDMYPEFSIVCR | 1522306 | Cell/Virus Enclosed | NO |
| VDNDLHPFVSDS | 1522308 | Cell/Virus Enclosed | NO |
| VDNDLNPFVSDS | 1522309 | Cell/Virus Enclosed | NO |
| VDNDLYPFVSDSVATY | 1702234 | Cell/Virus Enclosed | NO |
| VDNGSSINRRQLDVVK | 1702238 | Cell/Virus Enclosed | NO |
| VDNIYVTYAGNV | 1522315 | Cell/Virus Enclosed | NO |
| VDNLVFTNKTTLPTNV | 1702241 | Cell/Virus Enclosed | NO |
| VDNPHLMGWDYPKCDR | 1702244 | Cell/Virus Enclosed | NO |
| VDNPVLMGWDYPKCDR | 1702245 | Cell/Virus Enclosed | NO |
| VDNRECISMDYKQTQLCLIG | 111890 | Capsid | UN |
| VDNTLYSEFSTI | 1522322 | Undetermined | UN |
| VDNVKLPRWSAS | 1522325 | Ectodomain | YES |
| VDNVYMAKAGDKYYPV | 1702249 | Cell/Virus Enclosed | NO |
| VDNVYMAKVGDK | 1522330 | Cell/Virus Enclosed | NO |
| VDPAAAYLDAVKQGAK | 1702252 | Cell/Virus Enclosed | NO |
| VDPATTYLEAVKHGAK | 1702254 | Cell/Virus Enclosed | NO |
| VDPDDVNKNTLQAR | 68034 | Capsid | UN |
| VDPDLSAFYVKDGKYF | 1702255 | Cell/Virus Enclosed | NO |
| VDPKKTYLDFIQQGGT | 1702256 | Cell/Virus Enclosed | NO |
| VDPKKTYLDYIQQGGV | 1702257 | Cell/Virus Enclosed | NO |
| VDPKYQVIVLKDNVLS | 1702258 | Cell/Virus Enclosed | NO |
| VDPLSNFEHKDI | 1522343 | Cell/Virus Enclosed | NO |
| VDPNLSGFYVKD | 1522346 | Cell/Virus Enclosed | NO |
| VDPQIQLTITRM | 1522351 | Cell/Virus Enclosed | NO |
| VDPQIQLTTTRM | 1522352 | Cell/Virus Enclosed | NO |
| VDPQKAYLDFVNAGGA | 1702261 | Cell/Virus Enclosed | NO |
| VDPTTVLNDPVFTGDL | 1702263 | Cell/Virus Enclosed | NO |
| VDPTTVLNGPVF | 1522359 | Cell/Virus Enclosed | NO |
| VDQFERSHQTYN | 1522362 | Cell/Virus Enclosed | NO |
| VDQFNSSYFKLS | 1522363 | Ectodomain | YES |
| VDQLKHLLEQWN | 1522364 | Undetermined | UN |
| VDQLNSSYFKLSIPTN | 1702264 | Ectodomain | YES |

| | | | |
|---|---|---|---|
| VDQLNSSYKLSI | 1522365 | Ectodomain | YES |
| VDQLNSTYFKLS | 1522366 | Ectodomain | YES |
| VDQSERSHQTYN | 1522369 | Cell/Virus Enclosed | NO |
| VDQSRKWIGHCT | 1522370 | Cell/Virus Enclosed | NO |
| VDQSRKWISHCT | 1522371 | Cell/Virus Enclosed | NO |
| VDQVEEVNSIFDIETV | 1702266 | Cell/Virus Enclosed | NO |
| VDQYMCGVDGKPISAY | 1702270 | Cell/Virus Enclosed | NO |
| VDRDPSNDEAIP | 1522375 | Cell/Virus Enclosed | NO |
| VDRELTSTHDGP | 1522380 | Cell/Virus Enclosed | NO |
| VDRESTLTHYGL | 1522381 | Cell/Virus Enclosed | NO |
| VDRESTSIHDGP | 1522382 | Cell/Virus Enclosed | NO |
| VDRIRFCLMLNS | 1522393 | Cell/Virus Enclosed | NO |
| VDRQTAQAAGAD | 1522394 | Cell/Virus Enclosed | NO |
| VDRYPANAIVCR | 1522397 | Cell/Virus Enclosed | NO |
| VDSAQGSEYDYVIFAQ | 1702281 | Cell/Virus Enclosed | NO |
| VDSAQGSEYDYVIYSQ | 1702282 | Cell/Virus Enclosed | NO |
| VDSCLQGDRCNI | 1522400 | Ectodomain | YES |
| VDSDLNEFVSDA | 1522402 | Cell/Virus Enclosed | NO |
| VDSDQPWKVIPR | 1522404 | Cell/Virus Enclosed | NO |
| VDSEQPCLVIPR | 1522407 | Cell/Virus Enclosed | NO |
| VDSEQPWKVIPRNLVP | 1702287 | Capsid | UN |
| VDSEQPWKVVPR | 1522409 | Cell/Virus Enclosed | NO |
| VDSEQSWKVIPR | 1522410 | Capsid | UN |
| VDSFEGVLPINTDTVL | 1702288 | Cell/Virus Enclosed | NO |
| VDSFTCNNIDAA | 1522413 | Ectodomain | YES |
| VDSFTNGAIVGI | 1522414 | Cell/Virus Enclosed | NO |
| VDSFTNGAVIGT | 1522416 | Cell/Virus Enclosed | NO |
| VDSFTNGAVVGT | 1522417 | Cell/Virus Enclosed | NO |
| VDSFVVKGDDVR | 1522418 | Ectodomain | YES |
| VDSIVQRCYELS | 1522423 | Cell/Virus Enclosed | NO |
| VDSKCLTSIEGV | 1522425 | Cell/Virus Enclosed | NO |
| VDSKYQVIVLKD | 1522427 | Cell/Virus Enclosed | NO |
| VDSLCAKAVTAYSVDK | 1702292 | Cell/Virus Enclosed | NO |
| VDSNENLTWPLG | 1522430 | Cell/Virus Enclosed | NO |
| VDSNLNDFVSDA | 1522431 | Cell/Virus Enclosed | NO |
| VDSNWPLVIIANRYNE | 1702294 | Cell/Virus Enclosed | NO |
| VDSRWNNARHSD | 1522433 | Ectodomain | YES |
| VDSRWNNARLSD | 1522434 | Ectodomain | YES |
| VDSRWNNARQFD | 1522435 | Ectodomain | YES |
| VDSRWNNARQSDNMTA | 1702295 | Ectodomain | YES |
| VDSRWNNERASDISLS | 1702296 | Underneath Membrane | NO |
| VDSRWNNLRASD | 1522438 | Underneath Membrane | NO |
| VDSSQGSEYDYVIYAQ | 1702299 | Cell/Virus Enclosed | NO |
| VDSSQGSEYQYVIFCQ | 1390290 | Cell/Virus Enclosed | NO |
| VDSSYECDIPIG | 1522442 | Ectodomain | YES |
| VDSTPVEPPTVD | 1522443 | Cell/Virus Enclosed | NO |
| VDSVQYAHKHDIQITN | 1702304 | Cell/Virus Enclosed | NO |
| VDSVTVKETVAQ | 1522449 | Cell/Virus Enclosed | NO |
| VDSVTVKETVVQ | 1522450 | Cell/Virus Enclosed | NO |
| VDSVVRSFQPLL | 1522453 | Ectodomain | YES |
| VDSYFENDSILQSVAS | 1702308 | Cell/Virus Enclosed | NO |
| VDSYYSYLMPAL | 1522456 | Cell/Virus Enclosed | NO |
| VDTFLSMFDVDKKSLN | 1702318 | Cell/Virus Enclosed | NO |
| VDTFMSHFDVDRKSFN | 1702319 | Cell/Virus Enclosed | NO |
| VDTGPPSISTAT | 1522476 | Ectodomain | YES |
| VDTGVPPISTDT | 1522477 | Ectodomain | YES |
| VDTGVPSTSTDI | 1522478 | Ectodomain | YES |
| VDTHRYCLSLKD | 1522480 | Cell/Virus Enclosed | NO |
| VDTHRYRLSLKD | 1522481 | Cell/Virus Enclosed | NO |
| VDTILNMARNGVVPLS | 1702322 | Cell/Virus Enclosed | NO |
| VDTKDVIECLKL | 1522483 | Cell/Virus Enclosed | NO |
| VDTLQETPVVSD | 1522486 | Cell/Virus Enclosed | NO |
| VDTLVNKIPANIVLPQ | 1702329 | Cell/Virus Enclosed | NO |
| VDTNCLADSVMS | 1522489 | Cell/Virus Enclosed | NO |
| VDTSHFTTQFFG | 1522493 | Ectodomain | YES |
| VDTSHFTTQFVS | 1522494 | Ectodomain | YES |
| VDTSYDCDIPIG | 1522500 | Ectodomain | YES |
| VDTSYECDIPVG | 1522502 | Ectodomain | YES |
| VDTTPKEEFVVKEKLN | 1702338 | Cell/Virus Enclosed | NO |
| VDTTVESAAQAD | 1522507 | Cell/Virus Enclosed | NO |
| VDTVIFKNNTSFPTNI | 1702339 | Cell/Virus Enclosed | NO |
| VDTVKEAQSANP | 1522508 | Cell/Virus Enclosed | NO |
| VDTVKGLDYKTF | 1522509 | Cell/Virus Enclosed | NO |
| VDTVSALVYENKLKAK | 1702343 | Cell/Virus Enclosed | NO |
| VDTVVNGVDFDFIVNA | 1702344 | Cell/Virus Enclosed | NO |
| VDTVVNGVEFDF | 1522513 | Cell/Virus Enclosed | NO |

| | | | |
|---|---|---|---|
| VDVAIQEVIDDMHWGD | 1702345 | Cell/Virus Enclosed | NO |
| VDVANPSTPAYTFTTV | 1702346 | Cell/Virus Enclosed | NO |
| VDVANPSTPVYT | 1522514 | Cell/Virus Enclosed | NO |
| VDVDFHSVEVEA | 1522518 | Cell/Virus Enclosed | NO |
| VDVDFHSVKPFA | 1522519 | Cell/Virus Enclosed | NO |
| VDVDYKSCTKGLSIAD | 1702349 | Ectodomain | YES |
| VDVFHETSDDEW | 1522525 | Cell/Virus Enclosed | NO |
| VDVGPDSAKSVC | 1522531 | Ectodomain | YES |
| VDVGPDSFKSAC | 1522532 | Ectodomain | YES |
| VDVGPDSIKSAC | 1522533 | Ectodomain | YES |
| VDVGPDSLKSAC | 1522534 | Ectodomain | YES |
| VDVGPDSVKPAC | 1522535 | Ectodomain | YES |
| VDVGPDSVKSACIEVD | 1702352 | Ectodomain | YES |
| VDVIDNEIIVKP | 1522539 | Cell/Virus Enclosed | NO |
| VDVLKDMPVLKTINGL | 1702356 | Cell/Virus Enclosed | NO |
| VDVLPEGWHVVN | 1522545 | Cell/Virus Enclosed | NO |
| VDVLPEGWRVVN | 1522546 | Cell/Virus Enclosed | NO |
| VDVMRYNLNFSANSLD | 1702358 | Ectodomain | YES |
| VDVNDCFKNDNV | 1522548 | Cell/Virus Enclosed | NO |
| VDVNDSHSMAVVDGKQ | 1702359 | Cell/Virus Enclosed | NO |
| VDVNKTLQEFAK | 1522549 | Ectodomain | YES |
| VDVNKTLQEFAQNLPK | 1702360 | Ectodomain | YES |
| VDVPPQVVELPS | 1522554 | Cell/Virus Enclosed | NO |
| VDVPPQVVKLPS | 1522555 | Cell/Virus Enclosed | NO |
| VDVQQWGYVGNL | 1522558 | Cell/Virus Enclosed | NO |
| VDVQQWGYVSNL | 1522559 | Cell/Virus Enclosed | NO |
| VDVSHDVSDDEW | 1522562 | Cell/Virus Enclosed | NO |
| VDVSHETSDDDW | 1522564 | Cell/Virus Enclosed | NO |
| VDVSHETSDDEWAAAV | 1702364 | Cell/Virus Enclosed | NO |
| VDVSHEVFDDEW | 1522566 | Cell/Virus Enclosed | NO |
| VDVSHGLGTYYI | 1522568 | Ectodomain | YES |
| VDVTNGLGTYYI | 1522570 | Ectodomain | YES |
| VDVTNGLGTYYVLDRV | 1702367 | Ectodomain | YES |
| VDVVALQQEMTIVKCK | 1702368 | Cell/Virus Enclosed | NO |
| VDVVPEGWRVVNKFYQ | 1702370 | Cell/Virus Enclosed | NO |
| VDVWDMYLTGCD | 1522580 | Cell/Virus Enclosed | NO |
| VDVYDMWLTGCK | 1522581 | Cell/Virus Enclosed | NO |
| VDWDIEYPYISHEKKL | 1702374 | Cell/Virus Enclosed | NO |
| VDWRSCNYAVSS | 1522598 | Cell/Virus Enclosed | NO |
| VDWSITYPMIANENAI | 1702378 | Cell/Virus Enclosed | NO |
| VDYKSCTKGLSIADLA | 1702391 | Ectodomain | YES |
| VDYNSPSSSSSRRKRR | 1702393 | Ectodomain | YES |
| VDYNVVTQRMCA | 1522623 | Cell/Virus Enclosed | NO |
| VDYSNLLHSKVKSVPN | 1702396 | Cell/Virus Enclosed | NO |
| VDYSTKRRSRRA | 1522636 | Ectodomain | YES |
| VDYTFVNEYYEFLCKH | 1702397 | Cell/Virus Enclosed | NO |
| VDYTKEGSVINF | 1522640 | Cell/Virus Enclosed | NO |
| VDYTKKSLNVDG | 1522646 | Cell/Virus Enclosed | NO |
| VEAAFDLSKELK | 1522651 | Cell/Virus Enclosed | NO |
| VEAAIELSKELKRPVN | 1702403 | Cell/Virus Enclosed | NO |
| VEAALDLSKELKRPIQ | 1702404 | Cell/Virus Enclosed | NO |
| VEACIVQVTCGS | 1522652 | Cell/Virus Enclosed | NO |
| VEACMVQVTCGG | 1522653 | Cell/Virus Enclosed | NO |
| VEACVVQVTCGS | 1522656 | Cell/Virus Enclosed | NO |
| VEADFHLSGCDE | 1522657 | Ectodomain | YES |
| VEADVAARADYE | 1522662 | Cell/Virus Enclosed | NO |
| VEAGEVKPFAVY | 1522666 | Cell/Virus Enclosed | NO |
| VEAHFYLSGCDE | 1522667 | Ectodomain | YES |
| VEAKLSAGIEFL | 1522668 | Cell/Virus Enclosed | NO |
| VEASDLEEGESE | 1522675 | Cell/Virus Enclosed | NO |
| VEASTDFHQLEC | 1522678 | Cell/Virus Enclosed | NO |
| VEASTDFHQLEF | 1522679 | Cell/Virus Enclosed | NO |
| VEASTDFHQLEY | 1522680 | Cell/Virus Enclosed | NO |
| VEASTVVCGGKR | 1522681 | Cell/Virus Enclosed | NO |
| VEATGLFADRDG | 1522682 | Cell/Virus Enclosed | NO |
| VEATGLFAERDC | 1522683 | Cell/Virus Enclosed | NO |
| VEAVNFNVTTAETTDF | 1702421 | Ectodomain | YES |
| VEAWSGLCVDGI | 1522687 | Ectodomain | YES |
| VEAWSGLCVDGR | 1522688 | Ectodomain | YES |
| VEAYNNCTGGAG | 1522692 | Ectodomain | YES |
| VEAYNSILFENGIPLM | 1702425 | Cell/Virus Enclosed | NO |
| VEAYNSILVENG | 1522695 | Cell/Virus Enclosed | NO |
| VECDEECSEVEA | 1522700 | Cell/Virus Enclosed | NO |
| VECDFSPLLFGT | 1522707 | Ectodomain | YES |
| VECGSCGNDSWL | 1522713 | Cell/Virus Enclosed | NO |
| VECLSKEDLHTL | 1522716 | Cell/Virus Enclosed | NO |

| | | | |
|---|---|---|---|
| VECTTIVNGMKR | 1522718 | Cell/Virus Enclosed | NO |
| VECYDRFKVNETNSQY | 1702438 | Cell/Virus Enclosed | NO |
| VECYSGFKPNNNSAQY | 1702439 | Cell/Virus Enclosed | NO |
| VECYSGFKPNNTSAQY | 1702440 | Cell/Virus Enclosed | NO |
| VECYSILAAKTGVSVE | 1702441 | Cell/Virus Enclosed | NO |
| VEDAYAEVHAEPKGKY | 1702443 | Cell/Virus Enclosed | NO |
| VEDAYDLVHDEP | 1522723 | Cell/Virus Enclosed | NO |
| VEDAYDQVHDEPKGKF | 1702444 | Cell/Virus Enclosed | NO |
| VEDCKLPLGQSLCALP | 1702445 | Ectodomain | YES |
| VEDCKLPLGRSL | 1522727 | Ectodomain | YES |
| VEDCNLPLGQSL | 1522728 | Ectodomain | YES |
| VEDIAQVVIADTLQET | 1702448 | Cell/Virus Enclosed | NO |
| VEDLCARMTYVC | 1522736 | Cell/Virus Enclosed | NO |
| VEDSTSVTAVVT | 1522741 | Ectodomain | YES |
| VEDSVIVNDATS | 1522743 | Cell/Virus Enclosed | NO |
| VEDSVIVNDDTG | 1522744 | Cell/Virus Enclosed | NO |
| VEDSVIVNNDTS | 1522745 | Cell/Virus Enclosed | NO |
| VEDTIDGVVEDTINDD | 1702452 | Cell/Virus Enclosed | NO |
| VEDTINDDEDVVTGDN | 1702453 | Cell/Virus Enclosed | NO |
| VEEDDVEGEETDLTVT | 1702464 | Cell/Virus Enclosed | NO |
| VEEDLTKGPHEFCSQH | 1702465 | Cell/Virus Enclosed | NO |
| VEEDLTKGPYEF | 1522761 | Cell/Virus Enclosed | NO |
| VEEDVEMSDFVDLESV | 1702466 | Cell/Virus Enclosed | NO |
| VEEEHVNRMDTFCFNV | 1702468 | Cell/Virus Enclosed | NO |
| VEELHDDLVNYIRDKG | 1702469 | Cell/Virus Enclosed | NO |
| VEELKKLLEQWNLVIG | 1702470 | Undetermined | UN |
| VEEQEQEQEEQEEQELE | 68167 | Cell/Virus Enclosed | NO |
| VEEQEQEQEEQELEEVE | 68168 | Cell/Virus Enclosed | NO |
| VEESFIDDCYGY | 1522775 | Cell/Virus Enclosed | NO |
| VEESFIDDYYGY | 1522776 | Cell/Virus Enclosed | NO |
| VEETSITIKKPN | 1522777 | Cell/Virus Enclosed | NO |
| VEEVNSIFDIETVDVK | 1702475 | Cell/Virus Enclosed | NO |
| VEFDFIVNAANE | 1522783 | Cell/Virus Enclosed | NO |
| VEFEDRSNNSSRASSR | 1702478 | Capsid | UN |
| VEFHLDGEVLPL | 1522785 | Cell/Virus Enclosed | NO |
| VEFMLWCKDGKV | 1522793 | Cell/Virus Enclosed | NO |
| VEFNRLATCDWTESGD | 1702486 | Cell/Virus Enclosed | NO |
| VEFVEGQGVPIA | 1522799 | Cell/Virus Enclosed | NO |
| VEFWFAMRKDGDDVIF | 1702488 | Cell/Virus Enclosed | NO |
| VEFYEDRILDFKVLDN | 1702489 | Cell/Virus Enclosed | NO |
| VEGAHAFRNACG | 1522803 | Cell/Virus Enclosed | NO |
| VEGAHASRNACG | 1522805 | Cell/Virus Enclosed | NO |
| VEGAHATRENIGTNFP | 1702492 | Cell/Virus Enclosed | NO |
| VEGAHVTGDNVGTNVP | 1702493 | Cell/Virus Enclosed | NO |
| VEGCHATRDAVG | 1522808 | Cell/Virus Enclosed | NO |
| VEGCMVQVTCGT | 1522810 | Cell/Virus Enclosed | NO |
| VEGELLVQDLRF | 1522813 | Cell/Virus Enclosed | NO |
| VEGELLVQDLRL | 1522814 | Cell/Virus Enclosed | NO |
| VEGELLVQNQSF | 1522815 | Cell/Virus Enclosed | NO |
| VEGELPVAVVNDKIFT | 1702500 | Cell/Virus Enclosed | NO |
| VEGGLLVQDLRL | 1522818 | Cell/Virus Enclosed | NO |
| VEGKDGFVSVEL | 1522820 | Cell/Virus Enclosed | NO |
| VEGNFSFFDCGV | 1522822 | Cell/Virus Enclosed | NO |
| VEGQGPPIAPGV | 1522826 | Cell/Virus Enclosed | NO |
| VEGSGRSAPNSR | 1522829 | Cell/Virus Enclosed | NO |
| VEGSGRSASNSRPGSR | 1702506 | Capsid | UN |
| VEGVSSFMNVTL | 1522833 | Ectodomain | YES |
| VEGVVWVAVDGA | 1522834 | Cell/Virus Enclosed | NO |
| VEHALSIKQPFS | 1522836 | Cell/Virus Enclosed | NO |
| VEHGLLHSSQLP | 1522837 | Cell/Virus Enclosed | NO |
| VEHLELKETVFVEPKD | 1702512 | Cell/Virus Enclosed | NO |
| VEHSLPIKQPFS | 1522841 | Cell/Virus Enclosed | NO |
| VEHSLTIKQPFS | 1522842 | Cell/Virus Enclosed | NO |
| VEIDGNVMHANYLFWR | 1702520 | Cell/Virus Enclosed | NO |
| VEIEAGEVKPFAVYKN | 1702521 | Cell/Virus Enclosed | NO |
| VEIEVGEVKPFA | 1522847 | Cell/Virus Enclosed | NO |
| VEIFESKTTLPV | 1522849 | Cell/Virus Enclosed | NO |
| VEIIKSQDLSVI | 1522850 | Cell/Virus Enclosed | NO |
| VEILGPLSAQTG | 1522853 | Cell/Virus Enclosed | NO |
| VEINRLQEAIKV | 1522854 | Ectodomain | YES |
| VEISVTPIKNTVDTTP | 1702526 | Cell/Virus Enclosed | NO |
| VEKCVVRVCYGN | 1522858 | Cell/Virus Enclosed | NO |
| VEKFNFVTDWLK | 1522863 | Cell/Virus Enclosed | NO |
| VEKFTFVTDWLK | 1522864 | Cell/Virus Enclosed | NO |
| VEKICIIDNMYM | 1522867 | Cell/Virus Enclosed | NO |
| VEKLKQLLEQWN | 1522871 | Undetermined | UN |

| | | | |
|---|---|---|---|
| VEKLSPWFMDGENAYE | 1702538 | Cell/Virus Enclosed | NO |
| VEKLVSWYIYVR | 1522878 | Cell/Virus Enclosed | NO |
| VEKSFQLDTAPK | 1522881 | Ectodomain | YES |
| VELFAKRNVRHH | 1522887 | Cell/Virus Enclosed | NO |
| VELGSKHVDCWT | 1522892 | Cell/Virus Enclosed | NO |
| VELGTDPIEDYE | 1522893 | Cell/Virus Enclosed | NO |
| VELHTVIPDYVD | 1522896 | Ectodomain | YES |
| VELHTVYPDYVD | 1522897 | Ectodomain | YES |
| VELKIKIASGFGPLI | 68272 | Ectodomain | YES |
| VELLSEPQTIQP | 1522902 | Cell/Virus Enclosed | NO |
| VELPKFFYASNGTYNC | 1702555 | Ectodomain | YES |
| VELYELMGKFAW | 1522924 | Cell/Virus Enclosed | NO |
| VEMHNKINLCDDPETA | 1702566 | Cell/Virus Enclosed | NO |
| VEMNRLQDAIKV | 1522928 | Ectodomain | YES |
| VEMNRLQEAIKV | 1522929 | Ectodomain | YES |
| VEMVAYNPDLSQYYCD | 1702569 | Cell/Virus Enclosed | NO |
| VENDNISVAVPK | 1522934 | Cell/Virus Enclosed | NO |
| VENGIPLMPLLS | 1522936 | Cell/Virus Enclosed | NO |
| VENKAEAEWIEL | 1522937 | Cell/Virus Enclosed | NO |
| VENKTEADWIEL | 1522938 | Cell/Virus Enclosed | NO |
| VENPDIINIYKKLGPI | 1702577 | Cell/Virus Enclosed | NO |
| VENPDIINVYKKLGPI | 1702578 | Cell/Virus Enclosed | NO |
| VENPMLMGWDYPKCDR | 1702581 | Cell/Virus Enclosed | NO |
| VENTPPHYFTLG | 1522955 | Cell/Virus Enclosed | NO |
| VEPKNGGQFFVS | 1522963 | Cell/Virus Enclosed | NO |
| VEPVRDEVSTET | 1522971 | Cell/Virus Enclosed | NO |
| VEQADGVECDFS | 1522972 | Ectodomain | YES |
| VEQAEGFECDFS | 1522973 | Ectodomain | YES |
| VEQKPVDVKPKI | 1522979 | Cell/Virus Enclosed | NO |
| VEQLEPCVQSRL | 1522983 | Cell/Virus Enclosed | NO |
| VEQPKIEPKPVSVIKV | 1702602 | Cell/Virus Enclosed | NO |
| VEQSFQLDTAPQ | 1522989 | Ectodomain | YES |
| VEQYNQIILNLT | 1522992 | Ectodomain | YES |
| VEQYNQTILNLTSEIS | 1702606 | Ectodomain | YES |
| VERGLFHSSQLP | 1522995 | Cell/Virus Enclosed | NO |
| VERGLLHSSQLP | 1522996 | Cell/Virus Enclosed | NO |
| VERLIACENPFM | 1523001 | Cell/Virus Enclosed | NO |
| VERLIACENSFM | 1523002 | Cell/Virus Enclosed | NO |
| VERSVSLAIDAY | 1523006 | Cell/Virus Enclosed | NO |
| VESAAQADLQHE | 1523007 | Cell/Virus Enclosed | NO |
| VESAHVCGDNIGTNVP | 1702615 | Cell/Virus Enclosed | NO |
| VESANLMLSDNVVAFL | 1702616 | Cell/Virus Enclosed | NO |
| VESANQMLTVNVVAFL | 1702617 | Cell/Virus Enclosed | NO |
| VESCGNFKVTRG | 1523012 | Cell/Virus Enclosed | NO |
| VESCGNYKVTKG | 1523013 | Cell/Virus Enclosed | NO |
| VESGGYIPPNFA | 1523021 | Ectodomain | YES |
| VESGGYIPSDFAFNNW | 1702629 | Ectodomain | YES |
| VESGGYIPSNFA | 1523023 | Ectodomain | YES |
| VESGTEVTEFAC | 1523025 | Cell/Virus Enclosed | NO |
| VESGVYSVSSFEAKPS | 1702631 | Ectodomain | YES |
| VESNVVDVNDCF | 1523033 | Cell/Virus Enclosed | NO |
| VESPTVDVVALQ | 1523035 | Cell/Virus Enclosed | NO |
| VESQSKRYGFGG | 1523036 | Ectodomain | YES |
| VESSKSLGKQLGVVSD | 1702634 | Cell/Virus Enclosed | NO |
| VESTPVELPTVD | 1523040 | Cell/Virus Enclosed | NO |
| VESTPVEPPTVD | 1523041 | Cell/Virus Enclosed | NO |
| VESTPVEPSTVD | 1523042 | Cell/Virus Enclosed | NO |
| VESTSVEPPTVD | 1523044 | Cell/Virus Enclosed | NO |
| VESYNIFTQAGF | 1523045 | Cell/Virus Enclosed | NO |
| VETCTIAGKAFDKVFD | 1702638 | Cell/Virus Enclosed | NO |
| VETDLKKGPHEF | 1523050 | Cell/Virus Enclosed | NO |
| VETETYFKVNGLWSPT | 1702640 | Cell/Virus Enclosed | NO |
| VETKVQCSTNLF | 1523058 | Cell/Virus Enclosed | NO |
| VETLNEDLMEIL | 1523060 | Cell/Virus Enclosed | NO |
| VETLTQENLELE | 1523062 | Cell/Virus Enclosed | NO |
| VETLVGQLEQTN | 1523063 | Cell/Virus Enclosed | NO |
| VETNEIVDSVQY | 1523064 | Cell/Virus Enclosed | NO |
| VETSHFTTKYVA | 1523074 | Ectodomain | YES |
| VETVGGLFEIQI | 1523081 | Ectodomain | YES |
| VETVLNLARDGVVPLS | 1702662 | Cell/Virus Enclosed | NO |
| VETVSALVYDNKLKAK | 1702663 | Cell/Virus Enclosed | NO |
| VETVSLAGSYRD | 1523088 | Cell/Virus Enclosed | NO |
| VETYIKWPWWVWLCIS | 1702665 | Undetermined | UN |
| VEVFNRLATSDWTDVR | 1702667 | Cell/Virus Enclosed | NO |
| VEVFTDGKLNCSFQDA | 1702668 | Cell/Virus Enclosed | NO |
| VEVFTDGRLNCS | 1523098 | Cell/Virus Enclosed | NO |

| | | | |
|---|---|---|---|
| VEVHVEDIVQVV | 1523100 | Cell/Virus Enclosed | NO |
| VEVNAPYYNSWQ | 1523106 | Ectodomain | YES |
| VEVNATYYNSWQNLLY | 1702673 | Ectodomain | YES |
| VEVPPQLVELPS | 1523113 | Cell/Virus Enclosed | NO |
| VEVPPQVMKLPS | 1523114 | Cell/Virus Enclosed | NO |
| VEVPPQVVELLS | 1523115 | Cell/Virus Enclosed | NO |
| VEVPPQVVELPS | 1523116 | Cell/Virus Enclosed | NO |
| VEVPPQVVELPY | 1523117 | Cell/Virus Enclosed | NO |
| VEVQSSGIIKPN | 1523127 | Cell/Virus Enclosed | NO |
| VEVQSSGMIKPN | 1523129 | Cell/Virus Enclosed | NO |
| VEVQSSGMINPN | 1523130 | Cell/Virus Enclosed | NO |
| VEVSPQVVKLPS | 1523132 | Cell/Virus Enclosed | NO |
| VEVTPQVVKLPS | 1523134 | Cell/Virus Enclosed | NO |
| VEVVCEDVTQLA | 1523135 | Cell/Virus Enclosed | NO |
| VEWKFYDVQPCS | 1523141 | Cell/Virus Enclosed | NO |
| VEYAYDQVHDEP | 1523143 | Cell/Virus Enclosed | NO |
| VEYEVAKKNLADAKNS | 1702686 | Cell/Virus Enclosed | NO |
| VEYKITQWPTAT | 1523151 | Cell/Virus Enclosed | NO |
| VEYLQITSKPIV | 1523152 | Ectodomain | YES |
| VEYLQITSTPIV | 1523154 | Ectodomain | YES |
| VEYNIHAVLDTL | 1523158 | Cell/Virus Enclosed | NO |
| VEYPIIGDELKI | 1523161 | Cell/Virus Enclosed | NO |
| VEYPIISNELSINTSC | 1702697 | Cell/Virus Enclosed | NO |
| VEYSIHAVLDTL | 1523164 | Cell/Virus Enclosed | NO |
| VEYSKNRRSRRA | 1523165 | Ectodomain | YES |
| VEYSLPIKQPFS | 1523166 | Cell/Virus Enclosed | NO |
| VEYSLYGISGRG | 1523167 | Ectodomain | YES |
| VEYSLYGVLGRG | 1523168 | Ectodomain | YES |
| VEYSLYGVSGRG | 1523169 | Ectodomain | YES |
| VEYTYTDEVIYG | 1523170 | Cell/Virus Enclosed | NO |
| VEYTYTDEVIYS | 1523171 | Cell/Virus Enclosed | NO |
| VFAVESGGYIPP | 1523183 | Ectodomain | YES |
| VFAVESGGYIPSDFAF | 1702707 | Ectodomain | YES |
| VFCCLSTGCCGC | 1523189 | Undetermined | UN |
| VFCDGVNVTKHKCDIN | 1702711 | Cell/Virus Enclosed | NO |
| VFDDEWAAAVDE | 1523201 | Cell/Virus Enclosed | NO |
| VFDGKSKCEESA | 1523205 | Cell/Virus Enclosed | NO |
| VFDIKNSSVNTVCFVD | 1702726 | Cell/Virus Enclosed | NO |
| VFDITDAAVKAA | 1523208 | Cell/Virus Enclosed | NO |
| VFDKVVDNVYVTYAGS | 1702731 | Cell/Virus Enclosed | NO |
| VFDNFNQWHEAK | 1523214 | Cell/Virus Enclosed | NO |
| VFDSSKEFVEYK | 1523215 | Cell/Virus Enclosed | NO |
| VFDVDKTKTPHI | 1523217 | Cell/Virus Enclosed | NO |
| VFDVESGGYIPS | 1523219 | Ectodomain | YES |
| VFDVKGVEELHD | 1523220 | Cell/Virus Enclosed | NO |
| VFENFEDWHYAKFGFT | 1702738 | Cell/Virus Enclosed | NO |
| VFEVDDTVDMEEFYAV | 1702741 | Cell/Virus Enclosed | NO |
| VFFADNNDITNT | 1523232 | Cell/Virus Enclosed | NO |
| VFFKGNVQVDNGSSIN | 1702747 | Cell/Virus Enclosed | NO |
| VFFNAFVKCNCG | 1523241 | Cell/Virus Enclosed | NO |
| VFFNGADISDTI | 1523242 | Cell/Virus Enclosed | NO |
| VFFNGVDISDTI | 1523243 | Cell/Virus Enclosed | NO |
| VFFQFDNLSSEDLKAV | 1702750 | Cell/Virus Enclosed | NO |
| VFFQFENLSSED | 1523245 | Cell/Virus Enclosed | NO |
| VFFSPESVFSVFGDVK | 1702752 | Cell/Virus Enclosed | NO |
| VFGKRGGGNVTY | 1523261 | Cell/Virus Enclosed | NO |
| VFGSTMDNKSQS | 1523266 | Ectodomain | YES |
| VFGYVQQGIFNKATVA | 1702771 | Cell/Virus Enclosed | NO |
| VFGYVQQGVFNK | 1523273 | Cell/Virus Enclosed | NO |
| VFHETSDDEWTA | 1523274 | Cell/Virus Enclosed | NO |
| VFIKRSDALSTN | 1523292 | Cell/Virus Enclosed | NO |
| VFIKRSDAQGTN | 1523294 | Cell/Virus Enclosed | NO |
| VFINKYGLEDFNFEHV | 1702779 | Cell/Virus Enclosed | NO |
| VFIQKYGLEDFNFEHV | 1702781 | Cell/Virus Enclosed | NO |
| VFISPYNSQNFAAKRV | 1702787 | Cell/Virus Enclosed | NO |
| VFISPYNSQNYVAKRV | 1702789 | Cell/Virus Enclosed | NO |
| VFKADGIIYPQG | 1523312 | Ectodomain | YES |
| VFKDGCDVGFCP | 1523313 | Cell/Virus Enclosed | NO |
| VFKEGVMYSEFP | 1523315 | Ectodomain | YES |
| VFKHGVMYSEFP | 1523316 | Ectodomain | YES |
| VFKIGDVKFKRLGDYV | 1702795 | Cell/Virus Enclosed | NO |
| VFKNKDGFPYVY | 1523329 | Ectodomain | YES |
| VFKPQPAGVFTD | 1523333 | Ectodomain | YES |
| VFKPQPAGVLTD | 1523335 | Ectodomain | YES |
| VFKTLKPGDSFNILAC | 1702804 | Cell/Virus Enclosed | NO |
| VFLLVILAILTA | 1523376 | Undetermined | UN |

| | | | |
|---|---|---|---|
| VFLLVTLAILTALRL | 68513 | Undetermined | UN |
| VFMSEAKCWVETDLKK | 1702838 | Cell/Virus Enclosed | NO |
| VFMSESKCWVEHDINN | 1702839 | Cell/Virus Enclosed | NO |
| VFMSESKCWVENDITN | 1702840 | Cell/Virus Enclosed | NO |
| VFMSTSKCWVEEDLTK | 1702842 | Cell/Virus Enclosed | NO |
| VFNASISESKPL | 1523396 | Cell/Virus Enclosed | NO |
| VFNASRFPNVYA | 1523398 | Ectodomain | YES |
| VFNDVYNDALFVSYGS | 1702848 | Cell/Virus Enclosed | NO |
| VFNICQAVSANVCALM | 1702852 | Cell/Virus Enclosed | NO |
| VFNKATVAQIKA | 1523414 | Cell/Virus Enclosed | NO |
| VFNLLHFTNDKT | 1523416 | Cell/Virus Enclosed | NO |
| VFNNNGTNVTQVKNAS | 1702860 | Cell/Virus Enclosed | NO |
| VFNNTLSRQLQP | 1523419 | Ectodomain | YES |
| VFNVVKQGHFIG | 1523430 | Cell/Virus Enclosed | NO |
| VFPLDEAEDVTE | 1523435 | Cell/Virus Enclosed | NO |
| VFPYDMDFGVSS | 1523448 | Cell/Virus Enclosed | NO |
| VFPYDMDSGVSSFSEL | 1702874 | Cell/Virus Enclosed | NO |
| VFQNCSAVGVRQ | 1523458 | Ectodomain | YES |
| VFQSATKIIALN | 1523466 | Secreted | YES |
| VFRAFDICNYKAKVAG | 1702886 | Cell/Virus Enclosed | NO |
| VFRMPMGVYNYK | 1523485 | Cell/Virus Enclosed | NO |
| VFSAVGNICYTP | 1523496 | Cell/Virus Enclosed | NO |
| VFSLLHFTNDKT | 1523502 | Cell/Virus Enclosed | NO |
| VFSNMVETVVGQ | 1523510 | Cell/Virus Enclosed | NO |
| VFSPAASSCHNASGKEAKVC | 68567 | Capsid | UN |
| VFSTVNALPEVNADIV | 1702904 | Cell/Virus Enclosed | NO |
| VFSYCKKLGTTV | 1523518 | Cell/Virus Enclosed | NO |
| VFTACSHAAVDSLCAK | 1702911 | Cell/Virus Enclosed | NO |
| VFTDGKLNCSFQDAAS | 1702914 | Cell/Virus Enclosed | NO |
| VFTGYHITKNSKVQLG | 1702919 | Cell/Virus Enclosed | NO |
| VFTKSGVNDICI | 1523543 | Cell/Virus Enclosed | NO |
| VFTNKTTLPTNVAFEL | 1702927 | Cell/Virus Enclosed | NO |
| VFTTINALPEMVTDIV | 1702931 | Cell/Virus Enclosed | NO |
| VFVCTDNSANTKVLRN | 1702938 | Cell/Virus Enclosed | NO |
| VFVDDITKTDAVILLE | 1702940 | Cell/Virus Enclosed | NO |
| VFVDDVVKTDAVVLLE | 1702941 | Cell/Virus Enclosed | NO |
| VFVDKYMCGFDGKPVL | 1702942 | Cell/Virus Enclosed | NO |
| VFVEPKDGGHFF | 1523575 | Cell/Virus Enclosed | NO |
| VFVEPKDGGQFF | 1523576 | Cell/Virus Enclosed | NO |
| VFVEPKNGGQFF | 1523577 | Cell/Virus Enclosed | NO |
| VFVLGGCRHKLVCAPAPCNF | 68600 | Cell/Virus Enclosed | NO |
| VFVNKTSLPTNIAFEL | 1702953 | Cell/Virus Enclosed | NO |
| VFVPQPTGVFTN | 1523595 | Ectodomain | YES |
| VFVRYGDVDNLV | 1523601 | Cell/Virus Enclosed | NO |
| VFVVESGGYIPS | 1523617 | Ectodomain | YES |
| VFVYTDTEVCKV | 1523621 | Cell/Virus Enclosed | NO |
| VFWVANHQADTSTPSD | 1702971 | Capsid | UN |
| VFYCTNNTLVSGDAHI | 1702977 | Ectodomain | YES |
| VFYELDKDFNTI | 1523630 | Cell/Virus Enclosed | NO |
| VFYELDNDFNTILNTA | 1702978 | Cell/Virus Enclosed | NO |
| VFYFAVRKEGQDVIFS | 1702979 | Cell/Virus Enclosed | NO |
| VFYVKDGKYFTS | 1523647 | Cell/Virus Enclosed | NO |
| VFYVRVKHGDFSAVSS | 1702989 | Underneath Membrane | NO |
| VGADGELPVAISGDKV | 1702995 | Cell/Virus Enclosed | NO |
| VGAESCGRPFSGSLG | 68618 | Cell/Virus Enclosed | NO |
| VGALDASASVVA | 1523661 | Cell/Virus Enclosed | NO |
| VGALDVSASAVA | 1523662 | Cell/Virus Enclosed | NO |
| VGAMLSENFTSYGFSN | 1703003 | Ectodomain | YES |
| VGAPGGDLLYLD | 1523667 | Cell/Virus Enclosed | NO |
| VGAWDGYLSSCCGTPA | 1703006 | Cell/Virus Enclosed | NO |
| VGCNNADCVACS | 1523681 | Cell/Virus Enclosed | NO |
| VGDFQLNEEVAI | 1523698 | Cell/Virus Enclosed | NO |
| VGDLNCPLDPKL | 1523704 | Ectodomain | YES |
| VGDPAQLPTPRT | 1523708 | Cell/Virus Enclosed | NO |
| VGDPQQLPAPRV | 1523709 | Cell/Virus Enclosed | NO |
| VGEFVVNDVVIA | 1523734 | Cell/Virus Enclosed | NO |
| VGENVPDDLYIKGSG | 110733 | Capsid | UN |
| VGEPTISQPSKL | 1523752 | Cell/Virus Enclosed | NO |
| VGFDAEGAHATL | 1523773 | Cell/Virus Enclosed | NO |
| VGFGHVYNFKVE | 1523777 | Cell/Virus Enclosed | NO |
| VGFKNQYCNESI | 1523783 | Cell/Virus Enclosed | NO |
| VGFSNGVDFVAQPEGC | 1703077 | Cell/Virus Enclosed | NO |
| VGFVEAYNNCTGGAEI | 1703082 | Ectodomain | YES |
| VGFVEAYNNCTGGSEI | 1703083 | Ectodomain | YES |
| VGGCVTSNVPPI | 1523810 | Cell/Virus Enclosed | NO |
| VGGCVTSNVSPI | 1523812 | Cell/Virus Enclosed | NO |

| | | | |
|---|---|---|---|
| VGGMIRYYDVMANGGT | 1703097 | Cell/Virus Enclosed | NO |
| VGGTIQILASVP | 1523842 | Cell/Virus Enclosed | NO |
| VGGVYDNSITCPLTVA | 1703108 | Cell/Virus Enclosed | NO |
| VGHDICAQLNAK | 1523847 | Cell/Virus Enclosed | NO |
| VGHDICAQLNDK | 1523848 | Cell/Virus Enclosed | NO |
| VGHTVCDSLNAK | 1523858 | Cell/Virus Enclosed | NO |
| VGHYDGRTGELPCAIM | 1703115 | Cell/Virus Enclosed | NO |
| VGIADDTYVMGL | 1523870 | Cell/Virus Enclosed | NO |
| VGIGEHCPGLGI | 1523873 | Ectodomain | YES |
| VGIGEHCSGLAI | 1523874 | Ectodomain | YES |
| VGINYLGDFAQGPFID | 1703127 | Cell/Virus Enclosed | NO |
| VGITQQVLSENQKLIA | 1703128 | Ectodomain | YES |
| VGIVYSSQLYEDVPSI | 1703130 | Cell/Virus Enclosed | NO |
| VGKDGVMYSEFP | 1523890 | Ectodomain | YES |
| VGKIAQWIKNTF | 1523896 | Cell/Virus Enclosed | NO |
| VGKYDINHGDNG | 1523907 | Underneath Membrane | NO |
| VGLLDQAWRFPC | 1523916 | Cell/Virus Enclosed | NO |
| VGLMWLSYFIASFRL | 2001367 | Undetermined | UN |
| VGLRTPQRFTDLVKF | 68723 | Ectodomain | YES |
| VGLVFVTLRSIN | 1523924 | Cell/Virus Enclosed | NO |
| VGLVGVLTLDNQDLNG | 1703145 | Cell/Virus Enclosed | NO |
| VGMPSFVAYETARQEY | 1703147 | Cell/Virus Enclosed | NO |
| VGNCVKMLTNGSGSGQ | 1703151 | Cell/Virus Enclosed | NO |
| VGNFSDGKMGLF | 1523931 | Ectodomain | YES |
| VGNGPGIDAGYK | 1523936 | Ectodomain | YES |
| VGNKFYNVSTGLIPGF | 1703156 | Cell/Virus Enclosed | NO |
| VGNLKYSTFMLN | 1523943 | Ectodomain | YES |
| VGNVILKPSDEG | 1523953 | Cell/Virus Enclosed | NO |
| VGNVKFDRWSAS | 1523956 | Ectodomain | YES |
| VGNVKLDRWSAS | 1523957 | Ectodomain | YES |
| VGNYRLPSSKPSGMDT | 1703160 | Underneath Membrane | NO |
| VGNYRLPSTQKGSGMD | 1703161 | Ectodomain | YES |
| VGPDARAKQDVSLLSK | 1703162 | Cell/Virus Enclosed | NO |
| VGPDARTQGKQSYVLL | 1703164 | Cell/Virus Enclosed | NO |
| VGPDSIKSACIE | 1523975 | Ectodomain | YES |
| VGPKELLLLWESGKAK | 1703169 | Cell/Virus Enclosed | NO |
| VGPRAGKHEHSL | 1523994 | Cell/Virus Enclosed | NO |
| VGQEEFIQTNSPKVTI | 1703179 | Ectodomain | YES |
| VGRPIVYETDAN | 1524018 | Cell/Virus Enclosed | NO |
| VGRSVNETSQTGWAFY | 1703190 | Ectodomain | YES |
| VGRSVNSQNCTG | 1524021 | Underneath Membrane | NO |
| VGRSVNSQNSTGWVFY | 1703191 | Underneath Membrane | NO |
| VGRWSASINTGNCPFS | 1703193 | Ectodomain | YES |
| VGSANIFIGDKVGHYV | 1703194 | Cell/Virus Enclosed | NO |
| VGSATNASLFVLGVALSANG | 742127 | Undetermined | UN |
| VGSGYCVDYSTK | 1524047 | Ectodomain | YES |
| VGSPIQFHLDGE | 1524056 | Cell/Virus Enclosed | NO |
| VGSSFDGVMYGGFEDQ | 1703205 | Cell/Virus Enclosed | NO |
| VGSSVAVQSKDLNFLN | 1703206 | Cell/Virus Enclosed | NO |
| VGTAEEKLKKKSSF | 68800 | Capsid | UN |
| VGTCPAGTNSLT | 1524064 | Ectodomain | YES |
| VGTGIIRSSNQSLAGG | 1703210 | Ectodomain | YES |
| VGTNVPLQVGFSNGVD | 1703216 | Cell/Virus Enclosed | NO |
| VGTQSVDMLAVKTGVA | 1703219 | Cell/Virus Enclosed | NO |
| VGVIADKDKDLSGAVP | 1703229 | Cell/Virus Enclosed | NO |
| VGVIRISNETFL | 1524105 | Ectodomain | YES |
| VGVQTIEDLRAR | 1524113 | Cell/Virus Enclosed | NO |
| VGVQTLEDLHAR | 1524114 | Cell/Virus Enclosed | NO |
| VGVQTVEDLCAR | 1524115 | Cell/Virus Enclosed | NO |
| VGVQTVEDLRVR | 1524116 | Cell/Virus Enclosed | NO |
| VGVRQQRFVYDAYQNL | 1703236 | Ectodomain | YES |
| VGVSERINVVPGVPTN | 1703238 | Cell/Virus Enclosed | NO |
| VGVVDTEWGNMLTGIA | 1703240 | Cell/Virus Enclosed | NO |
| VGVYDINHGDAGFTSI | 1703241 | Ectodomain | YES |
| VGYQVRAFLNKLNENV | 1703253 | Cell/Virus Enclosed | NO |
| VGYTKDGSVINF | 1524156 | Cell/Virus Enclosed | NO |
| VHAEPKGKYSQKAYAL | 1703262 | Cell/Virus Enclosed | NO |
| VHAIATIPLQAS | 1524176 | Secreted | YES |
| VHARLKGGLILKFDSG | 1703267 | Cell/Virus Enclosed | NO |
| VHARLKGGLILR | 1524181 | Cell/Virus Enclosed | NO |
| VHATAMIPLQAS | 1524185 | Secreted | YES |
| VHDNVAFYQGDVDTVV | 1703273 | Cell/Virus Enclosed | NO |
| VHDSDLEHVLTPSTSWTTKILKFI | 68842 | Ectodomain | YES |
| VHDVEPTHKVKL | 1524199 | Cell/Virus Enclosed | NO |
| VHEPTTPISSAL | 1524205 | Cell/Virus Enclosed | NO |
| VHEWKMVVCDKV | 1524213 | Cell/Virus Enclosed | NO |

| | | | |
|---|---|---|---|
| VHGNAHVASCDA | 1524230 | Cell/Virus Enclosed | NO |
| VHGNGSMIEDMD | 1524232 | Cell/Virus Enclosed | NO |
| VHGNGSMIEDVD | 1524233 | Cell/Virus Enclosed | NO |
| VHGNVAFYQGDV | 1524234 | Cell/Virus Enclosed | NO |
| VHGQSNADPKVY | 1524235 | Cell/Virus Enclosed | NO |
| VHKDNFKDGLCMFWNC | 1703298 | Cell/Virus Enclosed | NO |
| VHLPYPDPSRIL | 1524261 | Cell/Virus Enclosed | NO |
| VHNLSGNPDEPQ | 1524278 | Cell/Virus Enclosed | NO |
| VHQVQLIDKYCS | 1524296 | Cell/Virus Enclosed | NO |
| VHRNEHVASGDAIMTR | 1703317 | Cell/Virus Enclosed | NO |
| VHSLSVHDGYTN | 1524315 | Cell/Virus Enclosed | NO |
| VHSVFLLMFLLT | 1524321 | Ectodomain | YES |
| VHTGFGAMDFTTLQANKSEV | 111915 | Capsid | UN |
| VHTPKHVFKTLKPGDS | 1703326 | Cell/Virus Enclosed | NO |
| VHVDSFSCNNLD | 1524341 | Ectodomain | YES |
| VHVGGVYDNSIT | 1524343 | Cell/Virus Enclosed | NO |
| VHVKCEQSYQLYDASN | 1703327 | Cell/Virus Enclosed | NO |
| VHVYVDDSIILR | 1524357 | Cell/Virus Enclosed | NO |
| VHYYQNNVFMSE | 1524372 | Cell/Virus Enclosed | NO |
| VIADKDKDLSGAVPSD | 1703339 | Cell/Virus Enclosed | NO |
| VIADTLQEIPVV | 1524380 | Cell/Virus Enclosed | NO |
| VIADTLQETPVVPDTV | 1703341 | Cell/Virus Enclosed | NO |
| VIADYNYKLPDDFMGC | 1703342 | Ectodomain | YES |
| VIALTGTNQLCA | 1524403 | Cell/Virus Enclosed | NO |
| VIANVPAIDVLKKLLS | 1703351 | Cell/Virus Enclosed | NO |
| VIASNTTSAIDYDHEY | 1703354 | Cell/Virus Enclosed | NO |
| VIAWNTAKQDQG | 1524412 | Ectodomain | YES |
| VIAWNTAQQDQG | 1524415 | Ectodomain | YES |
| VICSASDMTNPDYTNL | 1703366 | Cell/Virus Enclosed | NO |
| VICSSSNMNEPDYSAL | 1703367 | Cell/Virus Enclosed | NO |
| VIDDGDGVKPFR | 1524425 | Cell/Virus Enclosed | NO |
| VIDDVDVVKPFRVEGN | 1703373 | Cell/Virus Enclosed | NO |
| VIDGVTYQLAWD | 1524431 | Cell/Virus Enclosed | NO |
| VIDLLLNDFVEI | 1524435 | Cell/Virus Enclosed | NO |
| VIDLRLDDFVMI | 1524437 | Cell/Virus Enclosed | NO |
| VIDMRSYEKLANSISP | 1703384 | Cell/Virus Enclosed | NO |
| VIDQDFGSTVFNVPTK | 1703387 | Cell/Virus Enclosed | NO |
| VIDVKQVGCSMR | 1524443 | Cell/Virus Enclosed | NO |
| VIEVSQIQSKLTDVKC | 1703398 | Cell/Virus Enclosed | NO |
| VIFAQTSDTAHACNAN | 1703400 | Cell/Virus Enclosed | NO |
| VIFCQTADTAHANNIN | 1703401 | Cell/Virus Enclosed | NO |
| VIFINNTTYPTNVAVE | 1703404 | Cell/Virus Enclosed | NO |
| VIFKNNTSFPTNIAVE | 1703405 | Cell/Virus Enclosed | NO |
| VIFNAVDCKSDF | 1524463 | Ectodomain | YES |
| VIFPGQKYSSDC | 1524464 | Cell/Virus Enclosed | NO |
| VIFSDFYLPKIG | 1524466 | Cell/Virus Enclosed | NO |
| VIFSQFDSLGVSSNQS | 1703409 | Cell/Virus Enclosed | NO |
| VIFSQSDSLGVS | 1524471 | Cell/Virus Enclosed | NO |
| VIGAVILRGHLRIAG | 1543118 | Underneath Membrane | NO |
| VIGDFNCTNFAINDLN | 1703415 | Ectodomain | YES |
| VIGDFNCTNSFI | 1524483 | Ectodomain | YES |
| VIGDLKCPLDSR | 1524484 | Ectodomain | YES |
| VIGDLKCPLESR | 1524487 | Ectodomain | YES |
| VIGDLKCTSDNINDKD | 1703416 | Ectodomain | YES |
| VIGDLKCTSDTS | 1524488 | Ectodomain | YES |
| VIGDLNCPLDPK | 1524491 | Ectodomain | YES |
| VIGDLNCPLDSR | 1524493 | Ectodomain | YES |
| VIGDLNCPLDTR | 1524494 | Ectodomain | YES |
| VIGDLNCTLDPR | 1524495 | Ectodomain | YES |
| VIGDLNCTLHPR | 1524496 | Ectodomain | YES |
| VIGFLFLTWICLLQF | 1543119 | Transmembrane | NO |
| VIGIINNTVYDPLQPEL | 68971 | Ectodomain | YES |
| VIGTTKFCGGWN | 1524518 | Cell/Virus Enclosed | NO |
| VIGTTKFYGGWG | 1524519 | Cell/Virus Enclosed | NO |
| VIHYNINFEENL | 1524531 | Ectodomain | YES |
| VIIANRYNEVANAVMQ | 1703444 | Cell/Virus Enclosed | NO |
| VIIANRYNEVPA | 1524537 | Cell/Virus Enclosed | NO |
| VIIANRYNEVSA | 1524538 | Cell/Virus Enclosed | NO |
| VIIINNSTNVVIRACN | 1703449 | Ectodomain | YES |
| VIKDGVMYSEFP | 1524576 | Ectodomain | YES |
| VIKFETDAYKLF | 1524583 | Cell/Virus Enclosed | NO |
| VIKHGVMYSEFP | 1524586 | Ectodomain | YES |
| VIKNLTPIKLNFGMLN | 1703463 | Cell/Virus Enclosed | NO |
| VIKNVNSVRDWLKSLK | 1703464 | Cell/Virus Enclosed | NO |
| VIKPVCAKSPPGEQFR | 1703465 | Cell/Virus Enclosed | NO |
| VIKVAPKPYRVDGKFS | 1703470 | Cell/Virus Enclosed | NO |

| | | | |
|---|---|---|---|
| VILCWETGKVKPPLNK | 1703474 | Cell/Virus Enclosed | NO |
| VILLERYVSLAIDAYP | 1703480 | Cell/Virus Enclosed | NO |
| VILPGEFSKKQLGLLQ | 1703487 | Cell/Virus Enclosed | NO |
| VILQFGYTSRSM | 1524629 | Undetermined | UN |
| VILSTCDGQKFT | 1524634 | Cell/Virus Enclosed | NO |
| VILVSNNKEDFDIIQK | 1703497 | Cell/Virus Enclosed | NO |
| VINARIRAKHYVYIGD | 1703503 | Cell/Virus Enclosed | NO |
| VINITLANFNET | 1524663 | Ectodomain | YES |
| VINLRADQINDMVYSL | 1703513 | Cell/Virus Enclosed | NO |
| VINQLDEKAQKFFDFG | 1703516 | Cell/Virus Enclosed | NO |
| VIPATSASKLTIVSPD | 1703523 | Cell/Virus Enclosed | NO |
| VIPDFTVWNQVVTYPS | 1703524 | Cell/Virus Enclosed | NO |
| VIPDKQVFDKVVDNVY | 1703525 | Cell/Virus Enclosed | NO |
| VIPDYVDVNKTLQEFA | 1703527 | Ectodomain | YES |
| VIPFKDGIYFAATEKS | 1703529 | Ectodomain | YES |
| VIPLAFMPVQSW | 1524696 | Cell/Virus Enclosed | NO |
| VIPRNSVPVNKK | 1524700 | Cell/Virus Enclosed | NO |
| VIPSERLHYRNQGWRSVETSGVAEE | 69074 | Undetermined | UN |
| VIQDVKDGTVFS | 1524714 | Cell/Virus Enclosed | NO |
| VIQDVKDGTVTFNLLA | 1703542 | Cell/Virus Enclosed | NO |
| VIQDYENVCFDF | 1524716 | Cell/Virus Enclosed | NO |
| VIRDVSVEDVCFANKF | 1703548 | Cell/Virus Enclosed | NO |
| VIRGHLYIQGVKLGTG | 1703553 | Underneath Membrane | NO |
| VIRKDLSYEQQNVLAI | 1703554 | Cell/Virus Enclosed | NO |
| VIRYNLNFEENLRRGT | 1703558 | Ectodomain | YES |
| VISATSNYSGVV | 1524757 | Cell/Virus Enclosed | NO |
| VISEDLWQFVDH | 1524763 | Cell/Virus Enclosed | NO |
| VISQLDENAQKF | 1524771 | Cell/Virus Enclosed | NO |
| VISRLGFKLDAT | 1524773 | Cell/Virus Enclosed | NO |
| VISRMGFKLDAI | 1524774 | Cell/Virus Enclosed | NO |
| VISRMGFKLDATVPGY | 1703571 | Cell/Virus Enclosed | NO |
| VITGVPKPVEGV | 1524798 | Ectodomain | YES |
| VITKKEAAAAKN | 1524803 | Cell/Virus Enclosed | NO |
| VITKRDAAAAKN | 1524805 | Cell/Virus Enclosed | NO |
| VITNVGEPIISQPSKL | 1703590 | Cell/Virus Enclosed | NO |
| VITPGTNASSEF | 1524810 | Ectodomain | YES |
| VITPGTNASSEVAVLY | 69129 | Ectodomain | YES |
| VIVAWHVVRDPRYVMR | 1703602 | Cell/Virus Enclosed | NO |
| VIVDTLQETPVV | 1524831 | Cell/Virus Enclosed | NO |
| VIVEDDVIENVKSSLS | 1703604 | Cell/Virus Enclosed | NO |
| VIVGDYVIISEK | 1524835 | Cell/Virus Enclosed | NO |
| VIVKETNLICKA | 1524838 | Cell/Virus Enclosed | NO |
| VIVKRDATFATH | 1524840 | Cell/Virus Enclosed | NO |
| VIVNAANTHLKH | 1524849 | Cell/Virus Enclosed | NO |
| VIVNDDTSDTKY | 1524852 | Cell/Virus Enclosed | NO |
| VIVNDDTSETKYVKSL | 1703614 | Cell/Virus Enclosed | NO |
| VIVNNDTSETKY | 1524853 | Cell/Virus Enclosed | NO |
| VIVNNYDKSAGYPFNK | 1703616 | Cell/Virus Enclosed | NO |
| VIVQPTHYVRVT | 1524858 | Cell/Virus Enclosed | NO |
| VIVRRDATFATH | 1524861 | Cell/Virus Enclosed | NO |
| VIWLSHEKASLS | 1524881 | Cell/Virus Enclosed | NO |
| VIWLSHEQASLN | 1524882 | Cell/Virus Enclosed | NO |
| VIYDKETKTHATLFGS | 1703631 | Ectodomain | YES |
| VIYDKKTKTHAT | 1524886 | Ectodomain | YES |
| VIYGFNSTETIT | 1524889 | Ectodomain | YES |
| VIYGLNFTETIT | 1524890 | Ectodomain | YES |
| VIYQLDENAQKF | 1524906 | Cell/Virus Enclosed | NO |
| VIYSQTAETAHSVNVN | 1703640 | Cell/Virus Enclosed | NO |
| VKAAESKAFVDTIVPP | 1703642 | Cell/Virus Enclosed | NO |
| VKAKGYPQFAELVPST | 1703649 | Capsid | UN |
| VKALNESYIDLKELGN | 1703650 | Ectodomain | YES |
| VKAQSSGVIPENPVLF | 1703653 | Cell/Virus Enclosed | NO |
| VKATKGEGDGGITSEG | 1703655 | Cell/Virus Enclosed | NO |
| VKAVRNKTSACF | 1524933 | Cell/Virus Enclosed | NO |
| VKAVTAYSVDKC | 1524934 | Cell/Virus Enclosed | NO |
| VKAWSGICVDGI | 1524936 | Ectodomain | YES |
| VKCEQAYQLYDA | 1524937 | Cell/Virus Enclosed | NO |
| VKCEQFYQLYDA | 1524938 | Cell/Virus Enclosed | NO |
| VKCEQSYQLYDASNVK | 1703659 | Cell/Virus Enclosed | NO |
| VKCKGLNKPFVKDNVS | 1703662 | Cell/Virus Enclosed | NO |
| VKDDYCDGSLGCKMCL | 1703672 | Cell/Virus Enclosed | NO |
| VKDFDFKFYDAQPIVK | 1703673 | Cell/Virus Enclosed | NO |
| VKDFDFYRYNKPTILD | 1703674 | Cell/Virus Enclosed | NO |
| VKDFNTLSQEGKKYLV | 1703676 | Cell/Virus Enclosed | NO |
| VKDFVSGLLNVQ | 1524963 | Cell/Virus Enclosed | NO |
| VKDFVSGLVNVQKVEQ | 1703677 | Cell/Virus Enclosed | NO |

| | | | |
|---|---|---|---|
| VKDGKYFTSEPPVTYS | 1703679 | Cell/Virus Enclosed | NO |
| VKDNDGKNVHLKDVTK | 1703684 | Cell/Virus Enclosed | NO |
| VKDNVSFVADDSGTPV | 1703686 | Cell/Virus Enclosed | NO |
| VKDNVSFVVDDS | 1524987 | Cell/Virus Enclosed | NO |
| VKDPPNVQIHTIDGSS | 1703687 | Secreted | YES |
| VKDVDHVQIHTI | 1524998 | Secreted | YES |
| VKDVFVESSKSL | 1525000 | Cell/Virus Enclosed | NO |
| VKDYIQTVNVIA | 1525002 | Cell/Virus Enclosed | NO |
| VKEAQNANPTVI | 1525009 | Cell/Virus Enclosed | NO |
| VKEIAPVYEADT | 1525017 | Cell/Virus Enclosed | NO |
| VKEIISIPNIIK | 1525019 | Cell/Virus Enclosed | NO |
| VKEITRENVETL | 1525025 | Cell/Virus Enclosed | NO |
| VKEKLNAFLVHG | 1525029 | Cell/Virus Enclosed | NO |
| VKEQVVDLLVKL | 1525034 | Cell/Virus Enclosed | NO |
| VKETVAQFNYRR | 1525040 | Cell/Virus Enclosed | NO |
| VKFGAALAMEKVNECV | 1703712 | Ectodomain | YES |
| VKFKARIEEVNT | 1525064 | Cell/Virus Enclosed | NO |
| VKFQDSKPPLPPDEWV | 1703719 | Cell/Virus Enclosed | NO |
| VKFQTVKPGNFNQDFY | 1703720 | Cell/Virus Enclosed | NO |
| VKFSATQAMEKV | 1525075 | Ectodomain | YES |
| VKFVVESNVVDV | 1525082 | Cell/Virus Enclosed | NO |
| VKGDSCSSNCKH | 1525096 | Cell/Virus Enclosed | NO |
| VKGQIQVASDNL | 1525108 | Cell/Virus Enclosed | NO |
| VKGWVGFDVEGA | 1525115 | Cell/Virus Enclosed | NO |
| VKHDVSPFEMPFEELN | 1703749 | Cell/Virus Enclosed | NO |
| VKHGDFSAVSSPMSNM | 1703751 | Underneath Membrane | NO |
| VKHITNITGVSL | 1525121 | Cell/Virus Enclosed | NO |
| VKHLNKNLNEGVLESF | 1703758 | Cell/Virus Enclosed | NO |
| VKHLNKTLNEGVLESF | 1703759 | Cell/Virus Enclosed | NO |
| VKIDGDYVYLPYPDPS | 1703763 | Cell/Virus Enclosed | NO |
| VKIDNASHASTV | 1525130 | Secreted | YES |
| VKIDNASPASTA | 1525131 | Secreted | YES |
| VKIDNASPASTVHATA | 1703764 | Secreted | YES |
| VKIEDAIIVNDENSSI | 1703765 | Cell/Virus Enclosed | NO |
| VKIGAVKHCQCGTVAT | 1703767 | Cell/Virus Enclosed | NO |
| VKIGGLTFKTYAFDKP | 1703768 | Cell/Virus Enclosed | NO |
| VKIGPIKYCYCGNSAT | 1703771 | Cell/Virus Enclosed | NO |
| VKINNVDTVIFKNNTS | 1703774 | Cell/Virus Enclosed | NO |
| VKIQAVVNANAEALNN | 1703775 | Ectodomain | YES |
| VKKASYDTNLNK | 1525155 | Cell/Virus Enclosed | NO |
| VKKGCFTGVDGELPVA | 1703782 | Cell/Virus Enclosed | NO |
| VKKPFKVEDSVI | 1525158 | Cell/Virus Enclosed | NO |
| VKKVAFCGDQVH | 1525161 | Cell/Virus Enclosed | NO |
| VKKVAFGGDQVHEVAA | 1703787 | Cell/Virus Enclosed | NO |
| VKKVIDVTGKLS | 1525164 | Cell/Virus Enclosed | NO |
| VKLGAALAMEKV | 1525175 | Ectodomain | YES |
| VKLGAEKKDGFYETIV | 1703793 | Cell/Virus Enclosed | NO |
| VKLGAENKDGFY | 1525176 | Cell/Virus Enclosed | NO |
| VKLGTGYTLSDLPVYV | 1703794 | Underneath Membrane | NO |
| VKLNPSEEFIKH | 1525193 | Cell/Virus Enclosed | NO |
| VKLPFEPQTIQP | 1525194 | Cell/Virus Enclosed | NO |
| VKLPRGVGSANI | 1525195 | Cell/Virus Enclosed | NO |
| VKLPRWSASIDT | 1525196 | Ectodomain | YES |
| VKLPSAPQTIQPEVKE | 1703806 | Cell/Virus Enclosed | NO |
| VKLSDVGFVEAYNNCT | 1703812 | Ectodomain | YES |
| VKMDGDDVYLPYPNPS | 1703815 | Cell/Virus Enclosed | NO |
| VKMVNPTSKVEP | 1525231 | Cell/Virus Enclosed | NO |
| VKNHKNLLKIAQYAAQNRRGLDL | 69279 | Ectodomain | YES |
| VKNIPCYASAVA | 1525242 | Cell/Virus Enclosed | NO |
| VKNLNSSEGVSD | 1525250 | Underneath Membrane | NO |
| VKNLNSSQGIPD | 1525251 | Ectodomain | YES |
| VKNNKGFKKGVM | 1525255 | Ectodomain | YES |
| VKNSQSSPIIPG | 1525260 | Ectodomain | YES |
| VKNSTAFPFETLSACR | 1703840 | Cell/Virus Enclosed | NO |
| VKNTKLYVNKTLYSEF | 1703841 | Ectodomain | YES |
| VKNTKLYVNNIL | 1525266 | Ectodomain | YES |
| VKNTKVFKDGVM | 1525269 | Ectodomain | YES |
| VKNVTDVTGNLS | 1525283 | Cell/Virus Enclosed | NO |
| VKNYFITDAKTG | 1525286 | Cell/Virus Enclosed | NO |
| VKPDMADEIANL | 1525296 | Cell/Virus Enclosed | NO |
| VKPDMADEIASL | 1525297 | Cell/Virus Enclosed | NO |
| VKPFLGQAAITT | 1525306 | Cell/Virus Enclosed | NO |
| VKPFLGQAAVTT | 1525308 | Cell/Virus Enclosed | NO |
| VKPGDAVITTQQAGAG | 1703858 | Cell/Virus Enclosed | NO |
| VKPGGTSSGDATTAFA | 1703861 | Cell/Virus Enclosed | NO |
| VKPGHFNEEFYNFLRL | 1703863 | Cell/Virus Enclosed | NO |

| | | | |
|---|---|---|---|
| VKPGHFNKEFYDFLRS | 1703864 | Cell/Virus Enclosed | NO |
| VKPGNFNQDFYDFVLS | 1703866 | Cell/Virus Enclosed | NO |
| VKPGNFNQDFYEFVKS | 1703867 | Cell/Virus Enclosed | NO |
| VKPKPVINQLDEKAQK | 1703871 | Cell/Virus Enclosed | NO |
| VKPKRLRKKRNVDPLS | 1703872 | Cell/Virus Enclosed | NO |
| VKPNRTSVVSFNEWAL | 1703876 | Cell/Virus Enclosed | NO |
| VKPTVYVHSRVK | 1525341 | Ectodomain | YES |
| VKPVATVHRRIPDLPD | 1703884 | Ectodomain | YES |
| VKPVFFNAFVKC | 1525346 | Cell/Virus Enclosed | NO |
| VKPVRARAPPGEQFTH | 1703887 | Cell/Virus Enclosed | NO |
| VKQEQRIGLDAV | 1525354 | Cell/Virus Enclosed | NO |
| VKQGAKPVGNCVKMLT | 1703893 | Cell/Virus Enclosed | NO |
| VKQGHFIGVEGELPVA | 1703895 | Cell/Virus Enclosed | NO |
| VKQLFYVYDVHKDNFK | 1703896 | Cell/Virus Enclosed | NO |
| VKQLPEQFDSLNLSAGTQ | 1525367 | Cell/Virus Enclosed | NO |
| VKQMYKTPAIKD | 1525373 | Ectodomain | YES |
| VKQMYKTPTLKYFGGFN | 69299 | Ectodomain | YES |
| VKQSPELFDSLN | 1525382 | Cell/Virus Enclosed | NO |
| VKQSPELLDPLN | 1525383 | Cell/Virus Enclosed | NO |
| VKQSPESFDSLN | 1525384 | Cell/Virus Enclosed | NO |
| VKRDGVQVGYCVHGIK | 1703908 | Cell/Virus Enclosed | NO |
| VKRRFSLLRPEE | 1525410 | Cell/Virus Enclosed | NO |
| VKRTDLTIYNREMKCY | 1703920 | Cell/Virus Enclosed | NO |
| VKRTNLEVYNKEKTYY | 1703922 | Cell/Virus Enclosed | NO |
| VKRVRAWVGFDAEGAH | 1703925 | Cell/Virus Enclosed | NO |
| VKSACIKVDIQQ | 1525422 | Ectodomain | YES |
| VKSALLADKFSV | 1525424 | Cell/Virus Enclosed | NO |
| VKSDIFGSDFKTFDLL | 1703931 | Cell/Virus Enclosed | NO |
| VKSDYCRGNSCI | 1525433 | Ectodomain | YES |
| VKSFEIDKGIYQTSNF | 1703935 | Ectodomain | YES |
| VKSFKIDKGIYQ | 1525440 | Ectodomain | YES |
| VKSGTAILIKSP | 1525444 | Cell/Virus Enclosed | NO |
| VKSKGLFKEGST | 1525446 | Cell/Virus Enclosed | NO |
| VKSKVGNYRLPSSKPS | 1703943 | Underneath Membrane | NO |
| VKSKVGNYRLPSTQKG | 1703944 | Ectodomain | YES |
| VKSLSMIRGPPRAELN | 1703949 | Cell/Virus Enclosed | NO |
| VKSSDVVDSNENLTWP | 1703958 | Cell/Virus Enclosed | NO |
| VKSSLSSYEYCQPPKS | 1703960 | Cell/Virus Enclosed | NO |
| VKSSQSSPIIPGFGGD | 1703961 | Ectodomain | YES |
| VKSVSPGDSSWH | 1525479 | Ectodomain | YES |
| VKTCDLTVGSGY | 1525484 | Ectodomain | YES |
| VKTDAVVLLERYVSLA | 1703972 | Cell/Virus Enclosed | NO |
| VKTDGTLMVERS | 1525487 | Cell/Virus Enclosed | NO |
| VKTGHYTGQAGEMPCA | 1703977 | Cell/Virus Enclosed | NO |
| VKVGTGVMVECDSLRI | 1703998 | Cell/Virus Enclosed | NO |
| VKVPIDLIMIEF | 1525521 | Cell/Virus Enclosed | NO |
| VKVVRNKTSACF | 1525533 | Cell/Virus Enclosed | NO |
| VKWADAPEPQRG | 1525534 | Cell/Virus Enclosed | NO |
| VKWADASEPQRGRQGR | 1704009 | Capsid | UN |
| VKWADESEPQRG | 1525536 | Cell/Virus Enclosed | NO |
| VKWEYEGGCNTI | 1525537 | Cell/Virus Enclosed | NO |
| VKYFECPNSIYI | 1525548 | Cell/Virus Enclosed | NO |
| VKYYSRVKSVRG | 1525558 | Cell/Virus Enclosed | NO |
| VLAAYNGKPQGA | 1525572 | Cell/Virus Enclosed | NO |
| VLAAYNGRPQGA | 1525574 | Cell/Virus Enclosed | NO |
| VLAEFDTYDPIY | 1525578 | Cell/Virus Enclosed | NO |
| VLAIESIHYLGTTGHT | 1704037 | Cell/Virus Enclosed | NO |
| VLAKGGFKFGDPADSR | 1704040 | Cell/Virus Enclosed | NO |
| VLAKGHFKFDEPSDAT | 1704041 | Cell/Virus Enclosed | NO |
| VLAKLGKDATKPQQVT | 1704042 | Cell/Virus Enclosed | NO |
| VLARKHETSCSQ | 1525605 | Cell/Virus Enclosed | NO |
| VLASDDLYVKRYFKGC | 1704055 | Cell/Virus Enclosed | NO |
| VLATDDLYVKRYERGC | 1704056 | Cell/Virus Enclosed | NO |
| VLAVVNSQDVYK | 1525618 | Cell/Virus Enclosed | NO |
| VLAYFSKKHVEVFTDG | 1704062 | Cell/Virus Enclosed | NO |
| VLDDFIVFNNNGTNVT | 1704077 | Cell/Virus Enclosed | NO |
| VLDILLASSSLR | 1525655 | Cell/Virus Enclosed | NO |
| VLDITDAAVKAAESKA | 1704080 | Cell/Virus Enclosed | NO |
| VLDNGIIRDINP | 1525667 | Cell/Virus Enclosed | NO |
| VLDQSDNNCWISTTLV | 1704091 | Cell/Virus Enclosed | NO |
| VLDQSENNCWIS | 1525672 | Cell/Virus Enclosed | NO |
| VLDTTSYREAAC | 1525682 | Cell/Virus Enclosed | NO |
| VLDVKGLEELHY | 1525684 | Cell/Virus Enclosed | NO |
| VLDVKGVEELHDDLVN | 1704098 | Cell/Virus Enclosed | NO |
| VLECTRASTSAV | 1525693 | Cell/Virus Enclosed | NO |
| VLEDTLDDGPSVETSD | 1704103 | Cell/Virus Enclosed | NO |

| | | | |
|---|---|---|---|
| VLEENLNAYVNSNQNT | 1704104 | Capsid | UN |
| VLEMAKEVKTSSKVVL | 1704110 | Cell/Virus Enclosed | NO |
| VLENFIVYNNSG | 1525705 | Cell/Virus Enclosed | NO |
| VLEQDAQIDRLINGRL | 1704112 | Ectodomain | YES |
| VLESFSVTLLDNQEDK | 1704114 | Cell/Virus Enclosed | NO |
| VLETFQNNVVNYQHIG | 1704115 | Cell/Virus Enclosed | NO |
| VLETTDNNCWVN | 1525713 | Cell/Virus Enclosed | NO |
| VLFDGRDNGALEAFKR | 1704122 | Cell/Virus Enclosed | NO |
| VLFSGQKYSSDC | 1525753 | Cell/Virus Enclosed | NO |
| VLFSTVVIKNLTPIKL | 1704136 | Cell/Virus Enclosed | NO |
| VLFTNSTDTVNHDSFN | 1704139 | Cell/Virus Enclosed | NO |
| VLFVMFLSLIGLLAI | 69496 | Transmembrane | NO |
| VLFVNDKITEFQ | 1525765 | Cell/Virus Enclosed | NO |
| VLFYCSNSSSGVLDTT | 1704143 | Ectodomain | YES |
| VLGCLGSECNKV | 1525775 | Ectodomain | YES |
| VLGDAIQVAKCY | 1525780 | Cell/Virus Enclosed | NO |
| VLGEYVFDKSELTNGV | 1704149 | Cell/Virus Enclosed | NO |
| VLGLQTQTVDSAQGSE | 1704159 | Cell/Virus Enclosed | NO |
| VLGLYVPIATRN | 1525799 | Cell/Virus Enclosed | NO |
| VLGLYVPKATRN | 1525800 | Cell/Virus Enclosed | NO |
| VLGRGVFQNCTA | 1525805 | Ectodomain | YES |
| VLGYLDSLKPKV | 1525812 | Cell/Virus Enclosed | NO |
| VLHLGAGSDKEVAPGS | 1704178 | Cell/Virus Enclosed | NO |
| VLHTKSAAGYGG | 1525821 | Cell/Virus Enclosed | NO |
| VLHTKSASGYGG | 1525822 | Cell/Virus Enclosed | NO |
| VLHYGAGSDKGVAPGT | 1704186 | Cell/Virus Enclosed | NO |
| VLIDYDHAYSTMRLHN | 1704189 | Cell/Virus Enclosed | NO |
| VLIQNGAKHVQGNVAK | 1704196 | Cell/Virus Enclosed | NO |
| VLIVLLCLGLVT | 1525855 | Ectodomain | YES |
| VLKITEDGINVK | 1525869 | Cell/Virus Enclosed | NO |
| VLKLLTLFFYVP | 1525876 | Cell/Virus Enclosed | NO |
| VLKLPSLMKVMTHMDD | 1704221 | Cell/Virus Enclosed | NO |
| VLKQWLPSGSIL | 1525887 | Cell/Virus Enclosed | NO |
| VLLDWVKHLNKNLNEG | 1704247 | Cell/Virus Enclosed | NO |
| VLLERVYKHFNN | 1525924 | Cell/Virus Enclosed | NO |
| VLLERYVSLAIDAYPL | 1704250 | Cell/Virus Enclosed | NO |
| VLLIERFVSLAIDAYP | 1704260 | Cell/Virus Enclosed | NO |
| VLLLYFMVDNGT | 1525955 | Undetermined | UN |
| VLLNCLQHLHVASNSK | 1704269 | Cell/Virus Enclosed | NO |
| VLLSKGSLEPRHFNSI | 1704282 | Cell/Virus Enclosed | NO |
| VLMDGSIIQFPN | 1525995 | Cell/Virus Enclosed | NO |
| VLMGILSNMNIASNSK | 1704298 | Cell/Virus Enclosed | NO |
| VLMGWDYPKCDRAMPN | 1704299 | Cell/Virus Enclosed | NO |
| VLMTARTVYDDA | 1525998 | Cell/Virus Enclosed | NO |
| VLNASEIYCANI | 1526011 | Cell/Virus Enclosed | NO |
| VLNASEIYCANL | 1526012 | Cell/Virus Enclosed | NO |
| VLNASTPESKPL | 1526016 | Cell/Virus Enclosed | NO |
| VLNDPVFTGELF | 1526024 | Cell/Virus Enclosed | NO |
| VLNDVLSRLDKV | 1526027 | Ectodomain | YES |
| VLNENLNAYQQQDGMM | 1704311 | Cell/Virus Enclosed | NO |
| VLNKALEMCIDQ | 1526046 | Cell/Virus Enclosed | NO |
| VLNNCTKYNIYDYVGT | 1704320 | Ectodomain | YES |
| VLNNISLTTQPYFDSY | 1704326 | Ectodomain | YES |
| VLNNKSTYRSQL | 1526059 | Cell/Virus Enclosed | NO |
| VLNNLNLPGCNGGSLY | 1704328 | Cell/Virus Enclosed | NO |
| VLNNPAYLAPQA | 1526062 | Ectodomain | YES |
| VLNQSYINLKDIGTYE | 1704330 | Ectodomain | YES |
| VLNVVGLDARTQ | 1526077 | Cell/Virus Enclosed | NO |
| VLNVVGPDARTQGKQS | 1704335 | Cell/Virus Enclosed | NO |
| VLPETYVALPIYYQHT | 1704338 | Ectodomain | YES |
| VLPGAFAYSQMR | 1526094 | Cell/Virus Enclosed | NO |
| VLPINTDTVLSVAPEV | 1704343 | Cell/Virus Enclosed | NO |
| VLPLTCNAISSNTDNE | 1704346 | Ectodomain | YES |
| VLPPLLSDNQIS | 1526118 | Ectodomain | YES |
| VLPPLLSENQIN | 1526119 | Ectodomain | YES |
| VLPPLLSENQISGYTL | 1704353 | Ectodomain | YES |
| VLPPLLTDDMIAAYTAA | 69699 | Ectodomain | YES |
| VLPPLMDVNMEAAYTS | 1704356 | Ectodomain | YES |
| VLPQGYYIEGSG | 1526127 | Cell/Virus Enclosed | NO |
| VLPQRNIRSSRIAGRS | 1704362 | Ectodomain | YES |
| VLQEDQKILAAS | 1526149 | Ectodomain | YES |
| VLQENQKILAASFNKA | 1704378 | Ectodomain | YES |
| VLQNNELMPAKL | 1526162 | Cell/Virus Enclosed | NO |
| VLQYGHYMYSRF | 1526185 | Undetermined | UN |
| VLQYGYPSRSMT | 1526186 | Undetermined | UN |
| VLRCGDCLRKPM | 1526189 | Cell/Virus Enclosed | NO |

| | | | |
|---|---|---|---|
| VLRDNLKPVTTLSTIT | 1704397 | Cell/Virus Enclosed | NO |
| VLRIGAAANSTG | 1526196 | Ectodomain | YES |
| VLRNSFGKDLNANMSL | 1704403 | Cell/Virus Enclosed | NO |
| VLRNSNGVCIWN | 1526203 | Cell/Virus Enclosed | NO |
| VLRPILKLPFGI | 1526205 | Ectodomain | YES |
| VLRQWLPTDAIV | 1526210 | Cell/Virus Enclosed | NO |
| VLRQWLPTGTLL | 1526212 | Cell/Virus Enclosed | NO |
| VLRSTGQKSIVA | 1526213 | Ectodomain | YES |
| VLRTSPPHRPGVRMRRV | 69746 | Cell/Virus Enclosed | NO |
| VLSAAIKDCVDINKAI | 1704412 | Cell/Virus Enclosed | NO |
| VLSDNQKLIANK | 1526226 | Ectodomain | YES |
| VLSDVDGLKYTKIMKD | 1704417 | Cell/Virus Enclosed | NO |
| VLSDVIRYNLNFEENL | 1704418 | Ectodomain | YES |
| VLSENQKLIANKFNQA | 1704423 | Ectodomain | YES |
| VLSFELLNAPATVCGPK | 69760 | Ectodomain | YES |
| VLSKSILLVDSS | 1526267 | Cell/Virus Enclosed | NO |
| VLSLDKLKSLLT | 1526269 | Cell/Virus Enclosed | NO |
| VLSLRHDIALDDDART | 1704440 | Cell/Virus Enclosed | NO |
| VLSLVNFTVDPQKAYL | 1704442 | Cell/Virus Enclosed | NO |
| VLSMTDCLAAETHRDC | 1704445 | Cell/Virus Enclosed | NO |
| VLSSAVMFEDAQ | 1526289 | Cell/Virus Enclosed | NO |
| VLSSDVMFEDDQ | 1526290 | Cell/Virus Enclosed | NO |
| VLSSLCTMFKRGDGTP | 1704453 | Cell/Virus Enclosed | NO |
| VLSVDSVSEESQGNVV | 1704459 | Cell/Virus Enclosed | NO |
| VLSVNSSNCNNFNVKK | 1704462 | Cell/Virus Enclosed | NO |
| VLSYELLNAPAT | 1526307 | Ectodomain | YES |
| VLTDVDVPFTAH | 1526316 | Cell/Virus Enclosed | NO |
| VLTENALVRLTTEVVR | 1704470 | Cell/Virus Enclosed | NO |
| VLTENLNAYQQQ | 1526323 | Cell/Virus Enclosed | NO |
| VLTFTTKVTVPK | 1526328 | Cell/Virus Enclosed | NO |
| VLTGCCDNSYFDSKGW | 1704473 | Cell/Virus Enclosed | NO |
| VLTGDSVKFVYMHQLE | 1704475 | Cell/Virus Enclosed | NO |
| VLTIANNYSKLFDEGY | 1704479 | Cell/Virus Enclosed | NO |
| VLTIANNYSKLS | 1526333 | Cell/Virus Enclosed | NO |
| VLTLDNQDLNGKWYDF | 1704483 | Cell/Virus Enclosed | NO |
| VLTPLLTDDMIA | 1526347 | Ectodomain | YES |
| VLTPSSKRFQPFQQFGRD | 1073900 | Ectodomain | YES |
| VLTTFIDAARGPAGVE | 1704502 | Cell/Virus Enclosed | NO |
| VLTVTLQNSRTP | 1526363 | Cell/Virus Enclosed | NO |
| VLTYSNFGVCAD | 1526366 | Ectodomain | YES |
| VLVKLLEVYNGF | 1526373 | Cell/Virus Enclosed | NO |
| VLVNGLHYPTGS | 1526375 | Cell/Virus Enclosed | NO |
| VLVNGVHHPTGS | 1526377 | Cell/Virus Enclosed | NO |
| VLVNGVHYPTGI | 1526379 | Cell/Virus Enclosed | NO |
| VLVNGVHYPTGS | 1526380 | Cell/Virus Enclosed | NO |
| VLVNGVRYPTGS | 1526381 | Cell/Virus Enclosed | NO |
| VLVSNNQEDFDLISKC | 1704532 | Cell/Virus Enclosed | NO |
| VLVSPGLCISGDVGIA | 1704533 | Ectodomain | YES |
| VLWANGVKAKGY | 1526403 | Cell/Virus Enclosed | NO |
| VLWDYEAERPFT | 1526405 | Cell/Virus Enclosed | NO |
| VLWDYEAERPLTSFTK | 1704545 | Cell/Virus Enclosed | NO |
| VLYDNQKQIANQ | 1526420 | Ectodomain | YES |
| VLYENQKQIANQFNKAI | 69865 | Ectodomain | YES |
| VLYQGVNCTDVS | 1526446 | Ectodomain | YES |
| VLYSPNVSVPSSSSTPLLYPSLA | 69909 | Ectodomain | YES |
| VLYVDGHRLASG | 1526454 | Underneath Membrane | NO |
| VMAAPAAPRAVS | 1526461 | Cell/Virus Enclosed | NO |
| VMCPADQLSDPN | 1526477 | Cell/Virus Enclosed | NO |
| VMDVNDCFKNDNVVLK | 1704589 | Cell/Virus Enclosed | NO |
| VMDYLDSLKPRV | 1526486 | Cell/Virus Enclosed | NO |
| VMEPIDYNVVTQ | 1526491 | Cell/Virus Enclosed | NO |
| VMESQISTKEVKLKGV | 1704596 | Cell/Virus Enclosed | NO |
| VMFDLDSTFDDI | 1526492 | Cell/Virus Enclosed | NO |
| VMFEDAQGLKKF | 1526494 | Cell/Virus Enclosed | NO |
| VMFEDAQSLKKF | 1526495 | Cell/Virus Enclosed | NO |
| VMFEGAQGLKKF | 1526497 | Cell/Virus Enclosed | NO |
| VMFENAQGLKKF | 1526498 | Cell/Virus Enclosed | NO |
| VMFQPRLPVLSD | 1526505 | Ectodomain | YES |
| VMGLHGNQTLFCNIMK | 1704607 | Cell/Virus Enclosed | NO |
| VMILKSQDLGAV | 1526524 | Cell/Virus Enclosed | NO |
| VMISEWPLFDQQ | 1526525 | Cell/Virus Enclosed | NO |
| VMISEWPLSESD | 1526527 | Cell/Virus Enclosed | NO |
| VMISEWPLSVQQAQQE | 1704618 | Cell/Virus Enclosed | NO |
| VMISQWPISNDSNGCV | 1704619 | Cell/Virus Enclosed | NO |
| VMKLPSEPQTIQ | 1526536 | Cell/Virus Enclosed | NO |
| VMLKAAMLCNRY | 1526542 | Cell/Virus Enclosed | NO |

| | | | |
|---|---|---|---|
| VMLLVKHKHTFL | 1526544 | Cell/Virus Enclosed | NO |
| VMLNTSIPNLPDFKEE | 1704628 | Ectodomain | YES |
| VMLNTSTSNLPD | 1526553 | Ectodomain | YES |
| VMLPGEFSKKQL | 1526554 | Cell/Virus Enclosed | NO |
| VMMSAPPSEYTL | 1526565 | Cell/Virus Enclosed | NO |
| VMNDKNICLLDH | 1526566 | Cell/Virus Enclosed | NO |
| VMNFDVDTHRYR | 1526568 | Cell/Virus Enclosed | NO |
| VMNMDVSLHRHR | 1526572 | Cell/Virus Enclosed | NO |
| VMPLTAPTLVPQ | 1526580 | Cell/Virus Enclosed | NO |
| VMPTYTVHKSDF | 1526594 | Cell/Virus Enclosed | NO |
| VMPVHTLSIHDG | 1526596 | Cell/Virus Enclosed | NO |
| VMPVSMAKTSVD | 1526598 | Ectodomain | YES |
| VMQFGYIDCEQDSCDF | 1704647 | Cell/Virus Enclosed | NO |
| VMQFGYIDCEQT | 1526601 | Cell/Virus Enclosed | NO |
| VMQKLPFKFKDLAIEN | 1704649 | Cell/Virus Enclosed | NO |
| VMQMDIQGFVNI | 1526606 | Cell/Virus Enclosed | NO |
| VMQNNELMPHKL | 1526607 | Cell/Virus Enclosed | NO |
| VMRIMTWLELAD | 1526615 | Cell/Virus Enclosed | NO |
| VMRYNLNFSANSLDNL | 1704660 | Ectodomain | YES |
| VMSEDLWQFVDH | 1526628 | Cell/Virus Enclosed | NO |
| VMSLSYNSVLDLSKFN | 1704666 | Cell/Virus Enclosed | NO |
| VMTFYPRLQAASDWKP | 1704671 | Cell/Virus Enclosed | NO |
| VMVDCRRLLKQD | 1526650 | Cell/Virus Enclosed | NO |
| VMVDCRRLLKQE | 1526651 | Cell/Virus Enclosed | NO |
| VMWFYTYSIGEA | 1526666 | Cell/Virus Enclosed | NO |
| VMWLYTYSIGEASSPI | 1704687 | Cell/Virus Enclosed | NO |
| VMYGGFEDQPNLQVES | 1704694 | Cell/Virus Enclosed | NO |
| VMYMGTLSYEEL | 1526687 | Cell/Virus Enclosed | NO |
| VMYSEFPAIVIG | 1526692 | Ectodomain | YES |
| VNAANTHLKHGGGIAG | 1704705 | Cell/Virus Enclosed | NO |
| VNADIVVVDEVSMCTN | 1704710 | Cell/Virus Enclosed | NO |
| VNAGGAPLTNCVKMLT | 1704713 | Cell/Virus Enclosed | NO |
| VNAITTKISLKG | 1526709 | Cell/Virus Enclosed | NO |
| VNANGGSKFCKKHRFF | 1704725 | Cell/Virus Enclosed | NO |
| VNANGGTCFCSK | 1526720 | Cell/Virus Enclosed | NO |
| VNANHYSPCVSI | 1526721 | Ectodomain | YES |
| VNANQYSPCVPI | 1526722 | Ectodomain | YES |
| VNANQYSPCVST | 1526724 | Ectodomain | YES |
| VNAPYYNSWQNL | 1526727 | Ectodomain | YES |
| VNAQPKCNPNLHYWTTQD | 1058471 | Secreted | YES |
| VNAYNSTAISVK | 1526743 | Ectodomain | YES |
| VNCCRFQRFDEN | 1526748 | Cell/Virus Enclosed | NO |
| VNCCRFQRIDDDGNKL | 1704742 | Cell/Virus Enclosed | NO |
| VNCCRFQRVDENGDKL | 1704743 | Cell/Virus Enclosed | NO |
| VNCDSFGPGNTFINGD | 1704745 | Cell/Virus Enclosed | NO |
| VNCNGYQHHSFA | 1526762 | Ectodomain | YES |
| VNCNGYQHHSVA | 1526764 | Ectodomain | YES |
| VNCNGYQHNSFA | 1526765 | Ectodomain | YES |
| VNCNGYQHNSVA | 1526766 | Ectodomain | YES |
| VNCNGYQHNSVV | 1526767 | Ectodomain | YES |
| VNCTDVPTAIRA | 1526770 | Ectodomain | YES |
| VNCTDVPTSIHA | 1526772 | Ectodomain | YES |
| VNDAITQTSQALQTVA | 1704755 | Ectodomain | YES |
| VNDCFKKDHVVL | 1526785 | Cell/Virus Enclosed | NO |
| VNDDTGETKYVK | 1526788 | Cell/Virus Enclosed | NO |
| VNDDTSETKYVKSLSI | 1704757 | Cell/Virus Enclosed | NO |
| VNDFTCRQISPA | 1526791 | Ectodomain | YES |
| VNDFTCSQISPP | 1526792 | Ectodomain | YES |
| VNDICMFENKTTLPTN | 1704762 | Cell/Virus Enclosed | NO |
| VNDKALYEQVVM | 1526799 | Cell/Virus Enclosed | NO |
| VNDKIFTKSGFN | 1526800 | Cell/Virus Enclosed | NO |
| VNDKITEFQLDYSIDV | 1704766 | Cell/Virus Enclosed | NO |
| VNDLCFANKFFQFDQW | 1704768 | Cell/Virus Enclosed | NO |
| VNDNAQALNTLV | 1526812 | Ectodomain | YES |
| VNDSLKPVGGLY | 1526820 | Ectodomain | YES |
| VNDSYRQFDLVQYDFT | 1704773 | Cell/Virus Enclosed | NO |
| VNECVESQSKRY | 1526826 | Ectodomain | YES |
| VNELLDTTQLQVANSL | 1704785 | Ectodomain | YES |
| VNESRYGLHNLL | 1526844 | Ectodomain | YES |
| VNESRYGLQNLLQLPN | 1704787 | Ectodomain | YES |
| VNETNSQYLFSTINAL | 1704788 | Cell/Virus Enclosed | NO |
| VNETSQTGWALY | 1526847 | Ectodomain | YES |
| VNETTSQYLFST | 1526848 | Cell/Virus Enclosed | NO |
| VNFELRRGPGKC | 1526855 | Ectodomain | YES |
| VNFKSISLTVGEVFGK | 1704792 | Cell/Virus Enclosed | NO |
| VNFNFNGLRGTG | 1526867 | Ectodomain | YES |

| | | | |
|---|---|---|---|
| VNFNVDDINFSPVLGC | 1704799 | Ectodomain | YES |
| VNFNVDDINFSS | 1526874 | Ectodomain | YES |
| VNFNVDDVNFSP | 1526875 | Ectodomain | YES |
| VNFNVTNAATIV | 1526876 | Ectodomain | YES |
| VNFNVTTASATD | 1526877 | Ectodomain | YES |
| VNFTKALAKQDA | 1526884 | Cell/Virus Enclosed | NO |
| VNFTVDPQKAYLDFVN | 1704807 | Cell/Virus Enclosed | NO |
| VNFVVQPVGVVDTEWG | 1704810 | Cell/Virus Enclosed | NO |
| VNGDWFLFGDSRSDCN | 1704812 | Ectodomain | YES |
| VNGFYRLYSGDN | 1526901 | Cell/Virus Enclosed | NO |
| VNGLHYPTGSCL | 1526908 | Cell/Virus Enclosed | NO |
| VNGLLDTTQLHVADTL | 1704819 | Ectodomain | YES |
| VNGVDFDFIVNAANEN | 1704828 | Cell/Virus Enclosed | NO |
| VNGVHYPTGGCL | 1526928 | Cell/Virus Enclosed | NO |
| VNGVTLIGESVK | 1526933 | Cell/Virus Enclosed | NO |
| VNGYAKSNFVKDDYCD | 1704834 | Cell/Virus Enclosed | NO |
| VNGYFIKTNNTRIVDE | 1704835 | Ectodomain | YES |
| VNHNVLIKESIP | 1526948 | Cell/Virus Enclosed | NO |
| VNIAKNAYEKDKAVAR | 1704843 | Cell/Virus Enclosed | NO |
| VNINIVGDFQLN | 1526958 | Cell/Virus Enclosed | NO |
| VNINIVGDFQLT | 1526959 | Cell/Virus Enclosed | NO |
| VNINPHNYSYMDLNPV | 1704850 | Ectodomain | YES |
| VNIPADIVDRDP | 1526961 | Cell/Virus Enclosed | NO |
| VNISLNGNTSVCVRTS | 1704854 | Ectodomain | YES |
| VNITLANFNETKGPLC | 1704859 | Ectodomain | YES |
| VNKASYDTNLNKFNRA | 1704861 | Cell/Virus Enclosed | NO |
| VNKGSFVGADGELPVA | 1704869 | Cell/Virus Enclosed | NO |
| VNKHAFHKPAYD | 1526985 | Cell/Virus Enclosed | NO |
| VNKIQPWTNDALNTIC | 1704877 | Cell/Virus Enclosed | NO |
| VNKMVNWFNTMLDASA | 1704879 | Cell/Virus Enclosed | NO |
| VNKTLQEFAQNLPKYV | 1704881 | Ectodomain | YES |
| VNKTLQELSYKLPNYT | 1704882 | Ectodomain | YES |
| VNKTLYSEFSTIVIGS | 1704883 | Ectodomain | YES |
| VNKYFEIYDGGCLNAS | 1704885 | Cell/Virus Enclosed | NO |
| VNLALKNLGFDN | 1527006 | Cell/Virus Enclosed | NO |
| VNLNLPANIPYS | 1527018 | Cell/Virus Enclosed | NO |
| VNLNLPANVPYS | 1527019 | Cell/Virus Enclosed | NO |
| VNLPRGFPAGFS | 1527022 | Ectodomain | YES |
| VNLRLVDCAPVV | 1527025 | Cell/Virus Enclosed | NO |
| VNLVAVPTGYVD | 1527031 | Cell/Virus Enclosed | NO |
| VNMEAAYTSSLV | 1527037 | Ectodomain | YES |
| VNMQLMGVVMQS | 1527043 | Cell/Virus Enclosed | NO |
| VNMQQIYLINKFLKAN | 1704912 | Cell/Virus Enclosed | NO |
| VNMRVLHLGAGSDKEV | 1704914 | Cell/Virus Enclosed | NO |
| VNMRVLHLGAGSEKGV | 1704915 | Cell/Virus Enclosed | NO |
| VNNDYCCFKHAL | 1527053 | Cell/Virus Enclosed | NO |
| VNNGLTCSTDLQ | 1527059 | Ectodomain | YES |
| VNNHAFHTPAYDKRAM | 1704921 | Cell/Virus Enclosed | NO |
| VNNNAQALAKLA | 1527066 | Ectodomain | YES |
| VNNTWMFTGSRY | 1527080 | Ectodomain | YES |
| VNNYDKSAGYPFNKFG | 1704934 | Cell/Virus Enclosed | NO |
| VNNYEKSAGYPF | 1527085 | Cell/Virus Enclosed | NO |
| VNPDDKCFANKHRSFT | 1704939 | Cell/Virus Enclosed | NO |
| VNPTNCIAPVNG | 1527103 | Ectodomain | YES |
| VNQIANMFLEQT | 1527115 | Cell/Virus Enclosed | NO |
| VNQIVLRNSNGV | 1527118 | Cell/Virus Enclosed | NO |
| VNQLAYSSSANGSLVG | 1704951 | Cell/Virus Enclosed | NO |
| VNQQGNLLNHLN | 1527123 | Ectodomain | YES |
| VNQQGNSLNHLTSQLR | 1704953 | Ectodomain | YES |
| VNQQGNSLSHLT | 1527124 | Ectodomain | YES |
| VNQQGSALNHLTSQLR | 1704954 | Ectodomain | YES |
| VNQSLSNAKVSAEVSR | 1704957 | Cell/Virus Enclosed | NO |
| VNQTVLRNSNGA | 1527131 | Cell/Virus Enclosed | NO |
| VNSDFCPCADRS | 1527145 | Ectodomain | YES |
| VNSDTVGWSWPDGAELPFTID | 97748 | Ectodomain | YES |
| VNSGLLTACAFS | 1527153 | Cell/Virus Enclosed | NO |
| VNSIFDIETVDVKHDV | 1704981 | Cell/Virus Enclosed | NO |
| VNSINDSLEPVG | 1527160 | Ectodomain | YES |
| VNSKNDNLEPVS | 1527163 | Ectodomain | YES |
| VNSNAQALNSLLQQLF | 1704984 | Ectodomain | YES |
| VNSNQNTDSDSLSSKP | 1704985 | Capsid | UN |
| VNSNQYSPCVSI | 1527167 | Ectodomain | YES |
| VNSNVKLLDYDYATHG | 1704986 | Cell/Virus Enclosed | NO |
| VNSQDVYKSLTT | 1527172 | Cell/Virus Enclosed | NO |
| VNSSLFVEDCNL | 1527180 | Ectodomain | YES |
| VNSSNVTTMDFC | 1527186 | Cell/Virus Enclosed | NO |

| | | | |
|---|---|---|---|
| VNSVFNICQAVS | 1527191 | Cell/Virus Enclosed | NO |
| VNSVNDSLEPVGGLYE | 1704996 | Ectodomain | YES |
| VNSVNDSLKPVG | 1527200 | Ectodomain | YES |
| VNTAIPTIEDWL | 1527207 | Cell/Virus Enclosed | NO |
| VNTHSIRLTITE | 1527214 | Cell/Virus Enclosed | NO |
| VNTLLSNQLRLA | 1527217 | Cell/Virus Enclosed | NO |
| VNTPADILDRDP | 1527222 | Cell/Virus Enclosed | NO |
| VNTPADIVDRDP | 1527223 | Cell/Virus Enclosed | NO |
| VNTPADIVDRDQ | 1527224 | Capsid | UN |
| VNTSLGNRKRNQKPLE | 1705009 | Capsid | UN |
| VNTTIGNETTSAFVGA | 1705013 | Ectodomain | YES |
| VNTTIGNETTSS | 1527237 | Ectodomain | YES |
| VNTVAGIPSNVYLVGK | 1705014 | Cell/Virus Enclosed | NO |
| VNTVCFVDVDFHSVEI | 1705015 | Cell/Virus Enclosed | NO |
| VNTVKPKPVINQLDEK | 1705016 | Cell/Virus Enclosed | NO |
| VNTVSELVYENKFVPV | 1705017 | Cell/Virus Enclosed | NO |
| VNVAAWLYAAIL | 1527241 | Cell/Virus Enclosed | NO |
| VNVDGFNEWAMANGYT | 1705024 | Cell/Virus Enclosed | NO |
| VNVGHYDGRTGELPCA | 1705025 | Cell/Virus Enclosed | NO |
| VNVHDVTVTTDKSFEQ | 1705026 | Cell/Virus Enclosed | NO |
| VNVIGTRPEKFP | 1527255 | Cell/Virus Enclosed | NO |
| VNVITTKISLKG | 1527256 | Cell/Virus Enclosed | NO |
| VNVKPKRLRKKR | 1527259 | Cell/Virus Enclosed | NO |
| VNVMFDLDSTFD | 1527265 | Cell/Virus Enclosed | NO |
| VNVNNTWMFTGS | 1527267 | Ectodomain | YES |
| VNVNNTWMYTGSGYYY | 1705036 | Ectodomain | YES |
| VNVNRFNVAITRAKKG | 1705037 | Cell/Virus Enclosed | NO |
| VNVNVDFKDFQFMLWC | 1705038 | Cell/Virus Enclosed | NO |
| VNVPTIRKFIKF | 1527270 | Cell/Virus Enclosed | NO |
| VNVPYLYNGSAQ | 1527271 | Ectodomain | YES |
| VNVQKVEQPKIEPKPV | 1705043 | Cell/Virus Enclosed | NO |
| VNVSATNIQNLLYCDS | 1705045 | Ectodomain | YES |
| VNVSHVYNGSAQ | 1527278 | Ectodomain | YES |
| VNVSYVYNGSAQ | 1527286 | Ectodomain | YES |
| VNVTKHKCDINYKGKV | 1705050 | Cell/Virus Enclosed | NO |
| VNVTKHKCDISY | 1527289 | Cell/Virus Enclosed | NO |
| VNVTTLNGRIVN | 1527292 | Ectodomain | YES |
| VNVYRSTSPDPKFVDK | 1705057 | Cell/Virus Enclosed | NO |
| VNWADDRAARKKFPPP | 1705058 | Capsid | UN |
| VNWFNTMLDASAPATG | 1705059 | Cell/Virus Enclosed | NO |
| VNWNLDYPIISN | 1527307 | Cell/Virus Enclosed | NO |
| VNWNLEYPIISNEVSI | 1705060 | Cell/Virus Enclosed | NO |
| VNYTKAPDVMLN | 1527319 | Ectodomain | YES |
| VNYTVCDDCNGYTDNI | 1705070 | Ectodomain | YES |
| VPAAYKLTKLSV | 1527323 | Ectodomain | YES |
| VPANSAVLSFCA | 1527337 | Cell/Virus Enclosed | NO |
| VPANSTVLSFCA | 1527339 | Cell/Virus Enclosed | NO |
| VPARARVECFDK | 1527342 | Cell/Virus Enclosed | NO |
| VPATRDSVPLHIIAPS | 1705083 | Cell/Virus Enclosed | NO |
| VPDGFYSTSPIQSVEL | 1705094 | Ectodomain | YES |
| VPDHDSFVKMMVDGFV | 1705095 | Cell/Virus Enclosed | NO |
| VPDIVDVPPQVV | 1527375 | Cell/Virus Enclosed | NO |
| VPDIVEVPPQVV | 1527376 | Cell/Virus Enclosed | NO |
| VPDKSVYDQVVDNVYV | 1705096 | Cell/Virus Enclosed | NO |
| VPDLVVEQYNQI | 1527377 | Ectodomain | YES |
| VPDLVVEQYNQTILNL | 1705097 | Ectodomain | YES |
| VPDTLGGDPKFRSLTHEDHA | 70275 | Capsid | UN |
| VPDTVEVPPQVVKLPS | 1705099 | Cell/Virus Enclosed | NO |
| VPDTVGVPPQVV | 1527382 | Cell/Virus Enclosed | NO |
| VPEEFASHVANFQKSG | 1705102 | Cell/Virus Enclosed | NO |
| VPEHCQNYSGFM | 1527389 | Cell/Virus Enclosed | NO |
| VPEHCQNYSGSM | 1527390 | Cell/Virus Enclosed | NO |
| VPEKFLNAFDVFVTAI | 1705106 | Cell/Virus Enclosed | NO |
| VPESPFSRTGEGLDIRGNQGFP | 244302 | Cell/Virus Enclosed | NO |
| VPEYIDVNKTLQELSY | 1705108 | Ectodomain | YES |
| VPFGGTTVVRPLVEDS | 1705111 | Ectodomain | YES |
| VPFLICSNTPAS | 1527407 | Cell/Virus Enclosed | NO |
| VPFSPNGKPCTP | 1527413 | Ectodomain | YES |
| VPFSSDGKPCTP | 1527414 | Ectodomain | YES |
| VPGDVIITSTGA | 1527428 | Cell/Virus Enclosed | NO |
| VPGNVKPGDAVITTQQ | 1705134 | Cell/Virus Enclosed | NO |
| VPGNVVPGDVII | 1527445 | Cell/Virus Enclosed | NO |
| VPGQSKRVDFCG | 1527447 | Ectodomain | YES |
| VPGYPKLFITCE | 1527458 | Cell/Virus Enclosed | NO |
| VPHIVRKDLTKYTMLD | 1705140 | Cell/Virus Enclosed | NO |
| VPHNLTTITKPF | 1527463 | Ectodomain | YES |

| | | | |
|---|---|---|---|
| VPHNLTTITKPL | 1527464 | Ectodomain | YES |
| VPHNMRVLHYGAGSDK | 1705144 | Cell/Virus Enclosed | NO |
| VPHTRYSLADSN | 1527467 | Cell/Virus Enclosed | NO |
| VPHTRYSLANSN | 1527468 | Cell/Virus Enclosed | NO |
| VPIAFGIPPSEA | 1527476 | Cell/Virus Enclosed | NO |
| VPIAFGVPLSEA | 1527477 | Cell/Virus Enclosed | NO |
| VPIAFGVPPSEAKGYW | 1705151 | Capsid | UN |
| VPIDEYITTYPG | 1527486 | Cell/Virus Enclosed | NO |
| VPIEYQATRVVV | 1527490 | Ectodomain | YES |
| VPIGKGNKDEQIGYWN | 1705155 | Capsid | UN |
| VPIGKGSKDEQI | 1527496 | Cell/Virus Enclosed | NO |
| VPIGTNDPIRFCLENT | 1705156 | Cell/Virus Enclosed | NO |
| VPIGTNDPIRSC | 1527497 | Cell/Virus Enclosed | NO |
| VPIINCVKMLCDHAGT | 1705158 | Cell/Virus Enclosed | NO |
| VPINTNSGKDDQ | 1527503 | Cell/Virus Enclosed | NO |
| VPINTNSSKDDQ | 1527505 | Cell/Virus Enclosed | NO |
| VPINTNSTKDDQIGYY | 1705163 | Cell/Virus Enclosed | NO |
| VPIRLAGHIRHL | 1527507 | Cell/Virus Enclosed | NO |
| VPKIFKARAIEC | 1527518 | Cell/Virus Enclosed | NO |
| VPKIFKARAIEF | 1527519 | Cell/Virus Enclosed | NO |
| VPKNFCPCKLNG | 1527520 | Ectodomain | YES |
| VPLAFGVPLSEA | 1527527 | Cell/Virus Enclosed | NO |
| VPLCGSGNLVEK | 1527529 | Cell/Virus Enclosed | NO |
| VPLCIFNGKFLSNTKY | 1705174 | Ectodomain | YES |
| VPLGIKDPILYV | 1527538 | Cell/Virus Enclosed | NO |
| VPLKSATCITRC | 1527546 | Cell/Virus Enclosed | NO |
| VPLLEVYVHLRNWNFS | 1705182 | Undetermined | UN |
| VPLMHKGAAWPI | 1527555 | Cell/Virus Enclosed | NO |
| VPLNANSTPAQNAGYW | 1705188 | Cell/Virus Enclosed | NO |
| VPLNANSTPSQN | 1527558 | Cell/Virus Enclosed | NO |
| VPLNIVPLTTAA | 1527560 | Cell/Virus Enclosed | NO |
| VPLNSNSTPAQN | 1527562 | Cell/Virus Enclosed | NO |
| VPLQTIINGMHK | 1527566 | Cell/Virus Enclosed | NO |
| VPLRASSCVTRCNIGG | 1705195 | Cell/Virus Enclosed | NO |
| VPLSAIPALAANTLTI | 1705199 | Cell/Virus Enclosed | NO |
| VPLSEAKGYWYR | 1527572 | Cell/Virus Enclosed | NO |
| VPLSEVYVQLRN | 1527574 | Undetermined | UN |
| VPLSVIPATSASKLTI | 1705201 | Cell/Virus Enclosed | NO |
| VPLTFSPGQGVP | 1527580 | Cell/Virus Enclosed | NO |
| VPLTTAAKLMVV | 1527581 | Cell/Virus Enclosed | NO |
| VPLYVRDYVDKWDDFC | 1705206 | Cell/Virus Enclosed | NO |
| VPNAVGGYAISISFWPQTTTTPTSV | 70378 | Undetermined | UN |
| VPNLQGVPESPFSRT | 70384 | Cell/Virus Enclosed | NO |
| VPPALHLVNPQI | 1527601 | Cell/Virus Enclosed | NO |
| VPPIISEKISVM | 1527606 | Cell/Virus Enclosed | NO |
| VPPQVMKLPSEP | 1527609 | Cell/Virus Enclosed | NO |
| VPPQVVELPYEP | 1527612 | Cell/Virus Enclosed | NO |
| VPQLVNSNQYSP | 1527637 | Ectodomain | YES |
| VPQSVETKVQCS | 1527642 | Cell/Virus Enclosed | NO |
| VPRASADIGSGH | 1527649 | Cell/Virus Enclosed | NO |
| VPRASANIGASH | 1527650 | Cell/Virus Enclosed | NO |
| VPRNAELGDRKGKIHIPF | 177356 | Undetermined | UN |
| VPRNAELGDRQGKIHIPF | 177357 | Undetermined | UN |
| VPSAELTGTQSV | 1527665 | Cell/Virus Enclosed | NO |
| VPSDSCNNVNVR | 1527669 | Cell/Virus Enclosed | NO |
| VPSINDLSFDKNLGRC | 1705241 | Cell/Virus Enclosed | NO |
| VPSIVSEKISVM | 1527679 | Cell/Virus Enclosed | NO |
| VPSLPTSGSRVA | 1527681 | Ectodomain | YES |
| VPSPDPNNSPARP | 70481 | Ectodomain | YES |
| VPSTLWEDGDYY | 1527699 | Ectodomain | YES |
| VPSTSGTQVYSR | 1527703 | Cell/Virus Enclosed | NO |
| VPSTVWEDGDYYRKQL | 1705248 | Ectodomain | YES |
| VPSYVKPDSVSTSDLG | 1705250 | Cell/Virus Enclosed | NO |
| VPTAIRADQLTP | 1527716 | Ectodomain | YES |
| VPTIAHGDGRRPSKQRTFI | 94962 | Capsid | UN |
| VPTKYATAKVSP | 1527739 | Ectodomain | YES |
| VPTMADFVQIEN | 1527745 | Ectodomain | YES |
| VPTNVSSTTQPQL | 70496 | Undetermined | UN |
| VPTNVYLVGKTL | 1527749 | Cell/Virus Enclosed | NO |
| VPTREENVMQCF | 1527753 | Cell/Virus Enclosed | NO |
| VPTSIHADQLTP | 1527754 | Ectodomain | YES |
| VPTTLAWVNNQI | 1527758 | Cell/Virus Enclosed | NO |
| VPTTTGALVRFV | 1527760 | Cell/Virus Enclosed | NO |
| VPTTTGILVRFV | 1527762 | Cell/Virus Enclosed | NO |
| VPTYLKSDNIVAADLG | 1705271 | Cell/Virus Enclosed | NO |
| VPVAEYITTYPG | 1527767 | Cell/Virus Enclosed | NO |

| | | | |
|---|---|---|---|
| VPVEDIAQVVIADTLQ | 1705273 | Cell/Virus Enclosed | NO |
| VPVEDIVQVVIV | 1527773 | Cell/Virus Enclosed | NO |
| VPVEFYEDRILD | 1527775 | Cell/Virus Enclosed | NO |
| VPVGDSFGKSLG | 1527778 | Cell/Virus Enclosed | NO |
| VPVGESFGKSLG | 1527779 | Cell/Virus Enclosed | NO |
| VPVGIRDPVSYV | 1527782 | Cell/Virus Enclosed | NO |
| VPVIEVSQIQSKLTDV | 1705276 | Cell/Virus Enclosed | NO |
| VPVKMFDAYVDT | 1527786 | Cell/Virus Enclosed | NO |
| VPVQVVDSRWNNERAS | 1705282 | Underneath Membrane | NO |
| VPVSEYITTYPG | 1527787 | Cell/Virus Enclosed | NO |
| VPVSPAFHCIES | 1527790 | Cell/Virus Enclosed | NO |
| VPVSPAFHRIESTDSI | 1705284 | Cell/Virus Enclosed | NO |
| VPVSPALHRIES | 1527793 | Cell/Virus Enclosed | NO |
| VPVSPVFHRIES | 1527795 | Cell/Virus Enclosed | NO |
| VPVSVIYDKETKTHAT | 1705285 | Ectodomain | YES |
| VPVTRDSVPLHI | 1527802 | Cell/Virus Enclosed | NO |
| VPYCYDTNLLEG | 1527813 | Cell/Virus Enclosed | NO |
| VPYLYNGSAQAT | 1527818 | Ectodomain | YES |
| VPYNMRVIHFGA | 1527820 | Cell/Virus Enclosed | NO |
| VPYPKQSIFTINSVVQ | 1705299 | Cell/Virus Enclosed | NO |
| VPYQRNNTSFPE | 1527822 | Cell/Virus Enclosed | NO |
| VPYTALYYYATV | 1527829 | Cell/Virus Enclosed | NO |
| VPYVYNGSAQST | 1527835 | Ectodomain | YES |
| VPYVYNGSVQAR | 1527836 | Ectodomain | YES |
| VQADTAGLYTNF | 1527841 | Cell/Virus Enclosed | NO |
| VQALESKVLLER | 1527849 | Cell/Virus Enclosed | NO |
| VQAPNYTQHTSSMRGVY | 70527 | Ectodomain | YES |
| VQASESKVLLERFLPK | 1705314 | Cell/Virus Enclosed | NO |
| VQCMTGFNTLLV | 1527863 | Cell/Virus Enclosed | NO |
| VQDADGINKQLTDISV | 1705321 | Cell/Virus Enclosed | NO |
| VQDAKGLKIKYLYFVK | 1705322 | Cell/Virus Enclosed | NO |
| VQDAVNNNAQALSKLA | 1705323 | Ectodomain | YES |
| VQDPAPIDIDAFPVKP | 1705327 | Cell/Virus Enclosed | NO |
| VQEFVSYDFSIH | 1527900 | Cell/Virus Enclosed | NO |
| VQFCDAMRNAGI | 1527918 | Cell/Virus Enclosed | NO |
| VQFNYCRDGQPF | 1527925 | Cell/Virus Enclosed | NO |
| VQGGITQRTRTR | 1527942 | Cell/Virus Enclosed | NO |
| VQGNVAKVANIS | 1527952 | Cell/Virus Enclosed | NO |
| VQGSITQHTRTR | 1527958 | Cell/Virus Enclosed | NO |
| VQGSITQRIRTR | 1527960 | Cell/Virus Enclosed | NO |
| VQGSITQRSKSR | 1527961 | Cell/Virus Enclosed | NO |
| VQGSITQRTRTRPSVQ | 1705355 | Cell/Virus Enclosed | NO |
| VQIGNNNVCTYEHVIS | 1705365 | Cell/Virus Enclosed | NO |
| VQIGSNNVCTYEHVIS | 1705366 | Cell/Virus Enclosed | NO |
| VQIHTIDGSSGVANPA | 1705367 | Secreted | YES |
| VQITYTYKMLVAKDNK | 1705374 | Capsid | UN |
| VQKAKQKPIYLK | 1527998 | Cell/Virus Enclosed | NO |
| VQKESDEYILAKGPLQ | 1705379 | Cell/Virus Enclosed | NO |
| VQKHIGAPANLRVVGH | 1705380 | Cell/Virus Enclosed | NO |
| VQKLQILIDNIN | 1528014 | Ectodomain | YES |
| VQKLQSLIDNIN | 1528015 | Ectodomain | YES |
| VQKLQTLIDNINSTLV | 1705383 | Ectodomain | YES |
| VQKRFRTRKGKR | 1528018 | Cell/Virus Enclosed | NO |
| VQKVEQPKIEPKPVSV | 1705386 | Cell/Virus Enclosed | NO |
| VQLEHYSSCVEC | 1528028 | Cell/Virus Enclosed | NO |
| VQLFEKYFKYWDQTYH | 1705389 | Cell/Virus Enclosed | NO |
| VQLLIQDYIQSV | 1528034 | Cell/Virus Enclosed | NO |
| VQLPVKDYIQTV | 1528041 | Cell/Virus Enclosed | NO |
| VQLPVKDYVQTVNVIA | 1705394 | Cell/Virus Enclosed | NO |
| VQLQLTKLLDDS | 1528043 | Cell/Virus Enclosed | NO |
| VQLRAFDICNTNRAGI | 1705396 | Cell/Virus Enclosed | NO |
| VQLSEINMDNSQ | 1528052 | Cell/Virus Enclosed | NO |
| VQLTDKYCSVNV | 1528058 | Cell/Virus Enclosed | NO |
| VQMGSNNVCTYE | 1528068 | Cell/Virus Enclosed | NO |
| VQMIADYLAGSS | 1528070 | Cell/Virus Enclosed | NO |
| VQMLSDTLKGLS | 1528071 | Cell/Virus Enclosed | NO |
| VQMNIAKYMQLC | 1528075 | Cell/Virus Enclosed | NO |
| VQMTKLASTEEL | 1528076 | Cell/Virus Enclosed | NO |
| VQNDVCSTEDFNVWAM | 1705420 | Cell/Virus Enclosed | NO |
| VQNGVDAKGFPL | 1528082 | Cell/Virus Enclosed | NO |
| VQNGVDAKGFPQLAEL | 1705421 | Capsid | UN |
| VQNLFGNPDEPQ | 1528084 | Capsid | UN |
| VQNLSGNPDEPH | 1528085 | Cell/Virus Enclosed | NO |
| VQNLSGNPDEPQKDVY | 1705422 | Cell/Virus Enclosed | NO |
| VQNTVFNASISE | 1528089 | Cell/Virus Enclosed | NO |
| VQNTVLNASIPE | 1528090 | Cell/Virus Enclosed | NO |

| | | | |
|---|---|---|---|
| VQNTVLNASTPE | 1528091 | Cell/Virus Enclosed | NO |
| VQNVNLERCELANYKQ | 1705423 | Cell/Virus Enclosed | NO |
| VQPRNVLYDSVS | 1528115 | Ectodomain | YES |
| VQPRTIDSTQDG | 1528116 | Ectodomain | YES |
| VQPRTINATQDG | 1528119 | Ectodomain | YES |
| VQPRTINLDNKL | 1528120 | Ectodomain | YES |
| VQPRTINLTQDG | 1528121 | Ectodomain | YES |
| VQPRTINSQDGV | 1528123 | Ectodomain | YES |
| VQPRTINSTQDGDNKL | 1705431 | Ectodomain | YES |
| VQPVGALDVSAS | 1528139 | Cell/Virus Enclosed | NO |
| VQPVGVVDTEWGNMLT | 1705439 | Cell/Virus Enclosed | NO |
| VQQAQQEATHIA | 1528141 | Cell/Virus Enclosed | NO |
| VQQAQQEATLPDIAED | 1705440 | Cell/Virus Enclosed | NO |
| VQQAQQEVTHIA | 1528143 | Cell/Virus Enclosed | NO |
| VQQCFGKRGPNQNFGG | 1705441 | Cell/Virus Enclosed | NO |
| VQQCFGKRGPSQNFGN | 1705442 | Capsid | UN |
| VQQDFNVILPGE | 1528151 | Cell/Virus Enclosed | NO |
| VQQDGRIPNGFS | 1528153 | Ectodomain | YES |
| VQQDLDVILPGE | 1528155 | Cell/Virus Enclosed | NO |
| VQQDLDVMLPGE | 1528156 | Cell/Virus Enclosed | NO |
| VQQGIFNKATVAQIKA | 1705448 | Cell/Virus Enclosed | NO |
| VQQGVFNKATVA | 1528164 | Cell/Virus Enclosed | NO |
| VQQLFYTEDIAS | 1528167 | Cell/Virus Enclosed | NO |
| VQQLFYTEDMASRFAD | 1705450 | Cell/Virus Enclosed | NO |
| VQQNKIRELTAE | 1528172 | Cell/Virus Enclosed | NO |
| VQQWGFTGNLQS | 1528182 | Cell/Virus Enclosed | NO |
| VQQWGYVSNLAT | 1528185 | Cell/Virus Enclosed | NO |
| VQRCTRKLNRLM | 1528189 | Cell/Virus Enclosed | NO |
| VQRPPTLLLLFS | 1528194 | Cell/Virus Enclosed | NO |
| VQSNLIRDIFVH | 1528209 | Cell/Virus Enclosed | NO |
| VQSNVEVLPEGW | 1528212 | Cell/Virus Enclosed | NO |
| VQSSGIIKPNAL | 1528217 | Cell/Virus Enclosed | NO |
| VQSSGMIKPNAI | 1528219 | Cell/Virus Enclosed | NO |
| VQSSGMIKPNAL | 1528220 | Cell/Virus Enclosed | NO |
| VQTFYPRFQASA | 1528233 | Cell/Virus Enclosed | NO |
| VQTFYPRLQASADWKP | 1705489 | Cell/Virus Enclosed | NO |
| VQTIEDLRARMT | 1528239 | Cell/Virus Enclosed | NO |
| VQTLEDLHARMT | 1528241 | Cell/Virus Enclosed | NO |
| VQTVEDLCARMT | 1528247 | Cell/Virus Enclosed | NO |
| VQTVEDLRARMTYVCQC | 1390824 | Cell/Virus Enclosed | NO |
| VQTVEDLRVRMT | 1528248 | Cell/Virus Enclosed | NO |
| VQTVNVIAWLYAAILN | 1705495 | Cell/Virus Enclosed | NO |
| VQTYLKSDNIVA | 1528252 | Cell/Virus Enclosed | NO |
| VQVDNGSSINRKQLEI | 1705498 | Cell/Virus Enclosed | NO |
| VQVDNGSSINRRQLDV | 1705499 | Cell/Virus Enclosed | NO |
| VQVGQLPKYVIV | 1528260 | Ectodomain | YES |
| VQVHNLPEYMAV | 1528264 | Underneath Membrane | NO |
| VQVHNLPEYMTVAVPS | 1705503 | Underneath Membrane | NO |
| VQVKQMYKTPTL | 1528267 | Ectodomain | YES |
| VQVPIGCLDPIR | 1528270 | Cell/Virus Enclosed | NO |
| VQVVDSRWNNLR | 1528283 | Underneath Membrane | NO |
| VQVVIADTLQET | 1528285 | Cell/Virus Enclosed | NO |
| VQYAHKHDIQITNESY | 1705515 | Cell/Virus Enclosed | NO |
| VQYAVGIMTVTMTFKLGPRK | 70677 | Capsid | UN |
| VQYDFTDYKLELFNKY | 1705516 | Cell/Virus Enclosed | NO |
| VQYGTDTISVCP | 1528304 | Ectodomain | YES |
| VQYGTDTNSACP | 1528305 | Ectodomain | YES |
| VQYGTDTNSVCPKLEF | 1705520 | Ectodomain | YES |
| VQYGTDTNSVSP | 1528307 | Ectodomain | YES |
| VQYRINGLGVTMDVLN | 1705521 | Ectodomain | YES |
| VQYRINGLGVTMDVLS | 1705522 | Ectodomain | YES |
| VRAFDVYNKDASFIGK | 1705527 | Cell/Virus Enclosed | NO |
| VRASRQLAQQKVNECV | 1705533 | Ectodomain | YES |
| VRATATIPLQAS | 1528330 | Secreted | YES |
| VRAWVGFDAEGAHATR | 1705536 | Cell/Virus Enclosed | NO |
| VRAYSEAAAQGF | 1528338 | Cell/Virus Enclosed | NO |
| VRCAVTDAKWYC | 1528340 | Cell/Virus Enclosed | NO |
| VRDEVSIETDIIDEVN | 1705544 | Capsid | UN |
| VRDGNTDNLVFVNKTS | 1705545 | Cell/Virus Enclosed | NO |
| VRDPKTSEILDISPCA | 1705552 | Ectodomain | YES |
| VRDPRYVMRLQSAATI | 1705553 | Cell/Virus Enclosed | NO |
| VRDSRFVMRLQTIATI | 1705555 | Cell/Virus Enclosed | NO |
| VRDTNKEVFVGD | 1528374 | Cell/Virus Enclosed | NO |
| VRDYKLANDVKDTLRL | 1705560 | Cell/Virus Enclosed | NO |
| VRDYLLSKMTSLPKDW | 1705561 | Cell/Virus Enclosed | NO |
| VRDYVDKWDDFCRQYS | 1705562 | Cell/Virus Enclosed | NO |

| | | | |
|---|---|---|---|
| VREEVDIIEQPFEEVE | 1705563 | Cell/Virus Enclosed | NO |
| VRELYLSSSNVT | 1528387 | Cell/Virus Enclosed | NO |
| VRESYVNSSNVT | 1528390 | Cell/Virus Enclosed | NO |
| VREYIRYGIKPI | 1528399 | Cell/Virus Enclosed | NO |
| VRFGPFNFCSET | 1528409 | Cell/Virus Enclosed | NO |
| VRFYRALYSNMA | 1528425 | Ectodomain | YES |
| VRGFGDSVEEAL | 1528429 | Cell/Virus Enclosed | NO |
| VRGGDGKMKDLSPRWY | 1390837 | Cell/Virus Enclosed | NO |
| VRGGDGKMKELSPRWY | 1390838 | Capsid | UN |
| VRGRAIIVSVEQLEPC | 1705595 | Cell/Virus Enclosed | NO |
| VRGSIVNARIEPCSSG | 1705596 | Cell/Virus Enclosed | NO |
| VRGSRRLAQQKINECV | 1705597 | Ectodomain | YES |
| VRGSSAARLEPCNGTD | 1705598 | Cell/Virus Enclosed | NO |
| VRGWLGMDVESAHVCG | 1705606 | Cell/Virus Enclosed | NO |
| VRGWVFGSTMDN | 1528446 | Ectodomain | YES |
| VRGWVGFDVEGAHATR | 1705608 | Cell/Virus Enclosed | NO |
| VRHHPELKLFRNLNID | 1705609 | Cell/Virus Enclosed | NO |
| VRIGAAANATGT | 1528465 | Ectodomain | YES |
| VRIGASANSTGT | 1528469 | Ectodomain | YES |
| VRILQDKTGDFM | 1528476 | Cell/Virus Enclosed | NO |
| VRKDGKPVDHYDGFYT | 1705629 | Cell/Virus Enclosed | NO |
| VRKEGQDVIFSQFDSL | 1705631 | Cell/Virus Enclosed | NO |
| VRKLDNQALNSILDNA | 1705635 | Cell/Virus Enclosed | NO |
| VRKNGHFQDHYD | 1528496 | Cell/Virus Enclosed | NO |
| VRKNGQFQDHYDGFYT | 1705637 | Cell/Virus Enclosed | NO |
| VRLLQDKAGDFM | 1528511 | Cell/Virus Enclosed | NO |
| VRLLQDKTGDFT | 1528514 | Cell/Virus Enclosed | NO |
| VRLLQGKTGDFM | 1528515 | Cell/Virus Enclosed | NO |
| VRLQAGKQTEFVSNSH | 1705641 | Cell/Virus Enclosed | NO |
| VRLQAGSNTEFASNSS | 1705643 | Cell/Virus Enclosed | NO |
| VRLQAGTATEYASNSS | 1705644 | Cell/Virus Enclosed | NO |
| VRLSKMGILKAEEFVA | 1705645 | Cell/Virus Enclosed | NO |
| VRMYNLLACLWL | 1528532 | Cell/Virus Enclosed | NO |
| VRNDGTVEFCYLHQIE | 1705652 | Cell/Virus Enclosed | NO |
| VRNLFASVKNSQ | 1528541 | Ectodomain | YES |
| VRNLFASVKSSQ | 1528542 | Ectodomain | YES |
| VRPIIEDYHTLTVTII | 1705660 | Ectodomain | YES |
| VRPRIVQMFADH | 1528560 | Cell/Virus Enclosed | NO |
| VRPRNSSDNGISAIIT | 1705663 | Ectodomain | YES |
| VRPVIEDYHTLTATVI | 1705664 | Underneath Membrane | NO |
| VRQCSGVTFQGK | 1528564 | Cell/Virus Enclosed | NO |
| VRQPARPLGSAWRDQAQRPAVASRR | 70808 | Undetermined | UN |
| VRQQRFVYDAYQNLVG | 1705675 | Ectodomain | YES |
| VRQVAPGETGVI | 1528579 | Ectodomain | YES |
| VRQVRSWIGFDVEGAH | 1705678 | Cell/Virus Enclosed | NO |
| VRRGDNFHNVLT | 1528584 | Cell/Virus Enclosed | NO |
| VRRNLDSLEARA | 1528590 | Cell/Virus Enclosed | NO |
| VRRVNVPYVCLLANKE | 1705688 | Cell/Virus Enclosed | NO |
| VRSESAAISAQL | 1528596 | Ectodomain | YES |
| VRSHPDFKLLHNLQAD | 1705694 | Cell/Virus Enclosed | NO |
| VRSLKLSDNNCYLNAV | 1705698 | Cell/Virus Enclosed | NO |
| VRSSFNFDQKELLAYY | 1705699 | Cell/Virus Enclosed | NO |
| VRSVRGRAIIVSVEQL | 1705701 | Cell/Virus Enclosed | NO |
| VRSWIGFDVEGAHASR | 1705703 | Cell/Virus Enclosed | NO |
| VRTANALSRAVNVPTI | 1705704 | Cell/Virus Enclosed | NO |
| VRTNGGYNSWHLVEGE | 1705707 | Cell/Virus Enclosed | NO |
| VRTYSEAAAQGF | 1528655 | Cell/Virus Enclosed | NO |
| VRVCYGSTVLNG | 1528657 | Cell/Virus Enclosed | NO |
| VRVKHGDFSAVSSPMS | 1705723 | Underneath Membrane | NO |
| VRYGDVDNLVFTNKTT | 1705731 | Cell/Virus Enclosed | NO |
| VRYKCGKNRETIEKLLSKLL | 111921 | Cell/Virus Enclosed | NO |
| VSAAEFKYMVANGLHA | 1705735 | Cell/Virus Enclosed | NO |
| VSAAFHANSSEPALLF | 1705736 | Ectodomain | YES |
| VSAAFHQNASSL | 1528694 | Ectodomain | YES |
| VSAAYHANSSEP | 1528704 | Ectodomain | YES |
| VSAEVSRQVIQDVKDG | 1705740 | Cell/Virus Enclosed | NO |
| VSAGQEQTNCNI | 1528725 | Cell/Virus Enclosed | NO |
| VSALIGANGNKI | 1528730 | Cell/Virus Enclosed | NO |
| VSALKTAGVVTC | 1528731 | Cell/Virus Enclosed | NO |
| VSALKTAGVVTGNVVK | 1705748 | Cell/Virus Enclosed | NO |
| VSALMGANGNKIVDKE | 1705749 | Cell/Virus Enclosed | NO |
| VSALVYENKLKAKNES | 1705754 | Cell/Virus Enclosed | NO |
| VSALVYNNKLLAKKEL | 1705755 | Cell/Virus Enclosed | NO |
| VSAMQTMLFGMI | 1528740 | Cell/Virus Enclosed | NO |
| VSANVCALMSCNGNKI | 1705760 | Cell/Virus Enclosed | NO |
| VSARDPTIQEAI | 1528747 | Cell/Virus Enclosed | NO |

| | | | |
|---|---|---|---|
| VSATKLNDLCFSNVYA | 1705764 | Ectodomain | YES |
| VSATVTANLSIP | 1528755 | Ectodomain | YES |
| VSAVNQSLSNAKVSAE | 1705768 | Cell/Virus Enclosed | NO |
| VSAYGLCDSANP | 1528767 | Ectodomain | YES |
| VSAYGLCDVANP | 1528768 | Ectodomain | YES |
| VSCGHYQTNIYSQNLC | 1705771 | Cell/Virus Enclosed | NO |
| VSCIIGSLGFYN | 1528782 | Ectodomain | YES |
| VSCLYKRNFTYD | 1528788 | Ectodomain | YES |
| VSDAYANLVPYYQLIG | 1705788 | Cell/Virus Enclosed | NO |
| VSDDDFVSAVANAHRY | 1705789 | Cell/Virus Enclosed | NO |
| VSDDEWVAAVDE | 1528802 | Cell/Virus Enclosed | NO |
| VSDNVRALMSCN | 1528813 | Cell/Virus Enclosed | NO |
| VSDREVILCWETGKVK | 1705800 | Cell/Virus Enclosed | NO |
| VSDSLVDAGAVGPQGFVEED | 742160 | Ectodomain | YES |
| VSDTVEVPPHEA | 1528821 | Cell/Virus Enclosed | NO |
| VSDTVEVPPHVV | 1528822 | Cell/Virus Enclosed | NO |
| VSDTVEVPPQVV | 1528824 | Cell/Virus Enclosed | NO |
| VSDVDYVPLKSA | 1528826 | Cell/Virus Enclosed | NO |
| VSDVWHKQLLIN | 1528831 | Cell/Virus Enclosed | NO |
| VSEESQGNVVTSVMES | 1705813 | Cell/Virus Enclosed | NO |
| VSEETGTLIVNSVLL | 1313919 | Undetermined | UN |
| VSEKISVMDKLDTGAQ | 1705818 | Cell/Virus Enclosed | NO |
| VSELVYENKFVPVKEA | 1705824 | Cell/Virus Enclosed | NO |
| VSELVYENKFVPVKPA | 1705825 | Cell/Virus Enclosed | NO |
| VSEMLTFDKKAFTLAN | 1705826 | Ectodomain | YES |
| VSERINVVPGVPTNVY | 1705828 | Cell/Virus Enclosed | NO |
| VSERVNVVPGVP | 1528855 | Cell/Virus Enclosed | NO |
| VSFADNNDITNTNLSR | 1705830 | Cell/Virus Enclosed | NO |
| VSFADNNDTTNT | 1528862 | Cell/Virus Enclosed | NO |
| VSFDDLYVAIRG | 1528868 | Cell/Virus Enclosed | NO |
| VSFDYLIREIET | 1528870 | Cell/Virus Enclosed | NO |
| VSFSPAFHRIES | 1528886 | Cell/Virus Enclosed | NO |
| VSFVGELKAAEAKVIT | 1705846 | Cell/Virus Enclosed | NO |
| VSFVNDSIESVGGLYE | 1705847 | Ectodomain | YES |
| VSFVVDDSGTPV | 1528893 | Cell/Virus Enclosed | NO |
| VSGCETSNVPPI | 1528898 | Cell/Virus Enclosed | NO |
| VSGDAHIPFGTVLGNF | 1705851 | Ectodomain | YES |
| VSGDYLWYVVDD | 1528904 | Cell/Virus Enclosed | NO |
| VSGGLCSTRLHRHALLATCP | 70950 | Cell/Virus Enclosed | NO |
| VSGLVNVQKVAQ | 1528917 | Cell/Virus Enclosed | NO |
| VSGLVNVQKVEQPKIE | 1705857 | Cell/Virus Enclosed | NO |
| VSGMDTALLRNN | 1528919 | Underneath Membrane | NO |
| VSGTATAGWTFGAGAA | 1705864 | Ectodomain | YES |
| VSGYEKSTFNKDIYCG | 1705869 | Cell/Virus Enclosed | NO |
| VSHEVSDDEWAA | 1528955 | Cell/Virus Enclosed | NO |
| VSHVCKCGHNMTLIAA | 1705877 | Cell/Virus Enclosed | NO |
| VSIAPSTVWEDG | 1528982 | Ectodomain | YES |
| VSIPIDLLNLREIKPA | 1705891 | Cell/Virus Enclosed | NO |
| VSIPQGLTFSYD | 1529002 | Cell/Virus Enclosed | NO |
| VSIRTSNQGNKQ | 1529006 | Cell/Virus Enclosed | NO |
| VSISTGSRSARSAIED | 1705893 | Ectodomain | YES |
| VSIVPSTVWEDGDYYR | 1705896 | Ectodomain | YES |
| VSIVPSTVWKDG | 1529018 | Ectodomain | YES |
| VSKADGIIYPQGRTYS | 1705899 | Ectodomain | YES |
| VSKDGFFTYICH | 1529025 | Cell/Virus Enclosed | NO |
| VSKESGNTVVLTFTTR | 1705904 | Capsid | UN |
| VSKGNCSGPTTIRGQ | 71009 | Ectodomain | YES |
| VSKGTQQSSWVY | 1529033 | Ectodomain | YES |
| VSKIGDSIFTGC | 1529038 | Cell/Virus Enclosed | NO |
| VSKTSDWKCKVTDVLF | 1705914 | Cell/Virus Enclosed | NO |
| VSKVVNINVDFKDFQF | 1705921 | Cell/Virus Enclosed | NO |
| VSLAIDAYPLIK | 1529072 | Cell/Virus Enclosed | NO |
| VSLAIDAYPLSKHPNS | 1705925 | Cell/Virus Enclosed | NO |
| VSLAIDAYPLTK | 1529073 | Cell/Virus Enclosed | NO |
| VSLDSLGAAIHYLNSK | 1705932 | Cell/Virus Enclosed | NO |
| VSLEKFMSTPNA | 1529077 | Cell/Virus Enclosed | NO |
| VSLETLLAAIKRLKNG | 1705933 | Cell/Virus Enclosed | NO |
| VSLLRNTSQKSI | 1529092 | Ectodomain | YES |
| VSLTQSEMTIVK | 1529113 | Cell/Virus Enclosed | NO |
| VSLTYLLGTAKKQVVL | 1705952 | Cell/Virus Enclosed | NO |
| VSLVKPSFYVYSRVK | 1313920 | Ectodomain | YES |
| VSLYNVTRDKLE | 1529135 | Cell/Virus Enclosed | NO |
| VSMKGTGVFVQDPAPI | 1705965 | Cell/Virus Enclosed | NO |
| VSNCIKMLSNGAGNGQ | 1705971 | Cell/Virus Enclosed | NO |
| VSNGGNNCTTAVMTYS | 1705975 | Ectodomain | YES |
| VSNGSNNCTTAV | 1529161 | Ectodomain | YES |

| | | | |
|---|---|---|---|
| VSNLSLKQKLLY | 1529170 | Cell/Virus Enclosed | NO |
| VSNNKEDFDIIQKCQI | 1705980 | Cell/Virus Enclosed | NO |
| VSNNQEDFDLISKCQI | 1705981 | Cell/Virus Enclosed | NO |
| VSNSGNLLGFKNVSTG | 1705983 | Ectodomain | YES |
| VSNVKLDRWSAS | 1529181 | Ectodomain | YES |
| VSPALHRIESTD | 1529188 | Cell/Virus Enclosed | NO |
| VSPCIKVAAMQS | 1529190 | Cell/Virus Enclosed | NO |
| VSPFEMPFEELNGLKI | 1705989 | Cell/Virus Enclosed | NO |
| VSPTSKIEPCIV | 1529217 | Cell/Virus Enclosed | NO |
| VSPVFHRIESID | 1529220 | Cell/Virus Enclosed | NO |
| VSPYVCNAPGCDVNDV | 1705996 | Cell/Virus Enclosed | NO |
| VSQDSITHYPAL | 1529224 | Cell/Virus Enclosed | NO |
| VSQIQSKLTDVKCANV | 1706002 | Cell/Virus Enclosed | NO |
| VSQQVDLVISDM | 1529235 | Cell/Virus Enclosed | NO |
| VSRIGDSIFTGC | 1529251 | Cell/Virus Enclosed | NO |
| VSRIGDSVFTGC | 1529252 | Cell/Virus Enclosed | NO |
| VSRINWVTGGIA | 1529253 | Undetermined | UN |
| VSRNSSRSSSQG | 1529259 | Cell/Virus Enclosed | NO |
| VSRQVIQDVKDGTVTF | 1706017 | Cell/Virus Enclosed | NO |
| VSSAFWLFTHIPMAGL | 1706022 | Cell/Virus Enclosed | NO |
| VSSCDLRMGSGF | 1529283 | Ectodomain | YES |
| VSSFEAIPSGSV | 1529293 | Ectodomain | YES |
| VSSFEGYDFKYIENGQ | 1706029 | Cell/Virus Enclosed | NO |
| VSSIKGDKGVEK | 1529315 | Cell/Virus Enclosed | NO |
| VSSNGQPGGDAI | 1529325 | Cell/Virus Enclosed | NO |
| VSSNINCVLSVNSSNC | 1706039 | Cell/Virus Enclosed | NO |
| VSSNINMLLSVP | 1529326 | Cell/Virus Enclosed | NO |
| VSSNINRILGVN | 1529327 | Cell/Virus Enclosed | NO |
| VSSNINRLLSVPSDSC | 1706040 | Cell/Virus Enclosed | NO |
| VSSNINRVLSVP | 1529328 | Cell/Virus Enclosed | NO |
| VSSNMVETLVGQ | 1529329 | Cell/Virus Enclosed | NO |
| VSSNMVETVVGQLEQT | 1706041 | Cell/Virus Enclosed | NO |
| VSSNQSPQGNLGSNGK | 1706042 | Cell/Virus Enclosed | NO |
| VSSPMSNMTENERLLH | 1706045 | Underneath Membrane | NO |
| VSSQIPPNWHIPSIQDVAPH | 102215 | Ectodomain | YES |
| VSSSRRGVYYND | 1529343 | Ectodomain | YES |
| VSSVECYSILAAKTGV | 1706049 | Cell/Virus Enclosed | NO |
| VSSYAKPEEKLSAYDL | 1706051 | Cell/Virus Enclosed | NO |
| VSTDDCFANKHA | 1529359 | Cell/Virus Enclosed | NO |
| VSTDEEGDNVQI | 1529360 | Cell/Virus Enclosed | NO |
| VSTDFLYFHFYQ | 1529361 | Ectodomain | YES |
| VSTDTTVQSKDTNFLN | 1706055 | Cell/Virus Enclosed | NO |
| VSTFNVLQDVLSLRHD | 1706058 | Cell/Virus Enclosed | NO |
| VSTNFGDVNKPV | 1529390 | Cell/Virus Enclosed | NO |
| VSTNLFEGDKFVGTFE | 1706072 | Cell/Virus Enclosed | NO |
| VSTQEFRFMTANNLTA | 1706073 | Cell/Virus Enclosed | NO |
| VSTQPQENTIPH | 1529400 | Cell/Virus Enclosed | NO |
| VSTQPQGNSIPH | 1529403 | Cell/Virus Enclosed | NO |
| VSTSIPNFGSLTQINT | 1706080 | Ectodomain | YES |
| VSTSISNFGSLT | 1529412 | Ectodomain | YES |
| VSTVFVPAMRDS | 1529417 | Cell/Virus Enclosed | NO |
| VSTVPSTVWEDG | 1529424 | Ectodomain | YES |
| VSTVSSHGFPYG | 1529425 | Cell/Virus Enclosed | NO |
| VSTVSSHGMVPV | 1529427 | Cell/Virus Enclosed | NO |
| VSVCQFNMCEYP | 1529432 | Ectodomain | YES |
| VSVCQYNMCDYP | 1529433 | Ectodomain | YES |
| VSVDDKYKTIDE | 1529436 | Cell/Virus Enclosed | NO |
| VSVDRESTSTHD | 1529442 | Cell/Virus Enclosed | NO |
| VSVEDVCFANKFEQFD | 1706093 | Cell/Virus Enclosed | NO |
| VSVEQLEPCAQSRLLS | 1390917 | Cell/Virus Enclosed | NO |
| VSVEQLLASIQHLHEG | 1390918 | Cell/Virus Enclosed | NO |
| VSVGESFGKSLG | 1529447 | Cell/Virus Enclosed | NO |
| VSVIKVAPKPYRVDGK | 1706095 | Cell/Virus Enclosed | NO |
| VSVITPGTNASSEVAVL | 71189 | Ectodomain | YES |
| VSVPVSVIYDKETKTH | 1706105 | Ectodomain | YES |
| VSVPVSVIYDKK | 1529471 | Ectodomain | YES |
| VSVQELVQNTMD | 1529473 | Cell/Virus Enclosed | NO |
| VSVSRFDPSTWN | 1529474 | Ectodomain | YES |
| VSVSRFNPSTWN | 1529476 | Ectodomain | YES |
| VSVTDFSLNGTV | 1529484 | Cell/Virus Enclosed | NO |
| VSVTVFSNMVET | 1529486 | Cell/Virus Enclosed | NO |
| VSVTVSSNMVETVVGQ | 1706110 | Cell/Virus Enclosed | NO |
| VSYDSVSATVTA | 1529514 | Ectodomain | YES |
| VSYFYNGSAQAT | 1529521 | Ectodomain | YES |
| VSYGSSVETETYFKVN | 1706127 | Cell/Virus Enclosed | NO |
| VSYLSERCNVIADSKS | 1706132 | Cell/Virus Enclosed | NO |

| | | | |
|---|---|---|---|
| VSYSSYHATYSLYLFPHWTKKPNR | 71235 | Ectodomain | YES |
| VSYVNTNMGLKFRQL | 79677 | Capsid | UN |
| VSYVYNGSAQAT | 1529539 | Ectodomain | YES |
| VTAFSKPIYVPE | 1529552 | Cell/Virus Enclosed | NO |
| VTAFSKPLYVPE | 1529553 | Cell/Virus Enclosed | NO |
| VTAGPFMDDAIN | 1529555 | Cell/Virus Enclosed | NO |
| VTAGPFMGNAIN | 1529557 | Cell/Virus Enclosed | NO |
| VTAKGYPQFAEL | 1529563 | Cell/Virus Enclosed | NO |
| VTANVCSPMACN | 1529574 | Cell/Virus Enclosed | NO |
| VTAQDGCGHIST | 1529577 | Ectodomain | YES |
| VTASDTTLRCCTVTYL | 1706148 | Cell/Virus Enclosed | NO |
| VTASSKPLYVPEHCQN | 1706150 | Cell/Virus Enclosed | NO |
| VTAVVTNGYLKM | 1529587 | Ectodomain | YES |
| VTCCTASDKFYRLSNE | 1706157 | Cell/Virus Enclosed | NO |
| VTDAKWYCYDKNPINS | 1706165 | Cell/Virus Enclosed | NO |
| VTDAKWYCYDKQPVNS | 1706166 | Cell/Virus Enclosed | NO |
| VTDECRLAMLGAEMTS | 1706167 | Cell/Virus Enclosed | NO |
| VTDFSLNGTVST | 1529608 | Cell/Virus Enclosed | NO |
| VTDIVVVDEVSMLTNY | 1706171 | Cell/Virus Enclosed | NO |
| VTDPSLIISSSP | 1529610 | Cell/Virus Enclosed | NO |
| VTDTIVVTEGDGISTP | 1706174 | Secreted | YES |
| VTDTPKGPKVKY | 1529616 | Cell/Virus Enclosed | NO |
| VTDVKQVGCFMR | 1529621 | Cell/Virus Enclosed | NO |
| VTDVKQVGCSMR | 1529622 | Cell/Virus Enclosed | NO |
| VTDVLFPGQKYSSDCN | 1706179 | Cell/Virus Enclosed | NO |
| VTEDVKFAASTGVIDI | 1706184 | Cell/Virus Enclosed | NO |
| VTEFACVIAETV | 1529629 | Cell/Virus Enclosed | NO |
| VTESVQEEAQLV | 1529634 | Cell/Virus Enclosed | NO |
| VTESVQEKAQPV | 1529637 | Cell/Virus Enclosed | NO |
| VTEYSWNKKLYELVQR | 1706190 | Cell/Virus Enclosed | NO |
| VTEYYEFLNKHFSMMI | 1706191 | Cell/Virus Enclosed | NO |
| VTFCCKCDSTLRLCVQ | 110456 | Cell/Virus Enclosed | NO |
| VTFCDHMVKAGL | 1529641 | Cell/Virus Enclosed | NO |
| VTFEPFNVSFVNDSIE | 1706194 | Ectodomain | YES |
| VTFKEQPTVKEI | 1529656 | Cell/Virus Enclosed | NO |
| VTFLDLQVEMNRLQEA | 1706204 | Ectodomain | YES |
| VTFLNLQVEMNR | 1529664 | Ectodomain | YES |
| VTFSDSNMWYLR | 1529677 | Cell/Virus Enclosed | NO |
| VTFVNISRVKLH | 1529682 | Ectodomain | YES |
| VTGDDVDDIESIYDFD | 1706217 | Cell/Virus Enclosed | NO |
| VTGDNDDEDNKD | 1529685 | Cell/Virus Enclosed | NO |
| VTGDNDDEDVVTGDND | 1706218 | Cell/Virus Enclosed | NO |
| VTGDNDDEDVVTGDNN | 1706219 | Cell/Virus Enclosed | NO |
| VTGDNNDEEIVTGDND | 1706221 | Cell/Virus Enclosed | NO |
| VTGDSCYNFMLT | 1529692 | Cell/Virus Enclosed | NO |
| VTGINDDDYVIALTGT | 1706231 | Cell/Virus Enclosed | NO |
| VTGNVVQQCCTA | 1529718 | Cell/Virus Enclosed | NO |
| VTIADPGYMQGY | 1529737 | Ectodomain | YES |
| VTIALNKIQDVVNQQG | 1706252 | Ectodomain | YES |
| VTICDFYSKLGVGFTS | 1706254 | Cell/Virus Enclosed | NO |
| VTICDLYSIHDV | 1529748 | Cell/Virus Enclosed | NO |
| VTICDLYSIQDV | 1529749 | Cell/Virus Enclosed | NO |
| VTICDLYSMQDV | 1529750 | Cell/Virus Enclosed | NO |
| VTIDCAAFVCGD | 1529753 | Ectodomain | YES |
| VTIDCAAFVRGD | 1529754 | Ectodomain | YES |
| VTIDCVAFVCGD | 1529756 | Ectodomain | YES |
| VTIDKLCDYLAD | 1529757 | Cell/Virus Enclosed | NO |
| VTIDKLRDYLAD | 1529758 | Cell/Virus Enclosed | NO |
| VTILDGIFEQSL | 1529767 | Cell/Virus Enclosed | NO |
| VTINNFNPSSWN | 1529774 | Ectodomain | YES |
| VTINNYNPSFWN | 1529775 | Ectodomain | YES |
| VTISSYKLDLTS | 1529785 | Cell/Virus Enclosed | NO |
| VTKFSGDKFDFIVGYG | 1706277 | Cell/Virus Enclosed | NO |
| VTKHTAKEVRQK | 1529809 | Cell/Virus Enclosed | NO |
| VTKKSAAEASKKPRQKRTATKA | 1870827 | Capsid | UN |
| VTKKYTYSFLPKEDGD | 1706289 | Cell/Virus Enclosed | NO |
| VTKNGLMYSEFP | 1529831 | Ectodomain | YES |
| VTKVLAEKFDYNVSVM | 1706300 | Cell/Virus Enclosed | NO |
| VTLADAGFMKQYGECL | 1706305 | Ectodomain | YES |
| VTLDKCTKYNIYDVSG | 1706313 | Ectodomain | YES |
| VTLDKLRDYLADYDVA | 1390947 | Cell/Virus Enclosed | NO |
| VTLEKLRDYLAD | 1529851 | Cell/Virus Enclosed | NO |
| VTLEQYSTCDICKSTV | 1390951 | Cell/Virus Enclosed | NO |
| VTLEQYSTCDMC | 1529853 | Cell/Virus Enclosed | NO |
| VTLGDFTIMSGRMSLT | 1390952 | Cell/Virus Enclosed | NO |
| VTLKHFFFAQDG | 1529859 | Cell/Virus Enclosed | NO |

| | | | |
|---|---|---|---|
| VTLLDEHEFKFW | 1529871 | Cell/Virus Enclosed | NO |
| VTLLDEHESKFWDESF | 1706326 | Cell/Virus Enclosed | NO |
| VTLLDEQESKFW | 1529873 | Cell/Virus Enclosed | NO |
| VTLRSINHDLFS | 1529891 | Cell/Virus Enclosed | NO |
| VTLSDAGFMKQY | 1529892 | Ectodomain | YES |
| VTLSSNLNTNLHFDVD | 1706333 | Ectodomain | YES |
| VTLSTKLKDGVNFNVD | 1706334 | Ectodomain | YES |
| VTMDVLNKNQKLIATA | 1706339 | Ectodomain | YES |
| VTMDVLSQNQKLIANA | 1706340 | Ectodomain | YES |
| VTMTDLQSESSC | 1529911 | Cell/Virus Enclosed | NO |
| VTNAETTDFFTV | 1529914 | Ectodomain | YES |
| VTNEIGLNASVTLKIC | 1706346 | Ectodomain | YES |
| VTNLNKSAGWPLNKFG | 1706351 | Cell/Virus Enclosed | NO |
| VTPCNQPDQVAVYQQS | 1706357 | Ectodomain | YES |
| VTPDMADEIASL | 1529934 | Cell/Virus Enclosed | NO |
| VTPDMADHIASL | 1529935 | Cell/Virus Enclosed | NO |
| VTPDMADQIASL | 1529936 | Cell/Virus Enclosed | NO |
| VTPEANMDQGSF | 1529938 | Cell/Virus Enclosed | NO |
| VTPGGTSSGDAT | 1529940 | Cell/Virus Enclosed | NO |
| VTPIKNTVDTTPKEEF | 1706361 | Cell/Virus Enclosed | NO |
| VTPLPAADMDDF | 1529945 | Capsid | UN |
| VTPLTPRQYLLA | 1529950 | Ectodomain | YES |
| VTPVDQYMCGVD | 1529967 | Cell/Virus Enclosed | NO |
| VTQEYIQTTIQKVTVD | 1706374 | Ectodomain | YES |
| VTQNVLYDNQKQ | 1529988 | Ectodomain | YES |
| VTQNVLYENQKQIANQ | 1706384 | Ectodomain | YES |
| VTQRMCAIGPDVFLHK | 1706386 | Cell/Virus Enclosed | NO |
| VTRDIVSTDDCF | 1530007 | Cell/Virus Enclosed | NO |
| VTRDKLEKLISTARDG | 1706395 | Cell/Virus Enclosed | NO |
| VTREKLEKLIST | 1530012 | Cell/Virus Enclosed | NO |
| VTRGHRWDVVRP | 1530017 | Cell/Virus Enclosed | NO |
| VTRPLMESELVI | 1530024 | Underneath Membrane | NO |
| VTSAGEPCVASE | 1530028 | Cell/Virus Enclosed | NO |
| VTSAGQPYVASE | 1530031 | Cell/Virus Enclosed | NO |
| VTSAMQTMLFTM | 1530032 | Cell/Virus Enclosed | NO |
| VTSDKFDFIIGH | 1530036 | Cell/Virus Enclosed | NO |
| VTSDSCNNFMLT | 1530038 | Cell/Virus Enclosed | NO |
| VTSGNLFDVGYS | 1530043 | Cell/Virus Enclosed | NO |
| VTSNVPPIVSEK | 1530054 | Cell/Virus Enclosed | NO |
| VTSVMESQISTKEVKL | 1706422 | Cell/Virus Enclosed | NO |
| VTSVNFFDVGYS | 1530075 | Cell/Virus Enclosed | NO |
| VTTEVKVSALKT | 1530093 | Cell/Virus Enclosed | NO |
| VTTGDVKIMAKA | 1530099 | Cell/Virus Enclosed | NO |
| VTTKAKSVQQDLDVIL | 1706440 | Cell/Virus Enclosed | NO |
| VTTLNGRLVNYT | 1530115 | Ectodomain | YES |
| VTTMDFCEGSFL | 1530120 | Cell/Virus Enclosed | NO |
| VTTMDFCEGSFP | 1530121 | Cell/Virus Enclosed | NO |
| VTTNSDKCPIVVGVSE | 1706451 | Cell/Virus Enclosed | NO |
| VTTSMAPDFVAF | 1530130 | Cell/Virus Enclosed | NO |
| VTTSTEPDFVAF | 1530134 | Cell/Virus Enclosed | NO |
| VTVADEPDSRAP | 1530140 | Cell/Virus Enclosed | NO |
| VTVAEEPDSRAP | 1530141 | Cell/Virus Enclosed | NO |
| VTVAEEPDSRVP | 1530142 | Cell/Virus Enclosed | NO |
| VTVDCKQYICNG | 1530147 | Ectodomain | YES |
| VTVDGVNFRTVVLNNK | 1706465 | Cell/Virus Enclosed | NO |
| VTVDTTVESAAQ | 1530154 | Cell/Virus Enclosed | NO |
| VTVDVLKDMPVLKTIN | 1390978 | Cell/Virus Enclosed | NO |
| VTVELEPPCRFVIDTP | 1390979 | Cell/Virus Enclosed | NO |
| VTVKETVVQFNYRRDG | 1706468 | Cell/Virus Enclosed | NO |
| VTVLRDNLKPVTTLST | 1706471 | Cell/Virus Enclosed | NO |
| VTVPHNLTTITK | 1530176 | Ectodomain | YES |
| VTVQGPPGTGKS | 1530180 | Cell/Virus Enclosed | NO |
| VTVSSNMVETVVGQLE | 1706478 | Cell/Virus Enclosed | NO |
| VTYAGNVWQIQTIQDS | 1706488 | Cell/Virus Enclosed | NO |
| VTYAGSVWHIQTVQDA | 1706489 | Cell/Virus Enclosed | NO |
| VTYGNMTLNGLWLDDK | 1706494 | Cell/Virus Enclosed | NO |
| VTYGSMTLNGLWLDDK | 1706495 | Cell/Virus Enclosed | NO |
| VTYKSTATTKLVPGML | 1706499 | Cell/Virus Enclosed | NO |
| VTYLNELSSKVVCTYM | 1706502 | Cell/Virus Enclosed | NO |
| VTYPFIANEKFINGCG | 1706505 | Cell/Virus Enclosed | NO |
| VTYQNISINLPP | 1530212 | Ectodomain | YES |
| VTYQNISTNLPPPLLG | 1706508 | Ectodomain | YES |
| VTYTDQYLCGADGKPV | 1706511 | Cell/Virus Enclosed | NO |
| VTYVPSQEKNFT | 1530223 | Ectodomain | YES |
| VTYVPSQERNFTTAPAI | 71589 | Ectodomain | YES |
| VVANGVKAKGYPQFAE | 1706538 | Capsid | UN |

| | | | |
|---|---|---|---|
| VVANGVTAKGYP | 1530258 | Cell/Virus Enclosed | NO |
| VVAQCSGVTFQL | 1530259 | Cell/Virus Enclosed | NO |
| VVARTGQFYING | 1530263 | Ectodomain | YES |
| VVAVIDQDFGSTVFNV | 1706543 | Cell/Virus Enclosed | NO |
| VVAWHVVRDSRFVMRL | 1706546 | Cell/Virus Enclosed | NO |
| VVCASVFKDGCD | 1530275 | Cell/Virus Enclosed | NO |
| VVCATLINNISY | 1530277 | Cell/Virus Enclosed | NO |
| VVCDAIENALNSCKEH | 1706550 | Cell/Virus Enclosed | NO |
| VVCDKVRSLKLSDNNC | 1706551 | Cell/Virus Enclosed | NO |
| VVCNNSDYASKG | 1530295 | Cell/Virus Enclosed | NO |
| VVCYNSDYASKGYIAN | 1706567 | Cell/Virus Enclosed | NO |
| VVDCATYVCNGN | 1530306 | Ectodomain | YES |
| VVDDHVGLLDLA | 1530307 | Cell/Virus Enclosed | NO |
| VVDDHVGLLDQAWRVP | 1706574 | Cell/Virus Enclosed | NO |
| VVDDIYYPASCNGVLP | 1706575 | Cell/Virus Enclosed | NO |
| VVDEVSMCTNYDLSII | 1706577 | Cell/Virus Enclosed | NO |
| VVDEVSMCTNYDLSVI | 1706578 | Cell/Virus Enclosed | NO |
| VVDEVSMLTNYELSVI | 1706579 | Cell/Virus Enclosed | NO |
| VVDGFVRSFQPL | 1530322 | Ectodomain | YES |
| VVDGKQIDDYRI | 1530324 | Cell/Virus Enclosed | NO |
| VVDNDVVDYVSDADFS | 1706591 | Cell/Virus Enclosed | NO |
| VVDQVEEVNSIFDIET | 1706594 | Cell/Virus Enclosed | NO |
| VVDSRWNNARHS | 1530337 | Ectodomain | YES |
| VVDSRWNNARLS | 1530338 | Ectodomain | YES |
| VVDSRWNNLRAS | 1530342 | Underneath Membrane | NO |
| VVDSVAVKNGAL | 1530343 | Cell/Virus Enclosed | NO |
| VVDVMRYNLNFSANSL | 1706602 | Ectodomain | YES |
| VVDVNDSHSMAVVDGK | 1706603 | Cell/Virus Enclosed | NO |
| VVDYVSDADFSVTGDC | 1706608 | Cell/Virus Enclosed | NO |
| VVEDTINDDEDVVTGD | 1706614 | Cell/Virus Enclosed | NO |
| VVEEPDSRAPSRSQSRSQSR | 2060909 | Capsid | UN |
| VVEFEDRSNNSSRASS | 1706616 | Capsid | UN |
| VVELPKFFYASNGTYN | 1706617 | Ectodomain | YES |
| VVENDADKANFLNAAV | 1706619 | Cell/Virus Enclosed | NO |
| VVEQAEGFECDF | 1530393 | Ectodomain | YES |
| VVEQYNQTILNLTSEI | 1706625 | Ectodomain | YES |
| VVESGGYIPSNF | 1530401 | Ectodomain | YES |
| VVEVVTTSLTPCGYSE | 1706635 | Cell/Virus Enclosed | NO |
| VVEYLSKEDLHTLYVD | 1706636 | Cell/Virus Enclosed | NO |
| VVFDTVEVPPQV | 1530416 | Cell/Virus Enclosed | NO |
| VVFGKRGGGSVT | 1530425 | Cell/Virus Enclosed | NO |
| VVFVDKYMCGFDGKPV | 1706652 | Cell/Virus Enclosed | NO |
| VVGAMLSENFTSYGFS | 1706659 | Ectodomain | YES |
| VVGGCVASNVPS | 1530466 | Cell/Virus Enclosed | NO |
| VVGGCVTSNVPP | 1530467 | Cell/Virus Enclosed | NO |
| VVGGCVTSNVSP | 1530469 | Cell/Virus Enclosed | NO |
| VVGGTIQIFASV | 1530472 | Cell/Virus Enclosed | NO |
| VVGIVYSSQLYEDVPS | 1706672 | Cell/Virus Enclosed | NO |
| VVGKDGVMYSEF | 1530476 | Ectodomain | YES |
| VVGNVILKPSDE | 1530479 | Cell/Virus Enclosed | NO |
| VVGPDARAKQDVSLLS | 1706674 | Cell/Virus Enclosed | NO |
| VVGPDARTQGKQSYVL | 1706675 | Cell/Virus Enclosed | NO |
| VVGPKELLLSWESGKV | 1706676 | Cell/Virus Enclosed | NO |
| VVGPRTGKHEHS | 1530492 | Cell/Virus Enclosed | NO |
| VVGTKALAVRLT | 1530500 | Cell/Virus Enclosed | NO |
| VVGTLSSTIRLQAGVA | 1706688 | Cell/Virus Enclosed | NO |
| VVGVSERINVVPGVPT | 1706691 | Cell/Virus Enclosed | NO |
| VVIADTLQEIPV | 1530521 | Cell/Virus Enclosed | NO |
| VVIADTLQETPVVPDT | 1706700 | Cell/Virus Enclosed | NO |
| VVIADTLQETSV | 1530522 | Cell/Virus Enclosed | NO |
| VVIFINNTTYPTNVAV | 1706703 | Cell/Virus Enclosed | NO |
| VVIGDDVDDIDP | 1530529 | Cell/Virus Enclosed | NO |
| VVIKNLTPIKLNFGML | 1706713 | Cell/Virus Enclosed | NO |
| VVIMTLDLLKDI | 1530544 | Cell/Virus Enclosed | NO |
| VVITNVGEPIISQPSK | 1706726 | Cell/Virus Enclosed | NO |
| VVKALNESYIDLKELG | 1706728 | Ectodomain | YES |
| VVKAMLFKKEPL | 1530563 | Cell/Virus Enclosed | NO |
| VVKAMLLKKEPL | 1530564 | Cell/Virus Enclosed | NO |
| VVKEQVVDLLVK | 1530571 | Cell/Virus Enclosed | NO |
| VVKFPIDLTMIE | 1530572 | Cell/Virus Enclosed | NO |
| VVKHIINITGVS | 1530576 | Cell/Virus Enclosed | NO |
| VVKLPSAPQTIQPEVK | 1706742 | Cell/Virus Enclosed | NO |
| VVKLQFEPQTIQ | 1530583 | Cell/Virus Enclosed | NO |
| VVKLQNNEIMPG | 1530584 | Cell/Virus Enclosed | NO |
| VVKNSTAFPFETLSAC | 1706746 | Cell/Virus Enclosed | NO |
| VVKPGHFNEEFYNFLR | 1706749 | Cell/Virus Enclosed | NO |

| | | | |
|---|---|---|---|
| VVKPVRARAPPGEQFT | 1706751 | Cell/Virus Enclosed | NO |
| VVKQGHFIGVEGELPV | 1706753 | Cell/Virus Enclosed | NO |
| VVKQGHFTGVEG | 1530605 | Cell/Virus Enclosed | NO |
| VVKSALLADKFP | 1530621 | Cell/Virus Enclosed | NO |
| VVKTDAVVLLERYVSL | 1706767 | Cell/Virus Enclosed | NO |
| VVKVPIDLTMIE | 1530630 | Cell/Virus Enclosed | NO |
| VVLAEFDTYDPI | 1530635 | Cell/Virus Enclosed | NO |
| VVLASDDLYVKRYFKG | 1706776 | Cell/Virus Enclosed | NO |
| VVLATDDLYVKRYERG | 1706777 | Cell/Virus Enclosed | NO |
| VVLELDPPCKFS | 1530640 | Cell/Virus Enclosed | NO |
| VVLLERYVSLAIDAYP | 1706792 | Cell/Virus Enclosed | NO |
| VVLNNKNIYRSQ | 1530666 | Cell/Virus Enclosed | NO |
| VVLNNKNTYISQ | 1530668 | Cell/Virus Enclosed | NO |
| VVLNNKNTYRSQ | 1530669 | Cell/Virus Enclosed | NO |
| VVLSDALDKLYKVFGS | 1706804 | Cell/Virus Enclosed | NO |
| VVLSYELLNAPA | 1530690 | Ectodomain | YES |
| VVLVSNNQEDFDLISK | 1706813 | Cell/Virus Enclosed | NO |
| VVMEIPSLMTVK | 1530698 | Cell/Virus Enclosed | NO |
| VVMSMFSRTTSN | 1530708 | Ectodomain | YES |
| VVNAANENLLHGGGVA | 1706824 | Cell/Virus Enclosed | NO |
| VVNANAEALNNLLQQL | 1706828 | Ectodomain | YES |
| VVNAYNSTAIFV | 1530722 | Ectodomain | YES |
| VVNDKIFTKSGT | 1530725 | Cell/Virus Enclosed | NO |
| VVNDKIFTKSGVNDIC | 1706834 | Cell/Virus Enclosed | NO |
| VVNDKVFVRYGDVDNL | 1706835 | Cell/Virus Enclosed | NO |
| VVNDNAQALNTL | 1530729 | Ectodomain | YES |
| VVNGKYFTFKQANNNC | 1706839 | Cell/Virus Enclosed | NO |
| VVNGVDFDFIVNAANE | 1706841 | Cell/Virus Enclosed | NO |
| VVNINVDFKDFQFMLW | 1706843 | Cell/Virus Enclosed | NO |
| VVNLNVDFKDFQ | 1530754 | Cell/Virus Enclosed | NO |
| VVNNLDKSAGHPFNKF | 1706852 | Cell/Virus Enclosed | NO |
| VVNPANGHMAHGGGVA | 1706855 | Cell/Virus Enclosed | NO |
| VVNQQGNSLNHLTSQL | 1706857 | Ectodomain | YES |
| VVNQQGSALNHLTSQL | 1706858 | Ectodomain | YES |
| VVNSGSDINCNI | 1530771 | Cell/Virus Enclosed | NO |
| VVNSNAQALNSLLQQL | 1706861 | Ectodomain | YES |
| VVNTNPRNYSYM | 1530774 | Ectodomain | YES |
| VVNVNVDFKDFQFMLW | 1706865 | Cell/Virus Enclosed | NO |
| VVNVVKAVRNKT | 1530779 | Cell/Virus Enclosed | NO |
| VVNYTVCDDCNGYTDN | 1706869 | Ectodomain | YES |
| VVPDHDSFVKMMVDGF | 1706870 | Cell/Virus Enclosed | NO |
| VVPDIVEVPPQV | 1530785 | Cell/Virus Enclosed | NO |
| VVPDTVEVPPQVVKLP | 1706871 | Cell/Virus Enclosed | NO |
| VVPDTVGVPPQV | 1530787 | Cell/Virus Enclosed | NO |
| VVPGNVKPGDAVITTQ | 1706878 | Cell/Virus Enclosed | NO |
| VVPGSCNFPLEA | 1530794 | Ectodomain | YES |
| VVPGVADAERMA | 1530795 | Ectodomain | YES |
| VVPLENDADKAN | 1530797 | Cell/Virus Enclosed | NO |
| VVPLSVIPATSASKLT | 1706882 | Cell/Virus Enclosed | NO |
| VVPSINDLSFDKNLGR | 1706886 | Cell/Virus Enclosed | NO |
| VVQFNYCRDGQP | 1530816 | Cell/Virus Enclosed | NO |
| VVQKDGVEDTPP | 1530821 | Cell/Virus Enclosed | NO |
| VVQLPVKDYIQT | 1530829 | Cell/Virus Enclosed | NO |
| VVQLPVKDYVQTVNVI | 1706897 | Cell/Virus Enclosed | NO |
| VVQNTVLNASTP | 1530836 | Cell/Virus Enclosed | NO |
| VVQPRTIDSTQD | 1530840 | Ectodomain | YES |
| VVQPRTIISTQD | 1530841 | Ectodomain | YES |
| VVQPRTINSTQDGDNK | 1706902 | Ectodomain | YES |
| VVQPVGVVDTEWGNML | 1706903 | Cell/Virus Enclosed | NO |
| VVQTCDLTVGSG | 1530851 | Ectodomain | YES |
| VVQYEADRRAFVDYTG | 1706909 | Cell/Virus Enclosed | NO |
| VVRDISVNDLCFANKF | 1706914 | Cell/Virus Enclosed | NO |
| VVRDPRYVMRLQSAAT | 1706916 | Cell/Virus Enclosed | NO |
| VVRDSRFVMRLQTIAT | 1706917 | Cell/Virus Enclosed | NO |
| VVRIGAAANSTGTVII | 1706924 | Ectodomain | YES |
| VVRKDLEEGDLP | 1530884 | Cell/Virus Enclosed | NO |
| VVRNKTSACFNF | 1530889 | Cell/Virus Enclosed | NO |
| VVRQCSGVTFQG | 1530891 | Cell/Virus Enclosed | NO |
| VVRTANALSRAVNVPT | 1706937 | Cell/Virus Enclosed | NO |
| VVSAYGLCDASN | 1530909 | Ectodomain | YES |
| VVSAYGLCDLAN | 1530911 | Ectodomain | YES |
| VVSAYGLCDSAN | 1530912 | Ectodomain | YES |
| VVSDTVDVPPQV | 1530919 | Cell/Virus Enclosed | NO |
| VVSDTVEVPPHE | 1530920 | Cell/Virus Enclosed | NO |
| VVSDTVEVPPQV | 1530921 | Cell/Virus Enclosed | NO |
| VVSFNEWALANQFTEF | 1706950 | Cell/Virus Enclosed | NO |

| | | | |
|---|---|---|---|
| VVSGCETSNVPP | 1530932 | Cell/Virus Enclosed | NO |
| VVSHICKCGESMVLID | 1706951 | Cell/Virus Enclosed | NO |
| VVSIGYHYKELGVVMN | 1706955 | Cell/Virus Enclosed | NO |
| VVSIRSSNRGNK | 1530941 | Cell/Virus Enclosed | NO |
| VVSIRTSNQGNK | 1530942 | Cell/Virus Enclosed | NO |
| VVSKIGDSIFTG | 1530944 | Cell/Virus Enclosed | NO |
| VVSMLLLCCCST | 1530956 | Undetermined | UN |
| VVSTIGDSIFTG | 1530969 | Cell/Virus Enclosed | NO |
| VVSTVKIGGLTFKTYA | 1706970 | Cell/Virus Enclosed | NO |
| VVTDIKQVGCMM | 1530984 | Cell/Virus Enclosed | NO |
| VVTGDDVDDIESIYDF | 1706978 | Cell/Virus Enclosed | NO |
| VVTGDNDDEDNK | 1530988 | Cell/Virus Enclosed | NO |
| VVTGDNDDEDVVTGDN | 1706979 | Cell/Virus Enclosed | NO |
| VVTGDNDQIVVT | 1530989 | Cell/Virus Enclosed | NO |
| VVTGDNNDEEIVTGDN | 1706980 | Cell/Virus Enclosed | NO |
| VVTGNVVKQCCI | 1530993 | Cell/Virus Enclosed | NO |
| VVTGNVVKQCCTAAVD | 1706981 | Cell/Virus Enclosed | NO |
| VVTGNVVQQCCT | 1530995 | Cell/Virus Enclosed | NO |
| VVTQRMCAIGPDVFLH | 1706988 | Cell/Virus Enclosed | NO |
| VVTSVMESQISTKEVK | 1706992 | Cell/Virus Enclosed | NO |
| VVTTNSDKCPIVVGVS | 1706999 | Cell/Virus Enclosed | NO |
| VVTYPSLNYAGALWDI | 1707003 | Cell/Virus Enclosed | NO |
| VVVDEVSMCTNYDLSV | 1707006 | Cell/Virus Enclosed | NO |
| VVVDEVSMLTNYELSV | 1707007 | Cell/Virus Enclosed | NO |
| VVVGGCVASNVP | 1531037 | Cell/Virus Enclosed | NO |
| VVVGGCVTSNVP | 1531038 | Cell/Virus Enclosed | NO |
| VVVGGCVTSNVS | 1531039 | Cell/Virus Enclosed | NO |
| VVVMSTCAVNYTKAPY | 1707023 | Ectodomain | YES |
| VVVNGKYFTFKQANNN | 1707026 | Cell/Virus Enclosed | NO |
| VVVNGPFFSFEQSHNN | 1707027 | Cell/Virus Enclosed | NO |
| VVVNNLDKSAGHPFNK | 1707028 | Cell/Virus Enclosed | NO |
| VVVNSQDVYKSLTIVD | 1707030 | Cell/Virus Enclosed | NO |
| VVVPDHDSFVKMMVDG | 1707031 | Cell/Virus Enclosed | NO |
| VVVQPRTINYGV | 1531073 | Ectodomain | YES |
| VVVSGCETSNVP | 1531075 | Cell/Virus Enclosed | NO |
| VVVTAGPFMDNA | 1531079 | Cell/Virus Enclosed | NO |
| VVVVENDADKANFLNA | 1707039 | Cell/Virus Enclosed | NO |
| VVVVESDADRANFLNA | 1707040 | Cell/Virus Enclosed | NO |
| VVVVPDHDSFVKMMVD | 1707041 | Cell/Virus Enclosed | NO |
| VVWTLNDVKDNDGRPV | 1707046 | Cell/Virus Enclosed | NO |
| VVWTLQEVKDNDGKNV | 1707047 | Cell/Virus Enclosed | NO |
| VVWVAKEGAKTVNTSL | 1707048 | Capsid | UN |
| VVWVAVDGAKTEPTGY | 1707050 | Capsid | UN |
| VVYHTVGSPIQF | 1531113 | Cell/Virus Enclosed | NO |
| VVYQQAVVGAMLSENF | 1707068 | Ectodomain | YES |
| VVYSNGGFYFKPGGTT | 1707073 | Cell/Virus Enclosed | NO |
| VVYVNATGSDVN | 1531135 | Ectodomain | YES |
| VWEAGKSKPPLNRNYV | 1707084 | Cell/Virus Enclosed | NO |
| VWEEGDYYRKQL | 1531147 | Ectodomain | YES |
| VWGVKDFNTLSQEGKK | 1707090 | Cell/Virus Enclosed | NO |
| VWHIQTVQDADGINKQ | 1707093 | Cell/Virus Enclosed | NO |
| VWHVERKDVPYPKQSI | 1707095 | Cell/Virus Enclosed | NO |
| VWLLISFSFIIF | 1531183 | Transmembrane | NO |
| VWNGGAYSLFDMAKFP | 1707112 | Cell/Virus Enclosed | NO |
| VWPFYPYGRCPT | 1531195 | Ectodomain | YES |
| VWQIQTIQDSDGTNKQ | 1707120 | Cell/Virus Enclosed | NO |
| VWRVIALQSVDC | 1531210 | Cell/Virus Enclosed | NO |
| VWSYNAELLVAMENQHTIDLADSEM | 1179659 | Ectodomain | YES |
| VWTLNDVKDNDGRPVH | 1707126 | Cell/Virus Enclosed | NO |
| VWTLQEVKDNDGKNVH | 1707127 | Cell/Virus Enclosed | NO |
| VWVAKEGAKTVNTSLG | 1707128 | Capsid | UN |
| VWVANELSRLVKSPTV | 1707129 | Cell/Virus Enclosed | NO |
| VWVAVDGAKTEPTGYG | 1707131 | Capsid | UN |
| VWVHEDGATDAPSTFG | 1707132 | Cell/Virus Enclosed | NO |
| VWVHEHGATDAP | 1531223 | Cell/Virus Enclosed | NO |
| VWVHEYGATDAP | 1531224 | Cell/Virus Enclosed | NO |
| VYADDSIILRQYNLVD | 1707138 | Cell/Virus Enclosed | NO |
| VYAGCEQSYFTK | 1531240 | Cell/Virus Enclosed | NO |
| VYAIRNFDEQNC | 1531243 | Cell/Virus Enclosed | NO |
| VYALRHFDQNSEVLKA | 1707148 | Cell/Virus Enclosed | NO |
| VYAMRNFDEQNCDVLK | 1707149 | Cell/Virus Enclosed | NO |
| VYANDSIILRQY | 1531252 | Cell/Virus Enclosed | NO |
| VYANLGERVRQA | 1531257 | Cell/Virus Enclosed | NO |
| VYANLGERVRQS | 1531258 | Cell/Virus Enclosed | NO |
| VYANVGRWSASINTGN | 1707154 | Ectodomain | YES |
| VYAQHCFKAPKN | 1531263 | Ectodomain | YES |

| | | | |
|---|---|---|---|
| VYAQHCFKAPTN | 1531265 | Ectodomain | YES |
| VYAQHCFKVPKN | 1531267 | Ectodomain | YES |
| VYAVLSDVDGLKYTKI | 1707158 | Cell/Virus Enclosed | NO |
| VYCPRHVIASNTTSAI | 1707168 | Cell/Virus Enclosed | NO |
| VYCPRHVICSSSNMNE | 1707170 | Cell/Virus Enclosed | NO |
| VYCYNTDLIEGSKPYS | 1707174 | Cell/Virus Enclosed | NO |
| VYDAANNAVYDGARLF | 1707175 | Cell/Virus Enclosed | NO |
| VYDAYHNLVGYY | 1531309 | Ectodomain | YES |
| VYDAYQNLVGYYSDDG | 1707176 | Ectodomain | YES |
| VYDCFCNNINWN | 1531312 | Cell/Virus Enclosed | NO |
| VYDDIFAESKPR | 1531316 | Cell/Virus Enclosed | NO |
| VYDDVDASLFVDINNL | 1707182 | Cell/Virus Enclosed | NO |
| VYDFCVSPAEFKYMVA | 1707183 | Cell/Virus Enclosed | NO |
| VYDFKVEDAYDQVHDE | 1707184 | Cell/Virus Enclosed | NO |
| VYDFKVSAAEFKYMVA | 1707185 | Cell/Virus Enclosed | NO |
| VYDGARLFSSDL | 1531323 | Cell/Virus Enclosed | NO |
| VYDIACCMRAGSKVVN | 1707189 | Cell/Virus Enclosed | NO |
| VYDIGNPKGIPIVDDP | 1707190 | Cell/Virus Enclosed | NO |
| VYDMWLTGCKYV | 1531338 | Cell/Virus Enclosed | NO |
| VYDQVVDNVYVTYAGN | 1707202 | Cell/Virus Enclosed | NO |
| VYDSSEVKAQSSGVIP | 1707203 | Cell/Virus Enclosed | NO |
| VYDTAKKSMYDGDRFV | 1707204 | Cell/Virus Enclosed | NO |
| VYDTNHGDAGFT | 1531354 | Ectodomain | YES |
| VYDVEPTHKVKL | 1531355 | Cell/Virus Enclosed | NO |
| VYDVHKDNFKDGLCMF | 1707206 | Cell/Virus Enclosed | NO |
| VYEADTEQTQNVTVKP | 1707208 | Cell/Virus Enclosed | NO |
| VYEADTEQTQSV | 1531362 | Cell/Virus Enclosed | NO |
| VYELNGYTVQPV | 1531368 | Ectodomain | YES |
| VYENKFVPVKEASKQC | 1707216 | Cell/Virus Enclosed | NO |
| VYENKFVPVKPASKQC | 1707217 | Cell/Virus Enclosed | NO |
| VYERDKAVARKLERMA | 1707221 | Cell/Virus Enclosed | NO |
| VYEVGDLIAQSS | 1531378 | Cell/Virus Enclosed | NO |
| VYEVGDLMAQSS | 1531379 | Cell/Virus Enclosed | NO |
| VYFAVNDQDLYD | 1531384 | Cell/Virus Enclosed | NO |
| VYFDNPTLPFGD | 1531385 | Ectodomain | YES |
| VYFIKMIILWLM | 1531389 | Transmembrane | NO |
| VYFNATGSDVNCNGYQ | 1707232 | Ectodomain | YES |
| VYFNGTGRGDCQ | 1531397 | Ectodomain | YES |
| VYFSDSKPSTTE | 1531401 | Cell/Virus Enclosed | NO |
| VYGDFSHGQLGG | 1531413 | Cell/Virus Enclosed | NO |
| VYGFSNVVYPDL | 1531416 | Underneath Membrane | NO |
| VYGNFDDQPSLQVESA | 1707247 | Cell/Virus Enclosed | NO |
| VYHKLGERIRQA | 1531428 | Cell/Virus Enclosed | NO |
| VYHRYRIGNYKL | 1531436 | Underneath Membrane | NO |
| VYHTVGSPIQFH | 1531441 | Cell/Virus Enclosed | NO |
| VYIGDPVQLPAP | 1531445 | Cell/Virus Enclosed | NO |
| VYIWTADEAIKFLKEW | 1707265 | Undetermined | UN |
| VYKLDISEATQVEQH | 227687 | Secreted | YES |
| VYKSDIHLPAYASFKV | 1707282 | Cell/Virus Enclosed | NO |
| VYLEDKFDLLISDMYD | 1707289 | Cell/Virus Enclosed | NO |
| VYLEYIKKLYND | 1531482 | Cell/Virus Enclosed | NO |
| VYLNHSVPKLSDFESE | 1707299 | Ectodomain | YES |
| VYMAKAGDKYYPVVVD | 1707308 | Cell/Virus Enclosed | NO |
| VYMAKVGDKYYP | 1531514 | Cell/Virus Enclosed | NO |
| VYNDALFVSYGSSVET | 1707315 | Cell/Virus Enclosed | NO |
| VYNGSAQAISFC | 1531532 | Ectodomain | YES |
| VYNGSAQSTALC | 1531535 | Ectodomain | YES |
| VYNKDASFIGKNLKSN | 1707319 | Cell/Virus Enclosed | NO |
| VYNLVKTGHYTGQAGE | 1707321 | Cell/Virus Enclosed | NO |
| VYNLVNVGHYDG | 1531542 | Cell/Virus Enclosed | NO |
| VYNNSGSNITQIKNAC | 1707324 | Cell/Virus Enclosed | NO |
| VYPDYVDVNKTL | 1531557 | Ectodomain | YES |
| VYQNAYNCTYDR | 1531577 | Ectodomain | YES |
| VYQQAVVGAMLSENFT | 1707343 | Ectodomain | YES |
| VYQQSIIGAMTAVNES | 1707346 | Ectodomain | YES |
| VYRAFDIYNERV | 1531587 | Cell/Virus Enclosed | NO |
| VYRINWITGGIAIAM | 2001384 | Undetermined | UN |
| VYRINWVTGGIA | 1531592 | Undetermined | UN |
| VYRRKPDPPNCN | 1531601 | Underneath Membrane | NO |
| VYRRKPNLPNCNIEAW | 1707356 | Ectodomain | YES |
| VYRSTSPDPKFVDKYY | 1707359 | Cell/Virus Enclosed | NO |
| VYRVNTAIPTIE | 1531614 | Cell/Virus Enclosed | NO |
| VYSAGHATGITP | 1531617 | Ectodomain | YES |
| VYSAGHATGSTP | 1531618 | Ectodomain | YES |
| VYSAGHATGTTPQKLF | 1707365 | Ectodomain | YES |
| VYSAGYATGTTP | 1531623 | Ectodomain | YES |

| | | | |
|---|---|---|---|
| VYSNGGFYFKPGGTTS | 1707377 | Cell/Virus Enclosed | NO |
| VYSRKARNRSHS | 1531652 | Cell/Virus Enclosed | NO |
| VYSSGHATGTTP | 1531657 | Ectodomain | YES |
| VYSSNEEQAVLKFLDG | 1707384 | Cell/Virus Enclosed | NO |
| VYSVLETFQNNVVNYQ | 1707389 | Cell/Virus Enclosed | NO |
| VYSVSSFEAKPSGSVV | 1707391 | Ectodomain | YES |
| VYTDTEVCKVKDFVSG | 1707395 | Cell/Virus Enclosed | NO |
| VYTFTTVKPGAA | 1531674 | Cell/Virus Enclosed | NO |
| VYTNSCLVSSDGQPGG | 1707402 | Cell/Virus Enclosed | NO |
| VYTNSCLVSSNG | 1531683 | Cell/Virus Enclosed | NO |
| VYTWTADEAIKF | 1531695 | Ectodomain | YES |
| VYVDGLESKQVDYVPL | 1707409 | Cell/Virus Enclosed | NO |
| VYVDTFMSHFDVDRKS | 1707411 | Cell/Virus Enclosed | NO |
| VYVGDPAQLPAPRTLL | 1707413 | Cell/Virus Enclosed | NO |
| VYVGDPAQLPTP | 1531705 | Cell/Virus Enclosed | NO |
| VYVIKMIILWLM | 1531711 | Transmembrane | NO |
| VYVKFQDSKPPLPPDE | 1707420 | Cell/Virus Enclosed | NO |
| VYVKSKVGNYRLPSTQ | 1707421 | Ectodomain | YES |
| VYVRKDGKPVDHYDGF | 1707424 | Cell/Virus Enclosed | NO |
| VYWMRRHTQKAPKRIRLPHI | 742182 | Undetermined | UN |
| VYYGNSLDQAIS | 1531737 | Cell/Virus Enclosed | NO |
| VYYKGVTTHESSSAVN | 1707436 | Cell/Virus Enclosed | NO |
| VYYNSWQNLLYDSNGN | 1707437 | Ectodomain | YES |
| VYYPDEIFRSDTLYLTQD | 72205 | Ectodomain | YES |
| VYYVARLMKGDKGDAE | 1707451 | Cell/Virus Enclosed | NO |
| WAAAVDEAFPLDEAED | 1707453 | Cell/Virus Enclosed | NO |
| WAAAVDEAFPLN | 1531759 | Cell/Virus Enclosed | NO |
| WAAFYVYKLQPL | 1531766 | Ectodomain | YES |
| WADASEPQRGRQGRIPYSLY | 2060912 | Capsid | UN |
| WADESEPQRGRQ | 1531774 | Cell/Virus Enclosed | NO |
| WADQSDQFRNFQ | 1531780 | Cell/Virus Enclosed | NO |
| WADQSDQFRSVQ | 1531782 | Cell/Virus Enclosed | NO |
| WADQSDQVRNVQ | 1531783 | Cell/Virus Enclosed | NO |
| WADQSERNYQTFNRGR | 1707461 | Capsid | UN |
| WAFGAGAALQIP | 1531786 | Ectodomain | YES |
| WAFYVRAKHGDFSGVA | 1707467 | Ectodomain | YES |
| WAGSKINAVIYN | 1531798 | Cell/Virus Enclosed | NO |
| WAIFNDHHLNRPGVYC | 1707474 | Cell/Virus Enclosed | NO |
| WAPEFPWMFEDAEEKL | 1707485 | Cell/Virus Enclosed | NO |
| WARFPKSDGTGT | 1531834 | Cell/Virus Enclosed | NO |
| WARVEGKDGFVS | 1531835 | Cell/Virus Enclosed | NO |
| WAWNPEVNAITV | 1531844 | Underneath Membrane | NO |
| WAYCVDLHNKINLCDD | 1707492 | Cell/Virus Enclosed | NO |
| WCKDGKVQTFYP | 1531854 | Cell/Virus Enclosed | NO |
| WCKDGQVQTFYPRLQA | 1707503 | Cell/Virus Enclosed | NO |
| WCKDNAVATFYPQLQS | 1707504 | Cell/Virus Enclosed | NO |
| WCPRHVICPADQ | 1531870 | Cell/Virus Enclosed | NO |
| WCPRHVMCPADQ | 1531871 | Cell/Virus Enclosed | NO |
| WCRCSCLPDPIS | 1531873 | Ectodomain | YES |
| WCTYLFSLIAVL | 1531880 | Cell/Virus Enclosed | NO |
| WCVVLIVILLVC | 1531888 | Cell/Virus Enclosed | NO |
| WDDFCRQYSNESWFED | 1707526 | Cell/Virus Enclosed | NO |
| WDDMLRHLIKDVDNPV | 1707527 | Cell/Virus Enclosed | NO |
| WDDMLRRLIKDVDNPV | 1707528 | Cell/Virus Enclosed | NO |
| WDEGIIKSTIVEAAK | 72295 | Cell/Virus Enclosed | NO |
| WDFMLKTLYKDVDNPH | 1707531 | Cell/Virus Enclosed | NO |
| WDIEYPYISHEKKLNS | 1707533 | Cell/Virus Enclosed | NO |
| WDITVINNVDNEIVKS | 1707534 | Cell/Virus Enclosed | NO |
| WDQKLWCRHFCVLAD | 72324 | Ectodomain | YES |
| WDVIRKDLSYEQQNVL | 1707555 | Cell/Virus Enclosed | NO |
| WDYHVTKVLAEKFDYN | 1707561 | Cell/Virus Enclosed | NO |
| WDYPKCDRALPNMIRM | 1707564 | Cell/Virus Enclosed | NO |
| WDYPKCDRAMPNMCRI | 1707567 | Cell/Virus Enclosed | NO |
| WDYPKFDRAMPN | 1531953 | Cell/Virus Enclosed | NO |
| WEAGKSKPPLNRNYVF | 1707571 | Cell/Virus Enclosed | NO |
| WEDGDYYRKQLSPLEG | 1707573 | Ectodomain | YES |
| WEEGDYYRKQLS | 1531967 | Ectodomain | YES |
| WEFRDYFDDNTD | 1531971 | Cell/Virus Enclosed | NO |
| WEFRDYFDNNTD | 1531972 | Cell/Virus Enclosed | NO |
| WEFRDYFNDNTD | 1531973 | Cell/Virus Enclosed | NO |
| WEGLHEVLTSAM | 1531975 | Cell/Virus Enclosed | NO |
| WERKKISNCVVD | 1531991 | Ectodomain | YES |
| WERKTFSNCNFN | 1531993 | Ectodomain | YES |
| WESGKAKPPLNRNSVF | 1707588 | Cell/Virus Enclosed | NO |
| WESGKVKPPLNRNSVF | 1707589 | Cell/Virus Enclosed | NO |
| WFAMRKDGDDVIFSRT | 1707600 | Cell/Virus Enclosed | NO |

| | | | |
|---|---|---|---|
| WFDFIENPSVIG | 1532015 | Cell/Virus Enclosed | NO |
| WFDPIENEDIHRVYAA | 1707606 | Cell/Virus Enclosed | NO |
| WFKLSYHYENNS | 1532048 | Cell/Virus Enclosed | NO |
| WFKNHTSMAPNL | 1532049 | Ectodomain | YES |
| WFKNQTSVALDL | 1532052 | Ectodomain | YES |
| WFKNQTSVAPDF | 1532053 | Ectodomain | YES |
| WFKNQTSVAPDLSLDY | 1707622 | Ectodomain | YES |
| WFKNQTSVAPLL | 1532056 | Ectodomain | YES |
| WFKNQTSVAPVL | 1532057 | Ectodomain | YES |
| WFKNQTSVARDL | 1532058 | Ectodomain | YES |
| WFLLTNTSSVVDGVVR | 1707634 | Ectodomain | YES |
| WFMDGENAYEVVKAML | 1707640 | Cell/Virus Enclosed | NO |
| WFNTMLDASAPATGWL | 1707646 | Cell/Virus Enclosed | NO |
| WFYTYSIGEASS | 1532127 | Cell/Virus Enclosed | NO |
| WGLNWDLGLSQWAREALQTGITL | 72481 | Undetermined | UN |
| WGMKYHPNTVDCDNDR | 1707682 | Cell/Virus Enclosed | NO |
| WGNMLTGIAAHP | 1532138 | Cell/Virus Enclosed | NO |
| WGTSAAAYFVGYLKPTT | 72519 | Ectodomain | YES |
| WGVKDFNTLSQEGKKY | 1707687 | Cell/Virus Enclosed | NO |
| WHEGRTIYGNVSRQDL | 1707696 | Cell/Virus Enclosed | NO |
| WHINQTTNKTTY | 1532163 | Cell/Virus Enclosed | NO |
| WHIQTVQDADGINKQL | 1707702 | Cell/Virus Enclosed | NO |
| WHVERKDVPYPKQSIF | 1707714 | Cell/Virus Enclosed | NO |
| WHVLYTPNISIPQQTSSRTILFP | 72558 | Ectodomain | YES |
| WHVVRDPRYVMRLQSA | 1707718 | Cell/Virus Enclosed | NO |
| WHYFDAQPLTRK | 1532203 | Cell/Virus Enclosed | NO |
| WHYTSYTIVGALYVT | 1327764 | Ectodomain | YES |
| WICLLQFAYANRNRF | 1310941 | Undetermined | UN |
| WIELFPHNDRIKSFST | 1707726 | Cell/Virus Enclosed | NO |
| WIFDSSKEFVEY | 1532220 | Cell/Virus Enclosed | NO |
| WIGFDVEGAHAF | 1532230 | Cell/Virus Enclosed | NO |
| WIGFDVEGAHASRNAC | 1707729 | Cell/Virus Enclosed | NO |
| WILQEVKDNDGK | 1532245 | Cell/Virus Enclosed | NO |
| WINPVILTPVFC | 1532253 | Cell/Virus Enclosed | NO |
| WIPAGIGVTGVIIAV | 72638 | Secreted | YES |
| WIQFMMSRRRLL | 1532258 | Cell/Virus Enclosed | NO |
| WKCKSKNPLLYE | 1532280 | Secreted | YES |
| WKMVVCDKVRSF | 1532313 | Cell/Virus Enclosed | NO |
| WKMVVCDKVRSL | 1532314 | Cell/Virus Enclosed | NO |
| WKRQPNDDATSN | 1532320 | Cell/Virus Enclosed | NO |
| WKRQPNDDVTSNVTQC | 1707784 | Capsid | UN |
| WKRVPTREENVI | 1532323 | Cell/Virus Enclosed | NO |
| WLAPTNAYMRTTHTDI | 1707794 | Cell/Virus Enclosed | NO |
| WLGDTVTCPRHVIAPS | 1707824 | Cell/Virus Enclosed | NO |
| WLKGDKLFVEHY | 1532412 | Cell/Virus Enclosed | NO |
| WLKNQTSVAPDL | 1532415 | Ectodomain | YES |
| WLKPFGGQPVSE | 1532416 | Cell/Virus Enclosed | NO |
| WLLHKDGGIPDG | 1532426 | Cell/Virus Enclosed | NO |
| WLLHKDGGVPDG | 1532427 | Cell/Virus Enclosed | NO |
| WLLWPVTLACFVLAA | 1543360 | Transmembrane | NO |
| WLNDKAVPSPLN | 1532469 | Ectodomain | YES |
| WLNDKLVPSPLN | 1532470 | Ectodomain | YES |
| WLNDKSVPSPLNWERK | 1707869 | Ectodomain | YES |
| WLNNFNVPSPLNWERKI | 1532477 | Ectodomain | YES |
| WLPELFMFSTLH | 1532485 | Cell/Virus Enclosed | NO |
| WLPHDAIVVDNDVVDY | 1707877 | Cell/Virus Enclosed | NO |
| WLPPDAIIIDNDINDY | 1707878 | Cell/Virus Enclosed | NO |
| WLPTDAIIIDNDLNEF | 1707880 | Cell/Virus Enclosed | NO |
| WLPTDAIVIDND | 1532494 | Cell/Virus Enclosed | NO |
| WLSVIWMMWYWGPSL | 956994 | Undetermined | UN |
| WLTGNAIQGFAC | 1532525 | Cell/Virus Enclosed | NO |
| WLTIPPMKNLALGVI | 72808 | Ectodomain | YES |
| WLTYHGAIKLDDKDPQF | 72816 | Capsid | UN |
| WLTYTGAIKLDDKDPN | 1707901 | Cell/Virus Enclosed | NO |
| WLTYTGAIKLDDKDPS | 1707902 | Cell/Virus Enclosed | NO |
| WLYTYSIGEASSPIAY | 1707915 | Cell/Virus Enclosed | NO |
| WMDDFEADPKGKYAQN | 1707916 | Cell/Virus Enclosed | NO |
| WMFEDAEEKLDNPSSS | 1707918 | Cell/Virus Enclosed | NO |
| WMGIRFEYTSPTDKLA | 1707920 | Cell/Virus Enclosed | NO |
| WNCNVDMYPEFS | 1532608 | Cell/Virus Enclosed | NO |
| WNCNVDRYPANA | 1532609 | Cell/Virus Enclosed | NO |
| WNCQLSFAGYTESAVN | 1707944 | Cell/Virus Enclosed | NO |
| WNFSWNLILTLF | 1532625 | Undetermined | UN |
| WNGGAYSLFDMAKFPL | 1707952 | Cell/Virus Enclosed | NO |
| WNGGAYSLFDMTKFSL | 1707953 | Cell/Virus Enclosed | NO |
| WNGVSDLSLKQK | 1532632 | Cell/Virus Enclosed | NO |

| | | | |
|---|---|---|---|
| WNKDVNTHSTRLTITE | 1707961 | Cell/Virus Enclosed | NO |
| WNKKFGFIEDSV | 1532650 | Ectodomain | YES |
| WNKRFGFIEDSA | 1532652 | Ectodomain | YES |
| WNKRFGFIEDSVFKPR | 1707966 | Ectodomain | YES |
| WNKVTQIANMFLEQTQ | 1707967 | Cell/Virus Enclosed | NO |
| WNKYLYELIQKF | 1532659 | Cell/Virus Enclosed | NO |
| WNKYLYELIQRF | 1532660 | Cell/Virus Enclosed | NO |
| WNLDYPIISNEV | 1532666 | Cell/Virus Enclosed | NO |
| WNLEYPIISNEVSINT | 1707971 | Cell/Virus Enclosed | NO |
| WNNARLSDNMTA | 1532681 | Ectodomain | YES |
| WNNLRASDTSLS | 1532686 | Underneath Membrane | NO |
| WNNMLRTLIDGV | 1532687 | Cell/Virus Enclosed | NO |
| WNTAKQDQGQYY | 1532701 | Ectodomain | YES |
| WNTAQQDQGQYY | 1532705 | Ectodomain | YES |
| WNTRNIDATSTGNYNY | 1708000 | Ectodomain | YES |
| WNVEYPIISNELSINT | 1708001 | Cell/Virus Enclosed | NO |
| WNYGKPVTLPTG | 1532727 | Cell/Virus Enclosed | NO |
| WNYLFETIPTQL | 1532729 | Cell/Virus Enclosed | NO |
| WPFYPYGRCPTA | 1532733 | Ectodomain | YES |
| WPISNDSNCCVV | 1532737 | Cell/Virus Enclosed | NO |
| WPISNDSNGCVVEAST | 1708011 | Cell/Virus Enclosed | NO |
| WPLFESDKEEDL | 1532743 | Cell/Virus Enclosed | NO |
| WPLFESDKEEDV | 1532744 | Cell/Virus Enclosed | NO |
| WPLNDYGFYTTS | 1532752 | Ectodomain | YES |
| WPLSVQQAQQEATLPD | 1708018 | Cell/Virus Enclosed | NO |
| WPLTIILTIFNC | 1532763 | Undetermined | UN |
| WPLYPYGRCPTAADIN | 1708027 | Ectodomain | YES |
| WPQIAELVPTAS | 1532780 | Cell/Virus Enclosed | NO |
| WPQIAQFAPSASAFFGMSRI | 1392517 | Capsid | UN |
| WPRPIDVFKADG | 1532784 | Ectodomain | YES |
| WPRPIDVSKADGIIYP | 1708031 | Ectodomain | YES |
| WPTATGDVVLAK | 1532790 | Cell/Virus Enclosed | NO |
| WPVATGDVVLASDDLY | 1708034 | Cell/Virus Enclosed | NO |
| WPVTLACFVLAAVYR | 2001388 | Transmembrane | NO |
| WQFVDHFGENEEIIIN | 1708049 | Cell/Virus Enclosed | NO |
| WQIQTIQDSDGTNKQL | 1708051 | Cell/Virus Enclosed | NO |
| WQNLLYDFNGNI | 1532835 | Ectodomain | YES |
| WQNLLYDSNGNIIGFK | 1708056 | Ectodomain | YES |
| WRFPCAGRKVNFNEKP | 1708070 | Cell/Virus Enclosed | NO |
| WRIYSTGNNVFQTQAGC | 73036 | Ectodomain | YES |
| WRLINKFDVINGVKTV | 1708076 | Cell/Virus Enclosed | NO |
| WRMRRGQRVDLPPKVH | 1708077 | Capsid | UN |
| WRRQDRKINTGNGIKQ | 1708087 | Cell/Virus Enclosed | NO |
| WRVTAVHSDGIF | 1532902 | Cell/Virus Enclosed | NO |
| WRYDFDITESKYSCKE | 1708097 | Cell/Virus Enclosed | NO |
| WRYNFDITESKYSCKE | 1708098 | Cell/Virus Enclosed | NO |
| WSADSCLQGDKCNIFA | 1708100 | Ectodomain | YES |
| WSADSCLQGNKC | 1532916 | Ectodomain | YES |
| WSASINTGNCPFSFGK | 1708103 | Ectodomain | YES |
| WSDIRDYKLANDAKES | 1708105 | Cell/Virus Enclosed | NO |
| WSDQSDQFRNVQ | 1532924 | Cell/Virus Enclosed | NO |
| WSEGNSITGVPYPVSG | 1708107 | Ectodomain | YES |
| WSFDYHPNCSDCYDDM | 1708108 | Cell/Virus Enclosed | NO |
| WSFNPETNNLMCIDMK | 1708112 | Ectodomain | YES |
| WSGISDLSLKQK | 1532946 | Cell/Virus Enclosed | NO |
| WSGLCVDGRNGY | 1532948 | Ectodomain | YES |
| WSGVSDLSLKQK | 1532952 | Cell/Virus Enclosed | NO |
| WSITYPMIANENAINK | 1708118 | Cell/Virus Enclosed | NO |
| WSMPYHPNTVDCQDDR | 1708127 | Cell/Virus Enclosed | NO |
| WSNFTKVQGLEN | 1532973 | Cell/Virus Enclosed | NO |
| WSPQGNLGGNCAGNVI | 1708131 | Cell/Virus Enclosed | NO |
| WSVDAFNQFSSD | 1532996 | Cell/Virus Enclosed | NO |
| WSYTGSSFYAPEPITS | 1708154 | Ectodomain | YES |
| WTAAEGVPFYLN | 1533020 | Ectodomain | YES |
| WTAAVDEAFPLD | 1533021 | Cell/Virus Enclosed | NO |
| WTADEAIKFLKEWNFS | 1391194 | Undetermined | UN |
| WTDVRDYKLANDVKDT | 1708160 | Cell/Virus Enclosed | NO |
| WTESGDYTLANTTTEP | 1708162 | Cell/Virus Enclosed | NO |
| WTHCFDPQIQAIVSSPCHNSLIL | 73133 | Ectodomain | YES |
| WTHCYQPRLQAITTDNCNNSIIL | 73134 | Ectodomain | YES |
| WTLNDVKDNDGRPVHV | 1708174 | Cell/Virus Enclosed | NO |
| WTLQEVKDNDGKNVHL | 1708175 | Cell/Virus Enclosed | NO |
| WTLVEPGDKITFEATGNLVVPRYAF | 1179982 | Ectodomain | YES |
| WTMFCTSVNTSSSEAF | 1708176 | Cell/Virus Enclosed | NO |
| WTNDALNTICIKDADY | 1708180 | Cell/Virus Enclosed | NO |
| WTTVLSMAVAKV | 1533062 | Cell/Virus Enclosed | NO |

| | | | |
|---|---|---|---|
| WTVFCTNANASSSEGF | 1708186 | Cell/Virus Enclosed | NO |
| WTVFCTNVNASSSEGF | 1708187 | Cell/Virus Enclosed | NO |
| WTVTYPFIANEKFING | 1708188 | Cell/Virus Enclosed | NO |
| WTWETKGDLAEN | 73171 | Ectodomain | YES |
| WVAAVDEAFPLD | 1533065 | Cell/Virus Enclosed | NO |
| WVAKEGAKTVNTSLGN | 1708190 | Capsid | UN |
| WVANHQADTSTPSDVS | 1708192 | Capsid | UN |
| WVASNQADVNIP | 1533072 | Cell/Virus Enclosed | NO |
| WVASNQADVNTPADIV | 1708193 | Cell/Virus Enclosed | NO |
| WVATEGALNTPKDHIG | 1708194 | Cell/Virus Enclosed | NO |
| WVAVDGAKTEPTGYGV | 1708195 | Capsid | UN |
| WVDQFERSHQTY | 1533076 | Cell/Virus Enclosed | NO |
| WVEEDLTKGPHEFCSQ | 1708197 | Cell/Virus Enclosed | NO |
| WVENDITNGPHEFCSQ | 1708200 | Cell/Virus Enclosed | NO |
| WVFYVRVKHGDFSAVS | 1708206 | Underneath Membrane | NO |
| WVGFDAEGAHATRDSI | 1708207 | Cell/Virus Enclosed | NO |
| WVGFDVEGAHATRENI | 1708208 | Cell/Virus Enclosed | NO |
| WVGVNLEDPASRDLVVSYVN | 97104 | Capsid | UN |
| WVHEHGATDAPS | 1533100 | Cell/Virus Enclosed | NO |
| WVHEYGATDAPS | 1533101 | Cell/Virus Enclosed | NO |
| WVKHLNKNLNEGVLES | 1708212 | Cell/Virus Enclosed | NO |
| WVKHLNKTLNEGVLES | 1708213 | Cell/Virus Enclosed | NO |
| WVNATCIILQYL | 1533134 | Cell/Virus Enclosed | NO |
| WVNKASYDTNLNKFNR | 1708229 | Cell/Virus Enclosed | NO |
| WVSGNMMDGFSCPLCC | 1708239 | Cell/Virus Enclosed | NO |
| WWLKGDKLSVEH | 1533184 | Cell/Virus Enclosed | NO |
| WWLLAKFTKLLL | 1533186 | Cell/Virus Enclosed | NO |
| WWTVFCTNANASSSEG | 1708260 | Cell/Virus Enclosed | NO |
| WYCYDKNPINSNVKTL | 1708263 | Cell/Virus Enclosed | NO |
| WYDFVENPDIINVYKK | 1708271 | Cell/Virus Enclosed | NO |
| WYDPVENEDIHRVYAS | 1708273 | Cell/Virus Enclosed | NO |
| WYEFRAGRPHRL | 1533209 | Cell/Virus Enclosed | NO |
| WYESTFGLSYYS | 1533211 | Cell/Virus Enclosed | NO |
| WYGYHHQNEQGSGYAADLKSTQNAI | 1180001 | Ectodomain | YES |
| WYHEHIGGVYDN | 1533222 | Cell/Virus Enclosed | NO |
| WYHEHVGGVYDS | 1533224 | Cell/Virus Enclosed | NO |
| WYKGSNLEEEILLMLA | 1708294 | Cell/Virus Enclosed | NO |
| WYKPPFLSDFNN | 1533243 | Ectodomain | YES |
| WYQKPFLSDFNN | 1533254 | Ectodomain | YES |
| WYRHSRRSFKTADGQQ | 1708300 | Capsid | UN |
| WYSGAIKLDPKN | 1533257 | Capsid | UN |
| WYVNDGRVDDDY | 1533268 | Cell/Virus Enclosed | NO |
| WYVNDGRVDDGYICGD | 1708306 | Cell/Virus Enclosed | NO |
| WYVWLLISLAGV | 1533278 | Transmembrane | NO |
| WYWGPSLYNILSPFL | 1708311 | Undetermined | UN |
| XRAPRITFGGPSDSTD | 1708315 | Cell/Virus Enclosed | NO |
| YAADLKSTQNAIDEITNKVNSVIEK | 1180011 | Ectodomain | YES |
| YAADPAMHIASSNAFL | 1708325 | Cell/Virus Enclosed | NO |
| YAAILNNCAWFVQNDV | 1708331 | Cell/Virus Enclosed | NO |
| YAAILNNCNWFVQSDK | 1708332 | Cell/Virus Enclosed | NO |
| YAAKGYIAGIQNFKET | 1708334 | Cell/Virus Enclosed | NO |
| YAAQNRRGLDLLFWEQGGL | 73327 | Ectodomain | YES |
| YAASYNRYKYYSGNAN | 1708341 | Cell/Virus Enclosed | NO |
| YADKPNHFINFPLTHF | 1708355 | Cell/Virus Enclosed | NO |
| YADKTLQEMKALFPTW | 1708356 | Cell/Virus Enclosed | NO |
| YADKTLQEMKTL | 1533320 | Cell/Virus Enclosed | NO |
| YAFATLKEVVGPKELL | 1708372 | Cell/Virus Enclosed | NO |
| YAGNVWQIQTIQDSDG | 1708388 | Cell/Virus Enclosed | NO |
| YAGSRSSSGNRSGILK | 1708390 | Capsid | UN |
| YAGSVWHIQTVQDADG | 1708391 | Cell/Virus Enclosed | NO |
| YAGVVWTLNDVKDNDG | 1708392 | Cell/Virus Enclosed | NO |
| YAHKHDIQITNESYNN | 1708394 | Cell/Virus Enclosed | NO |
| YAHKHDIQLTNE | 1533365 | Cell/Virus Enclosed | NO |
| YAIQEGFDATNSALVK | 1708403 | Ectodomain | YES |
| YAIQQGFDATNS | 1533384 | Ectodomain | YES |
| YAIRNFDEQNCD | 1533388 | Cell/Virus Enclosed | NO |
| YAISGKERARTVGGVS | 1708406 | Cell/Virus Enclosed | NO |
| YAISNEQHYKSW | 1533392 | Ectodomain | YES |
| YAKDQYGTDIDG | 1533400 | Cell/Virus Enclosed | NO |
| YAKPEEKLSAYDLACC | 1708413 | Cell/Virus Enclosed | NO |
| YAKRAVRSHPDFKLLH | 1708416 | Cell/Virus Enclosed | NO |
| YAKSNFVKDDYCDGSL | 1708417 | Cell/Virus Enclosed | NO |
| YALLDWVKHLNKTLNE | 1708420 | Cell/Virus Enclosed | NO |
| YALLHIAGCQTTNGLN | 1708421 | Ectodomain | YES |
| YALRHFDQNSEVLKAI | 1708427 | Cell/Virus Enclosed | NO |
| YANIVHEPTTPI | 1533461 | Cell/Virus Enclosed | NO |

| | | | |
|---|---|---|---|
| YANVGRWSASINTGNC | 1708450 | Ectodomain | YES |
| YAPEPITSFNTK | 1533483 | Ectodomain | YES |
| YAPEPITSLNTKYVAP | 1708453 | Ectodomain | YES |
| YAPEPITYLNTK | 1533487 | Ectodomain | YES |
| YAPTYVSVDDKYKTSD | 1708454 | Cell/Virus Enclosed | NO |
| YAQHCFKAPTNF | 1533496 | Ectodomain | YES |
| YAQHCYKAPKNF | 1533499 | Ectodomain | YES |
| YATGTTPQKLFV | 1533530 | Ectodomain | YES |
| YATLYFNNEPAF | 1533537 | Cell/Virus Enclosed | NO |
| YAYATLKEIVGPKELL | 1708501 | Cell/Virus Enclosed | NO |
| YAYDQVHDEPKG | 1533556 | Cell/Virus Enclosed | NO |
| YAYFNGAIIRDSDVVK | 1708503 | Cell/Virus Enclosed | NO |
| YAYFTDAGVVTK | 1533564 | Ectodomain | YES |
| YAYMMPVMGMTN | 1533571 | Cell/Virus Enclosed | NO |
| YCADVAAEELMNALV | 73483 | Ectodomain | YES |
| YCAFTAPEDDDFLEES | 1708512 | Cell/Virus Enclosed | NO |
| YCDGSLGCKMCLFGYQ | 1708525 | Cell/Virus Enclosed | NO |
| YCDNGKYYTKPIIKAQ | 1708526 | Cell/Virus Enclosed | NO |
| YCDSPFEKLQCEHLQF | 1708527 | Ectodomain | YES |
| YCFNAEGDASWS | 1533612 | Cell/Virus Enclosed | NO |
| YCFSVNNTFCPCAKPS | 1708535 | Ectodomain | YES |
| YCFVNTTIGNETTSAF | 1708536 | Ectodomain | YES |
| YCFVNTTIGTET | 1533621 | Ectodomain | YES |
| YCGNNSCTCQPQ | 1533631 | Ectodomain | YES |
| YCGNSATCYNSVSNEY | 1708538 | Cell/Virus Enclosed | NO |
| YCHDPTVLPGAF | 1533639 | Cell/Virus Enclosed | NO |
| YCILEPRPGNHC | 1533643 | Ectodomain | YES |
| YCILEPRSGNHCPAGN | 1708543 | Ectodomain | YES |
| YCILEPRSGNYC | 1533645 | Ectodomain | YES |
| YCIPYNSITDTI | 1533646 | Secreted | YES |
| YCPAGNSYTSFA | 1533675 | Ectodomain | YES |
| YCPRHVIASNTTSAID | 1708563 | Cell/Virus Enclosed | NO |
| YCPRHVICSSSNMNEP | 1708565 | Cell/Virus Enclosed | NO |
| YCQVHGNAHVAS | 1533687 | Cell/Virus Enclosed | NO |
| YCRKIGVNVCSDSTFE | 1708582 | Cell/Virus Enclosed | NO |
| YCRVGECVESNAGVCF | 1708587 | Cell/Virus Enclosed | NO |
| YCSNSSSGALDT | 1533718 | Ectodomain | YES |
| YCSNSSSGVLDTTIPF | 1708593 | Ectodomain | YES |
| YCTFAIEDVEDVISSE | 1708600 | Cell/Virus Enclosed | NO |
| YCTNNTLVSGDAHIPF | 1708607 | Ectodomain | YES |
| YCTNTTLVSGDA | 1533732 | Ectodomain | YES |
| YCTVQGPPGTGKSHLA | 1708610 | Cell/Virus Enclosed | NO |
| YCVDLHNKINLCDDPE | 1708612 | Cell/Virus Enclosed | NO |
| YCVDYSKNRLSR | 1533744 | Ectodomain | YES |
| YCVEMHNKINLCDDPE | 1708614 | Cell/Virus Enclosed | NO |
| YCVRAFDGYNKD | 1533755 | Cell/Virus Enclosed | NO |
| YCVRAFDVYNKDASFI | 1708616 | Cell/Virus Enclosed | NO |
| YCVTTKAKSVQQDLDV | 1708617 | Cell/Virus Enclosed | NO |
| YCYCGNFATCYN | 1533763 | Cell/Virus Enclosed | NO |
| YCYCGNSATCYNSVSN | 1708618 | Cell/Virus Enclosed | NO |
| YCYDKNPINSNVKTLE | 1708619 | Cell/Virus Enclosed | NO |
| YCYDKQPVNSNVKLLD | 1708620 | Cell/Virus Enclosed | NO |
| YCYFRNSSANYVGKYD | 1708624 | Underneath Membrane | NO |
| YCYFRNSTTNYVGVYD | 1708625 | Ectodomain | YES |
| YCYNTDLIEGSKPYST | 1708627 | Cell/Virus Enclosed | NO |
| YCYSDGVMKNASLYTS | 1708628 | Cell/Virus Enclosed | NO |
| YCYTARDALDTI | 1533777 | Cell/Virus Enclosed | NO |
| YCYTDGLMQNAS | 1533779 | Cell/Virus Enclosed | NO |
| YCYTEGLMQNAS | 1533780 | Cell/Virus Enclosed | NO |
| YDALLHRNSLDPFCFD | 1708635 | Cell/Virus Enclosed | NO |
| YDANYFVCWHTN | 1533794 | Secreted | YES |
| YDASNVKNVTDV | 1533800 | Cell/Virus Enclosed | NO |
| YDAYQNLVGYYSDDGN | 1708643 | Ectodomain | YES |
| YDCFCNNINWNV | 1533808 | Cell/Virus Enclosed | NO |
| YDDCMHQGPASA | 1533813 | Ectodomain | YES |
| YDDCMQQGPSSA | 1533819 | Ectodomain | YES |
| YDDFLEESDVEE | 1533820 | Cell/Virus Enclosed | NO |
| YDDGSCEVVHDQVNYV | 1708653 | Cell/Virus Enclosed | NO |
| YDDIFAESKPRF | 1533822 | Cell/Virus Enclosed | NO |
| YDDRILEFKVLD | 1533830 | Cell/Virus Enclosed | NO |
| YDDSQYYFNKDTGVIY | 1708658 | Ectodomain | YES |
| YDDVDASLFVDINNLL | 1708659 | Cell/Virus Enclosed | NO |
| YDEEGGYDVSKPVMIS | 1708662 | Cell/Virus Enclosed | NO |
| YDENGTITDAVDCSQN | 1708663 | Ectodomain | YES |
| YDFDGVTTSDKCIFNS | 1708666 | Cell/Virus Enclosed | NO |
| YDFGDFIQTAPGFGVA | 1708669 | Cell/Virus Enclosed | NO |

| | | | |
|---|---|---|---|
| YDFGDFVITQPGSGVA | 1708671 | Cell/Virus Enclosed | NO |
| YDFGDFVVRLPN | 1533857 | Cell/Virus Enclosed | NO |
| YDFGDFVVSLPNMGVP | 1708674 | Cell/Virus Enclosed | NO |
| YDFGDYVIAAPGCGVA | 1708675 | Cell/Virus Enclosed | NO |
| YDFKVEDAYDQVHDEP | 1708676 | Cell/Virus Enclosed | NO |
| YDFLNIDMGVFI | 1533864 | Cell/Virus Enclosed | NO |
| YDFLNMDMGVFINKYG | 1708681 | Cell/Virus Enclosed | NO |
| YDFLPIILQGIL | 1533866 | Undetermined | UN |
| YDFNGNIIGFKD | 1533867 | Ectodomain | YES |
| YDFTDYKLELFNKYFK | 1708685 | Cell/Virus Enclosed | NO |
| YDFVENPDIINVYKKL | 1708693 | Cell/Virus Enclosed | NO |
| YDFYPNVPIEYQ | 1533882 | Ectodomain | YES |
| YDGARLSASDLS | 1533885 | Cell/Virus Enclosed | NO |
| YDGCAQGVFGVNMRTN | 1708698 | Cell/Virus Enclosed | NO |
| YDGDRFVKHDLSLLSV | 1708699 | Cell/Virus Enclosed | NO |
| YDGGCLNASEVVVNNL | 1708704 | Cell/Virus Enclosed | NO |
| YDGLRGAIRKAKDYGF | 1708705 | Cell/Virus Enclosed | NO |
| YDGNMFIKFPEVVFES | 1708706 | Cell/Virus Enclosed | NO |
| YDGRTGELPCAIMNDK | 1708708 | Cell/Virus Enclosed | NO |
| YDGSGVGVIRIS | 1533900 | Ectodomain | YES |
| YDHSQYYFNKDT | 1533906 | Ectodomain | YES |
| YDHVFGTDHKFI | 1533908 | Cell/Virus Enclosed | NO |
| YDIEFVTGKQNI | 1533915 | Cell/Virus Enclosed | NO |
| YDIGNPKGIPIVDDPV | 1708721 | Cell/Virus Enclosed | NO |
| YDINLNKFNRAS | 1533929 | Cell/Virus Enclosed | NO |
| YDKFKINDTTCKYVFT | 1708728 | Cell/Virus Enclosed | NO |
| YDKFKINDTTRKYVFT | 1708729 | Cell/Virus Enclosed | NO |
| YDKNPINSNVKTLEYD | 1708733 | Cell/Virus Enclosed | NO |
| YDKQPVNSNVKLLDYD | 1708734 | Cell/Virus Enclosed | NO |
| YDLEVLLQDALQSREA | 1708740 | Cell/Virus Enclosed | NO |
| YDLGVELGTDPI | 1533958 | Cell/Virus Enclosed | NO |
| YDLVNYIRDKGC | 1533964 | Cell/Virus Enclosed | NO |
| YDLYDGNMFIKF | 1533965 | Cell/Virus Enclosed | NO |
| YDMDFGVSSFSE | 1533967 | Cell/Virus Enclosed | NO |
| YDNCYRKSNVDE | 1533976 | Cell/Virus Enclosed | NO |
| YDNCYRNSNVGE | 1533977 | Cell/Virus Enclosed | NO |
| YDNFTSIWPRFS | 1533983 | Undetermined | UN |
| YDNGNFIKLPEVIAQG | 1708761 | Cell/Virus Enclosed | NO |
| YDNISSVWPLHP | 1533986 | Ectodomain | YES |
| YDNKLKAHKEKS | 1533990 | Cell/Virus Enclosed | NO |
| YDNQKQIANQFN | 1533993 | Ectodomain | YES |
| YDNVSSVWPHYP | 1533996 | Ectodomain | YES |
| YDNVSSVWPLYL | 1533997 | Ectodomain | YES |
| YDPITKNIGEYNVSKD | 1708770 | Cell/Virus Enclosed | NO |
| YDPLTKNIGDYNVSKD | 1708776 | Cell/Virus Enclosed | NO |
| YDPTTKNLTGSN | 1534020 | Cell/Virus Enclosed | NO |
| YDPTTKNVTGSNESKA | 1708779 | Cell/Virus Enclosed | NO |
| YDPVENEDIHRVYASL | 1708780 | Cell/Virus Enclosed | NO |
| YDQVVDNVYVTYAGNV | 1708782 | Cell/Virus Enclosed | NO |
| YDRDGQRTYDDVNASL | 1708783 | Cell/Virus Enclosed | NO |
| YDRDGQRVYDDVDASL | 1708784 | Cell/Virus Enclosed | NO |
| YDRFKVNETNSQYLFS | 1708785 | Cell/Virus Enclosed | NO |
| YDRLDIIQADQQ | 1534035 | Ectodomain | YES |
| YDRLDSIQADQQVDRL | 1708786 | Ectodomain | YES |
| YDRLDTIQADQQVDRL | 1708787 | Ectodomain | YES |
| YDRLPNEVTVAKPNVL | 1708788 | Ectodomain | YES |
| YDRLSNEVTVAK | 1534043 | Underneath Membrane | NO |
| YDRVEQSFQLDT | 1534045 | Ectodomain | YES |
| YDSADRGQESLA | 1534046 | Cell/Virus Enclosed | NO |
| YDSMSYEDQDVL | 1534056 | Cell/Virus Enclosed | NO |
| YDSNGNIIGFKDFVTN | 1708791 | Ectodomain | YES |
| YDSNGNLYGLRD | 1534059 | Ectodomain | YES |
| YDSPCFSQQGVF | 1534060 | Ectodomain | YES |
| YDSSDHGQESLA | 1534062 | Cell/Virus Enclosed | NO |
| YDSSEVKAQSSGVIPE | 1708794 | Cell/Virus Enclosed | NO |
| YDSSGNLYGFRD | 1534066 | Ectodomain | YES |
| YDSVSAIGTANL | 1534069 | Ectodomain | YES |
| YDTAKKSMYDGDRFVK | 1708798 | Cell/Virus Enclosed | NO |
| YDTNHGDAGFTS | 1534077 | Ectodomain | YES |
| YDTNLNKFNGAS | 1534079 | Cell/Virus Enclosed | NO |
| YDVANNAVYDGA | 1534083 | Cell/Virus Enclosed | NO |
| YDVAVTAGPFMDNAIN | 1708805 | Cell/Virus Enclosed | NO |
| YDVAVTAGPFMG | 1534085 | Cell/Virus Enclosed | NO |
| YDVHKDNFKDGLCMFW | 1708806 | Cell/Virus Enclosed | NO |
| YDVLFYCSNSSSGVLD | 1708807 | Ectodomain | YES |
| YDVVVTAGPFMD | 1534109 | Cell/Virus Enclosed | NO |

| | | | |
|---|---|---|---|
| YDYSQYYFNKDT | 1534119 | Ectodomain | YES |
| YDYTQYYFNKDT | 1534121 | Ectodomain | YES |
| YDYVGTGIIRSSNQSL | 1708820 | Ectodomain | YES |
| YEADTEQTQNVTVKPK | 1708829 | Cell/Virus Enclosed | NO |
| YEADTEQTQSVI | 1534130 | Cell/Virus Enclosed | NO |
| YEADTEQTQSVN | 1534131 | Cell/Virus Enclosed | NO |
| YEADTEQTQSVT | 1534132 | Cell/Virus Enclosed | NO |
| YEAMYTPHTVLQ | 1534139 | Cell/Virus Enclosed | NO |
| YEDAIANGSSSQLIKQ | 1708840 | Cell/Virus Enclosed | NO |
| YEDEEIPEETC | 1534150 | Cell/Virus Enclosed | NO |
| YEEQDAIFSLTKRNIL | 1708850 | Cell/Virus Enclosed | NO |
| YEEYDLEPHKVH | 1534167 | Underneath Membrane | NO |
| YEFLNKHFNMMI | 1534178 | Cell/Virus Enclosed | NO |
| YEFRAGRPHRLV | 1534179 | Cell/Virus Enclosed | NO |
| YEFYNDIKPPVL | 1534180 | Cell/Virus Enclosed | NO |
| YEGFITLKNNLY | 1534181 | Cell/Virus Enclosed | NO |
| YEGGCIKACEVVVTNL | 1708860 | Cell/Virus Enclosed | NO |
| YEGGCITSREVVVTNL | 1708861 | Cell/Virus Enclosed | NO |
| YEGVNFSPHHRF | 1534191 | Ectodomain | YES |
| YEGVNFTPYHAF | 1534192 | Ectodomain | YES |
| YEIQIPSKFTIG | 1534200 | Ectodomain | YES |
| YEKDKAVARKLERMAD | 1708872 | Cell/Virus Enclosed | NO |
| YEKSTFNKDIYCGNSM | 1708876 | Cell/Virus Enclosed | NO |
| YELDNDFNTILNTACG | 1708878 | Cell/Virus Enclosed | NO |
| YELLNAPATVCG | 1534219 | Ectodomain | YES |
| YELNGYTVQPVA | 1534226 | Ectodomain | YES |
| YELRYNGAIRFD | 1534228 | Cell/Virus Enclosed | NO |
| YELSQLISGSLG | 1534232 | Cell/Virus Enclosed | NO |
| YEMLSLQQVVKALNES | 1708893 | Ectodomain | YES |
| YEMYVKWPWYVWLLIS | 1708895 | Undetermined | UN |
| YENARQAYEDAIANGS | 1708899 | Cell/Virus Enclosed | NO |
| YENAVANGSSPQIIKQ | 1708900 | Cell/Virus Enclosed | NO |
| YENKFVPVKEASKQCF | 1708905 | Cell/Virus Enclosed | NO |
| YENKFVPVKPASKQCF | 1708906 | Cell/Virus Enclosed | NO |
| YENLSLADISAVQSSF | 1708908 | Cell/Virus Enclosed | NO |
| YENQKQIANQFNKAIS | 1708911 | Ectodomain | YES |
| YENRFAAIYGGD | 1534256 | Cell/Virus Enclosed | NO |
| YENYTILQAAGL | 1534260 | Cell/Virus Enclosed | NO |
| YEPYYHSDHAESSWVNRGES | 73744 | Ectodomain | YES |
| YEQAVANGDSEA | 1534275 | Cell/Virus Enclosed | NO |
| YEQLNDSSEEEDEIDG | 110460 | Cell/Virus Enclosed | NO |
| YEQNWNTKHGGG | 1534281 | Cell/Virus Enclosed | NO |
| YEQVVMDYLDSL | 1534287 | Cell/Virus Enclosed | NO |
| YEQYIKWPWYVWLGFI | 1708929 | Undetermined | UN |
| YERDKAVARKLERMAD | 1708930 | Cell/Virus Enclosed | NO |
| YERDNTSCPEWMDDFE | 1708931 | Cell/Virus Enclosed | NO |
| YERDRAVAKKLERMAD | 1708932 | Cell/Virus Enclosed | NO |
| YERQTPFEIKSA | 1534308 | Cell/Virus Enclosed | NO |
| YERVKDCKFVAE | 1534311 | Cell/Virus Enclosed | NO |
| YESFDVESGVYS | 1534314 | Ectodomain | YES |
| YETARQEYENAVANGS | 1708950 | Cell/Virus Enclosed | NO |
| YEVAKKNLDEARFSGS | 1708954 | Cell/Virus Enclosed | NO |
| YEVGDLIAQSSG | 1534342 | Cell/Virus Enclosed | NO |
| YEVGDLMAQSSG | 1534343 | Cell/Virus Enclosed | NO |
| YFAKLGRELNCNVCSN | 1708967 | Cell/Virus Enclosed | NO |
| YFAKVGREISCNVCTK | 1708968 | Cell/Virus Enclosed | NO |
| YFASHFISNSWL | 1534363 | Cell/Virus Enclosed | NO |
| YFAVRKEGEDVI | 1534367 | Cell/Virus Enclosed | NO |
| YFAVRKEGQDVIFSQF | 1708975 | Cell/Virus Enclosed | NO |
| YFCYGFNEPLNV | 1534382 | Ectodomain | YES |
| YFDSKGWYDPVENEDI | 1708991 | Cell/Virus Enclosed | NO |
| YFEIYDGGCIPASQVI | 1708996 | Cell/Virus Enclosed | NO |
| YFEIYDGGCLNASEVV | 1708997 | Cell/Virus Enclosed | NO |
| YFGGFNFSQILPDPLK | 1709010 | Ectodomain | YES |
| YFIKGCNTLARG | 1534443 | Cell/Virus Enclosed | NO |
| YFIKTNNTRIVDEWSY | 1709022 | Ectodomain | YES |
| YFITMSSKGQKGDAEE | 1709027 | Cell/Virus Enclosed | NO |
| YFKDCLFKDWEELGEE | 73843 | Cell/Virus Enclosed | NO |
| YFKHWSFDYHPN | 1534470 | Cell/Virus Enclosed | NO |
| YFKHWSMPYHPNTVDC | 1709034 | Cell/Virus Enclosed | NO |
| YFKNHTSPDVDF | 1534477 | Ectodomain | YES |
| YFKPGGTTSGDASTAY | 1709038 | Cell/Virus Enclosed | NO |
| YFKYWGQDYHPDCVDC | 1709045 | Cell/Virus Enclosed | NO |
| YFLDDVEMVAYNPDLS | 1709047 | Cell/Virus Enclosed | NO |
| YFLVRLLQGKTG | 1534513 | Cell/Virus Enclosed | NO |
| YFMVDNVTRLHDA | 1709059 | Undetermined | UN |

| | | | |
|---|---|---|---|
| YFMYDIVSSFEGYDFK | 1709060 | Cell/Virus Enclosed | NO |
| YFNATGSDVNCNGYQH | 1709061 | Ectodomain | YES |
| YFNGAIIRDSDVVKQP | 1709062 | Cell/Virus Enclosed | NO |
| YFNGTGRGACKG | 1534534 | Ectodomain | YES |
| YFNGTGRGDCKGFSSD | 1709063 | Ectodomain | YES |
| YFNGTGRGDCQG | 1534536 | Ectodomain | YES |
| YFNNEPAFIMHV | 1534555 | Cell/Virus Enclosed | NO |
| YFNRPLLVDDNK | 1534560 | Cell/Virus Enclosed | NO |
| YFNRPSLVDDNKFDVL | 1709072 | Cell/Virus Enclosed | NO |
| YFPKSGANFTDL | 1534571 | Ectodomain | YES |
| YFRNSSANYVGKYDIN | 1709085 | Underneath Membrane | NO |
| YFRNSTTNYVGVYDIN | 1709086 | Ectodomain | YES |
| YFSDSKIGTTEV | 1534595 | Cell/Virus Enclosed | NO |
| YFSGAMETAAYREAAA | 1709090 | Cell/Virus Enclosed | NO |
| YFSKKDWYDFVENPDI | 1709091 | Cell/Virus Enclosed | NO |
| YFSKKHVEVFTDGKLN | 1709092 | Cell/Virus Enclosed | NO |
| YFSLGDVEAVNF | 1534609 | Ectodomain | YES |
| YFSQLLCRPIKLVDSE | 1709095 | Cell/Virus Enclosed | NO |
| YFSYPFSMKSDL | 1534620 | Ectodomain | YES |
| YFSYPLSMKSDL | 1534621 | Ectodomain | YES |
| YFTKKDWYDFVENPDI | 1709106 | Cell/Virus Enclosed | NO |
| YFTLGCKILTLT | 1534636 | Cell/Virus Enclosed | NO |
| YFTNEPAFIMHV | 1534644 | Cell/Virus Enclosed | NO |
| YFTSAPPVTYSP | 1534652 | Cell/Virus Enclosed | NO |
| YFTSEPPVTYSS | 1534657 | Cell/Virus Enclosed | NO |
| YFVNVNNTWMYTGSGY | 1709138 | Ectodomain | YES |
| YFYYLGTGPEAGLPY | 1313991 | Cell/Virus Enclosed | NO |
| YGAFMDKQVHQVQLTD | 1709158 | Cell/Virus Enclosed | NO |
| YGAGSDKGVAPGTTVL | 1709161 | Cell/Virus Enclosed | NO |
| YGCCDATYFENK | 1534751 | Cell/Virus Enclosed | NO |
| YGCDYSGKLADCLQAY | 1709168 | Cell/Virus Enclosed | NO |
| YGDVDNLVFTNKTTLP | 1709187 | Cell/Virus Enclosed | NO |
| YGEPVLVNAANT | 1534786 | Cell/Virus Enclosed | NO |
| YGFCGNGTHIFS | 1534790 | Ectodomain | YES |
| YGFNSTETITTG | 1534796 | Ectodomain | YES |
| YGFSNVVEMPKF | 1534803 | Ectodomain | YES |
| YGGFEDQPNLQVESAN | 1709216 | Cell/Virus Enclosed | NO |
| YGGFGCKDSFTL | 1534818 | Cell/Virus Enclosed | NO |
| YGGWDDMLRRLIKDVD | 1709222 | Cell/Virus Enclosed | NO |
| YGGWDFMLKTLYKDVD | 1709223 | Cell/Virus Enclosed | NO |
| YGGWDNMLKNLMADVD | 1709224 | Cell/Virus Enclosed | NO |
| YGIPLSEAKGYW | 1534842 | Cell/Virus Enclosed | NO |
| YGIPPSEAKGYW | 1534843 | Cell/Virus Enclosed | NO |
| YGKMGRFFNHTL | 1534859 | Ectodomain | YES |
| YGKPINLPTGCMMNVA | 1709238 | Cell/Virus Enclosed | NO |
| YGKPVTLPTGCMMNVA | 1709239 | Cell/Virus Enclosed | NO |
| YGKSTFNKDIYC | 1534863 | Cell/Virus Enclosed | NO |
| YGLCDSANPTNC | 1534868 | Ectodomain | YES |
| YGLEYYAFEHIV | 1534877 | Cell/Virus Enclosed | NO |
| YGLNSTETISTG | 1534889 | Ectodomain | YES |
| YGLNSTETITTGFDLN | 1709254 | Ectodomain | YES |
| YGLNSTETITTS | 1534891 | Ectodomain | YES |
| YGNCTFGSPDDA | 1534907 | Cell/Virus Enclosed | NO |
| YGNFDDQPSLQVESAN | 1709263 | Cell/Virus Enclosed | NO |
| YGNMTLNGLWLDDKVY | 1709265 | Cell/Virus Enclosed | NO |
| YGNVSRHNLTKYTMMD | 1709268 | Cell/Virus Enclosed | NO |
| YGPYKDAQVVQL | 1534931 | Cell/Virus Enclosed | NO |
| YGPYRDAQVVQLPVKD | 1709275 | Cell/Virus Enclosed | NO |
| YGQCLGDINARD | 1534934 | Ectodomain | YES |
| YGRCPTAADINIPDLP | 1709280 | Ectodomain | YES |
| YGRCPTAADINN | 1534944 | Ectodomain | YES |
| YGRCPTAADINT | 1534945 | Ectodomain | YES |
| YGRGGYHYTPFH | 1534949 | Cell/Virus Enclosed | NO |
| YGRSGITLGVLV | 1534954 | Cell/Virus Enclosed | NO |
| YGSRLGVPDTLGGDPKFRSL | 74084 | Capsid | UN |
| YGSSVETETYFKVNGL | 1709294 | Cell/Virus Enclosed | NO |
| YGSTVLNGVWLGDTVT | 1709297 | Cell/Virus Enclosed | NO |
| YGTCLNCNAPRM | 1534977 | Cell/Virus Enclosed | NO |
| YGTDIDGIYWVA | 1534979 | Cell/Virus Enclosed | NO |
| YGTDTISVCPKL | 1534983 | Ectodomain | YES |
| YGTDTNSVCPKLEFAN | 1709302 | Ectodomain | YES |
| YGTFCDNINSILDEVN | 1709307 | Ectodomain | YES |
| YGTYKNTCDGNT | 1535006 | Cell/Virus Enclosed | NO |
| YGVALKDCGFVM | 1535008 | Cell/Virus Enclosed | NO |
| YGVCKYTDLKFIENLN | 1709313 | Cell/Virus Enclosed | NO |
| YGVCMYTDLKFIDKLN | 1709314 | Cell/Virus Enclosed | NO |

| | | | |
|---|---|---|---|
| YGVRRKNSEPEIPHFN | 1709320 | Capsid | UN |
| YGWVSGNMMDGFSCPL | 1709325 | Cell/Virus Enclosed | NO |
| YGYFLPYPDPSR | 1535026 | Cell/Virus Enclosed | NO |
| YGYLMKRFSMMI | 1535032 | Cell/Virus Enclosed | NO |
| YGYSNFVSSDLD | 1535037 | Underneath Membrane | NO |
| YGYTSRSMFVYV | 1535039 | Undetermined | UN |
| YHAFKCNHSGSN | 1535044 | Ectodomain | YES |
| YHAFKCNRSGSNDIWM | 1709336 | Ectodomain | YES |
| YHEHIGGVYDNS | 1535061 | Cell/Virus Enclosed | NO |
| YHFKQLGLVWNK | 1535066 | Cell/Virus Enclosed | NO |
| YHNSQDDIMKAV | 1535109 | Capsid | UN |
| YHPAHAPSFLAVDDKY | 1709363 | Cell/Virus Enclosed | NO |
| YHQKHFKSIVNT | 1535125 | Cell/Virus Enclosed | NO |
| YHQKMLKSMAATRGAT | 1709371 | Cell/Virus Enclosed | NO |
| YHSGLPLHMSIS | 1535141 | Cell/Virus Enclosed | NO |
| YHTASTLRSVGQ | 1535146 | Ectodomain | YES |
| YHTASVLRSTGQ | 1535147 | Ectodomain | YES |
| YHTPATDCSDGNYNRN | 1709381 | Ectodomain | YES |
| YHTPATNCSDGN | 1535158 | Ectodomain | YES |
| YHTPTTDCSDGN | 1535160 | Ectodomain | YES |
| YHTVGSPIQFHL | 1535164 | Cell/Virus Enclosed | NO |
| YHVTKVLAEKFDYNVS | 1709388 | Cell/Virus Enclosed | NO |
| YHYKELGVVMNLDVDT | 1709392 | Cell/Virus Enclosed | NO |
| YHYNELGIEMNM | 1535180 | Cell/Virus Enclosed | NO |
| YHYTPFHYERDNTSCP | 1709395 | Cell/Virus Enclosed | NO |
| YHYTPFHYERNN | 1535185 | Cell/Virus Enclosed | NO |
| YHYTPVRYERDN | 1535188 | Cell/Virus Enclosed | NO |
| YIAPQANSGIIT | 1535195 | Ectodomain | YES |
| YIAPRANSGIIT | 1535196 | Ectodomain | YES |
| YIAYENARQAYEDAIA | 1709407 | Cell/Virus Enclosed | NO |
| YICGDSTECANLLLQYG | 74173 | Ectodomain | YES |
| YIDCEQTPCGFK | 1535221 | Cell/Virus Enclosed | NO |
| YIDCEQYSCDFK | 1535222 | Cell/Virus Enclosed | NO |
| YIDVLRNSFGKDLNAN | 1709423 | Cell/Virus Enclosed | NO |
| YIDVNKTLQELSYKLP | 1709424 | Ectodomain | YES |
| YIECTYIGRFSK | 1535239 | Cell/Virus Enclosed | NO |
| YIEGSGKSAPNS | 1535245 | Cell/Virus Enclosed | NO |
| YIEGSGRSASNS | 1535250 | Cell/Virus Enclosed | NO |
| YIEGSGRSVPNS | 1535251 | Cell/Virus Enclosed | NO |
| YIEKLYKDLTGHMLDS | 1709429 | Cell/Virus Enclosed | NO |
| YIFERIDYSDAVSYKS | 1709434 | Cell/Virus Enclosed | NO |
| YIFSTANALPEC | 1535271 | Cell/Virus Enclosed | NO |
| YIGATVRLQAGKQTEF | 1709444 | Cell/Virus Enclosed | NO |
| YIGDPAQLPAPRVLLS | 1709446 | Cell/Virus Enclosed | NO |
| YIGDPVQLPAPR | 1535284 | Cell/Virus Enclosed | NO |
| YIIKLIFLWLLWPVTL | 1334490 | Undetermined | UN |
| YIISCYWGLFSL | 1535313 | Cell/Virus Enclosed | NO |
| YIKDGDDGYFLPYPDP | 1709460 | Cell/Virus Enclosed | NO |
| YIKKLYNDLGNQILDS | 1709462 | Cell/Virus Enclosed | NO |
| YIKKLYNDLGTQILDS | 1709463 | Cell/Virus Enclosed | NO |
| YILANECTERLKLFAA | 1709473 | Cell/Virus Enclosed | NO |
| YIMFEPIDPIWM | 1535363 | Cell/Virus Enclosed | NO |
| YINDKDTGPPPI | 1535369 | Ectodomain | YES |
| YINGFICEKLAI | 1535373 | Cell/Virus Enclosed | NO |
| YINGYRYFSLGN | 1535380 | Ectodomain | YES |
| YINIPDLPICVY | 1535383 | Ectodomain | YES |
| YINIPDLPVCVY | 1535384 | Ectodomain | YES |
| YINNKDTGPPFI | 1535396 | Ectodomain | YES |
| YINTHMIGVTLC | 1535398 | Cell/Virus Enclosed | NO |
| YINVTFLNLQVE | 1535407 | Ectodomain | YES |
| YIPLVGTPLHPV | 1535423 | Cell/Virus Enclosed | NO |
| YIQGIKLGTGYSLADL | 1709499 | Ectodomain | YES |
| YIQGVKLGTGYTLSDL | 1709500 | Underneath Membrane | NO |
| YIRRSIDCGFND | 1535446 | Ectodomain | YES |
| YISDAFSLDVSEKSGNF | 74367 | Ectodomain | YES |
| YISHETKLNSCC | 1535457 | Cell/Virus Enclosed | NO |
| YISSNGPLKVGAGVML | 1709519 | Cell/Virus Enclosed | NO |
| YITANGGISFCR | 1535479 | Cell/Virus Enclosed | NO |
| YIVGDVISSGDI | 1535505 | Cell/Virus Enclosed | NO |
| YIVNISAFQQVL | 1535515 | Cell/Virus Enclosed | NO |
| YIWTADEAIKFLKEWN | 1709547 | Undetermined | UN |
| YIYDEEGGNDLSLPVM | 1709549 | Cell/Virus Enclosed | NO |
| YIYDEEGGYDVSKPVM | 1709550 | Cell/Virus Enclosed | NO |
| YIYNRVKSGSPGDSSW | 1709552 | Ectodomain | YES |
| YIYSRVKNLNSS | 1535557 | Ectodomain | YES |
| YIYVRKNGQFQDHYDG | 1709553 | Cell/Virus Enclosed | NO |

| | | | |
|---|---|---|---|
| YIYVTNEIGLNASVTL | 1709554 | Ectodomain | YES |
| YKAACVVDVNDSHSMA | 1709555 | Cell/Virus Enclosed | NO |
| YKAFPPTEPKKD | 1535567 | Cell/Virus Enclosed | NO |
| YKATGDLAVCLGIGDS | 1709565 | Cell/Virus Enclosed | NO |
| YKDLTGYMLDSY | 1535599 | Cell/Virus Enclosed | NO |
| YKDSVFYVVKNSTAFP | 1709571 | Cell/Virus Enclosed | NO |
| YKDVDNPHLMGWDYPK | 1709572 | Cell/Virus Enclosed | NO |
| YKDVILWFSFGASCFILLAIVMGLV | 1180540 | Undetermined | UN |
| YKEGFRGFTYVW | 1535610 | Cell/Virus Enclosed | NO |
| YKELGLVMNMDV | 1535615 | Cell/Virus Enclosed | NO |
| YKELGVVMNLDVDTHR | 1709584 | Cell/Virus Enclosed | NO |
| YKETNWFLHFIP | 1535618 | Cell/Virus Enclosed | NO |
| YKGKWVQVPIGTNDPI | 1709605 | Cell/Virus Enclosed | NO |
| YKGLPLEFGAST | 1535649 | Cell/Virus Enclosed | NO |
| YKGNVTHDASSAINRP | 1709610 | Cell/Virus Enclosed | NO |
| YKGYQPIDVVRDLPSG | 1709618 | Ectodomain | YES |
| YKIDTTSSSCQLYYSL | 1709627 | Ectodomain | YES |
| YKITQWPTATGD | 1535686 | Cell/Virus Enclosed | NO |
| YKKCTKGLSIADLACA | 1709636 | Ectodomain | YES |
| YKKDGVYYPSNGTNIL | 1709638 | Cell/Virus Enclosed | NO |
| YKKLGPIFNRAL | 1535695 | Cell/Virus Enclosed | NO |
| YKKQQEGHIIME | 1535699 | Cell/Virus Enclosed | NO |
| YKKVNNEFIDPTECIY | 1709650 | Cell/Virus Enclosed | NO |
| YKLANDAKESLRLFAA | 1709651 | Cell/Virus Enclosed | NO |
| YKLANDVKDTLRLFAA | 1709652 | Cell/Virus Enclosed | NO |
| YKLFLSGDDCFVSNSS | 1709659 | Cell/Virus Enclosed | NO |
| YKLHPAFNVSDAYANL | 1709661 | Cell/Virus Enclosed | NO |
| YKLHPSFNVSDAYANL | 1709662 | Cell/Virus Enclosed | NO |
| YKLNTDHSGSND | 1535739 | Underneath Membrane | NO |
| YKLNTDHSNSSDNIAL | 1709670 | Underneath Membrane | NO |
| YKLNTDHSSSSDNIAL | 1709671 | Underneath Membrane | NO |
| YKLPNYTVPDLVVEQY | 1709675 | Ectodomain | YES |
| YKMFLHTGAVIVLPEDDLPT | 74479 | Cell/Virus Enclosed | NO |
| YKMLVAKDNKNLPKFI | 1709685 | Capsid | UN |
| YKNMAVYRSLTF | 1535763 | Ectodomain | YES |
| YKNMCTGSPSIVEFNR | 1709693 | Cell/Virus Enclosed | NO |
| YKNMYLKSAVMQSVGA | 1709694 | Cell/Virus Enclosed | NO |
| YKNSATGSLDVEVFNR | 1709696 | Cell/Virus Enclosed | NO |
| YKNSGIGTCPAG | 1535769 | Ectodomain | YES |
| YKPLLENIDYFNMRRA | 1709709 | Cell/Virus Enclosed | NO |
| YKQARAEDKKAKIVSA | 1709713 | Cell/Virus Enclosed | NO |
| YKQARSEDKRAQ | 1535794 | Cell/Virus Enclosed | NO |
| YKQQYSQLFVDT | 1535798 | Cell/Virus Enclosed | NO |
| YKQSCTGSPYIDDFNK | 1709717 | Cell/Virus Enclosed | NO |
| YKQSCTGSPYIDDFNR | 1709718 | Cell/Virus Enclosed | NO |
| YKRAFLDKLDVNSGFA | 1709721 | Underneath Membrane | NO |
| YKRNFTYDVNAD | 1535814 | Ectodomain | YES |
| YKRNFTYDVNAT | 1535817 | Ectodomain | YES |
| YKRQNNLAIEEIEYVH | 1709731 | Cell/Virus Enclosed | NO |
| YKRQQEGHIIME | 1535822 | Cell/Virus Enclosed | NO |
| YKRRMADAVSRKKM | 74503 | Capsid | UN |
| YKSCTKGLSIADLACA | 1709732 | Ectodomain | YES |
| YKSSALGSMDIDVFNK | 1709738 | Cell/Virus Enclosed | NO |
| YKTIDELCVNLN | 1535847 | Cell/Virus Enclosed | NO |
| YKTPTLKYFDGF | 1535854 | Ectodomain | YES |
| YKVCYSVKDTLA | 1535867 | Cell/Virus Enclosed | NO |
| YKVLPPLMDVNMEAAY | 1709762 | Ectodomain | YES |
| YKVNCCRFQRIDDDGN | 1709763 | Cell/Virus Enclosed | NO |
| YKVNENLAVNLNICEP | 1709764 | Cell/Virus Enclosed | NO |
| YKVTKGKPVKGA | 1535880 | Cell/Virus Enclosed | NO |
| YKYDAKNYVRFPEILA | 1709772 | Cell/Virus Enclosed | NO |
| YKYDNGNFIKLPEVIA | 1709773 | Cell/Virus Enclosed | NO |
| YKYLNSPMERVSLWNY | 1709778 | Cell/Virus Enclosed | NO |
| YKYYSGNANEADYRCA | 1709784 | Cell/Virus Enclosed | NO |
| YLADNLNADETK | 1535914 | Cell/Virus Enclosed | NO |
| YLADYDVAVTAGPFMD | 1709788 | Cell/Virus Enclosed | NO |
| YLAKAMLDFSRDHNDI | 1709790 | Cell/Virus Enclosed | NO |
| YLAYIKKLYNDLGNQI | 1709799 | Cell/Virus Enclosed | NO |
| YLCGADGKPVMSEDLW | 1709802 | Cell/Virus Enclosed | NO |
| YLCNLINNNLAL | 1535947 | Cell/Virus Enclosed | NO |
| YLDAVKQGAKPVGNCV | 1709807 | Cell/Virus Enclosed | NO |
| YLDDVKKIEYKPDLSQ | 1709809 | Cell/Virus Enclosed | NO |
| YLDFIQQGGTPIANCV | 1709810 | Cell/Virus Enclosed | NO |
| YLDFNALNLFFSPLEFQHLI | 74601 | Capsid | UN |
| YLDFVNAGGAPLTNCV | 1709811 | Cell/Virus Enclosed | NO |
| YLDIHPSLCNNGKISS | 1709815 | Ectodomain | YES |

| | | | |
|---|---|---|---|
| YLDLLNRLQALESGKV | 1709816 | Cell/Virus Enclosed | NO |
| YLDSLKPRVEAP | 1535963 | Cell/Virus Enclosed | NO |
| YLEDKFDLLISDMYDG | 1709824 | Cell/Virus Enclosed | NO |
| YLEYIKKLYNDLGTQI | 1709830 | Cell/Virus Enclosed | NO |
| YLFDEAGDEAMA | 1535980 | Cell/Virus Enclosed | NO |
| YLFSTINALPEI | 1535992 | Cell/Virus Enclosed | NO |
| YLFSTINALPETSADI | 1709842 | Cell/Virus Enclosed | NO |
| YLGCVVNAYNST | 1535999 | Ectodomain | YES |
| YLGDFIQGPFIAGNTV | 1709848 | Cell/Virus Enclosed | NO |
| YLGDISHLINCV | 1536003 | Cell/Virus Enclosed | NO |
| YLGDISHLVNCV | 1536004 | Cell/Virus Enclosed | NO |
| YLGKLKEQIDGY | 1536017 | Cell/Virus Enclosed | NO |
| YLGTGPEASLPYGANK | 74684 | Capsid | UN |
| YLGTGPHAKDQY | 1536036 | Cell/Virus Enclosed | NO |
| YLGTGPHSKDQY | 1536041 | Cell/Virus Enclosed | NO |
| YLGTGPHSKEQY | 1536042 | Cell/Virus Enclosed | NO |
| YLGTGPYAKDQY | 1536043 | Cell/Virus Enclosed | NO |
| YLGTIKENIDGGAMHA | 1709868 | Cell/Virus Enclosed | NO |
| YLGTVLSHYYVM | 1536053 | Ectodomain | YES |
| YLIAYISFAPWW | 1536066 | Cell/Virus Enclosed | NO |
| YLISKFLKANPVWNSA | 1709883 | Cell/Virus Enclosed | NO |
| YLKDQQLLGIWGCSGKL | 94978 | Undetermined | UN |
| YLKGSGSDFSLADSVV | 1709885 | Cell/Virus Enclosed | NO |
| YLKNLTQTFKSM | 1536097 | Cell/Virus Enclosed | NO |
| YLKNLTQTFTSMLTNY | 1709893 | Cell/Virus Enclosed | NO |
| YLKPTTFMLKYDENGT | 1709895 | Ectodomain | YES |
| YLKRARGSSAAR | 1536105 | Cell/Virus Enclosed | NO |
| YLLAFNQDGIIFNAVD | 1709904 | Ectodomain | YES |
| YLLEEFDVVFGKRGGG | 1709906 | Cell/Virus Enclosed | NO |
| YLLKFDDRGVIT | 1536129 | Ectodomain | YES |
| YLLKFDNRGVITNAVD | 1709912 | Ectodomain | YES |
| YLNDPSSKTVCTYMDL | 1709918 | Cell/Virus Enclosed | NO |
| YLNHSVPKLSDFESEL | 1709922 | Ectodomain | YES |
| YLNLSSELKQLEAKTA | 1709924 | Ectodomain | YES |
| YLNNSIPNLSDF | 1536152 | Ectodomain | YES |
| YLNVQYRINGLGVTMD | 1709941 | Ectodomain | YES |
| YLQITSIPIVVD | 1536191 | Ectodomain | YES |
| YLRHGKLRPFERDISN | 1709963 | Ectodomain | YES |
| YLRHSSLSAQSE | 1536215 | Cell/Virus Enclosed | NO |
| YLRRMGGIQRRDHLQTLRDQ | 74902 | Capsid | UN |
| YLSERCNVIADP | 1536232 | Cell/Virus Enclosed | NO |
| YLSERCNVIADSKSFT | 1709973 | Cell/Virus Enclosed | NO |
| YLSFNMDSATKV | 1536235 | Ectodomain | YES |
| YLSLSDRFKTSGDLAV | 1709976 | Cell/Virus Enclosed | NO |
| YLSSSNVTTMDF | 1536241 | Cell/Virus Enclosed | NO |
| YLSSVLLALQQL | 1536243 | Cell/Virus Enclosed | NO |
| YLTCHNAAQCDC | 1536249 | Ectodomain | YES |
| YLTCHNAAQCNC | 1536250 | Ectodomain | YES |
| YLTCHNAVQCNC | 1536251 | Ectodomain | YES |
| YLTKMSEYKGPV | 1536259 | Cell/Virus Enclosed | NO |
| YLTNQIRLNASV | 1536269 | Ectodomain | YES |
| YLTQDLFLPFYS | 1536272 | Ectodomain | YES |
| YLVFVTTLTSDY | 1536295 | Cell/Virus Enclosed | NO |
| YLVLPFGNYLAI | 1536301 | Ectodomain | YES |
| YLVQKRHYLVFLIAHHYYEA | 74975 | Capsid | UN |
| YLYKKVNNEFIDPTEC | 1710025 | Cell/Virus Enclosed | NO |
| YLYNTGRSVYVKFQDS | 1710030 | Cell/Virus Enclosed | NO |
| YMAKAGDKYYPVVVDD | 1710035 | Cell/Virus Enclosed | NO |
| YMAKSGDKYYPV | 1536341 | Cell/Virus Enclosed | NO |
| YMCGFDGKPMLP | 1536344 | Cell/Virus Enclosed | NO |
| YMCVVPSINDLSFDKN | 1710039 | Cell/Virus Enclosed | NO |
| YMCVVSSINDLS | 1536352 | Cell/Virus Enclosed | NO |
| YMDGMDAKQVDYVPLK | 1710040 | Cell/Virus Enclosed | NO |
| YMDLNPALCDSG | 1536362 | Ectodomain | YES |
| YMDLNSALCDSG | 1536364 | Ectodomain | YES |
| YMEAYNFVSASG | 1536367 | Cell/Virus Enclosed | NO |
| YMGKCGDKFFPIVMND | 1710048 | Cell/Virus Enclosed | NO |
| YMHQMELANGTH | 1536378 | Cell/Virus Enclosed | NO |
| YMIDPAGVCFGA | 1536379 | Cell/Virus Enclosed | NO |
| YMIDPAGVCLGA | 1536380 | Cell/Virus Enclosed | NO |
| YMIRDKLALGGS | 1536384 | Cell/Virus Enclosed | NO |
| YMLFTKFFYLLG | 1536388 | Cell/Virus Enclosed | NO |
| YMLRKRRKRLPYFF | 111065 | Capsid | UN |
| YMMPIMGLTICL | 1536392 | Cell/Virus Enclosed | NO |
| YMNANGLRPPKNSFEA | 1710067 | Cell/Virus Enclosed | NO |
| YMQGYDDCMHQG | 1536412 | Ectodomain | YES |

| | | | |
|---|---|---|---|
| YMTHGQMEGLCL | 1536433 | Cell/Virus Enclosed | NO |
| YMVANGLNAPNGPFDA | 1710090 | Cell/Virus Enclosed | NO |
| YMVGDVITSGDI | 1536439 | Cell/Virus Enclosed | NO |
| YNAFLDTVVSTVKIGG | 1710092 | Cell/Virus Enclosed | NO |
| YNAKVEISVTPIKNTV | 1710094 | Cell/Virus Enclosed | NO |
| YNAPCFSQQGVF | 1536448 | Ectodomain | YES |
| YNDIKPPVLDVD | 1536466 | Cell/Virus Enclosed | NO |
| YNELGIEMNMDV | 1536471 | Cell/Virus Enclosed | NO |
| YNEVPATVLQNN | 1536474 | Cell/Virus Enclosed | NO |
| YNEWAQANGFTAMNGE | 1710111 | Cell/Virus Enclosed | NO |
| YNFIIYDSSDRGQESL | 1710113 | Cell/Virus Enclosed | NO |
| YNGAIRSDSTLS | 1536499 | Capsid | UN |
| YNGIMVLPGAAD | 1536509 | Ectodomain | YES |
| YNGKPQGAFHVT | 1536511 | Cell/Virus Enclosed | NO |
| YNGRPQGAFHVT | 1536515 | Cell/Virus Enclosed | NO |
| YNGSAQSTALCK | 1536524 | Ectodomain | YES |
| YNIDTGSFYGFS | 1536540 | Underneath Membrane | NO |
| YNKDASFIGKNLKSNC | 1710142 | Cell/Virus Enclosed | NO |
| YNKDHGCGGFGH | 1536566 | Cell/Virus Enclosed | NO |
| YNKDYAELGYIADISA | 1710144 | Cell/Virus Enclosed | NO |
| YNKQKANVSVLT | 1536582 | Cell/Virus Enclosed | NO |
| YNKRKRIHIGPGRAFYTTKNIIG | 604752 | Ectodomain | YES |
| YNKWLELLEQNIDAYK | 1710153 | Cell/Virus Enclosed | NO |
| YNLKNGEVEFVYMHQI | 1710162 | Cell/Virus Enclosed | NO |
| YNLLKGCNAVAKHDFF | 1710166 | Cell/Virus Enclosed | NO |
| YNLNFSANSLDNLKSG | 1710169 | Ectodomain | YES |
| YNLPTMCDIKQM | 1536612 | Cell/Virus Enclosed | NO |
| YNLPTMCDIRQL | 1536613 | Cell/Virus Enclosed | NO |
| YNLSSVIPSLPRSGSRVAGR | 2060916 | Ectodomain | YES |
| YNLSSVLPQRNIRSSR | 1710175 | Ectodomain | YES |
| YNLSSVVPSLPT | 1536619 | Ectodomain | YES |
| YNLTKYTMMDLV | 1536624 | Cell/Virus Enclosed | NO |
| YNLTMPSQVQKA | 1536625 | Cell/Virus Enclosed | NO |
| YNLVKTGHYTGQAGEM | 1710179 | Cell/Virus Enclosed | NO |
| YNLVNVGHYDGRTGEL | 1710180 | Cell/Virus Enclosed | NO |
| YNLWQIFIETNL | 1536633 | Cell/Virus Enclosed | NO |
| YNMCDYPHTICH | 1536640 | Ectodomain | YES |
| YNMCEYPHTICH | 1536643 | Ectodomain | YES |
| YNMCEYPQTICH | 1536646 | Ectodomain | YES |
| YNMDTGFAYGYS | 1536648 | Underneath Membrane | NO |
| YNMLVNCFKWQV | 1536651 | Cell/Virus Enclosed | NO |
| YNMLVNCSKWQV | 1536652 | Cell/Virus Enclosed | NO |
| YNNCTGGAGIRD | 1536657 | Ectodomain | YES |
| YNNEGGRAFMYA | 1536661 | Cell/Virus Enclosed | NO |
| YNNKLLAKKDLS | 1536662 | Cell/Virus Enclosed | NO |
| YNNLVPTYLKSDNIVA | 1710199 | Cell/Virus Enclosed | NO |
| YNNSGSNITQIKNACV | 1710200 | Cell/Virus Enclosed | NO |
| YNNSNNGKIVYAILSD | 1710202 | Cell/Virus Enclosed | NO |
| YNNVSSVWPLYP | 1536671 | Ectodomain | YES |
| YNNYVPSYVKPDSVST | 1710204 | Cell/Virus Enclosed | NO |
| YNQIILNLTSEI | 1536686 | Ectodomain | YES |
| YNRNASLNSFKEYFNL | 1710221 | Ectodomain | YES |
| YNRVKSGSPGDSSWHI | 1710223 | Ectodomain | YES |
| YNRYKYYSGNANEADY | 1710224 | Cell/Virus Enclosed | NO |
| YNSDYASKGYIANISA | 1710230 | Cell/Virus Enclosed | NO |
| YNSPCFSQQGVF | 1536720 | Ectodomain | YES |
| YNSPSSSSRRKRRSISASY | 2060917 | Ectodomain | YES |
| YNSQNAVARSML | 1536723 | Cell/Virus Enclosed | NO |
| YNSQNAVSRSMLGLTT | 1710239 | Cell/Virus Enclosed | NO |
| YNSQNFAAKRVLGLQT | 1710240 | Cell/Virus Enclosed | NO |
| YNSQNYVAKRVLGVQT | 1710242 | Cell/Virus Enclosed | NO |
| YNSSCFSHQGVF | 1536733 | Ectodomain | YES |
| YNSSCFSQQGVF | 1536734 | Ectodomain | YES |
| YNSTAISVQKCD | 1536738 | Ectodomain | YES |
| YNSVSNDYCCFKHALG | 1710252 | Cell/Virus Enclosed | NO |
| YNSVSNEYCCFKHALG | 1710253 | Cell/Virus Enclosed | NO |
| YNSWQNLLYDSNGNII | 1710257 | Ectodomain | YES |
| YNSWQNLLYDSNGNLY | 1710258 | Ectodomain | YES |
| YNTDLIEGSKPYSTLQ | 1710261 | Cell/Virus Enclosed | NO |
| YNTGRSVYVKFQDSKP | 1710264 | Cell/Virus Enclosed | NO |
| YNTPCFSQQGVF | 1536768 | Ectodomain | YES |
| YNVRNDGTVEFCYLHQ | 1710272 | Cell/Virus Enclosed | NO |
| YNVSLSKFNSCKSDIL | 1710277 | Ectodomain | YES |
| YNVSVMQMDIQG | 1536785 | Cell/Virus Enclosed | NO |
| YNVTQAFGRRGPEQTQGNFG | 2060918 | Capsid | UN |
| YNYDFPEYGAMKPGAFGD | 558349 | Undetermined | UN |

| | | | |
|---|---|---|---|
| YNYGAGIKLPSGIMLN | 1710291 | Cell/Virus Enclosed | NO |
| YNYVVSSAFWLF | 1536829 | Cell/Virus Enclosed | NO |
| YNYYKYNLPTMVDIKQ | 1710299 | Cell/Virus Enclosed | NO |
| YPACINGHAVPT | 1536832 | Cell/Virus Enclosed | NO |
| YPACTNGHALPT | 1536834 | Cell/Virus Enclosed | NO |
| YPACTNGHAVPT | 1536835 | Cell/Virus Enclosed | NO |
| YPAFMLGSSVGN | 1536838 | Ectodomain | YES |
| YPAGVNITLANFNETK | 1710303 | Ectodomain | YES |
| YPCMVGPDYAYFNGAI | 1710310 | Cell/Virus Enclosed | NO |
| YPDYVDVNKTLQ | 1536856 | Ectodomain | YES |
| YPEPINENNVVV | 1536859 | Ectodomain | YES |
| YPEPITGNNVVV | 1536862 | Ectodomain | YES |
| YPEWMDDFEADP | 1536863 | Cell/Virus Enclosed | NO |
| YPFASPQECNNM | 1536864 | Cell/Virus Enclosed | NO |
| YPFIANEKFINGCGRN | 1710323 | Cell/Virus Enclosed | NO |
| YPFRLPIKGVPIELQ | 75287 | Ectodomain | YES |
| YPHMFITREEAI | 1536881 | Cell/Virus Enclosed | NO |
| YPHSEGRKSRRPLTTFGSGS | 742228 | Underneath Membrane | NO |
| YPIIGDELKINS | 1536896 | Cell/Virus Enclosed | NO |
| YPIISNEVSINTSCRL | 1710339 | Cell/Virus Enclosed | NO |
| YPILSCVYAICY | 1536904 | Cell/Virus Enclosed | NO |
| YPKCDRVLPNMI | 1536913 | Cell/Virus Enclosed | NO |
| YPKQSIFTINSVVQKD | 1710351 | Cell/Virus Enclosed | NO |
| YPLDCIKDFLAR | 1536921 | Cell/Virus Enclosed | NO |
| YPLSMKSDLSVS | 1536934 | Ectodomain | YES |
| YPLVYHENEEYQKVFR | 1710367 | Cell/Virus Enclosed | NO |
| YPMIANENAINKGGRT | 1710369 | Cell/Virus Enclosed | NO |
| YPNNAIVCRFDT | 1536956 | Cell/Virus Enclosed | NO |
| YPNTICHPNLGN | 1536958 | Ectodomain | YES |
| YPNVPIEYQATR | 1536959 | Ectodomain | YES |
| YPPDDEDECDEY | 1536961 | Cell/Virus Enclosed | NO |
| YPQTICHPNLGN | 1536976 | Ectodomain | YES |
| YPRLQAASDWKPGYSM | 1710390 | Cell/Virus Enclosed | NO |
| YPRLQASADWKPGHAM | 1710391 | Cell/Virus Enclosed | NO |
| YPSNHIDVVSAY | 1536992 | Ectodomain | YES |
| YPSNHIEVASAY | 1536993 | Ectodomain | YES |
| YPSNQIEVVSAY | 1536995 | Ectodomain | YES |
| YPTGSCLMAFLV | 1537010 | Cell/Virus Enclosed | NO |
| YPTSSSTYRNMA | 1537017 | Ectodomain | YES |
| YPVSGIREFSNL | 1537028 | Ectodomain | YES |
| YPYGRCPTAADINIPD | 1710411 | Ectodomain | YES |
| YPYISHENKLNS | 1537034 | Cell/Virus Enclosed | NO |
| YQCGHYTHVTVK | 1537049 | Cell/Virus Enclosed | NO |
| YQDYMQIAPVPAEVLN | 1710423 | Cell/Virus Enclosed | NO |
| YQEFNDLAHTSL | 1537068 | Cell/Virus Enclosed | NO |
| YQEFNDLVHTSL | 1537070 | Cell/Virus Enclosed | NO |
| YQEFVIKTSHDD | 1537071 | Underneath Membrane | NO |
| YQGDHGDMYVYS | 1537086 | Ectodomain | YES |
| YQGDVDTVVNGVDFDF | 1710435 | Cell/Virus Enclosed | NO |
| YQGLPLEFGASA | 1537093 | Cell/Virus Enclosed | NO |
| YQHIGMKRYCTVQGPP | 1710444 | Cell/Virus Enclosed | NO |
| YQHVIKGCKDTA | 1537108 | Cell/Virus Enclosed | NO |
| YQLDENAQKFFD | 1537126 | Cell/Virus Enclosed | NO |
| YQLESTQITTDTGIEN | 1710459 | Secreted | YES |
| YQLESTQITTNT | 1537129 | Secreted | YES |
| YQLIGKQKITTIQGPP | 1710461 | Cell/Virus Enclosed | NO |
| YQLYMIDPAGVC | 1537142 | Cell/Virus Enclosed | NO |
| YQNADTYVFVGSSRYSKKFKPEIAI | 1180945 | Ectodomain | YES |
| YQNGVFMSTAKCWTEE | 1710470 | Cell/Virus Enclosed | NO |
| YQNISTNLPPPLLGNS | 1710471 | Ectodomain | YES |
| YQNNVFMSEAKCWVET | 1710474 | Cell/Virus Enclosed | NO |
| YQNNVFMSESKCWVEN | 1710476 | Cell/Virus Enclosed | NO |
| YQNSSSPALLYR | 1537157 | Ectodomain | YES |
| YQPIDVVRDLPSGFNTL | 75528 | Ectodomain | YES |
| YQPIQQAPTGIT | 1537163 | Underneath Membrane | NO |
| YQPPTASVSTSFLQSG | 1710485 | Cell/Virus Enclosed | NO |
| YQQAVVGAMLSENFTS | 1710489 | Ectodomain | YES |
| YQQQDGMVNMSP | 1537180 | Cell/Virus Enclosed | NO |
| YQQSIIGAMTAVNESR | 1710493 | Ectodomain | YES |
| YQRNNTSFPEWI | 1537182 | Cell/Virus Enclosed | NO |
| YQTNIYSQNLCVDGFG | 1710496 | Cell/Virus Enclosed | NO |
| YQTNLYSFNKAIDGFG | 1710497 | Cell/Virus Enclosed | NO |
| YRAALKSEKHQD | 1537222 | Cell/Virus Enclosed | NO |
| YRAFDIYNERVA | 1537228 | Cell/Virus Enclosed | NO |
| YRAFISVLDITDAAVK | 1710514 | Cell/Virus Enclosed | NO |
| YRCPKEIVDTVSALVY | 1710526 | Cell/Virus Enclosed | NO |

| | | | |
|---|---|---|---|
| YRCPKEIVETVSALVY | 1710527 | Cell/Virus Enclosed | NO |
| YRDLYSSPTTLQ | 1537252 | Cell/Virus Enclosed | NO |
| YREAACSQLAKAMDTF | 1710537 | Cell/Virus Enclosed | NO |
| YRELYSSPTTLQ | 1537261 | Cell/Virus Enclosed | NO |
| YREYLESYNTATTAGF | 1710541 | Cell/Virus Enclosed | NO |
| YREYMEAYNFVS | 1537263 | Cell/Virus Enclosed | NO |
| YRFTNFEPFTVNSVND | 1710545 | Ectodomain | YES |
| YRFVTFEPFNVSFVND | 1710546 | Ectodomain | YES |
| YRIGNYKLNTDHSNSS | 1710558 | Underneath Membrane | NO |
| YRINGLGVTMDVLNKN | 1710560 | Ectodomain | YES |
| YRINGLGVTMDVLSQN | 1710561 | Ectodomain | YES |
| YRINWITGGIAIAMA | 1328509 | Undetermined | UN |
| YRKQLSPLEGGGWLVA | 1710570 | Ectodomain | YES |
| YRLANECAQVLSEYVL | 1710577 | Cell/Virus Enclosed | NO |
| YRLKDCVMYASA | 1537325 | Cell/Virus Enclosed | NO |
| YRLPSSKPSGMDTALL | 1710586 | Underneath Membrane | NO |
| YRLPSTQKGSGMDTAL | 1710587 | Ectodomain | YES |
| YRLPSTQKVSGM | 1537330 | Underneath Membrane | NO |
| YRNSNVGESFVD | 1537351 | Cell/Virus Enclosed | NO |
| YRRATRRIRGGDGKMKDLSPRW | 1862769 | Capsid | UN |
| YRRIPNLPDCNI | 1537371 | Ectodomain | YES |
| YRRKPDPPNCNI | 1537374 | Underneath Membrane | NO |
| YRRQQTSNLVAQ | 1537379 | Cell/Virus Enclosed | NO |
| YRRQQTSNLVVQEFVS | 1710611 | Cell/Virus Enclosed | NO |
| YRSGNGILKWAD | 1537391 | Cell/Virus Enclosed | NO |
| YRSTSPDPKFVDKYYA | 1710621 | Cell/Virus Enclosed | NO |
| YRSTSVEESFID | 1537411 | Cell/Virus Enclosed | NO |
| YRVFPYDMDSGG | 1537419 | Cell/Virus Enclosed | NO |
| YSAGHATGTTPK | 1537456 | Ectodomain | YES |
| YSAGHATGTTPQKLFV | 1710642 | Ectodomain | YES |
| YSAGHATGTTSQ | 1537459 | Ectodomain | YES |
| YSAGHATVTTPQ | 1537460 | Ectodomain | YES |
| YSAGYATGTTPQ | 1537461 | Ectodomain | YES |
| YSALKMVQTHLS | 1537469 | Cell/Virus Enclosed | NO |
| YSANFLDDNVLPETYV | 1710644 | Ectodomain | YES |
| YSASFKKGAKIL | 1537477 | Cell/Virus Enclosed | NO |
| YSCDFKGWIPGN | 1537479 | Cell/Virus Enclosed | NO |
| YSCMVCVDRSSLSSWHVLYTP | 75707 | Ectodomain | YES |
| YSEAAAQGFQAC | 1537502 | Cell/Virus Enclosed | NO |
| YSEAASNGFRAC | 1537503 | Cell/Virus Enclosed | NO |
| YSENVFAVESGGYIPS | 1710668 | Ectodomain | YES |
| YSFDKAIDGFGV | 1537524 | Cell/Virus Enclosed | NO |
| YSFEAHKDFFKD | 1537526 | Cell/Virus Enclosed | NO |
| YSFLPKEDGDVLLAEF | 1710679 | Cell/Virus Enclosed | NO |
| YSFLPKEDGDVV | 1537536 | Cell/Virus Enclosed | NO |
| YSFLPKEEGDVL | 1537537 | Cell/Virus Enclosed | NO |
| YSGDTFWRYDFDITES | 1710685 | Cell/Virus Enclosed | NO |
| YSGFKPNNTSAQYIFS | 1710687 | Cell/Virus Enclosed | NO |
| YSGNANEADYRCACYA | 1710691 | Cell/Virus Enclosed | NO |
| YSGRVSAAFHANSSEP | 1710696 | Ectodomain | YES |
| YSGSASEADYRC | 1537578 | Cell/Virus Enclosed | NO |
| YSGSIRFDSTLP | 1537579 | Cell/Virus Enclosed | NO |
| YSGVVGCKSIVK | 1537582 | Cell/Virus Enclosed | NO |
| YSGYHPAHAPSFLAVD | 1710702 | Cell/Virus Enclosed | NO |
| YSIDVIDNEIIVKPNI | 1710709 | Cell/Virus Enclosed | NO |
| YSILAAKTGVSVEQLL | 1710712 | Cell/Virus Enclosed | NO |
| YSIQDVGFKNQH | 1537615 | Cell/Virus Enclosed | NO |
| YSIQDVGFKNQYCNGS | 1710714 | Cell/Virus Enclosed | NO |
| YSKLDDEEPMEL | 1537628 | Cell/Virus Enclosed | NO |
| YSKLFITKDEAIKRVR | 1710722 | Cell/Virus Enclosed | NO |
| YSKLVKDVDHVQ | 1537632 | Secreted | YES |
| YSKTRRSRRAIT | 1537644 | Ectodomain | YES |
| YSKYVTVQGPPG | 1537646 | Cell/Virus Enclosed | NO |
| YSLADLPAYMSV | 1537647 | Ectodomain | YES |
| YSLADSNGFIRF | 1537648 | Cell/Virus Enclosed | NO |
| YSLDGNFRTGVD | 1537653 | Cell/Virus Enclosed | NO |
| YSLFDMAKFPLKLAGT | 1710733 | Cell/Virus Enclosed | NO |
| YSLKAAVHGWKM | 1537662 | Cell/Virus Enclosed | NO |
| YSLKAAVHKWKM | 1537663 | Cell/Virus Enclosed | NO |
| YSLPAINVTINNYNPS | 1710743 | Ectodomain | YES |
| YSMQDVGFKNQY | 1537691 | Cell/Virus Enclosed | NO |
| YSNASSDVIHYN | 1537694 | Ectodomain | YES |
| YSNESWFEDDYK | 1537696 | Cell/Virus Enclosed | NO |
| YSNFYASGCVLSSACT | 1710756 | Cell/Virus Enclosed | NO |
| YSNSMDCPIVVA | 1537717 | Cell/Virus Enclosed | NO |
| YSNVYRTDYVDYTFVN | 1710767 | Cell/Virus Enclosed | NO |

| | | | |
|---|---|---|---|
| YSPCVSIVPSTA | 1537731 | Ectodomain | YES |
| YSPCVSIVPSTVWEDG | 1710769 | Ectodomain | YES |
| YSPEPITSLNTK | 1537734 | Ectodomain | YES |
| YSPIDIPRGLPV | 1537736 | Ectodomain | YES |
| YSPLLVDSDQPW | 1537738 | Cell/Virus Enclosed | NO |
| YSPQIITTDNTF | 1537745 | Ectodomain | YES |
| YSPSRSFSYFYPFRL | 75863 | Ectodomain | YES |
| YSQMRPHVRYDLYDGN | 1710782 | Cell/Virus Enclosed | NO |
| YSQNLCVDGFGVNKIQ | 1710783 | Cell/Virus Enclosed | NO |
| YSQVLSKSILLV | 1537768 | Cell/Virus Enclosed | NO |
| YSQYYLNKDTGV | 1537770 | Ectodomain | YES |
| YSRPVVSANGEPTVKLYTSVENAQQ | 75883 | Undetermined | UN |
| YSRSTRSMLKRRDSTY | 1710793 | Ectodomain | YES |
| YSRVISRMGFKLDATV | 1710795 | Cell/Virus Enclosed | NO |
| YSRVKNLNSSQG | 1537793 | Ectodomain | YES |
| YSRYCFSVNNTF | 1537798 | Ectodomain | YES |
| YSRYRIGNYKLNTDHS | 1391666 | Underneath Membrane | NO |
| YSSCVECDAKFKNSVA | 1710803 | Cell/Virus Enclosed | NO |
| YSSCVECEAKFK | 1537804 | Cell/Virus Enclosed | NO |
| YSSLCDEFTLAEVVKQ | 1710810 | Cell/Virus Enclosed | NO |
| YSSNEEQAVLKFLDGL | 1710814 | Cell/Virus Enclosed | NO |
| YSSNEEQAVRKF | 1537819 | Cell/Virus Enclosed | NO |
| YSSSANGILVGT | 1537830 | Cell/Virus Enclosed | NO |
| YSSSANGRLVGT | 1537831 | Cell/Virus Enclosed | NO |
| YSSSANGSLDGT | 1537832 | Cell/Virus Enclosed | NO |
| YSSSANGSLVCT | 1537833 | Cell/Virus Enclosed | NO |
| YSTGIDFQDELD | 1537845 | Ectodomain | YES |
| YSTGNNVFQTLA | 1537846 | Ectodomain | YES |
| YSTGNNVFQTQAGCLI | 1710822 | Ectodomain | YES |
| YSTPWRYLDFNALNLFFSPL | 75905 | Capsid | UN |
| YSTSPIQPVELP | 1537860 | Ectodomain | YES |
| YSTTPCELHGNG | 1537865 | Cell/Virus Enclosed | NO |
| YSTTPCEVHGNGSMIE | 1710830 | Cell/Virus Enclosed | NO |
| YSTTPCGVHGNG | 1537867 | Cell/Virus Enclosed | NO |
| YSTVASSLVLVVSLGAISFWMCSNG | 1181119 | Undetermined | UN |
| YSVILSTCDGLKFTEE | 1710835 | Cell/Virus Enclosed | NO |
| YSVLETFQNNVVNYQH | 1710838 | Cell/Virus Enclosed | NO |
| YSVLSVDSVSEESQGN | 1710839 | Cell/Virus Enclosed | NO |
| YSVLYNSTFFSTFKCYG | 75920 | Ectodomain | YES |
| YSVSSFEAIPSG | 1537894 | Ectodomain | YES |
| YSVSSFEAKPSGSVVE | 1710848 | Ectodomain | YES |
| YSWADLPAYMTV | 1537910 | Ectodomain | YES |
| YSWFSGITQFQK | 1537911 | Cell/Virus Enclosed | NO |
| YSYFNKCSRLLS | 1537919 | Ectodomain | YES |
| YSYLMPVLSMTDCLAA | 1710863 | Cell/Virus Enclosed | NO |
| YTAEQCIRGAIG | 1537951 | Cell/Virus Enclosed | NO |
| YTAEQCIRGAMGDVGL | 1710872 | Cell/Virus Enclosed | NO |
| YTANDVEVQSSGMIKP | 1710877 | Cell/Virus Enclosed | NO |
| YTARDALDTILQ | 1537964 | Cell/Virus Enclosed | NO |
| YTARVVYTAASHAAVD | 1710878 | Cell/Virus Enclosed | NO |
| YTAVNLPRGFPA | 1537965 | Ectodomain | YES |
| YTCIVCIDRASLSTWHVLYSP | 75984 | Ectodomain | YES |
| YTCSITFEMSCDCGKK | 1710882 | Cell/Virus Enclosed | NO |
| YTDFAEDVCTCYDNSI | 1710885 | Cell/Virus Enclosed | NO |
| YTDGLMQNASLY | 1537979 | Cell/Virus Enclosed | NO |
| YTDGVMKNASLY | 1537980 | Cell/Virus Enclosed | NO |
| YTDIDVNPALNI | 1537983 | Cell/Virus Enclosed | NO |
| YTDIDVNSALNM | 1537985 | Cell/Virus Enclosed | NO |
| YTDLKFIDKLNV | 1537990 | Cell/Virus Enclosed | NO |
| YTDQYLCGADGKPVMS | 1710892 | Cell/Virus Enclosed | NO |
| YTEIQPKLDEYY | 1538007 | Cell/Virus Enclosed | NO |
| YTESAVNFTKAL | 1538015 | Cell/Virus Enclosed | NO |
| YTESAVNSTKALAKQD | 1710904 | Cell/Virus Enclosed | NO |
| YTEVRGSRRLAQ | 1538018 | Ectodomain | YES |
| YTFTTVKPGAAF | 1538030 | Cell/Virus Enclosed | NO |
| YTFYGKESFENP | 1538036 | Cell/Virus Enclosed | NO |
| YTFYGKESLENP | 1538038 | Cell/Virus Enclosed | NO |
| YTGAIKLDDKDPSFKD | 1710916 | Cell/Virus Enclosed | NO |
| YTGFQGKQILGS | 1538046 | Cell/Virus Enclosed | NO |
| YTGHQELVIKTS | 1538047 | Underneath Membrane | NO |
| YTGPVSSPSWKFHSDV | 76065 | Ectodomain | YES |
| YTGQAGEMPCAIINDK | 1710924 | Cell/Virus Enclosed | NO |
| YTGSLADGLEAYADKT | 1710926 | Cell/Virus Enclosed | NO |
| YTGSSFYAPEPITSLN | 1710930 | Ectodomain | YES |
| YTGSSFYSPEPI | 1538062 | Ectodomain | YES |
| YTGYQEFVIKTS | 1538070 | Underneath Membrane | NO |

| | | | |
|---|---|---|---|
| YTGYQESVIKTS | 1538072 | Underneath Membrane | NO |
| YTIKFSGFKLDGFNHQ | 1710941 | Cell/Virus Enclosed | NO |
| YTILPCYSGRVS | 1538087 | Ectodomain | YES |
| YTKAPDVMLNIS | 1538097 | Ectodomain | YES |
| YTKIFKDDGNFV | 1538108 | Cell/Virus Enclosed | NO |
| YTLANTTTAPLK | 1538129 | Cell/Virus Enclosed | NO |
| YTLANTTTEPLKLFAA | 1710960 | Cell/Virus Enclosed | NO |
| YTLCYDIGNPRG | 1538133 | Cell/Virus Enclosed | NO |
| YTLEEAKTAFKK | 1538135 | Cell/Virus Enclosed | NO |
| YTLEEAKTALRK | 1538137 | Cell/Virus Enclosed | NO |
| YTLEEARTALKK | 1538138 | Cell/Virus Enclosed | NO |
| YTLSDLPVYVTVAKVQ | 1391685 | Underneath Membrane | NO |
| YTMCEYPNTICH | 1538156 | Ectodomain | YES |
| YTMLDLCYALRHFDRN | 1710968 | Cell/Virus Enclosed | NO |
| YTMMDFVYALRH | 1538160 | Cell/Virus Enclosed | NO |
| YTMMDLCFALRNFDEK | 1391688 | Cell/Virus Enclosed | NO |
| YTMMDLVYALRHFDQN | 1391689 | Cell/Virus Enclosed | NO |
| YTMMDLVYAMRNFDEQ | 1391690 | Cell/Virus Enclosed | NO |
| YTMSLGADSSIAYSNN | 1710969 | Ectodomain | YES |
| YTNFRIDVPFAE | 1538173 | Cell/Virus Enclosed | NO |
| YTNIPVSILFNKEEADDNGD | 76132 | Cell/Virus Enclosed | NO |
| YTNSCLVSSDGQPGGD | 1710978 | Cell/Virus Enclosed | NO |
| YTNSCLVSSNGQ | 1538189 | Cell/Virus Enclosed | NO |
| YTPATDCSDGNY | 1538194 | Ectodomain | YES |
| YTPFHYERDNTSCPEW | 1710982 | Cell/Virus Enclosed | NO |
| YTPGHYAGSRSSSGNR | 1710983 | Capsid | UN |
| YTPIDEIPYKSF | 1538206 | Cell/Virus Enclosed | NO |
| YTPLVGTSLHSV | 1538211 | Cell/Virus Enclosed | NO |
| YTPLVGTSVHPV | 1538212 | Cell/Virus Enclosed | NO |
| YTPVHYERDNTS | 1538221 | Cell/Virus Enclosed | NO |
| YTQCAVIAVVAAVLNS | 1710994 | Cell/Virus Enclosed | NO |
| YTQGRNLQDFLPRSTM | 1710995 | Cell/Virus Enclosed | NO |
| YTQGRNLSDFTPRSDM | 1710996 | Cell/Virus Enclosed | NO |
| YTQLCQYLNSTTMCVP | 1710999 | Cell/Virus Enclosed | NO |
| YTSACKTIEDALRNSA | 1711008 | Ectodomain | YES |
| YTSKDTLDDILQQANK | 1711013 | Cell/Virus Enclosed | NO |
| YTSSSLGSIAGA | 1538280 | Ectodomain | YES |
| YTSTSGNLLGFKDVTK | 1711025 | Ectodomain | YES |
| YTTEPEFVKVLG | 1538288 | Cell/Virus Enclosed | NO |
| YTTSEYPSQPPS | 76192 | Secreted | YES |
| YTTTIKPVSLEH | 1538298 | Cell/Virus Enclosed | NO |
| YTVCDDCNGYTDNIFS | 1711044 | Ectodomain | YES |
| YTVPDLVVEQYNQTIL | 1711052 | Ectodomain | YES |
| YTVQKLQSLIDN | 1538325 | Ectodomain | YES |
| YTVQKLQTLIDNINST | 1711053 | Ectodomain | YES |
| YTVSKGTQQSSW | 1538331 | Ectodomain | YES |
| YTVYDTAKKSMYDGDR | 1711059 | Cell/Virus Enclosed | NO |
| YTWTADEAIKFL | 1538340 | Ectodomain | YES |
| YTYKMLVAKDNKNLPK | 1711063 | Capsid | UN |
| YTYSFLPKEDGDVLLA | 1711065 | Cell/Virus Enclosed | NO |
| YTYSFLPKEEGD | 1538351 | Cell/Virus Enclosed | NO |
| YVAALAVNVCII | 1538363 | Cell/Virus Enclosed | NO |
| YVAGYKVLPPLMDVNM | 1711072 | Ectodomain | YES |
| YVAKRVLGVQTQTVDS | 1711074 | Cell/Virus Enclosed | NO |
| YVANPTEDLCNG | 1538381 | Cell/Virus Enclosed | NO |
| YVAPQVTYQNISTNLP | 1711080 | Ectodomain | YES |
| YVAPVNTVKPKPVINQ | 1711081 | Cell/Virus Enclosed | NO |
| YVARLMKGDKGDAEDT | 1711083 | Cell/Virus Enclosed | NO |
| YVARLVKGDKGD | 1538389 | Cell/Virus Enclosed | NO |
| YVASRFLGLQIQTVDS | 1711084 | Cell/Virus Enclosed | NO |
| YVCKPHQVNISLNGNT | 1711090 | Ectodomain | YES |
| YVCNAPGCDVNDVTKL | 1711093 | Cell/Virus Enclosed | NO |
| YVCNASGCGVSDVKKL | 1711097 | Cell/Virus Enclosed | NO |
| YVCNGNPRCKNL | 1538415 | Ectodomain | YES |
| YVCNGNVRCVELLKQY | 1711101 | Ectodomain | YES |
| YVCNTSGCNVNDVTKL | 1711102 | Cell/Virus Enclosed | NO |
| YVCQCGGERHQQ | 1538417 | Cell/Virus Enclosed | NO |
| YVDFKPQSGGGKCFNC | 1711106 | Ectodomain | YES |
| YVDFKPQSGGGR | 1538425 | Ectodomain | YES |
| YVDGHRLASGVQVHNL | 1711107 | Underneath Membrane | NO |
| YVDGLESKQVDYVPLR | 1711108 | Cell/Virus Enclosed | NO |
| YVDKWDDFCRQYSNES | 1711109 | Cell/Virus Enclosed | NO |
| YVDNTLYSEFST | 1538435 | Undetermined | UN |
| YVDQYGCDYTGG | 1538437 | Cell/Virus Enclosed | NO |
| YVDQYGCDYTGSLADG | 1711117 | Cell/Virus Enclosed | NO |
| YVDVGPDSAKSV | 1538459 | Ectodomain | YES |

| | | | |
|---|---|---|---|
| YVDVGPDSFKSA | 1538460 | Ectodomain | YES |
| YVDVGPDSVKPA | 1538463 | Ectodomain | YES |
| YVDVNKTLQEFAQNLP | 1711127 | Ectodomain | YES |
| YVDYTFVNEYYE | 1538469 | Cell/Virus Enclosed | NO |
| YVFNNSLTRQLR | 1538497 | Ectodomain | YES |
| YVFSTVNALPEVNADI | 1711144 | Cell/Virus Enclosed | NO |
| YVGDPAQLPTPR | 1538514 | Cell/Virus Enclosed | NO |
| YVGKYDINHGDSGFIS | 1711163 | Undetermined | UN |
| YVGMQTVEDLRA | 1538518 | Cell/Virus Enclosed | NO |
| YVGSLSQNHHTF | 1538523 | Cell/Virus Enclosed | NO |
| YVGTGIIRSSNQSLAG | 1711168 | Ectodomain | YES |
| YVGVQTIEDLRA | 1538527 | Cell/Virus Enclosed | NO |
| YVGVQTVEDLCA | 1538529 | Cell/Virus Enclosed | NO |
| YVGVQTVEDLRARMTY | 1711169 | Cell/Virus Enclosed | NO |
| YVGVYDTNHGDA | 1538532 | Ectodomain | YES |
| YVHARLKCGLIL | 1538535 | Cell/Virus Enclosed | NO |
| YVHARLKGGLILKFDS | 1711173 | Cell/Virus Enclosed | NO |
| YVHKWFQFCRLY | 1538543 | Cell/Virus Enclosed | NO |
| YVHLPYPDPSRI | 1538545 | Cell/Virus Enclosed | NO |
| YVKDGKYFISEP | 1538579 | Cell/Virus Enclosed | NO |
| YVKDGKYFTSEPPVTY | 1711198 | Cell/Virus Enclosed | NO |
| YVKPGGTGSGDA | 1538596 | Cell/Virus Enclosed | NO |
| YVKPGGTSSGDATTAF | 1711204 | Cell/Virus Enclosed | NO |
| YVKPGRTSSGDA | 1538599 | Cell/Virus Enclosed | NO |
| YVKSKVGNYRLPSTQK | 1711210 | Ectodomain | YES |
| YVLLDWVKHLNKNLNE | 1711226 | Cell/Virus Enclosed | NO |
| YVLNNISLTTQPYFDS | 1711230 | Ectodomain | YES |
| YVLPLTCNAISSNTDN | 1711231 | Ectodomain | YES |
| YVLPSEAPNAKE | 1538633 | Cell/Virus Enclosed | NO |
| YVLPSETPNAKE | 1538634 | Cell/Virus Enclosed | NO |
| YVLTGDSVKFVYMHQL | 1711243 | Cell/Virus Enclosed | NO |
| YVMLNTSIPNLPDFKE | 1711251 | Ectodomain | YES |
| YVMLNTSTSNLP | 1538650 | Ectodomain | YES |
| YVMPLTCIRRSK | 1538657 | Ectodomain | YES |
| YVMPLTCNAISP | 1538660 | Ectodomain | YES |
| YVNANGGTCFCN | 1538671 | Cell/Virus Enclosed | NO |
| YVNATGSDVNCN | 1538673 | Ectodomain | YES |
| YVNFEHQRGPGK | 1538675 | Ectodomain | YES |
| YVNNWCRCSCLP | 1538689 | Ectodomain | YES |
| YVNSNQNTDSDSLSSK | 1711267 | Capsid | UN |
| YVNSSNVTTMDF | 1538692 | Cell/Virus Enclosed | NO |
| YVNVYRSTSPDPKFVD | 1711271 | Cell/Virus Enclosed | NO |
| YVPEHCQNYSGC | 1538702 | Cell/Virus Enclosed | NO |
| YVPEHCQNYSGS | 1538704 | Cell/Virus Enclosed | NO |
| YVPSQEKNFTTA | 1538724 | Ectodomain | YES |
| YVPSYVKPDSVSTSDL | 1711287 | Cell/Virus Enclosed | NO |
| YVPTKYVTARVS | 1538732 | Ectodomain | YES |
| YVQQGIFNKATVAQIK | 1711293 | Cell/Virus Enclosed | NO |
| YVRDYVDKWDDFCRQY | 1711296 | Cell/Virus Enclosed | NO |
| YVRKNGQFQDHYDGFY | 1711301 | Cell/Virus Enclosed | NO |
| YVRPIIEDYHTLTVTI | 1711303 | Ectodomain | YES |
| YVRVKHGDFSAVSSPM | 1711304 | Underneath Membrane | NO |
| YVSHGLDLMQAG | 1538769 | Cell/Virus Enclosed | NO |
| YVSNSGNLLGFKNVST | 1711315 | Ectodomain | YES |
| YVTHGFNLEDAA | 1538806 | Cell/Virus Enclosed | NO |
| YVTNEIGLNASVTLKI | 1711330 | Ectodomain | YES |
| YVTTKPGMKYVKWEHD | 1711332 | Cell/Virus Enclosed | NO |
| YVTVQGPPGTGK | 1538819 | Cell/Virus Enclosed | NO |
| YVTYAGNVWQIQTIQD | 1711337 | Cell/Virus Enclosed | NO |
| YVTYAGSVWHIQTVQD | 1711338 | Cell/Virus Enclosed | NO |
| YVVDDIYYPASCNGVL | 1391749 | Cell/Virus Enclosed | NO |
| YVVDSVAVKNGA | 1538825 | Cell/Virus Enclosed | NO |
| YVVDVSHGLGTY | 1538827 | Ectodomain | YES |
| YVVKNSTAFPFETLSA | 1711344 | Cell/Virus Enclosed | NO |
| YVVKNSTTFPFE | 1538834 | Cell/Virus Enclosed | NO |
| YVVVVESDADRANFLN | 1711351 | Cell/Virus Enclosed | NO |
| YVVYDAANNAVYDGAR | 1711353 | Cell/Virus Enclosed | NO |
| YVYDVHKDNFKDGLCM | 1711364 | Cell/Virus Enclosed | NO |
| YVYIGDPVQLPA | 1538868 | Cell/Virus Enclosed | NO |
| YVYKGYQPIDVVRDLPS | 76446 | Ectodomain | YES |
| YVYNGSAQAISF | 1538872 | Ectodomain | YES |
| YVYNGSAQVTSF | 1538874 | Ectodomain | YES |
| YVYSAGHATGTTPQKL | 1711378 | Ectodomain | YES |
| YWDQTYHANCVNCTDD | 1711382 | Cell/Virus Enclosed | NO |
| YWGQDYHPDCVDCHDE | 1711395 | Cell/Virus Enclosed | NO |
| YWIGYGCNCDSLRQAA | 1711396 | Cell/Virus Enclosed | NO |

| | | | |
|---|---|---|---|
| YWRRQDRKINTGNGIK | 1711404 | Cell/Virus Enclosed | NO |
| YWSPQGNLGGNCAGNV | 1711406 | Cell/Virus Enclosed | NO |
| YWVASSQADVNT | 1538938 | Capsid | UN |
| YYADVGMPTTFL | 1538959 | Ectodomain | YES |
| YYCVTTKAKSIQ | 1538988 | Cell/Virus Enclosed | NO |
| YYCVTTKAKSVQQDLD | 1711434 | Cell/Virus Enclosed | NO |
| YYCYTSKDTLDD | 1538993 | Cell/Virus Enclosed | NO |
| YYDDGSCEVVHDQVNY | 1711437 | Cell/Virus Enclosed | NO |
| YYDDSQYYFNKDTGVI | 1711438 | Ectodomain | YES |
| YYDDSQYYLNKD | 1538996 | Ectodomain | YES |
| YYDHSQYYFNKD | 1538999 | Ectodomain | YES |
| YYDHSQYYLNKD | 1539000 | Ectodomain | YES |
| YYDYSQYYLNKD | 1539008 | Ectodomain | YES |
| YYEFLNKHFNMM | 1539014 | Cell/Virus Enclosed | NO |
| YYGVFLYDNFTS | 1539037 | Undetermined | UN |
| YYIYVTNEIGLNASVT | 1711474 | Ectodomain | YES |
| YYKGVTTHESSSAVNM | 1711475 | Cell/Virus Enclosed | NO |
| YYKVEADFYLSG | 1539085 | Ectodomain | YES |
| YYKVNCCRFQRIDDDG | 1711485 | Cell/Virus Enclosed | NO |
| YYKWPWYIWLGF | 1539087 | Undetermined | UN |
| YYKYDNGNFIKLPEVI | 1711487 | Cell/Virus Enclosed | NO |
| YYLDDVKKIEYKPDLS | 1711490 | Cell/Virus Enclosed | NO |
| YYLGTGPEAGLPYGANKDGI | 1392539 | Capsid | UN |
| YYLGTGPHKDLK | 1539105 | Cell/Virus Enclosed | NO |
| YYLGTGPYANASYGES | 1711498 | Capsid | UN |
| YYLPVMAAPTGI | 1539116 | Transmembrane | NO |
| YYLPVMAAPTGV | 1539117 | Transmembrane | NO |
| YYNGIMVLPGAA | 1539122 | Ectodomain | YES |
| YYNGIMVVPGVA | 1539124 | Ectodomain | YES |
| YYNIDTGSFYGF | 1539126 | Underneath Membrane | NO |
| YYNNSNNGKIVYAILS | 1711513 | Cell/Virus Enclosed | NO |
| YYNNWQNLLYDS | 1539143 | Ectodomain | YES |
| YYNSWQNLLYDSNGNI | 1711514 | Ectodomain | YES |
| YYNSWQNLLYDSNGNL | 1711515 | Ectodomain | YES |
| YYPEPISDKNVVFMNT | 1711523 | Ectodomain | YES |
| YYPEPITENNVVVMST | 1711524 | Ectodomain | YES |
| YYPEPITGNNVV | 1539158 | Ectodomain | YES |
| YYPGTGPHKDLK | 1539159 | Cell/Virus Enclosed | NO |
| YYQLESTQITTDTGIE | 1711538 | Secreted | YES |
| YYQNGVFMSTAKCWTE | 1711542 | Cell/Virus Enclosed | NO |
| YYQNNVFMSEAKCWVE | 1711544 | Cell/Virus Enclosed | NO |
| YYQNNVFMSESKCWVE | 1711545 | Cell/Virus Enclosed | NO |
| YYQPIQQAPTGI | 1539183 | Underneath Membrane | NO |
| YYRITSRVMFEP | 1539187 | Ectodomain | YES |
| YYSFGRCPTSSI | 1539212 | Ectodomain | YES |
| YYSGNANEADYRCACY | 1711568 | Cell/Virus Enclosed | NO |
| YYSLPAINVTINNYNP | 1711573 | Ectodomain | YES |
| YYSTTPCELHGN | 1539237 | Cell/Virus Enclosed | NO |
| YYSTTPCEVHGNGSMI | 1711578 | Cell/Virus Enclosed | NO |
| YYSYMMPVMGMT | 1539245 | Cell/Virus Enclosed | NO |
| YYVEGSGRSASN | 1539269 | Cell/Virus Enclosed | NO |
| YYVKPGGTSSGDATTA | 1711600 | Cell/Virus Enclosed | NO |
| YYVKPGGTSSRD | 1539275 | Cell/Virus Enclosed | NO |
| YYVKWPWYVWLL | 1539277 | Undetermined | UN |
| YYVLHLCLAATKYPLLKLLG | 110468 | Cell/Virus Enclosed | NO |
| YYYKVEADFYLS | 1539305 | Ectodomain | YES |
| YYYNMATGFAYG | 1539308 | Ectodomain | YES |
| YYYPEPIIENNV | 1539310 | Ectodomain | YES |

UD: Undetermined visibility
