## Supplementary Dataset 3 for "Analysis of virus-specific B cell epitopes reveals extensive antigen processing prior to recognition"

**Supplementary Dataset 3.** Virus-specific linear B cell epitopes mapping in ectodomains of viral envelope proteins with available tertiary structures

| B cell epitope | IEDB | PDB | eRSA (%) |
|---|---|---|---|
| AASFNKAMTNIVDAFT | 1643154 | 5ZUV | 29.88 |
| ADLIIERREGSDVCYPGKFVNEEAL | 1126060 | 6IDZ | 23.27 |
| AGGQLFYSRPVVSANGEPTVKLYTS | 1533 | 2ZZQ | 19.87 |
| AINVTINNYNPSSWNR | 1644448 | 5GNB | 27.5 |
| AITQTSQALQTVATAL | 1644512 | 5ZUV | 14.22 |
| AKWAVPTTRTDDKLR | 2322 | 3INB | 19.97 |
| ALGVINTLEWIPRFK | 2598 | 3INB | 21.95 |
| ALNKIQDVVNQQGNSL | 1645025 | 5ZUV | 19.41 |
| ALQTVATALNKIQDVV | 1645078 | 5ZUV | 19.05 |
| ANCASILCKCYTTGT | 3236 | 5YZD | 34.13 |
| APAHDTPPVPDVDSRGAILRRQYNL | 3476 | 2ZZQ | 69.43 |
| APLSPLLPLQDGTNTHIMATEASNY | 3648 | 2ZZQ | 18.76 |
| ARATIRYRPLVPNAVGGYAISISFW | 4121 | 2ZZQ | 24 |
| ARSLDWTKVTLDGRPLSTIQQYSKT | 4260 | 2ZZQ | 44.59 |
| ASCPIGTNYRSCESTT | 1646072 | 5GNB | 45.31 |
| ATALNKIQDVVNQQGN | 1646370 | 5ZUV | 22.35 |
| AVSNGTKVNTLTERGVEVVNATETV | 1130264 | 6IDZ | 36 |
| CDIDKWLNNFNVPSPL | 1647541 | 5GNB | 21.08 |
| CKSHKPPSASCPIGTN | 1648203 | 5GNB | 32.84 |
| CLPDPITAYDPRSCSQ | 1648351 | 5GNB | 41.04 |
| CNHINNLKIKNFDYLD | 1648488 | 6Y3Y | 41.5 |
| CPIGTNYRSCESTTVL | 1648624 | 5GNB | 51.87 |
| CSNDLLQPNTEVFTDV | 1648866 | 5GNB | 47.78 |
| CYPGKFVNEEALRQILRESGGIDKE | 1130598 | 6IDZ | 19.12 |
| DAITQTSQALQTVATA | 1649573 | 5ZUV | 17.41 |
| DCDIDKWLNNFNVPSP | 1649748 | 5GNB | 25.62 |
| DCHAPTYLPAEVDGD | 7704 | 3INB | 39.95 |
| DCNHINNLKIKNFDYL | 1649807 | 6Y3Y | 41.86 |
| DDKLRMETCFQQACK | 7812 | 3INB | 35.83 |
| DFALELEFRNLTPGNTNTRVSRYSS | 8233 | 2ZZQ | 23.63 |
| DHCPVVEVNGVTIQV | 8530 | 5YZD | 37.73 |
| DHTDWCRCSCLPDPITAYDP | 2060803 | 5GNB | 30.34 |
| DIDKWLNNFNVPSPLN | 1650841 | 5GNB | 21.11 |
| DILSLNNPIFINYSKE | 1650928 | 6Y3Y | 34.64 |
| DIPYNVSLSKFNSCKS | 1650965 | 6Y3Y | 52.14 |
| DKGIAIPHDIDLGESRVVIQDYDNQ | 8875 | 2ZZQ | 21.77 |
| DKWLNNFNVPSPLNWE | 1651244 | 5GNB | 19.88 |
| DLQLGSSGFLQSSNYK | 1651479 | 5GNB | 28.09 |
| DLSNCMVALGELKLA | 9281 | 3INB | 20.31 |
| DPITAYDPRSCSQKKS | 1651915 | 5GNB | 43.59 |
| DPRSCSQKKSLVGVGE | 1651955 | 5GNB | 40.1 |
| DPVIDRLYLSSHRGV | 9804 | 3INB | 19.55 |
| DSFSCNNFDESKIYGS | 1652214 | 5GNB | 35.45 |
| DSNGNIIGFKDFVTNK | 1652303 | 5GNB | 41.36 |
| DSRGAILRRQYNLSTSPLTSSVATG | 10221 | 2ZZQ | 41.4 |
| DSRSDCNHINNLKIKN | 1652323 | 6Y3Y | 36.42 |
| DVVNQQGNSLNHLTSQ | 1652926 | 5ZUV | 23.4 |
| DYLDIHPSLCNNGKIS | 1653120 | 6Y3Y | 32.44 |
| EGDCYHSGGTIISNLPFQNIDSRAV | 1136912 | 6IDZ | 19.81 |
| EGDQIIFYEGVNFNPY | 1654020 | 6Y3Y | 6.96 |
| ELKLAALCHGEDSIT | 13126 | 3INB | 25.45 |
| EMKWLLSNTDNAAFPQMTKSYKNTR | 1138304 | 6IDZ | 33.1 |
| ENQKILAASFNKAMTN | 1654952 | 5ZUV | 38.01 |
| ESKIYGSCFKSIVLDK | 1655374 | 5GNB | 22.67 |
| ESTTVLDHTDWCRCSC | 1655443 | 5GNB | 49.63 |
| FDYLDIHPSLCNNGKI | 1656569 | 6Y3Y | 35.04 |
| FGDSRSDCNHINNLKI | 1656911 | 6Y3Y | 30.35 |
| FLFGDSRSDCNHINNL | 1657695 | 6Y3Y | 20.71 |
| FLQSSNYKIDTTSSSC | 1657862 | 5GNB | 18.72 |
| FNSCKSDILSLNNPIF | 1658214 | 6Y3Y | 33.03 |
| FPQMTKSYKNTRKSPALIVWGIHHS | 1141159 | 6IDZ | 24.02 |
| FSCNNFDESKIYGSCF | 1658601 | 5GNB | 27.89 |
| FSVNNTFCPCAKPSFA | 1658840 | 5GNB | 25.44 |
| GFGVDEEKCGVLDGSY | 1660684 | 5GNB | 25.84 |
| GFKDFVTNKTYNIFPC | 1660708 | 5GNB | 32.08 |
| GFLQSSNYKIDTTSSS | 1660736 | 5GNB | 20.63 |
| GINSGTTCSNDLLQPN | 1661381 | 5GNB | 17.58 |
| GNIIGFKDFVTNKTYN | 1662124 | 5GNB | 31.26 |
| GPVYVSDSVTLVNVATGAQAVARSL | 21834 | 2ZZQ | 29.9 |
| GSSGFLQSSNYKIDTT | 1662861 | 5GNB | 29.55 |
| GTTCSNDLLQPNTEVF | 1663199 | 5GNB | 37.96 |
| GVLDGSYNVSCLCSTD | 1663451 | 5GNB | 44.48 |
| GVNDAITQTSQALQTV | 1663503 | 5ZUV | 15.89 |
| GVNFNPYHRFKCFPNG | 1663507 | 6Y3Y | 28.56 |

| | | | |
|---|---|---|---|
| GYNVSSIVTMTSQGM | 23422 | 3INB | 29.33 |
| HKPPSASCPIGTNYRS | 1664569 | 5GNB | 32.88 |
| HPSLCNNGKISSSAGD | 1664829 | 6Y3Y | 33.83 |
| IAPDRASFLRGKSMGIQSGVQVDAN | 1147379 | 6IDZ | 27.92 |
| IGLDPVAPGDLTMVI | 227266 | 4LDI | 56.03 |
| IGTNYRSCESTTVLDH | 1666390 | 5GNB | 57.28 |
| ILAASFNKAMTNIVDA | 1666954 | 5ZUV | 31.83 |
| ILSLNNPIFINYSKEV | 1667197 | 6Y3Y | 29.31 |
| IPHDIDLGESRVVIQDYDNQHEQDR | 27861 | 2ZZQ | 17.5 |
| IPYNVSLSKFNSCKSD | 1667718 | 6Y3Y | 54.21 |
| IQDVVNQQGNSLNHLT | 1667736 | 5ZUV | 20.59 |
| ITQTSQALQTVATALN | 1668449 | 5ZUV | 16.79 |
| IVLDKFAIPNSRRSDL | 1668692 | 5GNB | 15.19 |
| KDNRIPSYGVLSVDL | 30220 | 3INB | 18.31 |
| KFNSCKSDILSLNNPI | 1669953 | 6Y3Y | 30.64 |
| KIDTTSSSCQLYYSLP | 1670454 | 5GNB | 18.97 |
| KILAASFNKAMTNIVD | 1670511 | 5ZUV | 35.17 |
| KIQDVVNQQGNSLNHL | 1670550 | 5ZUV | 22.49 |
| KPPSASCPIGTNYRSC | 1671614 | 5GNB | 30.77 |
| KPVATVHRRIPDLPDC | 1671663 | 5GNB | 40.08 |
| KSDILSLNNPIFINYS | 1672036 | 6Y3Y | 33.48 |
| KSHKPPSASCPIGTNY | 1672089 | 5GNB | 34.14 |
| LCENPEWAPLKDNRI | 35050 | 3INB | 41.42 |
| LCNNGKISSSAGDSIF | 1673512 | 6Y3Y | 17.45 |
| LDIHPSLCNNGKISSS | 1673697 | 6Y3Y | 30.69 |
| LFGDSRSDCNHINNLK | 1674247 | 6Y3Y | 12.55 |
| LGSSGFLQSSNYKIDT | 1674723 | 5GNB | 25.61 |
| LHYRNQGWRSVETSGVAEEEATSGL | 36443 | 2ZZQ | 22.96 |
| LKIKIASGFGPLITH | 36890 | 3INB | 14.6 |
| LLYDSNGNIIGFKDFV | 1676298 | 5GNB | 34.49 |
| LNKIQDVVNQQGNSLN | 1676619 | 5ZUV | 22.23 |
| LPAINVTINNYNPSSW | 1676821 | 5GNB | 30.38 |
| LPKYIGLDPVAPGDL | 227357 | 4LDI | 47.35 |
| LQENQKILAASFNKAM | 1677148 | 5ZUV | 42.11 |
| LQLGSSGFLQSSNYKI | 1677206 | 5GNB | 27.64 |
| LQSSNYKIDTTSSSCQ | 1677306 | 5GNB | 17.25 |
| LQTVATALNKIQDVVN | 1677325 | 5ZUV | 20.28 |
| LSLLDLYLGRGYNVS | 39483 | 3INB | 25.09 |
| LSVDLSLTVELKIKI | 39732 | 3INB | 28.89 |
| LYDSNGNIIGFKDFVT | 1679050 | 5GNB | 37.33 |
| LYKSNHNNVYWLTIP | 40826 | 3INB | 22.95 |
| MGIQSGVQVDANCEGDCYHSGGTII | 1160208 | 6IDZ | 25.53 |
| MQSWVPLSTDDPVID | 42413 | 3INB | 33.73 |
| NDAITQTSQALQTVAT | 1681148 | 5ZUV | 18.26 |
| NDKHSNGTIKDRSPY | 215815 | 6G02 | 45.54 |
| NKIQDVVNQQGNSLNH | 1682361 | 5ZUV | 25.29 |
| NLLYDSNGNIIGFKDF | 1682652 | 5GNB | 33.14 |
| NLPFQNIDSRAVGKCPRYVKQRSLL | 1161651 | 6IDZ | 23.77 |
| NLVILPGQDLQYVLA | 44915 | 3INB | 21.91 |
| NNLKIKNFDYLDIHPS | 1683027 | 6Y3Y | 38.68 |
| NPYHRFKCFPNGSNDV | 1683251 | 6Y3Y | 34.99 |
| NQKILAASFNKAMTNI | 1683305 | 5ZUV | 33.79 |
| NQQGNSLNHLTSQLRQ | 1683336 | 5ZUV | 25.53 |
| NSWQNLLYDSNGNIIG | 1683786 | 5GNB | 48.34 |
| NYKIDTTSSSCQLYYS | 1684489 | 5GNB | 16.9 |
| PAINVTINNYNPSSWN | 1684659 | 5GNB | 24.92 |
| PDLPDCDIDKWLNNFN | 1684914 | 5GNB | 31.82 |
| PGLGAPVFHMTNYLE | 47694 | 3INB | 10.33 |
| PIELQVECFTWDQKL | 47890 | 3INB | 28.85 |
| PIGTNYRSCESTTVLD | 1685475 | 5GNB | 57.39 |
| PLITHGSGMDLYKSN | 48370 | 3INB | 16.11 |
| PNDTVTFSFNGAFIAPDRASFLRGK | 1163297 | 6IDZ | 21.18 |
| PPSASCPIGTNYRSCE | 1686285 | 5GNB | 33.43 |
| PSASCPIGTNYRSCES | 1686512 | 5GNB | 36.02 |
| PSFASSCKSHKPPSAS | 1686541 | 5GNB | 27.02 |
| PSLCNNGKISSSAGDS | 1686590 | 6Y3Y | 31.01 |
| PTVKLYTSVENAQQDKGIAIPHDID | 49810 | 2ZZQ | 30.71 |
| PYHRFKCFPNGSNDVW | 1687141 | 6Y3Y | 32.93 |
| PYNVSLSKFNSCKSDI | 1687165 | 6Y3Y | 57.74 |
| QALQTVATALNKIQDV | 1687296 | 5ZUV | 21.89 |
| QDGTNTHIMATEASNYAQYRVARAT | 50464 | 2ZZQ | 25.12 |
| QDYDNQHEQDRPTPSPAPSRPFSVL | 50539 | 2ZZQ | 41.01 |
| QENQKILAASFNKAMT | 1687601 | 5ZUV | 41.67 |
| QILRESGGIDKEAMGFTYSGIRTNG | 1164116 | 6IDZ | 30.22 |
| QKILAASFNKAMTNIV | 1688113 | 5ZUV | 31.44 |
| QLGSSGFLQSSNYKID | 1688264 | 5GNB | 28.3 |

| | | | |
|---|---|---|---|
| QNLLYDSNGNIIGFKD | 1688526 | 5GNB | 35.96 |
| QQGNSLNHLTSQLRQN | 1688729 | 5ZUV | 26.78 |
| QSSNYKIDTTSSSCQL | 1689023 | 5GNB | 17.26 |
| QTVATALNKIQDVVNQ | 1689191 | 5ZUV | 22.04 |
| QYVLATYDTSRVEHA | 53046 | 3INB | 26.95 |
| RGADGTAELTTTAATRFMKDLYFTS | 53816 | 2ZZQ | 31.78 |
| RGVEVVNATETVERTNIPRICSKGK | 1165884 | 6IDZ | 31.79 |
| RHRLRRGADGTAELTTTAATRFMKD | 54090 | 2ZZQ | 28.66 |
| RNLTPGNTNTRVSRYSSTARHRLRR | 55026 | 2ZZQ | 31.79 |
| RQYNLSTSPLTSSVATGTNLVLYAA | 55519 | 2ZZQ | 23.18 |
| RSDLQLGSSGFLQSSN | 1691272 | 5GNB | 31.85 |
| RTNIPRICSKGKRTVDLGQCGLLGT | 1167503 | 6IDZ | 27.83 |
| RVFEVGVIRNPGLGA | 56283 | 3INB | 12.03 |
| RYSSTARHRLRRGADGTAELTTTAA | 56660 | 2ZZQ | 29.36 |
| SAGGQLFYSRPVVSANGE | 56796 | 2ZZQ | 21.64 |
| SANGEPTVKLYTSVENAQQDKGIAI | 56908 | 2ZZQ | 24.21 |
| SDCNHINNLKIKNFDY | 1692391 | 6Y3Y | 46.94 |
| SDILSLNNPIFINYSK | 1692472 | 6Y3Y | 33.59 |
| SFASSCKSHKPPSASC | 1692881 | 5GNB | 29.1 |
| SGFLQSSNYKIDTTSS | 1693229 | 5GNB | 26.13 |
| SGTTCSNDLLQPNTEV | 1693396 | 5GNB | 31.91 |
| SHRGVIADNQAKWAV | 58403 | 3INB | 15.46 |
| SISFWPQTTTTPTSVDMNSITSTDV | 58700 | 2ZZQ | 30.74 |
| SKFNSCKSDILSLNNP | 1693883 | 6Y3Y | 38.89 |
| SKIYGSCFKSIVLDKF | 1693953 | 5GNB | 15.69 |
| SLCNNGKISSSAGDSI | 1694147 | 6Y3Y | 33.25 |
| SLPAINVTINNYNPSS | 1694387 | 5GNB | 30.52 |
| SNDLLQPNTEVFTDVC | 1694719 | 5GNB | 50.28 |
| SNGNIIGFKDFVTNKT | 1694765 | 5GNB | 39.8 |
| SNYKIDTTSSSCQLYY | 1694926 | 5GNB | 16.6 |
| SPALIVWGIHHSVSTAEQTKLYGSG | 1170135 | 6IDZ | 22.26 |
| SRVVIQDYDNQHEQDRPTPSPAPSR | 60833 | 2ZZQ | 37.55 |
| SSGFLQSSNYKIDTTS | 1695540 | 5GNB | 26.35 |
| SSIQAIYDRLDTIQAD | 1695578 | 5ZUV | 44.75 |
| SSNYKIDTTSSSCQLY | 1695663 | 5GNB | 17.74 |
| STAEQTKLYGSGNKLVTVGSSNYQQ | 1171846 | 6IDZ | 39.75 |
| SWQNLLYDSNGNIIGF | 1696681 | 5GNB | 42.64 |
| TCSNDLLQPNTEVFTD | 1697298 | 5GNB | 43.69 |
| TDSFSCNNFDESKIYG | 1697467 | 5GNB | 36.52 |
| TEASNYAQYRVARATIRYRPLVPNA | 63307 | 2ZZQ | 26.76 |
| TGPPQCDQFLEFSADLIIERREGSD | 1173378 | 6IDZ | 22.38 |
| TIRGQFSNMSLSLLD | 64444 | 3INB | 33.5 |
| TNTRVSRYSSTARHRLRRGADGTAE | 65464 | 2ZZQ | 33.93 |
| TNYLEQPVSNDLSNC | 65486 | 3INB | 34.81 |
| TQTSQALQTVATALNK | 1699949 | 5ZUV | 18.71 |
| TSQALQTVATALNKIQ | 1700354 | 5ZUV | 17.39 |
| TSQGMYGGTYLVEKP | 66385 | 3INB | 20.7 |
| TSVENAQQDKGIAIPHDIDLGESRV | 66481 | 2ZZQ | 34.82 |
| TTAATRFMKDLYFTSTNGVGEIGRG | 66511 | 2ZZQ | 36.84 |
| TVATALNKIQDVVNQQ | 1700818 | 5ZUV | 19.75 |
| TVDLGQCGLLGTITGPPQCDQFLEF | 1175353 | 6IDZ | 17.48 |
| VATALNKIQDVVNQQG | 1701730 | 5ZUV | 18.9 |
| VDMNSITSTDVRILVQPGIASELVI | 68025 | 2ZZQ | 38.81 |
| VIPSERLHYRNQGWRSVETSGVAEE | 69074 | 2ZZQ | 31.24 |
| VKPVATVHRRIPDLPD | 1703884 | 5GNB | 35.82 |
| VNDAITQTSQALQTVA | 1704755 | 5ZUV | 15.9 |
| VNQQGNSLNHLTSQLR | 1704953 | 5ZUV | 23.48 |
| VPNAVGGYAISISFWPQTTTTPTSV | 70378 | 2ZZQ | 26.63 |
| VYYNSWQNLLYDSNGN | 1707437 | 5GNB | 55.64 |
| WLTIPPMKNLALGVI | 72808 | 3INB | 7 |
| YCFSVNNTFCPCAKPS | 1708535 | 5GNB | 27.15 |
| YKIDTTSSSCQLYYSL | 1709627 | 5GNB | 17.22 |
| YLDIHPSLCNNGKISS | 1709815 | 6Y3Y | 30.16 |
| YNSWQNLLYDSNGNII | 1710257 | 5GNB | 48.77 |
| YNVSLSKFNSCKSDIL | 1710277 | 6Y3Y | 54.04 |
| YPFRLPIKGVPIELQ | 75287 | 3INB | 36.41 |
| YSLPAINVTINNYNPS | 1710743 | 5GNB | 26.72 |
| YSRPVVSANGEPTVKLYTSVENAQQ | 75883 | 2ZZQ | 28.07 |
| YYNSWQNLLYDSNGNI | 1711514 | 5GNB | 50.51 |
| YYSLPAINVTINNYNP | 1711573 | 5GNB | 27.27 |

eRSA: Relative solvent accesibility (RSA) of the entire B cell epitope computed averaging the relevant residue RSAs
