## Supplementary Dataset 4 for "Analysis of virus-specific B cell epitopes reveals extensive antigen processing prior to recognition"

**Supplementary Dataset 4.** Discontinuous B cell epitopes extracted from antibody-antigen tertiary structures

| B cell epitope | PDB | eRSA (%) |
|---|---|---|
| 74-ASN,75-ASP,78-ALA,81-GLY,84-LYS,86-ILE,87-ASN,88-TRP,89-PHE,90-ASP,91-ILE,92-SER,95-LEU,96-TRP | 3LHP | 52.2 |
| 75-PRO,76-GLU,79-HIS,107-GLN,108-PRO,110-MET,111-LYS,112-LEU,113-GLY,114-THR,115-GLN,116-THR,117-VAL,118-PRO,119-CYS,148-THR,150-LEU,194-ARG,195-ARG,196-PHE,198-GLU,199-ALA,201-CYS,233-GLU,234-LYS | 4CMH | 40.99 |
| 472-ILE,473-SER,474-HIS,514-VAL,540-ASN,541-ASN,542-THR,543-ARG,545-PRO,546-LEU,547-GLY,548-ASN,549-TRP,567-PRO,591-GLU,595-SER,596-ARG,634-GLY | 6MEJ | 45.47 |
| 18-GLU,65-ASN,66-SER,67-ARG,96-MET,97-TYR,98-ASP,99-THR,123-ASP,124-THR,177-PHE,180-GLU | 6IAP | 54.38 |
| 29-ARG,51-GLU,141-ARG,157-TRP,183-TRP,185-ASN,201-ALA,225-PRO,226-HIS,242-TRP,243-SER,266-PHE | 3SOB | 34.73 |
| 36-LYS,54-ARG,61-VAL,62-LEU,63-LYS,64-GLU,65-ASP,66-ALA,67-LEU,68-PRO,69-GLY,73-GLU,75-LYS | 5X0T | 48.96 |
| 18-VAL,19-ASP,20-GLY,21-TRP,36-ALA,38-LYS,41-THR,42-GLN,45-ILE,46-ASP,48-VAL,49-THR | 4FQI | 46.91 |
| 314-LYS,317-LYS,318-GLU,320-LEU,321-ASN,324-GLN,327-LEU,355-LYS,356-ASP,357-GLU,358-LEU,359-ASP,360-TYR,364-ILE | 6B0S | 45.67 |
| 71-LYS,124-ASP,170-GLU,171-TRP,172-THR,173-ARG,174-GLU,175-PRO,176-ALA,177-ARG,197-THR,198-VAL,199-ASP,200-SER | 6D6U | 50.66 |
| 111-SER,112-PRO,116-ARG,138-GLN,168-ARG,169-ALA,170-CYS,171-HIS,172-PRO,173-CYS,175-PRO,180-SER,185-GLU | 3WLW | 54.61 |
| 91-SER,92-LYS,94-PHE,101-ASP,103-PRO,105-TYR,219-SER,220-ARG,221-PRO,222-TRP,223-VAL,231-SER,269-ARG,271-ASP | 6E56 | 53.19 |
| 18-ILE,19-ASP,20-GLY,21-TRP,37-ASP,38-LEU,41-THR,42-GLN,45-ILE,46-ASP,48-ILE,49-ASN,52-LEU,53-ASN,56-ILE | 4FQY | 47.01 |
| 202-PRO,203-ASP,226-ASN,229-SER,230-ARG,231-CYS,232-LEU,233-PHE,235-GLU,248-VAL,250-TYR,251-VAL,252-GLN,253-GLU,289-LYS,290-THR,291-ASN,292-LYS,293-LEU,294-CYS,295-TYR,296-GLU | 4HWB | 50.62 |
| 89-LYS,115-SER,116-ASP,117-LYS,120-ALA,121-PHE,139-LYS,140-ASN,141-ASN,144-VAL,145-LYS,166-THR,167-GLN,188-SER,189-GLU,192-ASP | 5GS0 | 44.61 |
| 72-THR,74-ASN,75-GLU,76-GLU,77-ALA,78-ALA,79-GLU,81-ASP,82-ARG,83-LEU,85-PRO,100-ARG,101-GLY,102-SER | 1AFV | 40.84 |
| 38-LEU,39-VAL,40-TRP,71-GLN,73-ARG,75-ILE,77-ILE,79-ARG,80-LYS,81-LYS,82-PRO,83-ILE,84-PHE,85-LYS,86-LYS | 4QCI | 55.82 |
| 18-ASP,19-ASN,22-GLY,23-TYR,24-SER,27-ASN,102-GLY,103-ASN,116-LYS,117-GLY,118-THR,119-ASP,120-VAL,121-GLN,124-ILE,125-ARG | 1G7H | 49.96 |
| 302-TYR,306-SER,307-LYS,308-ALA,309-PHE,330-THR,331-GLY,332-THR,333-ASP,365-ALA,366-THR,367-ALA,368-ASN,389-GLY,390-GLU,391-GLN | 1ZTX | 50.44 |
| 15-GLU,16-SER,17-HIS,18-VAL,19-ALA,20-ARG,21-ALA,22-ASN,42-LEU,43-LYS,44-ASN,45-ASN,46-ASN,47-ARG,49-VAL,57-TRP | 4LMQ | 49.89 |
| 408-ILE,428-GLN,429-GLN,460-ASN,464-ILE,465-THR,466-PHE,467-PRO,468-ASN,473-LYS,477-ASN,479-PRO,481-ASP,510-TYR,512-TYR,518-TRP | 5JQ6 | 52.86 |
| 309-THR,310-ALA,311-ALA,312-PHE,313-THR,314-PHE,331-GLN,332-TYR,333-ALA,334-GLY,335-THR,336-ASP,337-GLY,368-SER,370-GLU,371-ASN,393-GLU,394-LYS,395-LYS,396-ILE,397-THR | 5KVG | 42.56 |
| 346-GLY,347-ASP,349-ILE,351-ASP,355-GLN,364-LEU,365-GLU,366-PRO,367-SER,369-ILE,371-TYR,390-ASP,392-LYS,394-LYS,411-ILE,413-LYS,414-LYS,415-ASP,416-LYS | 6E62 | 49.13 |
| 96-THR,97-SER,98-LEU,99-ASP,159-ASP,161-ARG,162-LYS,164-TYR,235-GLU,236-SER,237-THR,239-GLY | 5XS7 | 62.35 |
| 20-PRO,21-GLN,23-GLU,65-LYS,66-GLY,67-GLN,110-GLU,111-ALA,113-PRO,115-TYR,140-ASP,141-TYR,143-ASP,144-PHE,145-ALA,146-GLU,147-SER | 3WD5 | 56.02 |
| 70-GLN,71-PHE,72-LEU,73-TYR,74-PHE,95-PHE,96-LYS,98-LYS,99-ASP,102-ASP,103-LYS,105-LYS,107-LYS,125-LYS,245-ASN | 3W2D | 42.99 |
| 275-LYS,276-ASN,278-ALA,279-ASP,280-ASN,282-LYS,356-LYS,365-SER,366-GLY,367-GLY,368-ASP,371-ILE,428-GLN,455-LEU,456-ARG,457-ASP,458-GLY,459-CYS,460-ASN,461-THR,465-ALA,469-ARG | 6MYY | 49.3 |
| 47-SER,48-ASN,57-LEU,103-ASN,104-ARG,152-GLN,153-ALA,154-GLN,155-ASN,156-GLN,157-TRP,158-LEU,159-GLN,160-ASP | 4CNI | 48.46 |
| 144-TRP,145-GLU,147-ARG,148-TYR,151-GLU,176-HIS,179-VAL,180-ASN,196-ASN,198-THR,199-GLU,200-THR,202-VAL,203-LYS,206-GLU,210-GLU | 6AQ7 | 48.48 |
| 12-LEU,23-GLU,25-GLU,37-ASP,58-ASP,59-SER,60-ILE,61-LEU,63-GLY,64-PRO,70-ASN,71-LYS,72-PHE,73-VAL | 5EII | 45.75 |
| 8-HIS,28-HIS,29-SER,30-VAL,40-THR,66-VAL,67-ASP,68-GLY,69-TRP,86-GLN,87-LYS,89-THR,90-GLN,91-ASN,93-ILE,94-ASN,97-THR,100-VAL,101-ASN,104-ILE | 5C0R | 41.02 |
| 25-THR,26-ARG,27-ASP,28-GLY,29-GLY,30-VAL,31-SER,32-ASN,35-THR,37-ILE,39-ARG,42-GLY,43-GLY,44-ASP,79-ASP,81-ARG,82-ASP,83-ASN,84-ALA,85-LYS,138-SER,139-GLY,140-GLY,141-ASP,144-PHE | 4JPK | 49.92 |
| 194-GLY,197-LEU,198-ASN,201-TRP,202-ARG,203-TRP,225-ILE,226-GLU,227-ASP,228-SER,229-GLU,230-ILE,255-ASN,453-SER,454-ILE,455-HIS,456-LEU,460-LYS | 5X2N | 47.76 |
| 627-PRO,628-GLN,629-ARG,631-SER,632-ARG,634-PHE,635-VAL,636-ARG,660-LYS,661-GLN,664-LEU | 1FNS | 49.95 |
| 266-THR,267-MET,268-GLY,270-GLN,273-GLU,276-TYR,277-ILE,278-LYS,282-ILE,283-ARG,285-ASN | 5Y9F | 73.88 |
| 123-PRO,124-HIS,126-ILE,127-GLN,130-ASN,131-SER,132-VAL,133-ASP,134-MET,142-THR,143-ASP,144-ALA,147-VAL,148-LYS,149-ASP,150-PRO,151-GLN | 3CX5 | 47.67 |
| 50-LYS,59-LEU,60-ASP,62-ILE,63-ASP,74-PRO,75-HIS,78-VAL,82-GLU,90-ARG,92-LYS,94-PHE,271-ASP,273-PRO | 1EO8 | 43.88 |
| 22-VAL,23-LYS,24-SER,25-ILE,26-SER,27-LYS,28-LEU,51-TRP,53-SER,54-SER,57-VAL,71-LEU,72-GLU,73-SER,74-SER,75-ASP,76-SER,77-ARG,78-ARG | 3B9K | 47.98 |
| 88-TYR,121-ARG,123-ASN,124-GLY,125-ALA,126-THR,127-SER,130-ARG,132-SER,133-GLY,134-SER,142-TRP,144-LEU,148-ASP,149-ASN,177-VAL,178-SER,180-ALA,184-LYS,185-LEU,213-GLN,216-GLY,217-LEU | 6II9 | 43.44 |
| 7-ARG,210-HIS,211-GLY,213-PRO,275-THR,276-PRO,277-PHE,278-LEU,279-ALA,282-ARG,297-ASP,298-THR,299-ASN,300-LEU,353-HIS,354-PRO,358-GLU,361-PRO | 4JQI | 44.51 |
| 6-GLU,8-ARG,9-ILE,10-PHE,11-PRO,12-LYS,14-MET,15-ASP,20-GLN,21-LYS,22-VAL,23-ASP,25-VAL,27-GLU,29-LEU,30-GLY,84-LYS,114-SER | 2ARJ | 43.9 |
| 60-PHE,61-PRO,62-THR,75-HIS,100-ALA,101-GLY,119-ASN,120-TRP,121-PHE,123-ILE,124-THR,127-LEU | 4M8Q | 51.52 |
| 34-GLY,35-LEU,36-THR,37-SER,38-PRO,39-CYS,40-LYS,41-ASP,85-GLY,86-GLY,87-SER,88-PRO,89-TRP,90-PRO,91-PRO,93-GLN | 1H0D | 52.83 |
| 48-LYS,79-GLN,80-ILE,81-MET,82-ARG,83-ILE,86-HIS,87-GLN,88-GLY,89-GLN,90-HIS,91-ILE,92-GLY,93-GLU,94-MET | 1BJ1 | 50.79 |
| 359-THR,362-SER,391-GLY,392-ASP,424-ASN,426-ARG,427-ASN,486-THR,487-THR,488-GLY,489-ILE,490-GLY,491-TYR,492-GLN,494-TYR | 3BGF | 49.71 |
| 81-ASN,83-VAL,95-GLY,148-LYS,149-ALA,150-VAL,151-ASP,152-GLY,155-LYS,157-ASP,163-ILE | 6B0E | 57.96 |
| 122-LEU,123-THR,124-GLY,198-GLY,200-VAL,279-ASN,280-ASN,367-GLY,368-ASP,371-ILE,429-GLY,430-THR,431-GLY,473-GLY,474-ASN,476-LYS | 5UEM | 55.62 |
| 125-GLN,129-GLN,132-GLN,135-VAL,136-VAL,144-LYS,161-THR,163-THR,164-ALA,165-LEU,167-THR,168-SER,171-LYS,172-ASN,181-ILE,182-ILE | 5DFV | 52.08 |
| 276-ASN,279-ASP,280-ASN,281-ALA,282-LYS,364-SER,365-GLY,366-GLY,367-ASP,370-ILE,455-LEU,456-ARG,457-ASP,458-GLY,459-GLY,460-ASP,461-THR,462-THR,465-THR,467-ILE,469-ARG | 5FEC | 49.53 |
| 72-ASN,89-GLU,90-ASN,91-SER,94-GLU,95-ILE,96-PHE,97-GLY,98-ASP,99-SER,116-TYR,117-ASN,133-TYR,138-ARG,139-GLN,140-LEU,145-LYS,241-ASP | 4FFW | 52.78 |
| 613-LYS,614-ARG,615-HIS,616-GLY,619-ASN,620-TYR,621-ARG,663-TRP,673-ARG,675-LEU,689-THR,693-ASN,696-ALA,697-LYS,698-ILE,700-SER,701-ASP,702-GLY,703-ASN | 1NBY | 48.99 |
| 25-GLU,27-ASN,28-ALA,29-SER,31-GLN,32-THR,33-LYS,35-ASP,58-SER,59-ALA,60-ASP,61-ALA,62-ASP,125-LYS,127-LYS | 4U6H | 56.84 |
| 129-GLY,130-VAL,131-THR,132-GLN,133-ASN,134-GLY,135-GLY,136-SER,137-ASN,153-TRP,155-ILE,156-LYS,157-SER,158-GLY,159-SER,193-SER,194-LEU | 2VIT | 41.09 |
| 170-LYS,172-GLY,173-ASN,174-SER,176-PRO,177-LYS,206-GLN,210-GLN,211-ASN,213-ASP,262-THR | 6A4K | 49.42 |
| 663-SER,665-LEU,666-SER,667-ILE,668-HIS,669-SER,670-ALA,671-LYS,673-LYS,702-THR,705-THR,707-VAL | 6MW9 | 49.43 |
| 416-LYS,417-ASN,418-LYS,440-LEU,441-ASN,442-GLU,465-TYR,466-ASN,467-LYS,468-TYR,489-ARG,491-ALA,492-LEU,493-LYS,515-ASN,516-ASN,517-ASN,519-ALA,539-HIS,541-ASN,571-SER,595-LEU | 3ULU | 37.01 |
| 251-LEU,252-MET,253-ILE,254-SER,255-ARG,384-ASN,385-GLY,386-GLN,424-SER,428-MET,433-HIS,434-ASN,435-HIS,436-TYR,438-GLN,440-SER | 1ADQ | 52.27 |
| 307-LEU,340-LYS,344-GLN,347-VAL,348-ASP,350-GLN,351-THR,352-LEU,353-THR,354-PRO,355-VAL,384-ASP,386-TYR | 5KVE | 49.26 |
| 305-THR,306-MET,307-CYS,308-ASP,309-LYS,310-THR,311-LYS,333-SER,334-GLY,335-THR,336-LYS,337-PRO,339-ARG,366-ASN,387-GLU | 5O6V | 50.01 |
| 297-MET,298-SER,299-TYR,300-SER,301-MET,303-THR,328-GLY,329-ASP,330-GLY,331-SER,332-PRO,334-LYS,361-LYS,382-VAL,383-GLU,384-PRO | 6FLC | 56.55 |
| 10-GLU,11-GLU,12-ASP,13-PRO,17-ARG,19-TYR,21-THR,23-TYR,28-GLN,31-GLN,32-CYS,33-GLU,34-ARG,36-LYS,50-LEU | 4DTG | 47.45 |
| 398-ASN,400-LYS,495-LEU,496-LYS,527-VAL,528-SER,529-ILE,530-VAL,531-PRO,532-SER,535-TRP,536-GLU,539-ASP,540-TYR,542-ARG,543-LYS,544-GLN | 5DO2 | 48.28 |
| 240-ARG,241-GLN,242-ASP,243-GLY,244-VAL,245-SER,247-SER,249-ASP,250-THR,279-ARG,283-TYR,290-HIS,291-SER,292-THR | 6DDM | 50.39 |
| 16-THR,18-ARG,19-LYS,20-ILE,21-SER,23-GLN,24-ARG,45-THR,46-ILE,47-VAL,49-LYS | 4DN4 | 52.84 |
| 71-ASN,73-TYR,74-TRP,206-ASN,207-LEU,208-TRP,210-LYS,211-ASP,262-TRP,264-ARG,267-PHE | 5K59 | 57.17 |
| 301-MET,334-LYS,336-PRO,337-PHE,338-SER,339-SER,340-GLN,343-LYS,344-GLY,345-VAL,346-THR,349-GLY,350-ARG,351-LEU,370-GLU,372-PRO,379-VAL,382-ALA,383-GLY,384-GLU | 4FFZ | 46.82 |
| 72-VAL,75-VAL,76-SER,79-ARG,80-ASP,83-GLU,84-LEU,145-LEU,148-LYS,149-VAL,150-LEU,153-ASP,155-GLY | 3UBX | 52.35 |
| 36-PHE,37-GLY,58-THR,60-LYS,61-GLU,62-GLU,96-ALA,99-LYS,100-LYS,103-ASN,104-GLU | 1WEJ | 51.36 |
| 19-LEU,20-LYS,33-GLU,34-GLY,282-PRO,284-TRP,348-ARG,349-GLU,352-ILE,353-ASN,358-ASN,411-THR,414-ASP | 4DAG | 51.18 |
| 199-ARG,221-HIS,222-GLN,223-LYS,224-GLU,225-PRO,226-PRO,227-PHE,229-TRP,230-MET,358-ARG | 1R0A | 52.86 |
| 222-PRO,223-THR,233-LEU,234-ASP,236-ARG,238-VAL,239-GLU,240-THR,242-PRO,243-PRO,244-PRO,245-TYR | 5J3H | 48.11 |
| 33-LEU,59-LEU,60-GLU,62-GLU,88-ALA,91-SER,92-SER,93-PHE,94-VAL,95-PRO,149-PRO,150-MET,153-HIS,202-GLU,203-PRO,204-SER,205-PHE | 4Y5Y | 46.58 |
| 137-MET,139-HIS,140-PHE,141-GLY,142-ASN,143-ASP,144-TRP,145-GLU,146-ASP,203-LYS,207-ARG,211-GLN | 4H88 | 52.22 |
| 8-HIS,28-HIS,30-VAL,31-ASN,32-LEU,40-THR,67-ASP,68-GLY,69-TRP,86-GLN,89-THR,90-GLN,93-ILE,94-ASN,97-THR,100-VAL,101-ASN,104-ILE,105-GLU | 5C0S | 37.24 |
| 232-TRP,235-GLY,236-ILE,237-LEU,239-SER,240-ASP,253-ASN,256-ARG,257-ARG,258-PRO,260-TYR,261-LYS,262-THR,264-ASN,265-GLN,289-GLU,293-HIS,294-ARG,314-ARG,315-SER,351-ASN | 4OII | 42.71 |
| 18-GLY,20-THR,21-GLN,22-GLU,23-ASP,24-ALA,25-THR,27-LYS,120-PRO,138-SER,139-LEU,141-LYS | 6IEA | 51.54 |
| 15-LEU,16-GLY,19-ASN,20-TYR,21-ARG,62-TRP,63-TRP,73-LYS,75-ALA,89-THR,93-ARG,96-LYS,97-ARG,100-SER,101-ASP,102-GLY | 5VJQ | 48.01 |
| 153-PRO,154-PRO,155-ASN,156-PRO,157-LEU,159-TRP,286-THR,287-SER,288-ASP,290-THR,291-TRP,293-GLN,294-LEU | 4XHJ | 52.34 |
| 32-GLU,34-ARG,39-ASP,40-GLU,41-TRP,43-ALA,65-ASN,67-ARG,68-PHE,75-LYS,76-ASN,77-VAL,78-PHE,79-ASP,80-ASP | 2VXQ | 43.52 |
| 305-TYR,306-SER,307-LEU,309-THR,310-ALA,311-ALA,334-GLY,335-THR,336-ASP,340-LYS,350-GLN,351-THR,352-LEU,391-VAL,392-GLY,393-GLU,394-LYS,395-LYS | 5VIG | 50.48 |
| 54-ASN,294-ASN,334-THR,366-TRP,383-LEU,384-LYS,387-GLN,390-ILE,391-ASP,393-ILE,394-ASN,397-LEU,398-ASN,401-ILE,403-LYS,404-THR,405-ASN | 4KVN | 48.48 |
| 83-LYS,145-LYS,148-ALA,149-PRO,150-CYS,151-TYR,152-GLN,154-TYR,155-VAL,160-GLU,161-ASN,162-PHE,163-ALA,193-PHE,198-THR,199-THR,200-ASN,201-LYS,202-TYR,206-LEU,208-LYS,255-GLU,256-LYS,257-GLU | 4IRZ | 46.85 |
| 126-GLU,128-SER,129-LEU,131-VAL,132-SER,133-SER,139-ARG,140-LYS,141-SER,152-LYS,153-LYS,154-ASN,155-SER,189-LYS,218-LYS | 6A67 | 57.71 |
| 201-LYS,202-ASN,204-TRP,205-ASN,206-THR,207-GLY,208-ASN,209-CYS,211-ASN,216-ASP,234-ARG,235-HIS,238-GLN,241-ARG | 6DZZ | 46.07 |
| 627-PRO,628-GLN,629-ARG,631-SER,632-ARG,634-PHE,635-VAL,636-ARG,658-ASN,660-LYS,661-GLN,664-LEU | 1OAK | 44.33 |
| 400-SER,403-PRO,405-ILE,406-GLY,407-GLY,408-HIS,409-GLY,410-SER,411-LYS,412-LYS,424-GLU,425-CYS,426-THR,427-ALA,428-GLN,429-TYR,431-ASN | 6I9I | 56.17 |
| 51-THR,52-LEU,53-PHE,54-CYS,71-THR,72-HIS,73-ALA,74-CYS,75-VAL,76-PRO,77-THR,78-ASP,103-GLN,106-GLU,107-ASP,217-TYR,221-ALA | 5KJR | 45.32 |
| 429-GLU,432-ARG,433-ASN,477-SER,480-GLU,481-THR,482-ASN,483-GLY,484-LEU,485-GLU,655-LEU,656-ARG,657-SER,658-GLU,660-PRO,662-TRP | 4YZF | 55.58 |
| 116-PRO,117-ASP,143-GLY,221-GLN,223-THR,233-LEU,268-ASN,269-THR,270-THR,272-LYS,273-LEU,274-ILE,305-GLU,307-SER,308-PHE | 5KEN | 59.53 |
| 362-LYS,363-MET,364-ARG,365-TYR,366-GLU,367-HIS,395-HIS,416-THR,418-ARG,419-THR,421-TYR | 1EGJ | 60.95 |
| 21-ARG,22-GLY,23-TYR,102-GLY,103-ASN,104-GLY,106-ASN,109-VAL,111-TRP,112-ARG,113-ASN,114-ARG,116-LYS,117-GLY,118-THR | 4TSB | 57.34 |
| 37-ASN,38-GLU,39-THR,41-GLU,44-SER,45-GLU,46-MET,96-CYS,97-ALA,98-THR,99-GLN,100-ILE,101-ILE,102-THR,104-GLU,105-SER,108-GLU,109-ASN,112-ASP | 6BFQ | 53.44 |
| 29-GLU,31-GLN,32-ASN,34-THR,35-GLU,36-VAL,37-TYR,39-LYS,46-ASP,51-ASP,53-ALA,54-LEU,97-GLU,99-THR,100-GLU,101-LEU,102-THR,103-ARG,104-GLU | 5TZU | 46.17 |
| 30-ARG,33-LEU,34-ASP,37-SER,40-ARG,67-MET,69-GLU,74-PHE,75-GLN,76-SER,164-HIS,168-ARG,171-LYS,172-GLU,175-GLN,176-SER,178-LEU,179-ARG,182-ARG,183-GLN | 4ZS7 | 48.24 |
| 22-ARG,24-ILE,25-SER,26-LEU,27-PHE,30-LEU,33-ARG,34-HIS,35-ASP,36-PHE,37-GLY,40-GLN,41-GLU,125-ARG,141-GLU,144-ARG,149-ARG,153-LEU | 4Z5R | 47.59 |
| 15-GLN,15-ARG,15-ASN,15-THR,15-ASP,15-GLY,15-SER,15-TYR,15-PRO,15-LEU | 1P2C | 53.3 |
| 1-SER,2-TYR,3-ASN,4-ASP,6-LYS,7-LYS,10-SER,11-ASN,13-ILE,14-ALA,21-GLU,24-ARG,25-SER,28-LEU,29-SER,31-ILE,32-ASN,33-ASP | 3QWO | 53.81 |
| 305-LYS,306-PHE,307-LYS,308-VAL,309-VAL,310-LYS,311-GLU,312-ILE,325-GLN,327-GLU,362-ASP,364-PRO,387-LEU,388-LYS,389-LEU,390-ASN | 2R29 | 44.66 |
| 97-LYS,276-ASN,278-THR,279-ASN,280-ASN,281-ALA,365-SER,366-GLY,367-GLY,368-ASP,371-ILE,430-THR,455-THR,456-ARG,457-ASP,458-GLY,459-GLY,461-ASN,469-ARG,474-ASN,476-LYS | 4XVT | 58.48 |
| 277-LEU,279-ALA,281-GLN,282-ALA,284-GLN,307-ARG,314-GLU,320-HIS,335-SER,336-VAL,337-PRO,339-GLY,340-GLY,341-ARG,342-VAL,365-ILE,367-ASP,369-ALA | 4AG4 | 41.09 |
| 47-LYS,49-TYR,50-GLY,51-VAL,52-LYS,53-ASN,54-SER,55-GLU,56-TRP,76-ASP,78-SER,79-ASN,82-TRP,84-ARG,98-LYS | 1JRH | 51.11 |
| 163-ASN,184-ARG,186-THR,187-GLU,188-LYS,192-HIS,239-HIS,240-ASN,241-TYR,242-PHE,243-VAL,245-ASN,248-ASP,280-GLU,282-GLU,283-LEU,284-ASP,285-THR,286-GLU,502-ASP,504-ALA,505-ALA,508-ASN | 5W0K | 45.59 |
| 163-TYR,205-THR,206-TYR,208-MET,209-GLY,215-LYS,221-GLY,222-ASP,224-LEU,225-SER,226-LEU,231-ARG,233-ILE,264-PRO,265-ARG,266-GLU | 5Y9J | 52.53 |
| 21-PHE,21-THR,21-ASN,21-MET,22-PHE,22-ARG,22-GLN,22-VAL,22-SER,22-LEU,22-THR,22-MET,23-HIS,23-GLN | 1IQD | 49.29 |
| 12-GLY,15-ILE,16-THR,17-ASP,19-ARG,24-ALA,25-PHE,27-GLN,28-GLN,29-ARG,32-ARG | 4JR9 | 60.61 |
| 194-ARG,195-GLU,238-ASP,239-ALA,243-LYS,341-ALA,342-GLN,364-PRO,366-GLU,367-ASP,368-ILE,369-ILE,375-CYS,376-SER,377-THR,378-CYS,379-PHE,381-SER | 5VL7 | 48.57 |
| 615-ARG,618-GLY,619-GLU,620-ARG,621-GLY,622-THR,656-ILE,658-PRO,659-CYS,660-THR,661-GLY,662-ASN,678-ASP | 5ZS0 | 58.42 |
| 210-TYR,211-THR,212-GLU,220-TYR,241-TYR,268-ARG,269-LEU,270-PRO,271-ARG,272-LEU,345-ALA,346-ARG,347-MET,348-ALA,349-PRO,350-GLU,351-GLU | 6I8S | 52.17 |
| 72-TRP,73-CYS,74-GLY,75-HIS,114-ARG,117-ASN,118-ILE,119-ALA,120-GLY,121-PHE,122-PRO,125-ARG,135-SER,136-GLY,137-ALA,138-THR,139-LEU,140-PRO,142-ALA,143-GLY | 5D96 | 49.74 |
| 20-MET,21-SER,22-GLY,23-PRO,24-TYR,25-GLU,27-LYS,64-GLU,65-LYS,66-ASN,67-LEU,74-LYS,81-GLN,82-LEU,84-SER,85-VAL,86-ASP,87-PRO | 5MVZ | 44.48 |
| 93-ALA,94-PHE,101-ASP,102-VAL,186-SER,190-GLU,219-SER,220-ARG,221-PRO,222-TRP,223-VAL,224-ARG,225-GLY,226-LEU,227-SER | 6N5E | 43.52 |
| 476-THR,477-ALA,479-GLU,484-THR,485-TYR,497-SER,498-VAL,499-THR,510-VAL,512-ARG,513-SER,514-LEU,515-ASP,534-LYS,573-ASN,574-ALA,575-ALA,577-HIS,578-ARG | 3RKD | 39.49 |
| 409-HIS,411-GLN,412-PHE,417-VAL,436-ASP,438-ILE,440-SER,453-TRP,454-LYS,457-PHE,460-SER,461-GLY,462-GLN,463-LYS,464-THR,465-LYS,467-ILE | 3P0Y | 42.12 |

| | | |
|---|---|---|
| 9-LYS,57-GLU,60-ALA,61-PHE,64-LYS,68-ASN,70-LYS,95-ASP,96-GLY,97-LYS,98-MET,105-ARG,106-GLN,120-THR,121-HIS,124-HIS,127-LYS,135-GLU | 1NSN | 48 |
| 478-SER,480-GLY,527-GLU,528-LYS,553-GLN,554-ASP,555-TYR,598-TRP,599-GLY,600-GLU,604-ARG | 4OGY | 60.02 |
| 32-ASP,34-LEU,35-TRP,36-VAL,37-THR,38-VAL,39-TYR,40-TYR,43-PRO,82-GLN,84-ILE,86-LEU,87-GLU,224-VAL,244-SER,246-GLN | 5W4L | 48.39 |
| 72-VAL,73-LEU,74-LYS,75-ASP,81-GLN,83-GLU,84-SER,86-ASP,89-ASN,90-TYR,92-LYS,94-LYS,96-GLU,97-LYS,98-ARG,115-ALA,116-GLN,117-PHE,118-PRO | 4G6M | 45.68 |
| 25-GLU,56-SER,58-GLN,60-GLU,61-ASP,64-TRP,95-PRO,97-GLU,99-ARG,101-THR,105-GLY,107-PRO,110-HIS,111-ARG,112-VAL,114-HIS | 2JIX | 44.06 |
| 98-TYR,131-THR,134-GLY,135-GLY,136-SER,137-SER,153-TRP,155-TYR,156-LYS,157-SER,158-GLY,159-SER,189-LYS,190-GLU,193-ASN,194-LEU,196-VAL,226-LEU | 4GMS | 38.59 |
| 81-LEU,84-HIS,85-GLN,87-TYR,88-PRO,89-LEU,91-LYS,134-LEU,137-LYS,138-PHE,139-GLY,140-PHE,141-GLN | 6O3A | 61.27 |
| 46-LYS,47-LEU,48-VAL,49-SER,50-ASP,51-CYS,52-THR,60-LEU,61-PRO,63-GLY,64-GLU,66-GLU,73-ARG,75-THR,76-HIS,77-CYS | 5DMI | 45.59 |
| 269-ASP,271-VAL,315-GLU,316-GLU,317-ILE,462-TYR,463-GLN,467-PRO,468-ALA,469-GLN,470-SER,493-LYS | 6N81 | 48.21 |
| 15-LEU,16-GLY,19-ASN,20-TYR,21-ARG,62-TRP,63-TRP,73-LYS,75-ALA,93-ARG,96-LYS,97-ARG,98-ILE,100-SER,101-ASP,102-GLY | 5VJO | 47.83 |
| 217-ARG,220-ASP,224-THR,225-LYS,295-GLN,298-LYS,302-GLU,305-ASP,306-LEU,308-GLU,309-LYS,311-LYS,312-CYS,313-SER,416-LYS | 6BPE | 48.13 |
| 53-PHE,58-ALA,59-LYS,60-ALA,61-TYR,71-THR,72-HIS,74-CYS,75-VAL,76-PRO,77-THR,78-ASP,79-PRO,218-CYS,219-VAL,220-PRO,246-GLN | 4R4H | 49.01 |
| 25-GLU,26-GLY,27-LEU,34-ILE,37-ASP,38-GLY,53-HIS,54-TRP,56-ASP,57-LEU,58-LEU,59-PHE,61-LEU,62-ARG | 1ZA3 | 57.03 |
| 476-ALA,477-ALA,478-ALA,479-LEU,480-PRO,481-GLY,482-GLN,483-PRO,497-PRO,498-ASP,499-ILE,500-PRO,501-GLY | 6JMR | 58.06 |
| 503-ALA,506-ASN,507-ALA,510-LYS,511-CYS,512-ASN,513-PRO,514-ASN,516-HIS,545-GLU,546-GLY,547-LEU,549-HIS,550-ASN,551-GLN,552-ASP,553-GLY,556-CYS,557-GLY | 5KEN | 54.54 |
| 45-TYR,48-LYS,81-MET,84-LYS,86-HIS,87-GLN,88-GLY,89-GLN,90-HIS,91-ILE,92-GLY,93-GLU | 3BDY | 52.69 |
| 114-LYS,115-LYS,116-PRO,117-ASP,118-GLY,120-GLU,142-SER,143-GLY,144-THR,145-GLY,146-PRO,221-GLN,223-THR,224-GLY,227-THR,231-GLU,233-LEU,241-TYR,269-THR,309-SER | 5FHC | 47.59 |
| 232-PRO,233-GLU,235-VAL,236-GLN,237-THR,255-GLN,263-GLU,270-ASN,271-ILE,272-GLN,273-PHE,274-SER,275-CYS | 3RAJ | 39.33 |
| 16-LYS,17-PHE,18-MET,19-ASP,21-TYR,22-GLN,23-ARG,25-TYR,61-CYS,66-LEU,101-LYS | 2FJH | 53.71 |
| 57-GLU,58-THR,61-GLN,68-THR,71-LEU,75-MET,98-ARG,102-GLY,105-GLN,106-SER,109-GLY,110-THR,111-GLN,112-LEU,113-PRO,114-PRO,115-GLN | 1V7M | 48.14 |
| 5-LEU,39-THR,40-GLY,41-LYS,44-ARG,174-ILE,175-ASN,176-HIS,177-SER,178-ALA,179-ASP,204-SER,205-LEU,215-ILE,217-GLU,219-TYR,220-ARG,221-ASN,223-TYR,224-LYS,226-ASN,227-ASP | 4ETQ | 48.29 |
| 308-ARG,309-ILE,310-GLN,314-GLY,315-ARG,316-ALA,317-PHE,318-VAL,319-THR,320-ILE,322-LYS,323-ILE,324-GLY,327-ARG,419-ARG,421-LYS,422-GLN,423-ILE,434-MET | 3J70 | 57.29 |
| 48-THR,49-GLY,50-LYS,60-ASP,62-ILE,63-ASP,74-PRO,75-HIS,78-VAL,90-ARG,92-LYS,94-PHE,143-PRO,271-ASP,272-ALA,273-PRO,274-ILE | 1QFU | 49.05 |
| 432-LYS,433-CYS,434-ILE,435-CYS,441-LEU,448-ASP,449-ALA,450-CYS,451-GLY,469-PHE,475-TYR,480-SER | 4ZFG | 47.43 |
| 238-VAL,241-LYS,243-VAL,266-LYS,267-HIS,270-LYS,271-LYS,272-LEU,275-ARG,293-THR,294-ILE,295-ASP,298-THR,299-ARG,326-GLU,327-LYS,328-PRO,329-PHE | 3S35 | 53.66 |
| 58-ARG,89-ARG,90-SER,91-ARG,92-TYR,94-GLU,112-SER,113-LEU,114-GLN,116-ARG,118-PRO,174-THR | 4QTI | 43.57 |
| 92-HIS,94-ASP,96-PHE,97-PRO,98-CYS,99-GLN,100-ASP,103-SER,104-LYS,105-ALA,107-LEU,160-CYS,162-PHE,163-SER,164-PRO,165-GLN,166-ILE,167-GLU,169-PRO,222-THR,224-ASP | 6I04 | 52.14 |
| 244-GLN,346-PHE,348-PRO,349-GLY,350-ILE,351-ARG,352-PRO,354-GLU,355-PRO,358-LYS,359-PHE | 2ZJS | 53.79 |
| 21-GLU,22-PRO,23-PRO,27-ARG,30-GLN,35-SER,36-GLN,37-CYS,39-SER,42-GLN,45-GLN,53-GLU,54-PHE,55-THR,56-GLU,57-THR,59-CYS,71-TRP | 6FAX | 52.26 |
| 58-TRP,65-ASP,66-SER,151-ARG,156-LYS,157-ASP,158-LYS,159-PRO,161-VAL,176-ARG,177-SER,180-ASN,200-ILE,201-GLY,202-TYR | 5OCC | 50.23 |
| 63-ASN,65-LYS,67-ASN,68-LYS,69-CYS,168-LYS,197-ASN,201-LYS,202-GLN,204-LEU,205-PRO,207-VAL,208-ASN,209-LYS,212-CYS,294-GLU,295-GLU | 5W23 | 46.4 |
| 6-PRO,18-VAL,21-THR,22-GLN,23-ASN,24-GLN,97-LYS,100-LEU,101-LEU,104-LYS,105-LYS,108-ARG,109-GLU | 3G6D | 53.15 |
| 64-ASN,65-ASN,66-THR,67-VAL,68-SER,69-CYS,70-SER,71-ASN,72-ARG,74-HIS,75-CYS,78-GLU,81-SER,150-ARG,151-PHE,153-ARG | 5J13 | 42.25 |
| 77-ARG,78-ASN,79-VAL,80-GLY,81-SER,82-GLN,84-LEU,149-ASN,154-ILE,158-ASN,205-SER,206-SER,208-ARG,210-GLY,211-ILE,213-LEU,214-TRP,215-HIS | 3T3P | 48.44 |
| 129-GLU,130-ALA,132-SER,141-ALA,142-GLU,143-LYS,144-GLY,146-TYR,178-CYS,179-SER,217-PRO,218-CYS,245-SER,246-GLN,248-SER,249-HIS,250-GLY,251-THR | 1I9R | 47.65 |
| 105-HIS,109-ILE,113-ASP,273-GLY,274-GLY,275-ASP,276-PRO,277-GLU,278-ILE,318-ARG,320-LYS,324-ASN,325-MET,372-GLY,373-ASP,375-ARG | 5UKR | 59.99 |
| 186-THR,187-GLU,188-PRO,189-LEU,190-MET,194-THR,196-ASP,197-GLU,199-ARG,200-HIS,201-PHE,202-TYR,203-LYS,204-ASP,205-ASN,208-VAL,222-GLY,223-ASN,224-MET,225-ILE,235-LYS | 2Q8A | 49.21 |
| 63-ASN,64-ILE,65-LYS,66-GLU,83-LEU,84-ASP,86-TYR,87-LYS,94-GLN,161-GLU,162-GLY,165-ASN,168-LYS,169-SER,172-LEU,204-LEU,208-ASN,294-GLU,295-GLU | 6OE4 | 50.33 |
| 430-LYS,431-GLU,433-SER,434-ASP,436-ASP,449-ASN,452-ASN,453-TRP,454-LYS,455-LYS,456-LEU,457-PHE,458-GLY,459-THR,460-SER,461-GLY,463-LYS,464-THR,466-ILE | 3C09 | 46.14 |
| 8-VAL,9-LYS,10-LYS,11-GLN,13-PRO,23-ASP,24-GLU,25-ARG,26-GLU,28-CYS,38-GLY,39-ASP | 1OB1 | 60.42 |
| 649-GLU,652-TYR,653-HIS,656-LEU,657-ARG,658-THR,659-GLN,660-GLY,661-PRO,665-ILE,668-TYR,676-VAL,678-PHE,679-SER,680-THR,681-ASP,682-ASP,728-ARG,735-LEU,736-GLN,739-GLN | 5W2B | 41.05 |
| 256-ARG,257-MET,258-ALA,259-TRP,260-PRO,315-TYR,317-VAL,318-GLY,342-GLU,343-ASP,350-THR,351-ARG,354-GLU,355-LEU,357-GLY,358-LEU,360-GLN,367-VAL,369-LEU,371-PRO,373-HIS | 5BV7 | 32.15 |
| 14-GLN,15-ALA,131-GLN,132-GLU,133-SER,134-PHE,135-GLU,192-MET,193-PRO,194-GLU,195-THR,197-THR,199-ILE,200-GLY,201-GLU,202-LYS,336-ARG,340-PHE | 6GV1 | 46.91 |
| 60-SER,62-SER,63-PHE,64-VAL,66-ASN,68-TYR,76-THR,78-LYS,83-PRO,85-ASP,86-ARG,87-SER,88-GLN,89-PRO,90-GLY,128-LEU,131-LYS,132-ALA,134-ILE | 5GGS | 49.4 |
| 119-GLU,120-ILE,125-SER,126-SER,128-PRO,129-ASN,165-SER,166-LYS,167-SER,168-TYR,169-ALA,170-ASN,171-ASN,173-GLU,240-GLY,242-THR | 5W6G | 49.06 |
| 96-TRP,275-GLU,276-ASN,277-LEU,278-THR,279-ASN,280-ASN,281-ALA,282-LYS,365-SER,366-GLY,367-GLY,368-ASP,371-ILE,429-GLY,455-THR,456-ARG,457-ASP,458-GLY,459-GLY,469-ARG,472-GLY,480-ARG | 5F96 | 54.78 |
| 141-TYR,143-ASP,144-THR,146-LEU,162-ILE,164-THR,166-ARG,167-SER,168-ARG,169-ALA,170-CYS,171-HIS,172-PRO,173-CYS,179-GLY,180-SER,181-ARG | 5O4G | 45.18 |
| 193-ILE,202-LYS,205-ALA,209-PHE,212-LYS,213-ILE,216-ALA,328-ILE,331-ASP,335-TYR,338-ASN,339-LEU,341-GLU,342-GLN,344-SER | 4U0R | 38.22 |
| 329-LEU,330-VAL,332-PHE,333-LYS,335-MET,337-TYR,338-GLU,345-LYS,396-LYS,399-ILE,400-ASP,403-THR,407-ASP | 6BPA | 44.48 |
| 12-ASN,14-LYS,15-GLY,16-PRO,17-PHE,18-PRO,48-VAL,52-ILE,63-THR,64-GLU,65-PRO,66-LEU,67-ASP,70-ARG | 6CXY | 51.06 |
| 18-THR,19-GLY,20-LEU,22-GLU,23-ASN,26-VAL,29-ASP,30-ASP,31-ASN,58-LEU,59-GLY,60-ASN,88-ASN,90-TYR,110-ARG,177-PHE,178-ASN | 4RGM | 45.71 |
| 353-PHE,354-PRO,355-ASN,357-LYS,358-ALA,360-LYS,456-ARG,462-ASP,463-SER,464-ASN,465-THR,466-GLU | 5F6J | 47.45 |
| 20-MET,21-SER,22-GLY,23-PRO,25-GLU,27-LYS,31-LEU,34-GLN,35-ASP,37-GLU,38-GLN,39-GLN,64-GLU,65-LYS,66-ASN,86-ASP,87-PRO,88-LYS,129-ASN | 4G6J | 52.65 |
| 142-GLY,143-SER,144-ASP,145-TYR,147-ASP,148-ARG,149-TYR,151-ARG,152-GLU,153-ASN,155-HIS,156-ARG,197-ASN,199-THR,204-LYS | 2W9E | 53.22 |
| 122-TYR,125-GLN,146-ARG,150-GLY,152-ASN,176-HIS,197-LYS,198-ARG,199-LEU,200-TYR,201-ARG | 4OQT | 45.27 |
| 152-ILE,154-THR,156-TYR,164-GLY,165-LYS,166-LYS,167-THR,168-ALA,169-LYS,170-THR,171-ASN,190-GLN,192-VAL,194-PRO,197-THR,198-VAL,199-ASN,200-ARG,201-LYS,203-THR,204-ASP | 1JPS | 46.79 |
| 9-ASP,10-VAL,11-ASN,13-THR,15-PHE,41-PRO,42-TYR,56-ARG,59-GLY,60-PRO,61-GLY,62-ARG,63-CYS,64-HIS | 4AEI | 60.98 |
| 97-LYS,276-ASN,278-THR,279-ASN,280-ASN,365-SER,366-GLY,367-GLY,368-ASP,371-ILE,428-GLN,429-GLY,456-ARG,457-ASP,458-GLY,459-GLY,460-ALA,465-ASN,466-GLU,471-GLY,473-GLY | 4S1S | 57.56 |
| 21-GLN,22-GLU,23-ASP,24-ALA,25-THR,30-THR,120-PRO,121-LYS,137-ALA,138-SER,141-LYS | 6IEB | 57.08 |
| 13-HIS,20-THR,21-GLN,23-ASP,24-ALA,27-LYS,28-PRO,120-PRO,138-SER,139-LEU,140-LYS,141-LYS | 6IEK | 50.23 |
| 144-HIS,145-ILE,179-GLU,180-GLY,213-HIS,214-GLY,215-LYS,216-ILE,232-ASP,233-THR,234-VAL,235-GLN,236-LEU,237-ASN,245-GLU,247-LYS,266-GLU,267-VAL,268-SER,269-SER,271-ASN,272-PRO,273-GLU,274-TYR | 5VL3 | 42.42 |
| 276-ASN,278-THR,280-ASN,281-ALA,365-SER,366-GLY,367-GLY,368-ASP,371-ILE,429-GLY,456-ARG,457-ASP,458-GLY,459-GLY,460-ALA,461-ASN,465-ASN,466-GLU,467-THR,469-ARG,473-GLY,476-LYS | 4OLW | 52.59 |
| 123-GLY,125-GLN,126-LEU,127-ASP,128-LYS,129-TYR,130-ARG,131-SER,149-GLN,153-LYS,226-LYS | 4RGN | 56.13 |
| 349-PRO,353-ARG,382-LEU,384-GLN,408-GLN,409-HIS,411-GLN,412-PHE,417-VAL,418-SER,438-ILE,440-SER,443-LYS,465-LYS,466-ILE,467-ILE,468-SER,469-ASN,471-GLY,473-ASN | 6ARU | 43.57 |
| 336-GLU,367-TYR,372-ASP,373-GLU,374-ARG,375-ASP,377-PRO,379-ASN,380-TRP,389-THR,391-MET,393-GLN,394-ALA,395-TRP,396-GLY,399-LEU,414-ILE,415-GLY | 2OZ4 | 46.33 |
| 22-ASP,23-ALA,26-TYR,45-LEU,46-GLU,48-GLN,49-VAL,52-LEU,53-GLU,55-GLY,64-GLU,67-ILE,68-ILE,71-ASN,72-ASN,75-SER,93-GLU | 2XQB | 54.98 |
| 1-GLU,2-LEU,4-ASP,5-ASP,6-ASP,25-MET,27-ASN,42-LEU,43-TYR,118-ILE,120-HIS,149-MET,150-THR,151-HIS,152-GLY,153-LYS,154-THR,155-ARG | 3NFP | 59.39 |
| 10-THR,11-ARG,68-TYR,270-ARG,271-LYS,272-ASP,274-ASP,431-LYS,434-GLU,435-GLU,438-ARG,439-GLY,443-LYS | 3TT1 | 45.02 |
| 150-HIS,197-HIS,198-ASP,199-GLU,218-TRP,219-SER,220-LYS,221-LYS,222-ILE,224-ARG,244-GLY,245-SER,246-ALA,248-GLY,249-ARG,250-ALA,251-ASP | 2AEP | 39.91 |
| 43-HIS,60-PHE,62-ILE,64-GLU,67-ASP,74-ALA,77-CYS,78-MET,79-LYS,80-TYR,81-GLU,84-ASP,86-GLN,90-SER,91-PRO,92-LYS,93-ALA,94-GLN,95-LEU,96-ARG,97-ARG | 3NH7 | 50.95 |
| 116-ILE,120-SER,121-SER,161-LYS,162-ARG,163-SER,164-TYR,165-ASN,167-THR,201-GLY,202-THR,203-SER,238-ALA,240-ASN | 6IUV | 46.26 |
| 35-SER,37-GLN,40-GLY,41-LEU,42-ASP,43-THR,44-SER,45-LEU,46-GLY,47-ALA,48-VAL,49-GLN,50-SER,51-ASP,52-THR | 4HC1 | 62.38 |
| 5-ASP,6-PRO,8-PRO,135-ARG,137-PHE,139-ASP,140-GLY,141-LYS,142-ASN,145-SER,186-TYR,188-LYS,190-ILE,229-LYS,231-GLU,233-TYR,235-THR,238-LYS | 4RGO | 51.57 |
| 524-LYS,525-ARG,526-SER,527-GLY,528-TYR,529-GLY,530-GLN,571-TRP,630-ARG,631-THR,632-ASN | 4F2M | 62.32 |
| 502-LYS,506-PHE,510-ASP,511-ARG,513-GLU,535-TRP,536-GLU,540-TYR,542-ARG,553-TRP,555-VAL,557-SER | 6C6Z | 51.38 |
| 557-PRO,558-GLU,560-ASP,561-GLN,569-LYS,570-ASP,571-PRO,572-PRO,573-PHE,588-TYR,589-MET,591-ILE,593-LYS,596-ASP,600-ALA,602-GLN,603-PRO,604-CYS | 6BGT | 49.68 |
| 320-ASN,322-GLU,337-ARG,339-ARG,340-PHE,343-LYS,352-ASP,353-GLY,354-ILE,356-ASP,360-ASP,361-LEU,362-HIS,363-MET,364-GLY,365-THR,366-GLN,367-LYS,383-GLU,425-GLU | 5O1R | 46.27 |
| 614-ARG,615-HIS,616-GLY,619-ASN,620-TYR,621-ARG,662-TRP,663-TRP,672-SER,673-ARG,674-ASN,675-LEU,677-ASN,689-THR,693-ASN,696-LYS,697-LYS,698-ILE,700-SER,701-ASP,702-GLY | 1NDG | 47.76 |
| 7-ASP,31-LYS,32-ALA,33-GLY,34-PRO,35-HIS,37-PRO,38-THR,39-ALA,56-GLN,63-ILE,68-LEU | 4RAU | 60.77 |
| 68-ASP,69-GLU,71-ILE,72-ASN,110-HIS,112-GLU,113-LYS,114-ILE,115-GLN,119-LYS,137-TYR,138-GLN,145-ARG,167-THR,168-ASN,169-GLN,170-GLU,252-TYR,255-LYS,257-VAL,258-LYS | 5DUR | 43.8 |
| 31-VAL,32-ASP,55-ILE,57-GLU,59-ILE,81-TYR,87-LEU,89-ASP,90-GLY,91-VAL,92-LEU,97-LEU,126-GLY,130-ARG,132-ARG,134-LEU,135-PRO,136-TRP | 4HT1 | 44.54 |
| 2-PRO,3-GLN,4-GLN,18-ASN,20-ARG,21-LEU,56-PHE,58-PRO,61-TYR,100-LYS,109-LYS,110-HIS,111-ASP,112-ASP,115-ILE,168-THR,170-LEU,171-GLY,179-ASP,181-VAL,228-ASP,229-THR,232-TYR | 5USH | 41.15 |
| 294-SER,295-SER,296-GLN,297-TYR,298-SER,299-THR,302-SER,318-GLY,319-GLU,321-SER,323-ARG,324-GLU | 2QQN | 45.83 |
| 301-MET,303-THR,305-LYS,327-GLU,328-GLY,336-PRO,337-PHE,338-GLU,345-ARG,381-GLY,382-VAL,383-GLU,384-PRO,385-GLY,386-GLN | 6FLB | 46.42 |
| 127-GLN,128-HIS,129-PRO,130-ALA,131-ASP,133-SER,134-TRP,137-LYS,139-ASP,142-ILE,143-TRP,146-GLN,156-TRP,157-TYR,167-ARG | 5W1K | 47.41 |
| 186-THR,187-GLU,188-PRO,189-LEU,190-MET,193-MET,194-THR,196-ASP,197-GLU,200-HIS,201-PHE,202-TYR,203-LYS,204-ASP,205-ASN,208-VAL,223-ASN,224-MET,225-ILE,228-ASN,230-LYS | 2Q8B | 51.53 |
| 21-GLU,21-SER,21-LYS,21-ALA,21-ASP,22-ARG,22-HIS,22-GLN,22-GLY | 4KI5 | 51.56 |
| 20-TYR,20-GLU,20-ASN,20-THR,20-ASP,20-GLY,20-PHE,20-SER,20-ALA,20-VAL | 4NP4 | 42.1 |
| 30-THR,30-ALA,30-LEU,30-GLU,30-PRO,30-TYR,30-ASN,30-GLN,30-CYS,30-SER,30-VAL,31-ILE,31-LYS,31-GLY | 1YNT | 41.79 |
| 35-ASN,38-LEU,39-GLU,52-GLU,54-ASP,55-LYS,60-TYR,62-SER,74-LYS,77-CYS,78-TRP,79-LEU,80-ASP,81-ASP,82-PHE,83-ASN,85-TYR,94-GLU,99-VAL,101-PHE | 5NGV | 45.66 |
| 1-GLU,2-LEU,4-ASP,5-ASP,6-ASP,22-GLU,23-GLY,25-MET,27-ASN,29-GLU,36-ARG,38-LYS,39-SER,40-GLY,42-LEU,43-TYR,45-LEU,47-THR,48-GLY,56-ASP,57-ASN,118-ILE,120-HIS | 3IU3 | 53.53 |
| 14-ARG,15-HIS,16-GLY,18-ASP,19-ASN,20-TYR,21-ARG,62-TRP,63-TRP,73-ARG,75-LEU,77-ASN,89-THR,93-ASN,96-LYS,97-LYS,100-SER,101-ASP,102-GLY,103-ASN | 1XGU | 49.82 |
| 301-LYS,315-THR,316-LYS,319-ALA,320-GLU,321-THR,322-LEU,327-THR,329-GLU,362-ASN,364-VAL,365-ILE,366-THR,367-GLU,372-SER,373-LYS,374-MET,375-MET,377-GLU | 5KVD | 38.82 |
| 100-TYR,101-ASP,216-ASN,217-ILE,218-GLY,219-SER,220-ARG,221-PRO,222-TRP,223-VAL,224-ARG,229-ARG | 6OCB | 46.77 |
| 149-LYS,152-ILE,154-THR,156-TYR,165-LYS,166-LYS,167-THR,168-ALA,169-LYS,170-THR,171-ASN,178-ASP,190-GLN,192-VAL,194-PRO,196-ARG,197-THR,198-VAL,200-ARG,201-LYS,204-ASP | 1AHW | 42.83 |
| 7-THR,9-CYS,10-GLY,11-GLY,12-GLU,14-VAL,15-ASP,17-LEU,26-PHE,34-ARG,35-VAL,40-ARG,43-VAL,44-GLU,45-GLU,47-CYS,48-PHE,49-ARG,59-TYR | 3KR3 | 44.98 |
| 48-THR,49-ASP,51-GLU,56-ALA,57-ASP,58-TYR,59-SER,60-MET,61-PRO,84-ARG,85-VAL,86-ALA,87-ASN,89-PRO,90-PHE,91-SER | 4JZJ | 51.97 |
| 15-GLU,16-CYS,17-LEU,18-LYS,19-LEU,22-ASN,24-TYR,27-ARG,42-TYR,43-ALA,44-PHE,45-ALA | 2YC1 | 48.37 |
| 505-ARG,507-LEU,514-VAL,516-GLN,519-ASN,521-ASN,522-GLN,523-TYR,525-PRO,528-SER,543-LYS,546-SER,547-PRO,548-LEU,549-GLU,550-GLY,554-LEU | 5YY5 | 49.95 |
| 25-GLU,26-ALA,31-GLN,32-THR,33-LYS,35-ASP,58-SER,59-ALA,60-ASP,61-ALA,63-ALA,125-LYS,127-LYS | 2I9L | 51.05 |
| 31-SER,32-PRO,33-PRO,55-GLU,56-LYS,57-ASP,58-ASP,59-PHE,60-HIS,61-HIS,65-VAL,67-ARG,92-ARG | 5B3J | 58.7 |
| 73-GLU,77-ARG,160-LEU,161-LYS,164-GLN,391-LYS,393-ASP,394-ASP,395-PHE,396-VAL,397-GLU | 5XBM | 58.23 |
| 36-LEU,39-ASP,40-GLY,41-GLN,42-PHE,43-CYS,44-HIS,45-LYS,46-PRO,47-CYS,48-PRO,49-PRO,59-VAL,60-ASN,78-LYS,79-ALA,81-PHE,86-ARG | 3THM | 56.28 |
| 448-THR,449-GLY,450-GLN,453-ARG,472-THR,473-ARG,474-ASN,477-ALA,478-LEU,498-SER,499-PRO,503-GLN,504-PRO,506-ARG,522-ASN,523-ASP,524-ASP,527-GLU | 3ULU | 39.6 |
| 502-LYS,506-PHE,510-ASP,513-GLU,535-TRP,536-GLU,538-GLY,540-TYR,542-ARG,553-TRP,555-VAL,556-ALA,557-SER | 4XAK | 47.34 |
| 16-VAL,20-GLN,23-ARG,49-LEU,50-GLN,83-HIS,86-GLN,87-HIS,88-CYS,89-PRO,114-LEU,115-LEU,116-VAL,117-ILE,118-PRO,119-PHE,120-ASP,121-CYS,122-TRP,123-GLU,124-PRO | 6BFS | 40.91 |
| 321-ASP,322-SER,345-PRO,346-GLY,347-ASP,349-ILE,351-ASP,354-PHE,355-GLN,369-ILE,371-TYR,392-LYS,394-LYS,411-ILE,413-LYS,414-LYS,415-ASP,416-LYS | 6E63 | 44.02 |
| 314-SER,316-GLU,317-LYS,318-PHE,319-PRO,327-GLN,328-LEU,329-VAL,330-CYS,331-TRP,332-GLN,333-ALA,335-THR,337-PRO,340-ILE,375-THR,378-ASP,380-CYS,426-PHE | 3R1G | 46.72 |
| 226-SER,228-GLU,229-GLU,230-CYS,231-SER,232-LYS,233-ASP,234-LYS,235-ASN,236-ASN,239-CYS,242-GLU | 4OKV | 56.12 |
| 8-LYS,9-THR,11-LYS,226-GLU,230-GLU,234-GLN,235-THR,238-GLY,239-TRP,240-LYS,241-ASP,242-VAL,243-ASP,244-MET,246-MET,247-GLU,250-ARG | 4YBQ | 47.85 |
| 32-GLU,34-ARG,49-VAL,51-TYR,59-GLN,61-TYR,62-SER,63-LYS,64-THR,65-GLY,97-GLU,99-MET,100-TYR,104-TYR | 1YJD | 43.04 |
| 63-PRO,64-LEU,68-LYS,70-TRP,71-ALA,72-PHE,100-PRO,123-HIS,125-ILE,126-GLN,128-GLN,129-ASN,130-PRO,131-HIS | 5UQY | 53.23 |
| 56-ARG,60-PHE,75-HIS,100-ALA,101-GLY,119-ASN,120-TRP,121-PHE,122-ASP,123-ILE,124-THR,127-LEU | 4M62 | 50.83 |
| 41-ASN,67-TRP,72-PHE,74-LYS,75-GLU,76-VAL,87-GLU,90-ARG,91-PRO,92-GLY,110-PRO,111-GLY,113-GLN,115-GLU,116-GLU,203-MET,204-ASP | 4ZSO | 52.67 |

| | | |
|---|---|---|
| 281-ALA,283-THR,365-SER,366-GLY,367-GLY,368-ASP,371-ILE,372-VAL,373-THR,425-ASN,474-ASP | 3IDX | 53.54 |
| 19-ASP,20-GLY,21-TRP,38-LYS,41-THR,42-GLN,45-ILE,46-ASP,48-VAL,49-THR,52-VAL,53-ASN | 3GBM | 47.56 |
| 25-GLN,27-HIS,29-VAL,31-GLU,32-TYR,33-ASP,34-PRO,36-ILE,37-GLU,38-ASP,39-SER,40-TYR,61-GLN,63-GLU,64-TYR | 2UZI | 51.22 |
| 310-LYS,313-ALA,314-GLU,315-THR,316-GLN,317-HIS,321-LEU,352-ILE,354-ALA,364-PRO,368-GLU | 4AL8 | 51.01 |
| 39-MET,71-PHE,74-GLU,75-GLU,78-PRO,79-GLN,82-ASN,83-GLN,125-LYS,128-PHE,129-ASN,130-LYS,131-LEU,132-GLN,133-GLU,134-LYS,136-ILE,137-TYR | 1LK3 | 42.61 |
| 112-TRP,281-ALA,365-SER,366-GLY,367-GLY,368-ASP,370-GLU,371-ILE,426-MET,429-GLU,473-GLY,474-ASP,476-ARG | 3HI1 | 57.65 |
| 45-TYR,49-LEU,50-ALA,52-ARG,53-GLY,54-ALA,55-PRO,56-GLY,57-ALA,58-GLN,60-ILE,61-THR,62-TYR,64-ARG | 2IH3 | 55.63 |
| 47-TYR,51-ARG,53-GLN,54-SER,55-LEU,83-ILE,84-GLN,85-HIS,86-ASN,87-GLY,90-GLN,93-TYR,107-ASN,108-ALA,109-GLN,110-ARG,111-PHE,112-GLY,113-ILE,114-SER,221-ILE,222-LEU | 5VCN | 39.89 |
| 251-LEU,252-MET,253-ILE,254-SER,310-HIS,311-GLN,314-LEU,382-GLU,384-ASN,385-GLY,424-SER,431-ALA,432-LEU,433-HIS,434-ASN,435-HIS,436-TYR,437-THR,438-GLN,440-SER | 5XMH | 46.05 |
| 64-SER,65-ILE,66-SER,67-ASP,68-MET,69-ALA,84-LYS,87-ASP,89-GLN,90-TYR,118-LYS,119-PHE,120-ALA,233-THR,252-ARG | 5GZO | 41.79 |
| 76-GLY,87-ASN,88-GLU,90-LYS,91-GLN,92-VAL,102-ASP,105-ASN,106-PRO,107-VAL,108-LYS,109-THR,110-GLY,111-VAL,122-VAL,123-GLN,125-GLN,126-ASN | 6AZZ | 59.68 |
| 41-ARG,126-LYS,128-GLN,130-ARG,214-TYR,215-ASN,217-ALA,218-GLU,219-LYS,220-GLY,221-SER,239-GLU,241-LYS,243-VAL,244-ASN,245-GLY,246-ILE,248-HIS | 2YPV | 52.69 |
| 32-SER,52-ILE,53-ASN,54-VAL,55-SER,56-GLY,57-CYS,58-SER,59-ALA,60-ILE,61-GLU,62-LYS,64-GLN,65-ARG,66-MET,68-SER,69-GLY,74-LYS,76-SER,77-ALA | 4I77 | 49.99 |
| 24-GLU,25-PRO,27-THR,28-PHE,30-VAL,42-CYS,43-ASP,44-PRO,45-CYS,46-ILE,47-PRO,49-VAL,63-SER,64-CYS | 5TLK | 43.45 |
| 363-VAL,365-SER,367-ARG,387-GLU,390-LEU,391-LYS,393-LEU,394-HIS,395-GLY,396-ARG,397-LYS,401-ASP,408-ASP,410-ILE,454-LYS | 4LVN | 45.85 |
| 102-PHE,103-ARG,104-VAL,148-MET,167-ASP,195-HIS,199-TYR,201-ARG,202-ASN,203-THR,204-SER,205-ASN,257-ARG,259-ARG,261-LEU,262-ASP,265-LYS,266-ILE,269-LEU | 6CW3 | 46.29 |
| 276-ASN,278-THR,279-ASN,280-ASN,281-ALA,365-SER,366-GLY,367-GLY,368-ASP,370-GLU,371-ILE,425-ASN,427-TRP,429-GLY,457-ASP,458-GLY,459-GLY,460-ALA,461-ASN,465-ASN,466-GLU,467-THR,469-ARG,473-GLY | 4OLZ | 52.12 |
| 34-THR,35-THR,37-GLN,46-ASN,47-PHE,48-LYS,96-TRP,98-LYS,102-SER,103-SER,104-TYR,140-SER,145-ASN,147-SER,148-ALA,149-PRO,151-ASP | 5USL | 46.74 |
| 256-SER,280-ASN,364-HIS,365-SER,366-GLY,367-GLY,368-ASP,369-LEU,370-GLU,371-ILE,455-THR,457-ASP,458-GLY,459-GLY,460-ASN,461-ASP,462-ASP,463-ASN,469-ARG,470-PRO,471-GLY,472-GLY | 4JAN | 51.36 |
| 20-MET,21-SER,22-GLY,23-PRO,27-LYS,32-GLN,34-GLN,35-ASP,37-GLU,38-GLN,39-GLN,41-VAL,64-GLU,65-LYS,66-ASN,86-ASP,87-PRO,88-LYS,129-ASN | 5BVP | 51.6 |
| 8-HIS,9-ARG,10-GLY,11-GLU,12-PHE,13-SER,17-SER,19-SER,20-VAL,21-TRP,23-GLY,52-TYR,54-PHE,56-THR,106-THR,107-ALA,108-CYS | 4EDW | 57.26 |
| 89-GLU,91-SER,92-THR,94-GLU,95-ILE,96-PHE,97-GLY,98-ASP,99-SER,116-TYR,117-ASN,138-ARG,139-GLN,140-LEU,241-ASP,243-SER,747-THR | 5VTA | 56.4 |
| 451-VAL,557-ARG,589-ASP,591-LYS,628-PRO,638-TYR,639-CYS,640-ASP,641-VAL,642-PHE,644-ARG,646-ARG | 5L0Q | 59.58 |
| 144-PRO,147-PHE,148-GLN,151-LEU,155-LYS,191-ASP,193-ASP,194-ALA,197-LYS,198-HIS,199-VAL,200-LYS,201-HIS,203-LEU | 3EOA | 42.28 |
| 308-ILE,309-ASP,310-LYS,311-GLU,312-MET,313-ALA,314-GLU,315-THR,316-GLN,323-LYS,344-LYS,364-VAL,377-TYR,390-HIS,391-TRP,392-PHE,393-ARG,394-LYS,396-HIS | 4BZ2 | 47.58 |
| 373-THR,374-TRP,375-SER,376-ARG,377-ALA,378-SER,379-GLY,380-LYS,412-GLU,414-GLU,417-GLN,419-ARG,430-MET | 5HYS | 46.79 |
| 305-SER,306-PHE,307-LYS,309-GLU,310-LYS,325-LYS,326-TYR,327-GLU,328-GLY,329-THR,330-ASP,361-LYS,362-GLU | 4L5F | 47.18 |
| 21-ARG,23-TYR,103-ASP,106-ASN,112-ARG,113-LYS,116-LYS,117-GLY,118-THR,119-ASP,121-ASN | 1JHL | 58.19 |
| 60-ASN,62-ASP,63-ARG,64-TYR,65-PHE,93-ASP,94-ASN,95-ILE,96-ASN,132-PHE,134-LYS,135-LYS,136-VAL | 5IKC | 52.03 |
| 359-THR,363-THR,365-LYS,390-LYS,391-GLY,392-ASP,395-ARG,426-ARG,436-TYR,482-GLY,484-TYR,485-THR,486-THR,487-THR,488-GLY,489-ILE,491-TYR,492-GLN,494-TYR | 2DD8 | 43.14 |
| 407-ALA,408-LYS,409-GLN,427-LEU,428-ASN,429-CYS,430-ASN,431-ASP,432-SER,433-LEU,434-ASN,435-THR,436-GLY,437-TRP,438-LEU,439-ALA,443-TYR,446-LYS,447-PHE,448-ASN | 6MEJ | 55.8 |
| 116-ILE,121-SER,123-SER,124-ASP,159-THR,161-LYS,162-ARG,163-SER,164-TYR,165-ASN,240-ASN,241-PHE,242-GLU | 5DUM | 42.51 |
| 41-GLY,42-GLU,45-GLU,63-ASP,66-LYS,67-GLU,68-THR,69-ASP,70-LEU,71-THR,91-GLU,99-LYS,103-THR,116-ASN,117-ASN,118-ALA,119-GLU | 5TIH | 52.67 |
| 7-LEU,8-ARG,9-CYS,12-LEU,13-GLN,32-GLY,33-PRO,34-HIS,35-CYS,36-ALA,37-GLN | 5OB5 | 59.49 |
| 206-SER,207-SER,208-ALA,209-ALA,211-LYS,213-THR,214-ALA,215-ALA,226-THR,228-ASN,229-SER,230-LYS,249-ASP,250-SER,251-ASN,252-GLY,253-THR | 1FJ1 | 54.35 |
| 314-THR,315-VAL,316-GLY,317-GLU,318-GLY,319-LEU,320-ASN,367-SER,369-PRO,370-ARG,372-LYS,373-PRO,375-GLU,401-PRO,402-PRO,493-SER | 4LIQ | 45.61 |
| 196-LYS,197-SER,198-SER,344-SER,345-CYS,346-TYR,348-ASN,349-ASN,350-PHE,352-ASN,354-ASN,451-GLN,452-LYS | 4U1G | 57 |
| 122-LEU,124-GLY,276-ASN,278-THR,280-ASN,281-ALA,365-SER,366-GLY,367-GLY,368-ASP,370-GLU,371-ILE,425-ASN,427-TRP,431-GLY,432-GLN,456-ARG,457-ASP,458-GLY,459-GLY,467-THR,469-ARG,473-GLY,476-LYS | 4JKP | 49.12 |
| 250-PHE,253-PRO,255-GLY,256-ALA,257-PHE,396-HIS,397-GLN,398-ASN,401-GLN,435-ARG,446-ASN,502-THR,503-GLY,504-GLN,505-HIS,506-ASP | 6N8D | 50.52 |
| 57-HIS,62-GLU,64-GLY,65-LEU,66-GLU,68-HIS,69-GLN,70-PHE,71-TRP,72-PRO,73-LEU,120-MET,123-TYR,125-PHE,128-PRO,130-ARG | 6O39 | 37.04 |
| 156-ILE,157-PRO,159-GLU,160-HIS,161-SER,166-SER,181-ARG,190-PHE,194-PRO,197-ARG,198-TYR | 4D3C | 54.58 |
| 70-ASN,71-GLY,201-LYS,202-GLN,205-PRO,206-ILE,208-ASN,209-LYS,210-GLN,211-SER,213-SER | 6APD | 63.2 |
| 15-HIS,16-GLY,20-TYR,21-ARG,61-ARG,62-TRP,73-ARG,75-LEU,77-ASN,93-ASN,96-LYS,97-LYS,100-SER,101-ASP | 1FBI | 47.36 |
| 45-THR,46-TYR,47-SER,48-LYS,49-PRO,72-SER,73-GLY,74-PHE,75-GLN,76-ARG,78-SER,92-TYR,115-SER,118-ASN,120-THR,123-GLN,137-VAL,138-GLN,139-ILE,140-SER,142-GLU,143-ASP,146-ILE | 5VOB | 57.67 |
| 23-GLU,24-GLY,25-GLN,65-LYS,66-GLY,67-GLN,69-CYS,70-PRO,104-GLU,105-THR,107-GLU,108-GLY,109-ALA,110-GLU,111-ALA,138-ARG,139-PRO,140-ASP,141-TYR | 5YOY | 51.58 |
| 226-ARG,255-ASP,256-VAL,257-LEU,258-PRO,259-ASP,260-GLY,261-ASN,265-GLN,266-THR,267-TRP | 6DDV | 54.39 |
| 14-SER,14-ARG,14-THR,14-CYS,14-ASN,14-PRO,14-VAL,14-TYR,14-GLU,14-GLN,14-GLY,16-ARG,16-ASP | 5CZX | 61.46 |
| 32-ILE,33-LYS,36-GLN,143-ARG,146-ASN,150-SER,153-HIS,199-HIS,202-GLN,203-LEU,204-THR,206-TRP,207-GLY,209-LYS,210-GLN,211-LEU,213-ALA,214-ARG | 2CMR | 45.05 |
| 30-VAL,33-ASP,34-LEU,35-PRO,42-VAL,43-SER,44-LYS,45-GLU,46-LYS,52-TYR,69-LYS,70-ASN,71-ASN,92-ASP,93-ASP,94-LEU,95-GLY,117-LYS,118-ASP,119-LYS | 1OSP | 44.19 |
| 57-GLN,77-ASP,78-ASP,80-GLN,81-ASN,82-LYS,83-GLU,119-GLU,120-PHE,121-THR,122-GLN,124-SER,133-ASP,141-ARG,144-VAL,172-ASP,257-TYR,259-LYS | 5W42 | 50.13 |
| 18-SER,19-GLY,87-ASN,88-GLU,91-GLN,92-VAL,93-THR,102-ASP,104-SER,105-ASN,106-PRO,107-VAL,108-LYS,111-VAL,112-CYS,120-PRO,123-GLN,125-GLN,126-ASN,127-LYS | 6B0A | 55.52 |
| 15-HIS,16-GLY,20-TYR,21-ARG,62-TRP,63-TRP,73-ARG,74-ASN,75-LEU,77-ASN,89-THR,93-ASN,96-LYS,97-LYS,98-ILE,100-SER,101-ASP,102-GLY,103-ASN | 3A6C | 46.04 |
| 95-TYR,134-GLY,135-VAL,136-SER,137-ALA,143-GLY,144-GLU,145-SER,153-TRP,155-THR,157-LYS,158-ASN,159-GLY,190-ASP,194-LEU,196-HIS,226-ARG | 5UG0 | 42.99 |
| 615-HIS,616-GLY,618-ASP,619-ASN,620-TYR,621-ARG,663-TRP,673-ARG,675-LEU,689-THR,693-ASN,696-LYS,697-LYS,698-ILE,700-SER,701-ASP,702-GLY,703-ASN | 1NDM | 45.41 |
| 16-HIS,19-LEU,20-ASP,23-MET,76-LYS,77-ASN,78-PHE,79-HIS,80-LEU,81-ARG,84-ASP,85-LEU,88-ASN | 5UTZ | 47.34 |
| 251-SER,252-SER,253-SER,256-PHE,275-LYS,276-THR,277-SER,278-GLY,284-MET,286-PHE,287-CYS,288-LYS,289-VAL,290-ALA,291-GLY,292-CYS,293-GLU,294-HIS | 5Y11 | 41.23 |
| 283-THR,365-SER,366-GLY,367-GLY,368-ASP,369-LEU,371-ILE,372-THR,373-MET,384-TYR,419-LYS,420-ILE,421-LYS,423-ILE,455-THR,469-ARG,470-PRO,471-GLY,472-GLY,473-GLY,474-ASN | 4YDJ | 41.6 |
| 433-VAL,434-PHE,435-PRO,436-GLY,496-GLU,529-VAL,530-ASN,531-GLN,532-PHE,533-TYR,534-THR,535-LEU,536-ALA,537-PRO | 3V7A | 50.63 |
| 45-LEU,50-THR,51-GLY,178-VAL,264-MET,265-PRO,266-ILE,268-ASN,269-ASP,272-LYS,273-LEU,305-LEU,309-ILE,310-ASP,311-THR,312-PRO,344-ASP,345-ASN,346-ALA,347-GLY,364-ARG | 6APD | 43.6 |
| 25-LEU,26-ASP,27-GLN,38-TYR,39-HIS,139-ASP,140-SER,193-LEU,196-ARG,215-ASP,216-SER,218-GLY,220-LEU,221-PRO,222-ARG,223-PHE,224-ILE,227-ASN,231-VAL,234-TYR,235-SER,238-ILE,239-ALA | 3W9E | 51.18 |
| 190-TYR,201-ARG,202-PRO,203-ARG,204-ASP,205-ASP,206-PHE,207-PHE,209-HIS,211-ALA,221-MET,222-GLU,224-TRP,246-LYS,247-LEU,250-ASP,252-ARG | 5BO1 | 55.08 |
| 109-LYS,111-SER,113-ASP,114-ASP,115-ILE,116-ALA,117-VAL,119-LEU,122-TYR,169-LYS,170-THR,171-GLU,210-GLY,211-LYS,212-LEU,216-LYS,218-GLN,219-PRO,220-THR | 5EN2 | 49.97 |
| 408-VAL,412-LYS,476-ASP,484-GLU,486-ASN,487-GLU,488-VAL,489-ALA,491-GLU,492-ASN,493-GLU,497-ARG,498-ASP,499-GLY,500-ALA,502-ILE,503-GLU | 6R2S | 46.98 |
| 164-HIS,167-PRO,168-ALA,169-ALA,170-ASN,171-THR,172-VAL,173-LYS,175-ARG,205-LYS,207-ARG,210-HIS,214-ILE,216-GLU,248-ARG | 4WV1 | 49.86 |
| 62-SER,63-PRO,64-LEU,98-THR,99-ASP,100-PRO,101-SER,102-GLY,121-THR,122-VAL,123-HIS,124-HIS,126-GLN,127-GLY,128-GLN,130-PRO | 6BP2 | 53.65 |
| 51-THR,52-LEU,53-PHE,58-ALA,60-ALA,71-THR,72-HIS,73-ALA,74-CYS,75-VAL,76-PRO,77-THR,78-ASP,79-PRO,103-GLN,106-GLU,107-ASP,114-GLN | 4H8W | 49.48 |
| 85-GLU,90-VAL,101-TYR,133-ASN,215-SER,216-ARG,217-PRO,218-TRP,219-VAL,220-ARG,221-GLY,266-ASN | 6N5B | 52.62 |
| 82-PRO,83-LYS,84-ASN,281-TRP,282-HIS,285-ASN,289-ARG,293-LYS,295-TRP,364-ARG,365-GLY,367-ALA,368-ASP,408-ASN,409-ASN,410-PRO,411-ARG,448-ARG,451-GLU,453-ARG | 5WI9 | 46.05 |
| 30-ASN,63-GLU,64-PRO,65-LEU,66-HIS,67-ALA,68-PRO,69-ASN,70-VAL,72-ASP,74-TYR | 3KS0 | 61.98 |
| 12-ASN,14-ASP,15-LYS,42-LYS,44-GLU,45-GLU,49-GLN,62-TRP,65-ASP,66-ARG,152-GLN,153-GLU,154-PRO,155-TYR,207-ASP,209-ASP,211-SER,330-MET,334-PRO,337-SER,340-TRP,341-TYR,344-ARG | 5BK1 | 41.68 |
| 410-VAL,412-LEU,414-ALA,415-GLN,416-ARG,418-LEU,419-CYS,434-HIS,435-CYS,437-ASP,438-GLY,477-VAL,578-SER | 4OGY | 39.9 |
| 19-GLU,22-GLN,23-HIS,26-GLN,27-LEU,28-THR,30-TRP,31-GLY,33-LYS,34-GLN,35-LEU,38-ARG,50-TRP,51-MET,53-TRP,54-ASP,58-ASN,62-SER,65-HIS,68-ILE,205-VAL,208-ILE,209-LYS,212-GLN | 3MA9 | 35.89 |
| 24-ASN,27-PRO,29-VAL,155-THR,156-LYS,159-GLU,161-LEU,190-TYR,192-THR,194-VAL,195-THR,196-ASN,199-ILE | 4TNW | 56.42 |
| 59-THR,60-SER,61-GLU,62-SER,63-PHE,64-VAL,66-ASN,68-TYR,76-THR,78-LYS,82-PHE,83-PRO,85-ASP,86-ARG,87-SER,88-GLN,89-PRO,90-GLY,128-LEU,129-ALA,130-PRO | 5B8C | 48.63 |
| 149-GLU,152-TRP,153-GLU,156-LEU,160-LYS,161-ASN,162-ASN,163-ILE,164-ASN,167-LYS,168-ASN,169-ILE | 4K2U | 49.95 |
| 49-ASP,51-ALA,54-ILE,56-TYR,66-GLN,68-VAL,69-HIS,115-MET,117-SER,119-GLY,120-GLY,121-ALA | 5GGT | 44.67 |
| 28-TYR,30-ARG,31-ILE,32-THR,34-SER,37-PRO,38-LYS,39-GLU,40-ALA,41-VAL,55-PRO,56-LYS,61-GLN,64-MET,65-ASP,68-ASP,69-LYS | 2BDN | 49.18 |
| 246-PHE,249-LEU,250-TYR,253-GLU,254-ARG,256-ILE,257-GLN,260-ILE,301-GLN,302-ILE,304-GLU,305-LEU,306-TYR,307-GLU,308-ASP,309-PHE | 4XTR | 43.48 |
| 234-HIS,243-LYS,250-SER,251-GLU,254-LYS,255-LYS,256-ASP,257-ILE,258-PHE,259-GLY,260-GLU,272-LYS | 5HDQ | 45.68 |
| 56-ARG,60-PHE,75-HIS,78-SER,100-ALA,101-GLY,119-ASN,120-TRP,121-PHE,122-ASP,123-ILE,124-THR,127-LEU | 4ODX | 48.47 |
| 365-ASN,369-LEU,370-LEU,372-ARG,373-GLN,374-ALA,375-SER,376-PRO,377-PRO,379-PRO,383-ASP,385-GLN,386-ASN,388-TRP,389-PHE,397-THR,449-PRO | 5TQQ | 45.69 |
| 345-PRO,346-GLY,347-ASP,348-ILE,349-ILE,351-ASP,354-PHE,355-GLN,369-ILE,371-TYR,392-LYS,394-LYS,413-LYS,415-ASP,416-LYS,418-SER | 6H5N | 42.14 |
| 38-ASN,318-THR,344-GLU,345-GLY,347-ILE,348-ASP,349-GLY,350-TRP,354-ARG,363-GLN,364-ALA,365-ALA,367-LEU,370-THR,371-GLN,374-ILE,377-ILE | 5K9K | 38.15 |
| 47-THR,48-GLU,49-ALA,51-ASP,52-ALA,53-THR,65-SER,70-HIS,72-HIS,74-SER,76-GLU,107-VAL,108-SER,109-LYS | 5B71 | 52.22 |
| 34-ILE,35-SER,36-GLU,37-ASP,38-GLY,53-HIS,56-ASP,57-LEU,58-LEU,59-PHE,61-LEU,62-ARG,64-THR | 2H9G | 53.02 |
| 45-LYS,65-ARG,66-CYS,67-ASP,68-SER,70-GLU,71-VAL,72-GLU,87-GLU,88-GLU,90-THR,99-MET,102-LYS | 4OD2 | 59.97 |
| 348-LEU,349-PRO,384-GLN,409-HIS,417-VAL,418-SER,420-ASP,440-SER,441-GLY,443-LYS,464-THR,465-LYS,466-ILE,467-ILE,468-ARG,469-ASN,471-GLY,473-ASN | 5SX5 | 44.53 |
| 41-GLN,43-THR,44-ASN,45-ARG,46-ASN,47-THR,48-ASP,49-GLY,53-TYR,67-GLY,68-ARG,69-THR,70-PRO,84-LEU | 1YQV | 51.72 |
| 15-TRP,17-PRO,18-ASP,20-PRO,23-MET,40-LEU,42-GLN,43-SER,45-GLU,47-LEU,54-THR,55-ILE,56-GLN,58-LYS,59-GLU,61-GLY,62-ASP,66-TYR | 3HMX | 52.66 |
| 156-SER,157-ASN,158-SER,160-TYR,192-GLU,218-GLN,219-ARG,220-GLY,221-GLY,222-ARG,223-GLN,224-THR,259-GLU,260-SER,261-HIS,263-ASN,264-TYR,294-LEU | 2B2X | 44.22 |
| 65-LYS,66-GLU,68-LYS,197-SER,201-ASN,202-GLN,204-LEU,206-ILE,208-ASN,209-GLN,210-GLN | 5UDD | 65.65 |
| 24-GLY,44-ARG,45-ASP,72-THR,73-HIS,77-THR,83-ILE,85-VAL,86-SER,87-TYR,88-GLN,89-THR,90-LYS,131-ARG,135-GLU,136-ILE,137-ASN,138-ARG,140-ASP | 5WUX | 54.31 |
| 80-THR,81-GLU,89-GLY,90-ILE,91-ASP,92-ASN,94-ARG,95-GLU,96-ALA,97-LYS,100-ASN | 1RVF | 43.58 |
| 317-PRO,318-MET,319-ILE,320-ASN,329-GLU,330-ASN,331-VAL,332-ASP,334-ILE,336-GLU,358-LYS,360-GLU,362-TYR,364-LYS,365-SER,366-GLU,372-ARG,376-GLU,378-HIS,380-THR,381-ARG | 4K94 | 45.83 |
| 20-GLY,22-THR,25-LEU,28-PHE,122-ARG,123-GLU,126-LEU,127-GLU,130-VAL,131-SER,134-VAL,137-ARG,138-THR,139-PRO,142-TYR,144-PRO,149-ILE | 3V6Z | 51.59 |
| 300-VAL,301-MET,302-CYS,303-THR,304-GLY,305-SER,326-TYR,327-GLU,328-GLY,329-THR,330-ASP,336-PRO,340-GLN,361-LYS,382-ALA,383-GLY,384-GLU,385-LYS | 5VIC | 50.64 |
| 498-PRO,500-VAL,501-ILE,502-ASN,505-ILE,513-GLN,517-LYS,520-ASN,523-CYS,524-ASN,550-CYS,551-PRO,552-LEU,553-GLN,556-ASP | 6H3U | 46.17 |
| 71-SER,73-ALA,74-SER,111-PHE,112-GLU,113-ARG,115-GLU,118-PRO,120-THR,121-SER,162-SER,163-LYS,164-SER,169-LYS,171-LYS,253-TYR | 4LVH | 43.77 |
| 45-ASN,46-VAL,47-PRO,66-ASP,68-LEU,74-GLU,76-HIS,78-TYR,87-TRP,88-ILE,89-THR,90-LEU,91-ASN,92-ASP,95-LYS,96-GLU,136-PHE,144-TYR,145-ASN,146-ASN | 5EZO | 42.02 |
| 25-THR,26-ARG,27-GLN,28-GLY,29-GLY,30-TYR,37-ILE,39-ARG,42-GLY,43-GLY,44-ASP,79-ASP,81-ARG,82-ASP,83-ASN,84-ALA,85-LYS,138-SER,139-GLY,140-GLY,141-ASP,144-PHE | 5IES | 53.6 |
| 54-PRO,55-LYS,57-THR,58-ARG,61-THR,62-PHE,63-LYS,91-ASN,92-LEU,93-ALA,94-GLN,96-LYS,97-ASN | 5LQB | 59.03 |
| 6-LEU,7-LEU,8-GLY,9-ILE,47-SER,48-SER,49-ARG,51-LEU,52-ASP,94-VAL,96-LEU,110-GLY,112-TYR,113-VAL,114-ASN,138-ASN,140-LEU,142-GLU,143-LYS,144-PRO,145-ARG,146-VAL,147-THR,148-ARG | 6AYZ | 45.29 |
| 216-ILE,219-TRP,220-ARG,221-ASN,250-GLN,251-ALA,252-SER,254-LYS,263-ILE,264-VAL,266-SER,267-VAL,268-GLU,270-ASN | 4QNP | 36.26 |
| 127-GLY,128-LEU,156-ASN,158-TYR,159-ARG,185-ILE,188-LYS,189-GLN,191-THR,192-VAL,193-THR,194-THR,195-THR,196-THR,197-LYS,198-GLY,199-GLU,200-ASN | 1TQB | 52.53 |
| 232-SER,233-ASN,250-ALA,251-ASP,252-SER,253-LYS,272-ASN,273-THR,274-ALA,276-THR,277-SER | 1RJL | 59.94 |
| 27-HIS,29-LYS,31-SER,32-ASN,33-GLN,34-ILE,35-LYS,46-LYS,47-GLY,48-PRO,49-SER,50-LYS,52-ASN,53-ASP,77-GLU,90-LYS | 4Q6I | 50.01 |
| 121-ARG,123-ASN,124-GLY,125-ALA,134-SER,146-ASN,147-THR,148-ASP,149-ASN,150-ALA,178-SER,179-THR,180-ALA,181-GLU,183-THR,184-LYS | 6II4 | 51.09 |
| 101-ILE,102-ASN,103-ASP,104-PRO,119-SER,120-GLN,121-LEU,122-ASP,125-THR,154-GLN,156-ASN,157-GLY,186-PRO,188-LEU,190-GLU,192-ILE | 3AB0 | 40.15 |
| 172-ASN,176-ASP,180-SER,330-GLY,333-ASP,334-LEU,335-THR,338-SER,405-SER,406-LEU,407-PRO,408-LEU | 6CSE | 57.73 |
| 65-SER,68-LEU,69-SER,72-ASN,73-ASP,78-ASN,79-ASP,82-LYS,83-LEU,85-SER,86-ASN,89-LEU | 4JLR | 60.18 |
| 14-VAL,14-CYS,14-SER,14-LEU,14-GLN,14-ASN,14-TRP,14-LYS,15-TYR,15-PHE,15-SER,15-ASP,15-HIS,17-ALA,17-SER,17-LEU,17-GLY,17-ASN,17-ILE,17-PRO,17-LYS | 3L95 | 33.48 |
| 470-GLU,482-GLU,485-PHE,488-ASN,489-GLN,490-PRO,491-GLU,492-GLY,493-ARG,496-LYS,498-LYS,499-PRO,500-GLY | 6BDZ | 57.92 |
| 19-GLU,22-GLN,23-HIS,24-LEU,26-GLN,27-LEU,28-THR,30-TRP,31-GLY,33-LYS,34-GLN,35-LEU,38-ARG,51-MET,54-ASP,65-HIS,68-ILE,208-ILE,209-LYS,212-GLN | 3MAC | 36.73 |
| 116-VAL,117-PRO,161-ARG,162-ASP,165-ILE,195-VAL,198-GLY,200-GLU,215-LYS,216-LYS,250-ARG,276-LYS,277-ASP,278-PRO,279-LYS,280-THR,281-GLY,283-TYR,288-PHE,289-PRO | 3WKM | 59.11 |

| | | |
|---|---|---|
| 90-TYR,91-PRO,92-PHE,95-ASP,98-ARG,171-TYR,256-LYS,305-LYS,308-GLU,311-LEU,312-ALA,318-ALA,319-ALA,321-MET,322-GLU,325-GLN,328-GLU | 5BJZ | 48.28 |
| 68-ASP,89-GLU,92-LYS,95-SER,96-ASN,100-TYR,105-TYR | 6N5E | 29.74 |
| 44-LYS,47-LEU,48-GLY,56-GLN,57-CYS,58-ILE,59-GLU,60-ASN,61-PRO,62-ASP,63-PRO,65-GLN,66-VAL,67-ASN,68-MET,69-TYR,72-GLY,73-CYS,75-GLU | 1Z3G | 49.42 |
| 332-VAL,333-ALA,334-PRO,335-ALA,337-GLN,350-PRO,351-ARG,353-ASP,354-LEU,356-ARG,357-ALA,358-THR,361-GLU,362-ALA,365-ARG,370-LYS,371-PRO,372-PHE,373-ASP | 2XQY | 46.69 |
| 280-ASN,281-ALA,362-GLN,365-SER,366-GLY,459-GLY,460-ASN,461-ASN,462-ASN,463-LYS,471-ARG | 4YE4 | 70.65 |
| 161-ASP,163-GLY,180-LYS,181-ILE,182-GLU,183-HIS,184-LEU,185-LYS,187-PRO,190-ASN,191-VAL,192-ASP,213-LEU,214-TYR,215-ASN,243-VAL | 5O14 | 43.03 |
| 349-PRO,384-GLN,408-GLN,409-HIS,411-GLN,412-PHE,417-VAL,418-SER,438-ILE,440-SER,441-GLY,443-LYS,464-THR,465-LYS,466-ILE,467-ILE,468-SER,469-ASN,473-ASN | 3B2U | 44.08 |
| 28-GLN,29-ARG,31-GLU,79-HIS,146-LYS,148-VAL,149-THR,150-LYS,169-TYR,170-ASP,171-SER,172-ASN,173-MET,174-ARG,175-SER,176-GLY,177-LYS,178-PHE,190-LEU,191-THR,193-ASP,194-ILE,195-SER,196-ARG | 3MJ9 | 51.28 |
| 582-TYR,584-ASP,585-PRO,590-HIS,591-PRO,592-CYS,593-HIS,603-PRO,604-THR,605-SER,606-HIS,607-ASP | 3U9U | 47 |
| 317-PRO,318-MET,319-ILE,320-ASN,323-VAL,324-PHE,325-VAL,326-ASN,329-GLU,330-ASN,331-VAL,332-ASP,334-ILE,336-GLU,358-LYS,360-GLU,362-TYR,363-PRO,364-LYS,365-SER,366-GLU,372-ARG,376-GLU,378-HIS,381-ARG | 4K9E | 41.42 |
| 504-ILE,505-VAL,506-ASN,507-ALA,508-GLN,510-LYS,511-CYS,512-ASN,513-PRO,514-ASN,549-HIS,550-ASN,551-GLN,552-ASP,553-GLY,556-CYS,557-GLY | 5KEL | 53.32 |
| 54-TYR,55-LEU,58-LEU,59-TYR,107-GLU,111-ALA,112-GLY,113-LEU,114-GLY,115-LEU,116-PHE,117-ARG,119-VAL,123-ARG | 1ORQ | 55.68 |
| 100-TYR,101-ASP,216-ASN,218-GLY,219-PHE,220-ARG,221-PRO,222-ARG,223-ILE,224-ARG,229-ARG | 6E4X | 47.44 |
| 135-VAL,158-ASN,159-GLY,160-LEU,187-ASN,188-ILE,189-GLY,190-ASP,193-ALA,196-HIS,222-LYS,225-ASP,226-ARG,227-GLU | 5UGY | 54.5 |
| 139-SER,140-MET,141-SER,142-LEU,143-GLN,144-PRO,200-LYS,201-HIS,202-MET,203-LEU,204-LEU,205-LEU,206-THR,239-ASP,241-GLU,242-ALA,243-THR,263-LYS,264-HIS,268-LYS,269-GLU,273-THR | 3HI6 | 47.32 |
| 322-LYS,323-ASP,325-LEU,346-HIS,348-LEU,349-PRO,350-VAL,353-ARG,355-ASP,357-PHE,380-PHE,382-LEU,384-GLN,407-LYS,408-GLN,409-HIS,417-VAL,418-SER,438-ILE,440-SER,441-GLY,443-LYS,465-LYS,467-ILE,468-SER | 5XWD | 42.76 |
| 3-PRO,4-ALA,5-PRO,6-SER,7-LEU,8-LEU,9-THR,83-SER,84-ARG,87-GLN,100-GLN,101-SER,103-ASN,120-ARG,121-LYS,122-THR,127-GLN | 3U9P | 41.84 |
| 53-PHE,58-ALA,59-LYS,60-ALA,61-TYR,62-ASP,71-THR,72-HIS,73-ALA,74-CYS,75-VAL,76-PRO,77-THR,78-ASP,79-PRO,80-SER,219-ALA,220-PRO,221-ALA,246-GLN | 4R4N | 57.12 |
| 195-ASN,197-ASP,198-CYS,200-THR,201-ILE,203-ARG,204-ALA,205-LEU,206-GLY,207-PRO,208-GLY,213-GLU,217-ALA,218-CYS,219-GLN,221-VAL | 3VRL | 58.02 |
| 155-PRO,192-ASP,194-ARG,197-GLU,237-ARG,238-ASP,239-ALA,376-SER,377-THR,378-CYS,379-PHE | 2XTJ | 54.15 |
| 36-GLN,37-GLU,40-SER,43-GLU,44-ASN,45-TYR,46-ARG,49-LYS,52-ARG,53-MET,93-GLN,130-VAL,134-ARG | 4YQX | 47.74 |
| 32-VAL,33-GLU,35-GLN,37-HIS,42-LEU,47-ARG,48-PRO,49-LYS,51-PRO,75-ILE,77-ARG,78-VAL,79-GLY,80-GLU,81-LEU,82-TRP | 6IAP | 40.63 |
| 27-VAL,29-GLY,177-ASP,181-ALA,184-GLU,185-GLU,188-ALA,234-MET,235-GLY,236-PRO,237-TRP,238-PRO | 6MLK | 54.4 |
| 116-TYR,117-GLN,118-MET,155-ASP,157-ASN,168-ASP,170-ASP,171-VAL,172-SER,173-GLN,174-GLU,175-VAL | 4LU5 | 56.72 |
| 279-ASN,280-ASN,281-VAL,282-LYS,357-LYS,365-SER,366-GLY,367-GLY,455-THR,456-ARG,457-ASP,458-GLY,459-GLY,460-VAL,461-ASP,462-THR,465-LYS | 5T33 | 55.23 |
| 150-HIS,152-ARG,153-THR,154-PRO,197-HIS,198-ASP,199-GLU,218-TRP,219-SER,220-LYS,221-LYS,222-ILE,224-ARG,244-GLY,246-ALA,248-GLY,249-ARG,250-ALA,251-ASP | 2AEQ | 39.08 |
| 80-TYR,117-GLU,118-MET,145-GLN,146-ASP,150-ASP,151-GLY,177-PHE,198-ASN,199-VAL,200-TYR,201-SER,203-LYS,204-TYR,205-ASN,207-GLN,212-THR | 4O02 | 49.21 |
| 53-ILE,56-ARG,92-PHE,98-GLN,99-GLU,101-ALA,102-GLU,103-ALA,105-THR,106-HIS,109-THR,112-GLN,113-ASN,114-ARG,116-THR | 4KUC | 42.14 |
| 82-ILE,84-VAL,85-ASP,89-ILE,91-LEU,115-GLN,117-SER,118-GLU,119-HIS,121-GLY,122-LYS,123-MET | 5T5F | 58.6 |
| 25-GLU,30-ASP,31-TRP,33-ILE,34-ALA,35-PRO,36-LYS,83-ASN,85-LEU,87-PHE,91-GLU,92-GLN,93-ILE,94-ILE,95-TYR | 5F3B | 53.73 |
| 82-LEU,83-GLY,84-GLY,85-LYS,86-GLY,88-TYR,112-LEU,113-ASN,130-ASP,131-TRP,132-PRO,133-GLU,135-LEU | 5WHK | 55.95 |
| 123-THR,276-ASN,278-THR,279-ASN,280-ASN,281-ALA,282-LYS,365-SER,366-GLY,367-GLY,368-ASP,371-ILE,431-GLY,456-ARG,457-ASP,458-GLY,459-GLY,461-ASN,469-ARG,472-GLY,473-GLY,474-ASN | 5WB9 | 53.03 |
| 47-TYR,55-LYS,61-ASN,63-LYS,73-ARG,74-ILE,158-LEU,160-ARG,162-LYS,166-ASP,167-ASN,170-ARG | 4DKF | 43.77 |
| 32-ALA,33-GLY,36-TYR,37-GLN,39-TRP,40-GLU,43-ARG,47-GLN,65-PHE,67-MET,68-TYR,87-TYR,89-PRO,90-TRP,113-ARG,115-HIS,119-GLU | 4HJ0 | 39.46 |
| 72-THR,110-SER,114-PHE,146-LYS,166-ILE,256-ALA | 4LVH | 47.1 |
| 217-ARG,220-ASP,221-ILE,223-LYS,224-THR,225-LYS,295-GLN,298-LYS,302-GLU,305-ASP,306-LEU,308-GLU,309-LYS,311-LYS,416-LYS | 6BPC | 47.55 |
| 204-ASP,205-GLU,207-VAL,208-GLN,211-GLY,212-ALA,213-ASP,214-GLY,215-LEU,216-GLY,217-GLU | 4G7V | 60.17 |
| 68-TYR,71-VAL,73-THR,74-SER,75-TYR,76-PRO,77-GLY,78-PHE,79-GLN,82-GLU,86-PRO,89-GLY,90-MET,272-GLN,341-LYS,342-ASP,431-LEU | 4QWW | 45.11 |
| 32-LYS,105-GLU,143-SER,146-THR | 5USL | 52.05 |
| 96-LEU,103-SER,120-GLU,155-LYS,157-TYR,159-TRP,162-ASP,163-ARG,164-PRO,166-ILE,217-VAL,218-HIS,219-PRO,220-LYS,221-GLY | 6AL5 | 38.23 |
| 94-PHE,136-SER,137-ASN,138-ALA,219-SER,220-ARG,221-PRO,222-TRP,223-VAL,224-ARG,225-GLY,226-LEU,229-ARG | 6N5D | 46.85 |
| 51-THR,52-LEU,53-PHE,60-ALA,71-THR,72-HIS,73-ALA,74-CYS,75-VAL,76-PRO,103-GLN,106-GLU,107-ASP,113-ASP,114-GLN | 4YBL | 50.64 |
| 47-ALA,48-ASN,54-LEU,56-ASP,57-GLY,59-SER,60-LEU,61-SER,62-ASN,66-LEU,68-PRO,69-THR,70-SER,106-SER,142-THR,143-GLN,144-GLY,145-ASP,146-GLN | 4MXV | 49.68 |
| 17-ASP,18-GLY,21-GLN,70-LYS,72-ASP,73-PRO,74-PRO,75-SER,77-ASP,79-GLN,117-LYS,118-CYS,119-PRO,120-PRO,121-LYS,133-GLN | 6IEC | 52.06 |
| 58-THR,59-HIS,60-THR,74-ASN,75-ARG,76-GLN,77-ASN,78-GLU,202-THR,203-GLY,204-GLN,206-TYR | 5W3M | 64.7 |
| 194-ARG,195-HIS,198-PRO,199-THR,200-ARG,201-LEU,202-GLU,203-ARG,204-CYS,205-GLN,229-ARG,231-SER | 3V4V | 48.9 |
| 31-THR,32-PHE,33-SER,34-ALA,55-THR,57-MET,58-PHE,59-GLY,60-THR,61-ALA,62-ASN,63-TYR,68-GLN,75-ALA,77-GLU,105-TYR,106-TYR | 5JO4 | 45.53 |
| 70-GLY,71-ALA,72-GLN,73-GLY,75-GLY,86-ARG,188-SER,265-TYR,267-SER,268-ALA,270-ARG,361-GLN,364-ILE,368-GLU,369-ASP,373-LEU,374-SER,375-ARG,434-GLU,435-GLU,438-ARG,511-GLU | 3TT3 | 49.8 |
| 33-GLY,34-PRO,35-ILE,37-GLU,41-ARG,43-TYR,46-VAL,47-GLY,48-PRO,87-ARG,88-GLY,89-ILE,106-GLY,107-ALA,108-LEU,109-SER,112-GLN | 1OAZ | 50.9 |
| 2-ALA,3-MET,33-GLU,39-LEU,45-GLN,95-LYS,99-MET,100-TYR,102-PRO,103-PRO,104-TYR,105-TYR,106-LEU,107-GLY,108-ILE | 5TRU | 49.87 |
| 160-VAL,161-THR,162-ALA,163-MET,168-SER,170-SER,171-VAL,172-GLU,173-VAL,174-LYS,176-PRO,177-ASP,180-GLU,291-LYS,293-ARG | 3UC0 | 48.41 |
| 535-SER,536-VAL,537-HIS,538-THR,540-PRO,541-PRO,542-ALA,543-GLU,544-ALA,546-MET,547-GLY,548-THR,549-ARG,553-HIS,554-GLN,557-HIS,567-GLU,568-VAL | 5VLP | 47.39 |
| 15-PRO,16-ASN,18-GLU,19-THR,21-THR,53-ASP,55-ILE,56-THR,59-PRO,63-HIS,66-LYS,104-ASP,105-PRO,107-LEU,132-LEU,196-PRO | 4I18 | 43.43 |
| 90-HIS,337-TYR,498-TYR,502-ARG,505-GLU,506-ASP,508-ARG,509-ASP,511-ILE,512-GLY,513-PHE,514-PRO,598-ARG,599-ASP | 4XP4 | 41.59 |
| 327-ALA,328-GLN,331-ARG,332-GLN,333-ILE,334-GLY,335-ALA,336-SER,337-LEU,338-ASN,339-ASP,368-LYS,369-TYR,426-ARG,448-GLY,449-ASP,469-ARG | 4K3J | 52.36 |
| 333-LYS,336-TYR,337-LYS,338-LEU,339-SER,340-THR,341-LYS,419-LEU,422-ARG,423-LYS,439-LYS,545-GLU | 4QEX | 57.25 |
| 26-LEU,27-ASN,28-ILE,30-ASN,35-THR,36-ASN,39-ARG,55-ARG,57-GLU,59-PRO,60-GLU,61-ARG,62-TYR,64-SER,65-VAL,67-TRP,99-LEU,100-ARG,101-ARG,102-GLU,103-PRO,104-PRO,110-PHE | 4QHU | 50.09 |
| 27-THR,28-PHE,41-GLN,43-ASP,44-PRO,45-CYS,46-ILE,47-PRO,48-GLY,49-VAL,60-HIS,63-SER,64-CYS,66-HIS,78-ILE | 5TLJ | 50.76 |
| 21-ASN,22-GLU,23-LEU,24-LEU,26-ASP,27-LEU,30-THR,31-LYS,36-GLU,39-ARG,43-LYS,47-ASP | 5VYF | 51.23 |
| 258-LEU,262-ASN,263-ASP,265-PRO,266-ILE,267-THR,268-ASN,269-ASP,271-LYS,272-LYS,275-SER | 5J3D | 58.56 |
| 43-GLU,49-ASP,68-GLU,69-TYR,73-LYS,98-ASP,113-ASP,114-HIS,115-ARG,116-ILE,117-LYS,119-GLU,120-ALA | 6GKU | 50.24 |
| 520-ALA,545-LEU | 5YY5 | 30.05 |
| 34-THR,35-THR,37-GLN,39-THR,44-ARG,46-ASN,64-SER,65-SER,104-TYR,105-GLU,108-LYS,142-ARG,143-SER,144-ALA,145-ASN,146-THR,147-SER,175-ASN,203-LEU,204-SER | 6B9J | 37.61 |
| 1-VAL,2-LYS,3-PRO,27-LEU,32-ARG,35-SER,36-VAL,37-GLN,39-ARG,41-LEU,71-VAL,73-VAL,74-GLY,75-SER,77-GLY,78-GLY,79-ARG,80-THR,81-PHE,82-GLN,83-HIS,84-THR | 4HCR | 51.73 |
| 18-VAL,19-ASP,20-GLY,21-TRP,38-LYS,41-THR,42-GLN,45-ILE,49-THR,52-VAL,53-ASN | 6B3M | 50.5 |
| 13-GLN,23-ASP,24-GLY,25-THR,27-VAL,29-SER,60-GLU,61-ASN,63-VAL,65-GLN,67-ARG,68-TYR | 6A77 | 50 |
| 474-LYS,475-PHE,476-SER,477-TYR,478-ILE,479-ARG,480-THR,481-SER,486-LEU,488-ARG,490-GLU,544-LYS,546-GLN,548-HIS,552-LEU,591-ASP,596-GLU | 5KQV | 42.73 |
| 67-GLN,70-PRO,71-SER,73-HIS,75-LEU,77-THR,97-ILE,105-THR,107-GLU,108-GLY,109-ALA,110-GLU,111-ALA,137-ASN,138-ARG,139-PRO,140-ASP,141-TYR | 4G3Y | 51.85 |
| 1-LYS,49-GLN,52-ALA,53-THR,73-SER,75-LEU,341-TYR,342-ALA,345-THR,349-ASN,354-ARG,355-GLN,359-GLU,362-LYS,363-ASP,365-GLN,366-THR,367-ASN | 5BK2 | 52.3 |
| 153-SER,155-PRO,194-ARG,195-GLU,197-GLU,237-ARG,238-ASP,369-ILE,374-ASP,375-CYS,377-THR,378-CYS,379-PHE,381-SER | 3SQO | 52.15 |
| 266-ASN,267-LYS,268-LEU,269-THR,270-PHE,271-GLN,273-GLU,274-PRO,276-PRO,277-HIS,315-ASN,571-ASP,583-PRO,584-HIS,596-ALA,597-LYS,602-LYS,611-ARG,612-PRO,613-CYS,614-HIS,615-GLU | 4P59 | 54.32 |
| 141-GLN,164-SER,165-GLU,166-VAL,167-ALA,168-SER,169-LEU,170-ASP,172-PHE,179-ASP,180-ILE,181-VAL,182-ALA,184-PRO,202-ASP,204-ILE,206-ARG | 6CNJ | 42.54 |
| 108-GLN,109-THR,110-PHE,111-GLU,124-TRP,126-GLN,130-GLU,160-TYR,161-SER,162-ARG,163-ASP,167-VAL,169-GLN,174-GLU,176-GLY,177-ASP,178-GLY,179-TYR,196-PRO,197-GLY,198-ILE,199-GLN,201-ASP | 5TH9 | 50.23 |
| 503-ASP,520-GLY,522-GLN,523-PHE,524-SER,525-PHE,526-SER,527-LEU,529-PRO,532-ALA,533-SER,539-ILE,540-GLN,541-ASP,542-ASN,543-LYS,552-ARG,571-SER,572-ASP,573-ASN,574-ASP,575-TYR,576-PRO,577-VAL | 5VEB | 53.15 |
| 3-VAL,31-SER,80-ASP,94-GLN,96-LEU,120-SER,122-PRO,123-GLY,124-SER,125-SER,126-PRO,127-SER,140-GLY,141-GLY,142-LYS,162-LEU,163-GLN,164-ASN,165-GLN,166-LYS | 3O2D | 49.23 |
| 566-TYR,567-TRP,578-GLU,579-ALA,580-LEU,581-HIS,582-SER,586-THR,587-ALA,588-VAL,590-LYS,592-ILE,593-ASP,594-TYR,595-GLY,596-ASN,597-TYR,599-GLU | 3PNW | 48.78 |
| 1-MET,2-TYR,5-LEU,6-GLU,9-ASN,55-ILE,57-PRO,59-GLN,62-LYS,107-LEU,110-SER,111-LEU,113-TYR,114-SER,115-TYR,164-TYR,168-THR,169-PRO,172-HIS,173-HIS,176-LEU | 5H35 | 42.25 |
| 8-LYS,9-ASN,31-GLN,32-GLN,33-TYR,34-GLY,35-LYS,52-TYR,54-GLN,56-ILE,62-ASN | 4V1D | 67.08 |
| 25-SER,26-PRO,29-HIS,30-ASP,33-TRP,166-VAL,167-GLY,205-SER,206-VAL,207-ASP,208-GLN,209-GLU,235-SER,236-GLN,237-GLU,238-LYS,240-GLU | 3HB3 | 37.45 |
| 44-TYR,45-LYS,87-LYS,123-ARG,125-THR,126-GLU,131-ASP,132-GLY,133-HIS,134-HIS,135-SER,136-GLU,147-ASP,149-THR,153-ARG,155-ARG,174-TYR,176-GLU,177-SER,178-LYS,179-ALA,180-HIS | 3MXW | 39.4 |
| 126-GLU,128-SER,130-GLY,131-VAL,132-SER,149-TRP,151-ILE,152-LYS,153-LYS,154-ASN,155-ASN,189-LYS,190-LEU,222-GLN | 5DUP | 39.82 |
| 6-LEU,7-LEU,8-GLY,47-SER,49-ARG,50-SER,51-LEU,52-ASP,94-VAL,96-LEU,110-GLY,112-TYR,113-VAL,114-ASN,138-ASN,140-LEU,142-GLU,143-LYS,144-PRO,145-ARG,146-VAL,147-THR,148-ARG,149-PHE | 5UEK | 48.17 |
| 14-GLN,14-GLU,14-CYS,14-PRO,14-ALA,14-SER,14-TYR,14-GLY,15-ARG,15-SER,15-PRO,15-GLU,15-ASN,15-ASP,15-HIS,16-PRO,16-SER | 5CZV | 55.65 |
| 557-PRO,558-GLU,560-ASP,561-GLN,569-LYS,570-ASP,571-PRO,572-PRO,573-PHE,591-ILE,593-LYS,596-ASP,598-GLU,600-ALA,602-GLN,603-PRO,604-CYS | 1N8Z | 47.66 |
| 81-GLU,83-GLY,84-ASN,128-PHE,129-PRO,132-ASP,135-ILE,146-ARG,147-TYR,149-GLU,169-PHE,171-ILE,172-ALA,173-GLU,174-ASP,175-GLN,176-ASN,196-TYR,315-ASN,316-ASN,317-ARG,318-ASP,321-LYS | 4UAO | 51.95 |
| 21-SER,22-SER,23-GLU,26-ASP,27-LYS,30-ARG,73-CYS,74-PHE,75-GLN,78-PHE,80-GLU,129-LYS,179-ARG,182-ARG,183-GLN,184-MET | 4O9H | 51.85 |
| 68-LEU,69-GLU,156-LYS,159-LYS,160-LYS,161-ILE,162-ILE,163-GLY,164-GLN,165-VAL,166-ARG,168-GLN,172-LEU,182-ILE,188-LYS,189-GLY,190-GLY | 5EU7 | 46.67 |
| 51-LEU,93-PRO,140-ARG,143-PRO,144-GLY,145-HIS,148-LYS,157-GLU,160-PHE,176-LYS,177-GLU,178-ASP,179-GLU,180-LEU,181-GLY,182-ASP,183-ARG,186-MET | 2VXT | 53.84 |
| 1-ASP,2-ASP,3-ALA,4-GLU,5-ASN,24-ASP,26-ASP,27-GLU,28-ASN,30-LYS,63-TYR,64-THR,65-PRO,66-ALA,68-ASN,70-ASN | 3R08 | 66.84 |
| 5-SER,8-VAL,9-ALA,12-LEU,13-ARG,16-THR,23-ARG,27-ARG,327-LEU,328-GLU,331-ASP | 6C9U | 51.86 |
| 19-ASP,20-GLY,38-LEU,39-LYS,41-THR,42-GLN,45-ILE,53-ASN,56-ILE,58-LYS,59-THR | 5K9Q | 55.56 |
| 193-GLN,195-TYR,207-SER,211-PHE,212-LEU,214-ASP,268-LYS,269-SER,270-GLY,271-CYS,273-HIS,275-ASP | 4K24 | 51.12 |
| 64-ILE,65-LYS,66-GLU,67-ASN,68-LYS,69-CYS,201-LYS,202-GLN,204-LEU,206-ILE,208-ASN,209-LYS,210-GLN,211-SER,216-ASN | 4JHW | 56.14 |
| 115-GLN,116-ILE,118-PRO,120-SER,121-SER,122-TRP,123-SER,124-ASP,158-PRO,159-THR,160-ILE,161-LYS,162-ARG,163-SER,164-TYR,165-ASN,167-THR,168-ASN,240-ASN | 6IUT | 43.74 |
| 1-MET,3-LEU,4-LYS,307-GLU,309-PRO,310-GLU,311-PHE,312-PHE,313-GLU,316-VAL,318-PHE,319-LYS,430-ASN,435-SER,436-LEU,437-GLU | 3GI9 | 52.21 |
| 173-ASN,179-TRP,181-PRO,182-TYR,183-ASP,185-ASP,186-SER,187-TRP,188-ASN,189-PRO,190-VAL,191-TYR,200-ARG,263-THR,264-ASN,266-LYS | 4U6V | 43.79 |
| 65-LYS,66-ASP,67-VAL,68-LYS,69-GLN,70-THR,101-THR,103-TYR,104-LEU,129-ASP,136-ARG,141-LEU,142-SER,144-ARG,145-ASP,146-VAL,148-GLY,149-LYS,150-ASP,172-THR,194-PRO,195-SER,197-THR | 5W06 | 44.95 |
| 31-THR,32-ALA,33-GLY,35-LEU,36-TYR,39-TRP,66-ASP,67-MET,87-TYR,90-TRP,113-ARG,115-HIS,123-LYS,125-GLU,128-LEU,129-ASP | 6DKJ | 50.11 |
| 48-LEU,49-GLN,50-ALA,51-VAL,52-PRO,53-THR,54-GLY,56-PRO,57-ALA,70-SER,71-TYR,78-GLN,79-SER,81-ARG,94-ALA | 3KJ4 | 49.58 |
| 111-SER,113-ASP,114-ASP,115-ILE,116-ALA,117-VAL,119-LEU,165-ARG,171-GLU,174-ILE,210-GLY,211-LYS,212-LEU,216-LYS,218-GLN | 5NUZ | 41.51 |
| 51-SER,52-GLN,54-ALA,121-CYS,122-THR,123-SER,124-LYS,128-ARG,131-GLN,133-GLU,134-ASN,136-LYS,207-GLY,208-SER,230-SER,274-TYR,275-SER,276-SER | 5YWP | 49.92 |
| 28-PRO,55-ILE,73-PRO,76-SER | 3KJ4 | 26.98 |
| 29-THR,30-VAL,31-SER,32-LEU,33-TRP,35-THR,36-VAL,39-TRP,40-ARG,43-ARG,68-GLU | 5E94 | 55.77 |
| 576-THR,580-CYS,581-ASN,582-THR,584-PRO,585-HIS,590-ARG,610-SER,611-ILE,612-TYR,614-ASP,615-GLY,616-HIS,618-ASP,619-ARG | 6H3T | 48.75 |
| 64-HIS,87-ASP,90-TYR,91-PRO,92-PHE,93-THR,95-ASP,98-ARG,110-VAL,259-VAL,301-ALA,302-VAL,305-LYS,308-GLU,309-GLU,321-MET,324-ALA,325-GLN,327-GLY,328-GLU,329-ILE,330-MET,331-PRO,332-ASN | 3PGF | 34.95 |
| 422-ILE,427-LEU,429-CYS,430-ASN,431-ASP,434-GLN,439-THR,441-LEU,442-PHE,529-TRP,531-GLU,613-TYR | 6BKC | 53.77 |
| 34-PRO,36-THR,37-PHE,38-SER,39-PRO,40-ALA,41-LEU,42-LEU,43-VAL,136-GLU,137-SER,138-LEU,139-ARG,143-ARG | 6HIG | 47.29 |
| 128-LYS,156-GLN,236-PHE,245-HIS,246-CYS,248-ALA,252-TYR,254-THR,255-ASP,256-THR,257-PHE,268-THR,286-VAL,287-GLY,288-SER,290-THR,292-VAL,294-PRO,295-LEU,296-HIS,311-LYS,315-PRO | 1S78 | 58.6 |
| 41-GLN,45-ARG,46-ASN,47-THR,48-ASP,49-GLY,50-SER,51-THR,53-TYR,66-ASP,67-GLY,68-ARG,70-PRO,81-SER,84-LEU | 1MLC | 48.3 |
| 11-LYS,18-GLU,19-SER,21-ILE,22-PHE,24-LYS,25-LYS,26-TRP,27-GLU,45-ASN,46-ALA,47-ILE,48-PRO,49-PHE,50-THR,51-TYR,52-GLU | 4F3F | 53.87 |
| 308-ASP,309-LYS,310-SER,311-LYS,312-PHE,313-ALA,332-PHE,333-SER,334-GLY,335-SER,336-LYS,387-GLU | 6J5D | 47.25 |
| 298-SER,299-TYR,300-VAL,301-MET,303-THR,304-GLY,326-TYR,328-GLY,329-THR,330-ASP,331-ALA,332-PRO,361-LYS,362-GLU,382-ALA,383-GLY,384-GLU,385-LYS | 6DFJ | 53.38 |
| 220-GLY,221-TYR,222-ARG,251-GLU,255-GLY,256-ILE,257-ASP,259-ASN,261-GLU,308-ALA,310-SER,311-SER,312-GLY,313-LEU,314-MET,315-THR,316-LYS,317-LYS | 3S37 | 50.01 |
| 15-HIS,16-GLY,19-ASN,20-TYR,21-ARG,62-TRP,75-LEU,93-ASN,96-LYS,97-LYS,100-SER,101-ASP,102-GLY | 2DQF | 48.46 |

| | | |
|---|---|---|
| 67-HIS,69-LYS,70-ILE,71-ASN,72-ASP,73-LYS,81-HIS,84-GLU,99-ARG,101-TRP,103-ASN,104-GLY,240-ALA,241-THR,243-ARG | 6IVZ | 42.14 |
| 45-TYR,79-GLN,82-ARG,83-ILE,84-LYS,85-PRO,86-HIS,87-GLN,88-GLY,89-GLN,90-HIS,91-ILE,92-GLY | 1TZH | 50.1 |
| 25-ILE,25-TYR,25-PRO,25-ASN,25-THR,25-ASP,25-ALA,25-ARG,25-GLN,25-PHE,25-LEU | 5UMI | 42 |
| 505-ARG,507-LEU,509-ASP,519-ASN,521-ASN,522-GLN,523-TYR,524-SER,525-PRO,528-SER,543-LYS,545-LEU,546-SER,547-PRO,548-LEU,549-GLU,550-GLY | 5ZXV | 48.59 |
| 11-ASP,14-LYS,15-ALA,18-LEU,21-PHE,22-LEU,48-ARG,51-GLN,55-ILE,58-ARG,59-ILE,62-THR,63-TYR | 5DHV | 44.89 |
| 174-LEU,205-ALA,208-ARG,308-LYS,309-GLN,310-ARG,311-GLY,313-ASP,317-GLU,318-PHE,320-HIS,322-GLU | 5MES | 53.62 |
| 506-LEU,510-ASP,529-ILE,535-TRP,536-GLU,540-TYR,541-TYR,542-ARG,543-LYS,544-GLN,553-TRP | 5GMQ | 47.81 |
| 55-THR,56-PHE,57-LYS,59-TYR,71-GLN,75-ASP,76-GLU,77-LEU,78-GLY,79-PRO,82-HIS,105-ARG,117-THR,119-GLU | 4YUE | 47.64 |
| 51-THR,52-LEU,53-PHE,54-CYS,55-ALA,72-HIS,73-ALA,74-CYS,75-VAL,76-PRO,77-THR,78-ASP,79-PRO,80-ASN,107-ASP,217-TYR,220-PRO,221-ALA,246-GLN,247-CYS | 5FCU | 49.62 |
| 69-TRP,70-CYS,71-GLY,72-HIS,76-PHE,114-ASN,115-ILE,116-PRO,117-GLY,118-PHE,119-PRO,120-THR,122-ARG,132-SER,133-GLY,134-ALA,135-VAL,137-PRO,139-ALA,149-ARG | 4IJ3 | 45.05 |
| 38-ARG,56-THR,58-LYS,80-ARG,82-TYR,105-HIS,107-GLU,109-ARG,110-ASN,129-LYS,130-PHE,151-ASP,152-ILE,153-PHE,155-ILE,157-GLU,183-LYS,185-TYR,208-ASN,209-LYS,235-GLN,255-ARG,256-ASN | 3G04 | 36.92 |
| 18-GLY,19-THR,49-THR,50-ARG,51-TYR,52-HIS,53-ILE,54-ASN,55-LYS,56-THR,68-PHE,69-LEU,70-VAL,71-PRO,75-TYR | 3WIH | 47.67 |
| 187-GLU,204-ALA,205-LEU,206-GLY,207-PRO,208-ALA,209-ALA,210-THR,212-GLU,213-GLU,216-THR,217-ALA | 1E6J | 57.84 |
| 123-THR,124-GLY,280-ASN,281-ALA,282-LYS,283-THR,365-SER,366-GLY,367-GLY,368-ASP,369-LEU,425-ASN,426-MET,429-GLY,430-THR,431-GLY,432-GLN,455-THR,456-ARG,457-ASP,458-GLY,459-GLY,460-ALA,469-ARG | 4YDK | 51.73 |
| 22-VAL,22-ASN,22-PRO,22-LYS,22-GLU,22-TRP,22-LEU,22-SER,22-PHE,23-ARG,23-HIS,23-GLN | 4KI5 | 31.92 |
| 35-GLU,37-TYR,39-LYS,46-ASP,49-THR,53-ALA,54-LEU,56-LYS,58-THR,59-VAL,99-THR,101-LEU,102-THR | 5TZ2 | 47.96 |
| 1-MET,2-PHE,3-GLN,4-GLN,34-THR,66-GLU,67-GLY,68-GLU,70-GLU,71-GLN,72-LYS,75-GLU,76-HIS | 2JEL | 50.92 |
| 427-LEU,429-CYS,430-ASN,431-GLU,432-SER,433-LEU,436-GLY,438-LEU,439-ALA,441-LEU,442-PHE,443-TYR,446-LYS,529-TRP | 4MWF | 48.58 |
| 30-LYS,31-GLU,64-GLN,66-ARG,68-TYR,69-SER,70-GLU,72-GLY,79-ALA,205-LYS,253-ARG,274-TRP | 4IDJ | 48.8 |
| 169-SER,170-ALA,172-LEU,173-SER,174-THR,175-ASN,176-LYS,177-ALA,178-VAL,188-LEU,191-LYS,194-ASP,197-ASN,201-LYS,226-LYS,263-ASP | 5TPN | 47.88 |
| 465-ARG,466-TYR,467-HIS,468-ARG,469-SER,470-SER,471-LEU,480-HIS,481-PRO,482-ILE,484-GLU,502-PRO,504-PHE | 3V6O | 53.5 |
| 349-PRO,353-ARG,382-LEU,384-GLN,408-GLN,409-HIS,411-GLN,412-PHE,417-VAL,418-SER,438-ILE,440-SER,443-LYS,465-LYS,467-ILE,468-SER,469-ASN,471-GLY,473-ASN | 1YY9 | 44.72 |
| 27-SER,28-PRO,29-ASP,30-ARG,60-SER,61-GLU,128-LEU,129-ALA,130-PRO,131-LYS,132-ALA | 5GGR | 78.56 |
| 42-GLU,43-ASN,44-ILE,45-GLU,46-GLY,47-ASN,48-GLY,49-GLY,50-PRO,51-GLY,52-THR,53-ILE,70-ARG,72-ASP,76-HIS,87-GLU,97-LYS | 1FSK | 47.21 |
| 74-CYS,75-PRO,76-THR,77-MET,78-GLY,79-GLU,104-GLY,105-CYS,106-GLY,107-LEU,109-GLY,110-LYS | 3I50 | 54.01 |
| 31-SER,37-SER,38-GLY,40-TYR,117-SER,118-GLY,180-ASN,181-ASN,183-TYR,186-GLU,187-ILE,188-PHE,189-ASN,190-GLN,191-SER,194-TYR,234-GLN,235-GLU,238-ALA,239-ILE,240-THR,241-GLN,242-ASP | 4CAD | 41.37 |
| 87-ASN,88-GLU,90-LYS,91-GLN,92-VAL,102-ASP,105-ASN,106-PRO,107-VAL,108-LYS,109-THR,110-GLY,111-VAL,112-CYS,120-PRO,122-VAL,125-GLN,126-ASN | 6B0H | 54.2 |
| 483-SER | 3V6O | 20.3 |
| 12-ALA,13-GLU,17-ARG,18-GLN,20-ARG,156-ARG,157-THR,158-ILE,178-SER,179-ASN,180-ALA,181-GLN,182-GLY,185-TYR,198-ASP,199-ASN,203-TYR | 4PP2 | 53.22 |
| 1-GLY,2-GLN,30-LYS,34-GLY,40-TYR,45-SER,53-ASN,101-ARG,102-ASP,103-HIS,104-SER,105-TYR,106-GLN,107-GLU,108-GLU | 1PKQ | 59.2 |
| 64-ILE,65-LYS,66-GLU,68-LYS,69-CYS,197-ASN,201-LYS,202-GLN,204-LEU,206-ILE,208-ASN,209-LYS,210-GLN,212-CYS | 5UDC | 52.27 |
| 233-ILE,234-ASP,235-LYS,236-SER,238-GLY,239-ARG,240-PHE,241-HIS,242-VAL,266-LEU,270-LYS | 2R4R | 65.6 |
| 60-PHE,73-ILE,75-HIS,78-SER,100-ALA,101-GLY,119-ASN,120-TRP,121-PHE,122-ASP,123-ILE,124-THR,125-LYS,127-LEU,131-LYS | 6MTQ | 46.48 |
| 316-GLN,318-PHE,320-ASN,321-GLY,322-ALA,323-ILE,325-ASN,358-ASP,360-THR,362-ILE,364-LYS,369-LYS,370-ASP,371-GLU,372-LYS,373-GLN | 5MO9 | 48.85 |
| 427-LEU,429-CYS,430-ASN,431-ASP,432-SER,434-HIS,435-THR,436-GLY,437-PHE,438-LEU,439-ALA,440-ALA,442-PHE,443-TYR,445-HIS,446-LYS,447-PHE,448-ASN | 6MEH | 52.22 |
| 77-ASN,78-ARG,79-SER,80-LYS,81-GLY,82-THR,83-ALA,84-GLU,85-LYS,119-GLU,120-ASP,151-ASN,154-ARG,155-ARG,156-ILE,157-THR,158-SER,399-SER,400-ILE | 3VI3 | 57.12 |
| 19-TRP,20-TYR,43-VAL,44-GLU,45-GLU,47-LYS,57-LEU,59-GLN,65-GLU,66-CYS,67-ALA,68-GLN,70-LYS,125-THR,126-PRO,127-GLU,154-THR,156-LEU,157-GLU,158-GLU,162-ILE | 2R56 | 40.43 |
| 18-ARG,19-SER,21-ARG,156-GLY,157-ARG,158-THR,159-ILE,181-THR,182-GLN,186-TYR,199-ASP,202-TYR,204-TYR | 5VPL | 44.94 |
| 37-VAL,38-LEU,71-SER,72-TRP,74-GLY,75-ALA,77-ILE,78-ALA,81-PRO,82-THR,84-LYS,85-GLU,142-ALA,143-GLY,144-ALA,145-ASN,146-VAL,148-THR | 6HF1 | 54.84 |
| 105-VAL,106-PRO,107-THR,108-ALA,109-LYS,115-GLU,116-GLU,117-PHE,118-MET,119-ASN,120-PHE,121-ASP,122-LEU,123-LYS,124-GLN,129-GLY,130-ASP,131-TRP,132-PRO,133-GLU | 6FGB | 41.38 |
| 72-MET,93-ASP,96-PHE,99-PHE,101-MET,102-LEU,104-TYR,106-THR,129-TYR,132-LYS,154-GLU,156-VAL,158-LYS,159-ASP,162-ARG | 4LEO | 43.69 |
| 31-THR,32-LYS,34-ARG,35-ALA,38-LEU,39-ASP,41-GLN,42-LYS,65-PHE,66-ASP,69-VAL,70-GLY,73-ASP,76-LEU,77-LYS,80-ASN | 6CBV | 44.18 |

eRSA: Relative solvent accesibility (RSA) of the entire B cell epitope computed averaging the relevant residue RSAs
